# DIAlignR provides precise retention time alignment across distant runs in DIA and targeted proteomics

**DOI:** 10.1101/438309

**Authors:** Shubham Gupta, Sara Ahadi, Wenyu Zhou, Hannes Röst

## Abstract

SWATH-MS has been widely used for proteomics analysis given its high-throughput and reproducibility but ensuring consistent quantification of analytes across large-scale studies of heterogeneous samples such as human-plasma remains challenging. Heterogeneity in large-scale studies can be caused by large time intervals between data-acquisition, acquisition by different operators or instruments, intermittent repair or replacement of parts, such as the liquid chromatography column, all of which affect retention time (RT) reproducibility and successively performance of SWATH-MS data analysis. Here, we present a novel algorithm for retention time alignment of SWATH-MS data based on direct alignment of raw MS2 chromatograms using a hybrid dynamic programming approach. The algorithm does not impose a chronological order of elution and allows for alignment of elution-order swapped peaks. Furthermore, allowing RT-mapping in a certain window around coarse global fit makes it robust against noise. On a manually validated dataset, this strategy outperforms the current state-of-the-art approaches. In addition, on a real-world clinical data, our approach outperforms global alignment methods by mapping 98% of peaks compared to 67% cumulatively and DIAlignR can reduce alignment error up to 30-fold for extremely distant runs. The robustness of technical parameters used in this pairwise alignment strategy has also been demonstrated. The source code is released under the BSD license at https://github.com/Roestlab/DIAlignR.

**Abbreviations:**

AUC: Area Under the Curve
DIA: Data-independent acquisition
LC: Liquid chromatography
LOESS: Local weighted regression
RSE: Residual Standard Error
RT: Retention time
XIC: Extracted ion chromatograms

**Data Availability:** Raw chromatograms and features extracted by OpenSWATH are available on PeptideAtlas.

Servername: ftp.peptideatlas.org

Username: PASS01280

Password: KQ2592b

## Introduction

In translational research, protein biomarkers and therapeutic targets are usually discovered by data-driven methods such as by linking protein abundance patterns with disease conditions. A large sample cohort is essential in these studies as substantial biological variability exists in the population and enough statistical power is required to identify disease specific events^1,2^. Blood plasma is a good source of clinical information of a patient as it can be obtained noninvasively and proteins from affected tissue can potentially leak into the blood. Plasma samples, unfortunately, are highly challenging for proteomic analysis due to the diversity of peptides within the samples and high dynamic range of plasma proteins^3^. Therefore, quantification of plasma proteins requires a highly reproducible reduction of complexity and measurement within a wide dynamic range. The situation is exacerbated across large-scale studies which makes development of plasma biomarker challenging^2,3^.

In the past two decades, mass spectrometry (MS) based proteomics has made rapid advances and high degree of innovation in obtaining identification and quantification of proteins in various biological samples^2,4^. Targeted proteomics methods, specifically selected reaction monitoring (SRM), can provide high reproducibility across multiple runs. However, it is limited by low throughput and can measure abundance of only a few tens to low hundreds of proteins per study^1,5^.

Recently, we developed SWATH-MS, an approach for targeted analysis of data-independent acquisition (DIA) data, which can reproducibly quantify larger sets of peptides in large-scale clinical studies^5,6^. Implementing this method in the clinical field could provide comprehensive characterization of sample across various conditions. It has allowed to reproducibly quantify about 2000 proteins in a biomarker study on tumorous kidney and healthy tissues^1,7^ and has the potential to record a molecular inventory of samples comprising a large number of proteotypes, thus making longitudinal monitoring of a patient possible^1^.

In DIA mode, precursors in MS1 are selected for a predetermined m/z range and fragmented non-specifically. This produces multiplexed MS2 spectra of fragment-ions of all selected precursors. The DIA data can be analyzed by using either a library-based approach^5,8^ or a library-free approach^9^. Library-based approaches have shown to be capable of accurate peptide and protein quantification in complex samples^5,10,11^. Nonetheless, obtaining reproducible and robust analysis of clinical plasma samples has been challenging even with SWATH-MS, as large variations in number of proteins in individual runs were observed^5,10,11^. One of the major factors driving variability is the retention time deviation between assay library and plasma peptides’ elution profiles. In experiments carried out by Nigjeh and coworkers, most of the peptides had RT variation of about 10 minutes between technical replicates, affecting the robustness of peptide quantification^3^. This variation, if left uncorrected, may also result into incorrect and inconsistent identification of the peptides^3^.

Current DIA data analysis software use iRT peptides to calculate a monotonic retention time function (linear regression^12^ or segmented regression^13^) with respect to a library. Using this mapping, extracted ion chromatograms (XICs) from MS2 spectra are obtained for peak-picking. Software usually finds multiple potential peak-groups in XICs, which makes downstream analysis challenging. By establishing peak correspondence among runs, correct peptide elution time could be determined for each MS run^5,14^. A shift in retention time (RT) is often considered as a system-level variation which is modelled using monotonic functions between two runs^14^. However, this assumption may not always be accurate, and specifically among distant runs, singularities specific to a single peptide are common that produces relative peak switching where the elution order of two peptides is swapped across two runs^14–16^. This phenomenon is increasingly likely in larger studies and very probable in large-scale clinical studies in which data-acquisition happens over a span of years.

There are many methods in the literature for establishing correspondence in retention times. Current RT alignment algorithms in metabolomics and proteomics were mostly developed before the development of SWATH-MS^14^ and, therefore, rely on either MS1 chromatograms^17–23^, picked features^24–27^, or a combination of both ^28,29^. These algorithms usually find a global pairwise alignment function using dynamic programming on raw MS1 chromatograms^20–23,28^ or on feature-lists^24,25^ which include methods using band constraints. For complex samples, so-called “landmark peaks”^28,29^ have been used to improve RT alignment accuracy. However, most of these approaches rely on MS1 data and the resulting alignment functions are influenced by all constituent peptides. In SWATH-MS runs, MS2 data has high signal-to-noise ratio and is reproducible across multiple runs. Previous research on RT alignment of SWATH runs has relied on MS2 feature finding software. These either align MS2 features using bipartite matching^16^ or use the features to calculate a global function by local weighted regression (LOESS)^30^ or by a kernel density approach^5,31^ (see Supplemental Section S1). These approaches, however, provide suboptimal results in case of high noise, missing features or when feature detection algorithms malfunction. Furthermore, the global monotone functions do not account for peptide switching as a monotone function disallows retention time reversal between any two peptides^16^.

Here we present DIAlignR, a retention time alignment algorithm that addresses these shortcomings of previous methods. Our algorithm does not require features and is capable of directly aligning the raw multiplexed MS2 chromatographic traces from targeted proteomics data. Our approach uses dynamic programming to obtain an optimal mapping between chromatograms which contain local information as multiple, close-by peaks around the eluted peak-group. Independent RT alignment of each precursor facilitates the alignment of elution-order swapped peaks. Our method is also capable of using a global whole-run alignment for guidance, making it robust against noise. Thus, DIAlignR can flexibly handles user preference of selecting between extremes of global and local alignment.

We provide free-access to our source-code and our R-package at https://github.com/Roestlab/DIAlignR. We have tested our algorithm on a manually validated dataset of over 7000 chromatograms and demonstrate improved performance over existing methods. We have also tested our algorithm on 24 randomly selected blood plasma runs, selected from a heterogeneous cohort measured across many months. For both datasets, our algorithm outperforms global alignment methods and is capable of correcting mis-annotations introduced by feature detection algorithms. For very distant runs, it could also precisely align switched peaks which is not possible using global alignment methods^8,12^.

## Material and Methods

Alignment algorithms are useful for mapping signals, which are close to noise, over several runs. Few datasets exist that can be used for benchmarking as often the ground truth is unknown.

### Validation dataset

For benchmarking, we have used a previously published dataset^8^ of 16 SWATH runs from *Streptococcus pyogenes* bacterial strains. In these runs, 452 randomly selected precursors were manually annotated^5^. Out of these, eight precursors have annotation for less than two runs, and were thus removed. Seven other precursors from the remaining set have annotated peaks outside of the XICs, making them inapplicable and, hence, were removed from benchmarking (see the supplemental Section S2). Therefore, 437 annotated precursors are considered for performance testing of the developed DIAlignR tool against global alignment approaches. Annotated retention times of the precursors are available in the Supplemental Table 1. Since, annotated peaks were selected randomly from the validation dataset, this dataset has 4.9% peaks with signal-to-noise ratio less than 1 (Supplemental Figure 1*a*).

**Table I.**
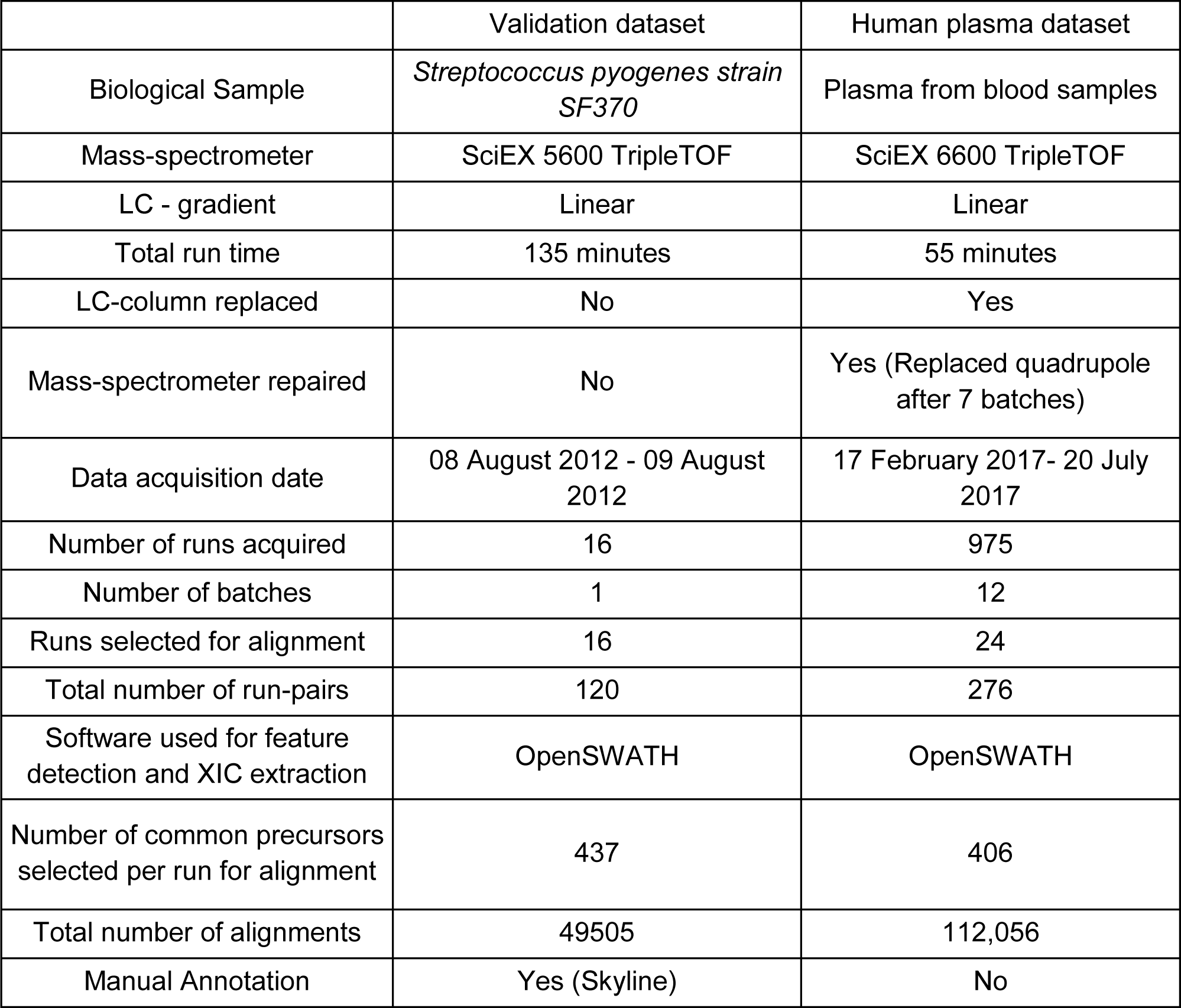
Summary description of validation and human plasma datasets

**FIG. 1.**
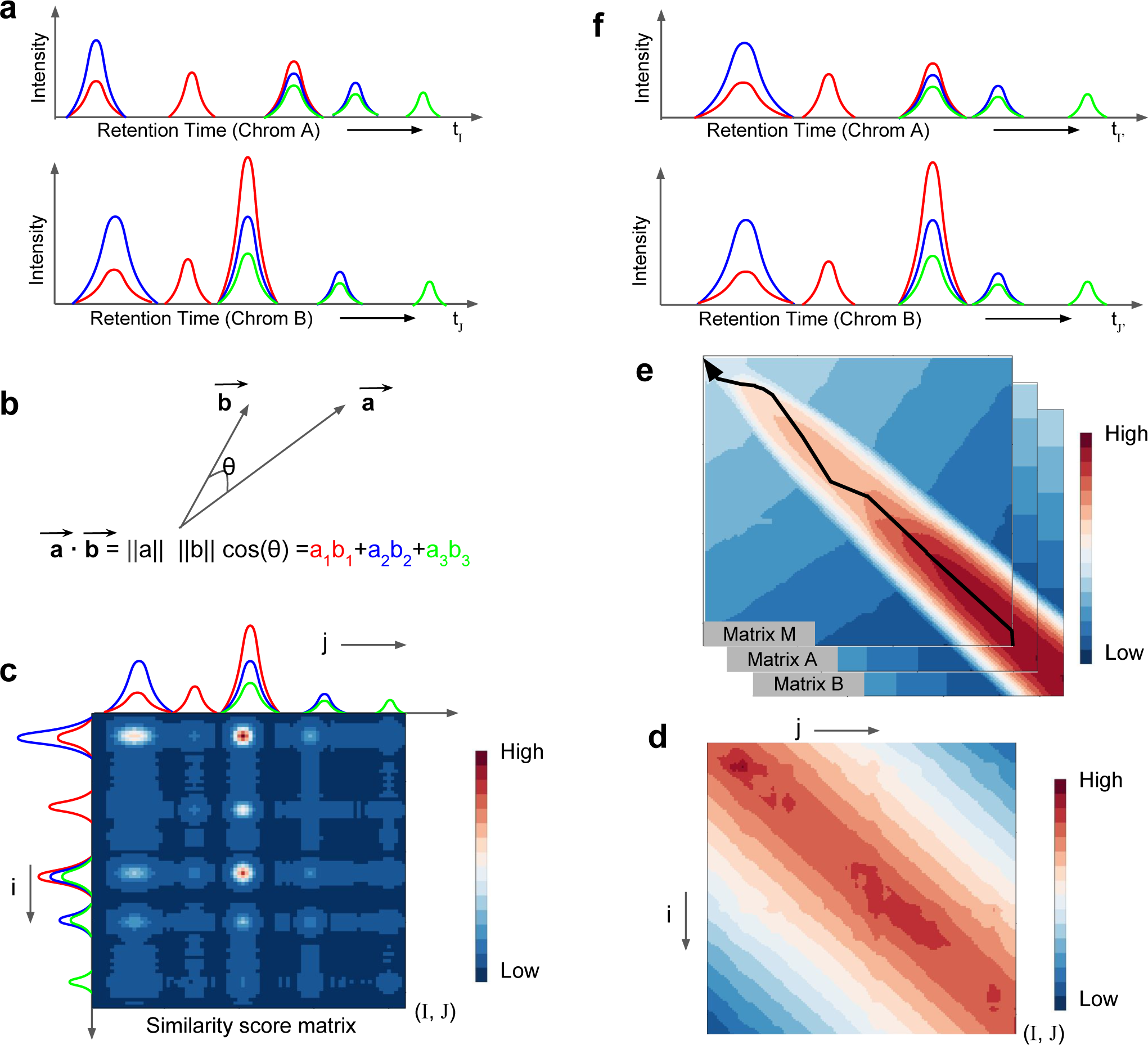
Alignment algorithm for targeted proteomics MS2 chromatograms. *a*, Fragment-ions chromatograms of a peptide for two runs; *run A* at top and *run B* at bottom. Correct peak, typically, has all library fragment-ions (*n* = 3) coeluting. *b*, similarity between chromatograms of both runs is calculated by dot-product of intensity vector; defined in *n* dimensional space. *c*, outer dot-product of chromatograms provides an *I* x *J* similarity score matrix (*S*). *d*, feature-based complete run alignment is used as an approximate path for alignment. Time points farther from an allowed window in similarity score matrix are penalized by adding negative score. *e*, Affine gap penalty based overlap alignment strategy is employed for calculating best scoring path through the similarity matrix. This dynamic programming based strategy utilizes three matrices for recursively calculating multiple gap length scores. Calculated alignment path is indicated using black arrow. *f*, Chromatograms recreated by mapping intensity back to aligned time path.

### Large-scale human plasma dataset

We have performed SWATH-MS on 975 human plasma samples in 12 batches from 17 February 2017 to 20 July 2017 (IRB 23602). Tryptic peptides of plasma samples were separated on a NanoLC 425 System (SCIEX). 5ul/min flow was used with trap-elute setting using a 0.5 x 10 mm ChromXP (SCIEX). LC gradient was set to a 43-minute gradient from 4-32% B with 1-hour total run. Mobile phase A was 100% water with 0.1% formic acid. Mobile phase B was 100% acetonitrile with 0.1% formic acid. 8ug load of undepleted plasma on 15cm ChromXP column. MS analysis was performed using SWATH Acquisition on a TripleTOF 6600 System equipped with a DuoSpray Source and 25μm I.D. electrode (SCIEX). Variable Q1 window SWATH Acquisition methods (100 windows) were built in high sensitivity MS/MS mode with Analyst TF Software 1.7.

To reduce the number of pairwise alignment, randomly two runs from each batch were selected; their metadata and OpenSWATH output files are described in the supplemental Table 2. Since peaks were not visually validated, only peaks with low FDR score were considered for performance evaluation. Therefore, peaks of target precursors with a q-value less than 10^-3^ (m-score < 1e-03, peak-group rank = 1) were selected and precursors were required to be present in all 24 runs. Successively, fragment-ion chromatograms of selected 406 precursors were extracted and parsed using OpenSWATH^8,32^ and “mzR” package^33^ with default parameters. The retention time of the peptides in all 24 runs is provided in the Supplemental Table 3. The chromatograms are available on PeptideAtlas (ftp.peptideatlas.org PASS01280:KQ2592b).

**Table II.**
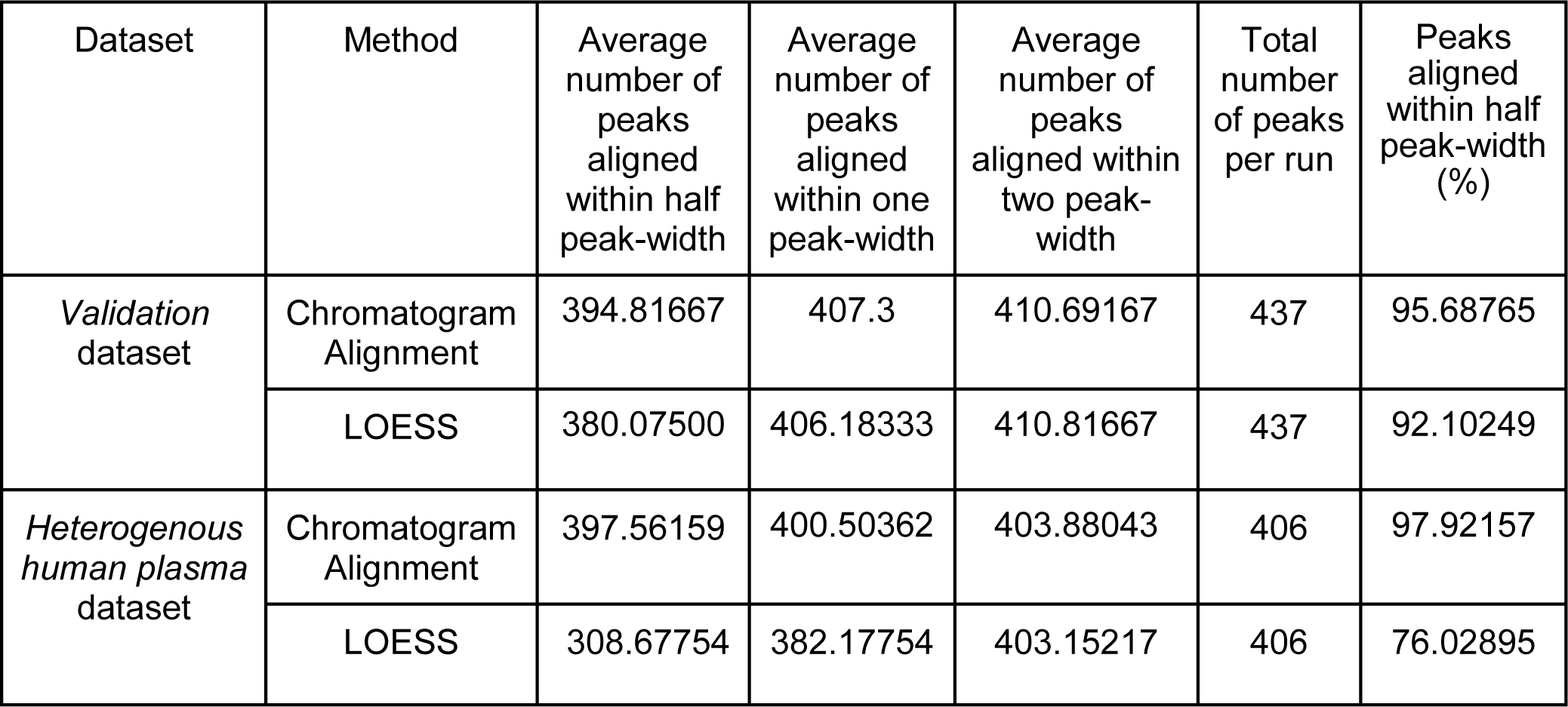
Below is presented average number of peptides aligned from manually validated *S. Pyogenes* dataset and from *heterogenous human plasma measurements*. For plasma data, peaks with OpenSWATH m-score < 0.001 are used for evaluation of the algorithm.

A tabular description of both datasets is provided as Table 1.

## Chromatogram Alignment Algorithm

In targeted proteomics or SWATH-MS experiments, each precursor is measured using one or more fragment ions (transitions). In general, we recommend for DIA / SWATH-MS analysis to use at least six fragment ions^4^. For each fragment-ion an extracted-ion chromatogram (XIC or chromatogram) is obtained. A collection of one or more chromatograms is called a “chromatogram group” which maps to the given precursor. If a precursor is measured using *n* transitions, then for each run the respective “chromatogram group” consists of *n* XICs, which is the raw data for our alignment procedure.

A chromatogram group can be considered a collection of time-series signals. The similarity of the time-series signals between chromatogram groups from runA (ChromA) and runB (ChromB) can be calculated. If a precursor has *n* fragment-ions, and each XIC has *I* and *J* time-points in *ChromA* and *ChromB*, respectively as shown in Fig. 1*a*, the similarity between all time-points can be represented as a similarity matrix *s* (Fig. 1*b* and *c*). Thus,

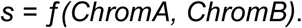

The function *ƒ* is termed as a similarity measure and can be selected by the user (see below).

### a. Similarity measure

In our R package DIAlignR, we have implemented several similarity measures which have been suggested in previous literature for chromatograms such as covariance, dot-product, Pearson’s correlation, spectral angle and euclidean distance^8,28^. We have observed that the dot-product between all *I* and *J* data-points provide information about both magnitude and angle between two time-points, hence segregating elution signal from the background. If each data point of chromatogram is represented by a vector in *n* dimensional space (*n* = 3 in Fig. 1*a*), the resulting dot-product of the two vectors will be as shown in Fig. 1*b*. Thus, with dot-product similarity, the matrix *s*from all vectors of both chromatogram-groups is defined as,

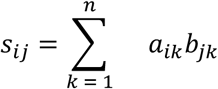

Where *i* ∈ {1,…, *I*}and *j* ∈ {1,…, *J*}represents index of vectors in *ChromA* and *ChromB*, respectively. A color-coded similarity matrix of size *I x J* is shown in Fig. 1*c*. However, to reduce the impact of noise peaks, a modified dot-product, termed as “Masked dot-product”, is used where higher similarity scores are checked again for spectral angle similarity (see the Supplemental Section S5). A path in the resulting similarity matrix is calculated using dynamic programming which directly translates to a retention time alignment that maps indices/time from *ChromA* to *ChromB* and vice-versa.

### b. Penalizing similarity matrix with global alignment

While dynamic programming will find a path of the highest cumulative score, in some instances the score is driven by alignment to noise and can lead to a solution where the alignment is highly divergent from a global linear or non-linear alignment. To make the alignment robust against noise and in order to incorporate information from the global context, we have added an option in our algorithm to modify the similarity matrix *s* (Fig. 1*d*) using feature-based global alignment such as LOESS. Residual standard error of the fit is utilized to define a region of non-interference in the similarity matrix and values outside of it punished with negative score (see the Supplemental Section S5). This allows us to find an alignment path within a reasonable time window relative to global prediction and avoid large deviations.

### c. Overlap Alignment with affine gap penalty

The optimal alignment path is found by recursively calculating all possible optimal paths from the start of the similarity matrix (1,1) to the end of it (I, J) using dynamic programming^34^. Chromatogram-groups *ChromA* and *ChromB* may not have end-to-end mapping as these could be partial chromatograms which were extracted around the expected peptide elution (as determined by iRT peptides for example). Therefore, overlap alignment instead of a global alignment of MS2 chromatogram groups is employed. This approach allows free end-gaps and thus, allows to slide chromatograms freely without incurring any gap-penalty for it.

To widen or shrink chromatogram peaks, a gap of unit length is a reasonable choice as it will distribute gaps along the complete peak. Therefore, an affine gap penalty scheme is utilized with higher gap penalty for gap length of more than one. In this approach, three matrices (Matrix M, A, B) are defined which recursively calculates score for gaps of more than unit length^34^. The overlap alignment path using affine gap-penalty is presented in Fig. 1*e*. The running time of such alignment is O(*max*(*I, J*)^3^). A heuristic data-driven approach is employed to obtain suitable affine gap penalties from the similarity matrix (see the Supplemental Section S5). Mapping the alignment path to the initial time values provides aligned chromatograms as depicted in Fig. 1*f*.

### d. Running time for alignment

Alignment of MS2 chromatograms of each peptide/precursor has running time of order O(max(I, J)^3); however, chromatograms of different precursors can be aligned independently. Therefore, we employ parallelization for different peptides to obtain much faster speed for complete run time-mapping.

### e. Optimization of algorithm parameters

There are various parameters used in DIAlignR. A description of these parameters is available in the Supplemental Section S5. We have employed validation dataset for parameter optimization and have used the number of peaks aligned within half chromatographic peak-width and cumulative RT alignment error as optimization target.

## Performance metrics for comparison with current algorithms

We have used the manually validated dataset^5^ to compare DIAlignR to the current state-of-the-art method (e.g. TRIC^5^) which utilizes a set of high confidence peaks (“anchor peptides”) to compute a linear or non-linear alignment function that transforms RT values from run1 to run2. We have chosen LOESS (local regression) as well as linear regression for evaluation. For LOESS, both optimized spanvalue from cross-validation (as used in TRIC) and default spanvalue (= 0.75) of the R software environment are tested^30^. For LOESS fit, ⅓ cross-validation is performed to obtain the optimum span value between two runs^5,30^. Steps to obtain a global fit (monotone mapping function) are detailed in the Supplemental Section S3.

Retention time error is calculated by comparing against the manual annotation of the *S. pyogenes* dataset^5^ and the resulting distribution of the number of peptides aligned within a certain RT tolerance is used as a measure of overall accuracy of the alignment algorithm. Manual annotations are not available for the human plasma dataset, therefore, the high-quality results (peaks with low FDR cutoff) of OpenSWATH are used for benchmarking.

## Results

### Parameter optimization

Here, we present an algorithm for multi-trace chromatographic alignment that only uses raw MS2 data from targeted proteomics or DIA experiments for retention time alignment. To optimize the performance of our algorithm, we have investigated the effect of algorithmic parameters on the accuracy of the alignment of runs from validation dataset^5^. First, we evaluate the performance for different similarity measures of chromatogram groups. The dot product masked with spectral angle (see the Supplemental Section S5) as a similarity measure provides the highest fraction of peptides aligned for all 120 possible run-pairs (Fig. 2*a*). Within an RT error tolerance of half peak-width (here: 15.3 sec), this similarity measure aligns 94.33% of annotated peaks with the highest area under the curve of all approaches (see the supplemental Table 7 and 8).

**FIG. 2.**
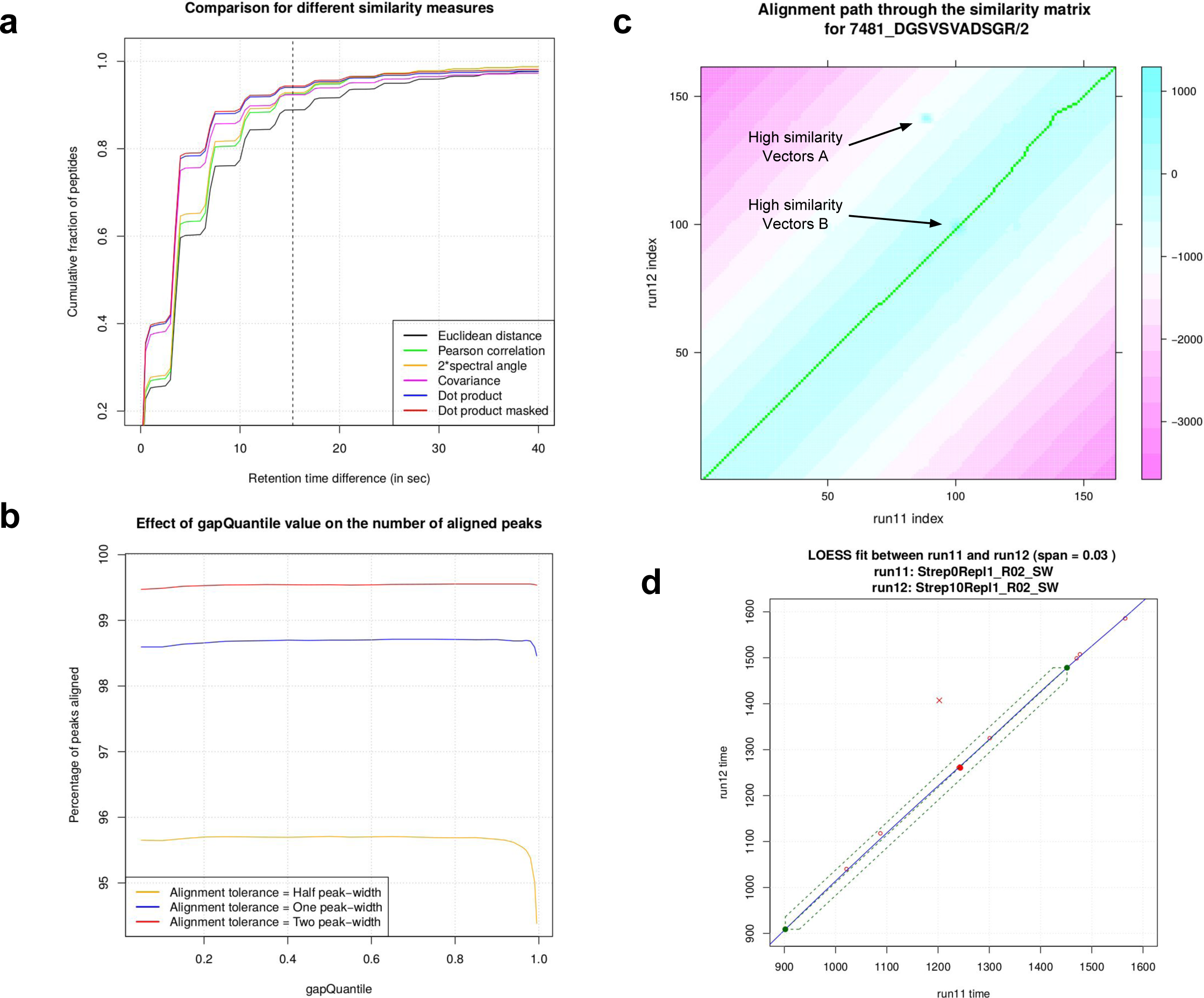
Comparison of different similarity measurements, technical parameters and effect of penalizing similarity using global prior on the accuracy of alignment in *S. Pyogenes* dataset. *a*, Performance of various similarity measures as the cumulative fraction of peptides having error less than RT difference is plotted. *b*, the effect of gap penalty selection using gapQuantile on the percentage of peaks aligned within certain RT difference tolerance is depicted. *c*, The penalized similarity matrix for peptide DGSVSVADSGR/2 between run11 and run12 is presented. From available two high similarity vectors, alignment path passes through high similarity vectors B. *d*, The end-points of extracted ion chromatograms (XICs) for the peptide are shown as green dots. Penalizing similarity gives preference to alignment within a certain window around LOESS fit, depicted as dashed green lines. Here, alignment of high similarity vectors B (solid red circle 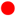) is preferred over high similarity vectors A (red cross 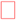).

We then investigated the effect of gap penalty used in dynamic programming. In DIAlignR, the gap penalty was calculated heuristically as a fixed quantile value of the distribution of similarity scores. We find that the selection of quantile value does not have a considerable impact on the percentage of peaks aligned within certain RT tolerance (Fig. 2*b*). From the figure, 20^th^ to 90^th^ quantile values yield approximate 95.6% of aligned peaks within half peak-width. The effect of gapQuantile is less pronounced for wider RT tolerance. For further analysis, the 65^th^ quantile is selected as the base gap penalty for chromatogram alignment. For the affine gap penalty, a gap opening factor of 0.125 was used, while a gap extension factor of 40 was considered (see the Supplemental Section S5).

We next investigated the impact of the number of transitions on the alignment(see the Supplemental Section S5 and supplemental Figure 10*a*). We find that as number of fragment-ions increases, alignment accuracy improves with best alignment 94.3% from using all library fragment-ions compared to 88.7% with only one transition (Supplemental Table 15). However, the largest improvement was going from 1 transition (88.7%) to two transitions (92.8%) with smaller improvements for using all transitions. This indicates single transition is not sufficient for this algorithm, but the relative gain is more modest after two transitions.

Our algorithm can constrain the similarity matrix using a global alignment function. Constraining the alignment in a certain window (given by RSEdistFactor) around the global fit derived from “anchor peptides” improves the alignment accuracy. We have observed that with a constrained similarity matrix 95.4% peaks get aligned compared to 94.3% with non-constrained one (see the supplemental Figure 10*b*). An example of such alignment is shown in Fig. 2*c* and 2*d*, in which the similarity matrix has two high similarity hot-spots. By constraining the alignment inside the dashed region in Fig. 2*d*, the alignment path goes through the correct hot spot. With the unconstrained similarity matrix, an incorrect alignment results as shown in the supplemental Figure 11.

### Validation using “gold standard” reference dataset

Using validation dataset, we have compared DIAlignR to current alignment methods. In terms of number of peptides aligned and alignment precision, chromatogram alignment outperforms LOESS and linear regression methods (see the Fig. 3*a* and Table I). On the *S. pyogenes* benchmark dataset, DIAlignR decreases error rates by 1.8-fold compared to the state-of-the-art methods. Cumulatively, chromatographic alignment only mis-aligns 4.3% of all peaks within 15.3 seconds (half peak width) of the true RT compared to 7.9% for LOESS (while LOESS with default parameters mis-aligns 22.8% of all peaks and linear regression mis-aligns 44.8%; see Fig. 3*a*).

**FIG. 3.**
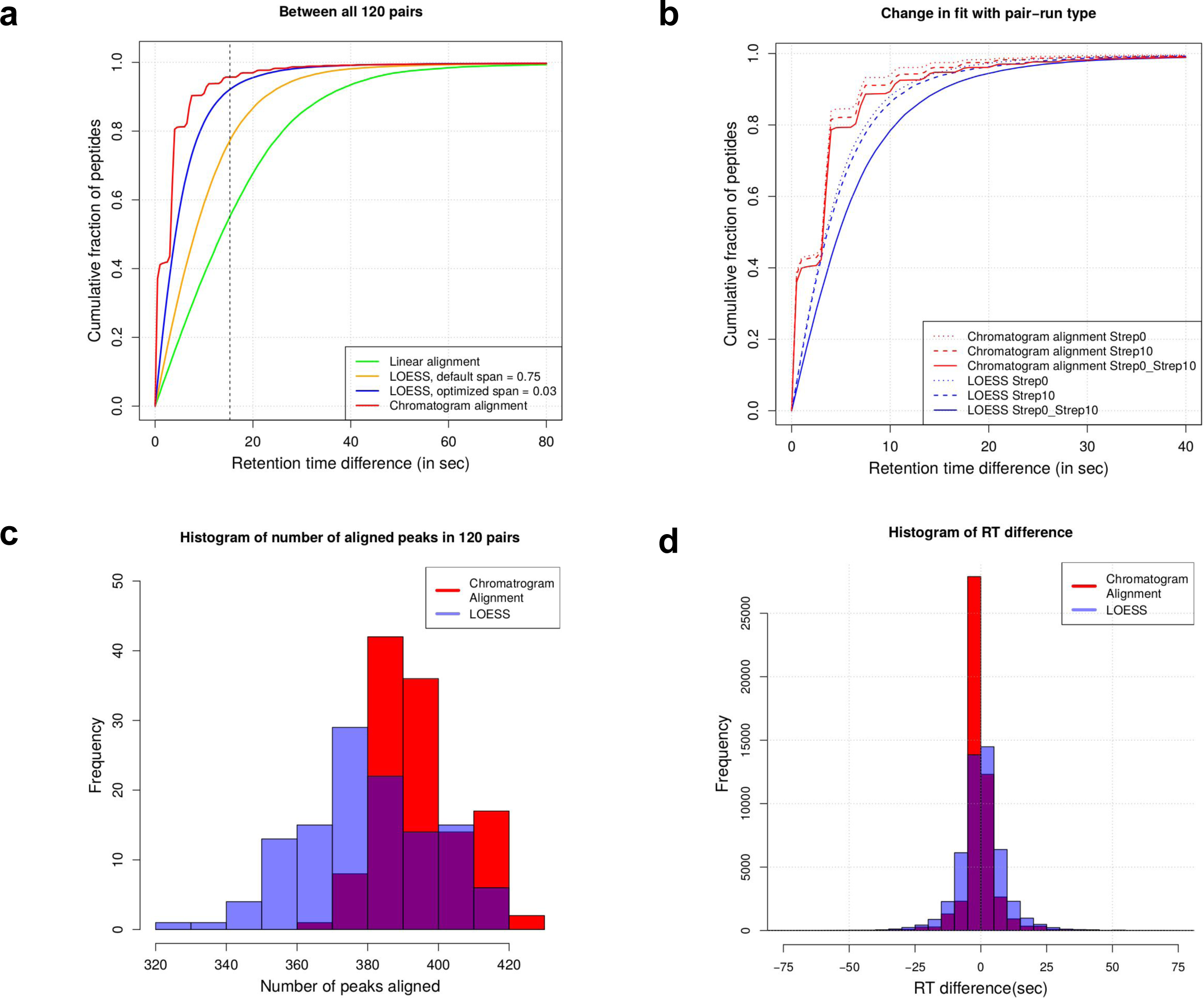
Alignment accuracy of MS2 chromatogram alignment on a validation dataset of 16 runs with manually annotated 437 peak groups in each run. *a*, cumulative fraction of peptides having error less than RT difference is plotted for all possible C(16,2) = 120 pairs for chromatogram alignment, linear fit and k-nearest neighbor smoothing (LOESS) with and without optimum span. *b*, cumulative fraction of peptides with alignment accuracy is plotted for chromatogram alignment and LOESS for pairs with different biological conditions. Strep0 pair constitutes both 0% plasma runs, Strep10 pair is composed of both 10% plasma runs and Strep0_Strep10 pair have one run with 0% plasma and other with 10% plasma. There are 28 Strep0 pairs, 28 Strep10 pairs and 64 pairs for Strep0_Strep10 case in the validation dataset. *c*, histogram of number of peptides matched within half peak-width for LOESS and chromatogram alignment. *d*, Histogram of retention time (RT) prediction error is plotted for chromatogram alignment and LOESS. RT difference standard deviation for both approaches is 9.56 sec and 10.98 sec, respectively.

We next investigated the effect of experimental perturbation on the performance of the alignment method. We compare within-condition alignments with between-condition alignments (in the validation dataset, the conditions are 0% and 10% human plasma added to *S. pyogenes* during growth). For both alignment methods, we observed decreased performance for between-condition alignments compared to within-condition alignments (Fig. 3*b*). However, the performance drop of the LOESS method (4.93%) is substantially larger than the corresponding performance drop of DIAlignR (2.7%), indicating increased robustness to sample heterogeneity for DIAlignR.

To evaluate the consistency of alignment approaches across multiple run-pairs, we have computed the number of correct aligned peaks (defined as instances where the alignment is correct within half peak-width) for each run-pair. This distribution is shifted towards the right with low standard deviation for chromatogram alignment method compared to LOESS, indicating that the former is consistent in its performance (Fig. 3*c*). In terms of the precision of the alignment, chromatogram alignment consistently performs better than global alignment methods as the former has higher area under the cumulative peptide frequency curve for each run-pair (higher AUC for 120 out of 120 pairs, see the supplemental Figure 12*c*). Similarly, we observe a larger RT variation (standard deviation = 18.45 sec) with the LOESS approach which chromatogram alignment corrects satisfactorily with the standard deviation being 11.68 sec (Fig. 3*d* and supplemental Fig. 12*a, b*). We conclude that on the validation dataset, DIAlignR performs consistently better in terms of accuracy of alignment and number of correctly aligned peaks across a range of different RT cutoffs and LC-MS/MS runs.

Next, we were interested in how the global differences between the two methods translate to individual alignments. We, therefore, computed the alignment error for each pairwise alignment of each peptide (total 49,505 alignments) and found that chromatographic alignment outperforms LOESS in 4.7% of all cases, whereas, LOESS achieves better results in 1.1% cases, while comparable performance was achieved in the remaining 94.2% cases (see the Supplemental Fig. 13*b*). On average, DIAlignR has reduced the RT error by 2.3 seconds with a median of 1.7 seconds (see the supplemental Fig. 13*c*). Overall, our method aligns 47.3k peaks correctly compared to 45.6k by an optimized LOESS within half peak-width (15.3 sec). However, in general we observed that on the validation dataset both methods perform with similar consistency which may be due to the low complexity of a bacterial sample and the high homogeneity of the data as it was acquired within two consecutive days on the same LC column.

### Application to large-scale heterogeneous human plasma measurements

After demonstrating consistently improved performance on the *S. pyogenes* validation dataset, we investigated the performance of our algorithm on a large-scale SWATH-MS experiments on human plasma. These experiments provided a more challenging dataset as the data was acquired over the period of six months with an intermittent repair of the instrument and replacement of the old column (column1) with a new column (column2). Two LC-MS/MS runs from each of the 12 batches were selected at random and 406 peptides were used for testing our algorithm. Since we did not have manually validate peaks, high confidence peak groups (q-value < 10^-3^) were used instead as validation set.

Comparing our chromatogram alignment algorithm (DIAlignR) with the LOESS method on a highly heterogeneous human plasma dataset, we found that our approach aligns 97.92% of peaks compared to 76.03% by LOESS with a maximal error of 20 seconds (half chromatographic peak-width) as depicted in Fig. 4*a*. All tested 276 pairwise alignments shown improved performance using chromatographic alignment (see Supplemental Fig. 15). Next, we were interested in the performance of our method on the alignment of runs acquired on the two different columns. We find that for runs acquired on different columns, chromatogram alignment method aligns 97.7% (compared to 97.84% within-column alignment) of peaks compared to 63.38% (89.06% for within-column alignment) by LOESS method (Fig. 4*b*), suggesting that DIAlignR retains performance even for highly heterogeneous datasets while the LOESS approach does not. Specifically, we find not only that DIAlignR outperforms LOESS on between-column alignments, but that the performance loss for DIAlignR is much less pronounced than for LOESS compared to within-column alignments. After validating the performance of chromatogram alignment cumulatively, we decided to investigate its consistency across individual run-pair alignments. Fig. 4*c* presents the distribution of the number of peaks aligned in all 276 pairs. DIAlignR is capable of aligning 400 peaks on average within half-peak width (while LOESS aligns 309 peaks on average), a 29% improvement. This indicates substantial alignment error when using LOESS, which could be reduced drastically by DIAlignR.

**FIG. 4.**
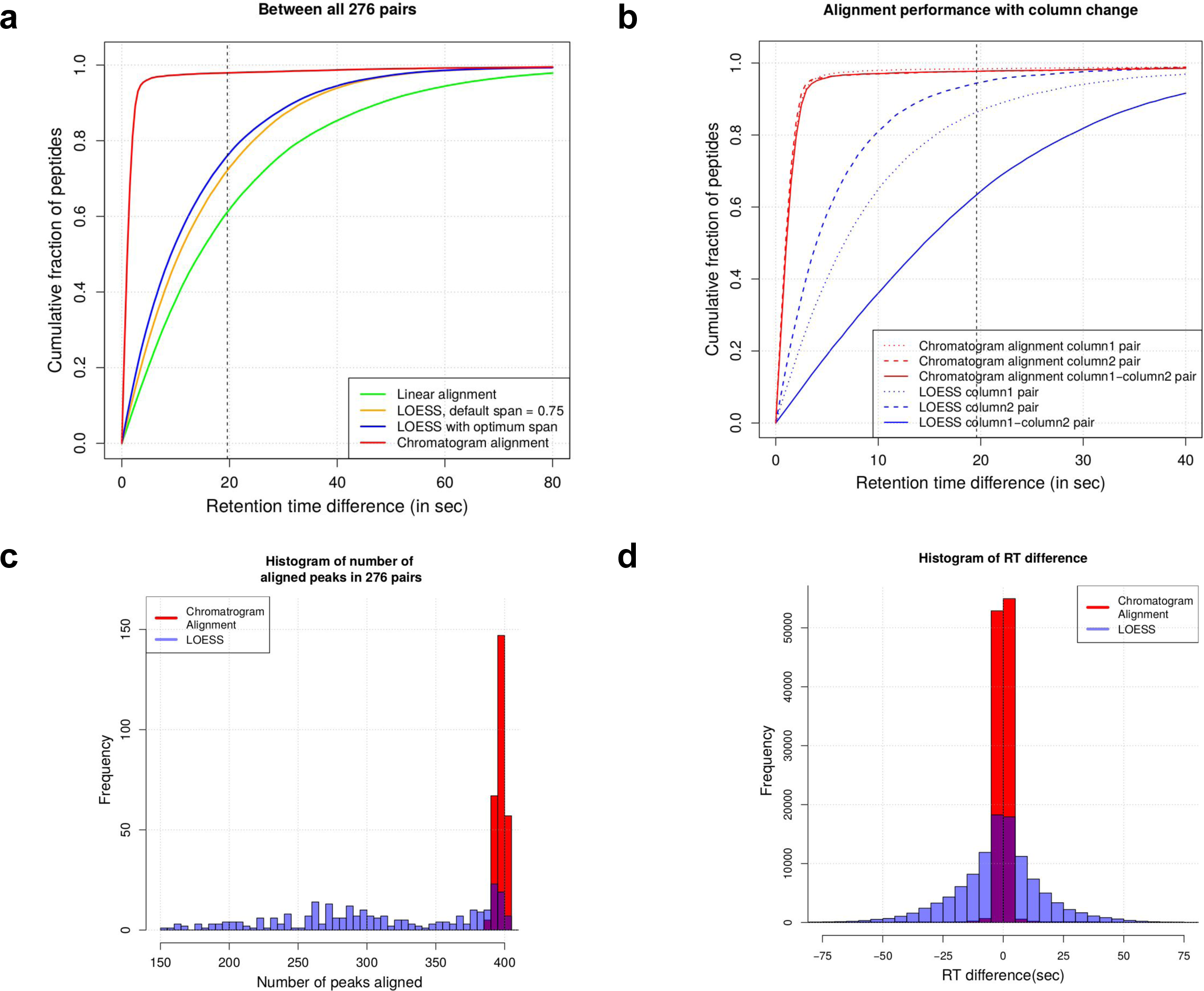
Alignment accuracy of MS2 chromatogram alignment on 24 runs of clinical plasma measurement dataset annotated with OpenSWATH. 406 peak groups are selected in each run with m-score < 0.001. *a*, cumulative fraction of peptides having error less than RT difference is plotted for all possible C(24,2) = 276 pairs for chromatogram alignment, linear fit and k-nearest neighbor smoothing (LOESS) with and without optimum span. *b*, cumulative fraction of peptides with alignment accuracy is plotted for chromatogram alignment and LOESS for pairs with different data acquisition conditions. LC column was changed together with quadrupole replacement. 14 runs were acquired on column1 which makes 91 pairs, labeled as “column1”. 10 runs were acquired after quadrupole replacement on column2 which results into 45 pairs, labelled as “column2”. There are 140 pairs composed of “column1” and “column2” labelled runs; these pairs are labelled as “column1-column2”. *c*, histogram of number of peptides matched within half peak-width for LOESS and chromatogram alignment. *d*, Histogram of retention time (RT) prediction error is plotted for chromatogram alignment and LOESS. RT difference standard deviation for both approaches is 22.91 sec and 13.7 sec, respectively.

To validate the performance on individual alignment, we further computed the alignment error for each pairwise alignment for every peptide. The standard deviation of alignment error for LOESS was 22.91 sec compared to 13.7 sec for DIAlignR (Fig. 4*d*). This indicates the higher precision of RT alignment with our approach. Out of 112,056 alignments, we found that DIAlignR outperforms LOESS in 23% of all cases and performs similarly in 76% cases (see the Supplemental Figure 14 - while performing worse in 1.13% cases). Upon manual validation, several of these worse-performing peaks were found to be due to wrong annotation by OpenSWATH (Supplemental Figure 18). Thus, testing of chromatogram alignment approach on the heterogeneous human plasma dataset again validates its consistent and improved RT alignment performance.

### Switching of peptide elution order

In liquid chromatography, retention time drift is often observed from one run to another run. However, we were interested whether this drift may be variable for different peptides and thus could result in reversal of retention order^15^. In such a scenario, two peptides which are eluting in order in one run may reverse their elution order in another run. Since our approach does not make an assumption of order preservation of peptide elution and facilitate independent alignment, we hypothesized that DIAlignR would be capable of uncovering instances of non-order preserving chromatographic alignment. Specifically, we analyzed peptide pairs that switch elution order from the heterogeneous and distant plasma runs.

To confirm the alignment for such peak-switching cases by chromatogram alignment algorithm, we have specifically looked at the alignment of the pair “run4_run23” as it has the highest number of peak switching pairs. run4 is part of batch V4, acquired on February 28^th^, 2017 whereas run23 is from batch M3, acquired on July 20^th^, 2017. The LOESS fitting from common high scoring training peptides for this pair is presented in Fig. 5*a*. Most of the test peptides are scattered around the global fit line, instead of being directly on the line. This graph quickly suggests 407 peptide pairs (one from either side of the line) comprising of 237 peptides that have switched their elution order out of 406 peptides (see the supplemental Section S6). We thus found that overall, 58.4% of peptides were involved in at least one event of non-order preserving elution.

**FIG. 5.**
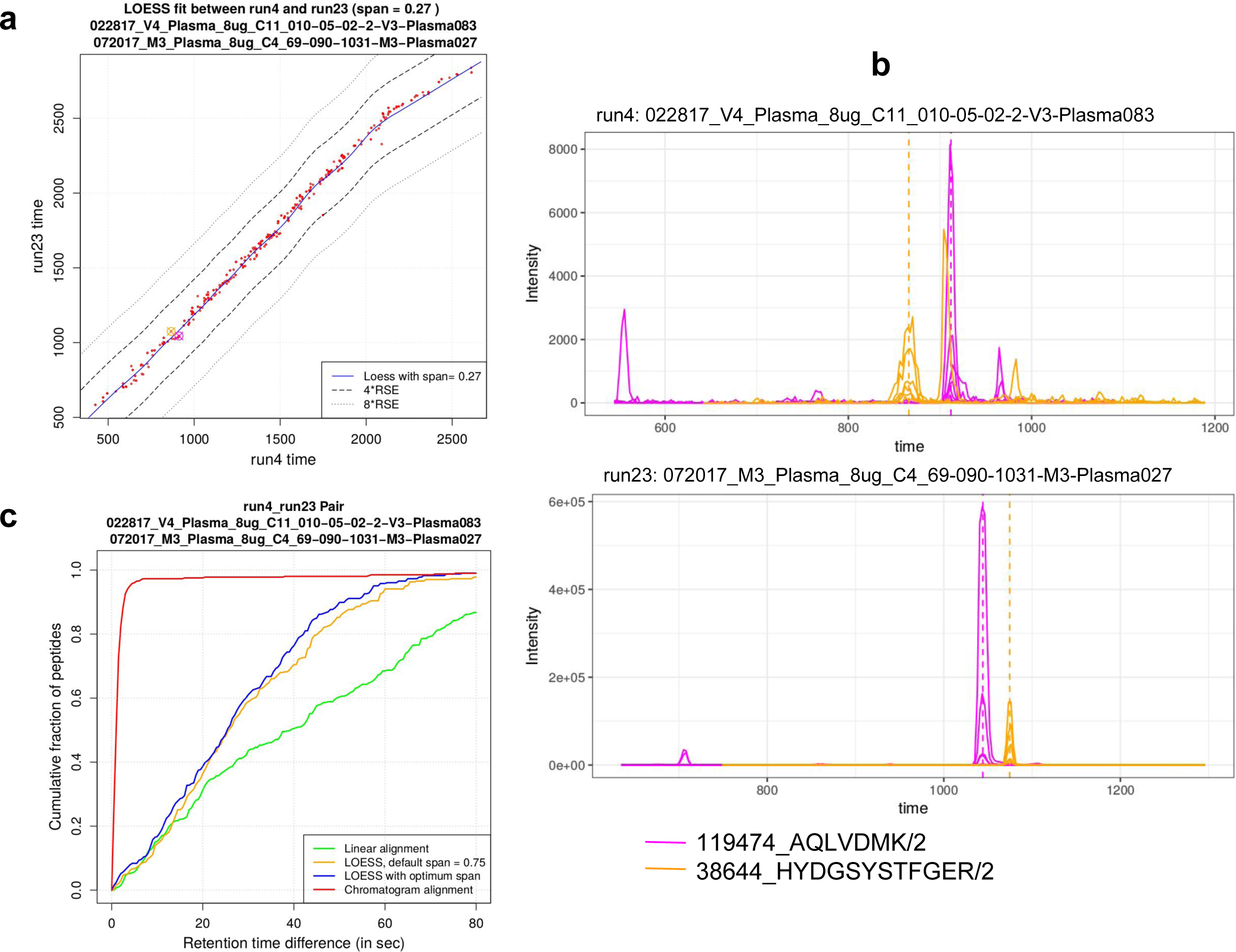
Alignment of 406 peptides in pair *run4 and run23* from clinical plasma measurement dataset. run4 “022817_V4_Plasma_8ug_C11_010-05-02-2-V3-Plasma083” was acquired on February 28^th^, 2017 whereas run23 “072017_M3_Plasma_8ug_C4_69-090-1031-M3-Plasma027” was acquired on July 20^th^, 2017. *a*, LOESS fit between two runs is obtained using confident peaks. Test peptides are shown in red color around the fit line. Span value = 0.27 for fit is obtained by ⅓ cross-validation. Precursors AQLVDMK/2 and HYDGSYSTFGER/2 are shown in magenta and orange circle-cross symbols, respectively. *b*, Two peptides AQLVDMK/2 and HYDGSYSTFGER/2 have their elution order reversed in these runs. This phenomenon makes alignment of peaks theoretically impossible for global monotonic methods. Chromatogram alignment uses fragment-ions as additional dimensions and hence can align them precisely. *c*, fraction of peptides having error less than RT difference is plotted for pair *run4 and run23* for chromatogram alignment, linear fit, k-nearest neighbor smoothing (LOESS) with and without optimum span and without any alignment.

One of the peak switching cases is presented in Fig 5*b*. In run4 peptide AQLVDMK/2 elutes after HYDGSYSTFGER/2, whereas in run23 the elution order has been reversed. Both peptides have seen positive RT drift in run23 from run4, however, HYDGSYSTFGER/2 shifts of 1070-850 = 270 seconds whereas AQLVDMK/2 drifts of only 1050-900 = 150 seconds. This varying RT drift between two runs has caused the peptides to elute in different order. The peptide-pair cannot be aligned with a global alignment approach, which in the best-case scenario will be off by 120 seconds -- however, our chromatogram alignment method has mapped the peaks correctly from run4 to run23 (see the supplemental Figure 17).

We, further, calculate the cumulative fraction of peptides aligned for pair “run4_run23” (Fig. 5*c*). Chromatogram alignment correctly aligns 98% peaks compared to LOESS which is able to align only 37.93%, thus DIAlignR is capable of decreasing the error by up to 30-fold. Eight peaks, which are not correctly aligned by chromatogram alignment, are further inspected visually by the authors and are found to be the cases of mis-annotation by OpenSWATH, mainly due to the post-translational modifications (see the supplemental Section S7 and supplemental Figure 18).

## Discussion

Correcting for retention time drift and aligning retention times between LC-MS/MS runs has been a long-standing problem in proteomics and it has become of particular importance as proteomics moves towards large-scale analysis of human cohorts. However, most efforts so far have focussed on MS1 data and few algorithms are available that can exploit the full information present in MS2 spectra produced by targeted methods or DIA / SWATH-MS.

In this paper, we have presented a novel algorithm that uses the raw fragment-ion chromatograms directly to perform retention time alignment for targeted proteomics and DIA data. Our algorithm uses XICs to map peaks across a pair of runs and improves accuracy compared to current state-of-the-art methods. We have, furthermore, extended the algorithm and implemented a hybrid approach, which uses a feature-based global alignment to condition the similarity matrix *s* that led to further gains in accuracy (see the Supplemental Fig. 10*b*). This hybrid approach provides the best of both worlds with a flexible “knob” which allows the user to either put more focus on global features or rely more on local information. To our knowledge, researchers have not yet explored dynamic programming-based alignment on raw fragment-ion-chromatograms. The dynamic programming approach is essential for obtaining a non-linear (or gapped) alignment as distant runs also have varying drift even for local peaks. Using LOESS to partially constrain the alignment makes our algorithm more stable and provides the robustness of global alignment methods.

We have shown that on a “gold-standard” validation dataset, DIAlignR consistently outperforms a global alignment method (using either linear or non-linear approaches), the current state-of-the-art (Supplemental Figure 12*c*). We have observed that alignment accuracy increases almost 4% if a precursor has two transitions instead of one (Supplemental Figure 10*a*), We get additional 1.5% aligned precursors by using all six fragment-ions which also corresponds to recommended guidelines^4^. Since, the DIAlignR approach puts more emphasis on local data, we also observe instances of over-fitting where the global alignment function produces better alignment (Supplemental Figure 13*b*). Overall, however, we see increased performance and the DIAlignR algorithm can decrease error rates from 7.9% to 4.3%. Interestingly, we find that our method is less sensitive to changes in chromatographic condition or sample matrix than global alignment approaches (Fig. 3*b*).

This finding led us to speculate that the novel chromatographic alignment would be less sensitive to heterogeneity in sample composition and chromatographic condition in large-scale studies. We have tested our algorithm on a large-scale SWATH-MS experiment of human plasma acquired over several months. On this heterogeneous dataset, DIAlignR reduces RT alignment error from 24% to 2%, which is a significant improvement over current state-of-the-art methods. Our approach has outperformed other methods and has consistently mapped the highest number of peaks within half-peak width irrespective of acquisition time interval, column change or instrument repair between two runs (Fig. 4*b*). DIAlignR improves retention time alignment accuracy which has the potential to improve peak-group identification and quantification through downstream tools. We have manually identified an example of wrongly aligned peak-group as shown in Supplemental Figure 19. We have also observed that in the case of a peak being outside of an extracted chromatogram, our method is able to map retention time outside of it as our hybrid approach also uses global alignment (see Supplemental Figure 18). Chromatograms can then be re-extracted and be used to correctly annotate peaks. Thus, this method can further be employed to extract chromatograms by OpenSWATH and other tools.

The improvement in retention time alignment comes with non-negligible computation cost. For an alignment of 10-min wide XICs with 3.4 sec cycle time, DIAlignR takes on average 0.16 seconds. Thus, pairwise alignment of selected 437 peptides takes approx 1 minute per run-pair. The highest cost in the alignment of each peptide is the calculation of the alignment path using dynamic programming. However, this problem scales linearly with the number of peptides in the library and can be easily parallelized on a computing cluster.

We believe that our approach is most useful for large-scale heterogeneous targeted proteomics studies where runs are acquired by different personnel and data is collected over several months or even years. Applying a single mapping function in such experiments becomes a very challenging task considering the switching of elution order of peptides. Global alignment functions, being monotonic in nature, assume chronological order of peptide elution and, therefore, cannot align switched peptides. Hence, we have observed substantial underfitting with the global function and an overall reduction of error using DIAlignR. Our hybrid approach aligns these peptides accurately as it mostly relies on additional dimensions of fragment-ion m/z to align peaks. It is possible that switching peptides may share many fragment-ions, however, this is very rare scenario if library is designed carefully^4^ and in such cases our method will perform no worse than global alignment methods.

Accurate RT alignment has multiple uses in the application of mass spectrometry-based proteomics for large-scale systems biology studies. Correct identification and improved quantitation of large number of analytes are few of them. This seems intuitive as most quantitative approaches currently available at least to some degree rely on accurate retention time alignment. We present a tool that can align retention times of DIA data by establishing correspondence between analytes across large number of samples, making DIA amenable for multi-center and longitudinal studies. We also expect that this tool can be utilized by existing proteomics software to streamline analyte identification and improve the quantification.

## Supporting information

Comparison to previous version

Supplemental Figures and Table titles

## Acknowledgements

We are grateful to Michael Snyder for supervising data-acquisition and providing access to the heterogenous plasma dataset. We also thank Michael Brudno for valuable discussion on chromatogram alignment using dynamic programming.

## Author contributions

S.G. designed and wrote code, performed data analysis and produced the figures. S.A. and W.Z. performed human plasma related experiments, acquired MS data. H.R. designed and supervised the study. S.G. and H.R. contributed to writing the manuscript. All authors have contributed to the final manuscript.

## Competing Financial Interest

The authors declare no competing financial interests.

## Data Availability

Raw chromatograms and features extracted by OpenSWATH are available on PeptideAtlas under accession code PASS01280.

## Notes

#### Summary of Updates

Main manuscript and Supplemental text are updated. Effect of the number of transitions on RT-alignment is included. Titles of Supplementary Figures and Tables are added. Supplementary Figure 1a and Supplementary Figure 10a are added. Figure 1c and Supplementary Figure 10b are corrected. Table 1, Supplemental Table 4 and Supplemental Table 15 are added. R-Package name is changed to DIAlignR.

