## Supplementary material for "DIAlignR provides precise retention time alignment across distant runs in DIA and targeted proteomics": Comparison to previous version

**Title of Paper**

**Abbreviations:**

AUC Area Under the Curve

DIA Data-independent acquisition

LC Liquid chromatography

LOESS Local weighted regression

RSE Residual Standard Error

RT Retention time

XIC Extracted ion chromatograms

**Running Title:**

~~DIAlign~~DIAlignR: Mapping retention time using MS2 chromatograms

**Data Availability:**

Raw chromatograms and features extracted by OpenSWATH are available on PeptideAtlas.

Servername: ftp.peptideatlas.org

Username: PASS01280

Password: KQ2592b

**Abstract**

SWATH-MS has been widely used for proteomics analysis given its high-throughput and ~~quantitative~~ reproducibility~~,~~ but ensuring consistent quantification of analytes across large-scale studies of heterogeneous samples such as human~~-~~-plasma remains challenging. Heterogeneity in large-scale studies can be ~~due to~~caused by large time intervals between data-acquisition, acquisition by different operators or instruments~~ or~~, intermittent repair or replacement of parts, such as the liquid chromatography column, all of which affect retention time (RT) reproducibility and successively ~~quantitative~~ performance of SWATH-MS~~.~~ data analysis. Here, we present a novel algorithm for retention time alignment of SWATH-MS data based on direct alignment of raw MS2 chromatograms using a hybrid dynamic programming approach. The algorithm does not impose a chronological order of elution and allows for ~~aligning~~alignment of elution-order swapped peaks. Furthermore, allowing RT~~-~~-mapping in a certain window around coarse global fit makes it robust against noise. On a manually validated dataset, this strategy outperforms the current state-of-the-art approaches. In addition, on a real-world clinical data, our approach outperforms global alignment methods by mapping 98% of peaks compared to 67% cumulatively~~, is capable of reducing~~ and DIAlignR can reduce alignment error up to 30-fold for extremely distant runs. The robustness of technical parameters used in this pairwise alignment strategy has also been demonstrated. The source code is released under the BSD license at https://github.com/Roestlab/~~DIAlign~~DIAlignR.

**Introduction:**

In translational research, protein biomarkers and therapeutic targets are usually discovered by data-driven methods such as by linking protein abundance patterns with disease conditions. A large sample cohort is essential in these studies as ~~huge~~substantial biological variability exists in the population and enough statistical power is required to identify disease specific events~~ (Uzozie and Aebersold 2018; Surinova et al. 2011)~~[1,2]. ~~Plasma~~Blood plasma is a good source of clinical information of a patient as it is noninvasivecan be obtained noninvasively and proteins from affected tissue can potentially leak into the blood. Plasma samples, unfortunately, are highly challenging for proteomic analysis due to the diversity of peptides within the samples and high dynamic range of plasma proteins(Nigjeh et al. 2017)[3]. Therefore, quantification of plasma proteins requires a highly reproducible reduction of complexity and measurement within a wide dynamic range. The situation exacerbatesis exacerbated across large-scale studies and makewhich makes development of plasma biomarker challenging(Nigjeh et al. 2017; Surinova et al. 2011)[2,3].

In the past two decades, mass spectrometry (MS) based proteomics has made rapid advances to obtain near-exhaustiveand high degree of innovation in obtaining identification and quantification of proteins in various biological samples(Surinova et al. 2011; Schubert et al. 2017)[2,4]. Targeted proteomics methods, specifically selected reaction monitoring (SRM), can provide reproducible protein quantificationhigh reproducibility across multiple runs. However, they areit is limited by low throughput and can measure abundance of only a few tens to low hundreds of proteins per study(Röst et al. 2016; Uzozie and Aebersold 2018)[1,5].

Recently, we developed SWATH-MS, an approach for targeted analysis of data-independent acquisition (DIA) data, which has potential tocan reproducibly quantify largelarger sets of peptides in large-scale clinical studystudies(Röst et al. 2016; Gillet et al. 2012)[5,6]. Implementing this method in the clinical field could provide comprehensive characterization of sample across various clinical conditions. It also facilitates creatinghas allowed to reproducibly quantify about 2000 proteins in a digital inventory of a tissue proteomebiomarker study on tumorous kidney and healthy tissues(Uzozie and Aebersold 2018; Guo et al. 2015)[1,7] and opens uphas the potential for clinics to record a molecular inventory of samples through whichcomprising a large number of proteotypes, thus making longitudinal monitoring of a patient is possible(Uzozie and Aebersold 2018)[1].

In DIA mode, after MS1 precursors in MS1 are selected for a predetermined m/z range and fragmented non-specifically. This produces a multiplexed MS2 spectra of fragment-ions of all selected precursors. The DIA data can then be analyzed by using either using a library-based approach (Röst et al. 2014; Röst et al. 2016)[5,8] or a library-free approach (Tsou et al. 2015)[9]. Library-based approaches have shown to be capable of accurate peptide and protein quantification in complex samples (Navarro et al. 2016; Röst et al. 2016; Röst et al. 2014; Liu et al. 2015).[5,10,11]. Nonetheless, obtaining reproducible protein quantification fromand robust analysis of clinical plasma samplesamples has been challenging even with SWATH-MS, as large variations in number of proteins in individual runs were observed (Nigjeh et al. 2017, (Röst et al. 2016; Liu et al. 2015; Navarro et al. 2016).[5,10,11]. One of the major factors driving variability is the retention time deviation between the assay library and plasma peptides' DIA elution profiles. In experiments carried out by Nigjeh and coworkers, most of the peptides had RT variation of about 10 minutes between technical replicates, affecting the robustness of peptide quantification (Nigjeh et al. 2017)[3] . RT variations. This variation, if left uncorrected, may also produceresult into incorrect and inconsistent identification of the peptides(Nigjeh et al. 2017)[3].

Current DIA data analysis softwares are capable of finding multiple peak-groups in MS2 software use iRT peptides to calculate a monotonic retention time function (linear regression[12] or segmented regression[13]) with respect to a library. Using this mapping, extracted ion chromatograms (XICs), however it is often) from MS2 spectra are obtained for peak-picking. Software usually finds multiple potential peak-groups in XICs, which makes downstream analysis challenging to efficiently integrate this information across multiple runs.. By establishing peak correspondence among runs, correct peptide elution time could be determined infor each MS run (Smith et al. 2015; Röst et al. 2016)[5,14]. A shift in retention time (RT) is often considered as a system-level variation which is then modelled using monotonic functions between two runs (Smith et al. 2015)[14]. However, this assumption may not always be accurate, and specifically among distant runs, singularities specific to a single peptide are also common that produces relative ~~peptide~~peak switching ~~(peak switching)~~ where the elution order of two peptides is swapped across two runs ~~(Smith et al. 2015; Spicer et al. 2010; Wu et al. 2016)~~[14–16]~~]~~. This phenomenon is increasingly likely in larger studies and very probable in large-scale clinical studies in which data ~~-~~acquisition happens over a span of years.

There are many methods in the literature for establishing correspondence in retention times. Current RT alignment ~~-~~algorithms in metabolomics and proteomics were mostly developed before the ~~arrival~~development of SWATH-MS ~~technique (Smith et al. 2015)~~[14] and, therefore ~~mostly~~, rely on either MS1 chromatograms ~~or~~ [17–23], picked features ~~or~~ [24–27], or a combination of both ~~(Listgarten et al. 2005; Prince and Marcotte 2006; Sandin et al. 2013).~~[28,29]. These algorithms usually find a global pairwise alignment function using dynamic programming on raw MS1 chromatograms[20–23,28] or on feature-lists[24,25] which include methods using band constraints. For complex samples, so-called “landmark peaks”[28,29] have been used to improve RT alignment accuracy. However, most of these approaches rely on MS1 data and the resulting alignment functions are influenced by all constituent peptides. In SWATH-MS runs, MS2 data has high signal-to-noise ratio and is reproducible across multiple runs. Previous research on RT alignment ~~in DIA data~~of SWATH runs has ~~also~~ relied on MS2 feature finding~~, using~~ software. These either ~~a~~align MS2 features using bipartite matching ~~approach (Wu et al. 2016)~~[16] ~~to match MS2~~or use the features ~~or using a global function calculated either by~~to calculate a global function by local weighted regression (LOESS~~)~~ )~~(Chambers et al. 1992)~~[30] or by a kernel density approach ~~to match features between two runs (Röst et al. 2016; Searle et al. 2018)~~[5,31]~~.~~ (see Supplemental Section S1). These approaches, however, provide suboptimal results in case of high noise, missing features or when feature detection algorithms malfunction. Furthermore, the global monotone functions do not account for peptide switching as a monotone function disallows retention time reversal between any two peptides ~~(Wu et al. 2016)~~[16].

Here~~,~~ we present ~~an~~DIAlignR, a retention time alignment algorithm ~~which~~that addresses these shortcomings of previous methods. Our algorithm does not require features and is capable of directly aligning the raw multiplexed MS2 chromatographic traces from targeted ~~proteomic~~proteomics data. Our approach uses dynamic programming to obtain an optimal mapping between ~~two~~ chromatograms which contain local information ~~in the form of~~ as multiple, close-by peaks around the ~~elution~~eluted peak-group. Independent RT alignment of each precursor facilitates the alignment ~~for~~of elution-order swapped peaks. Our method is also capable of using a global whole-run alignment for guidance, making it robust against noise. ~~With this flexibility, our DIAlign tool provides a knob to select~~Thus, DIAlignR can flexibly handles user preference of selecting between extremes of global and local alignment ~~fit as user decides~~.

We provide free-access to our source-code and our R-package ~~can be downloaded from CRAN.~~at https://github.com/Roestlab/DIAlignR. We have tested our algorithm on a manually validated dataset ~~which shows~~of over 7000 chromatograms and demonstrate improved performance over existing methods. We have also ~~have~~ tested our algorithm on 24 randomly selected ~~distant~~blood plasma runs~~. We observed that~~, selected from a heterogeneous cohort measured across many months. For both datasets, our algorithm outperforms global alignment methods and is capable of correcting mis-annotations introduced by feature detection algorithms. For very distant runs, it could also precisely align switched peaks ~~precisely~~ which is not possible using global alignment methods ~~(Escher et al. 2012; Röst et al. 2014)~~[8,12].

**Material and Methods**:

Alignment algorithms are useful for mapping signals, which are close to noise, over several runs. Few datasets exist that can be used for benchmarking as often the ground truth is unknown.

Validation dataset

For benchmarking ~~of the developed algorithm,~~, we have used a previously published ~~and manually validated~~ dataset ~~of~~ [8] of 16 SWATH runs from *Streptococcus* ~~*Pyogenes*~~*pyogenes* bacterial ~~strain is used (Röst et al. 2016). Out of 452 transition IDs from (Röst et al. 2016), eight IDs has~~ strains. In these runs, 452 randomly selected precursors were manually annotated ~~peak~~[5]. Out of these, eight precursors have annotation for less than two runs ~~out of 16 runs, making them unsuitable for benchmarking.~~, and were thus removed. Seven ~~transition IDs~~other precursors from the remaining set ~~had extracted fragment-ion chromatograms (XICs)~~have annotated peaks outside of ~~annotated peak~~the XICs, making them inapplicable and, hence, were removed from benchmarking (see the supplemental Section ~~S1). Total~~S2). Therefore, 437 ~~transition IDs~~ annotated ~~in 16 runs were~~ precursors are considered for ~~testing the~~ performance testing of the developed ~~DIAlign~~DIAlignR tool against global alignment approaches. ~~Retention time of these transition IDs in all 16 runs and run names~~Annotated retention times of the precursors are available in the ~~supplemental~~Supplemental Table 1 ~~and supplemental Section S2, respectively. For global alignment, LOESS fit with optimum span value is obtained between two runs (Chambers et al. 1992; Röst et al. 2016). ⅓ cross-~~. Since, annotated peaks were selected randomly from the validation ~~is performed~~ dataset, this dataset has 4.9% peaks with signal-to ~~obtain the optimum span value. Steps to obtain a global fit (monotone mapping function) are detailed in the supplemental Section S3.~~-noise ratio less than 1 (Supplemental Figure 1*a*).

### Large-scale human plasma dataset

We have performed SWATH-MS on 975 human plasma samples in 12 batches from 17 February 2017 to 20 July 2017 (-IRB 23602). Tryptic peptides of plasma samples were separated on a NanoLC™ 425 System (SCIEX). 5ul/min flow was used with trap-elute setting using a 0.5 x 10 mm ChromXP™ (SCIEX). LC gradient was set to a 43-minute gradient from 4-32% B with 1-hour total run. Mobile phase A was 100% water with 0.1% formic acid. Mobile phase B was 100% acetonitrile with 0.1% formic acid. 8ug load of undepleted plasma on 15cm ChromXP column. MS analysis ~~were~~was performed using SWATH® Acquisition on a

TripleTOF~~®~~ 6600 System equipped with a DuoSpray~~™~~ Source and 25µm I.D. electrode (SCIEX). Variable Q1 window SWATH Acquisition methods (100 windows) were built in high sensitivity MS/MS mode with Analyst~~®~~ TF Software 1.7.

To reduce the number of pairwise alignment, randomly two runs from each batch ~~are~~were selected; their metadata and OpenSWATH output files are described in the supplemental Table ~~3 and SelectedOSW.tar.gz, respectively. In the absence of manually annotated~~2. Since peaks~~,~~ were not visually validated, only peaks with low FDR score ~~and highest peak-group rank are~~were considered for performance evaluation. Therefore, ~~the best peak~~peaks of target precursors with a q-value less than $10^{-3}$ (m-score < 1e-03, peak-group rank = 1) ~~and common~~were selected and precursors were required to be present in all 24 runs~~ are selected, and successively, their~~. Successively, fragment-ion chromatograms ~~are~~of selected 406 precursors were extracted and parsed using OpenSWATH[8,32] and “mzR” package[33] with default parameters~~ (Rosenberger et al. 2017; Röst et al. 2014). Fragment ion chromatograms were parsed from OpenSWATH output using “mzR” package (Chambers et al. 2012).~~. The retention time of ~~these 406~~the peptides in all 24 runs is provided in the ~~supplemental~~Supplemental Table ~~4~~3. The chromatograms are available ~~in the file SelectedChroms.tar.gz. The global alignment function was fit~~on PeptideAtlas (ftp.peptideatlas.org PASS01280:KQ2592b).

A tabular description of both datasets is provided as ~~described above (see the Supplemental Section S4).~~Table 1.

### Chromatogram Alignment Algorithm

In targeted proteomics or SWATH-MS experiments, each precursor is measured using one or more fragment ions (transitions~~) which are measured using~~ ). In general, we recommend for DIA / SWATH-MS analysis to use at least six fragment ions[4]. For each fragment-ion an extracted~~ fragment~~-ion chromatogram (XIC or chromatogram~~).~~) is obtained. A collection of one or more ~~chromatogram~~chromatograms is called a "chromatogram group" which ~~all map~~maps to the ~~same~~given precursor~~ ion~~. If ~~the same~~a precursor is measured ~~across multiple runs,~~using $n$ transitions, then for each run ~~produces a~~the respective "chromatogram group" ~~for that precursor and this constitutes~~consists of $n$ XICs, which is the raw data for our alignment procedure.

A chromatogram group can be considered a collection of time-series signals. The similarity of the time-series signals between chromatogram groups from runA (ChromA) and runB (ChromB) can be calculated. If a precursor has $n$ fragment-ions, ~~therefore, $n$ XICs;~~ ~~where~~and each XIC has $I$ and $J$ time-points in *ChromA* and *ChromB*, respectively as shown in Fig. 1*a*, the similarity between all time-points ~~is~~can be represented as a similarity matrix $s$~~,~~ ~~where~~ (Fig. 1*b* and *c*). Thus,

$$s = f(ChromA, ChromB).$$

The function $f$ is termed as a similarity measure and can be selected by the user (see below).

**a. Similarity measure:**

In our R package ~~DIAlign~~DIAlignR, we have implemented several similarity measures which have been suggested in previous literature for chromatograms such as covariance, dot-product, Pearson's correlation, spectral angle and euclidean distance ~~(Röst et al. 2014; Prince and Marcotte 2006)~~[8,28]. We have observed that the dot-product between all $I$ and $J$ data~~-~~-points ~~provides~~provide information about both magnitude and angle between two ~~data-vector~~time-points, hence segregating elution signal from the background. If each data point of chromatogram is represented by a vector in $n$ dimensional space ($n$ = 3 in Fig. 1*a*), the resulting dot-product of the two vectors ~~is~~will be as shown in Fig. 1*b*. Thus, ~~in the case of the~~with dot-product~~, a~~ similarity, the matrix $s$ from all vectors of both chromatogram-groups is defined as,

$$s_{ij} = \sum_{k=1}^{n} a_{ik} b_{jk}$$

Where $i \in \{1, \ldots, I\}$ and $j \in \{1, \ldots, J\}$ represents index of vectors in *ChromA* and *ChromB*, respectively. A color-coded similarity matrix of size *I x J* is shown in Fig. 1*c*. However, to reduce the impact of noise peaks, a modified dot-product, termed as “Masked dot-product”, is used where higher similarity scores are checked again for spectral angle similarity (see the Supplemental Section S5). A path in the resulting similarity matrix is calculated using dynamic programming which directly translates to a retention time alignment that maps indices/time from *ChromA* to *ChromB* and vice-versa.

**b. Penalizing similarity matrix with global alignment:**

While dynamic programming will find a path ~~which results into~~of the highest cumulative score, in some instances the score is driven by alignment to noise and can lead to a solution where the alignment is highly divergent from a global linear or non-linear alignment. To make the alignment robust against noise and in order to incorporate information from ~~a~~the global context, we have added an option in our algorithm to modify the ~~the~~ similarity matrix *s* (Fig. 1*d*) using feature-based global alignment ~~(~~such as LOESS~~).~~. Residual ~~Sum~~standard error of ~~Error (RSE) of~~the fit is utilized to define a region of non-interference in the similarity matrix and values outside of it punished with negative score (see the Supplemental Section S5). This allows us to find an alignment path within a reasonable time window relative to global prediction and avoid large deviations~~ as shown in Fig. 1*d*~~.

**c. Overlap Alignment with affine gap penalty:**

The optimal alignment path is found by recursively calculating all possible optimal paths from the start of the similarity matrix (1,1) to the end of it (I, J) using dynamic programming ~~(Durbin et al. 1998)~~[34]. Chromatogram-groups *ChromA* and *ChromB* may not have end-to-end mapping as these ~~may only~~could be partial chromatograms which were extracted around the expected peptide elution (as determined by iRT peptides for example). Therefore, overlap alignment instead of a global alignment of MS2 chromatogram groups is employed. This approach allows free end-gaps and thus, allows to slide chromatograms freely without incurring any gap-penalty for it.

To widen or shrink chromatogram peaks, a gap of unit length is a reasonable choice as it will distribute gaps along the complete peak. Therefore, an affine gap penalty scheme is utilized with higher gap penalty for gap length of more than one. In this approach, three matrices (Matrix M, A, B) are defined which recursively calculates score for gaps of more than unit length ~~(Durbin et al. 1998)~~[34]. The overlap alignment path using affine gap-penalty is presented in Fig. 1*e*. The running time of such alignment is O($max(I, J)^3$). A heuristic data-driven approach is employed to obtain suitable affine gap penalties from the similarity matrix (see the Supplemental Section S5). Mapping the alignment path to the initial time values provides aligned chromatograms as depicted in Fig. 1*f*.

#### d. Running time for alignment:

Alignment of MS2 chromatograms of each peptide/precursor has running time of order O(max(I, J)^3); however, chromatograms of different precursors can be aligned independently. Therefore, we employ parallelization for different peptides to obtain much faster speed for complete run time-mapping.

#### e. Optimization of algorithm parameters:

There are various parameters used in ~~DIAlign~~DIAlignR. A description of these parameters is available in the Supplemental Section S5. We have ~~used a manually validated~~employed validation dataset ~~of 437 *S. pyogenes* peptides acquired with SWATH-MS across 16 LC-MS/MS runs~~ for parameter optimization~~, using~~ and have used the number of peaks aligned within half chromatographic peak-width and cumulative RT alignment error as ~~our~~ optimization target.

**Performance metrics for comparison with current algorithms:**

We have used the manually validated dataset (Röst et al. 2016)[5] to compare DIAlignDIAlignR to the current state-of-the-art method (e.g. TRIC[5]) which utilizes a set of high confidence peaks (“anchor peptides”) to compute a linear or non-linear alignment function that transforms RT values offrom run1 to RT values of run2. We chosehave chosen LOESS (local regression) as well as linear regression for our evaluation of a nonlinear alignment function. For LOESS, both optimized spanvalue from cross-validation (as used in TRIC) and default spanvalue (= 0.75) of the R software environment are tested (Chambers et al. 1992)[30].. For LOESS fit, ⅓ cross-validation is performed to obtain the optimum span value between two runs[5,30]. Steps to obtain a global fit (monotone mapping function) are detailed in the Supplemental Section S3.

Retention time error is calculated by comparing against the manual annotation of the *S. Pyogenespyogenes* dataset (Röst et al. 2016)[5] and the resulting distribution of the number of peptides aligned within a certain RT tolerance is used as a measure of overall accuracy of the alignment algorithm. Manual annotations are not available for the iPOPhuman plasma dataset, therefore, the high--quality results (peaks with low FDR cutoff) of the automated OpenSWATH tool isare used for benchmarking.

**Results**:

*Parameter optimization.*

Here, we present an algorithm for multi-trace chromatographic alignment that can directly useonly uses raw MS2 data from targeted proteomics or DIA experiments for retention time alignment. To optimize the performance of our algorithm, we used a manually validated dataset of 7,232 peakgroups (Röst et al. 2016) to investigatehave investigated the effect of algorithmic parameters on the accuracy of the results.alignment of runs from validation dataset[5]. First, we evaluatedevaluate the performance for different similarity measures of chromatogram groups. The dot product masked with spectral angle (see the Supplemental Section S5) as a similarity measure provides the highest fraction of peptides aligned for all 120 possible run~~-~~-pairs ~~on the validation dataset~~ (Fig. 2*a*~~) .~~). Within an RT error tolerance of half peak-width (here: 15.3 sec), this similarity measure aligns 94.33% of annotated peaks with the highest area under the curve of all approaches (see the supplemental Table ~~6~~7 and ~~7~~8).

We then investigated the effect of gap penalty used in dynamic programming. In ~~DIAlign~~DIAlignR, the gap penalty ~~is~~was calculated heuristically ~~from~~ as a fixed quantile value of the distribution of similarity scores ~~using a fixed quantile value.~~. We ~~found~~find that the selection of quantile value does not have a considerable impact on the percentage of peaks aligned within certain RT tolerance (Fig. 2*b*). From the figure, 20$^{th}$ to 90$^{th}$ quantile values yield approximate 95.6% of aligned peaks within half peak-width. The effect of gapQuantile is less pronounced for wider RT tolerance. For further analysis, the 65$^{th}$ quantile is selected as the base gap penalty for chromatogram alignment. For the affine gap penalty, a gap opening factor ~~is considered as~~of 0.125 was used, while a gap extension factor ~~is~~of 40 was considered (see the Supplemental Section S5).

We next investigated the impact of the number of transitions on the alignment(see the Supplemental Section S5 and supplemental Figure 10*a*). We find that as number of fragment-ions increases, alignment accuracy improves with best alignment 94.3% from using all library fragment-ions compared to 88.7% with only one transition (Supplemental Table 15). However, the largest improvement was going from 1 transition (88.7%) to two transitions (92.8%) with smaller improvements for using all transitions. This indicates single transition is not sufficient for this algorithm, but the relative gain is more modest after two transitions.

Our algorithm ~~is capable of constraining~~can constrain the similarity matrix using a global alignment function. Constraining the alignment in a certain window (given by RSEdistFactor) ~~about~~around the global fit derived from “anchor peptides” improves the alignment accuracy. We have observed that with a constrained similarity matrix 95.4% peaks get aligned compared to

94.3% with non-constrained one (see the supplemental Figure ~~10~~10*b*). An example of such alignment is shown in Fig. 2*c* and 2*d*, in which the similarity matrix has two high similarity hot-spots. ~~Constraining similarity outside of~~ By constraining the alignment inside the dashed region in Fig. 2*d*, the alignment path goes through the correct hot spot. With the unconstrained similarity matrix, an incorrect alignment ~~resulted~~results as shown in the supplemental Figure 11.

*Validation using "gold standard" reference dataset*

~~We then used the manually validated dataset to compare DIAlign to the current state-of-the-art method which utilizes a set of high confidence peaks ("anchor peptides") to compute a linear or non-linear alignment function that transforms RT values of run1 to RT values of run2.~~Using validation dataset, we have compared DIAlignR to current alignment methods. In terms of number of peptides aligned and alignment precision, chromatogram alignment outperforms LOESS and linear regression methods (see the Fig. 3*a* and Table I). On the *S. pyogenes* benchmark dataset, ~~DIAlign improves~~DIAlignR decreases error rates by 1.8-fold compared to the state-of-the-art methods. Cumulatively, chromatographic alignment only mis-aligns 4.3% of all peaks within 15.3 seconds (half peak width) of the true RT compared to 7.9% for LOESS (while LOESS with default parameters mis-aligns 22.8% of all peaks and linear regression mis-aligns 44.8%; see Fig. 3*a*).

We next investigated the effect of experimental perturbation on the performance of the alignment method. We ~~compared~~compare within-condition alignments with between-condition alignments (in the validation dataset, the conditions ~~were~~are 0% and 10% human plasma added to *S. pyogenes*~~). When human plasma (10% volume) is added to the sample, the performance of~~ during growth). For both alignment methods ~~degrades compared to samples without plasma~~, we observed decreased performance for between-condition alignments compared to within-condition alignments (Fig. 3*b*). ~~Additional drift in LC retention time is expected for a sample of increased complexity (Nigjeh et al. 2017).~~ However, the ~~LOESS~~ performance drop of the

LOESS method (4.93%) is substantially larger than the corresponding performance drop of ~~DIAlign~~DIAlignR (2.7~~%) (Fig 3*b*).~~%), indicating increased robustness to sample heterogeneity for DIAlignR.

To evaluate the consistency of alignment approaches across multiple run-pairs, we have computed the number of correct aligned~~-~~ peaks (~~time mapping falls~~defined as instances where the alignment is correct within half peak-width~~ from annotated RT~~) for each run-pair. This distribution is shifted towards the right with low standard deviation for chromatogram alignment method compared to ~~the same for~~ LOESS, indicating that the former is consistent in its performance (Fig. 3*c*). In terms of the precision of the alignment, chromatogram alignment consistently performs better than global alignment methods ~~such as LOESS~~ as the former has higher area under the cumulative peptide frequency curve for each run-pair (higher AUC for 120 out of 120 pairs, see the supplemental Figure 12*c*). Similarly, we ~~observed~~observe a~~-~~ larger RT variation (standard deviation = 18.45 sec) with the LOESS approach which chromatogram alignment ~~is able to correct~~corrects satisfactorily with the standard deviation being 11.68 sec (Fig. 3*d* and supplemental Fig. 12*a*, *b*). We conclude that on the validation dataset, ~~DIAlign~~DIAlignR performs consistently better in terms of accuracy of alignment and number of correctly aligned~~-~~ peaks across a range of different RT cutoffs and LC-MS/MS runs.

Next, we were interested in how the global differences between the two methods translate to individual alignments. We, therefore, computed the alignment error for each pairwise alignment of each peptide (total 49,505 alignments) and found that chromatographic alignment outperforms LOESS in 4.7% of all cases, whereas, LOESS achieves better results in 1.1% cases, while comparable performance was achieved in the remaining 94.2% cases (see the ~~supplemental~~Supplemental Fig. 13*b*). On average, ~~DIAlign reduces~~DIAlignR has reduced the RT error by 2.3 seconds with a median of 1.7 seconds (see the supplemental Fig. 13*c*). Overall, our method aligns 47.3k peaks correctly compared to 45.6k by an optimized ~~global lowess method~~LOESS within ~~15.3 seconds (~~half peak-width (15.3 sec). However, in general we observed that on the validation dataset both methods perform with similar consistency which may be due to the low complexity of a bacterial sample and the high homogeneity of the data ~~which~~as it was acquired within ~~a single week~~two consecutive days on the same LC column.

*Application to large-scale heterogeneous human plasma measurements*

After demonstrating consistently improved performance on the *S. pyogenes* validation dataset, we investigated the performance of our algorithm on a large-scale SWATH-MS experiments on human plasma. ~~This experiment~~These experiments provided a more challenging dataset as the data was acquired over the period of six months with an intermittent repair of the instrument and ~~change~~replacement of ~~LC~~the old column~~. We selected 2~~ (column1) with a new column (column2). Two LC-MS/MS runs from each of the 12 batches were selected at random and ~~used~~ 406 peptides were used for testing our ~~algorithms~~algorithm. Since we did not have ~~manual validations, we selected~~manually validate peaks, high confidence peak groups (q-value < $10^{-3}$) were used instead as ~~our~~ validation ~~peptide~~ set.

Comparing our chromatogram alignment algorithm (~~DIAlign~~DIAlignR) with the LOESS method on a highly heterogeneous human plasma dataset, we found that our approach aligns 97.92% of peaks compared to 76.03% ~~using the~~by LOESS ~~method~~ with a maximal error of 20 seconds (half chromatographic peak-width) as depicted in Fig. 4*a*. All tested 276 pairwise alignments shown improved performance using chromatographic alignment (see ~~supplemental~~Supplemental Fig. 15). Next, we were interested in the performance of our method on the alignment of runs acquired on the ~~same and~~two different columns. We ~~found~~find that for runs acquired on different columns, chromatogram alignment method aligns 97.7% (compared to 97.84% within-column alignment) of peaks compared to 63.38% (89.06% for within-column alignment) by LOESS method (Fig. 4*b*), suggesting that ~~DIAlign~~DIAlignR retains performance even for highly heterogeneous datasets~~. New column and instrument repair adds more features~~

~~in~~ while the ~~LC/MS-MS output (see supplemental Figure 2 and supplemental Table 5), therefore~~LOESS approach does not. Specifically, we ~~observed an improvement in LOESS'~~find not only that DIAlignR outperforms LOESS on between-column alignments, but that the performance ~~for "column2 pair". However, despite such changes our approach had steady response validating its robustness to such events.~~

loss for DIAlignR is much less pronounced than for LOESS compared to within-column alignments. After validating the performance of chromatogram alignment cumulatively, we decided to investigate its consistency across individual run-pair alignments. Fig. 4*c* presents the distribution of the number of peaks aligned in all 276 pairs. ~~DIAlign~~DIAlignR is capable of aligning 400 peaks on average within half-peak width (while LOESS ~~aligned~~aligns 309 peaks on average), a 29% improvement. This indicates ~~the inconsistency of~~substantial alignment error when using LOESS~~ approach~~, which ~~was not observed with DIAlign~~could be reduced drastically by DIAlignR.

To validate the performance on individual alignment, we further computed the alignment error for each pairwise alignment ~~of each~~for every peptide. The standard deviation of alignment error for LOESS was 22.91 sec compared to 13.7 sec for ~~DIAlign~~DIAlignR (Fig. 4*d*). This indicates the higher precision of RT alignment with our approach. Out of 112,056 alignments, we found that ~~DIAlign~~DIAlignR outperforms LOESS in 23% of all cases and performs similarly in 76% cases (see the ~~supplemental~~Supplemental Figure 14 - while performing worse in 1.13% cases). Upon manual validation, several of these worse-performing peaks were found to be due to wrong annotation by OpenSWATH (Supplemental Figure 18). Thus, testing of chromatogram alignment approach on the heterogeneous human plasma dataset again validates its consistent and improved RT alignment performance.

*Switching of peptide elution order*

In liquid chromatography, retention time drift is often observed from one run to another run. However, thewe were interested whether this drift canmay be variable for different peptides and thus willcould result in reversal of retention order (Spicer et al. 2010)[15]. In such a scenario, two peptides which are eluting in order in one run may reverse their elution order in otheranother run. Since our approach does not make an assumption of order preservation of peptide elution and facilitate independent alignment, we hypothesized that DIAlignDIAlignR would be capable of uncovering instances of non-order preserving chromatographic alignment. Specifically, we analyzed peptide pairs that switch elution order from the heterogeneous and distant blood plasma runs for peptide pairs that switch elution order.

To confirm the alignment for such peak-switching cases by chromatogram alignment algorithm, we have specifically looked at the alignment of the pair “run4_run23” as it hadhas the highest number of peak switching pairs. run4 wasis part of batch V4 and was, acquired on February 28th, 2017 whereas run23 wasis from batch M3 and was, acquired on July 20th, 2017. The LOESS fitting from common high scoring training peptides for this pair is presented in Fig. 5*a*. Most of the test peptides are scattered around the global fit line, instead of being directly on the line. This graph quickly suggests 407 peptide pairs (one from either side of the line) compromisingcomprising of 237 out of 406 peptides whichthat have switched their elution order out of 406 peptides (see the supplemental Section S6). We thus found that overall, 58.4% of peptides were involved in at least one event of non-order preserving elution.

One of the peak switching cases is presented in Fig 5*b*. In run4 peptide AQLVDMK/2 elutes after HYDGSYSTFGER/2, whereas in run23 the elution order has been reversed. Both peptides have seen positive RT drift in run23 from run4, however, HYDGSYSTFGER/2 had shiftshifts of 1070-850 = 270 seconds whereas AQLVDMK/2 had shiftdrifts of only 1050-900 = 150 seconds. This varying RT drift between two runs has caused the peptides to elute in different order. ThisThe peptide--pair cannot be aligned with a global alignment approach, which in the best-case scenario will be off by 120 seconds -- however, our chromatogram alignment method has mapped the peaks correctly from run4 to run23 (see the supplemental Figure 17).

~~To compare DIAlign against other state-of-the-art approaches, we calculated~~We, further, calculate the cumulative fraction of peptides aligned for pair “run4_run23” (Fig. 5*c*). Chromatogram alignment correctly ~~aligned~~aligns 98% peaks compared to LOESS which ~~was~~is able to align only 37.93%, thus DIAlignR is capable of decreasing the error by up to 30-fold. Eight peaks, which ~~were~~are not correctly aligned ~~were~~by chromatogram alignment, are further inspected visually by the authors and are found to be the cases of ~~incorrect~~ mis-annotation ~~of~~by OpenSWATH, mainly due to the~~ mis-annotations of peptides carrying of~~ post-translational modifications (see the supplemental Section S7 and supplemental Figure 18).

**Discussion:**

Correcting for retention time drift and aligning retention times between LC-MS/MS runs has been a long-standing problem in proteomics and it has become of particular importance as proteomics moves towards large-scale analysis of human cohorts. However, most efforts so far have focussed on MS1 data and few algorithms are available that can exploit the full information present in MS2 ~~information (such as~~spectra produced by targeted methods or DIA / SWATH-MS~~)~~..

In this paper, we have presented a novel algorithm that uses the raw fragment~~-~~-ion ~~chromatogram data~~chromatograms directly to perform retention time alignment for targeted proteomics and DIA data. Our algorithm uses ~~extracted-ion chromatograms~~XICs to map peaks across ~~multiple~~a pair of runs and improves accuracy compared to current state-of-the-art methods. We have, furthermore, extended the algorithm and implemented a hybrid approach ~~that also~~, which uses a feature-based global alignment to condition the similarity matrix *s* ~~which~~that led to further gains in accuracy (see the Supplemental Fig. ~~8~~10*b*). This hybrid approach provides the best of both worlds with a flexible “knob” which allows the user to either put more focus on global features or rely more on local information. To our knowledge, researchers have not yet explored dynamic programming--based alignment on raw fragment-ion-chromatograms. The dynamic programming approach is essential for obtaining a non-linear (or gapped) alignment as distant runs also have varying drift even for local peaks. Using a feature-based LOESS method to partially constrain the alignment makes our algorithm more stable and provideprovides the robustness of global alignment methods.

We have shown that on a “gold-standard” validation dataset, our methodDIAlignR consistently outperforms a global alignment method (using either linear or non-linear approaches), the current state-of-the-art (Supplemental Figure 12*c*). The DIAlign tool is able to decrease error rates from 7.9% to 4.3% overall.We have observed that alignment accuracy increases almost 4% if a precursor has two transitions instead of one (Supplemental Figure 10*a*), We get additional 1.5% aligned precursors by using all six fragment-ions which also corresponds to recommended guidelines[4]. Since, the DIAlignR approach puts more emphasis on local data, we also observe instances of over-fitting where the global alignment function produces better alignment (Supplemental Figure 13*b*). Overall, however, we see increased performance and the DIAlignR algorithm can decrease error rates from 7.9% to 4.3%. Interestingly, we find that our method is less sensitive to changes in chromatographic condition or sample matrix than global alignment approaches (Fig. 3*b*).

This finding led us to speculate that the novel chromatographic alignment would be less sensitive to heterogeneity in sample composition and chromatographic condition in large-scale studies. We have tested our algorithm on a large-scale SWATH-MS experiment of human blood plasma acquired over several months. On this heterogeneous dataset, DIAlignDIAlignR reduces RT alignment error from 24% to 2%, which is a significant improvement over current state-of-the-art methods. Our approach has outperformed other methods and has consistently mapped the highest number of peaks within half-peak width irrespective of acquisition time interval, column change or instrument repair between two runs (Fig. 4*b*). DIAlign picks correctDIAlignR improves retention time alignment accuracy which has the potential to improve peak-group ~~and reduces~~ identification ~~errors;~~and quantification through downstream tools. We have manually identified an example of ~~wrong~~wrongly aligned peak-group ~~picking by global alignment method is presented~~ as shown in ~~the~~ Supplemental Figure 19. We have also ~~shown~~observed that in the case of a peak being outside of ~~chromatograms~~an extracted chromatogram, our method is able to map retention time outside of it as ~~it~~our hybrid approach also uses global alignment (~~the~~see Supplemental Figure 18). Chromatograms can then be re-extracted and be used to correctly annotate peaks. Thus, this method can further be employed to extract chromatograms by OpenSWATH and other tools.

The improvement in retention time alignment comes with non-negligible computation cost. For an alignment of 10-min wide XICs with 3.4 sec cycle time, DIAlignR takes on average 0.16 seconds. Thus, pairwise alignment of selected 437 peptides takes approx 1 minute per run-pair. The highest cost in the alignment of each peptide is the calculation of the alignment path using dynamic programming. However, this problem scales linearly with the number of peptides in the library and can be easily parallelized on a computing cluster.

We believe that our approach is most useful for large-scale heterogeneous targeted proteomics studies where runs are acquired by different personnel and data is collected over several months or even years. Applying a single mapping function in such experiments becomes a very challenging task considering the switching of elution order of peptides. Global alignment functions, being monotonic in nature, assume chronological order of peptide elution and, therefore, cannot align switched peptides. ~~However~~Hence, we have ~~shown that our~~observed substantial underfitting with the global function and an overall reduction of error using DIAlignR. Our hybrid approach aligns these peptides accurately as it mostly relies on additional dimensions of fragment-ion m/z to align peaks. It is possible that switching peptides may share many fragment-~~ion m/z~~ions, however, this is very rare scenario if library is designed carefully[4] and in ~~that case~~such cases our method will perform no worse than global alignment methods.

~~Applying~~ Accurate RT alignment has multiple uses in the application of mass spectrometry-based proteomics ~~in~~for large-scale systems biology studies~~, high reproducibility~~. Correct identification and improved quantitation of large number of analytes ~~is imperative.~~are few of them. This seems intuitive as most quantitative approaches currently available at least to some degree rely on accurate retention time alignment. We present a tool that can ~~be used to establish~~align retention times of DIA data by establishing correspondence between analytes across large number of samples, making DIA amenable for multi-center and longitudinal studies. We also expect that this tool can be utilized by existing proteomics ~~softwares~~software to streamline analyte identification and improve the quantification.

**Acknowledgements:**

We are grateful to Michael Snyder for supervising data-acquisition and providing access to the heterogenous plasma dataset. We also thank Michael Brudno for valuable discussion on chromatogram alignment using dynamic programming.

**Author contributions:**

S.G. designed and wrote code, performed data analysis and produced the figures. S.A. and W.Z. performed human plasma related experiments, acquired MS data. H.R. designed and supervised the study. S.G. and H.R. contributed to writing the manuscript. All authors have contributed to the final manuscript.

**Competing Financial Interest:**

The authors declare no competing financial interests.

**Supplementary Material:**

**Data Availability:**

Raw chromatograms and features extracted by OpenSWATH are available on PeptideAtlas under accession code PASS01280.

34. Durbin, R., Eddy, S. R., Krogh, A. & Mitchison, G. *Biological sequence analysis:*

*probabilistic models of proteins and nucleic acids*. (Cambridge university press, 1998).

Table I

Summary description of validation and human plasma datasets

| | Validation dataset | Human plasma dataset |
|---|---|---|
| Biological Sample | *Streptococcus pyogenes strain SF370* | Plasma from blood samples |
| Mass-spectrometer | SciEX 5600 TripleTOF | SciEX 6600 TripleTOF |
| LC - gradient | Linear | Linear |
| Total run time | 135 minutes | 55 minutes |
| LC-column replaced | No | Yes |
| Mass-spectrometer repaired | No | Yes (Replaced quadrupole after 7 batches) |
| Data acquisition date | 08 August 2012 - 09 August 2012 | 17 February 2017- 20 July 2017 |
| Number of runs acquired | 16 | 975 |
| Number of batches | 1 | 12 |
| Runs selected for alignment | 16 | 24 |
| Total number of run-pairs | 120 | 276 |
| Software used for feature detection and XIC extraction | OpenSWATH | OpenSWATH |
| Number of common precursors selected per run for alignment | 437 | 406 |
| Total number of alignments | 49505 | 112,056 |
| Manual Annotation | Yes (Skyline) | No |

Table II

Below is presented average number of peptides aligned from manually validated *S. Pyogenes* dataset and from *clinicalheterogenous human plasma measurements* for peptides. For plasma data, peaks with OpenSWATH m-score < 0.001. For iPOP dataset, instead of manual annotation OpenSWATH annotation was are used for benchmarkingevaluation of the algorithm.

| Dataset | Method | Average number of peaks aligned within half peak-width | Average number of peaks aligned within one peak-width | Average number of peaks aligned within two peak-width | Total number of peaks per run | Peaks aligned within half peak-width (%) |
|---|---|---|---|---|---|---|
| *Validation* dataset | Chromatogram Alignment | 394.81667 | 407.3 | 410.69167 | 437 | 95.68765 |
| | LOESS | 380.07500 | 406.18333 | 410.81667 | 437 | 92.10249 |
| *Heterogenous human plasma* dataset | Chromatogram Alignment | 397.56159 | 400.50362 | 403.88043 | 406 | 97.92157 |
| | LOESS | 308.67754 | 382.17754 | 403.15217 | 406 | 76.02895 |
