## Supplemental Text for "DIAlignR provides precise retention time alignment across distant runs in DIA and targeted proteomics"

### S1: MS2 Chromatogram alignment in the context of current approaches

Most of the existing retention time alignment approaches in proteomics and metabolomics were developed for MS1 data. Most of these approaches uses dynamic programming to align either raw chromatograms[1–5] or feature(peak) lists[6,7]. While aligning raw chromatograms using dynamic programming, Sakoe-Chiba constraining[8–11] is used to limit the flexibility of algorithm. This constraining also reduces search space and make algorithm faster. Anchor-peptides have been suggested to improve the alignment accuracy[3,12]. These features act as landmarks in search space and guide the alignment path.

Some other concepts like linear time warping[13], LOESS[14–16], cross-correlation[17], parameter warping[18], correlation optimized warping[19,20] have been proposed in the literature for alignment of raw chromatogram. However, all aforementioned methods incorporate global alignment and fail to align peptides independently and to establish many-to-many RT mapping required to align inverted order peaks. For feature-based alignment, graph based bipartite matching[12,14,21,22] have been recommended, these approaches absolutely depend on efficiency of feature-finding software. A comparative discussion of current alignment strategies for proteomics is presented by Smith *et al*.[23].

With SWATH data, it is possible now to have local information about precursor and corresponding fragment ions. Because of multiplexing of m/z windows, we get addition neighboring signals in XICs which makes it advantageous for alignment. Exploiting the presence of additional signal in XICs, we have used dynamic programming to align MS2 fragment-ion chromatograms for each precursor and have established independent RT mapping for all peptides. Thus, we have developed a tool which only takes MS2 data for RT alignment compared to previous methods which relied on MS1 data.

### S2: Precursors excluded and run names of validation dataset

For four precursors, there was no annotated peaks in the validation dataset. These precursor IDs were:

1. 4410_QEVNELSR/2
2. 4571_AIEEDGSIEIVTPDHK/3
3. 4815_TINNIIK/2
4. 469_YADSVPLVYEIASIPEK/2

Another four precursors had annotation in one run out of 16 runs. The precursor IDs were:

1. 13115_GKGDDMIIER/2
2. 13289_QLLHAQLASTDKK/2
3. 8794_NAPYLPYLGNISR/2
4. 9809_LEDIGTDAYQLISDLR/3

Nine precursor did not have extracted fragment-ion chromatograms(XICs) covering manual annotations for few pairs. These retention time values were updated for not to be considered in the benchmarking.

| Precursor IDs | Number of runs where annotation was out of XICs | Number of runs where annotation was not out of XICs | Number of runs of unannotated peak |
|---|---|---|---|
| 10748_RVLESGQDYTM[147]DPMSNSER/3 | 16 | 0 | 0 |
| 13391_TPLFNLLK/2 | 16 | 0 | 0 |
| 19650_AAQALANTIIDHGPEAVKK/3 | 16 | 0 | 0 |
| 4920_VVSM[147]PSQNIFDEQSAEYKESILPAAVTK/3 | 16 | 0 | 0 |
| 7636_M[147]YYTSGNYEAFATPR/2 | 5 | 11 | 0 |
| 19612_LVLDDLTFSIPK/2 | 16 | 0 | 0 |
| 8834_EQGYDVIGVFMK/2 | 1 | 8 | 7 |
| 423_LLIIDHR/2 | 15 | 0 | 1 |
| 11349_ATGEVMAIGR/2 | 16 | 0 | 0 |

Hence, precursor IDs removed from the analysis are:

1. 4410_QEVNELSR/2
2. 4571_AIEEDGSIEIVTPDHK/3
3. 4815_TINNIIK/2
4. 469_YADSVPLVYEIASIPEK/2
5. 13115_GKGDDMIIER/2
6. 13289_QLLHAQLASTDKK/2
7. 8794_NAPYLPYLGNISR/2
8. 9809_LEDIGTDAYQLISDLR/3
9. 10748_RVLESGQDYTM[147]DPMSNSER/3
10. 13391_TPLFNLLK/2
11. 19650_AAQALANTIIDHGPEAVKK/3
12. 4920_VVSM[147]PSQNIFDEQSAEYKESILPAAVTK/3
13. 19612_LVLDDLTFSIPK/2
14. 423_LLIIDHR/2
15. 11349_ATGEVMAIGR/2

Incorrect annotation of precursor 4205_SEASVGFGAAR/2 for run14 is corrected with retention time = 1980.7 sec.

Out of 437 annotated precursors, five had doubtful manual annotation of peptide peaks. These precursor IDs are as follows:

1. 7118_MDELLDSLNANLDEM[147]SER/3
2. 6857_SEFPENELWDLTALYK/2

3. 1288_FILTSDELFELK/3
4. 20672_TGLLAETFGGQVMETVGIENMIGTLYTEGPK/3
5. 18091_Q[111]LDLLAHGER/2

However, these annotations are not modified as it will affect the performance of both global and local alignment approaches equally.

<u>Run names of validation dataset</u>

| **Run** | **Run name** |
|---|---|
| run0 | Strep10Repl1_R03 |
| run1 | Strep10Repl2_R04 |
| run2 | Strep0Repl2_R04 |
| run3 | Strep10Repl1_R04 |
| run4 | Strep0Repl2_R02 |
| run5 | Strep10Repl2_R03 |
| run6 | Strep10Repl2_R02 |
| run7 | Strep0Repl1_R04 |
| run8 | Strep10Repl2_R01 |
| run9 | Strep0Repl1_R01 |
| run10 | Strep10Repl1_R01 |
| run11 | Strep0Repl1_R02 |
| run12 | Strep10Repl1_R02 |
| run13 | Strep0Repl2_R03 |
| run14 | Strep0Repl2_R01 |
| run15 | Strep0Repl1_R03 |

### S3: Optimized LOESS fit between a run-pair in the validation dataset

Common precursors with m-score (FDR score) less than 1e-05 are chosen[15,24]. Precursors with higher cut-off value will add noise without providing any benefit results, whereas, with more strict cut-off there will be not-enough IDs to fit an optimum alignment. Post m-score cut-off, if multiple peak-groups are remained with the same precursor ID, peak-group with rank = 1 is considered[25]. The distribution of number of peak-groups for calculating LOESS fit is depicted in Supplemental Figure 1*b*. On average

4159 points are available to obtain global alignment fit. The number of common peak-groups, optimum span value and associated RSE is provided in the Supplemental Table 5.

### S4: Optimized LOESS fit between a run-pair in the clinical plasma dataset

Common peak-groups with m-score (FDR score) less than 1e-03 and highest rank are chosen to obtain a global alignment. m-score cutoff is same for training peptides and test peptides. A wider distribution of number of datapoints to obtain a global alignment fit is observed as shown in the supplemental Figure 2*a* with minimum being only 317 and maximum being 4532 common peak-groups. There are 406 test points, certainly for few pairs we have lesser number of training points available. This affects the residual standard error of the alignment as depicted in the supplemental Figure 2*b*. Certainly, higher RSE is observed specifically for pairs with a little number of common peak-groups. This is more pronounced for pairs which constitutes one run before quadrupole replacement and second run after it. This indicates that all peptides do not have consistent m-score (FDR cut-off) below the 1e-03. This is expected as quadrupole replacement has changed sensitivity of the instrument, therefore, probability of detection above noise threshold in OpenSWATH[25].

### S5: Parameters for chromatogram alignment

<u>Similarity measures</u>

For chromatogram alignment, similarity between indices of *ChromA* and *ChromB*, each of which has *n* fragment-ions, is calculated. Following similarity measures are tested in this manuscript:

1) Euclidean distance
2) Pearson's correlation
3) Covariance
4) Cosine similarity (with 2*spectral angle)
5) Dot-product
6) Masked dot-product

Euclidean distance is calculated between indices in *n* dimensional space. The similarity value is calculated as:

$$S_{ij} = \frac{1}{1+EuclDist_{ij}}$$

The euclidean distance does not necessarily distinguish distance among high-intensity signals and low-intensity background noise. Hence, generally provides high similarity values for background signals as well. This is evident from the distribution of similarity scores of chromatograms of an example peptide KALVETDGDMDK/2 from run11 and run12 of the validation dataset (Supplemental Figure 4*a*). The peak of the histogram at 0.5 indicates that euclidean distance based similarity does not provide zero (minimum) score for background region.

Similarly, cosine similarity and pearson correlation coefficient takes into account the relative intensity ratio of fragment-ion peaks and provides high similarity when it is same between two chromatograms, however, by nature it is unbiased towards intensity thus can't differentiate signal from the background. As presented in Supplemental Figure 4*b* and 4*c*, histograms of these similarity measures also indicate their inability in distinguishing signals from the background. Since, the intensity is always positive, the angle always remains between 0° and 90°. To use full-spectrum of cosine similarity, spectral angle is doubled, providing range of [-1, 1] for similarity score.

On the contrast, covariance and dot-product provides high similarity score when there is high joint variability of the two peak-groups and can also distinguish signal from noise as most of the scores are concentrated around zero or negative side and only few value attain maximum unit score as presented in the Supplemental Figure 5. However, since the covariance subtracts the expected/average intensity value of each fragment-ion, it distorts information present in terms of relative intensity ratio of fragment-ion peaks. Effect of this distortion on the chromatogram alignment is depicted in the Supplemental Figure 6*a* and 6*b*, in which noise peaks in run12 get aligned with peptide elution-peaks in run 11 because when relative intensity information is lost, noise peak being with larger area provides the high similarity hot-spot. On the contrary, with dot-product correct peak-groups are aligned in Supplemental Figure 6*c* and 6*d*.

Listing 1: Similarity matrix calculation pseudocode. A pseudocode representation of calculating similarity matrix for dot-product similarity is illustrated.

```
getSimilarityMatrix(XIC.run1.peptide, XIC.run2.peptide)
s1 = s2 = 0
for transition in peptide.transitions:
   s1 = s1 + sum(XIC.run1.peptide.transition.intensity)
   s2 = s2 + sum(XIC.run2.peptide.transition.intensity)
Initialize similarity_matrix of size (nrow = xic1_time_points, ncol = xic2_time_points)
select case (similarityMeasure)
case (dot-product):
   for transition in peptide.transitions:
        // intensity1 = XIC.run1.peptide.transition.intensity
        // intensity2 = XIC.run2.peptide.transition.intensity
        for i in xic1_time_points:
           for j in xic2_time_points:
              S[transition][i, j] = intensity1[i]/s1 * intensity2[i]/s2
              similarity_matrix[i, j] = similarity_matrix[i, j] + S[transition][i, j]
 case (dot-product-masked):
    // some other code see DIAlignR R-package
```

### Masked dot-product as similarity measure

The co-elution signal in both chromatograms provides high dot-product similarity, however, a very high intensity noise-peak in any chromatogram may also result in higher dot-product similarity and can drive the alignment. To reduce its impact, one way is to check for spectral angle at the high similarity hot-spots in the matrix and lower the similarity score if angle is higher than a threshold.

Hence, if $SpecAng$ is the spectral angle matrix and $S$ is dot-product similarity matrix, both of size *I x J,* then $mask$ for a certain similarity threshold $simThreshQuantile$ will be:

$$Mask = S > quantile(S, simThreshQuantile)$$
$$CosineSim_{new} = cos(2 \times SpecAng)$$
$$AllowDotProd = [Mask \times CosineSim_{new} + (1 - Mask)] > cosSimThresh$$
$$S_{new} = S \times AllowDotProd$$
$$S = S_{new}$$

Since, the intensity is always positive, the angle always remains between 0° and 90°. To use full-spectrum of cosine similarity, $SpecAng$ is doubled, providing range of [-1, 1] for $CosineSim_{new}$. The optimum values for $simThreshQuantile$ and $cosSimThresh$ are obtained by parameter sweep on a validation dataset (see the Supplemental Figure 7). From the supplemental Figures 7*a* and *b*, it is evident that 0.1 < $cosSimThresh$ < 0.8 provides optimal alignment. The optimal value for $simThreshQuantile$ falls between 0.96 and 0.985. For further analysis, $cosSimThresh$ = 0.3 and $simThreshQuantile$ = 0.96 are considered. Note that length of XIC and sensitivity of mass-spectrometer should also be considered while defining these parameters. The summary of number of peaks aligned, and AUC within a RT tolerance for the complete grid search are provided in the Supplemental Table 9 and 10, respectively.

### Effect of number of fragment-ions (transitions)

In the benchmarking dataset, 436 precursors have six fragment-ions and one precursor has five fragment-ions. Whereas, in human plasma dataset, number of transitions are more varied:

| | | | | | |
|---|---|---|---|---|---|
| Number of transitions per precursor | 2 | 3 | 4 | 5 | 6 |
| Number of precursors from plasma data | 1 | 4 | 11 | 10 | 380 |

To investigate the effect of number of transitions, we have ordered chromatograms from runA in the reverse-order of maximum intensity and aligned respective chromatograms from runB from benchmarking dataset using dynamic programming. As more transitions are included in the alignment, better RT alignment accuracy is obtained (Supplemental Figure 10*a*). Non-penalized similarity matrix is used for alignment as constraining with global alignment will introduce bias in the search space. Moving from one transition to two transitions results into 4.1% increase in number of aligned peaks (Supplemental Table 15). Adding more transitions improves alignment gradually with additional 1.5% increase with all library transitions.

In the gold-standard bacterial dataset we had six fragment-ions for each precursor, whereas, for human plasma dataset we have used 4, 5 or 6 fragment-ions for each precursor. It is possible that two precursors falling in the same SWATH window, may share one or more fragment ions and also have their retention times switched. In general, we recommend for DIA / SWATH-MS analysis to use at least 6 fragment ions (see best practices[26]). We have also cautioned readers about this in the manuscript. With one or two shared fragment ions, dot-product similarity and spectral angle similarity of switched-peaks would be very-low and most likely will not affect the alignment path and independence of alignment function will still remain. (Shown in ExampleFigure 1 below)

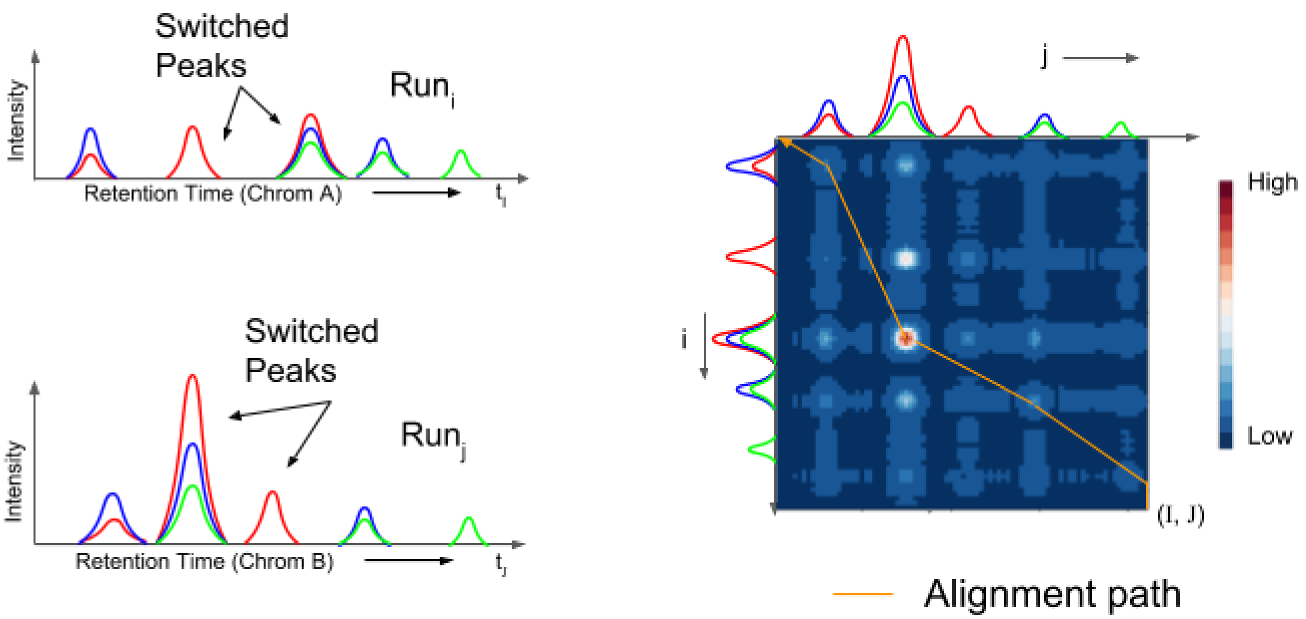


**ExampleFigure 1**. Alignment path when less than half library fragment-ions are shared between switched peaks. Library has three fragment-ions. Red fragment-ion is shared with other precursor which has switched its elution order with our peptide-of-interest. An approximate alignment path is directly overlaid on the similarity matrix (path for illustration only).

If there are half or more than half shared fragment-ions, then, it may not align both peptides correctly as second peptide's elution peak can substantially affect the similarity score and eventually the alignment path (Shown in ExampleFigure 2 below).

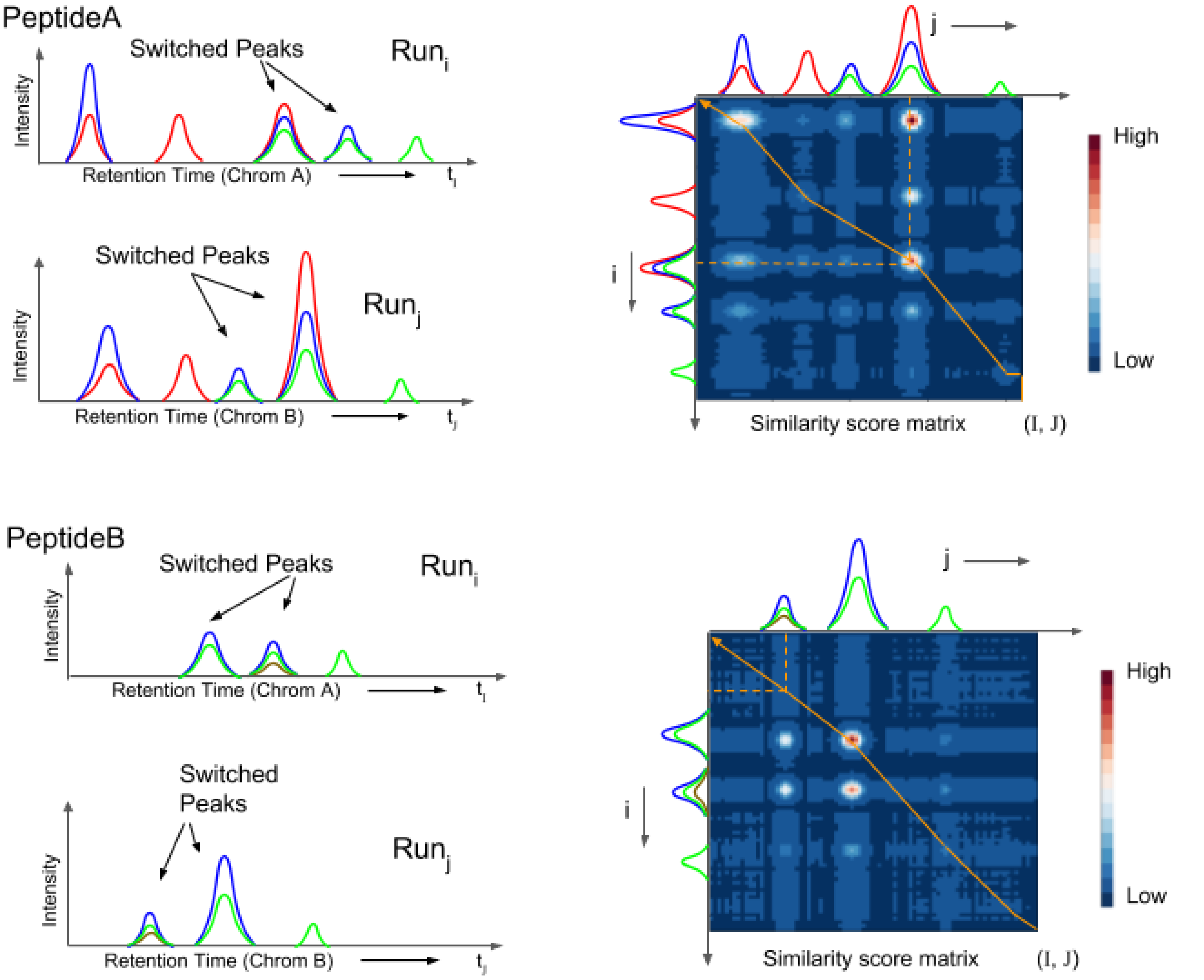


**ExampleFigure 2**. Alignment path when more than half library fragment-ions are shared between switched peaks. Library has three fragment-ions. Blue and green fragment-ions are shared between PeptideA and PeptideB which has switched their elution order. An approximate alignment path is directly overlaid on similarity matrix (path for illustration only).

PeptideA is aligned accurately, however, since more than half high-intensity fragment ions are shared, they also drive the alignment of XICs of PeptideB. This leads to incorrect alignment of PeptideB in a degenerate case where PeptideB does not have any unique fragment ions. However, we note that this situation is highly unlikely to occur within current spectral libraries that use 5-6 fragment ions due to the low probability of all fragment ions being shared between two peptides (see our theoretical analysis[27]).

Penalizing similarity matrix with global alignment

In the alignment of MS1 chromatogram, Sakoe-Chiba constraining have been suggested to reduce the search space[8–11]. It allows to search an optimal path only within a certain window along the diagonal of the similarity matrix by limiting number of consecutive gaps. However, this is a hard constraining. Instead by penalizing a similarity matrix, the probability of finding alignment path outside of the window is less, but possible.

Establishing RT correspondence using DIAlignR relies on extracted ion chromatograms. These chromatograms are crude, and therefore, restricting path along the diagonal using Sakoe-Chiba constraining will only be detrimental for the alignment. With inclusion of LOESS, additional information (feature-based) is imposed on the similarity matrix and high probable search space can be asymmetrical about the diagonal. High-confidence MS2 identification has been used as landmarks in alignment of MS1 chromatograms[3]. The LOESS constraint could also be thought as an approximate function derived from these landmarks. This provides soft-constraining instead of using hard landmarks directly.

The residual standard error (RSE) is the positive square root of the mean square error in the training data-set. Hence, RSE provides an approximate estimate of error that arises by applying regression fit on the test data. Considering this property, RSE is utilized to create a window outside of which alignment is not preferred. Supplemental Figure 3 indicates scattering of high scoring peak-groups within a certain window around the LOESS fit line.

Window of non-interference = RSEdistFactor * RSE

Slope of penalizing scores from the window = -2*max($S$)/samples4gradient

The optimum value for the parameters *RSEdistFactor* and *samples4gradient* can be estimated by grid search (see the Supplemental Figure 8). The optimal value for *RSEdistFactor* is between 3 and 4. And, the optimal value for *samples4gradient* is between 80 and 110. In this paper, *RSEdistFactor* = 3.5 and *samples4gradient* = 100 are chosen. A summary of the number of peaks aligned for *RSEdistFactor* and *samples4gradient* are provided as supplemental Table 13 and 14, respectively.

### Gap penalty calculation for overlap alignment

Base gap penalty is calculated heuristically from the distribution of scores in the similarity matrix $S$. A gapQuantile$^{th}$ value from $S$ is considered as base gap penalty. For affine gap opening and extension penalties, optimum gap opening factor(goFactor) and gap extension factor (geFactor) are determined by parameter sweep (see the Supplemental Figure 9). The optimal value for goFactor is < ¼ and for geFactor is > 10. Higher value for geFactor is recommended to force single gap effectively. Therefore, the optimal value for these parameters are chosen as ⅛ and 40, respectively. The summary matrix containing number of peaks aligned, and area under the curve within a RT tolerance for the complete grid search are provided in the Supplemental Table 11 and 12, respectively.

base gap penalty = quantile(s, gapQuantile)

Gap open penalty = goFactor * base gap penalty

Gap extension penalty = geFactor * base gap penalty

### Psuedocode of DIAlignR workflow

DIAlignR uses fragment-ion XICs which are usually of 10 min (+/- 5 minutes window about the library retention time) length from a LC-MS/MS run. Thus alignment path is restricted within this extracted time window. We are using overlap alignment, therefore, ‘start’ and ‘end’ region of XICs may not be aligned. Alignment path starts and stops with the end-point of XICs. The length of XICs can be changed using SWATH analysis softwares (e.g. OpenSWATH), however, 10 minute long extraction window was found to be optimal for feature extraction and we have carried on with the same.

Listing 2: DIAlignR Pseudocode. A pseudocode representation of the RT alignment algorithm which illustrates calculating similarity matrix and finding path using dynamic programming.

```
for (run1, run2) in pairs:
    for peptide in peptide_list:
        similarity_matrix = getSimilarityMatrix(XIC.run1.peptide, XIC.run2.peptide)

        IF alignment is to be constrained with LOESS:
            loess = LOESS fit between run1 and run2
            (p1, p2) = points where loess intersects the similarity_matrix
            similarity_matrix = penalizeMatrix(similarity_matrix, p1, p2, noPenaltyWidow, penalty)

        Perform dynamic programming to obtain affine_gap_alignment
        Follow the maximum scoring path
        Map retention-time back to the aligned path
```

### S6: Peaks with their elution order reversed in the clinical plasma dataset

To obtain peptide-pairs with their elution order reversed, the RT difference for each peptide-pair in each run is calculated. Pair "run4_run23" has maximum number of peak-swapped points. The peptide-pairs, which have product of this RT difference less than -300 sec$^2$, are considered for further analysis. On a visual verification following peptides were found incorrectly annotated by OpenSWATH:

1. 119603_M(UniMod:35)KPVPDLVPGNFK/3
2. 46713_M(UniMod:35)KPVPDLVPGNFK/3
3. 118295_N(UniMod:7)GFYPATR/2

These peptides and corresponding pairs are removed. Following this, 407 peptide pairs are plotted to visualize elution-order reversal cases (see the Supplemental Table 20 and Supplemental Figure 16). Out of 406 high m-scoring peptides, 237 peptides contributed towards it (Supplemental Table 21).

### S7: Incorrect annotation by OpenSWATH

Following peptides are not aligned within 19.6 sec for pair “run4_run23” out of 406 peptides:

1. 34143_FQNALIVR/2
2. 115879_DDNPNLPR/2
3. 118295_N(UniMod:7)GFYPATR/2
4. 46713_M(UniMod:35)KPVPDLVPGNFK/3
5. 119603_M(UniMod:35)KPVPDLVPGNFK/3
6. 116251_Q(UniMod:7)TALVELVK/2
7. 119597_MELERPGGN(UniMod:7)EITR/3
8. 117986_QQNAQGGFSSTQDTVVALHALSK/3

The chromatogram alignment for these peptides is presented in the Supplemental Figure 18.
