## Supplemental Figures and Table titles for "DIAlignR provides precise retention time alignment across distant runs in DIA and targeted proteomics"

| | |
|---|---|
| Supplemental Figure 1 | Distribution of signal-to-noise ratio of peaks and the number of training precursors used for LOESS fit across 120 pairs in benchmarking dataset |
| Supplemental Figure 2 | Effect of number of training point on Residual Standard Error of LOESS fit |
| Supplemental Figure 3 | An example of LOESS fit between two runs using common precursors |
| Supplemental Figure 4 | Distribution of Euclidean distance, cosine and correlation similarity scores for an example run-pair |
| Supplemental Figure 5 | Distribution of covariance and dot-product similarity scores for an example run-pair |
| Supplemental Figure 6 | Alignment path through chromatogram similarity matrix for covariance and dot-product similarity measure |
| Supplemental Figure 7 | Effect of masking parameters of dotProductMasked similarity on the proportion of aligned peaks |
| Supplemental Figure 8 | Effect of similarity-matrix penalizing parameters on the proportion of aligned peaks |
| Supplemental Figure 9 | Effect of gap-opening and gap-extension factors of dynamic programming on the proportion of aligned peaks |
| Supplemental Figure 10 | The fraction of peptides aligned with varying number of fragment-ions and without penalized similarity matrix |
| Supplemental Figure 11 | Effect of penalizing similarity matrix on the alignment of example chromatograms |
| Supplemental Figure 12 | Errors of reported RTs plotted along the runtime and boxplot of AUC of alignment of each run-pair |
| Supplemental Figure 13 | Difference of number of peptides aligned within certain RT error for each run-pair. Scatter plot and histogram of RT error for peptide |
| Supplemental Figure 14 | Errors of reported RTs plotted along the runtime and boxplot of AUC of alignment of each run-pair from human plasma dataset |
| Supplemental Figure 15 | The fraction of peptides aligned before certain RT error for each run-pair from human plasma dataset |
| Supplemental Figure 16 | Chromatograms of 407 peptide-pairs exhibiting peak-switching between highly heterogeneous plasma runs “run4” and “run23” |

| | |
|---|---|
| Supplemental Figure 17 | Chromatograms of 237 peptides which are aligned precisely with DIAlignR compared to LOESS |
| Supplemental Figure 18 | Chromatograms of eight peptides annotated incorrectly by OpenSWATH but aligned correctly by DIAlignR |
| Supplemental Figure 19 | An example alignment with LOESS and DIAlignR of peptide 42259_ALVQQMEQLR/2 from the human plasma dataset |

### Supplemental Tables

| | |
|---|---|
| Supplemental Table 1 | Annotation of the retention time of 437 precursors across 16 runs from the benchmarking dataset |
| Supplemental Table 2 | Description of 24 runs from human plasma dataset |
| Supplemental Table 3 | OpenSWATH annotation of the retention time of 406 precursors across 24 runs from the human plasma dataset |
| Supplemental Table 4 | Summary description of validation and human plasma datasets |
| Supplemental Table 5 | Number of common features, optimum span value and RSE for 120 pairs from benchmarking dataset |
| Supplemental Table 6 | Number of common features, optimum span value and RSE for 276 pairs from the human plasma dataset |
| Supplemental Table 7 | Percentage of aligned peaks cumulatively from benchmarking dataset with different similarity measures |
| Supplemental Table 8 | AUC of the fraction of aligned peaks from benchmarking dataset with different similarity measures |
| Supplemental Table 9 | Summary of number of peaks aligned for a combination of masking parameters |
| Supplemental Table 10 | Summary of AUC of a fraction of aligned peaks for a combination of masking parameters |
| Supplemental Table 11 | Summary of number of peaks aligned for a combination of multiple gap-opening and gap-extension factors |
| Supplemental Table 12 | AUC (of a fraction of peaks aligned within certain RT tolerance) for a combination of multiple gap-opening and gap-extension factors |
| Supplemental Table 13 | Summary of number of peaks aligned for multiple widths of a no-penalty window through similarity matrix |
| Supplemental Table 14 | Summary of number of peaks aligned for different penalizing gradients through similarity matrix |
| Supplemental Table 15 | Summary of number of peaks aligned for different number of transitions |
| Supplemental Table 16 | RT Alignment Error by LOESS for peaks from benchmarking dataset |
| Supplemental Table 17 | RT Alignment Error by DIAlignR for peaks from benchmarking dataset |

| | |
|---|---|
| Supplemental Table 18 | RT Alignment Error by LOESS for peaks from human plasma dataset |
| Supplemental Table 19 | RT Alignment Error by DIAlignR for peaks from human plasma dataset |
| Supplemental Table 20 | IDs of constituent precursors of 437 peptide-pairs that exhibit peak-switching |
| Supplemental Table 21 | IDs of 237 precursors which exhibit peak-switching |
