## Supplemental Figures for "DIAlignR provides precise retention time alignment across distant runs in DIA and targeted proteomics": Supplemental Figure15.pdf

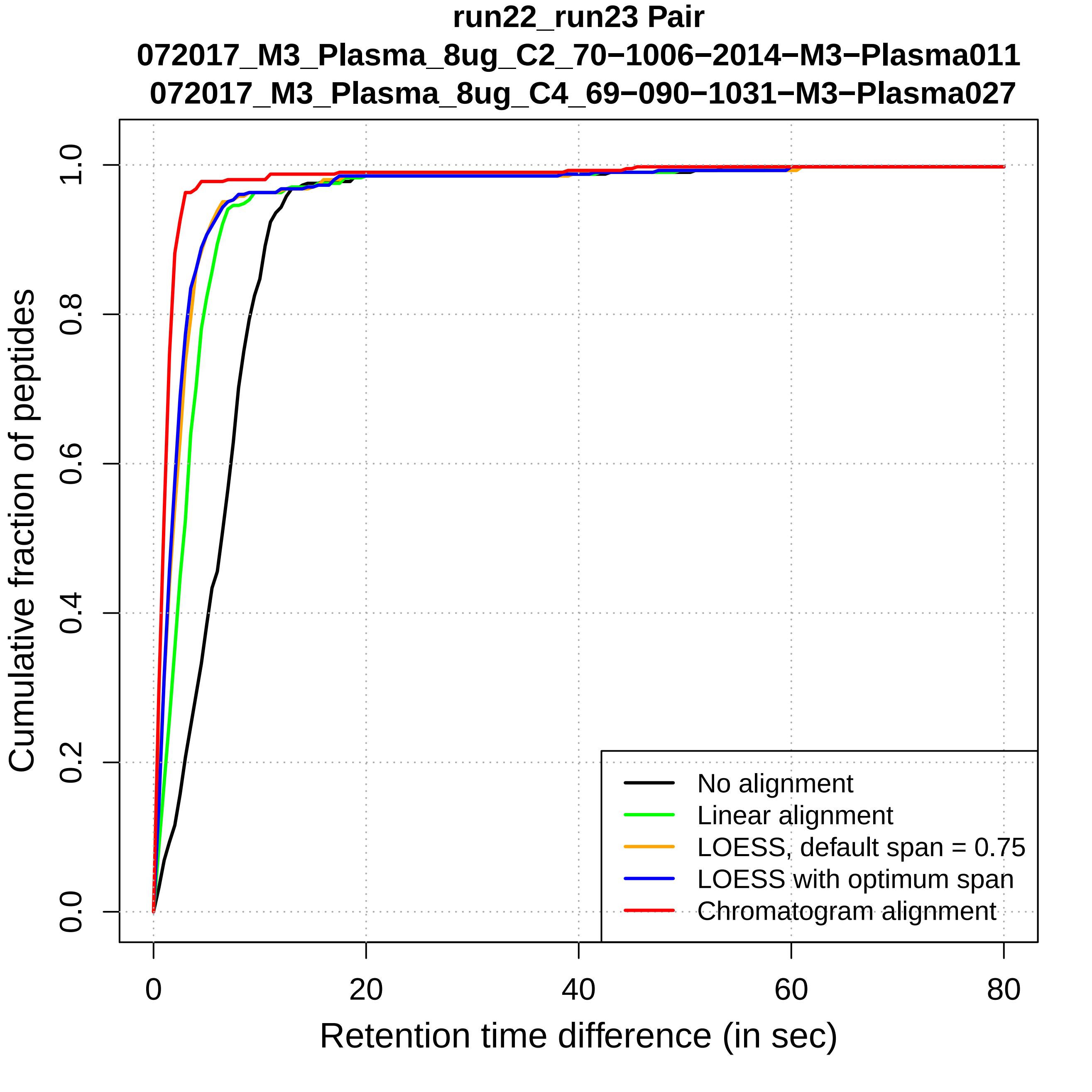
run22_run23 Pair
072017_M3_Plasma_8ug_C2_70−1006−2014−M3−Plasma011
072017_M3_Plasma_8ug_C4_69−090−1031−M3−Plasma027
Cumulative fraction of peptides
0.0
0.2
0.4
0.6
0.8
1.0
0
20
40
60
80
Retention time difference (in sec)
No alignment
Linear alignment
LOESS, default span = 0.75
LOESS with optimum span
Chromatogram alignment

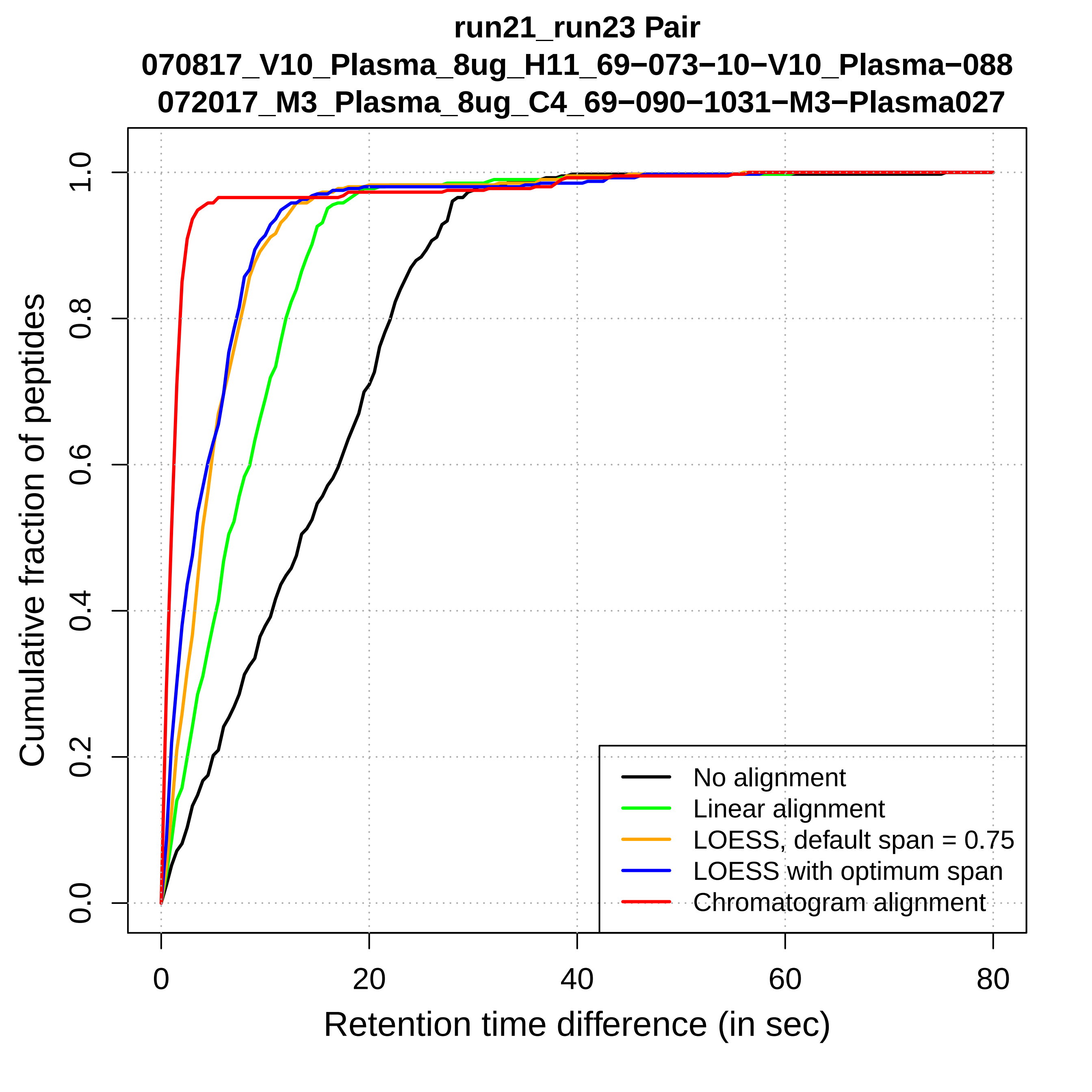
run21_run23 Pair
070817_V10_Plasma_8ug_H11_69−073−10−V10_Plasma−088
072017_M3_Plasma_8ug_C4_69−090−1031−M3−Plasma027
Cumulative fraction of peptides
Retention time difference (in sec)
0.0
0.2
0.4
0.6
0.8
1.0
0
20
40
60
80
No alignment
Linear alignment
LOESS, default span = 0.75
LOESS with optimum span
Chromatogram alignment

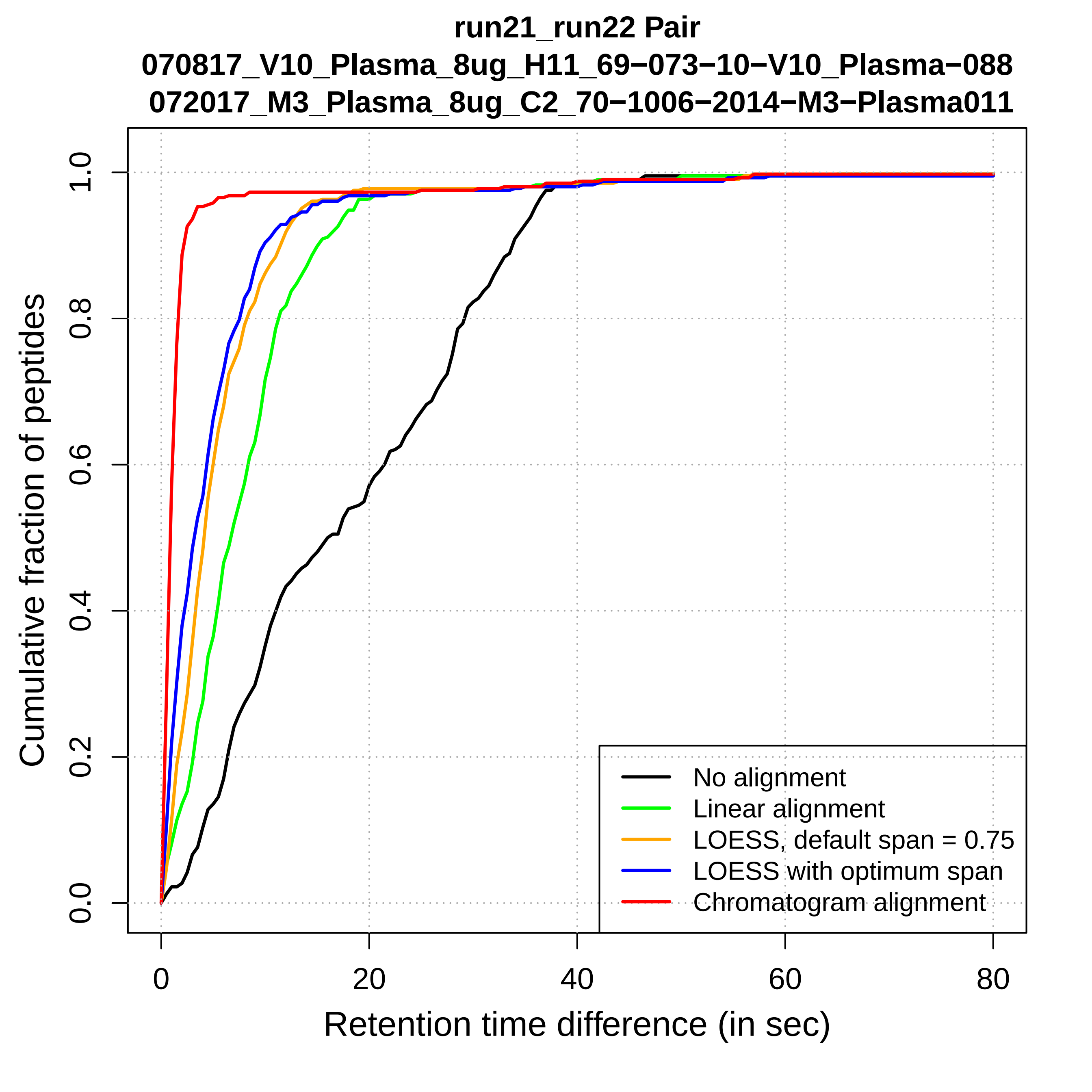

run21_run22 Pair
070817_V10_Plasma_8ug_H11_69−073−10−V10_Plasma−088
072017_M3_Plasma_8ug_C2_70−1006−2014−M3−Plasma011
Cumulative fraction of peptides
0.0
0.2
0.4
0.6
0.8
1.0
0
20
40
60
80
Retention time difference (in sec)
No alignment
Linear alignment
LOESS, default span = 0.75
LOESS with optimum span
Chromatogram alignment

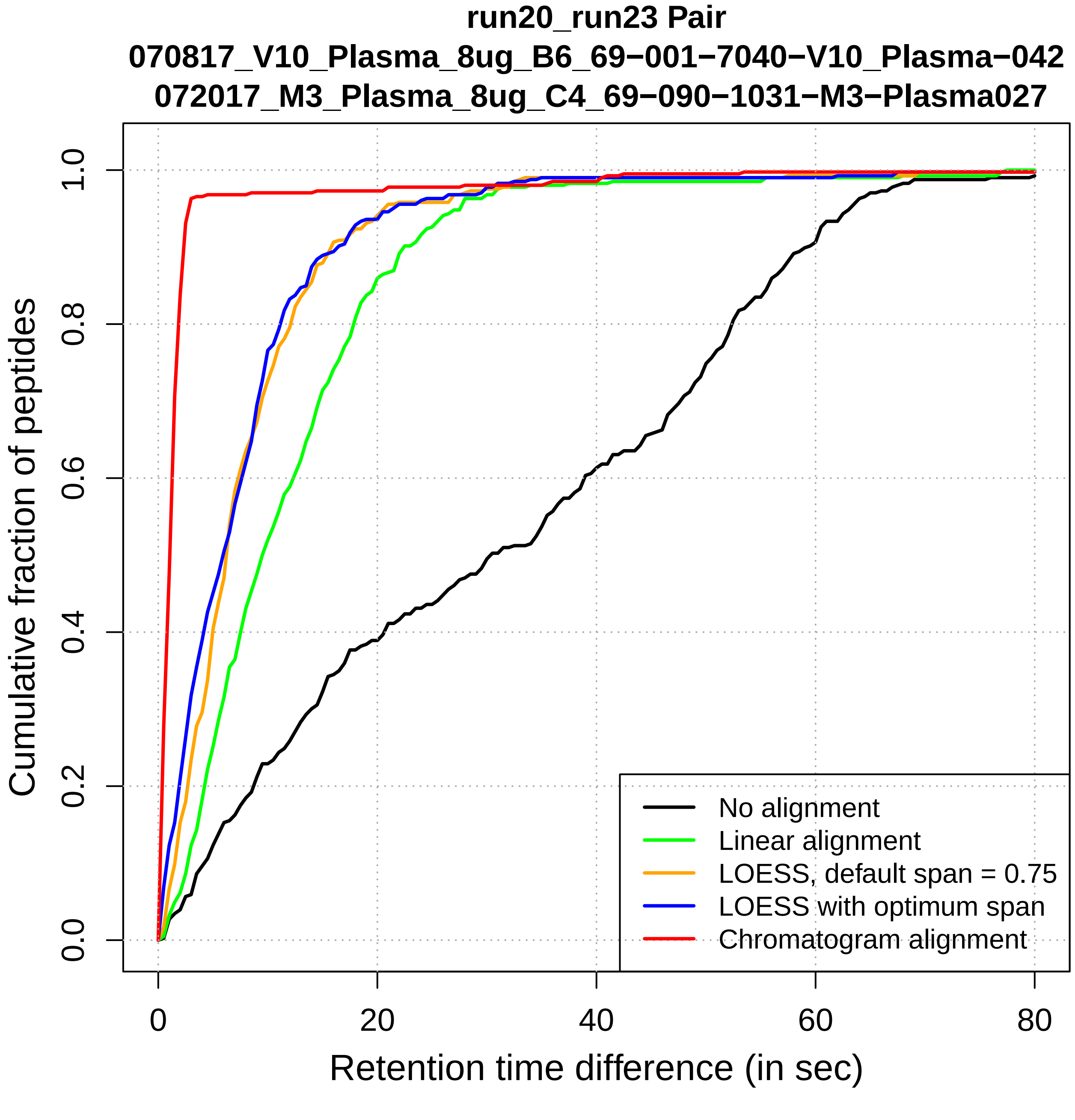
run20_run23 Pair
070817_V10_Plasma_8ug_B6_69−001−7040−V10_Plasma−042
072017_M3_Plasma_8ug_C4_69−090−1031−M3−Plasma027
Cumulative fraction of peptides
0.0
0.2
0.4
0.6
0.8
1.0
0
20
40
60
80
Retention time difference (in sec)
No alignment
Linear alignment
LOESS, default span = 0.75
LOESS with optimum span
Chromatogram alignment

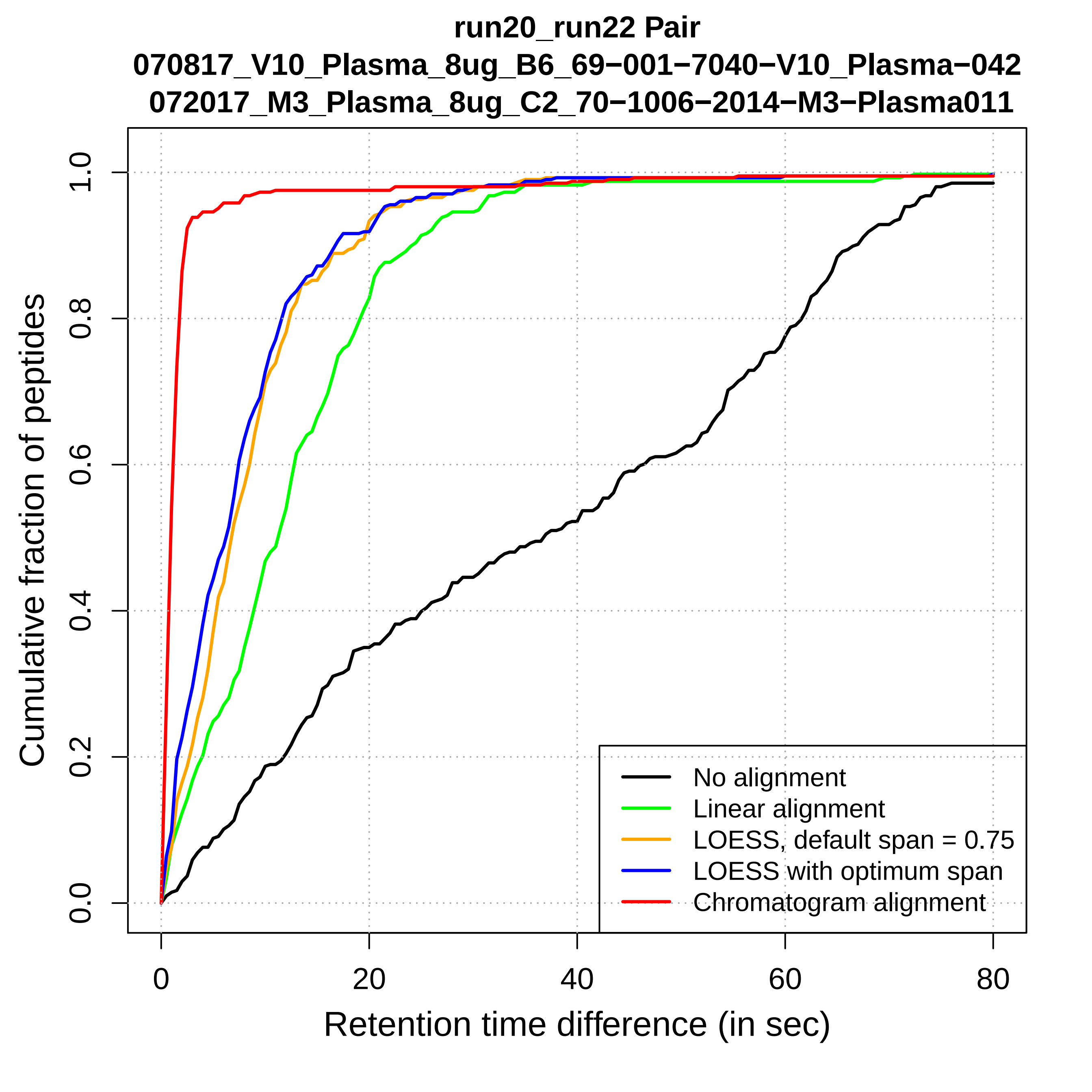
run20_run22 Pair
070817_V10_Plasma_8ug_B6_69−001−7040−V10_Plasma−042
072017_M3_Plasma_8ug_C2_70−1006−2014−M3−Plasma011
Cumulative fraction of peptides
Retention time difference (in sec)
0.0
0.2
0.4
0.6
0.8
1.0
0
20
40
60
80
No alignment
Linear alignment
LOESS, default span = 0.75
LOESS with optimum span
Chromatogram alignment

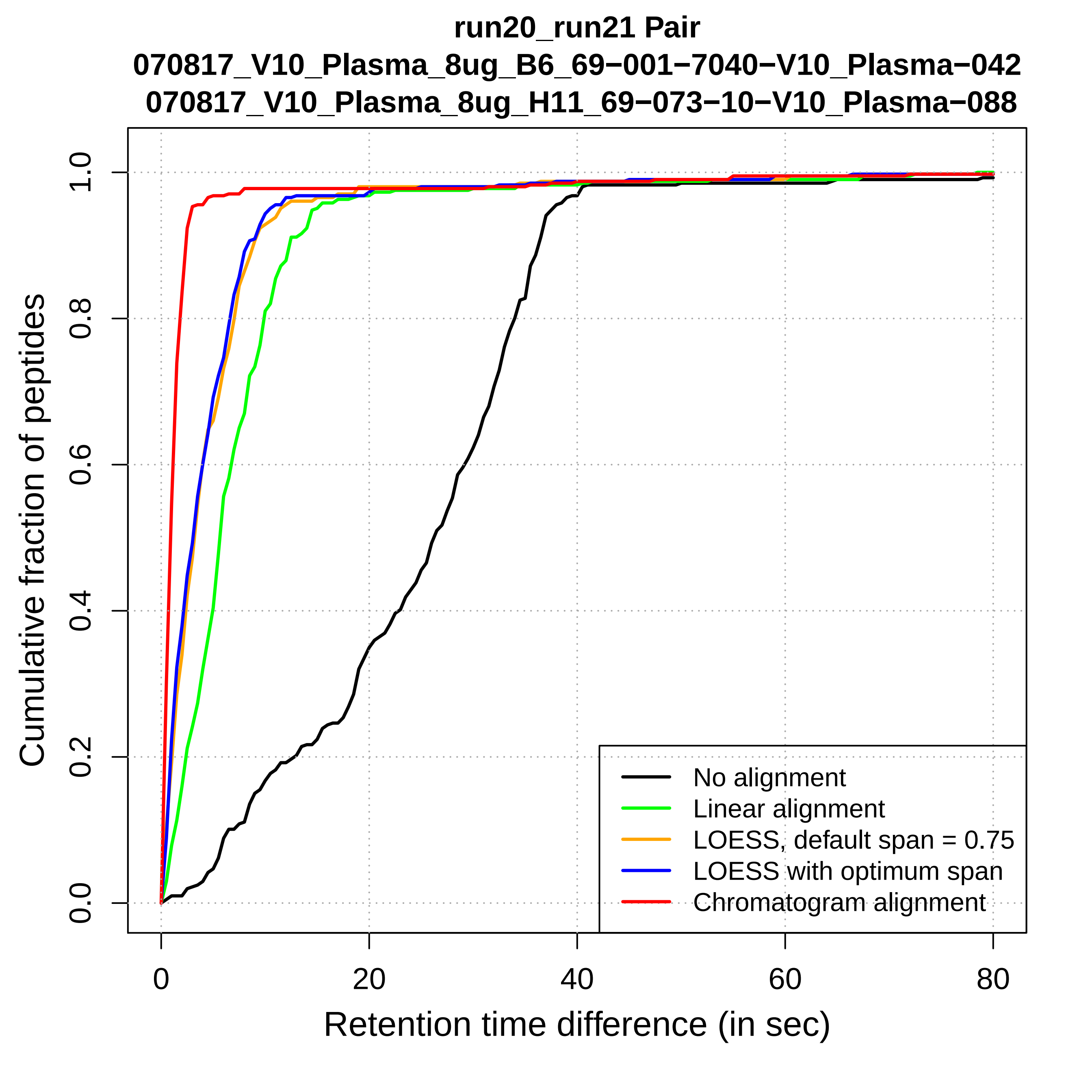

run20_run21 Pair
070817_V10_Plasma_8ug_B6_69−001−7040−V10_Plasma−042
070817_V10_Plasma_8ug_H11_69−073−10−V10_Plasma−088
Cumulative fraction of peptides
0.0
0.2
0.4
0.6
0.8
1.0
0
20
40
60
80
Retention time difference (in sec)
No alignment
Linear alignment
LOESS, default span = 0.75
LOESS with optimum span
Chromatogram alignment

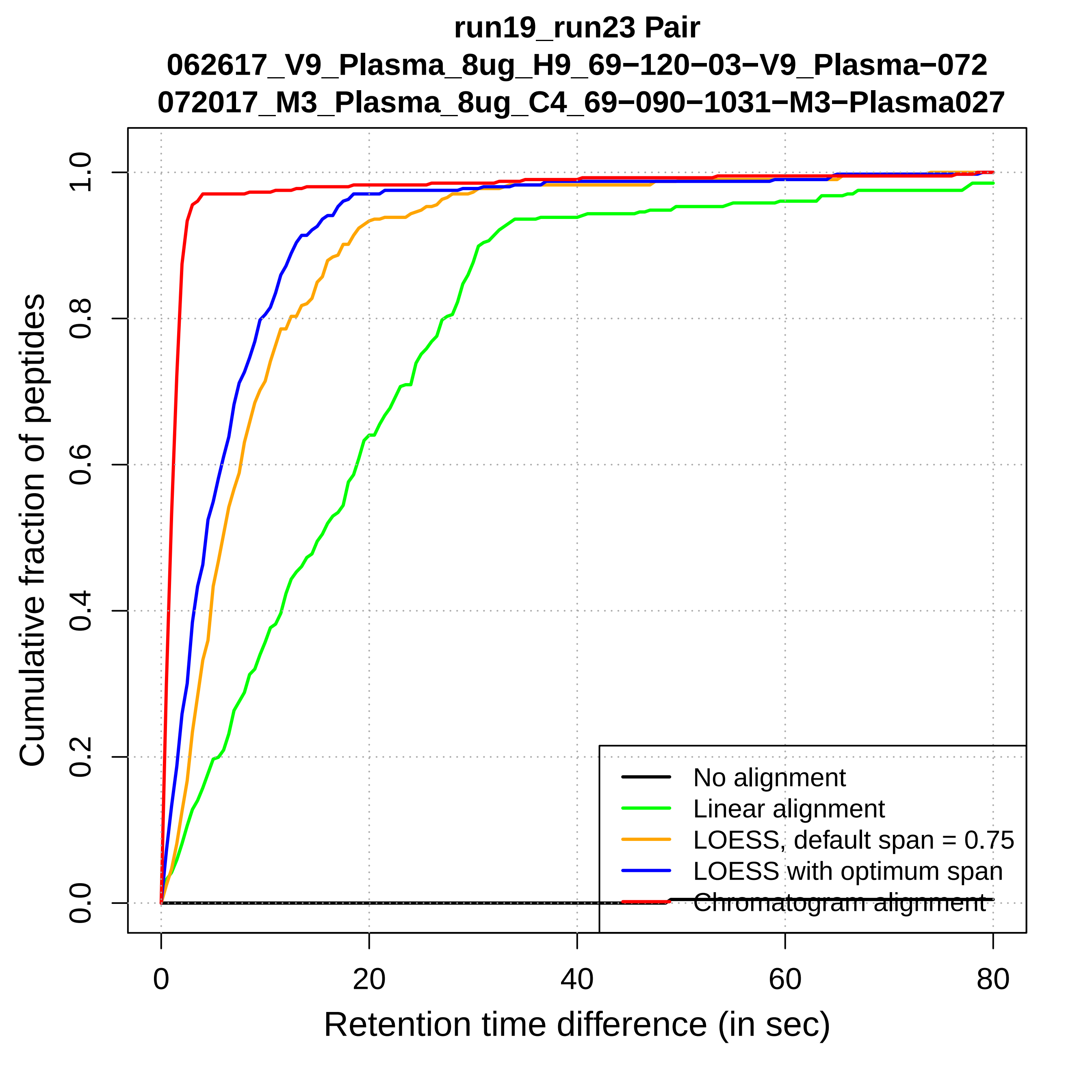

run19_run23 Pair
062617_V9_Plasma_8ug_H9_69−120−03−V9_Plasma−072
072017_M3_Plasma_8ug_C4_69−090−1031−M3−Plasma027
Cumulative fraction of peptides
0.0
0.2
0.4
0.6
0.8
1.0
0
20
40
60
80
Retention time difference (in sec)
No alignment
Linear alignment
LOESS, default span = 0.75
LOESS with optimum span
Chromatogram alignment

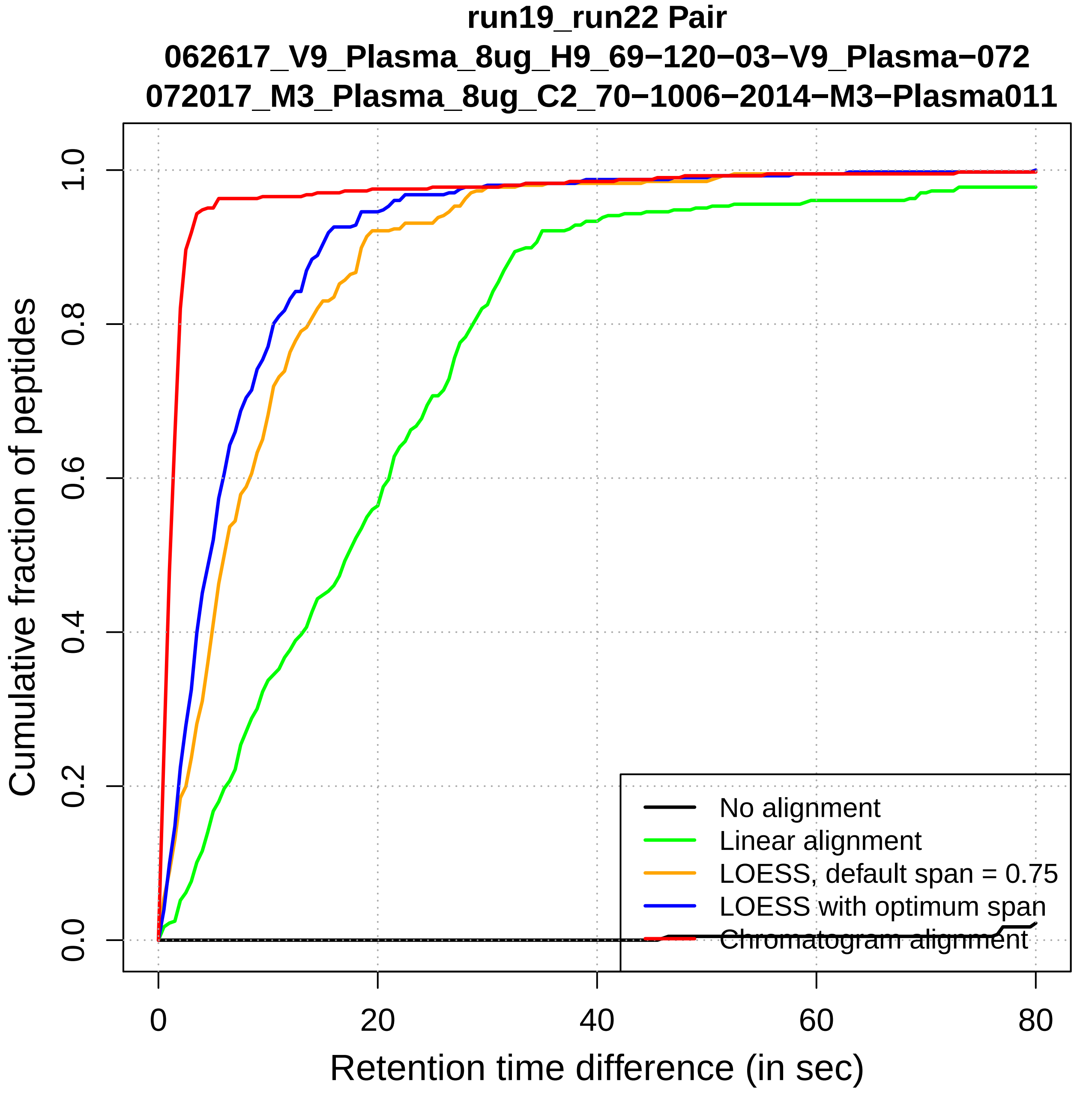

run19_run22 Pair
062617_V9_Plasma_8ug_H9_69−120−03−V9_Plasma−072
072017_M3_Plasma_8ug_C2_70−1006−2014−M3−Plasma011
Cumulative fraction of peptides
0.0
0.2
0.4
0.6
0.8
1.0
0
20
40
60
80
Retention time difference (in sec)
No alignment
Linear alignment
LOESS, default span = 0.75
LOESS with optimum span
Chromatogram alignment

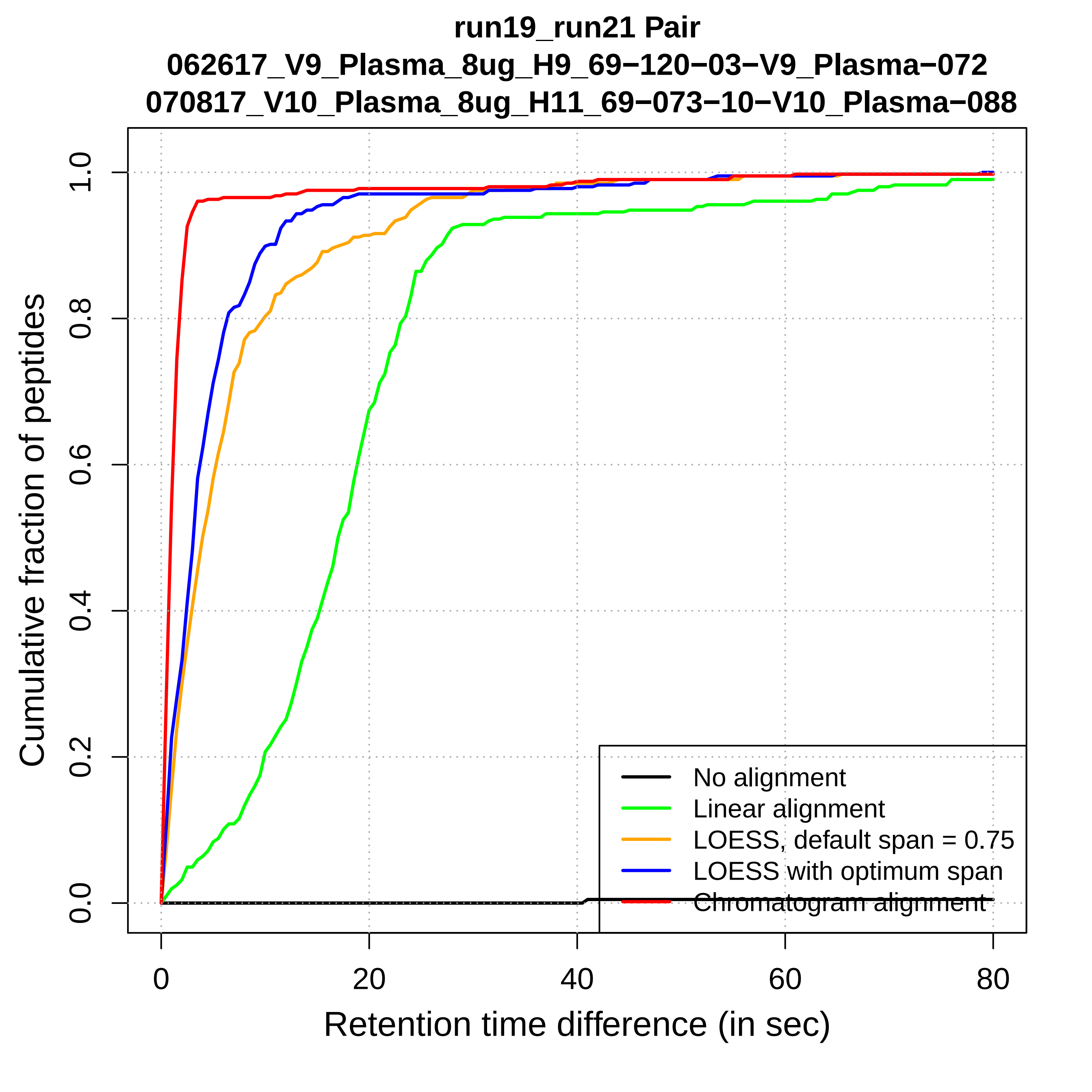
run19_run21 Pair
062617_V9_Plasma_8ug_H9_69−120−03−V9_Plasma−072
070817_V10_Plasma_8ug_H11_69−073−10−V10_Plasma−088
Cumulative fraction of peptides
0.0
0.2
0.4
0.6
0.8
1.0
0
20
40
60
80
Retention time difference (in sec)
No alignment
Linear alignment
LOESS, default span = 0.75
LOESS with optimum span
Chromatogram alignment

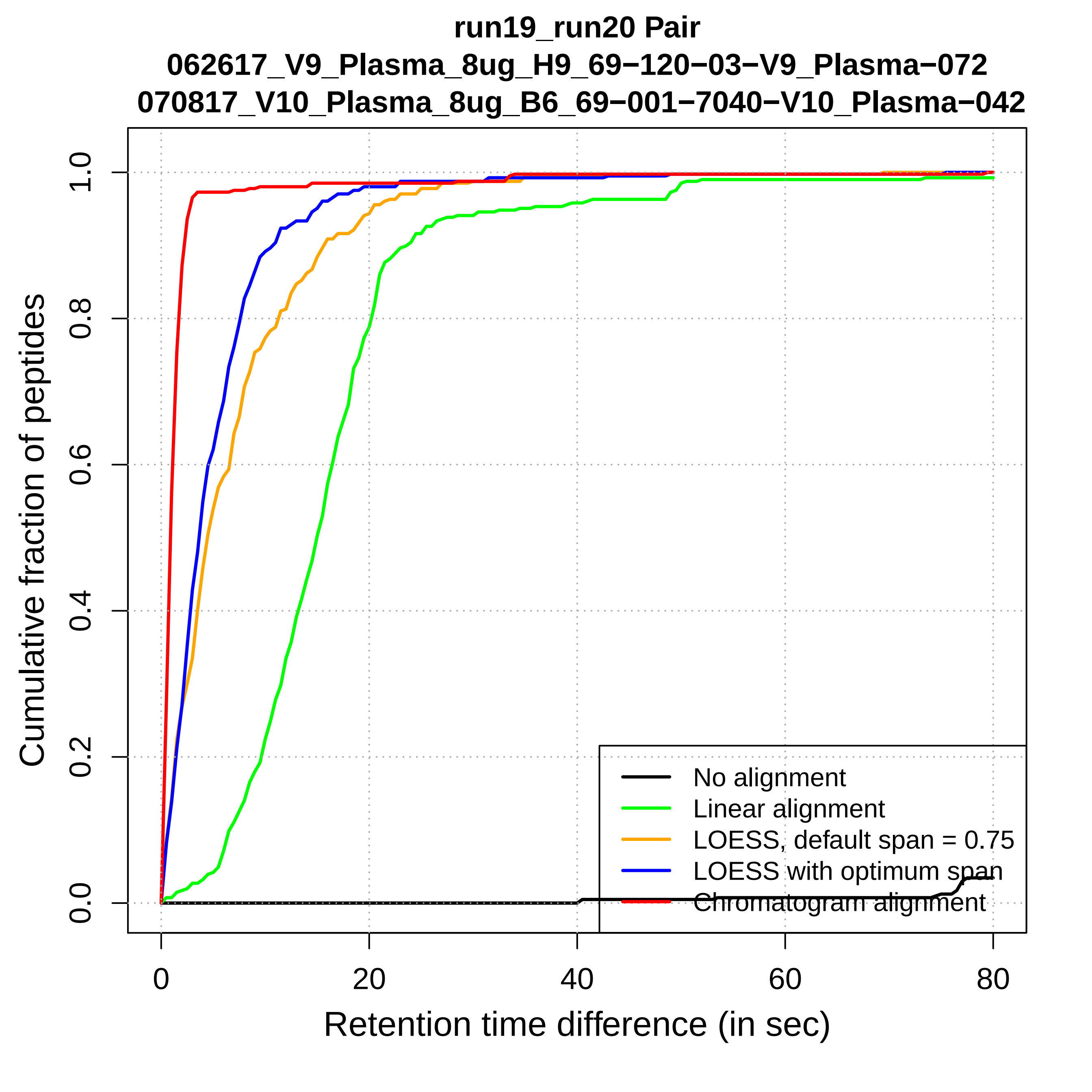

run19_run20 Pair
062617_V9_Plasma_8ug_H9_69−120−03−V9_Plasma−072
070817_V10_Plasma_8ug_B6_69−001−7040−V10_Plasma−042
Cumulative fraction of peptides
0.0
0.2
0.4
0.6
0.8
1.0
0
20
40
60
80
Retention time difference (in sec)
No alignment
Linear alignment
LOESS, default span = 0.75
LOESS with optimum span
Chromatogram alignment

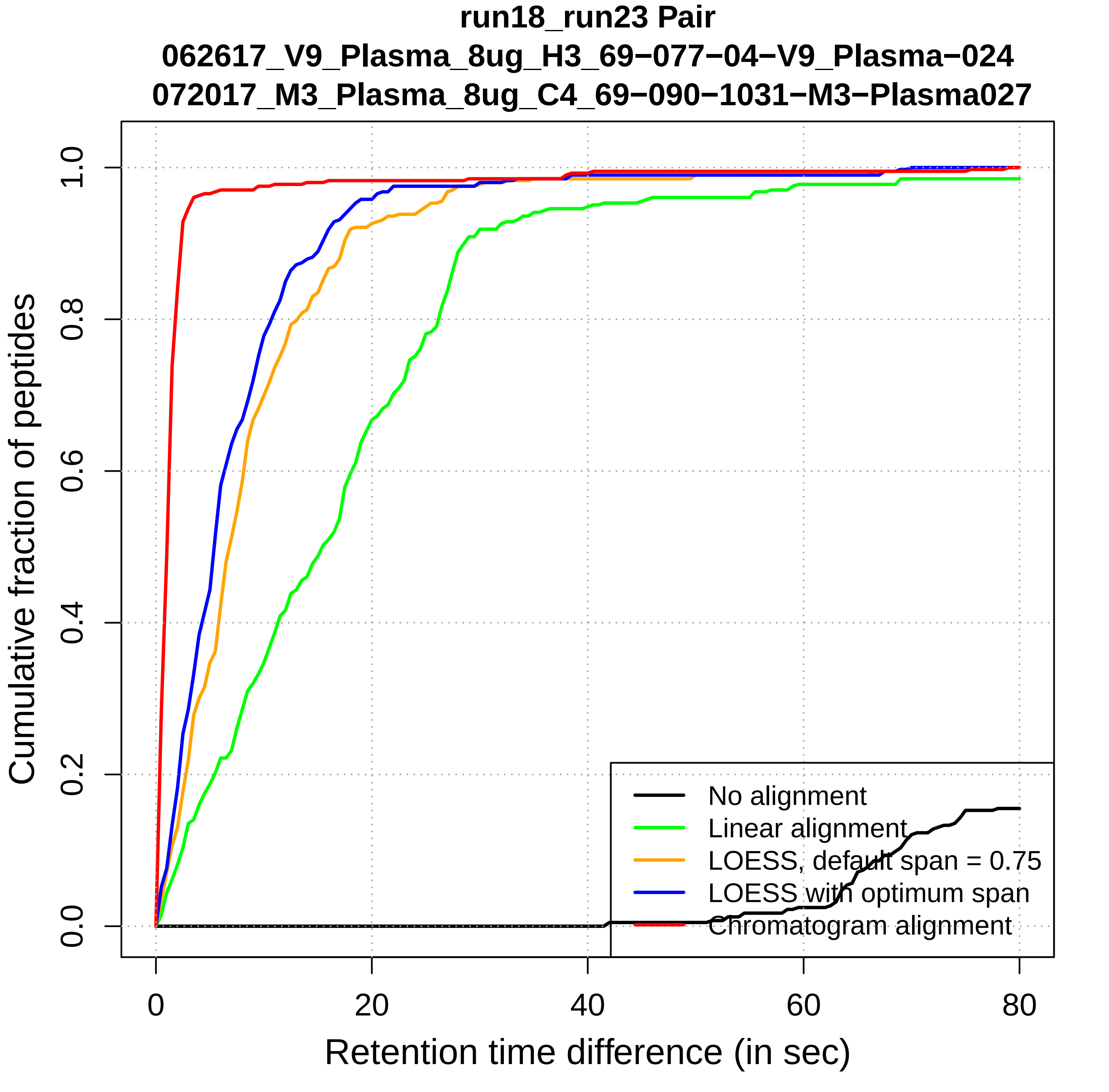
run18_run23 Pair
062617_V9_Plasma_8ug_H3_69−077−04−V9_Plasma−024
072017_M3_Plasma_8ug_C4_69−090−1031−M3−Plasma027
Cumulative fraction of peptides
0.0
0.2
0.4
0.6
0.8
1.0
0
20
40
60
80
Retention time difference (in sec)
No alignment
Linear alignment
LOESS, default span = 0.75
LOESS with optimum span
Chromatogram alignment

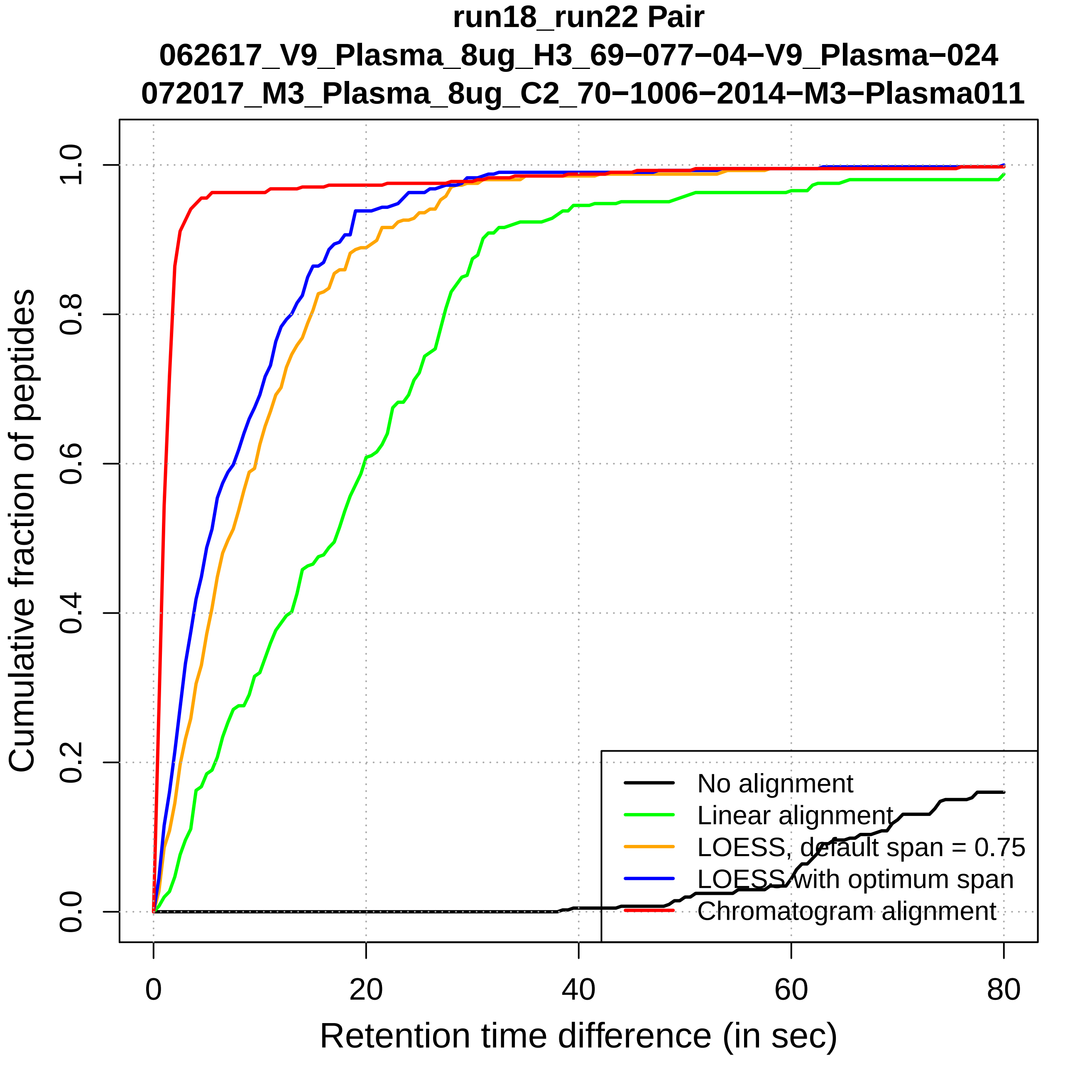
run18_run22 Pair
062617_V9_Plasma_8ug_H3_69−077−04−V9_Plasma−024
072017_M3_Plasma_8ug_C2_70−1006−2014−M3−Plasma011
Cumulative fraction of peptides
0.0
0.2
0.4
0.6
0.8
1.0
0
20
40
60
80
Retention time difference (in sec)
No alignment
Linear alignment
LOESS, default span = 0.75
LOESS with optimum span
Chromatogram alignment

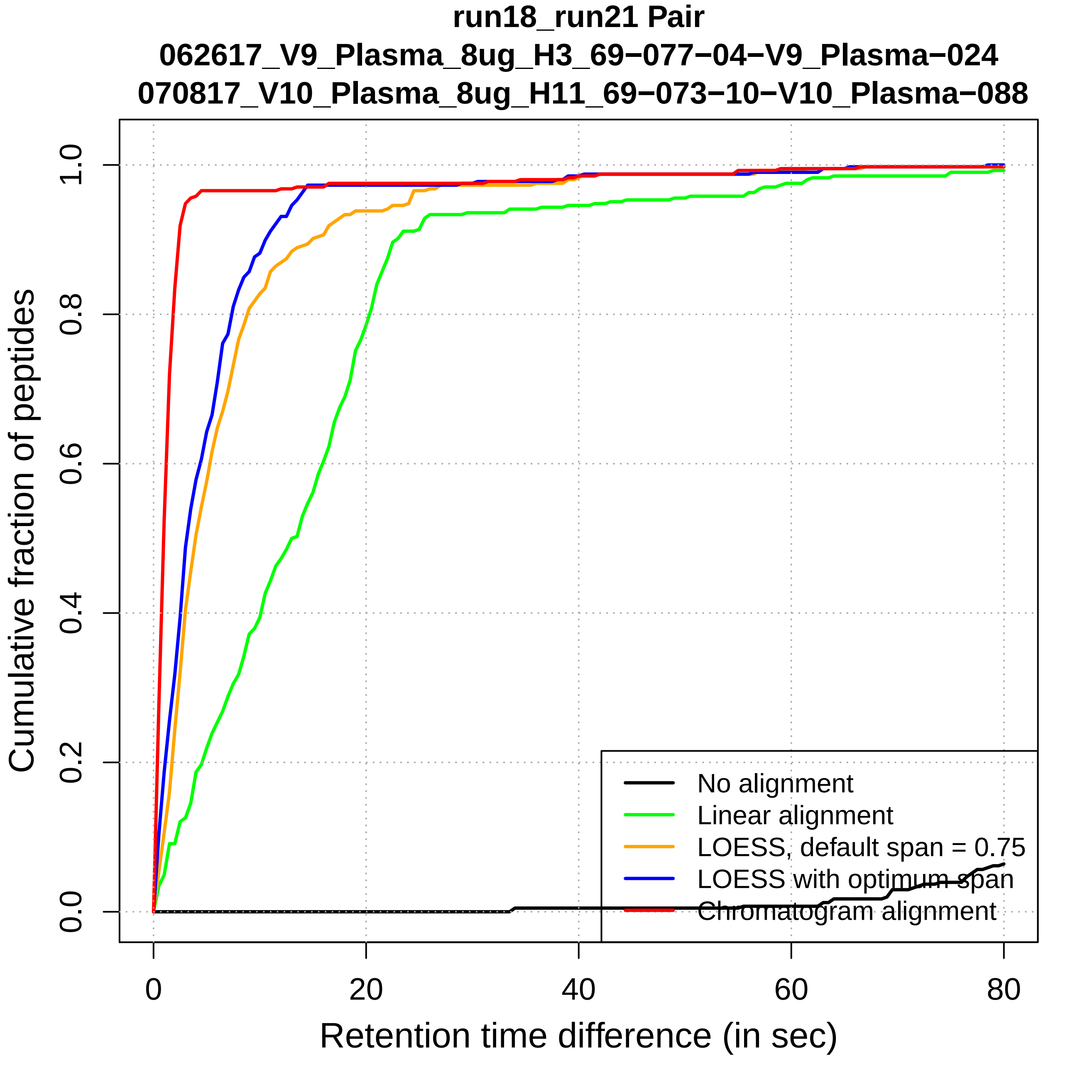

run18_run21 Pair
062617_V9_Plasma_8ug_H3_69−077−04−V9_Plasma−024
070817_V10_Plasma_8ug_H11_69−073−10−V10_Plasma−088
Cumulative fraction of peptides
0.0
0.2
0.4
0.6
0.8
1.0
0
20
40
60
80
Retention time difference (in sec)
No alignment
Linear alignment
LOESS, default span = 0.75
LOESS with optimum span
Chromatogram alignment

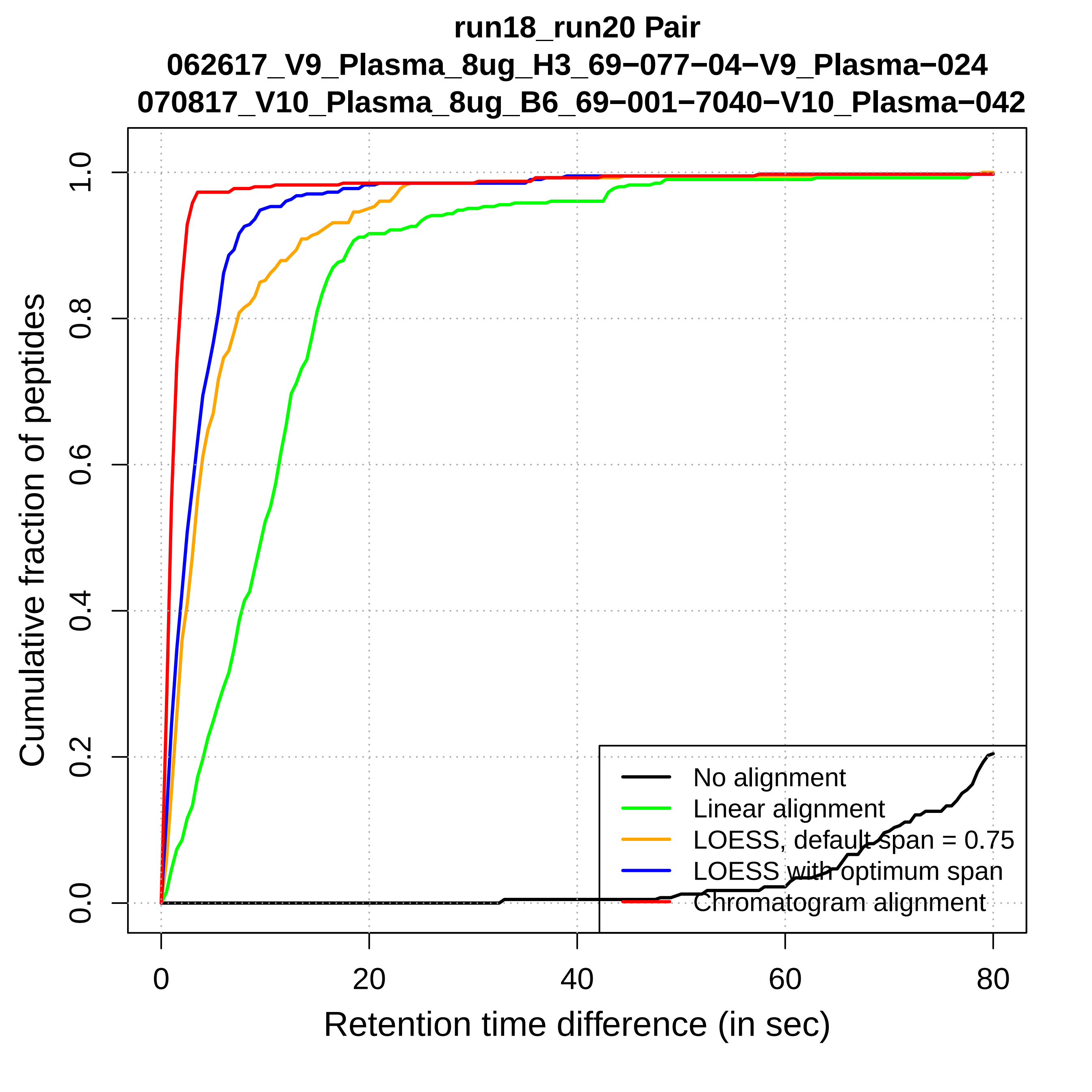

run18_run20 Pair
062617_V9_Plasma_8ug_H3_69−077−04−V9_Plasma−024
070817_V10_Plasma_8ug_B6_69−001−7040−V10_Plasma−042
Cumulative fraction of peptides
0.0
0.2
0.4
0.6
0.8
1.0
0
20
40
60
80
Retention time difference (in sec)
No alignment
Linear alignment
LOESS, default span = 0.75
LOESS with optimum span
Chromatogram alignment

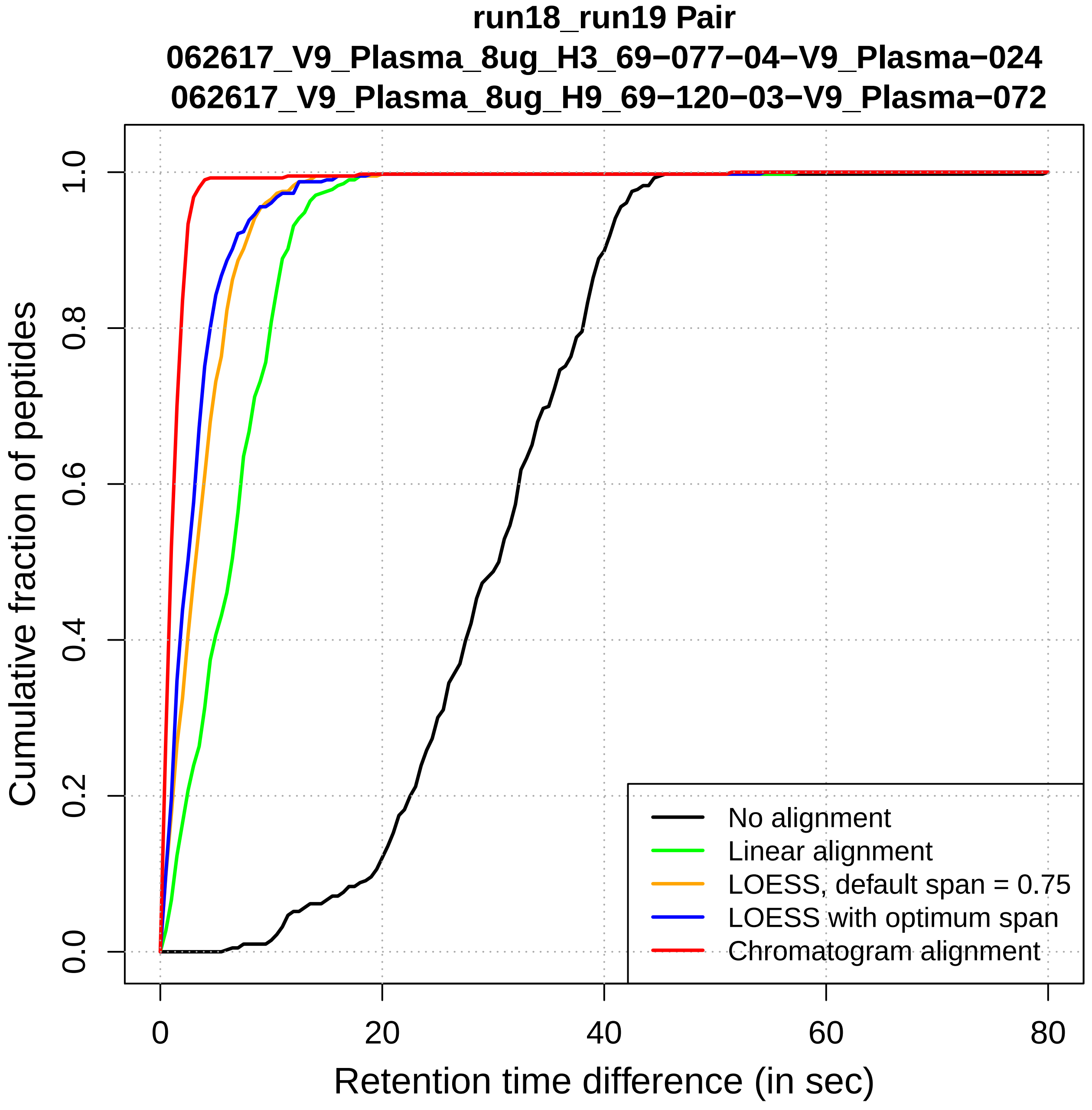
run18_run19 Pair
062617_V9_Plasma_8ug_H3_69−077−04−V9_Plasma−024
062617_V9_Plasma_8ug_H9_69−120−03−V9_Plasma−072
Cumulative fraction of peptides
0.0
0.2
0.4
0.6
0.8
1.0
0
20
40
60
80
Retention time difference (in sec)
No alignment
Linear alignment
LOESS, default span = 0.75
LOESS with optimum span
Chromatogram alignment

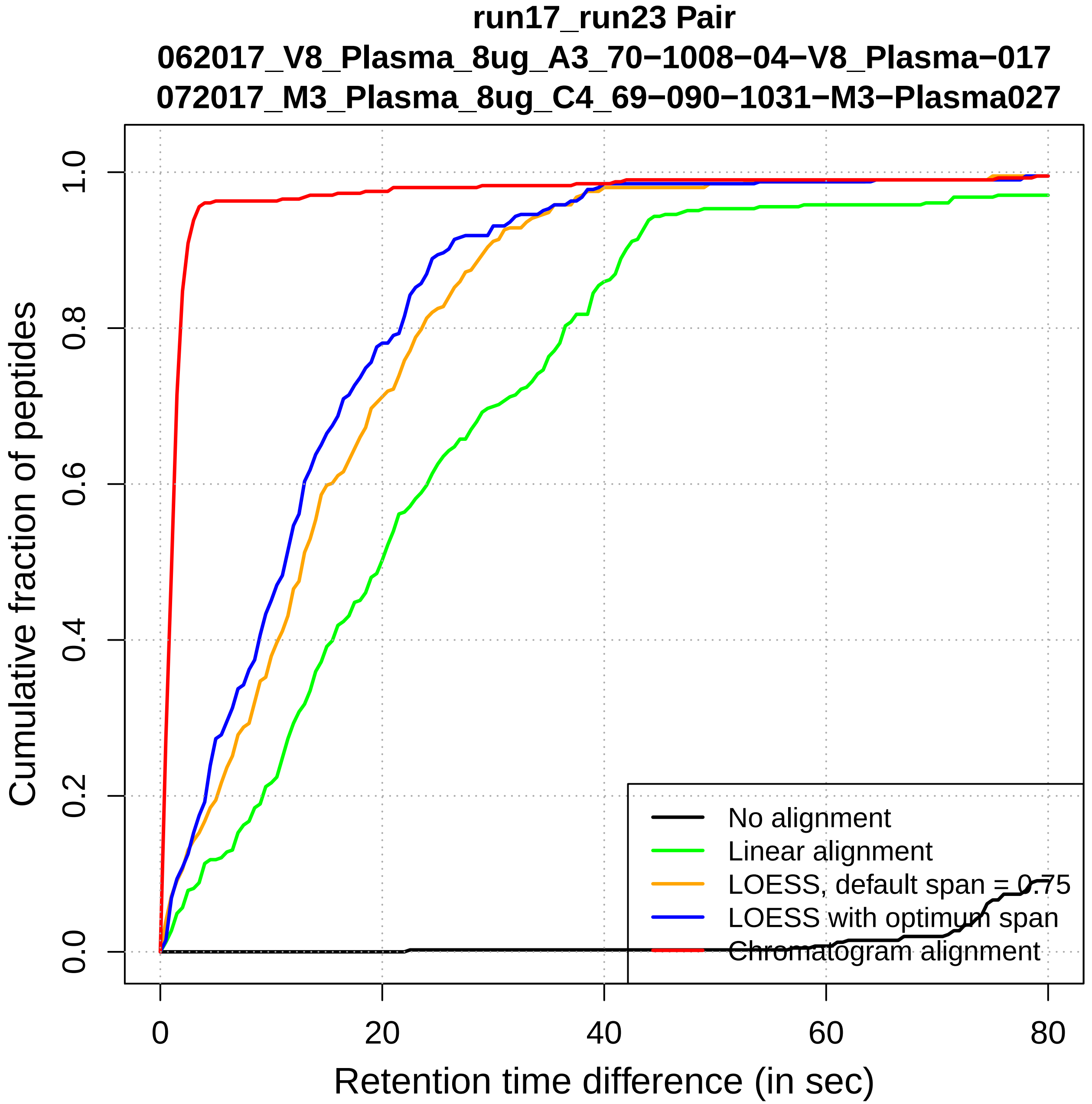
run17_run23 Pair
062017_V8_Plasma_8ug_A3_70−1008−04−V8_Plasma−017
072017_M3_Plasma_8ug_C4_69−090−1031−M3−Plasma027
Cumulative fraction of peptides
0.0
0.2
0.4
0.6
0.8
1.0
0
20
40
60
80
Retention time difference (in sec)
No alignment
Linear alignment
LOESS, default span = 0.75
LOESS with optimum span
Chromatogram alignment

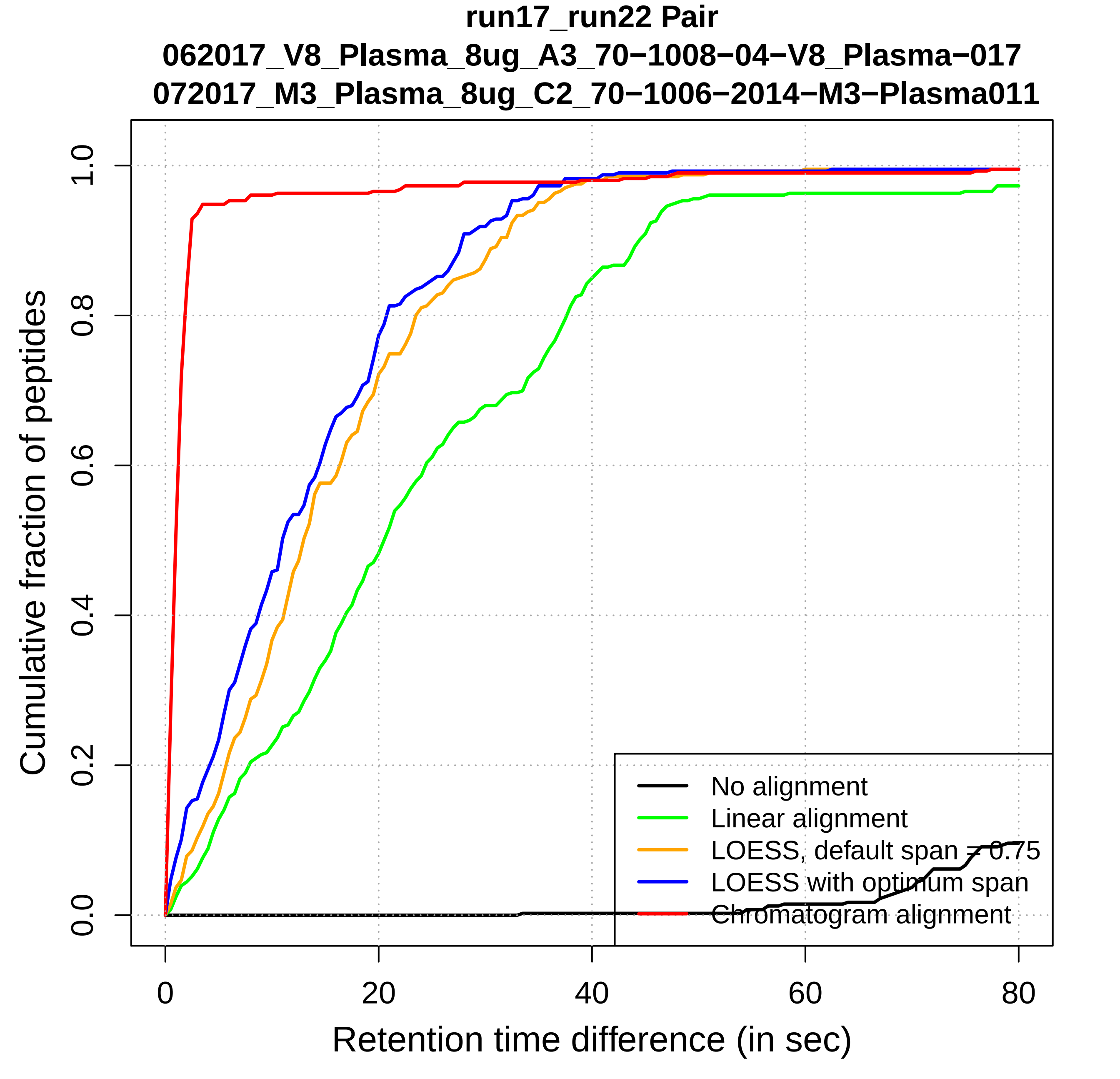
run17_run22 Pair
062017_V8_Plasma_8ug_A3_70−1008−04−V8_Plasma−017
072017_M3_Plasma_8ug_C2_70−1006−2014−M3−Plasma011
Cumulative fraction of peptides
0.0
0.2
0.4
0.6
0.8
1.0
0
20
40
60
80
Retention time difference (in sec)
No alignment
Linear alignment
LOESS, default span = 0.75
LOESS with optimum span
Chromatogram alignment

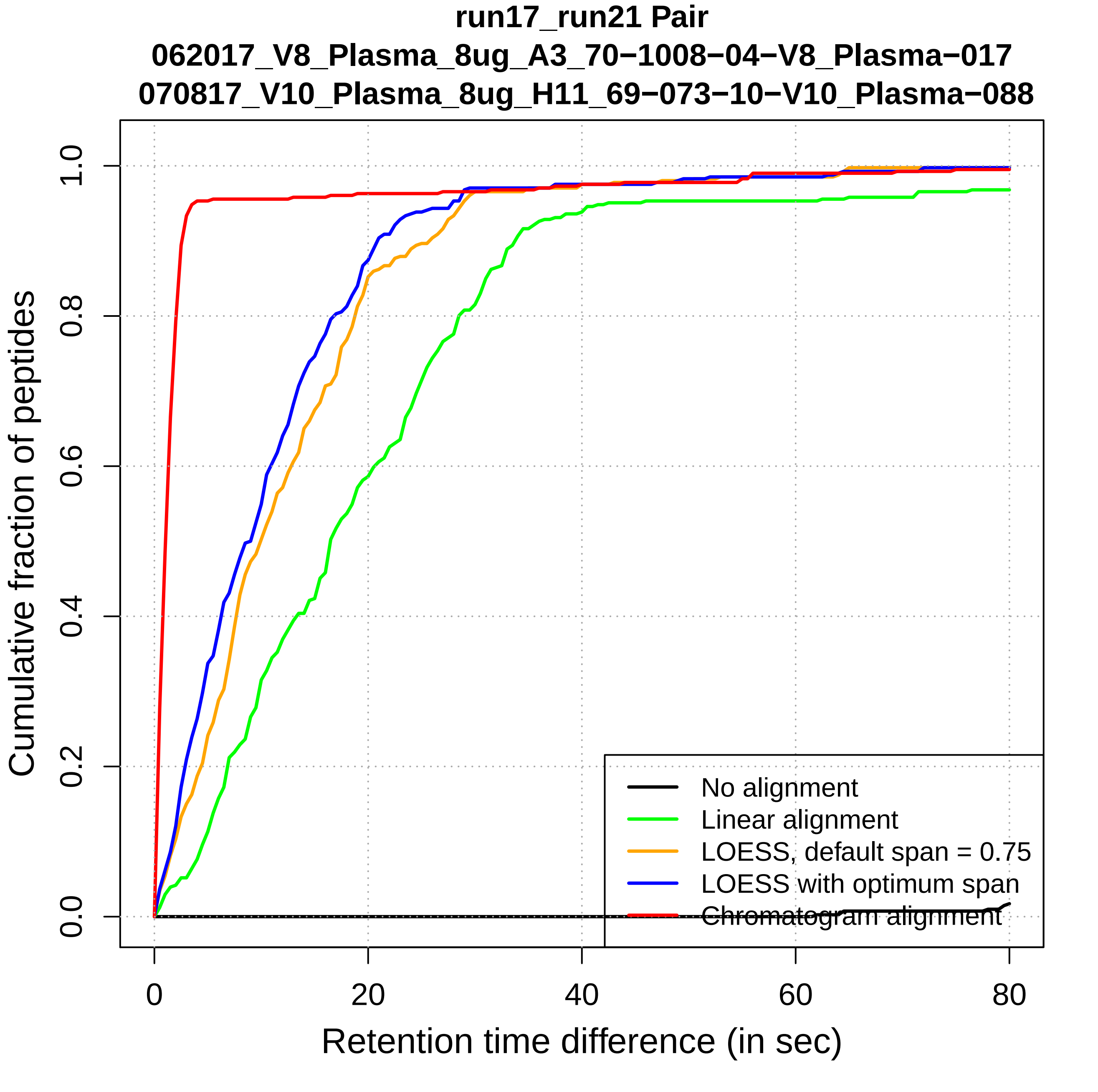
run17_run21 Pair
062017_V8_Plasma_8ug_A3_70−1008−04−V8_Plasma−017
070817_V10_Plasma_8ug_H11_69−073−10−V10_Plasma−088
Cumulative fraction of peptides
0.0
0.2
0.4
0.6
0.8
1.0
0
20
40
60
80
Retention time difference (in sec)
No alignment
Linear alignment
LOESS, default span = 0.75
LOESS with optimum span
Chromatogram alignment

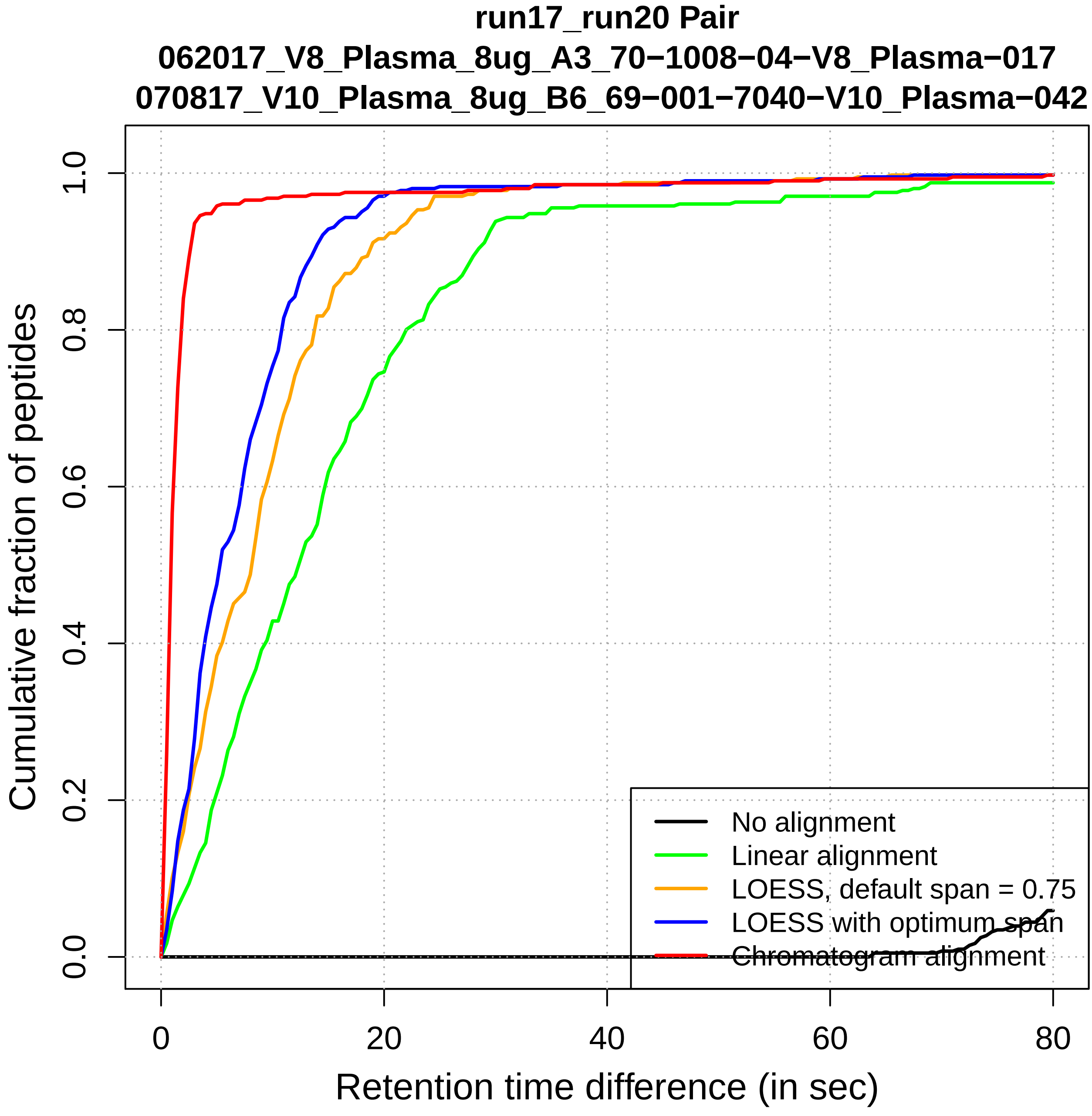

run17_run20 Pair
062017_V8_Plasma_8ug_A3_70−1008−04−V8_Plasma−017
070817_V10_Plasma_8ug_B6_69−001−7040−V10_Plasma−042
Cumulative fraction of peptides
0.0
0.2
0.4
0.6
0.8
1.0
0
20
40
60
80
Retention time difference (in sec)
No alignment
Linear alignment
LOESS, default span = 0.75
LOESS with optimum span
Chromatogram alignment

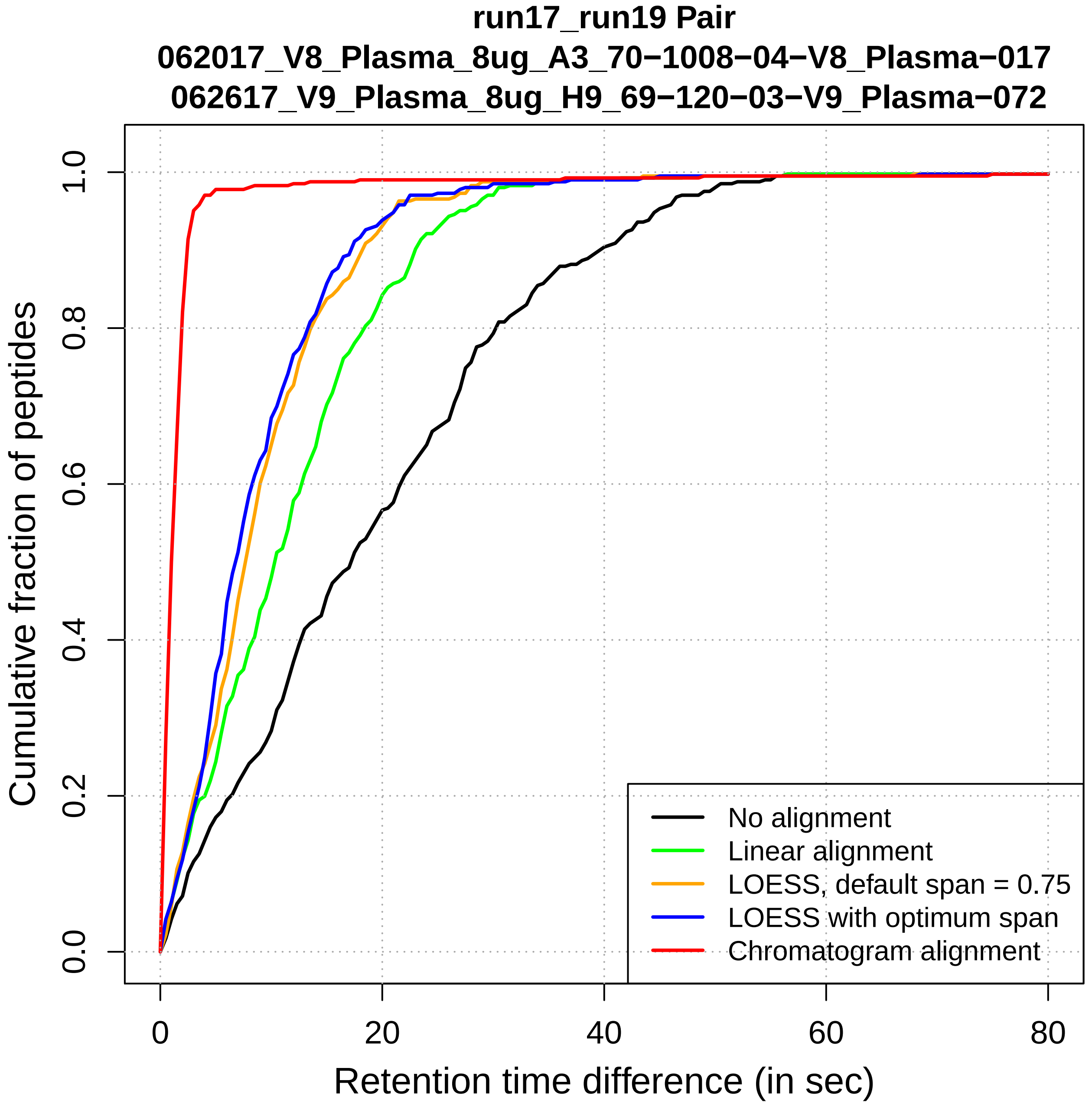
run17_run19 Pair
062017_V8_Plasma_8ug_A3_70−1008−04−V8_Plasma−017
062617_V9_Plasma_8ug_H9_69−120−03−V9_Plasma−072
Cumulative fraction of peptides
0.0
0.2
0.4
0.6
0.8
1.0
0
20
40
60
80
Retention time difference (in sec)
No alignment
Linear alignment
LOESS, default span = 0.75
LOESS with optimum span
Chromatogram alignment

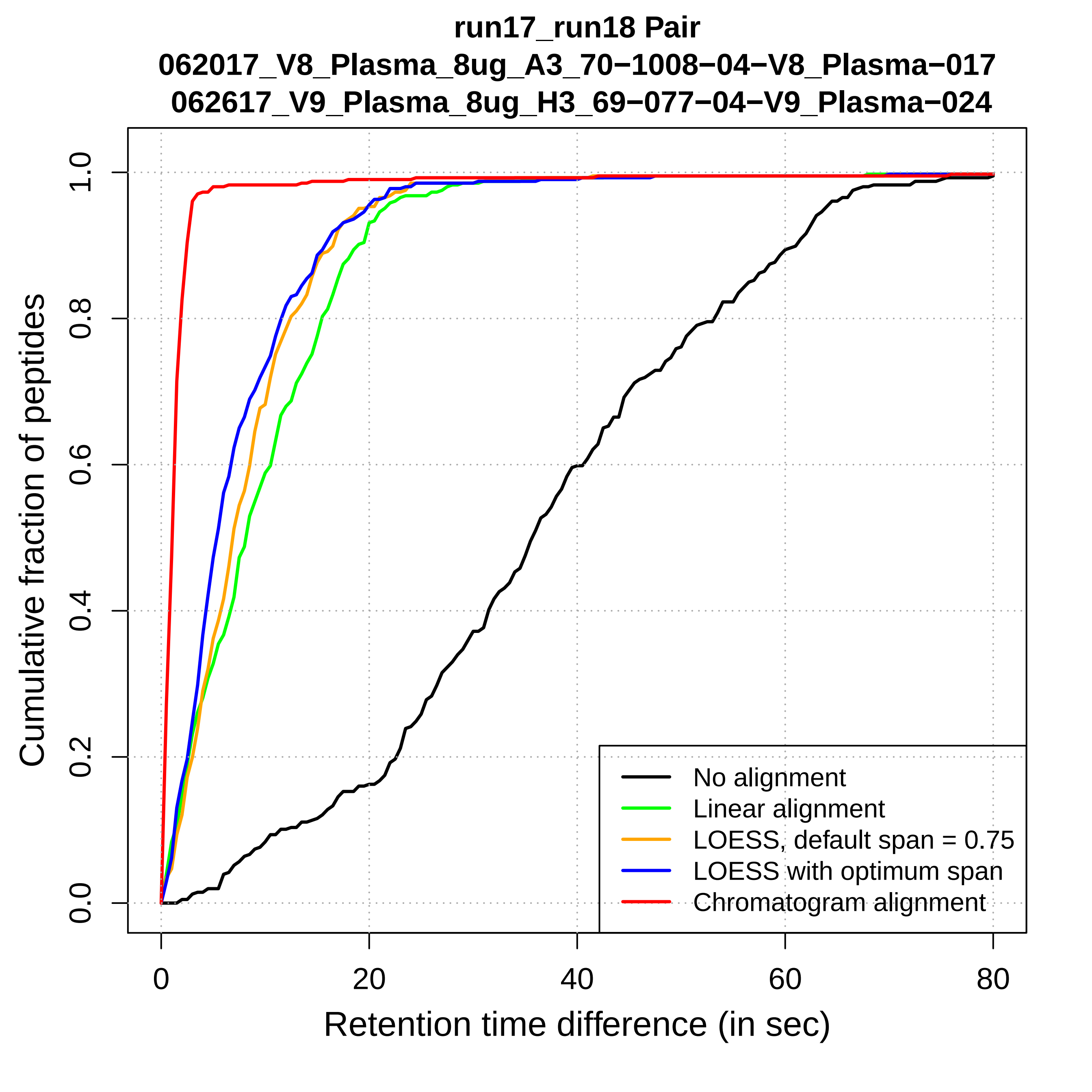

run17_run18 Pair
062017_V8_Plasma_8ug_A3_70−1008−04−V8_Plasma−017
062617_V9_Plasma_8ug_H3_69−077−04−V9_Plasma−024
Cumulative fraction of peptides
0.0
0.2
0.4
0.6
0.8
1.0
0
20
40
60
80
Retention time difference (in sec)
No alignment
Linear alignment
LOESS, default span = 0.75
LOESS with optimum span
Chromatogram alignment

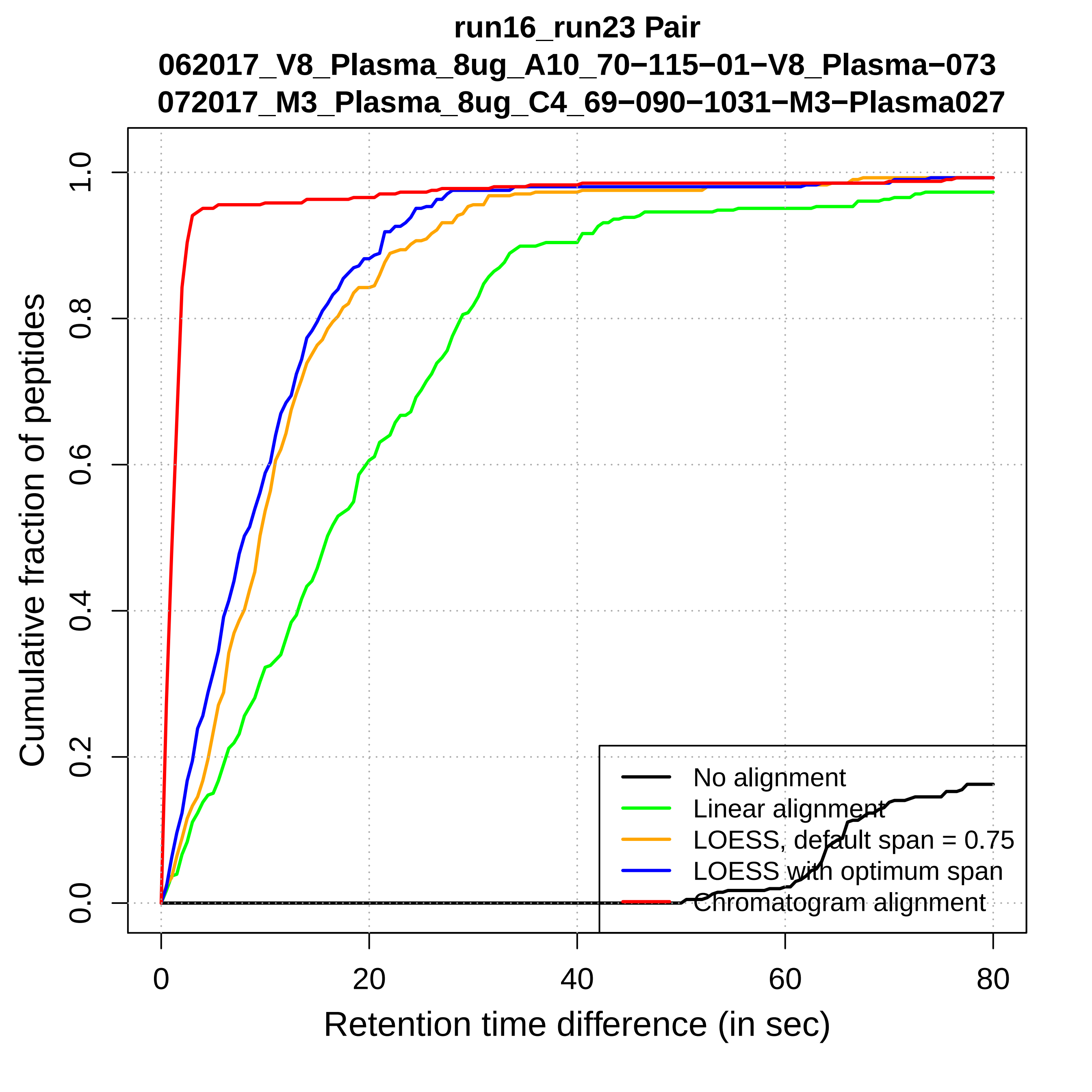

run16_run23 Pair
062017_V8_Plasma_8ug_A10_70−115−01−V8_Plasma−073
072017_M3_Plasma_8ug_C4_69−090−1031−M3−Plasma027
Cumulative fraction of peptides
0.0
0.2
0.4
0.6
0.8
1.0
0
20
40
60
80
Retention time difference (in sec)
No alignment
Linear alignment
LOESS, default span = 0.75
LOESS with optimum span
Chromatogram alignment

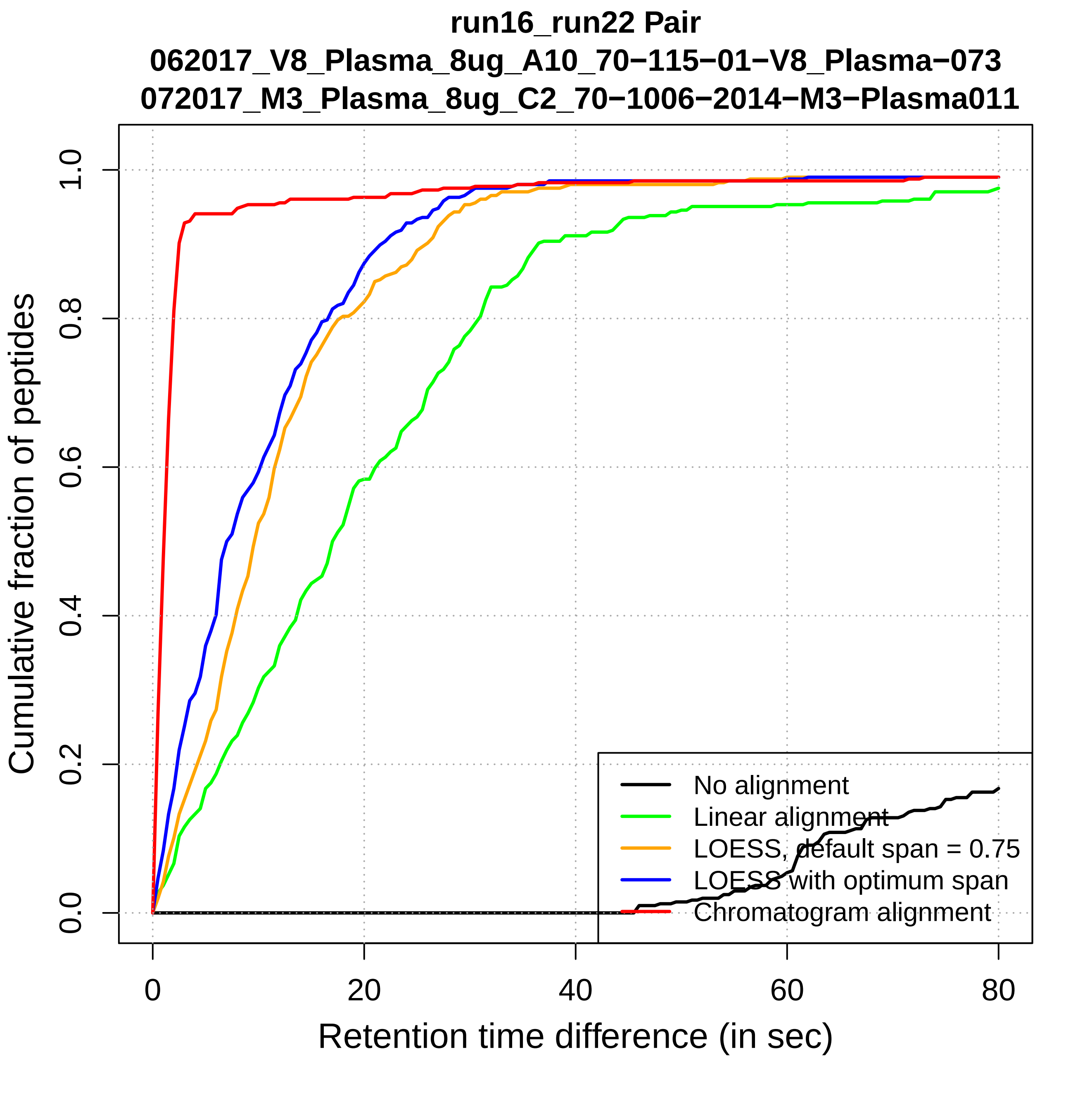

run16_run22 Pair
062017_V8_Plasma_8ug_A10_70−115−01−V8_Plasma−073
072017_M3_Plasma_8ug_C2_70−1006−2014−M3−Plasma011
Cumulative fraction of peptides
0.0
0.2
0.4
0.6
0.8
1.0
0
20
40
60
80
Retention time difference (in sec)
No alignment
Linear alignment
LOESS, default span = 0.75
LOESS with optimum span
Chromatogram alignment

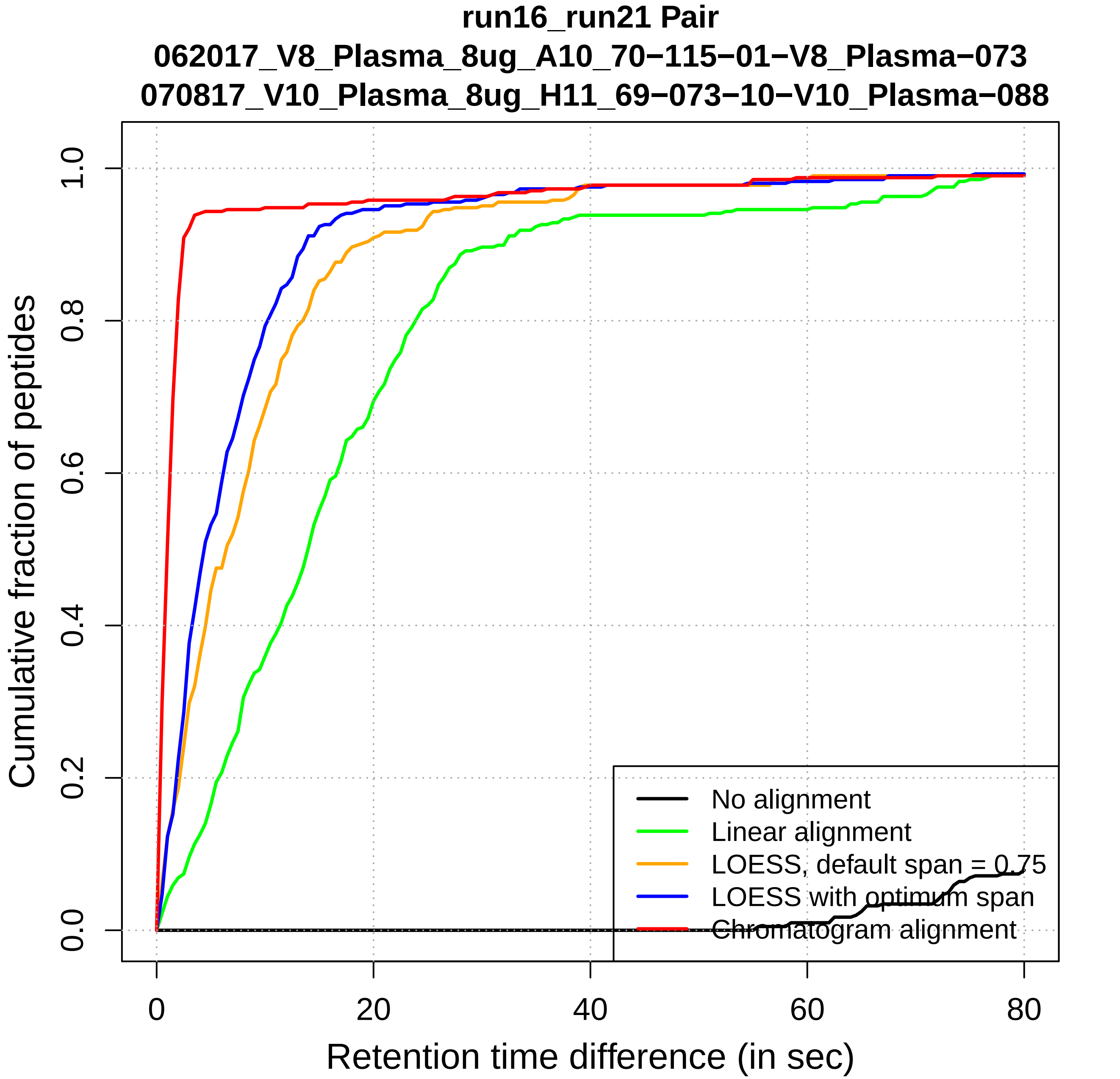
run16_run21 Pair
062017_V8_Plasma_8ug_A10_70−115−01−V8_Plasma−073
070817_V10_Plasma_8ug_H11_69−073−10−V10_Plasma−088
Cumulative fraction of peptides
Retention time difference (in sec)
0.0
0.2
0.4
0.6
0.8
1.0
0
20
40
60
80
No alignment
Linear alignment
LOESS, default span = 0.75
LOESS with optimum span
Chromatogram alignment

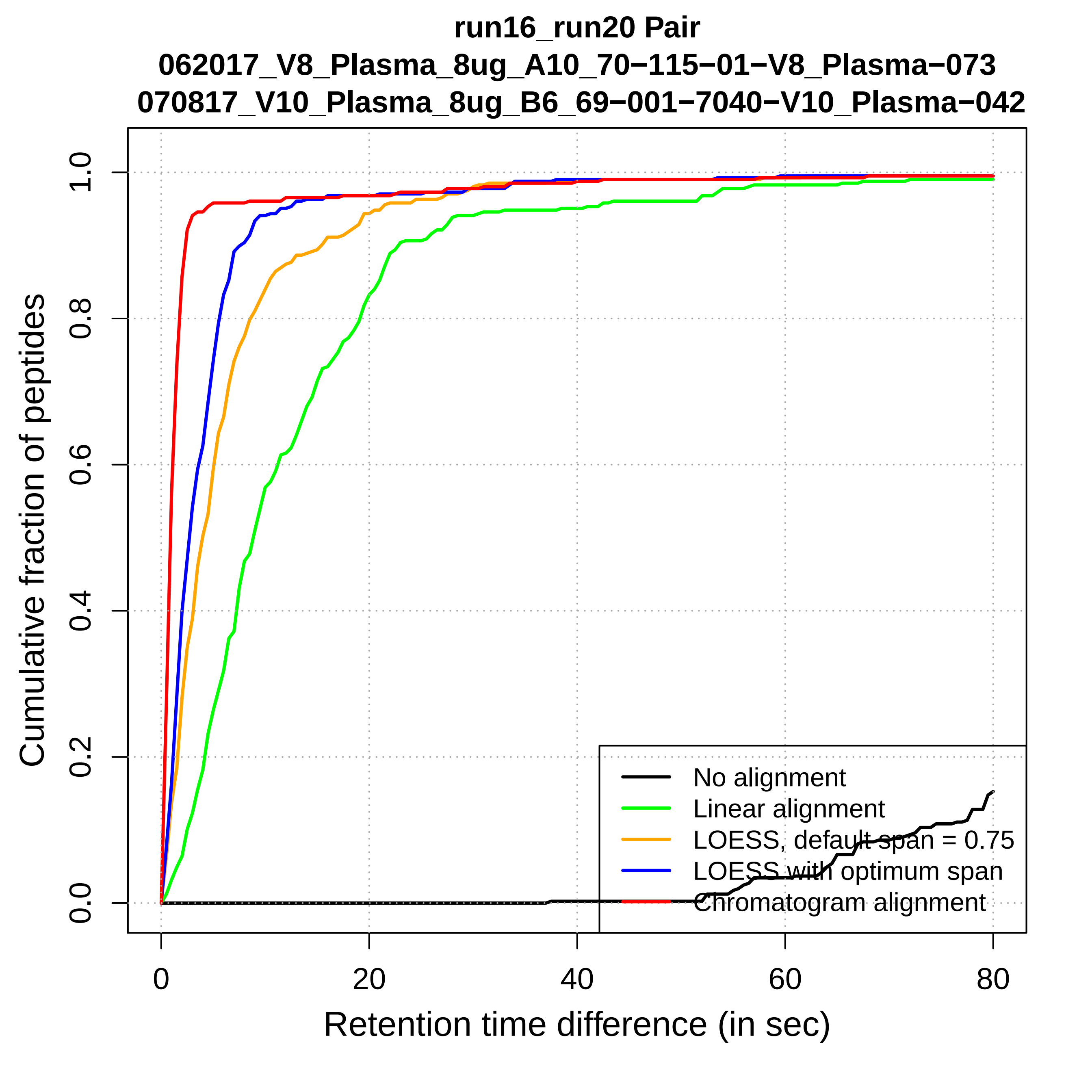
run16_run20 Pair
062017_V8_Plasma_8ug_A10_70−115−01−V8_Plasma−073
070817_V10_Plasma_8ug_B6_69−001−7040−V10_Plasma−042
Cumulative fraction of peptides
0.0
0.2
0.4
0.6
0.8
1.0
0
20
40
60
80
Retention time difference (in sec)
No alignment
Linear alignment
LOESS, default span = 0.75
LOESS with optimum span
Chromatogram alignment

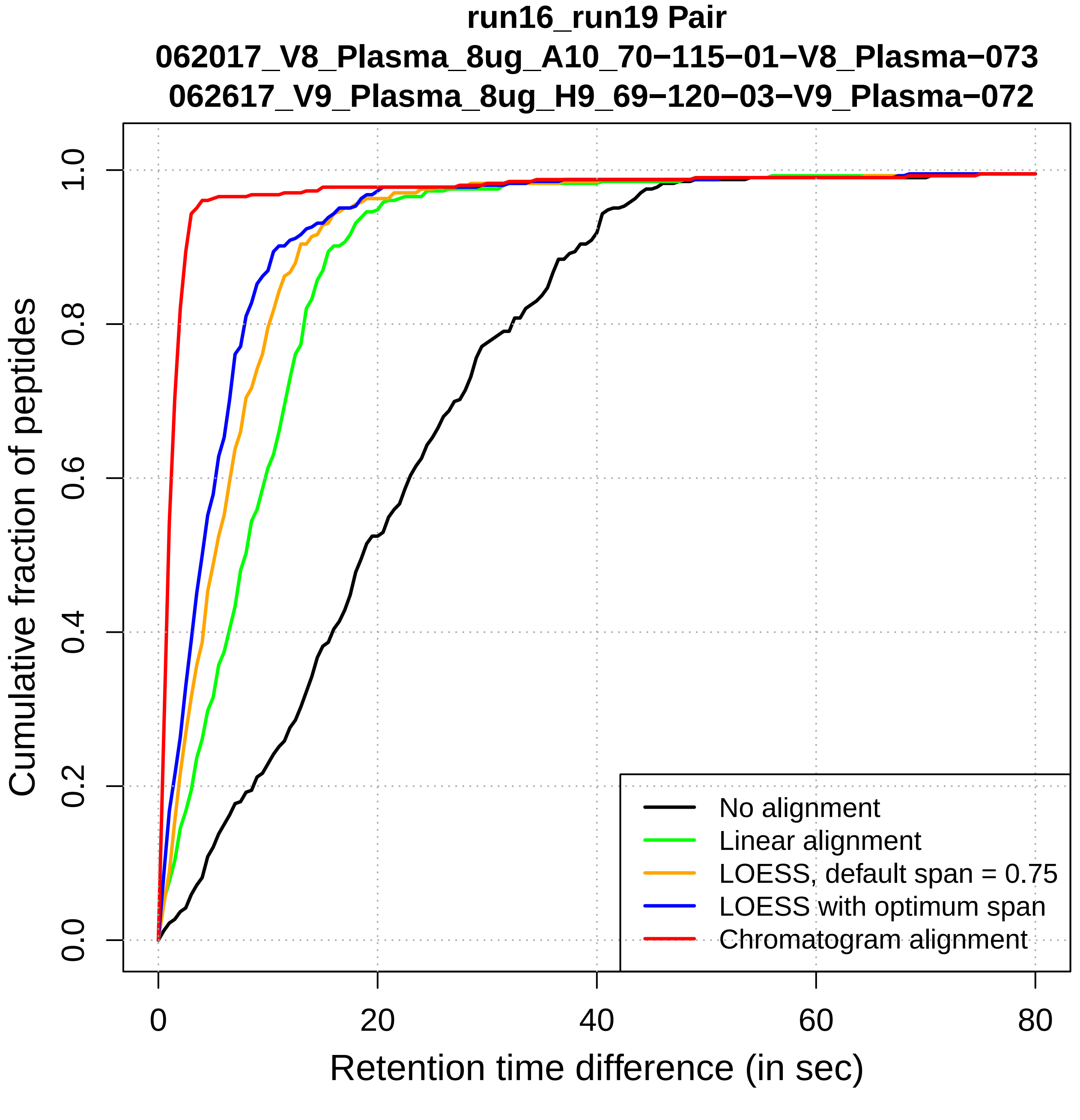

run16_run19 Pair
062017_V8_Plasma_8ug_A10_70−115−01−V8_Plasma−073
062617_V9_Plasma_8ug_H9_69−120−03−V9_Plasma−072
Cumulative fraction of peptides
0.0
0.2
0.4
0.6
0.8
1.0
0
20
40
60
80
Retention time difference (in sec)
No alignment
Linear alignment
LOESS, default span = 0.75
LOESS with optimum span
Chromatogram alignment

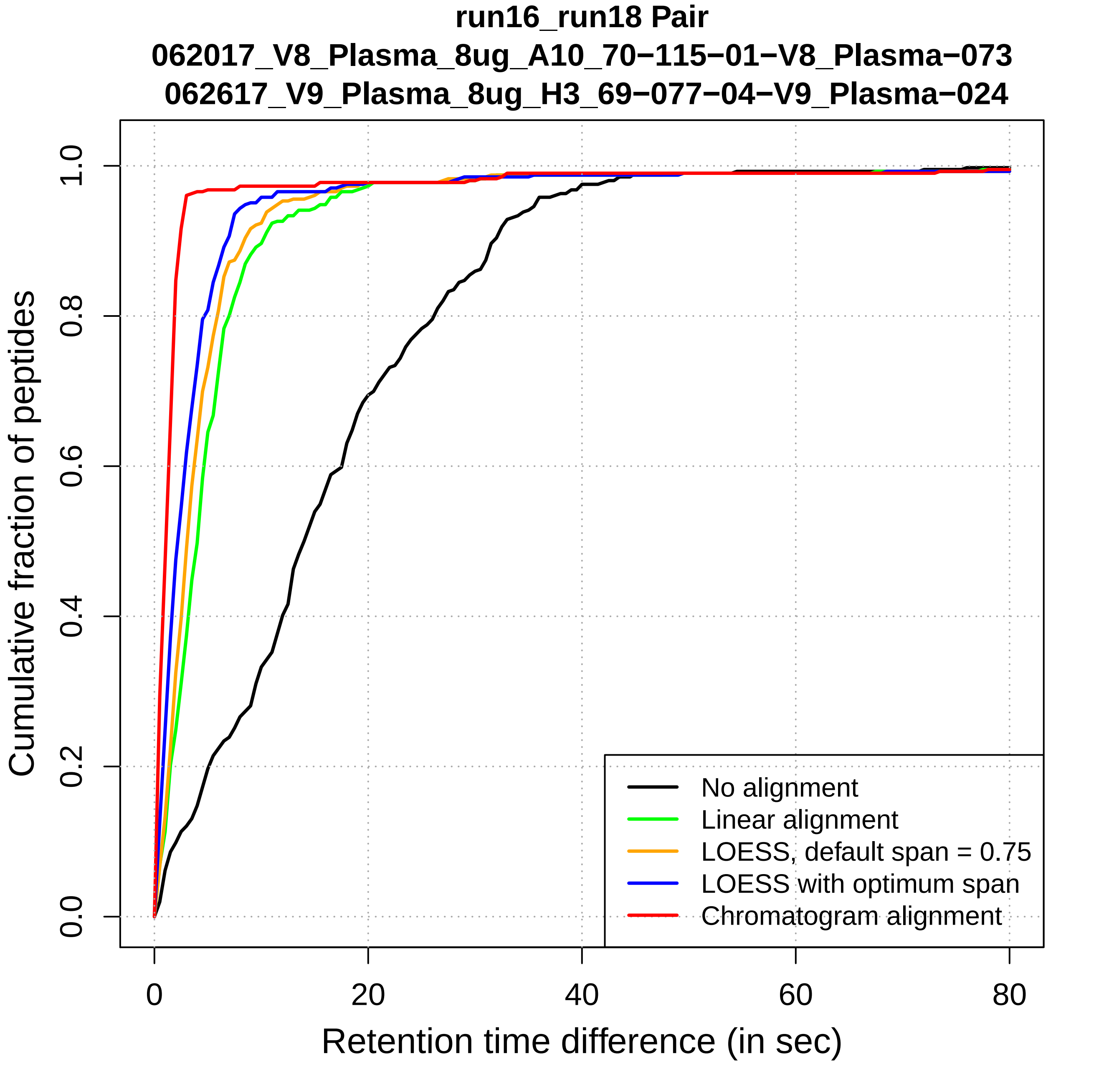

run16_run18 Pair
062017_V8_Plasma_8ug_A10_70−115−01−V8_Plasma−073
062617_V9_Plasma_8ug_H3_69−077−04−V9_Plasma−024
Cumulative fraction of peptides
0.0
0.2
0.4
0.6
0.8
1.0
0
20
40
60
80
Retention time difference (in sec)
No alignment
Linear alignment
LOESS, default span = 0.75
LOESS with optimum span
Chromatogram alignment

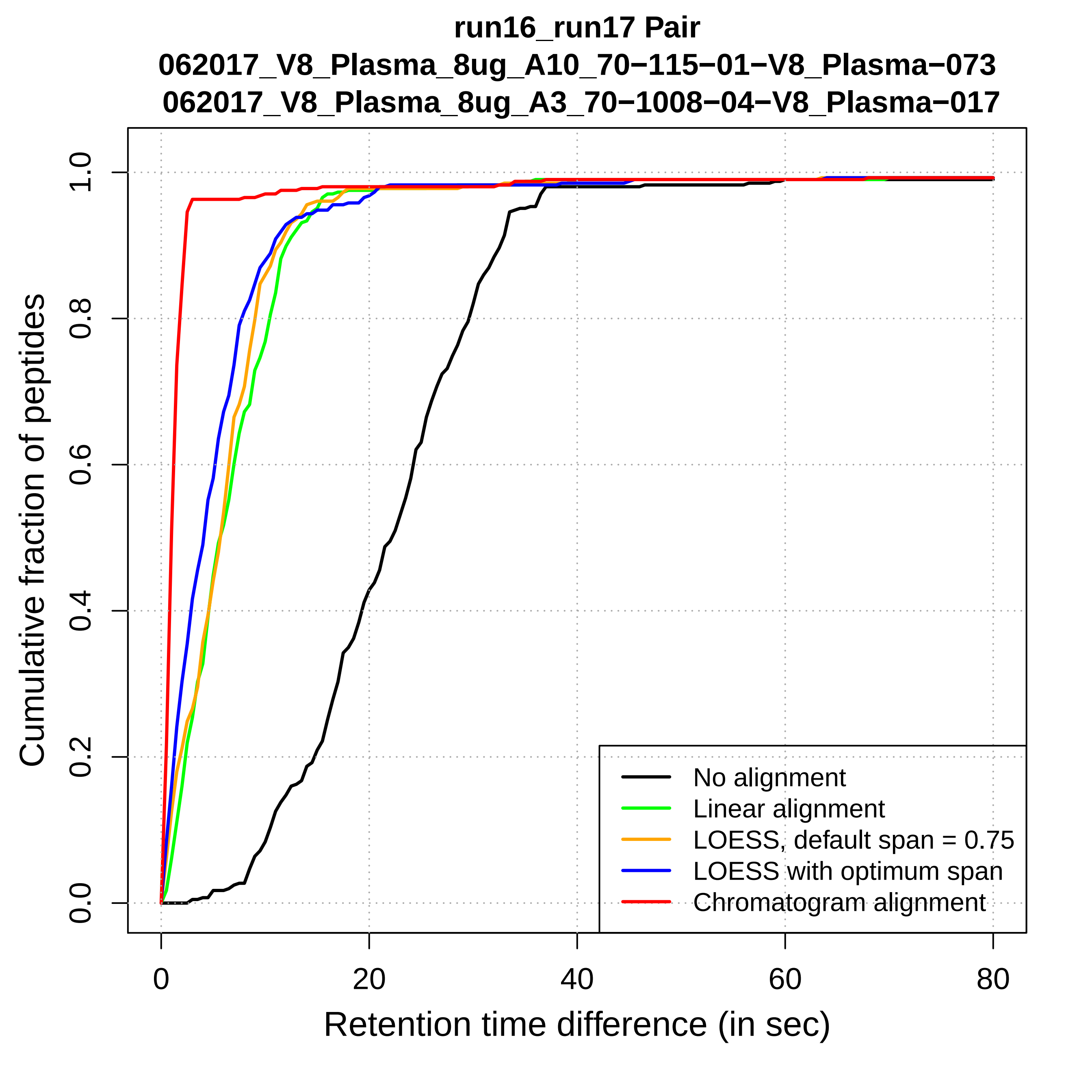
run16_run17 Pair
062017_V8_Plasma_8ug_A10_70−115−01−V8_Plasma−073
062017_V8_Plasma_8ug_A3_70−1008−04−V8_Plasma−017
Cumulative fraction of peptides
0.0
0.2
0.4
0.6
0.8
1.0
0
20
40
60
80
Retention time difference (in sec)
No alignment
Linear alignment
LOESS, default span = 0.75
LOESS with optimum span
Chromatogram alignment

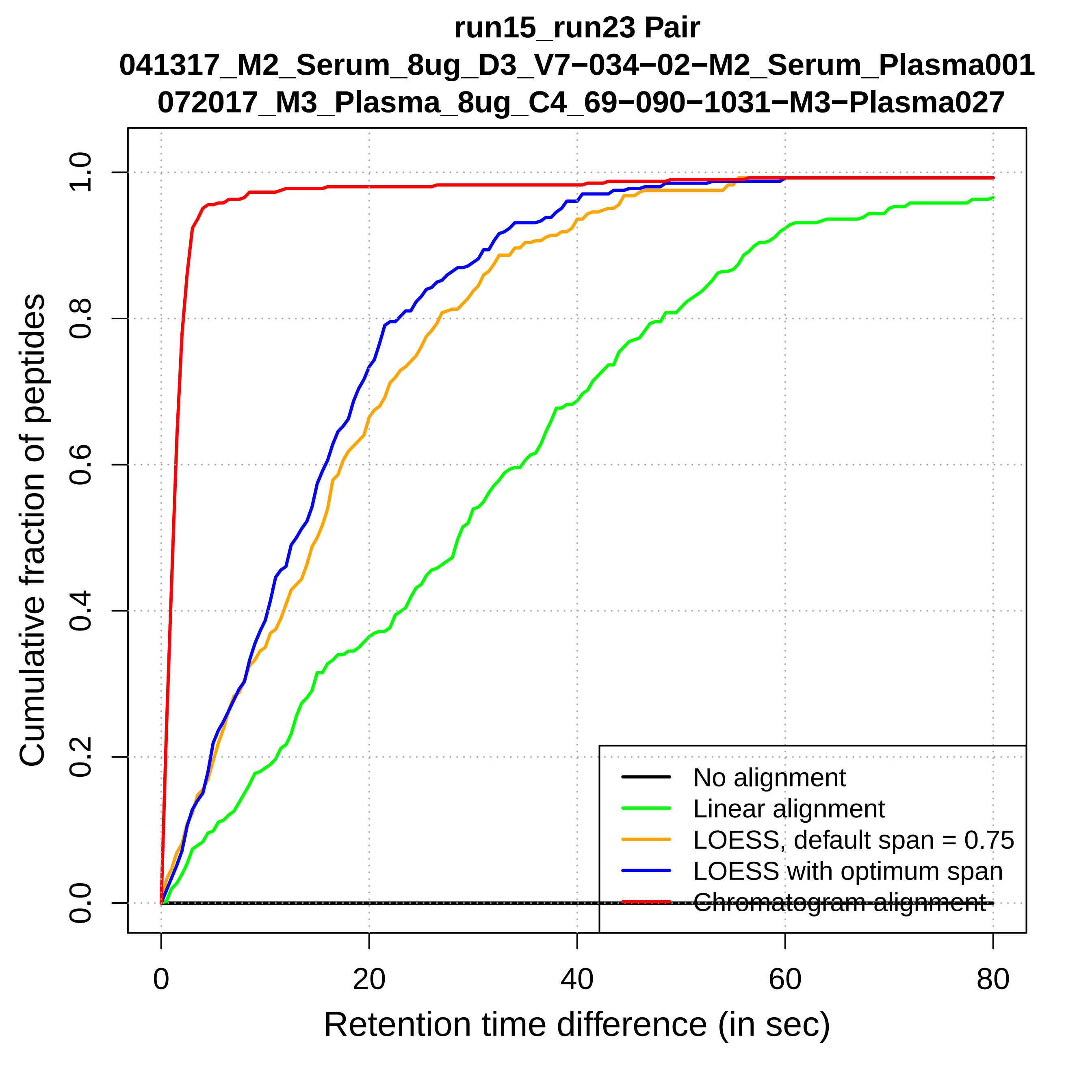

run15_run23 Pair
041317_M2_Serum_8ug_D3_V7−034−02−M2_Serum_Plasma001
072017_M3_Plasma_8ug_C4_69−090−1031−M3−Plasma027
Cumulative fraction of peptides
Retention time difference (in sec)
0.0
0.2
0.4
0.6
0.8
1.0
0
20
40
60
80
No alignment
Linear alignment
LOESS, default span = 0.75
LOESS with optimum span
Chromatogram alignment

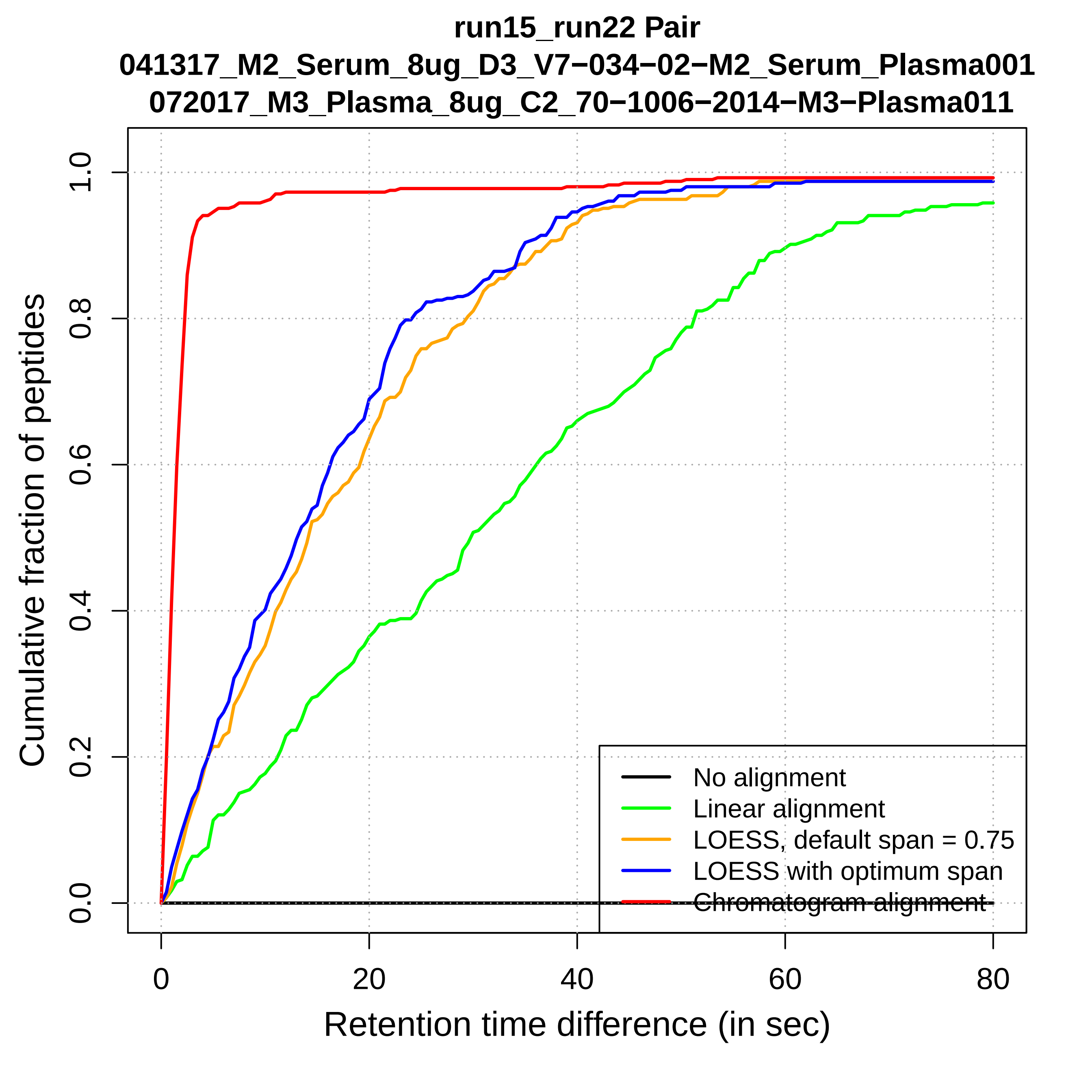
run15_run22 Pair
041317_M2_Serum_8ug_D3_V7−034−02−M2_Serum_Plasma001
072017_M3_Plasma_8ug_C2_70−1006−2014−M3−Plasma011
Cumulative fraction of peptides
0.0
0.2
0.4
0.6
0.8
1.0
0
20
40
60
80
Retention time difference (in sec)
No alignment
Linear alignment
LOESS, default span = 0.75
LOESS with optimum span
Chromatogram alignment

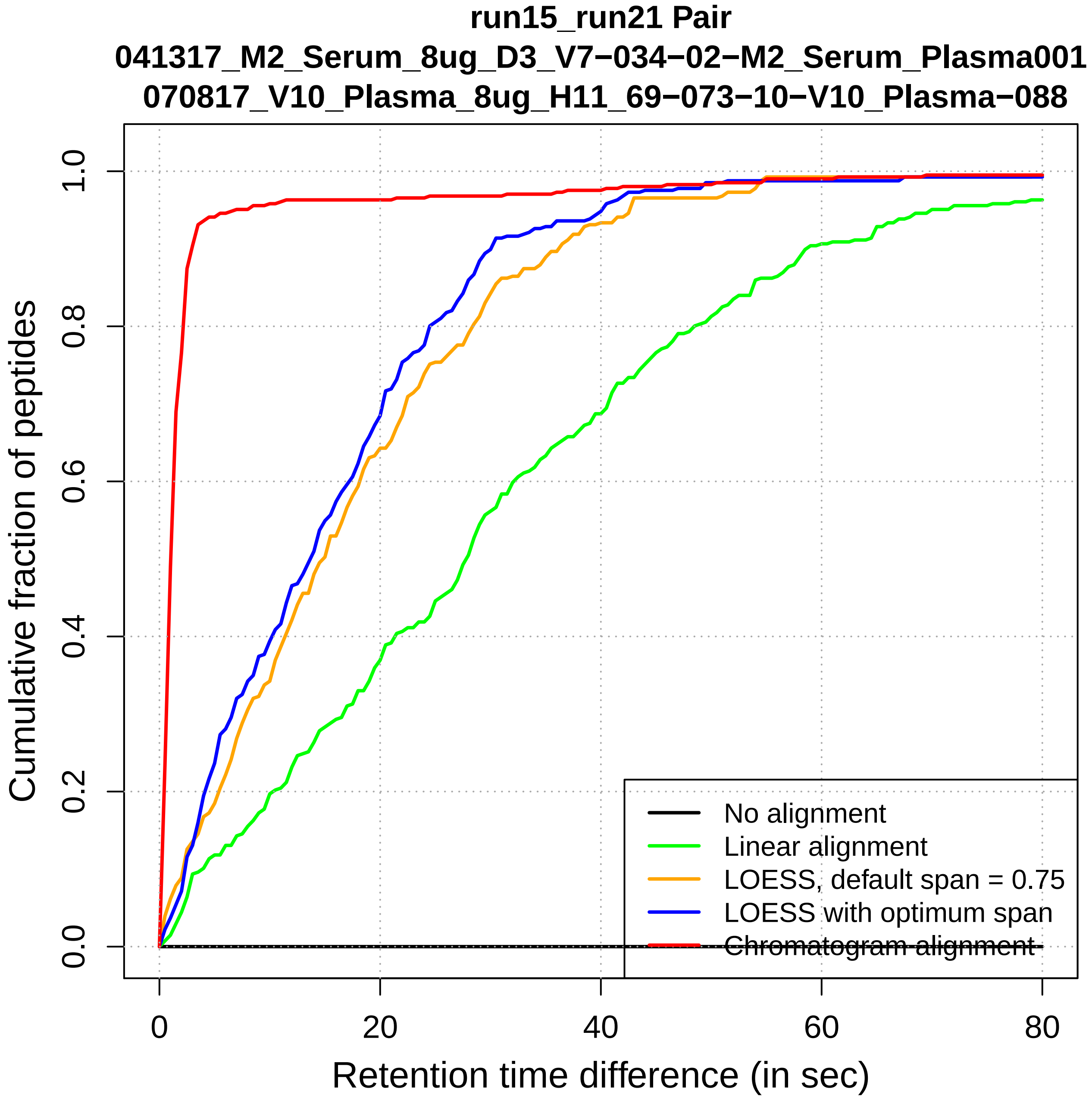

run15_run21 Pair
041317_M2_Serum_8ug_D3_V7−034−02−M2_Serum_Plasma001
070817_V10_Plasma_8ug_H11_69−073−10−V10_Plasma−088
Cumulative fraction of peptides
0.0
0.2
0.4
0.6
0.8
1.0
0
20
40
60
80
Retention time difference (in sec)
No alignment
Linear alignment
LOESS, default span = 0.75
LOESS with optimum span
Chromatogram alignment

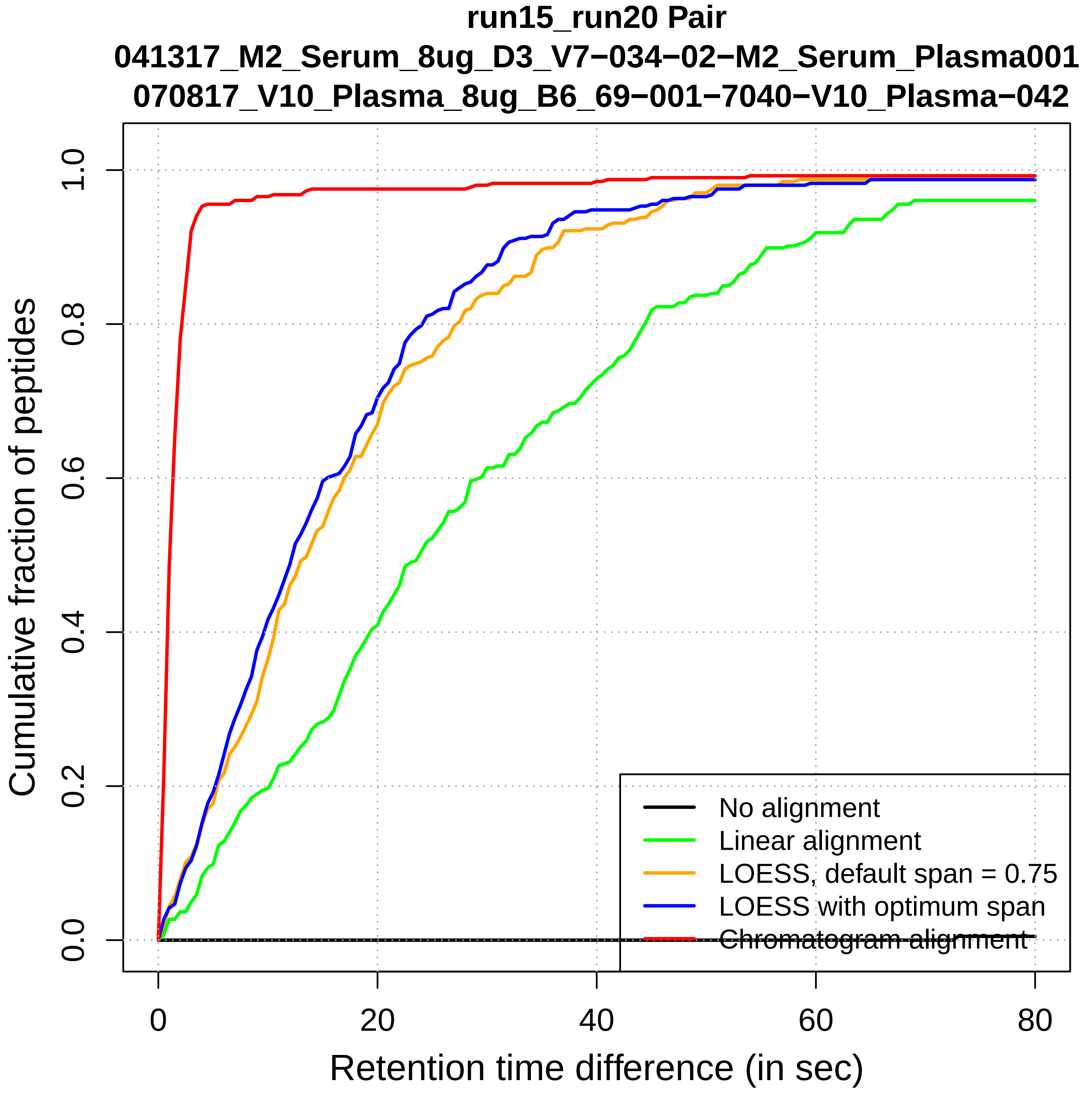
run15_run20 Pair
041317_M2_Serum_8ug_D3_V7−034−02−M2_Serum_Plasma001
070817_V10_Plasma_8ug_B6_69−001−7040−V10_Plasma−042
Cumulative fraction of peptides
0.0
0.2
0.4
0.6
0.8
1.0
0
20
40
60
80
Retention time difference (in sec)
No alignment
Linear alignment
LOESS, default span = 0.75
LOESS with optimum span
Chromatogram alignment

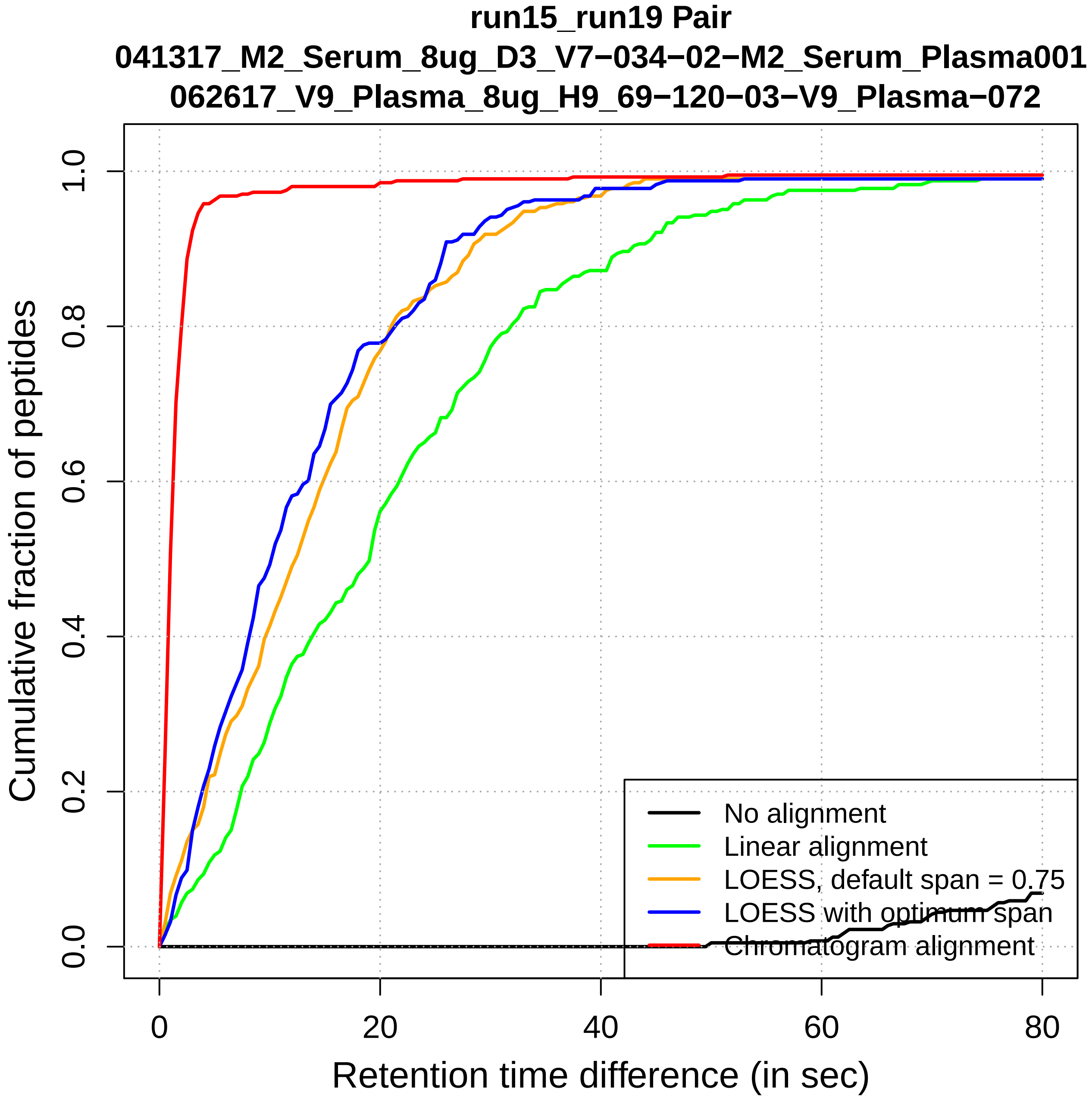

run15_run19 Pair
041317_M2_Serum_8ug_D3_V7−034−02−M2_Serum_Plasma001
062617_V9_Plasma_8ug_H9_69−120−03−V9_Plasma−072
Cumulative fraction of peptides
0.0
0.2
0.4
0.6
0.8
1.0
0
20
40
60
80
Retention time difference (in sec)
No alignment
Linear alignment
LOESS, default span = 0.75
LOESS with optimum span
Chromatogram alignment

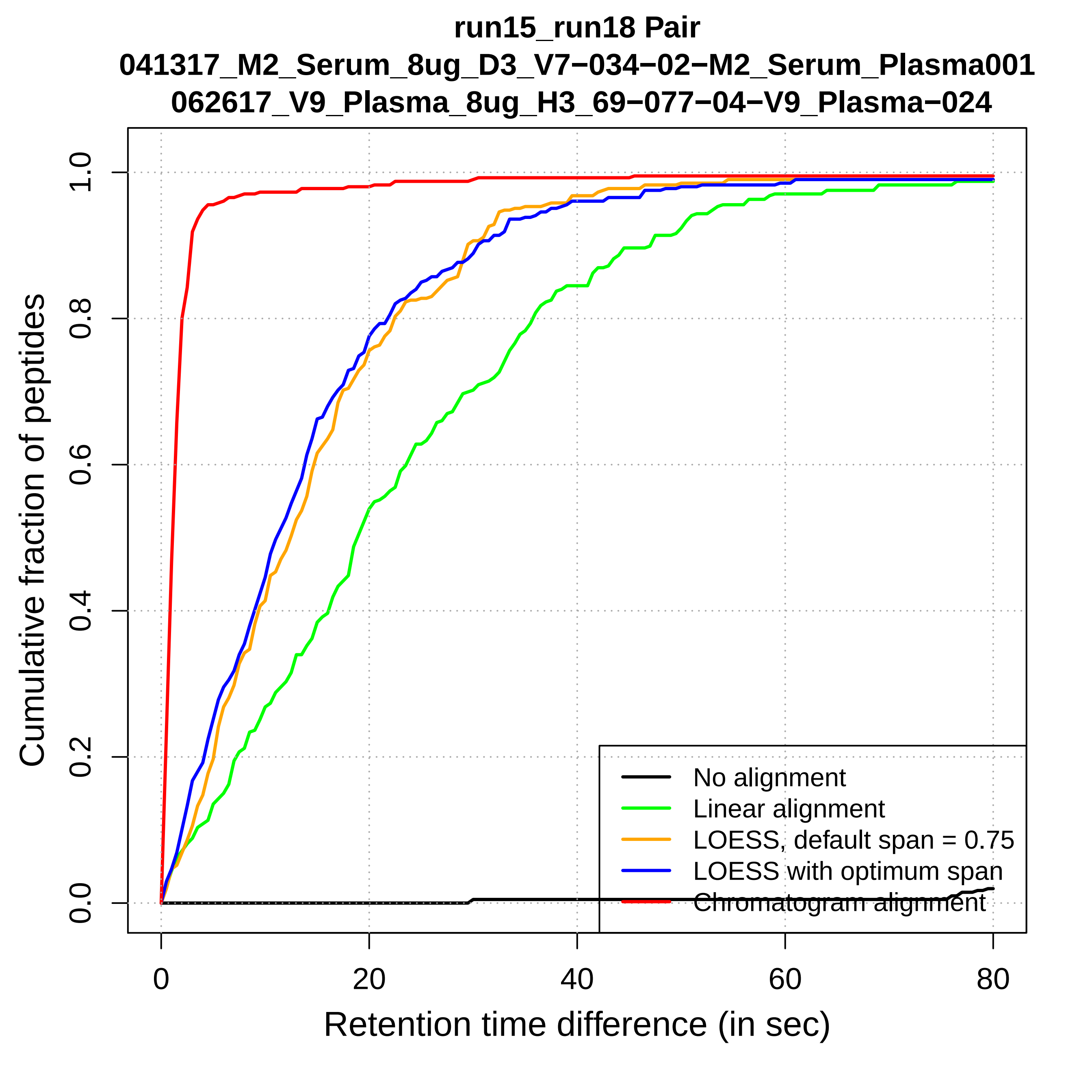
run15_run18 Pair
041317_M2_Serum_8ug_D3_V7−034−02−M2_Serum_Plasma001
062617_V9_Plasma_8ug_H3_69−077−04−V9_Plasma−024
Cumulative fraction of peptides
Retention time difference (in sec)
0.0
0.2
0.4
0.6
0.8
1.0
0
20
40
60
80
No alignment
Linear alignment
LOESS, default span = 0.75
LOESS with optimum span
Chromatogram alignment

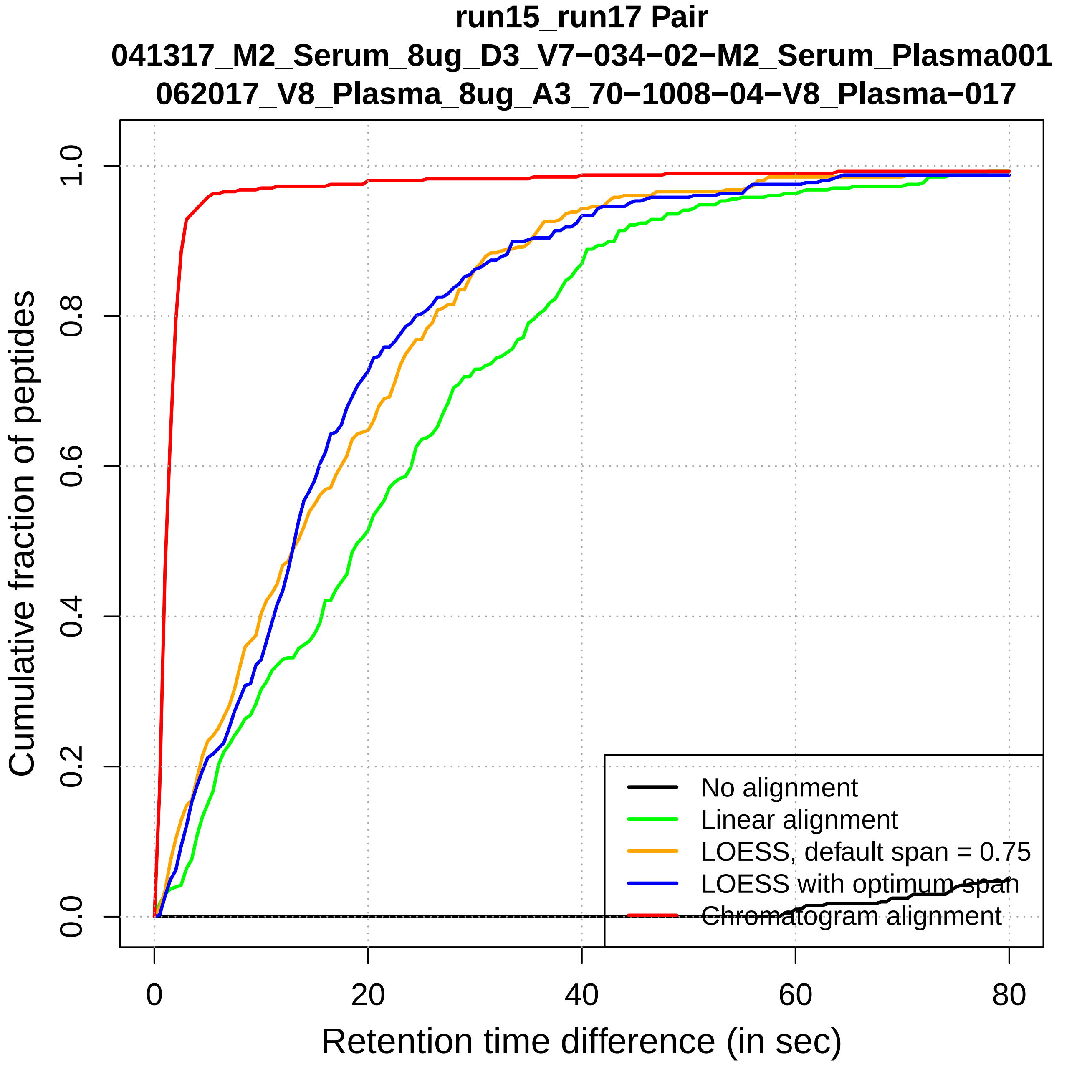
run15_run17 Pair
041317_M2_Serum_8ug_D3_V7−034−02−M2_Serum_Plasma001
062017_V8_Plasma_8ug_A3_70−1008−04−V8_Plasma−017
Cumulative fraction of peptides
0.0
0.2
0.4
0.6
0.8
1.0
0
20
40
60
80
Retention time difference (in sec)
No alignment
Linear alignment
LOESS, default span = 0.75
LOESS with optimum span
Chromatogram alignment

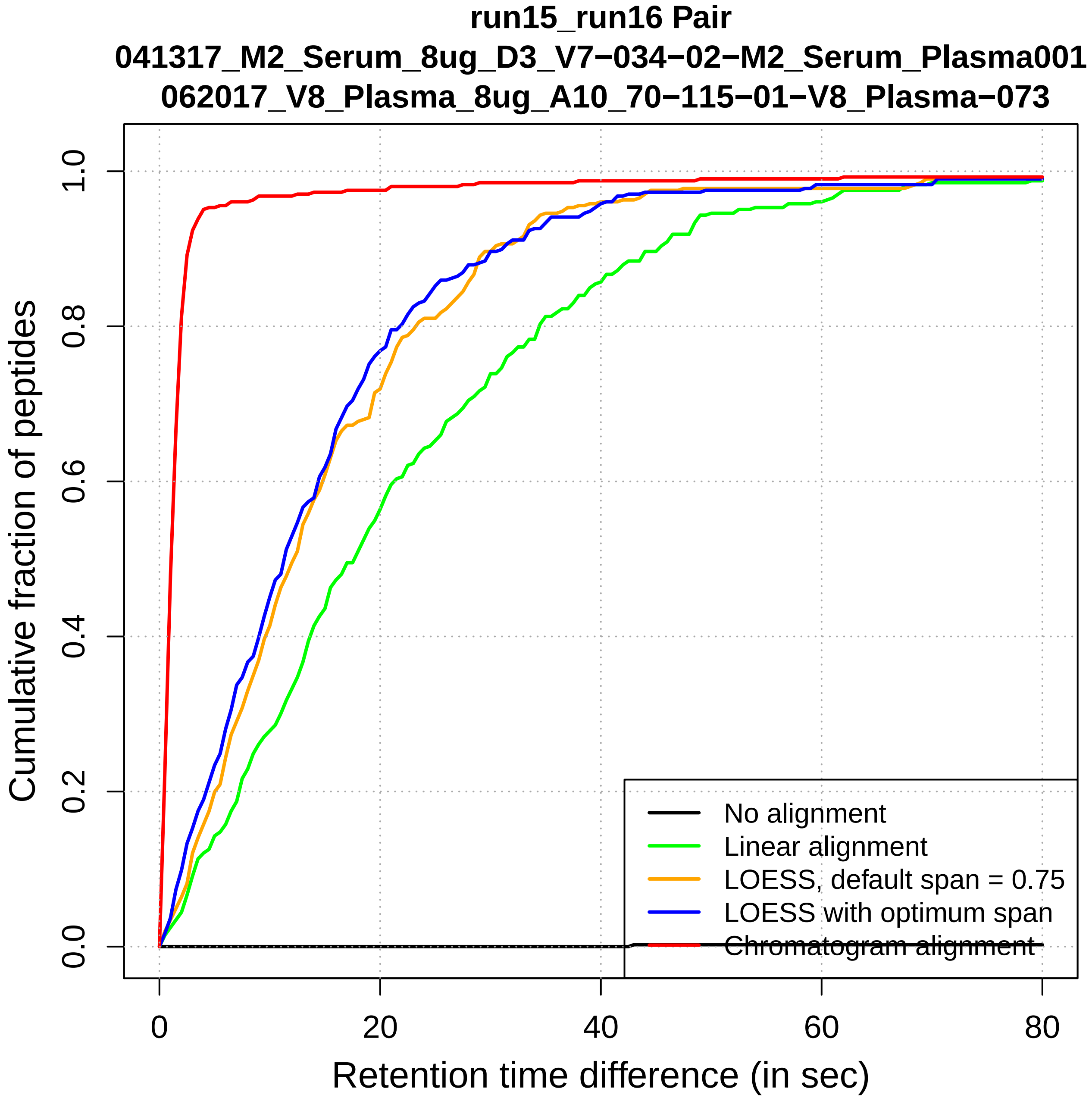

run15_run16 Pair
041317_M2_Serum_8ug_D3_V7−034−02−M2_Serum_Plasma001
062017_V8_Plasma_8ug_A10_70−115−01−V8_Plasma−073
Cumulative fraction of peptides
0.0
0.2
0.4
0.6
0.8
1.0
0
20
40
60
80
Retention time difference (in sec)
No alignment
Linear alignment
LOESS, default span = 0.75
LOESS with optimum span
Chromatogram alignment

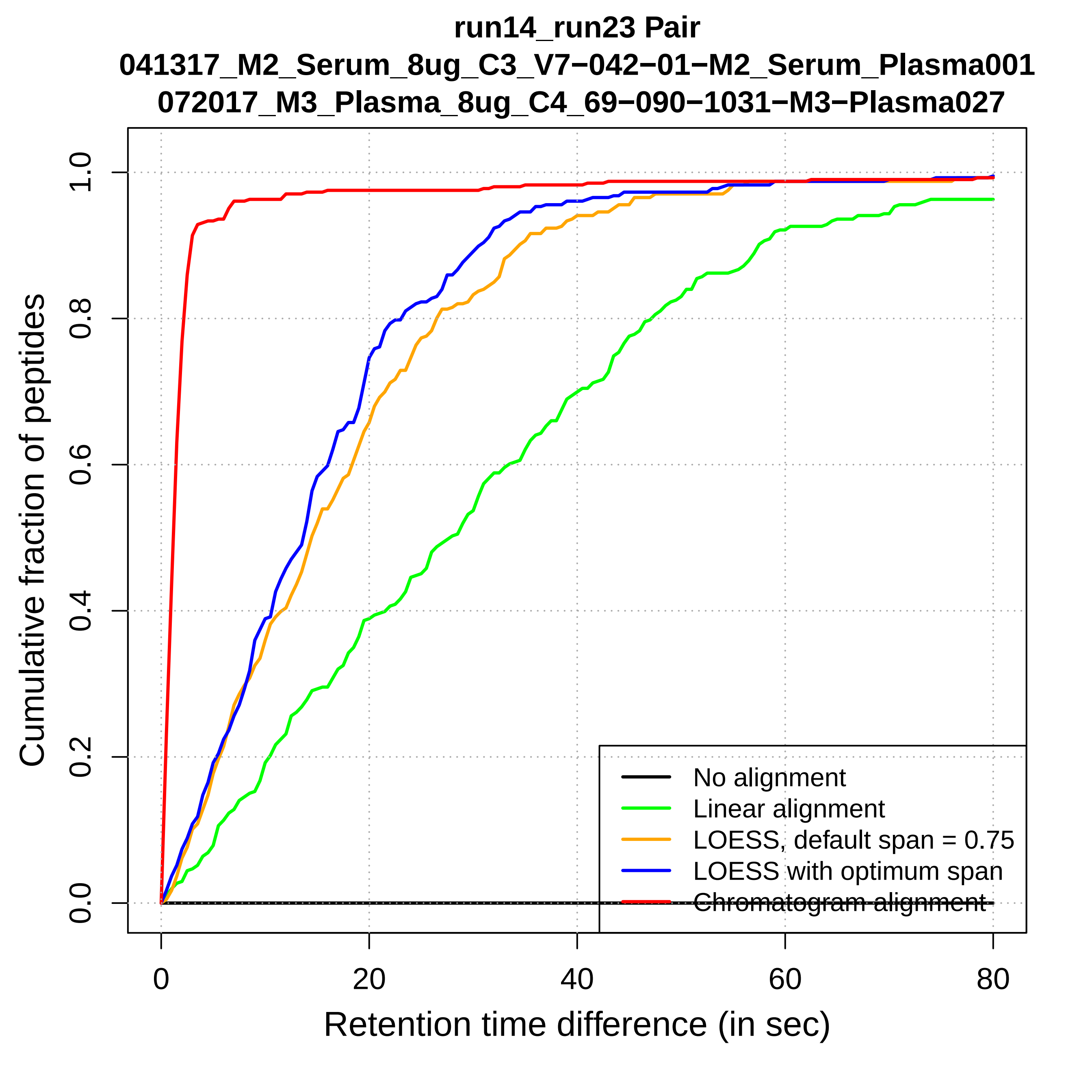
run14_run23 Pair
041317_M2_Serum_8ug_C3_V7−042−01−M2_Serum_Plasma001
072017_M3_Plasma_8ug_C4_69−090−1031−M3−Plasma027
Cumulative fraction of peptides
0.0
0.2
0.4
0.6
0.8
1.0
0
20
40
60
80
Retention time difference (in sec)
No alignment
Linear alignment
LOESS, default span = 0.75
LOESS with optimum span
Chromatogram alignment

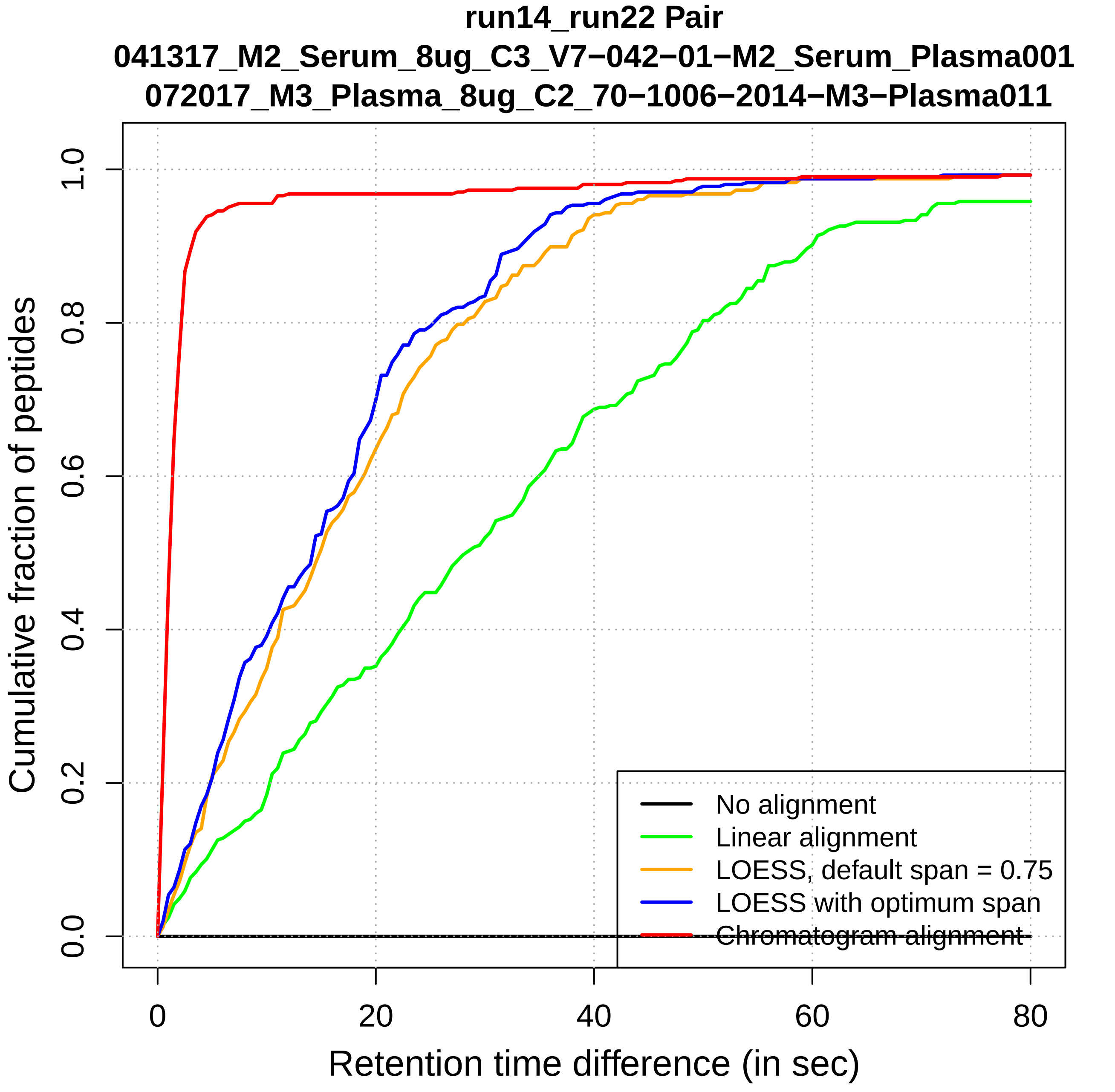

run14_run22 Pair
041317_M2_Serum_8ug_C3_V7−042−01−M2_Serum_Plasma001
072017_M3_Plasma_8ug_C2_70−1006−2014−M3−Plasma011
Cumulative fraction of peptides
0.0
0.2
0.4
0.6
0.8
1.0
0
20
40
60
80
Retention time difference (in sec)
No alignment
Linear alignment
LOESS, default span = 0.75
LOESS with optimum span
Chromatogram alignment

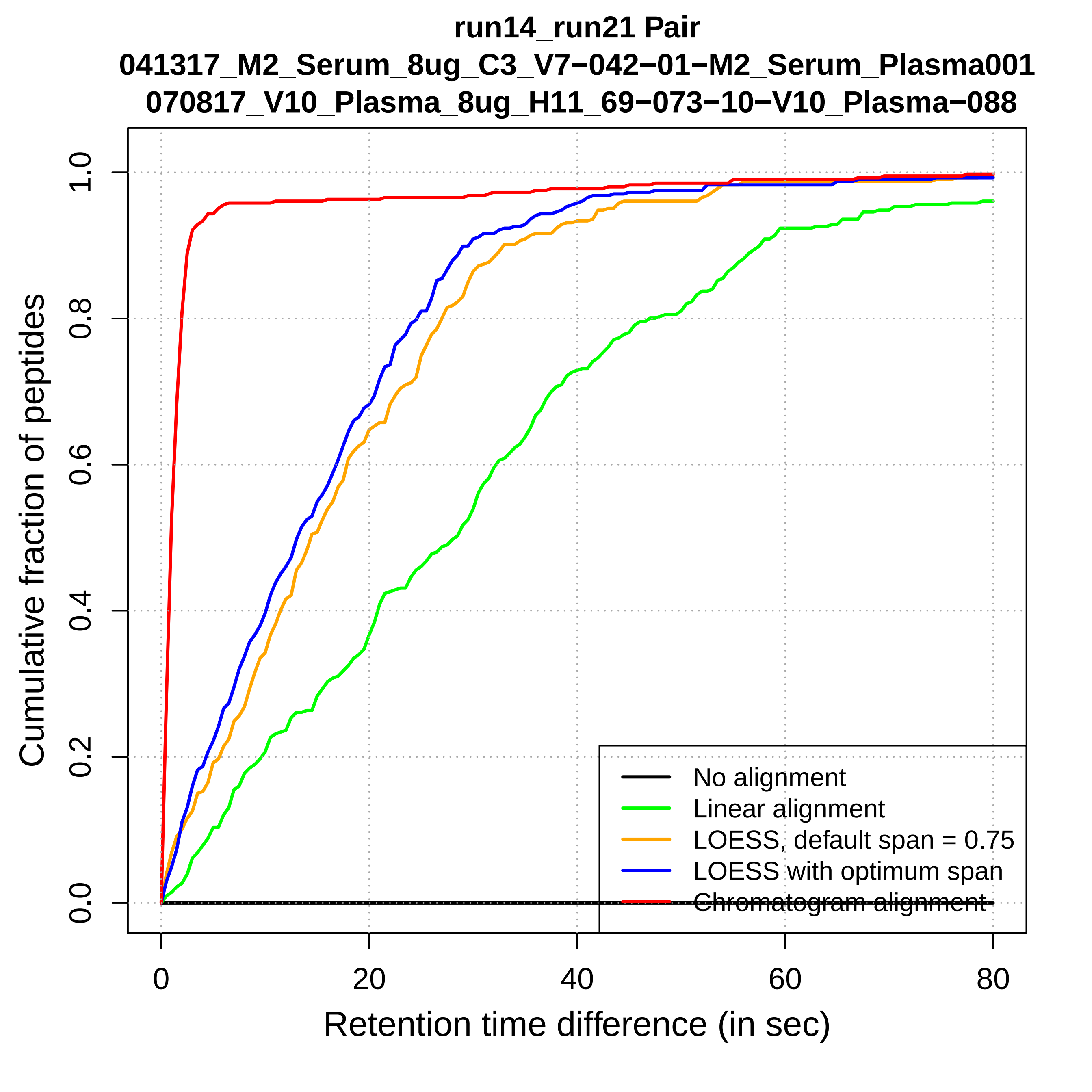

run14_run21 Pair
041317_M2_Serum_8ug_C3_V7−042−01−M2_Serum_Plasma001
070817_V10_Plasma_8ug_H11_69−073−10−V10_Plasma−088
Cumulative fraction of peptides
Retention time difference (in sec)
0.0
0.2
0.4
0.6
0.8
1.0
0
20
40
60
80
No alignment
Linear alignment
LOESS, default span = 0.75
LOESS with optimum span
Chromatogram alignment

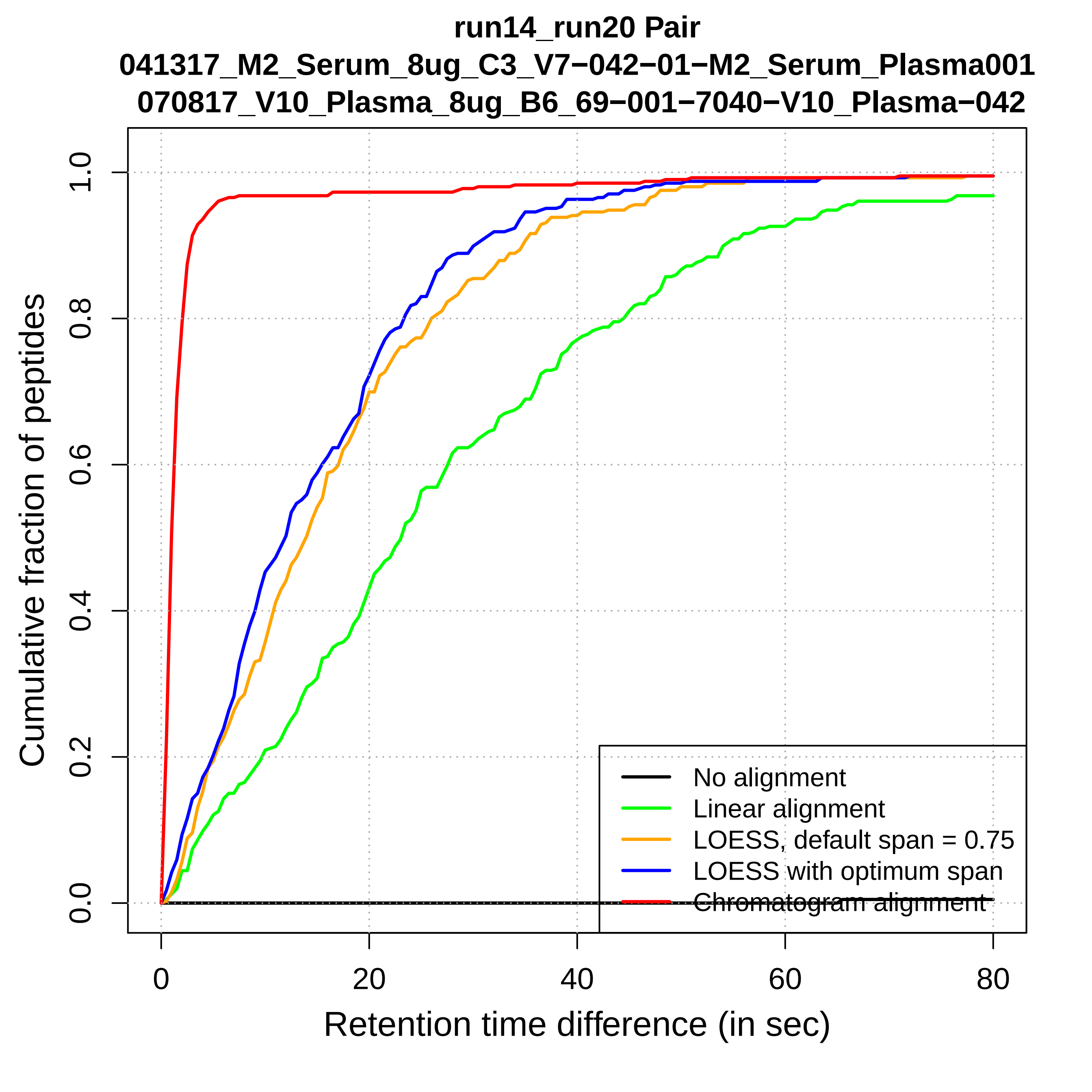
run14_run20 Pair
041317_M2_Serum_8ug_C3_V7−042−01−M2_Serum_Plasma001
070817_V10_Plasma_8ug_B6_69−001−7040−V10_Plasma−042
Cumulative fraction of peptides
0.0
0.2
0.4
0.6
0.8
1.0
0
20
40
60
80
Retention time difference (in sec)
No alignment
Linear alignment
LOESS, default span = 0.75
LOESS with optimum span
Chromatogram alignment

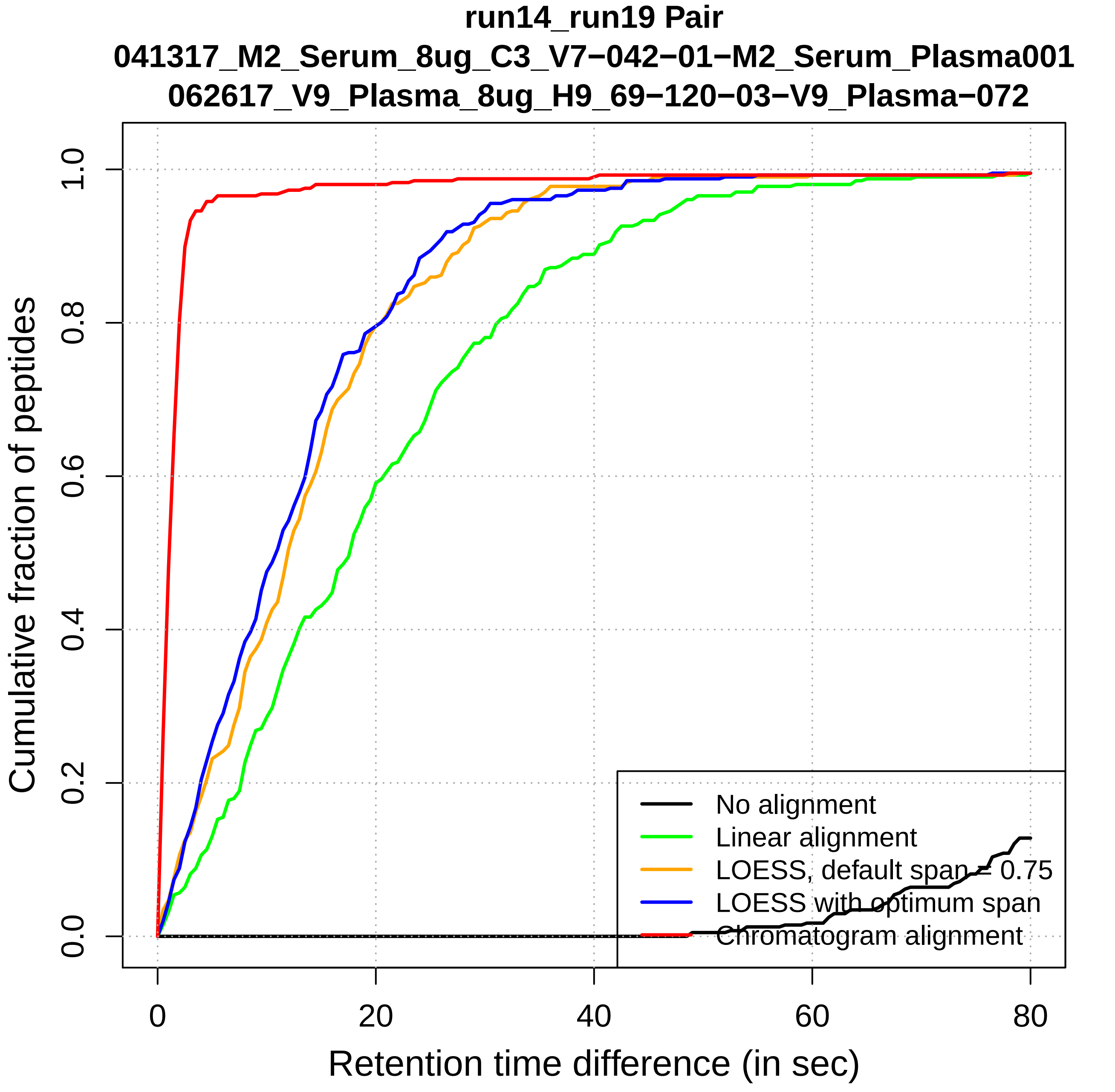

run14_run19 Pair
041317_M2_Serum_8ug_C3_V7−042−01−M2_Serum_Plasma001
062617_V9_Plasma_8ug_H9_69−120−03−V9_Plasma−072
Cumulative fraction of peptides
Retention time difference (in sec)
0.0
0.2
0.4
0.6
0.8
1.0
0
20
40
60
80
No alignment
Linear alignment
LOESS, default span = 0.75
LOESS with optimum span
Chromatogram alignment

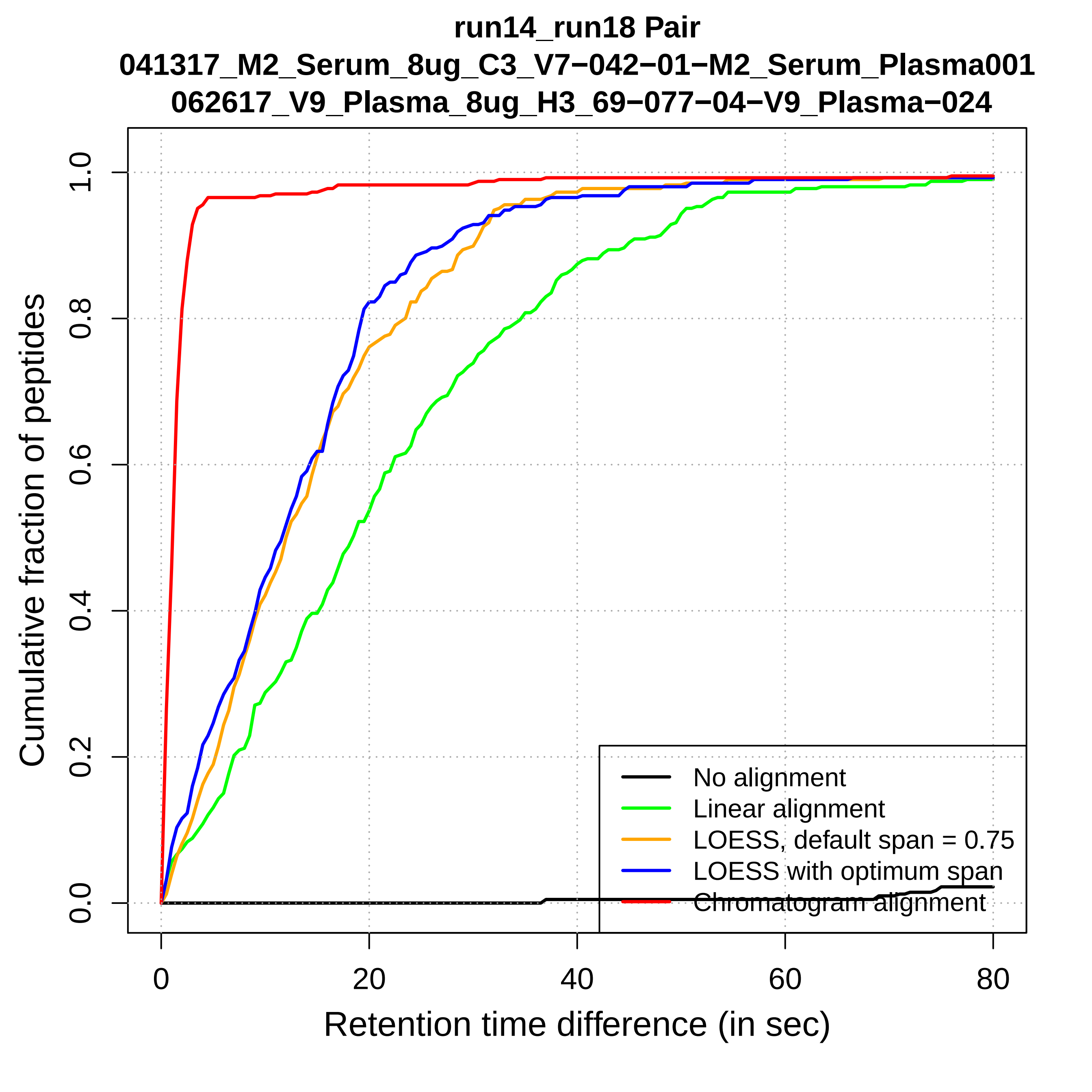
run14_run18 Pair
041317_M2_Serum_8ug_C3_V7−042−01−M2_Serum_Plasma001
062617_V9_Plasma_8ug_H3_69−077−04−V9_Plasma−024
Cumulative fraction of peptides
0.0
0.2
0.4
0.6
0.8
1.0
0
20
40
60
80
Retention time difference (in sec)
No alignment
Linear alignment
LOESS, default span = 0.75
LOESS with optimum span
Chromatogram alignment

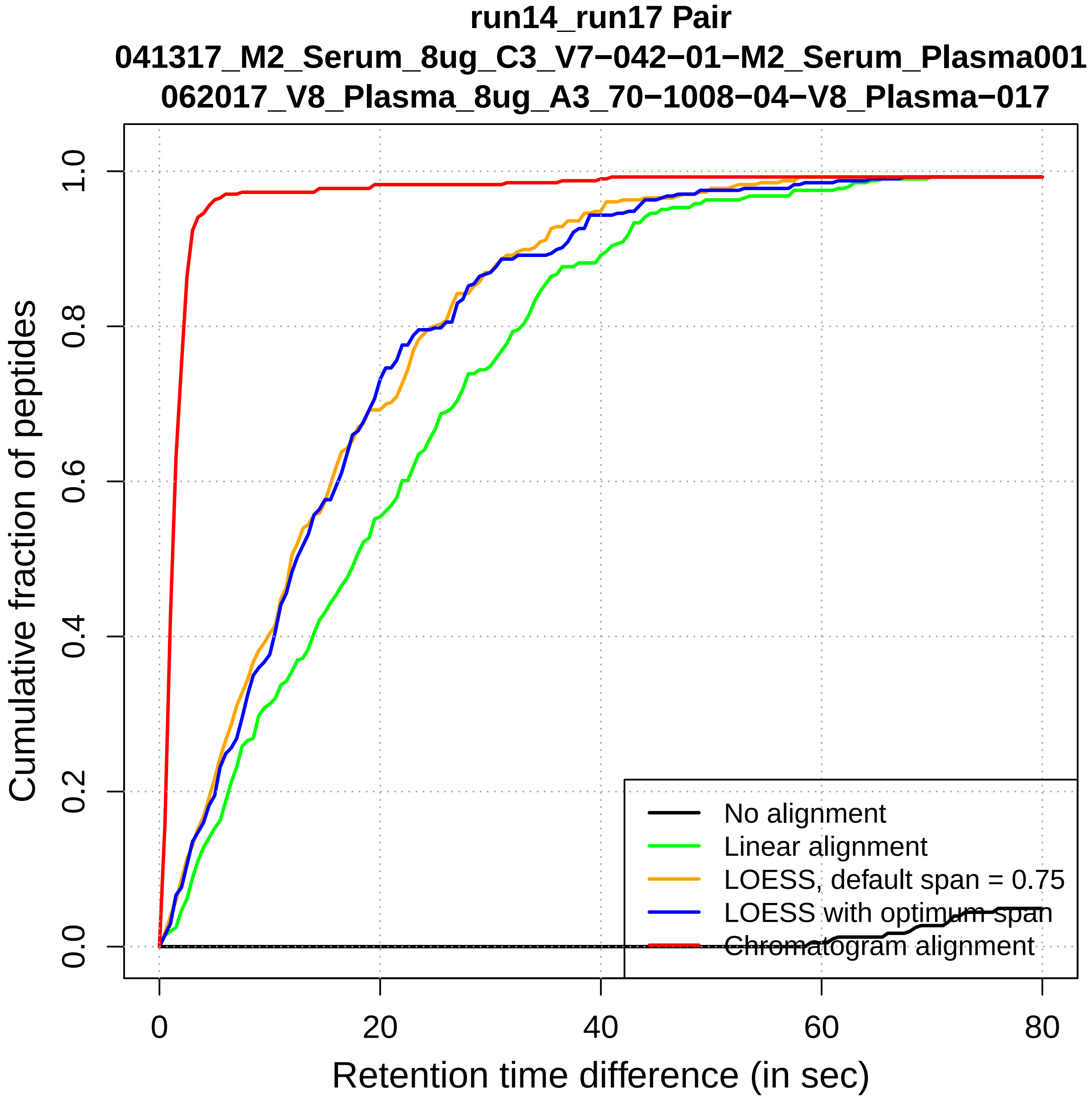
run14_run17 Pair
041317_M2_Serum_8ug_C3_V7−042−01−M2_Serum_Plasma001
062017_V8_Plasma_8ug_A3_70−1008−04−V8_Plasma−017
Cumulative fraction of peptides
Retention time difference (in sec)
0.0
0.2
0.4
0.6
0.8
1.0
0
20
40
60
80
No alignment
Linear alignment
LOESS, default span = 0.75
LOESS with optimum span
Chromatogram alignment

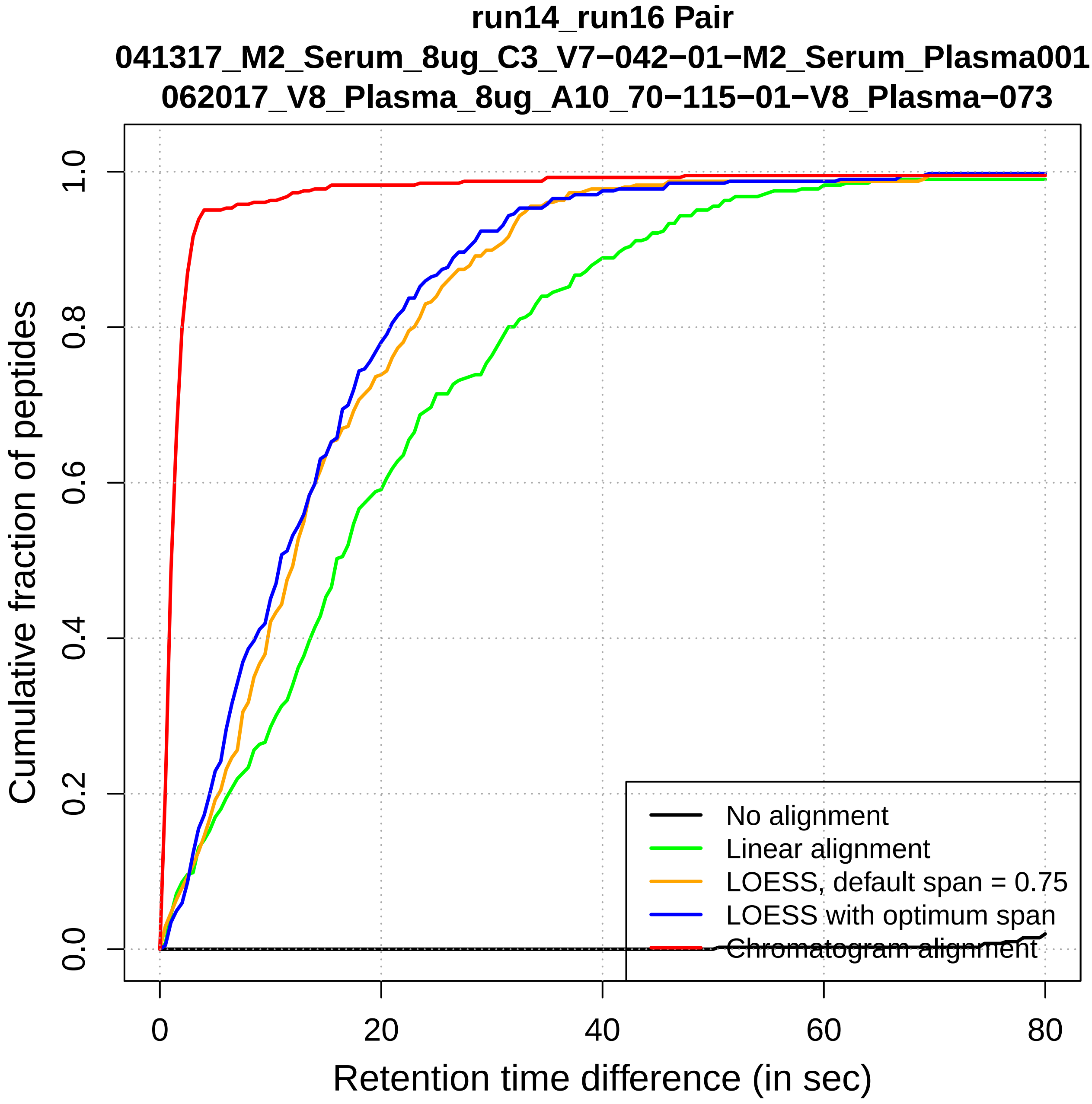

run14_run16 Pair
041317_M2_Serum_8ug_C3_V7−042−01−M2_Serum_Plasma001
062017_V8_Plasma_8ug_A10_70−115−01−V8_Plasma−073
Cumulative fraction of peptides
0.0
0.2
0.4
0.6
0.8
1.0
0
20
40
60
80
Retention time difference (in sec)
No alignment
Linear alignment
LOESS, default span = 0.75
LOESS with optimum span
Chromatogram alignment

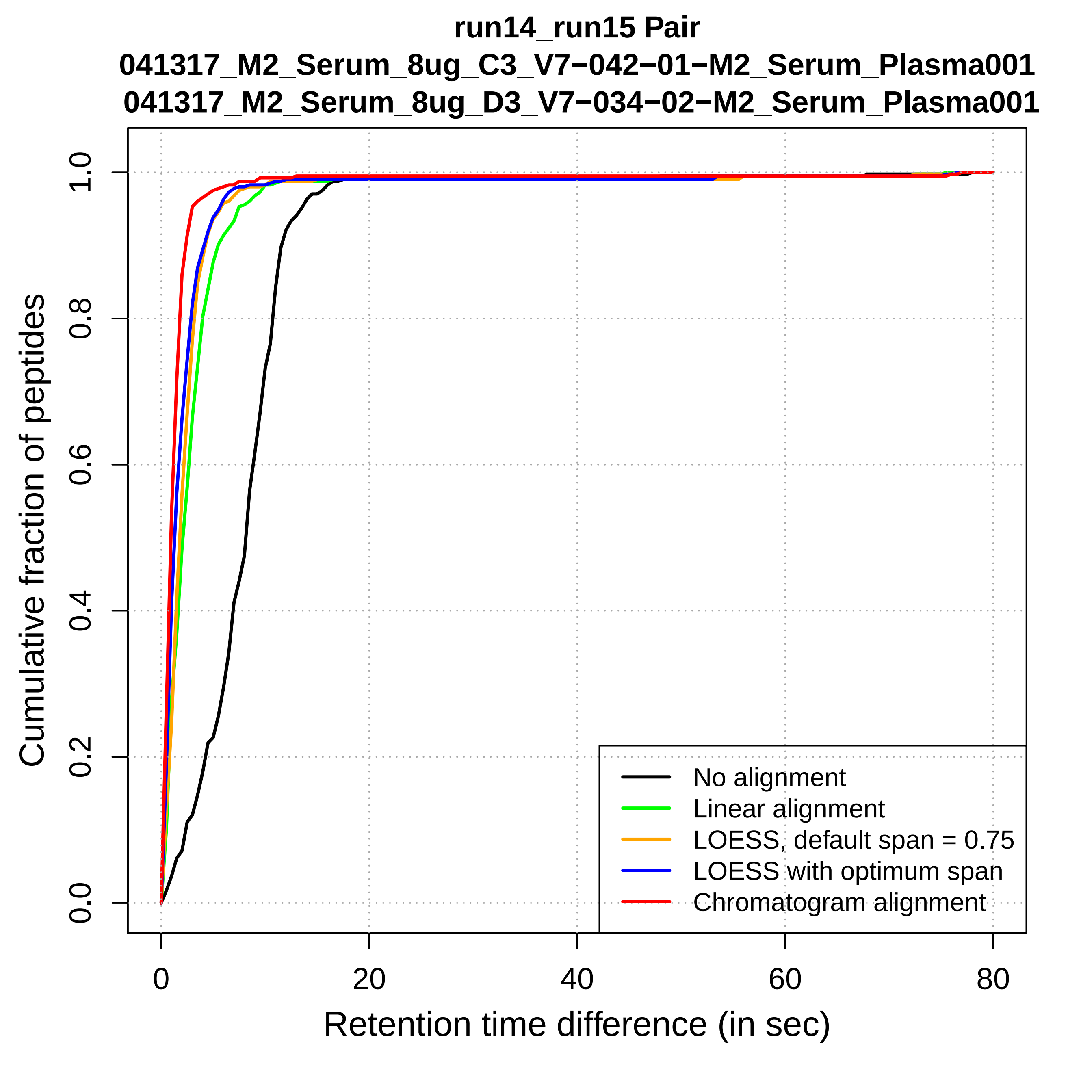

run14_run15 Pair
041317_M2_Serum_8ug_C3_V7−042−01−M2_Serum_Plasma001
041317_M2_Serum_8ug_D3_V7−034−02−M2_Serum_Plasma001
Cumulative fraction of peptides
Retention time difference (in sec)
0.0
0.2
0.4
0.6
0.8
1.0
0
20
40
60
80
No alignment
Linear alignment
LOESS, default span = 0.75
LOESS with optimum span
Chromatogram alignment

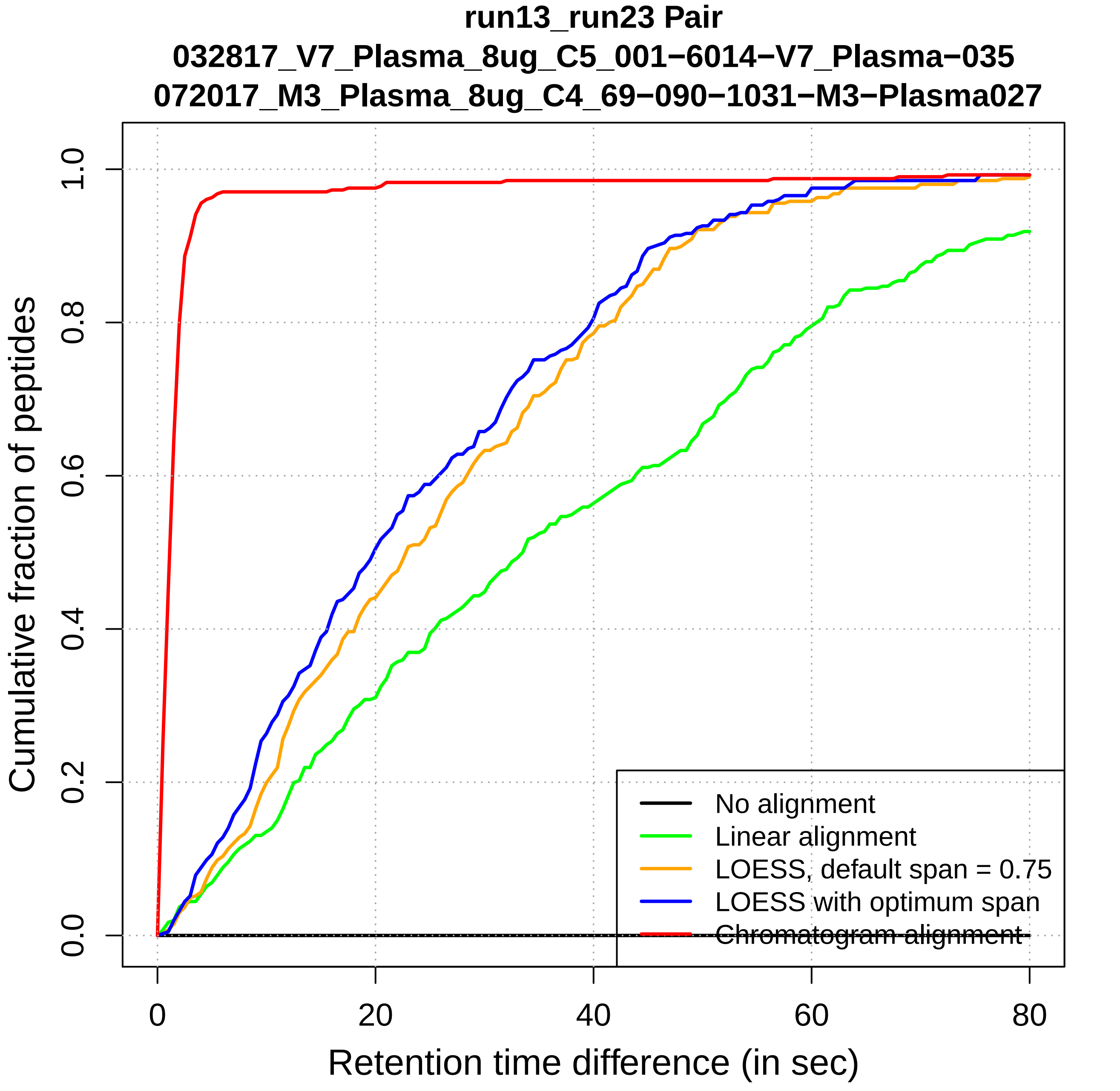

run13_run23 Pair
032817_V7_Plasma_8ug_C5_001−6014−V7_Plasma−035
072017_M3_Plasma_8ug_C4_69−090−1031−M3−Plasma027
Cumulative fraction of peptides
0.0
0.2
0.4
0.6
0.8
1.0
0
20
40
60
80
Retention time difference (in sec)
No alignment
Linear alignment
LOESS, default span = 0.75
LOESS with optimum span
Chromatogram alignment

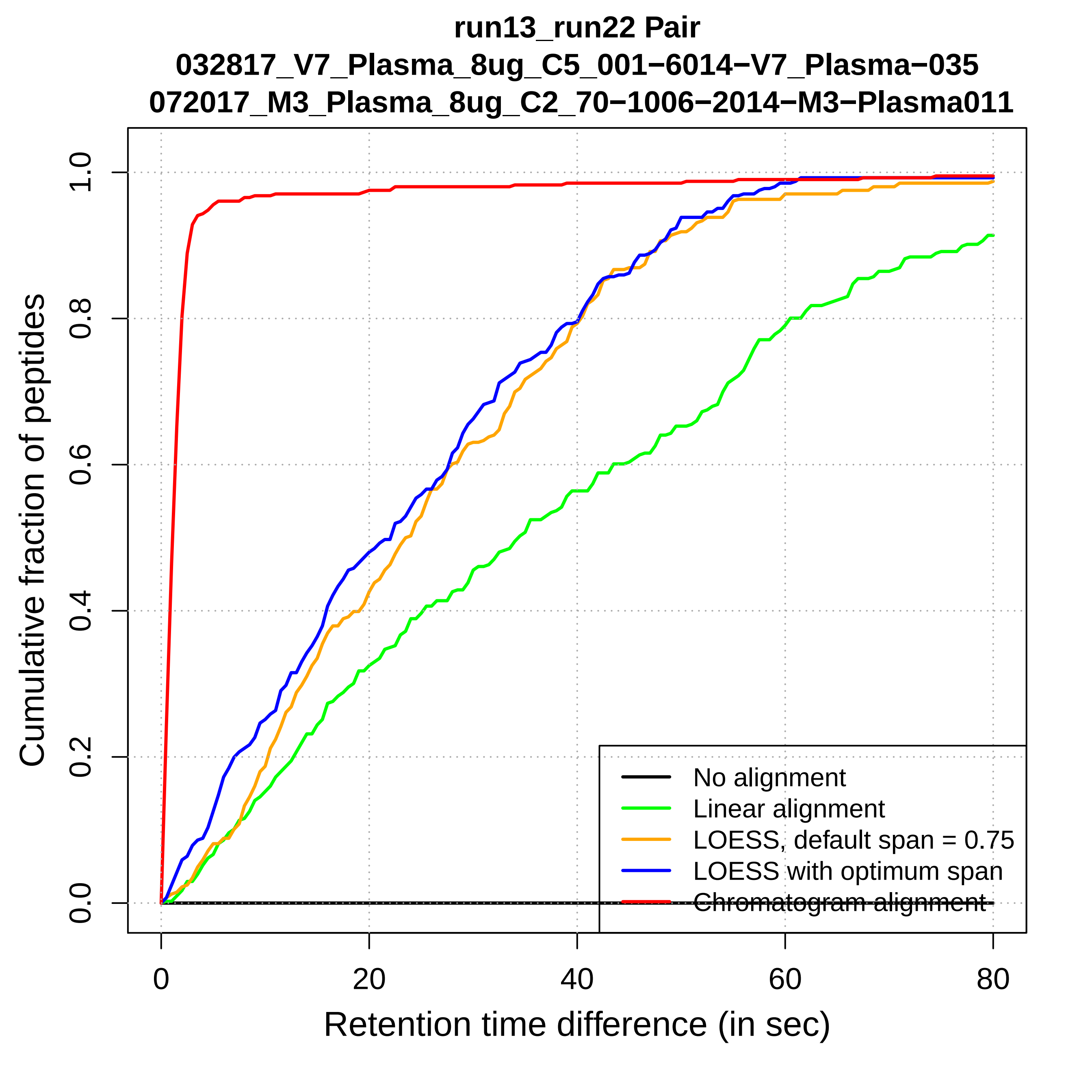
run13_run22 Pair
032817_V7_Plasma_8ug_C5_001−6014−V7_Plasma−035
072017_M3_Plasma_8ug_C2_70−1006−2014−M3−Plasma011
Cumulative fraction of peptides
0.0
0.2
0.4
0.6
0.8
1.0
0
20
40
60
80
Retention time difference (in sec)
No alignment
Linear alignment
LOESS, default span = 0.75
LOESS with optimum span
Chromatogram alignment

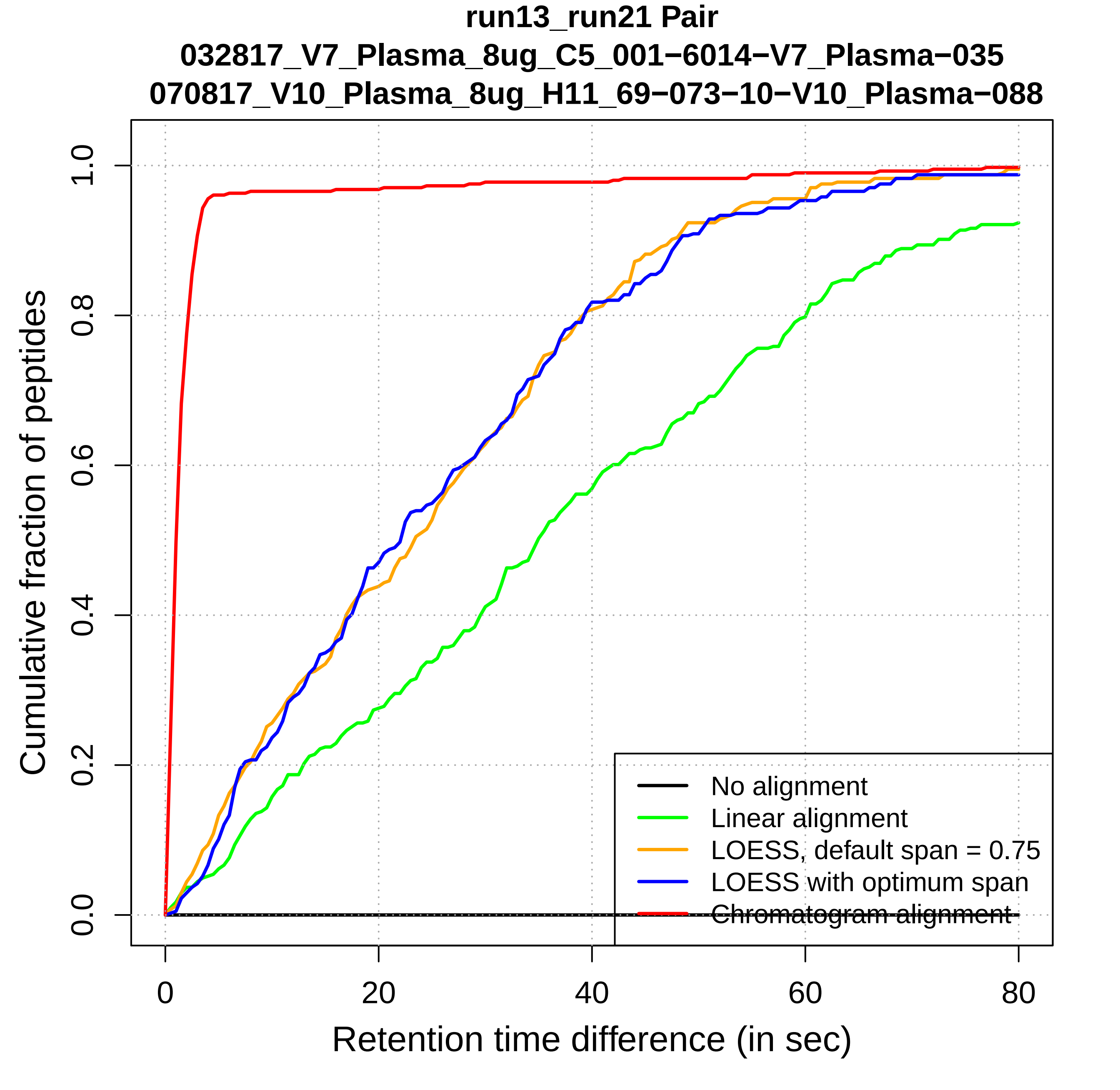
run13_run21 Pair
032817_V7_Plasma_8ug_C5_001−6014−V7_Plasma−035
070817_V10_Plasma_8ug_H11_69−073−10−V10_Plasma−088
Cumulative fraction of peptides
0.0
0.2
0.4
0.6
0.8
1.0
0
20
40
60
80
Retention time difference (in sec)
No alignment
Linear alignment
LOESS, default span = 0.75
LOESS with optimum span
Chromatogram alignment

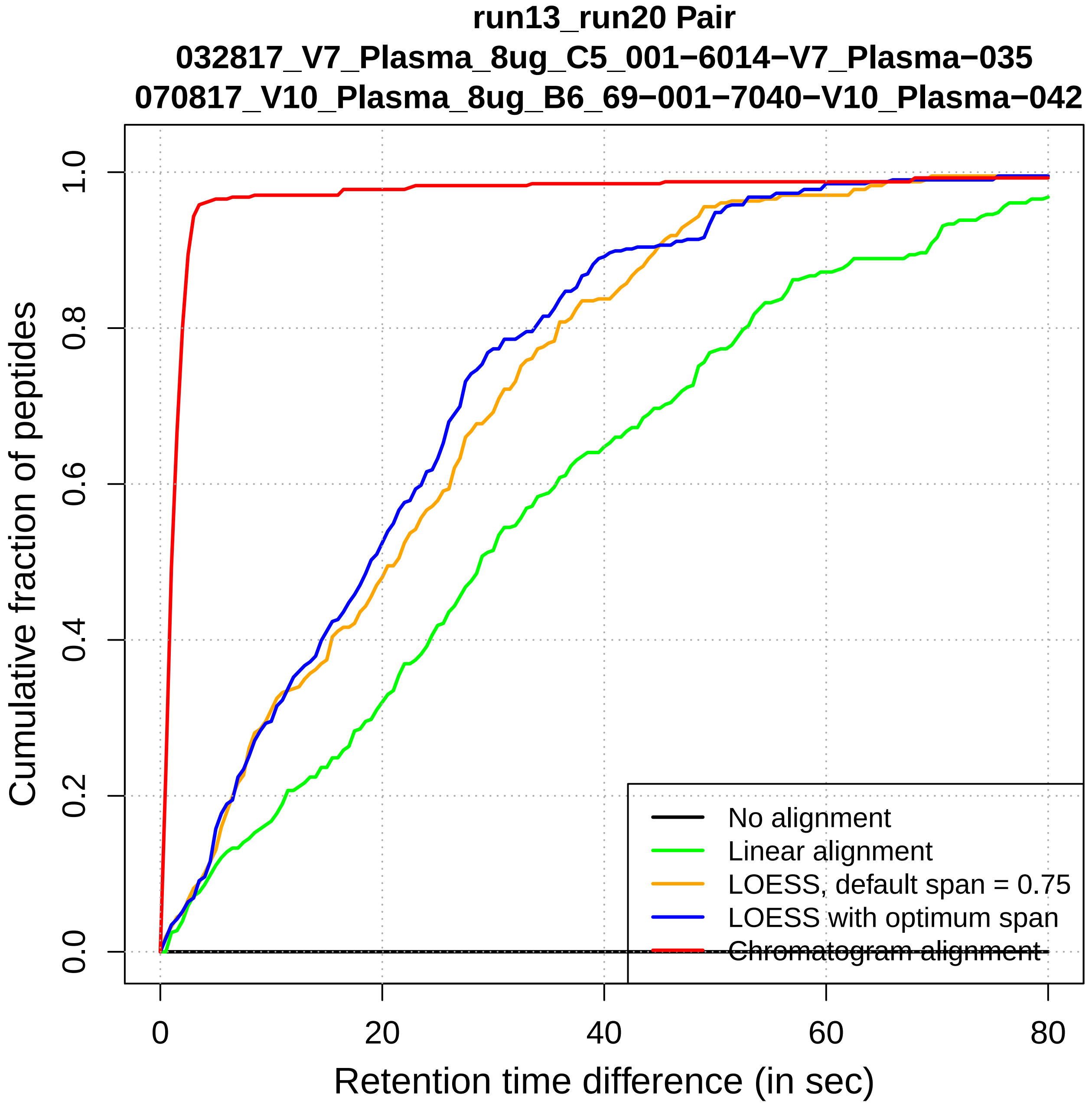

run13_run20 Pair
032817_V7_Plasma_8ug_C5_001−6014−V7_Plasma−035
070817_V10_Plasma_8ug_B6_69−001−7040−V10_Plasma−042
Cumulative fraction of peptides
Retention time difference (in sec)
0.0
0.2
0.4
0.6
0.8
1.0
0
20
40
60
80
No alignment
Linear alignment
LOESS, default span = 0.75
LOESS with optimum span
Chromatogram alignment

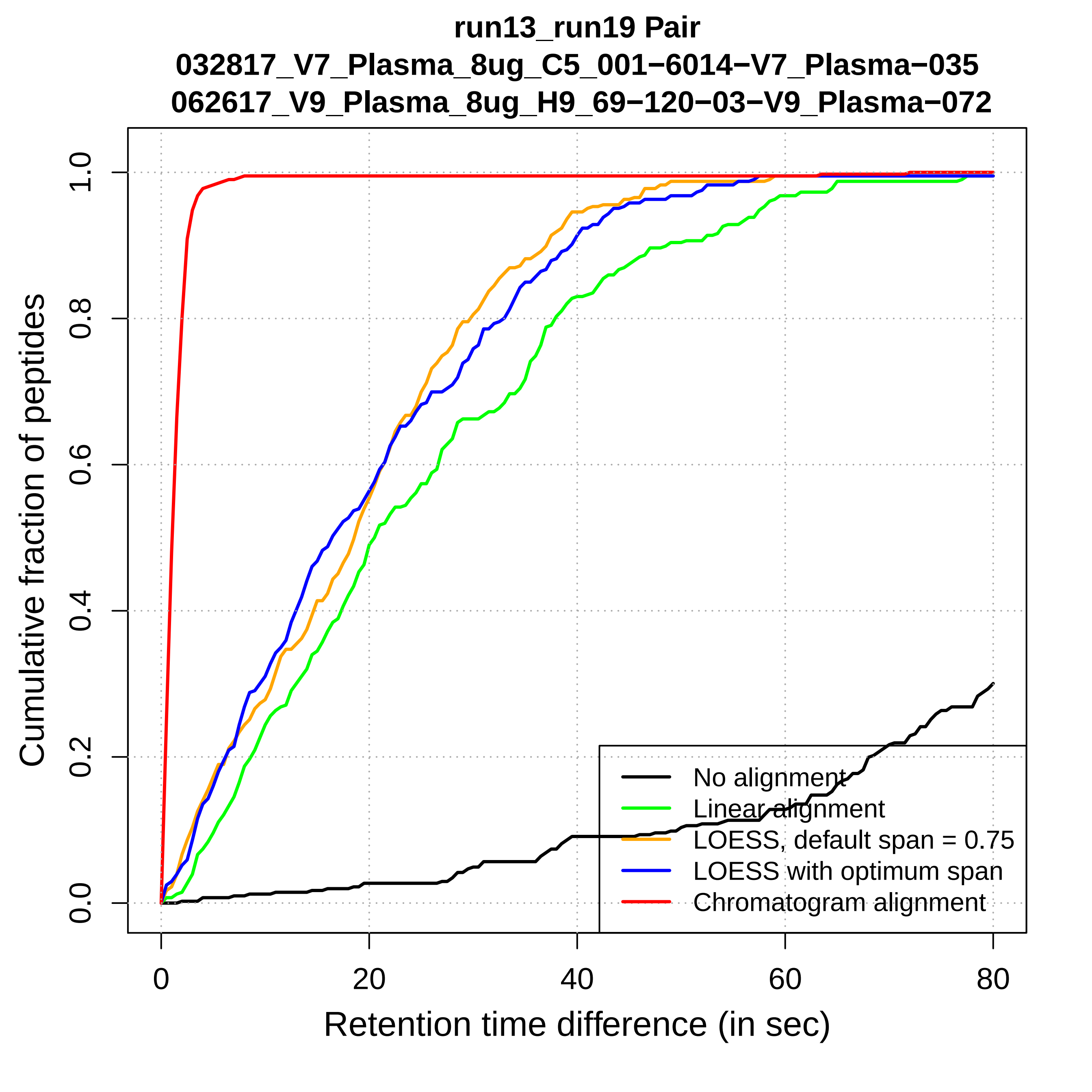

run13_run19 Pair
032817_V7_Plasma_8ug_C5_001−6014−V7_Plasma−035
062617_V9_Plasma_8ug_H9_69−120−03−V9_Plasma−072
Cumulative fraction of peptides
0.0
0.2
0.4
0.6
0.8
1.0
0
20
40
60
80
Retention time difference (in sec)
No alignment
Linear alignment
LOESS, default span = 0.75
LOESS with optimum span
Chromatogram alignment

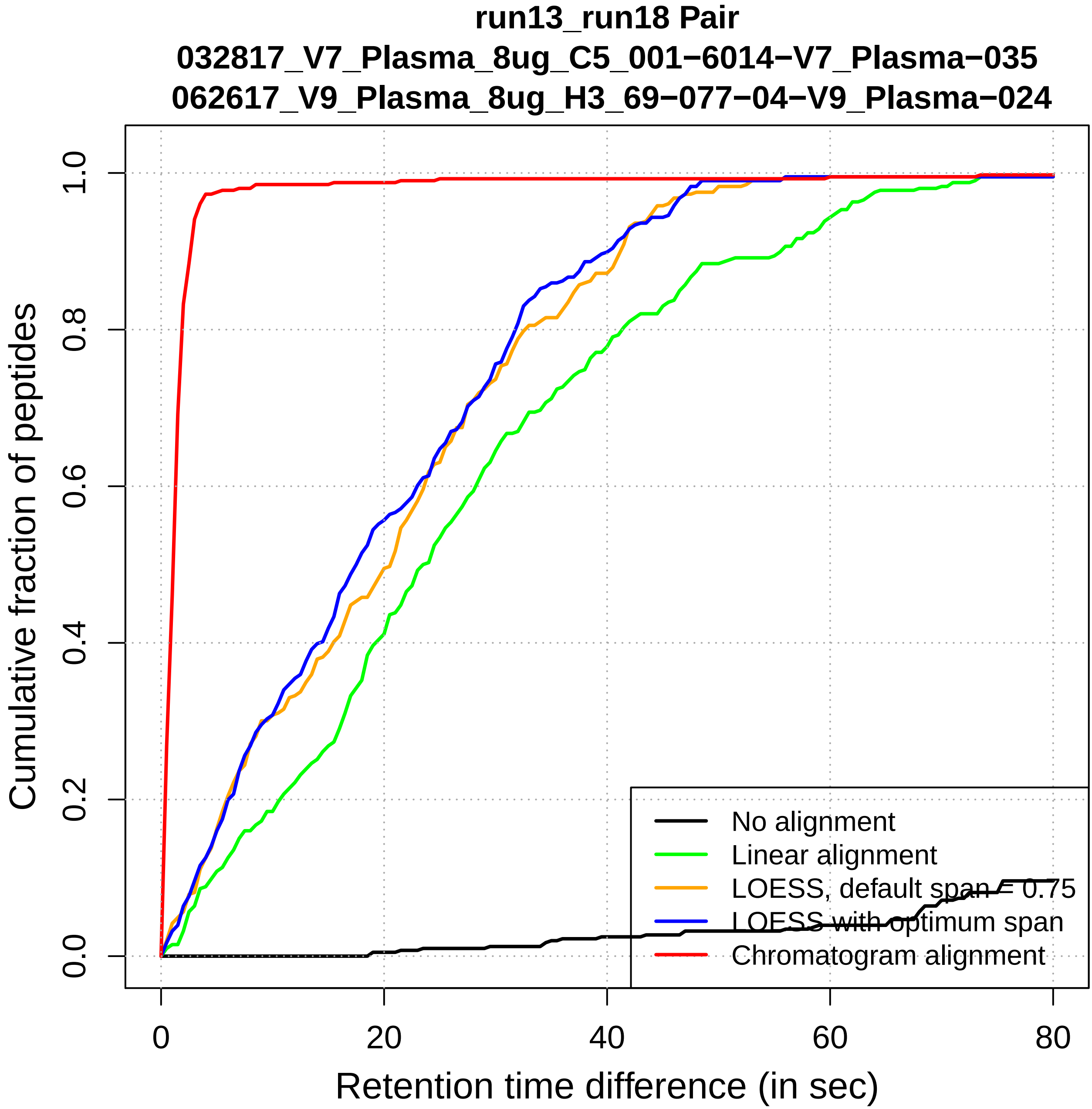
run13_run18 Pair
032817_V7_Plasma_8ug_C5_001−6014−V7_Plasma−035
062617_V9_Plasma_8ug_H3_69−077−04−V9_Plasma−024
Cumulative fraction of peptides
0.0
0.2
0.4
0.6
0.8
1.0
0
20
40
60
80
Retention time difference (in sec)
No alignment
Linear alignment
LOESS, default span = 0.75
LOESS with optimum span
Chromatogram alignment

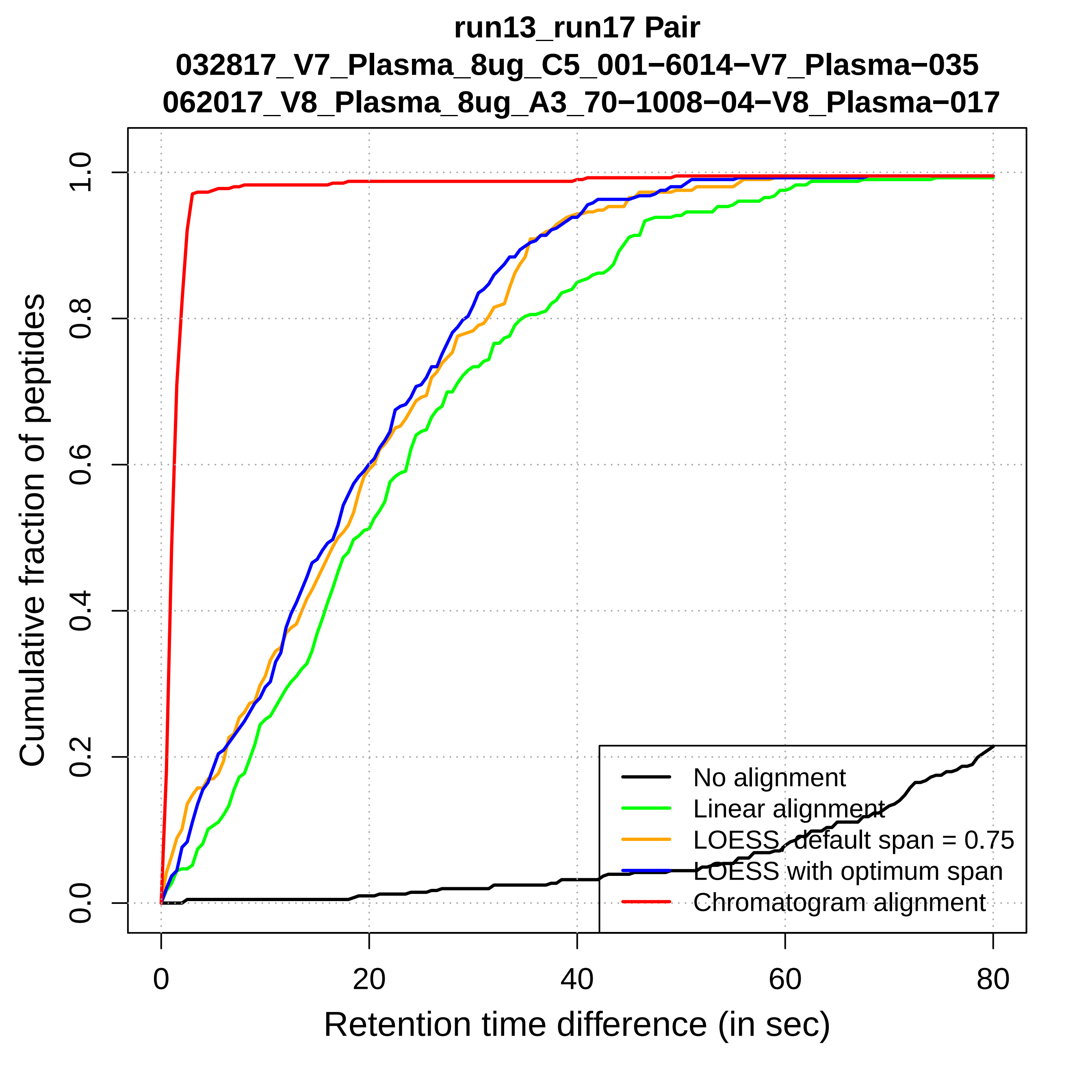
run13_run17 Pair
032817_V7_Plasma_8ug_C5_001−6014−V7_Plasma−035
062017_V8_Plasma_8ug_A3_70−1008−04−V8_Plasma−017
Cumulative fraction of peptides
0.0
0.2
0.4
0.6
0.8
1.0
0
20
40
60
80
Retention time difference (in sec)
No alignment
Linear alignment
LOESS, default span = 0.75
LOESS with optimum span
Chromatogram alignment

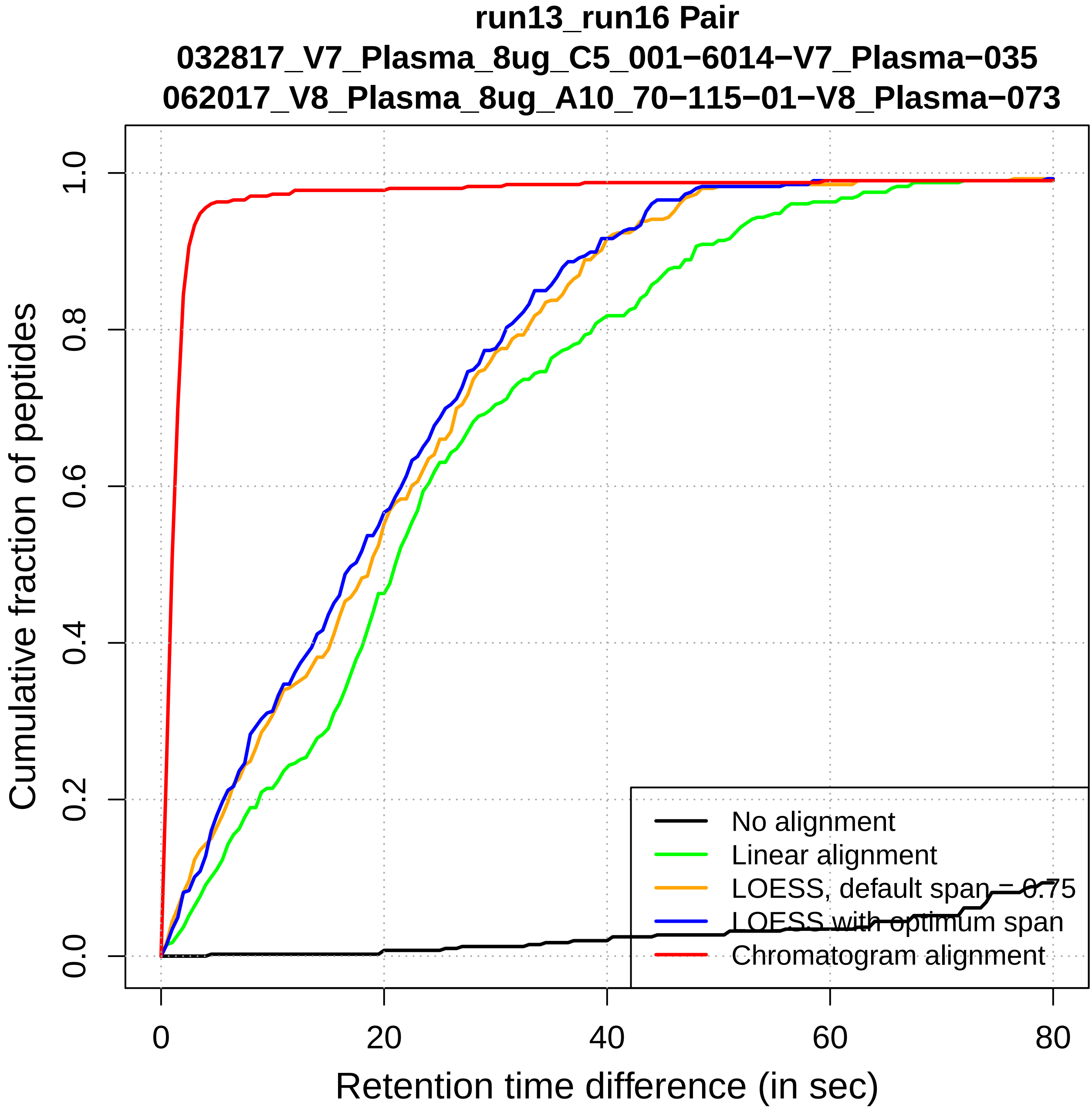

run13_run16 Pair
032817_V7_Plasma_8ug_C5_001−6014−V7_Plasma−035
062017_V8_Plasma_8ug_A10_70−115−01−V8_Plasma−073
Cumulative fraction of peptides
0.0
0.2
0.4
0.6
0.8
1.0
0
20
40
60
80
Retention time difference (in sec)
No alignment
Linear alignment
LOESS, default span = 0.75
LOESS with optimum span
Chromatogram alignment

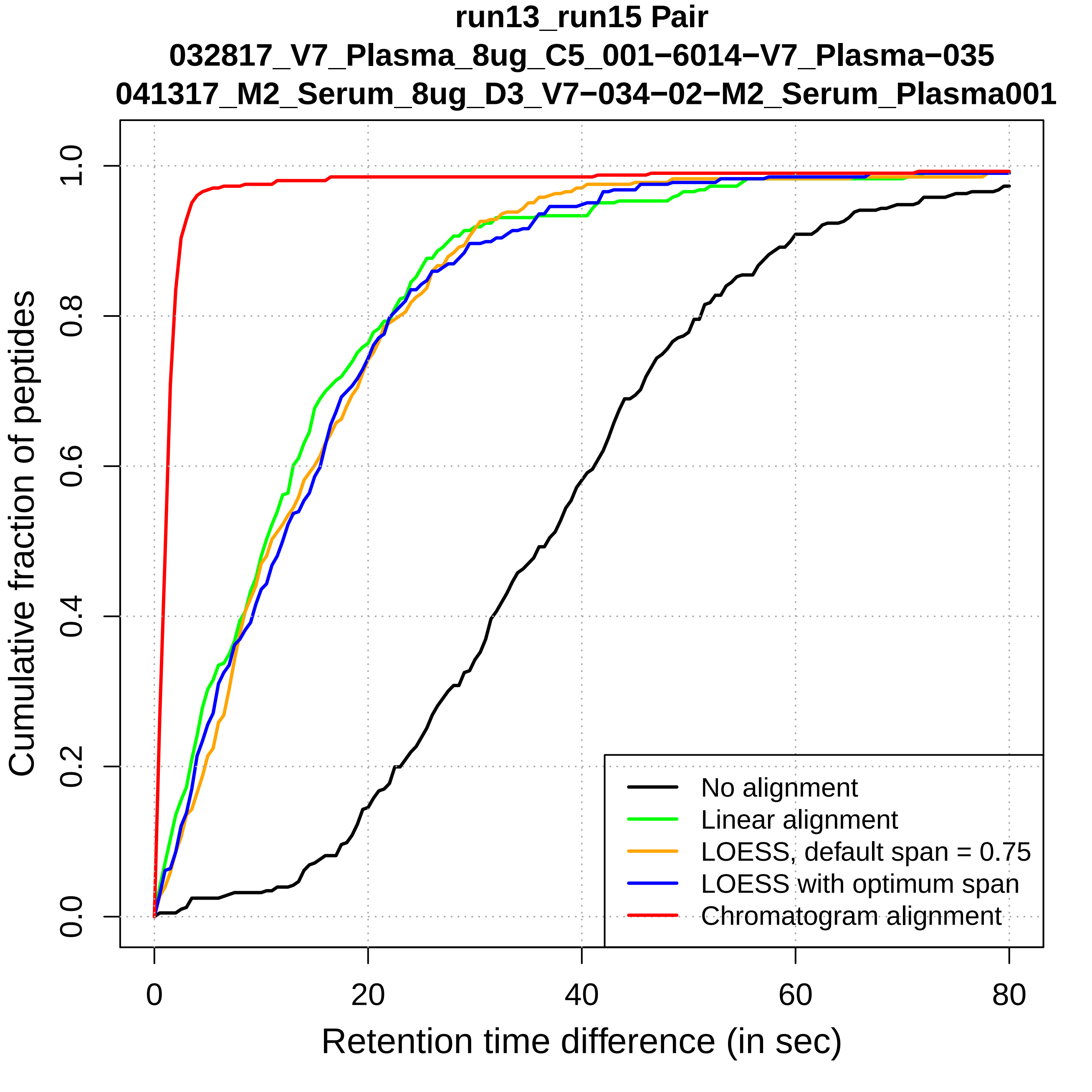
run13_run15 Pair
032817_V7_Plasma_8ug_C5_001−6014−V7_Plasma−035
041317_M2_Serum_8ug_D3_V7−034−02−M2_Serum_Plasma001
Cumulative fraction of peptides
0.0
0.2
0.4
0.6
0.8
1.0
0
20
40
60
80
Retention time difference (in sec)
No alignment
Linear alignment
LOESS, default span = 0.75
LOESS with optimum span
Chromatogram alignment

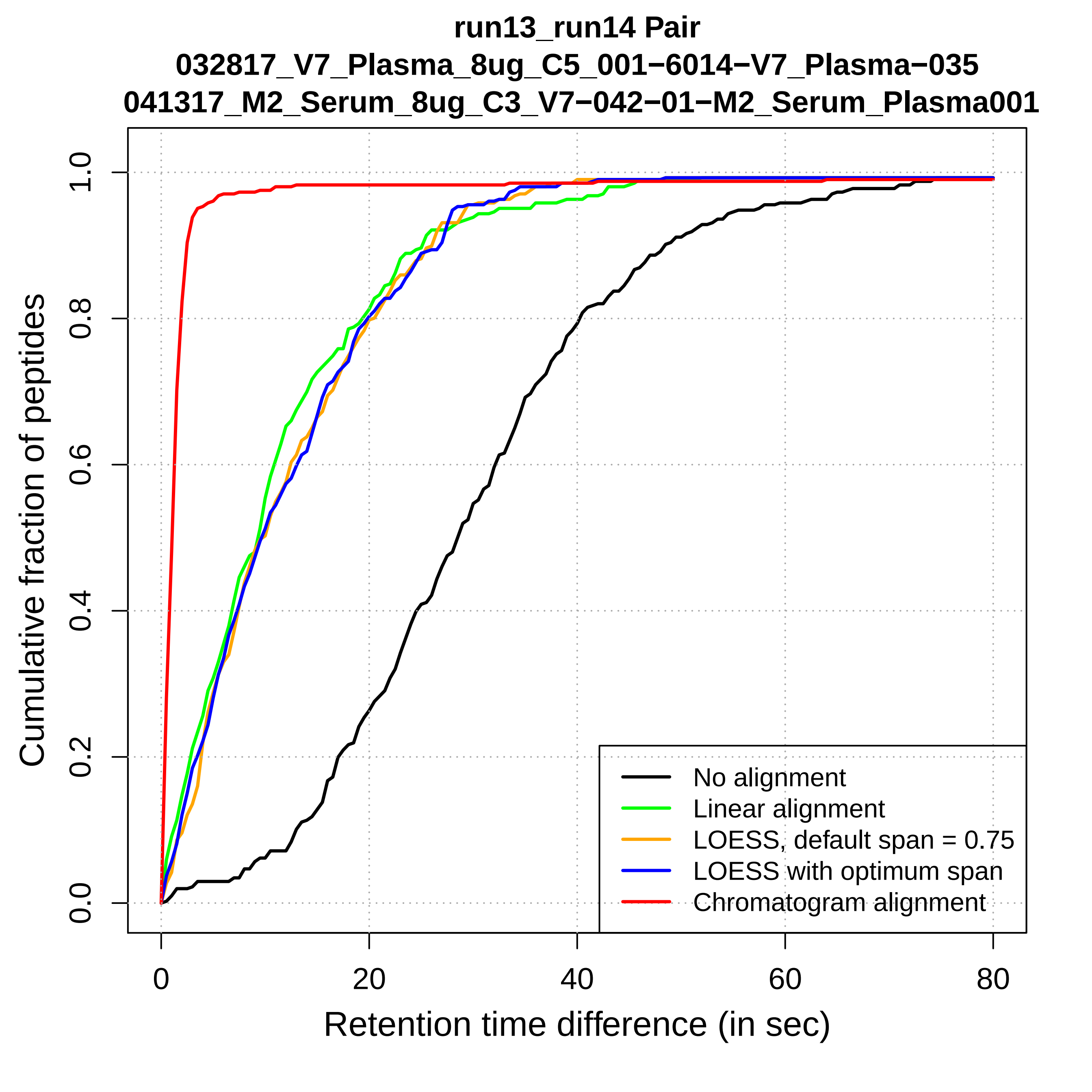
run13_run14 Pair
032817_V7_Plasma_8ug_C5_001−6014−V7_Plasma−035
041317_M2_Serum_8ug_C3_V7−042−01−M2_Serum_Plasma001
Cumulative fraction of peptides
0.0
0.2
0.4
0.6
0.8
1.0
0
20
40
60
80
Retention time difference (in sec)
No alignment
Linear alignment
LOESS, default span = 0.75
LOESS with optimum span
Chromatogram alignment

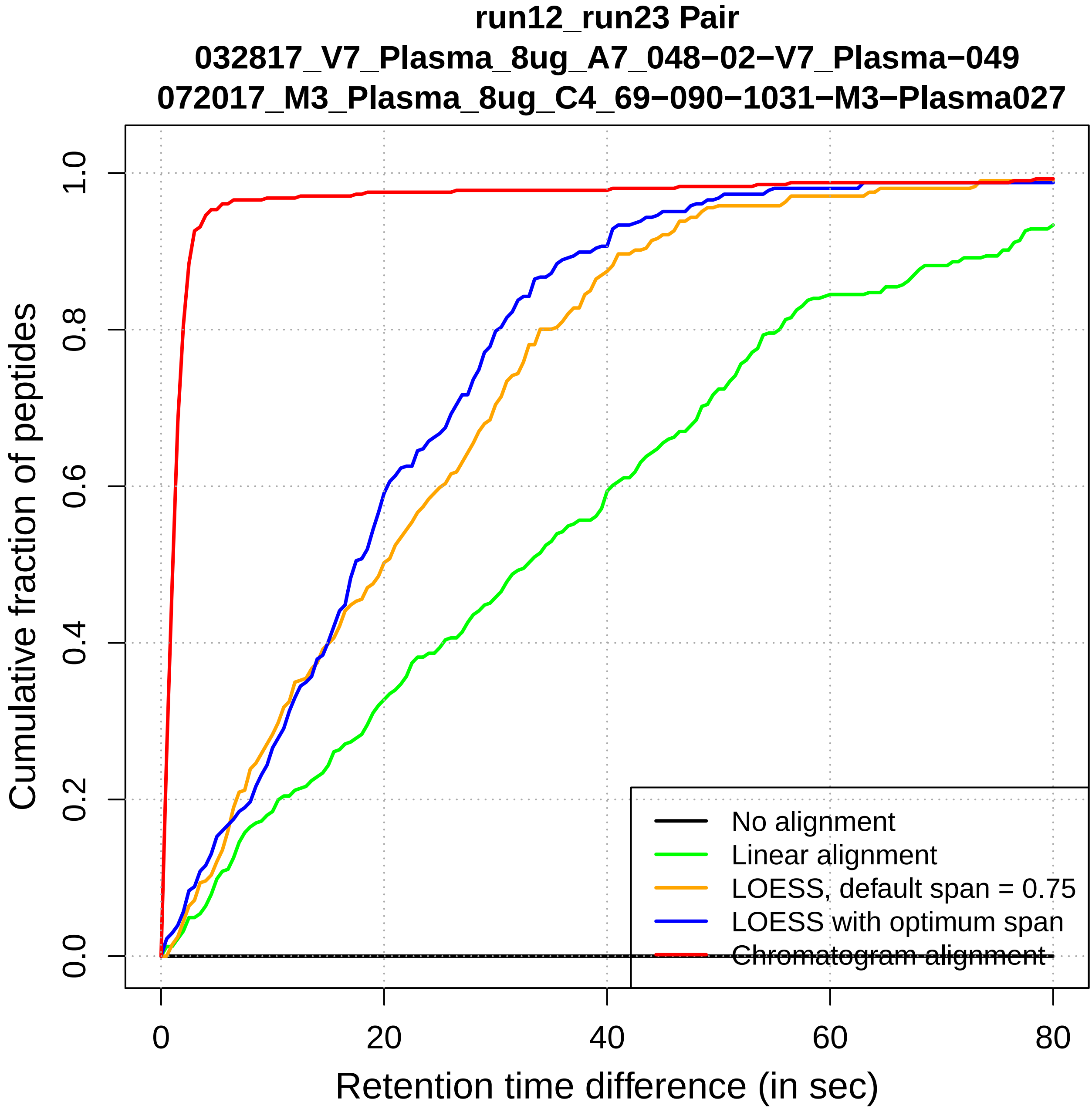
run12_run23 Pair
032817_V7_Plasma_8ug_A7_048−02−V7_Plasma−049
072017_M3_Plasma_8ug_C4_69−090−1031−M3−Plasma027
Cumulative fraction of peptides
0.0
0.2
0.4
0.6
0.8
1.0
0
20
40
60
80
Retention time difference (in sec)
No alignment
Linear alignment
LOESS, default span = 0.75
LOESS with optimum span
Chromatogram alignment

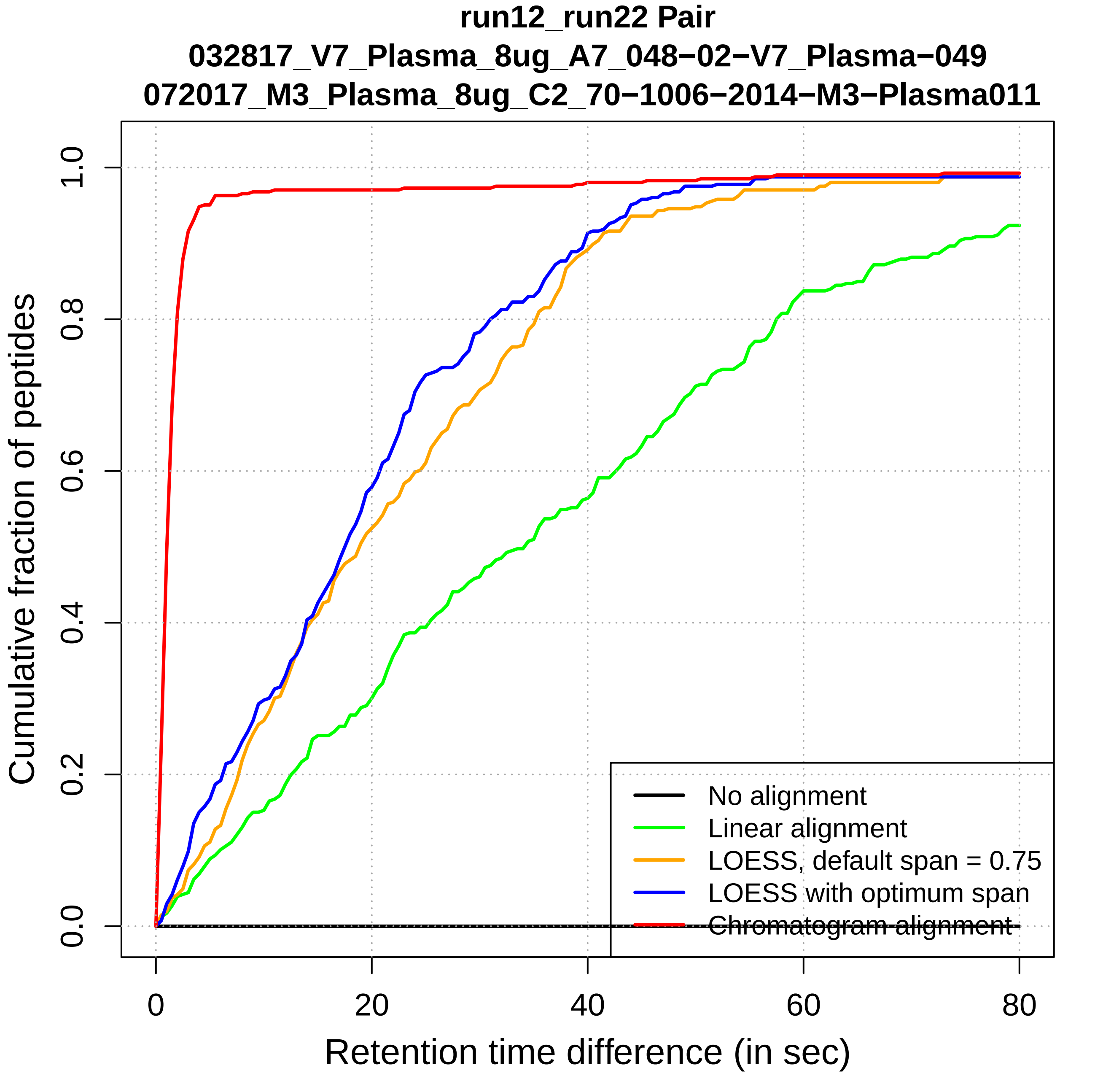
run12_run22 Pair
032817_V7_Plasma_8ug_A7_048−02−V7_Plasma−049
072017_M3_Plasma_8ug_C2_70−1006−2014−M3−Plasma011
Cumulative fraction of peptides
0.0
0.2
0.4
0.6
0.8
1.0
0
20
40
60
80
Retention time difference (in sec)
No alignment
Linear alignment
LOESS, default span = 0.75
LOESS with optimum span
Chromatogram alignment

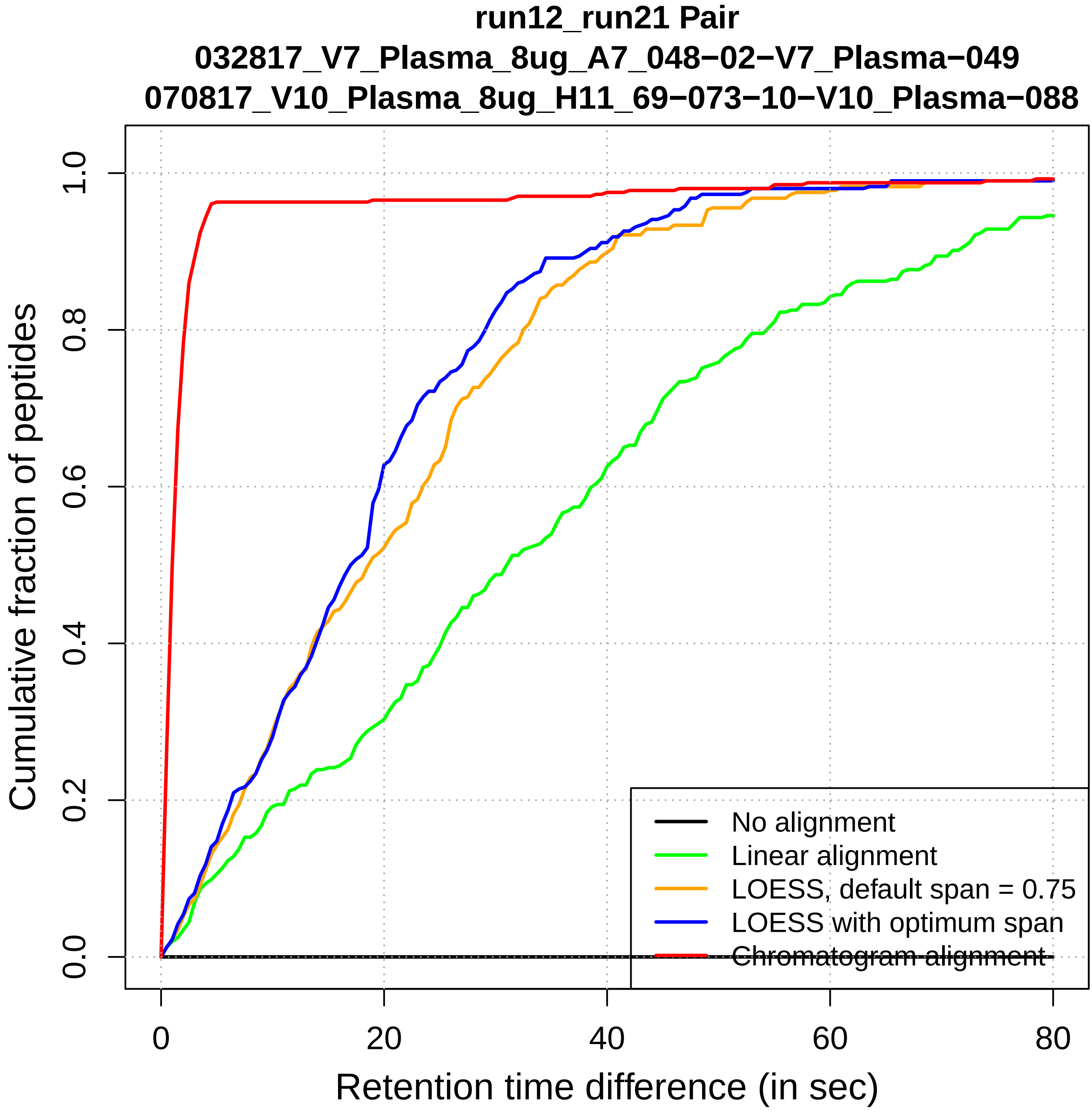
run12_run21 Pair
032817_V7_Plasma_8ug_A7_048−02−V7_Plasma−049
070817_V10_Plasma_8ug_H11_69−073−10−V10_Plasma−088
Cumulative fraction of peptides
0.0
0.2
0.4
0.6
0.8
1.0
0
20
40
60
80
Retention time difference (in sec)
No alignment
Linear alignment
LOESS, default span = 0.75
LOESS with optimum span
Chromatogram alignment

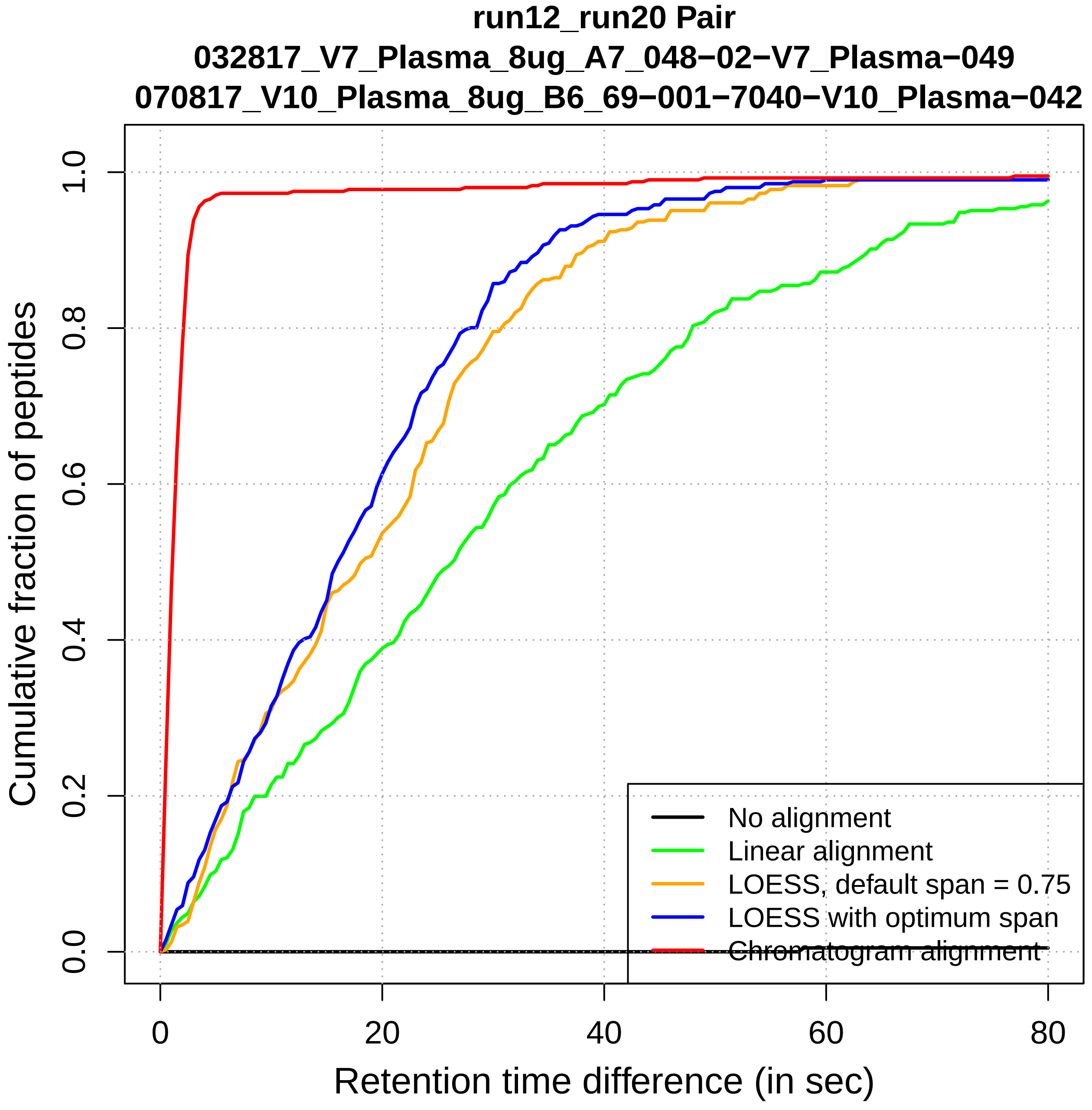
run12_run20 Pair
032817_V7_Plasma_8ug_A7_048−02−V7_Plasma−049
070817_V10_Plasma_8ug_B6_69−001−7040−V10_Plasma−042
Cumulative fraction of peptides
Retention time difference (in sec)
0.0
0.2
0.4
0.6
0.8
1.0
0
20
40
60
80
No alignment
Linear alignment
LOESS, default span = 0.75
LOESS with optimum span
Chromatogram alignment

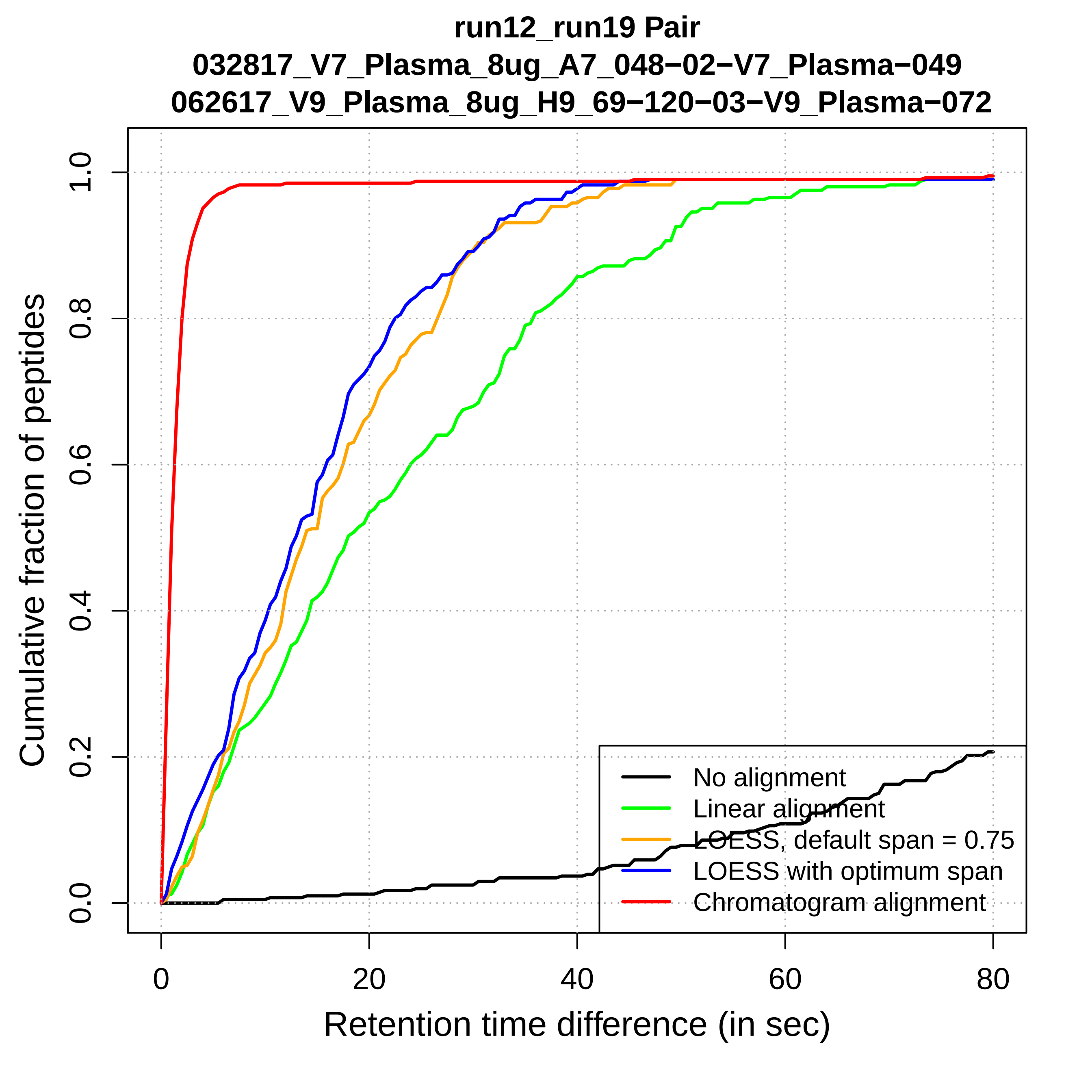

run12_run19 Pair
032817_V7_Plasma_8ug_A7_048−02−V7_Plasma−049
062617_V9_Plasma_8ug_H9_69−120−03−V9_Plasma−072
Cumulative fraction of peptides
0.0
0.2
0.4
0.6
0.8
1.0
0
20
40
60
80
Retention time difference (in sec)
No alignment
Linear alignment
LOESS, default span = 0.75
LOESS with optimum span
Chromatogram alignment

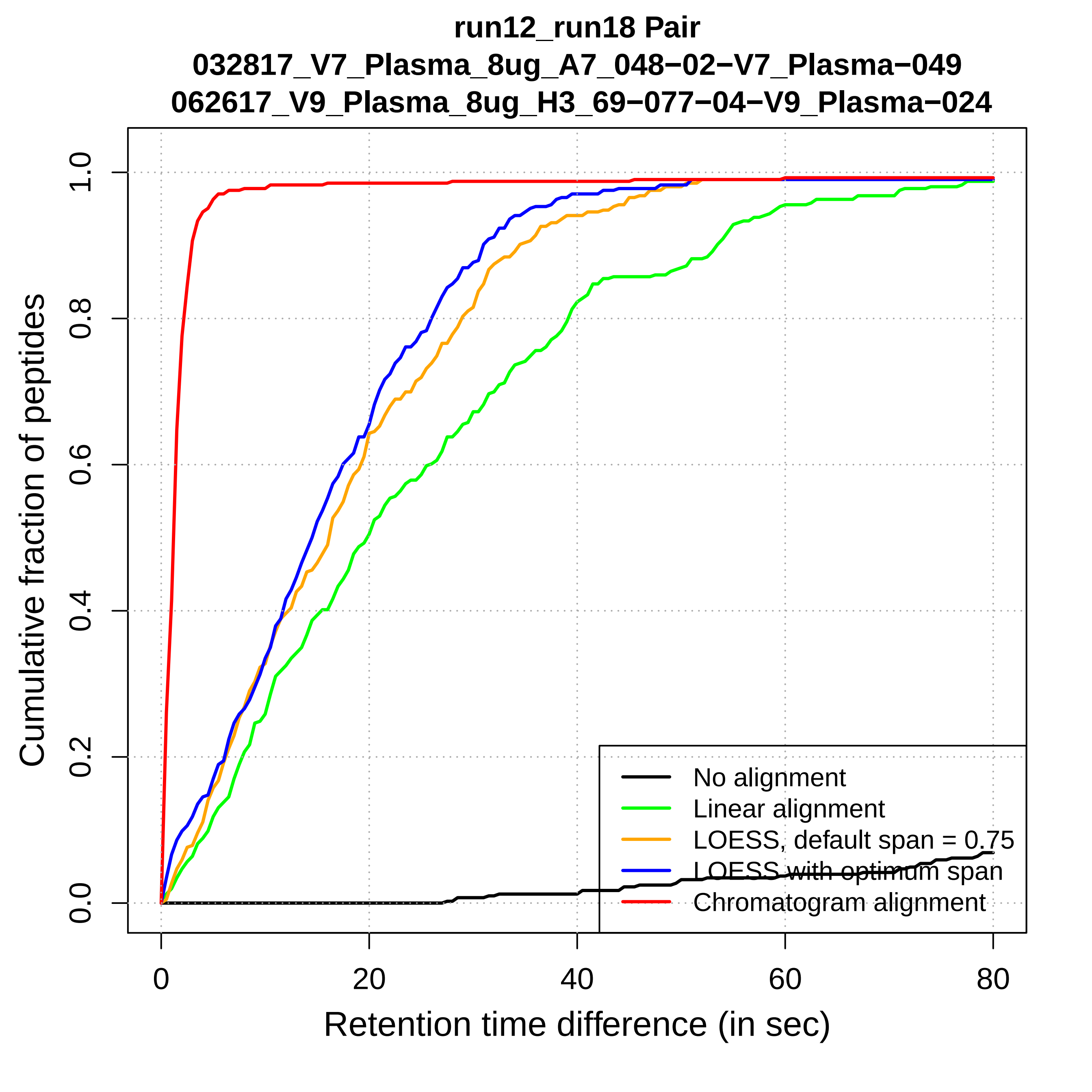
run12_run18 Pair
032817_V7_Plasma_8ug_A7_048−02−V7_Plasma−049
062617_V9_Plasma_8ug_H3_69−077−04−V9_Plasma−024
Cumulative fraction of peptides
0.0
0.2
0.4
0.6
0.8
1.0
0
20
40
60
80
Retention time difference (in sec)
No alignment
Linear alignment
LOESS, default span = 0.75
LOESS with optimum span
Chromatogram alignment

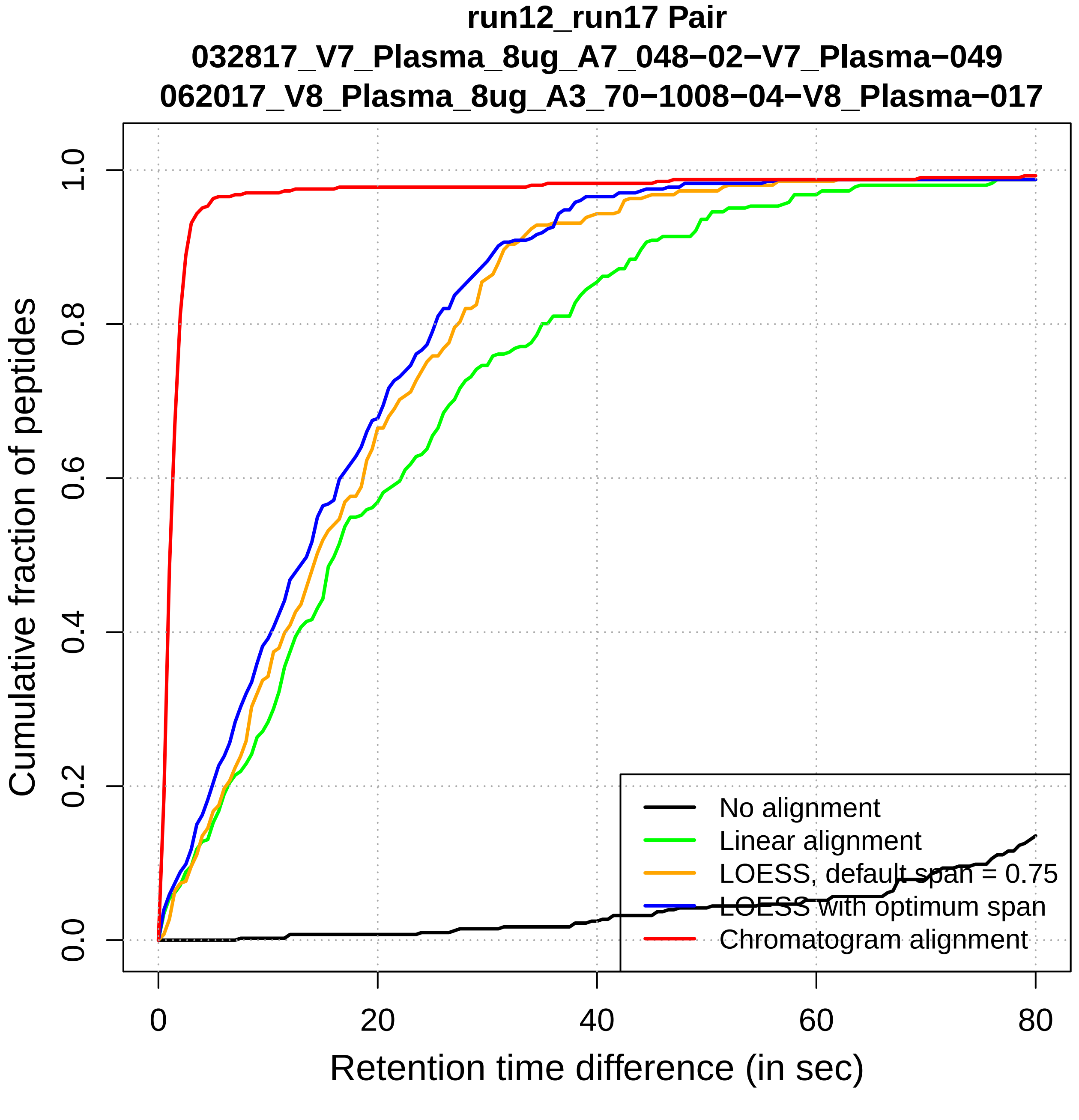
run12_run17 Pair
032817_V7_Plasma_8ug_A7_048−02−V7_Plasma−049
062017_V8_Plasma_8ug_A3_70−1008−04−V8_Plasma−017
Cumulative fraction of peptides
0.0
0.2
0.4
0.6
0.8
1.0
0
20
40
60
80
Retention time difference (in sec)
No alignment
Linear alignment
LOESS, default span = 0.75
LOESS with optimum span
Chromatogram alignment

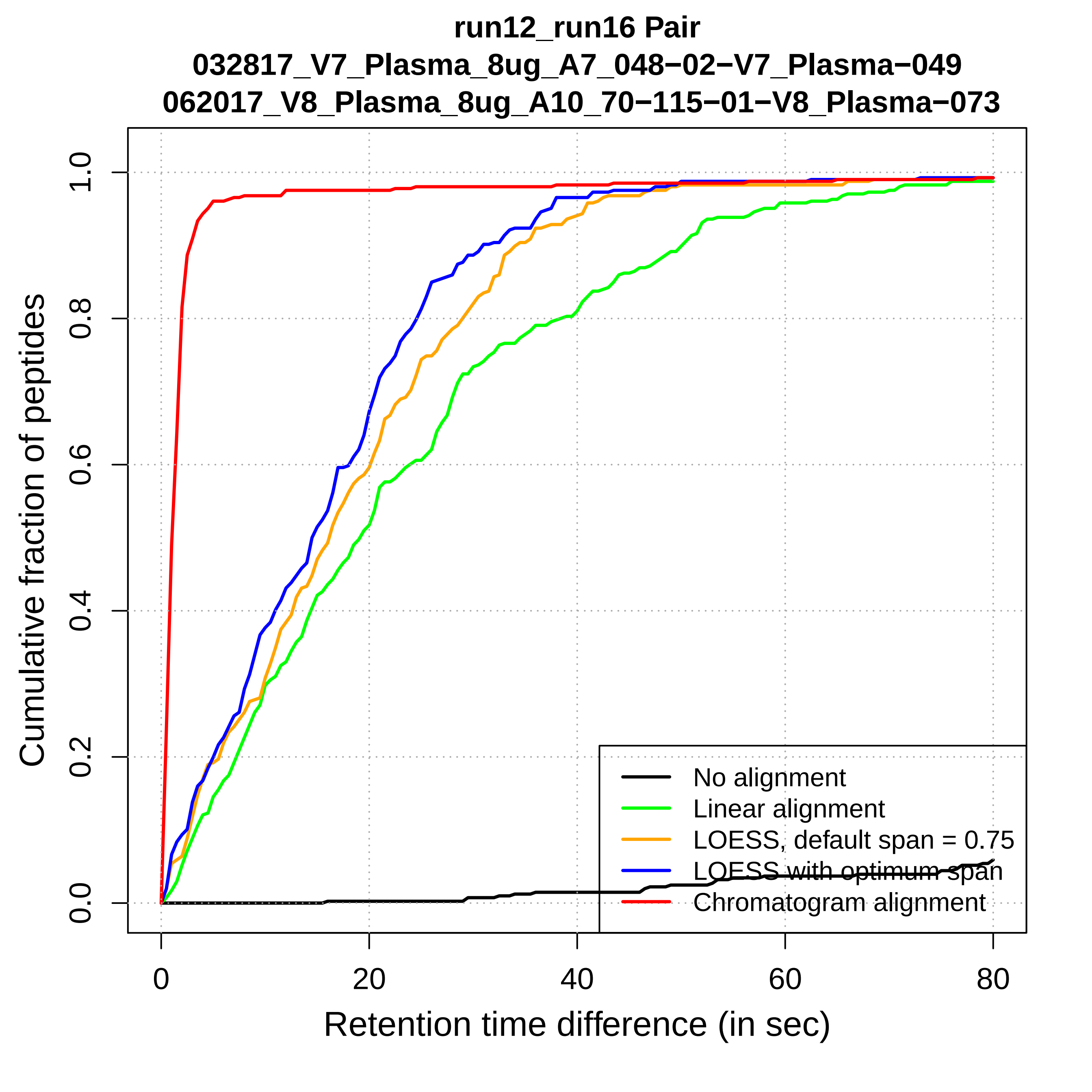
run12_run16 Pair
032817_V7_Plasma_8ug_A7_048−02−V7_Plasma−049
062017_V8_Plasma_8ug_A10_70−115−01−V8_Plasma−073
Cumulative fraction of peptides
0.0
0.2
0.4
0.6
0.8
1.0
0
20
40
60
80
Retention time difference (in sec)
No alignment
Linear alignment
LOESS, default span = 0.75
LOESS with optimum span
Chromatogram alignment

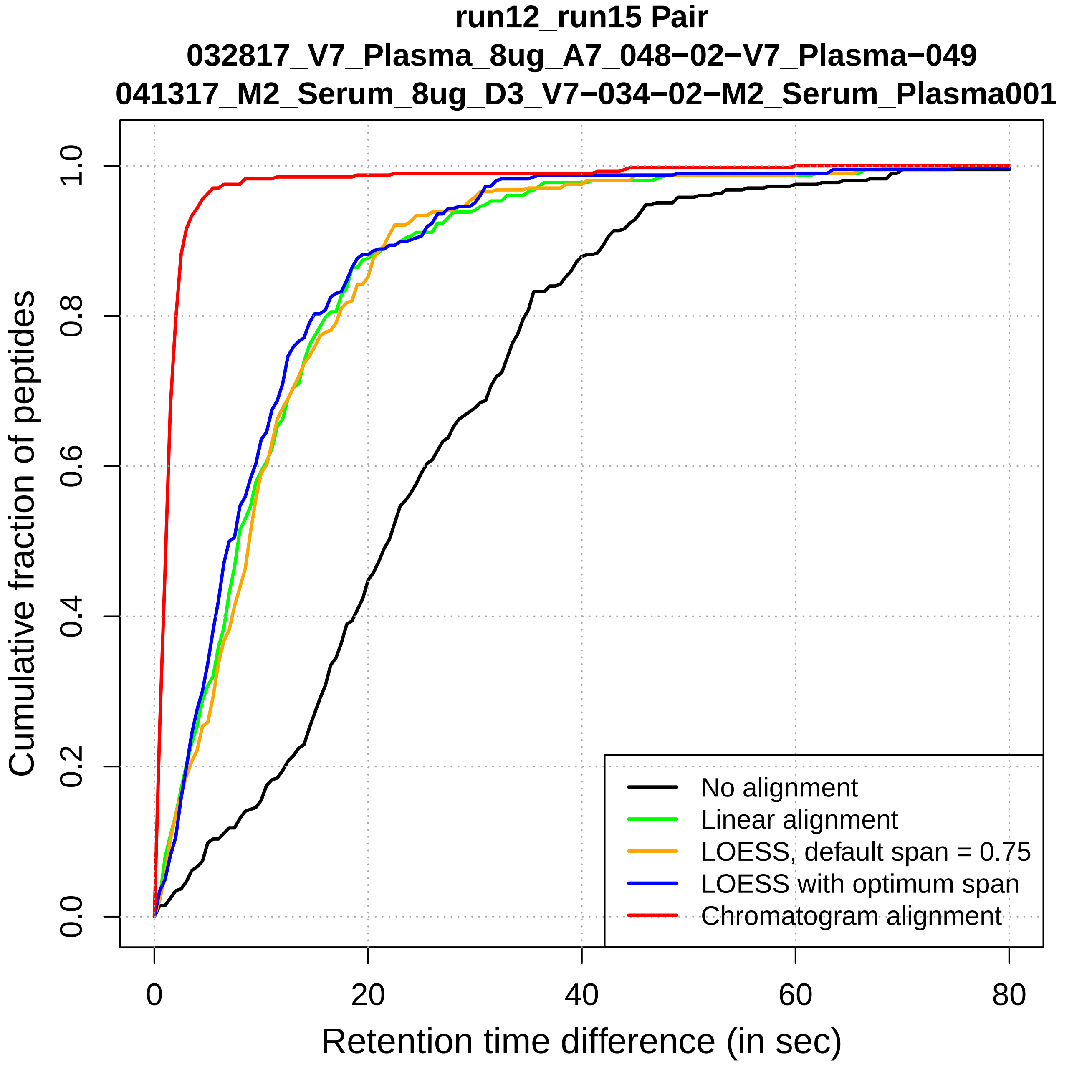

run12_run15 Pair
032817_V7_Plasma_8ug_A7_048−02−V7_Plasma−049
041317_M2_Serum_8ug_D3_V7−034−02−M2_Serum_Plasma001
Cumulative fraction of peptides
0.0
0.2
0.4
0.6
0.8
1.0
0
20
40
60
80
Retention time difference (in sec)
No alignment
Linear alignment
LOESS, default span = 0.75
LOESS with optimum span
Chromatogram alignment

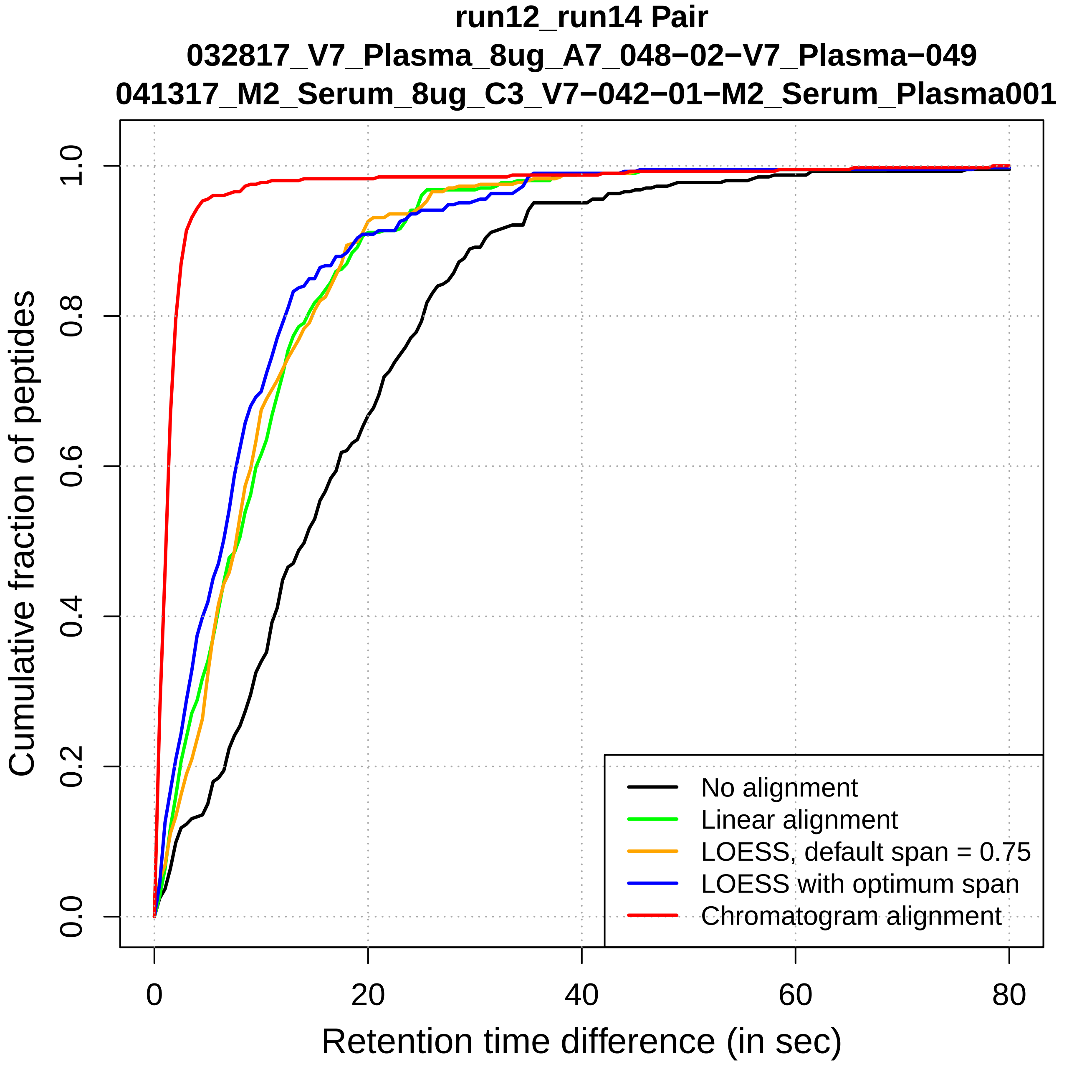

run12_run14 Pair
032817_V7_Plasma_8ug_A7_048−02−V7_Plasma−049
041317_M2_Serum_8ug_C3_V7−042−01−M2_Serum_Plasma001
Cumulative fraction of peptides
0.0
0.2
0.4
0.6
0.8
1.0
0
20
40
60
80
Retention time difference (in sec)
No alignment
Linear alignment
LOESS, default span = 0.75
LOESS with optimum span
Chromatogram alignment

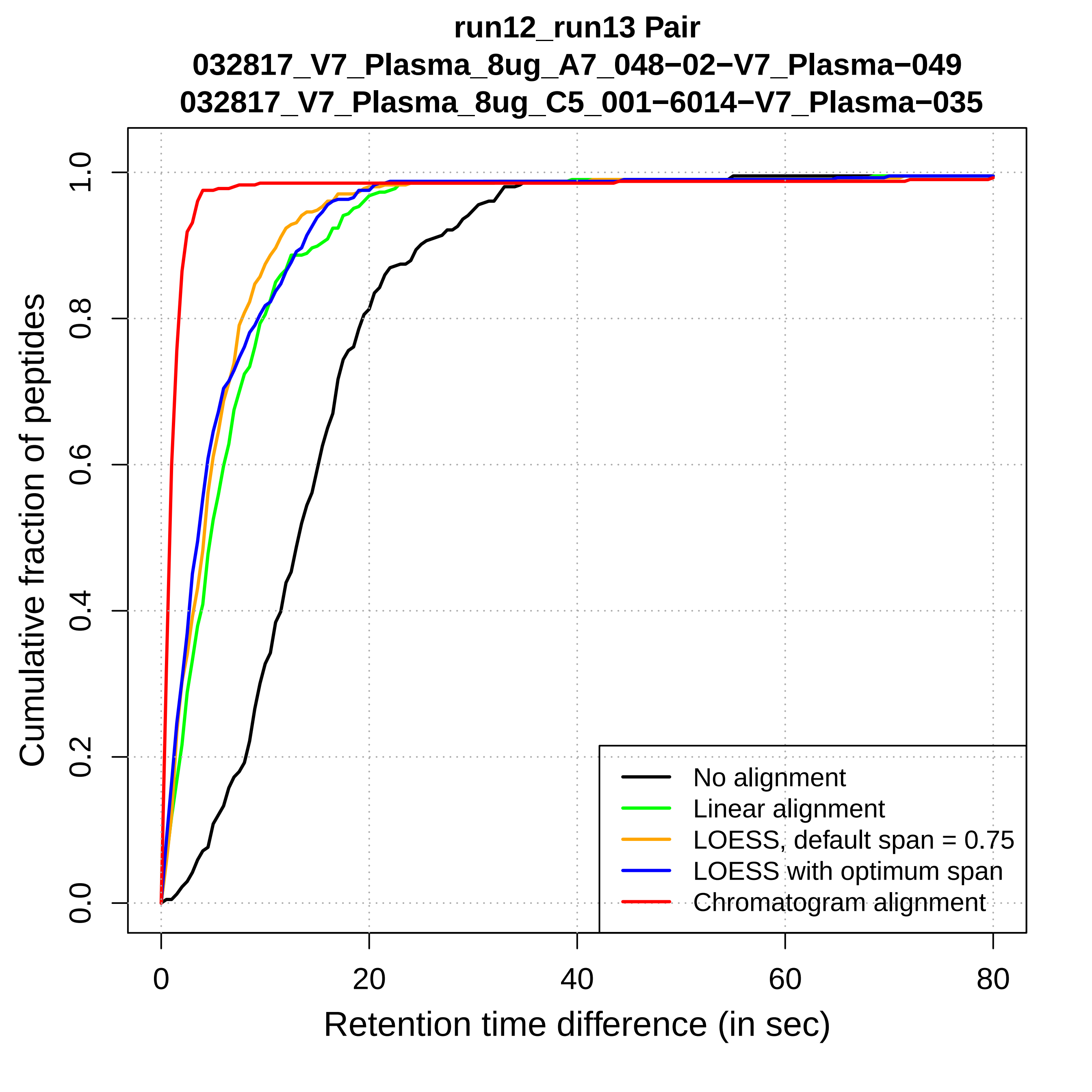

run12_run13 Pair
032817_V7_Plasma_8ug_A7_048−02−V7_Plasma−049
032817_V7_Plasma_8ug_C5_001−6014−V7_Plasma−035
Cumulative fraction of peptides
0.0
0.2
0.4
0.6
0.8
1.0
0
20
40
60
80
Retention time difference (in sec)
No alignment
Linear alignment
LOESS, default span = 0.75
LOESS with optimum span
Chromatogram alignment

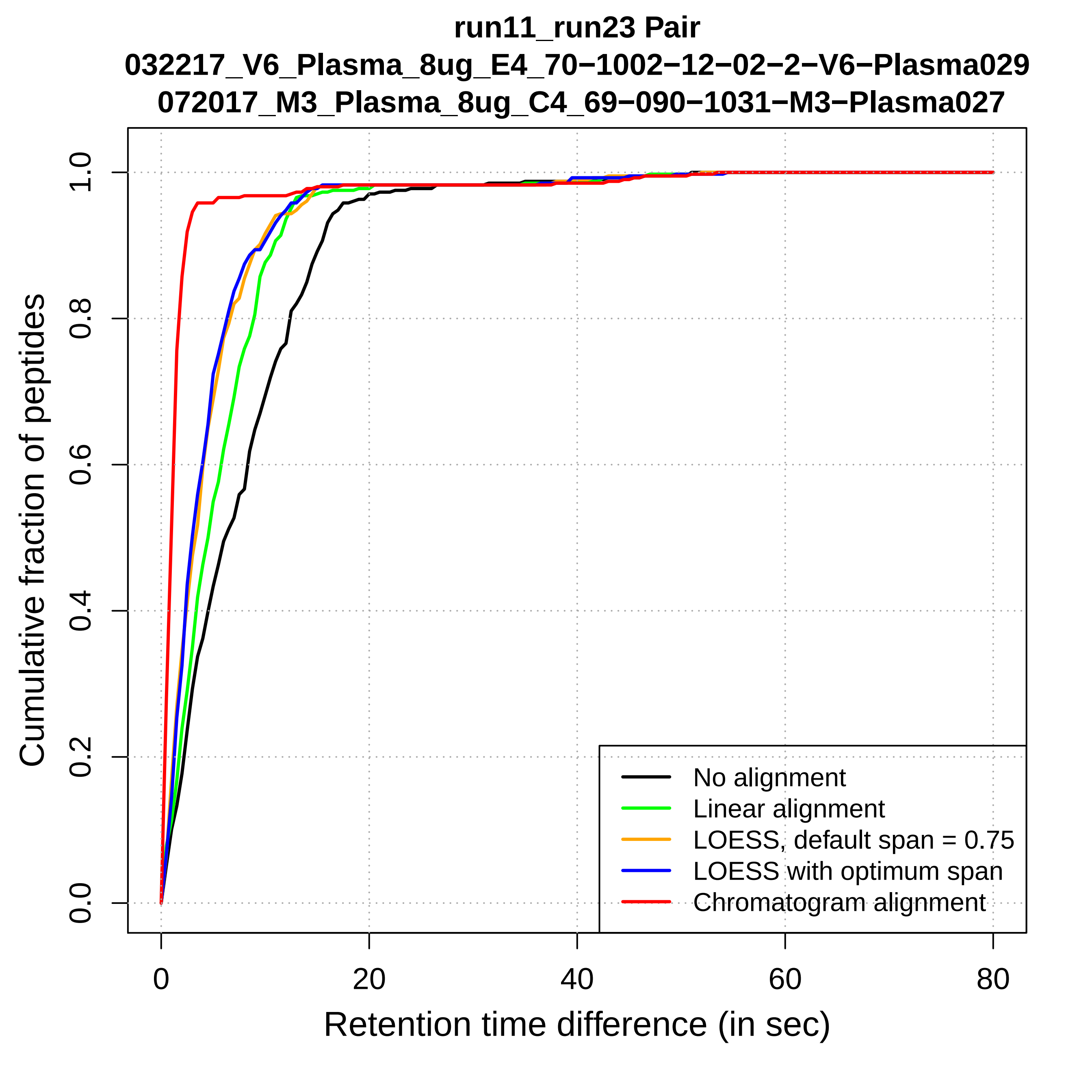
run11_run23 Pair
032217_V6_Plasma_8ug_E4_70−1002−12−02−2−V6−Plasma029
072017_M3_Plasma_8ug_C4_69−090−1031−M3−Plasma027
Cumulative fraction of peptides
Retention time difference (in sec)
0.0
0.2
0.4
0.6
0.8
1.0
0
20
40
60
80
No alignment
Linear alignment
LOESS, default span = 0.75
LOESS with optimum span
Chromatogram alignment

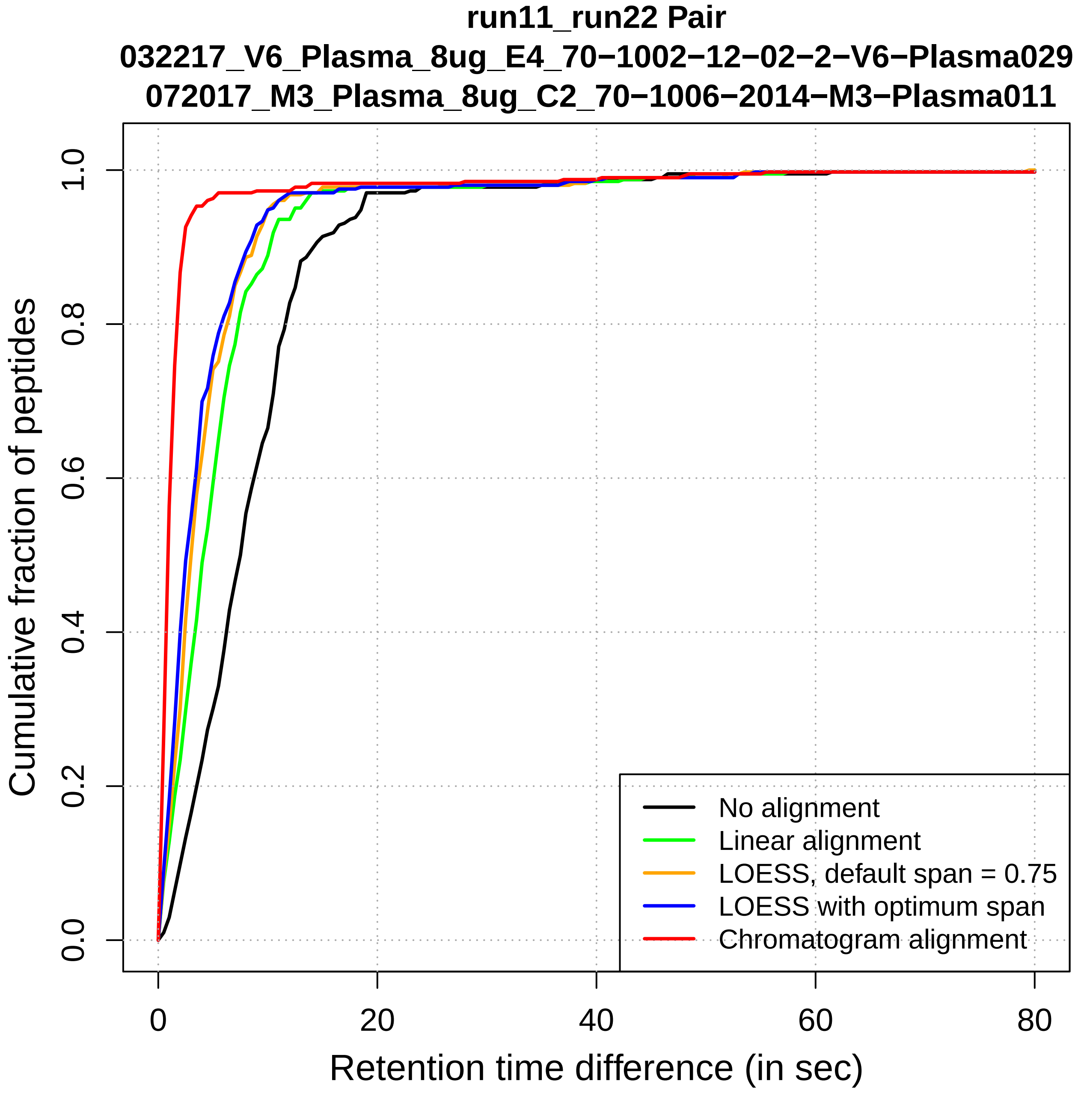

run11_run22 Pair
032217_V6_Plasma_8ug_E4_70−1002−12−02−2−V6−Plasma029
072017_M3_Plasma_8ug_C2_70−1006−2014−M3−Plasma011
Cumulative fraction of peptides
0.0
0.2
0.4
0.6
0.8
1.0
0
20
40
60
80
Retention time difference (in sec)
No alignment
Linear alignment
LOESS, default span = 0.75
LOESS with optimum span
Chromatogram alignment

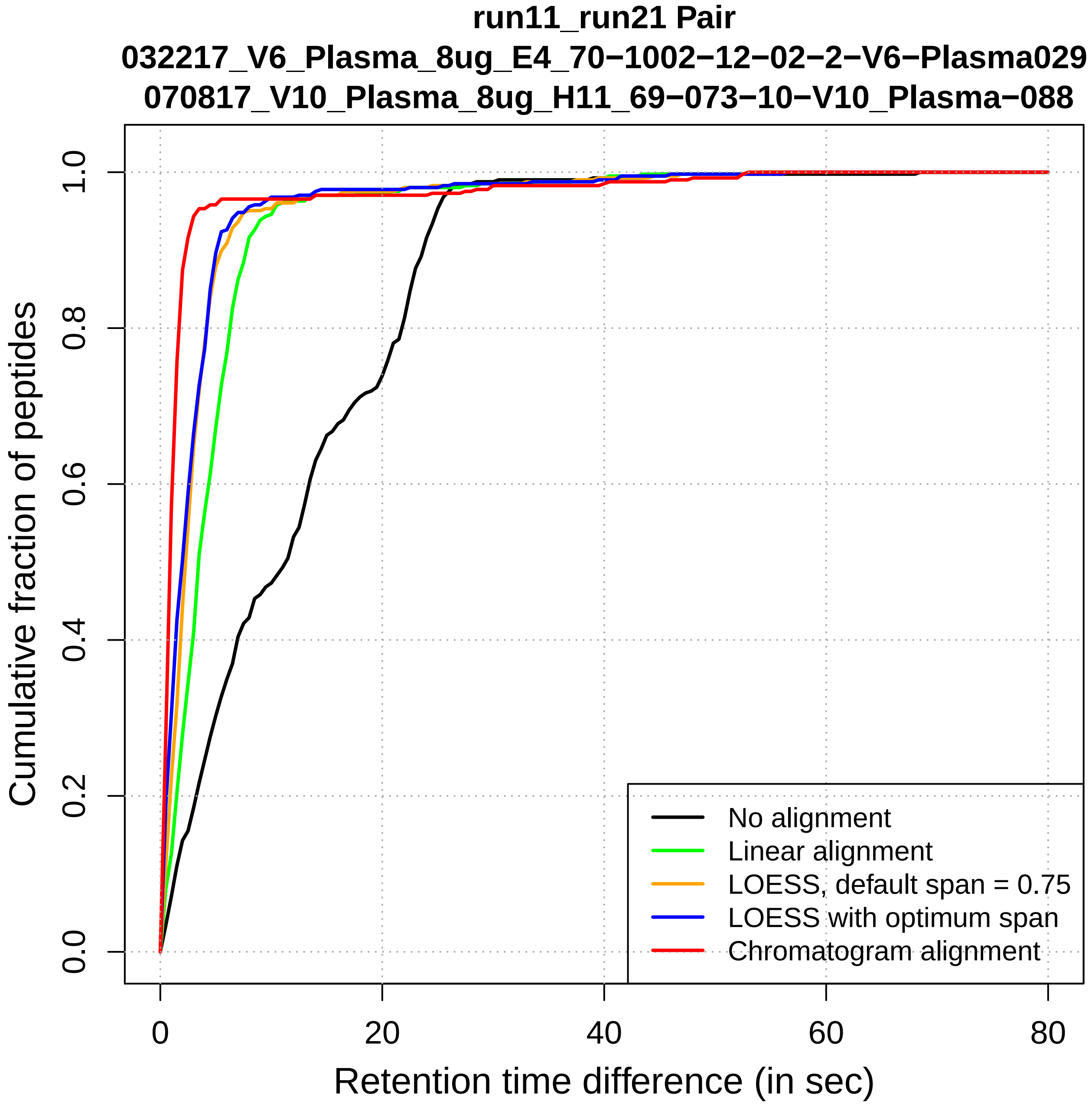
run11_run21 Pair
032217_V6_Plasma_8ug_E4_70−1002−12−02−2−V6−Plasma029
070817_V10_Plasma_8ug_H11_69−073−10−V10_Plasma−088
Cumulative fraction of peptides
0.0
0.2
0.4
0.6
0.8
1.0
0
20
40
60
80
Retention time difference (in sec)
No alignment
Linear alignment
LOESS, default span = 0.75
LOESS with optimum span
Chromatogram alignment

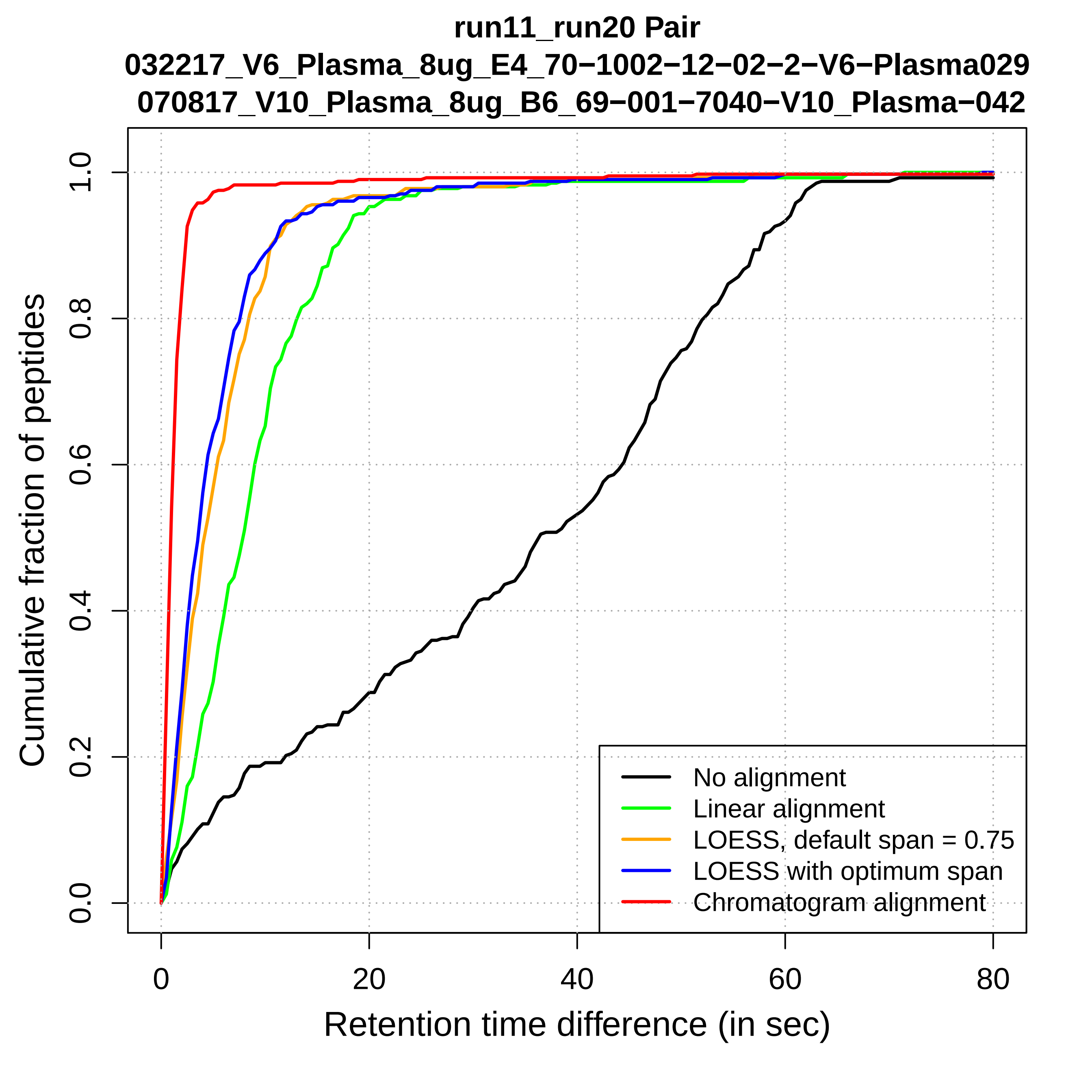

run11_run20 Pair
032217_V6_Plasma_8ug_E4_70−1002−12−02−2−V6−Plasma029
070817_V10_Plasma_8ug_B6_69−001−7040−V10_Plasma−042
Cumulative fraction of peptides
0.0
0.2
0.4
0.6
0.8
1.0
0
20
40
60
80
Retention time difference (in sec)
No alignment
Linear alignment
LOESS, default span = 0.75
LOESS with optimum span
Chromatogram alignment

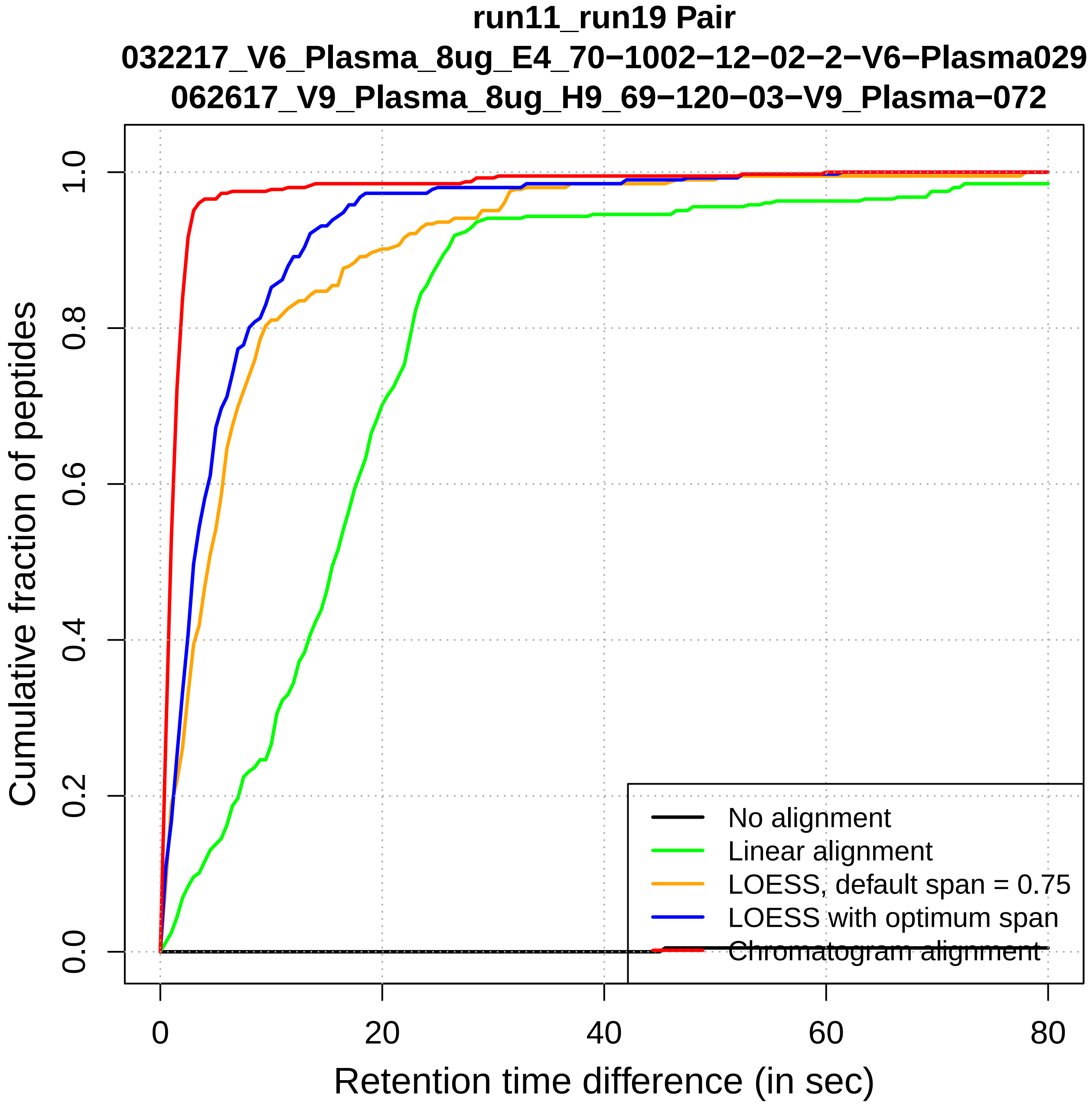

run11_run19 Pair
032217_V6_Plasma_8ug_E4_70−1002−12−02−2−V6−Plasma029
062617_V9_Plasma_8ug_H9_69−120−03−V9_Plasma−072
Cumulative fraction of peptides
0.0
0.2
0.4
0.6
0.8
1.0
0
20
40
60
80
Retention time difference (in sec)
No alignment
Linear alignment
LOESS, default span = 0.75
LOESS with optimum span
Chromatogram alignment

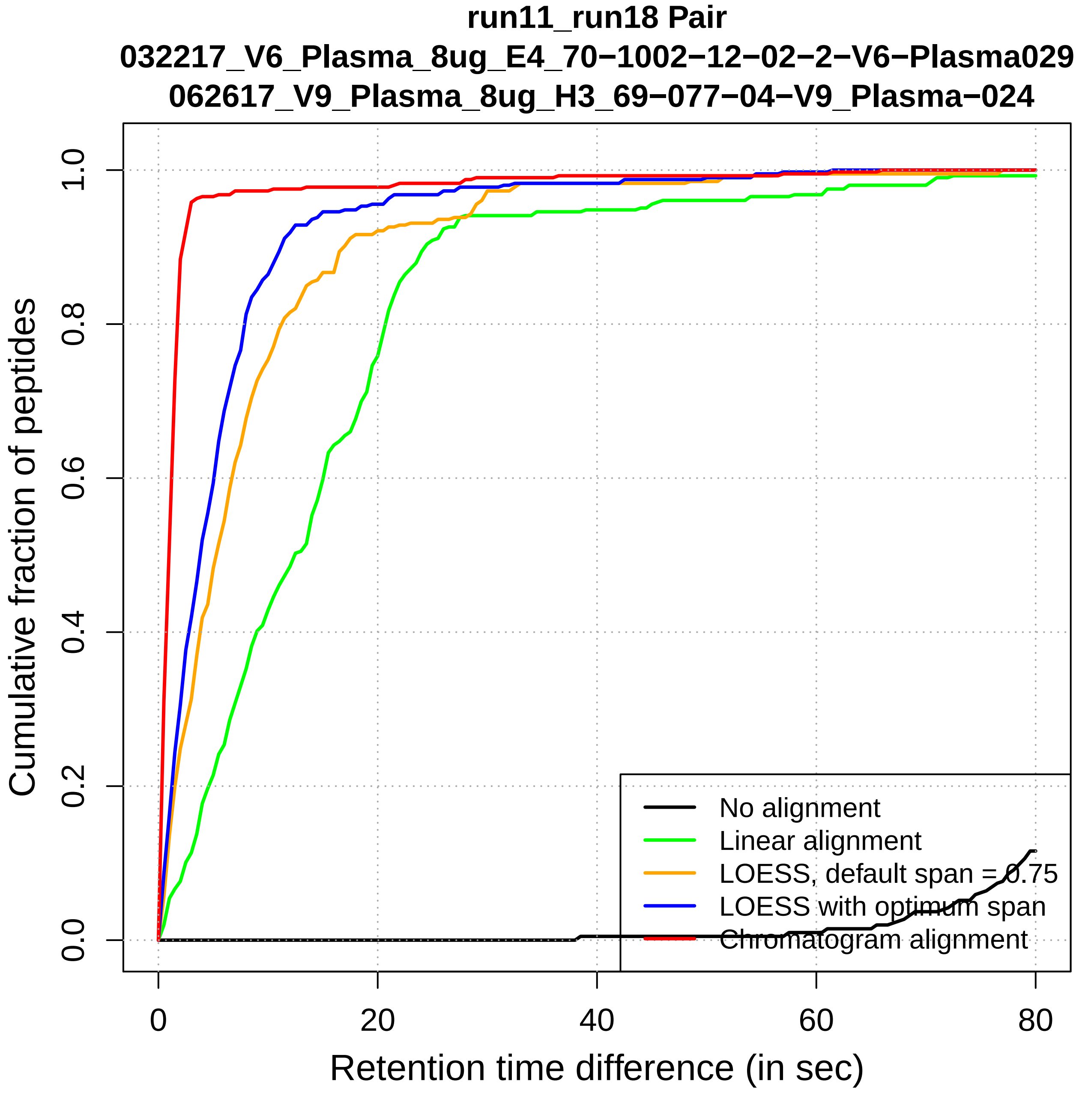
run11_run18 Pair
032217_V6_Plasma_8ug_E4_70−1002−12−02−2−V6−Plasma029
062617_V9_Plasma_8ug_H3_69−077−04−V9_Plasma−024
Cumulative fraction of peptides
0.0
0.2
0.4
0.6
0.8
1.0
0
20
40
60
80
Retention time difference (in sec)
No alignment
Linear alignment
LOESS, default span = 0.75
LOESS with optimum span
Chromatogram alignment

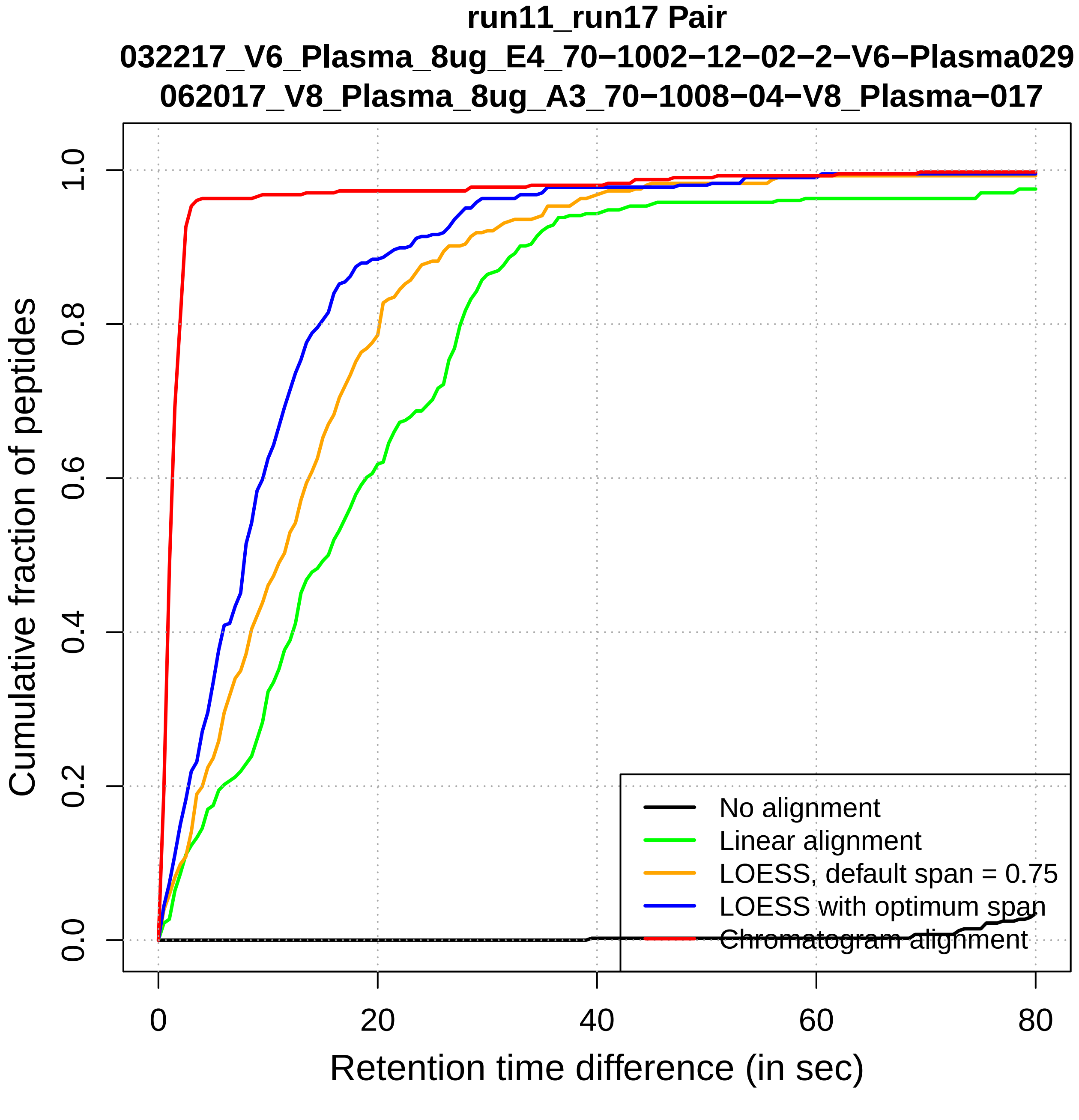
run11_run17 Pair
032217_V6_Plasma_8ug_E4_70−1002−12−02−2−V6−Plasma029
062017_V8_Plasma_8ug_A3_70−1008−04−V8_Plasma−017
Cumulative fraction of peptides
0.0
0.2
0.4
0.6
0.8
1.0
0
20
40
60
80
Retention time difference (in sec)
No alignment
Linear alignment
LOESS, default span = 0.75
LOESS with optimum span
Chromatogram alignment

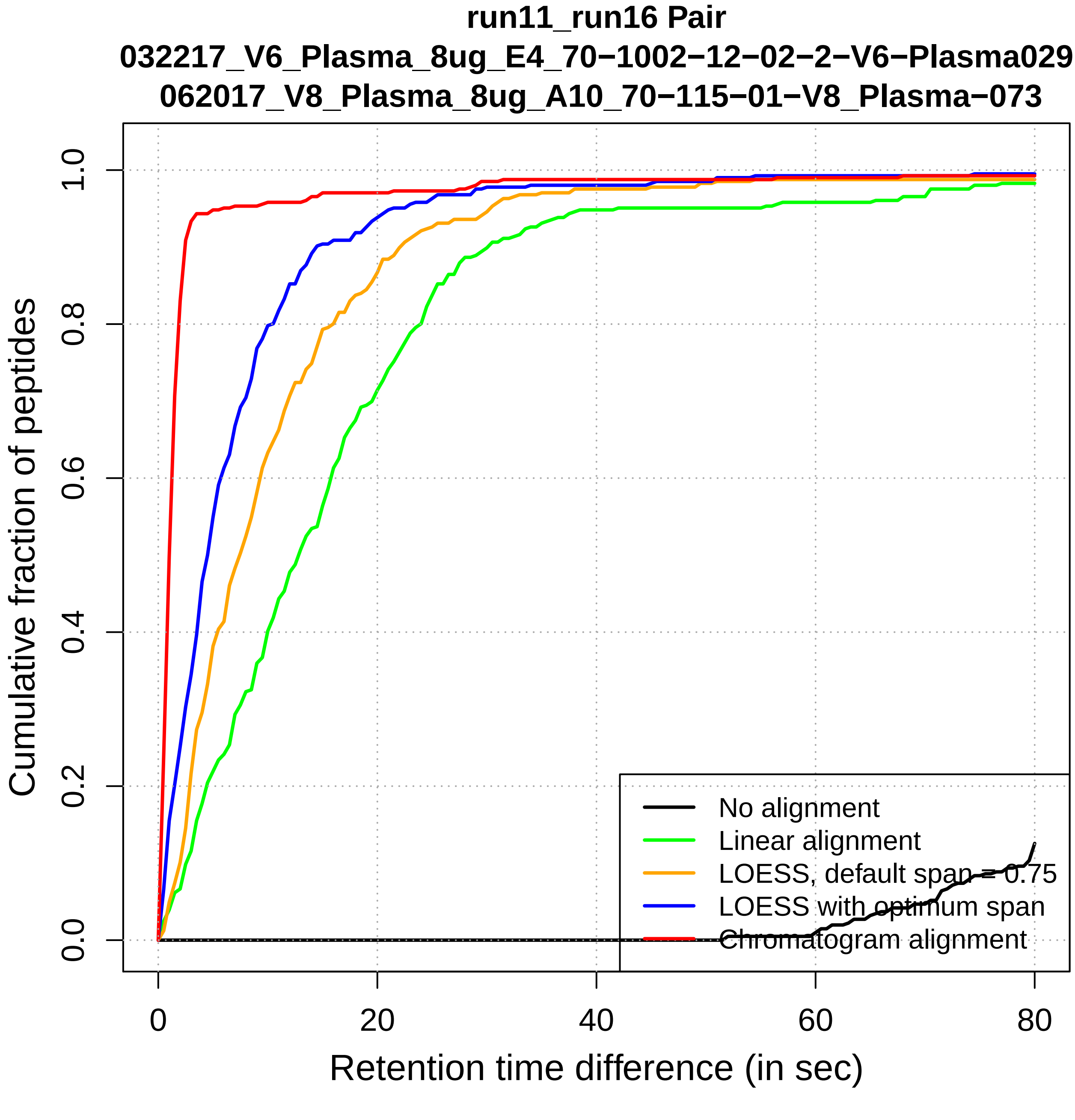

run11_run16 Pair
032217_V6_Plasma_8ug_E4_70−1002−12−02−2−V6−Plasma029
062017_V8_Plasma_8ug_A10_70−115−01−V8_Plasma−073
Cumulative fraction of peptides
0.0
0.2
0.4
0.6
0.8
1.0
0
20
40
60
80
Retention time difference (in sec)
No alignment
Linear alignment
LOESS, default span = 0.75
LOESS with optimum span
Chromatogram alignment

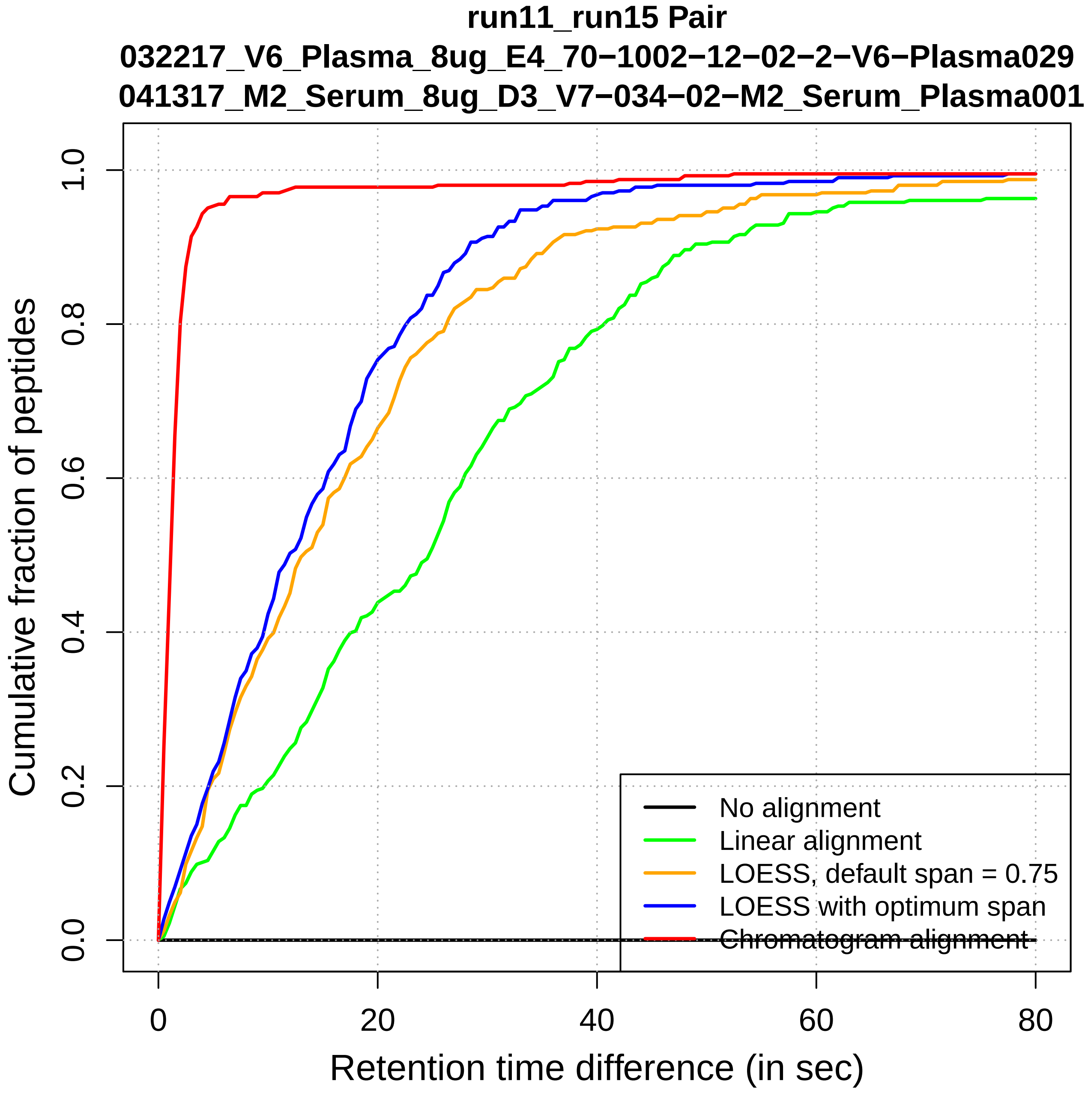

run11_run15 Pair
032217_V6_Plasma_8ug_E4_70−1002−12−02−2−V6−Plasma029
041317_M2_Serum_8ug_D3_V7−034−02−M2_Serum_Plasma001
Cumulative fraction of peptides
0.0
0.2
0.4
0.6
0.8
1.0
0
20
40
60
80
Retention time difference (in sec)
No alignment
Linear alignment
LOESS, default span = 0.75
LOESS with optimum span
Chromatogram alignment

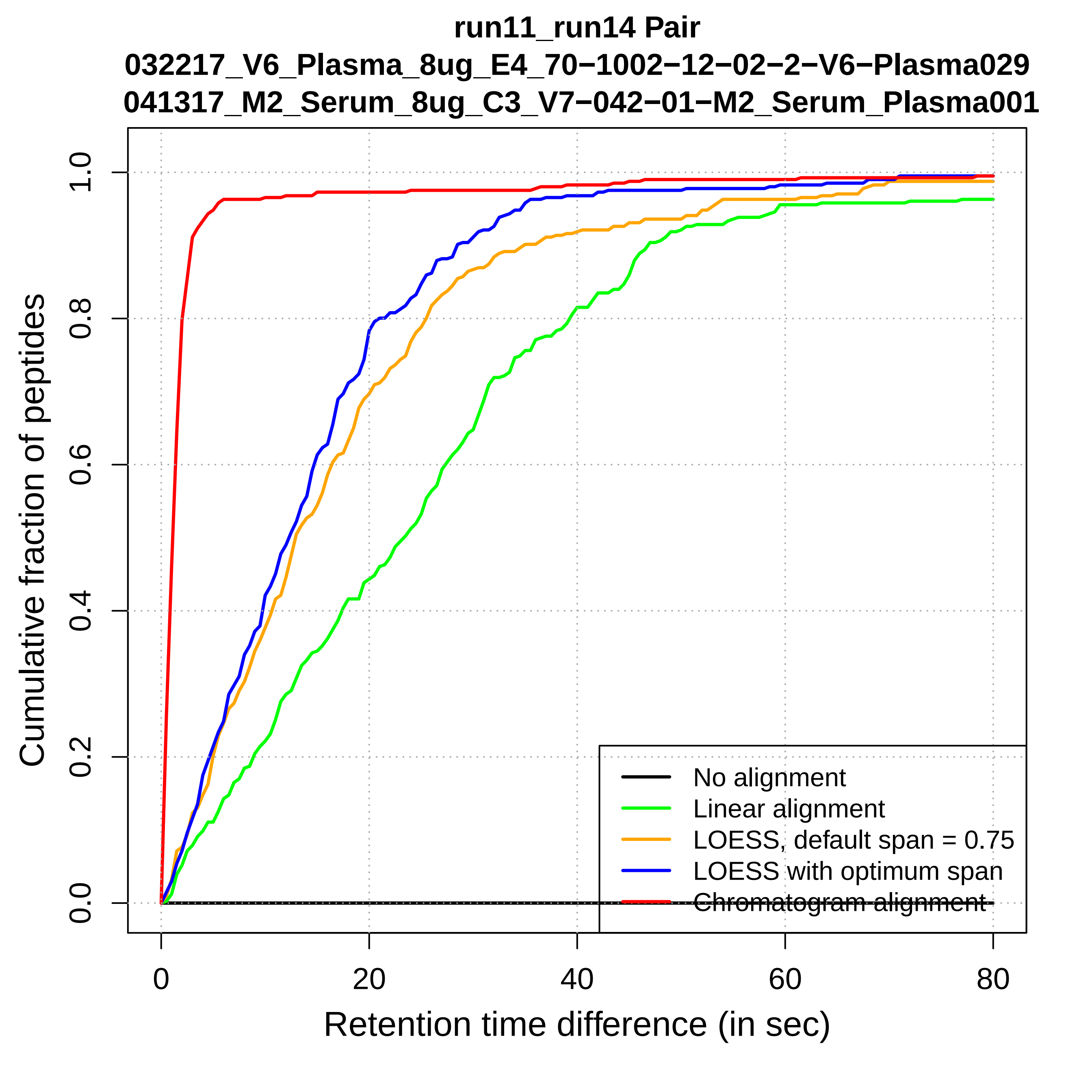
run11_run14 Pair
032217_V6_Plasma_8ug_E4_70−1002−12−02−2−V6−Plasma029
041317_M2_Serum_8ug_C3_V7−042−01−M2_Serum_Plasma001
Cumulative fraction of peptides
0.0
0.2
0.4
0.6
0.8
1.0
0
20
40
60
80
Retention time difference (in sec)
No alignment
Linear alignment
LOESS, default span = 0.75
LOESS with optimum span
Chromatogram alignment

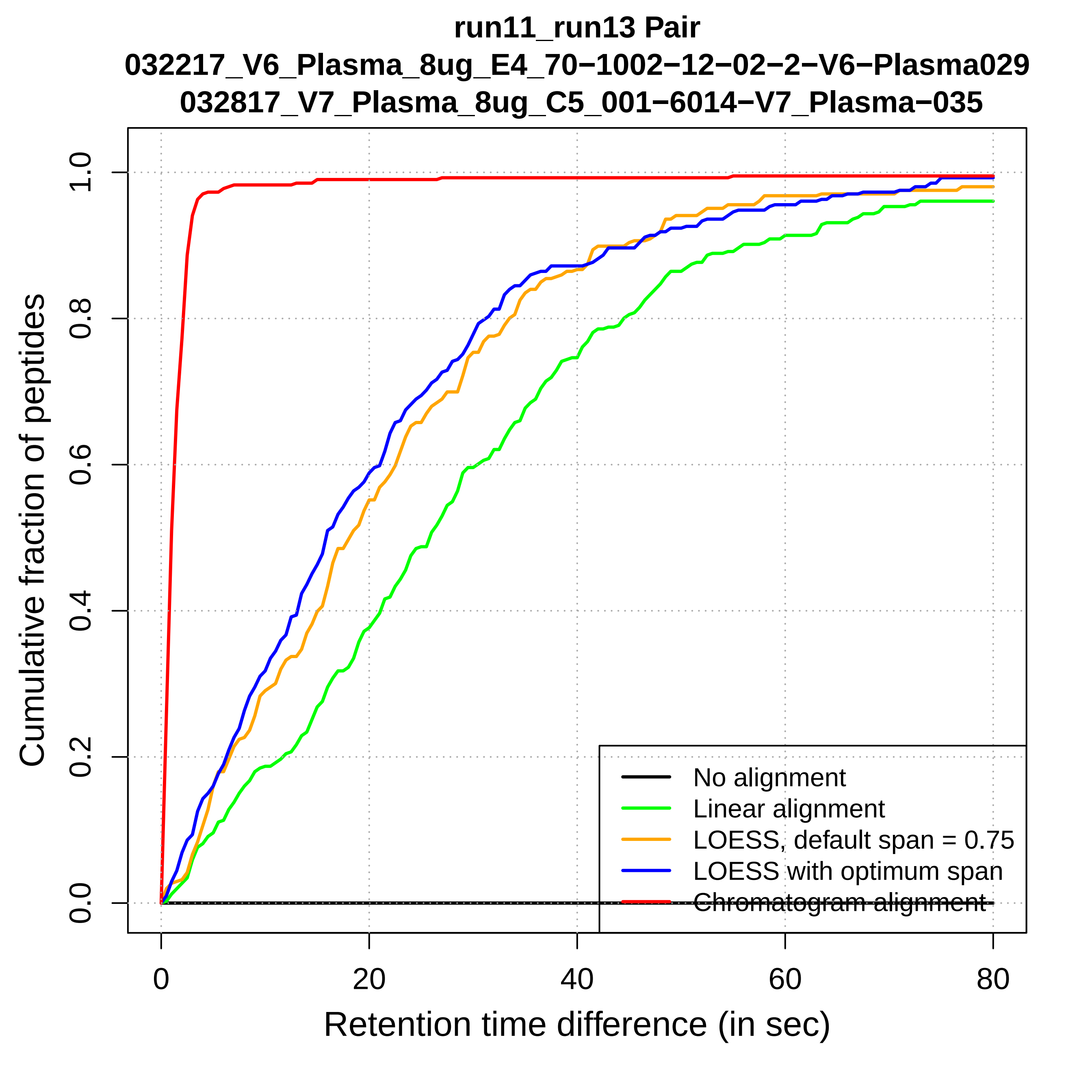

run11_run13 Pair
032217_V6_Plasma_8ug_E4_70−1002−12−02−2−V6−Plasma029
032817_V7_Plasma_8ug_C5_001−6014−V7_Plasma−035
Cumulative fraction of peptides
0.0
0.2
0.4
0.6
0.8
1.0
0
20
40
60
80
Retention time difference (in sec)
No alignment
Linear alignment
LOESS, default span = 0.75
LOESS with optimum span
Chromatogram alignment

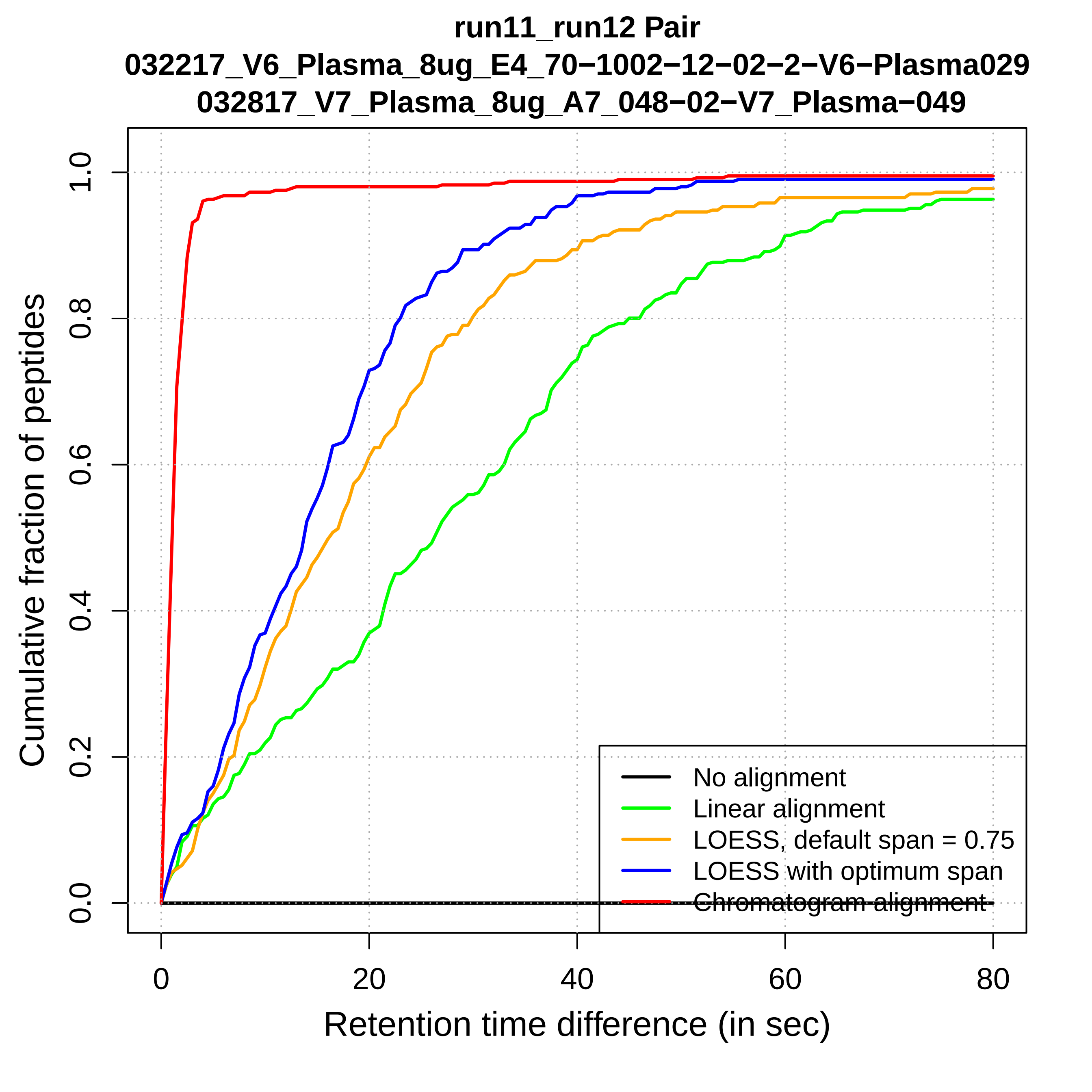
run11_run12 Pair
032217_V6_Plasma_8ug_E4_70−1002−12−02−2−V6−Plasma029
032817_V7_Plasma_8ug_A7_048−02−V7_Plasma−049
Cumulative fraction of peptides
0.0
0.2
0.4
0.6
0.8
1.0
0
20
40
60
80
Retention time difference (in sec)
No alignment
Linear alignment
LOESS, default span = 0.75
LOESS with optimum span
Chromatogram alignment

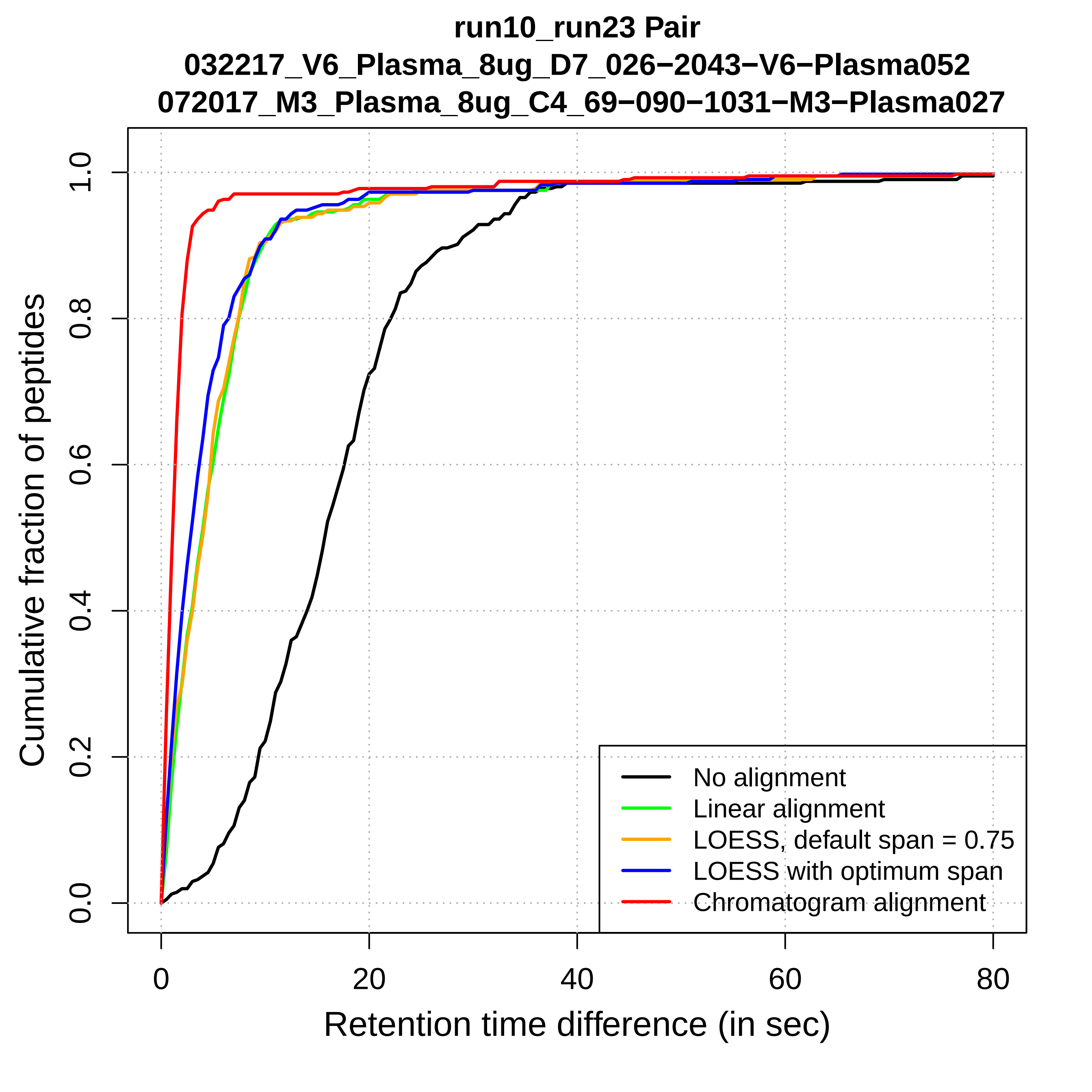
run10_run23 Pair
032217_V6_Plasma_8ug_D7_026−2043−V6−Plasma052
072017_M3_Plasma_8ug_C4_69−090−1031−M3−Plasma027
Cumulative fraction of peptides
Retention time difference (in sec)
0.0
0.2
0.4
0.6
0.8
1.0
0
20
40
60
80
No alignment
Linear alignment
LOESS, default span = 0.75
LOESS with optimum span
Chromatogram alignment

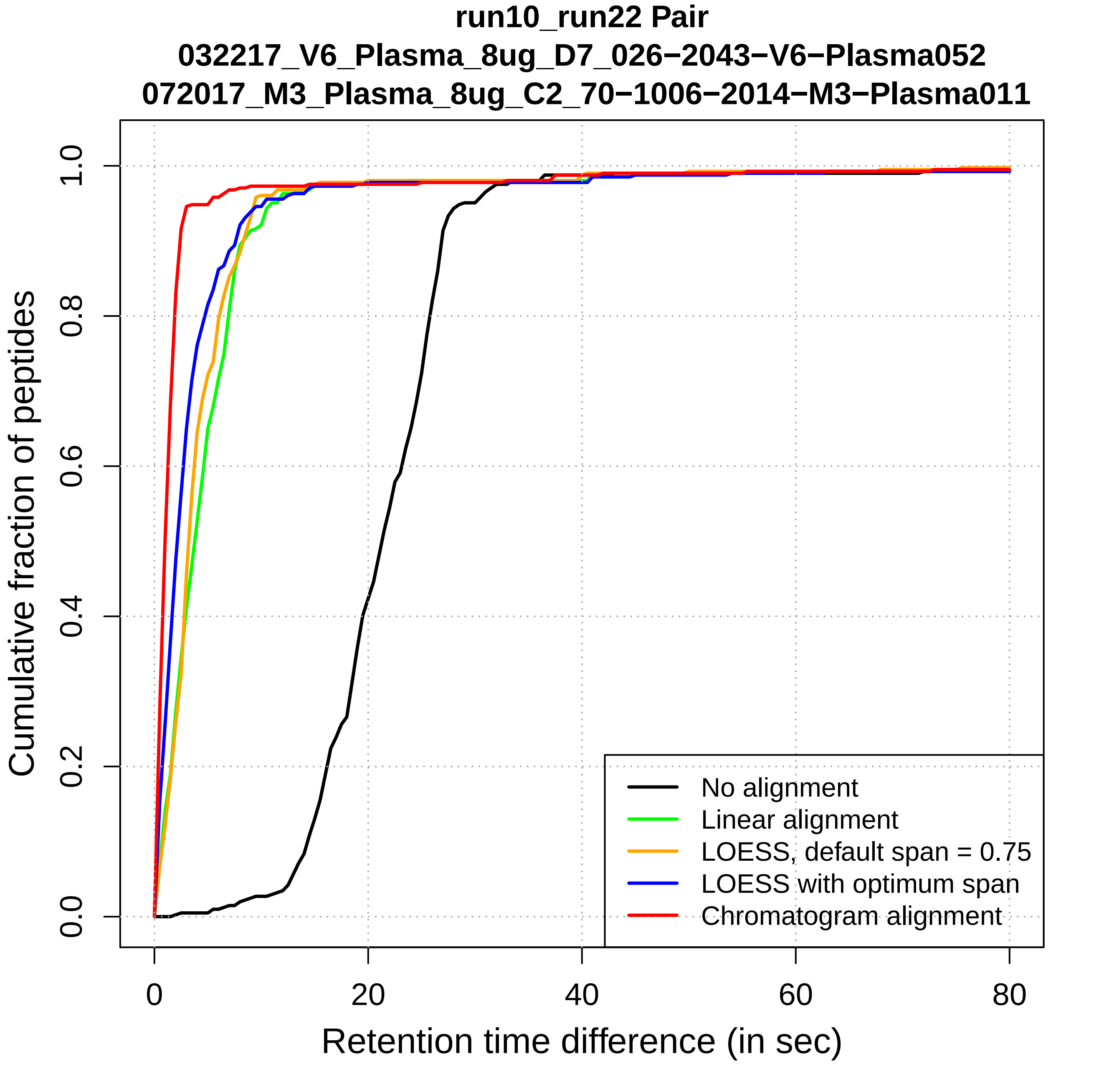
run10_run22 Pair
032217_V6_Plasma_8ug_D7_026−2043−V6−Plasma052
072017_M3_Plasma_8ug_C2_70−1006−2014−M3−Plasma011
Cumulative fraction of peptides
0.0
0.2
0.4
0.6
0.8
1.0
0
20
40
60
80
Retention time difference (in sec)
No alignment
Linear alignment
LOESS, default span = 0.75
LOESS with optimum span
Chromatogram alignment

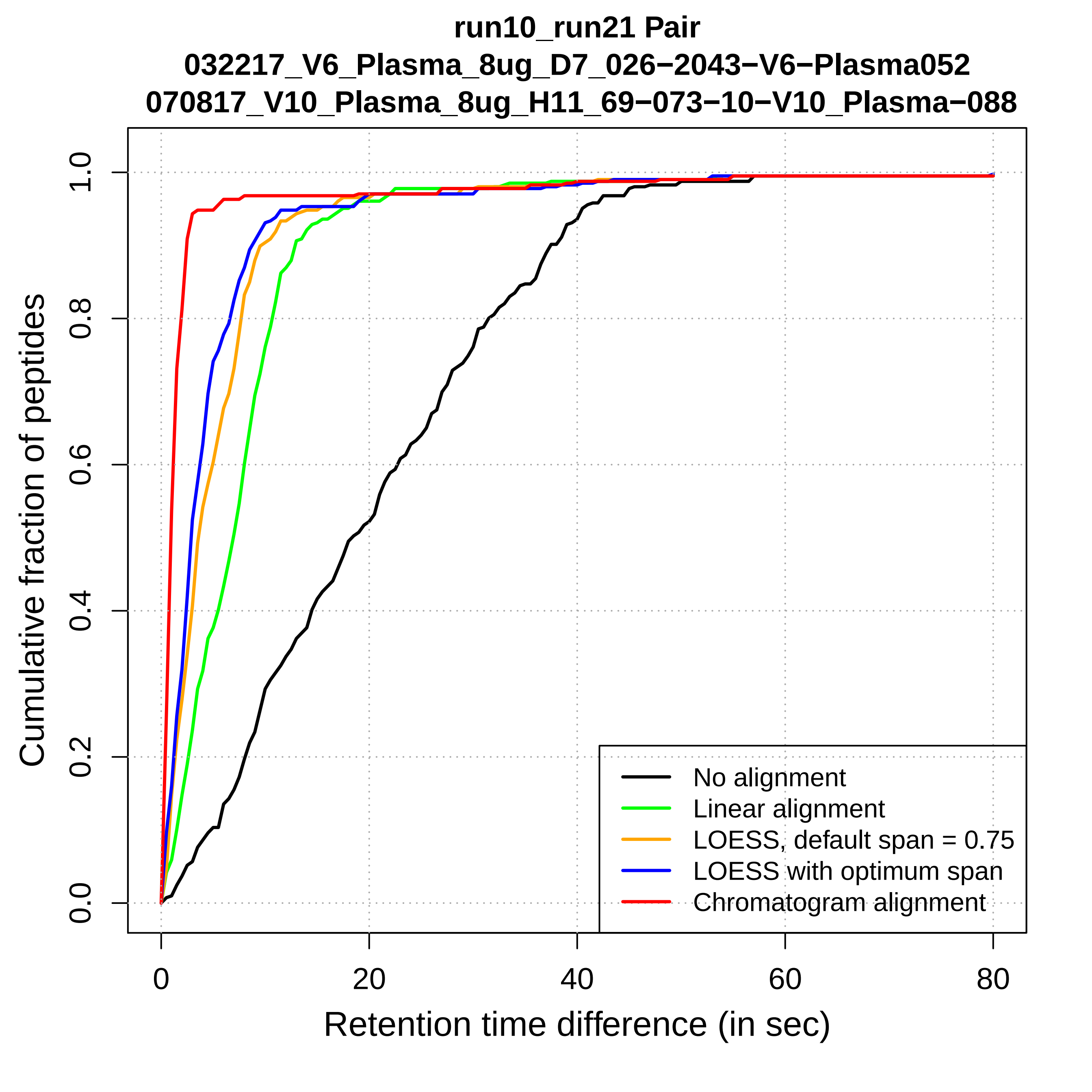
run10_run21 Pair
032217_V6_Plasma_8ug_D7_026−2043−V6−Plasma052
070817_V10_Plasma_8ug_H11_69−073−10−V10_Plasma−088
Cumulative fraction of peptides
0.0
0.2
0.4
0.6
0.8
1.0
0
20
40
60
80
Retention time difference (in sec)
No alignment
Linear alignment
LOESS, default span = 0.75
LOESS with optimum span
Chromatogram alignment

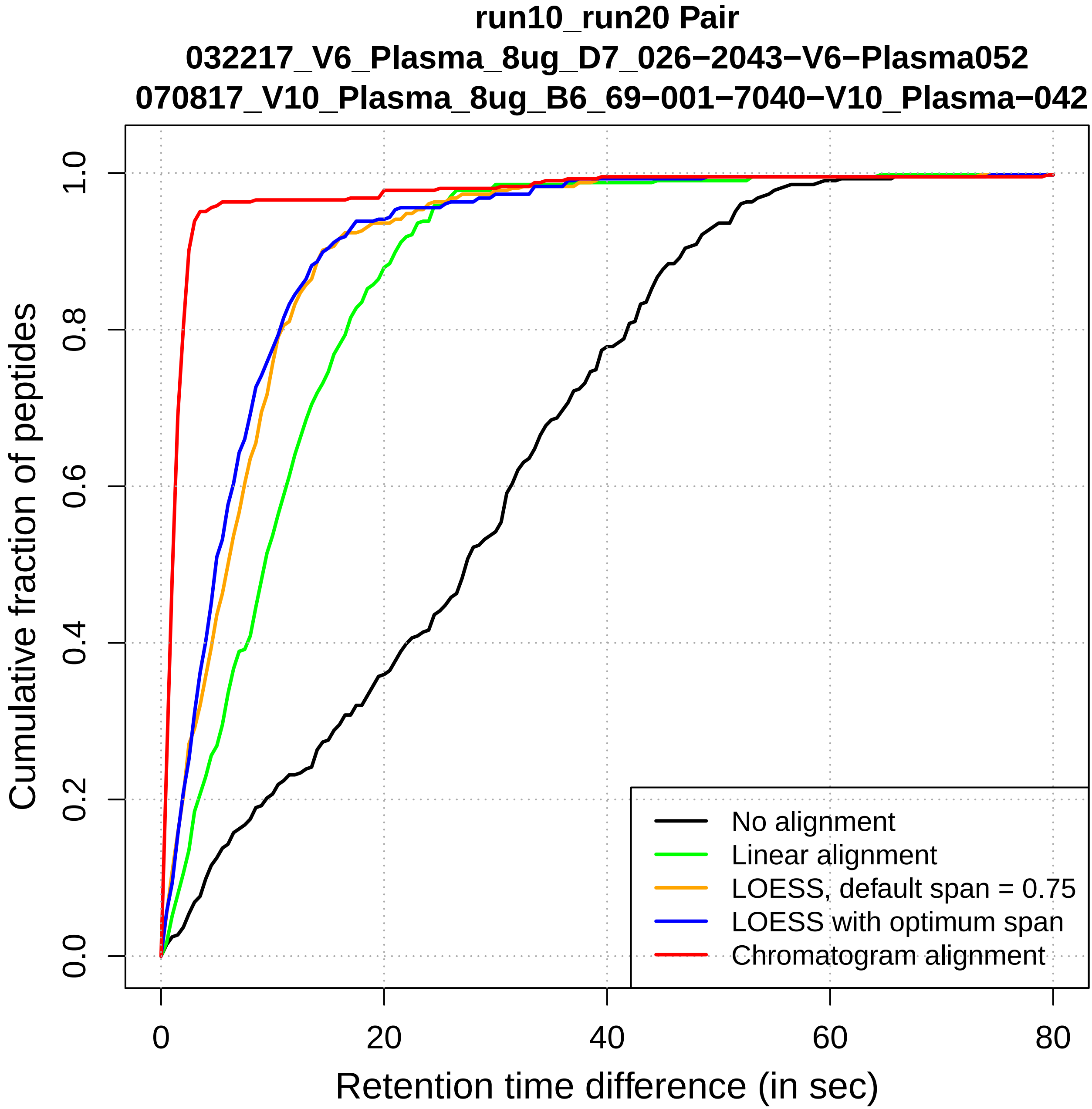

run10_run20 Pair
032217_V6_Plasma_8ug_D7_026−2043−V6−Plasma052
070817_V10_Plasma_8ug_B6_69−001−7040−V10_Plasma−042
Cumulative fraction of peptides
0.0
0.2
0.4
0.6
0.8
1.0
0
20
40
60
80
Retention time difference (in sec)
No alignment
Linear alignment
LOESS, default span = 0.75
LOESS with optimum span
Chromatogram alignment

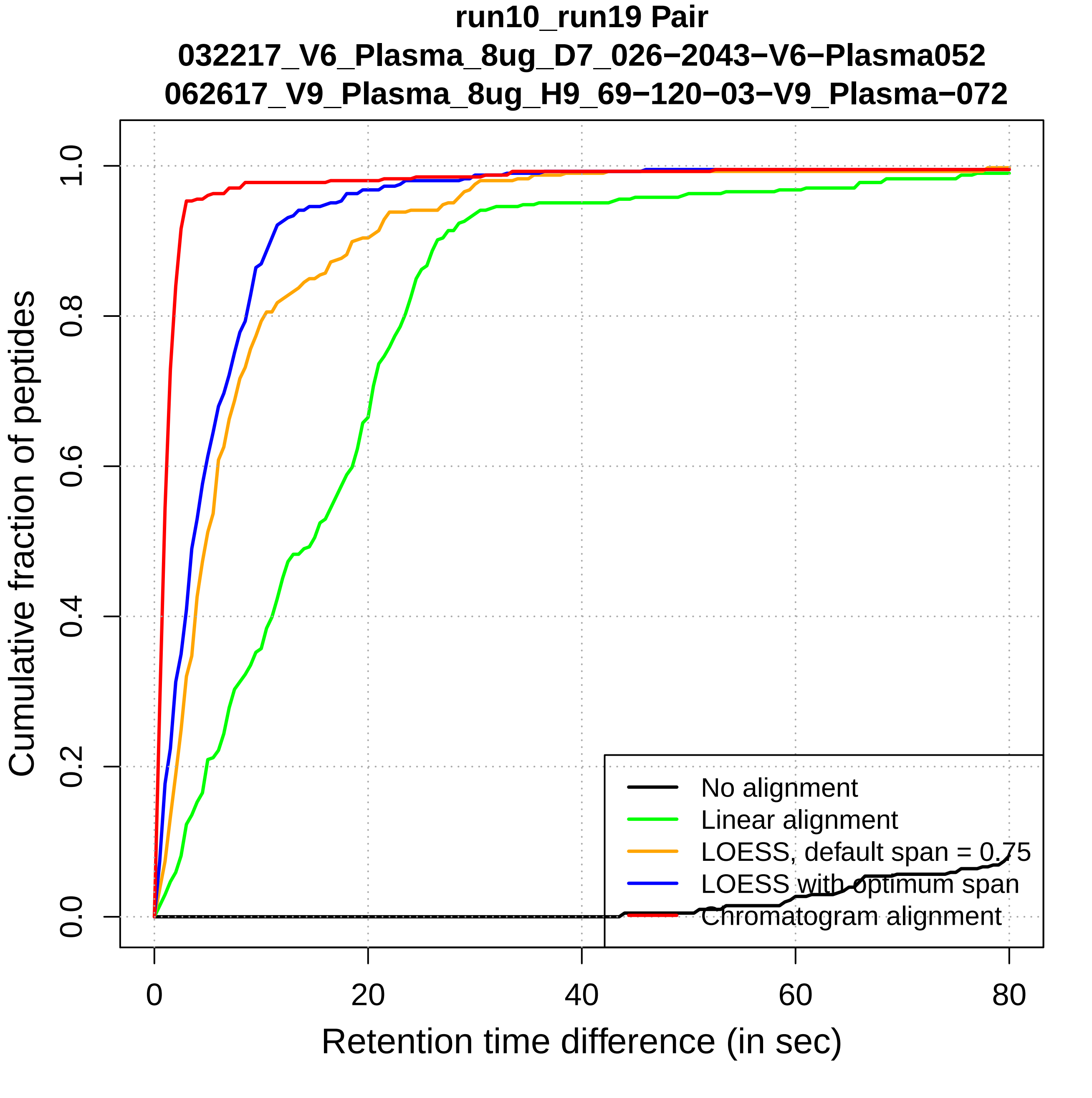

run10_run19 Pair
032217_V6_Plasma_8ug_D7_026−2043−V6−Plasma052
062617_V9_Plasma_8ug_H9_69−120−03−V9_Plasma−072
Cumulative fraction of peptides
0.0
0.2
0.4
0.6
0.8
1.0
0
20
40
60
80
Retention time difference (in sec)
No alignment
Linear alignment
LOESS, default span = 0.75
LOESS with optimum span
Chromatogram alignment

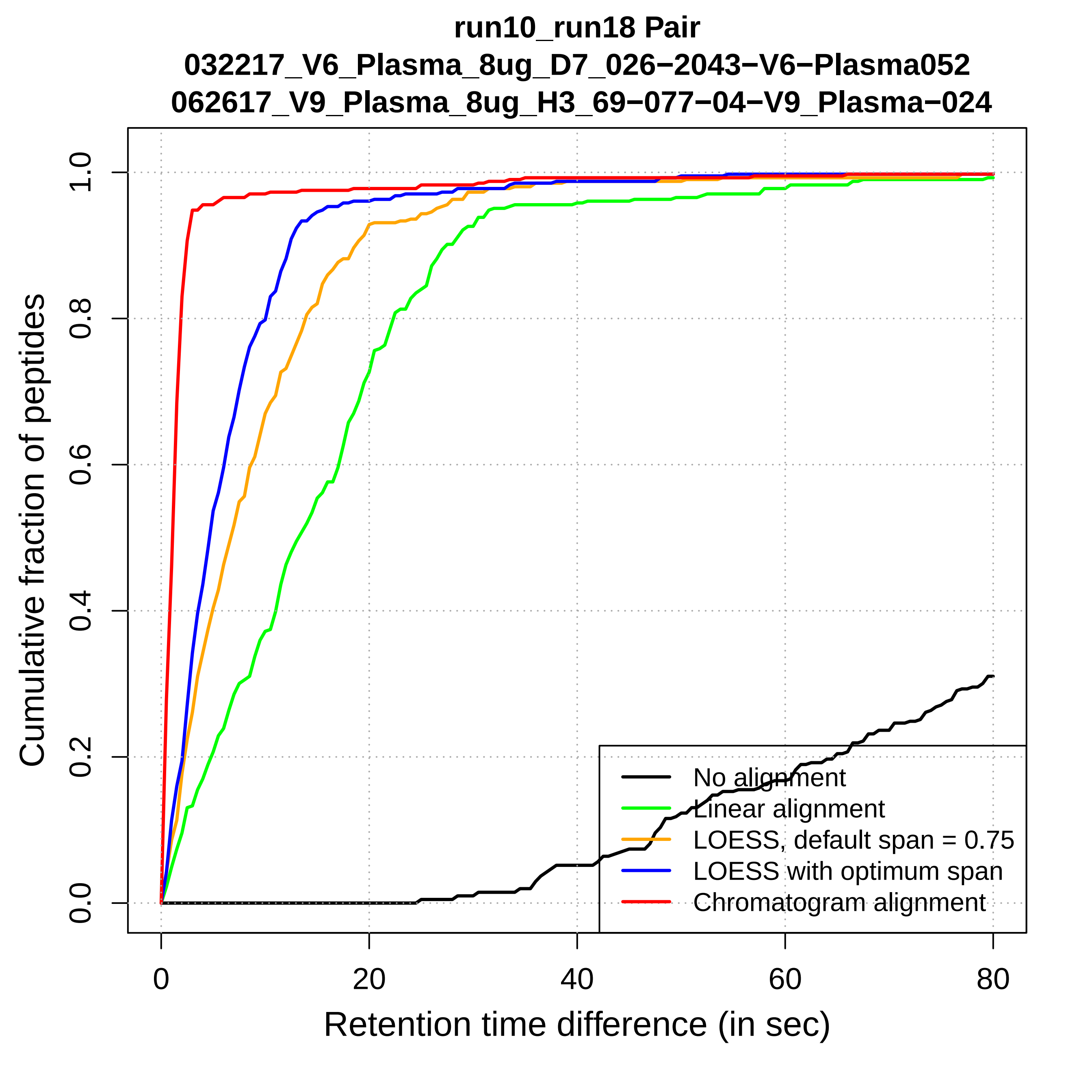

run10_run18 Pair
032217_V6_Plasma_8ug_D7_026−2043−V6−Plasma052
062617_V9_Plasma_8ug_H3_69−077−04−V9_Plasma−024
Cumulative fraction of peptides
0.0
0.2
0.4
0.6
0.8
1.0
0
20
40
60
80
Retention time difference (in sec)
No alignment
Linear alignment
LOESS, default span = 0.75
LOESS with optimum span
Chromatogram alignment

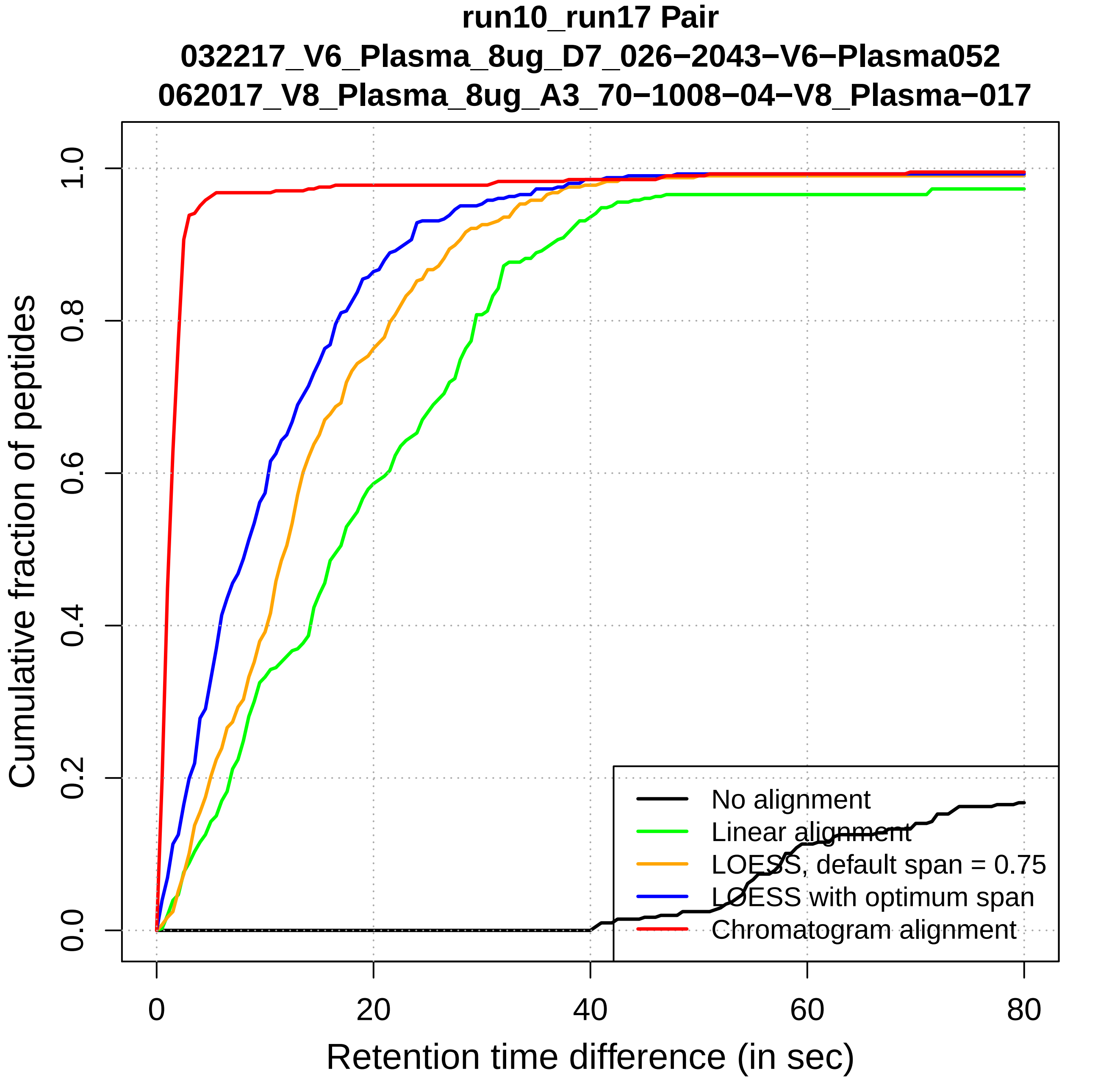

run10_run17 Pair
032217_V6_Plasma_8ug_D7_026−2043−V6−Plasma052
062017_V8_Plasma_8ug_A3_70−1008−04−V8_Plasma−017
Cumulative fraction of peptides
0.0
0.2
0.4
0.6
0.8
1.0
0
20
40
60
80
Retention time difference (in sec)
No alignment
Linear alignment
LOESS, default span = 0.75
LOESS with optimum span
Chromatogram alignment

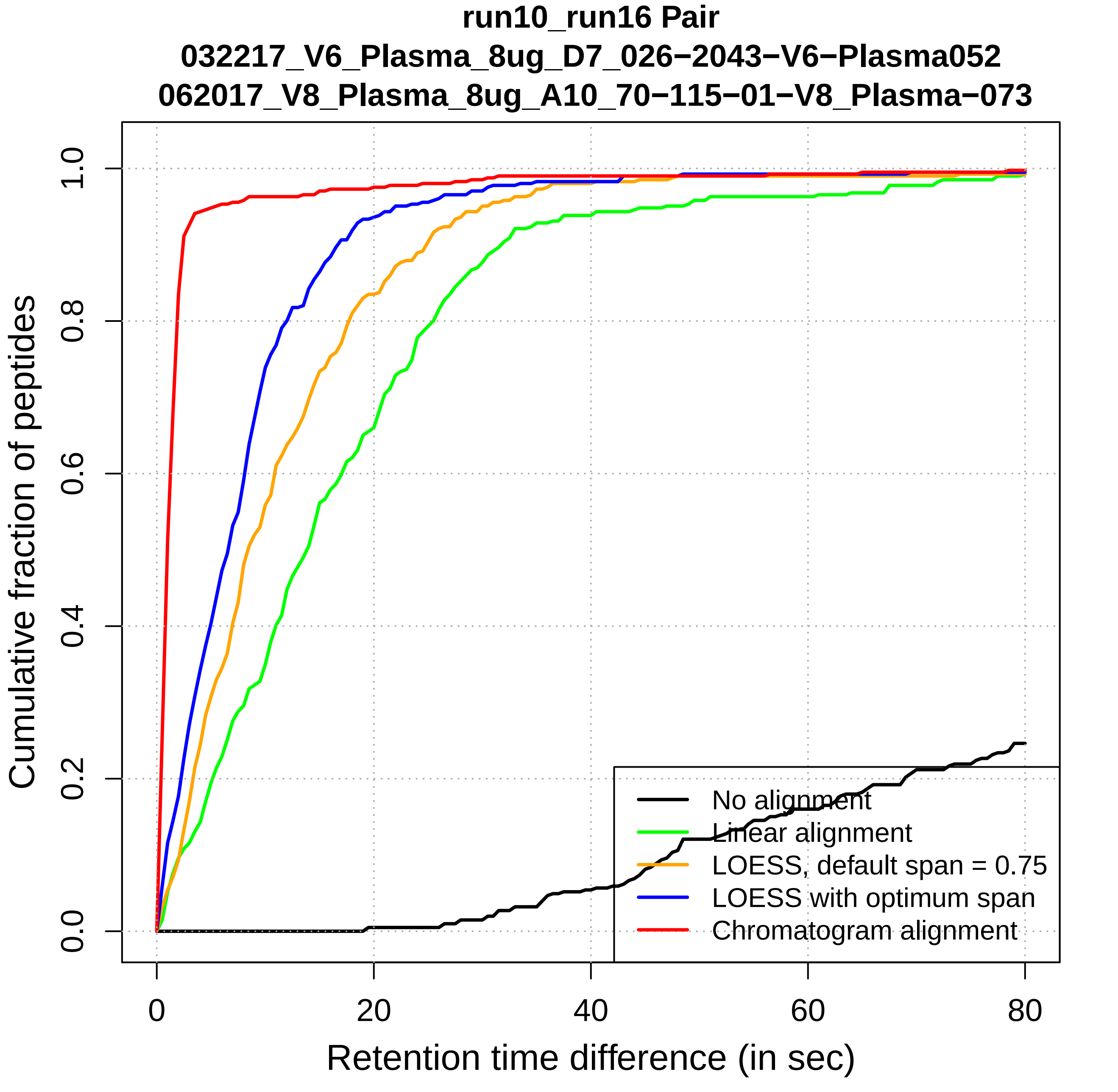
run10_run16 Pair
032217_V6_Plasma_8ug_D7_026−2043−V6−Plasma052
062017_V8_Plasma_8ug_A10_70−115−01−V8_Plasma−073
Cumulative fraction of peptides
Retention time difference (in sec)
0.0
0.2
0.4
0.6
0.8
1.0
0
20
40
60
80
No alignment
Linear alignment
LOESS, default span = 0.75
LOESS with optimum span
Chromatogram alignment

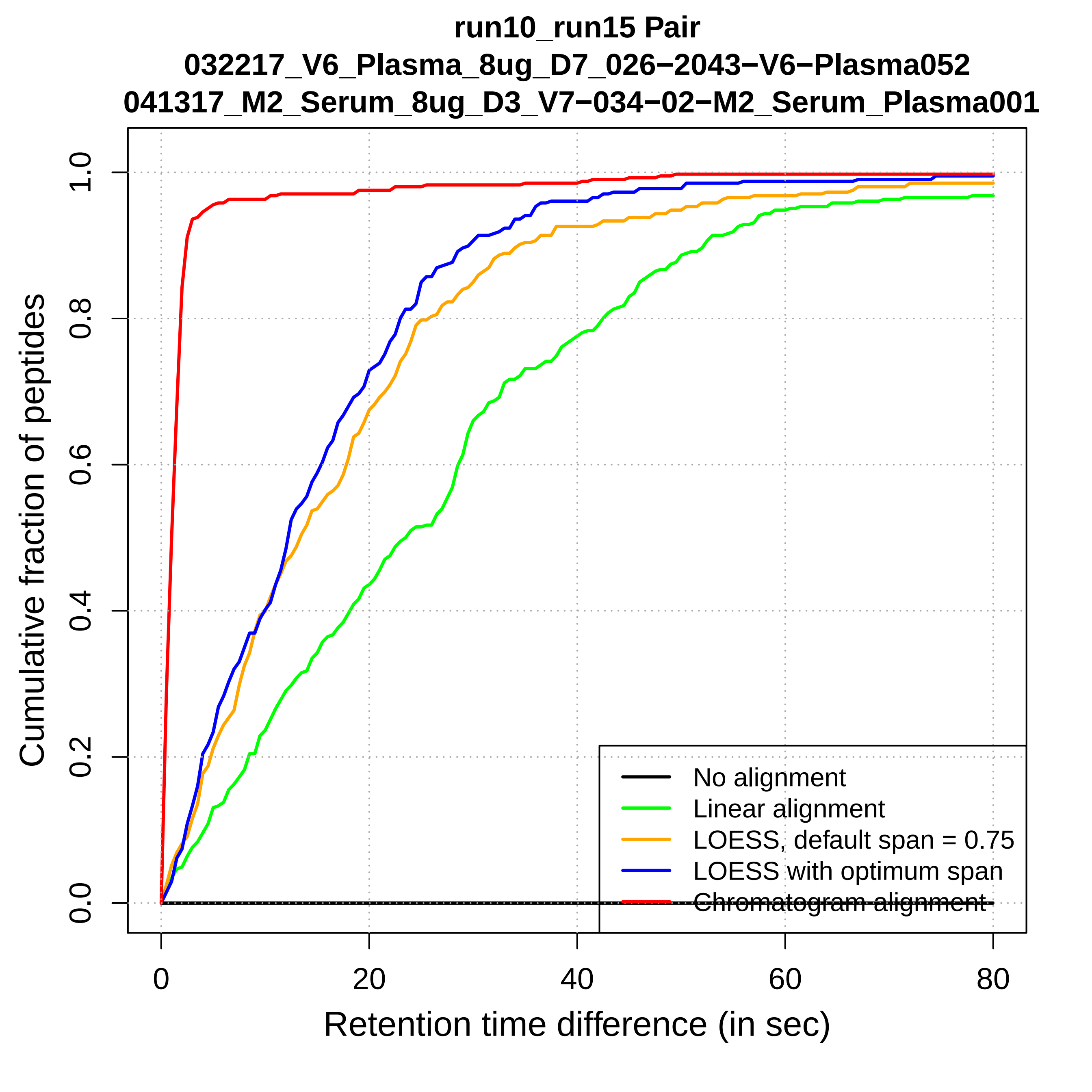
run10_run15 Pair
032217_V6_Plasma_8ug_D7_026−2043−V6−Plasma052
041317_M2_Serum_8ug_D3_V7−034−02−M2_Serum_Plasma001
Cumulative fraction of peptides
0.0
0.2
0.4
0.6
0.8
1.0
0
20
40
60
80
Retention time difference (in sec)
No alignment
Linear alignment
LOESS, default span = 0.75
LOESS with optimum span
Chromatogram alignment

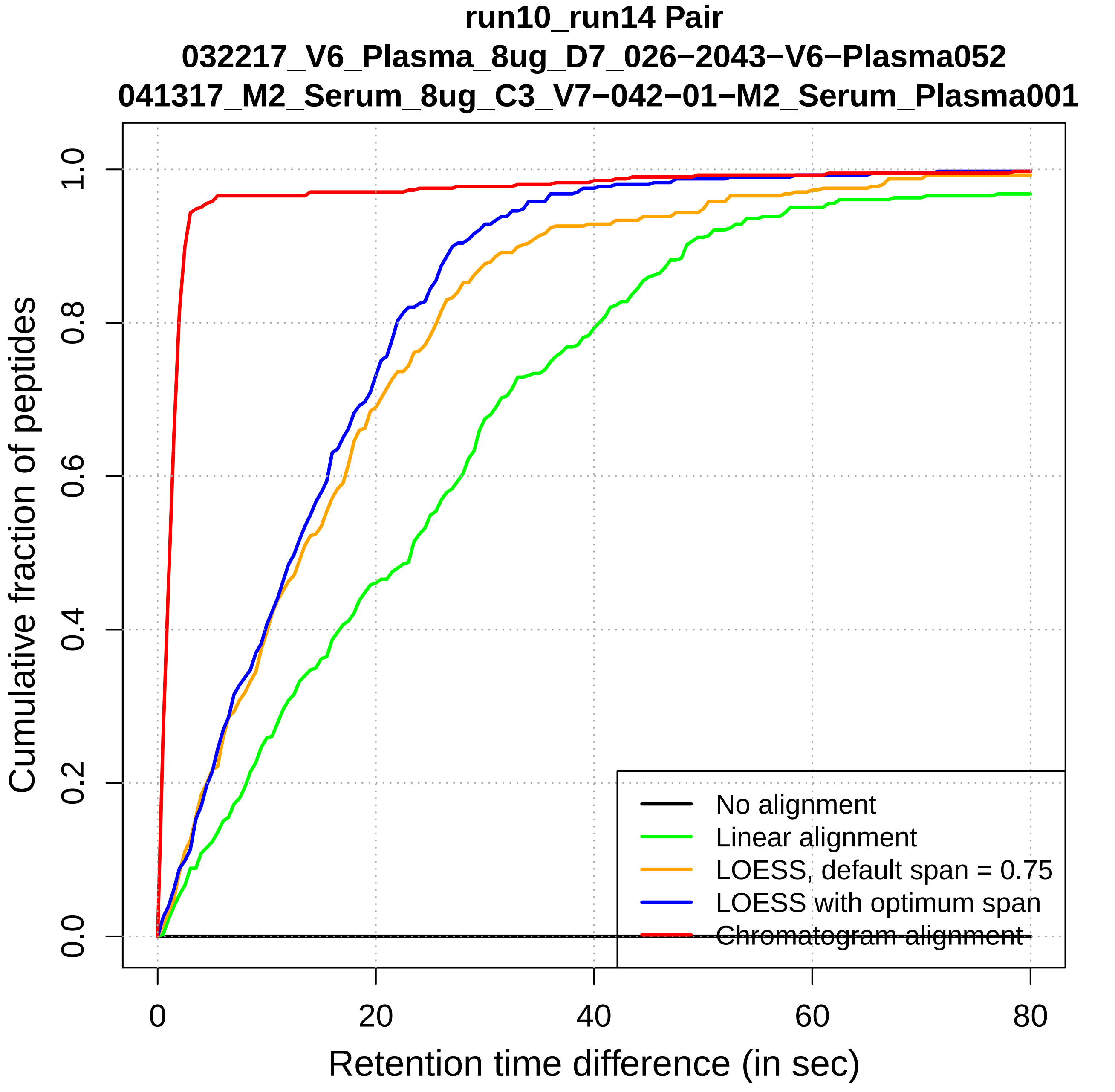
run10_run14 Pair
032217_V6_Plasma_8ug_D7_026−2043−V6−Plasma052
041317_M2_Serum_8ug_C3_V7−042−01−M2_Serum_Plasma001
Cumulative fraction of peptides
0.0
0.2
0.4
0.6
0.8
1.0
0
20
40
60
80
Retention time difference (in sec)
No alignment
Linear alignment
LOESS, default span = 0.75
LOESS with optimum span
Chromatogram alignment

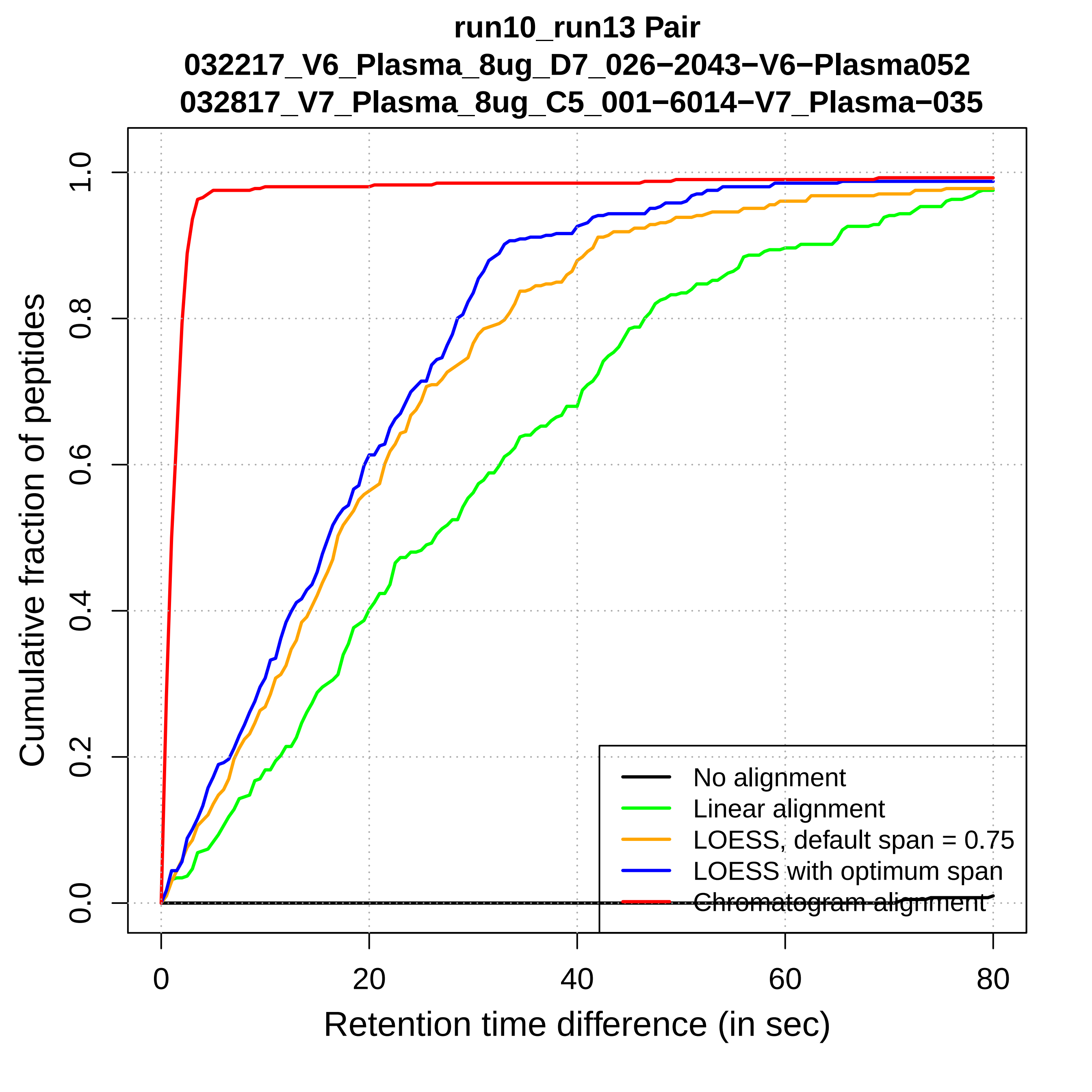
run10_run13 Pair
032217_V6_Plasma_8ug_D7_026−2043−V6−Plasma052
032817_V7_Plasma_8ug_C5_001−6014−V7_Plasma−035
Cumulative fraction of peptides
0.0
0.2
0.4
0.6
0.8
1.0
0
20
40
60
80
Retention time difference (in sec)
No alignment
Linear alignment
LOESS, default span = 0.75
LOESS with optimum span
Chromatogram alignment

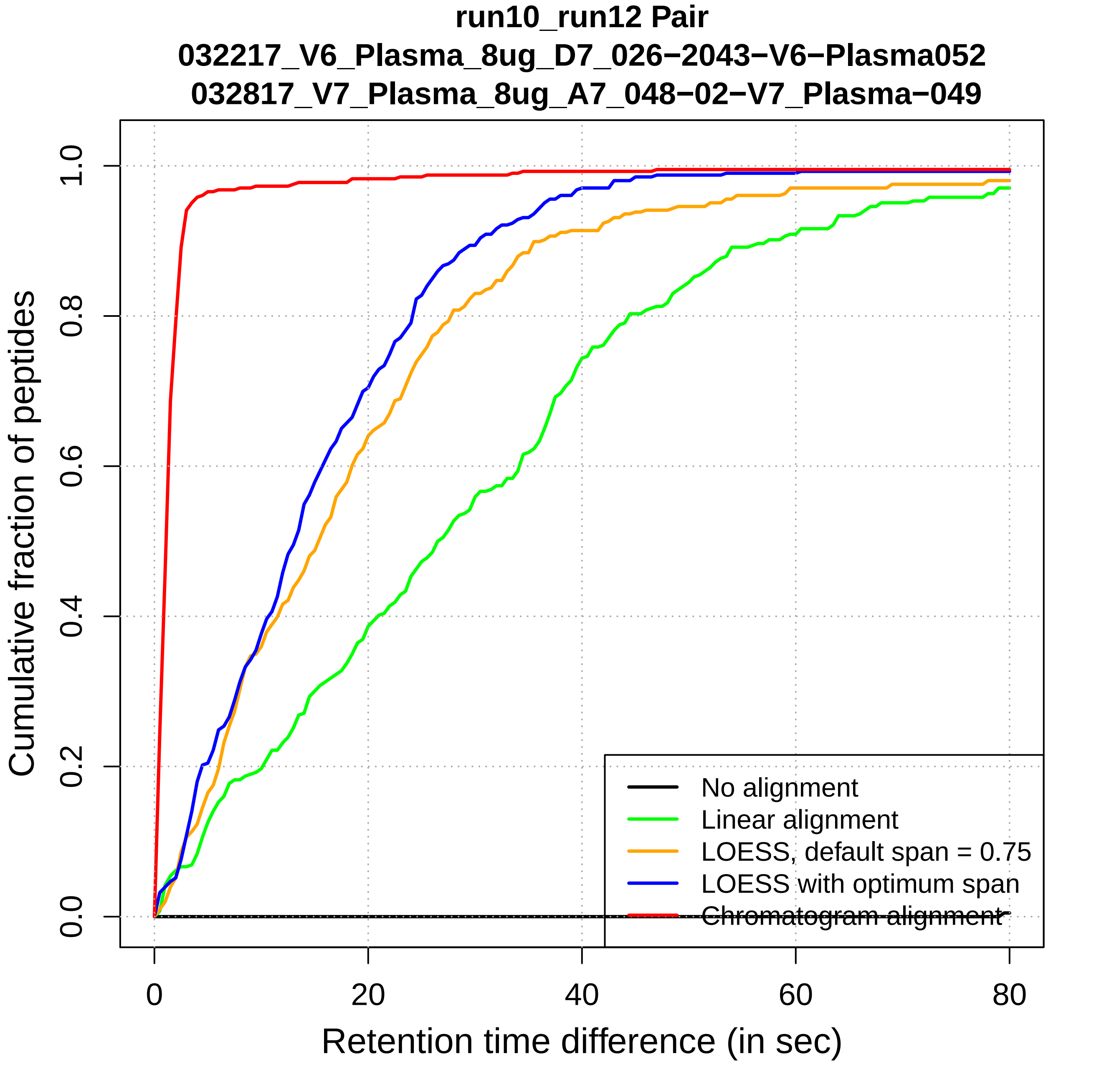
run10_run12 Pair
032217_V6_Plasma_8ug_D7_026−2043−V6−Plasma052
032817_V7_Plasma_8ug_A7_048−02−V7_Plasma−049
Cumulative fraction of peptides
0.0
0.2
0.4
0.6
0.8
1.0
0
20
40
60
80
Retention time difference (in sec)
No alignment
Linear alignment
LOESS, default span = 0.75
LOESS with optimum span
Chromatogram alignment

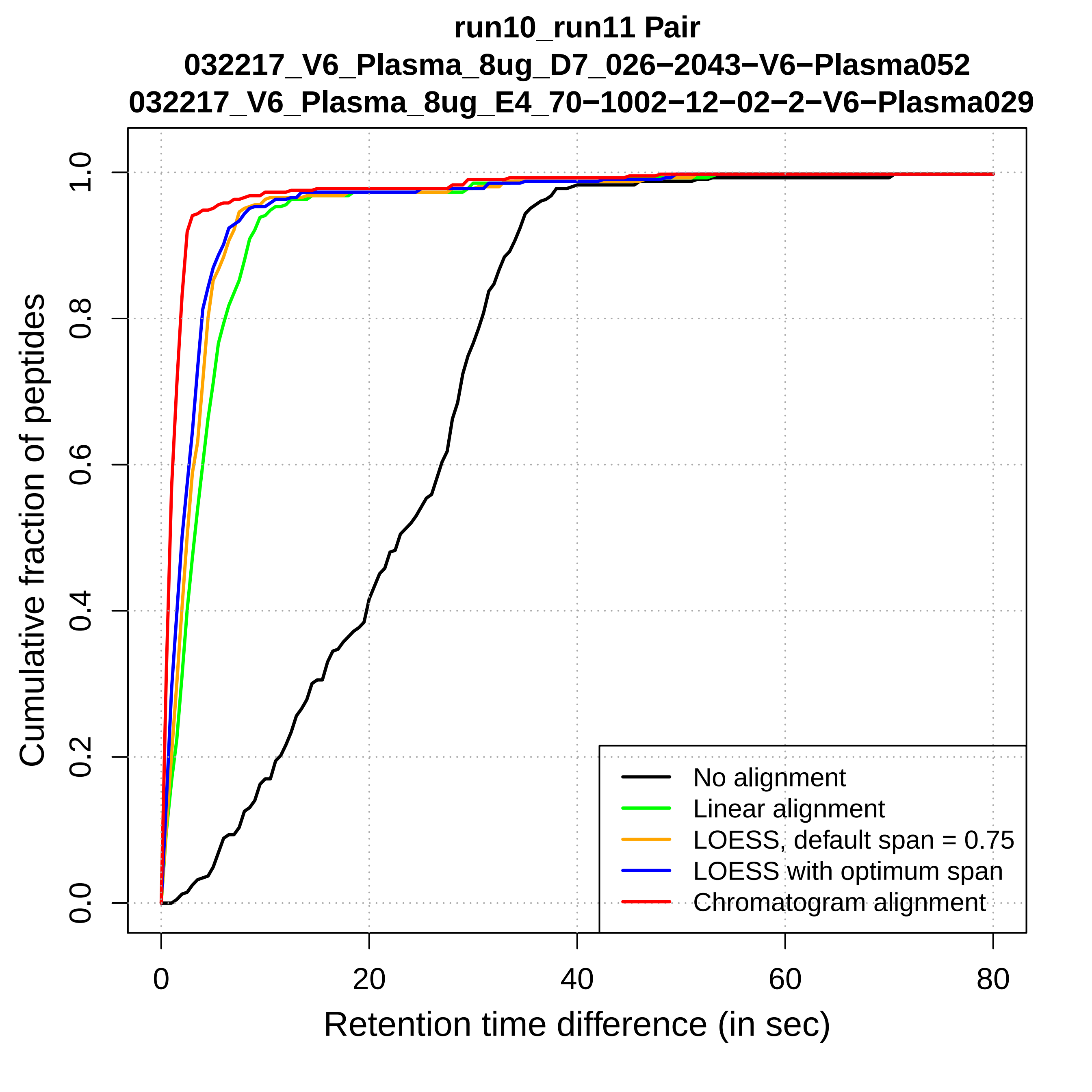
run10_run11 Pair
032217_V6_Plasma_8ug_D7_026−2043−V6−Plasma052
032217_V6_Plasma_8ug_E4_70−1002−12−02−2−V6−Plasma029
Cumulative fraction of peptides
0.0
0.2
0.4
0.6
0.8
1.0
0
20
40
60
80
Retention time difference (in sec)
No alignment
Linear alignment
LOESS, default span = 0.75
LOESS with optimum span
Chromatogram alignment

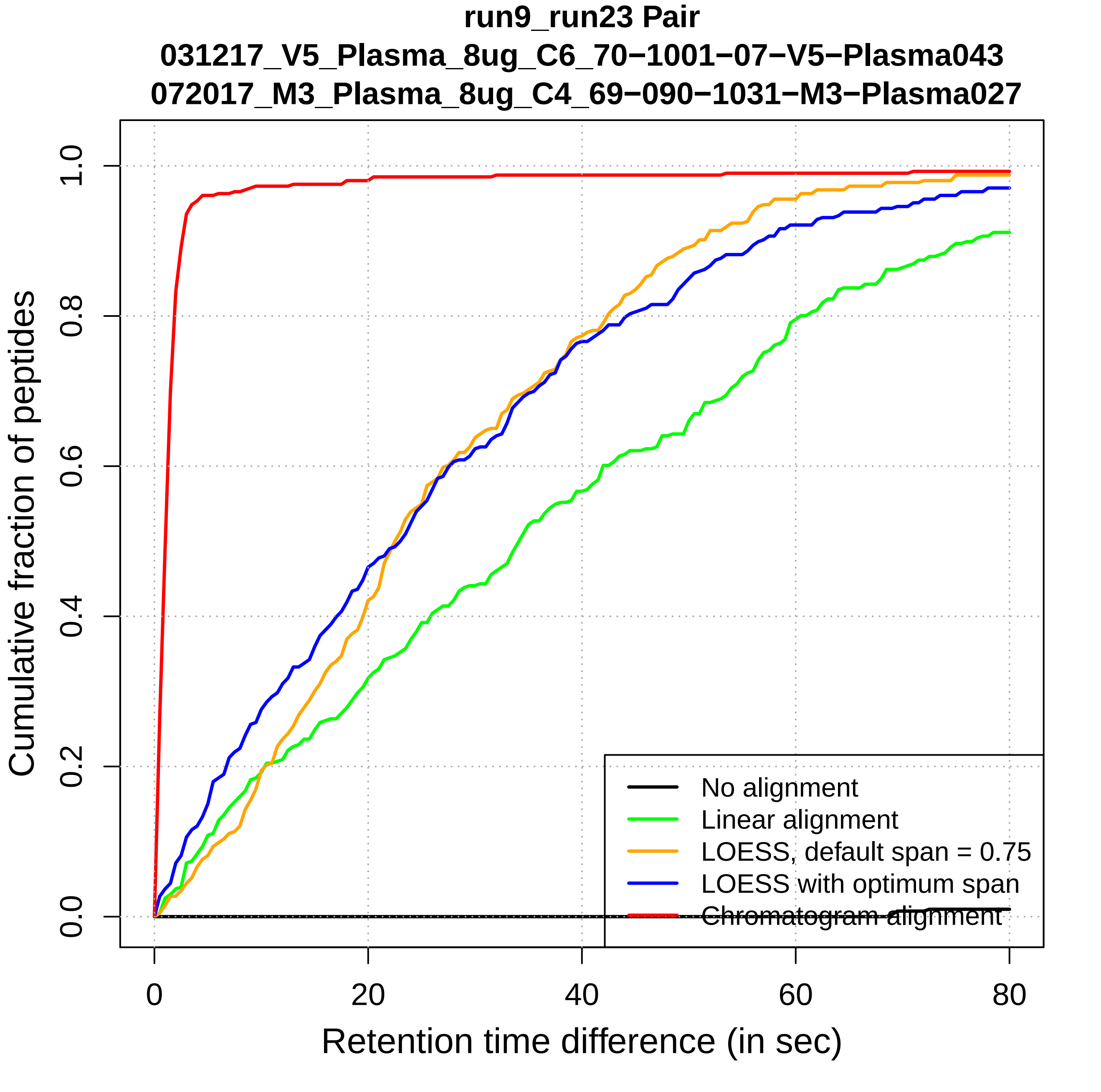
run9_run23 Pair
031217_V5_Plasma_8ug_C6_70−1001−07−V5−Plasma043
072017_M3_Plasma_8ug_C4_69−090−1031−M3−Plasma027
Cumulative fraction of peptides
0.0
0.2
0.4
0.6
0.8
1.0
0
20
40
60
80
Retention time difference (in sec)
No alignment
Linear alignment
LOESS, default span = 0.75
LOESS with optimum span
Chromatogram alignment

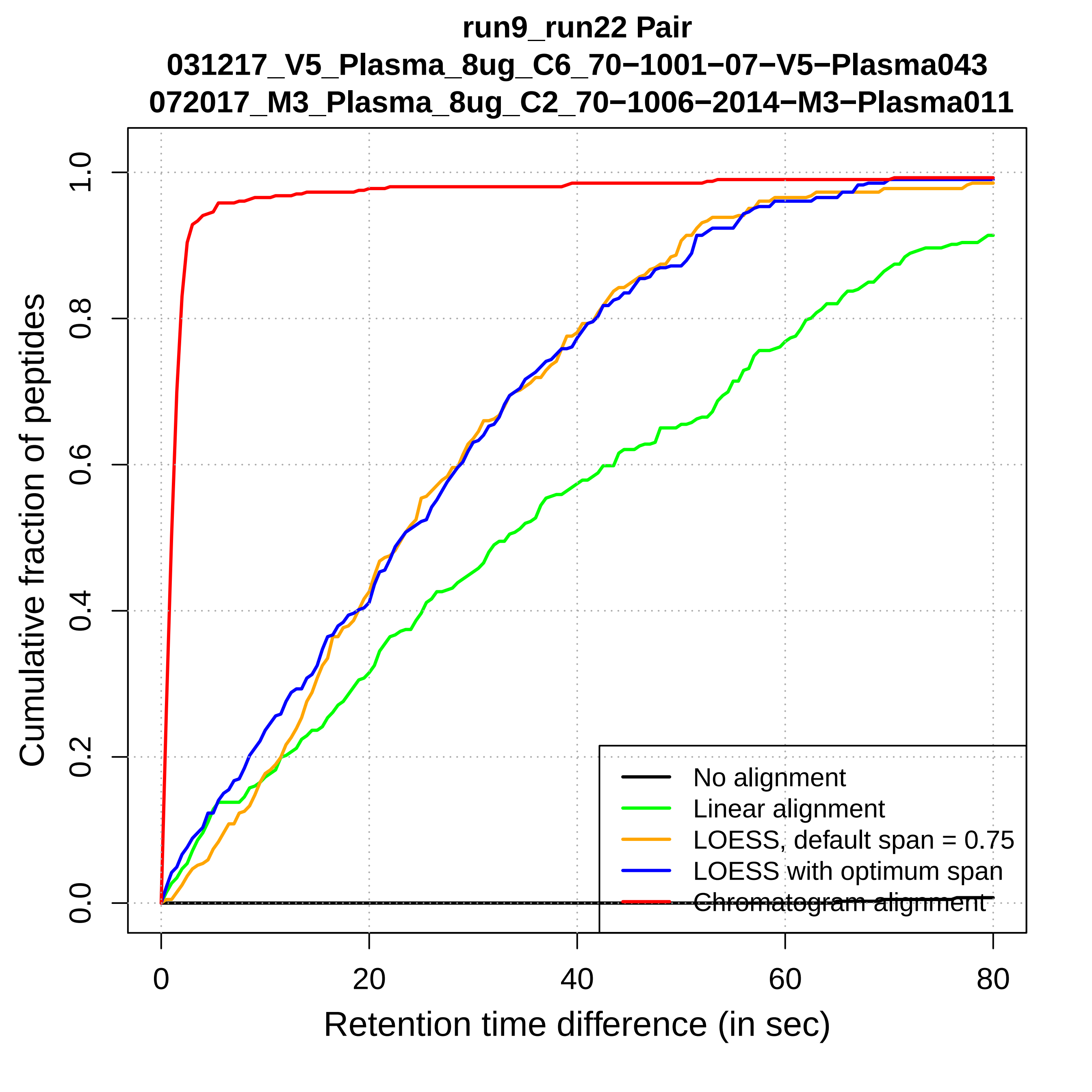

run9_run22 Pair
031217_V5_Plasma_8ug_C6_70−1001−07−V5−Plasma043
072017_M3_Plasma_8ug_C2_70−1006−2014−M3−Plasma011
Cumulative fraction of peptides
0.0
0.2
0.4
0.6
0.8
1.0
0
20
40
60
80
Retention time difference (in sec)
No alignment
Linear alignment
LOESS, default span = 0.75
LOESS with optimum span
Chromatogram alignment

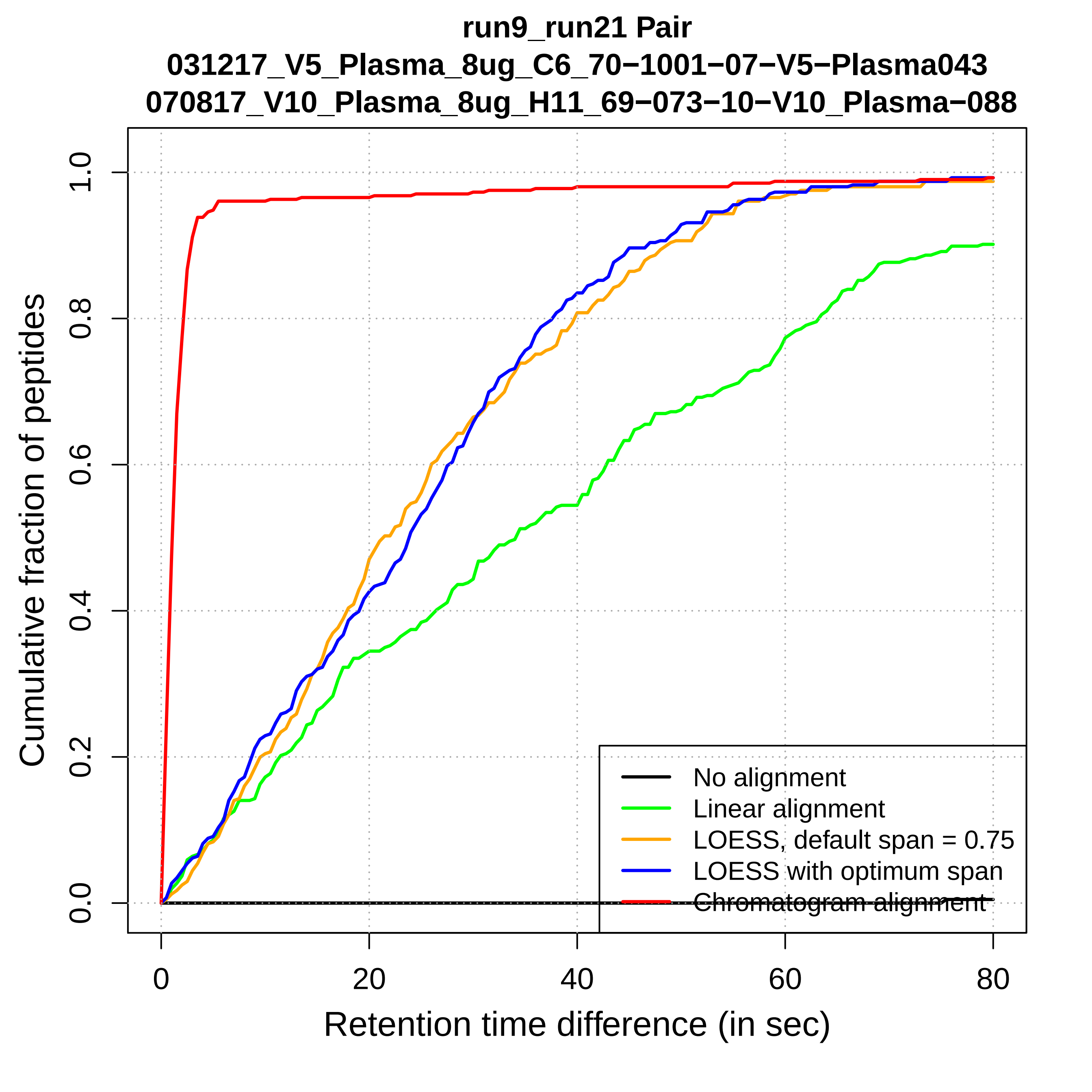
run9_run21 Pair
031217_V5_Plasma_8ug_C6_70−1001−07−V5−Plasma043
070817_V10_Plasma_8ug_H11_69−073−10−V10_Plasma−088
Cumulative fraction of peptides
0.0
0.2
0.4
0.6
0.8
1.0
0
20
40
60
80
Retention time difference (in sec)
No alignment
Linear alignment
LOESS, default span = 0.75
LOESS with optimum span
Chromatogram alignment

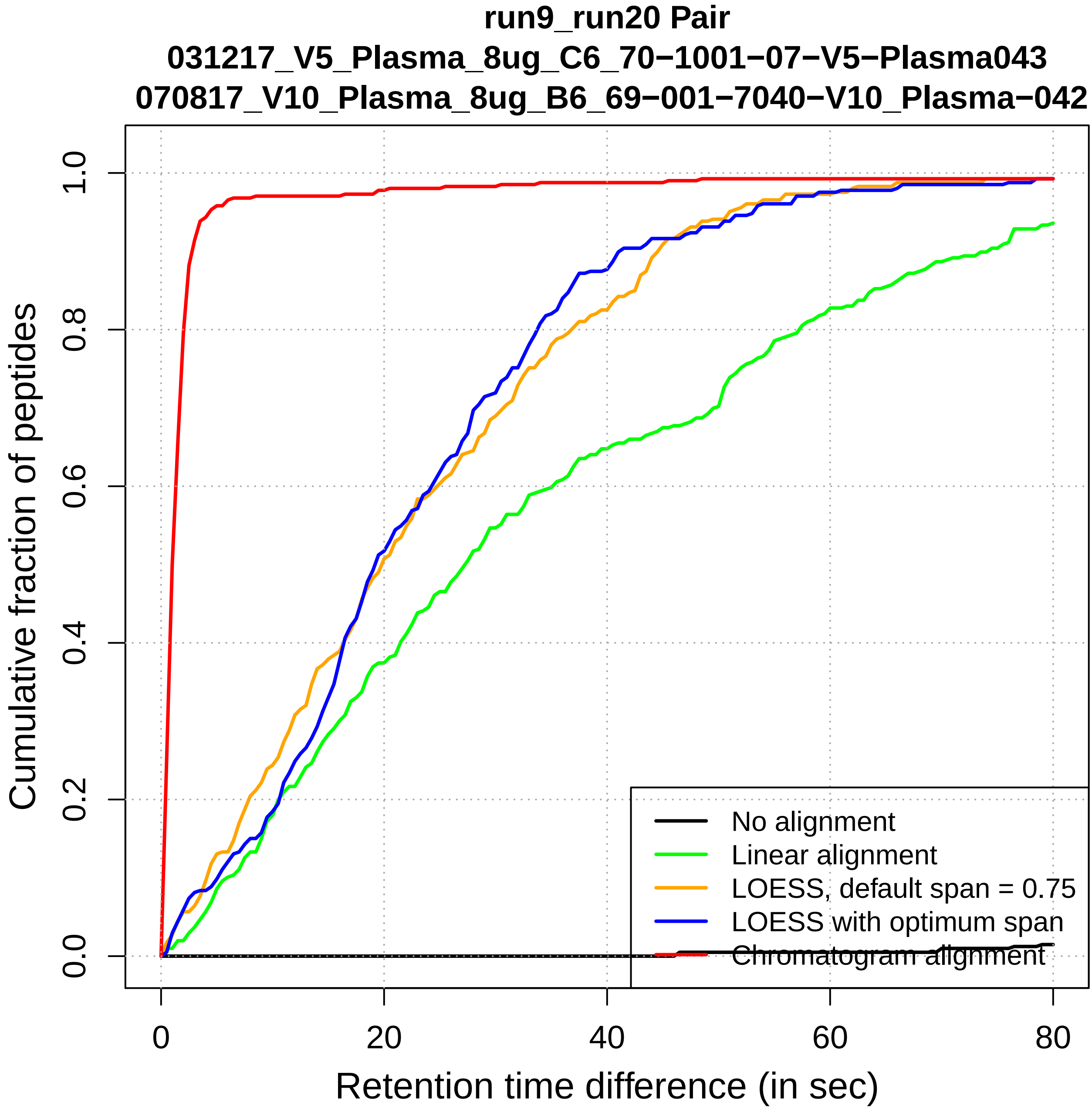

run9_run20 Pair
031217_V5_Plasma_8ug_C6_70−1001−07−V5−Plasma043
070817_V10_Plasma_8ug_B6_69−001−7040−V10_Plasma−042
Cumulative fraction of peptides
0.0
0.2
0.4
0.6
0.8
1.0
0
20
40
60
80
Retention time difference (in sec)
No alignment
Linear alignment
LOESS, default span = 0.75
LOESS with optimum span
Chromatogram alignment

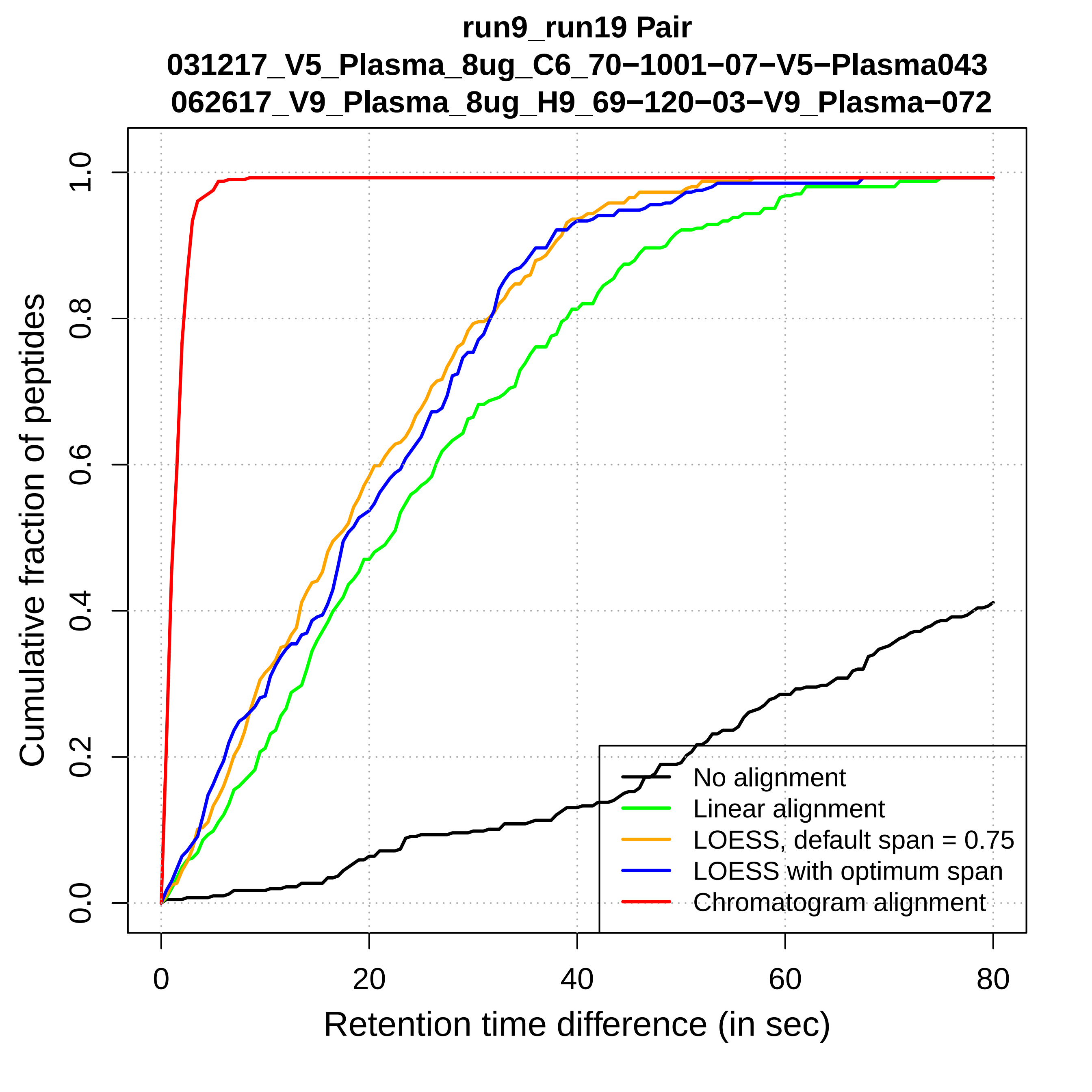

run9_run19 Pair
031217_V5_Plasma_8ug_C6_70−1001−07−V5−Plasma043
062617_V9_Plasma_8ug_H9_69−120−03−V9_Plasma−072
Cumulative fraction of peptides
0.0
0.2
0.4
0.6
0.8
1.0
0
20
40
60
80
Retention time difference (in sec)
No alignment
Linear alignment
LOESS, default span = 0.75
LOESS with optimum span
Chromatogram alignment

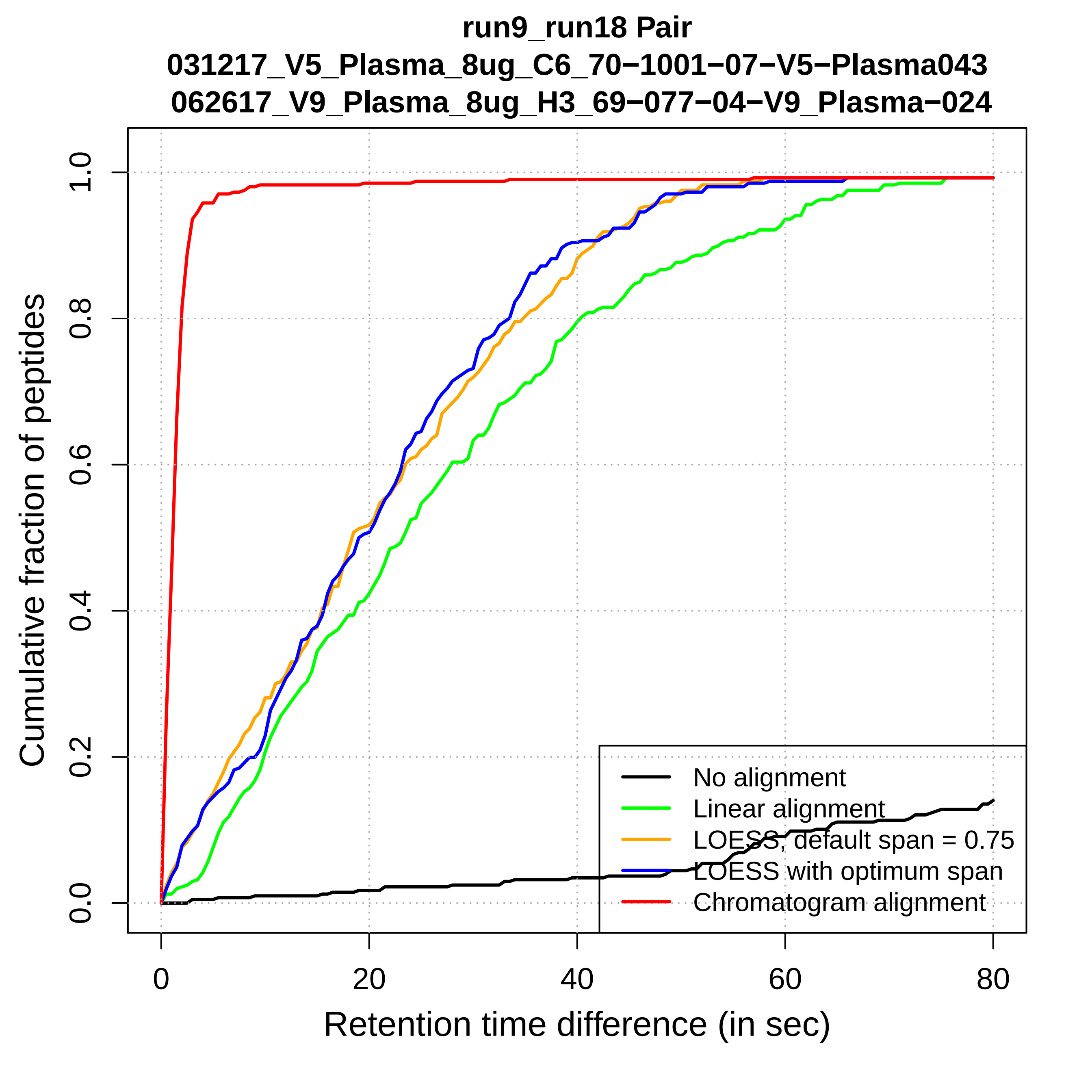

run9_run18 Pair
031217_V5_Plasma_8ug_C6_70−1001−07−V5−Plasma043
062617_V9_Plasma_8ug_H3_69−077−04−V9_Plasma−024
Cumulative fraction of peptides
Retention time difference (in sec)
0.0
0.2
0.4
0.6
0.8
1.0
0
20
40
60
80
No alignment
Linear alignment
LOESS, default span = 0.75
LOESS with optimum span
Chromatogram alignment

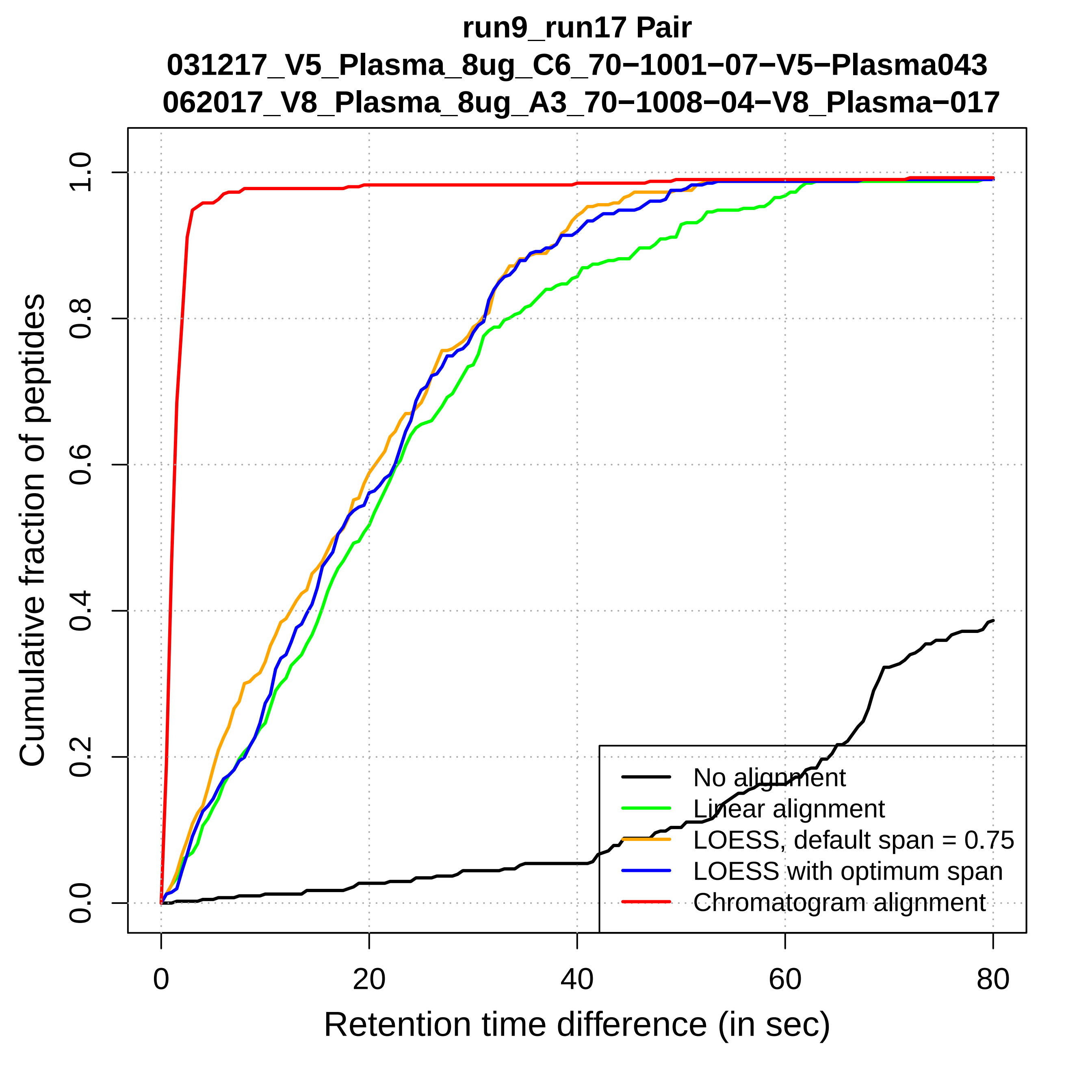
run9_run17 Pair
031217_V5_Plasma_8ug_C6_70−1001−07−V5−Plasma043
062017_V8_Plasma_8ug_A3_70−1008−04−V8_Plasma−017
Cumulative fraction of peptides
0.0
0.2
0.4
0.6
0.8
1.0
0
20
40
60
80
Retention time difference (in sec)
No alignment
Linear alignment
LOESS, default span = 0.75
LOESS with optimum span
Chromatogram alignment

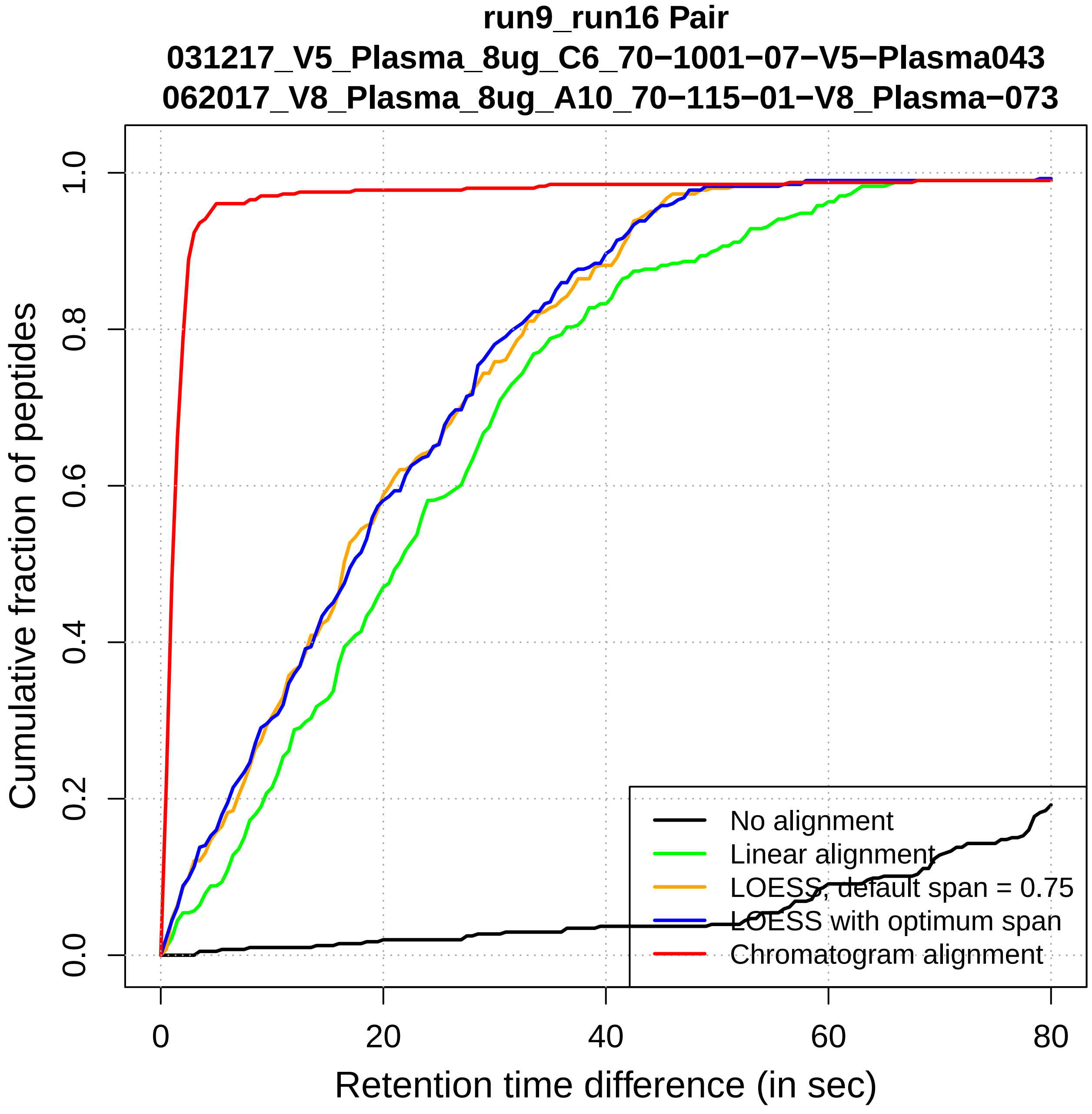

run9_run16 Pair
031217_V5_Plasma_8ug_C6_70−1001−07−V5−Plasma043
062017_V8_Plasma_8ug_A10_70−115−01−V8_Plasma−073
Cumulative fraction of peptides
0.0
0.2
0.4
0.6
0.8
1.0
0
20
40
60
80
Retention time difference (in sec)
No alignment
Linear alignment
LOESS, default span = 0.75
LOESS with optimum span
Chromatogram alignment

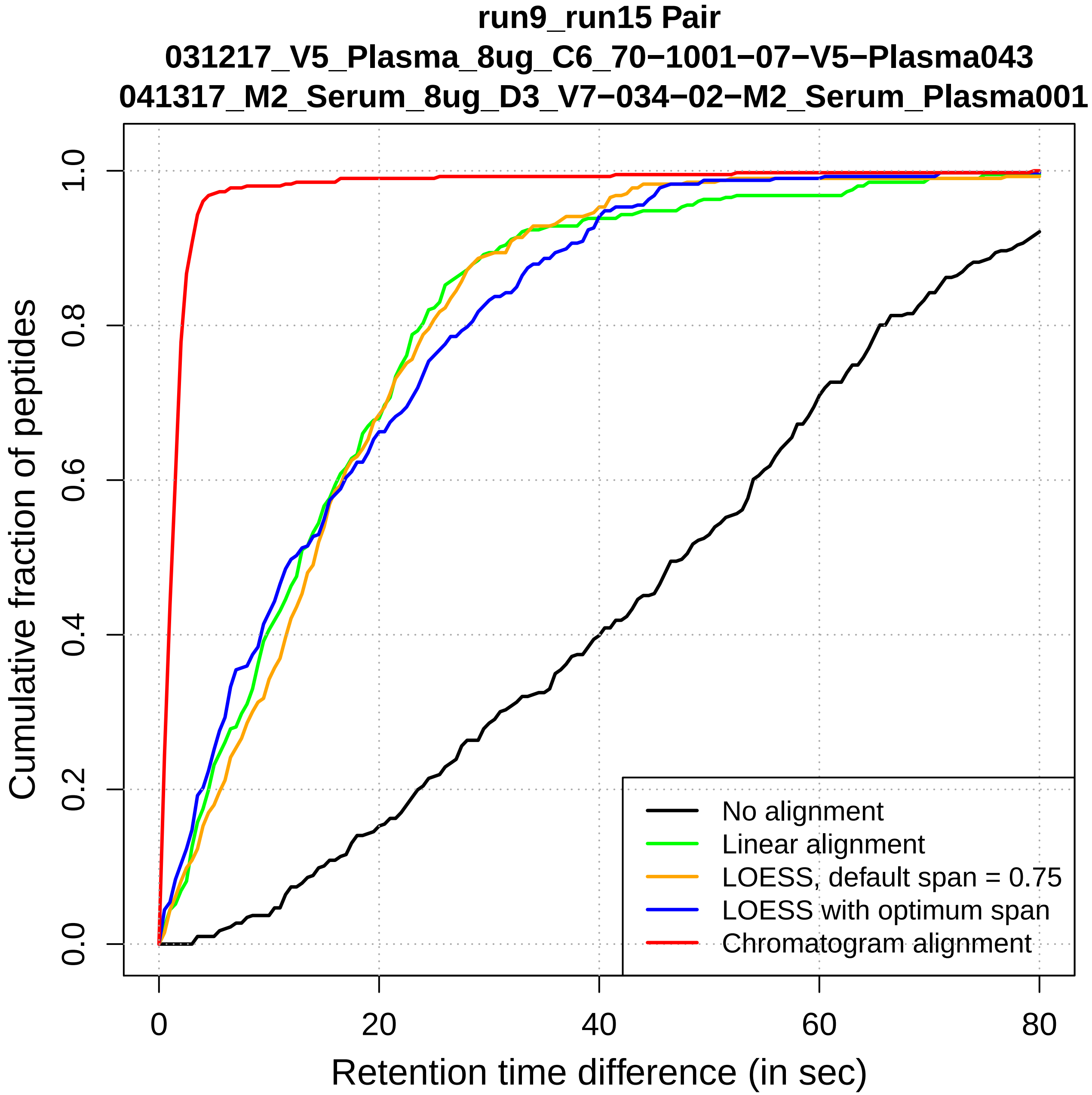

run9_run15 Pair
031217_V5_Plasma_8ug_C6_70−1001−07−V5−Plasma043
041317_M2_Serum_8ug_D3_V7−034−02−M2_Serum_Plasma001
Cumulative fraction of peptides
0.0
0.2
0.4
0.6
0.8
1.0
0
20
40
60
80
Retention time difference (in sec)
No alignment
Linear alignment
LOESS, default span = 0.75
LOESS with optimum span
Chromatogram alignment

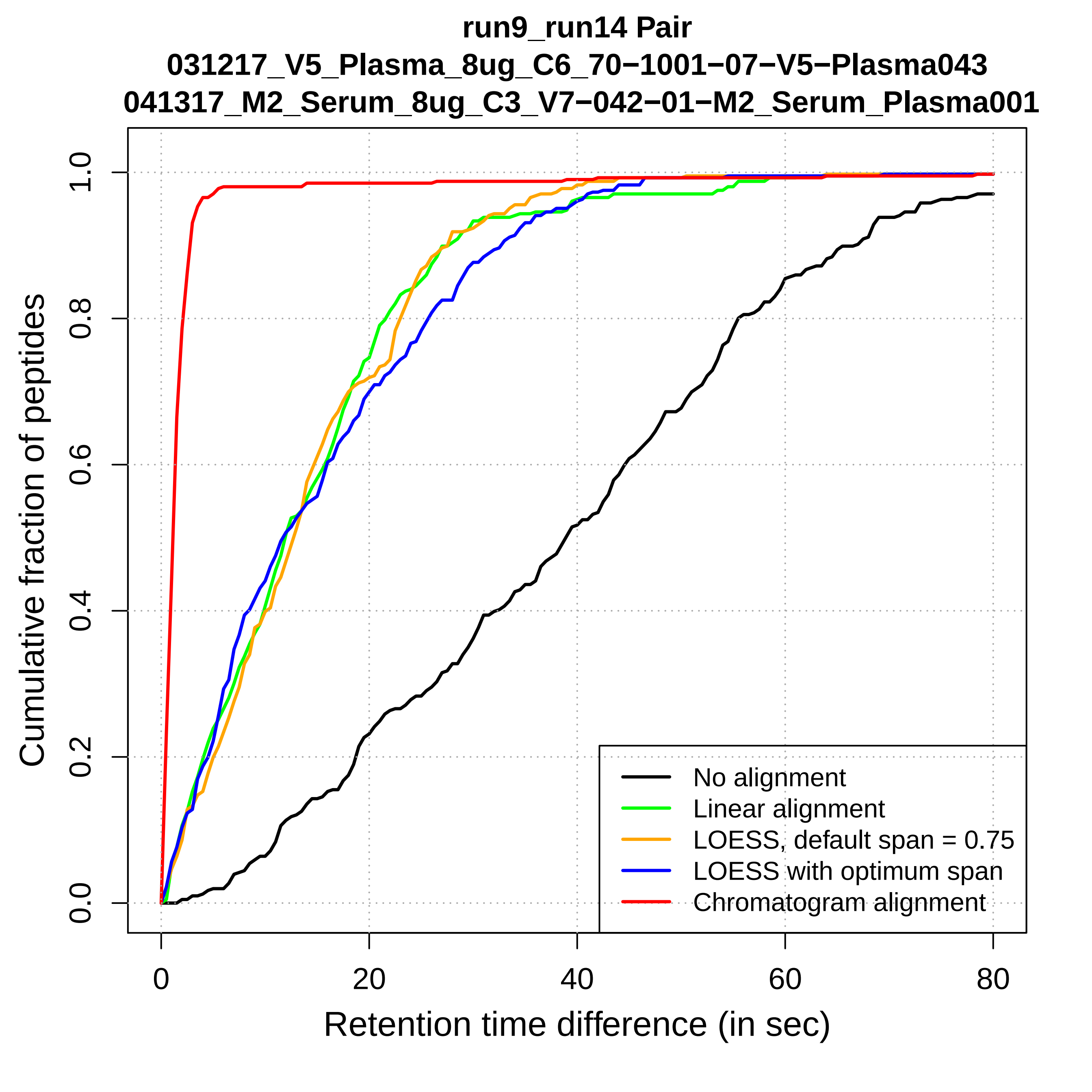

run9_run14 Pair
031217_V5_Plasma_8ug_C6_70−1001−07−V5−Plasma043
041317_M2_Serum_8ug_C3_V7−042−01−M2_Serum_Plasma001
Cumulative fraction of peptides
0.0
0.2
0.4
0.6
0.8
1.0
0
20
40
60
80
Retention time difference (in sec)
No alignment
Linear alignment
LOESS, default span = 0.75
LOESS with optimum span
Chromatogram alignment

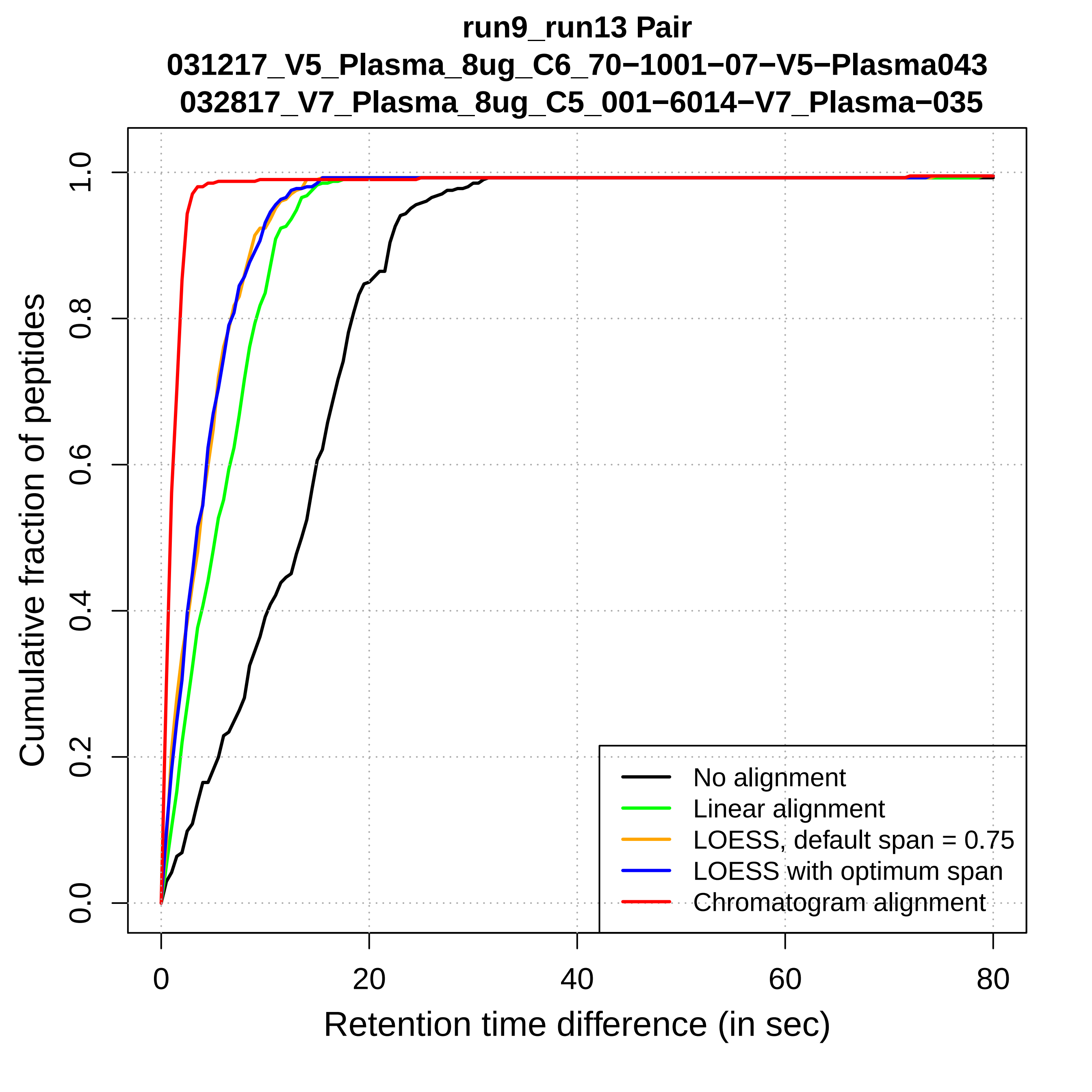
run9_run13 Pair
031217_V5_Plasma_8ug_C6_70−1001−07−V5−Plasma043
032817_V7_Plasma_8ug_C5_001−6014−V7_Plasma−035
Cumulative fraction of peptides
Retention time difference (in sec)
0.0
0.2
0.4
0.6
0.8
1.0
0
20
40
60
80
No alignment
Linear alignment
LOESS, default span = 0.75
LOESS with optimum span
Chromatogram alignment

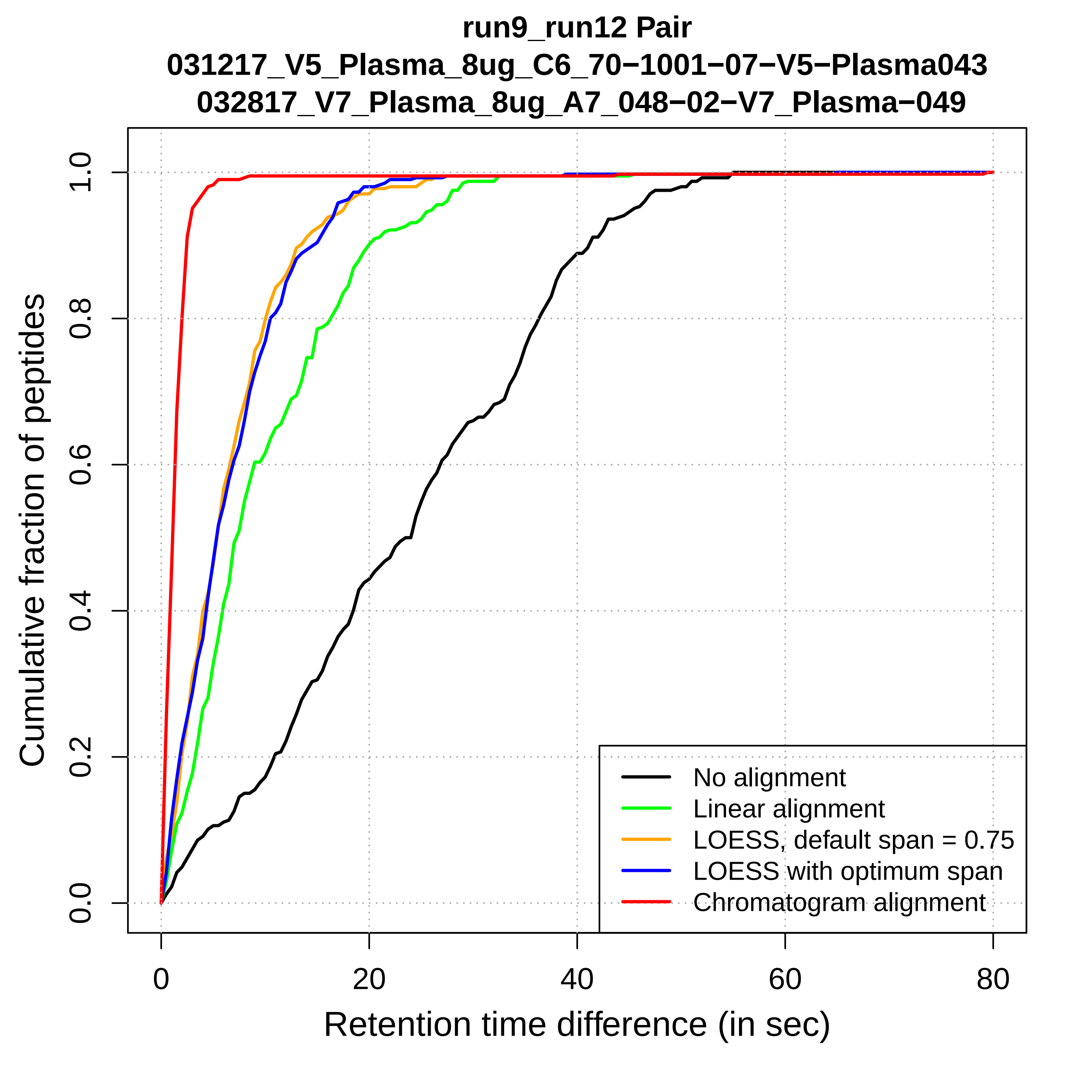

run9_run12 Pair
031217_V5_Plasma_8ug_C6_70−1001−07−V5−Plasma043
032817_V7_Plasma_8ug_A7_048−02−V7_Plasma−049
Cumulative fraction of peptides
0.0
0.2
0.4
0.6
0.8
1.0
0
20
40
60
80
Retention time difference (in sec)
No alignment
Linear alignment
LOESS, default span = 0.75
LOESS with optimum span
Chromatogram alignment

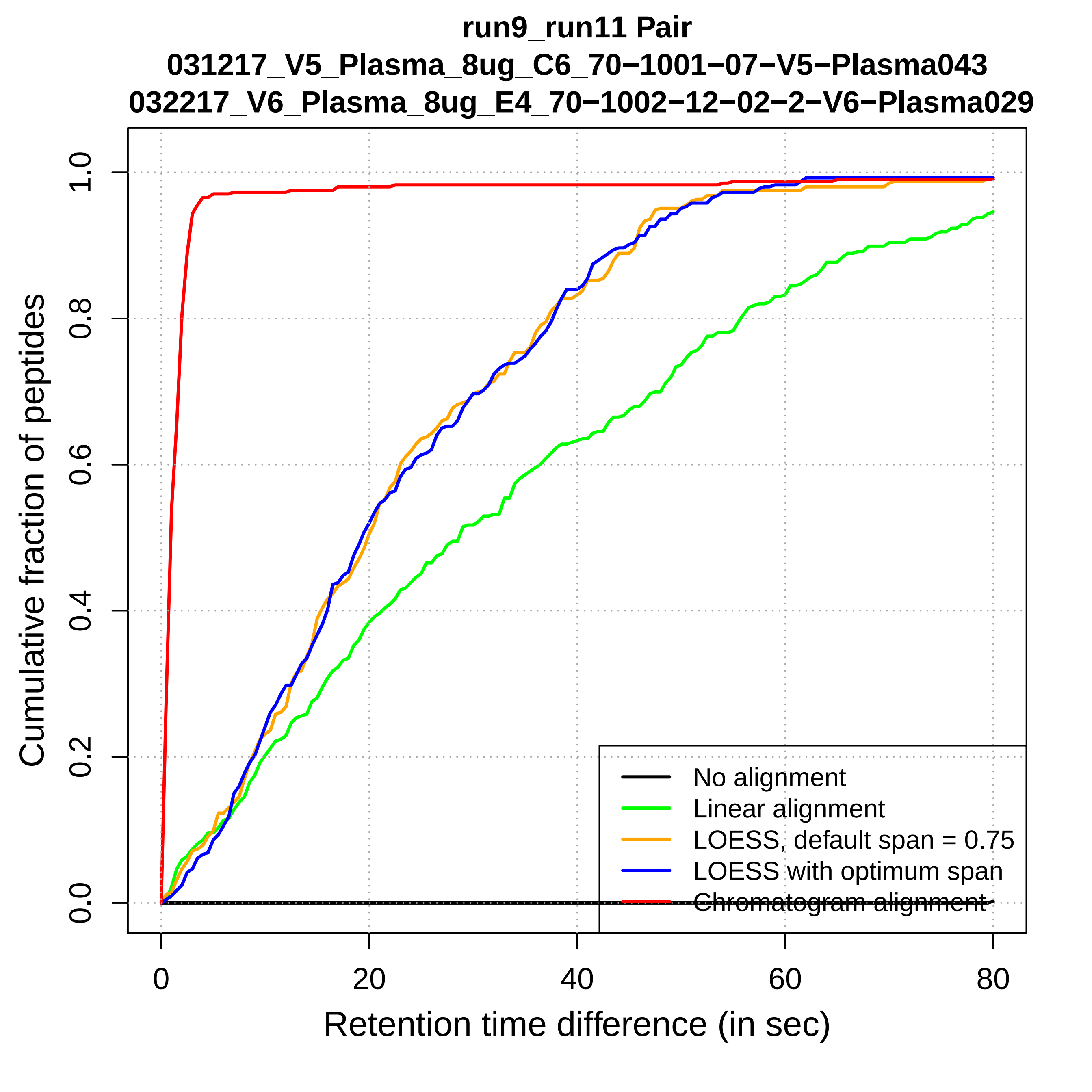
run9_run11 Pair
031217_V5_Plasma_8ug_C6_70−1001−07−V5−Plasma043
032217_V6_Plasma_8ug_E4_70−1002−12−02−2−V6−Plasma029
Cumulative fraction of peptides
0.0
0.2
0.4
0.6
0.8
1.0
0
20
40
60
80
Retention time difference (in sec)
No alignment
Linear alignment
LOESS, default span = 0.75
LOESS with optimum span
Chromatogram alignment

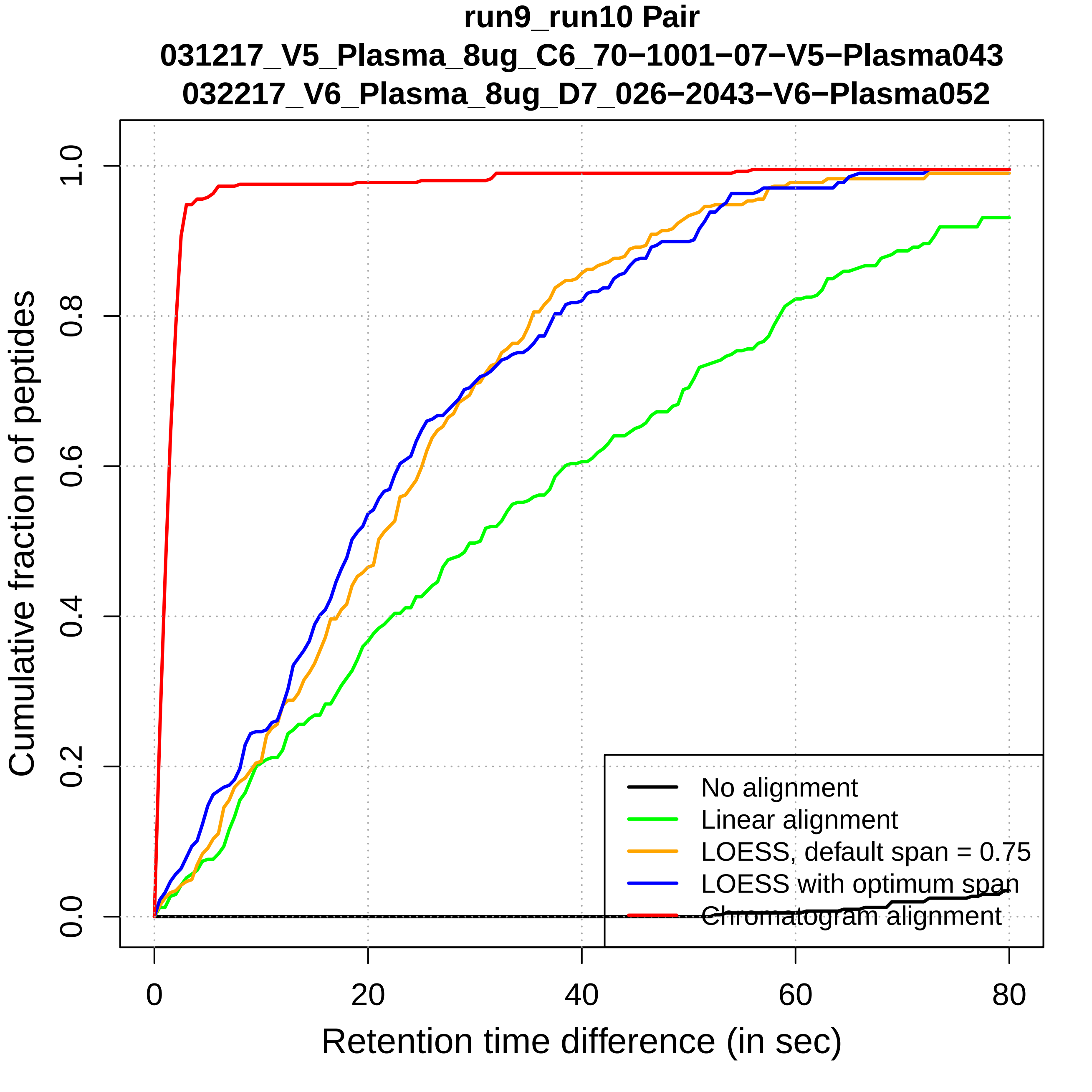

run9_run10 Pair
031217_V5_Plasma_8ug_C6_70−1001−07−V5−Plasma043
032217_V6_Plasma_8ug_D7_026−2043−V6−Plasma052
Cumulative fraction of peptides
0.0
0.2
0.4
0.6
0.8
1.0
0
20
40
60
80
Retention time difference (in sec)
No alignment
Linear alignment
LOESS, default span = 0.75
LOESS with optimum span
Chromatogram alignment

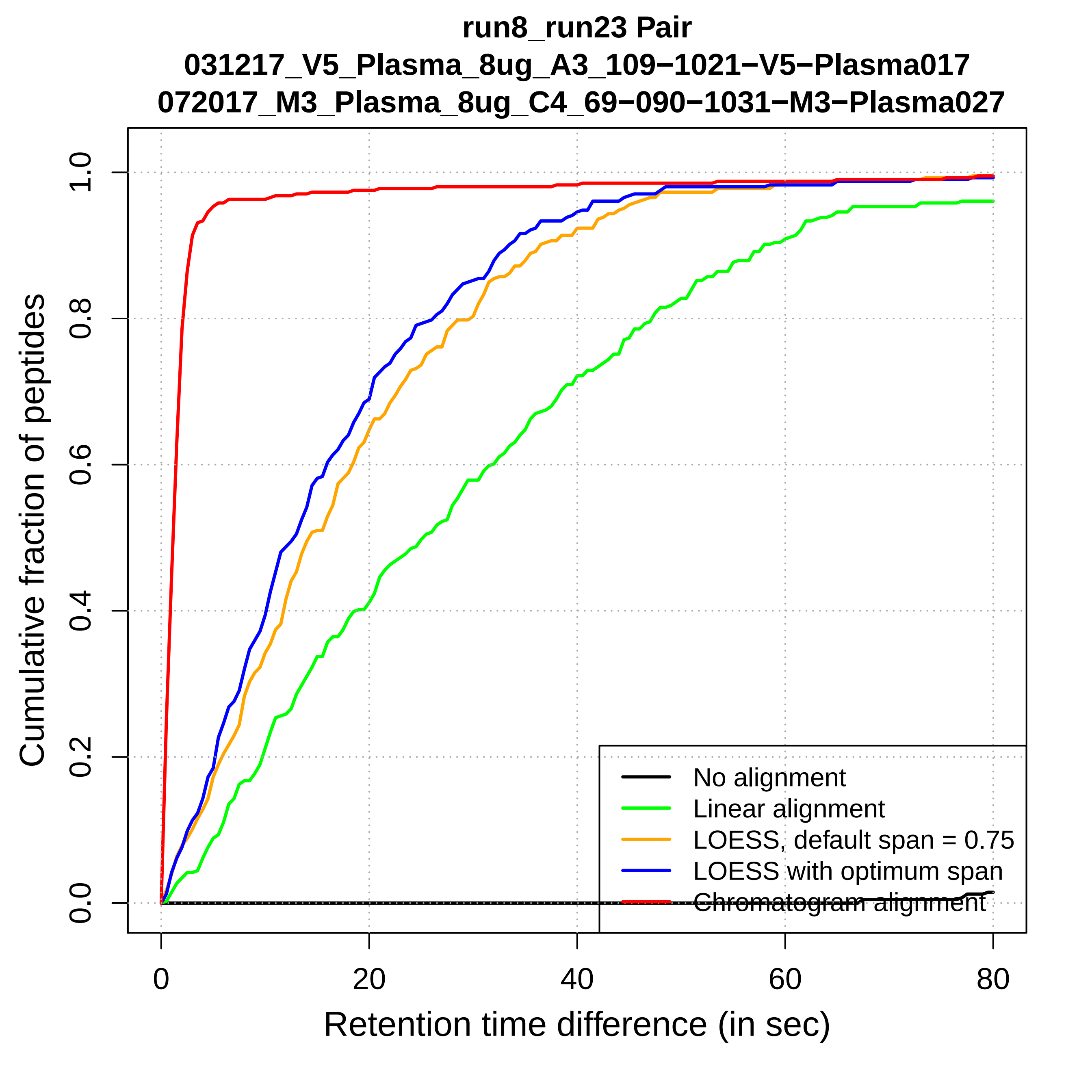
run8_run23 Pair
031217_V5_Plasma_8ug_A3_109−1021−V5−Plasma017
072017_M3_Plasma_8ug_C4_69−090−1031−M3−Plasma027
Cumulative fraction of peptides
0.0
0.2
0.4
0.6
0.8
1.0
0
20
40
60
80
Retention time difference (in sec)
No alignment
Linear alignment
LOESS, default span = 0.75
LOESS with optimum span
Chromatogram alignment

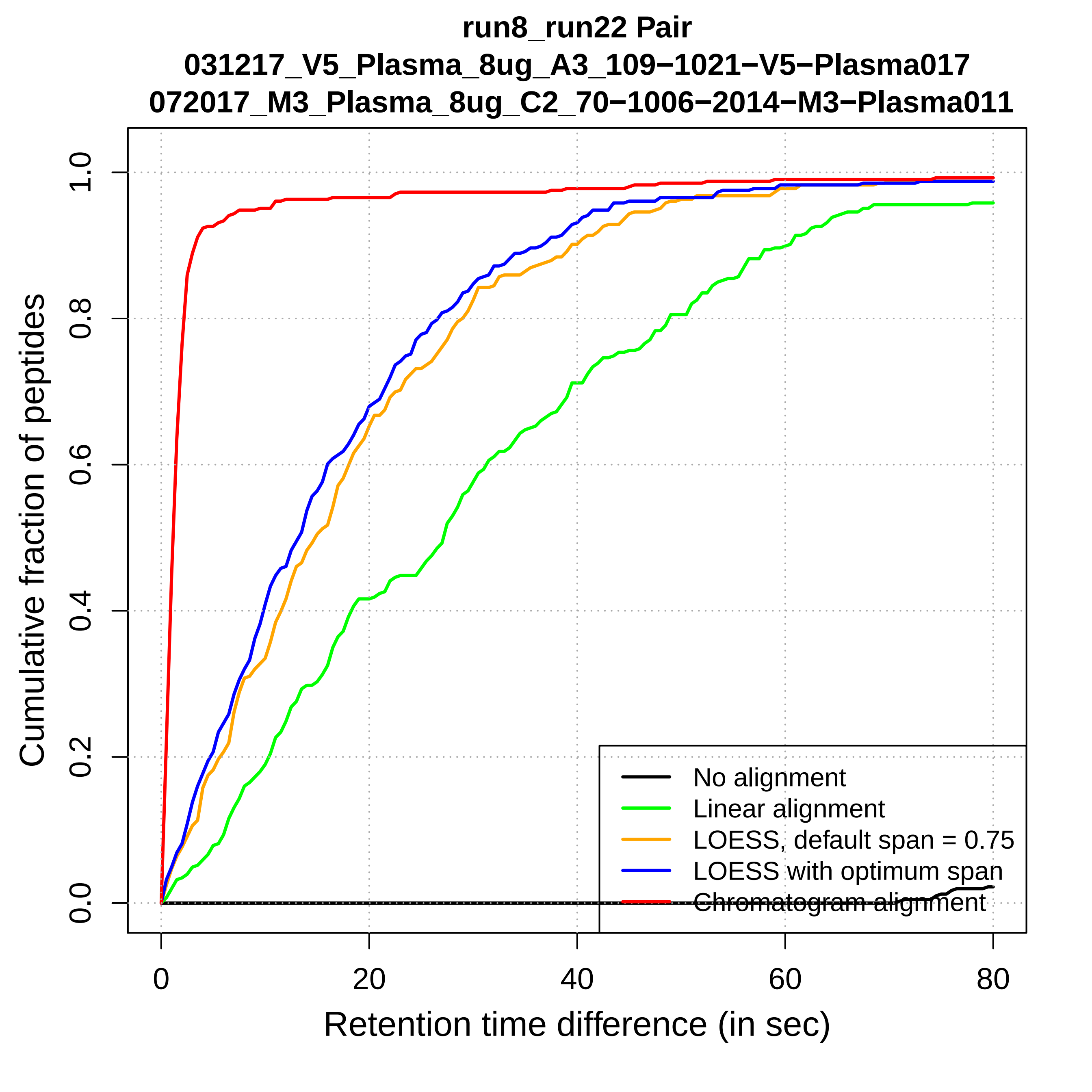
run8_run22 Pair
031217_V5_Plasma_8ug_A3_109−1021−V5−Plasma017
072017_M3_Plasma_8ug_C2_70−1006−2014−M3−Plasma011
Cumulative fraction of peptides
Retention time difference (in sec)
0.0
0.2
0.4
0.6
0.8
1.0
0
20
40
60
80
No alignment
Linear alignment
LOESS, default span = 0.75
LOESS with optimum span
Chromatogram alignment

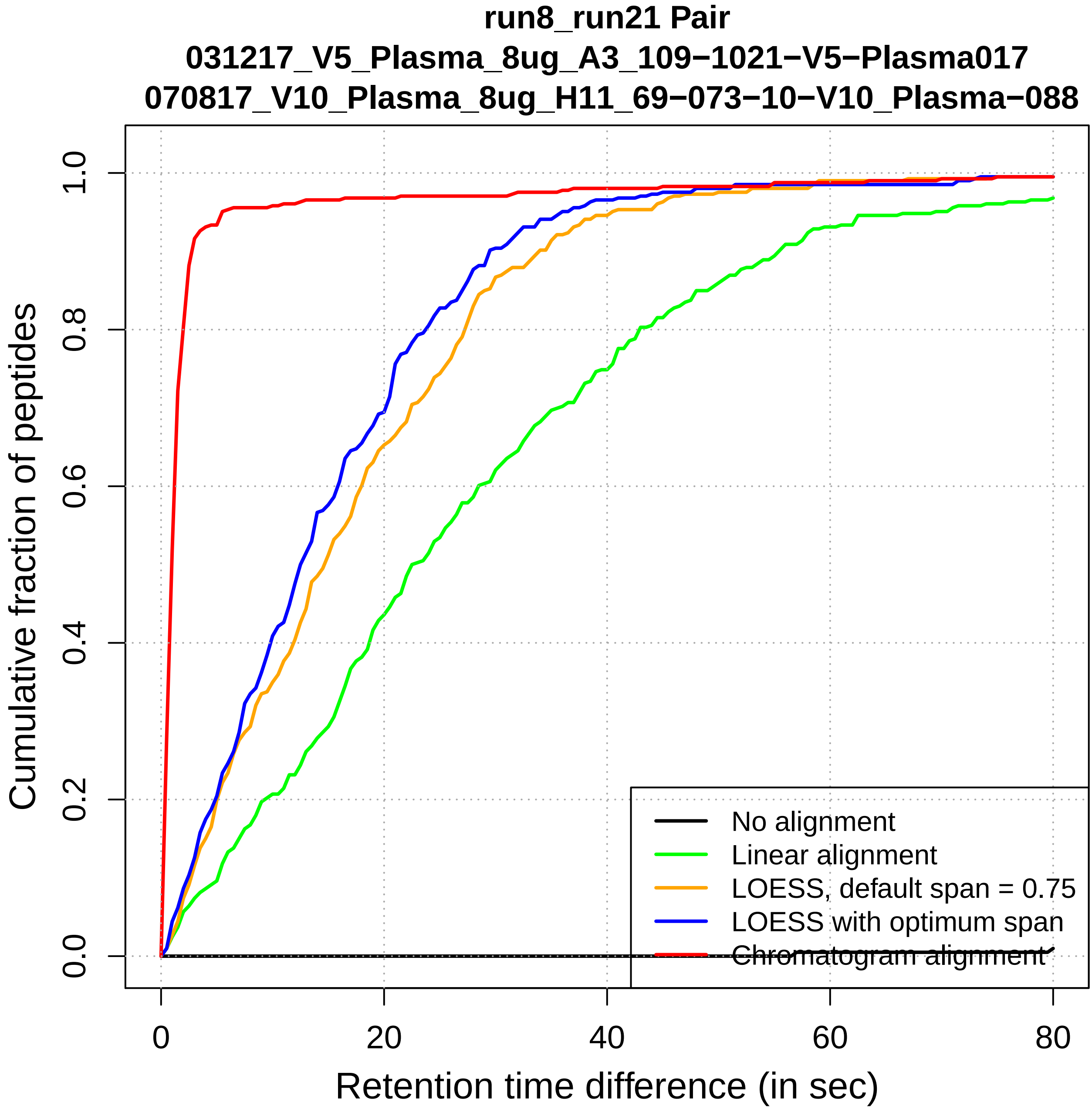

run8_run21 Pair
031217_V5_Plasma_8ug_A3_109−1021−V5−Plasma017
070817_V10_Plasma_8ug_H11_69−073−10−V10_Plasma−088
Cumulative fraction of peptides
0.0
0.2
0.4
0.6
0.8
1.0
0
20
40
60
80
Retention time difference (in sec)
No alignment
Linear alignment
LOESS, default span = 0.75
LOESS with optimum span
Chromatogram alignment

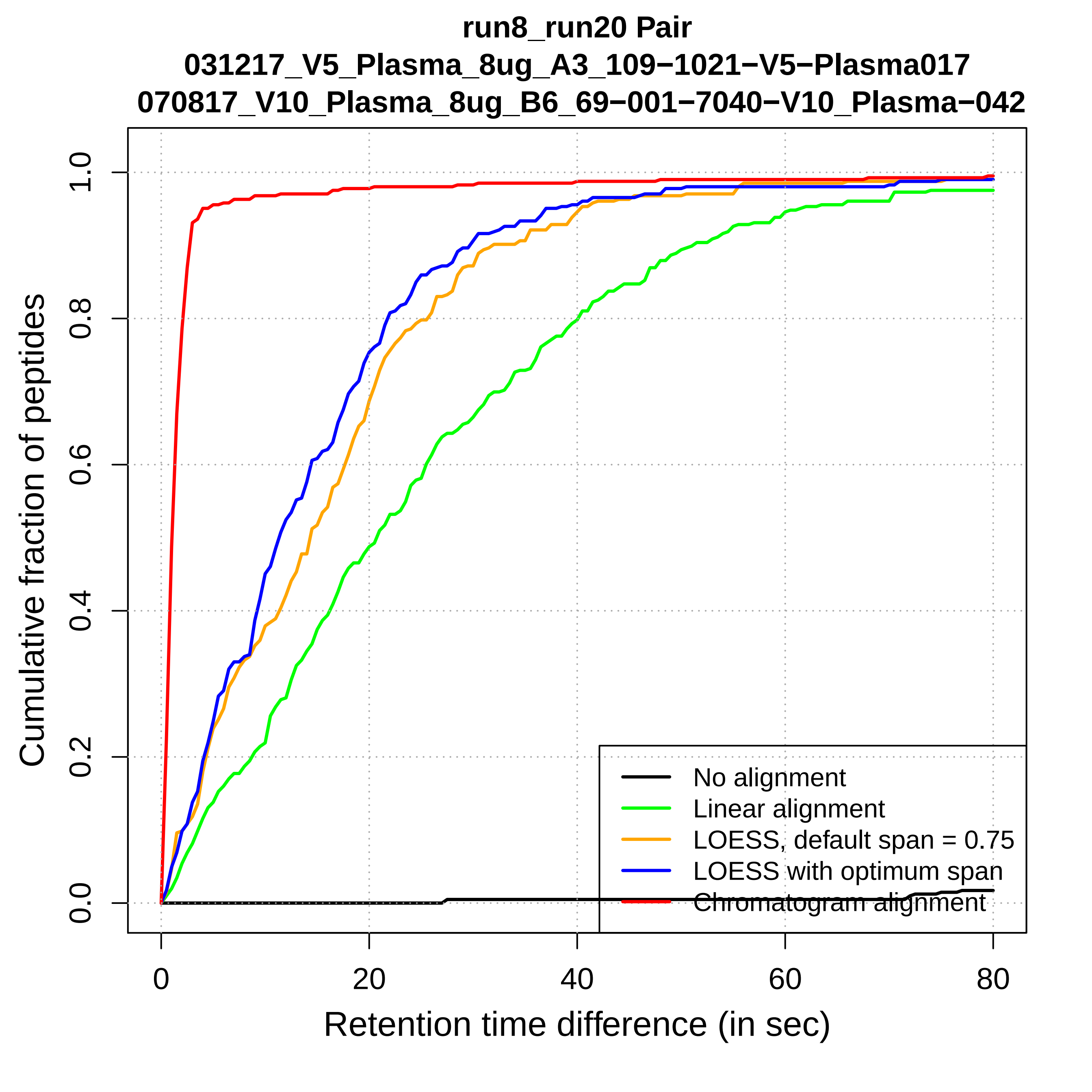
run8_run20 Pair
031217_V5_Plasma_8ug_A3_109−1021−V5−Plasma017
070817_V10_Plasma_8ug_B6_69−001−7040−V10_Plasma−042
Cumulative fraction of peptides
0.0
0.2
0.4
0.6
0.8
1.0
0
20
40
60
80
Retention time difference (in sec)
No alignment
Linear alignment
LOESS, default span = 0.75
LOESS with optimum span
Chromatogram alignment

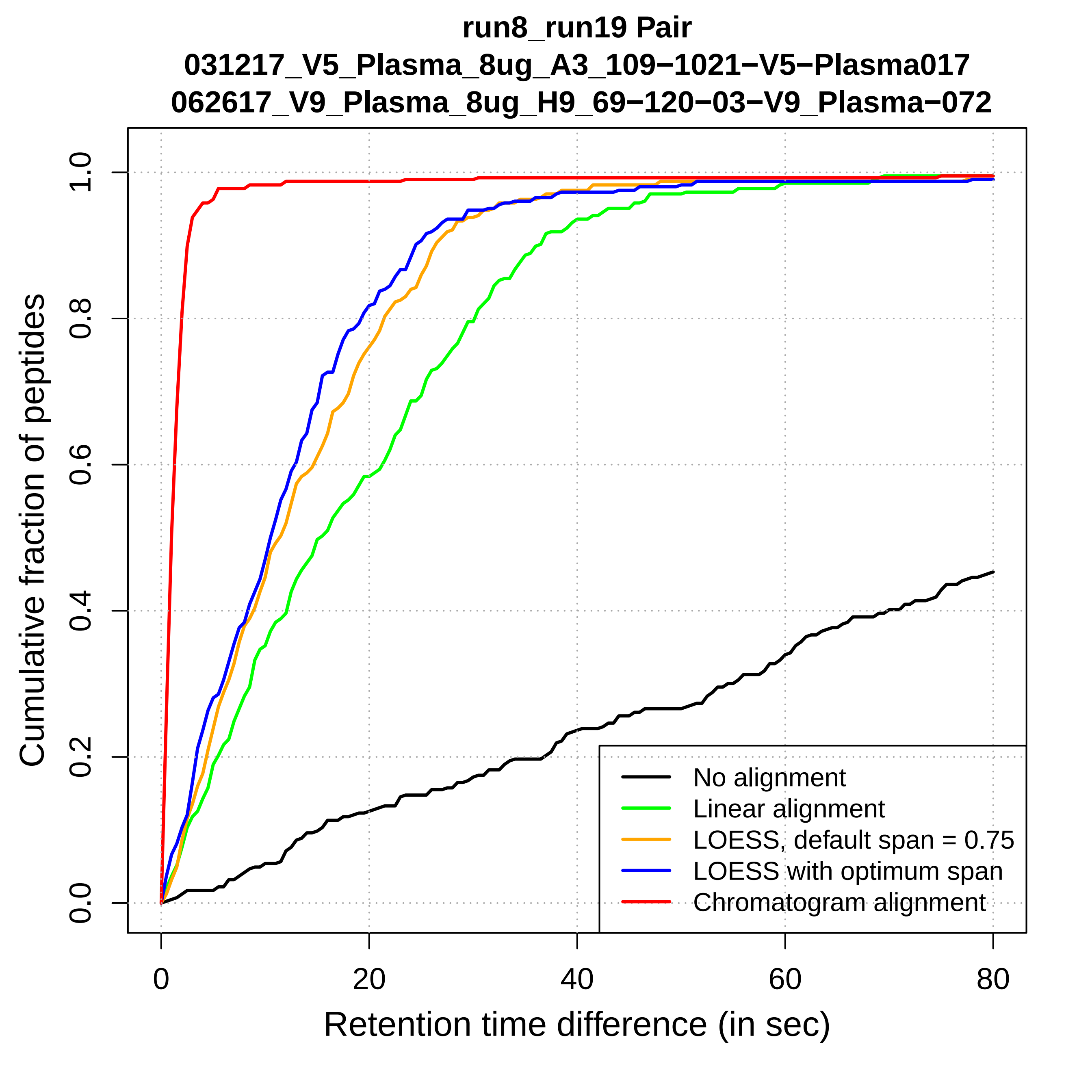

run8_run19 Pair
031217_V5_Plasma_8ug_A3_109−1021−V5−Plasma017
062617_V9_Plasma_8ug_H9_69−120−03−V9_Plasma−072
Cumulative fraction of peptides
0.0
0.2
0.4
0.6
0.8
1.0
0
20
40
60
80
Retention time difference (in sec)
No alignment
Linear alignment
LOESS, default span = 0.75
LOESS with optimum span
Chromatogram alignment

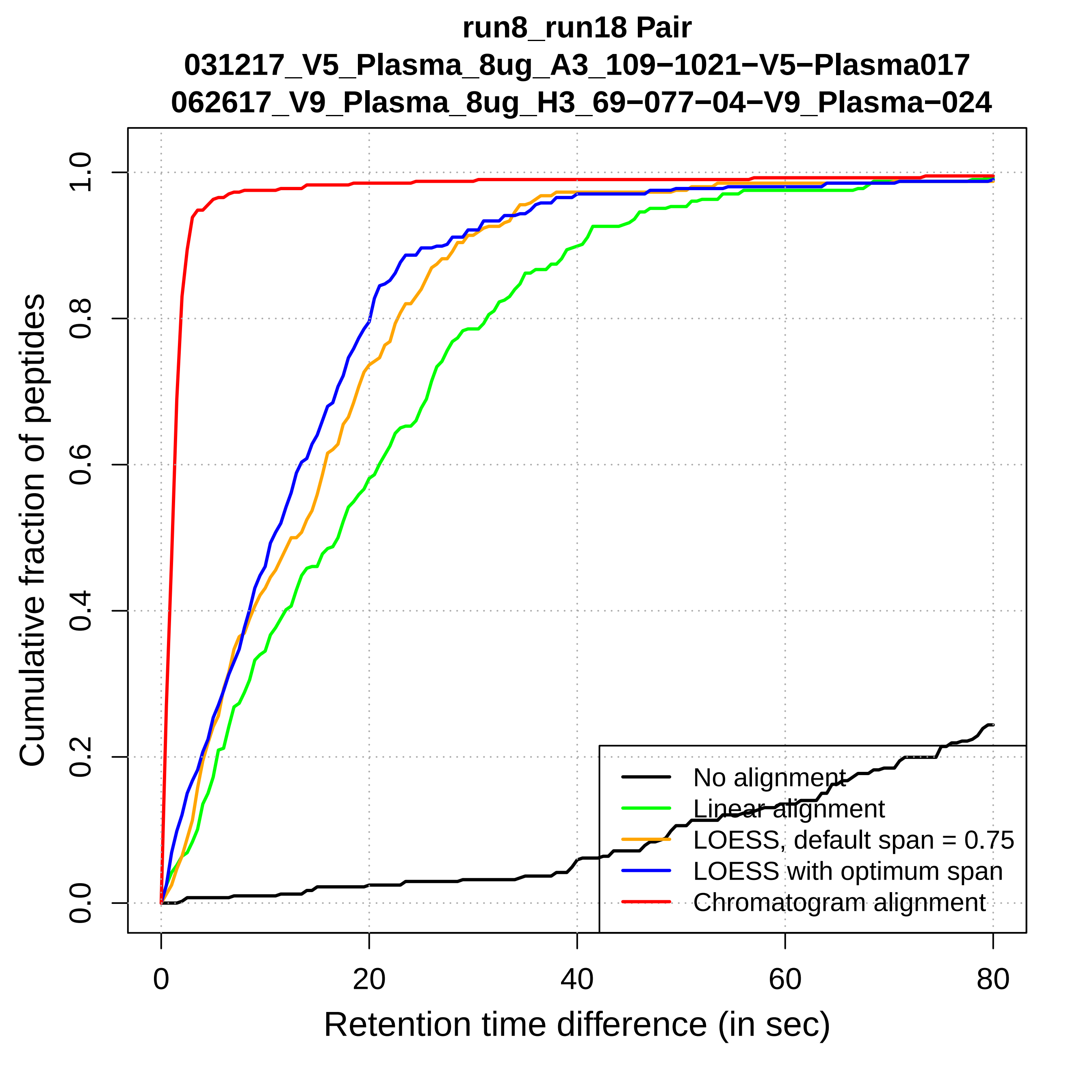

run8_run18 Pair
031217_V5_Plasma_8ug_A3_109−1021−V5−Plasma017
062617_V9_Plasma_8ug_H3_69−077−04−V9_Plasma−024
Cumulative fraction of peptides
0.0
0.2
0.4
0.6
0.8
1.0
0
20
40
60
80
Retention time difference (in sec)
No alignment
Linear alignment
LOESS, default span = 0.75
LOESS with optimum span
Chromatogram alignment

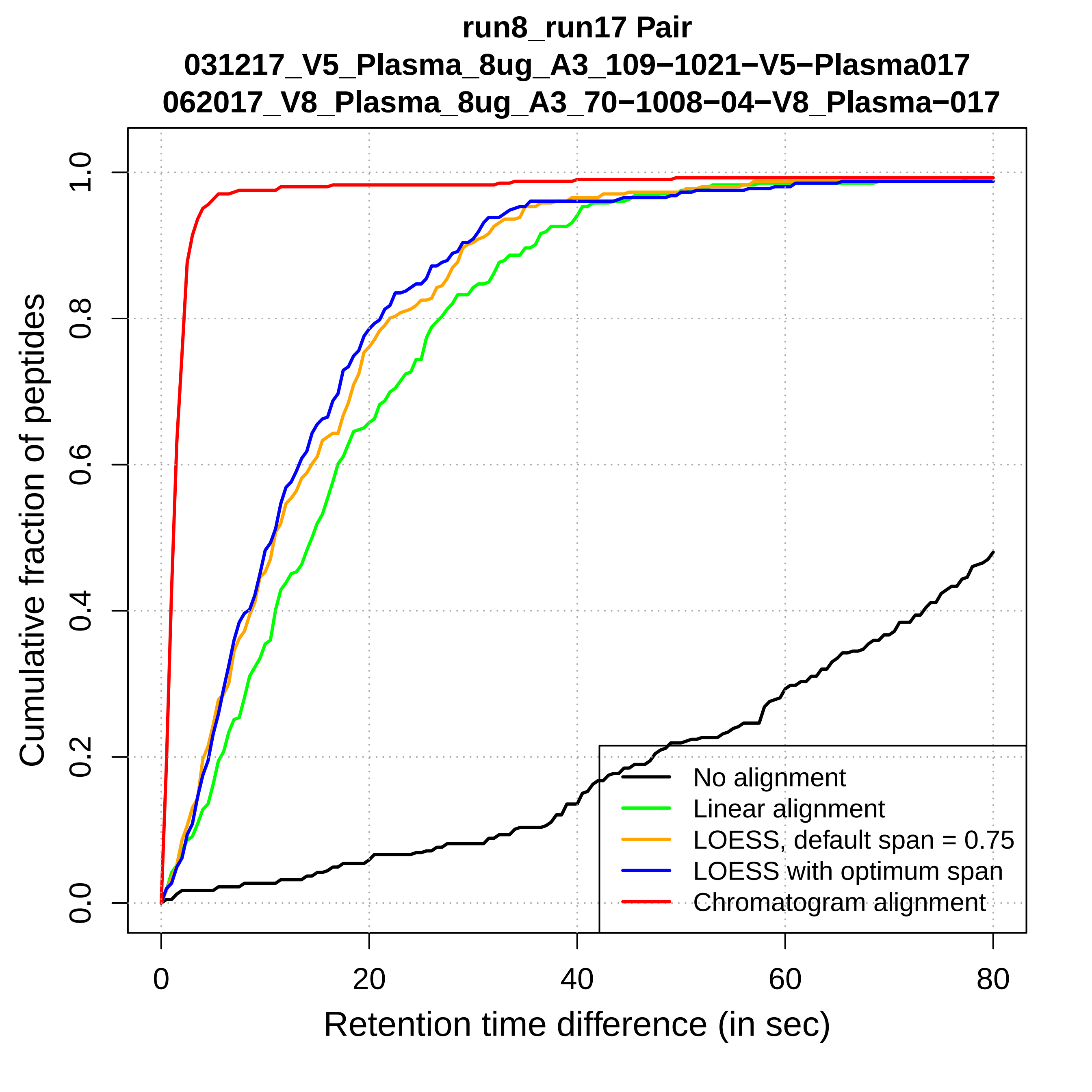
run8_run17 Pair
031217_V5_Plasma_8ug_A3_109−1021−V5−Plasma017
062017_V8_Plasma_8ug_A3_70−1008−04−V8_Plasma−017
Cumulative fraction of peptides
0.0
0.2
0.4
0.6
0.8
1.0
0
20
40
60
80
Retention time difference (in sec)
No alignment
Linear alignment
LOESS, default span = 0.75
LOESS with optimum span
Chromatogram alignment

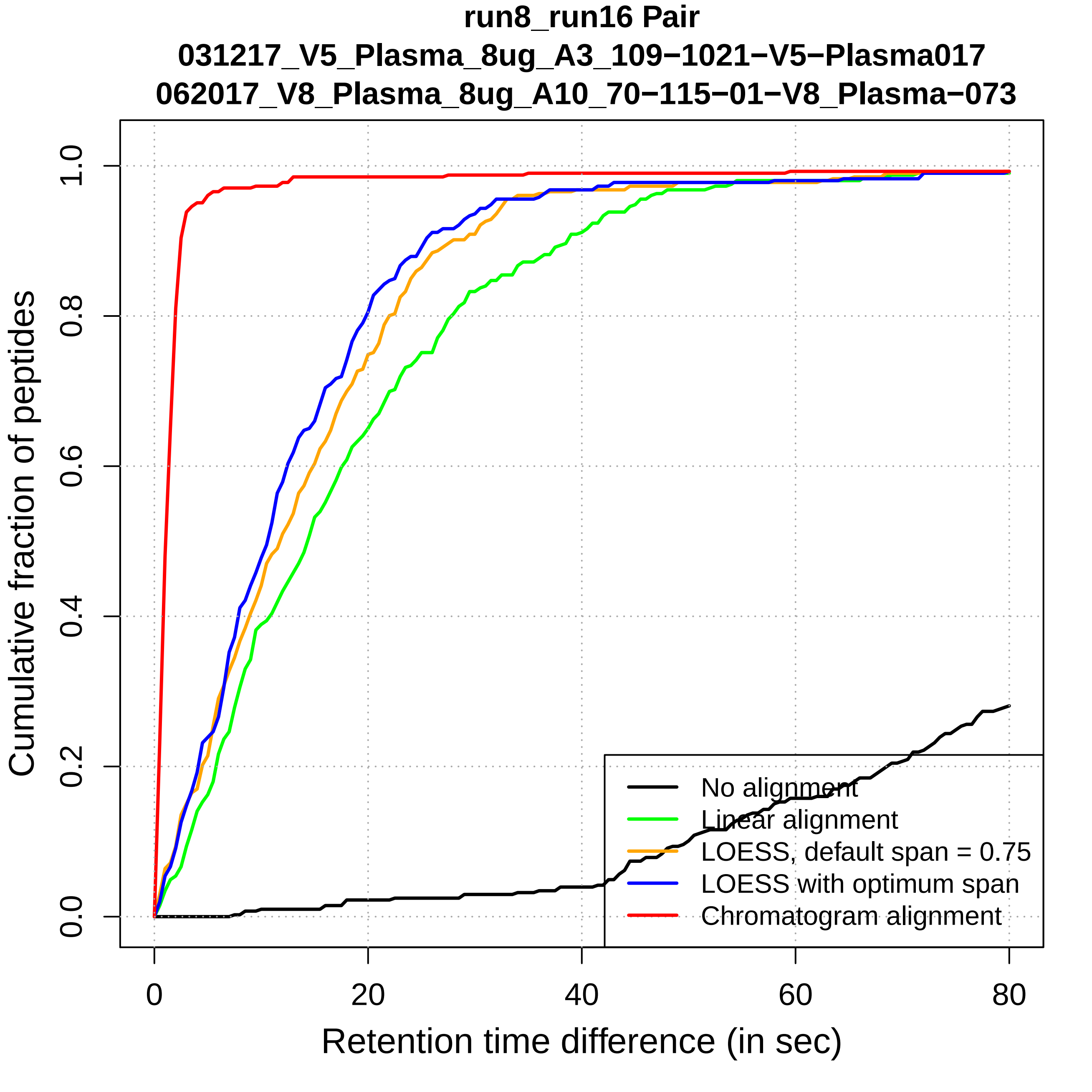
run8_run16 Pair
031217_V5_Plasma_8ug_A3_109−1021−V5−Plasma017
062017_V8_Plasma_8ug_A10_70−115−01−V8_Plasma−073
Cumulative fraction of peptides
Retention time difference (in sec)
0.0
0.2
0.4
0.6
0.8
1.0
0
20
40
60
80
No alignment
Linear alignment
LOESS, default span = 0.75
LOESS with optimum span
Chromatogram alignment

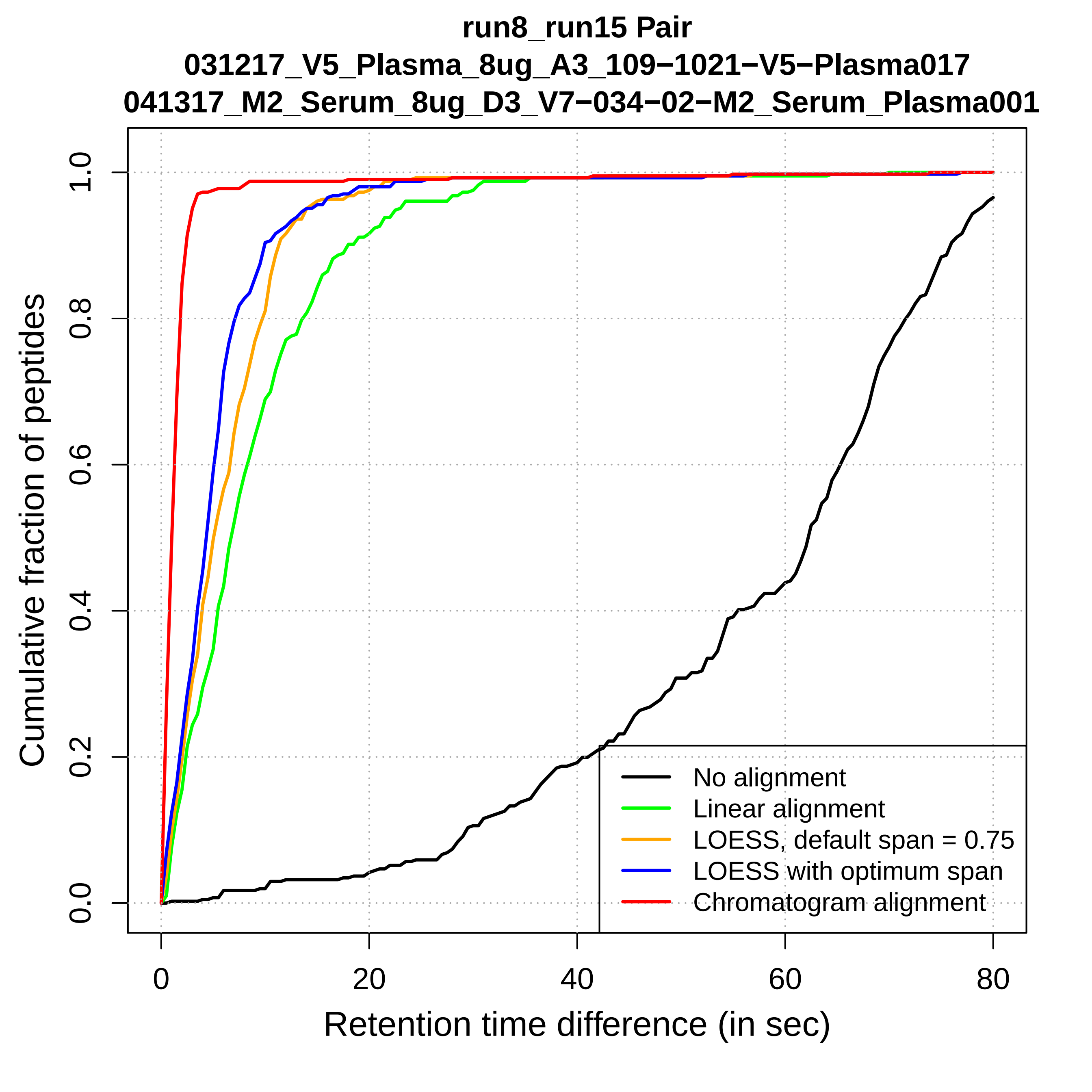
run8_run15 Pair
031217_V5_Plasma_8ug_A3_109−1021−V5−Plasma017
041317_M2_Serum_8ug_D3_V7−034−02−M2_Serum_Plasma001
Cumulative fraction of peptides
0.0
0.2
0.4
0.6
0.8
1.0
0
20
40
60
80
Retention time difference (in sec)
No alignment
Linear alignment
LOESS, default span = 0.75
LOESS with optimum span
Chromatogram alignment

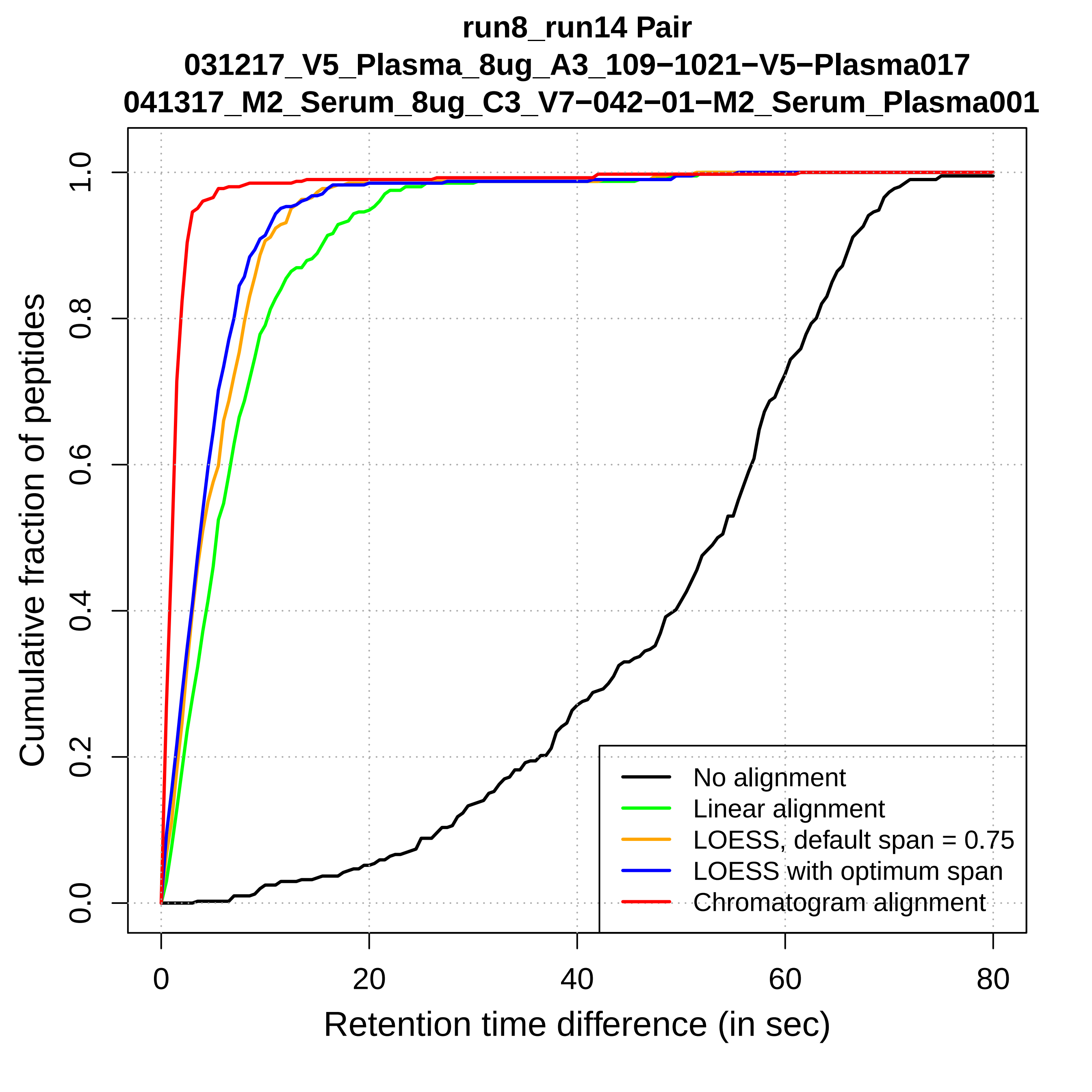

run8_run14 Pair
031217_V5_Plasma_8ug_A3_109−1021−V5−Plasma017
041317_M2_Serum_8ug_C3_V7−042−01−M2_Serum_Plasma001
Cumulative fraction of peptides
0.0
0.2
0.4
0.6
0.8
1.0
0
20
40
60
80
Retention time difference (in sec)
No alignment
Linear alignment
LOESS, default span = 0.75
LOESS with optimum span
Chromatogram alignment

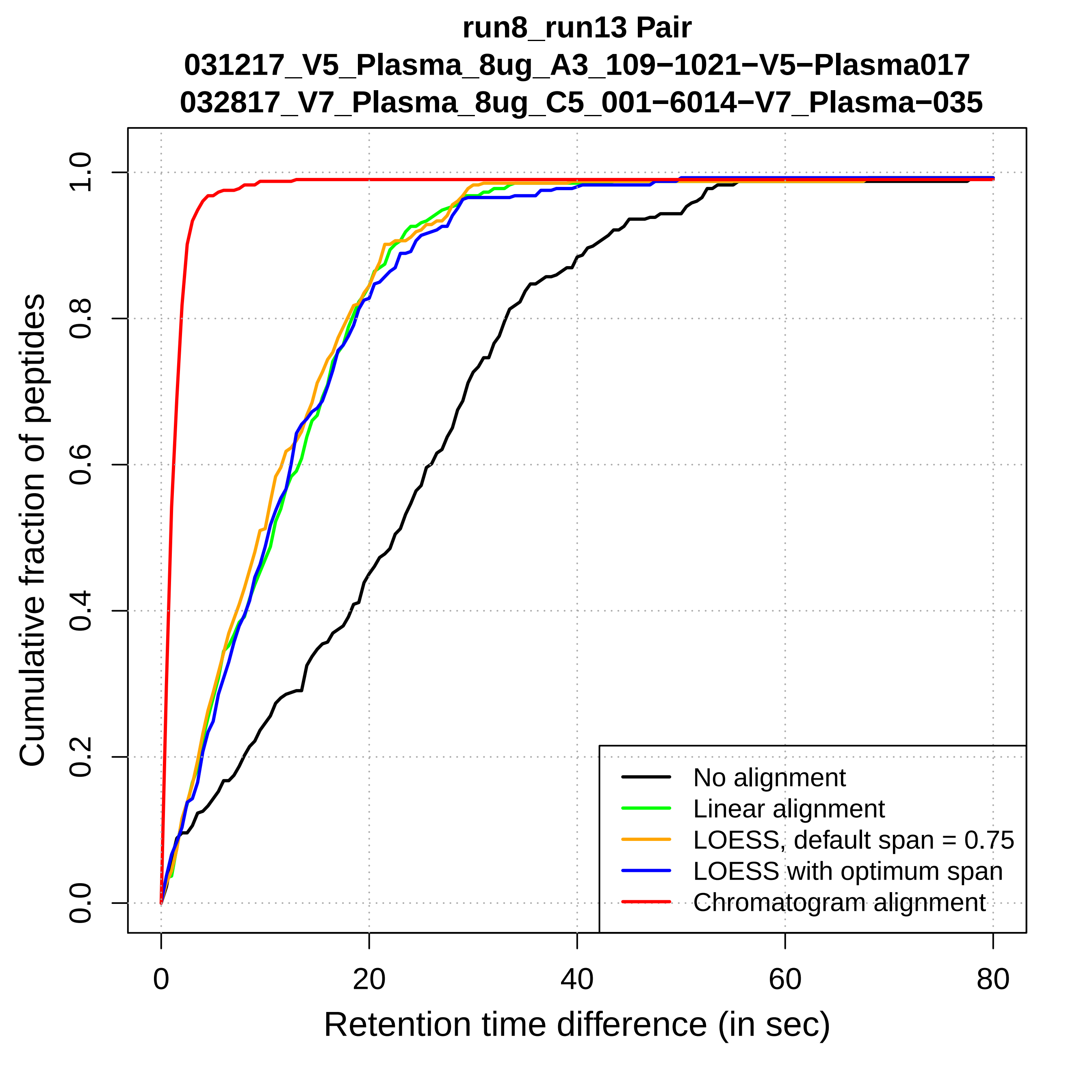

run8_run13 Pair
031217_V5_Plasma_8ug_A3_109−1021−V5−Plasma017
032817_V7_Plasma_8ug_C5_001−6014−V7_Plasma−035
Cumulative fraction of peptides
0.0
0.2
0.4
0.6
0.8
1.0
0
20
40
60
80
Retention time difference (in sec)
No alignment
Linear alignment
LOESS, default span = 0.75
LOESS with optimum span
Chromatogram alignment

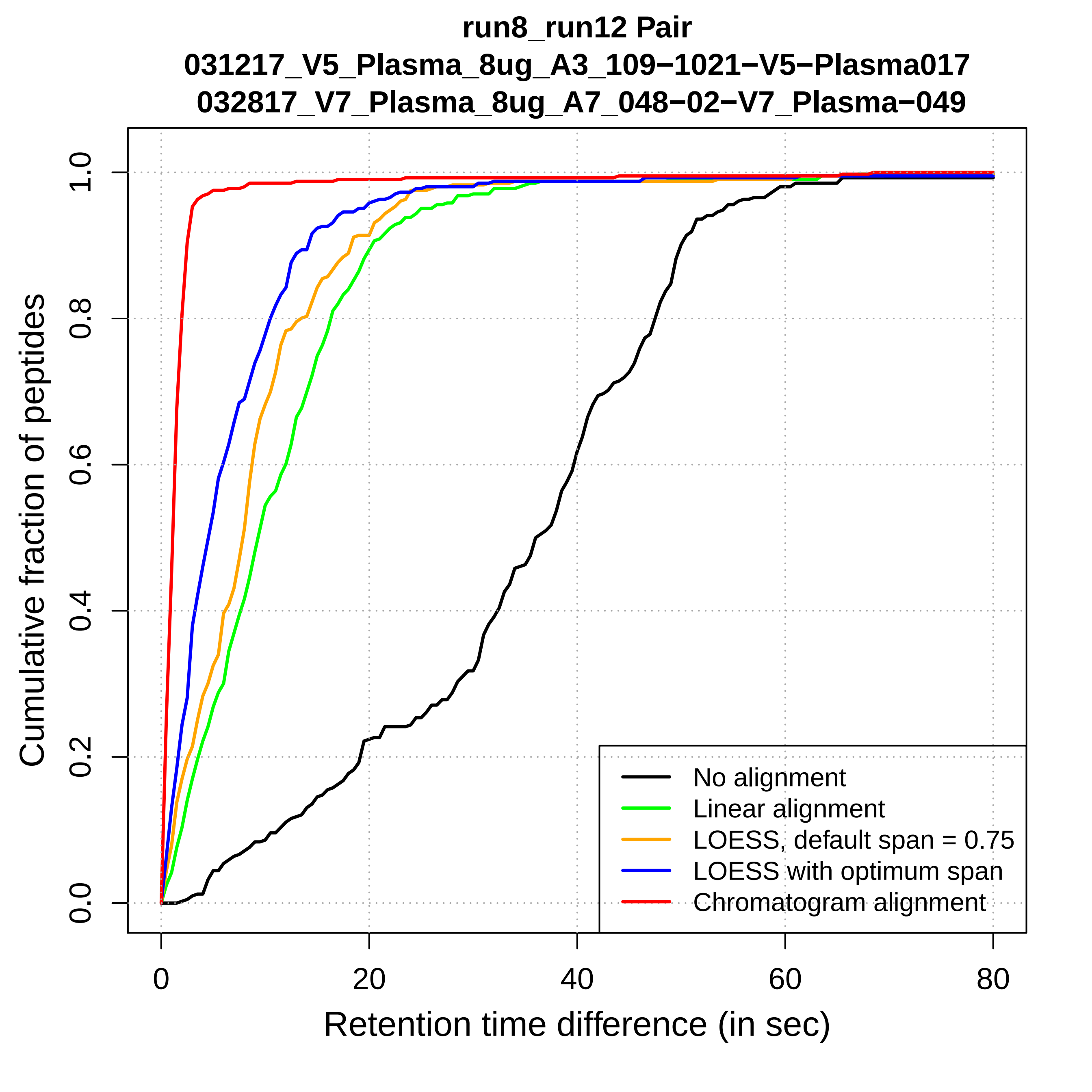
run8_run12 Pair
031217_V5_Plasma_8ug_A3_109−1021−V5−Plasma017
032817_V7_Plasma_8ug_A7_048−02−V7_Plasma−049
Cumulative fraction of peptides
0.0
0.2
0.4
0.6
0.8
1.0
0
20
40
60
80
Retention time difference (in sec)
No alignment
Linear alignment
LOESS, default span = 0.75
LOESS with optimum span
Chromatogram alignment

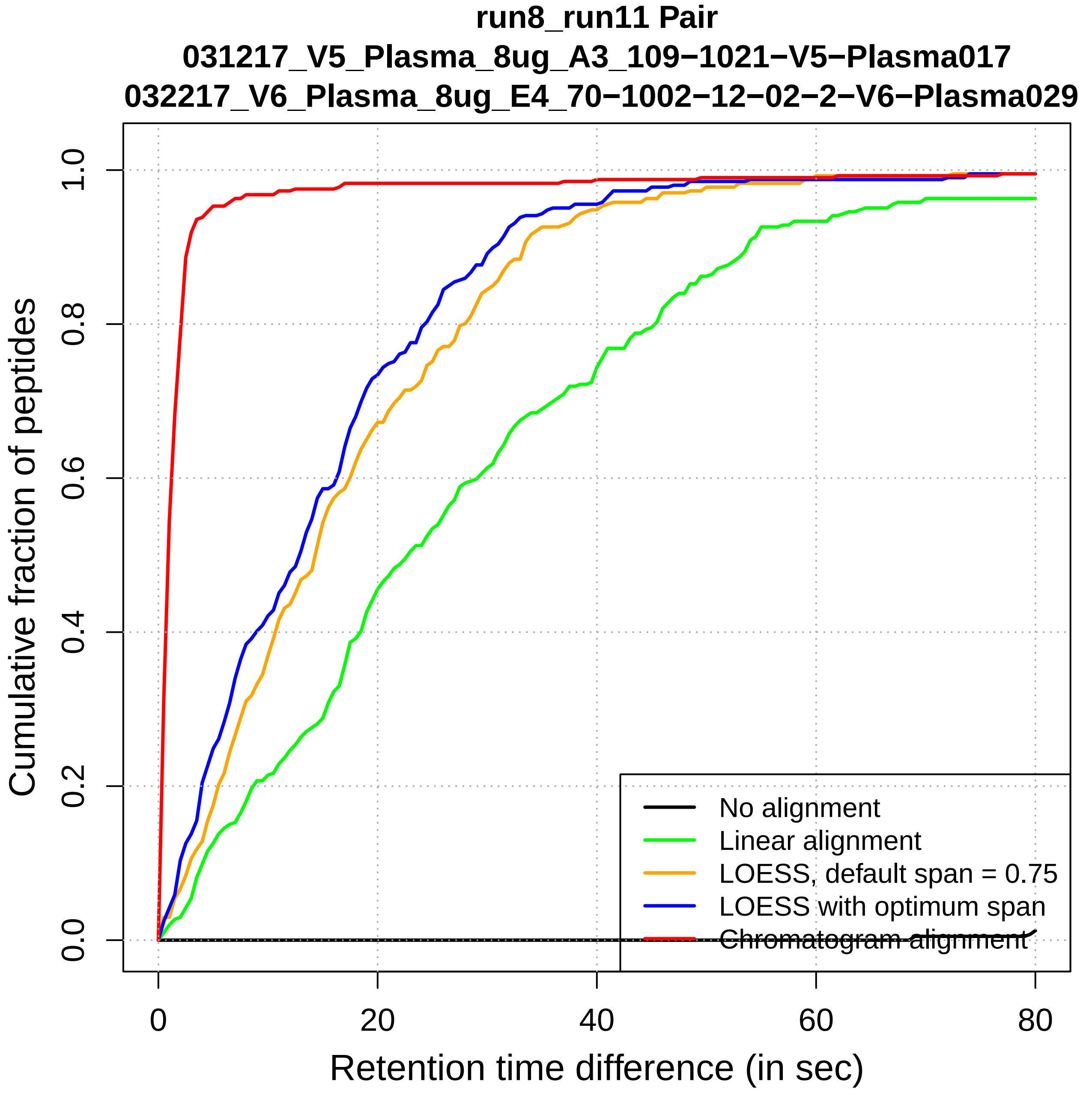

run8_run11 Pair
031217_V5_Plasma_8ug_A3_109−1021−V5−Plasma017
032217_V6_Plasma_8ug_E4_70−1002−12−02−2−V6−Plasma029
Cumulative fraction of peptides
Retention time difference (in sec)
0.0
0.2
0.4
0.6
0.8
1.0
0
20
40
60
80
No alignment
Linear alignment
LOESS, default span = 0.75
LOESS with optimum span
Chromatogram alignment

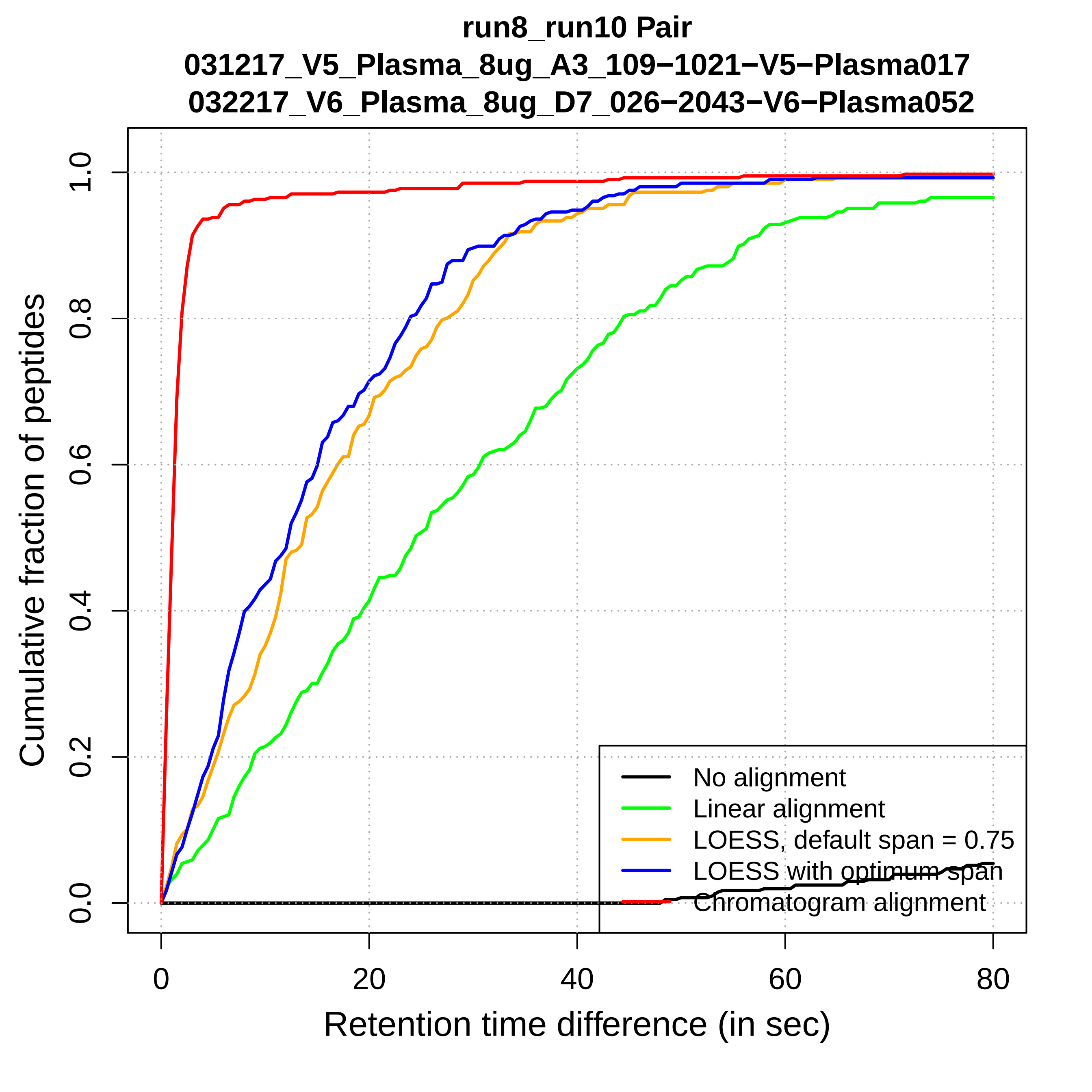

run8_run10 Pair
031217_V5_Plasma_8ug_A3_109−1021−V5−Plasma017
032217_V6_Plasma_8ug_D7_026−2043−V6−Plasma052
Cumulative fraction of peptides
0.0
0.2
0.4
0.6
0.8
1.0
0
20
40
60
80
Retention time difference (in sec)
No alignment
Linear alignment
LOESS, default span = 0.75
LOESS with optimum span
Chromatogram alignment

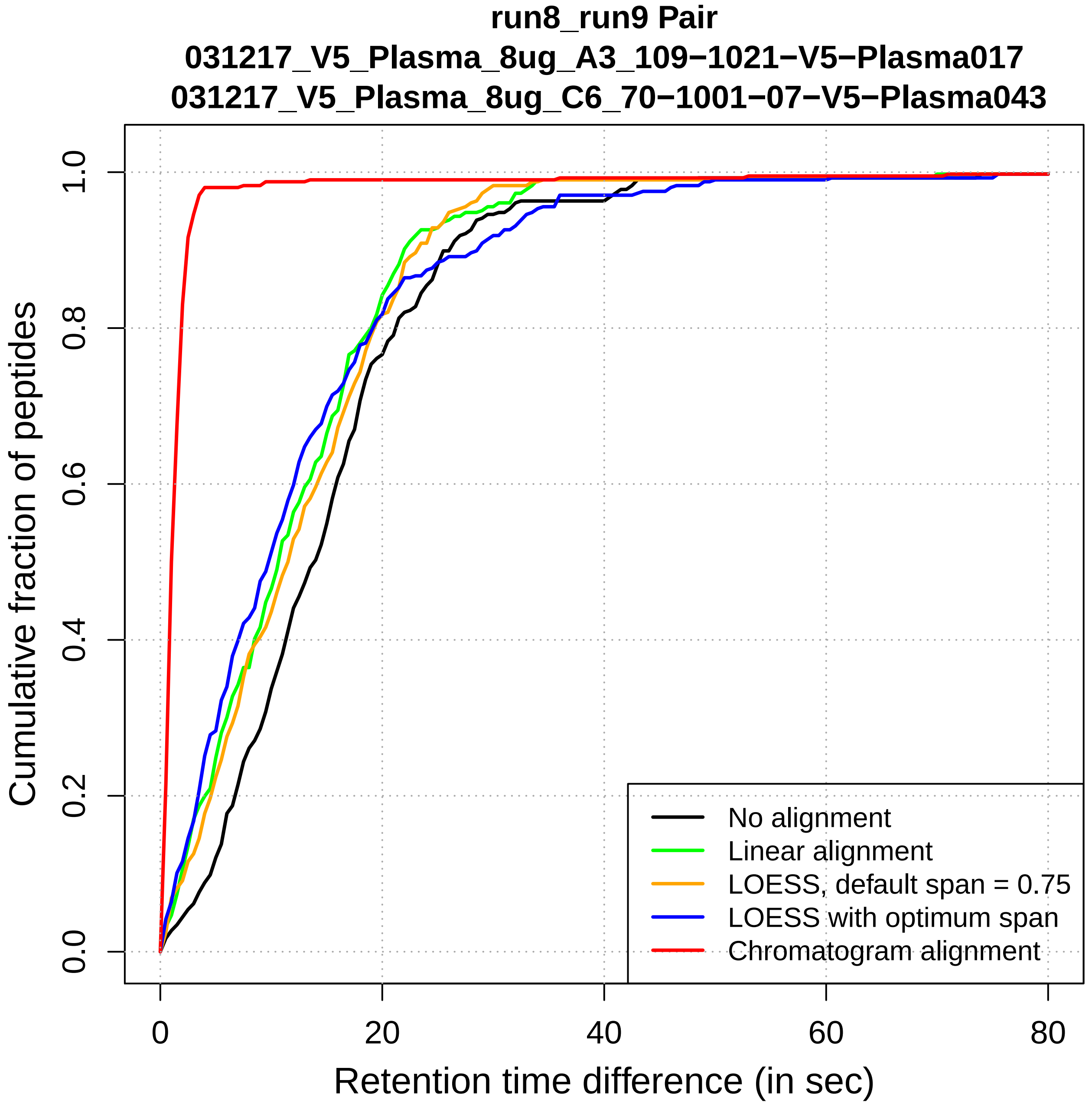
run8_run9 Pair
031217_V5_Plasma_8ug_A3_109−1021−V5−Plasma017
031217_V5_Plasma_8ug_C6_70−1001−07−V5−Plasma043
Cumulative fraction of peptides
0.0
0.2
0.4
0.6
0.8
1.0
0
20
40
60
80
Retention time difference (in sec)
No alignment
Linear alignment
LOESS, default span = 0.75
LOESS with optimum span
Chromatogram alignment

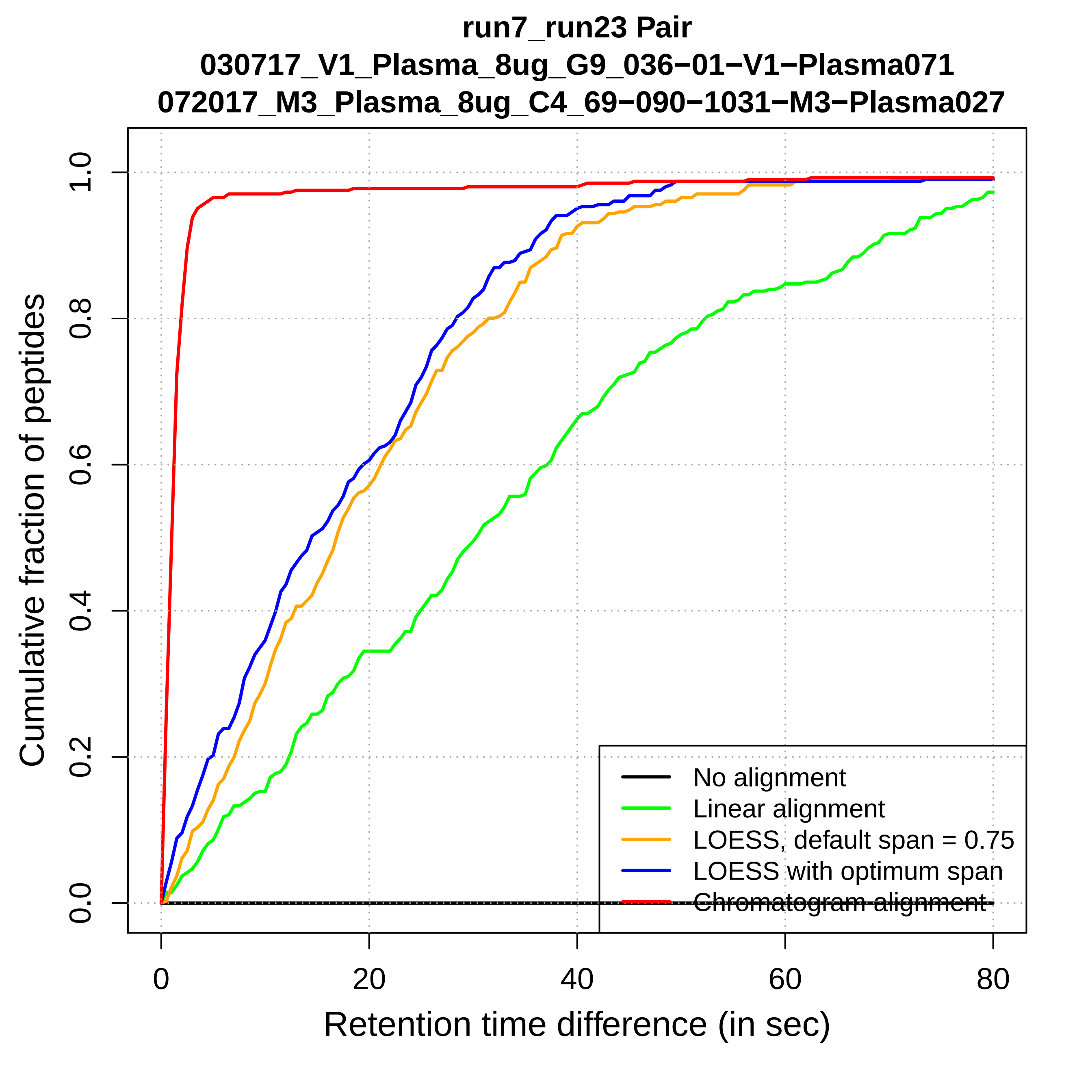
run7_run23 Pair
030717_V1_Plasma_8ug_G9_036−01−V1−Plasma071
072017_M3_Plasma_8ug_C4_69−090−1031−M3−Plasma027
Cumulative fraction of peptides
0.0
0.2
0.4
0.6
0.8
1.0
0
20
40
60
80
Retention time difference (in sec)
No alignment
Linear alignment
LOESS, default span = 0.75
LOESS with optimum span
Chromatogram alignment

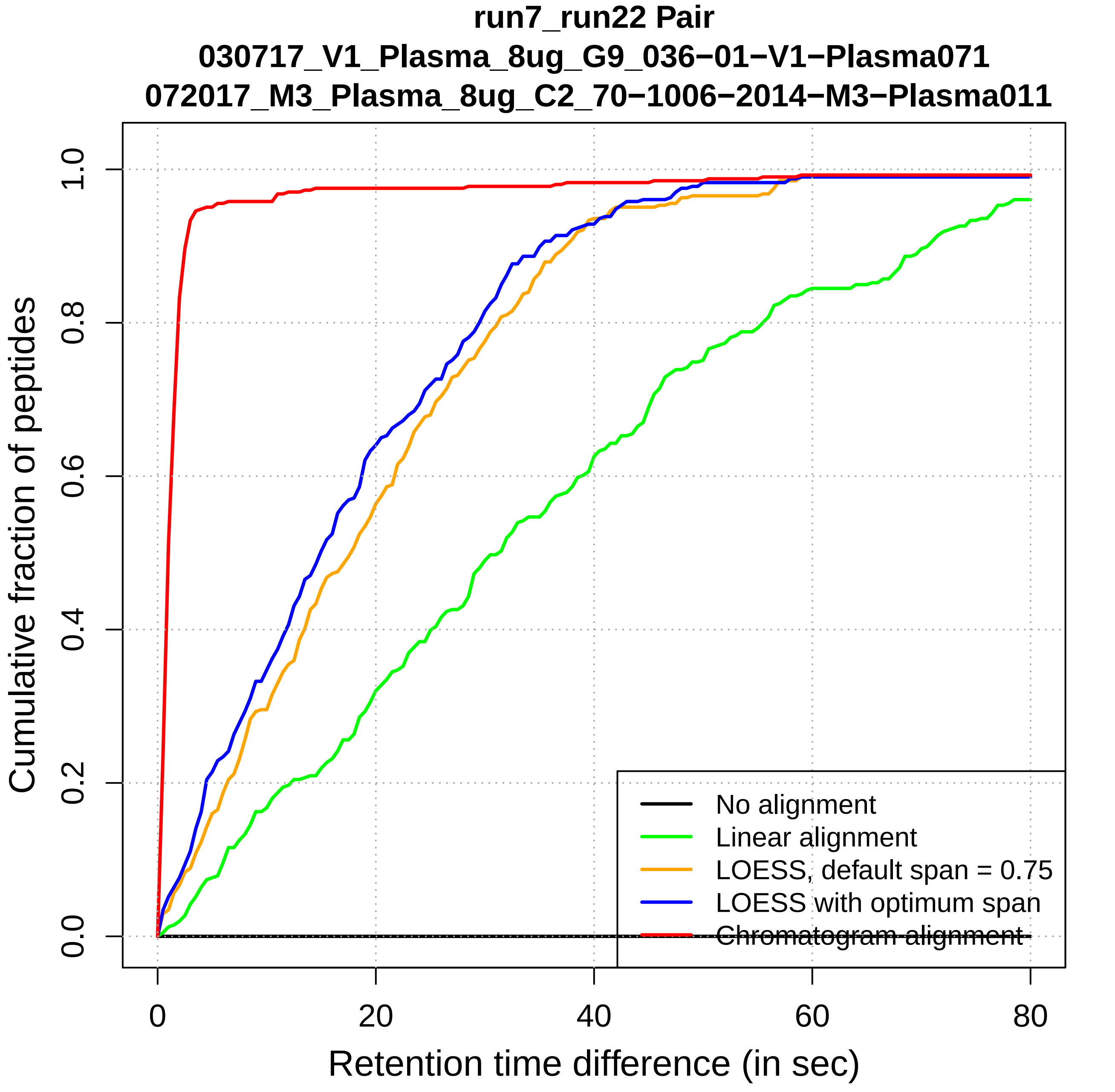

run7_run22 Pair
030717_V1_Plasma_8ug_G9_036−01−V1−Plasma071
072017_M3_Plasma_8ug_C2_70−1006−2014−M3−Plasma011
Cumulative fraction of peptides
0.0
0.2
0.4
0.6
0.8
1.0
0
20
40
60
80
Retention time difference (in sec)
No alignment
Linear alignment
LOESS, default span = 0.75
LOESS with optimum span
Chromatogram alignment

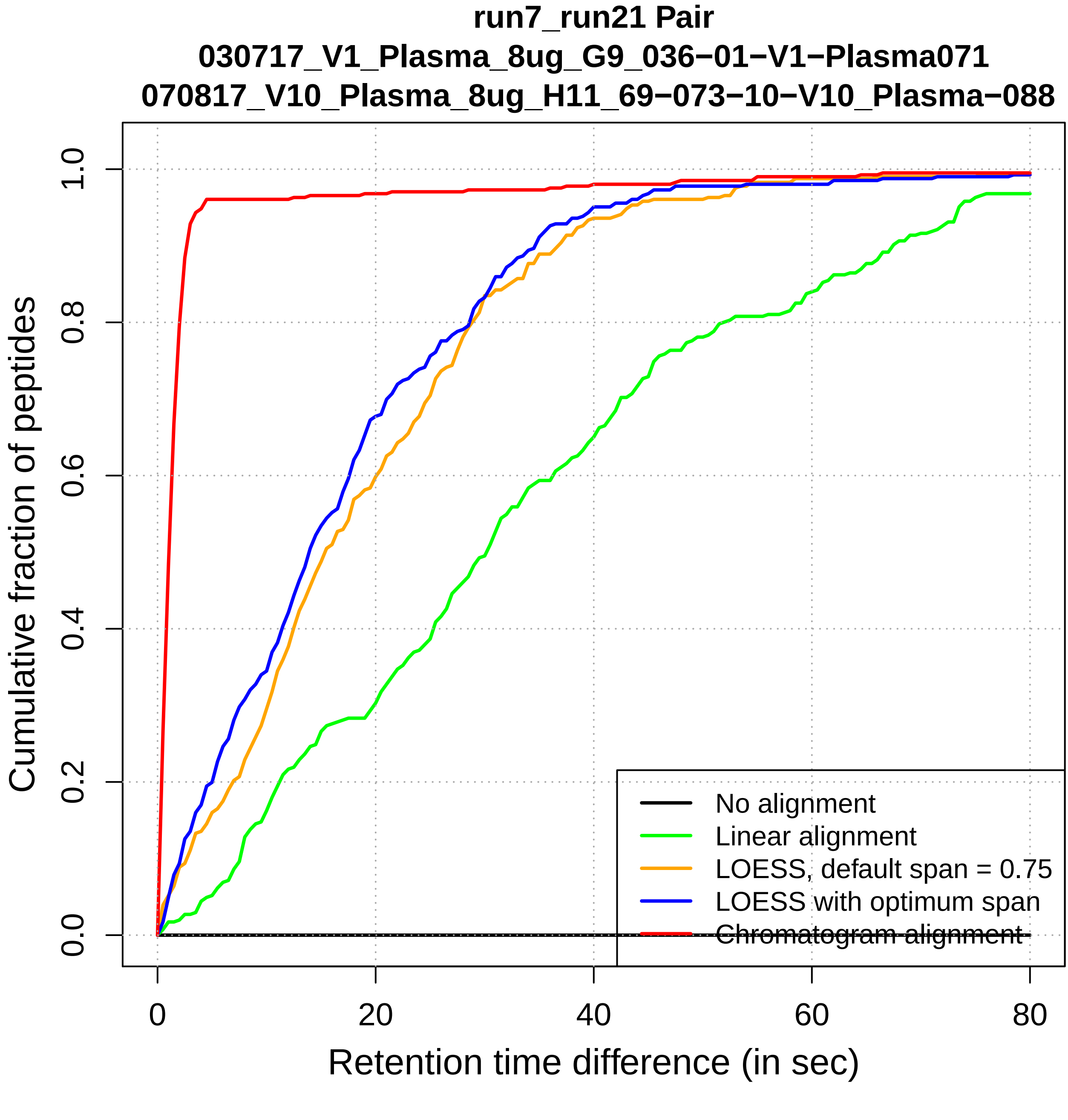

run7_run21 Pair
030717_V1_Plasma_8ug_G9_036−01−V1−Plasma071
070817_V10_Plasma_8ug_H11_69−073−10−V10_Plasma−088
Cumulative fraction of peptides
0.0
0.2
0.4
0.6
0.8
1.0
0
20
40
60
80
Retention time difference (in sec)
No alignment
Linear alignment
LOESS, default span = 0.75
LOESS with optimum span
Chromatogram alignment

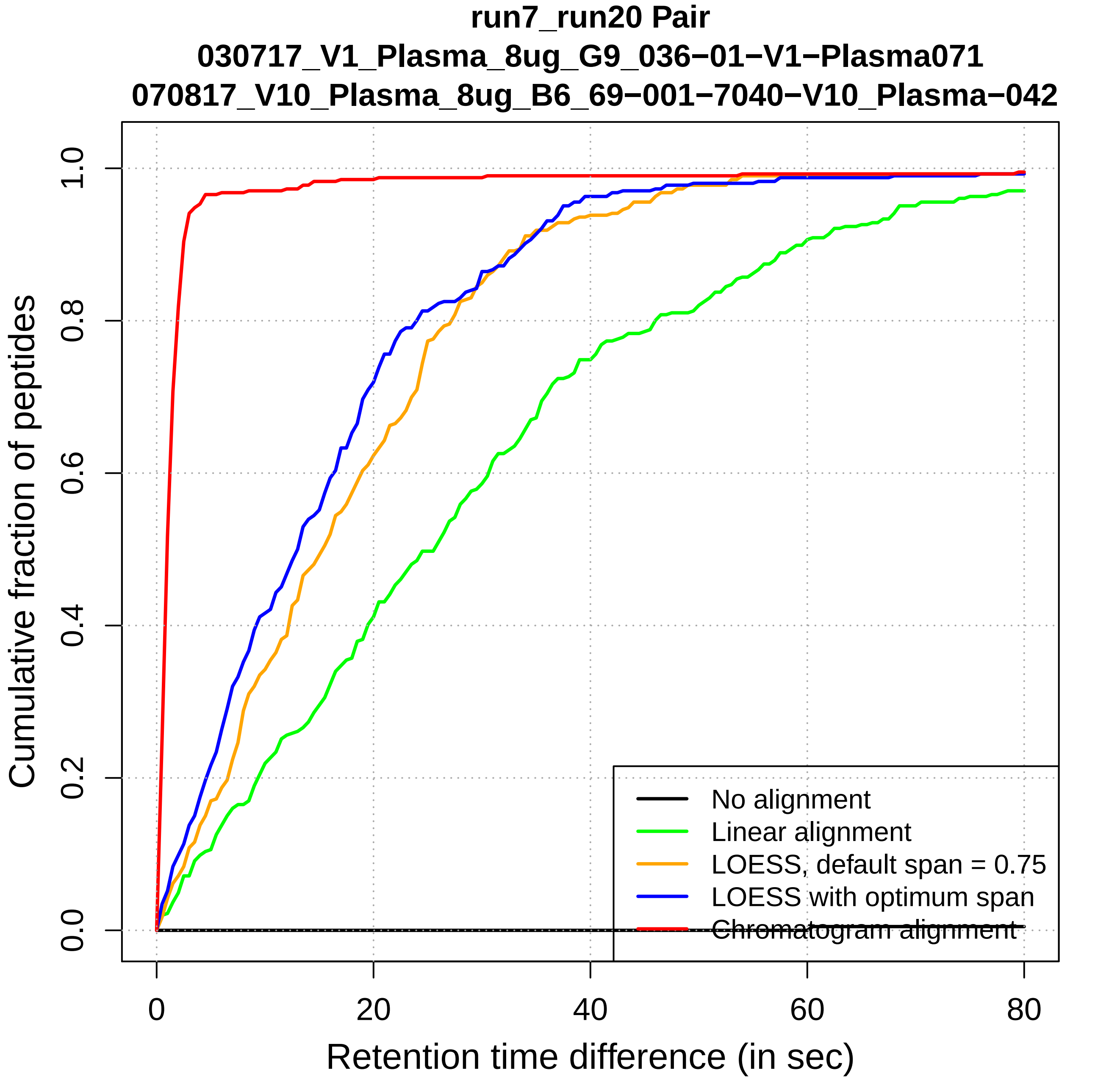
run7_run20 Pair
030717_V1_Plasma_8ug_G9_036−01−V1−Plasma071
070817_V10_Plasma_8ug_B6_69−001−7040−V10_Plasma−042
Cumulative fraction of peptides
Retention time difference (in sec)
0.0
0.2
0.4
0.6
0.8
1.0
0
20
40
60
80
No alignment
Linear alignment
LOESS, default span = 0.75
LOESS with optimum span
Chromatogram alignment

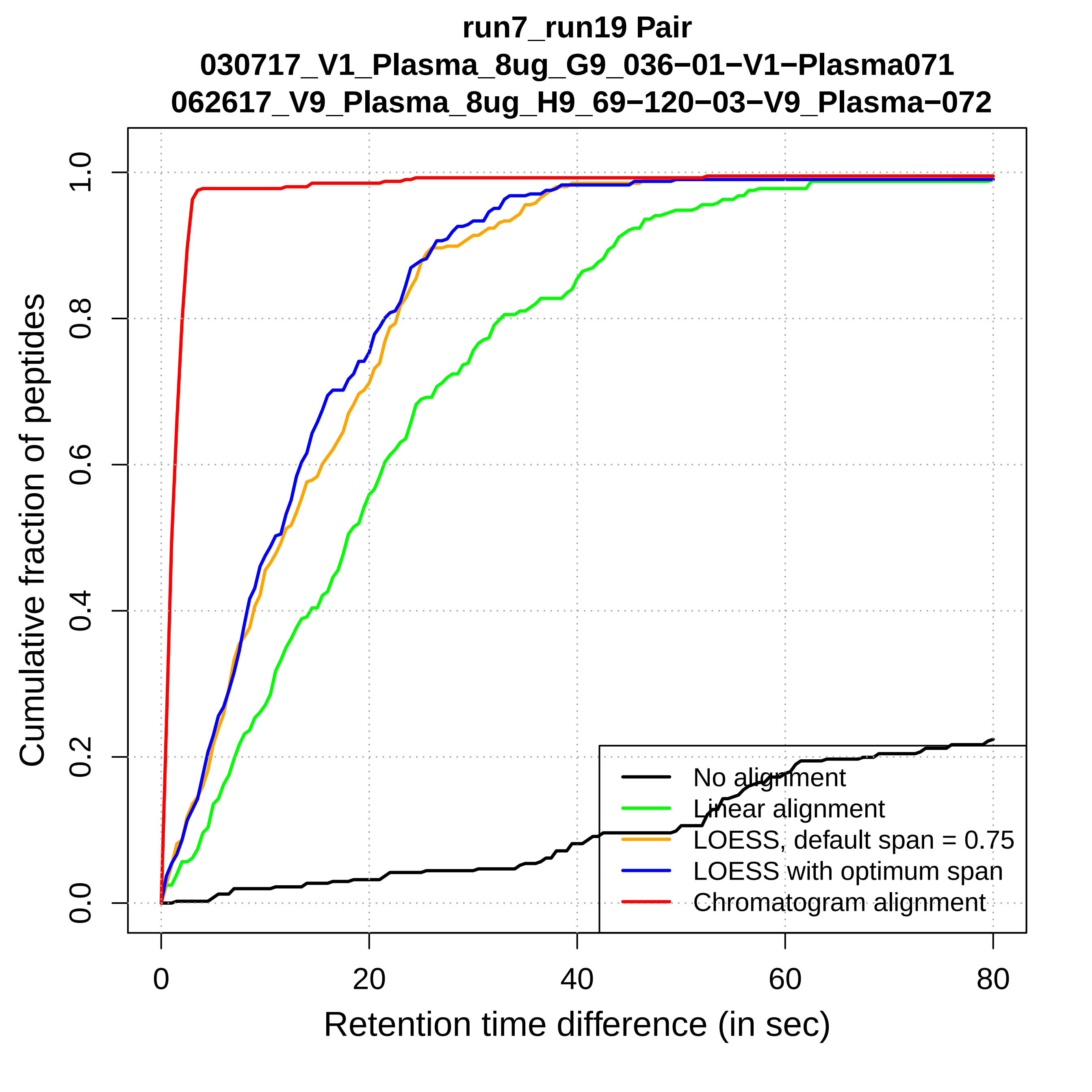
run7_run19 Pair
030717_V1_Plasma_8ug_G9_036−01−V1−Plasma071
062617_V9_Plasma_8ug_H9_69−120−03−V9_Plasma−072
Cumulative fraction of peptides
0.0
0.2
0.4
0.6
0.8
1.0
0
20
40
60
80
Retention time difference (in sec)
No alignment
Linear alignment
LOESS, default span = 0.75
LOESS with optimum span
Chromatogram alignment

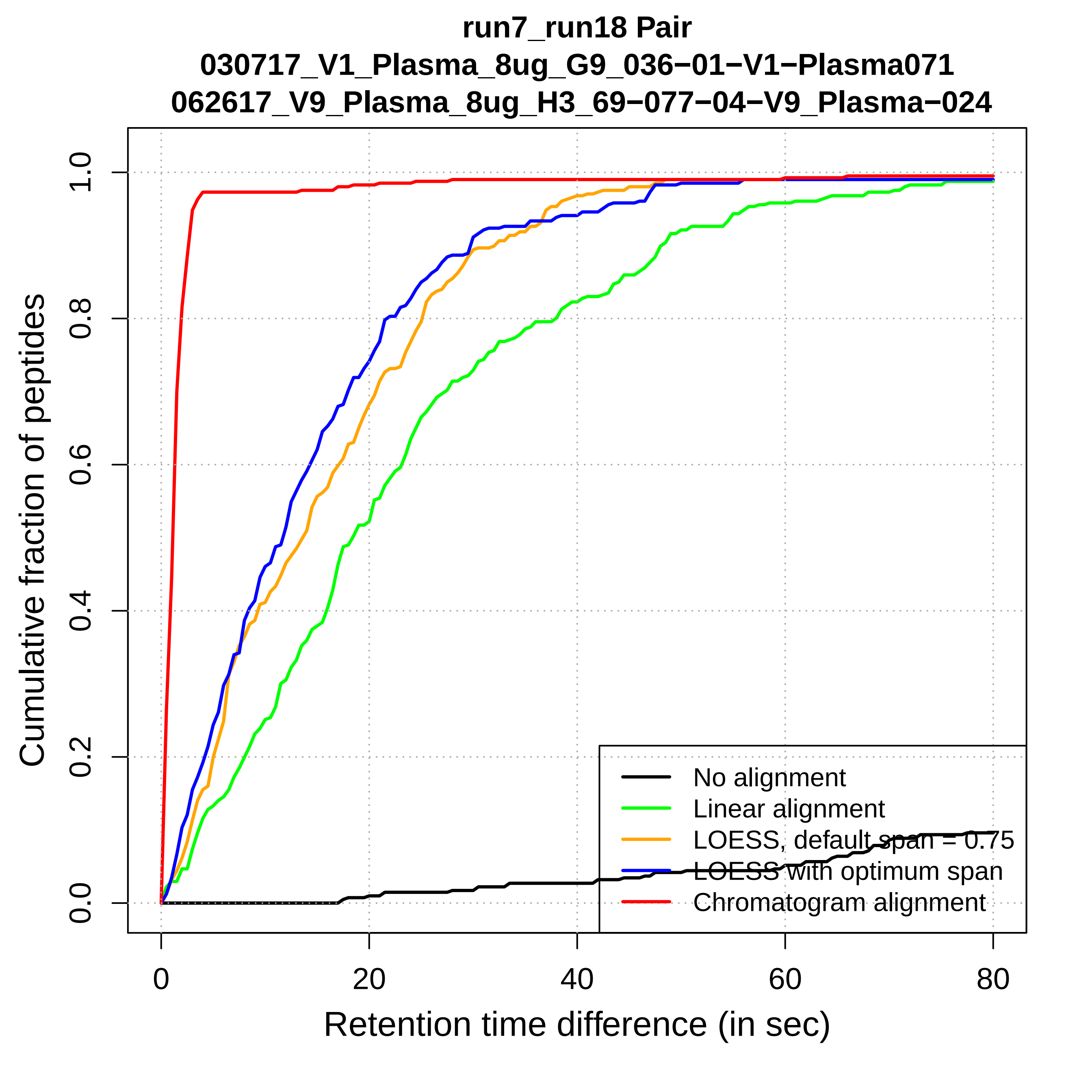

run7_run18 Pair
030717_V1_Plasma_8ug_G9_036−01−V1−Plasma071
062617_V9_Plasma_8ug_H3_69−077−04−V9_Plasma−024
Cumulative fraction of peptides
0.0
0.2
0.4
0.6
0.8
1.0
0
20
40
60
80
Retention time difference (in sec)
No alignment
Linear alignment
LOESS, default span = 0.75
LOESS with optimum span
Chromatogram alignment

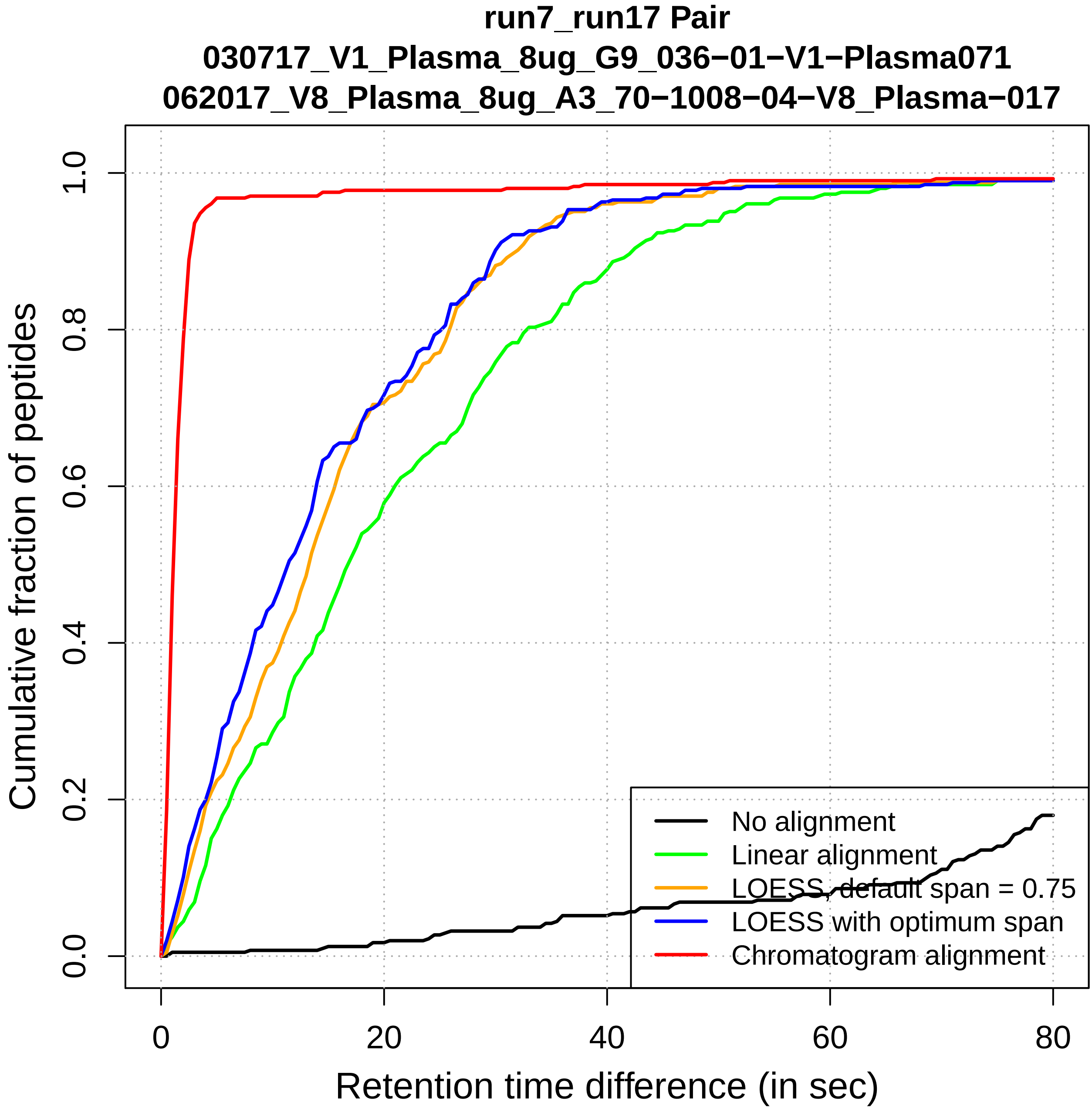
run7_run17 Pair
030717_V1_Plasma_8ug_G9_036−01−V1−Plasma071
062017_V8_Plasma_8ug_A3_70−1008−04−V8_Plasma−017
Cumulative fraction of peptides
0.0
0.2
0.4
0.6
0.8
1.0
0
20
40
60
80
Retention time difference (in sec)
No alignment
Linear alignment
LOESS, default span = 0.75
LOESS with optimum span
Chromatogram alignment

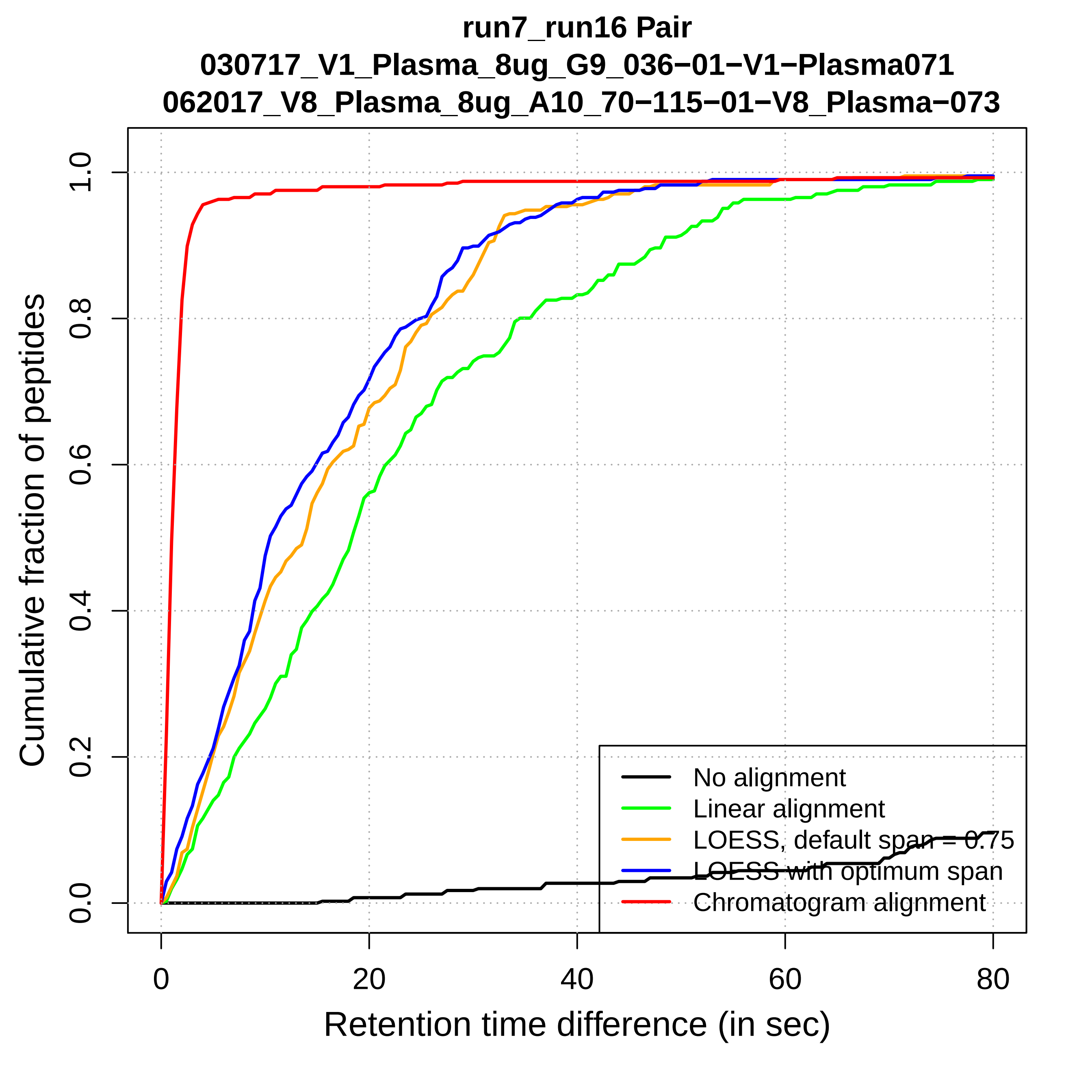
run7_run16 Pair
030717_V1_Plasma_8ug_G9_036−01−V1−Plasma071
062017_V8_Plasma_8ug_A10_70−115−01−V8_Plasma−073
Cumulative fraction of peptides
Retention time difference (in sec)
0.0
0.2
0.4
0.6
0.8
1.0
0
20
40
60
80
No alignment
Linear alignment
LOESS, default span = 0.75
LOESS with optimum span
Chromatogram alignment

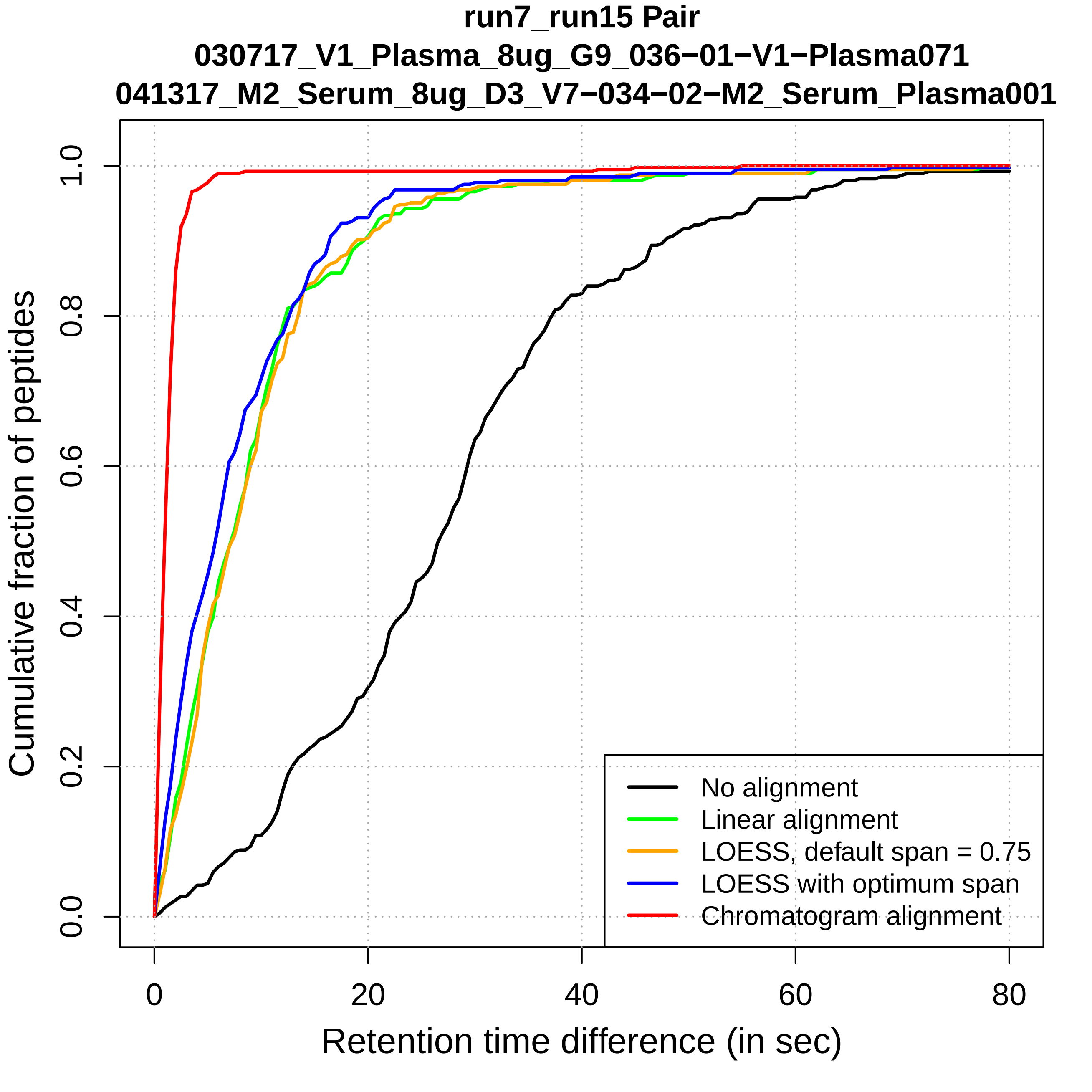

run7_run15 Pair
030717_V1_Plasma_8ug_G9_036−01−V1−Plasma071
041317_M2_Serum_8ug_D3_V7−034−02−M2_Serum_Plasma001
Cumulative fraction of peptides
0.0
0.2
0.4
0.6
0.8
1.0
0
20
40
60
80
Retention time difference (in sec)
No alignment
Linear alignment
LOESS, default span = 0.75
LOESS with optimum span
Chromatogram alignment

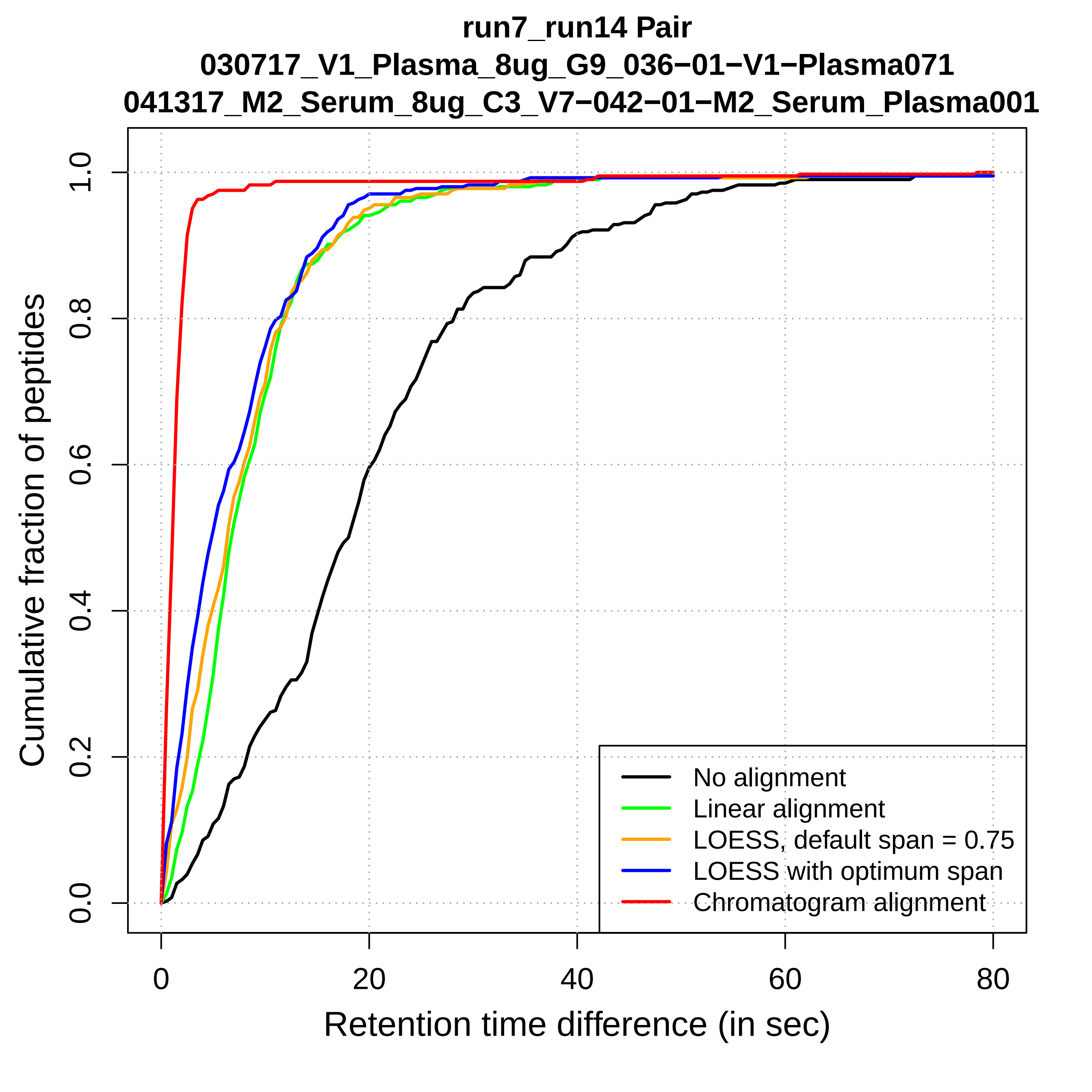
run7_run14 Pair
030717_V1_Plasma_8ug_G9_036−01−V1−Plasma071
041317_M2_Serum_8ug_C3_V7−042−01−M2_Serum_Plasma001
Cumulative fraction of peptides
0.0
0.2
0.4
0.6
0.8
1.0
0
20
40
60
80
Retention time difference (in sec)
No alignment
Linear alignment
LOESS, default span = 0.75
LOESS with optimum span
Chromatogram alignment

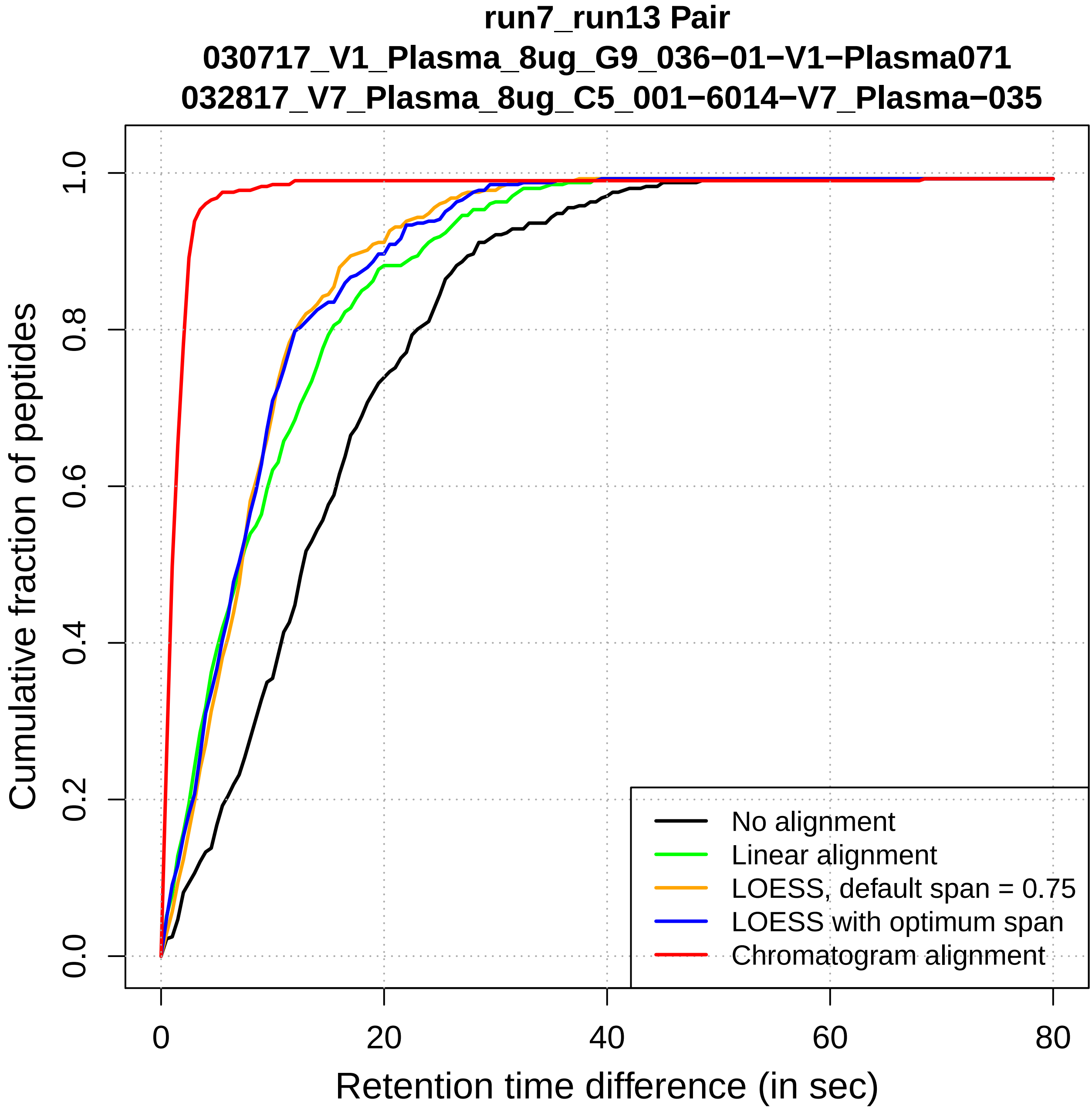
run7_run13 Pair
030717_V1_Plasma_8ug_G9_036−01−V1−Plasma071
032817_V7_Plasma_8ug_C5_001−6014−V7_Plasma−035
Cumulative fraction of peptides
0.0
0.2
0.4
0.6
0.8
1.0
0
20
40
60
80
Retention time difference (in sec)
No alignment
Linear alignment
LOESS, default span = 0.75
LOESS with optimum span
Chromatogram alignment

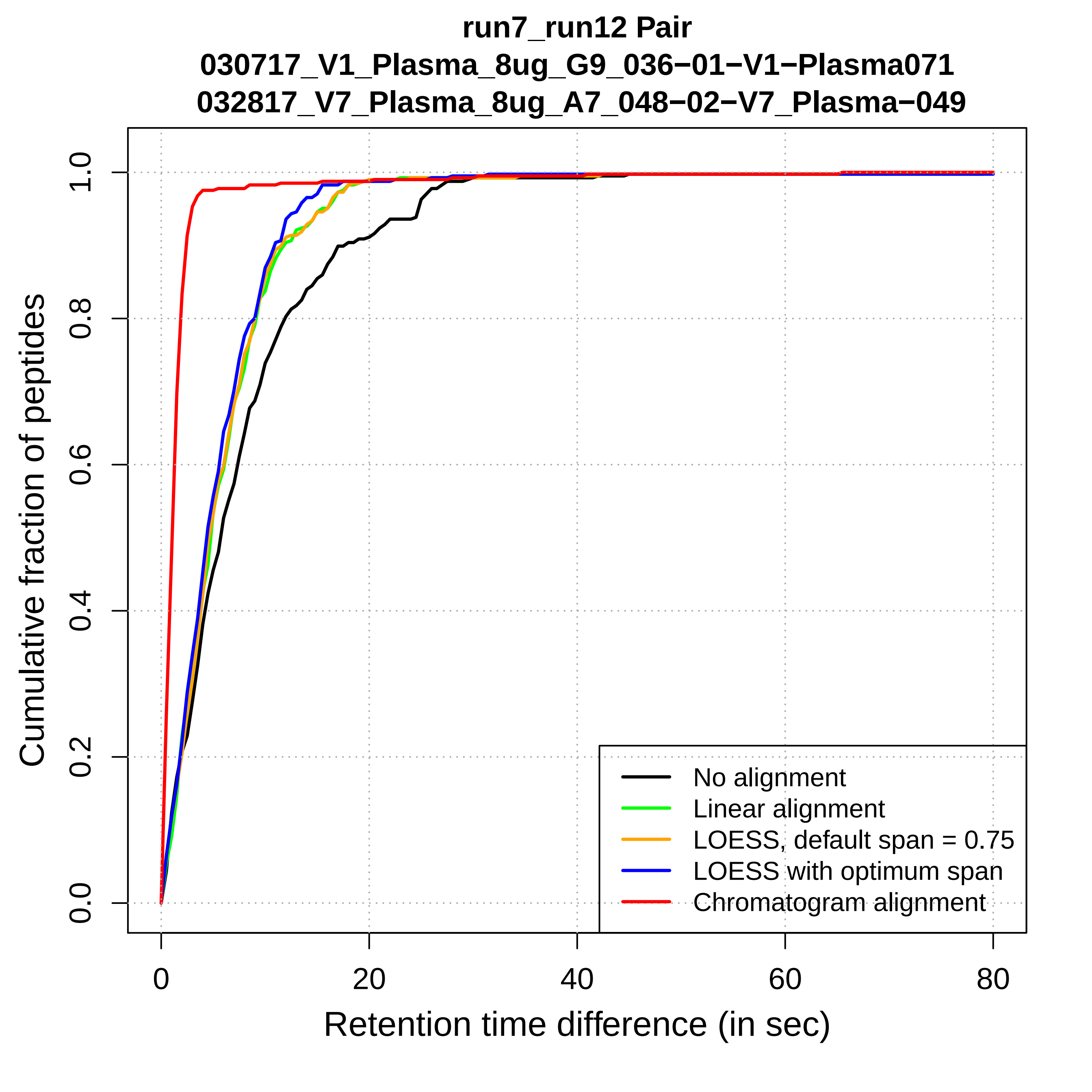
run7_run12 Pair
030717_V1_Plasma_8ug_G9_036−01−V1−Plasma071
032817_V7_Plasma_8ug_A7_048−02−V7_Plasma−049
Cumulative fraction of peptides
0.0
0.2
0.4
0.6
0.8
1.0
0
20
40
60
80
Retention time difference (in sec)
No alignment
Linear alignment
LOESS, default span = 0.75
LOESS with optimum span
Chromatogram alignment

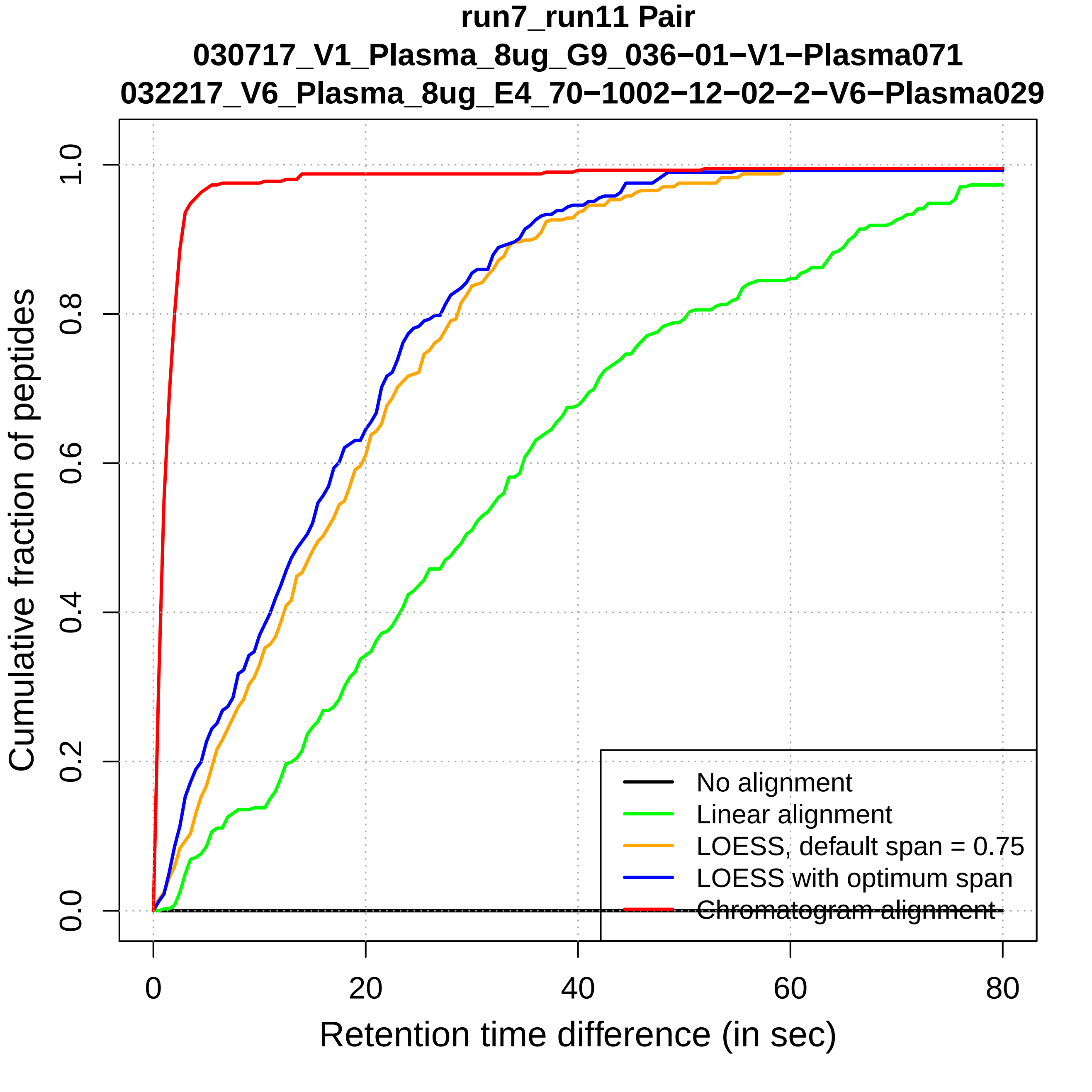
run7_run11 Pair
030717_V1_Plasma_8ug_G9_036−01−V1−Plasma071
032217_V6_Plasma_8ug_E4_70−1002−12−02−2−V6−Plasma029
Cumulative fraction of peptides
0.0
0.2
0.4
0.6
0.8
1.0
0
20
40
60
80
Retention time difference (in sec)
No alignment
Linear alignment
LOESS, default span = 0.75
LOESS with optimum span
Chromatogram alignment

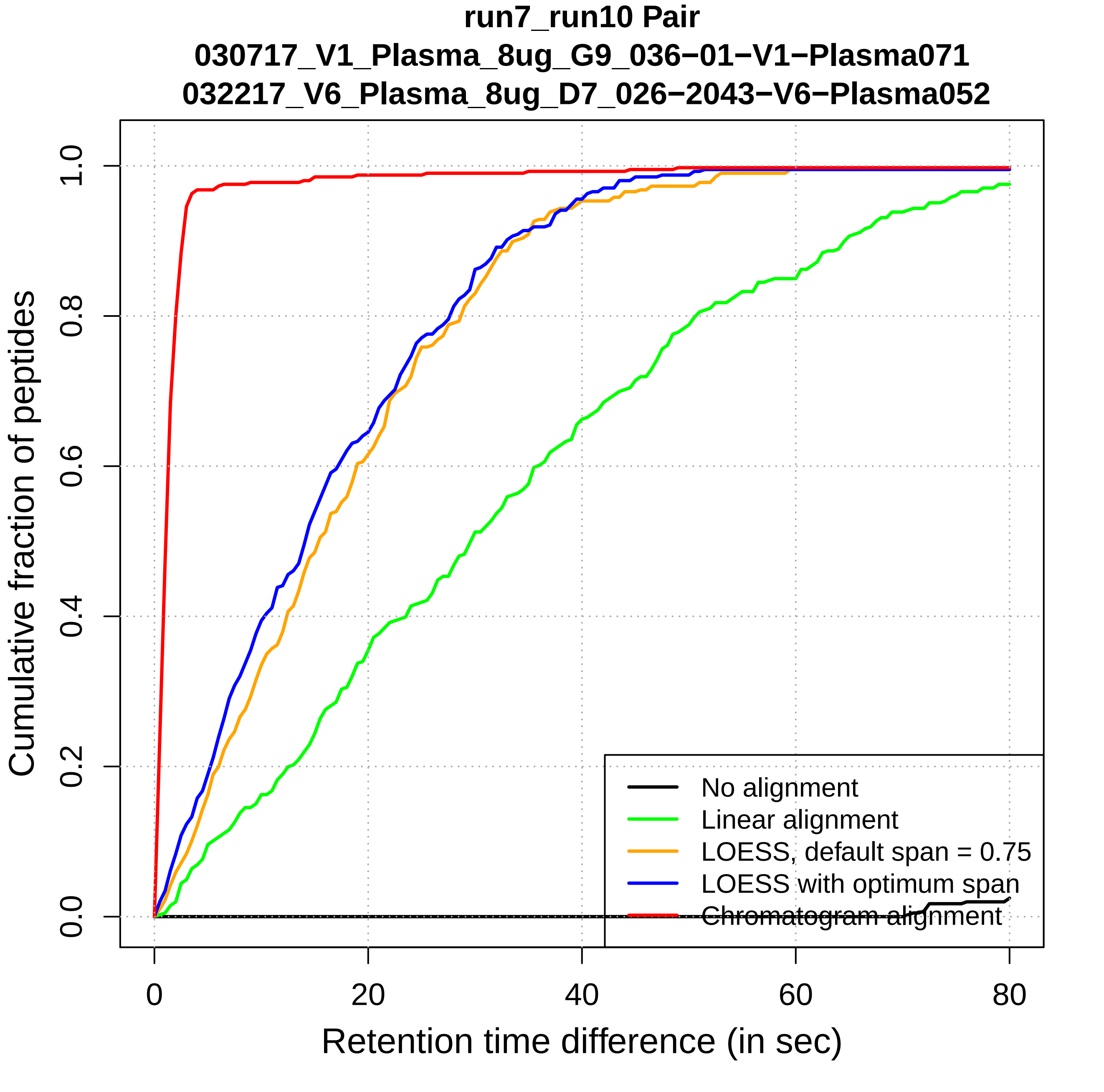
run7_run10 Pair
030717_V1_Plasma_8ug_G9_036−01−V1−Plasma071
032217_V6_Plasma_8ug_D7_026−2043−V6−Plasma052
Cumulative fraction of peptides
0.0
0.2
0.4
0.6
0.8
1.0
0
20
40
60
80
Retention time difference (in sec)
No alignment
Linear alignment
LOESS, default span = 0.75
LOESS with optimum span
Chromatogram alignment

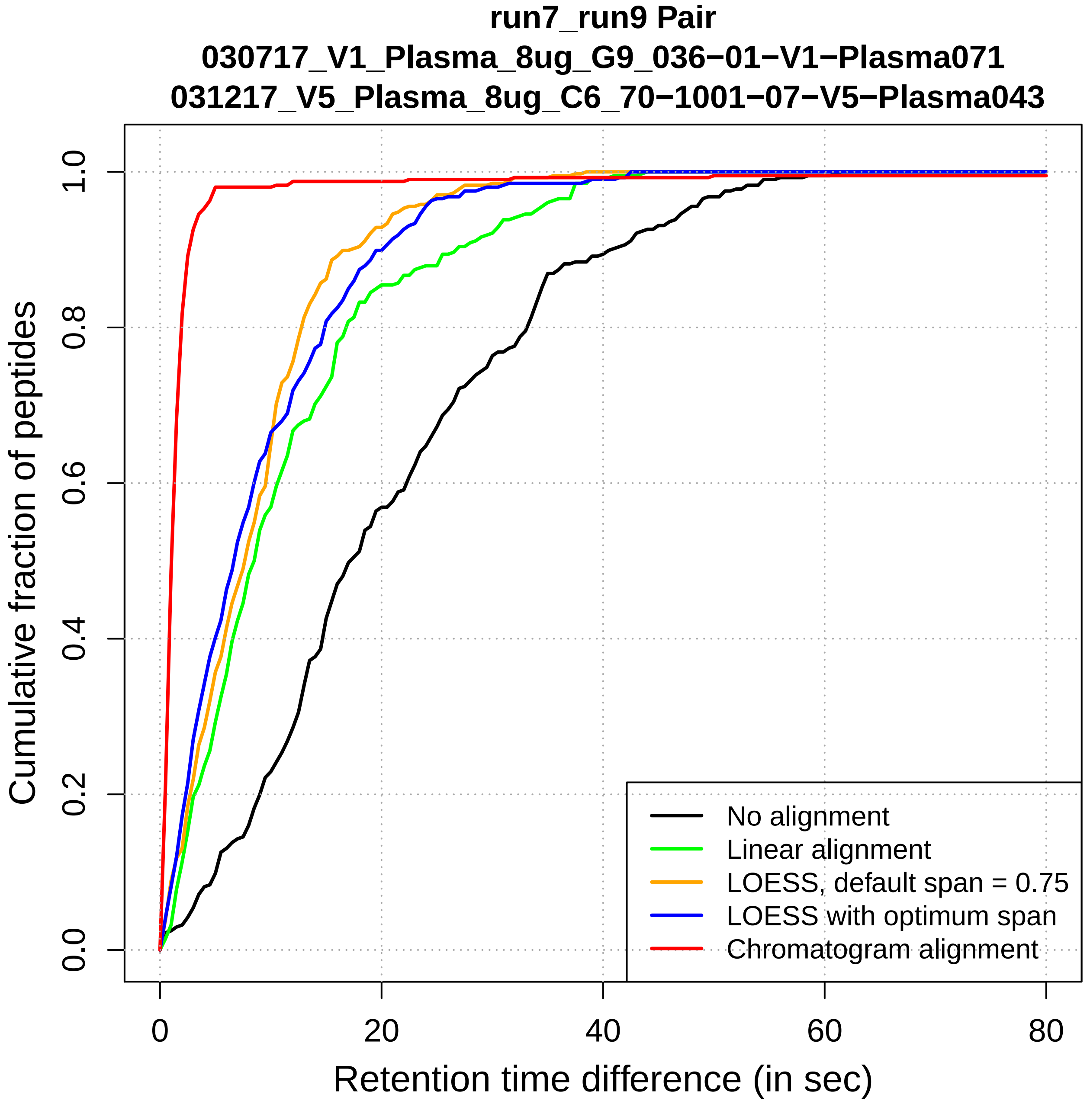
run7_run9 Pair
030717_V1_Plasma_8ug_G9_036−01−V1−Plasma071
031217_V5_Plasma_8ug_C6_70−1001−07−V5−Plasma043
Cumulative fraction of peptides
0.0
0.2
0.4
0.6
0.8
1.0
0
20
40
60
80
Retention time difference (in sec)
No alignment
Linear alignment
LOESS, default span = 0.75
LOESS with optimum span
Chromatogram alignment

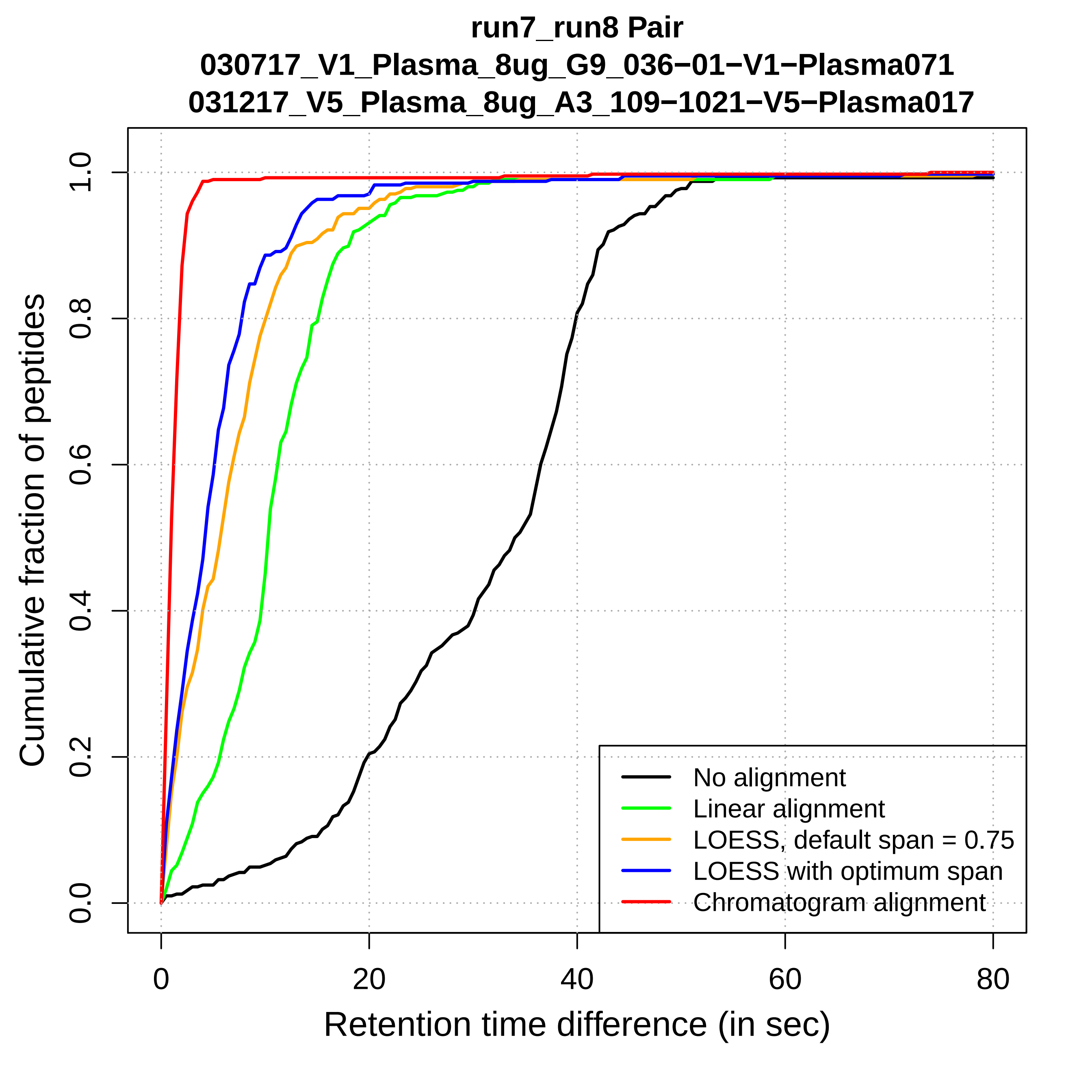
run7_run8 Pair
030717_V1_Plasma_8ug_G9_036−01−V1−Plasma071
031217_V5_Plasma_8ug_A3_109−1021−V5−Plasma017
Cumulative fraction of peptides
0.0
0.2
0.4
0.6
0.8
1.0
0
20
40
60
80
Retention time difference (in sec)
No alignment
Linear alignment
LOESS, default span = 0.75
LOESS with optimum span
Chromatogram alignment

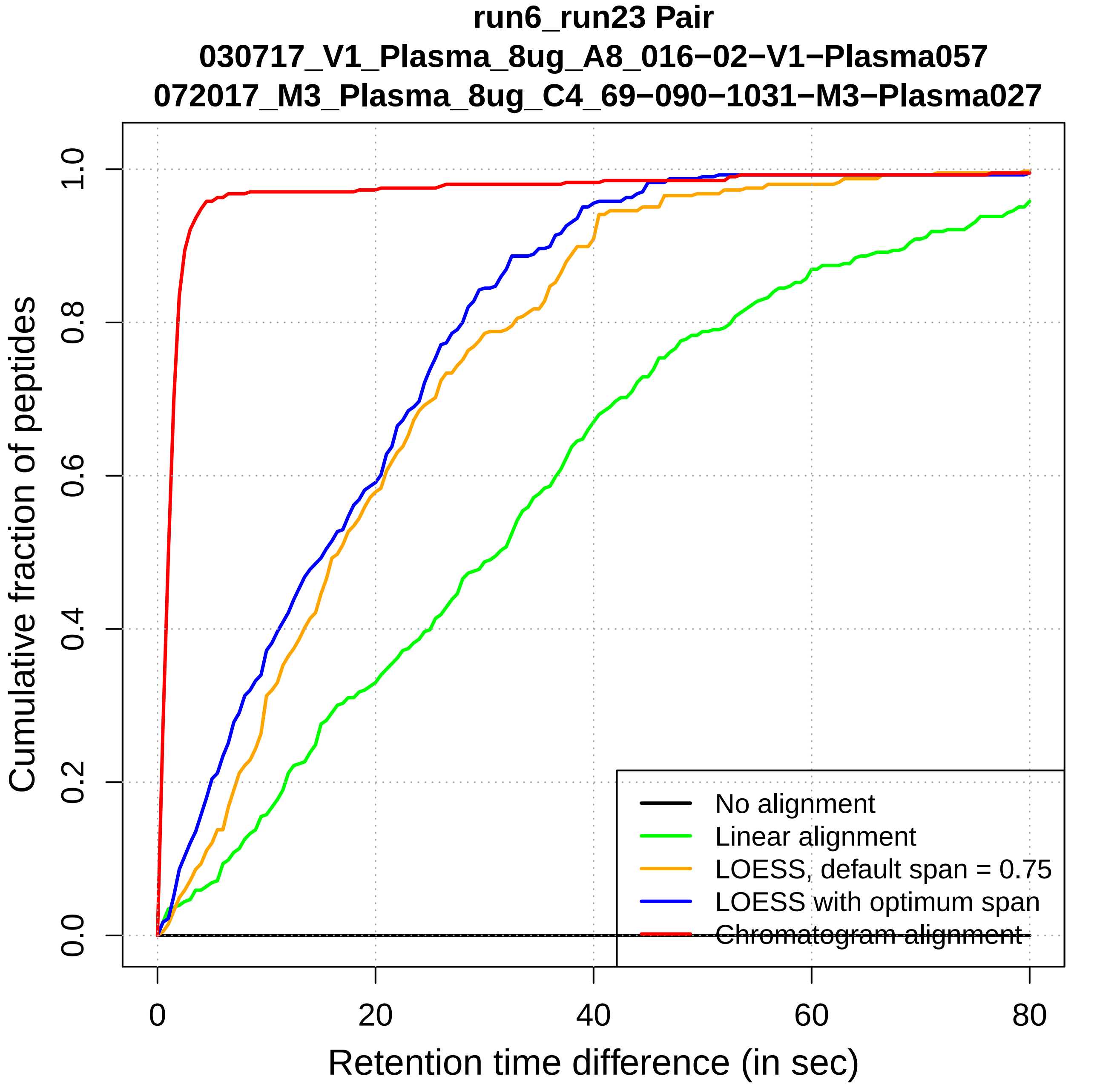

run6_run23 Pair
030717_V1_Plasma_8ug_A8_016−02−V1−Plasma057
072017_M3_Plasma_8ug_C4_69−090−1031−M3−Plasma027
Cumulative fraction of peptides
0.0
0.2
0.4
0.6
0.8
1.0
0
20
40
60
80
Retention time difference (in sec)
No alignment
Linear alignment
LOESS, default span = 0.75
LOESS with optimum span
Chromatogram alignment

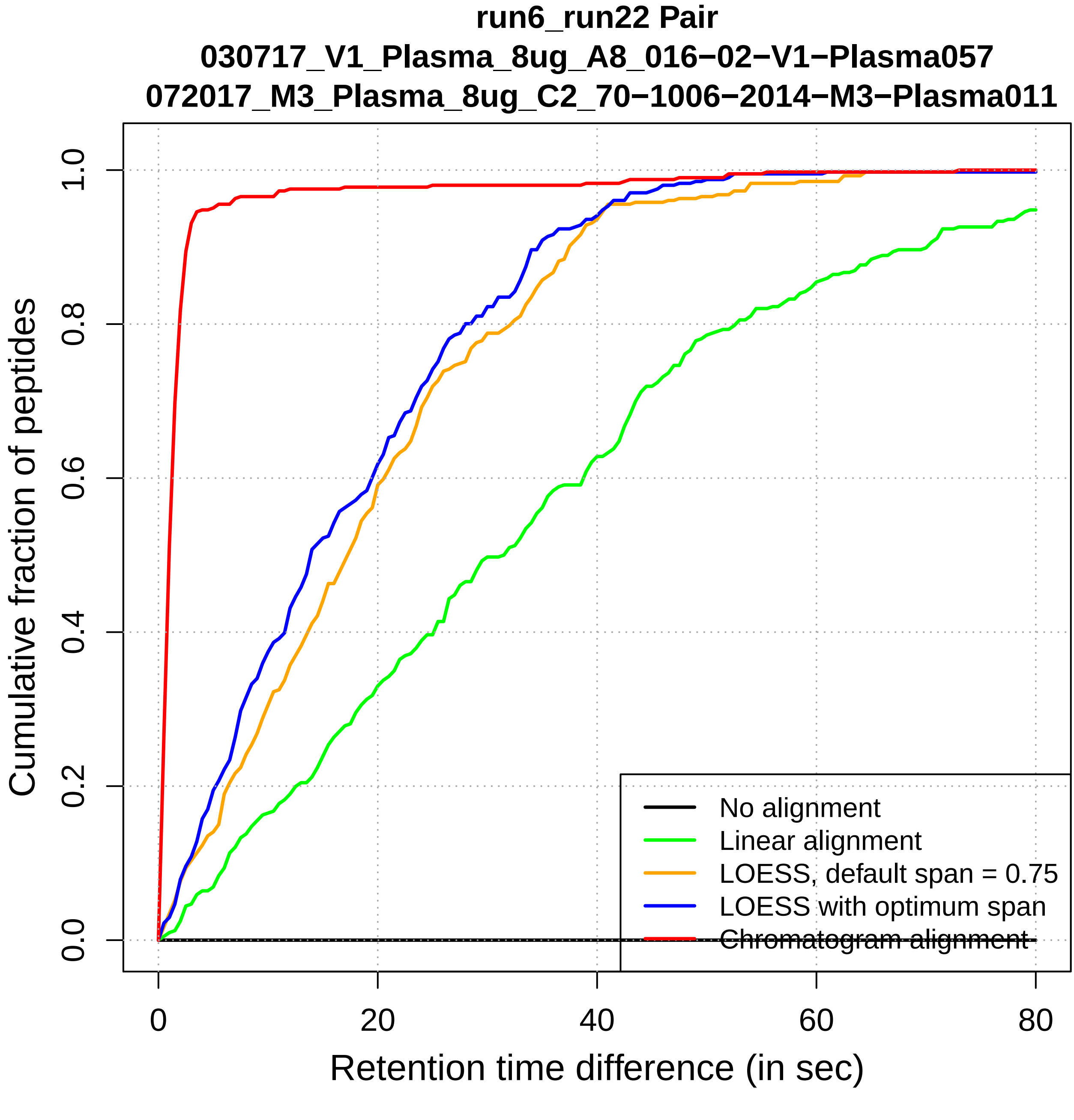

run6_run22 Pair
030717_V1_Plasma_8ug_A8_016−02−V1−Plasma057
072017_M3_Plasma_8ug_C2_70−1006−2014−M3−Plasma011
Cumulative fraction of peptides
0.0
0.2
0.4
0.6
0.8
1.0
0
20
40
60
80
Retention time difference (in sec)
No alignment
Linear alignment
LOESS, default span = 0.75
LOESS with optimum span
Chromatogram alignment

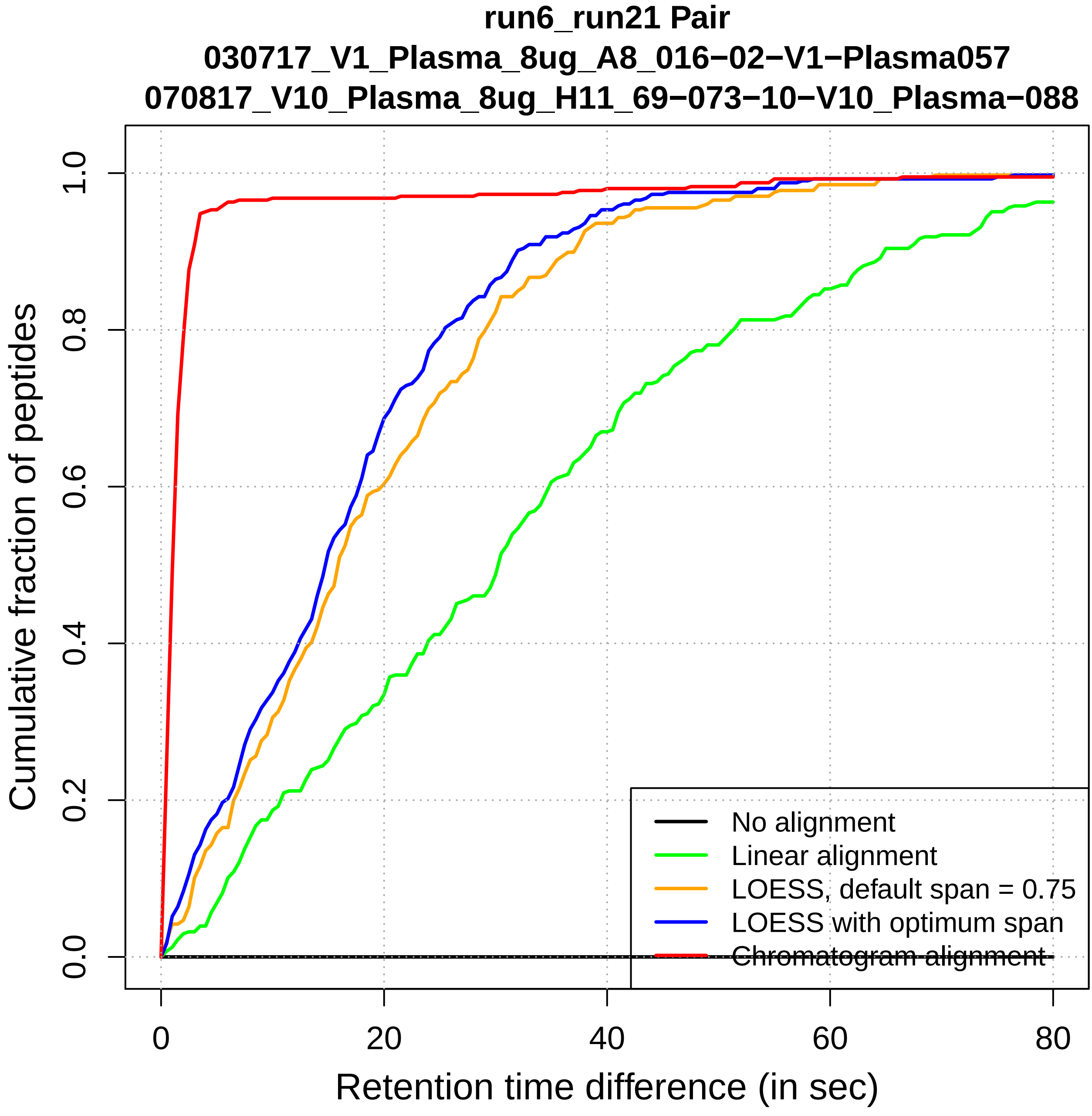

run6_run21 Pair
030717_V1_Plasma_8ug_A8_016−02−V1−Plasma057
070817_V10_Plasma_8ug_H11_69−073−10−V10_Plasma−088
Cumulative fraction of peptides
0.0
0.2
0.4
0.6
0.8
1.0
0
20
40
60
80
Retention time difference (in sec)
No alignment
Linear alignment
LOESS, default span = 0.75
LOESS with optimum span
Chromatogram alignment

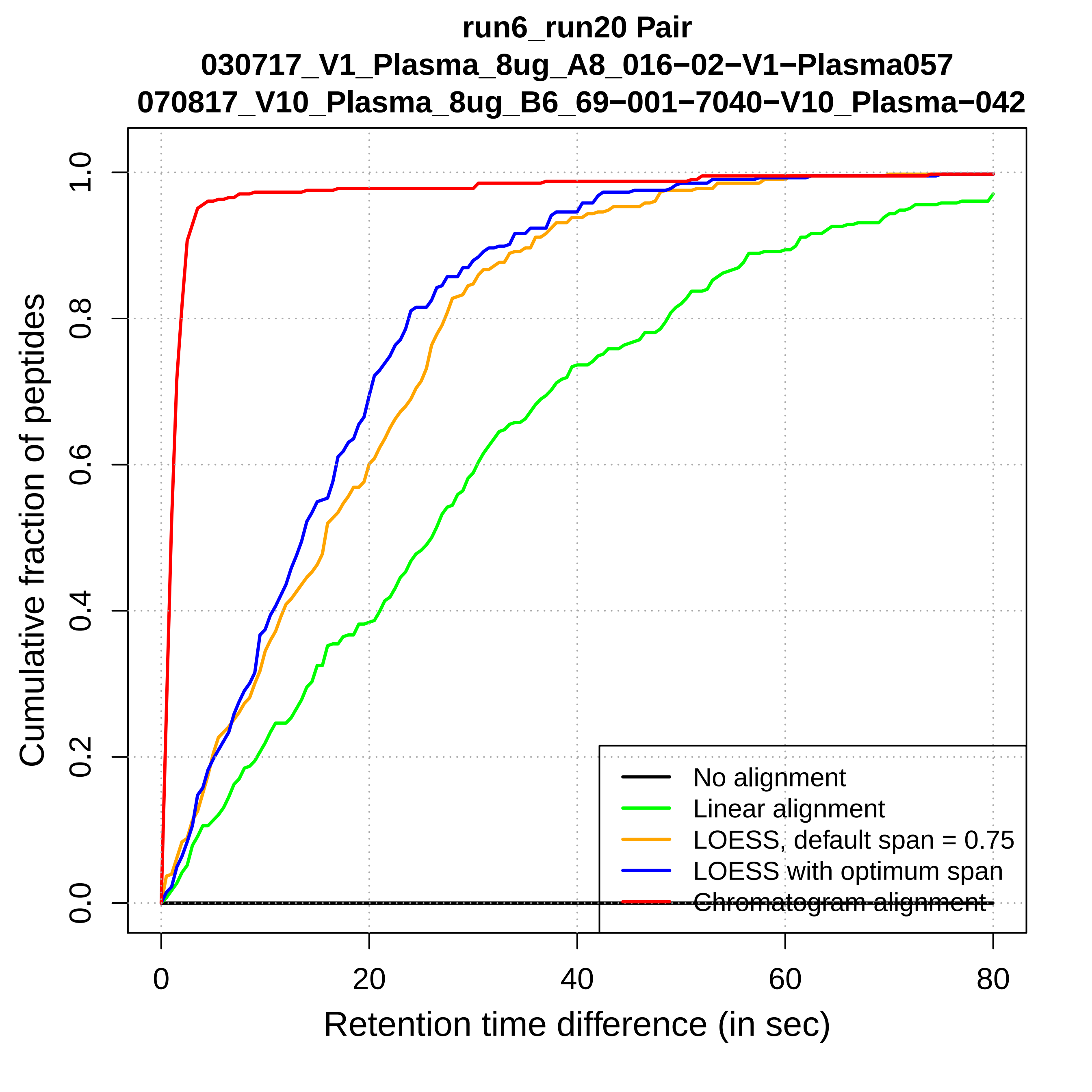
run6_run20 Pair
030717_V1_Plasma_8ug_A8_016−02−V1−Plasma057
070817_V10_Plasma_8ug_B6_69−001−7040−V10_Plasma−042
Cumulative fraction of peptides
0.0
0.2
0.4
0.6
0.8
1.0
0
20
40
60
80
Retention time difference (in sec)
No alignment
Linear alignment
LOESS, default span = 0.75
LOESS with optimum span
Chromatogram alignment

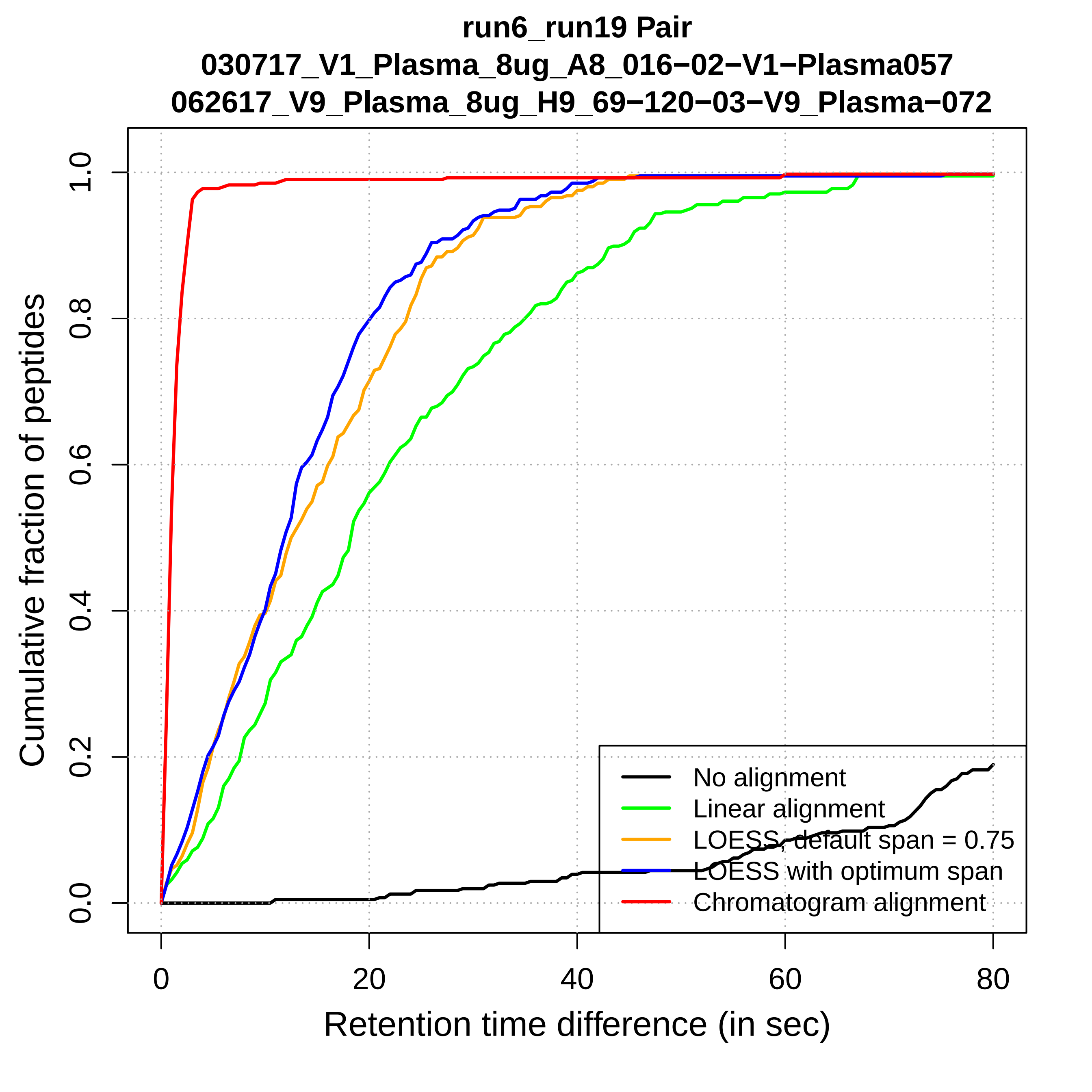

run6_run19 Pair
030717_V1_Plasma_8ug_A8_016−02−V1−Plasma057
062617_V9_Plasma_8ug_H9_69−120−03−V9_Plasma−072
Cumulative fraction of peptides
Retention time difference (in sec)
0.0
0.2
0.4
0.6
0.8
1.0
0
20
40
60
80
No alignment
Linear alignment
LOESS, default span = 0.75
LOESS with optimum span
Chromatogram alignment

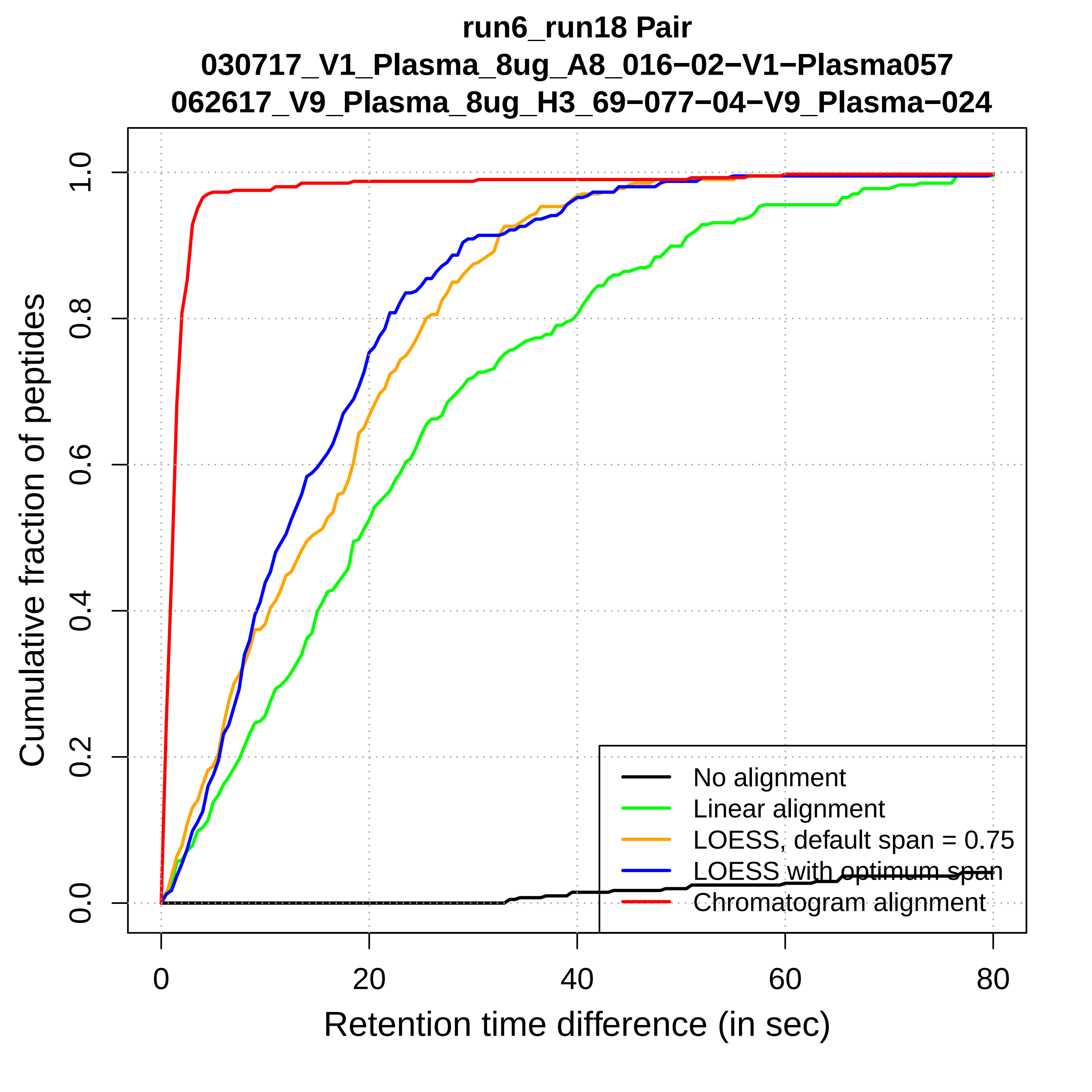
run6_run18 Pair
030717_V1_Plasma_8ug_A8_016−02−V1−Plasma057
062617_V9_Plasma_8ug_H3_69−077−04−V9_Plasma−024
Cumulative fraction of peptides
Retention time difference (in sec)
0.0
0.2
0.4
0.6
0.8
1.0
0
20
40
60
80
No alignment
Linear alignment
LOESS, default span = 0.75
LOESS with optimum span
Chromatogram alignment

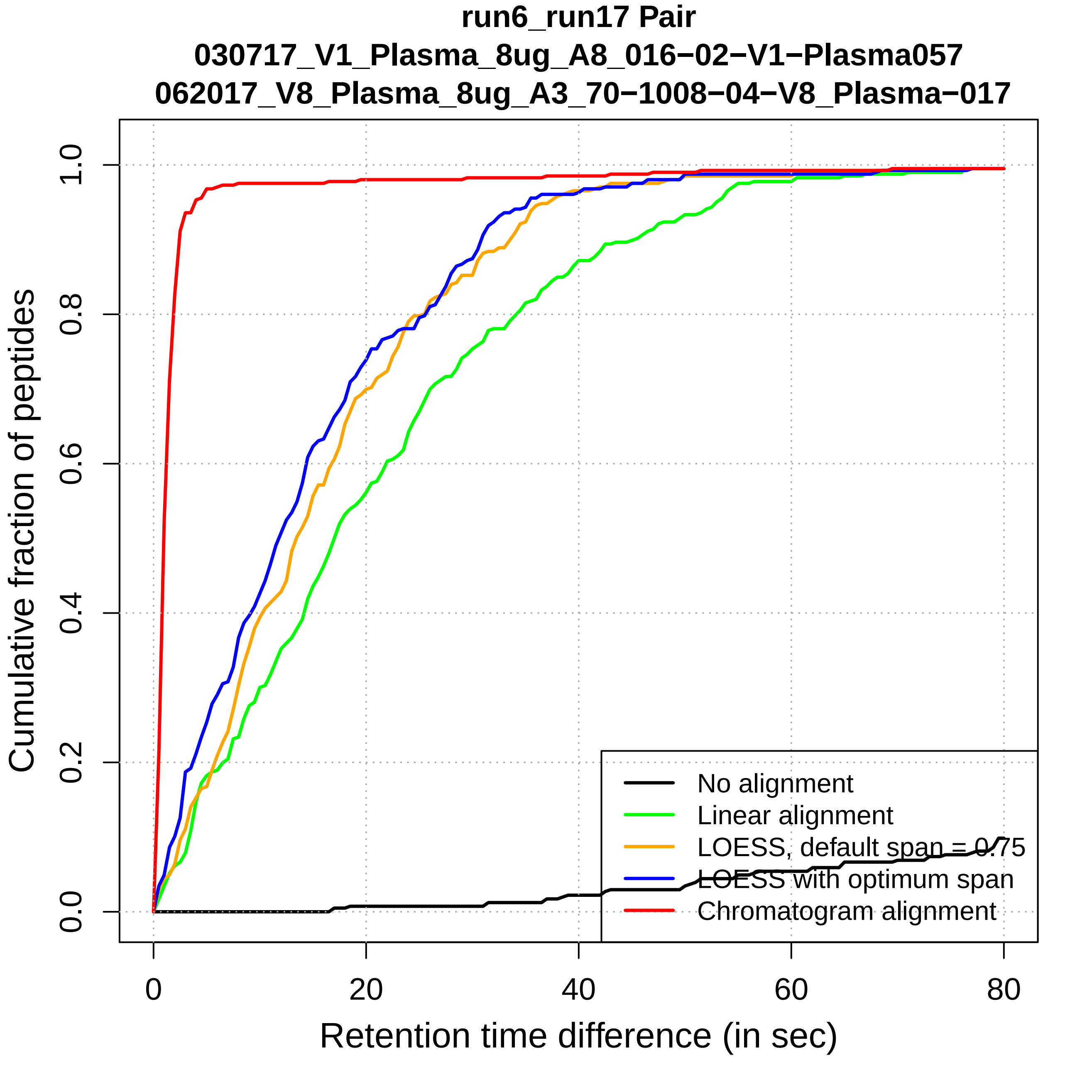

run6_run17 Pair
030717_V1_Plasma_8ug_A8_016−02−V1−Plasma057
062017_V8_Plasma_8ug_A3_70−1008−04−V8_Plasma−017
Cumulative fraction of peptides
0.0
0.2
0.4
0.6
0.8
1.0
0
20
40
60
80
Retention time difference (in sec)
No alignment
Linear alignment
LOESS, default span = 0.75
LOESS with optimum span
Chromatogram alignment

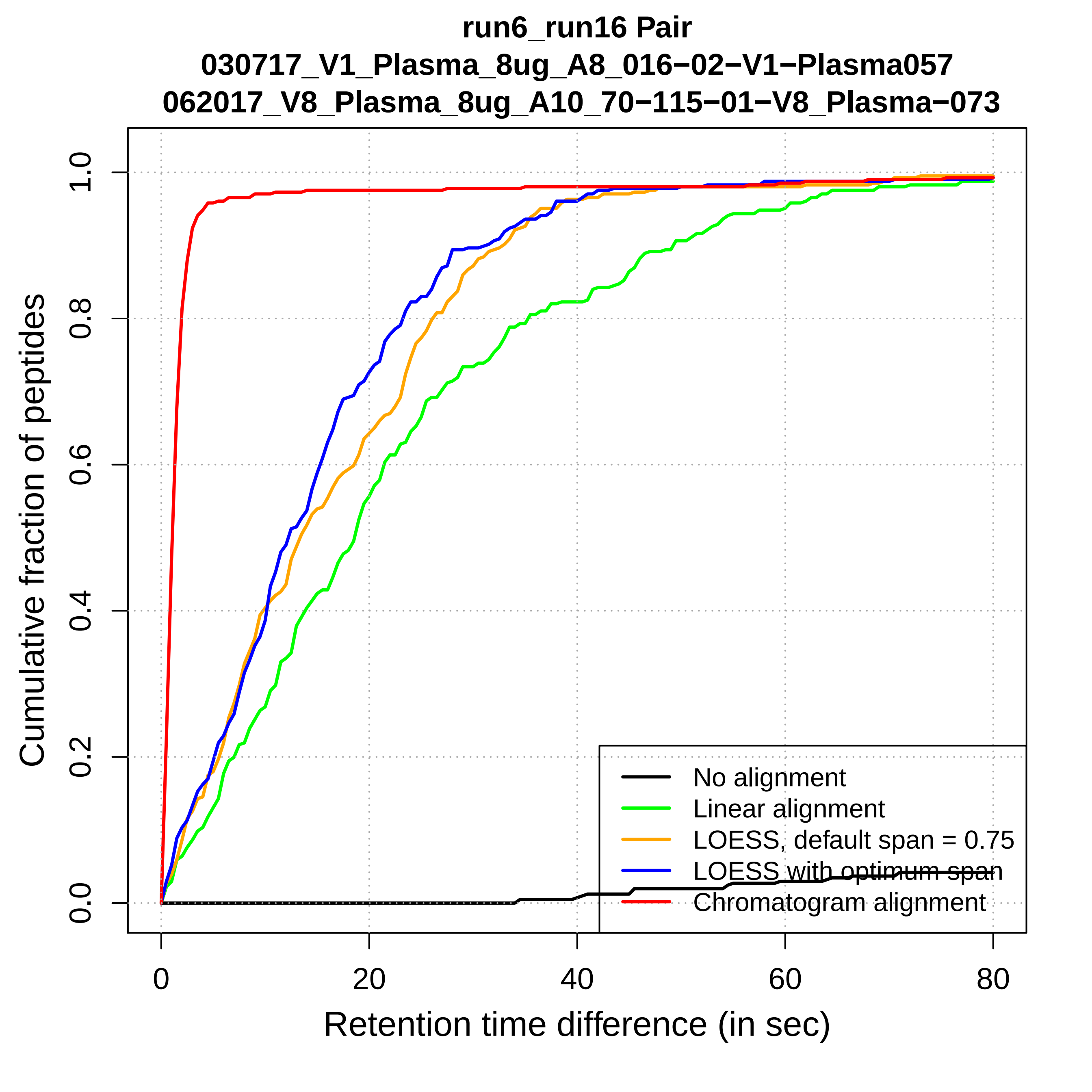

run6_run16 Pair
030717_V1_Plasma_8ug_A8_016−02−V1−Plasma057
062017_V8_Plasma_8ug_A10_70−115−01−V8_Plasma−073
Cumulative fraction of peptides
0.0
0.2
0.4
0.6
0.8
1.0
0
20
40
60
80
Retention time difference (in sec)
No alignment
Linear alignment
LOESS, default span = 0.75
LOESS with optimum span
Chromatogram alignment

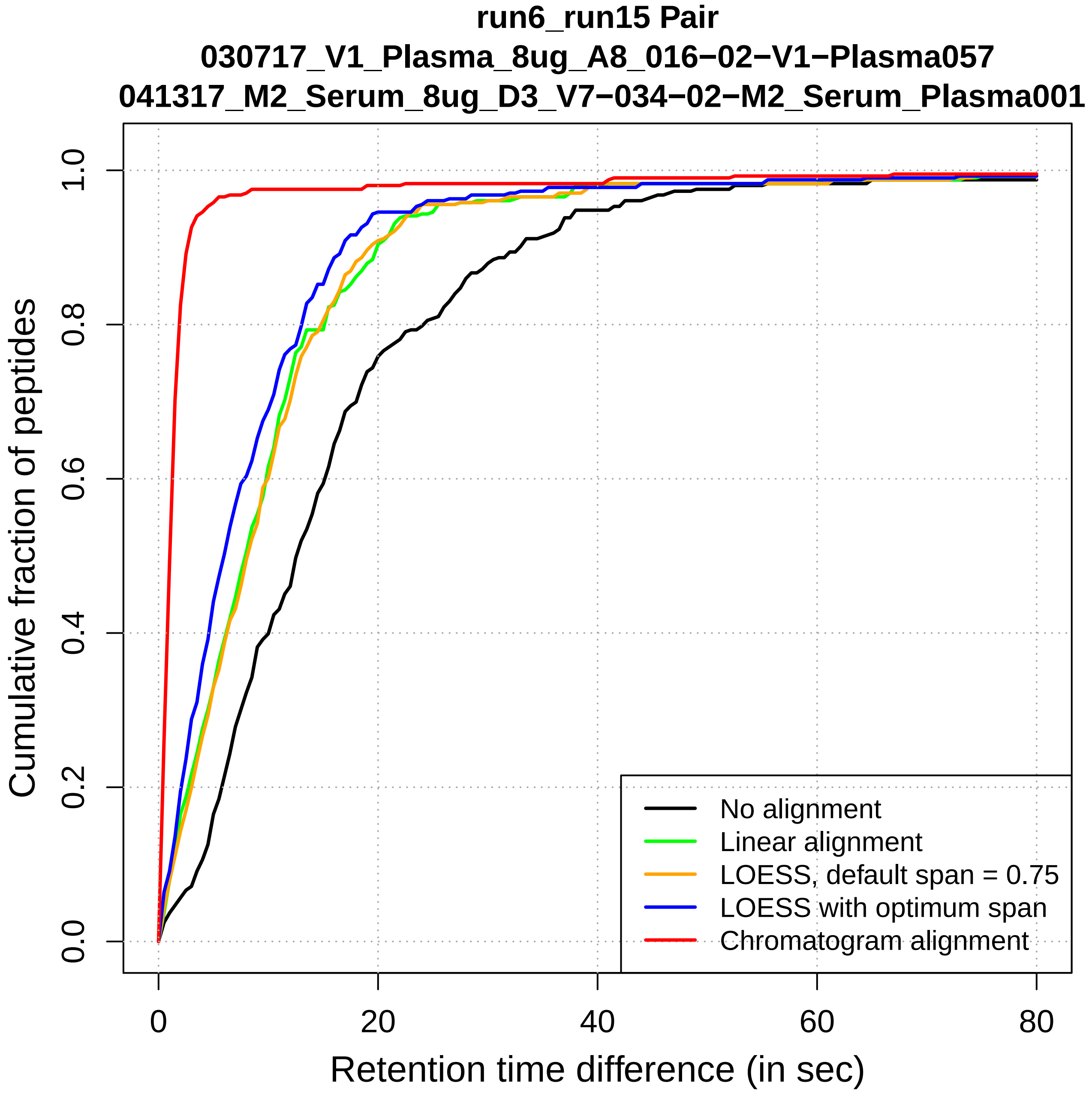
run6_run15 Pair
030717_V1_Plasma_8ug_A8_016−02−V1−Plasma057
041317_M2_Serum_8ug_D3_V7−034−02−M2_Serum_Plasma001
Cumulative fraction of peptides
0.0
0.2
0.4
0.6
0.8
1.0
0
20
40
60
80
Retention time difference (in sec)
No alignment
Linear alignment
LOESS, default span = 0.75
LOESS with optimum span
Chromatogram alignment

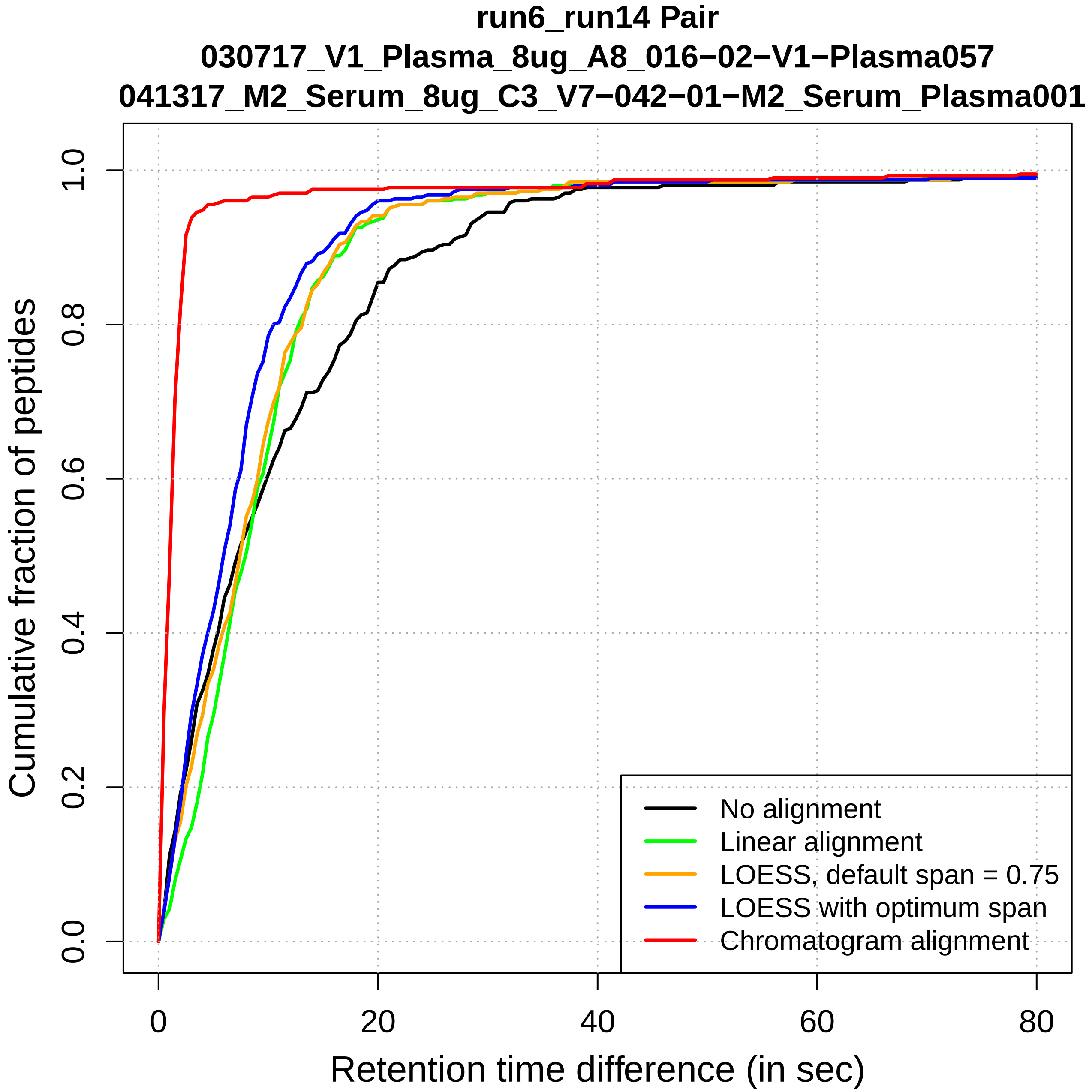

run6_run14 Pair
030717_V1_Plasma_8ug_A8_016−02−V1−Plasma057
041317_M2_Serum_8ug_C3_V7−042−01−M2_Serum_Plasma001
Cumulative fraction of peptides
Retention time difference (in sec)
0.0
0.2
0.4
0.6
0.8
1.0
0
20
40
60
80
No alignment
Linear alignment
LOESS, default span = 0.75
LOESS with optimum span
Chromatogram alignment

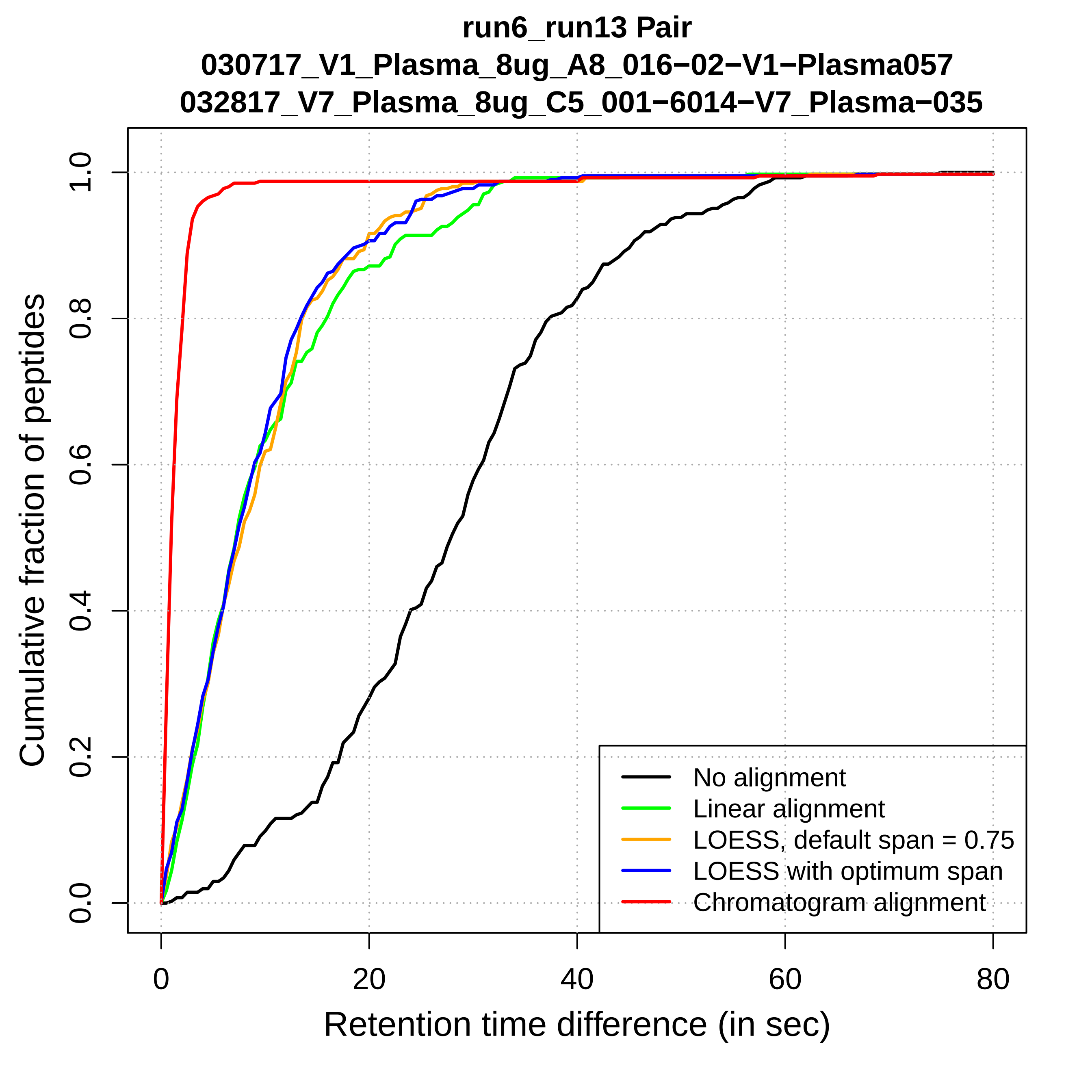
run6_run13 Pair
030717_V1_Plasma_8ug_A8_016−02−V1−Plasma057
032817_V7_Plasma_8ug_C5_001−6014−V7_Plasma−035
Cumulative fraction of peptides
0.0
0.2
0.4
0.6
0.8
1.0
0
20
40
60
80
Retention time difference (in sec)
No alignment
Linear alignment
LOESS, default span = 0.75
LOESS with optimum span
Chromatogram alignment

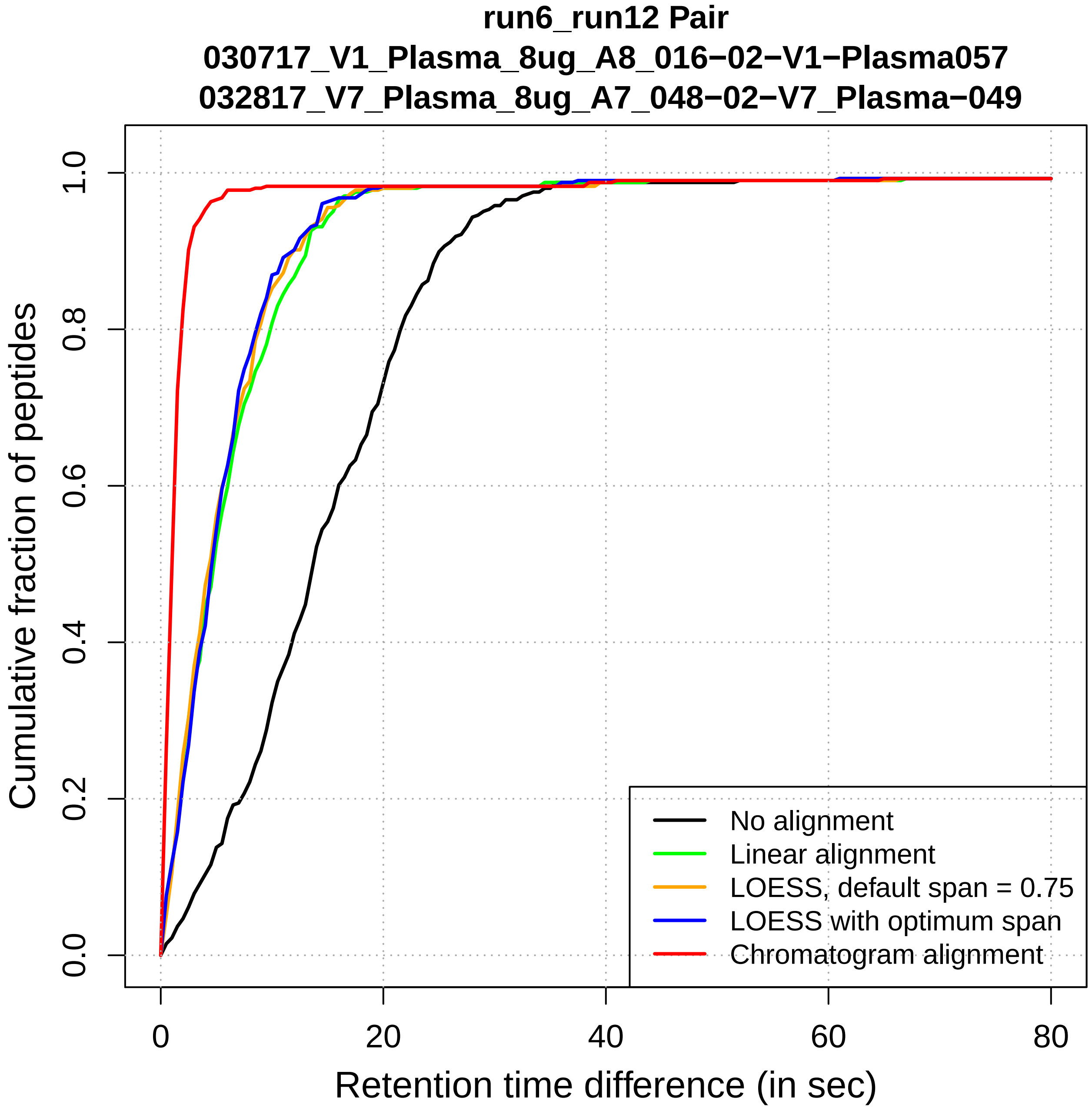
run6_run12 Pair
030717_V1_Plasma_8ug_A8_016−02−V1−Plasma057
032817_V7_Plasma_8ug_A7_048−02−V7_Plasma−049
Cumulative fraction of peptides
0.0
0.2
0.4
0.6
0.8
1.0
0
20
40
60
80
Retention time difference (in sec)
No alignment
Linear alignment
LOESS, default span = 0.75
LOESS with optimum span
Chromatogram alignment

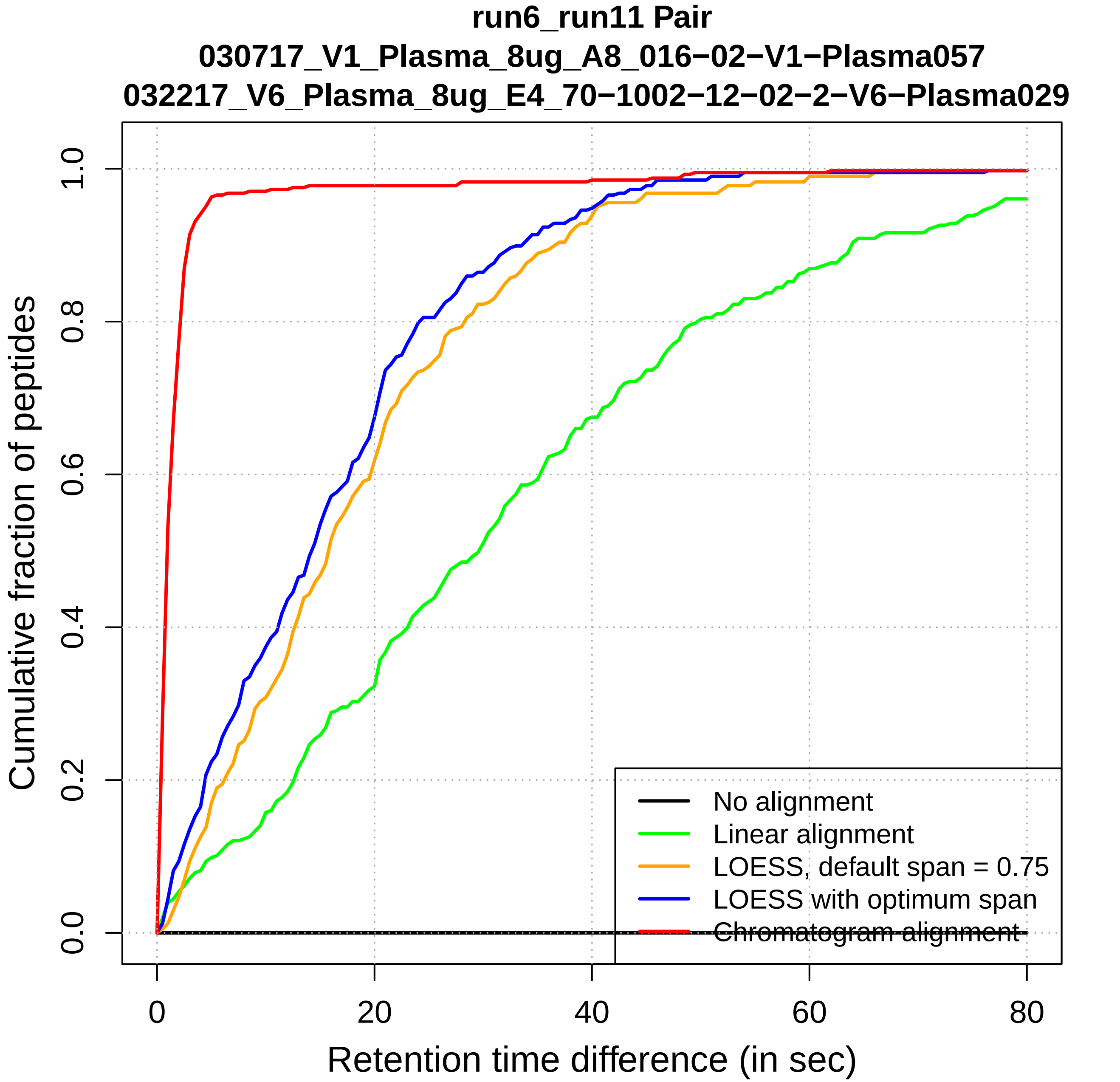

run6_run11 Pair
030717_V1_Plasma_8ug_A8_016−02−V1−Plasma057
032217_V6_Plasma_8ug_E4_70−1002−12−02−2−V6−Plasma029
Cumulative fraction of peptides
0.0
0.2
0.4
0.6
0.8
1.0
0
20
40
60
80
Retention time difference (in sec)
No alignment
Linear alignment
LOESS, default span = 0.75
LOESS with optimum span
Chromatogram alignment

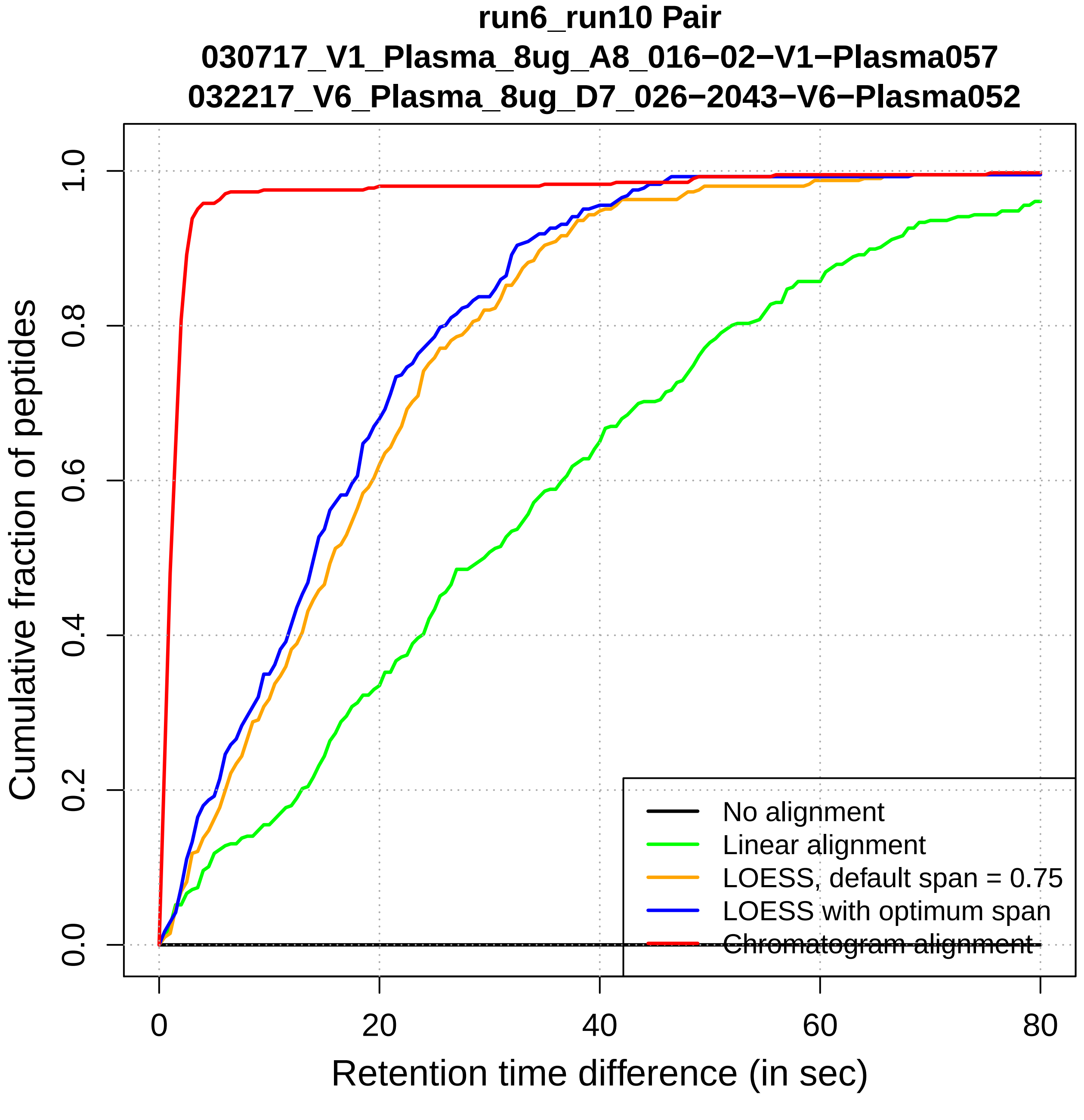
run6_run10 Pair
030717_V1_Plasma_8ug_A8_016−02−V1−Plasma057
032217_V6_Plasma_8ug_D7_026−2043−V6−Plasma052
Cumulative fraction of peptides
0.0
0.2
0.4
0.6
0.8
1.0
0
20
40
60
80
Retention time difference (in sec)
No alignment
Linear alignment
LOESS, default span = 0.75
LOESS with optimum span
Chromatogram alignment

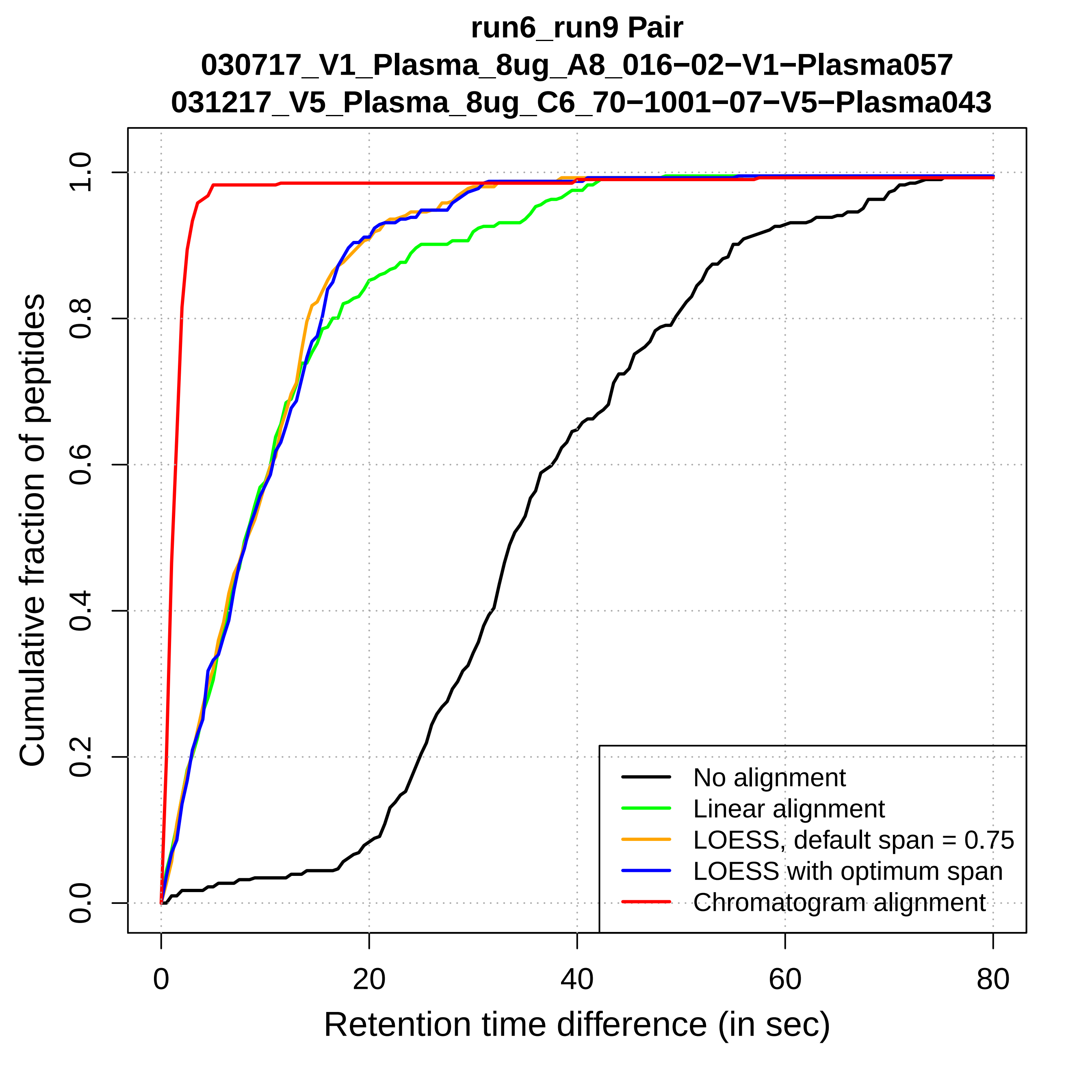
run6_run9 Pair
030717_V1_Plasma_8ug_A8_016−02−V1−Plasma057
031217_V5_Plasma_8ug_C6_70−1001−07−V5−Plasma043
Cumulative fraction of peptides
0.0
0.2
0.4
0.6
0.8
1.0
0
20
40
60
80
Retention time difference (in sec)
No alignment
Linear alignment
LOESS, default span = 0.75
LOESS with optimum span
Chromatogram alignment

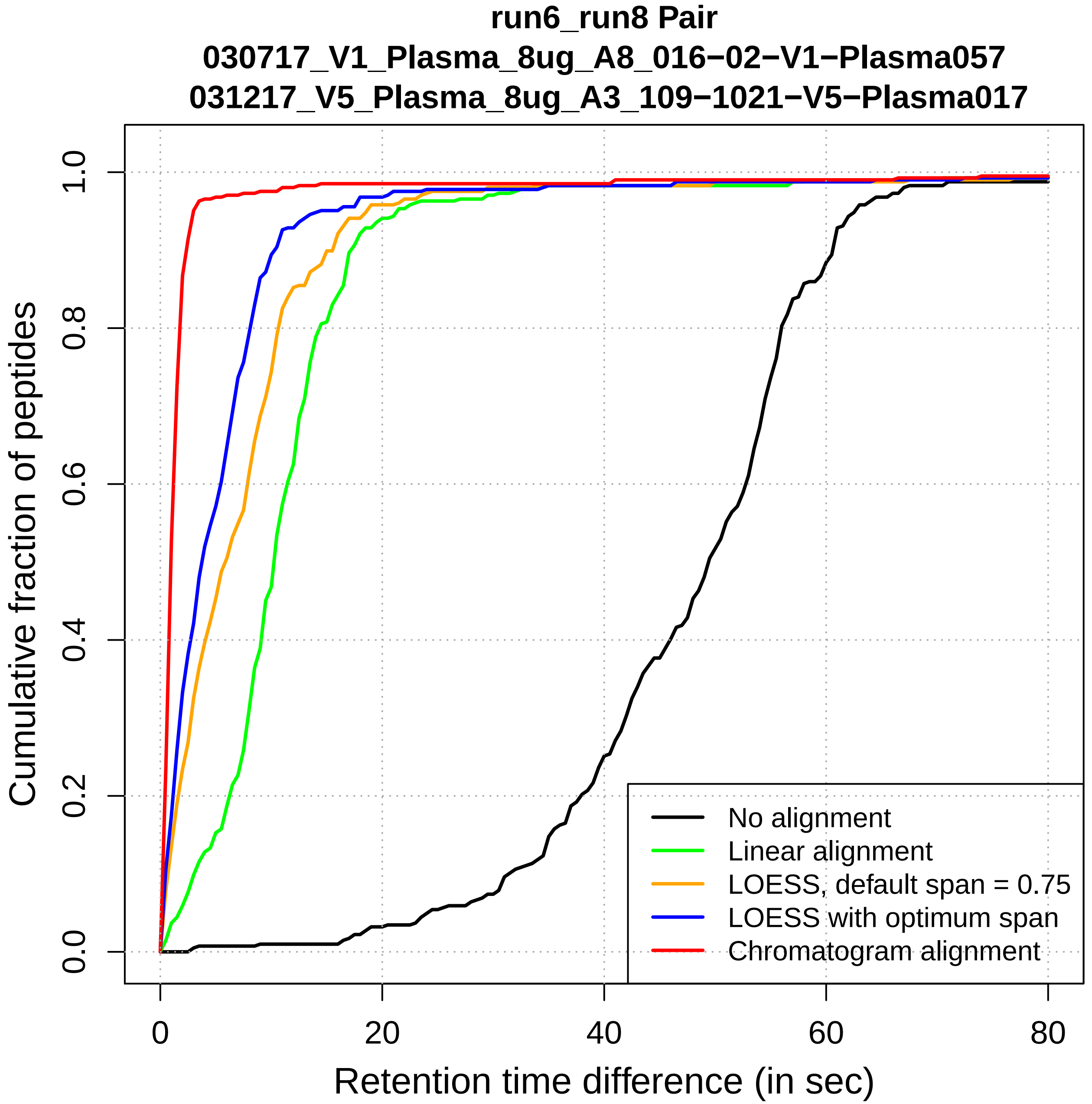
run6_run8 Pair
030717_V1_Plasma_8ug_A8_016−02−V1−Plasma057
031217_V5_Plasma_8ug_A3_109−1021−V5−Plasma017
Cumulative fraction of peptides
0.0
0.2
0.4
0.6
0.8
1.0
0
20
40
60
80
Retention time difference (in sec)
No alignment
Linear alignment
LOESS, default span = 0.75
LOESS with optimum span
Chromatogram alignment

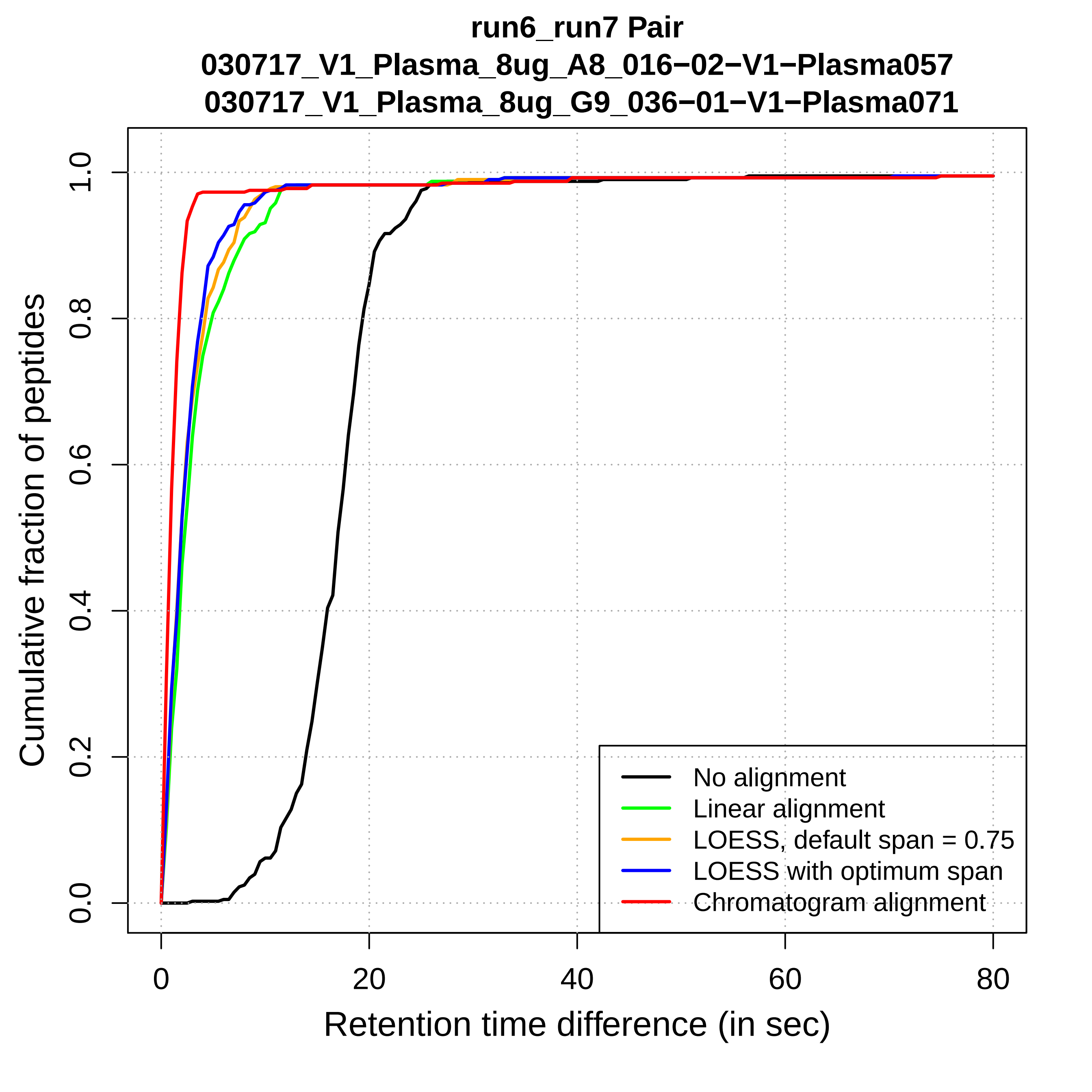
run6_run7 Pair
030717_V1_Plasma_8ug_A8_016−02−V1−Plasma057
030717_V1_Plasma_8ug_G9_036−01−V1−Plasma071
Cumulative fraction of peptides
0.0
0.2
0.4
0.6
0.8
1.0
0
20
40
60
80
Retention time difference (in sec)
No alignment
Linear alignment
LOESS, default span = 0.75
LOESS with optimum span
Chromatogram alignment

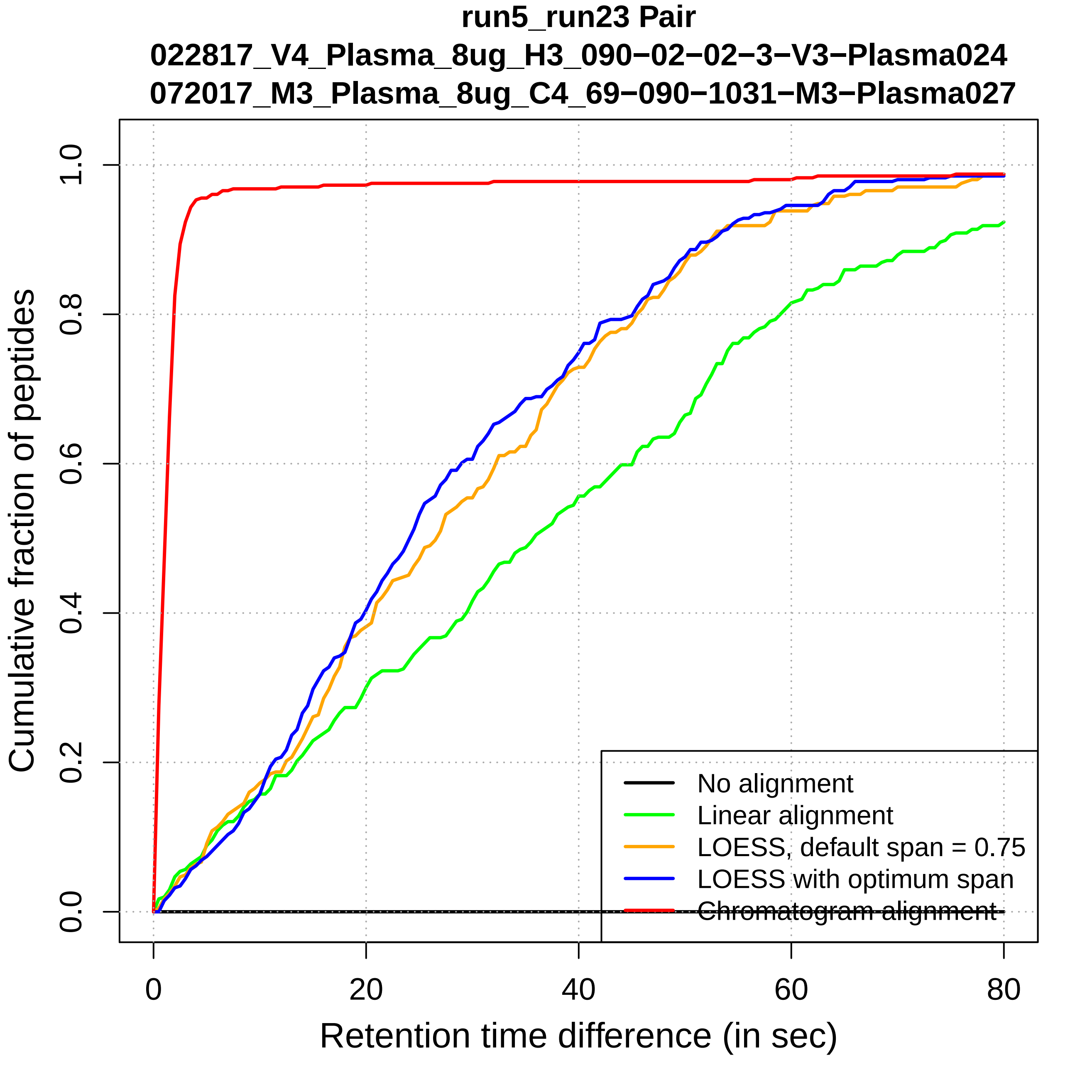
run5_run23 Pair
022817_V4_Plasma_8ug_H3_090−02−02−3−V3−Plasma024
072017_M3_Plasma_8ug_C4_69−090−1031−M3−Plasma027
Cumulative fraction of peptides
0.0
0.2
0.4
0.6
0.8
1.0
0
20
40
60
80
Retention time difference (in sec)
No alignment
Linear alignment
LOESS, default span = 0.75
LOESS with optimum span
Chromatogram alignment

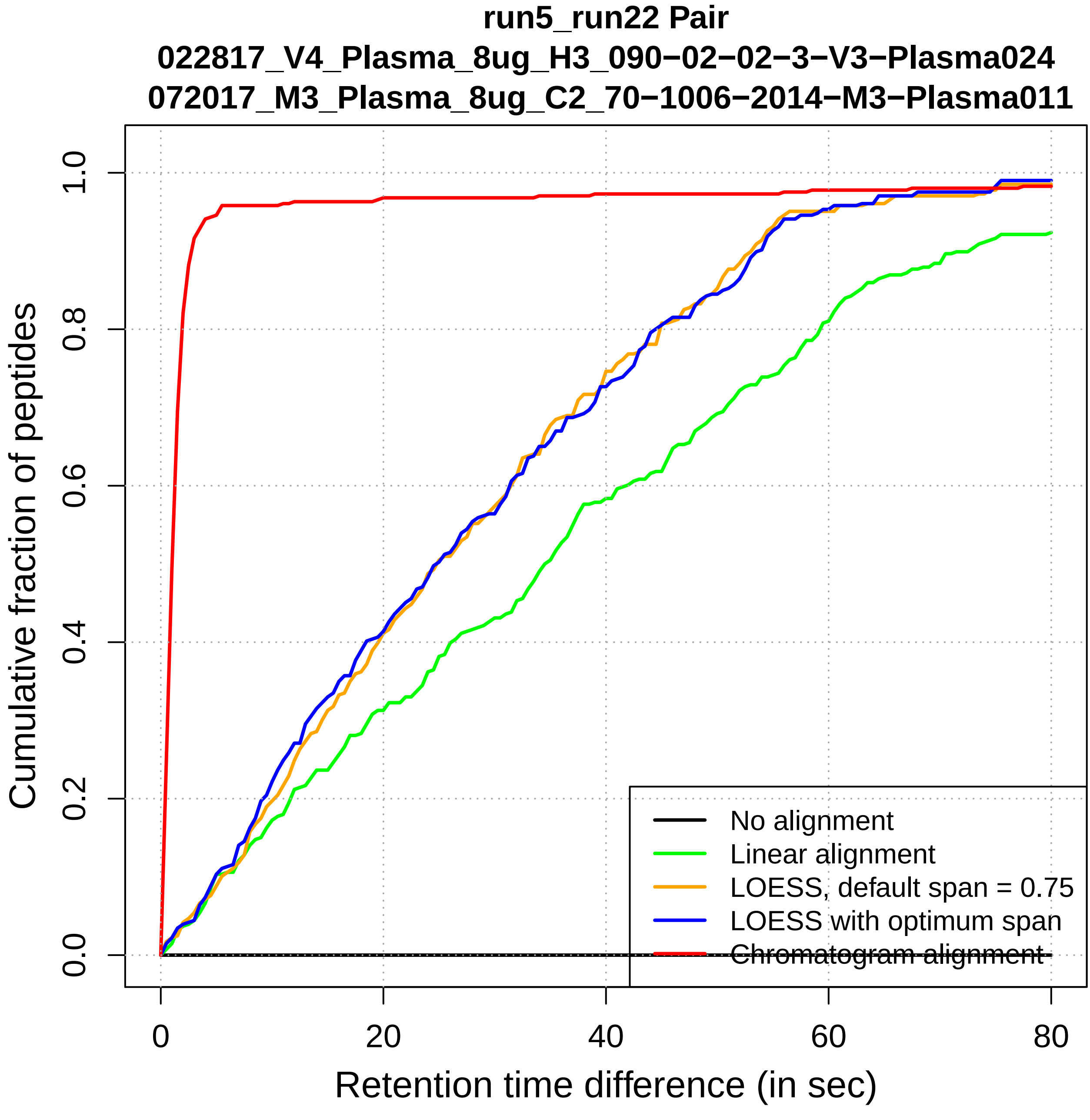

run5_run22 Pair
022817_V4_Plasma_8ug_H3_090−02−02−3−V3−Plasma024
072017_M3_Plasma_8ug_C2_70−1006−2014−M3−Plasma011
Cumulative fraction of peptides
Retention time difference (in sec)
0.0
0.2
0.4
0.6
0.8
1.0
0
20
40
60
80
No alignment
Linear alignment
LOESS, default span = 0.75
LOESS with optimum span
Chromatogram alignment

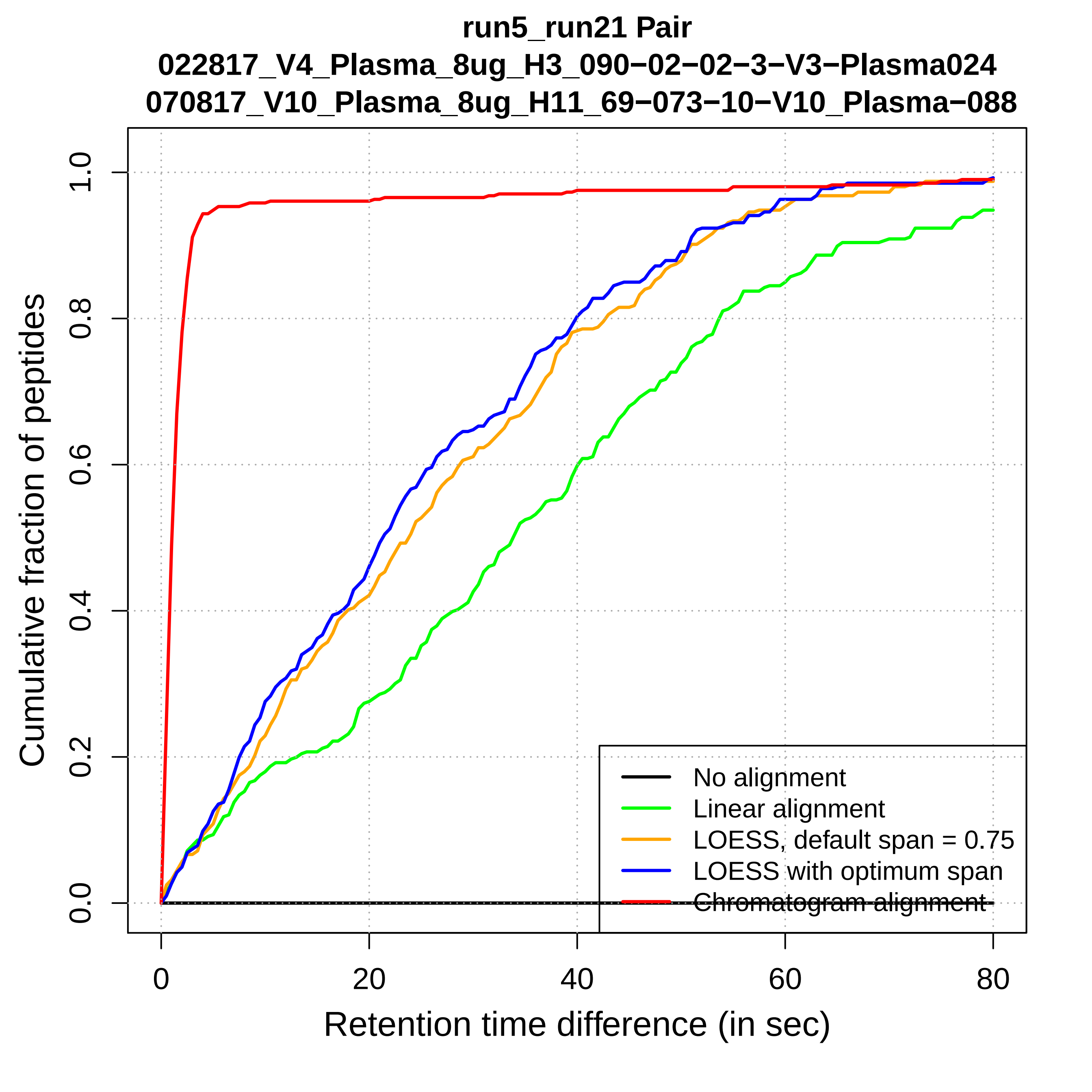
run5_run21 Pair
022817_V4_Plasma_8ug_H3_090−02−02−3−V3−Plasma024
070817_V10_Plasma_8ug_H11_69−073−10−V10_Plasma−088
Cumulative fraction of peptides
0.0
0.2
0.4
0.6
0.8
1.0
0
20
40
60
80
Retention time difference (in sec)
No alignment
Linear alignment
LOESS, default span = 0.75
LOESS with optimum span
Chromatogram alignment

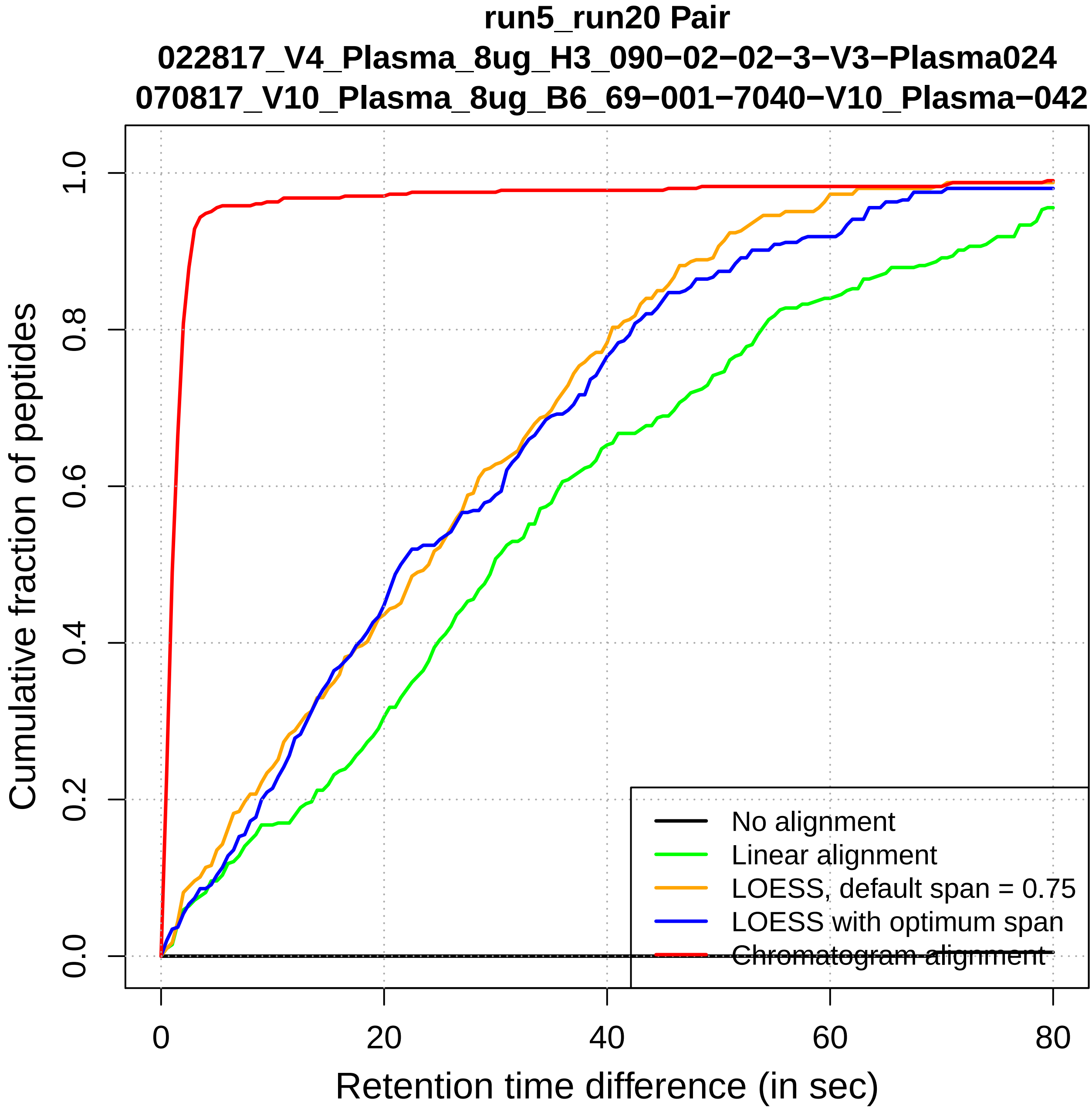

run5_run20 Pair
022817_V4_Plasma_8ug_H3_090−02−02−3−V3−Plasma024
070817_V10_Plasma_8ug_B6_69−001−7040−V10_Plasma−042
Cumulative fraction of peptides
0.0
0.2
0.4
0.6
0.8
1.0
0
20
40
60
80
Retention time difference (in sec)
No alignment
Linear alignment
LOESS, default span = 0.75
LOESS with optimum span
Chromatogram alignment

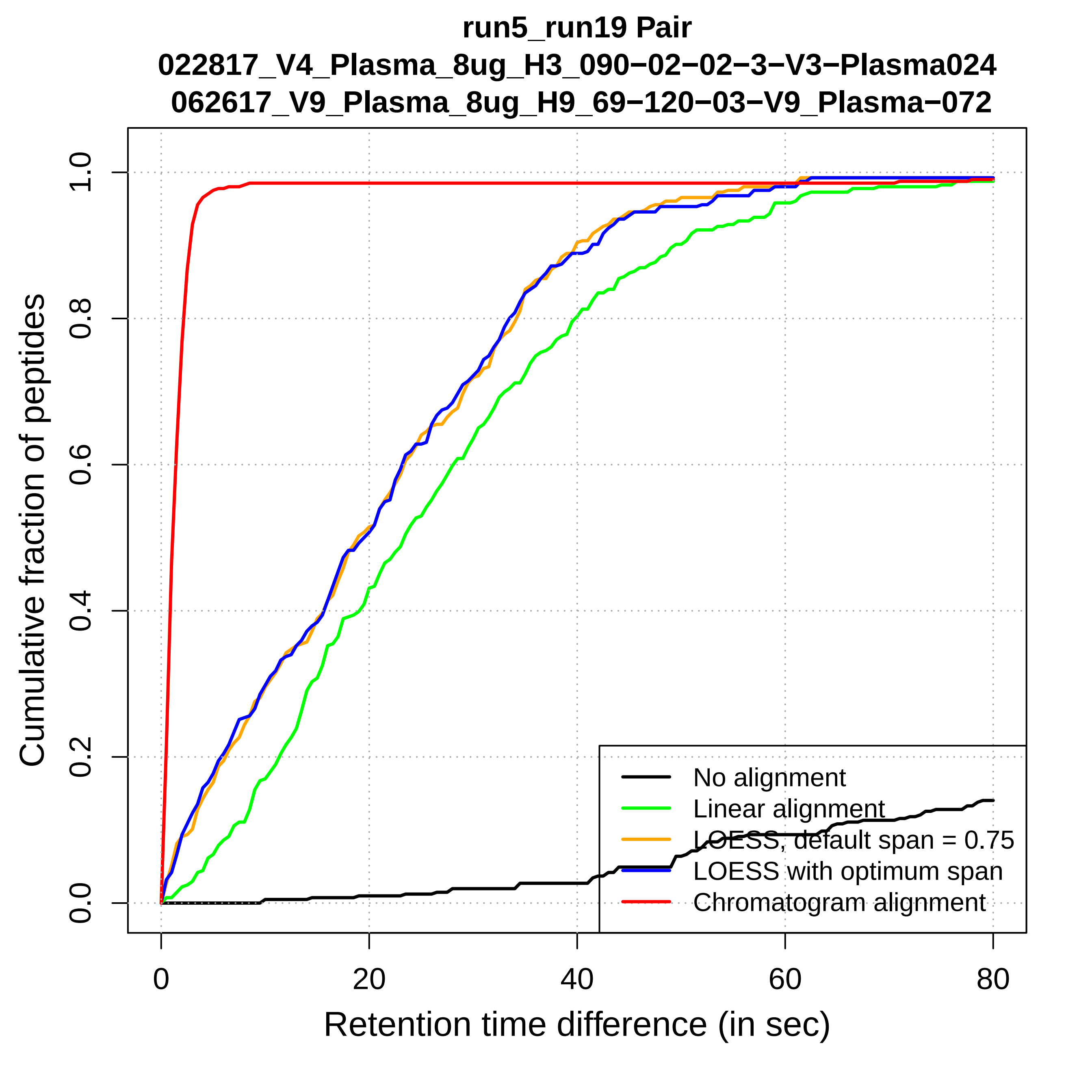
run5_run19 Pair
022817_V4_Plasma_8ug_H3_090−02−02−3−V3−Plasma024
062617_V9_Plasma_8ug_H9_69−120−03−V9_Plasma−072
Cumulative fraction of peptides
Retention time difference (in sec)
0.0
0.2
0.4
0.6
0.8
1.0
0
20
40
60
80
No alignment
Linear alignment
LOESS, default span = 0.75
LOESS with optimum span
Chromatogram alignment

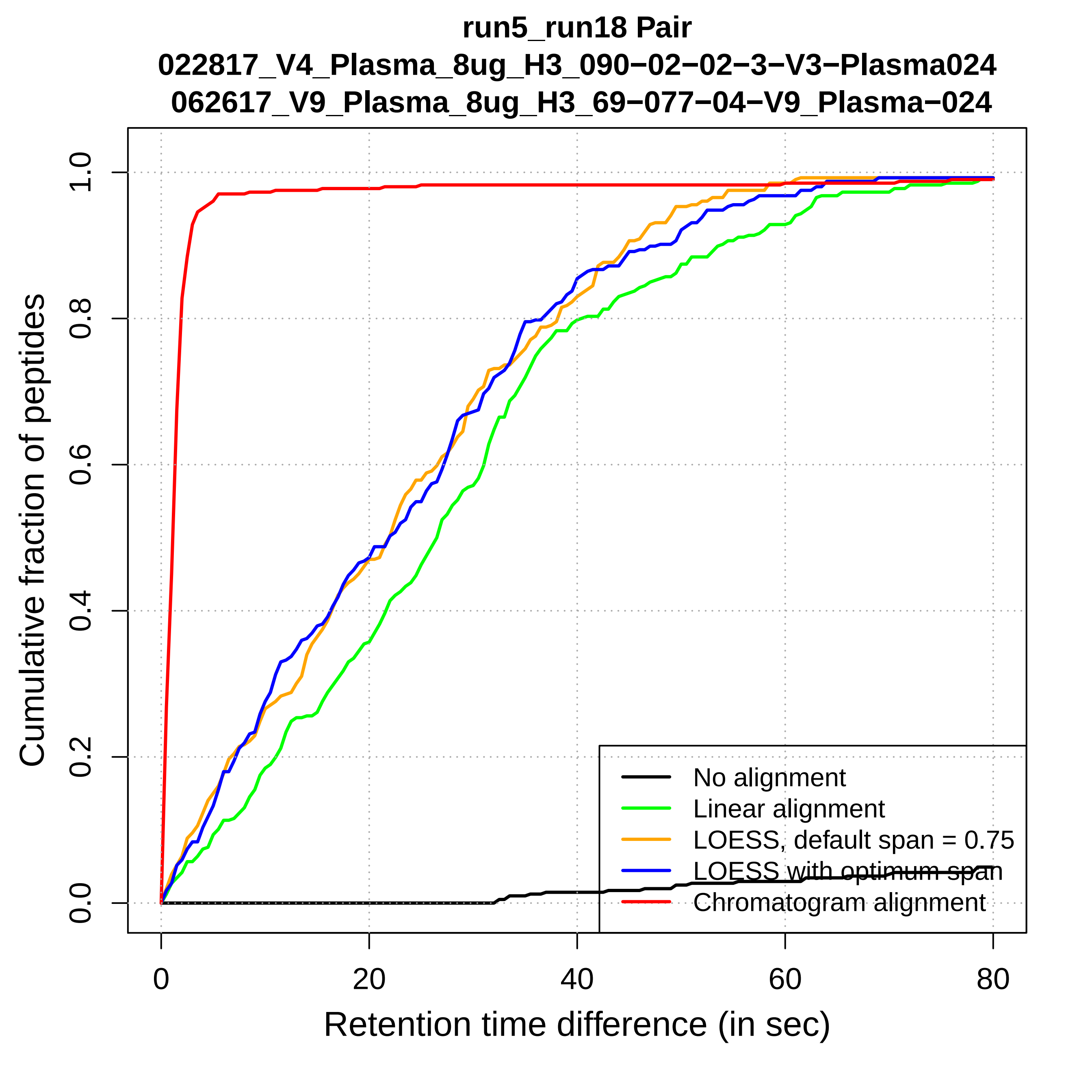

run5_run18 Pair
022817_V4_Plasma_8ug_H3_090−02−02−3−V3−Plasma024
062617_V9_Plasma_8ug_H3_69−077−04−V9_Plasma−024
Cumulative fraction of peptides
0.0
0.2
0.4
0.6
0.8
1.0
0
20
40
60
80
Retention time difference (in sec)
No alignment
Linear alignment
LOESS, default span = 0.75
LOESS with optimum span
Chromatogram alignment

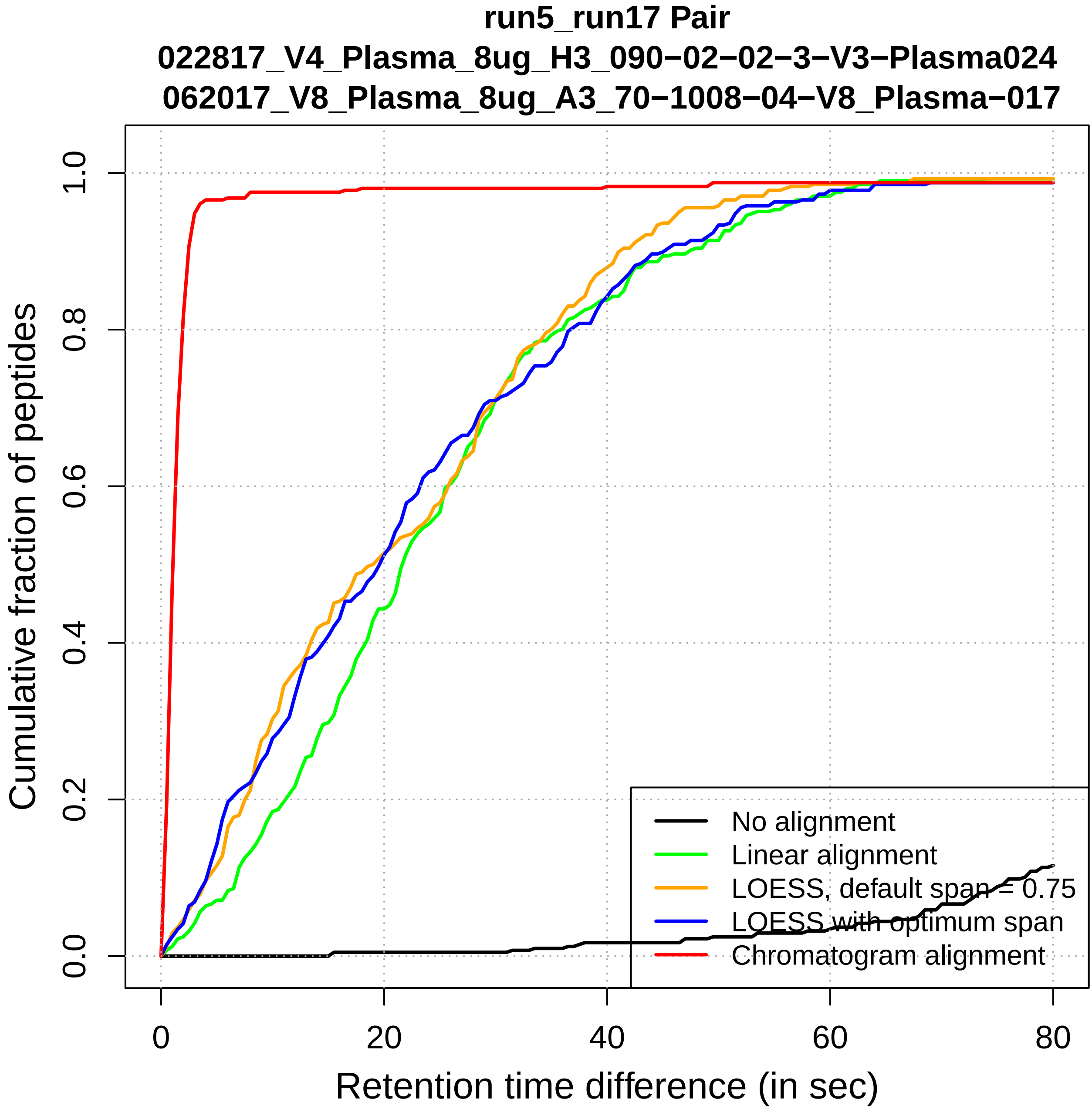

run5_run17 Pair
022817_V4_Plasma_8ug_H3_090−02−02−3−V3−Plasma024
062017_V8_Plasma_8ug_A3_70−1008−04−V8_Plasma−017
Cumulative fraction of peptides
0.0
0.2
0.4
0.6
0.8
1.0
0
20
40
60
80
Retention time difference (in sec)
No alignment
Linear alignment
LOESS, default span = 0.75
LOESS with optimum span
Chromatogram alignment

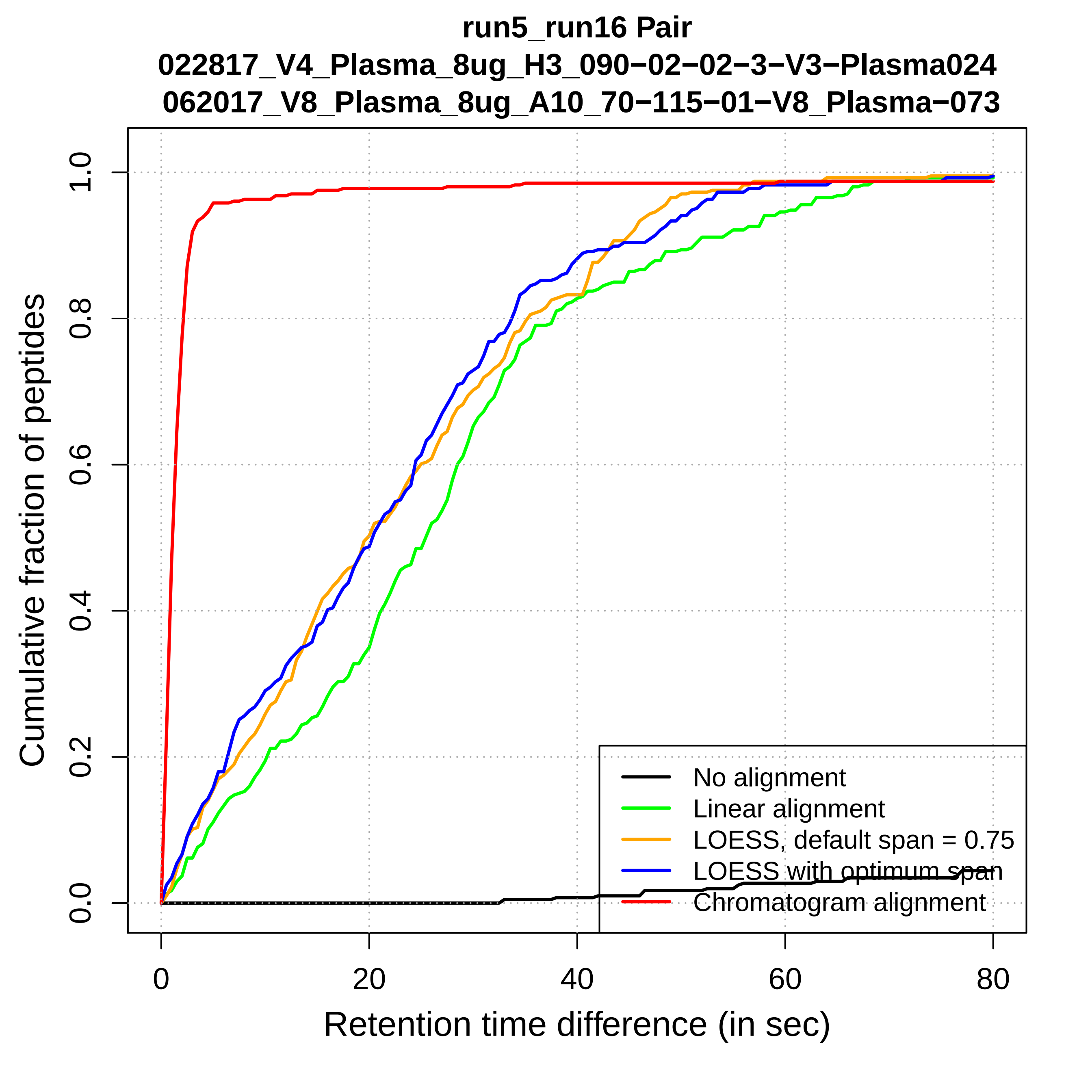

run5_run16 Pair
022817_V4_Plasma_8ug_H3_090−02−02−3−V3−Plasma024
062017_V8_Plasma_8ug_A10_70−115−01−V8_Plasma−073
Cumulative fraction of peptides
0.0
0.2
0.4
0.6
0.8
1.0
0
20
40
60
80
Retention time difference (in sec)
No alignment
Linear alignment
LOESS, default span = 0.75
LOESS with optimum span
Chromatogram alignment

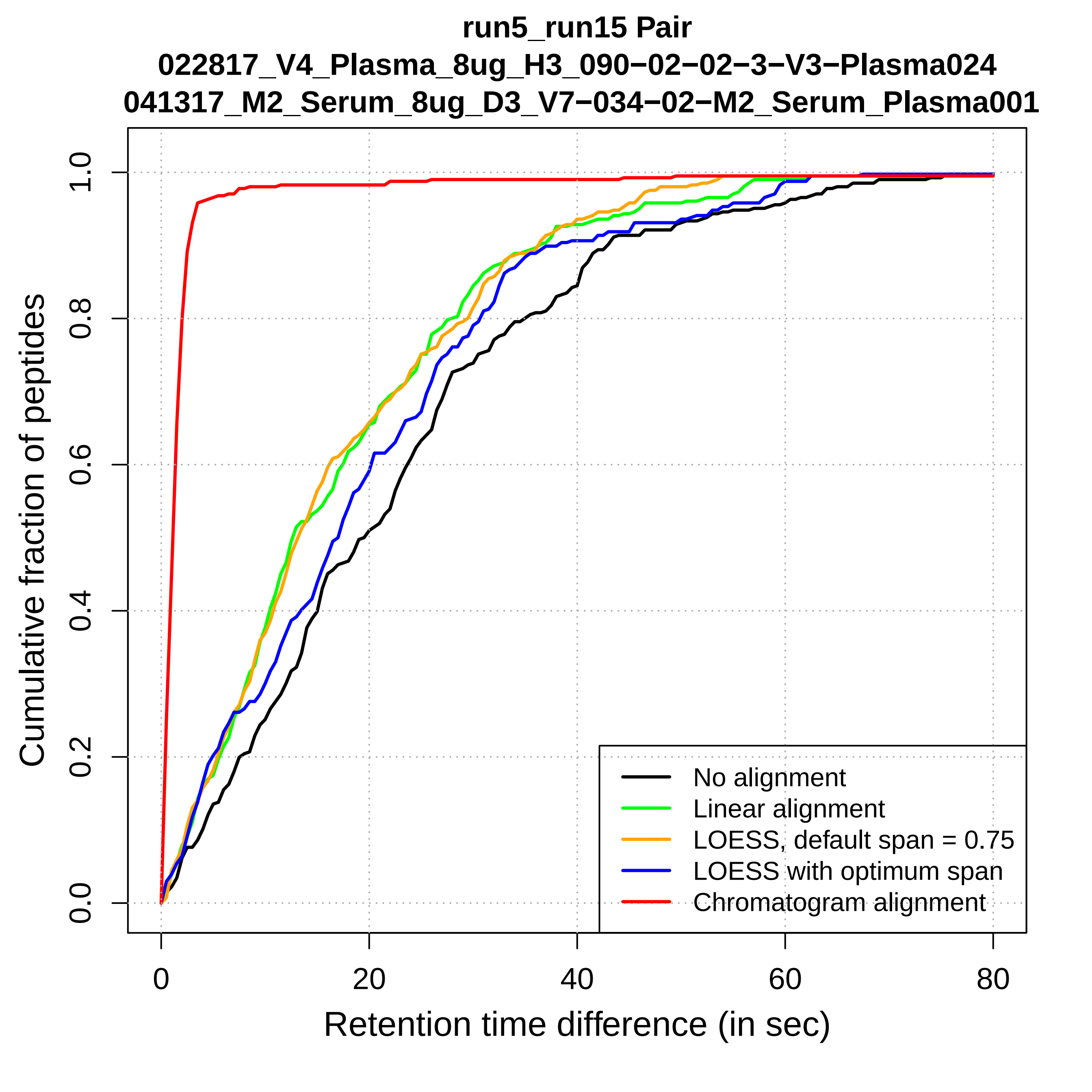
run5_run15 Pair
022817_V4_Plasma_8ug_H3_090−02−02−3−V3−Plasma024
041317_M2_Serum_8ug_D3_V7−034−02−M2_Serum_Plasma001
Cumulative fraction of peptides
0.0
0.2
0.4
0.6
0.8
1.0
0
20
40
60
80
Retention time difference (in sec)
No alignment
Linear alignment
LOESS, default span = 0.75
LOESS with optimum span
Chromatogram alignment

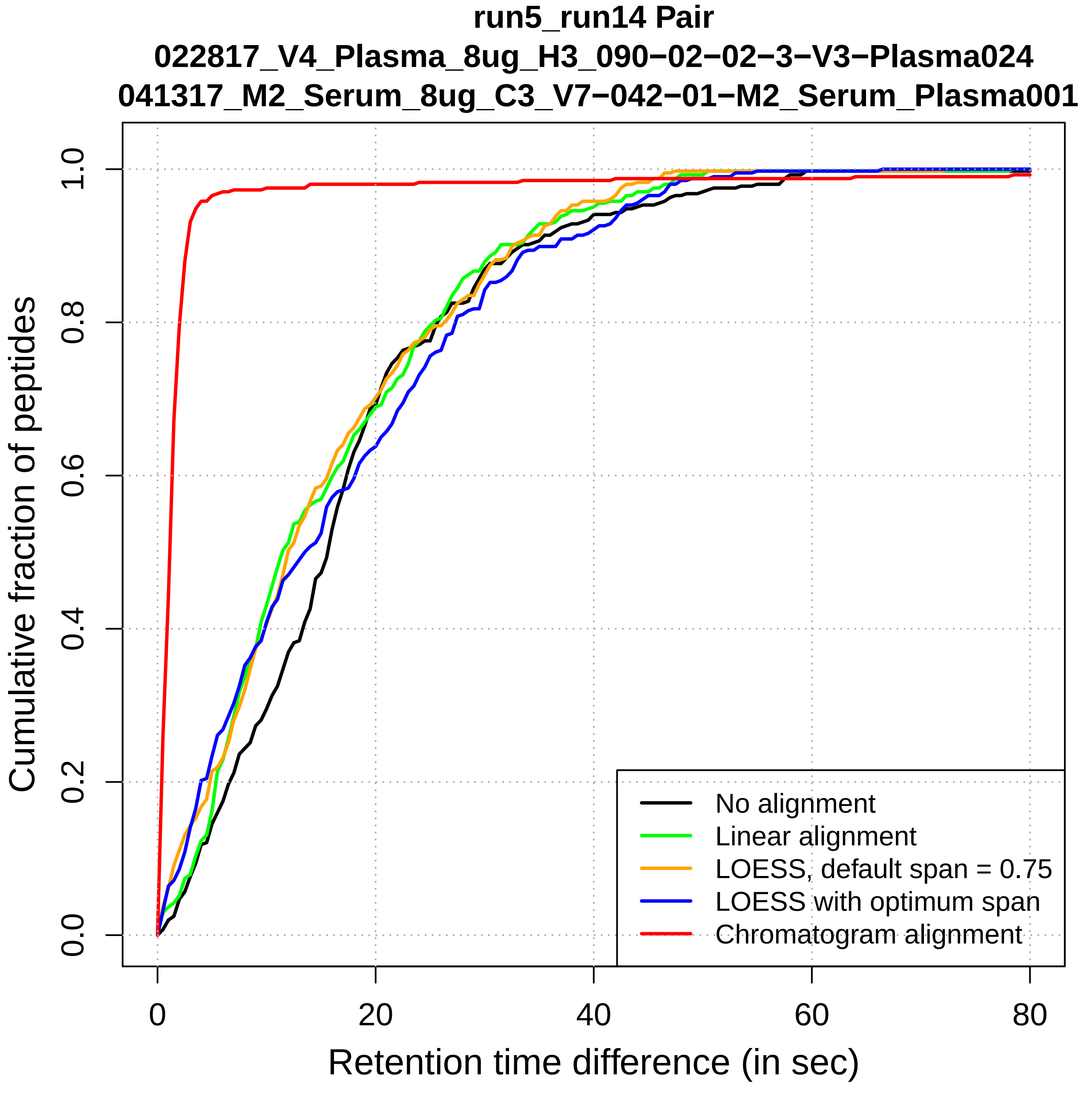

run5_run14 Pair
022817_V4_Plasma_8ug_H3_090−02−02−3−V3−Plasma024
041317_M2_Serum_8ug_C3_V7−042−01−M2_Serum_Plasma001
Cumulative fraction of peptides
0.0
0.2
0.4
0.6
0.8
1.0
0
20
40
60
80
Retention time difference (in sec)
No alignment
Linear alignment
LOESS, default span = 0.75
LOESS with optimum span
Chromatogram alignment

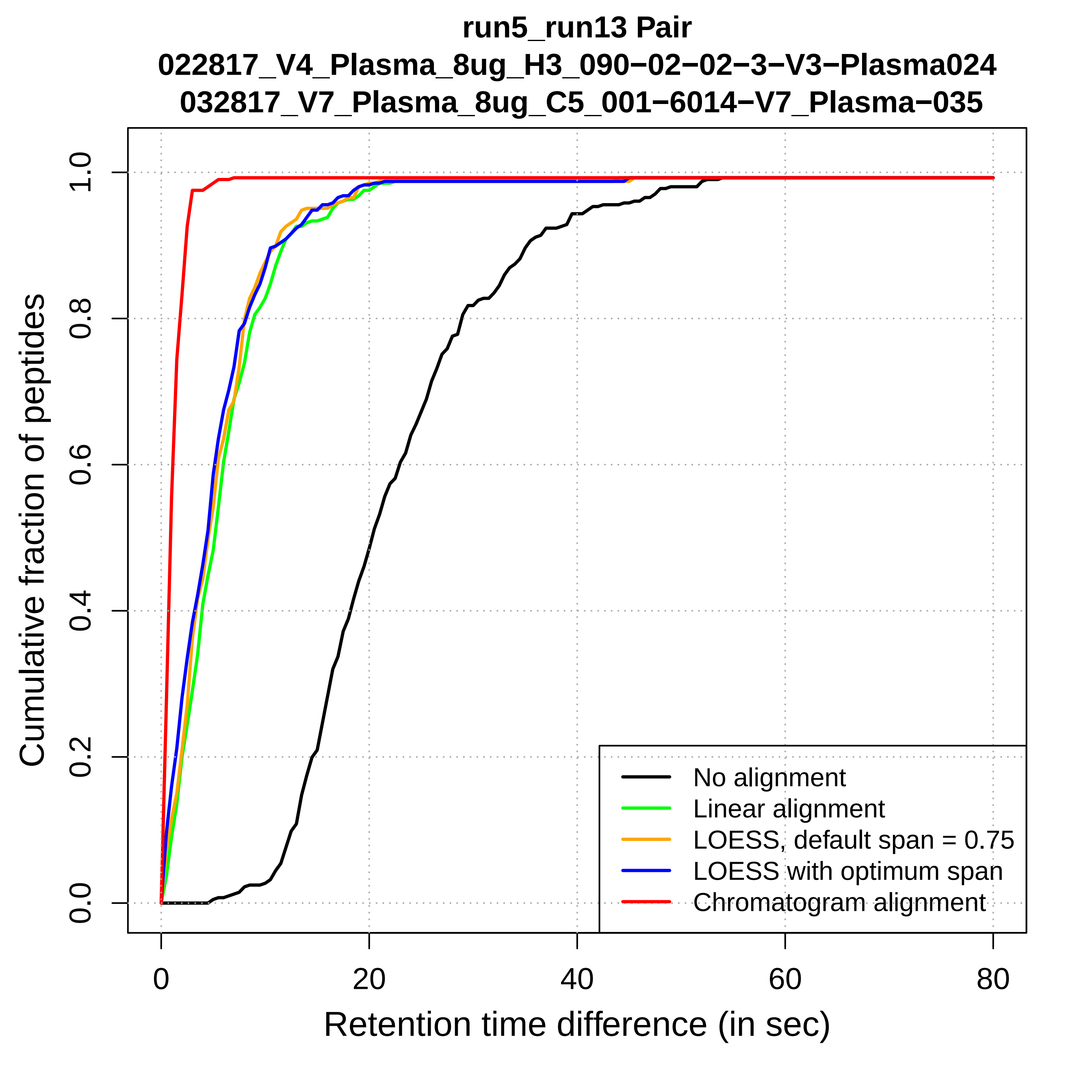
run5_run13 Pair
022817_V4_Plasma_8ug_H3_090−02−02−3−V3−Plasma024
032817_V7_Plasma_8ug_C5_001−6014−V7_Plasma−035
Cumulative fraction of peptides
0.0
0.2
0.4
0.6
0.8
1.0
0
20
40
60
80
Retention time difference (in sec)
No alignment
Linear alignment
LOESS, default span = 0.75
LOESS with optimum span
Chromatogram alignment

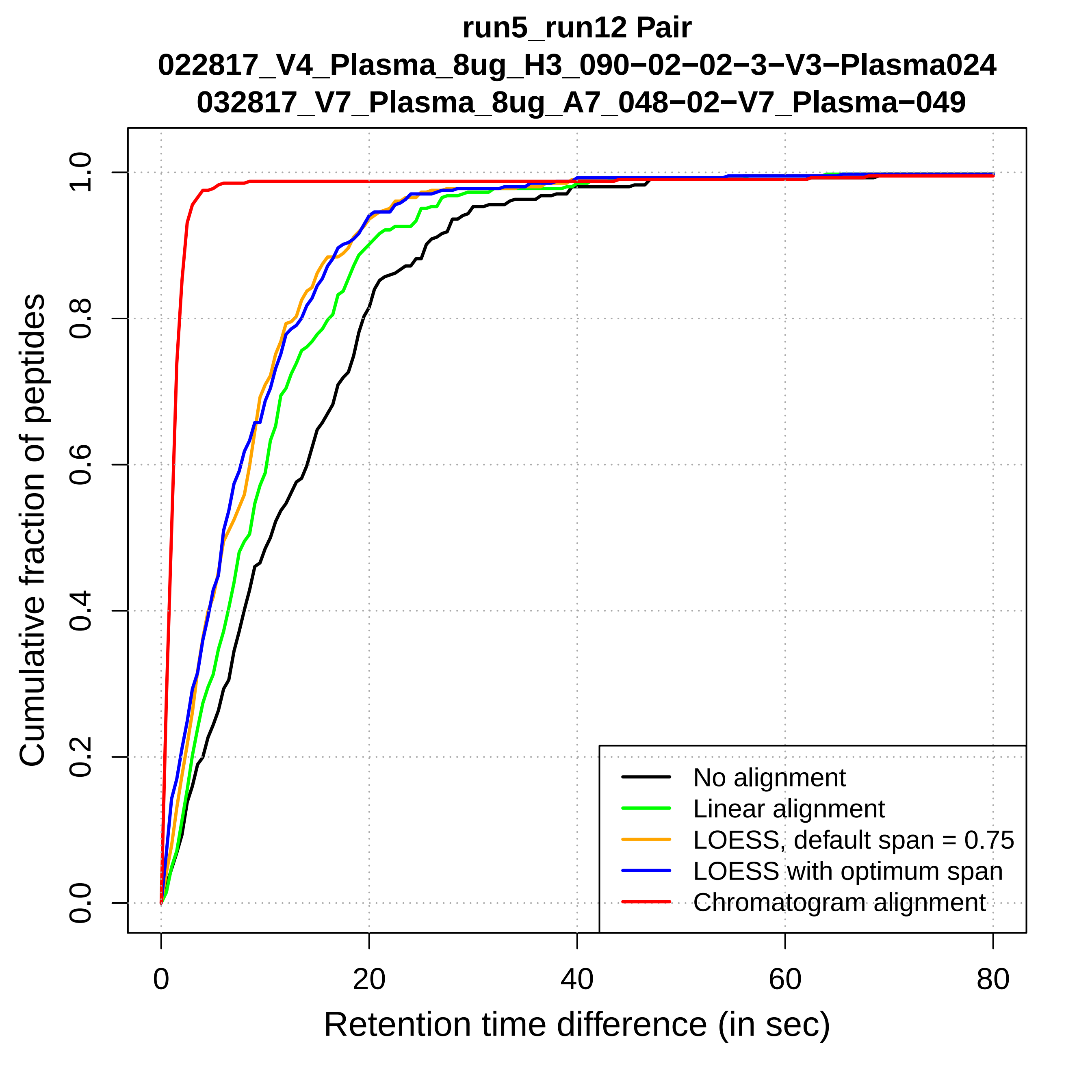

run5_run12 Pair
022817_V4_Plasma_8ug_H3_090−02−02−3−V3−Plasma024
032817_V7_Plasma_8ug_A7_048−02−V7_Plasma−049
Cumulative fraction of peptides
0.0
0.2
0.4
0.6
0.8
1.0
0
20
40
60
80
Retention time difference (in sec)
No alignment
Linear alignment
LOESS, default span = 0.75
LOESS with optimum span
Chromatogram alignment

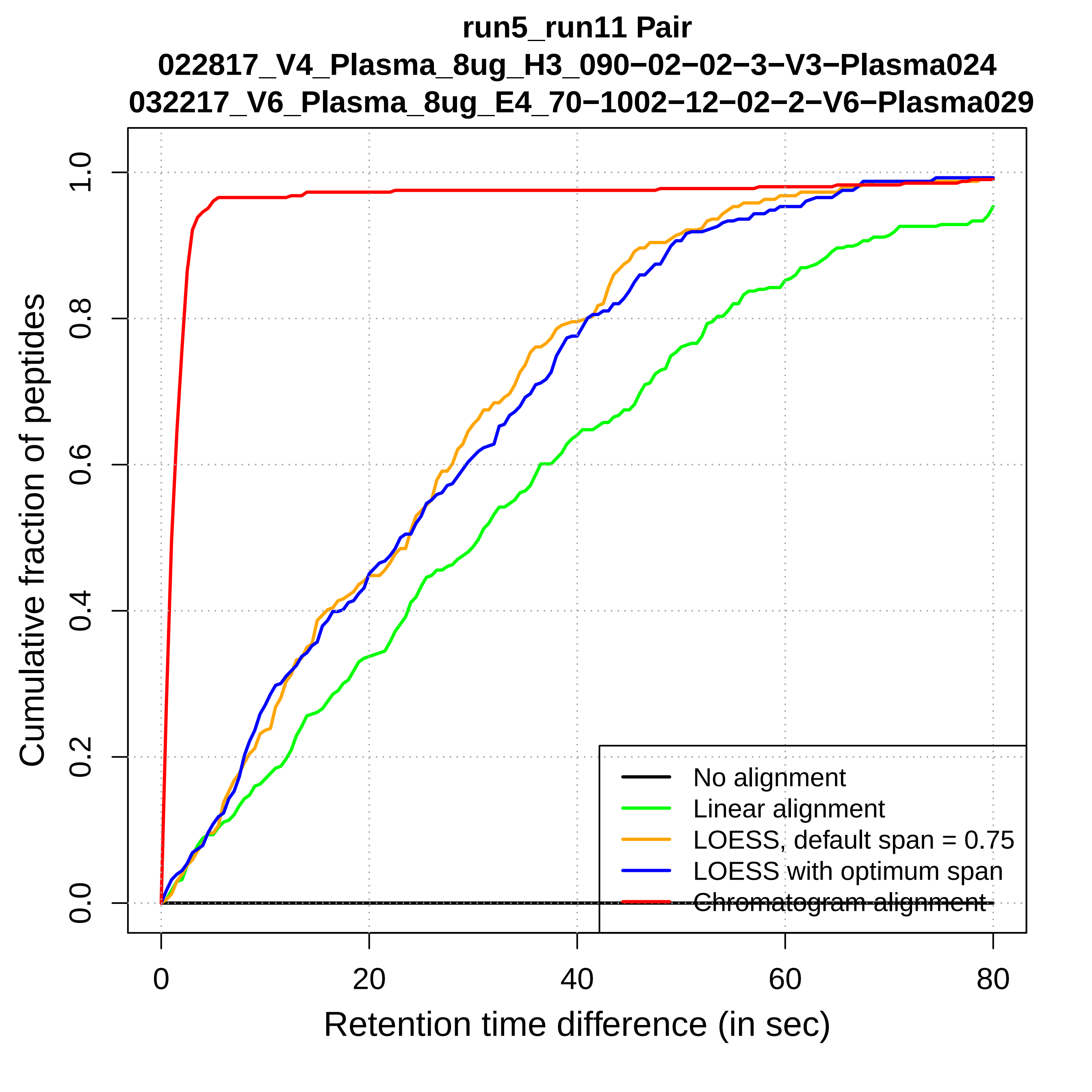

run5_run11 Pair
022817_V4_Plasma_8ug_H3_090−02−02−3−V3−Plasma024
032217_V6_Plasma_8ug_E4_70−1002−12−02−2−V6−Plasma029
Cumulative fraction of peptides
0.0
0.2
0.4
0.6
0.8
1.0
0
20
40
60
80
Retention time difference (in sec)
No alignment
Linear alignment
LOESS, default span = 0.75
LOESS with optimum span
Chromatogram alignment

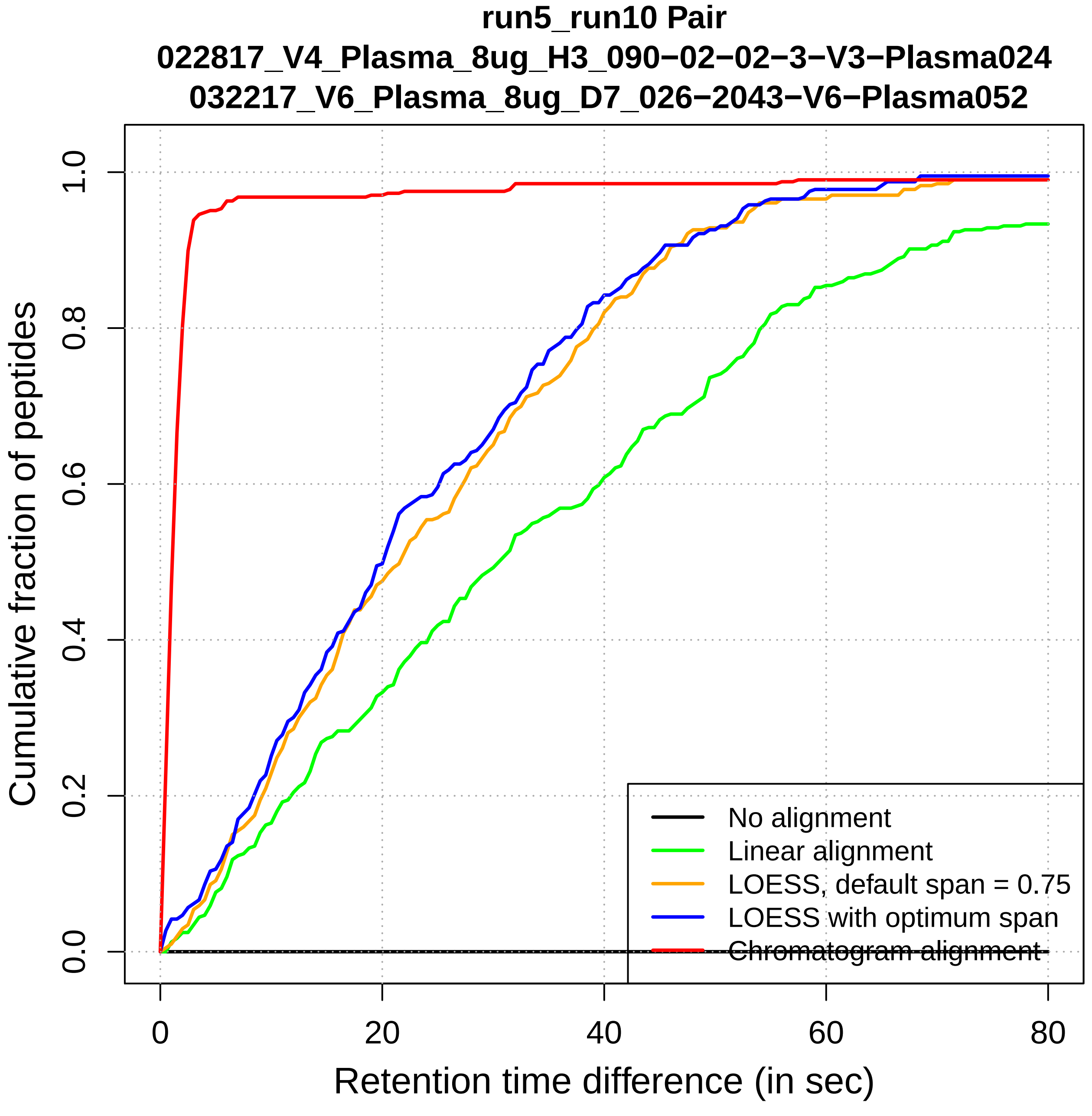

run5_run10 Pair
022817_V4_Plasma_8ug_H3_090−02−02−3−V3−Plasma024
032217_V6_Plasma_8ug_D7_026−2043−V6−Plasma052
Cumulative fraction of peptides
0.0
0.2
0.4
0.6
0.8
1.0
0
20
40
60
80
Retention time difference (in sec)
No alignment
Linear alignment
LOESS, default span = 0.75
LOESS with optimum span
Chromatogram alignment

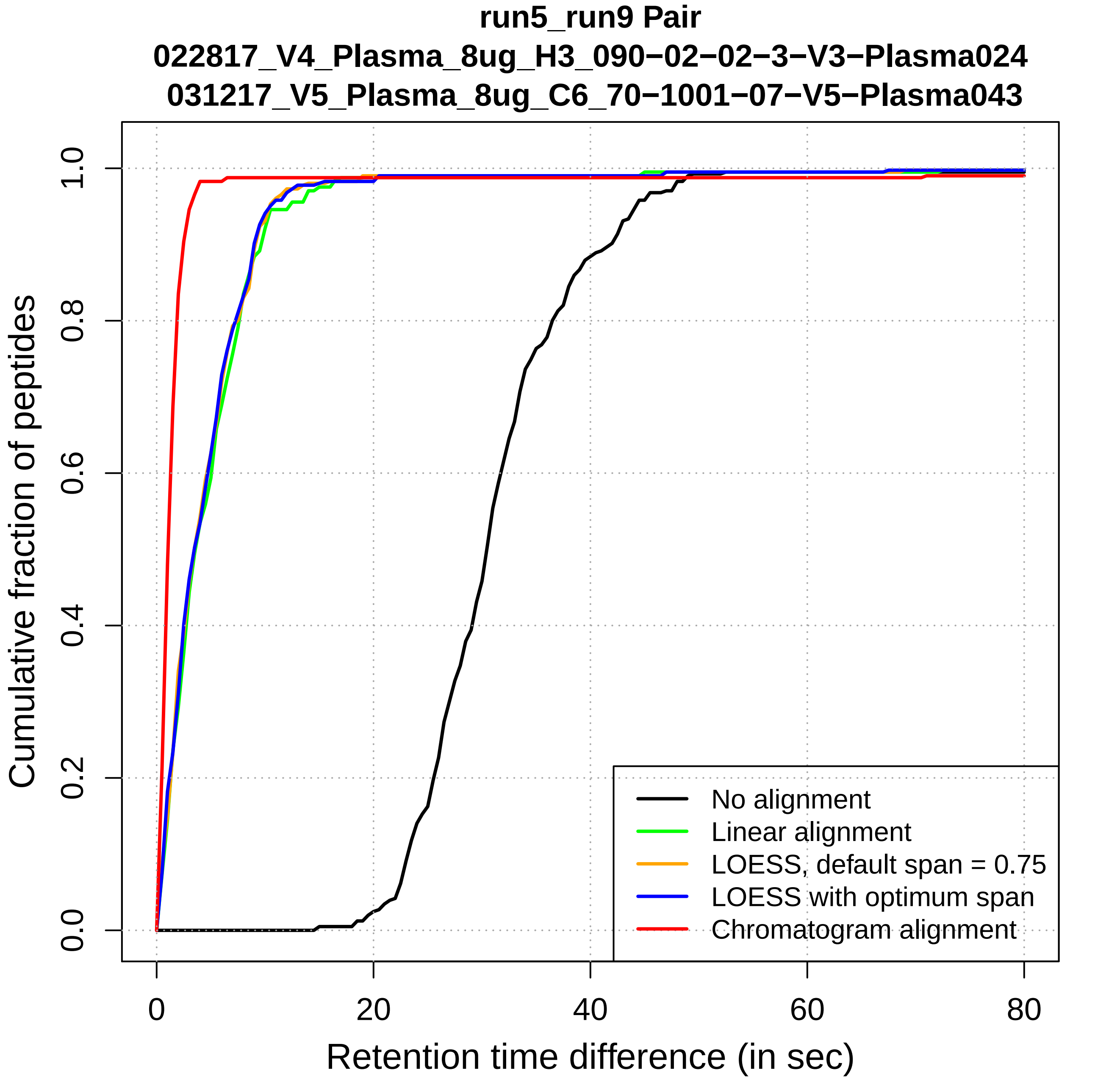
run5_run9 Pair
022817_V4_Plasma_8ug_H3_090−02−02−3−V3−Plasma024
031217_V5_Plasma_8ug_C6_70−1001−07−V5−Plasma043
Cumulative fraction of peptides
0.0
0.2
0.4
0.6
0.8
1.0
0
20
40
60
80
Retention time difference (in sec)
No alignment
Linear alignment
LOESS, default span = 0.75
LOESS with optimum span
Chromatogram alignment

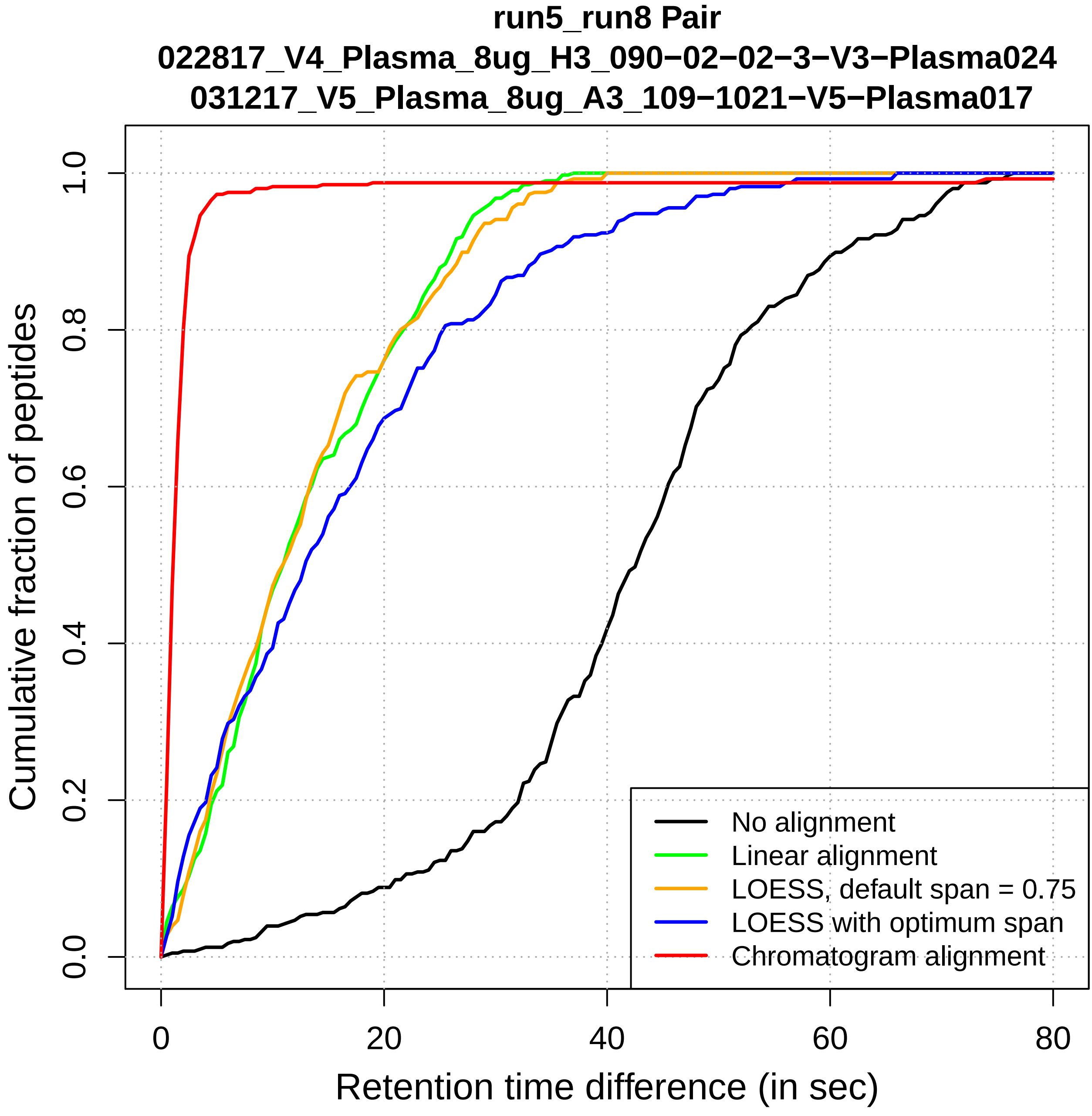
run5_run8 Pair
022817_V4_Plasma_8ug_H3_090−02−02−3−V3−Plasma024
031217_V5_Plasma_8ug_A3_109−1021−V5−Plasma017
Cumulative fraction of peptides
0.0
0.2
0.4
0.6
0.8
1.0
0
20
40
60
80
Retention time difference (in sec)
No alignment
Linear alignment
LOESS, default span = 0.75
LOESS with optimum span
Chromatogram alignment

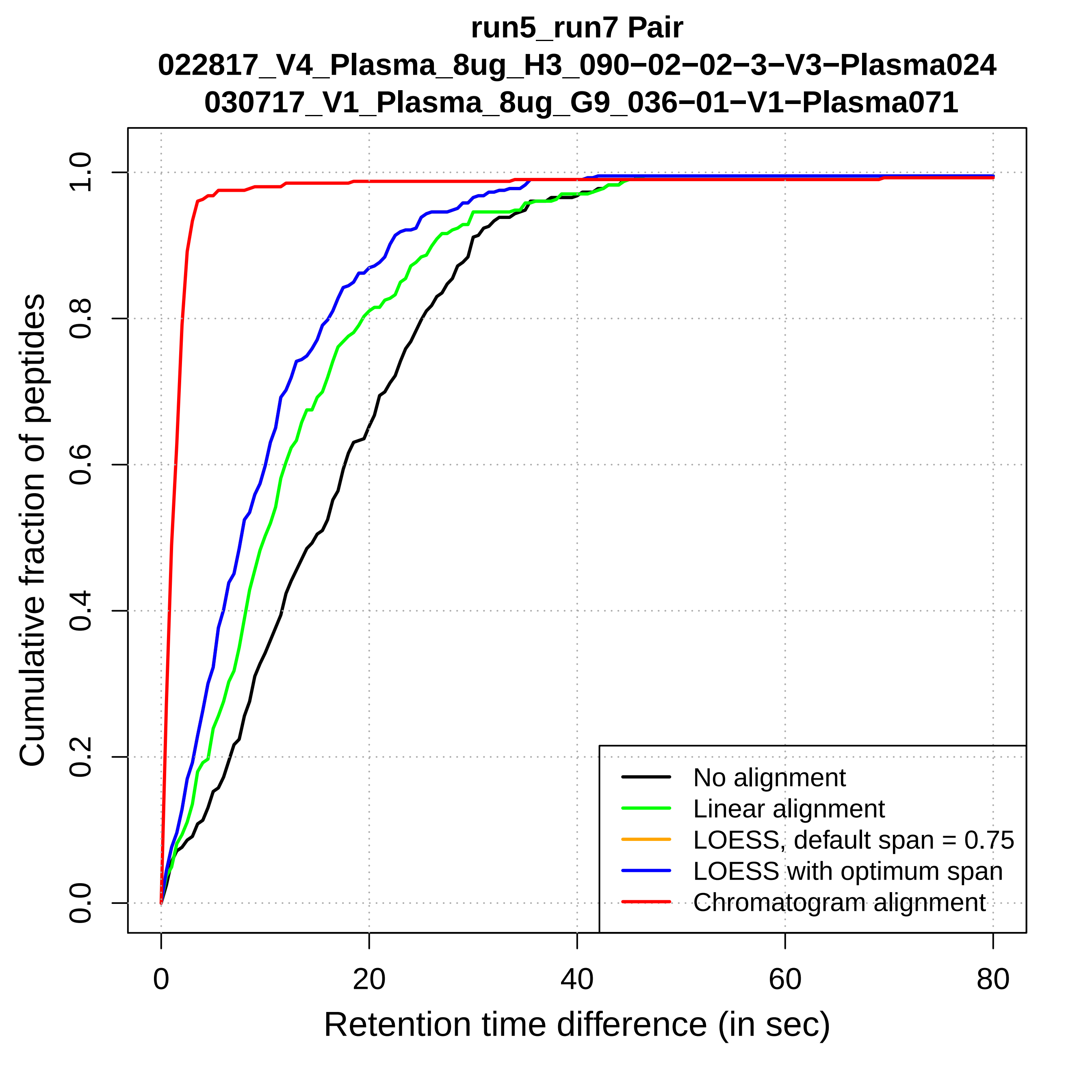

run5_run7 Pair
022817_V4_Plasma_8ug_H3_090−02−02−3−V3−Plasma024
030717_V1_Plasma_8ug_G9_036−01−V1−Plasma071
Cumulative fraction of peptides
0.0
0.2
0.4
0.6
0.8
1.0
0
20
40
60
80
Retention time difference (in sec)
No alignment
Linear alignment
LOESS, default span = 0.75
LOESS with optimum span
Chromatogram alignment

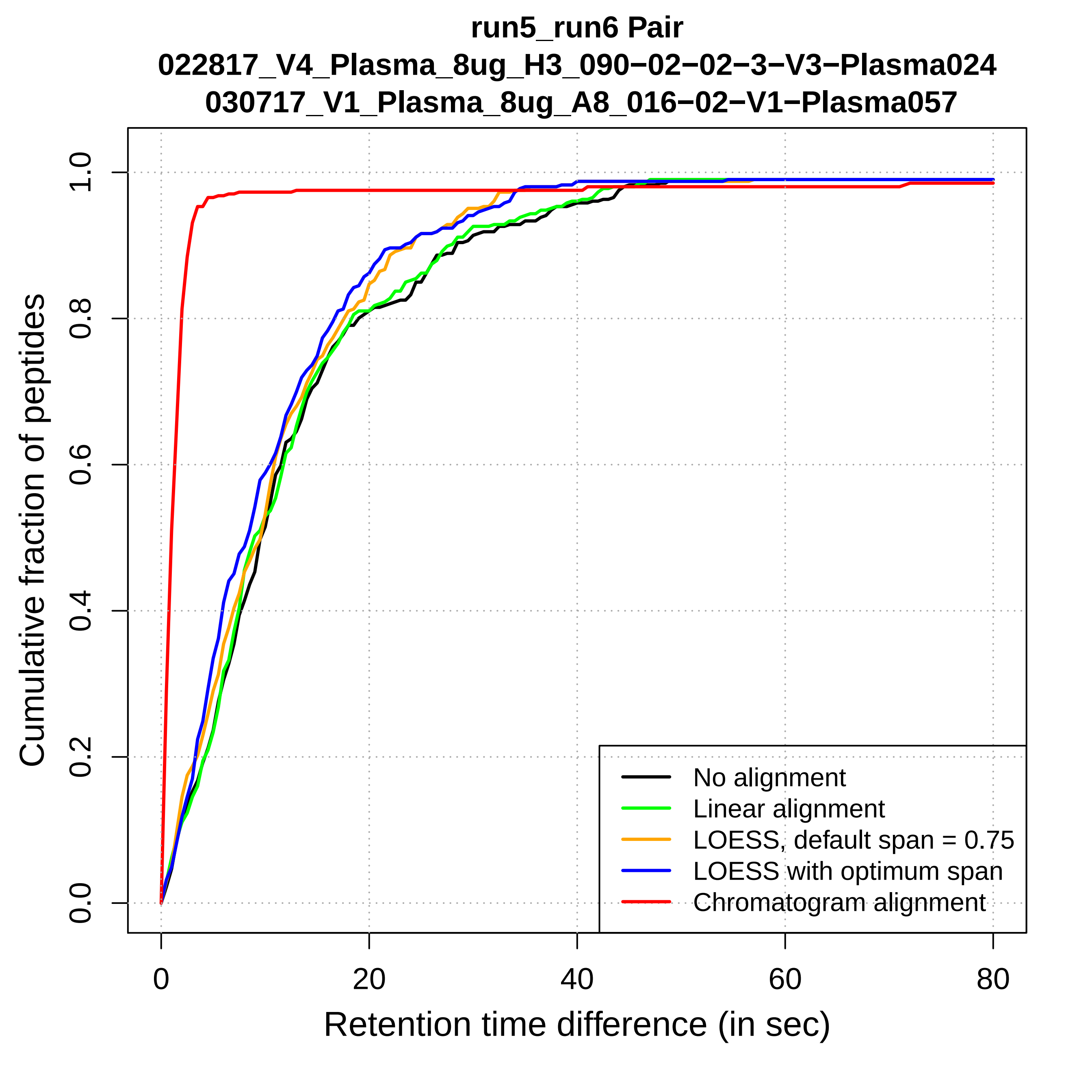

run5_run6 Pair
022817_V4_Plasma_8ug_H3_090−02−02−3−V3−Plasma024
030717_V1_Plasma_8ug_A8_016−02−V1−Plasma057
Cumulative fraction of peptides
0.0
0.2
0.4
0.6
0.8
1.0
0
20
40
60
80
Retention time difference (in sec)
No alignment
Linear alignment
LOESS, default span = 0.75
LOESS with optimum span
Chromatogram alignment

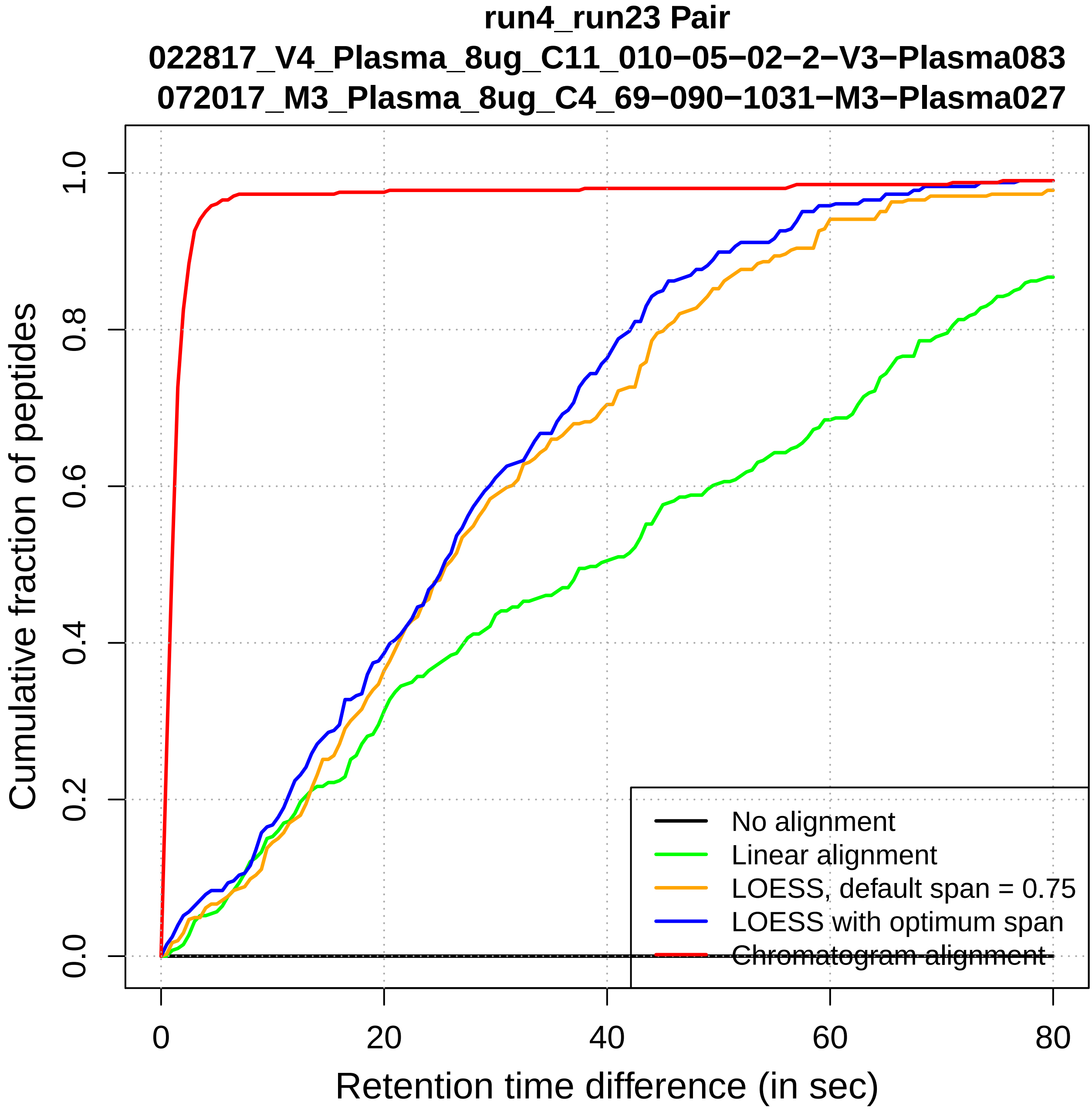

run4_run23 Pair
022817_V4_Plasma_8ug_C11_010−05−02−2−V3−Plasma083
072017_M3_Plasma_8ug_C4_69−090−1031−M3−Plasma027
Cumulative fraction of peptides
0.0
0.2
0.4
0.6
0.8
1.0
0
20
40
60
80
Retention time difference (in sec)
No alignment
Linear alignment
LOESS, default span = 0.75
LOESS with optimum span
Chromatogram alignment

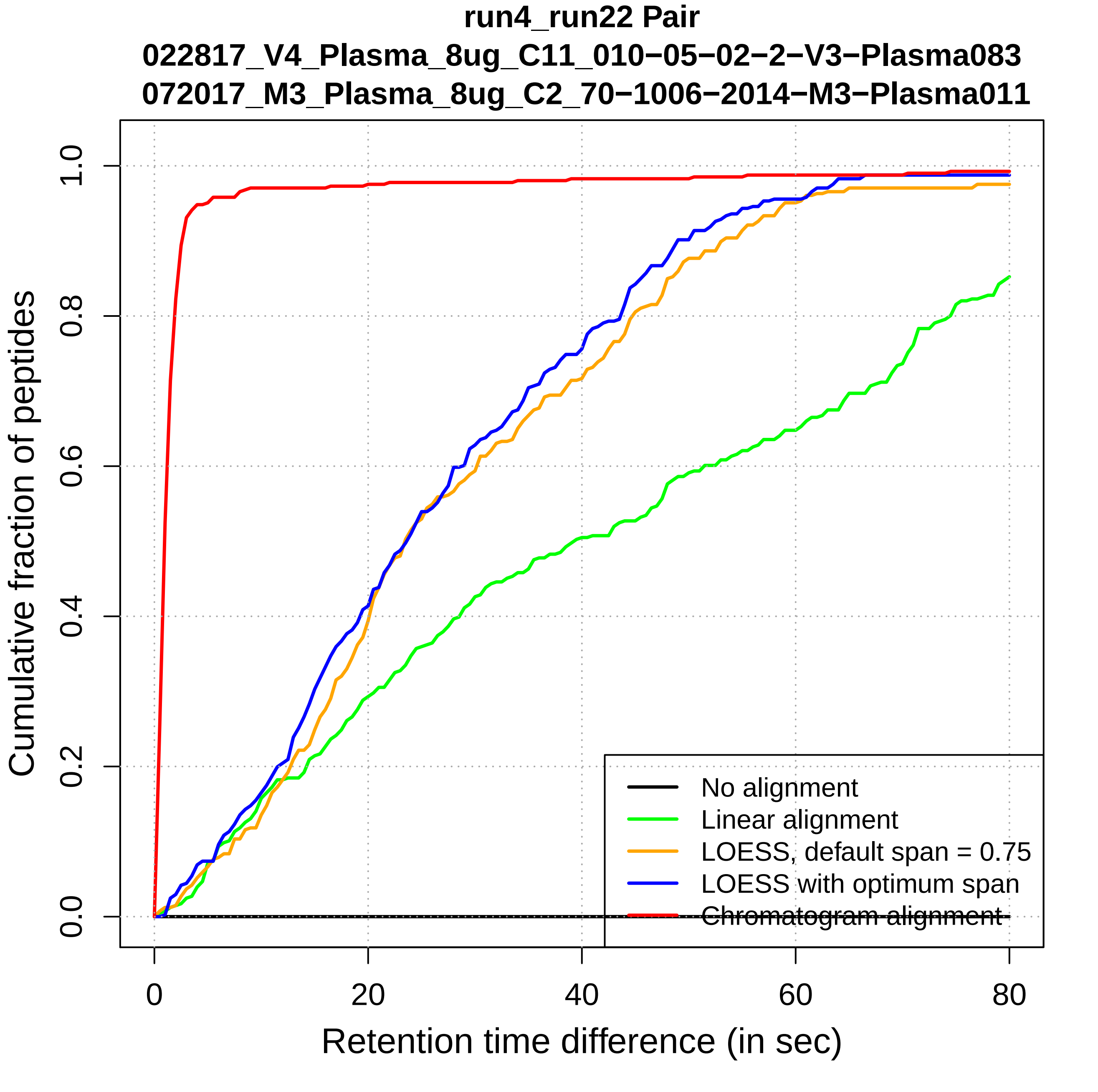

run4_run22 Pair
022817_V4_Plasma_8ug_C11_010−05−02−2−V3−Plasma083
072017_M3_Plasma_8ug_C2_70−1006−2014−M3−Plasma011
Cumulative fraction of peptides
0.0
0.2
0.4
0.6
0.8
1.0
0
20
40
60
80
Retention time difference (in sec)
No alignment
Linear alignment
LOESS, default span = 0.75
LOESS with optimum span
Chromatogram alignment

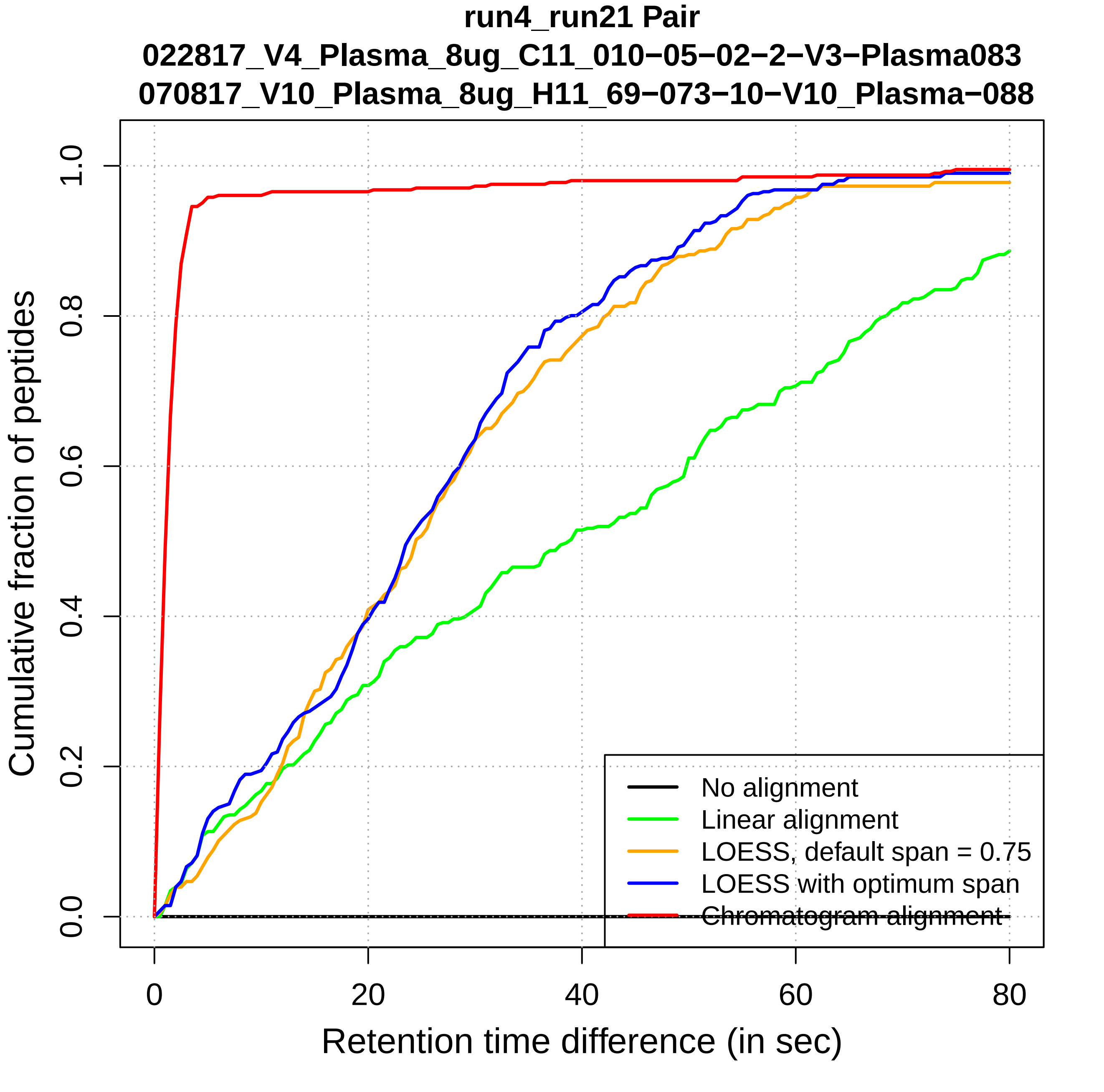

run4_run21 Pair
022817_V4_Plasma_8ug_C11_010−05−02−2−V3−Plasma083
070817_V10_Plasma_8ug_H11_69−073−10−V10_Plasma−088
Cumulative fraction of peptides
0.0
0.2
0.4
0.6
0.8
1.0
0
20
40
60
80
Retention time difference (in sec)
No alignment
Linear alignment
LOESS, default span = 0.75
LOESS with optimum span
Chromatogram alignment

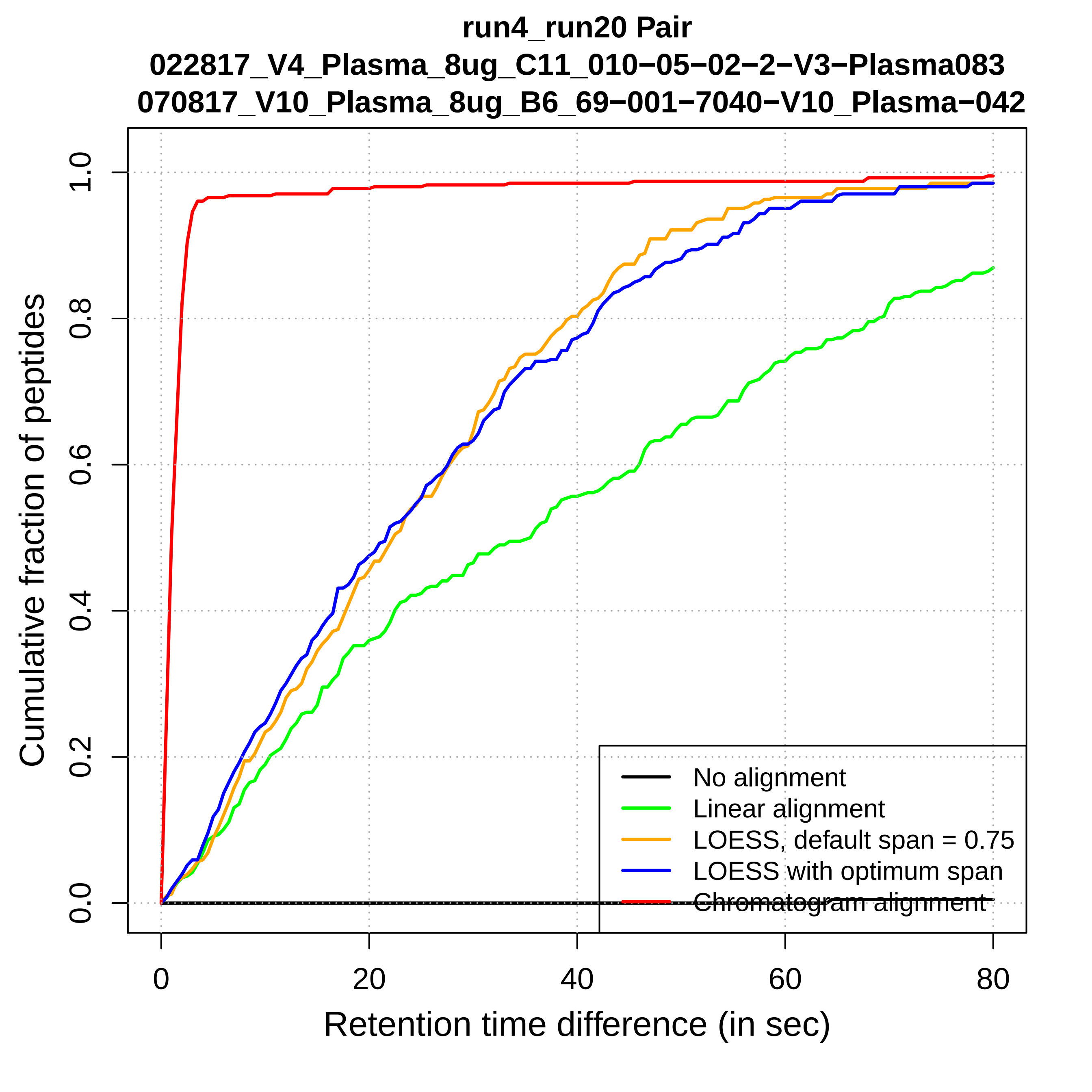

run4_run20 Pair
022817_V4_Plasma_8ug_C11_010−05−02−2−V3−Plasma083
070817_V10_Plasma_8ug_B6_69−001−7040−V10_Plasma−042
Cumulative fraction of peptides
0.0
0.2
0.4
0.6
0.8
1.0
0
20
40
60
80
Retention time difference (in sec)
No alignment
Linear alignment
LOESS, default span = 0.75
LOESS with optimum span
Chromatogram alignment

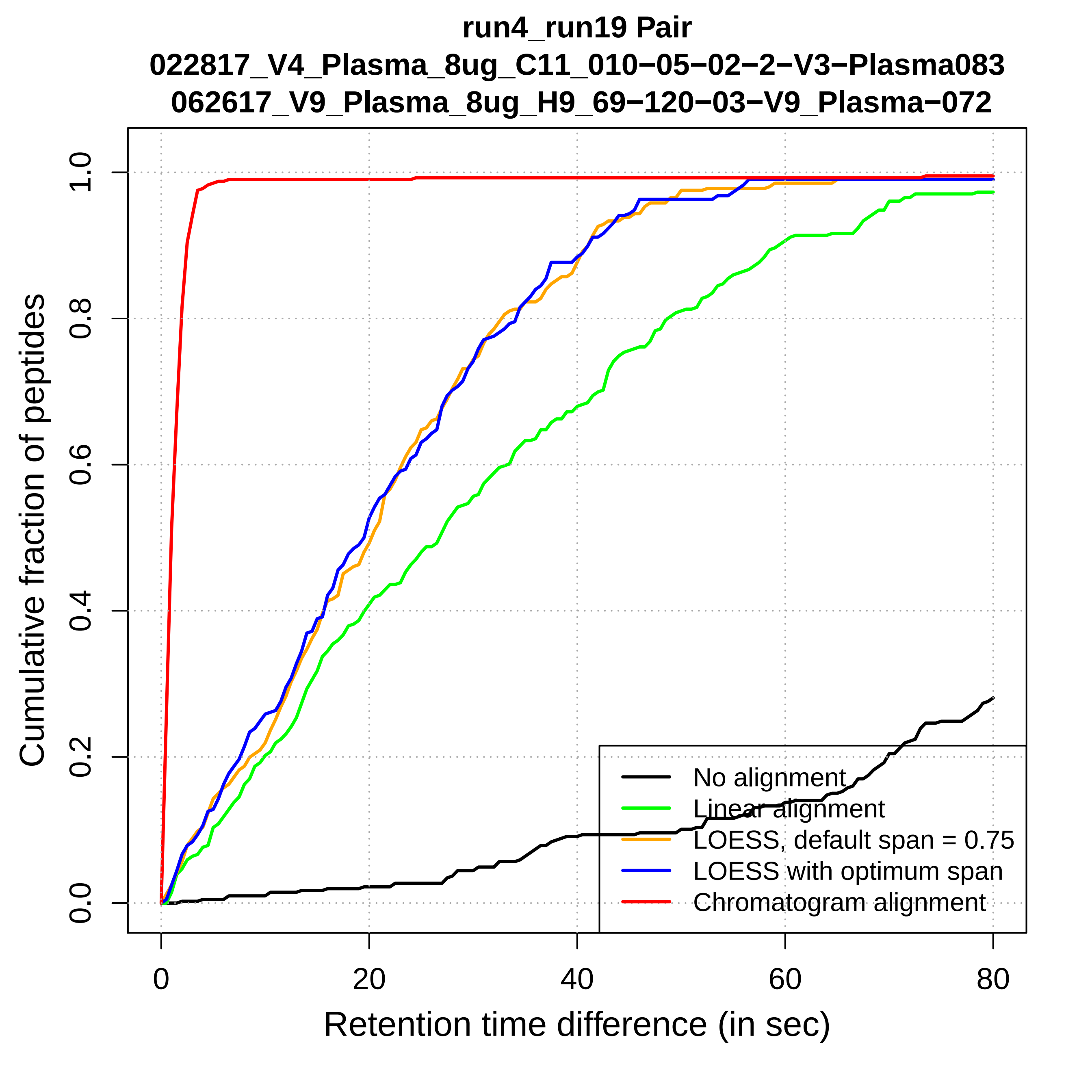

run4_run19 Pair
022817_V4_Plasma_8ug_C11_010−05−02−2−V3−Plasma083
062617_V9_Plasma_8ug_H9_69−120−03−V9_Plasma−072
Cumulative fraction of peptides
0.0
0.2
0.4
0.6
0.8
1.0
0
20
40
60
80
Retention time difference (in sec)
No alignment
Linear alignment
LOESS, default span = 0.75
LOESS with optimum span
Chromatogram alignment

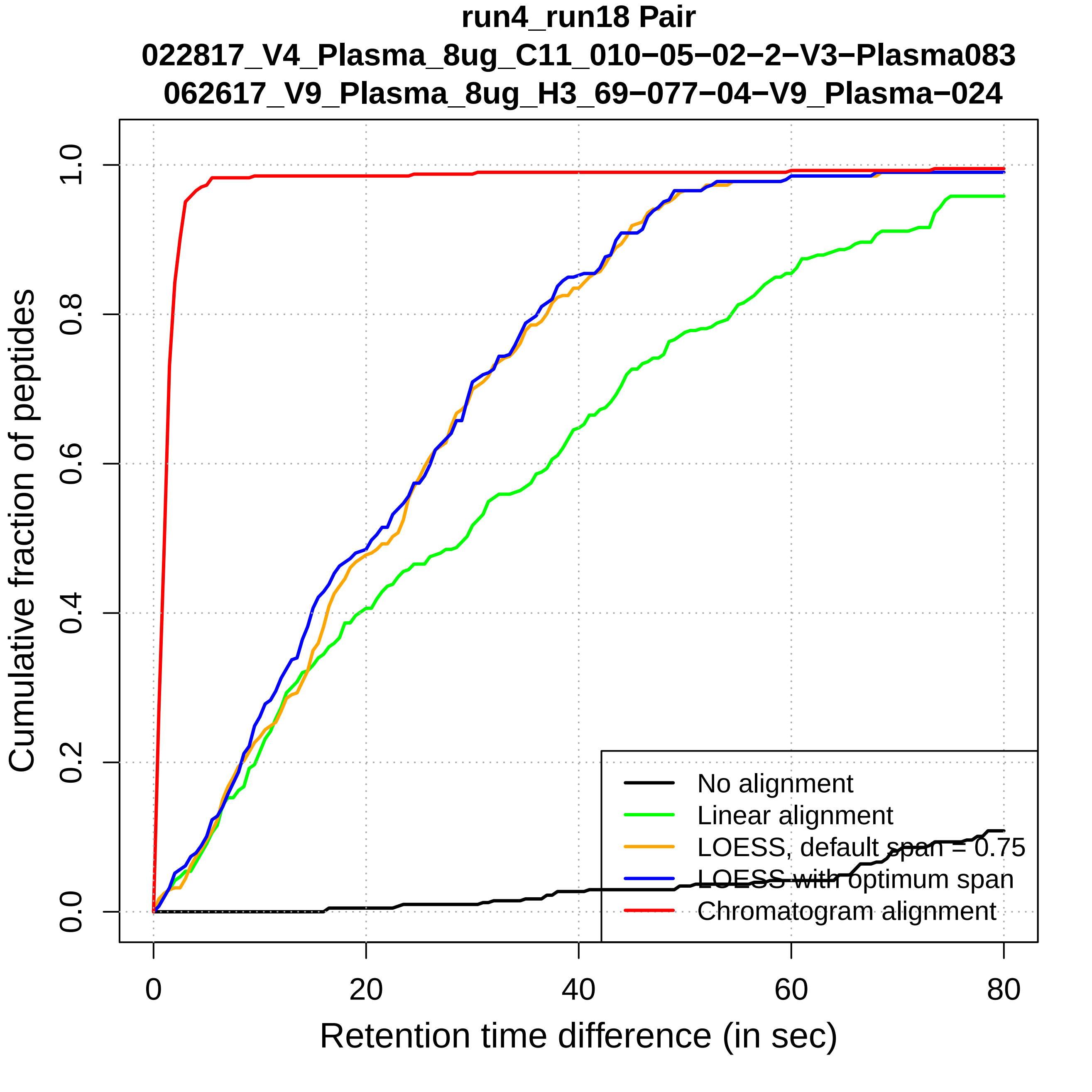

run4_run18 Pair
022817_V4_Plasma_8ug_C11_010−05−02−2−V3−Plasma083
062617_V9_Plasma_8ug_H3_69−077−04−V9_Plasma−024
Cumulative fraction of peptides
0.0
0.2
0.4
0.6
0.8
1.0
0
20
40
60
80
Retention time difference (in sec)
No alignment
Linear alignment
LOESS, default span = 0.75
LOESS with optimum span
Chromatogram alignment

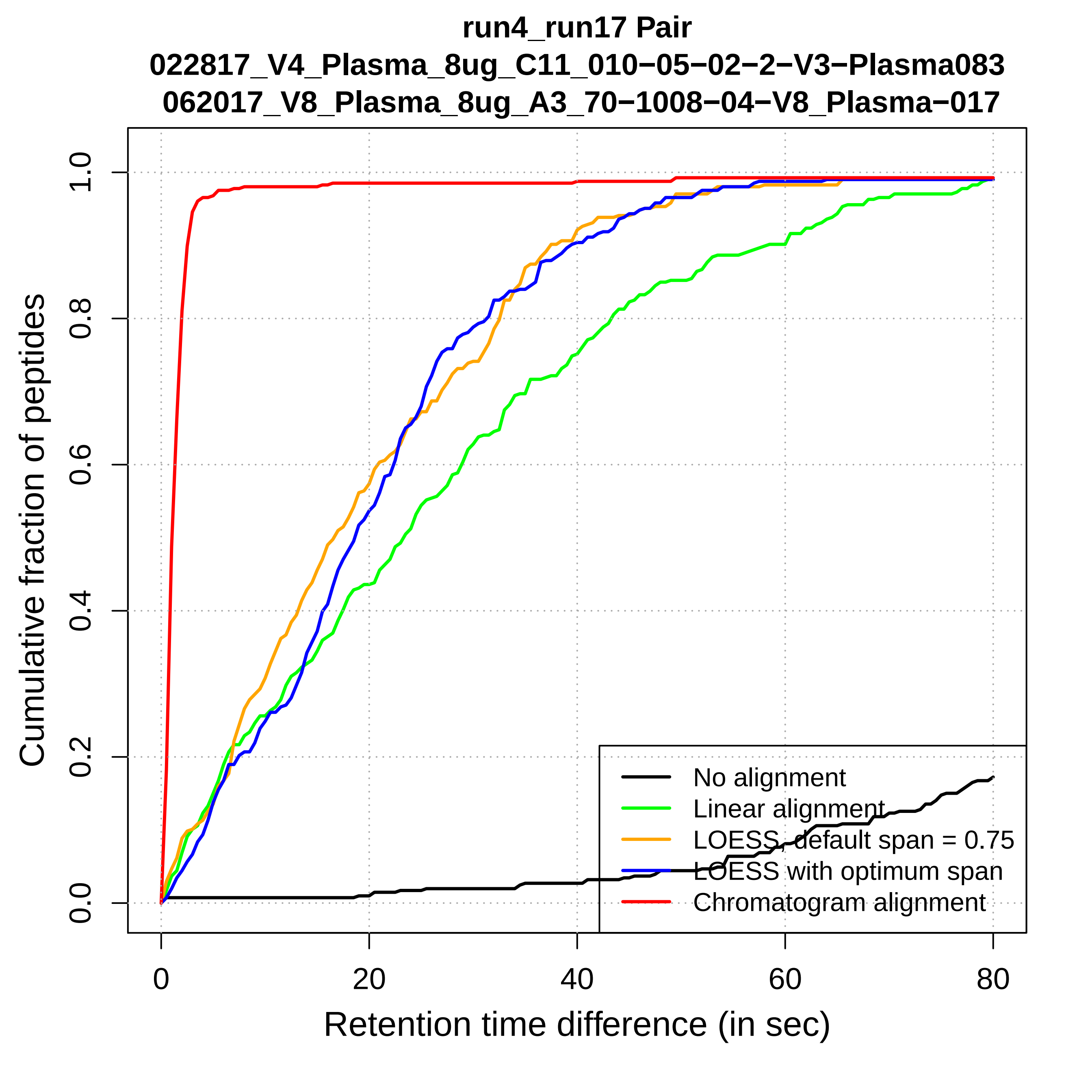
run4_run17 Pair
022817_V4_Plasma_8ug_C11_010−05−02−2−V3−Plasma083
062017_V8_Plasma_8ug_A3_70−1008−04−V8_Plasma−017
Cumulative fraction of peptides
Retention time difference (in sec)
0.0
0.2
0.4
0.6
0.8
1.0
0
20
40
60
80
No alignment
Linear alignment
LOESS, default span = 0.75
LOESS with optimum span
Chromatogram alignment

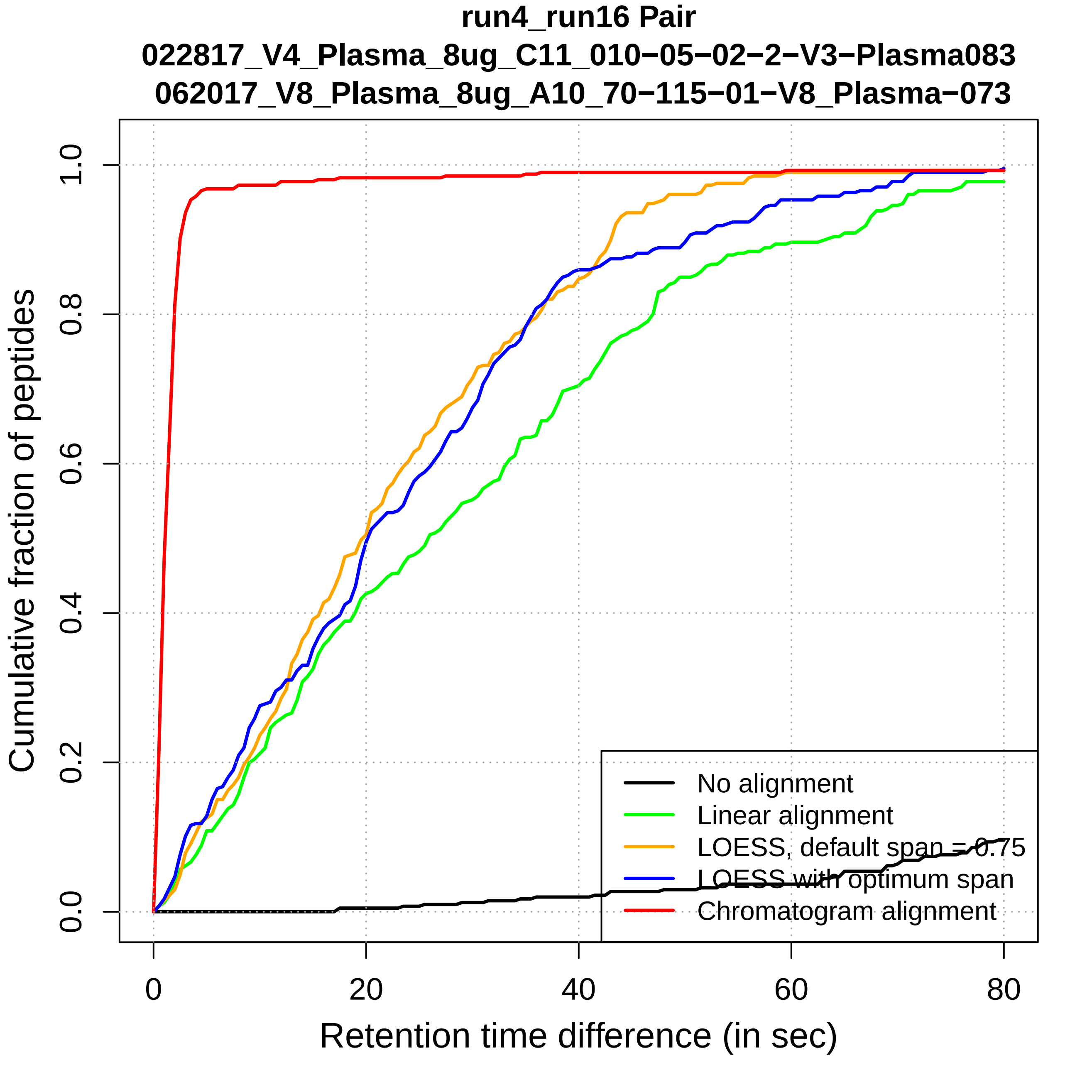

run4_run16 Pair
022817_V4_Plasma_8ug_C11_010−05−02−2−V3−Plasma083
062017_V8_Plasma_8ug_A10_70−115−01−V8_Plasma−073
Cumulative fraction of peptides
0.0
0.2
0.4
0.6
0.8
1.0
0
20
40
60
80
Retention time difference (in sec)
No alignment
Linear alignment
LOESS, default span = 0.75
LOESS with optimum span
Chromatogram alignment

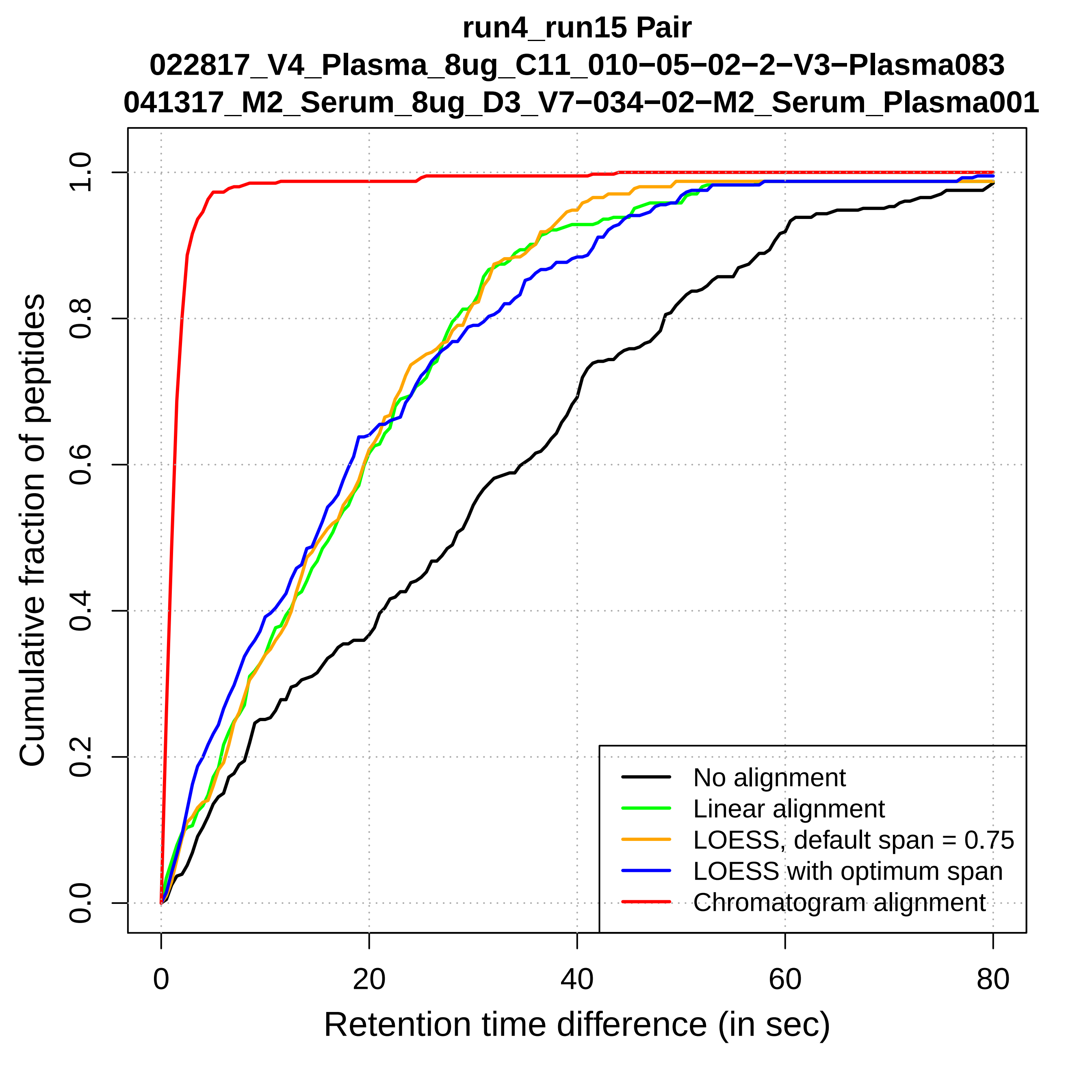
run4_run15 Pair
022817_V4_Plasma_8ug_C11_010−05−02−2−V3−Plasma083
041317_M2_Serum_8ug_D3_V7−034−02−M2_Serum_Plasma001
Cumulative fraction of peptides
0.0
0.2
0.4
0.6
0.8
1.0
0
20
40
60
80
Retention time difference (in sec)
No alignment
Linear alignment
LOESS, default span = 0.75
LOESS with optimum span
Chromatogram alignment

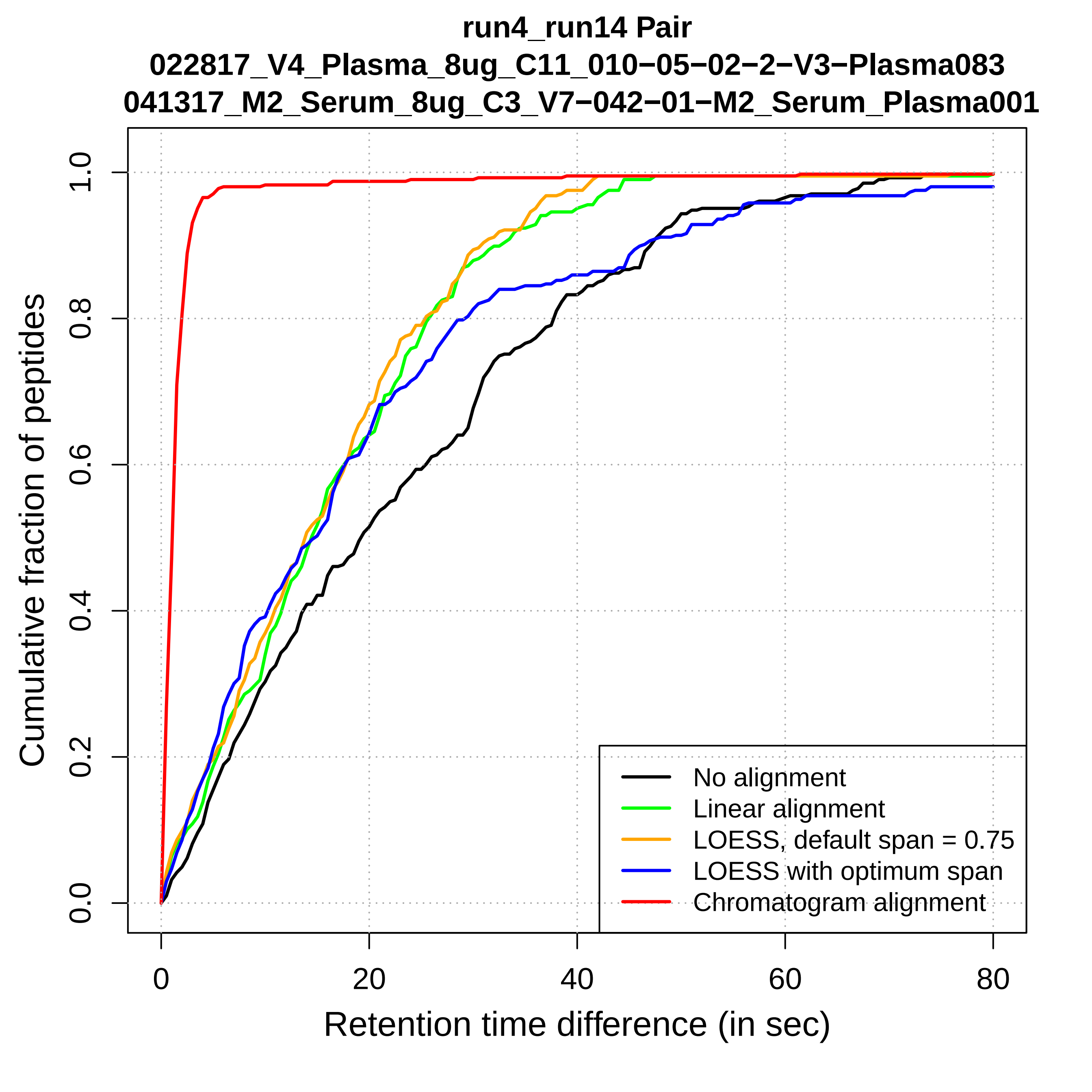
run4_run14 Pair
022817_V4_Plasma_8ug_C11_010−05−02−2−V3−Plasma083
041317_M2_Serum_8ug_C3_V7−042−01−M2_Serum_Plasma001
Cumulative fraction of peptides
0.0
0.2
0.4
0.6
0.8
1.0
0
20
40
60
80
Retention time difference (in sec)
No alignment
Linear alignment
LOESS, default span = 0.75
LOESS with optimum span
Chromatogram alignment

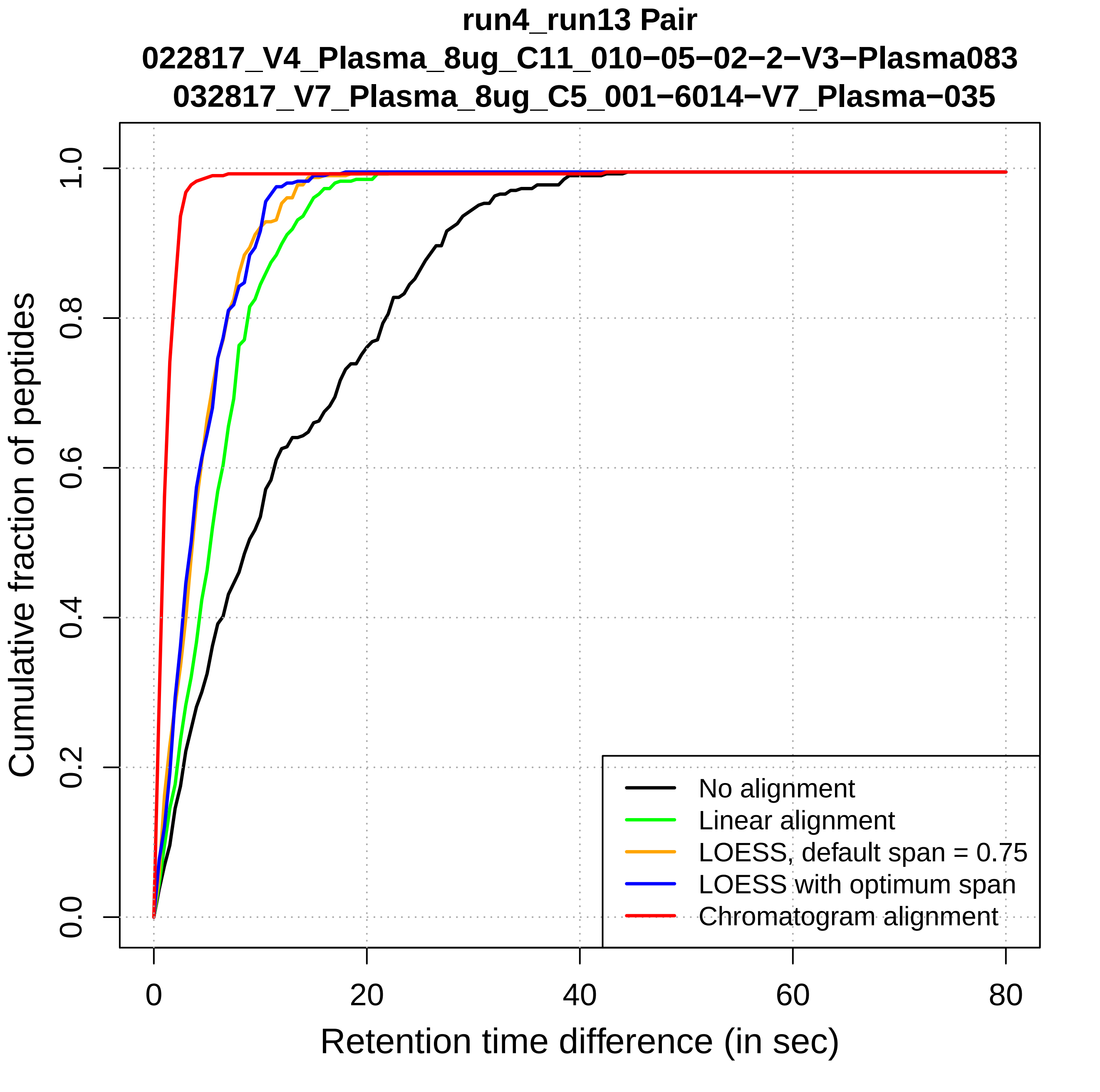
run4_run13 Pair
022817_V4_Plasma_8ug_C11_010−05−02−2−V3−Plasma083
032817_V7_Plasma_8ug_C5_001−6014−V7_Plasma−035
Cumulative fraction of peptides
0.0
0.2
0.4
0.6
0.8
1.0
0
20
40
60
80
Retention time difference (in sec)
No alignment
Linear alignment
LOESS, default span = 0.75
LOESS with optimum span
Chromatogram alignment

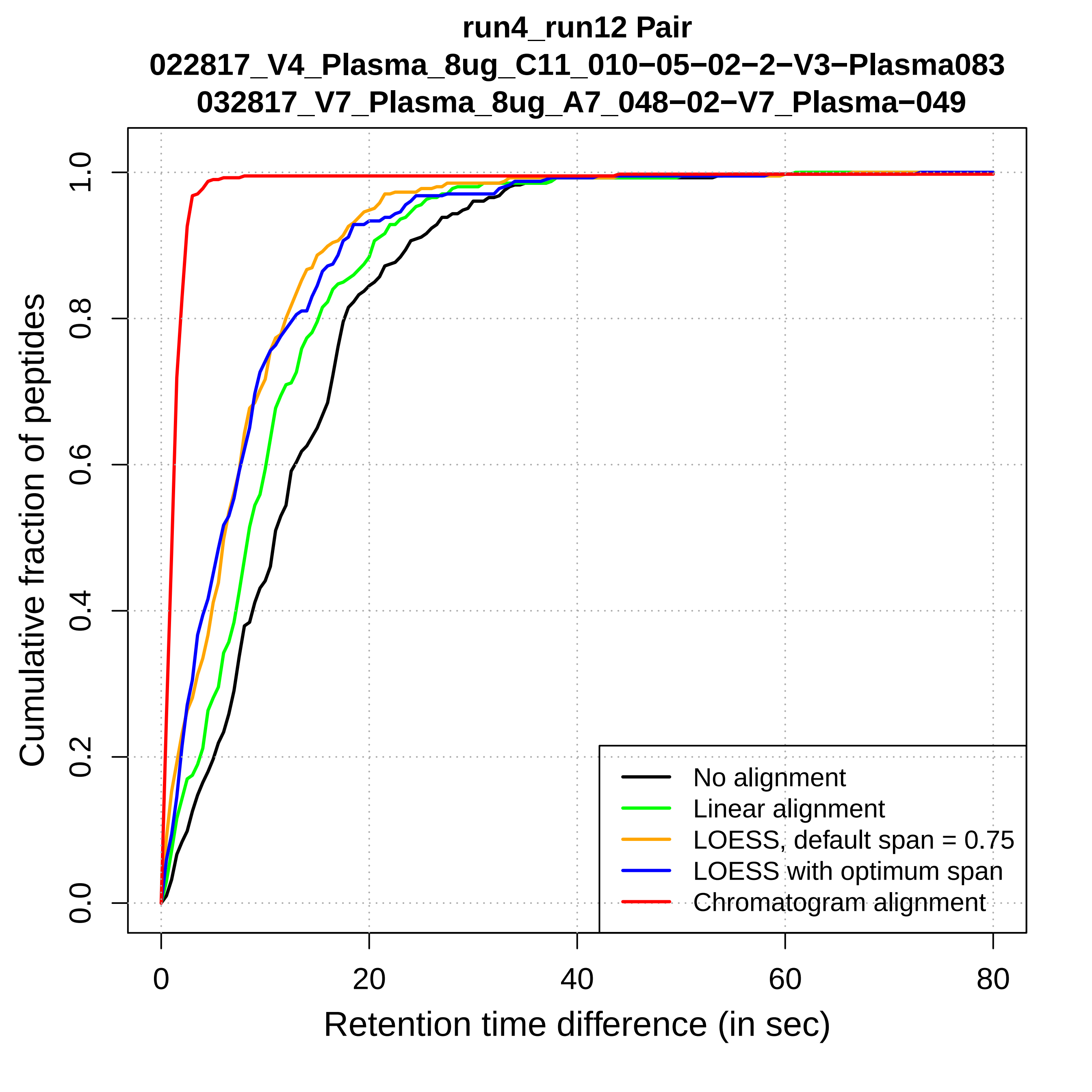
run4_run12 Pair
022817_V4_Plasma_8ug_C11_010−05−02−2−V3−Plasma083
032817_V7_Plasma_8ug_A7_048−02−V7_Plasma−049
Cumulative fraction of peptides
0.0
0.2
0.4
0.6
0.8
1.0
0
20
40
60
80
Retention time difference (in sec)
No alignment
Linear alignment
LOESS, default span = 0.75
LOESS with optimum span
Chromatogram alignment

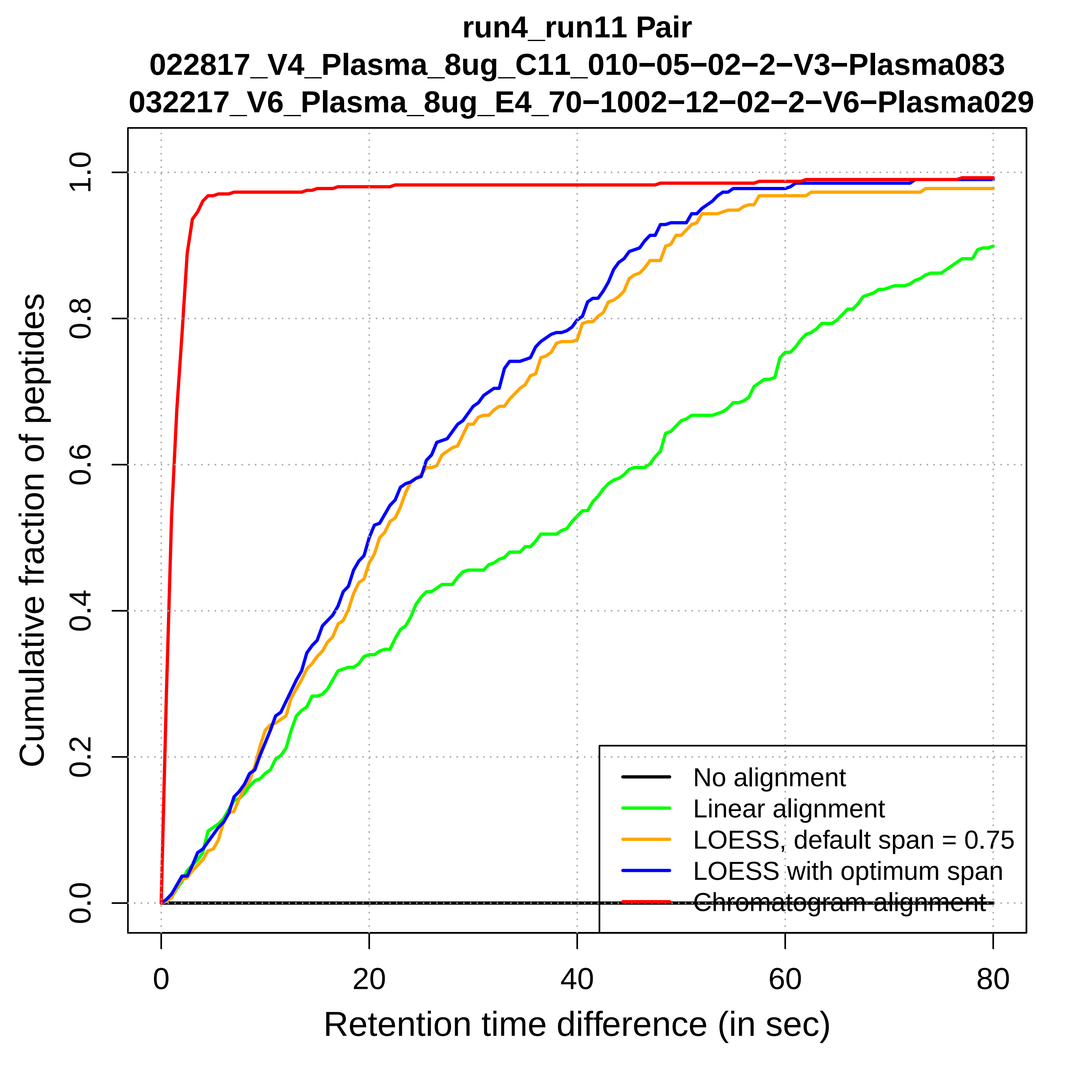
run4_run11 Pair
022817_V4_Plasma_8ug_C11_010−05−02−2−V3−Plasma083
032217_V6_Plasma_8ug_E4_70−1002−12−02−2−V6−Plasma029
Cumulative fraction of peptides
0.0
0.2
0.4
0.6
0.8
1.0
0
20
40
60
80
Retention time difference (in sec)
No alignment
Linear alignment
LOESS, default span = 0.75
LOESS with optimum span
Chromatogram alignment

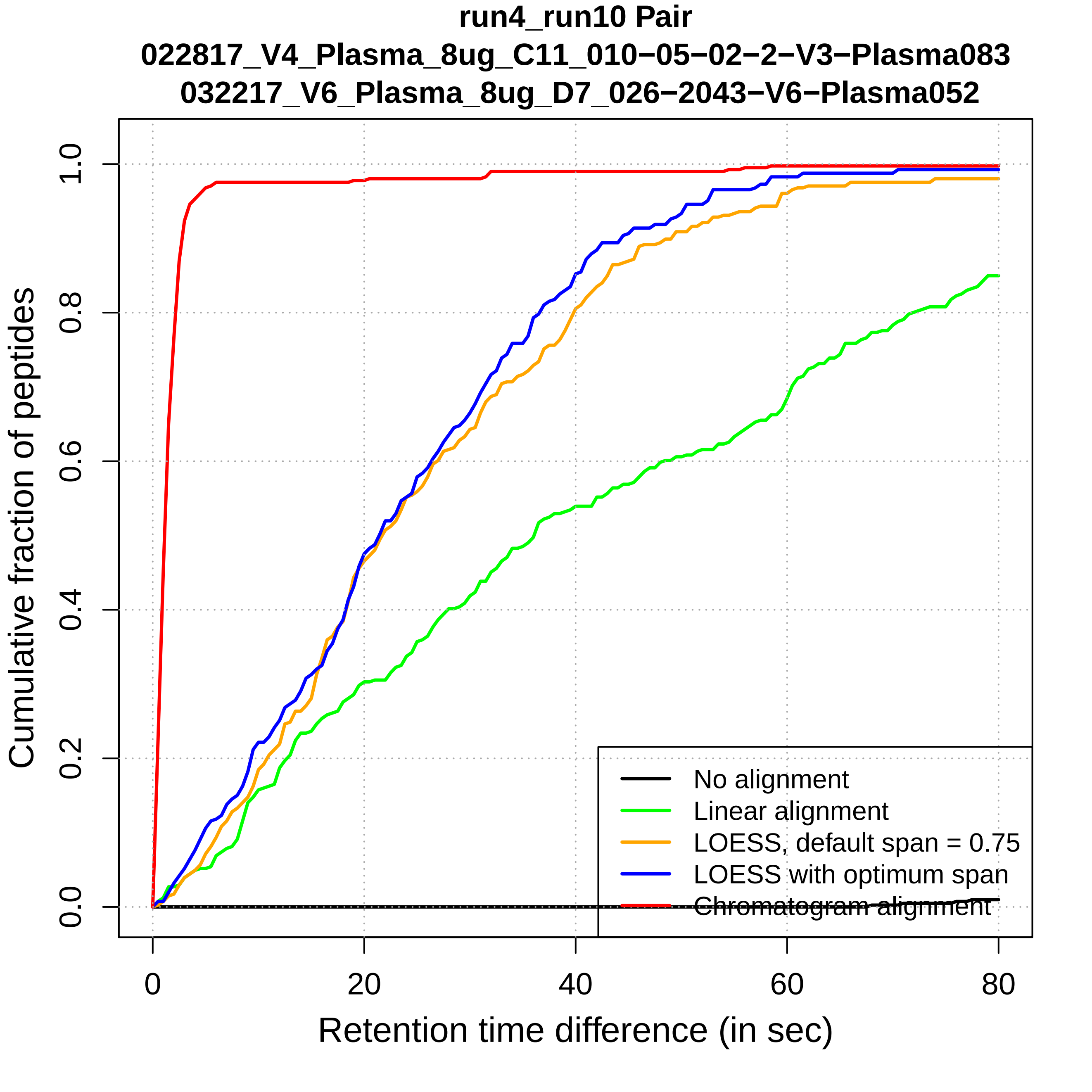

run4_run10 Pair
022817_V4_Plasma_8ug_C11_010−05−02−2−V3−Plasma083
032217_V6_Plasma_8ug_D7_026−2043−V6−Plasma052
Cumulative fraction of peptides
0.0
0.2
0.4
0.6
0.8
1.0
0
20
40
60
80
Retention time difference (in sec)
No alignment
Linear alignment
LOESS, default span = 0.75
LOESS with optimum span
Chromatogram alignment

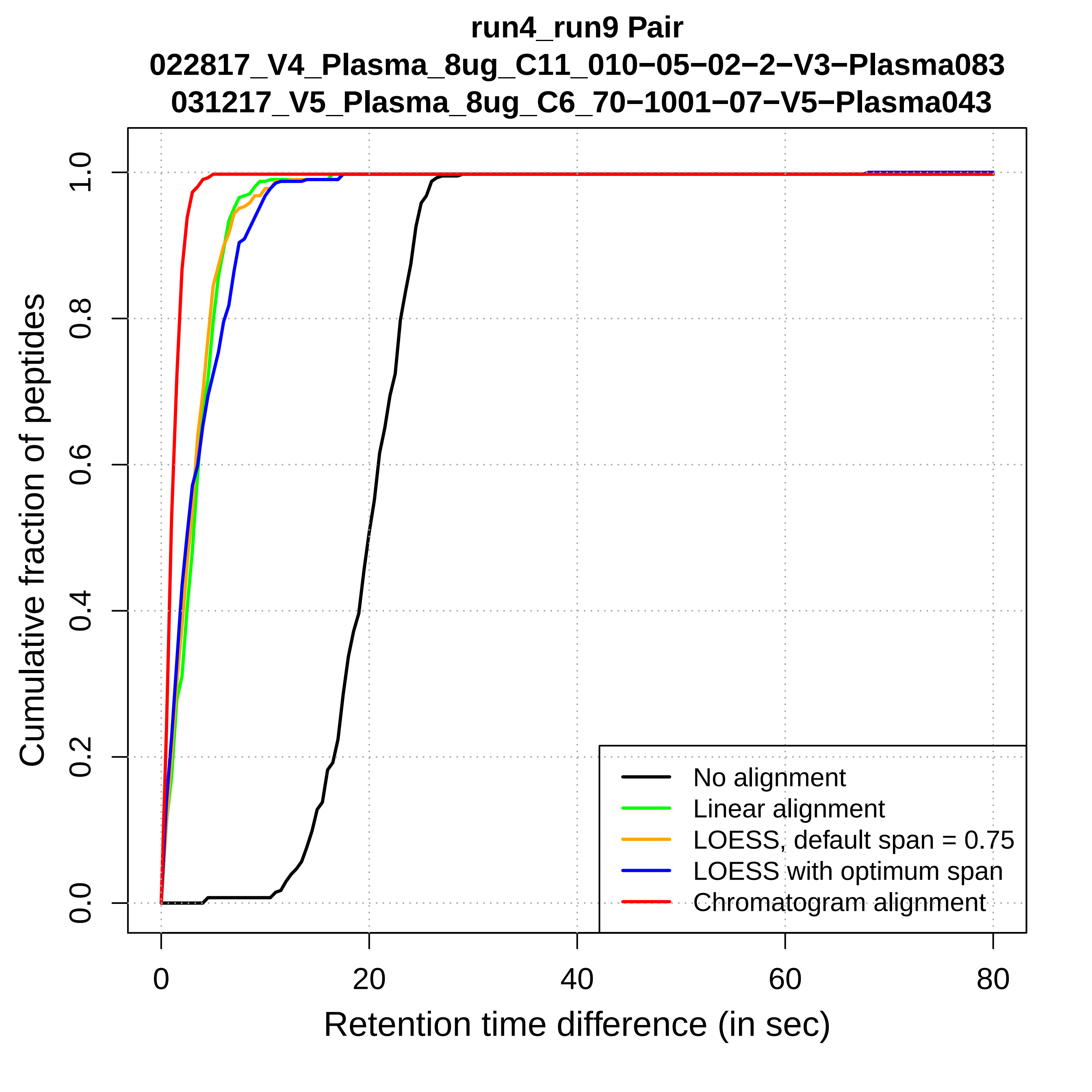

run4_run9 Pair
022817_V4_Plasma_8ug_C11_010−05−02−2−V3−Plasma083
031217_V5_Plasma_8ug_C6_70−1001−07−V5−Plasma043
Cumulative fraction of peptides
0.0
0.2
0.4
0.6
0.8
1.0
0
20
40
60
80
Retention time difference (in sec)
No alignment
Linear alignment
LOESS, default span = 0.75
LOESS with optimum span
Chromatogram alignment

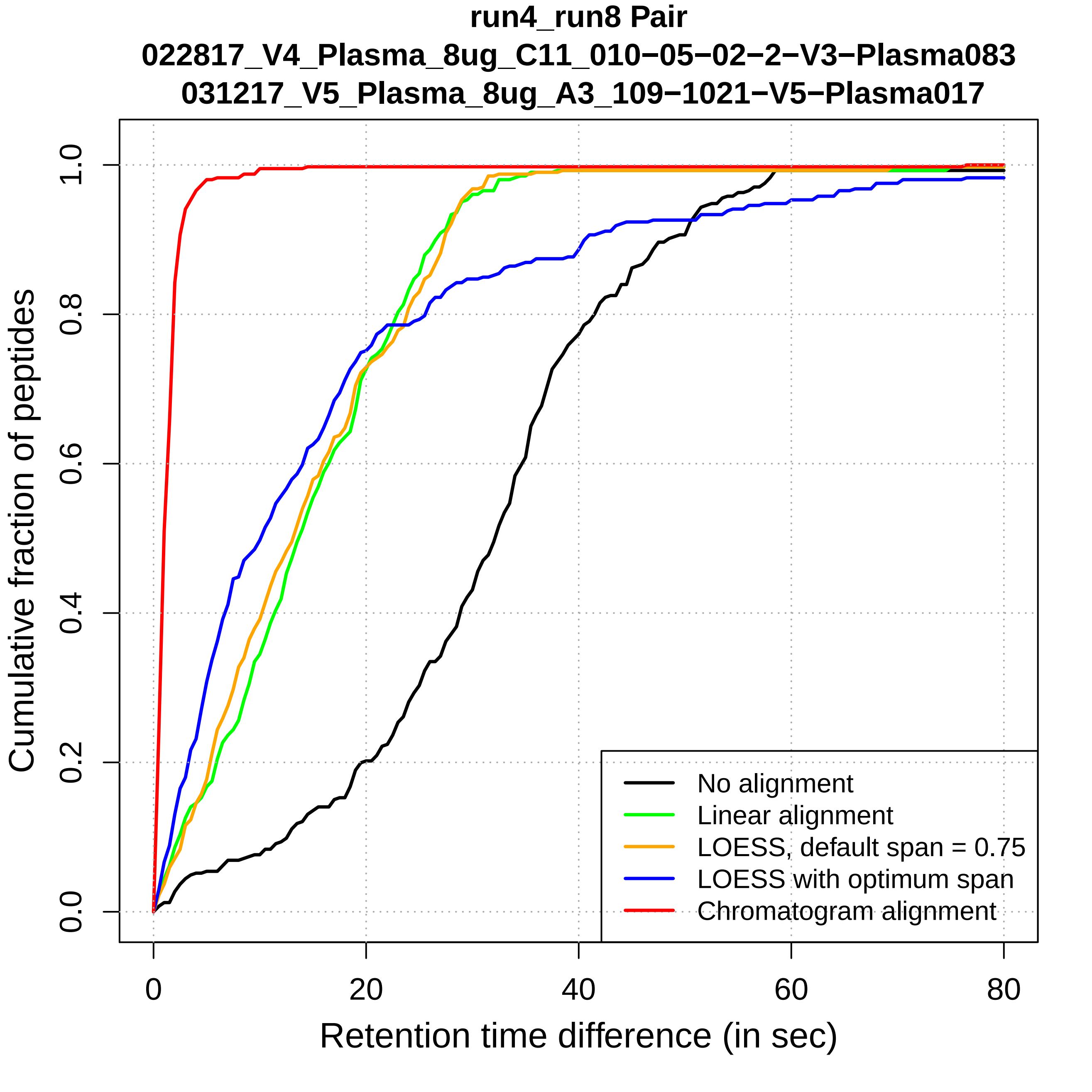
run4_run8 Pair
022817_V4_Plasma_8ug_C11_010−05−02−2−V3−Plasma083
031217_V5_Plasma_8ug_A3_109−1021−V5−Plasma017
Cumulative fraction of peptides
0.0
0.2
0.4
0.6
0.8
1.0
0
20
40
60
80
Retention time difference (in sec)
No alignment
Linear alignment
LOESS, default span = 0.75
LOESS with optimum span
Chromatogram alignment

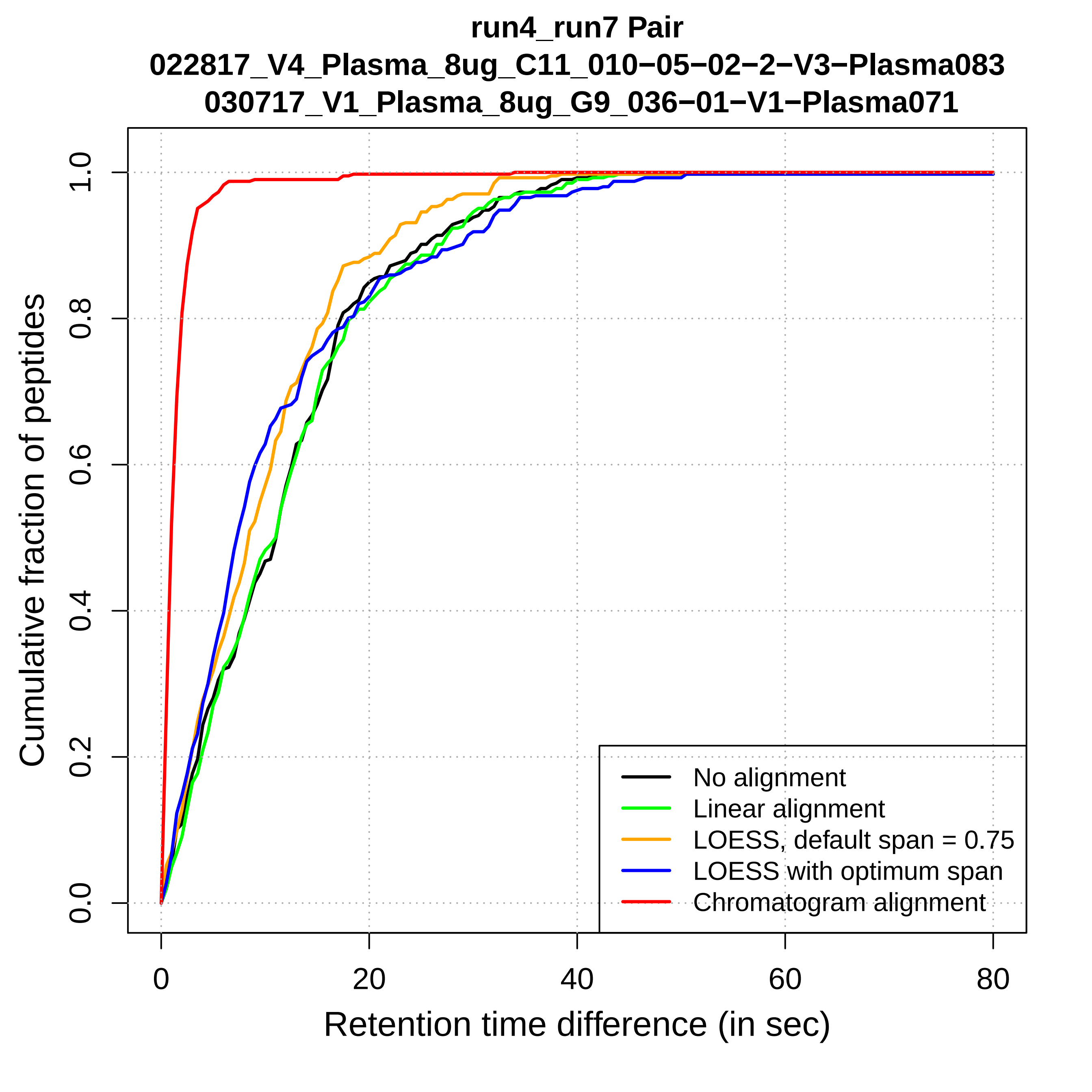

run4_run7 Pair
022817_V4_Plasma_8ug_C11_010−05−02−2−V3−Plasma083
030717_V1_Plasma_8ug_G9_036−01−V1−Plasma071
Cumulative fraction of peptides
0.0
0.2
0.4
0.6
0.8
1.0
0
20
40
60
80
Retention time difference (in sec)
No alignment
Linear alignment
LOESS, default span = 0.75
LOESS with optimum span
Chromatogram alignment

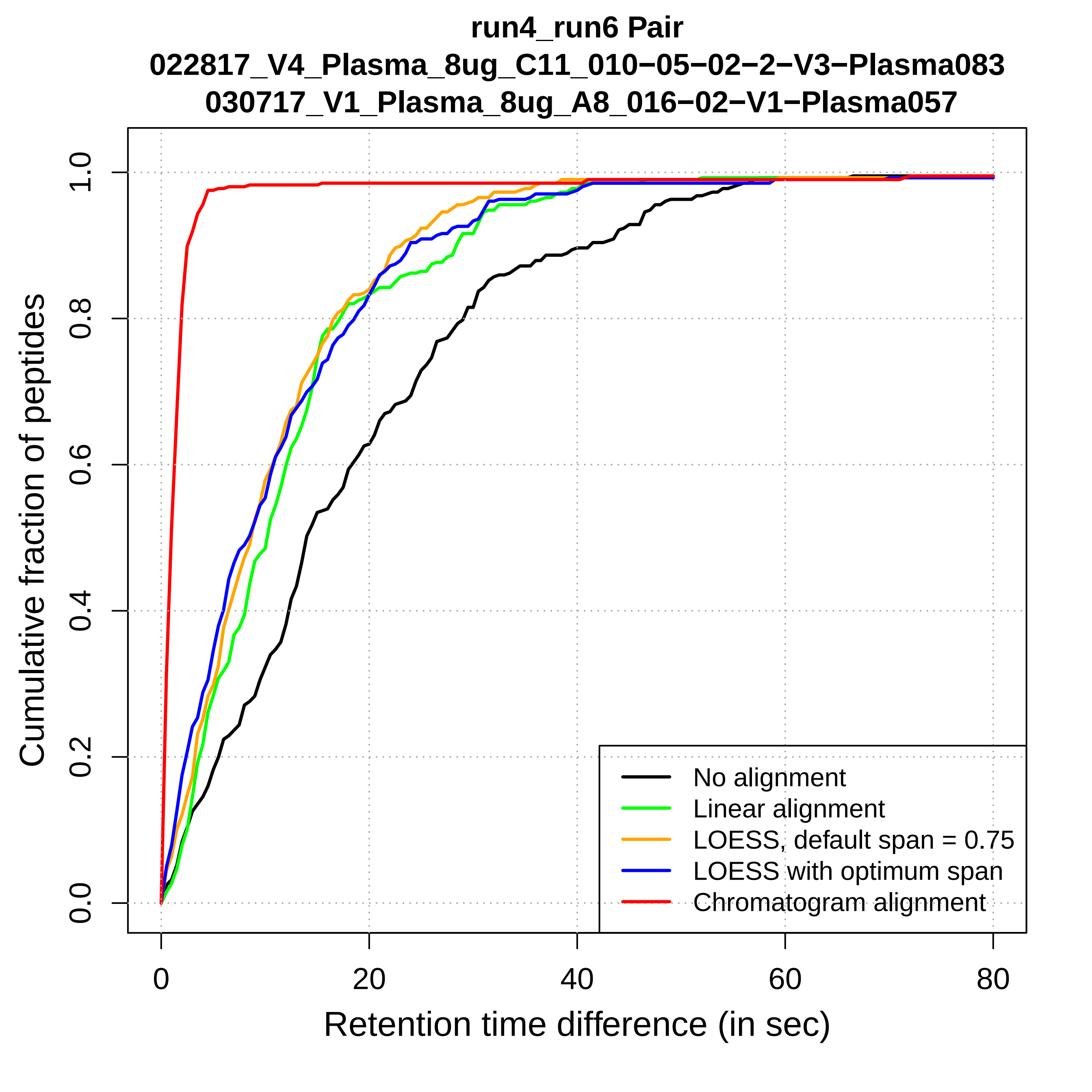
run4_run6 Pair
022817_V4_Plasma_8ug_C11_010−05−02−2−V3−Plasma083
030717_V1_Plasma_8ug_A8_016−02−V1−Plasma057
Cumulative fraction of peptides
0.0
0.2
0.4
0.6
0.8
1.0
0
20
40
60
80
Retention time difference (in sec)
No alignment
Linear alignment
LOESS, default span = 0.75
LOESS with optimum span
Chromatogram alignment

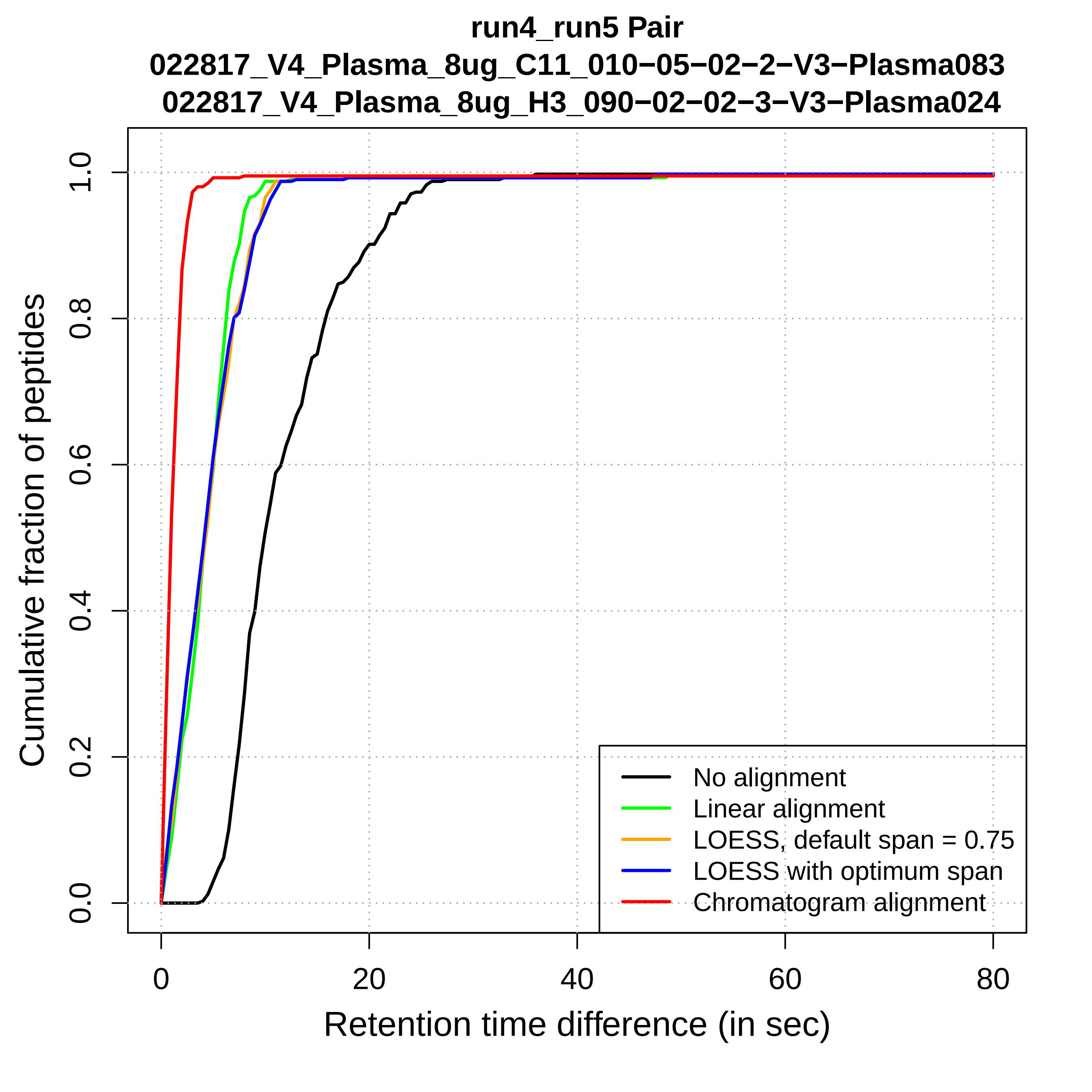
run4_run5 Pair
022817_V4_Plasma_8ug_C11_010−05−02−2−V3−Plasma083
022817_V4_Plasma_8ug_H3_090−02−02−3−V3−Plasma024
Cumulative fraction of peptides
0.0
0.2
0.4
0.6
0.8
1.0
0
20
40
60
80
Retention time difference (in sec)
No alignment
Linear alignment
LOESS, default span = 0.75
LOESS with optimum span
Chromatogram alignment

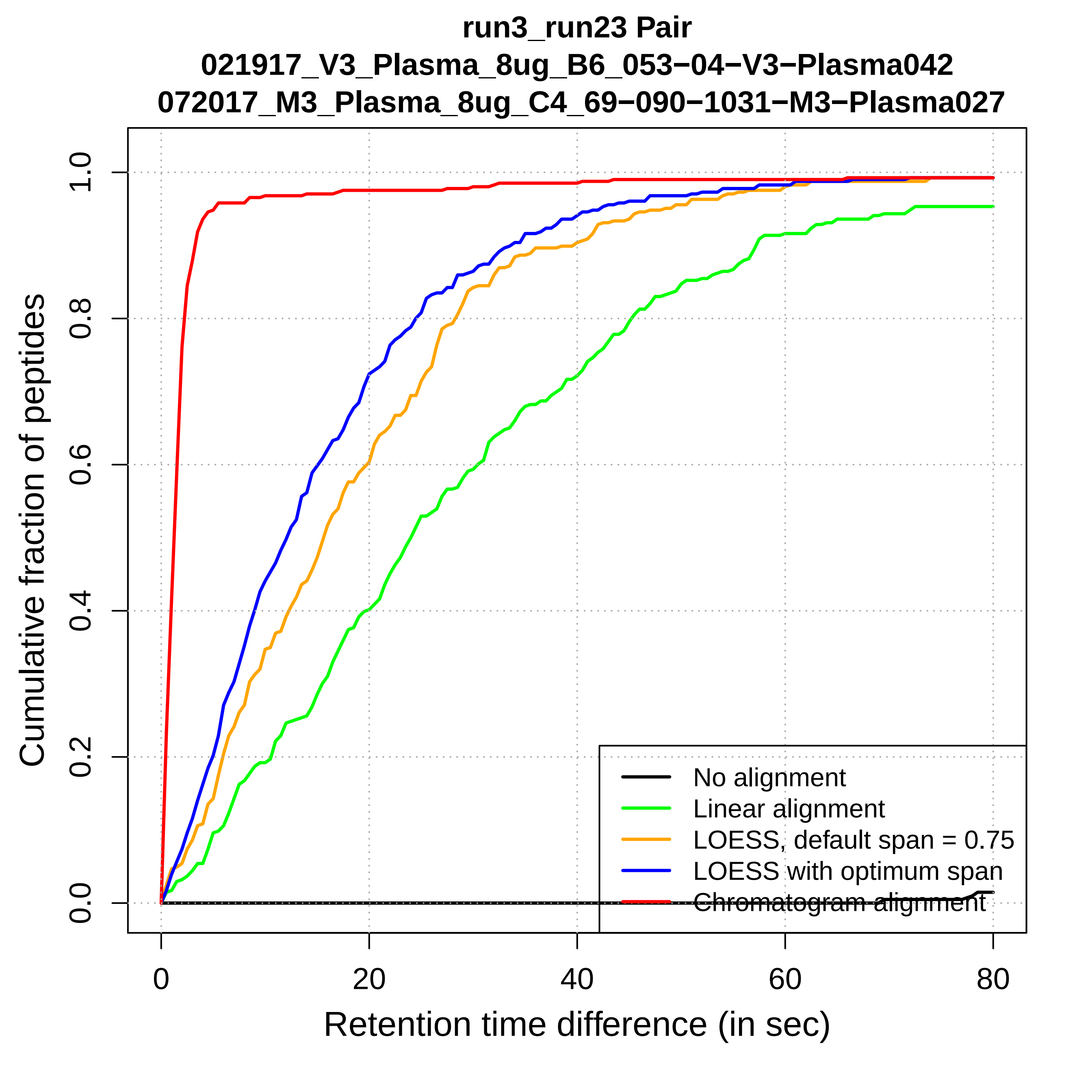

run3_run23 Pair
021917_V3_Plasma_8ug_B6_053−04−V3−Plasma042
072017_M3_Plasma_8ug_C4_69−090−1031−M3−Plasma027
Cumulative fraction of peptides
0.0
0.2
0.4
0.6
0.8
1.0
0
20
40
60
80
Retention time difference (in sec)
No alignment
Linear alignment
LOESS, default span = 0.75
LOESS with optimum span
Chromatogram alignment

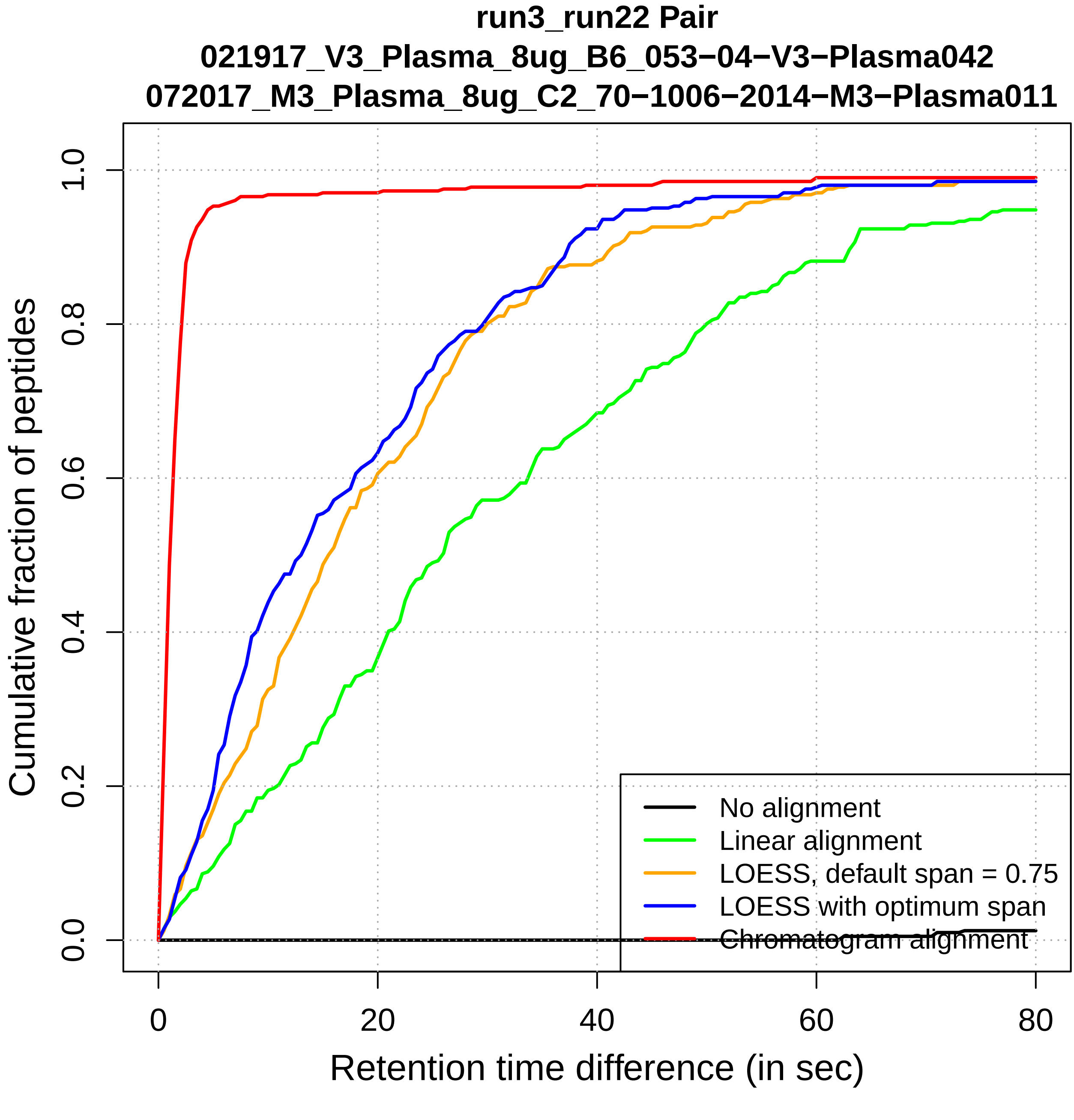
run3_run22 Pair
021917_V3_Plasma_8ug_B6_053−04−V3−Plasma042
072017_M3_Plasma_8ug_C2_70−1006−2014−M3−Plasma011
Cumulative fraction of peptides
0.0
0.2
0.4
0.6
0.8
1.0
0
20
40
60
80
Retention time difference (in sec)
No alignment
Linear alignment
LOESS, default span = 0.75
LOESS with optimum span
Chromatogram alignment

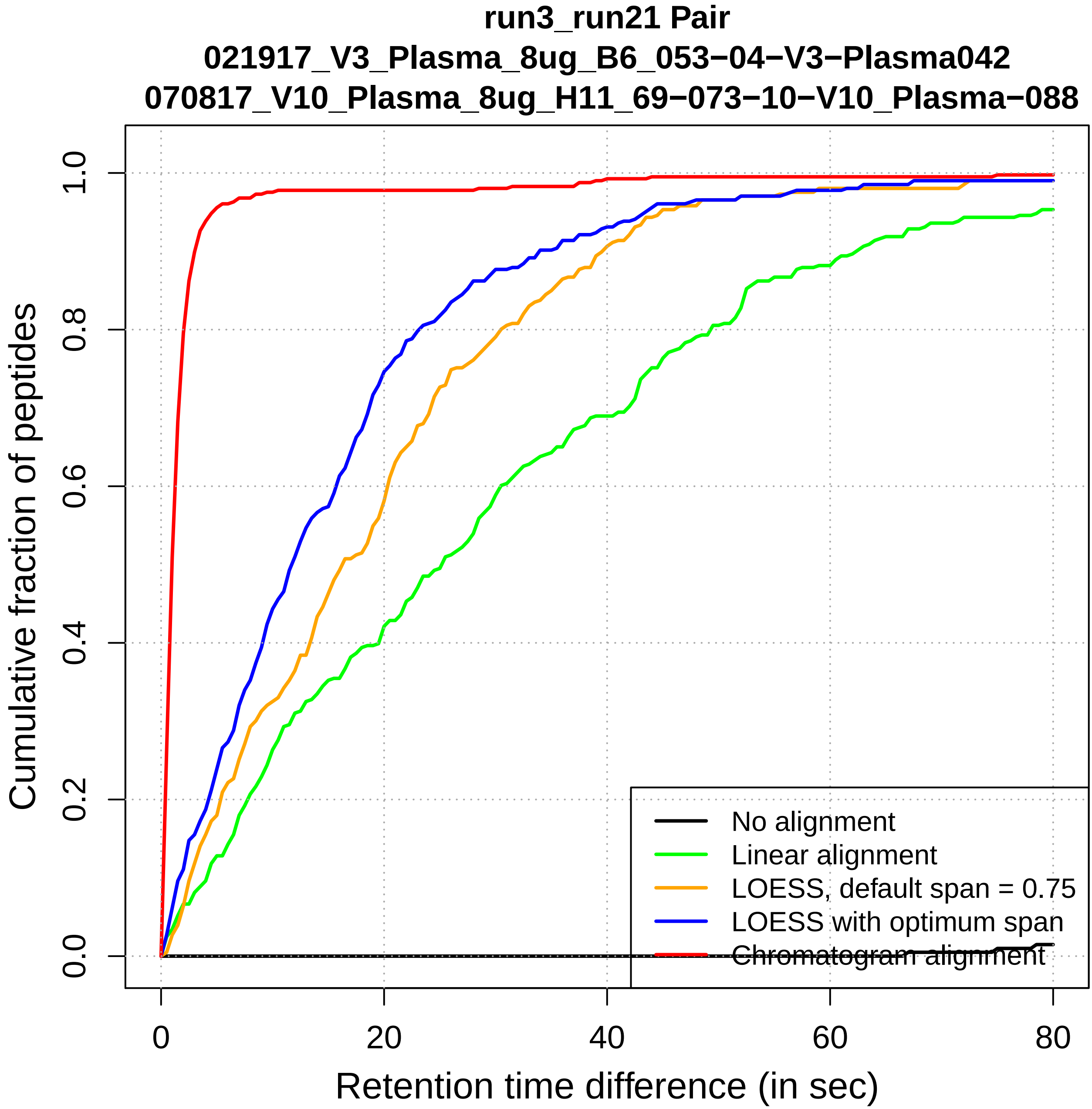
run3_run21 Pair
021917_V3_Plasma_8ug_B6_053−04−V3−Plasma042
070817_V10_Plasma_8ug_H11_69−073−10−V10_Plasma−088
Cumulative fraction of peptides
0.0
0.2
0.4
0.6
0.8
1.0
0
20
40
60
80
Retention time difference (in sec)
No alignment
Linear alignment
LOESS, default span = 0.75
LOESS with optimum span
Chromatogram alignment

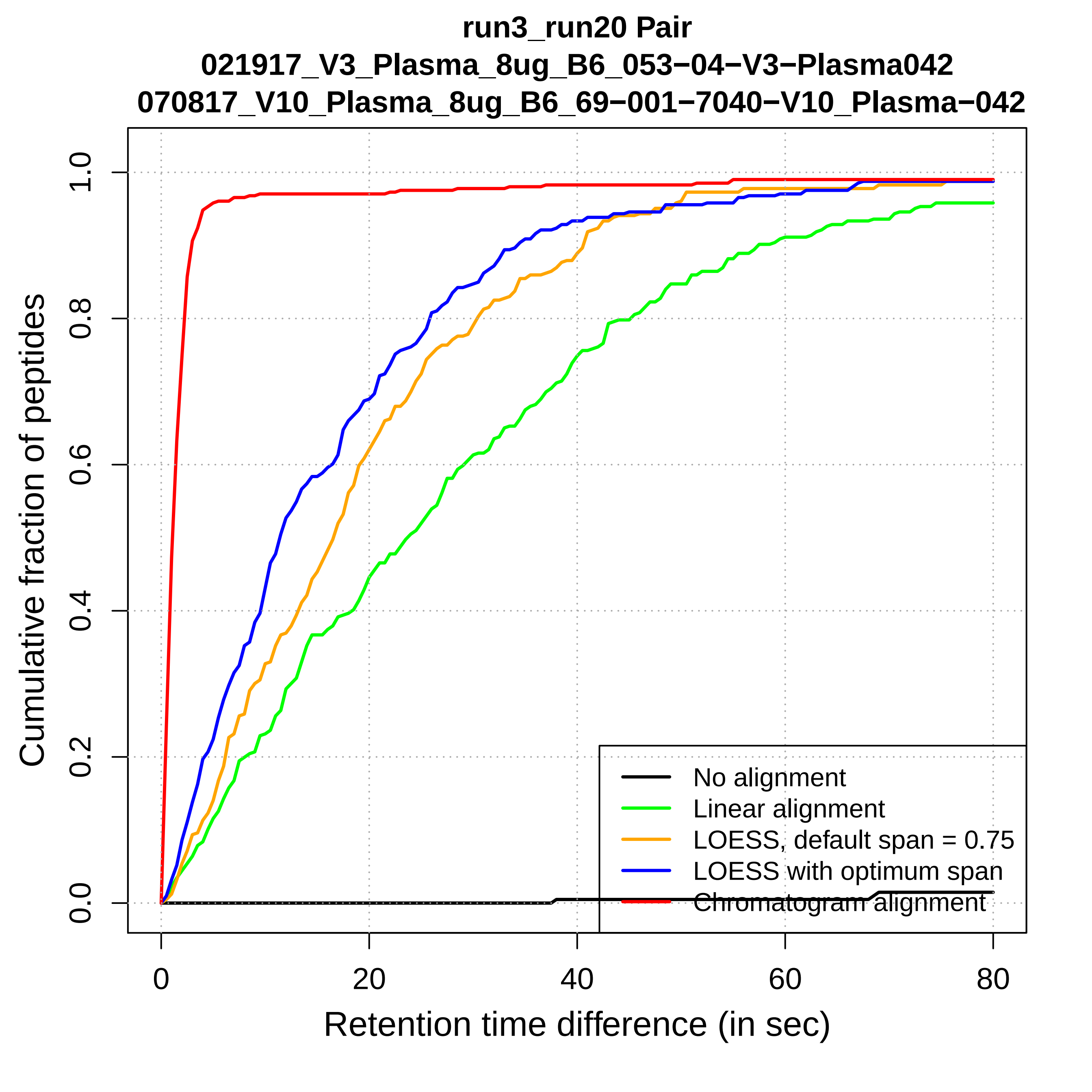
run3_run20 Pair
021917_V3_Plasma_8ug_B6_053−04−V3−Plasma042
070817_V10_Plasma_8ug_B6_69−001−7040−V10_Plasma−042
Cumulative fraction of peptides
Retention time difference (in sec)
0.0
0.2
0.4
0.6
0.8
1.0
0
20
40
60
80
No alignment
Linear alignment
LOESS, default span = 0.75
LOESS with optimum span
Chromatogram alignment

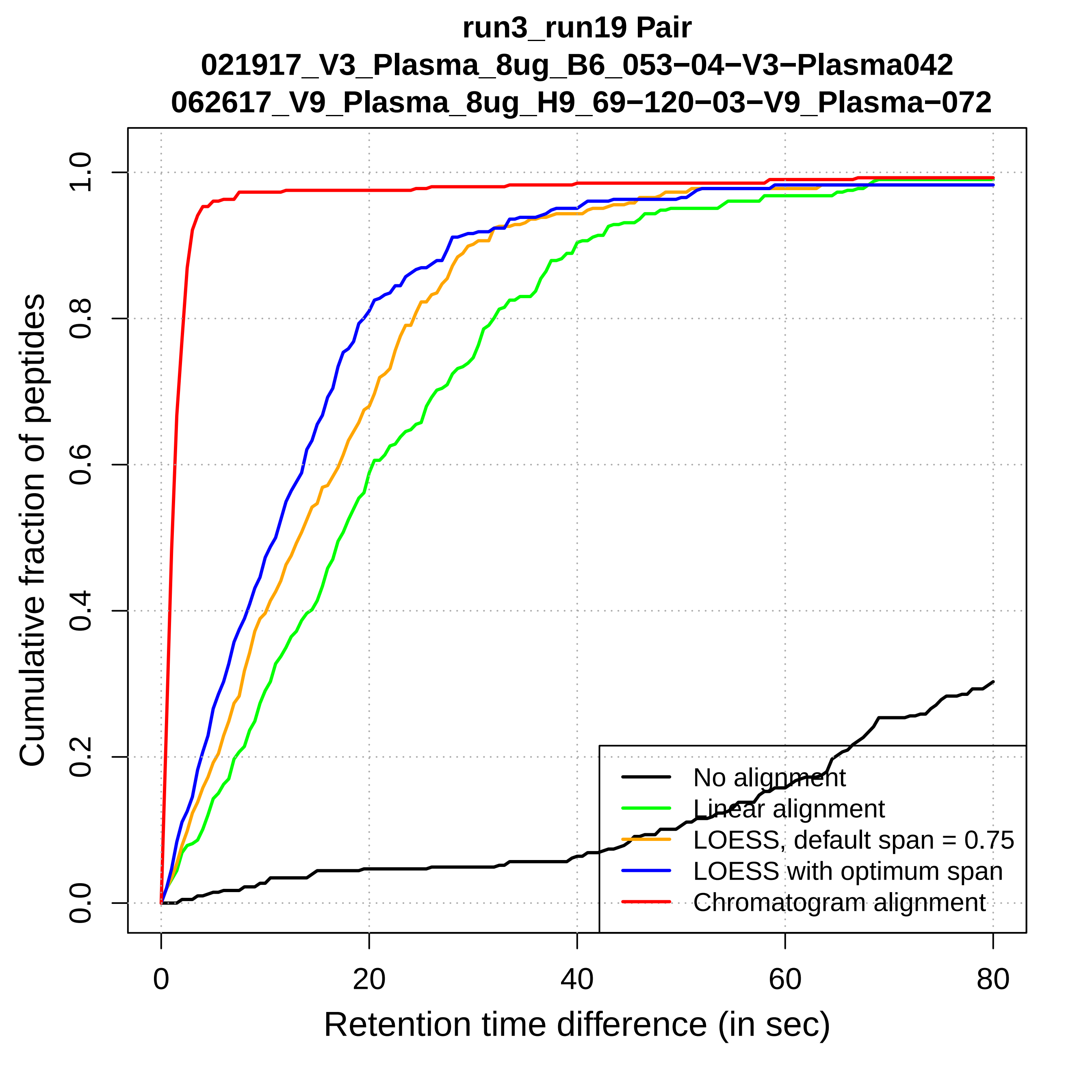
run3_run19 Pair
021917_V3_Plasma_8ug_B6_053−04−V3−Plasma042
062617_V9_Plasma_8ug_H9_69−120−03−V9_Plasma−072
Cumulative fraction of peptides
0.0
0.2
0.4
0.6
0.8
1.0
0
20
40
60
80
Retention time difference (in sec)
No alignment
Linear alignment
LOESS, default span = 0.75
LOESS with optimum span
Chromatogram alignment

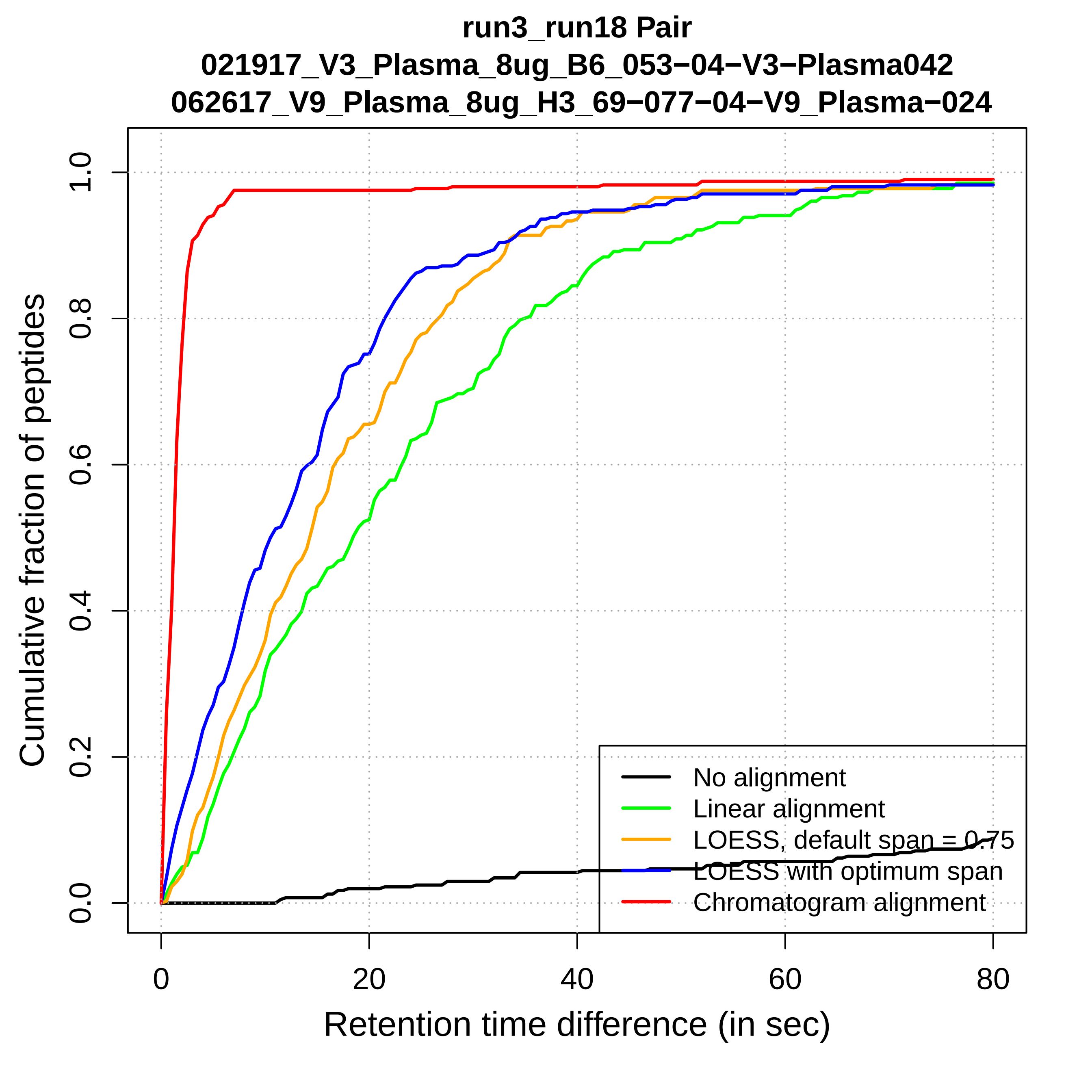

run3_run18 Pair
021917_V3_Plasma_8ug_B6_053−04−V3−Plasma042
062617_V9_Plasma_8ug_H3_69−077−04−V9_Plasma−024
Cumulative fraction of peptides
0.0
0.2
0.4
0.6
0.8
1.0
0
20
40
60
80
Retention time difference (in sec)
No alignment
Linear alignment
LOESS, default span = 0.75
LOESS with optimum span
Chromatogram alignment

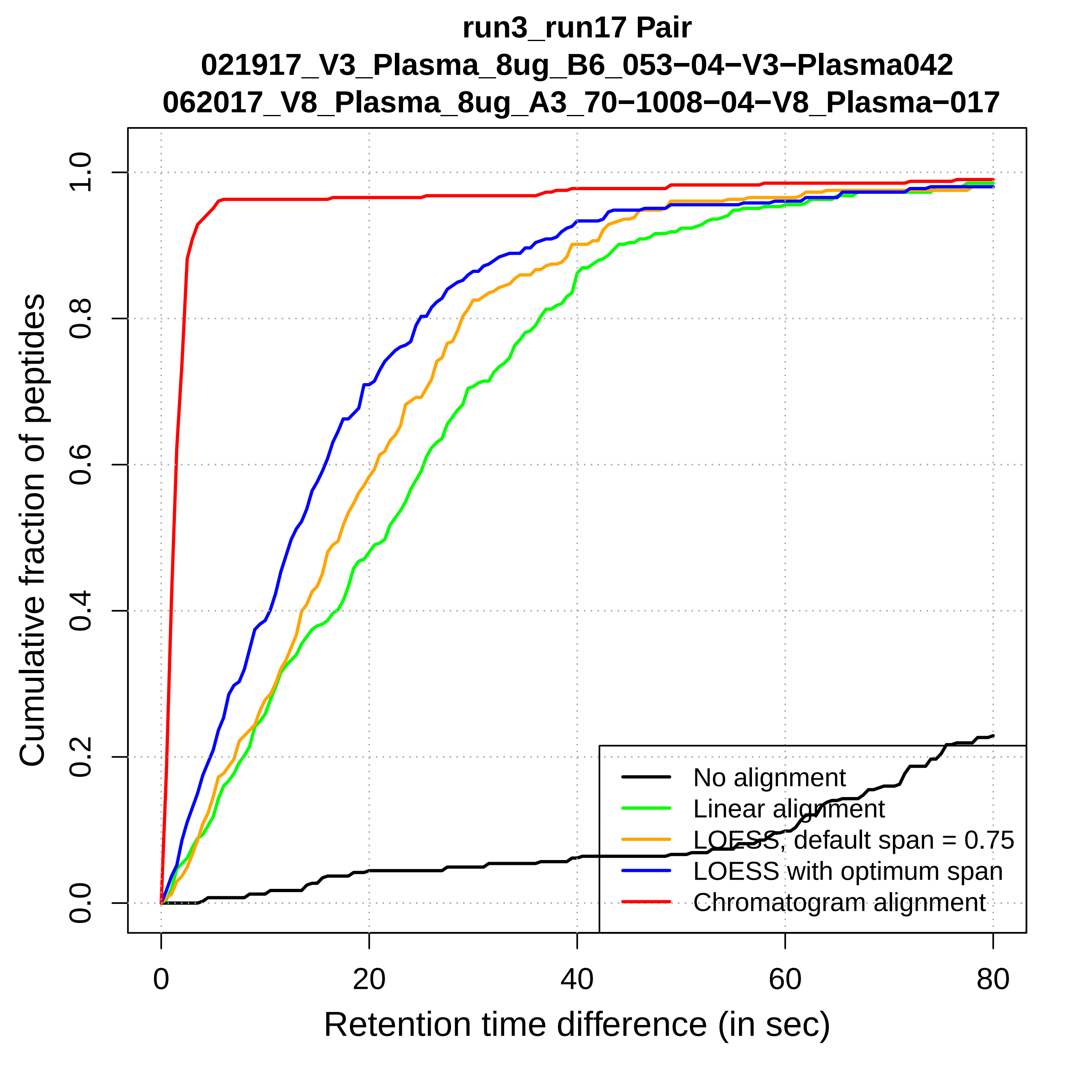

run3_run17 Pair
021917_V3_Plasma_8ug_B6_053−04−V3−Plasma042
062017_V8_Plasma_8ug_A3_70−1008−04−V8_Plasma−017
Cumulative fraction of peptides
0.0
0.2
0.4
0.6
0.8
1.0
0
20
40
60
80
Retention time difference (in sec)
No alignment
Linear alignment
LOESS, default span = 0.75
LOESS with optimum span
Chromatogram alignment

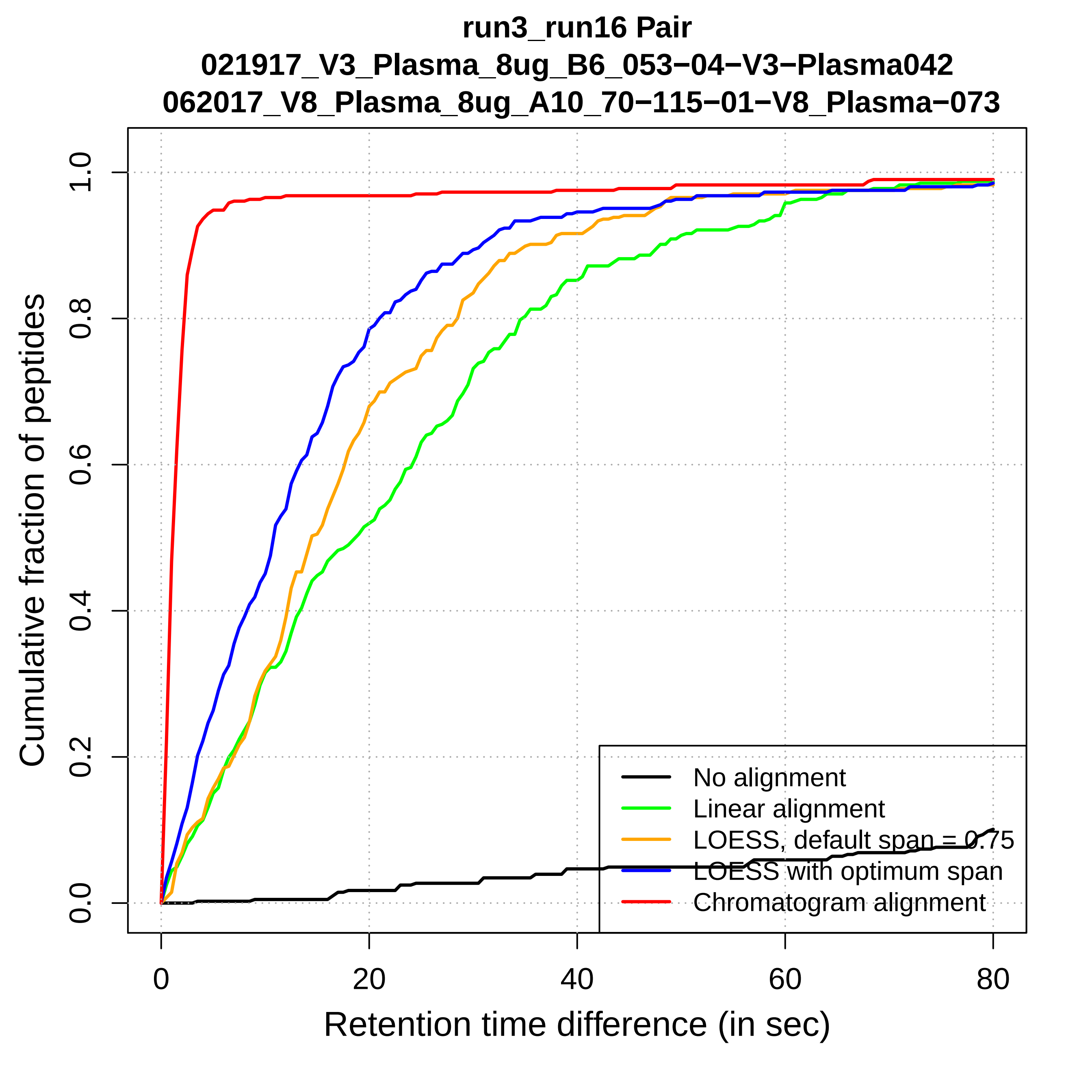
run3_run16 Pair
021917_V3_Plasma_8ug_B6_053−04−V3−Plasma042
062017_V8_Plasma_8ug_A10_70−115−01−V8_Plasma−073
Cumulative fraction of peptides
0.0
0.2
0.4
0.6
0.8
1.0
0
20
40
60
80
Retention time difference (in sec)
No alignment
Linear alignment
LOESS, default span = 0.75
LOESS with optimum span
Chromatogram alignment

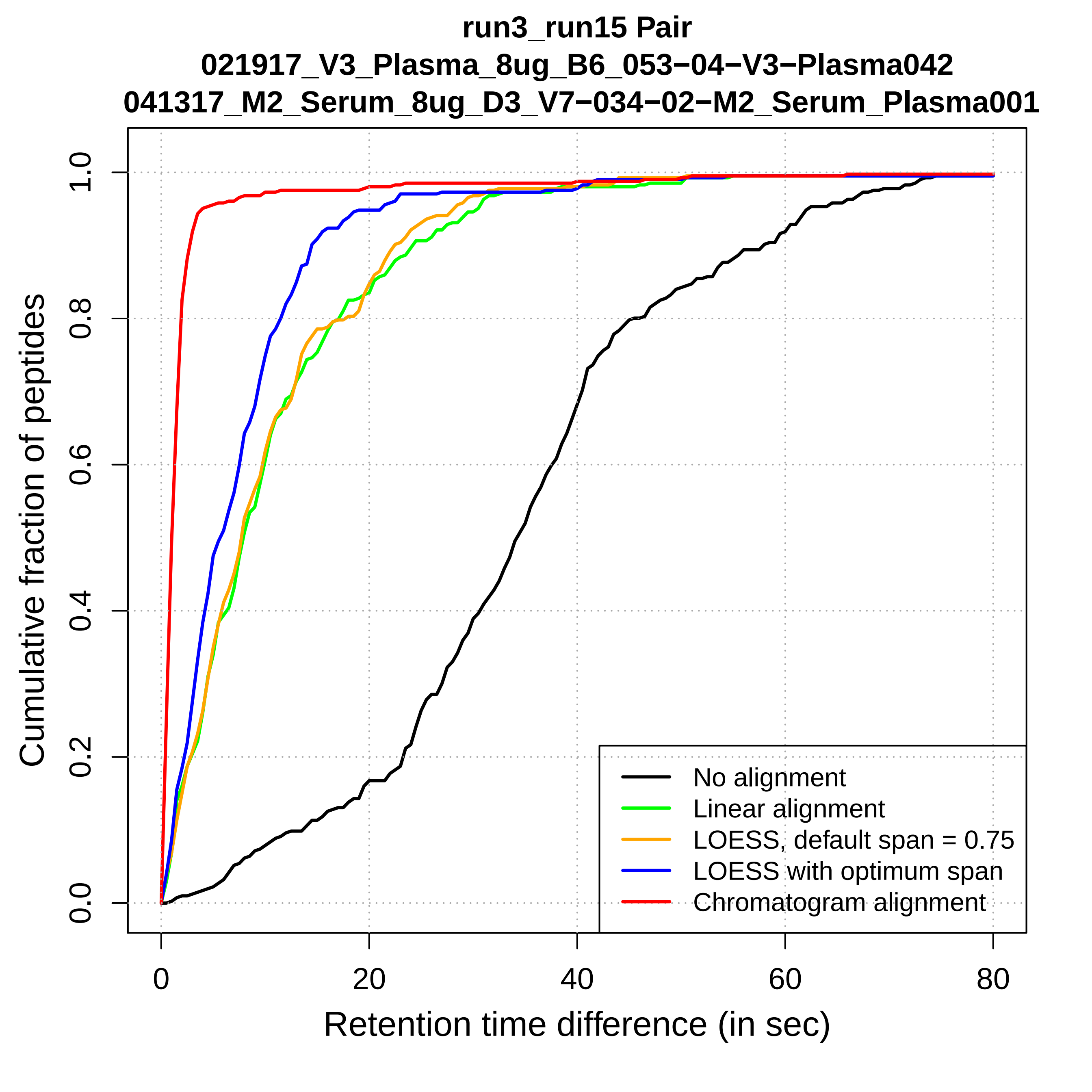
run3_run15 Pair
021917_V3_Plasma_8ug_B6_053−04−V3−Plasma042
041317_M2_Serum_8ug_D3_V7−034−02−M2_Serum_Plasma001
Cumulative fraction of peptides
0.0
0.2
0.4
0.6
0.8
1.0
0
20
40
60
80
Retention time difference (in sec)
No alignment
Linear alignment
LOESS, default span = 0.75
LOESS with optimum span
Chromatogram alignment

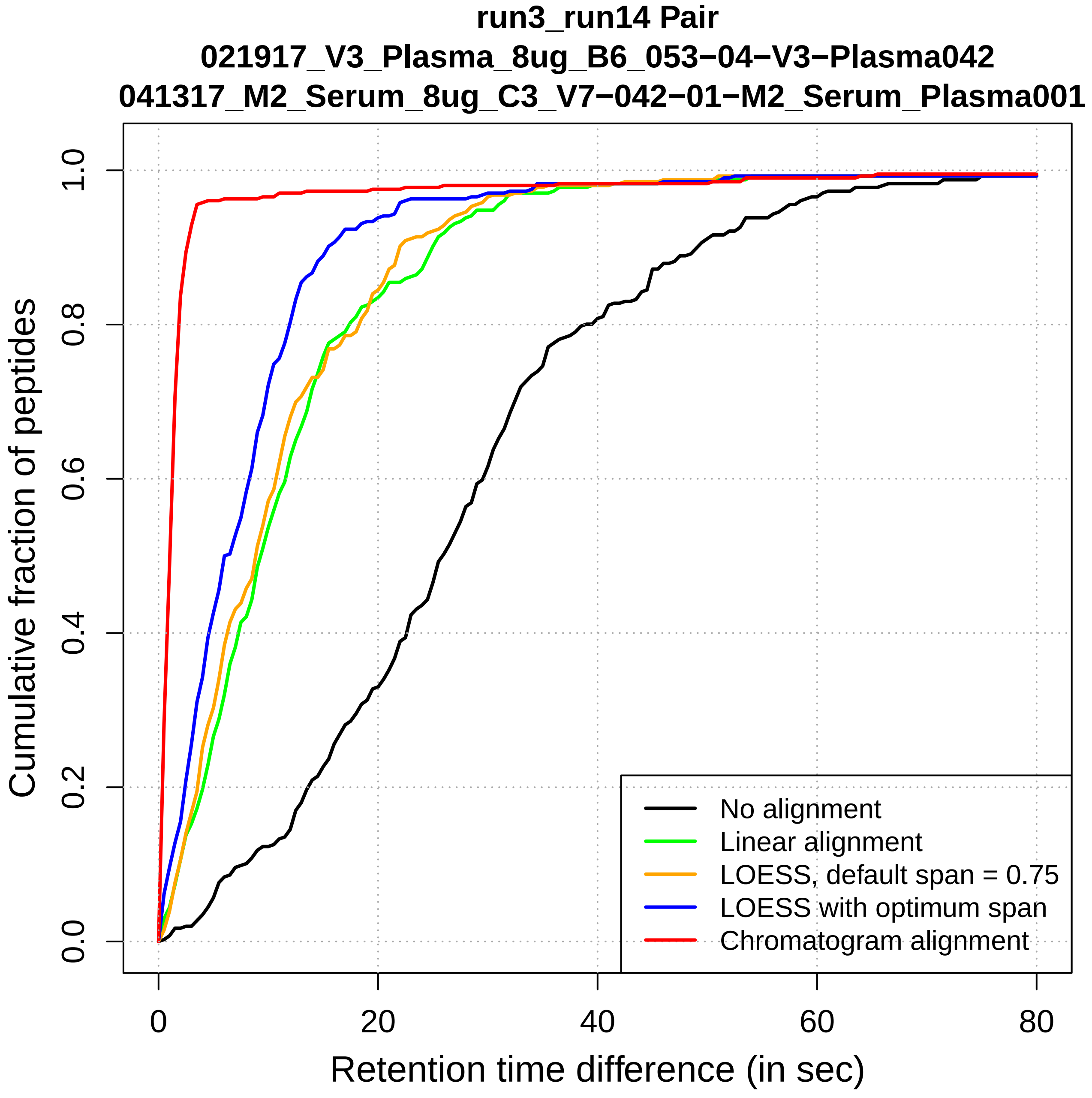
run3_run14 Pair
021917_V3_Plasma_8ug_B6_053−04−V3−Plasma042
041317_M2_Serum_8ug_C3_V7−042−01−M2_Serum_Plasma001
Cumulative fraction of peptides
0.0
0.2
0.4
0.6
0.8
1.0
0
20
40
60
80
Retention time difference (in sec)
No alignment
Linear alignment
LOESS, default span = 0.75
LOESS with optimum span
Chromatogram alignment

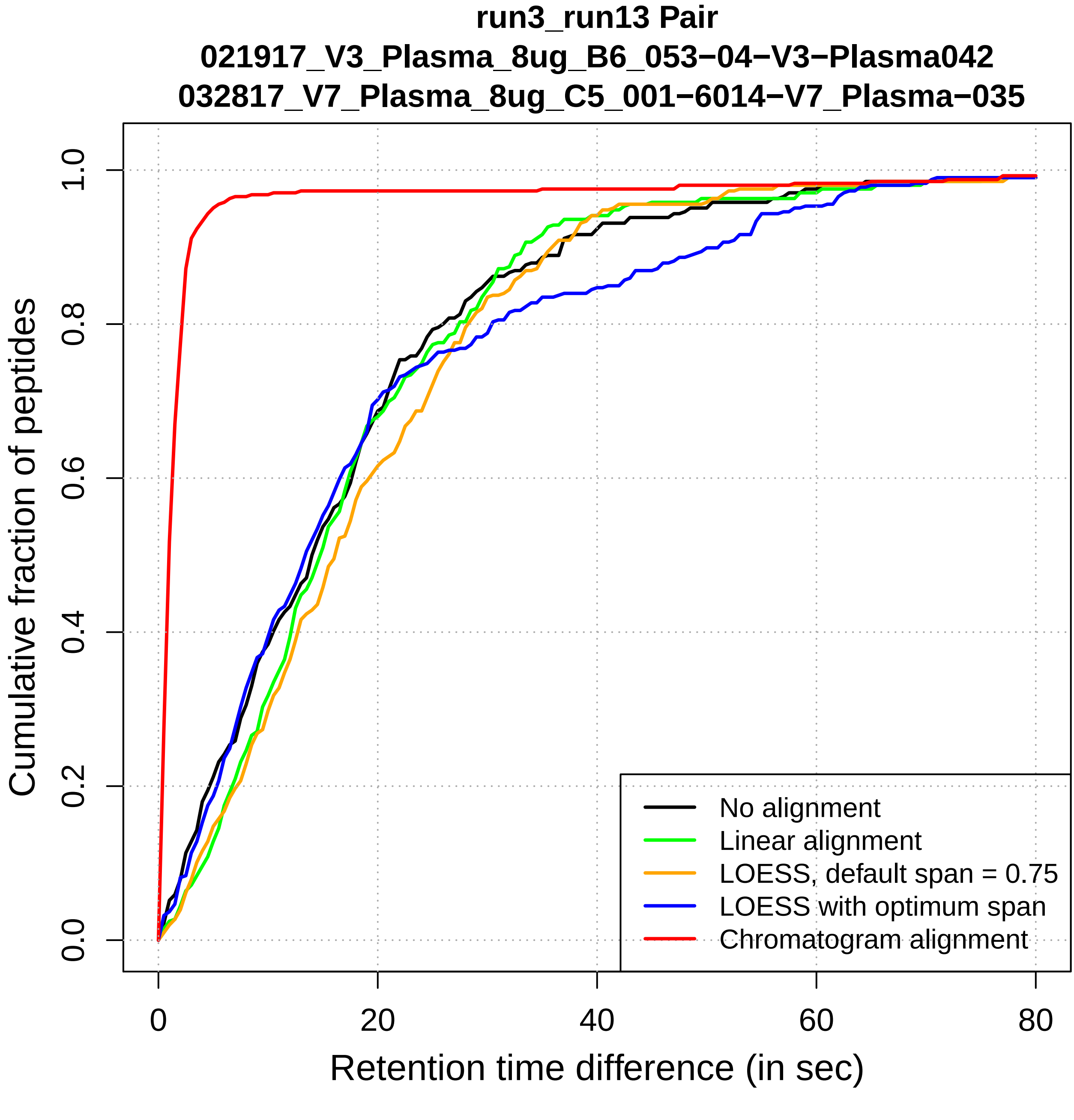
run3_run13 Pair
021917_V3_Plasma_8ug_B6_053−04−V3−Plasma042
032817_V7_Plasma_8ug_C5_001−6014−V7_Plasma−035
Cumulative fraction of peptides
0.0
0.2
0.4
0.6
0.8
1.0
0
20
40
60
80
Retention time difference (in sec)
No alignment
Linear alignment
LOESS, default span = 0.75
LOESS with optimum span
Chromatogram alignment

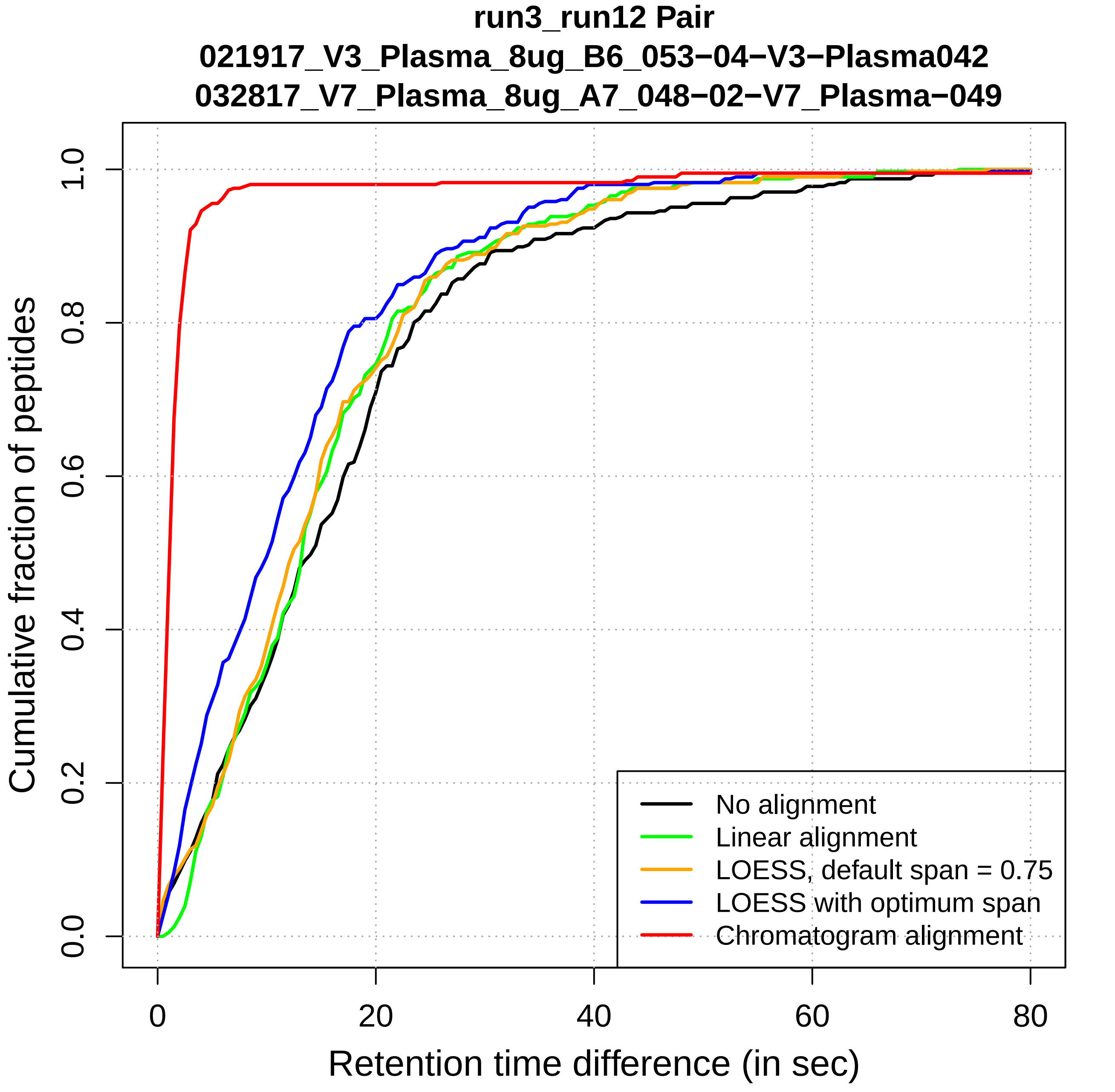
run3_run12 Pair
021917_V3_Plasma_8ug_B6_053−04−V3−Plasma042
032817_V7_Plasma_8ug_A7_048−02−V7_Plasma−049
Cumulative fraction of peptides
Retention time difference (in sec)
0.0
0.2
0.4
0.6
0.8
1.0
0
20
40
60
80
No alignment
Linear alignment
LOESS, default span = 0.75
LOESS with optimum span
Chromatogram alignment

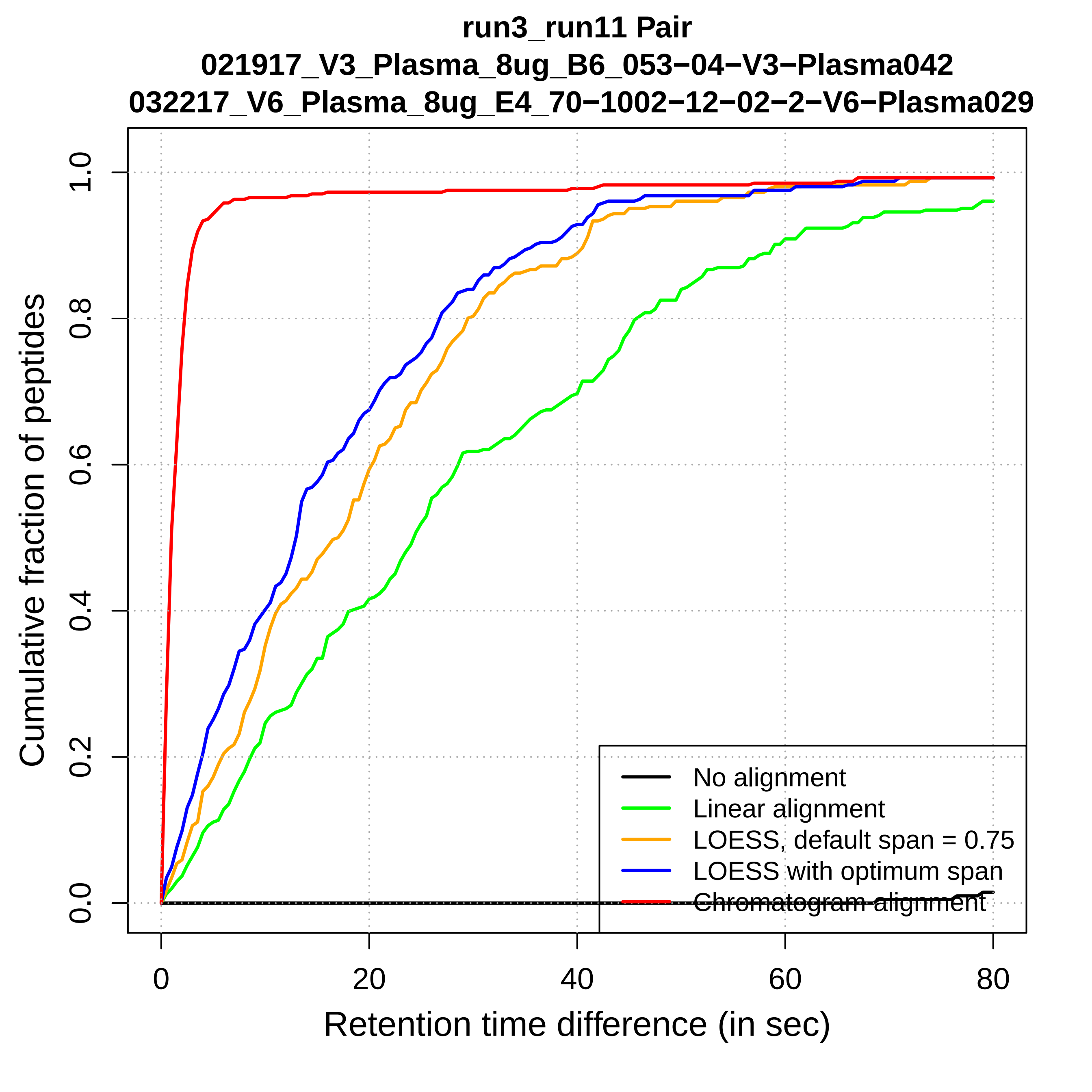
run3_run11 Pair
021917_V3_Plasma_8ug_B6_053−04−V3−Plasma042
032217_V6_Plasma_8ug_E4_70−1002−12−02−2−V6−Plasma029
Cumulative fraction of peptides
0.0
0.2
0.4
0.6
0.8
1.0
0
20
40
60
80
Retention time difference (in sec)
No alignment
Linear alignment
LOESS, default span = 0.75
LOESS with optimum span
Chromatogram alignment

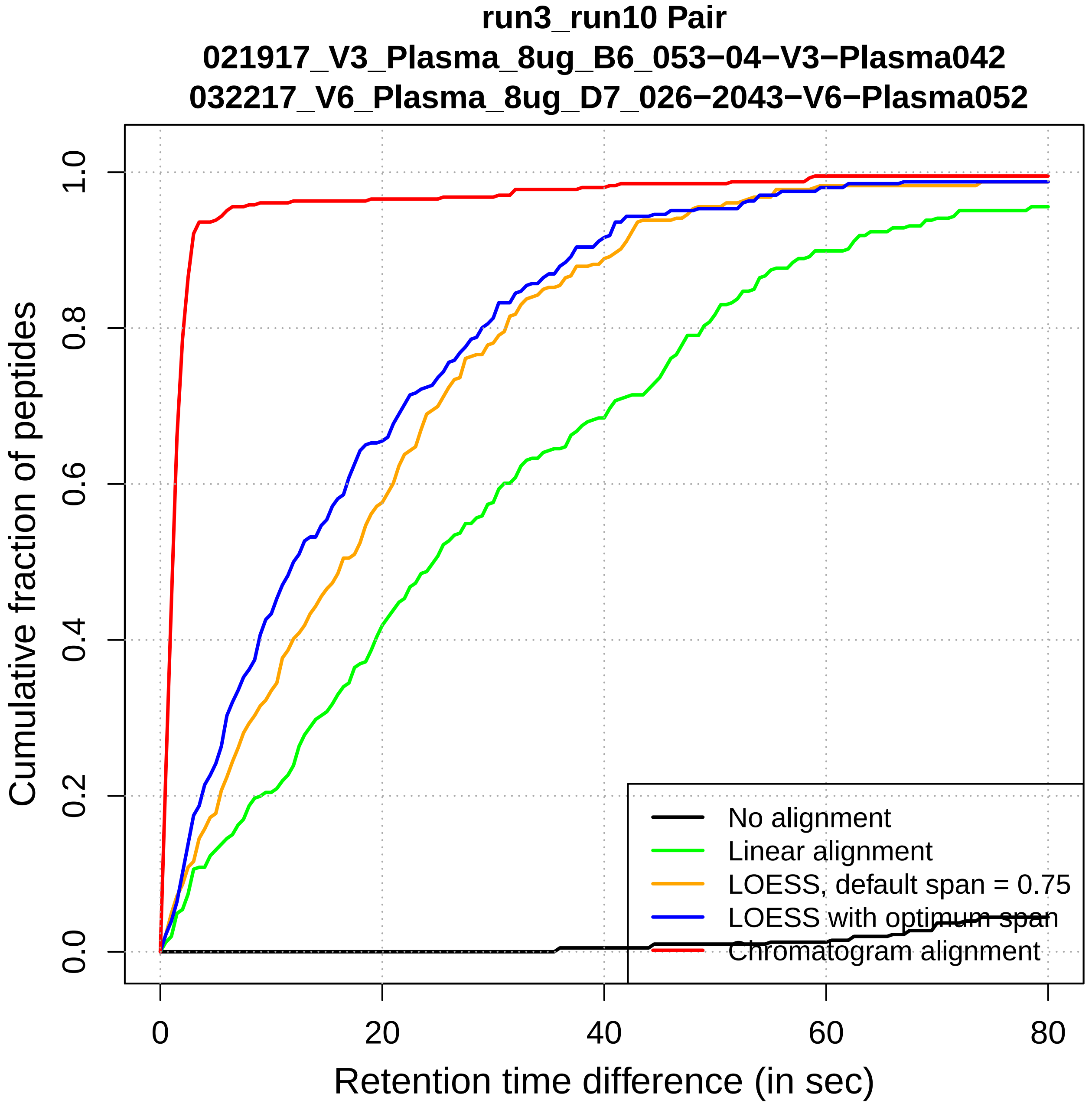

run3_run10 Pair
021917_V3_Plasma_8ug_B6_053−04−V3−Plasma042
032217_V6_Plasma_8ug_D7_026−2043−V6−Plasma052
Cumulative fraction of peptides
0.0
0.2
0.4
0.6
0.8
1.0
0
20
40
60
80
Retention time difference (in sec)
No alignment
Linear alignment
LOESS, default span = 0.75
LOESS with optimum span
Chromatogram alignment

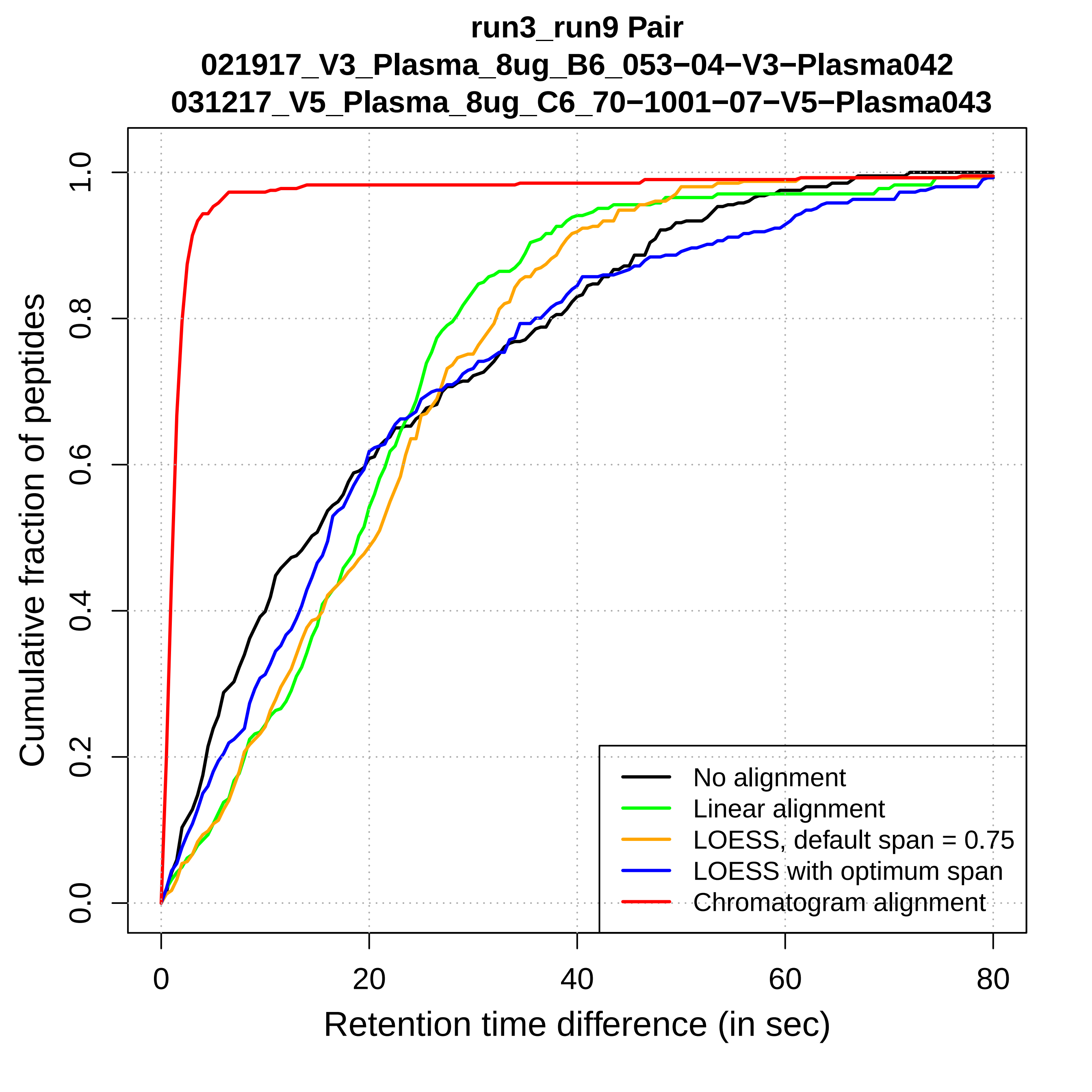

run3_run9 Pair
021917_V3_Plasma_8ug_B6_053−04−V3−Plasma042
031217_V5_Plasma_8ug_C6_70−1001−07−V5−Plasma043
Cumulative fraction of peptides
0.0
0.2
0.4
0.6
0.8
1.0
0
20
40
60
80
Retention time difference (in sec)
No alignment
Linear alignment
LOESS, default span = 0.75
LOESS with optimum span
Chromatogram alignment

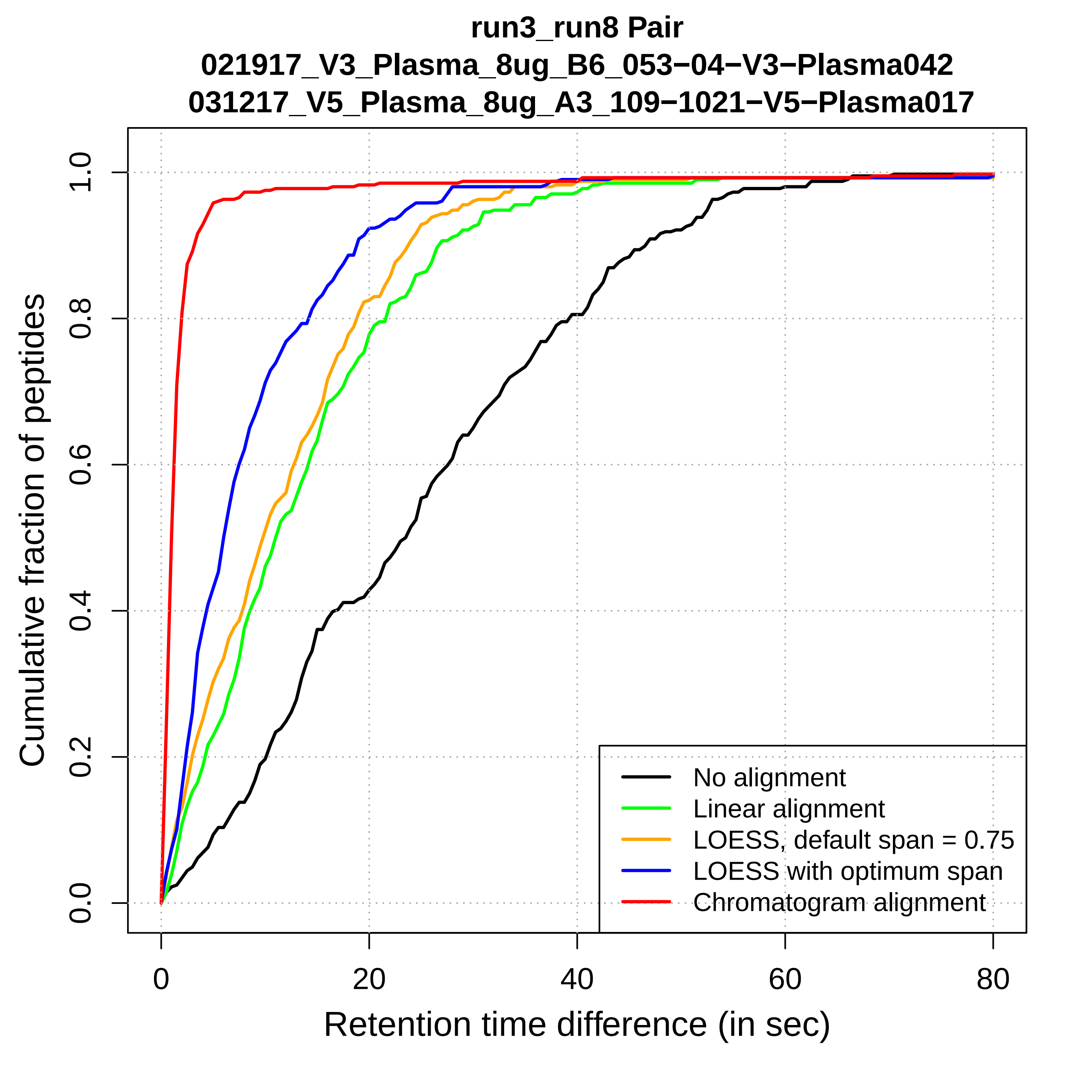

run3_run8 Pair
021917_V3_Plasma_8ug_B6_053−04−V3−Plasma042
031217_V5_Plasma_8ug_A3_109−1021−V5−Plasma017
Cumulative fraction of peptides
Retention time difference (in sec)
0.0
0.2
0.4
0.6
0.8
1.0
0
20
40
60
80
No alignment
Linear alignment
LOESS, default span = 0.75
LOESS with optimum span
Chromatogram alignment

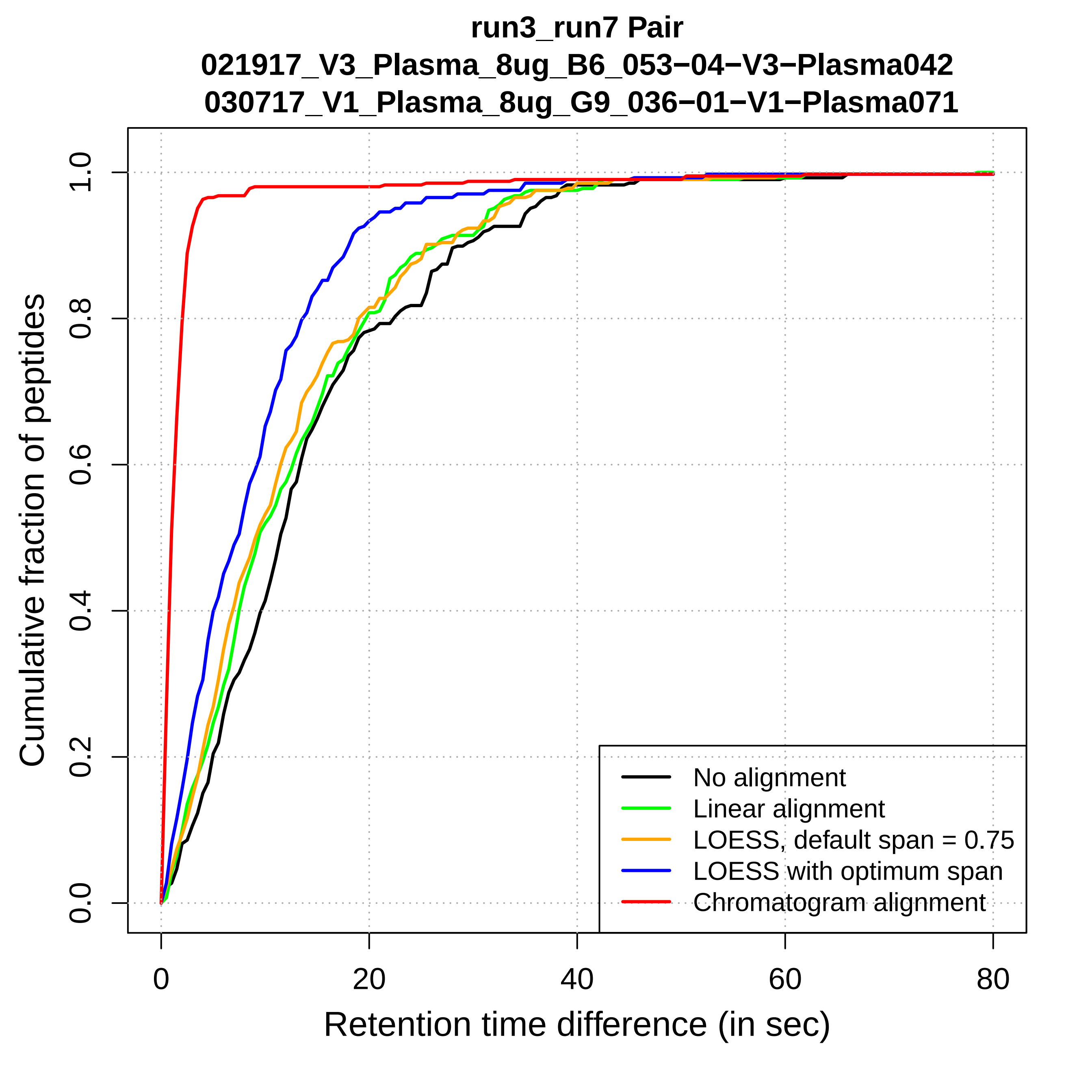

run3_run7 Pair
021917_V3_Plasma_8ug_B6_053−04−V3−Plasma042
030717_V1_Plasma_8ug_G9_036−01−V1−Plasma071
Cumulative fraction of peptides
0.0
0.2
0.4
0.6
0.8
1.0
0
20
40
60
80
Retention time difference (in sec)
No alignment
Linear alignment
LOESS, default span = 0.75
LOESS with optimum span
Chromatogram alignment

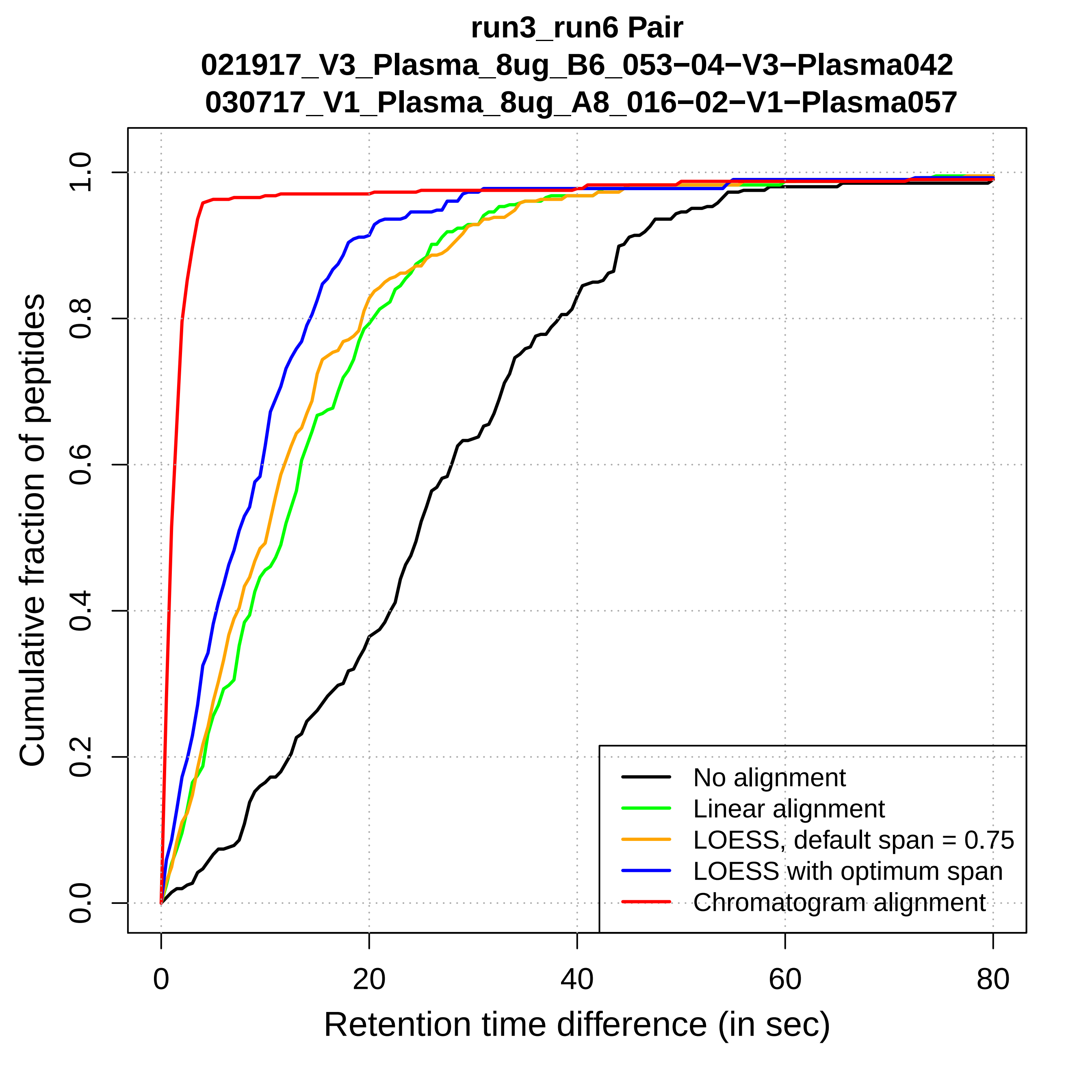

run3_run6 Pair
021917_V3_Plasma_8ug_B6_053−04−V3−Plasma042
030717_V1_Plasma_8ug_A8_016−02−V1−Plasma057
Cumulative fraction of peptides
0.0
0.2
0.4
0.6
0.8
1.0
0
20
40
60
80
Retention time difference (in sec)
No alignment
Linear alignment
LOESS, default span = 0.75
LOESS with optimum span
Chromatogram alignment

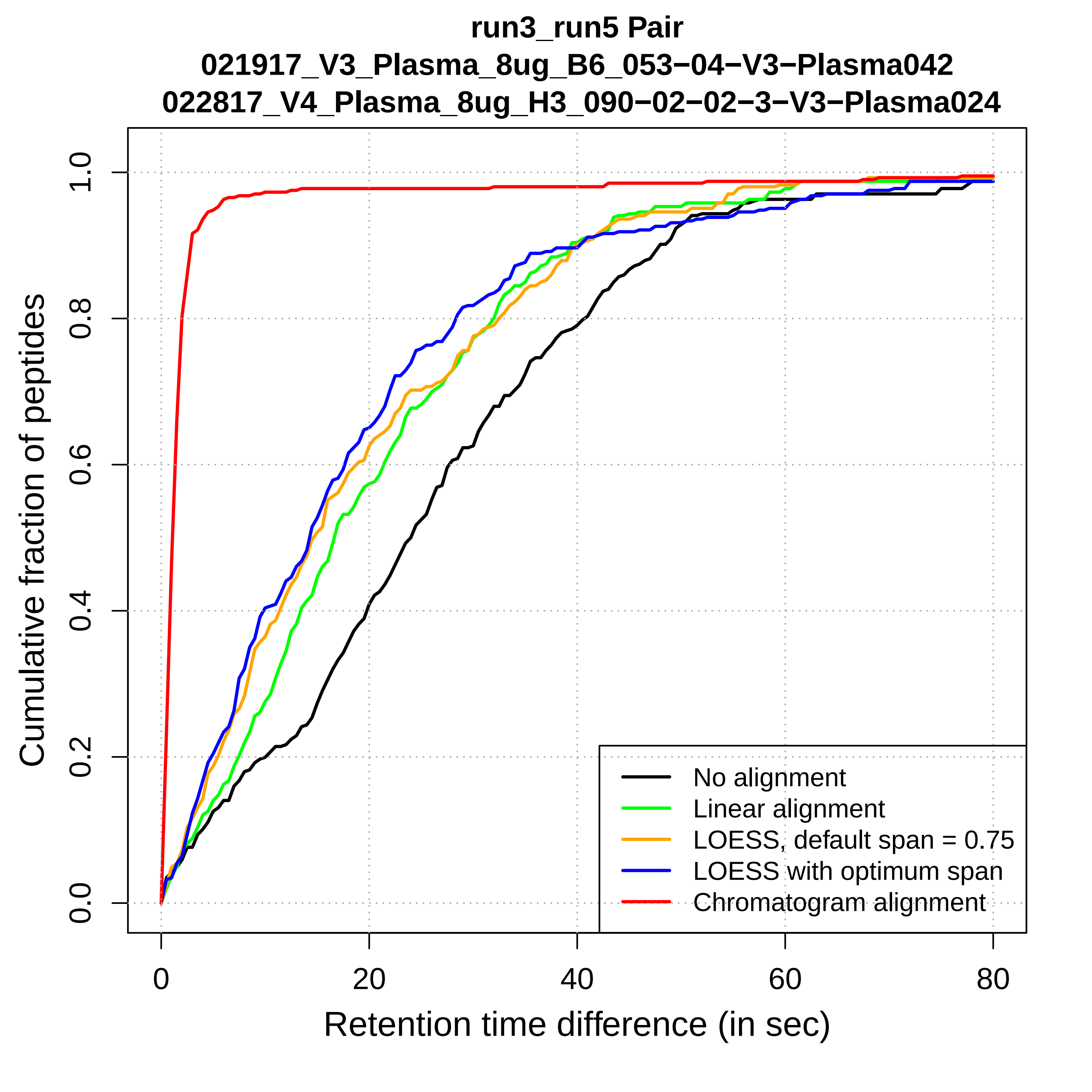
run3_run5 Pair
021917_V3_Plasma_8ug_B6_053−04−V3−Plasma042
022817_V4_Plasma_8ug_H3_090−02−02−3−V3−Plasma024
Cumulative fraction of peptides
0.0
0.2
0.4
0.6
0.8
1.0
0
20
40
60
80
Retention time difference (in sec)
No alignment
Linear alignment
LOESS, default span = 0.75
LOESS with optimum span
Chromatogram alignment

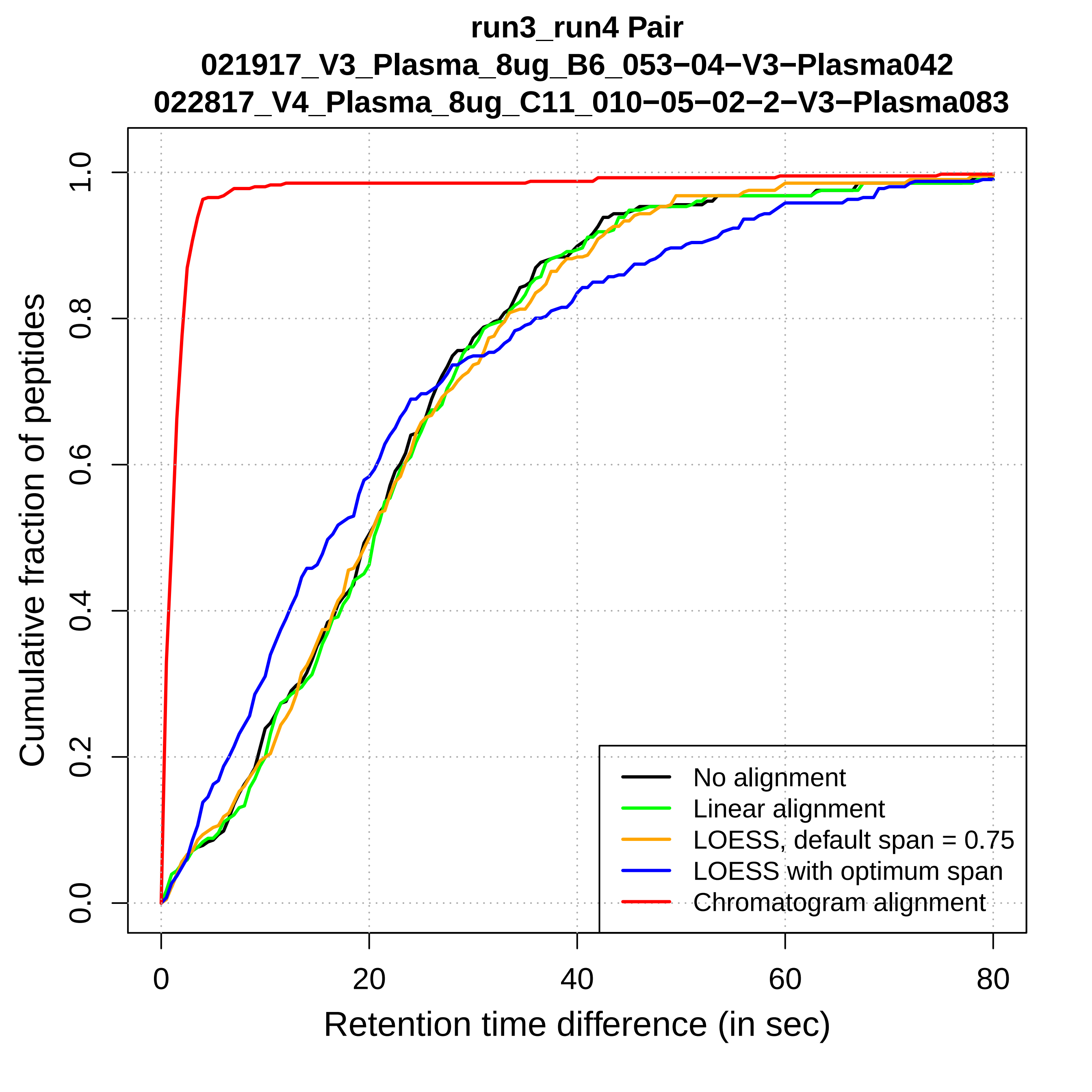

run3_run4 Pair
021917_V3_Plasma_8ug_B6_053−04−V3−Plasma042
022817_V4_Plasma_8ug_C11_010−05−02−2−V3−Plasma083
Cumulative fraction of peptides
Retention time difference (in sec)
0.0
0.2
0.4
0.6
0.8
1.0
0
20
40
60
80
No alignment
Linear alignment
LOESS, default span = 0.75
LOESS with optimum span
Chromatogram alignment

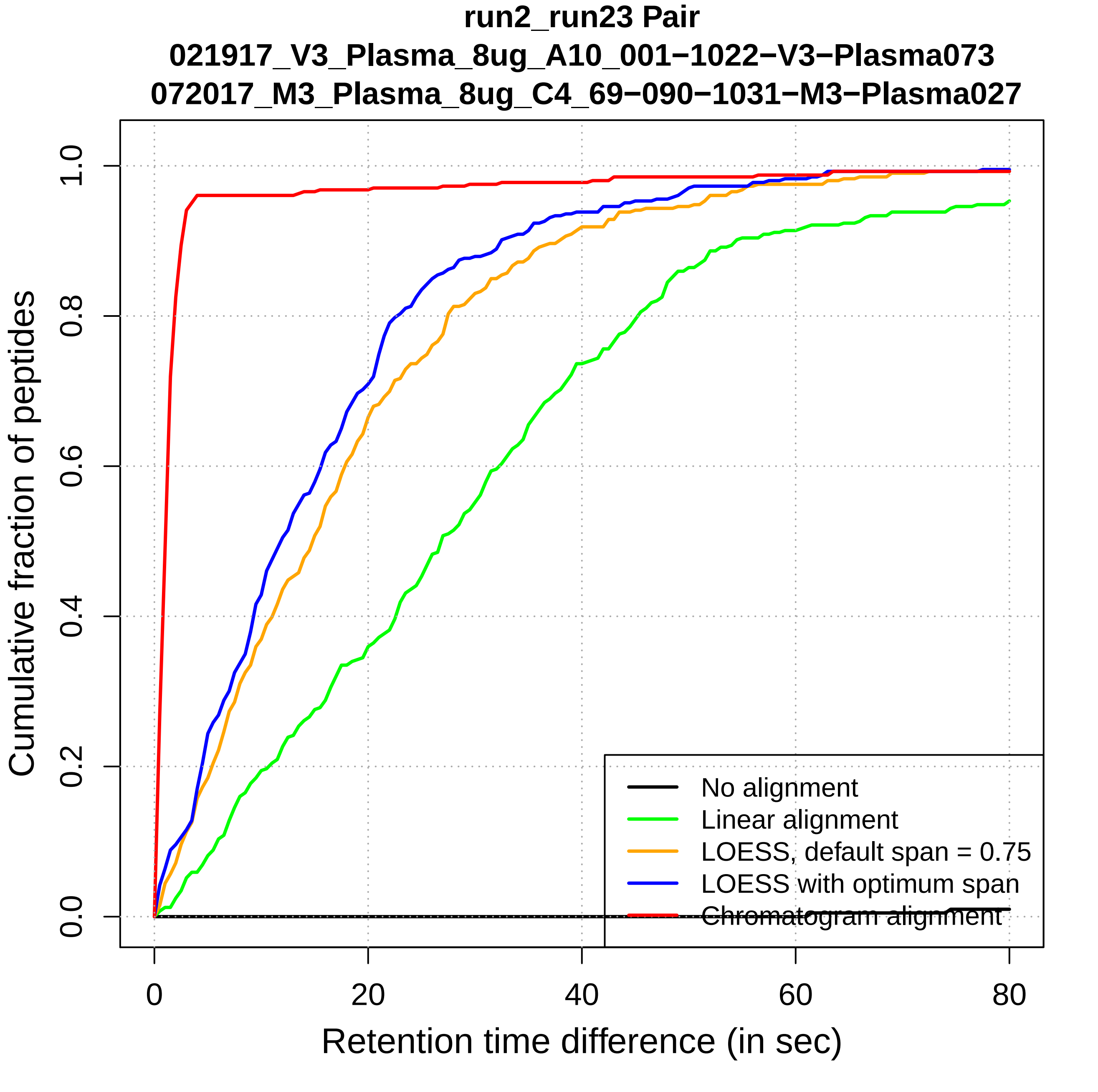
run2_run23 Pair
021917_V3_Plasma_8ug_A10_001−1022−V3−Plasma073
072017_M3_Plasma_8ug_C4_69−090−1031−M3−Plasma027
Cumulative fraction of peptides
0.0
0.2
0.4
0.6
0.8
1.0
0
20
40
60
80
Retention time difference (in sec)
No alignment
Linear alignment
LOESS, default span = 0.75
LOESS with optimum span
Chromatogram alignment

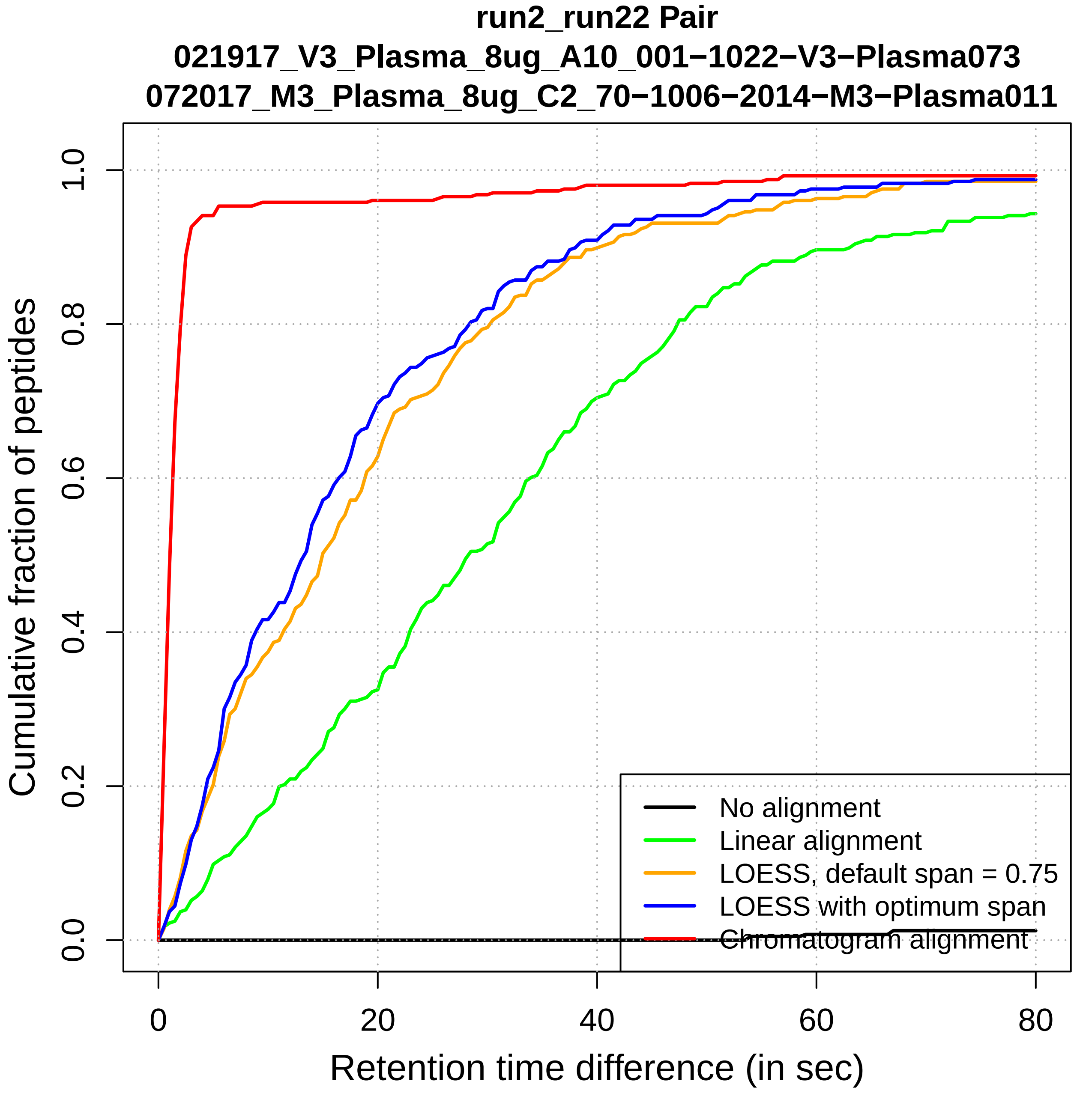
run2_run22 Pair
021917_V3_Plasma_8ug_A10_001−1022−V3−Plasma073
072017_M3_Plasma_8ug_C2_70−1006−2014−M3−Plasma011
Cumulative fraction of peptides
0.0
0.2
0.4
0.6
0.8
1.0
0
20
40
60
80
Retention time difference (in sec)
No alignment
Linear alignment
LOESS, default span = 0.75
LOESS with optimum span
Chromatogram alignment

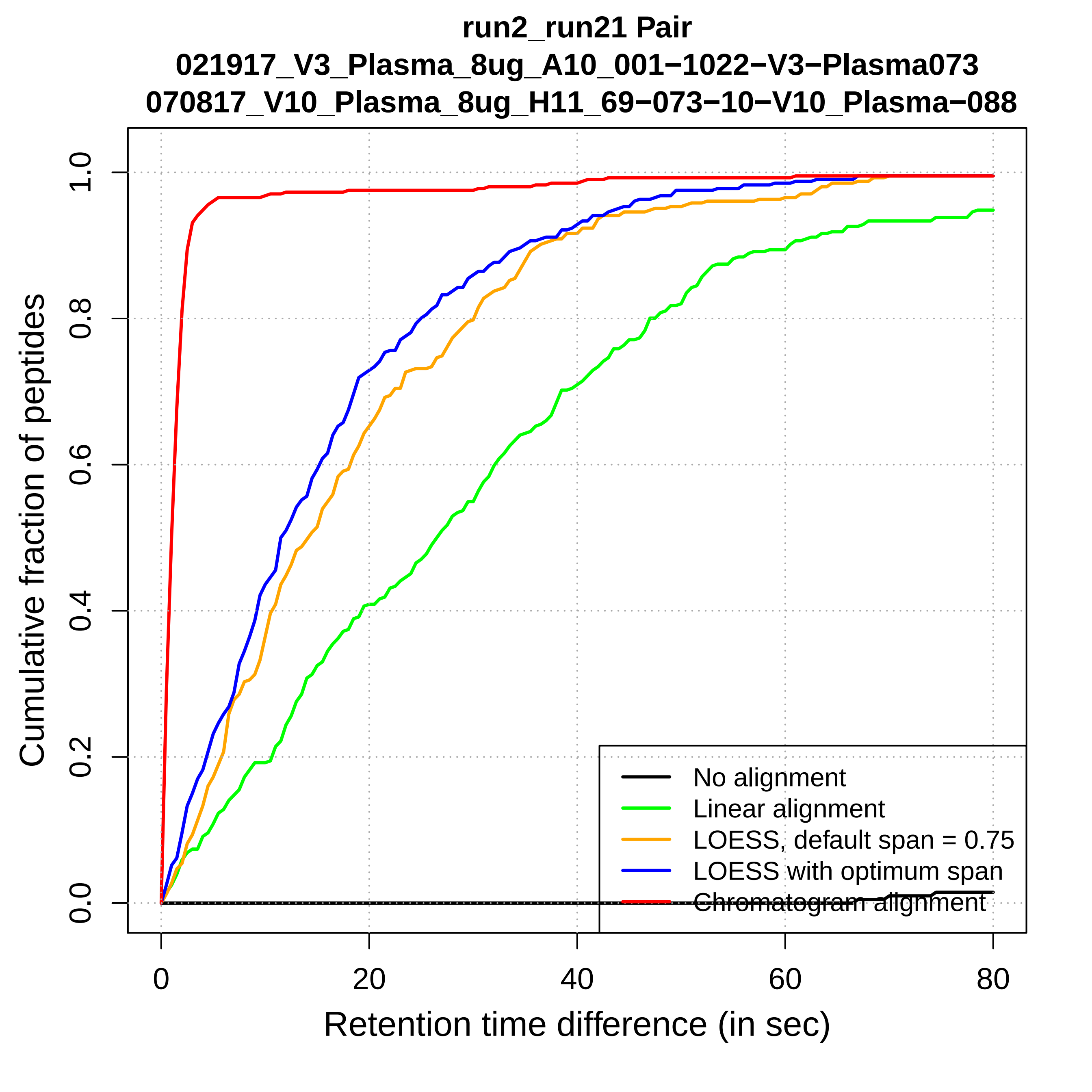
run2_run21 Pair
021917_V3_Plasma_8ug_A10_001−1022−V3−Plasma073
070817_V10_Plasma_8ug_H11_69−073−10−V10_Plasma−088
Cumulative fraction of peptides
Retention time difference (in sec)
0.0
0.2
0.4
0.6
0.8
1.0
0
20
40
60
80
No alignment
Linear alignment
LOESS, default span = 0.75
LOESS with optimum span
Chromatogram alignment

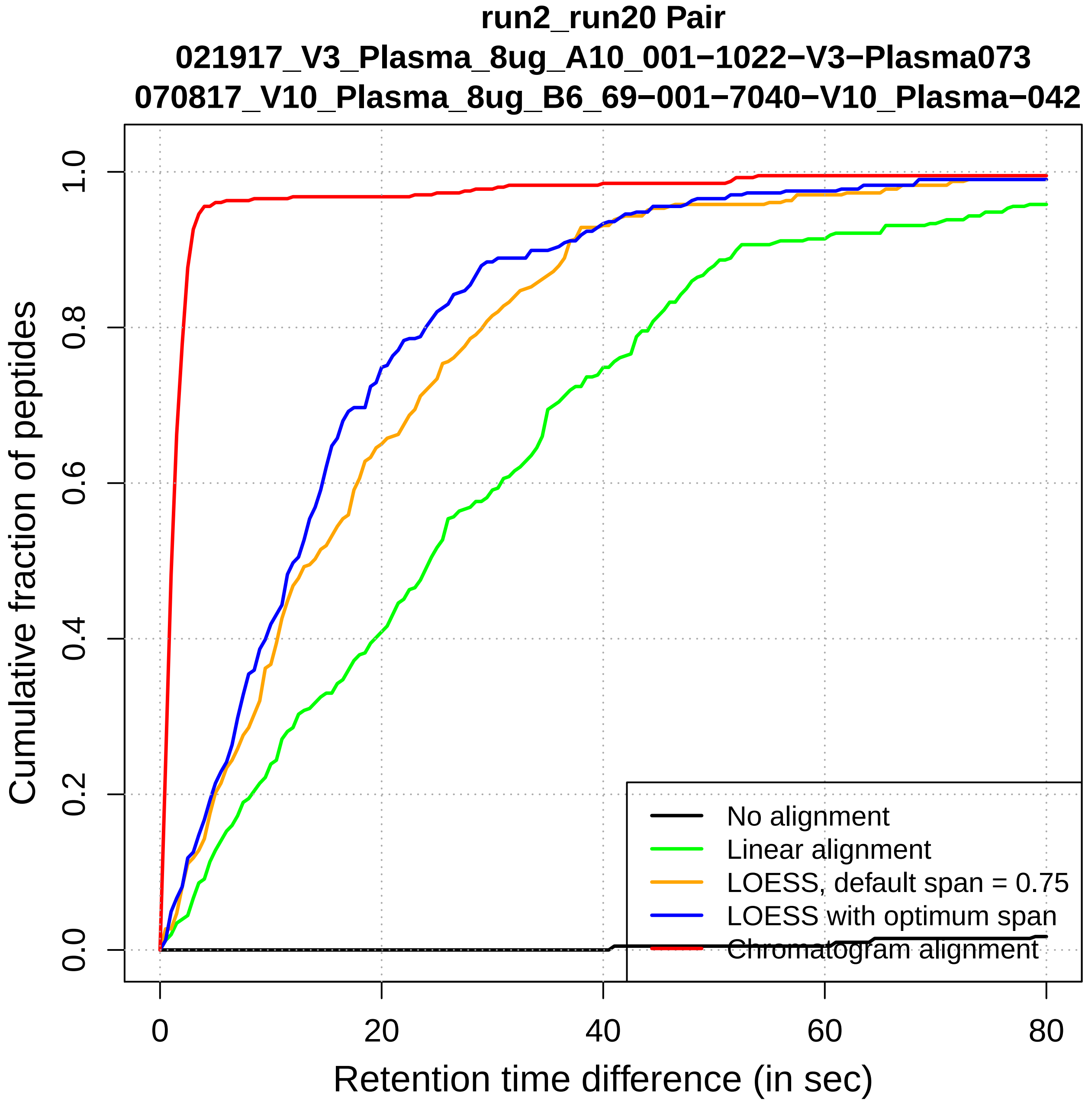

run2_run20 Pair
021917_V3_Plasma_8ug_A10_001−1022−V3−Plasma073
070817_V10_Plasma_8ug_B6_69−001−7040−V10_Plasma−042
Cumulative fraction of peptides
0.0
0.2
0.4
0.6
0.8
1.0
0
20
40
60
80
Retention time difference (in sec)
No alignment
Linear alignment
LOESS, default span = 0.75
LOESS with optimum span
Chromatogram alignment

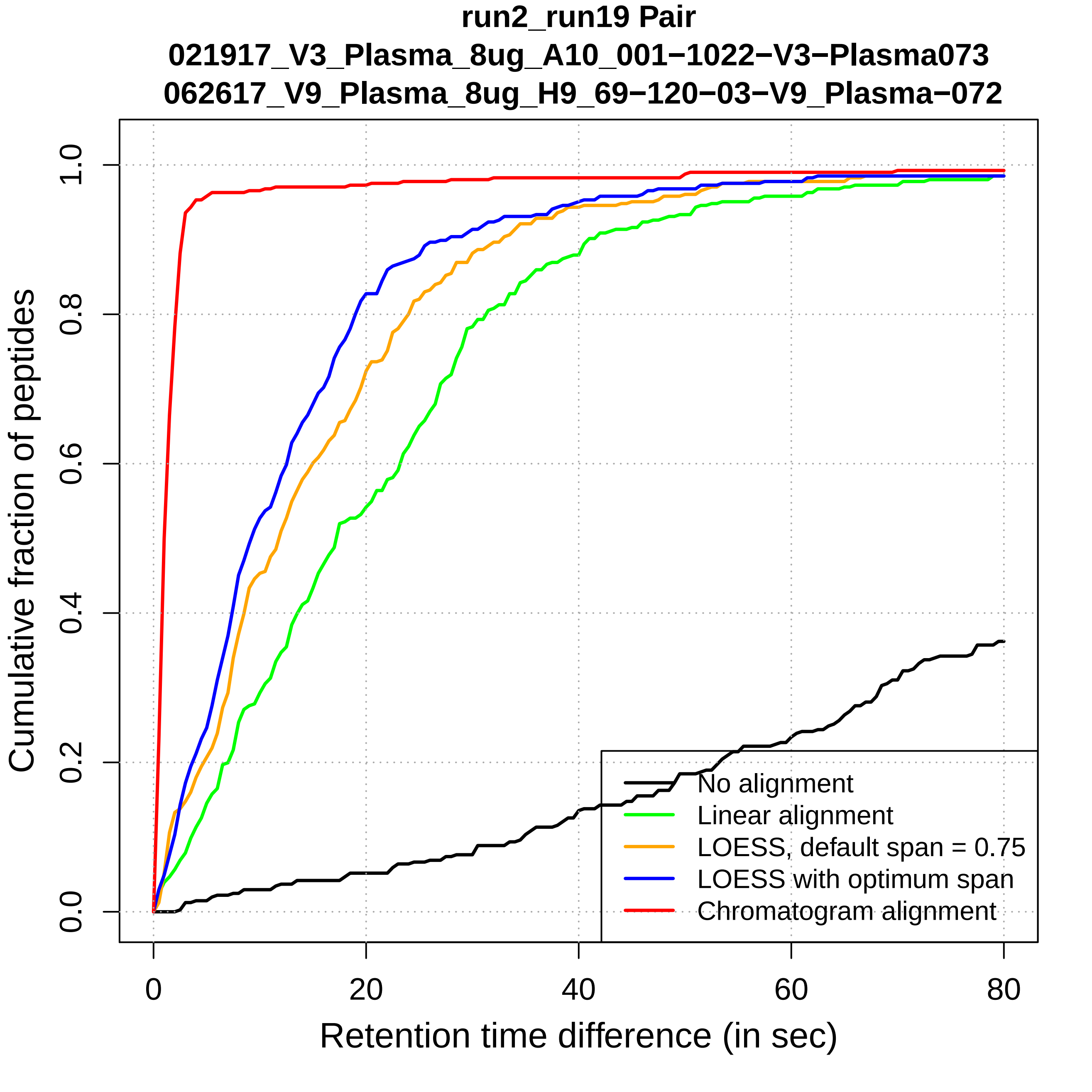

run2_run19 Pair
021917_V3_Plasma_8ug_A10_001−1022−V3−Plasma073
062617_V9_Plasma_8ug_H9_69−120−03−V9_Plasma−072
Cumulative fraction of peptides
0.0
0.2
0.4
0.6
0.8
1.0
0
20
40
60
80
Retention time difference (in sec)
No alignment
Linear alignment
LOESS, default span = 0.75
LOESS with optimum span
Chromatogram alignment

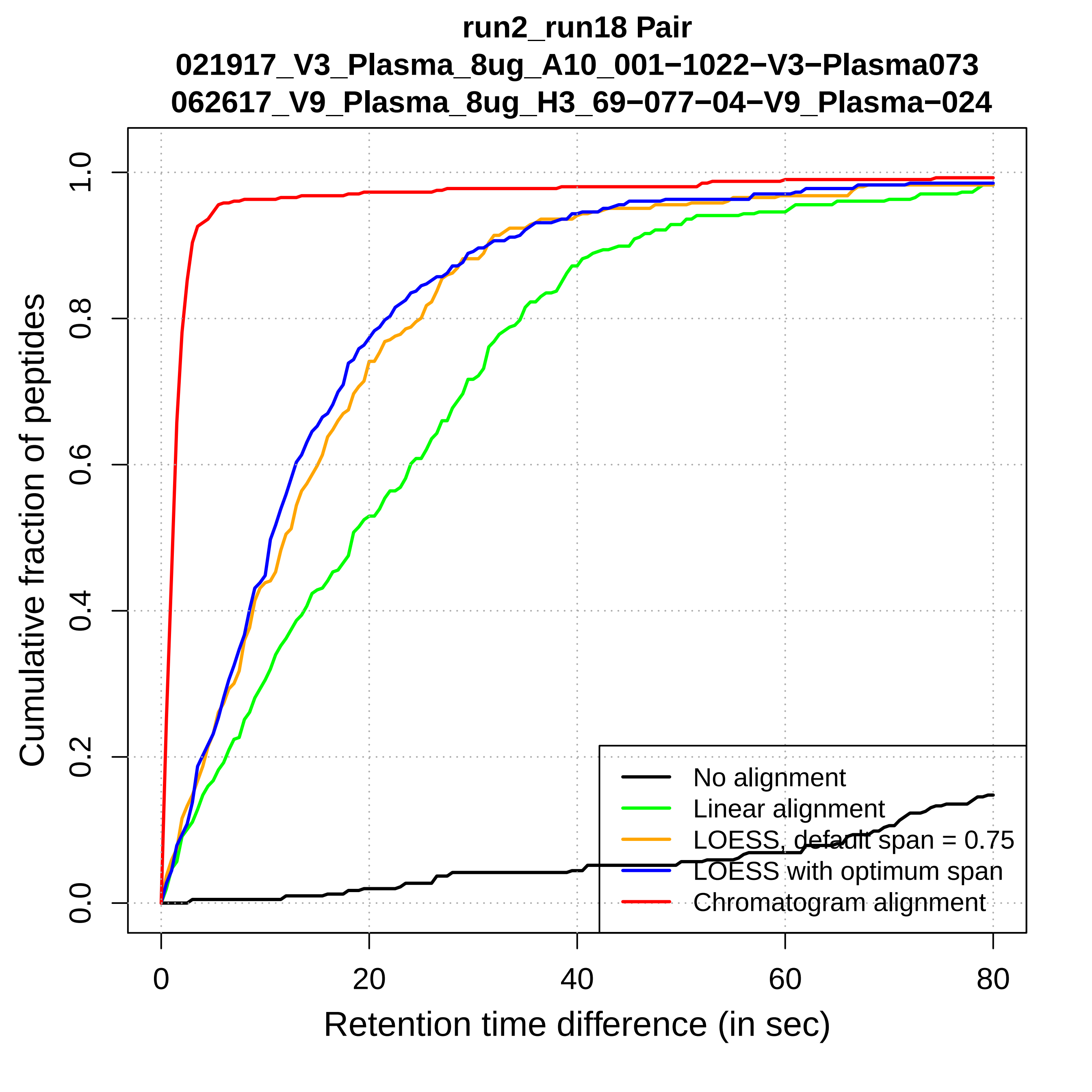

run2_run18 Pair
021917_V3_Plasma_8ug_A10_001−1022−V3−Plasma073
062617_V9_Plasma_8ug_H3_69−077−04−V9_Plasma−024
Cumulative fraction of peptides
0.0
0.2
0.4
0.6
0.8
1.0
0
20
40
60
80
Retention time difference (in sec)
No alignment
Linear alignment
LOESS, default span = 0.75
LOESS with optimum span
Chromatogram alignment

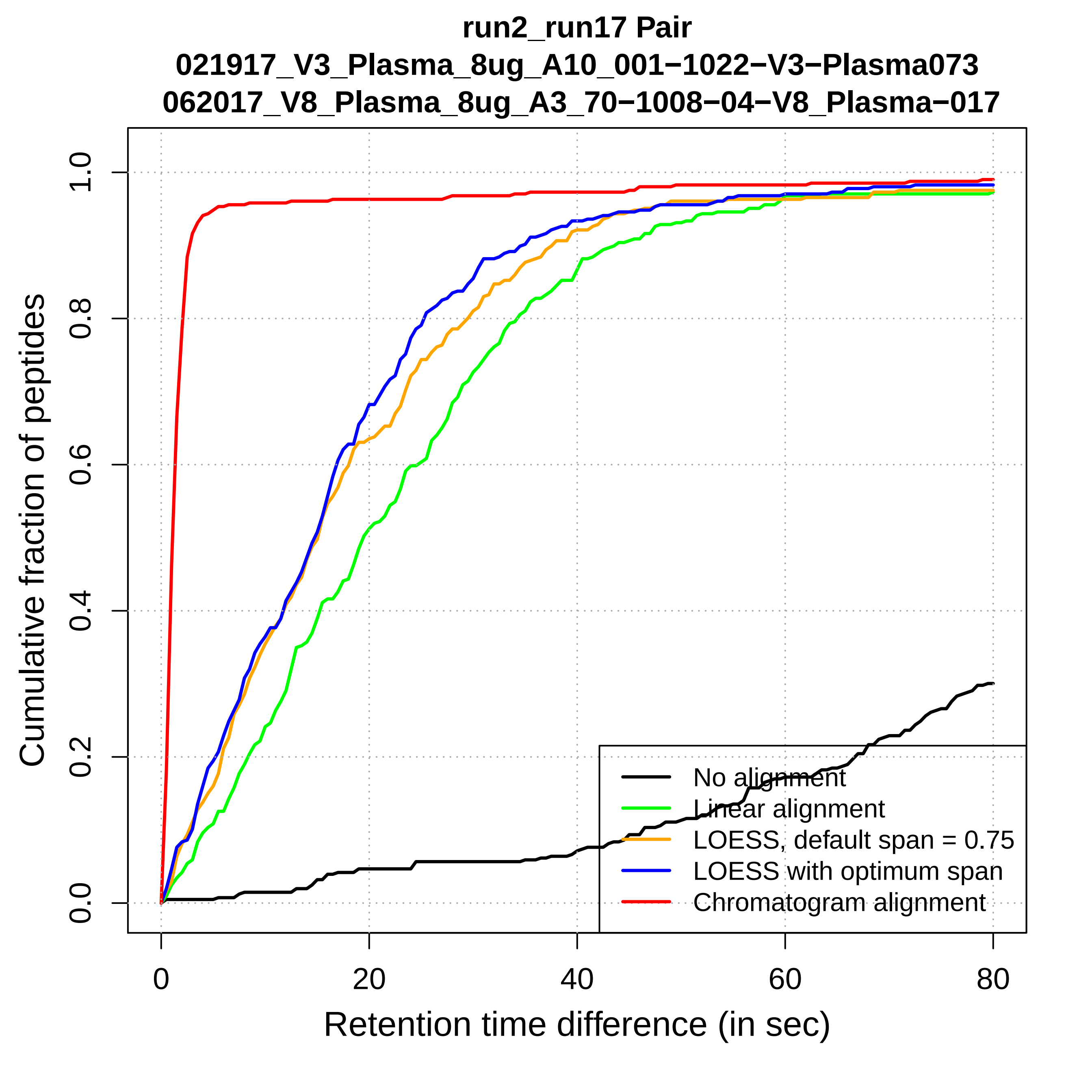
run2_run17 Pair
021917_V3_Plasma_8ug_A10_001−1022−V3−Plasma073
062017_V8_Plasma_8ug_A3_70−1008−04−V8_Plasma−017
Cumulative fraction of peptides
0.0
0.2
0.4
0.6
0.8
1.0
0
20
40
60
80
Retention time difference (in sec)
No alignment
Linear alignment
LOESS, default span = 0.75
LOESS with optimum span
Chromatogram alignment

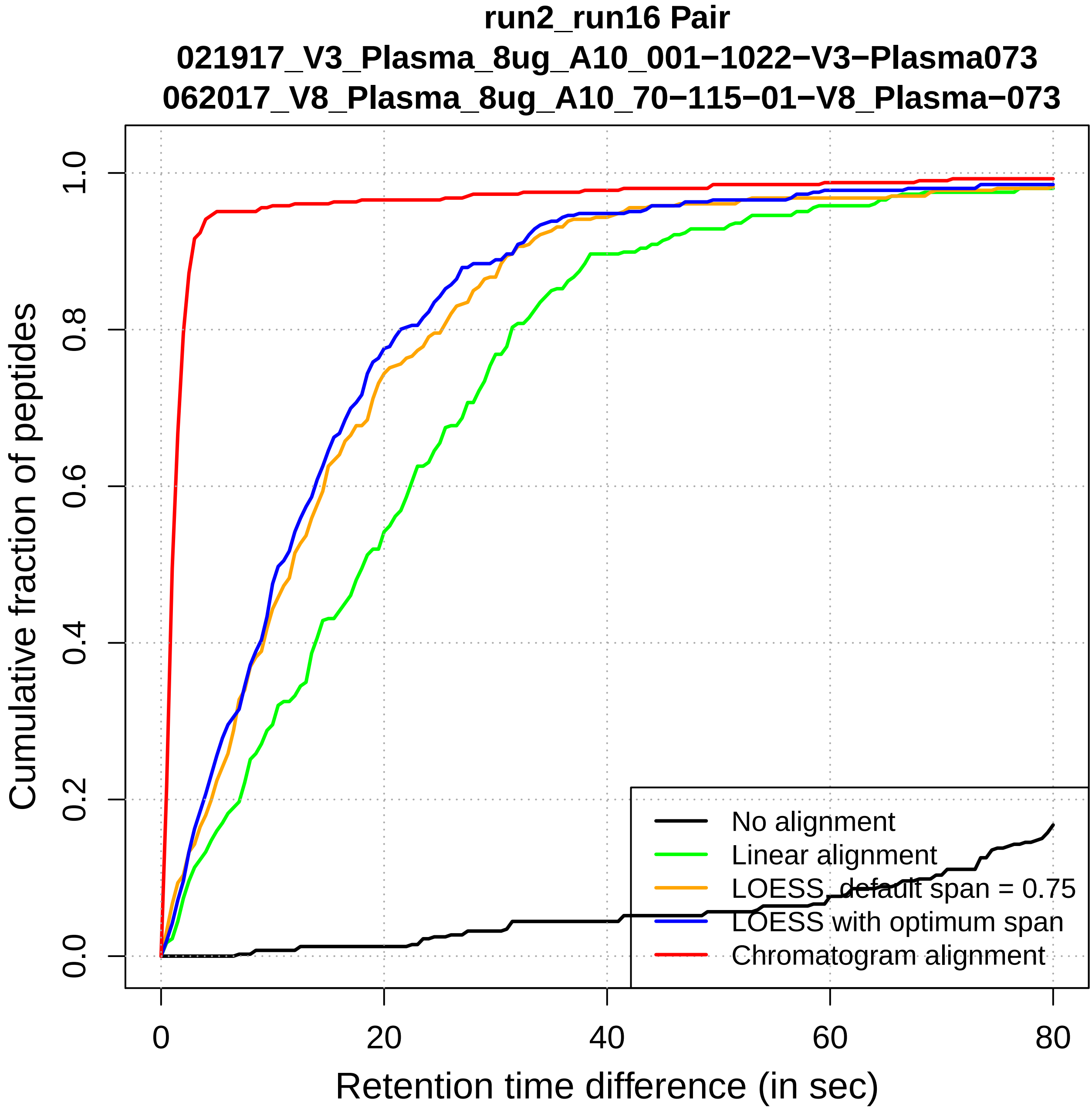
run2_run16 Pair
021917_V3_Plasma_8ug_A10_001−1022−V3−Plasma073
062017_V8_Plasma_8ug_A10_70−115−01−V8_Plasma−073
Cumulative fraction of peptides
0.0
0.2
0.4
0.6
0.8
1.0
0
20
40
60
80
Retention time difference (in sec)
No alignment
Linear alignment
LOESS, default span = 0.75
LOESS with optimum span
Chromatogram alignment

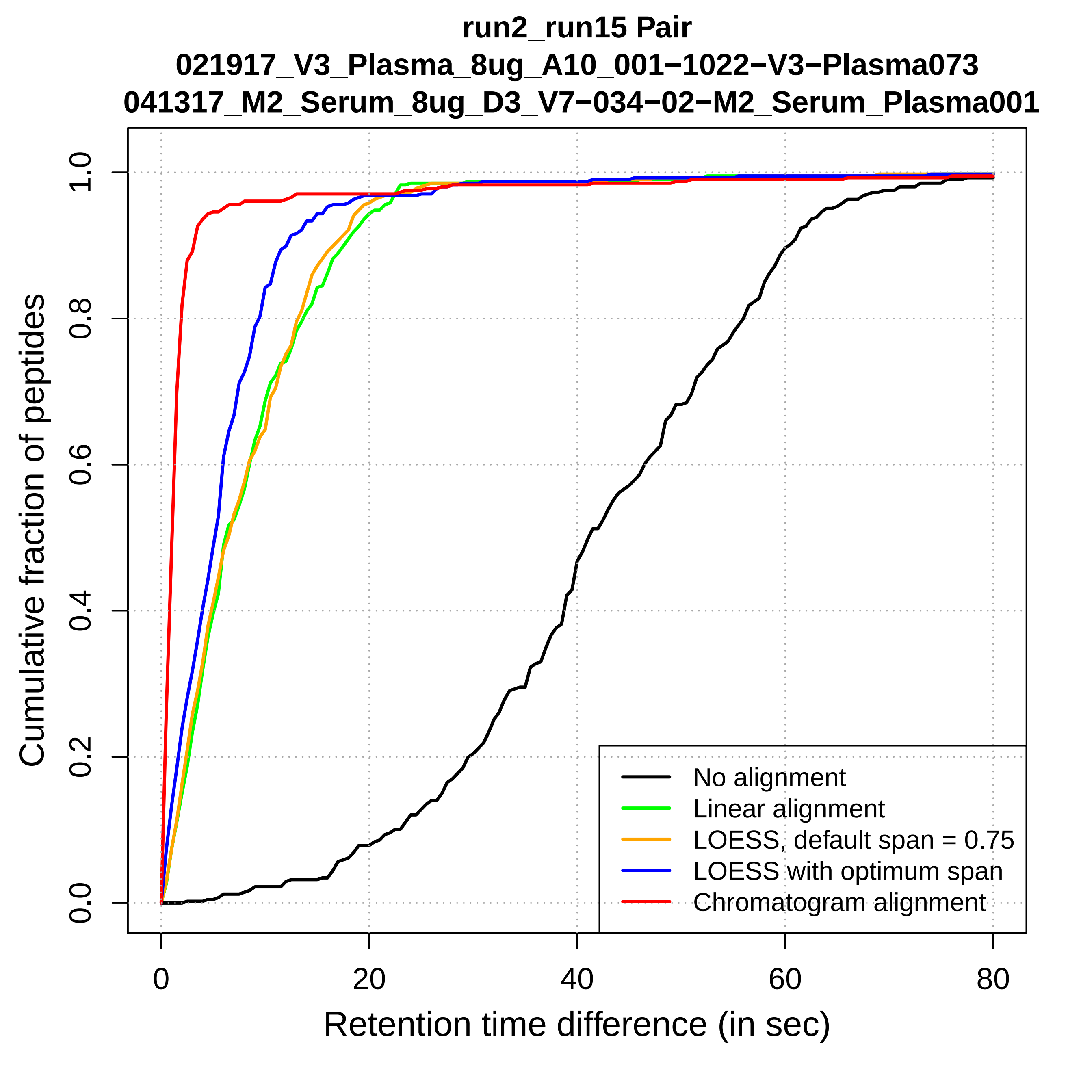

run2_run15 Pair
021917_V3_Plasma_8ug_A10_001−1022−V3−Plasma073
041317_M2_Serum_8ug_D3_V7−034−02−M2_Serum_Plasma001
Cumulative fraction of peptides
0.0
0.2
0.4
0.6
0.8
1.0
0
20
40
60
80
Retention time difference (in sec)
No alignment
Linear alignment
LOESS, default span = 0.75
LOESS with optimum span
Chromatogram alignment

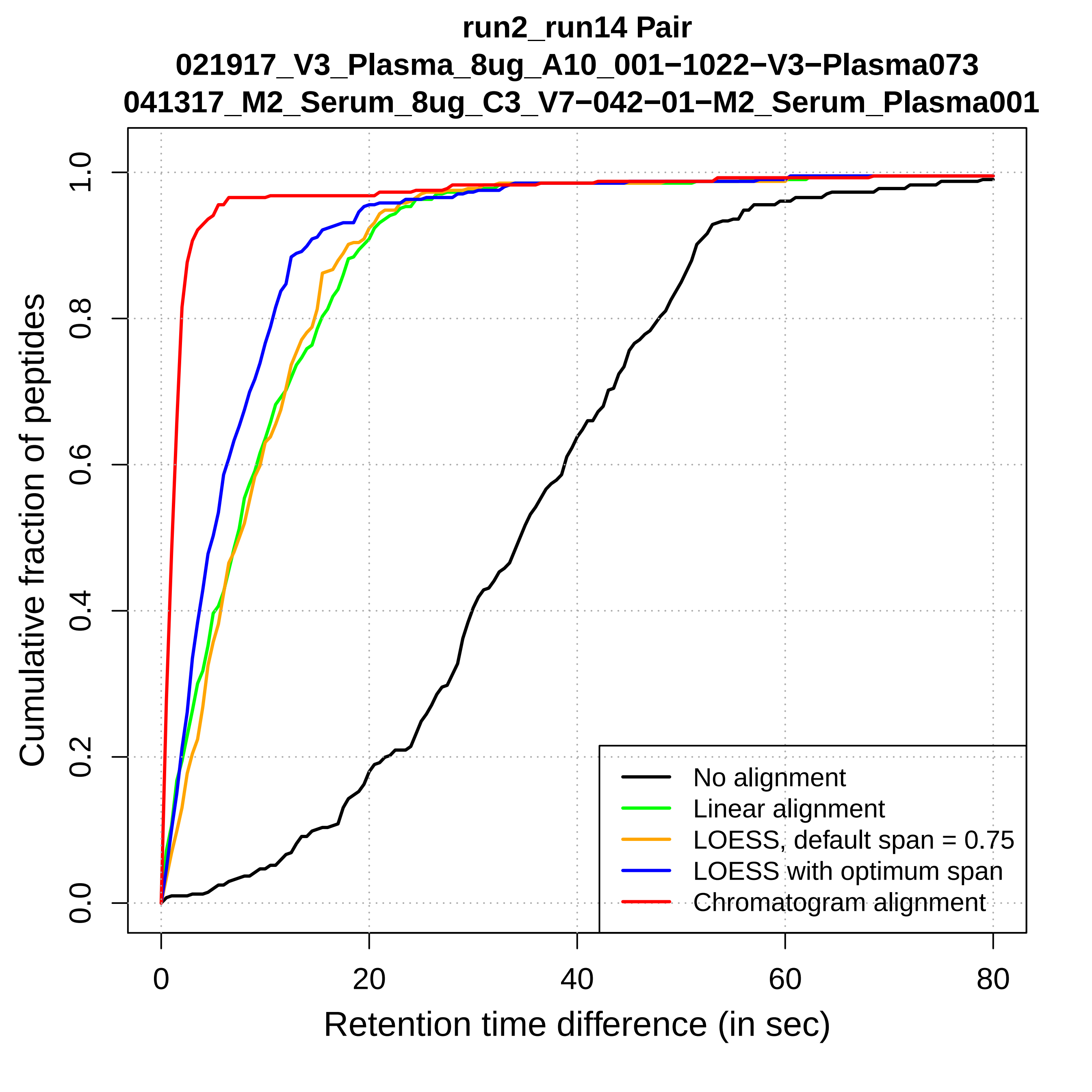
run2_run14 Pair
021917_V3_Plasma_8ug_A10_001−1022−V3−Plasma073
041317_M2_Serum_8ug_C3_V7−042−01−M2_Serum_Plasma001
Cumulative fraction of peptides
0.0
0.2
0.4
0.6
0.8
1.0
0
20
40
60
80
Retention time difference (in sec)
No alignment
Linear alignment
LOESS, default span = 0.75
LOESS with optimum span
Chromatogram alignment

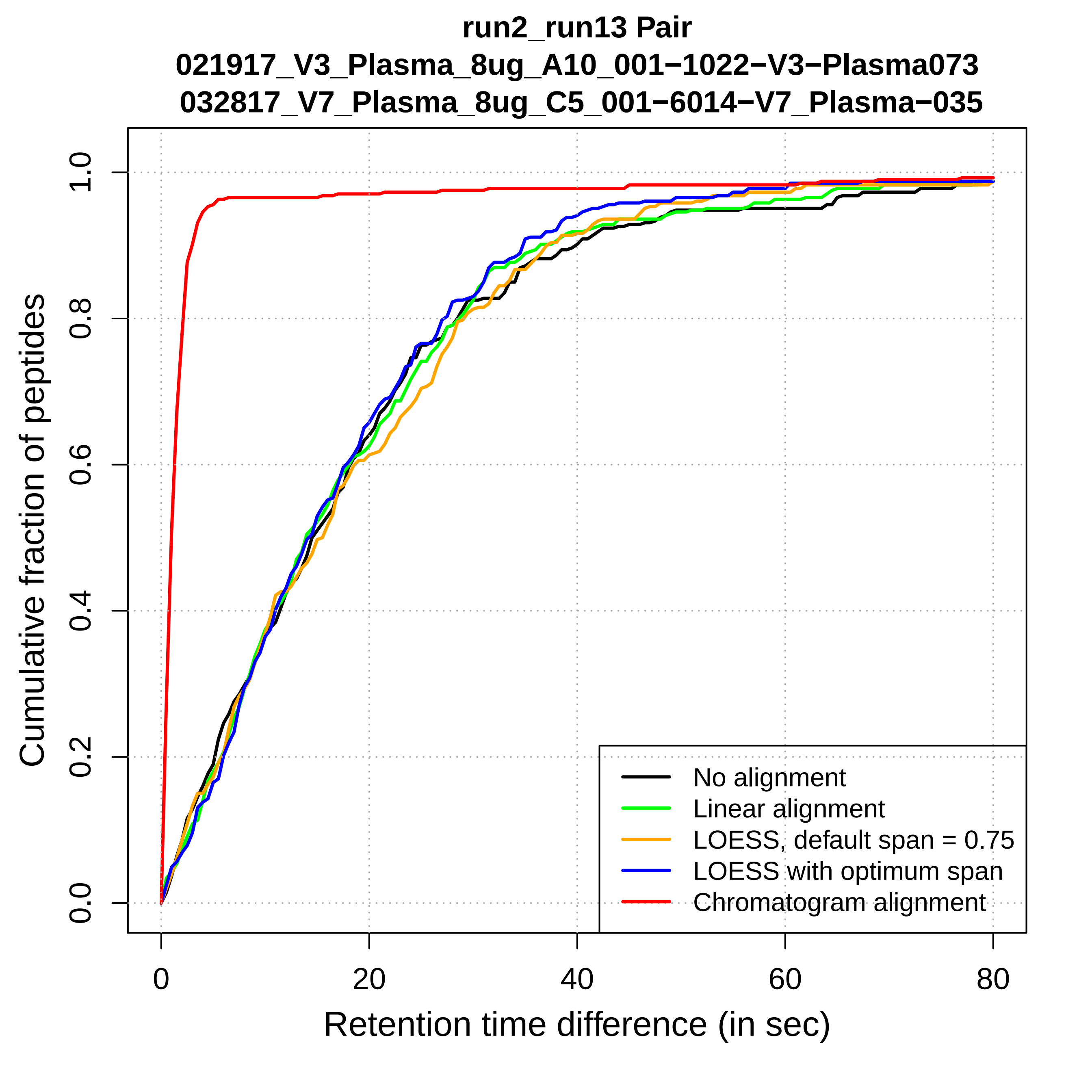

run2_run13 Pair
021917_V3_Plasma_8ug_A10_001−1022−V3−Plasma073
032817_V7_Plasma_8ug_C5_001−6014−V7_Plasma−035
Cumulative fraction of peptides
0.0
0.2
0.4
0.6
0.8
1.0
0
20
40
60
80
Retention time difference (in sec)
No alignment
Linear alignment
LOESS, default span = 0.75
LOESS with optimum span
Chromatogram alignment

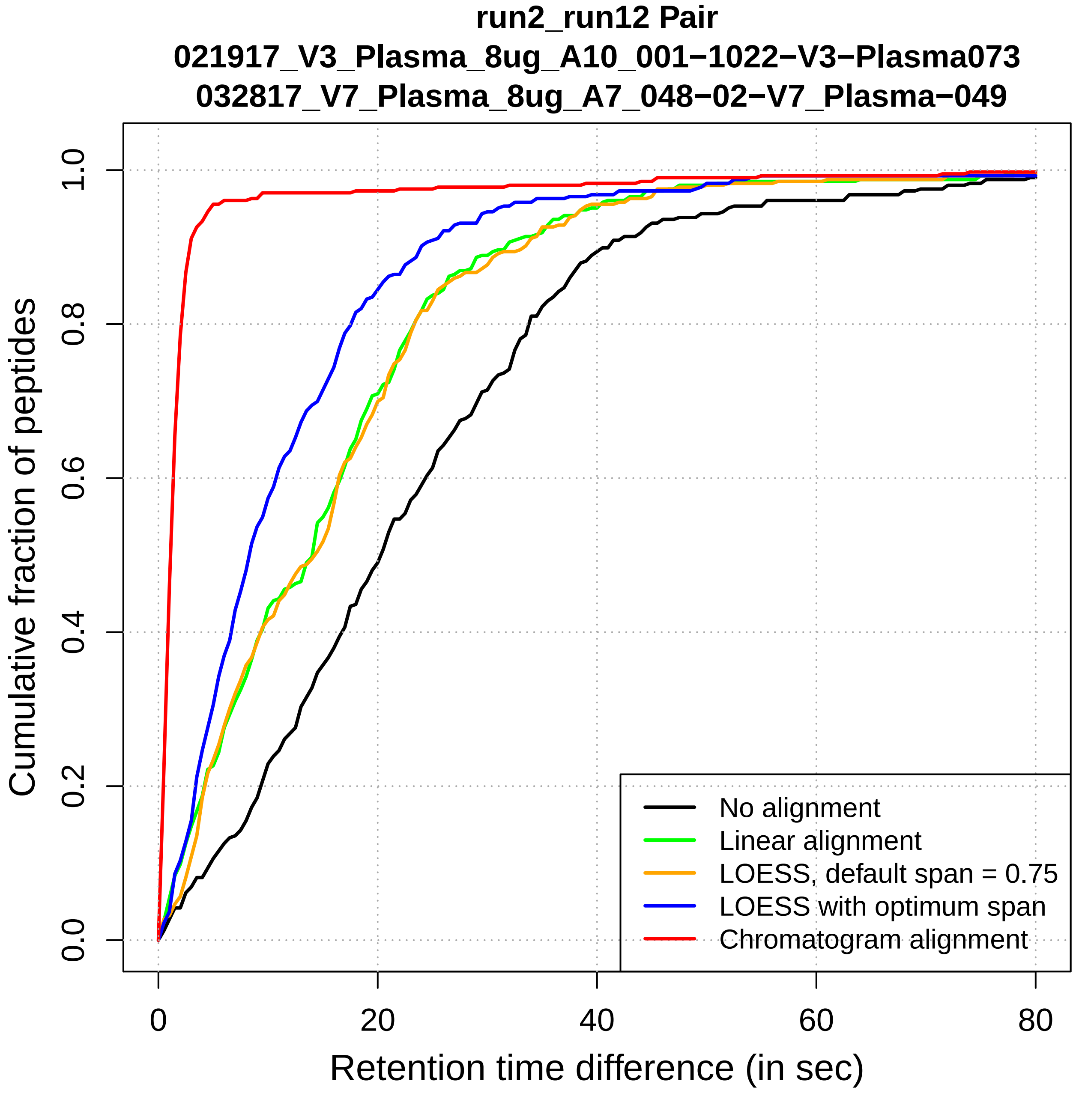
run2_run12 Pair
021917_V3_Plasma_8ug_A10_001−1022−V3−Plasma073
032817_V7_Plasma_8ug_A7_048−02−V7_Plasma−049
Cumulative fraction of peptides
0.0
0.2
0.4
0.6
0.8
1.0
0
20
40
60
80
Retention time difference (in sec)
No alignment
Linear alignment
LOESS, default span = 0.75
LOESS with optimum span
Chromatogram alignment

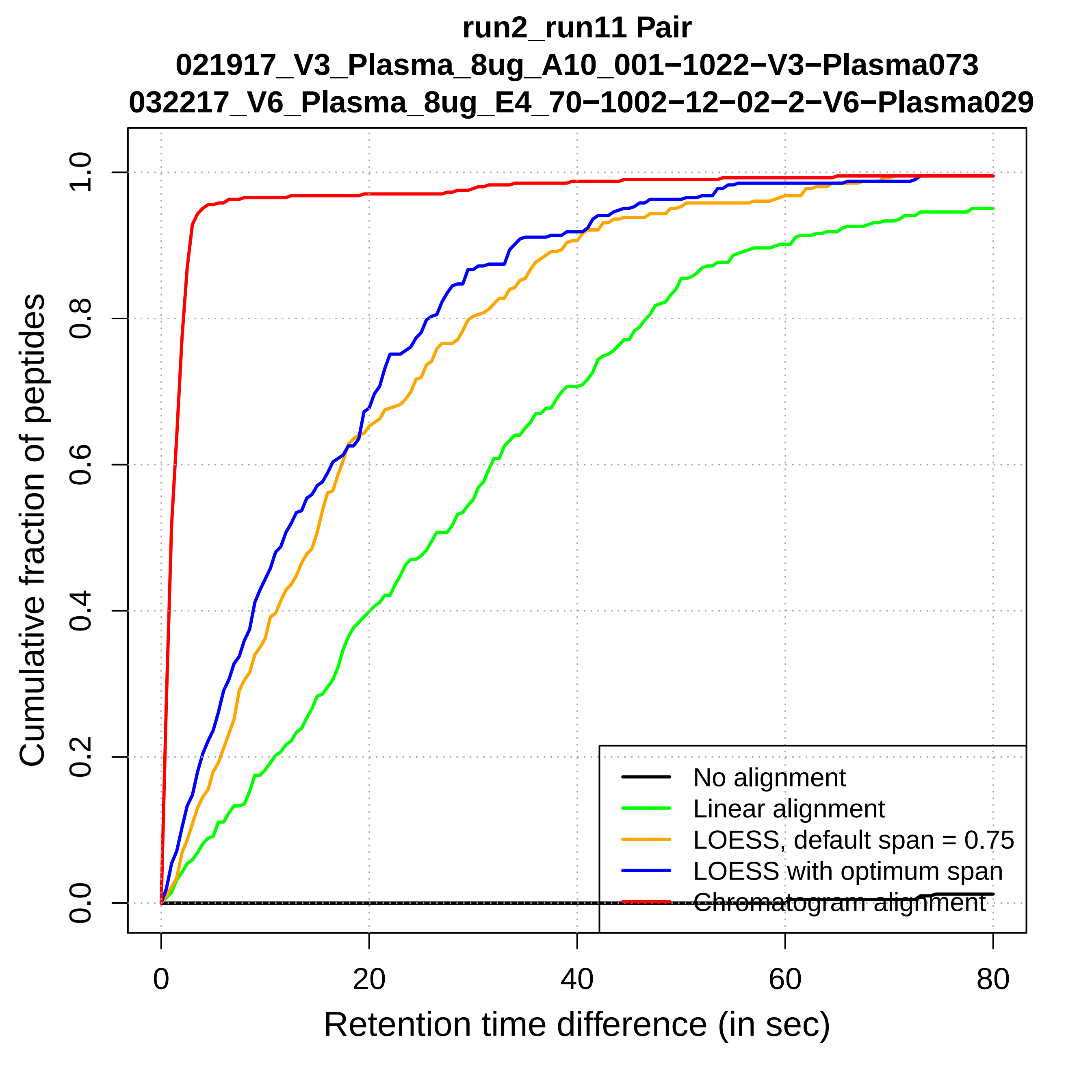
run2_run11 Pair
021917_V3_Plasma_8ug_A10_001−1022−V3−Plasma073
032217_V6_Plasma_8ug_E4_70−1002−12−02−2−V6−Plasma029
Cumulative fraction of peptides
0.0
0.2
0.4
0.6
0.8
1.0
0
20
40
60
80
Retention time difference (in sec)
No alignment
Linear alignment
LOESS, default span = 0.75
LOESS with optimum span
Chromatogram alignment

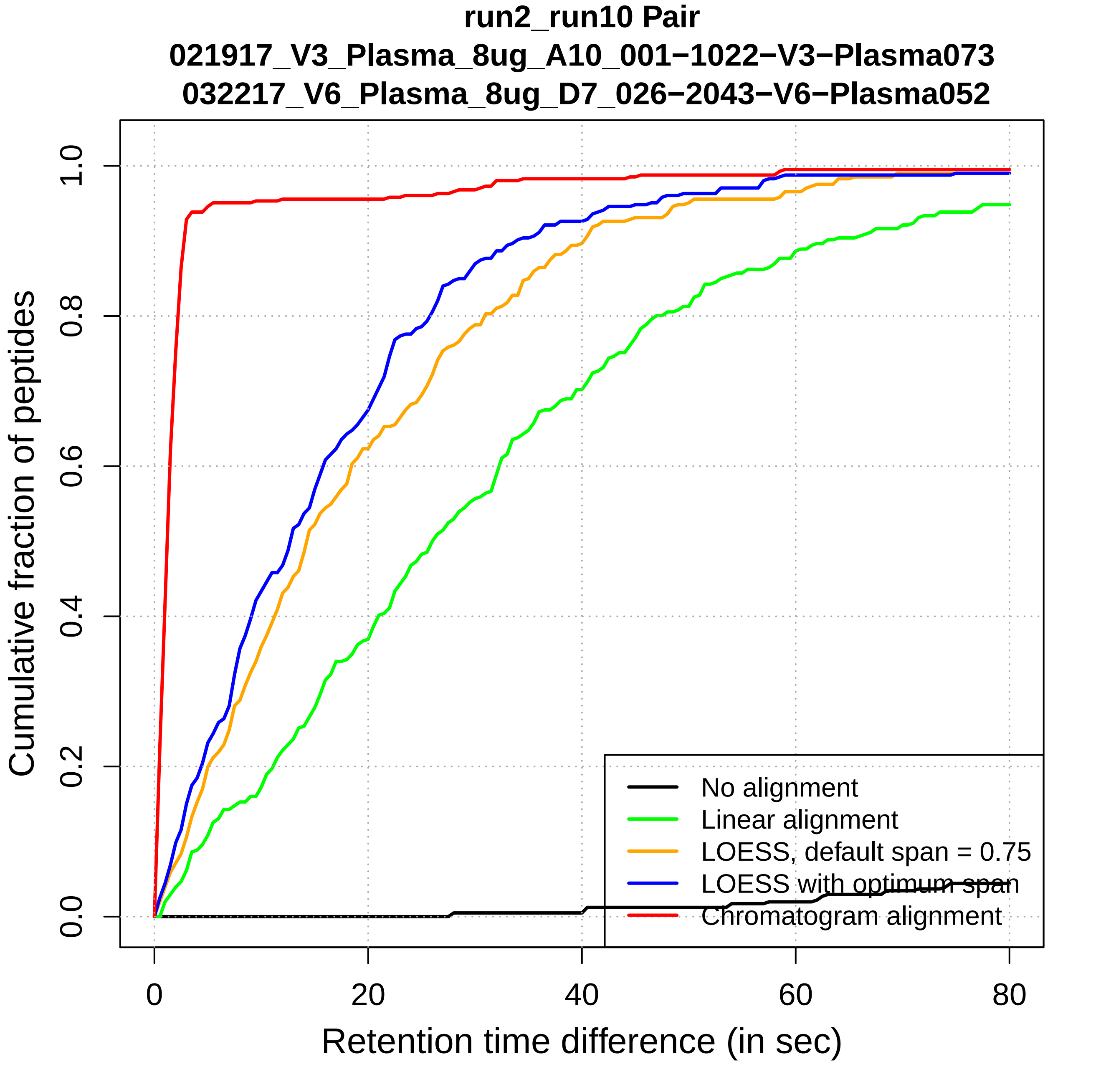

run2_run10 Pair
021917_V3_Plasma_8ug_A10_001−1022−V3−Plasma073
032217_V6_Plasma_8ug_D7_026−2043−V6−Plasma052
Cumulative fraction of peptides
0.0
0.2
0.4
0.6
0.8
1.0
0
20
40
60
80
Retention time difference (in sec)
No alignment
Linear alignment
LOESS, default span = 0.75
LOESS with optimum span
Chromatogram alignment

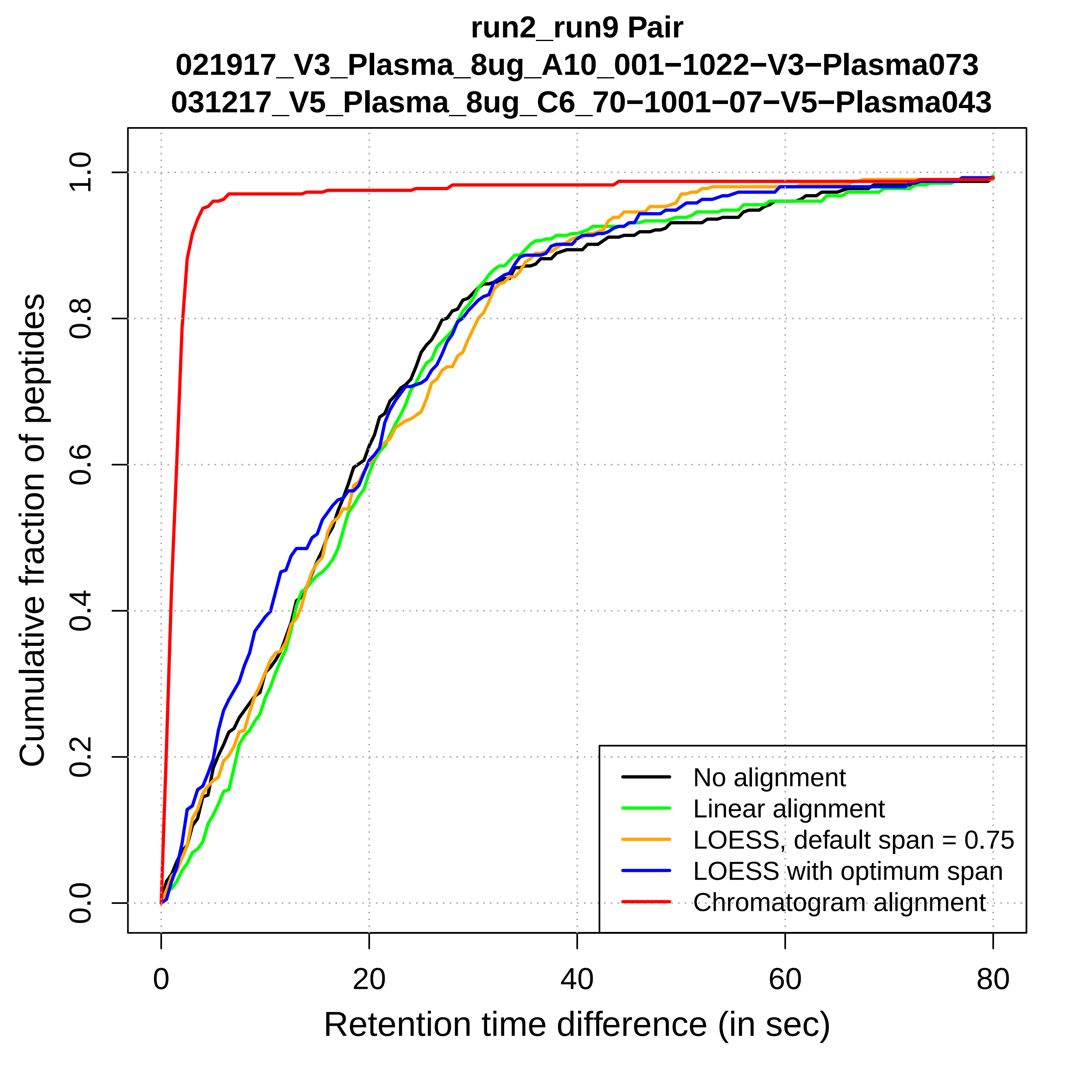
run2_run9 Pair
021917_V3_Plasma_8ug_A10_001−1022−V3−Plasma073
031217_V5_Plasma_8ug_C6_70−1001−07−V5−Plasma043
Cumulative fraction of peptides
0.0
0.2
0.4
0.6
0.8
1.0
0
20
40
60
80
Retention time difference (in sec)
No alignment
Linear alignment
LOESS, default span = 0.75
LOESS with optimum span
Chromatogram alignment

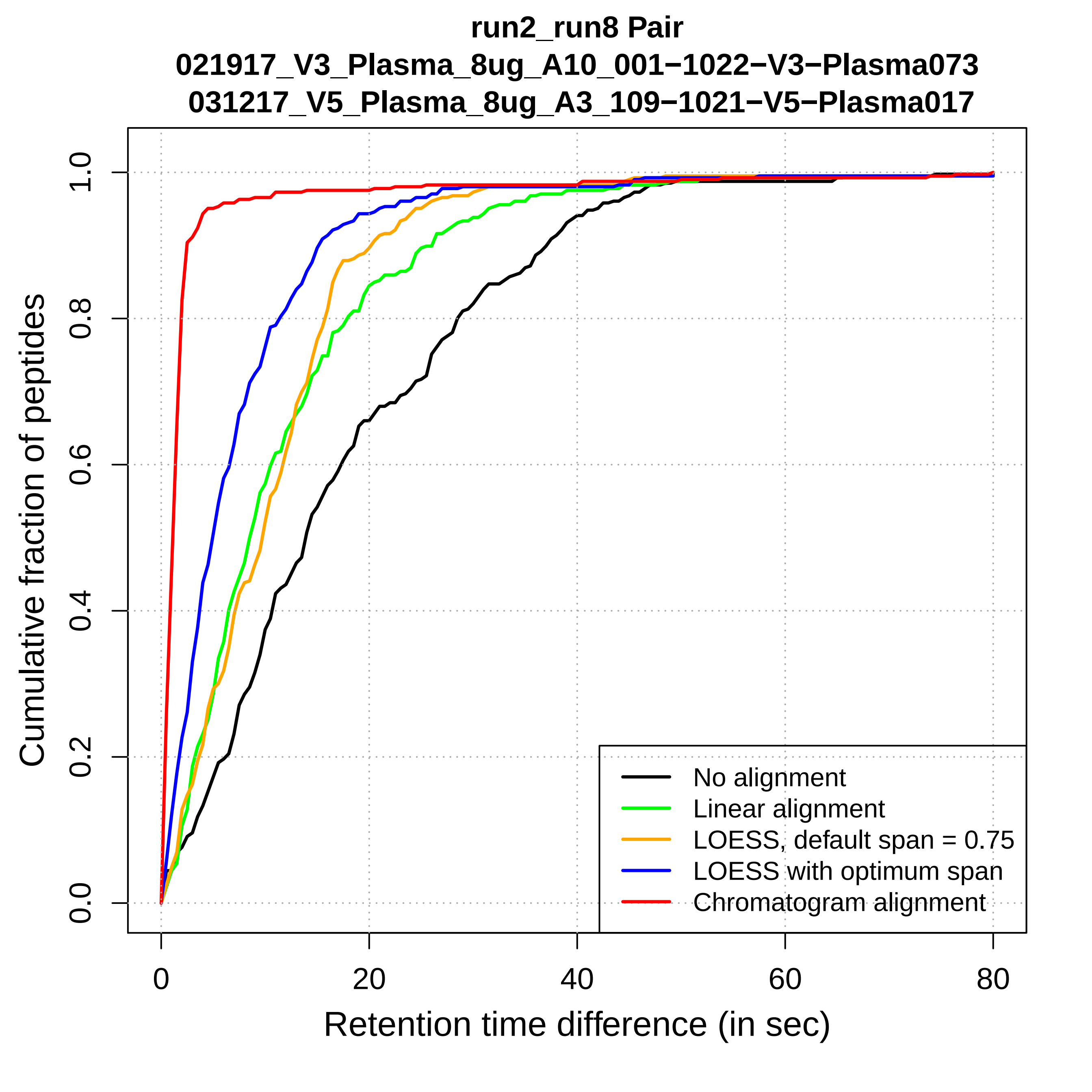
run2_run8 Pair
021917_V3_Plasma_8ug_A10_001−1022−V3−Plasma073
031217_V5_Plasma_8ug_A3_109−1021−V5−Plasma017
Cumulative fraction of peptides
0.0
0.2
0.4
0.6
0.8
1.0
0
20
40
60
80
Retention time difference (in sec)
No alignment
Linear alignment
LOESS, default span = 0.75
LOESS with optimum span
Chromatogram alignment

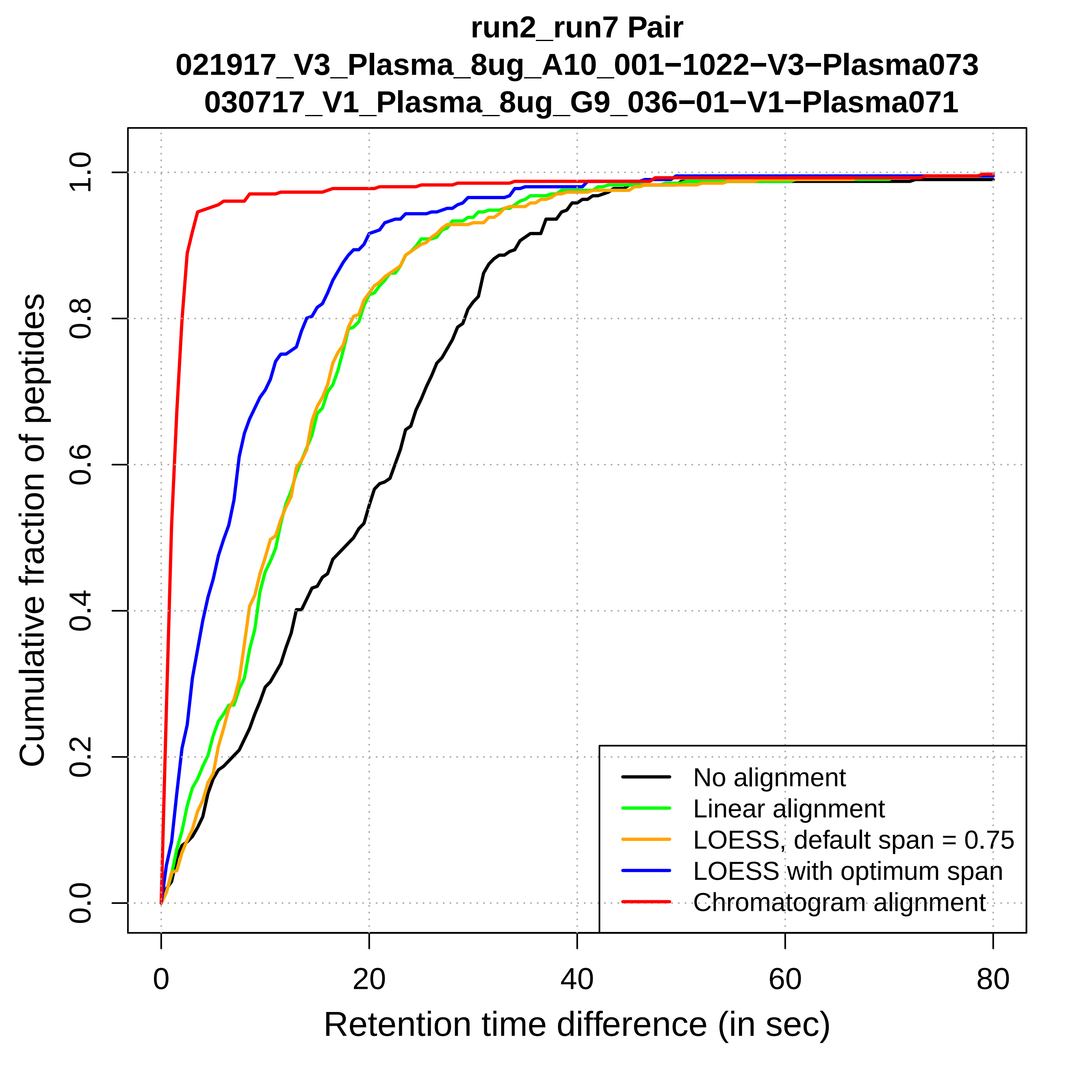
run2_run7 Pair
021917_V3_Plasma_8ug_A10_001−1022−V3−Plasma073
030717_V1_Plasma_8ug_G9_036−01−V1−Plasma071
Cumulative fraction of peptides
Retention time difference (in sec)
0.0
0.2
0.4
0.6
0.8
1.0
0
20
40
60
80
No alignment
Linear alignment
LOESS, default span = 0.75
LOESS with optimum span
Chromatogram alignment

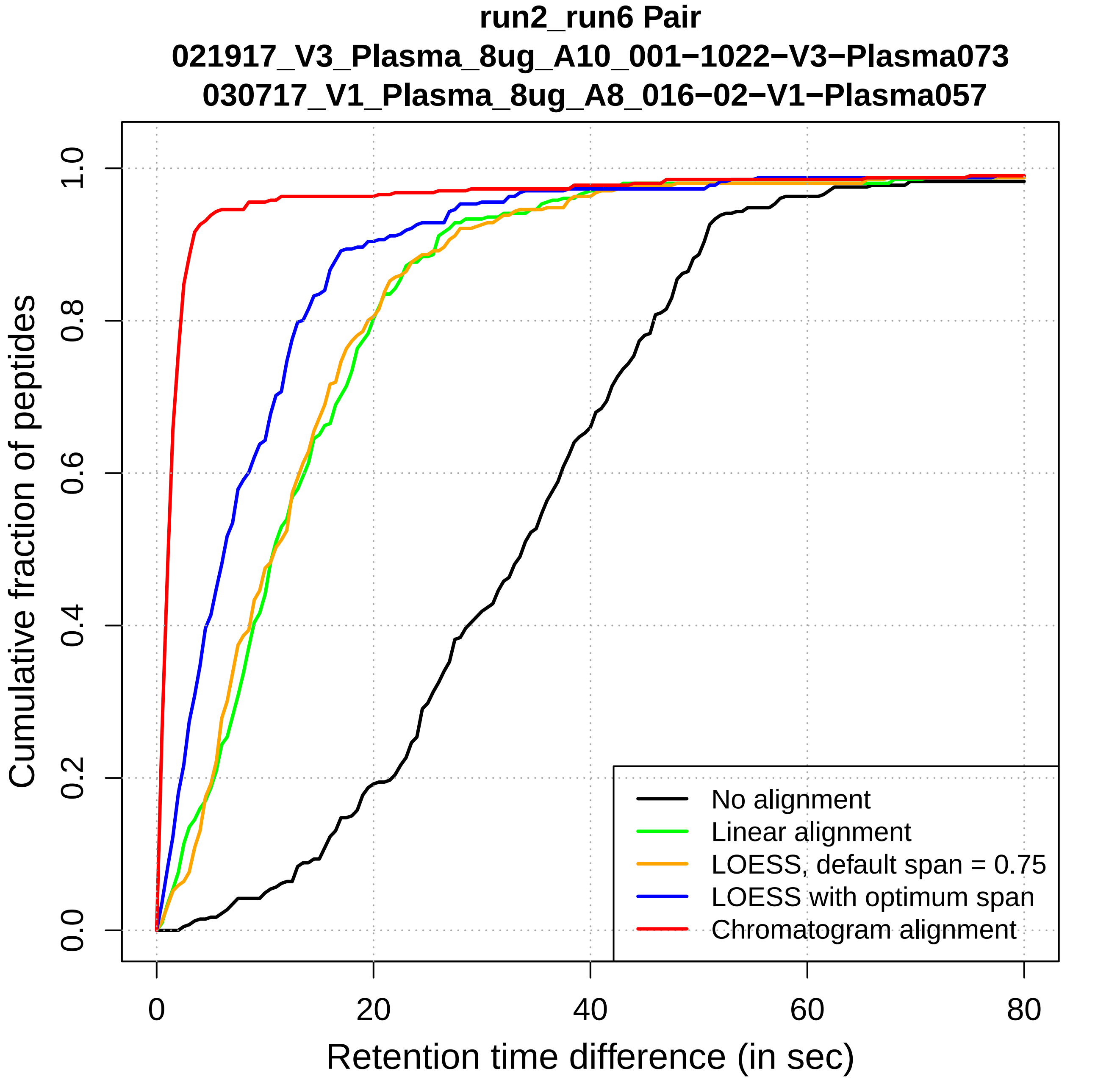

run2_run6 Pair
021917_V3_Plasma_8ug_A10_001−1022−V3−Plasma073
030717_V1_Plasma_8ug_A8_016−02−V1−Plasma057
Cumulative fraction of peptides
Retention time difference (in sec)
0.0
0.2
0.4
0.6
0.8
1.0
0
20
40
60
80
No alignment
Linear alignment
LOESS, default span = 0.75
LOESS with optimum span
Chromatogram alignment

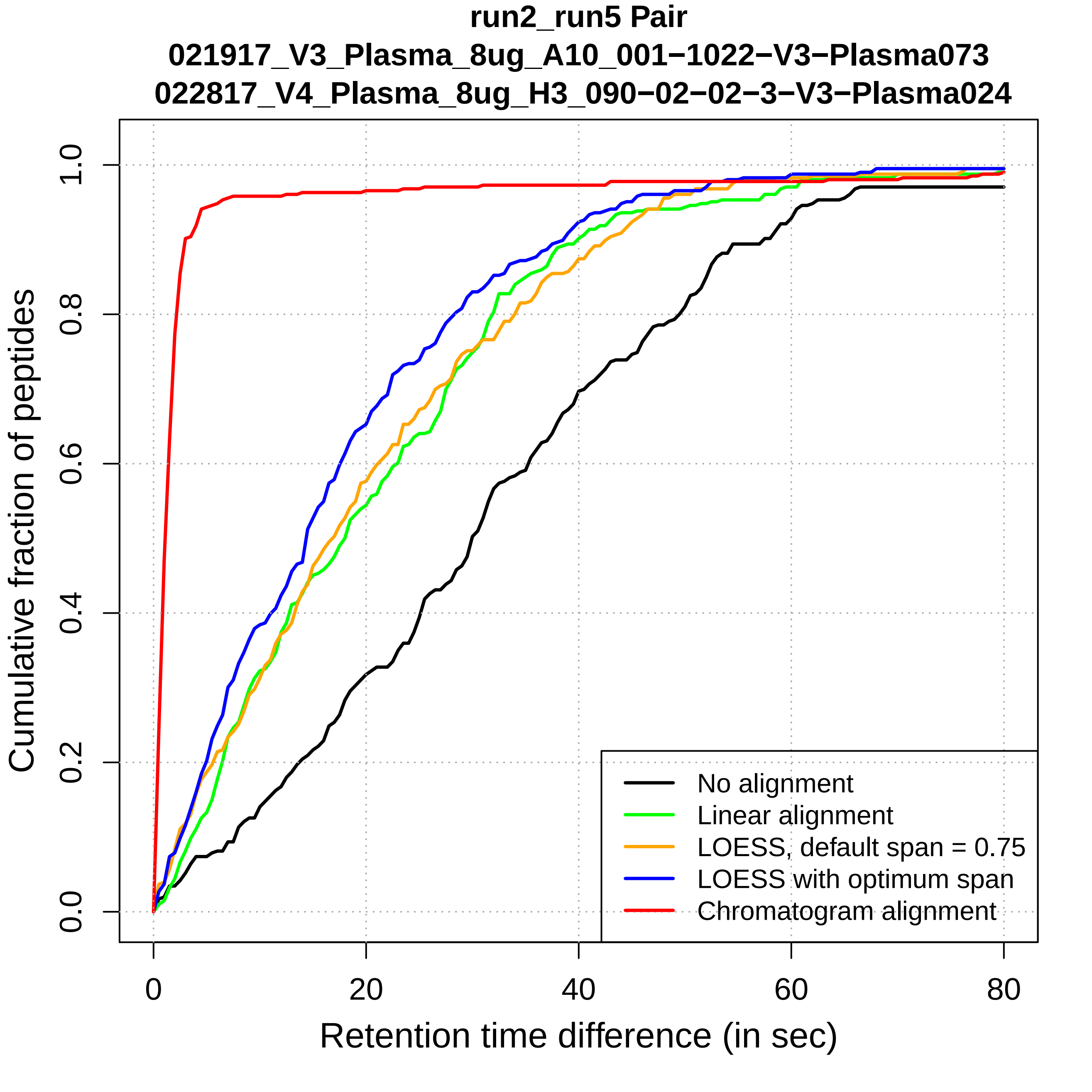
run2_run5 Pair
021917_V3_Plasma_8ug_A10_001−1022−V3−Plasma073
022817_V4_Plasma_8ug_H3_090−02−02−3−V3−Plasma024
Cumulative fraction of peptides
0.0
0.2
0.4
0.6
0.8
1.0
0
20
40
60
80
Retention time difference (in sec)
No alignment
Linear alignment
LOESS, default span = 0.75
LOESS with optimum span
Chromatogram alignment

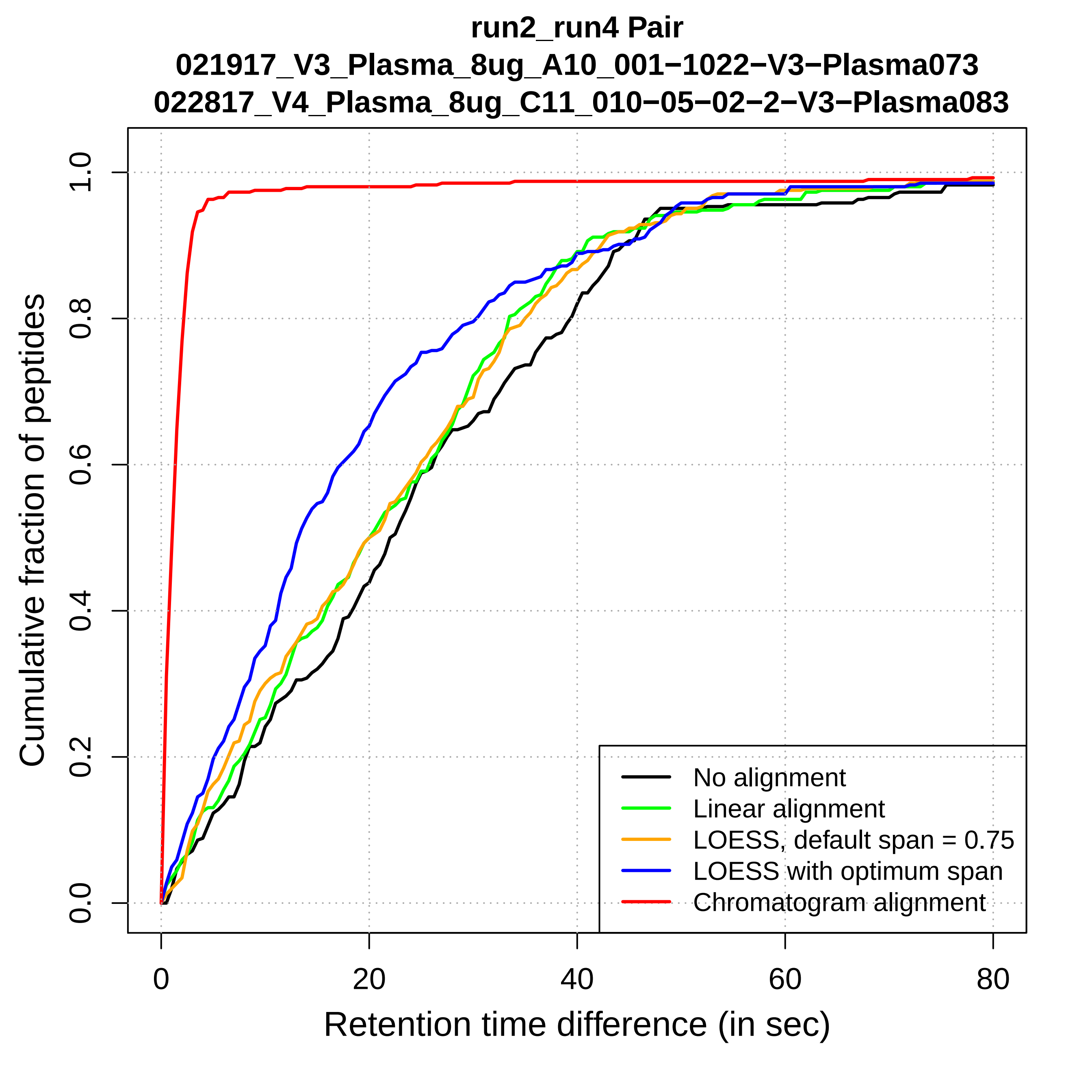
run2_run4 Pair
021917_V3_Plasma_8ug_A10_001−1022−V3−Plasma073
022817_V4_Plasma_8ug_C11_010−05−02−2−V3−Plasma083
Cumulative fraction of peptides
0.0
0.2
0.4
0.6
0.8
1.0
0
20
40
60
80
Retention time difference (in sec)
No alignment
Linear alignment
LOESS, default span = 0.75
LOESS with optimum span
Chromatogram alignment

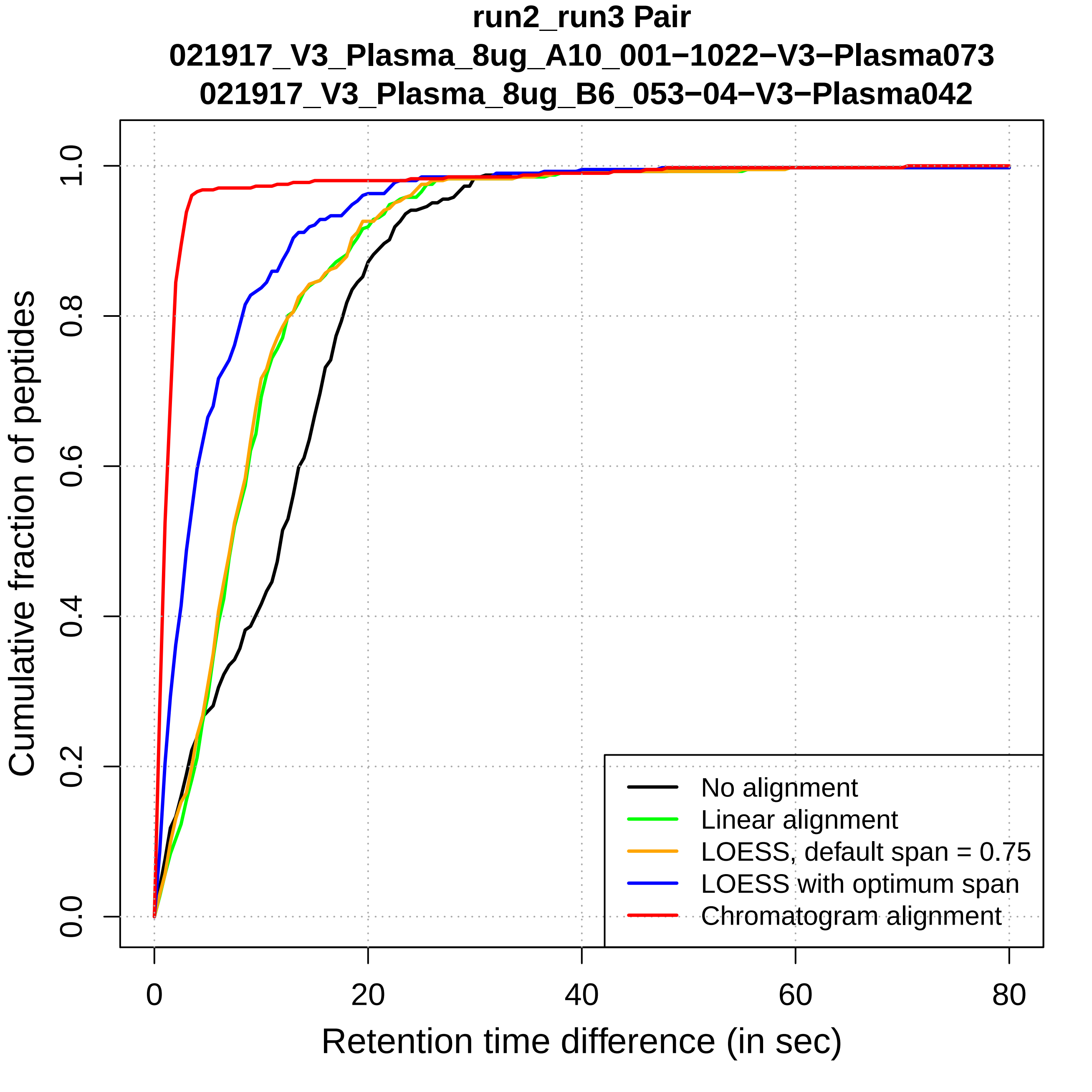

run2_run3 Pair
021917_V3_Plasma_8ug_A10_001−1022−V3−Plasma073
021917_V3_Plasma_8ug_B6_053−04−V3−Plasma042
Cumulative fraction of peptides
0.0
0.2
0.4
0.6
0.8
1.0
0
20
40
60
80
Retention time difference (in sec)
No alignment
Linear alignment
LOESS, default span = 0.75
LOESS with optimum span
Chromatogram alignment

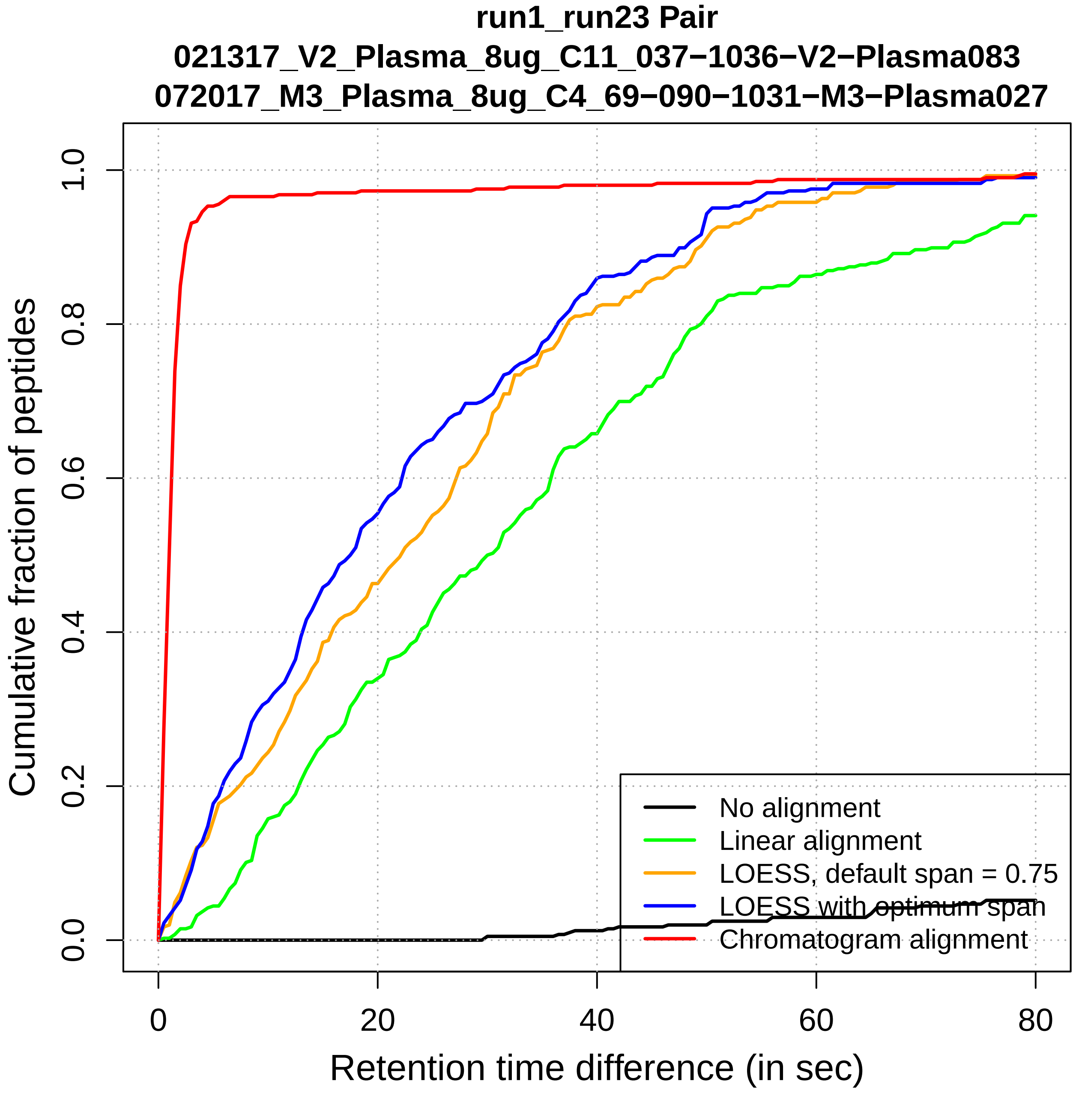

run1_run23 Pair
021317_V2_Plasma_8ug_C11_037−1036−V2−Plasma083
072017_M3_Plasma_8ug_C4_69−090−1031−M3−Plasma027
Cumulative fraction of peptides
0.0
0.2
0.4
0.6
0.8
1.0
0
20
40
60
80
Retention time difference (in sec)
No alignment
Linear alignment
LOESS, default span = 0.75
LOESS with optimum span
Chromatogram alignment

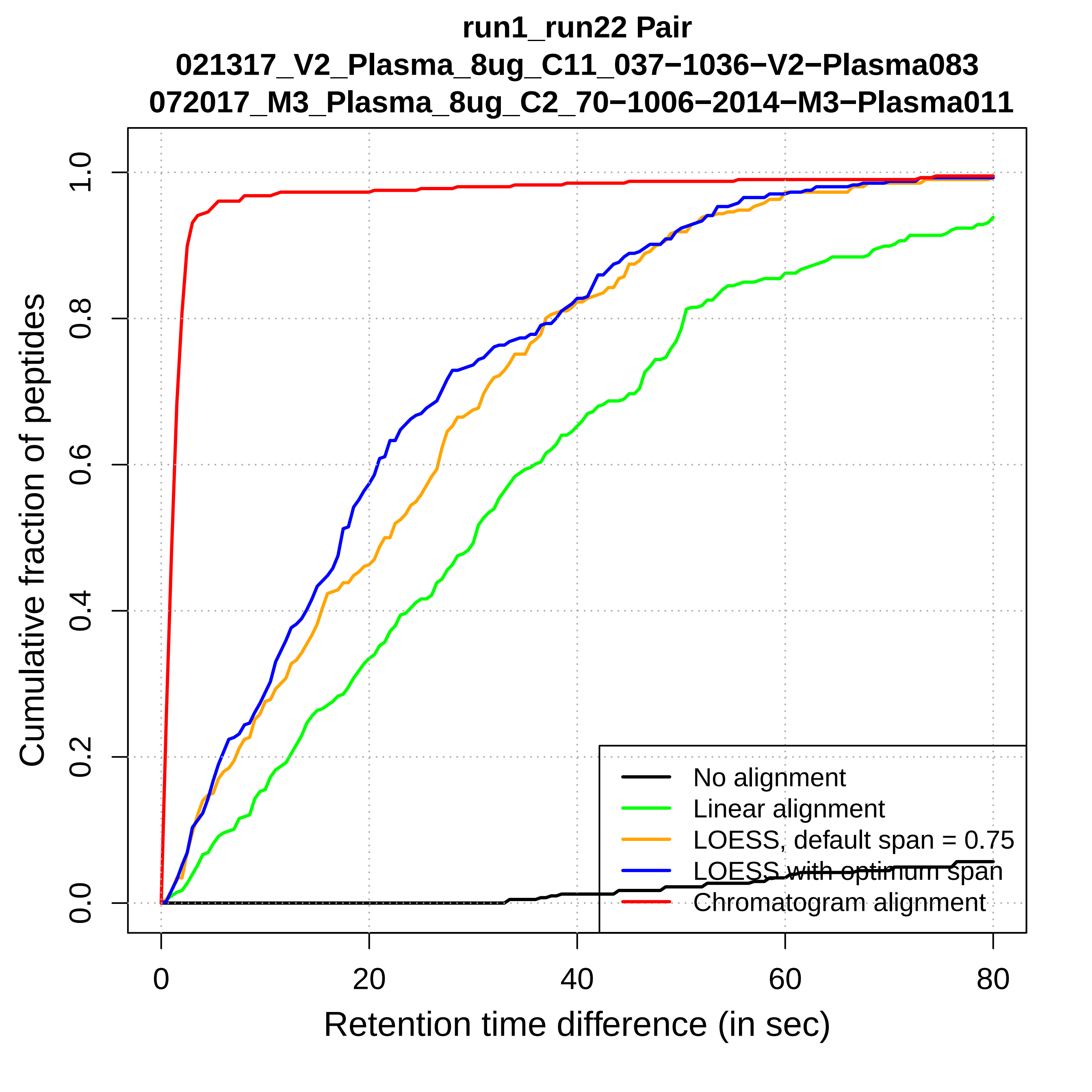
run1_run22 Pair
021317_V2_Plasma_8ug_C11_037−1036−V2−Plasma083
072017_M3_Plasma_8ug_C2_70−1006−2014−M3−Plasma011
Cumulative fraction of peptides
0.0
0.2
0.4
0.6
0.8
1.0
0
20
40
60
80
Retention time difference (in sec)
No alignment
Linear alignment
LOESS, default span = 0.75
LOESS with optimum span
Chromatogram alignment

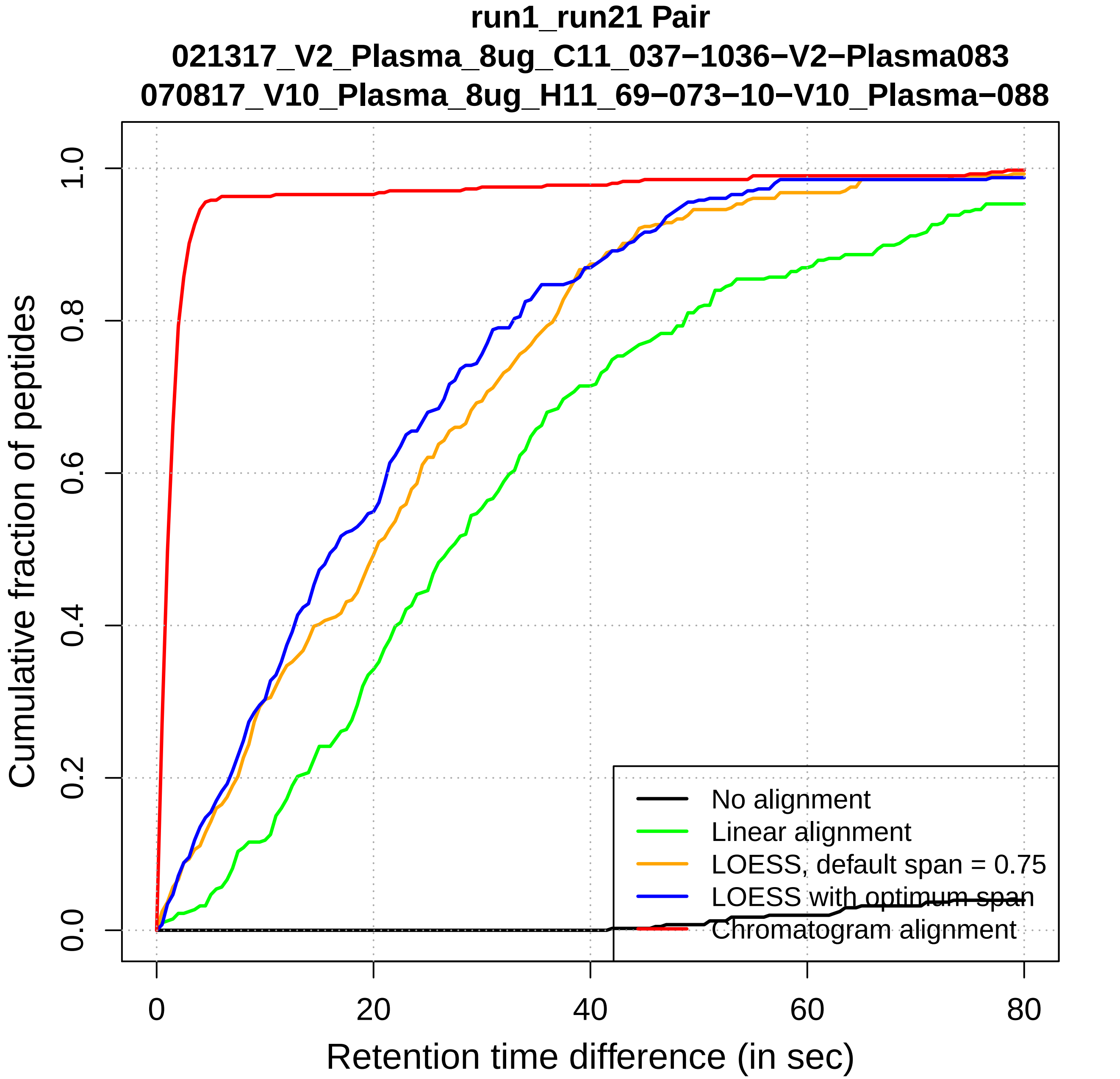
run1_run21 Pair
021317_V2_Plasma_8ug_C11_037−1036−V2−Plasma083
070817_V10_Plasma_8ug_H11_69−073−10−V10_Plasma−088
Cumulative fraction of peptides
0.0
0.2
0.4
0.6
0.8
1.0
0
20
40
60
80
Retention time difference (in sec)
No alignment
Linear alignment
LOESS, default span = 0.75
LOESS with optimum span
Chromatogram alignment

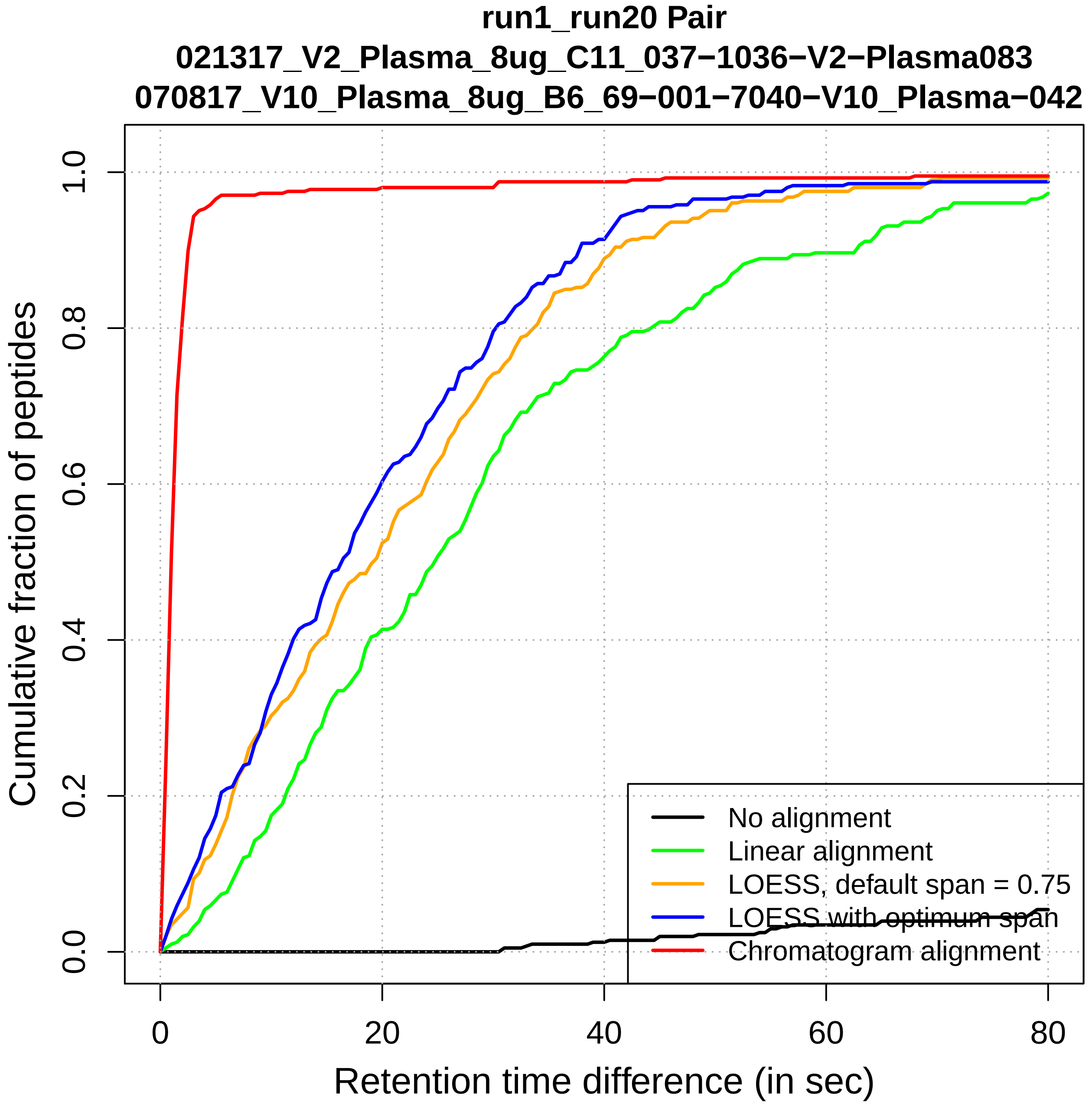

run1_run20 Pair
021317_V2_Plasma_8ug_C11_037−1036−V2−Plasma083
070817_V10_Plasma_8ug_B6_69−001−7040−V10_Plasma−042
Cumulative fraction of peptides
0.0
0.2
0.4
0.6
0.8
1.0
0
20
40
60
80
Retention time difference (in sec)
No alignment
Linear alignment
LOESS, default span = 0.75
LOESS with optimum span
Chromatogram alignment

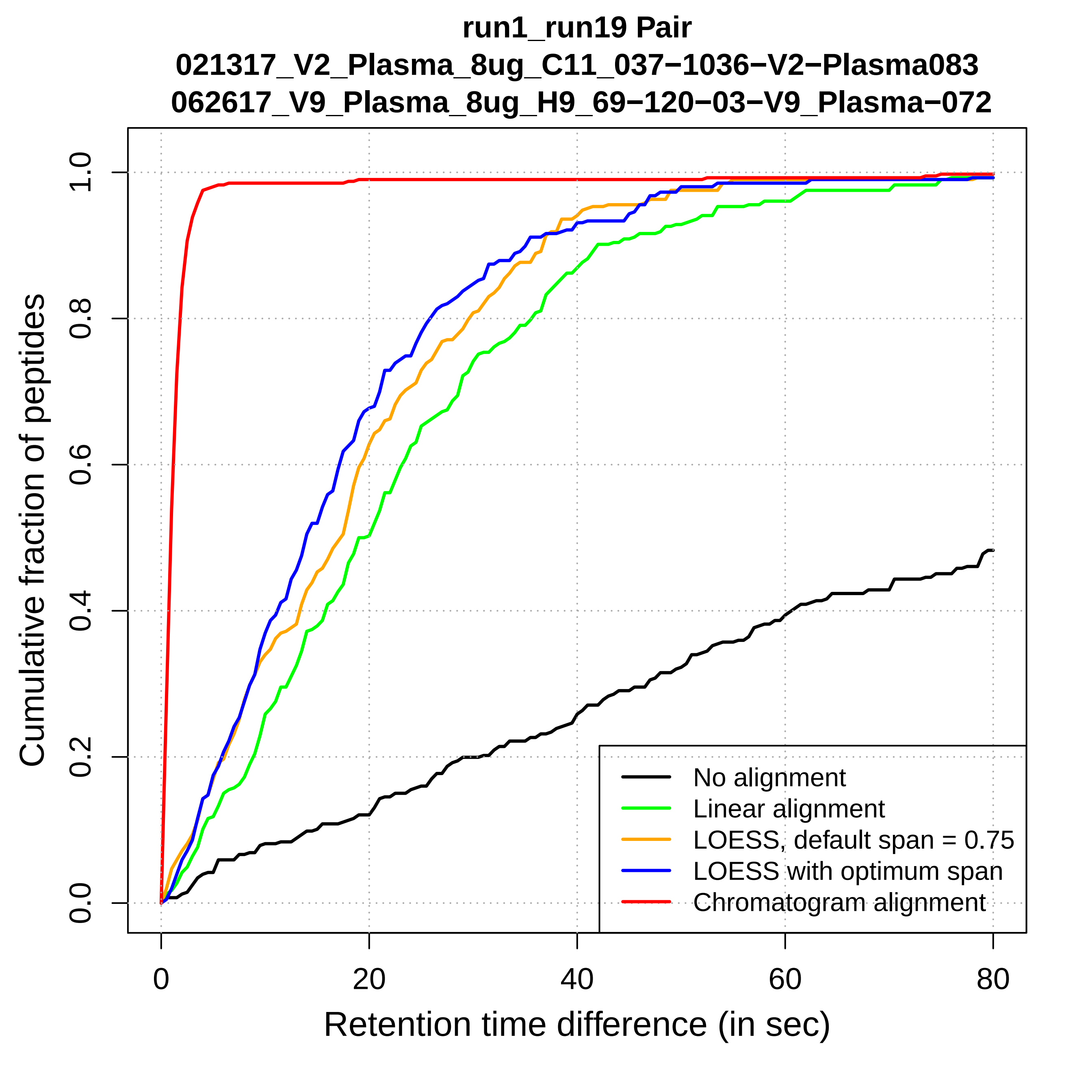

run1_run19 Pair
021317_V2_Plasma_8ug_C11_037−1036−V2−Plasma083
062617_V9_Plasma_8ug_H9_69−120−03−V9_Plasma−072
Cumulative fraction of peptides
0.0
0.2
0.4
0.6
0.8
1.0
0
20
40
60
80
Retention time difference (in sec)
No alignment
Linear alignment
LOESS, default span = 0.75
LOESS with optimum span
Chromatogram alignment

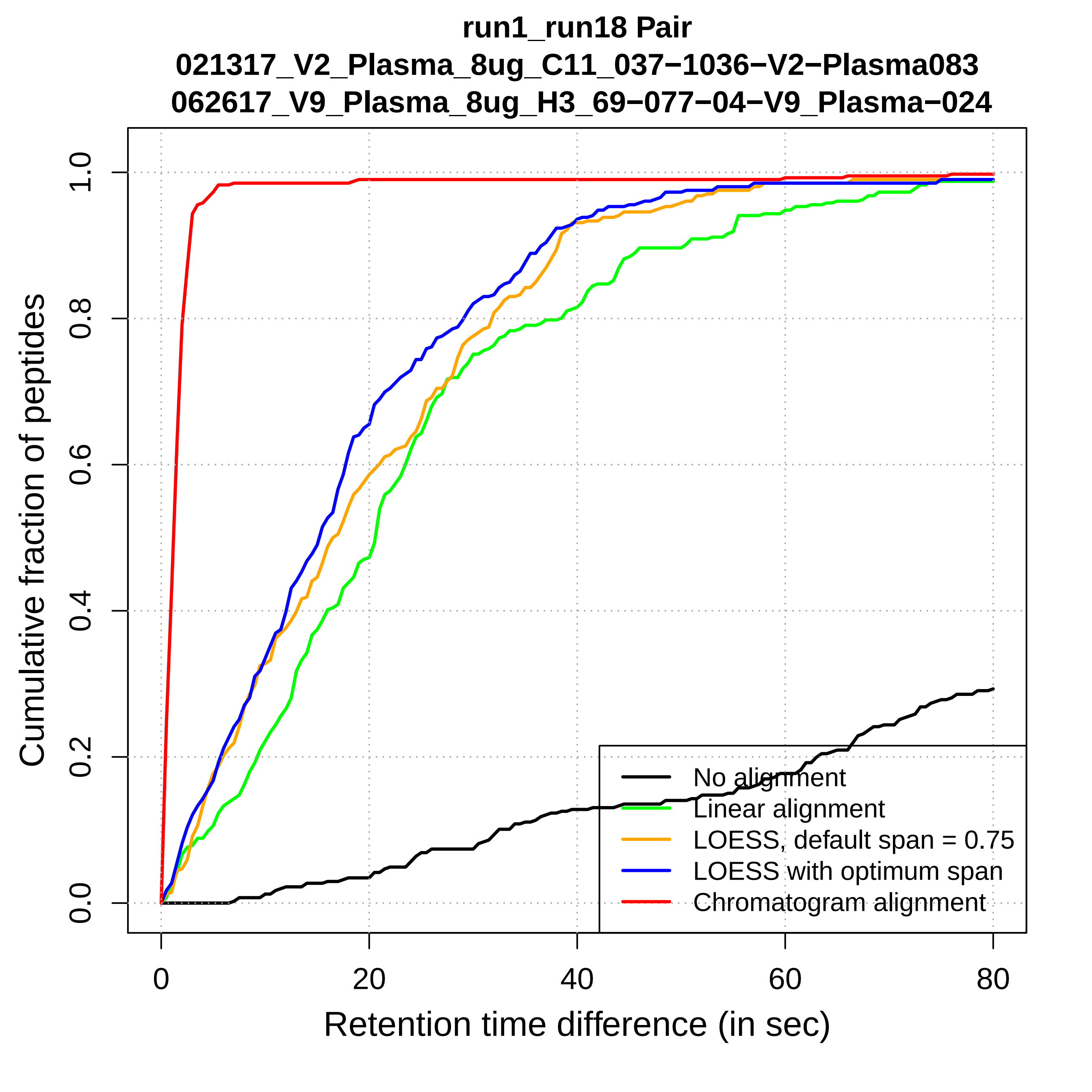
run1_run18 Pair
021317_V2_Plasma_8ug_C11_037−1036−V2−Plasma083
062617_V9_Plasma_8ug_H3_69−077−04−V9_Plasma−024
Cumulative fraction of peptides
0.0
0.2
0.4
0.6
0.8
1.0
0
20
40
60
80
Retention time difference (in sec)
No alignment
Linear alignment
LOESS, default span = 0.75
LOESS with optimum span
Chromatogram alignment

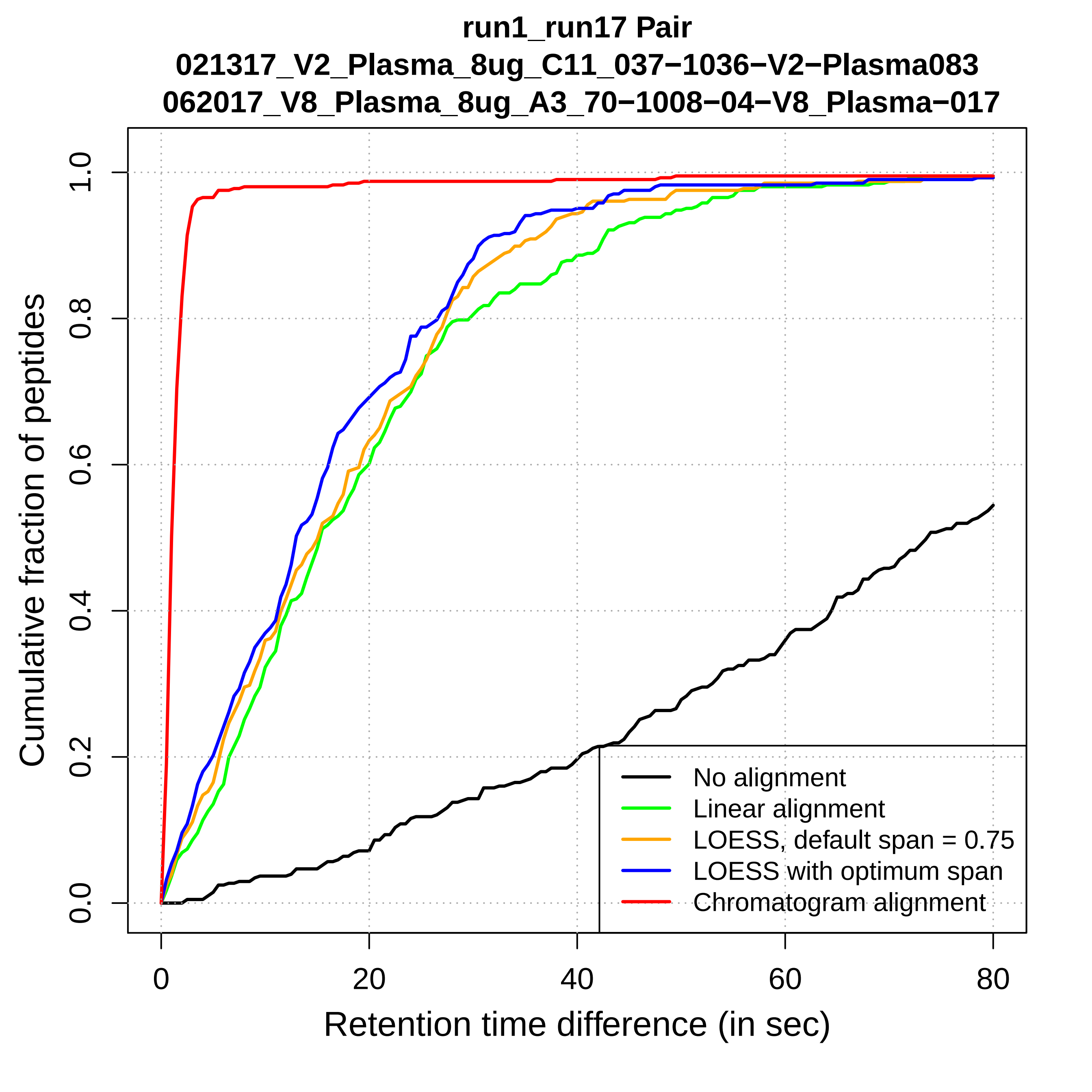
run1_run17 Pair
021317_V2_Plasma_8ug_C11_037−1036−V2−Plasma083
062017_V8_Plasma_8ug_A3_70−1008−04−V8_Plasma−017
Cumulative fraction of peptides
0.0
0.2
0.4
0.6
0.8
1.0
0
20
40
60
80
Retention time difference (in sec)
No alignment
Linear alignment
LOESS, default span = 0.75
LOESS with optimum span
Chromatogram alignment

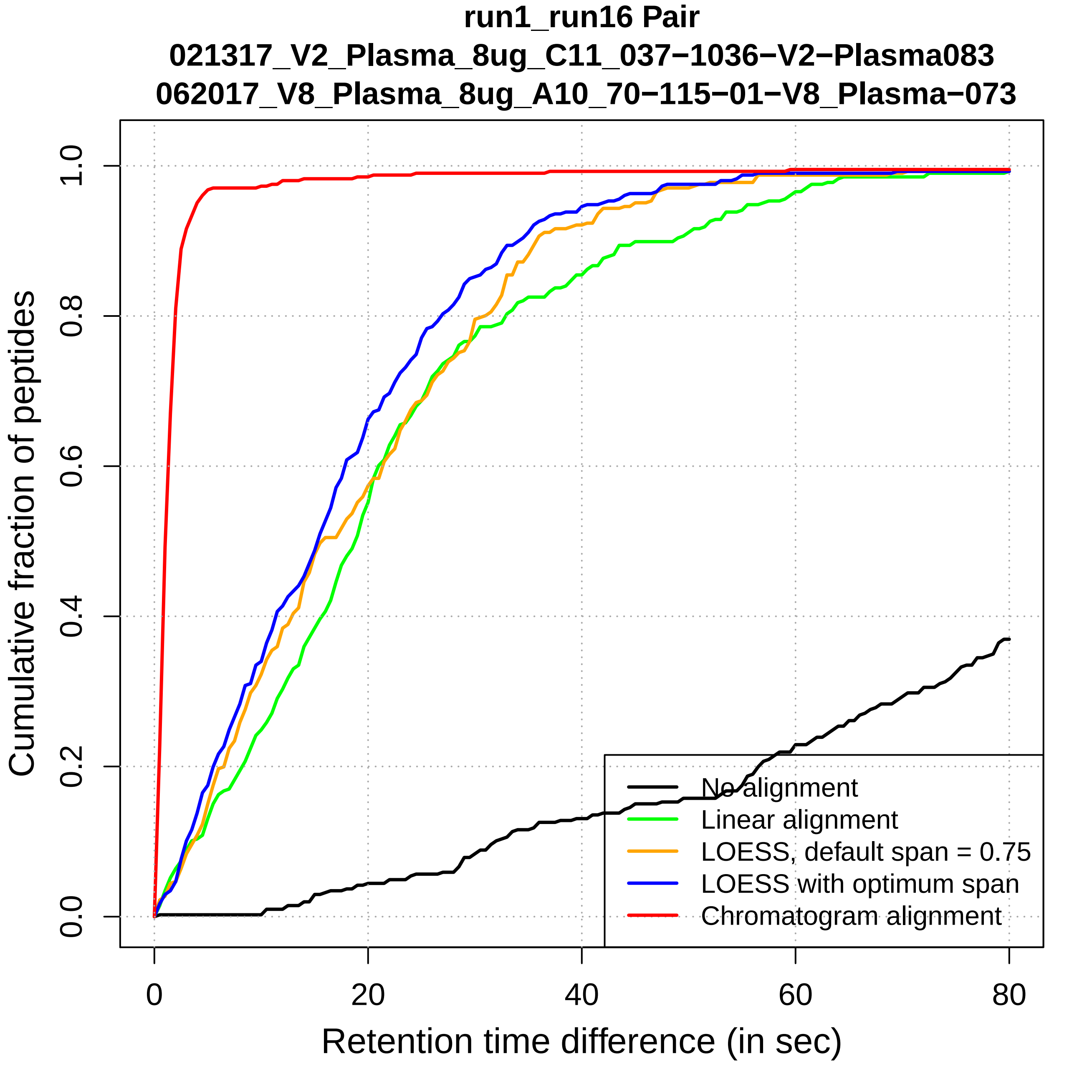

run1_run16 Pair
021317_V2_Plasma_8ug_C11_037−1036−V2−Plasma083
062017_V8_Plasma_8ug_A10_70−115−01−V8_Plasma−073
Cumulative fraction of peptides
Retention time difference (in sec)
0.0
0.2
0.4
0.6
0.8
1.0
0
20
40
60
80
No alignment
Linear alignment
LOESS, default span = 0.75
LOESS with optimum span
Chromatogram alignment

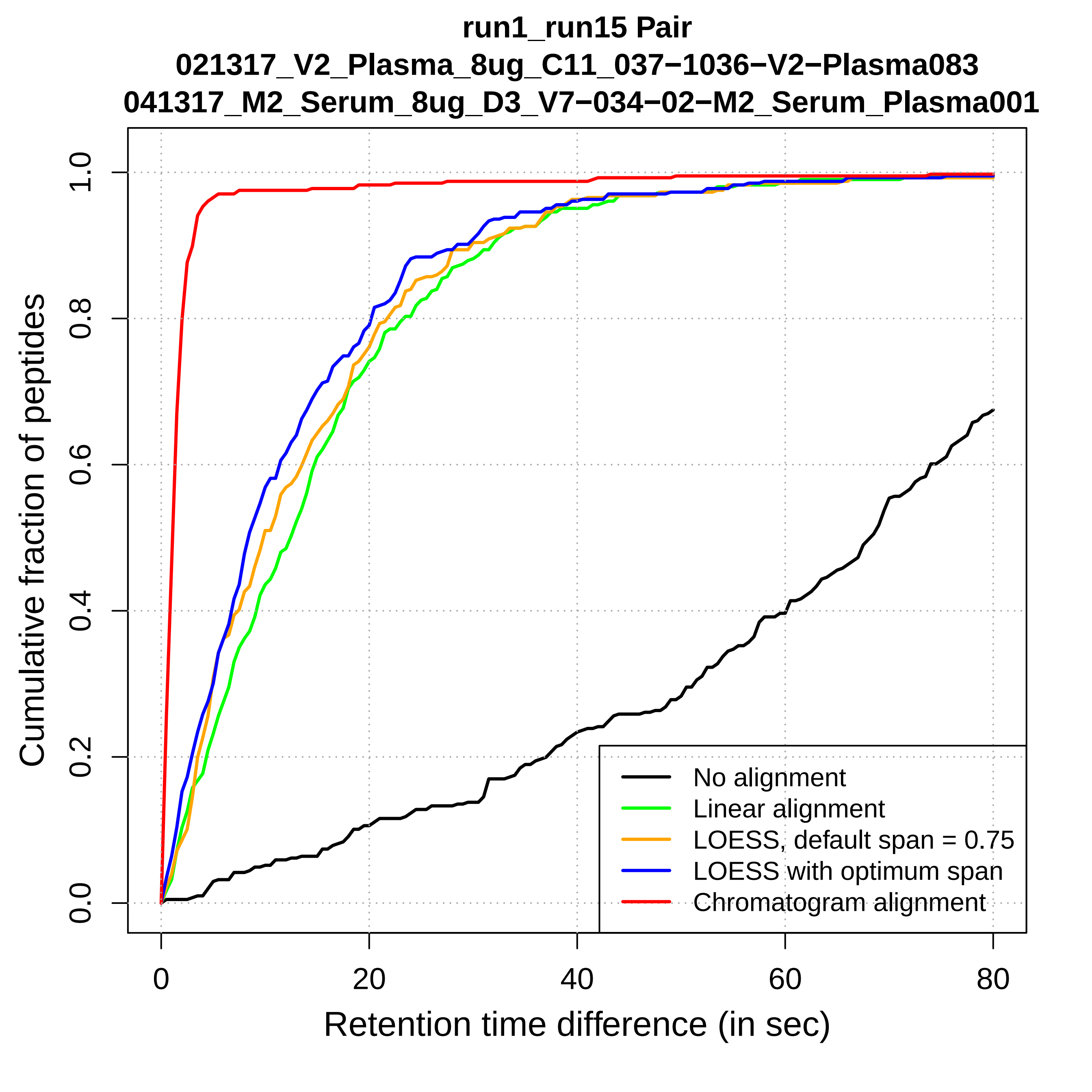

run1_run15 Pair
021317_V2_Plasma_8ug_C11_037−1036−V2−Plasma083
041317_M2_Serum_8ug_D3_V7−034−02−M2_Serum_Plasma001
Cumulative fraction of peptides
0.0
0.2
0.4
0.6
0.8
1.0
0
20
40
60
80
Retention time difference (in sec)
No alignment
Linear alignment
LOESS, default span = 0.75
LOESS with optimum span
Chromatogram alignment

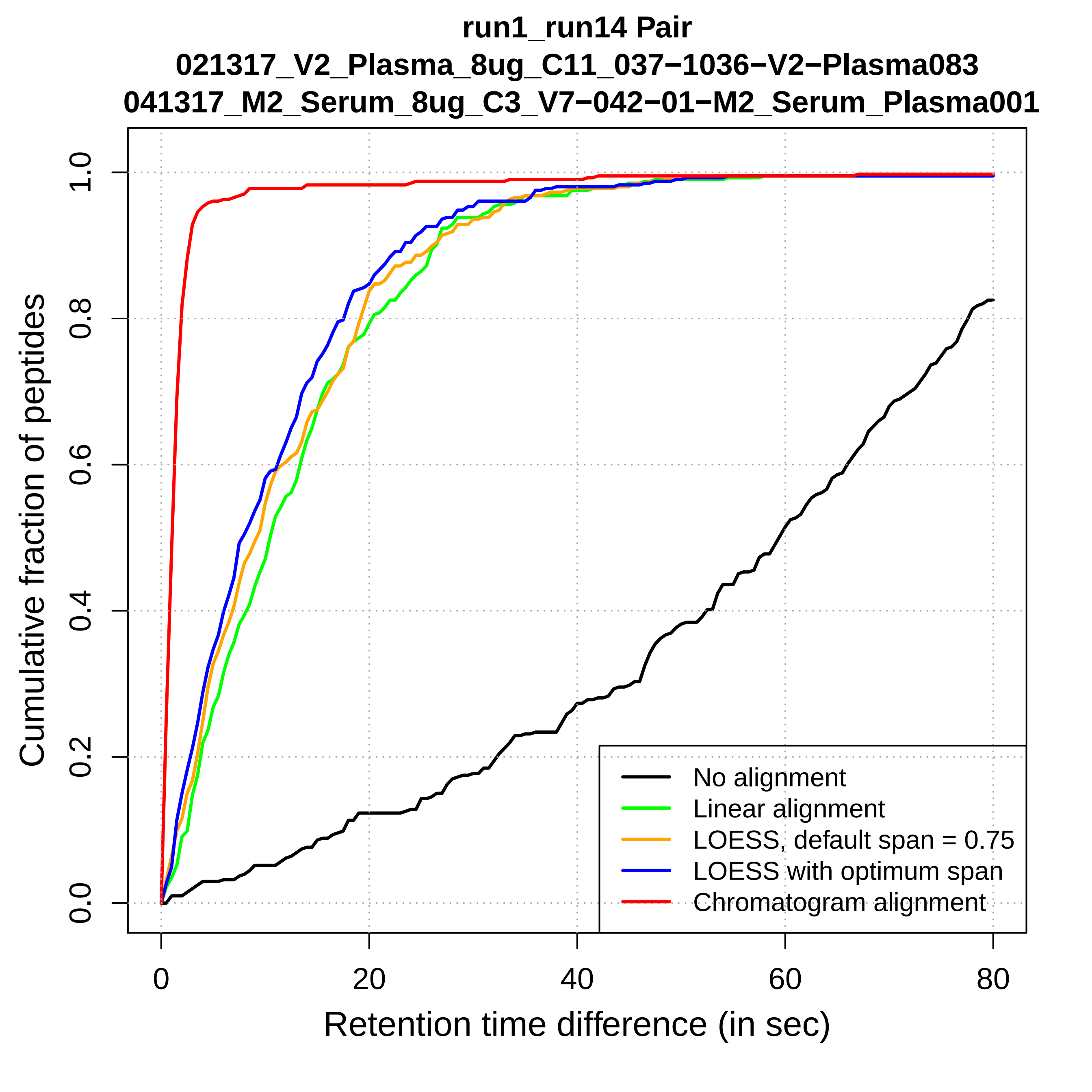

run1_run14 Pair
021317_V2_Plasma_8ug_C11_037−1036−V2−Plasma083
041317_M2_Serum_8ug_C3_V7−042−01−M2_Serum_Plasma001
Cumulative fraction of peptides
0.0
0.2
0.4
0.6
0.8
1.0
0
20
40
60
80
Retention time difference (in sec)
No alignment
Linear alignment
LOESS, default span = 0.75
LOESS with optimum span
Chromatogram alignment

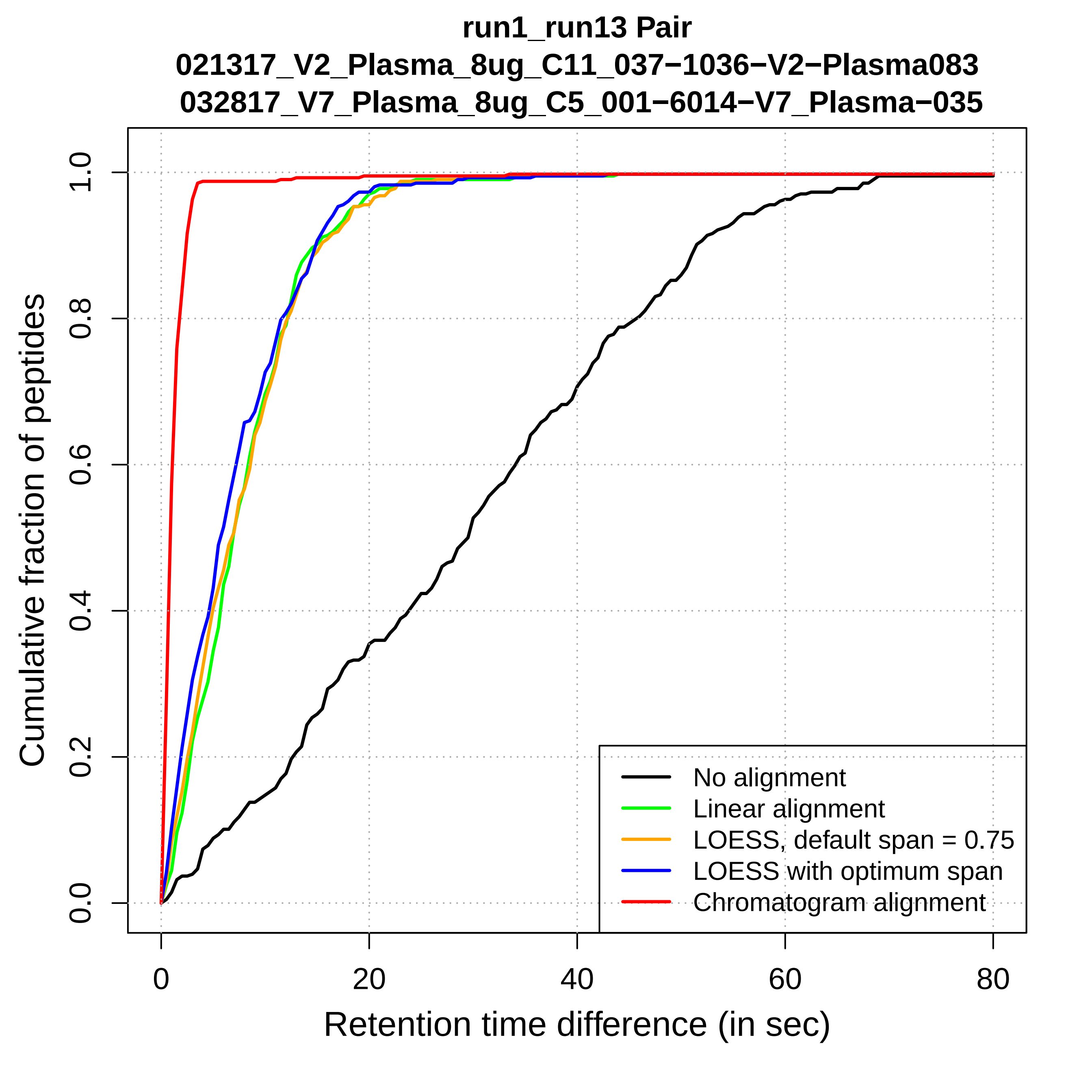
run1_run13 Pair
021317_V2_Plasma_8ug_C11_037−1036−V2−Plasma083
032817_V7_Plasma_8ug_C5_001−6014−V7_Plasma−035
Cumulative fraction of peptides
0.0
0.2
0.4
0.6
0.8
1.0
0
20
40
60
80
Retention time difference (in sec)
No alignment
Linear alignment
LOESS, default span = 0.75
LOESS with optimum span
Chromatogram alignment

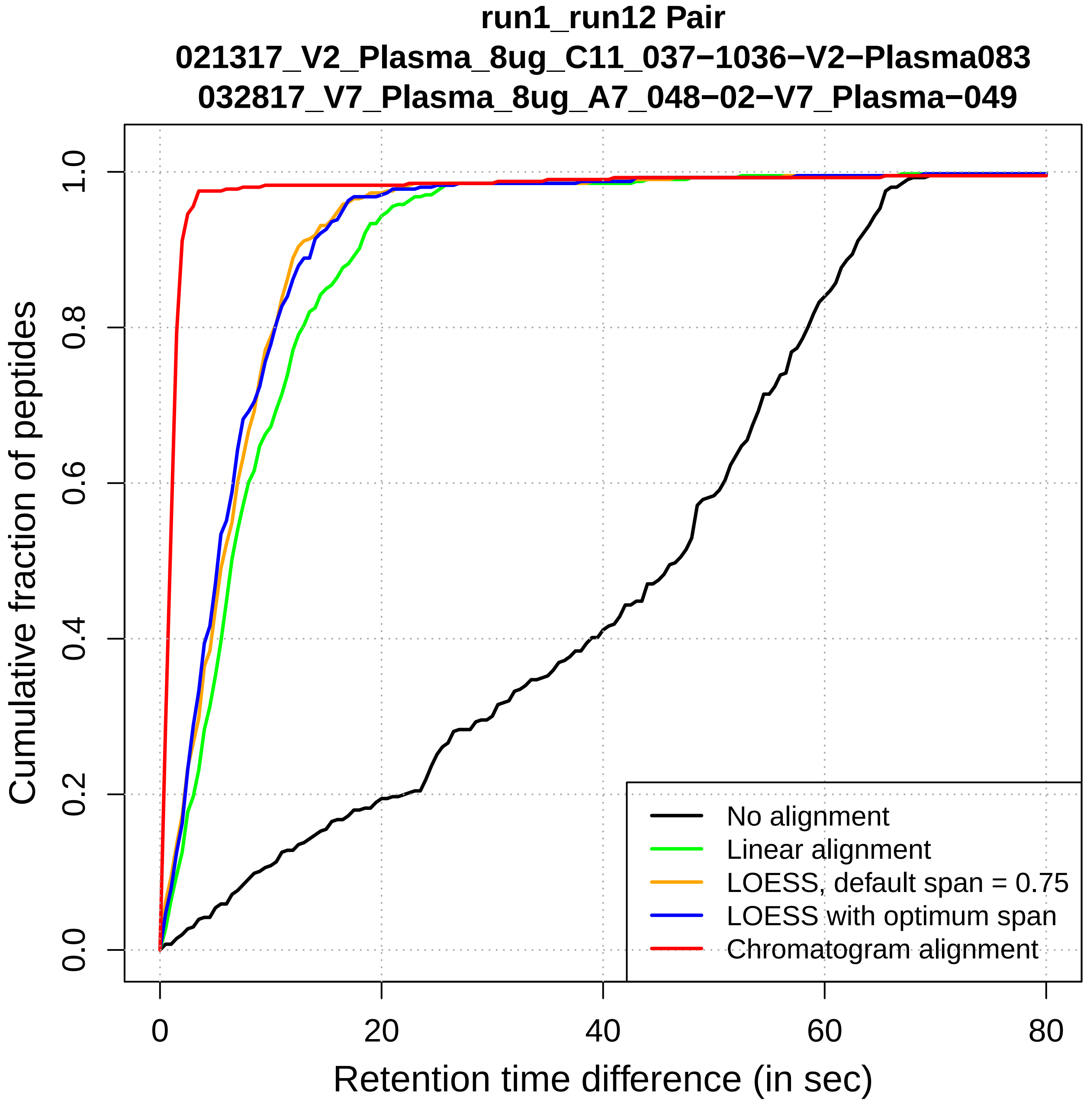

run1_run12 Pair
021317_V2_Plasma_8ug_C11_037−1036−V2−Plasma083
032817_V7_Plasma_8ug_A7_048−02−V7_Plasma−049
Cumulative fraction of peptides
0.0
0.2
0.4
0.6
0.8
1.0
0
20
40
60
80
Retention time difference (in sec)
No alignment
Linear alignment
LOESS, default span = 0.75
LOESS with optimum span
Chromatogram alignment

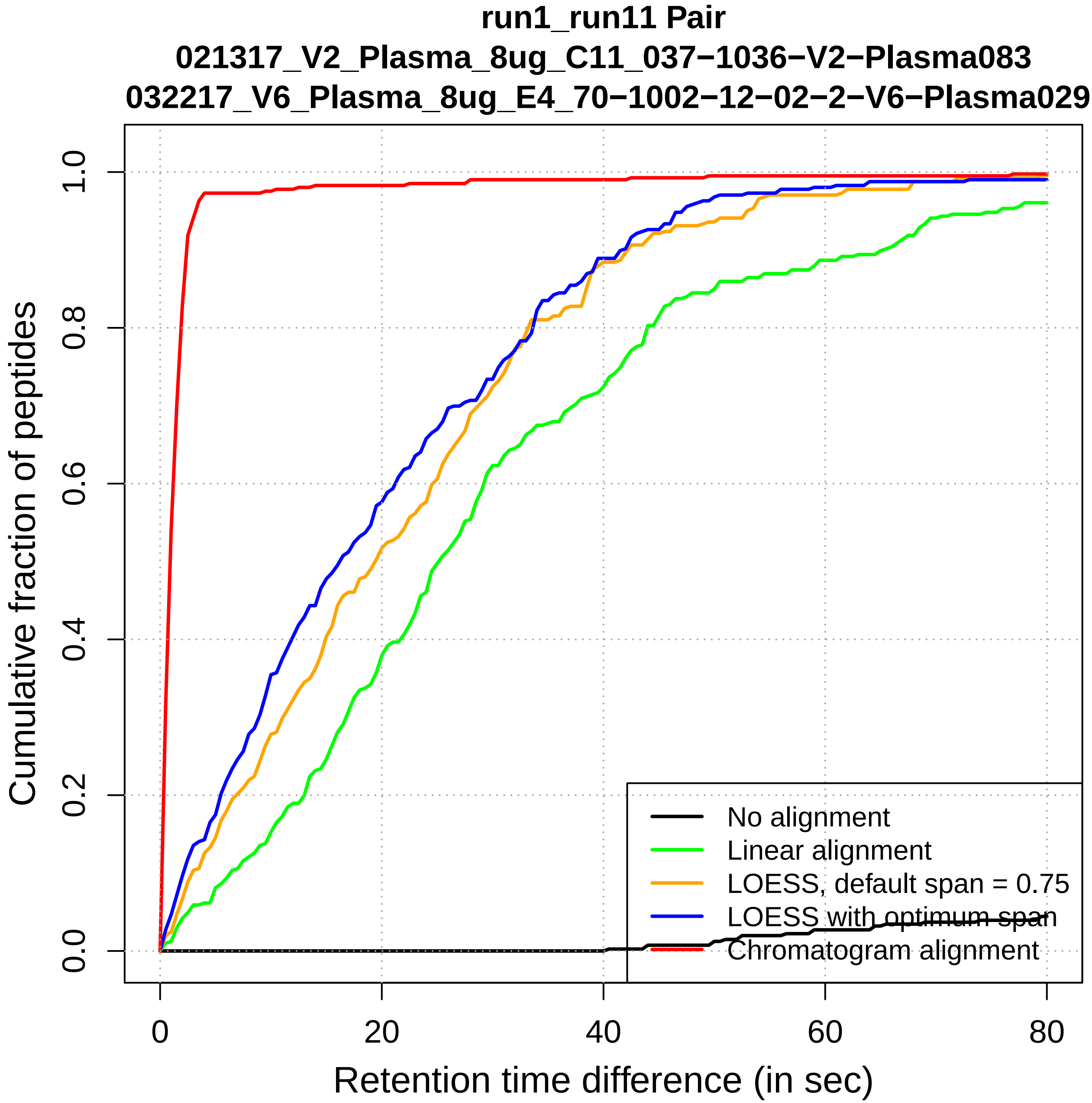

run1_run11 Pair
021317_V2_Plasma_8ug_C11_037−1036−V2−Plasma083
032217_V6_Plasma_8ug_E4_70−1002−12−02−2−V6−Plasma029
Cumulative fraction of peptides
0.0
0.2
0.4
0.6
0.8
1.0
0
20
40
60
80
Retention time difference (in sec)
No alignment
Linear alignment
LOESS, default span = 0.75
LOESS with optimum span
Chromatogram alignment

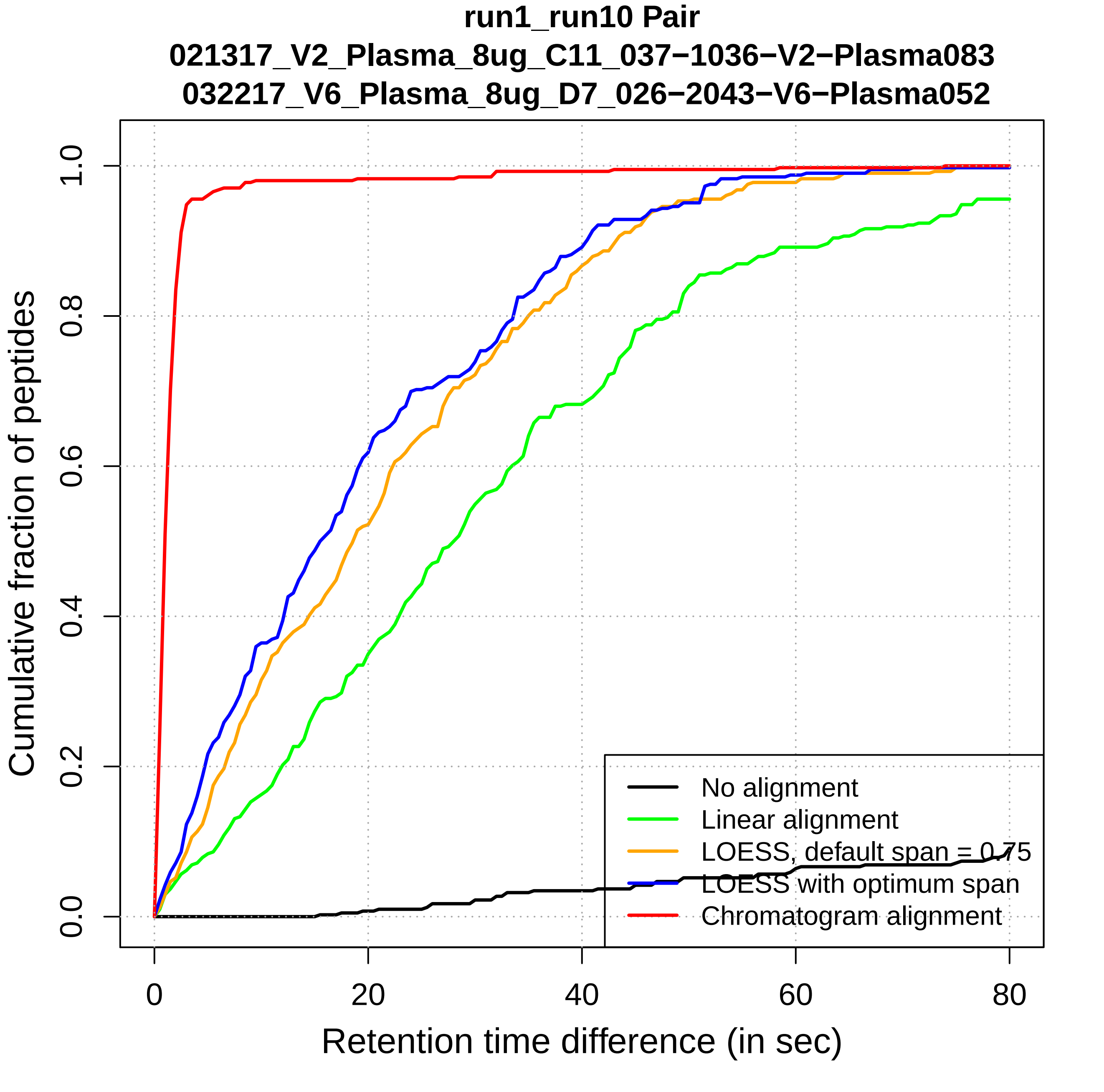

run1_run10 Pair
021317_V2_Plasma_8ug_C11_037−1036−V2−Plasma083
032217_V6_Plasma_8ug_D7_026−2043−V6−Plasma052
Cumulative fraction of peptides
0.0
0.2
0.4
0.6
0.8
1.0
0
20
40
60
80
Retention time difference (in sec)
No alignment
Linear alignment
LOESS, default span = 0.75
LOESS with optimum span
Chromatogram alignment

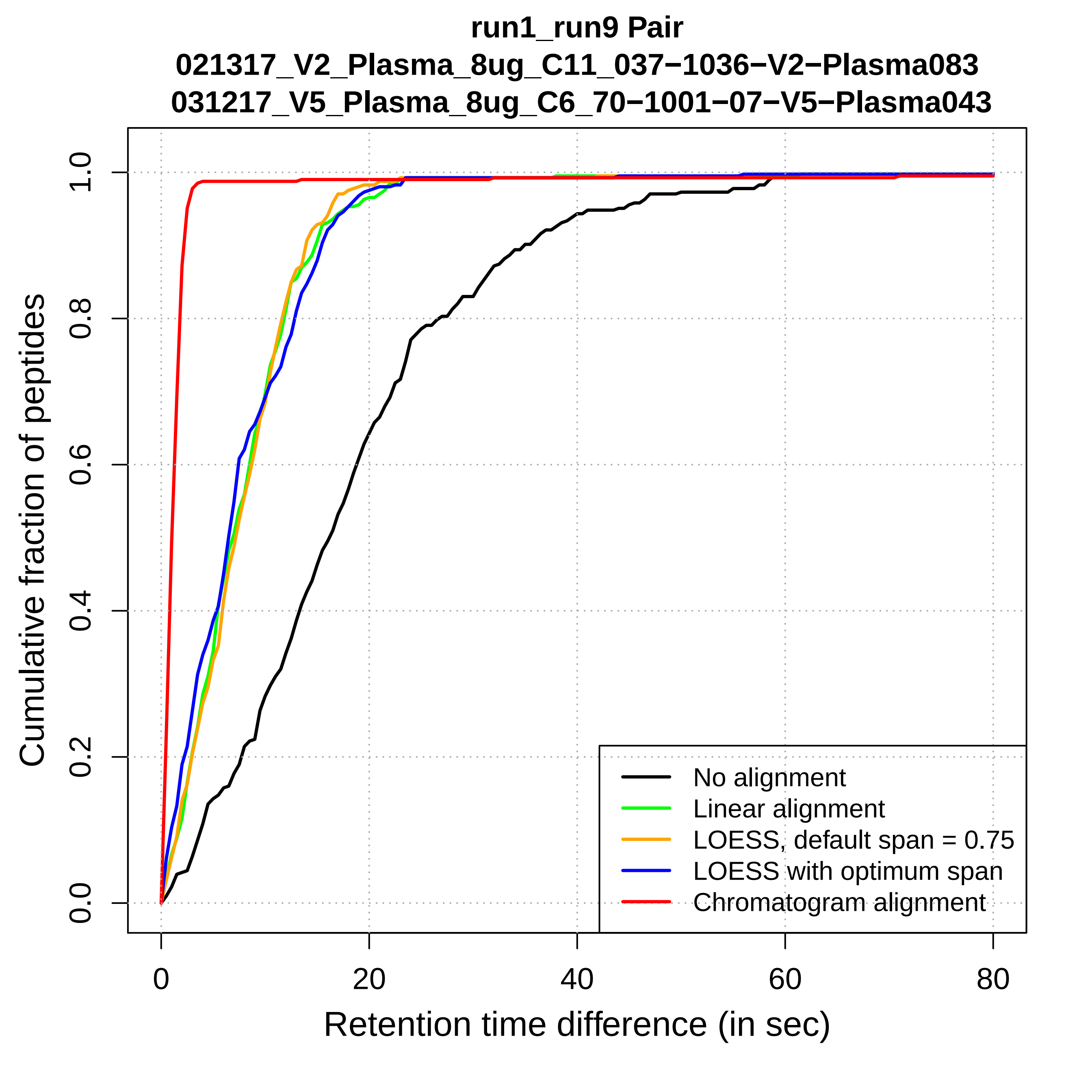
run1_run9 Pair
021317_V2_Plasma_8ug_C11_037−1036−V2−Plasma083
031217_V5_Plasma_8ug_C6_70−1001−07−V5−Plasma043
Cumulative fraction of peptides
Retention time difference (in sec)
0.0
0.2
0.4
0.6
0.8
1.0
0
20
40
60
80
No alignment
Linear alignment
LOESS, default span = 0.75
LOESS with optimum span
Chromatogram alignment

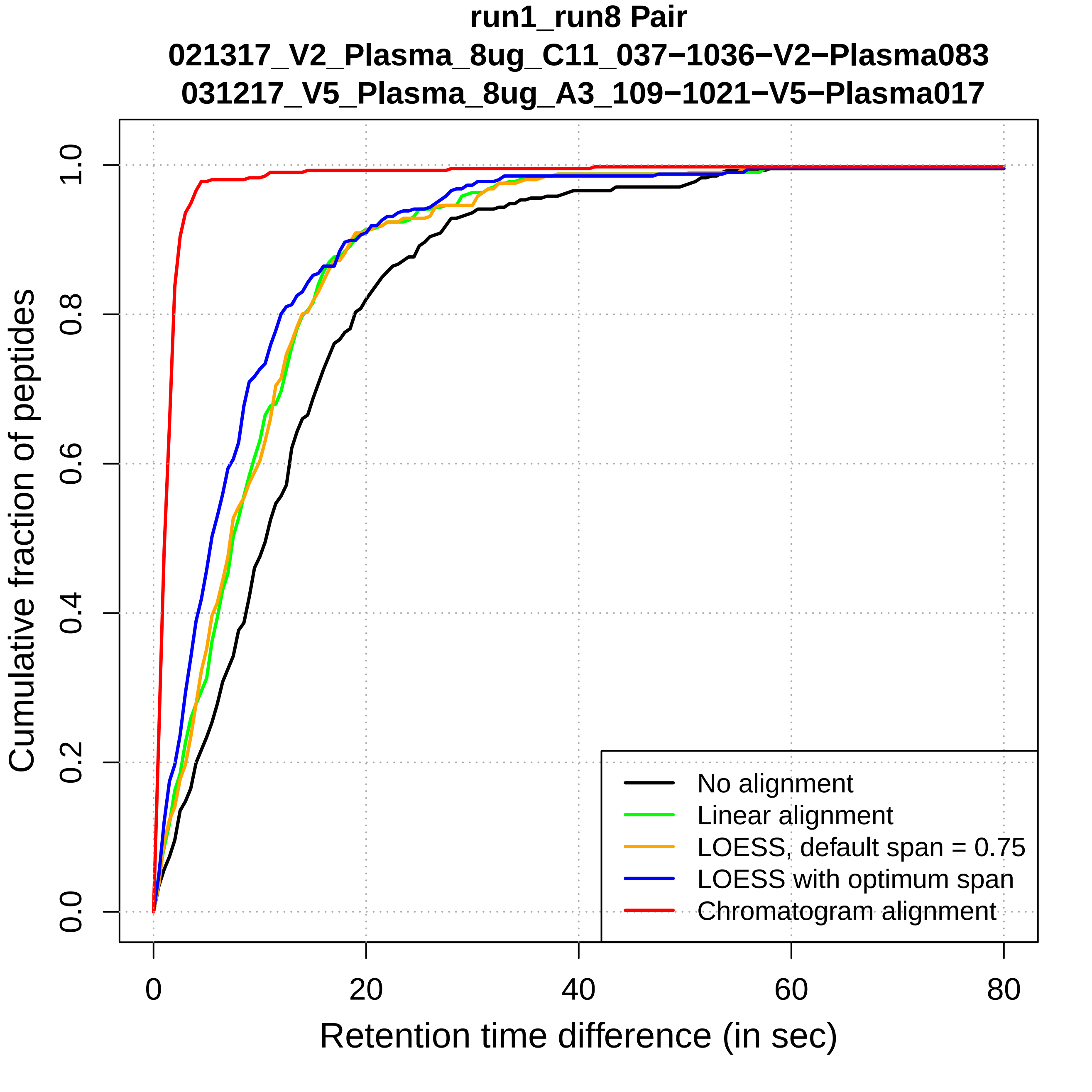
run1_run8 Pair
021317_V2_Plasma_8ug_C11_037−1036−V2−Plasma083
031217_V5_Plasma_8ug_A3_109−1021−V5−Plasma017
Cumulative fraction of peptides
0.0
0.2
0.4
0.6
0.8
1.0
0
20
40
60
80
Retention time difference (in sec)
No alignment
Linear alignment
LOESS, default span = 0.75
LOESS with optimum span
Chromatogram alignment

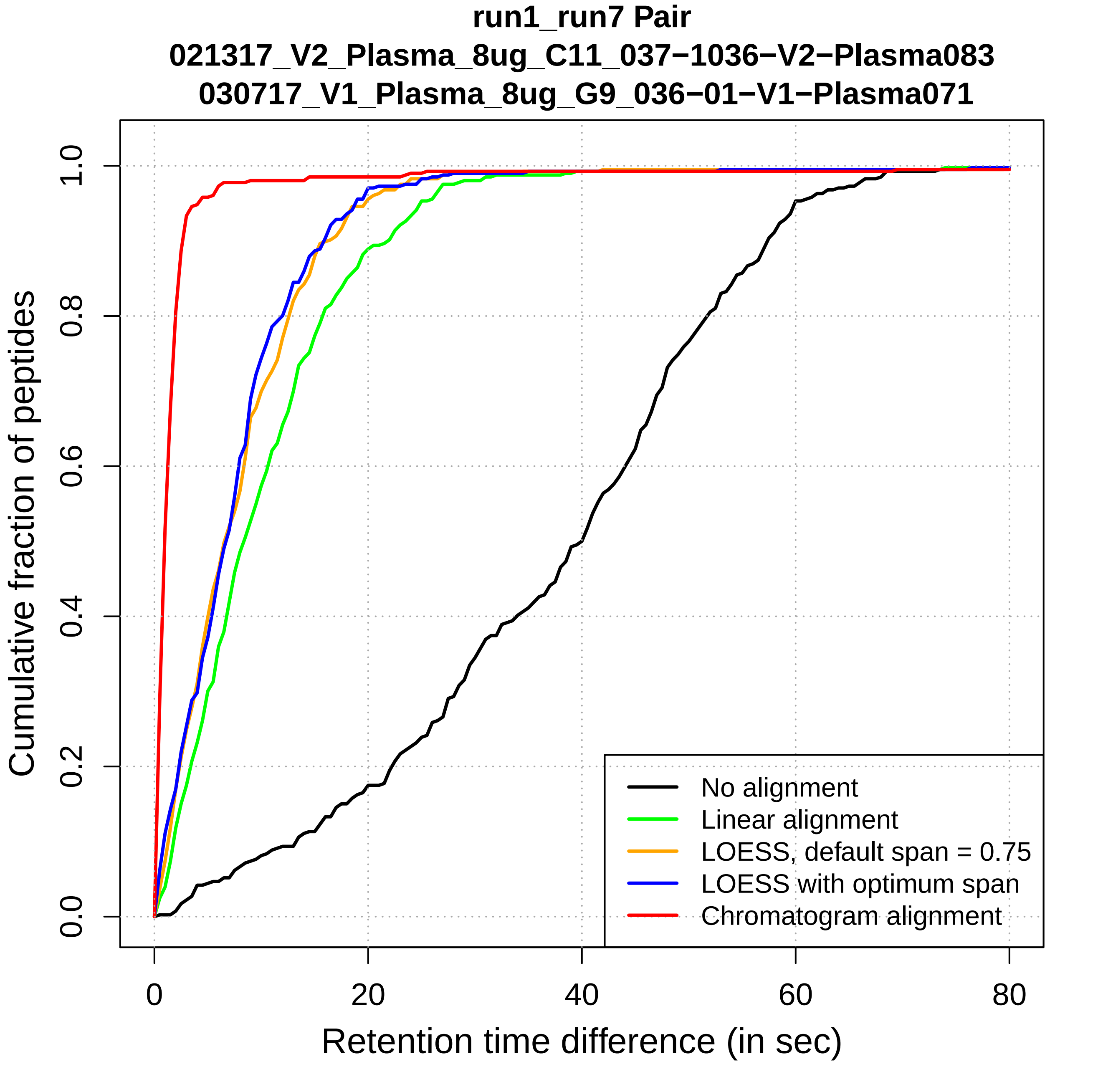
run1_run7 Pair
021317_V2_Plasma_8ug_C11_037−1036−V2−Plasma083
030717_V1_Plasma_8ug_G9_036−01−V1−Plasma071
Cumulative fraction of peptides
0.0
0.2
0.4
0.6
0.8
1.0
0
20
40
60
80
Retention time difference (in sec)
No alignment
Linear alignment
LOESS, default span = 0.75
LOESS with optimum span
Chromatogram alignment

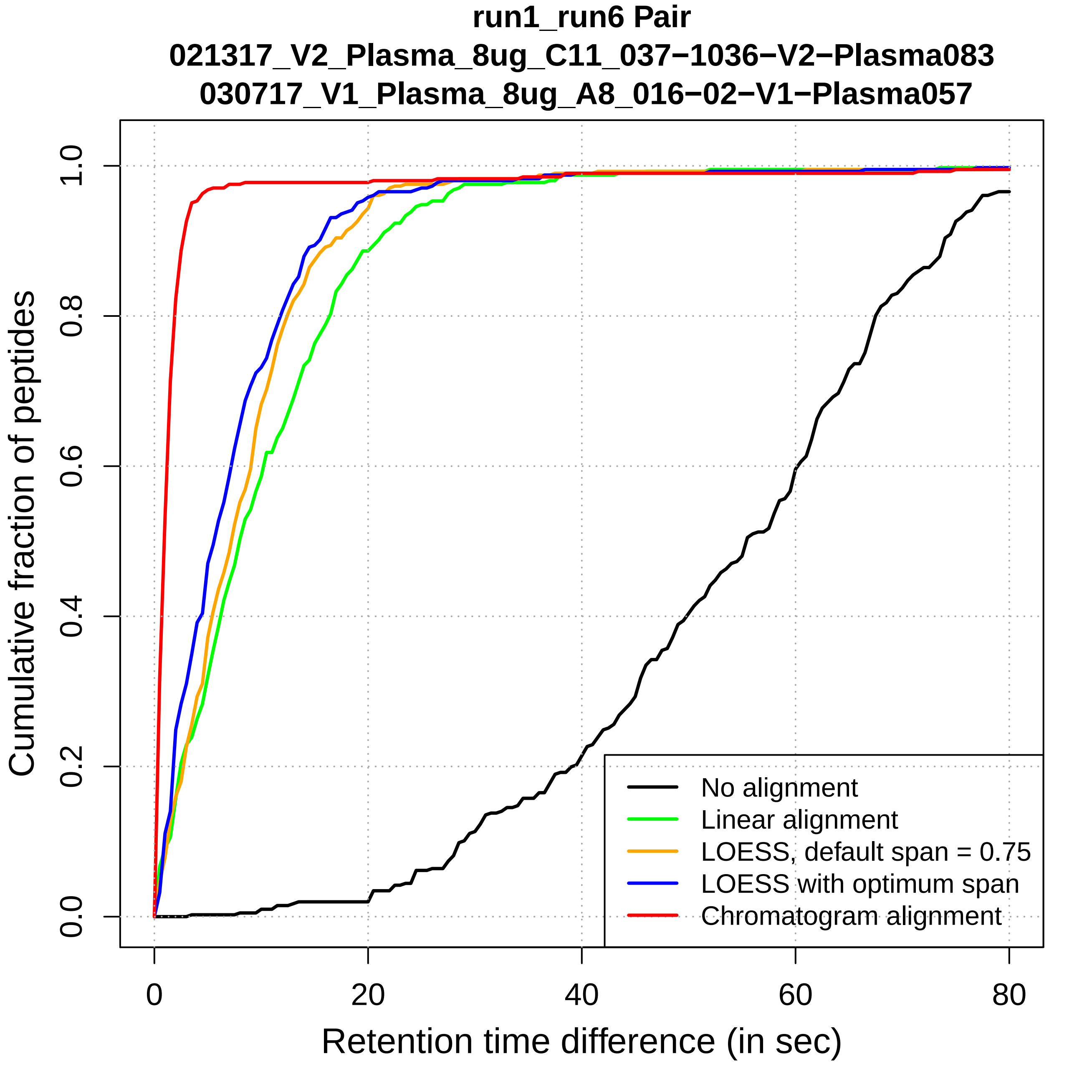
run1_run6 Pair
021317_V2_Plasma_8ug_C11_037−1036−V2−Plasma083
030717_V1_Plasma_8ug_A8_016−02−V1−Plasma057
Cumulative fraction of peptides
0.0
0.2
0.4
0.6
0.8
1.0
0
20
40
60
80
Retention time difference (in sec)
No alignment
Linear alignment
LOESS, default span = 0.75
LOESS with optimum span
Chromatogram alignment

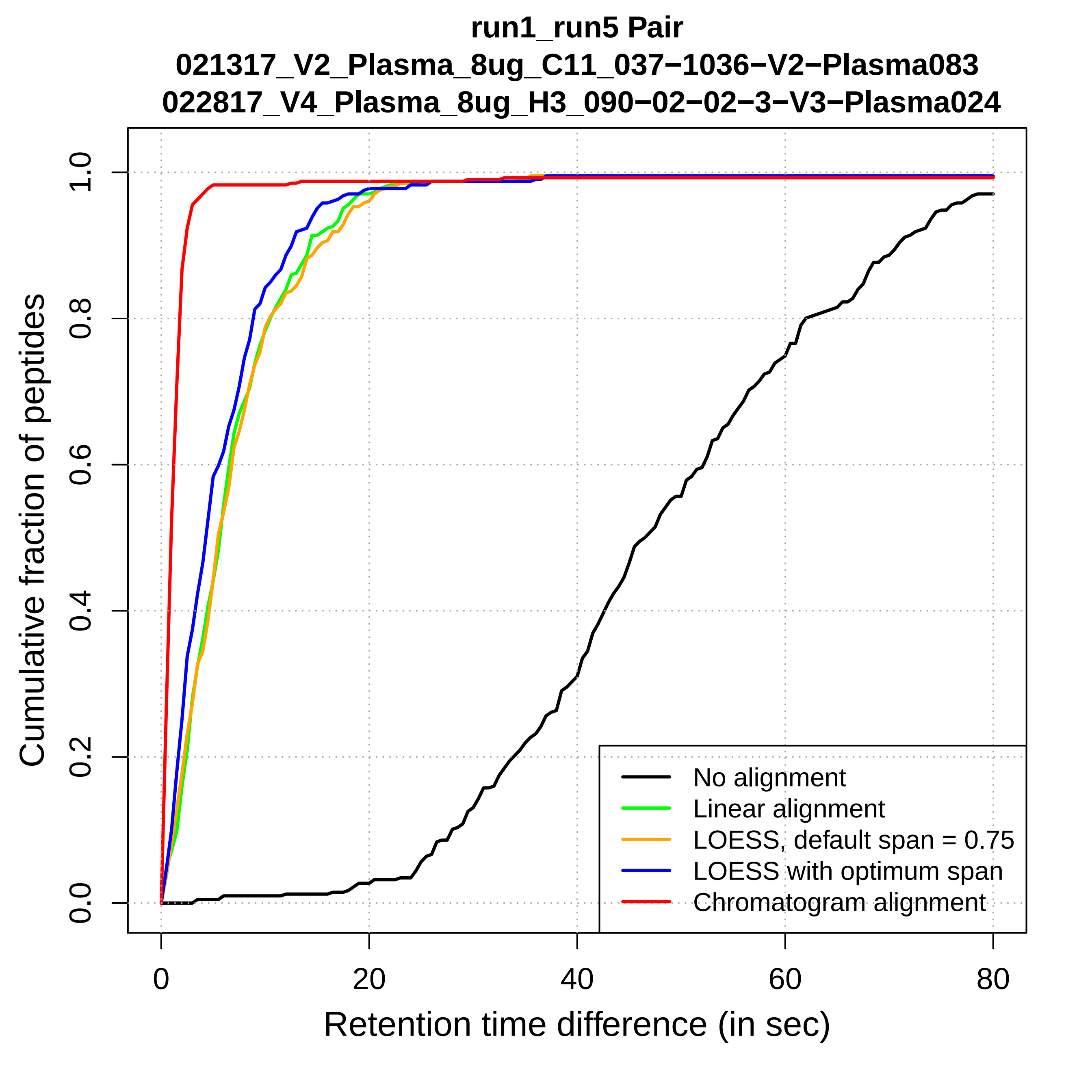
run1_run5 Pair
021317_V2_Plasma_8ug_C11_037−1036−V2−Plasma083
022817_V4_Plasma_8ug_H3_090−02−02−3−V3−Plasma024
Cumulative fraction of peptides
0.0
0.2
0.4
0.6
0.8
1.0
0
20
40
60
80
Retention time difference (in sec)
No alignment
Linear alignment
LOESS, default span = 0.75
LOESS with optimum span
Chromatogram alignment

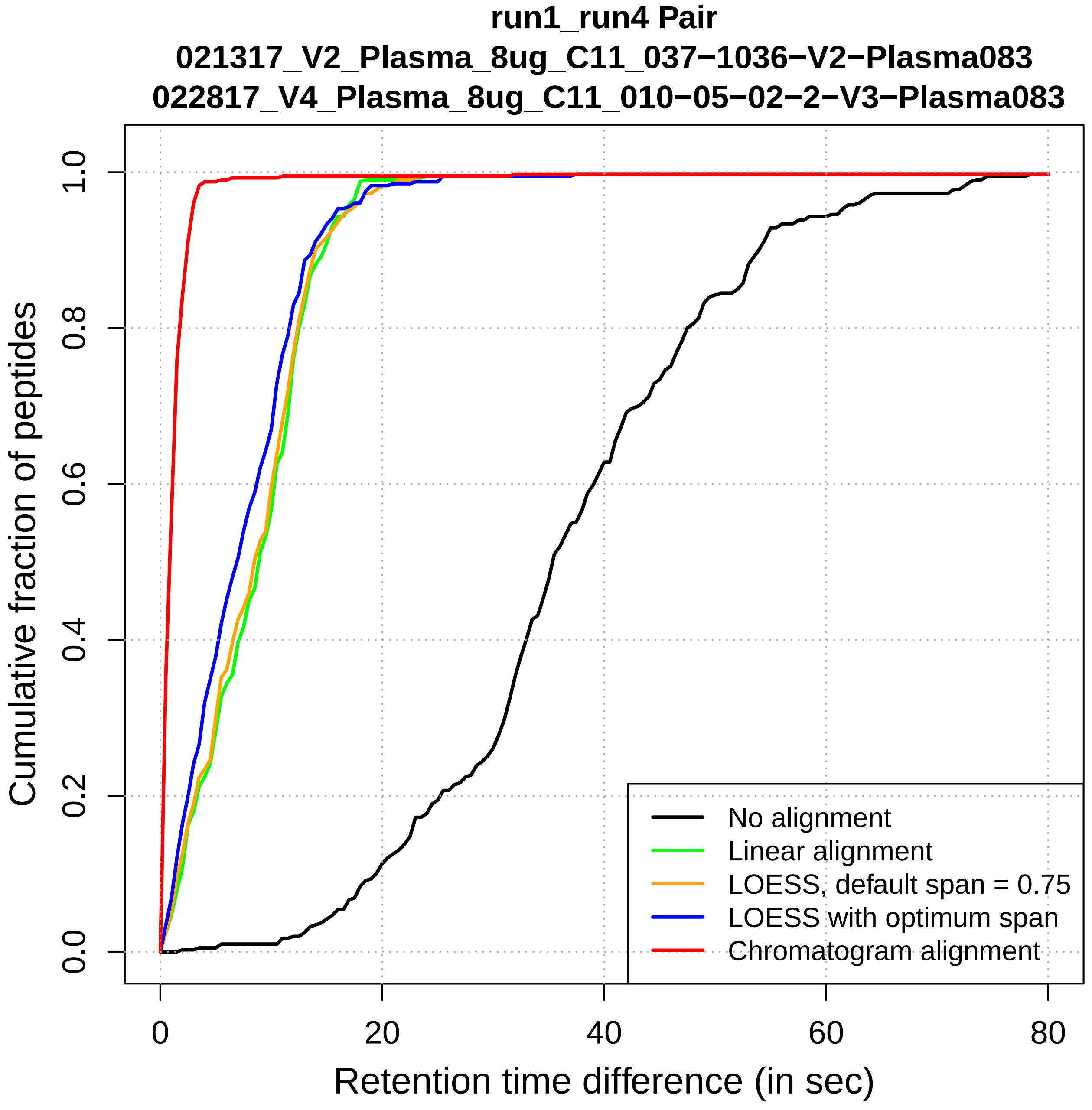
run1_run4 Pair
021317_V2_Plasma_8ug_C11_037−1036−V2−Plasma083
022817_V4_Plasma_8ug_C11_010−05−02−2−V3−Plasma083
Cumulative fraction of peptides
0.0
0.2
0.4
0.6
0.8
1.0
0
20
40
60
80
Retention time difference (in sec)
No alignment
Linear alignment
LOESS, default span = 0.75
LOESS with optimum span
Chromatogram alignment

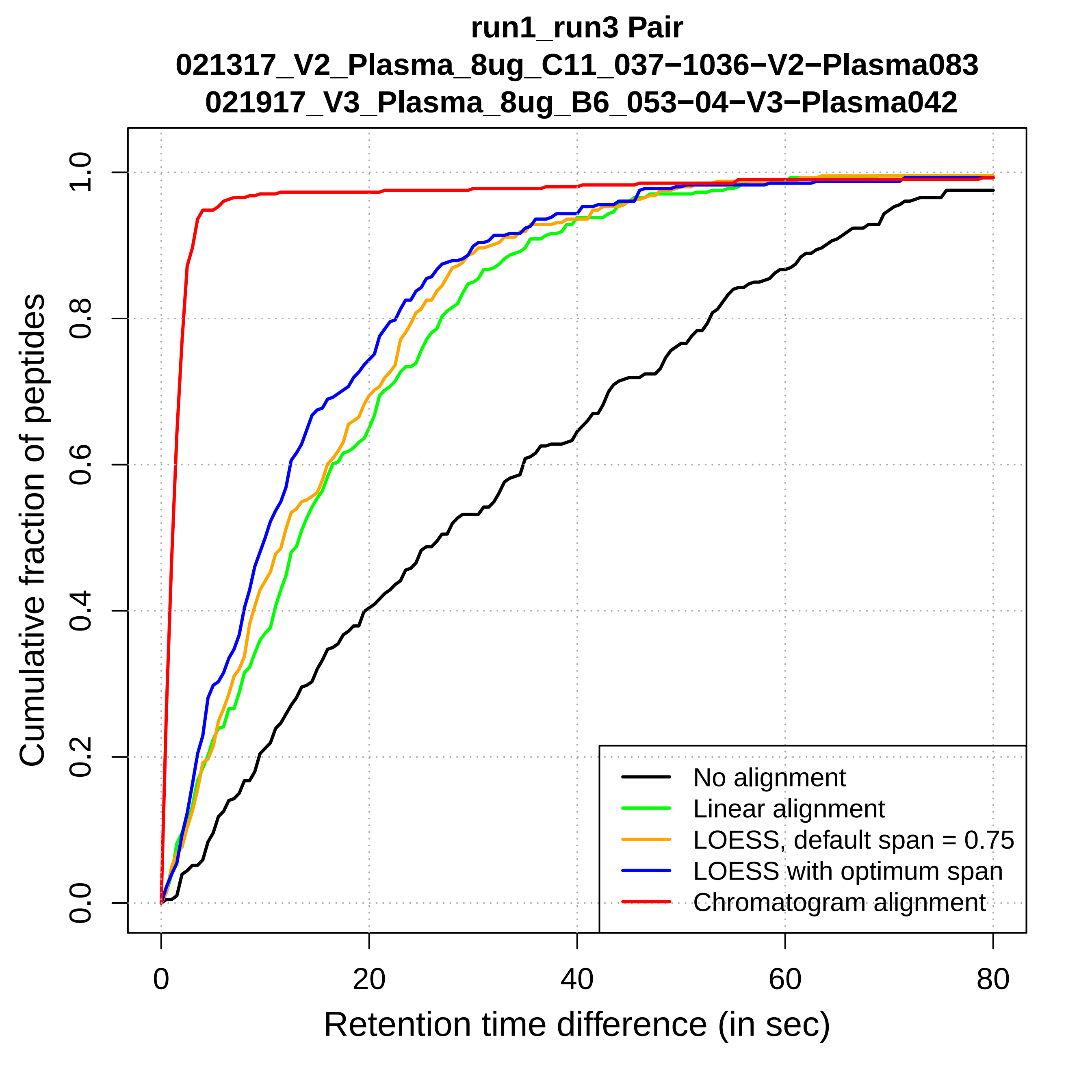

run1_run3 Pair
021317_V2_Plasma_8ug_C11_037−1036−V2−Plasma083
021917_V3_Plasma_8ug_B6_053−04−V3−Plasma042
Cumulative fraction of peptides
0.0
0.2
0.4
0.6
0.8
1.0
0
20
40
60
80
Retention time difference (in sec)
No alignment
Linear alignment
LOESS, default span = 0.75
LOESS with optimum span
Chromatogram alignment

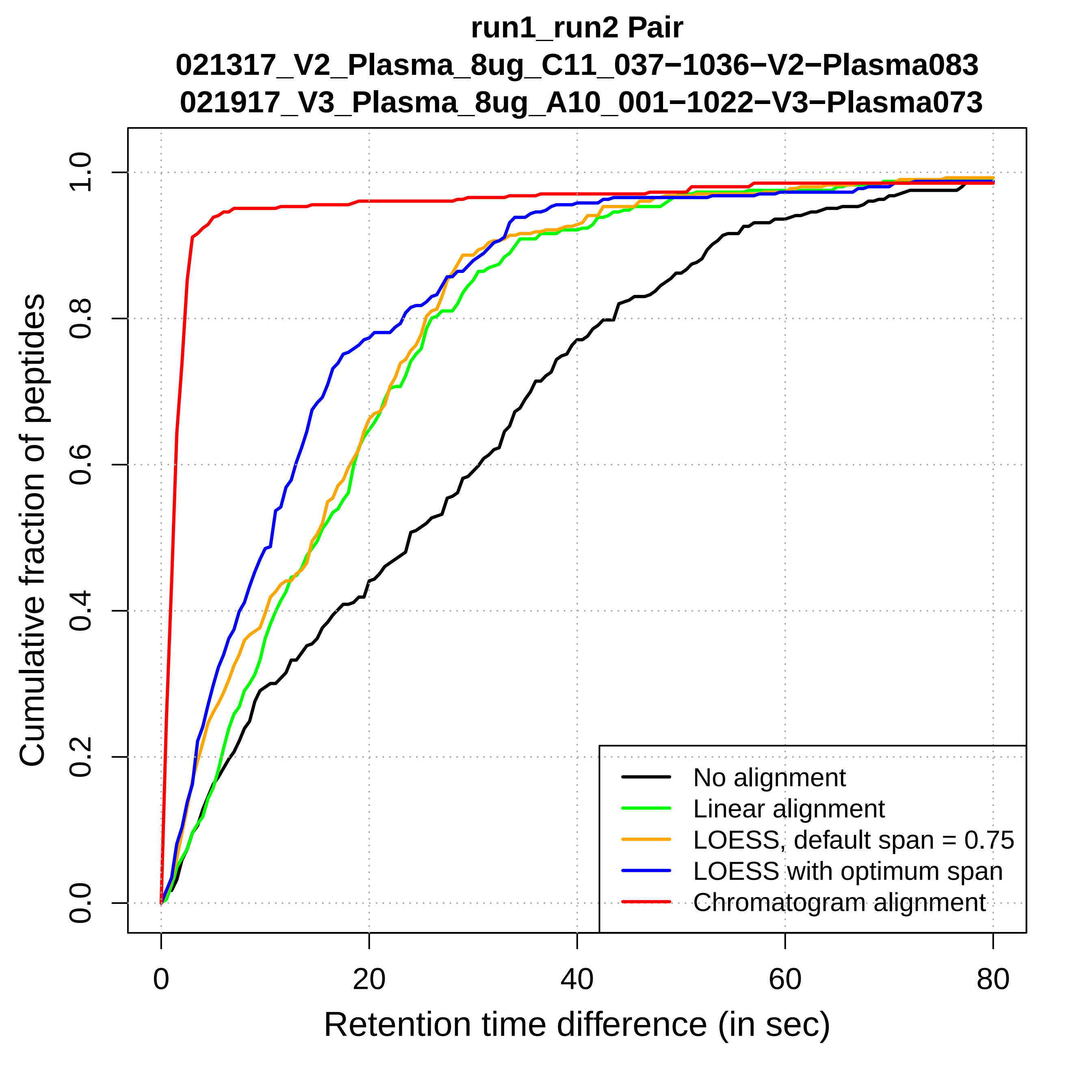

run1_run2 Pair
021317_V2_Plasma_8ug_C11_037−1036−V2−Plasma083
021917_V3_Plasma_8ug_A10_001−1022−V3−Plasma073
Cumulative fraction of peptides
0.0
0.2
0.4
0.6
0.8
1.0
0
20
40
60
80
Retention time difference (in sec)
No alignment
Linear alignment
LOESS, default span = 0.75
LOESS with optimum span
Chromatogram alignment

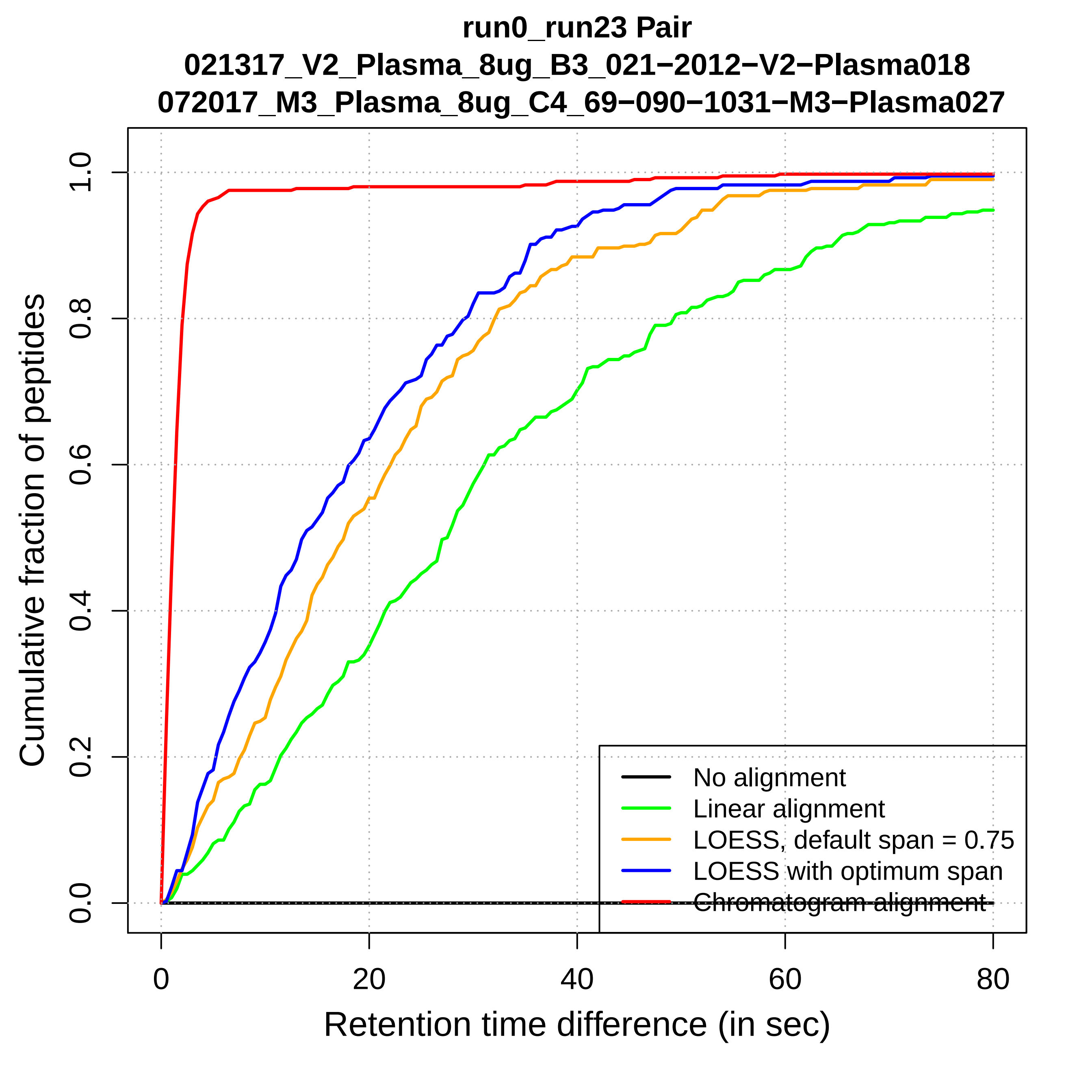

run0_run23 Pair
021317_V2_Plasma_8ug_B3_021−2012−V2−Plasma018
072017_M3_Plasma_8ug_C4_69−090−1031−M3−Plasma027
Cumulative fraction of peptides
0.0
0.2
0.4
0.6
0.8
1.0
0
20
40
60
80
Retention time difference (in sec)
No alignment
Linear alignment
LOESS, default span = 0.75
LOESS with optimum span
Chromatogram alignment

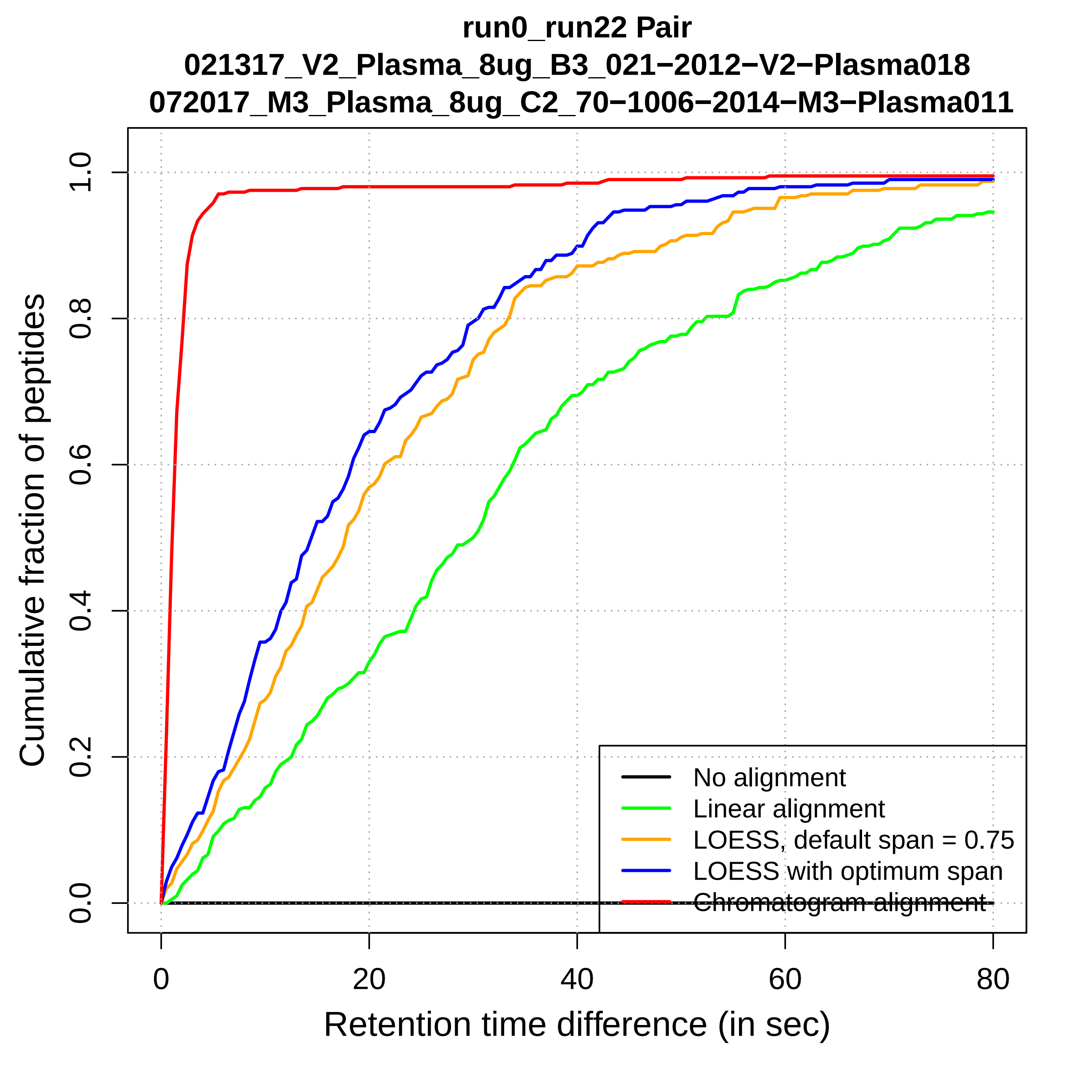
run0_run22 Pair
021317_V2_Plasma_8ug_B3_021−2012−V2−Plasma018
072017_M3_Plasma_8ug_C2_70−1006−2014−M3−Plasma011
Cumulative fraction of peptides
0.0
0.2
0.4
0.6
0.8
1.0
0
20
40
60
80
Retention time difference (in sec)
No alignment
Linear alignment
LOESS, default span = 0.75
LOESS with optimum span
Chromatogram alignment

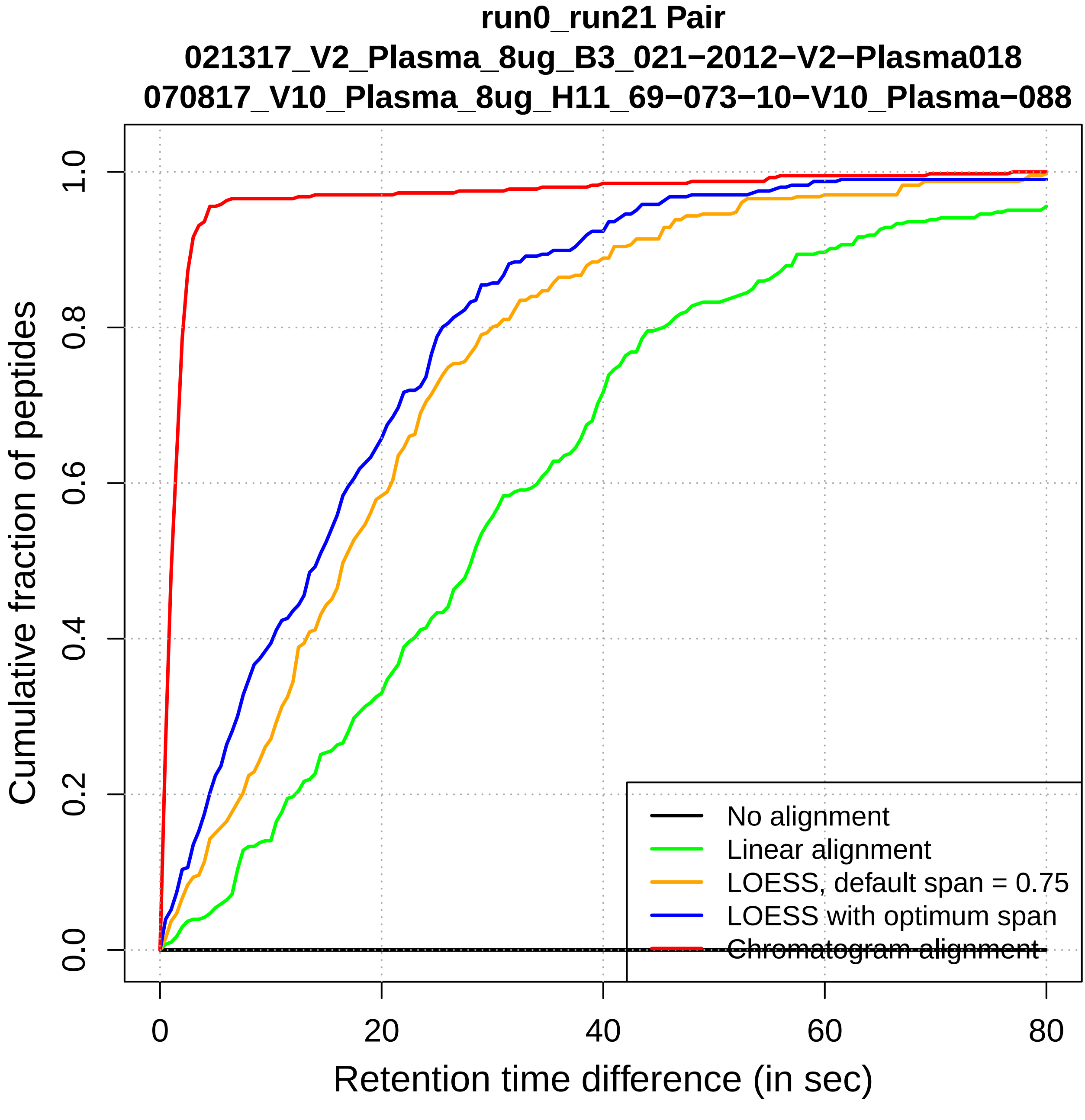
run0_run21 Pair
021317_V2_Plasma_8ug_B3_021−2012−V2−Plasma018
070817_V10_Plasma_8ug_H11_69−073−10−V10_Plasma−088
Cumulative fraction of peptides
0.0
0.2
0.4
0.6
0.8
1.0
0
20
40
60
80
Retention time difference (in sec)
No alignment
Linear alignment
LOESS, default span = 0.75
LOESS with optimum span
Chromatogram alignment

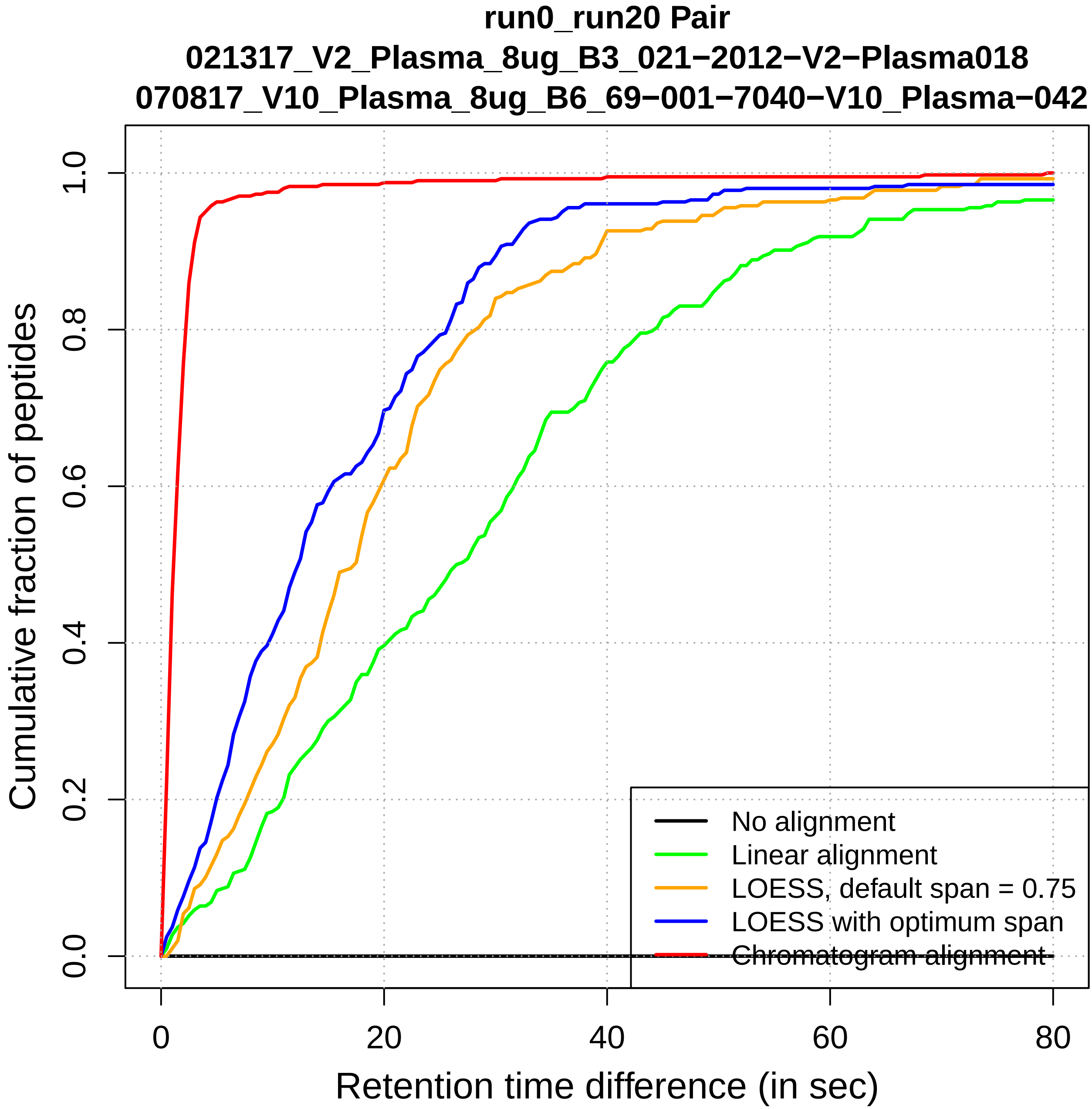

run0_run20 Pair
021317_V2_Plasma_8ug_B3_021−2012−V2−Plasma018
070817_V10_Plasma_8ug_B6_69−001−7040−V10_Plasma−042
Cumulative fraction of peptides
0.0
0.2
0.4
0.6
0.8
1.0
0
20
40
60
80
Retention time difference (in sec)
No alignment
Linear alignment
LOESS, default span = 0.75
LOESS with optimum span
Chromatogram alignment

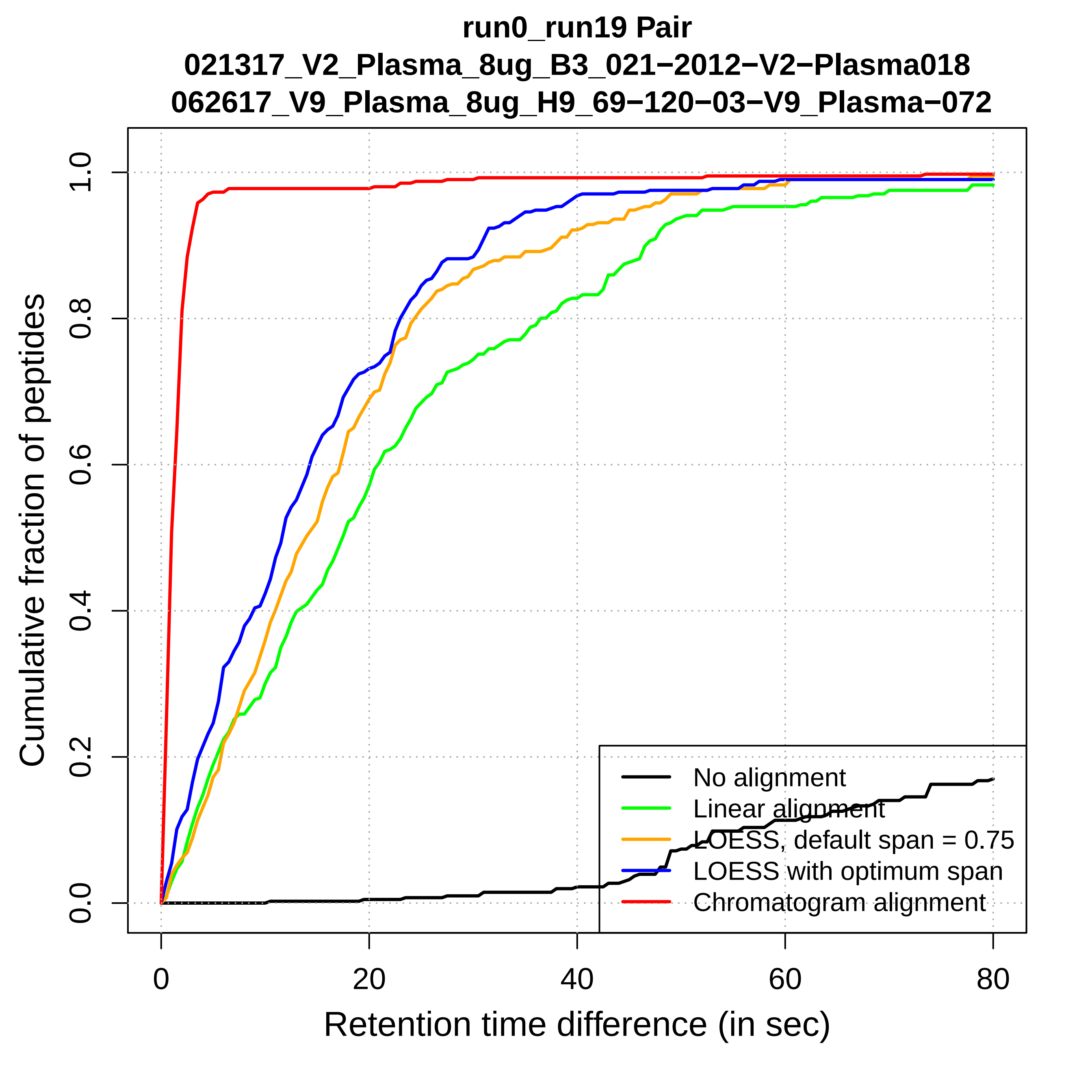

run0_run19 Pair
021317_V2_Plasma_8ug_B3_021−2012−V2−Plasma018
062617_V9_Plasma_8ug_H9_69−120−03−V9_Plasma−072
Cumulative fraction of peptides
0.0
0.2
0.4
0.6
0.8
1.0
0
20
40
60
80
Retention time difference (in sec)
No alignment
Linear alignment
LOESS, default span = 0.75
LOESS with optimum span
Chromatogram alignment

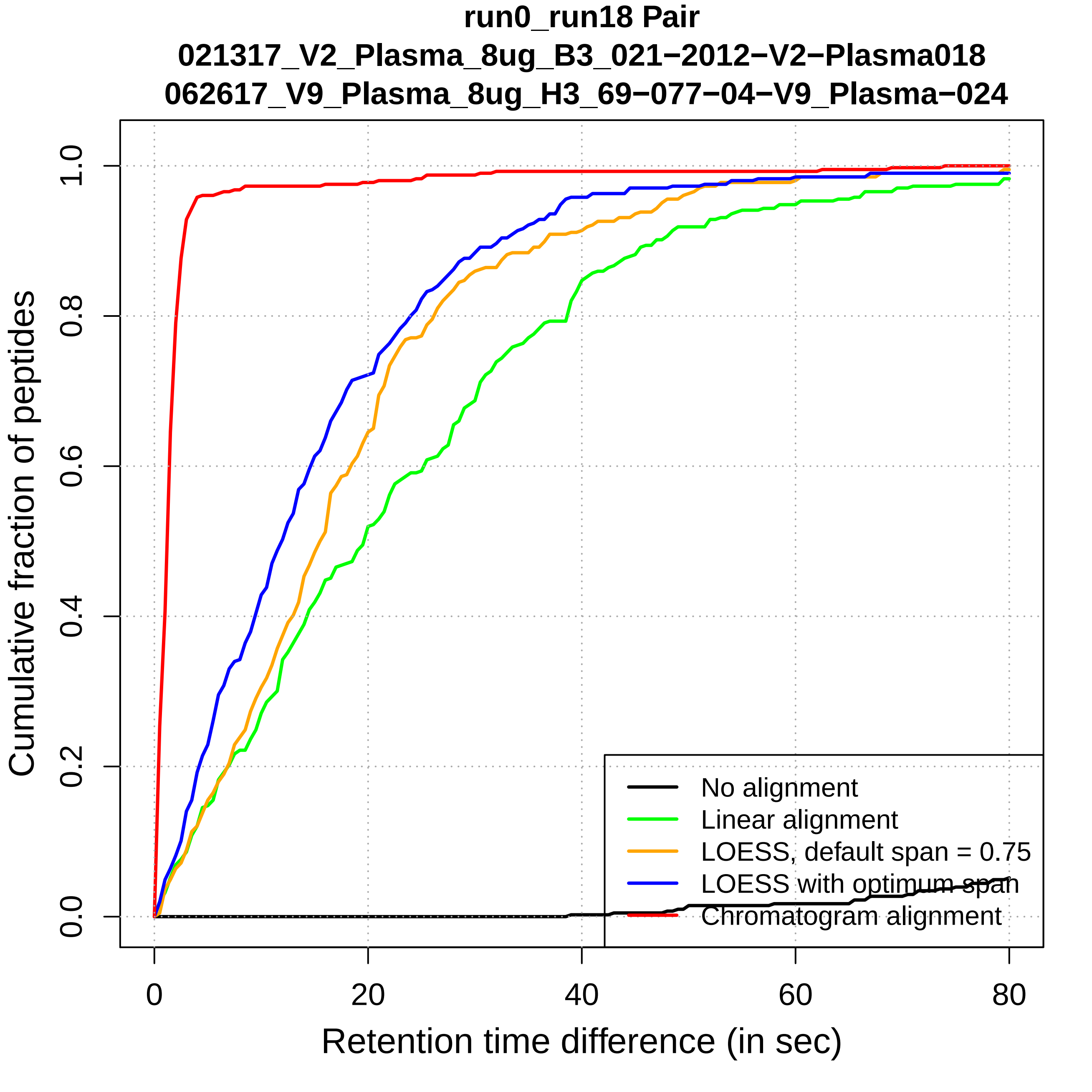
run0_run18 Pair
021317_V2_Plasma_8ug_B3_021−2012−V2−Plasma018
062617_V9_Plasma_8ug_H3_69−077−04−V9_Plasma−024
Cumulative fraction of peptides
0.0
0.2
0.4
0.6
0.8
1.0
0
20
40
60
80
Retention time difference (in sec)
No alignment
Linear alignment
LOESS, default span = 0.75
LOESS with optimum span
Chromatogram alignment

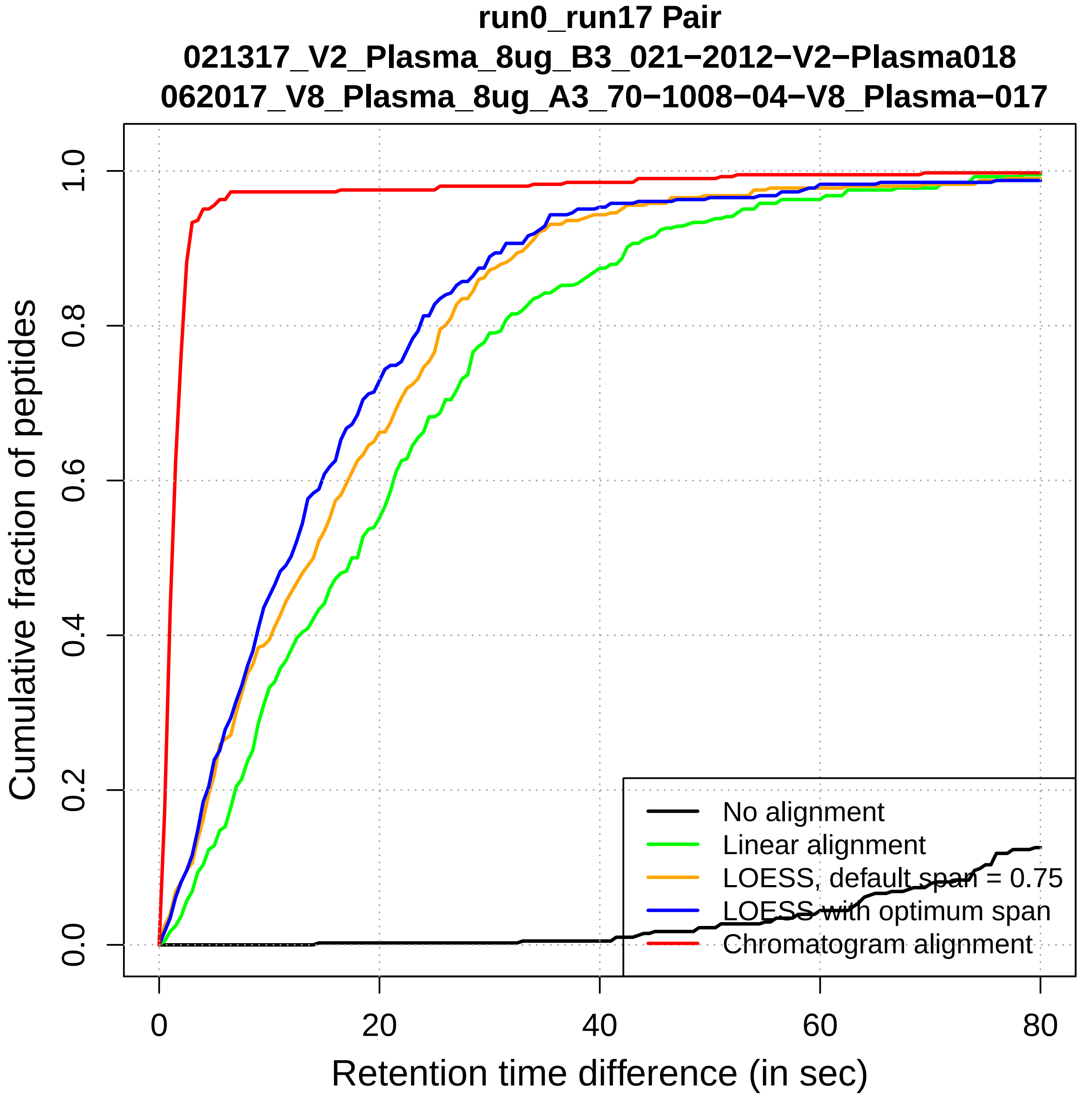
run0_run17 Pair
021317_V2_Plasma_8ug_B3_021−2012−V2−Plasma018
062017_V8_Plasma_8ug_A3_70−1008−04−V8_Plasma−017
Cumulative fraction of peptides
Retention time difference (in sec)
0.0
0.2
0.4
0.6
0.8
1.0
0
20
40
60
80
No alignment
Linear alignment
LOESS, default span = 0.75
LOESS with optimum span
Chromatogram alignment

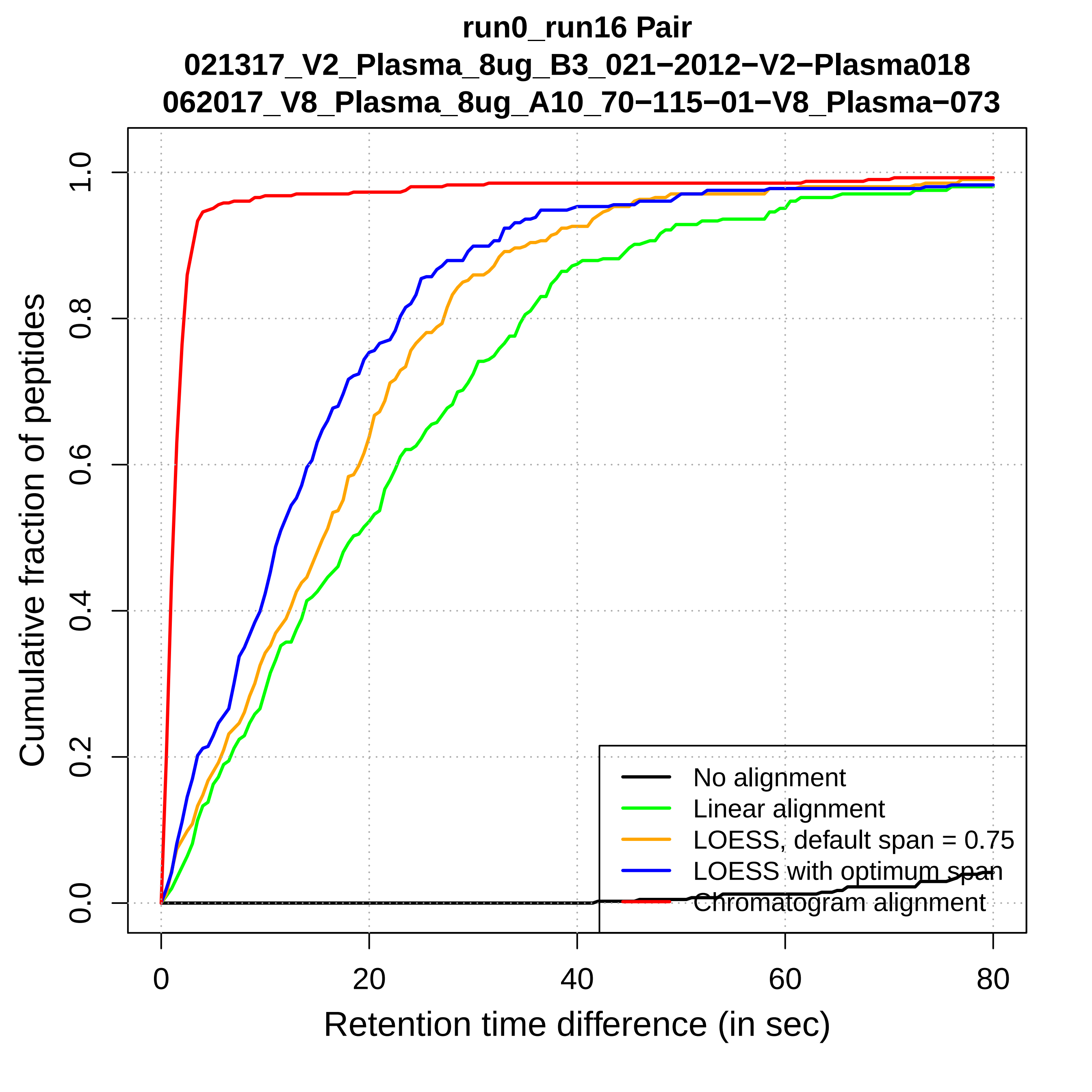

run0_run16 Pair
021317_V2_Plasma_8ug_B3_021−2012−V2−Plasma018
062017_V8_Plasma_8ug_A10_70−115−01−V8_Plasma−073
Cumulative fraction of peptides
0.0
0.2
0.4
0.6
0.8
1.0
0
20
40
60
80
Retention time difference (in sec)
No alignment
Linear alignment
LOESS, default span = 0.75
LOESS with optimum span
Chromatogram alignment

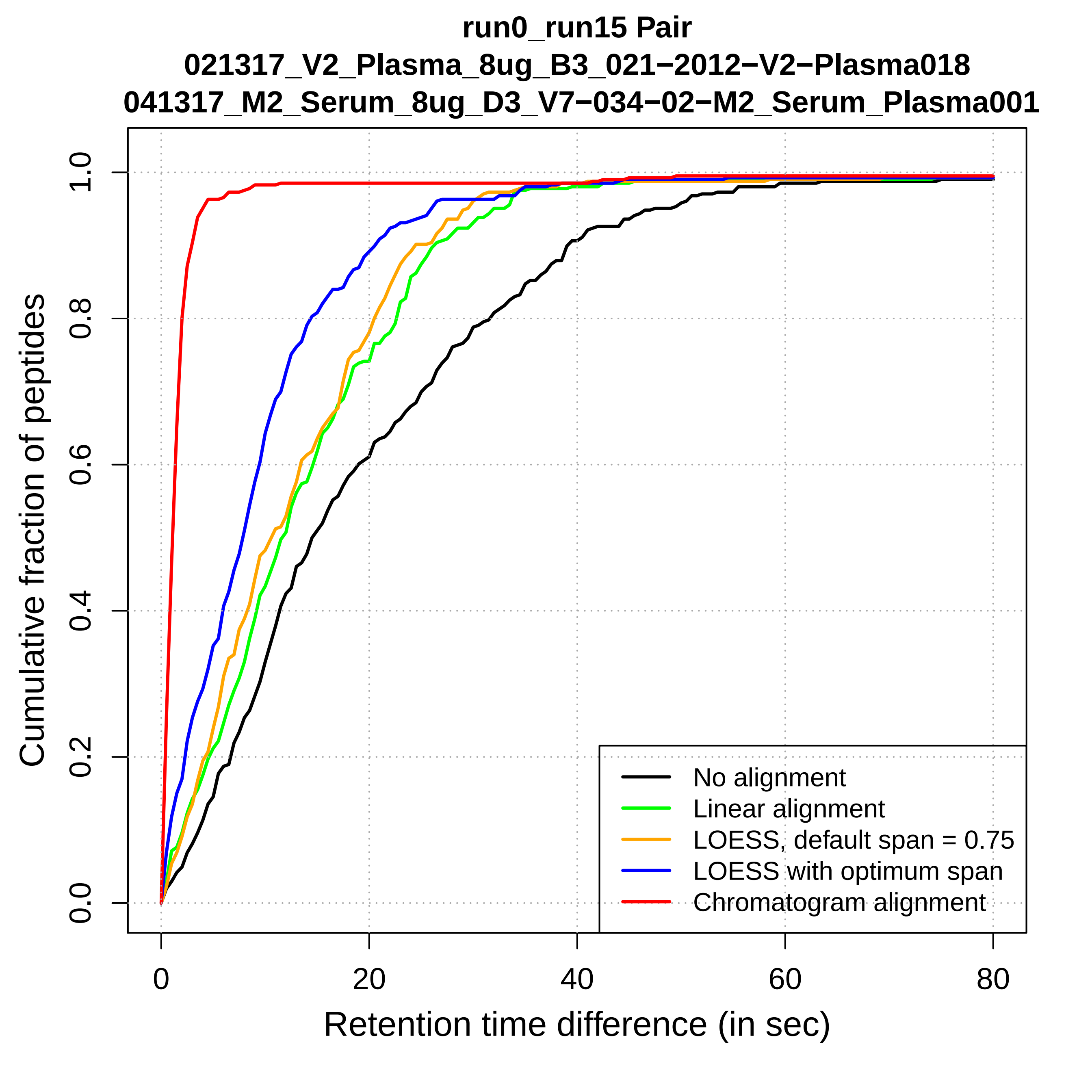
run0_run15 Pair
021317_V2_Plasma_8ug_B3_021−2012−V2−Plasma018
041317_M2_Serum_8ug_D3_V7−034−02−M2_Serum_Plasma001
Cumulative fraction of peptides
Retention time difference (in sec)
0.0
0.2
0.4
0.6
0.8
1.0
0
20
40
60
80
No alignment
Linear alignment
LOESS, default span = 0.75
LOESS with optimum span
Chromatogram alignment

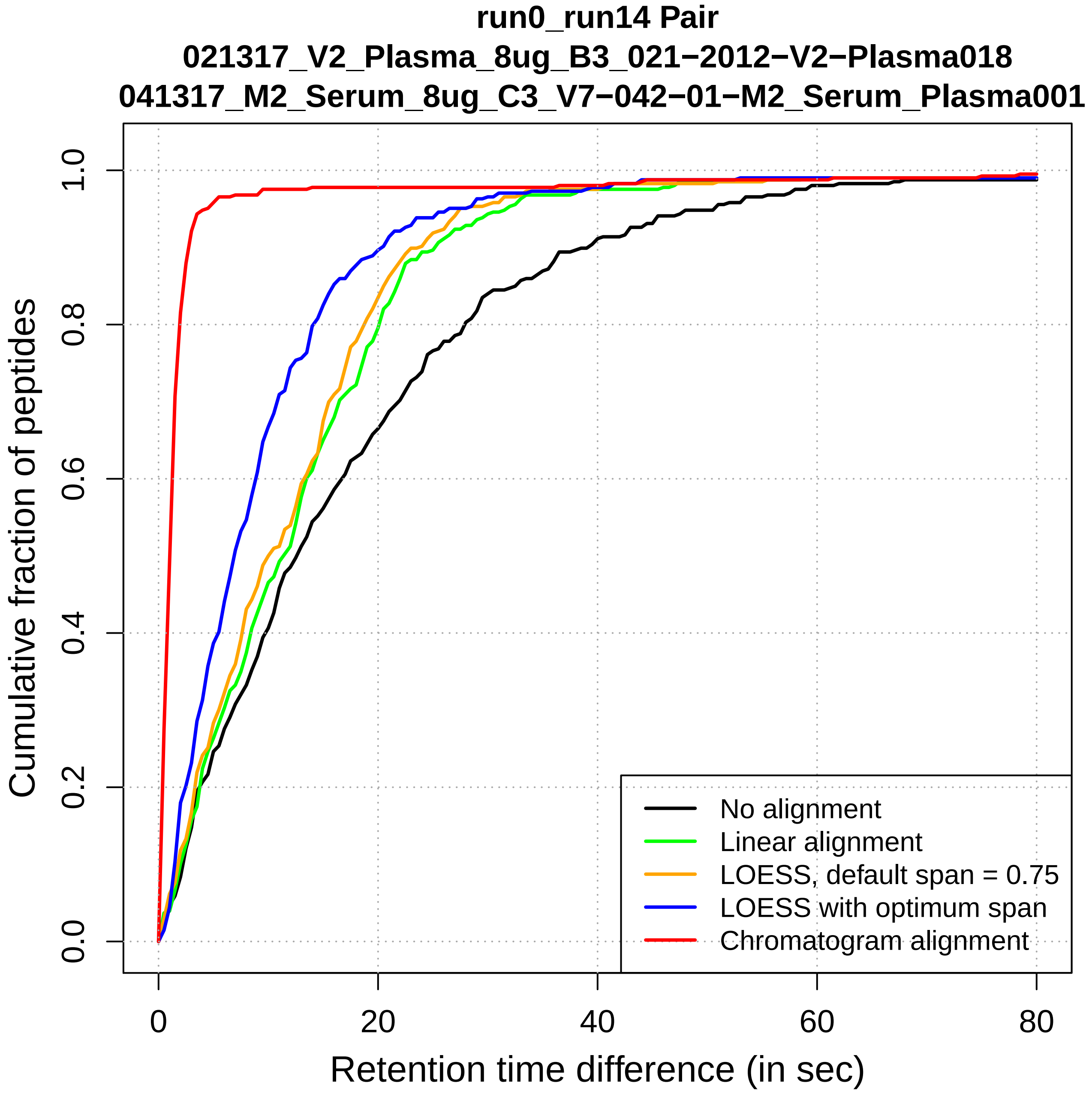
run0_run14 Pair
021317_V2_Plasma_8ug_B3_021−2012−V2−Plasma018
041317_M2_Serum_8ug_C3_V7−042−01−M2_Serum_Plasma001
Cumulative fraction of peptides
0.0
0.2
0.4
0.6
0.8
1.0
0
20
40
60
80
Retention time difference (in sec)
No alignment
Linear alignment
LOESS, default span = 0.75
LOESS with optimum span
Chromatogram alignment

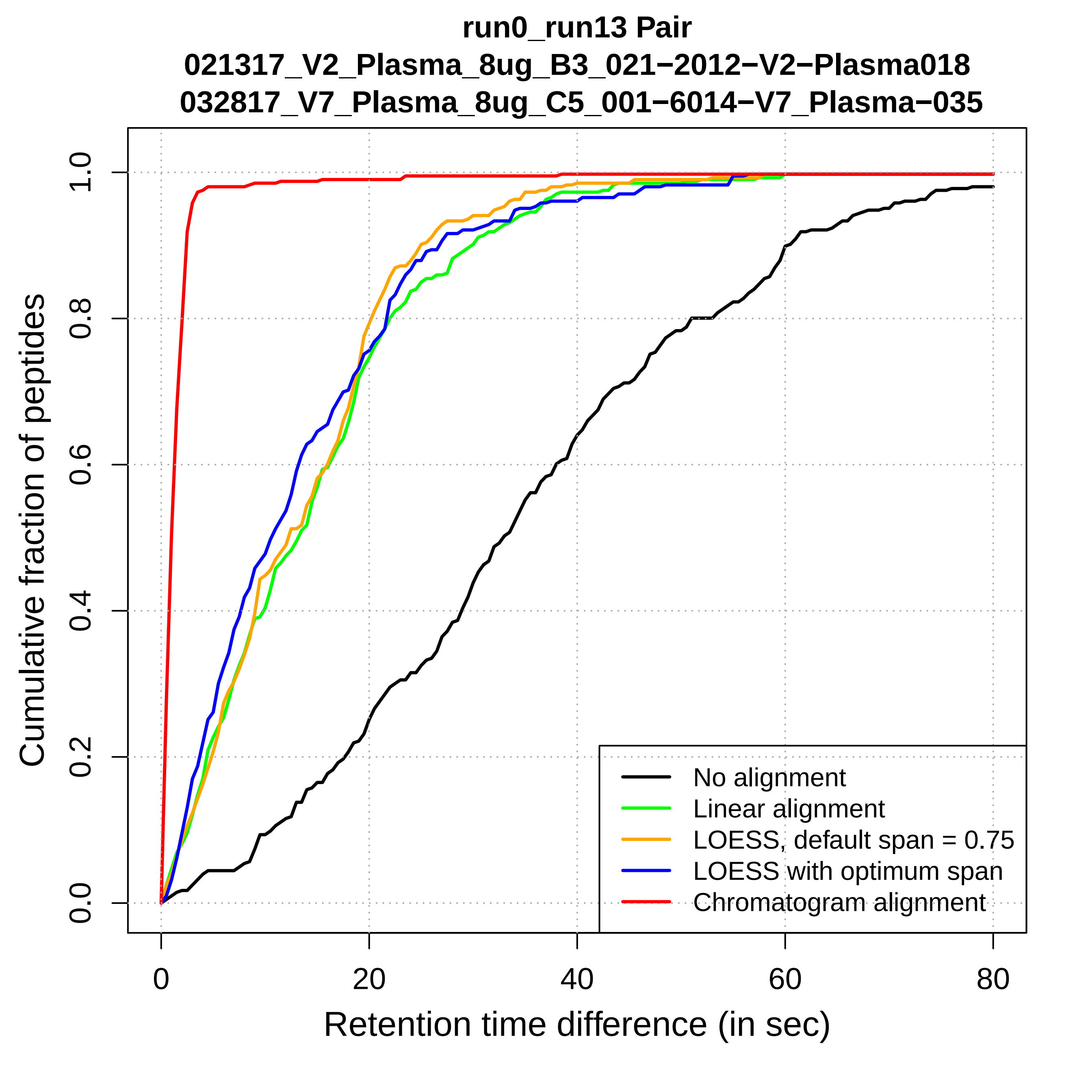
run0_run13 Pair
021317_V2_Plasma_8ug_B3_021−2012−V2−Plasma018
032817_V7_Plasma_8ug_C5_001−6014−V7_Plasma−035
Cumulative fraction of peptides
0.0
0.2
0.4
0.6
0.8
1.0
0
20
40
60
80
Retention time difference (in sec)
No alignment
Linear alignment
LOESS, default span = 0.75
LOESS with optimum span
Chromatogram alignment

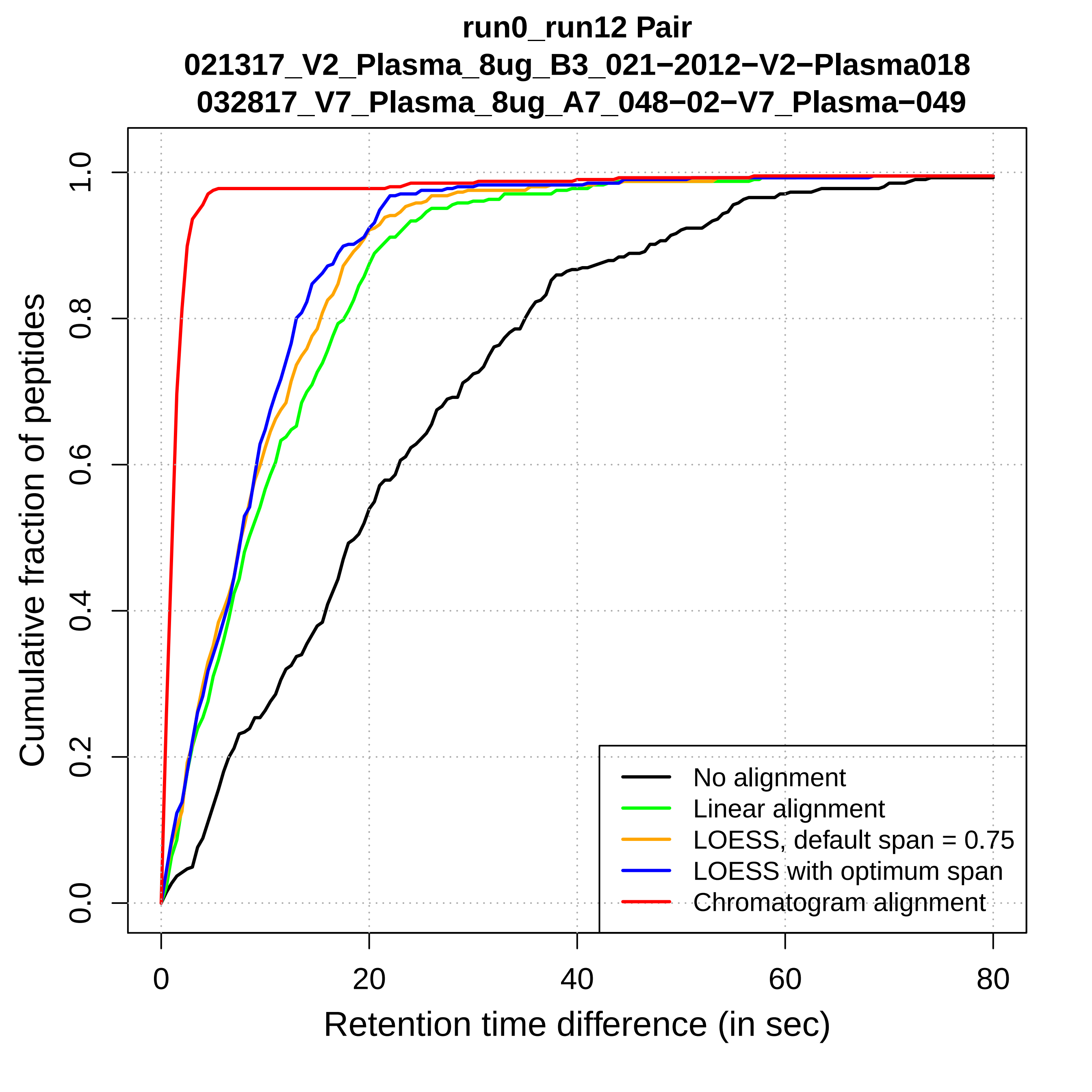
run0_run12 Pair
021317_V2_Plasma_8ug_B3_021−2012−V2−Plasma018
032817_V7_Plasma_8ug_A7_048−02−V7_Plasma−049
Cumulative fraction of peptides
0.0
0.2
0.4
0.6
0.8
1.0
0
20
40
60
80
Retention time difference (in sec)
No alignment
Linear alignment
LOESS, default span = 0.75
LOESS with optimum span
Chromatogram alignment

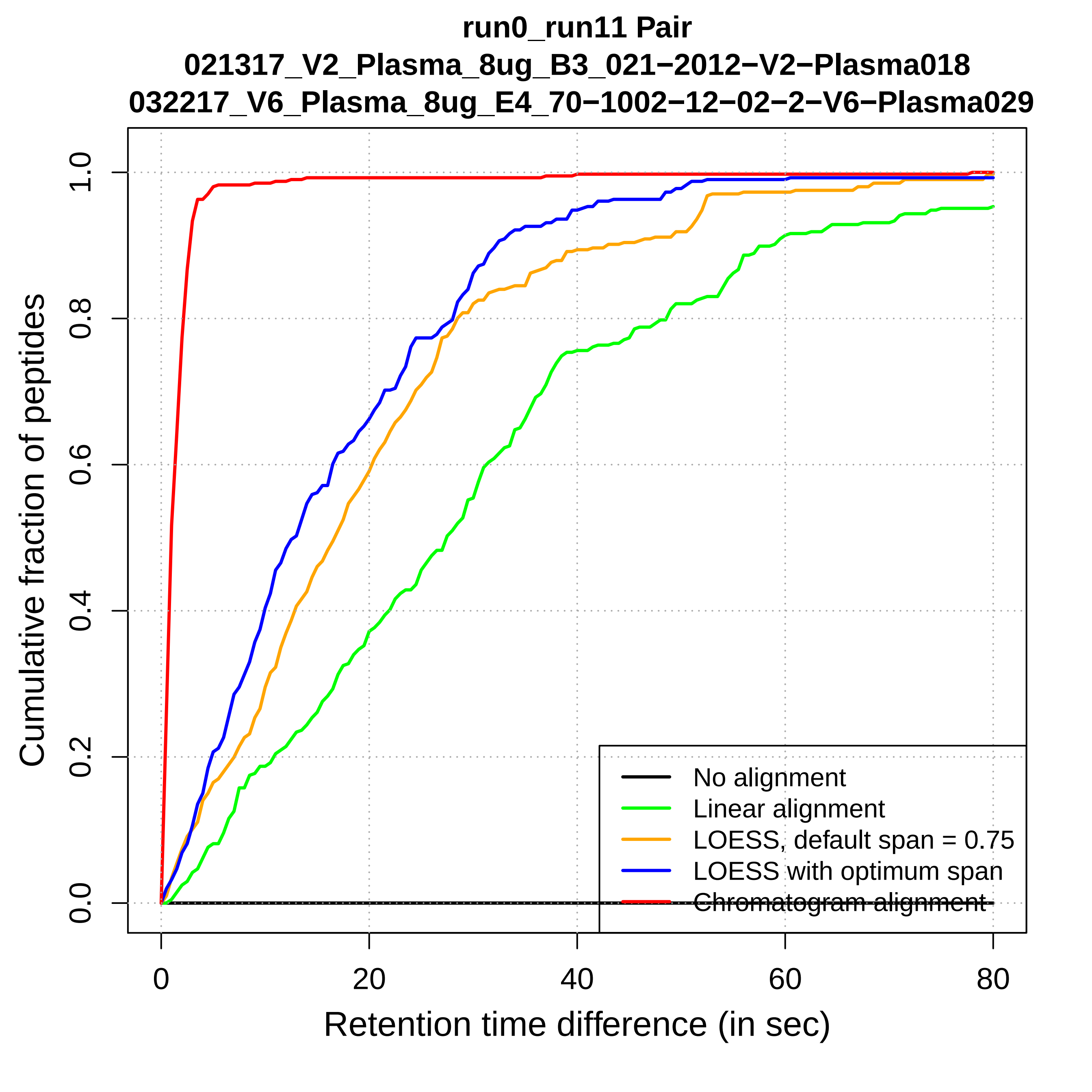

run0_run11 Pair
021317_V2_Plasma_8ug_B3_021−2012−V2−Plasma018
032217_V6_Plasma_8ug_E4_70−1002−12−02−2−V6−Plasma029
Cumulative fraction of peptides
0.0
0.2
0.4
0.6
0.8
1.0
0
20
40
60
80
Retention time difference (in sec)
No alignment
Linear alignment
LOESS, default span = 0.75
LOESS with optimum span
Chromatogram alignment

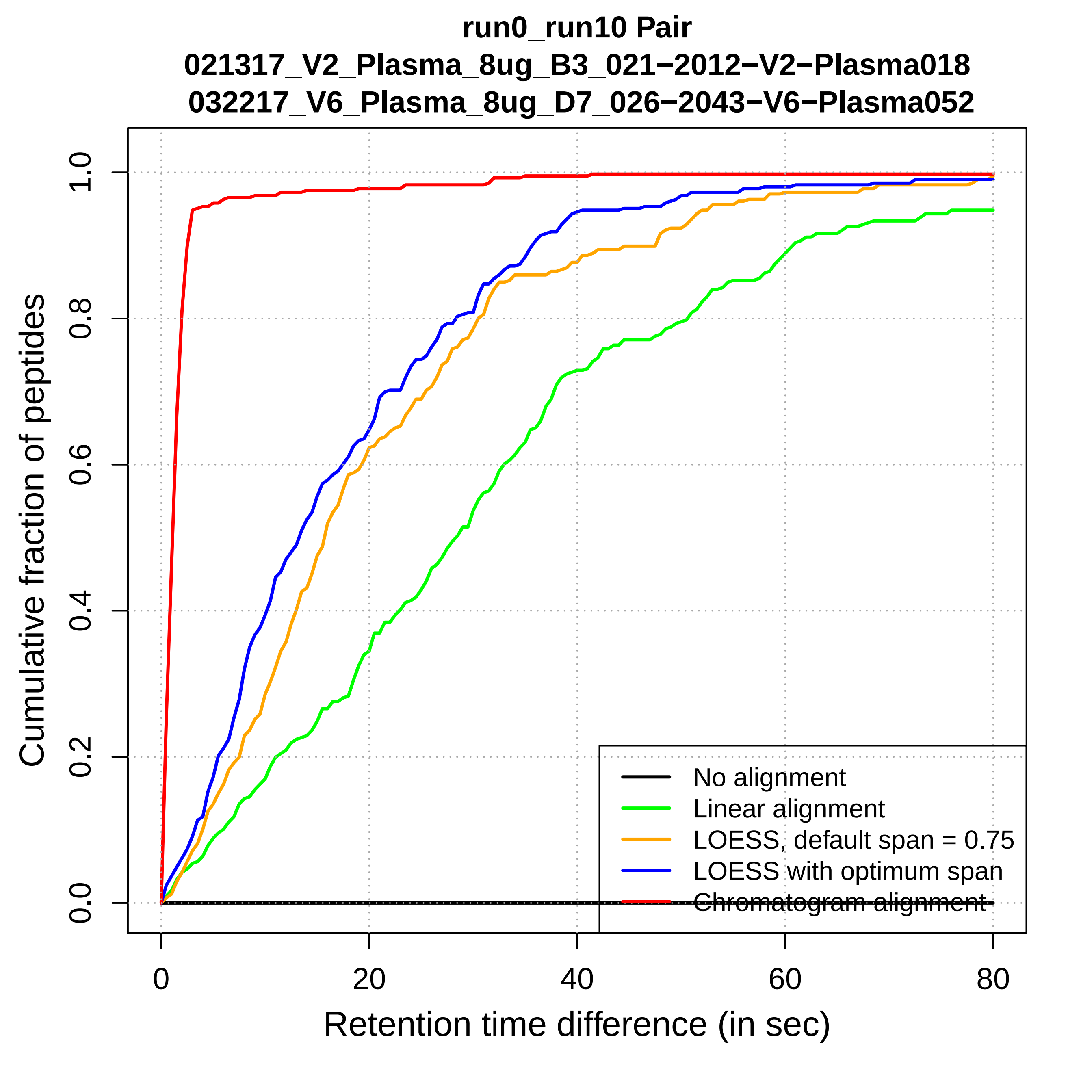
run0_run10 Pair
021317_V2_Plasma_8ug_B3_021−2012−V2−Plasma018
032217_V6_Plasma_8ug_D7_026−2043−V6−Plasma052
Cumulative fraction of peptides
Retention time difference (in sec)
0.0
0.2
0.4
0.6
0.8
1.0
0
20
40
60
80
No alignment
Linear alignment
LOESS, default span = 0.75
LOESS with optimum span
Chromatogram alignment

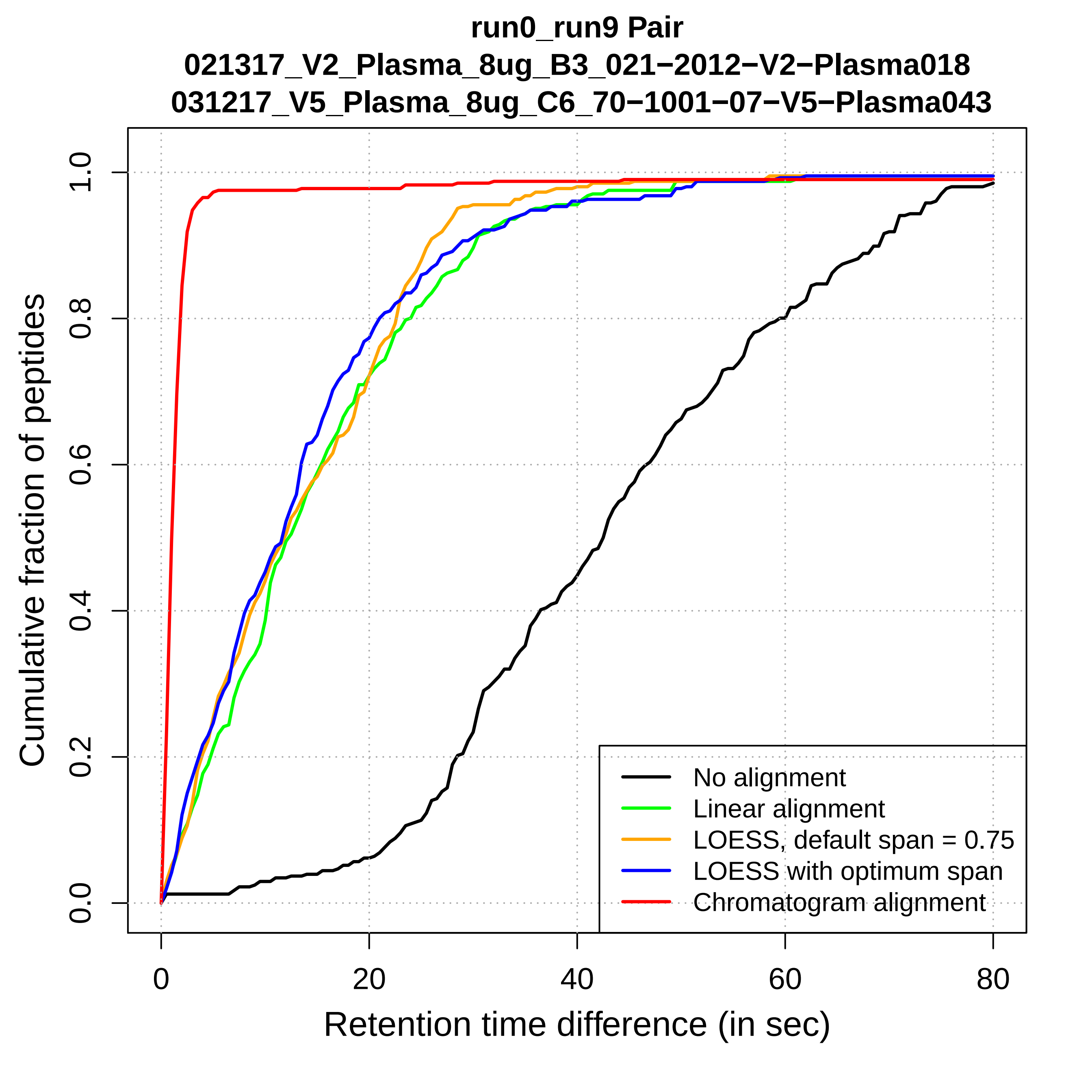
run0_run9 Pair
021317_V2_Plasma_8ug_B3_021−2012−V2−Plasma018
031217_V5_Plasma_8ug_C6_70−1001−07−V5−Plasma043
Cumulative fraction of peptides
0.0
0.2
0.4
0.6
0.8
1.0
0
20
40
60
80
Retention time difference (in sec)
No alignment
Linear alignment
LOESS, default span = 0.75
LOESS with optimum span
Chromatogram alignment

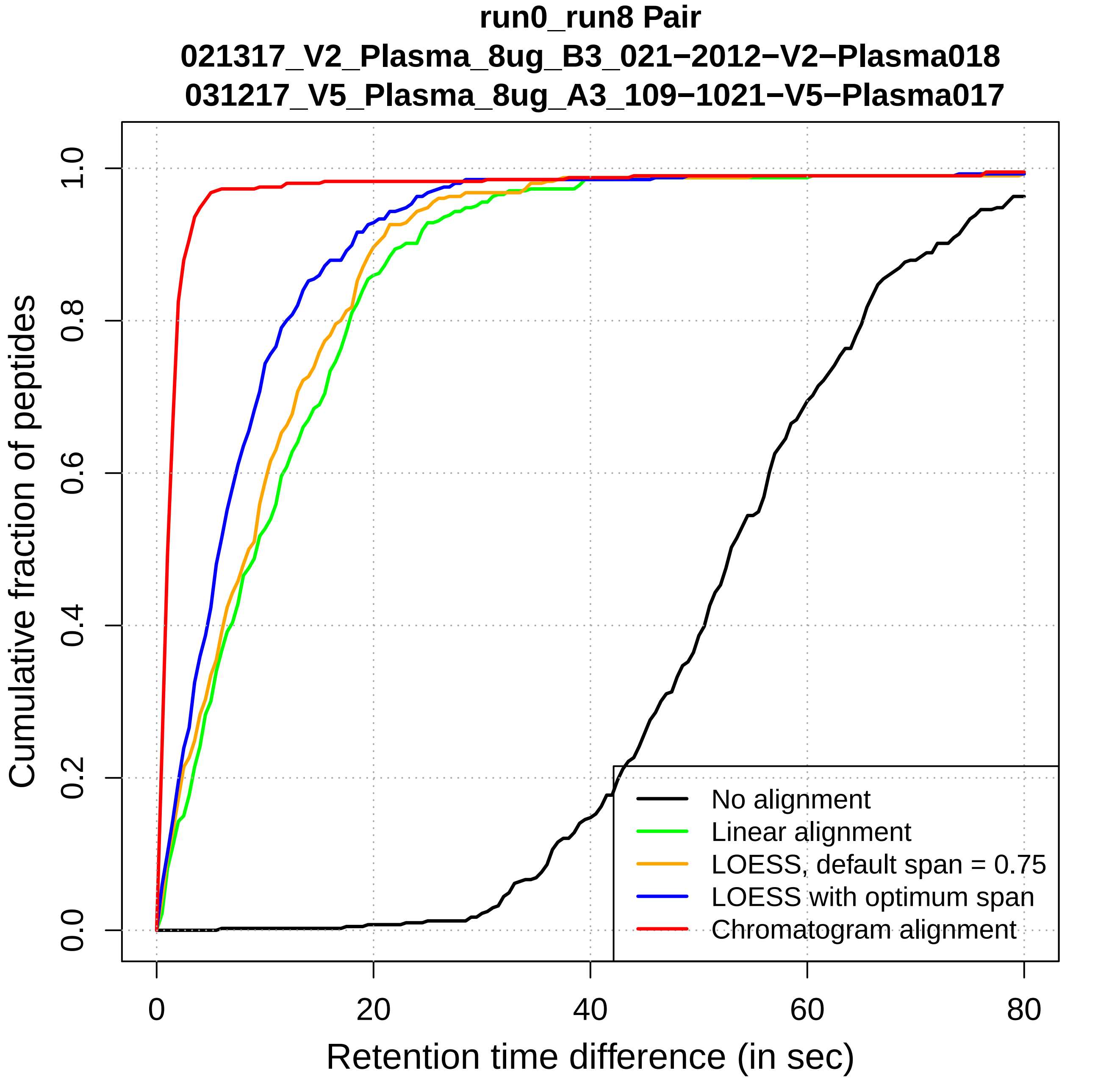

run0_run8 Pair
021317_V2_Plasma_8ug_B3_021−2012−V2−Plasma018
031217_V5_Plasma_8ug_A3_109−1021−V5−Plasma017
Cumulative fraction of peptides
0.0
0.2
0.4
0.6
0.8
1.0
0
20
40
60
80
Retention time difference (in sec)
No alignment
Linear alignment
LOESS, default span = 0.75
LOESS with optimum span
Chromatogram alignment

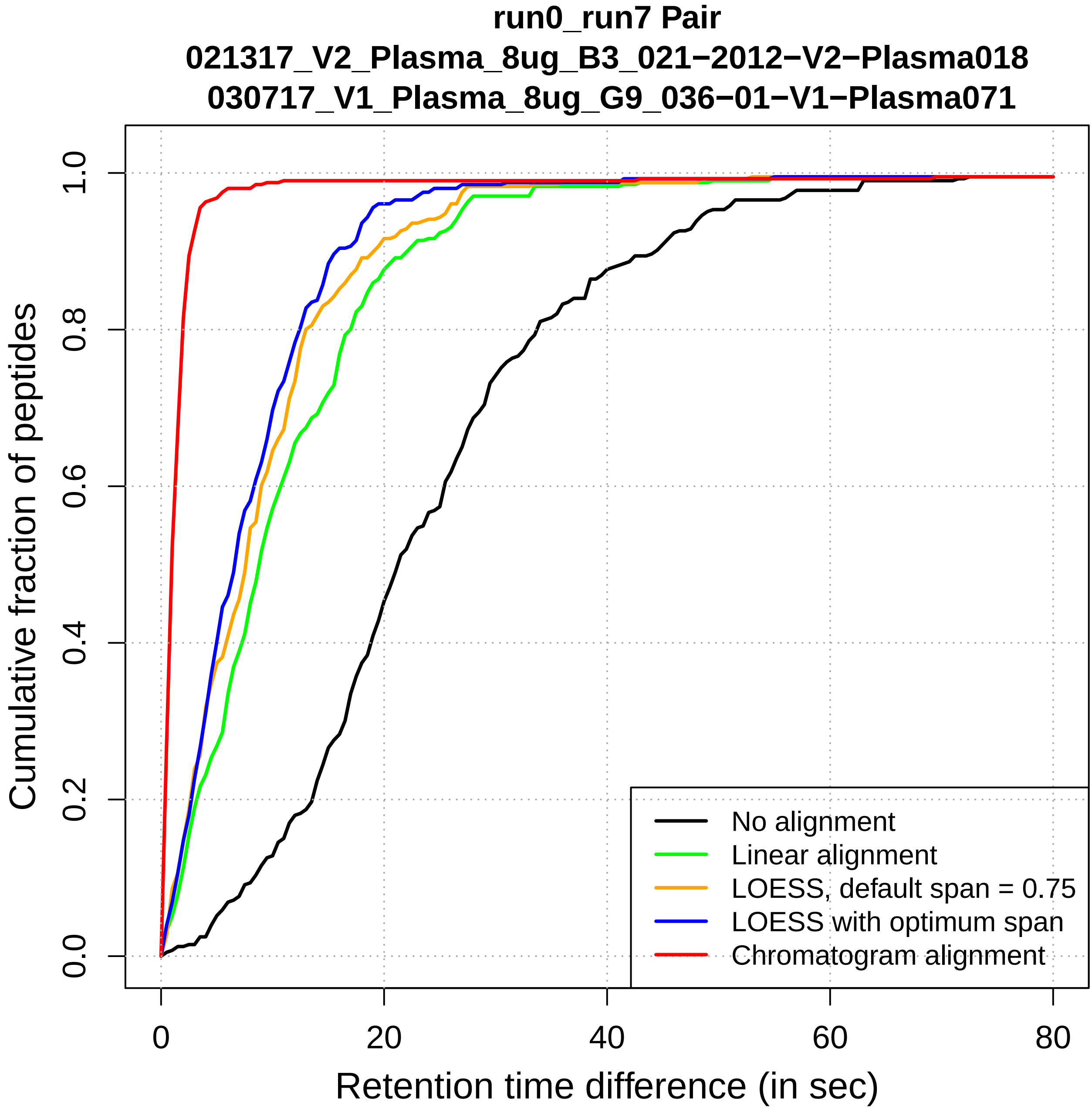
run0_run7 Pair
021317_V2_Plasma_8ug_B3_021−2012−V2−Plasma018
030717_V1_Plasma_8ug_G9_036−01−V1−Plasma071
Cumulative fraction of peptides
Retention time difference (in sec)
0.0
0.2
0.4
0.6
0.8
1.0
0
20
40
60
80
No alignment
Linear alignment
LOESS, default span = 0.75
LOESS with optimum span
Chromatogram alignment

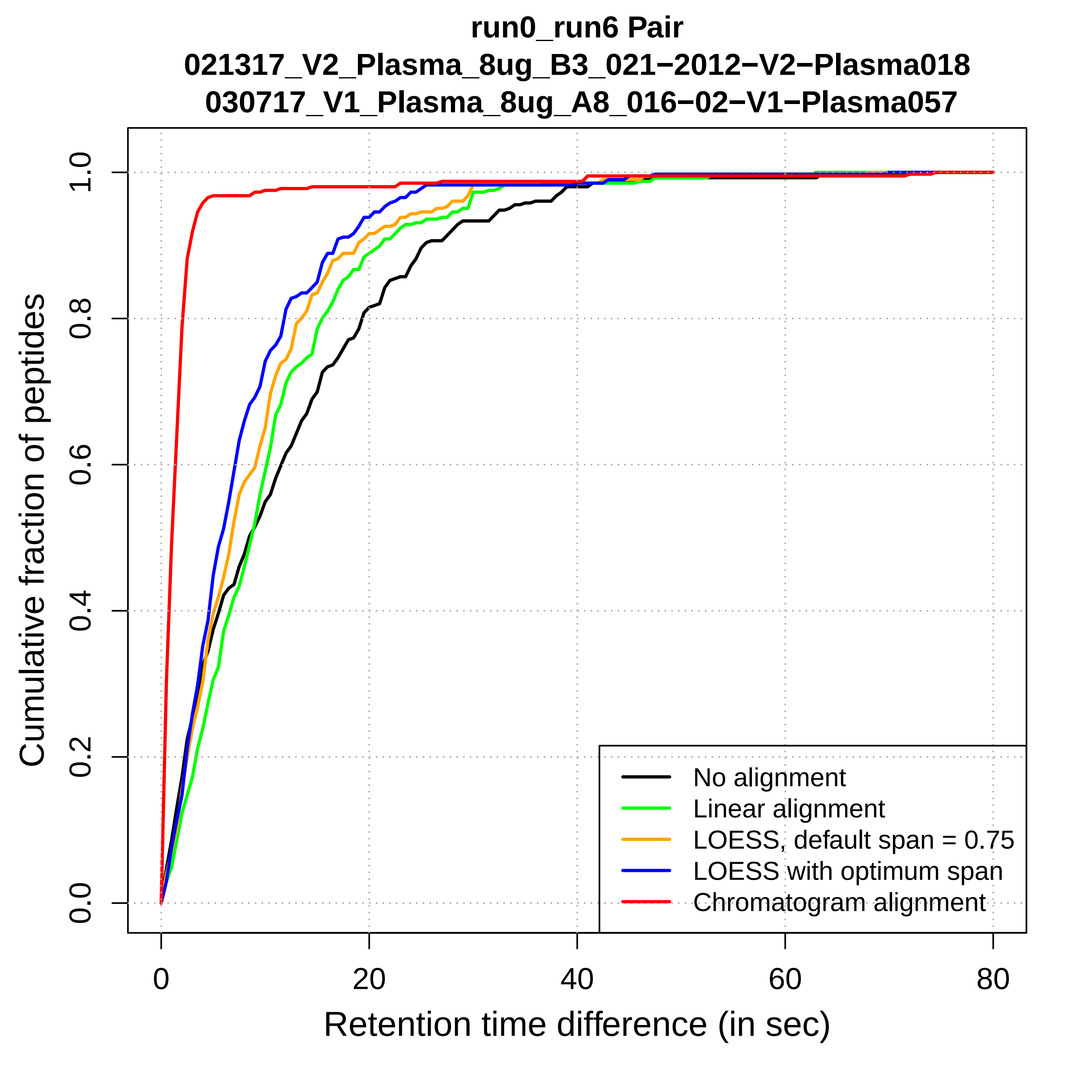
run0_run6 Pair
021317_V2_Plasma_8ug_B3_021−2012−V2−Plasma018
030717_V1_Plasma_8ug_A8_016−02−V1−Plasma057
Cumulative fraction of peptides
0.0
0.2
0.4
0.6
0.8
1.0
0
20
40
60
80
Retention time difference (in sec)
No alignment
Linear alignment
LOESS, default span = 0.75
LOESS with optimum span
Chromatogram alignment

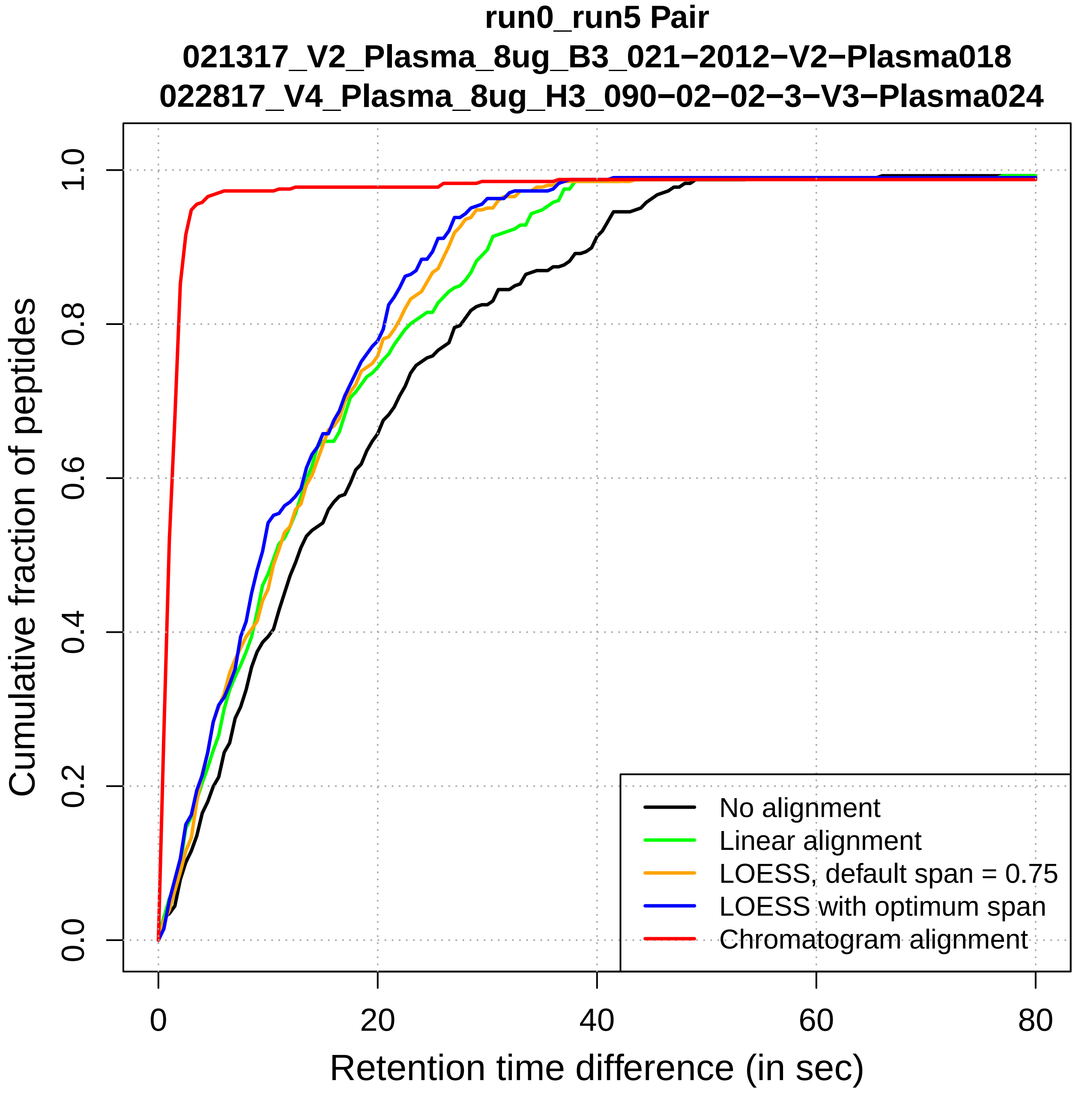
run0_run5 Pair
021317_V2_Plasma_8ug_B3_021−2012−V2−Plasma018
022817_V4_Plasma_8ug_H3_090−02−02−3−V3−Plasma024
Cumulative fraction of peptides
0.0
0.2
0.4
0.6
0.8
1.0
0
20
40
60
80
Retention time difference (in sec)
No alignment
Linear alignment
LOESS, default span = 0.75
LOESS with optimum span
Chromatogram alignment

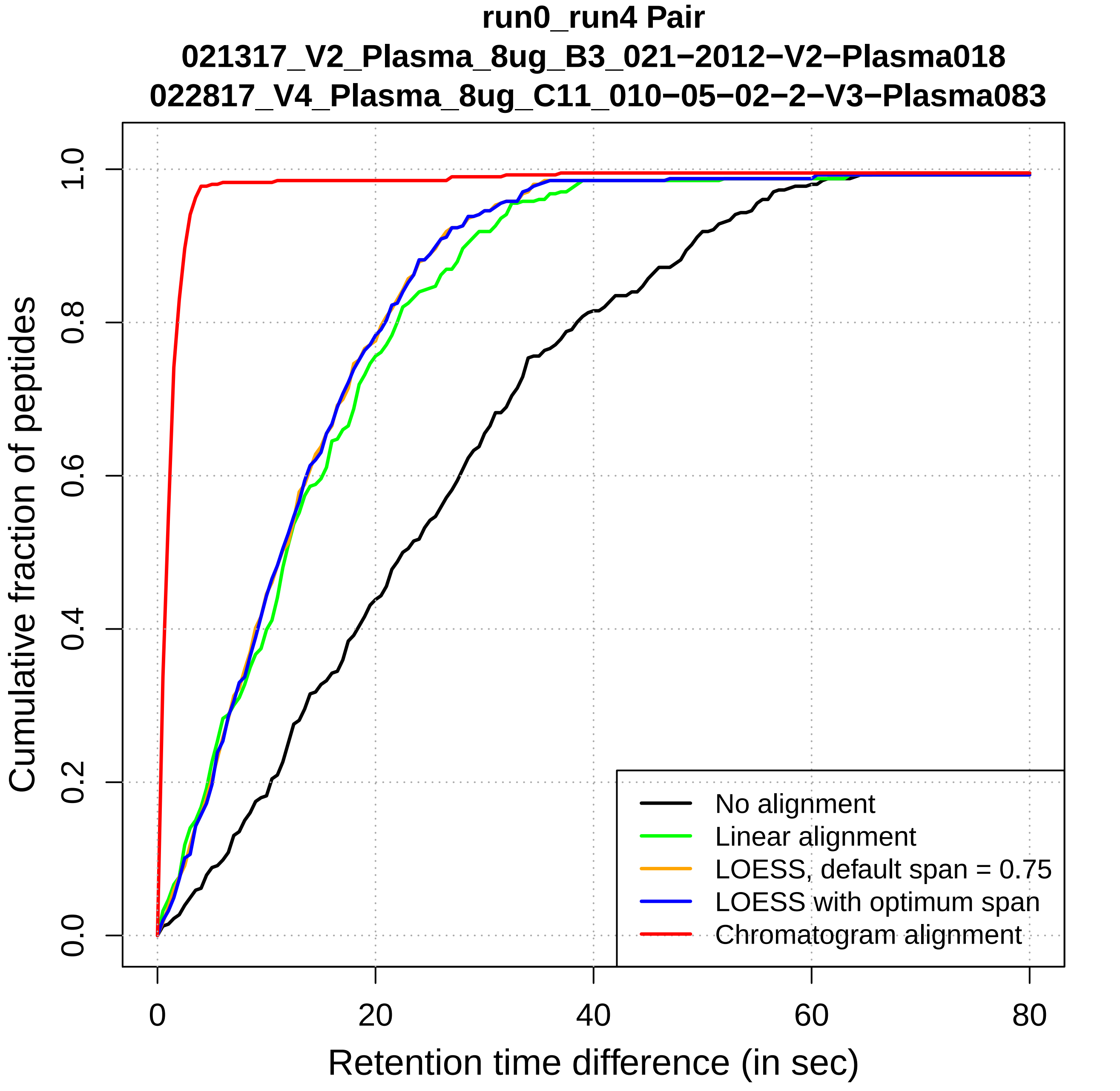

run0_run4 Pair
021317_V2_Plasma_8ug_B3_021−2012−V2−Plasma018
022817_V4_Plasma_8ug_C11_010−05−02−2−V3−Plasma083
Cumulative fraction of peptides
0.0
0.2
0.4
0.6
0.8
1.0
0
20
40
60
80
Retention time difference (in sec)
No alignment
Linear alignment
LOESS, default span = 0.75
LOESS with optimum span
Chromatogram alignment

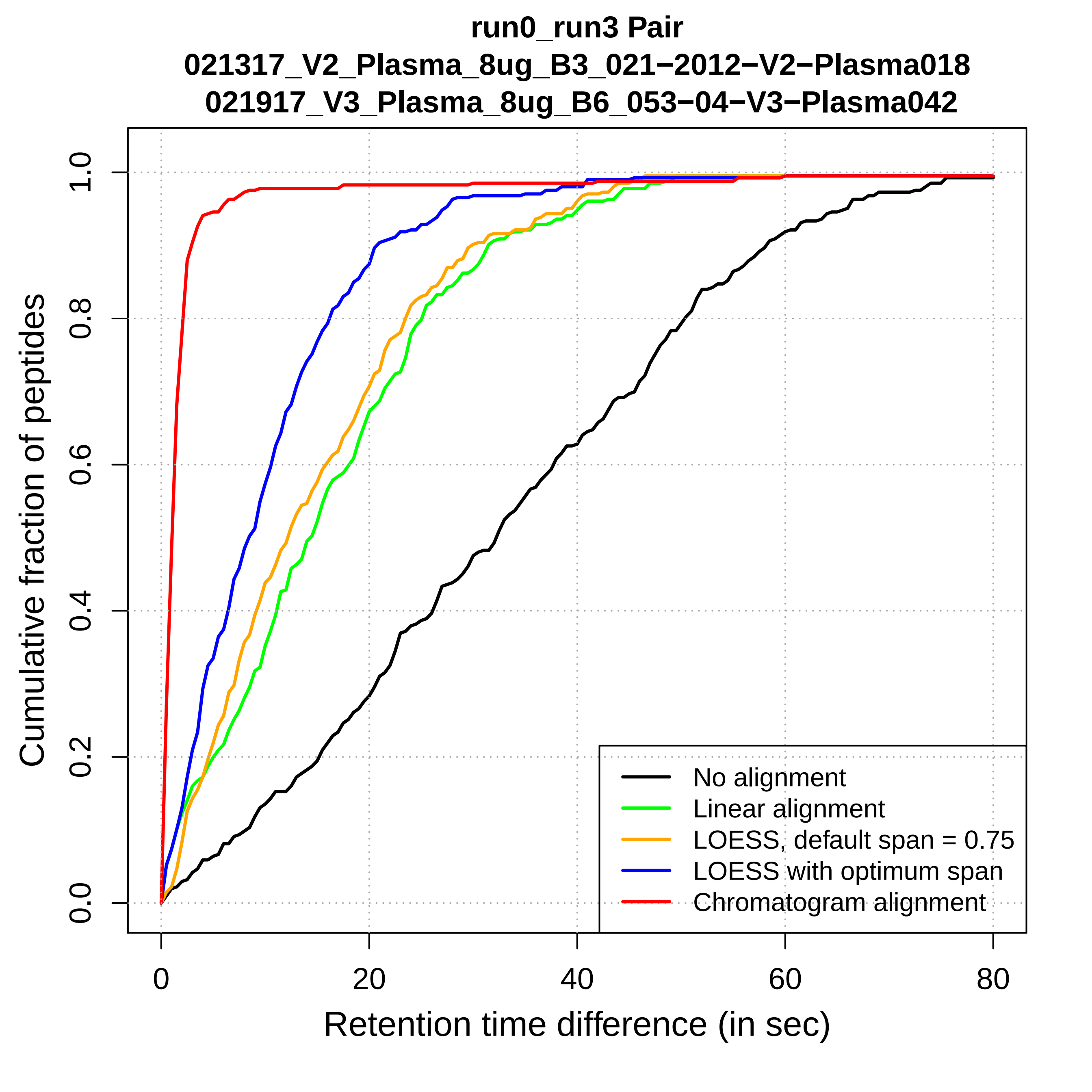

run0_run3 Pair
021317_V2_Plasma_8ug_B3_021−2012−V2−Plasma018
021917_V3_Plasma_8ug_B6_053−04−V3−Plasma042
Cumulative fraction of peptides
0.0
0.2
0.4
0.6
0.8
1.0
0
20
40
60
80
Retention time difference (in sec)
No alignment
Linear alignment
LOESS, default span = 0.75
LOESS with optimum span
Chromatogram alignment

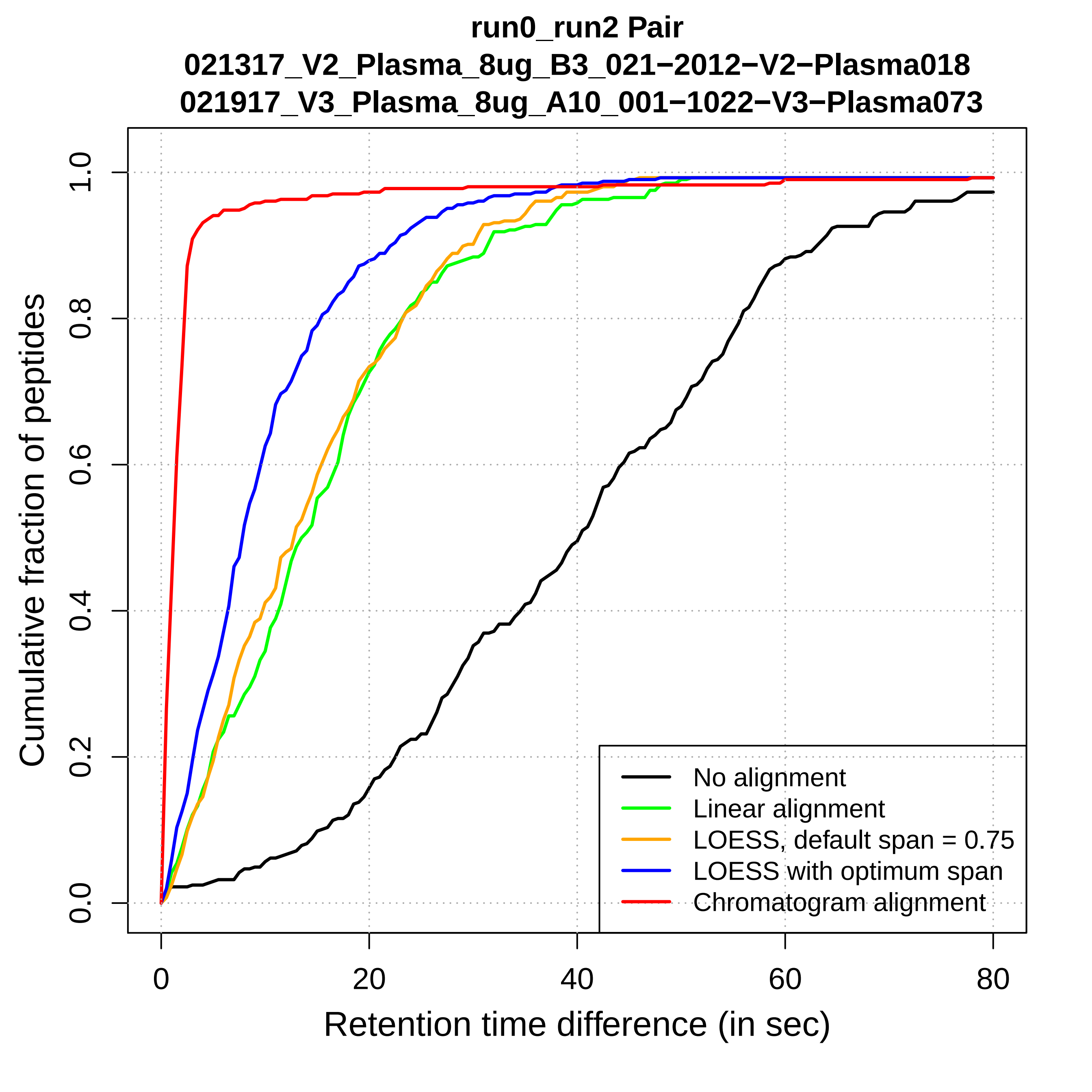
run0_run2 Pair
021317_V2_Plasma_8ug_B3_021−2012−V2−Plasma018
021917_V3_Plasma_8ug_A10_001−1022−V3−Plasma073
Cumulative fraction of peptides
0.0
0.2
0.4
0.6
0.8
1.0
0
20
40
60
80
Retention time difference (in sec)
No alignment
Linear alignment
LOESS, default span = 0.75
LOESS with optimum span
Chromatogram alignment

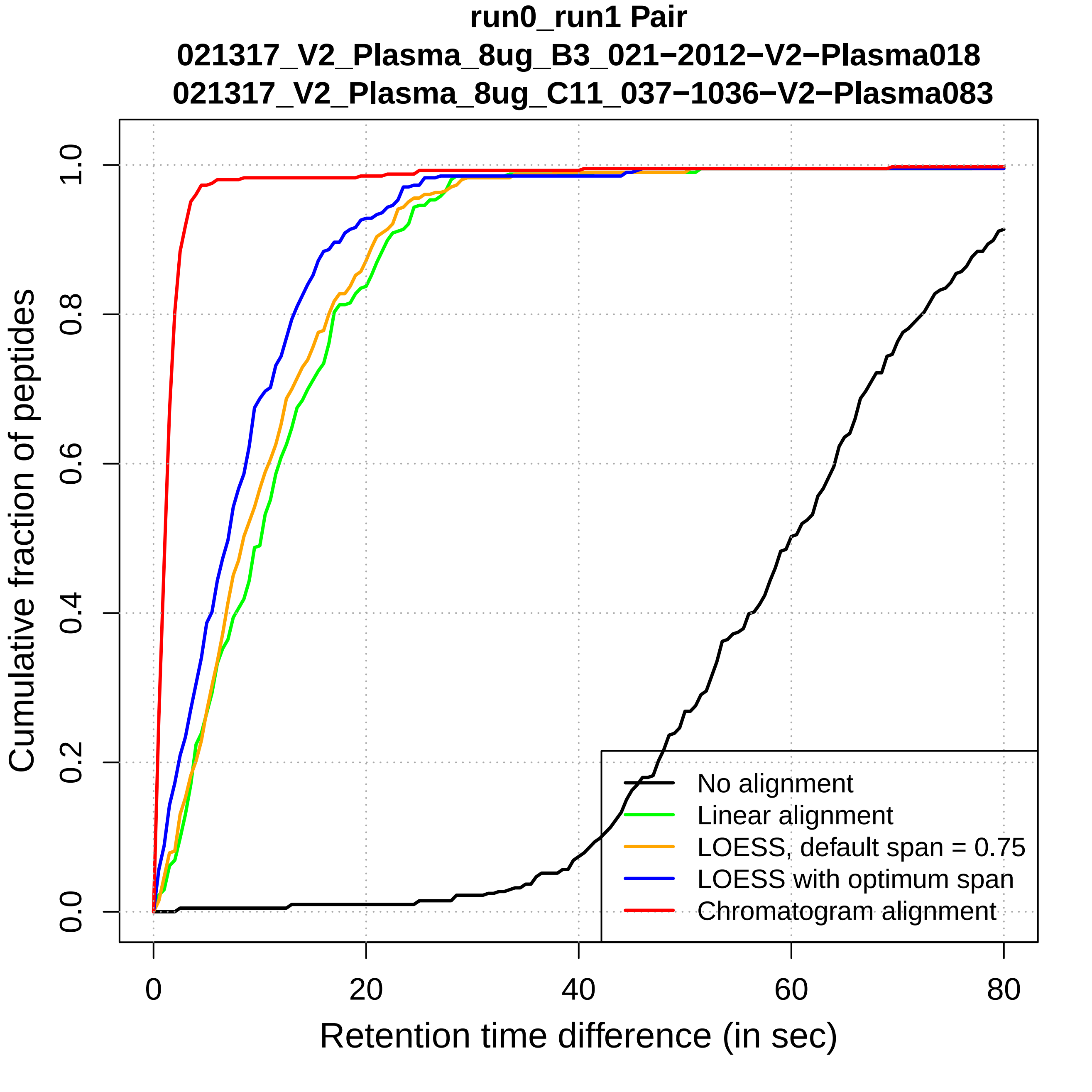
run0_run1 Pair
021317_V2_Plasma_8ug_B3_021−2012−V2−Plasma018
021317_V2_Plasma_8ug_C11_037−1036−V2−Plasma083
Cumulative fraction of peptides
0.0
0.2
0.4
0.6
0.8
1.0
0
20
40
60
80
Retention time difference (in sec)
No alignment
Linear alignment
LOESS, default span = 0.75
LOESS with optimum span
Chromatogram alignment
