## Supplemental Figures for "DIAlignR provides precise retention time alignment across distant runs in DIA and targeted proteomics": Supplemental Figure16.pdf

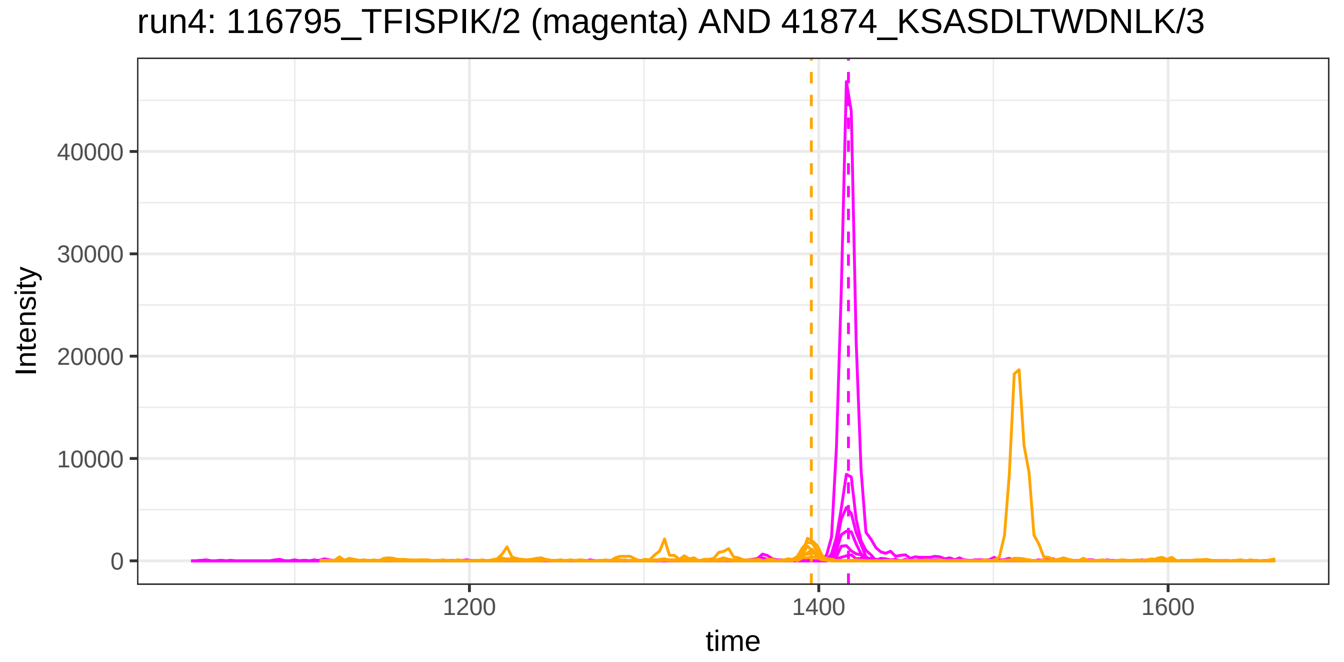
run4: 116795_TFISPIK/2 (magenta) AND 41874_KSASDLTWDNLK/3
Intensity
40000
30000
20000
10000
0
1200
1400
1600
time

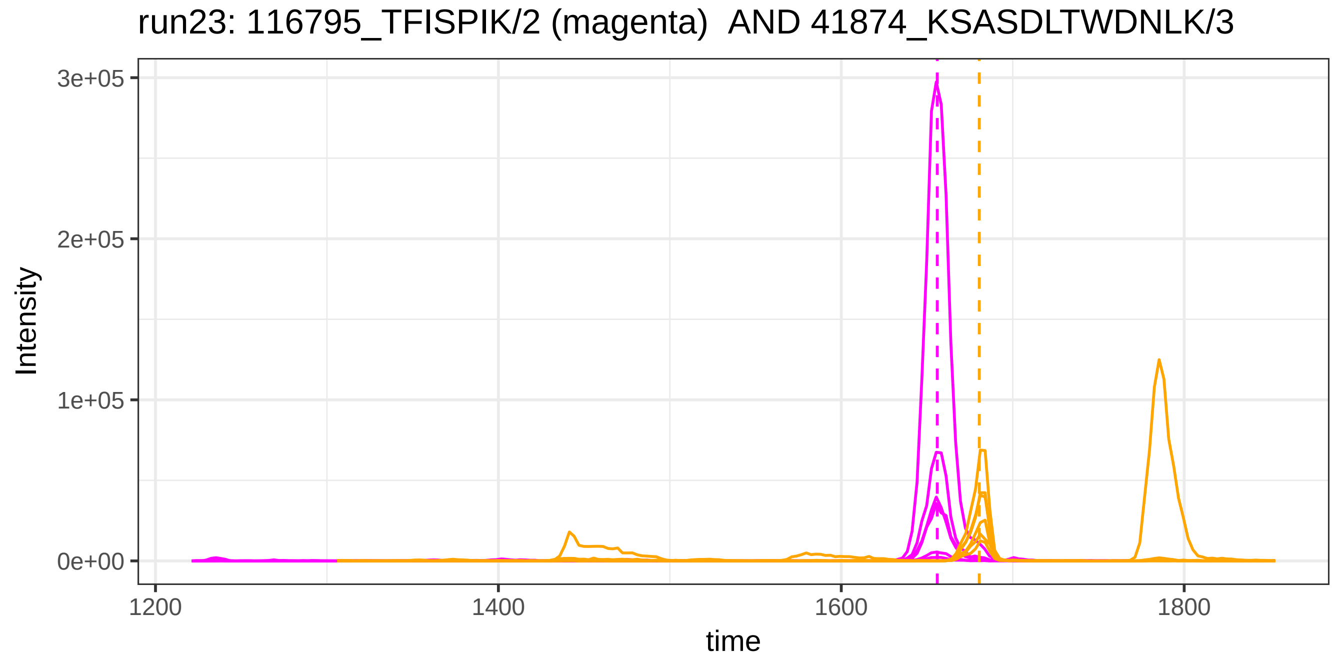
run23: 116795_TFISPIK/2 (magenta) AND 41874_KSASDLTWDNLK/3
Intensity
3e+05
2e+05
1e+05
0e+00
1200
1400
1600
1800
time

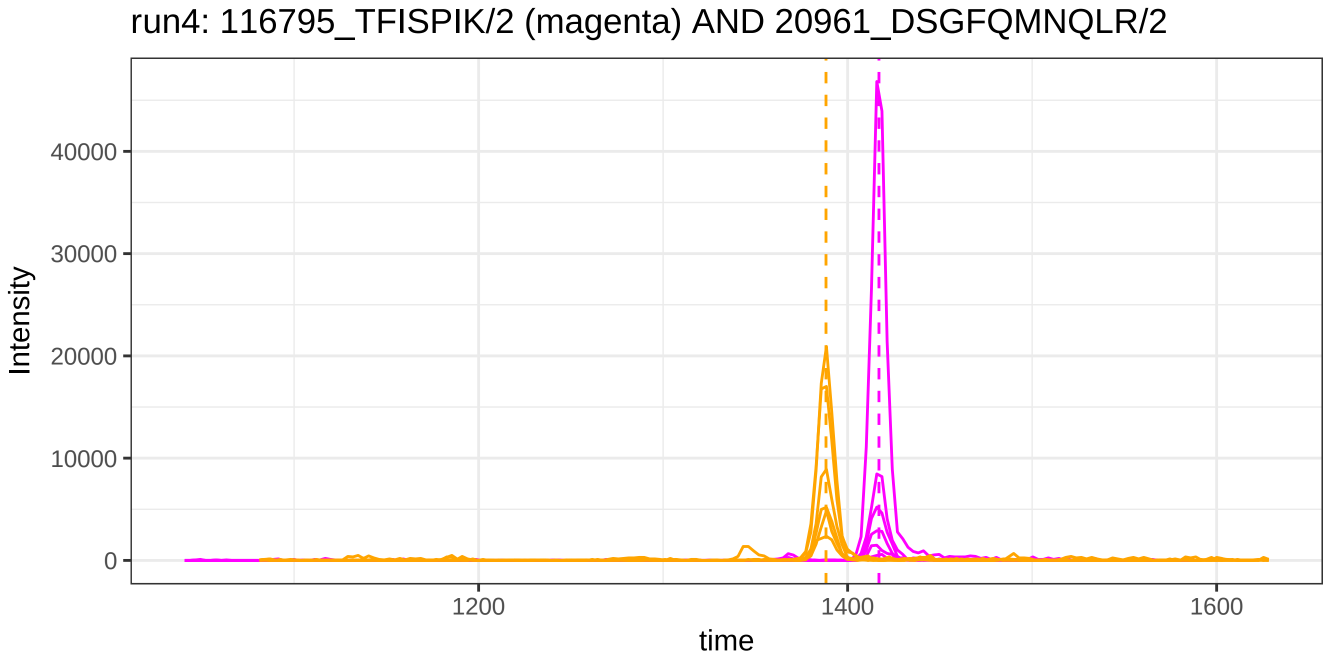

run4: 116795_TFISPIK/2 (magenta) AND 20961_DSGFQMNQLR/2
Intensity
40000
30000
20000
10000
0
1200
1400
1600
time


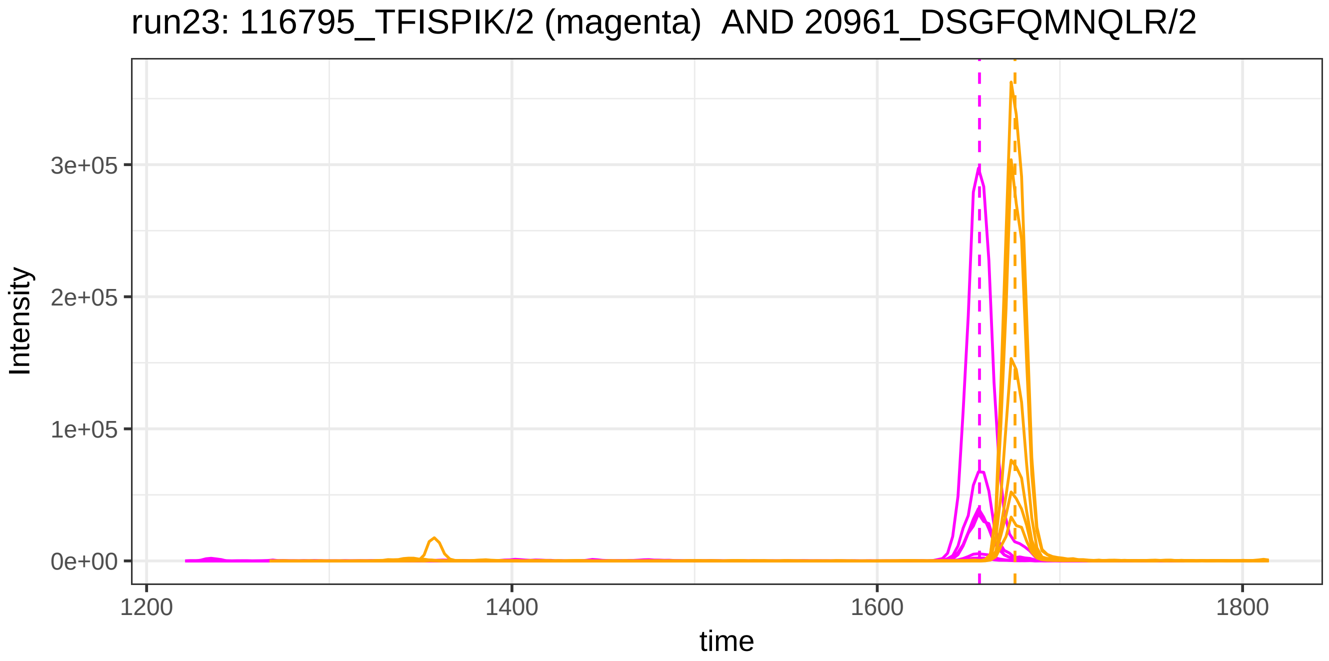

run23: 116795_TFISPIK/2 (magenta)  AND 20961_DSGFQMNQLR/2
Intensity
3e+05
2e+05
1e+05
0e+00
1200
1400
1600
1800
time

run4: 116795_TFISPIK/2 (magenta) AND 118454_DSGFQMNQLR/2

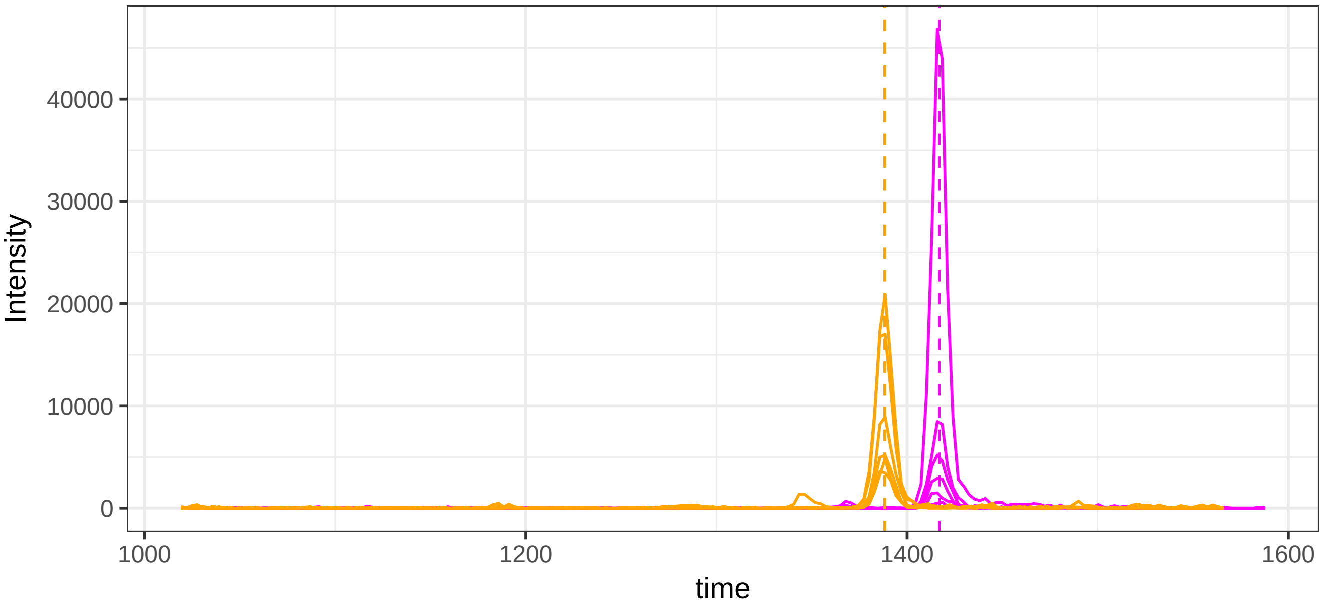


run23: 116795_TFISPIK/2 (magenta)  AND 118454_DSGFQMNQLR/2

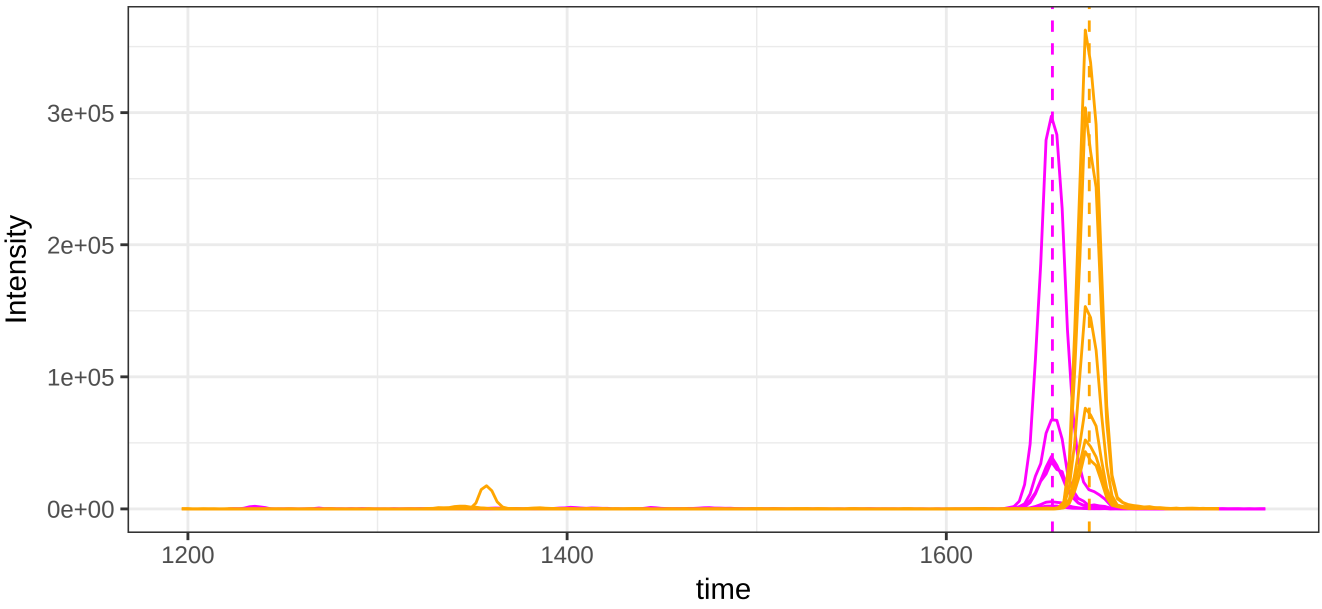

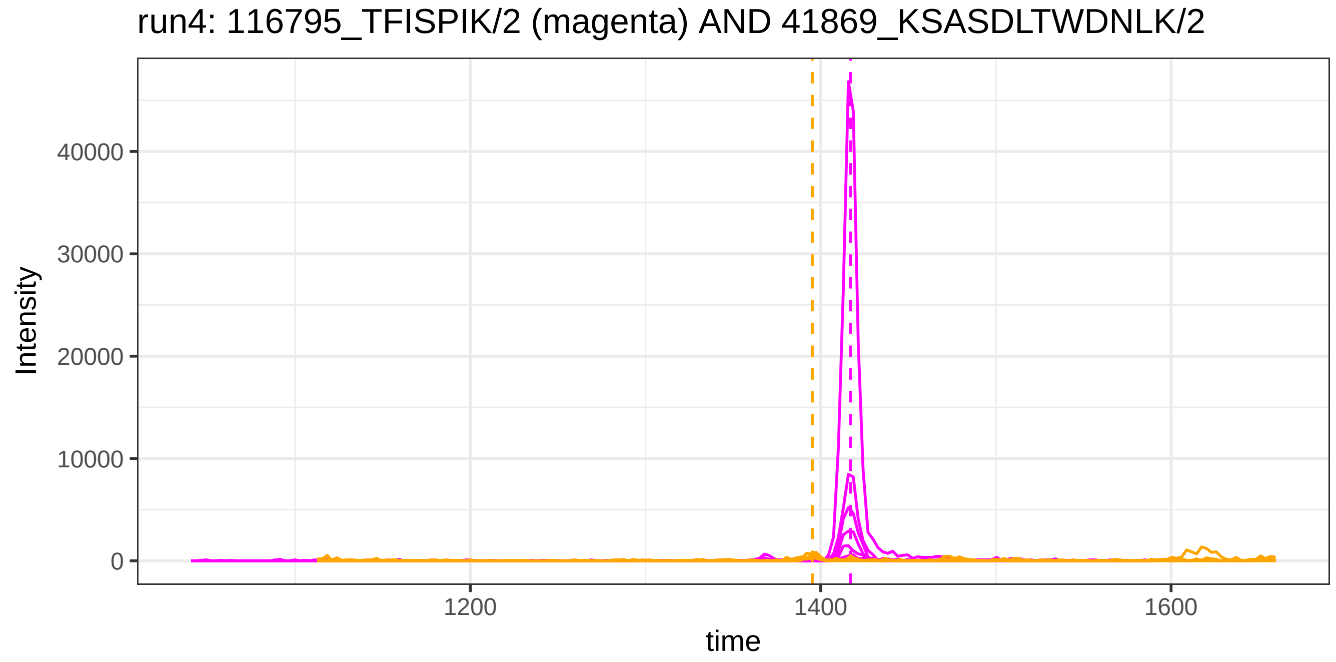
run4: 116795_TFISPIK/2 (magenta) AND 41869_KSASDLTWDNLK/2
Intensity
40000
30000
20000
10000
0
1200
1400
1600
time

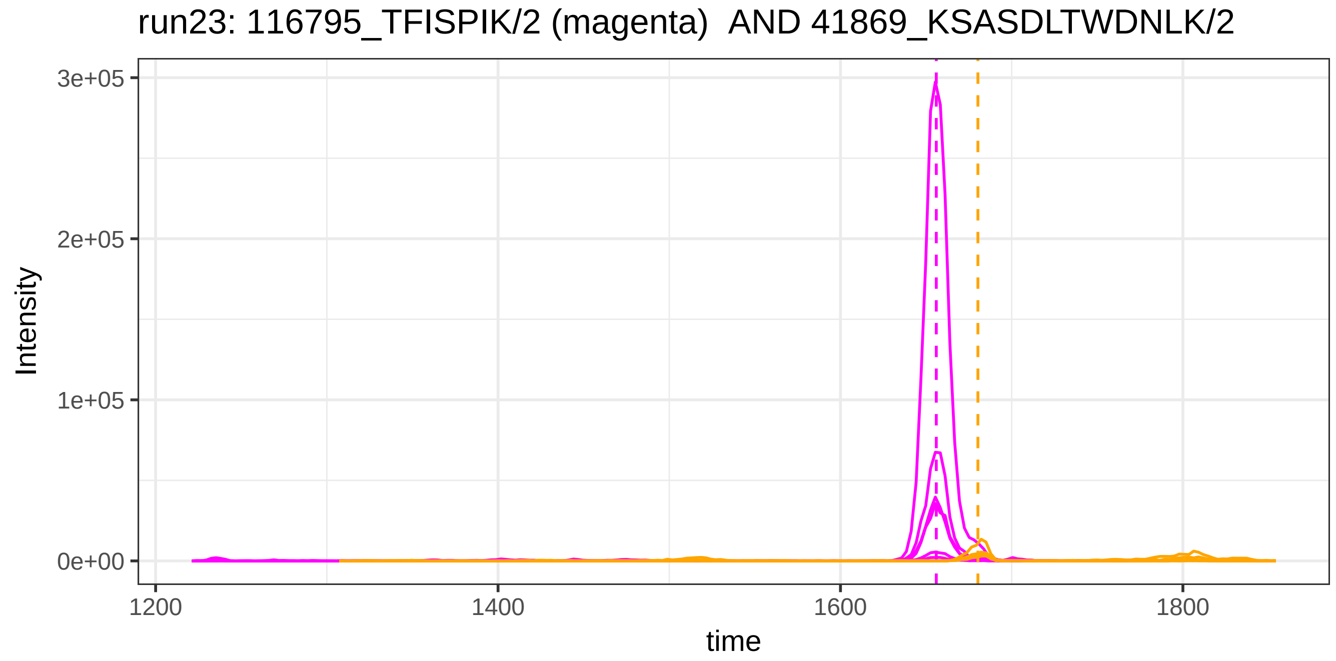
run23: 116795_TFISPIK/2 (magenta) AND 41869_KSASDLTWDNLK/2
Intensity
3e+05
2e+05
1e+05
0e+00
1200
1400
1600
1800
time

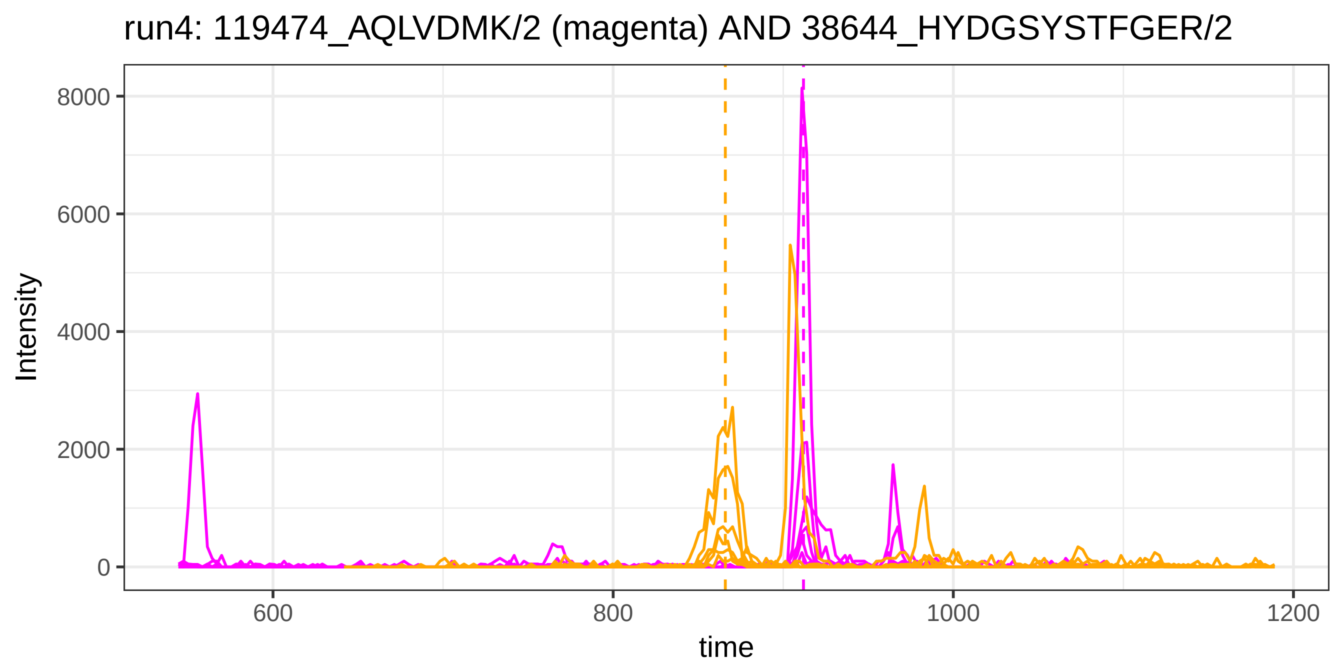

run4: 119474_AQLVDMK/2 (magenta) AND 38644_HYDGSYSTFGER/2
Intensity
8000
6000
4000
2000
0
600
800
1000
1200
time


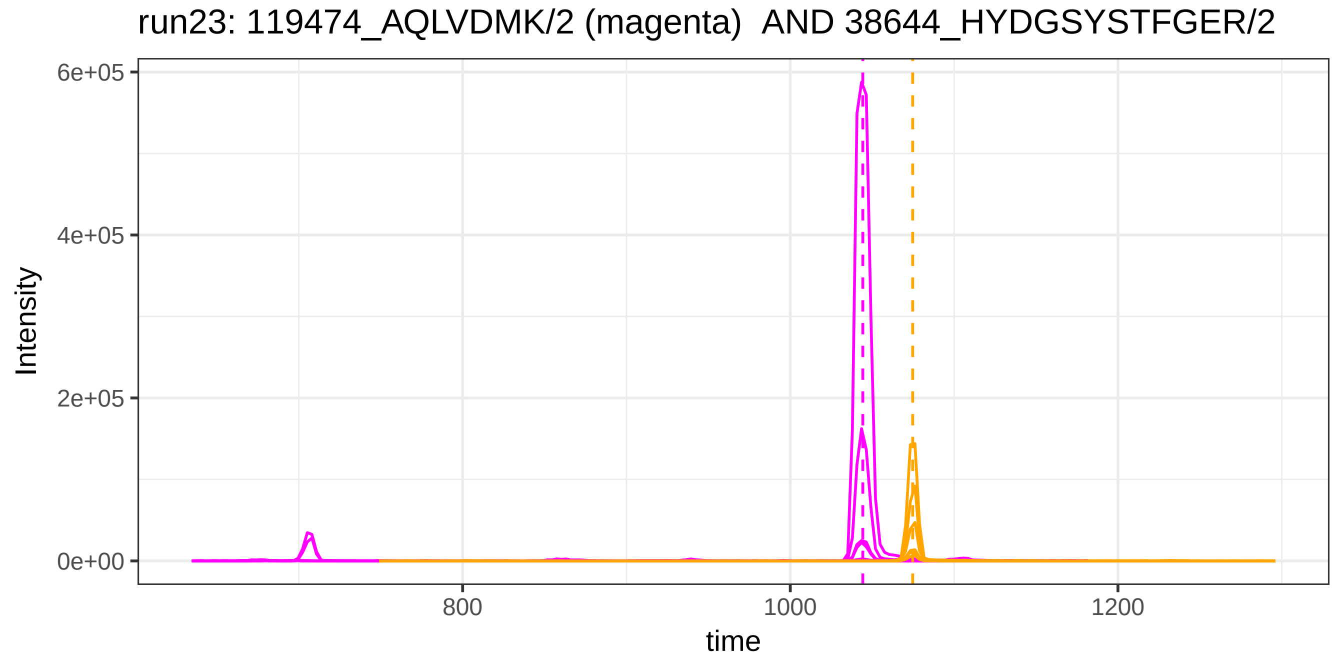

run23: 119474_AQLVDMK/2 (magenta) AND 38644_HYDGSYSTFGER/2
Intensity
6e+05
4e+05
2e+05
0e+00
800
1000
1200
time

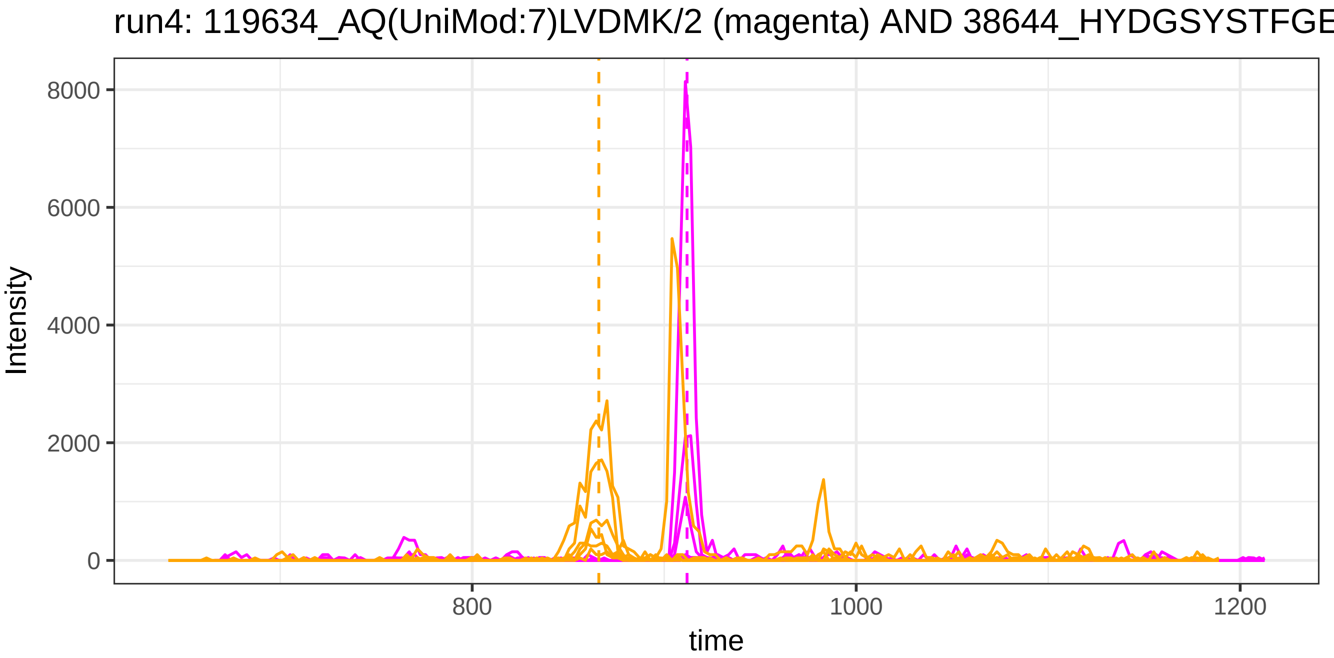
run4: 119634_AQ(UniMod:7)LVDMK/2 (magenta) AND 38644_HYDGSYSTFGE
Intensity
8000
6000
4000
2000
0
800
1000
1200
time

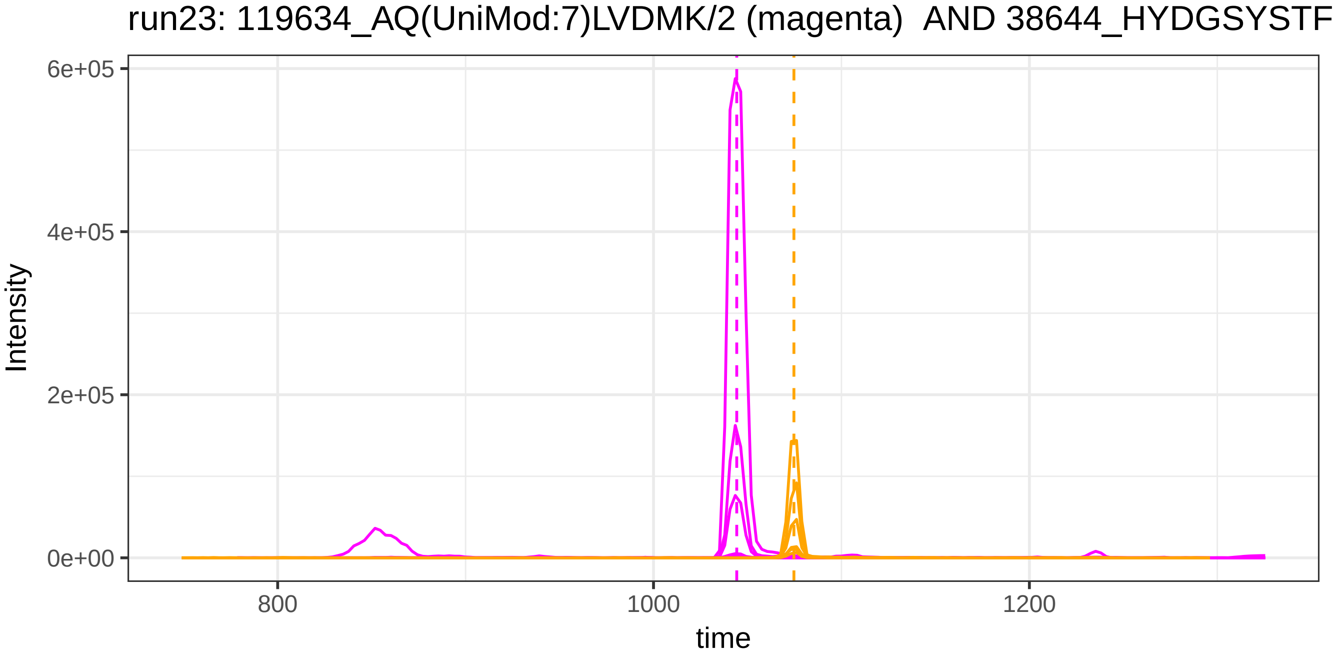
run23: 119634_AQ(UniMod:7)LVDMK/2 (magenta) AND 38644_HYDGSYSTF
Intensity
6e+05
4e+05
2e+05
0e+00
800
1000
1200
time

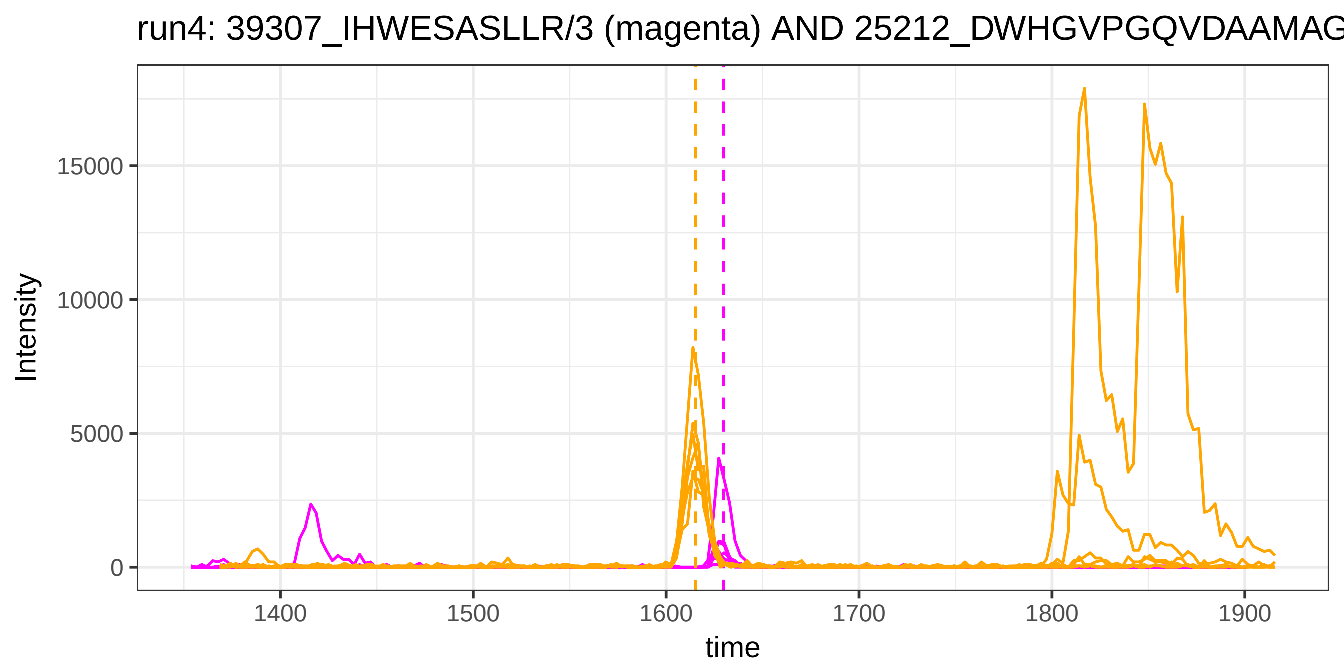
run4: 39307_IHWESASLLR/3 (magenta) AND 25212_DWHGVPGQVDAAMAG
Intensity
15000
10000
5000
0
1400
1500
1600
1700
1800
1900
time

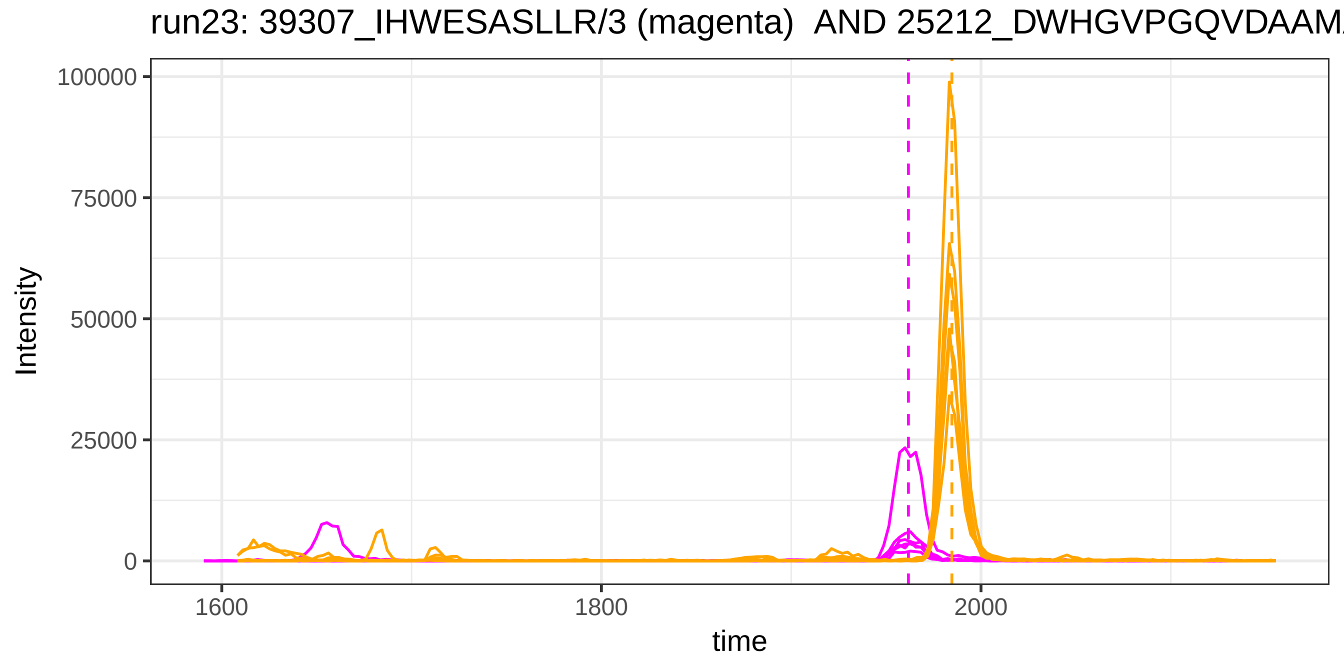
run23: 39307_IHWESASLLR/3 (magenta) AND 25212_DWHGVPGQVDAAM
Intensity
100000
75000
50000
25000
0
1600
1800
2000
time

run4: 39307_IHWESASLLR/3 (magenta) AND 125447_DWHGVPGQVDAAMA

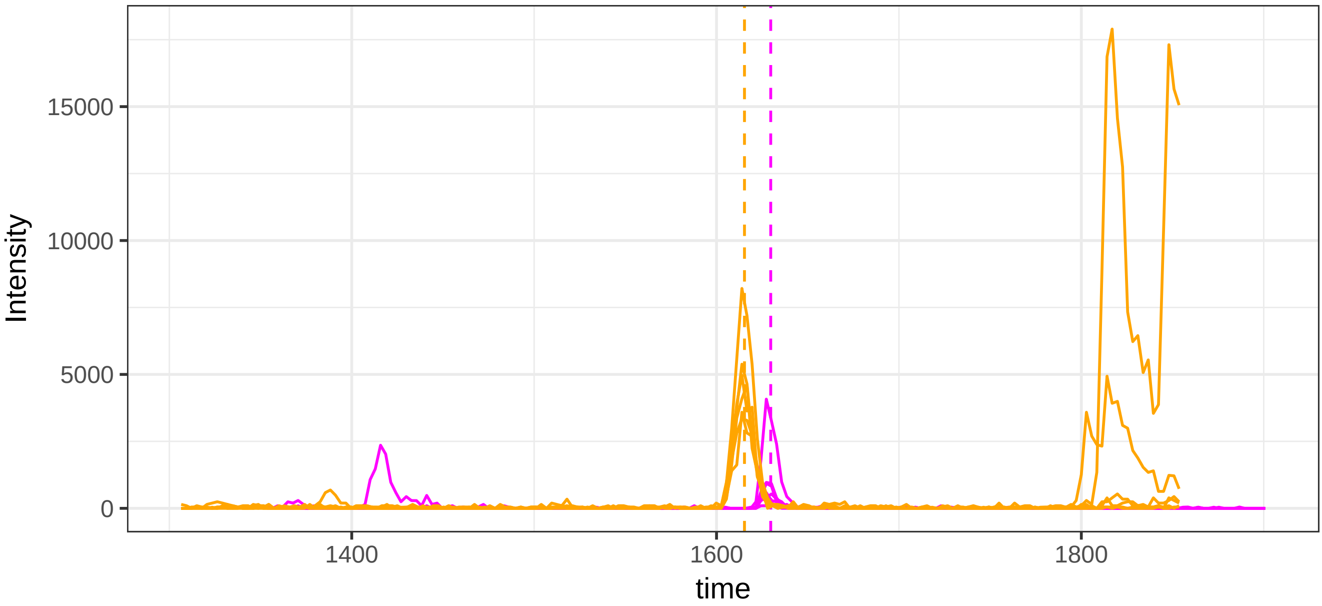


run23: 39307_IHWESASLLR/3 (magenta) AND 125447_DWHGVPGQVDAAM

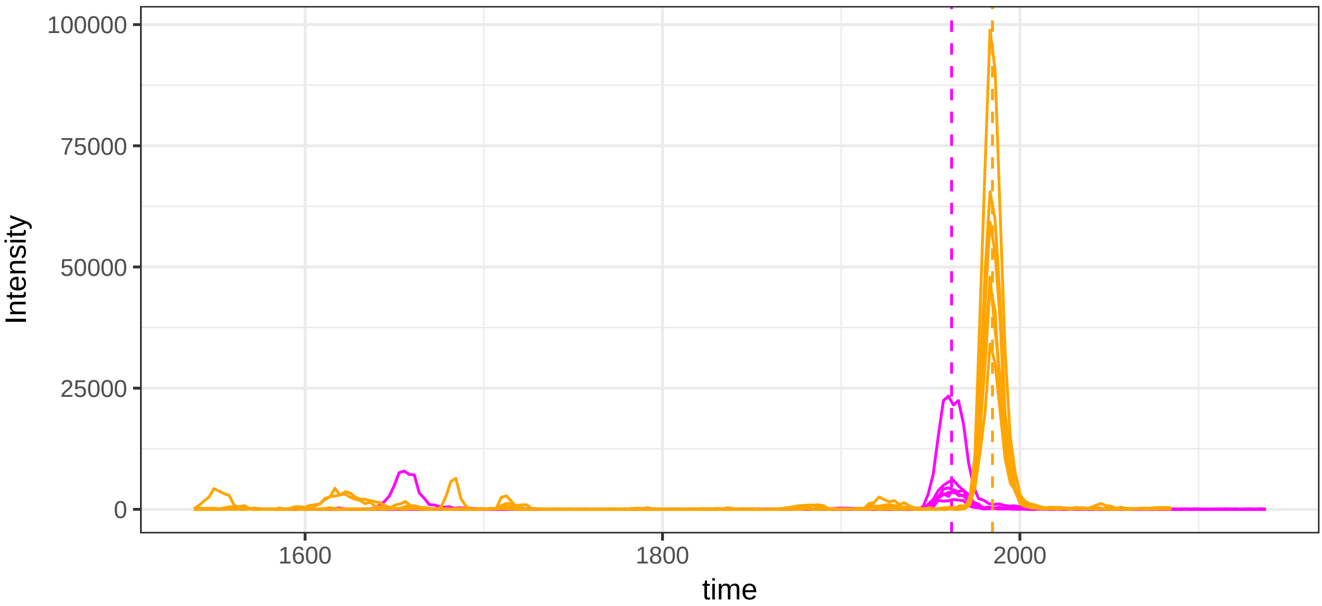

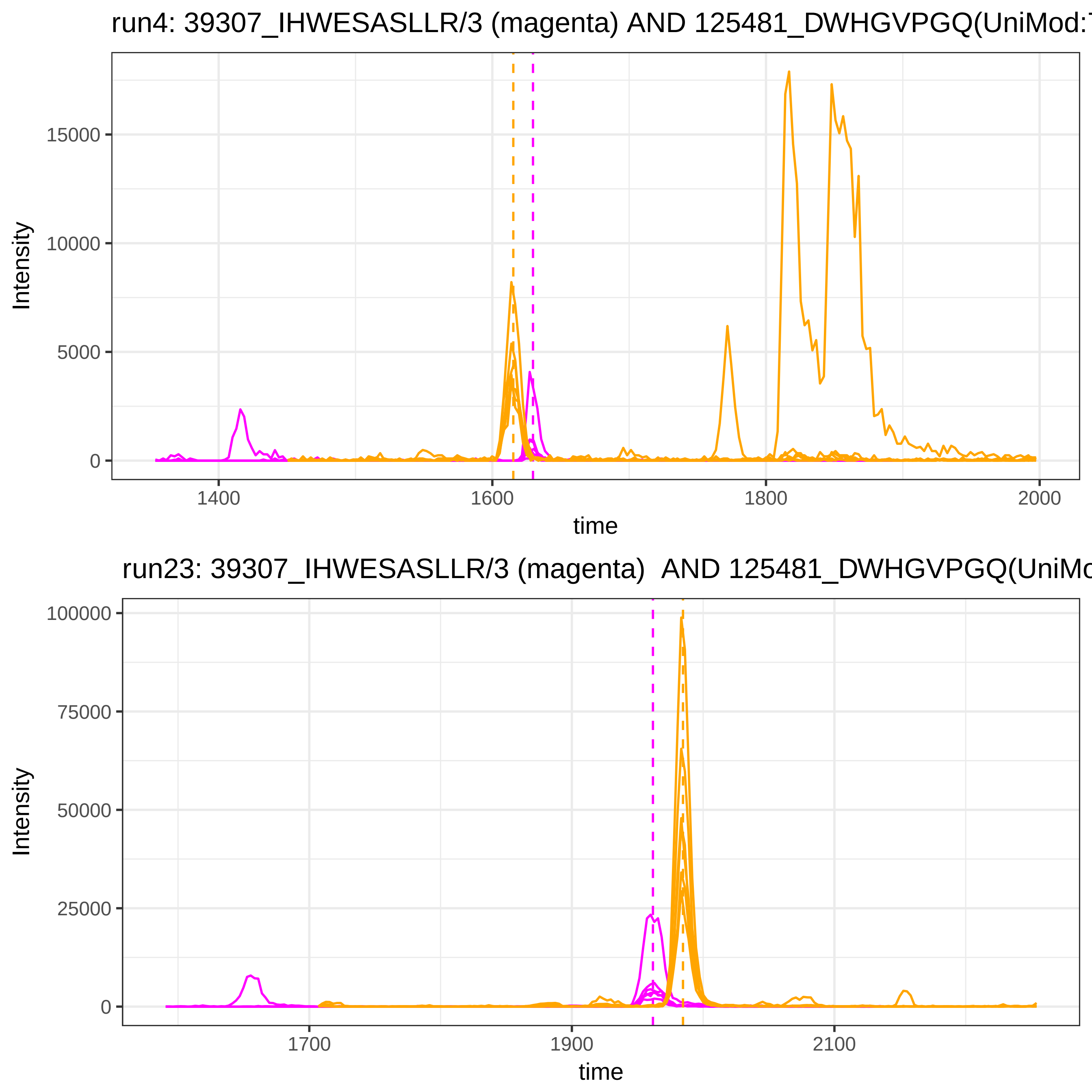
run4: 39307_IHWESASLLR/3 (magenta) AND 125481_DWHGVPGQ(UniMod:7
Intensity
15000
10000
5000
0
1400
1600
1800
2000
time
run23: 39307_IHWESASLLR/3 (magenta) AND 125481_DWHGVPGQ(UniMo
Intensity
100000
75000
50000
25000
0
1700
1900
2100
time

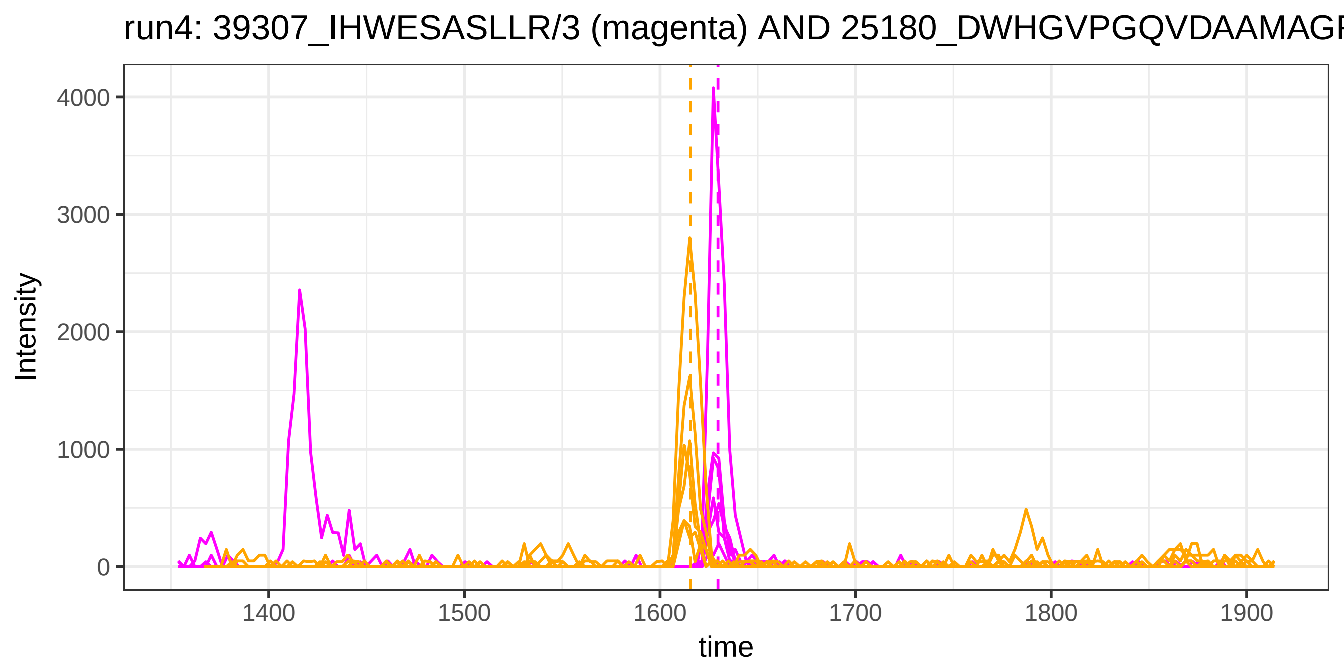
run4: 39307_IHWESASLLR/3 (magenta) AND 25180_DWHGVPGQVDAAMAGF
Intensity
4000
3000
2000
1000
0
1400
1500
1600
1700
1800
1900
time

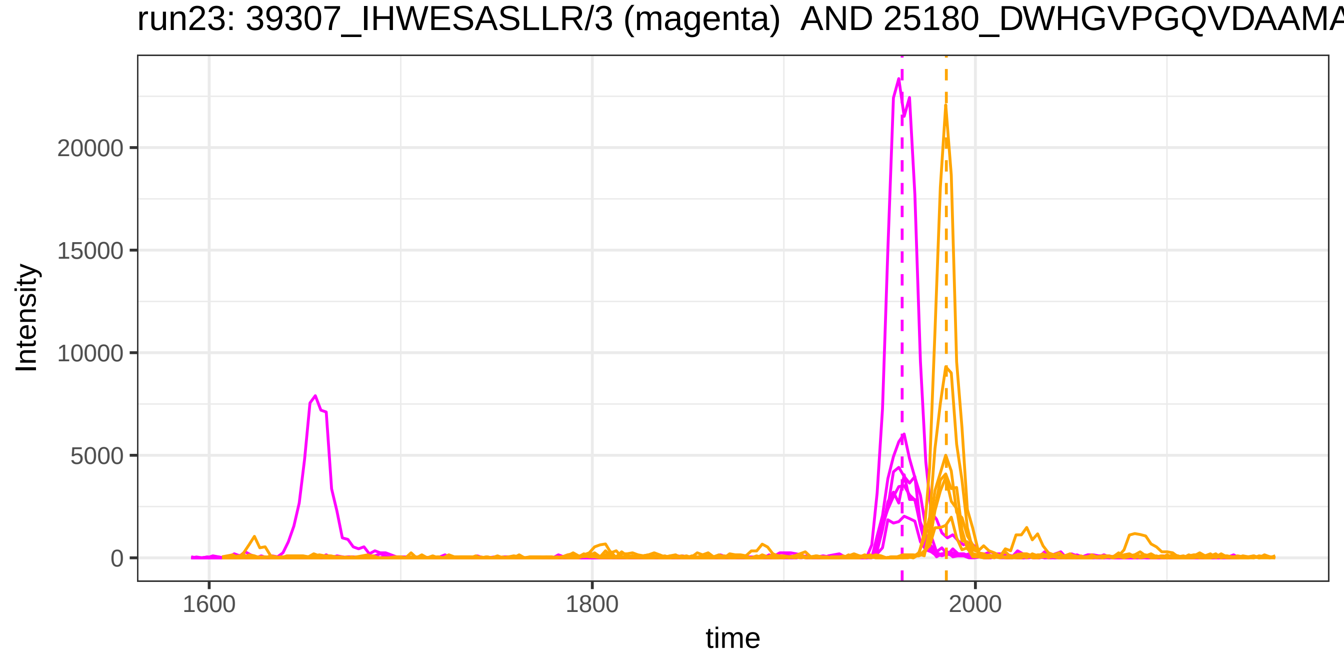
run23: 39307_IHWESASLLR/3 (magenta) AND 25180_DWHGVPGQVDAAMA
Intensity
20000
15000
10000
5000
0
1600
1800
2000
time

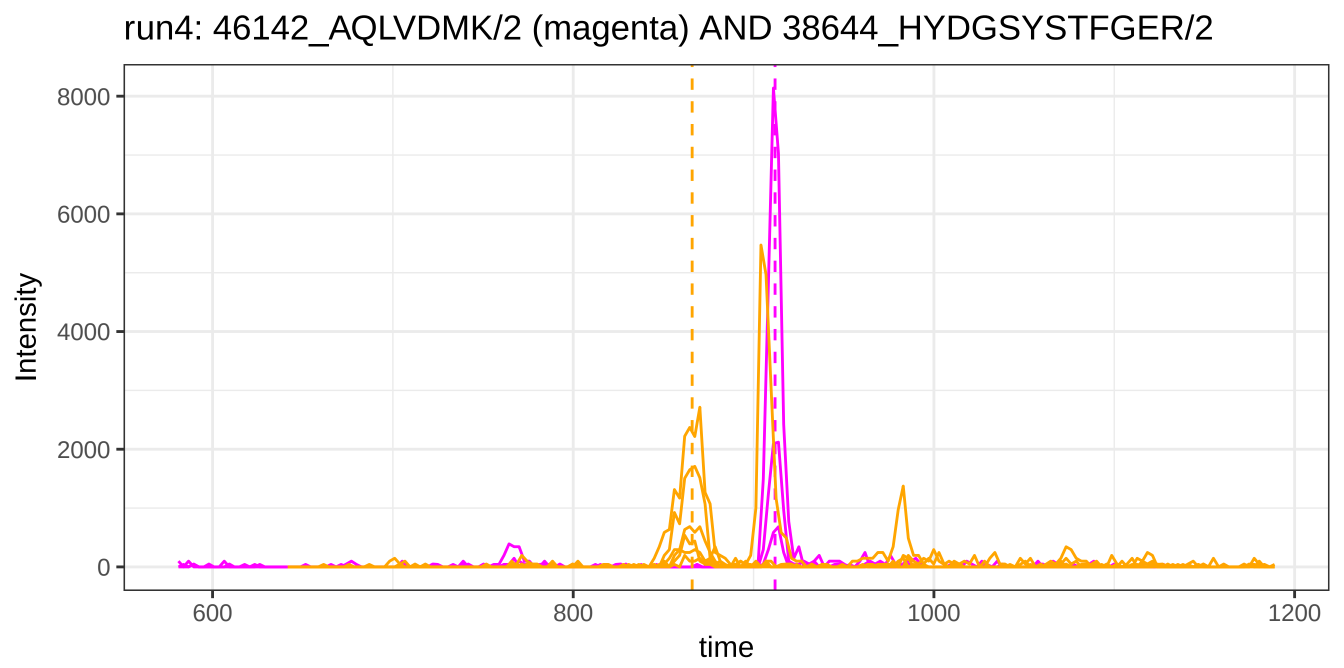

run4: 46142_AQLVDMK/2 (magenta) AND 38644_HYDGSYSTFGER/2
Intensity
8000
6000
4000
2000
0
600
800
1000
1200
time


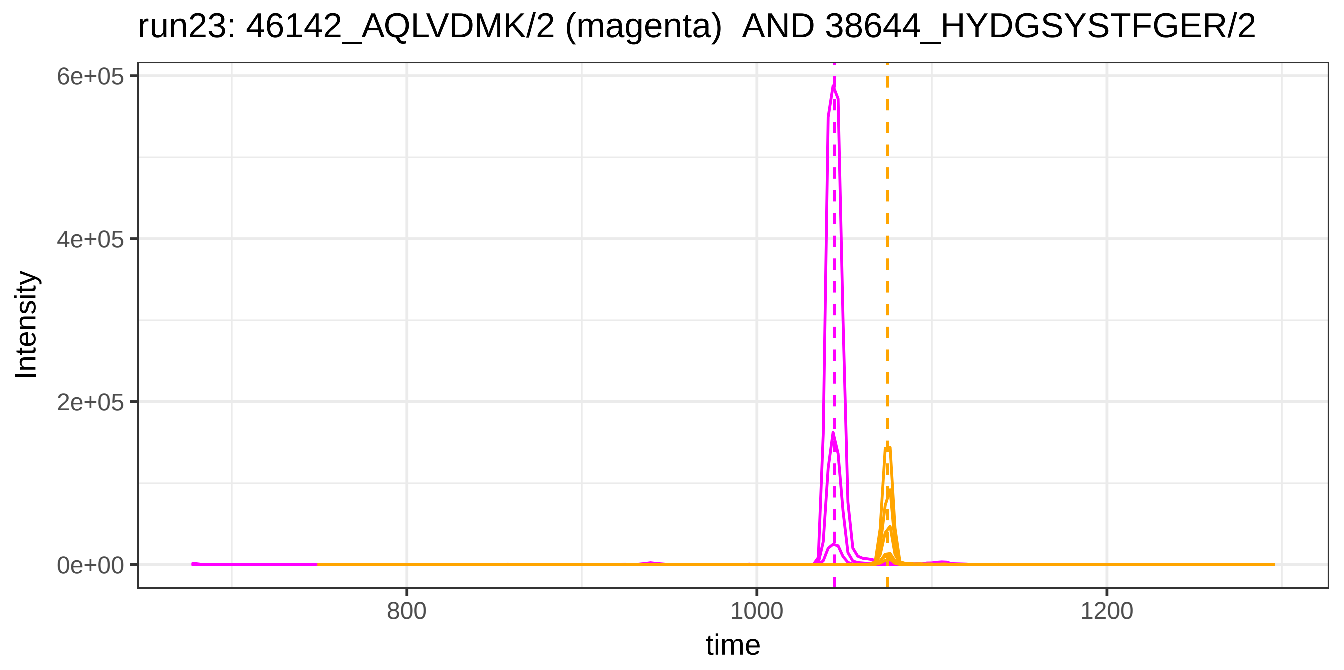

run23: 46142_AQLVDMK/2 (magenta) AND 38644_HYDGSYSTFGER/2
Intensity
6e+05
4e+05
2e+05
0e+00
800
1000
1200
time

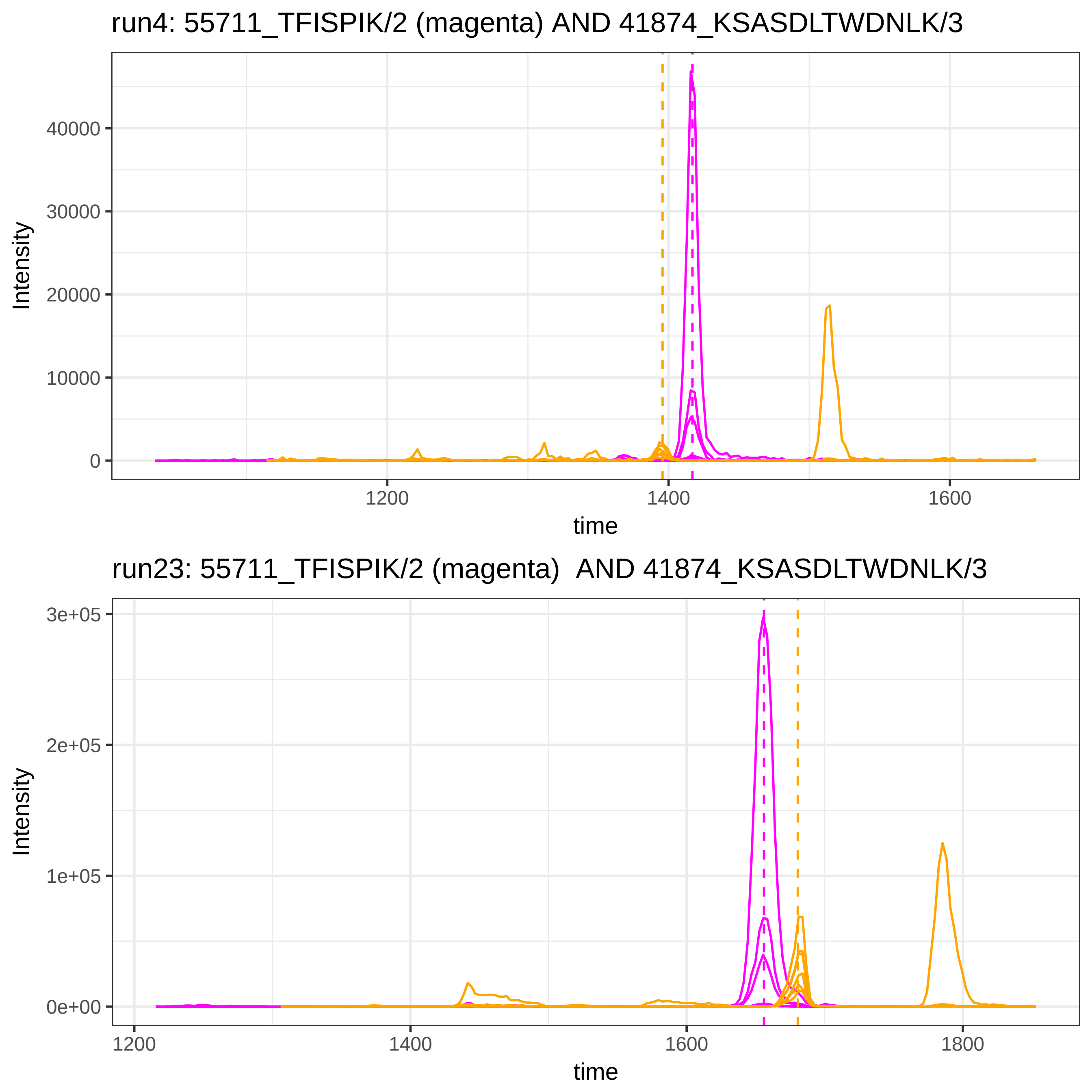

run4: 55711_TFISPIK/2 (magenta) AND 41874_KSASDLTWDNLK/3
Intensity
40000
30000
20000
10000
0
1200
1400
1600
time
run23: 55711_TFISPIK/2 (magenta)  AND 41874_KSASDLTWDNLK/3
Intensity
3e+05
2e+05
1e+05
0e+00
1200
1400
1600
1800
time

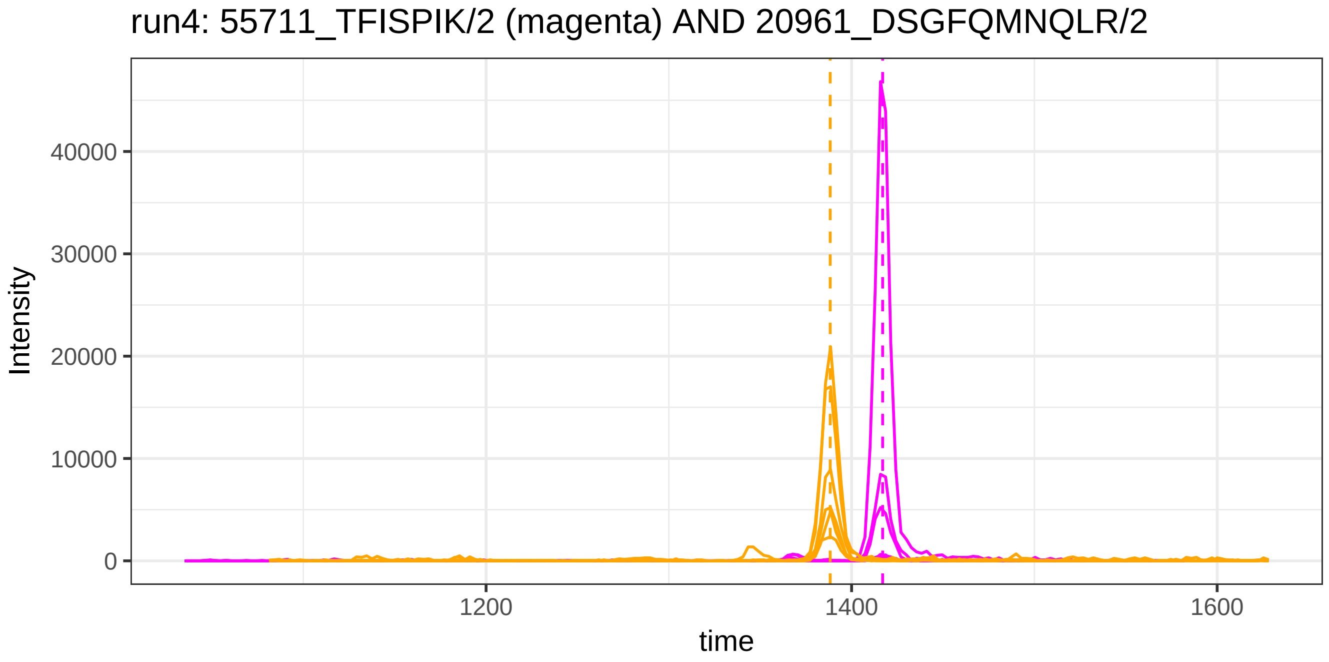
run4: 55711_TFISPIK/2 (magenta) AND 20961_DSGFQMNQLR/2
Intensity
40000
30000
20000
10000
0
1200
1400
1600
time

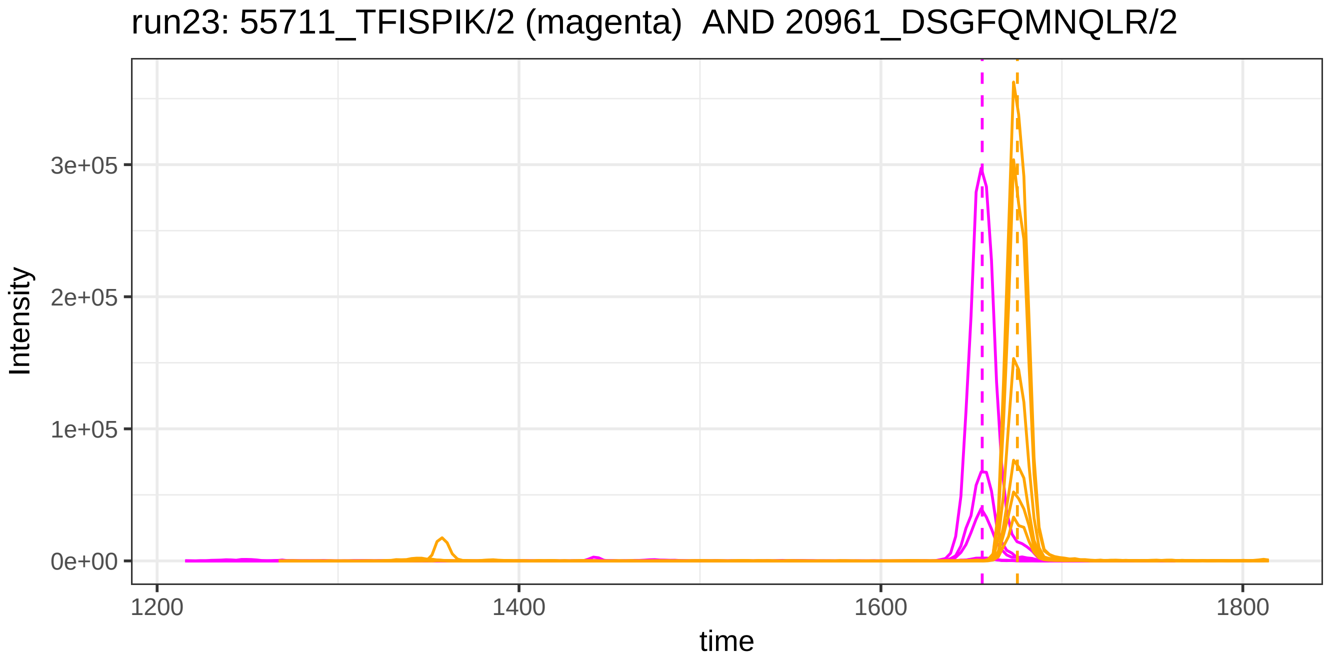
run23: 55711_TFISPIK/2 (magenta)  AND 20961_DSGFQMNQLR/2
Intensity
3e+05
2e+05
1e+05
0e+00
1200
1400
1600
1800
time

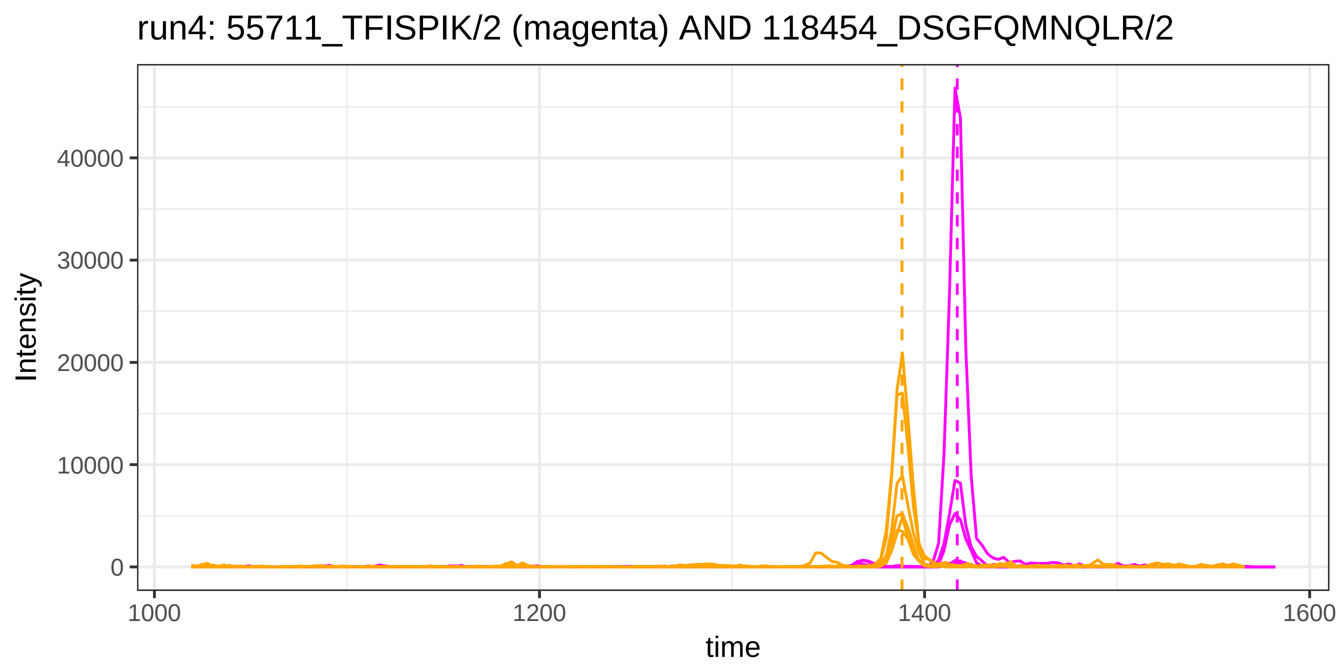
run4: 55711_TFISPIK/2 (magenta) AND 118454_DSGFQMNQLR/2
Intensity
40000
30000
20000
10000
0
1000
1200
1400
1600
time

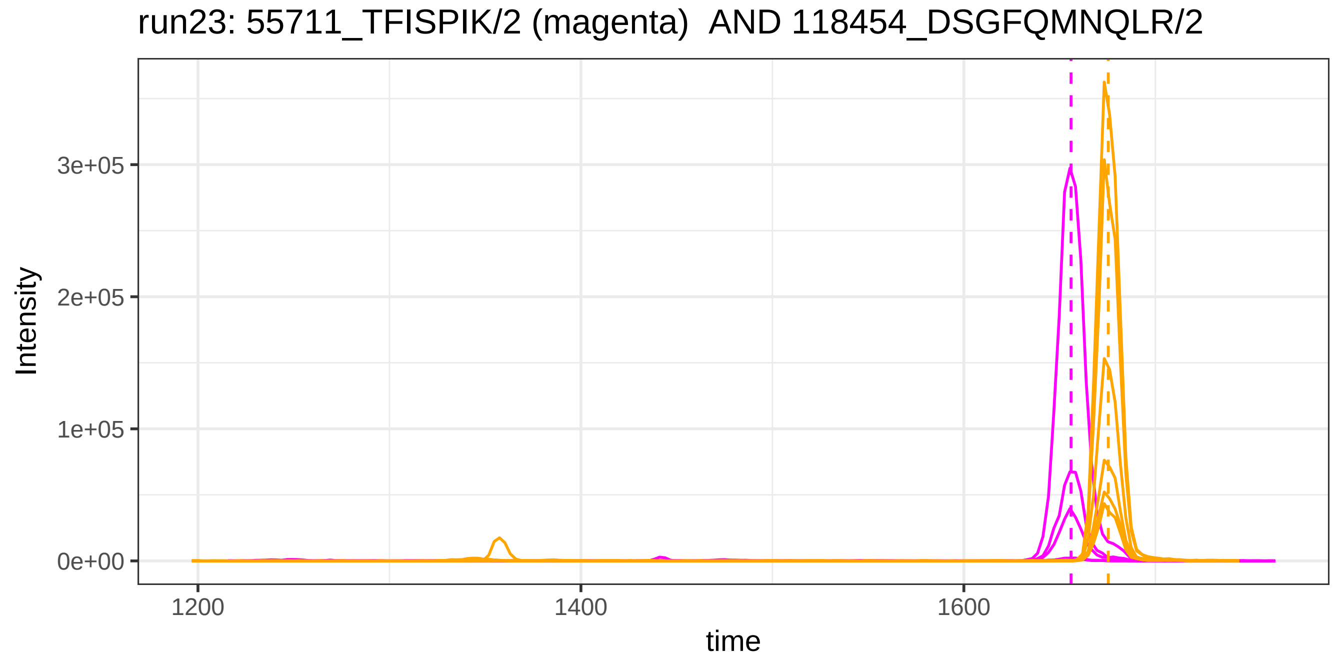
run23: 55711_TFISPIK/2 (magenta) AND 118454_DSGFQMNQLR/2
Intensity
3e+05
2e+05
1e+05
0e+00
1200
1400
1600
time

run4: 55711_TFISPIK/2 (magenta) AND 41869_KSASDLTWDNLK/2

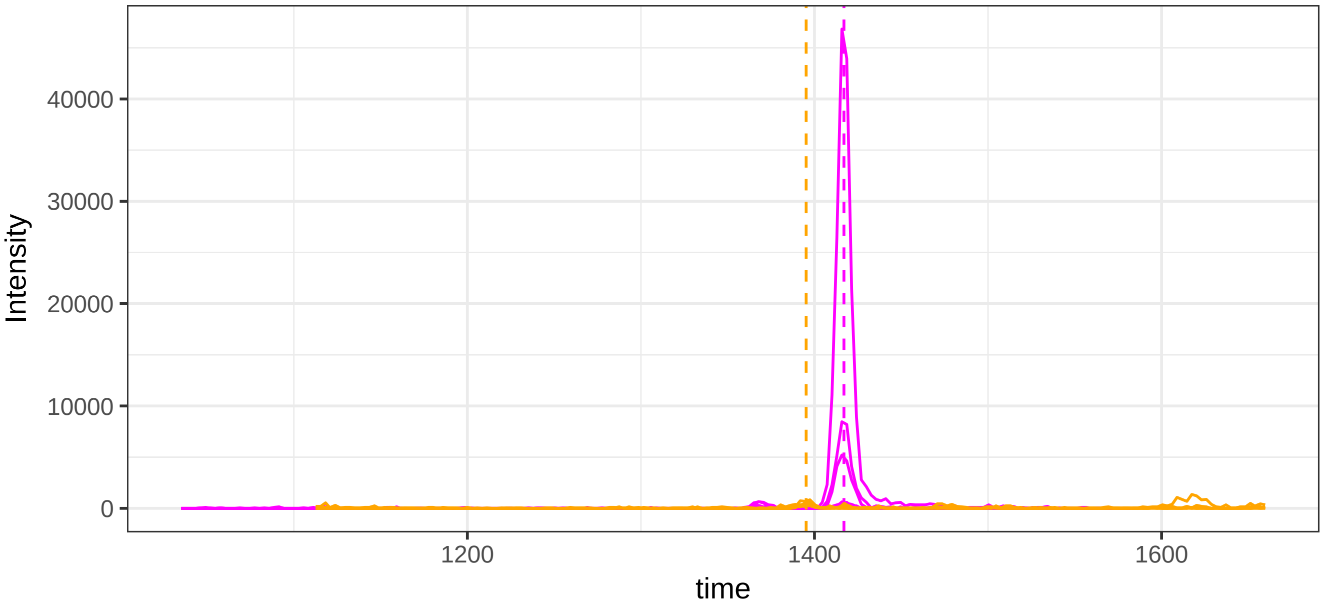


run23: 55711_TFISPIK/2 (magenta)  AND 41869_KSASDLTWDNLK/2

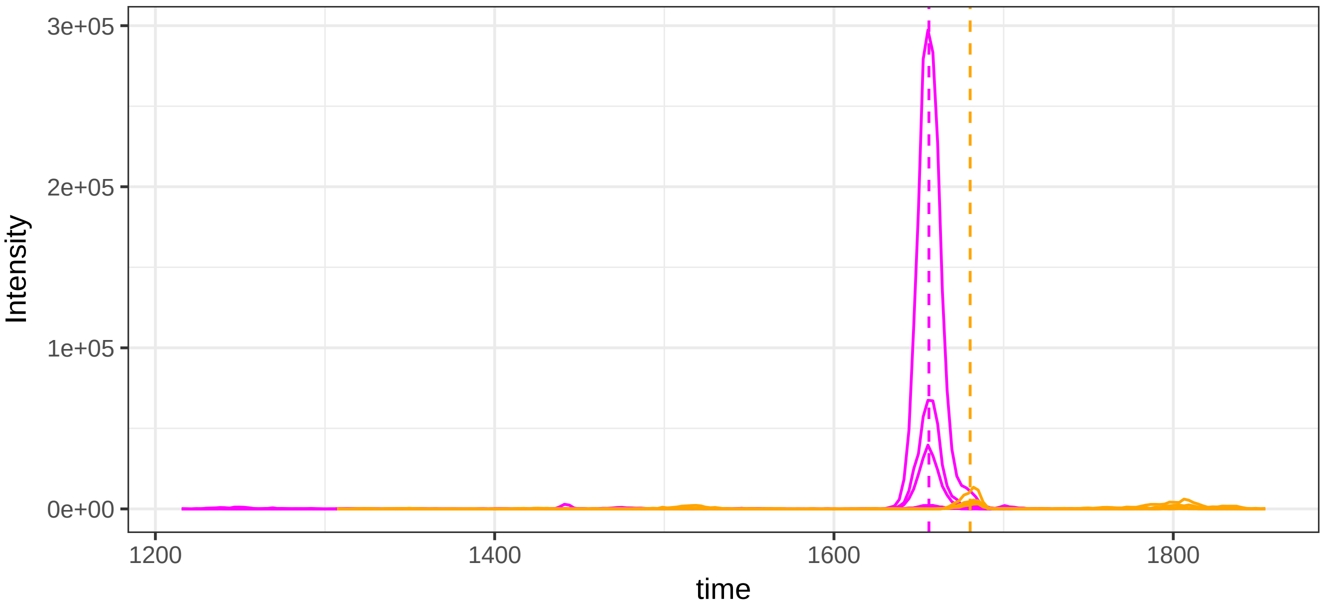

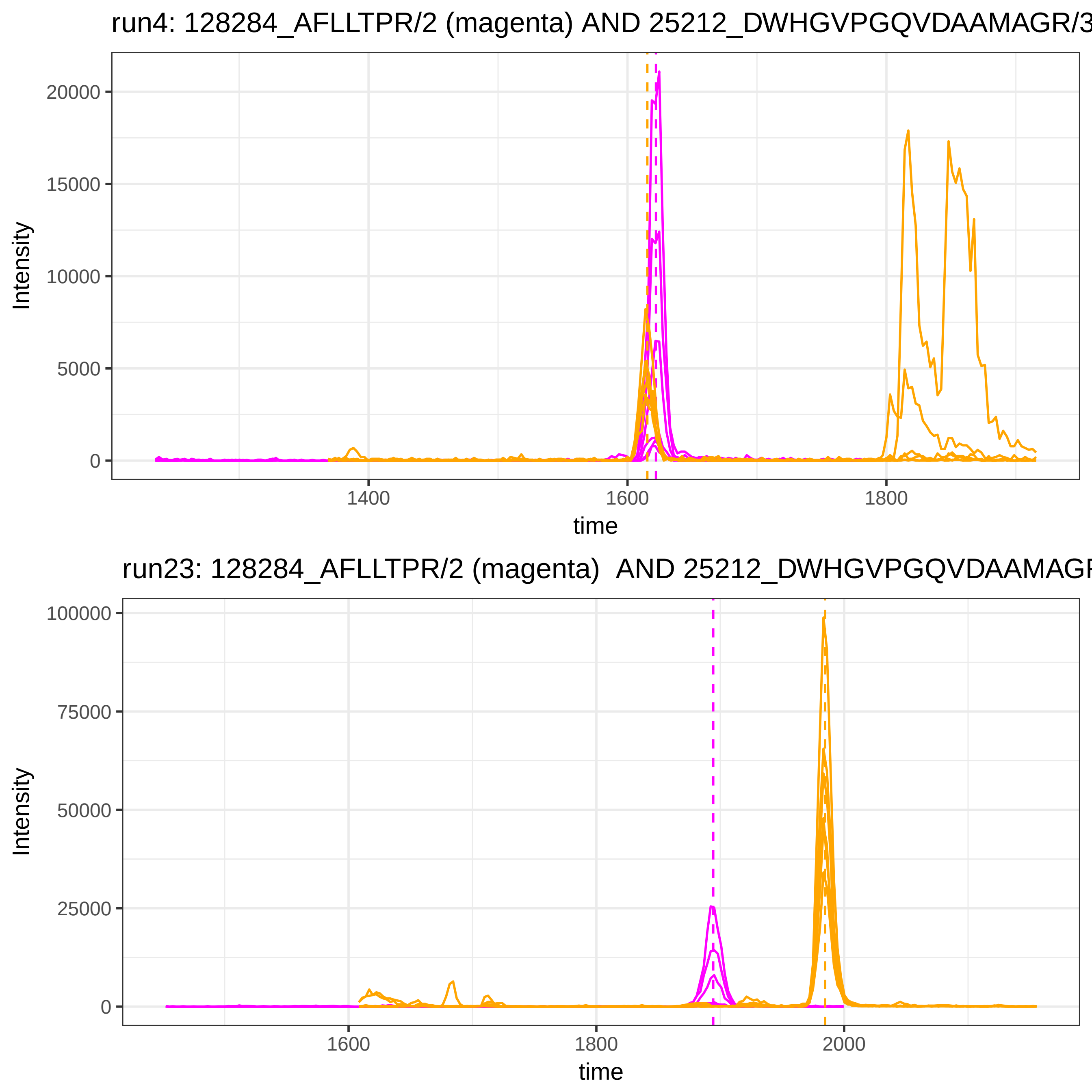
run4: 128284_AFLLTPR/2 (magenta) AND 25212_DWHGVPGQVDAAMAGR/3
Intensity
20000
15000
10000
5000
0
1400
1600
1800
time
run23: 128284_AFLLTPR/2 (magenta)  AND 25212_DWHGVPGQVDAAMAGR
Intensity
100000
75000
50000
25000
0
1600
1800
2000
time

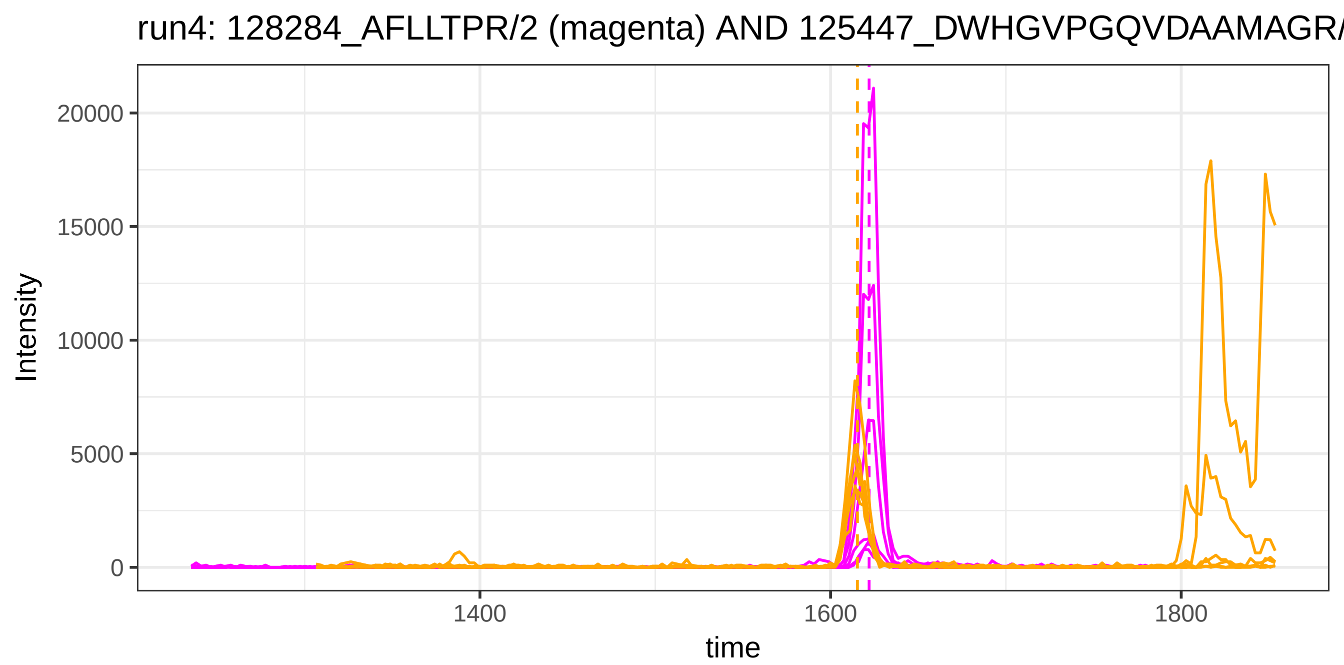

run4: 128284_AFLLTPR/2 (magenta) AND 125447_DWHGVPGQVDAAMAGR/
Intensity
20000
15000
10000
5000
0
1400
1600
1800
time


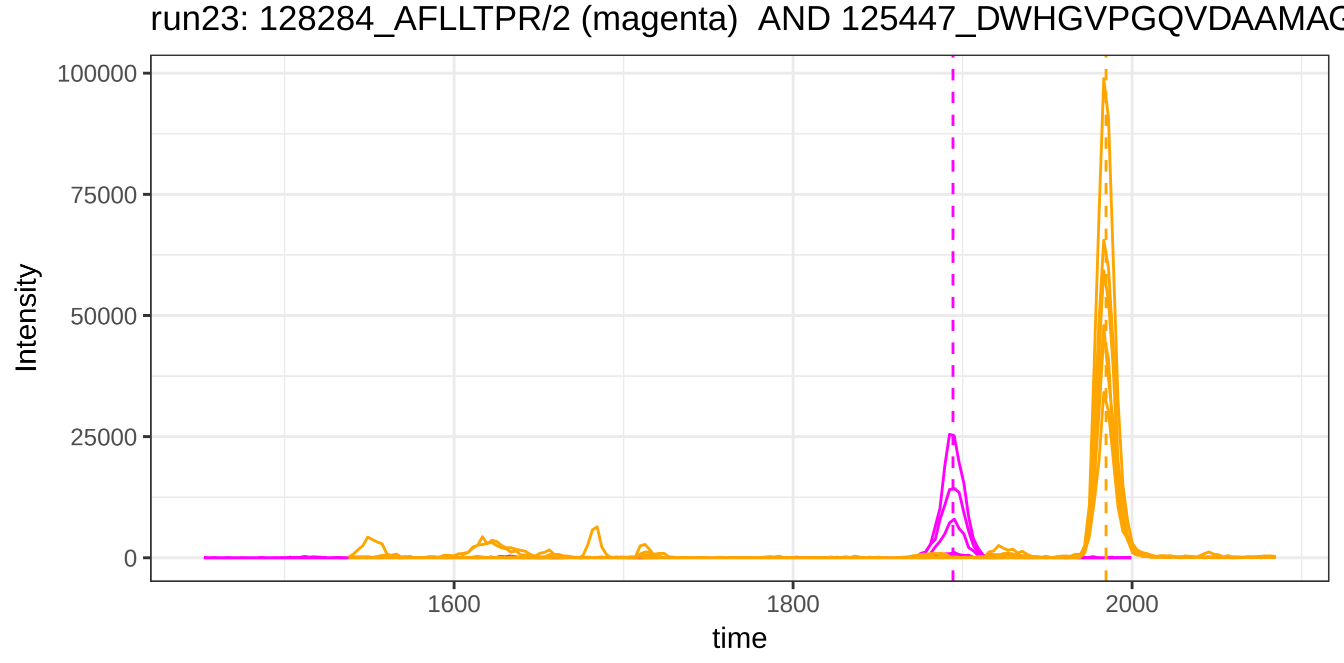

run23: 128284_AFLLTPR/2 (magenta) AND 125447_DWHGVPGQVDAAMAG
Intensity
100000
75000
50000
25000
0
1600
1800
2000
time

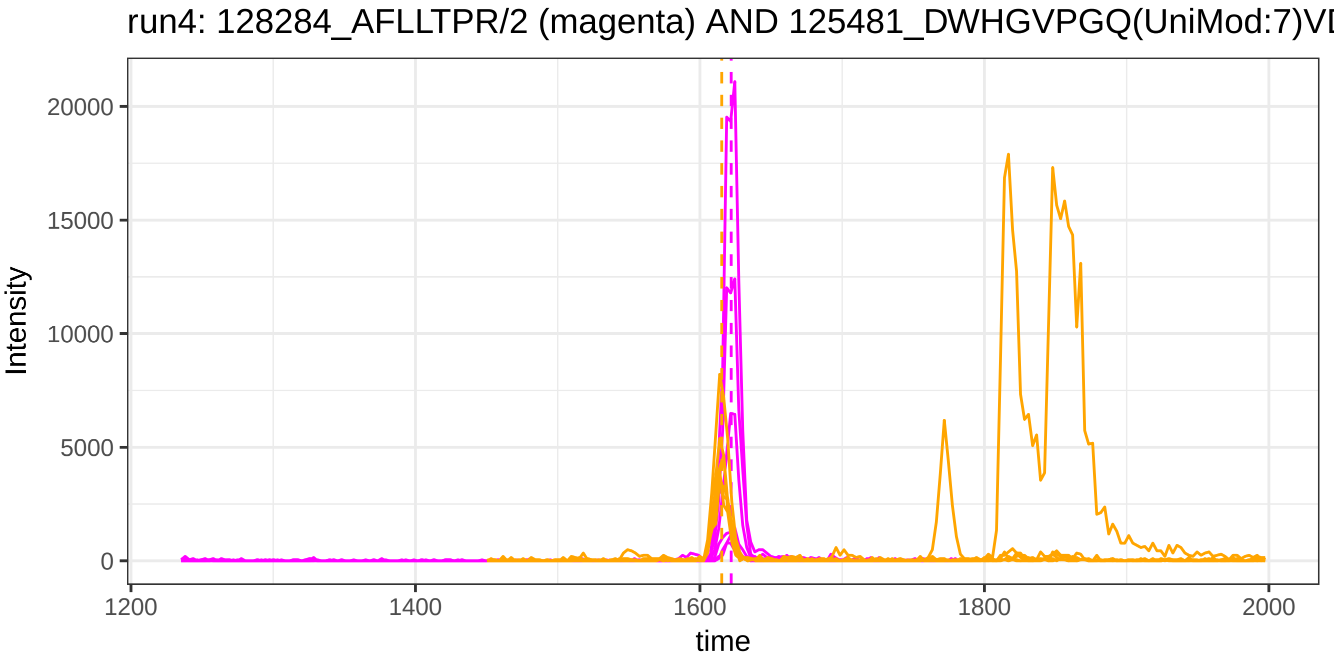
run4: 128284_AFLLTPR/2 (magenta) AND 125481_DWHGVPGQ(UniMod:7)VD
Intensity
20000
15000
10000
5000
0
1200
1400
1600
1800
2000
time

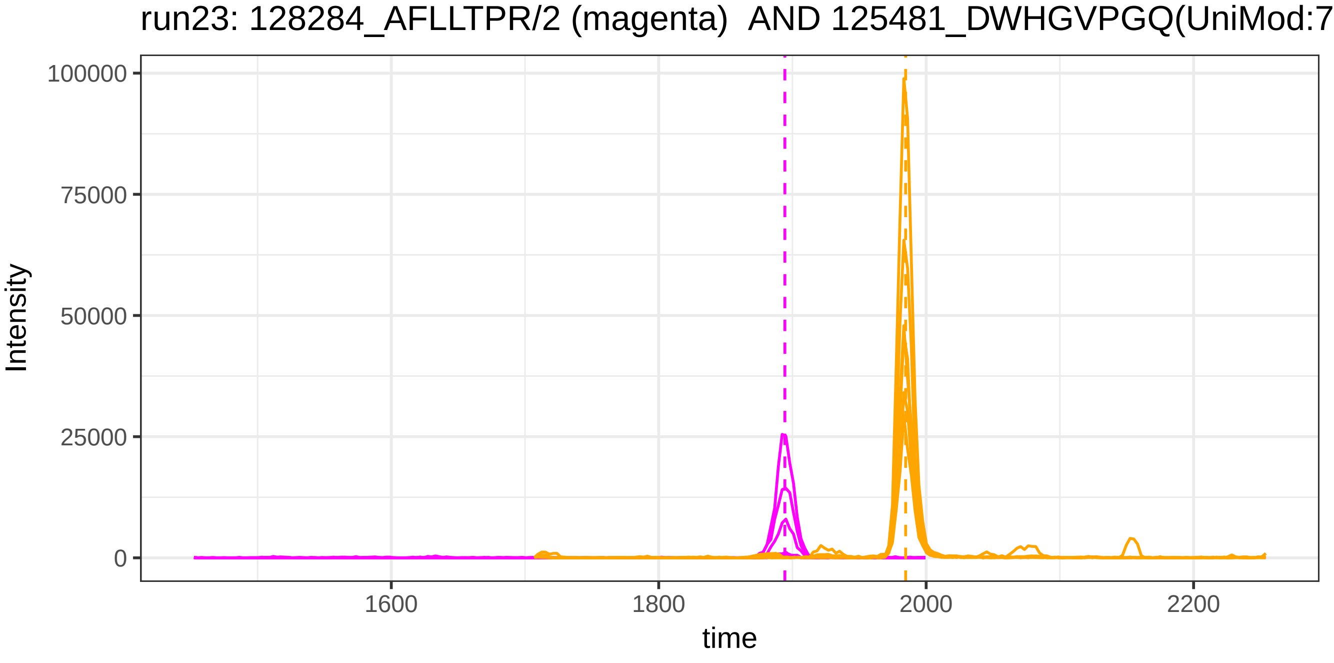
run23: 128284_AFLLTPR/2 (magenta) AND 125481_DWHGVPGQ(UniMod:7
Intensity
100000
75000
50000
25000
0
1600
1800
2000
2200
time

run4: 128284_AFLLTPR/2 (magenta) AND 2924_VTSIQDWVQK/2

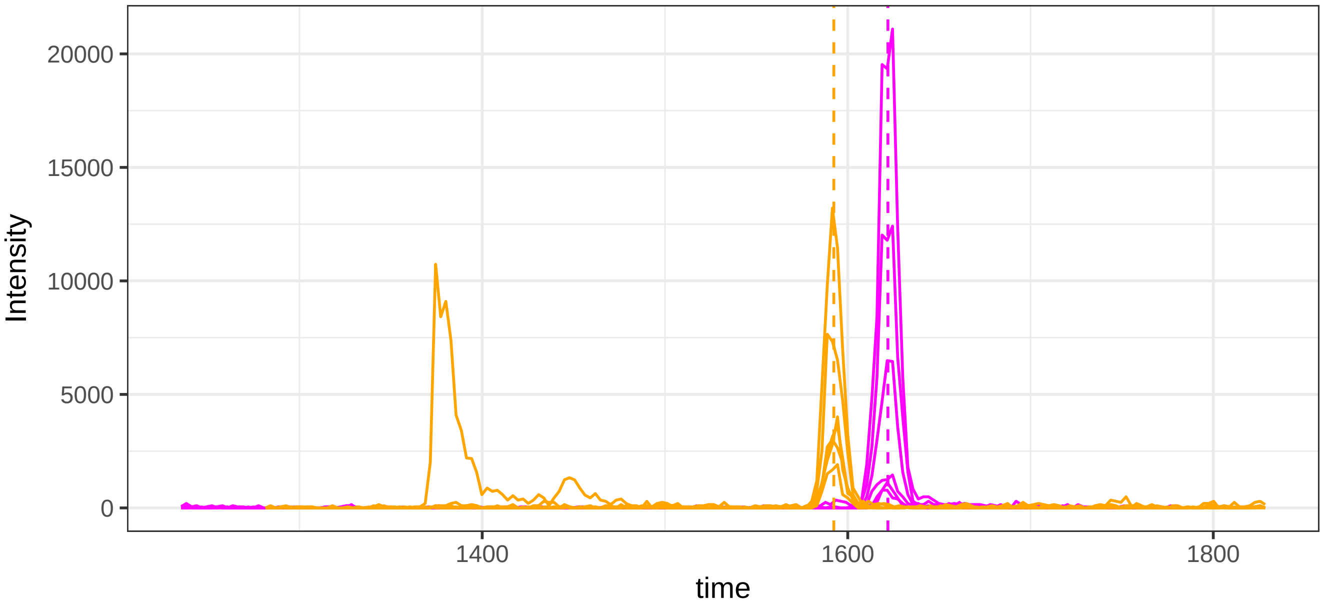


run23: 128284_AFLLTPR/2 (magenta)  AND 2924_VTSIQDWVQK/2

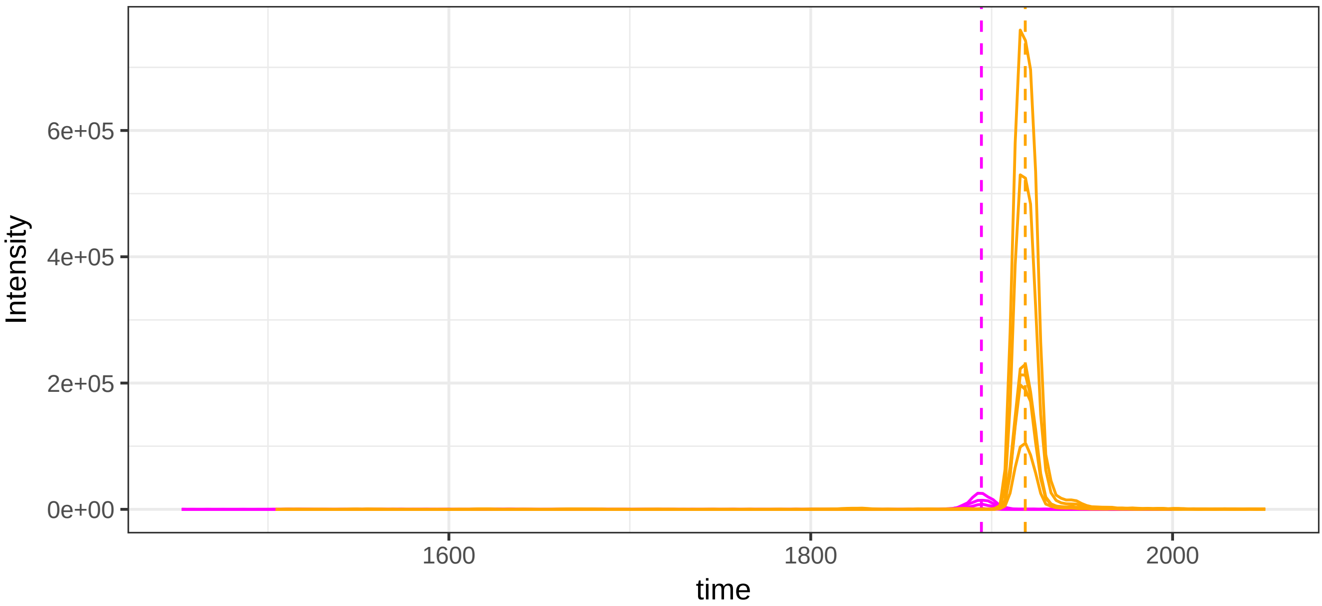

run4: 128284_AFLLTPR/2 (magenta) AND 42259_ALVQQMEQLR/2

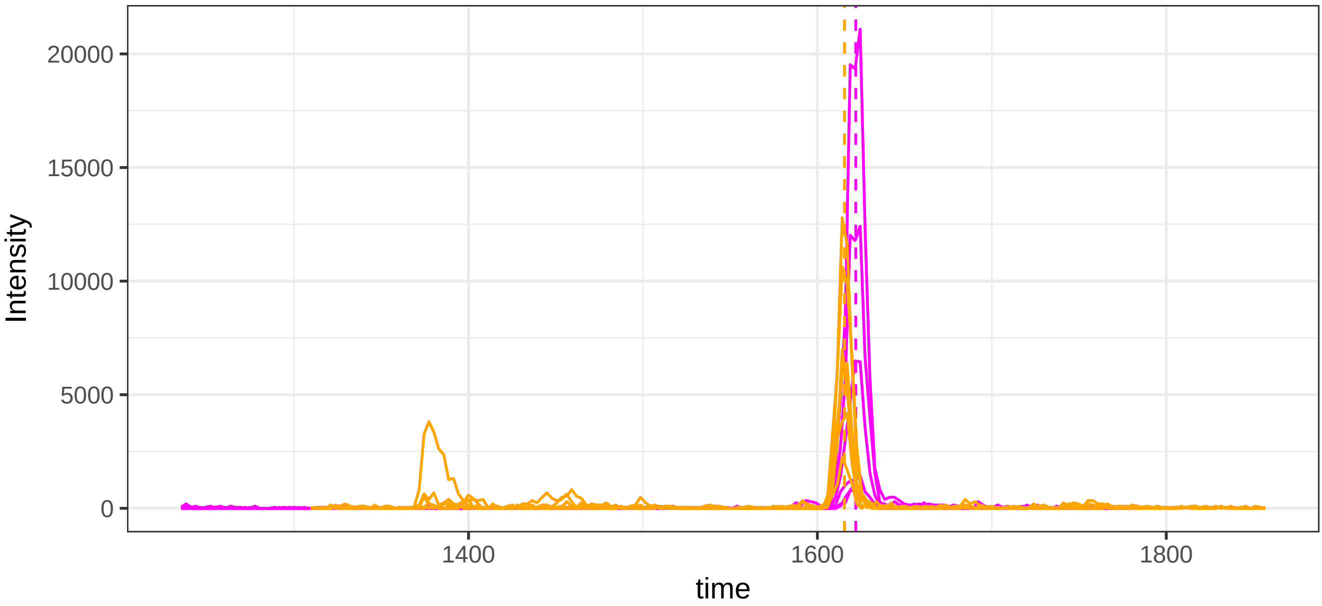


run23: 128284_AFLLTPR/2 (magenta)  AND 42259_ALVQQMEQLR/2

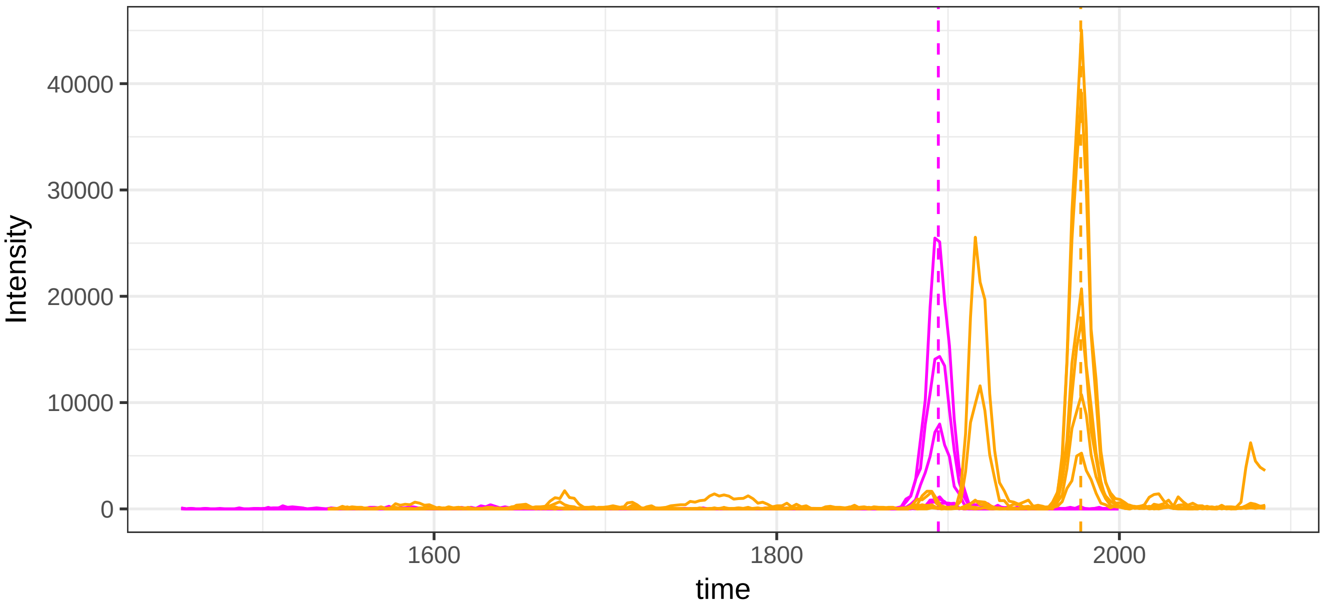

run4: 128284_AFLLTPR/2 (magenta) AND 1326_VGFYESDVMGR/2

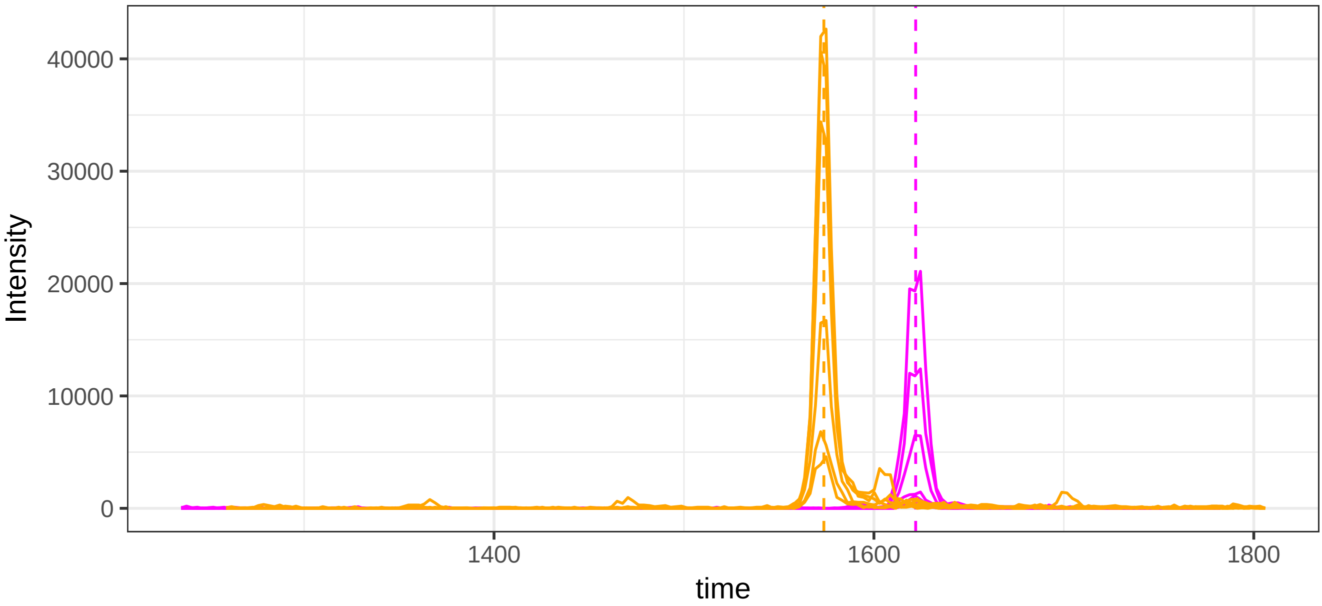


run23: 128284_AFLLTPR/2 (magenta)  AND 1326_VGFYESDVMGR/2

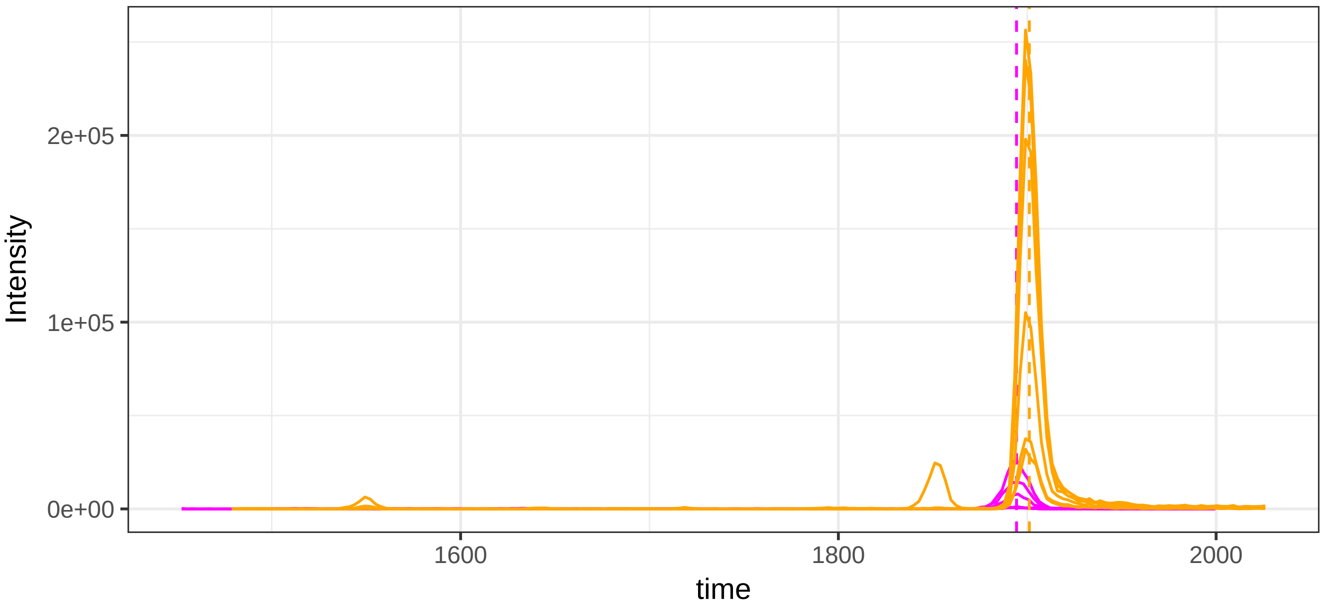

run4: 128284_AFLLTPR/2 (magenta) AND 52314_SGLSTGWTQLSK/2

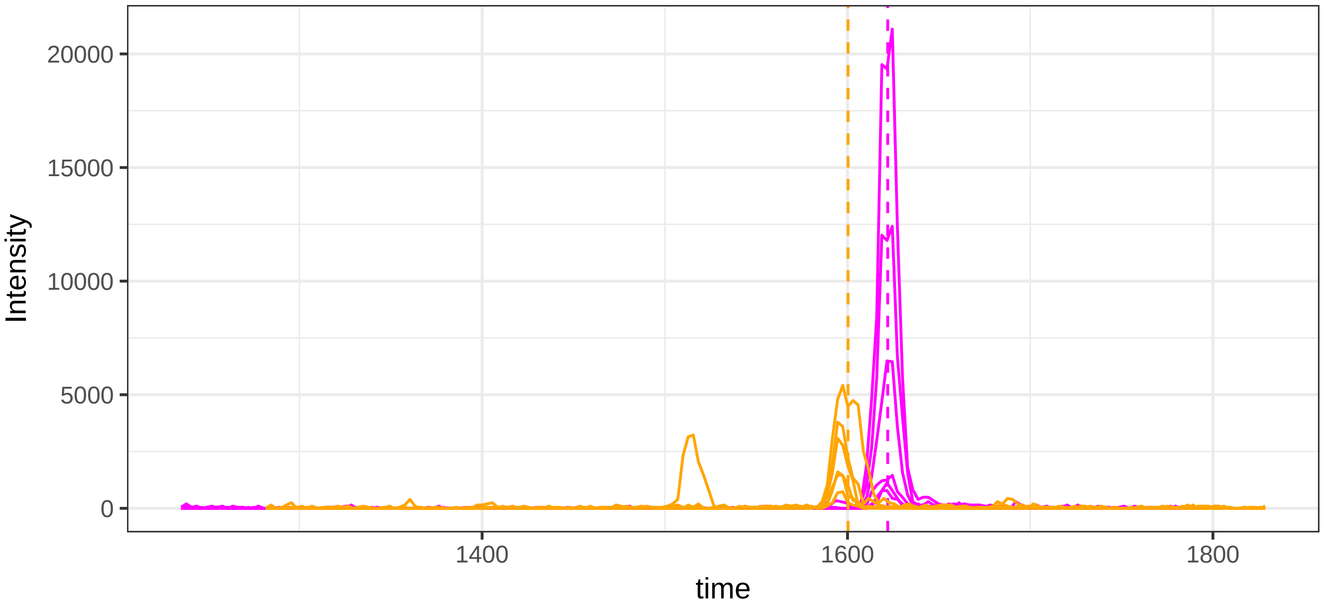


run23: 128284_AFLLTPR/2 (magenta)  AND 52314_SGLSTGWTQLSK/2

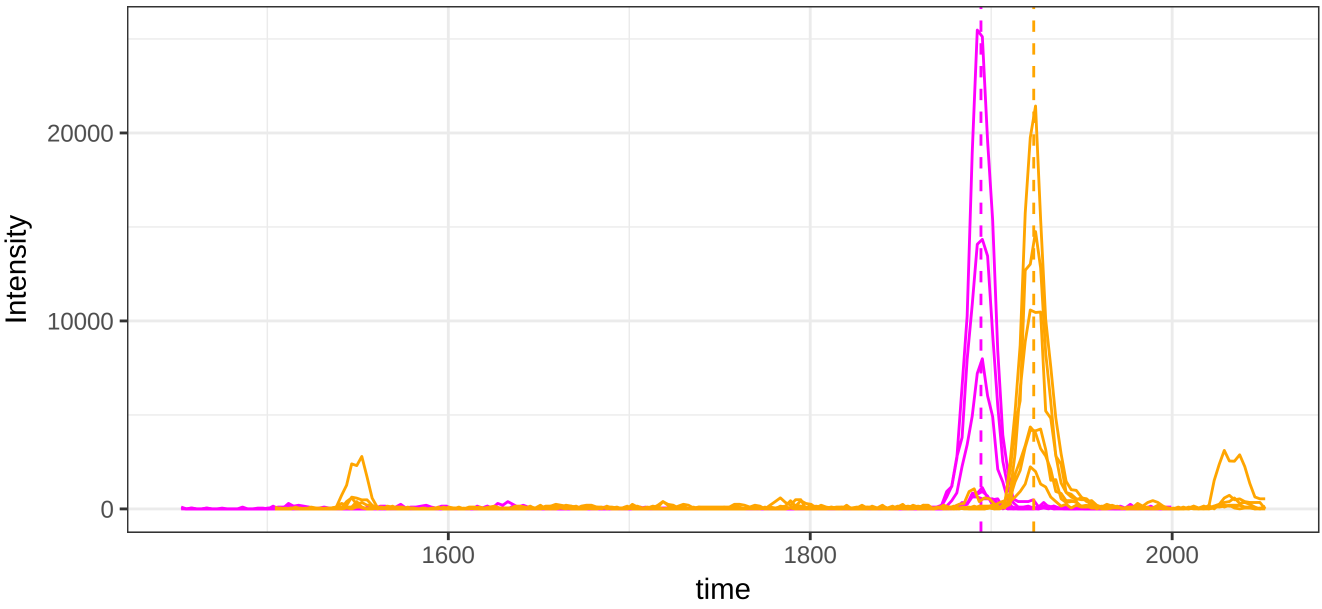

run4: 128284_AFLLTPR/2 (magenta) AND 119859_EDGGGWWYNR/2

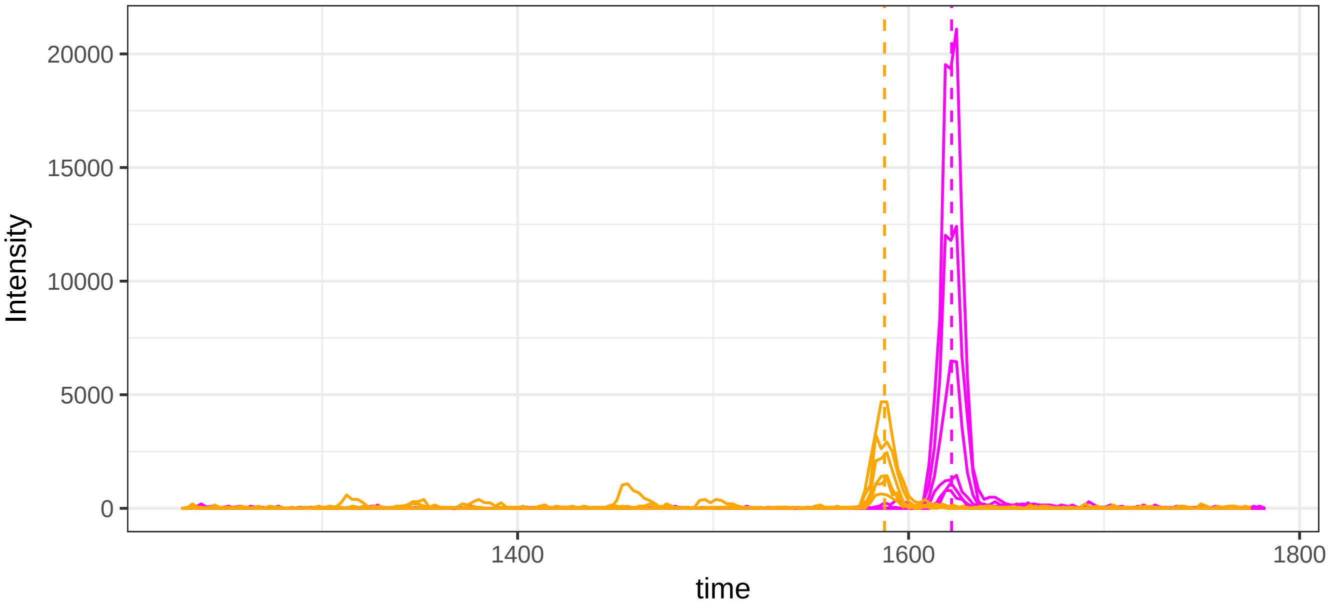


run23: 128284_AFLLTPR/2 (magenta)  AND 119859_EDGGGWWYNR/2

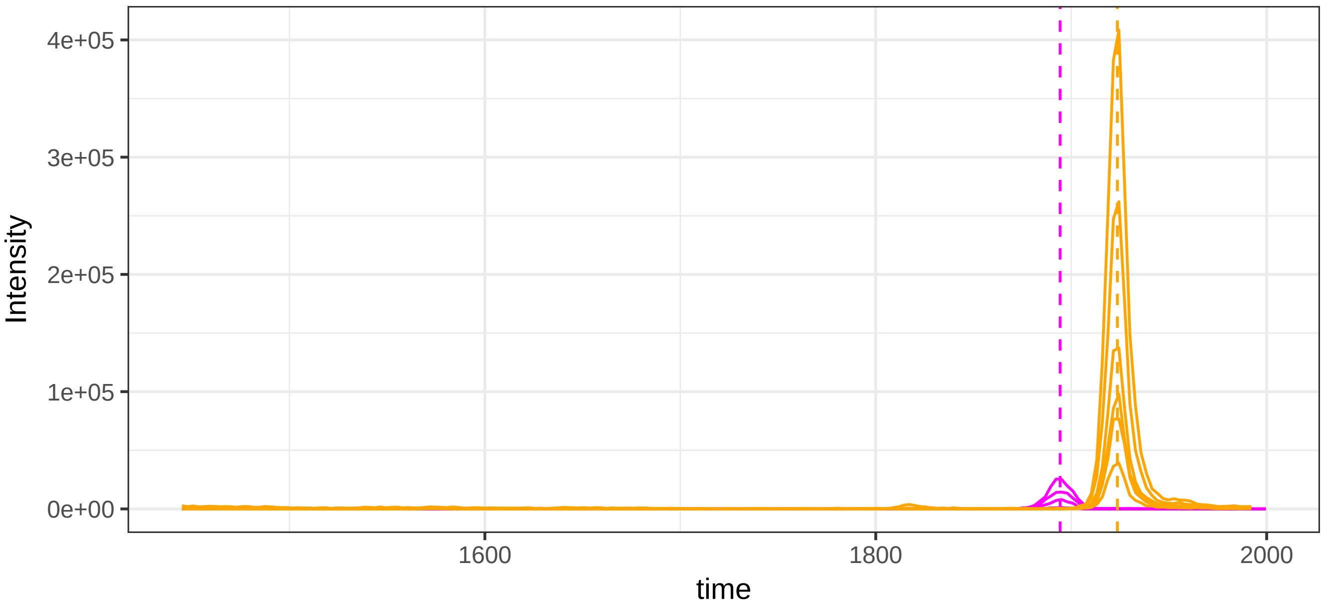

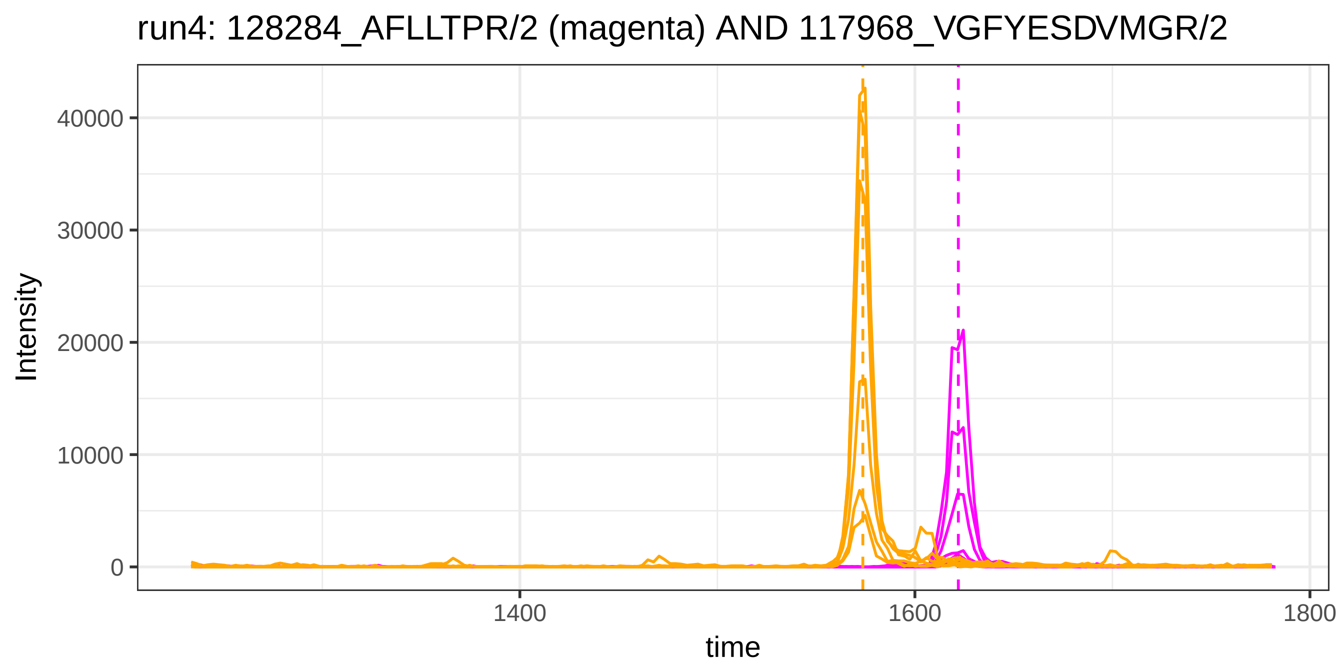

run4: 128284_AFLLTPR/2 (magenta) AND 117968_VGFYESDVMGR/2
Intensity
40000
30000
20000
10000
0
1400
1600
1800
time


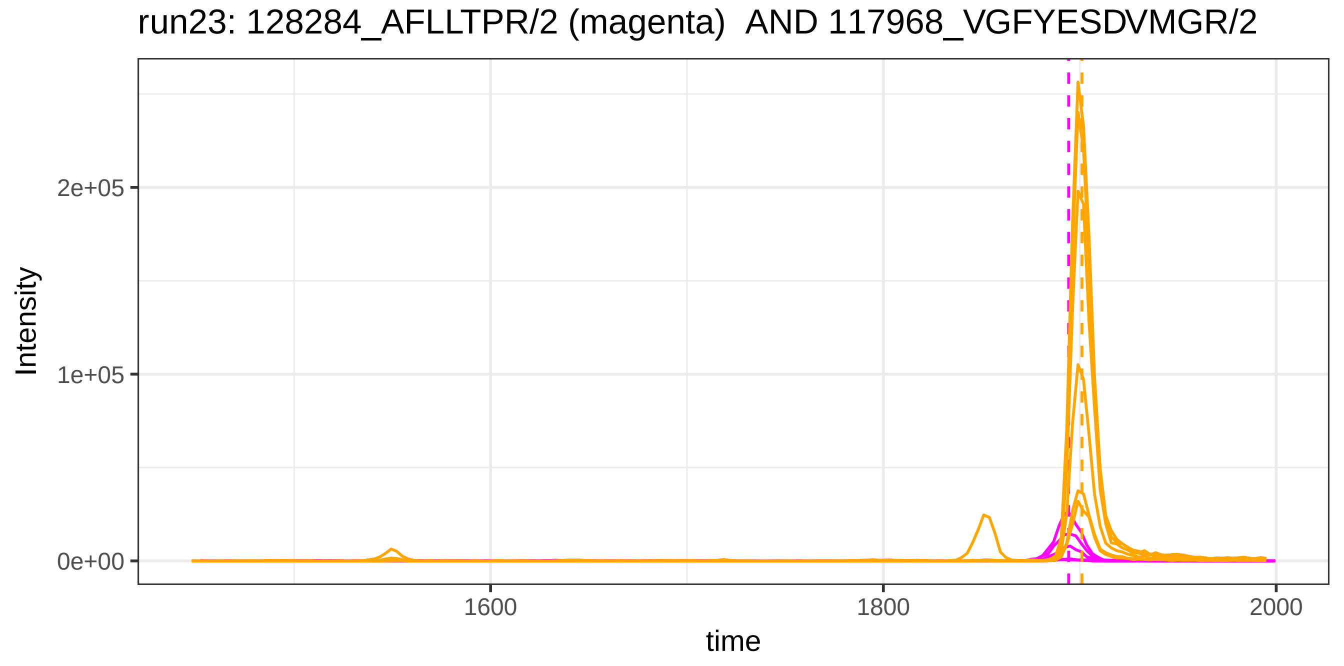

run23: 128284_AFLLTPR/2 (magenta)  AND 117968_VGFYESDVMGR/2
Intensity
2e+05
1e+05
0e+00
1600
1800
2000
time

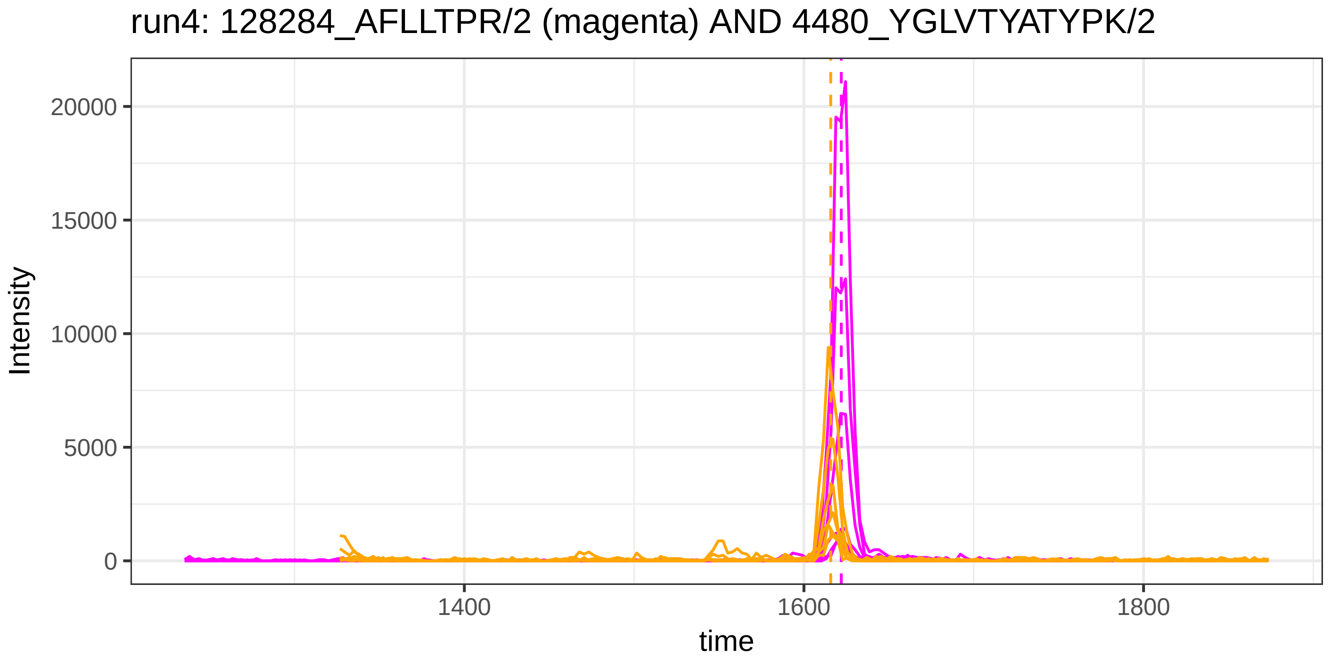
run4: 128284_AFLLTPR/2 (magenta) AND 4480_YGLVTYATYPK/2
Intensity
20000
15000
10000
5000
0
1400
1600
1800
time

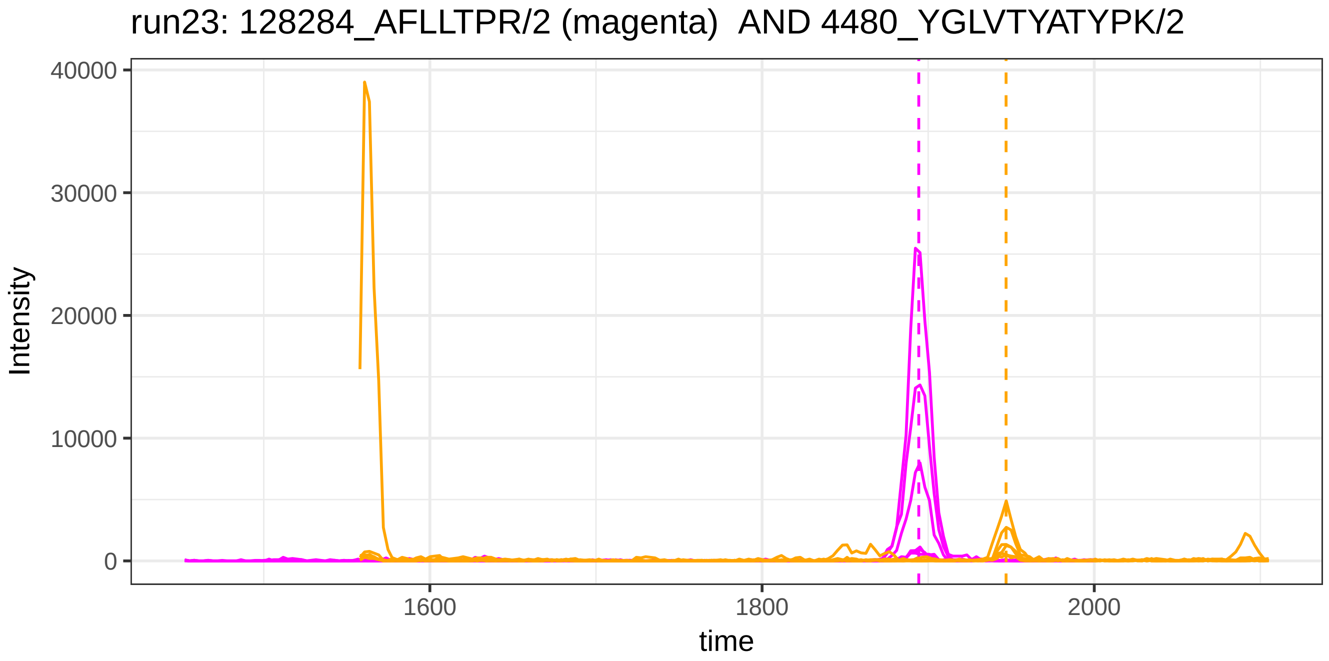
run23: 128284_AFLLTPR/2 (magenta) AND 4480_YGLVTYATYPK/2
Intensity
40000
30000
20000
10000
0
1600
1800
2000
time

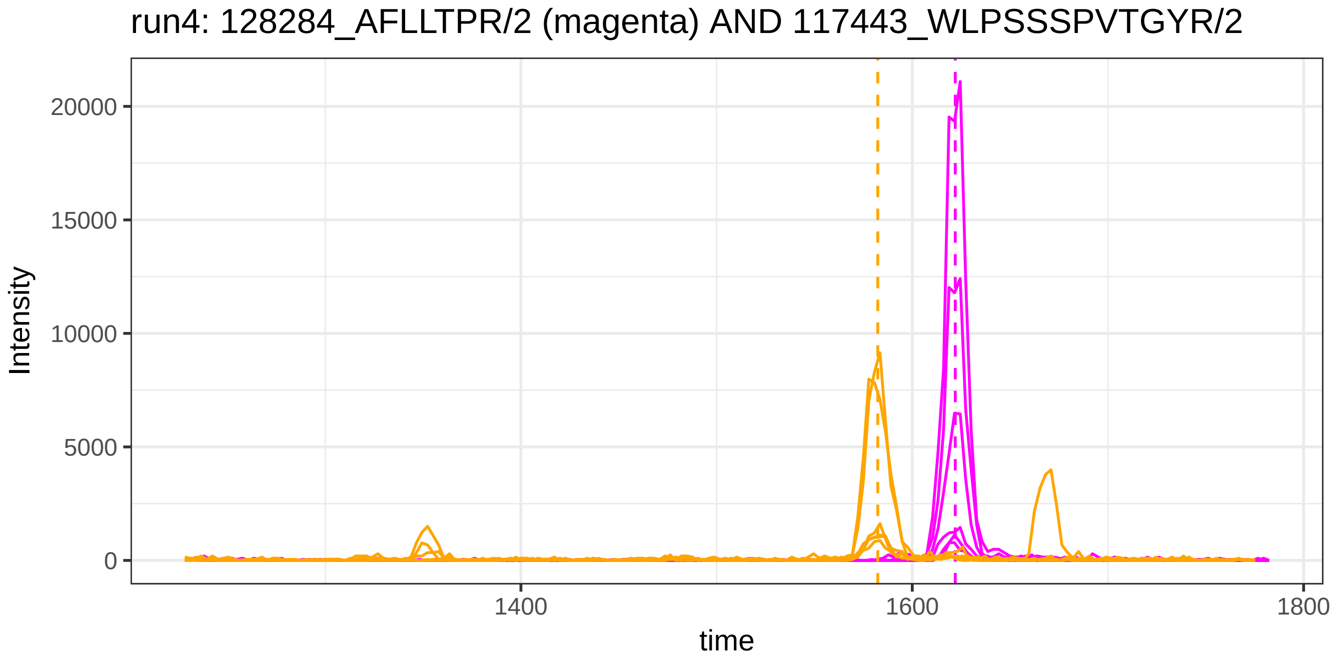
run4: 128284_AFLLTPR/2 (magenta) AND 117443_WLPSSSPVTGYR/2
Intensity
20000
15000
10000
5000
0
1400
1600
1800
time

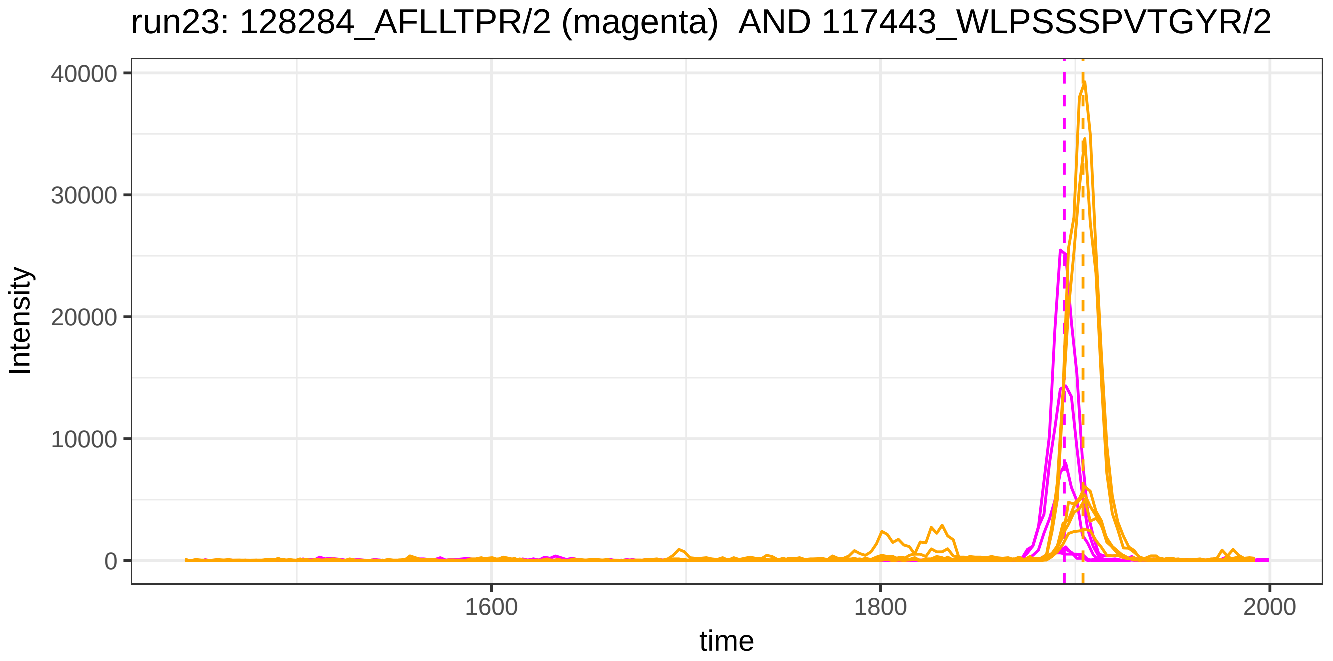
run23: 128284_AFLLTPR/2 (magenta) AND 117443_WLPSSSPVTGYR/2
Intensity
40000
30000
20000
10000
0
1600
1800
2000
time

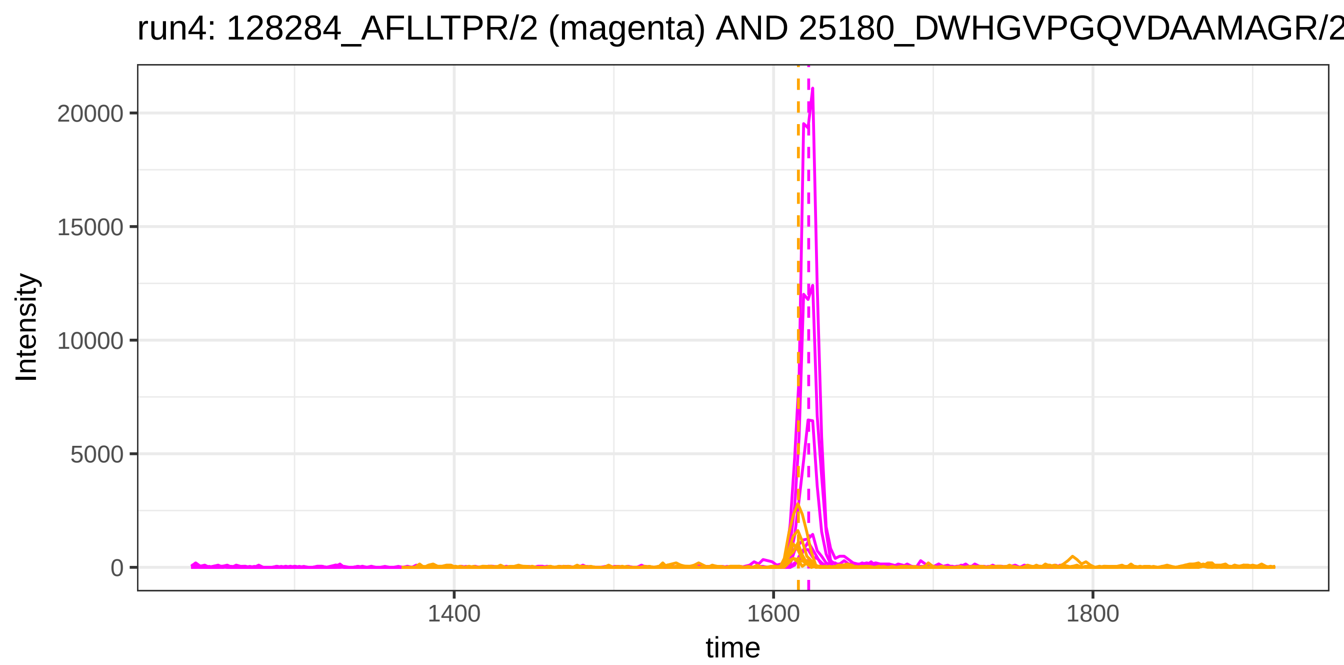
run4: 128284_AFLLTPR/2 (magenta) AND 25180_DWHGVPGQVDAAMAGR/2
Intensity
20000
15000
10000
5000
0
1400
1600
1800
time

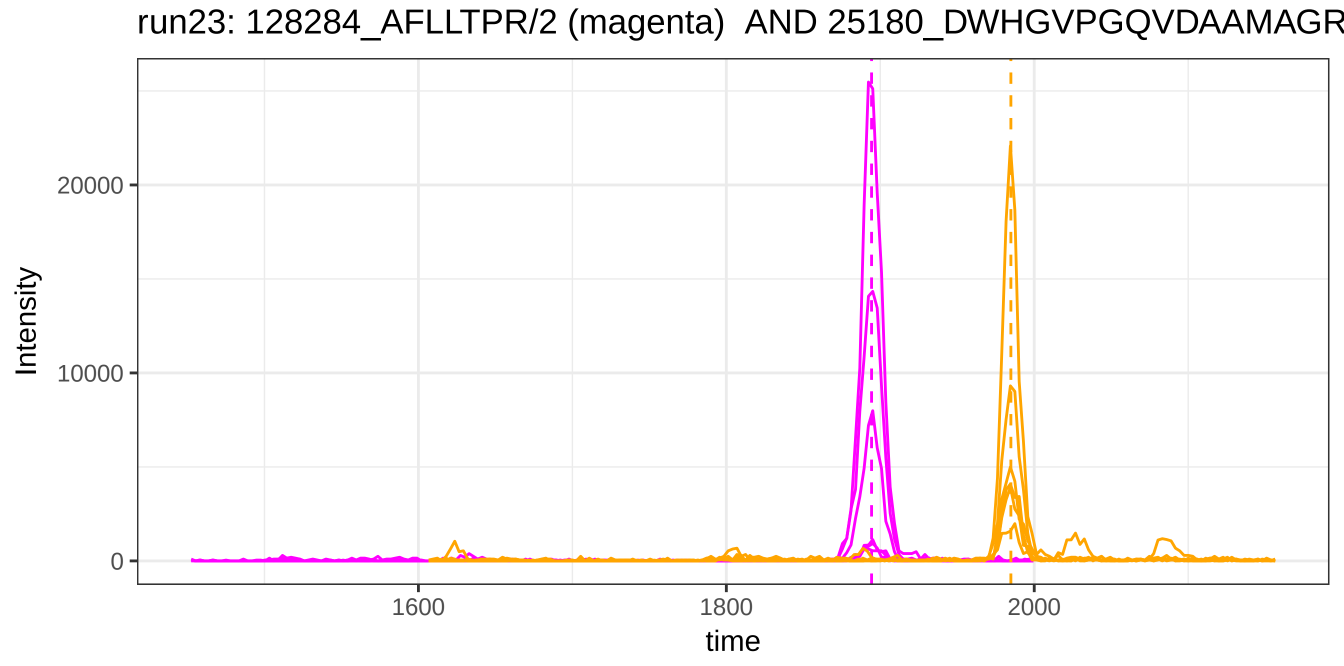
run23: 128284_AFLLTPR/2 (magenta) AND 25180_DWHGVPGQVDAAMAGR
Intensity
20000
10000
0
1600
1800
2000
time

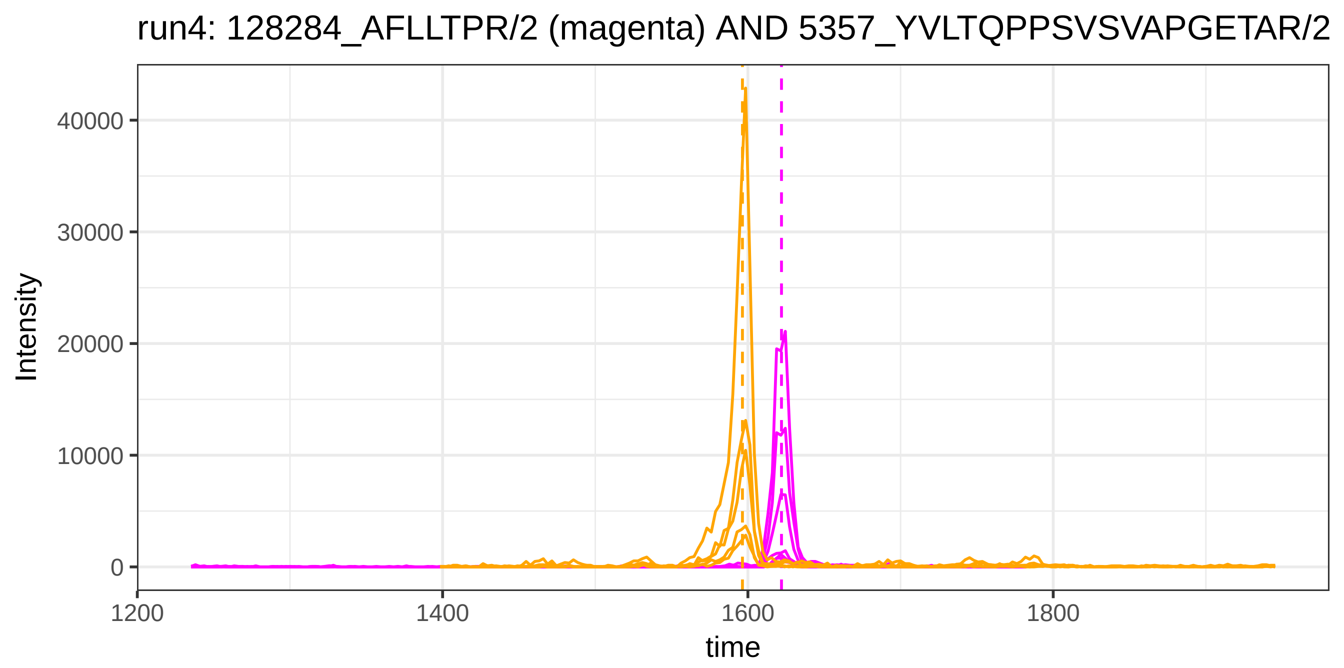
run4: 128284_AFLLTPR/2 (magenta) AND 5357_YVLTQPPSVSVAPGETAR/2
Intensity
40000
30000
20000
10000
0
1200
1400
1600
1800
time

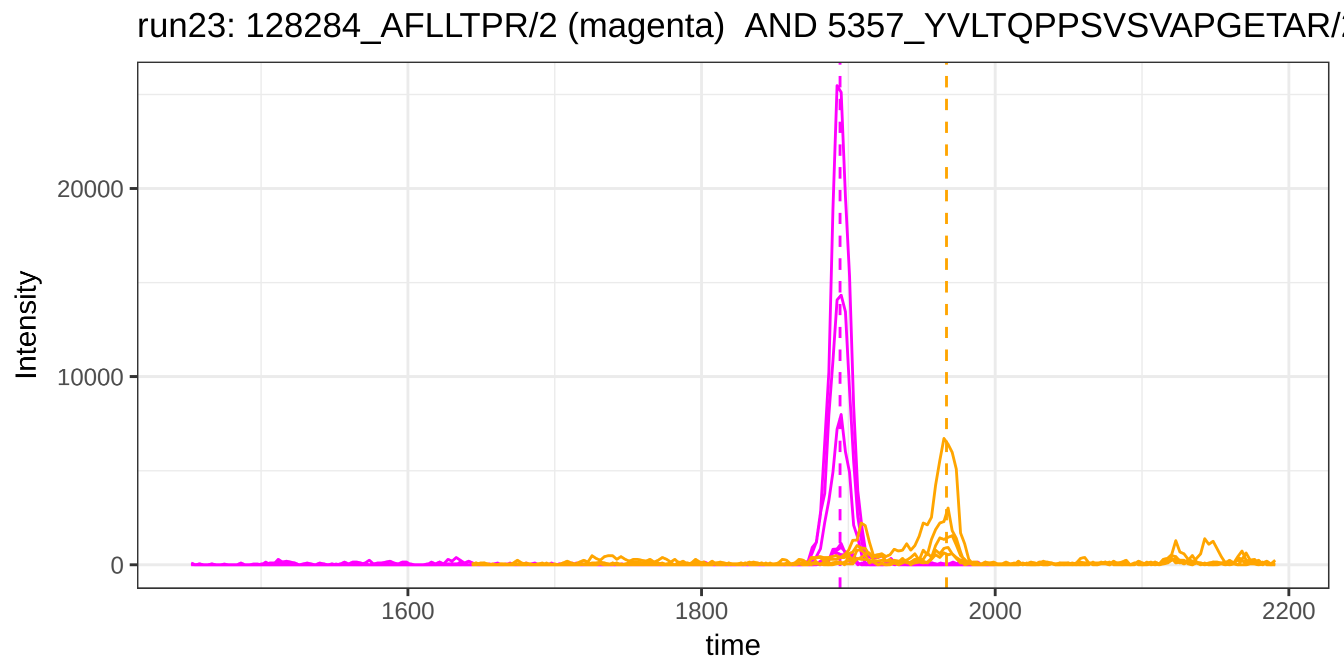
run23: 128284_AFLLTPR/2 (magenta) AND 5357_YVLTQPPSVSVAPGETAR/
Intensity
20000
10000
0
1600
1800
2000
2200
time

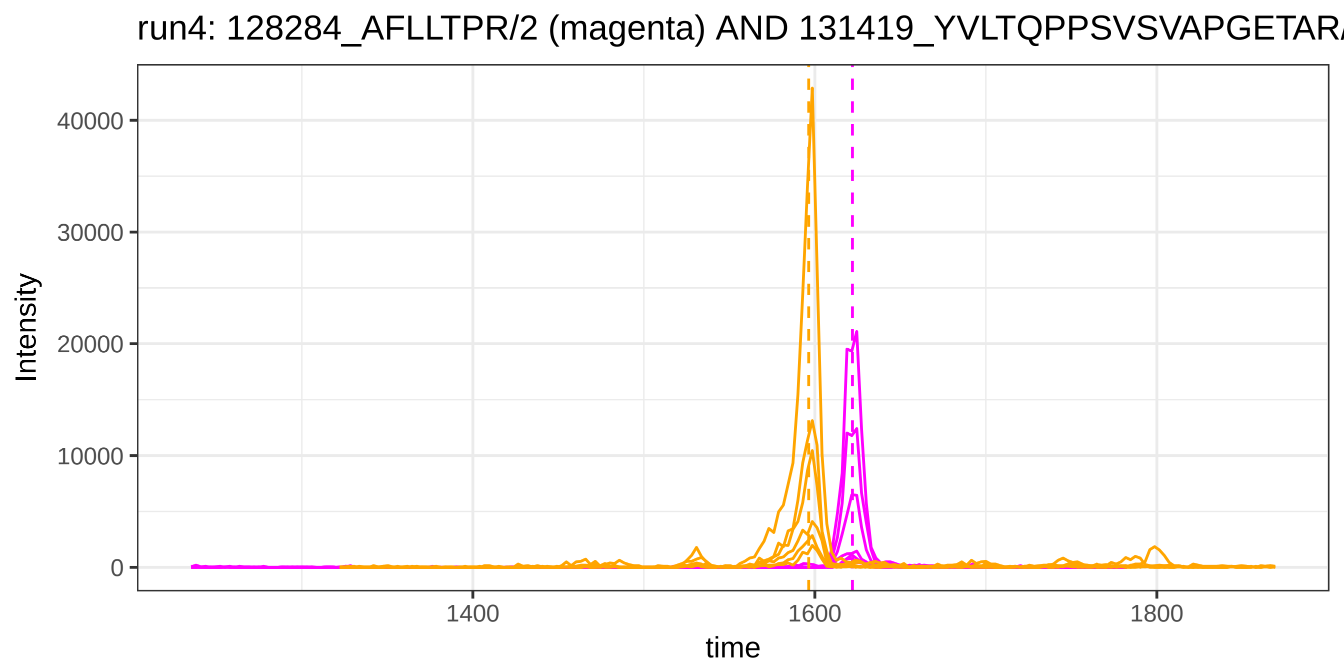
run4: 128284_AFLLTPR/2 (magenta) AND 131419_YVLTQPPSVSVAPGETAR/
Intensity
40000
30000
20000
10000
0
1400
1600
1800
time

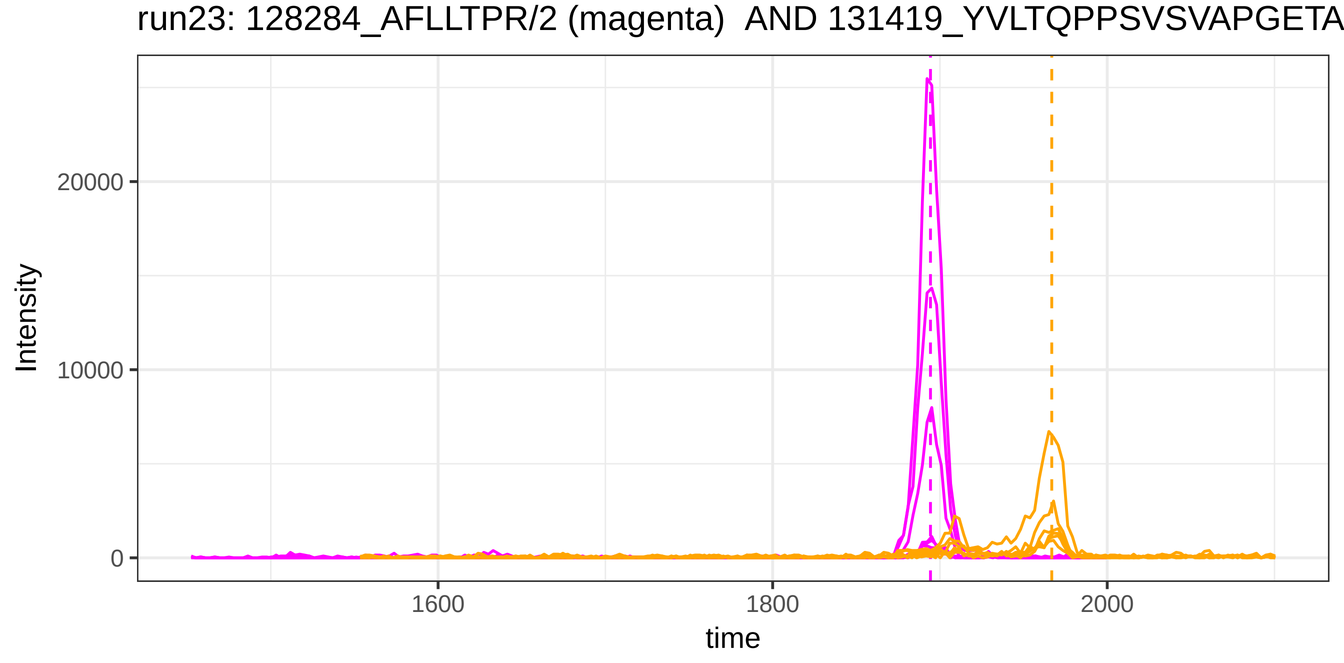
run23: 128284_AFLLTPR/2 (magenta) AND 131419_YVLTQPPSVSVAPGETA
Intensity
20000
10000
0
1600
1800
2000
time

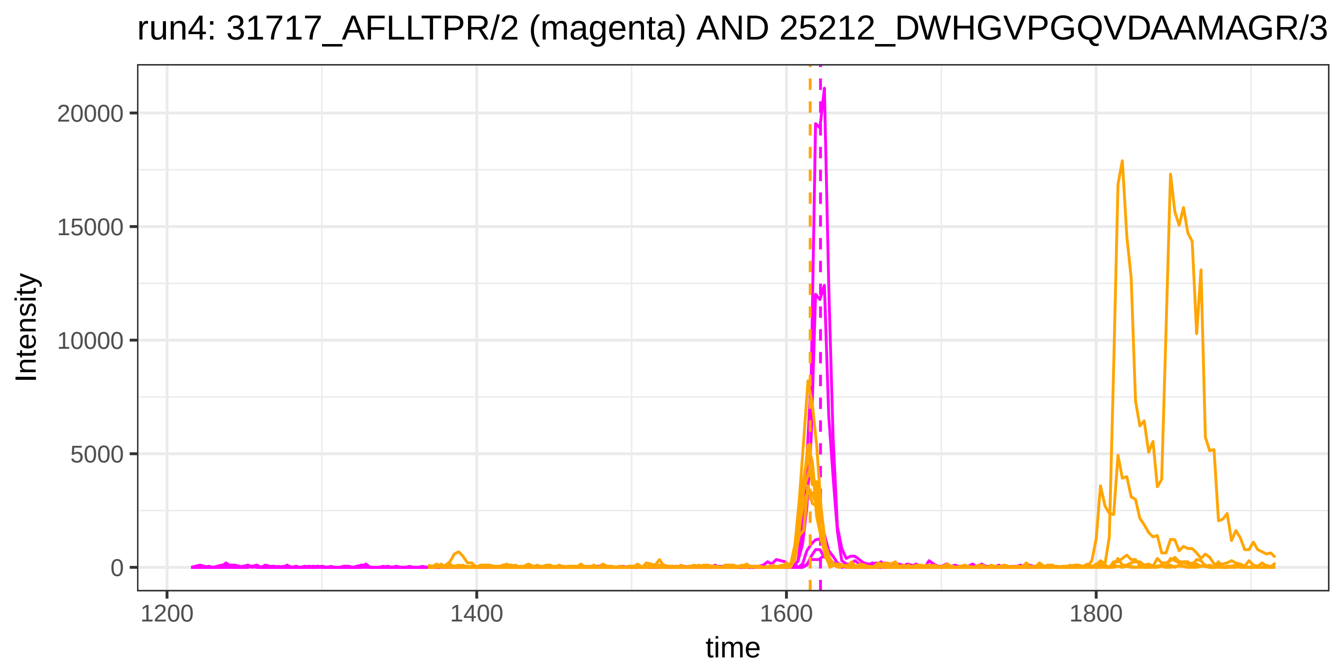

run4: 31717_AFLLTPR/2 (magenta) AND 25212_DWHGVPGQVDAAMAGR/3
Intensity
20000
15000
10000
5000
0
1200
1400
1600
1800
time


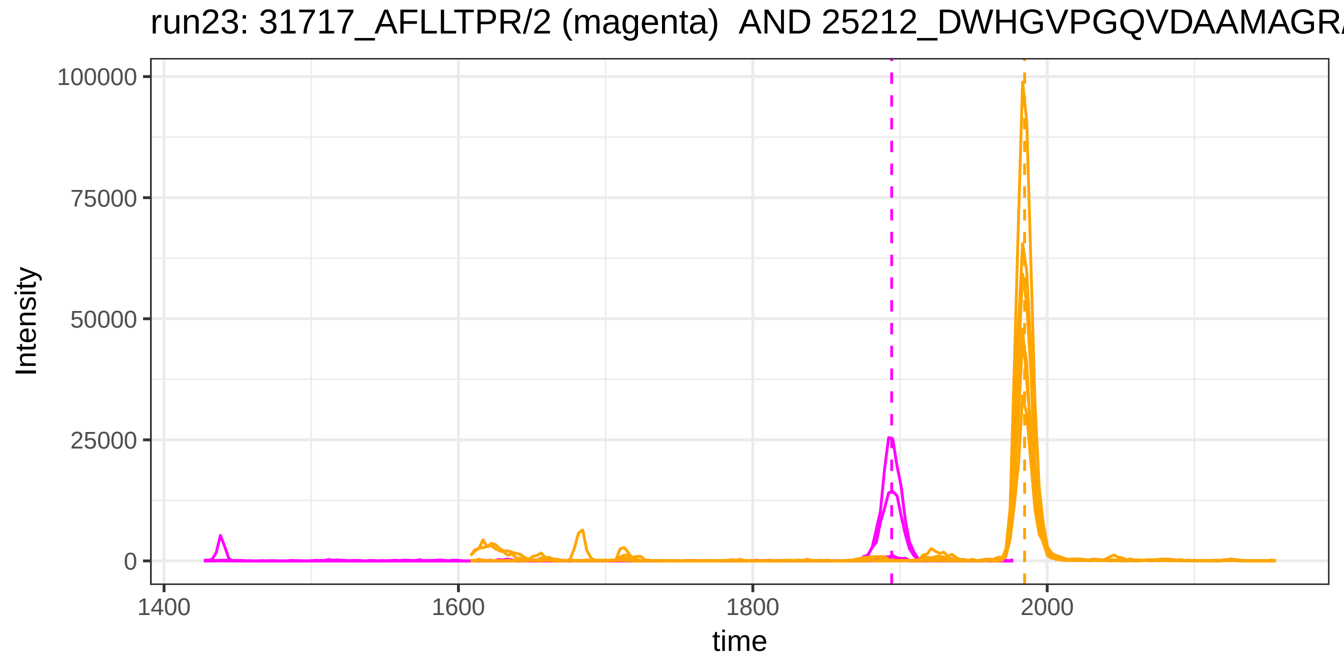

run23: 31717_AFLLTPR/2 (magenta) AND 25212_DWHGVPGQVDAAMAGR/
Intensity
100000
75000
50000
25000
0
1400
1600
1800
2000
time

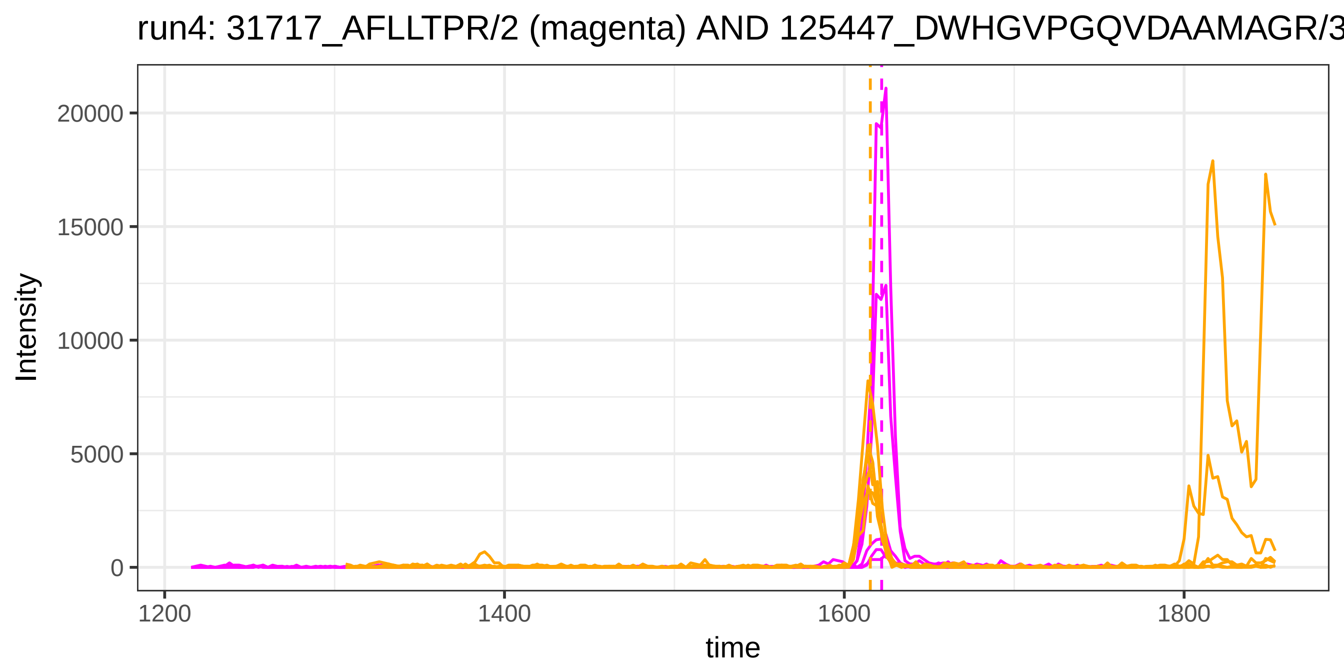
run4: 31717_AFLLTPR/2 (magenta) AND 125447_DWHGVPGQVDAAMAGR/3
Intensity
time
20000
15000
10000
5000
0
1200
1400
1600
1800

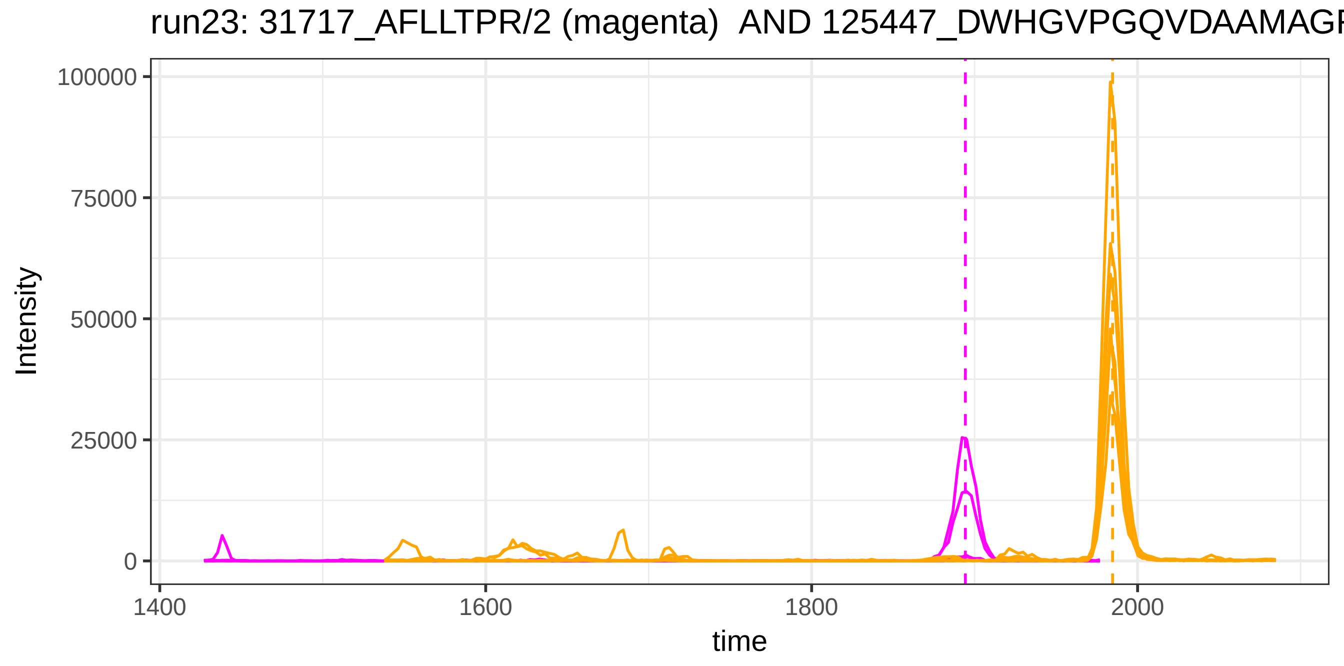
run23: 31717_AFLLTPR/2 (magenta) AND 125447_DWHGVPGQVDAAMAGR
Intensity
time
100000
75000
50000
25000
0
1400
1600
1800
2000

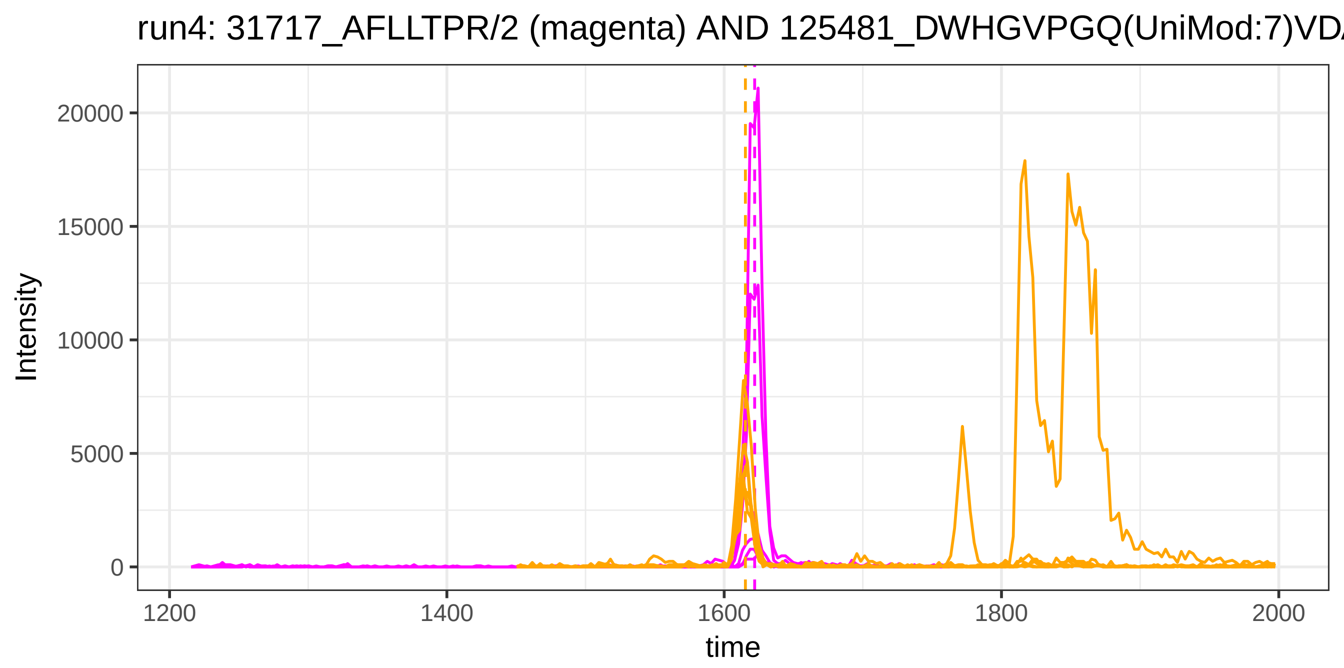

run4: 31717_AFLLTPR/2 (magenta) AND 125481_DWHGVPGQ(UniMod:7)VD
Intensity
20000
15000
10000
5000
0
1200
1400
1600
1800
2000
time


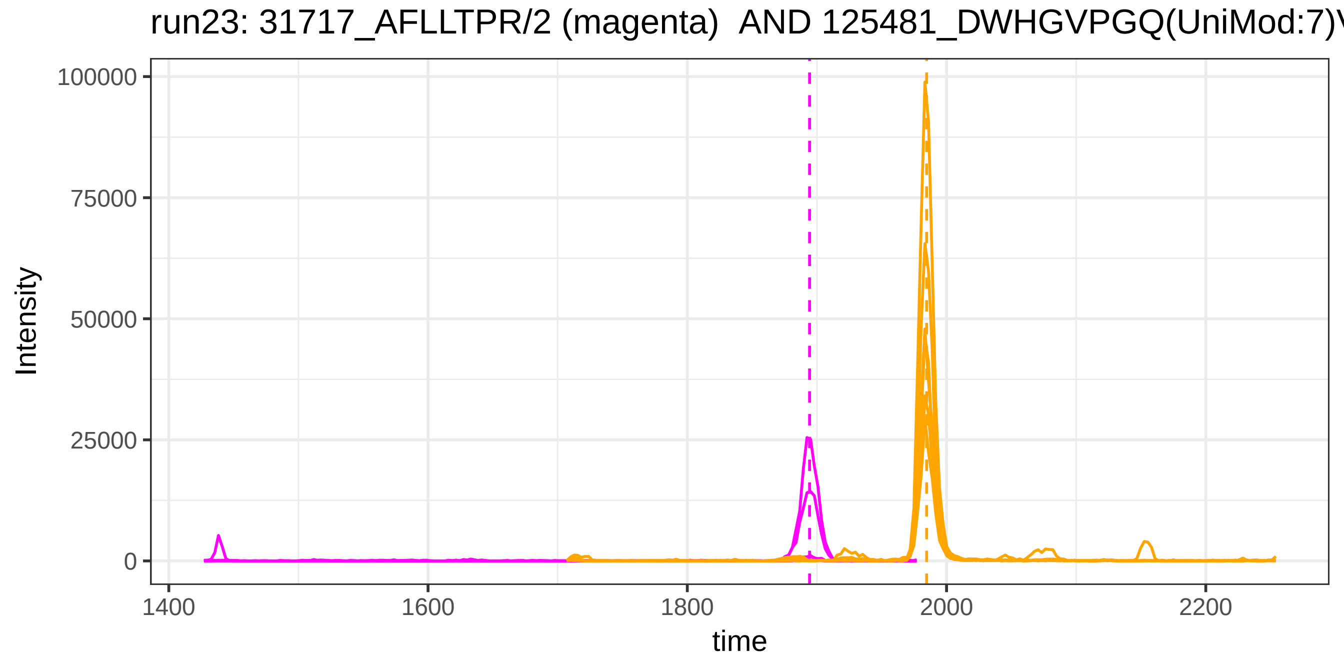

run23: 31717_AFLLTPR/2 (magenta) AND 125481_DWHGVPGQ(UniMod:7)V
Intensity
100000
75000
50000
25000
0
1400
1600
1800
2000
2200
time

run4: 31717_AFLLTPR/2 (magenta) AND 2924_VTSIQDWVQK/2

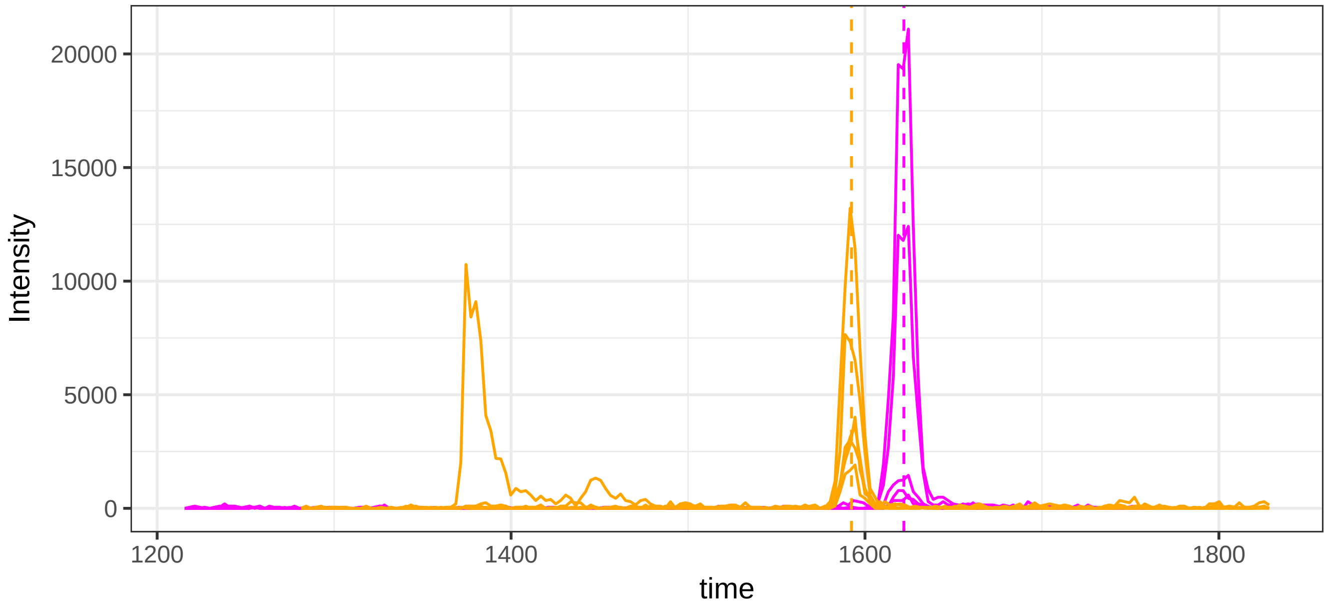


run23: 31717_AFLLTPR/2 (magenta)  AND 2924_VTSIQDWVQK/2

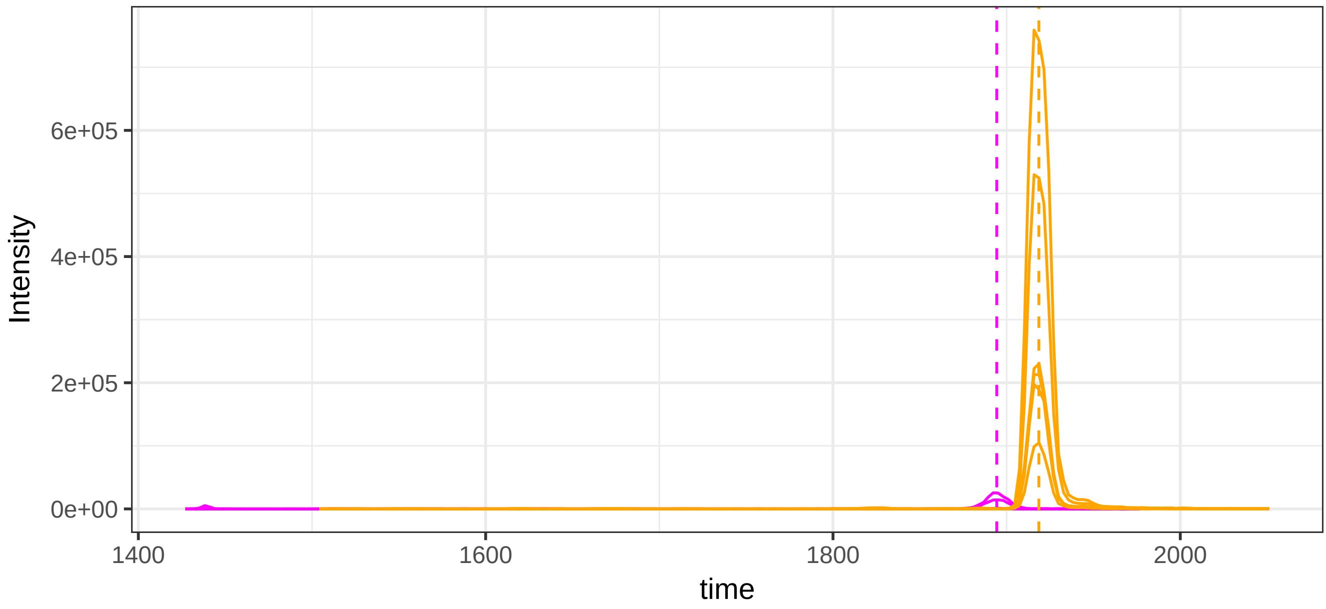

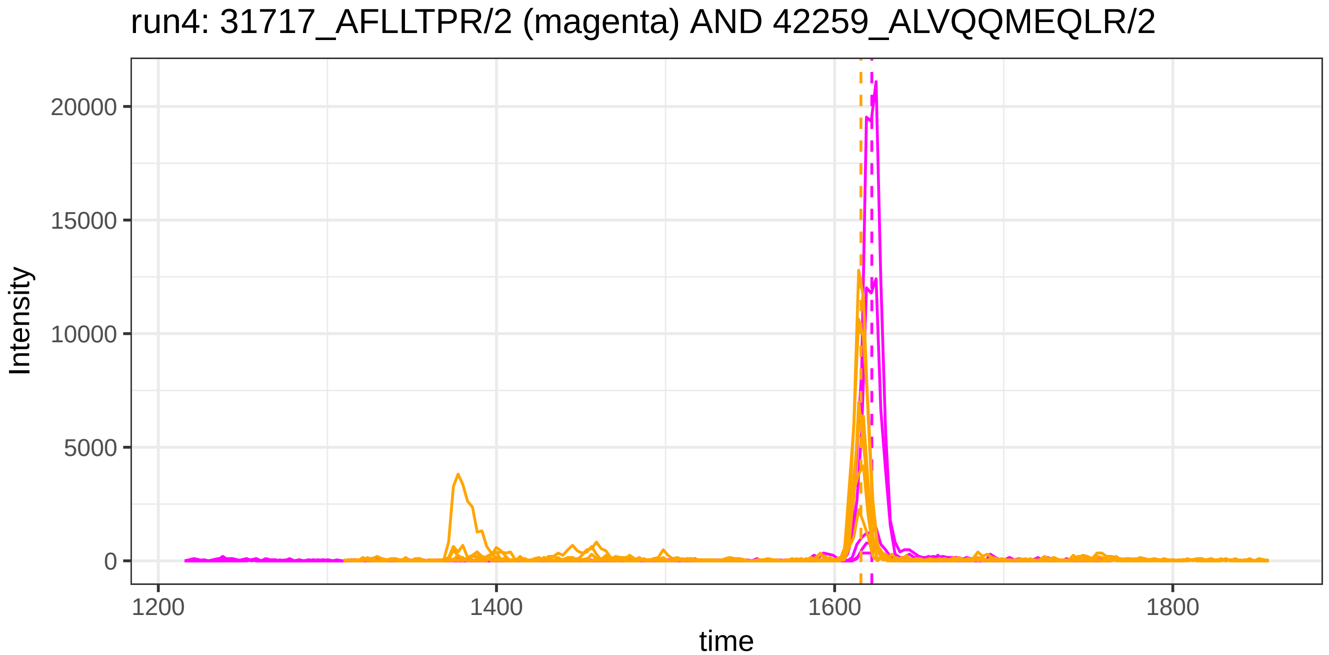
run4: 31717_AFLLTPR/2 (magenta) AND 42259_ALVQQMEQLR/2
Intensity
20000
15000
10000
5000
0
1200
1400
1600
1800
time

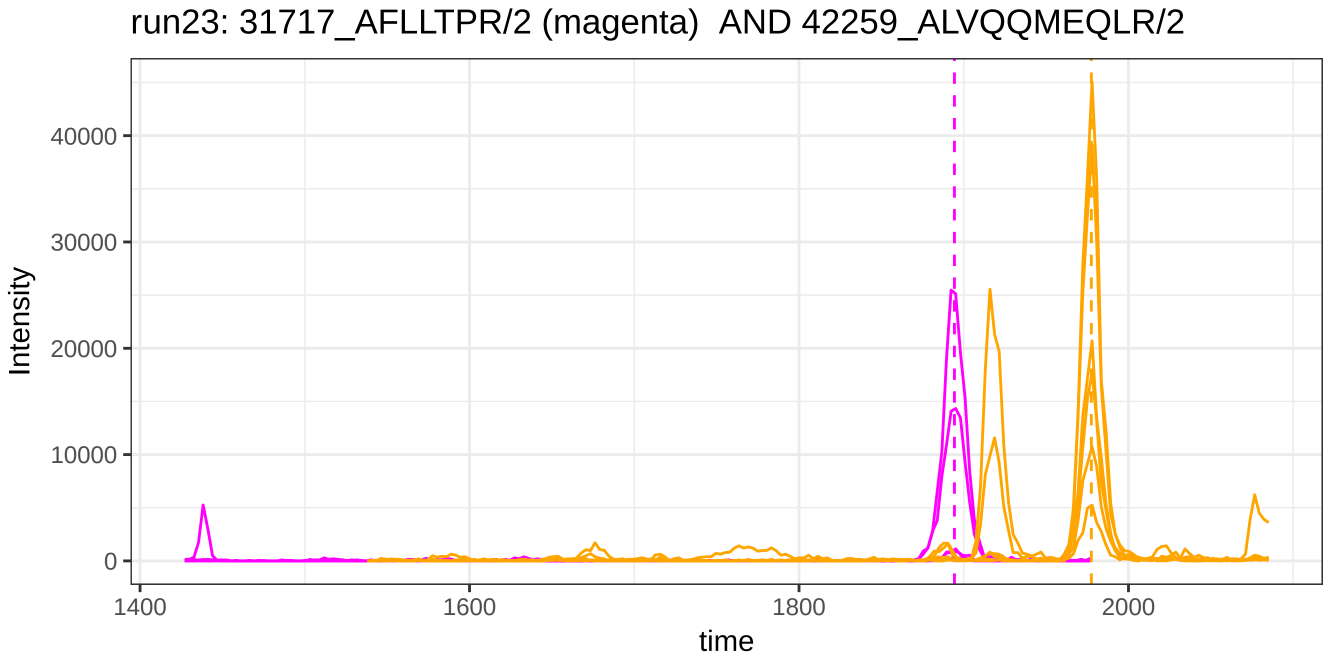
run23: 31717_AFLLTPR/2 (magenta)  AND 42259_ALVQQMEQLR/2
Intensity
40000
30000
20000
10000
0
1400
1600
1800
2000
time

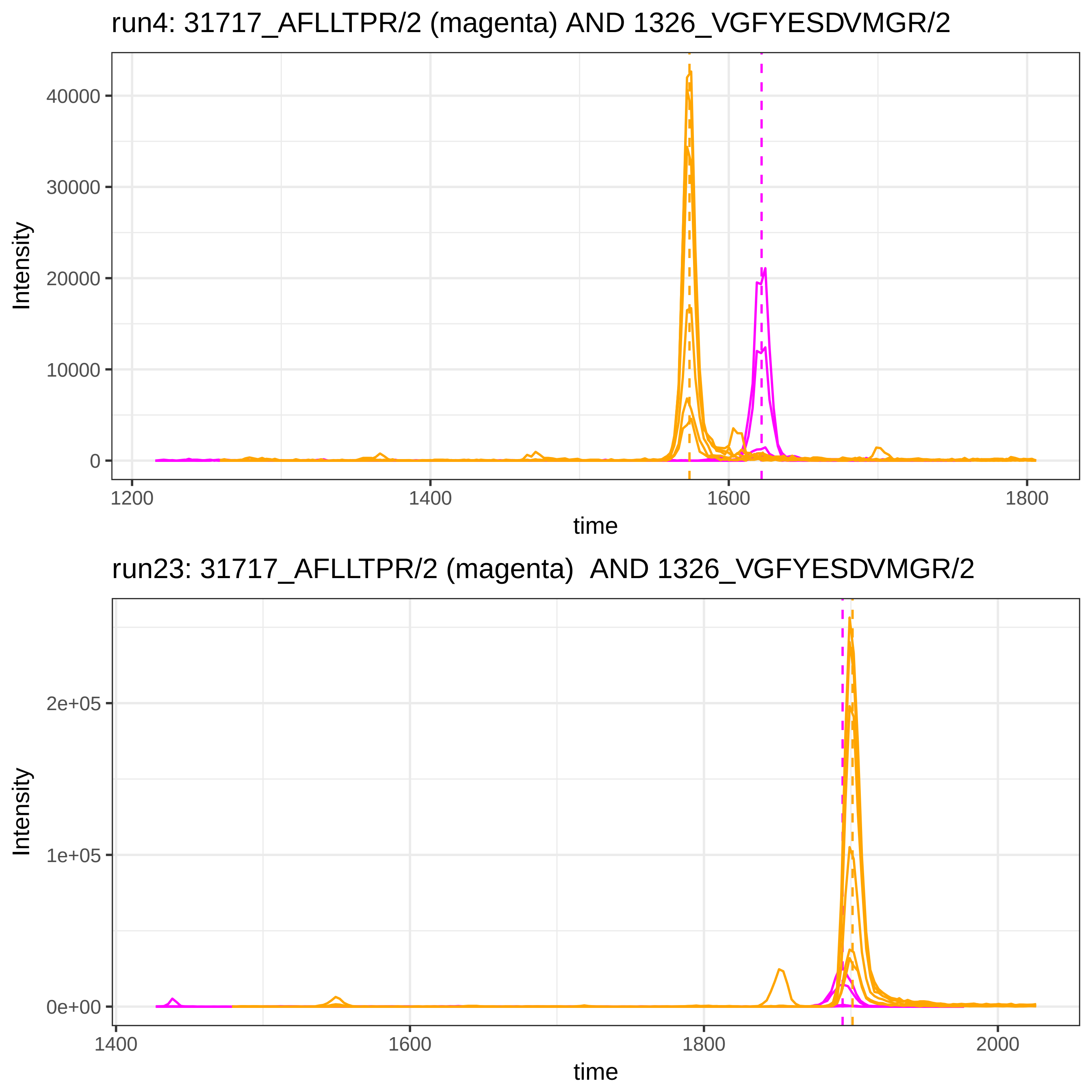
run4: 31717_AFLLTPR/2 (magenta) AND 1326_VGFYESDVMGR/2
Intensity
40000
30000
20000
10000
0
1200
1400
1600
1800
time
run23: 31717_AFLLTPR/2 (magenta)  AND 1326_VGFYESDVMGR/2
Intensity
2e+05
1e+05
0e+00
1400
1600
1800
2000
time

run4: 31717_AFLLTPR/2 (magenta) AND 52314_SGLSTGWTQLSK/2

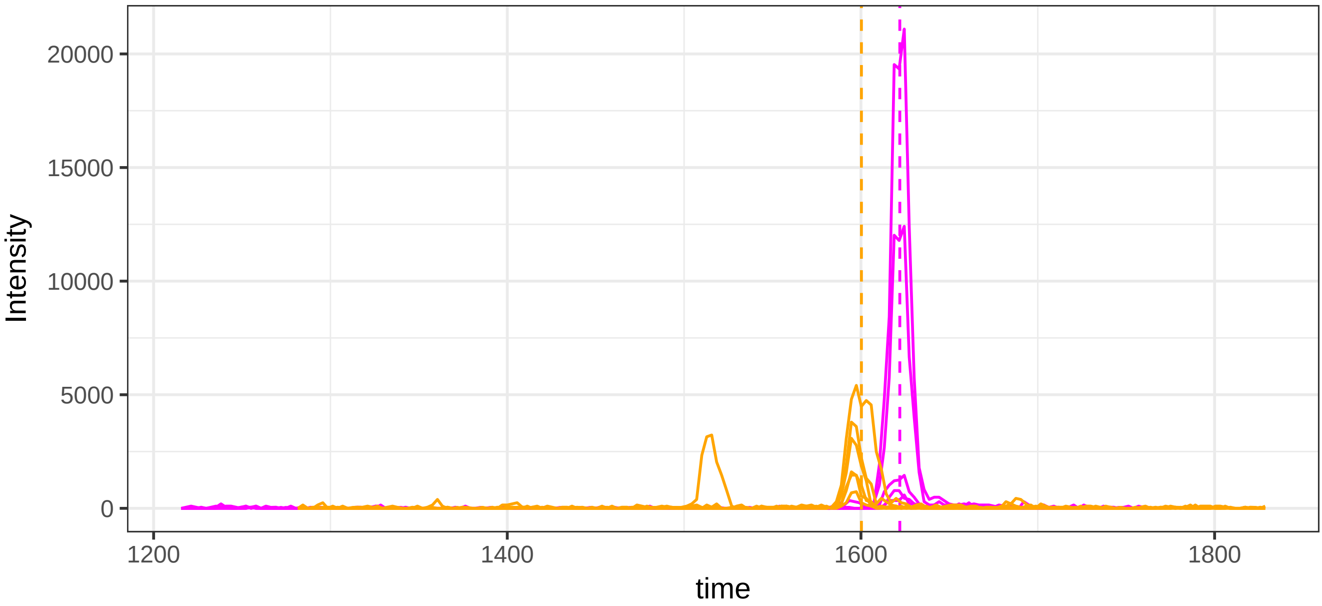


run23: 31717_AFLLTPR/2 (magenta)  AND 52314_SGLSTGWTQLSK/2

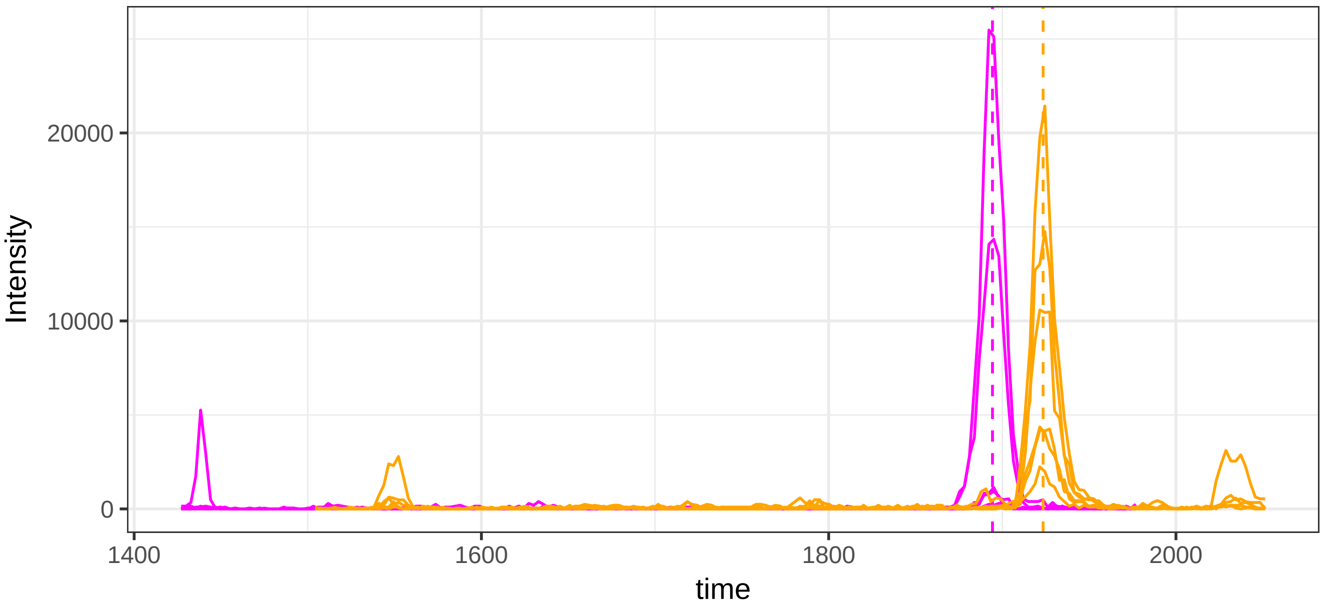

run4: 31717_AFLLTPR/2 (magenta) AND 119859_EDGGGWWYNR/2

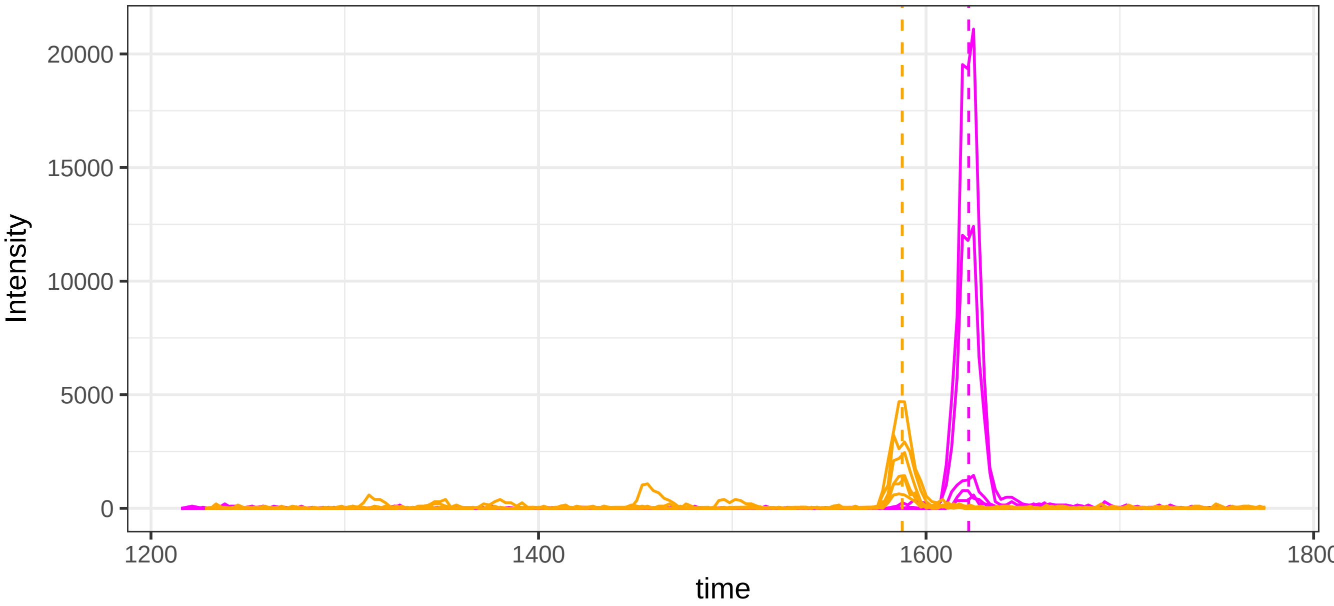


run23: 31717_AFLLTPR/2 (magenta)  AND 119859_EDGGGWWYNR/2

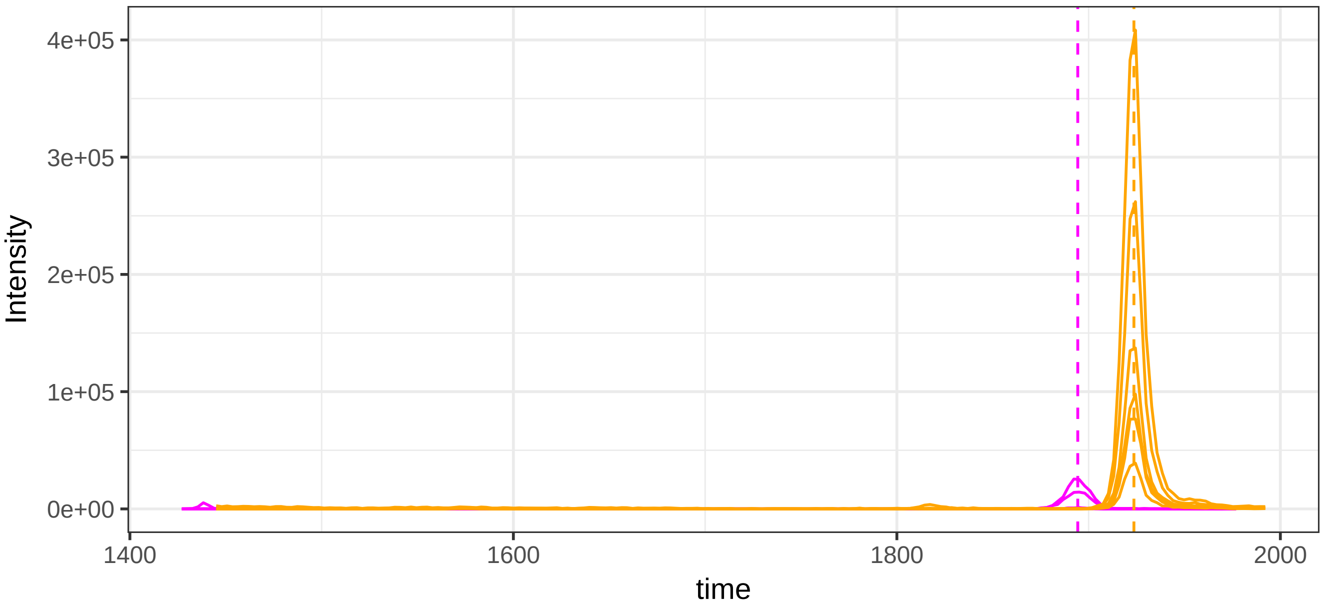

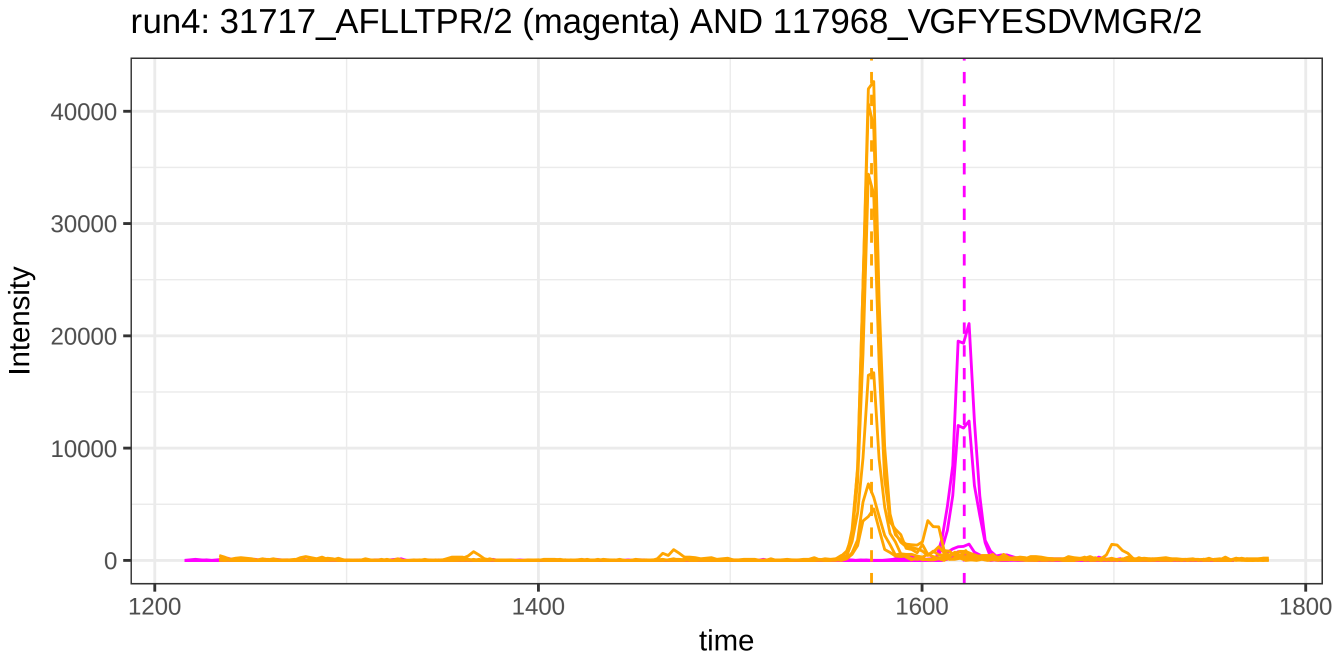
run4: 31717_AFLLTPR/2 (magenta) AND 117968_VGFYESDVMGR/2
Intensity
40000
30000
20000
10000
0
1200
1400
1600
1800
time

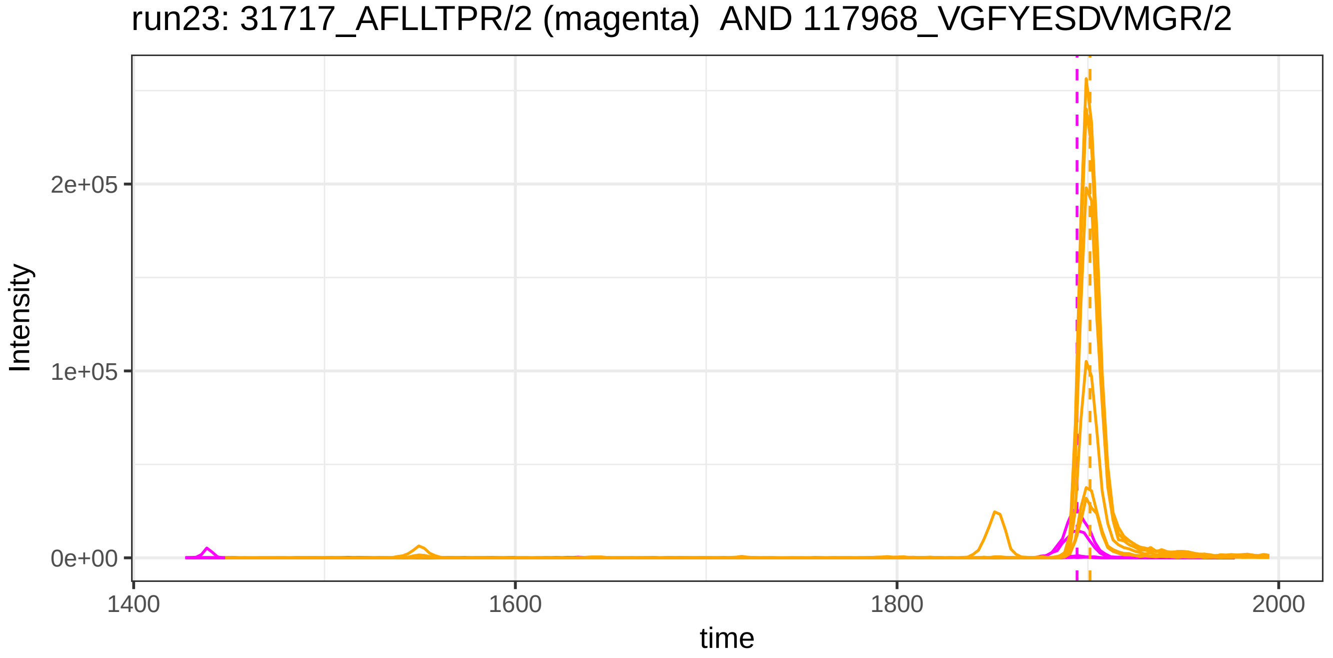
run23: 31717_AFLLTPR/2 (magenta)  AND 117968_VGFYESDVMGR/2
Intensity
2e+05
1e+05
0e+00
1400
1600
1800
2000
time

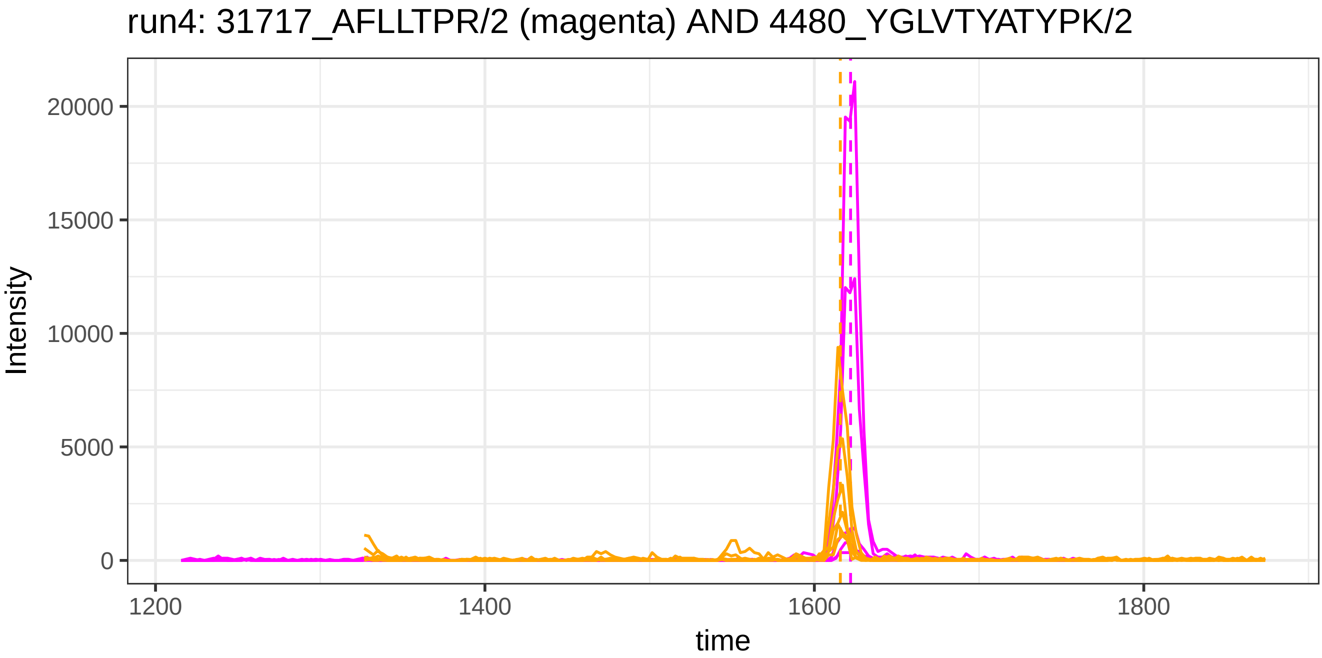
run4: 31717_AFLLTPR/2 (magenta) AND 4480_YGLVTYATYPK/2
Intensity
20000
15000
10000
5000
0
1200
1400
1600
1800
time

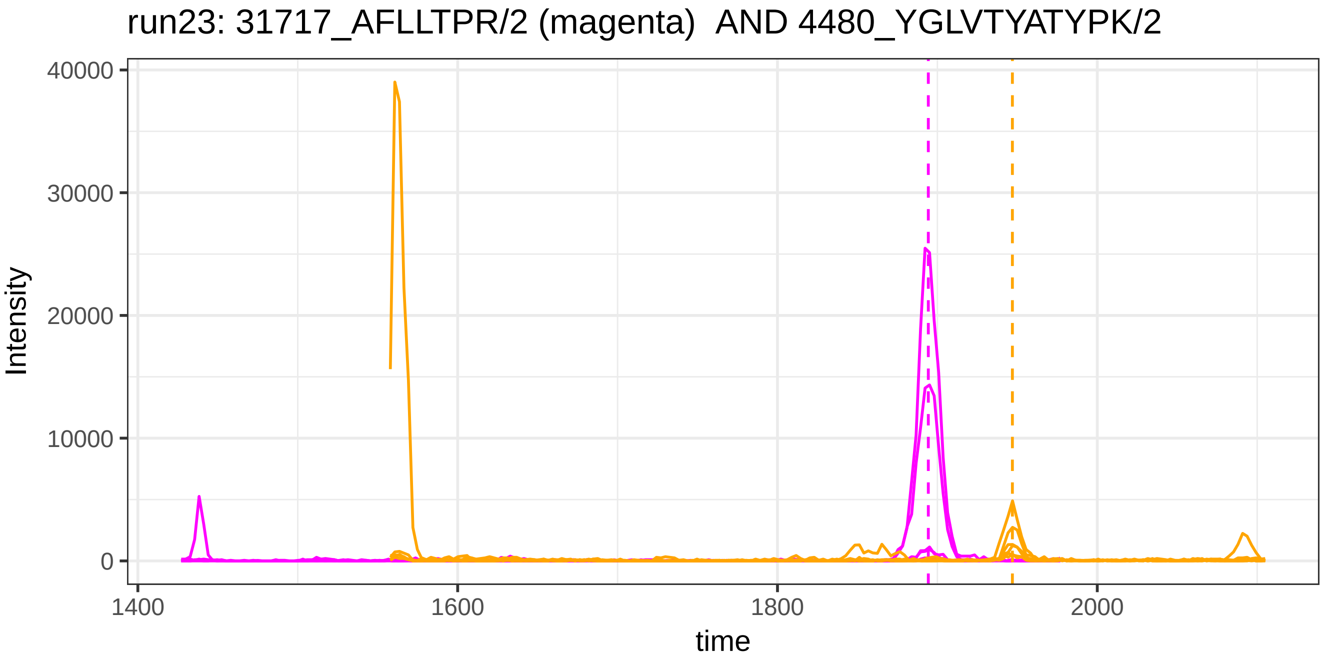
run23: 31717_AFLLTPR/2 (magenta) AND 4480_YGLVTYATYPK/2
Intensity
40000
30000
20000
10000
0
1400
1600
1800
2000
time

run4: 31717_AFLLTPR/2 (magenta) AND 117443_WLPSSSPVTGYR/2

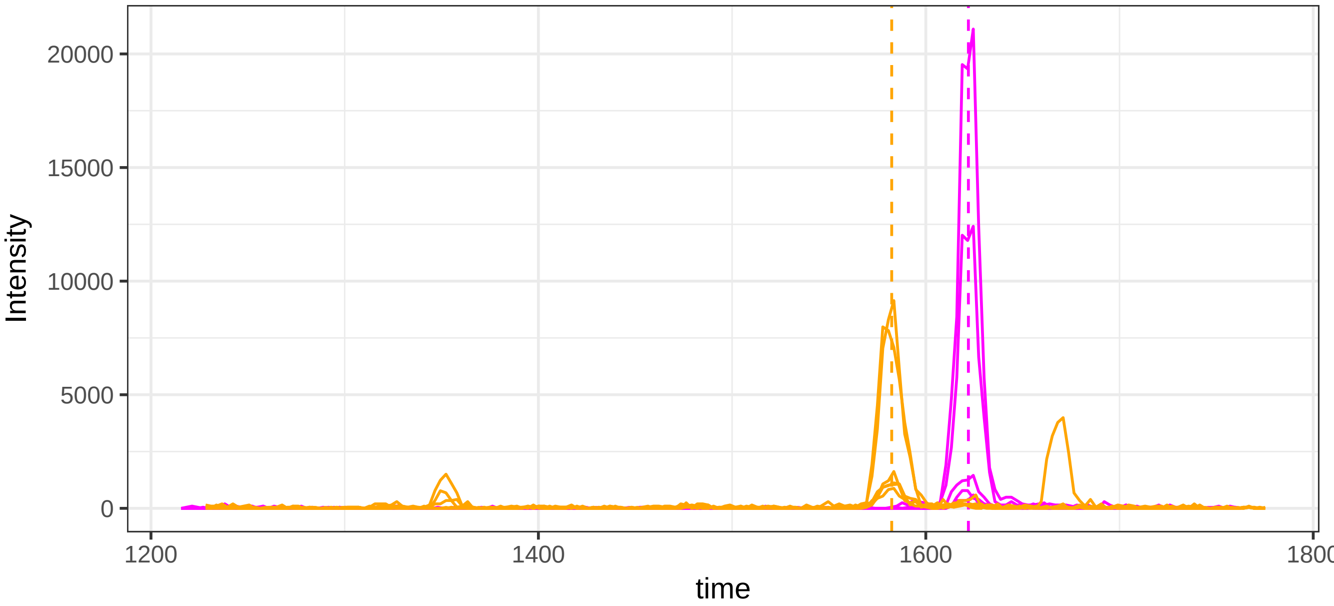


run23: 31717_AFLLTPR/2 (magenta)  AND 117443_WLPSSSPVTGYR/2

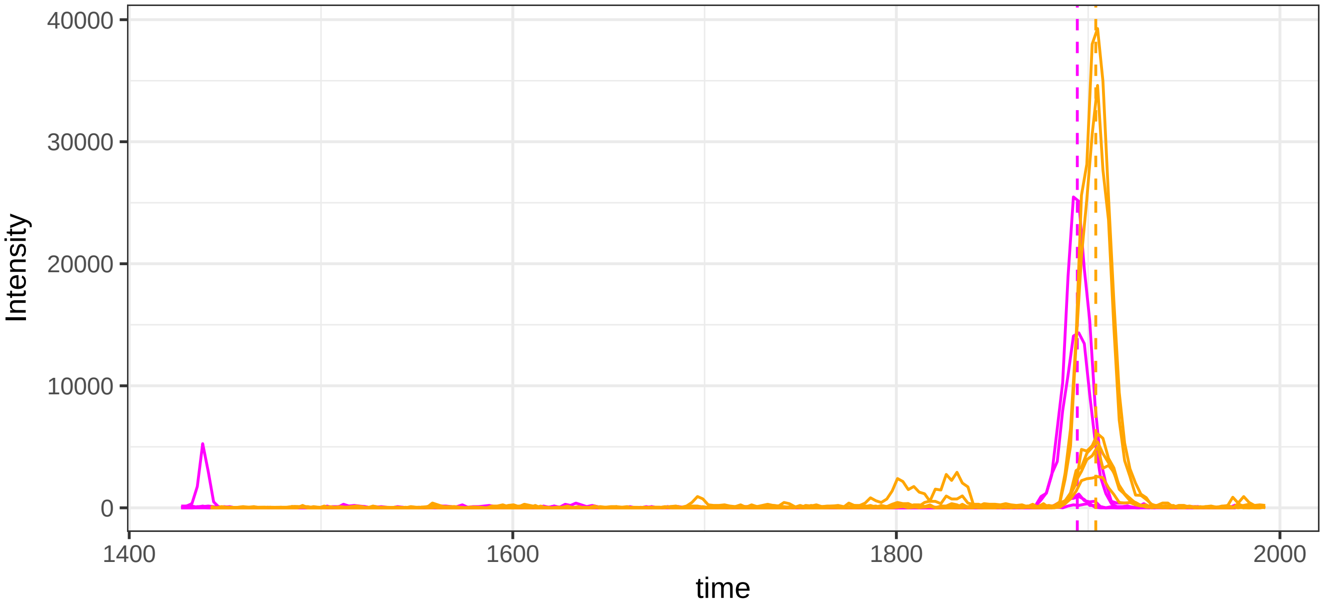

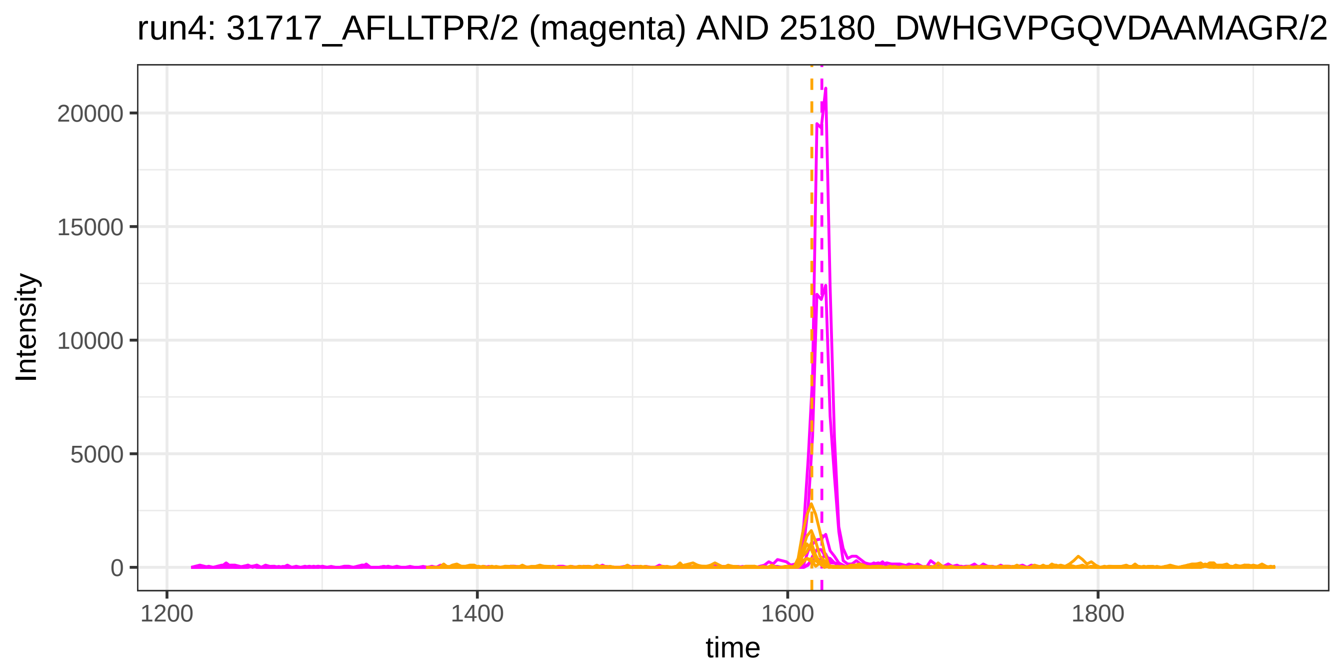

run4: 31717_AFLLTPR/2 (magenta) AND 25180_DWHGVPGQVDAAMAGR/2
Intensity
20000
15000
10000
5000
0
1200
1400
1600
1800
time


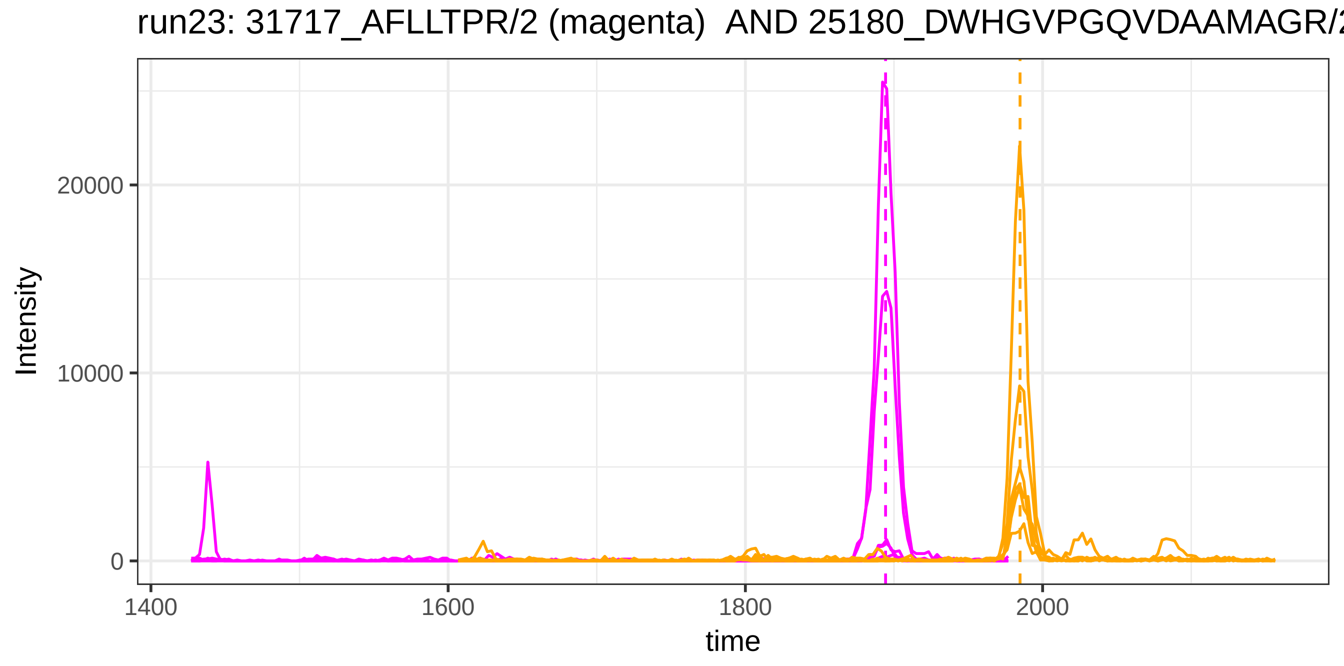

run23: 31717_AFLLTPR/2 (magenta) AND 25180_DWHGVPGQVDAAMAGR/2
Intensity
20000
10000
0
1400
1600
1800
2000
time

run4: 31717_AFLLTPR/2 (magenta) AND 5357_YVLTQPPSVSVAPGETAR/2

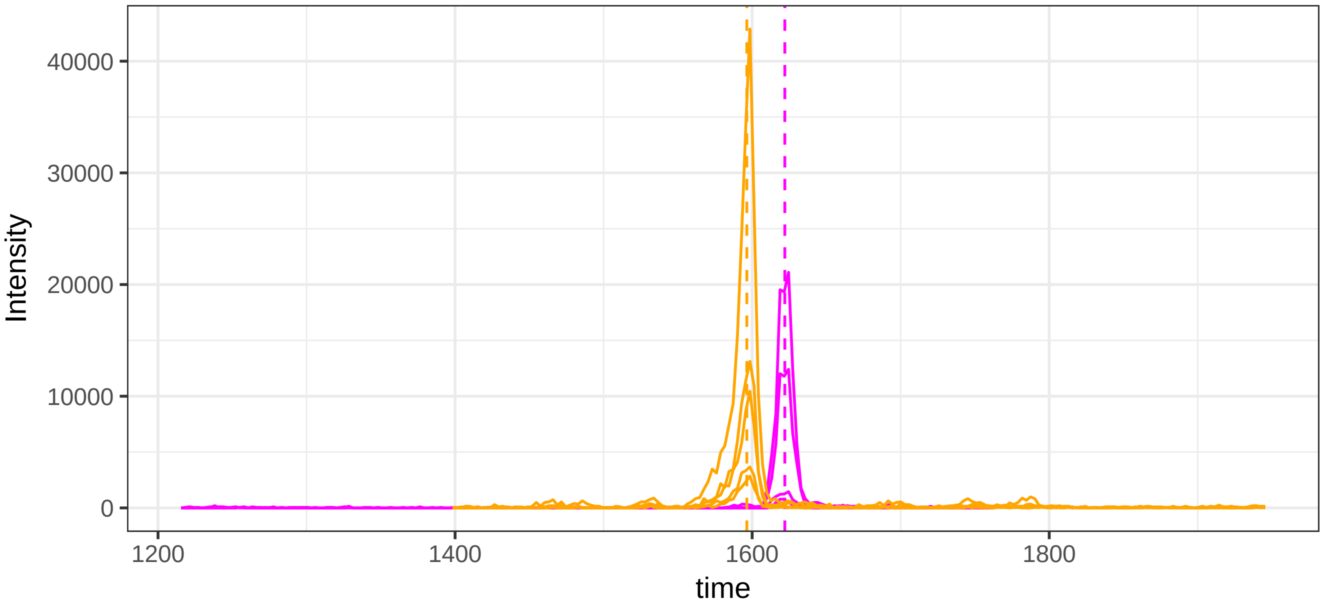


run23: 31717_AFLLTPR/2 (magenta)  AND 5357_YVLTQPPSVSVAPGETAR/2

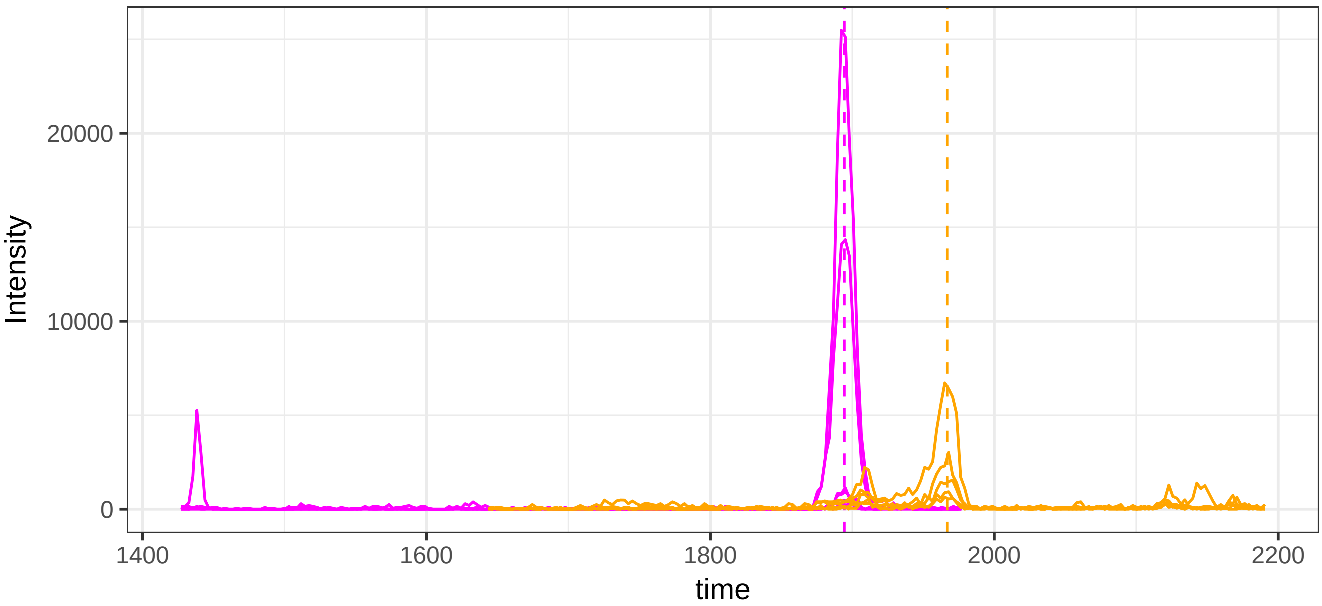

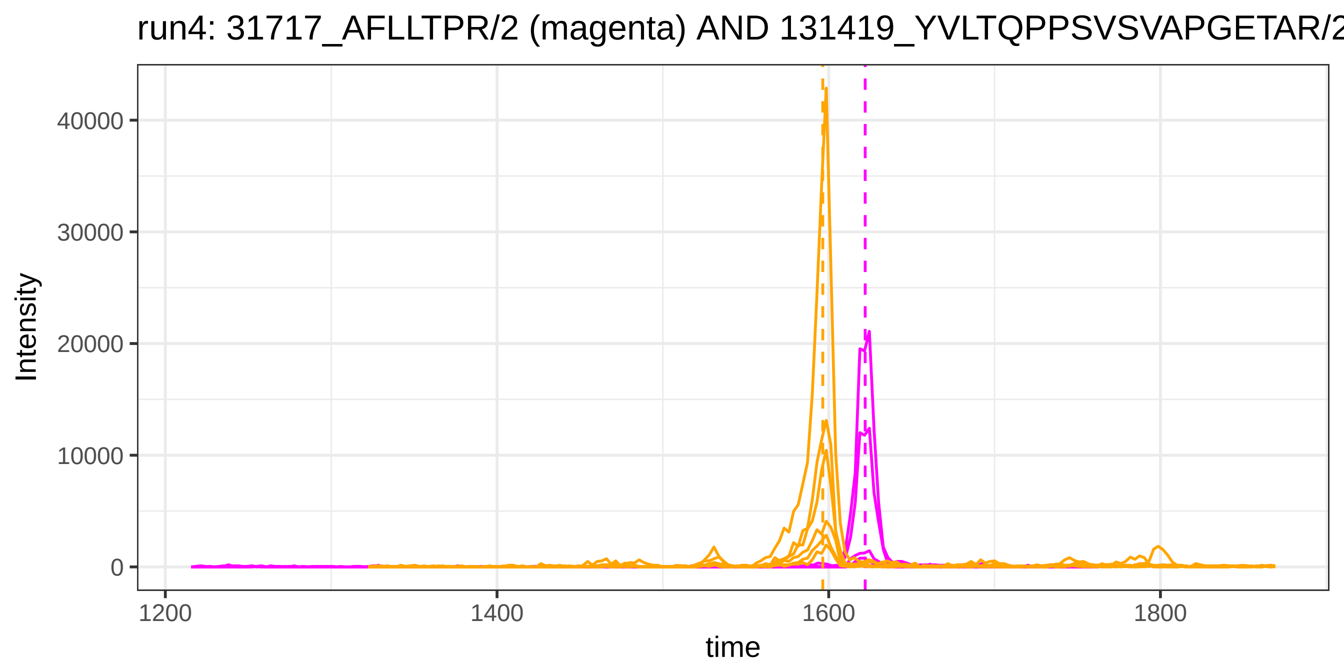
run4: 31717_AFLLTPR/2 (magenta) AND 131419_YVLTQPPSVSVAPGETAR/2
Intensity
40000
30000
20000
10000
0
1200
1400
1600
1800
time

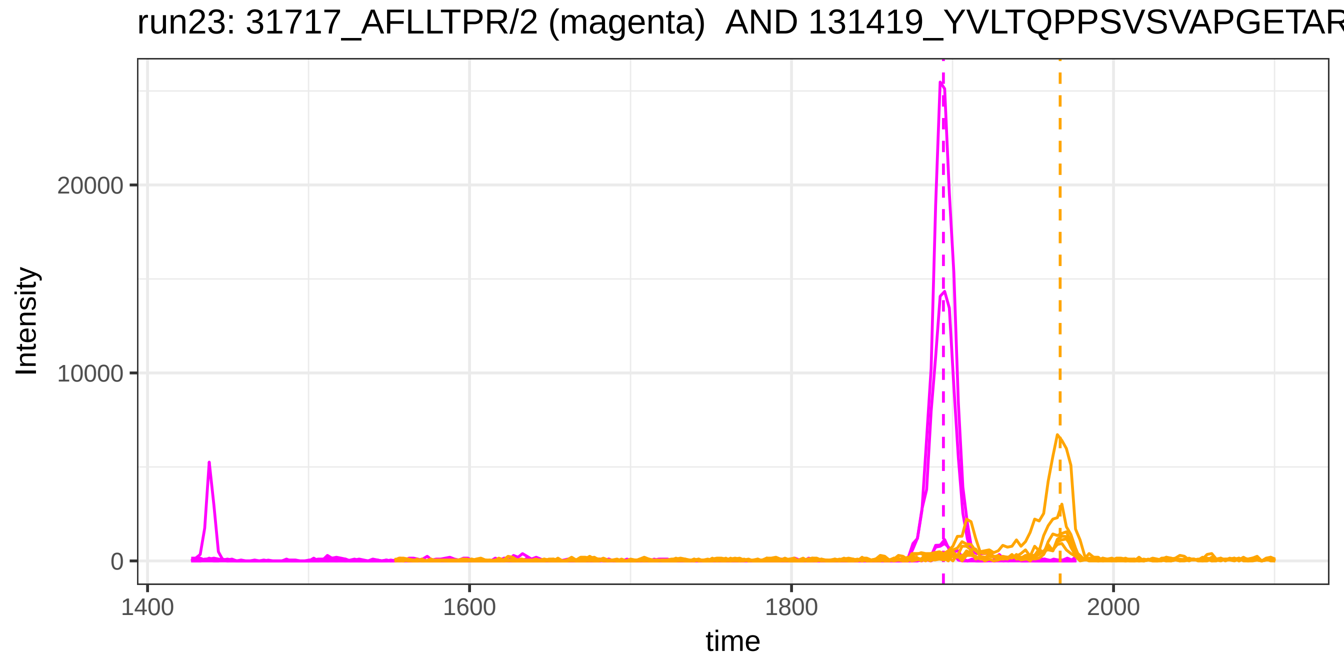
run23: 31717_AFLLTPR/2 (magenta) AND 131419_YVLTQPPSVSVAPGETAR
Intensity
20000
10000
0
1400
1600
1800
2000
time

run4: 116326_EVTEFAK/2 (magenta) AND 115879_DDNPNLPR/2

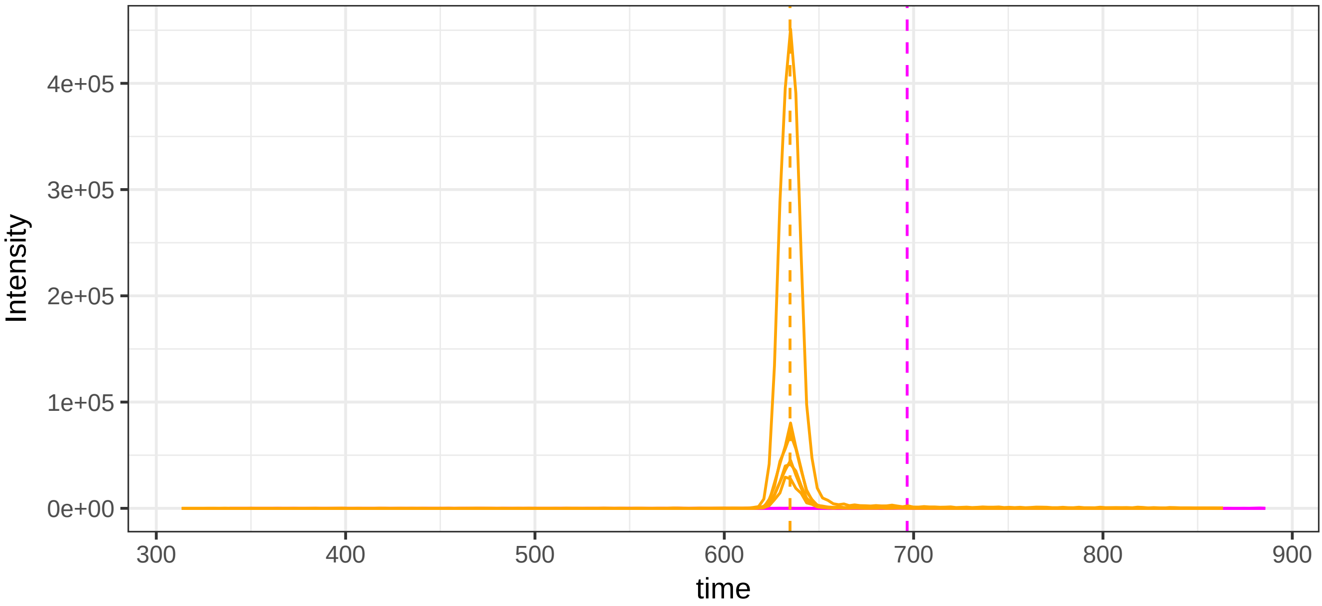


run23: 116326_EVTEFAK/2 (magenta)  AND 115879_DDNPNLPR/2

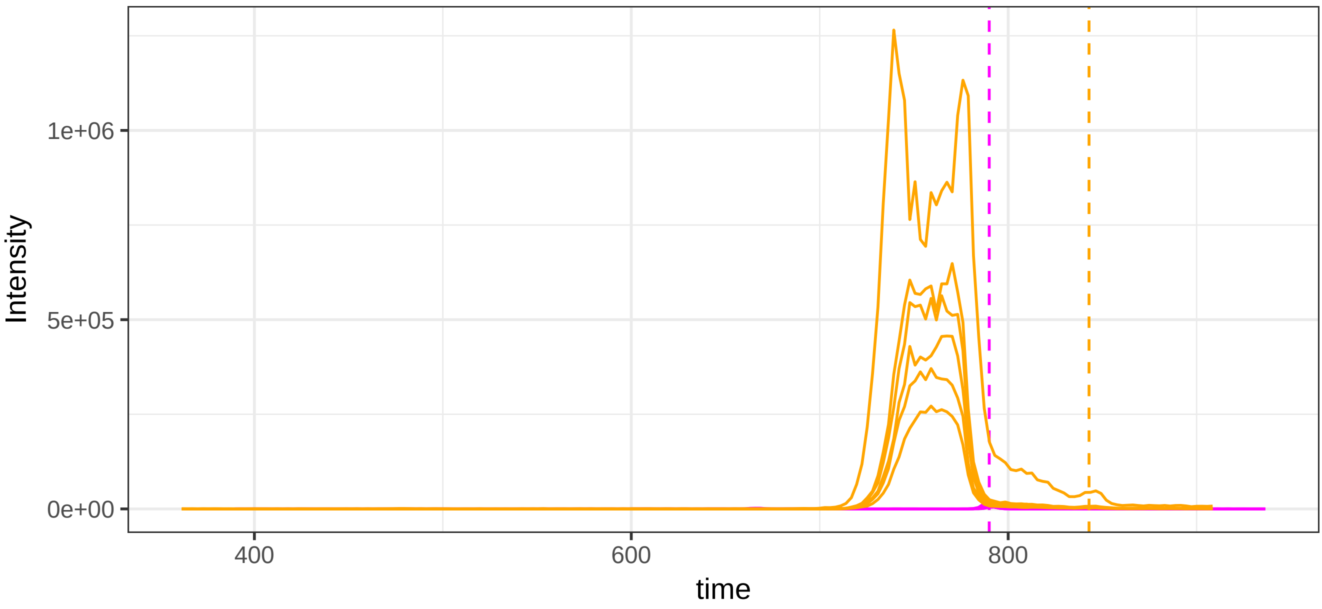

run4: 116326_EVTEFAK/2 (magenta) AND 49292_QELSEAEQATR/2

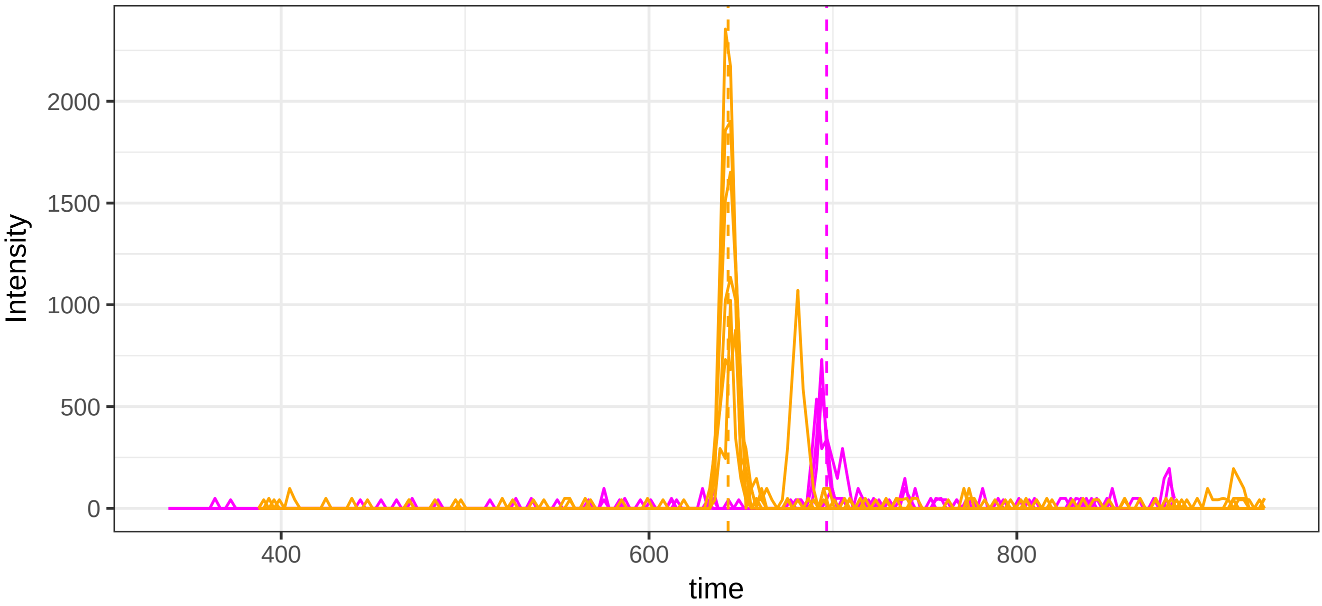


run23: 116326_EVTEFAK/2 (magenta)  AND 49292_QELSEAEQATR/2

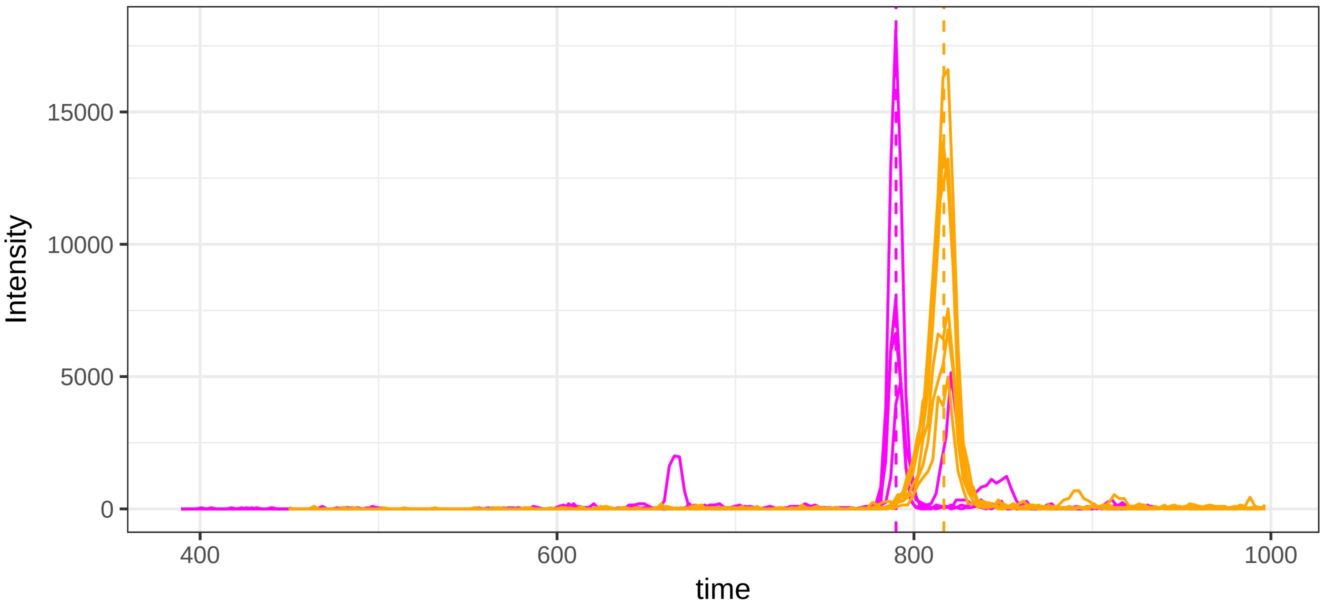

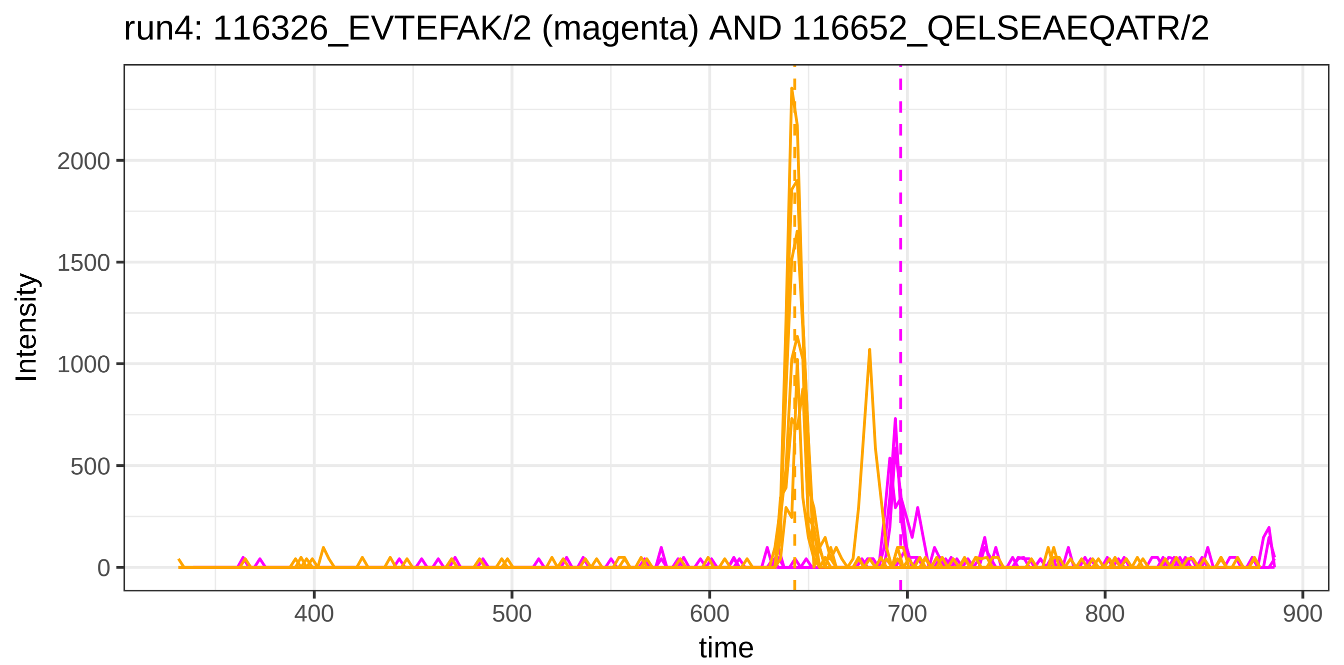
run4: 116326_EVTEFAK/2 (magenta) AND 116652_QELSEAEQATR/2
Intensity
2000
1500
1000
500
0
400
500
600
700
800
900
time

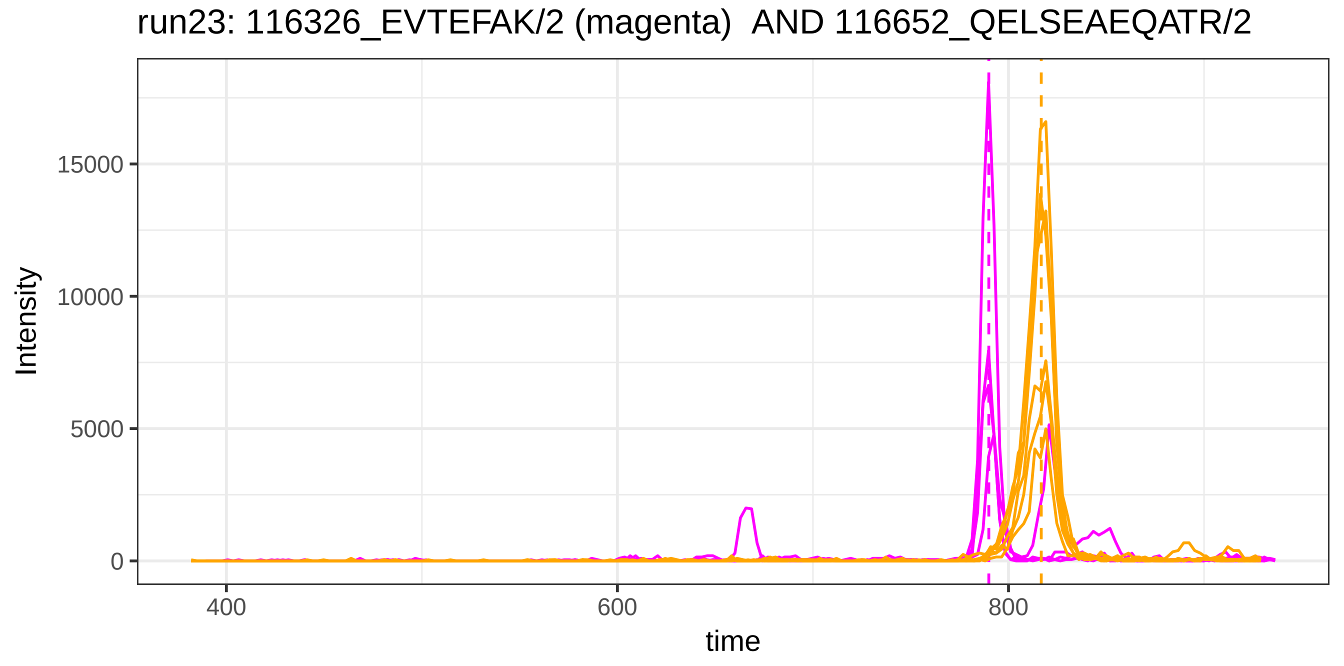
run23: 116326_EVTEFAK/2 (magenta) AND 116652_QELSEAEQATR/2
Intensity
15000
10000
5000
0
400
600
800
time

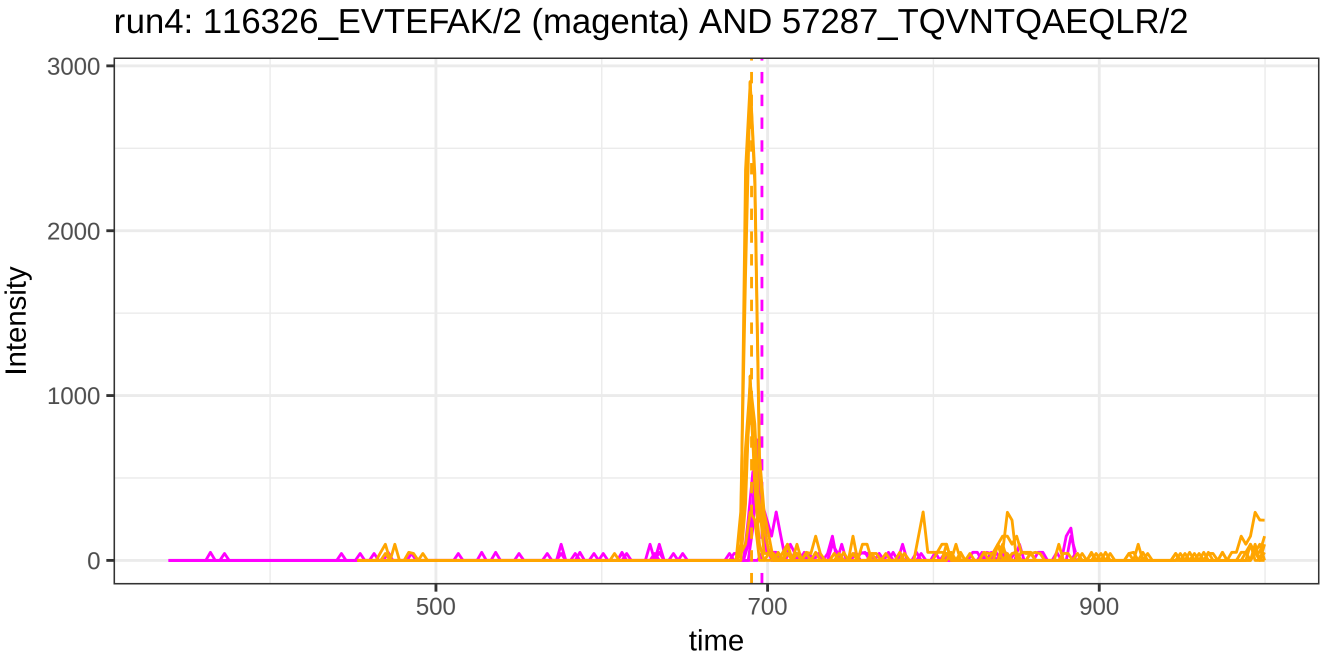
run4: 116326_EVTEFAK/2 (magenta) AND 57287_TQVNTQAEQLR/2
Intensity
3000
2000
1000
0
500
700
900
time

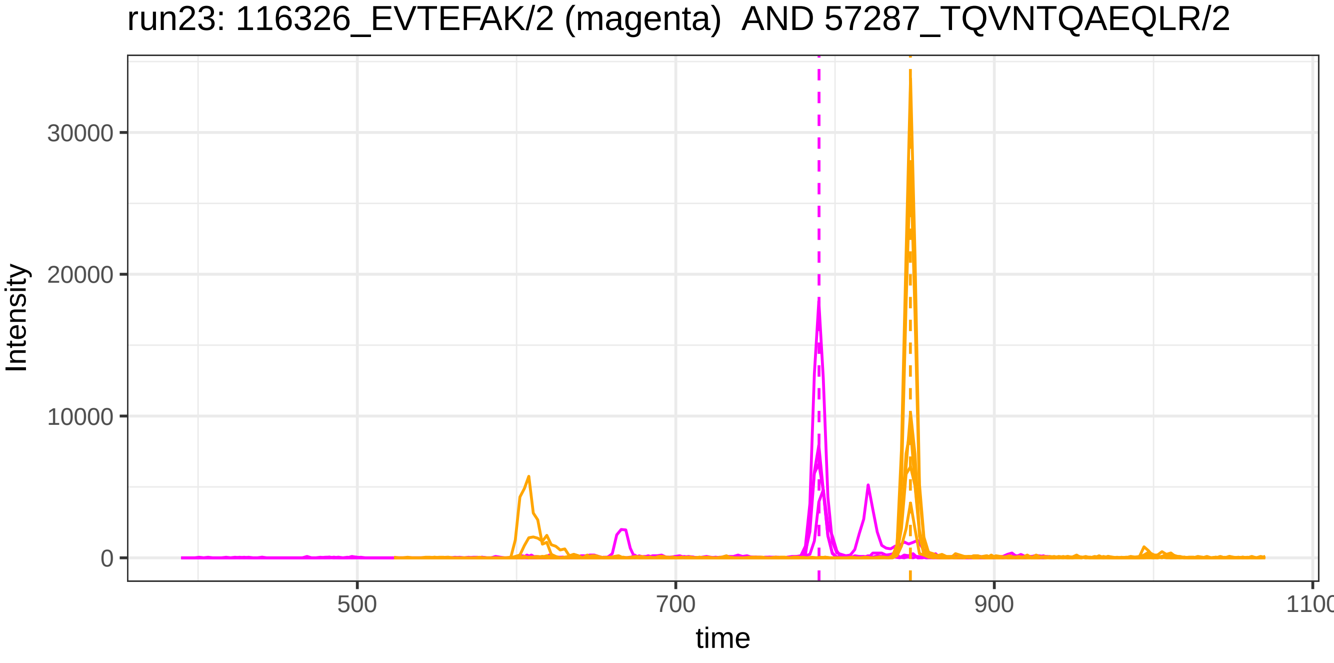
run23: 116326_EVTEFAK/2 (magenta) AND 57287_TQVNTQAEQLR/2
Intensity
30000
20000
10000
0
500
700
900
1100
time

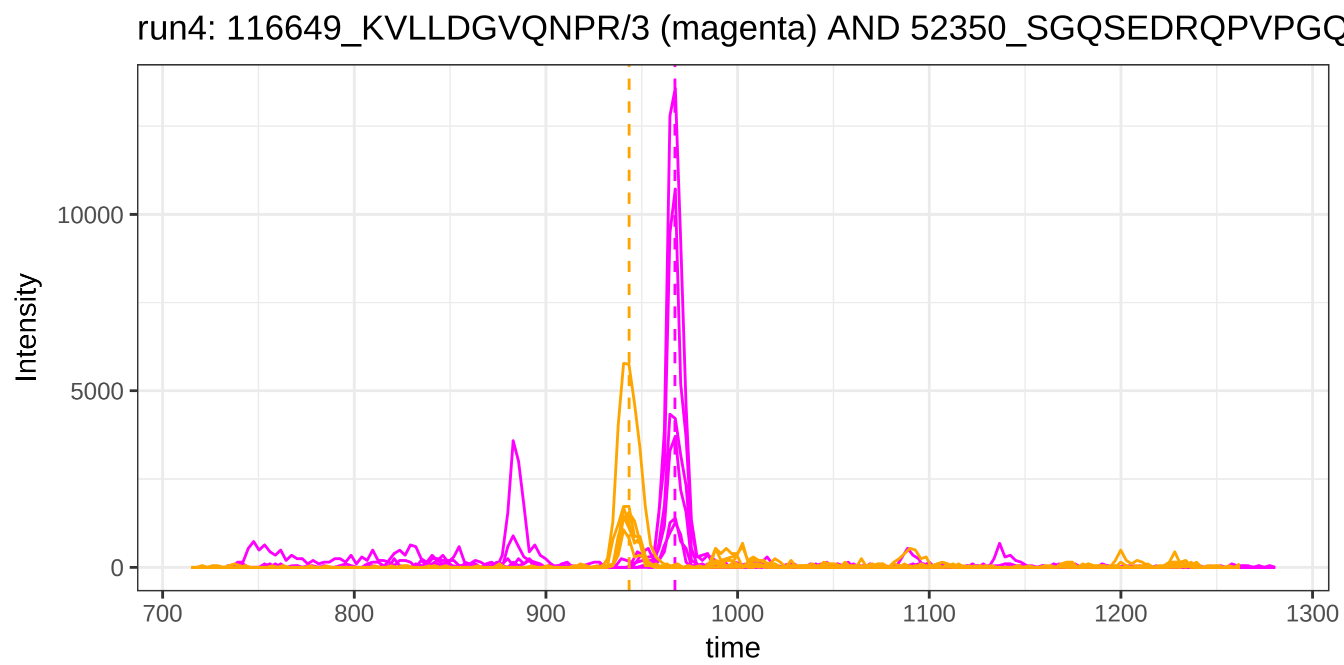
run4: 116649_KVLLDGVQNPR/3 (magenta) AND 52350_SGQSEDRQPVPGQ
Intensity
10000
5000
0
700
800
900
1000
1100
1200
1300
time

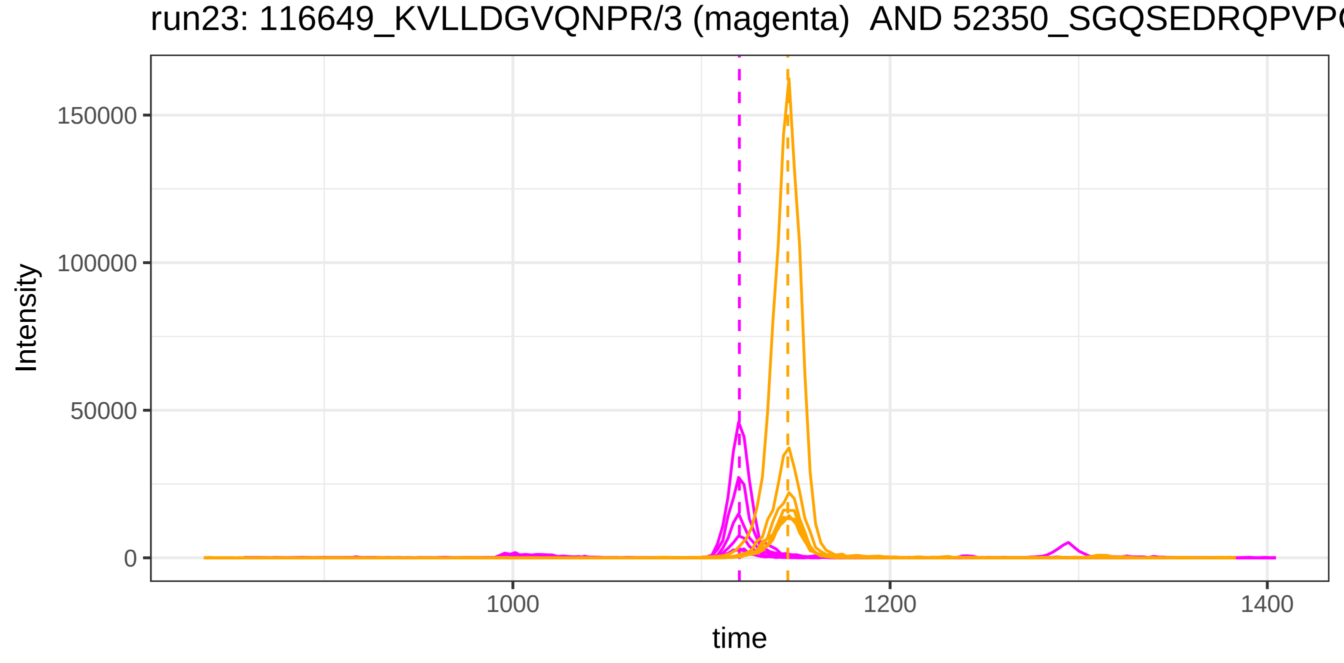
run23: 116649_KVLLDGVQNPR/3 (magenta) AND 52350_SGQSEDRQPVP
Intensity
150000
100000
50000
0
1000
1200
1400
time

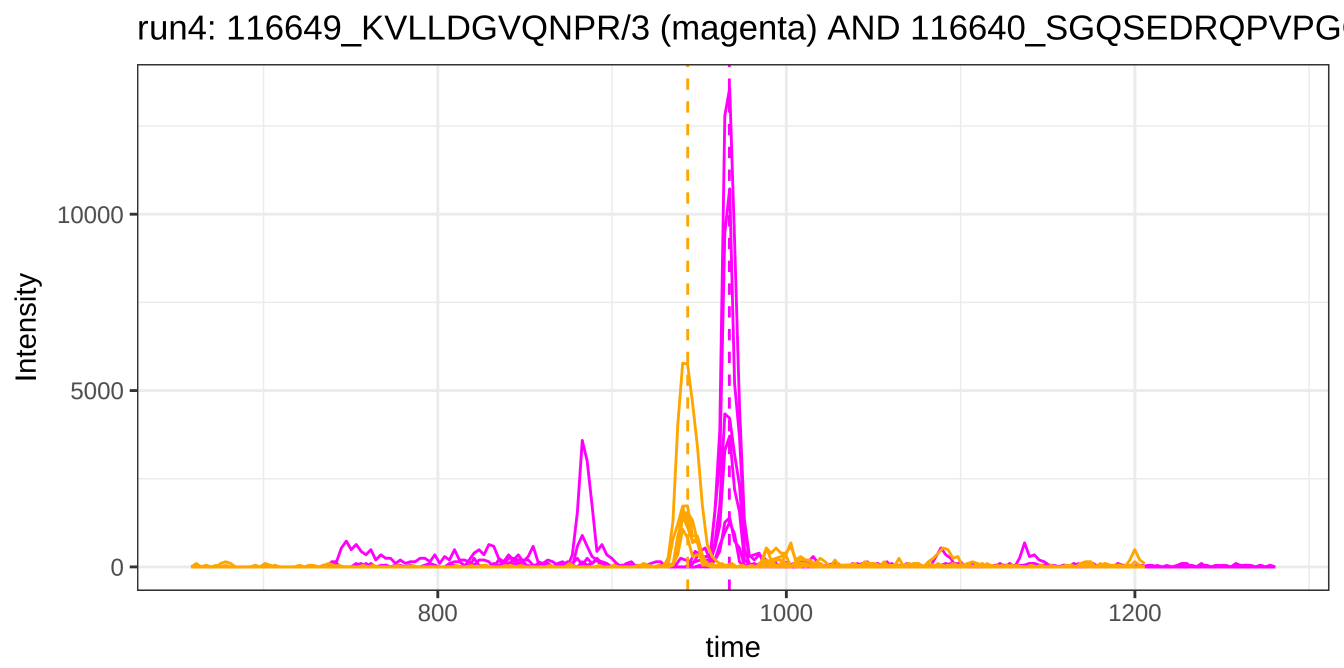
run4: 116649_KVLLDGVQNPR/3 (magenta) AND 116640_SGQSEDRQPVPG
Intensity
10000
5000
0
800
1000
1200
time

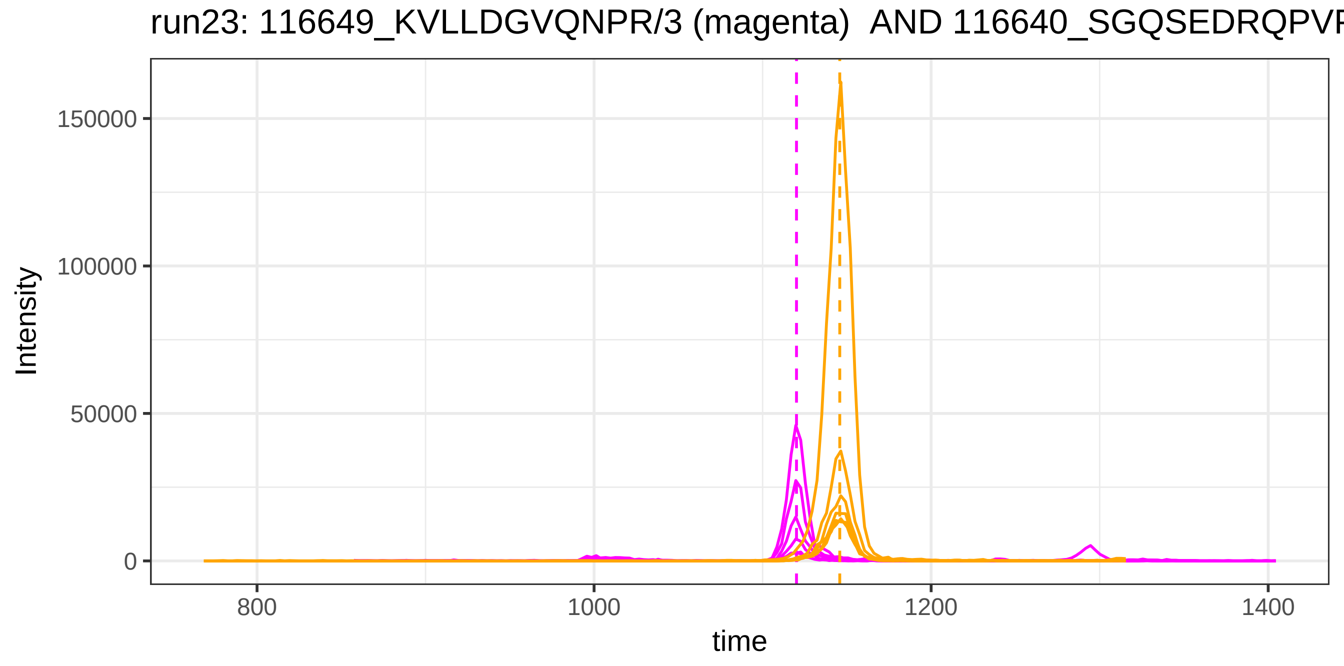
run23: 116649_KVLLDGVQNPR/3 (magenta) AND 116640_SGQSEDRQPVP
Intensity
150000
100000
50000
0
800
1000
1200
1400
time

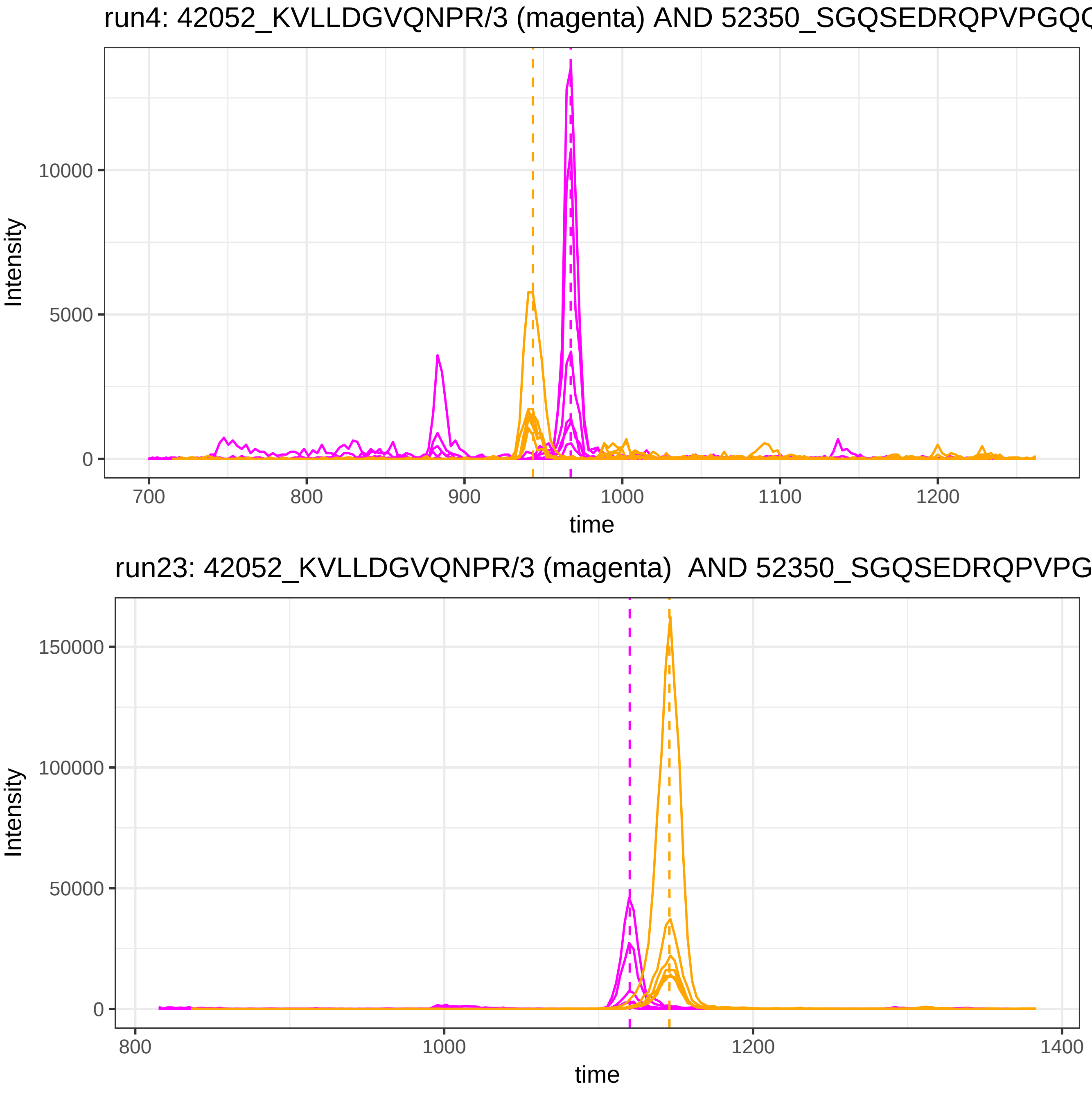

run4: 42052_KVLLDGVQNPR/3 (magenta) AND 52350_SGQSEDRQPVPGQ
Intensity
10000
5000
0
700
800
900
1000
1100
1200
time
run23: 42052_KVLLDGVQNPR/3 (magenta) AND 52350_SGQSEDRQPVPG
Intensity
150000
100000
50000
0
800
1000
1200
1400
time

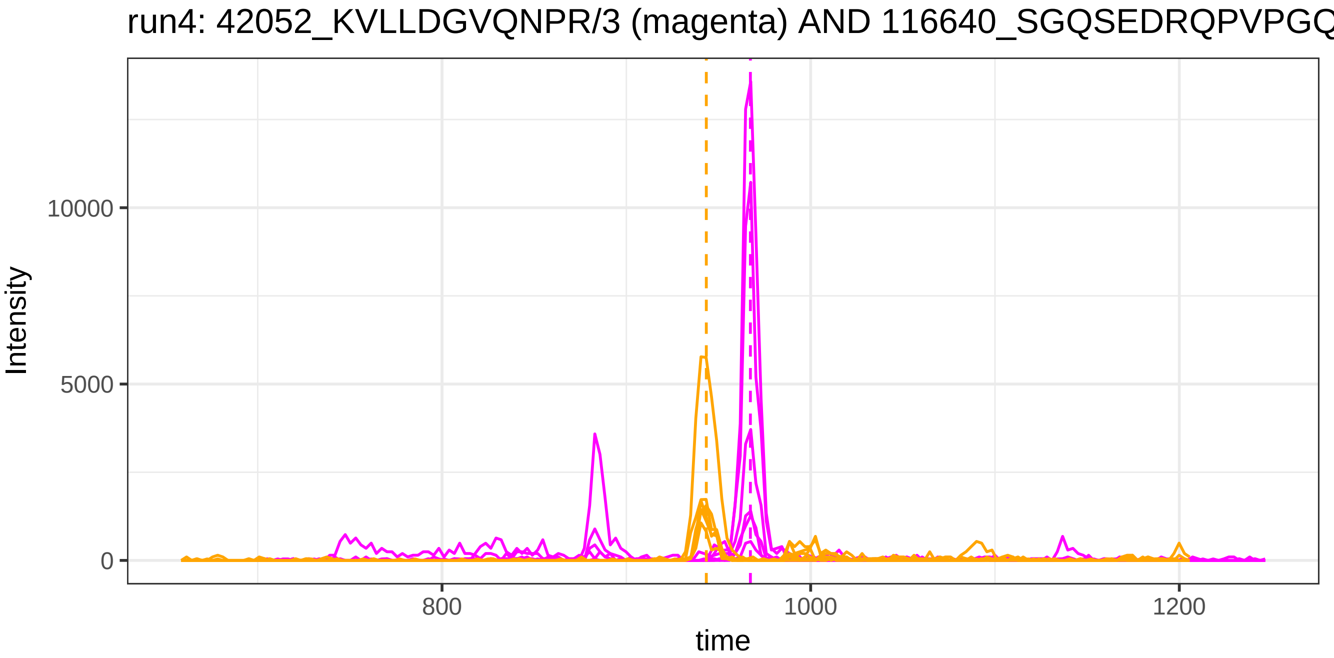
run4: 42052_KVLLDGVQNPR/3 (magenta) AND 116640_SGQSEDRQPVPGQ
Intensity
10000
5000
0
800
1000
1200
time

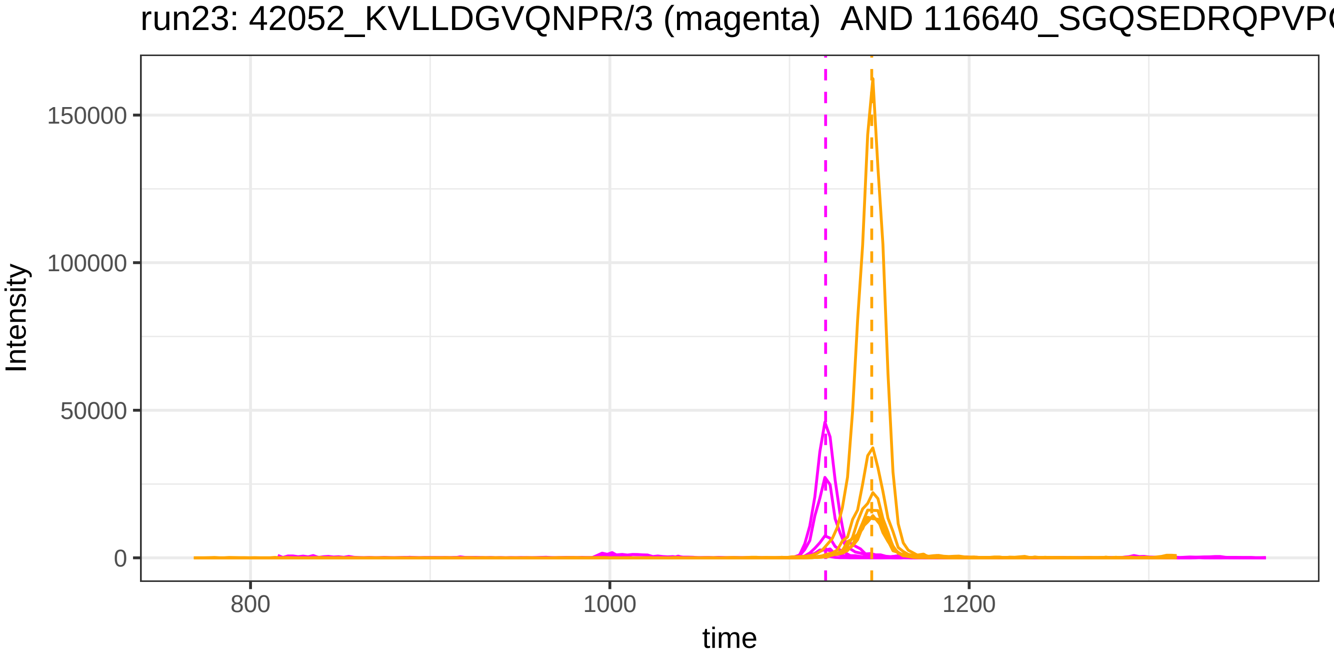
run23: 42052_KVLLDGVQNPR/3 (magenta) AND 116640_SGQSEDRQPVP
Intensity
150000
100000
50000
0
800
1000
1200
time

run4: 116796_WLNEQR/2 (magenta) AND 115879_DDNPNLPR/2

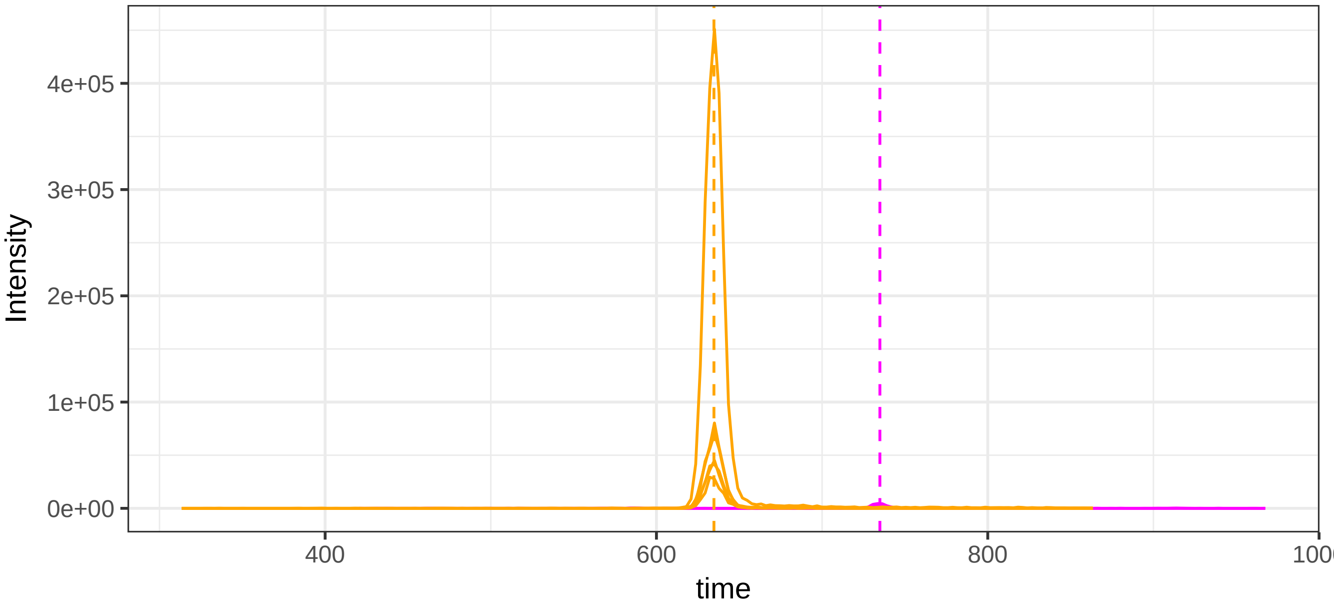


run23: 116796_WLNEQR/2 (magenta)  AND 115879_DDNPNLPR/2

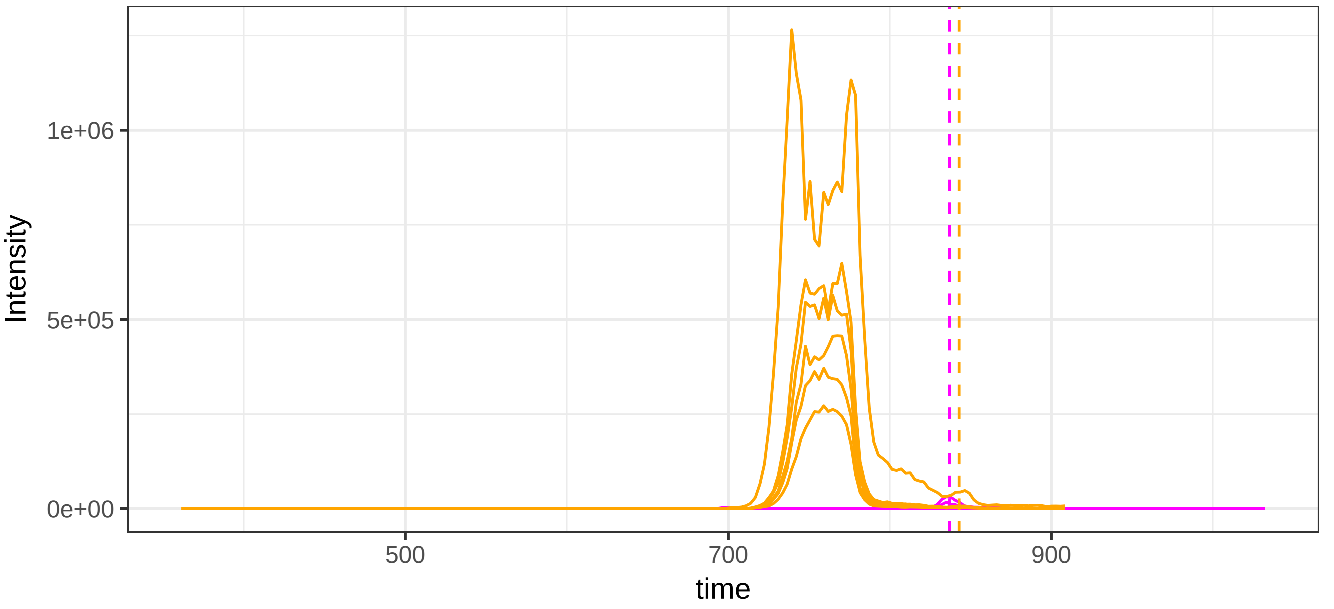

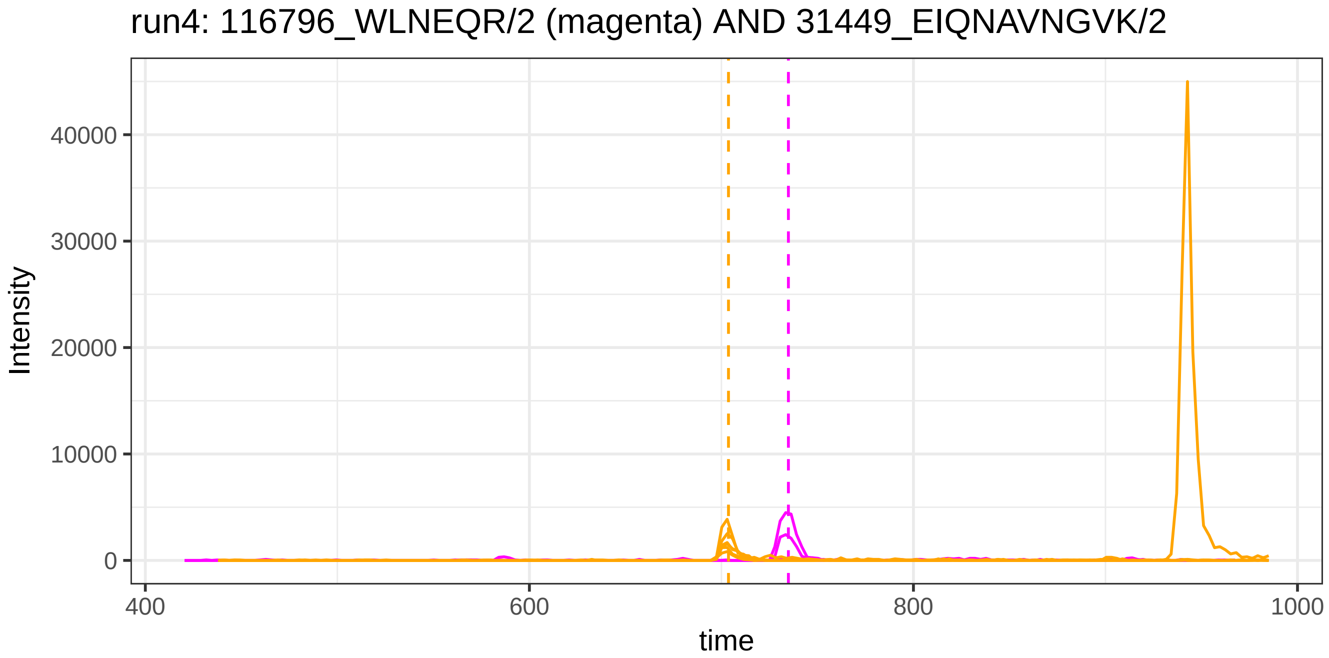
run4: 116796_WLNEQR/2 (magenta) AND 31449_EIQNAVNGVK/2
Intensity
0
10000
20000
30000
40000
400
600
800
1000
time

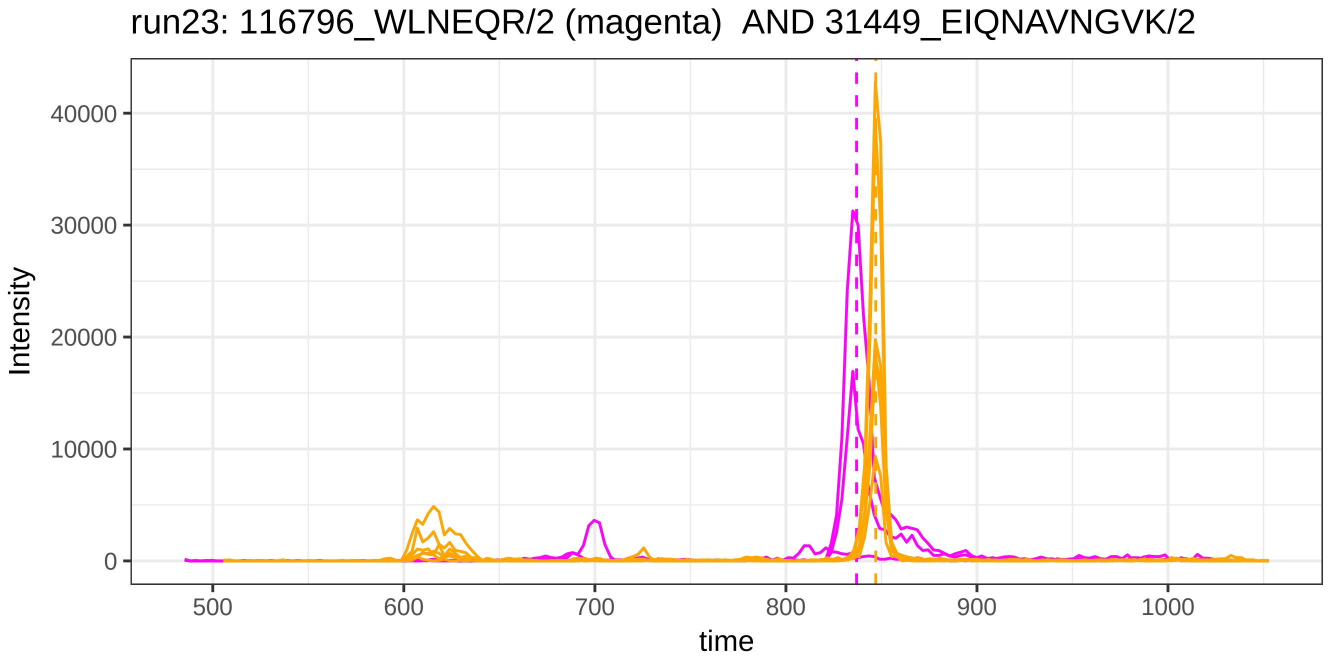
run23: 116796_WLNEQR/2 (magenta)  AND 31449_EIQNAVNGVK/2
Intensity
0
10000
20000
30000
40000
500
600
700
800
900
1000
time

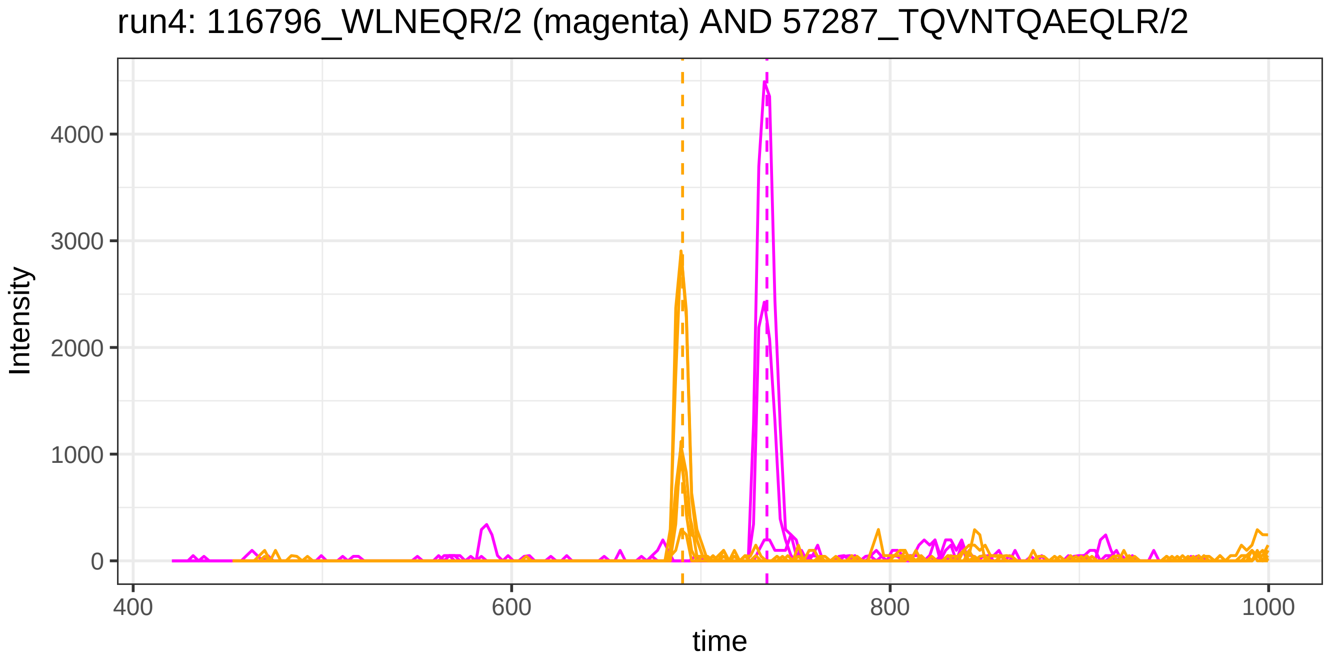
run4: 116796_WLNEQR/2 (magenta) AND 57287_TQVNTQAEQLR/2
Intensity
4000
3000
2000
1000
0
400
600
800
1000
time

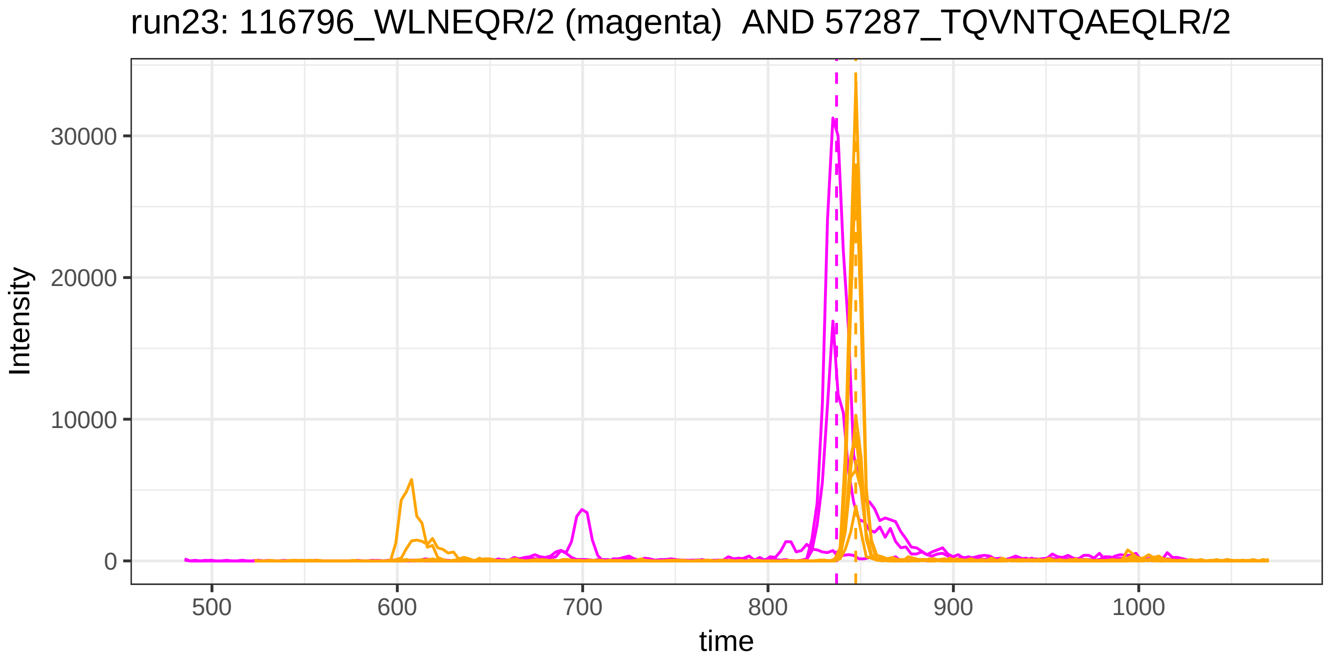
run23: 116796_WLNEQR/2 (magenta) AND 57287_TQVNTQAEQLR/2
Intensity
30000
20000
10000
0
500
600
700
800
900
1000
time

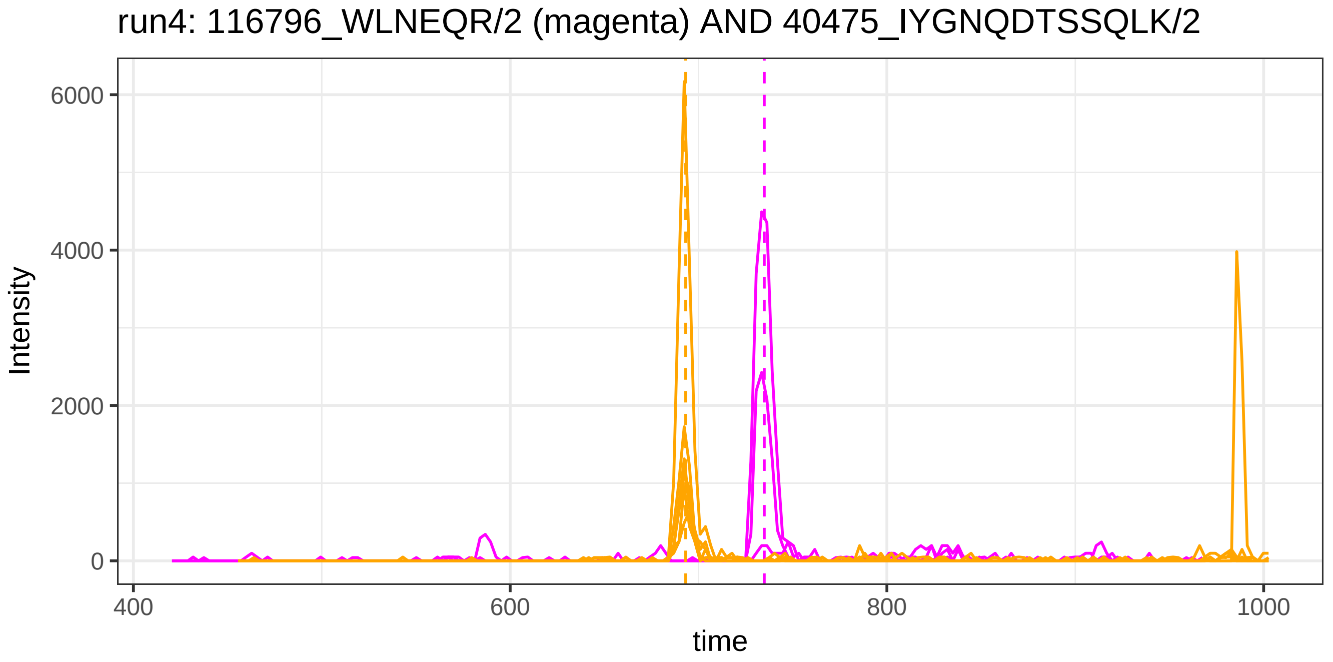
run4: 116796_WLNEQR/2 (magenta) AND 40475_IYGNQDTSSQLK/2
Intensity
6000
4000
2000
0
400
600
800
1000
time

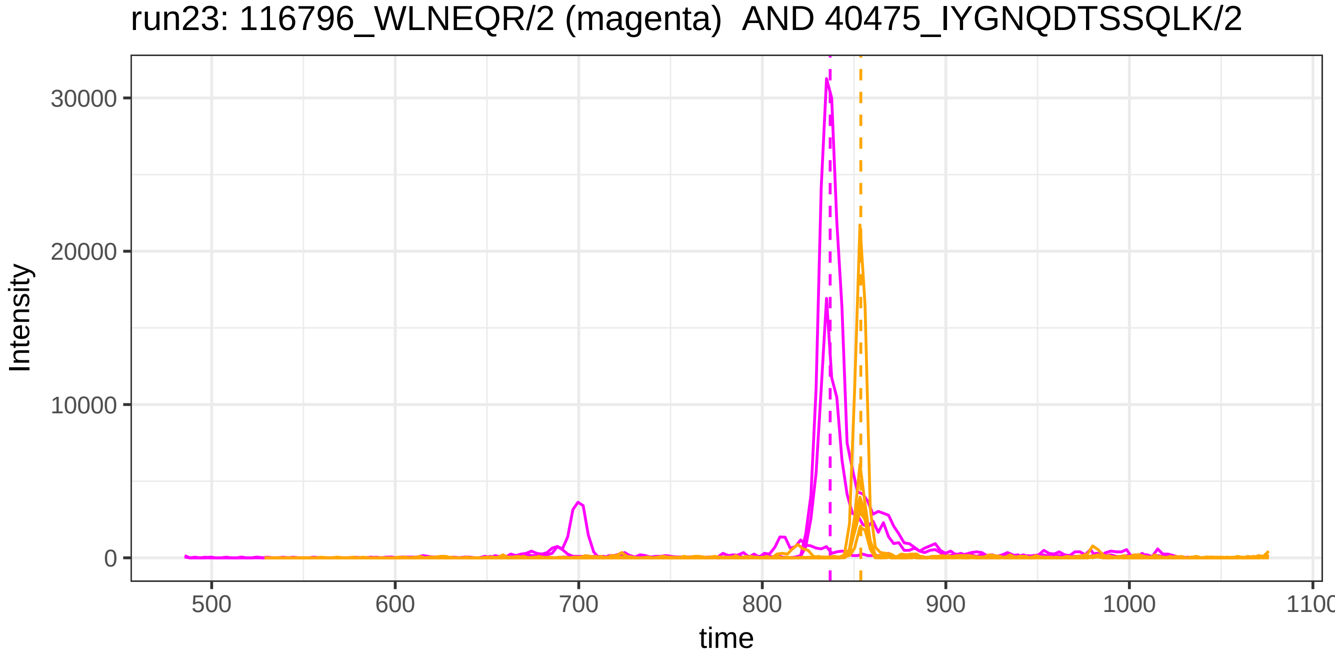
run23: 116796_WLNEQR/2 (magenta) AND 40475_IYGNQDTSSQLK/2
Intensity
30000
20000
10000
0
500
600
700
800
900
1000
1100
time

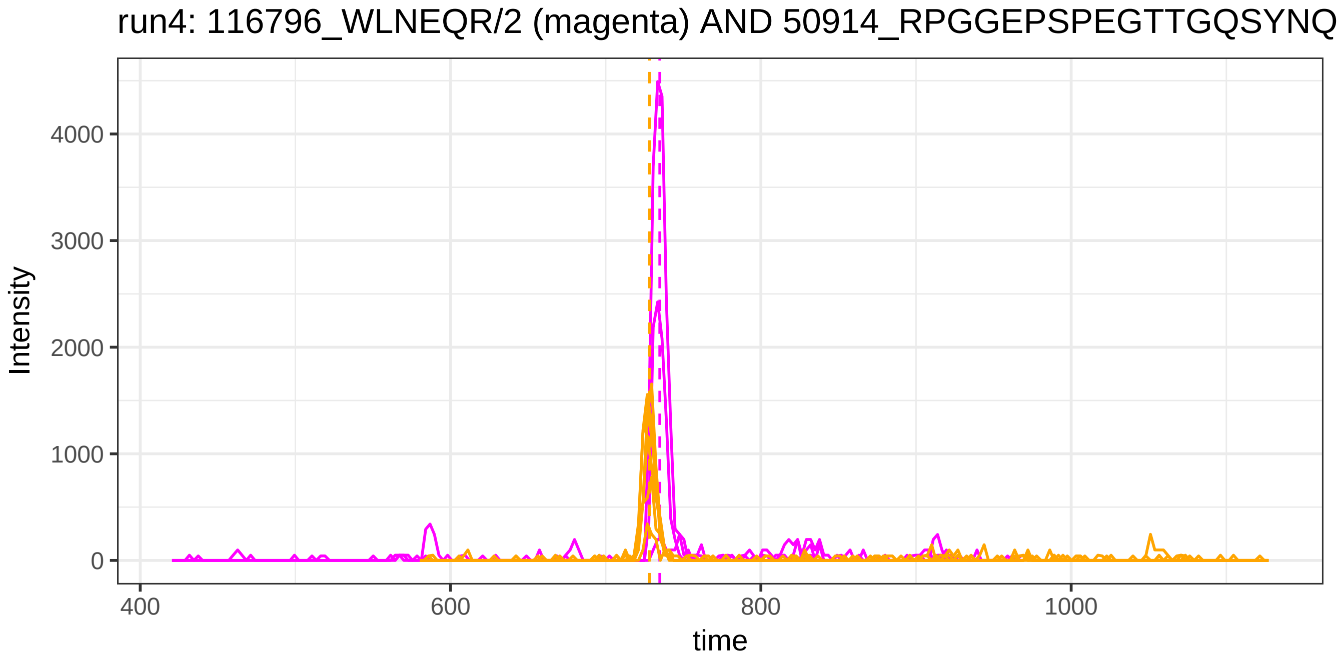
run4: 116796_WLNEQR/2 (magenta) AND 50914_RPGGEPSPEGTTGQSYNQ
Intensity
4000
3000
2000
1000
0
400
600
800
1000
time

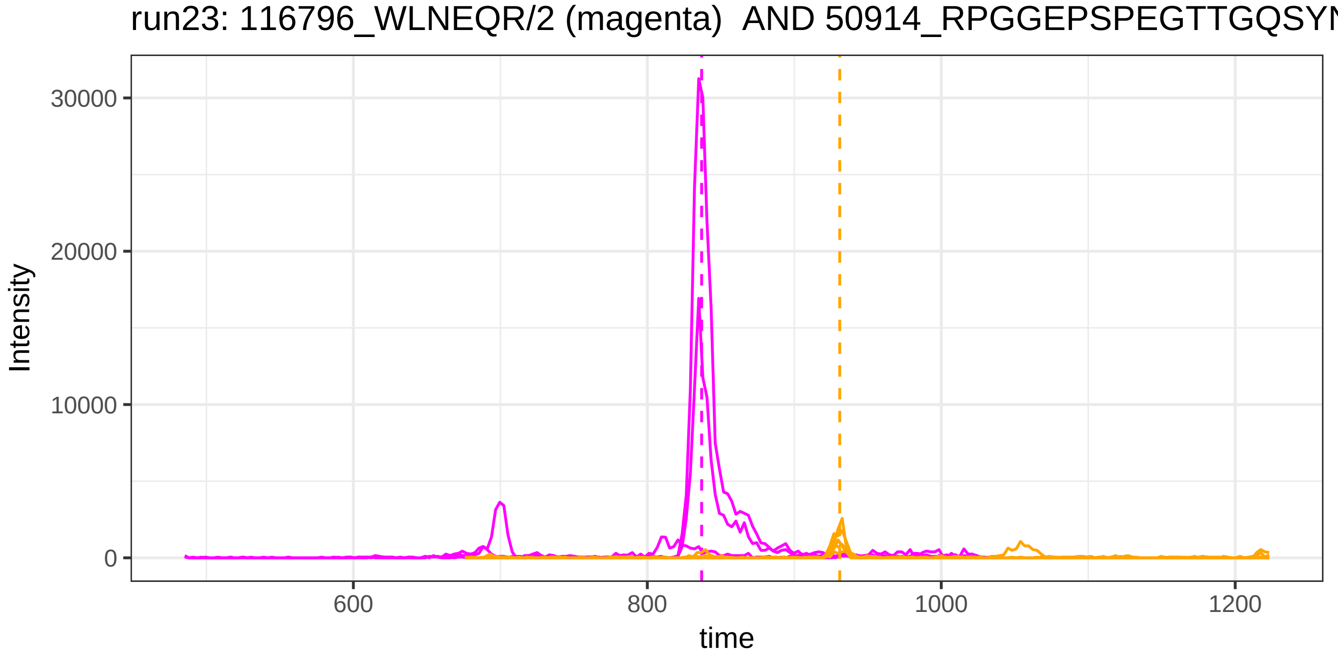
run23: 116796_WLNEQR/2 (magenta) AND 50914_RPGGEPSPEGTTGQSYN
Intensity
30000
20000
10000
0
600
800
1000
1200
time

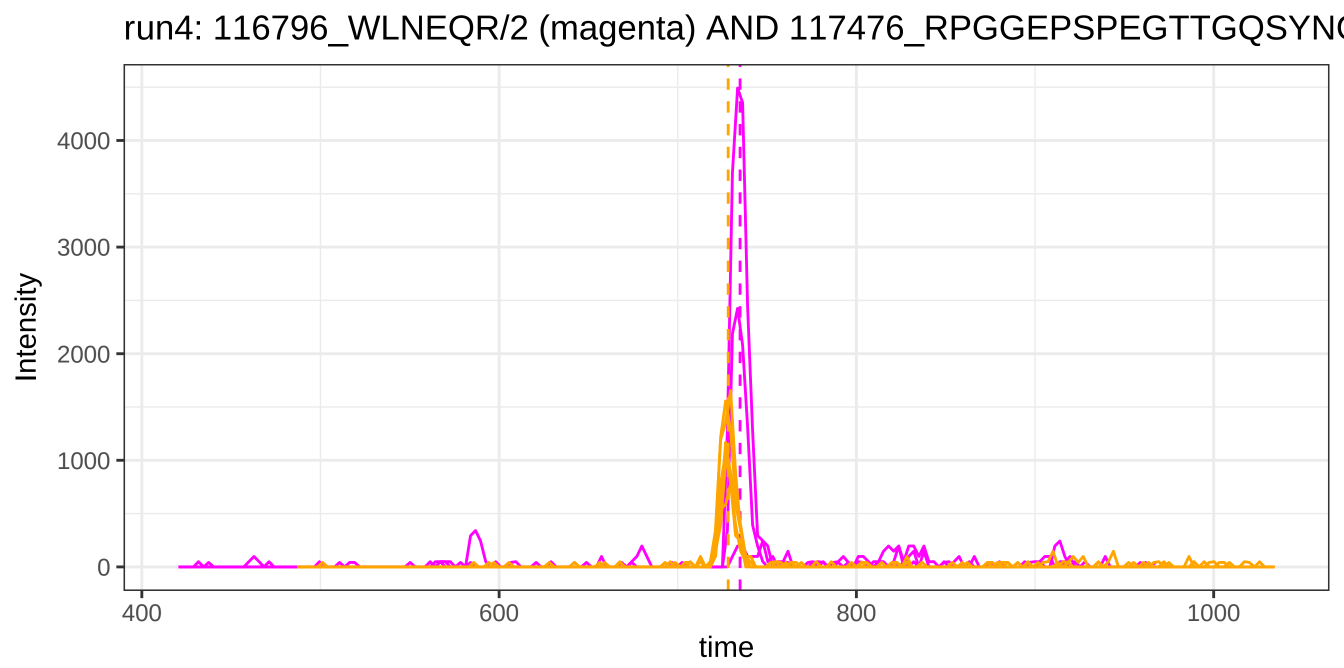
run4: 116796_WLNEQR/2 (magenta) AND 117476_RPGGEPSPEGTTGQSYNG
Intensity
0
1000
2000
3000
4000
time
400
600
800
1000

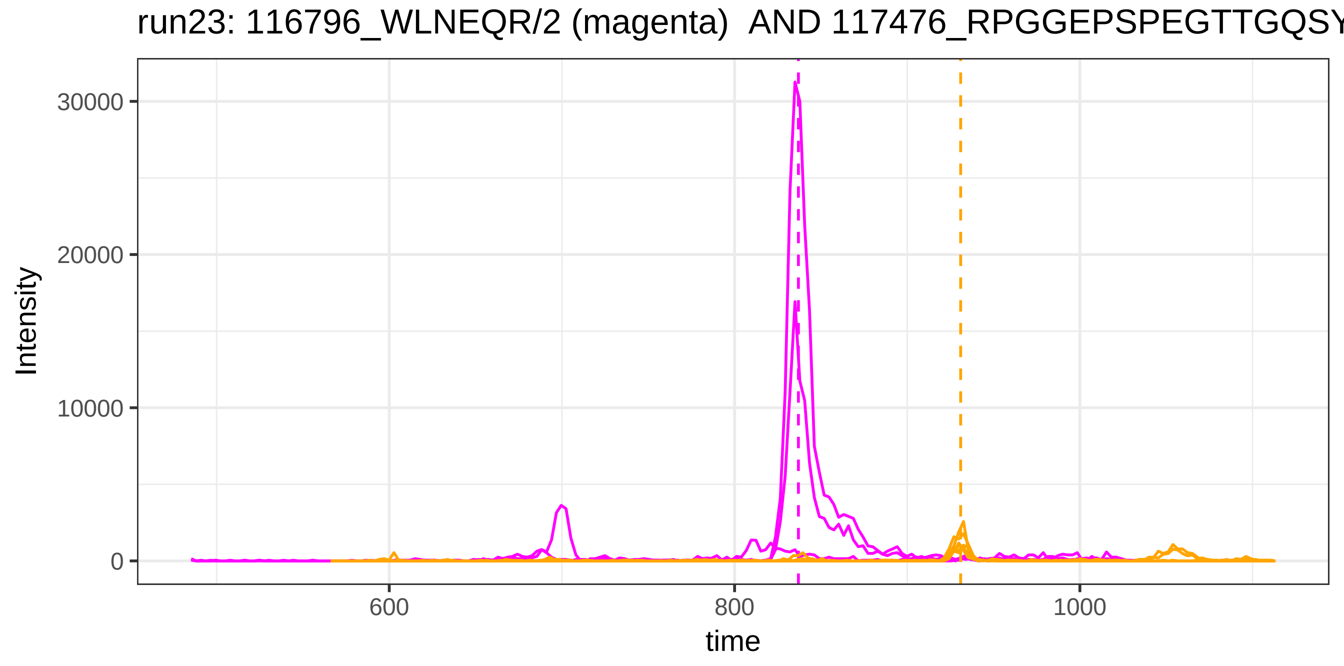
run23: 116796_WLNEQR/2 (magenta) AND 117476_RPGGEPSPEGTTGQSY
Intensity
0
10000
20000
30000
time
600
800
1000

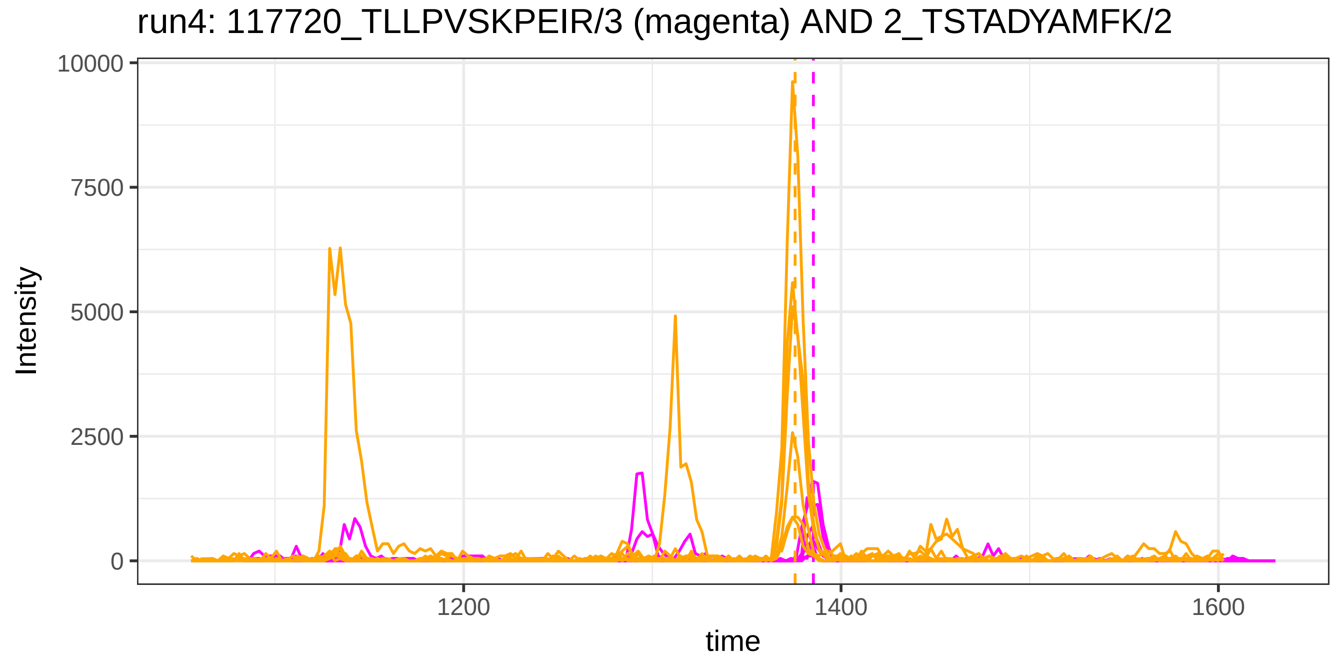

run4: 117720_TLLPVSKPEIR/3 (magenta) AND 2_TSTADYAMFK/2
Intensity
10000
7500
5000
2500
0
1200
1400
1600
time


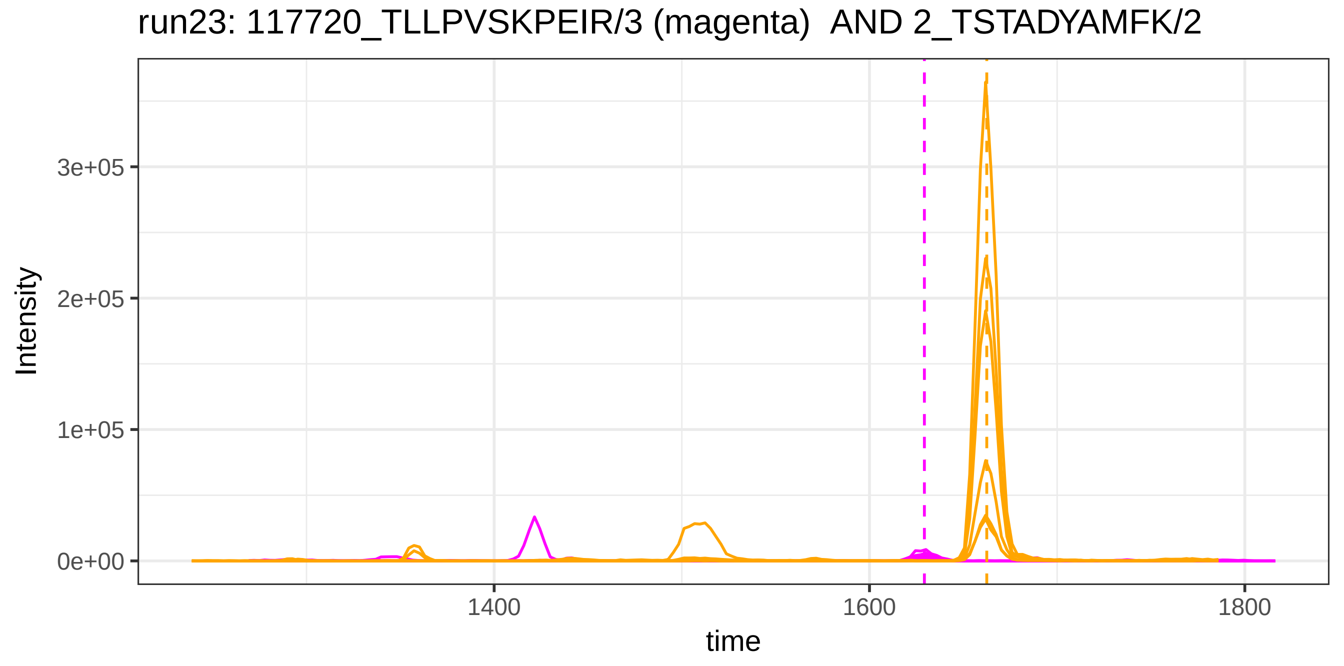

run23: 117720_TLLPVSKPEIR/3 (magenta) AND 2_TSTADYAMFK/2
Intensity
3e+05
2e+05
1e+05
0e+00
1400
1600
1800
time

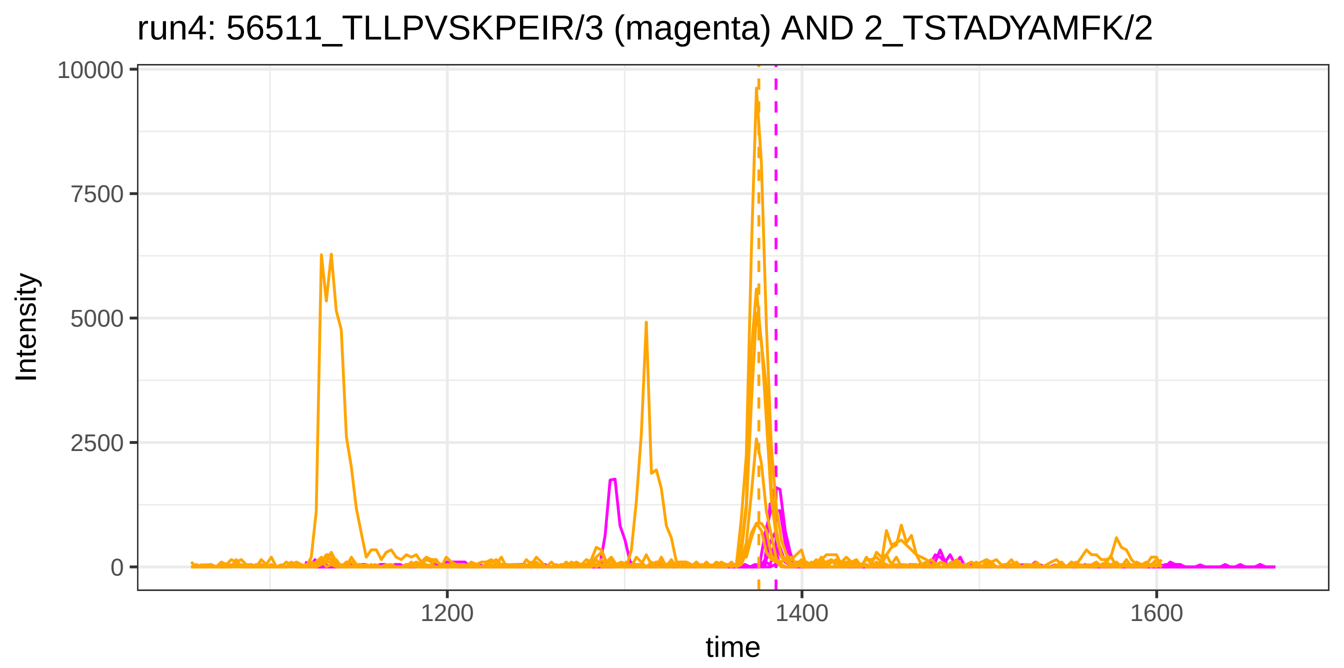
run4: 56511_TLLPVSKPEIR/3 (magenta) AND 2_TSTADYAMFK/2
Intensity
10000
7500
5000
2500
0
1200
1400
1600
time

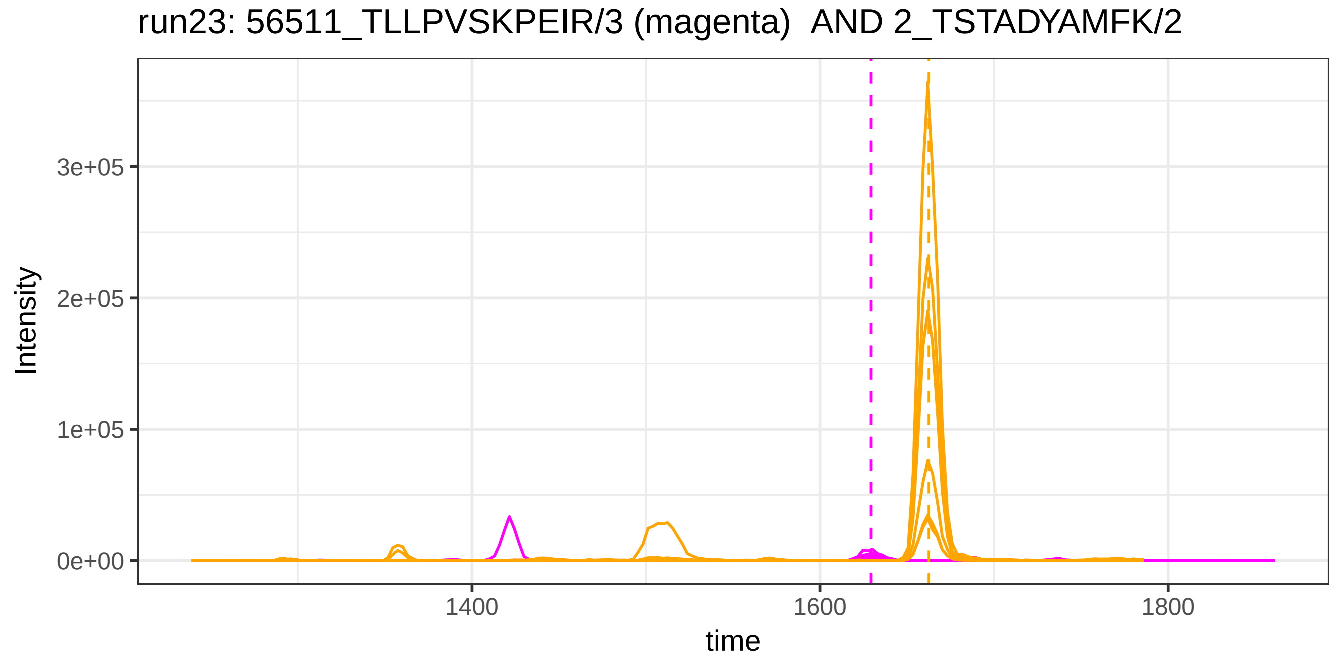
run23: 56511_TLLPVSKPEIR/3 (magenta) AND 2_TSTADYAMFK/2
Intensity
3e+05
2e+05
1e+05
0e+00
1400
1600
1800
time

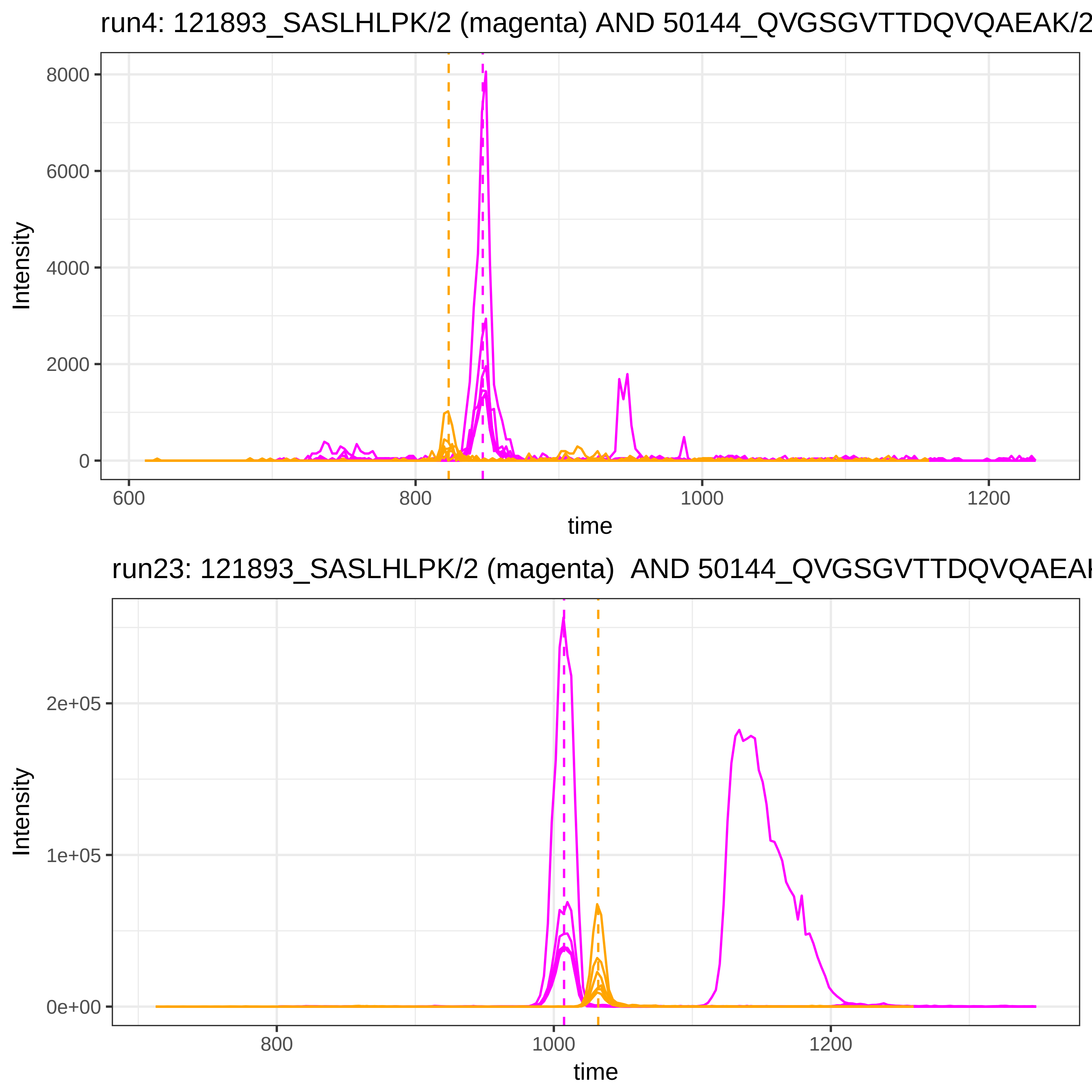
run4: 121893_SASLHLPK/2 (magenta) AND 50144_QVGSGVTTDQVQAEAK/2
Intensity
8000
6000
4000
2000
0
600
800
1000
1200
time
run23: 121893_SASLHLPK/2 (magenta) AND 50144_QVGSGVTTDQVQAEAK
Intensity
2e+05
1e+05
0e+00
800
1000
1200
time

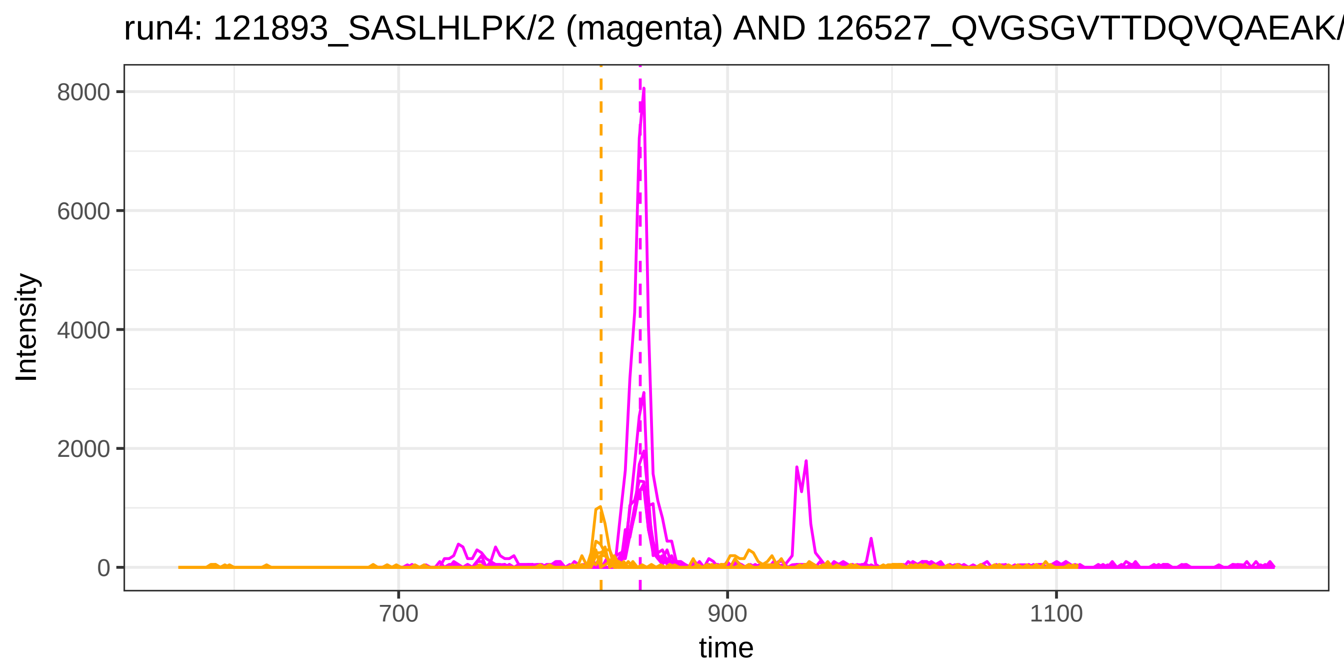
run4: 121893_SASLHLPK/2 (magenta) AND 126527_QVGSGVTTDQVQAEAK/
Intensity
8000
6000
4000
2000
0
700
900
1100
time

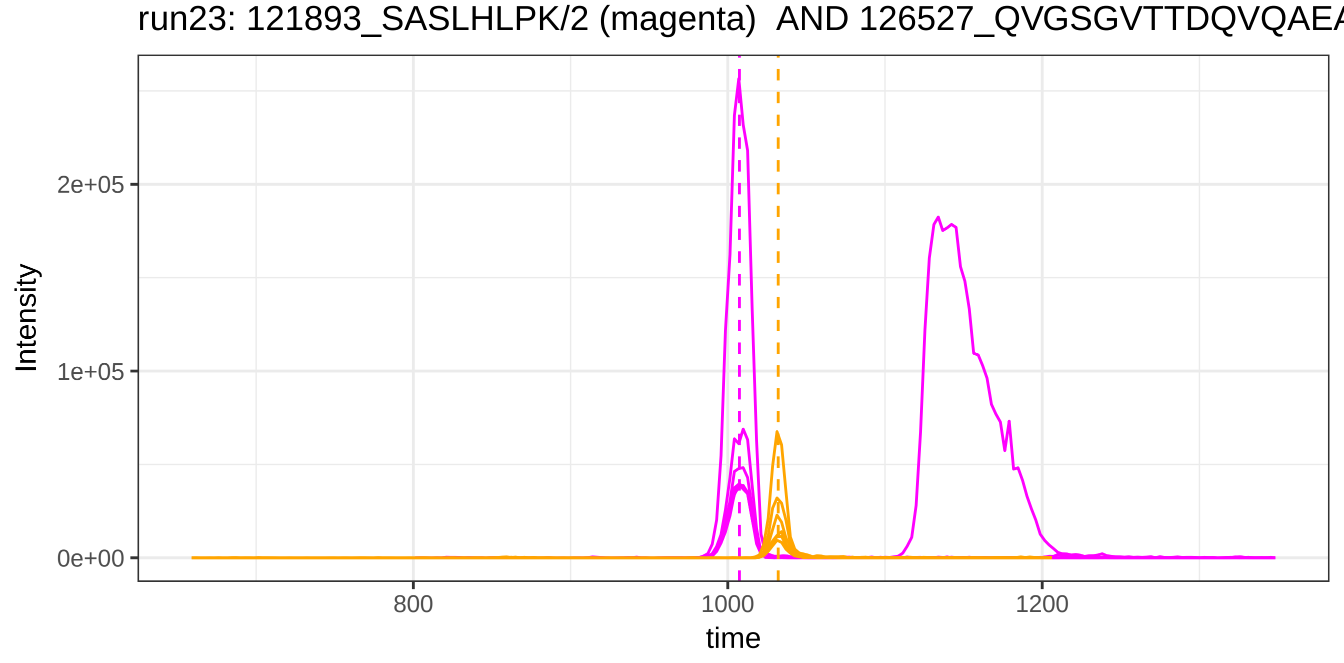
run23: 121893_SASLHLPK/2 (magenta) AND 126527_QVGSGVTTDQVQAEA
Intensity
2e+05
1e+05
0e+00
800
1000
1200
time

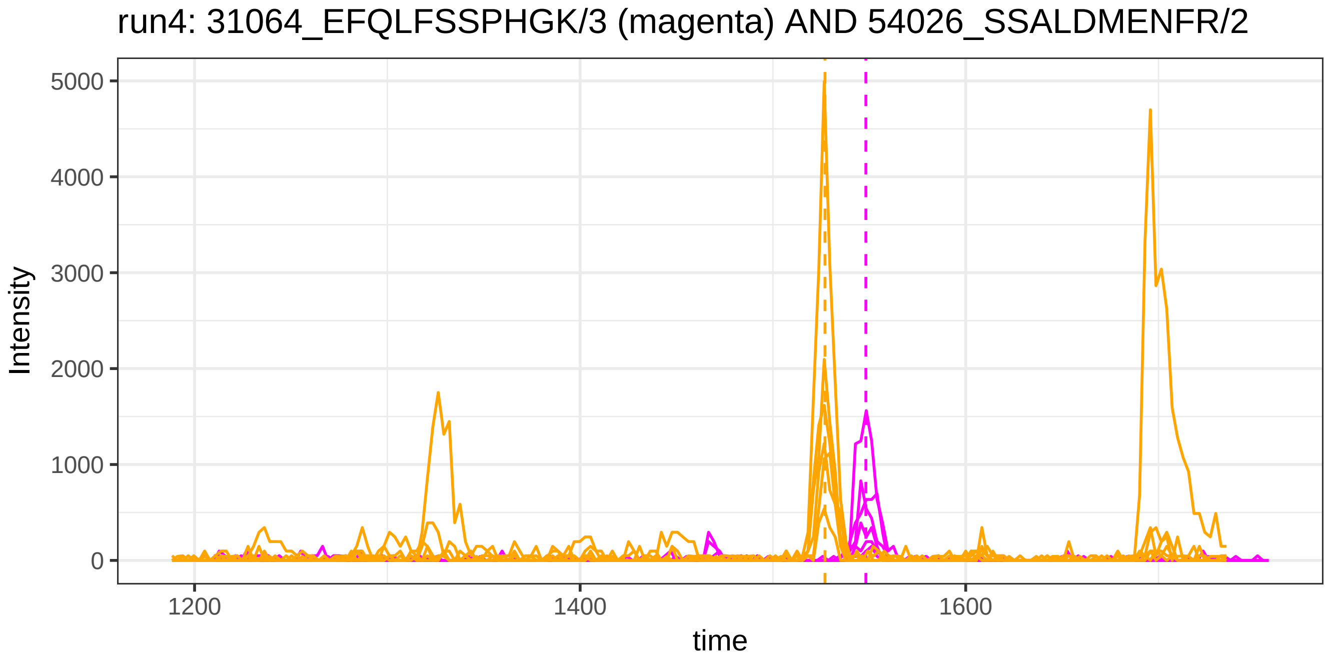
run4: 31064_EFQLFSSPHGK/3 (magenta) AND 54026_SSALDMENFR/2
Intensity
5000
4000
3000
2000
1000
0
1200
1400
1600
time

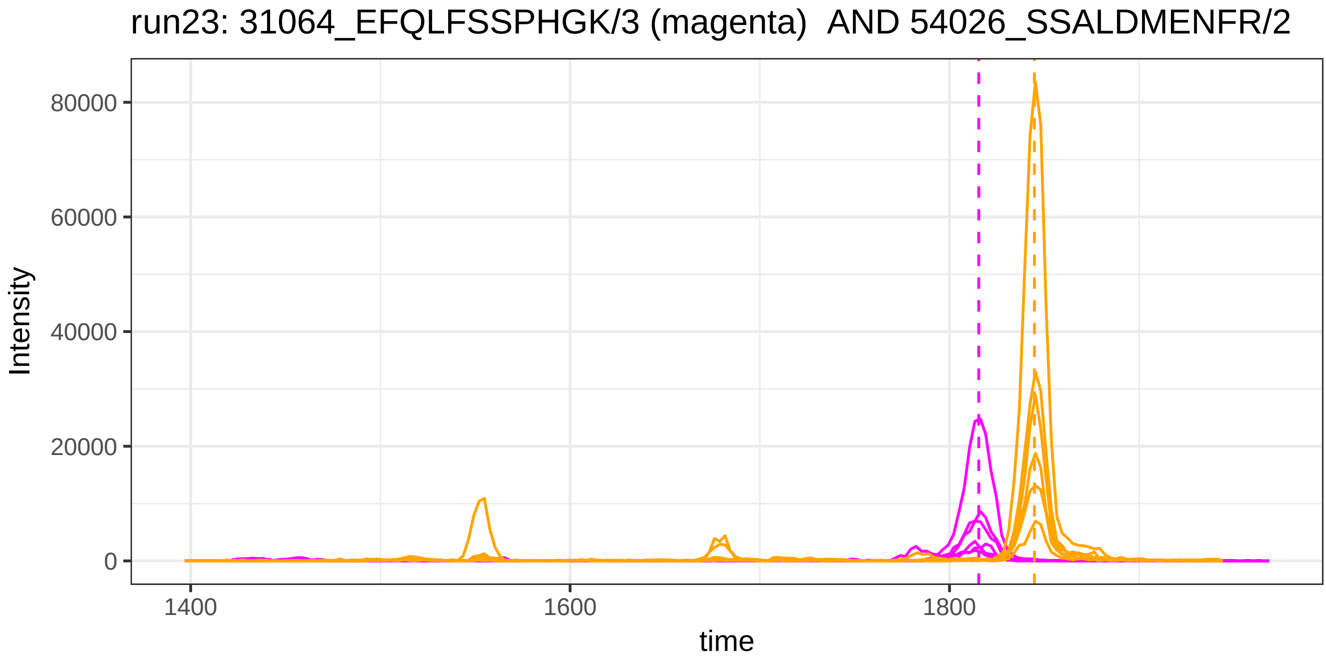
run23: 31064_EFQLFSSPHGK/3 (magenta)  AND 54026_SSALDMENFR/2
Intensity
80000
60000
40000
20000
0
1400
1600
1800
time

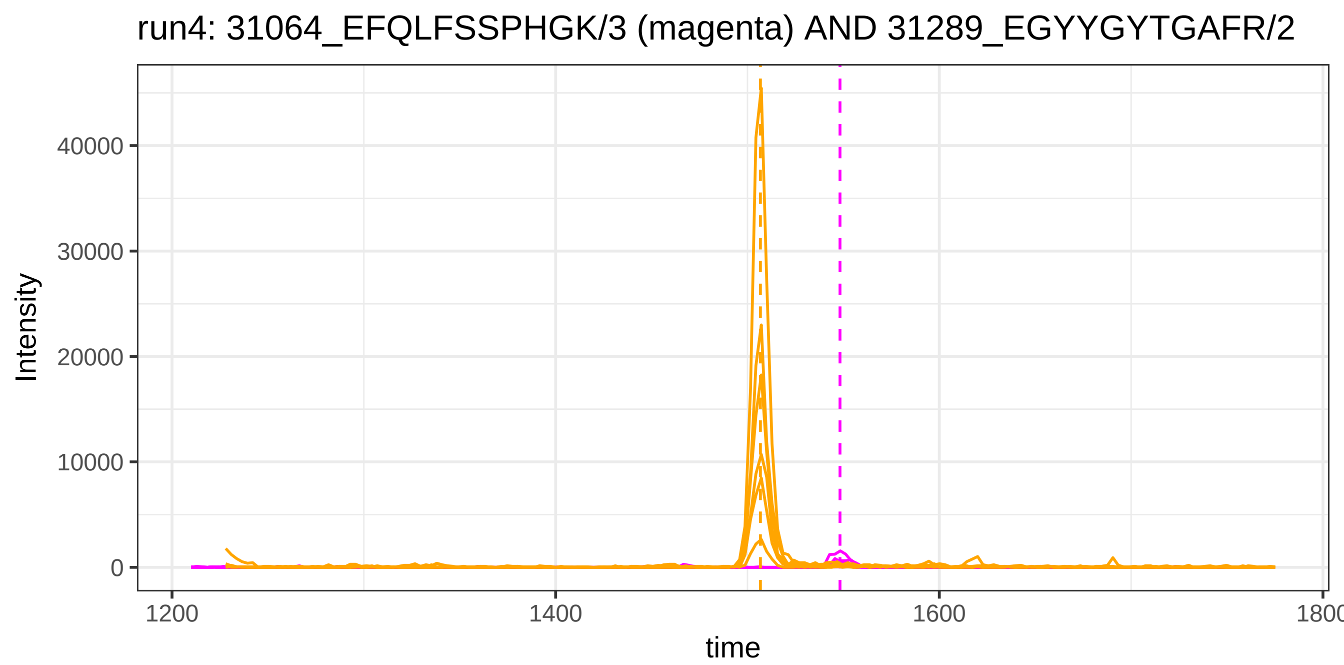
run4: 31064_EFQLFSSPHGK/3 (magenta) AND 31289_EGYYGYTGAFR/2
Intensity
40000
30000
20000
10000
0
1200
1400
1600
1800
time

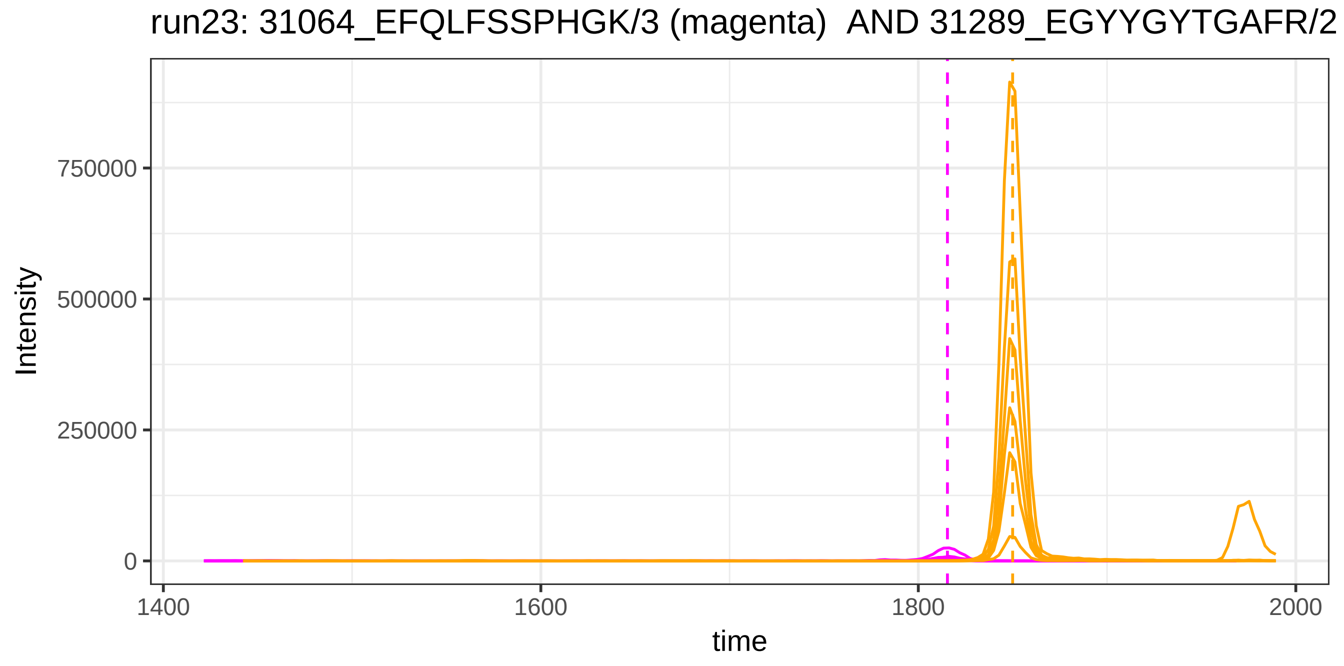
run23: 31064_EFQLFSSPHGK/3 (magenta) AND 31289_EGYYGYTGAFR/2
Intensity
750000
500000
250000
0
1400
1600
1800
2000
time

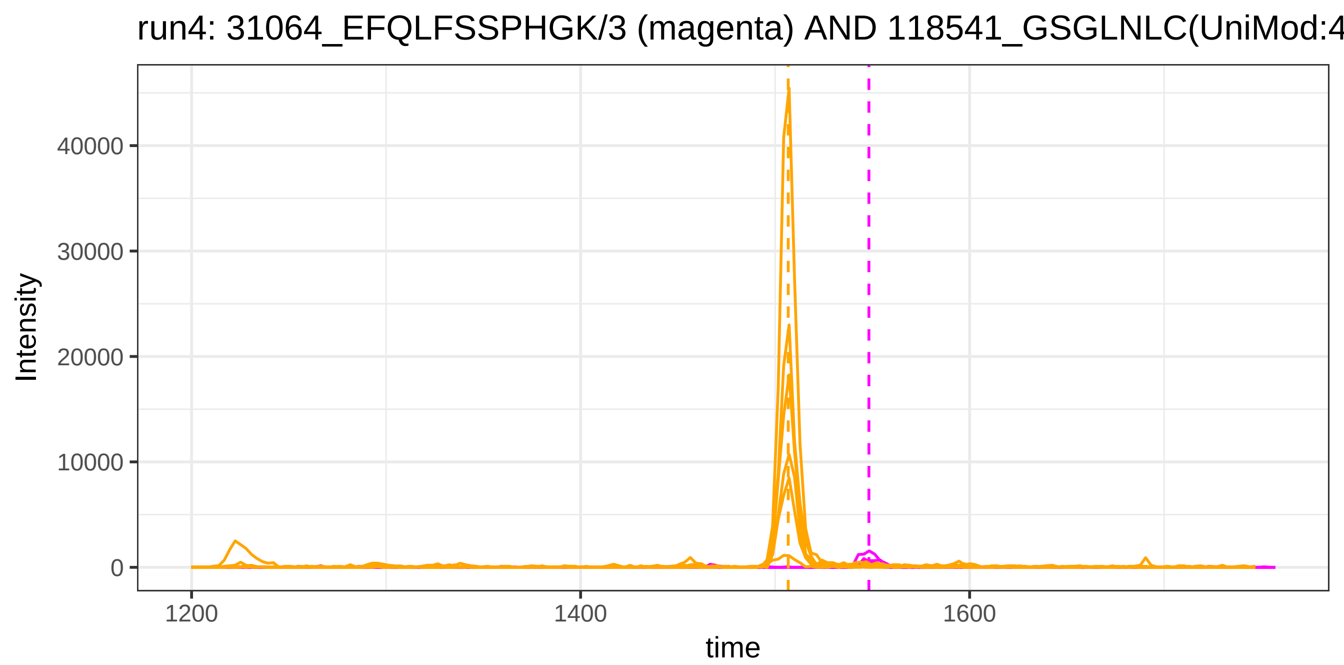
run4: 31064_EFQLFSSPHGK/3 (magenta) AND 118541_GSGLNLC(UniMod:4
Intensity
40000
30000
20000
10000
0
1200
1400
1600
time

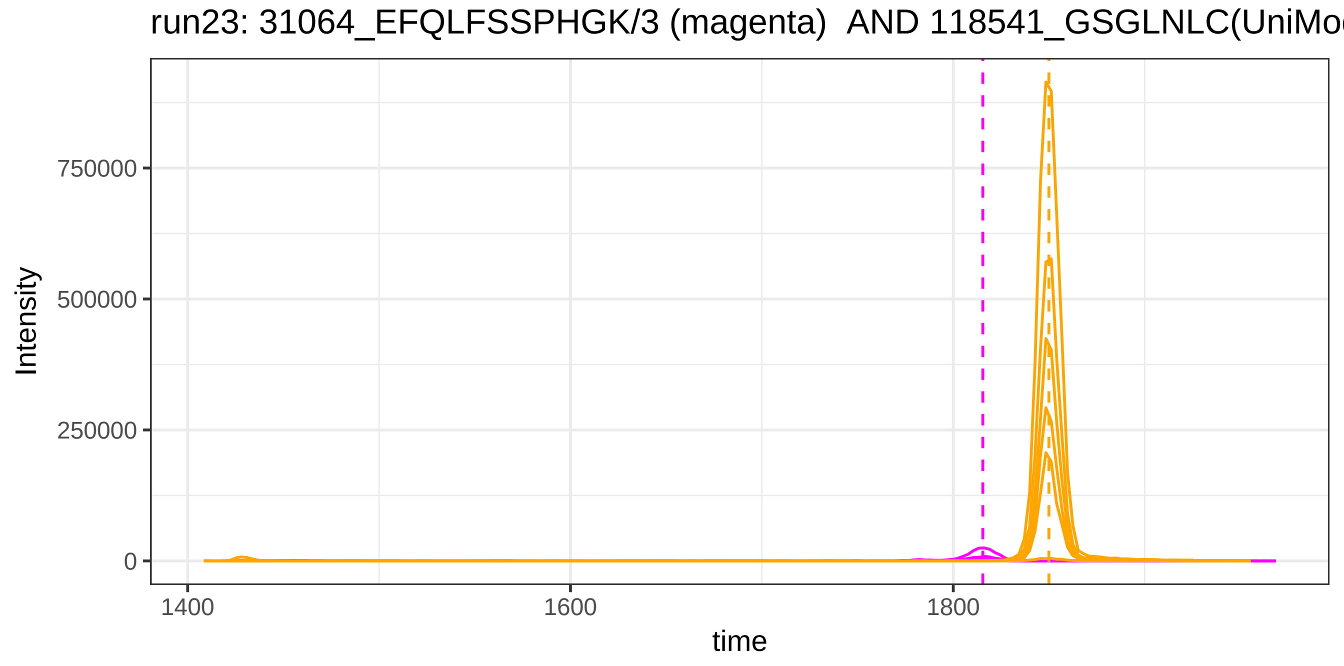
run23: 31064_EFQLFSSPHGK/3 (magenta) AND 118541_GSGLNLC(UniMo
Intensity
750000
500000
250000
0
1400
1600
1800
time

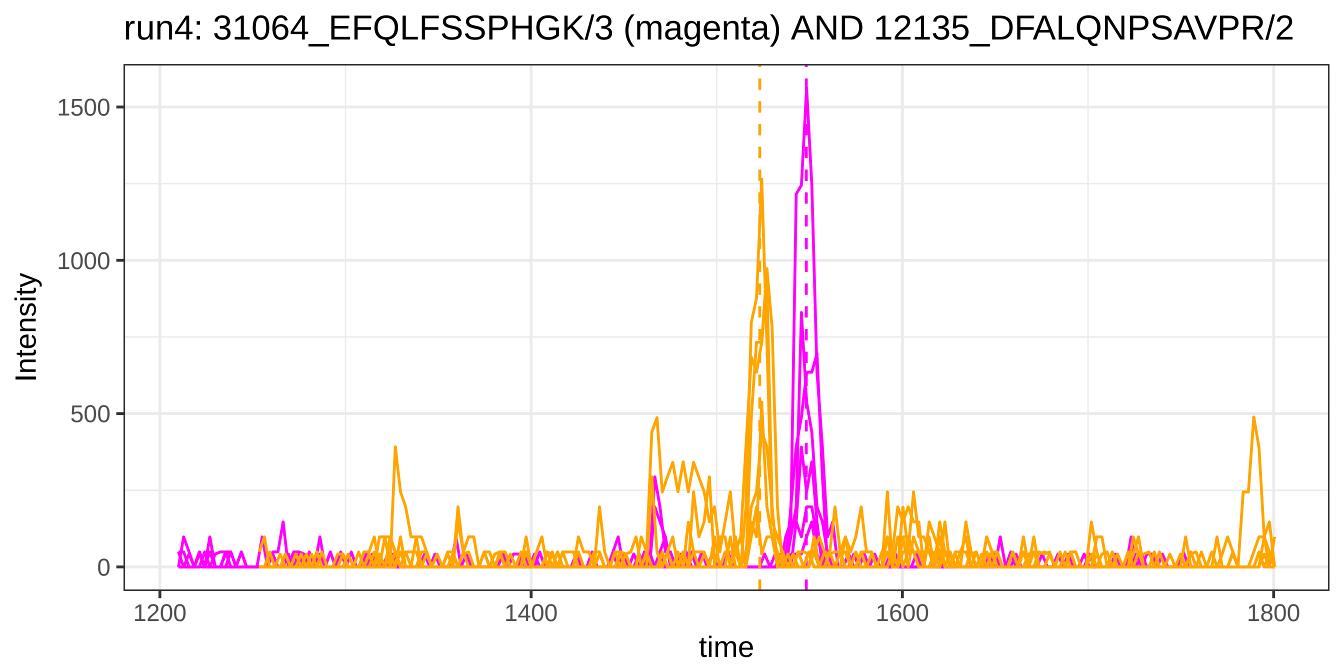
run4: 31064_EFQLFSSPHGK/3 (magenta) AND 12135_DFALQNPSAVPR/2
Intensity
1500
1000
500
0
1200
1400
1600
1800
time

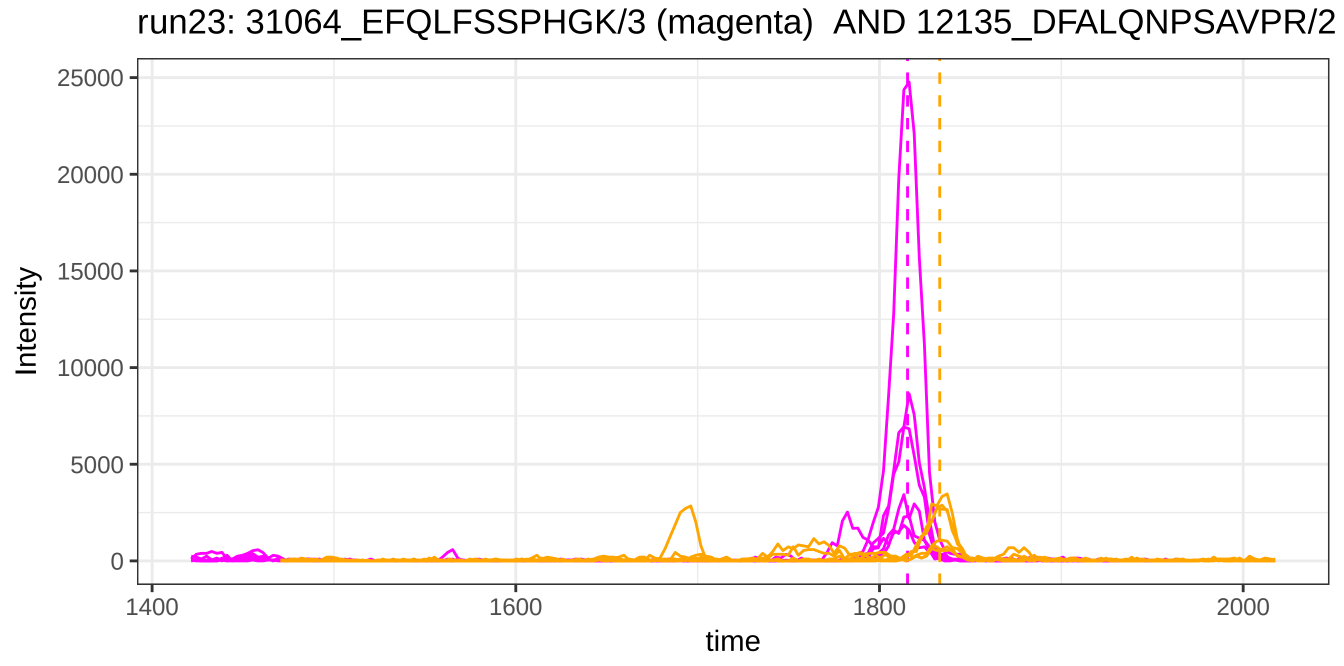
run23: 31064_EFQLFSSPHGK/3 (magenta) AND 12135_DFALQNPSAVPR/2
Intensity
25000
20000
15000
10000
5000
0
1400
1600
1800
2000
time

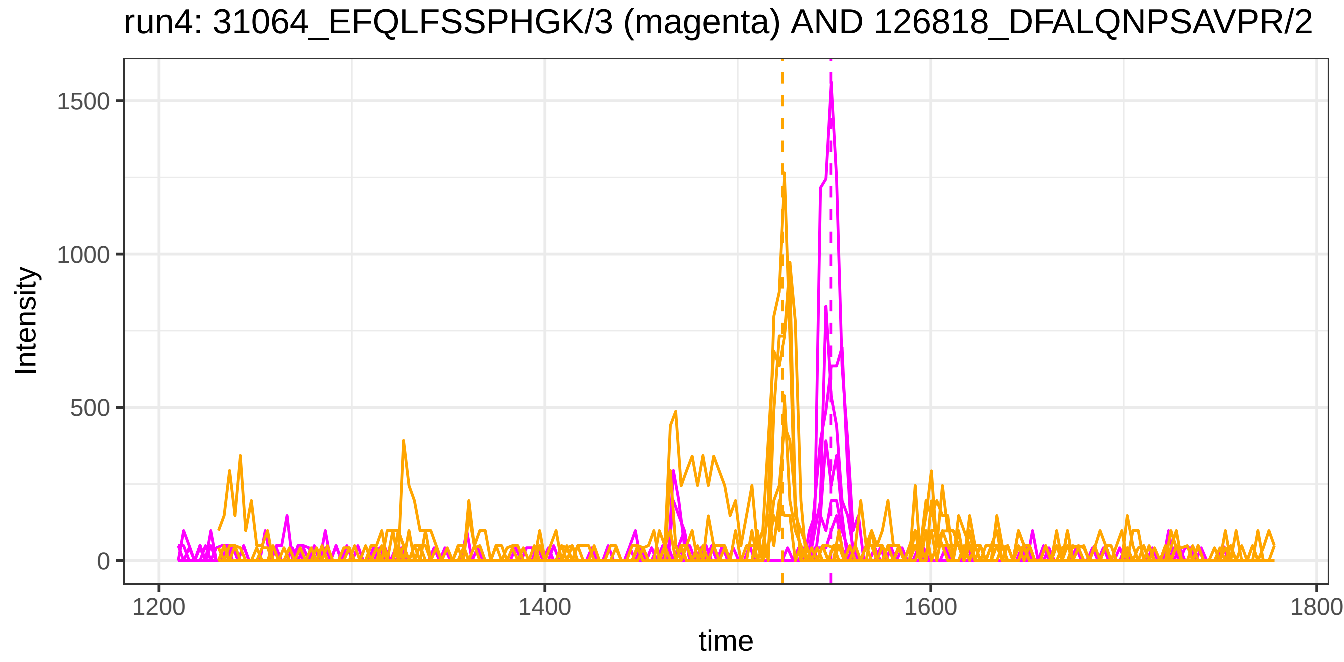
run4: 31064_EFQLFSSPHGK/3 (magenta) AND 126818_DFALQNPSAVPR/2
Intensity
1500
1000
500
0
1200
1400
1600
1800
time

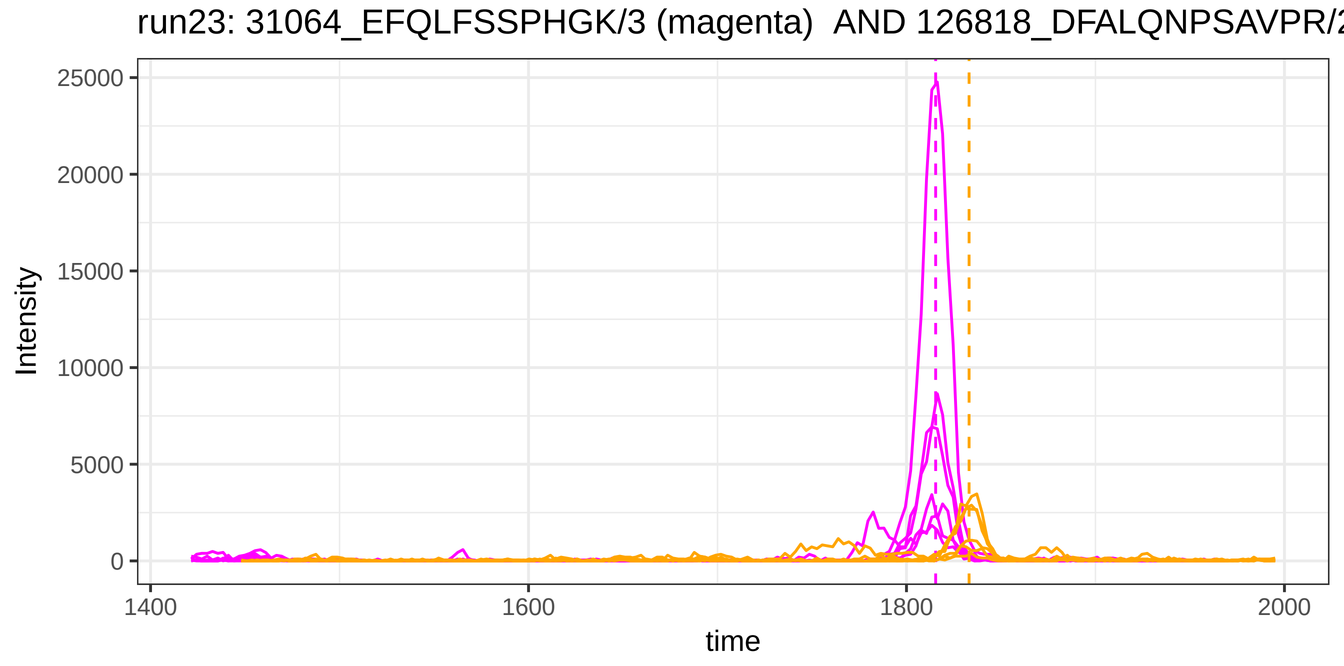
run23: 31064_EFQLFSSPHGK/3 (magenta) AND 126818_DFALQNPSAVPR/2
Intensity
25000
20000
15000
10000
5000
0
1400
1600
1800
2000
time

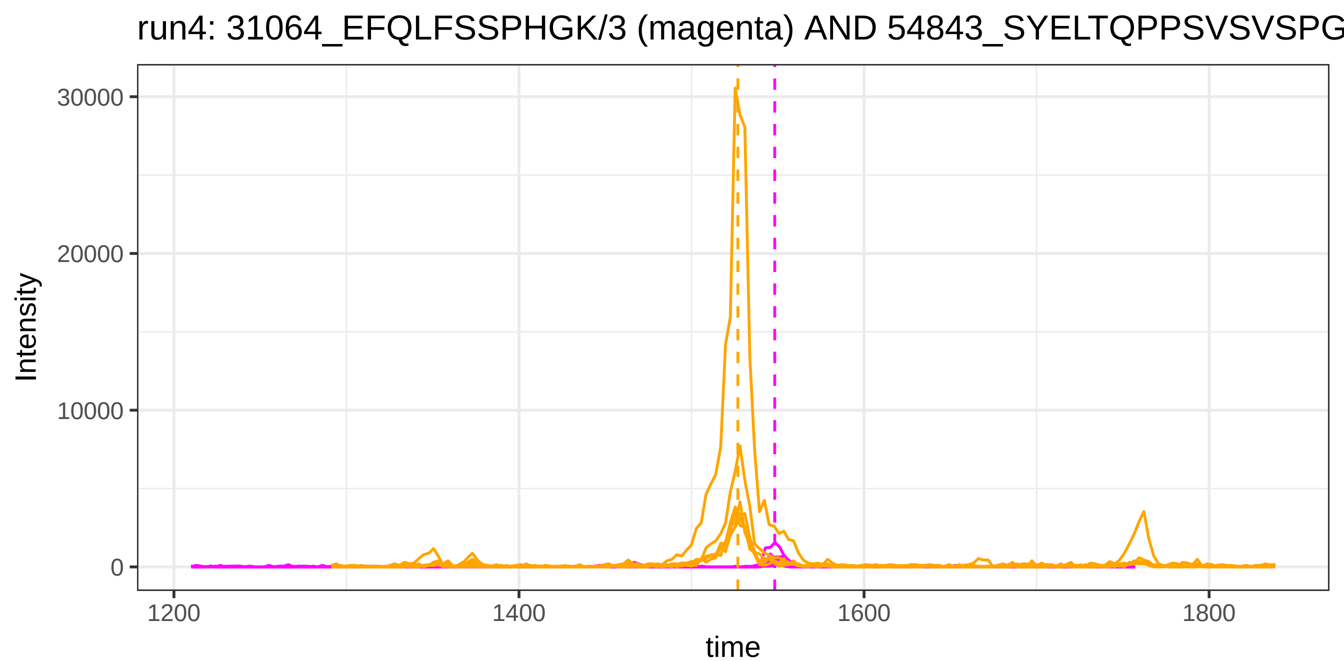
run4: 31064_EFQLFSSPHGK/3 (magenta) AND 54843_SYELTQPPSVSVSPG
Intensity
30000
20000
10000
0
1200
1400
1600
1800
time

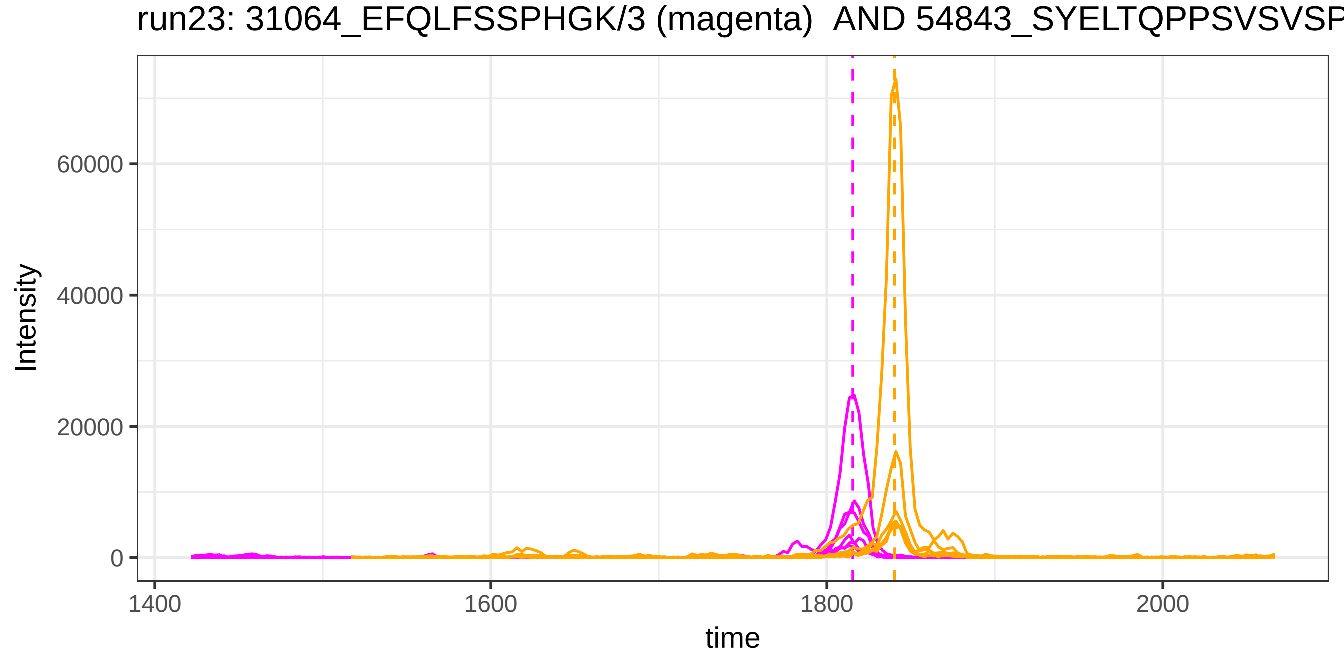
run23: 31064_EFQLFSSPHGK/3 (magenta) AND 54843_SYELTQPPSVSVSP
Intensity
60000
40000
20000
0
1400
1600
1800
2000
time

run4: 40510_IYLQPGR/2 (magenta) AND 38644_HYDGSYSTFGER/2

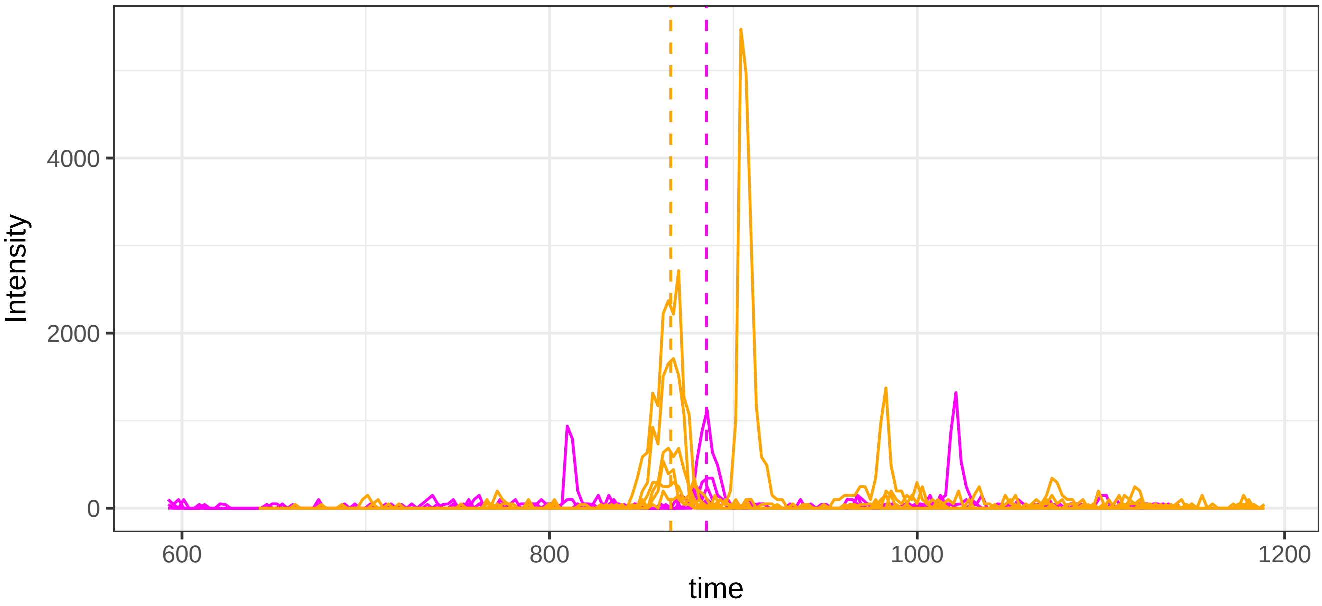


run23: 40510_IYLQPGR/2 (magenta)  AND 38644_HYDGSYSTFGER/2

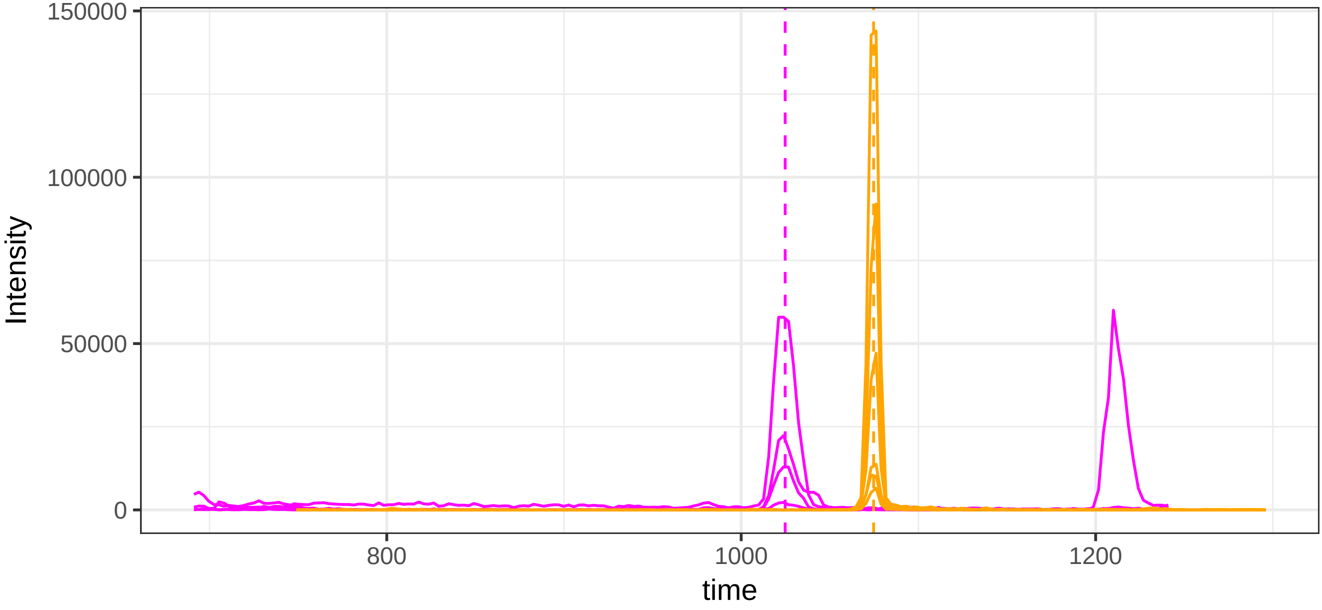

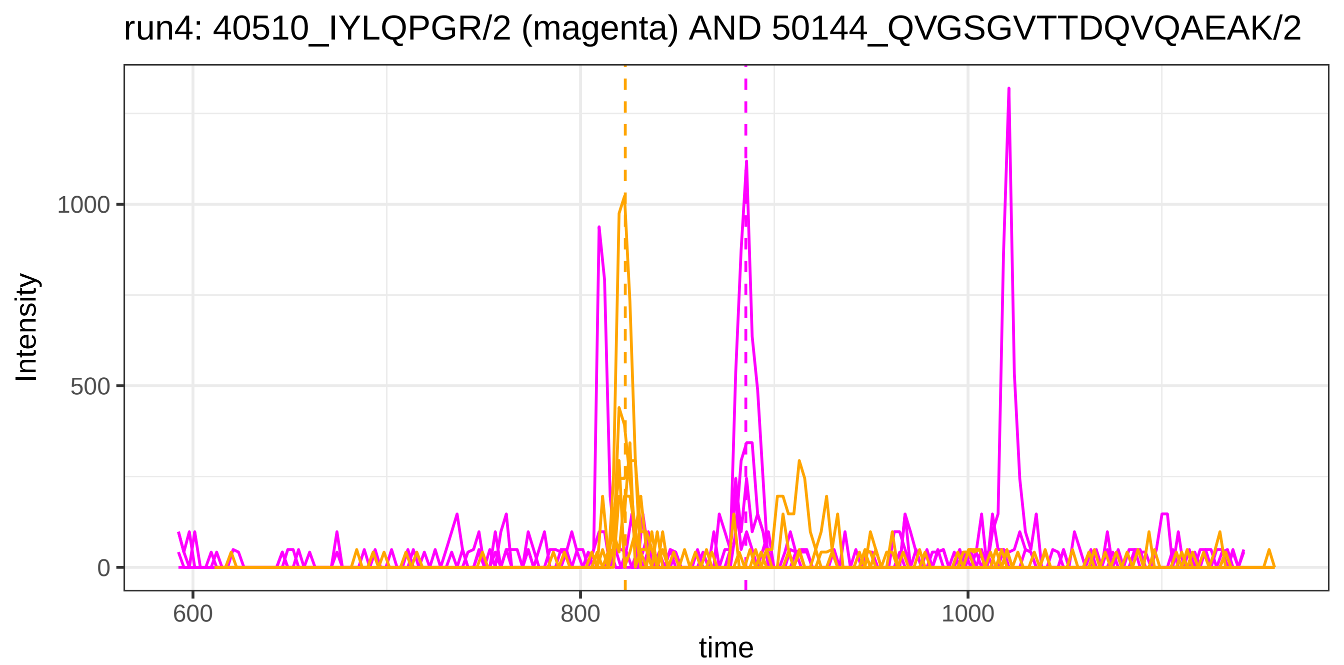
run4: 40510_IYLQPGR/2 (magenta) AND 50144_QVGSGVTTDQVQAEAK/2
Intensity
1000
500
0
600
800
1000
time

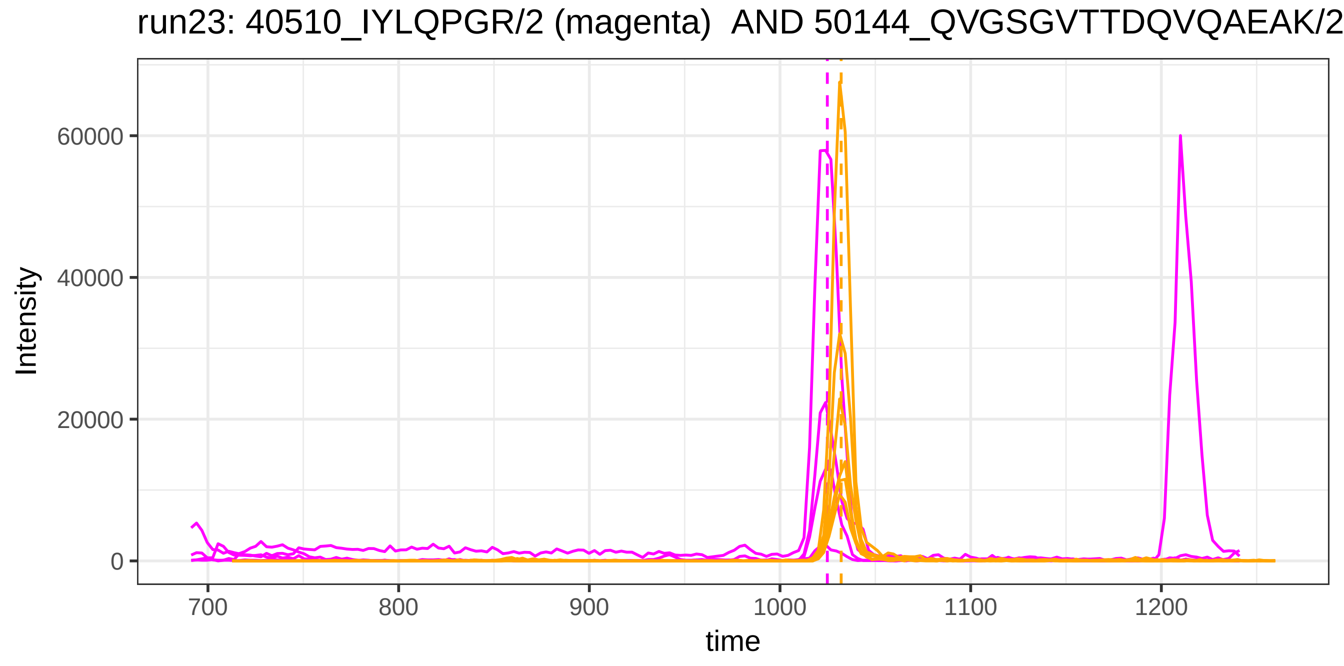
run23: 40510_IYLQPGR/2 (magenta) AND 50144_QVGSGVTTDQVQAEAK/2
Intensity
60000
40000
20000
0
700
800
900
1000
1100
1200
time

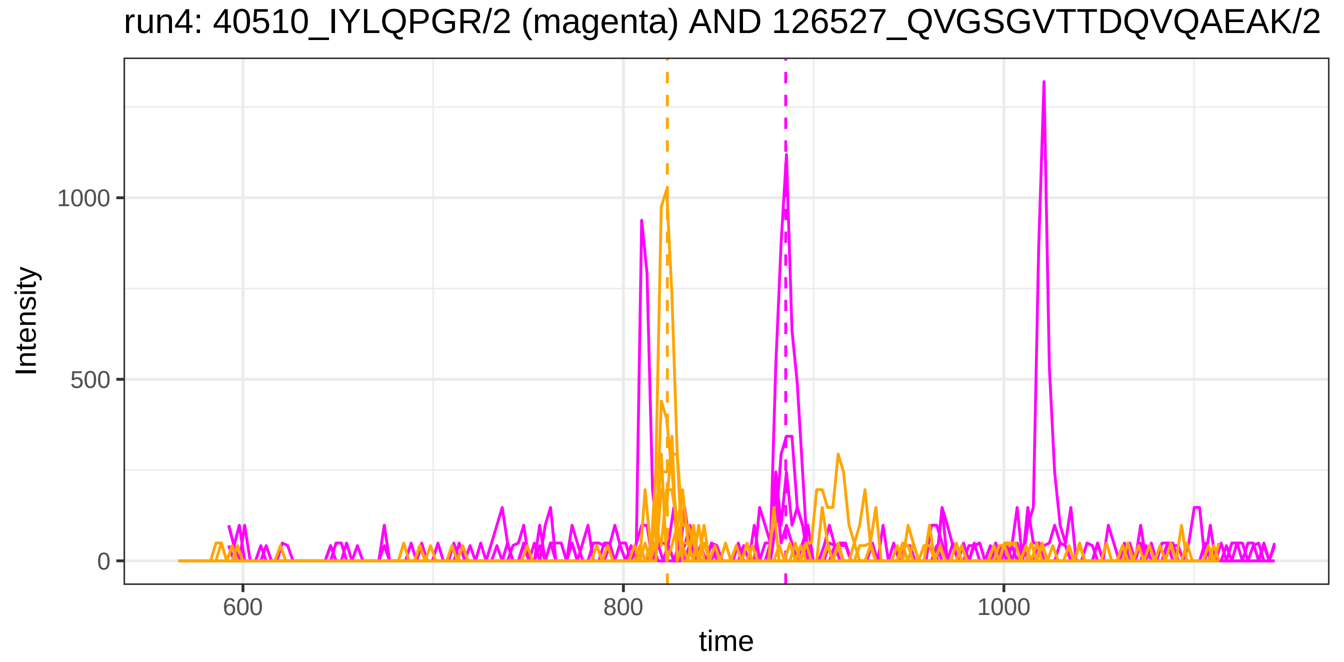
run4: 40510_IYLQPGR/2 (magenta) AND 126527_QVGSGVTTDQVQAEAK/2
Intensity
1000
500
0
600
800
1000
time

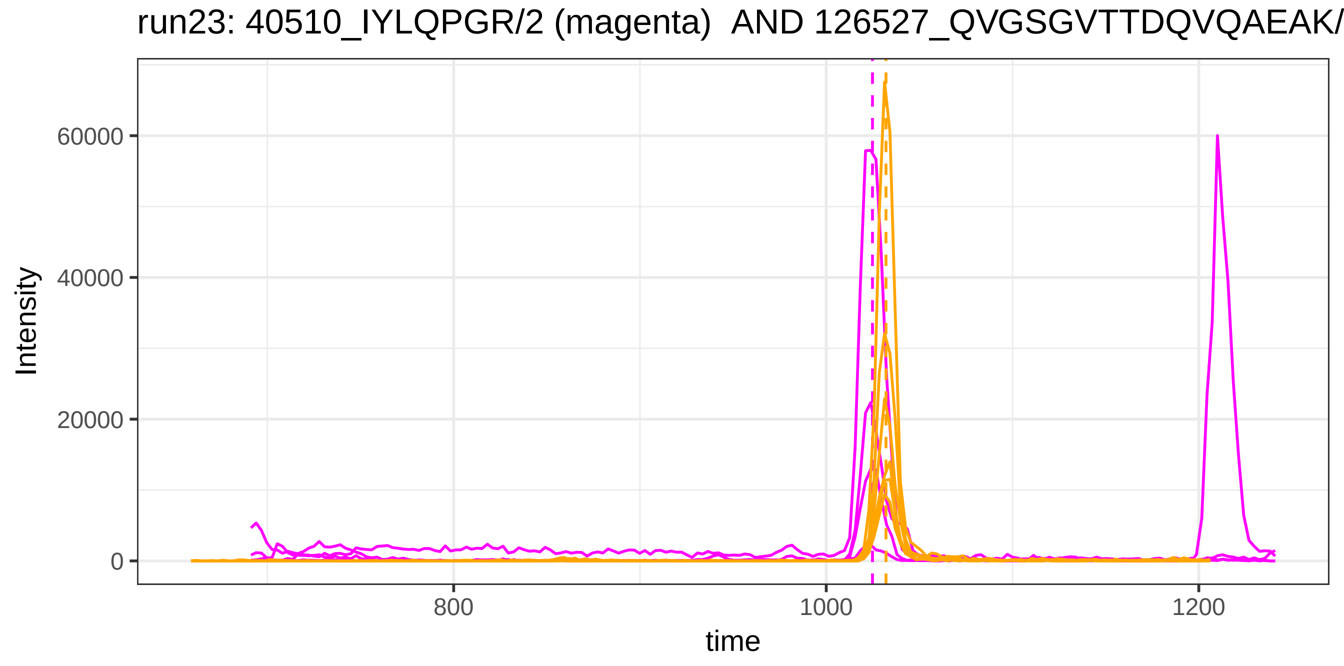
run23: 40510_IYLQPGR/2 (magenta) AND 126527_QVGSGVTTDQVQAEAK/
Intensity
60000
40000
20000
0
800
1000
1200
time

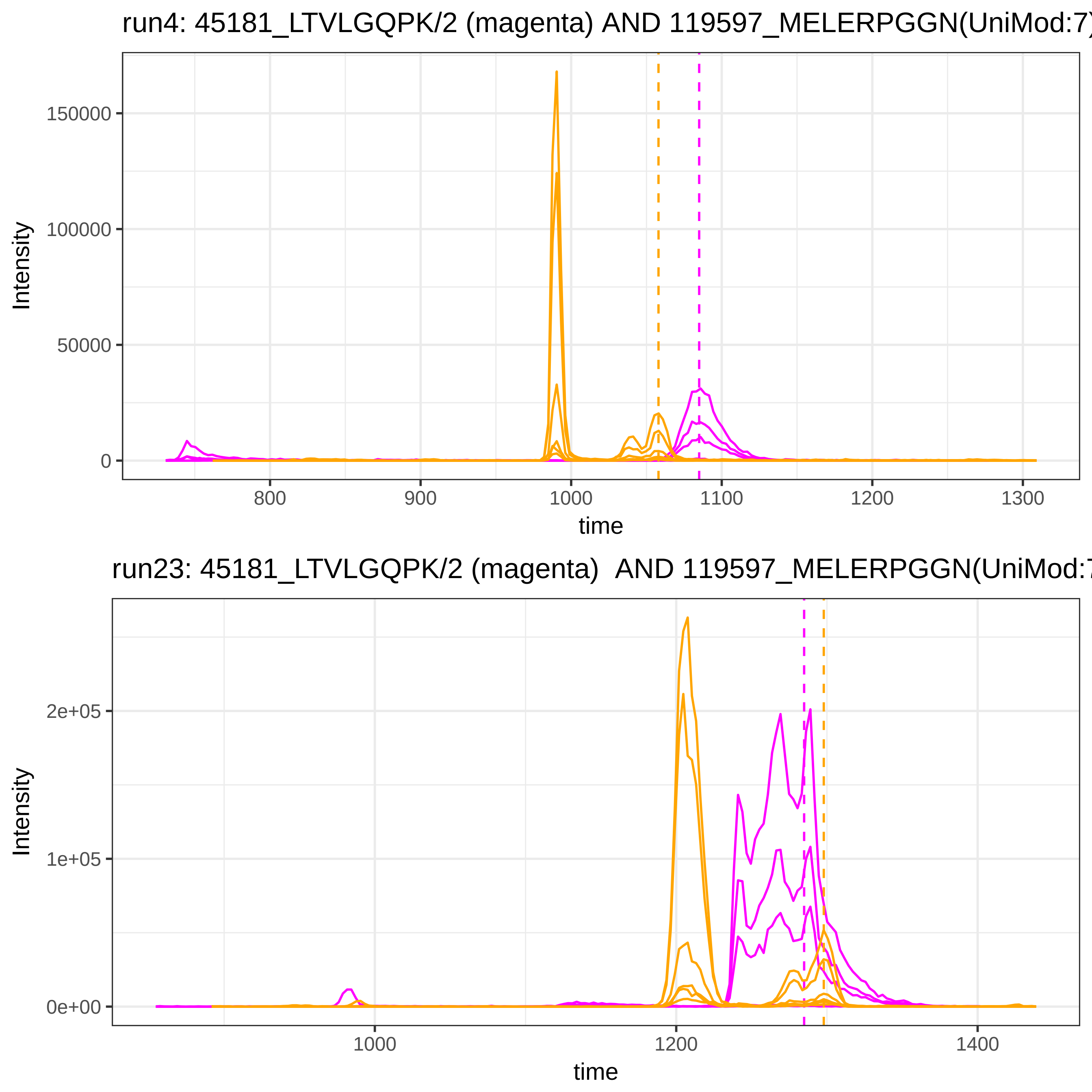
run4: 45181_LTVLGQPK/2 (magenta) AND 119597_MELERPGGN(UniMod:7)
Intensity
150000
100000
50000
0
800
900
1000
1100
1200
1300
time
run23: 45181_LTVLGQPK/2 (magenta) AND 119597_MELERPGGN(UniMod:7
Intensity
2e+05
1e+05
0e+00
1000
1200
1400
time

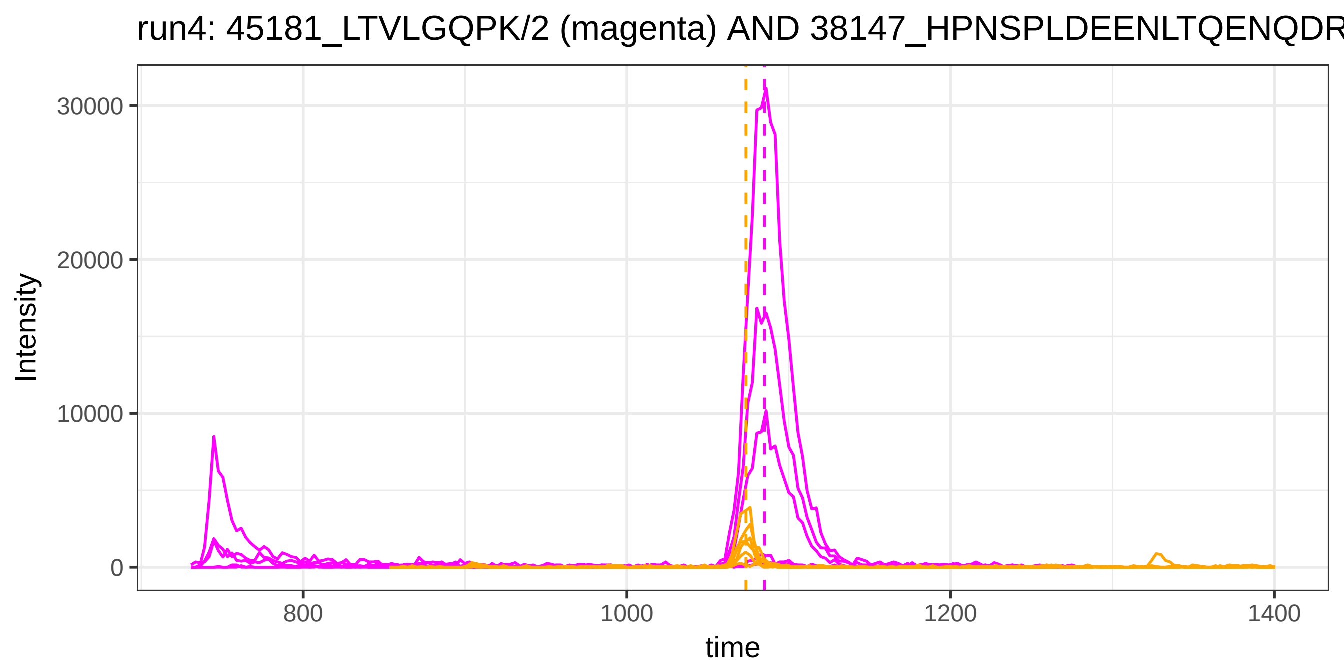
run4: 45181_LTVLGQPK/2 (magenta) AND 38147_HPNSPLDEENLTQENQDR
Intensity
30000
20000
10000
0
800
1000
1200
1400
time

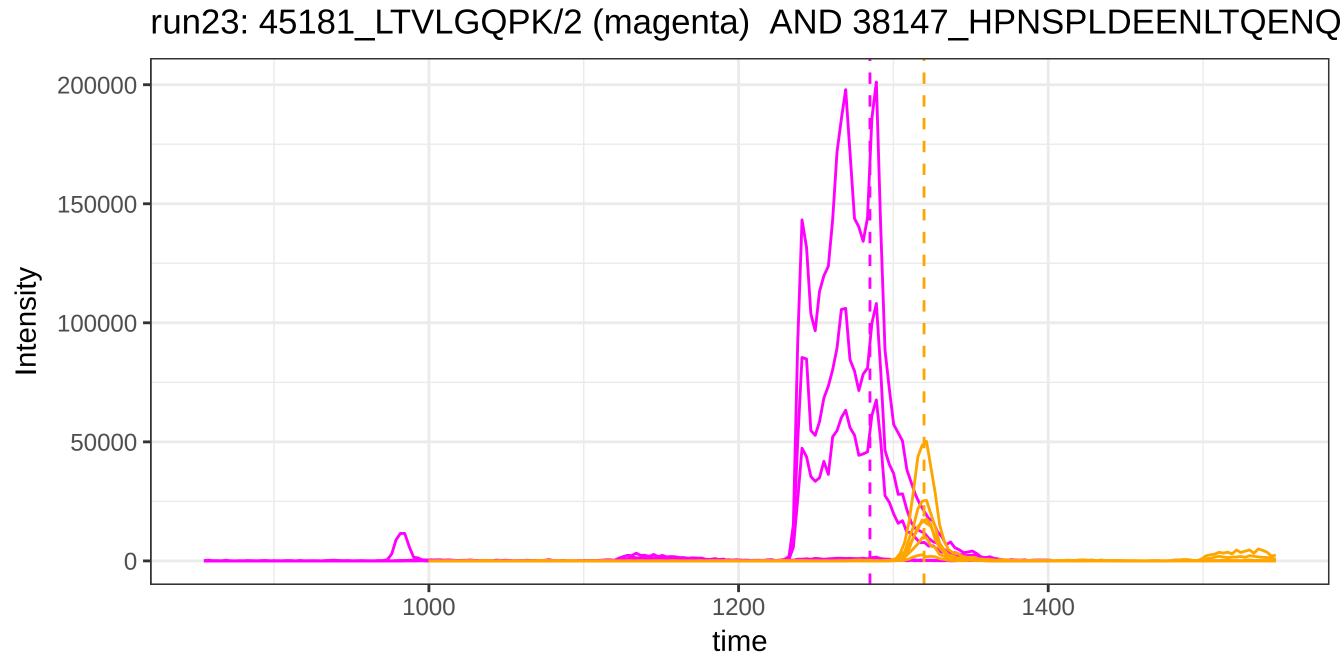
run23: 45181_LTVLGQPK/2 (magenta) AND 38147_HPNSPLDEENLTQENQ
Intensity
200000
150000
100000
50000
0
1000
1200
1400
time

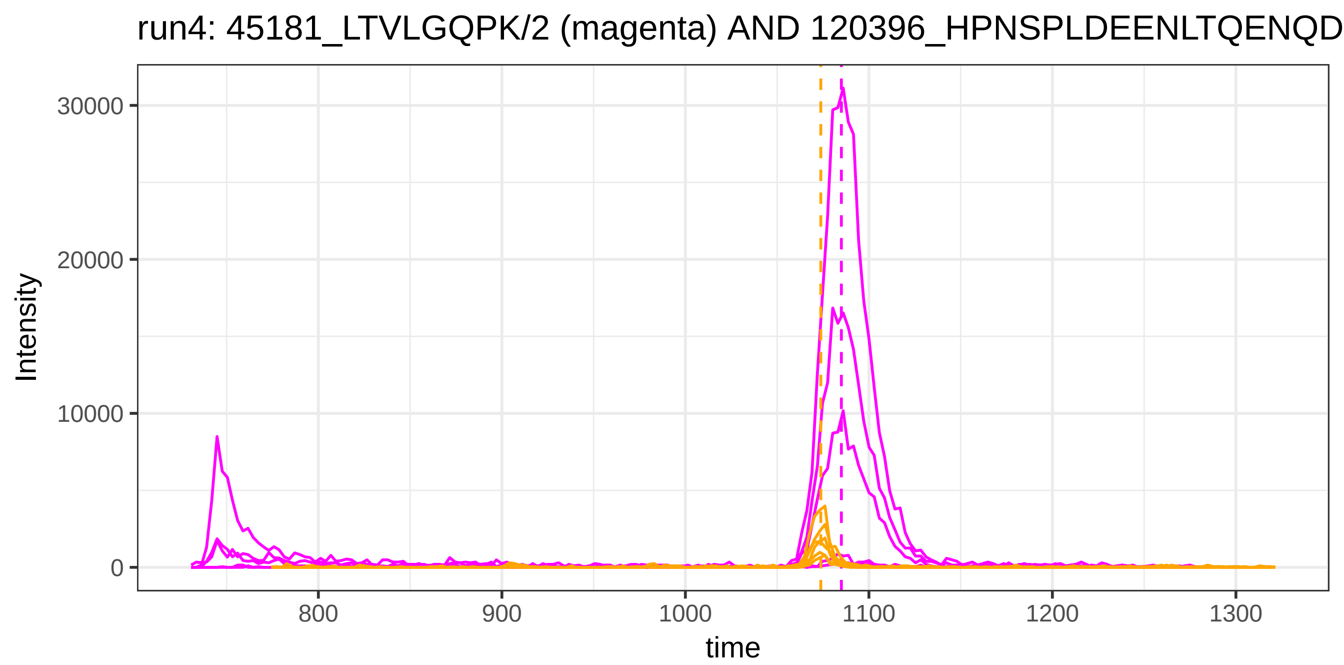
run4: 45181_LTVLGQPK/2 (magenta) AND 120396_HPNSPLDEENLTQENQD
Intensity
30000
20000
10000
0
800
900
1000
1100
1200
1300
time

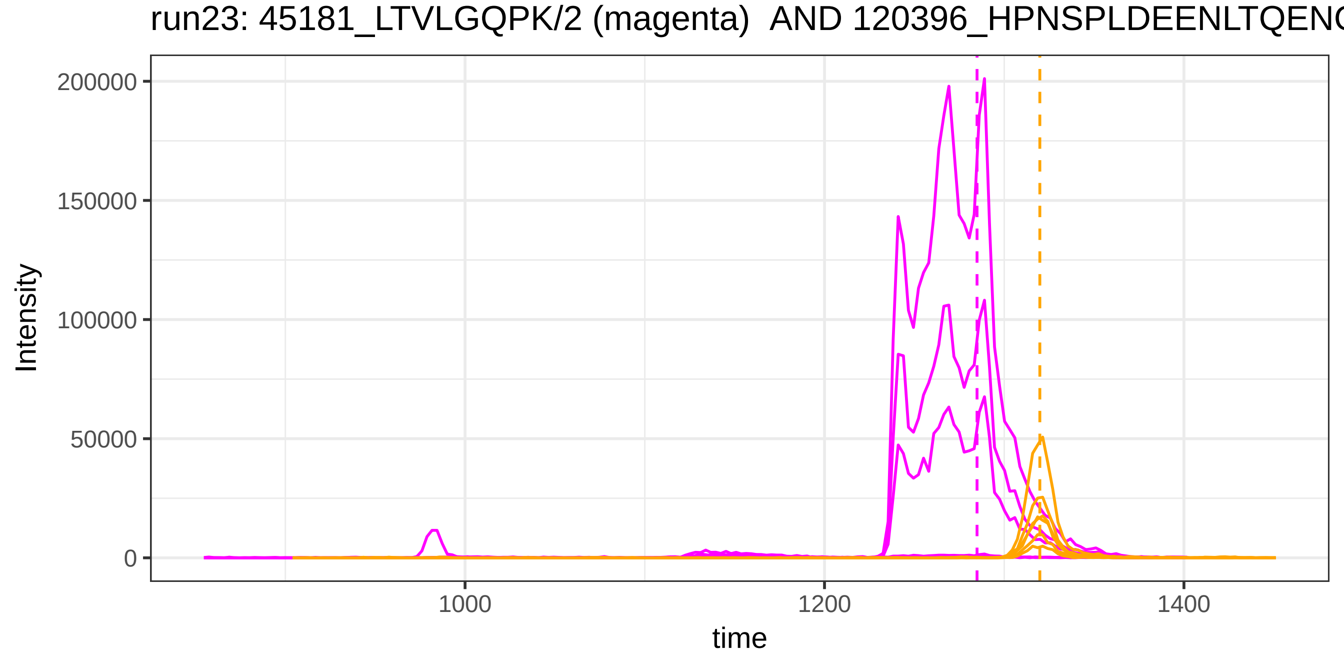
run23: 45181_LTVLGQPK/2 (magenta) AND 120396_HPNSPLDEENLTQENC
Intensity
200000
150000
100000
50000
0
1000
1200
1400
time

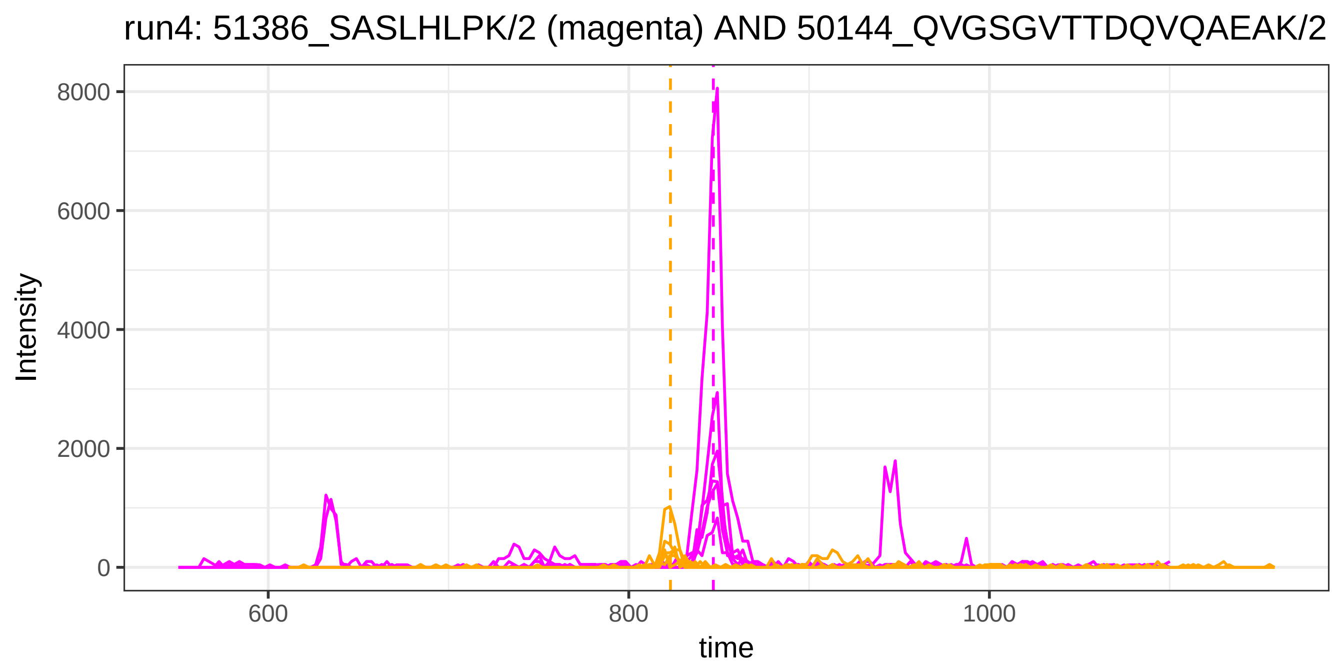
run4: 51386_SASLHLPK/2 (magenta) AND 50144_QVGSGVTTDQVQAEAK/2
Intensity
time
0
2000
4000
6000
8000
600
800
1000

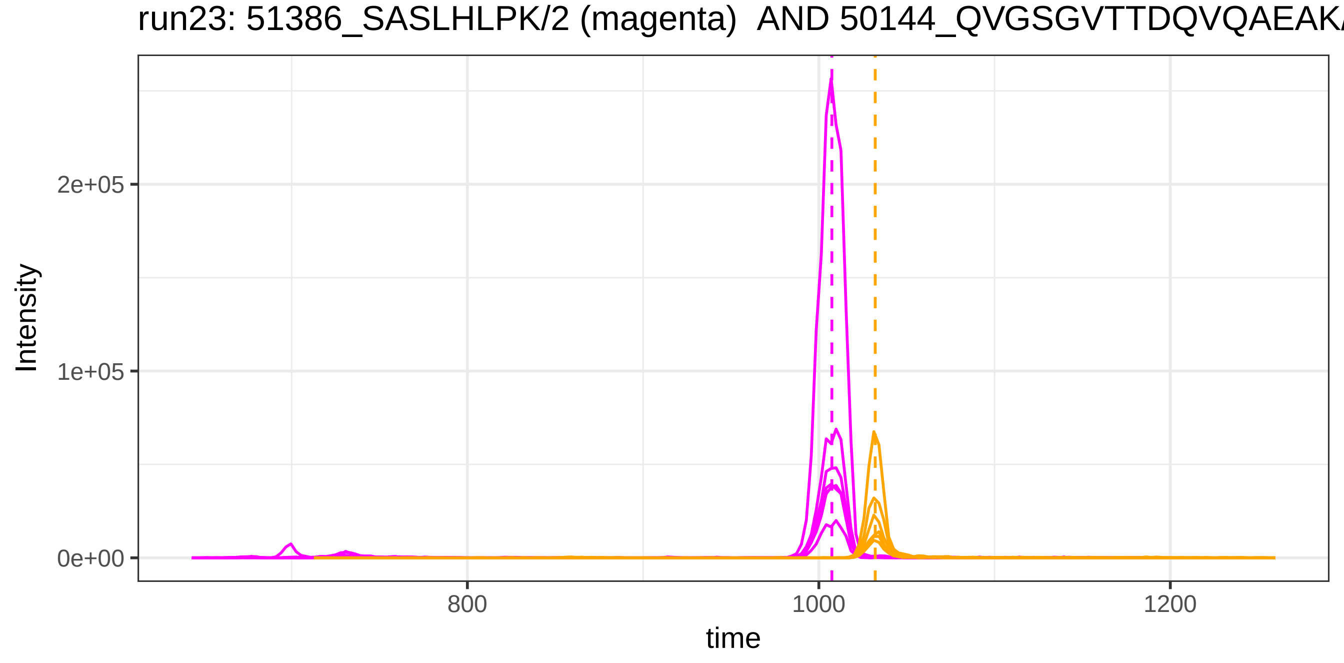
run23: 51386_SASLHLPK/2 (magenta) AND 50144_QVGSGVTTDQVQAEAK/
Intensity
time
0e+00
1e+05
2e+05
800
1000
1200

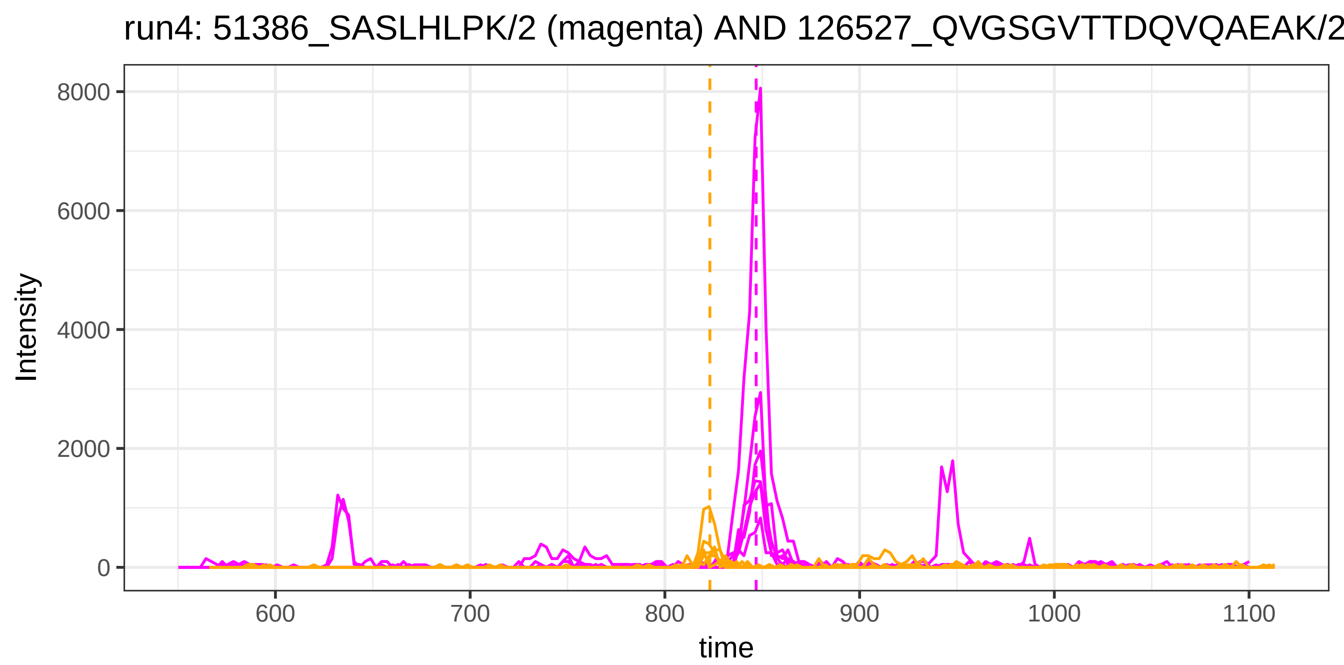
run4: 51386_SASLHLPK/2 (magenta) AND 126527_QVGSGVTTDQVQAEAK/2
Intensity
8000
6000
4000
2000
0
600
700
800
900
1000
1100
time

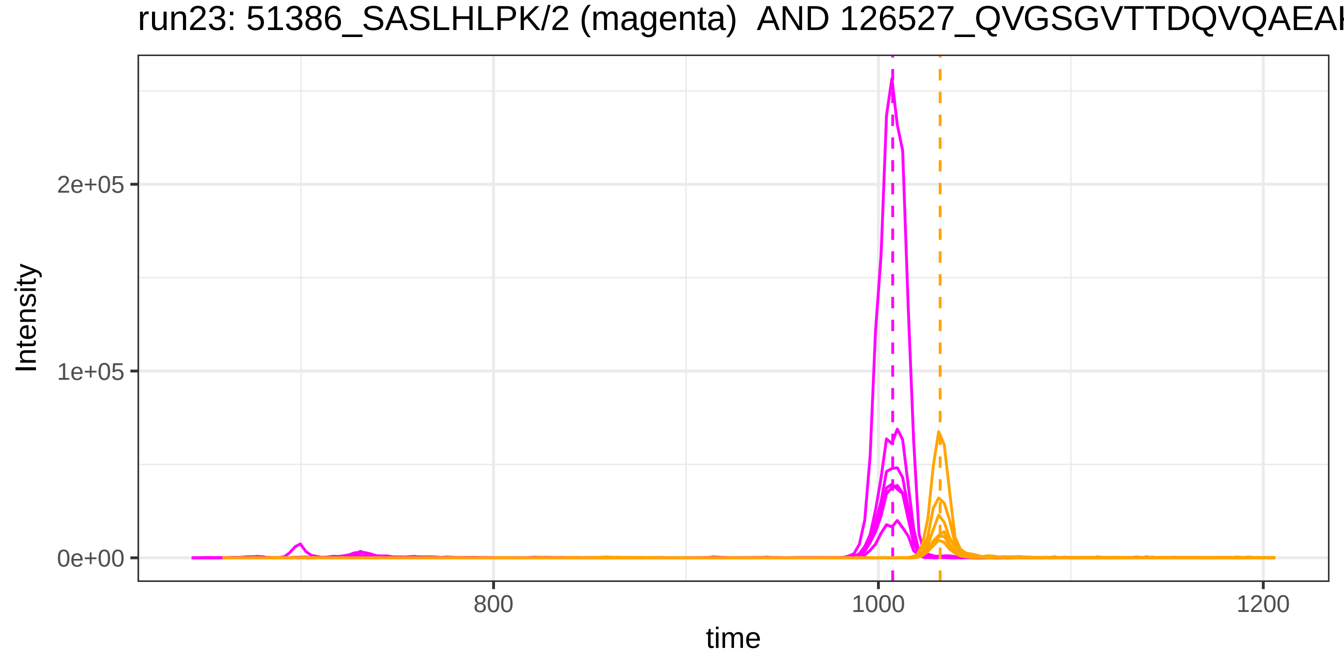
run23: 51386_SASLHLPK/2 (magenta) AND 126527_QVGSGVTTDQVQAEAK
Intensity
2e+05
1e+05
0e+00
800
1000
1200
time

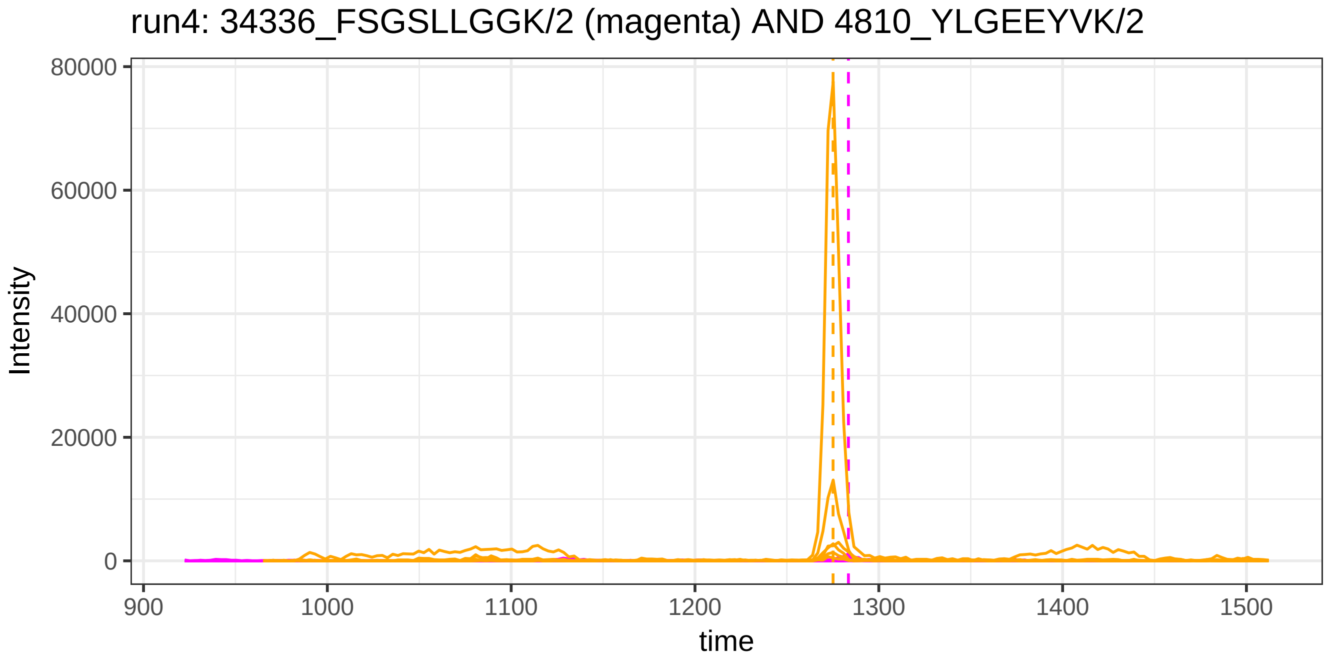
run4: 34336_FSGSLLGGK/2 (magenta) AND 4810_YLGEEYVK/2
Intensity
80000
60000
40000
20000
0
900
1000
1100
1200
1300
1400
1500
time

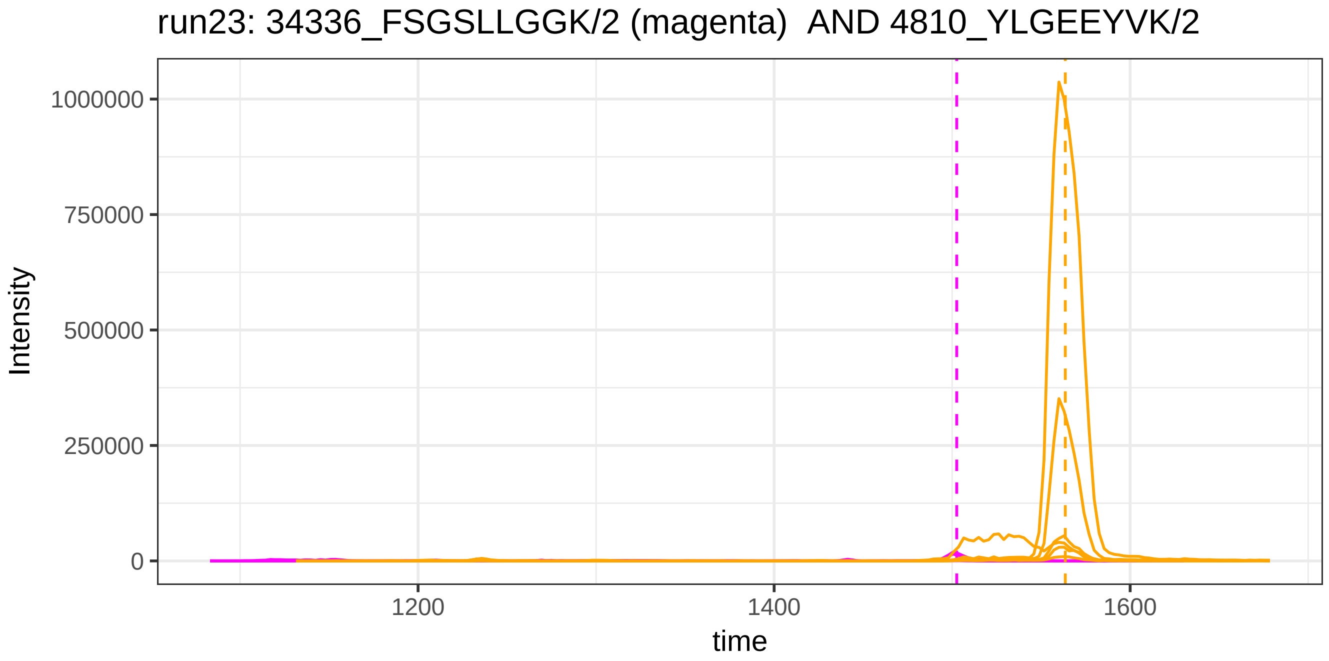
run23: 34336_FSGSLLGGK/2 (magenta) AND 4810_YLGEEYVK/2
Intensity
1000000
750000
500000
250000
0
1200
1400
1600
time

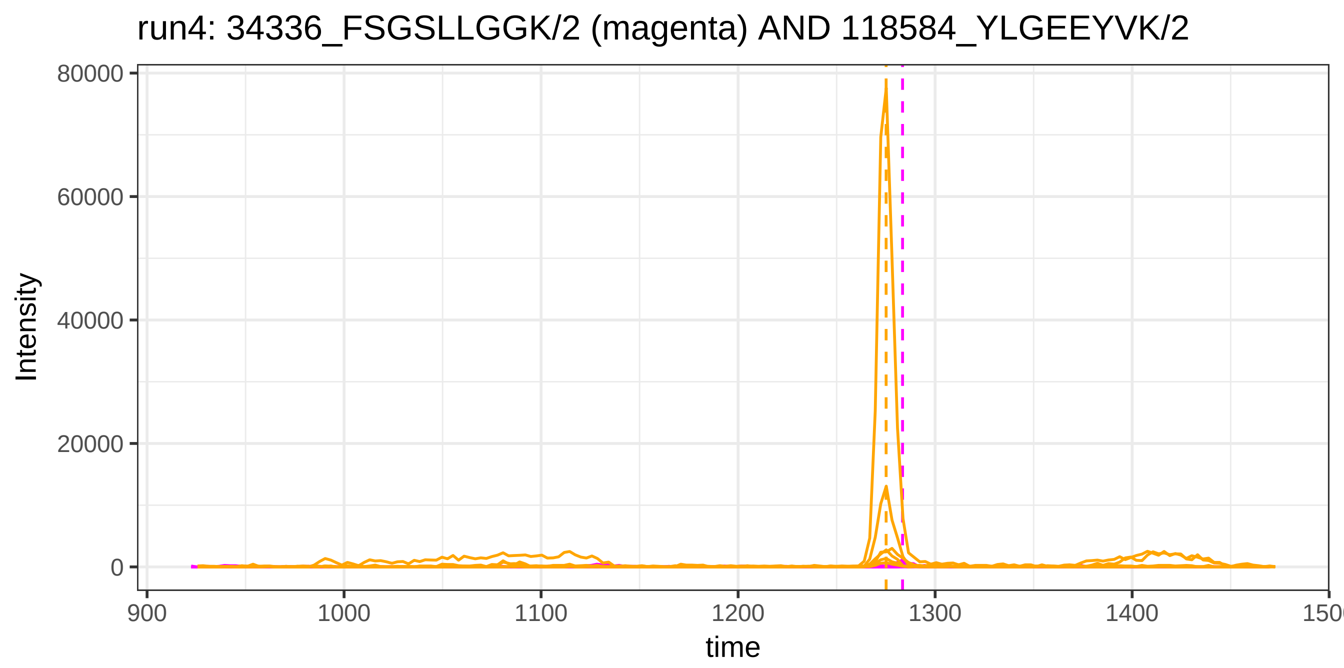
run4: 34336_FSGSLLGGK/2 (magenta) AND 118584_YLGEEYVK/2
Intensity
80000
60000
40000
20000
0
900
1000
1100
1200
1300
1400
150
time

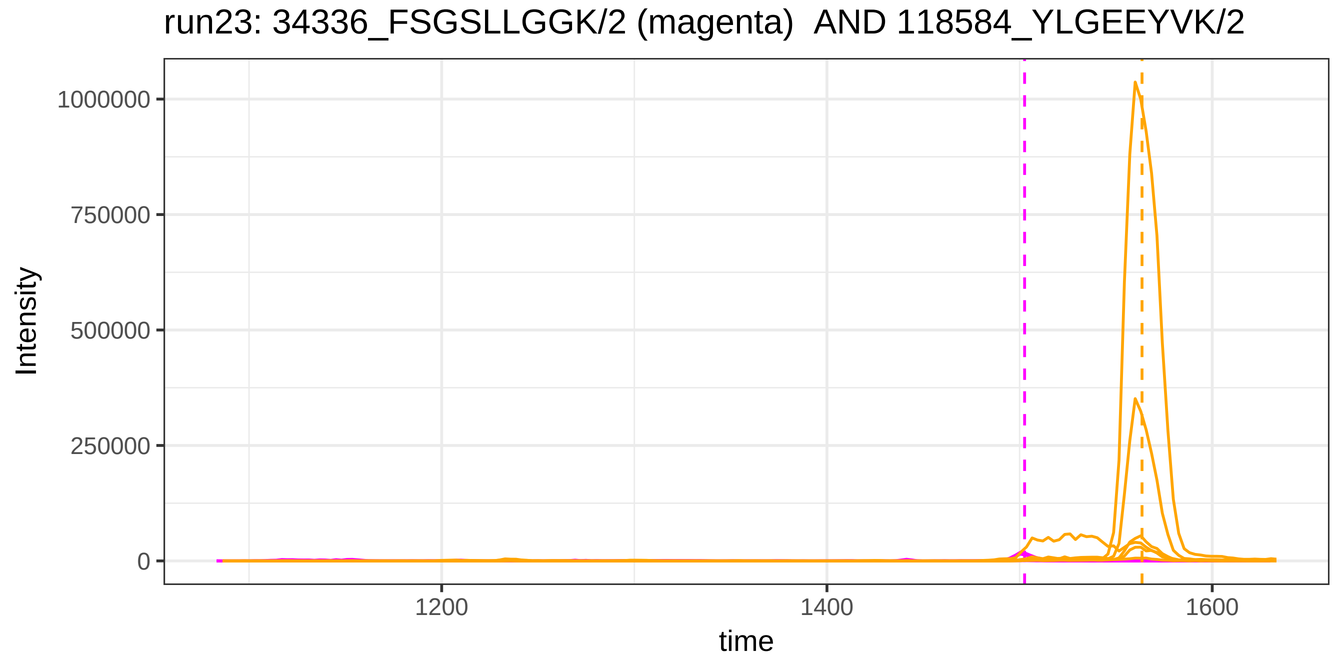
run23: 34336_FSGSLLGGK/2 (magenta) AND 118584_YLGEEYVK/2
Intensity
1000000
750000
500000
250000
0
1200
1400
1600
time

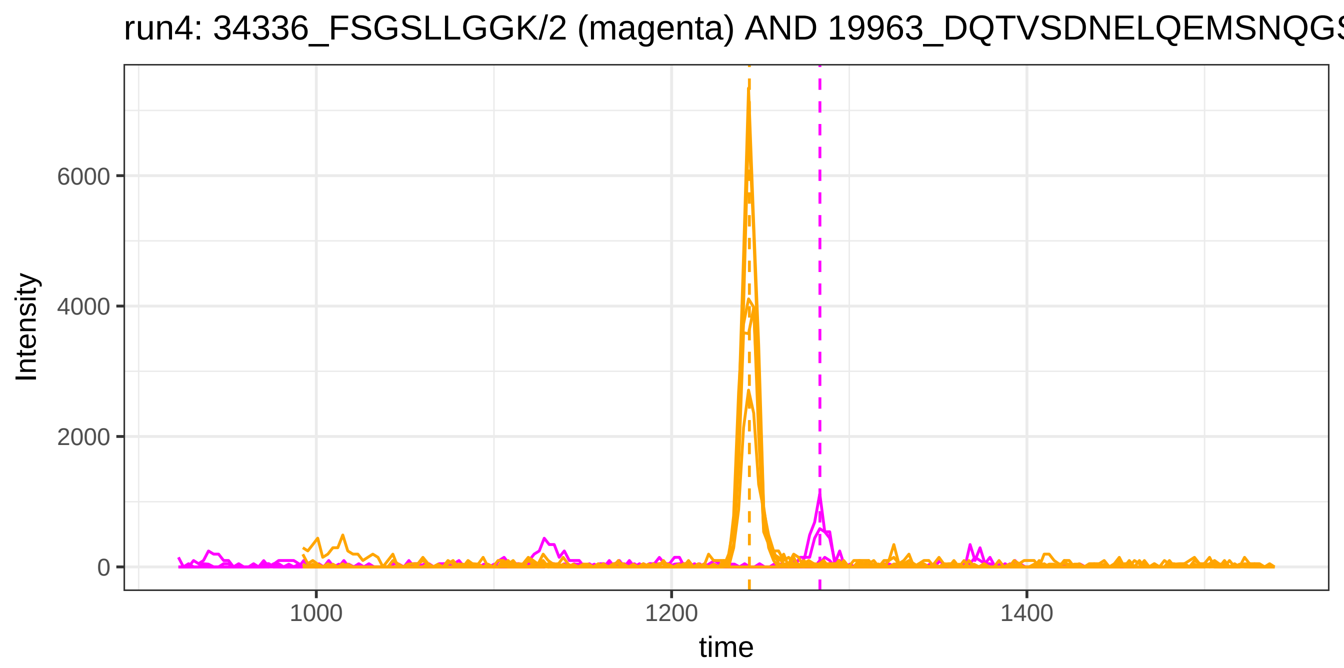
run4: 34336_FSGSLLGGK/2 (magenta) AND 19963_DQTVSDNELQEMSNQGS
Intensity
0
2000
4000
6000
1000
1200
1400
time

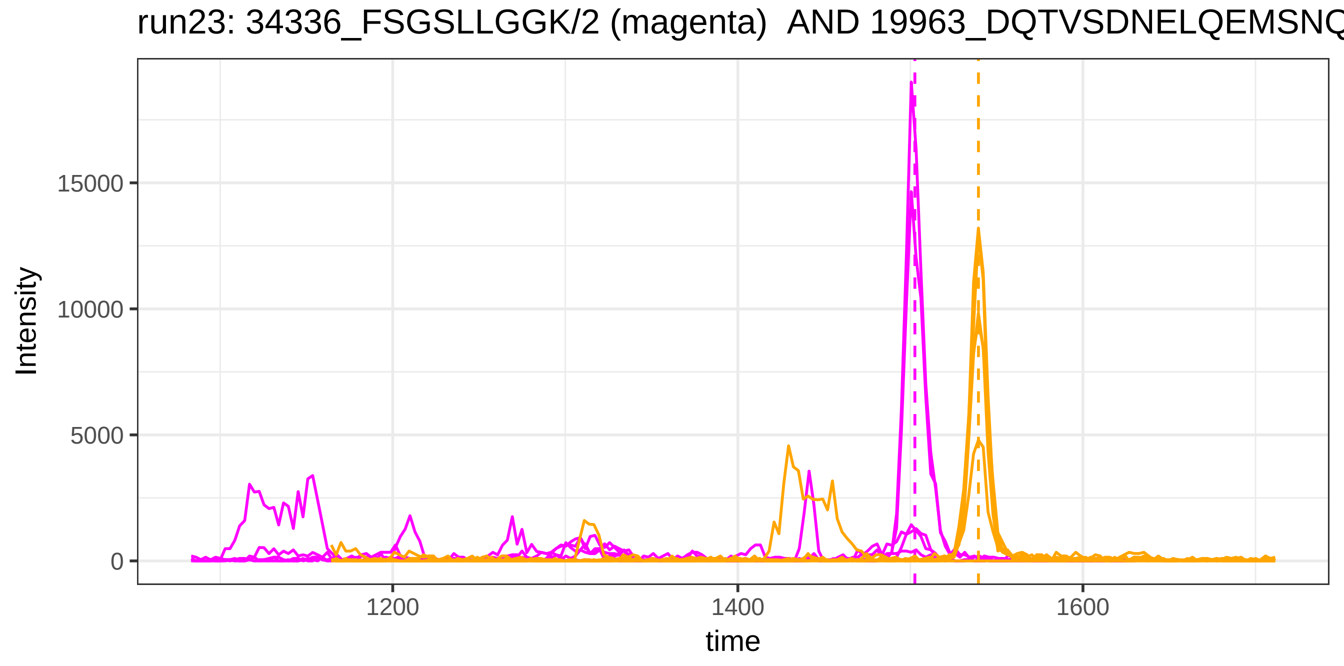
run23: 34336_FSGSLLGGK/2 (magenta) AND 19963_DQTVSDNELQEMSNQ
Intensity
0
5000
10000
15000
1200
1400
1600
time

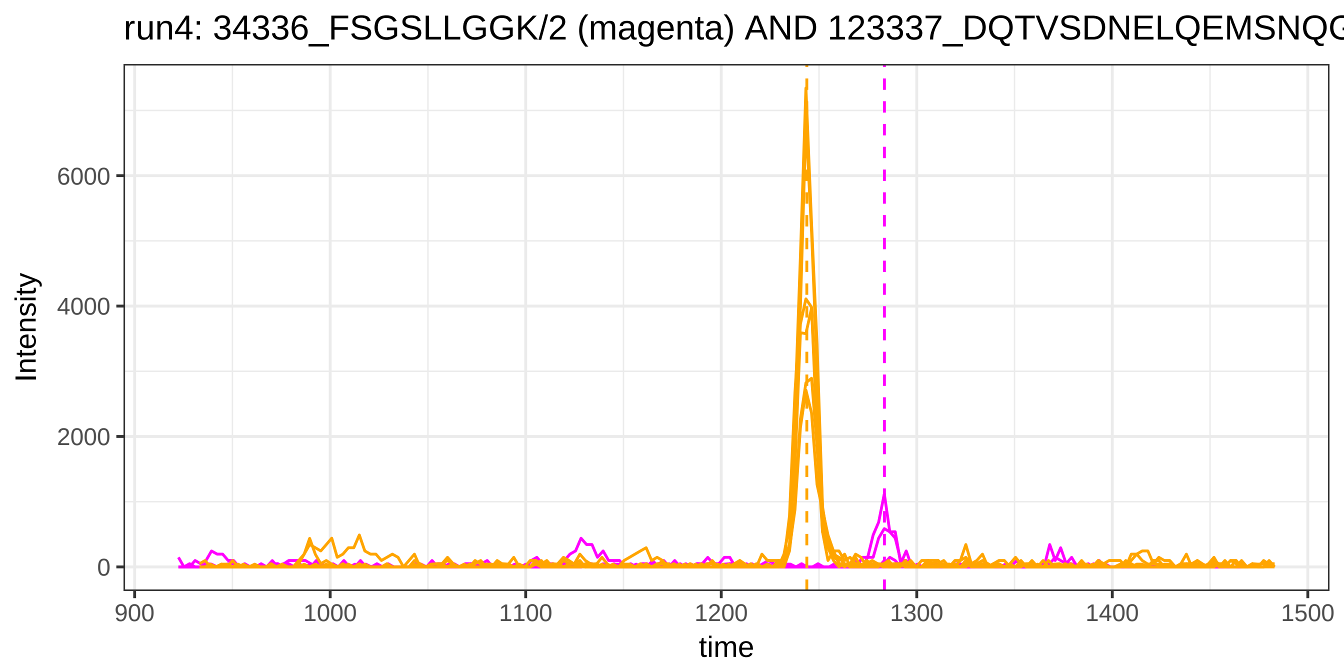
run4: 34336_FSGSLLGGK/2 (magenta) AND 123337_DQTVSDNELQEMSNQG
Intensity
6000
4000
2000
0
900
1000
1100
1200
1300
1400
1500
time

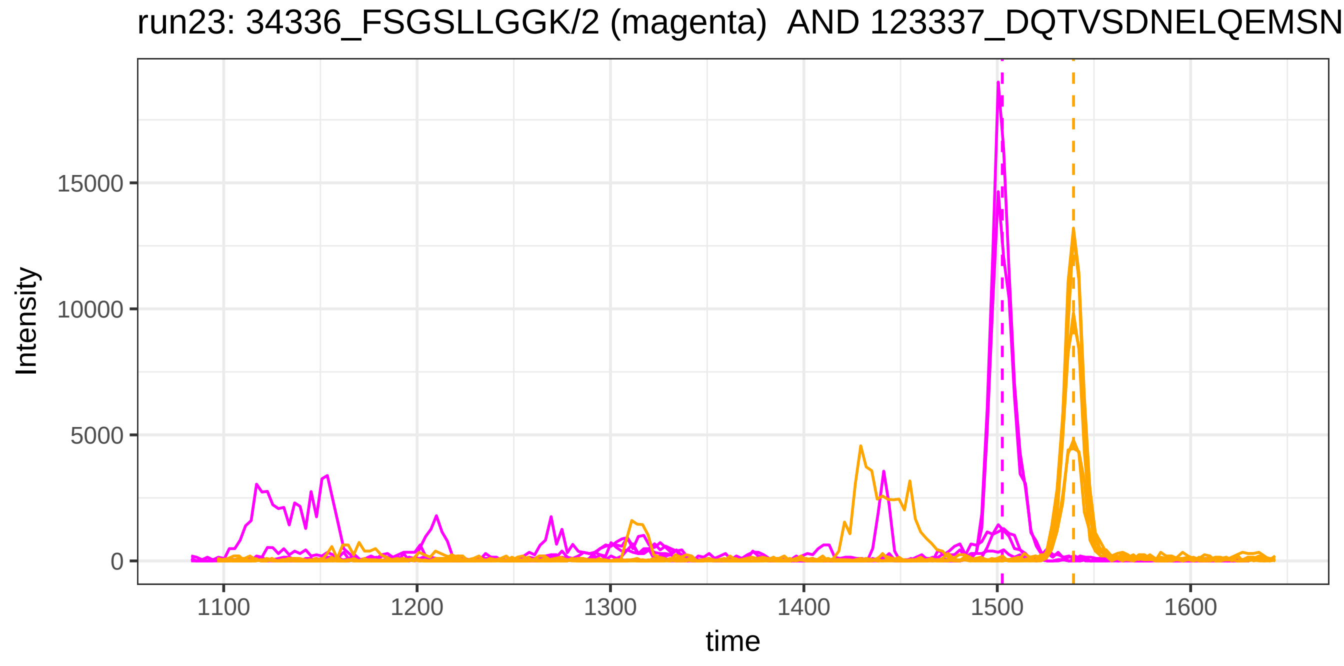
run23: 34336_FSGSLLGGK/2 (magenta) AND 123337_DQTVSDNELQEMSN
Intensity
15000
10000
5000
0
1100
1200
1300
1400
1500
1600
time

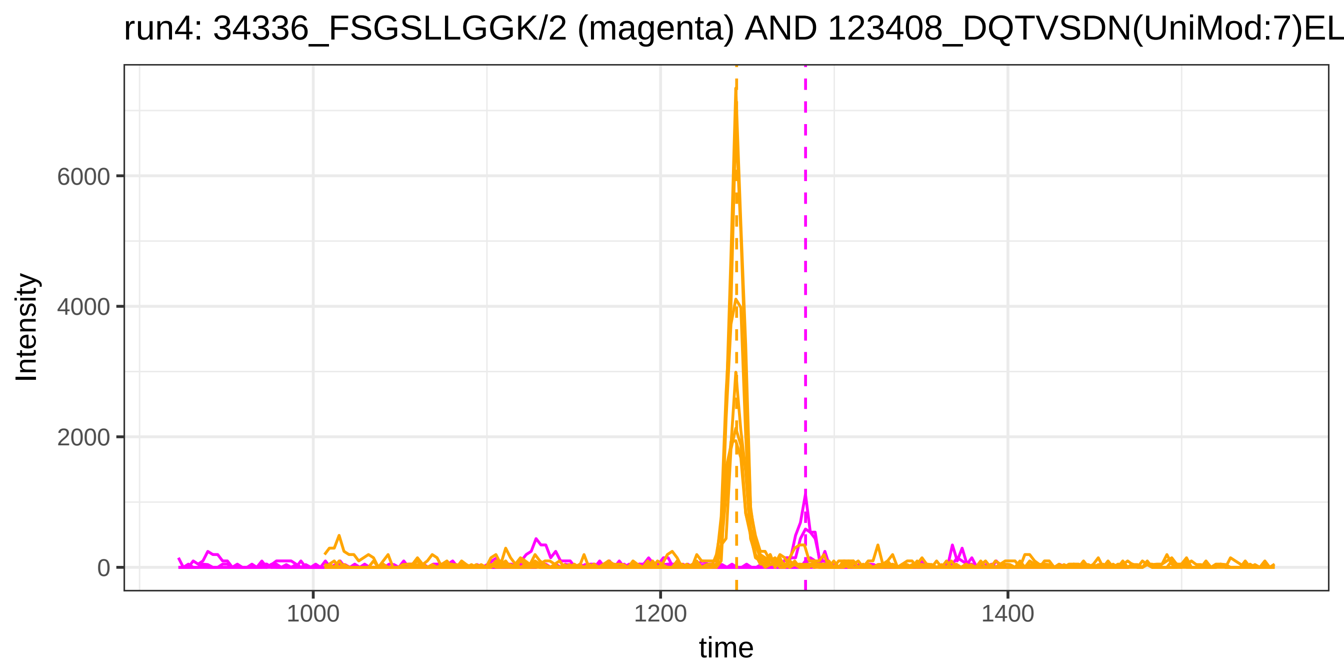

run4: 34336_FSGSLLGGK/2 (magenta) AND 123408_DQTVSDN(UniMod:7)EL
Intensity
6000
4000
2000
0
1000
1200
1400
time


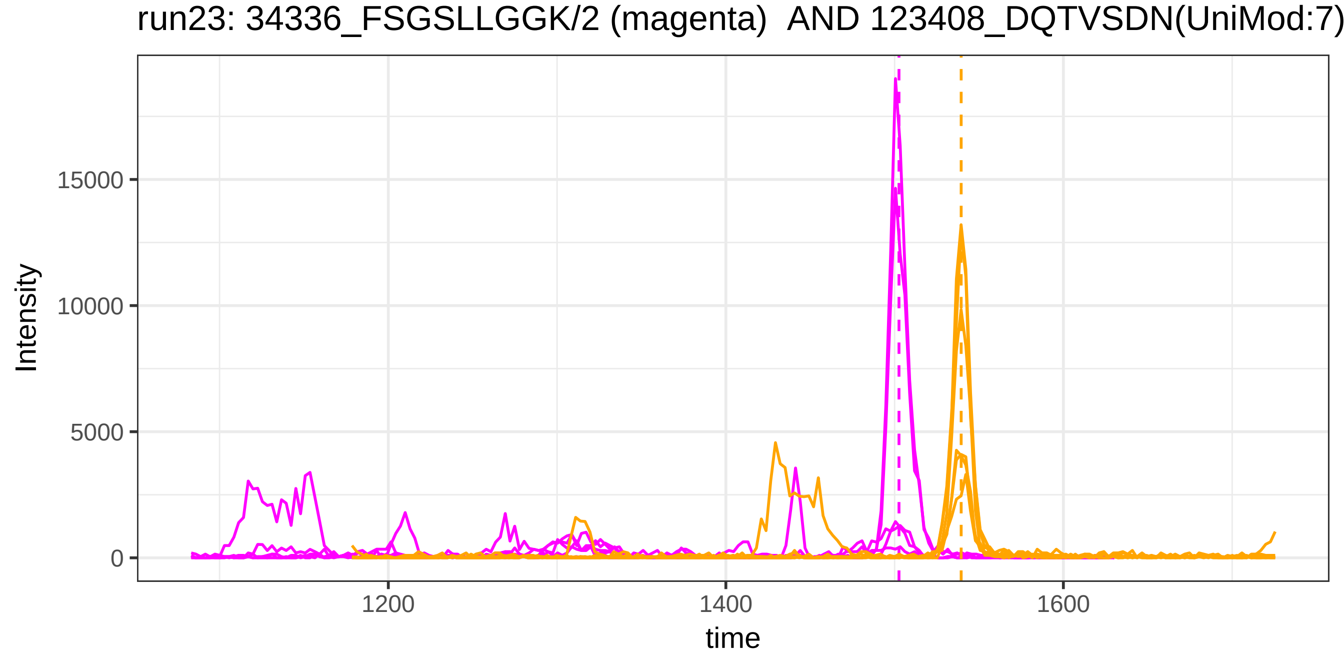

run23: 34336_FSGSLLGGK/2 (magenta) AND 123408_DQTVSDN(UniMod:7)
Intensity
15000
10000
5000
0
1200
1400
1600
time

run4: 43789_LLELTGPK/2 (magenta) AND 41874_KSASDLTWDNLK/3

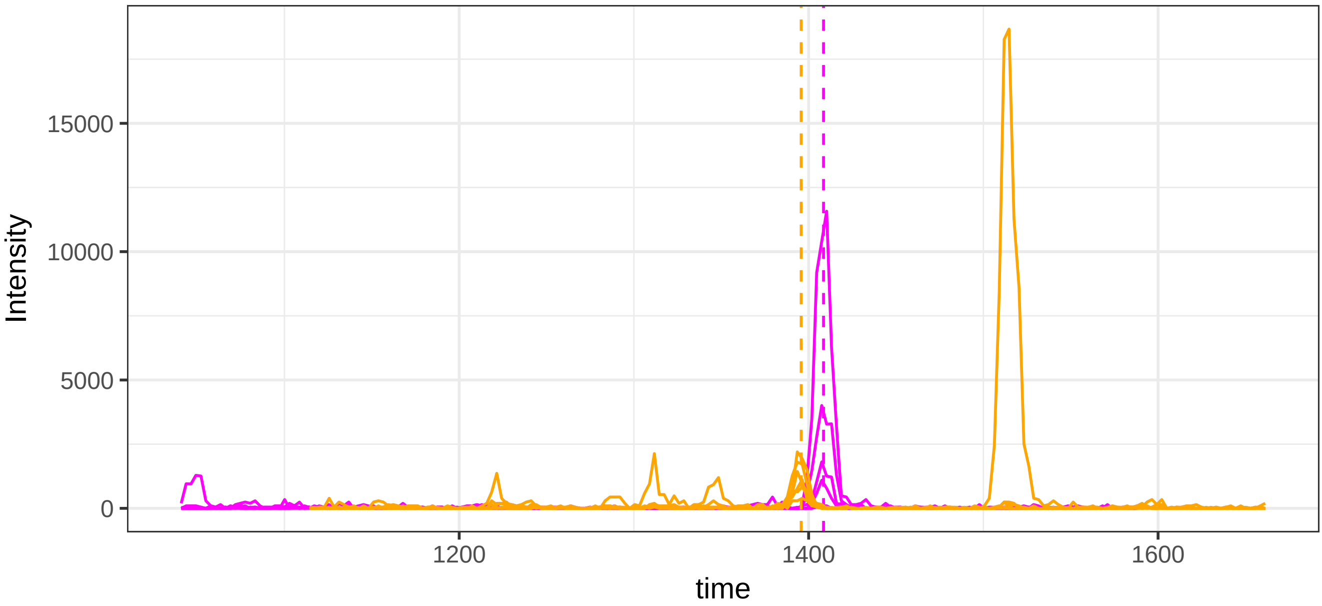


run23: 43789_LLELTGPK/2 (magenta)  AND 41874_KSASDLTWDNLK/3

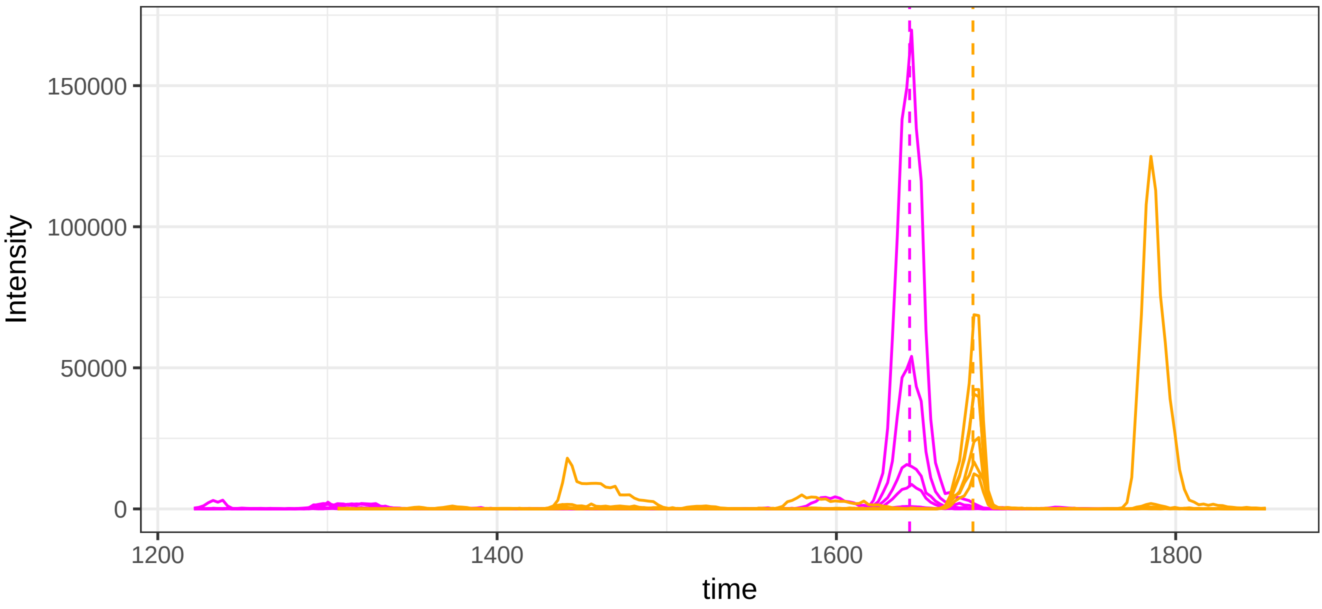

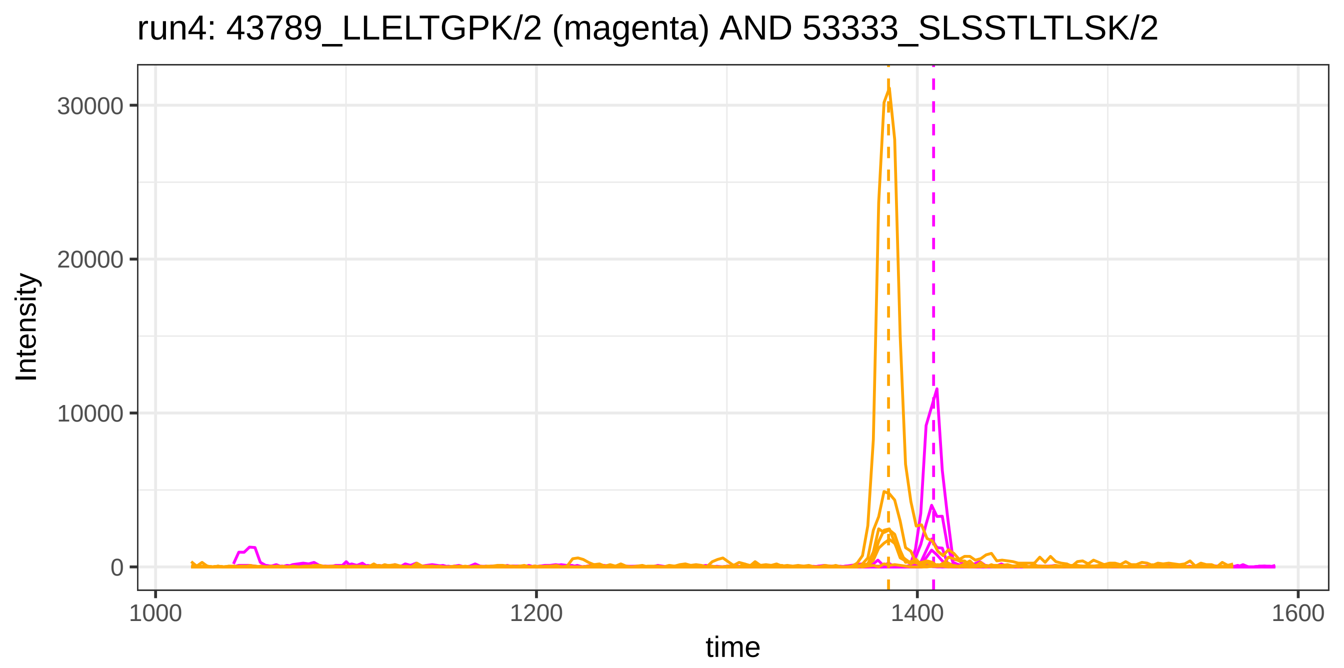
run4: 43789_LLELTGPK/2 (magenta) AND 53333_SLSSTLTLSK/2
Intensity
30000
20000
10000
0
1000
1200
1400
1600
time

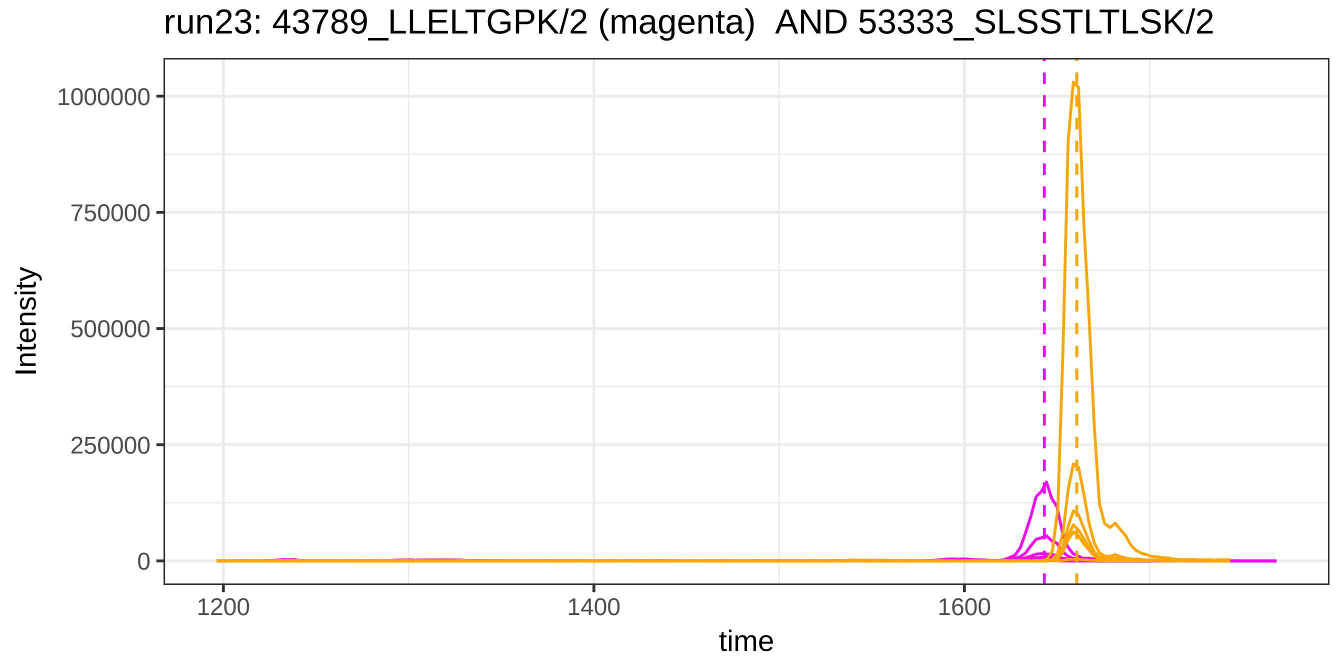
run23: 43789_LLELTGPK/2 (magenta) AND 53333_SLSSTLTLSK/2
Intensity
1000000
750000
500000
250000
0
1200
1400
1600
time

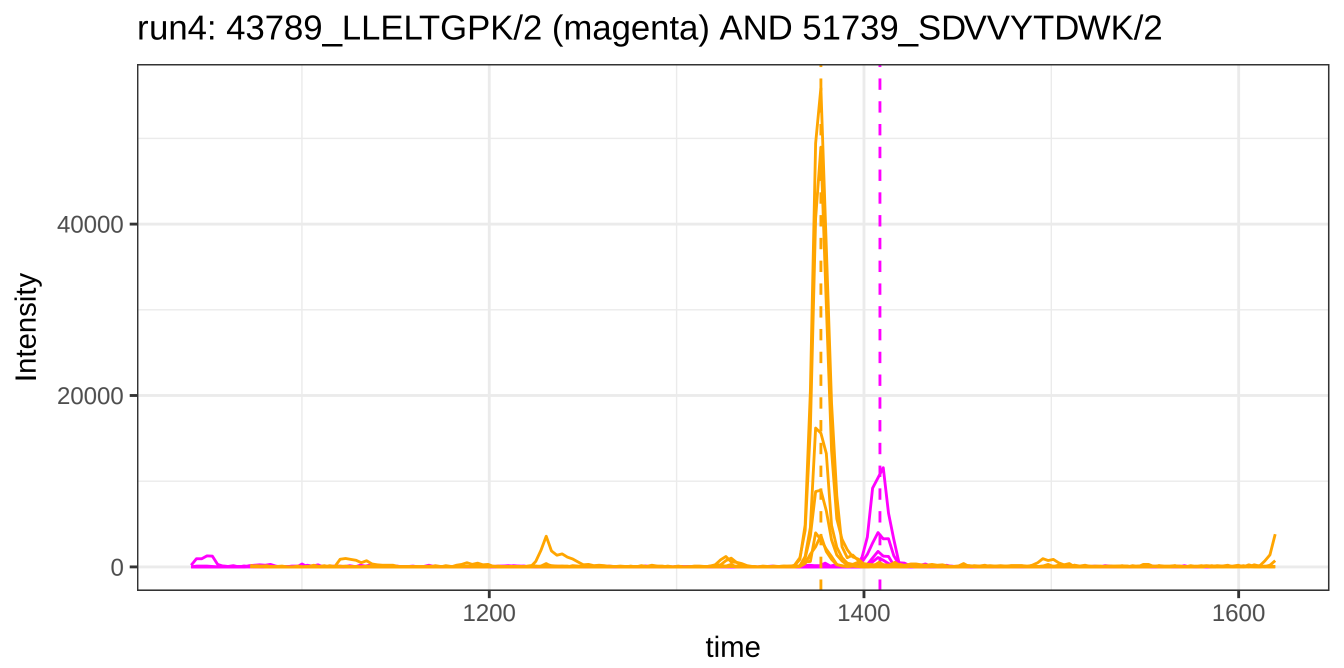

run4: 43789_LLELTGPK/2 (magenta) AND 51739_SDVVYTDWK/2
Intensity
40000
20000
0
1200
1400
1600
time


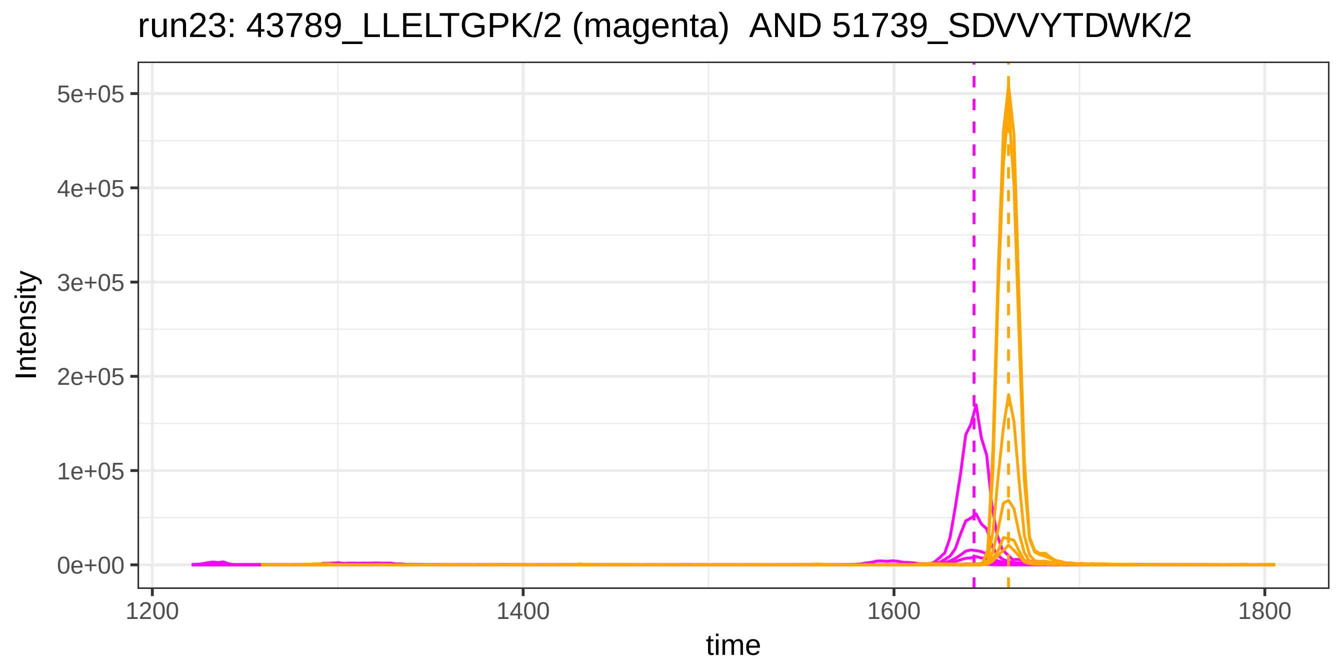

run23: 43789_LLELTGPK/2 (magenta)  AND 51739_SDVVYTDWK/2
Intensity
5e+05
4e+05
3e+05
2e+05
1e+05
0e+00
1200
1400
1600
1800
time

run4: 43789_LLELTGPK/2 (magenta) AND 127673_SDVVYTDWK/2

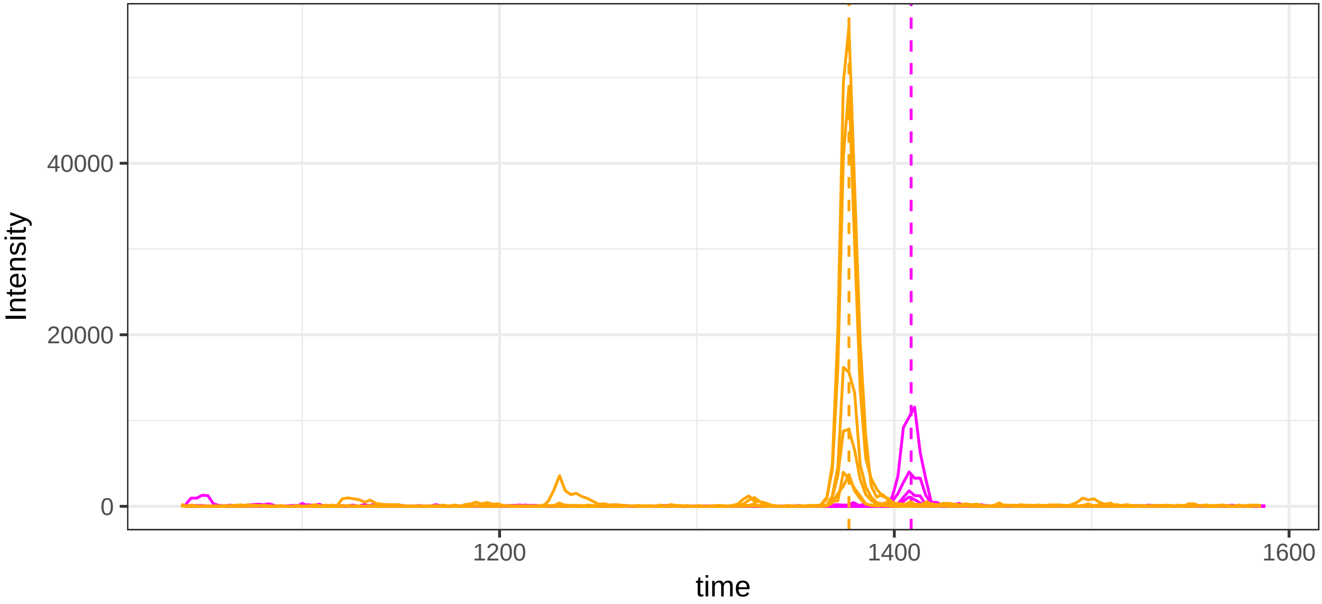


run23: 43789_LLELTGPK/2 (magenta)  AND 127673_SDVVYTDWK/2

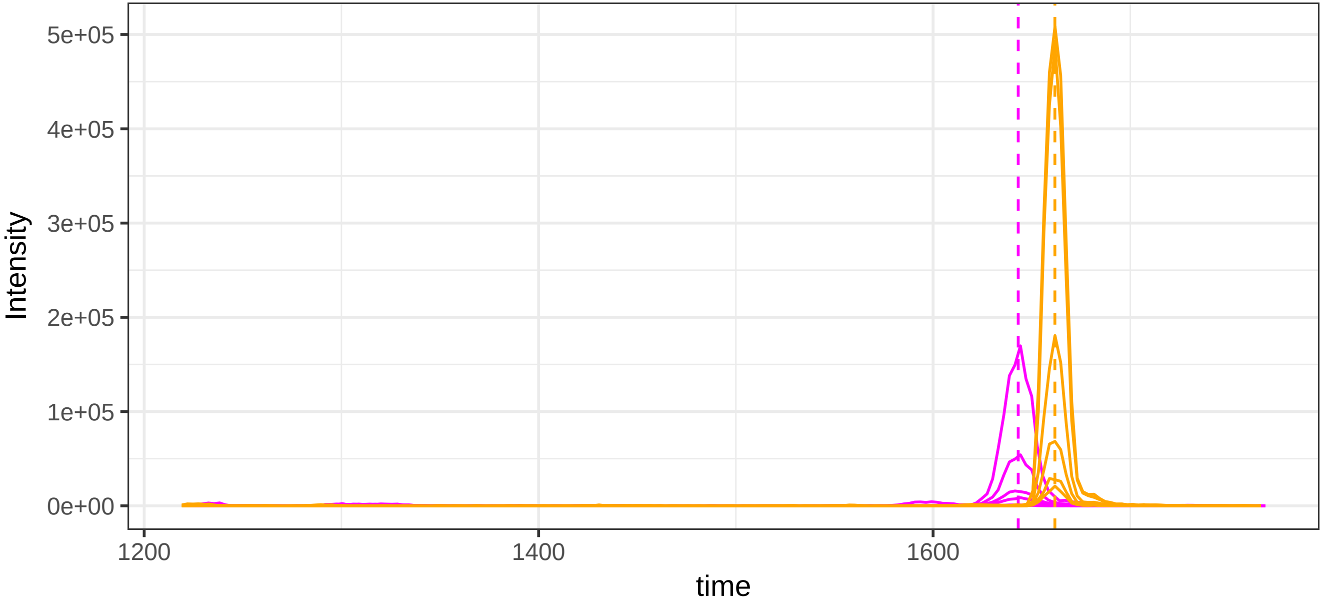

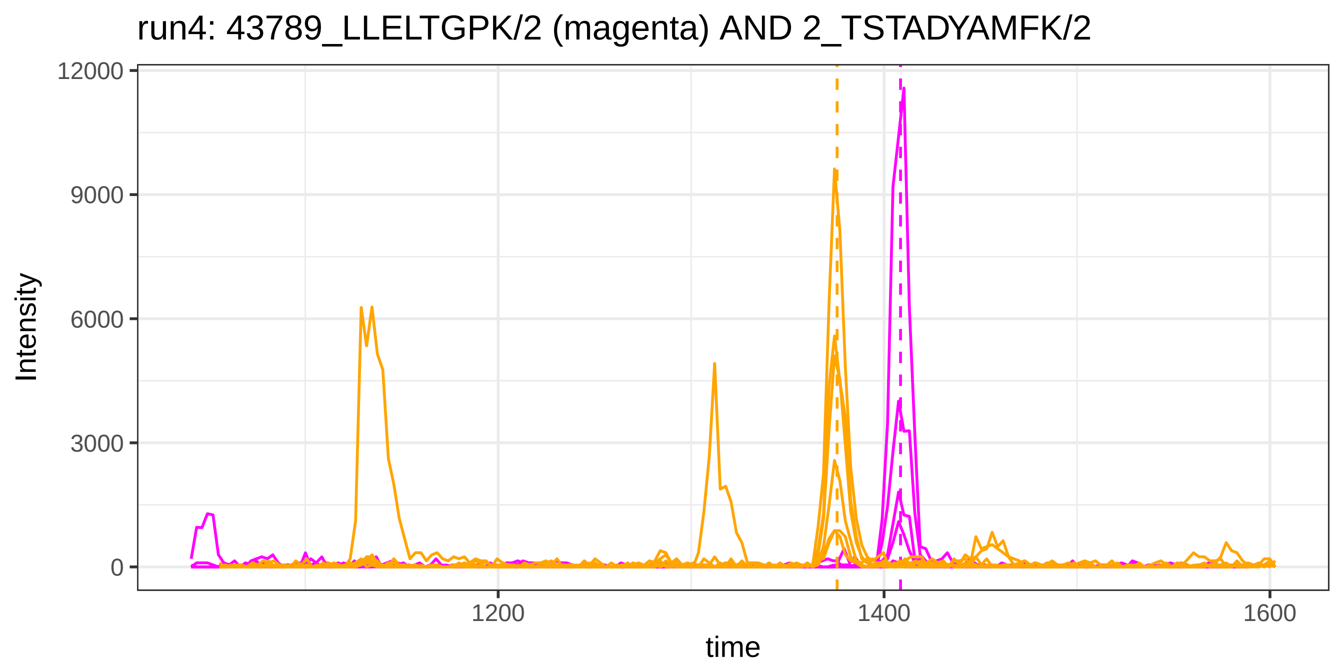
run4: 43789_LLELTGPK/2 (magenta) AND 2_TSTADYAMFK/2
Intensity
12000
9000
6000
3000
0
1200
1400
1600
time

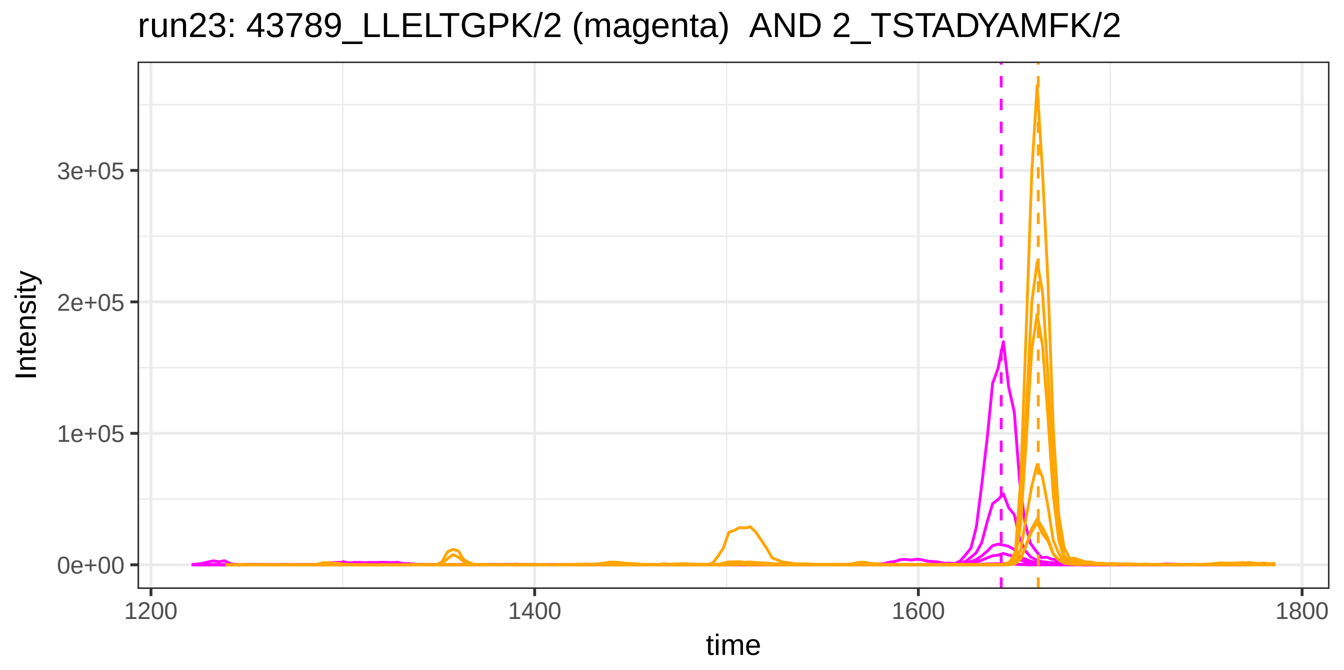
run23: 43789_LLELTGPK/2 (magenta) AND 2_TSTADYAMFK/2
Intensity
3e+05
2e+05
1e+05
0e+00
1200
1400
1600
1800
time

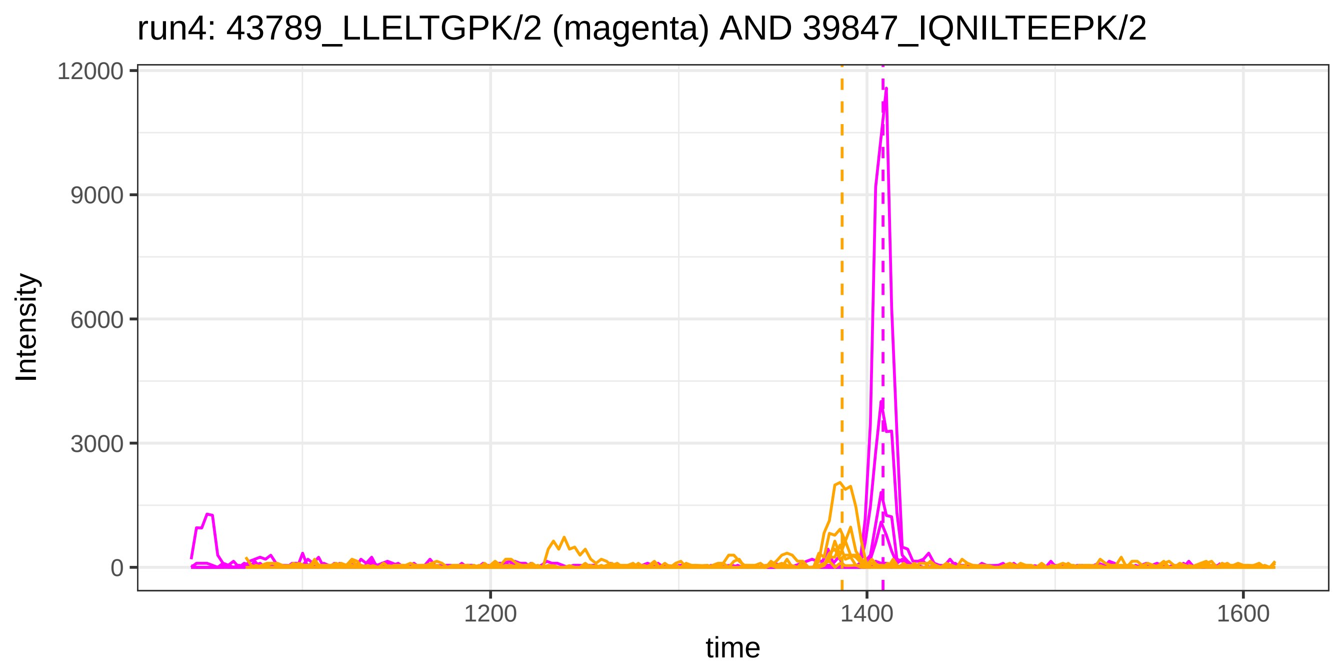
run4: 43789_LLELTGPK/2 (magenta) AND 39847_IQNILTEEPK/2
Intensity
12000
9000
6000
3000
0
1200
1400
1600
time

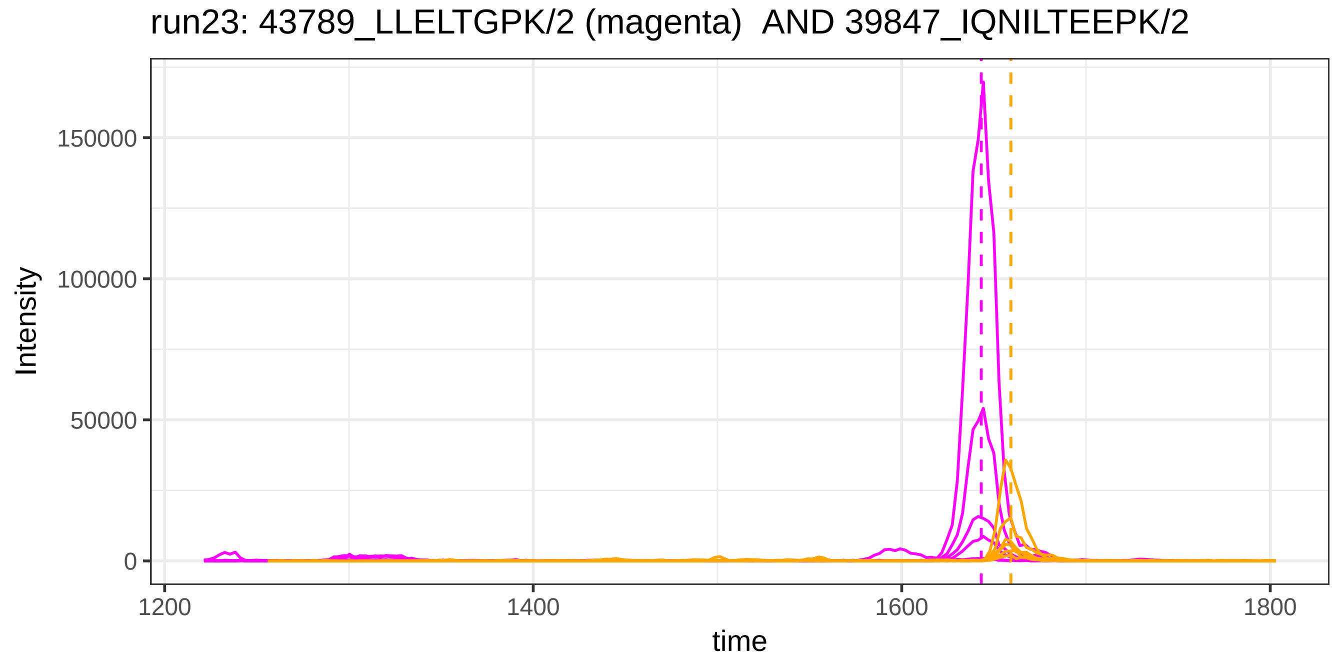
run23: 43789_LLELTGPK/2 (magenta)  AND 39847_IQNILTEEPK/2
Intensity
150000
100000
50000
0
1200
1400
1600
1800
time

run4: 43789_LLELTGPK/2 (magenta) AND 20961_DSGFQMNQLR/2

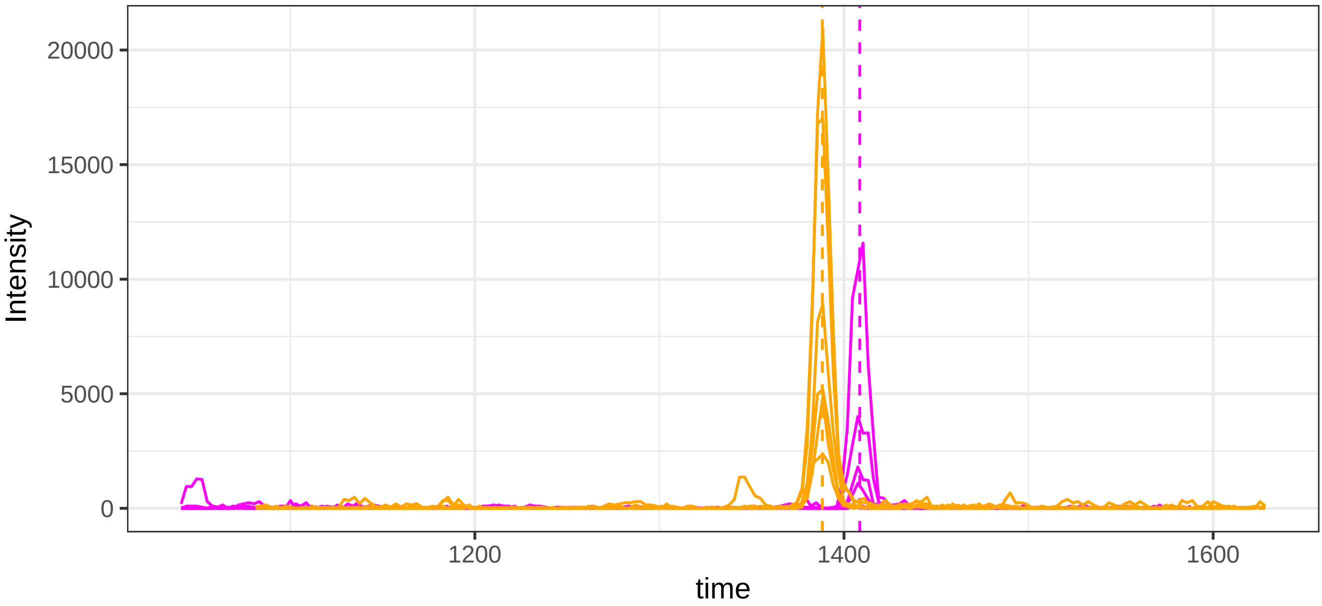


run23: 43789_LLELTGPK/2 (magenta) AND 20961_DSGFQMNQLR/2

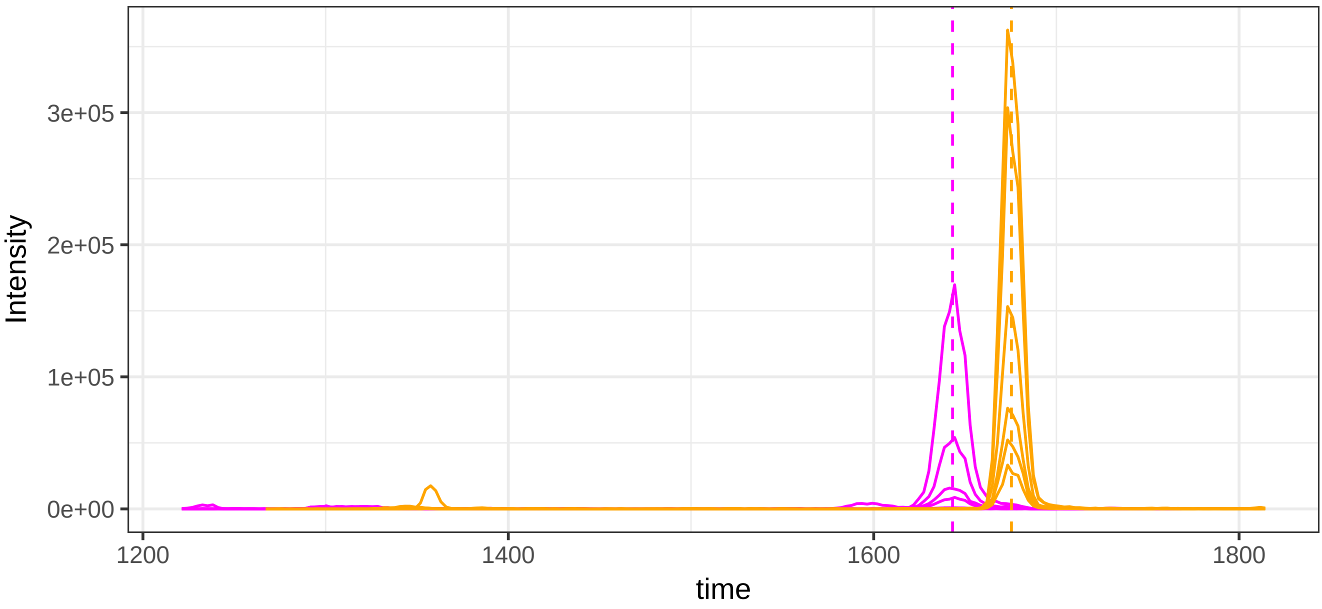

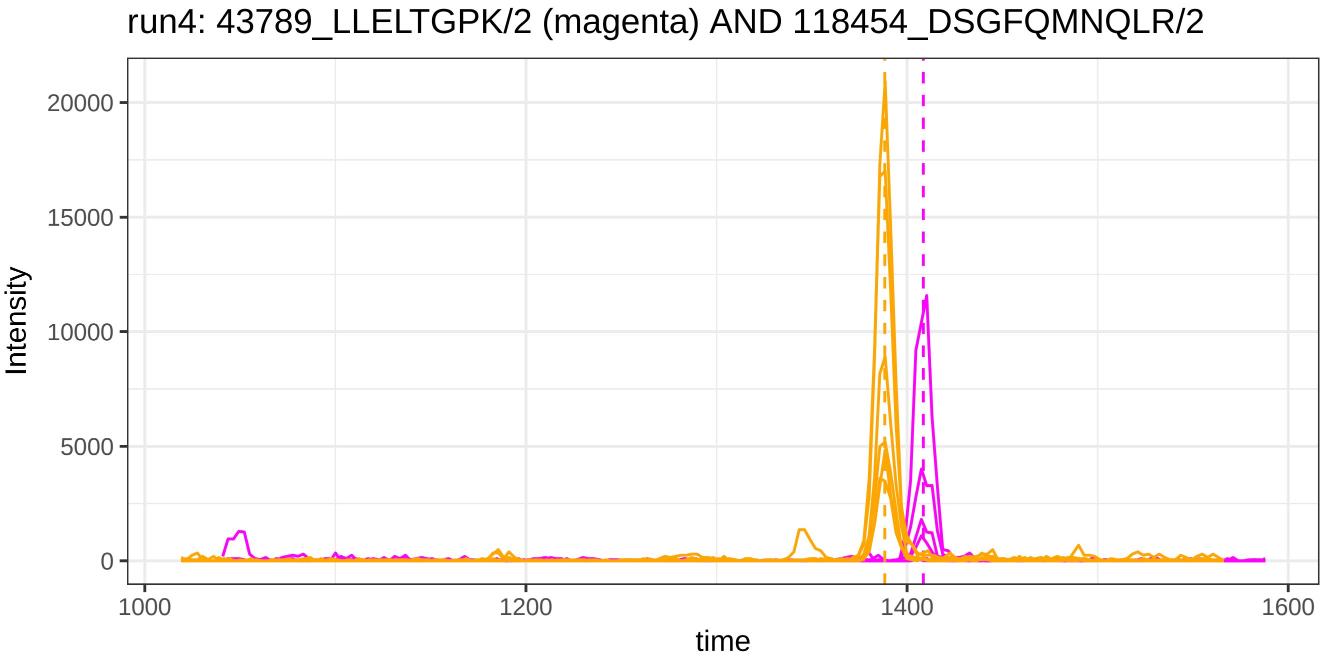
run4: 43789_LLELTGPK/2 (magenta) AND 118454_DSGFQMNQLR/2
Intensity
20000
15000
10000
5000
0
1000
1200
1400
1600
time

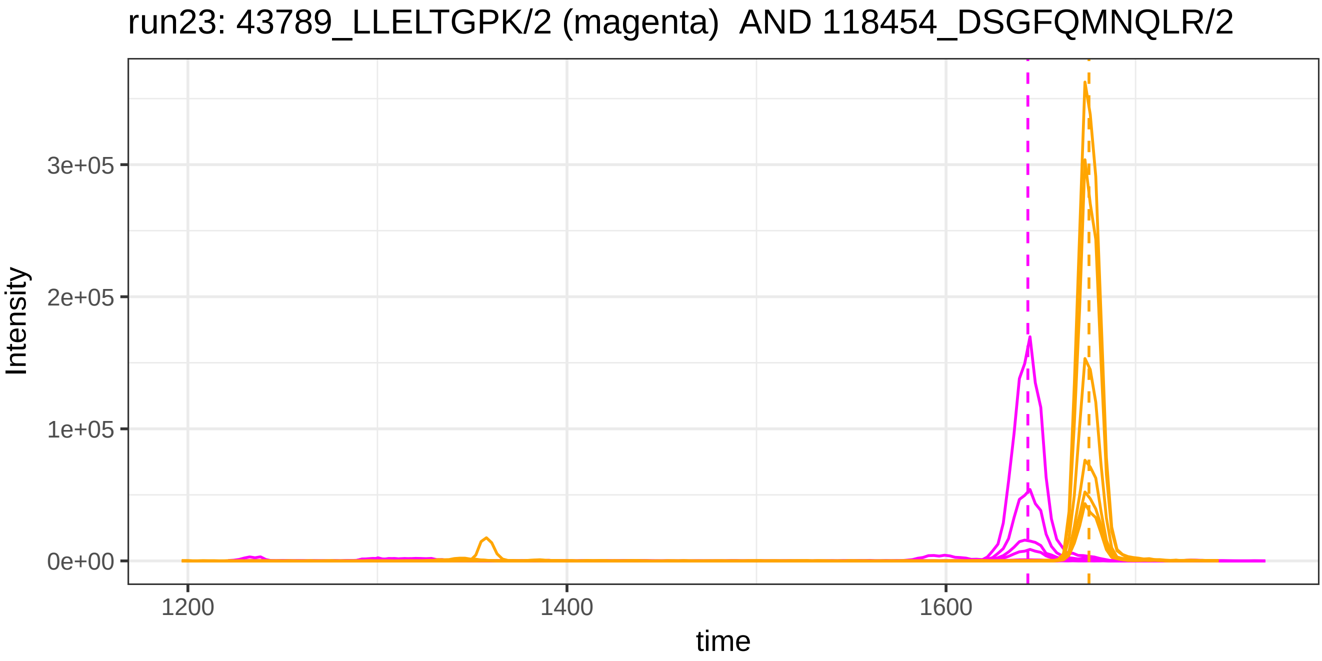
run23: 43789_LLELTGPK/2 (magenta)  AND 118454_DSGFQMNQLR/2
Intensity
3e+05
2e+05
1e+05
0e+00
1200
1400
1600
time

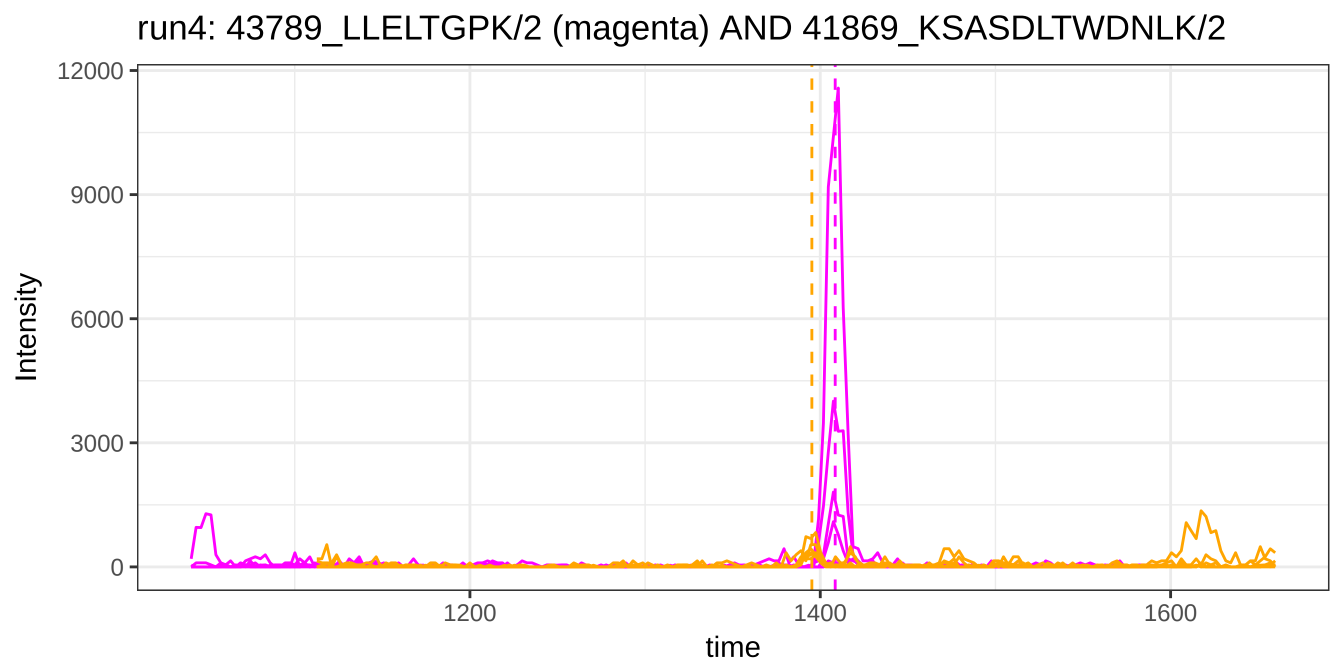
run4: 43789_LLELTGPK/2 (magenta) AND 41869_KSASDLTWDNLK/2
Intensity
12000
9000
6000
3000
0
1200
1400
1600
time

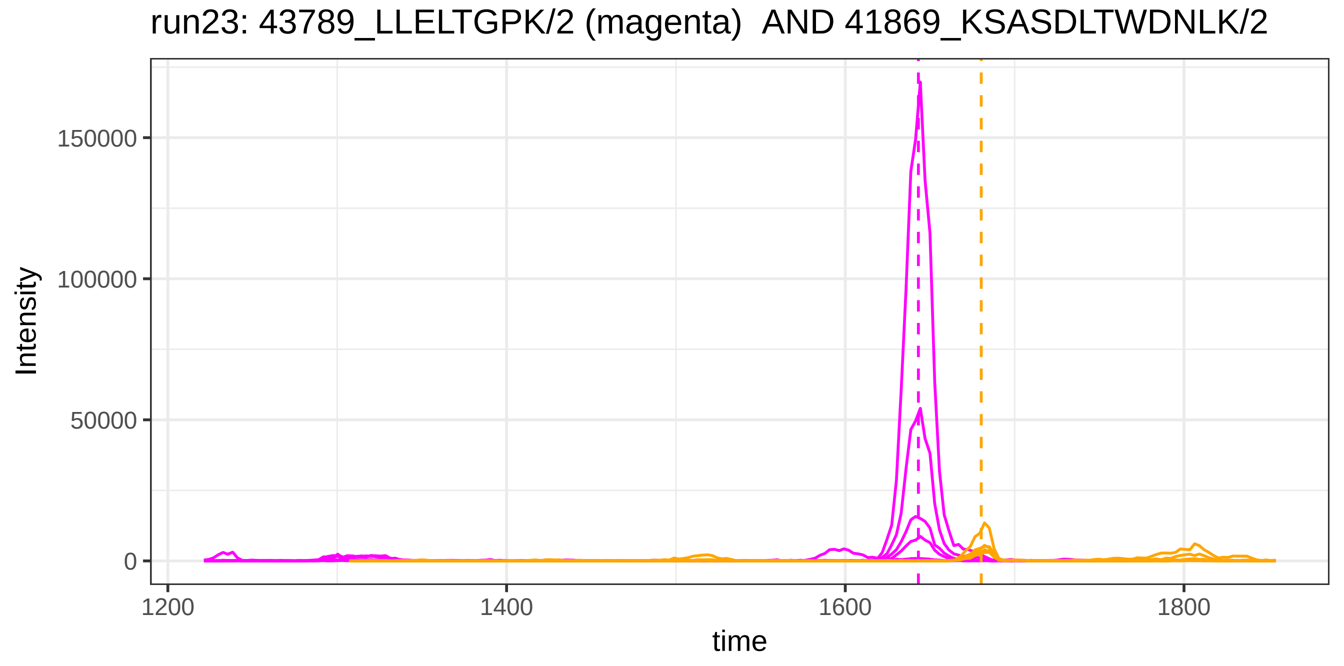
run23: 43789_LLELTGPK/2 (magenta)  AND 41869_KSASDLTWDNLK/2
Intensity
150000
100000
50000
0
1200
1400
1600
1800
time

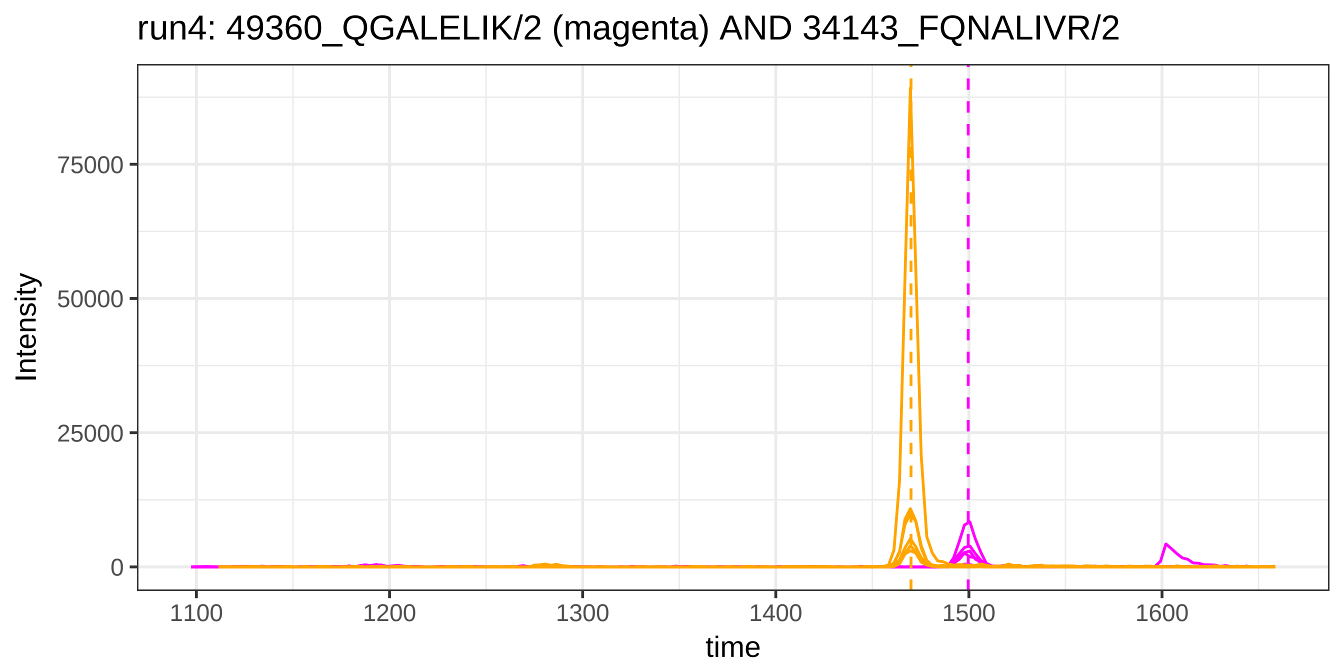

run4: 49360_QGALELIK/2 (magenta) AND 34143_FQNALIVR/2
Intensity
75000
50000
25000
0
1100
1200
1300
1400
1500
1600
time


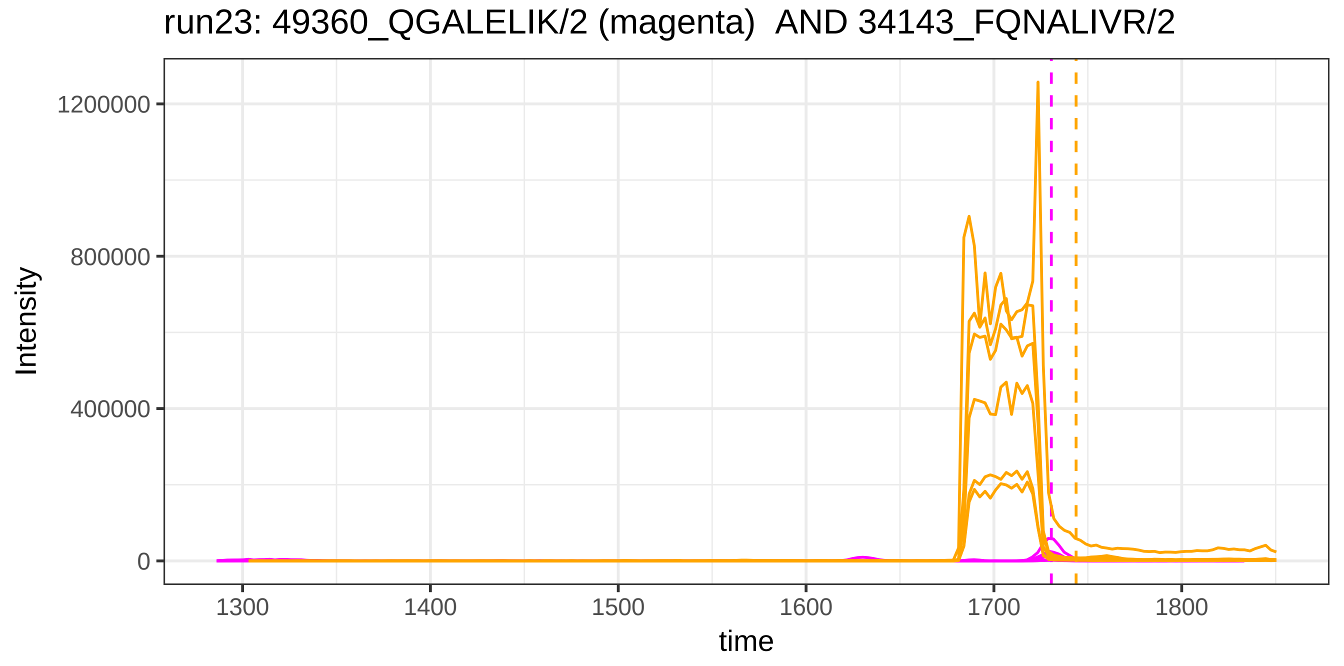

run23: 49360_QGALELIK/2 (magenta) AND 34143_FQNALIVR/2
Intensity
1200000
800000
400000
0
1300
1400
1500
1600
1700
1800
time

run4: 49360_QGALELIK/2 (magenta) AND 41242_KGGETSEMYLIQPDSSVKF

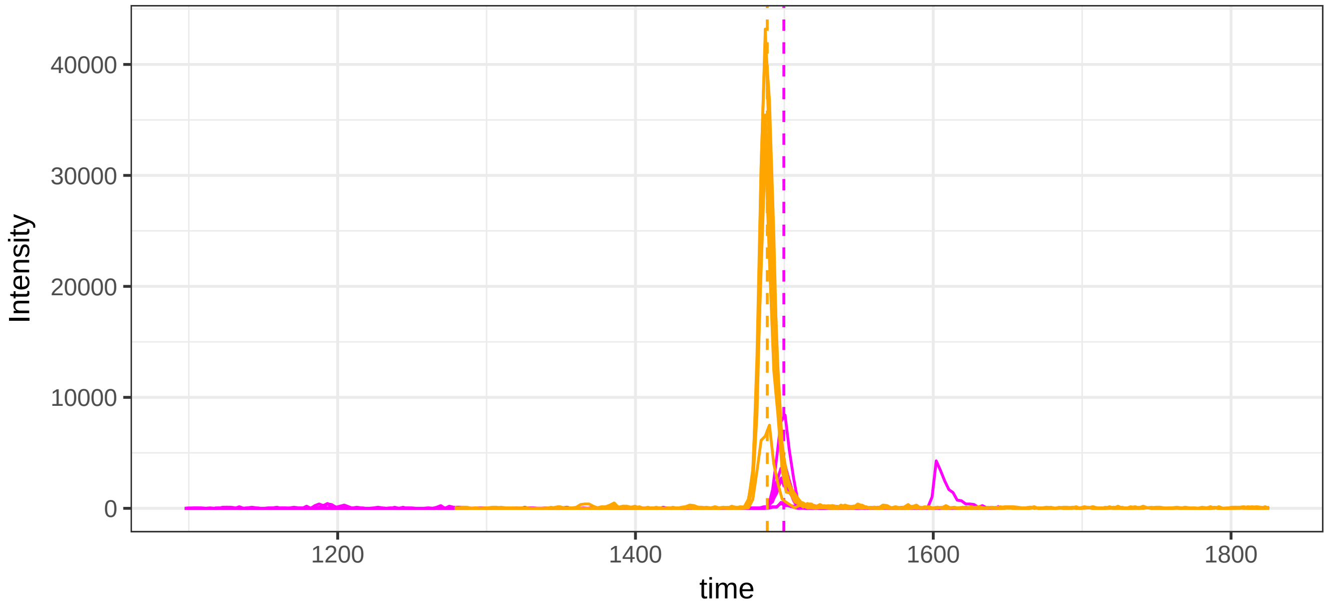


run23: 49360_QGALELIK/2 (magenta)  AND 41242_KGGETSEMYLIQPDSSVI

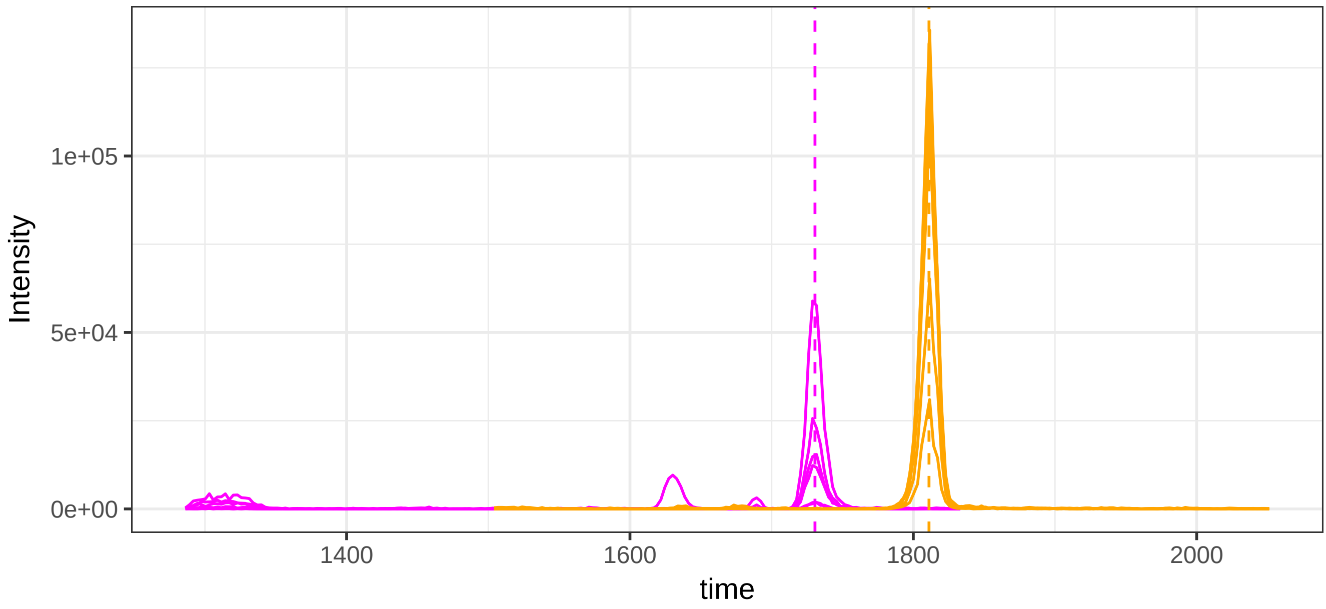

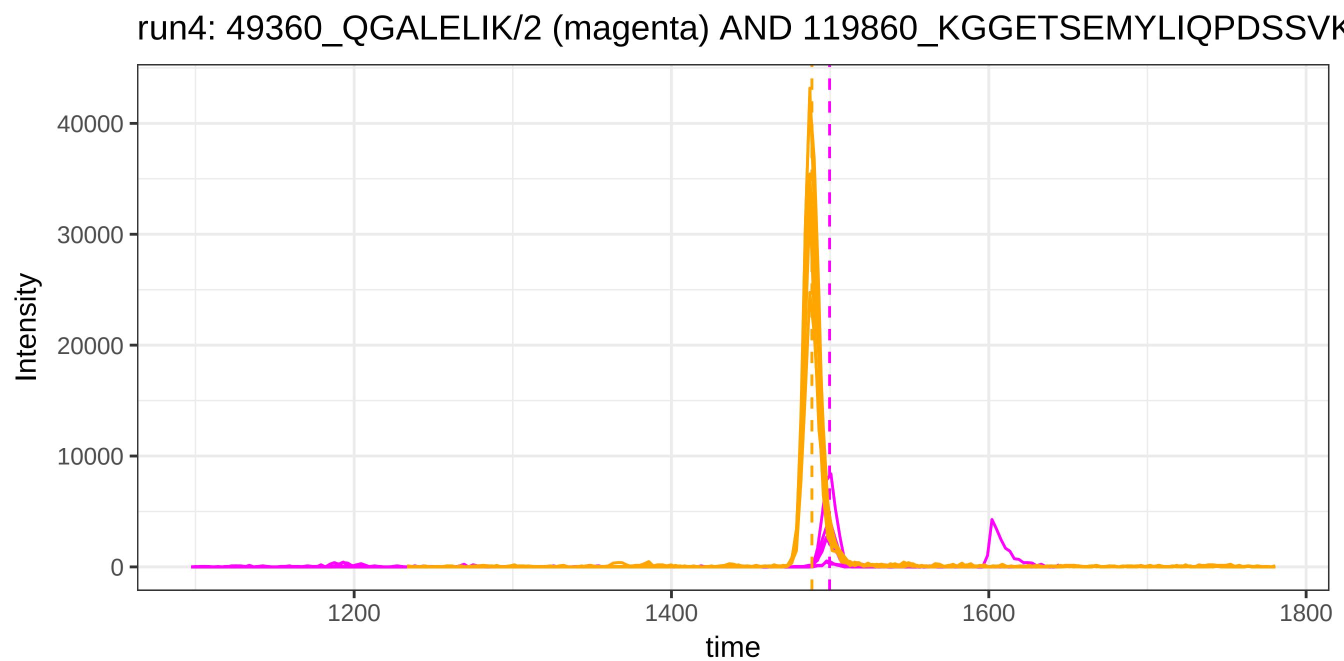

run4: 49360_QGALELIK/2 (magenta) AND 119860_KGGETSEMYLIQPDSSVK
Intensity
40000
30000
20000
10000
0
1200
1400
1600
1800
time


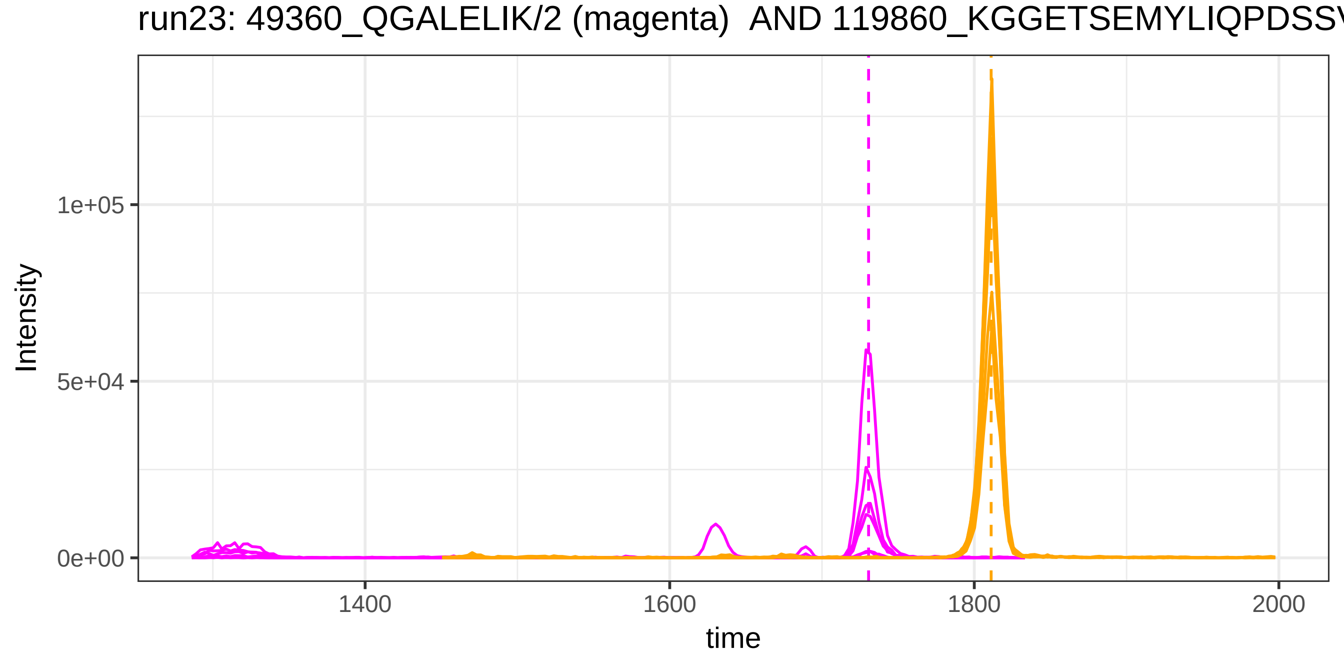

run23: 49360_QGALELIK/2 (magenta) AND 119860_KGGETSEMYLIQPDSSV
Intensity
1e+05
5e+04
0e+00
1400
1600
1800
2000
time

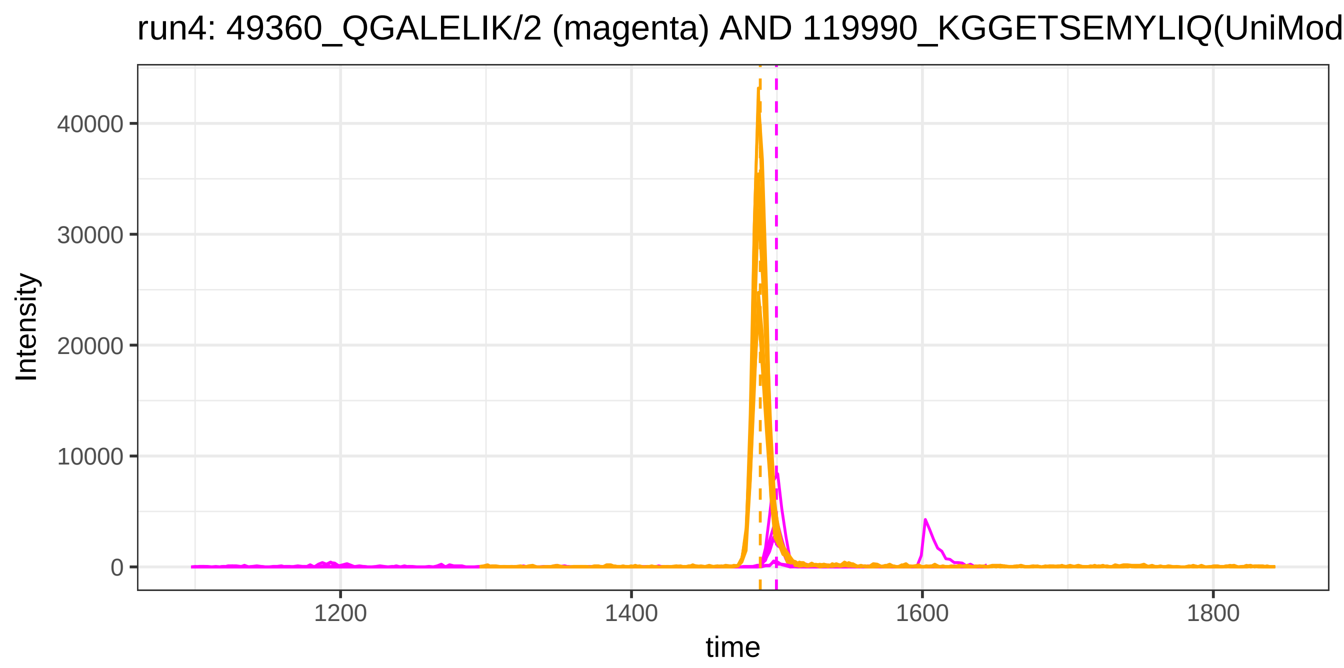
run4: 49360_QGALELIK/2 (magenta) AND 119990_KGGETSEMYLIQ(UniMod
Intensity
40000
30000
20000
10000
0
1200
1400
1600
1800
time

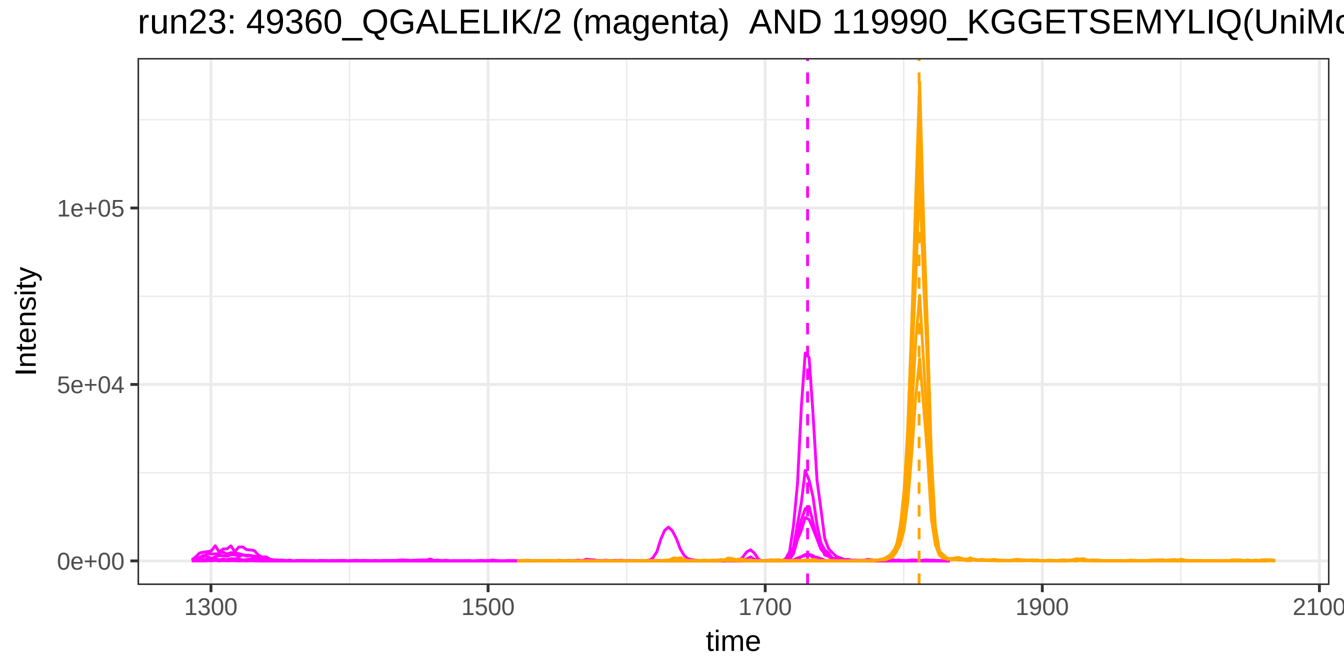
run23: 49360_QGALELIK/2 (magenta) AND 119990_KGGETSEMYLIQ(UniMo
Intensity
1e+05
5e+04
0e+00
1300
1500
1700
1900
2100
time

run4: 49360_QGALELIK/2 (magenta) AND 20944_DSGEGDFLAEGGGVR/2

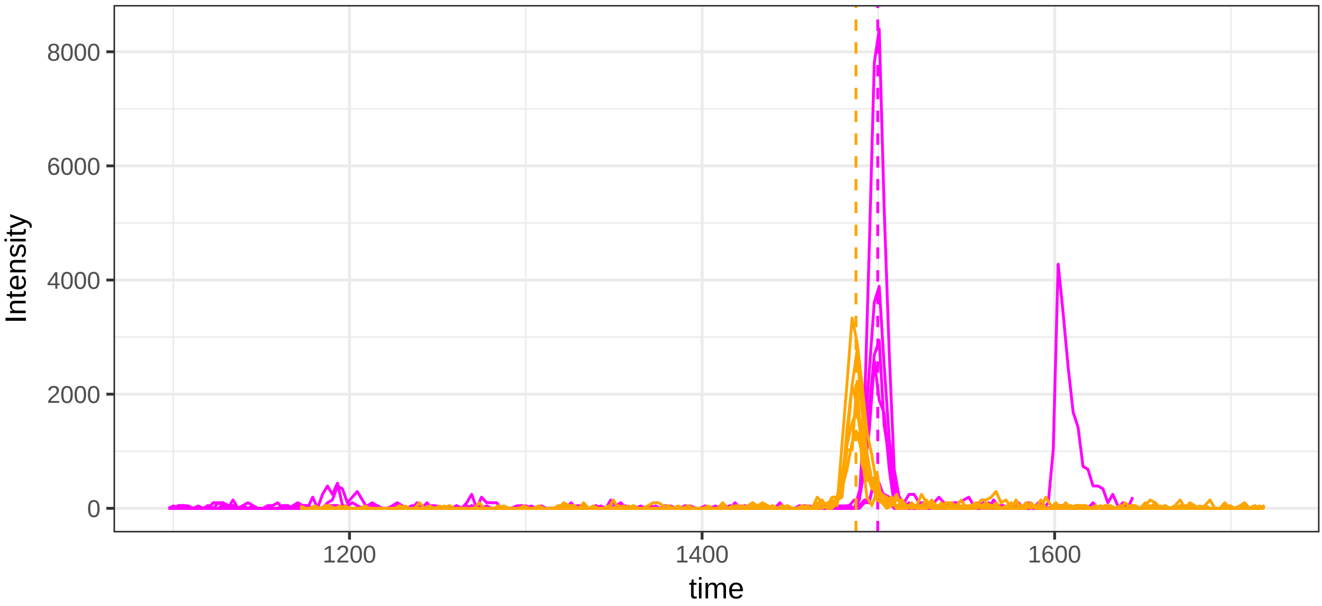


run23: 49360_QGALELIK/2 (magenta)  AND 20944_DSGEGDFLAEGGGVR/2

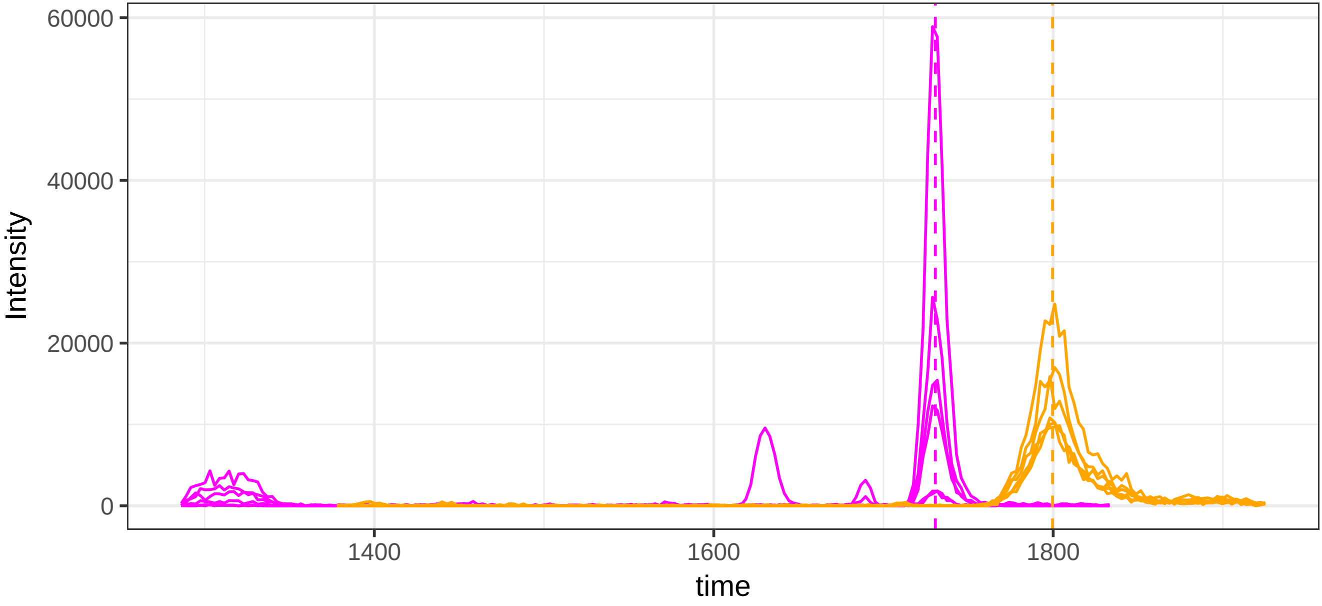

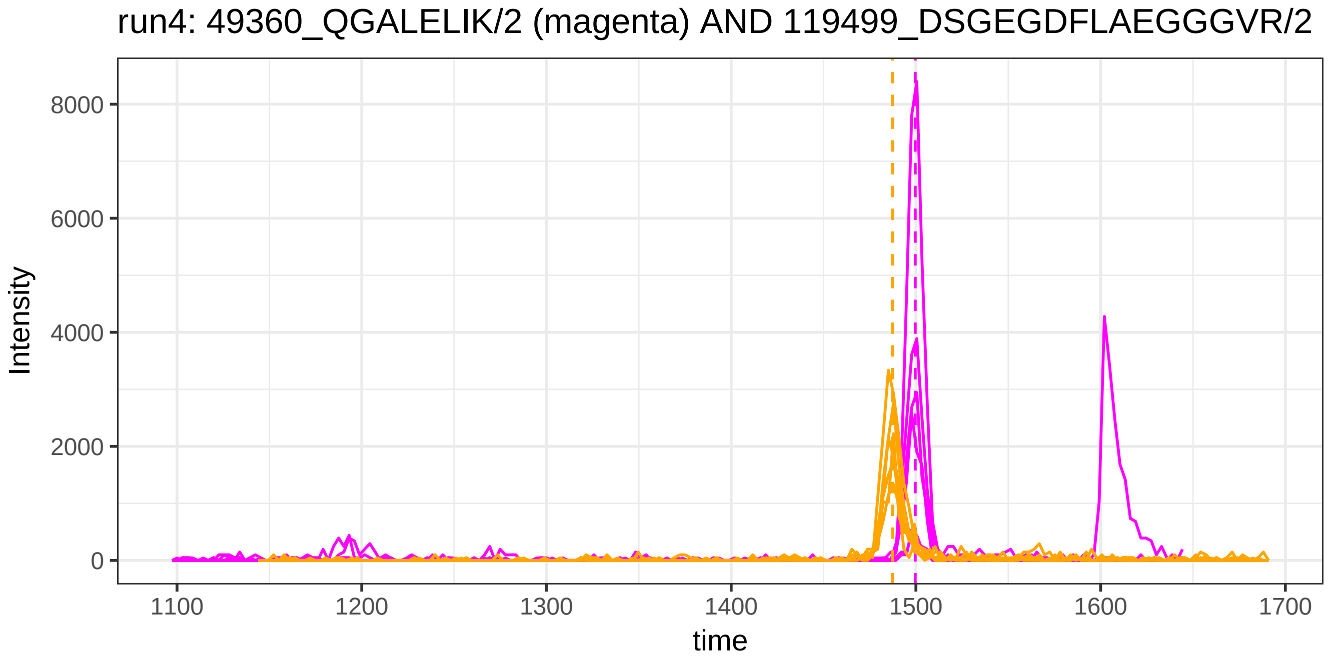
run4: 49360_QGALELIK/2 (magenta) AND 119499_DSGEGDFLAEGGGVR/2
Intensity
8000
6000
4000
2000
0
1100
1200
1300
1400
1500
1600
1700
time

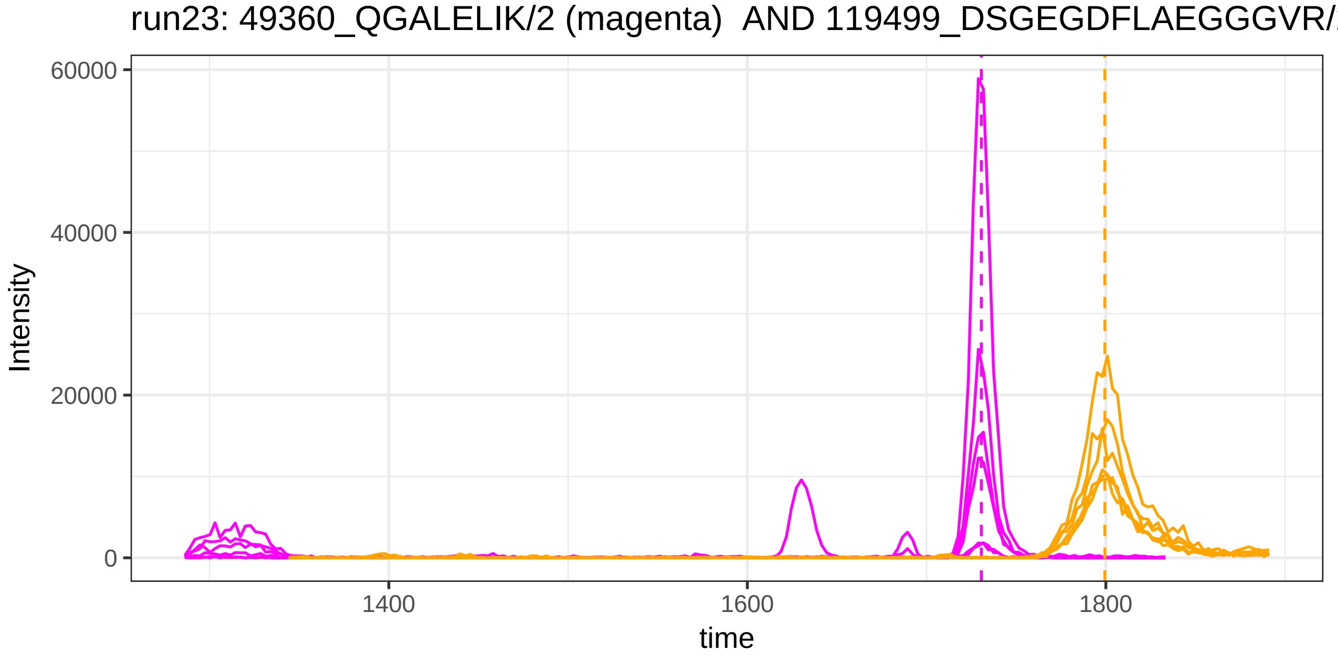
run23: 49360_QGALELIK/2 (magenta) AND 119499_DSGEGDFLAEGGGVR/
Intensity
60000
40000
20000
0
1400
1600
1800
time

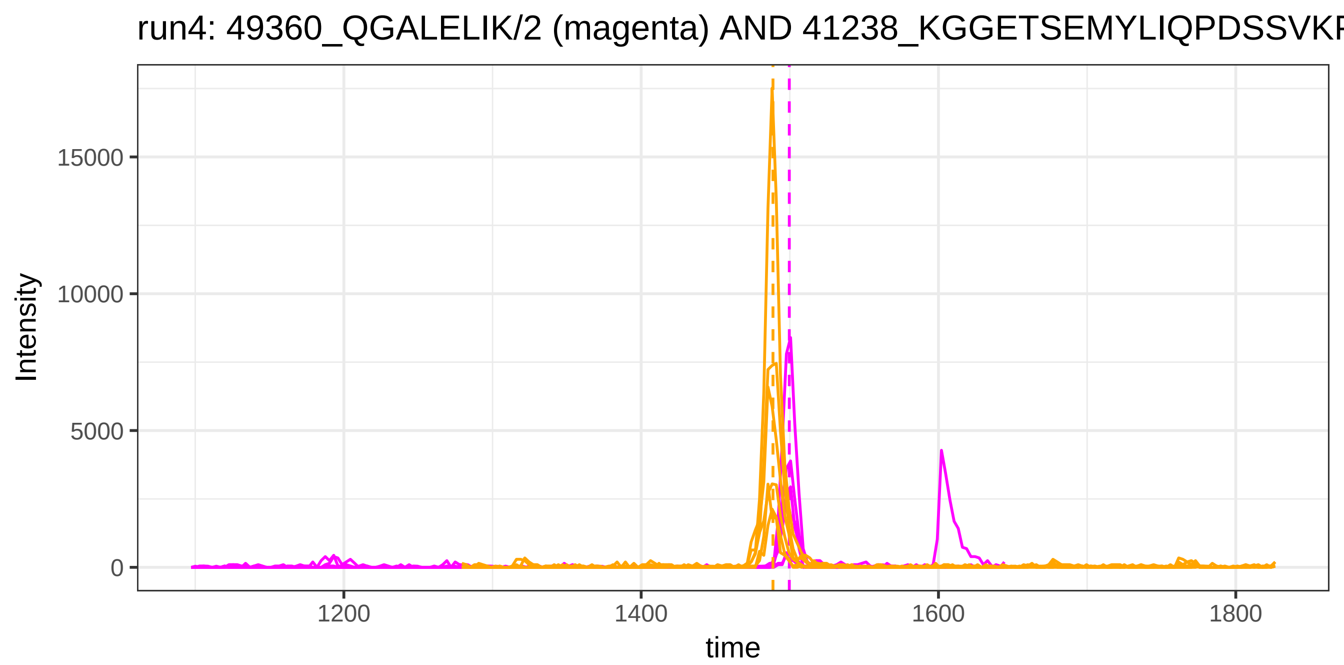
run4: 49360_QGALELIK/2 (magenta) AND 41238_KGGETSEMYLIQPDSSVKF
Intensity
15000
10000
5000
0
1200
1400
1600
1800
time

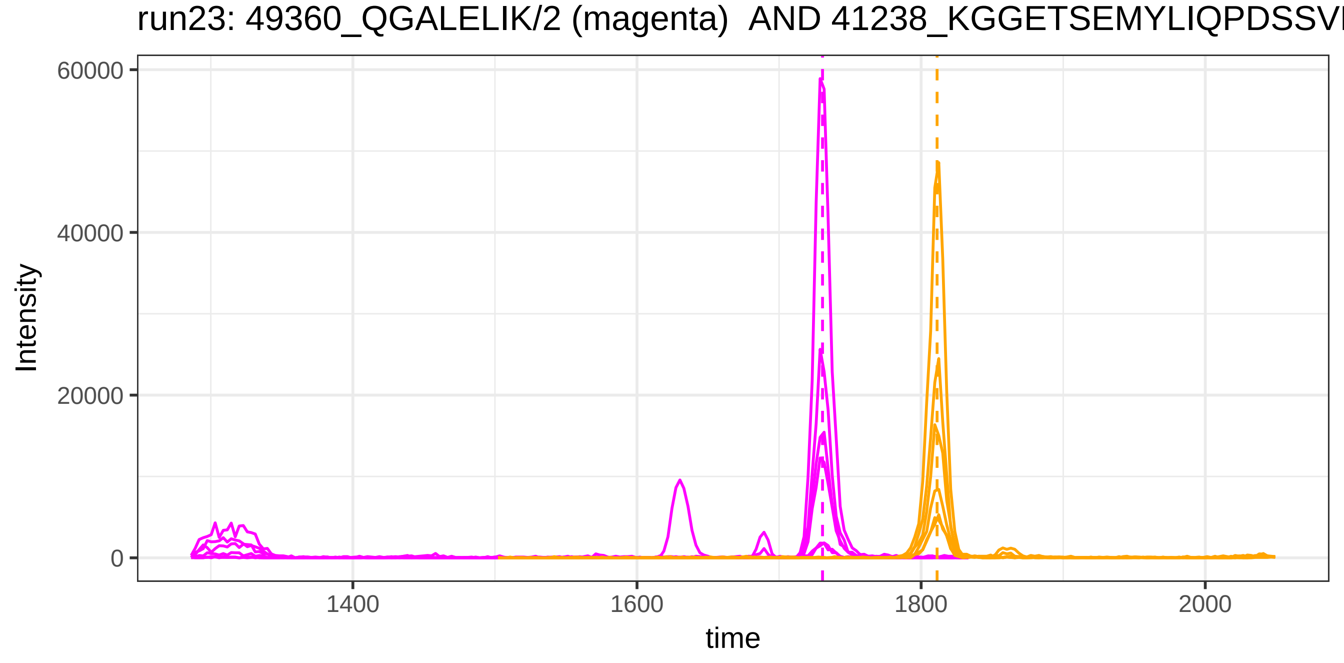
run23: 49360_QGALELIK/2 (magenta) AND 41238_KGGETSEMYLIQPDSSVI
Intensity
60000
40000
20000
0
1400
1600
1800
2000
time

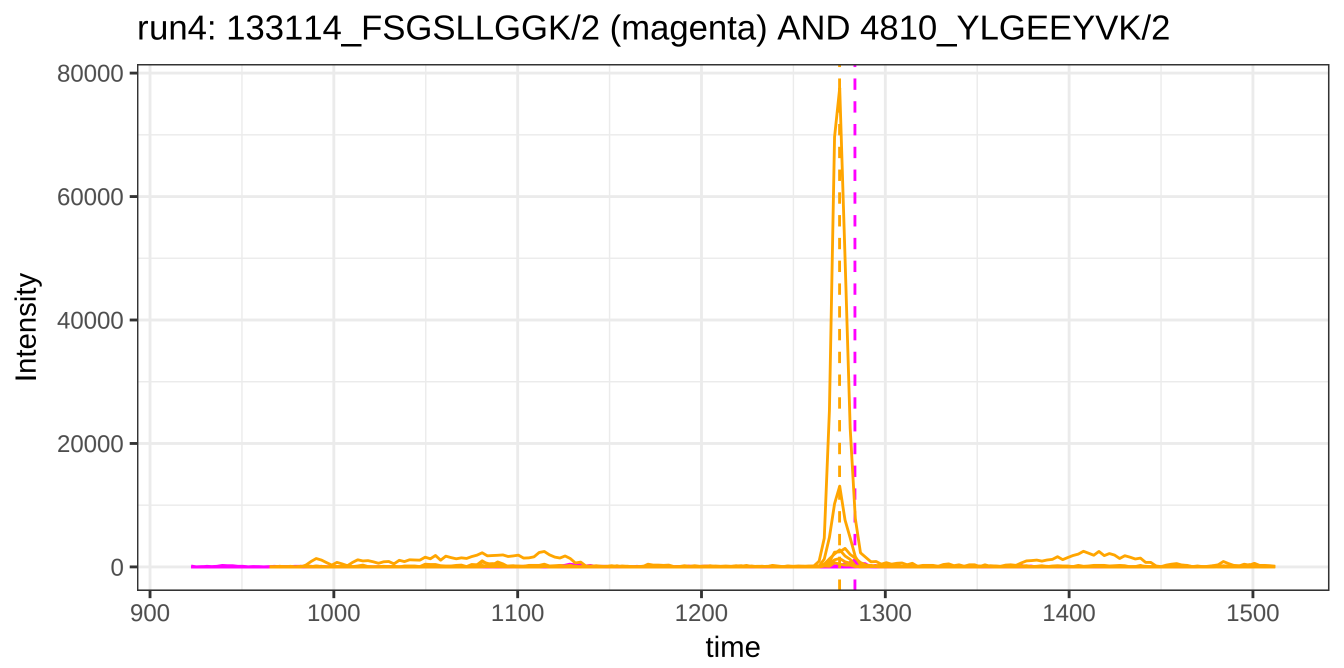

run4: 133114_FSGSLLGGK/2 (magenta) AND 4810_YLGEEYVK/2
Intensity
80000
60000
40000
20000
0
900
1000
1100
1200
1300
1400
1500
time


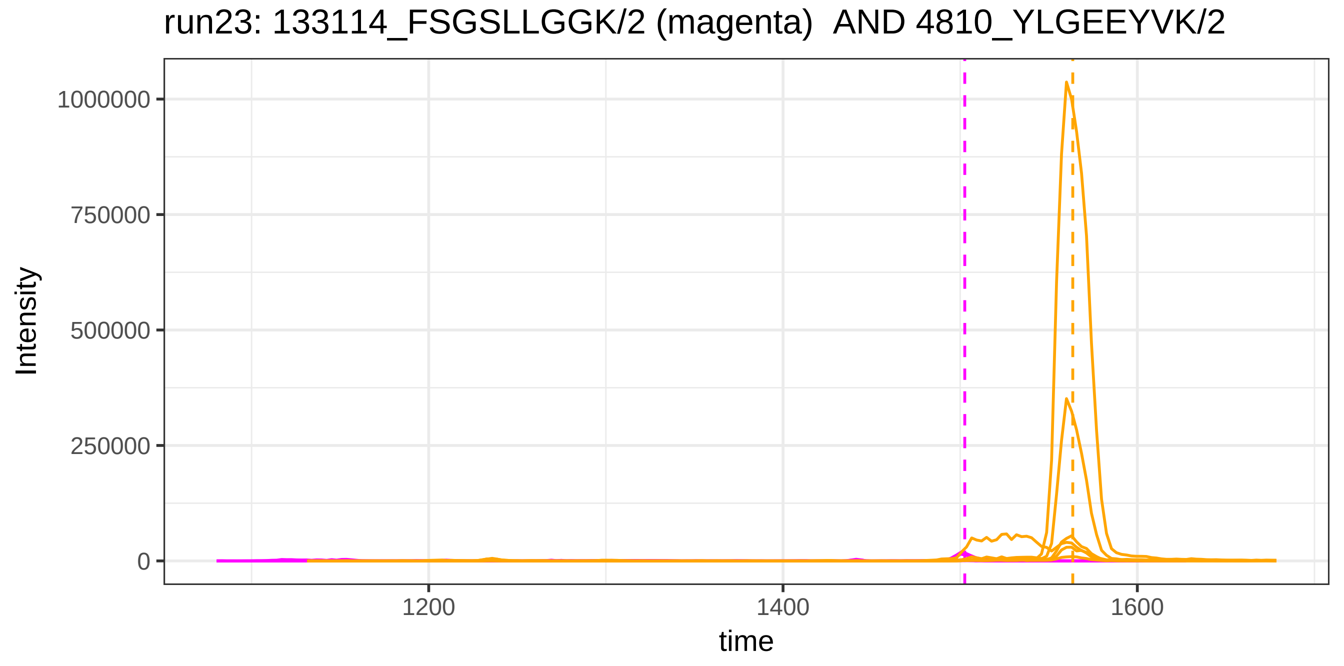

run23: 133114_FSGSLLGGK/2 (magenta) AND 4810_YLGEEYVK/2
Intensity
1000000
750000
500000
250000
0
1200
1400
1600
time

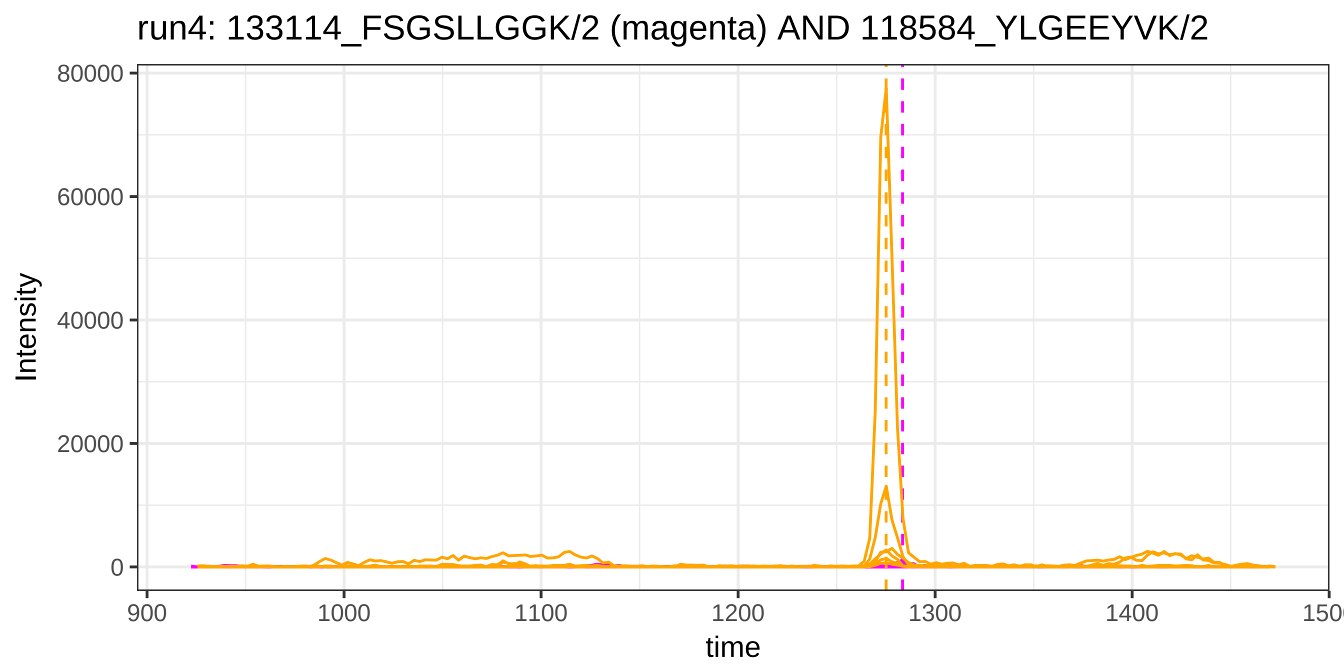
run4: 133114_FSGSLLGGK/2 (magenta) AND 118584_YLGEEYVK/2
Intensity
80000
60000
40000
20000
0
900
1000
1100
1200
1300
1400
time

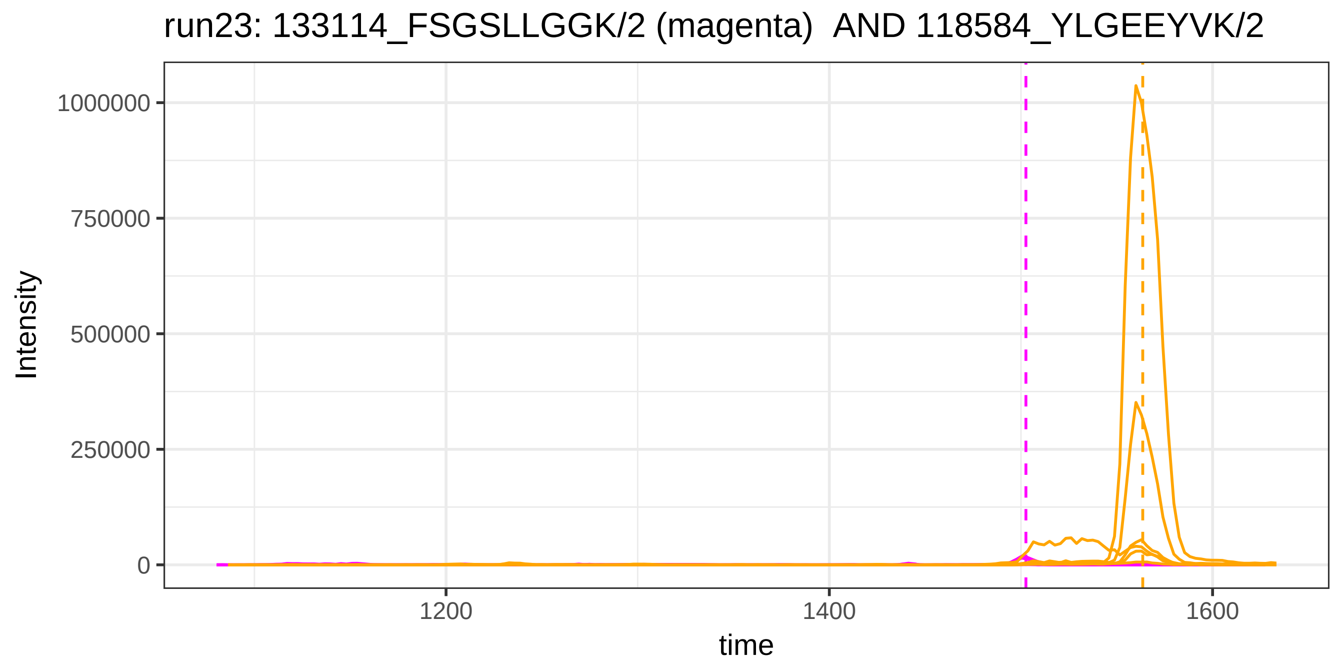
run23: 133114_FSGSLLGGK/2 (magenta) AND 118584_YLGEEYVK/2
Intensity
1000000
750000
500000
250000
0
1200
1400
1600
time

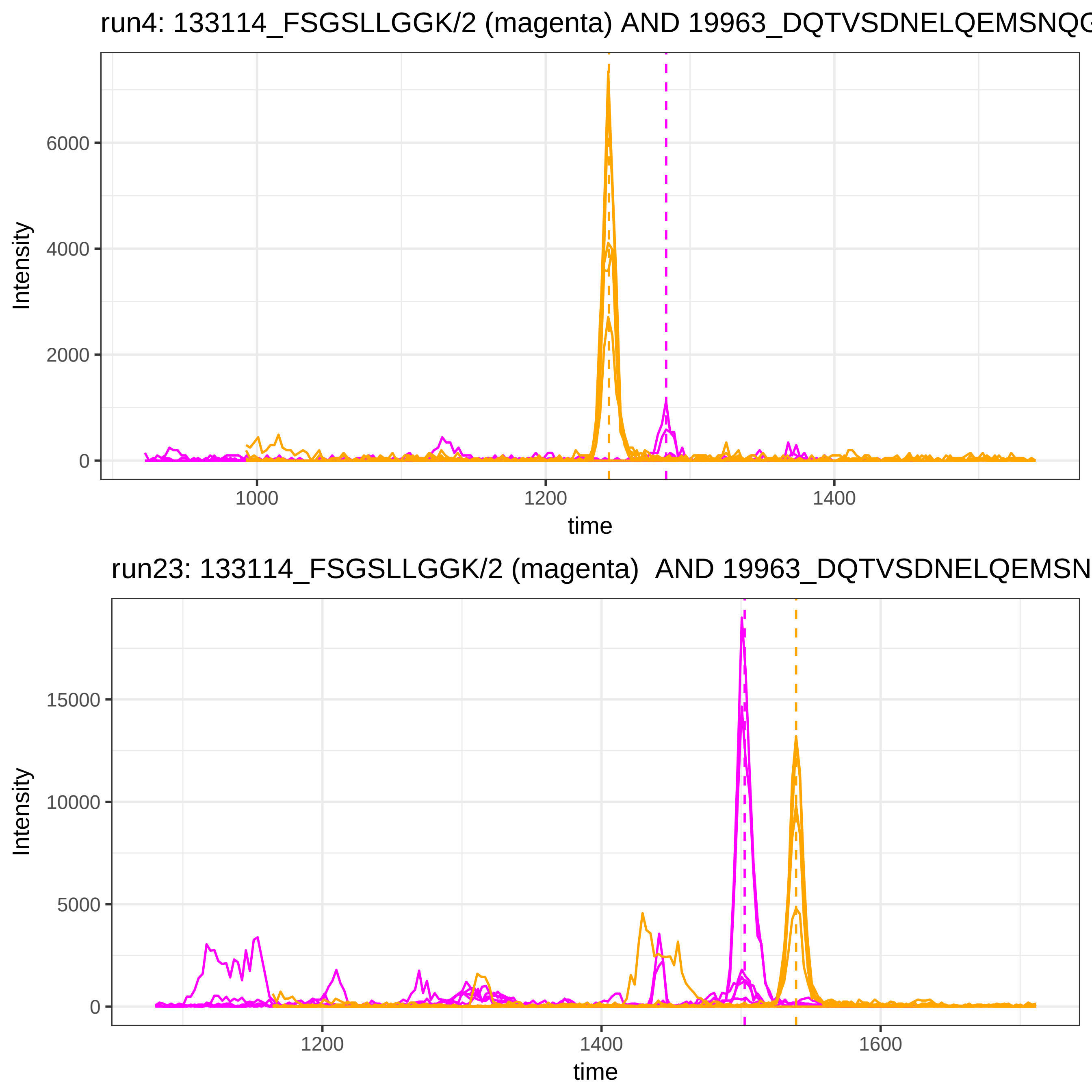
run4: 133114_FSGSLLGGK/2 (magenta) AND 19963_DQTVSDNELQEMSNQG
Intensity
6000
4000
2000
0
1000
1200
1400
time
run23: 133114_FSGSLLGGK/2 (magenta) AND 19963_DQTVSDNELQEMSN
Intensity
15000
10000
5000
0
1200
1400
1600
time

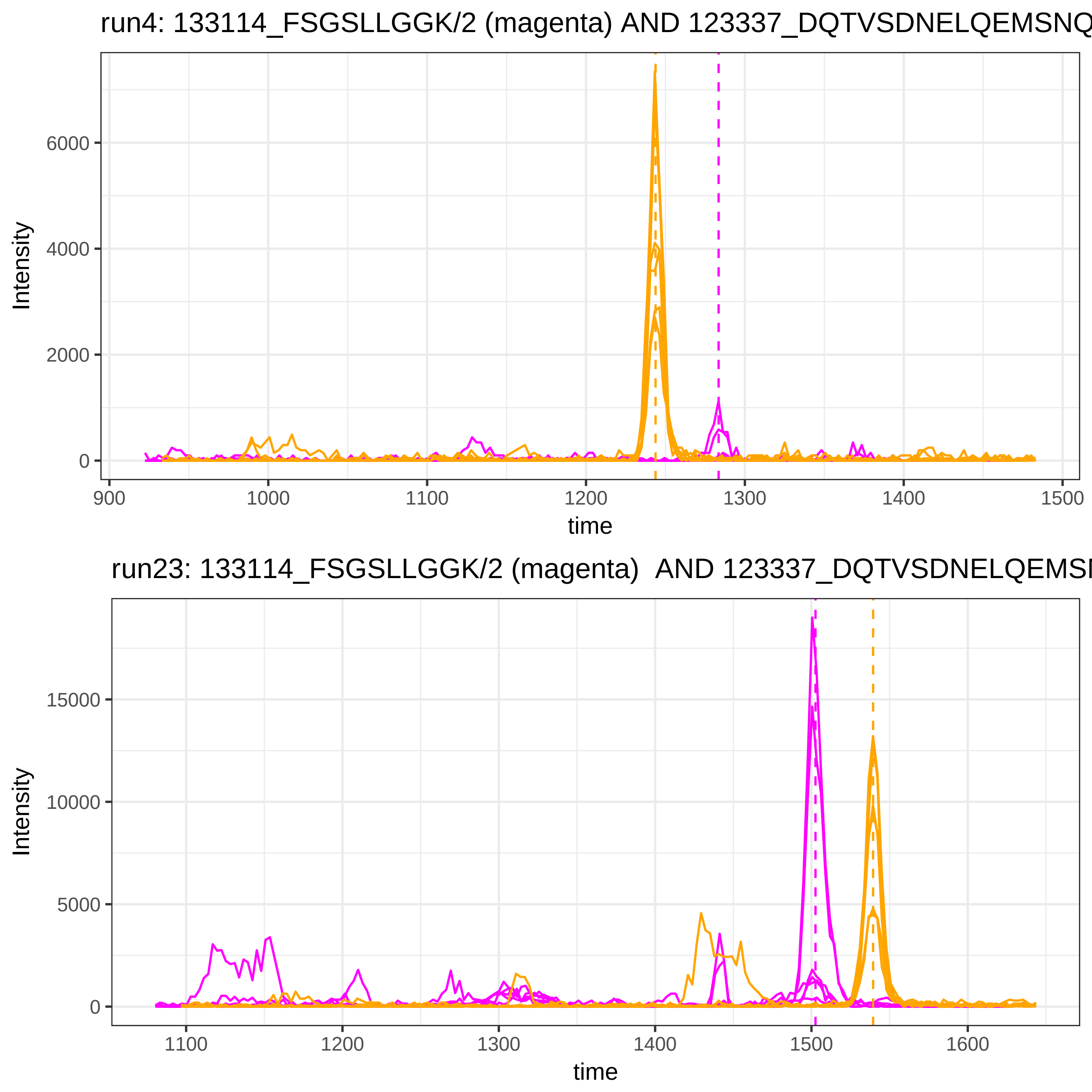
run4: 133114_FSGSLLGGK/2 (magenta) AND 123337_DQTVSDNELQEMSNQ
Intensity
6000
4000
2000
0
900
1000
1100
1200
1300
1400
1500
time
run23: 133114_FSGSLLGGK/2 (magenta)  AND 123337_DQTVSDNELQEMSN
Intensity
15000
10000
5000
0
1100
1200
1300
1400
1500
1600
time

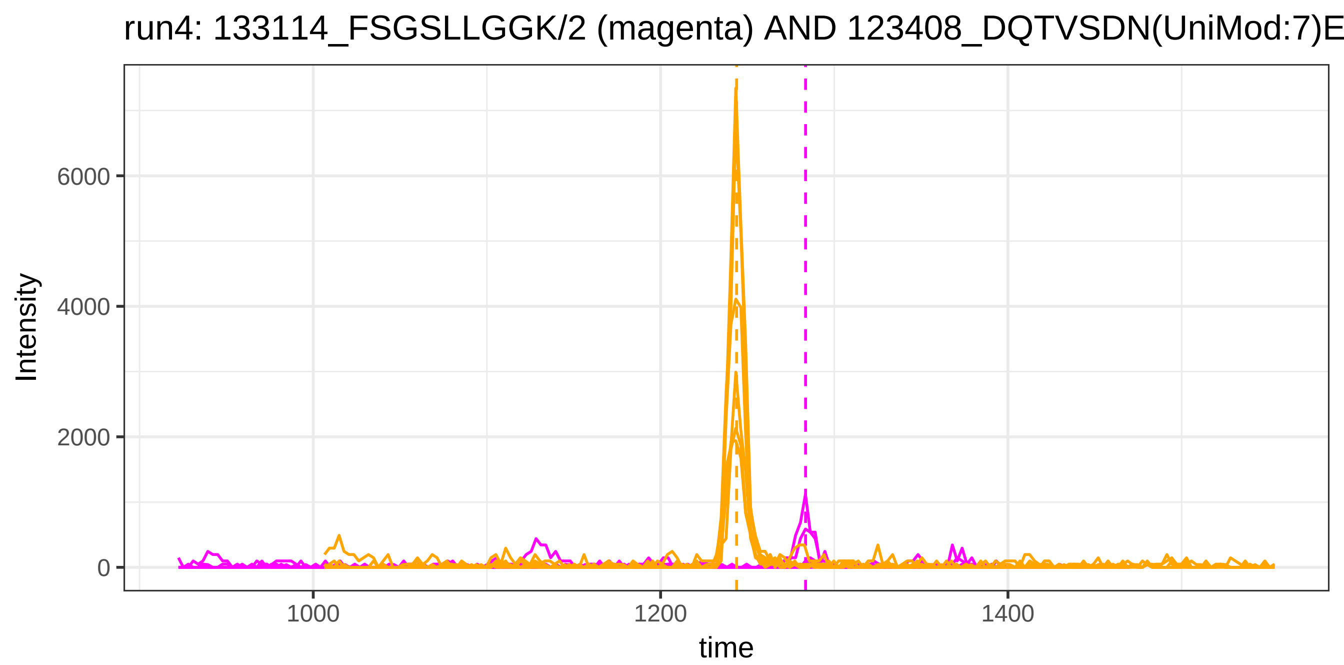

run4: 133114_FSGSLLGGK/2 (magenta) AND 123408_DQTVSDN(UniMod:7)E
Intensity
6000
4000
2000
0
1000
1200
1400
time


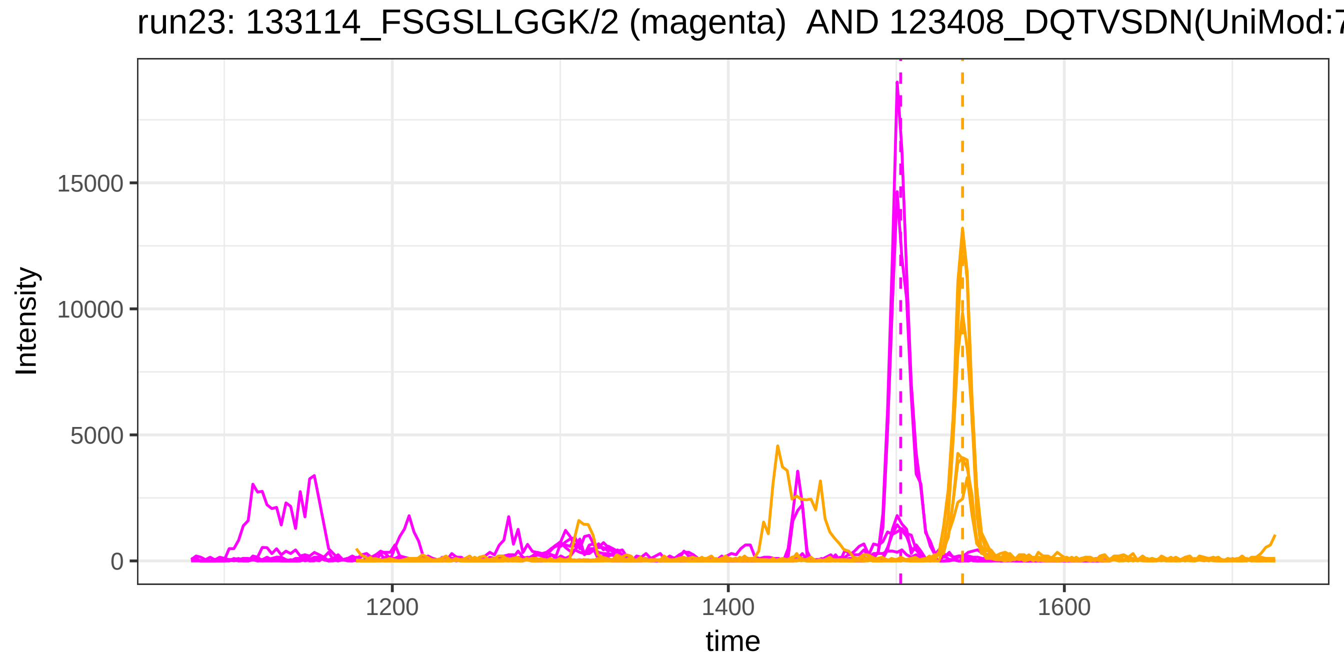

run23: 133114_FSGSLLGGK/2 (magenta) AND 123408_DQTVSDN(UniMod:7
Intensity
15000
10000
5000
0
1200
1400
1600
time

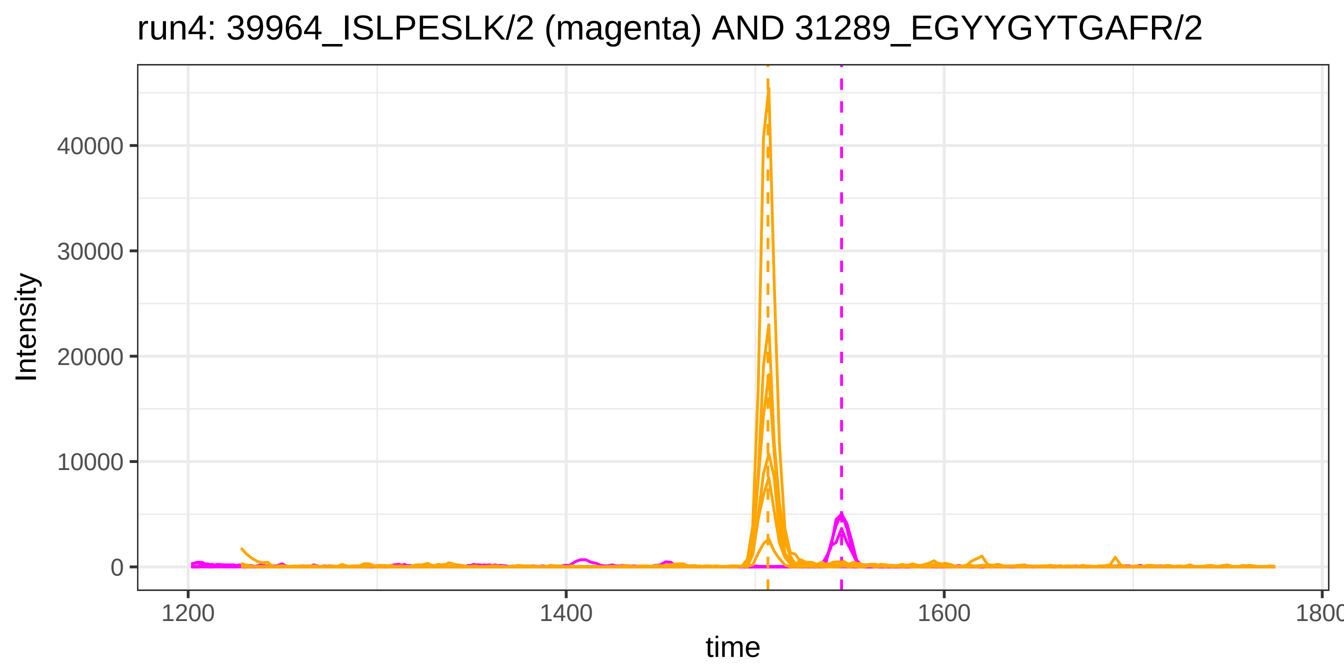
run4: 39964_ISLPESLK/2 (magenta) AND 31289_EGYYGYTGAFR/2
Intensity
40000
30000
20000
10000
0
1200
1400
1600
1800
time

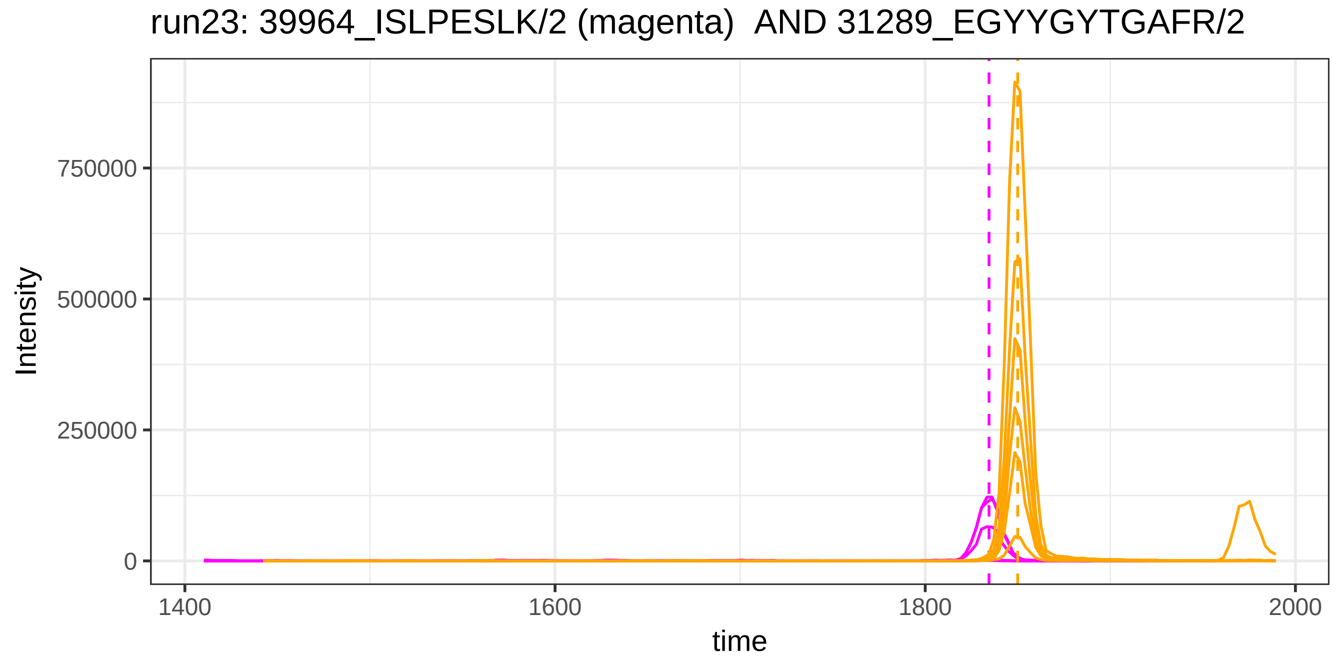
run23: 39964_ISLPESLK/2 (magenta) AND 31289_EGYYGYTGAFR/2
Intensity
750000
500000
250000
0
1400
1600
1800
2000
time

run4: 39964_ISLPESLK/2 (magenta) AND 118541_GSGLNLC(UniMod:4)EPNI

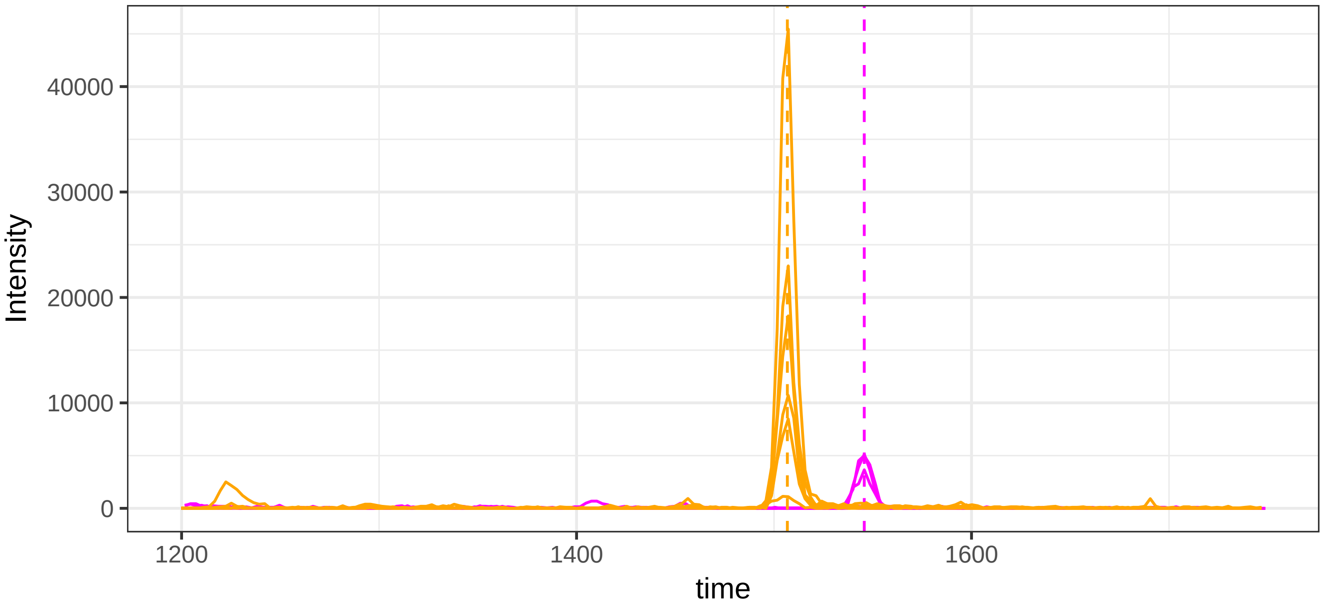


run23: 39964_ISLPESLK/2 (magenta)  AND 118541_GSGLNLC(UniMod:4)EF

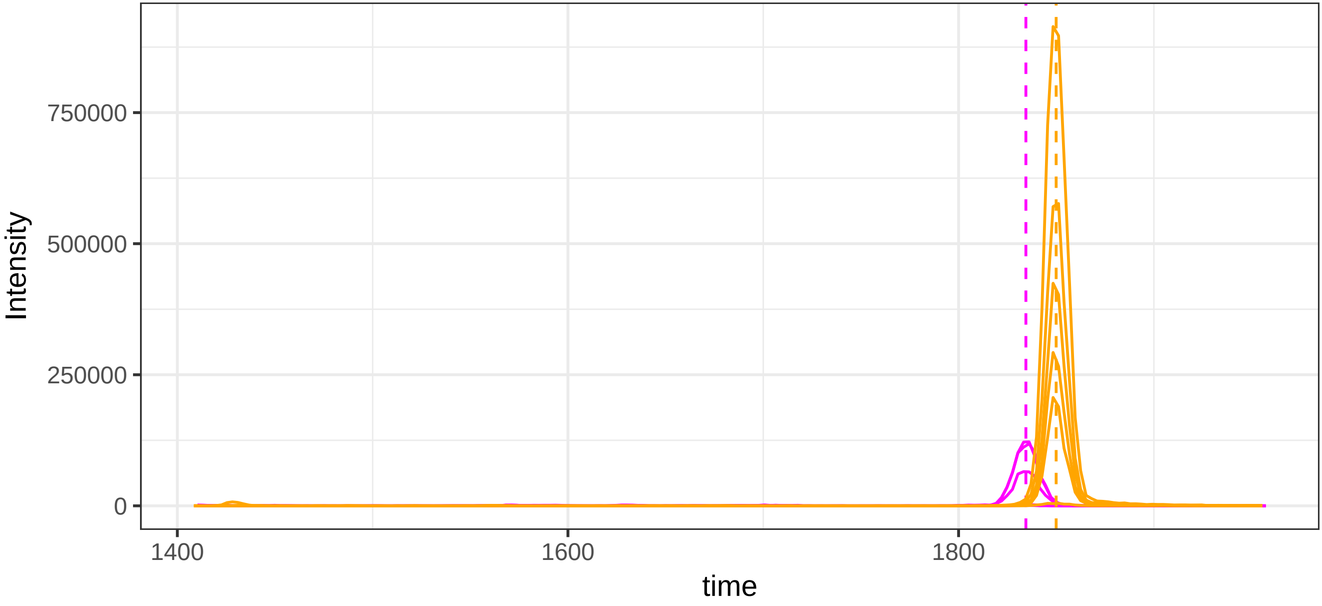

run4: 51131_RTHLPEVFLSK/3 (magenta) AND 41559_ALTDMPQMR/2

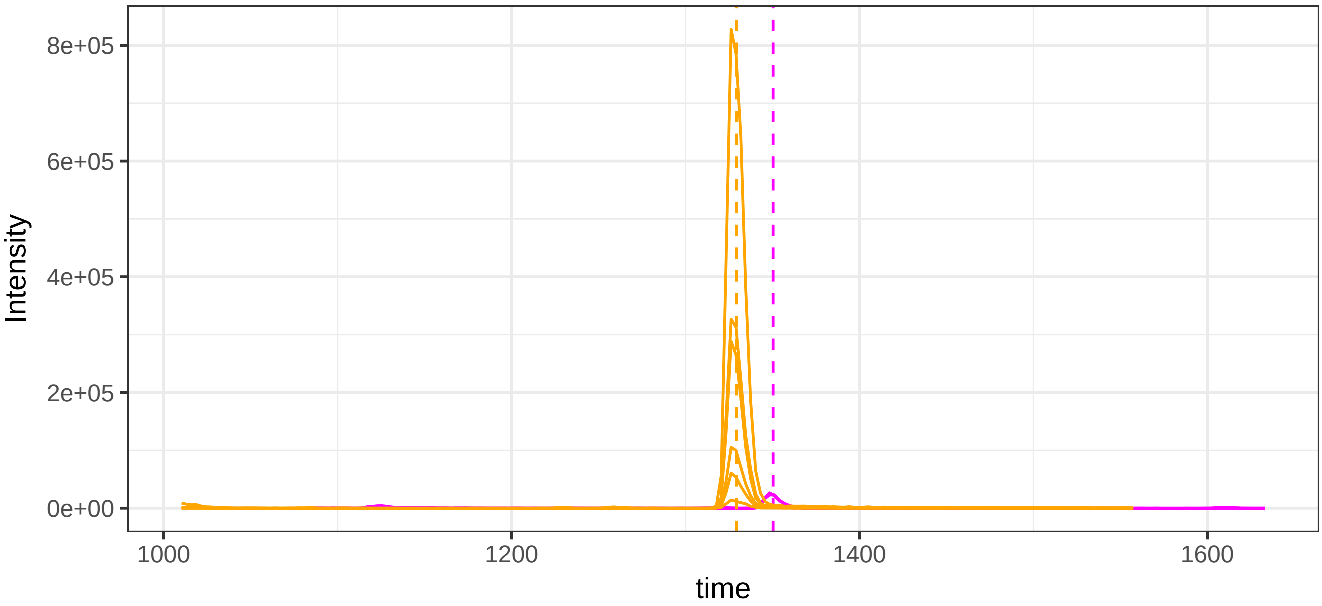


run23: 51131_RTHLPEVFLSK/3 (magenta)  AND 41559_ALTDMPQMR/2

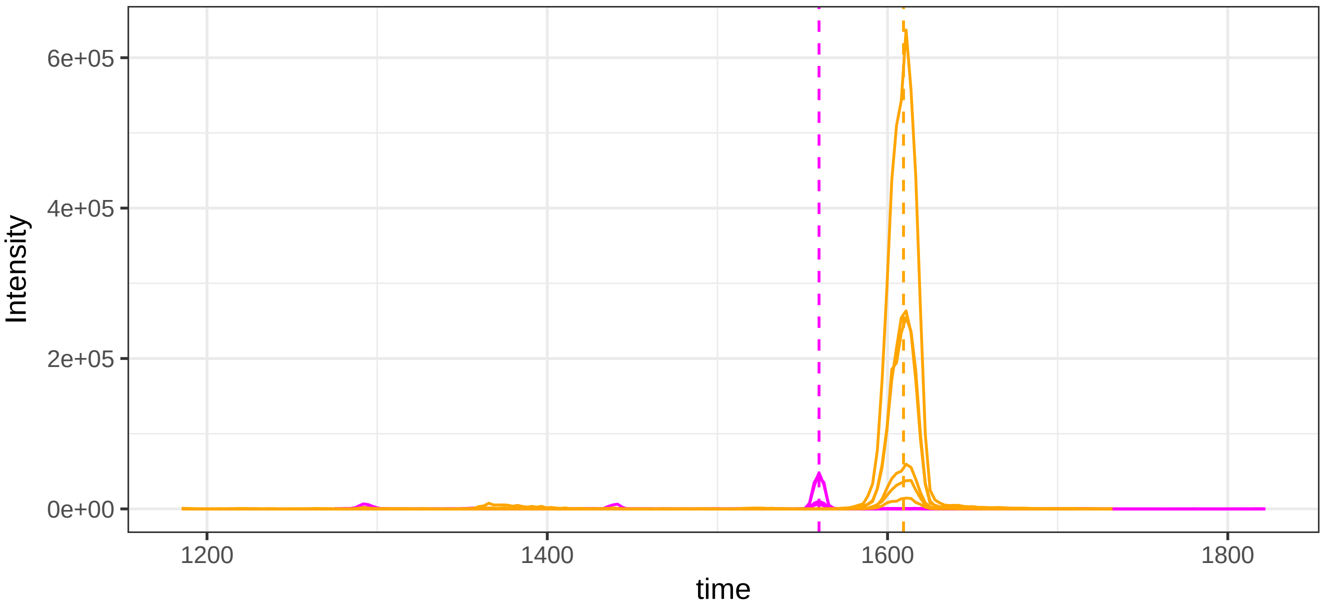

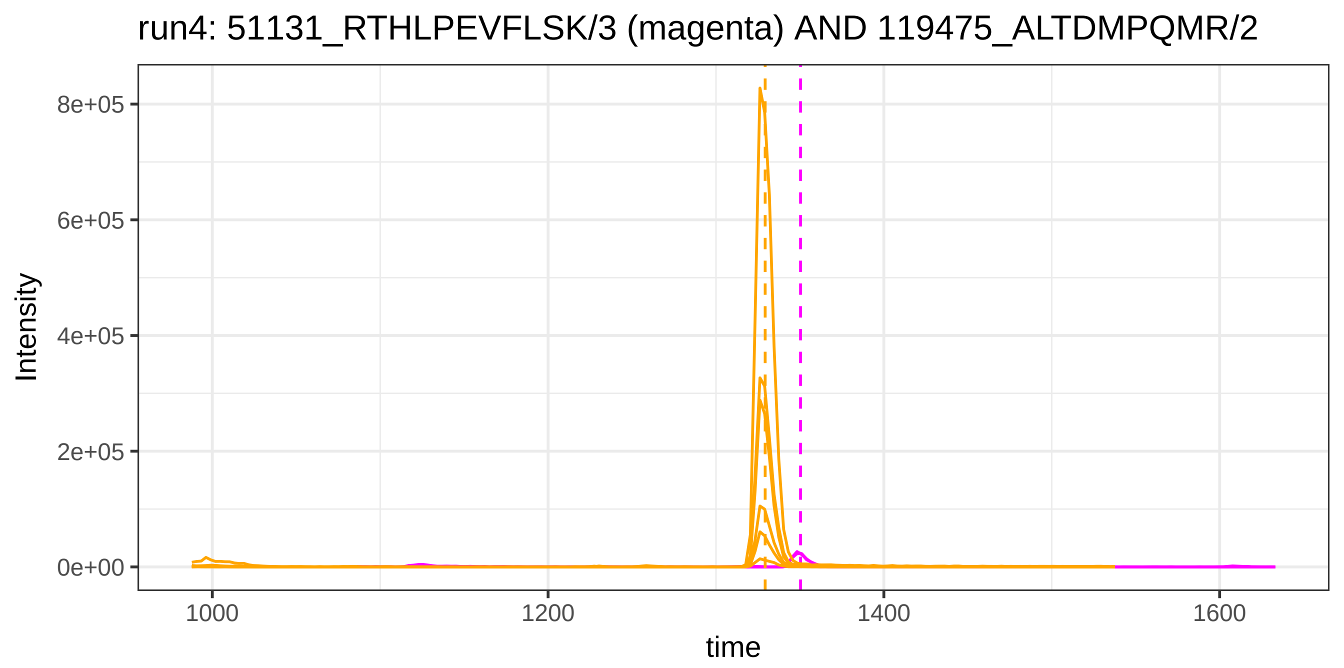
run4: 51131_RTHLPEVFLSK/3 (magenta) AND 119475_ALTDMPQMR/2
Intensity
8e+05
6e+05
4e+05
2e+05
0e+00
1000
1200
1400
1600
time

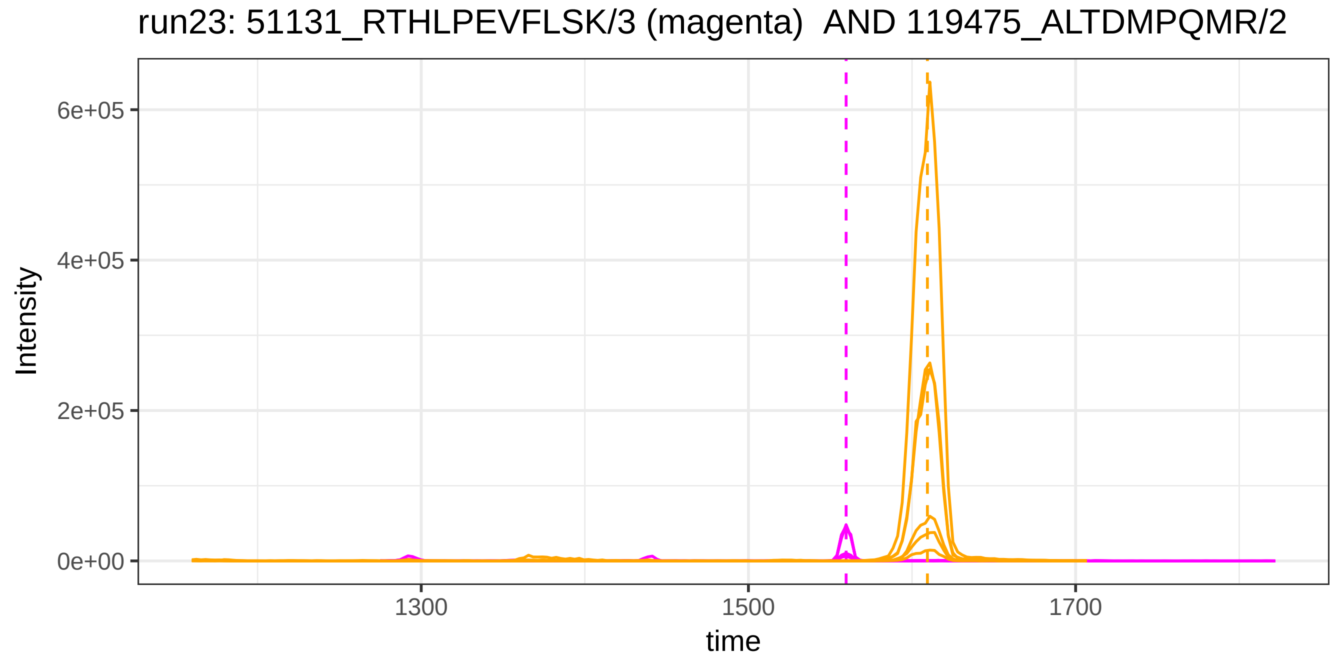
run23: 51131_RTHLPEVFLSK/3 (magenta) AND 119475_ALTDMPQMR/2
Intensity
6e+05
4e+05
2e+05
0e+00
1300
1500
1700
time

run4: 51131_RTHLPEVFLSK/3 (magenta) AND 1030_VELEDWNGR/2

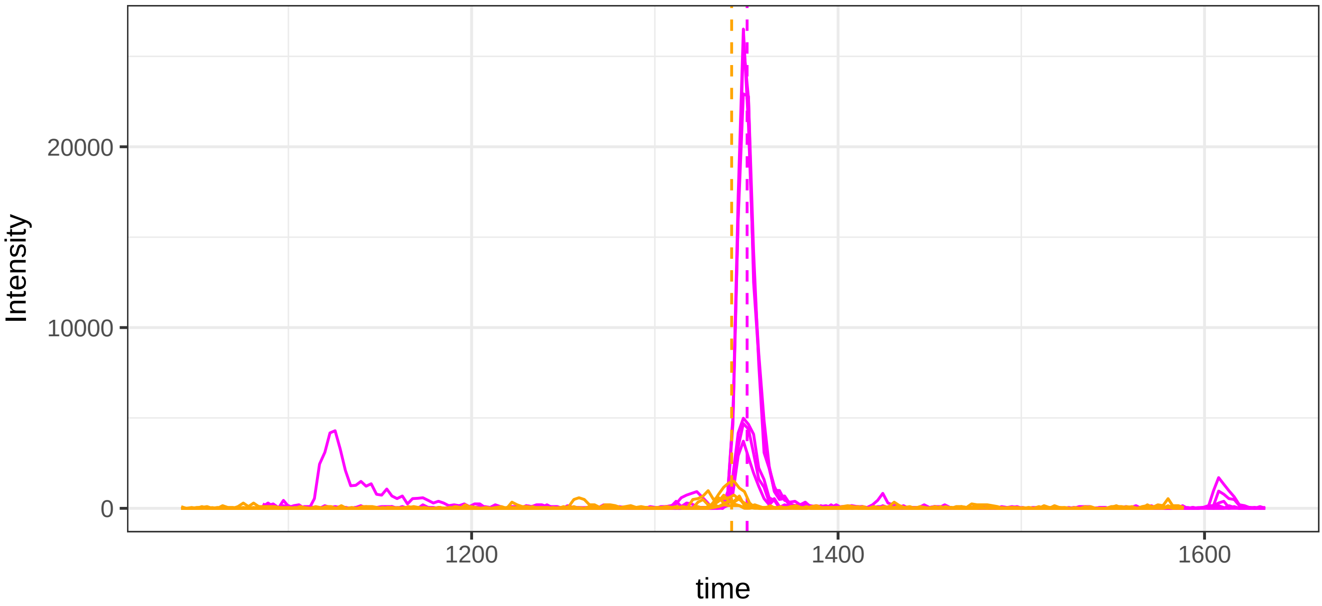


run23: 51131_RTHLPEVFLSK/3 (magenta)  AND 1030_VELEDWNGR/2

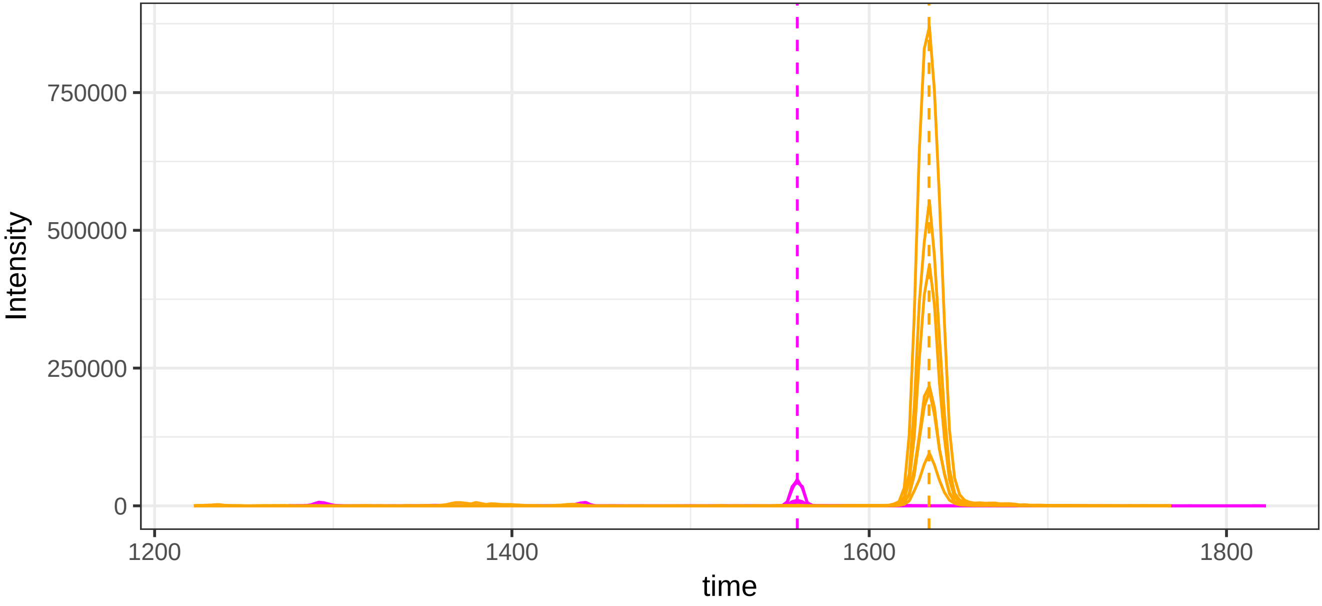

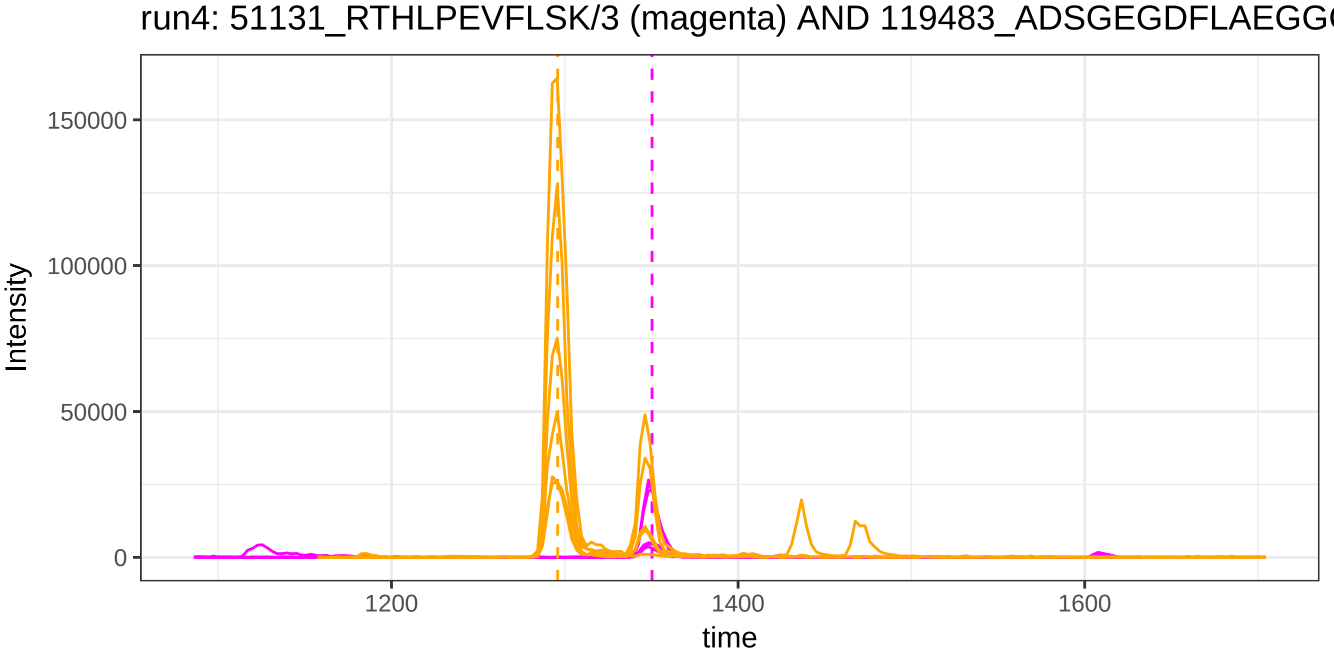

run4: 51131_RTHLPEVFLSK/3 (magenta) AND 119483_ADSGEGDFLAEGG
Intensity
150000
100000
50000
0
1200
1400
1600
time


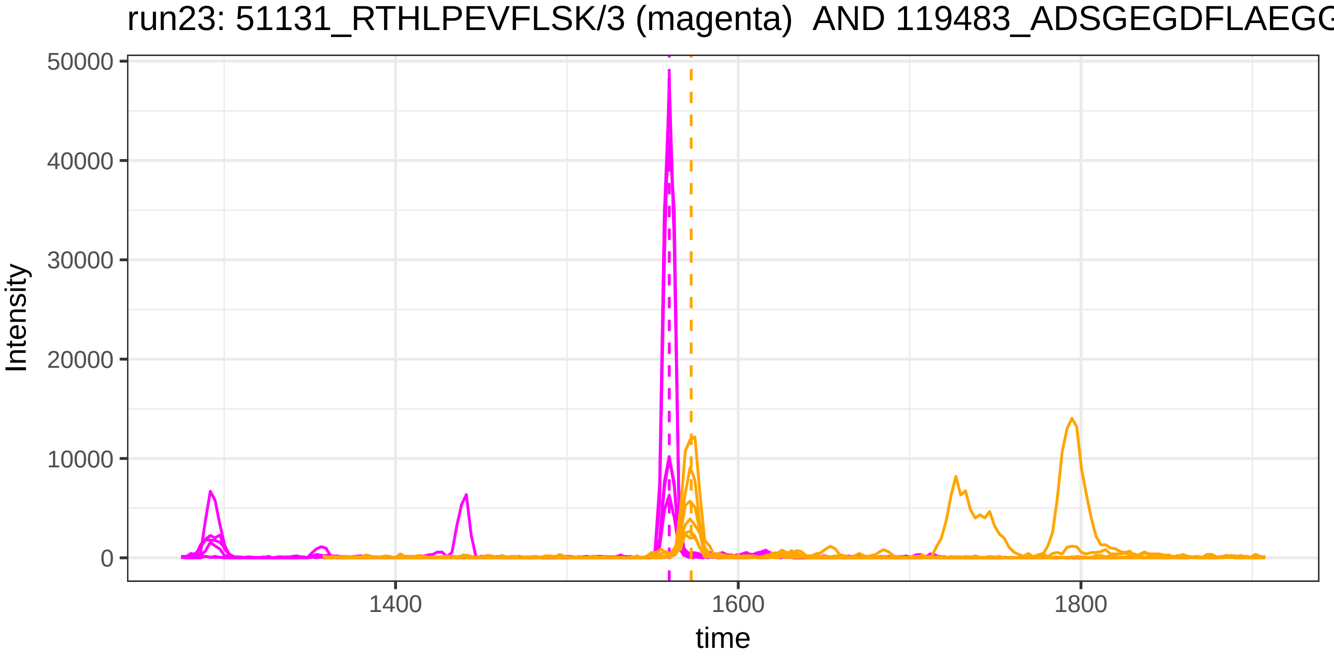

run23: 51131_RTHLPEVFLSK/3 (magenta) AND 119483_ADSGEGDFLAEGG
Intensity
50000
40000
30000
20000
10000
0
1400
1600
1800
time

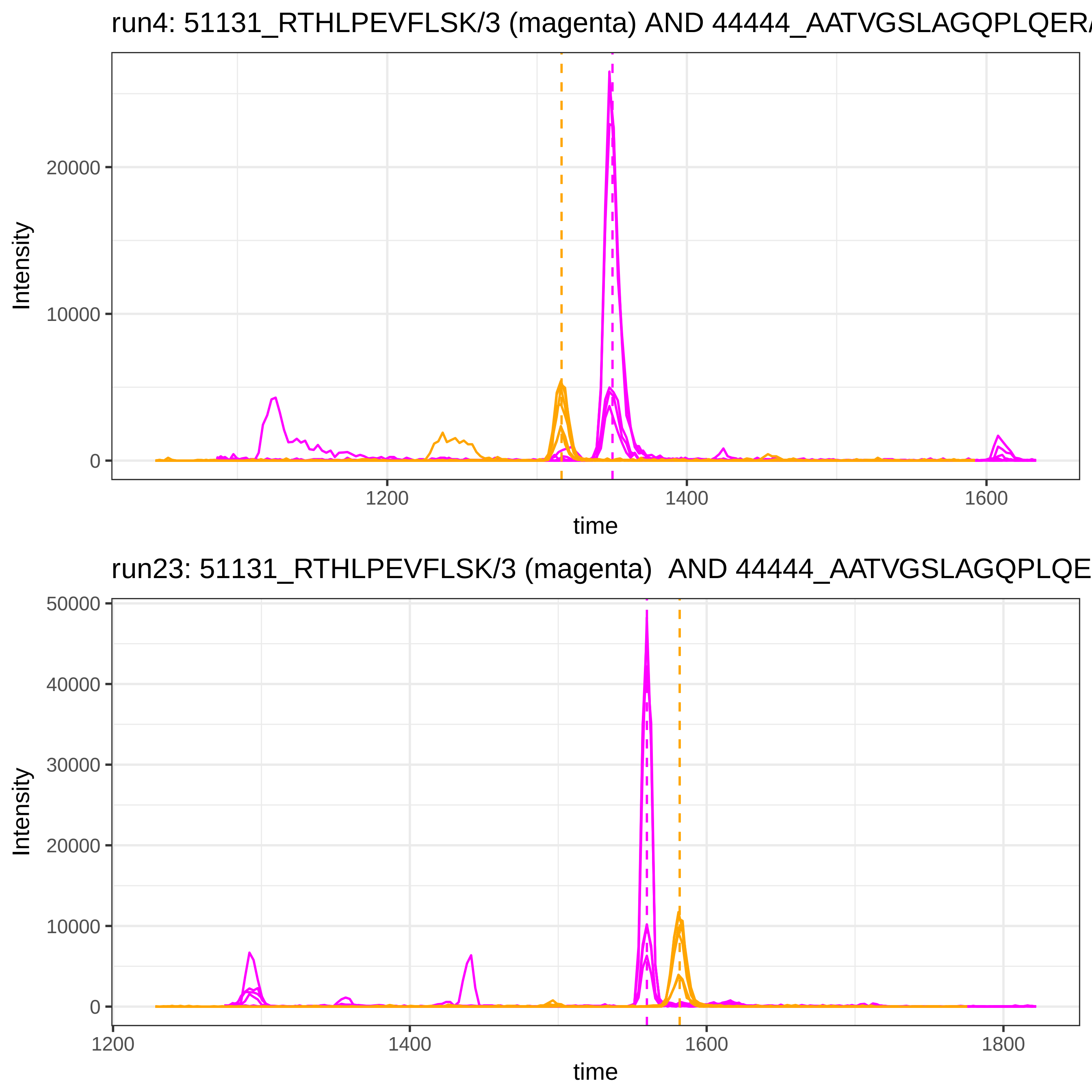

run4: 51131_RTHLPEVFLSK/3 (magenta) AND 44444_AATVGSLAGQPLQER/
Intensity
20000
10000
0
1200
1400
1600
time
run23: 51131_RTHLPEVFLSK/3 (magenta) AND 44444_AATVGSLAGQPLQE
Intensity
50000
40000
30000
20000
10000
0
1200
1400
1600
1800
time

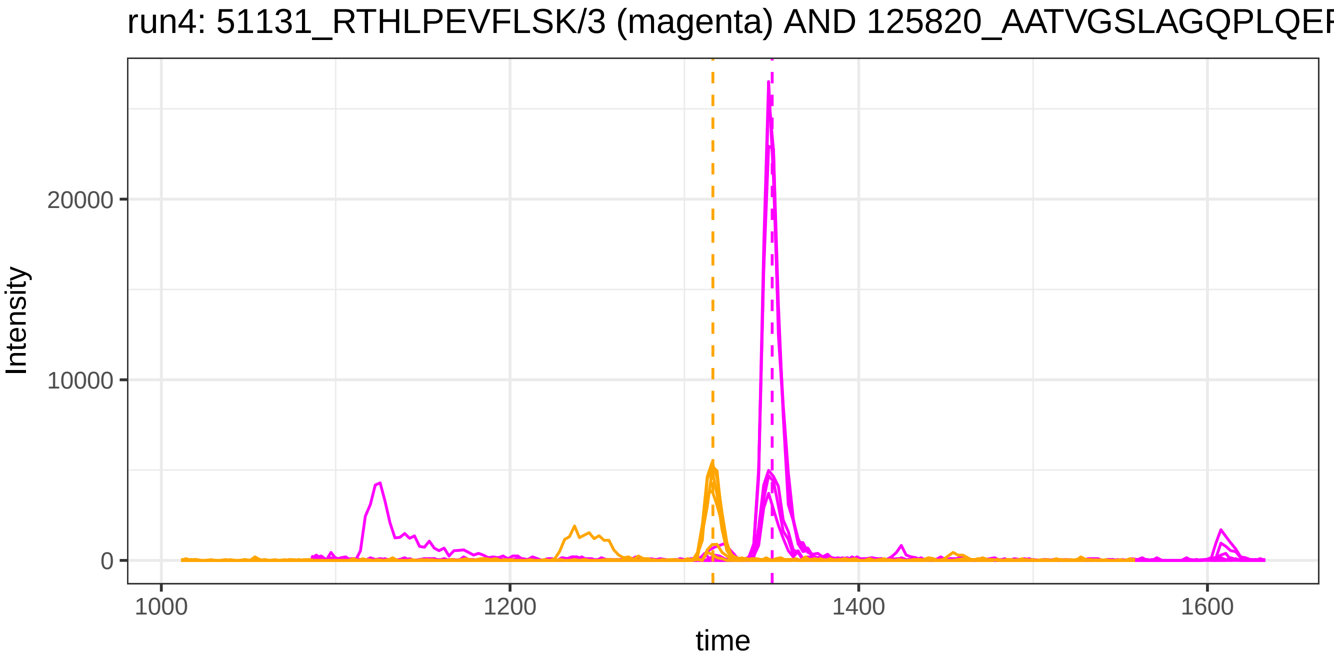
run4: 51131_RTHLPEVFLSK/3 (magenta) AND 125820_AATVGSLAGQPLQEF
Intensity
20000
10000
0
1000
1200
1400
1600
time

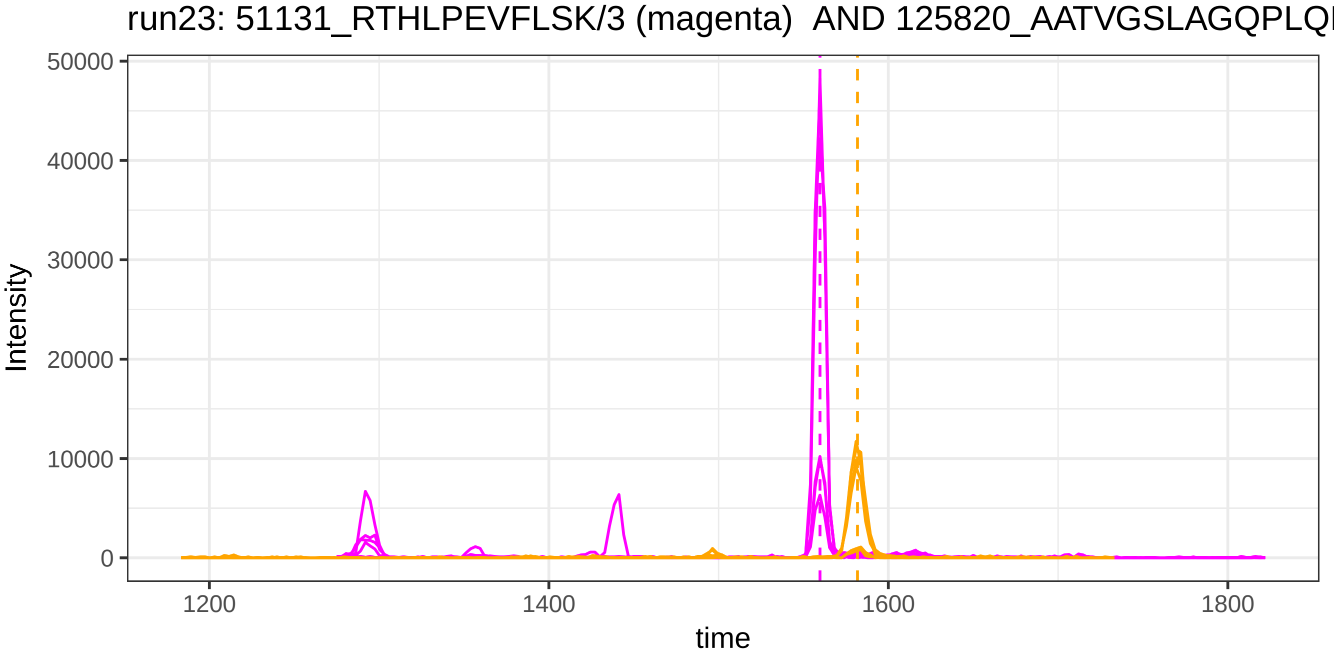
run23: 51131_RTHLPEVFLSK/3 (magenta) AND 125820_AATVGSLAGQPLQ
Intensity
50000
40000
30000
20000
10000
0
1200
1400
1600
1800
time

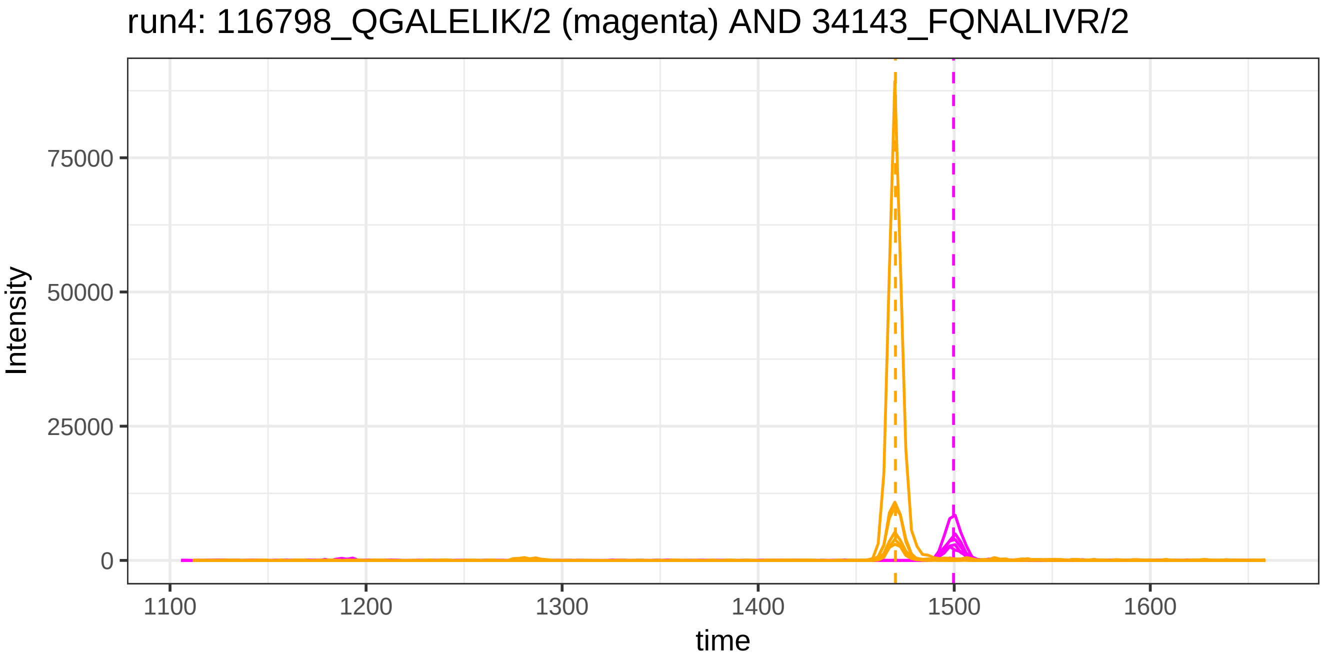
run4: 116798_QGALELIK/2 (magenta) AND 34143_FQNALIVR/2
Intensity
75000
50000
25000
0
1100
1200
1300
1400
1500
1600
time

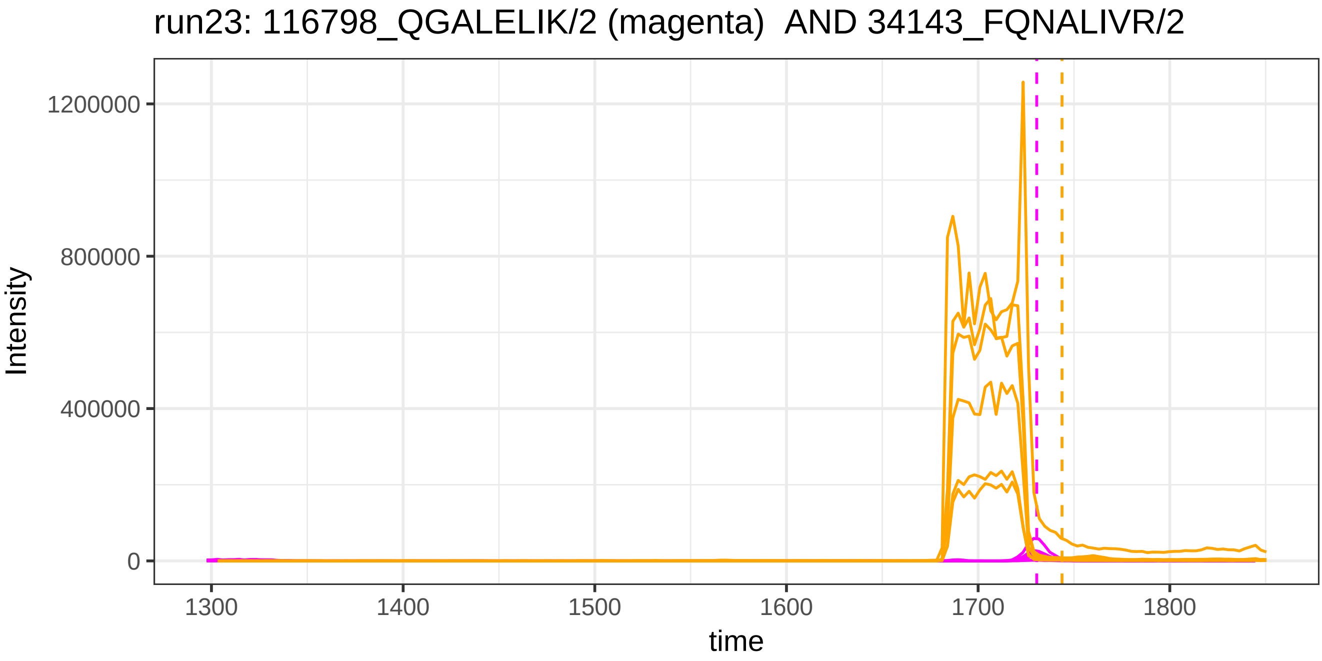
run23: 116798_QGALELIK/2 (magenta)  AND 34143_FQNALIVR/2
Intensity
1200000
800000
400000
0
1300
1400
1500
1600
1700
1800
time

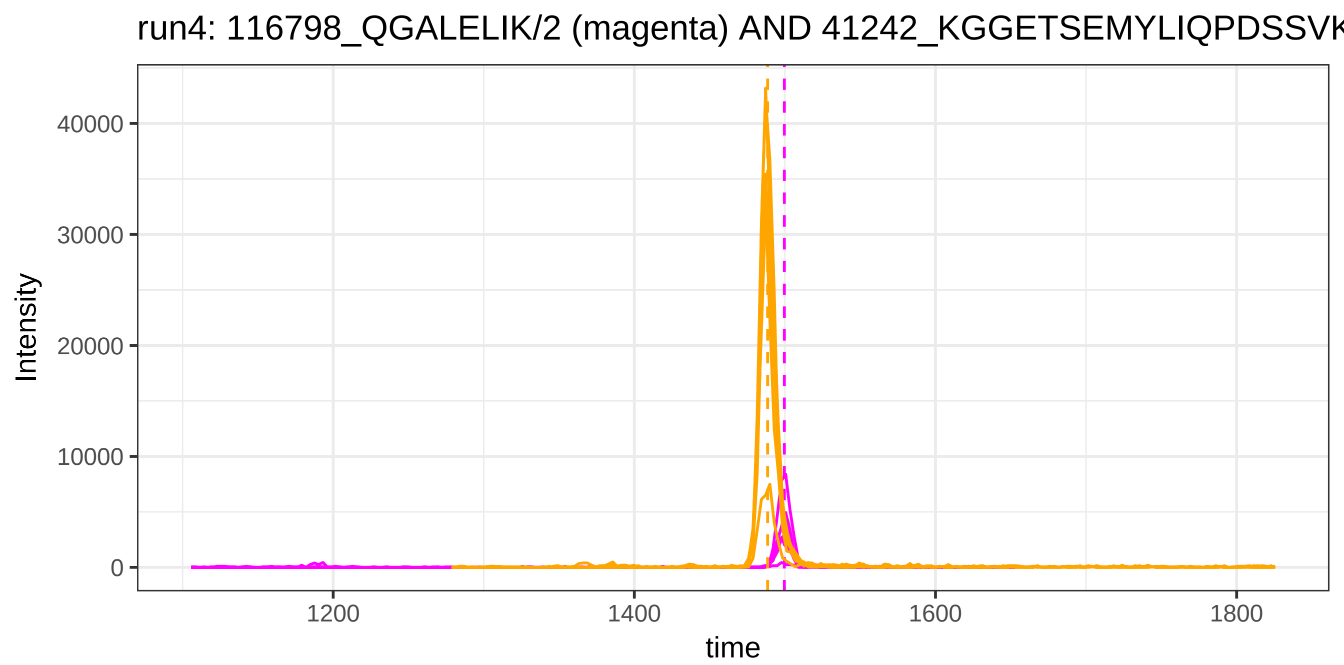

run4: 116798_QGALELIK/2 (magenta) AND 41242_KGGETSEMYLIQPDSSVK
Intensity
40000
30000
20000
10000
0
1200
1400
1600
1800
time


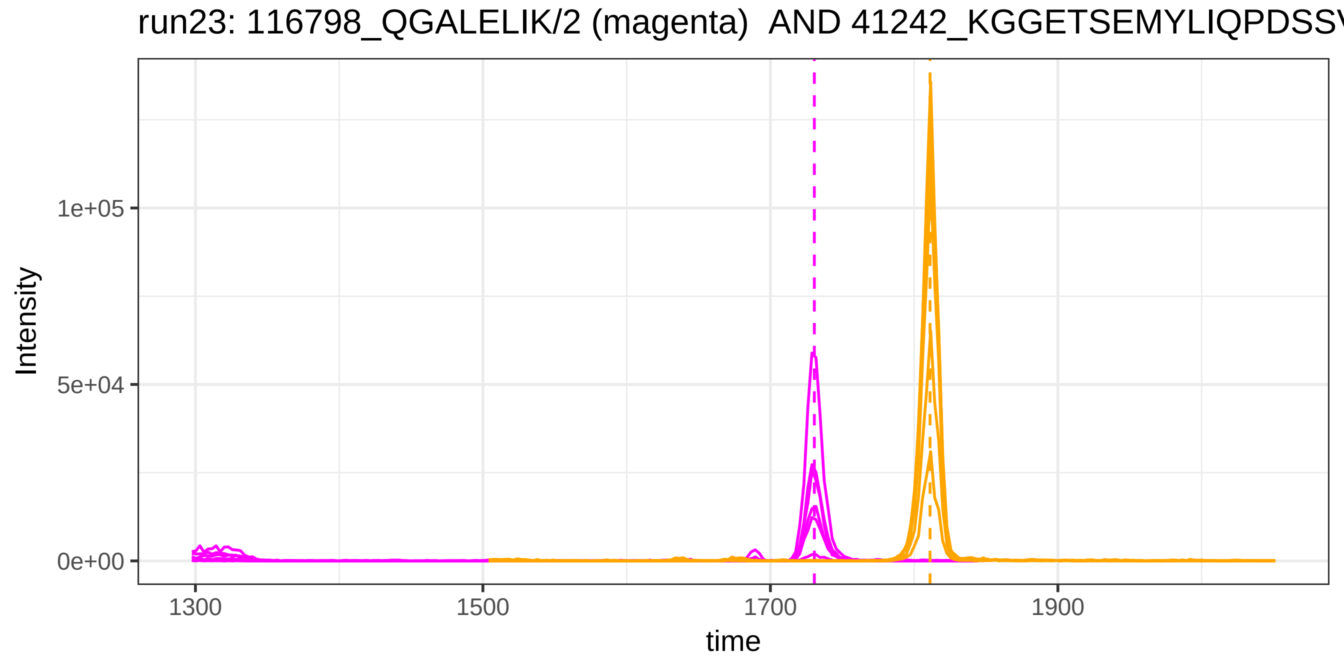

run23: 116798_QGALELIK/2 (magenta) AND 41242_KGGETSEMYLIQPDSSV
Intensity
1e+05
5e+04
0e+00
1300
1500
1700
1900
time

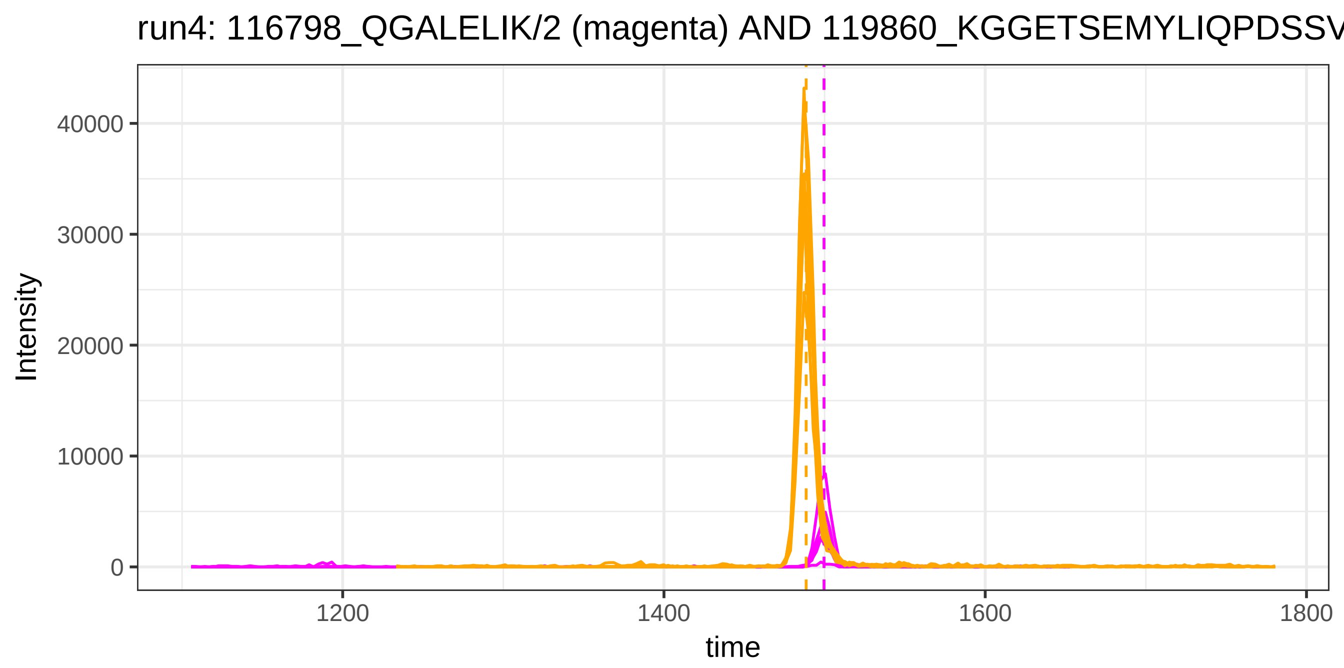
run4: 116798_QGALELIK/2 (magenta) AND 119860_KGGETSEMYLIQPDSSV
Intensity
40000
30000
20000
10000
0
1200
1400
1600
1800
time

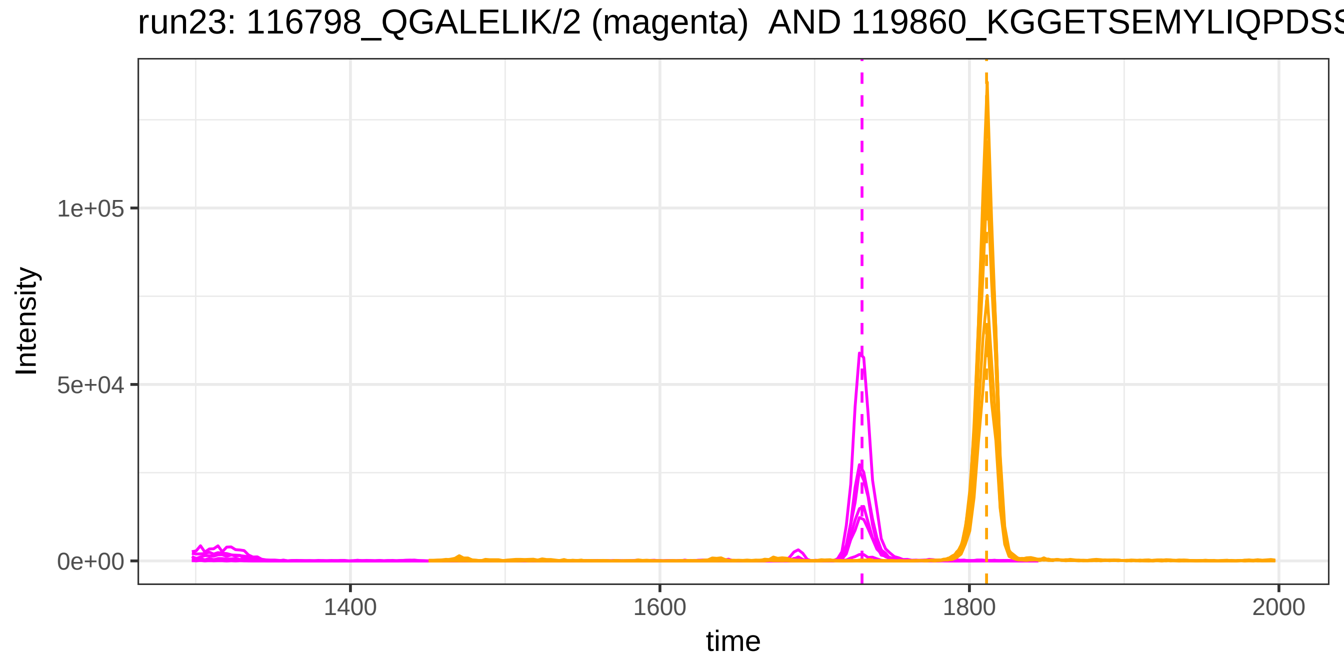
run23: 116798_QGALELIK/2 (magenta) AND 119860_KGGETSEMYLIQPDSS
Intensity
1e+05
5e+04
0e+00
1400
1600
1800
2000
time

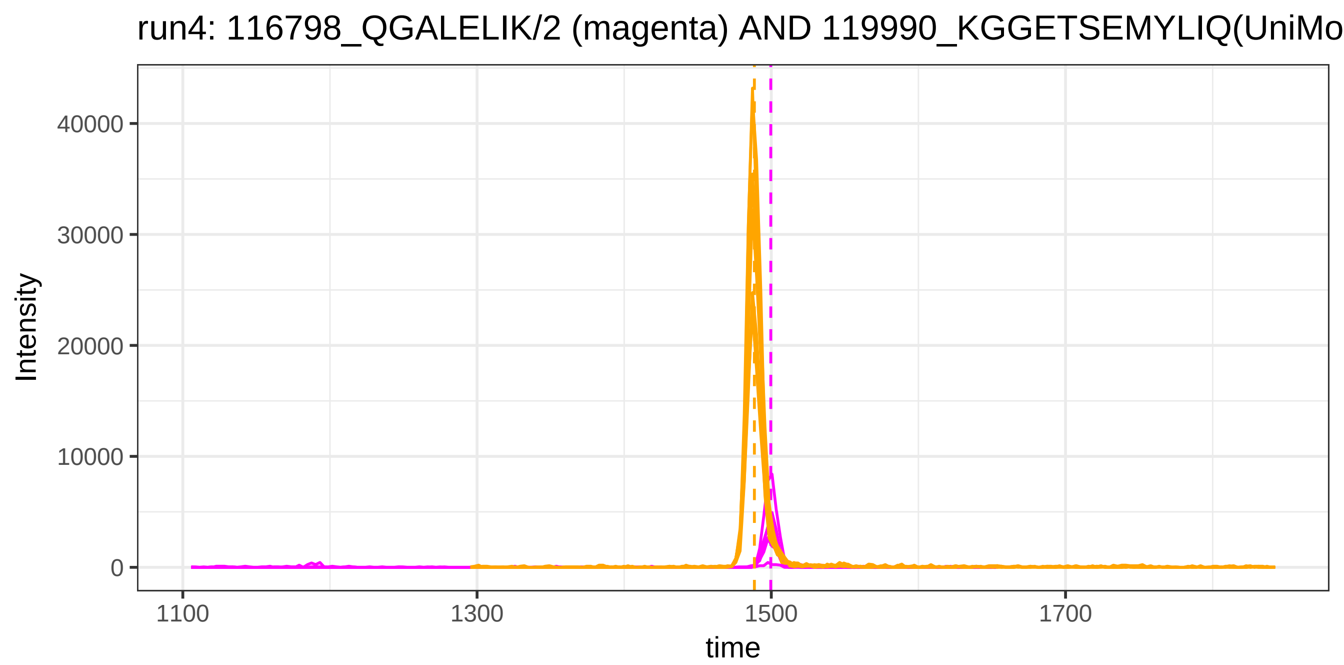
run4: 116798_QGALELIK/2 (magenta) AND 119990_KGGETSEMYLIQ(UniMo
Intensity
40000
30000
20000
10000
0
1100
1300
1500
1700
time

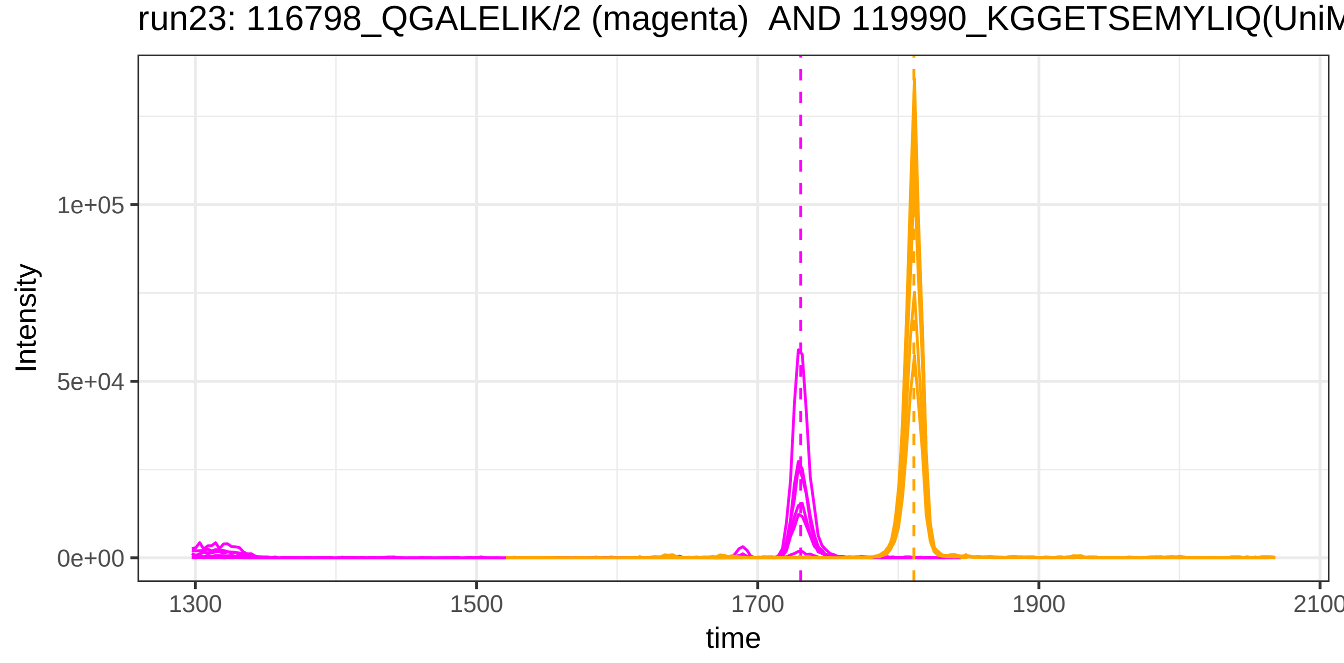
run23: 116798_QGALELIK/2 (magenta) AND 119990_KGGETSEMYLIQ(UniM
Intensity
1e+05
5e+04
0e+00
1300
1500
1700
1900
2100
time

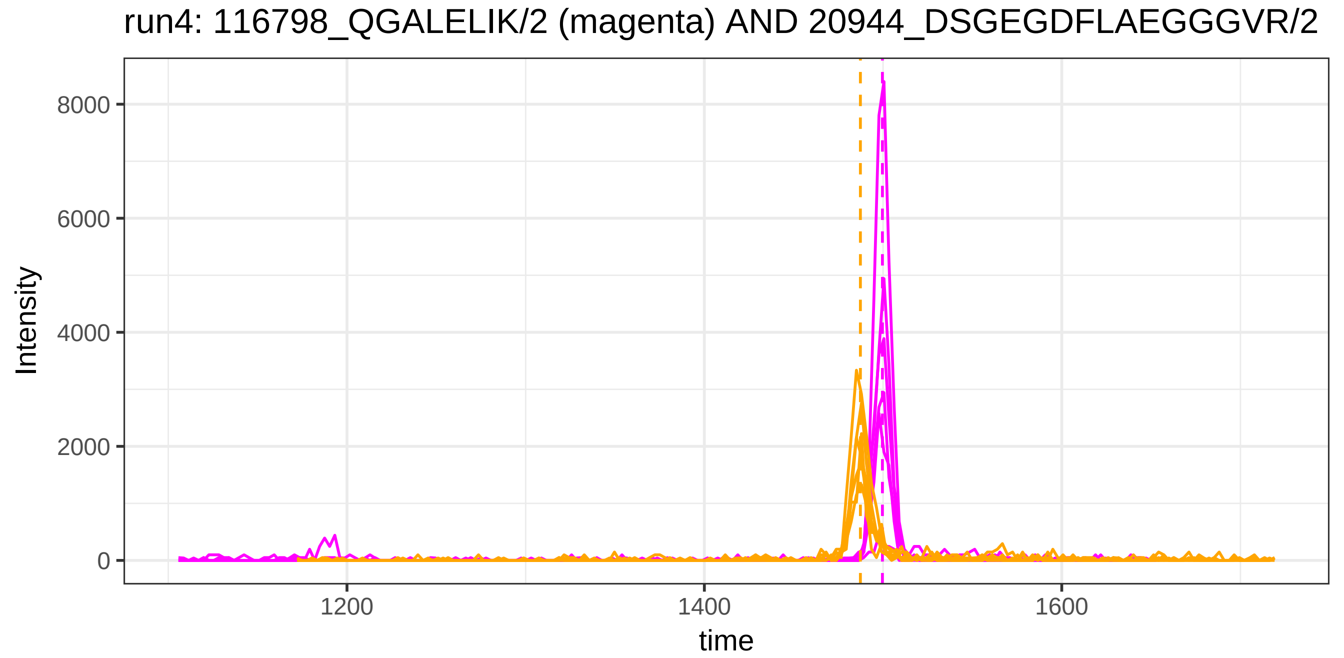

run4: 116798_QGALELIK/2 (magenta) AND 20944_DSGEGDFLAEGGGVR/2
Intensity
8000
6000
4000
2000
0
1200
1400
1600
time


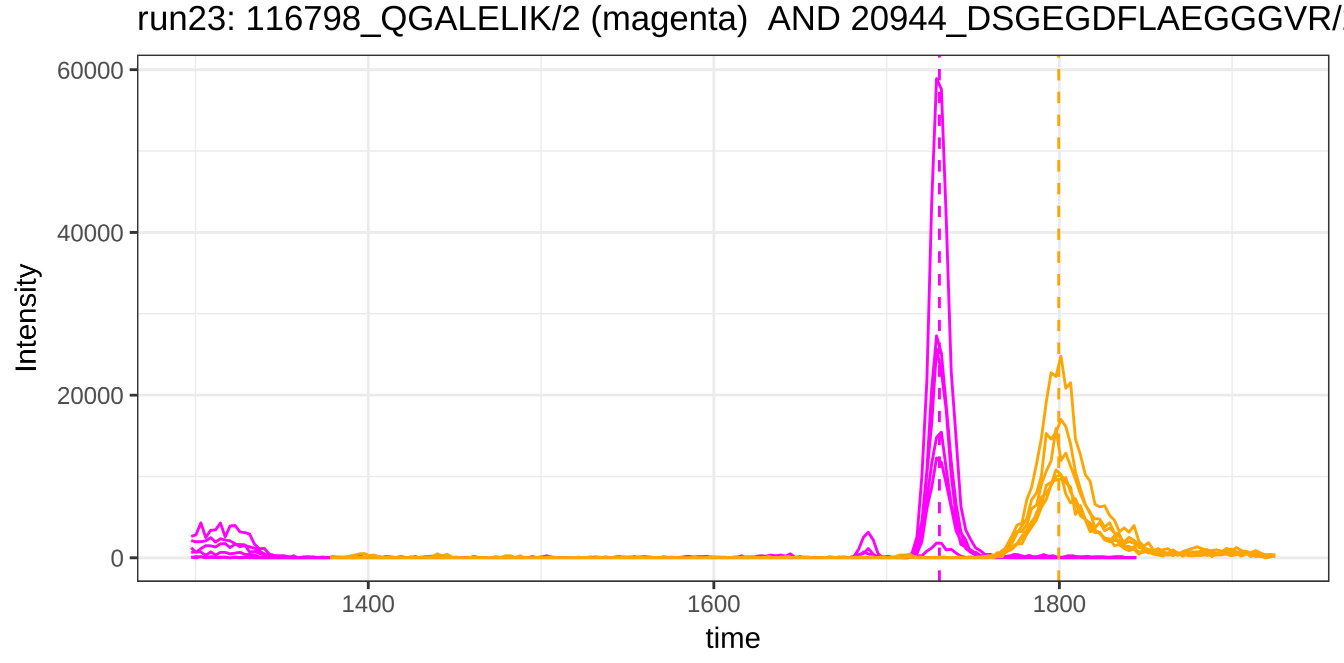

run23: 116798_QGALELIK/2 (magenta) AND 20944_DSGEGDFLAEGGGVR/
Intensity
60000
40000
20000
0
1400
1600
1800
time

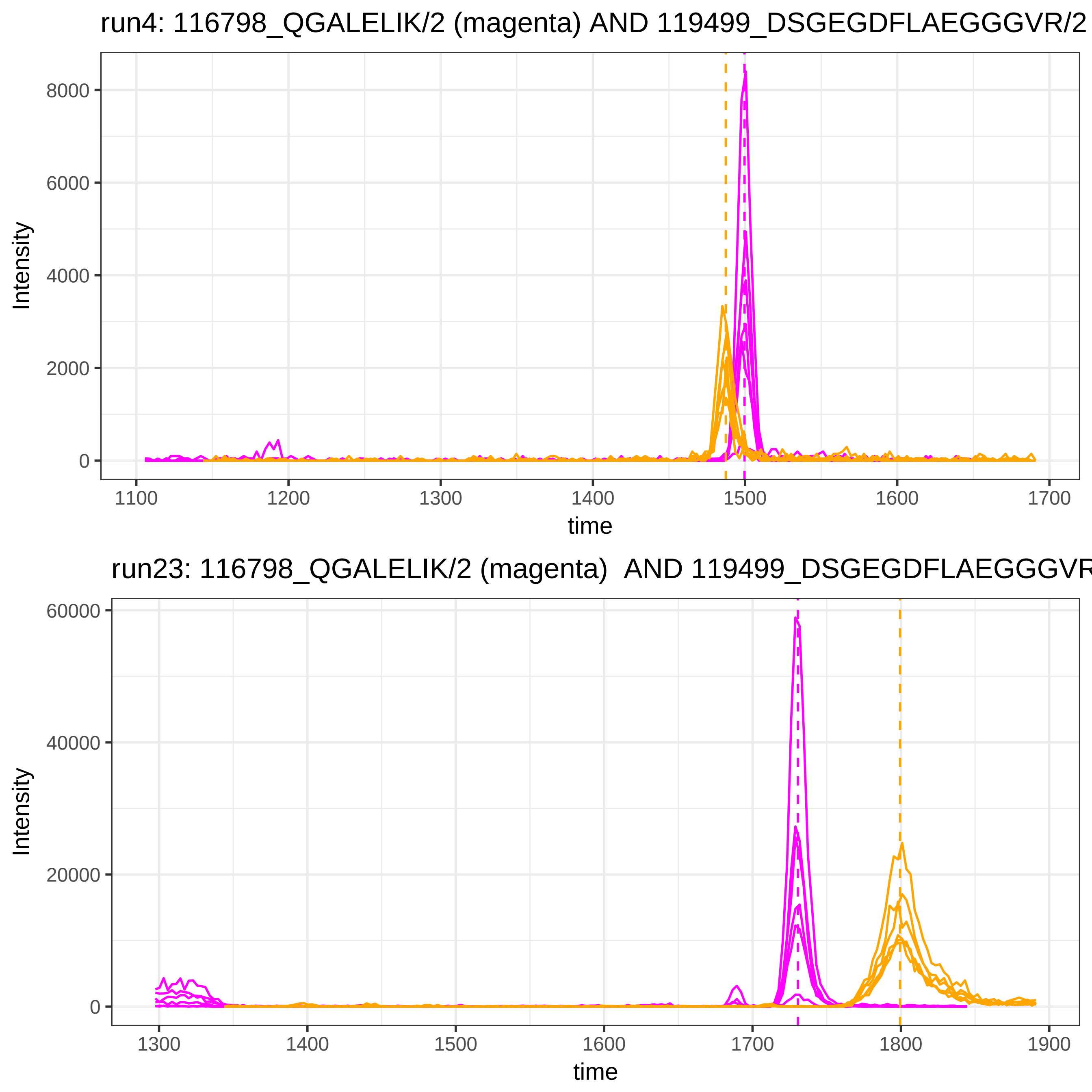
run4: 116798_QGALELIK/2 (magenta) AND 119499_DSGEGDFLAEGGGVR/2
Intensity
8000
6000
4000
2000
0
1100
1200
1300
1400
1500
1600
1700
time
run23: 116798_QGALELIK/2 (magenta) AND 119499_DSGEGDFLAEGGGVR
Intensity
60000
40000
20000
0
1300
1400
1500
1600
1700
1800
1900
time

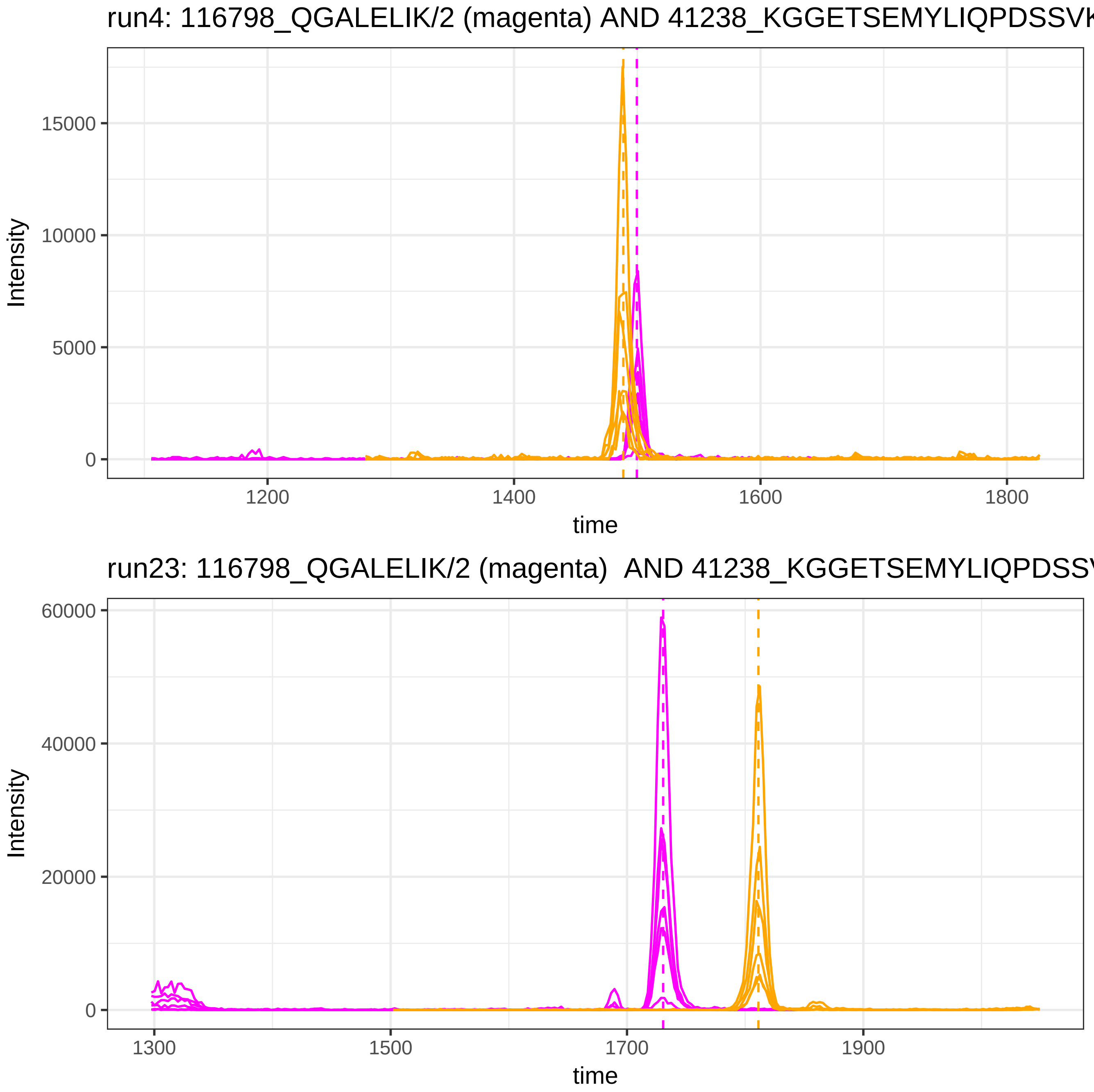
run4: 116798_QGALELIK/2 (magenta) AND 41238_KGGETSEMYLIQPDSSVK
Intensity
15000
10000
5000
0
1200
1400
1600
1800
time
run23: 116798_QGALELIK/2 (magenta) AND 41238_KGGETSEMYLIQPDSSV
Intensity
60000
40000
20000
0
1300
1500
1700
1900
time

run4: 120686_ELAAQTIK/2 (magenta) AND 115879_DDNPNLPR/2

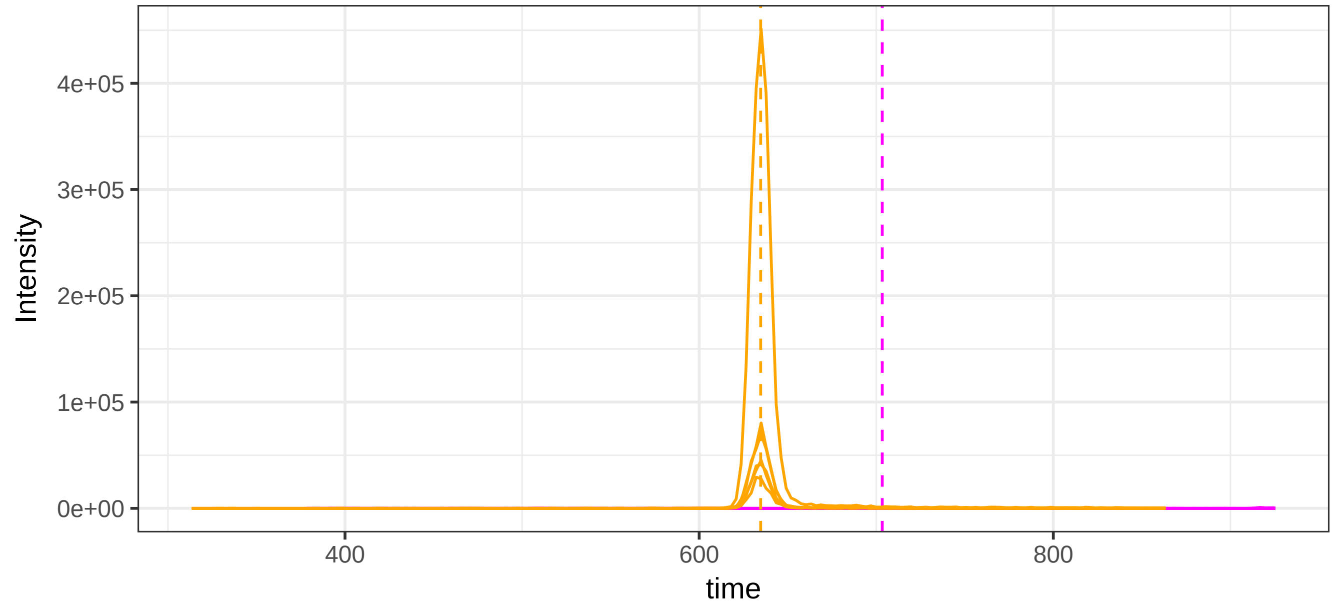


run23: 120686_ELAAQTIK/2 (magenta)  AND 115879_DDNPNLPR/2

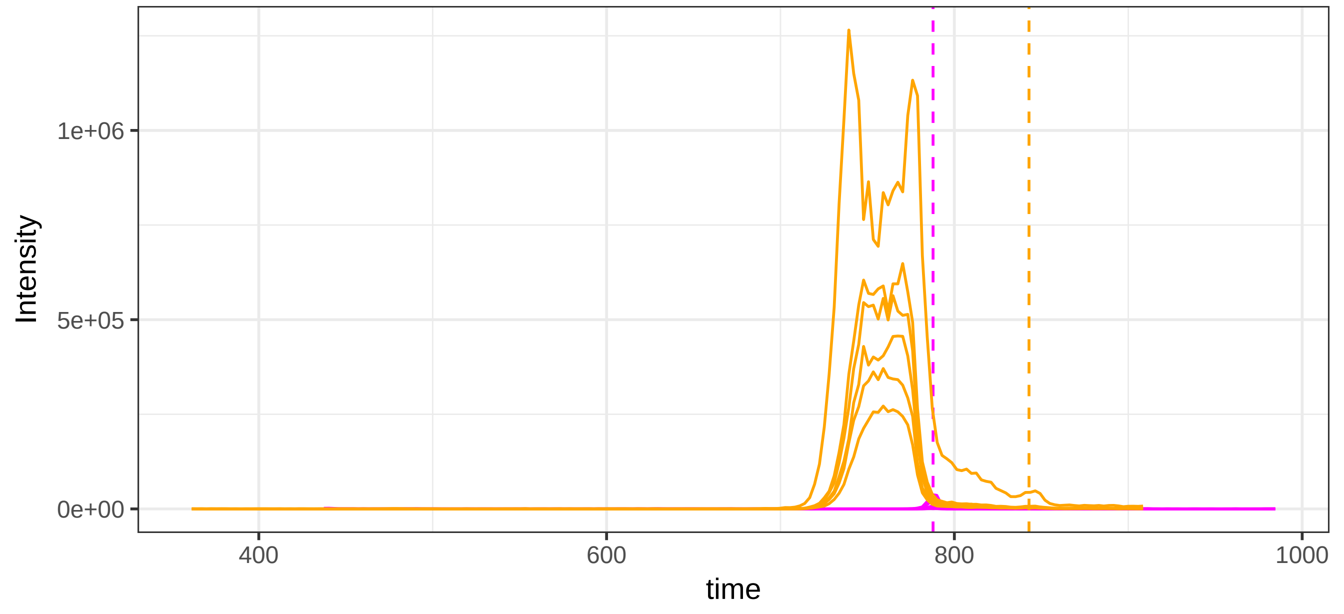

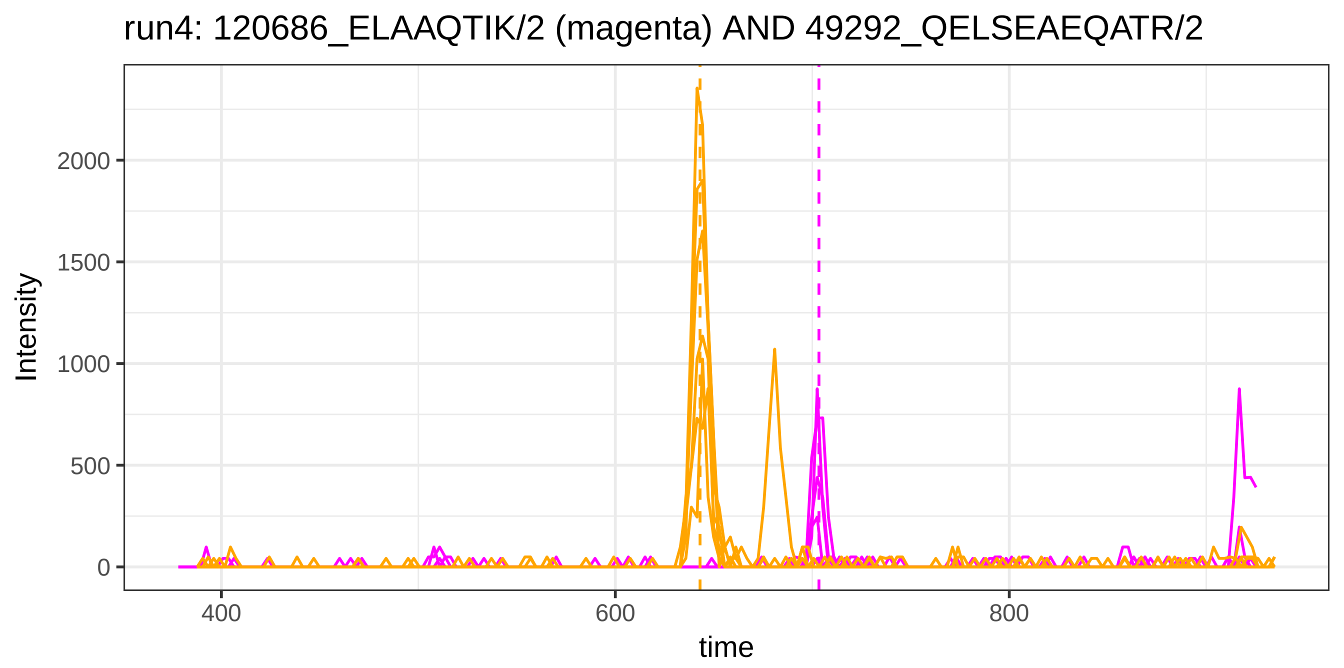
run4: 120686_ELAAQTIK/2 (magenta) AND 49292_QELSEAEQATR/2
Intensity
2000
1500
1000
500
0
400
600
800
time

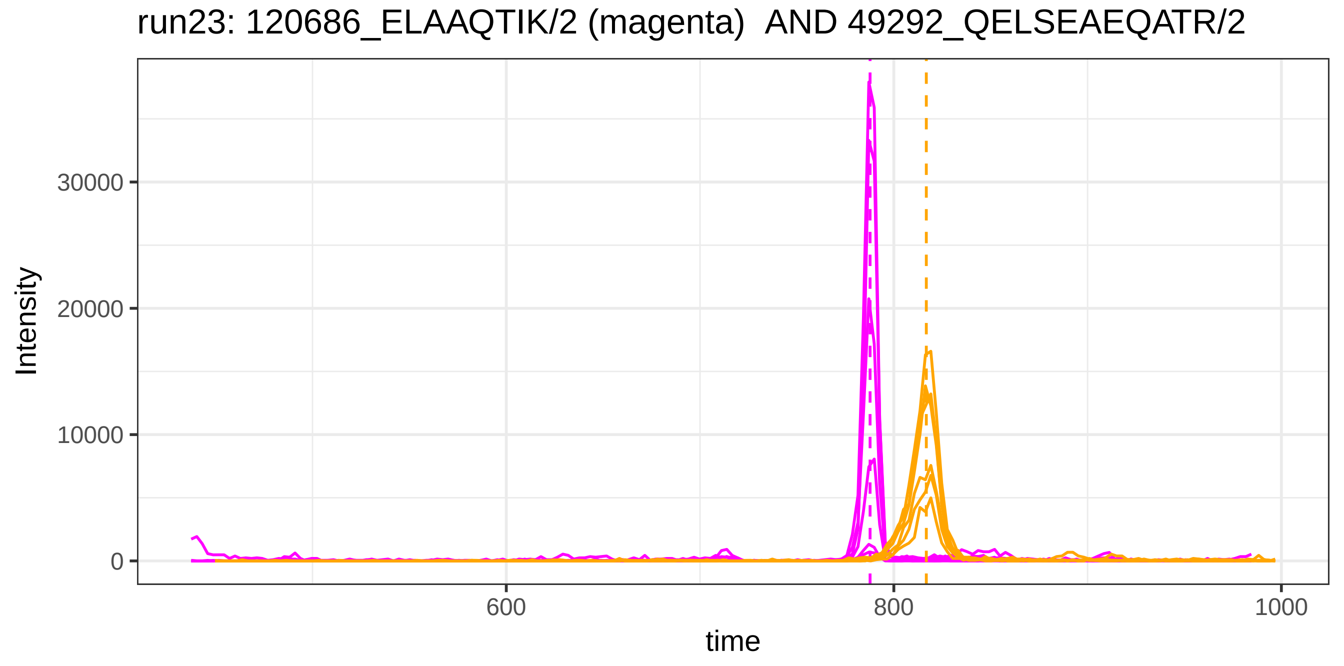
run23: 120686_ELAAQTIK/2 (magenta) AND 49292_QELSEAEQATR/2
Intensity
30000
20000
10000
0
600
800
1000
time

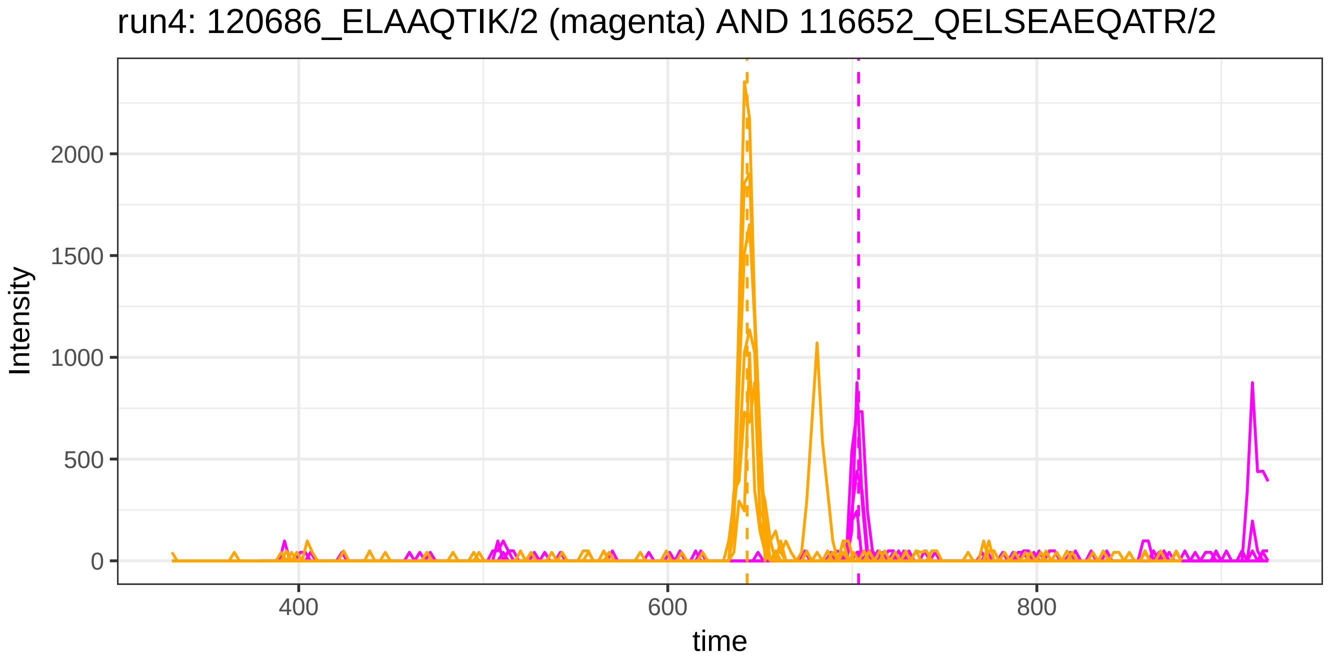
run4: 120686_ELAAQTIK/2 (magenta) AND 116652_QELSEAEQATR/2
Intensity
2000
1500
1000
500
0
400
600
800
time

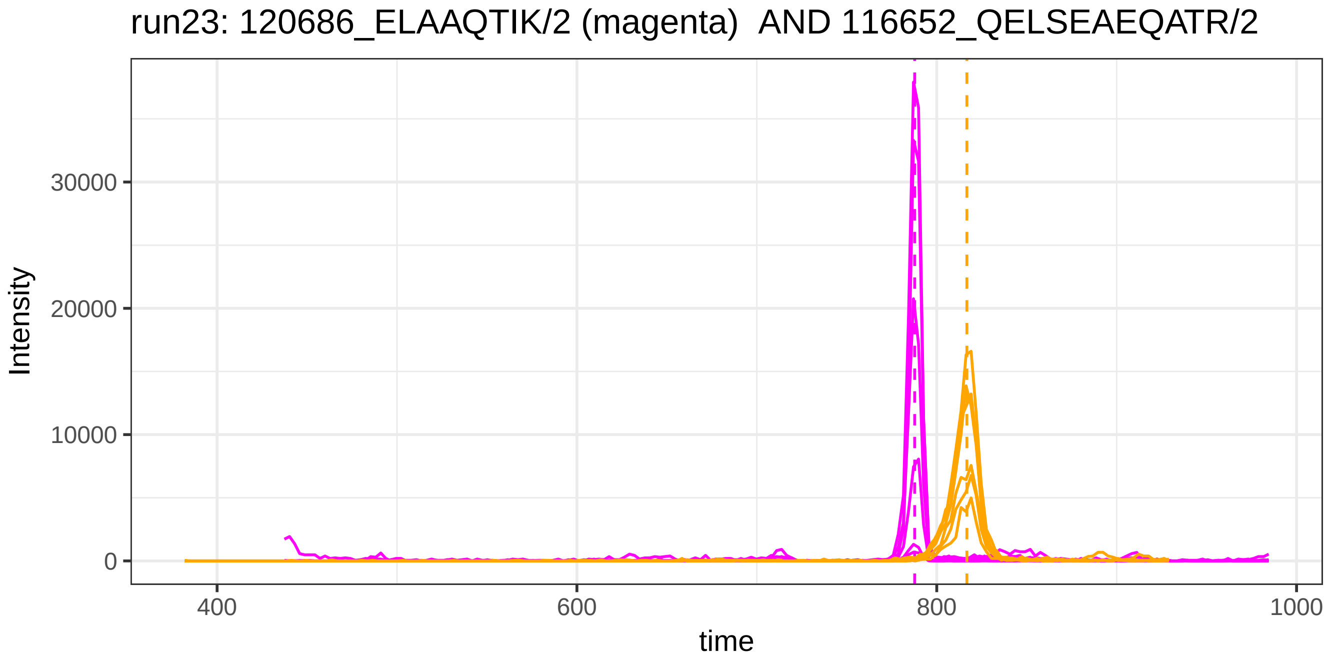
run23: 120686_ELAAQTIK/2 (magenta) AND 116652_QELSEAEQATR/2
Intensity
30000
20000
10000
0
400
600
800
1000
time

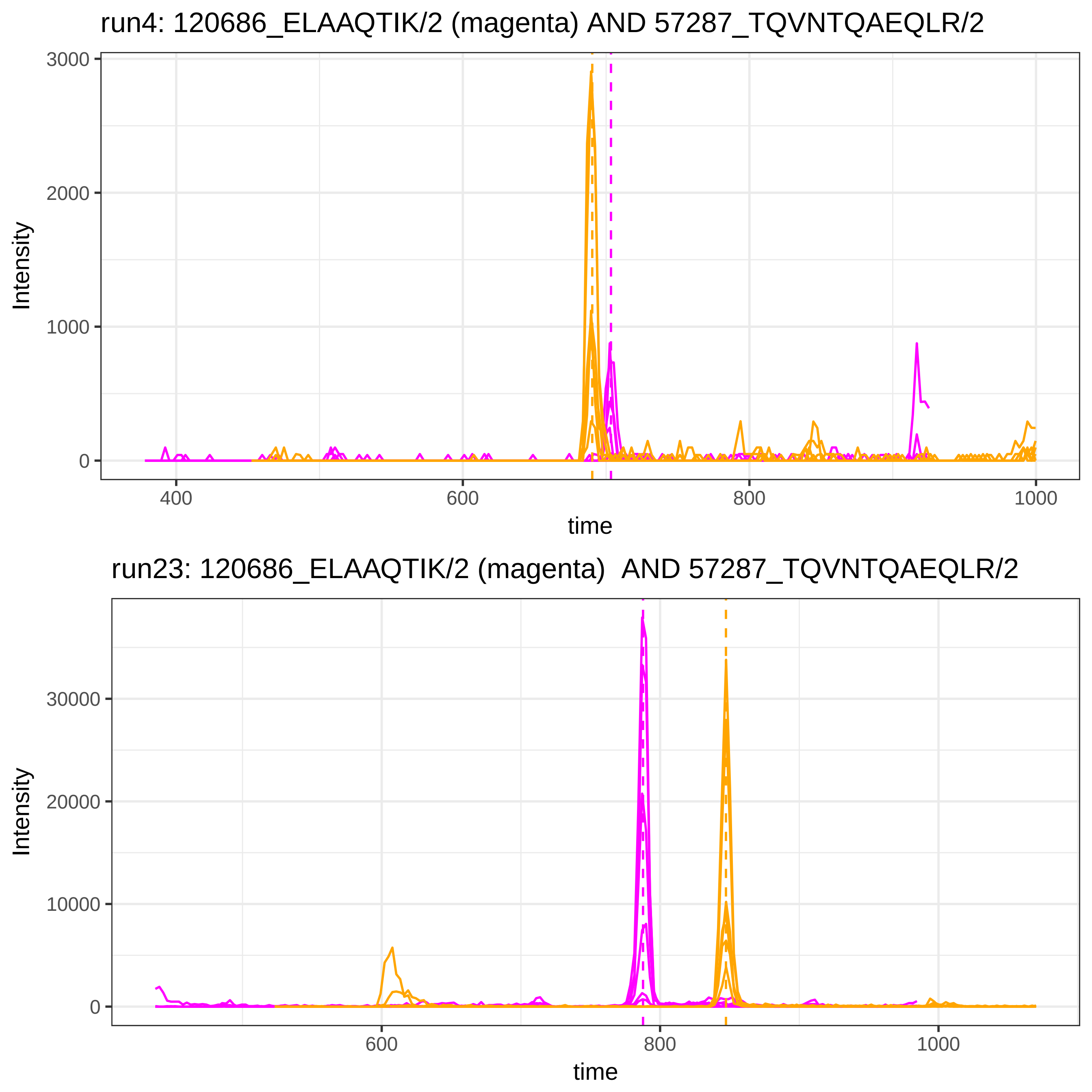
run4: 120686_ELAAQTIK/2 (magenta) AND 57287_TQVNTQAEQLR/2
Intensity
3000
2000
1000
0
400
600
800
1000
time
run23: 120686_ELAAQTIK/2 (magenta)  AND 57287_TQVNTQAEQLR/2
Intensity
30000
20000
10000
0
600
800
1000
time

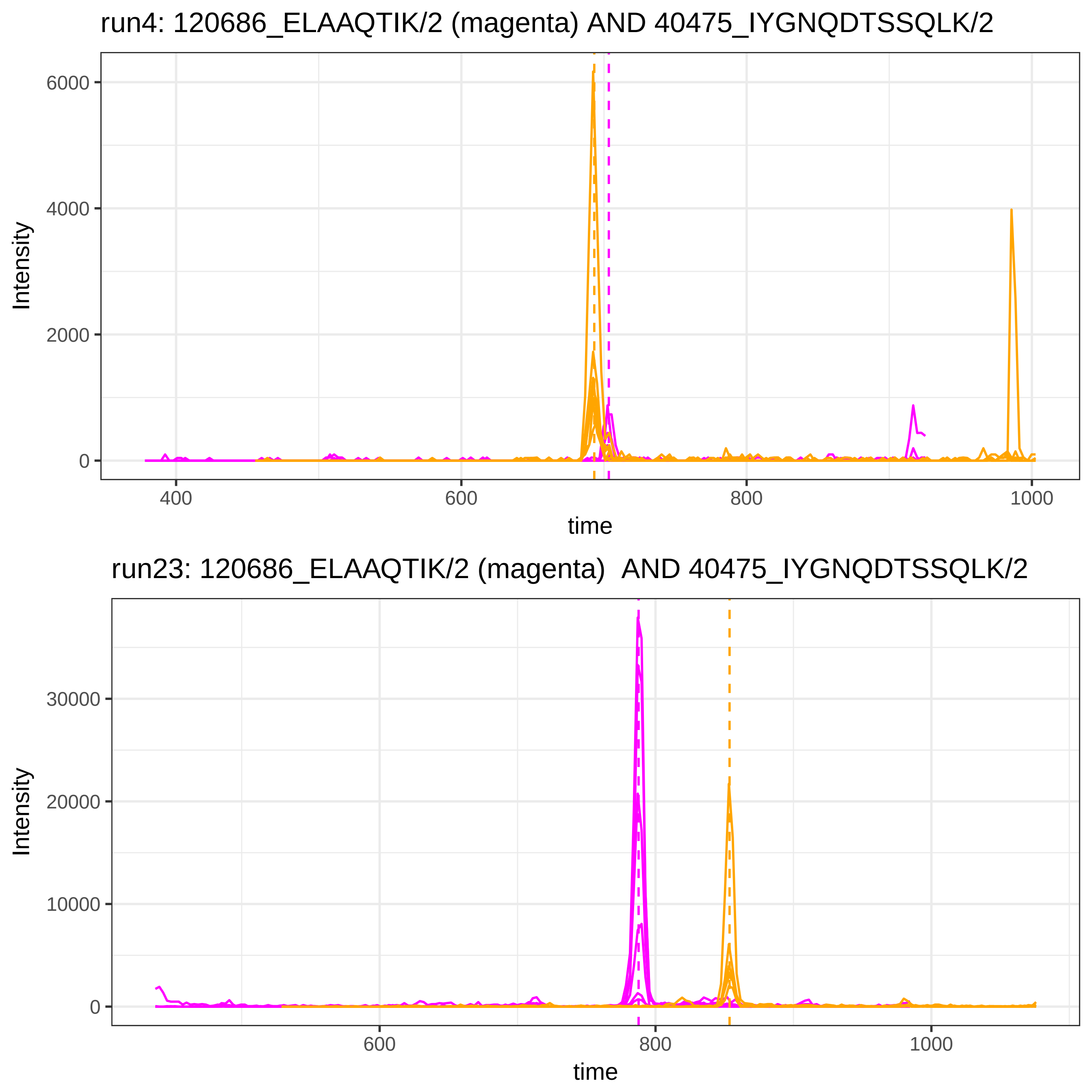
run4: 120686_ELAAQTIK/2 (magenta) AND 40475_IYGNQDTSSQLK/2
Intensity
6000
4000
2000
0
400
600
800
1000
time
run23: 120686_ELAAQTIK/2 (magenta) AND 40475_IYGNQDTSSQLK/2
Intensity
30000
20000
10000
0
600
800
1000
time

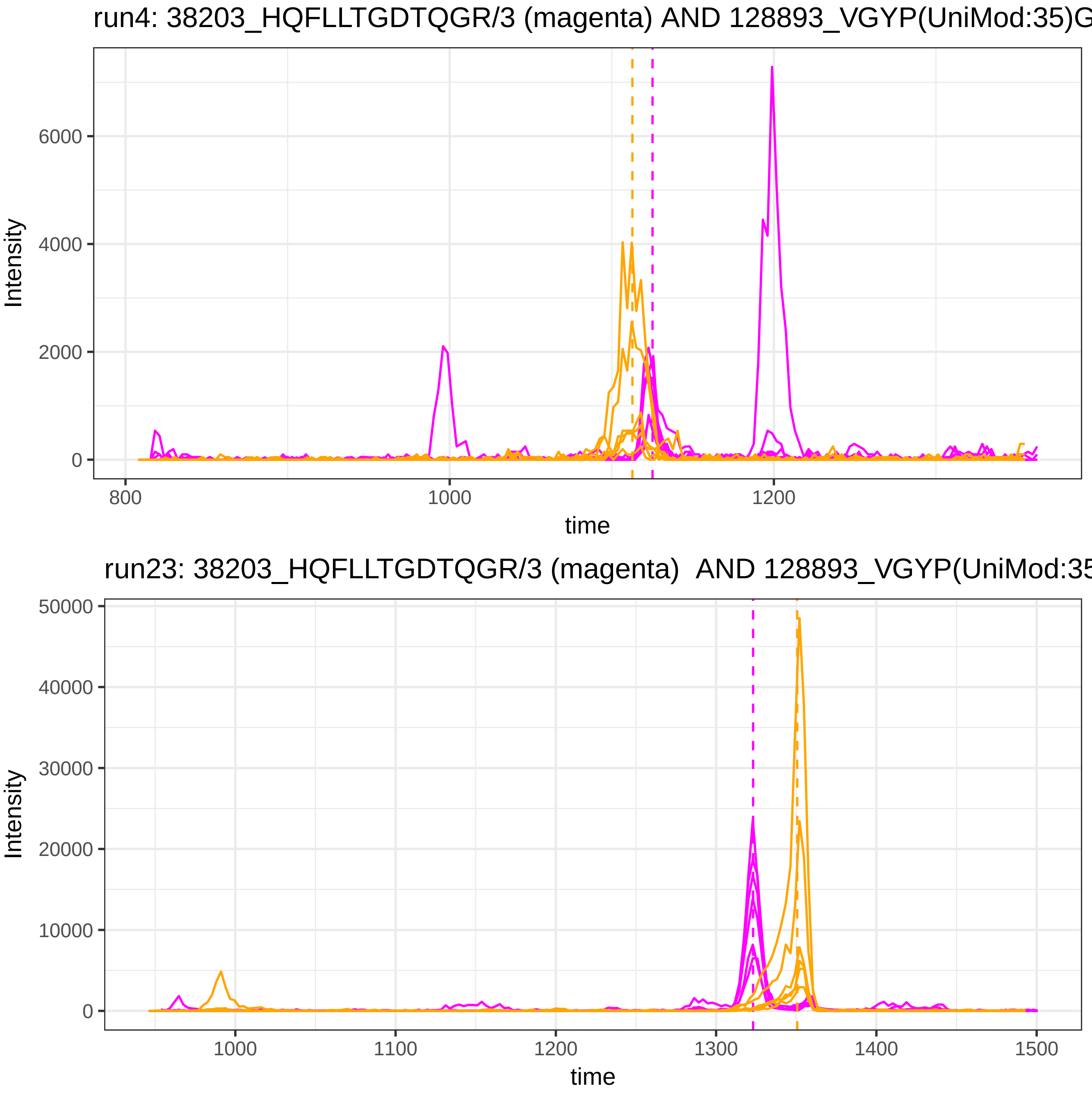

run4: 38203_HQFLLTGDTQGR/3 (magenta) AND 128893_VGYP(UniMod:35)G
Intensity
6000
4000
2000
0
800
1000
1200
time
run23: 38203_HQFLLTGDTQGR/3 (magenta) AND 128893_VGYP(UniMod:35
Intensity
50000
40000
30000
20000
10000
0
1000
1100
1200
1300
1400
1500
time

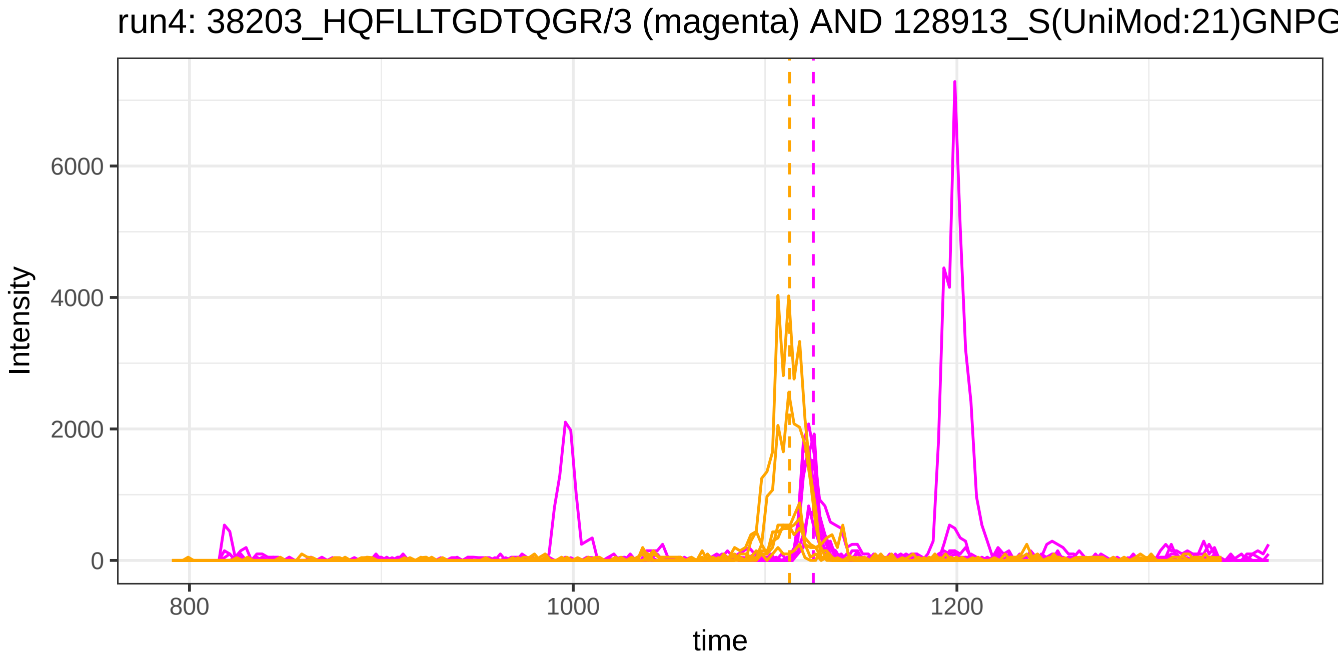
run4: 38203_HQFLLTGDTQGR/3 (magenta) AND 128913_S(UniMod:21)GNPG
Intensity
6000
4000
2000
0
800
1000
1200
time

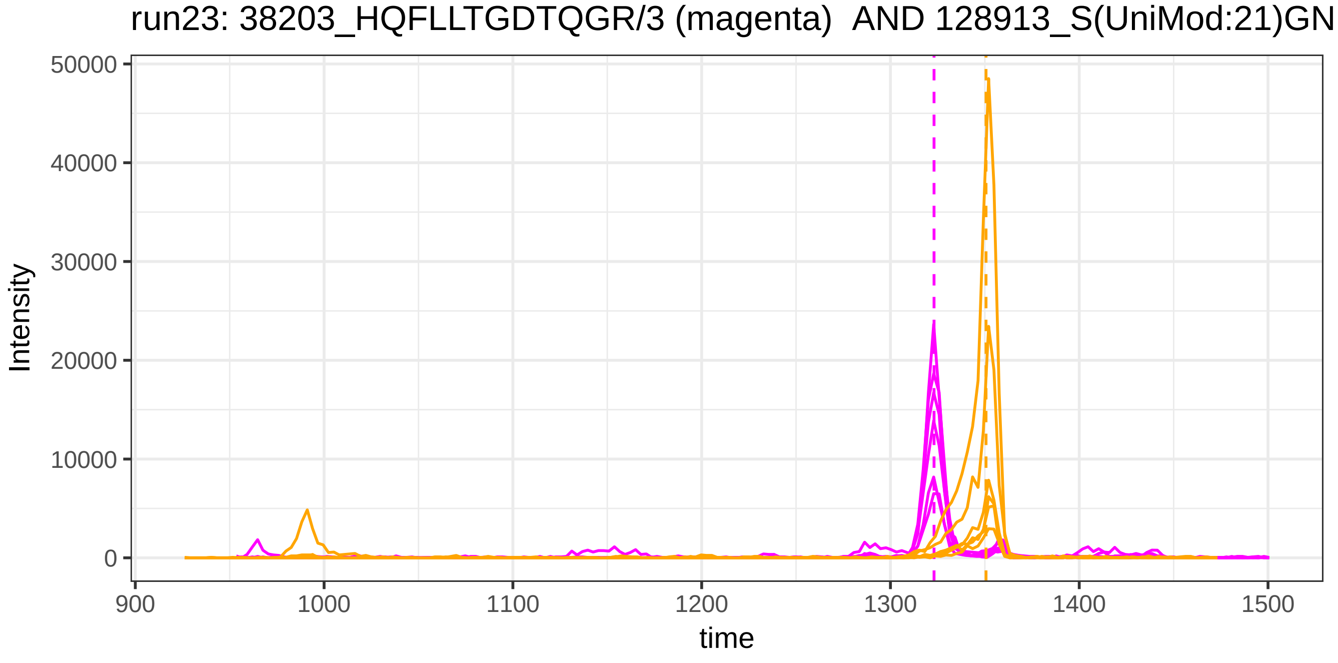
run23: 38203_HQFLLTGDTQGR/3 (magenta) AND 128913_S(UniMod:21)GN
Intensity
50000
40000
30000
20000
10000
0
900
1000
1100
1200
1300
1400
1500
time

run4: 38203_HQFLLTGDTQGR/3 (magenta) AND 4844_YLQEIYNSNNQK/2

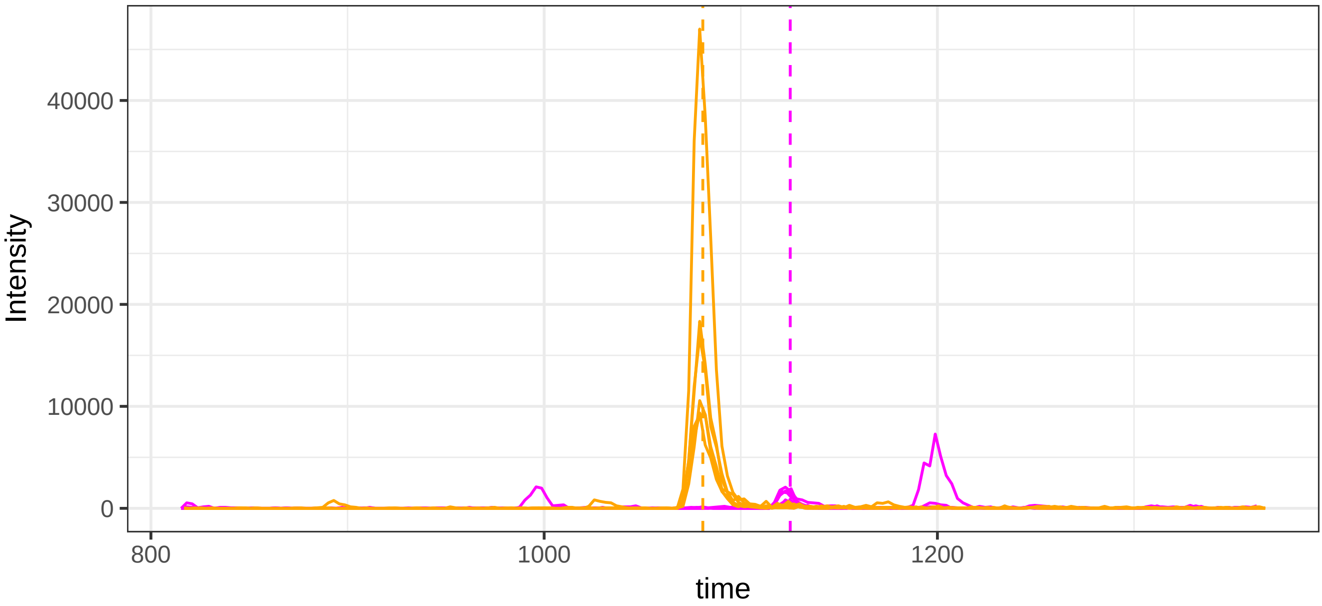


run23: 38203_HQFLLTGDTQGR/3 (magenta)  AND 4844_YLQEIYNSNNQK/2

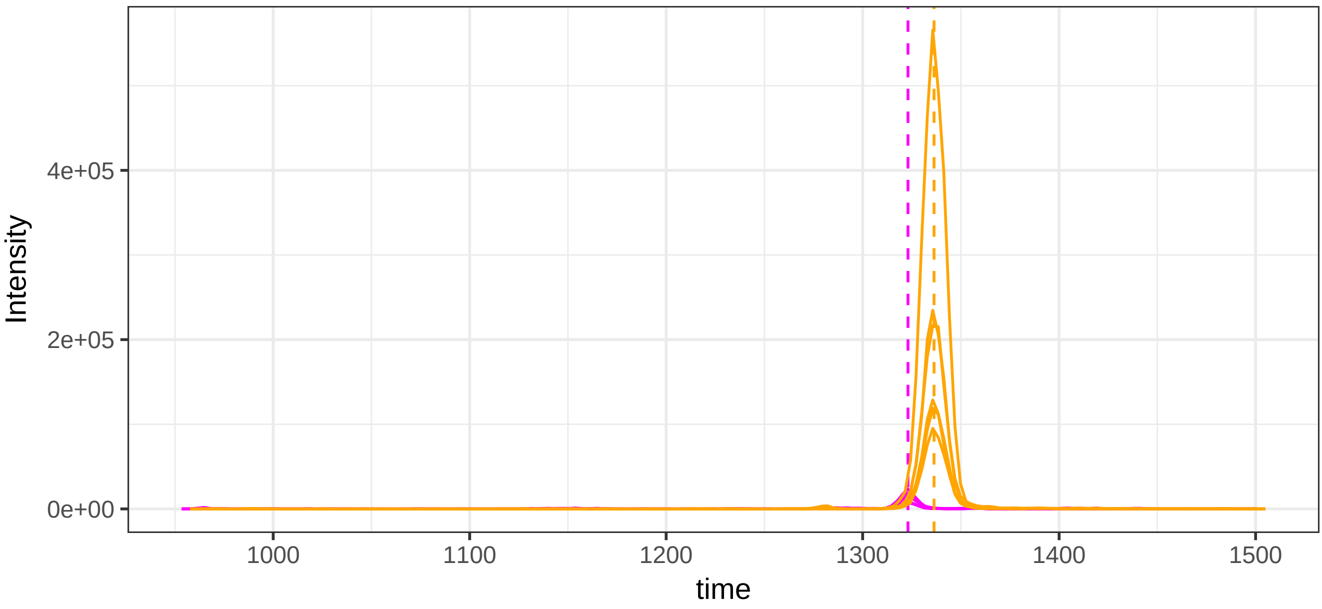

run4: 33823_FLENEDR/2 (magenta) AND 115879_DDNPNLPR/2

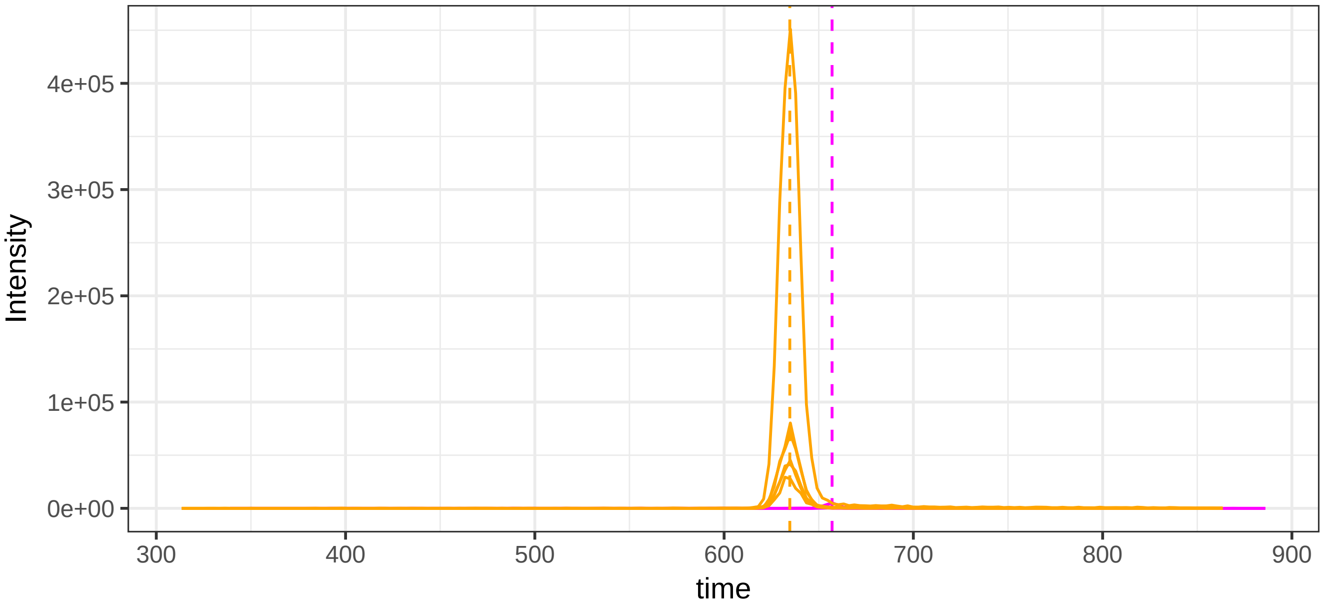


run23: 33823_FLENEDR/2 (magenta)  AND 115879_DDNPNLPR/2

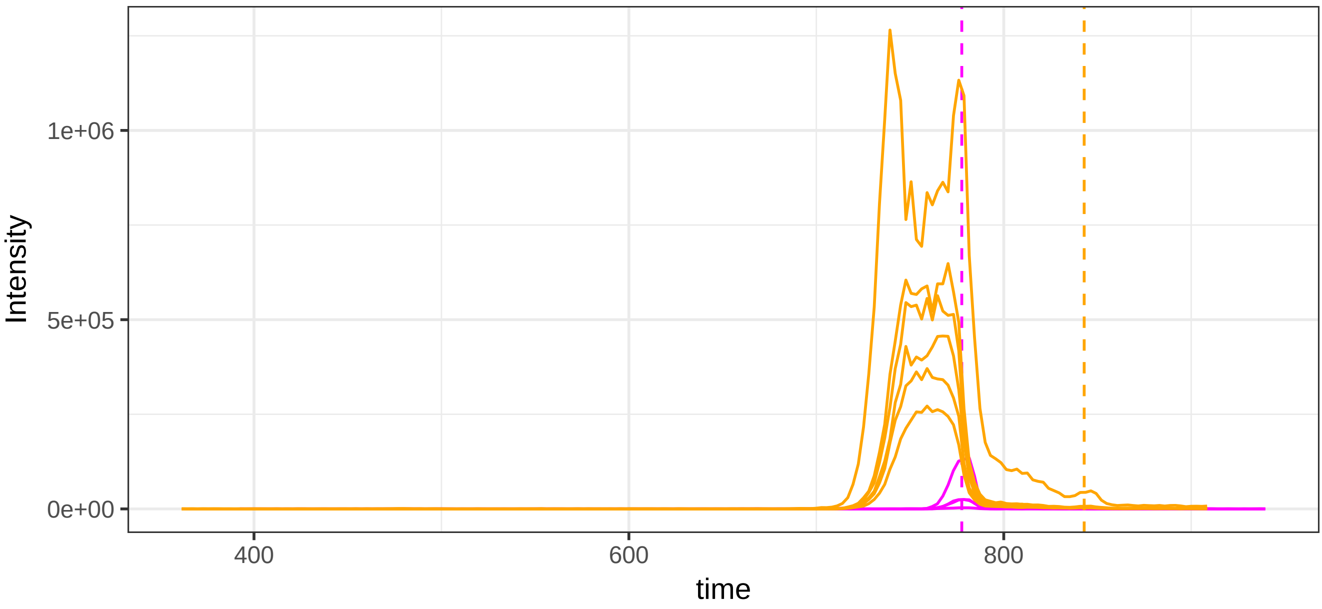

run4: 33823_FLENEDR/2 (magenta) AND 49292_QELSEAEQATR/2

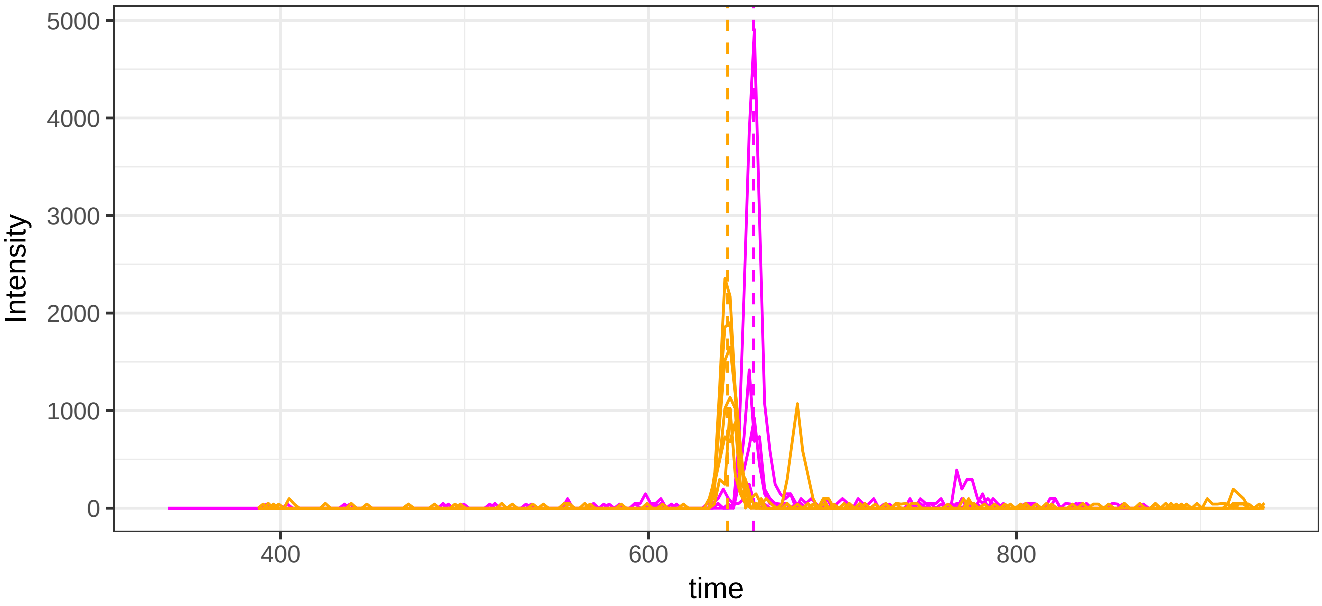


run23: 33823_FLENEDR/2 (magenta)  AND 49292_QELSEAEQATR/2

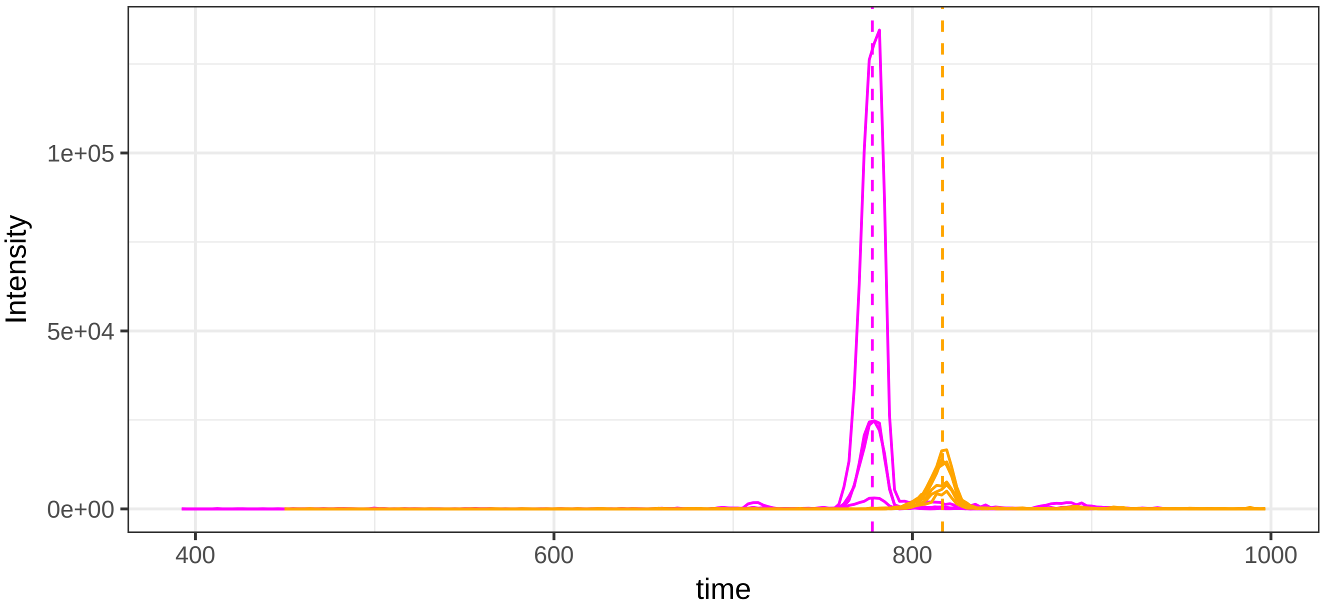

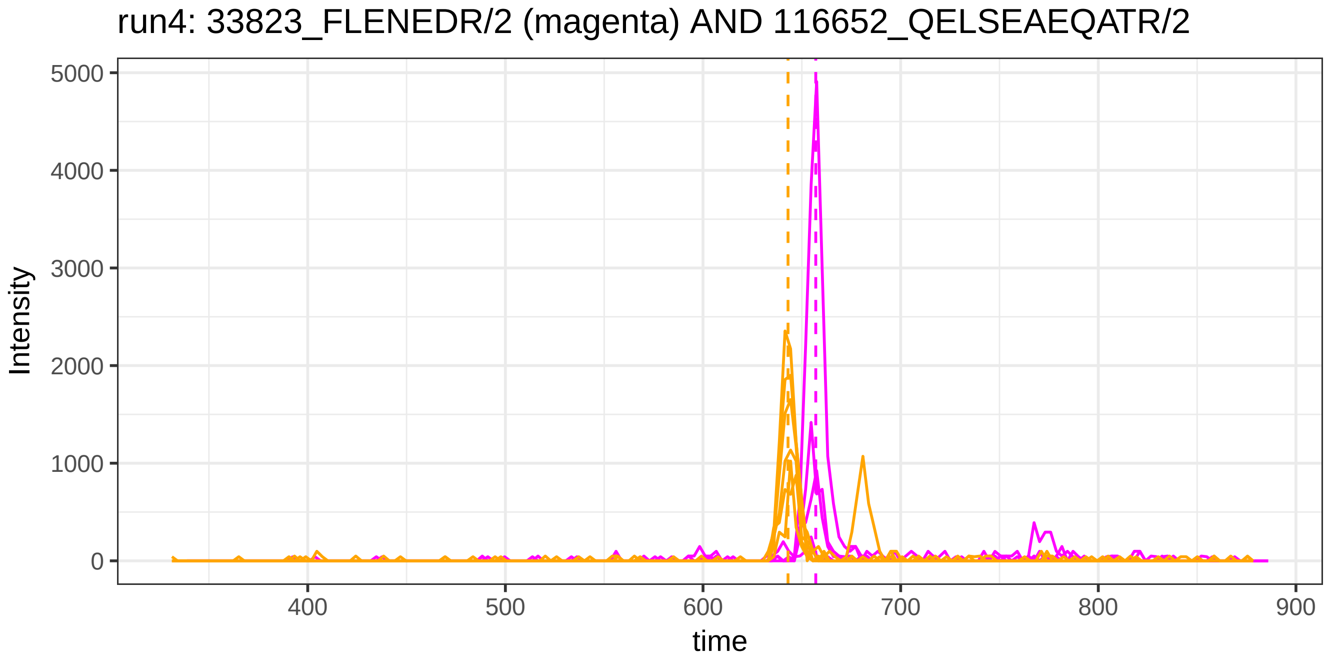
run4: 33823_FLENEDR/2 (magenta) AND 116652_QELSEAEQATR/2
Intensity
5000
4000
3000
2000
1000
0
400
500
600
700
800
900
time

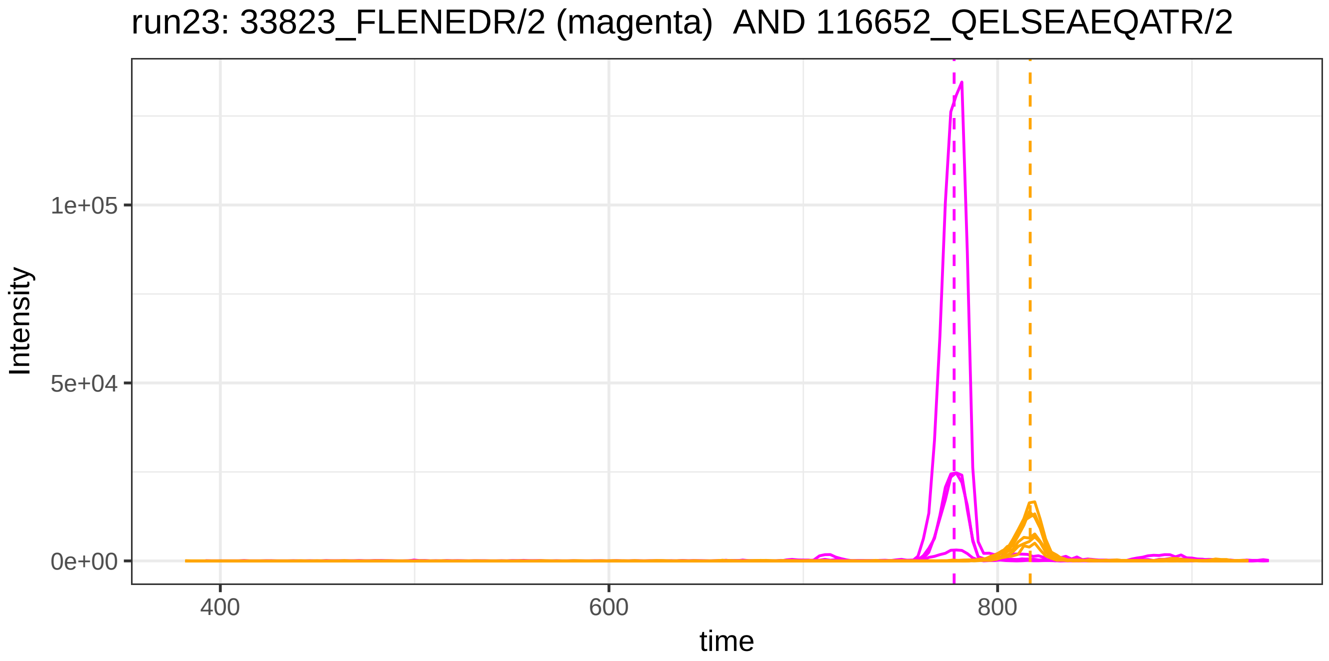
run23: 33823_FLENEDR/2 (magenta) AND 116652_QELSEAEQATR/2
Intensity
1e+05
5e+04
0e+00
400
600
800
time

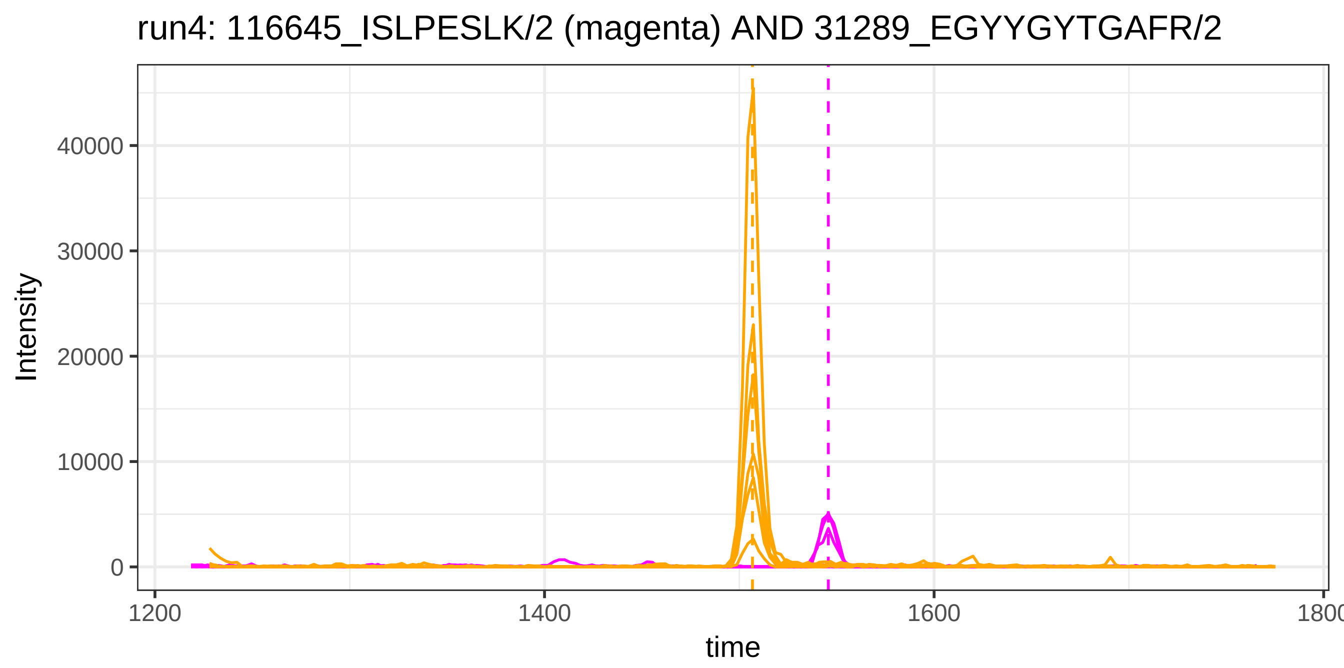

run4: 116645_ISLPESLK/2 (magenta) AND 31289_EGYYGYTGAFR/2
Intensity
40000
30000
20000
10000
0
1200
1400
1600
1800
time


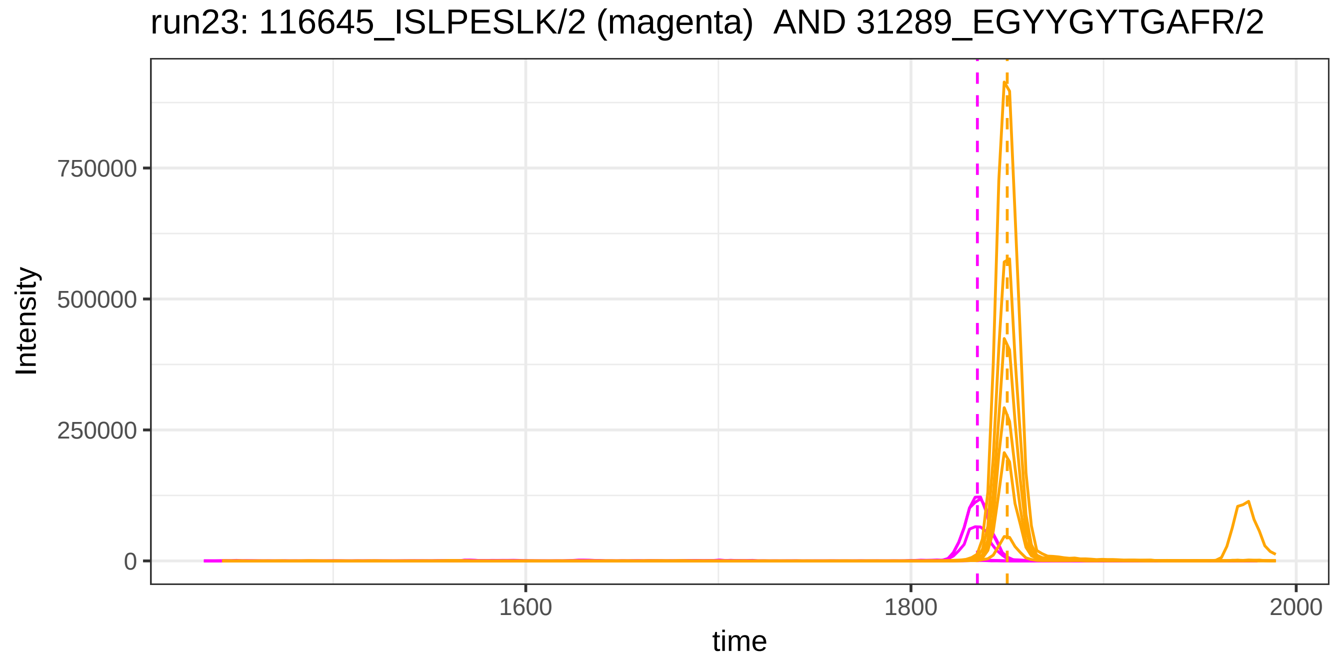

run23: 116645_ISLPESLK/2 (magenta) AND 31289_EGYYGYTGAFR/2
Intensity
750000
500000
250000
0
1600
1800
2000
time

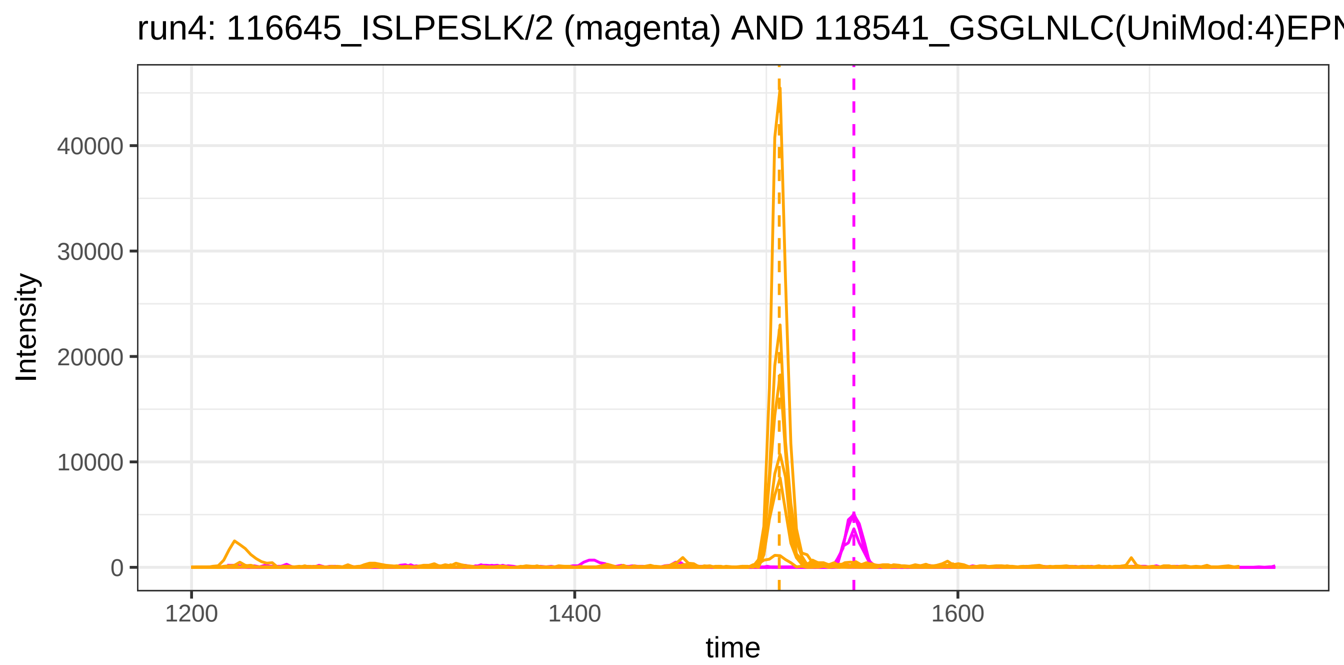
run4: 116645_ISLPESLK/2 (magenta) AND 118541_GSGLNLC(UniMod:4)EPN
Intensity
40000
30000
20000
10000
0
1200
1400
1600
time

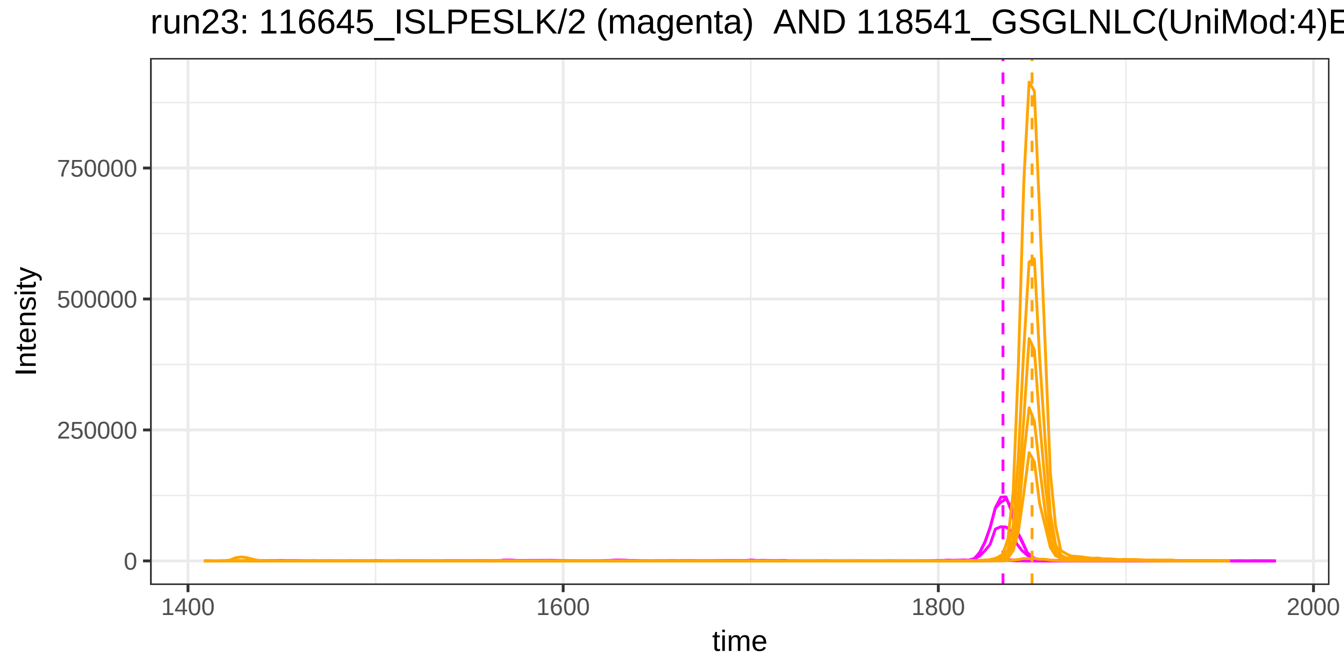
run23: 116645_ISLPESLK/2 (magenta) AND 118541_GSGLNLC(UniMod:4)E
Intensity
750000
500000
250000
0
1400
1600
1800
2000
time

run4: 120092_RTHLPEVFLSK/3 (magenta) AND 41559_ALTDMPQMR/2

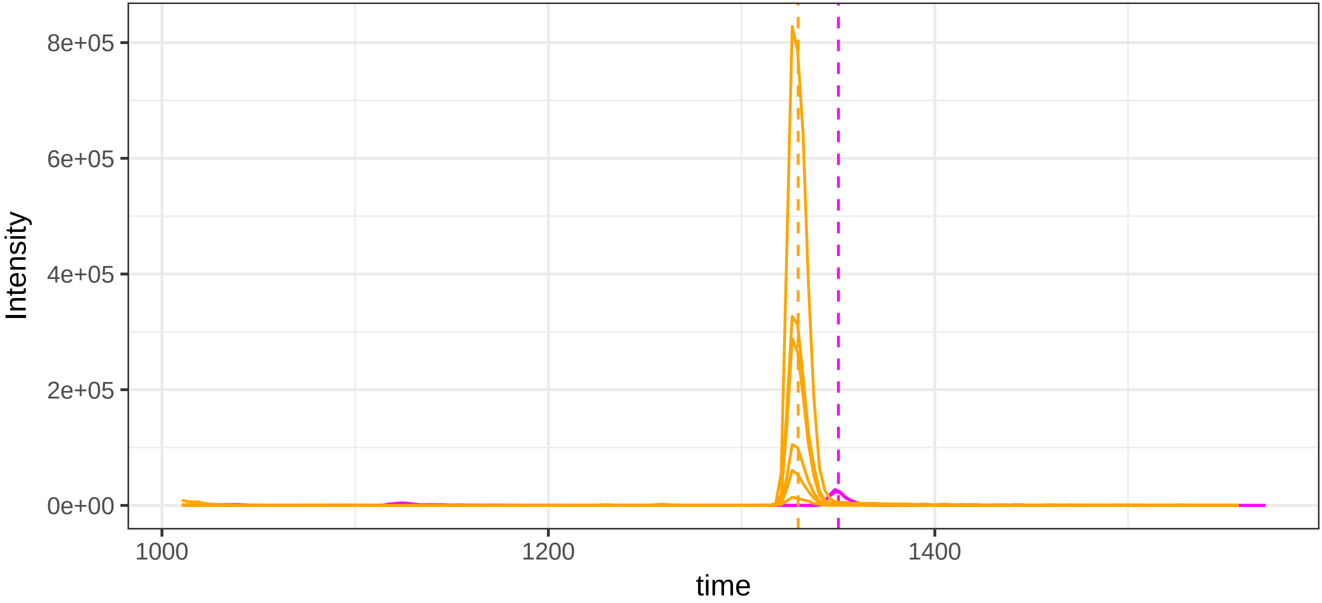


run23: 120092_RTHLPEVFLSK/3 (magenta) AND 41559_ALTDMPQMR/2

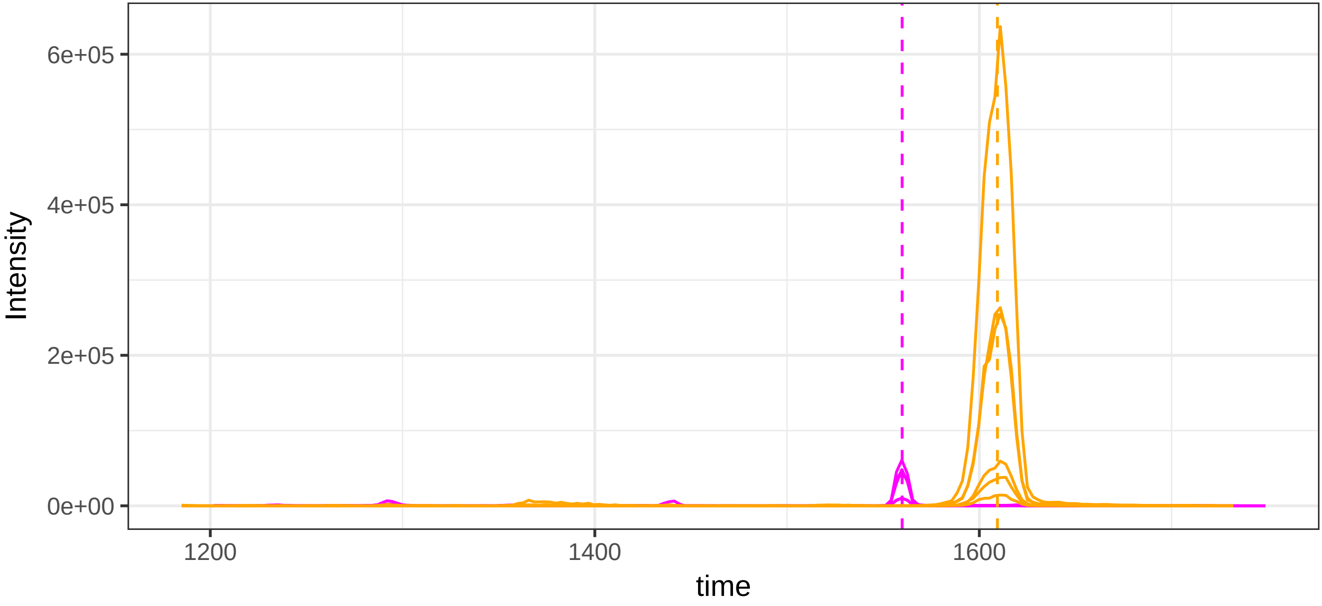

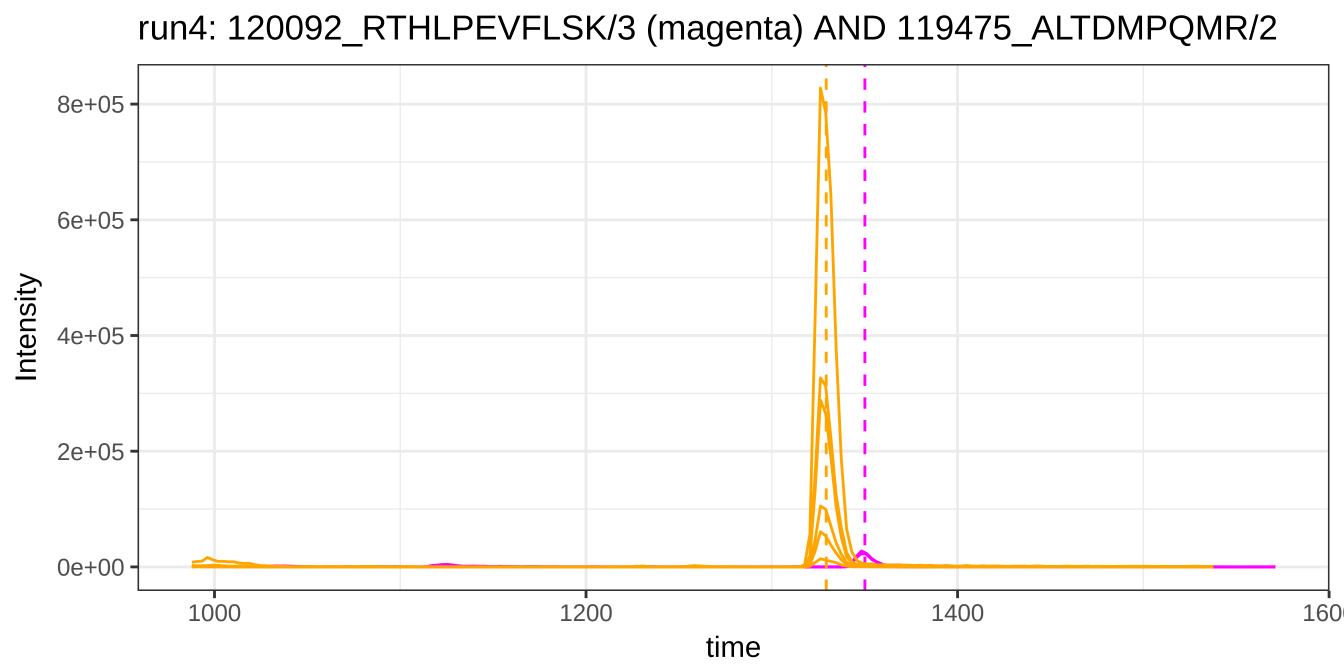
run4: 120092_RTHLPEVFLSK/3 (magenta) AND 119475_ALTDMPQMR/2
Intensity
8e+05
6e+05
4e+05
2e+05
0e+00
1000
1200
1400
1600
time

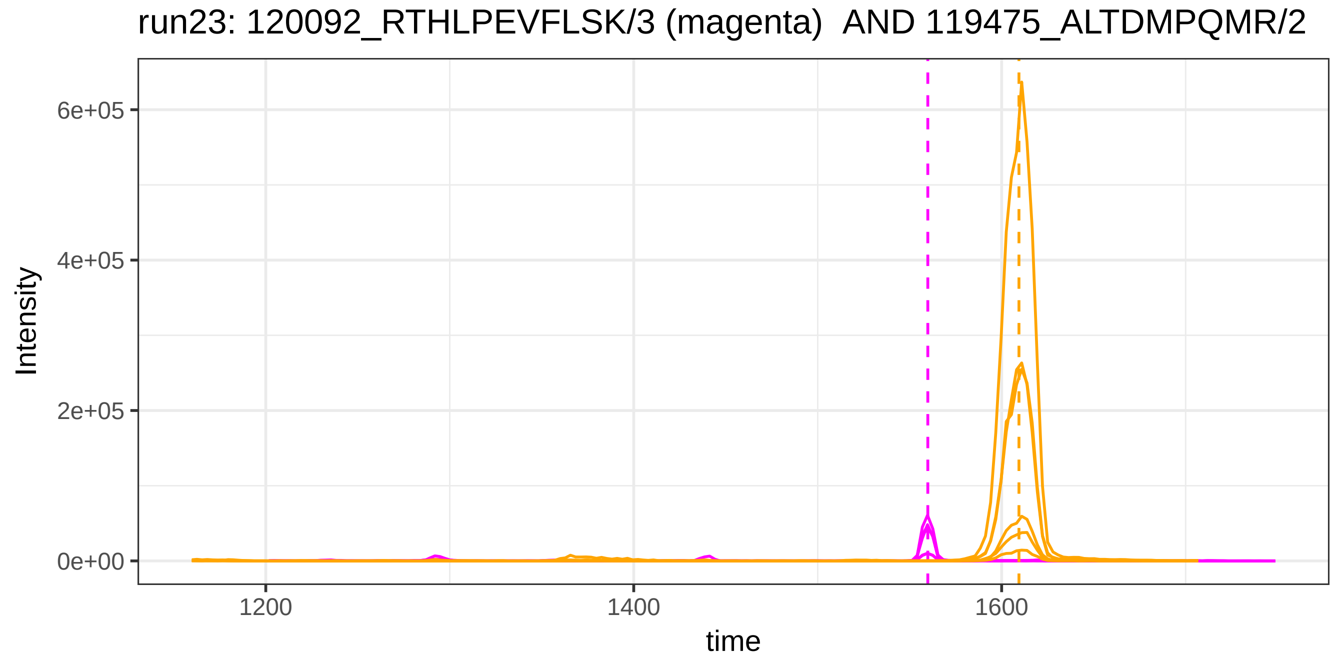
run23: 120092_RTHLPEVFLSK/3 (magenta)  AND 119475_ALTDMPQMR/2
Intensity
6e+05
4e+05
2e+05
0e+00
1200
1400
1600
time

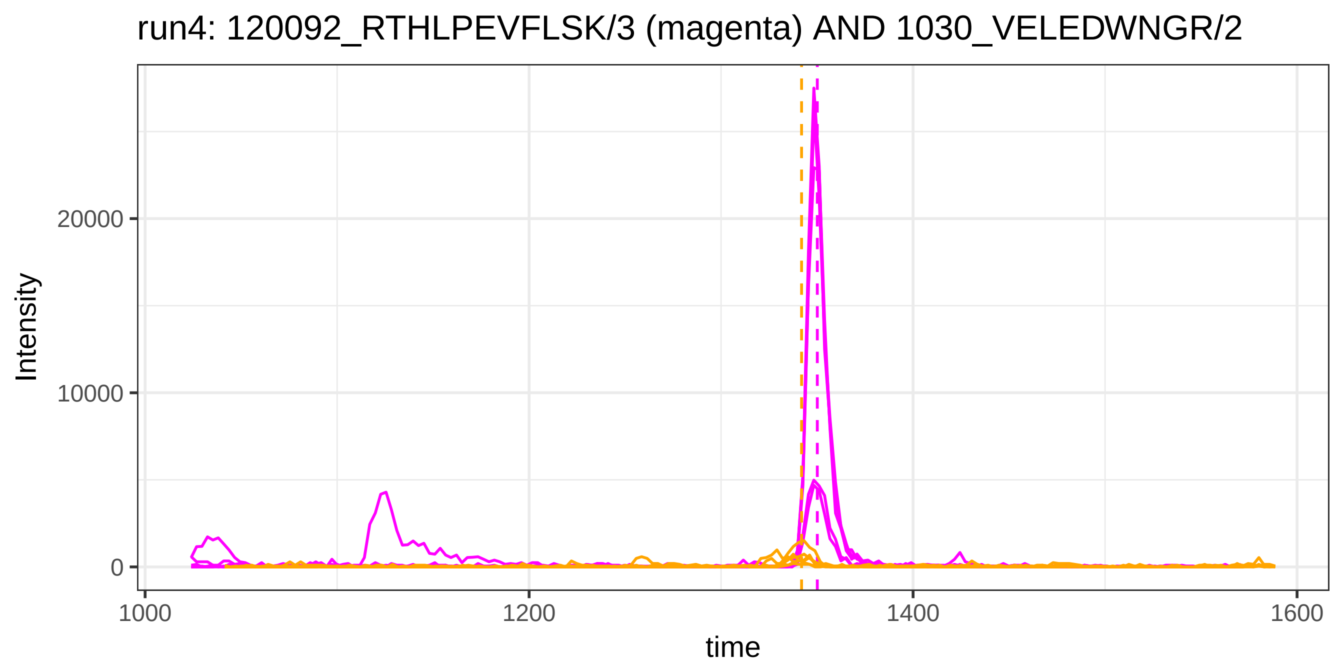
run4: 120092_RTHLPEVFLSK/3 (magenta) AND 1030_VELEDWNGR/2
Intensity
20000
10000
0
1000
1200
1400
1600
time

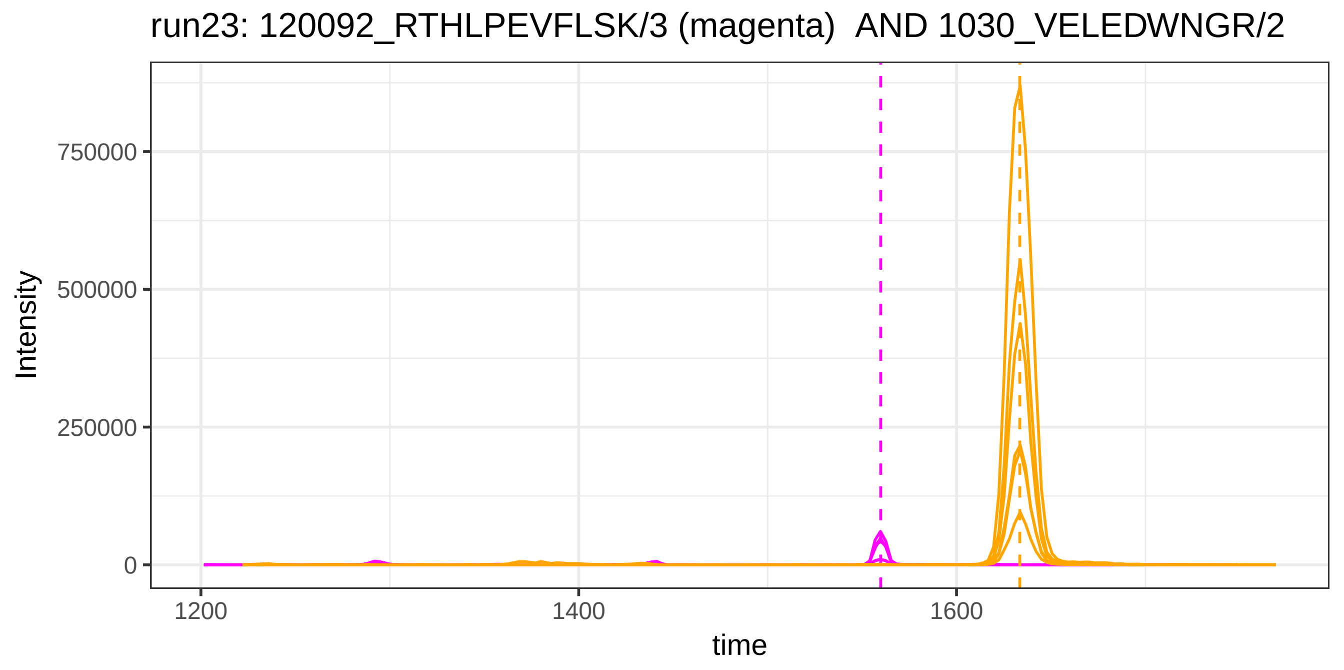
run23: 120092_RTHLPEVFLSK/3 (magenta) AND 1030_VELEDWNGR/2
Intensity
750000
500000
250000
0
1200
1400
1600
time

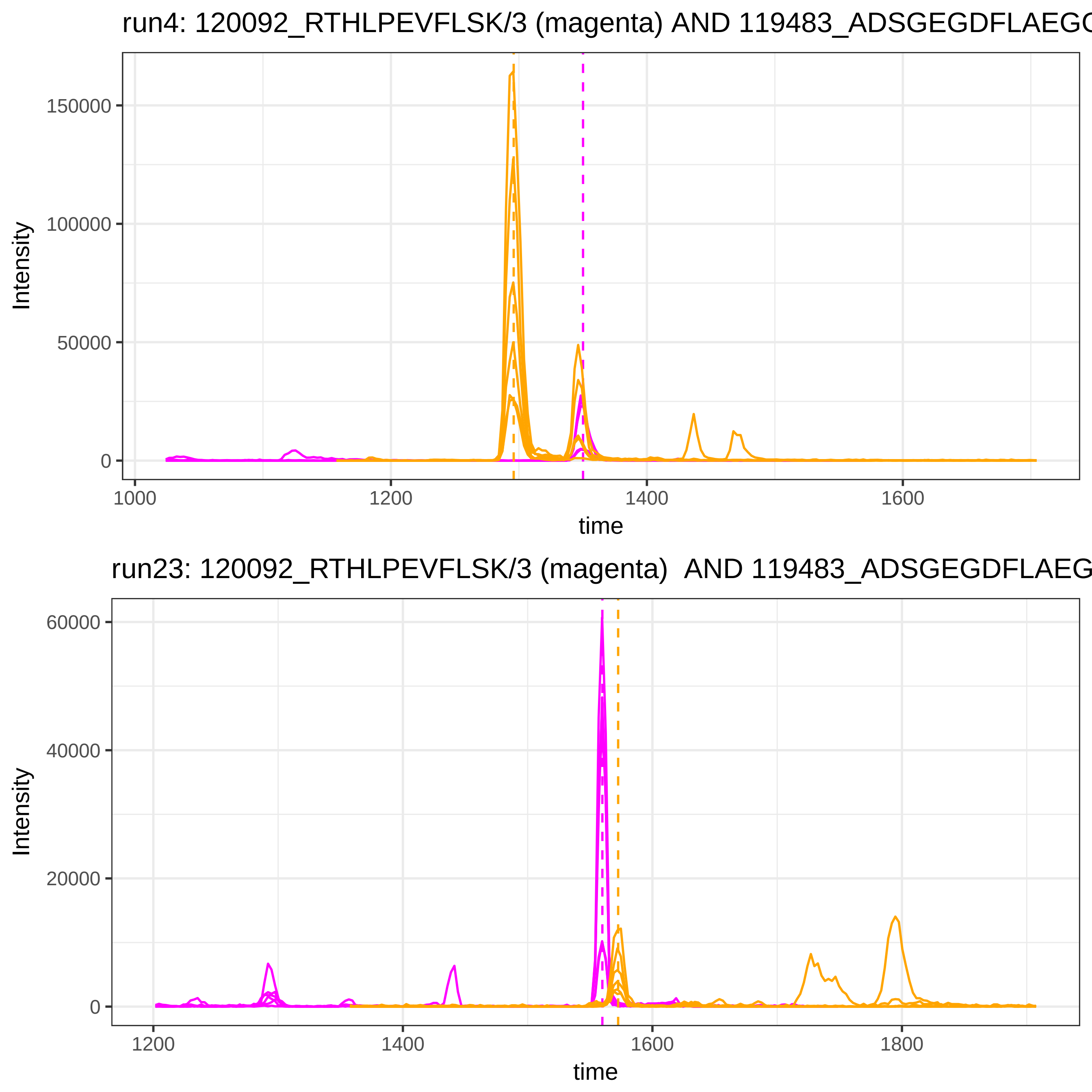
run4: 120092_RTHLPEVFLSK/3 (magenta) AND 119483_ADSGEGDFLAEGG
Intensity
150000
100000
50000
0
1000
1200
1400
1600
time
run23: 120092_RTHLPEVFLSK/3 (magenta) AND 119483_ADSGEGDFLAEG
Intensity
60000
40000
20000
0
1200
1400
1600
1800
time

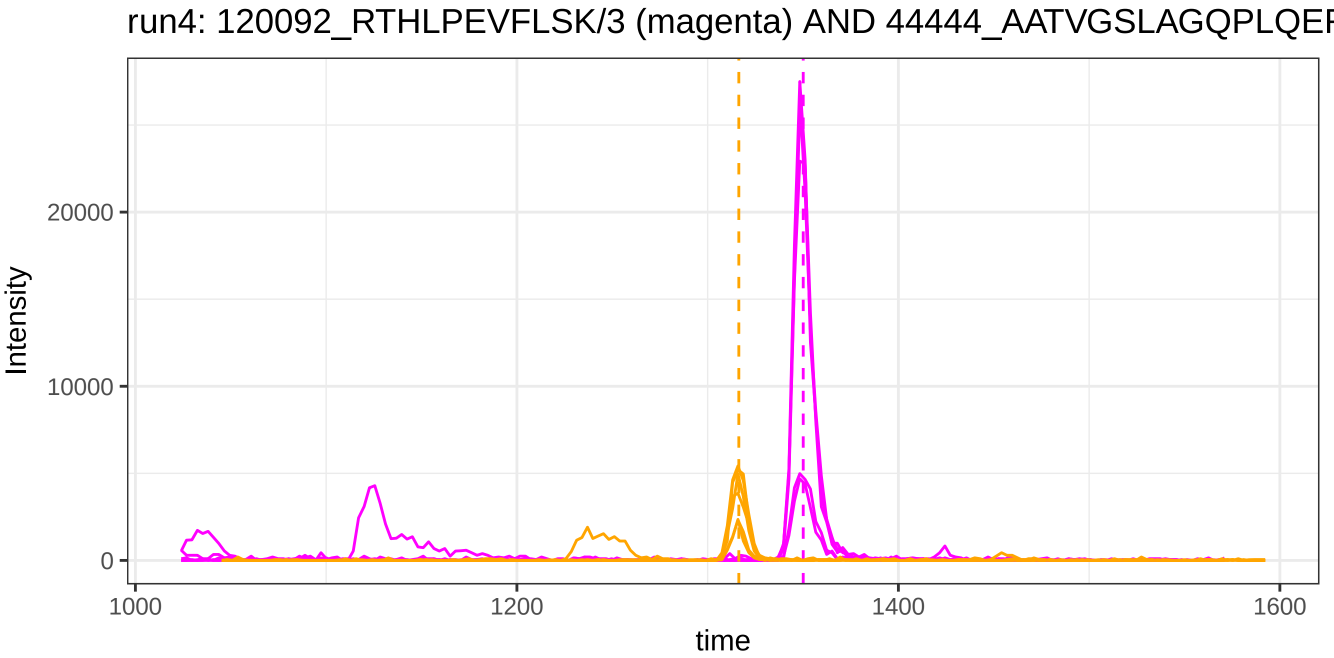
run4: 120092_RTHLPEVFLSK/3 (magenta) AND 44444_AATVGSLAGQPLQEF
Intensity
20000
10000
0
1000
1200
1400
1600
time

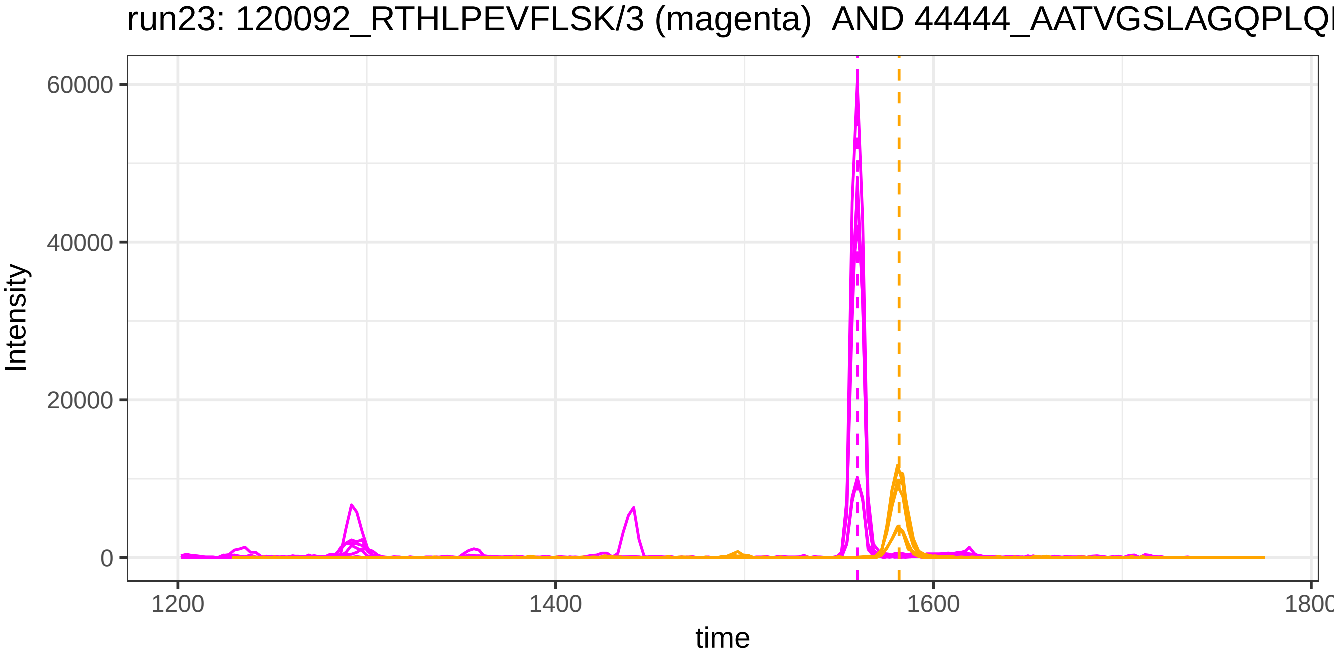
run23: 120092_RTHLPEVFLSK/3 (magenta) AND 44444_AATVGSLAGQPLQI
Intensity
60000
40000
20000
0
1200
1400
1600
1800
time

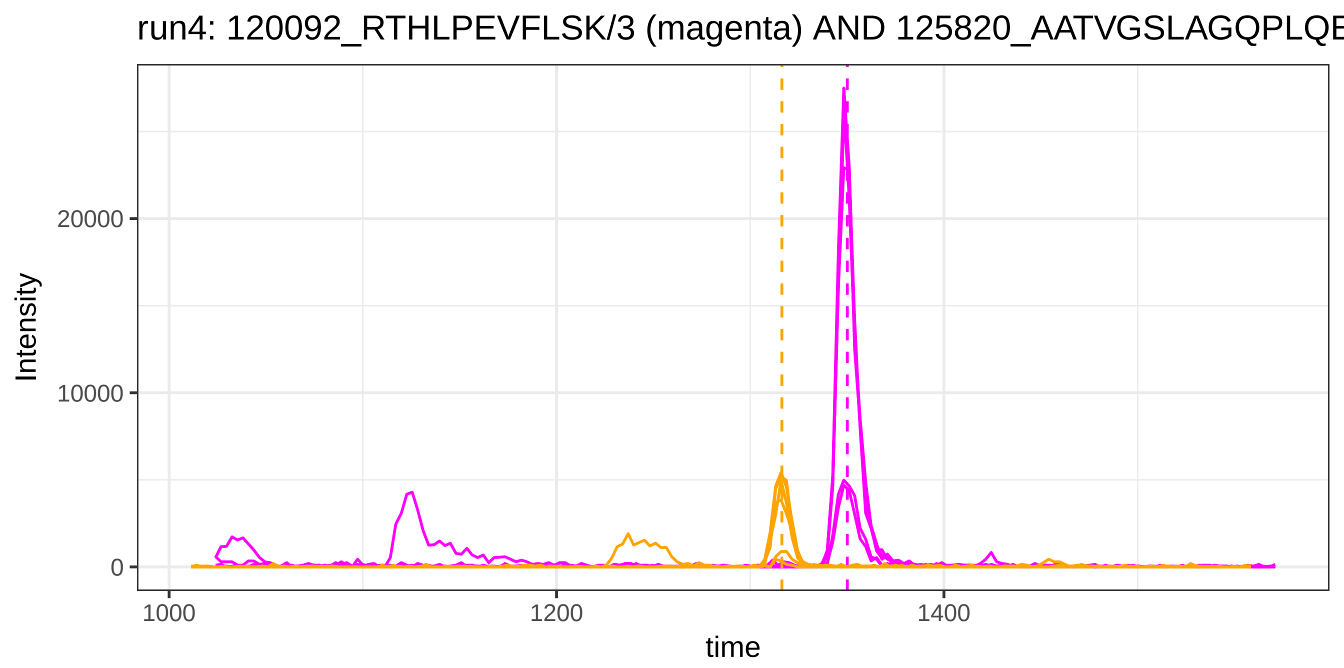
run4: 120092_RTHLPEVFLSK/3 (magenta) AND 125820_AATVGSLAGQPLQE
Intensity
20000
10000
0
time
1000
1200
1400

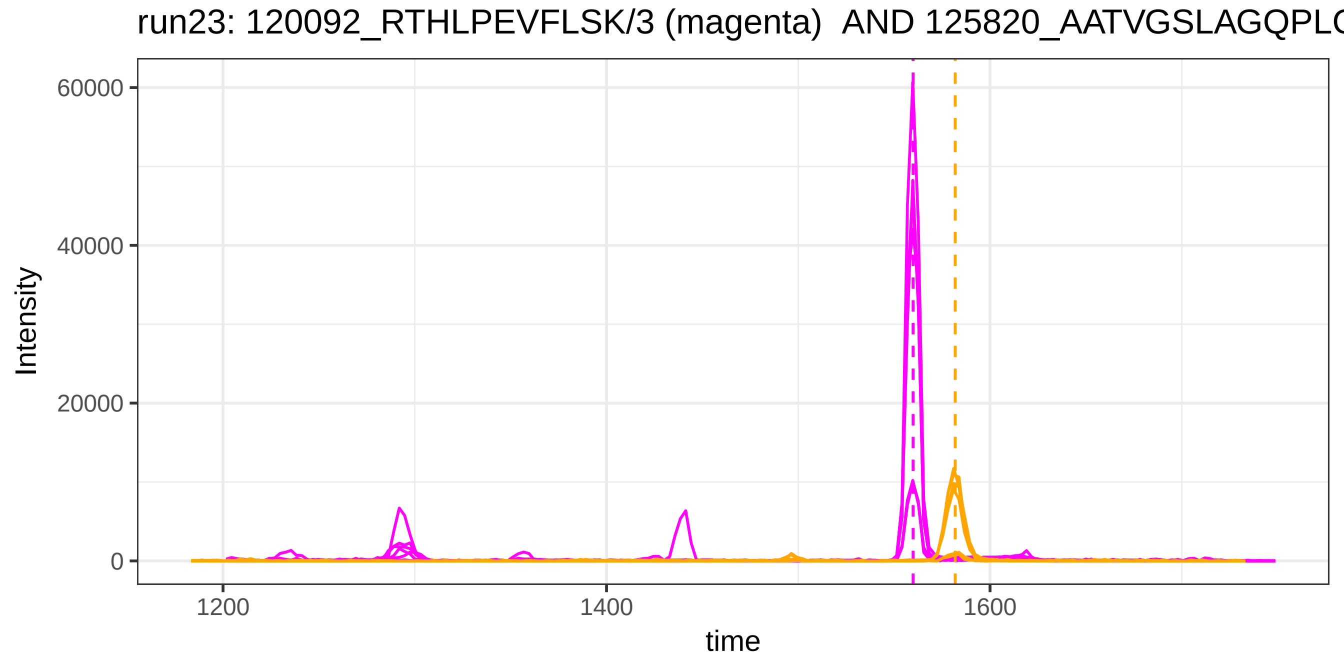
run23: 120092_RTHLPEVFLSK/3 (magenta) AND 125820_AATVGSLAGQPLQ
Intensity
60000
40000
20000
0
time
1200
1400
1600

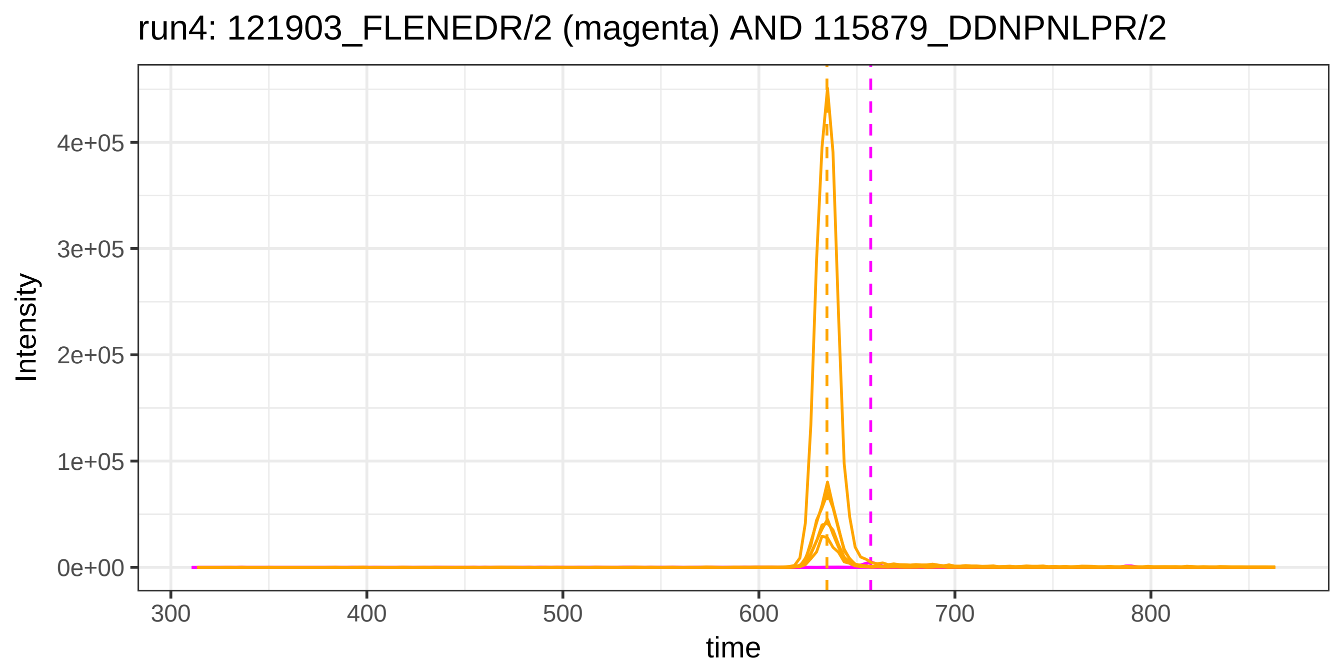
run4: 121903_FLENEDR/2 (magenta) AND 115879_DDNPNLPR/2
Intensity
4e+05
3e+05
2e+05
1e+05
0e+00
300
400
500
600
700
800
time

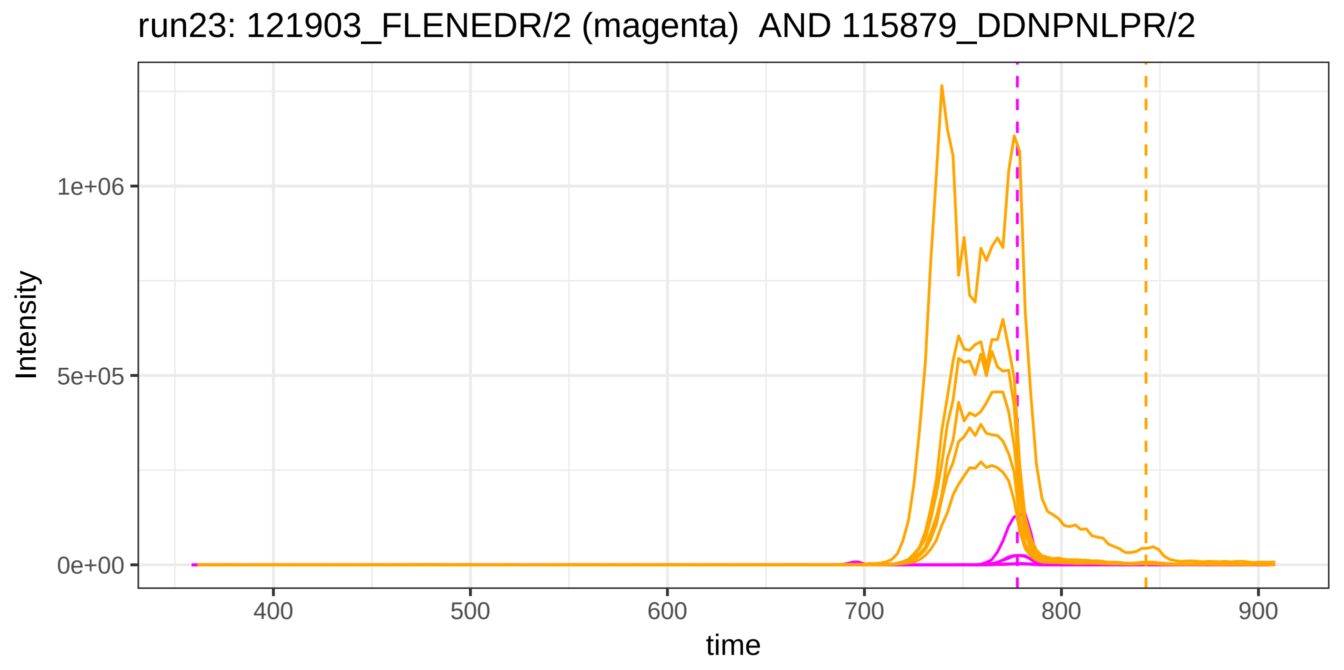
run23: 121903_FLENEDR/2 (magenta) AND 115879_DDNPNLPR/2
Intensity
1e+06
5e+05
0e+00
400
500
600
700
800
900
time

run4: 121903_FLENEDR/2 (magenta) AND 49292_QELSEAEQATR/2

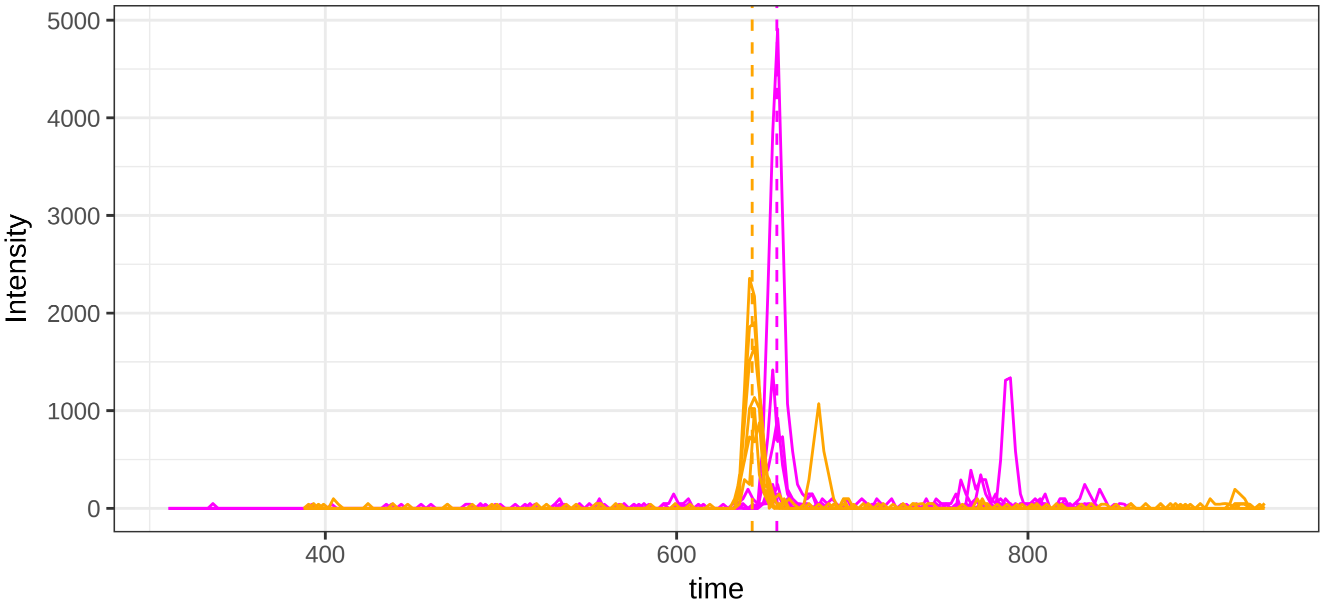


run23: 121903_FLENEDR/2 (magenta)  AND 49292_QELSEAEQATR/2

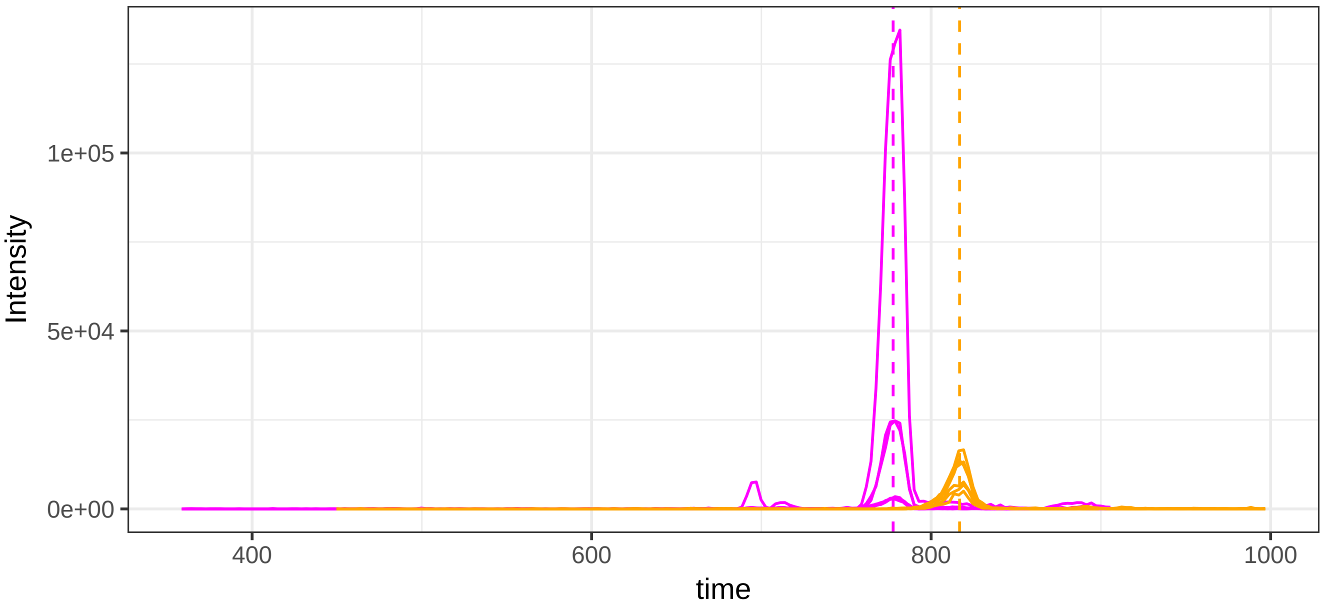

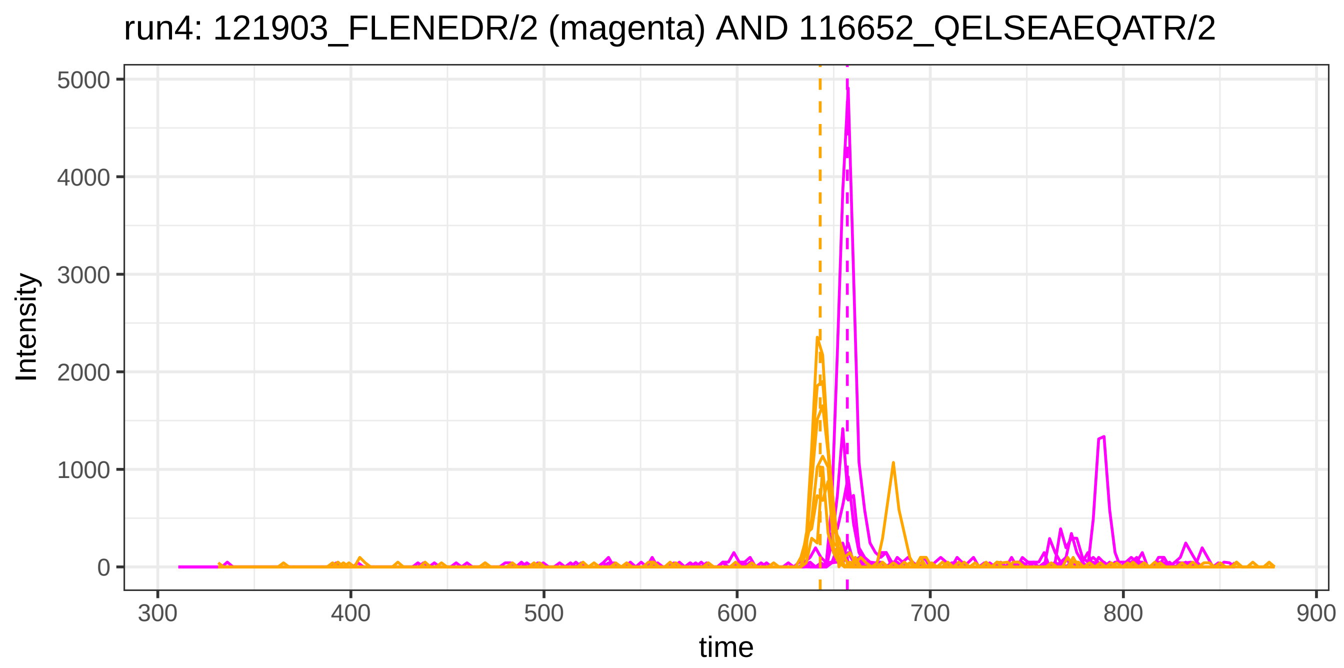

run4: 121903_FLENEDR/2 (magenta) AND 116652_QELSEAEQATR/2
Intensity
0
1000
2000
3000
4000
5000
300
400
500
600
700
800
900
time


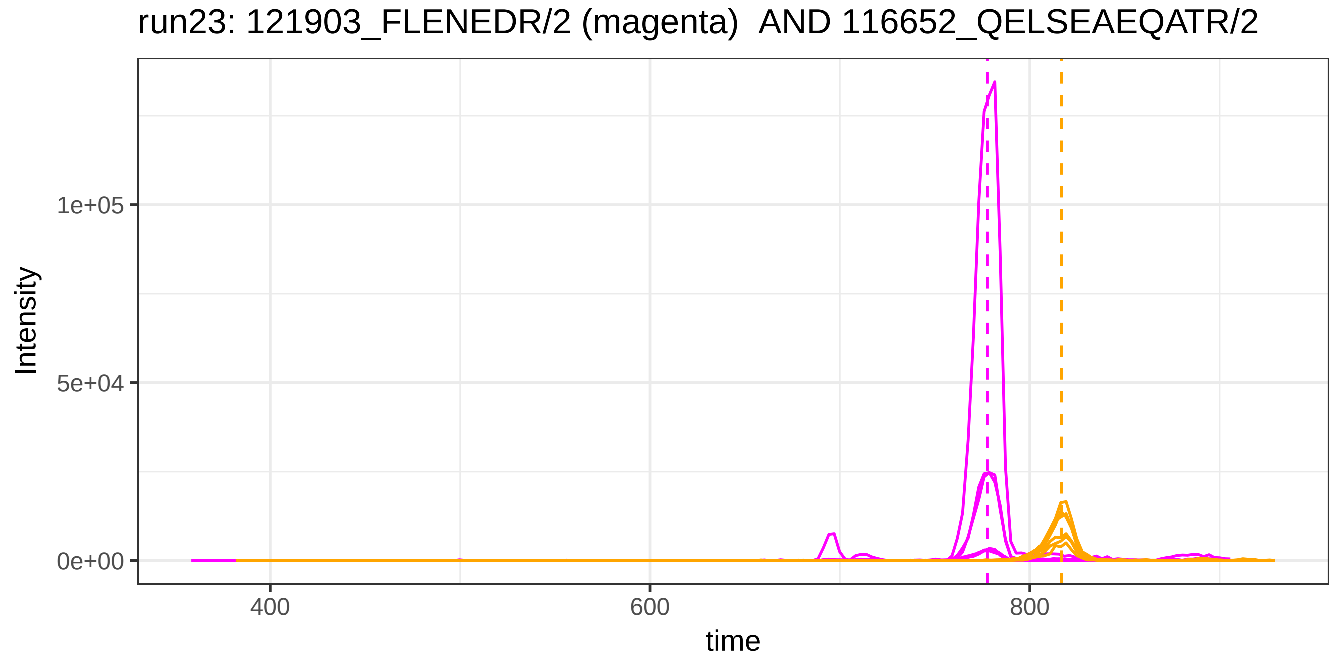

run23: 121903_FLENEDR/2 (magenta) AND 116652_QELSEAEQATR/2
Intensity
0e+00
5e+04
1e+05
400
600
800
time

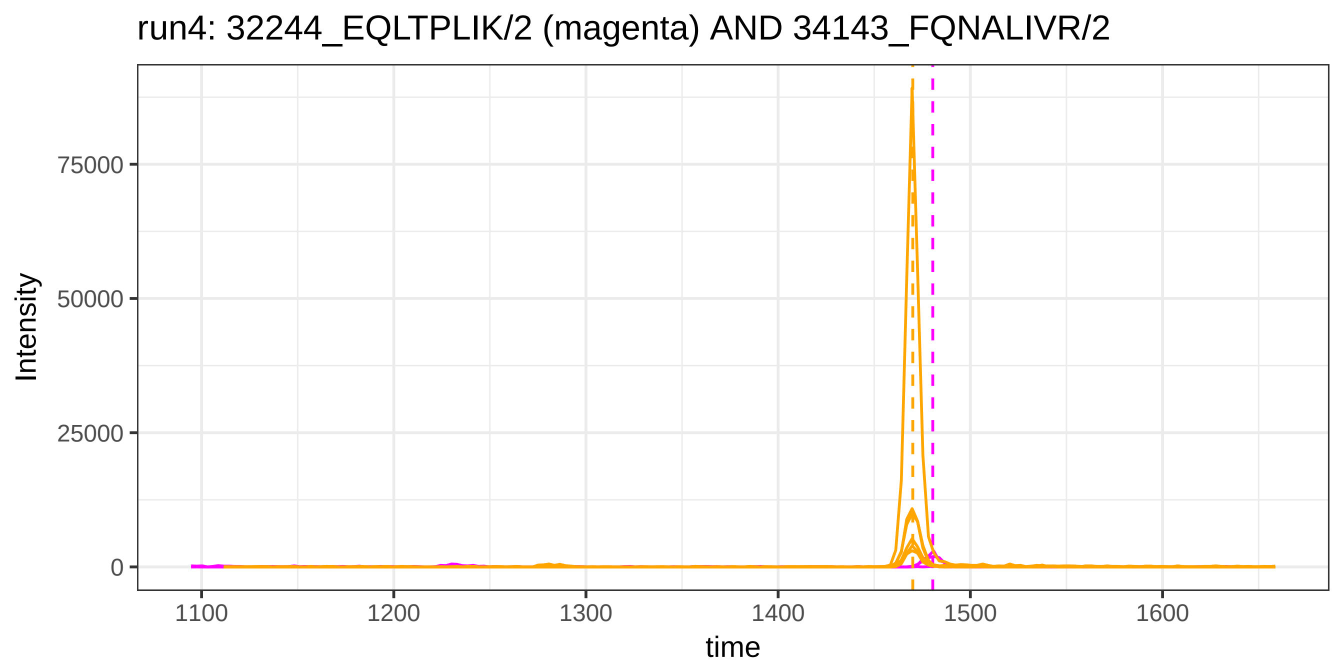
run4: 32244_EQLTPLIK/2 (magenta) AND 34143_FQNALIVR/2
Intensity
75000
50000
25000
0
1100
1200
1300
1400
1500
1600
time

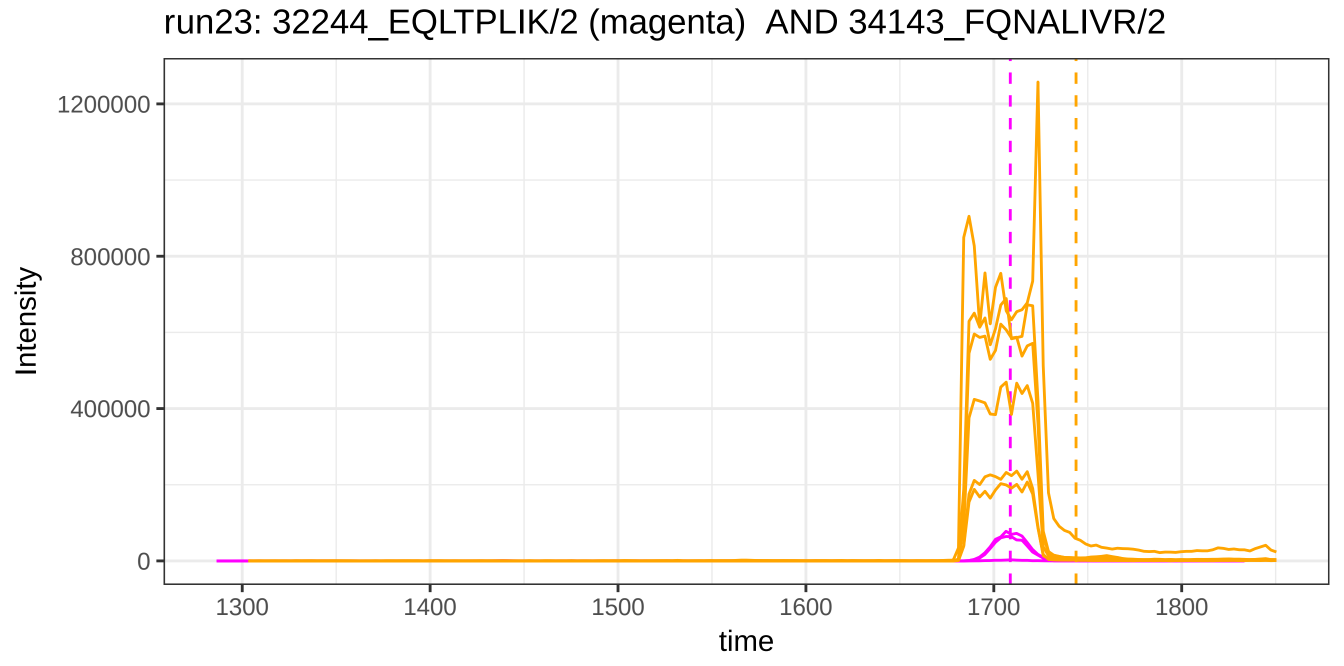
run23: 32244_EQLTPLIK/2 (magenta)  AND 34143_FQNALIVR/2
Intensity
1200000
800000
400000
0
1300
1400
1500
1600
1700
1800
time

run4: 32244_EQLTPLIK/2 (magenta) AND 51729_SDVMYTDWK/2

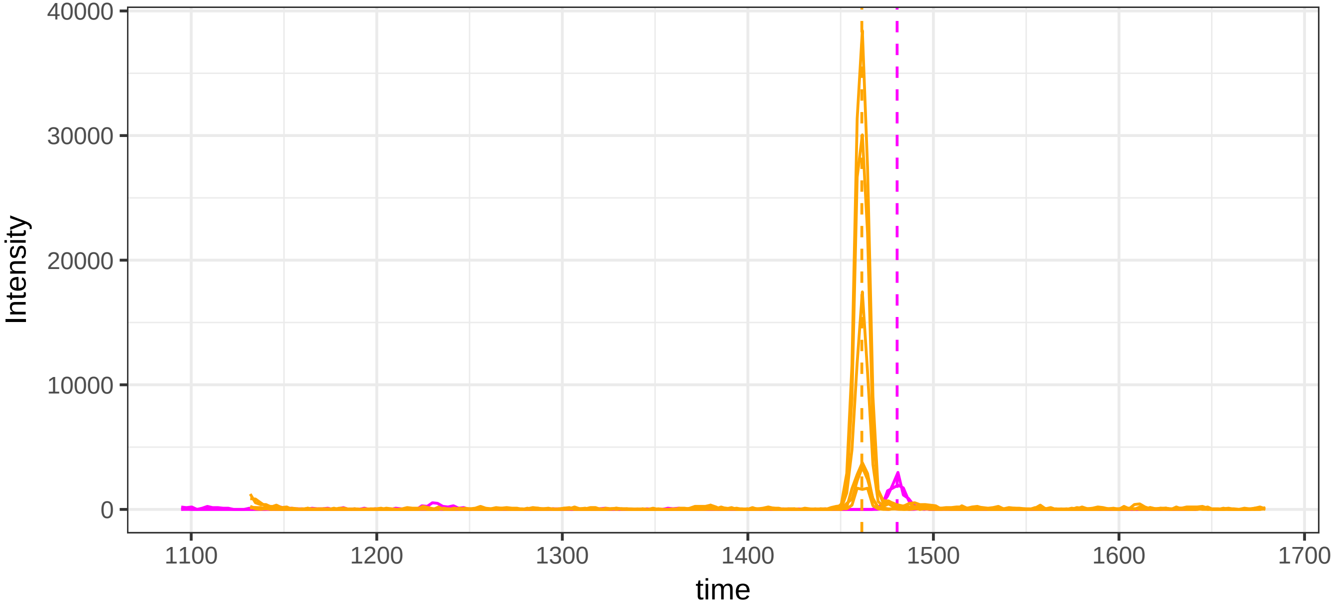


run23: 32244_EQLTPLIK/2 (magenta)  AND 51729_SDVMYTDWK/2

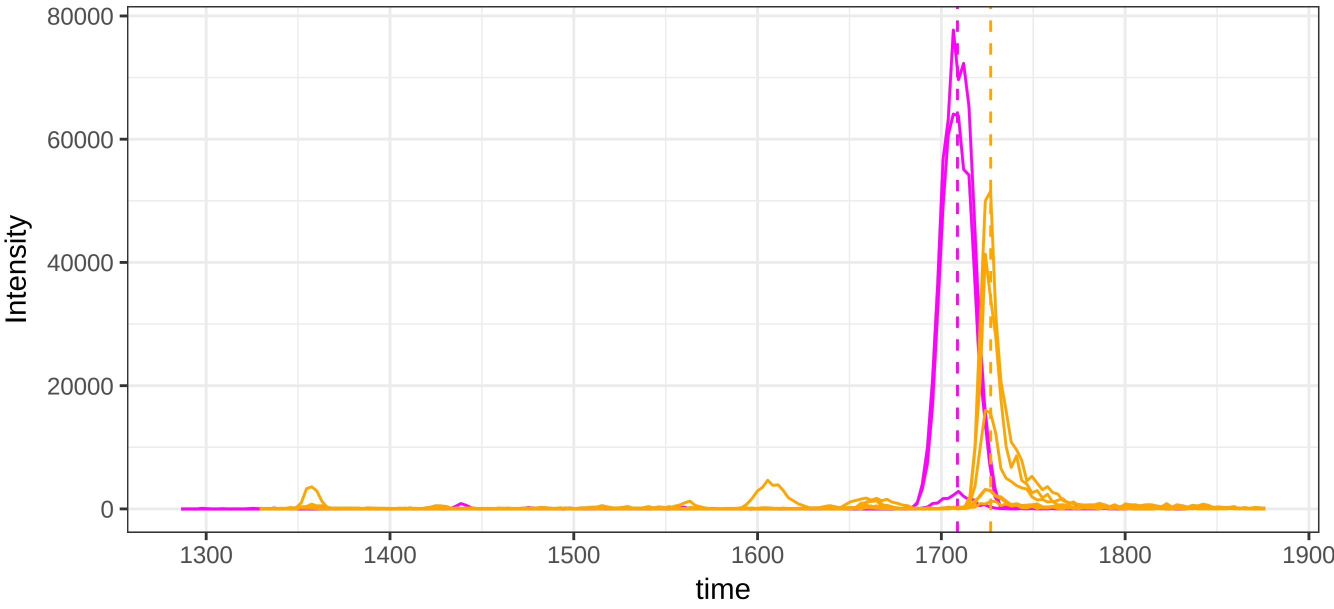

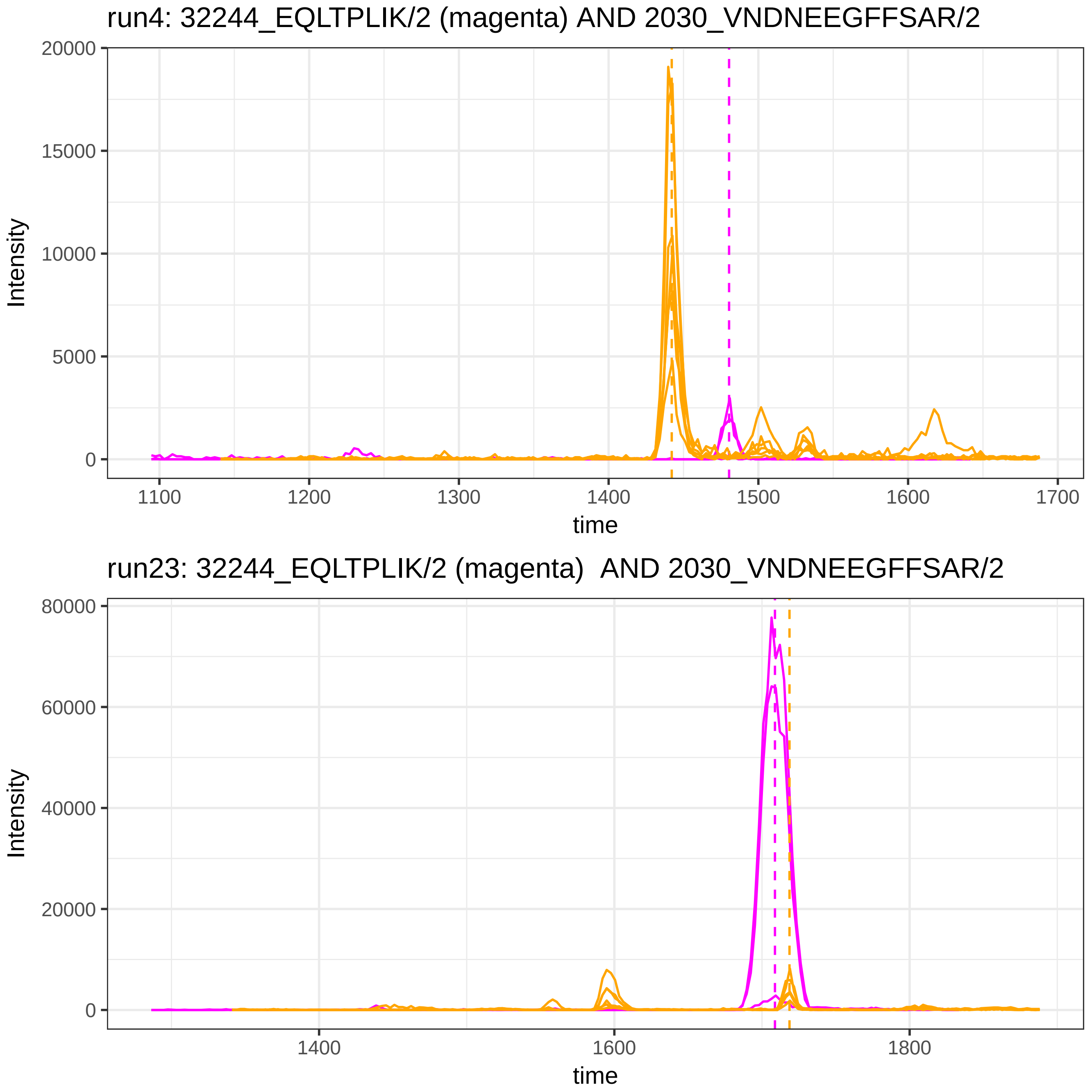

run4: 32244_EQLTPLIK/2 (magenta) AND 2030_VNDNEEGFFSAR/2
Intensity
time
run23: 32244_EQLTPLIK/2 (magenta)  AND 2030_VNDNEEGFFSAR/2
Intensity
time

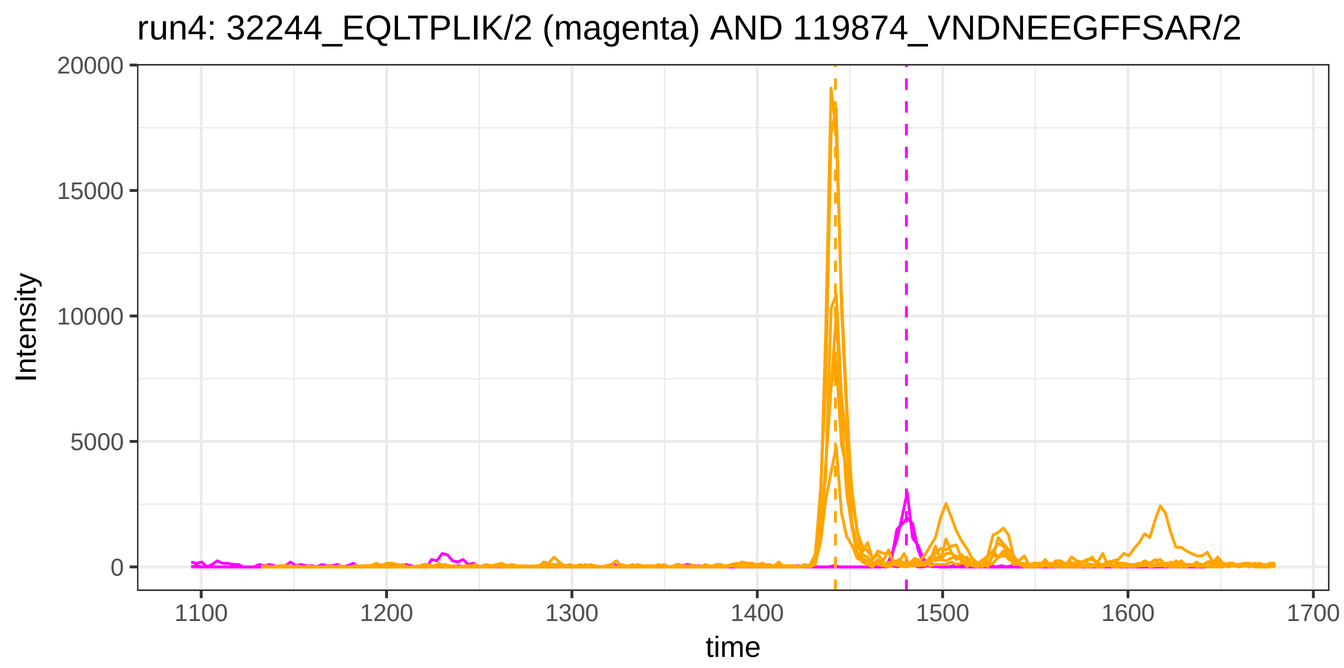
run4: 32244_EQLTPLIK/2 (magenta) AND 119874_VNDNEEGFFSAR/2
Intensity
20000
15000
10000
5000
0
1100
1200
1300
1400
1500
1600
1700
time

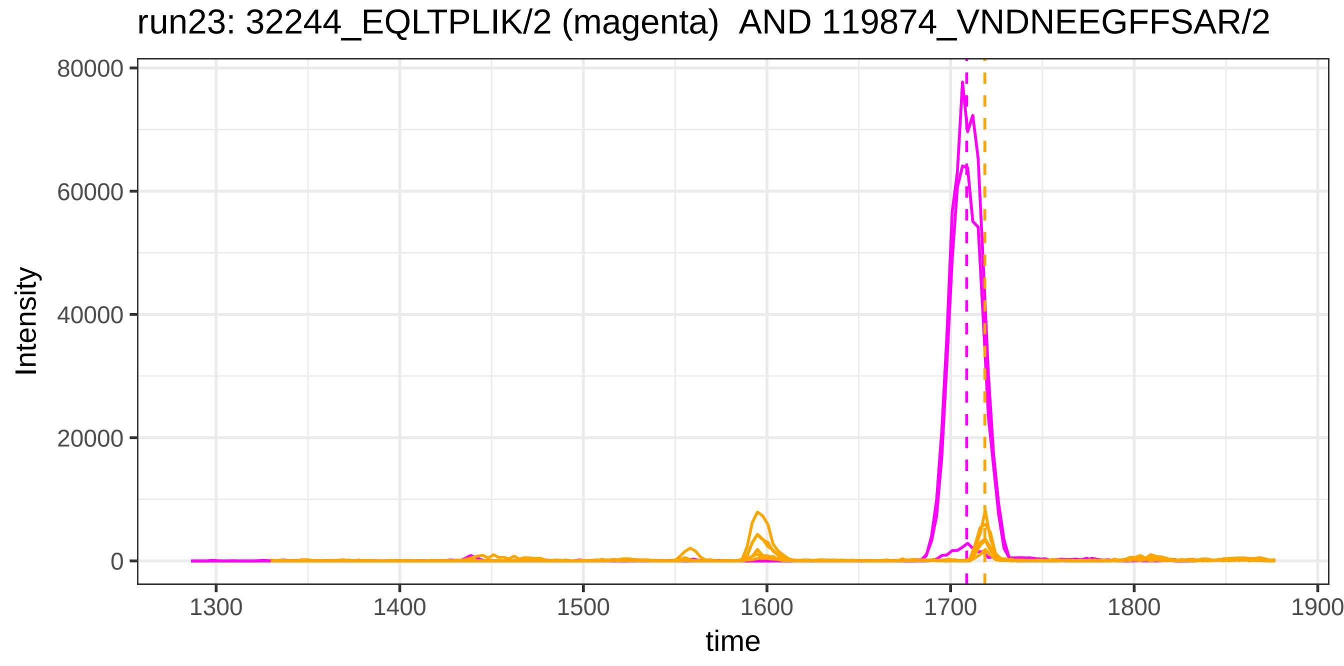
run23: 32244_EQLTPLIK/2 (magenta) AND 119874_VNDNEEGFFSAR/2
Intensity
80000
60000
40000
20000
0
1300
1400
1500
1600
1700
1800
1900
time

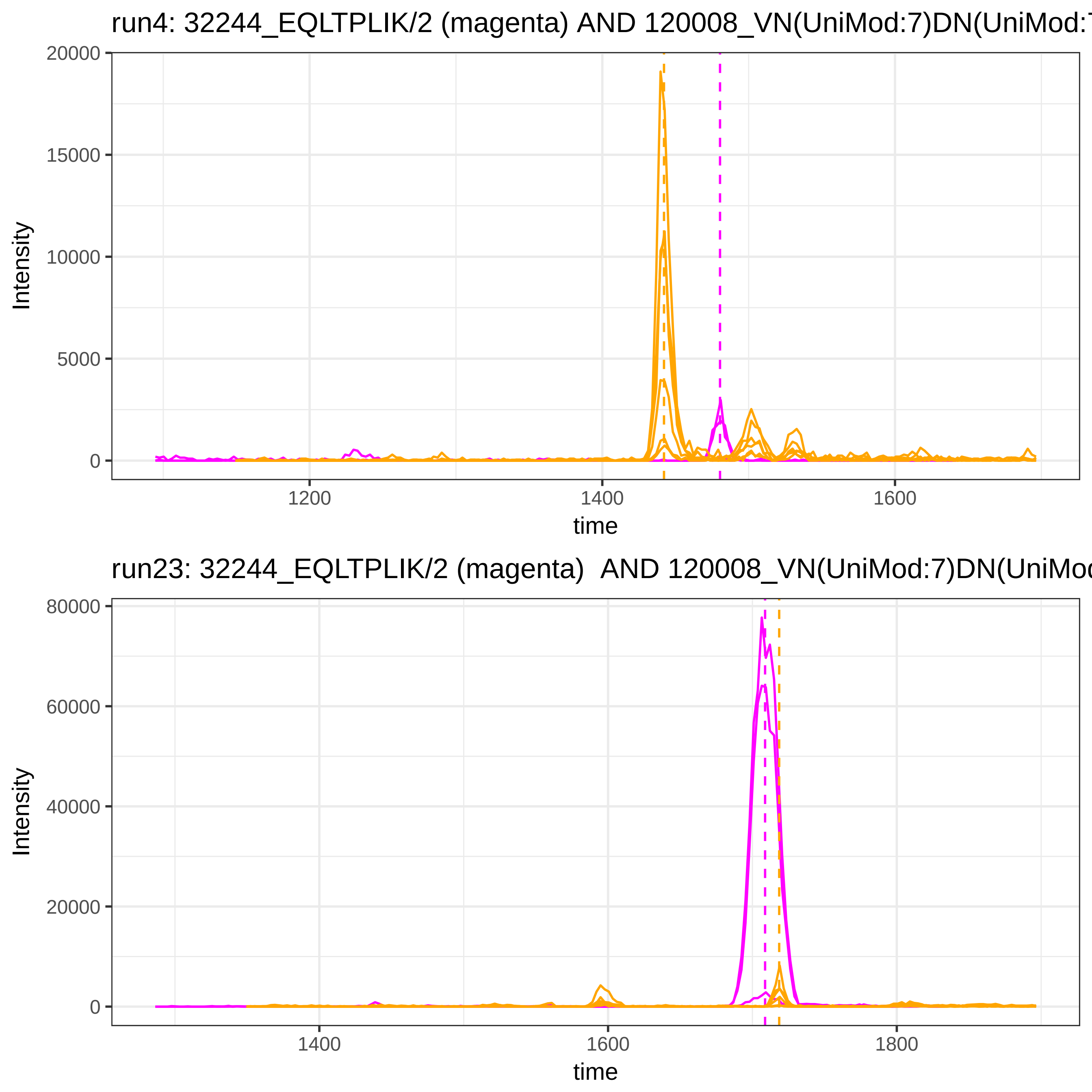

run4: 32244_EQLTPLIK/2 (magenta) AND 120008_VN(UniMod:7)DN(UniMod:7
Intensity
20000
15000
10000
5000
0
1200
1400
1600
time
run23: 32244_EQLTPLIK/2 (magenta)  AND 120008_VN(UniMod:7)DN(UniMod
Intensity
80000
60000
40000
20000
0
1400
1600
1800
time

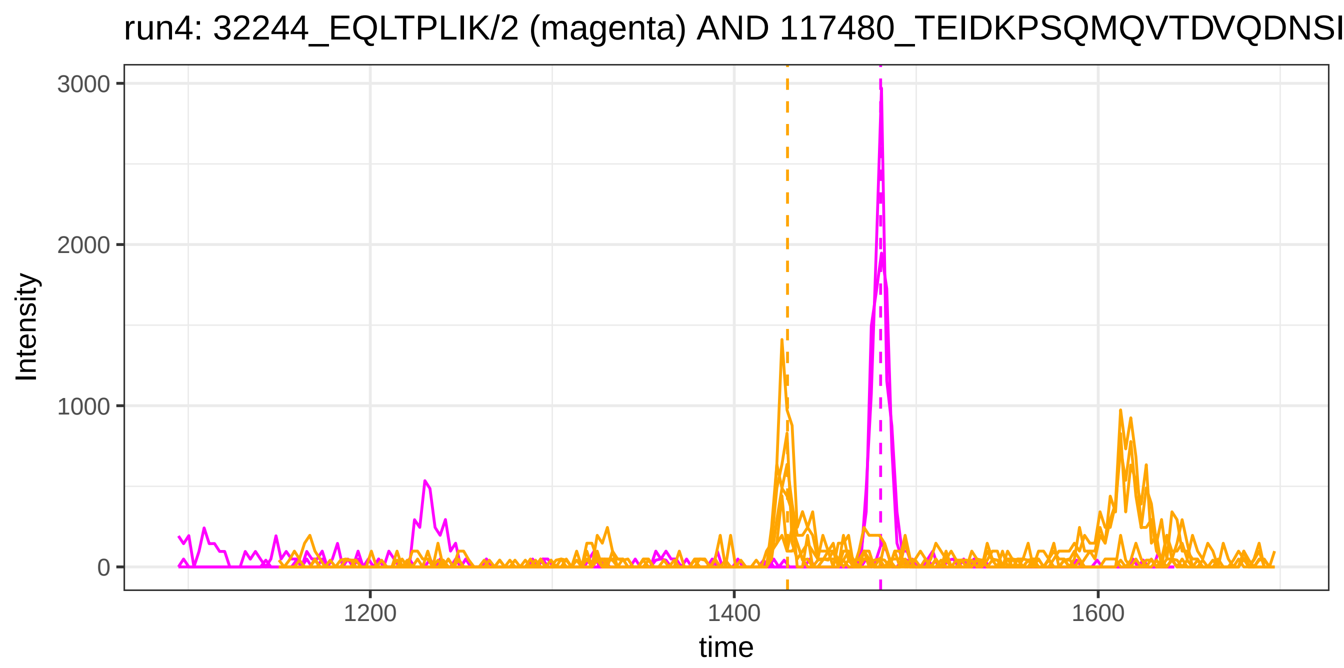

run4: 32244_EQLTPLIK/2 (magenta) AND 117480_TEIDKPSQMQVTDVQDNSI
Intensity
3000
2000
1000
0
1200
1400
1600
time


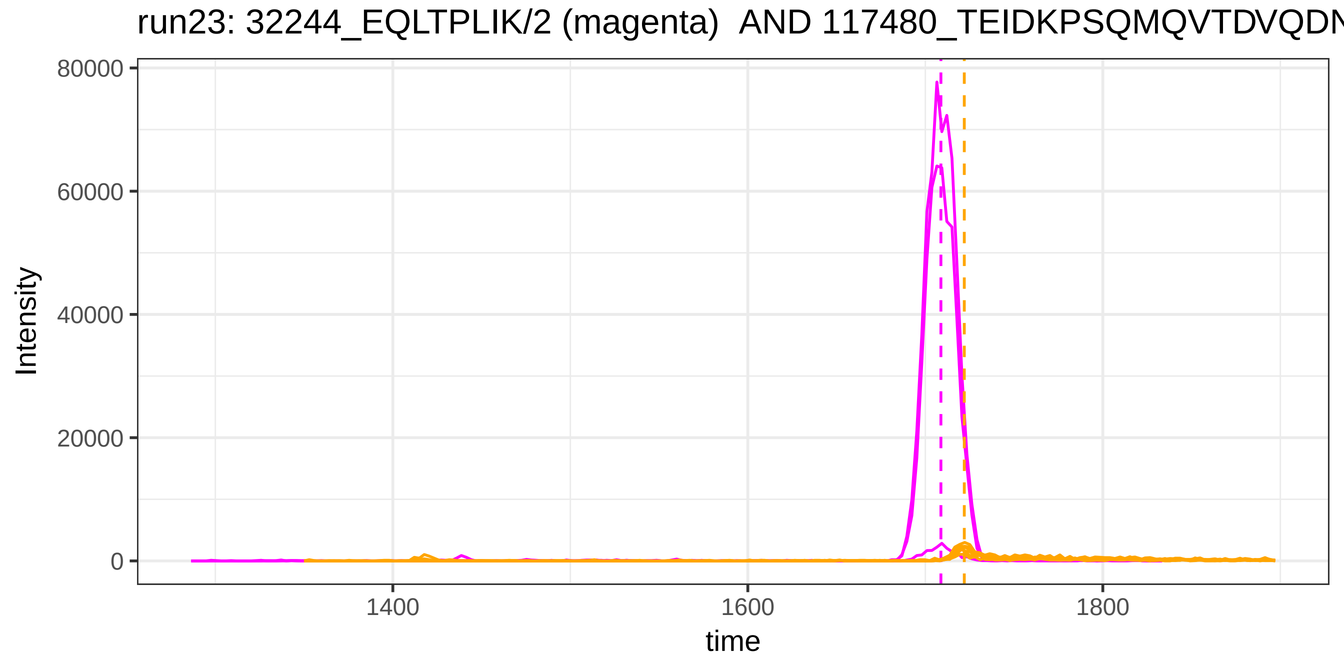

run23: 32244_EQLTPLIK/2 (magenta) AND 117480_TEIDKPSQMQVTDVQDN
Intensity
80000
60000
40000
20000
0
1400
1600
1800
time

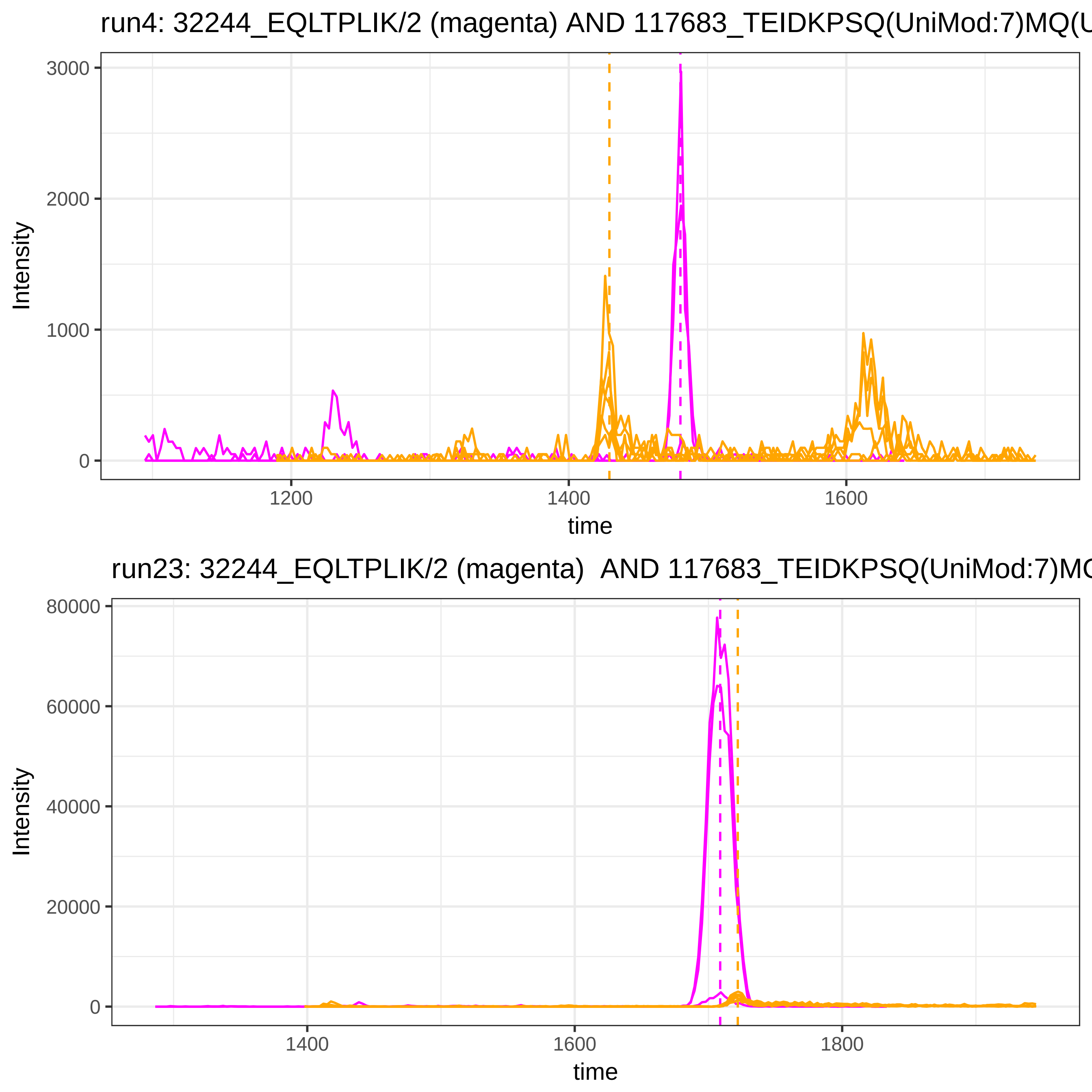
run4: 32244_EQLTPLIK/2 (magenta) AND 117683_TEIDKPSQ(UniMod:7)MQ(U
Intensity
3000
2000
1000
0
1200
1400
1600
time
run23: 32244_EQLTPLIK/2 (magenta) AND 117683_TEIDKPSQ(UniMod:7)MQ
Intensity
80000
60000
40000
20000
0
1400
1600
1800
time

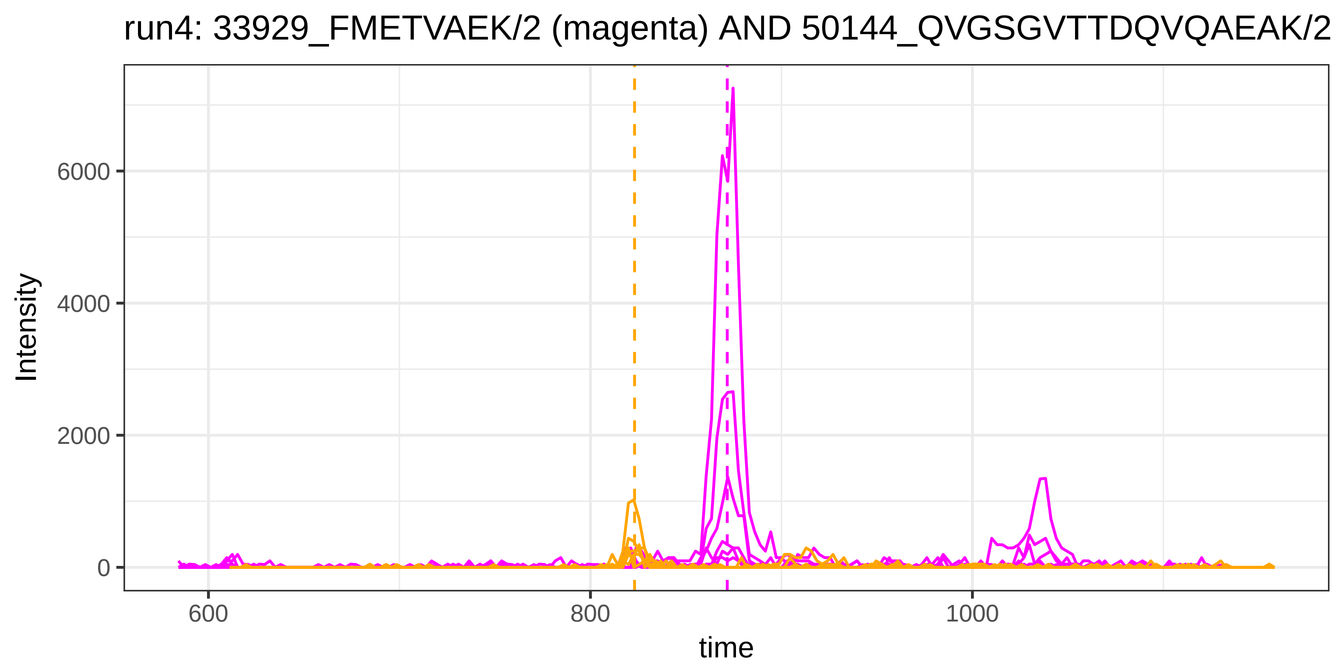
run4: 33929_FMETVAEK/2 (magenta) AND 50144_QVGSGVTTDQVQAEAK/2
Intensity
6000
4000
2000
0
600
800
1000
time

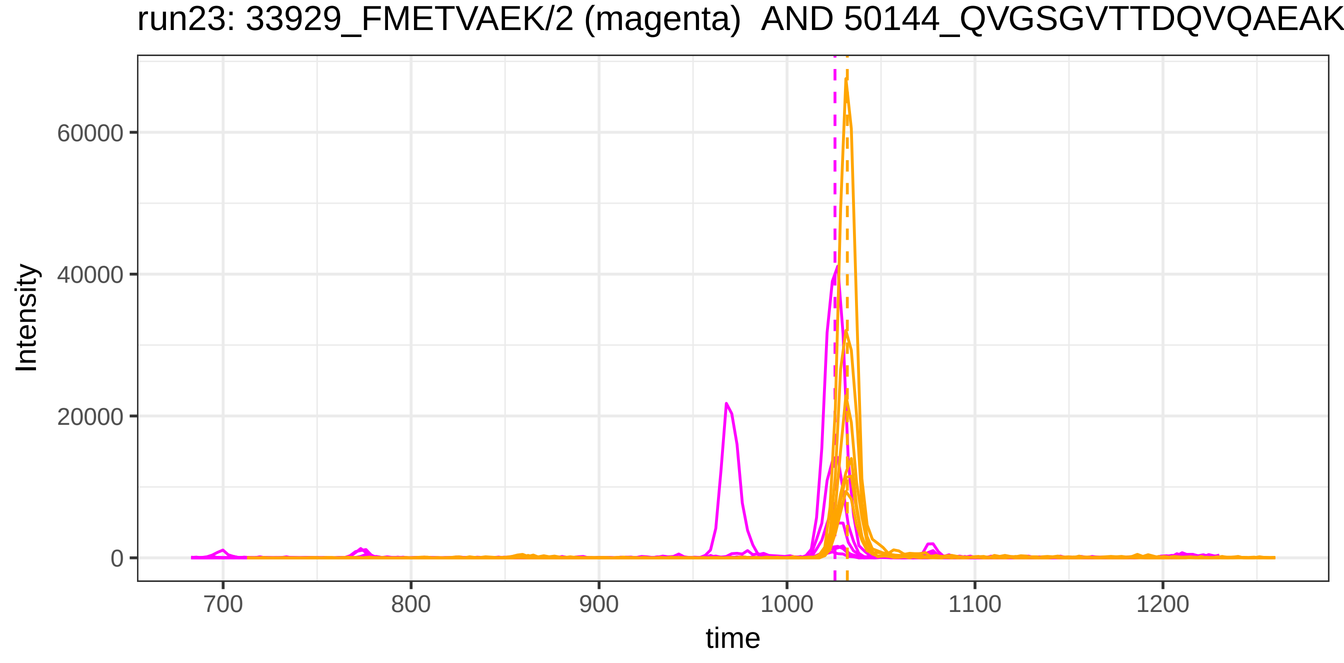
run23: 33929_FMETVAEK/2 (magenta) AND 50144_QVGSGVTTDQVQAEAK
Intensity
60000
40000
20000
0
700
800
900
1000
1100
1200
time

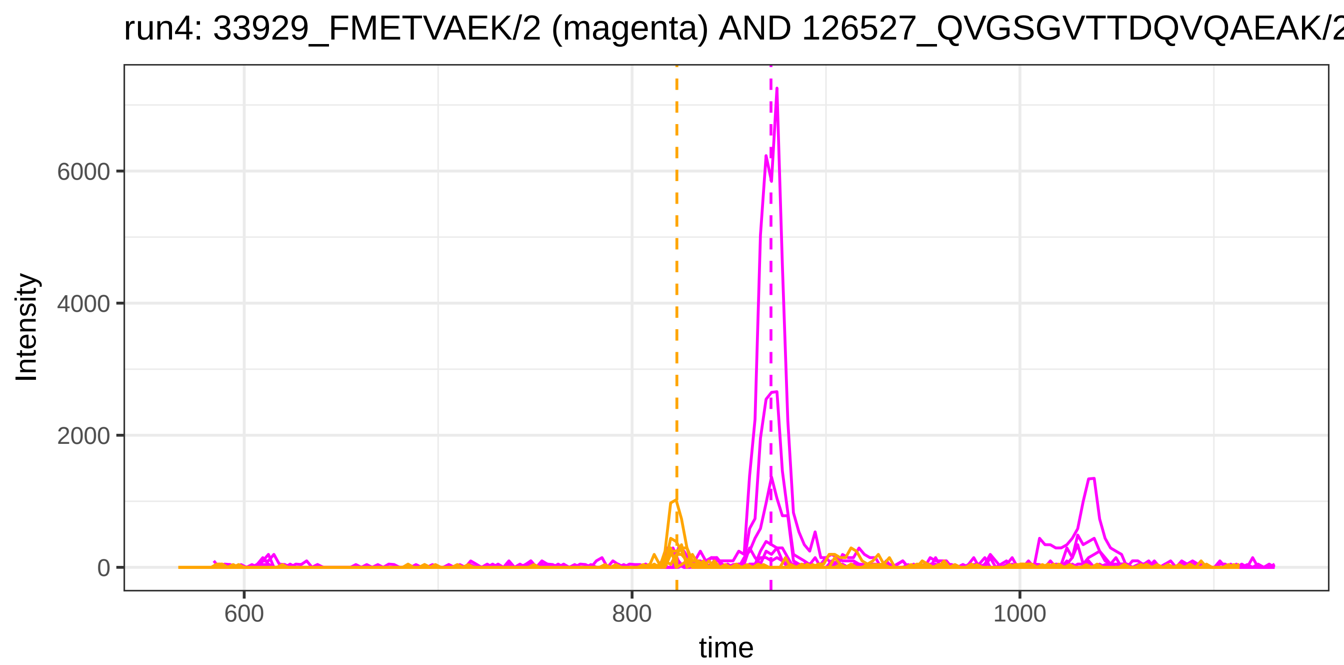
run4: 33929_FMETVAEK/2 (magenta) AND 126527_QVGSGVTTDQVQAEAK/2
Intensity
6000
4000
2000
0
600
800
1000
time

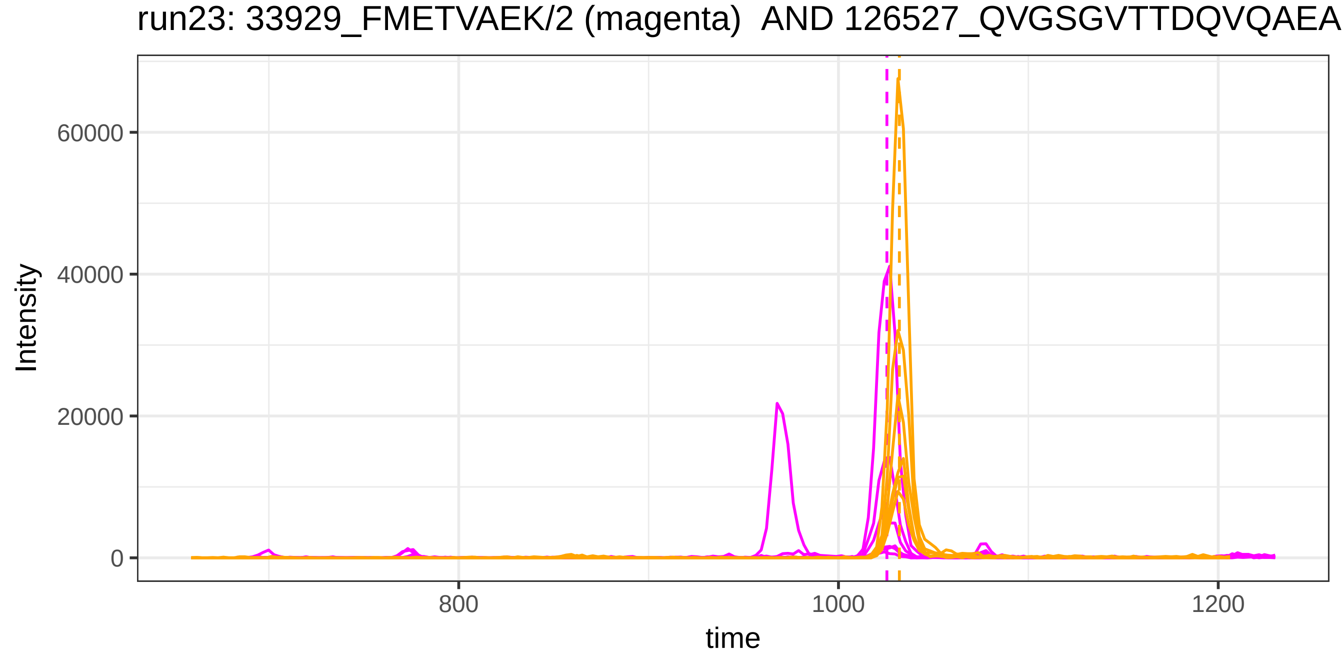
run23: 33929_FMETVAEK/2 (magenta) AND 126527_QVGSGVTTDQVQAEA
Intensity
60000
40000
20000
0
800
1000
1200
time

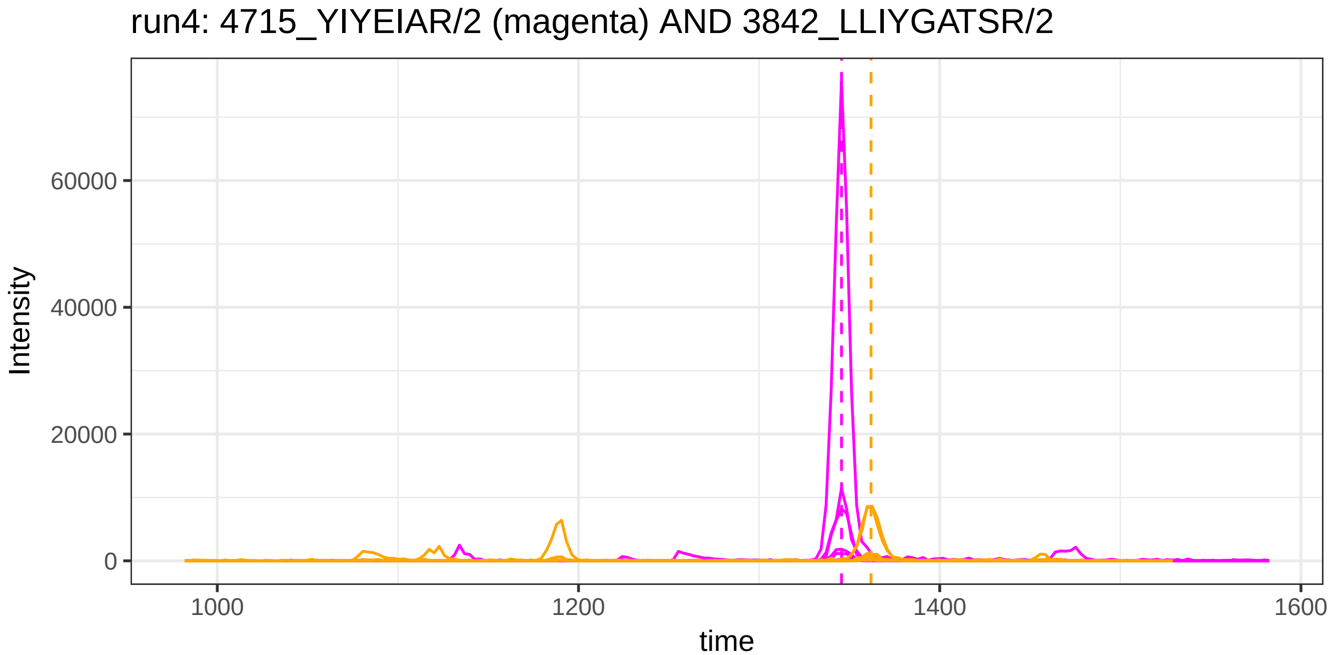

run4: 4715_YIYEIAR/2 (magenta) AND 3842_LLIYGATSR/2
Intensity
60000
40000
20000
0
1000
1200
1400
1600
time


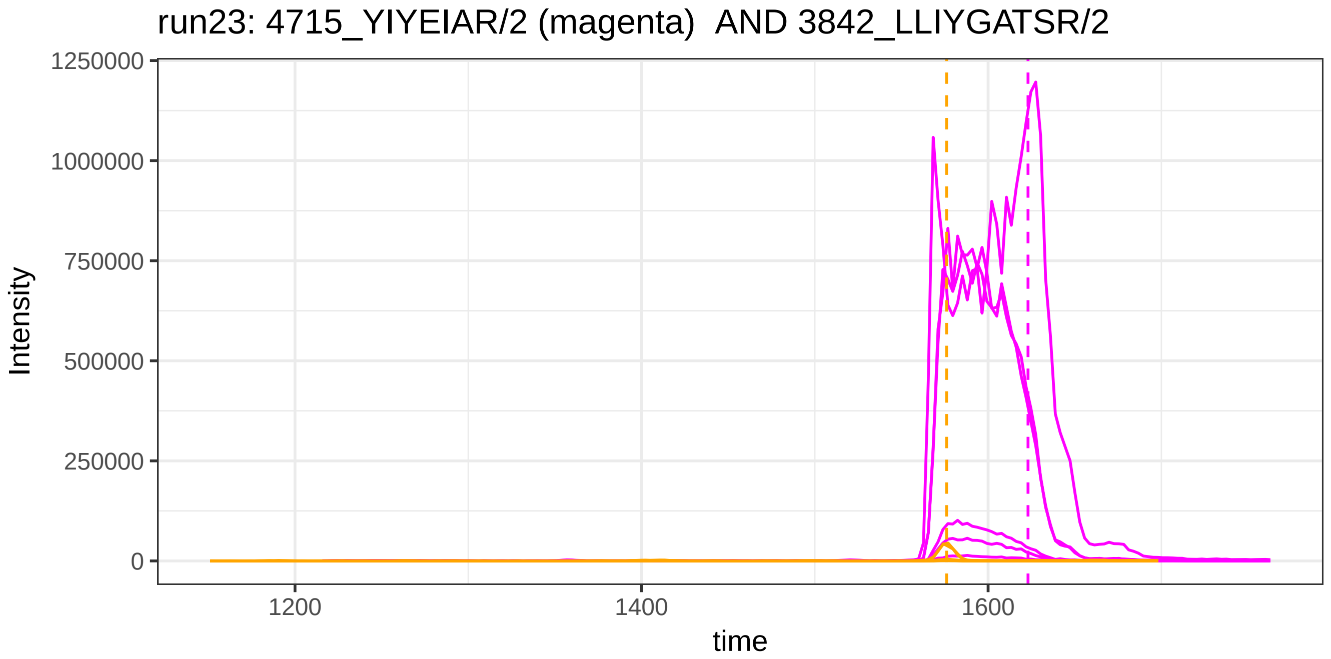

run23: 4715_YIYEIAR/2 (magenta) AND 3842_LLIYGATSR/2
Intensity
1250000
1000000
750000
500000
250000
0
1200
1400
1600
time

run4: 4715_YIYEIAR/2 (magenta) AND 43888_LLIYDASNR/2

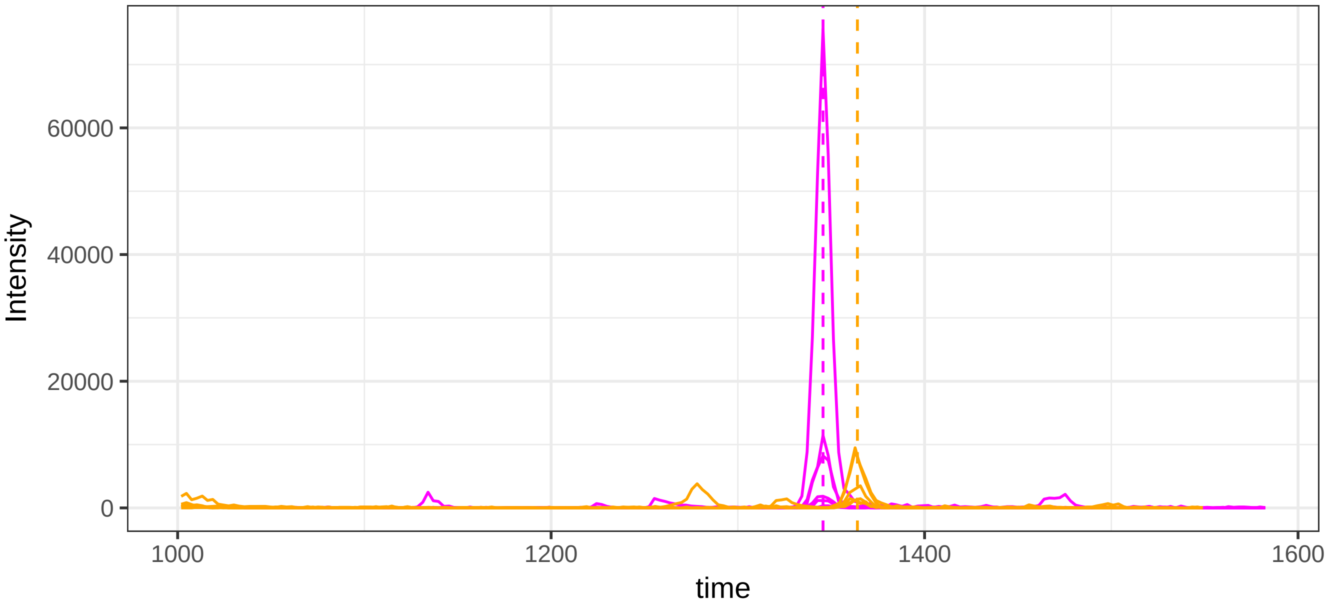


run23: 4715_YIYEIAR/2 (magenta) AND 43888_LLIYDASNR/2

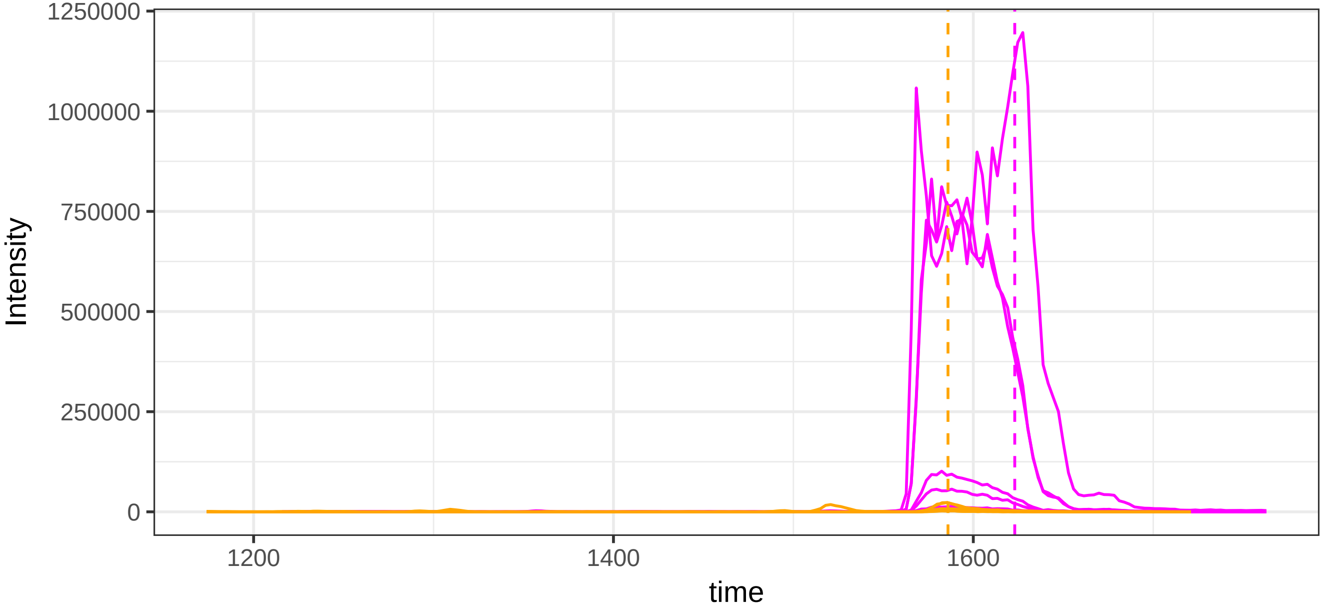

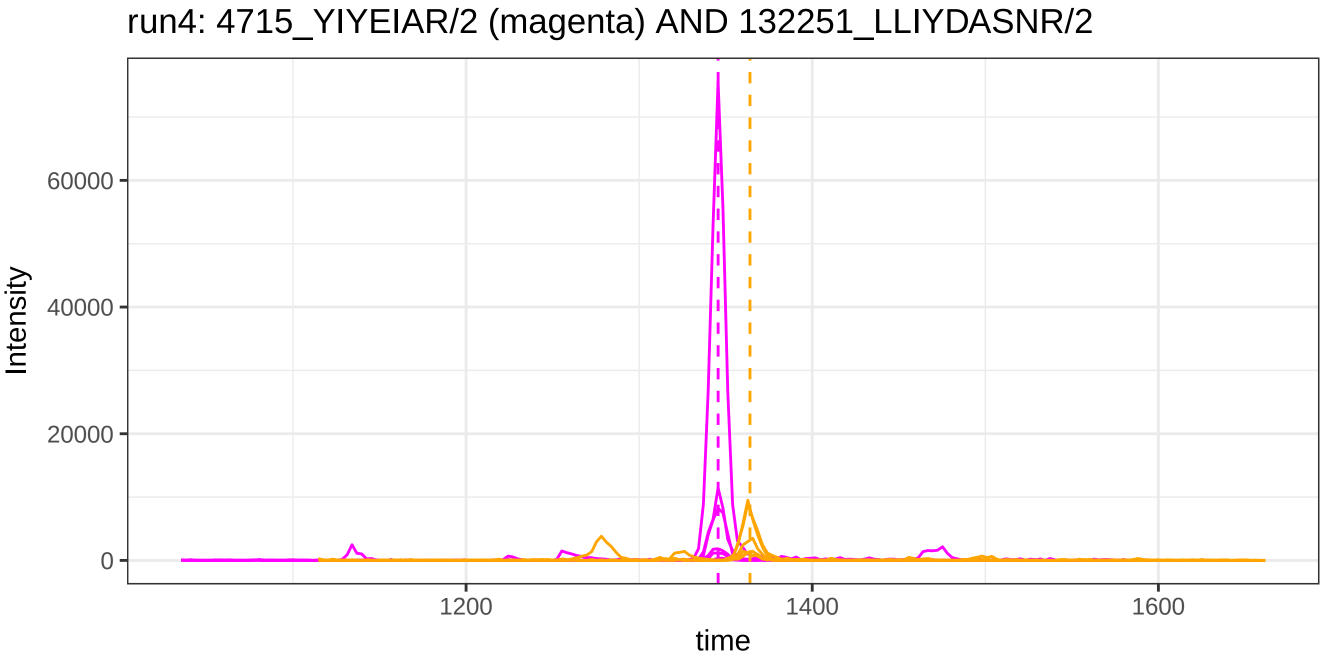

run4: 4715_YIYEIAR/2 (magenta) AND 132251_LLIYDASNR/2
Intensity
60000
40000
20000
0
1200
1400
1600
time


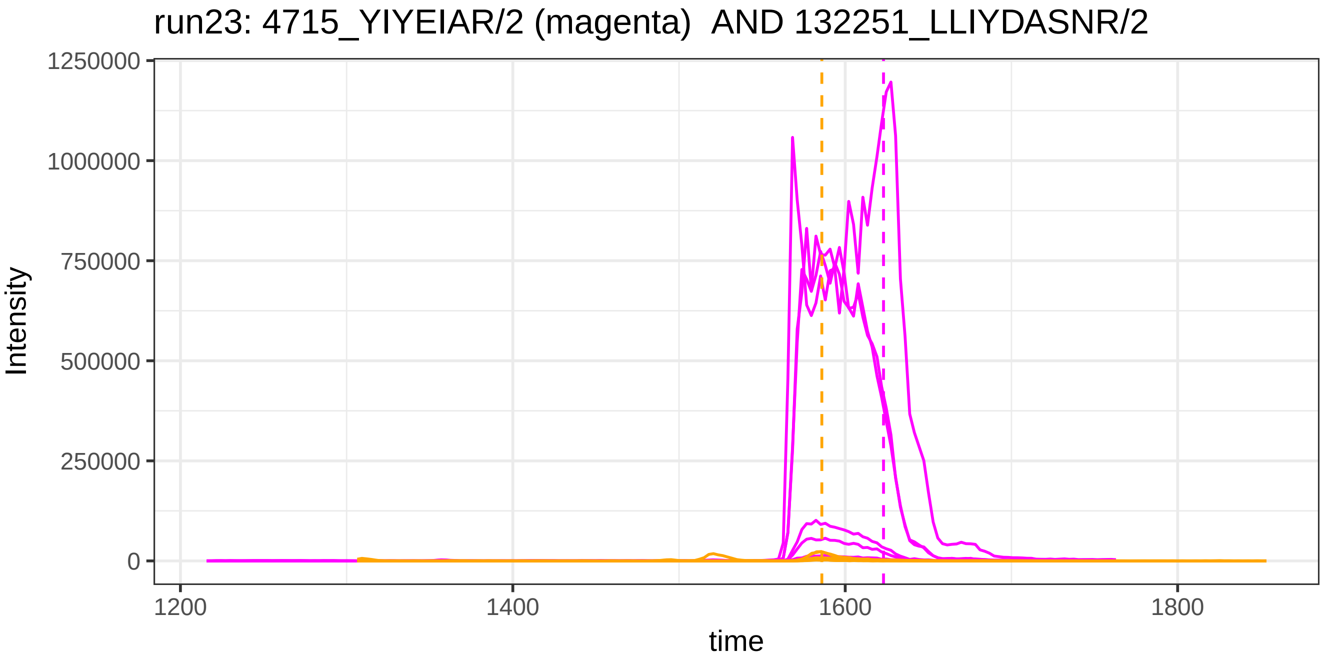

run23: 4715_YIYEIAR/2 (magenta) AND 132251_LLIYDASNR/2
Intensity
1250000
1000000
750000
500000
250000
0
1200
1400
1600
1800
time

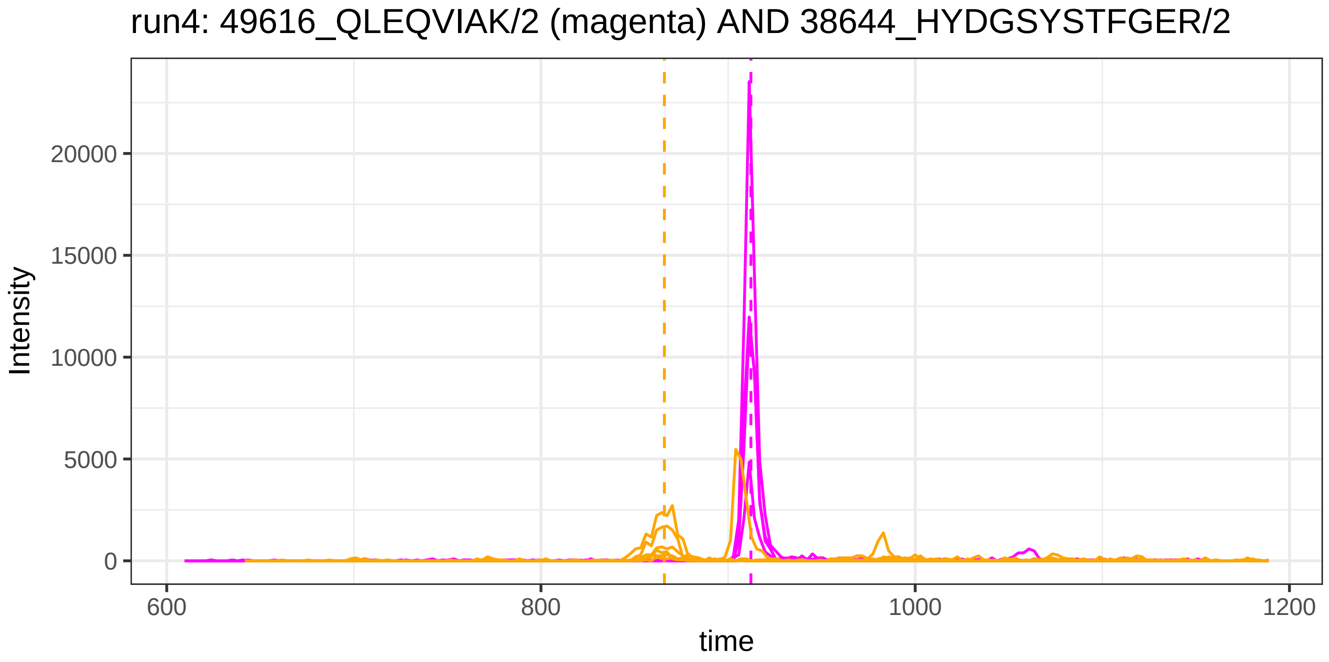

run4: 49616_QLEQVIAK/2 (magenta) AND 38644_HYDGSYSTFGER/2
Intensity
20000
15000
10000
5000
0
600
800
1000
1200
time


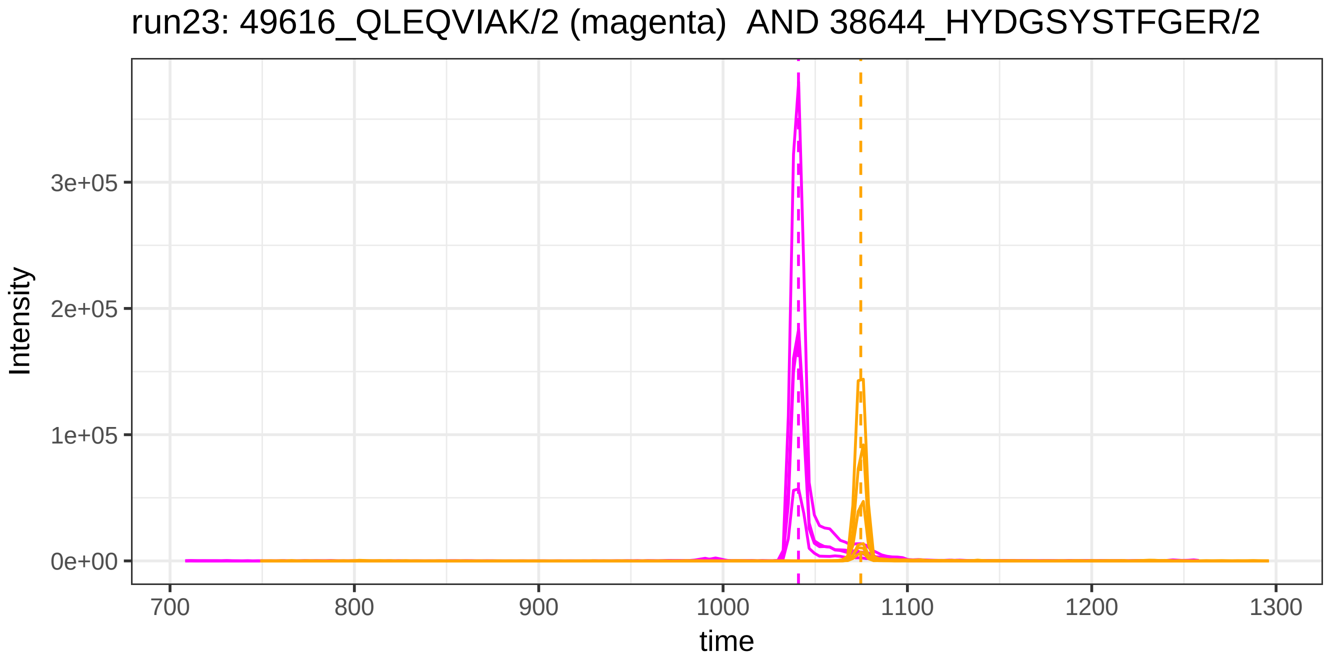

run23: 49616_QLEQVIAK/2 (magenta) AND 38644_HYDGSYSTFGER/2
Intensity
3e+05
2e+05
1e+05
0e+00
700
800
900
1000
1100
1200
1300
time

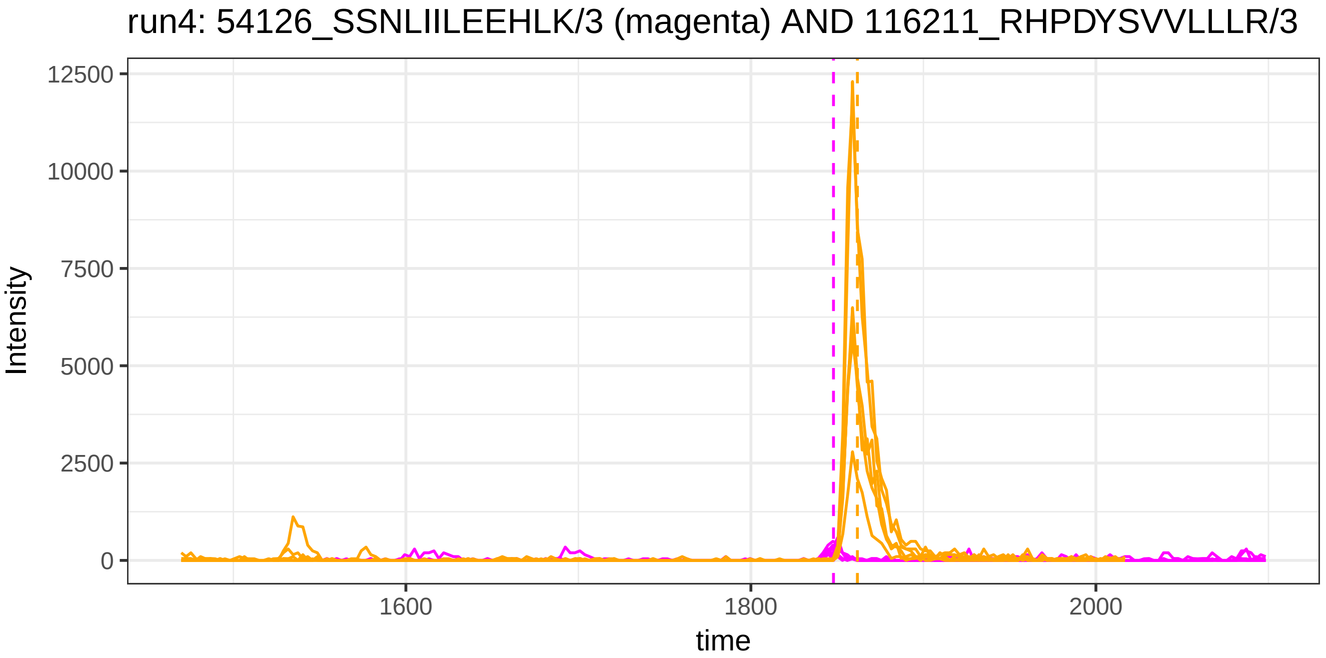

run4: 54126_SSNLIILEEHLK/3 (magenta) AND 116211_RHPDYSVVLLLR/3
Intensity
12500
10000
7500
5000
2500
0
1600
1800
2000
time


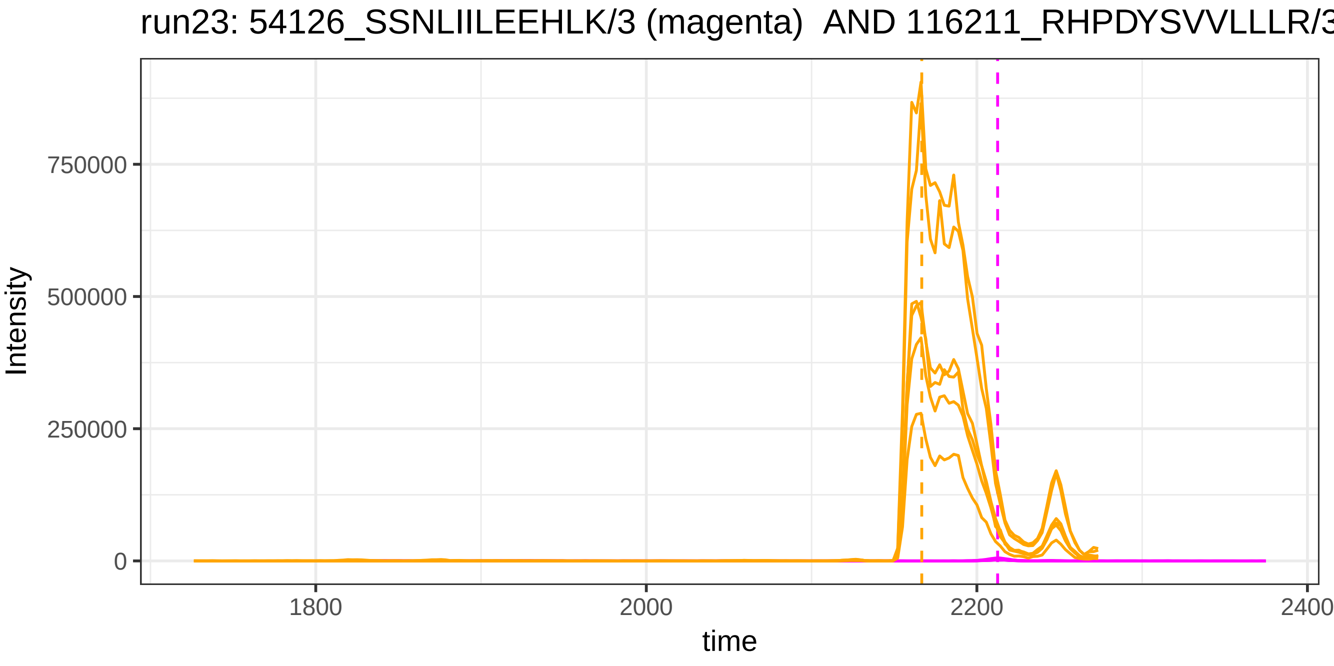

run23: 54126_SSNLIILEEHLK/3 (magenta) AND 116211_RHPDYSVVLLLR/3
Intensity
750000
500000
250000
0
1800
2000
2200
2400
time

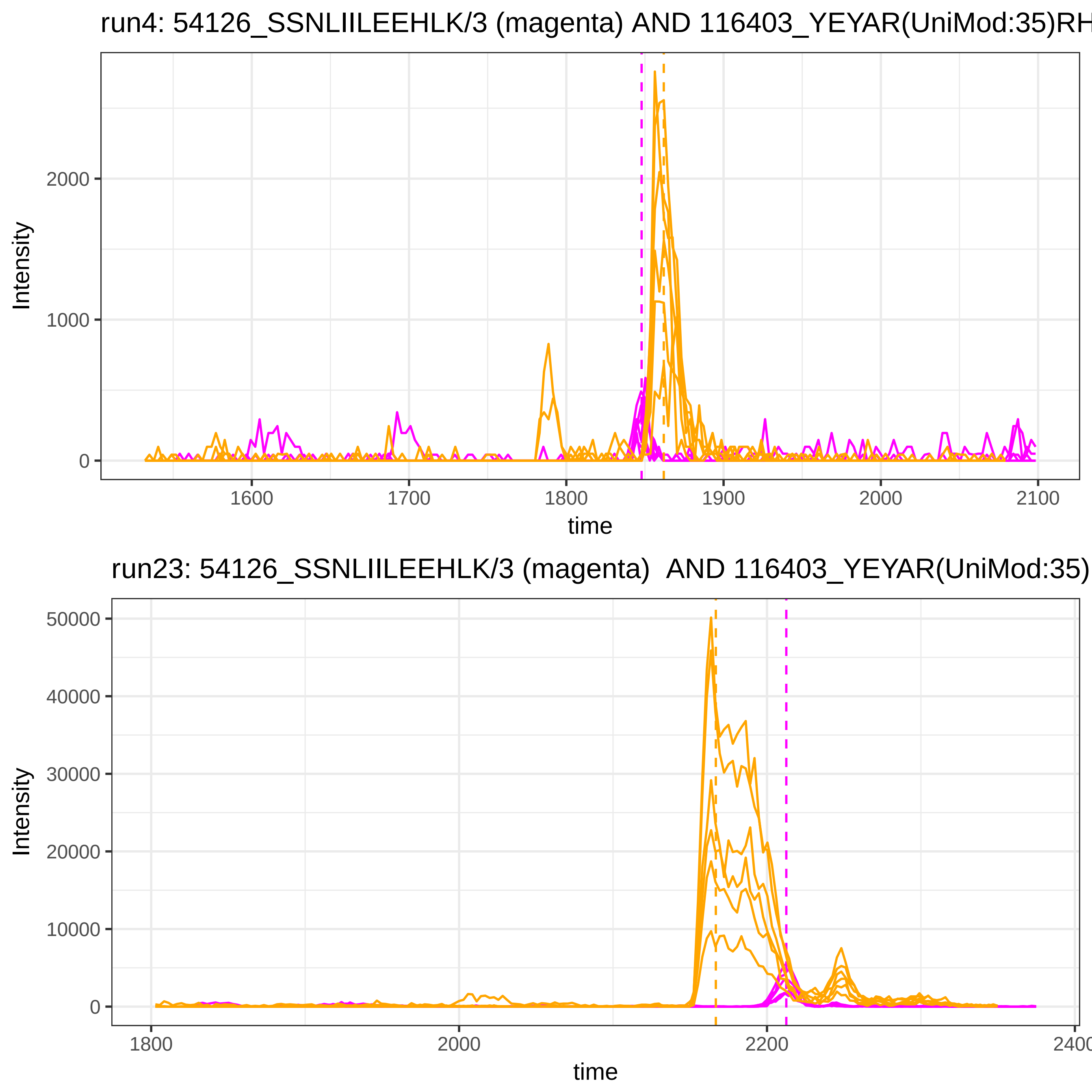
run4: 54126_SSNLIILEEHLK/3 (magenta) AND 116403_YEYAR(UniMod:35)RH
Intensity
2000
1000
0
1600
1700
1800
1900
2000
2100
time
run23: 54126_SSNLIILEEHLK/3 (magenta) AND 116403_YEYAR(UniMod:35)
Intensity
50000
40000
30000
20000
10000
0
1800
2000
2200
2400
time

run4: 54126_SSNLIILEEHLK/3 (magenta) AND 55601_TEVIPPLIENR/2

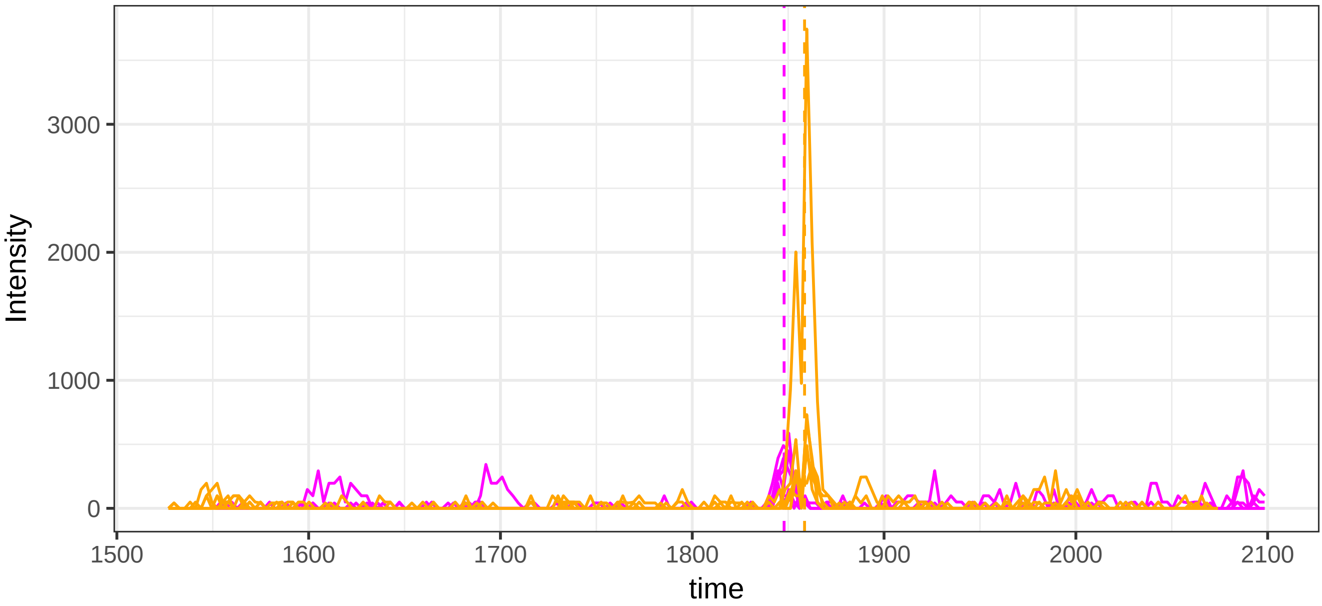


run23: 54126_SSNLIILEEHLK/3 (magenta)  AND 55601_TEVIPPLIENR/2

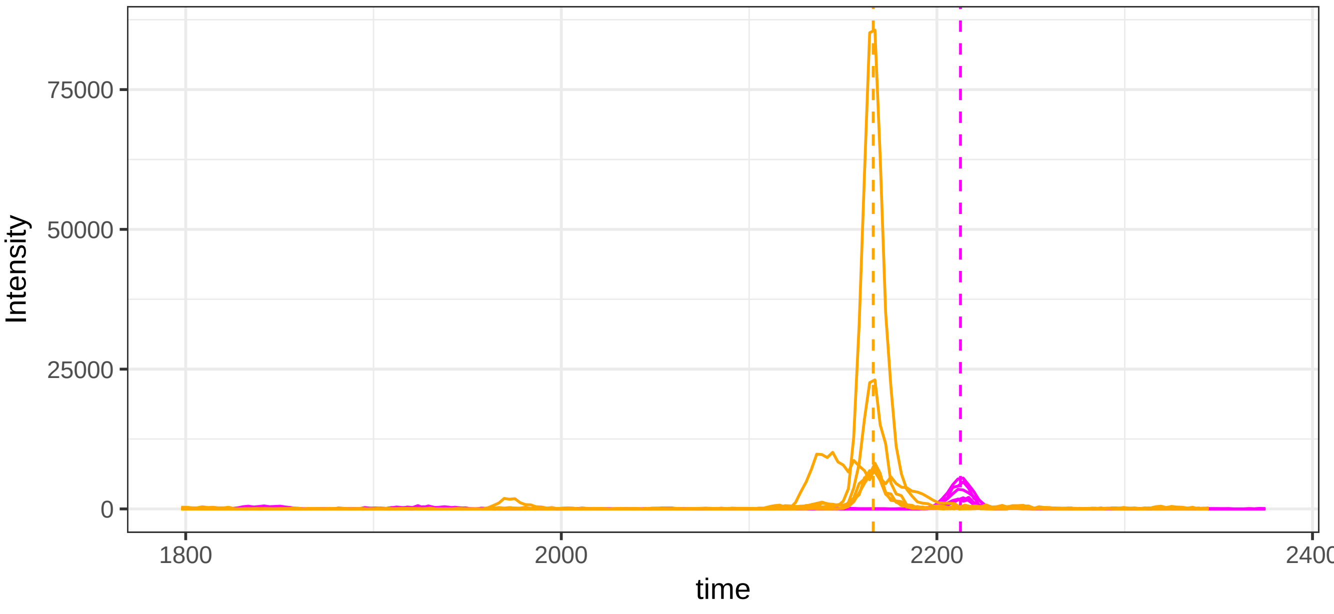

run4: 54126_SSNLIILEEHLK/3 (magenta) AND 114992_TEVIPPLIENR/2

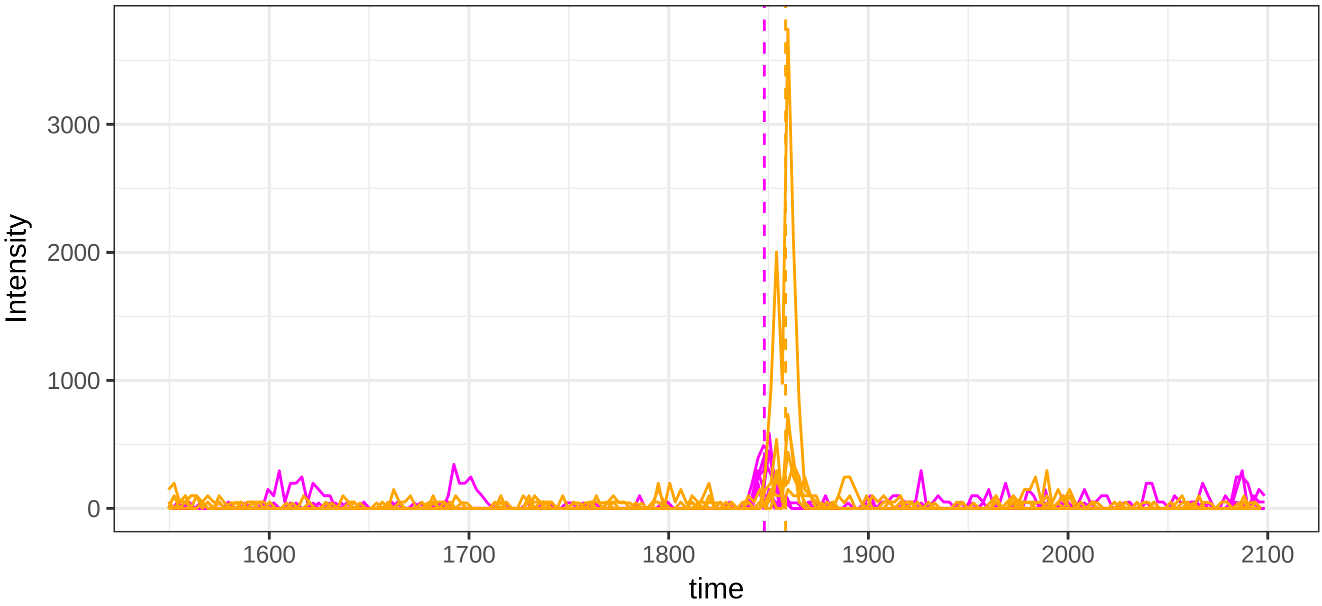


run23: 54126_SSNLIILEEHLK/3 (magenta) AND 114992_TEVIPPLIENR/2

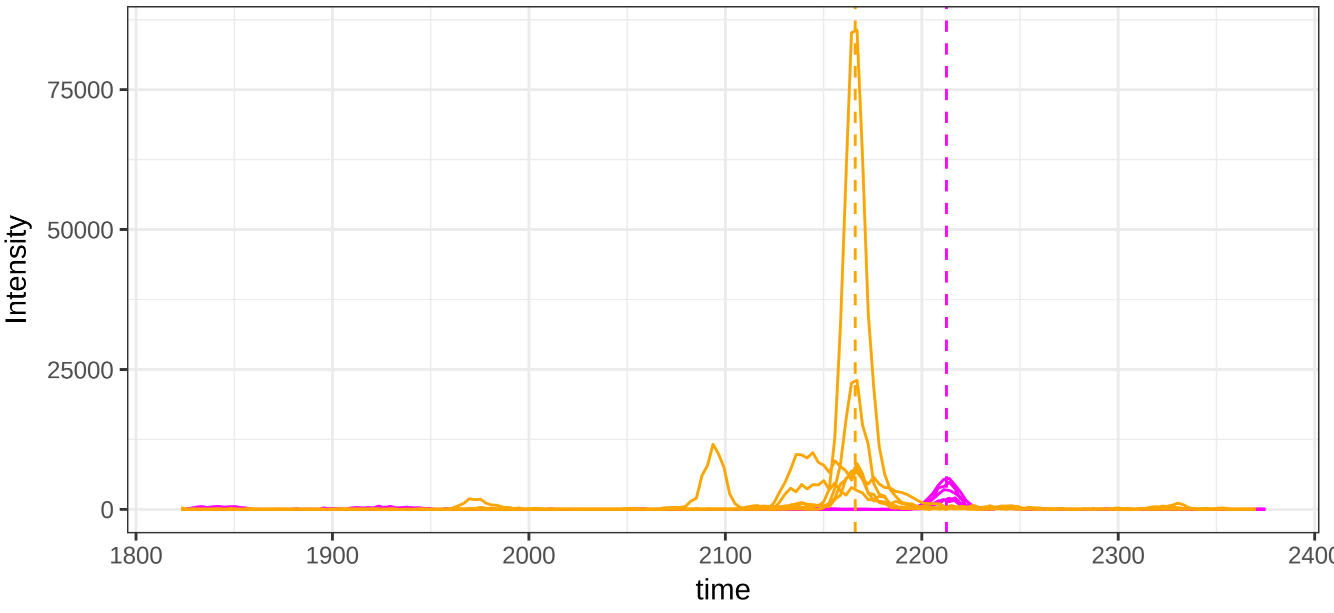

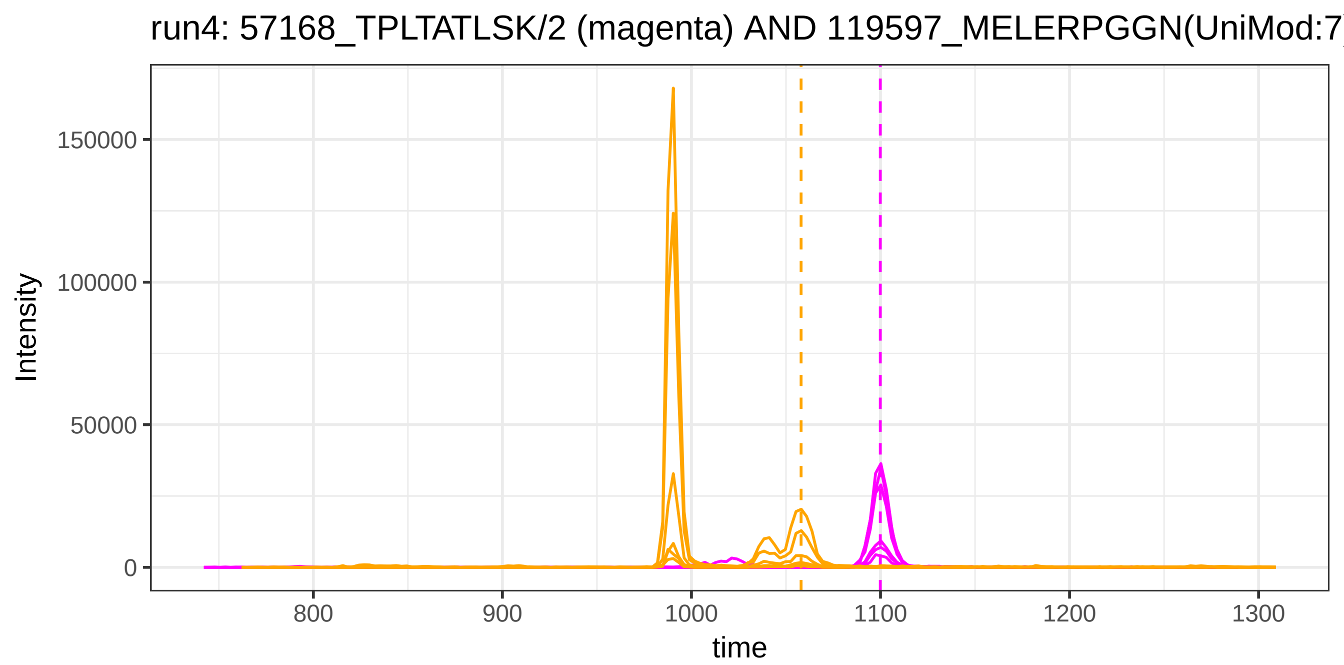
run4: 57168_TPLTATLSK/2 (magenta) AND 119597_MELERPGGN(UniMod:7
Intensity
150000
100000
50000
0
800
900
1000
1100
1200
1300
time

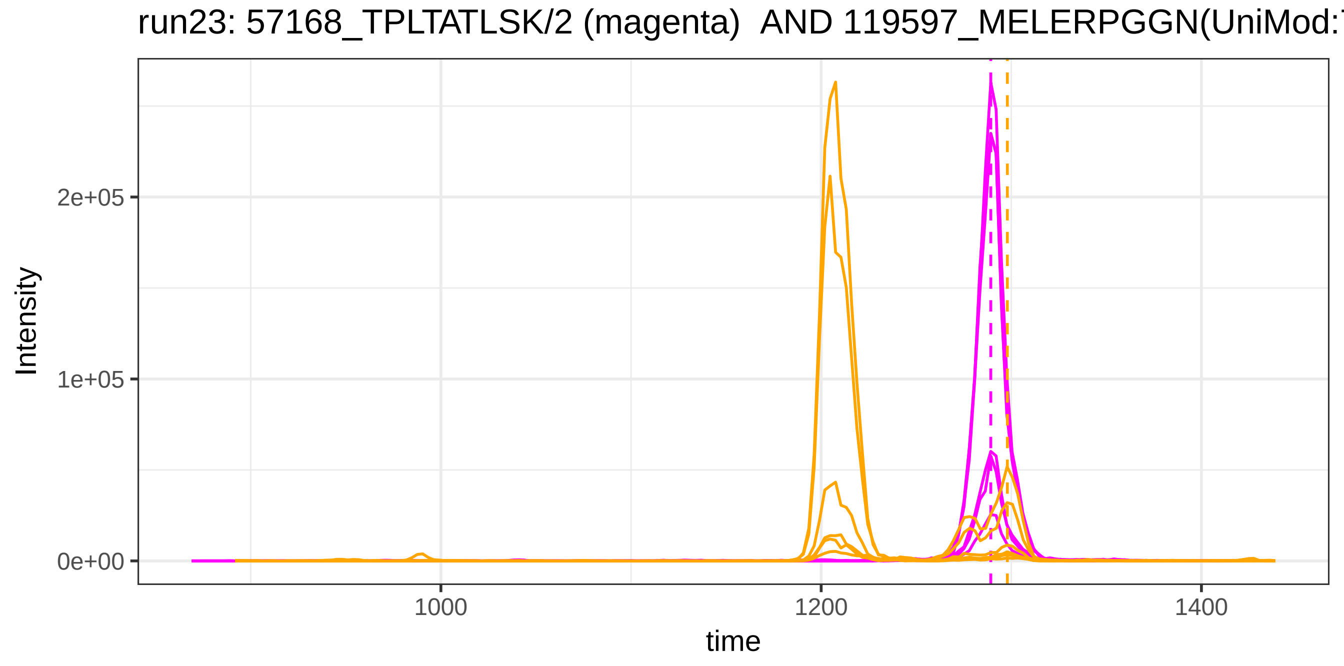
run23: 57168_TPLTATLSK/2 (magenta)  AND 119597_MELERPGGN(UniMod:
Intensity
2e+05
1e+05
0e+00
1000
1200
1400
time

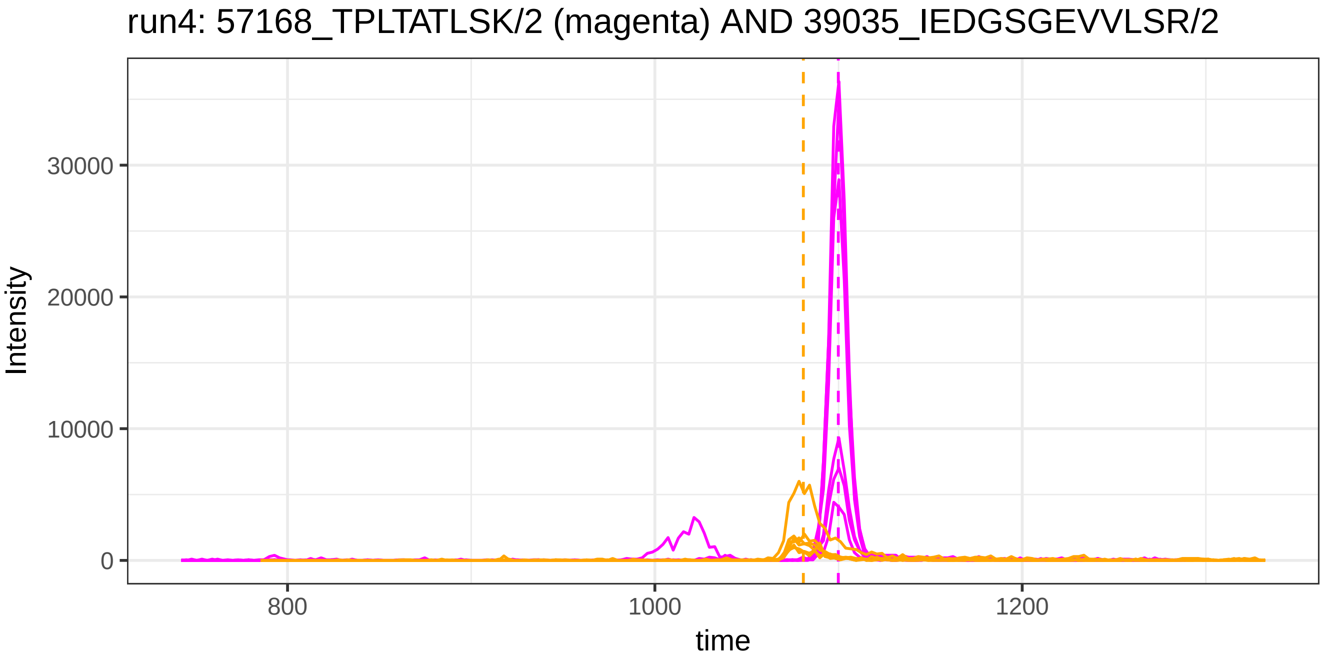
run4: 57168_TPLTATLSK/2 (magenta) AND 39035_IEDGSGEVVLSR/2
Intensity
30000
20000
10000
0
800
1000
1200
time

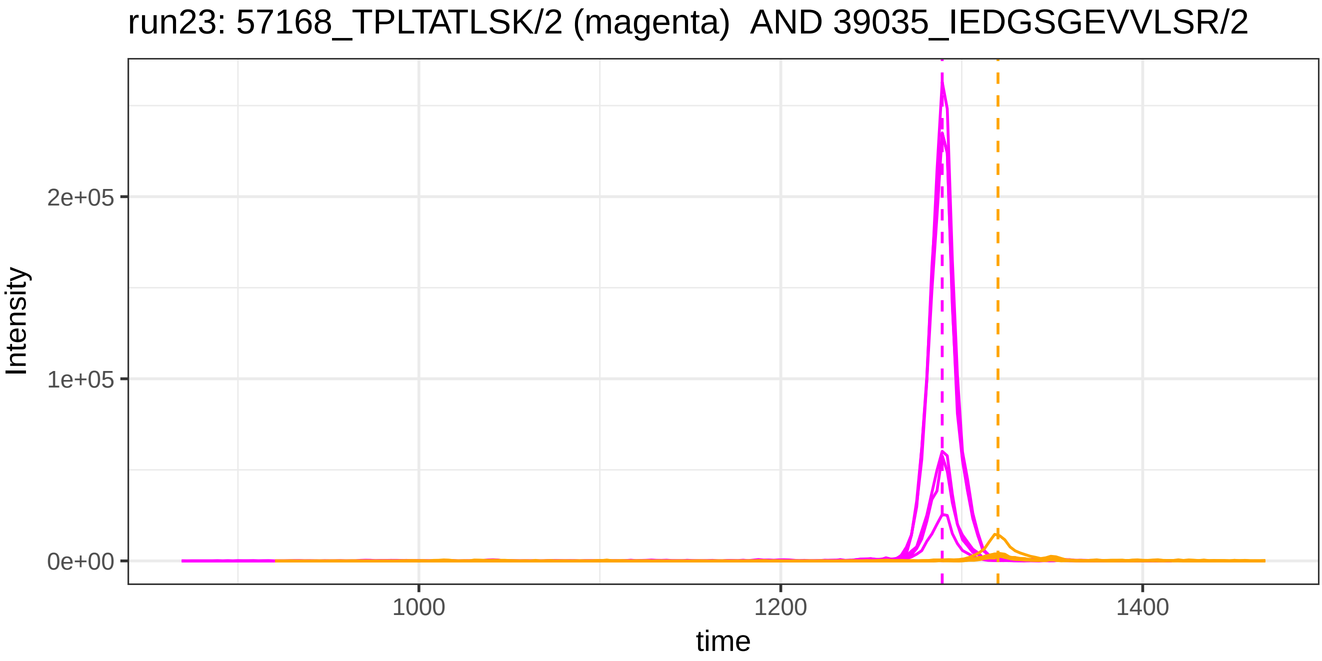
run23: 57168_TPLTATLSK/2 (magenta) AND 39035_IEDGSGEVVLSR/2
Intensity
2e+05
1e+05
0e+00
1000
1200
1400
time

run4: 57168_TPLTATLSK/2 (magenta) AND 116721_IEDGSGEVVLSR/2

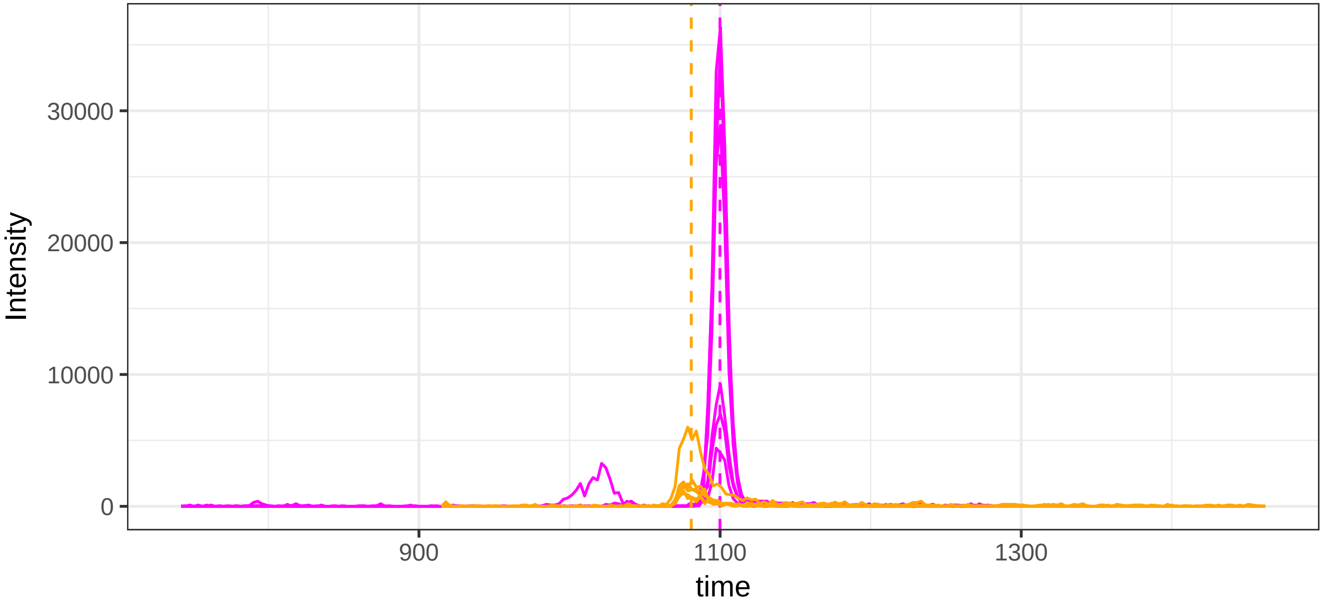


run23: 57168_TPLTATLSK/2 (magenta)  AND 116721_IEDGSGEVVLSR/2

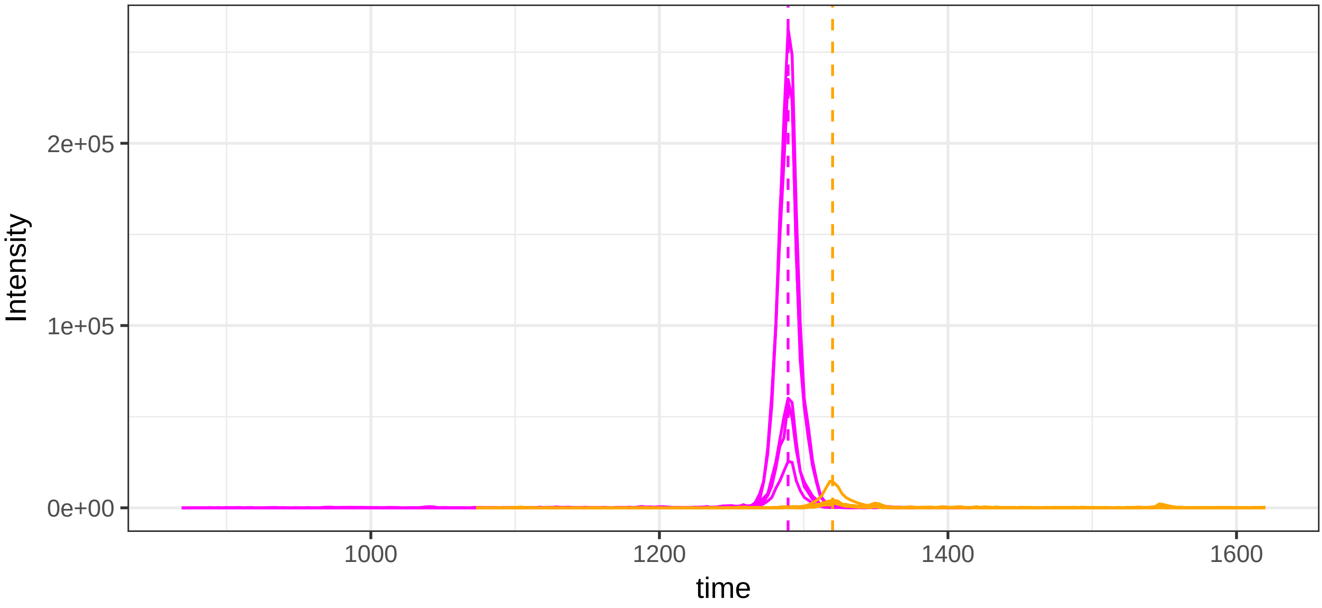

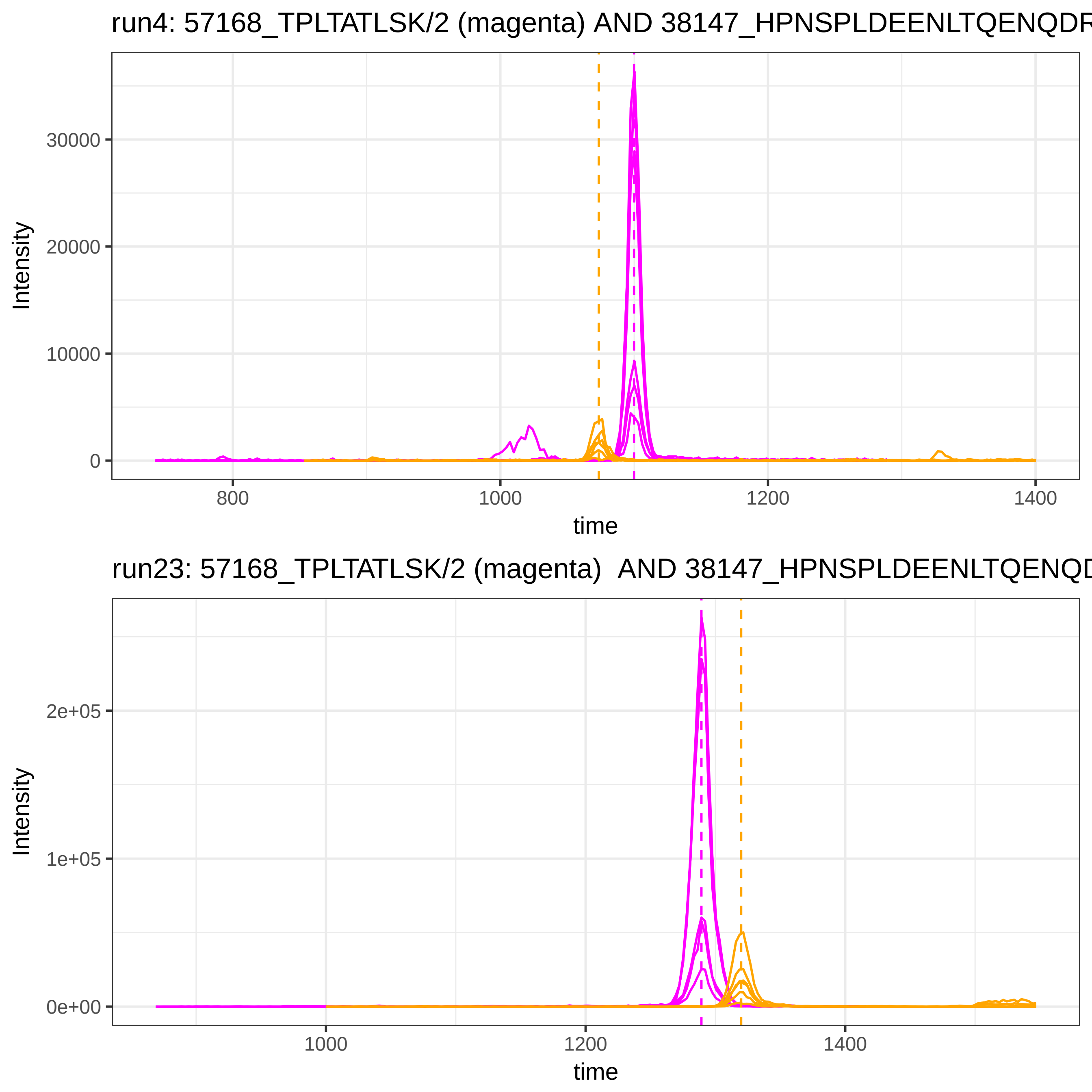

run4: 57168_TPLTATLSK/2 (magenta) AND 38147_HPNSPLDEENLTQENQDR
Intensity
30000
20000
10000
0
800
1000
1200
1400
time
run23: 57168_TPLTATLSK/2 (magenta) AND 38147_HPNSPLDEENLTQENQD
Intensity
2e+05
1e+05
0e+00
1000
1200
1400
time

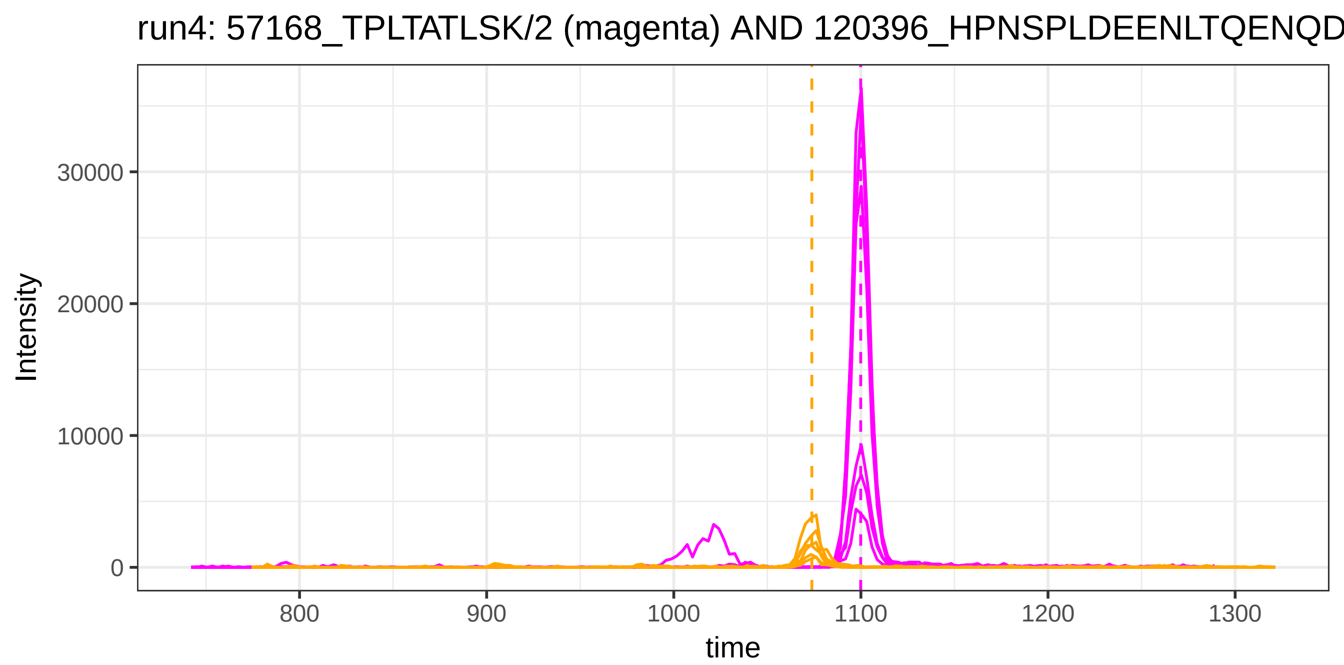
run4: 57168_TPLTATLSK/2 (magenta) AND 120396_HPNSPLDEENLTQENQD
Intensity
30000
20000
10000
0
800
900
1000
1100
1200
1300
time

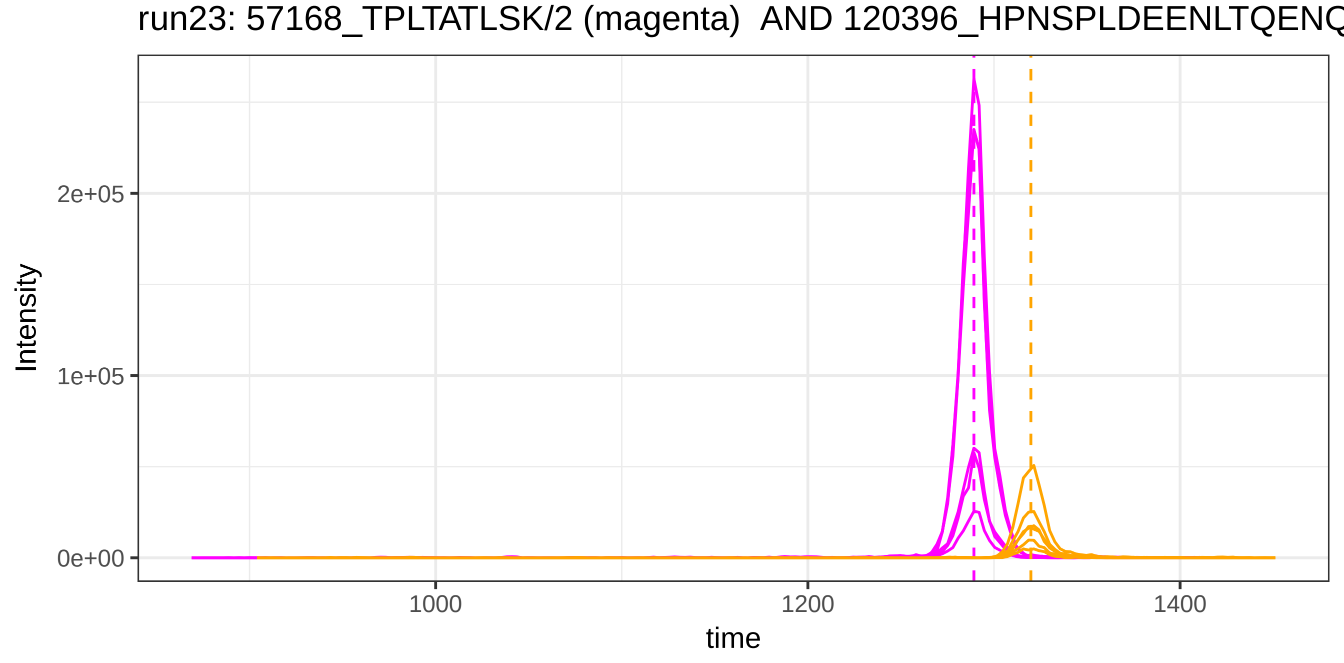
run23: 57168_TPLTATLSK/2 (magenta) AND 120396_HPNSPLDEENLTQENQ
Intensity
2e+05
1e+05
0e+00
1000
1200
1400
time

run4: 57168_TPLTATLSK/2 (magenta) AND 4844_YLQEIYNSNNQK/2

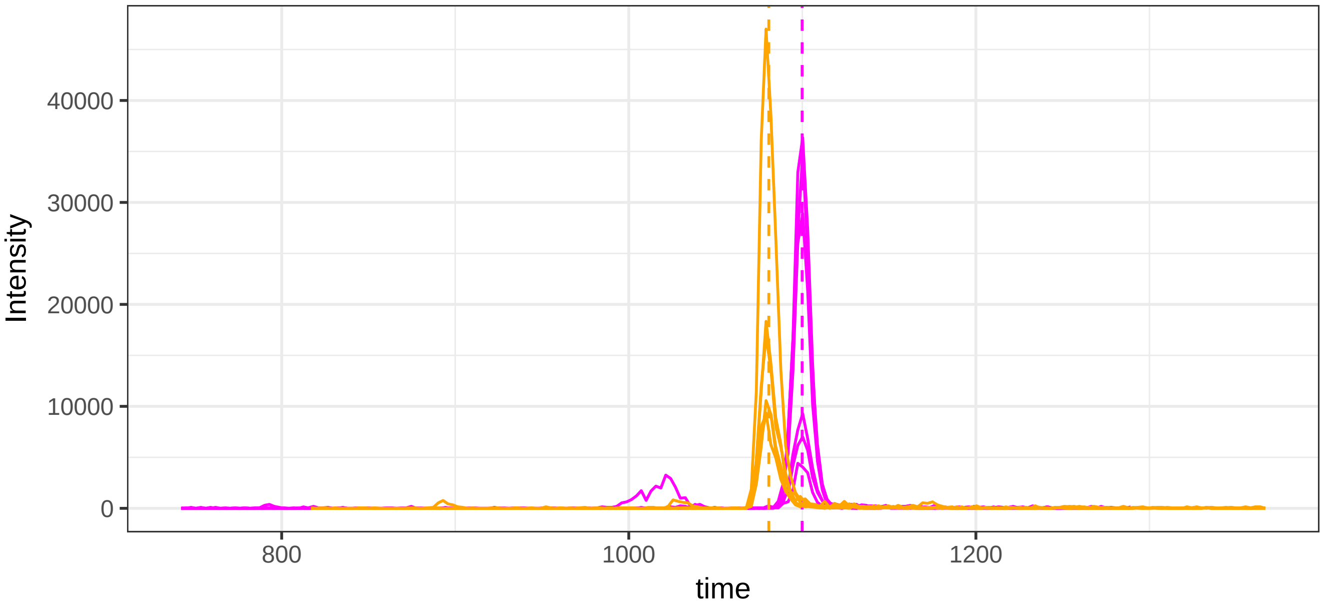


run23: 57168_TPLTATLSK/2 (magenta)  AND 4844_YLQEIYNSNNQK/2

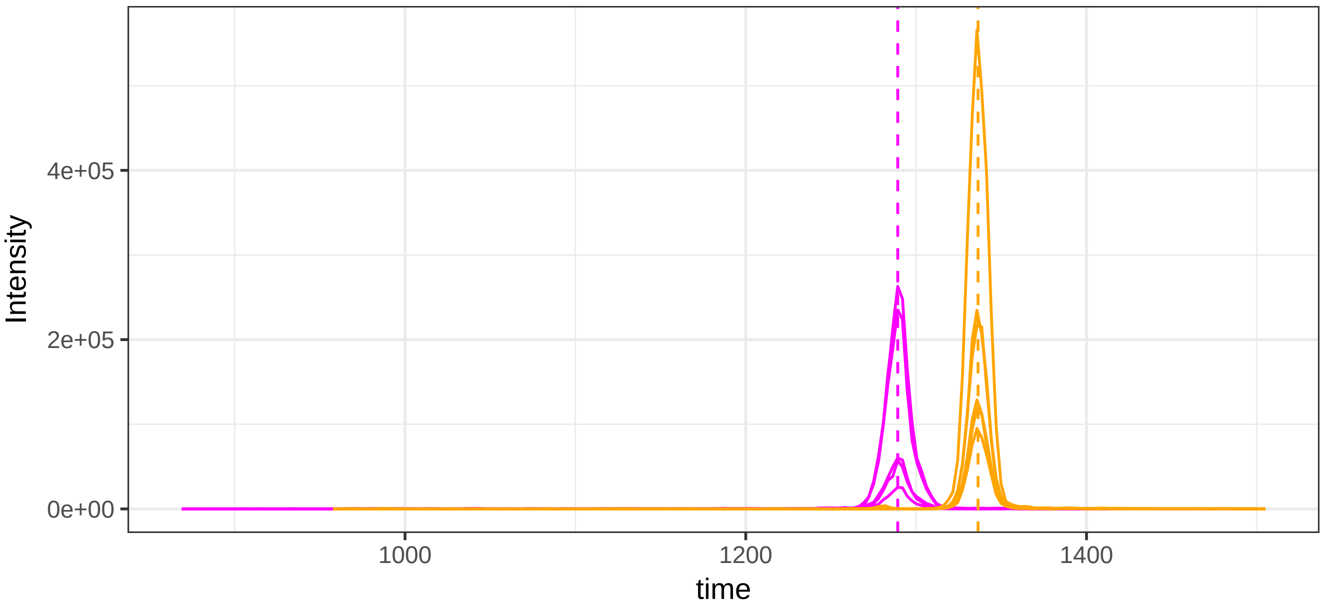

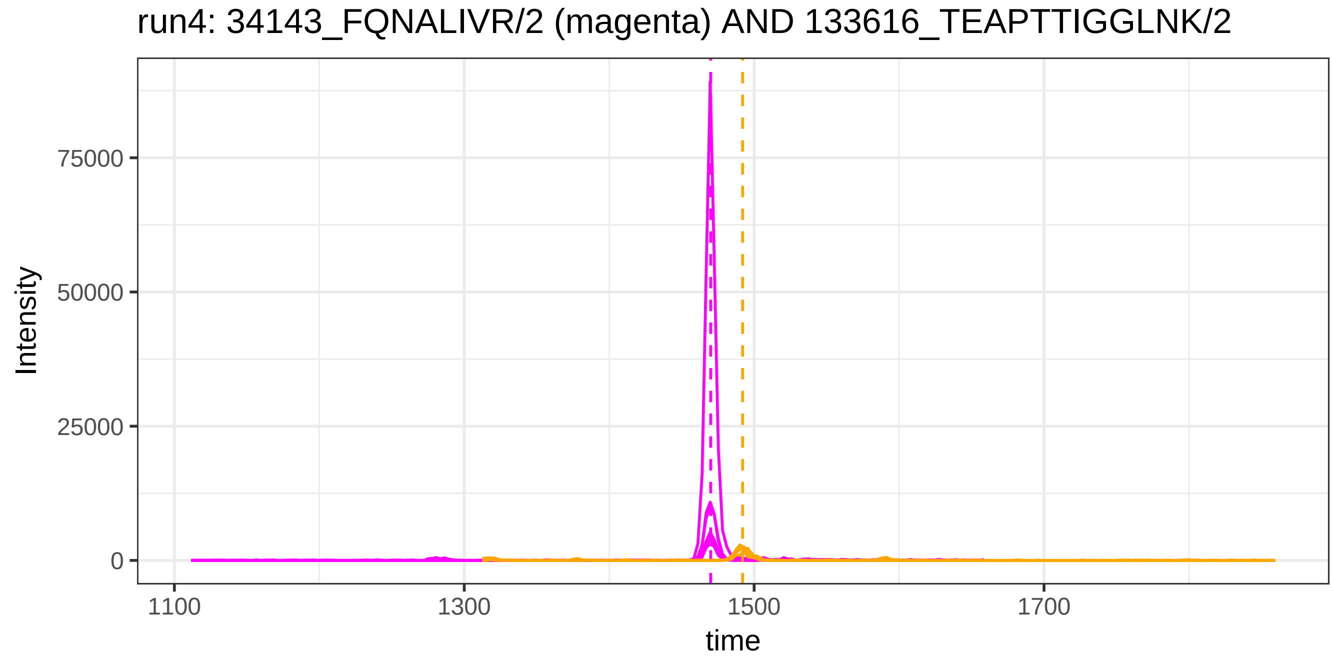
run4: 34143_FQNALIVR/2 (magenta) AND 133616_TEAPTTIGGLNK/2
Intensity
75000
50000
25000
0
1100
1300
1500
1700
time

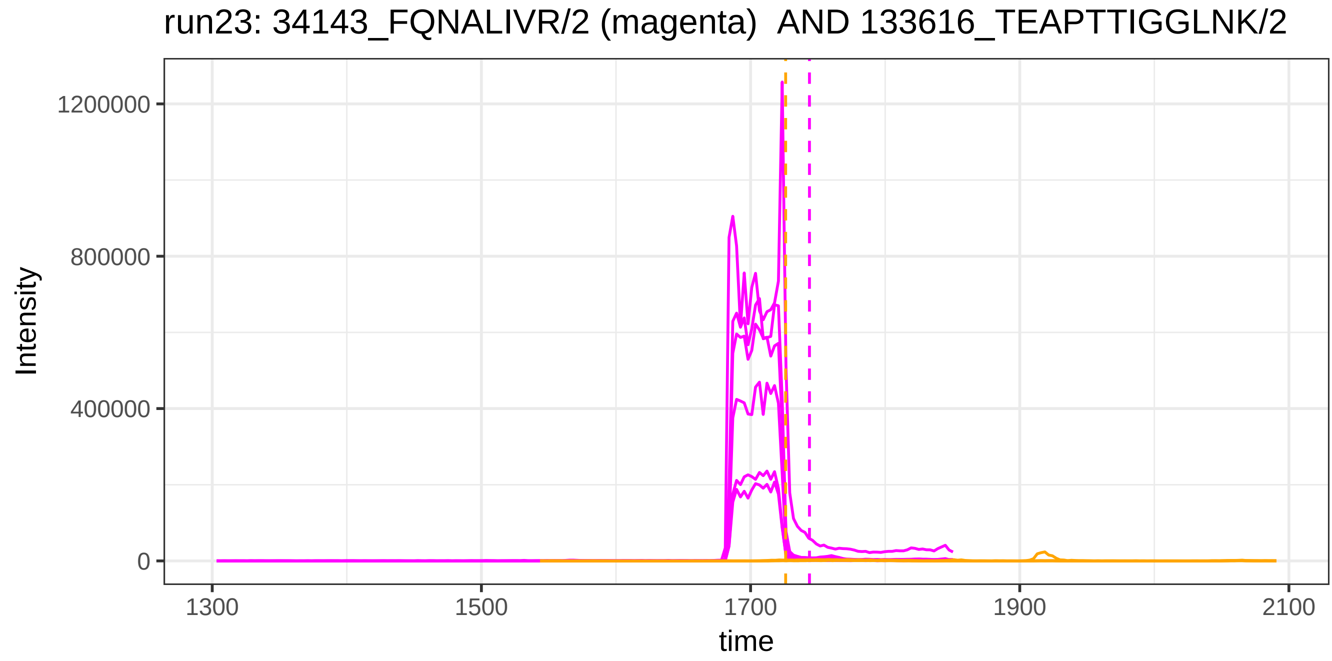
run23: 34143_FQNALIVR/2 (magenta)  AND 133616_TEAPTTIGGLNK/2
Intensity
1200000
800000
400000
0
1300
1500
1700
1900
2100
time

run4: 115879_DDNPNLPR/2 (magenta) AND 35386_AHVSFKPTVAQQR/3

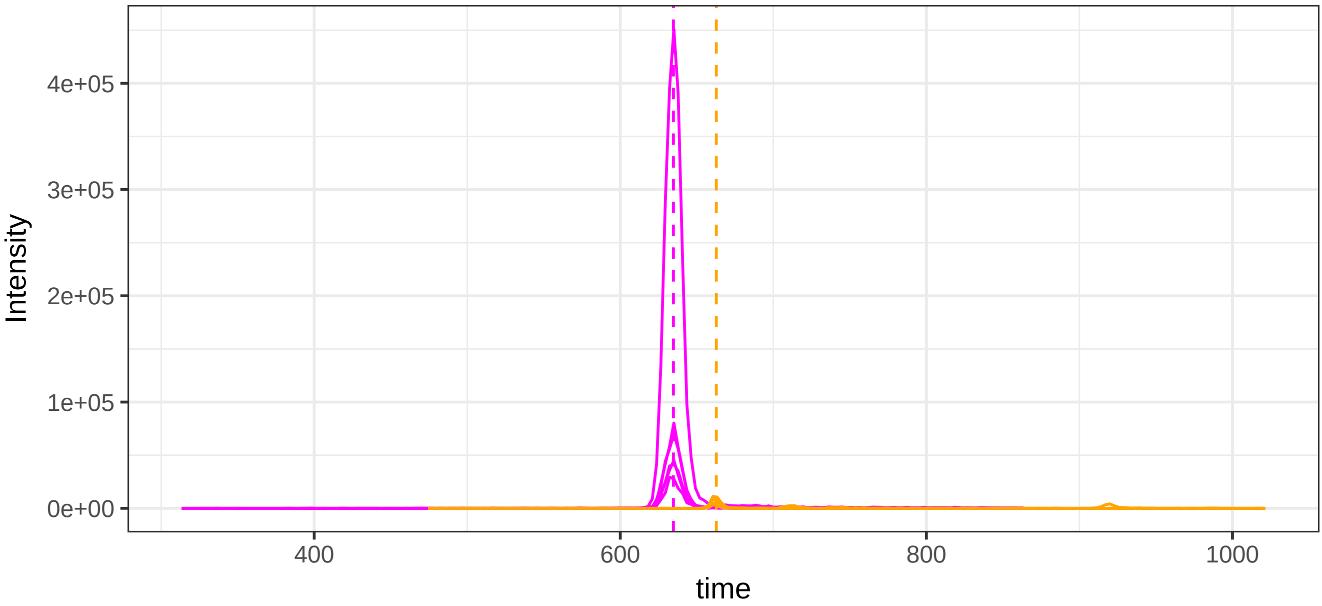


run23: 115879_DDNPNLPR/2 (magenta)  AND 35386_AHVSFKPTVAQQR/3

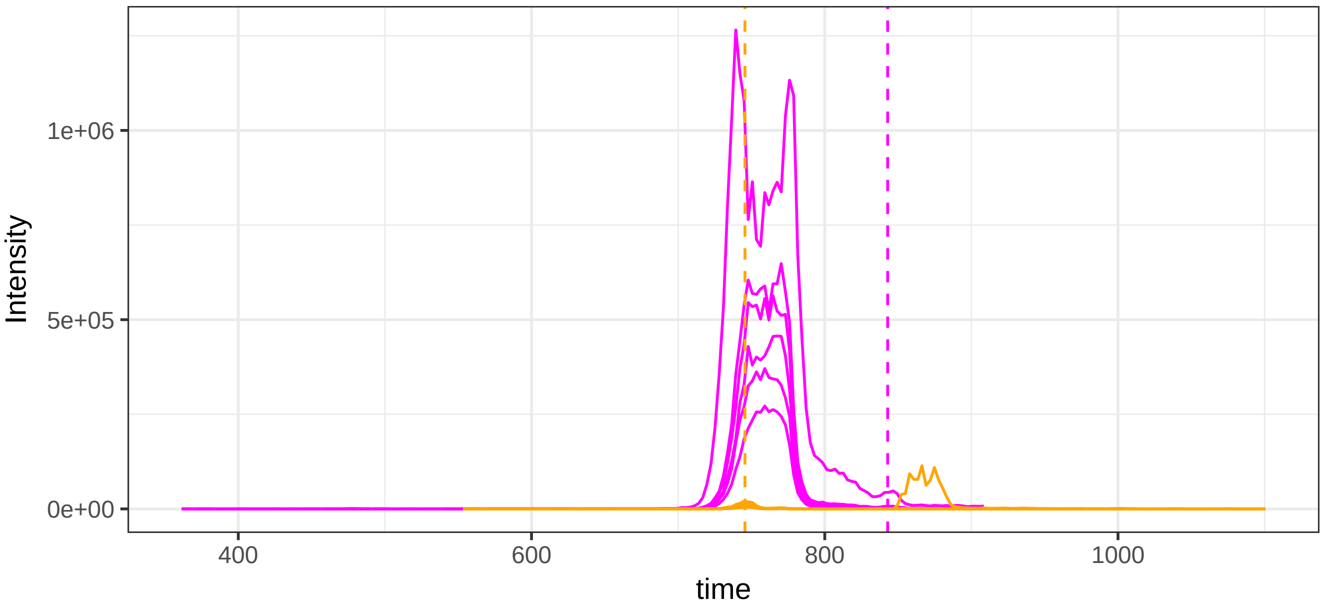

run4: 115879_DDNPNLPR/2 (magenta) AND 120914_AHVSFKPTVAQQR/3

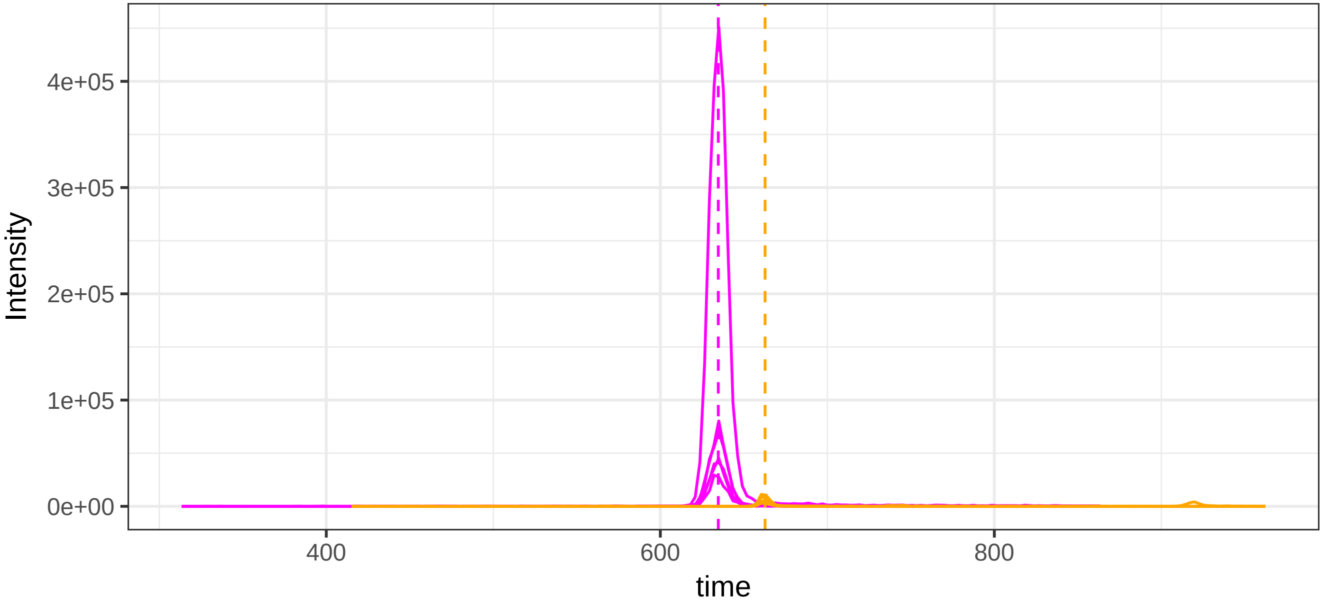


run23: 115879_DDNPNLPR/2 (magenta)  AND 120914_AHVSFKPTVAQQR/3

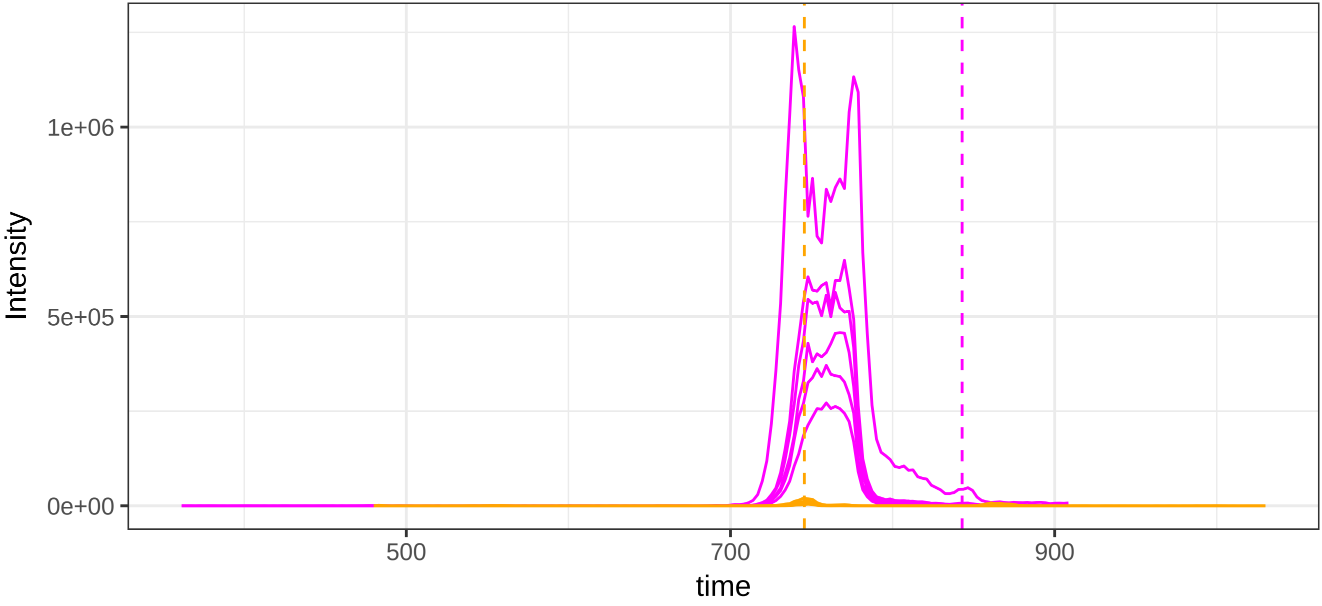

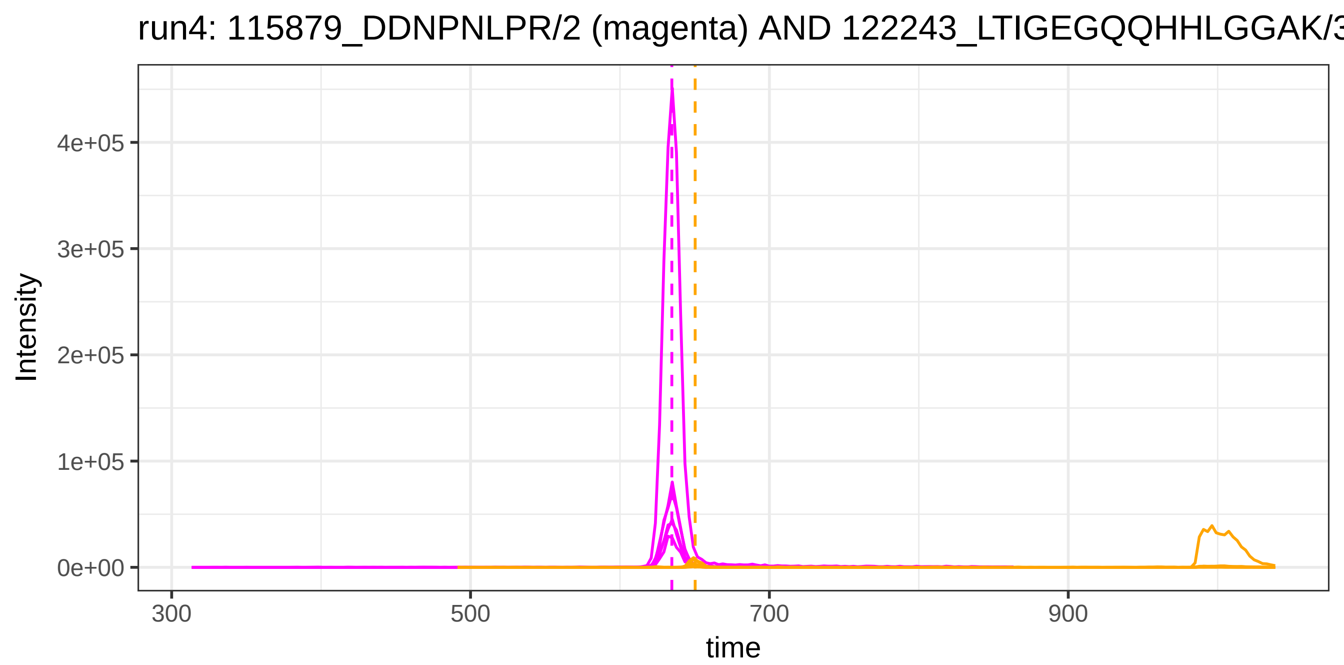
run4: 115879_DDNPNLPR/2 (magenta) AND 122243_LTIGEGQQHHLGGAK/3
Intensity
4e+05
3e+05
2e+05
1e+05
0e+00
300
500
700
900
time

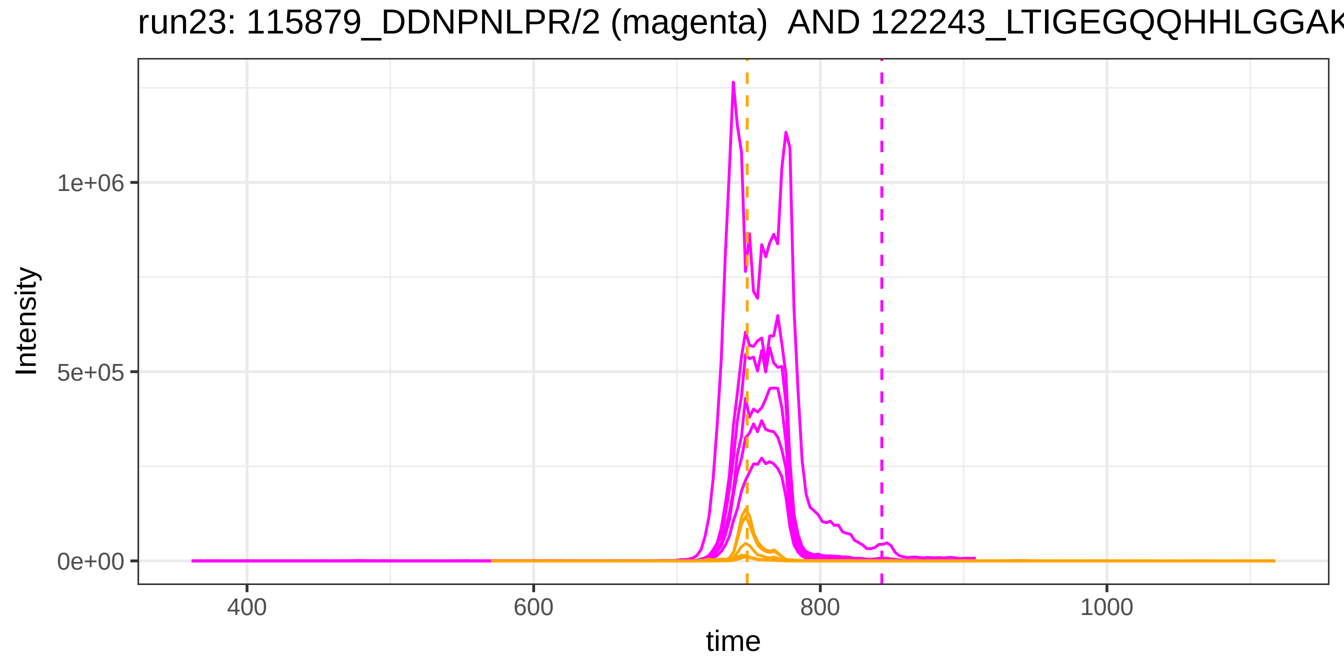
run23: 115879_DDNPNLPR/2 (magenta) AND 122243_LTIGEGQQHHLGGAK
Intensity
1e+06
5e+05
0e+00
400
600
800
1000
time

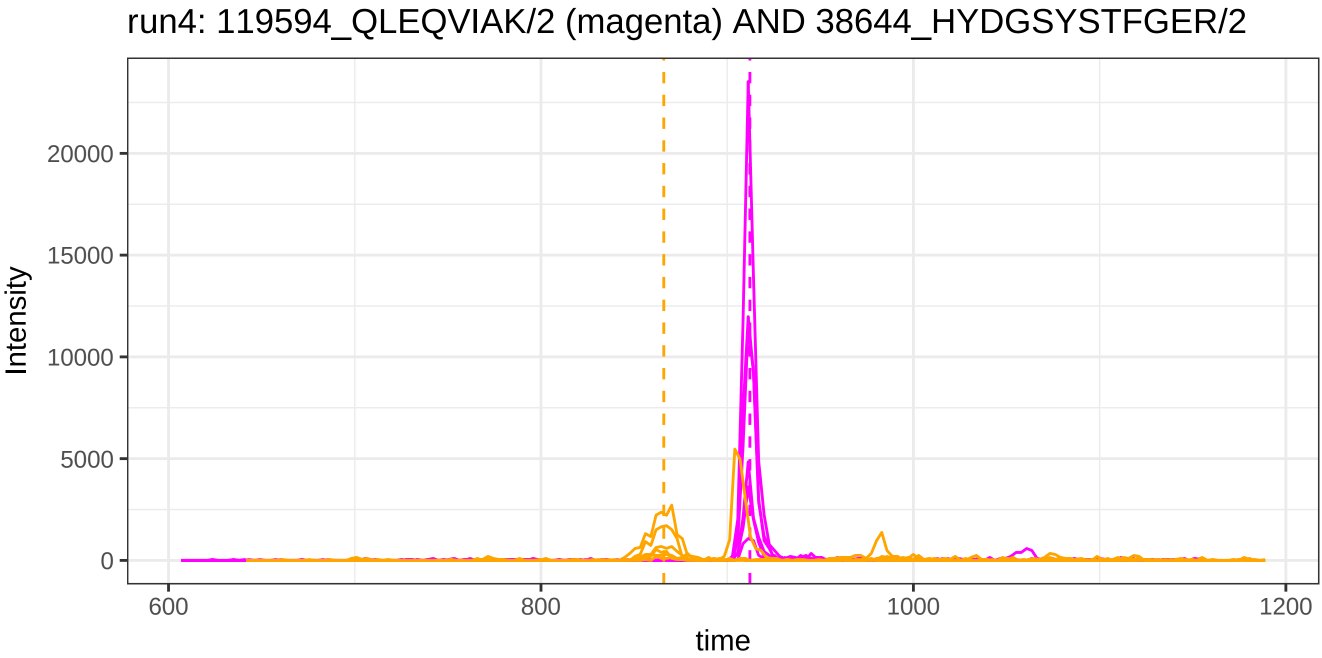
run4: 119594_QLEQVIAK/2 (magenta) AND 38644_HYDGSYSTFGER/2
Intensity
0
5000
10000
15000
20000
600
800
1000
1200
time

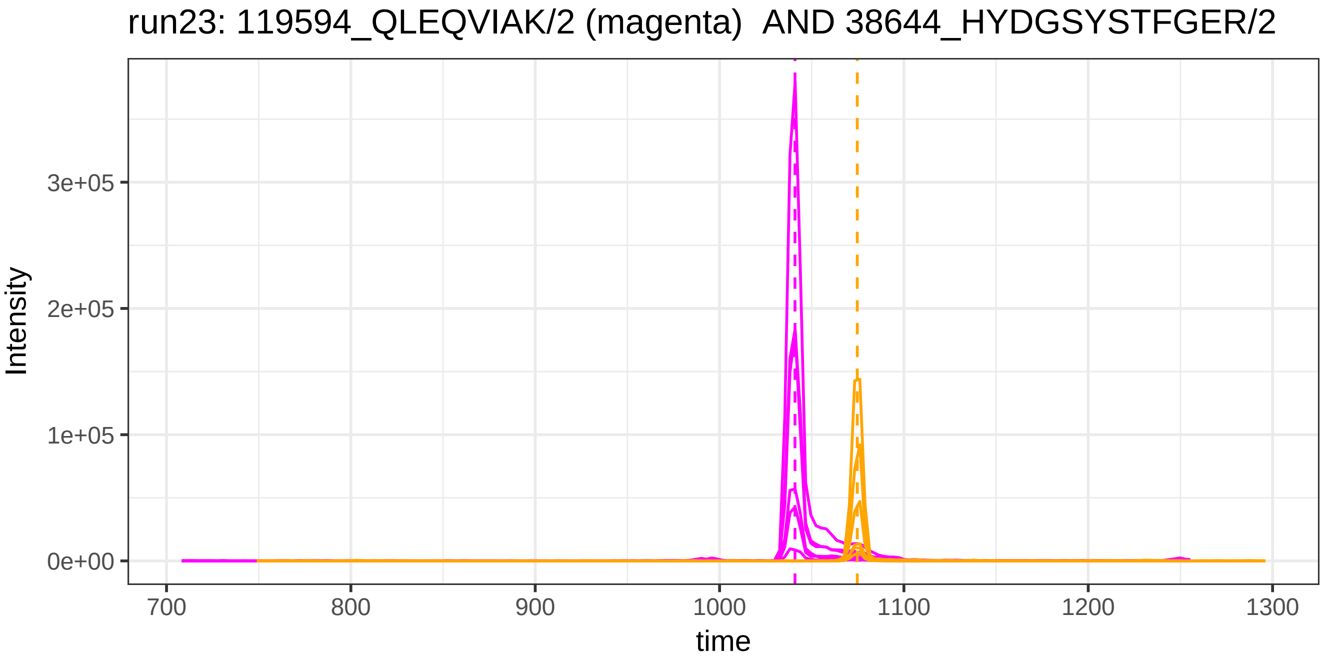
run23: 119594_QLEQVIAK/2 (magenta) AND 38644_HYDGSYSTFGER/2
Intensity
0e+00
1e+05
2e+05
3e+05
700
800
900
1000
1100
1200
1300
time

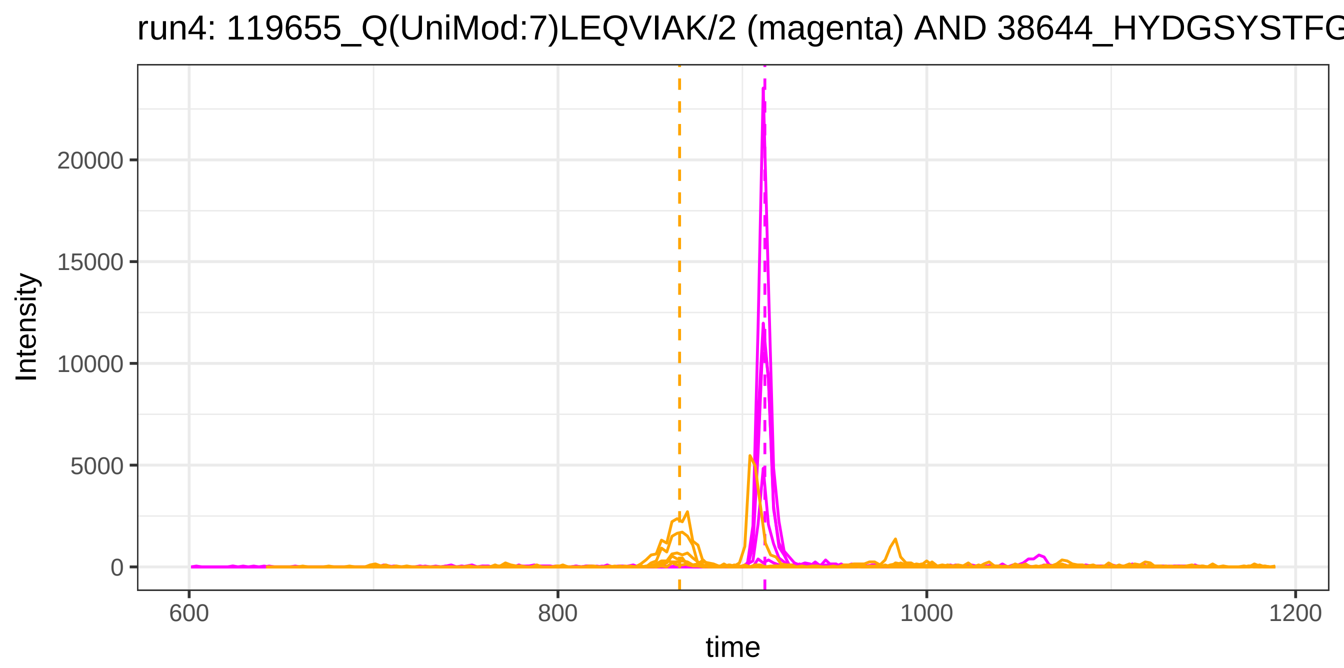

run4: 119655_Q(UniMod:7)LEQVIAK/2 (magenta) AND 38644_HYDGSYSTFG
Intensity
0
5000
10000
15000
20000
600
800
1000
1200
time


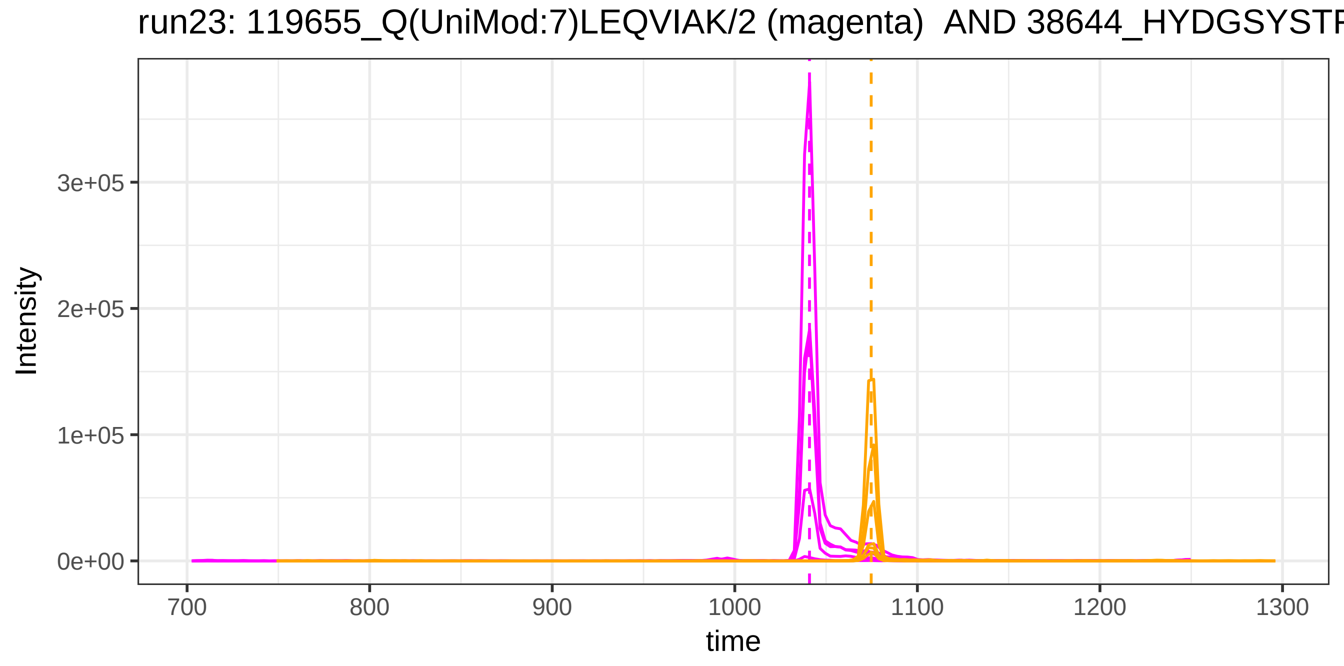

run23: 119655_Q(UniMod:7)LEQVIAK/2 (magenta) AND 38644_HYDGSYSTF
Intensity
0e+00
1e+05
2e+05
3e+05
700
800
900
1000
1100
1200
1300
time

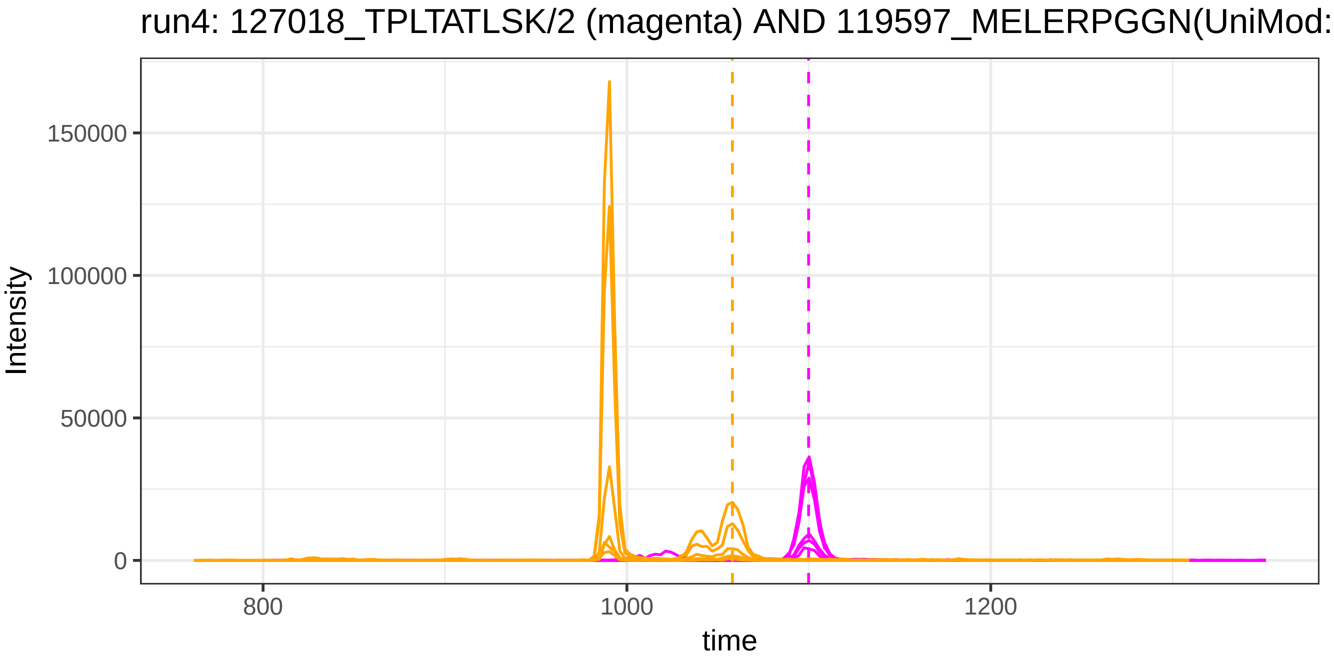

run4: 127018_TPLTATLSK/2 (magenta) AND 119597_MELERPGGN(UniMod:
Intensity
150000
100000
50000
0
800
1000
1200
time


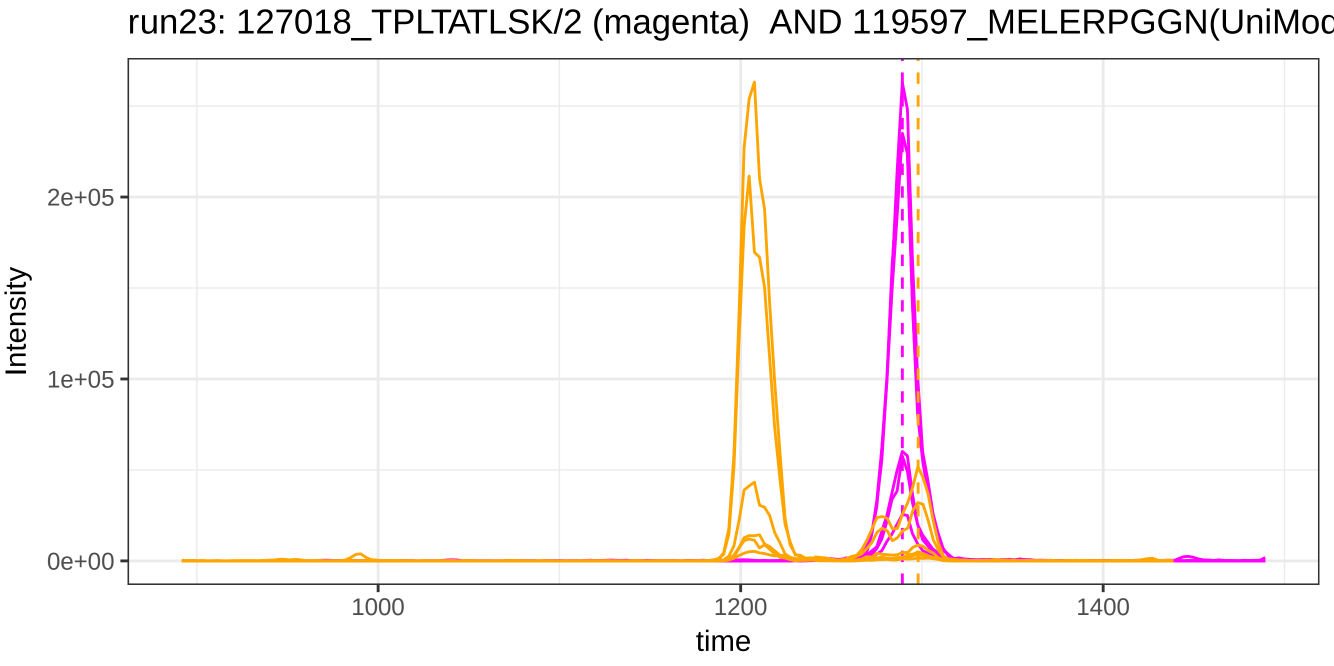

run23: 127018_TPLTATLSK/2 (magenta) AND 119597_MELERPGGN(UniMod
Intensity
2e+05
1e+05
0e+00
1000
1200
1400
time

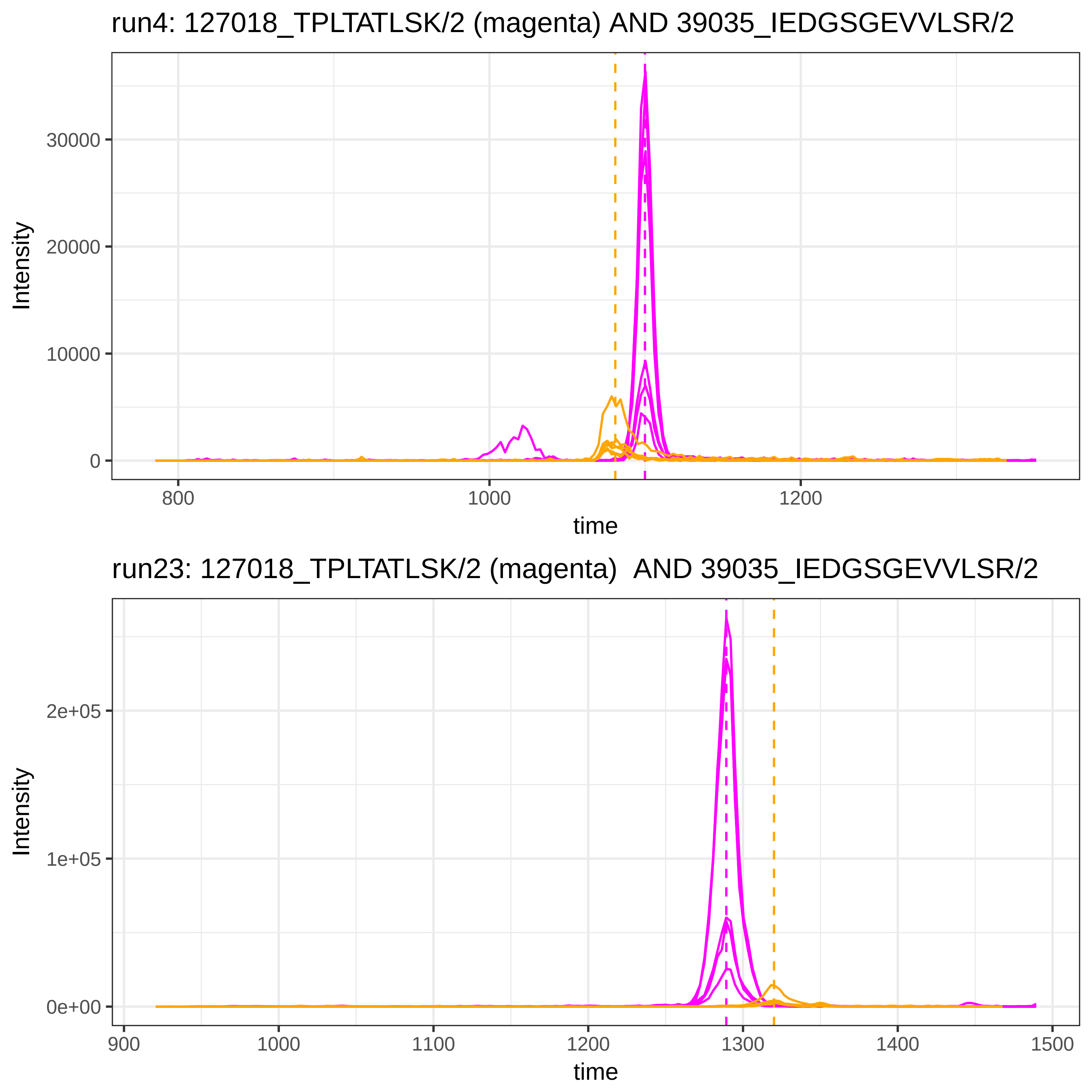

run4: 127018_TPLTATLSK/2 (magenta) AND 39035_IEDGSGEVVLSR/2
Intensity
30000
20000
10000
0
800
1000
1200
time
run23: 127018_TPLTATLSK/2 (magenta) AND 39035_IEDGSGEVVLSR/2
Intensity
2e+05
1e+05
0e+00
900
1000
1100
1200
1300
1400
1500
time

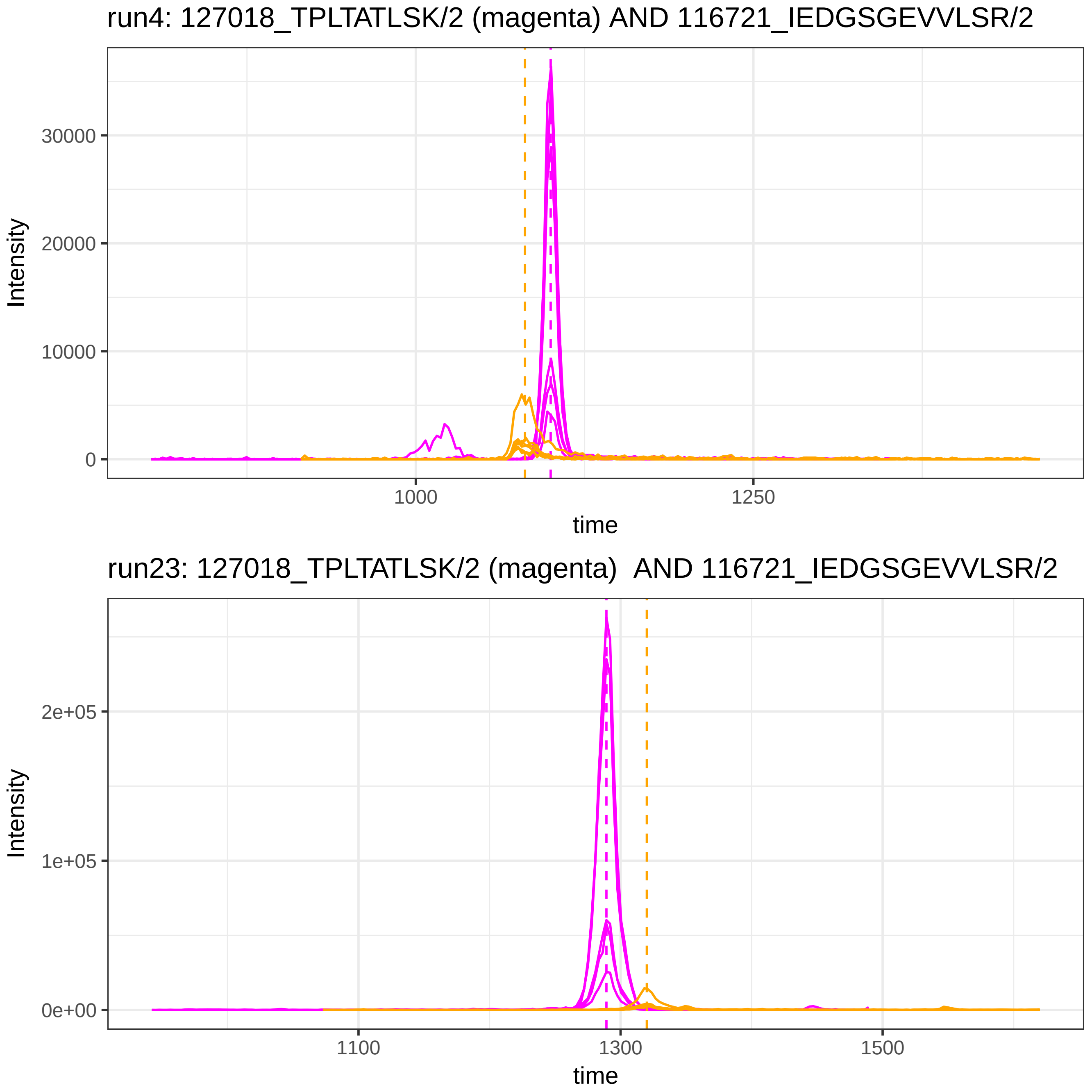
run4: 127018_TPLTATLSK/2 (magenta) AND 116721_IEDGSGEVVLSR/2
Intensity
30000
20000
10000
0
1000
1250
time
run23: 127018_TPLTATLSK/2 (magenta)  AND 116721_IEDGSGEVVLSR/2
Intensity
2e+05
1e+05
0e+00
1100
1300
1500
time

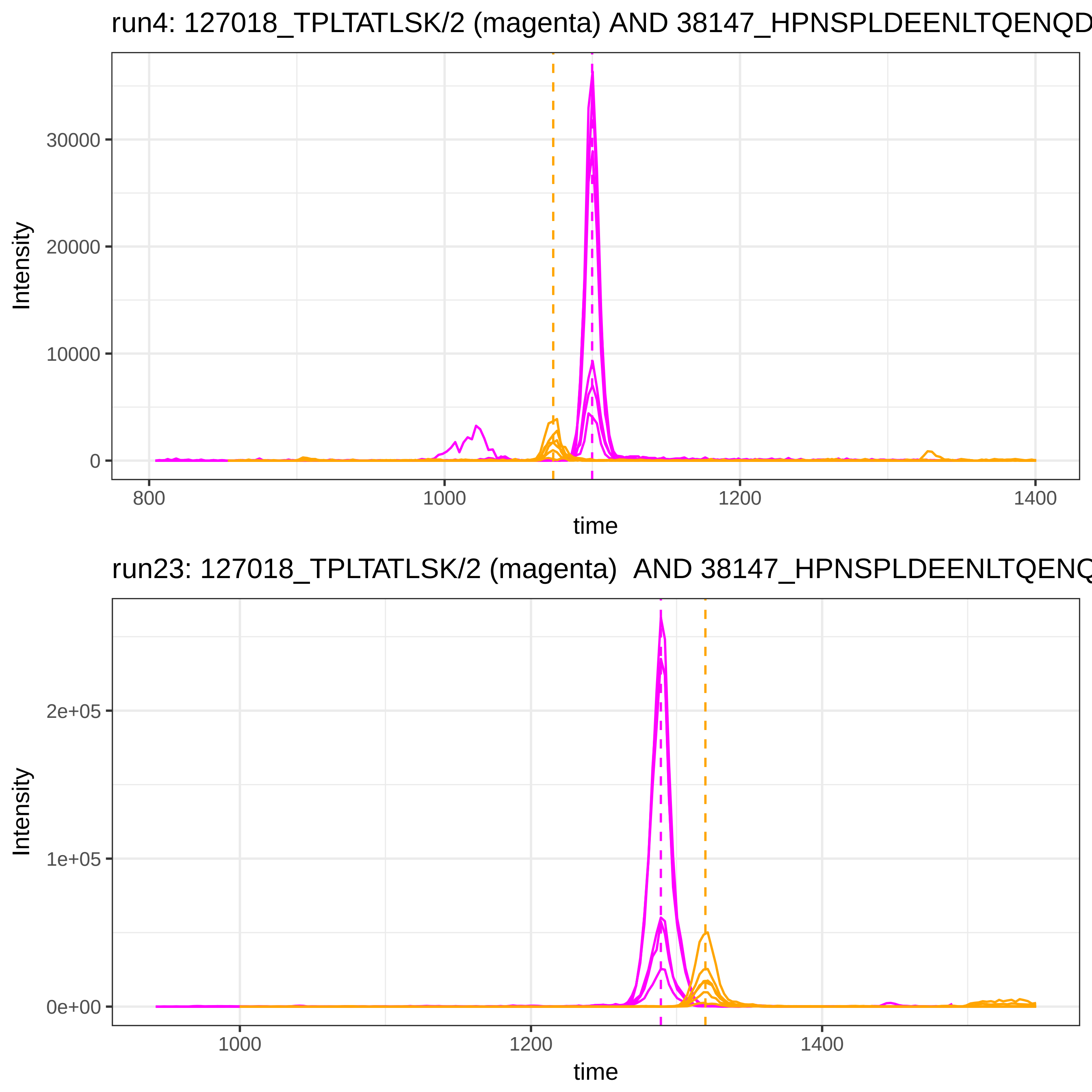
run4: 127018_TPLTATLSK/2 (magenta) AND 38147_HPNSPLDEENLTQENQD
Intensity
30000
20000
10000
0
800
1000
1200
1400
time
run23: 127018_TPLTATLSK/2 (magenta) AND 38147_HPNSPLDEENLTQENQ
Intensity
2e+05
1e+05
0e+00
1000
1200
1400
time

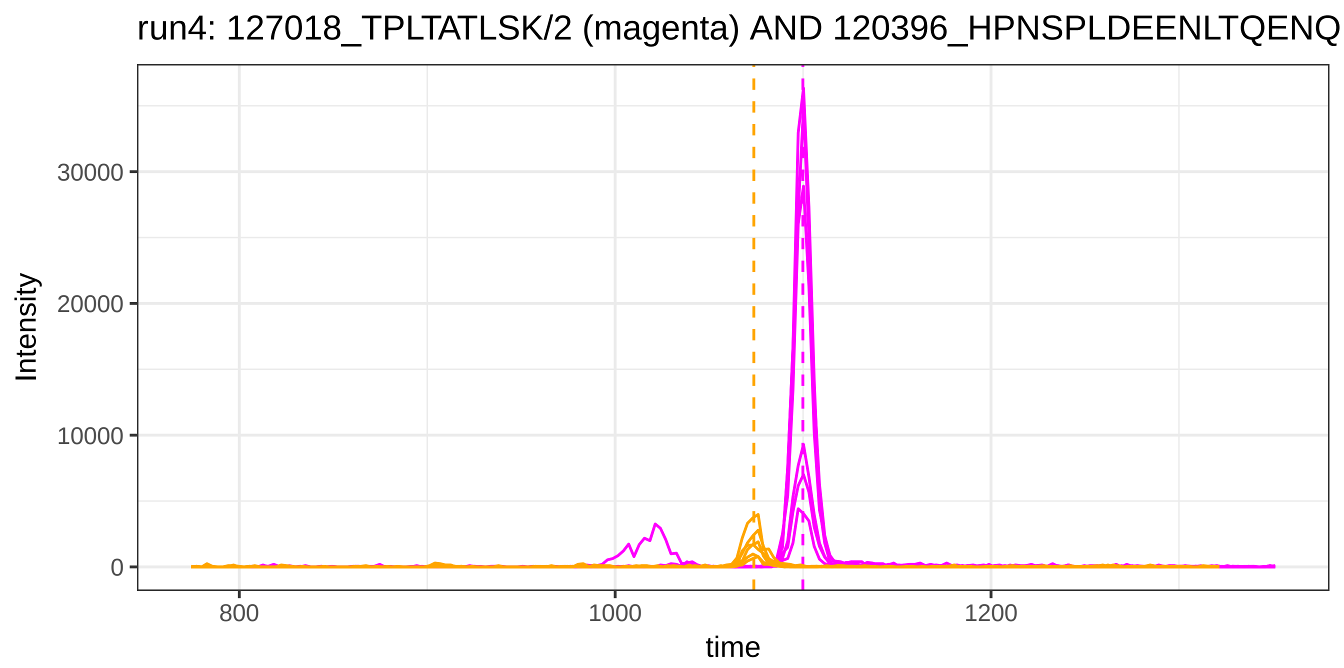

run4: 127018_TPLTATLSK/2 (magenta) AND 120396_HPNSPLDEENLTQENQ
Intensity
30000
20000
10000
0
800
1000
1200
time


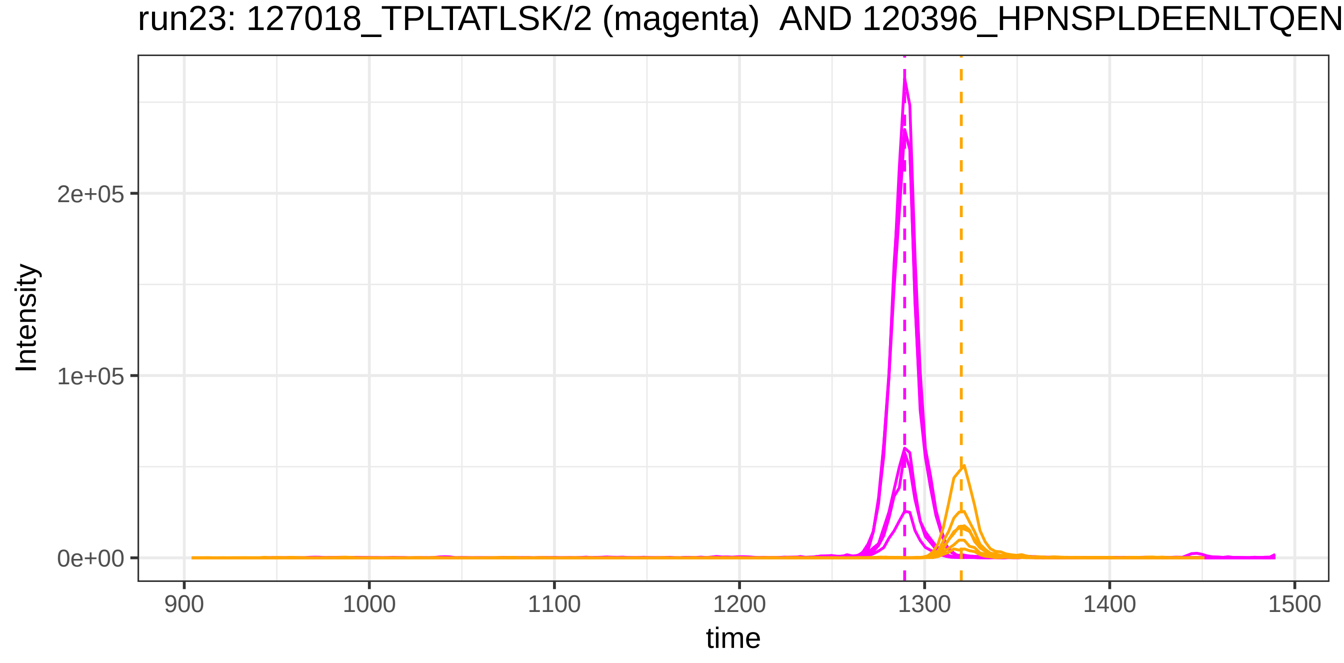

run23: 127018_TPLTATLSK/2 (magenta) AND 120396_HPNSPLDEENLTQEN
Intensity
2e+05
1e+05
0e+00
900
1000
1100
1200
1300
1400
1500
time

run4: 127018_TPLTATLSK/2 (magenta) AND 4844_YLQEIYNSNNQK/2

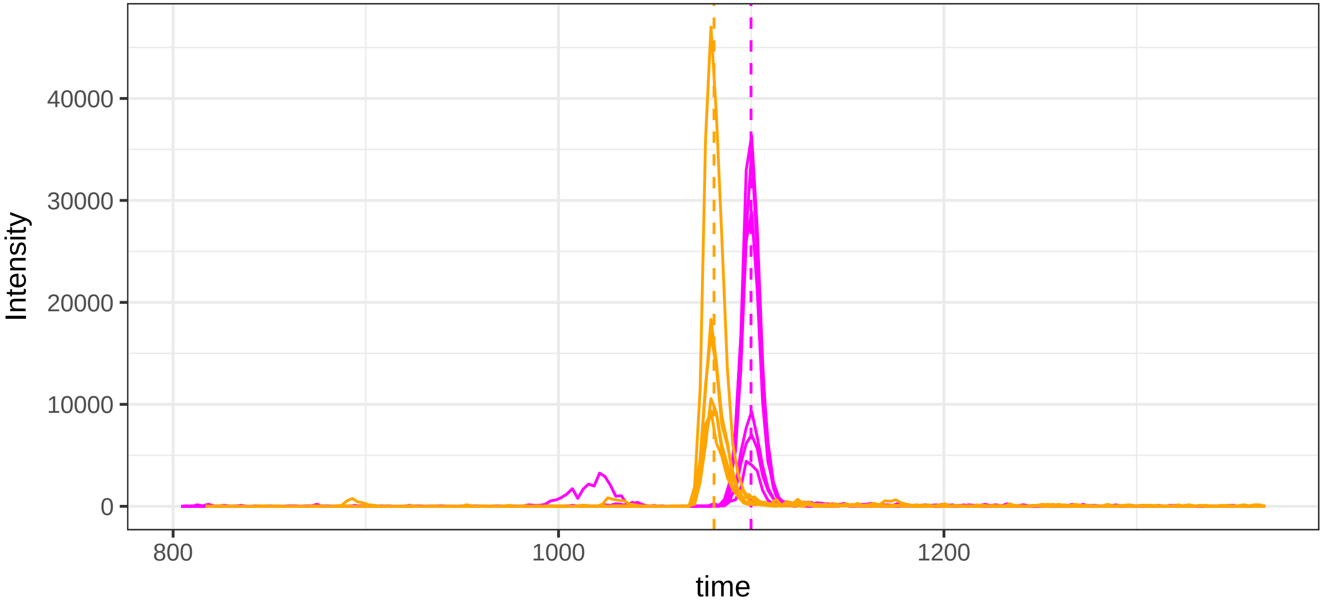


run23: 127018_TPLTATLSK/2 (magenta)  AND 4844_YLQEIYNSNNQK/2

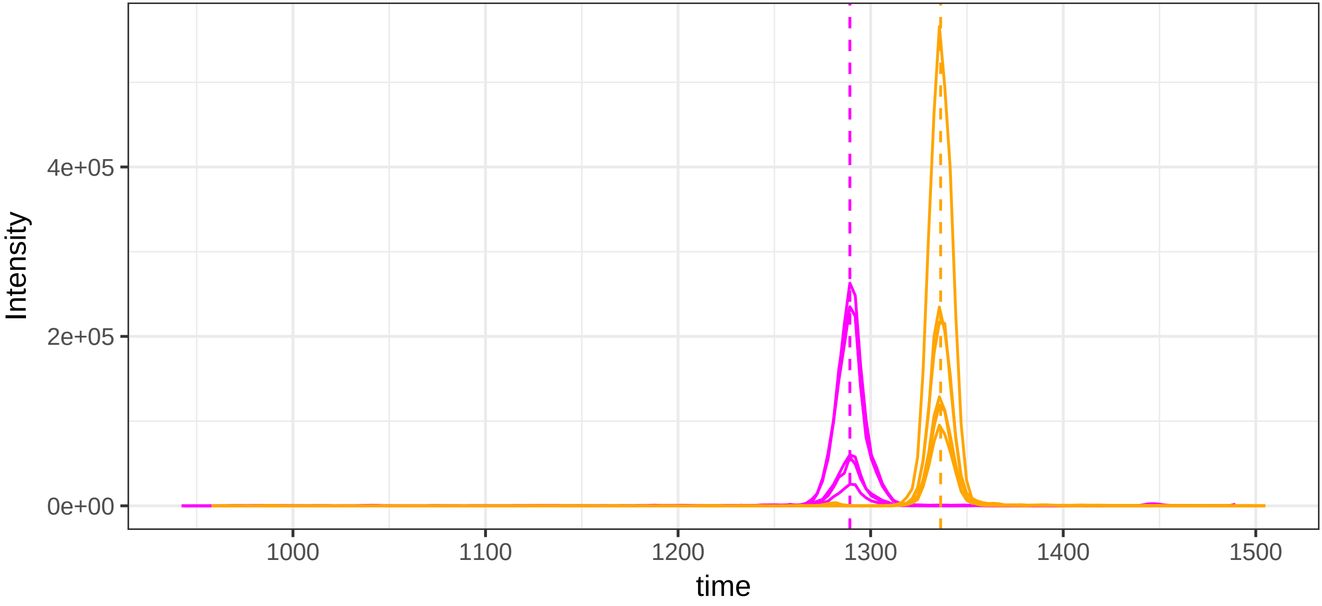

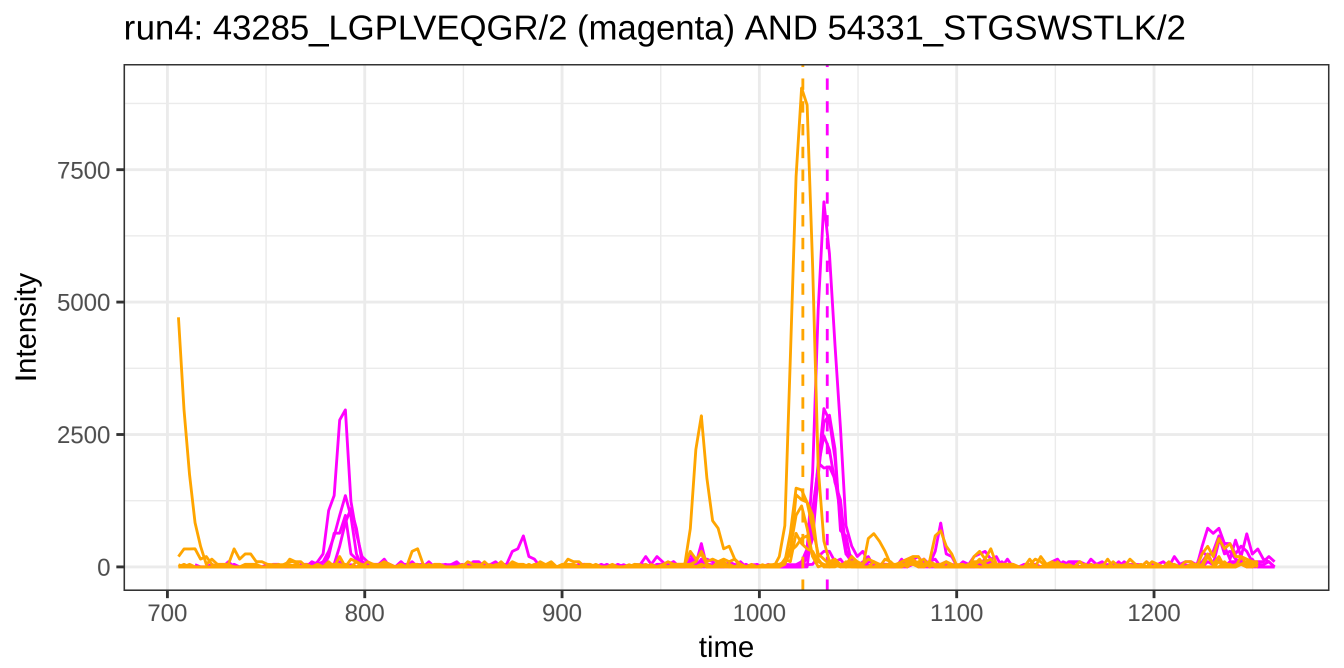

run4: 43285_LGPLVEQGR/2 (magenta) AND 54331_STGSWSTLK/2
Intensity
7500
5000
2500
0
700
800
900
1000
1100
1200
time


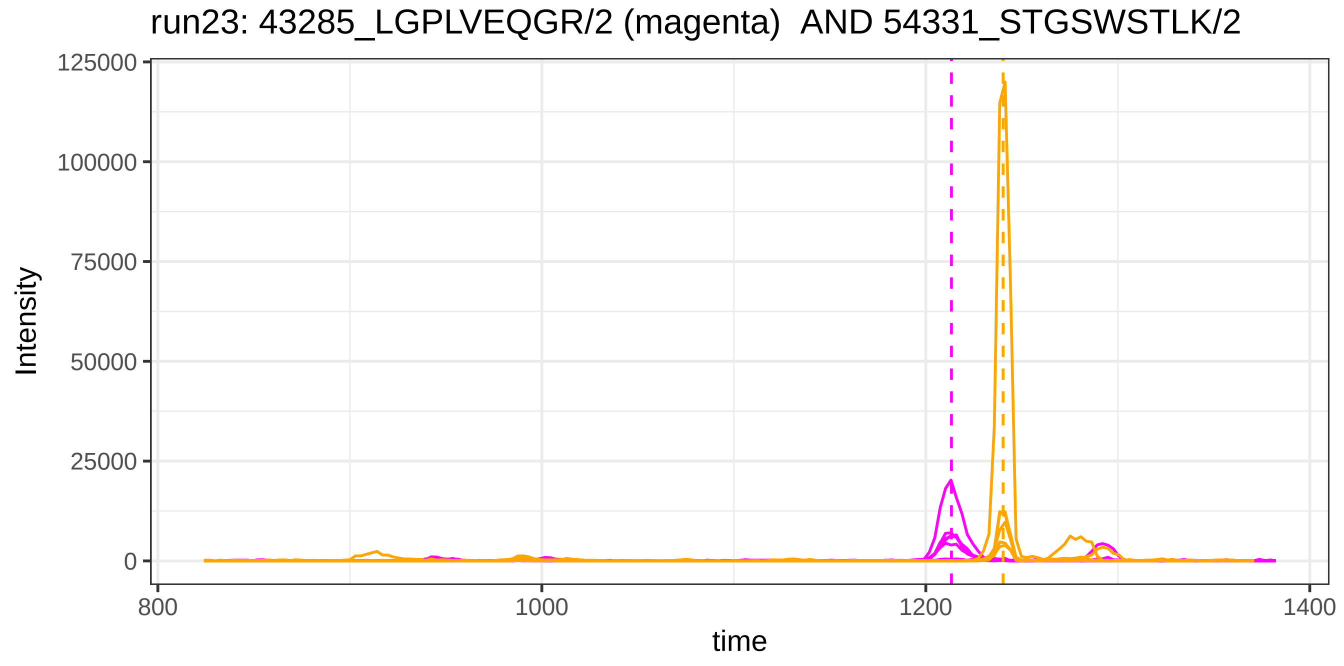

run23: 43285_LGPLVEQGR/2 (magenta) AND 54331_STGSWSTLK/2
Intensity
125000
100000
75000
50000
25000
0
800
1000
1200
1400
time

run4: 43285_LGPLVEQGR/2 (magenta) AND 119020_STGSWSTLK/2

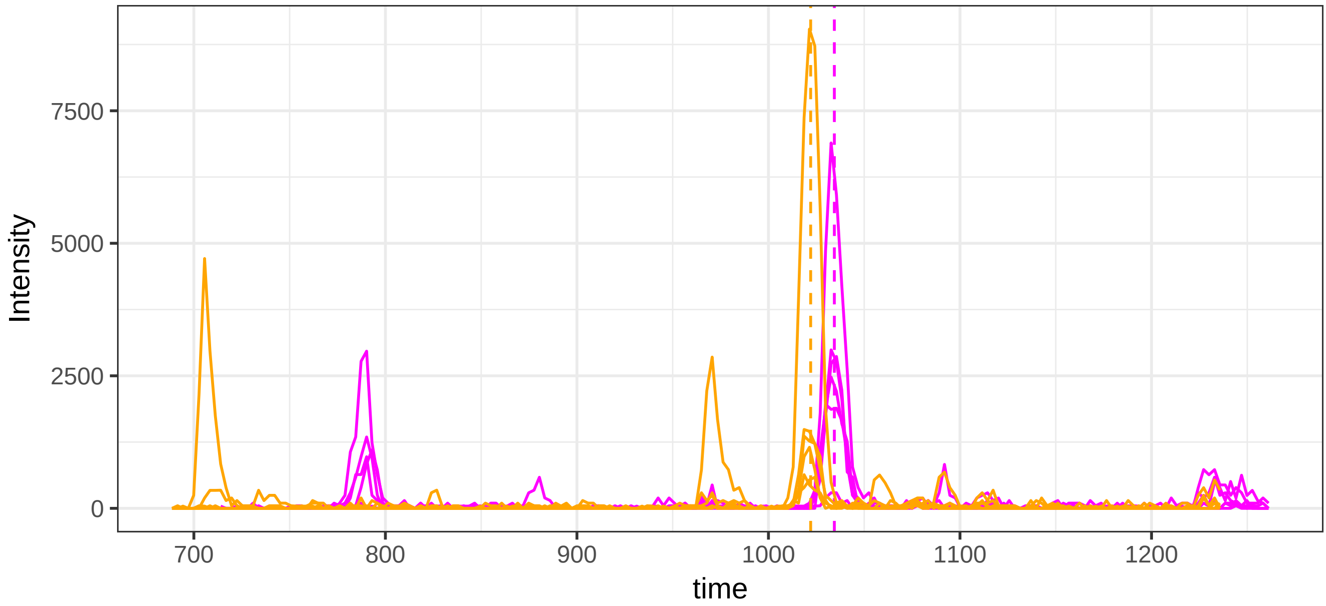


run23: 43285_LGPLVEQGR/2 (magenta)  AND 119020_STGSWSTLK/2

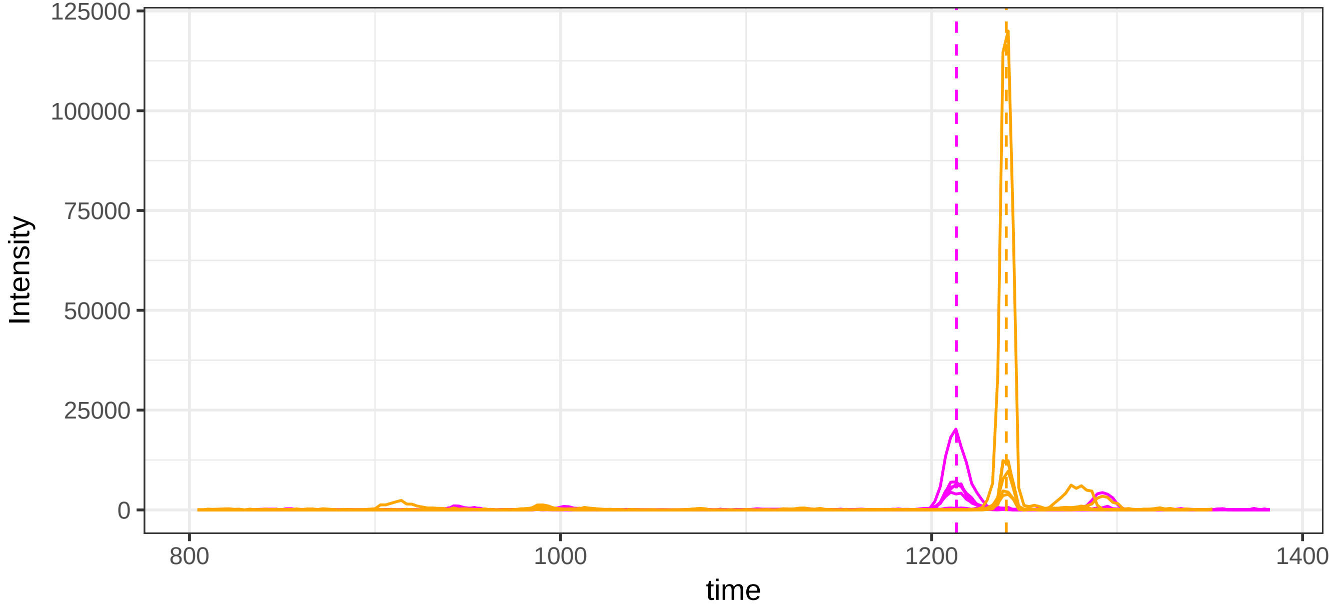

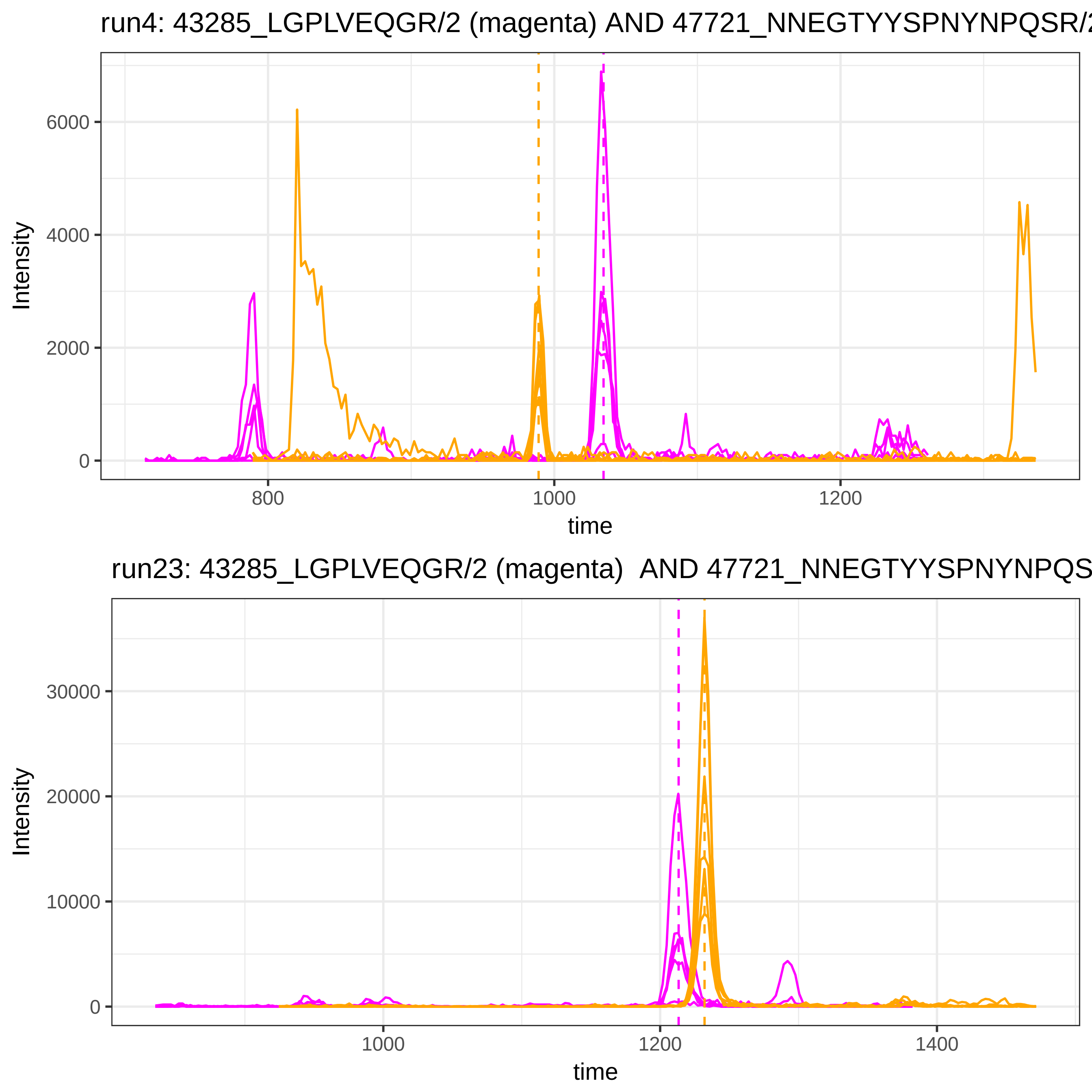

run4: 43285_LGPLVEQGR/2 (magenta) AND 47721_NNEGTYYSPNYNPQSR/2
Intensity
6000
4000
2000
0
800
1000
1200
time
run23: 43285_LGPLVEQGR/2 (magenta) AND 47721_NNEGTYYSPNYNPQS
Intensity
30000
20000
10000
0
1000
1200
1400
time

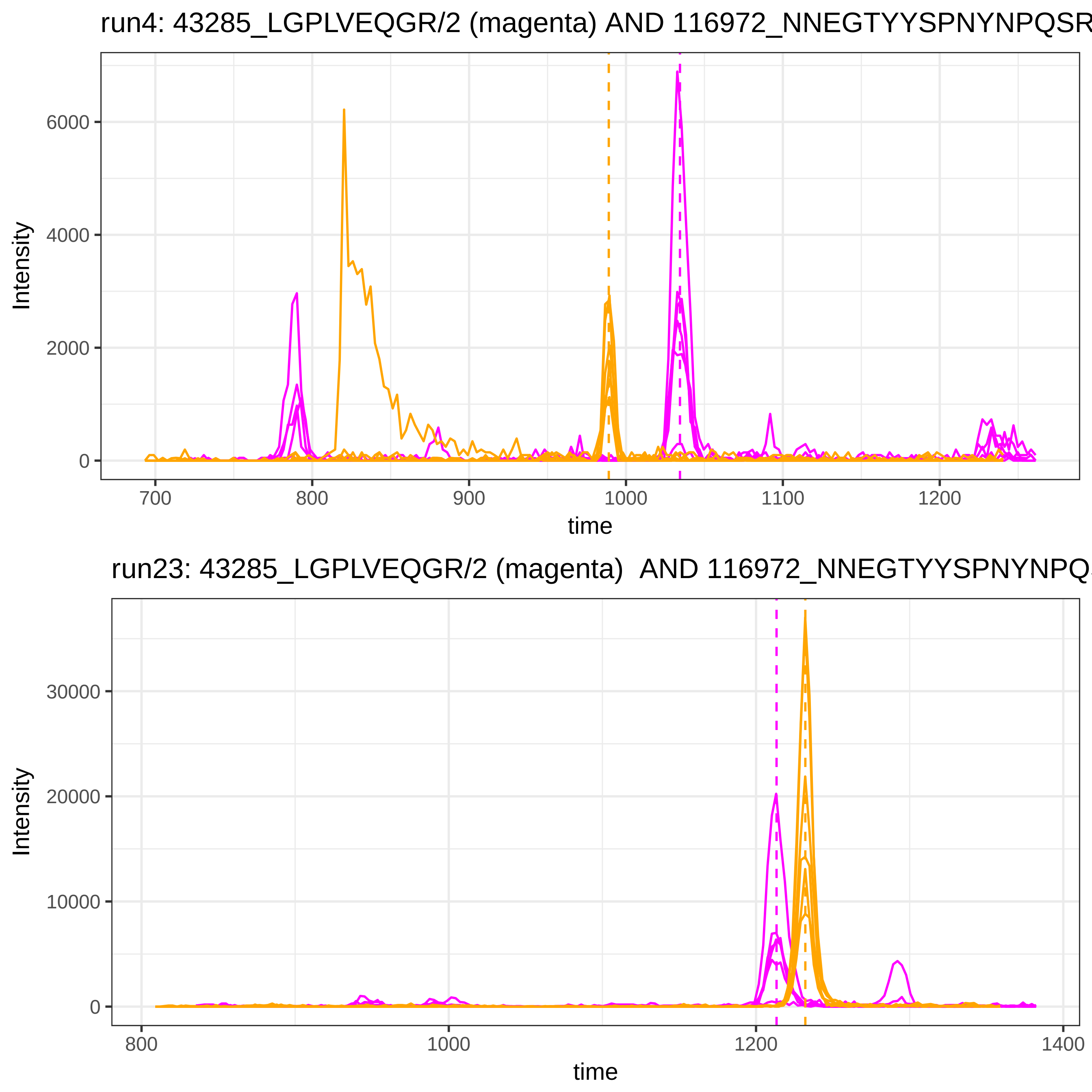

run4: 43285_LGPLVEQGR/2 (magenta) AND 116972_NNEGTYYSPNYNPQSR
Intensity
6000
4000
2000
0
700
800
900
1000
1100
1200
time
run23: 43285_LGPLVEQGR/2 (magenta) AND 116972_NNEGTYYSPNYNPQ
Intensity
30000
20000
10000
0
800
1000
1200
1400
time

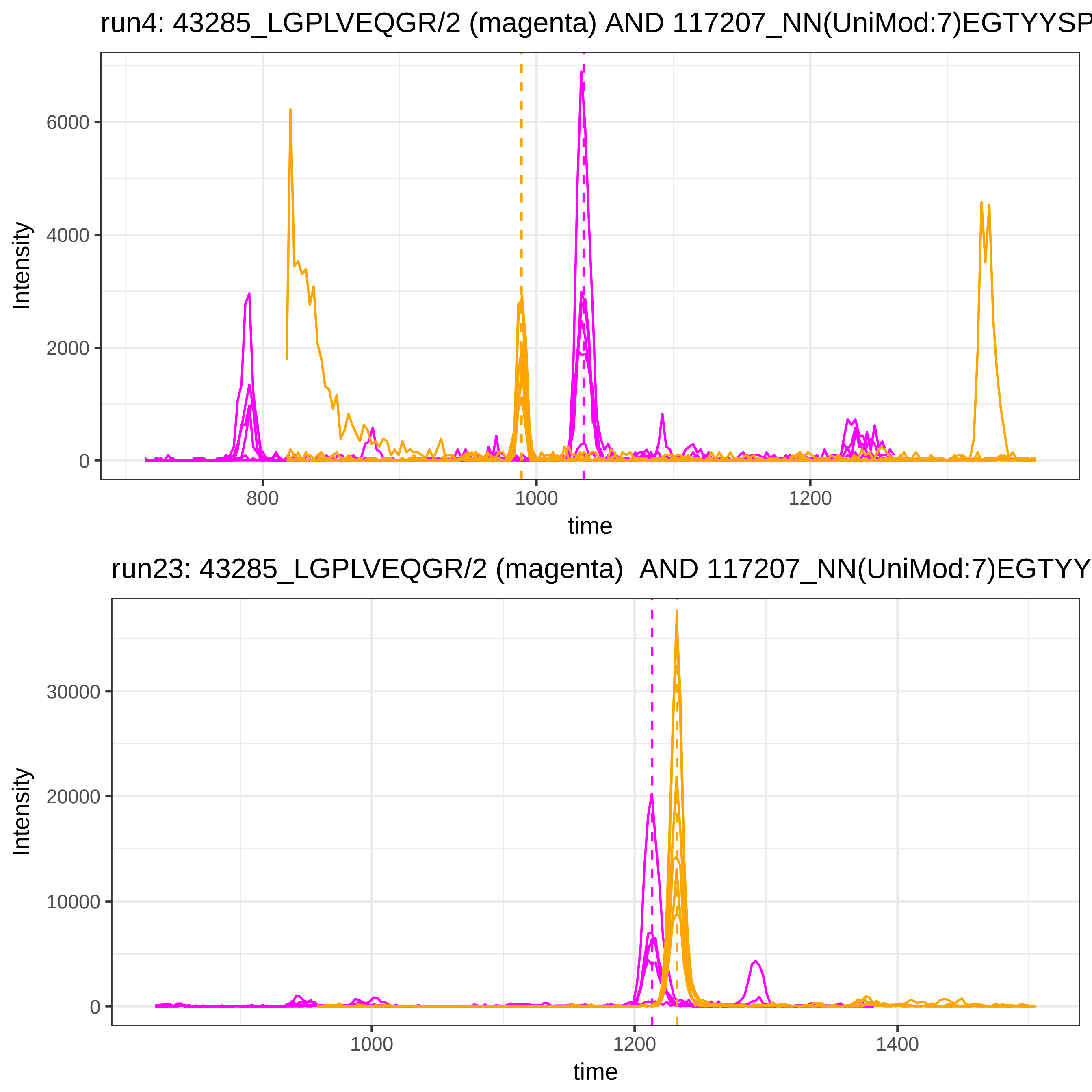

run4: 43285_LGPLVEQGR/2 (magenta) AND 117207_NN(UniMod:7)EGTYYSP
Intensity
6000
4000
2000
0
800
1000
1200
time
run23: 43285_LGPLVEQGR/2 (magenta) AND 117207_NN(UniMod:7)EGTYY
Intensity
30000
20000
10000
0
1000
1200
1400
time

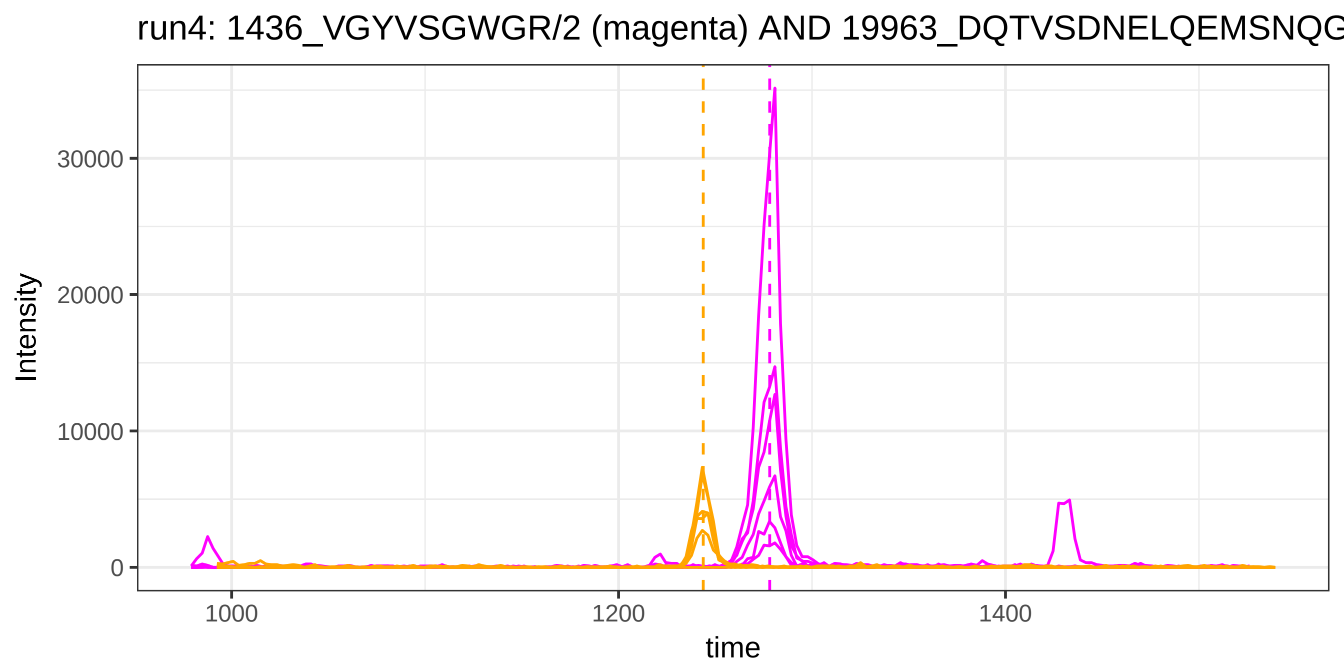
run4: 1436_VGYVSGWGR/2 (magenta) AND 19963_DQTVSDNELQEMSNQG
Intensity
30000
20000
10000
0
1000
1200
1400
time

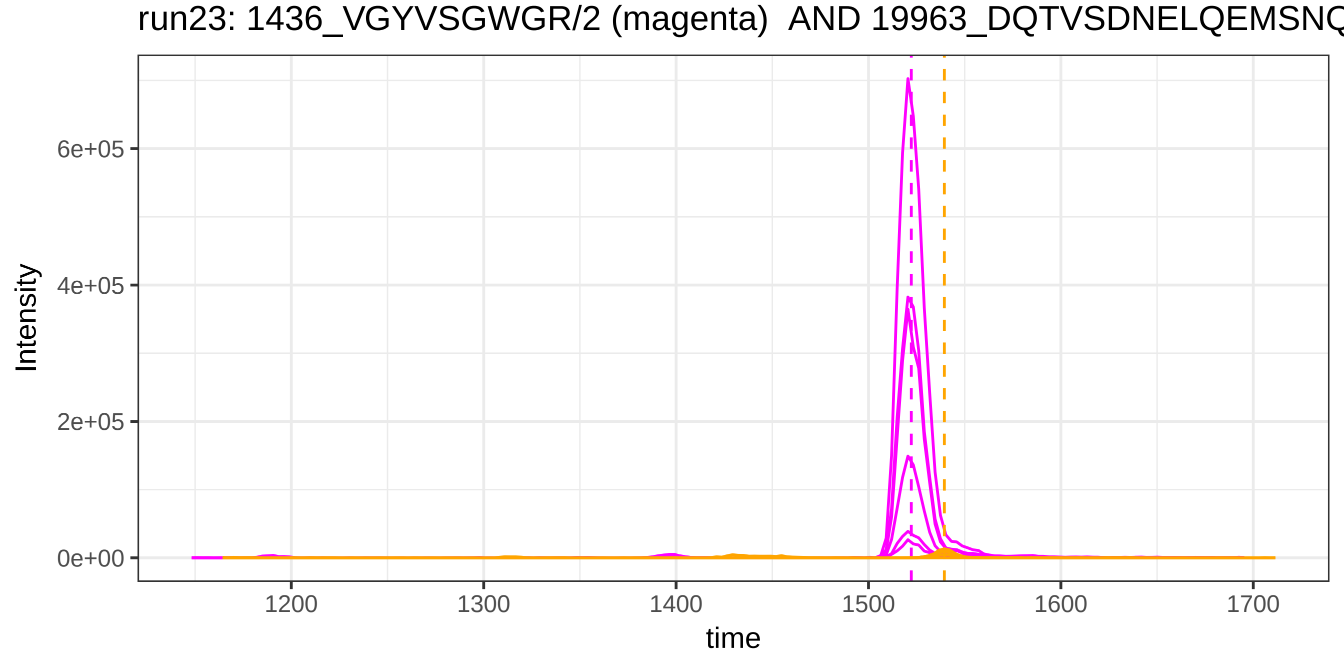
run23: 1436_VGYVSGWGR/2 (magenta) AND 19963_DQTVSDNELQEMSNQ
Intensity
6e+05
4e+05
2e+05
0e+00
1200
1300
1400
1500
1600
1700
time

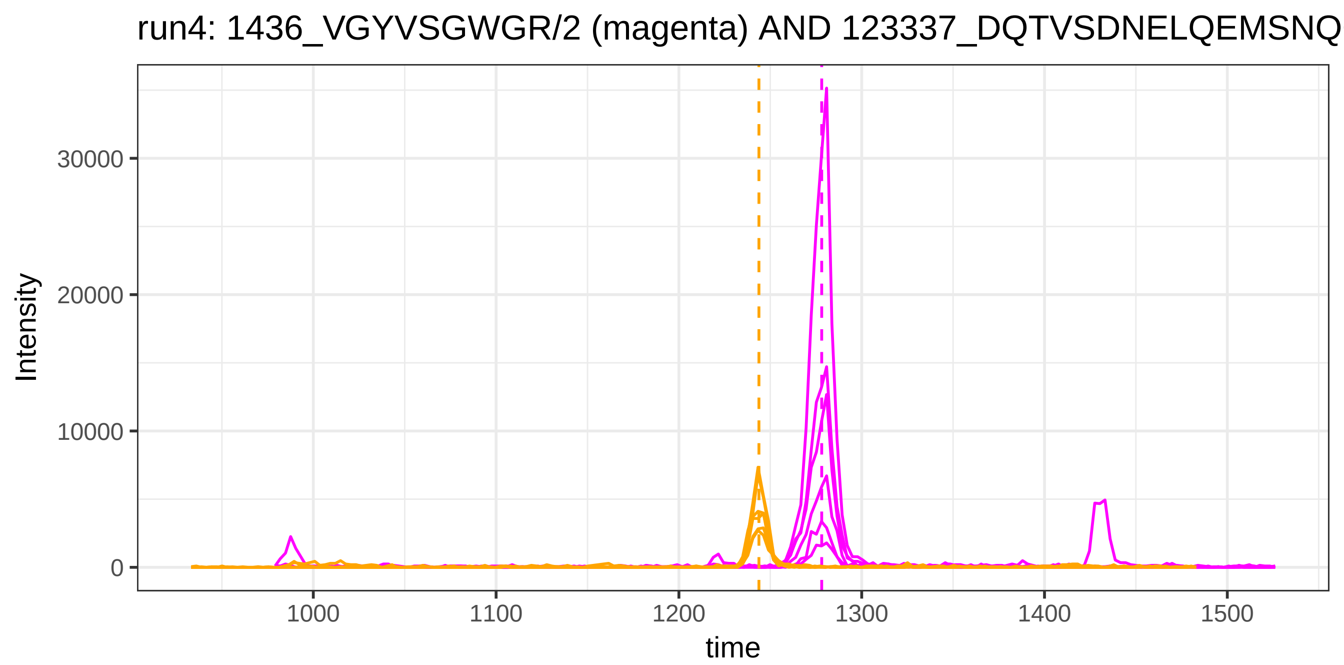

run4: 1436_VGYVSGWGR/2 (magenta) AND 123337_DQTVSDNELQEMSNQ
Intensity
30000
20000
10000
0
1000
1100
1200
1300
1400
1500
time


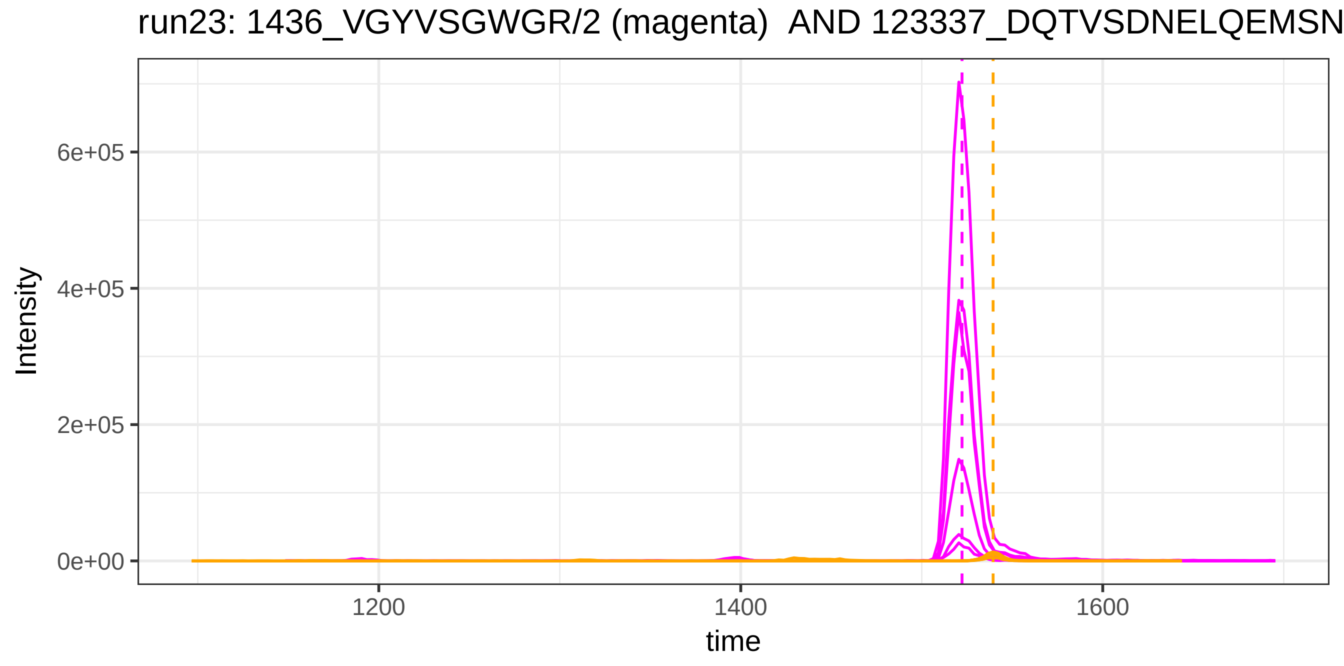

run23: 1436_VGYVSGWGR/2 (magenta) AND 123337_DQTVSDNELQEMSN
Intensity
6e+05
4e+05
2e+05
0e+00
1200
1400
1600
time

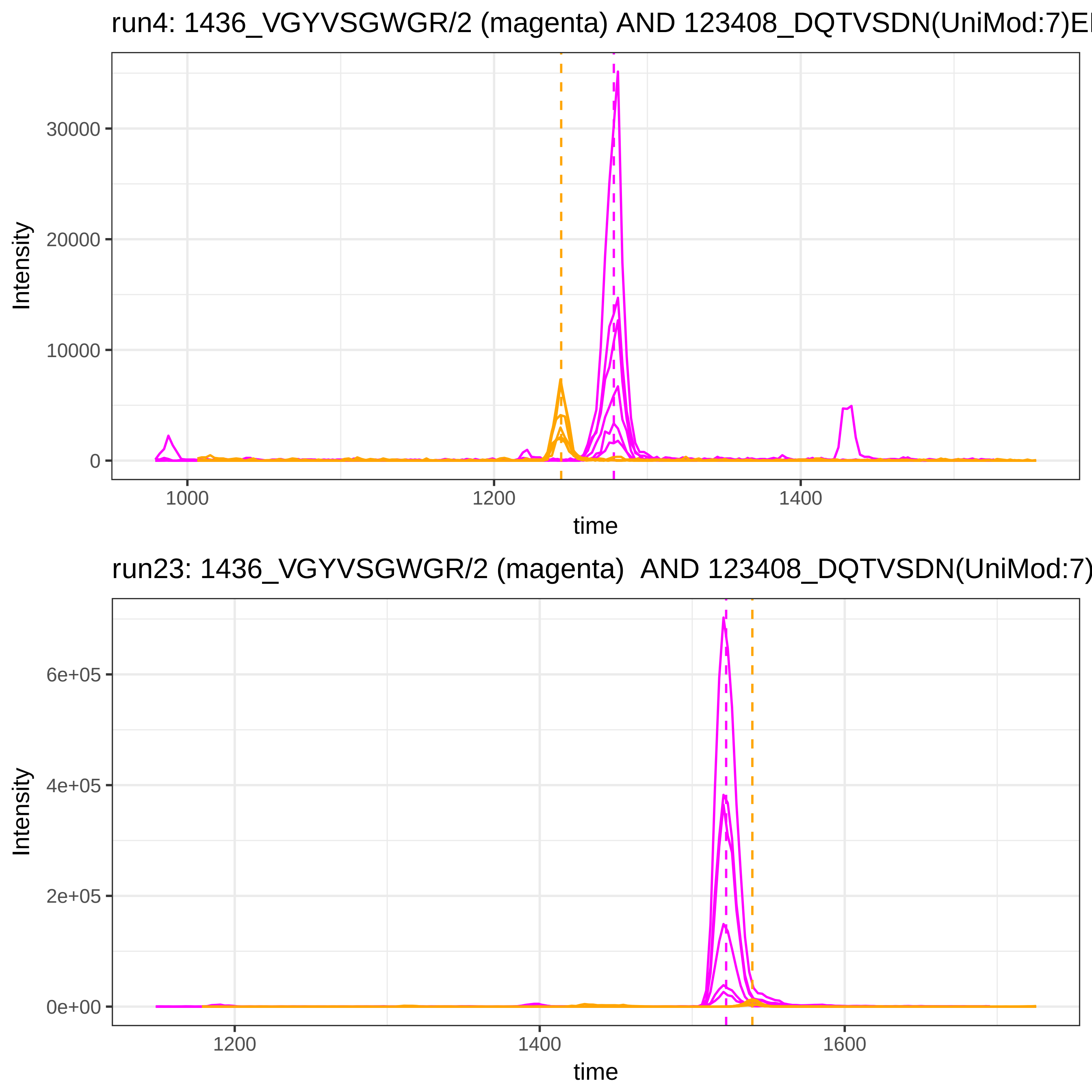
run4: 1436_VGYVSGWGR/2 (magenta) AND 123408_DQTVSDN(UniMod:7)E
30000
20000
10000
0
Intensity
1000
1200
1400
time
run23: 1436_VGYVSGWGR/2 (magenta) AND 123408_DQTVSDN(UniMod:7)
6e+05
4e+05
2e+05
0e+00
Intensity
1200
1400
1600
time

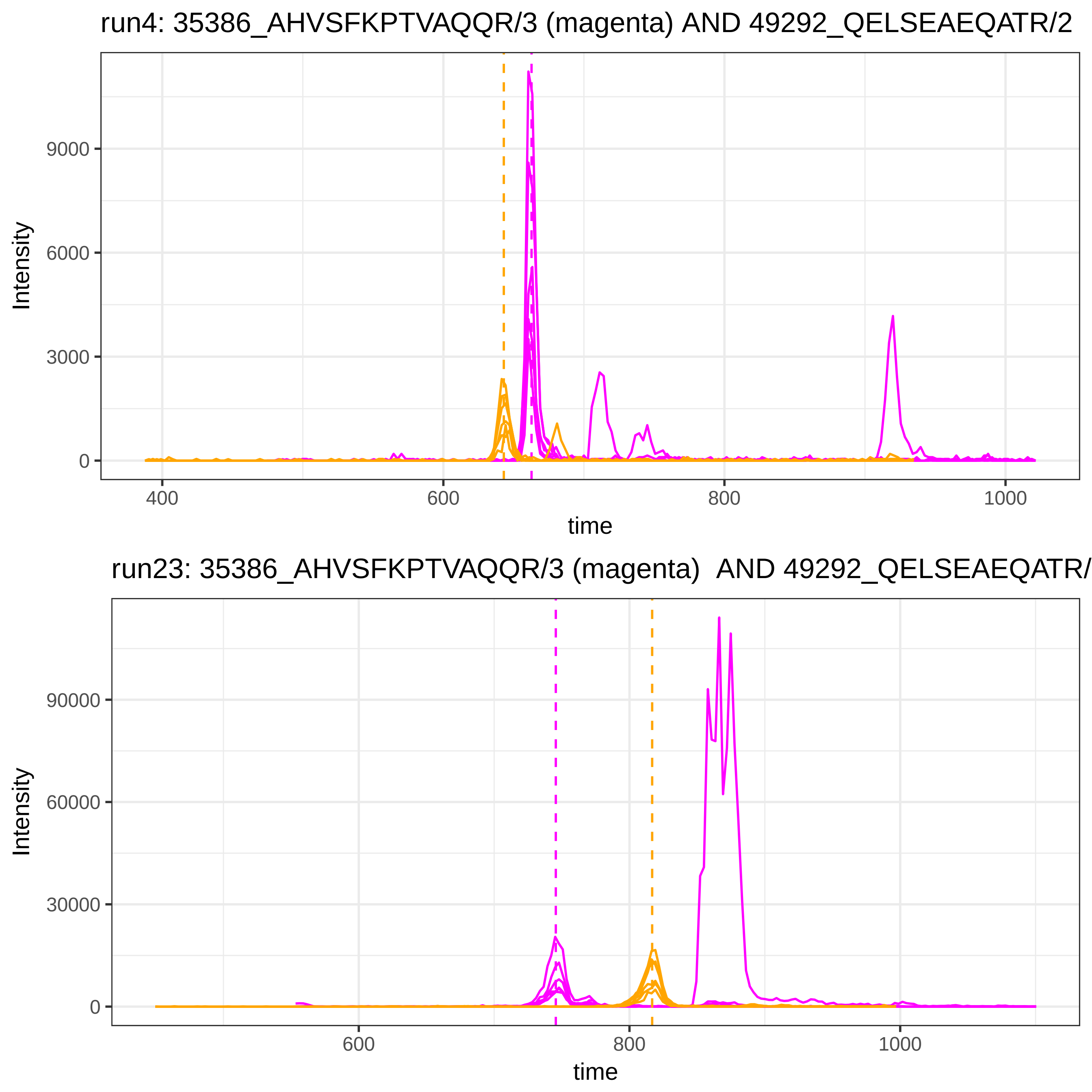
run4: 35386_AHVSFKPTVAQQR/3 (magenta) AND 49292_QELSEAEQATR/2
Intensity
0
3000
6000
9000
400
600
800
1000
time
run23: 35386_AHVSFKPTVAQQR/3 (magenta) AND 49292_QELSEAEQATR/
Intensity
0
30000
60000
90000
600
800
1000
time

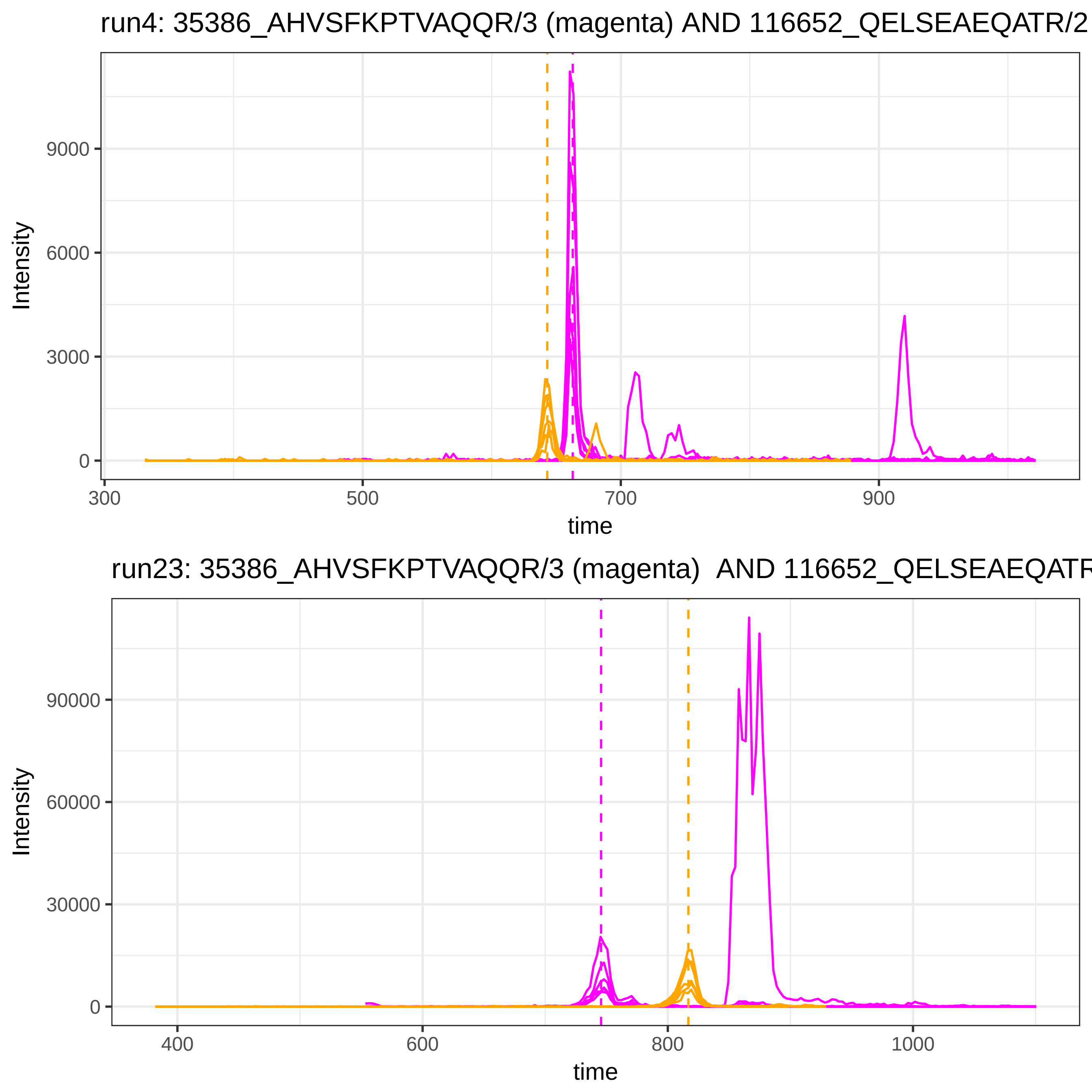
run4: 35386_AHVSFKPTVAQQR/3 (magenta) AND 116652_QELSEAEQATR/2
Intensity
9000
6000
3000
0
300
500
700
900
time
run23: 35386_AHVSFKPTVAQQR/3 (magenta) AND 116652_QELSEAEQATR
Intensity
90000
60000
30000
0
400
600
800
1000
time

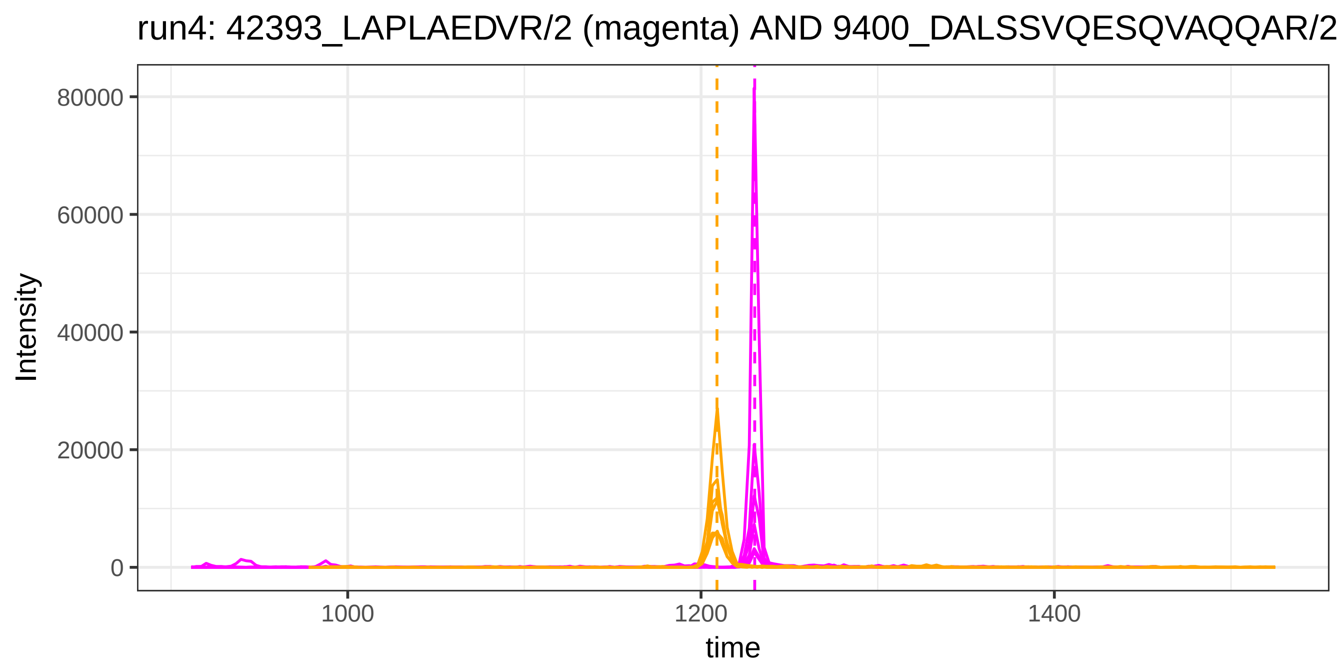

run4: 42393_LAPLAEDVR/2 (magenta) AND 9400_DALSSVQESQVAQQAR/2
Intensity
80000
60000
40000
20000
0
1000
1200
1400
time


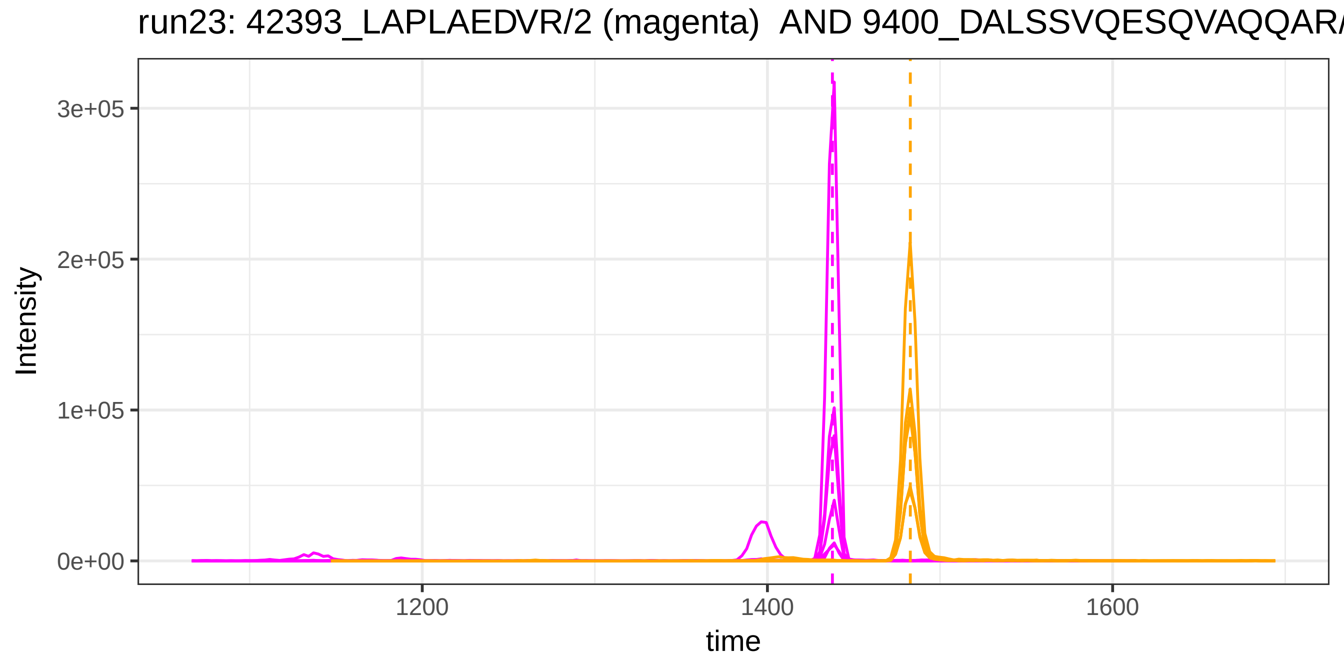

run23: 42393_LAPLAEDVR/2 (magenta) AND 9400_DALSSVQESQVAQQAR/
Intensity
3e+05
2e+05
1e+05
0e+00
1200
1400
1600
time

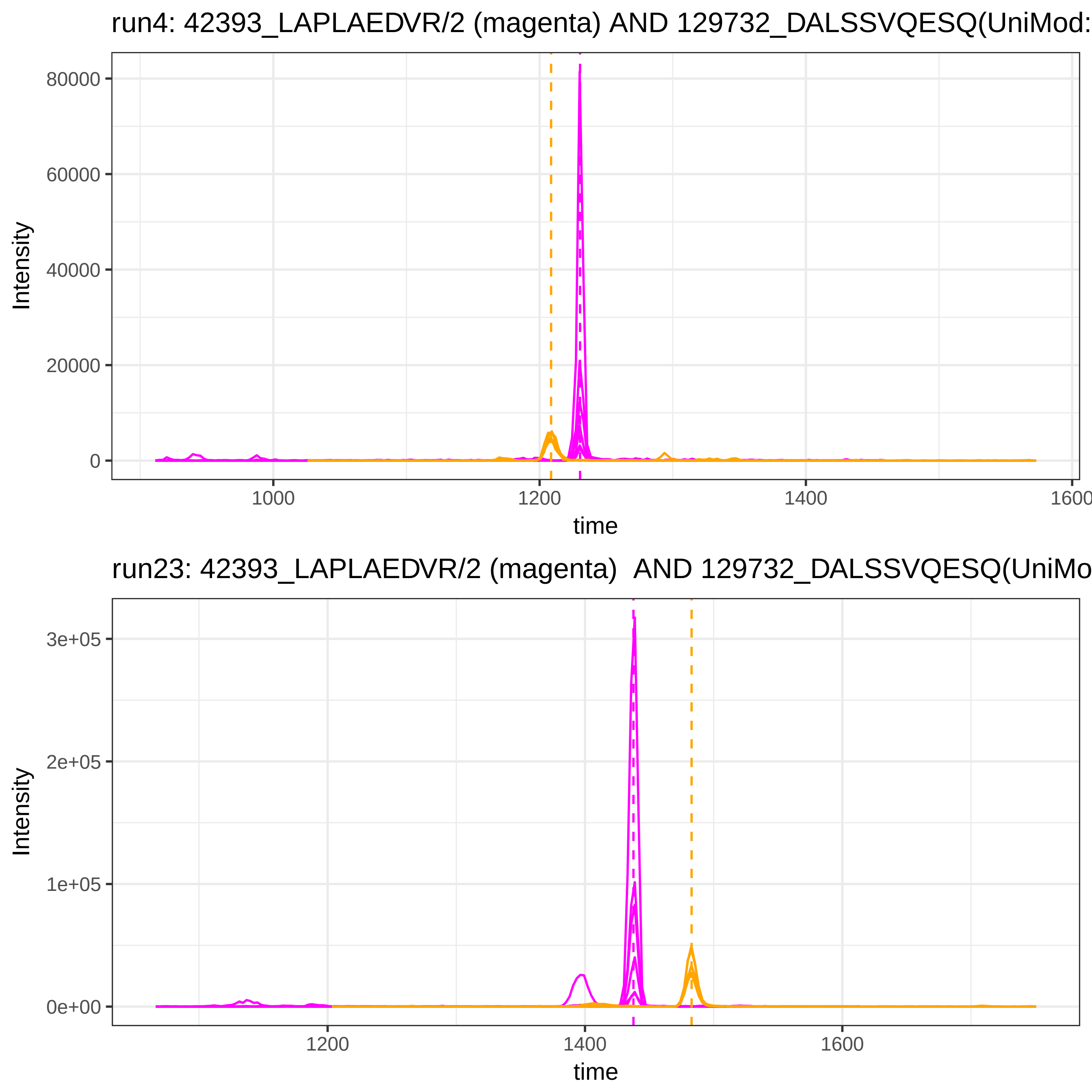
run4: 42393_LAPLAEDVR/2 (magenta) AND 129732_DALSSVQESQ(UniMod:
Intensity
80000
60000
40000
20000
0
1000
1200
1400
1600
time
run23: 42393_LAPLAEDVR/2 (magenta)  AND 129732_DALSSVQESQ(UniMo
Intensity
3e+05
2e+05
1e+05
0e+00
1200
1400
1600
time

run4: 42393_LAPLAEDVR/2 (magenta) AND 120401_NSPLDEENLTQENQDR

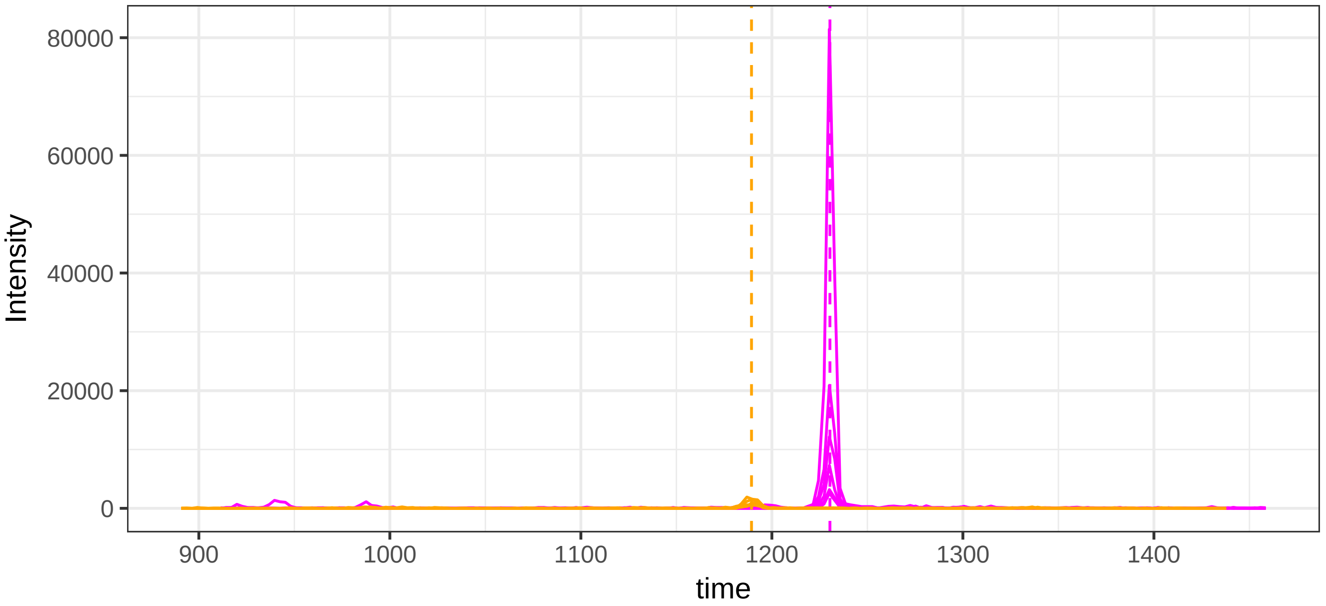


run23: 42393_LAPLAEDVR/2 (magenta) AND 120401_NSPLDEENLTQENQD

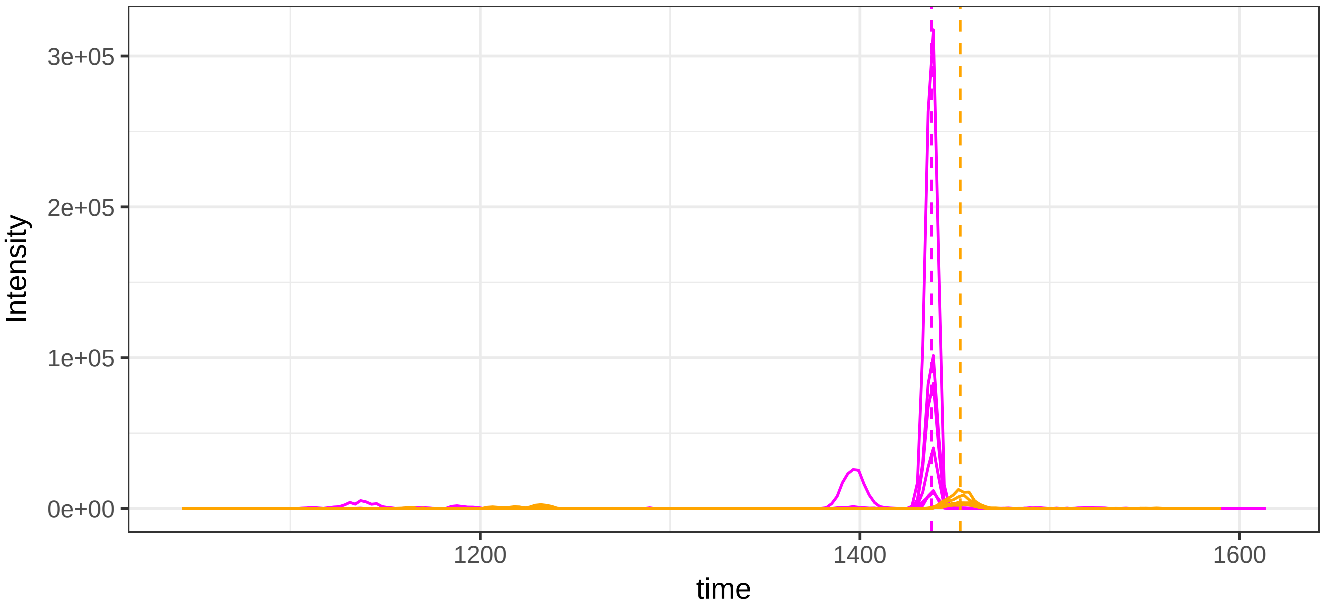

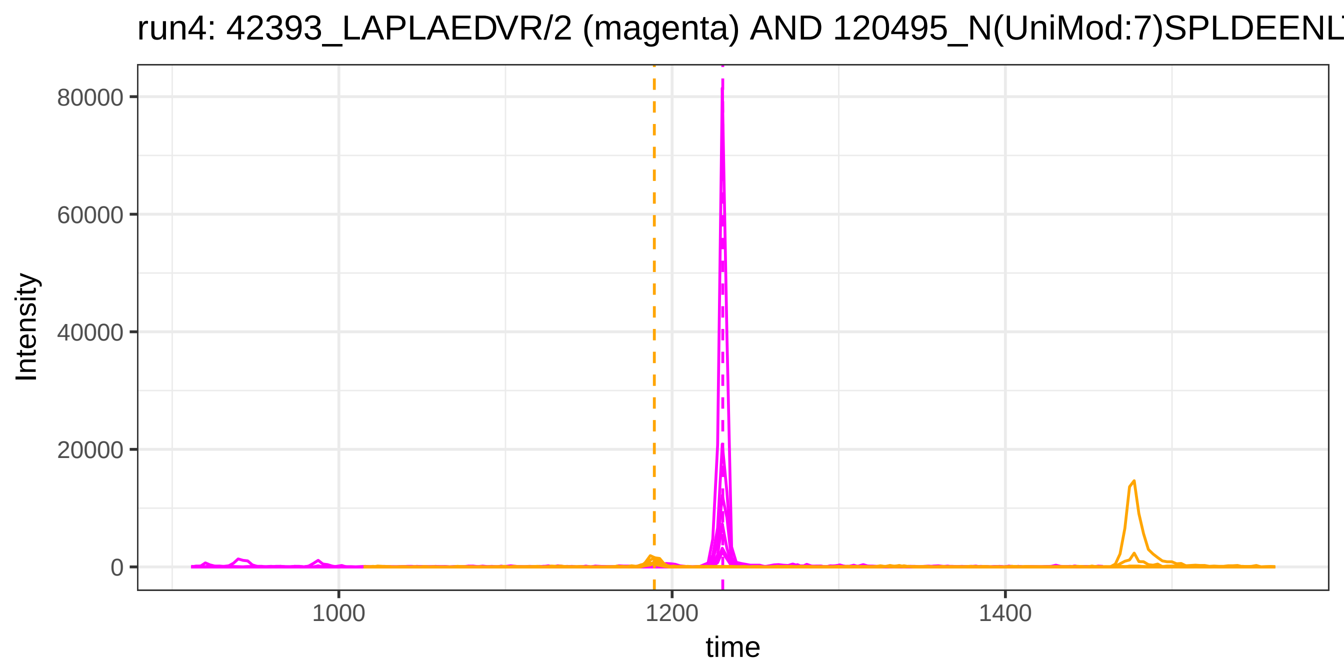
run4: 42393_LAPLAEDVR/2 (magenta) AND 120495_N(UniMod:7)SPLDEENL
80000
60000
40000
20000
0
Intensity
1000
1200
1400
time

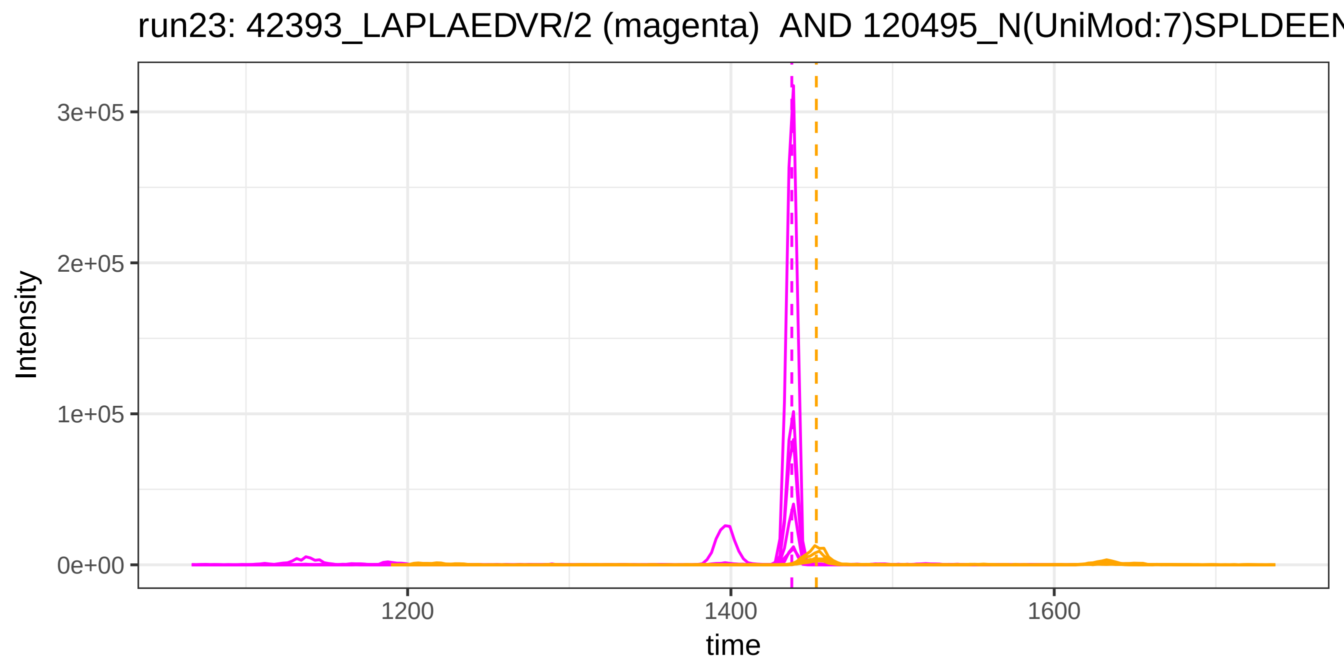
run23: 42393_LAPLAEDVR/2 (magenta) AND 120495_N(UniMod:7)SPLDEEN
3e+05
2e+05
1e+05
0e+00
Intensity
1200
1400
1600
time

run4: 3842_LLIYGATSR/2 (magenta) AND 41559_ALTDMPQMR/2

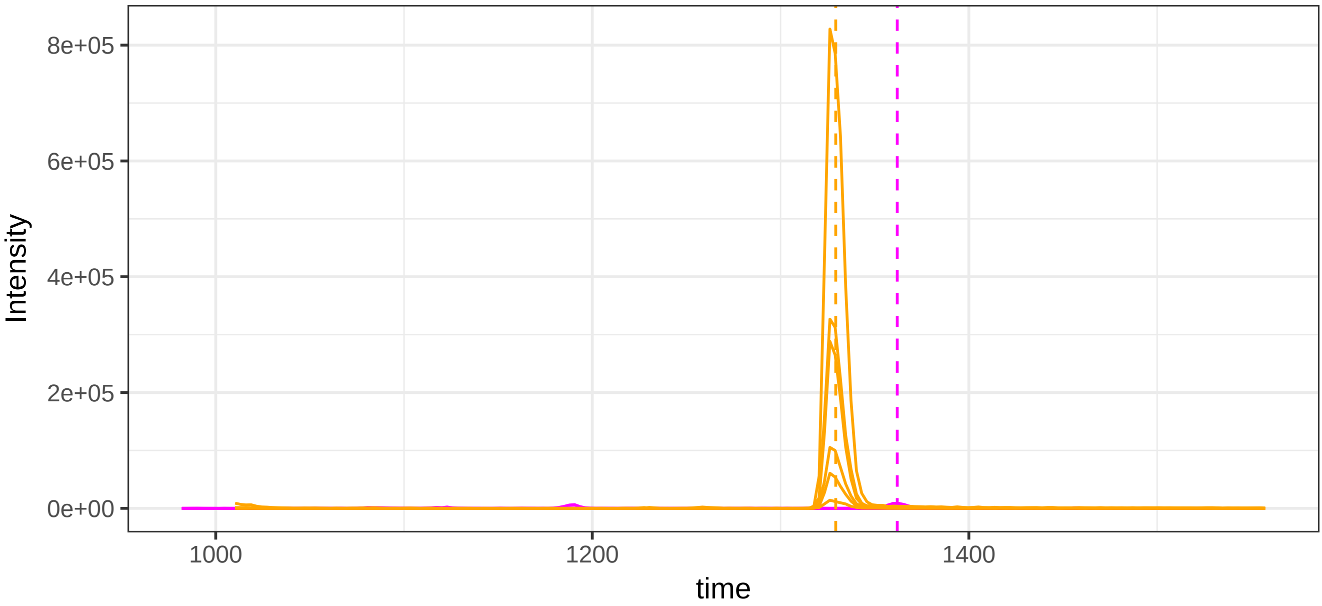


run23: 3842_LLIYGATSR/2 (magenta)  AND 41559_ALTDMPQMR/2

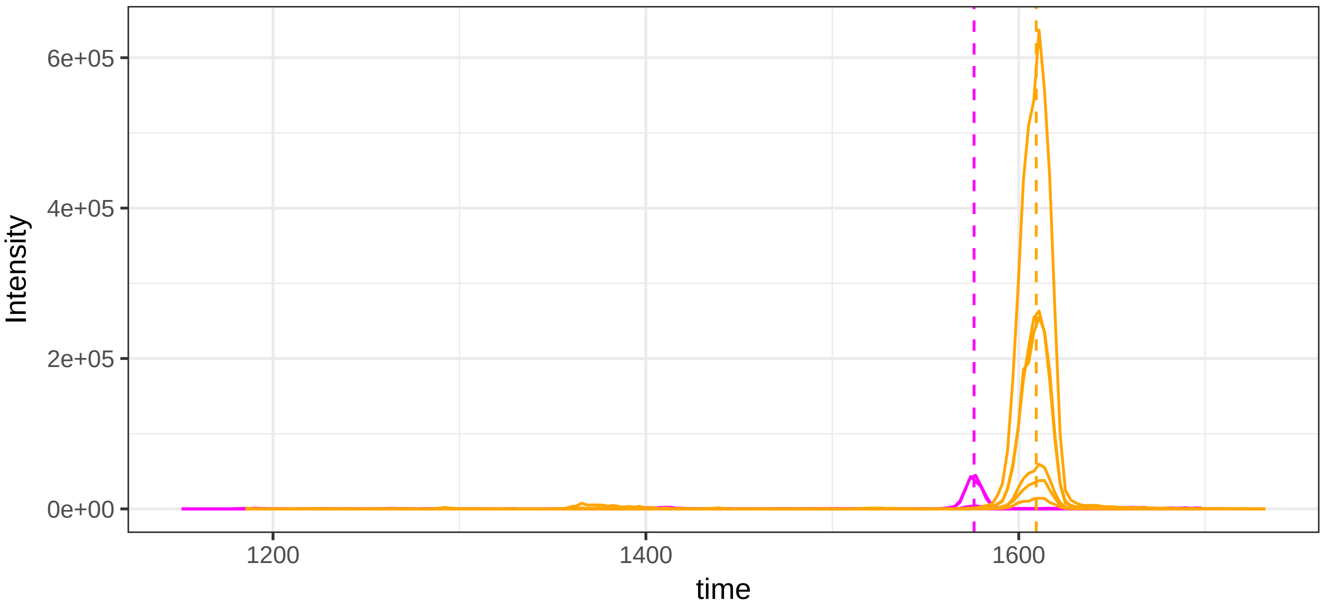

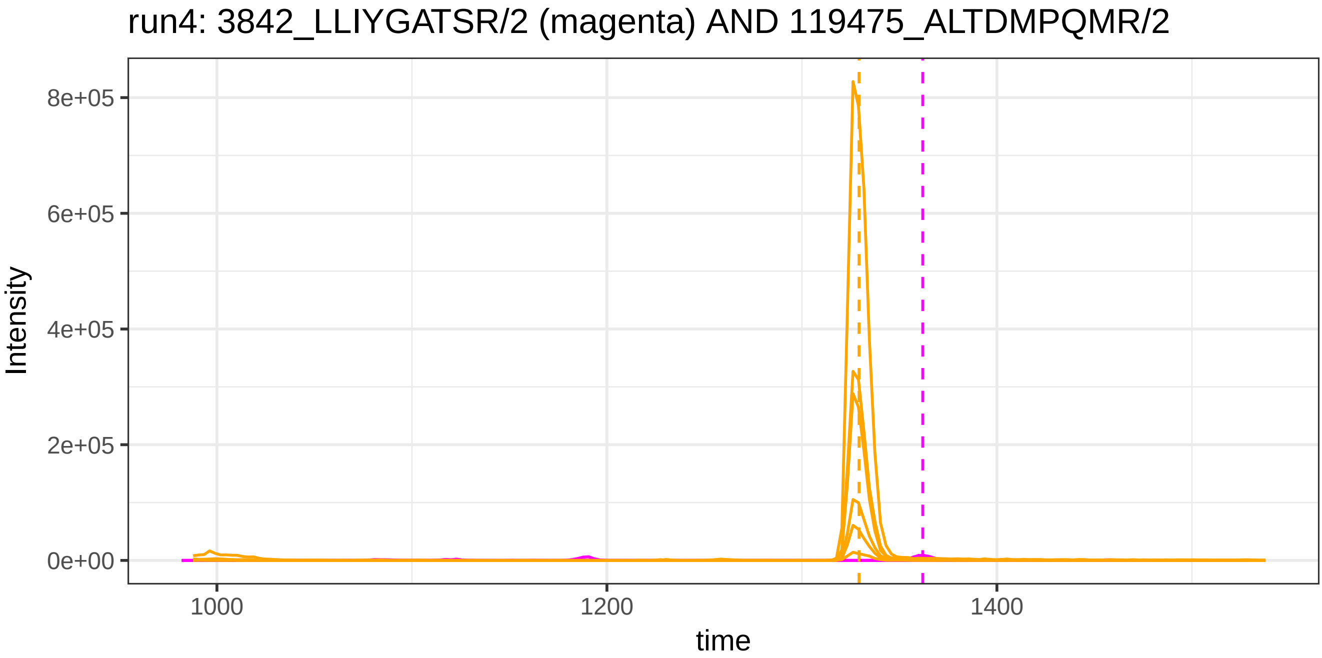
run4: 3842_LLIYGATSR/2 (magenta) AND 119475_ALTDMPQMR/2
Intensity
8e+05
6e+05
4e+05
2e+05
0e+00
1000
1200
1400
time

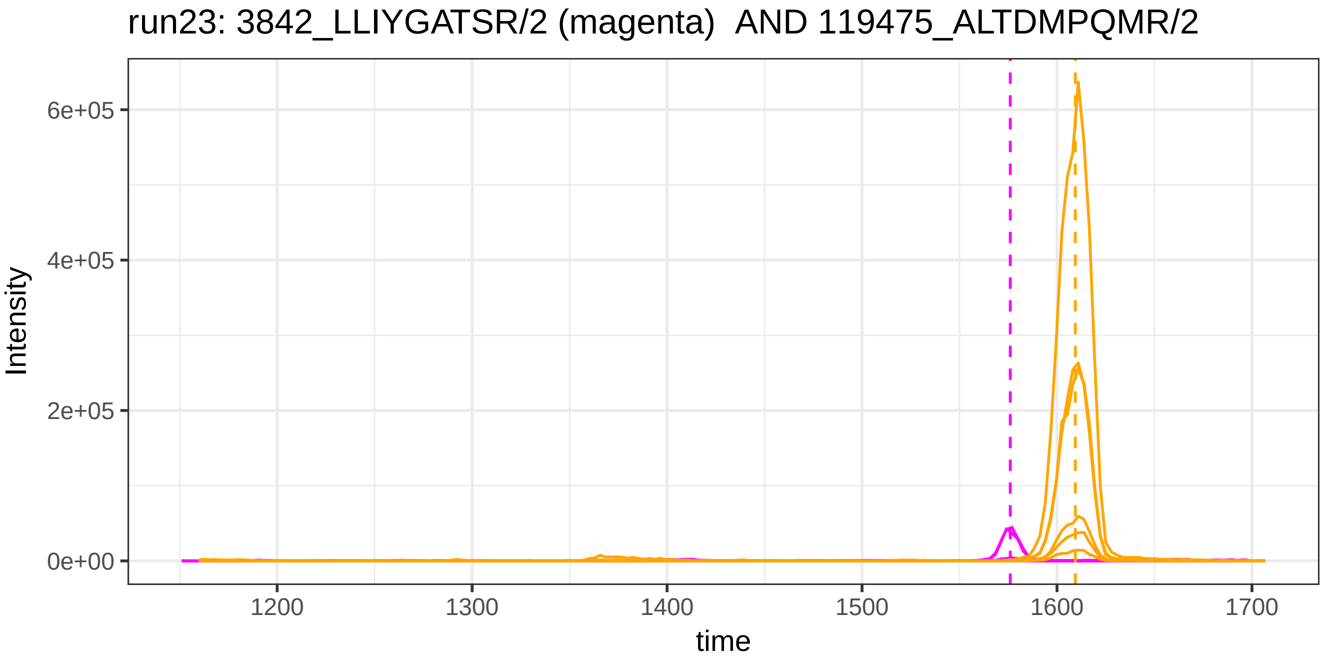
run23: 3842_LLIYGATSR/2 (magenta) AND 119475_ALTDMPQMR/2
Intensity
6e+05
4e+05
2e+05
0e+00
1200
1300
1400
1500
1600
1700
time

run4: 3842_LLIYGATSR/2 (magenta) AND 1030_VELEDWNGR/2

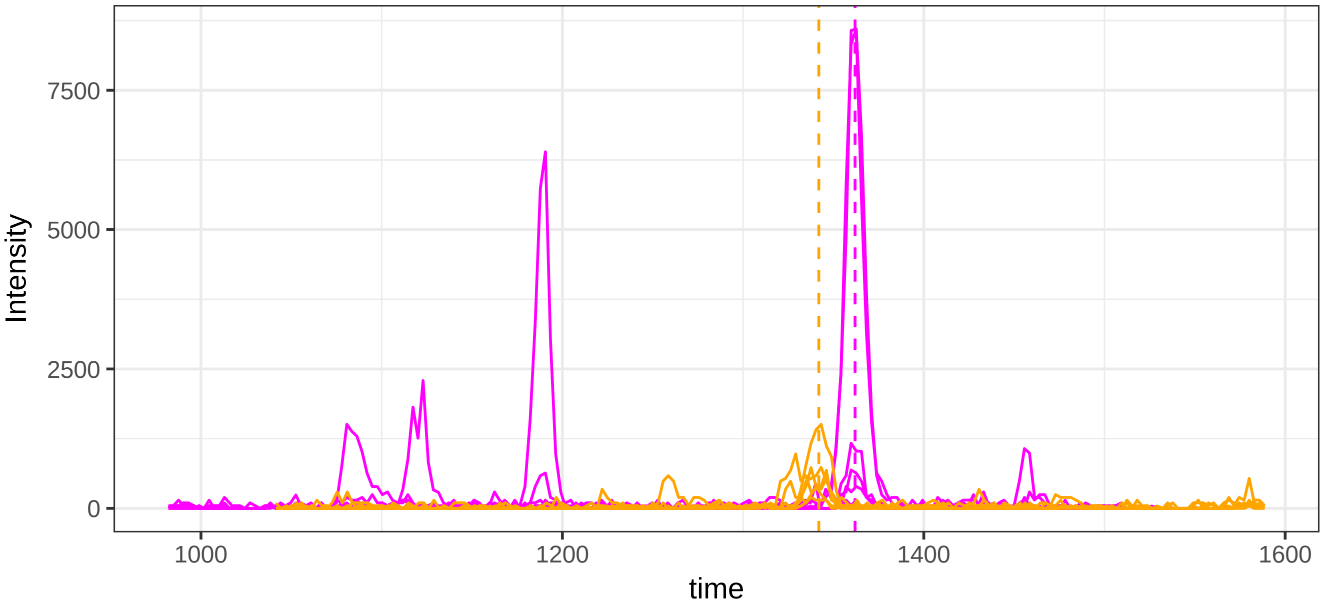


run23: 3842_LLIYGATSR/2 (magenta)  AND 1030_VELEDWNGR/2

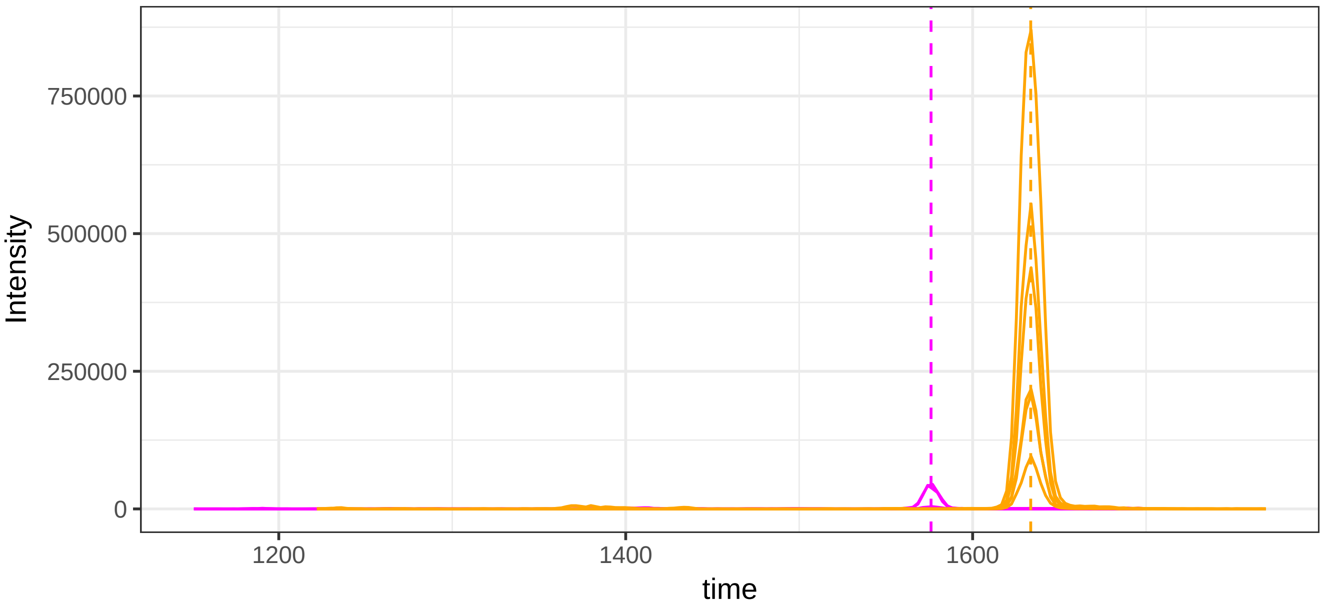

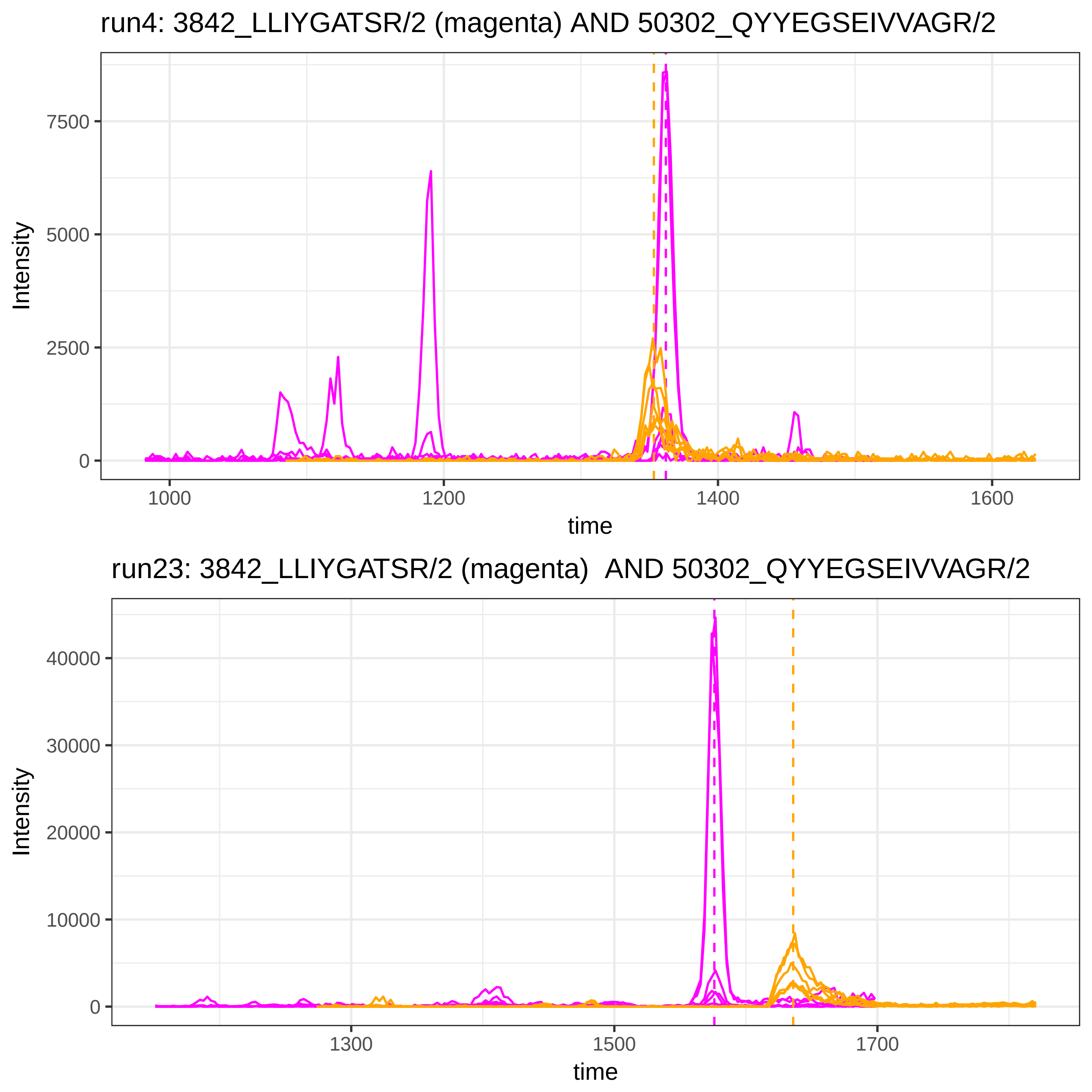

run4: 3842_LLIYGATSR/2 (magenta) AND 50302_QYYEGSEIVVAGR/2
Intensity
7500
5000
2500
0
1000
1200
1400
1600
time
run23: 3842_LLIYGATSR/2 (magenta) AND 50302_QYYEGSEIVVAGR/2
Intensity
40000
30000
20000
10000
0
1300
1500
1700
time

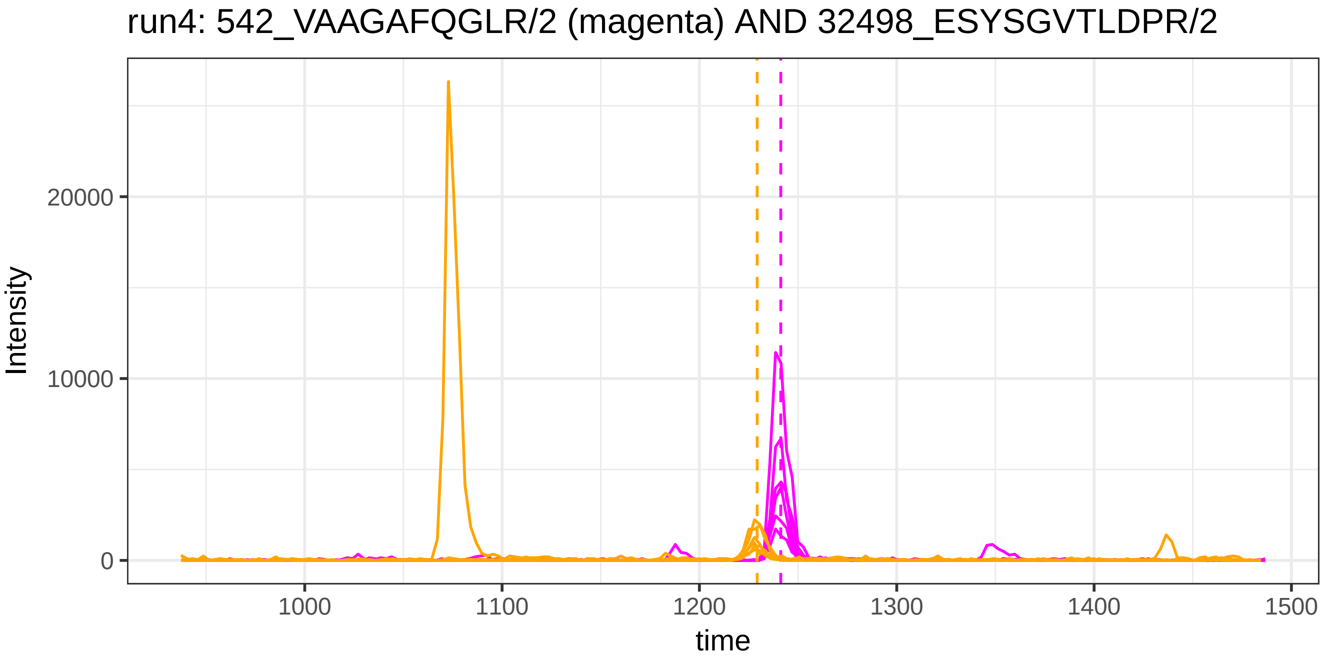
run4: 542_VAAGAFQGLR/2 (magenta) AND 32498_ESYSGVTLDPR/2
Intensity
20000
10000
0
1000
1100
1200
1300
1400
1500
time

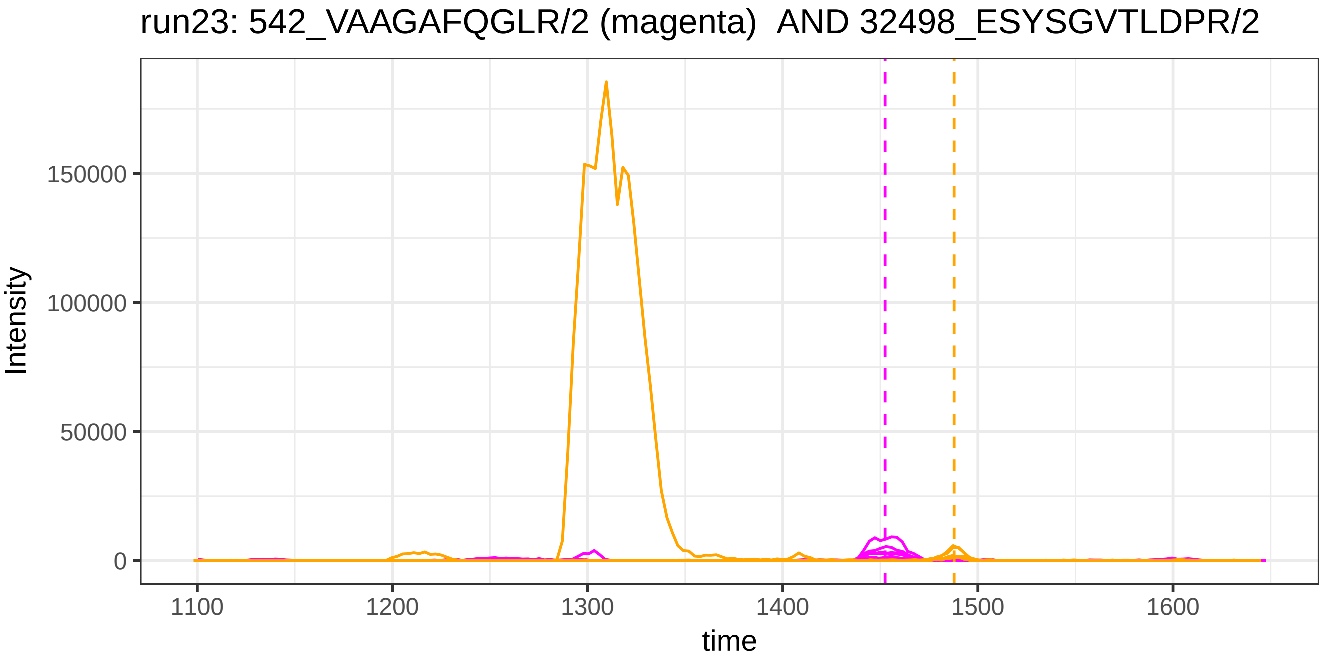
run23: 542_VAAGAFQGLR/2 (magenta)  AND 32498_ESYSGVTLDPR/2
Intensity
150000
100000
50000
0
1100
1200
1300
1400
1500
1600
time

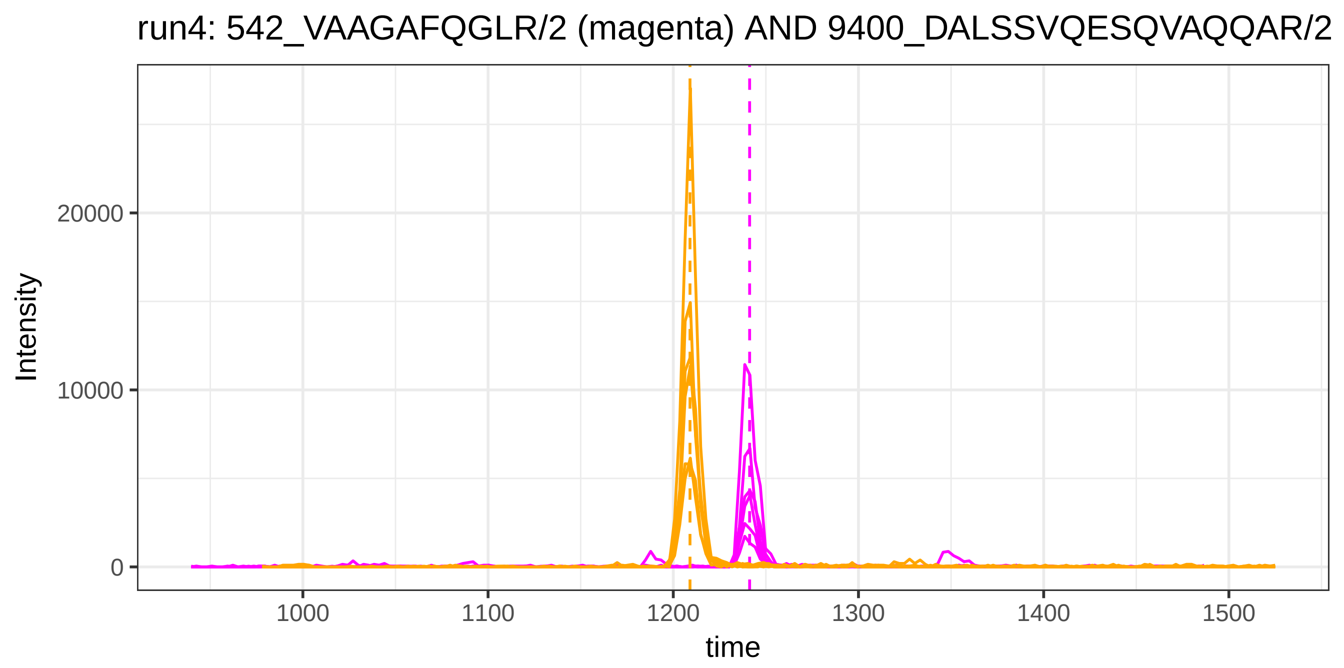
run4: 542_VAAGAFQGLR/2 (magenta) AND 9400_DALSSVQESQVAQQAR/2
Intensity
20000
10000
0
1000
1100
1200
1300
1400
1500
time

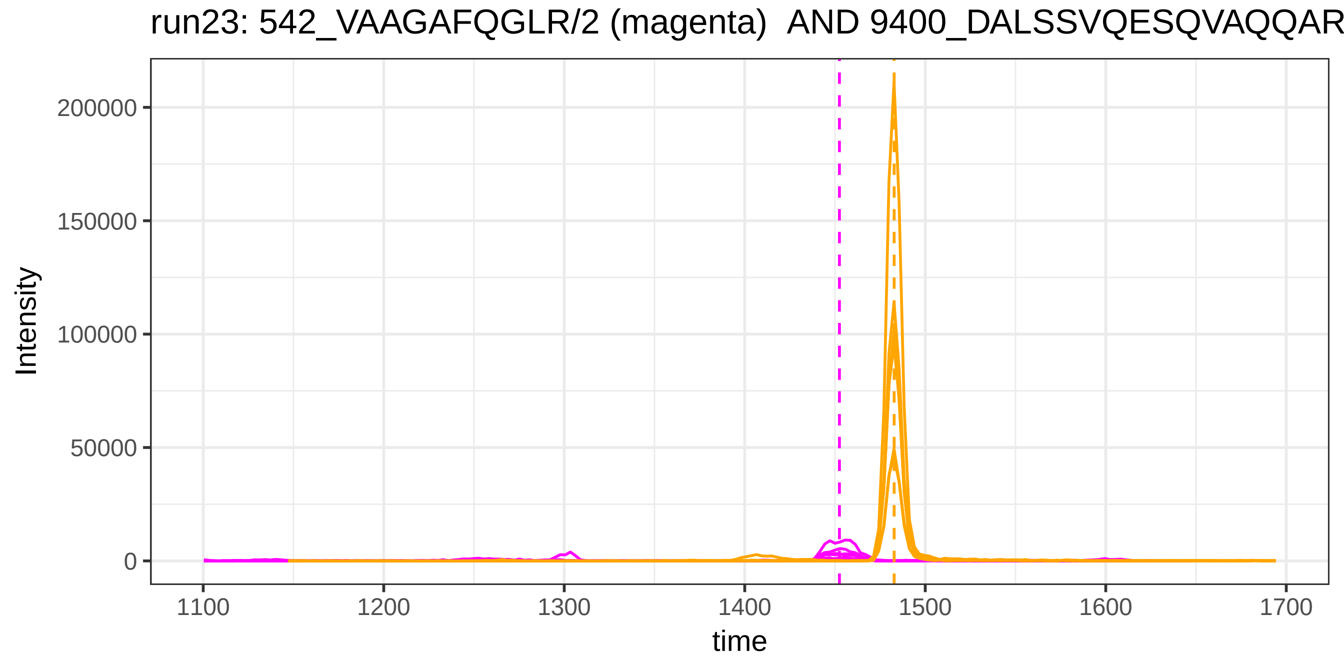
run23: 542_VAAGAFQGLR/2 (magenta) AND 9400_DALSSVQESQVAQQAR
Intensity
200000
150000
100000
50000
0
1100
1200
1300
1400
1500
1600
1700
time

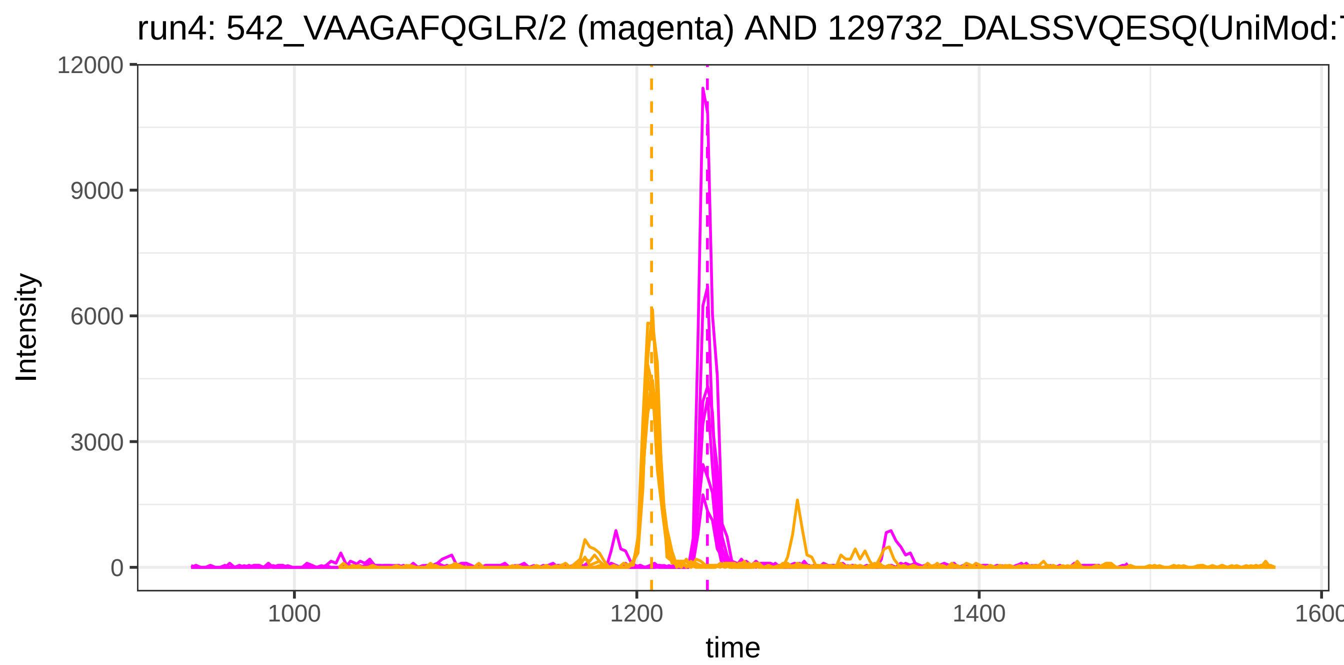
run4: 542_VAAGAFQGLR/2 (magenta) AND 129732_DALSSVQESQ(UniMod:
Intensity
12000
9000
6000
3000
0
1000
1200
1400
1600
time

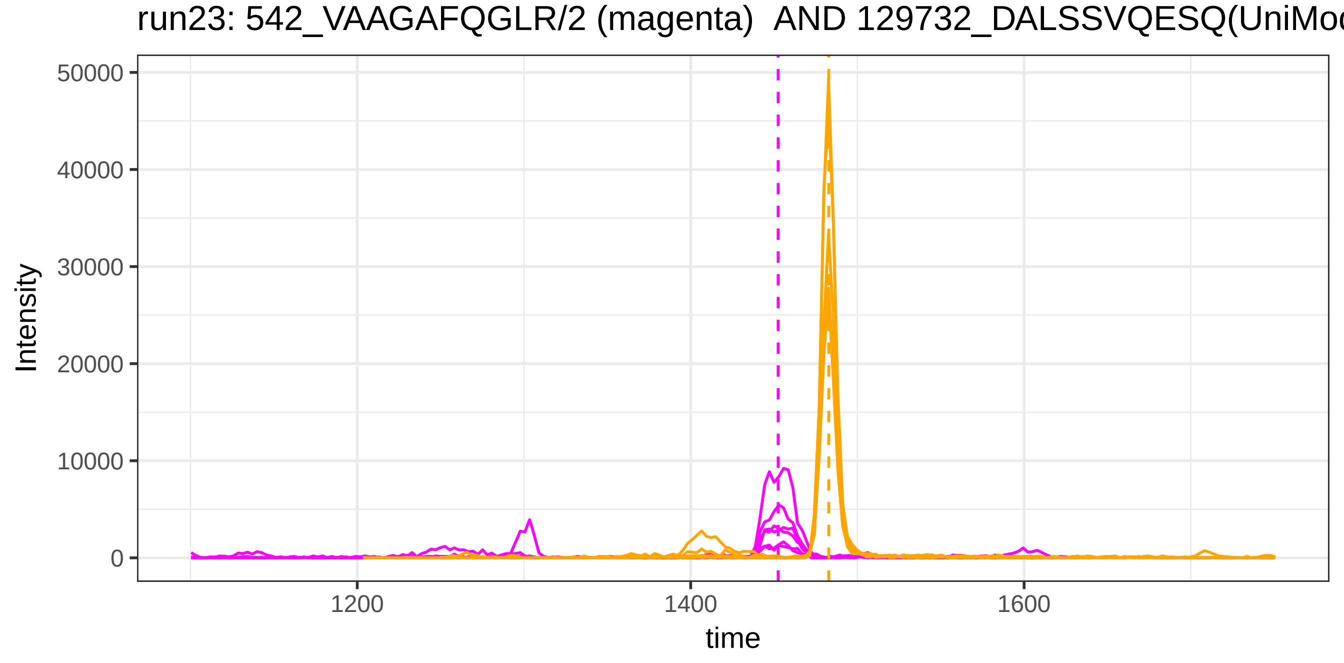
run23: 542_VAAGAFQGLR/2 (magenta)  AND 129732_DALSSVQESQ(UniMod
Intensity
50000
40000
30000
20000
10000
0
1200
1400
1600
time

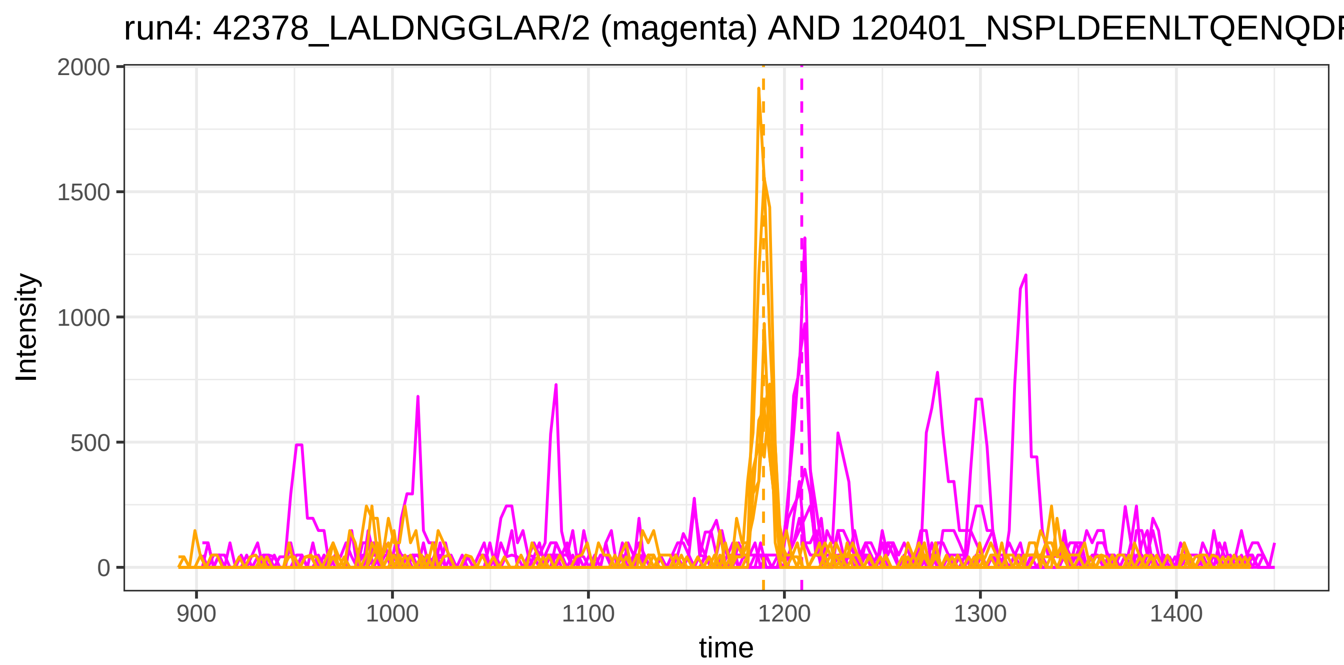
run4: 42378_LALDNGGLAR/2 (magenta) AND 120401_NSPLDEENLTQENQDF
Intensity
2000
1500
1000
500
0
900
1000
1100
1200
1300
1400
time

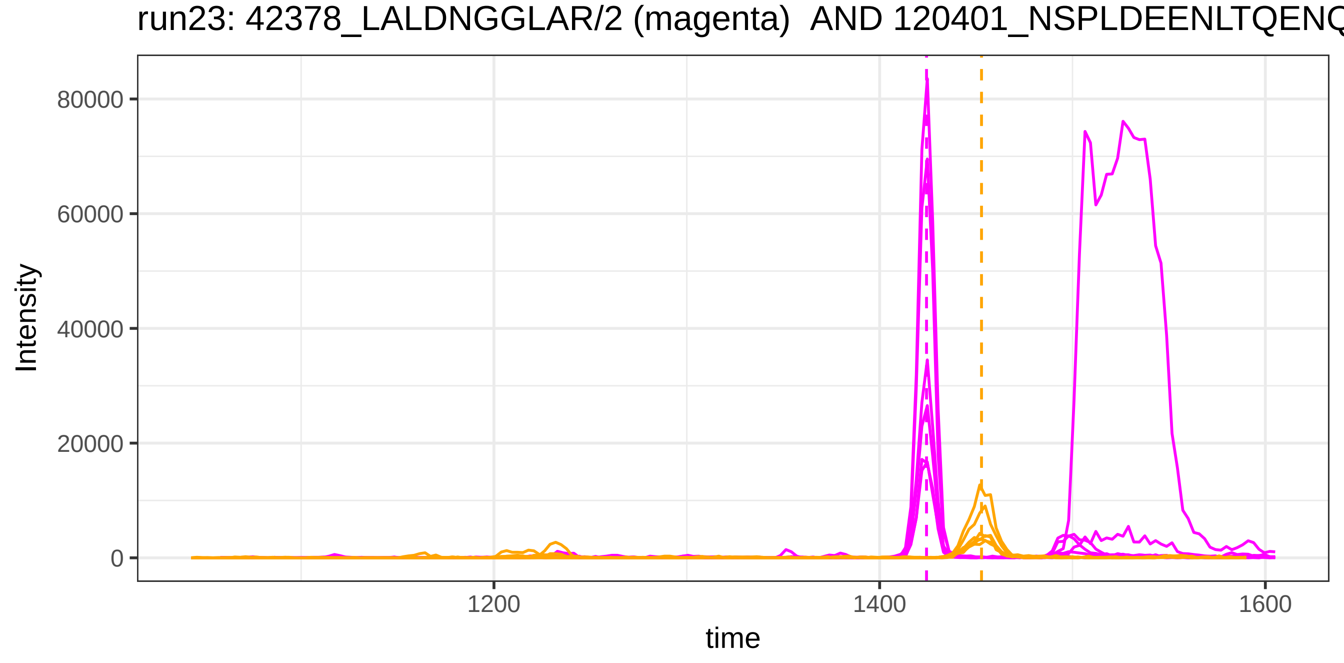
run23: 42378_LALDNGGLAR/2 (magenta) AND 120401_NSPLDEENLTQENQ
Intensity
80000
60000
40000
20000
0
1200
1400
1600
time

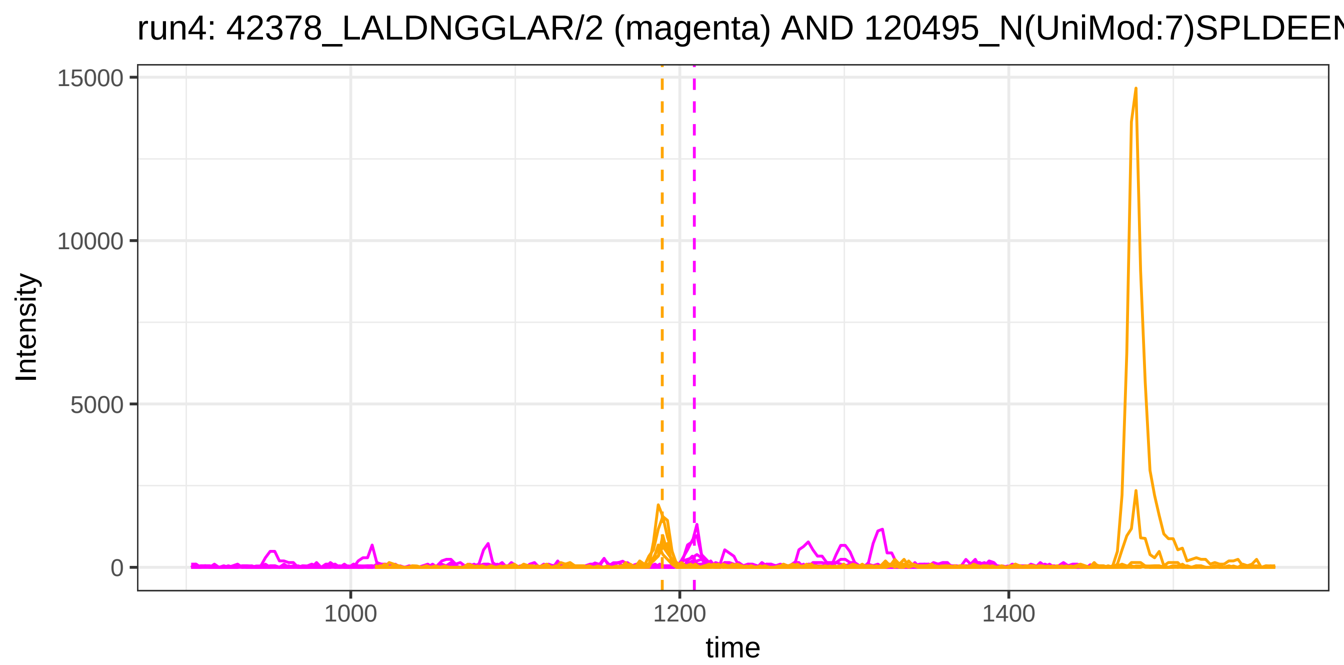

run4: 42378_LALDNGGLAR/2 (magenta) AND 120495_N(UniMod:7)SPLDEEN
15000
10000
5000
0
Intensity
1000
1200
1400
time


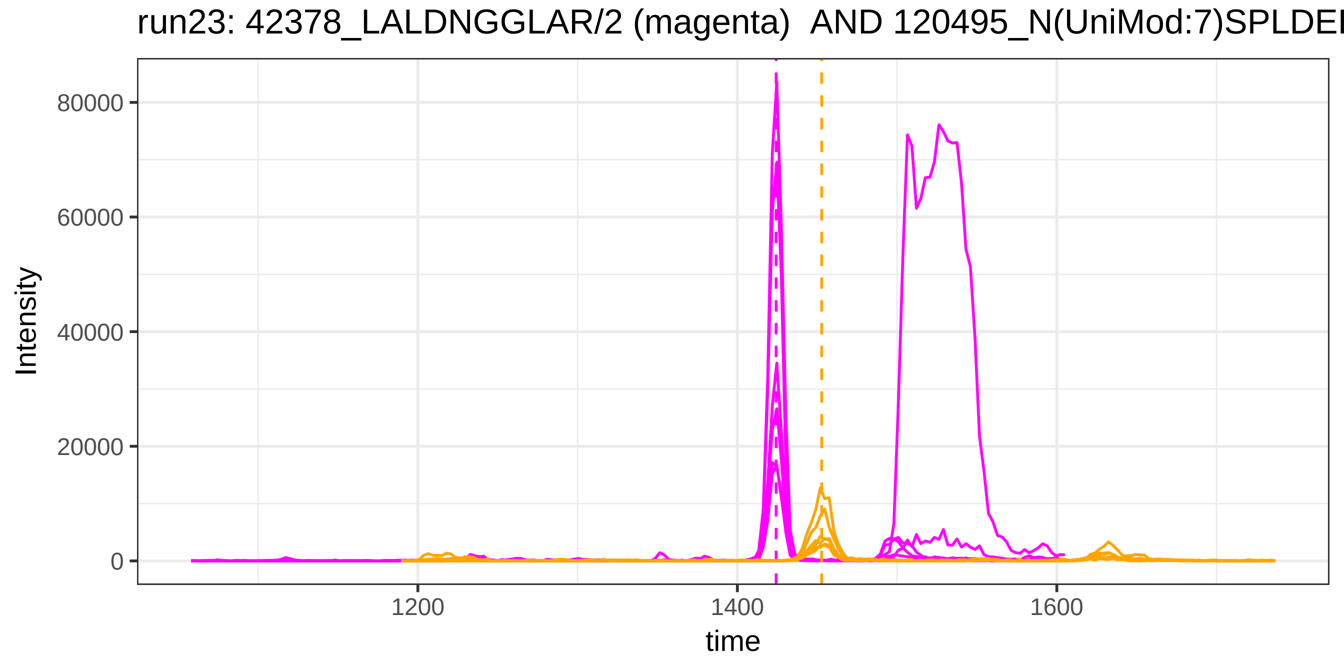

run23: 42378_LALDNGGLAR/2 (magenta) AND 120495_N(UniMod:7)SPLDEE
80000
60000
40000
20000
0
Intensity
1200
1400
1600
time

run4: 45875_MELERPGGNEITR/3 (magenta) AND 126974_EYKC(UniMod:4)K

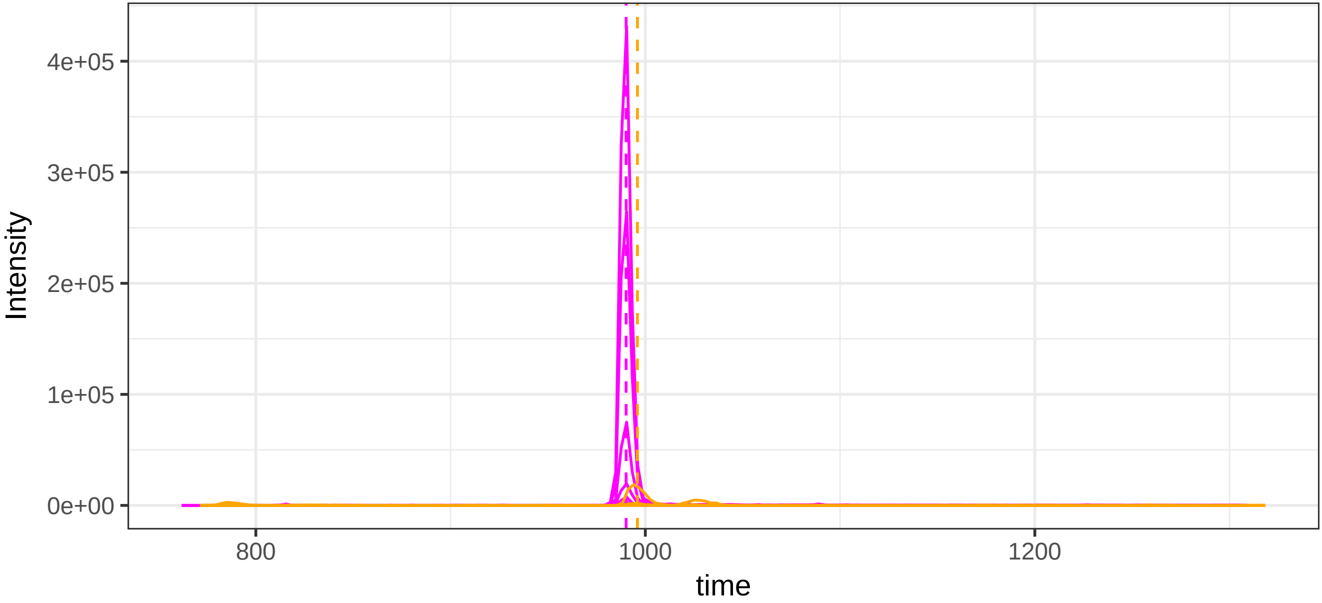


run23: 45875_MELERPGGNEITR/3 (magenta)  AND 126974_EYKC(UniMod:4

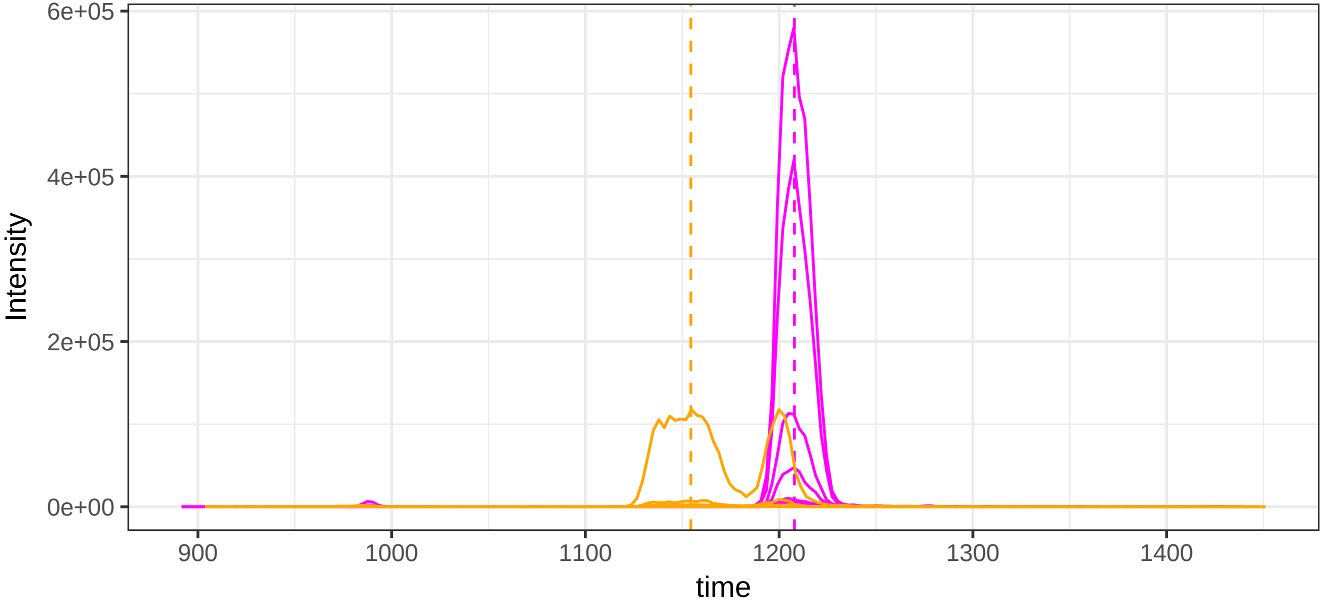

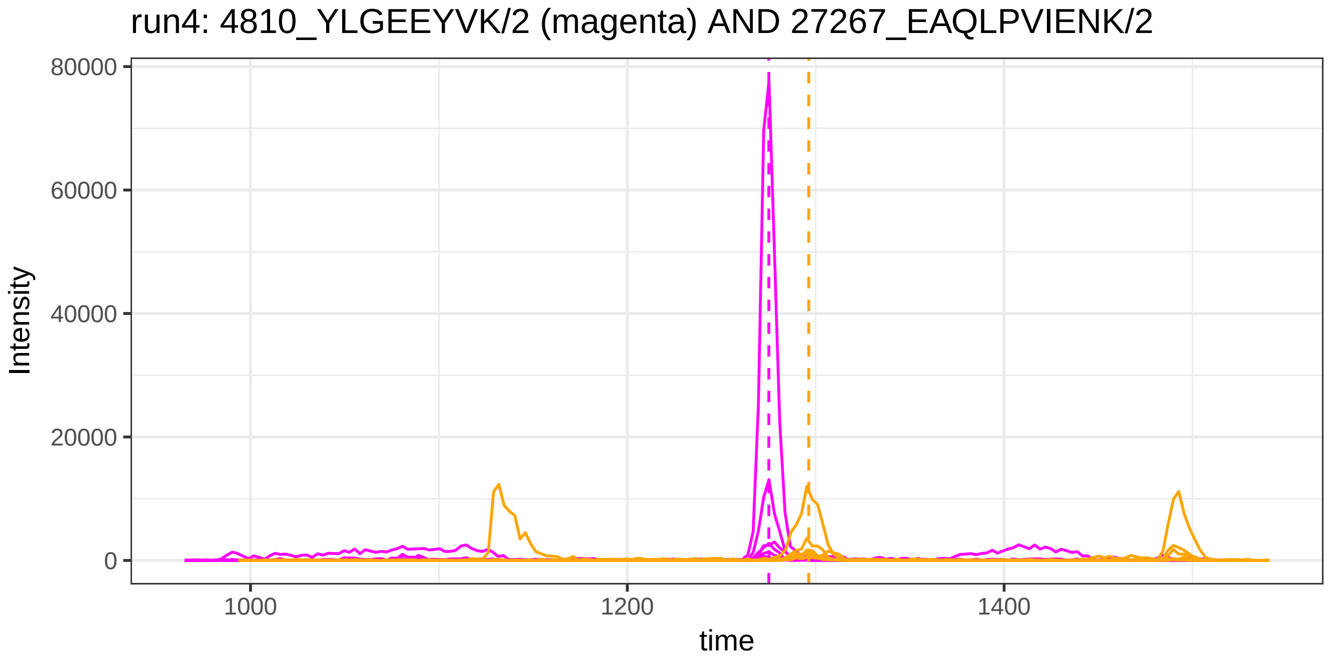
run4: 4810_YLGEEYVK/2 (magenta) AND 27267_EAQLPVIENK/2
Intensity
80000
60000
40000
20000
0
1000
1200
1400
time

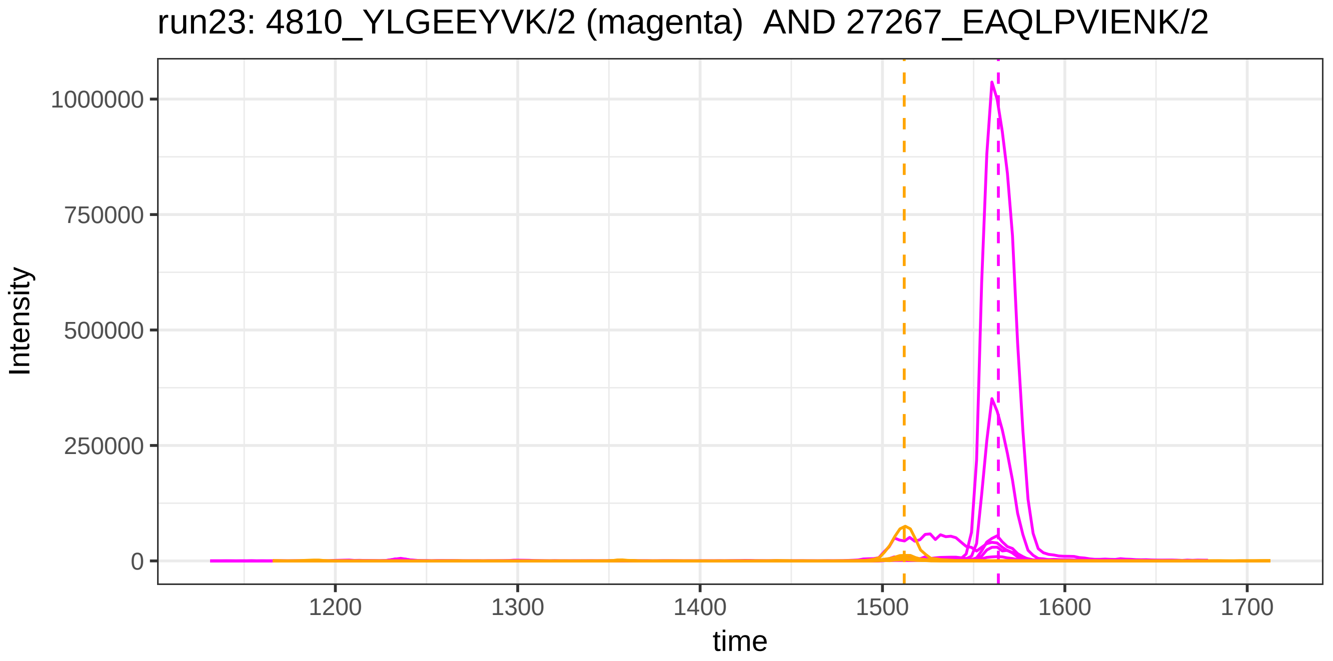
run23: 4810_YLGEEYVK/2 (magenta) AND 27267_EAQLPVIENK/2
Intensity
1000000
750000
500000
250000
0
1200
1300
1400
1500
1600
1700
time

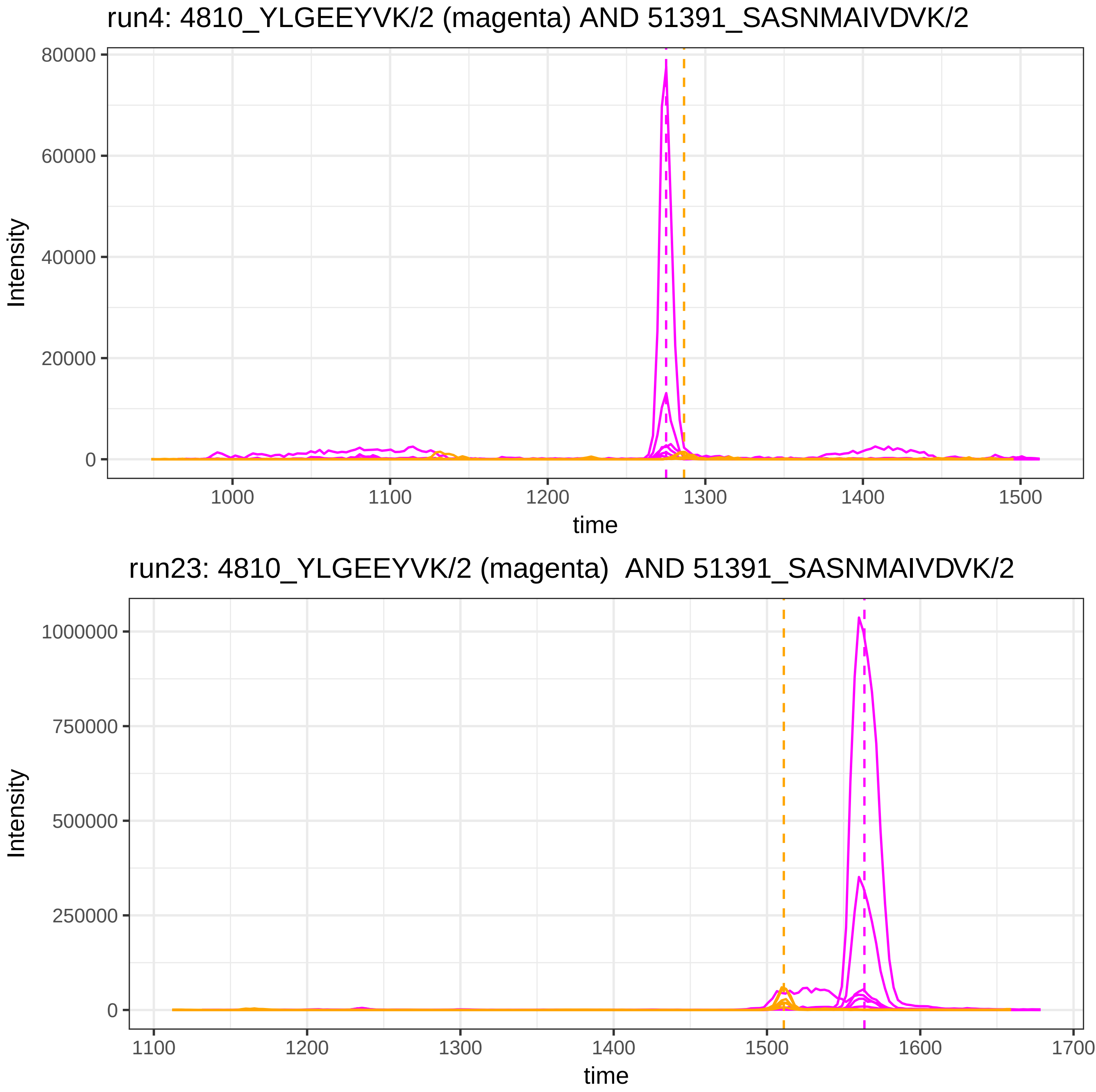
run4: 4810_YLGEEYVK/2 (magenta) AND 51391_SASNMAIVDVK/2
Intensity
80000
60000
40000
20000
0
1000
1100
1200
1300
1400
1500
time
run23: 4810_YLGEEYVK/2 (magenta) AND 51391_SASNMAIVDVK/2
Intensity
1000000
750000
500000
250000
0
1100
1200
1300
1400
1500
1600
1700
time

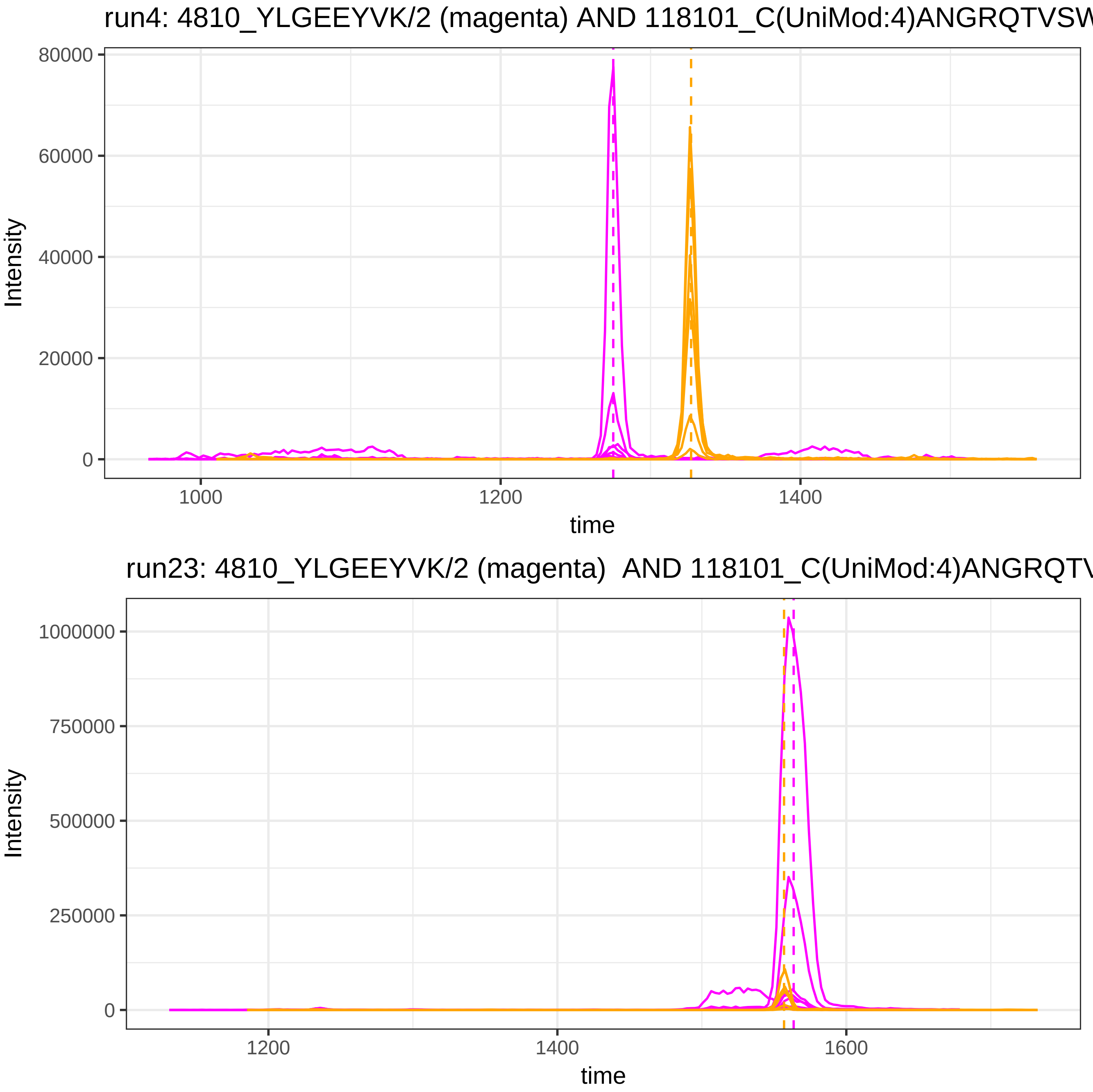

run4: 4810_YLGEEYVK/2 (magenta) AND 118101_C(UniMod:4)ANGRQTVSW
Intensity
80000
60000
40000
20000
0
1000
1200
1400
time
run23: 4810_YLGEEYVK/2 (magenta) AND 118101_C(UniMod:4)ANGRQTV
Intensity
1000000
750000
500000
250000
0
1200
1400
1600
time

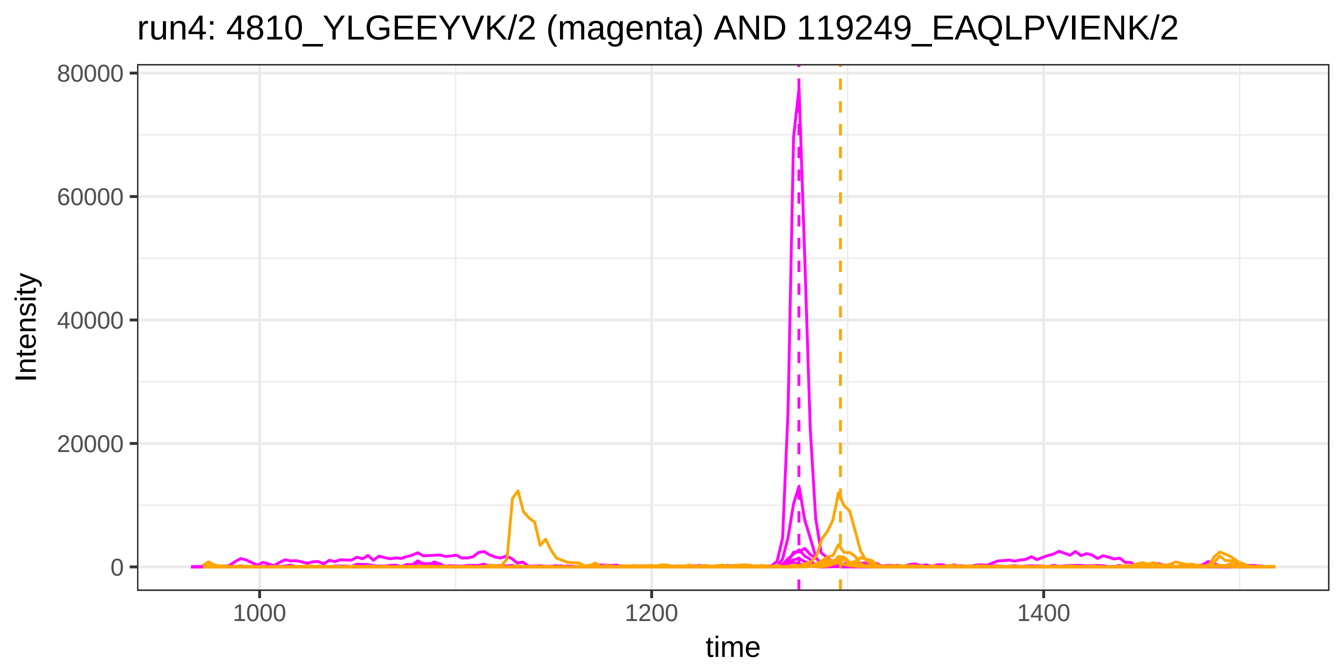
run4: 4810_YLGEEYVK/2 (magenta) AND 119249_EAQLPVIENK/2
Intensity
80000
60000
40000
20000
0
1000
1200
1400
time

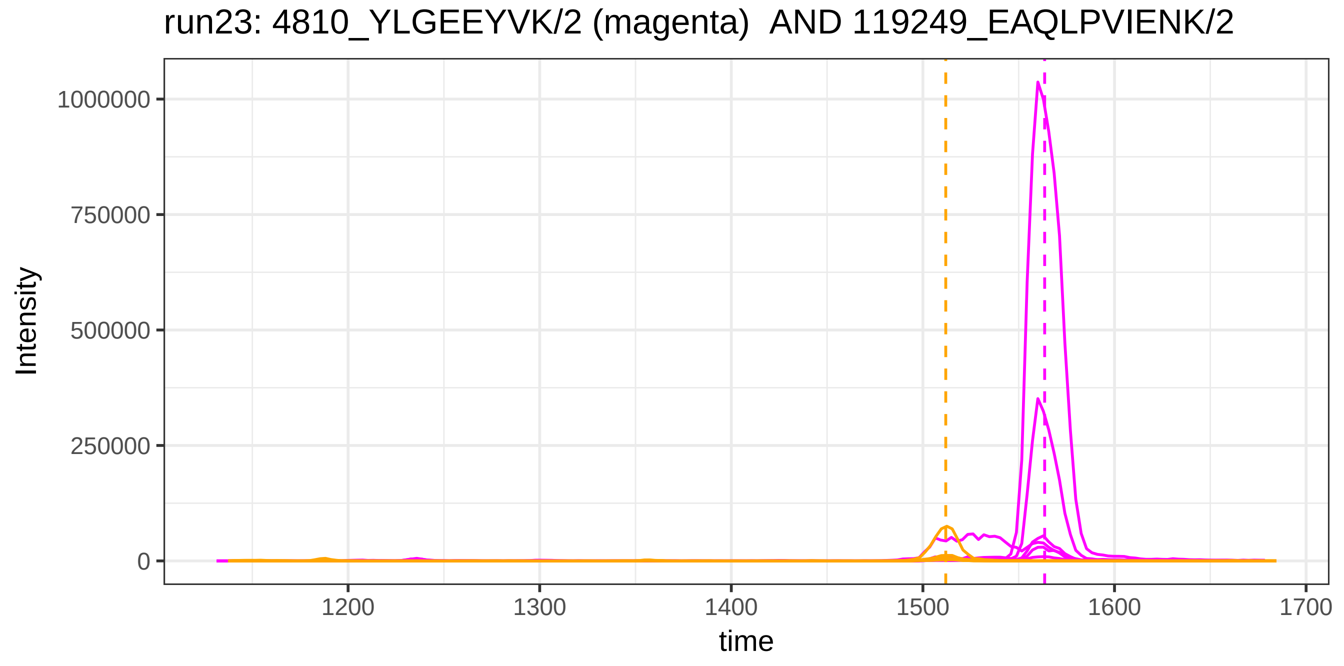
run23: 4810_YLGEEYVK/2 (magenta)  AND 119249_EAQLPVIENK/2
Intensity
1000000
750000
500000
250000
0
1200
1300
1400
1500
1600
1700
time

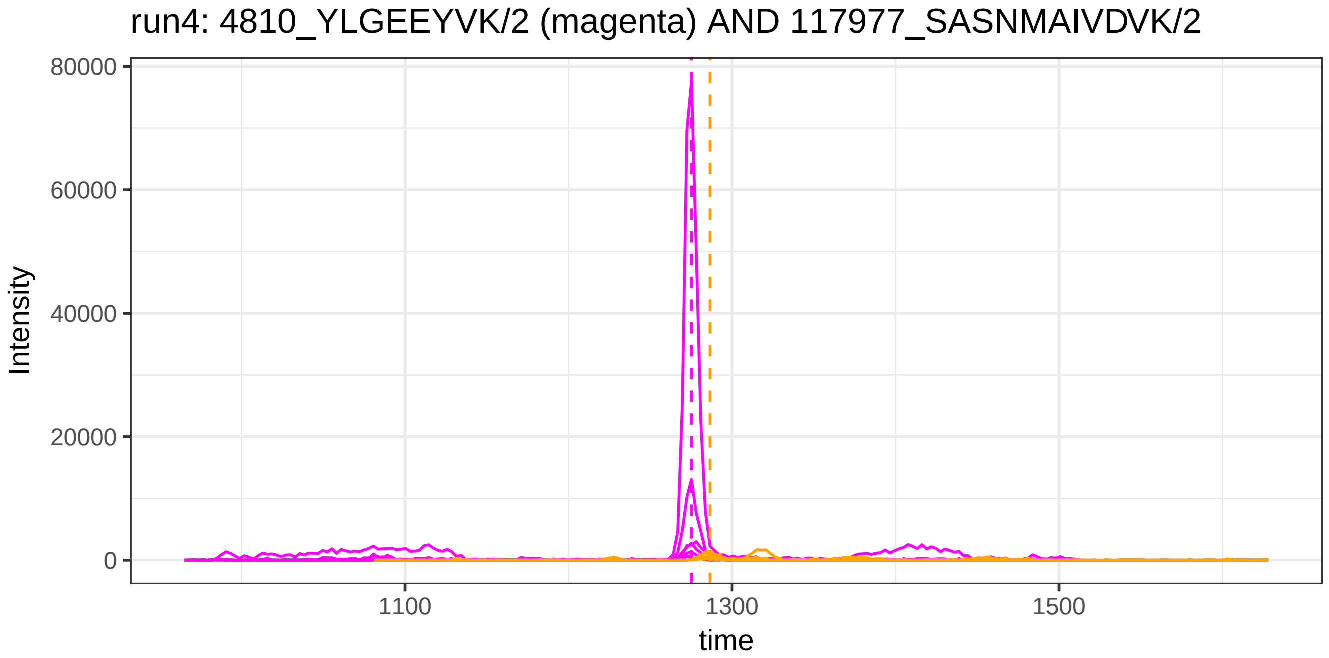
run4: 4810_YLGEEYVK/2 (magenta) AND 117977_SASNMAIVDVK/2
Intensity
80000
60000
40000
20000
0
1100
1300
1500
time

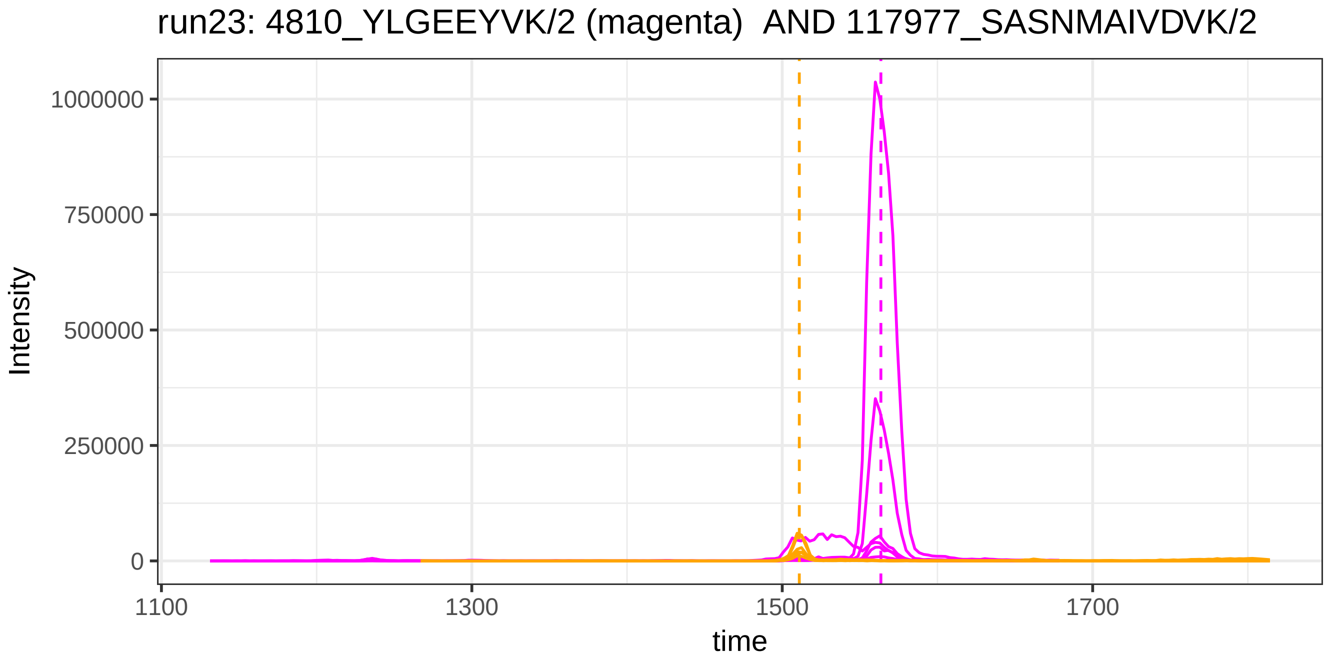
run23: 4810_YLGEEYVK/2 (magenta) AND 117977_SASNMAIVDVK/2
Intensity
1000000
750000
500000
250000
0
1100
1300
1500
1700
time

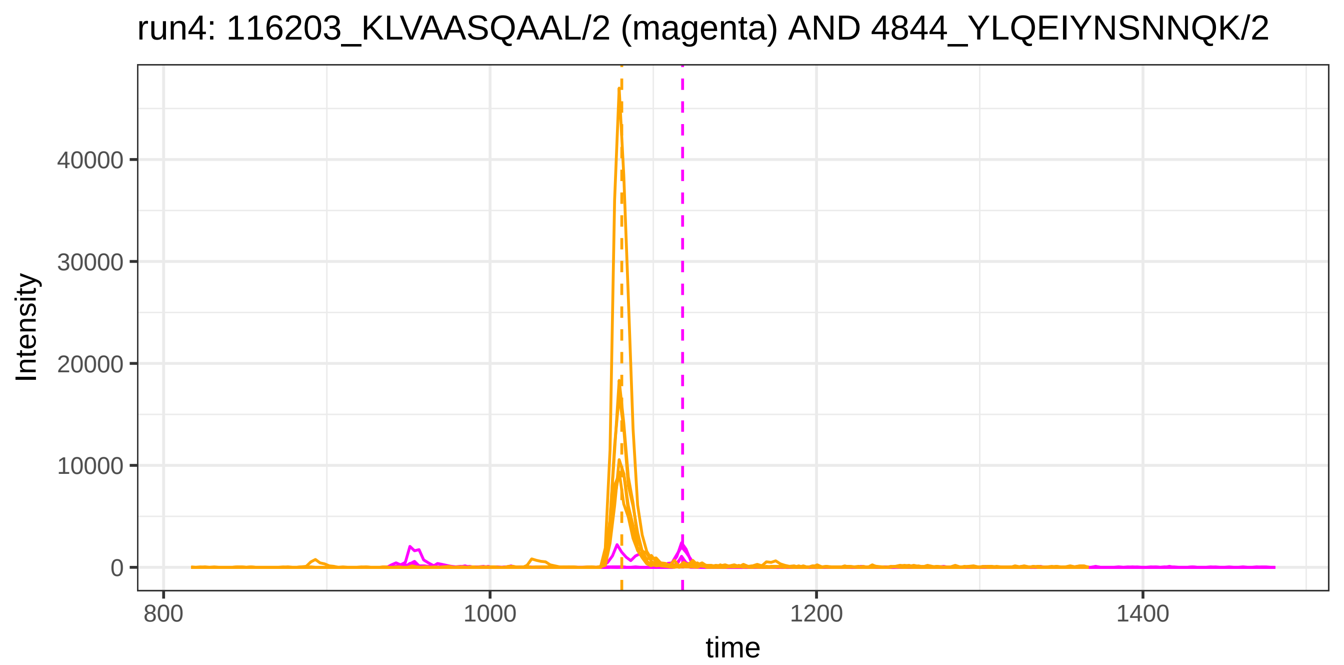
run4: 116203_KLVAASQAAL/2 (magenta) AND 4844_YLQEIYNSNNQK/2
Intensity
40000
30000
20000
10000
0
800
1000
1200
1400
time

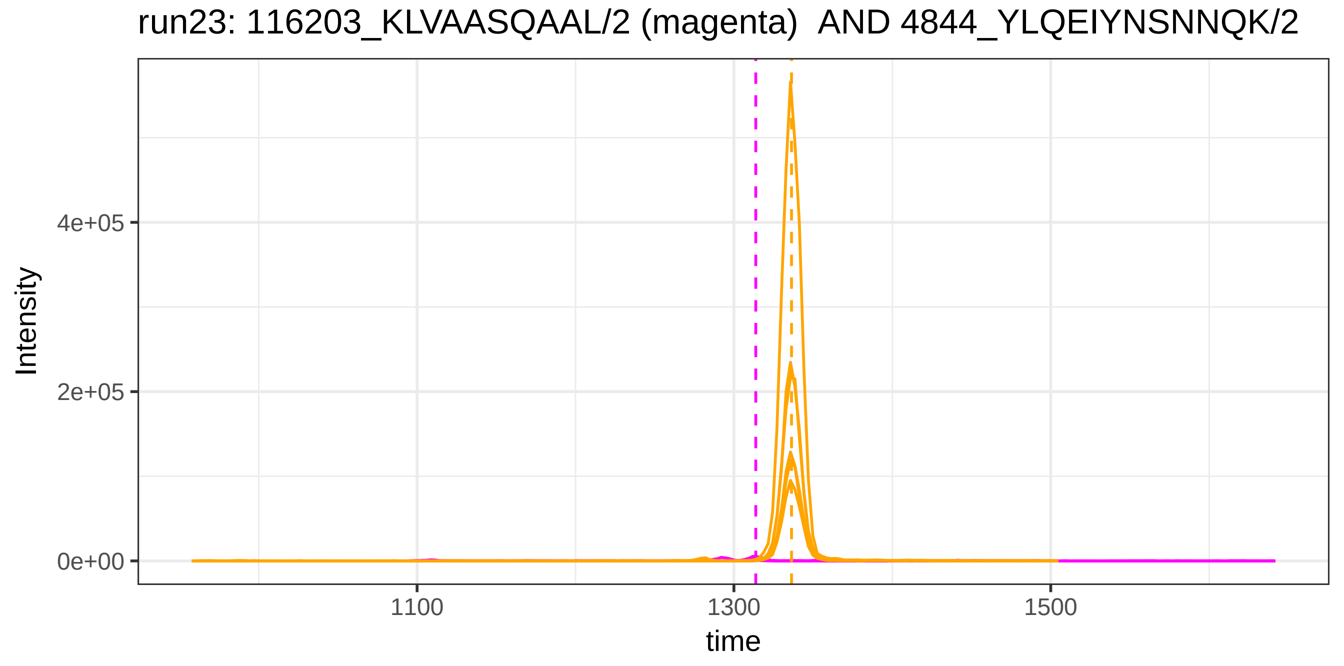
run23: 116203_KLVAASQAAL/2 (magenta)  AND 4844_YLQEIYNSNNQK/2
Intensity
4e+05
2e+05
0e+00
1100
1300
1500
time

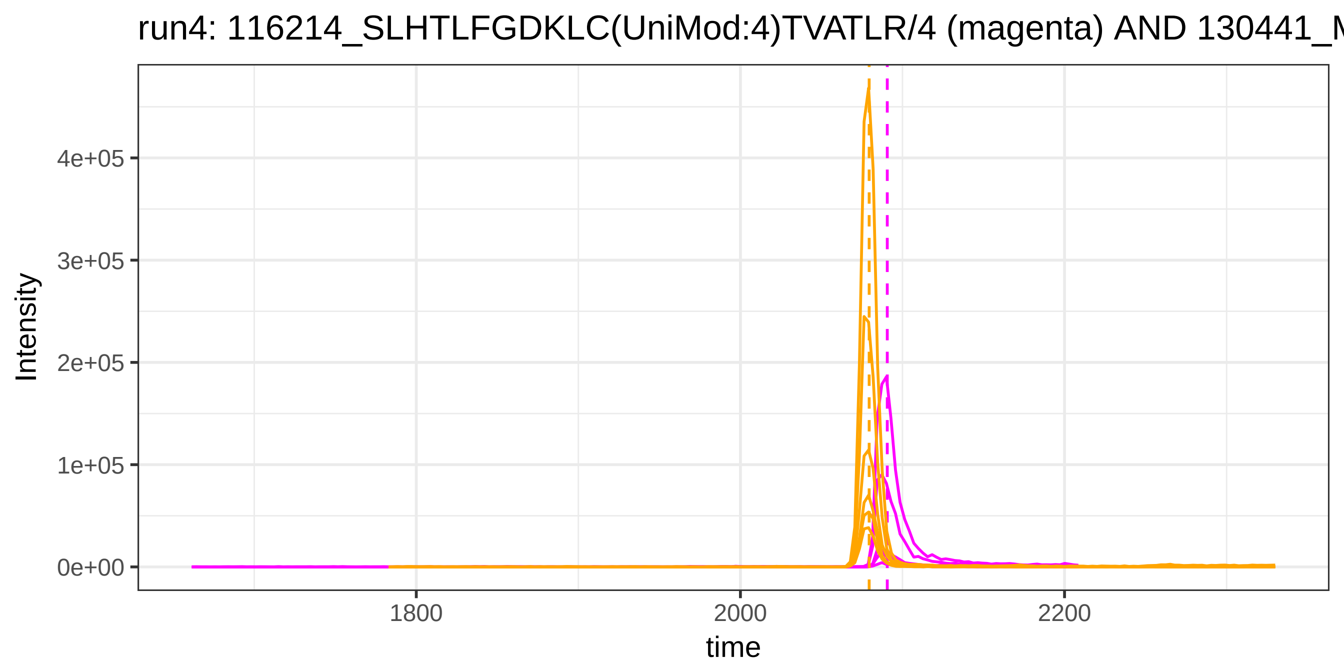

run4: 116214_SLHTLFGDKLC(UniMod:4)TVATLR/4 (magenta) AND 130441_M
Intensity
4e+05
3e+05
2e+05
1e+05
0e+00
1800
2000
2200
time


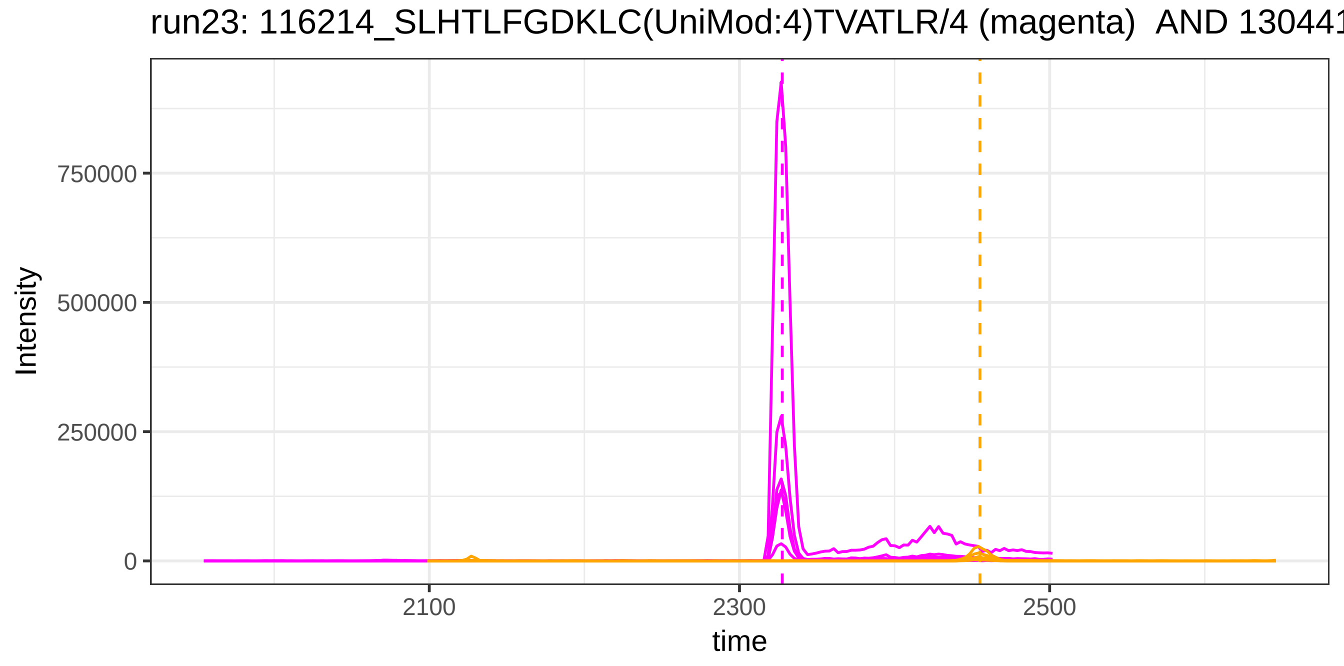

run23: 116214_SLHTLFGDKLC(UniMod:4)TVATLR/4 (magenta) AND 13044
Intensity
750000
500000
250000
0
2100
2300
2500
time

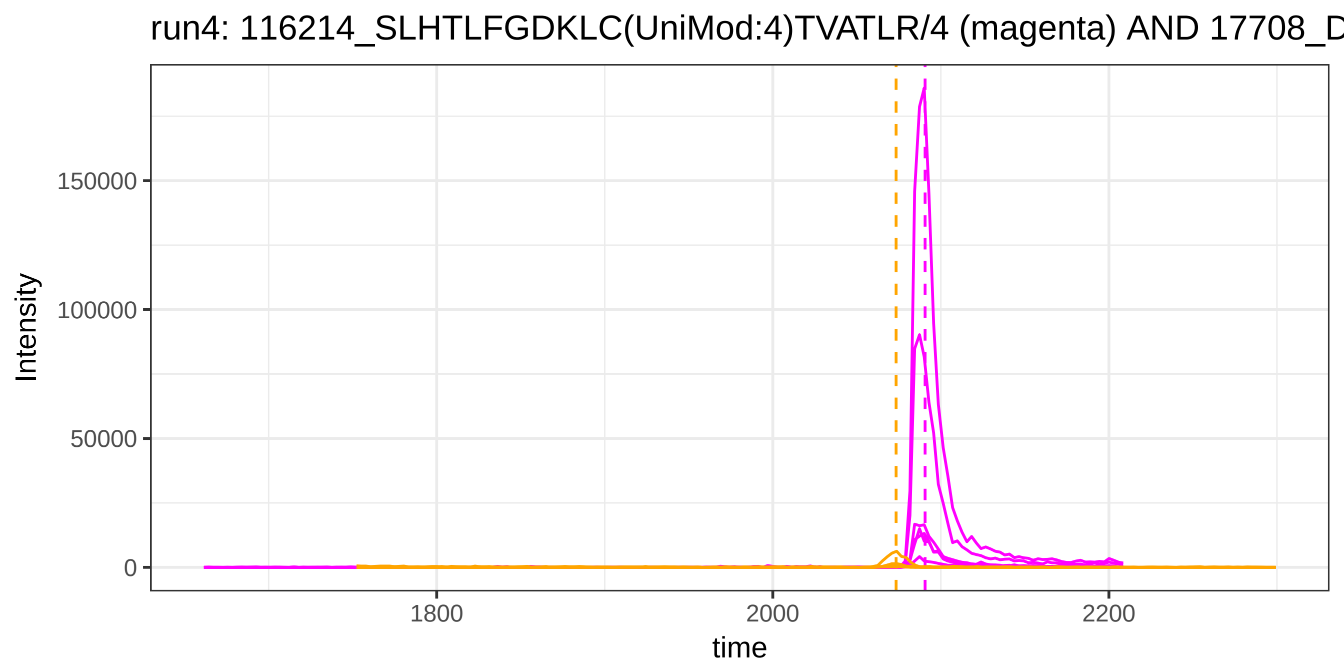

run4: 116214_SLHTLFGDKLC(UniMod:4)TVATLR/4 (magenta) AND 17708_D
Intensity
150000
100000
50000
0
1800
2000
2200
time


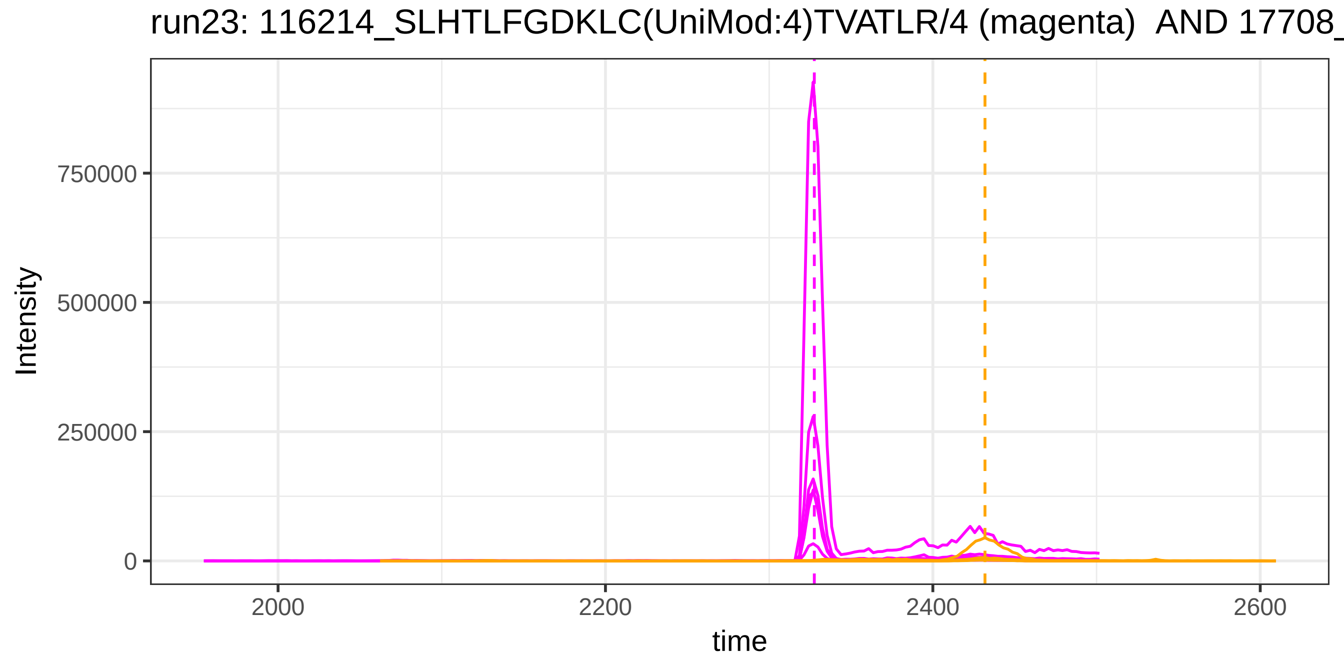

run23: 116214_SLHTLFGDKLC(UniMod:4)TVATLR/4 (magenta) AND 17708_
Intensity
750000
500000
250000
0
2000
2200
2400
2600
time

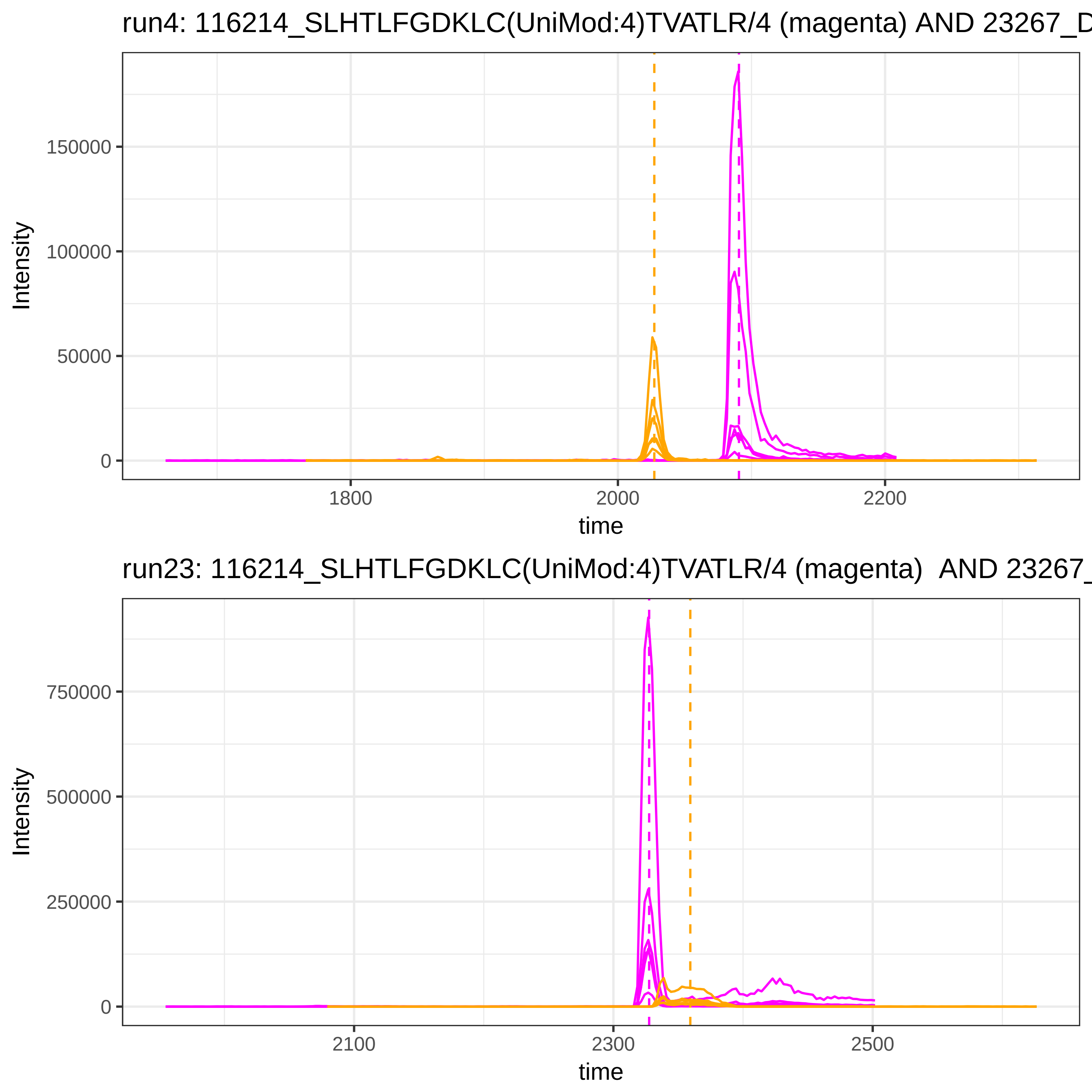

run4: 116214_SLHTLFGDKLC(UniMod:4)TVATLR/4 (magenta) AND 23267_D
Intensity
150000
100000
50000
0
1800
2000
2200
time
run23: 116214_SLHTLFGDKLC(UniMod:4)TVATLR/4 (magenta) AND 23267_
Intensity
750000
500000
250000
0
2100
2300
2500
time

run4: 116214_SLHTLFGDKLC(UniMod:4)TVATLR/4 (magenta) AND 117975_

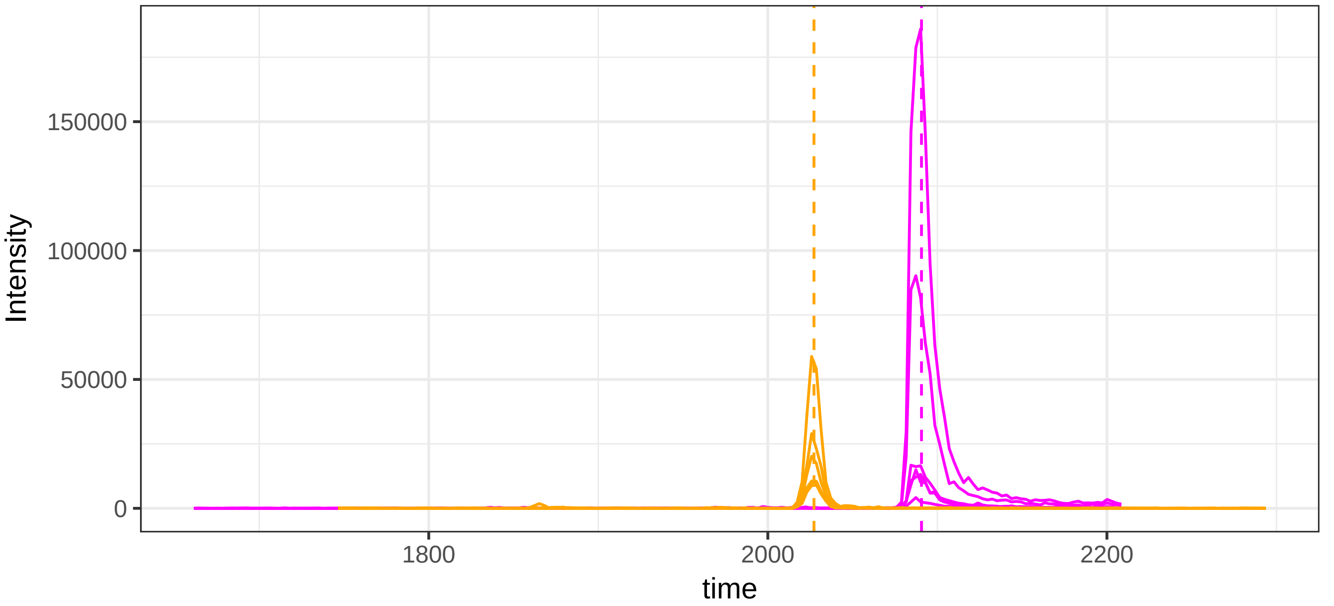


run23: 116214_SLHTLFGDKLC(UniMod:4)TVATLR/4 (magenta)  AND 11797

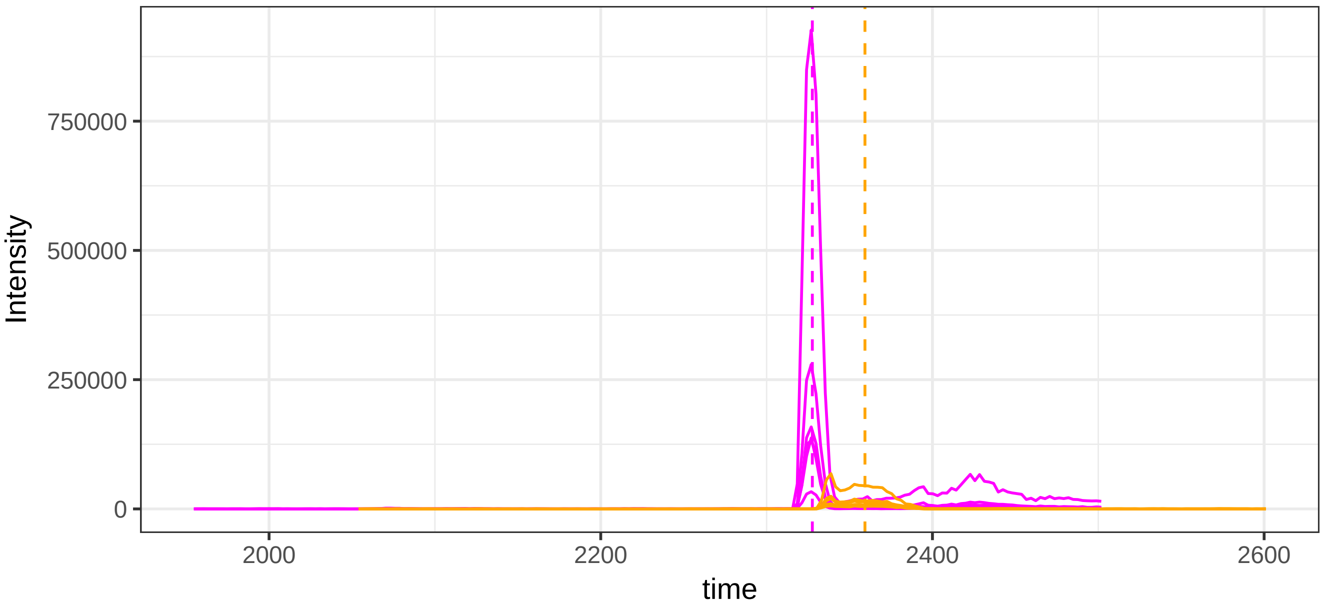

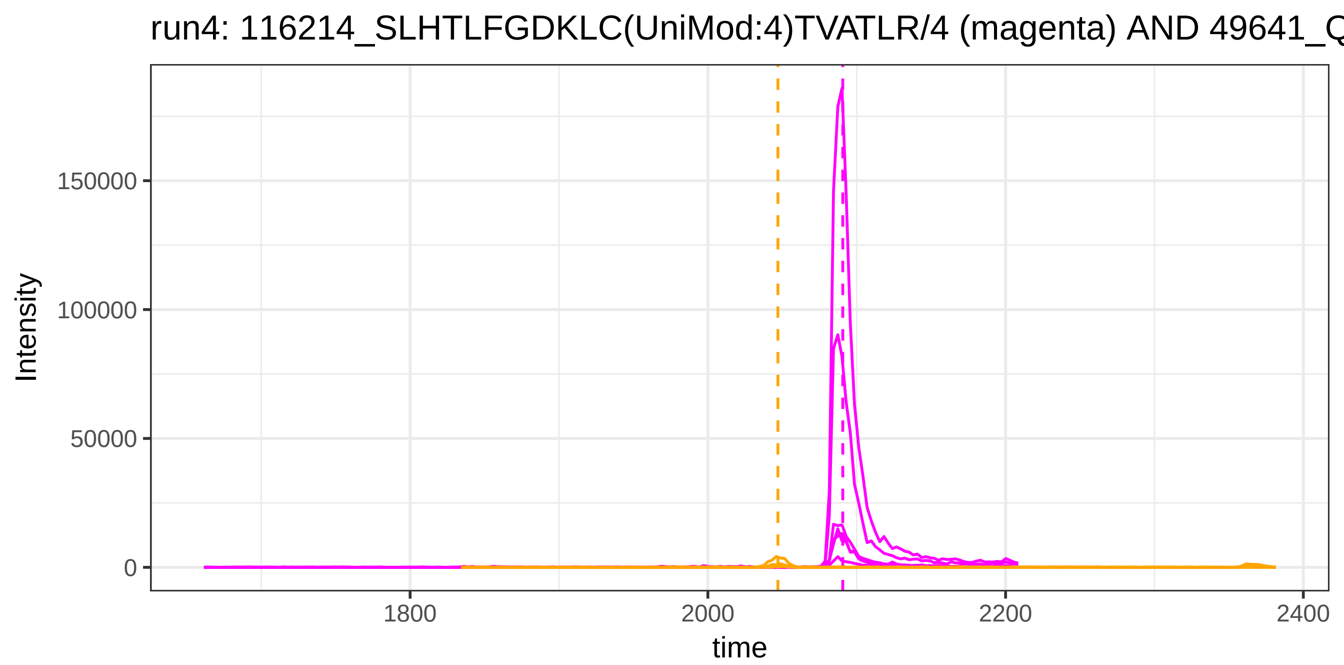
run4: 116214_SLHTLFGDKLC(UniMod:4)TVATLR/4 (magenta) AND 49641_Q
Intensity
150000
100000
50000
0
1800
2000
2200
2400
time

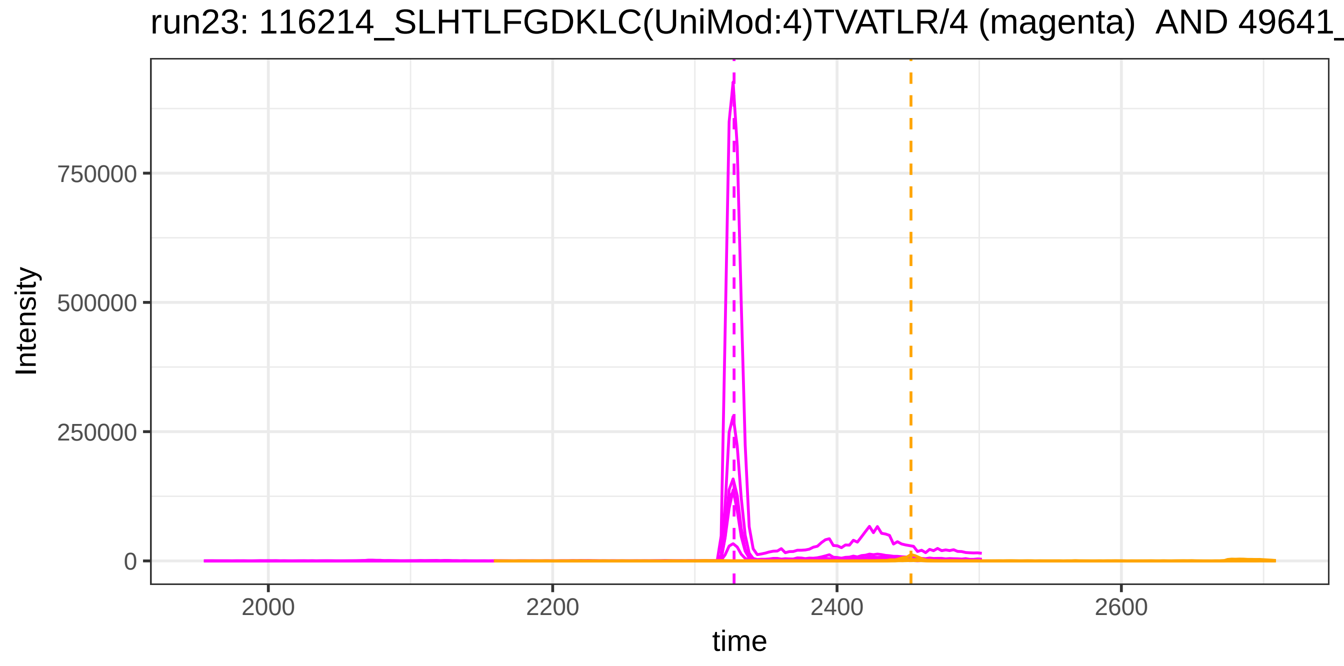
run23: 116214_SLHTLFGDKLC(UniMod:4)TVATLR/4 (magenta) AND 49641_
Intensity
750000
500000
250000
0
2000
2200
2400
2600
time

run4: 116214_SLHTLFGDKLC(UniMod:4)TVATLR/4 (magenta) AND 34881_G

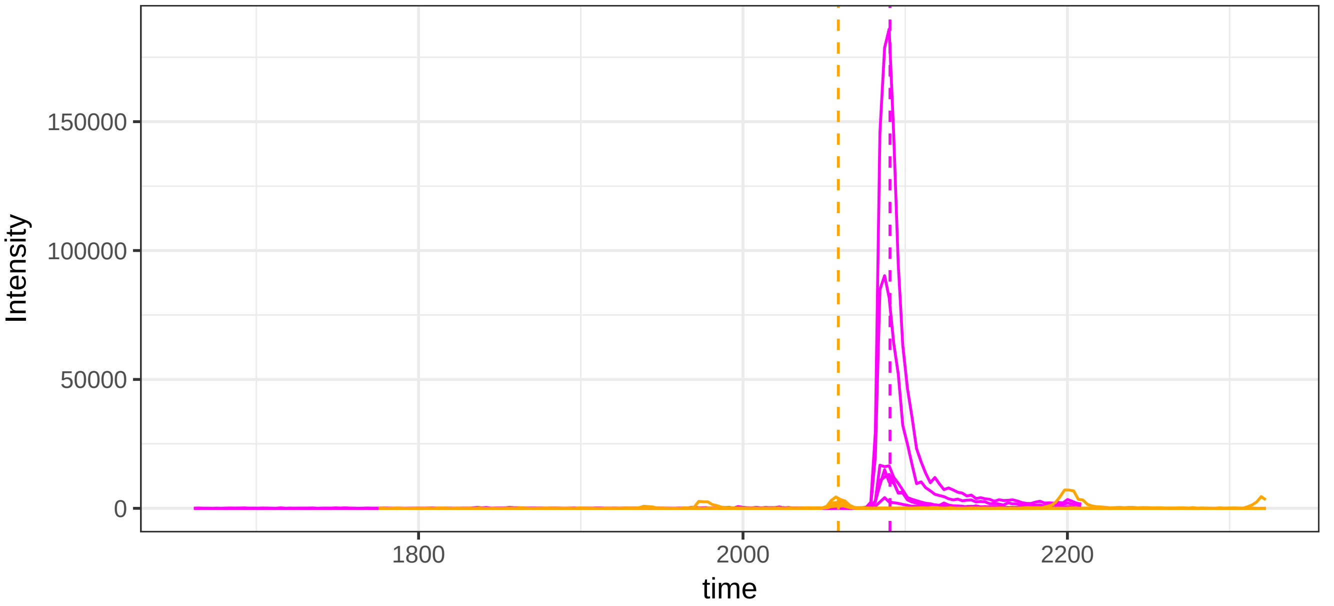


run23: 116214_SLHTLFGDKLC(UniMod:4)TVATLR/4 (magenta) AND 34881_

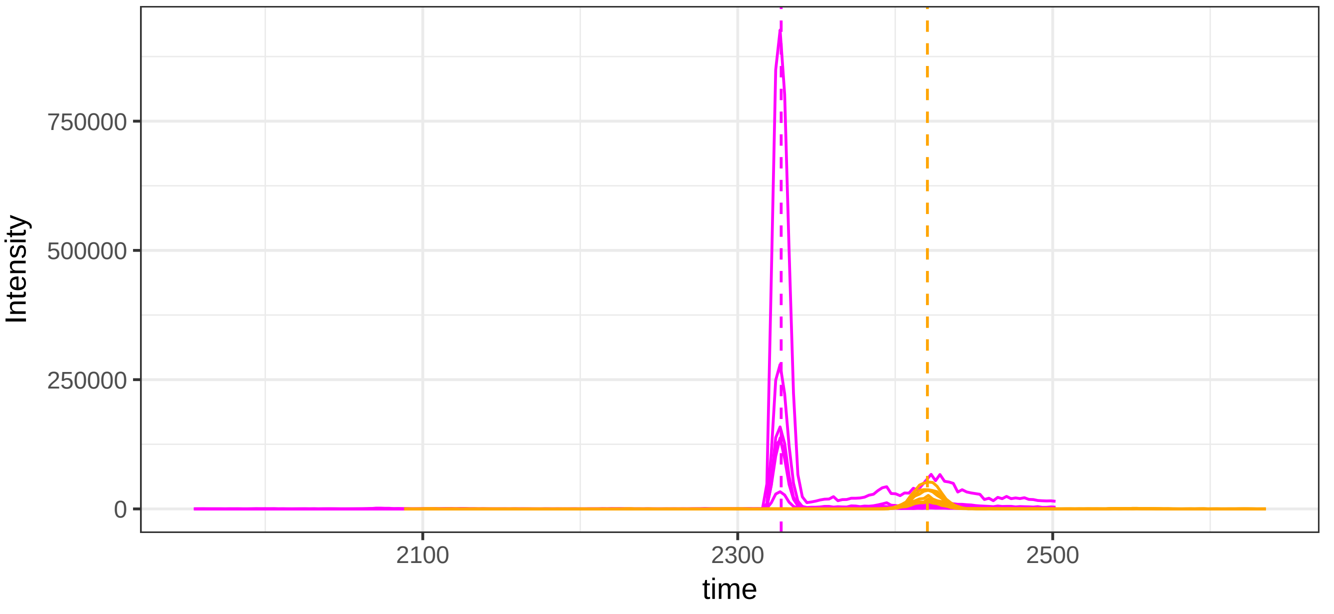

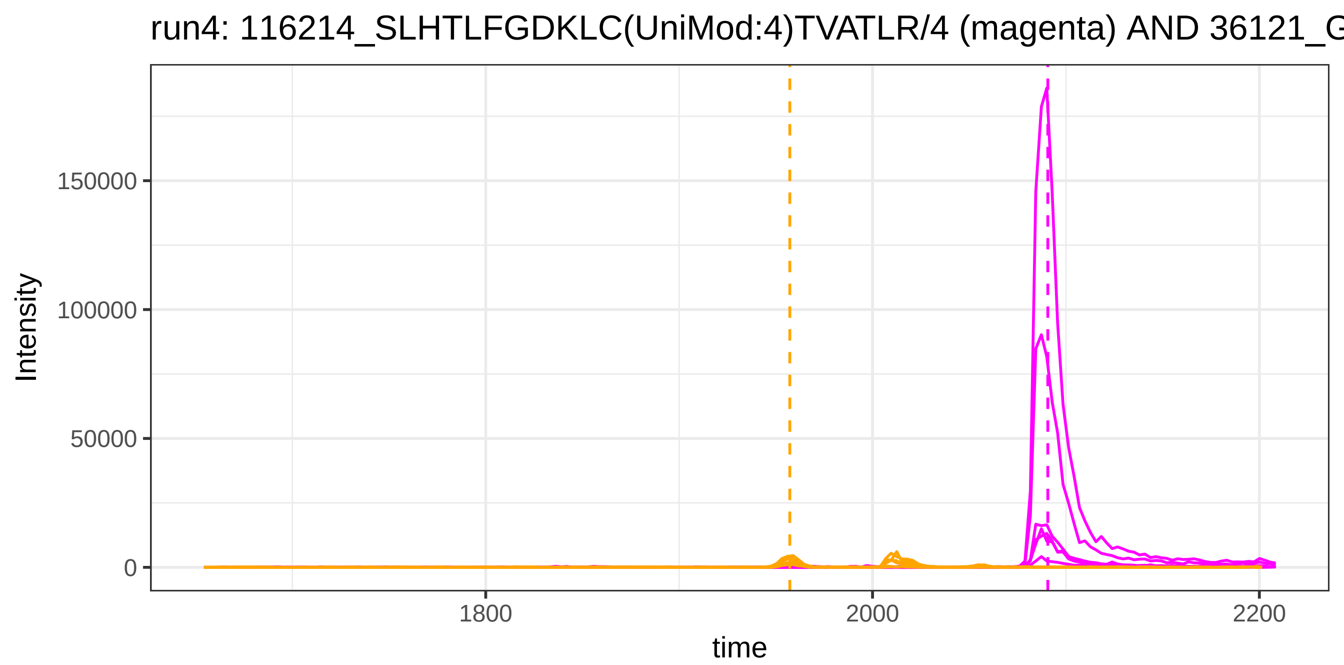
run4: 116214_SLHTLFGDKLC(UniMod:4)TVATLR/4 (magenta) AND 36121_G
Intensity
150000
100000
50000
0
1800
2000
2200
time

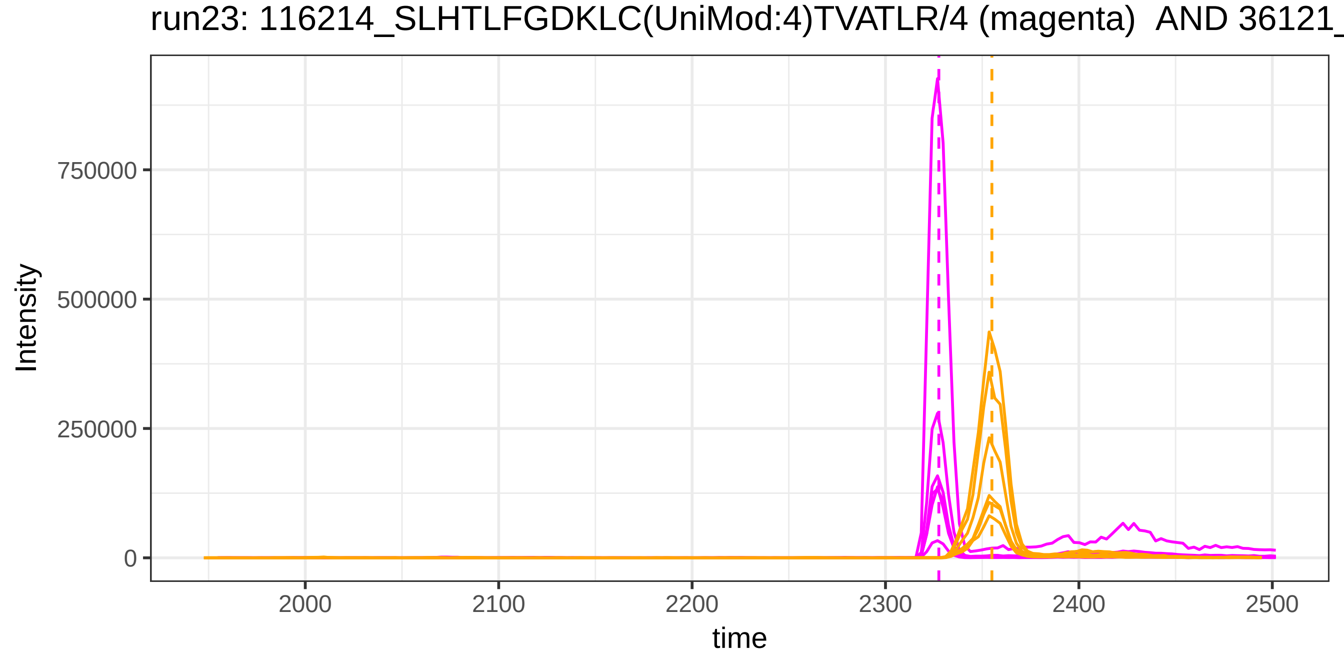
run23: 116214_SLHTLFGDKLC(UniMod:4)TVATLR/4 (magenta) AND 36121_
Intensity
750000
500000
250000
0
2000
2100
2200
2300
2400
2500
time

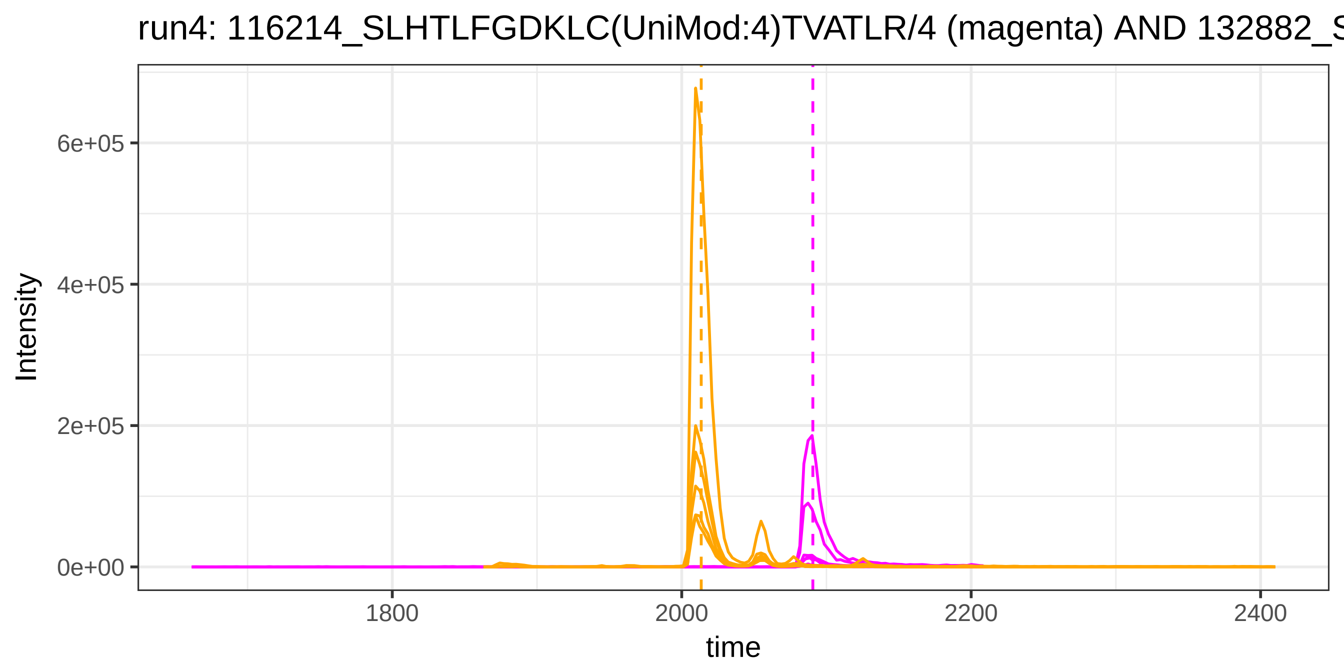

run4: 116214_SLHTLFGDKLC(UniMod:4)TVATLR/4 (magenta) AND 132882_S
Intensity
6e+05
4e+05
2e+05
0e+00
1800
2000
2200
2400
time


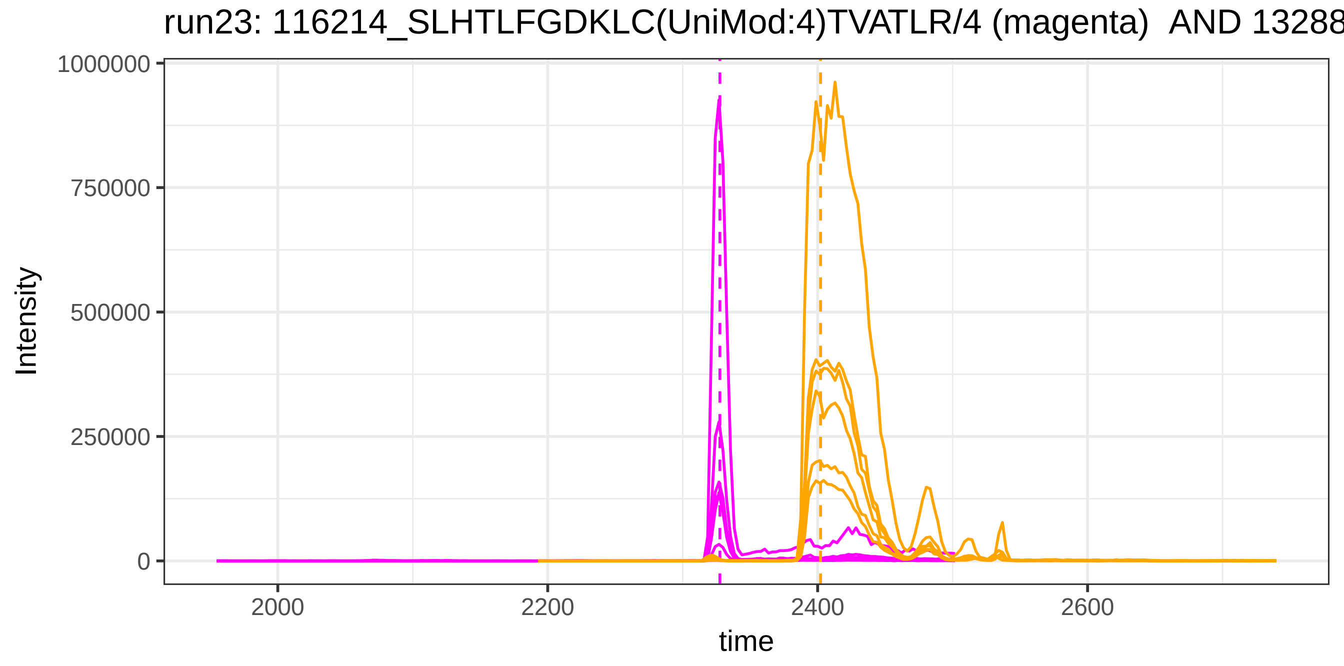

run23: 116214_SLHTLFGDKLC(UniMod:4)TVATLR/4 (magenta) AND 13288
Intensity
1000000
750000
500000
250000
0
2000
2200
2400
2600
time

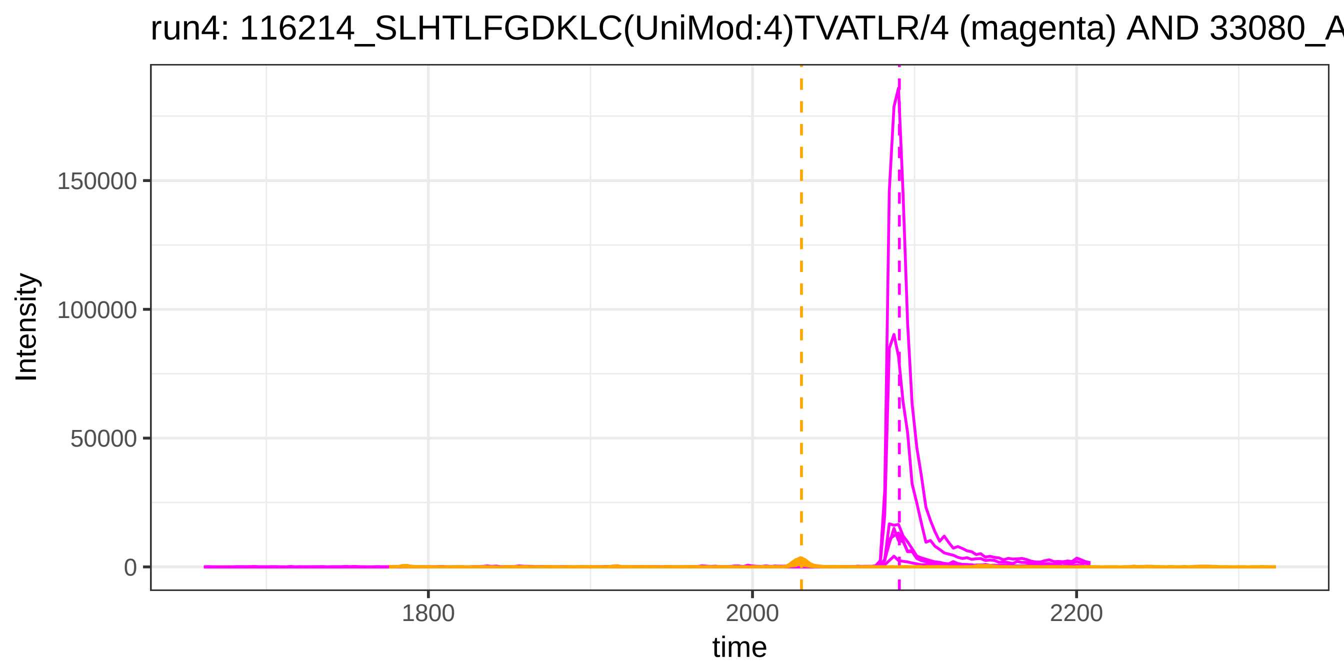

run4: 116214_SLHTLFGDKLC(UniMod:4)TVATLR/4 (magenta) AND 33080_A
Intensity
150000
100000
50000
0
1800
2000
2200
time


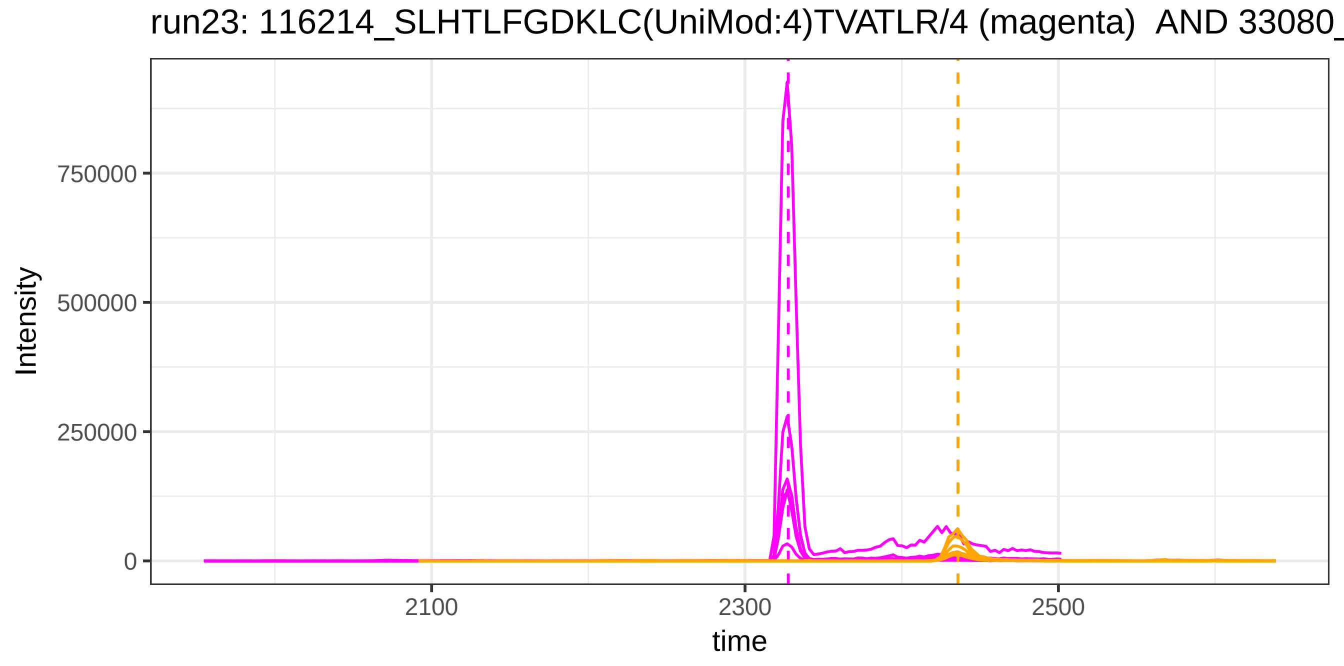

run23: 116214_SLHTLFGDKLC(UniMod:4)TVATLR/4 (magenta) AND 33080_
Intensity
750000
500000
250000
0
2100
2300
2500
time

run4: 116214_SLHTLFGDKLC(UniMod:4)TVATLR/4 (magenta) AND 116622_

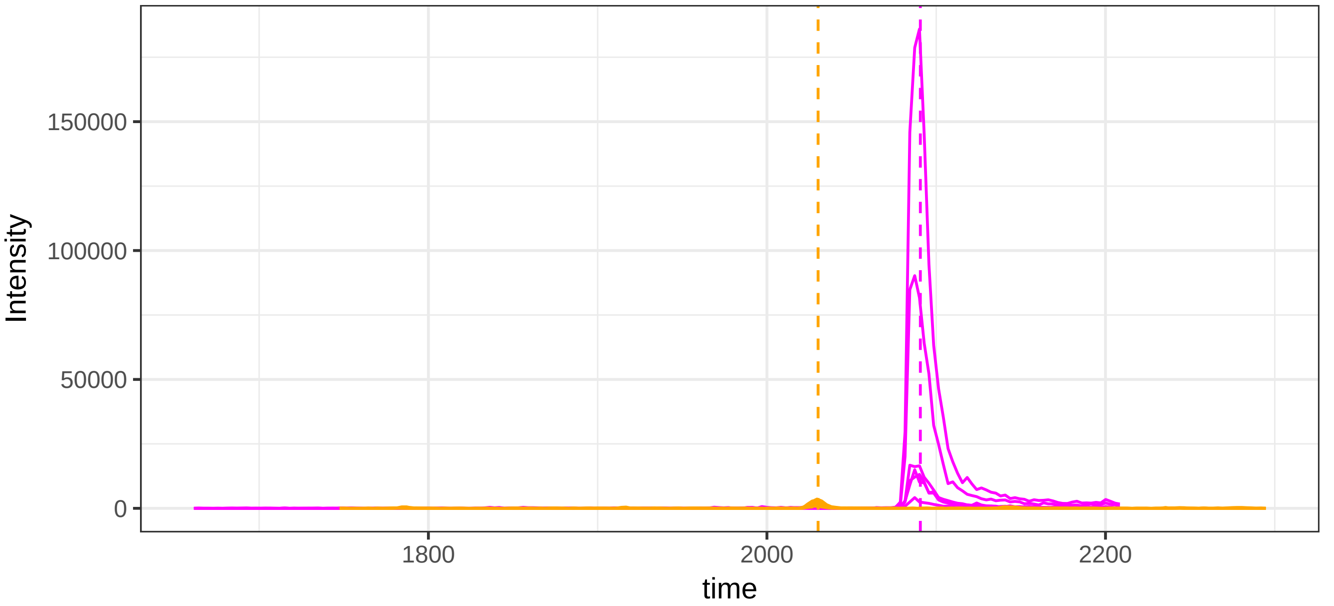


run23: 116214_SLHTLFGDKLC(UniMod:4)TVATLR/4 (magenta) AND 116622

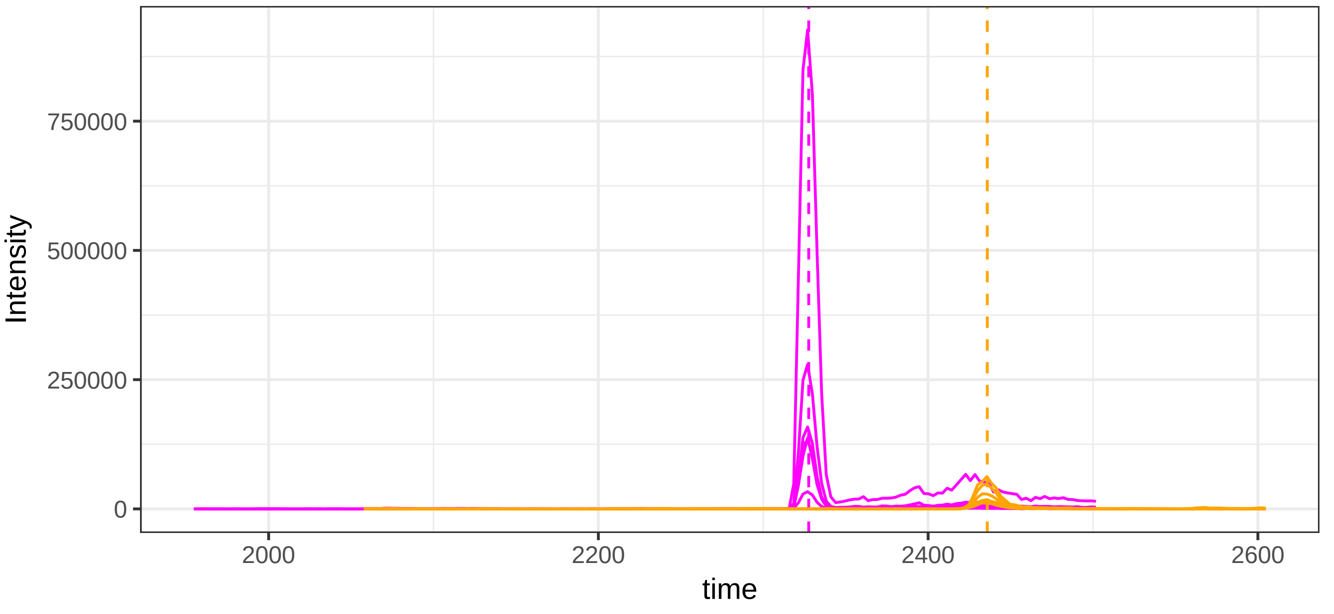

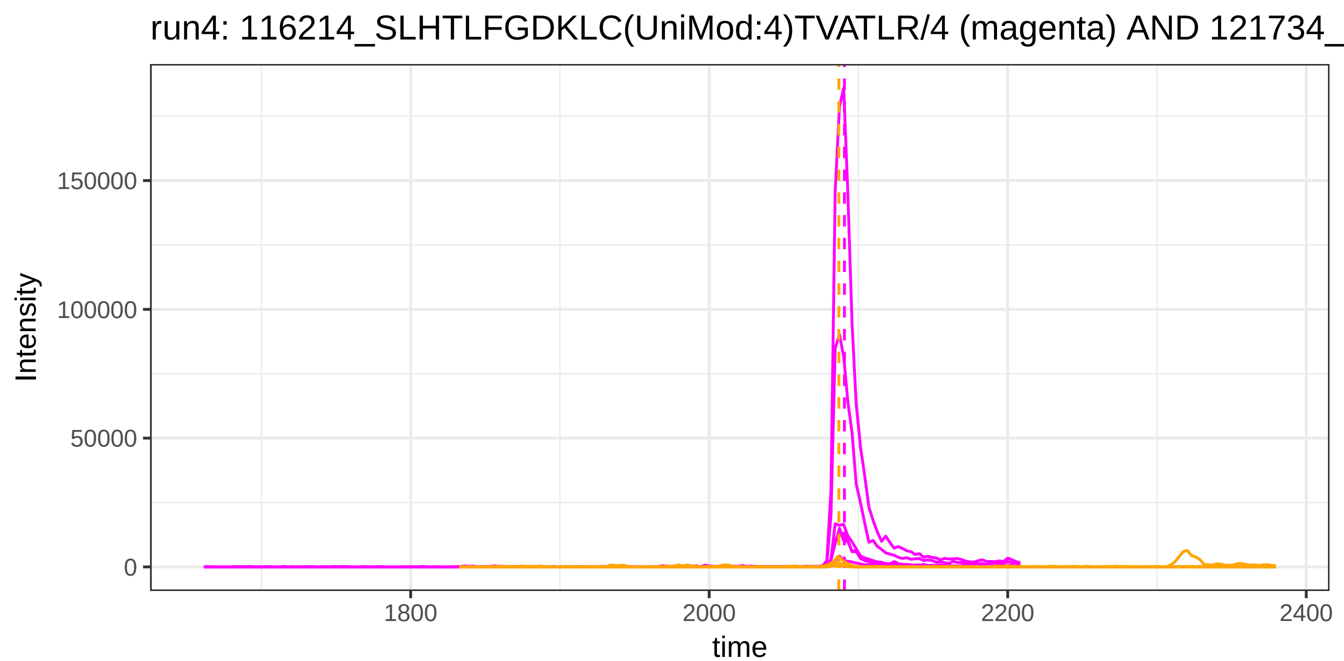
run4: 116214_SLHTLFGDKLC(UniMod:4)TVATLR/4 (magenta) AND 121734_
Intensity
150000
100000
50000
0
1800
2000
2200
2400
time

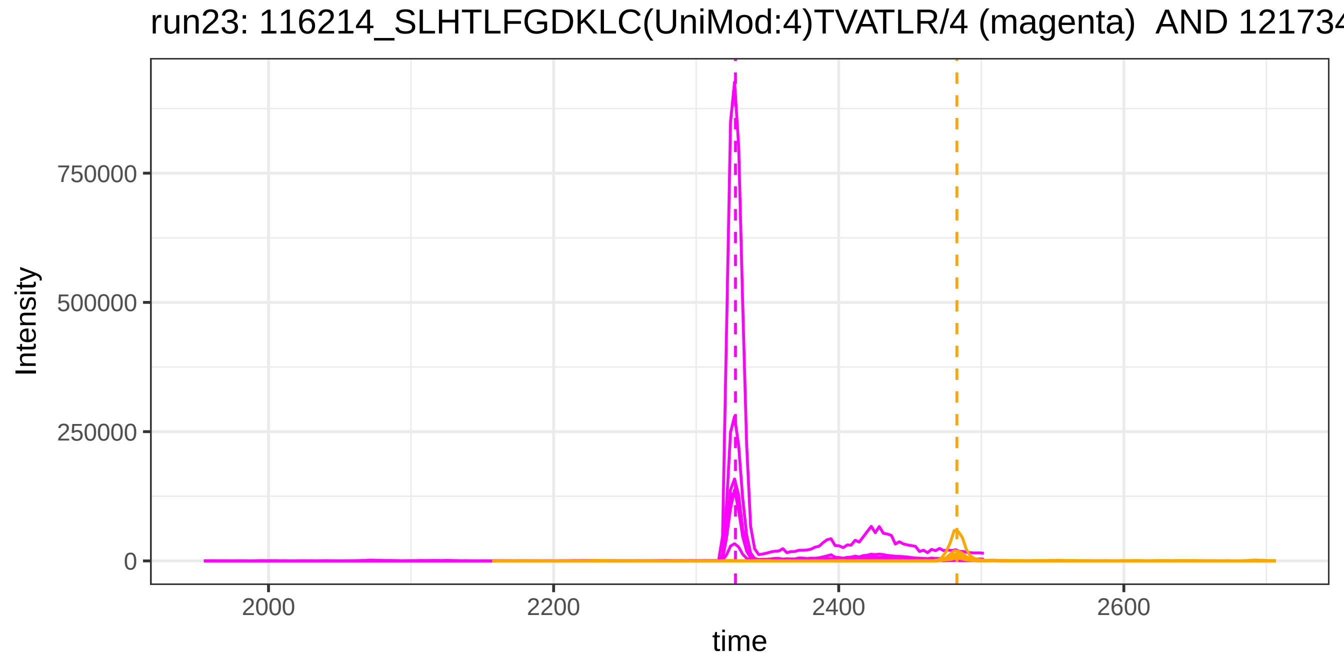
run23: 116214_SLHTLFGDKLC(UniMod:4)TVATLR/4 (magenta) AND 121734
Intensity
750000
500000
250000
0
2000
2200
2400
2600
time

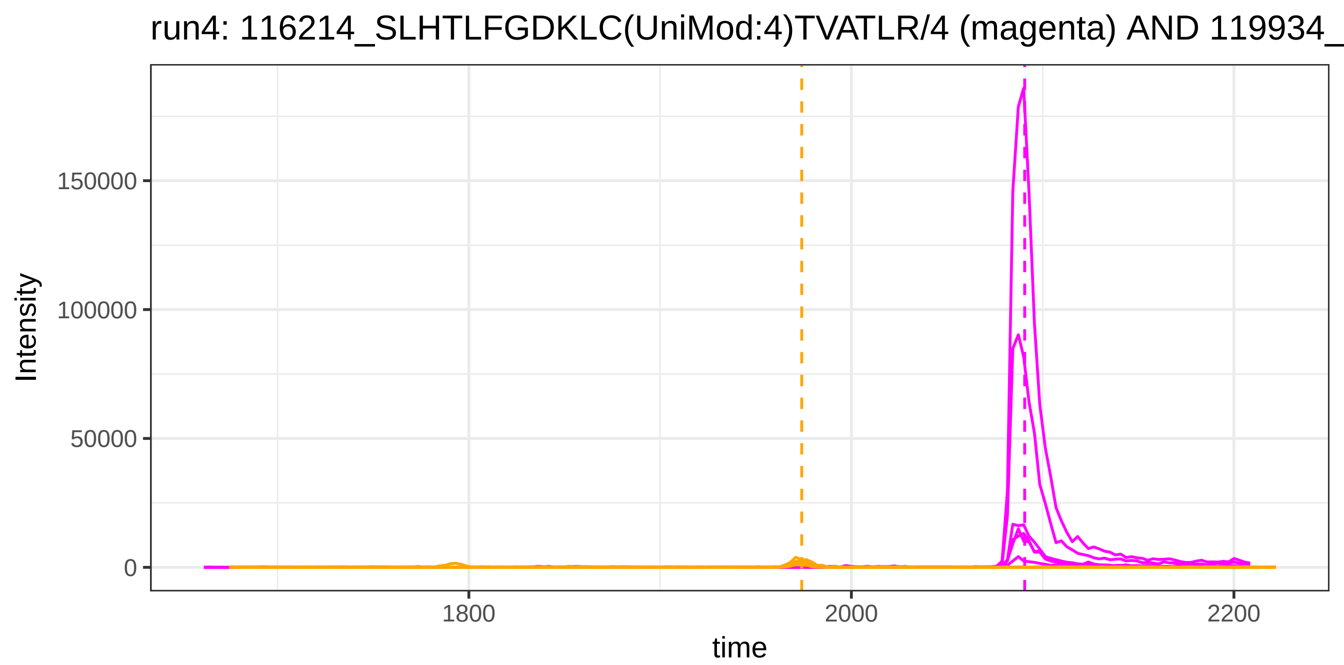
run4: 116214_SLHTLFGDKLC(UniMod:4)TVATLR/4 (magenta) AND 119934_
Intensity
150000
100000
50000
0
1800
2000
2200
time

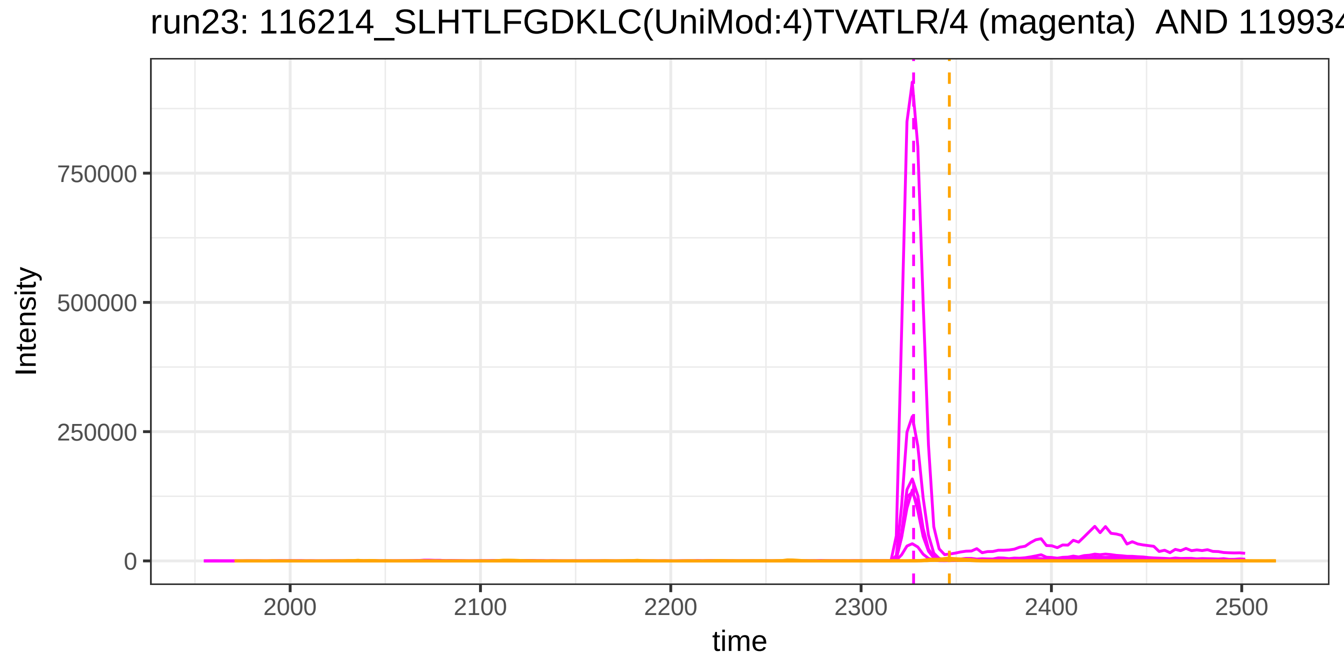
run23: 116214_SLHTLFGDKLC(UniMod:4)TVATLR/4 (magenta) AND 119934
Intensity
750000
500000
250000
0
2000
2100
2200
2300
2400
2500
time

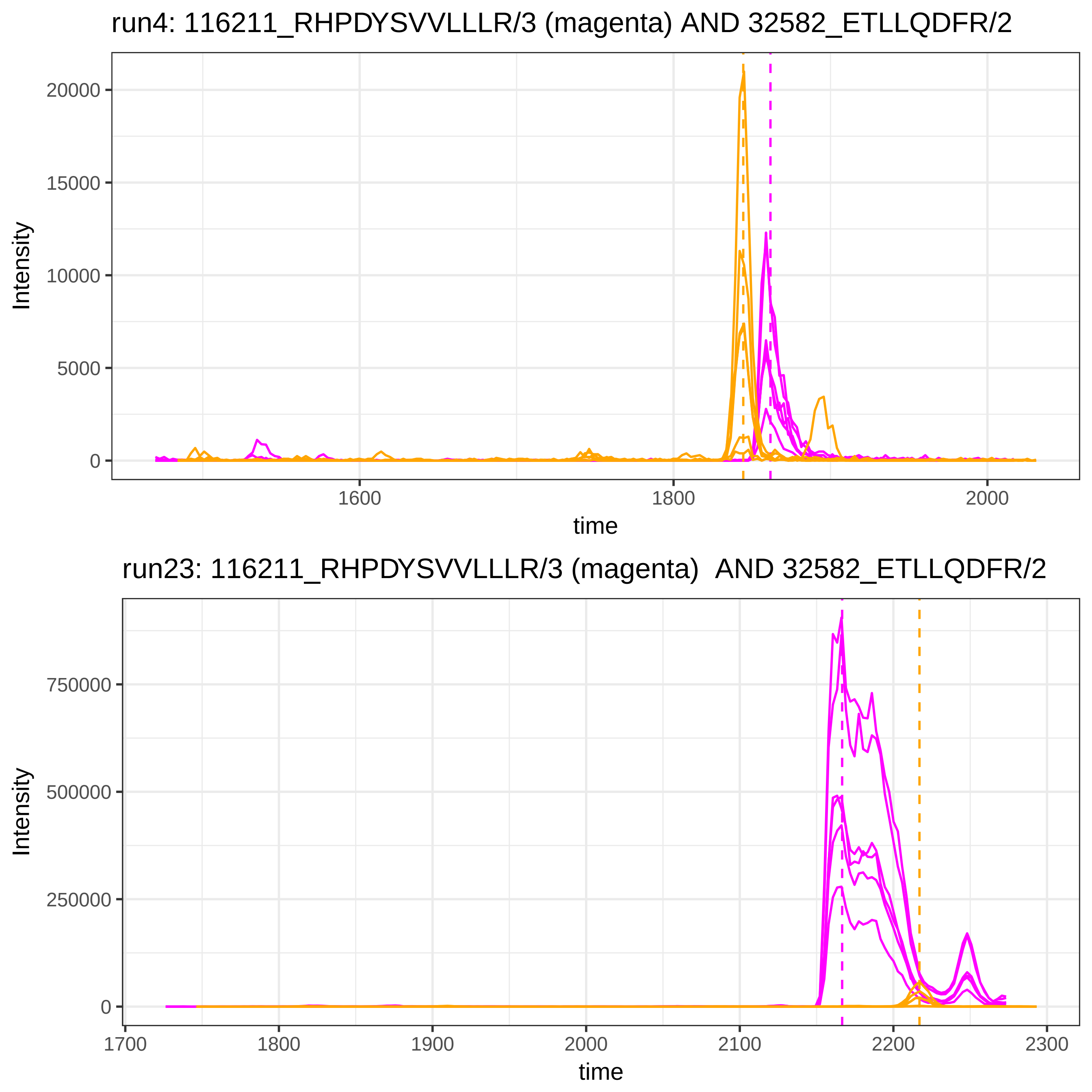

run4: 116211_RHPDYSVVLLLR/3 (magenta) AND 32582_ETLLQDFR/2
Intensity
20000
15000
10000
5000
0
1600
1800
2000
time
run23: 116211_RHPDYSVVLLLR/3 (magenta) AND 32582_ETLLQDFR/2
Intensity
750000
500000
250000
0
1700
1800
1900
2000
2100
2200
2300
time

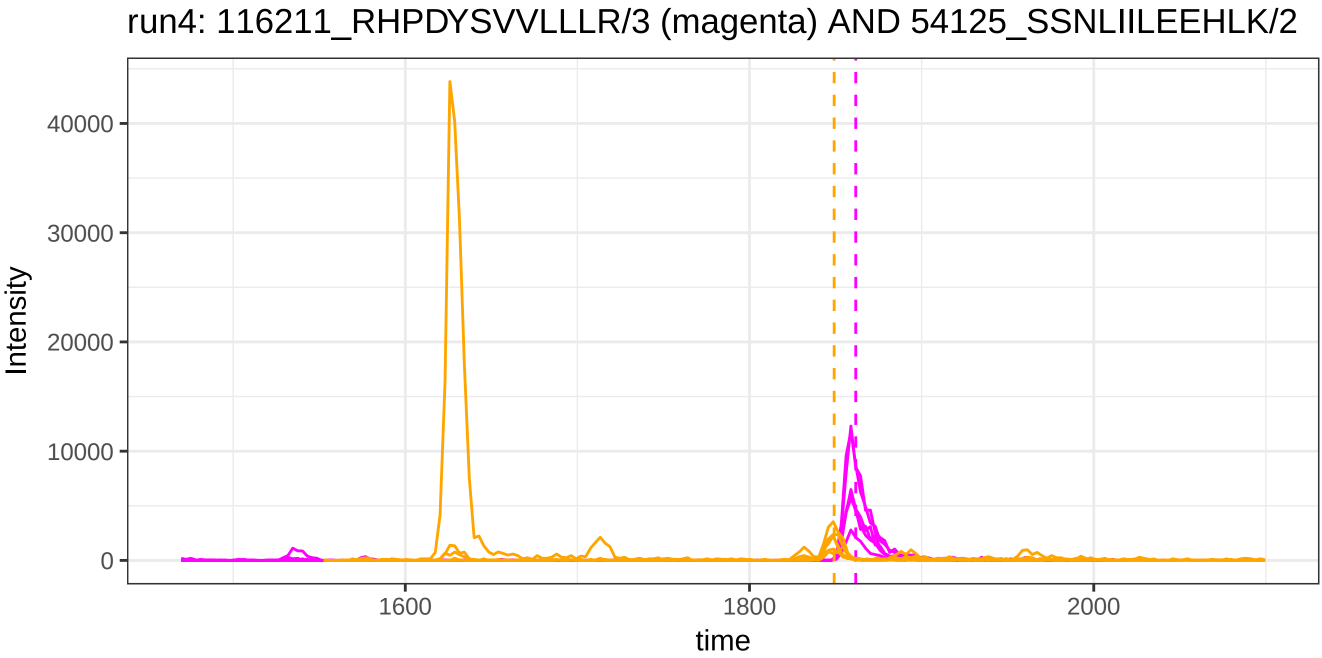
run4: 116211_RHPDYSVVLLLR/3 (magenta) AND 54125_SSNLIILEEHLK/2
Intensity
40000
30000
20000
10000
0
1600
1800
2000
time

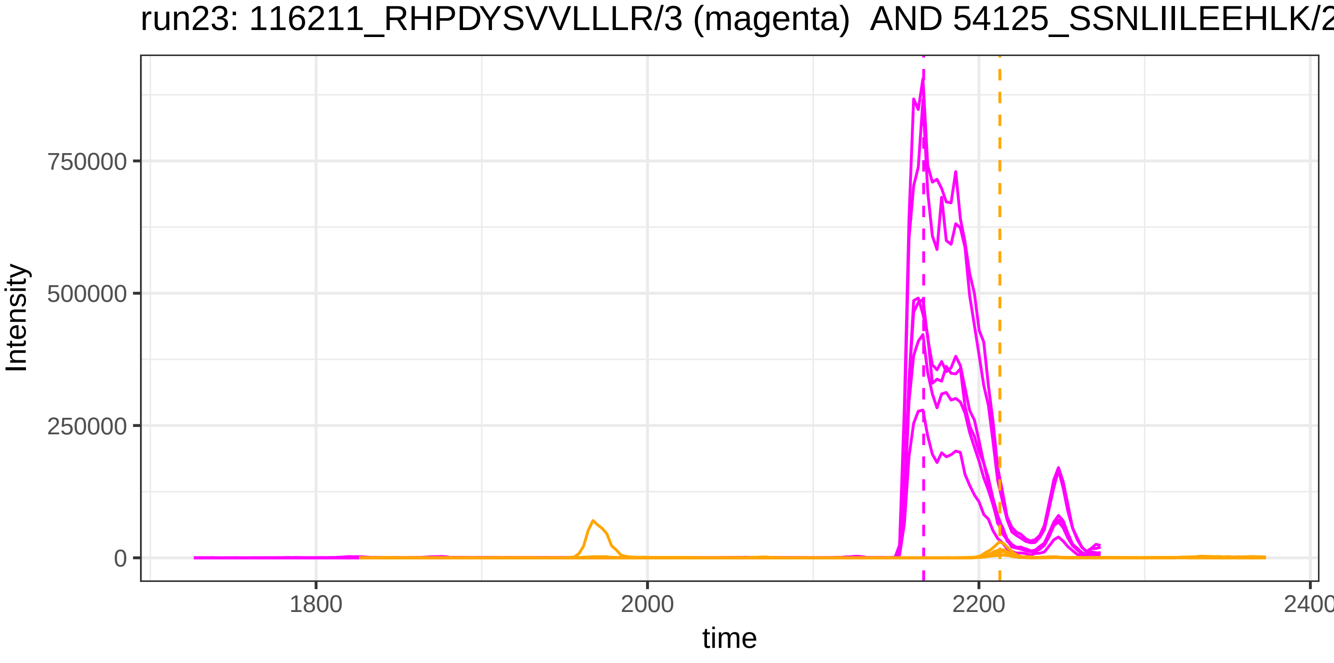
run23: 116211_RHPDYSVVLLLR/3 (magenta) AND 54125_SSNLIILEEHLK/2
Intensity
750000
500000
250000
0
1800
2000
2200
2400
time

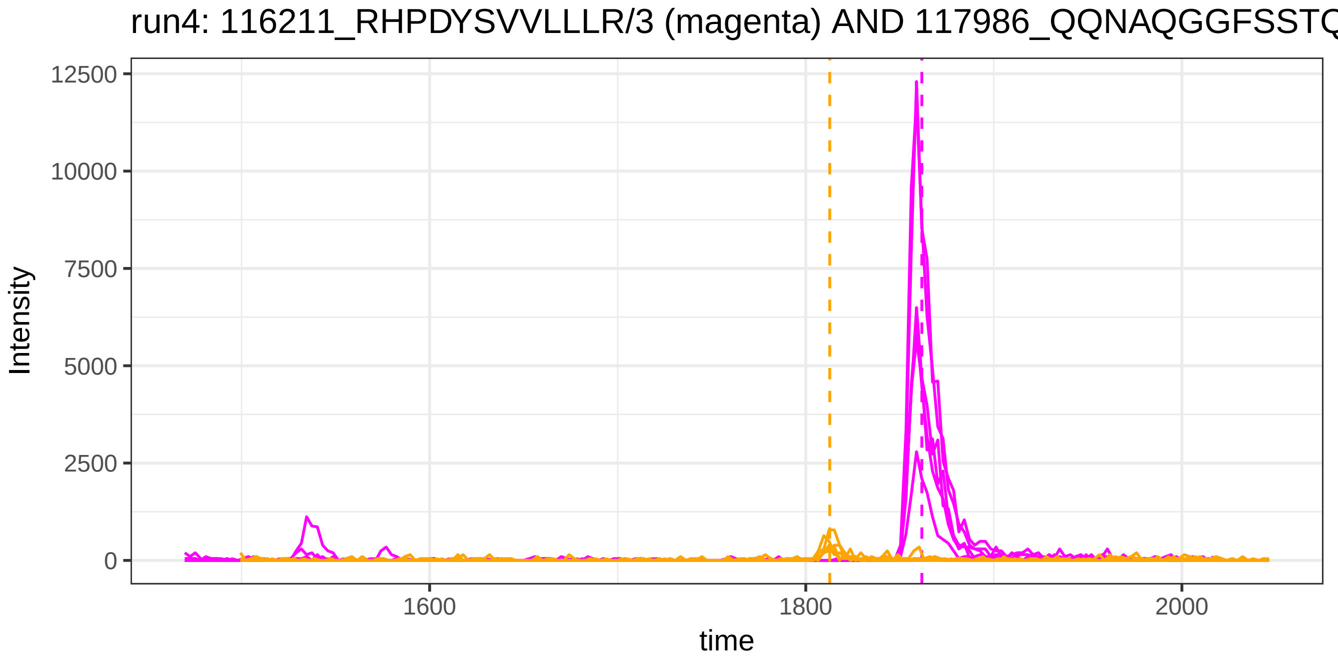
run4: 116211_RHPDYSVVLLLR/3 (magenta) AND 117986_QQNAQGGFSSTQ
Intensity
12500
10000
7500
5000
2500
0
time
1600
1800
2000

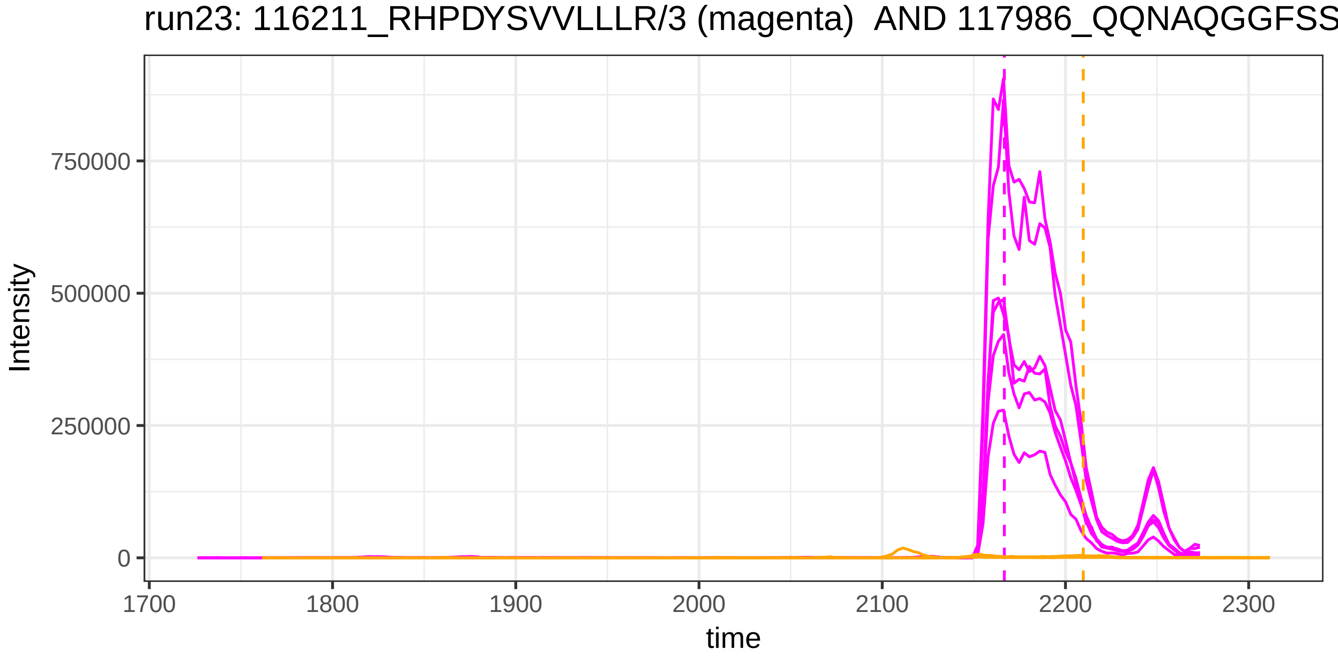
run23: 116211_RHPDYSVVLLLR/3 (magenta) AND 117986_QQNAQGGFSS
Intensity
750000
500000
250000
0
time
1700
1800
1900
2000
2100
2200
2300

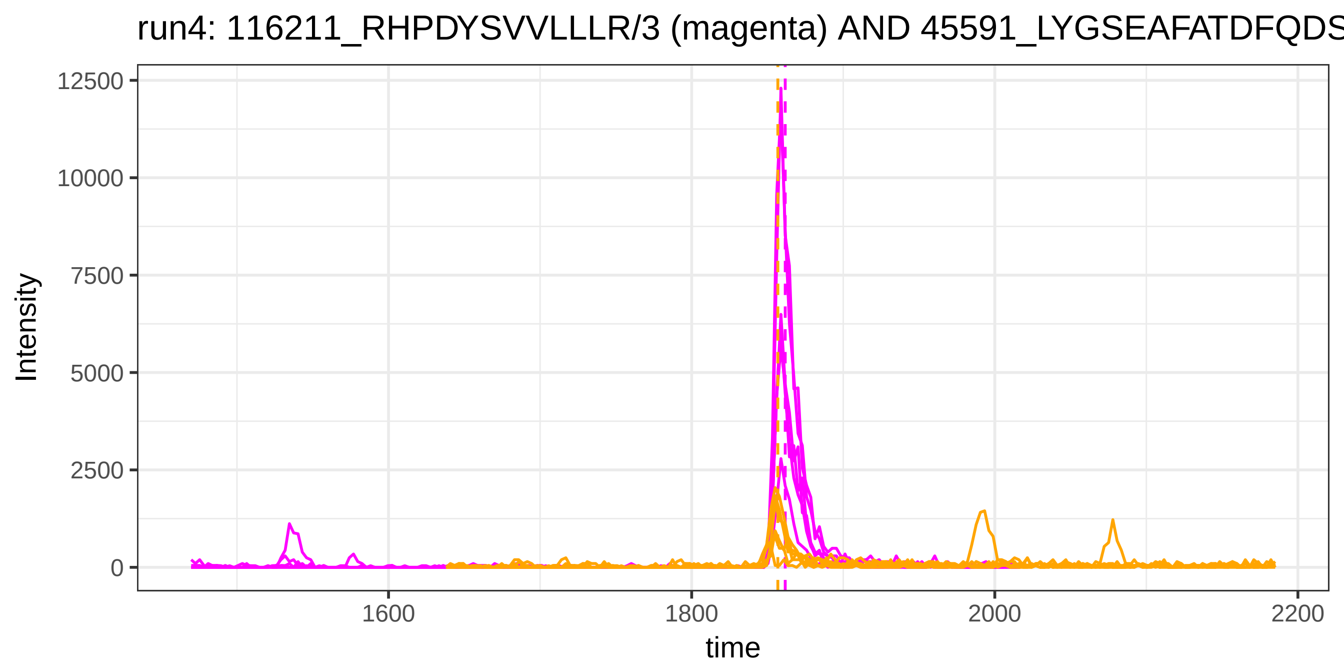
run4: 116211_RHPDYSVVLLLR/3 (magenta) AND 45591_LYGSEAFATDFQDS
Intensity
12500
10000
7500
5000
2500
0
1600
1800
2000
2200
time

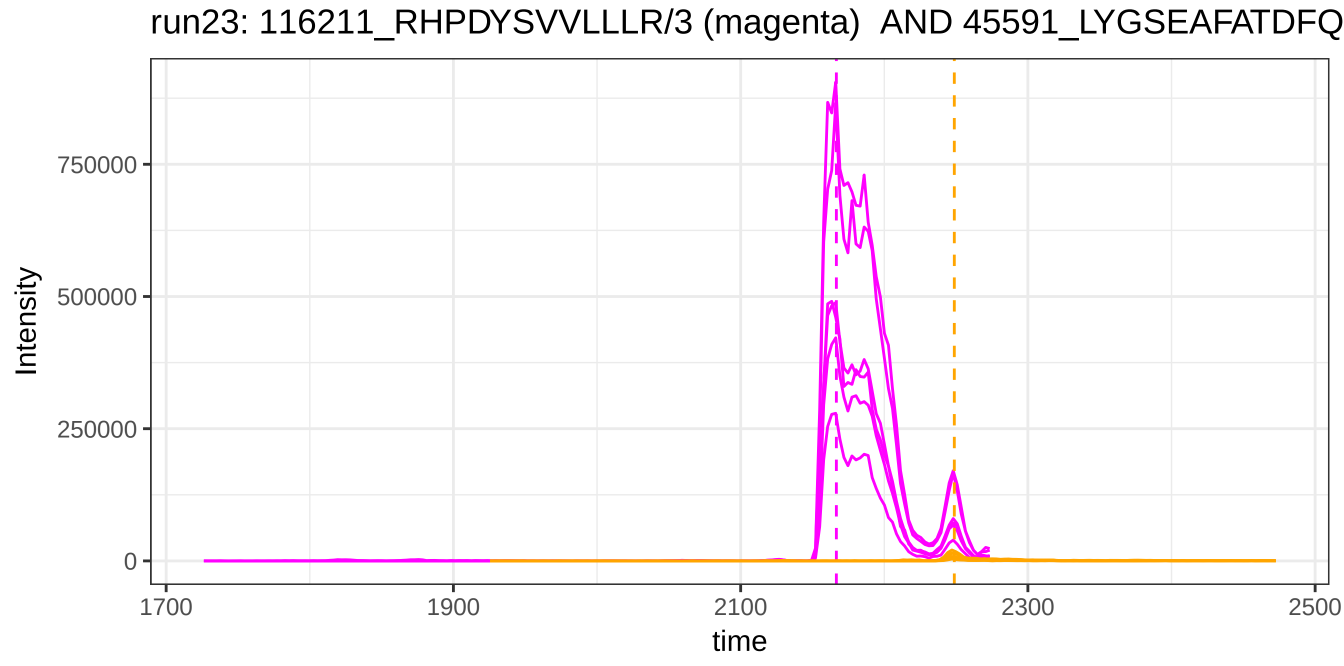
run23: 116211_RHPDYSVVLLLR/3 (magenta) AND 45591_LYGSEAFATDFQ
Intensity
750000
500000
250000
0
1700
1900
2100
2300
2500
time

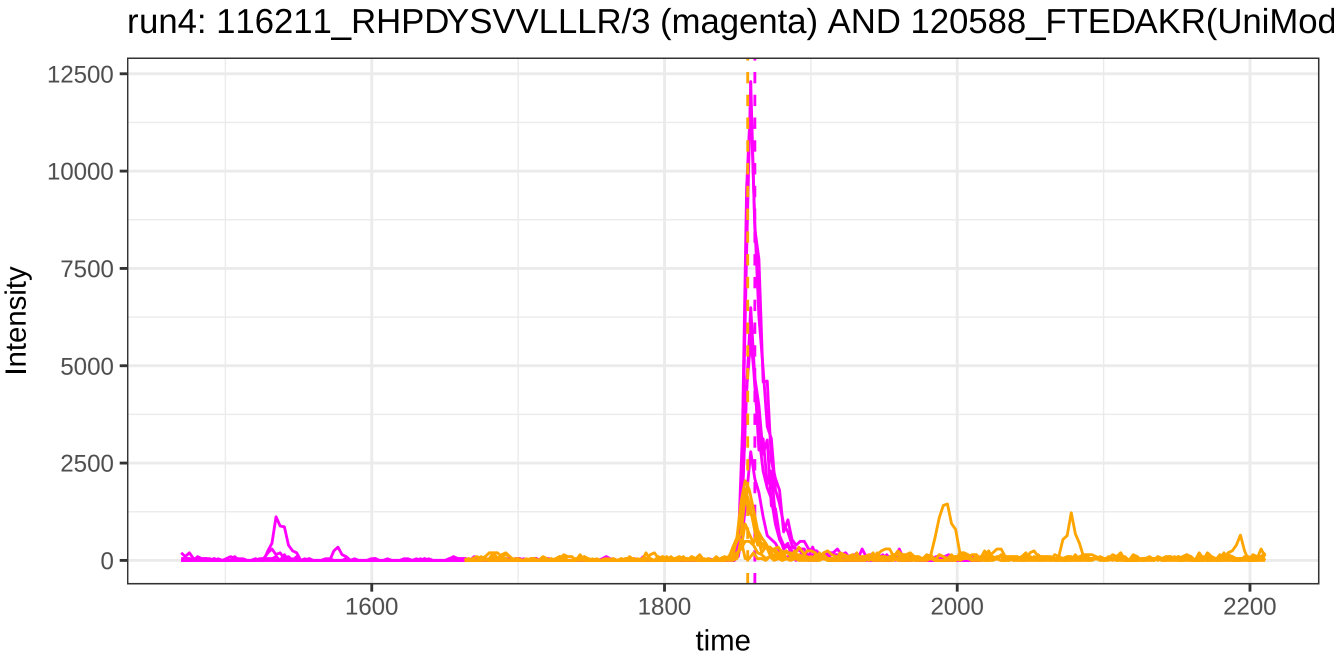

run4: 116211_RHPDYSVVLLLR/3 (magenta) AND 120588_FTEDAKR(UniMod
Intensity
12500
10000
7500
5000
2500
0
1600
1800
2000
2200
time


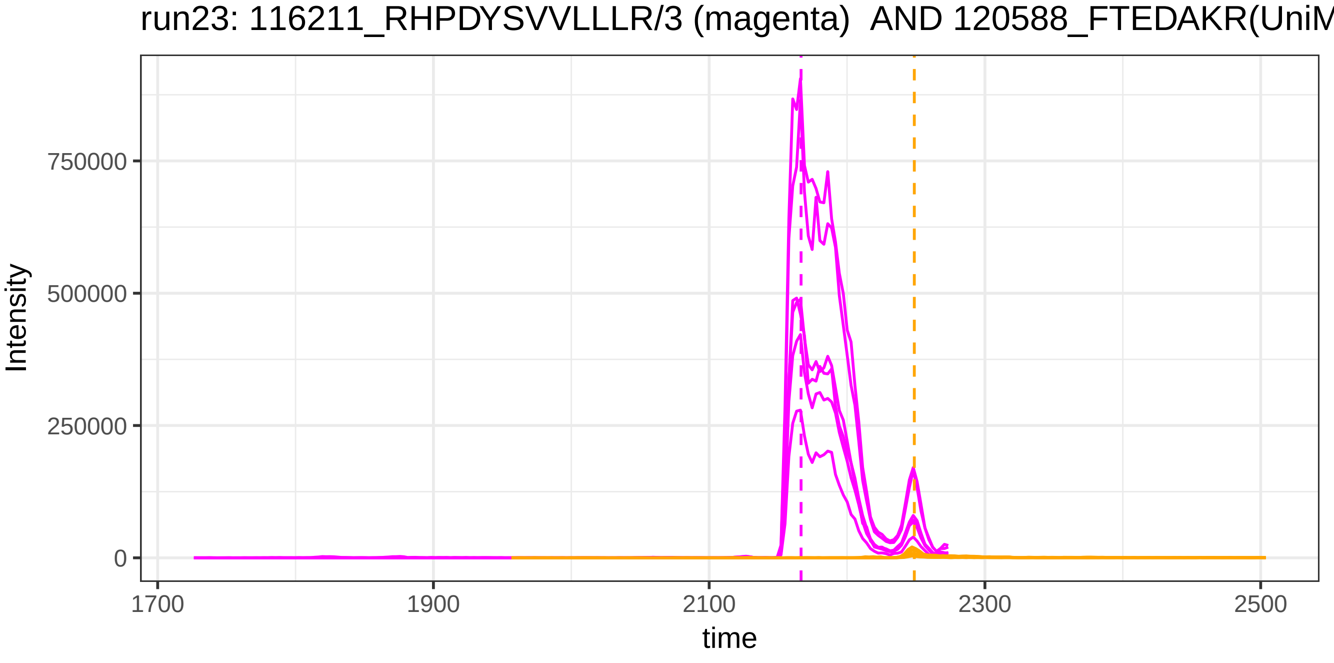

run23: 116211_RHPDYSVVLLLR/3 (magenta) AND 120588_FTEDAKR(UniM
Intensity
750000
500000
250000
0
1700
1900
2100
2300
2500
time

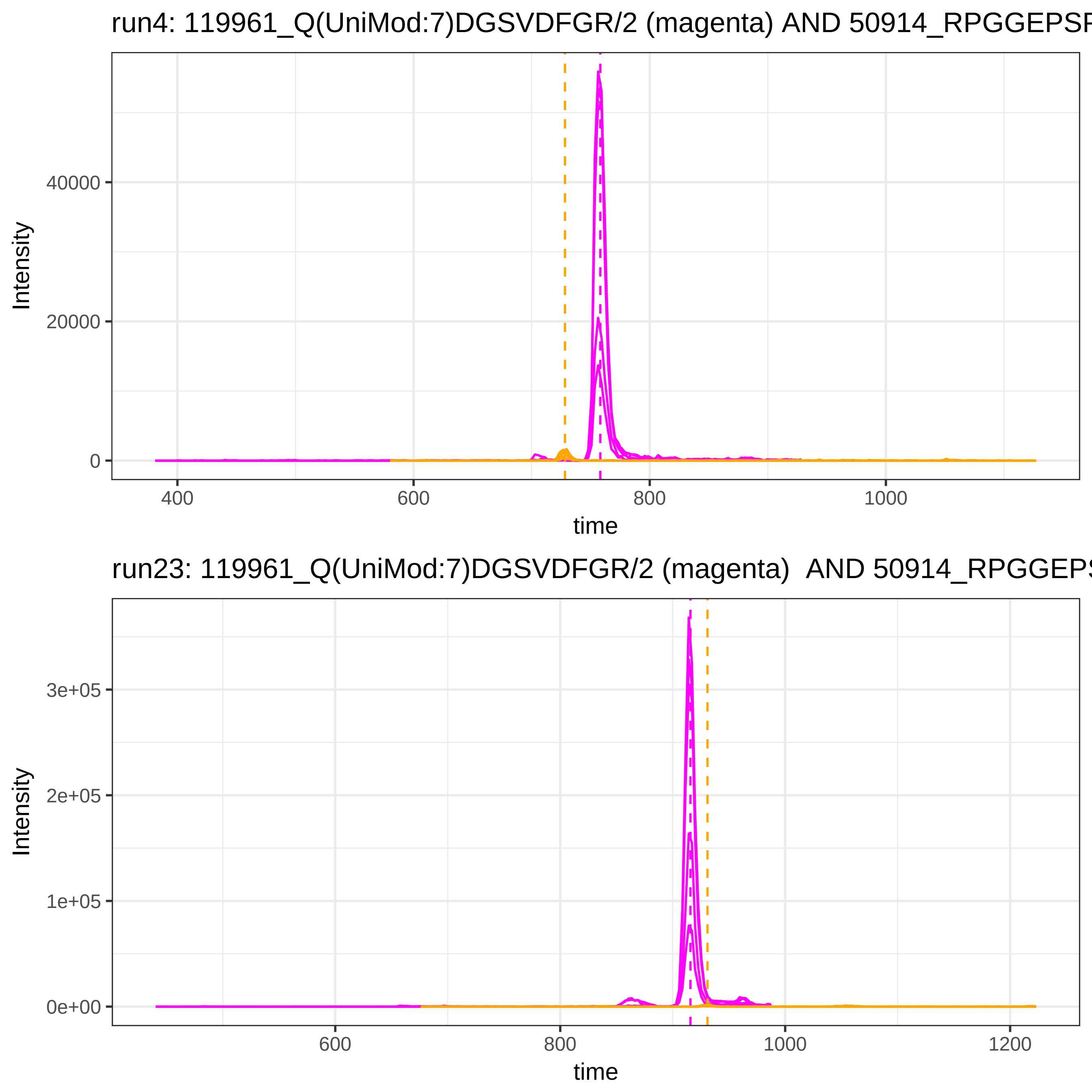
run4: 119961_Q(UniMod:7)DGSVDFGR/2 (magenta) AND 50914_RPGGEPSF
Intensity
40000
20000
0
400
600
800
1000
time
run23: 119961_Q(UniMod:7)DGSVDFGR/2 (magenta) AND 50914_RPGGEPS
Intensity
3e+05
2e+05
1e+05
0e+00
600
800
1000
1200
time

run4: 119961_Q(UniMod:7)DGSVDFGR/2 (magenta) AND 117476_RPGGEPS

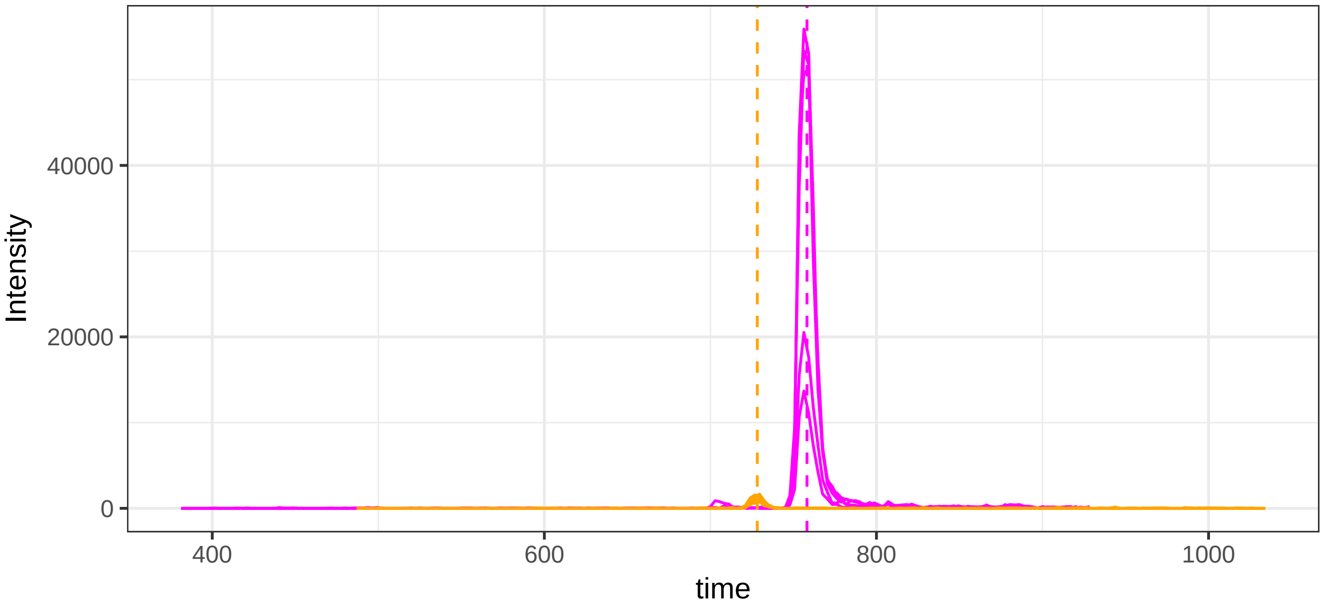


run23: 119961_Q(UniMod:7)DGSVDFGR/2 (magenta)  AND 117476_RPGGEF

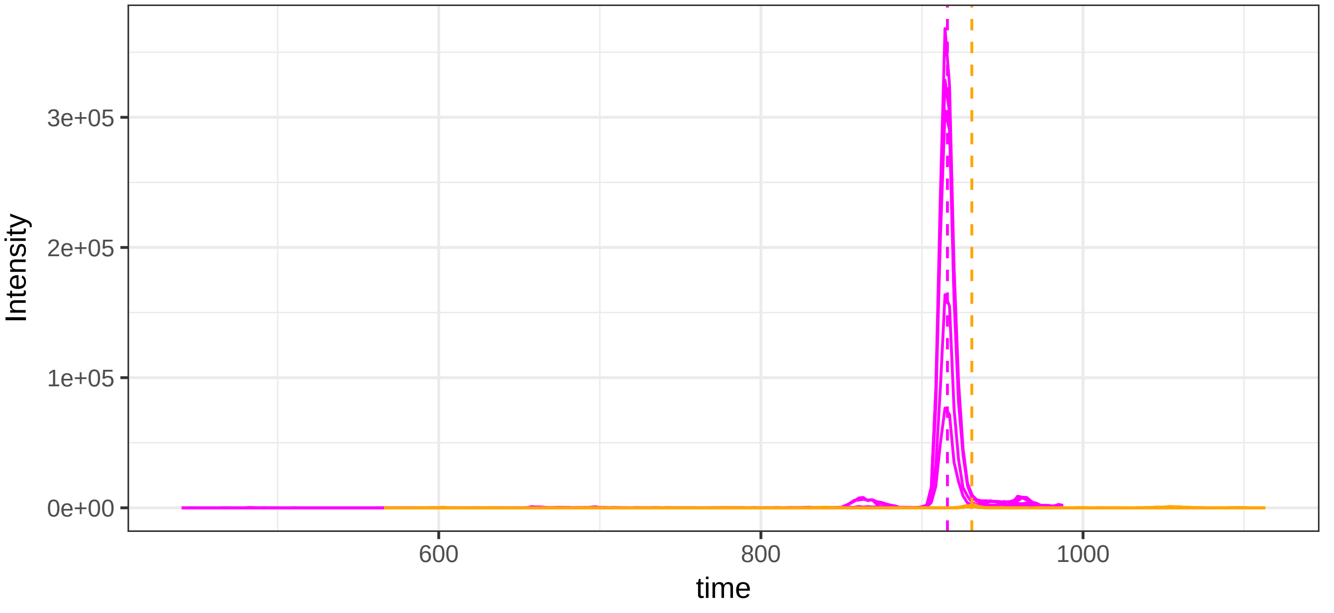

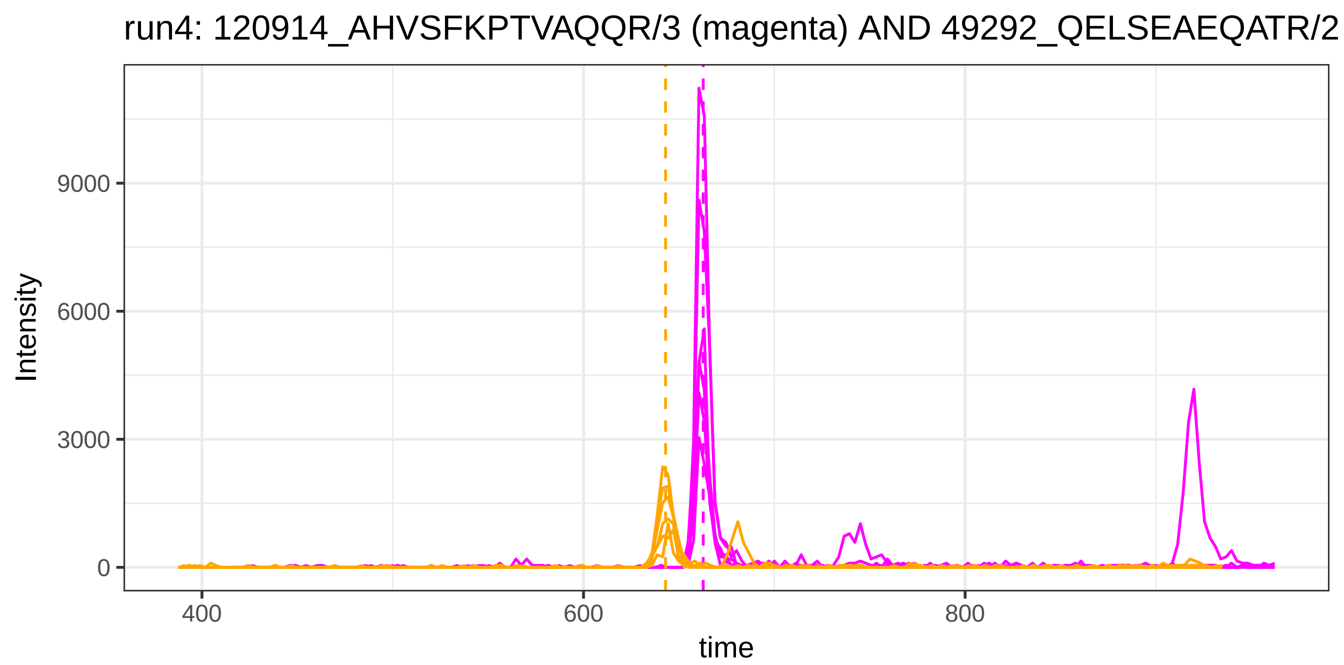
run4: 120914_AHVSFKPTVAQQR/3 (magenta) AND 49292_QELSEAEQATR/2
Intensity
9000
6000
3000
0
400
600
800
time

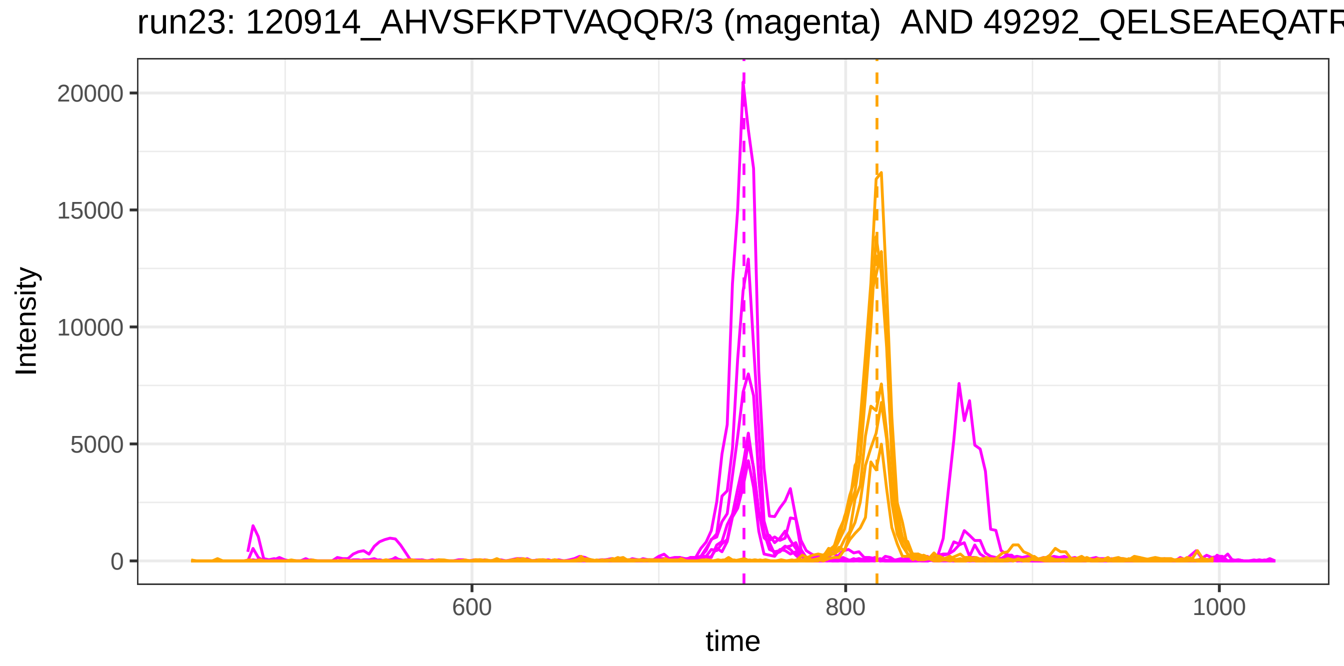
run23: 120914_AHVSFKPTVAQQR/3 (magenta)  AND 49292_QELSEAEQATR
Intensity
20000
15000
10000
5000
0
600
800
1000
time

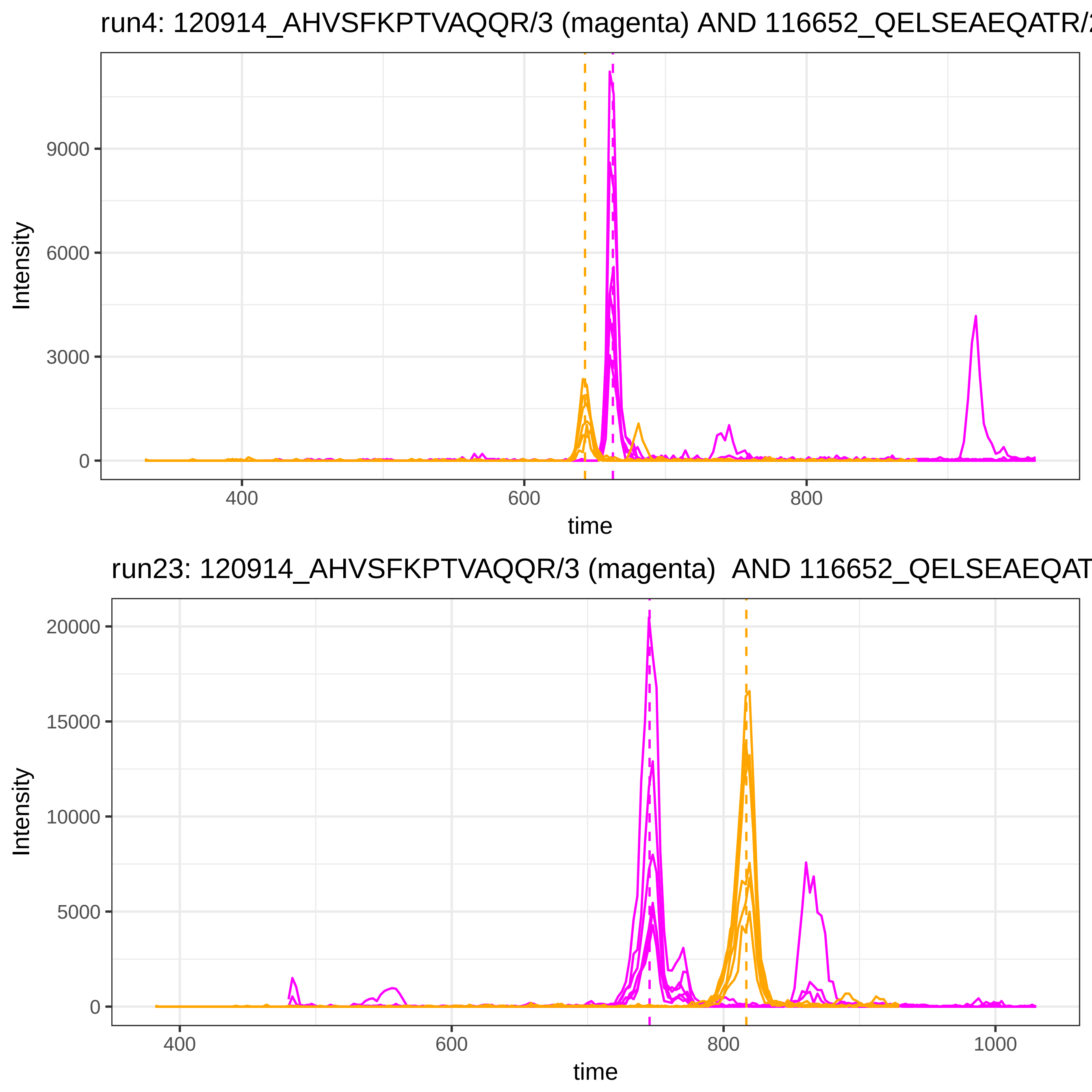

run4: 120914_AHVSFKPTVAQQR/3 (magenta) AND 116652_QELSEAEQATR/
Intensity
9000
6000
3000
0
400
600
800
time
run23: 120914_AHVSFKPTVAQQR/3 (magenta) AND 116652_QELSEAEQAT
Intensity
20000
15000
10000
5000
0
400
600
800
1000
time

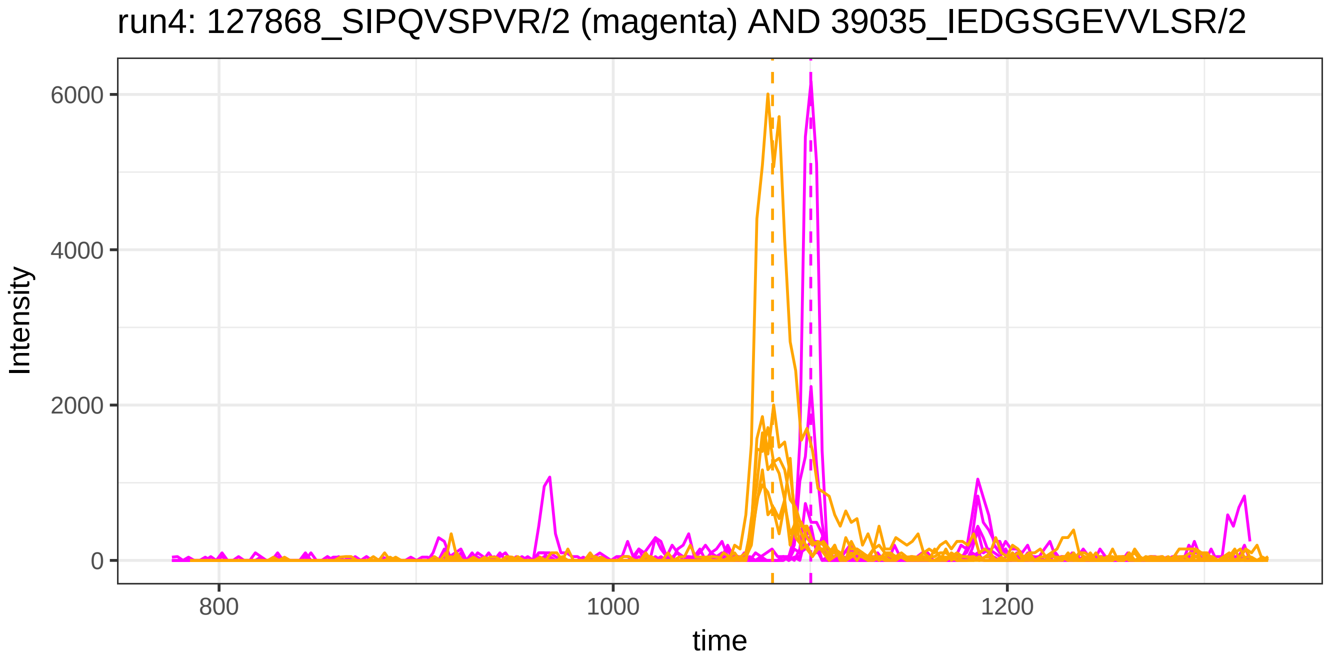
run4: 127868_SIPQVSPVR/2 (magenta) AND 39035_IEDGSGEVVLSR/2
Intensity
6000
4000
2000
0
800
1000
1200
time

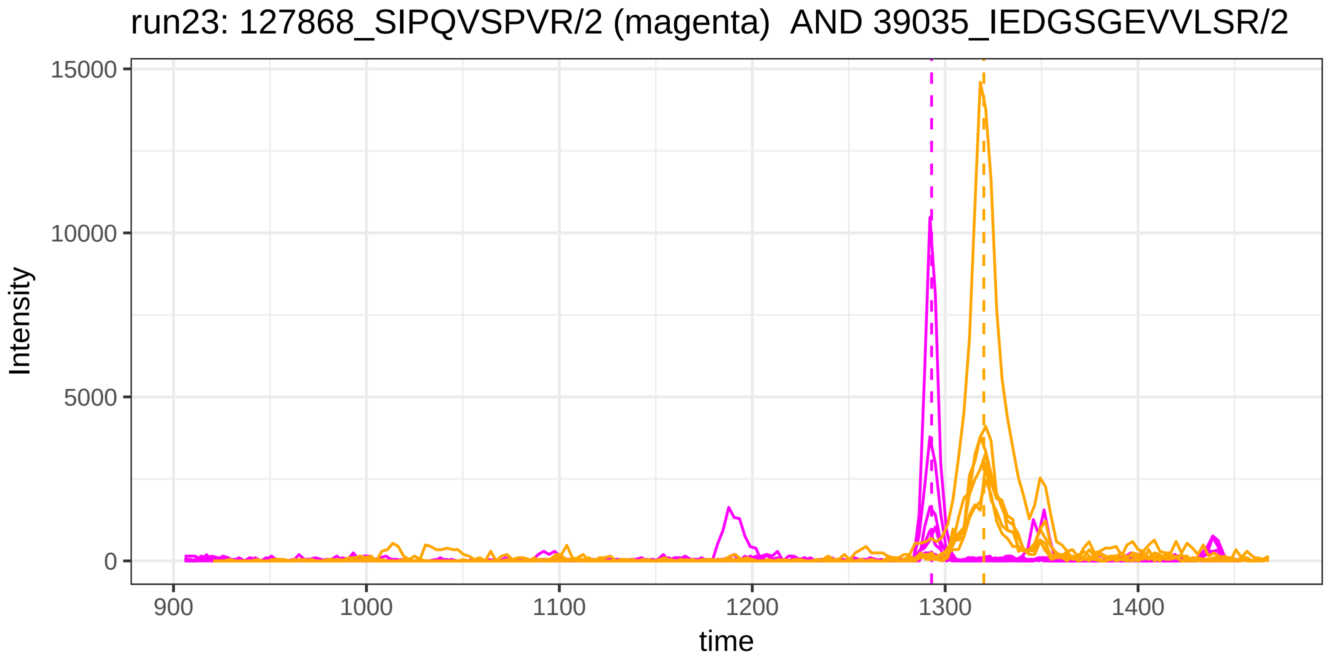
run23: 127868_SIPQVSPVR/2 (magenta) AND 39035_IEDGSGEVVLSR/2
Intensity
15000
10000
5000
0
900
1000
1100
1200
1300
1400
time

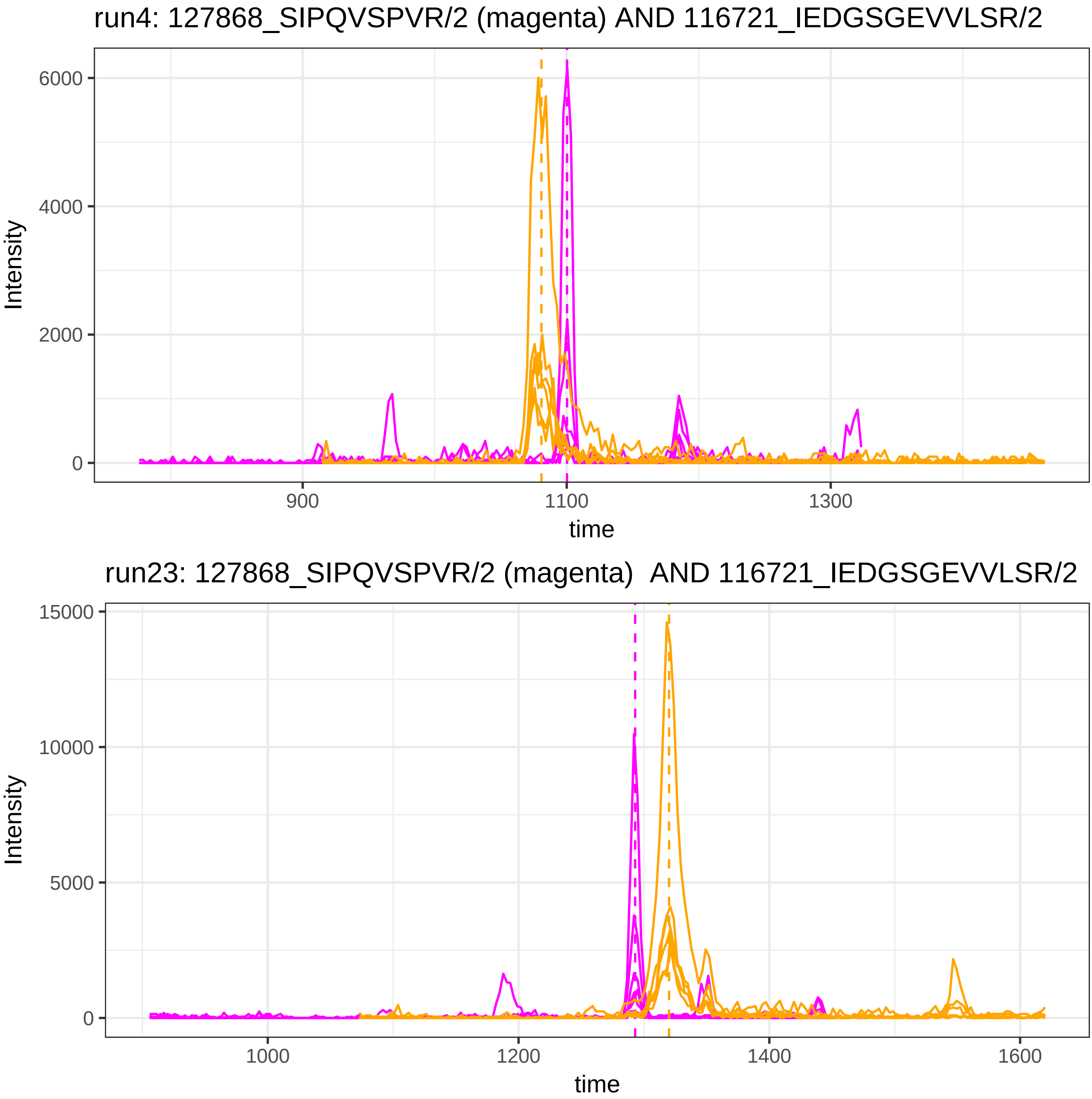

run4: 127868_SIPQVSPVR/2 (magenta) AND 116721_IEDGSGEVVLSR/2
Intensity
6000
4000
2000
0
900
1100
1300
time
run23: 127868_SIPQVSPVR/2 (magenta) AND 116721_IEDGSGEVVLSR/2
Intensity
15000
10000
5000
0
1000
1200
1400
1600
time

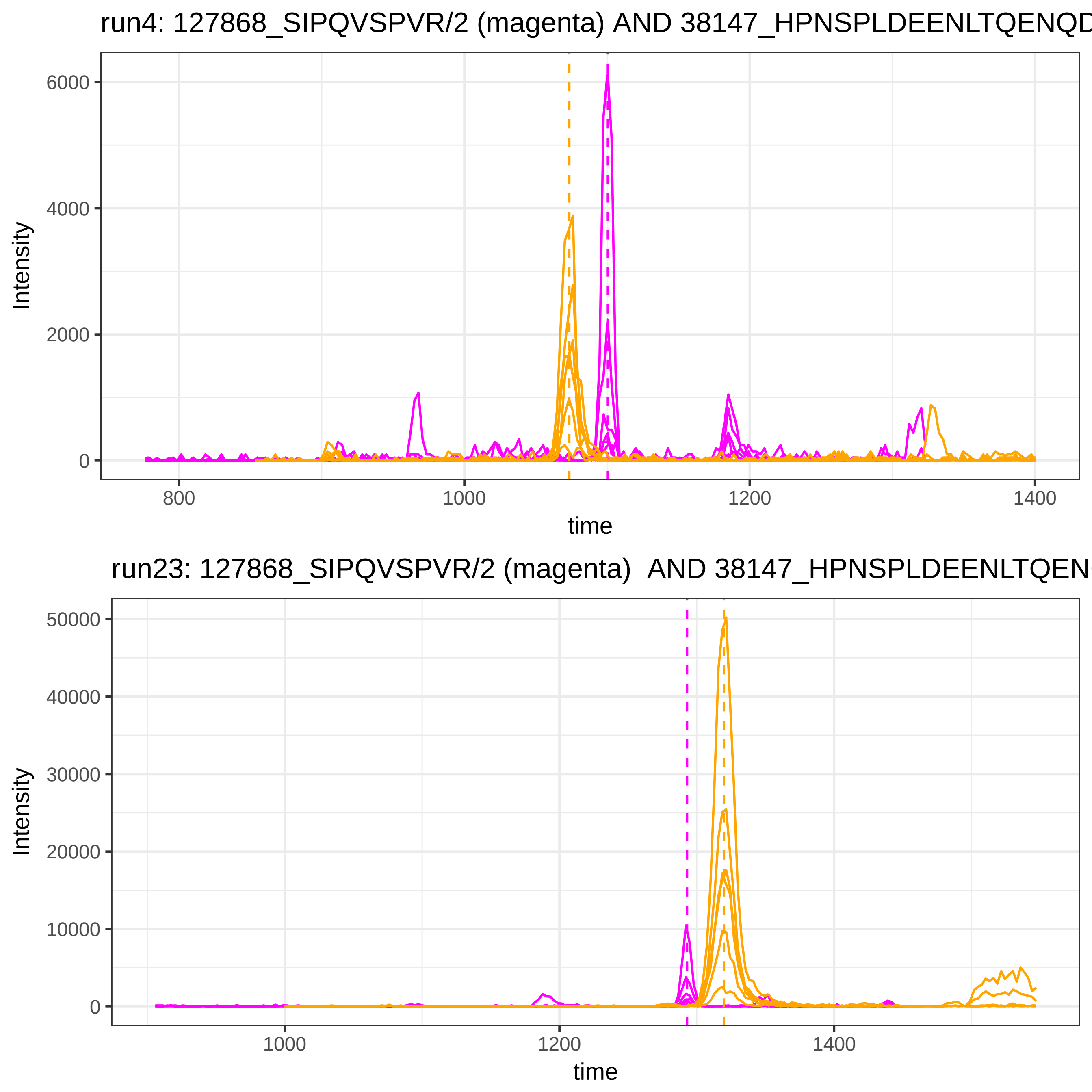
run4: 127868_SIPQVSPVR/2 (magenta) AND 38147_HPNSPLDEENLTQENQD
Intensity
6000
4000
2000
0
800
1000
1200
1400
time
run23: 127868_SIPQVSPVR/2 (magenta) AND 38147_HPNSPLDEENLTQEN
Intensity
50000
40000
30000
20000
10000
0
1000
1200
1400
time

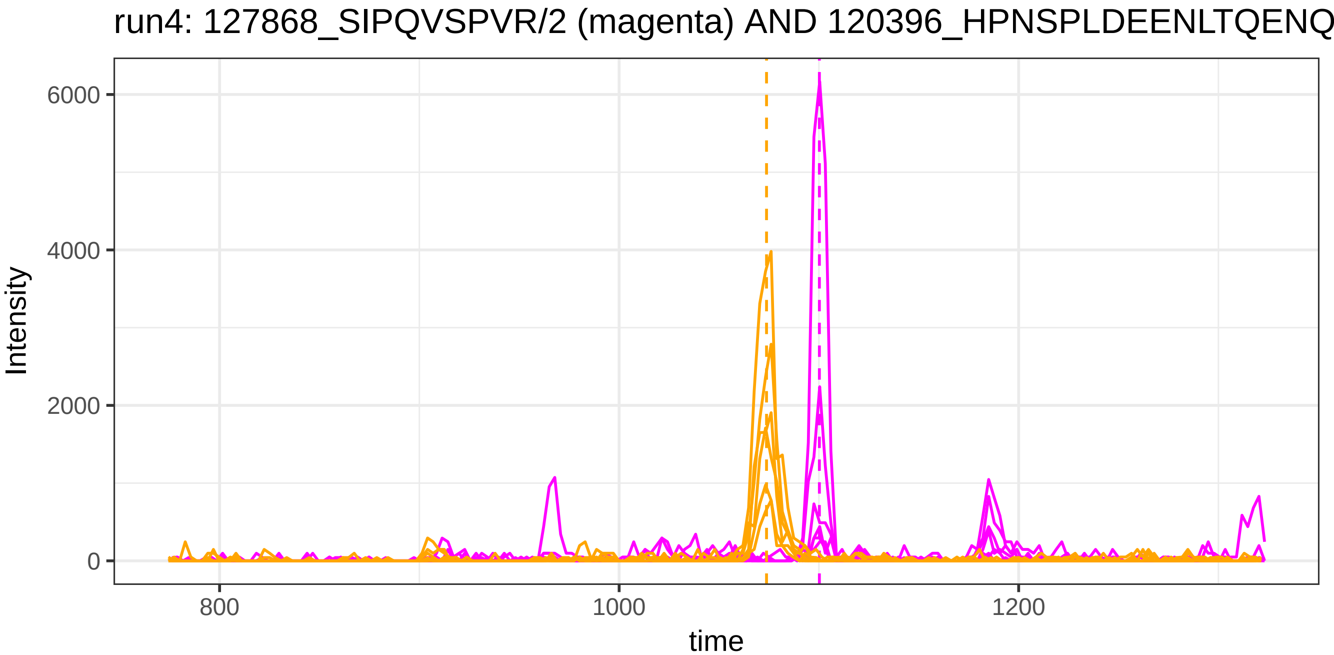
run4: 127868_SIPQVSPVR/2 (magenta) AND 120396_HPNSPLDEENLTQENQ
Intensity
6000
4000
2000
0
800
1000
1200
time

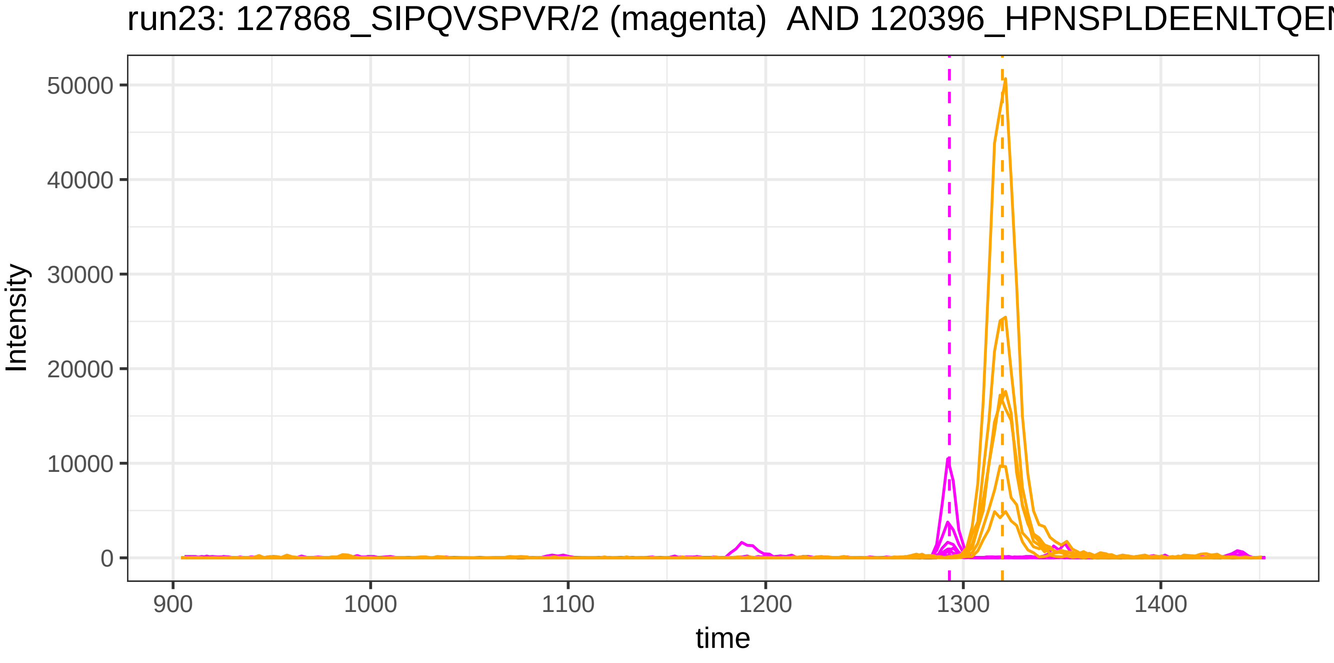
run23: 127868_SIPQVSPVR/2 (magenta) AND 120396_HPNSPLDEENLTQEN
Intensity
50000
40000
30000
20000
10000
0
900
1000
1100
1200
1300
1400
time

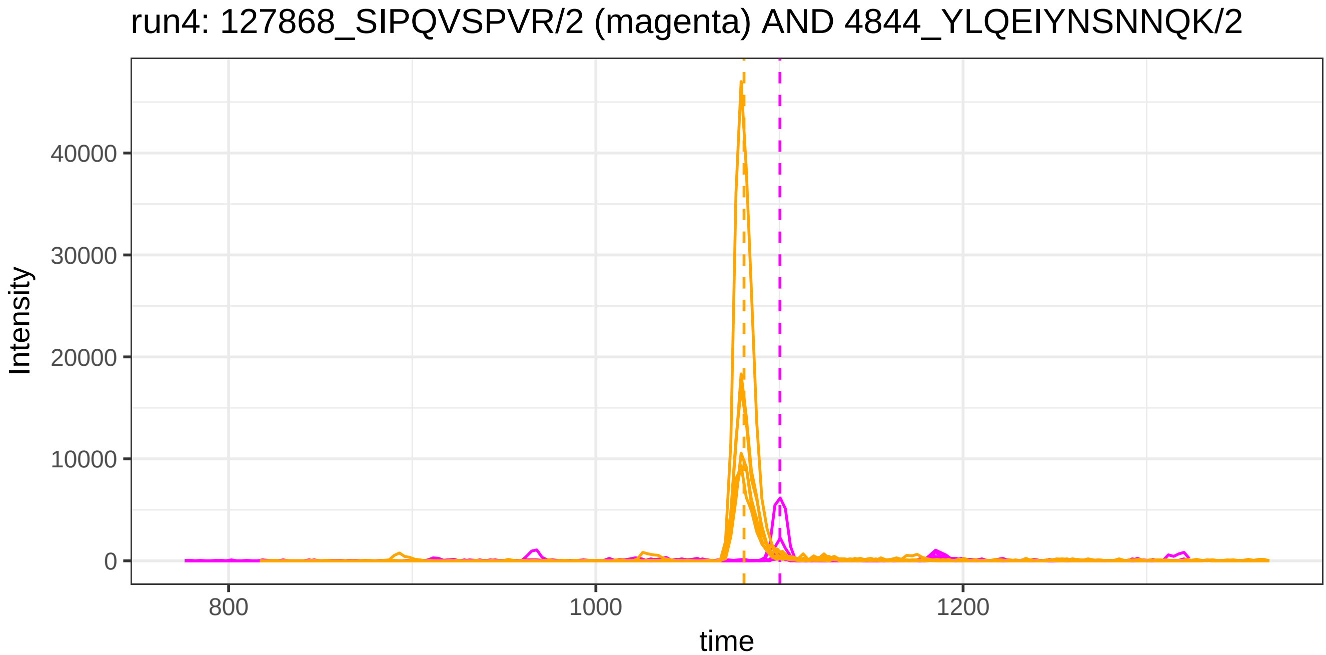
run4: 127868_SIPQVSPVR/2 (magenta) AND 4844_YLQEIYNSNNQK/2
Intensity
40000
30000
20000
10000
0
800
1000
1200
time

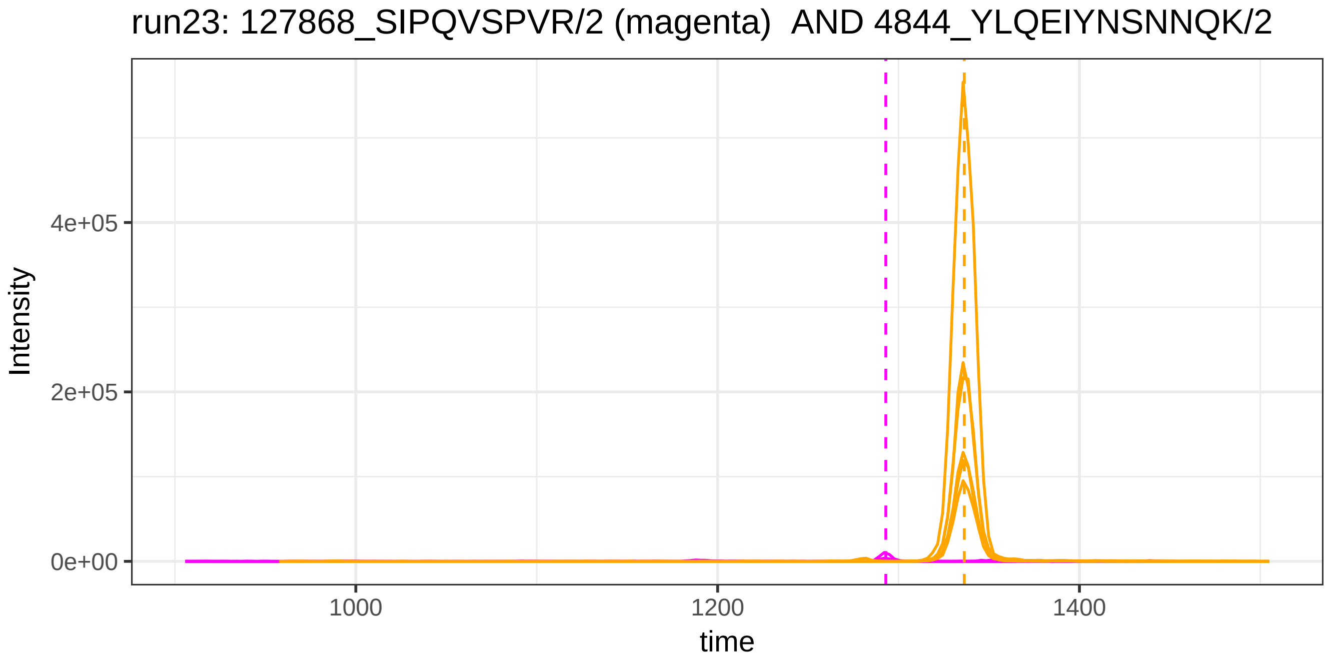
run23: 127868_SIPQVSPVR/2 (magenta)  AND 4844_YLQEIYNSNNQK/2
Intensity
4e+05
2e+05
0e+00
1000
1200
1400
time

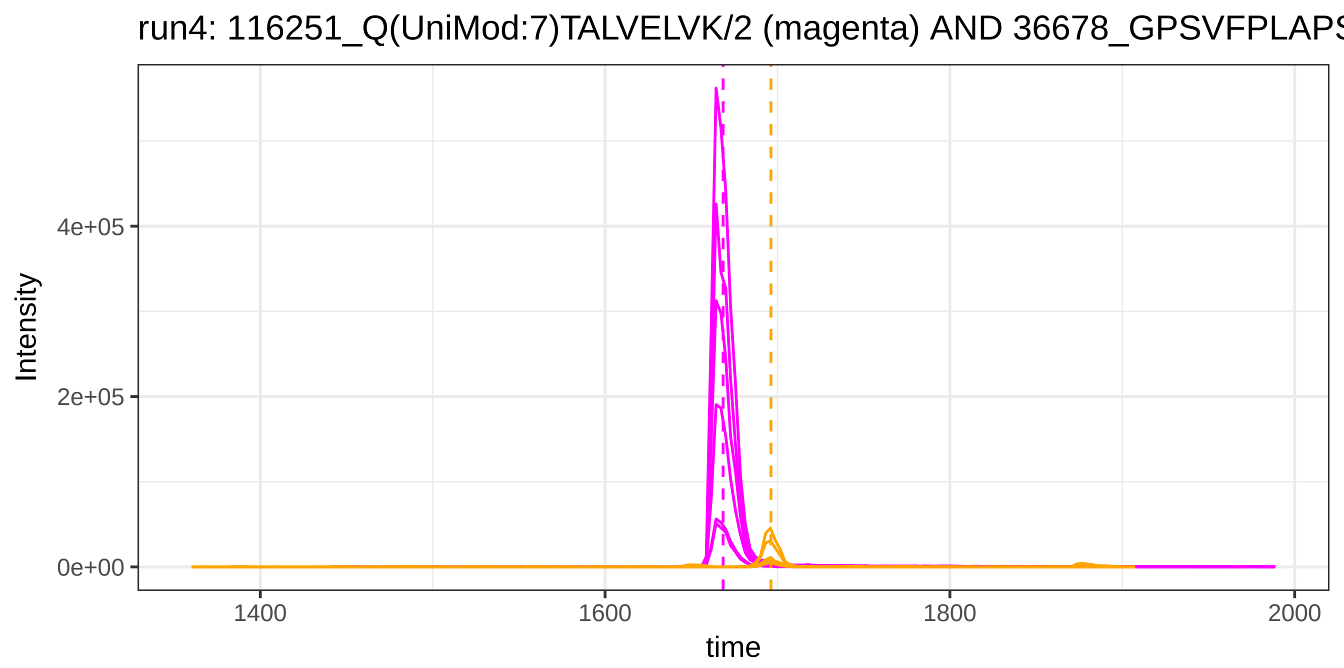

run4: 116251_Q(UniMod:7)TALVELVK/2 (magenta) AND 36678_GPSVFPLAPS
Intensity
4e+05
2e+05
0e+00
1400
1600
1800
2000
time


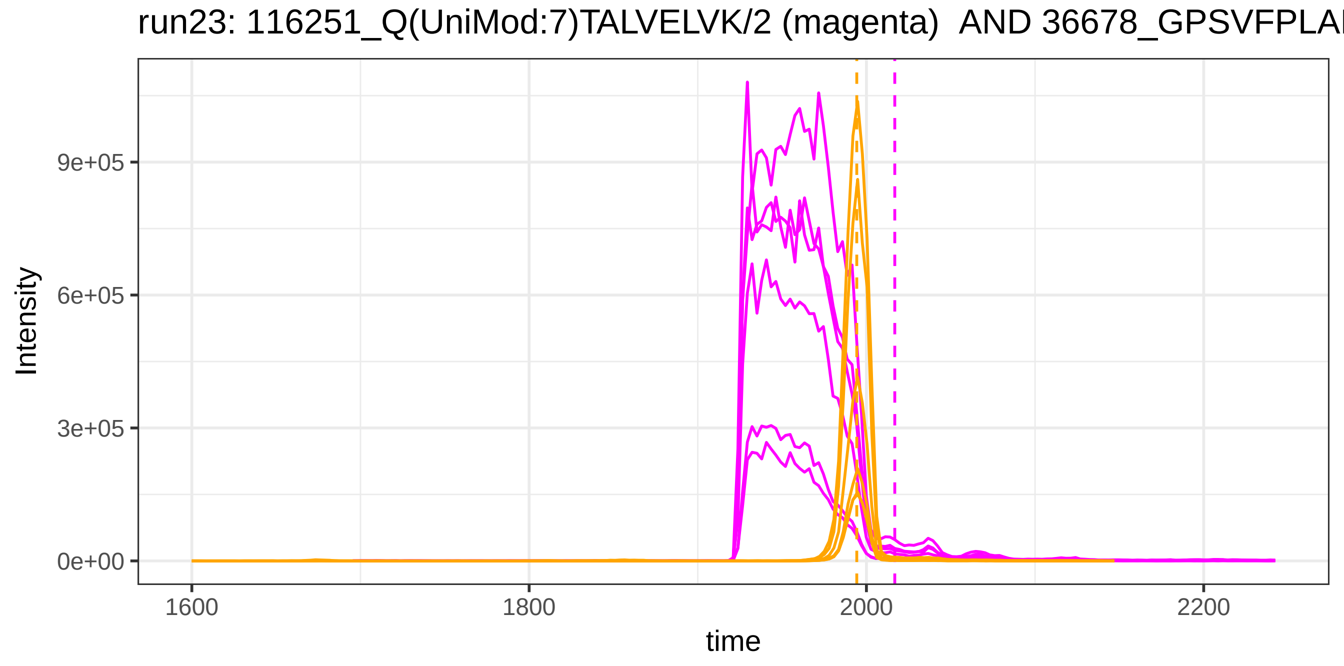

run23: 116251_Q(UniMod:7)TALVELVK/2 (magenta) AND 36678_GPSVFPLAI
Intensity
9e+05
6e+05
3e+05
0e+00
1600
1800
2000
2200
time

run4: 116480_Q(UniMod:7)IKKQTALVELVK/3 (magenta) AND 25212_DWHGV

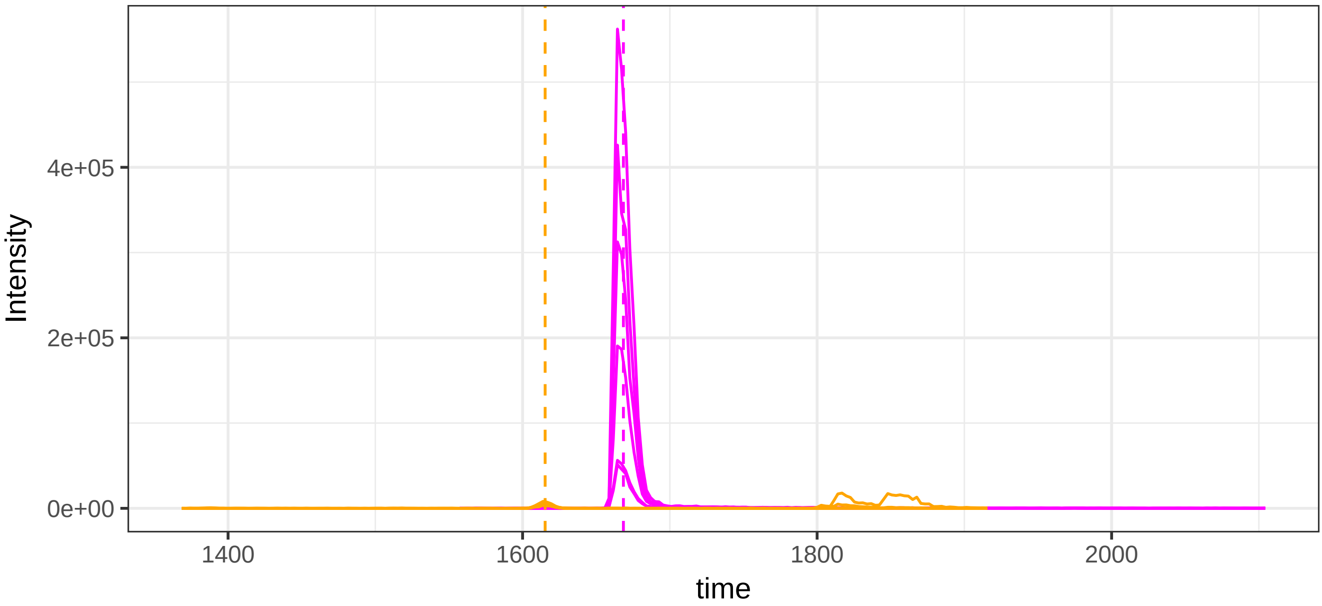


run23: 116480_Q(UniMod:7)IKKQTALVELVK/3 (magenta)  AND 25212_DWHG

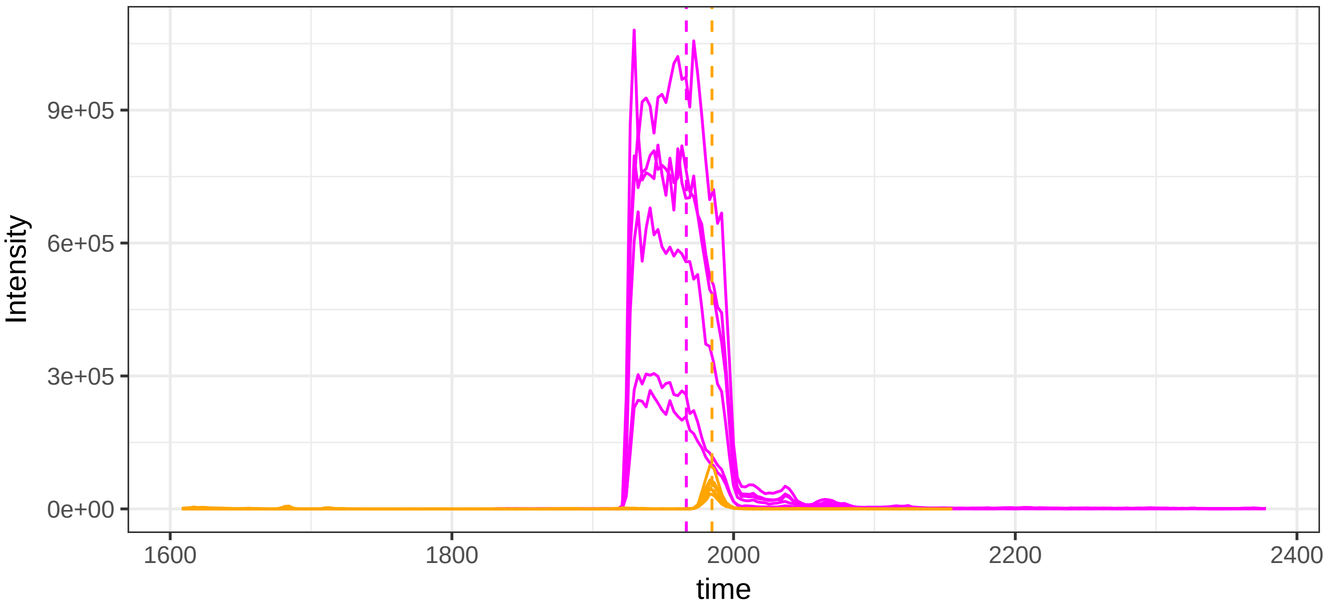

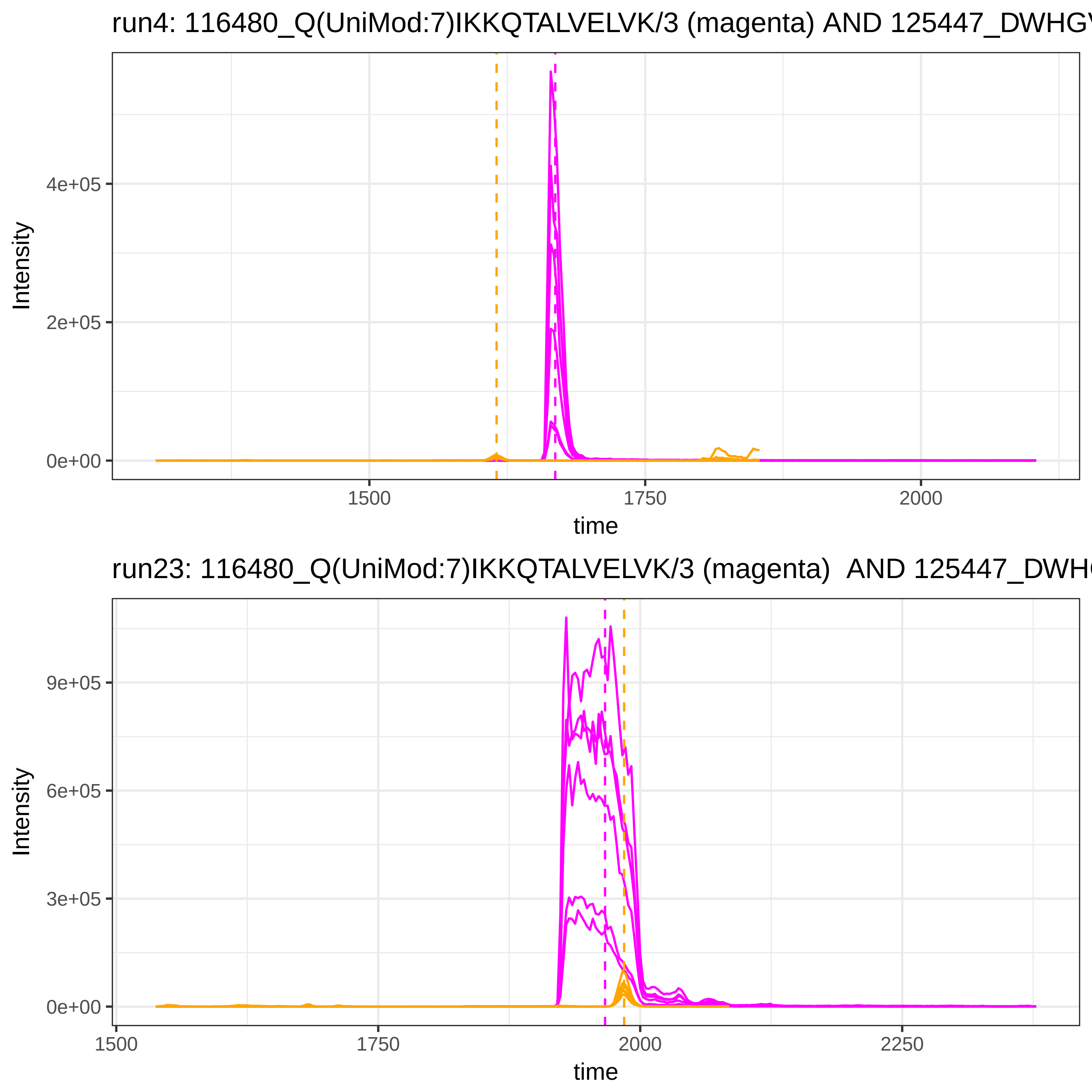

run4: 116480_Q(UniMod:7)IKKQTALVELVK/3 (magenta) AND 125447_DWHGV
Intensity
4e+05
2e+05
0e+00
1500
1750
2000
time
run23: 116480_Q(UniMod:7)IKKQTALVELVK/3 (magenta) AND 125447_DWH
Intensity
9e+05
6e+05
3e+05
0e+00
1500
1750
2000
2250
time

run4: 116480_Q(UniMod:7)IKKQTALVELVK/3 (magenta) AND 125481_DWHGV

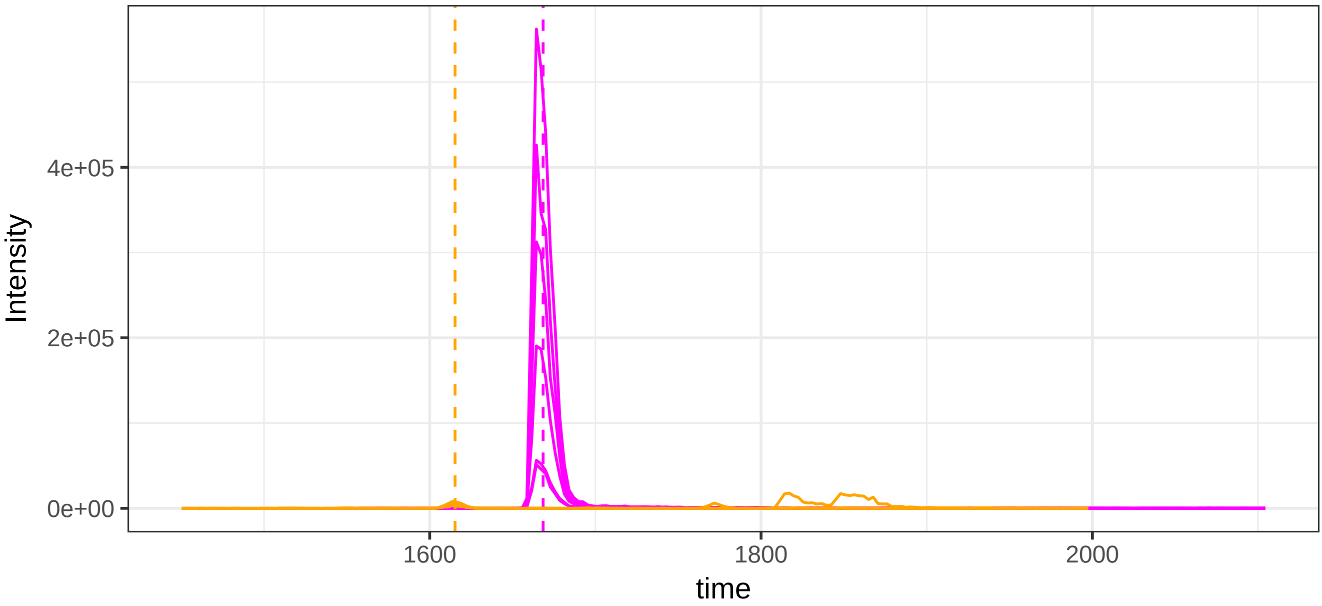


run23: 116480_Q(UniMod:7)IKKQTALVELVK/3 (magenta)  AND 125481_DWHG

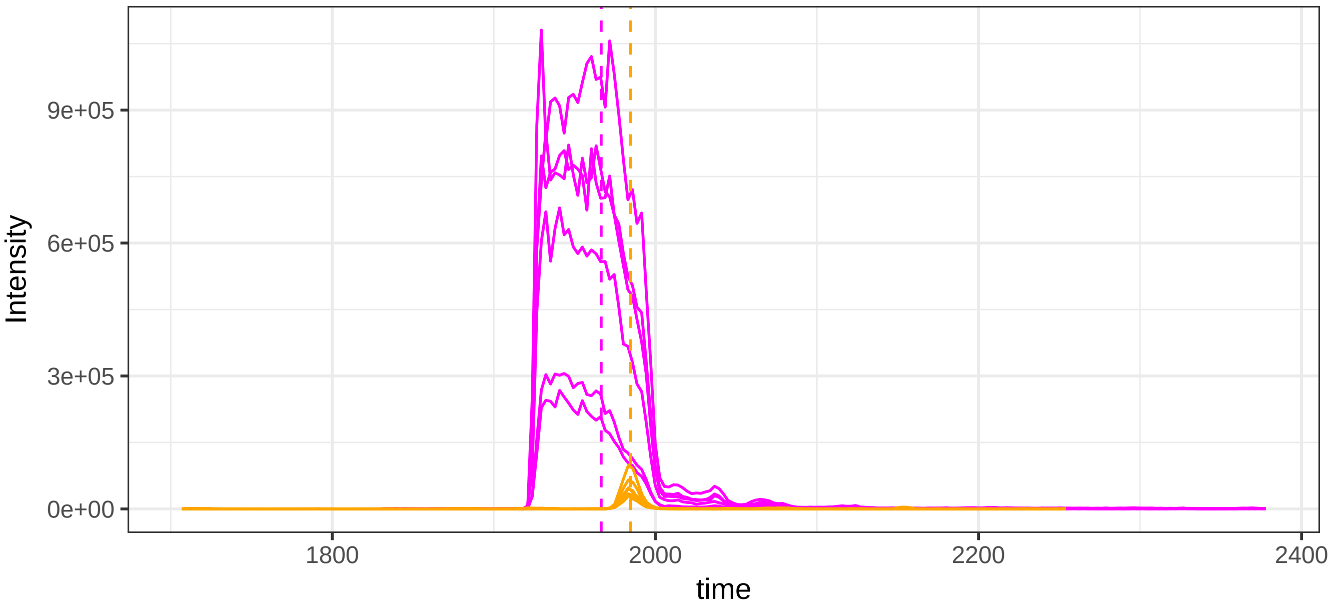

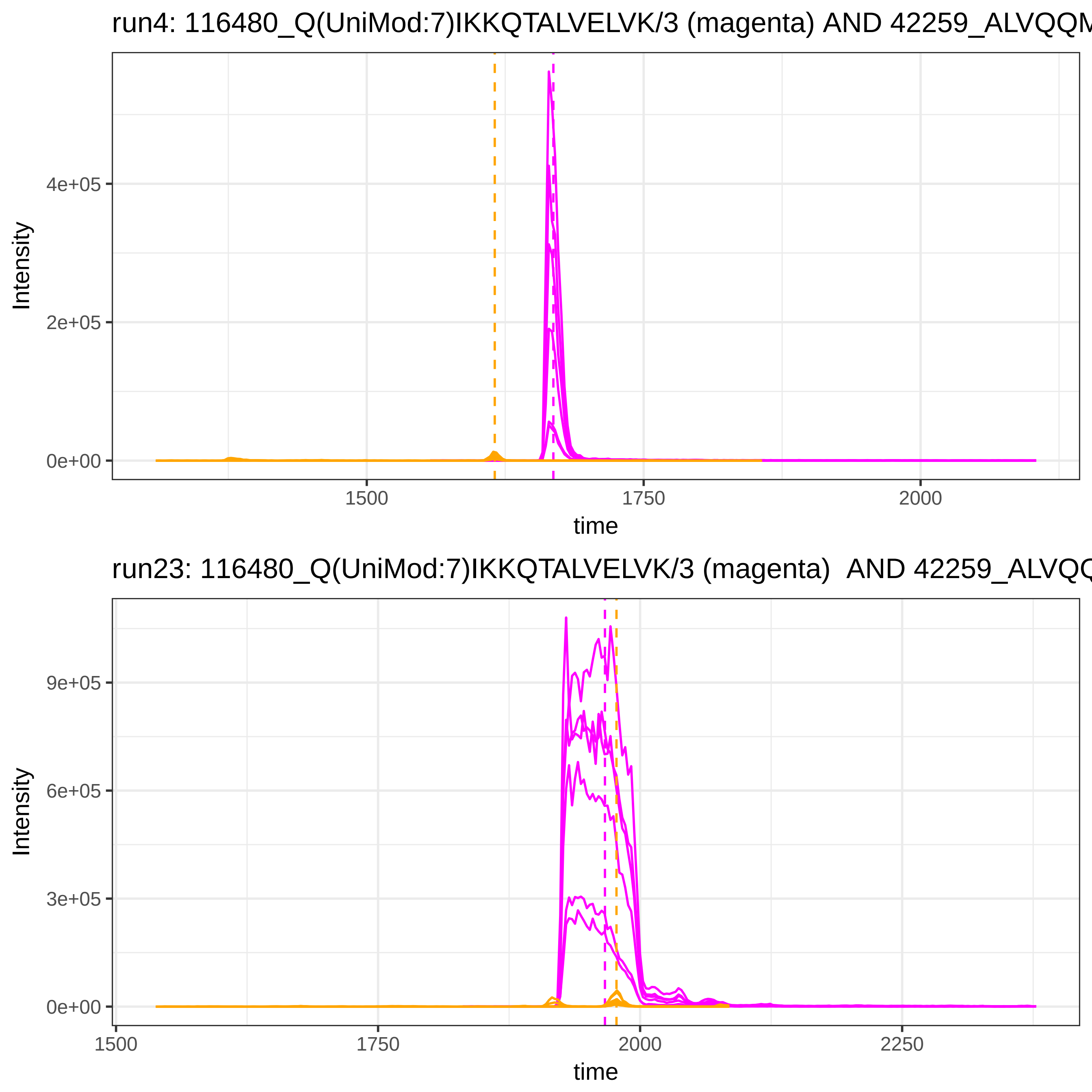

run4: 116480_Q(UniMod:7)IKKQTALVELVK/3 (magenta) AND 42259_ALVQQM
Intensity
4e+05
2e+05
0e+00
1500
1750
2000
time
run23: 116480_Q(UniMod:7)IKKQTALVELVK/3 (magenta) AND 42259_ALVQQ
Intensity
9e+05
6e+05
3e+05
0e+00
1500
1750
2000
2250
time

run4: 116480_Q(UniMod:7)IKKQTALVELVK/3 (magenta) AND 51375_SASDLTV

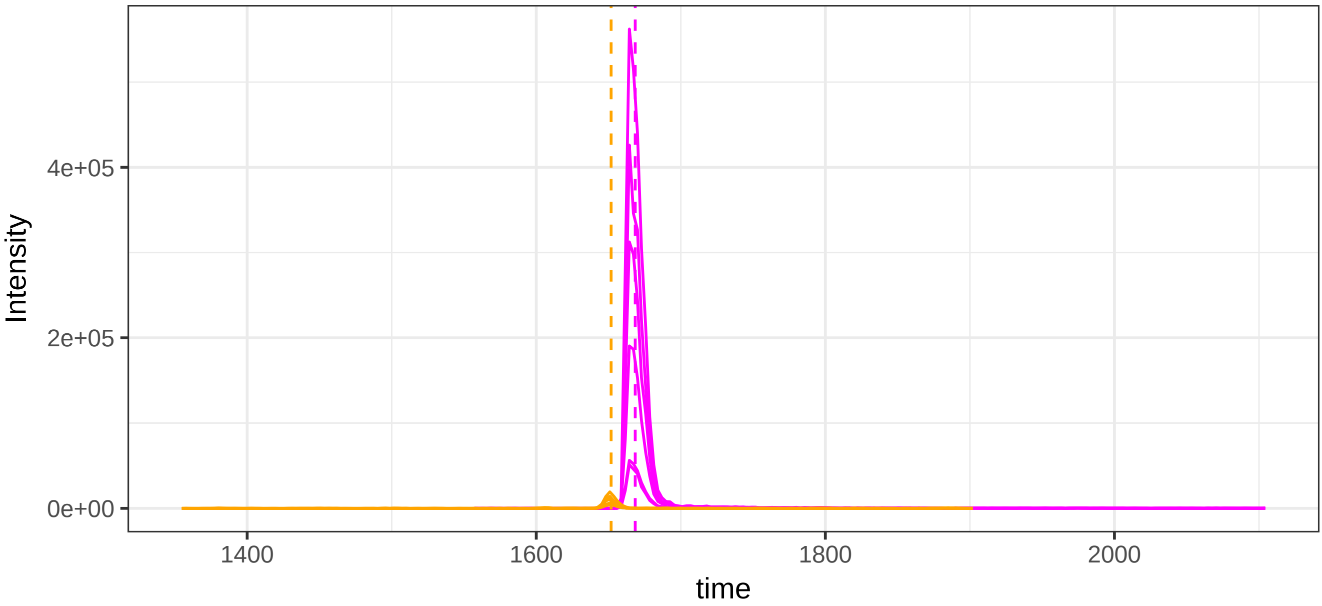


run23: 116480_Q(UniMod:7)IKKQTALVELVK/3 (magenta)  AND 51375_SASDL

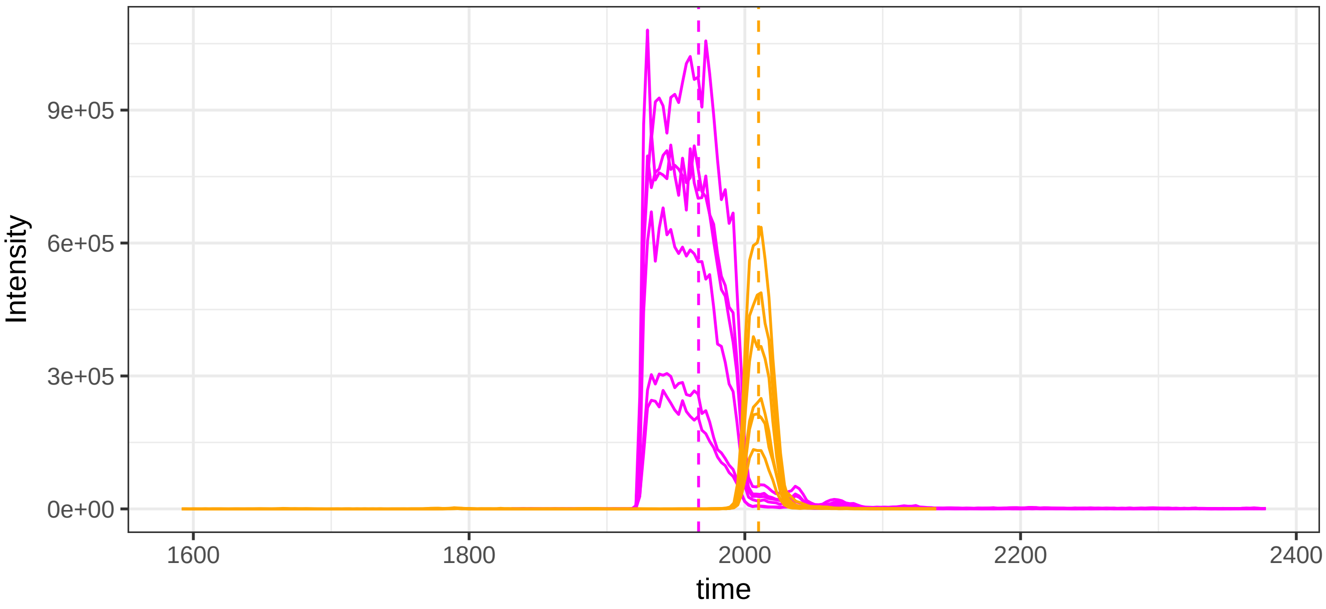

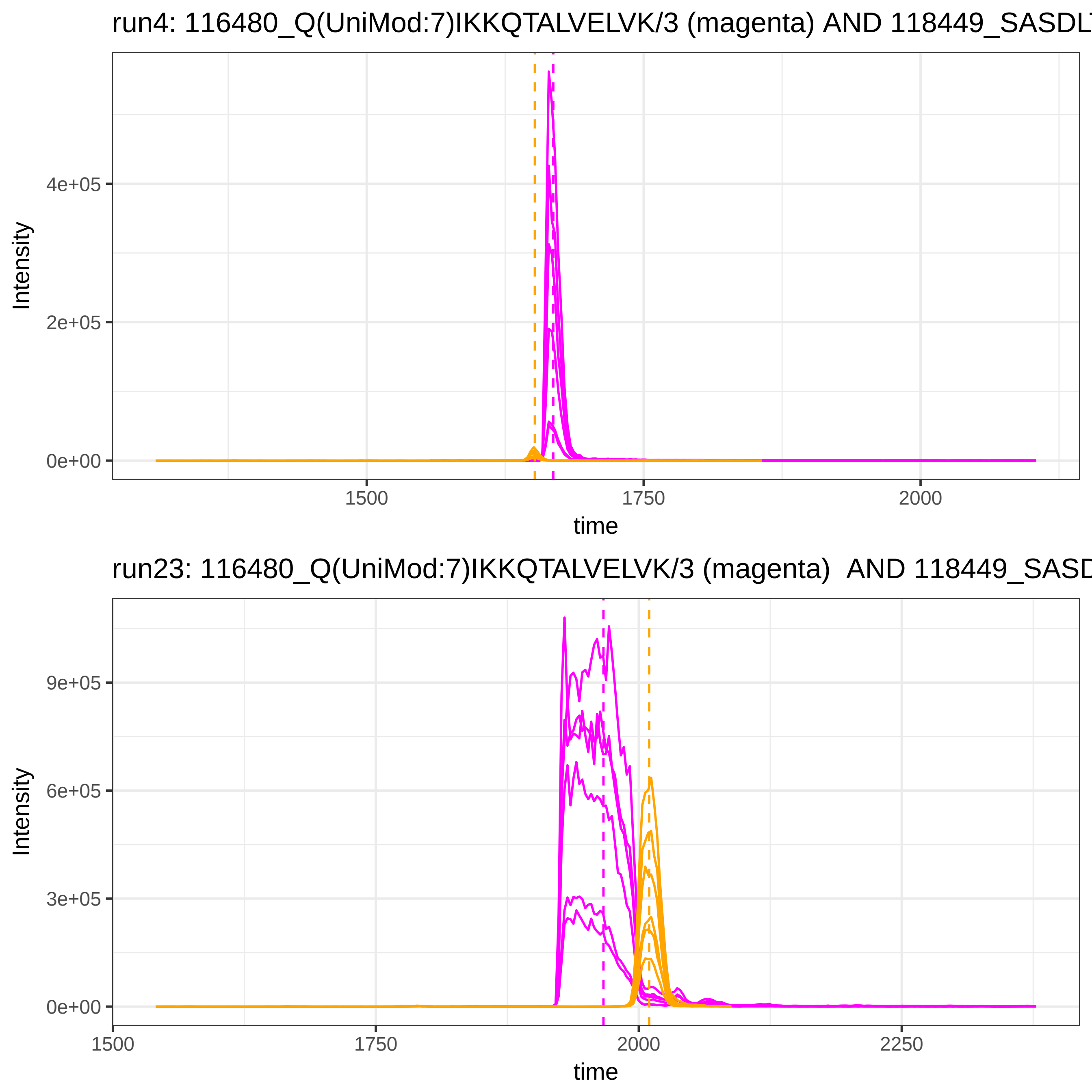
run4: 116480_Q(UniMod:7)IKKQTALVELVK/3 (magenta) AND 118449_SASDL
Intensity
4e+05
2e+05
0e+00
1500
1750
2000
time
run23: 116480_Q(UniMod:7)IKKQTALVELVK/3 (magenta) AND 118449_SASD
Intensity
9e+05
6e+05
3e+05
0e+00
1500
1750
2000
2250
time

run4: 116480_Q(UniMod:7)IKKQTALVELVK/3 (magenta) AND 25180_DWHGV

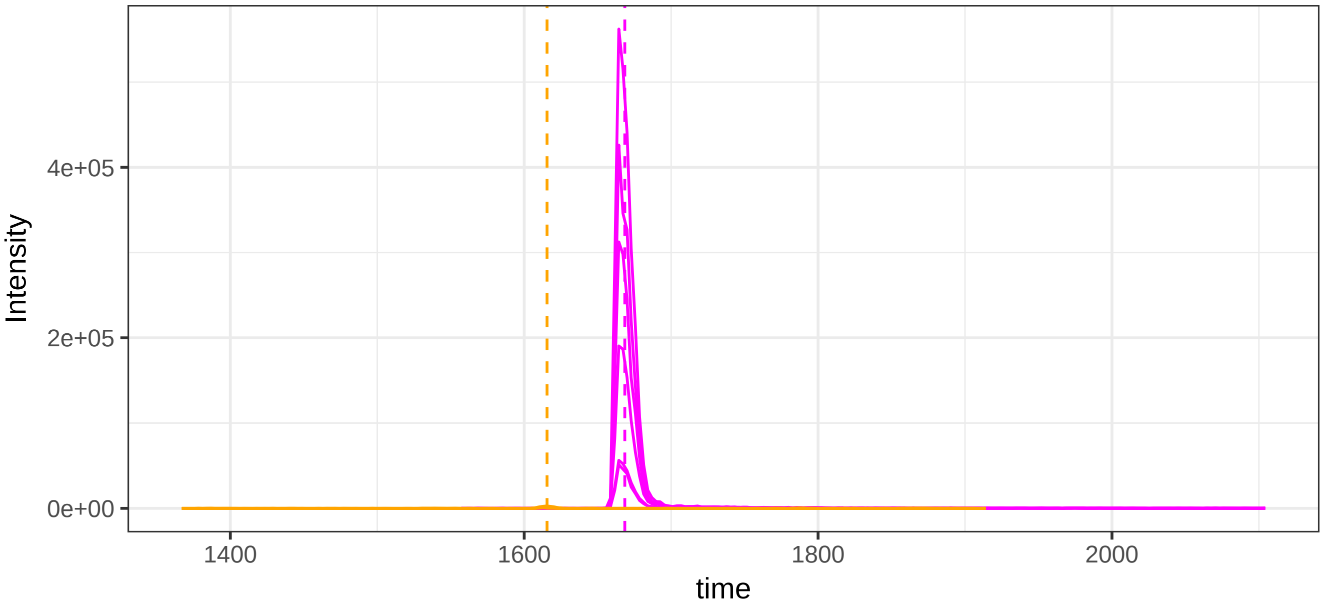


run23: 116480_Q(UniMod:7)IKKQTALVELVK/3 (magenta)  AND 25180_DWHG

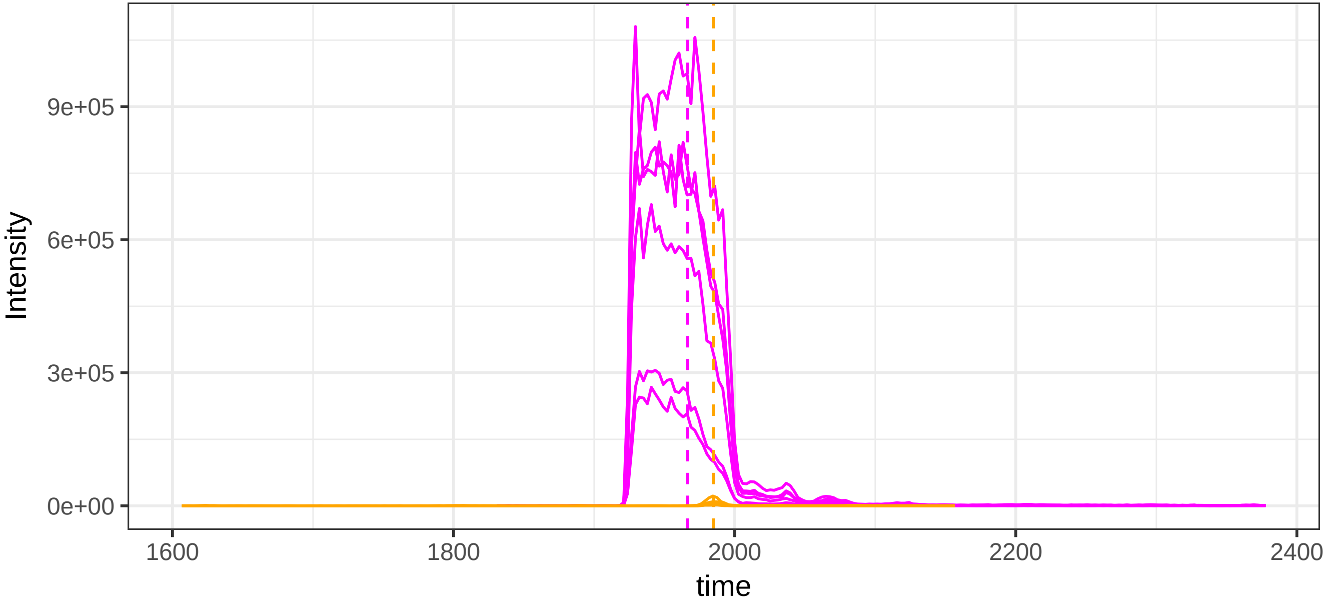

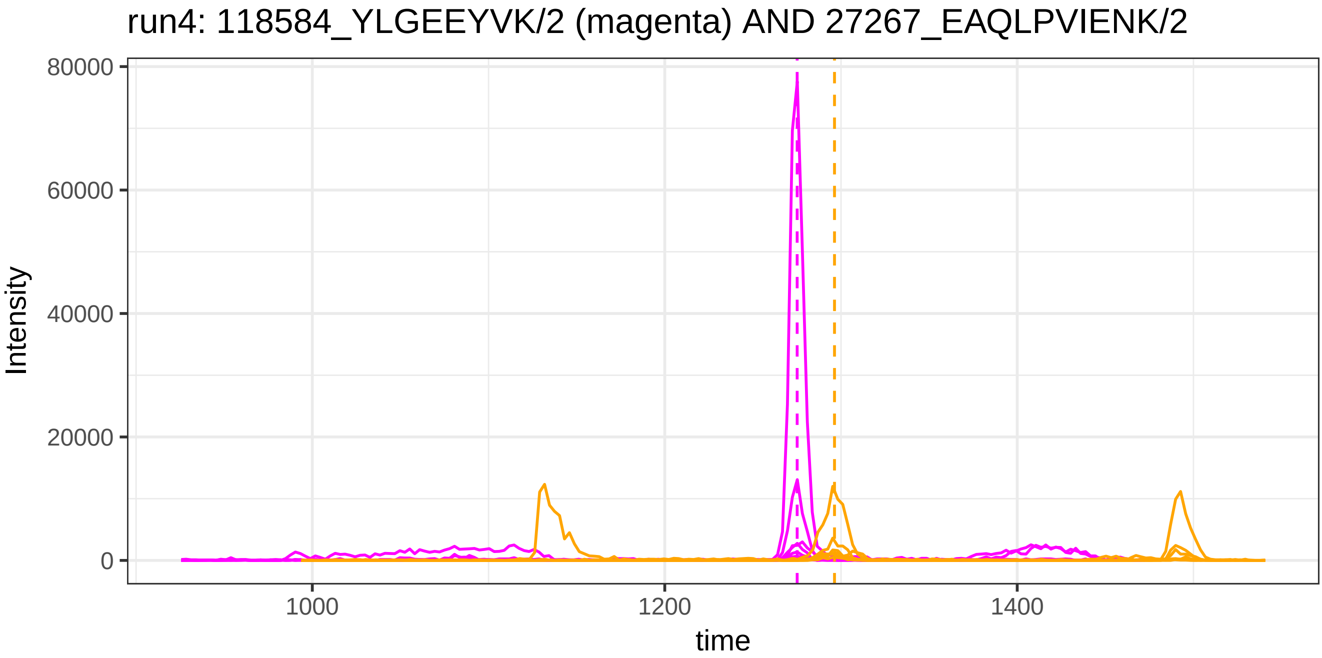

run4: 118584_YLGEEYVK/2 (magenta) AND 27267_EAQLPVIENK/2
Intensity
80000
60000
40000
20000
0
1000
1200
1400
time


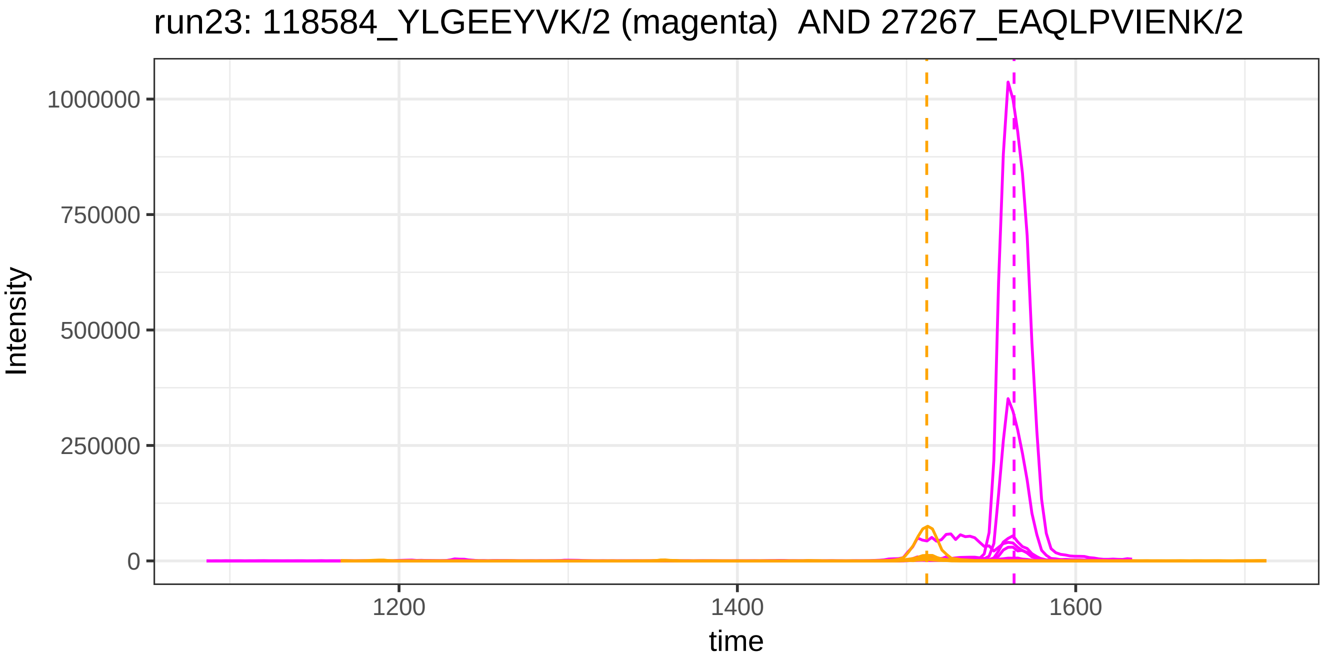

run23: 118584_YLGEEYVK/2 (magenta) AND 27267_EAQLPVIENK/2
Intensity
1000000
750000
500000
250000
0
1200
1400
1600
time

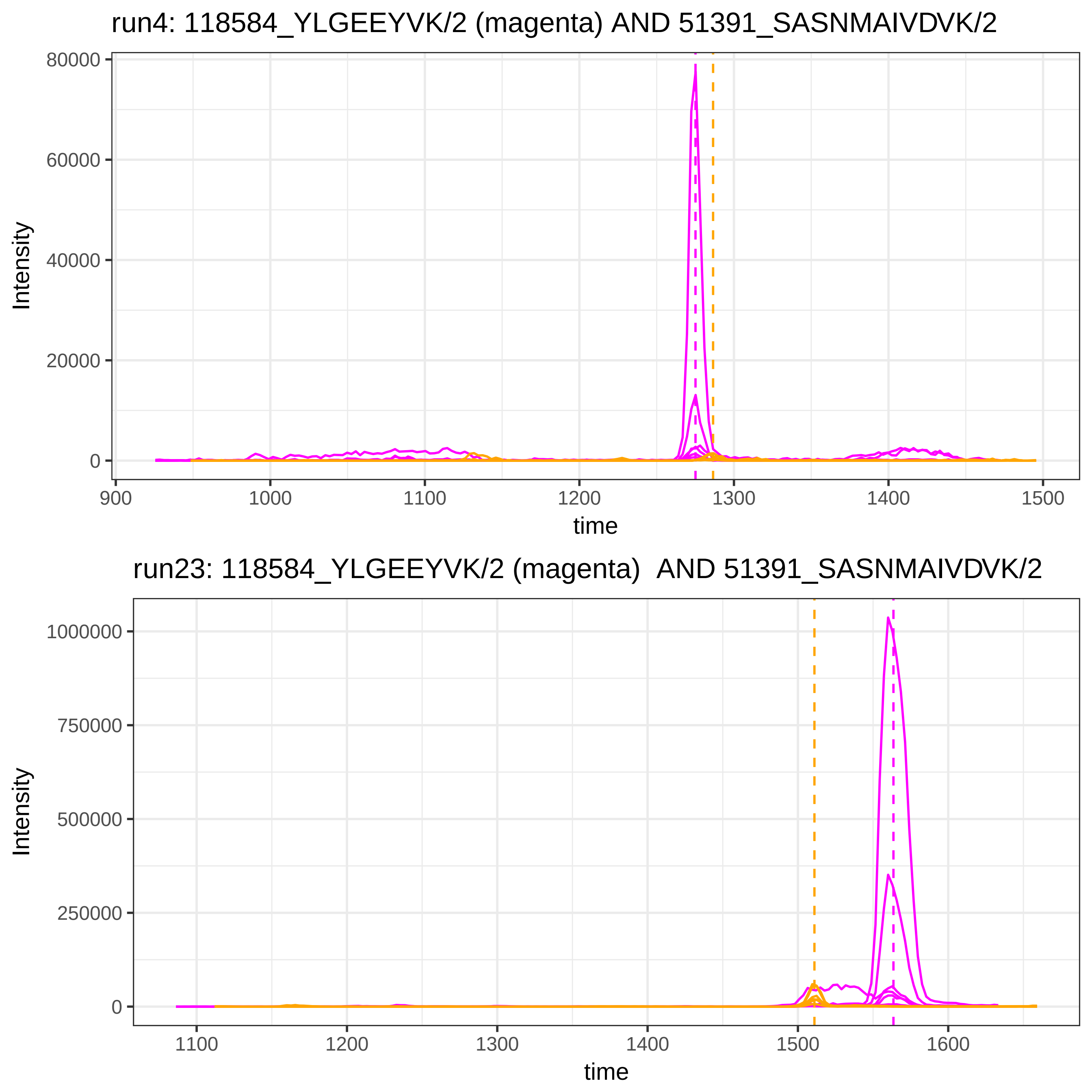

run4: 118584_YLGEEYVK/2 (magenta) AND 51391_SASNMAIVDVK/2
Intensity
80000
60000
40000
20000
0
900
1000
1100
1200
1300
1400
1500
time
run23: 118584_YLGEEYVK/2 (magenta) AND 51391_SASNMAIVDVK/2
Intensity
1000000
750000
500000
250000
0
1100
1200
1300
1400
1500
1600
time

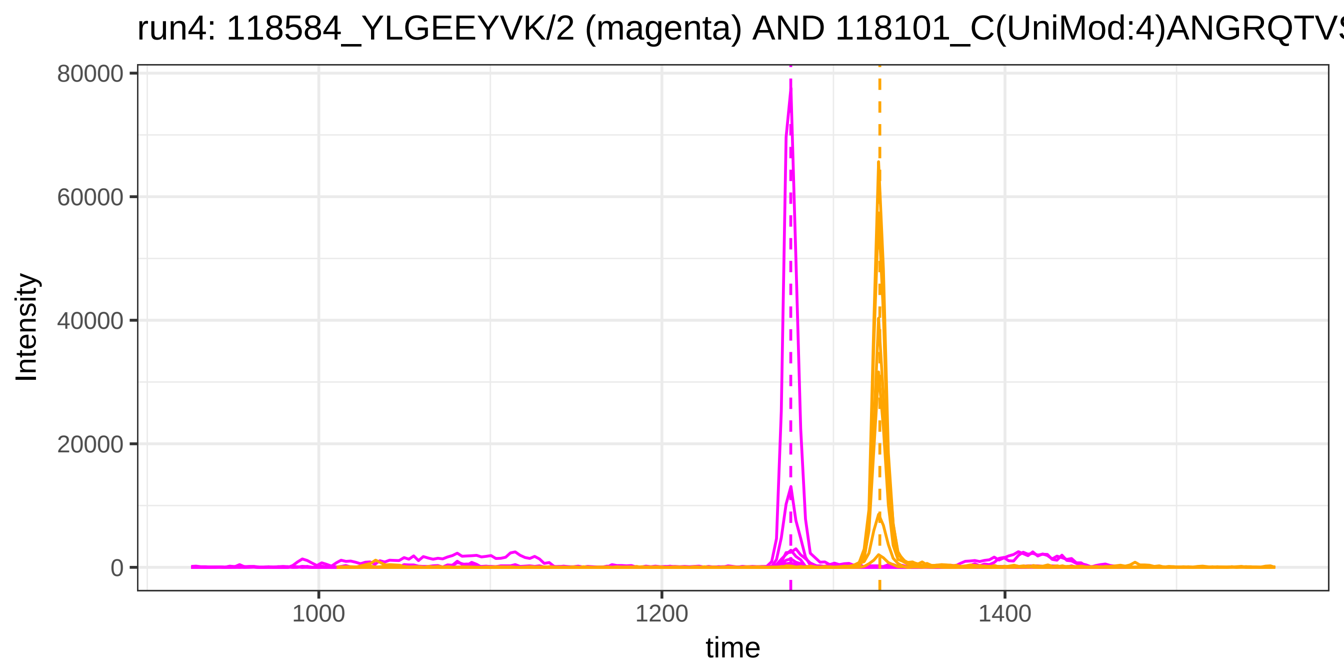
run4: 118584_YLGEEYVK/2 (magenta) AND 118101_C(UniMod:4)ANGRQTVS
Intensity
80000
60000
40000
20000
0
1000
1200
1400
time

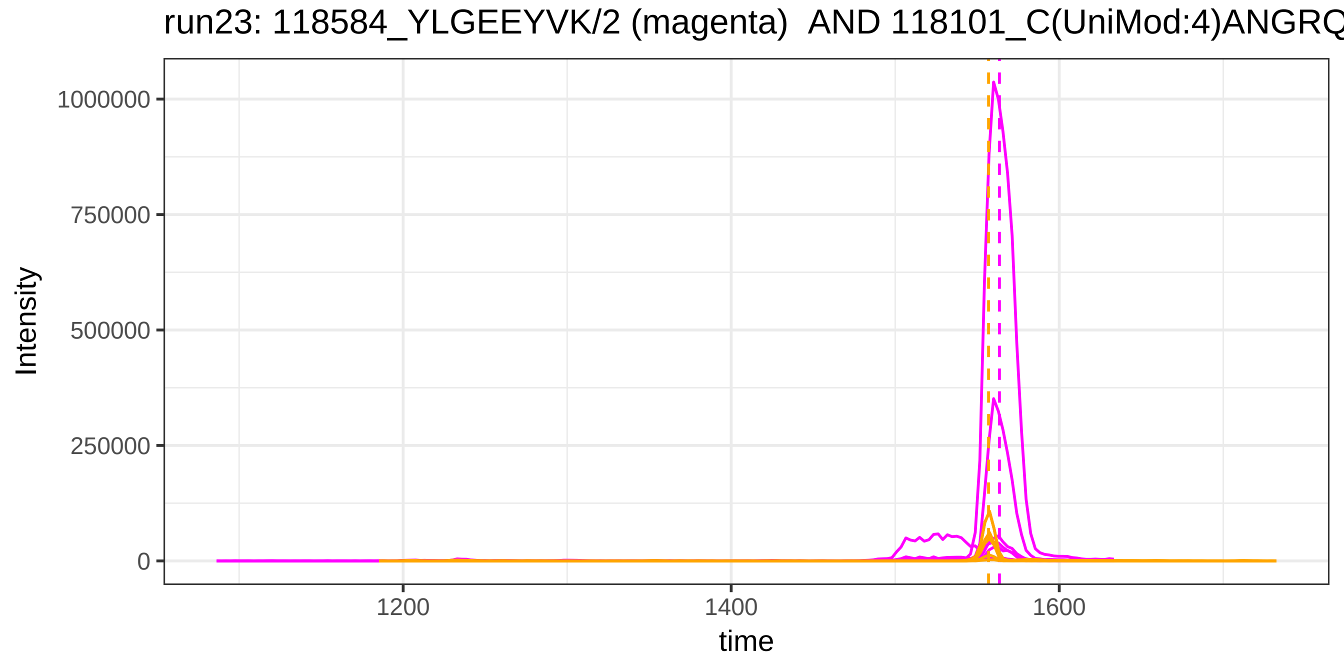
run23: 118584_YLGEEYVK/2 (magenta) AND 118101_C(UniMod:4)ANGRQ
Intensity
1000000
750000
500000
250000
0
1200
1400
1600
time

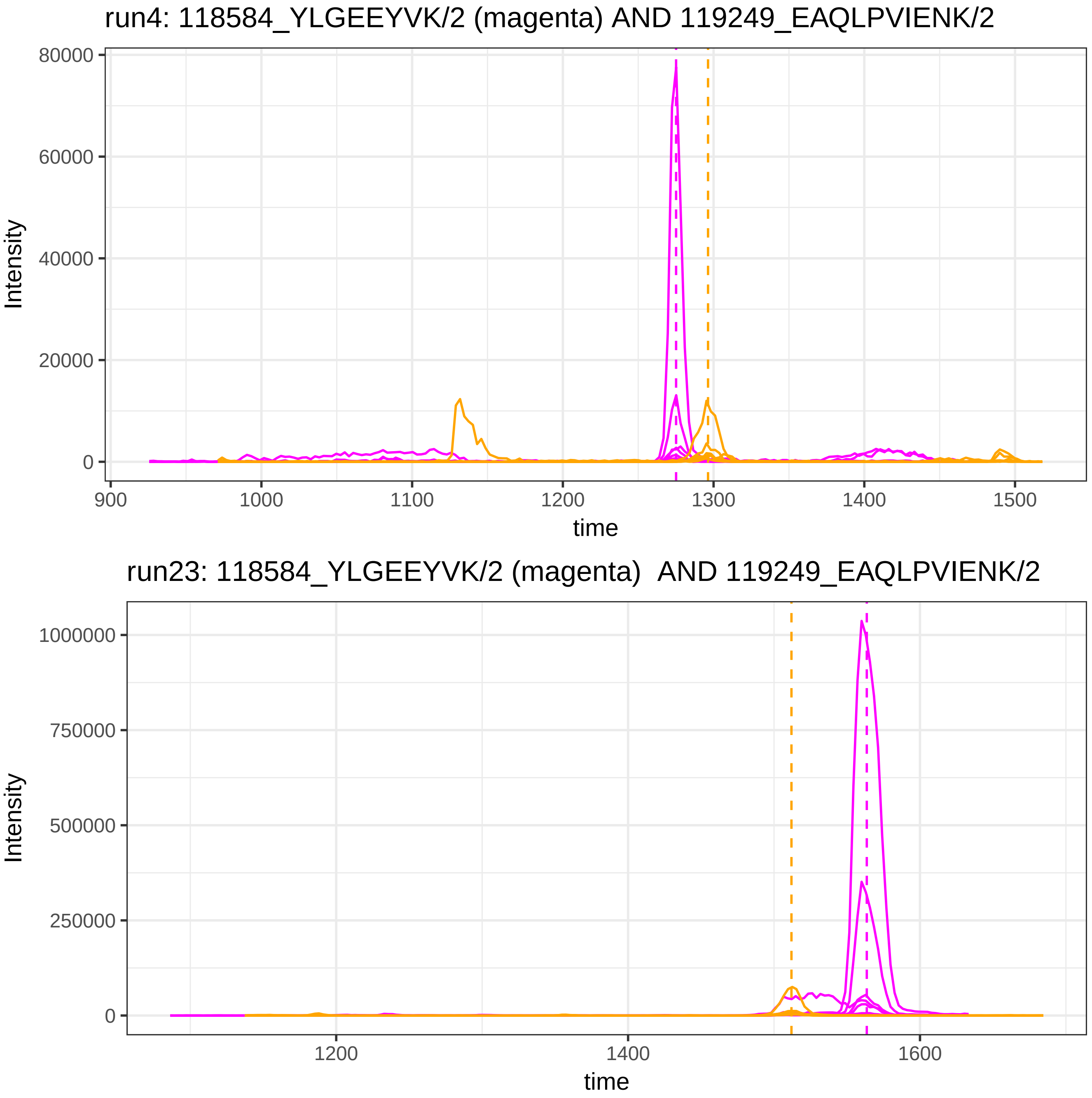
run4: 118584_YLGEEYVK/2 (magenta) AND 119249_EAQLPVIENK/2
Intensity
80000
60000
40000
20000
0
time
900
1000
1100
1200
1300
1400
1500
run23: 118584_YLGEEYVK/2 (magenta) AND 119249_EAQLPVIENK/2
Intensity
1000000
750000
500000
250000
0
time
1200
1400
1600

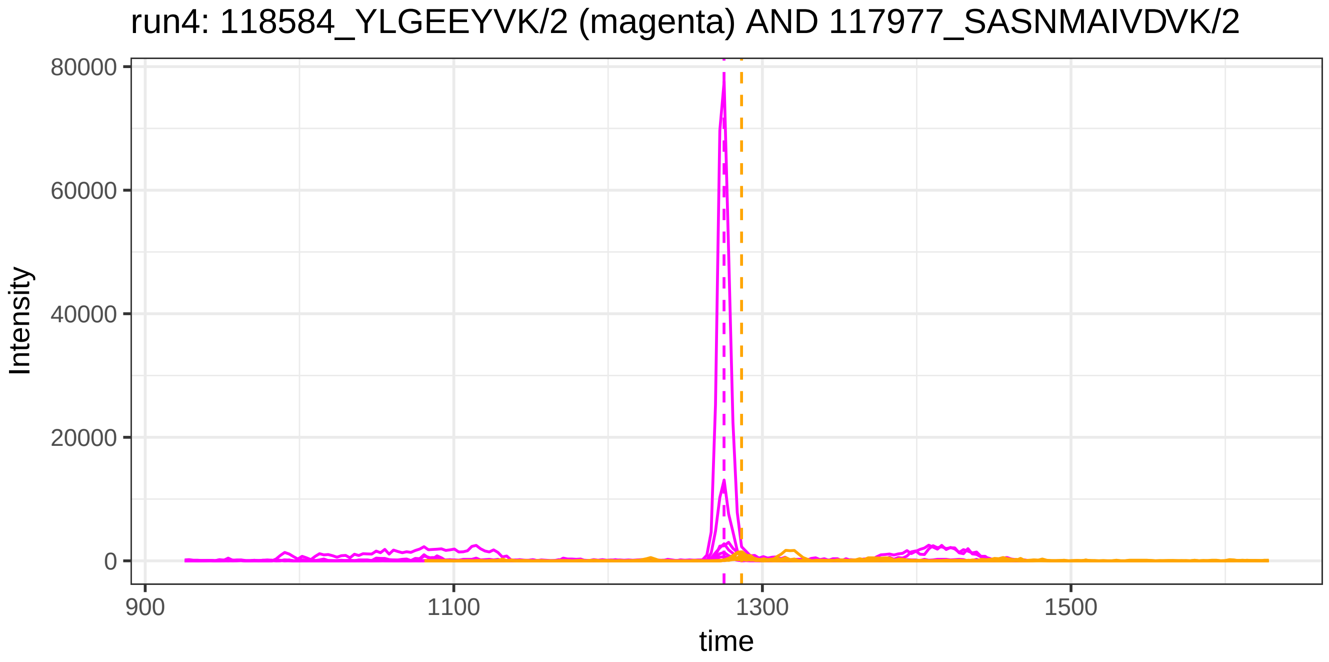
run4: 118584_YLGEEYVK/2 (magenta) AND 117977_SASNMAIVDVK/2
Intensity
80000
60000
40000
20000
0
900
1100
1300
1500
time

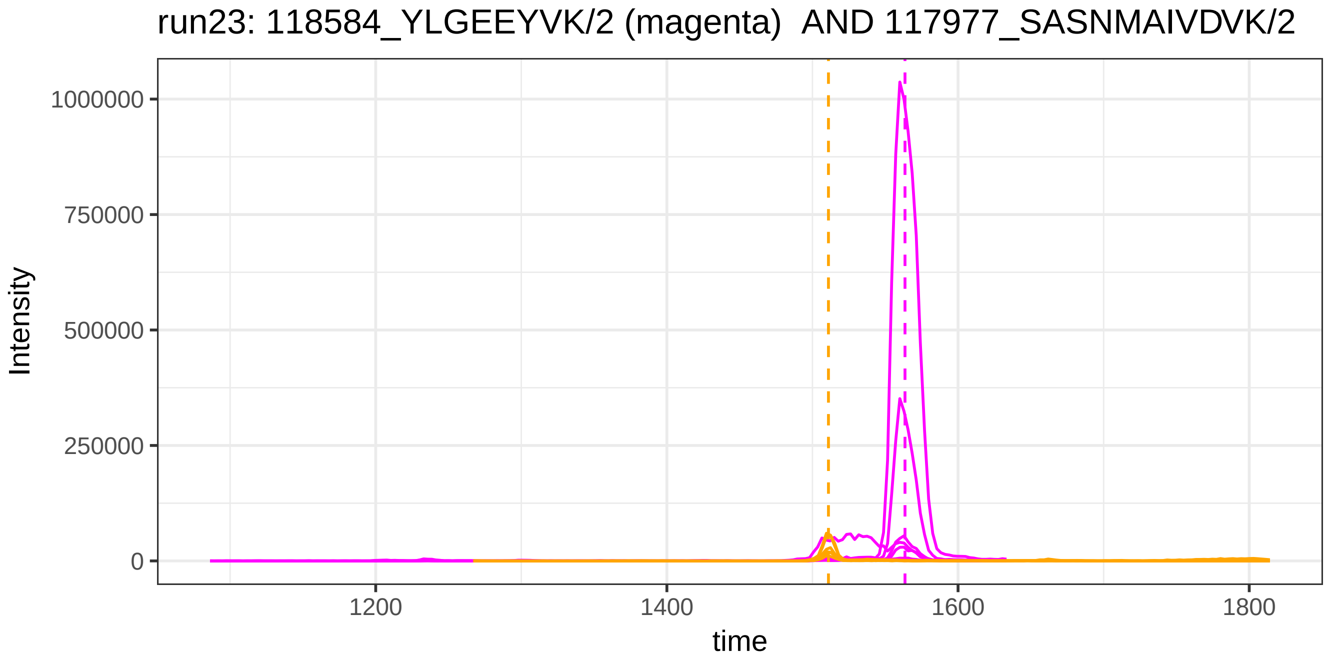
run23: 118584_YLGEEYVK/2 (magenta) AND 117977_SASNMAIVDVK/2
Intensity
1000000
750000
500000
250000
0
1200
1400
1600
1800
time

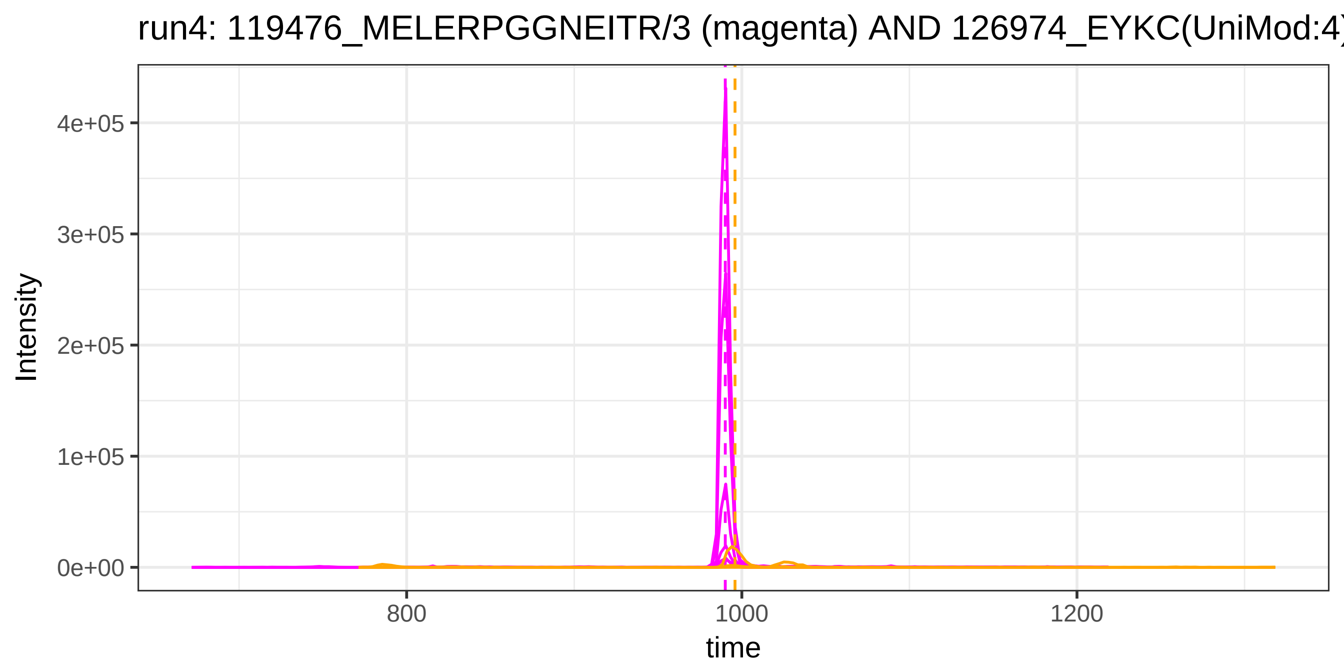
run4: 119476_MELERPGGNEITR/3 (magenta) AND 126974_EYKC(UniMod:4)
Intensity
4e+05
3e+05
2e+05
1e+05
0e+00
800
1000
1200
time

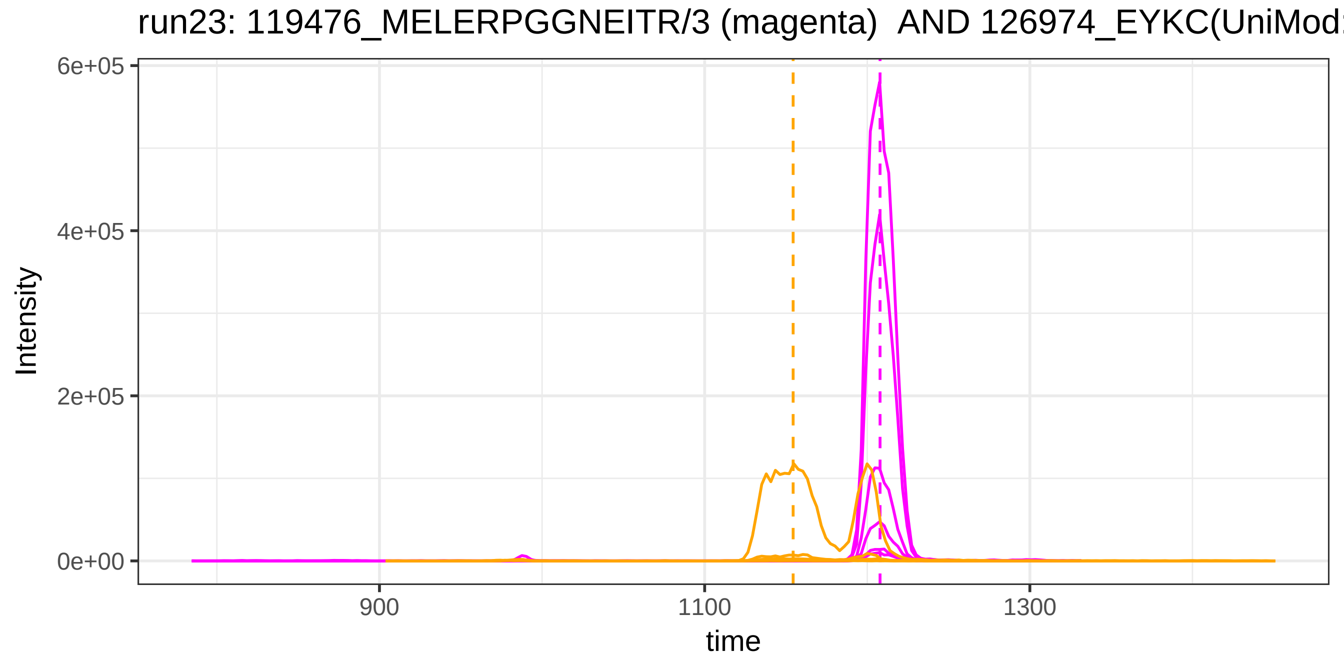
run23: 119476_MELERPGGNEITR/3 (magenta) AND 126974_EYKC(UniMod
Intensity
6e+05
4e+05
2e+05
0e+00
900
1100
1300
time

run4: 126775_VAAGAFQGLR/2 (magenta) AND 32498_ESYSGVTLDPR/2

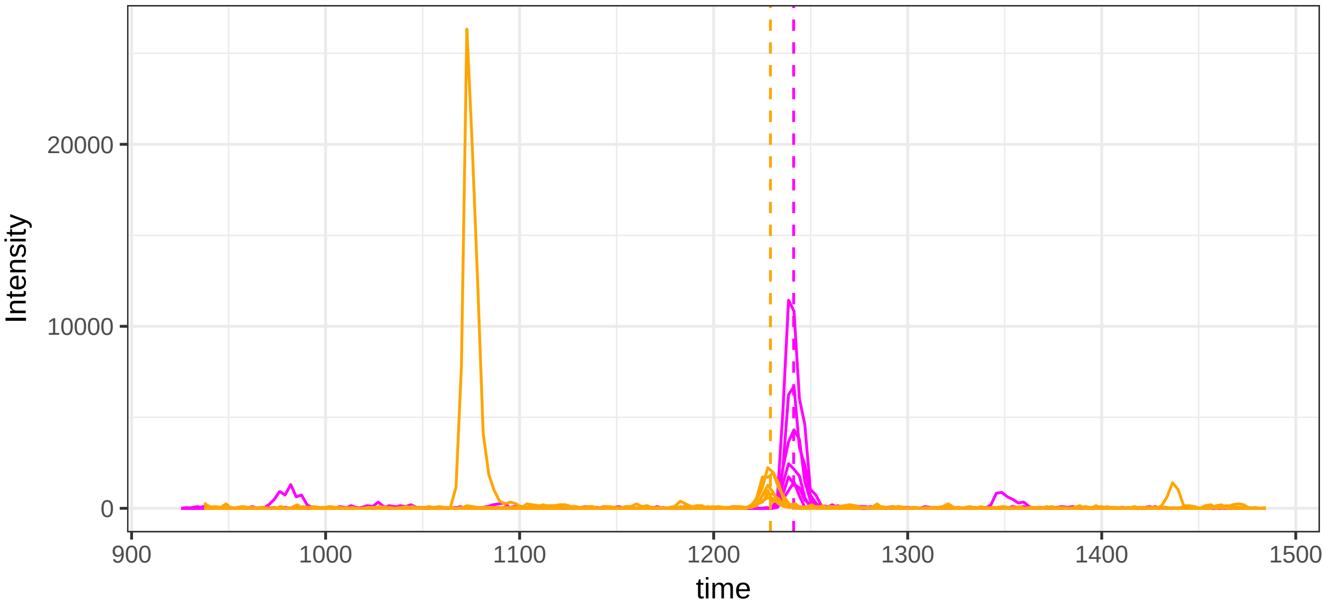


run23: 126775_VAAGAFQGLR/2 (magenta) AND 32498_ESYSGVTLDPR/2

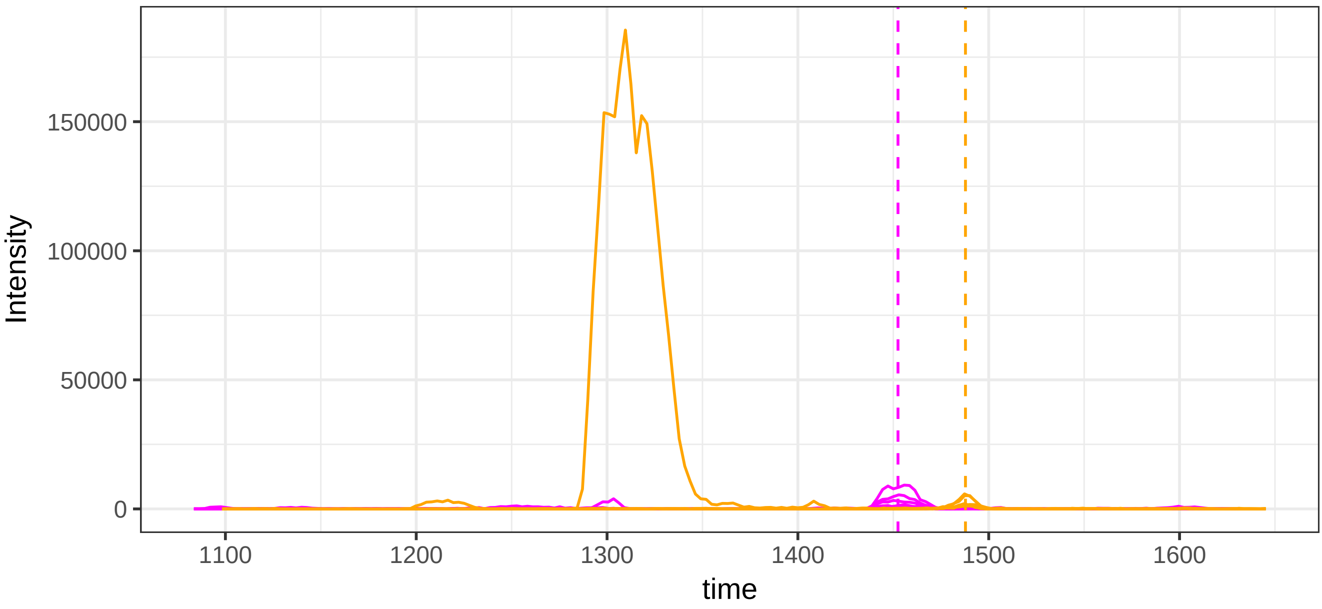

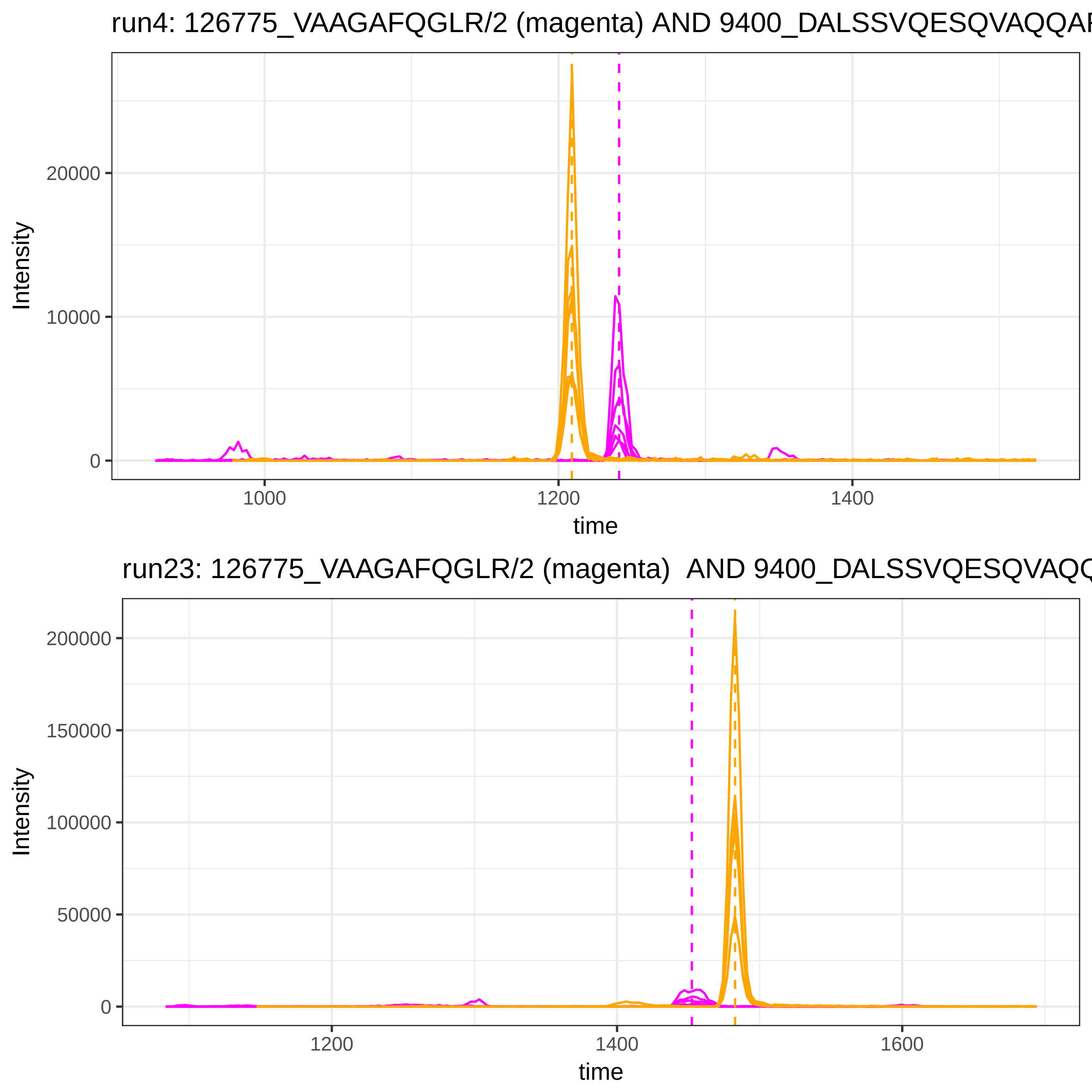
run4: 126775_VAAGAFQGLR/2 (magenta) AND 9400_DALSSVQESQVAQQAR
Intensity
20000
10000
0
1000
1200
1400
time
run23: 126775_VAAGAFQGLR/2 (magenta) AND 9400_DALSSVQESQVAQQ
Intensity
200000
150000
100000
50000
0
1200
1400
1600
time

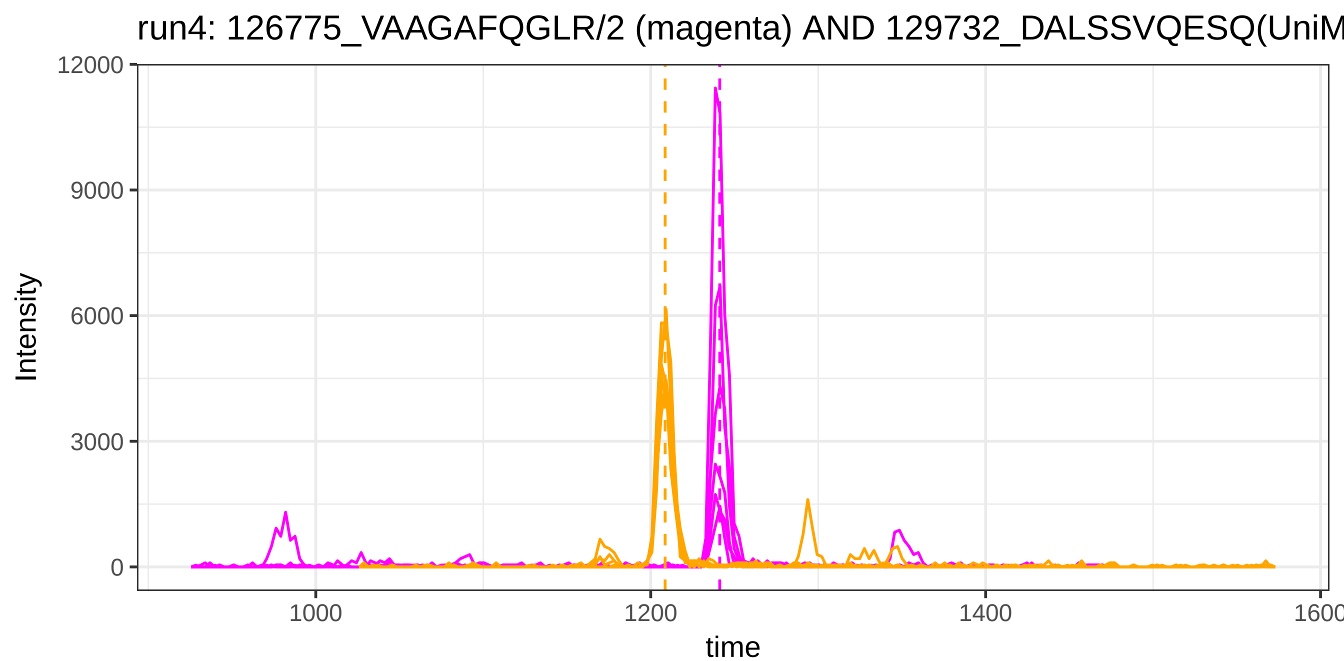

run4: 126775_VAAGAFQGLR/2 (magenta) AND 129732_DALSSVQESQ(UniM
Intensity
time
12000
9000
6000
3000
0
1000
1200
1400
1600


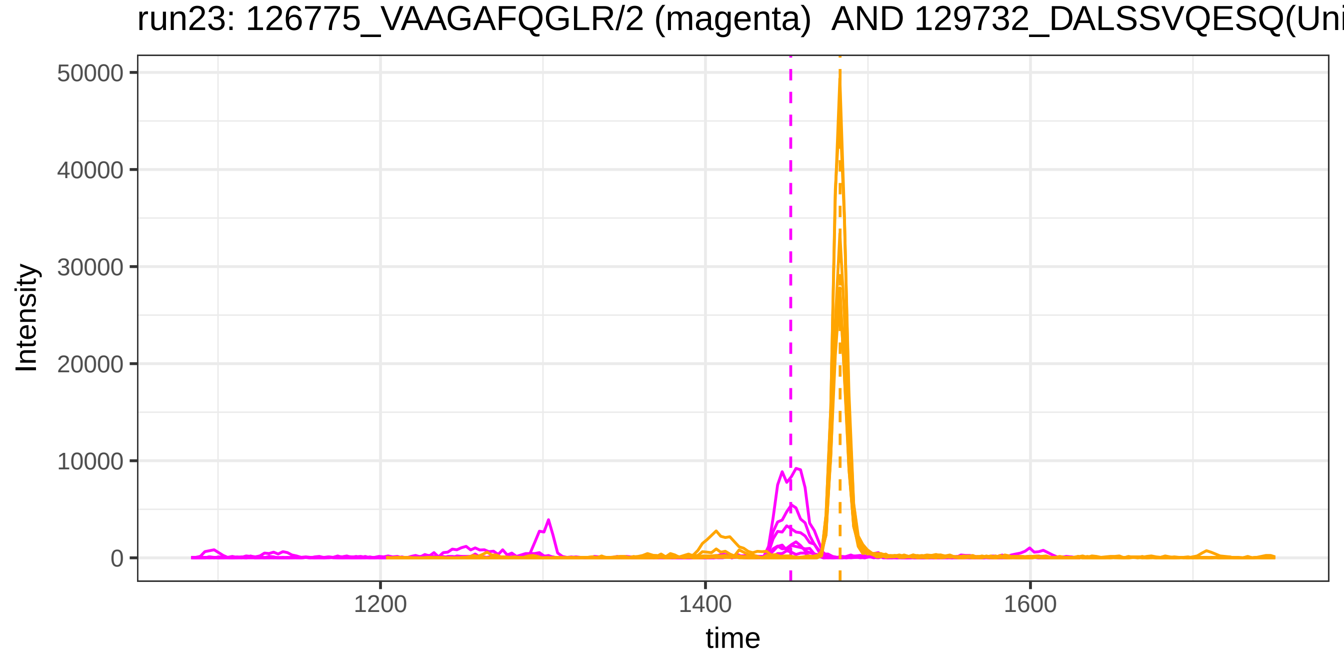

run23: 126775_VAAGAFQGLR/2 (magenta) AND 129732_DALSSVQESQ(Uni
Intensity
time
50000
40000
30000
20000
10000
0
1200
1400
1600

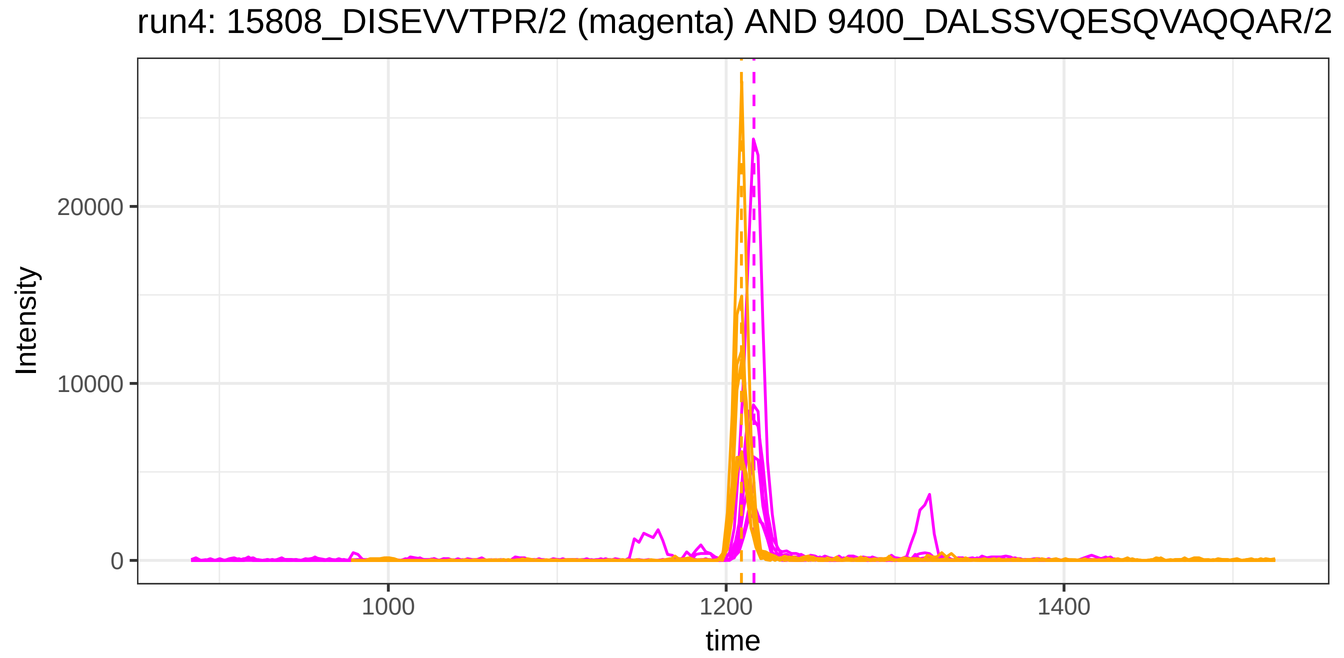
run4: 15808_DISEVVTPR/2 (magenta) AND 9400_DALSSVQESQVAQQAR/2
Intensity
20000
10000
0
1000
1200
1400
time

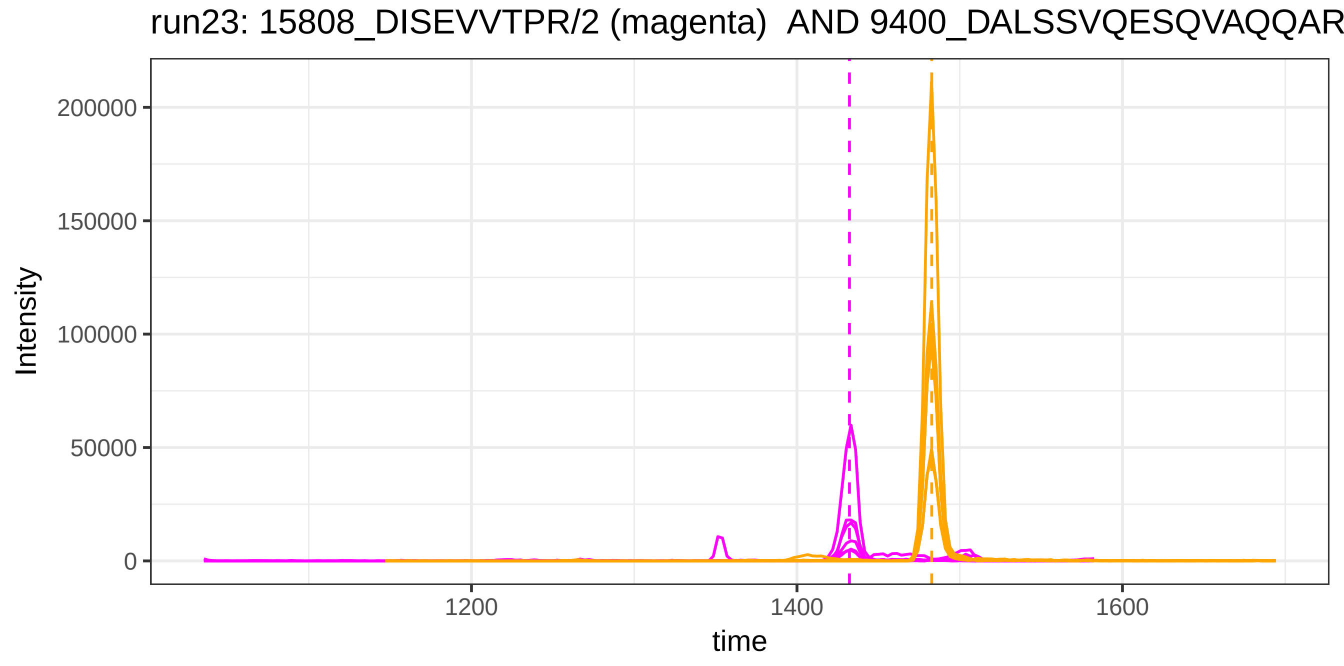
run23: 15808_DISEVVTPR/2 (magenta) AND 9400_DALSSVQESQVAQQAR
Intensity
200000
150000
100000
50000
0
1200
1400
1600
time

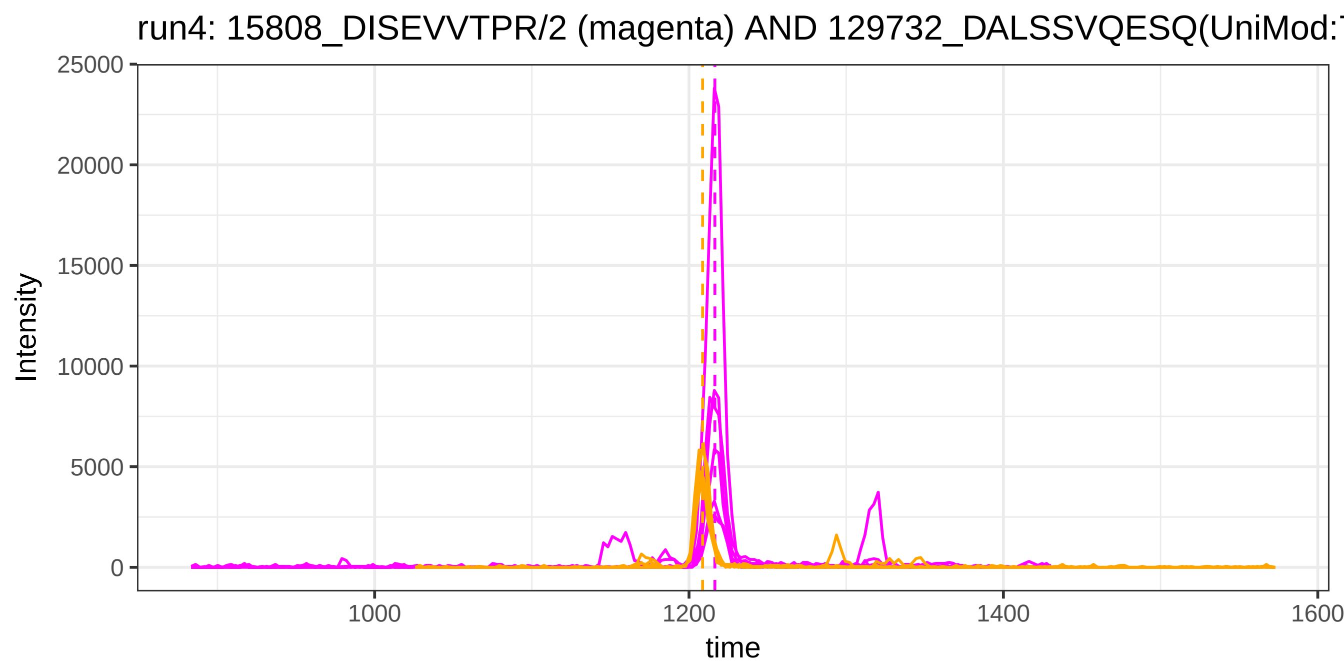
run4: 15808_DISEVVTPR/2 (magenta) AND 129732_DALSSVQESQ(UniMod:
Intensity
25000
20000
15000
10000
5000
0
1000
1200
1400
1600
time

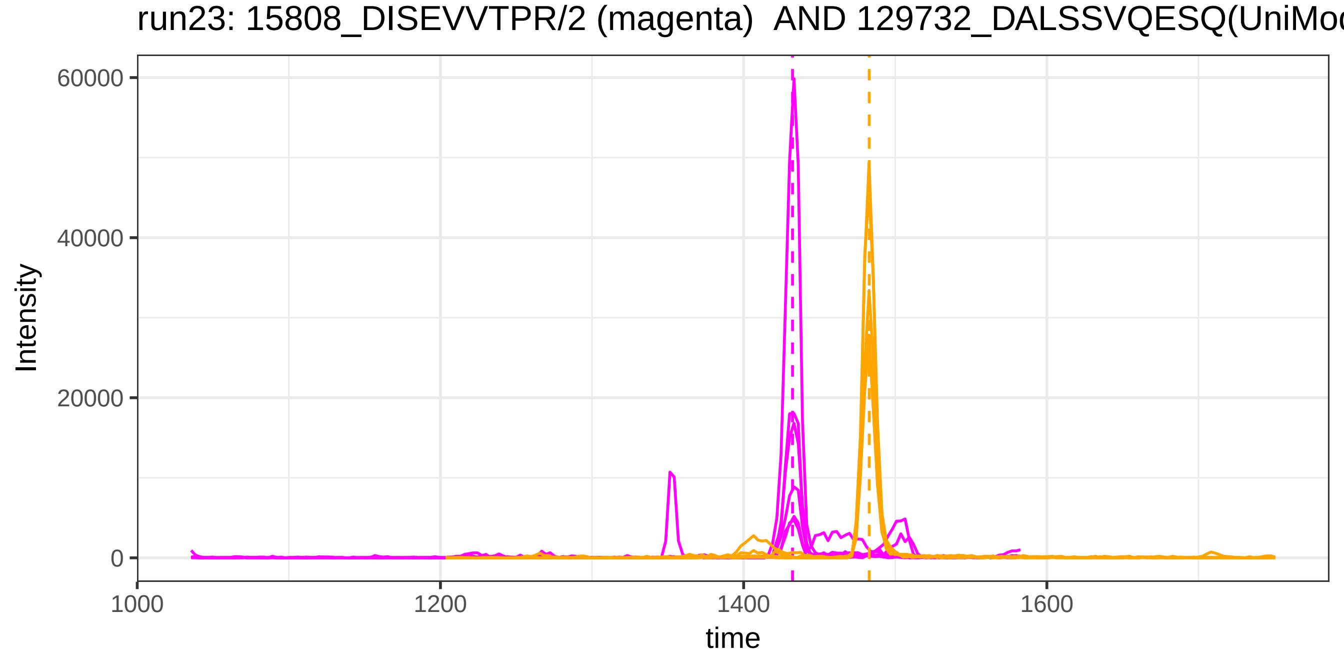
run23: 15808_DISEVVTPR/2 (magenta) AND 129732_DALSSVQESQ(UniMod
Intensity
60000
40000
20000
0
1000
1200
1400
1600
time

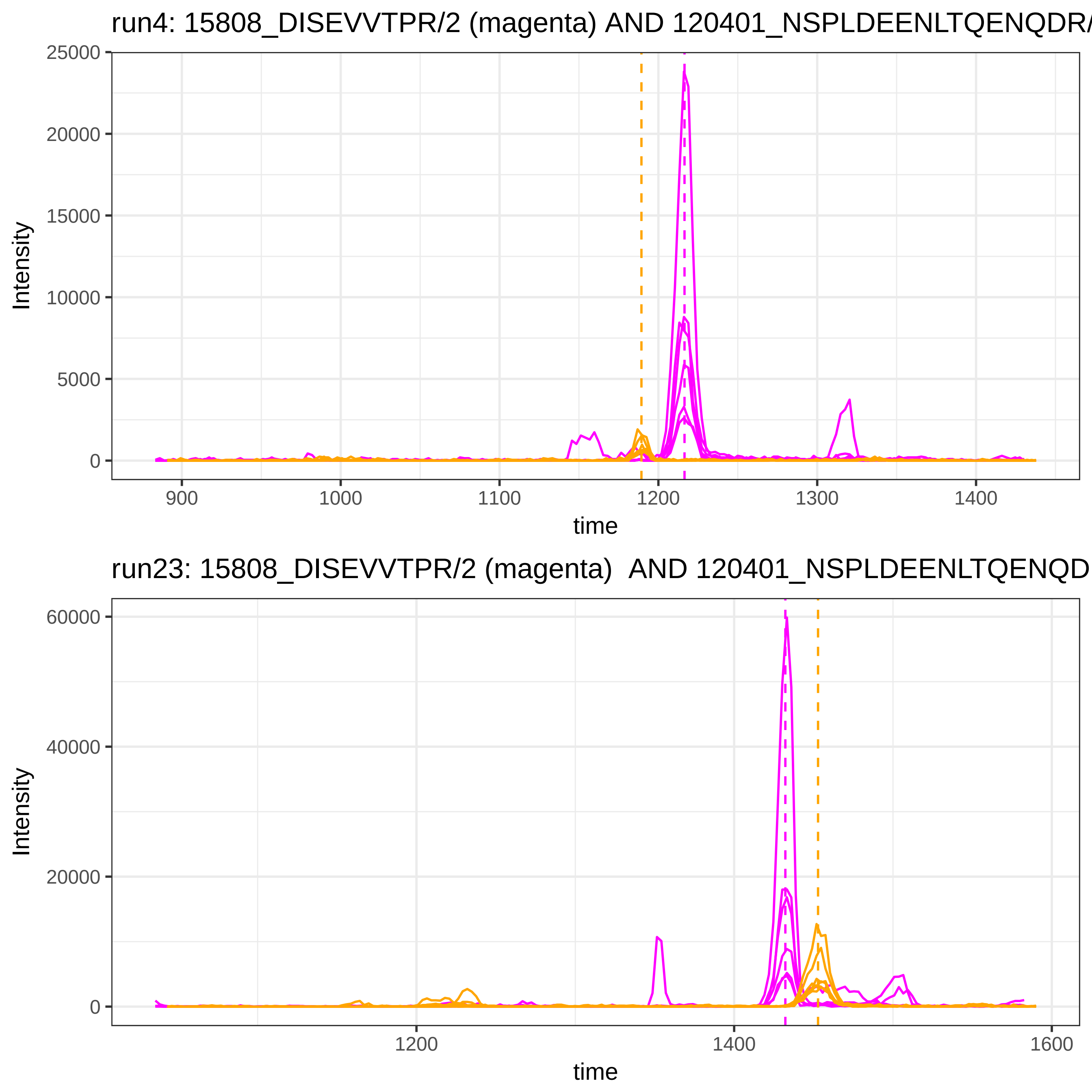
run4: 15808_DISEVVTPR/2 (magenta) AND 120401_NSPLDEENLTQENQDR/
Intensity
25000
20000
15000
10000
5000
0
900
1000
1100
1200
1300
1400
time
run23: 15808_DISEVVTPR/2 (magenta)  AND 120401_NSPLDEENLTQENQD
60000
40000
20000
0
1200
1400
1600

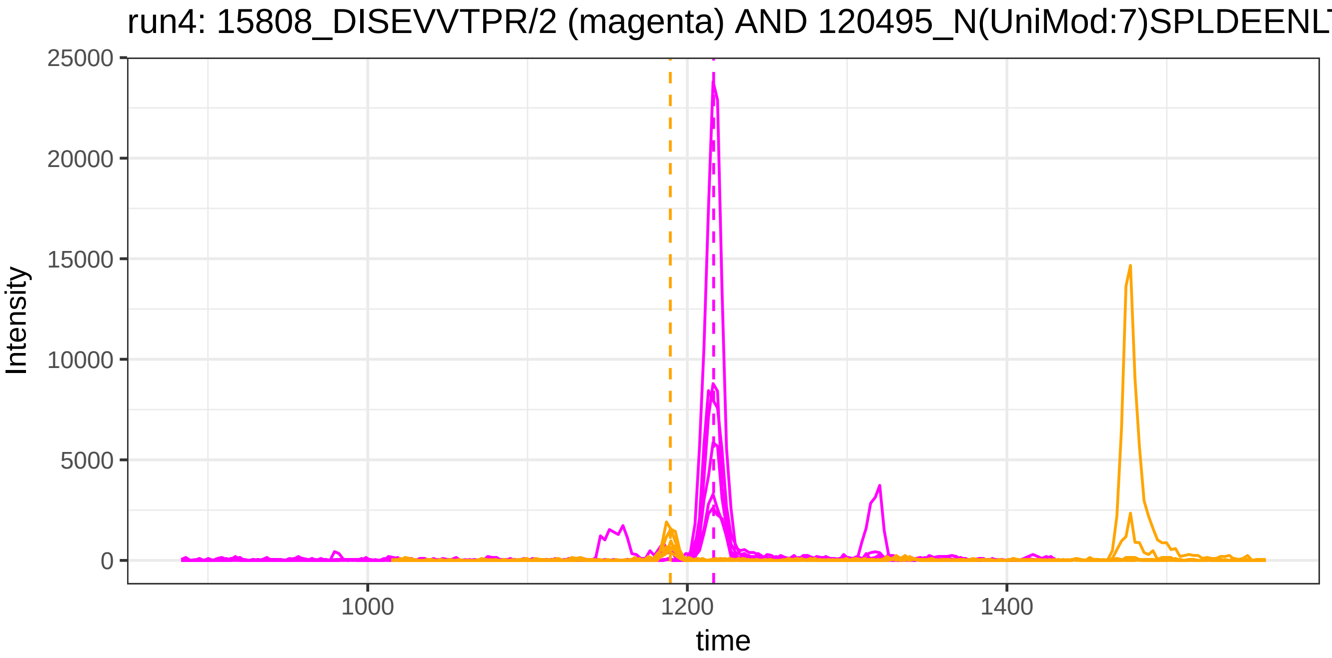
run4: 15808_DISEVVTPR/2 (magenta) AND 120495_N(UniMod:7)SPLDEENLT
Intensity
25000
20000
15000
10000
5000
0
1000
1200
1400
time

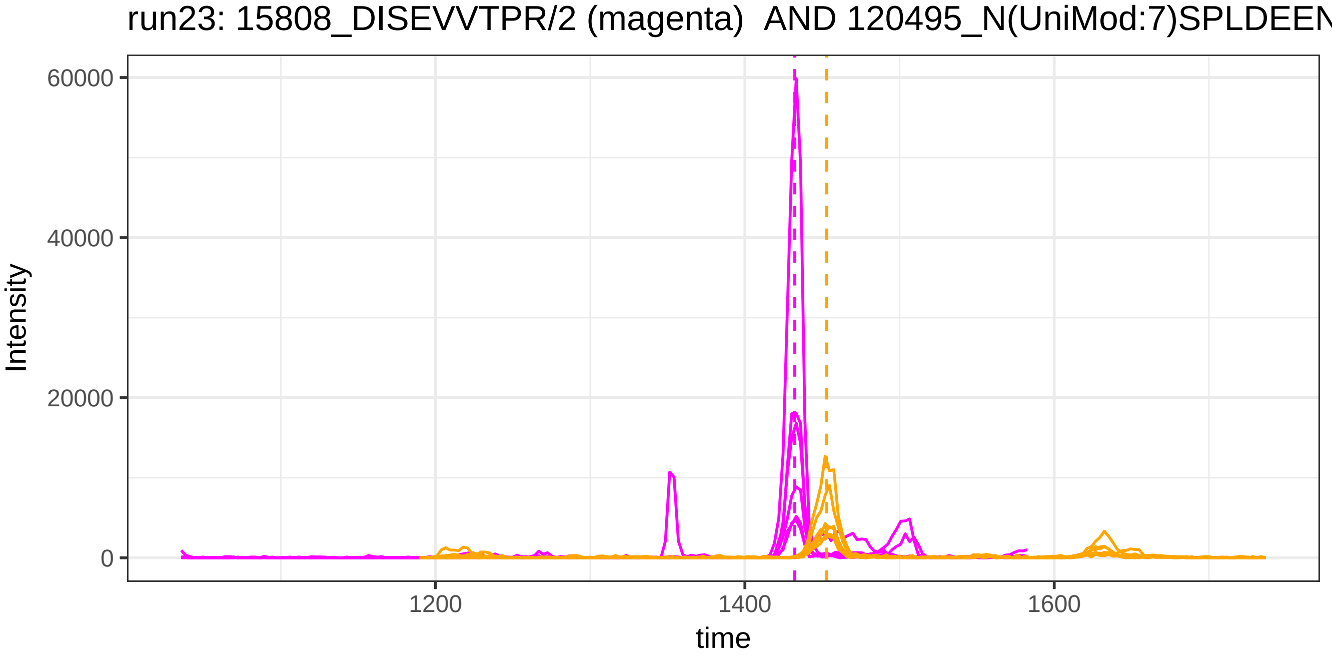
run23: 15808_DISEVVTPR/2 (magenta) AND 120495_N(UniMod:7)SPLDEEN
Intensity
60000
40000
20000
0
1200
1400
1600
time

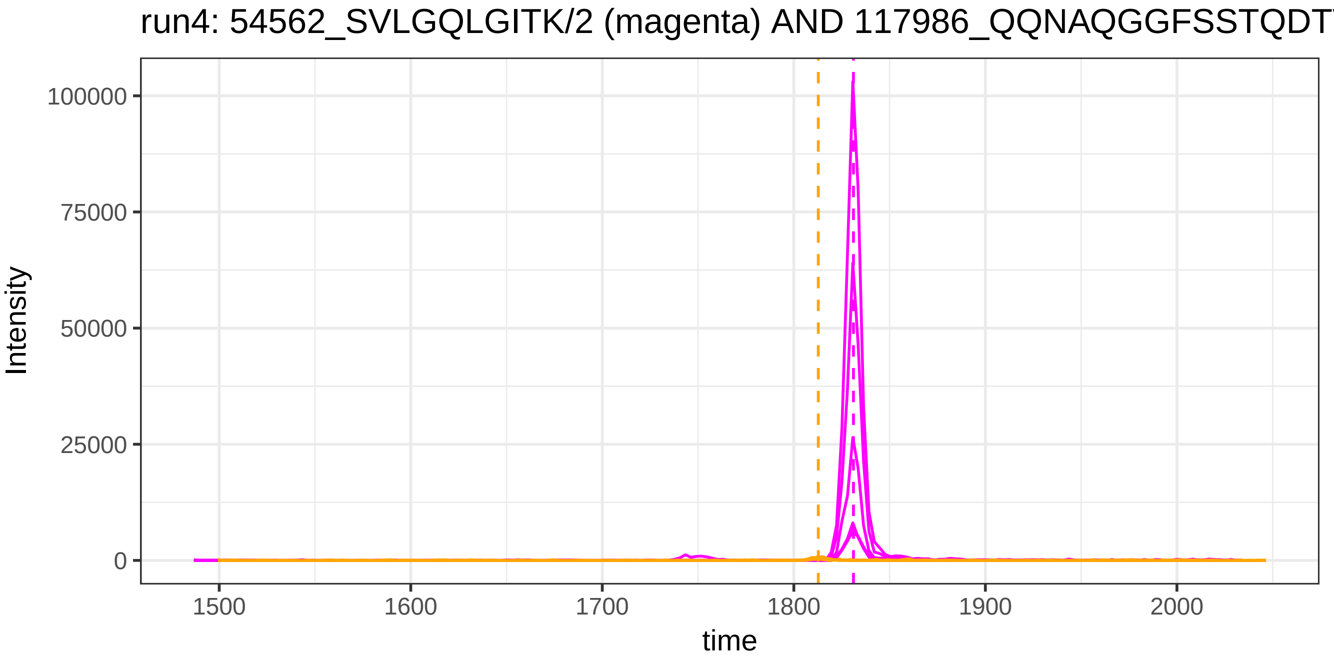

run4: 54562_SVLGQLGITK/2 (magenta) AND 117986_QQNAQGGFSSTQDT
Intensity
100000
75000
50000
25000
0
1500
1600
1700
1800
1900
2000
time


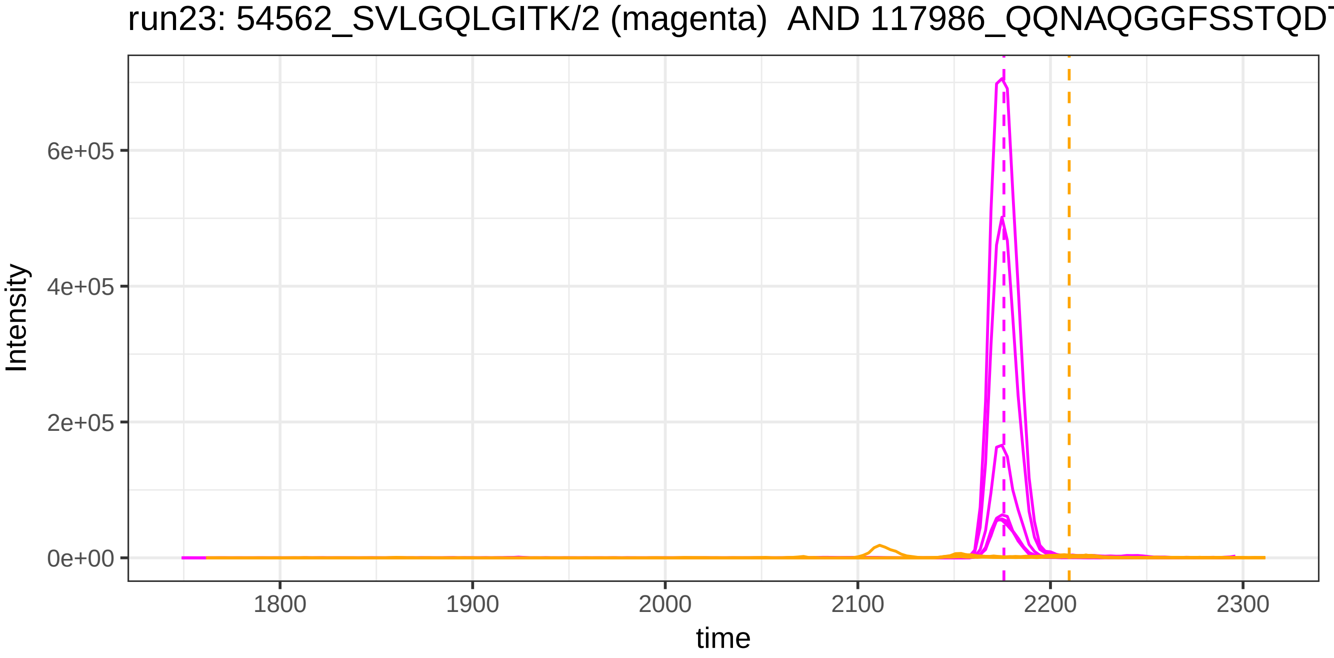

run23: 54562_SVLGQLGITK/2 (magenta) AND 117986_QQNAQGGFSSTQDT
Intensity
6e+05
4e+05
2e+05
0e+00
1800
1900
2000
2100
2200
2300
time

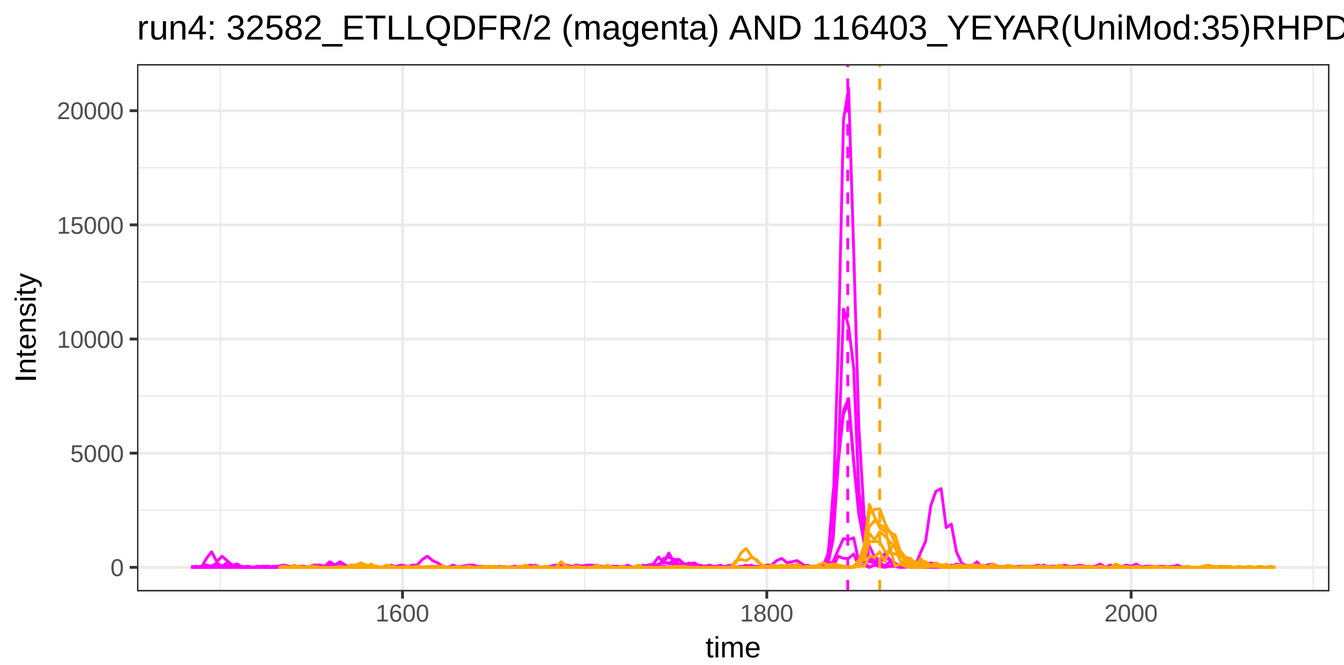

run4: 32582_ETLLQDFR/2 (magenta) AND 116403_YEYAR(UniMod:35)RHPD
Intensity
20000
15000
10000
5000
0
1600
1800
2000
time


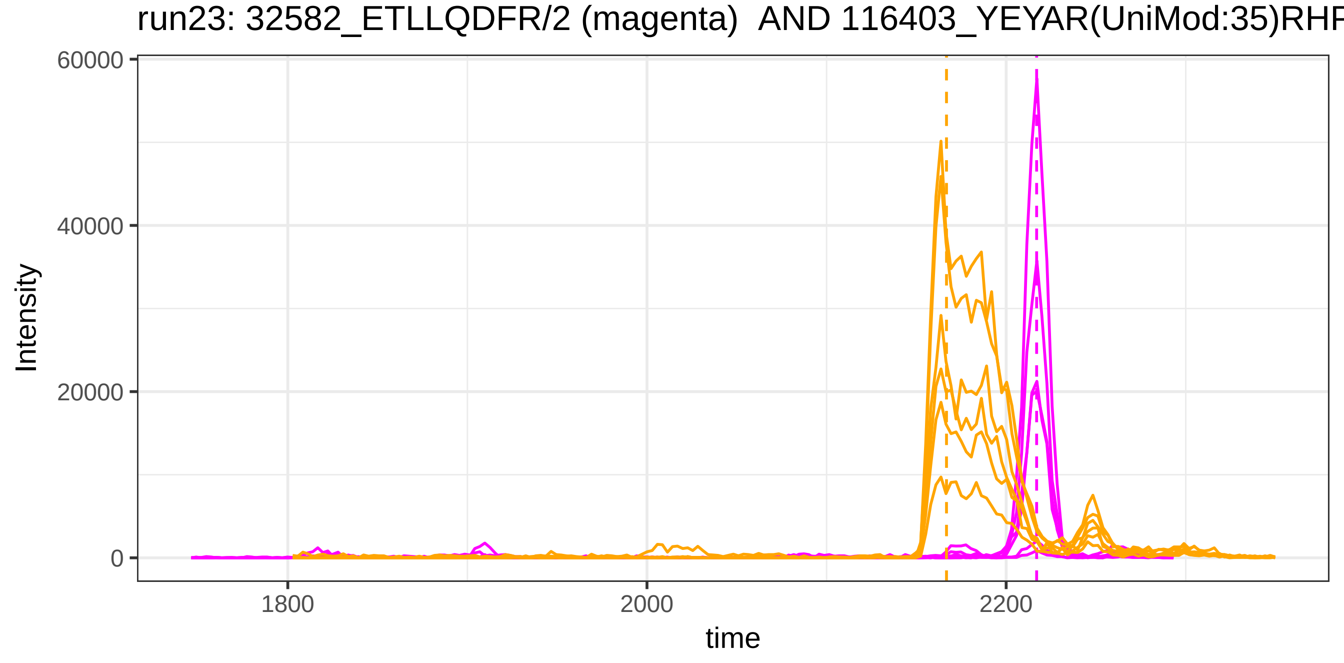

run23: 32582_ETLLQDFR/2 (magenta) AND 116403_YEYAR(UniMod:35)RHF
Intensity
60000
40000
20000
0
1800
2000
2200
time

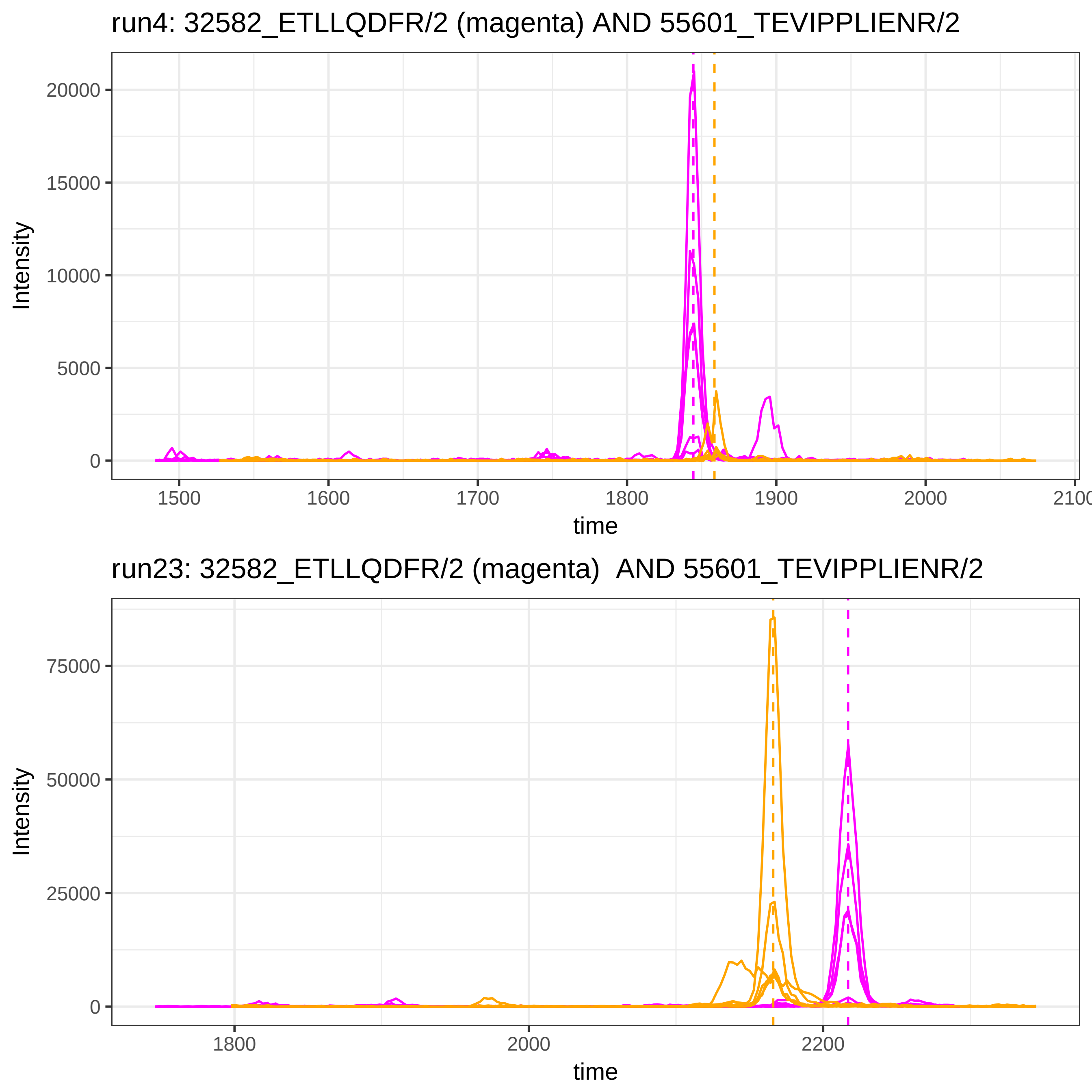
run4: 32582_ETLLQDFR/2 (magenta) AND 55601_TEVIPPLIENR/2
Intensity
20000
15000
10000
5000
0
1500
1600
1700
1800
1900
2000
2100
time
run23: 32582_ETLLQDFR/2 (magenta) AND 55601_TEVIPPLIENR/2
Intensity
75000
50000
25000
0
1800
2000
2200
time

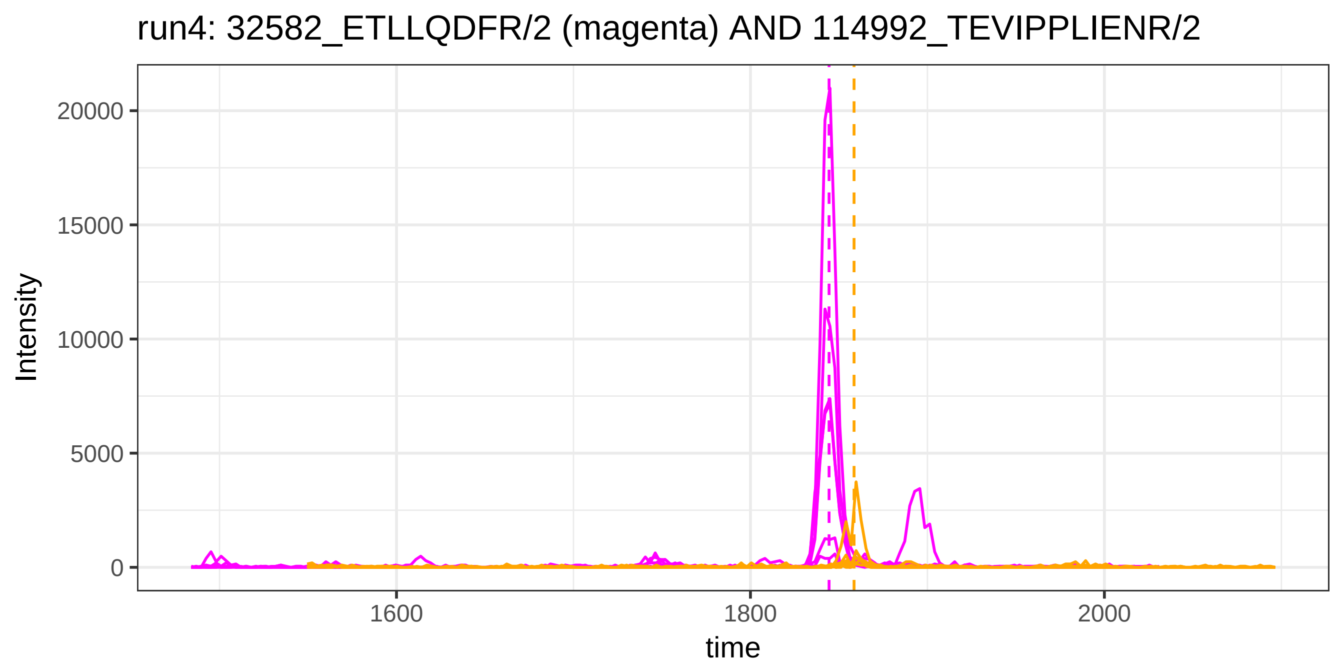
run4: 32582_ETLLQDFR/2 (magenta) AND 114992_TEVIPPLIENR/2
Intensity
20000
15000
10000
5000
0
1600
1800
2000
time

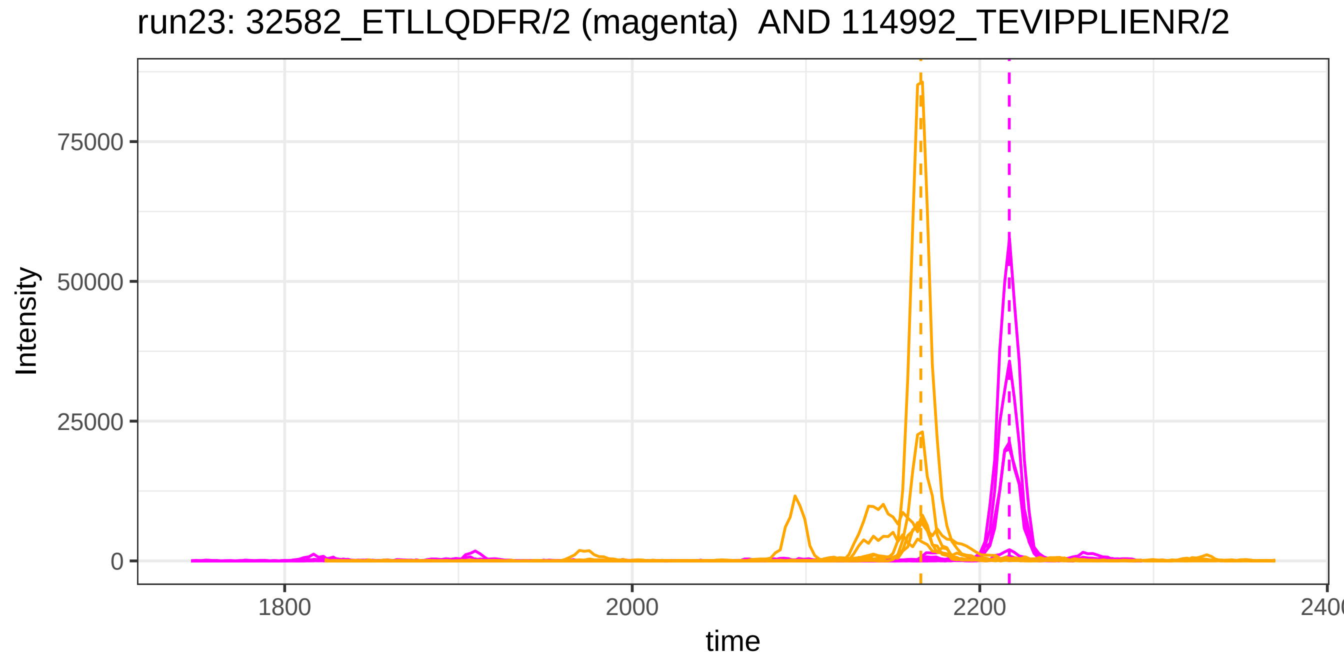
run23: 32582_ETLLQDFR/2 (magenta)  AND 114992_TEVIPPLIENR/2
Intensity
75000
50000
25000
0
1800
2000
2200
2400
time

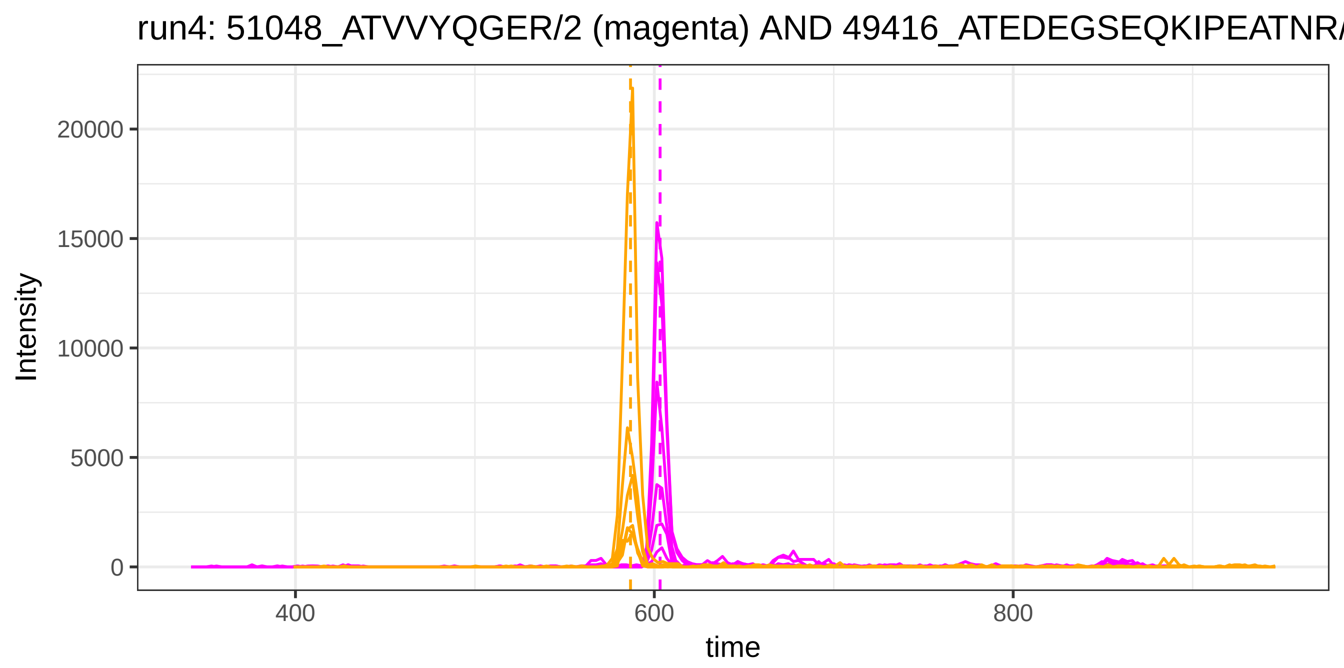
run4: 51048_ATVVYQGER/2 (magenta) AND 49416_ATEDEGSEQKIPEATNR/
Intensity
20000
15000
10000
5000
0
400
600
800
time

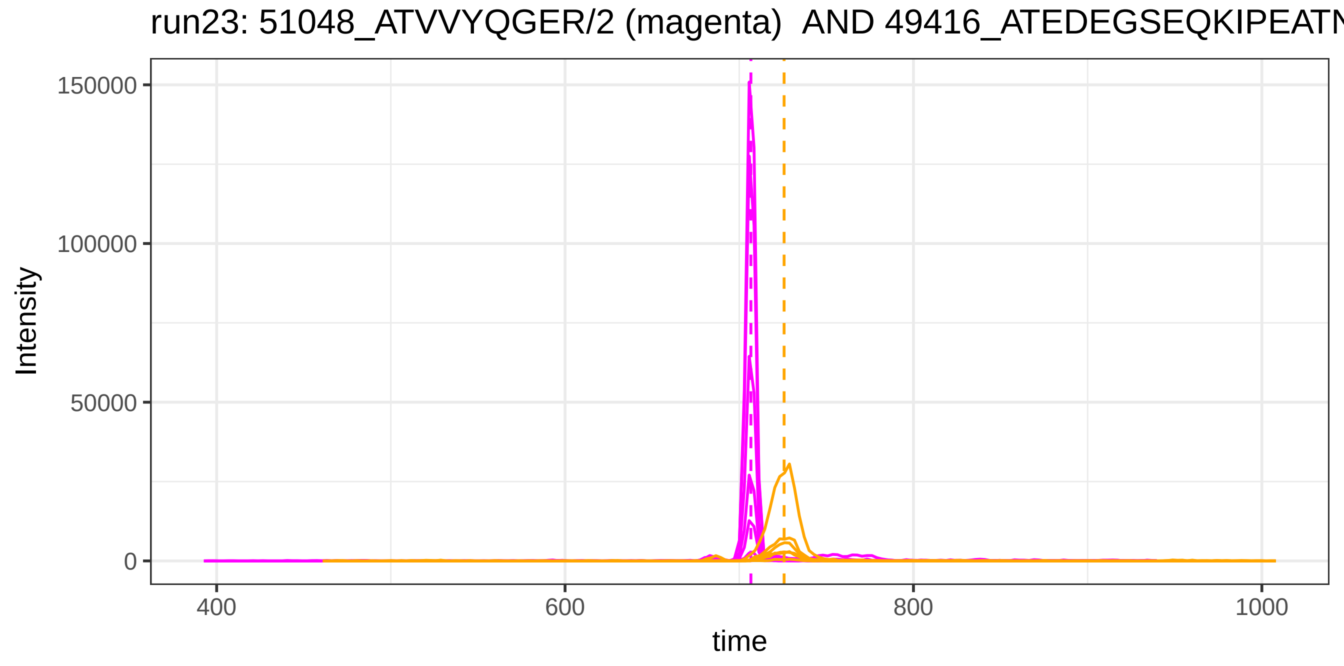
run23: 51048_ATVVYQGER/2 (magenta) AND 49416_ATEDEGSEQKIPEATN
Intensity
150000
100000
50000
0
400
600
800
1000
time

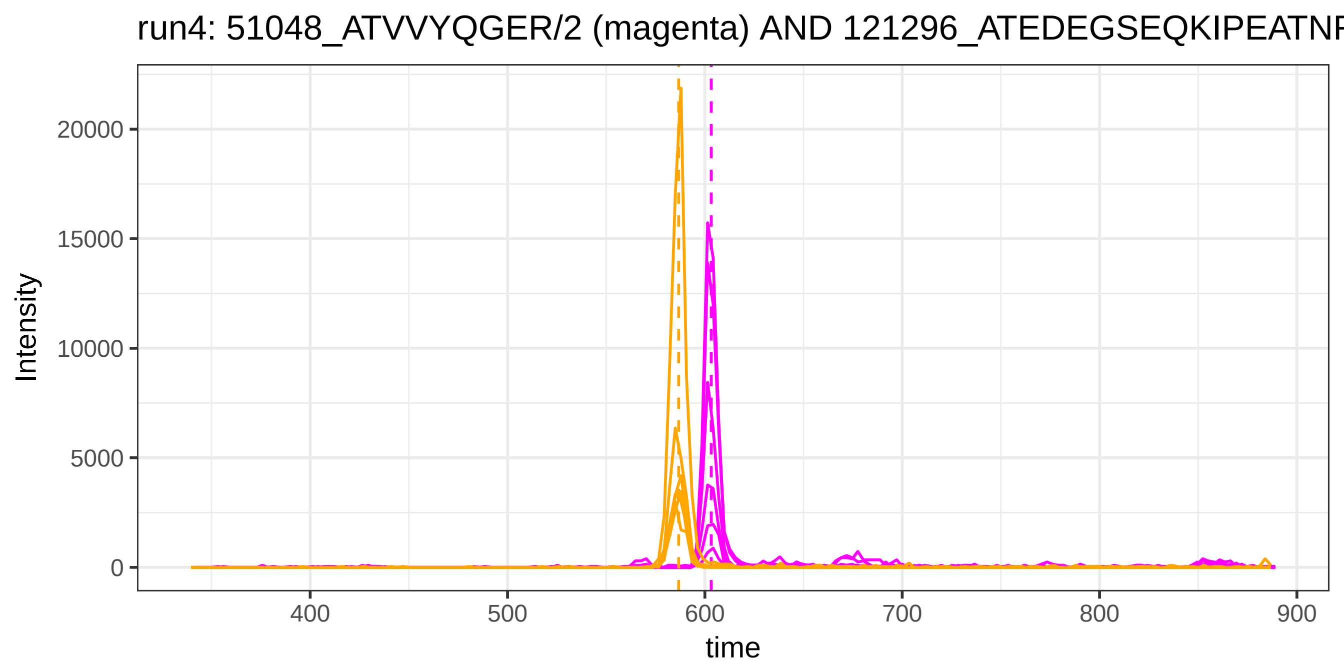
run4: 51048_ATVVYQGER/2 (magenta) AND 121296_ATEDEGSEQKIPEATNF
Intensity
20000
15000
10000
5000
0
400
500
600
700
800
900
time

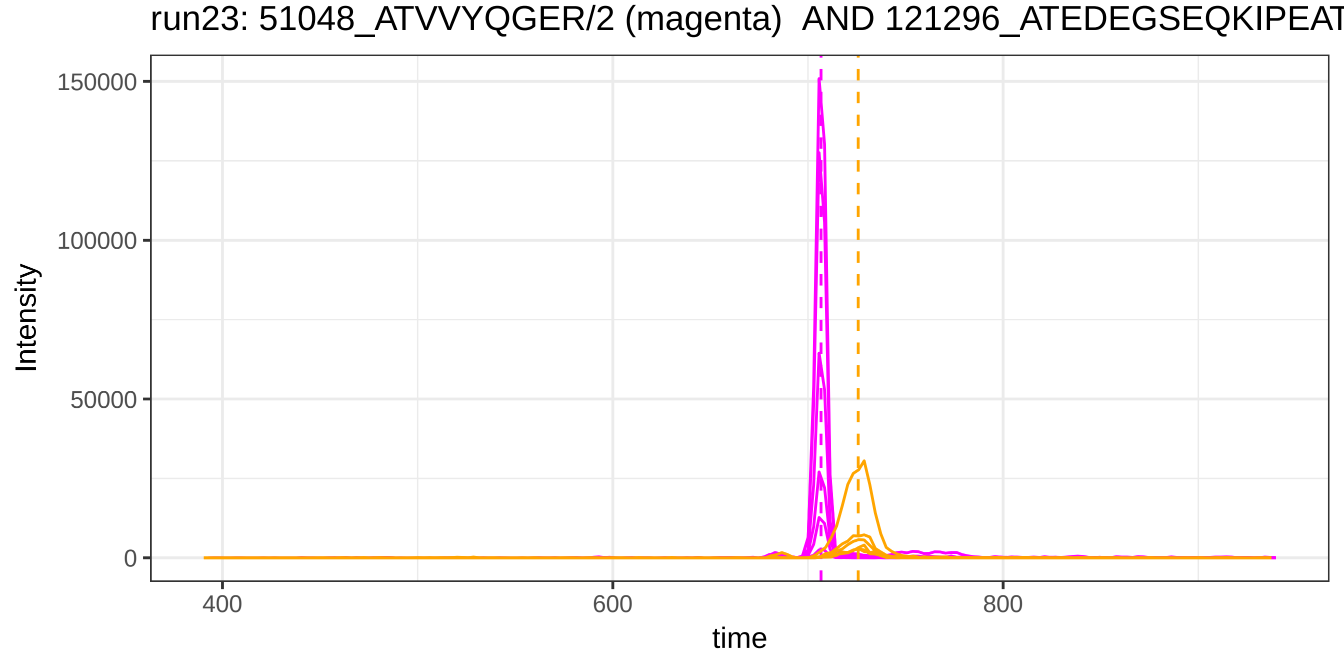
run23: 51048_ATVVYQGER/2 (magenta) AND 121296_ATEDEGSEQKIPEAT
Intensity
150000
100000
50000
0
400
600
800
time

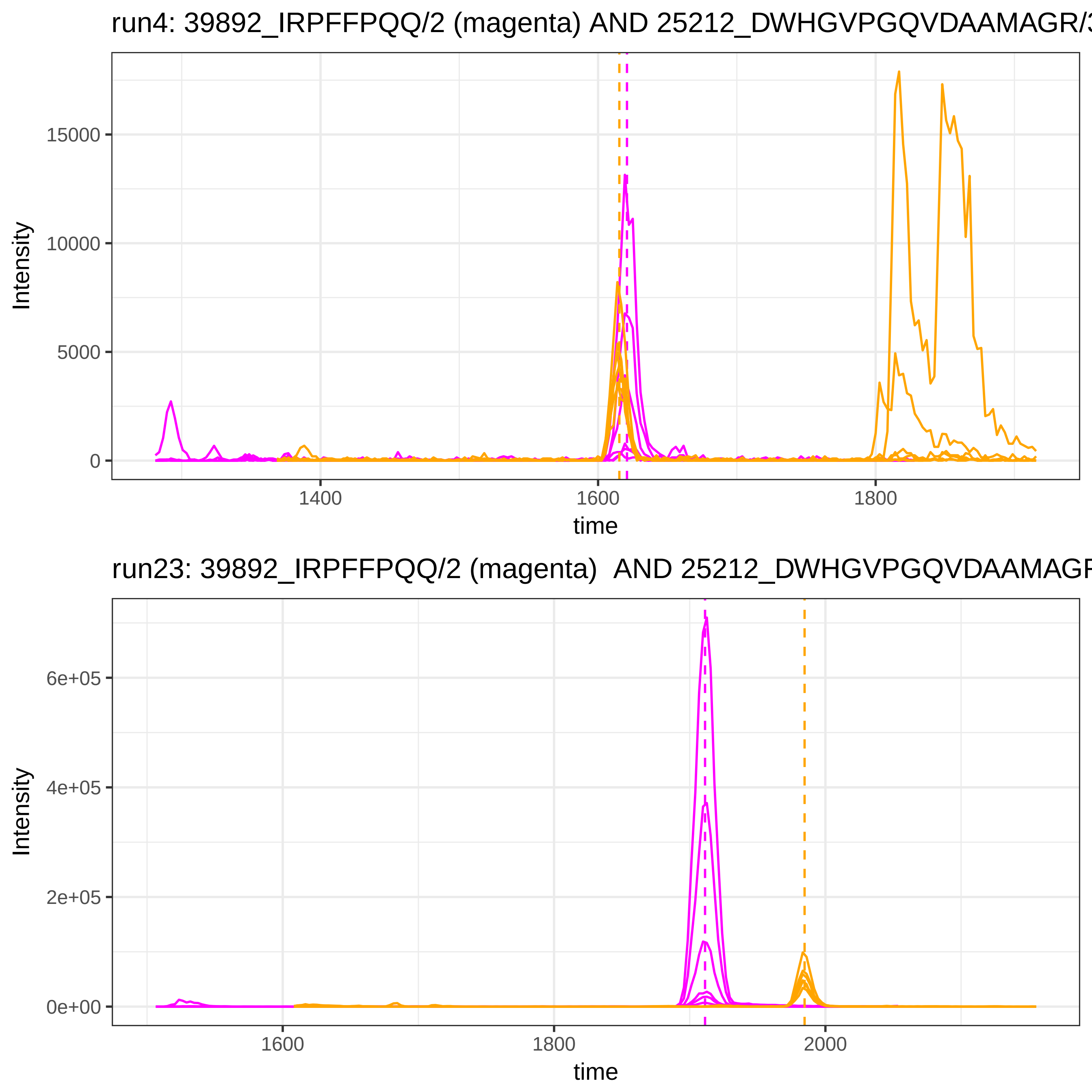

run4: 39892_IRPFFPQQ/2 (magenta) AND 25212_DWHGVPGQVDAAMAGR/3
Intensity
15000
10000
5000
0
1400
1600
1800
time
run23: 39892_IRPFFPQQ/2 (magenta) AND 25212_DWHGVPGQVDAAMAGR
Intensity
6e+05
4e+05
2e+05
0e+00
1600
1800
2000
time

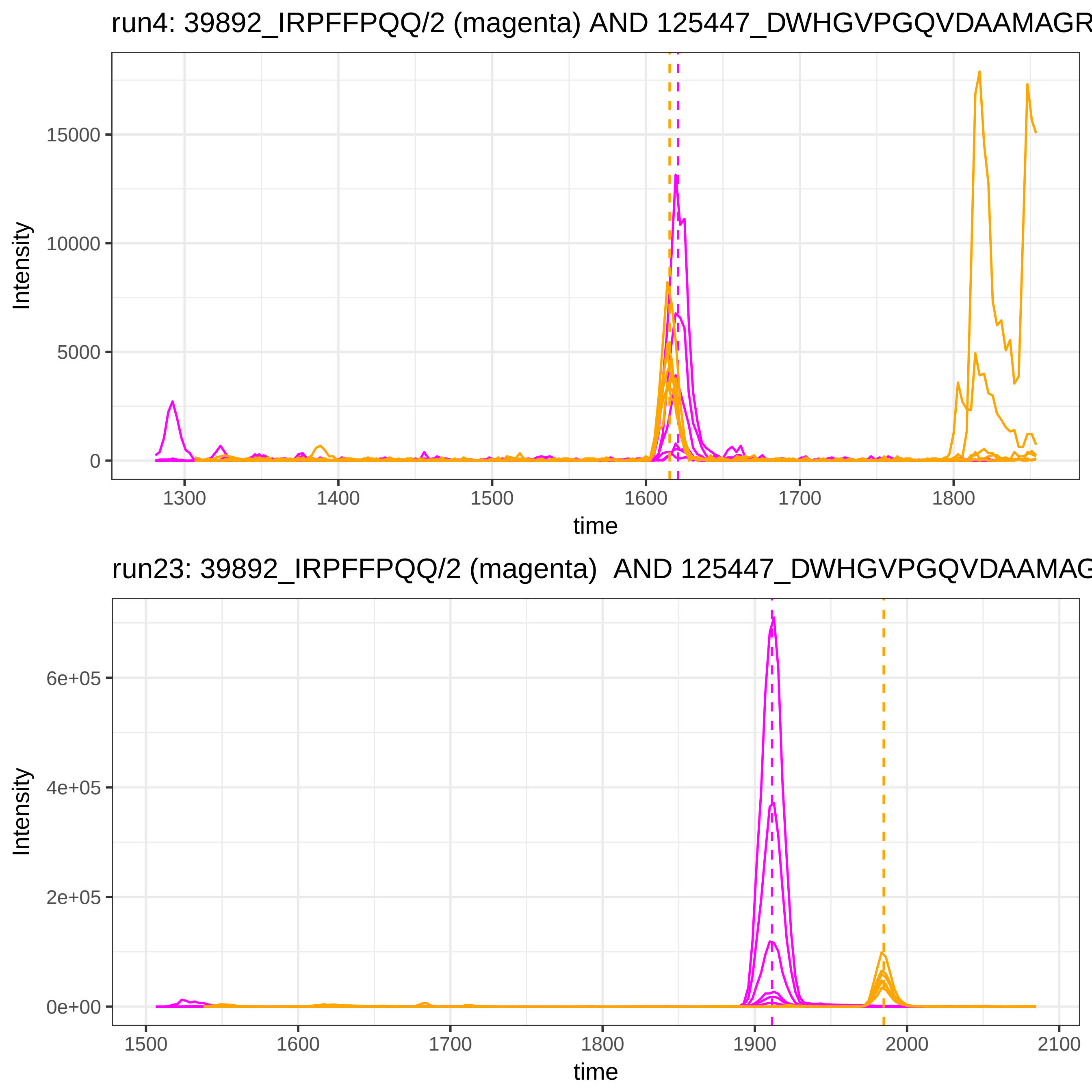
run4: 39892_IRPFFPQQ/2 (magenta) AND 125447_DWHGVPGQVDAAMAGR
15000
10000
5000
0
Intensity
1300
1400
1500
1600
1700
1800
time
run23: 39892_IRPFFPQQ/2 (magenta) AND 125447_DWHGVPGQVDAAMAG
6e+05
4e+05
2e+05
0e+00
Intensity
1500
1600
1700
1800
1900
2000
2100
time

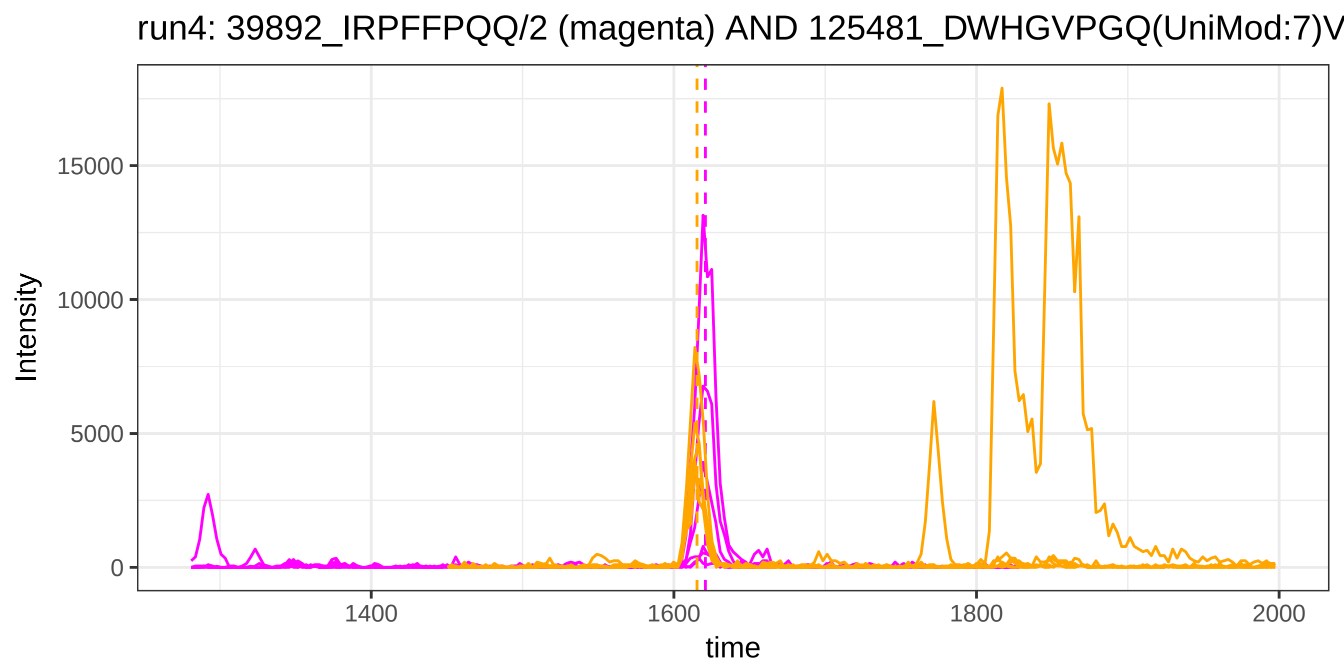
run4: 39892_IRPFFPQQ/2 (magenta) AND 125481_DWHGVPGQ(UniMod:7)V
Intensity
15000
10000
5000
0
1400
1600
1800
2000
time

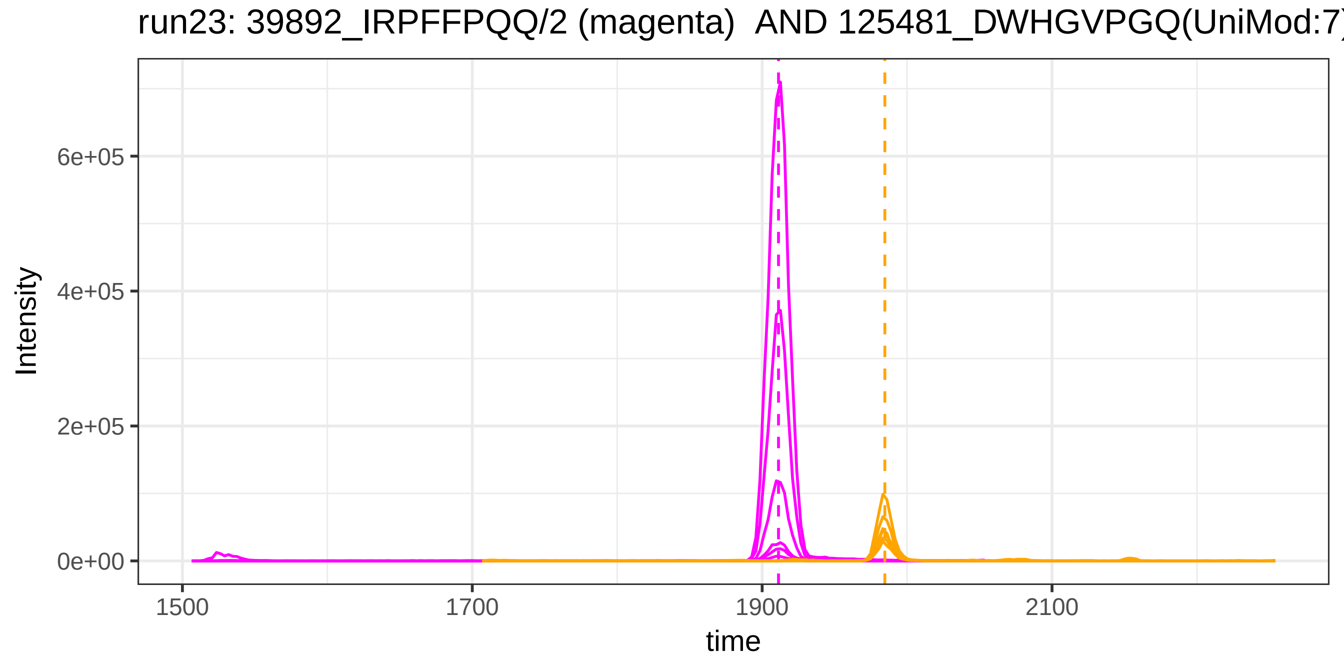
run23: 39892_IRPFFPQQ/2 (magenta) AND 125481_DWHGVPGQ(UniMod:7)
Intensity
6e+05
4e+05
2e+05
0e+00
1500
1700
1900
2100
time

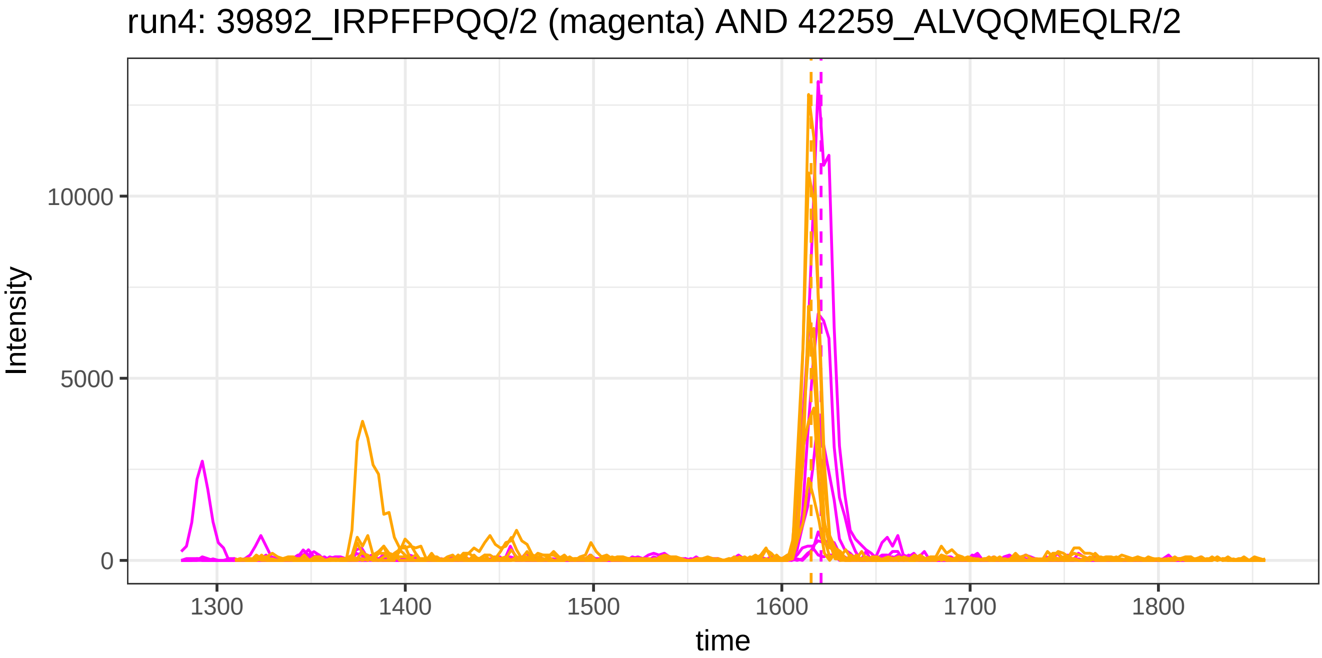
run4: 39892_IRPFFPQQ/2 (magenta) AND 42259_ALVQQMEQLR/2
Intensity
10000
5000
0
1300
1400
1500
1600
1700
1800
time

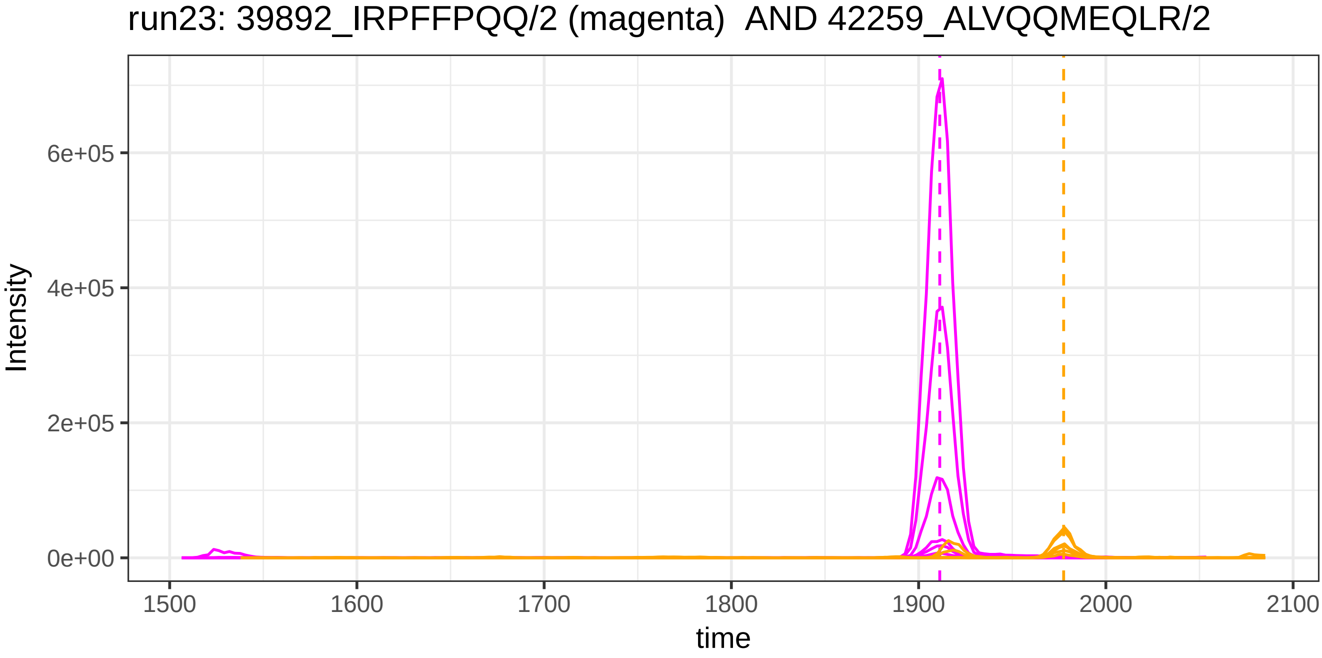
run23: 39892_IRPFFPQQ/2 (magenta) AND 42259_ALVQQMEQLR/2
Intensity
6e+05
4e+05
2e+05
0e+00
1500
1600
1700
1800
1900
2000
2100
time

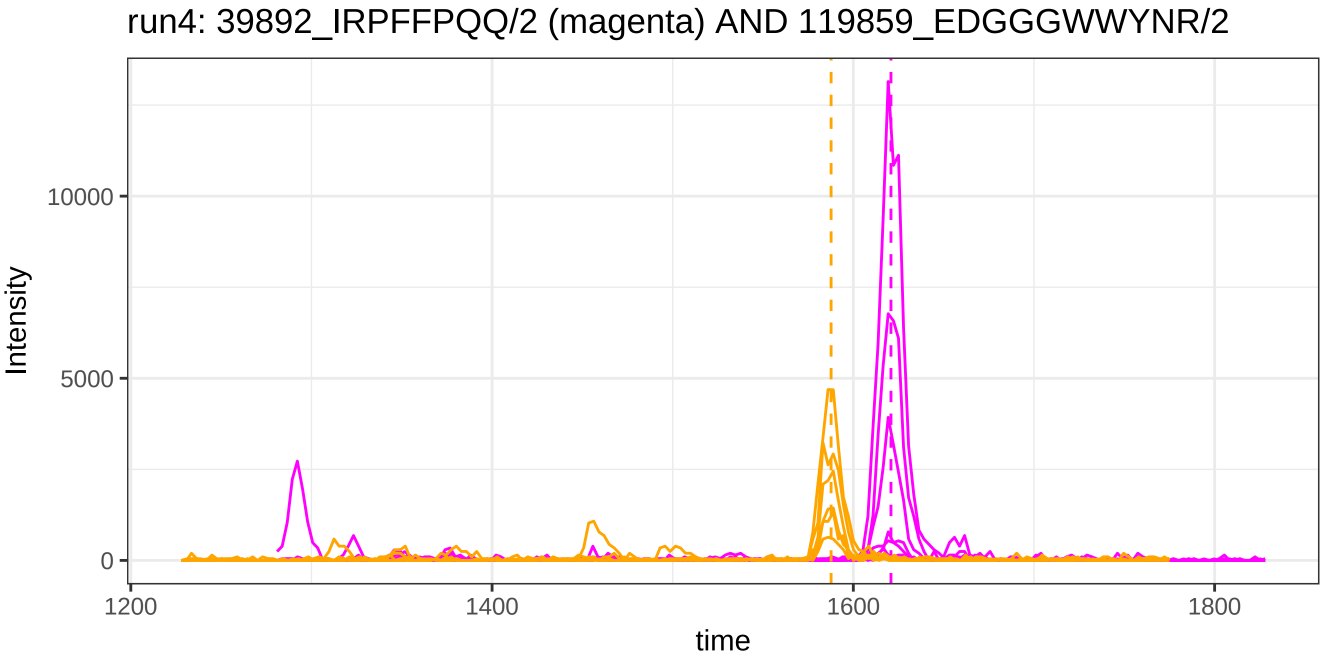
run4: 39892_IRPFFPQQ/2 (magenta) AND 119859_EDGGGWWYNR/2
Intensity
10000
5000
0
1200
1400
1600
1800
time

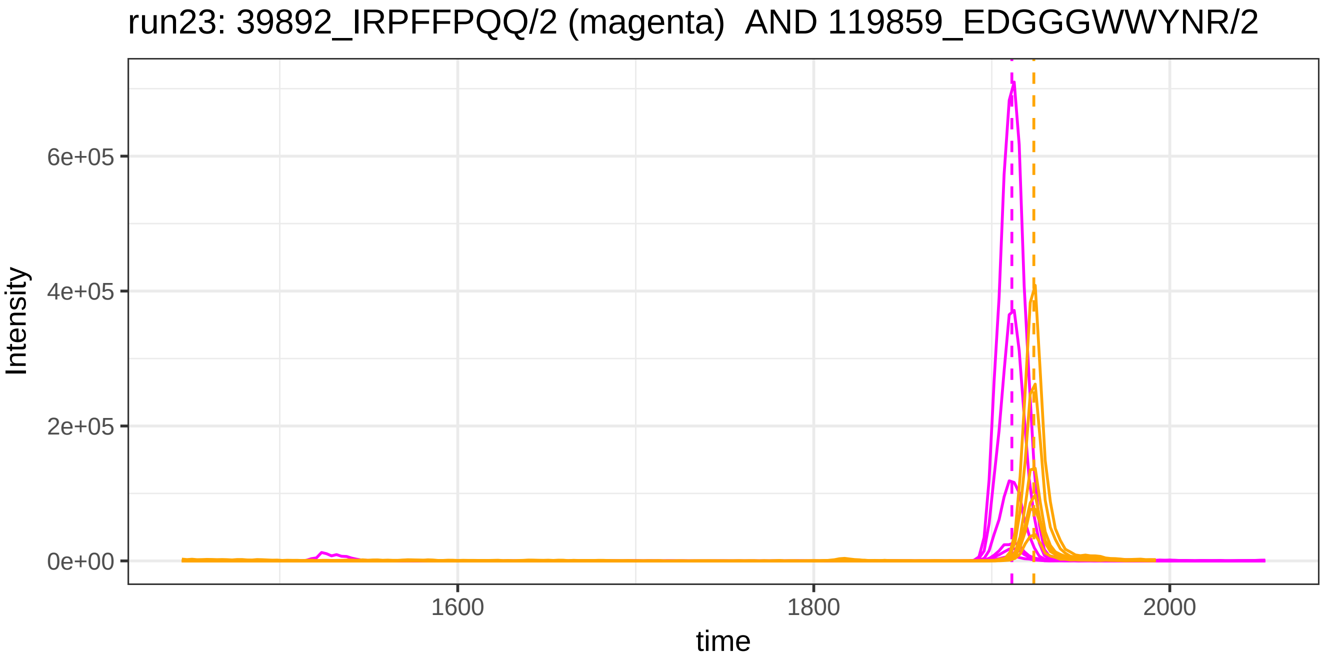
run23: 39892_IRPFFPQQ/2 (magenta) AND 119859_EDGGGWWYNR/2
Intensity
6e+05
4e+05
2e+05
0e+00
1600
1800
2000
time

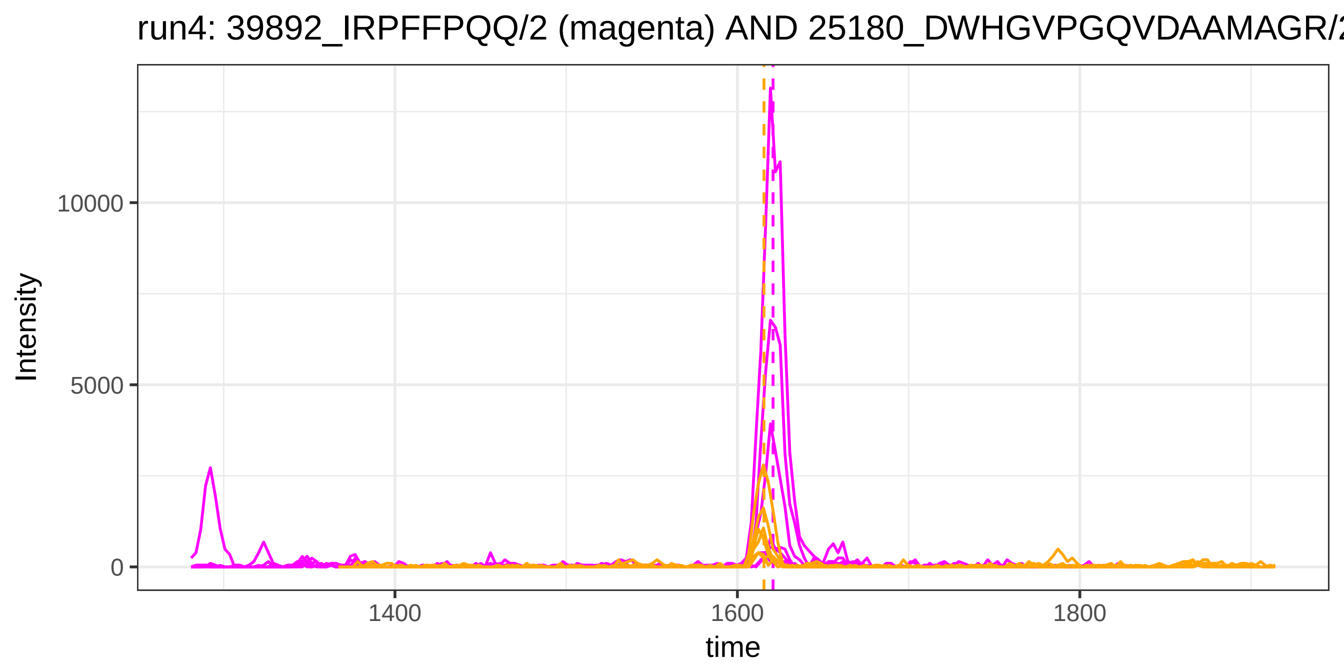
run4: 39892_IRPFFPQQ/2 (magenta) AND 25180_DWHGVPGQVDAAMAGR/2
Intensity
10000
5000
0
1400
1600
1800
time

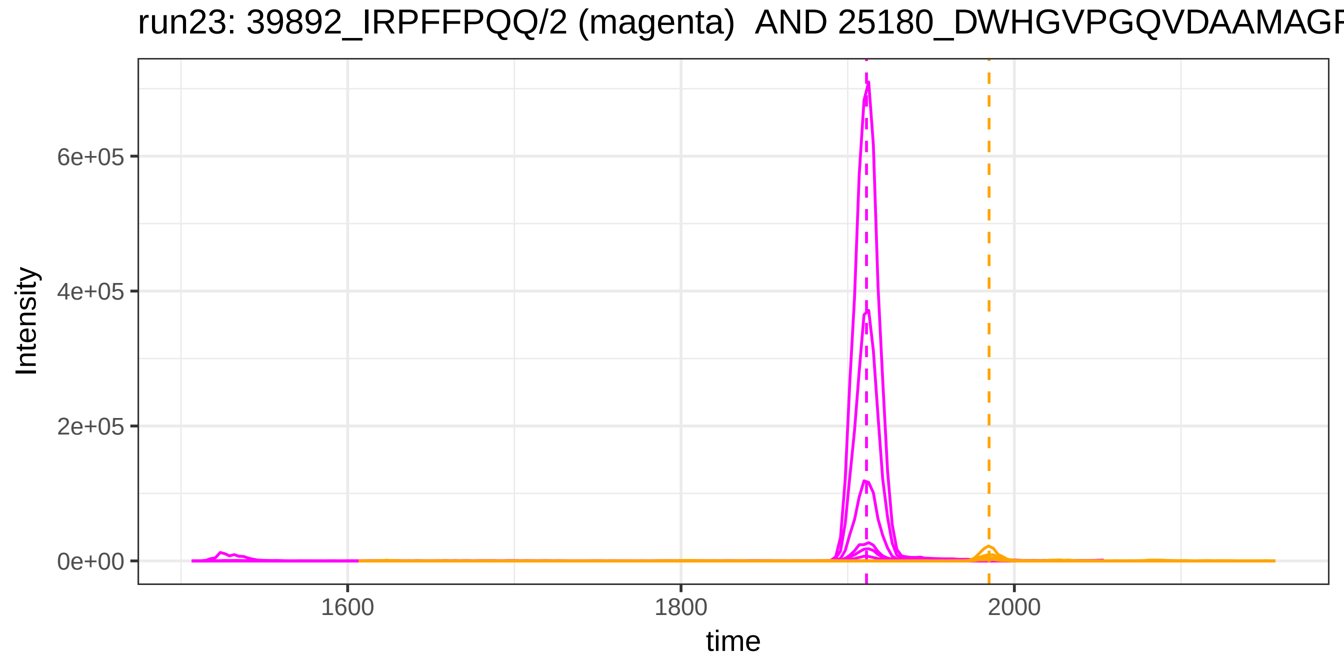
run23: 39892_IRPFFPQQ/2 (magenta) AND 25180_DWHGVPGQVDAAMAGR
Intensity
6e+05
4e+05
2e+05
0e+00
1600
1800
2000
time

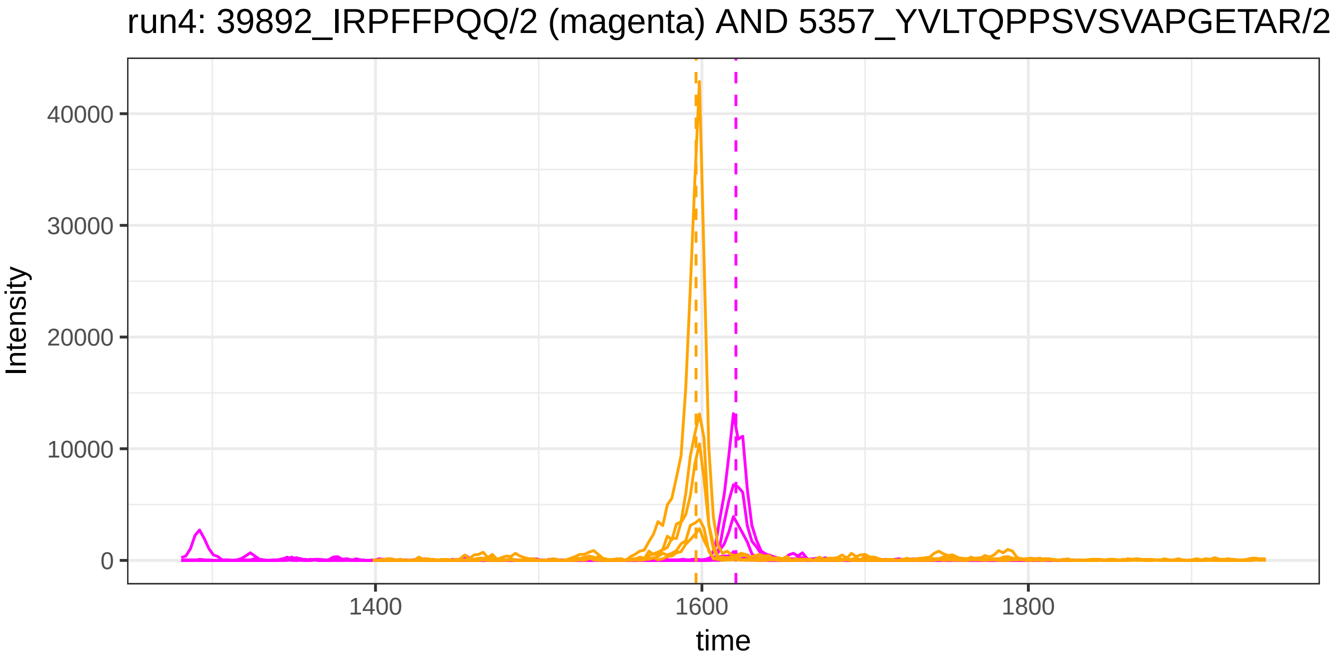

run4: 39892_IRPFFPQQ/2 (magenta) AND 5357_YVLTQPPSVSVAPGETAR/2
Intensity
40000
30000
20000
10000
0
1400
1600
1800
time


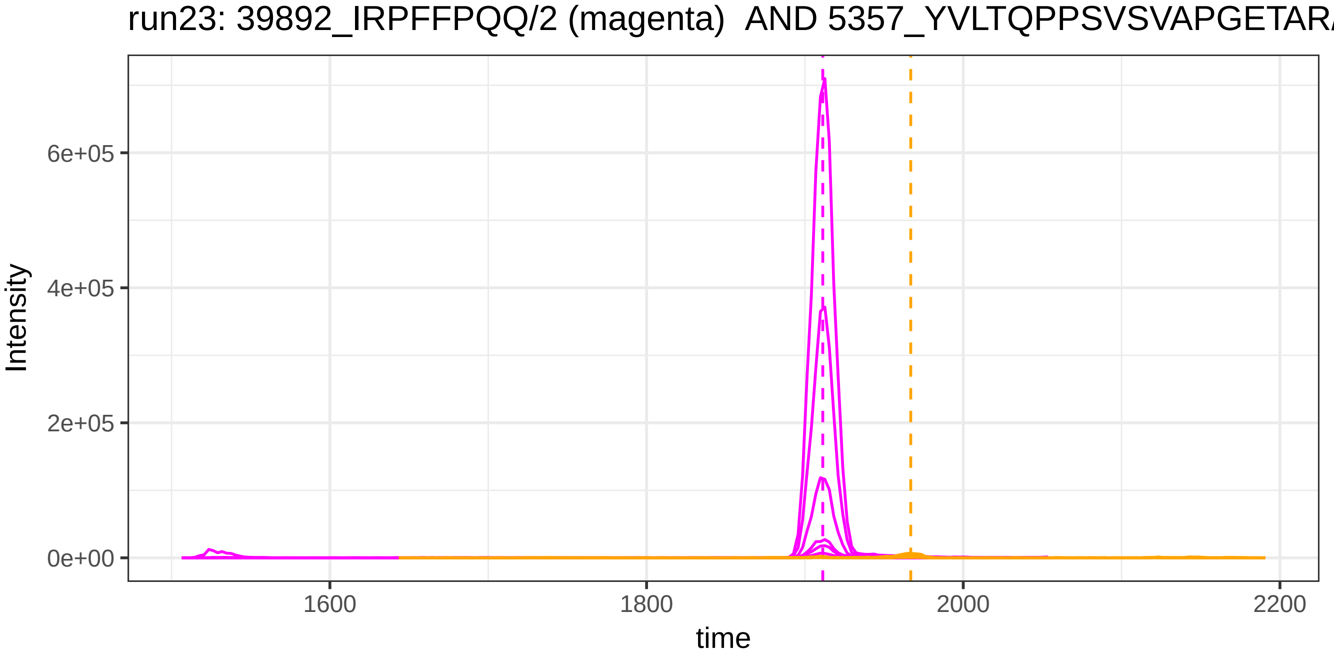

run23: 39892_IRPFFPQQ/2 (magenta) AND 5357_YVLTQPPSVSVAPGETAR/
Intensity
6e+05
4e+05
2e+05
0e+00
1600
1800
2000
2200
time

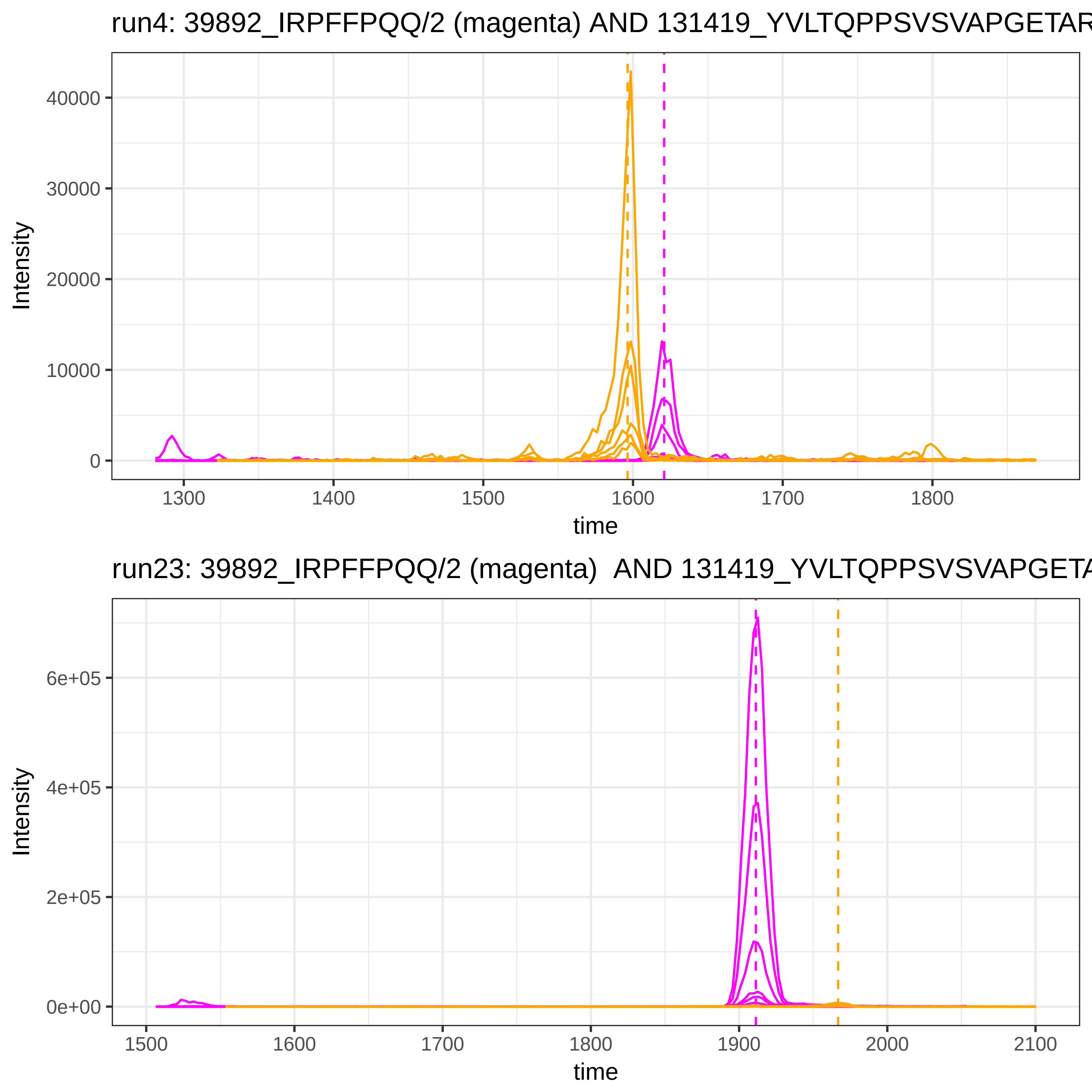

run4: 39892_IRPFFPQQ/2 (magenta) AND 131419_YVLTQPPSVSVAPGETAR
Intensity
40000
30000
20000
10000
0
1300
1400
1500
1600
1700
1800
time
run23: 39892_IRPFFPQQ/2 (magenta) AND 131419_YVLTQPPSVSVAPGETA
Intensity
6e+05
4e+05
2e+05
0e+00
1500
1600
1700
1800
1900
2000
2100
time

run4: 49520_QIQVSWLR/2 (magenta) AND 117986_QQNAQGGFSSTQDTVV

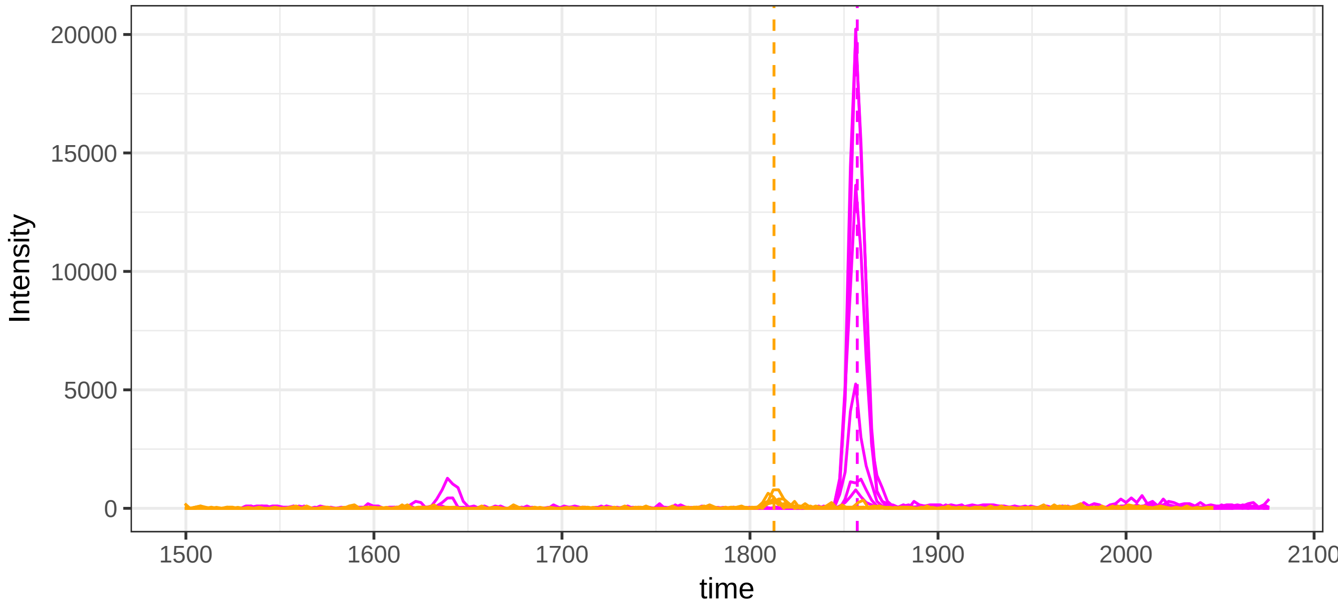


run23: 49520_QIQVSWLR/2 (magenta)  AND 117986_QQNAQGGFSSTQDTV

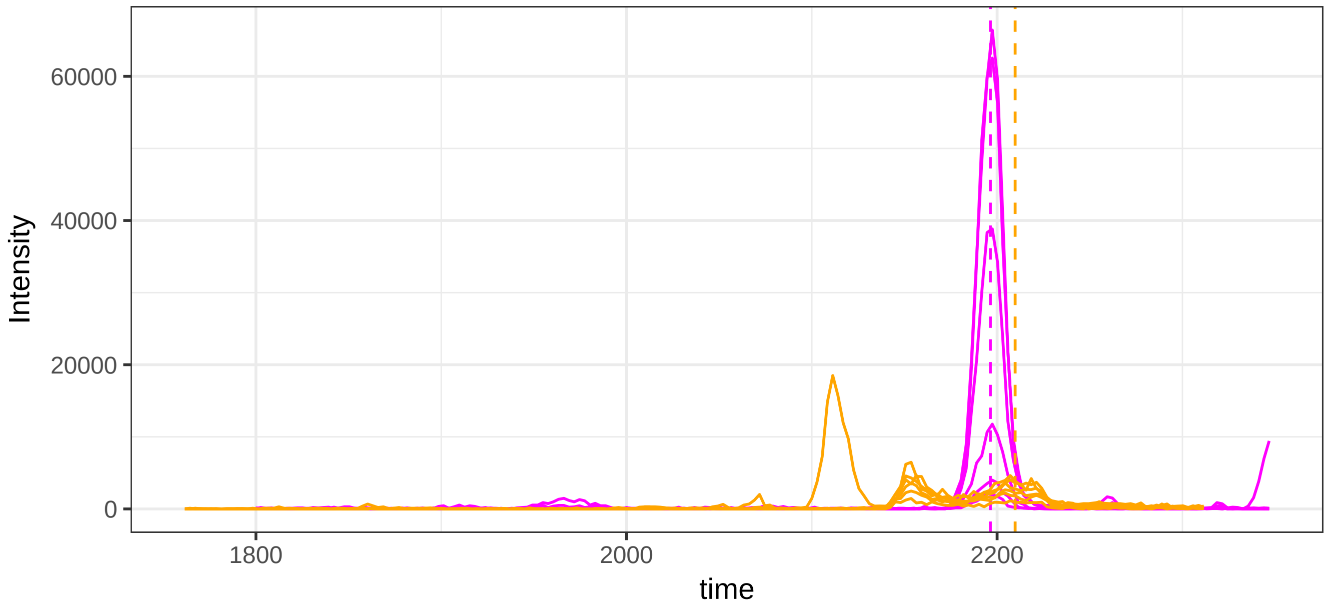

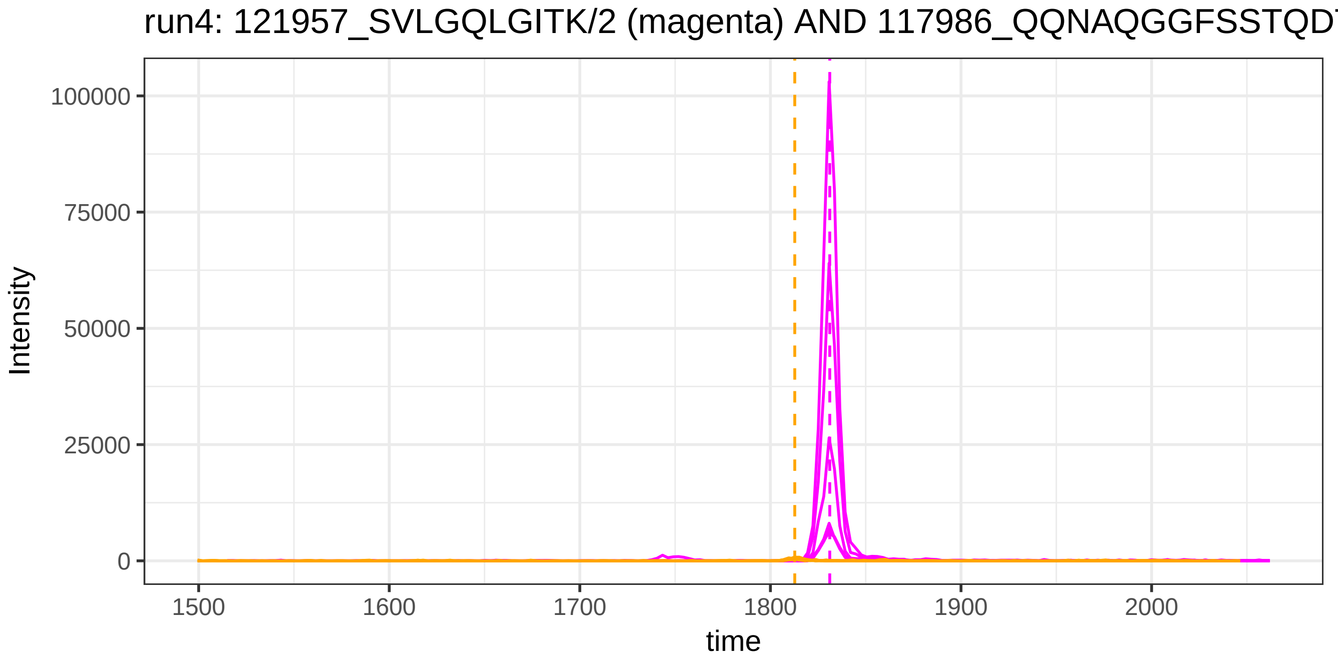
run4: 121957_SVLGQLGITK/2 (magenta) AND 117986_QQNAQGGFSSTQD
Intensity
100000
75000
50000
25000
0
1500
1600
1700
1800
1900
2000
time

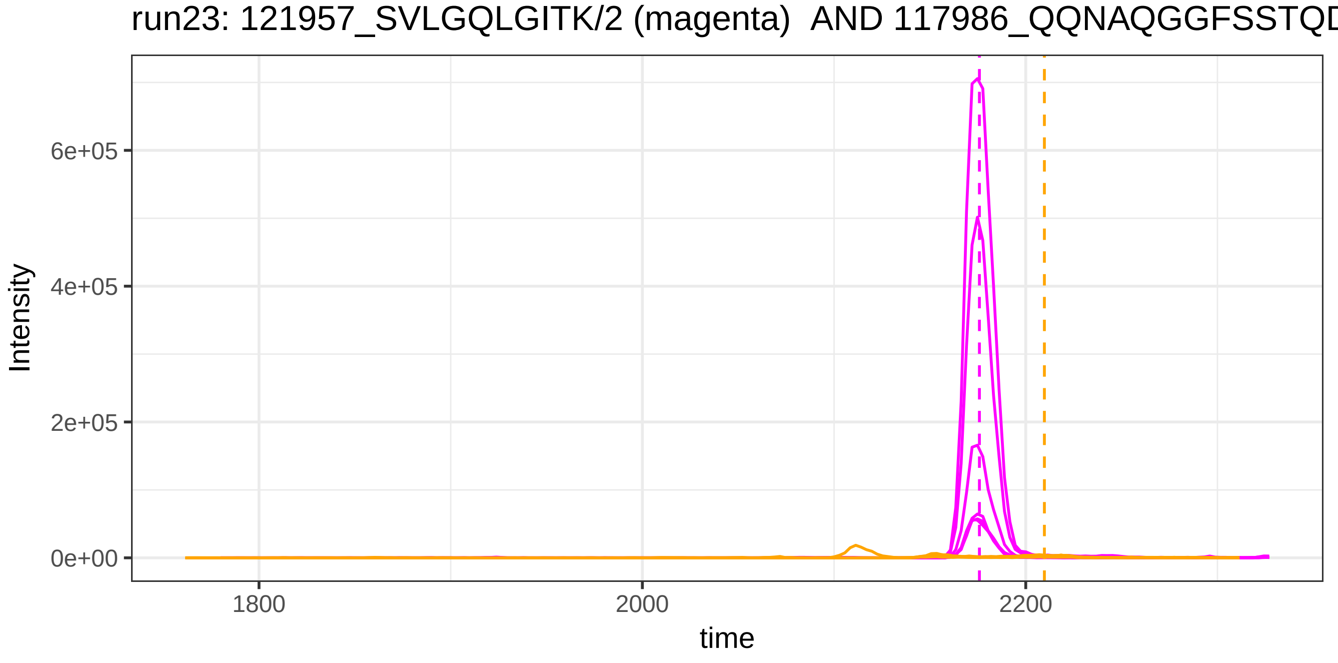
run23: 121957_SVLGQLGITK/2 (magenta) AND 117986_QQNAQGGFSSTQD
Intensity
6e+05
4e+05
2e+05
0e+00
1800
2000
2200
time

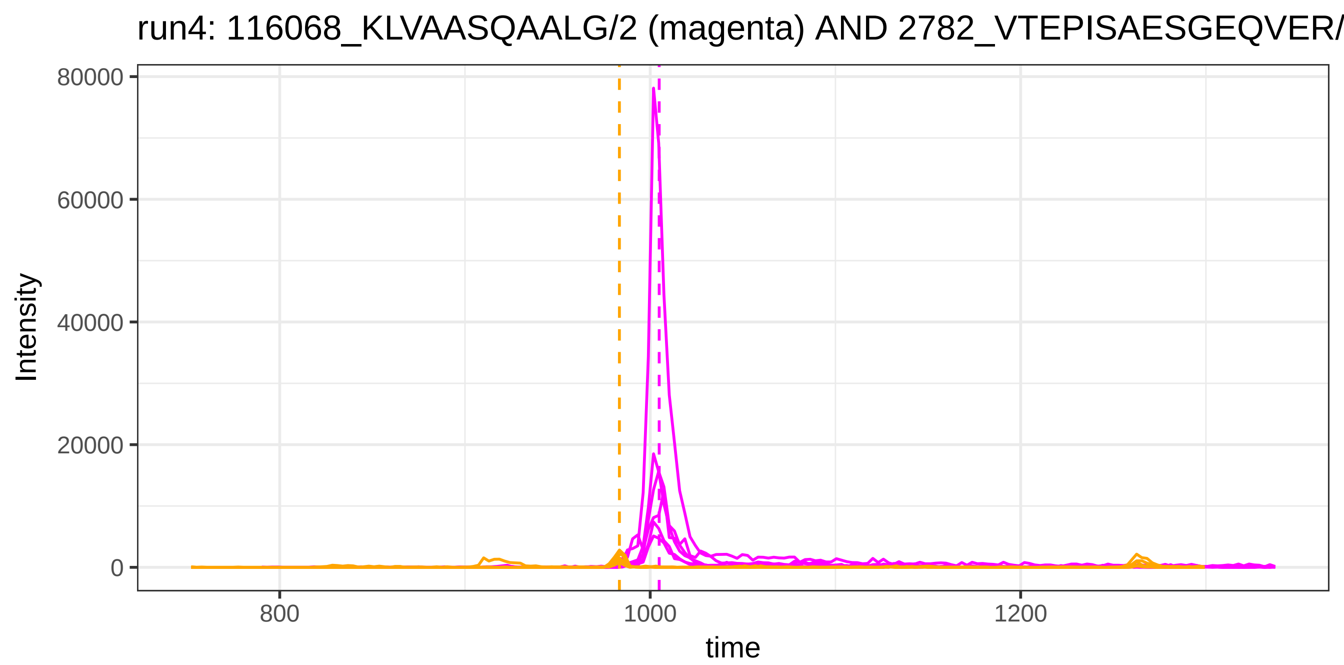

run4: 116068_KLVAASQAALG/2 (magenta) AND 2782_VTEPISAESGEQVER/
80000
60000
40000
20000
0
Intensity
800
1000
1200
time


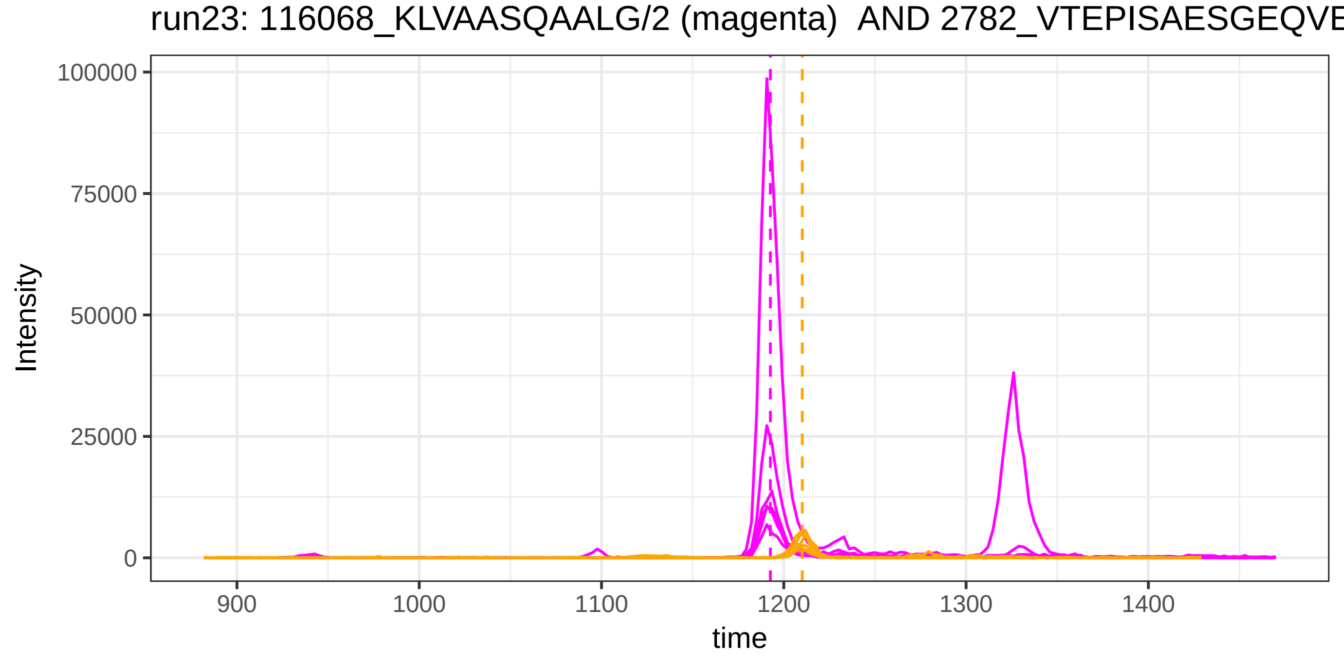

run23: 116068_KLVAASQAALG/2 (magenta) AND 2782_VTEPISAESGEQVE
100000
75000
50000
25000
0
Intensity
900
1000
1100
1200
1300
1400
time

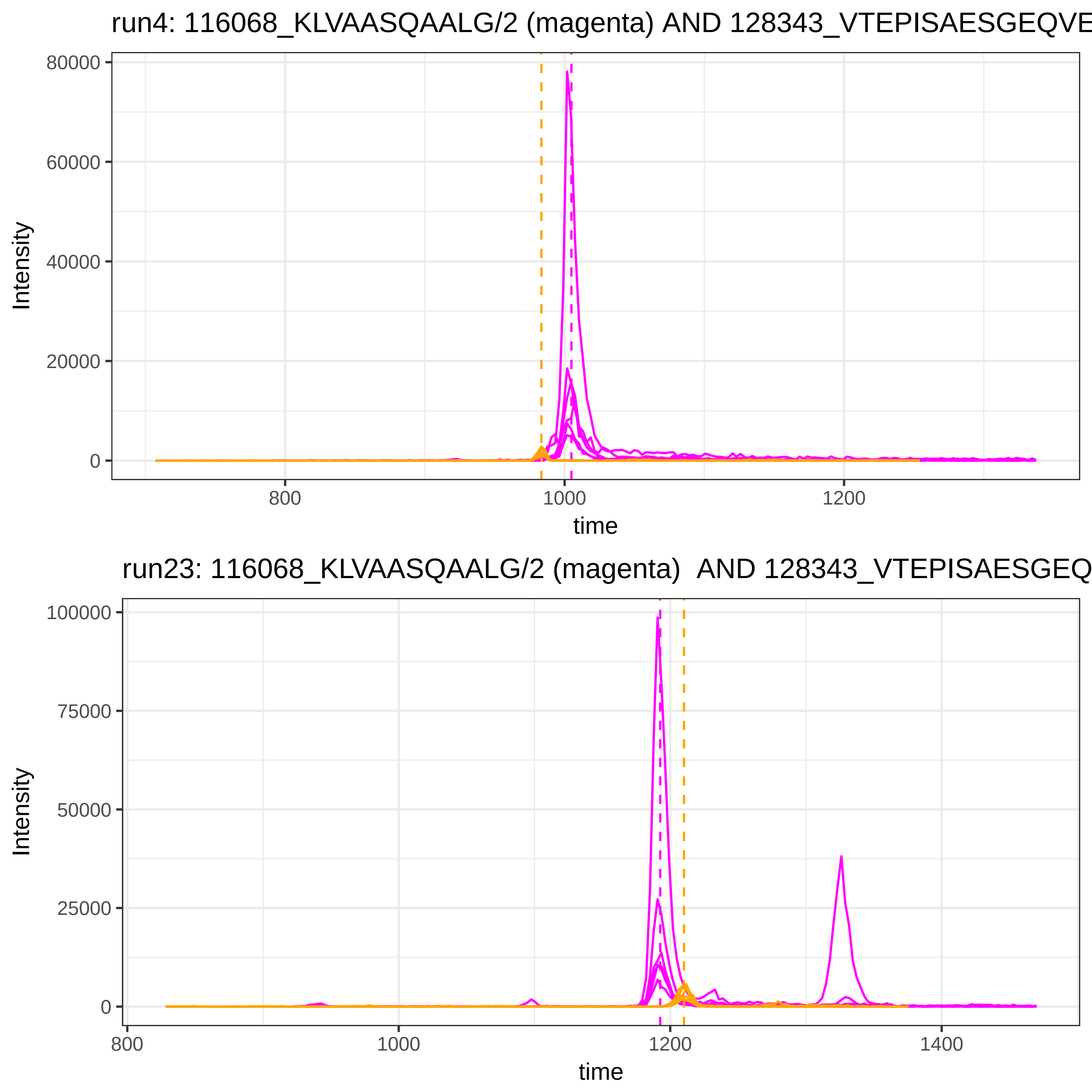
run4: 116068_KLVAASQAALG/2 (magenta) AND 128343_VTEPISAESGEQVE
Intensity
time
80000
60000
40000
20000
0
800
1000
1200
run23: 116068_KLVAASQAALG/2 (magenta) AND 128343_VTEPISAESGEQ
Intensity
time
100000
75000
50000
25000
0
800
1000
1200
1400

run4: 116068_KLVAASQAALG/2 (magenta) AND 47721_NNEGTYYSPNYNPQ

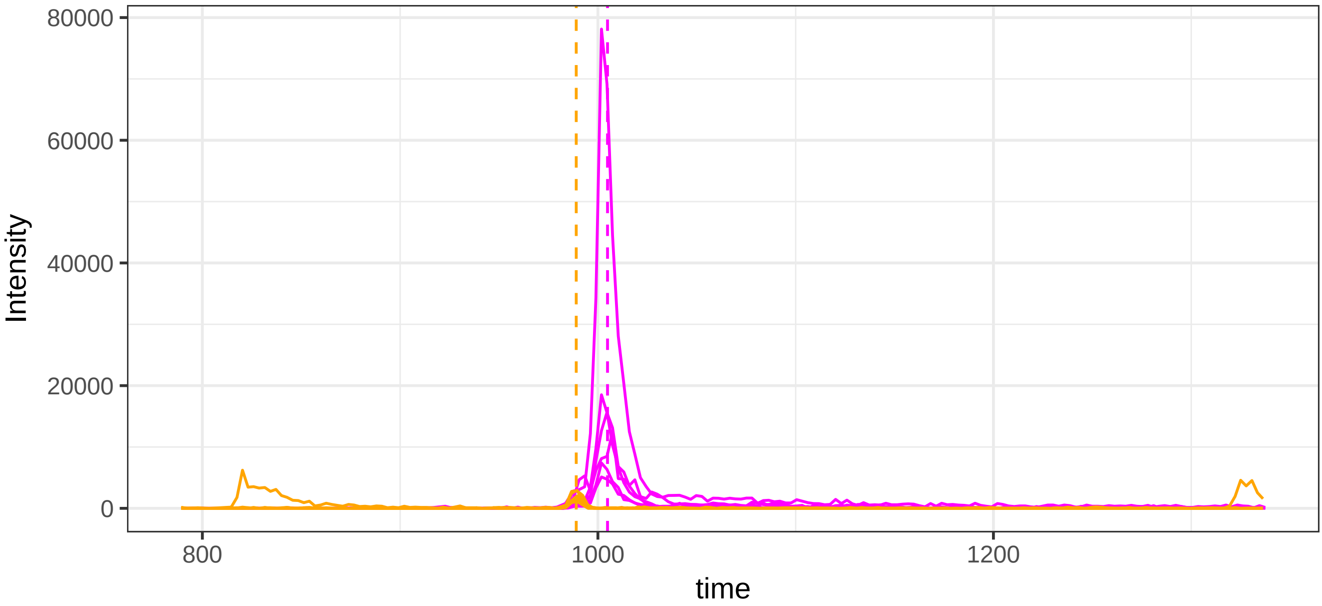


run23: 116068_KLVAASQAALG/2 (magenta)  AND 47721_NNEGTYYSPNYN

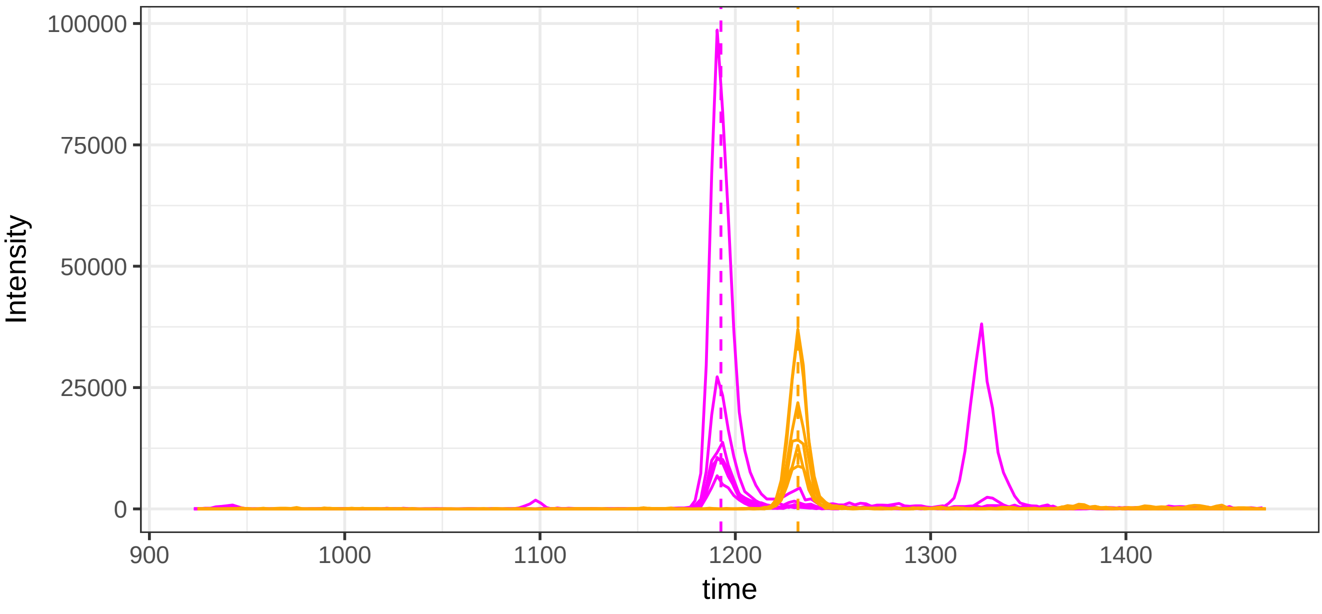

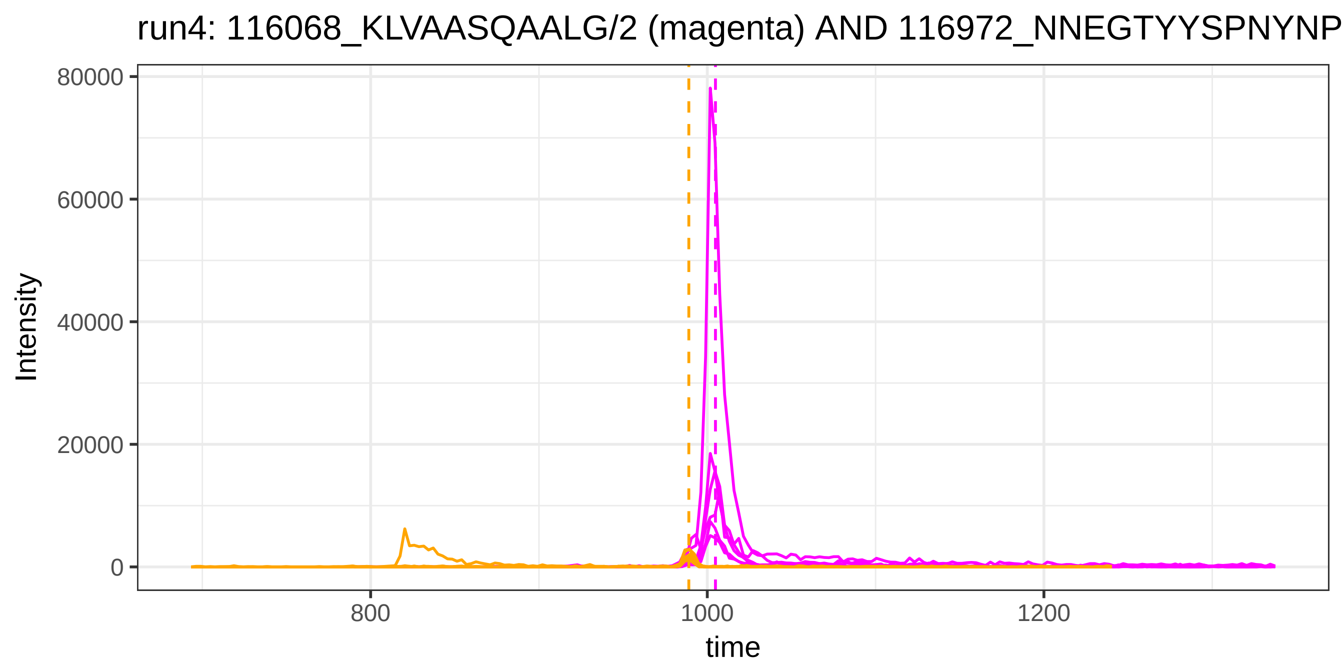

run4: 116068_KLVAASQAALG/2 (magenta) AND 116972_NNEGTYYSPNYNP
Intensity
80000
60000
40000
20000
0
800
1000
1200
time


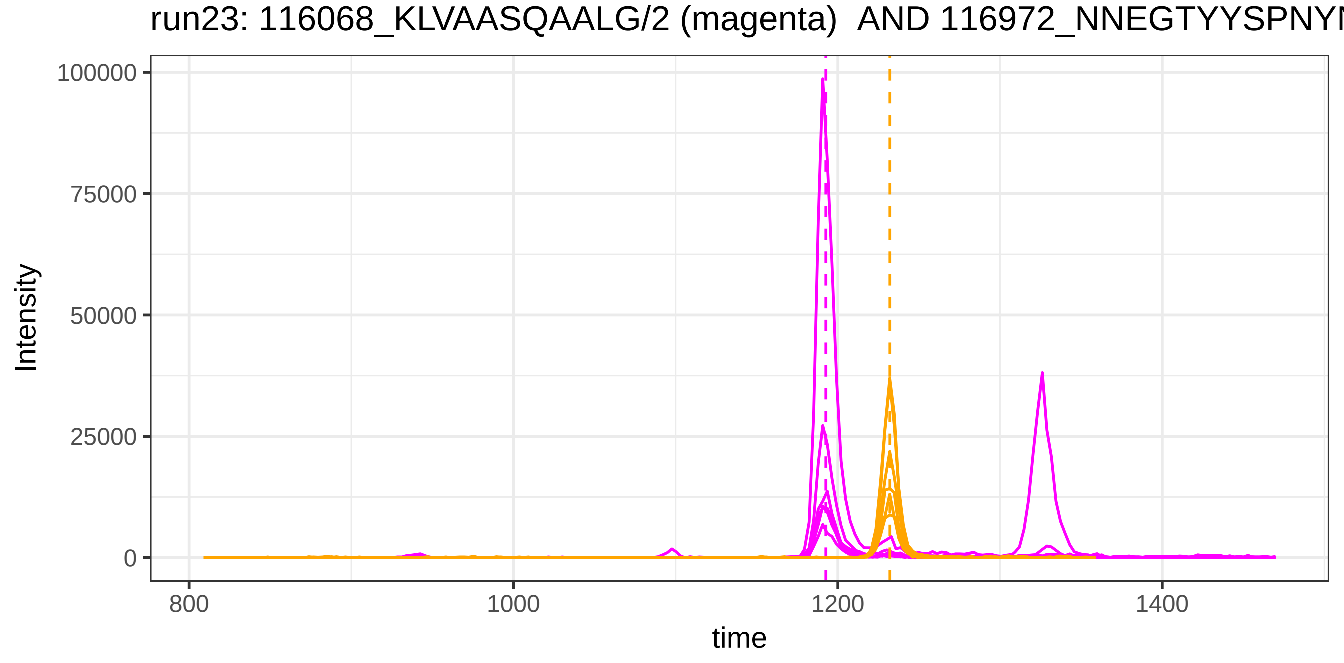

run23: 116068_KLVAASQAALG/2 (magenta) AND 116972_NNEGTYYSPNYN
Intensity
100000
75000
50000
25000
0
800
1000
1200
1400
time

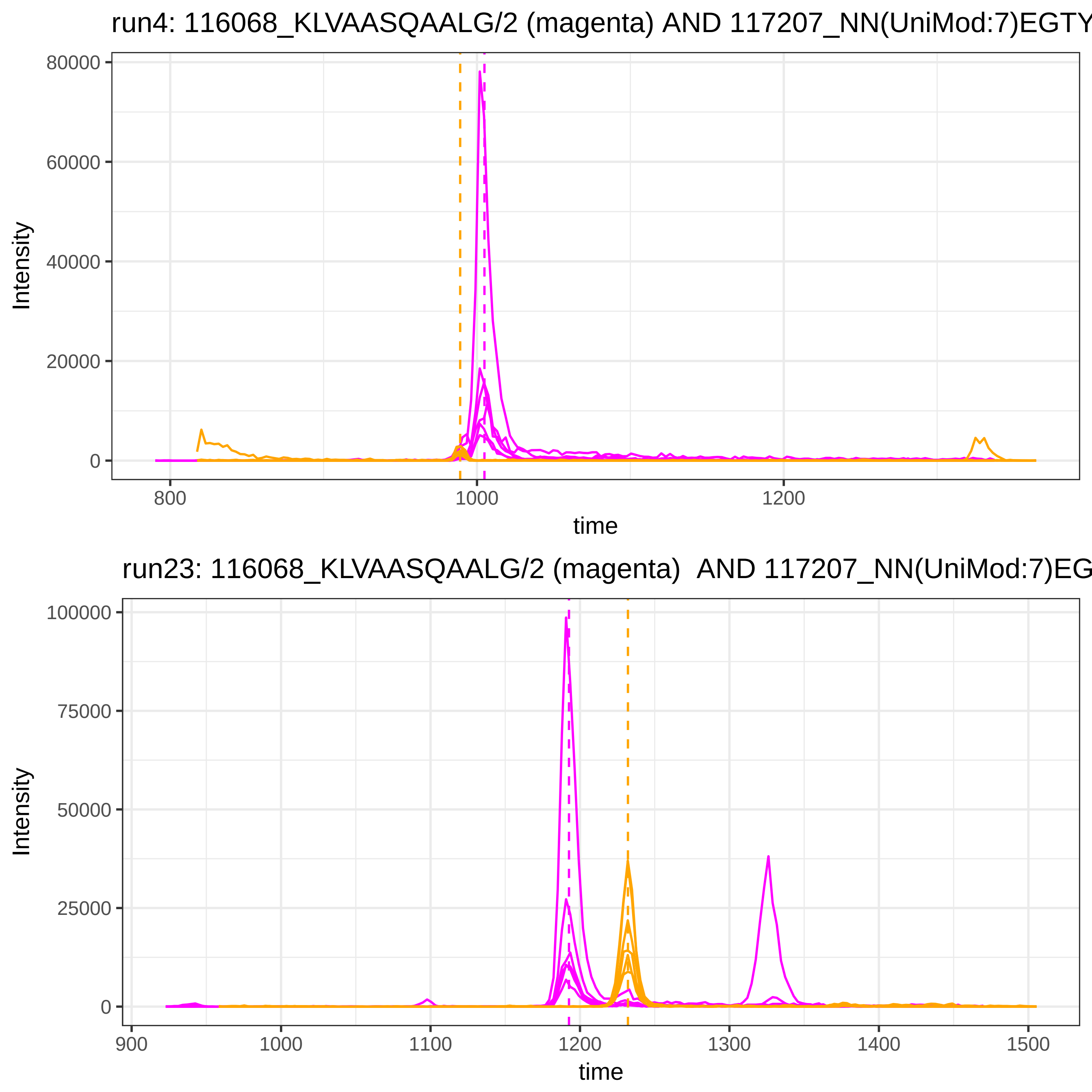

run4: 116068_KLVAASQAALG/2 (magenta) AND 117207_NN(UniMod:7)EGTY
80000
60000
40000
20000
0
Intensity
800
1000
1200
time
run23: 116068_KLVAASQAALG/2 (magenta) AND 117207_NN(UniMod:7)EG
100000
75000
50000
25000
0
Intensity
900
1000
1100
1200
1300
1400
1500
time

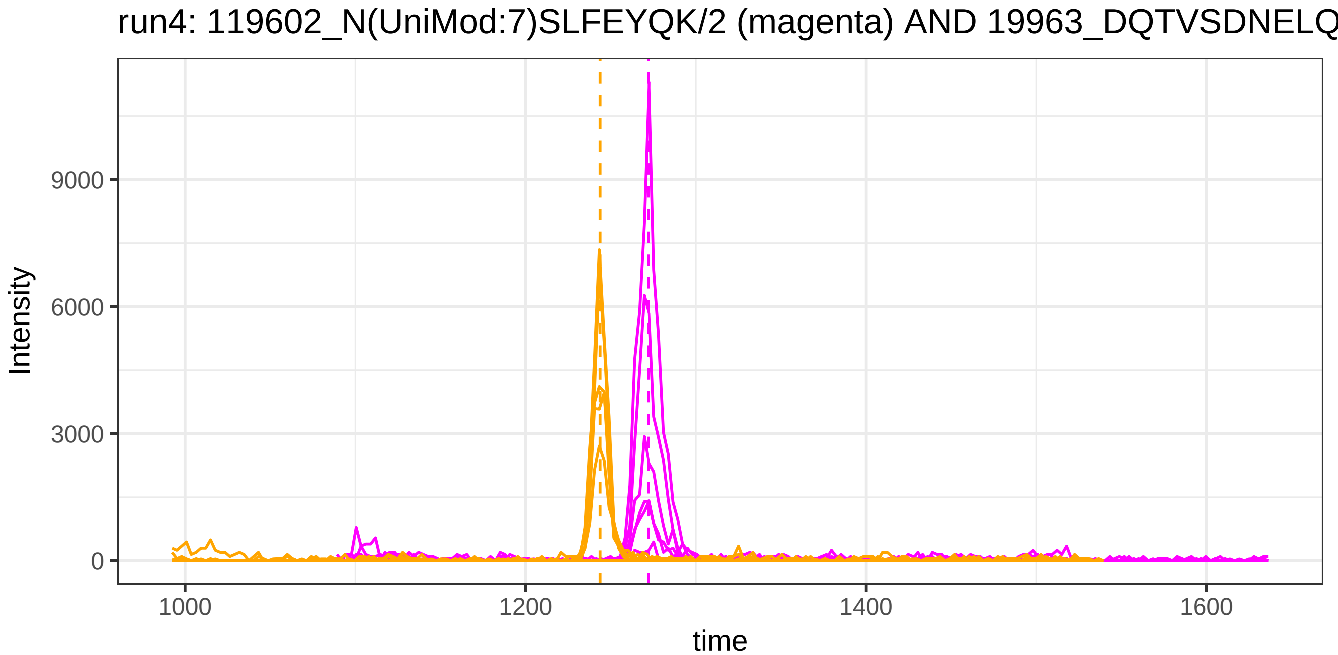
run4: 119602_N(UniMod:7)SLFEYQK/2 (magenta) AND 19963_DQTVSDNELQ
Intensity
9000
6000
3000
0
1000
1200
1400
1600
time

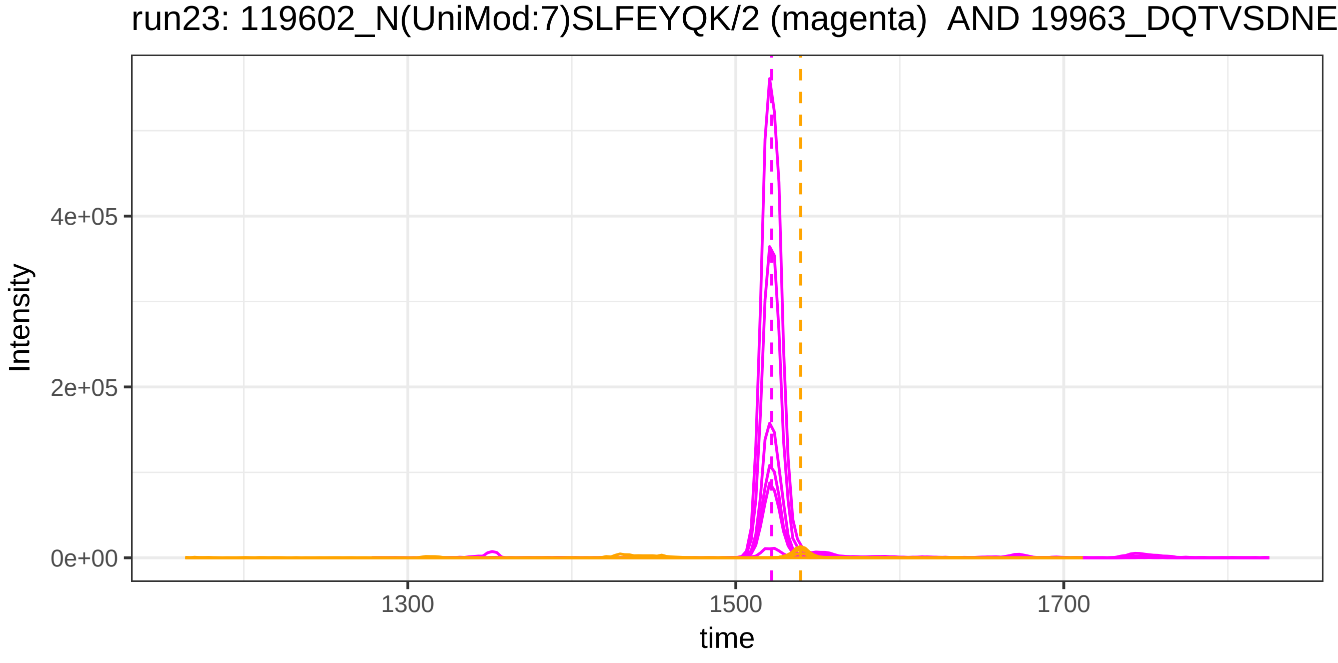
run23: 119602_N(UniMod:7)SLFEYQK/2 (magenta) AND 19963_DQTVSDNE
Intensity
4e+05
2e+05
0e+00
1300
1500
1700
time

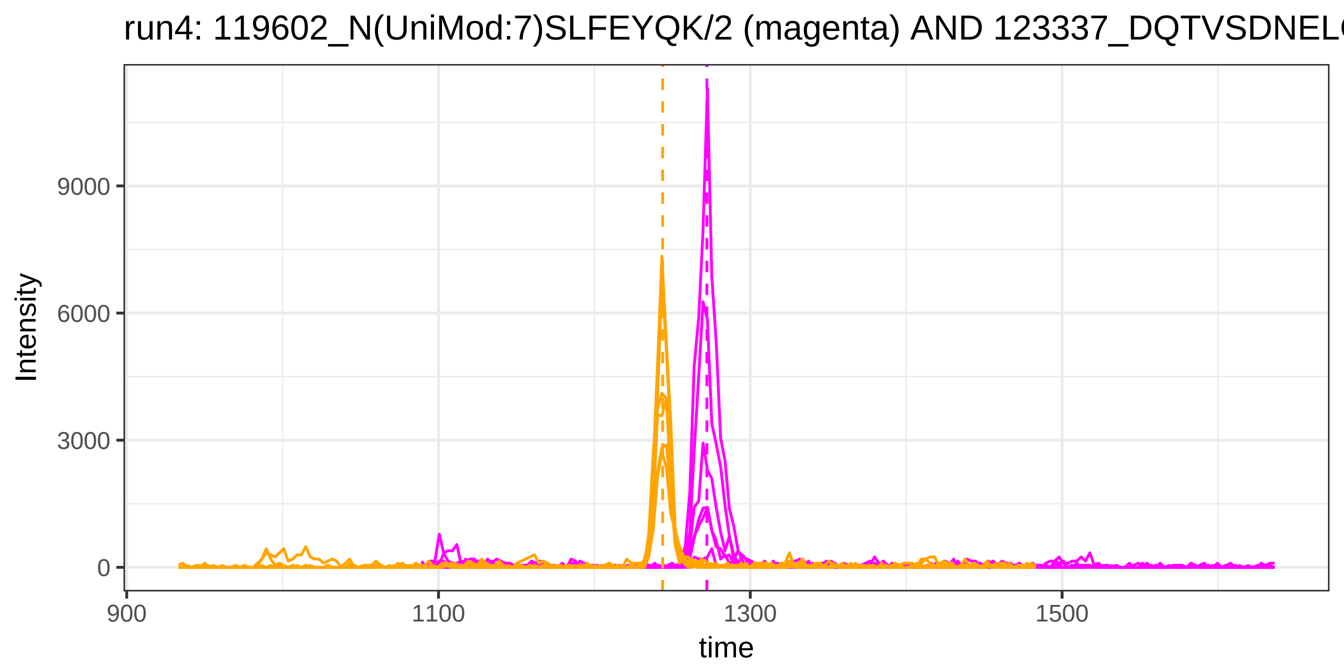

run4: 119602_N(UniMod:7)SLFEYQK/2 (magenta) AND 123337_DQTVSDNELQ
Intensity
9000
6000
3000
0
900
1100
1300
1500
time


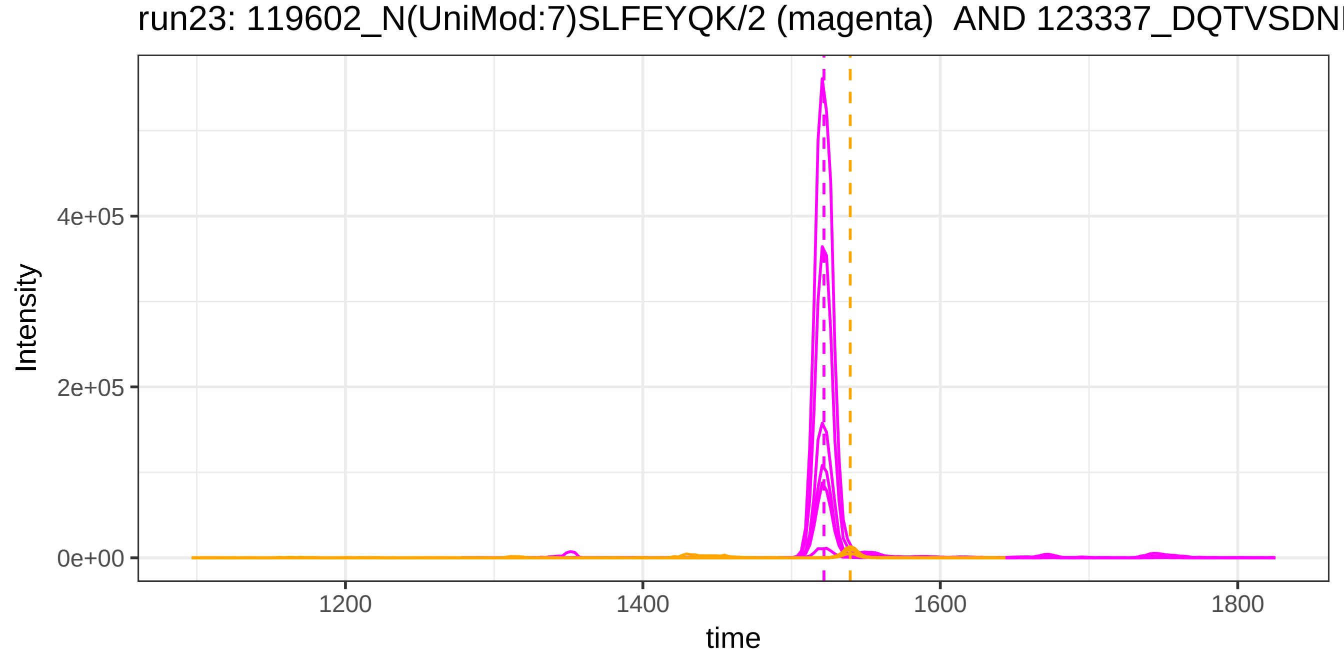

run23: 119602_N(UniMod:7)SLFEYQK/2 (magenta) AND 123337_DQTVSDNE
Intensity
4e+05
2e+05
0e+00
1200
1400
1600
1800
time

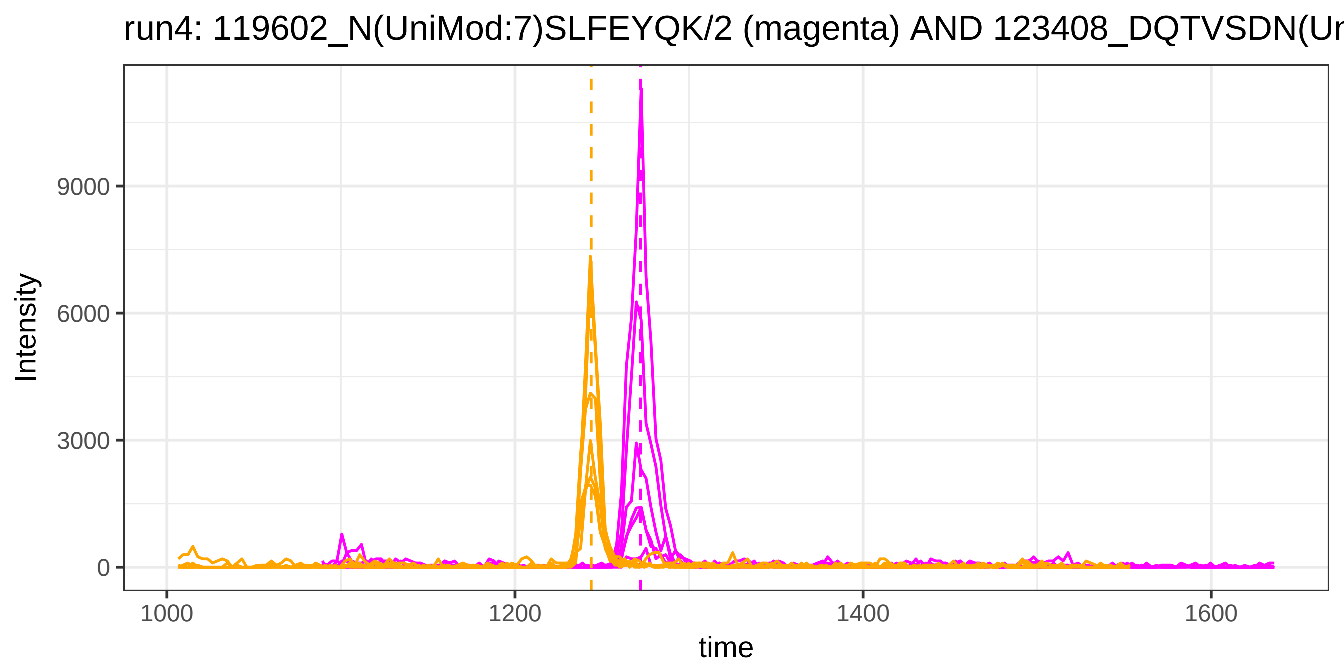
run4: 119602_N(UniMod:7)SLFEYQK/2 (magenta) AND 123408_DQTVSDN(Ur
Intensity
9000
6000
3000
0
1000
1200
1400
1600
time

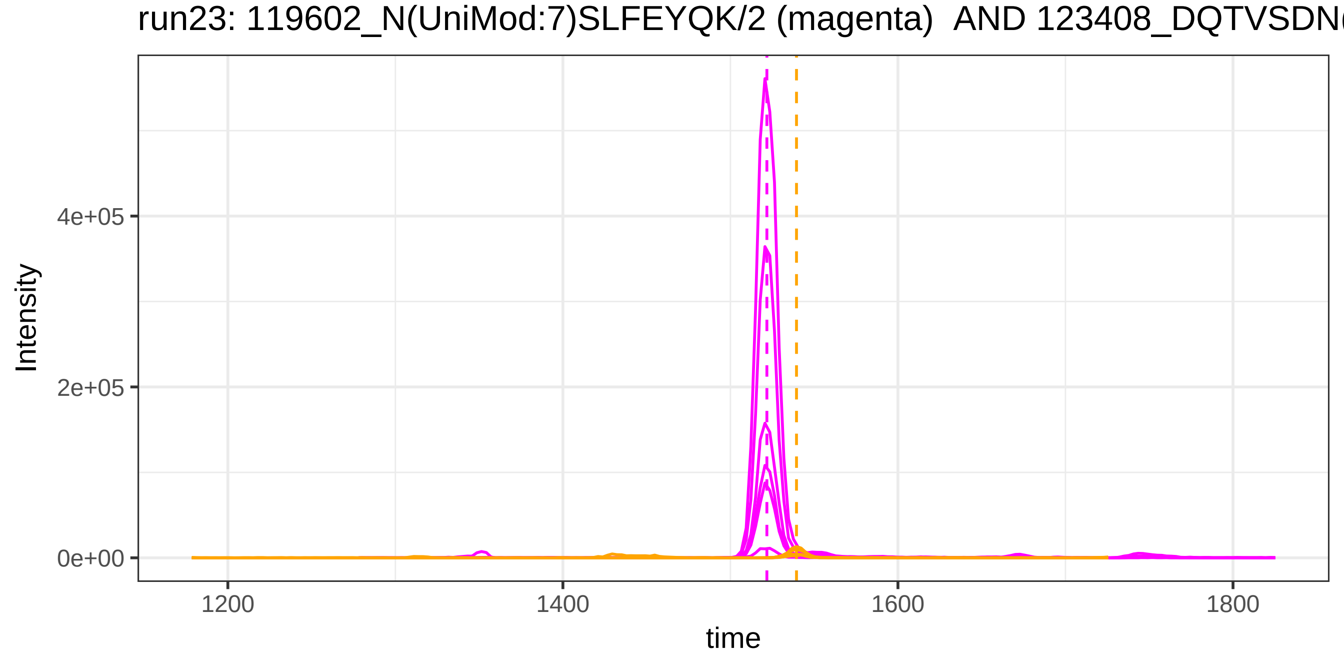
run23: 119602_N(UniMod:7)SLFEYQK/2 (magenta) AND 123408_DQTVSDN
Intensity
4e+05
2e+05
0e+00
1200
1400
1600
1800
time

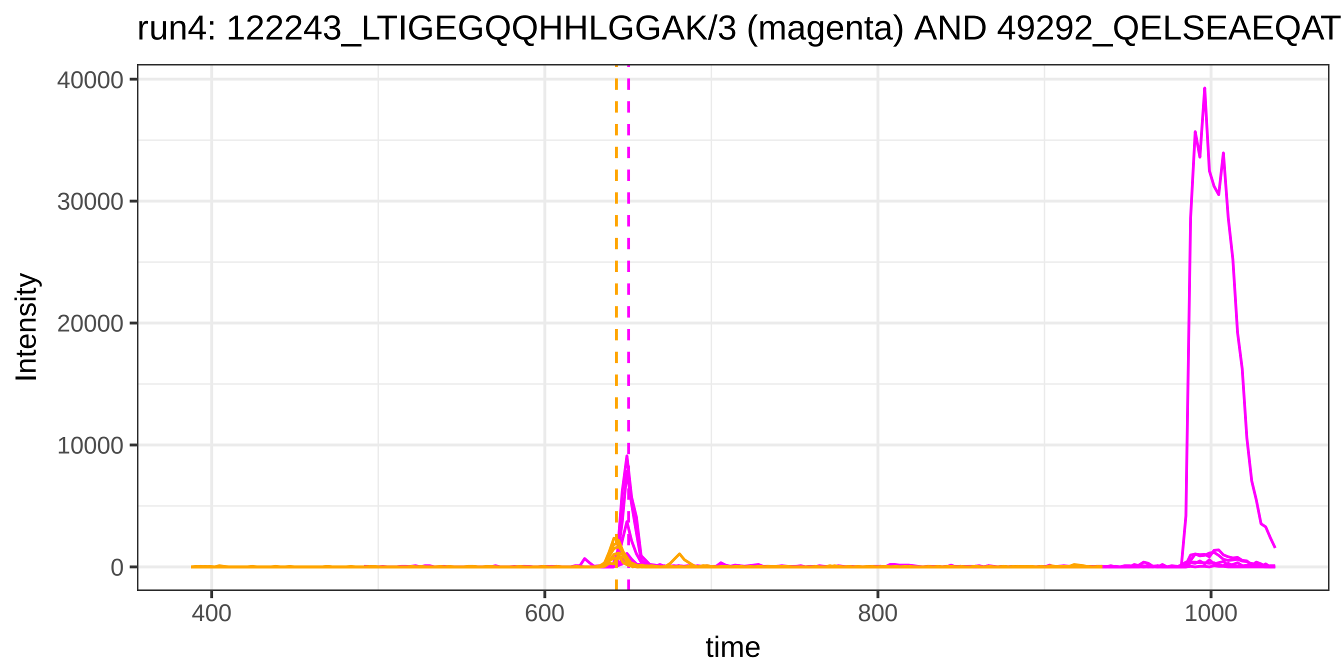
run4: 122243_LTIGEGQQHHLGGAK/3 (magenta) AND 49292_QELSEAEQAT
Intensity
40000
30000
20000
10000
0
400
600
800
1000
time

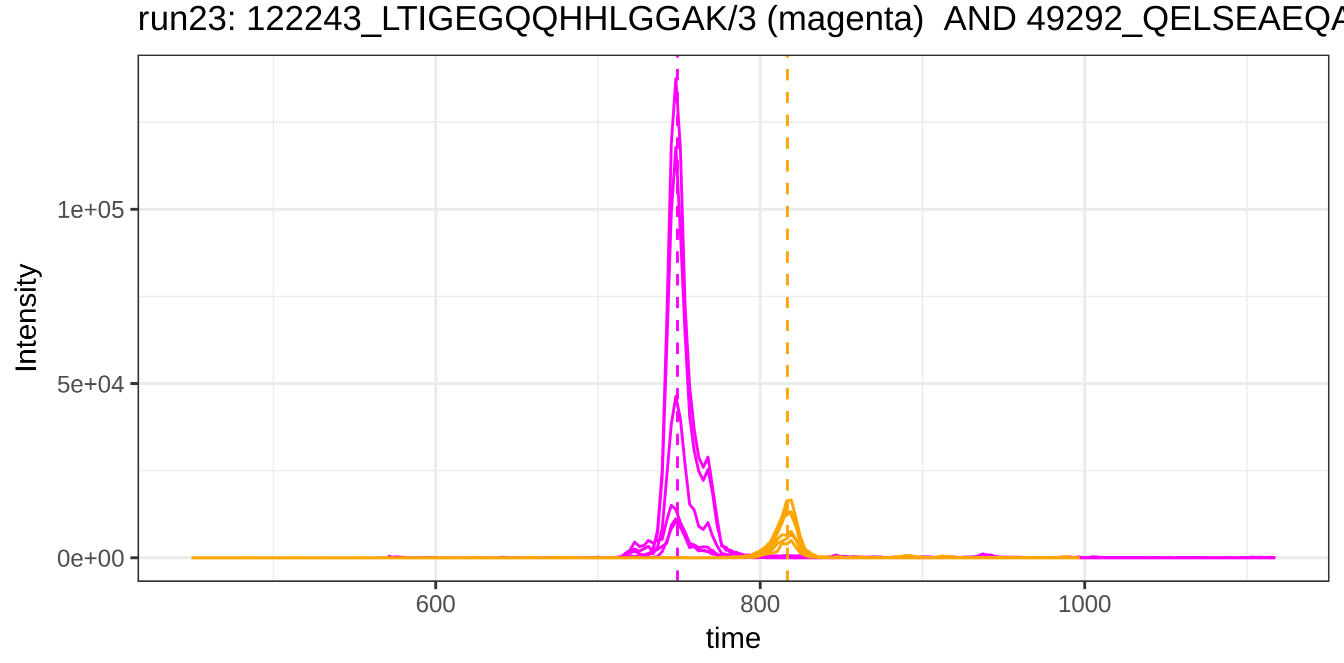
run23: 122243_LTIGEGQQHHLGGAK/3 (magenta) AND 49292_QELSEAEQA
Intensity
1e+05
5e+04
0e+00
600
800
1000
time

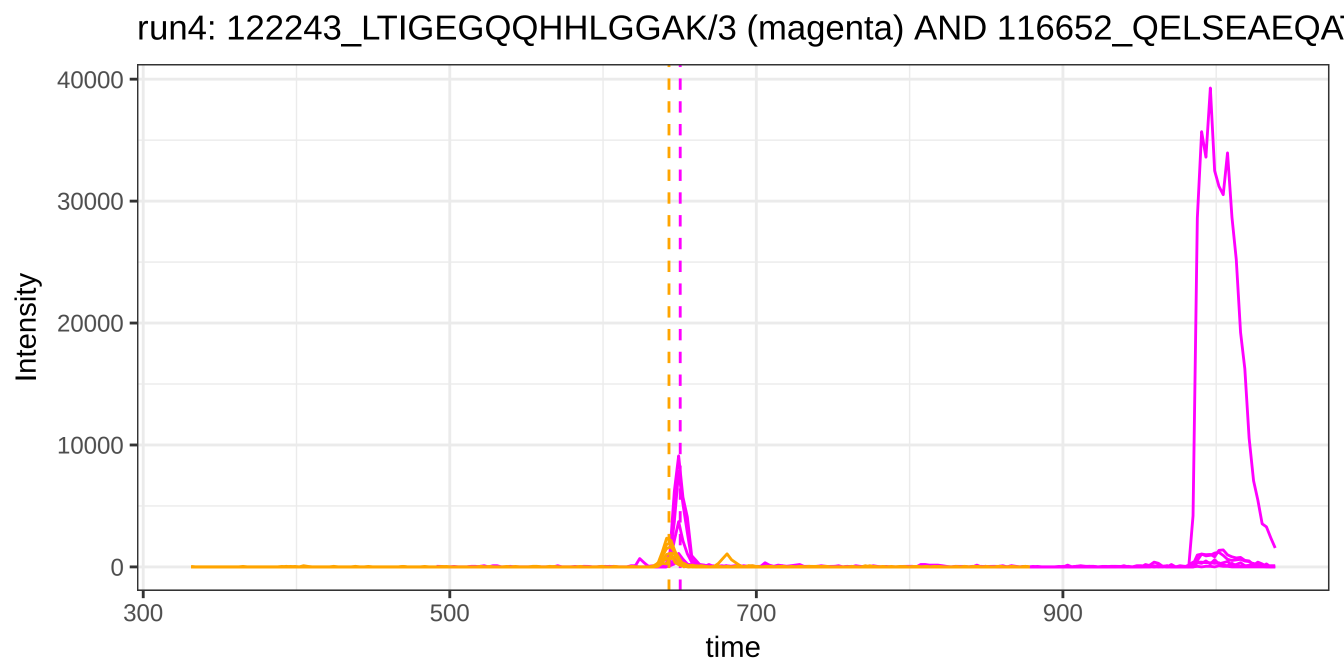
run4: 122243_LTIGEGQQHHLGGAK/3 (magenta) AND 116652_QELSEAEQA
Intensity
40000
30000
20000
10000
0
300
500
700
900
time

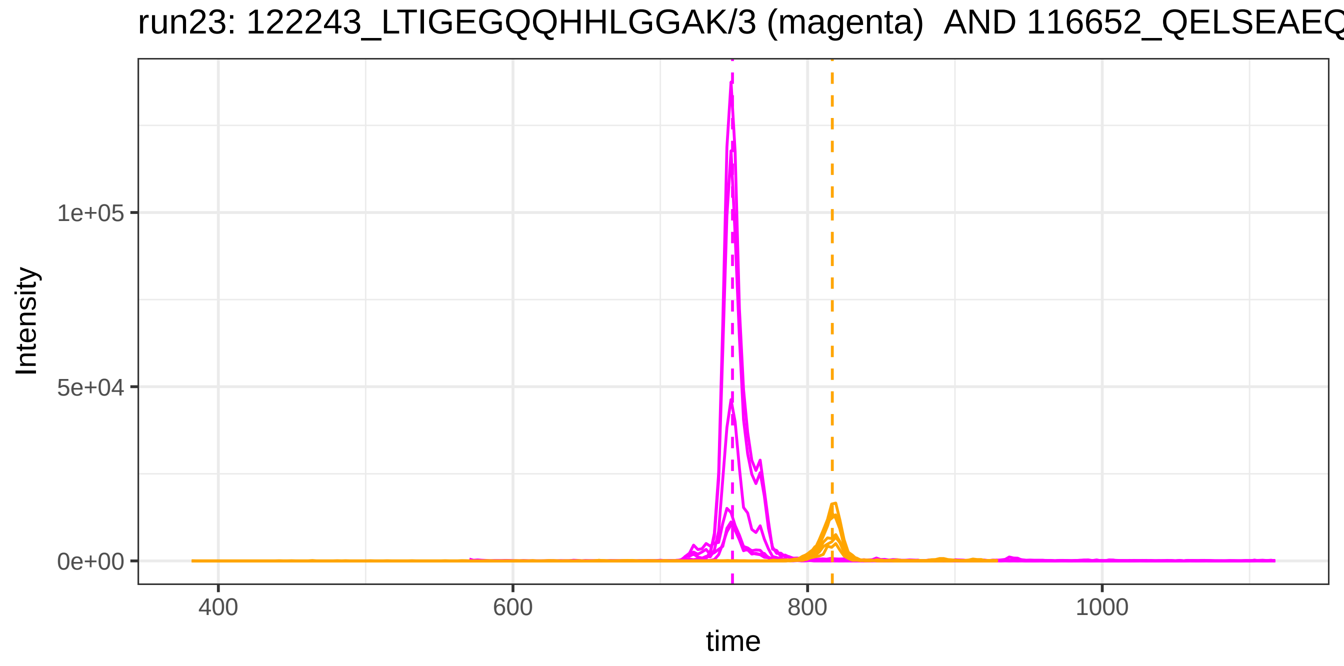
run23: 122243_LTIGEGQQHHLGGAK/3 (magenta) AND 116652_QELSEAEQ
Intensity
1e+05
5e+04
0e+00
400
600
800
1000
time

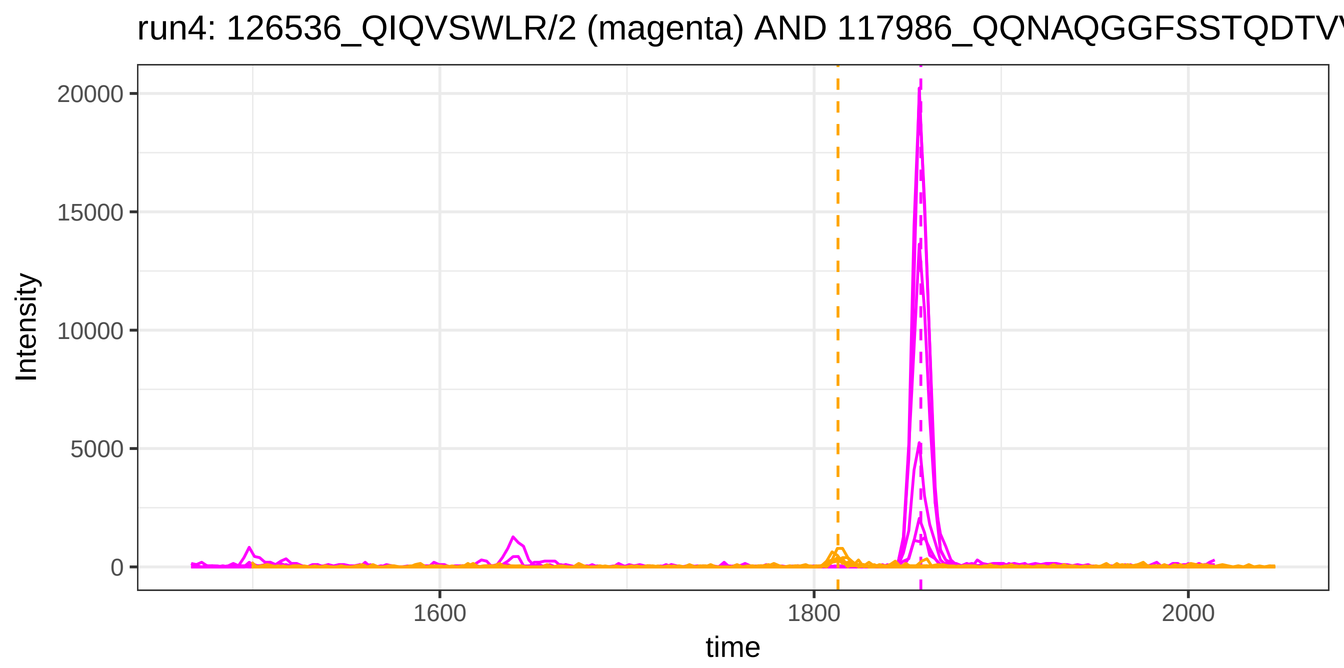
run4: 126536_QIQVSWLR/2 (magenta) AND 117986_QQNAQGGFSSTQDTV
Intensity
20000
15000
10000
5000
0
1600
1800
2000
time

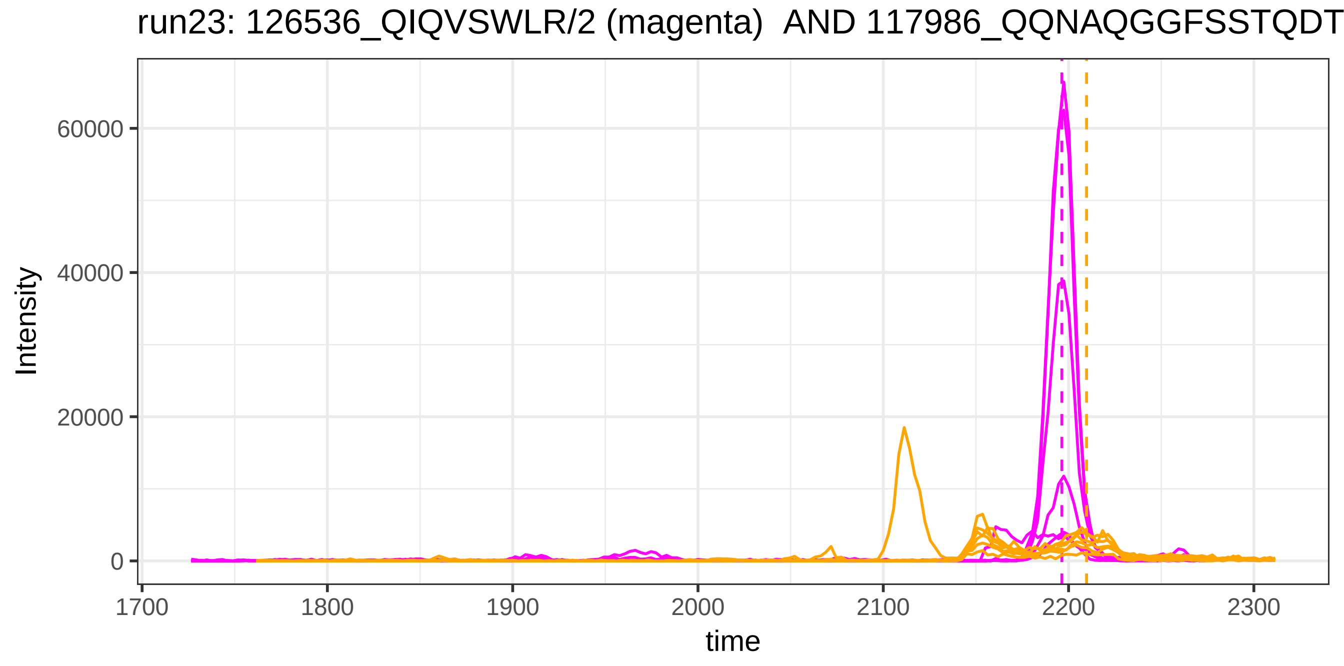
run23: 126536_QIQVSWLR/2 (magenta) AND 117986_QQNAQGGFSSTQDT
Intensity
60000
40000
20000
0
1700
1800
1900
2000
2100
2200
2300
time

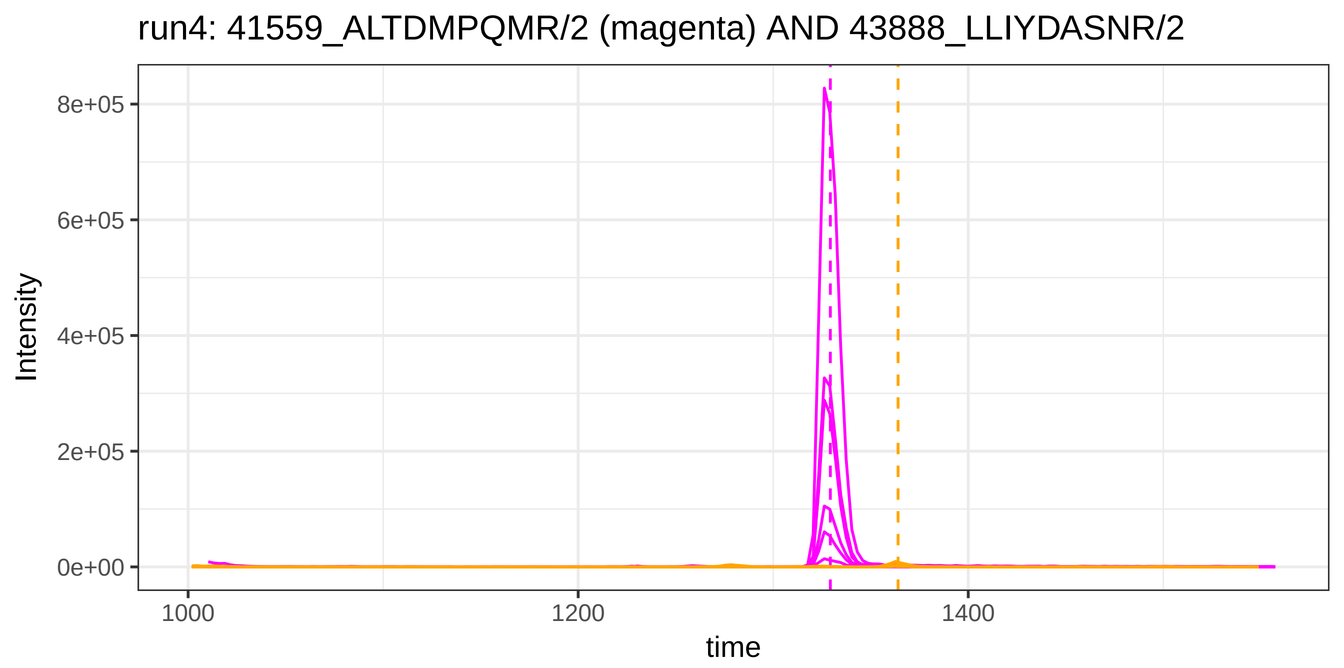

run4: 41559_ALTDMPQMR/2 (magenta) AND 43888_LLIYDASNR/2
Intensity
8e+05
6e+05
4e+05
2e+05
0e+00
1000
1200
1400
time


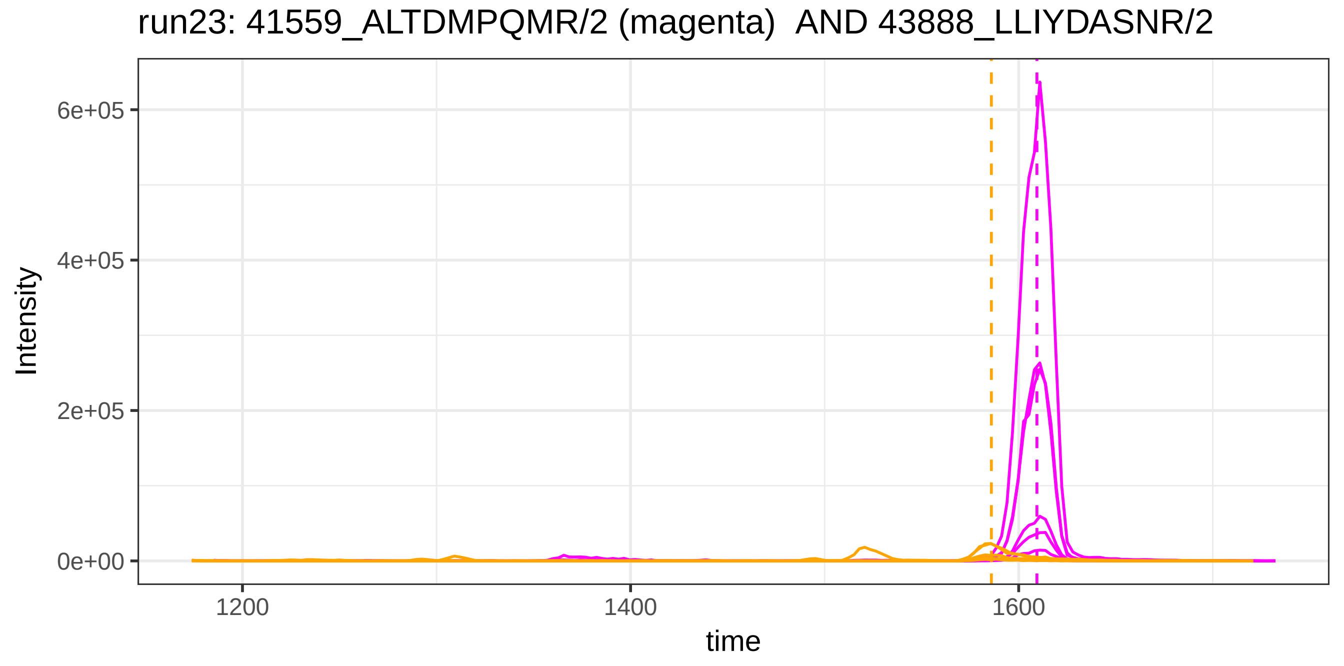

run23: 41559_ALTDMPQMR/2 (magenta)  AND 43888_LLIYDASNR/2
Intensity
6e+05
4e+05
2e+05
0e+00
1200
1400
1600
time

run4: 41559_ALTDMPQMR/2 (magenta) AND 132251_LLIYDASNR/2

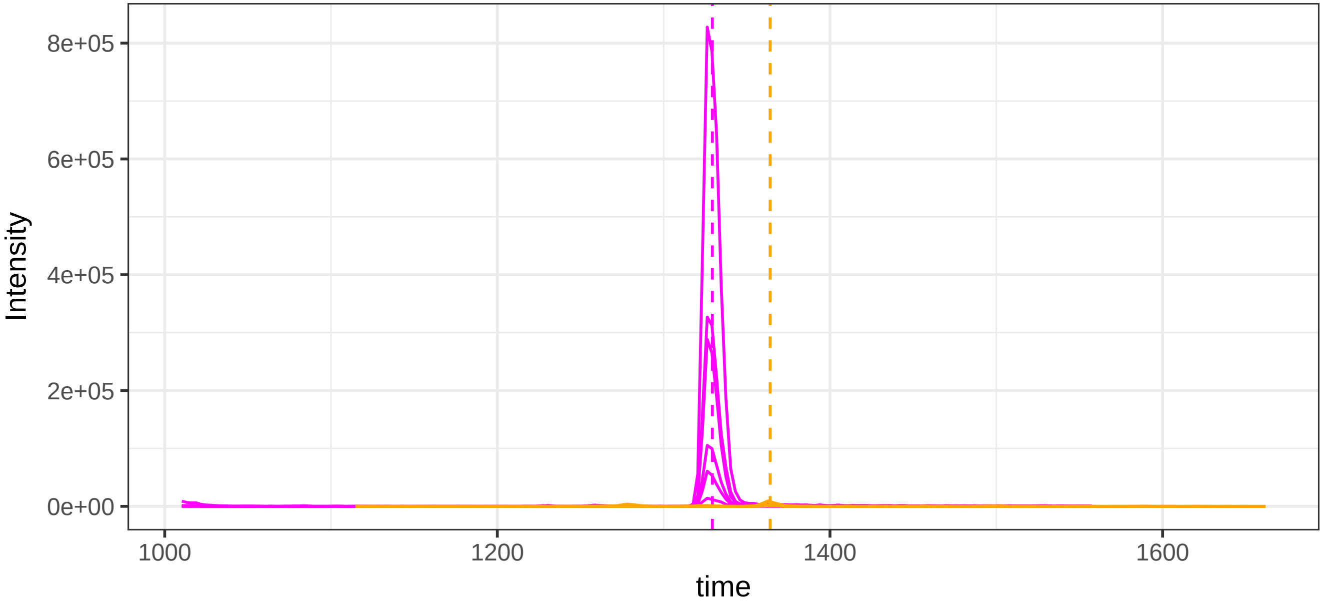


run23: 41559_ALTDMPQMR/2 (magenta)  AND 132251_LLIYDASNR/2

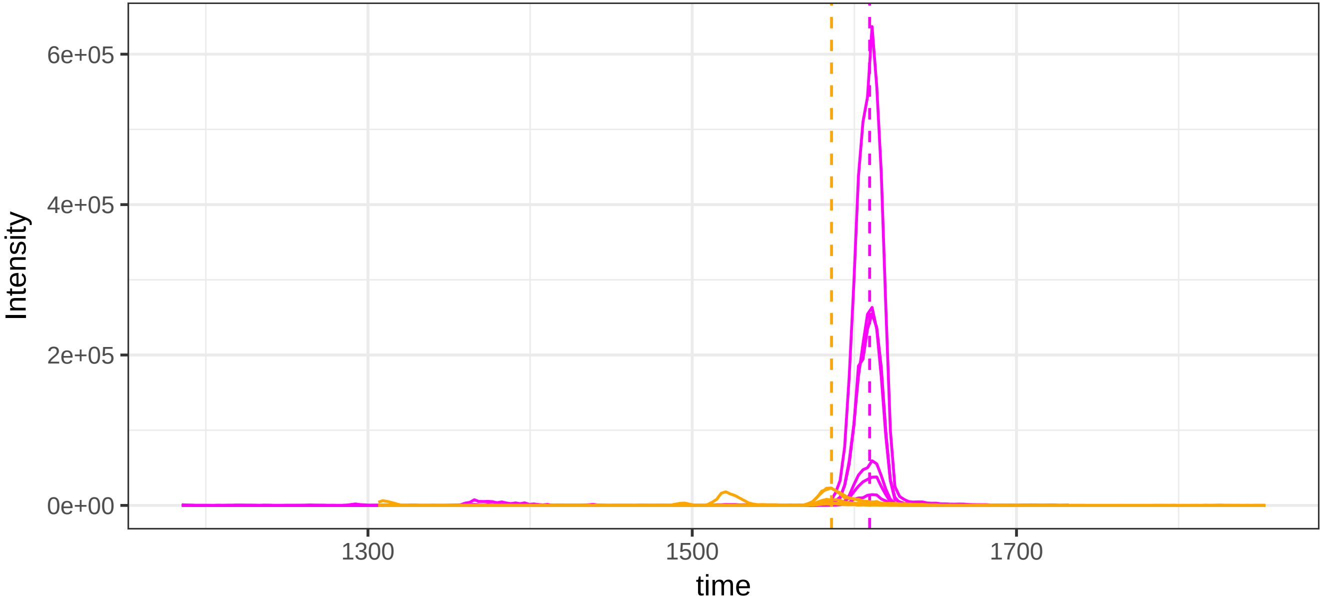

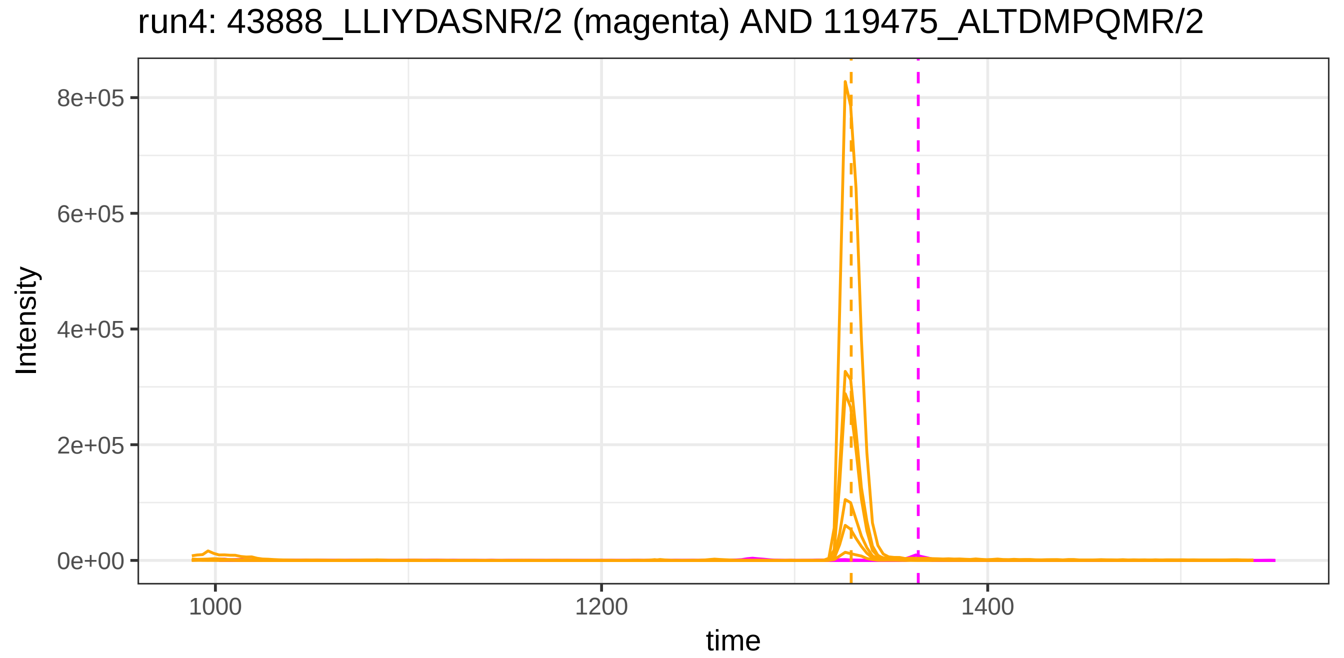

run4: 43888_LLIYDASNR/2 (magenta) AND 119475_ALTDMPQMR/2
Intensity
8e+05
6e+05
4e+05
2e+05
0e+00
1000
1200
1400
time


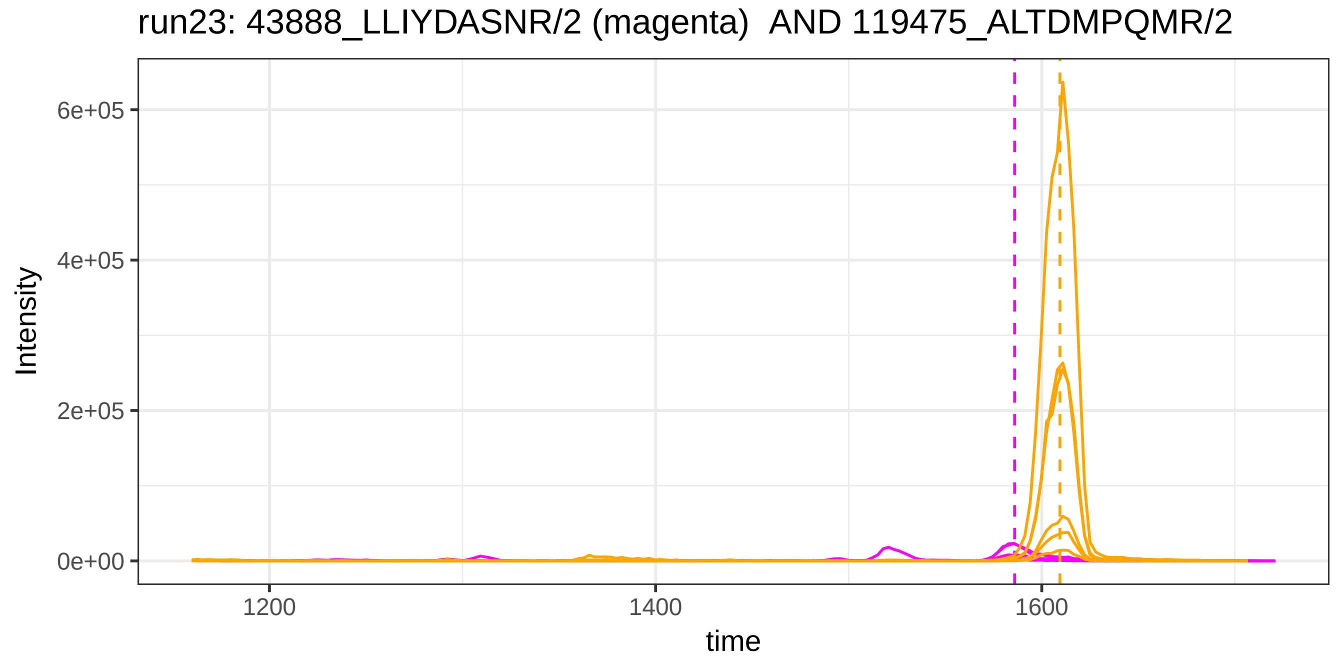

run23: 43888_LLIYDASNR/2 (magenta) AND 119475_ALTDMPQMR/2
Intensity
6e+05
4e+05
2e+05
0e+00
1200
1400
1600
time

run4: 43888_LLIYDASNR/2 (magenta) AND 1030_VELEDWNGR/2

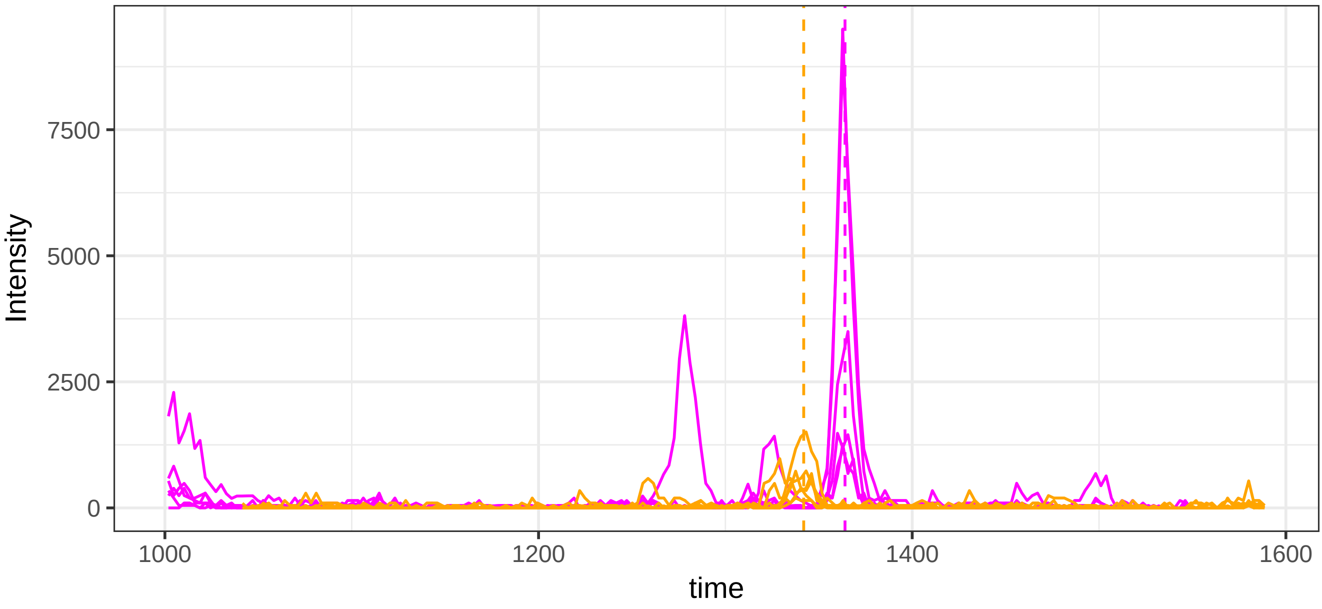


run23: 43888_LLIYDASNR/2 (magenta)  AND 1030_VELEDWNGR/2

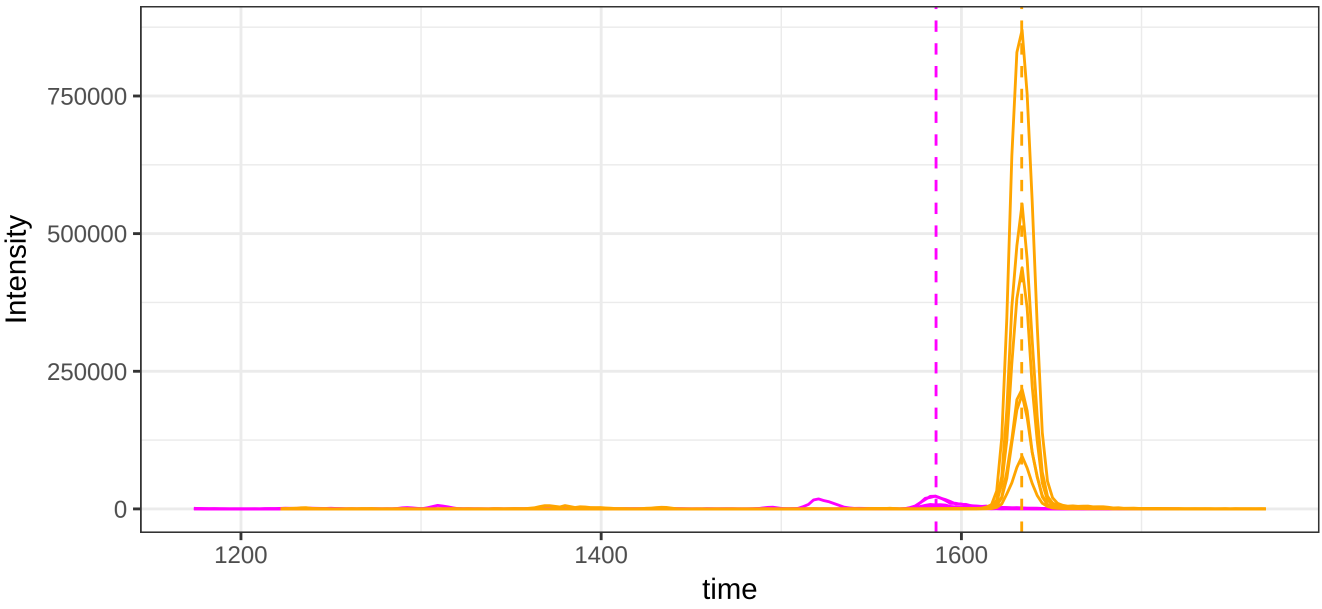

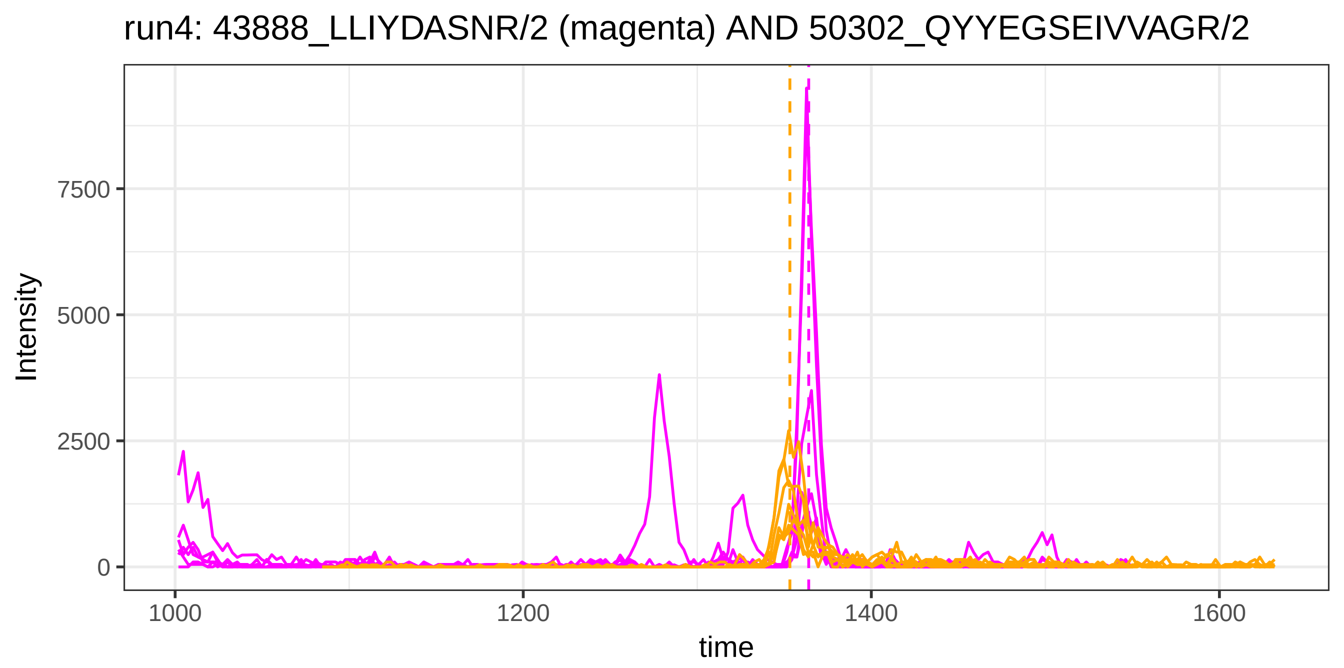

run4: 43888_LLIYDASNR/2 (magenta) AND 50302_QYYEGSEIVVAGR/2
Intensity
7500
5000
2500
0
1000
1200
1400
1600
time


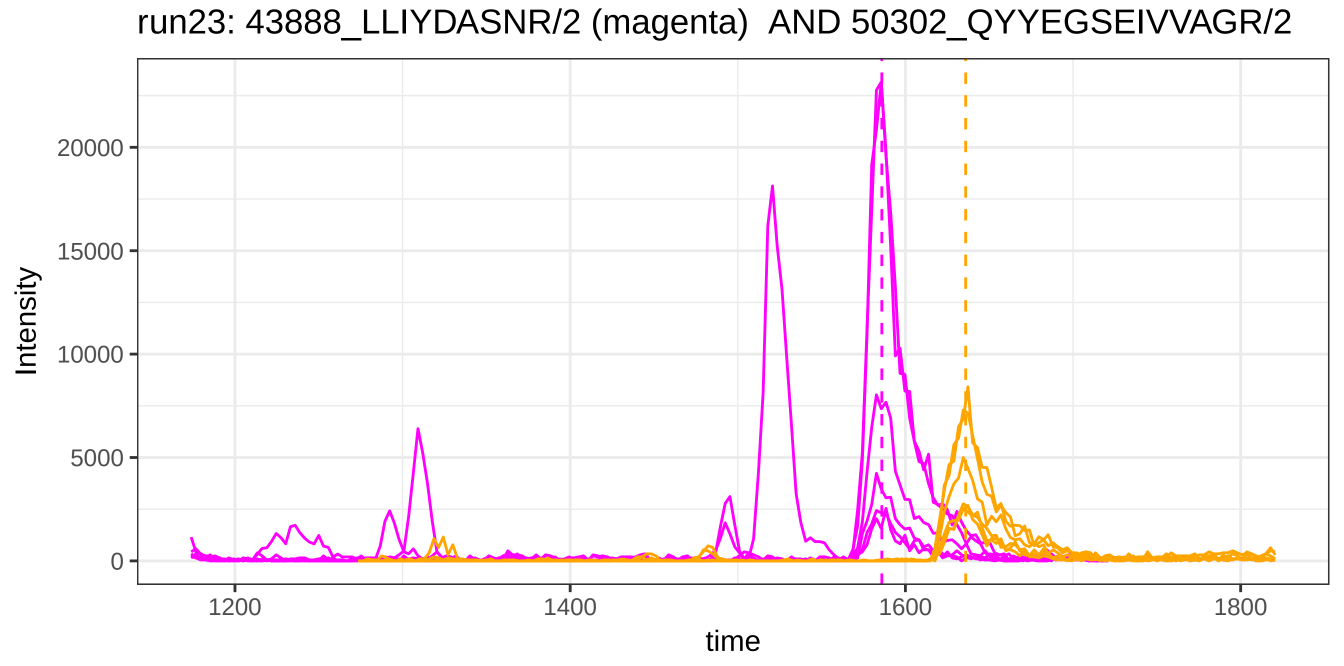

run23: 43888_LLIYDASNR/2 (magenta)  AND 50302_QYYEGSEIVVAGR/2
Intensity
20000
15000
10000
5000
0
1200
1400
1600
1800
time

run4: 115980_LVRPEVDVM/2 (magenta) AND 54026_SSALDMENFR/2

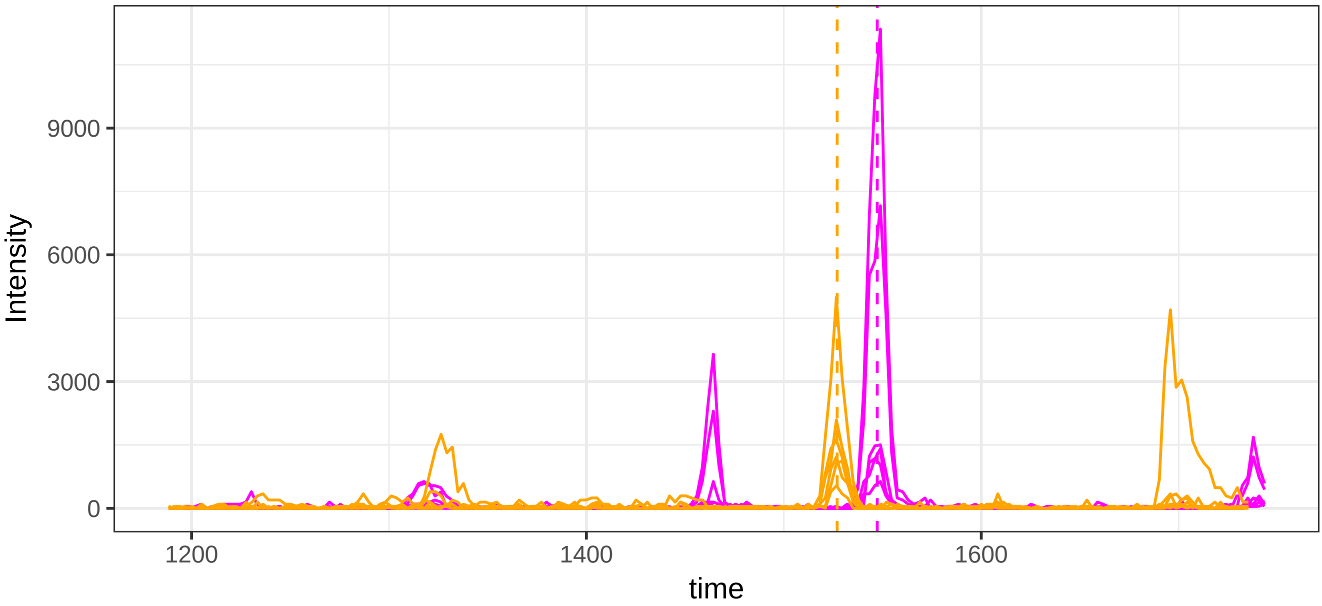


run23: 115980_LVRPEVDVM/2 (magenta)  AND 54026_SSALDMENFR/2

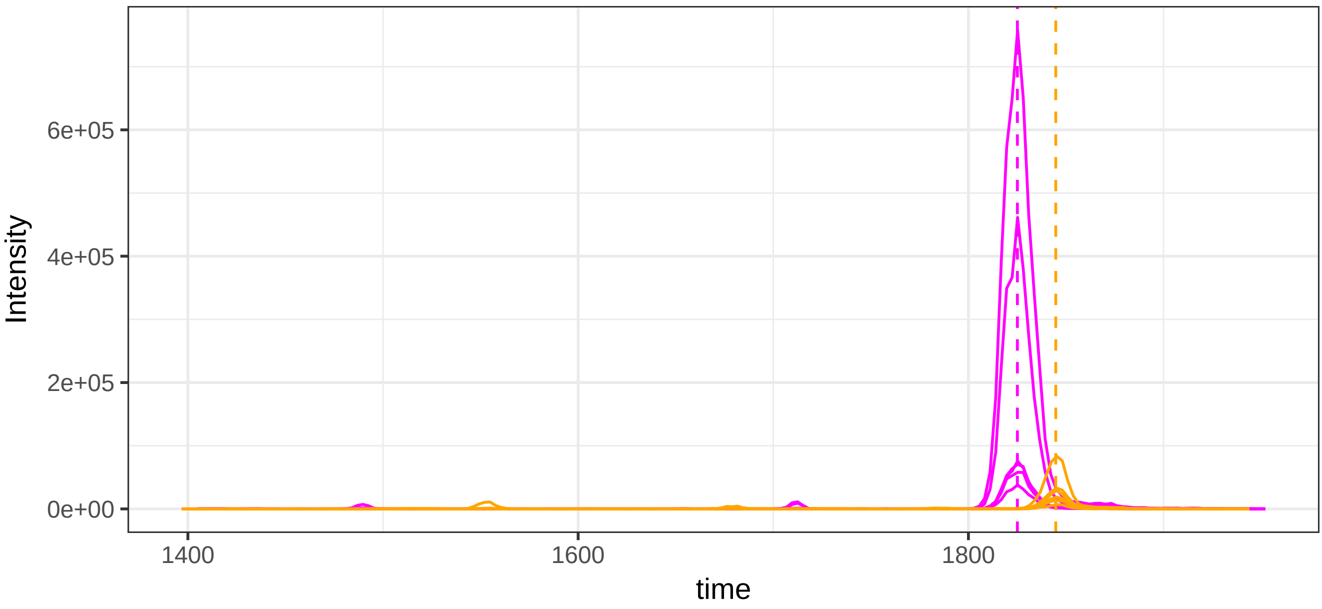

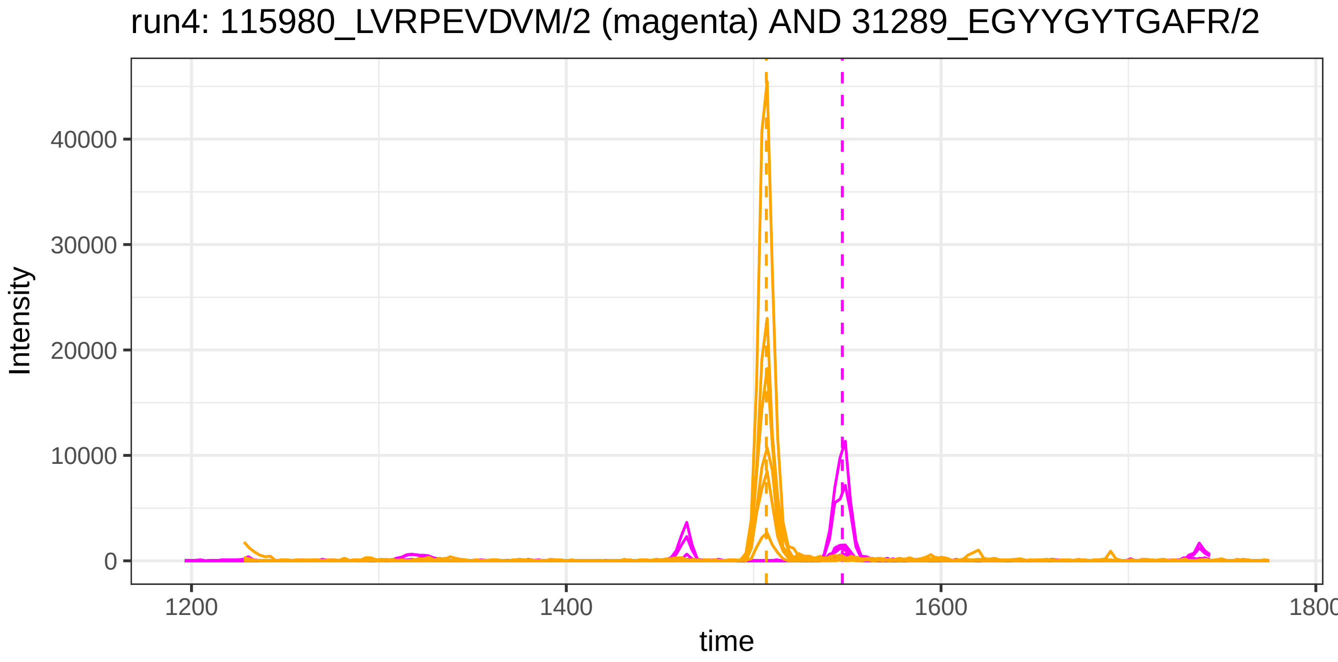
run4: 115980_LVRPEVDVM/2 (magenta) AND 31289_EGYYGYTGAFR/2
Intensity
40000
30000
20000
10000
0
1200
1400
1600
1800
time

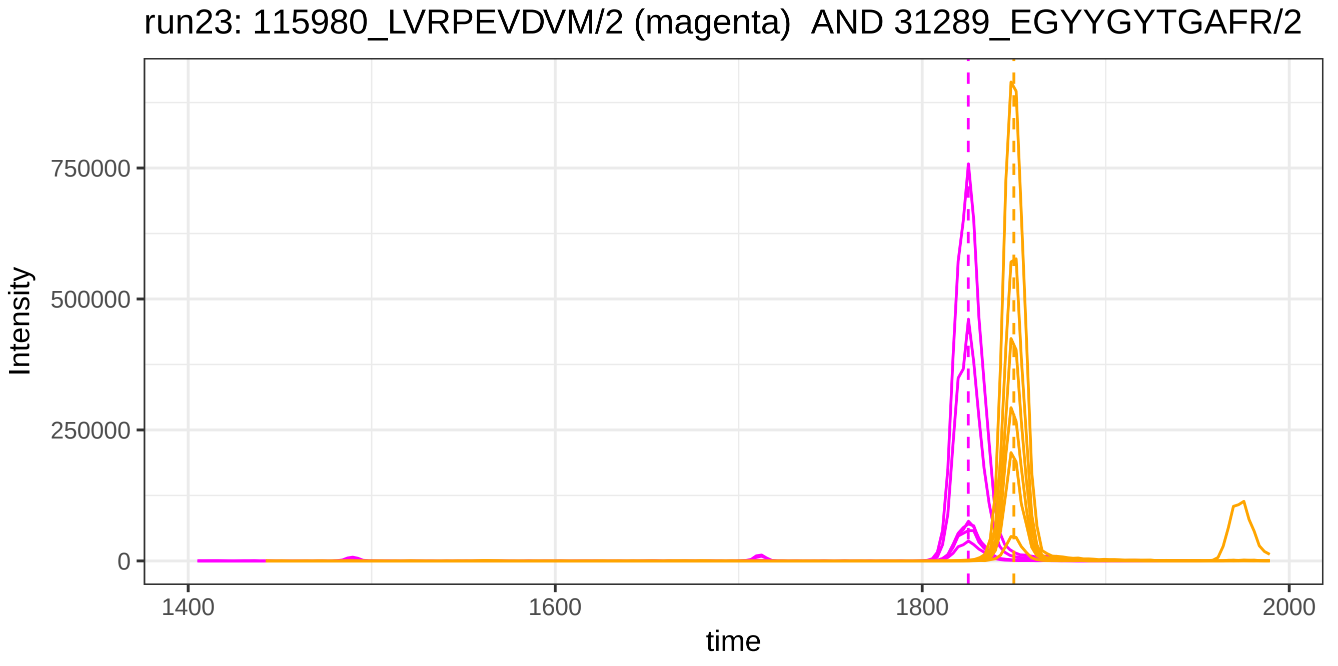
run23: 115980_LVRPEVDVM/2 (magenta) AND 31289_EGYYGYTGAFR/2
Intensity
750000
500000
250000
0
1400
1600
1800
2000
time

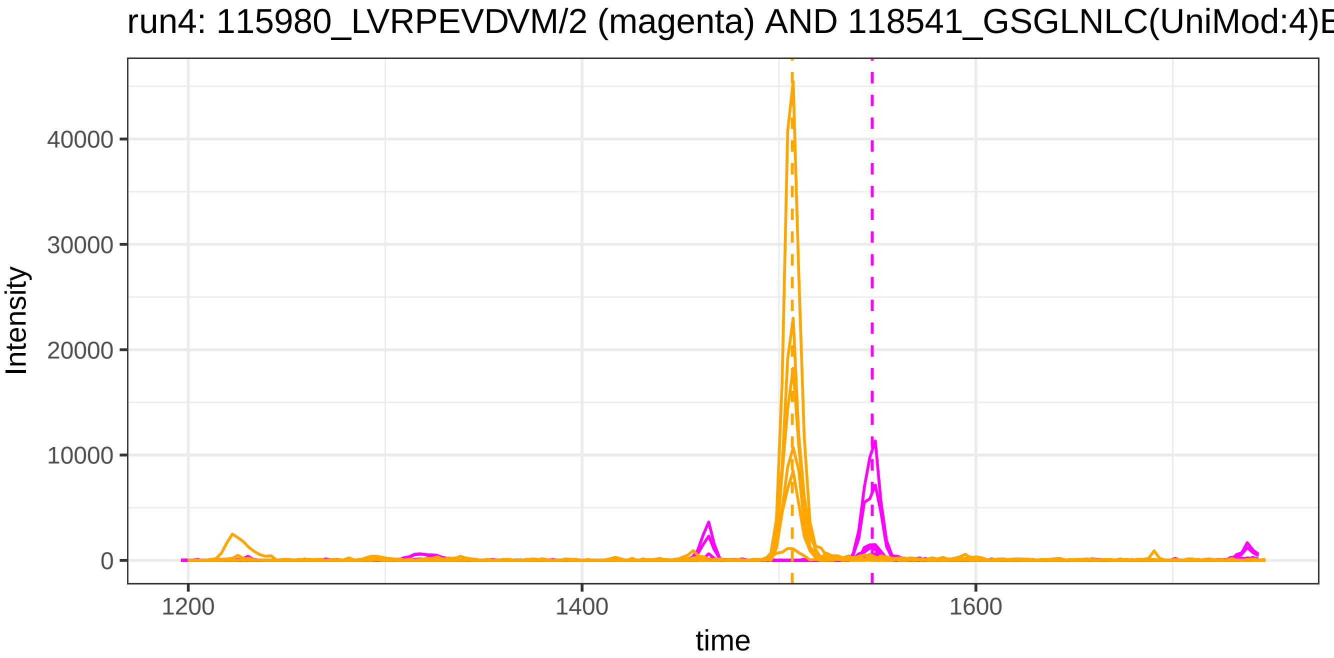
run4: 115980_LVRPEVDVM/2 (magenta) AND 118541_GSGLNLC(UniMod:4)E
Intensity
40000
30000
20000
10000
0
1200
1400
1600
time

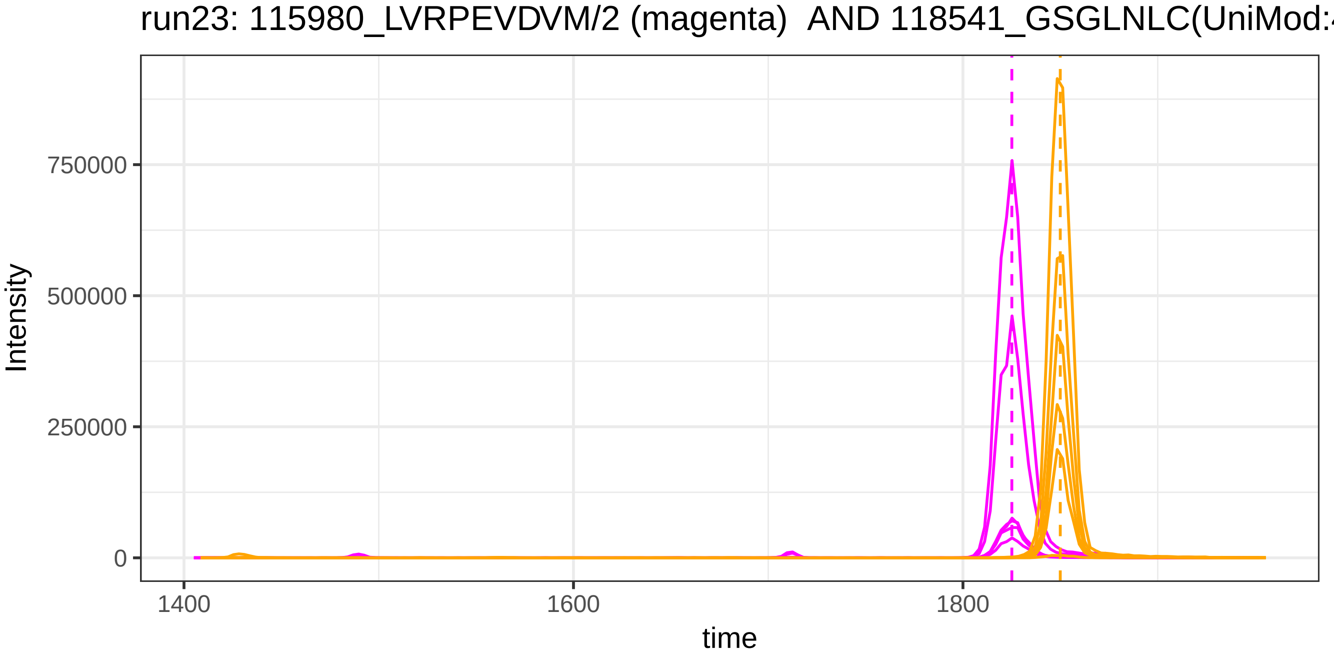
run23: 115980_LVRPEVDVM/2 (magenta) AND 118541_GSGLNLC(UniMod:4
Intensity
750000
500000
250000
0
1400
1600
1800
time

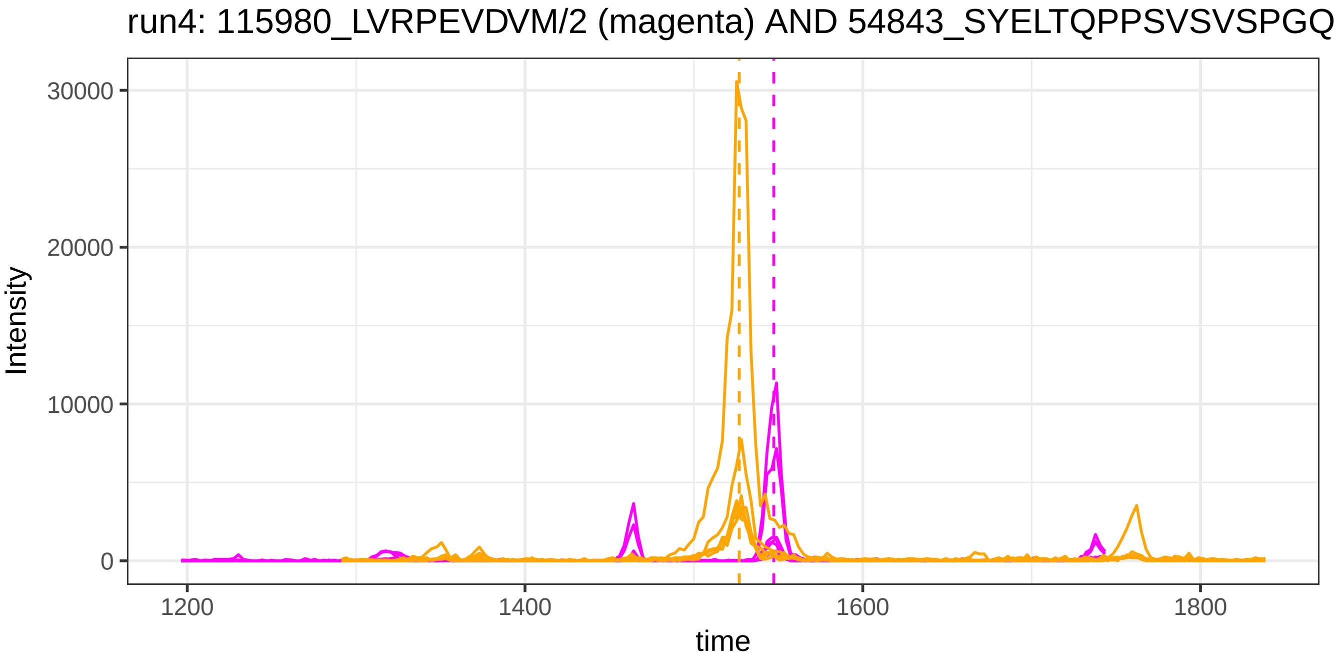
run4: 115980_LVRPEVDVM/2 (magenta) AND 54843_SYELTQPPSVSVSPGQ
Intensity
30000
20000
10000
0
1200
1400
1600
1800
time

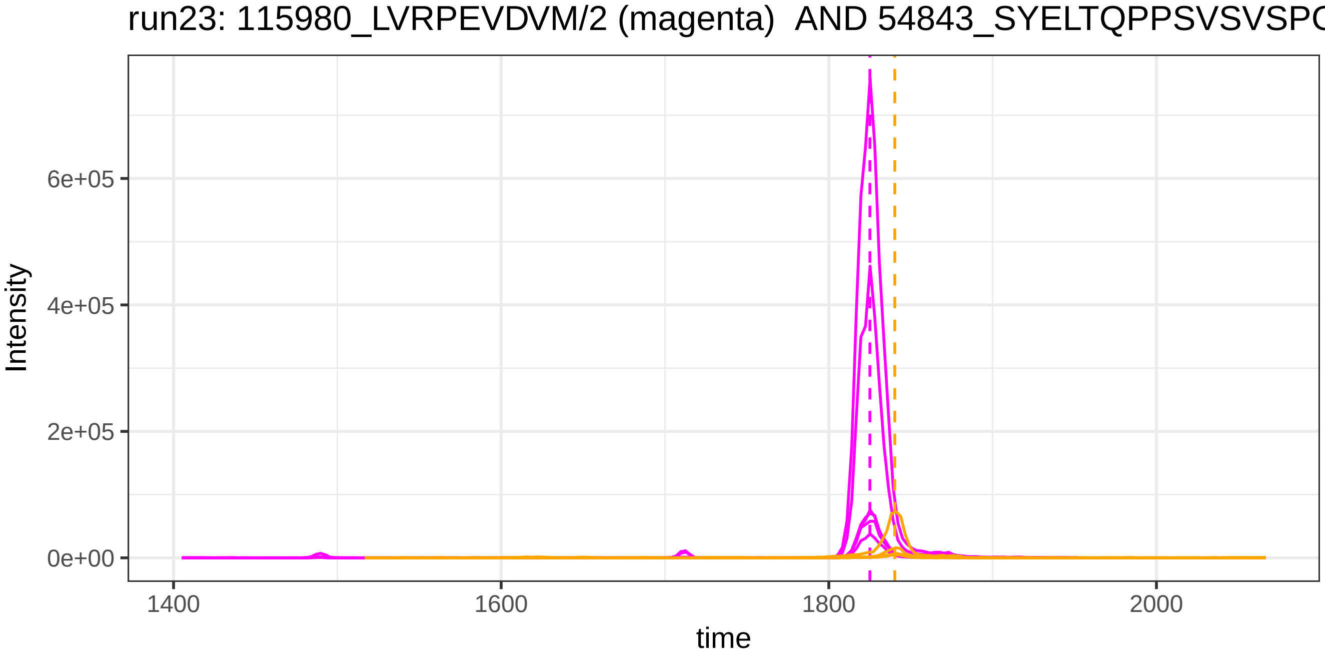
run23: 115980_LVRPEVDVM/2 (magenta) AND 54843_SYELTQPPSVSVSPG
Intensity
6e+05
4e+05
2e+05
0e+00
1400
1600
1800
2000
time

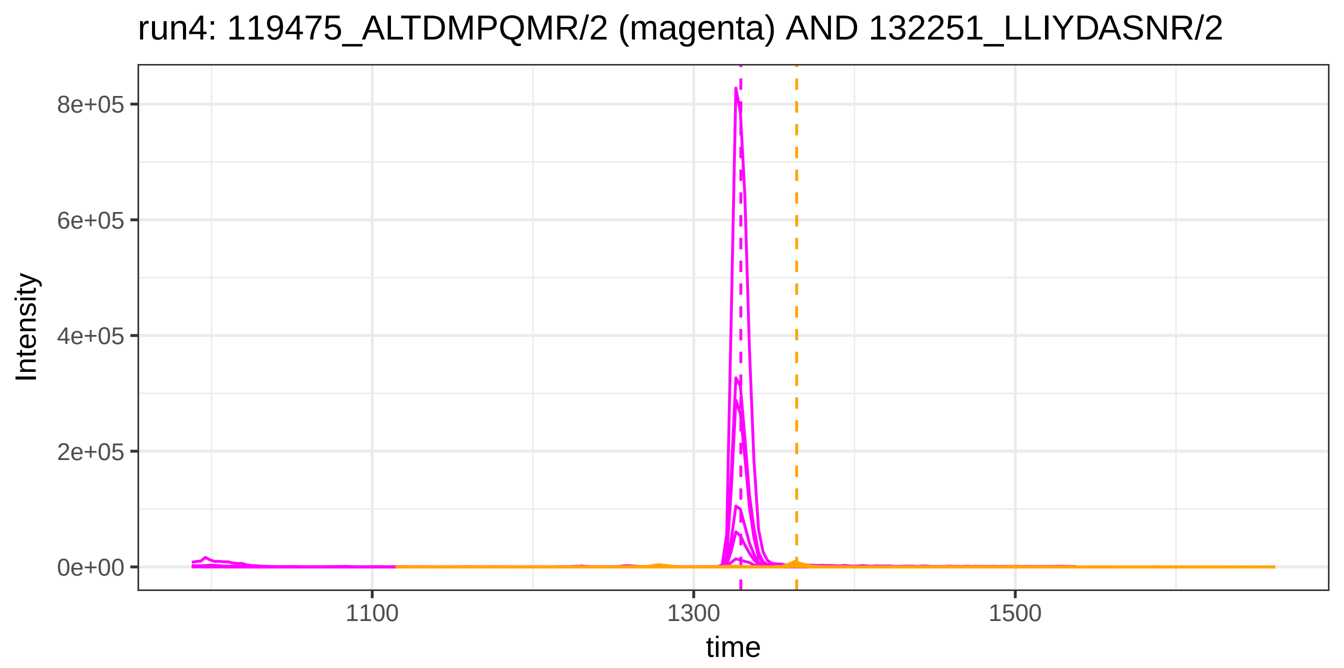
run4: 119475_ALTDMPQMR/2 (magenta) AND 132251_LLIYDASNR/2
Intensity
8e+05
6e+05
4e+05
2e+05
0e+00
1100
1300
1500
time

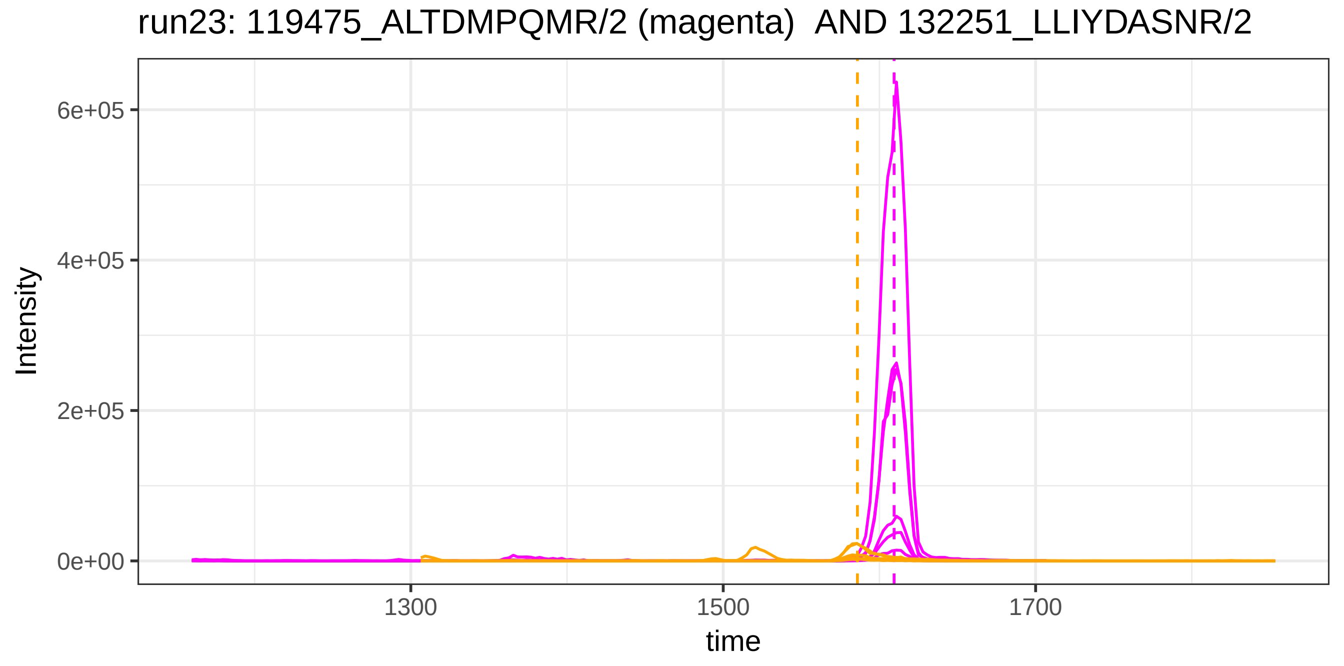
run23: 119475_ALTDMPQMR/2 (magenta)  AND 132251_LLIYDASNR/2
Intensity
6e+05
4e+05
2e+05
0e+00
1300
1500
1700
time

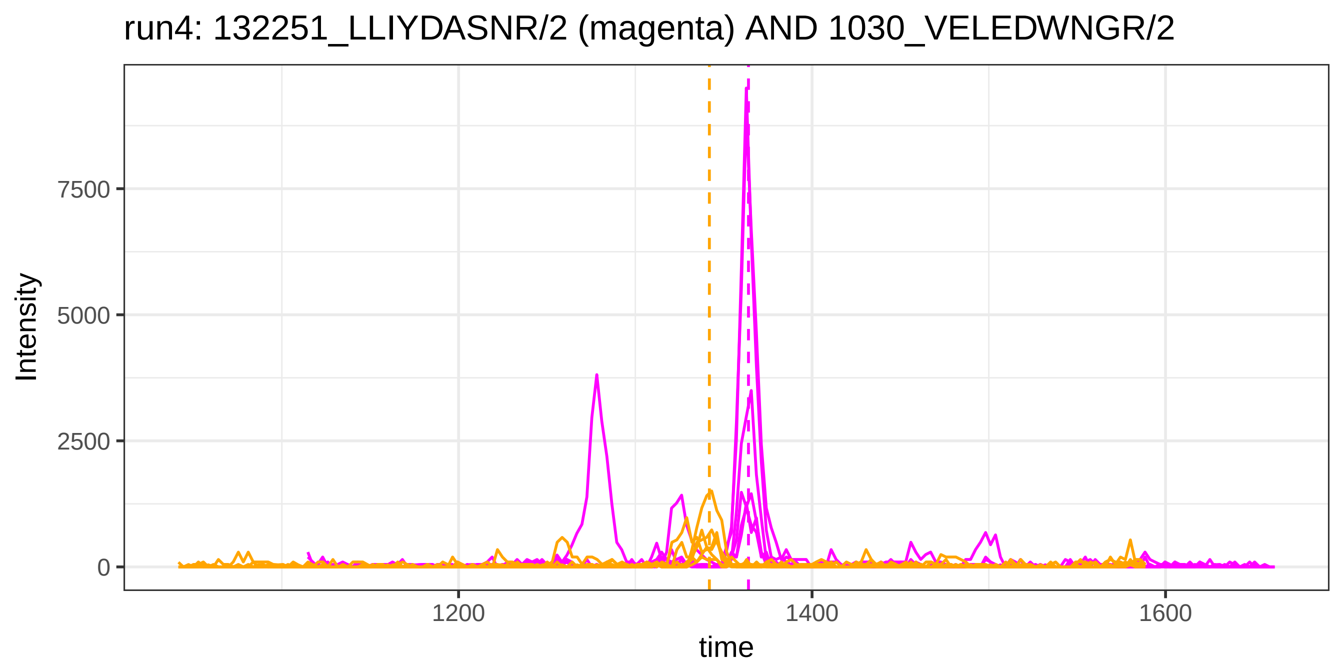
run4: 132251_LLIYDASNR/2 (magenta) AND 1030_VELEDWNGR/2
Intensity
7500
5000
2500
0
1200
1400
1600
time

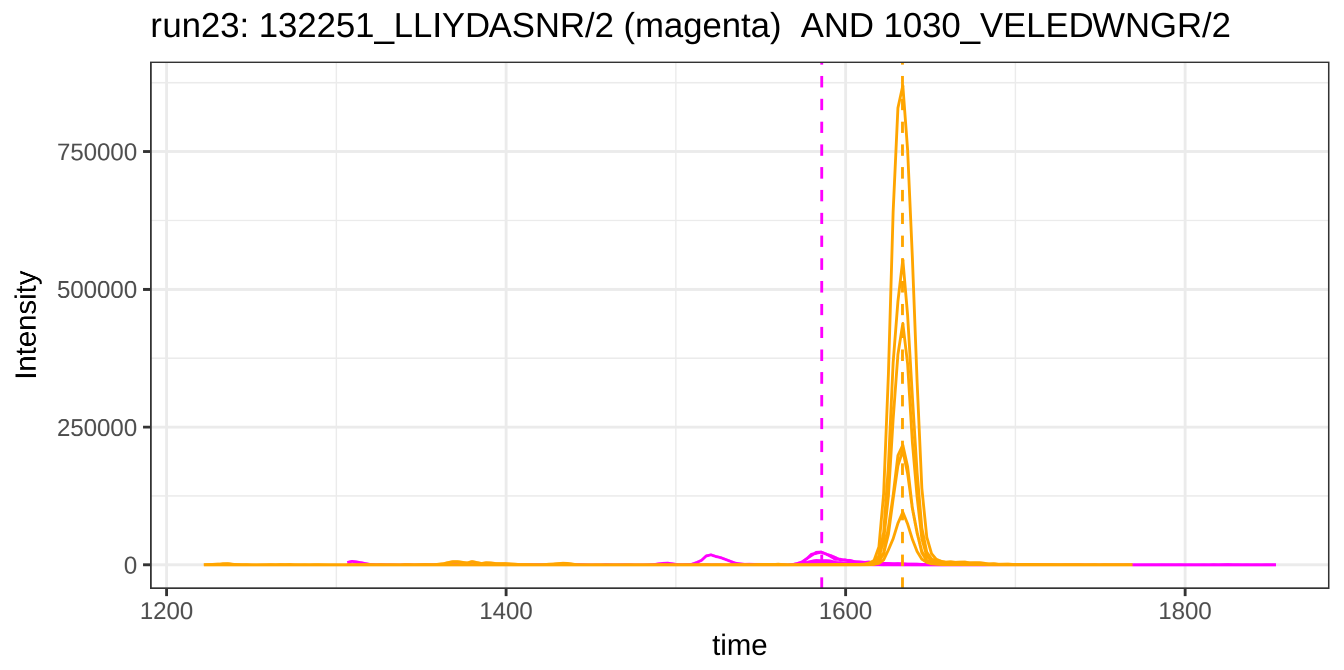
run23: 132251_LLIYDASNR/2 (magenta) AND 1030_VELEDWNGR/2
Intensity
750000
500000
250000
0
1200
1400
1600
1800
time

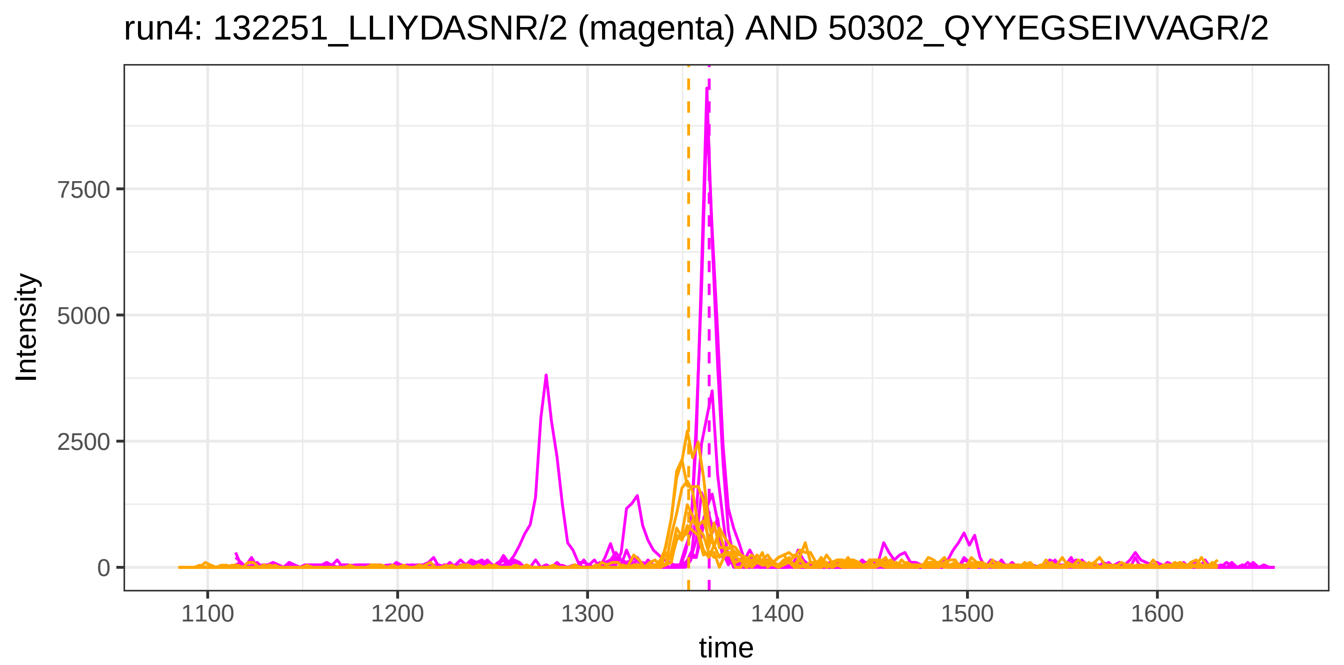
run4: 132251_LLIYDASNR/2 (magenta) AND 50302_QYYEGSEIVVAGR/2
Intensity
0
2500
5000
7500
1100
1200
1300
1400
1500
1600
time

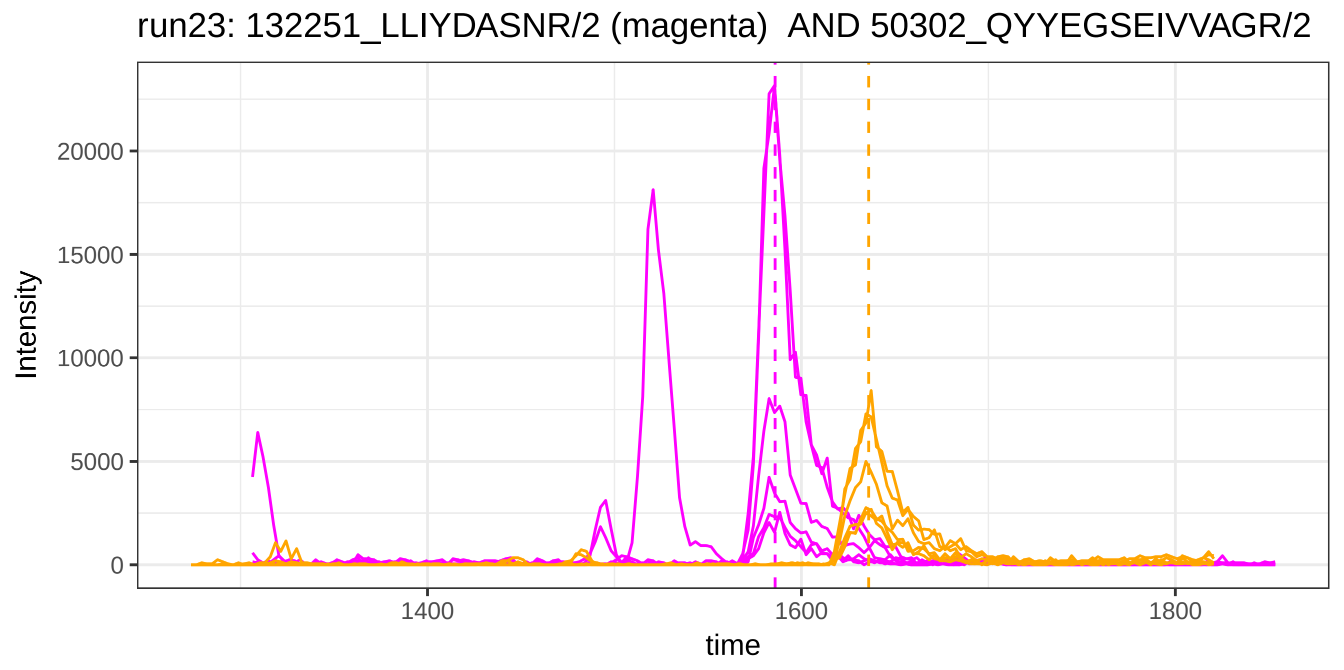
run23: 132251_LLIYDASNR/2 (magenta)  AND 50302_QYYEGSEIVVAGR/2
Intensity
0
5000
10000
15000
20000
1400
1600
1800
time

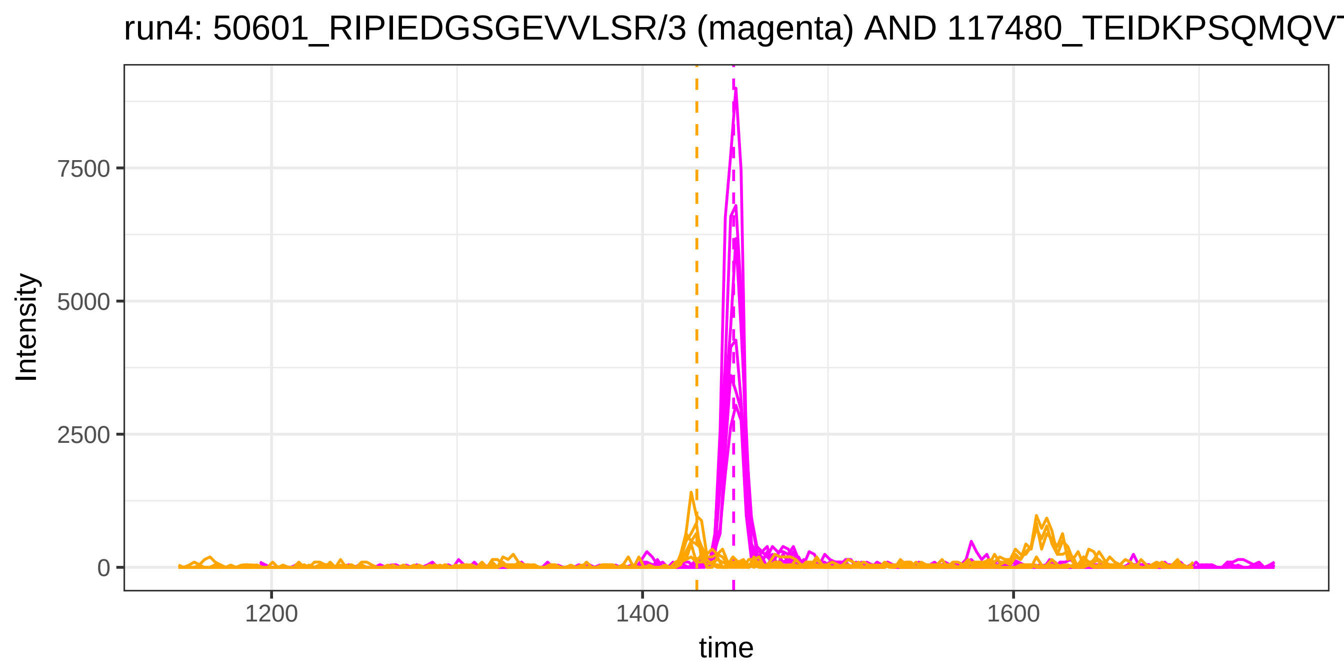
run4: 50601_RIPIEDGSGEVVLSR/3 (magenta) AND 117480_TEIDKPSQMQVT
Intensity
7500
5000
2500
0
1200
1400
1600
time

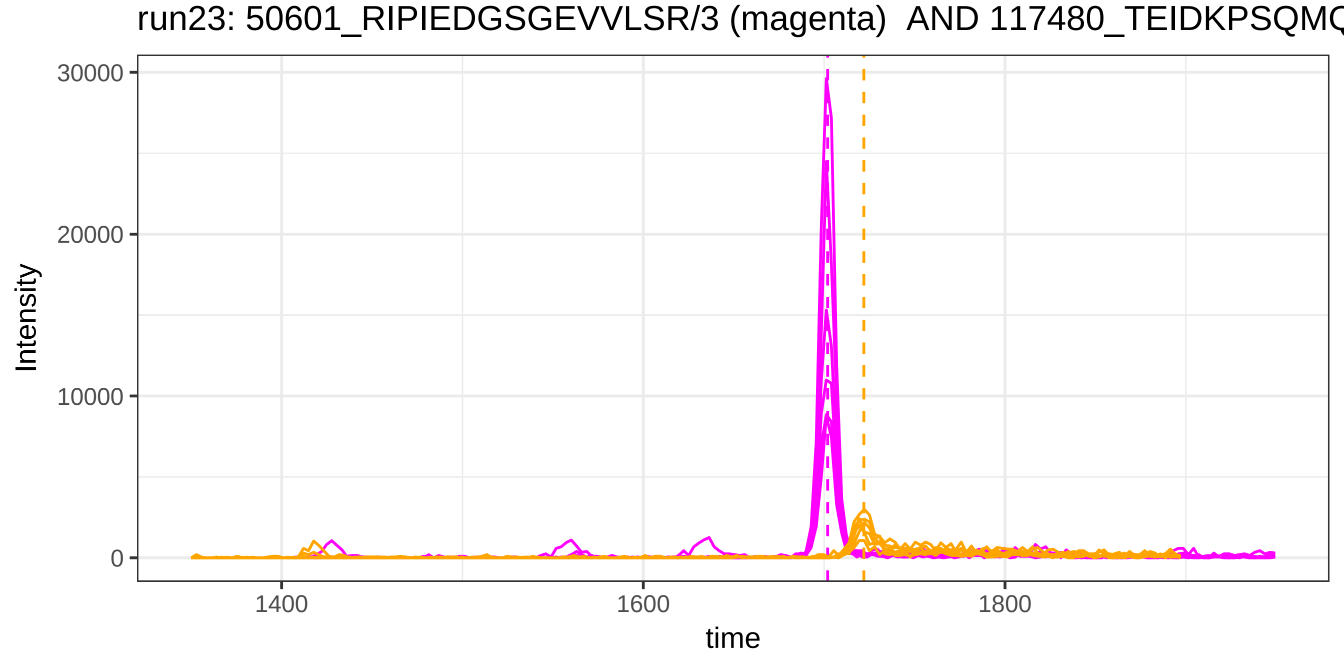
run23: 50601_RIPIEDGSGEVVLSR/3 (magenta) AND 117480_TEIDKPSQMQ
Intensity
30000
20000
10000
0
1400
1600
1800
time

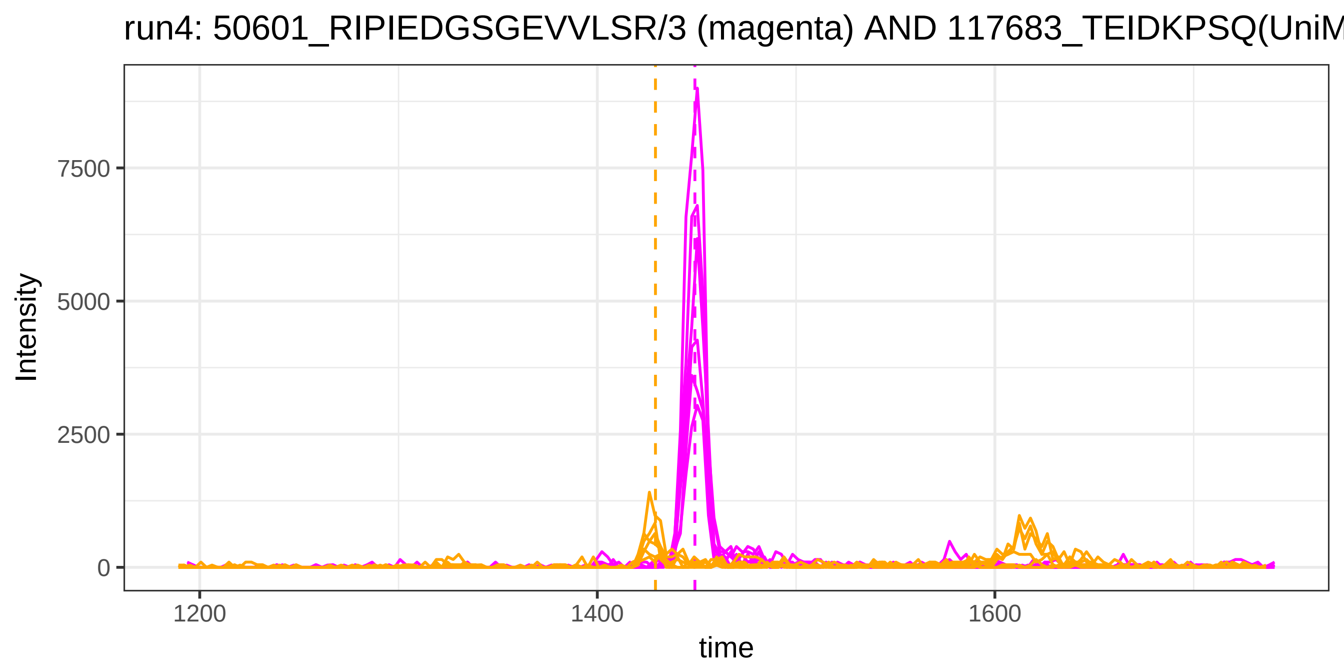
run4: 50601_RIPIEDGSGEVVLSR/3 (magenta) AND 117683_TEIDKPSQ(UniM
Intensity
7500
5000
2500
0
1200
1400
1600
time

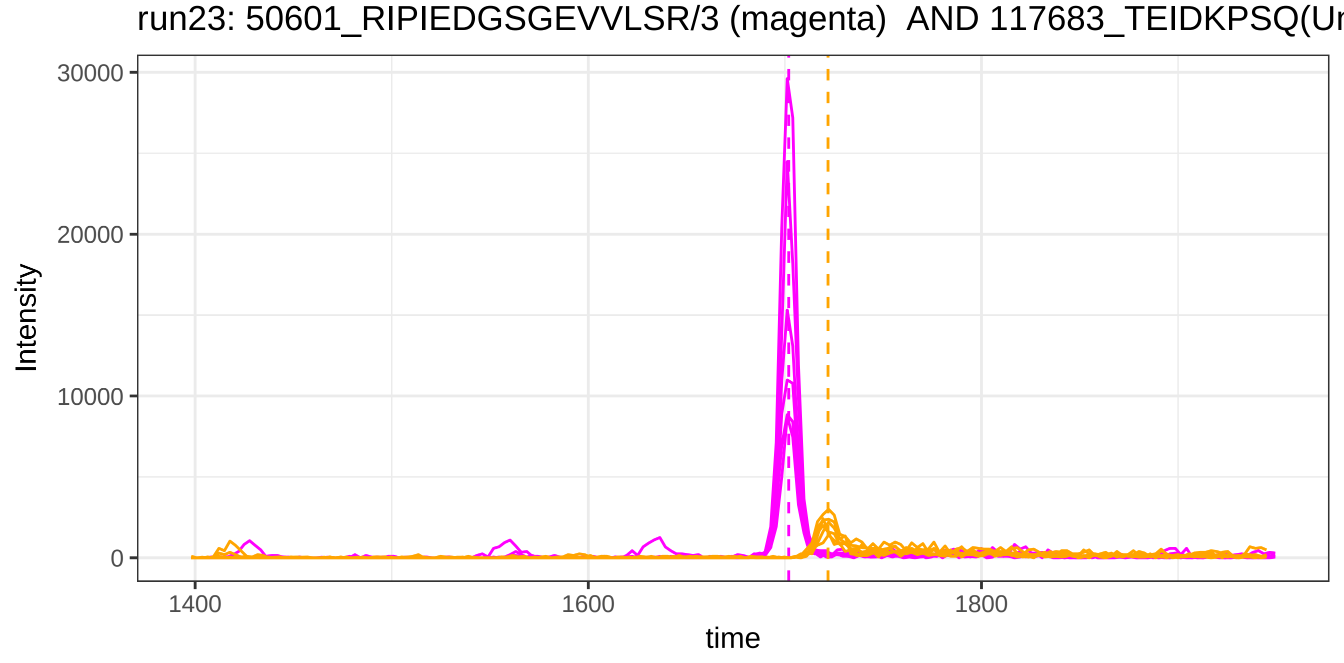
run23: 50601_RIPIEDGSGEVVLSR/3 (magenta) AND 117683_TEIDKPSQ(Ur
Intensity
30000
20000
10000
0
1400
1600
1800
time

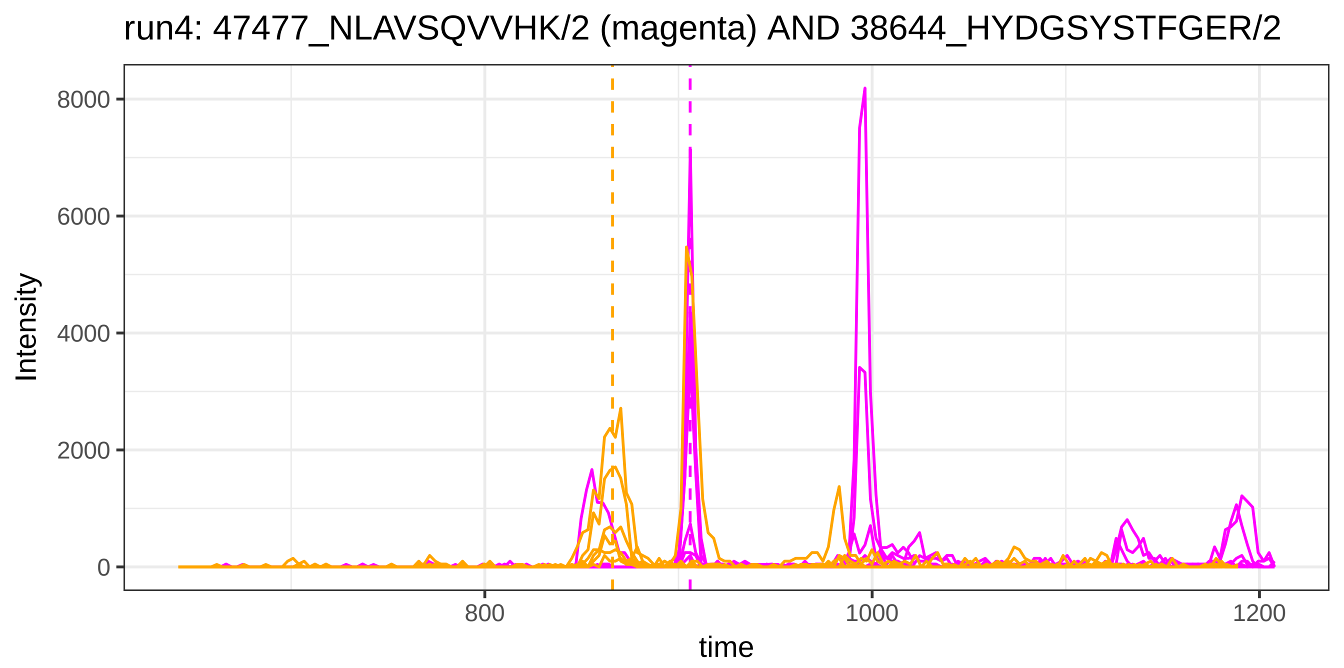

run4: 47477_NLAVSQVVHK/2 (magenta) AND 38644_HYDGSYSTFGER/2
Intensity
8000
6000
4000
2000
0
800
1000
1200
time


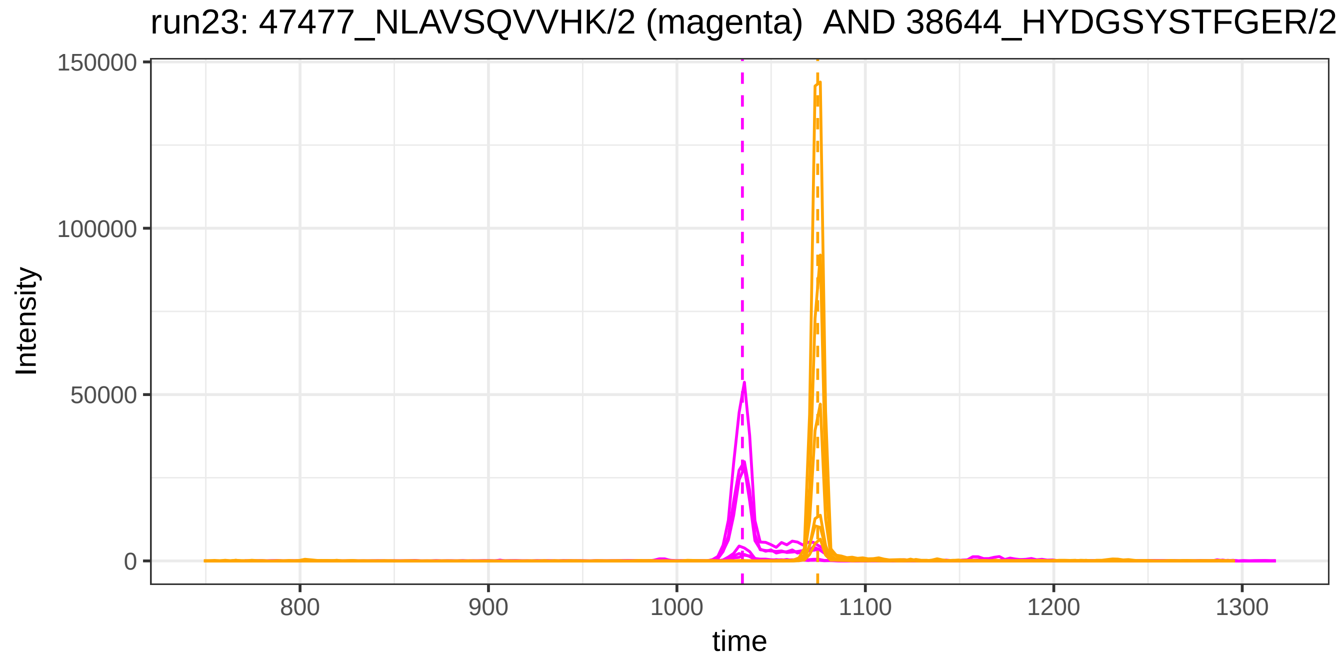

run23: 47477_NLAVSQVVHK/2 (magenta)  AND 38644_HYDGSYSTFGER/2
Intensity
150000
100000
50000
0
800
900
1000
1100
1200
1300
time

run4: 1809_VLLDGVQNPR/2 (magenta) AND 120401_NSPLDEENLTQENQDR

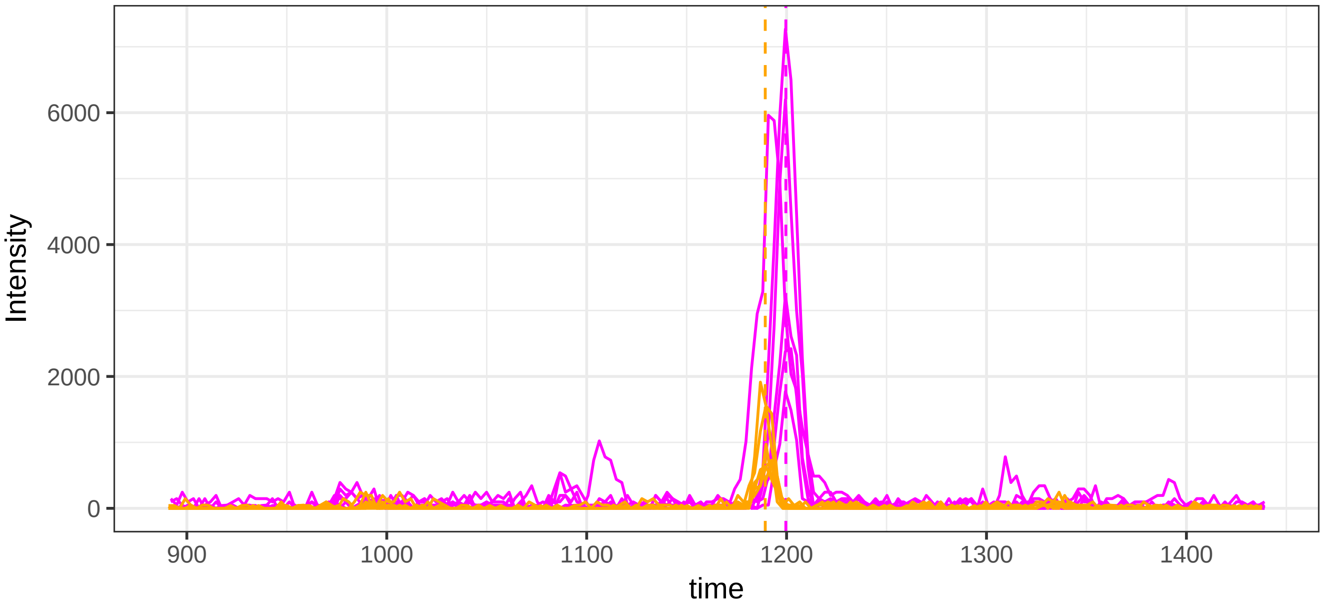


run23: 1809_VLLDGVQNPR/2 (magenta)  AND 120401_NSPLDEENLTQENQI

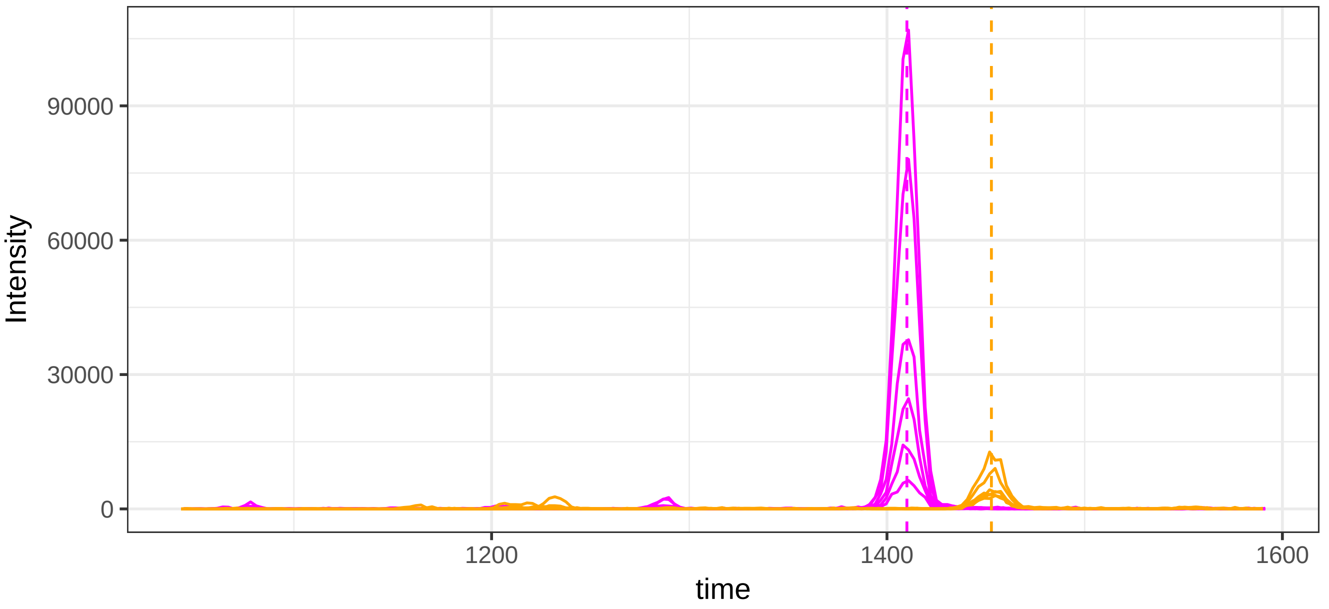

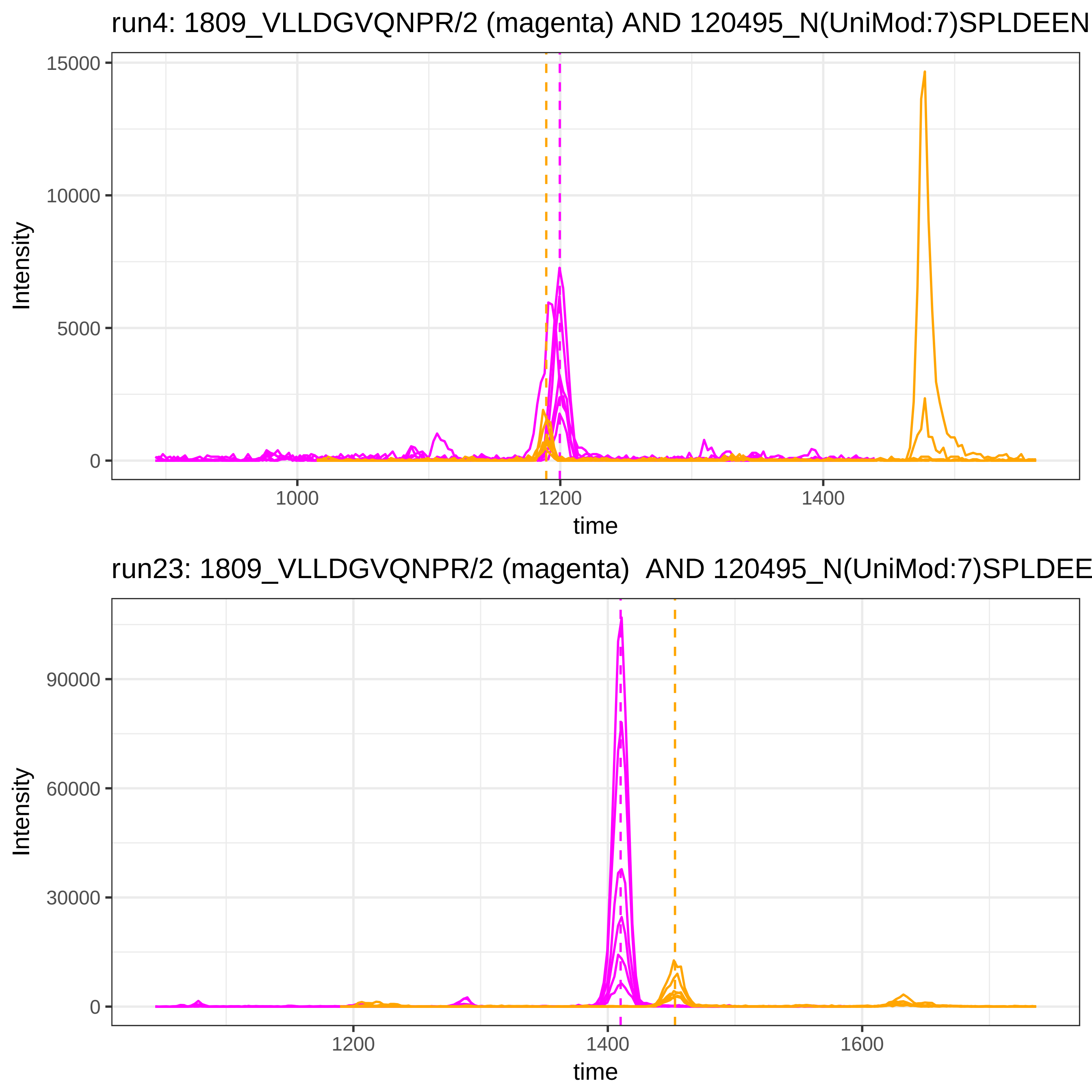
run4: 1809_VLLDGVQNPR/2 (magenta) AND 120495_N(UniMod:7)SPLDEEN
Intensity
15000
10000
5000
0
1000
1200
1400
time
run23: 1809_VLLDGVQNPR/2 (magenta) AND 120495_N(UniMod:7)SPLDEE
Intensity
90000
60000
30000
0
1200
1400
1600
time

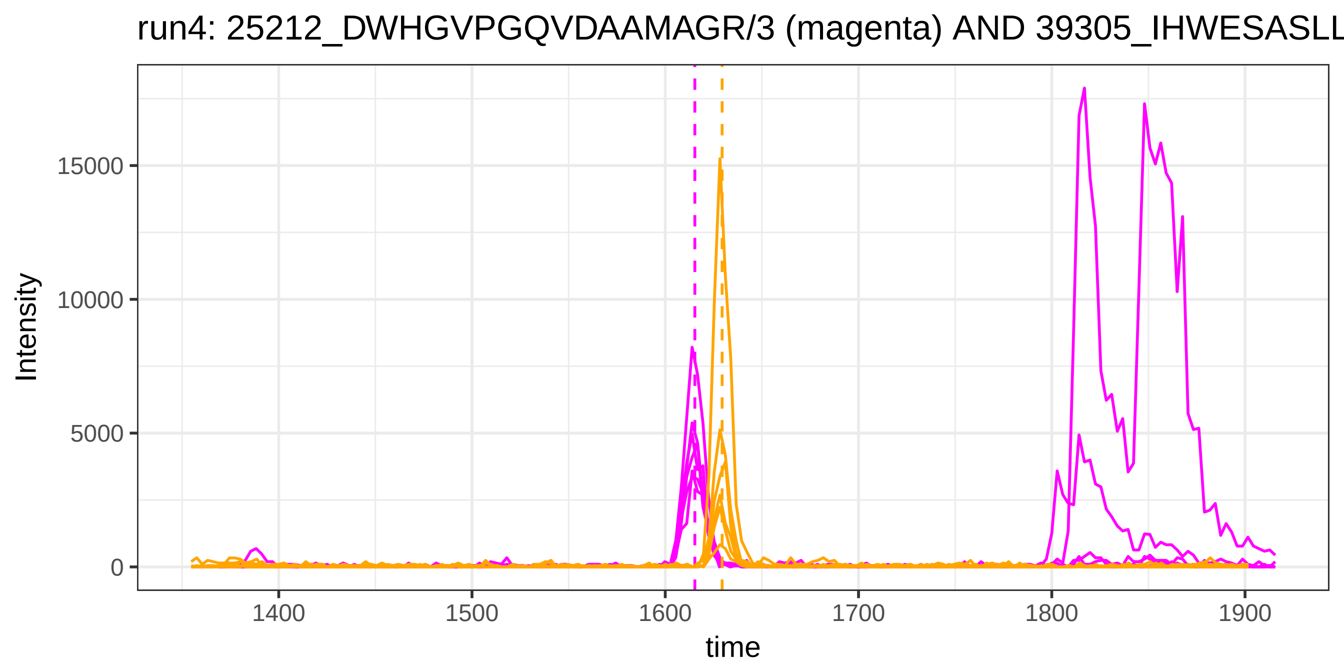
run4: 25212_DWHGVPGQVDAAMAGR/3 (magenta) AND 39305_IHWESASLL
Intensity
15000
10000
5000
0
1400
1500
1600
1700
1800
1900
time

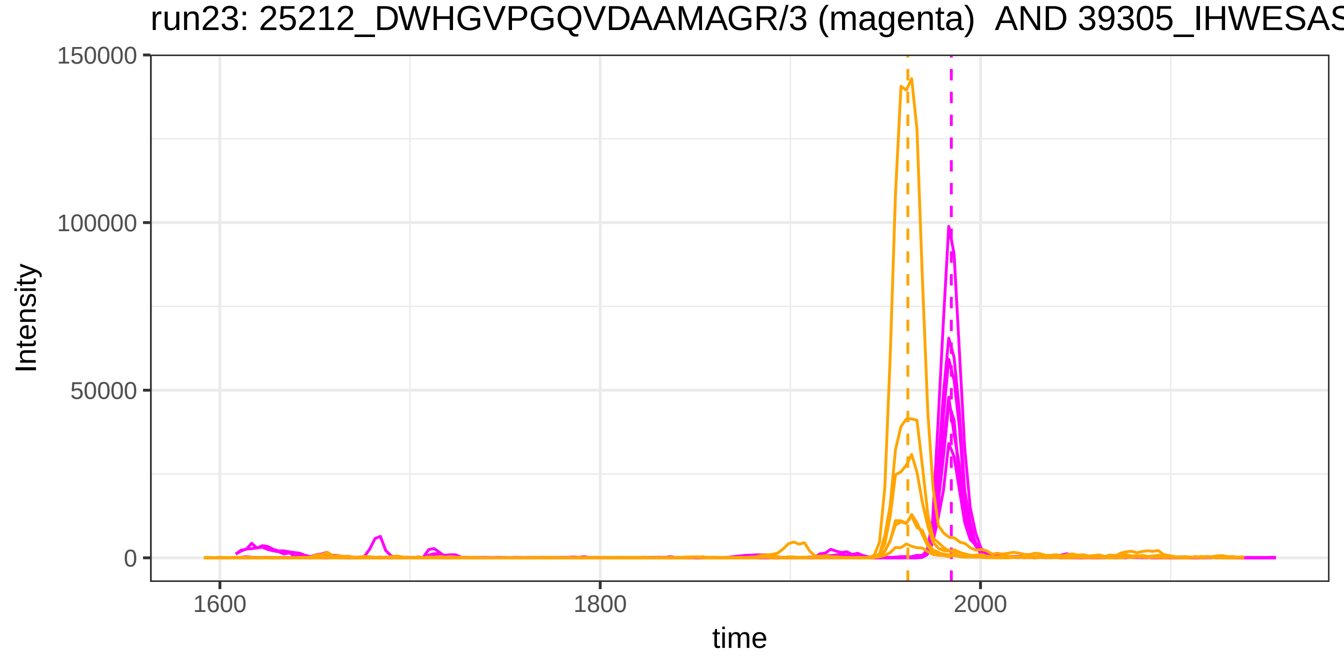
run23: 25212_DWHGVPGQVDAAMAGR/3 (magenta) AND 39305_IHWESAS
Intensity
150000
100000
50000
0
1600
1800
2000
time

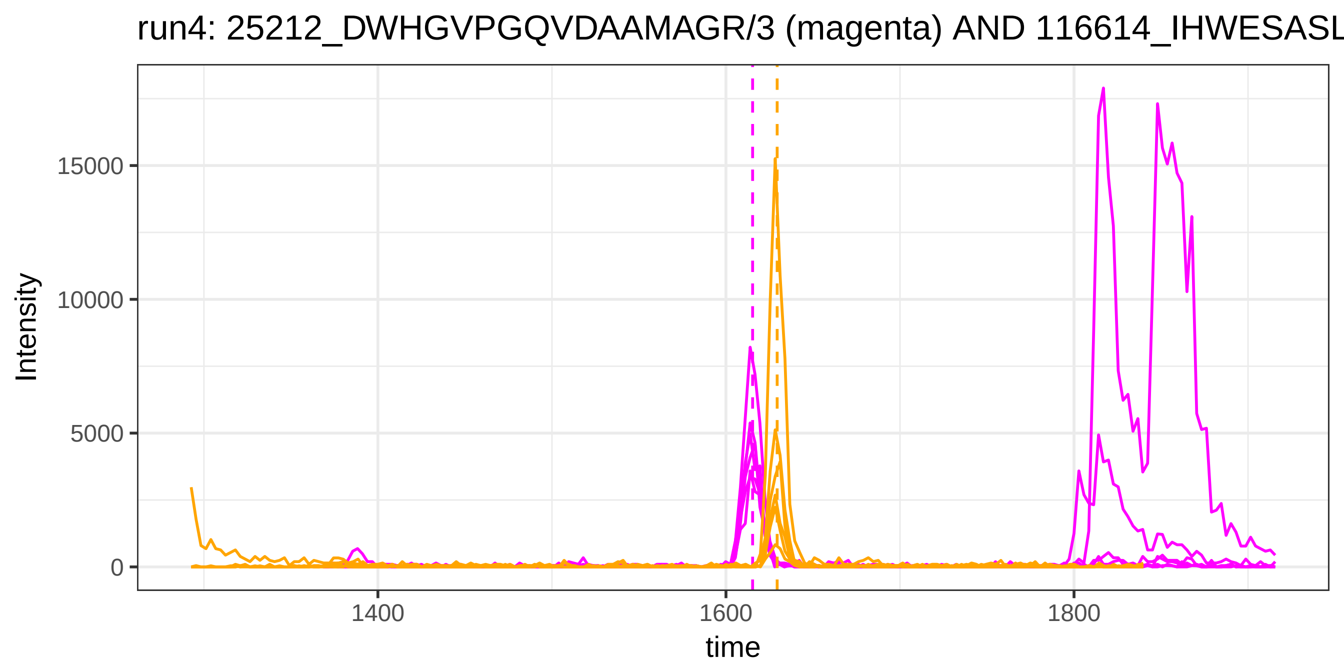
run4: 25212_DWHGVPGQVDAAMAGR/3 (magenta) AND 116614_IHWESASL
Intensity
15000
10000
5000
0
1400
1600
1800
time

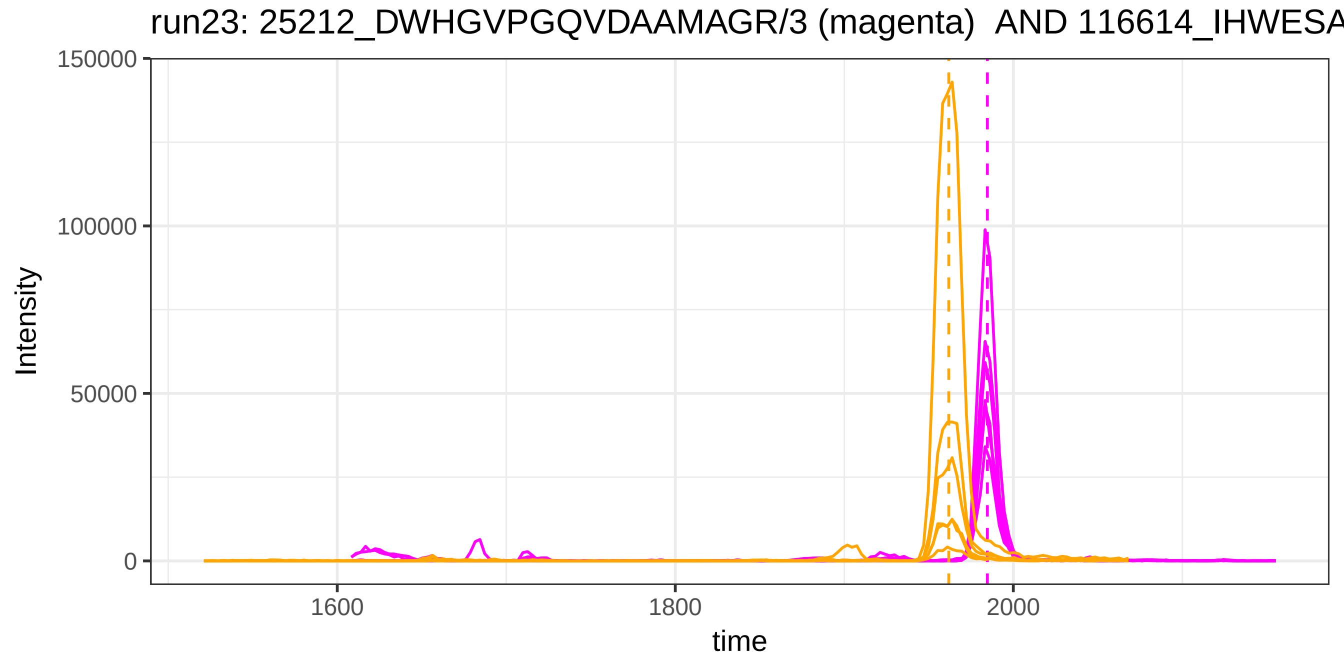
run23: 25212_DWHGVPGQVDAAMAGR/3 (magenta) AND 116614_IHWESA
Intensity
150000
100000
50000
0
1600
1800
2000
time

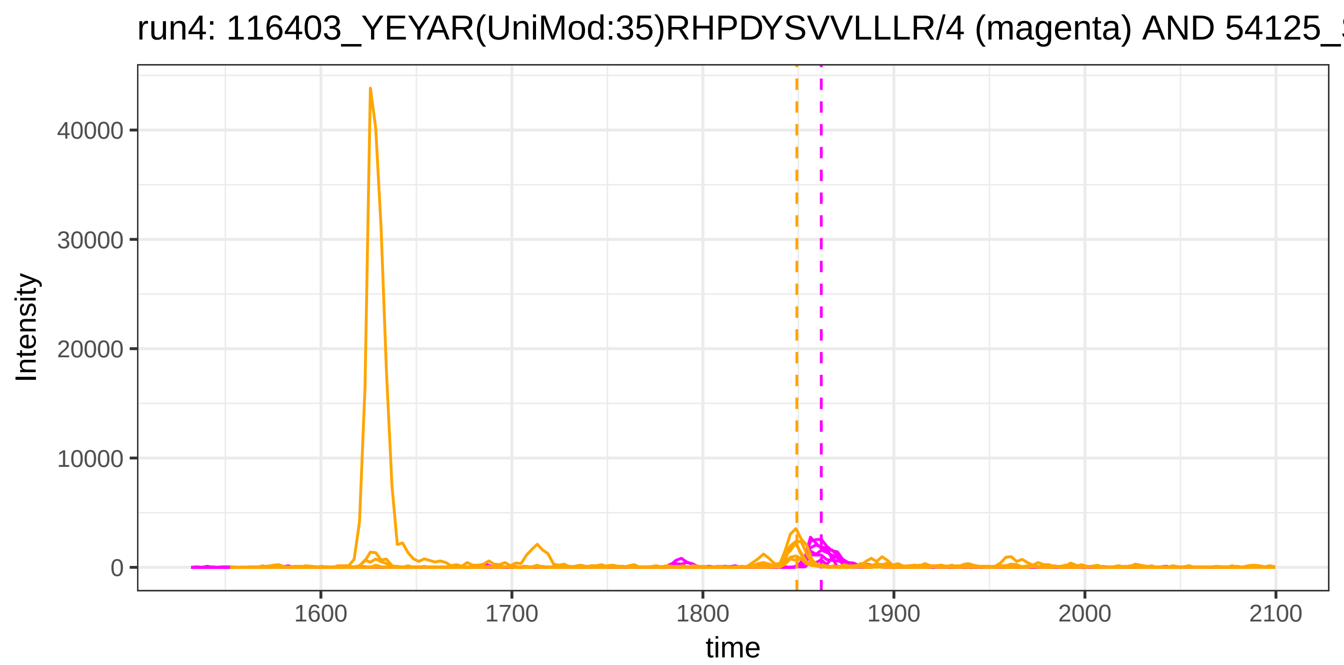
run4: 116403_YEYAR(UniMod:35)RHPDYSVVLLLR/4 (magenta) AND 54125_
Intensity
40000
30000
20000
10000
0
1600
1700
1800
1900
2000
2100
time

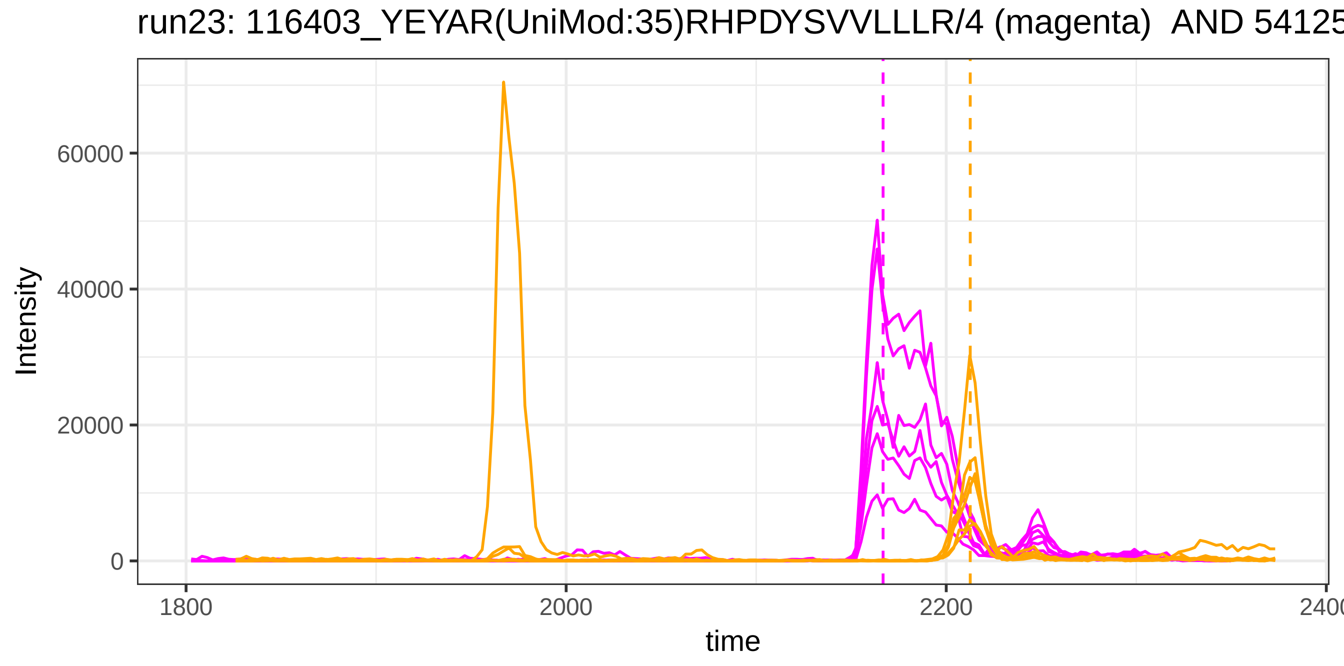
run23: 116403_YEYAR(UniMod:35)RHPDYSVVLLLR/4 (magenta) AND 54125
Intensity
60000
40000
20000
0
1800
2000
2200
time

run4: 116403_YEYAR(UniMod:35)RHPDYSVVLLLR/4 (magenta) AND 117986_

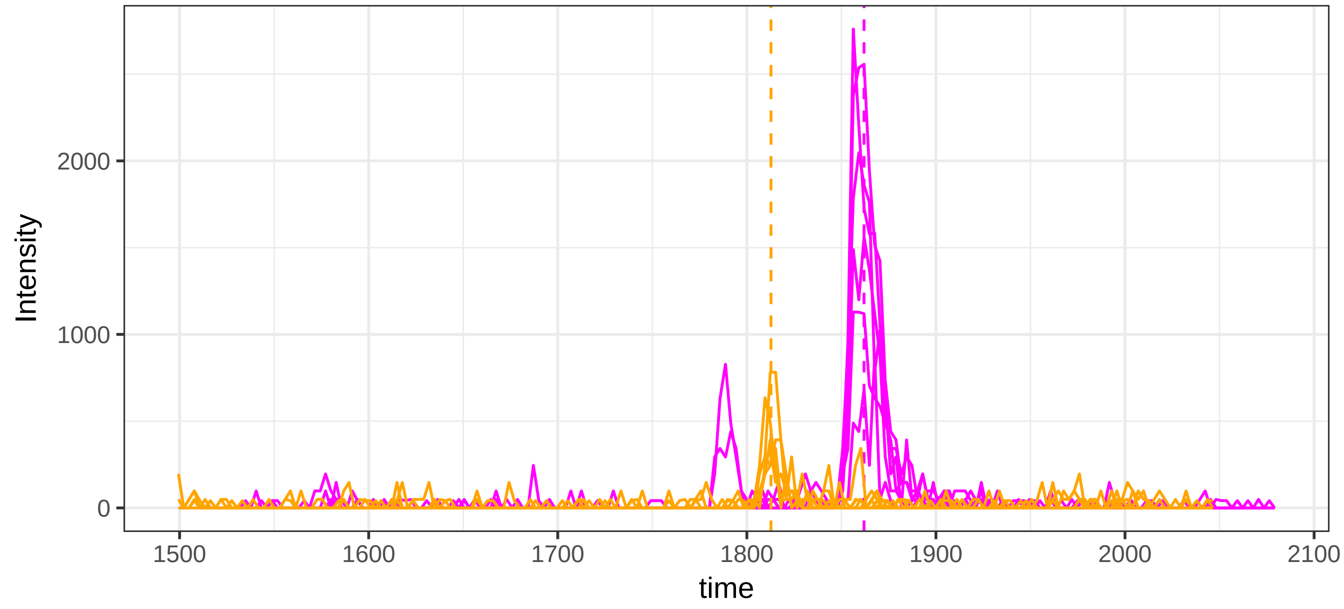


run23: 116403_YEYAR(UniMod:35)RHPDYSVVLLLR/4 (magenta) AND 11798

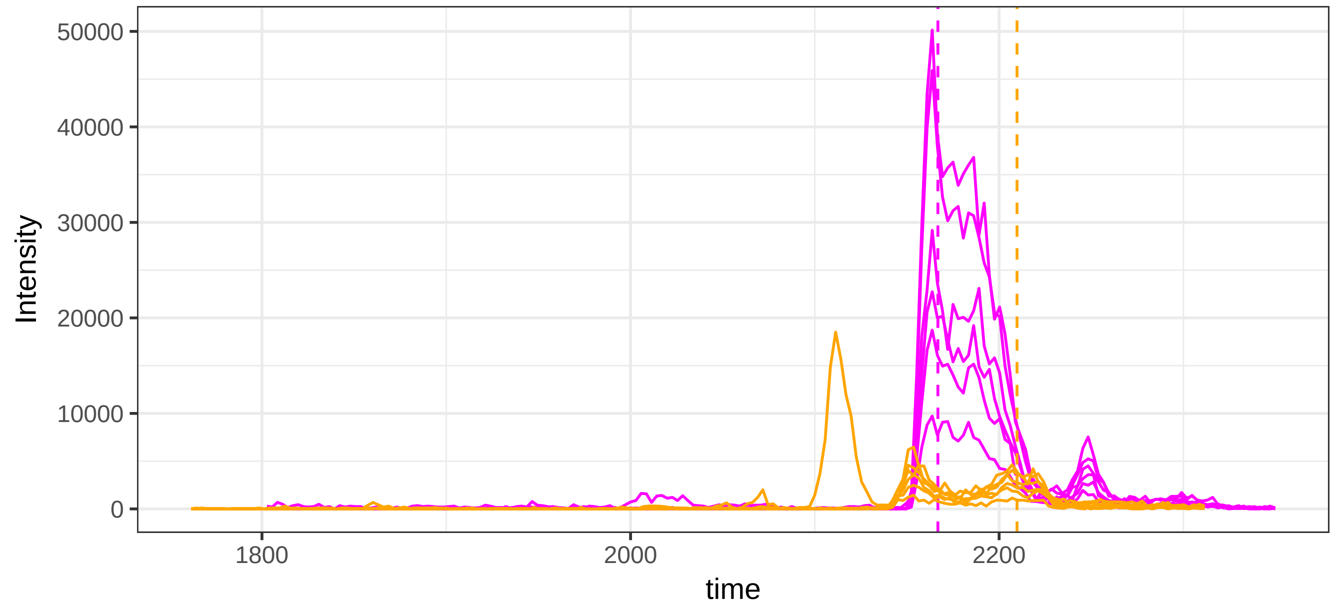

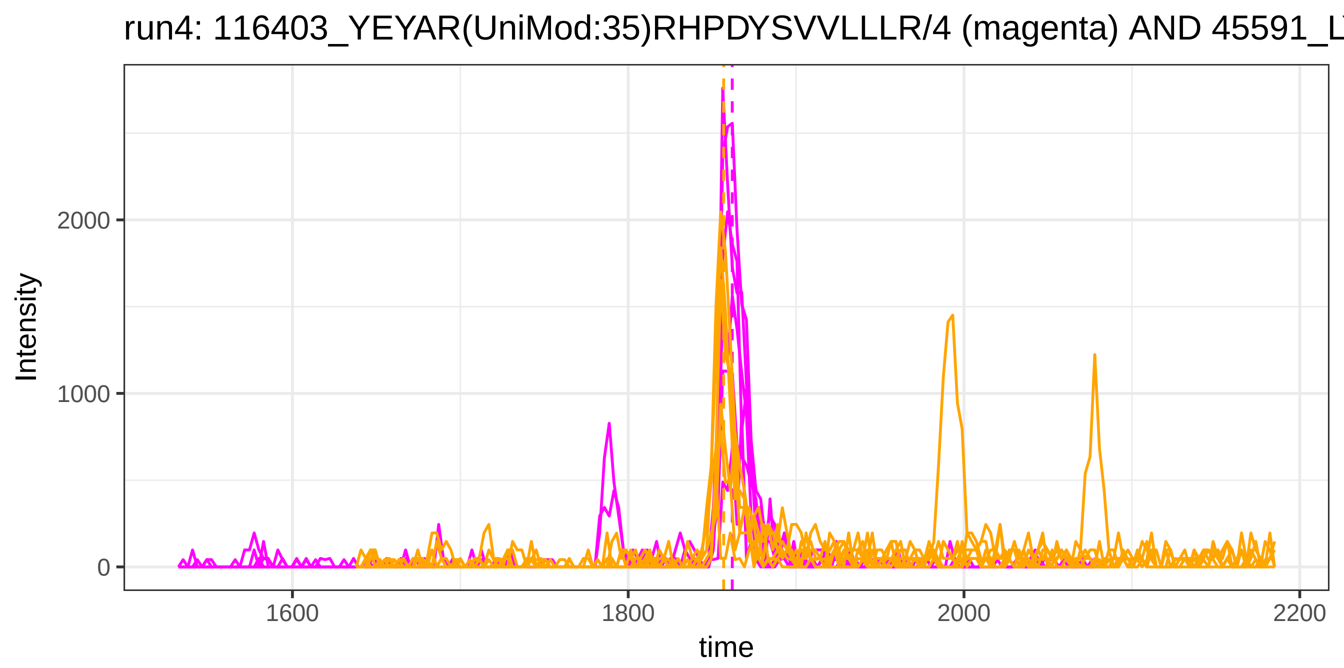
run4: 116403_YEYAR(UniMod:35)RHPDYSVVLLLR/4 (magenta) AND 45591_L
Intensity
2000
1000
0
1600
1800
2000
2200
time

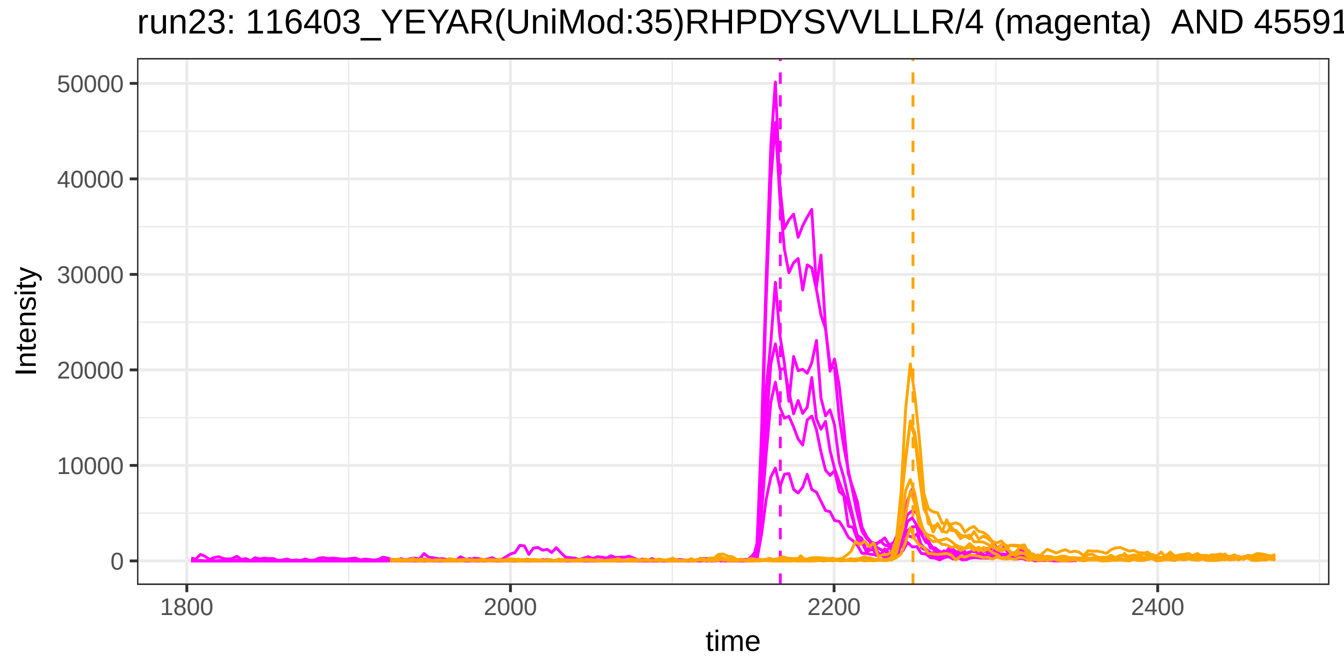
run23: 116403_YEYAR(UniMod:35)RHPDYSVVLLLR/4 (magenta) AND 45591
Intensity
50000
40000
30000
20000
10000
0
1800
2000
2200
2400
time

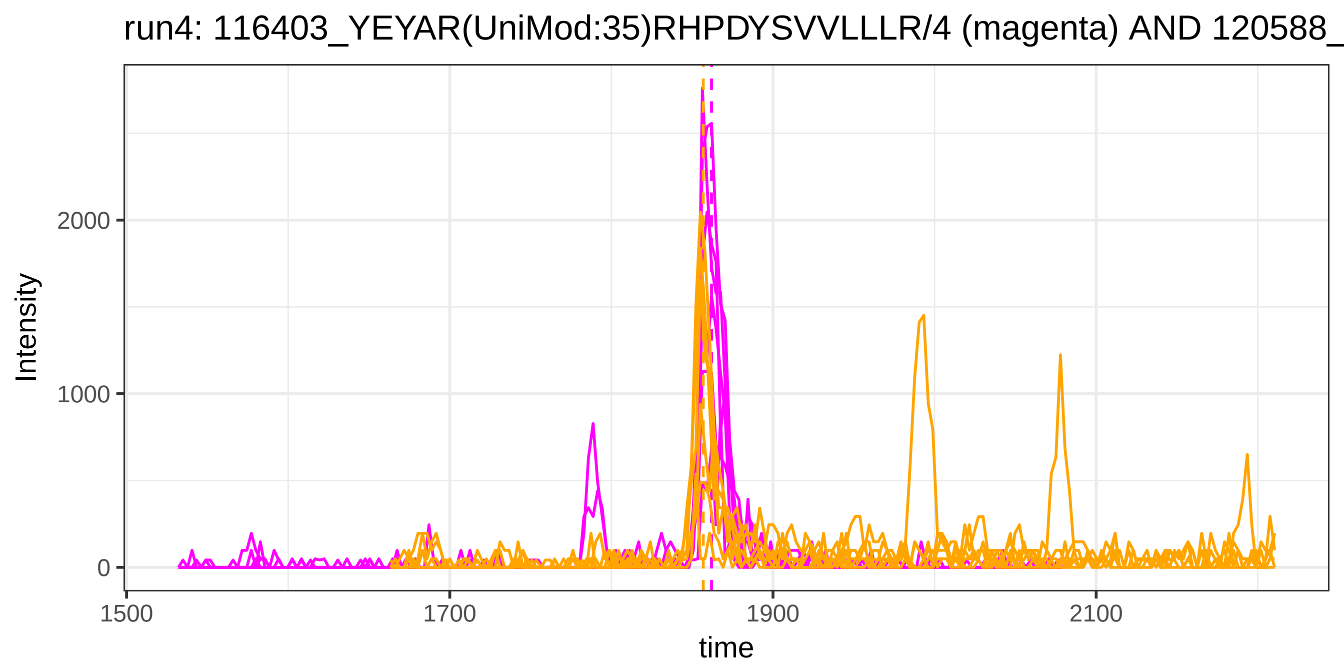
run4: 116403_YEYAR(UniMod:35)RHPDYSVVLLLR/4 (magenta) AND 120588_
Intensity
2000
1000
0
1500
1700
1900
2100
time

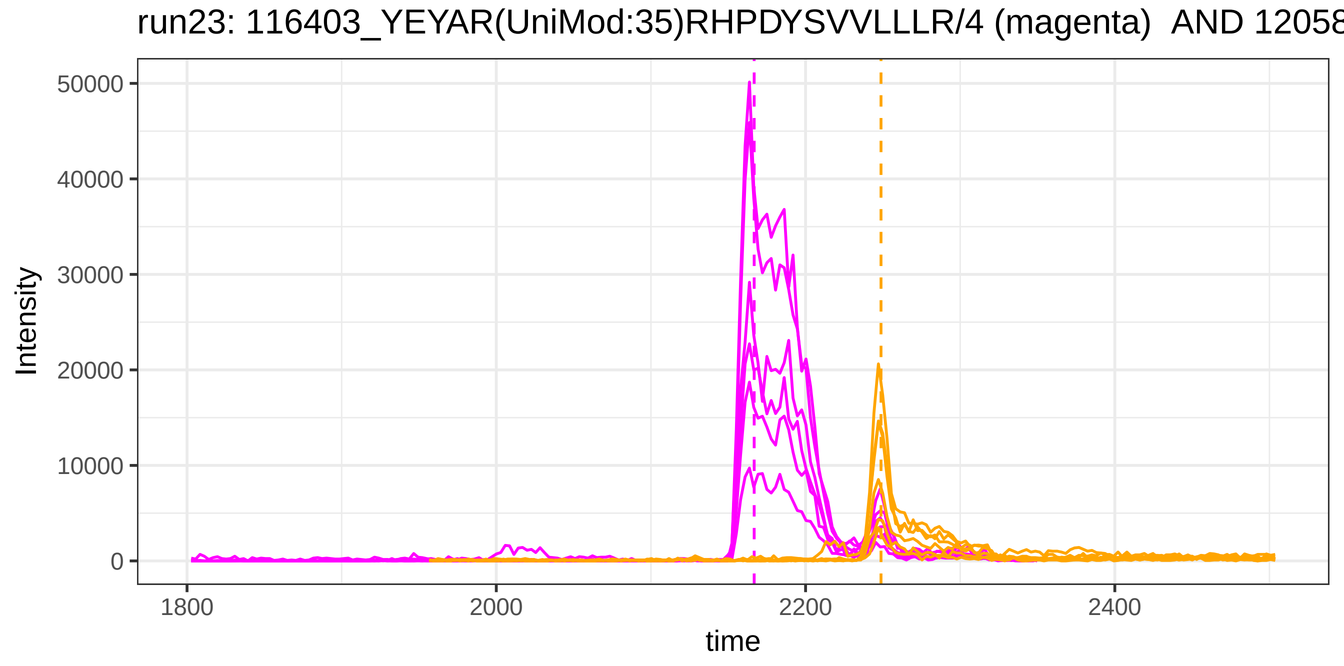
run23: 116403_YEYAR(UniMod:35)RHPDYSVVLLLR/4 (magenta) AND 12058
Intensity
50000
40000
30000
20000
10000
0
1800
2000
2200
2400
time

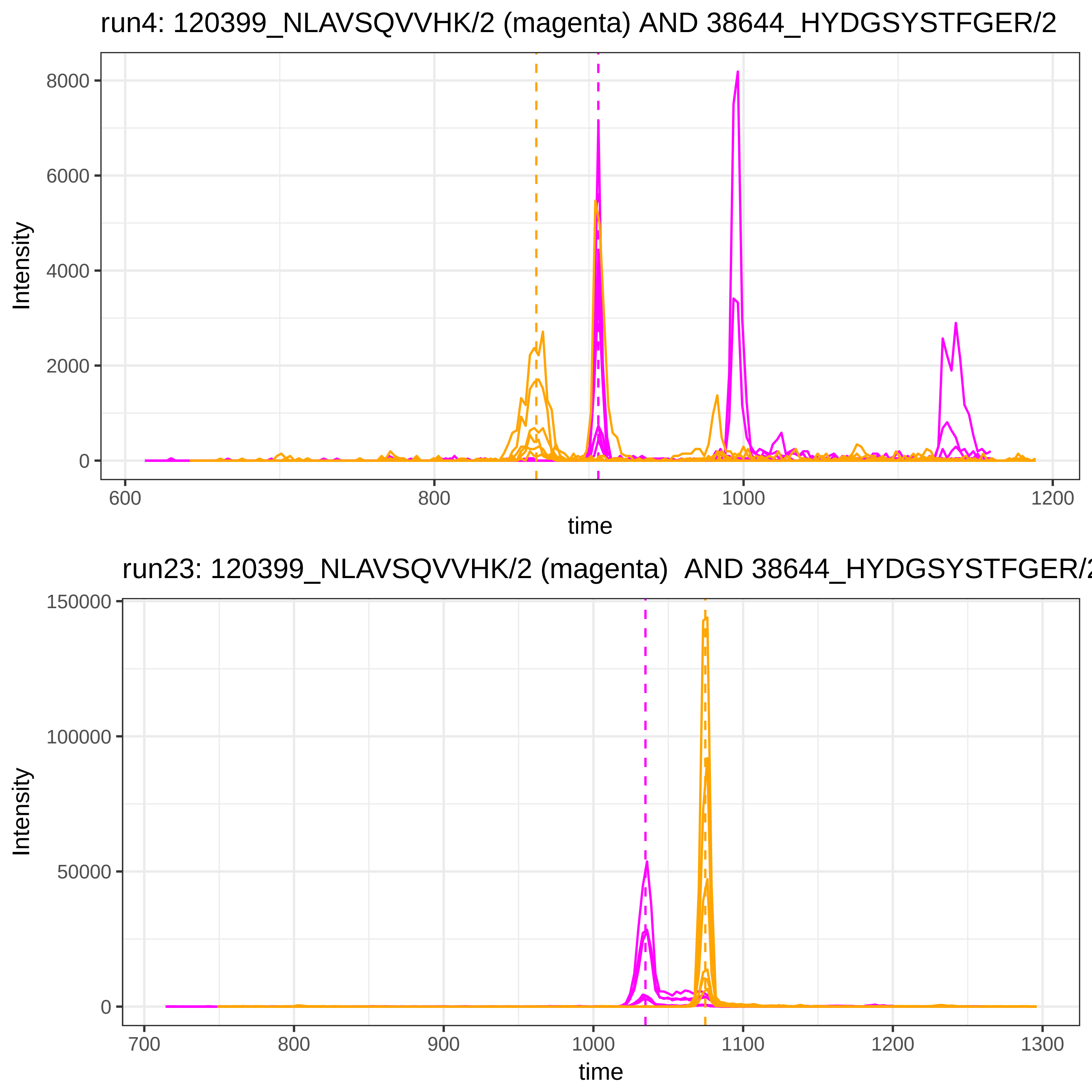
run4: 120399_NLAVSQVVHK/2 (magenta) AND 38644_HYDGSYSTFGER/2
Intensity
8000
6000
4000
2000
0
600
800
1000
1200
time
run23: 120399_NLAVSQVVHK/2 (magenta) AND 38644_HYDGSYSTFGER/2
Intensity
150000
100000
50000
0
700
800
900
1000
1100
1200
1300
time

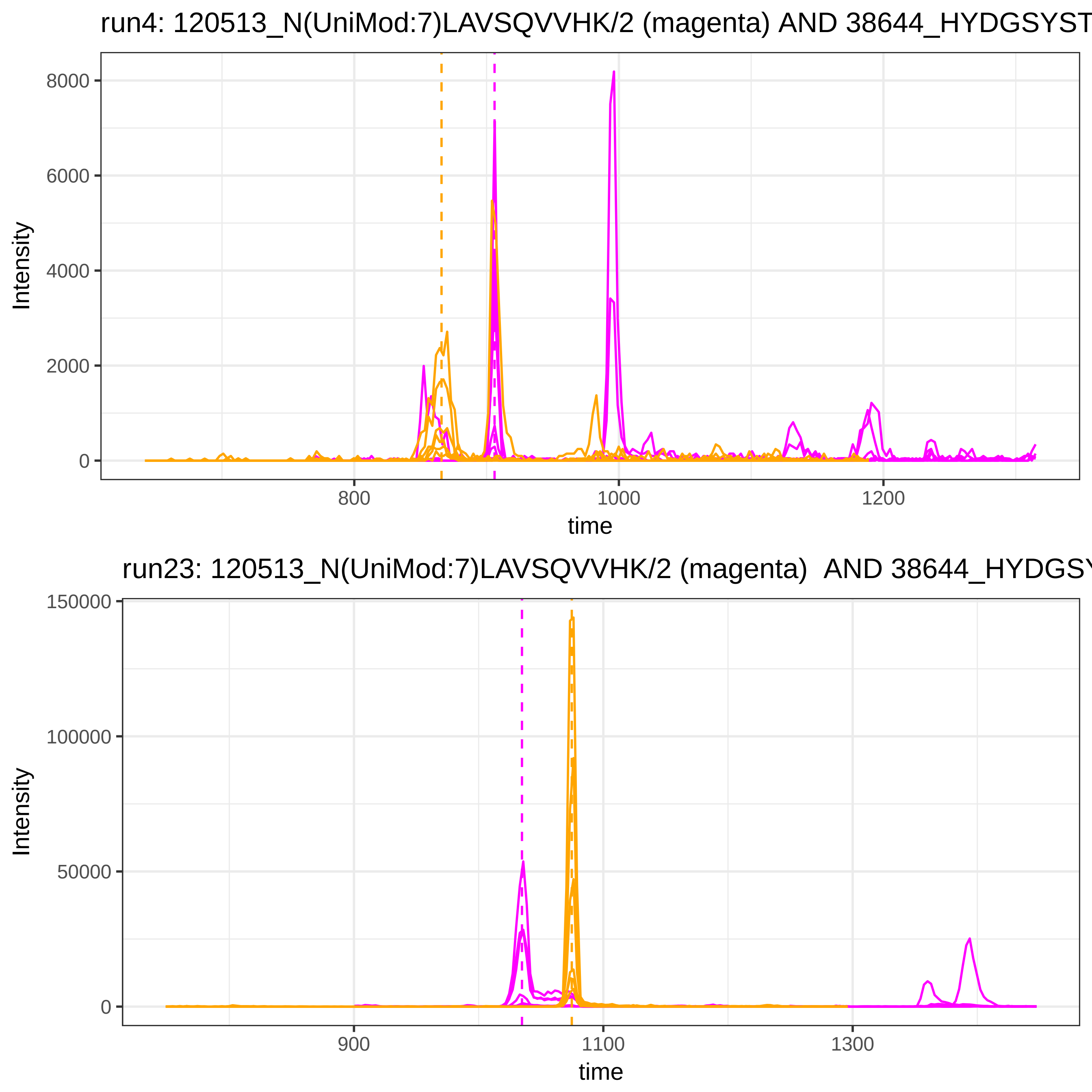
run4: 120513_N(UniMod:7)LAVSQVVHK/2 (magenta) AND 38644_HYDGSYST
Intensity
8000
6000
4000
2000
0
800
1000
1200
time
run23: 120513_N(UniMod:7)LAVSQVVHK/2 (magenta) AND 38644_HYDGSY
Intensity
150000
100000
50000
0
900
1100
1300
time

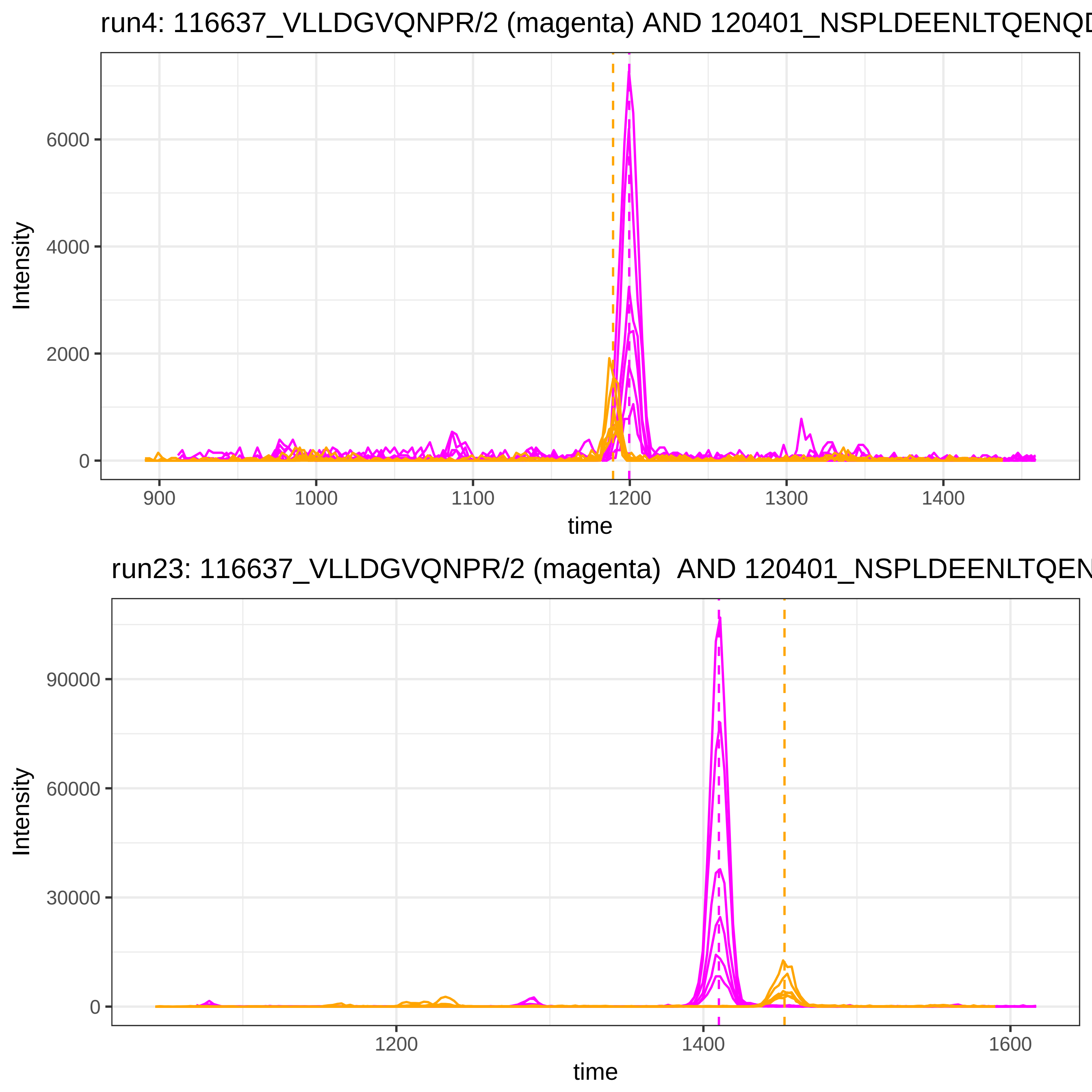
run4: 116637_VLLDGVQNPR/2 (magenta) AND 120401_NSPLDEENLTQENQD
6000
4000
2000
0
Intensity
900
1000
1100
1200
1300
1400
time
run23: 116637_VLLDGVQNPR/2 (magenta) AND 120401_NSPLDEENLTQEN
90000
60000
30000
0
Intensity
1200
1400
1600
time

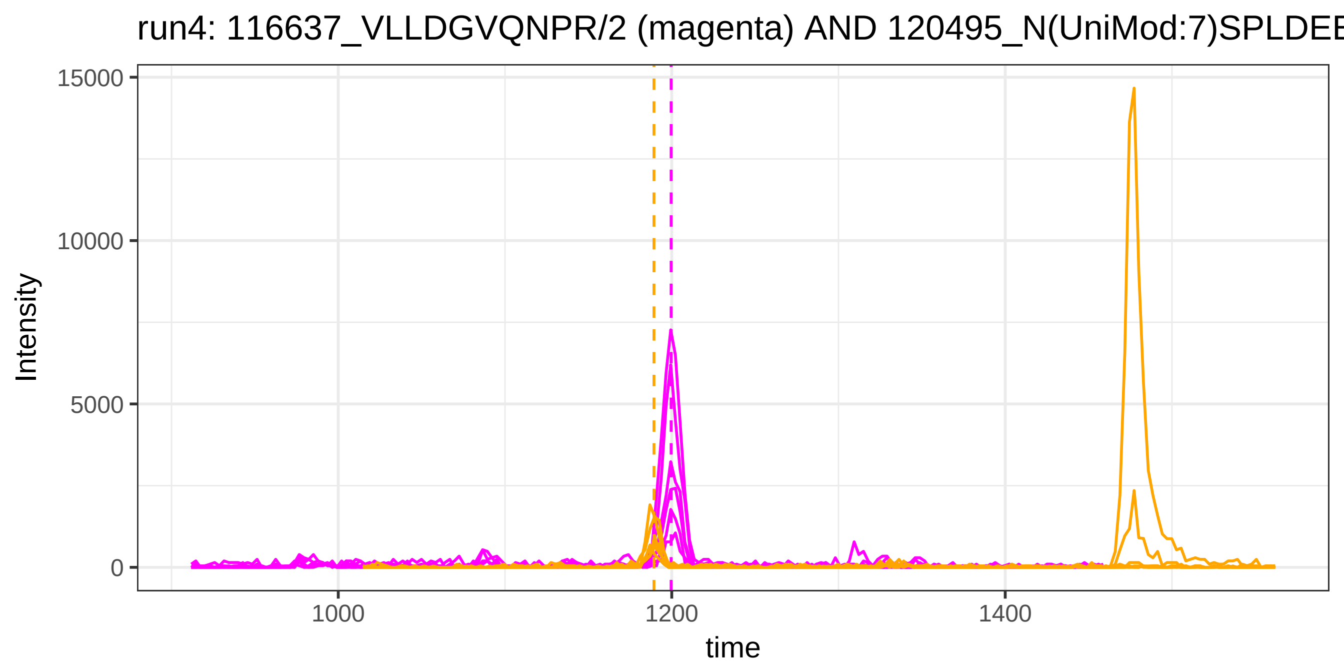

run4: 116637_VLLDGVQNPR/2 (magenta) AND 120495_N(UniMod:7)SPLDEE
15000
10000
5000
0
Intensity
1000
1200
1400
time


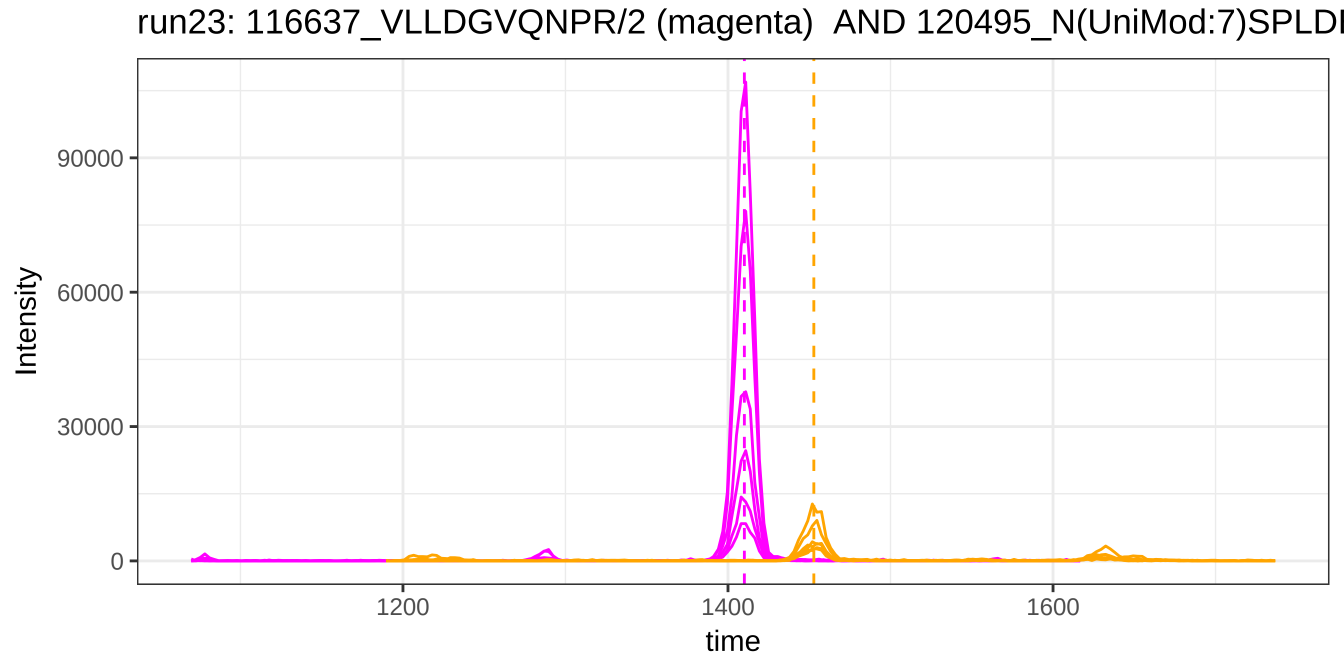

run23: 116637_VLLDGVQNPR/2 (magenta) AND 120495_N(UniMod:7)SPLDI
90000
60000
30000
0
Intensity
1200
1400
1600
time

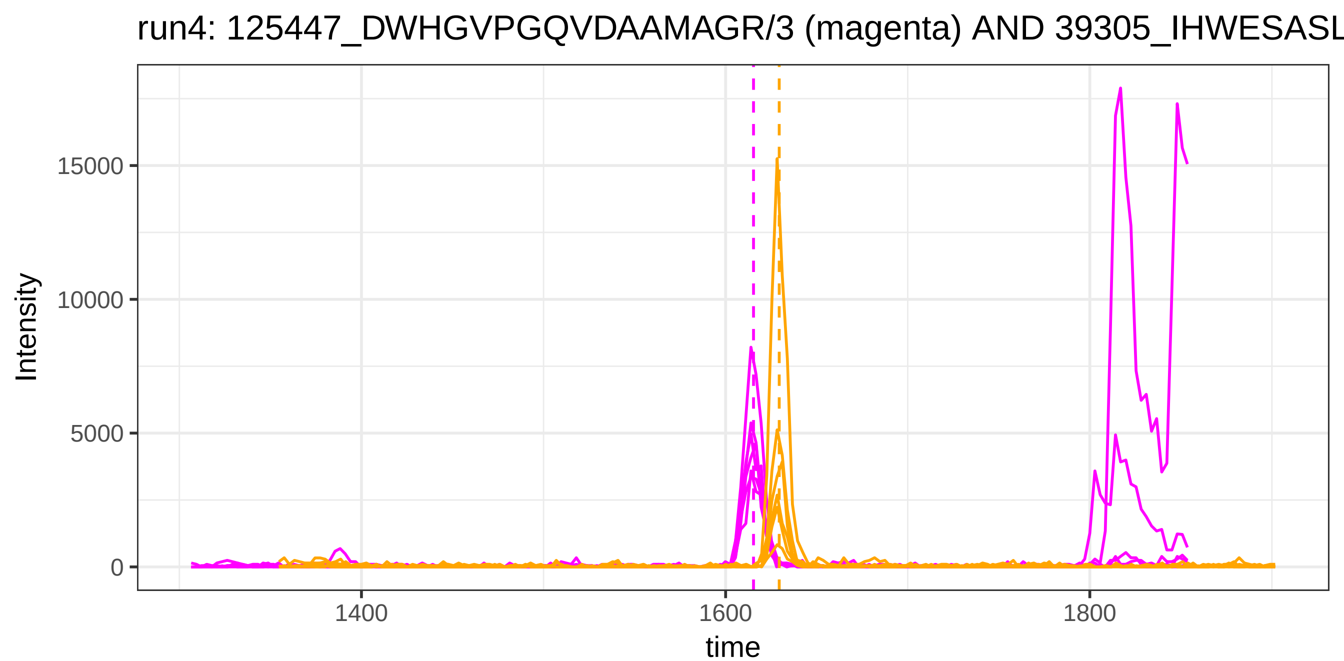

run4: 125447_DWHGVPGQVDAAMAGR/3 (magenta) AND 39305_IHWESASL
Intensity
15000
10000
5000
0
1400
1600
1800
time


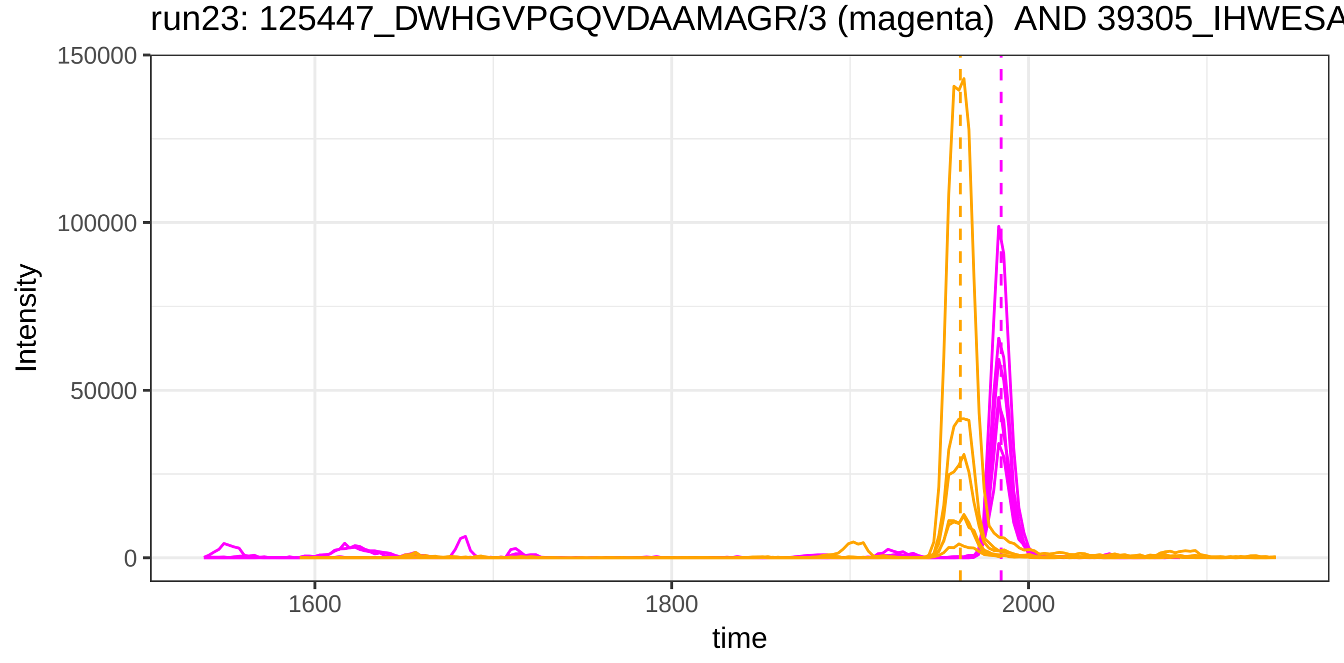

run23: 125447_DWHGVPGQVDAAMAGR/3 (magenta) AND 39305_IHWESA
Intensity
150000
100000
50000
0
1600
1800
2000
time

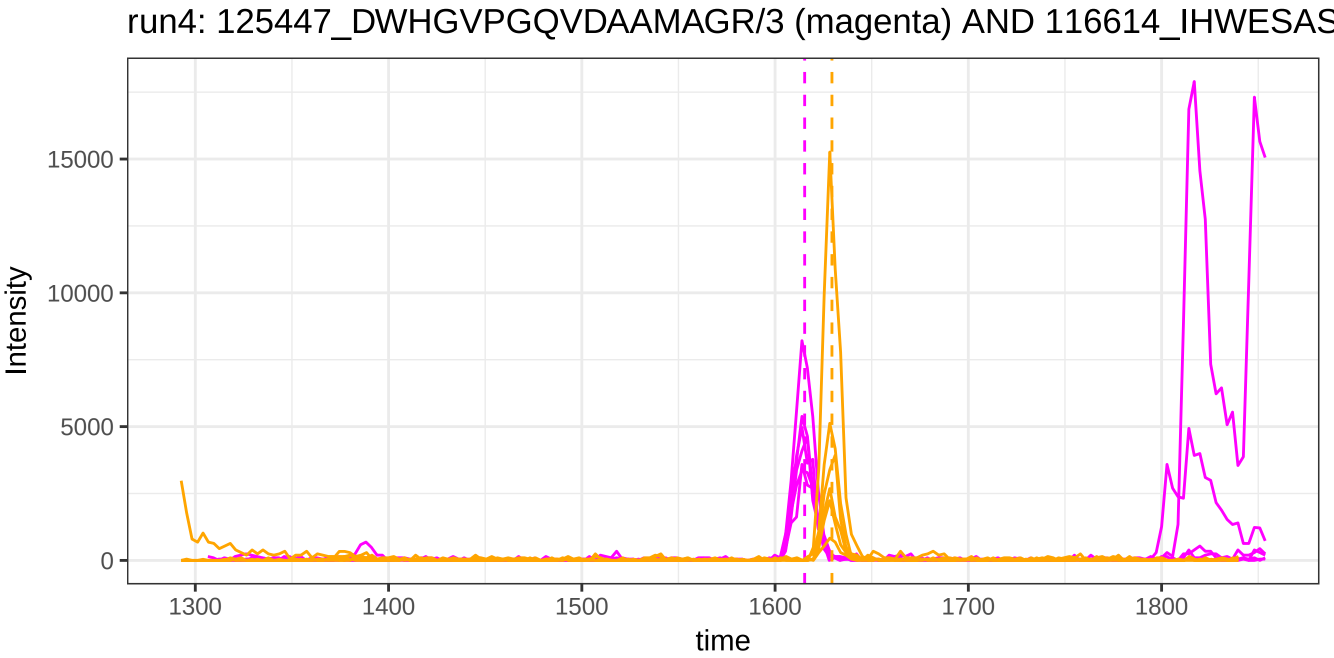

run4: 125447_DWHGVPGQVDAAMAGR/3 (magenta) AND 116614_IHWESAS
Intensity
15000
10000
5000
0
1300
1400
1500
1600
1700
1800
time


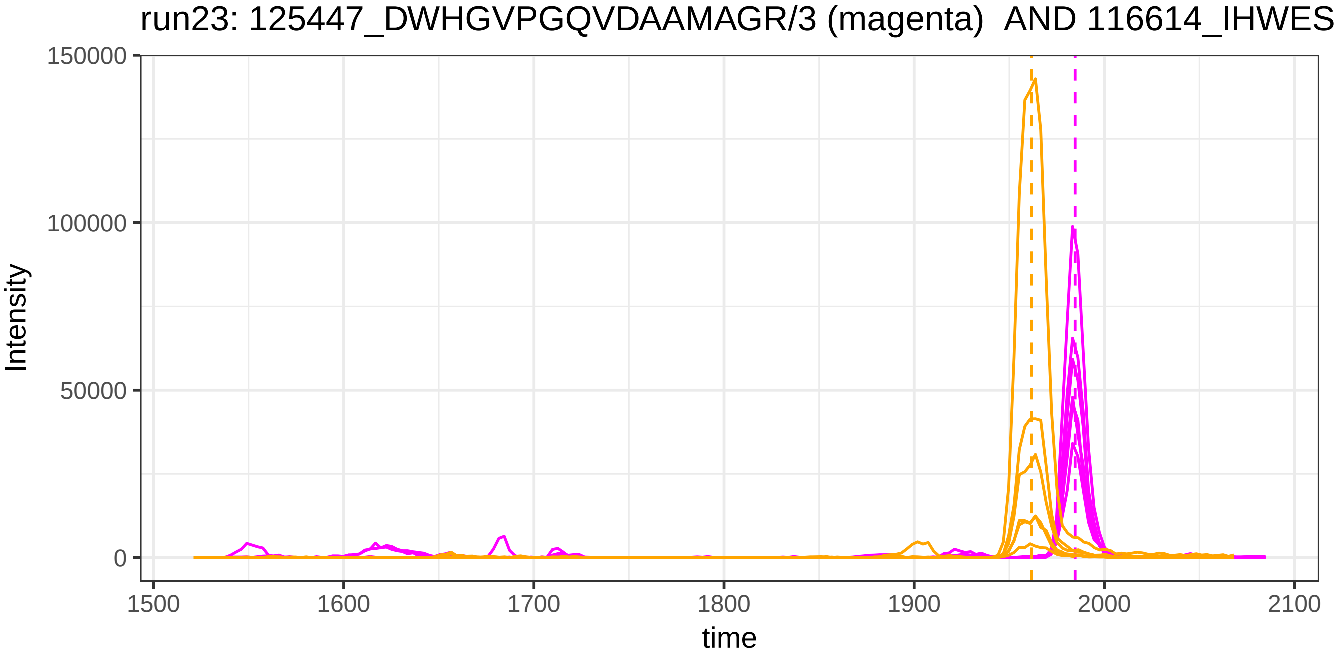

run23: 125447_DWHGVPGQVDAAMAGR/3 (magenta) AND 116614_IHWES
Intensity
150000
100000
50000
0
1500
1600
1700
1800
1900
2000
2100
time

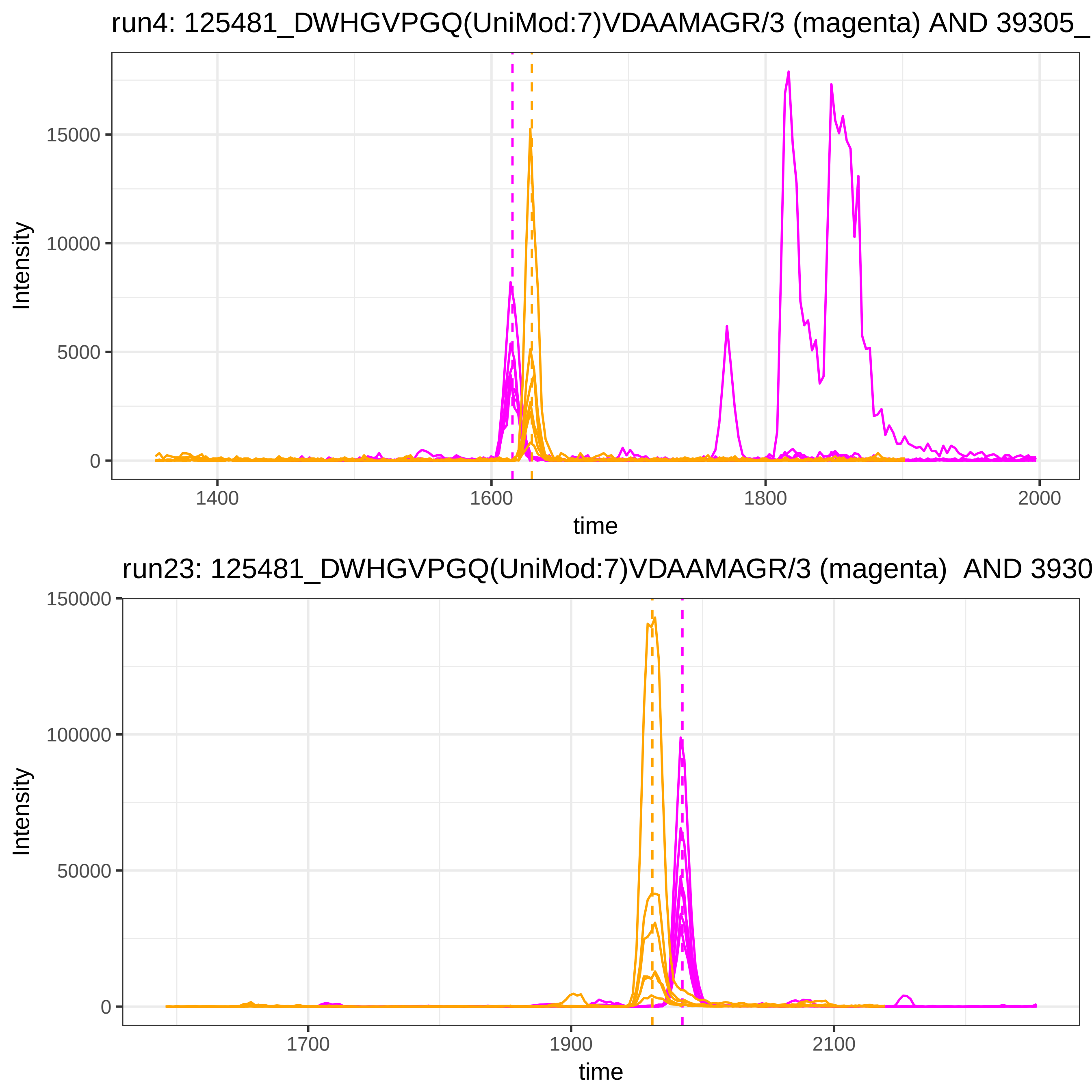
run4: 125481_DWHGVPGQ(UniMod:7)VDAAMAGR/3 (magenta) AND 39305_
Intensity
15000
10000
5000
0
1400
1600
1800
2000
time
run23: 125481_DWHGVPGQ(UniMod:7)VDAAMAGR/3 (magenta) AND 3930
Intensity
150000
100000
50000
0
1700
1900
2100
time

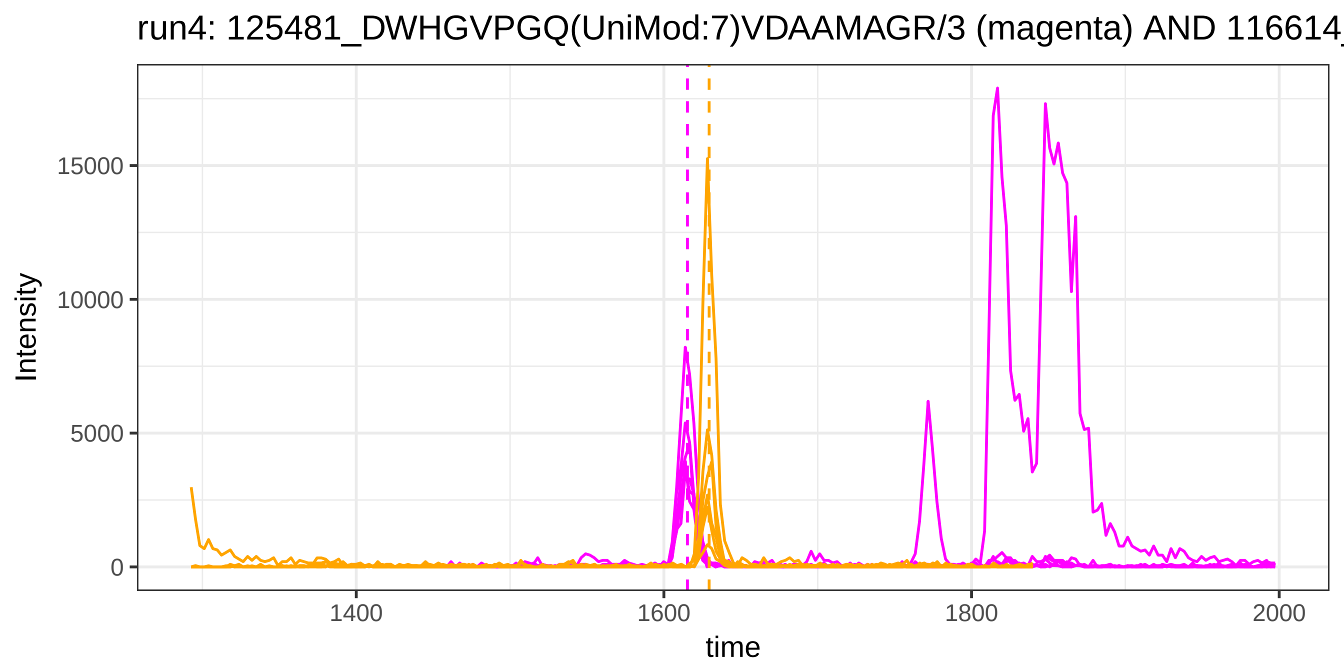

run4: 125481_DWHGVPGQ(UniMod:7)VDAAMAGR/3 (magenta) AND 116614_
Intensity
15000
10000
5000
0
1400
1600
1800
2000
time


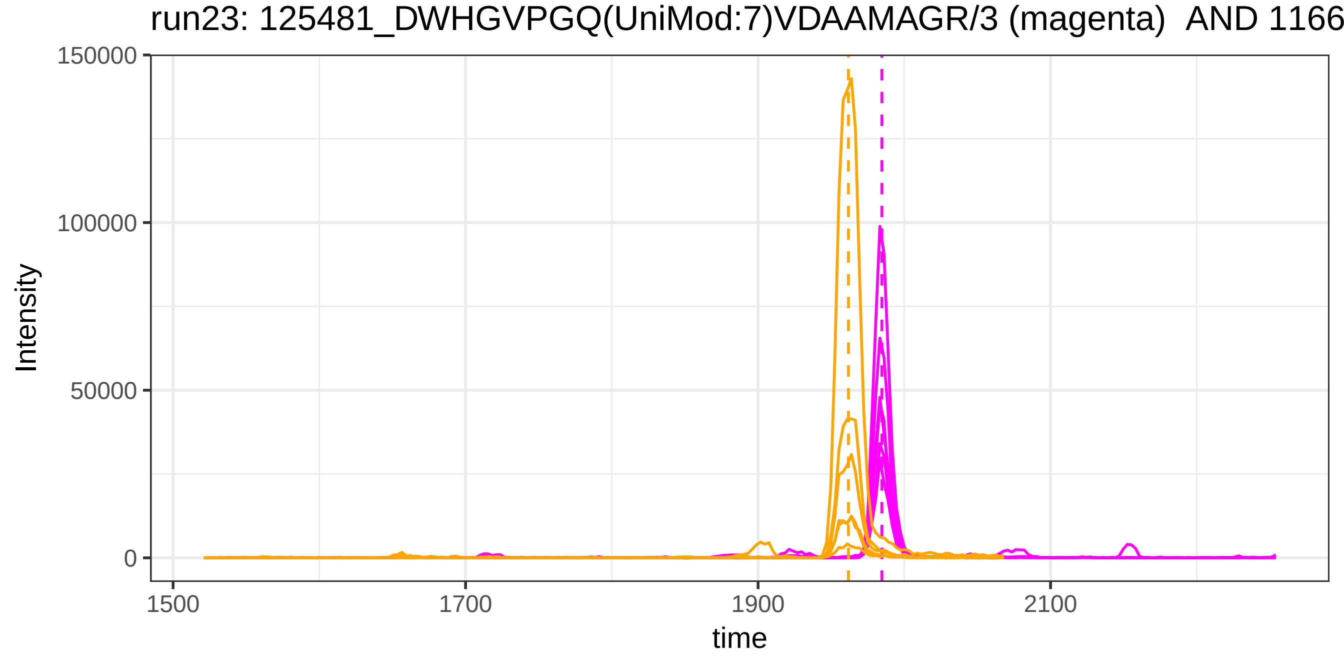

run23: 125481_DWHGVPGQ(UniMod:7)VDAAMAGR/3 (magenta) AND 1166
Intensity
150000
100000
50000
0
1500
1700
1900
2100
time

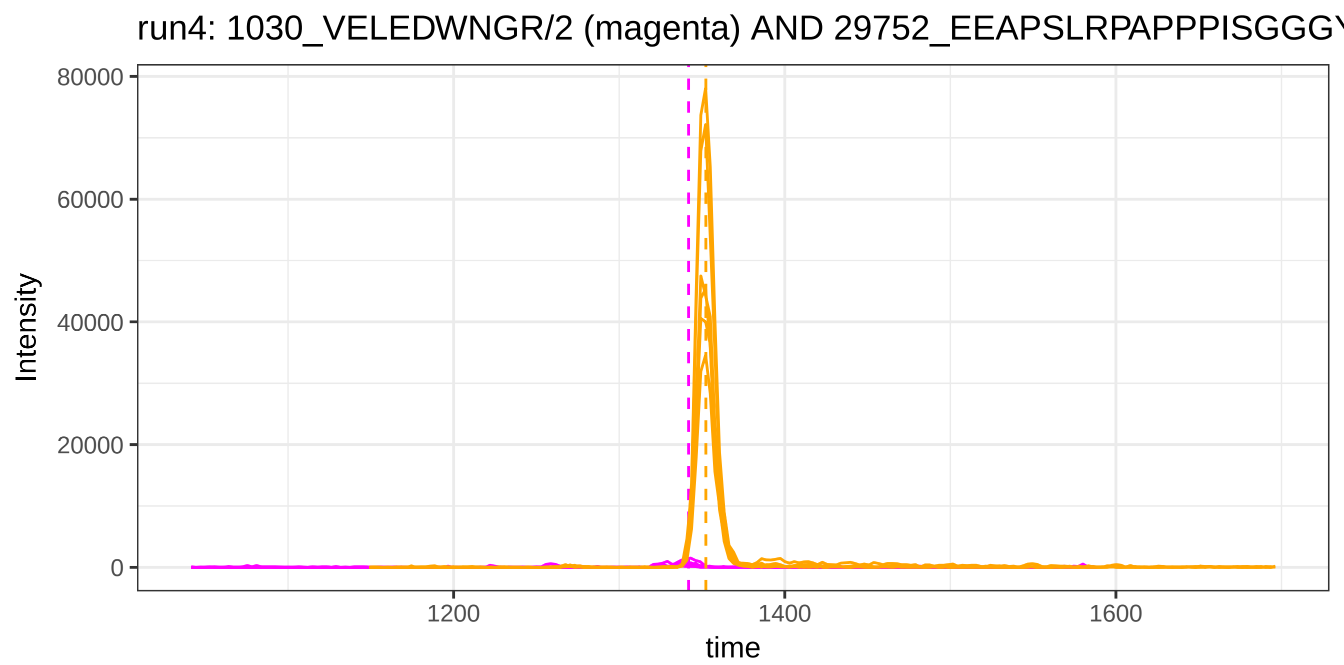

run4: 1030_VELEDWNGR/2 (magenta) AND 29752_EEAPSLRPAPPPISGGGY
Intensity
80000
60000
40000
20000
0
1200
1400
1600
time


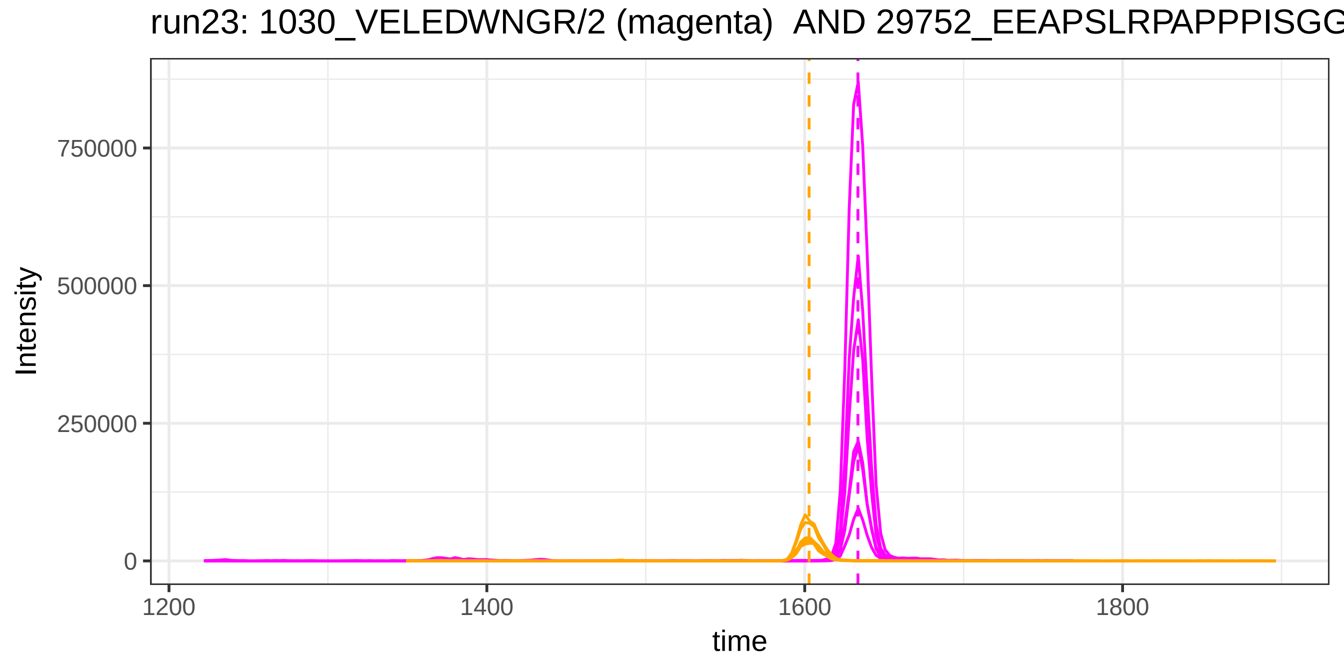

run23: 1030_VELEDWNGR/2 (magenta) AND 29752_EEAPSLRPAPPPISGG
Intensity
750000
500000
250000
0
1200
1400
1600
1800
time

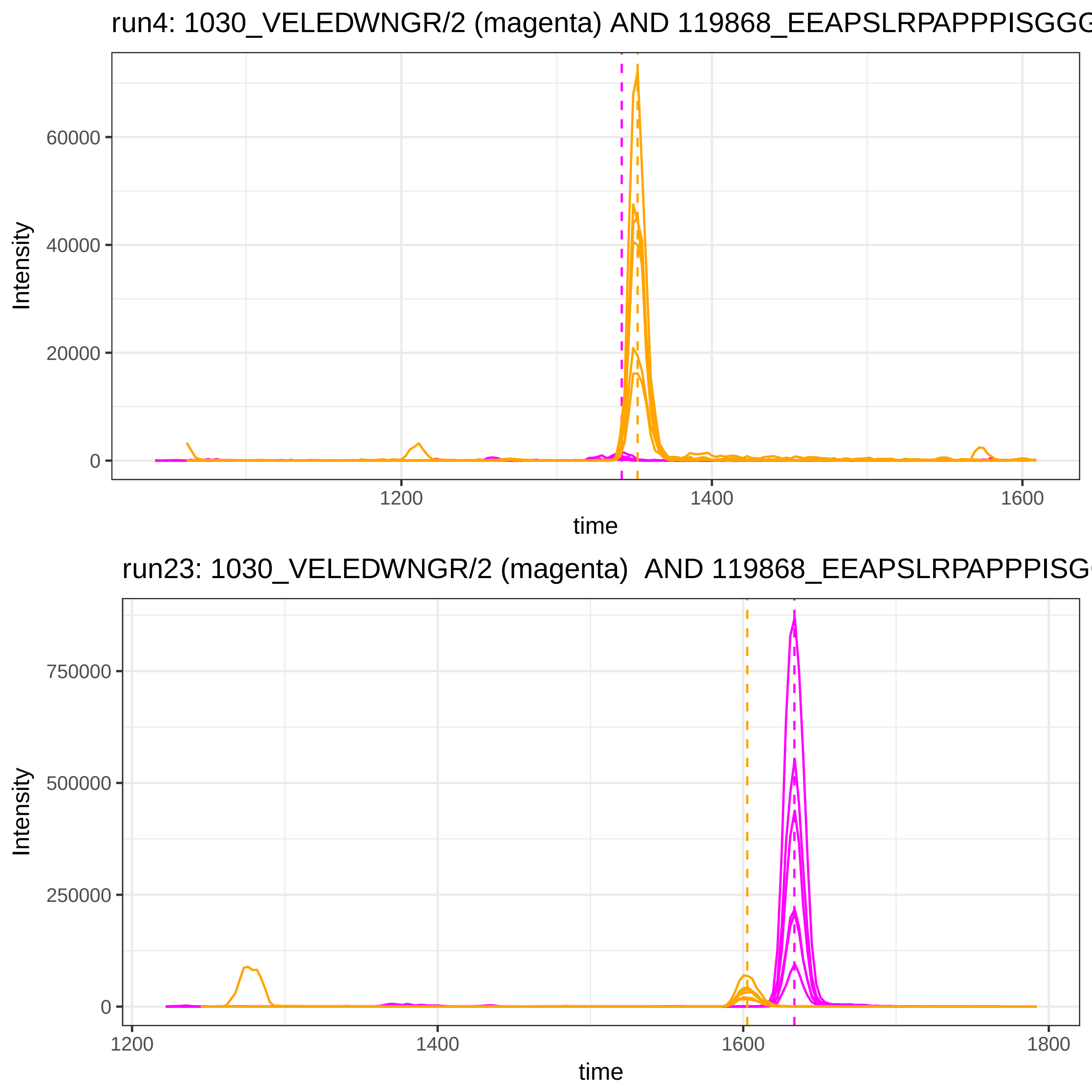
run4: 1030_VELEDWNGR/2 (magenta) AND 119868_EEAPSLRPAPPPISGGG
Intensity
60000
40000
20000
0
1200
1400
1600
time
run23: 1030_VELEDWNGR/2 (magenta) AND 119868_EEAPSLRPAPPPISG
Intensity
750000
500000
250000
0
1200
1400
1600
1800
time

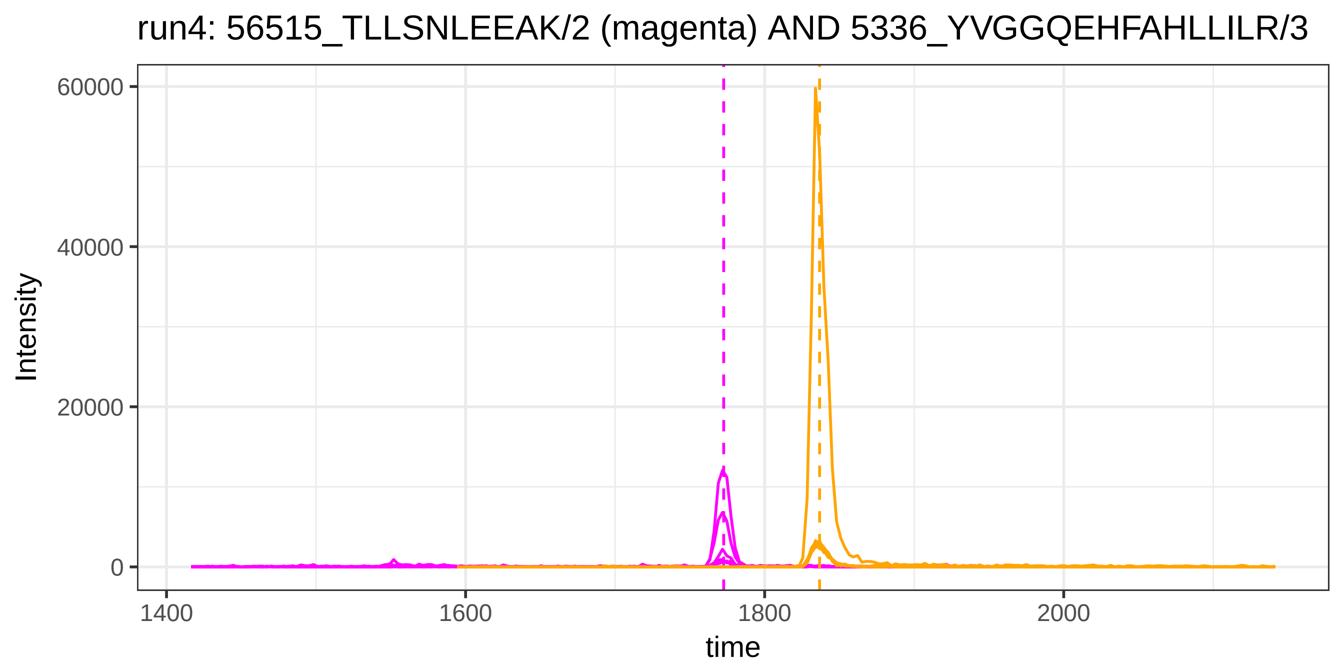

run4: 56515_TLLSNLEEAK/2 (magenta) AND 5336_YVGGQEHFAHLLILR/3
Intensity
60000
40000
20000
0
1400
1600
1800
2000
time


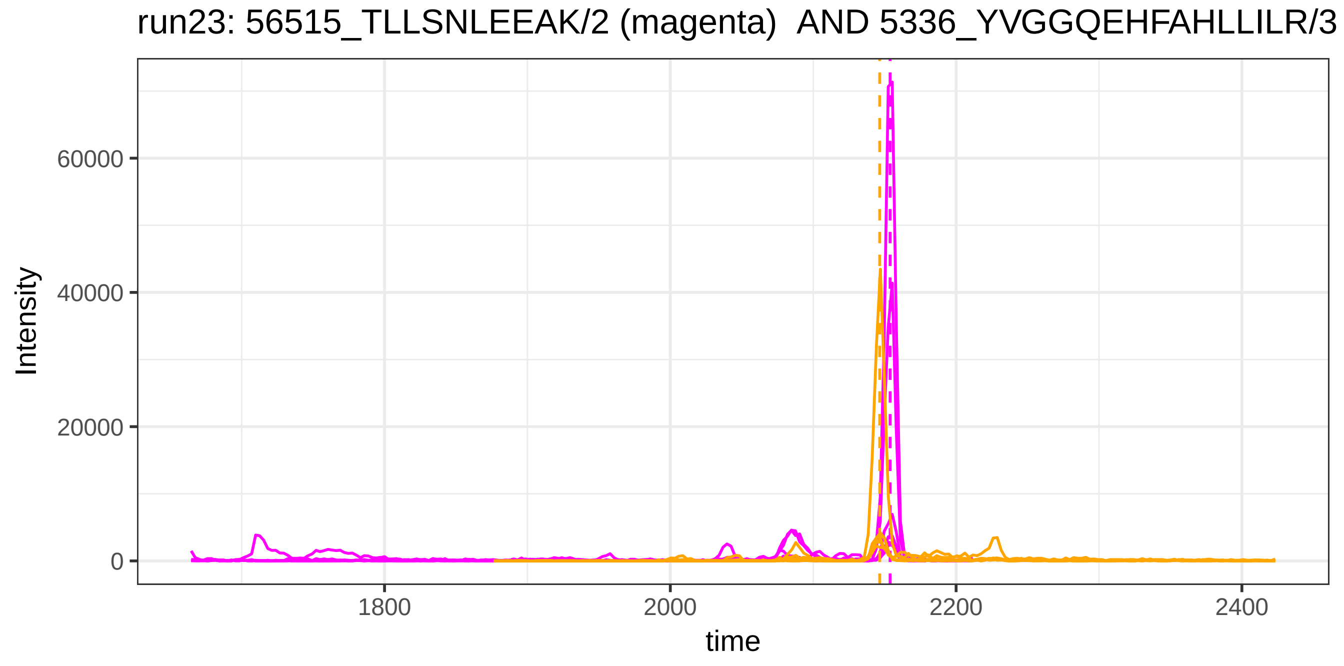

run23: 56515_TLLSNLEEAK/2 (magenta) AND 5336_YVGGQEHFAHLLILR/3
Intensity
60000
40000
20000
0
1800
2000
2200
2400
time

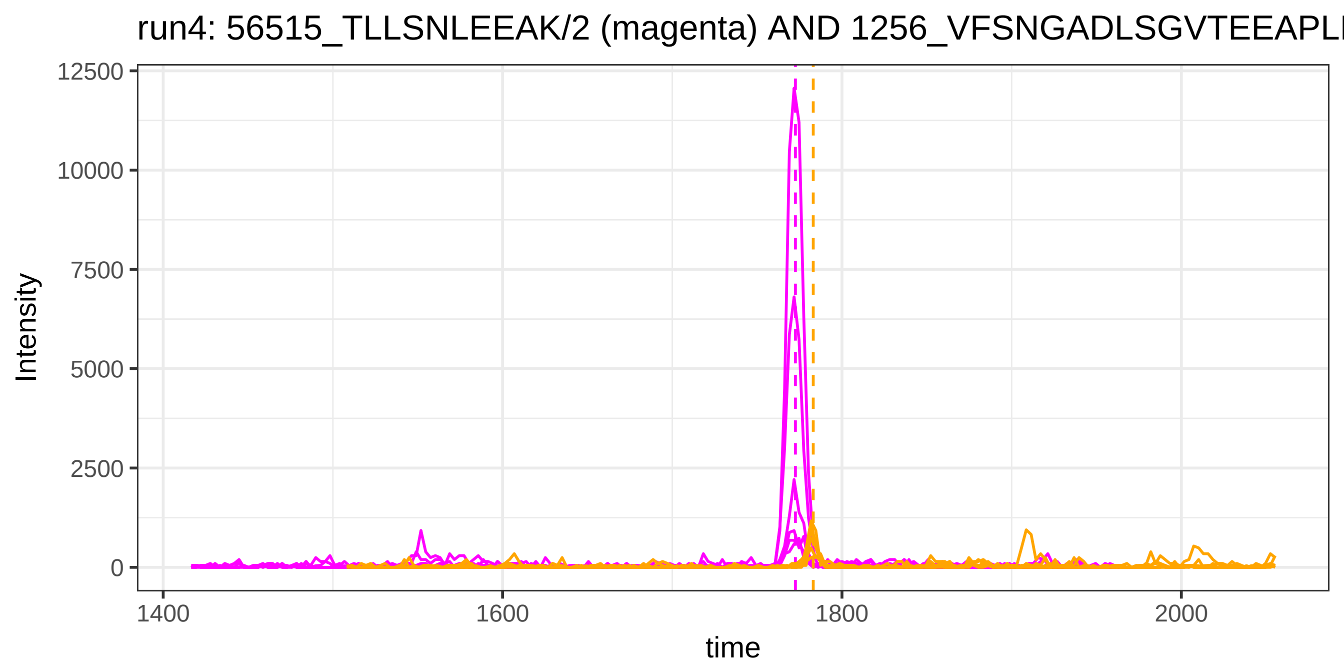

run4: 56515_TLLSNLEEAK/2 (magenta) AND 1256_VFSNGADLSGVTEEAPLI
Intensity
12500
10000
7500
5000
2500
0
1400
1600
1800
2000
time


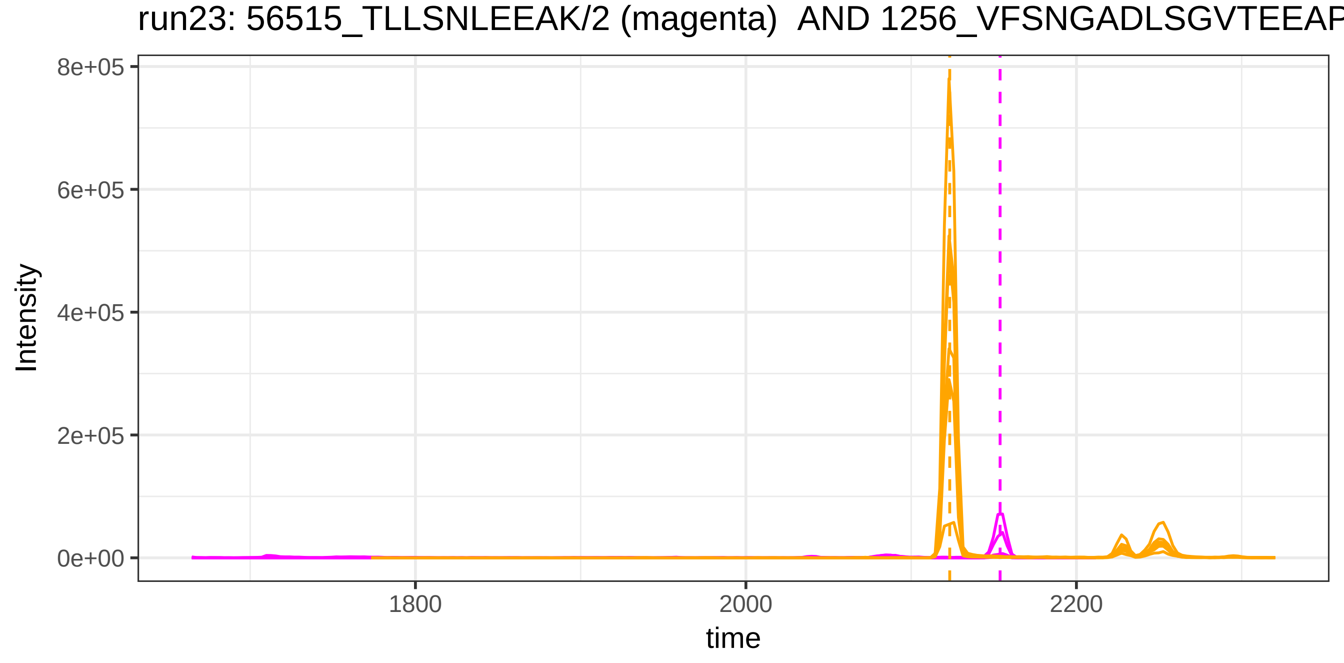

run23: 56515_TLLSNLEEAK/2 (magenta) AND 1256_VFSNGADLSGVTEEAP
Intensity
8e+05
6e+05
4e+05
2e+05
0e+00
1800
2000
2200
time

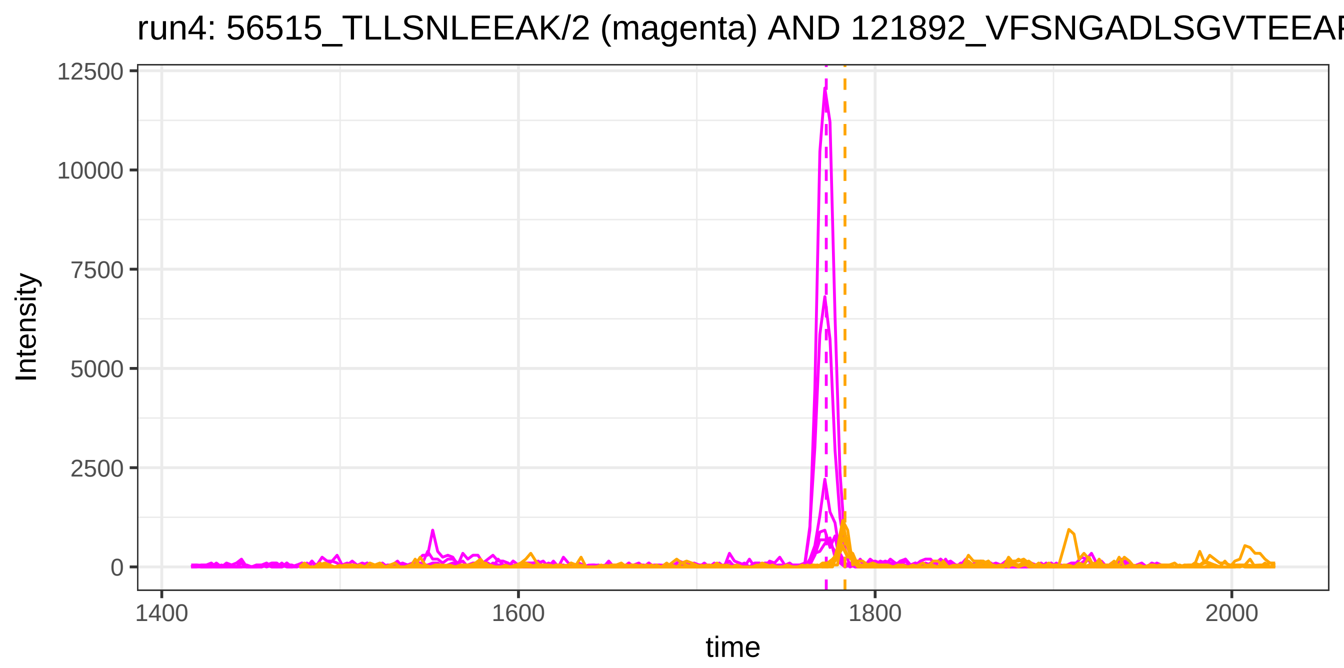
run4: 56515_TLLSNLEEAK/2 (magenta) AND 121892_VFSNGADLSGVTEEAF
Intensity
12500
10000
7500
5000
2500
0
1400
1600
1800
2000
time

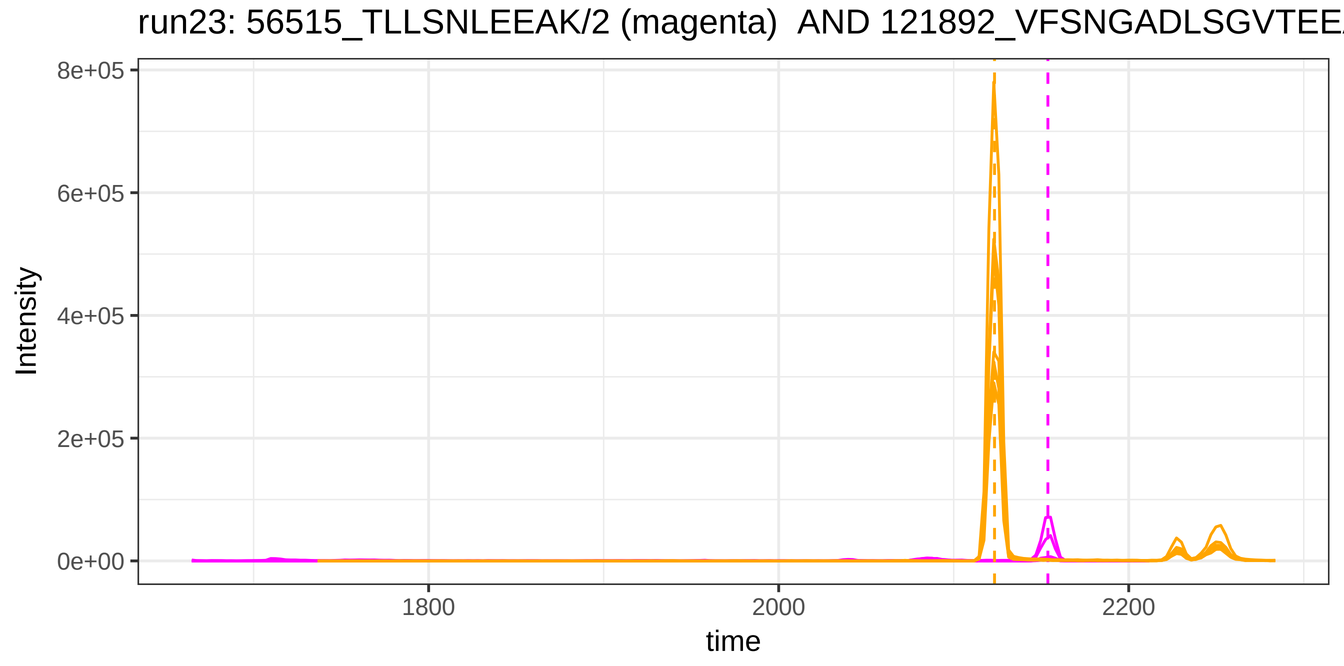
run23: 56515_TLLSNLEEAK/2 (magenta) AND 121892_VFSNGADLSGVTEE
Intensity
8e+05
6e+05
4e+05
2e+05
0e+00
1800
2000
2200
time

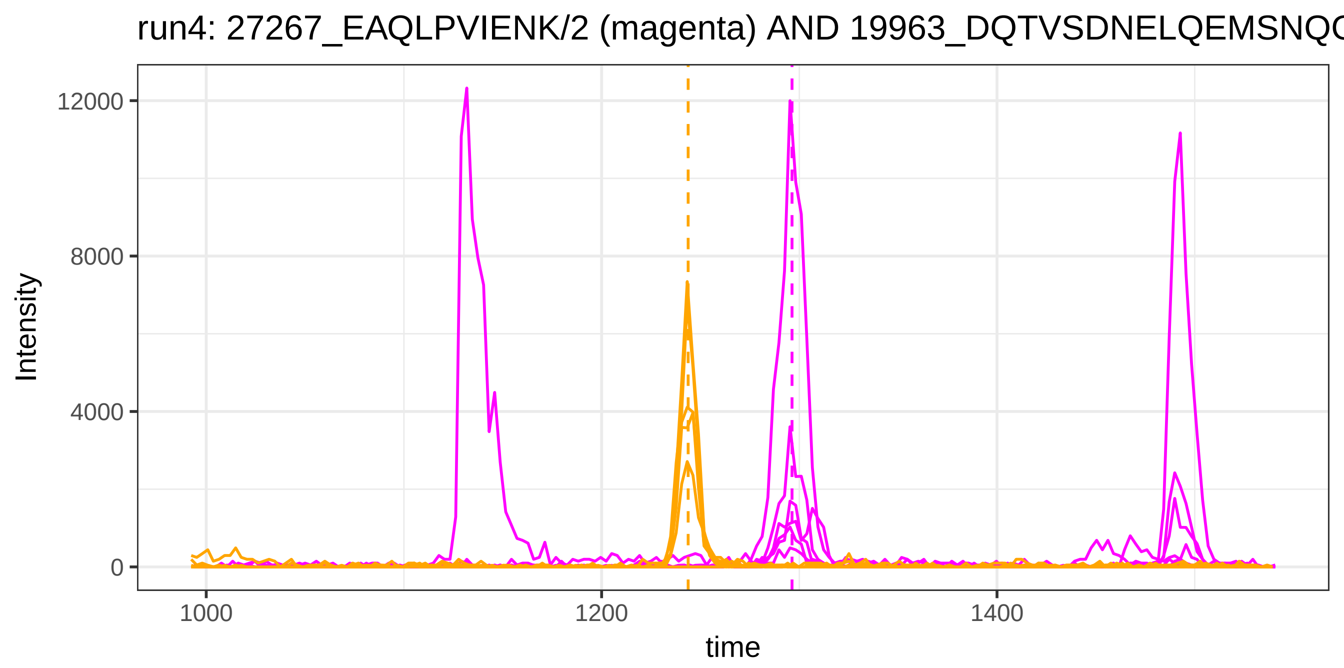
run4: 27267_EAQLPVIENK/2 (magenta) AND 19963_DQTVSDNELQEMSNQ
Intensity
12000
8000
4000
0
1000
1200
1400
time

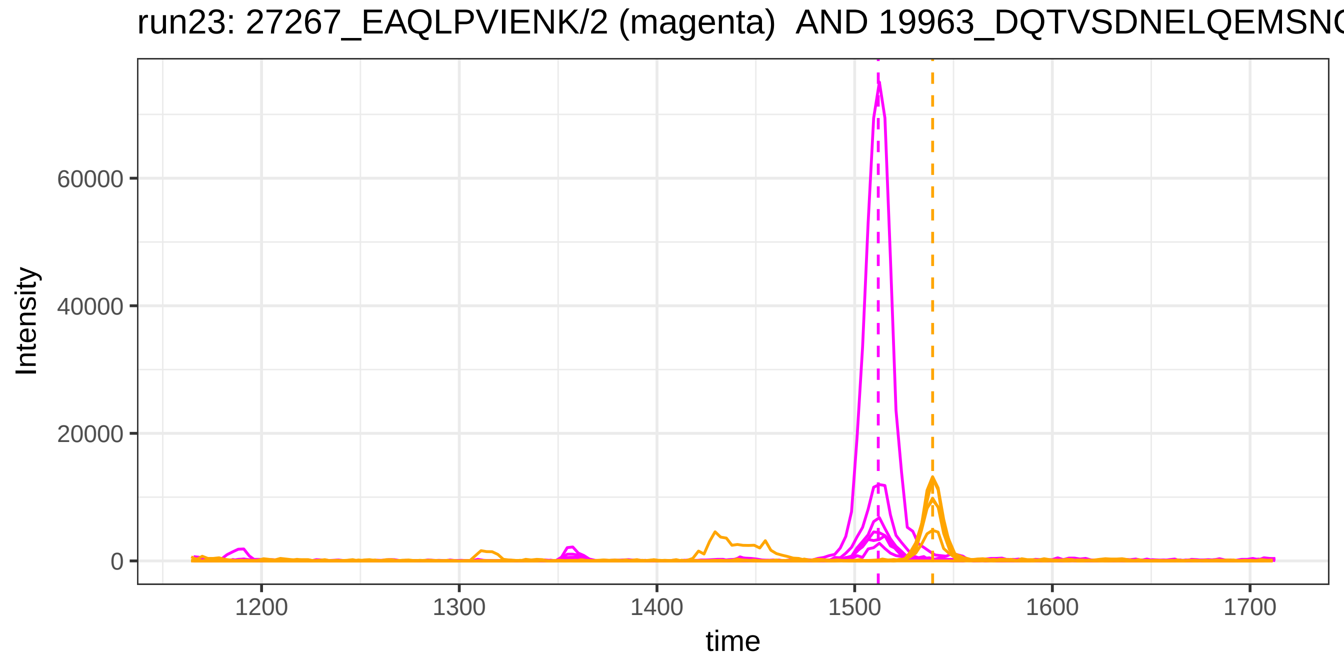
run23: 27267_EAQLPVIENK/2 (magenta) AND 19963_DQTVSDNELQEMSN
Intensity
60000
40000
20000
0
1200
1300
1400
1500
1600
1700
time

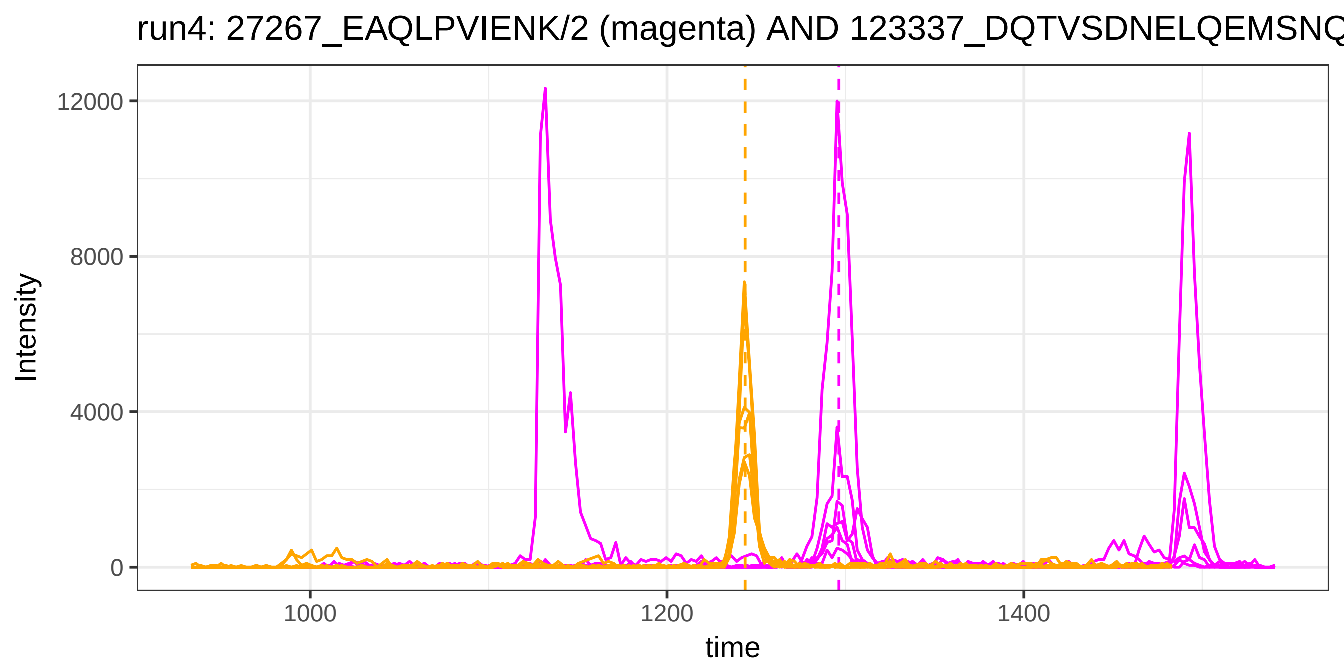
run4: 27267_EAQLPVIENK/2 (magenta) AND 123337_DQTVSDNELQEMSNQ
Intensity
12000
8000
4000
0
1000
1200
1400
time

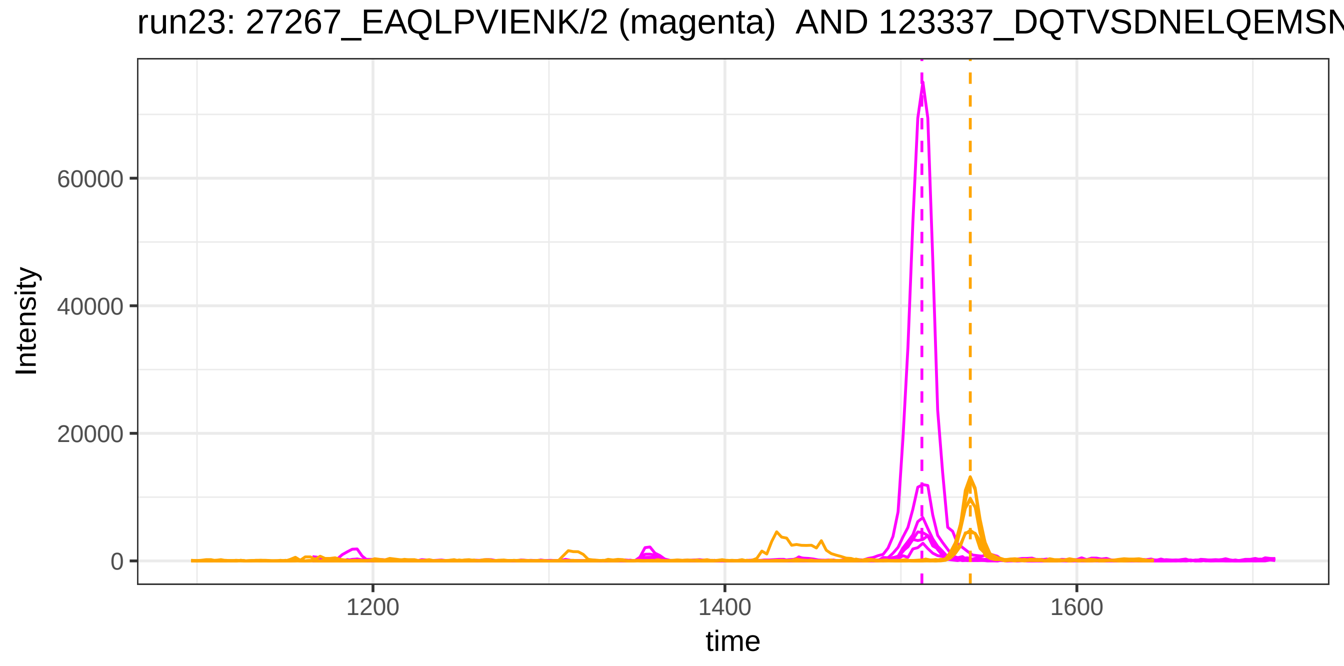
run23: 27267_EAQLPVIENK/2 (magenta)  AND 123337_DQTVSDNELQEMSN
Intensity
60000
40000
20000
0
1200
1400
1600
time

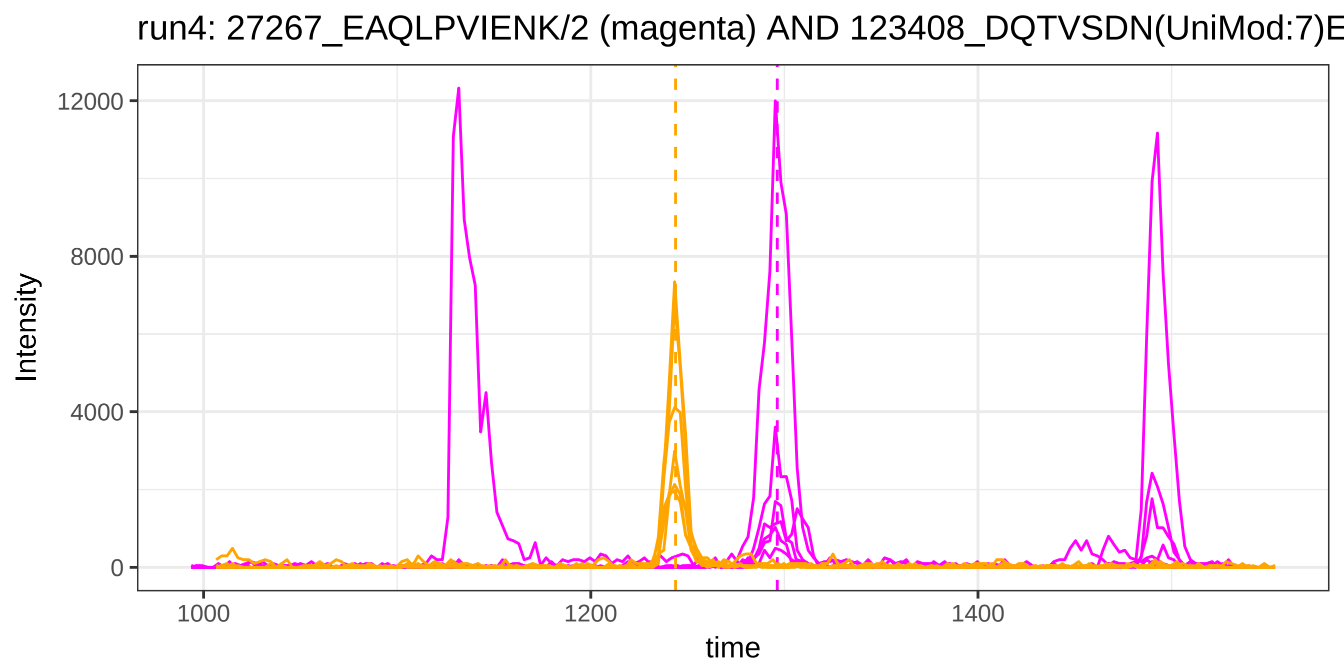
run4: 27267_EAQLPVIENK/2 (magenta) AND 123408_DQTVSDN(UniMod:7)E
Intensity
12000
8000
4000
0
1000
1200
1400
time

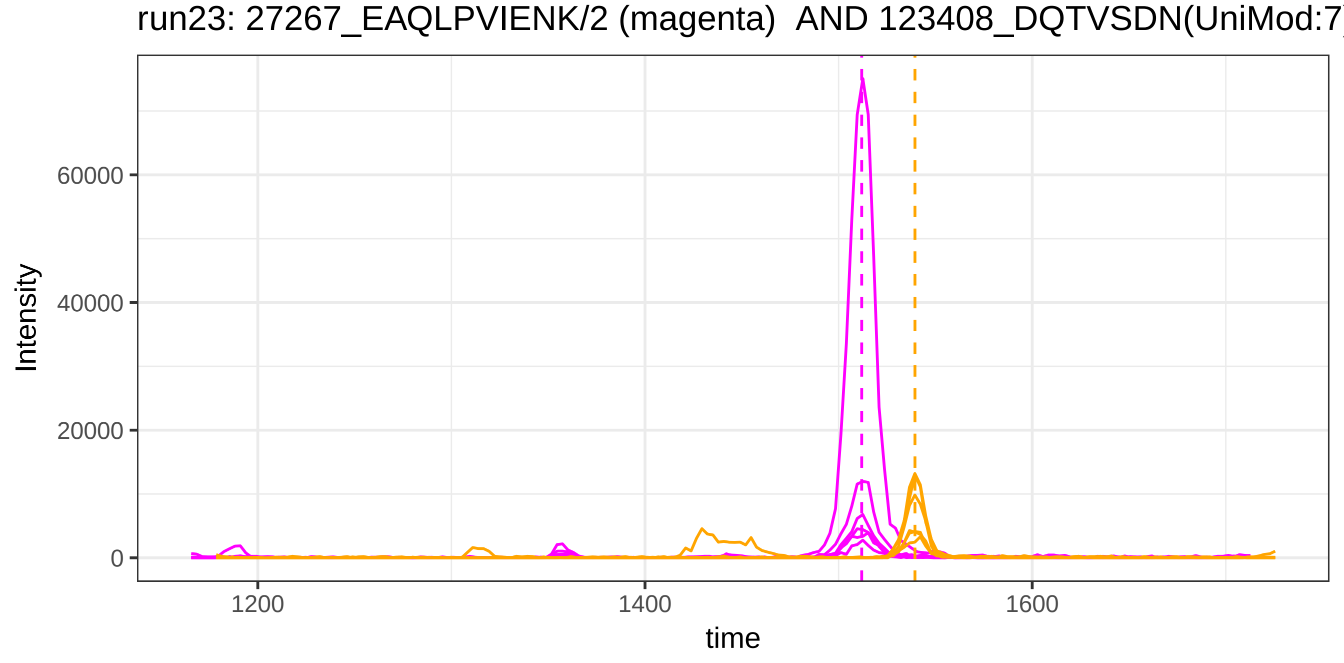
run23: 27267_EAQLPVIENK/2 (magenta)  AND 123408_DQTVSDN(UniMod:7)
Intensity
60000
40000
20000
0
1200
1400
1600
time

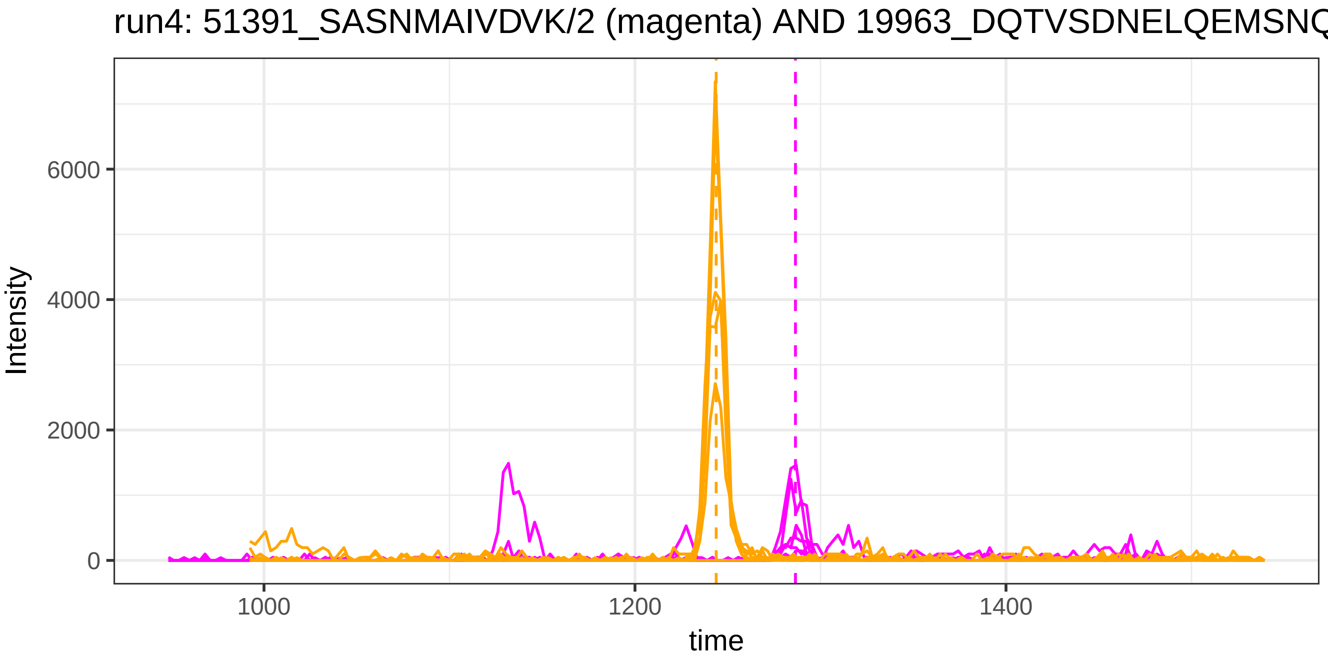
run4: 51391_SASNMAIVDVK/2 (magenta) AND 19963_DQTVSDNELQEMSNQ
Intensity
6000
4000
2000
0
1000
1200
1400
time

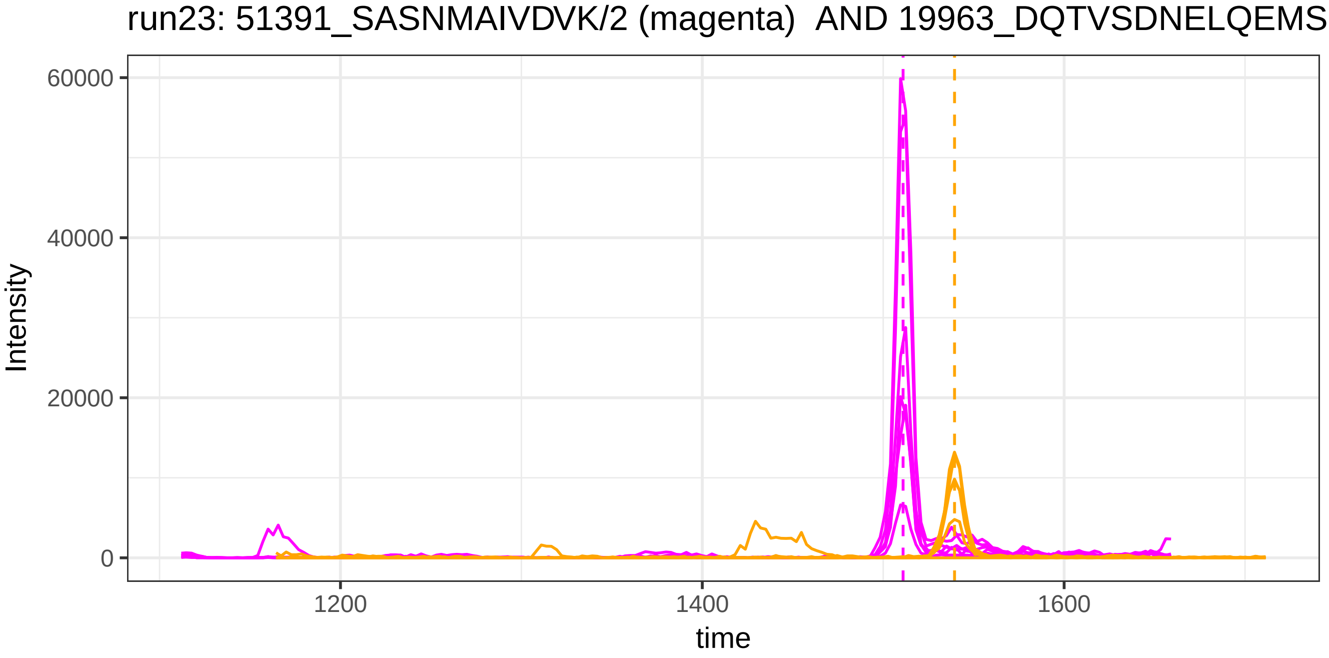
run23: 51391_SASNMAIVDVK/2 (magenta) AND 19963_DQTVSDNELQEMSN
Intensity
60000
40000
20000
0
1200
1400
1600
time

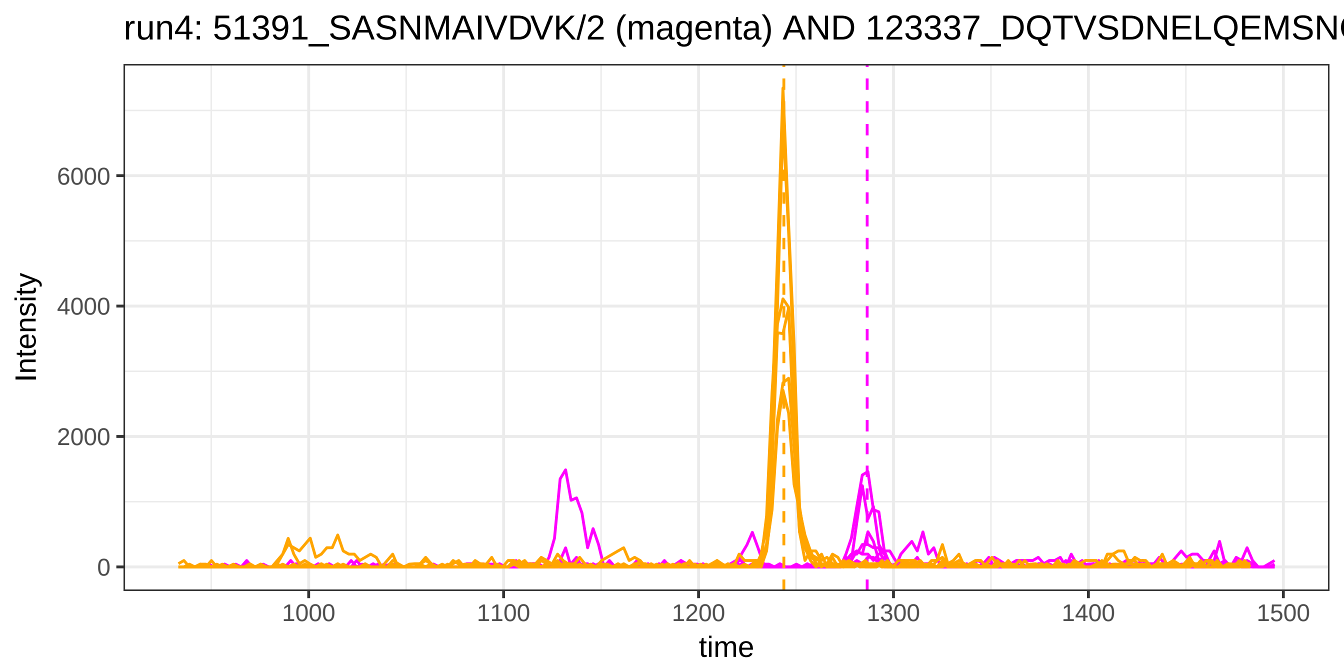
run4: 51391_SASNMAIVDVK/2 (magenta) AND 123337_DQTVSDNELQEMSN
Intensity
6000
4000
2000
0
1000
1100
1200
1300
1400
1500
time

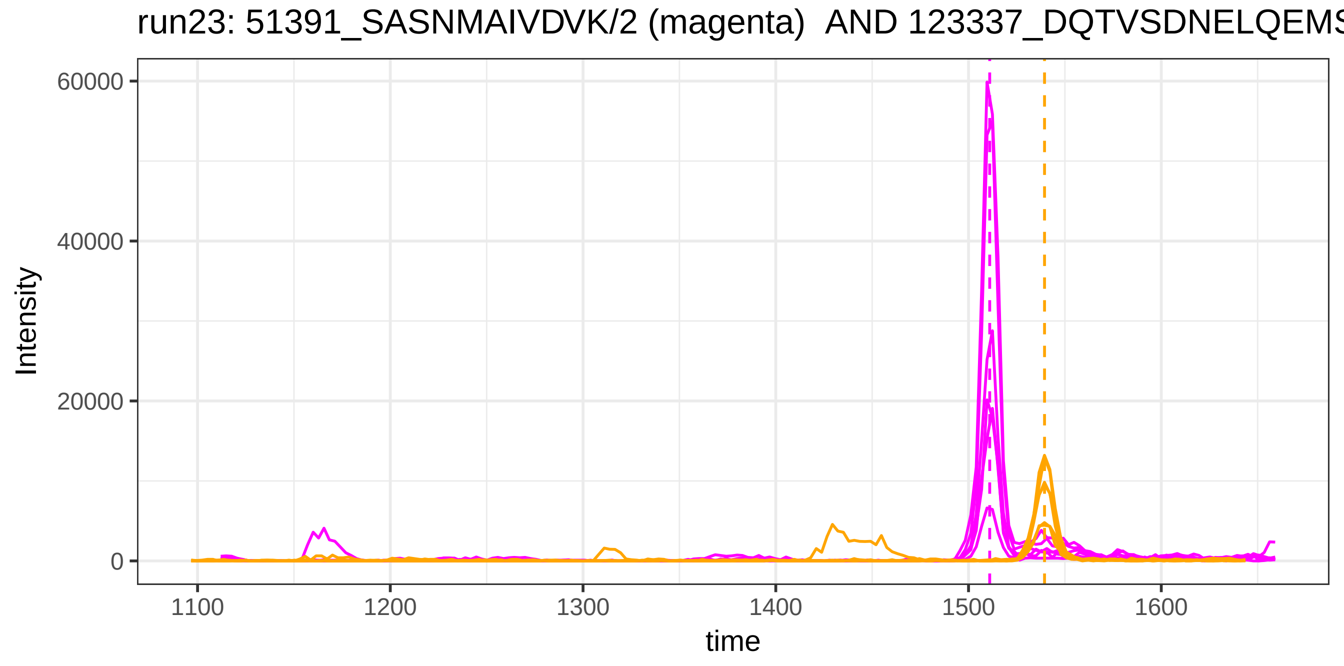
run23: 51391_SASNMAIVDVK/2 (magenta) AND 123337_DQTVSDNELQEMS
Intensity
60000
40000
20000
0
1100
1200
1300
1400
1500
1600
time

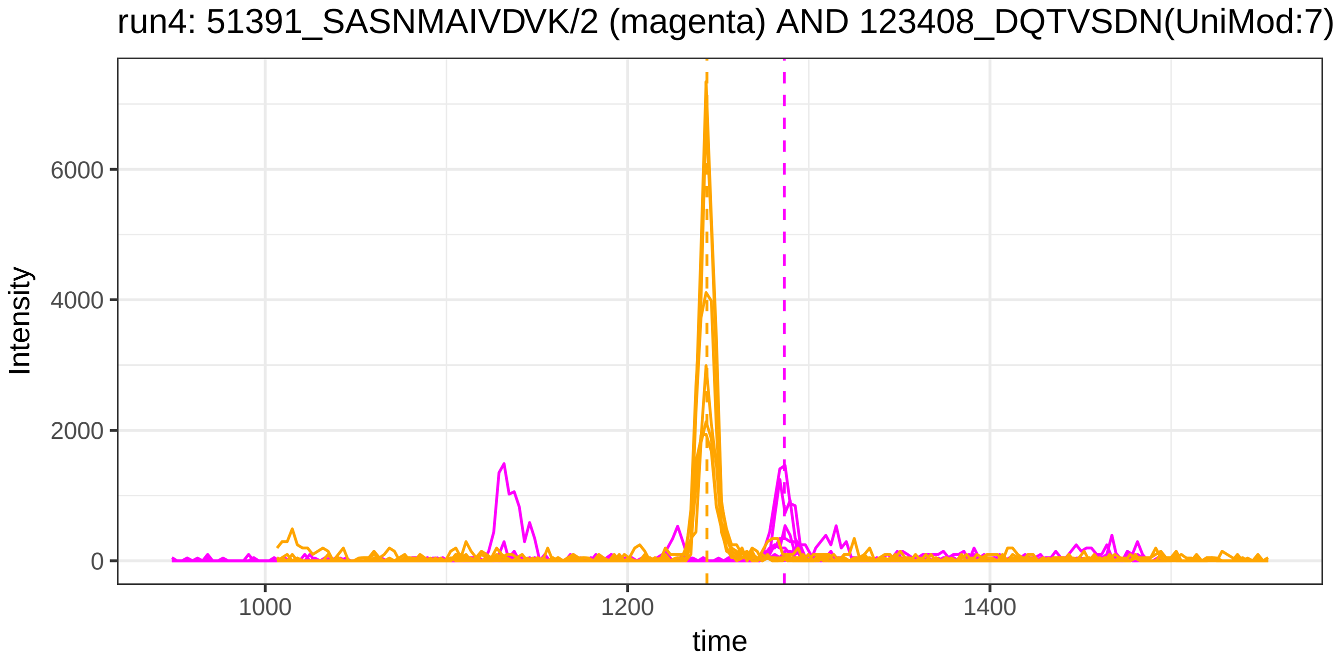
run4: 51391_SASNMAIVDVK/2 (magenta) AND 123408_DQTVSDN(UniMod:7)
Intensity
6000
4000
2000
0
1000
1200
1400
time

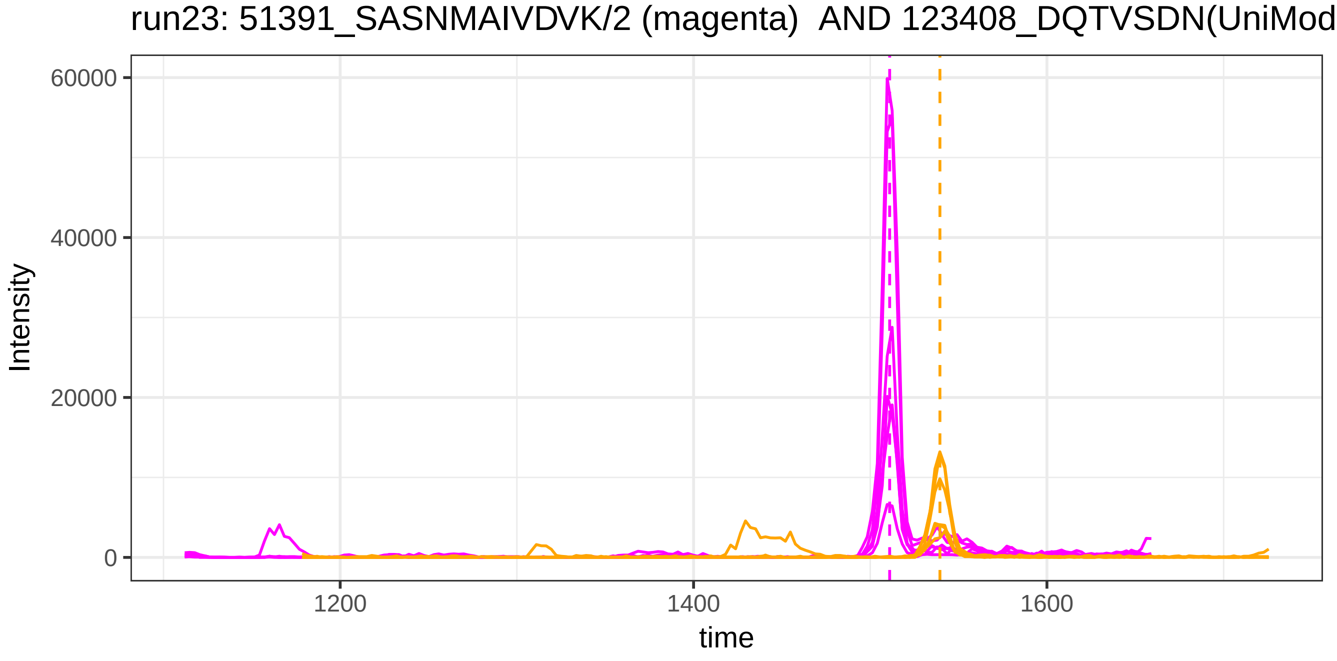
run23: 51391_SASNMAIVDVK/2 (magenta) AND 123408_DQTVSDN(UniMod
Intensity
60000
40000
20000
0
1200
1400
1600
time

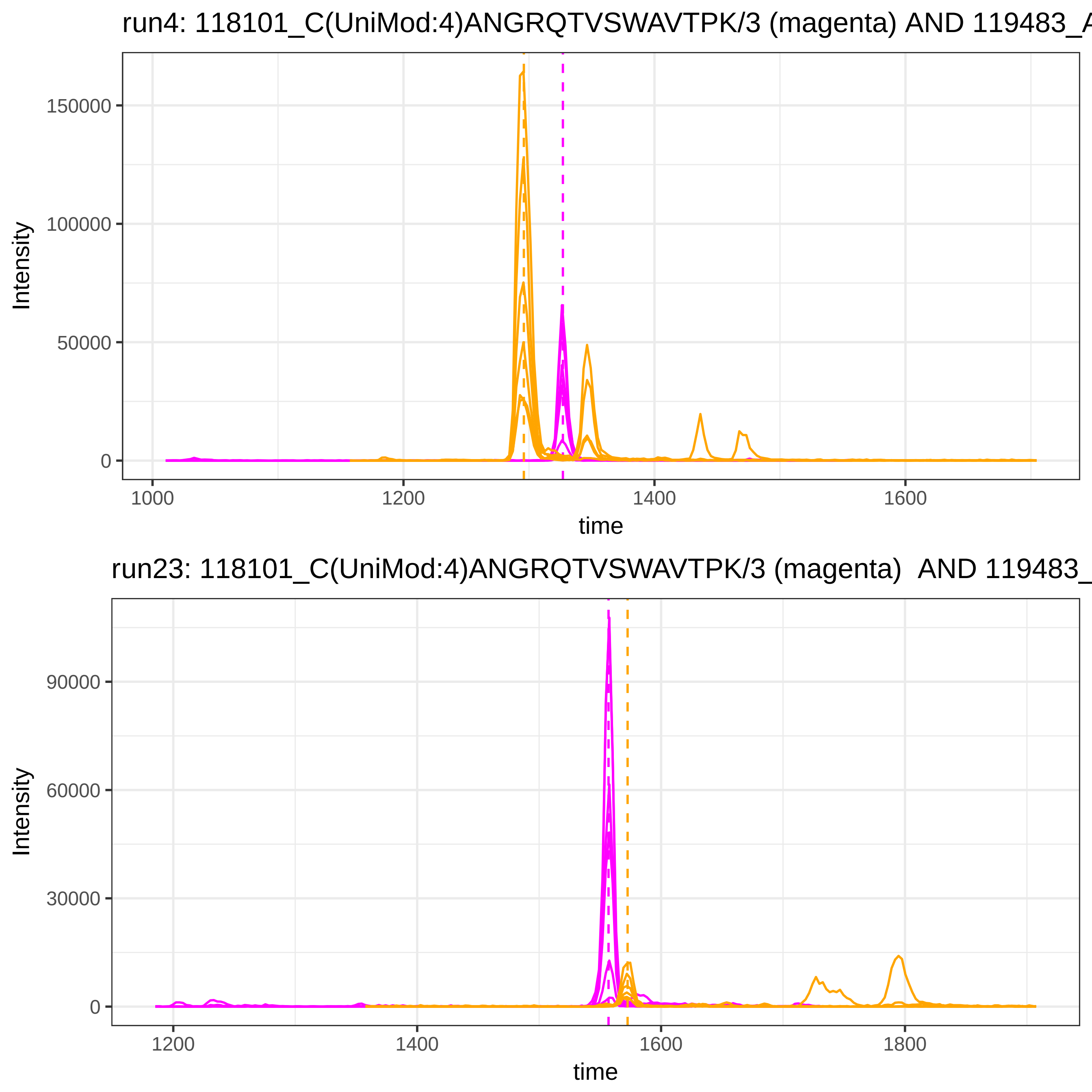

run4: 118101_C(UniMod:4)ANGRQTVSWAVTPK/3 (magenta) AND 119483_A
Intensity
150000
100000
50000
0
1000
1200
1400
1600
time
run23: 118101_C(UniMod:4)ANGRQTVSWAVTPK/3 (magenta) AND 119483_
Intensity
90000
60000
30000
0
1200
1400
1600
1800
time

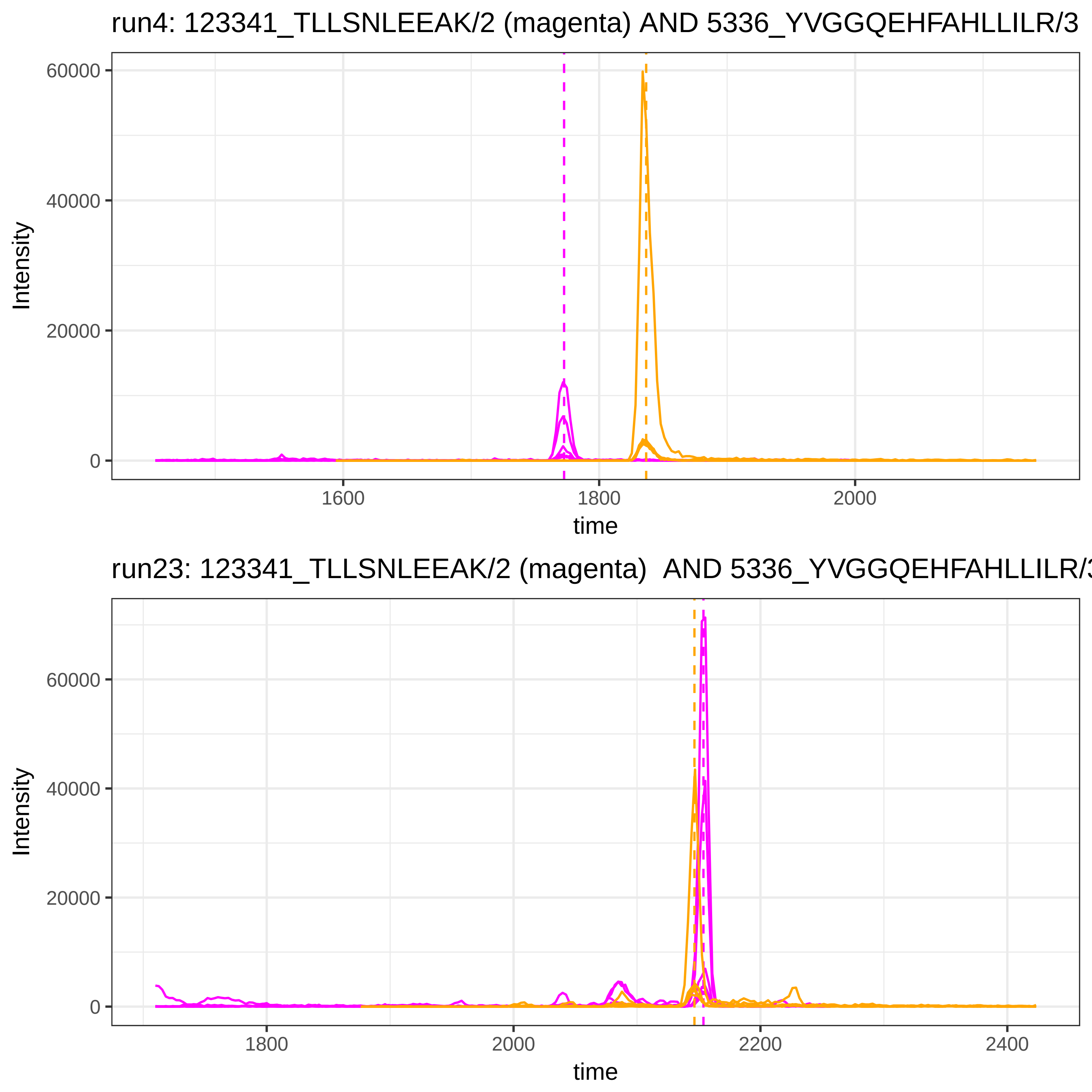

run4: 123341_TLLSNLEEAK/2 (magenta) AND 5336_YVGGQEHFAHLLILR/3
Intensity
60000
40000
20000
0
1600
1800
2000
time
run23: 123341_TLLSNLEEAK/2 (magenta) AND 5336_YVGGQEHFAHLLILR/3
Intensity
60000
40000
20000
0
1800
2000
2200
2400
time

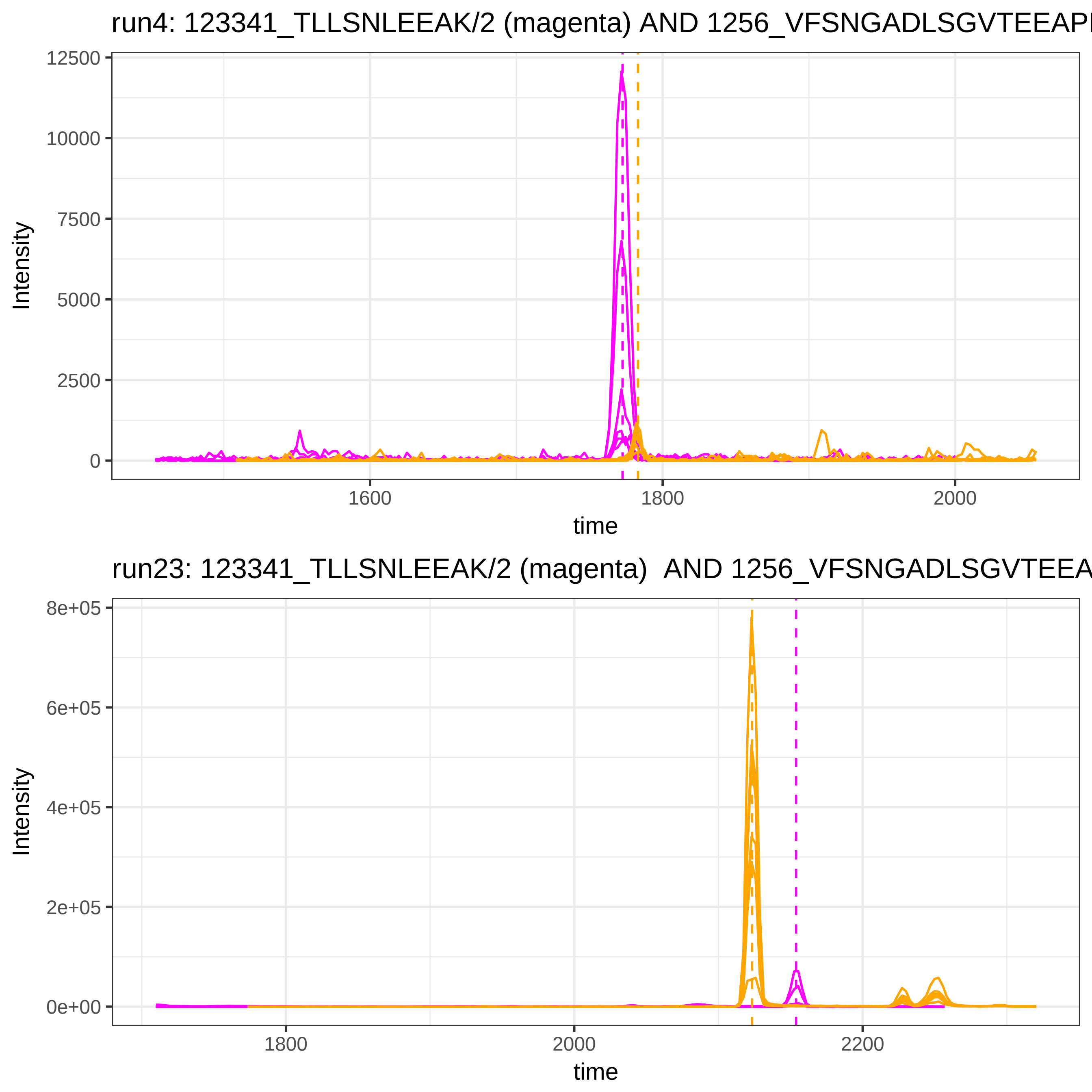

run4: 123341_TLLSNLEEAK/2 (magenta) AND 1256_VFSNGADLSGVTEEAP
Intensity
12500
10000
7500
5000
2500
0
1600
1800
2000
time
run23: 123341_TLLSNLEEAK/2 (magenta) AND 1256_VFSNGADLSGVTEEA
Intensity
8e+05
6e+05
4e+05
2e+05
0e+00
1800
2000
2200
time

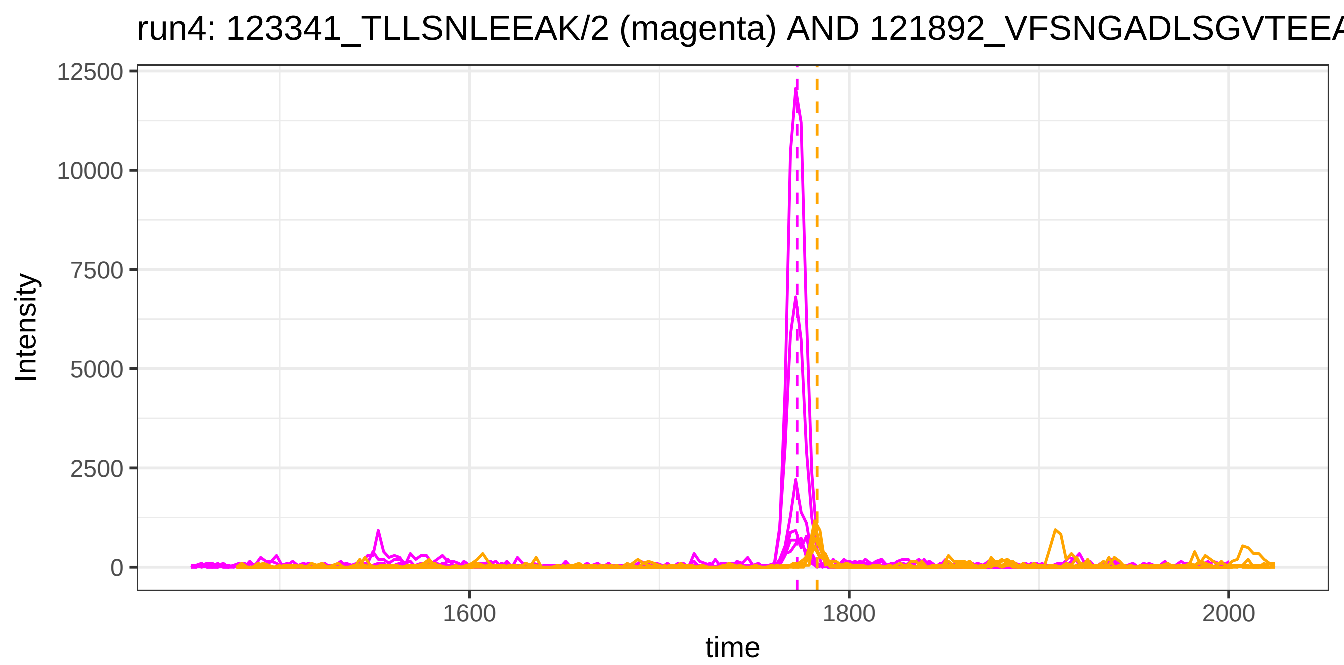

run4: 123341_TLLSNLEEAK/2 (magenta) AND 121892_VFSNGADLSGVTEEA
Intensity
12500
10000
7500
5000
2500
0
1600
1800
2000
time


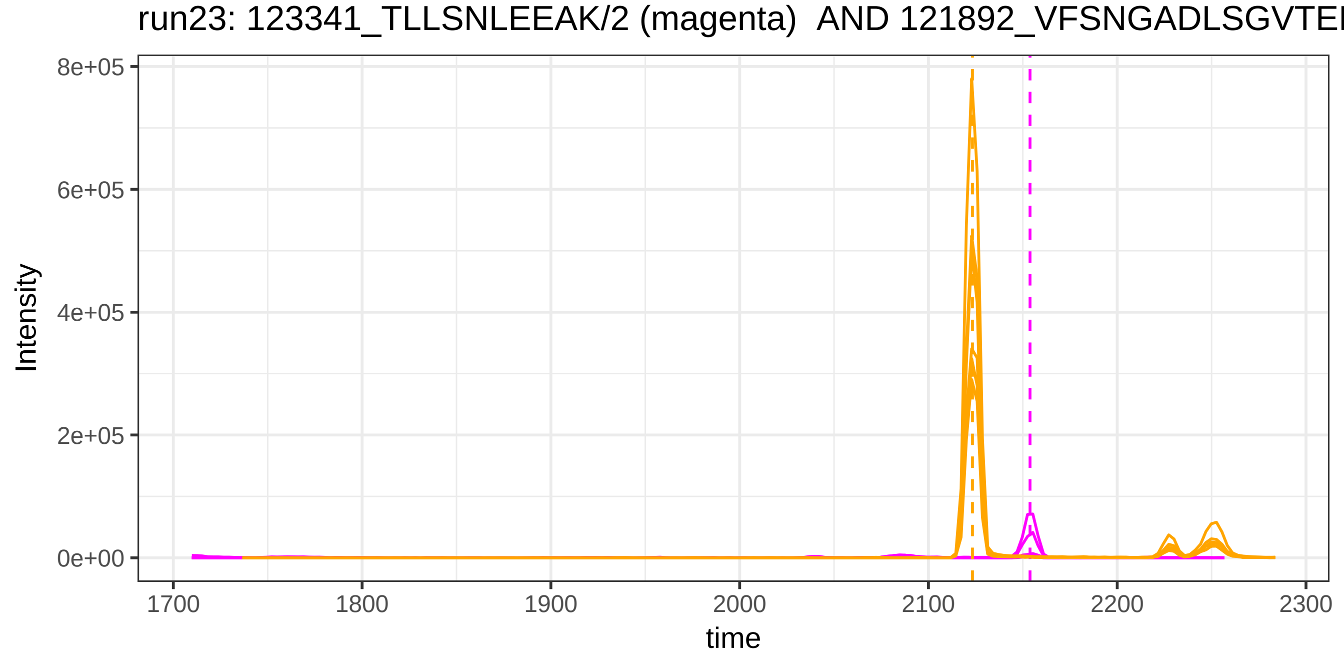

run23: 123341_TLLSNLEEAK/2 (magenta) AND 121892_VFSNGADLSGVTEE
Intensity
8e+05
6e+05
4e+05
2e+05
0e+00
1700
1800
1900
2000
2100
2200
2300
time

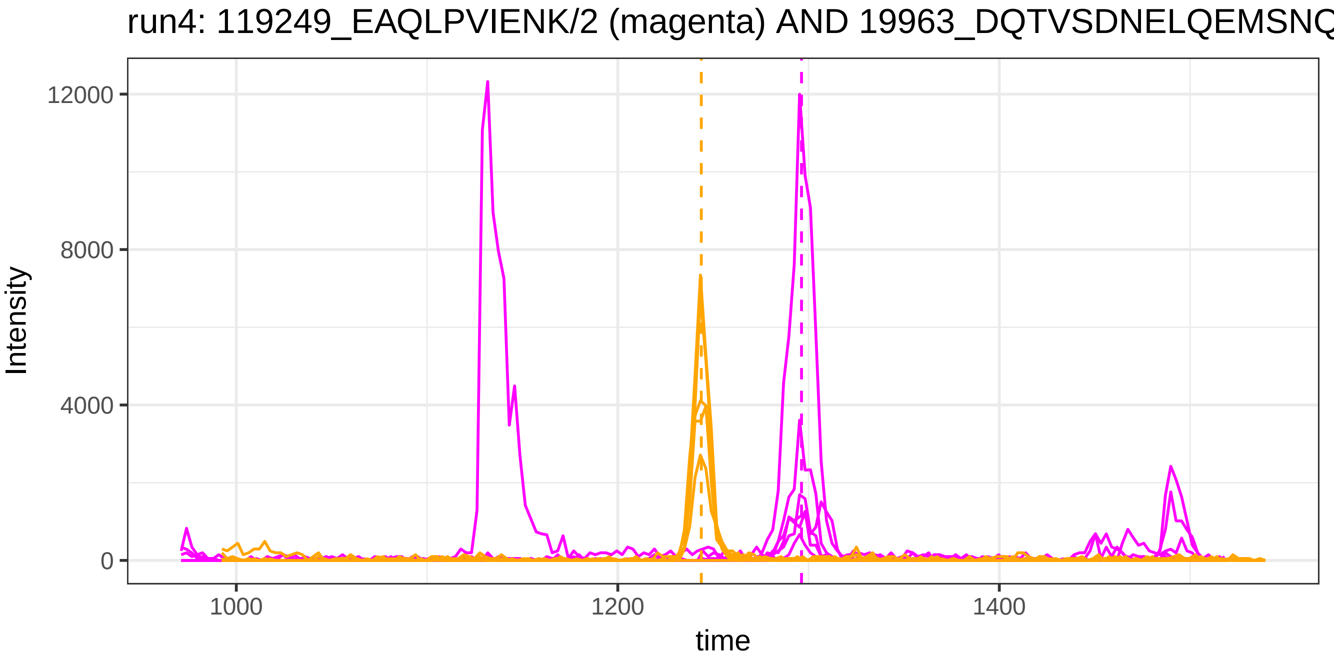

run4: 119249_EAQLPVIENK/2 (magenta) AND 19963_DQTVSDNELQEMSNQ
Intensity
12000
8000
4000
0
1000
1200
1400
time


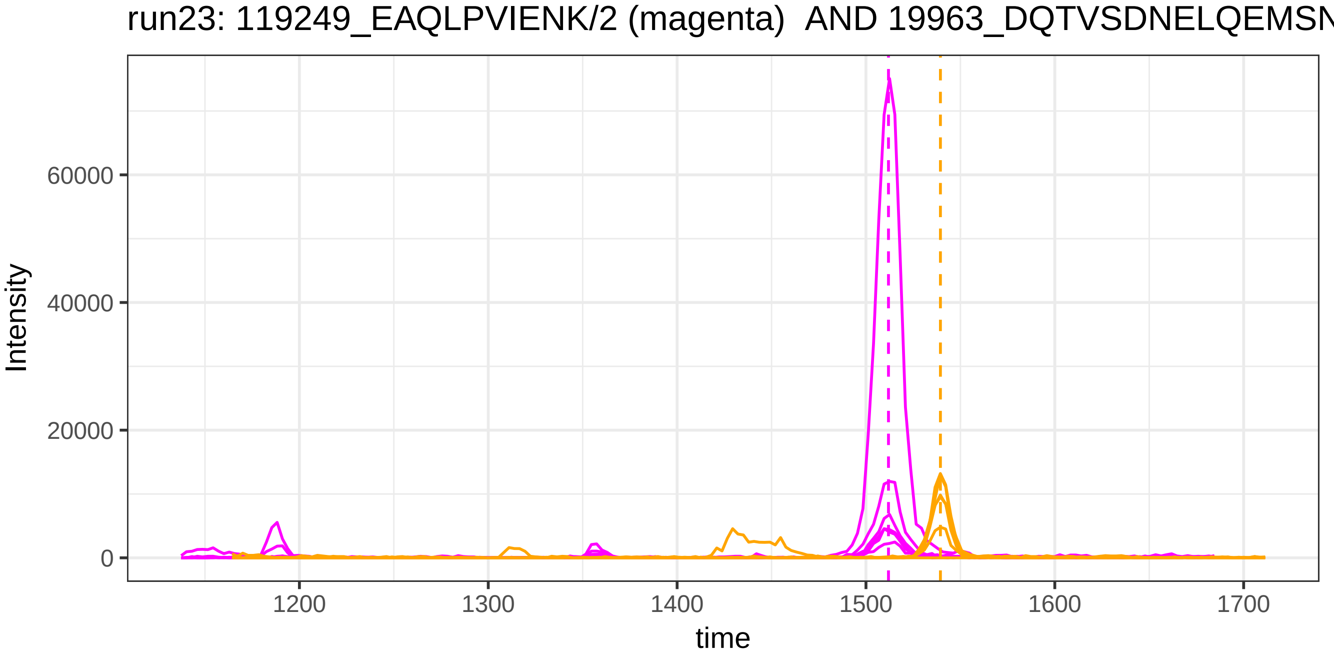

run23: 119249_EAQLPVIENK/2 (magenta) AND 19963_DQTVSDNELQEMSN
Intensity
60000
40000
20000
0
1200
1300
1400
1500
1600
1700
time

run4: 119249_EAQLPVIENK/2 (magenta) AND 123337_DQTVSDNELQEMSN

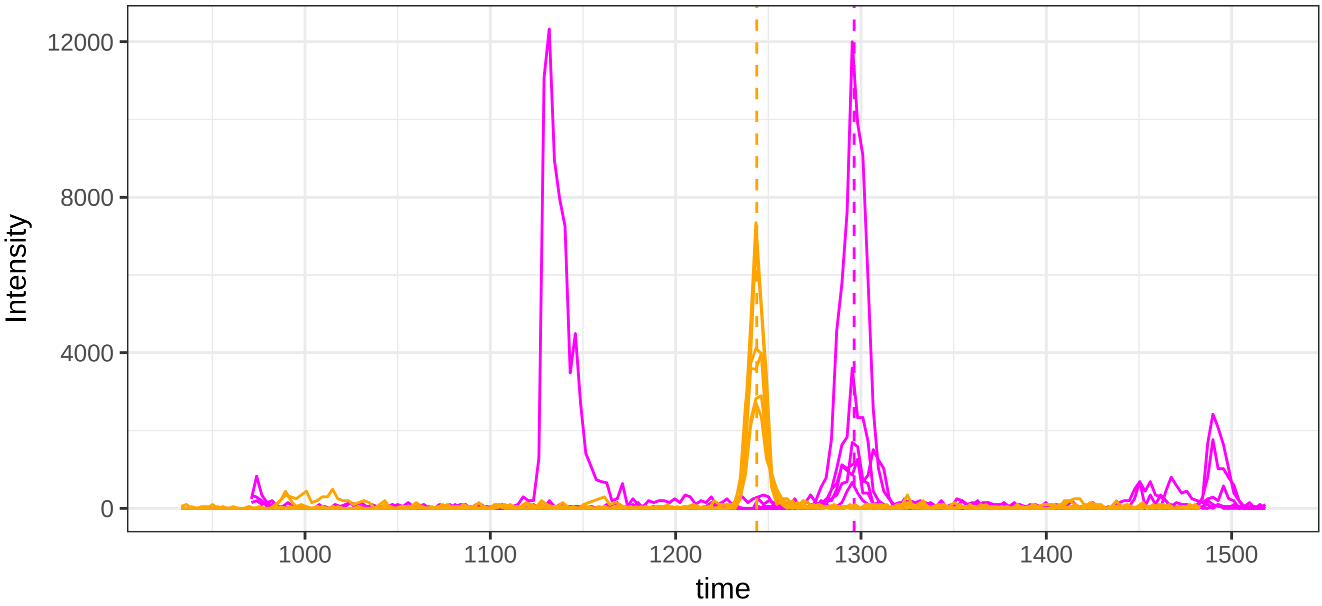


run23: 119249_EAQLPVIENK/2 (magenta)  AND 123337_DQTVSDNELQEMS

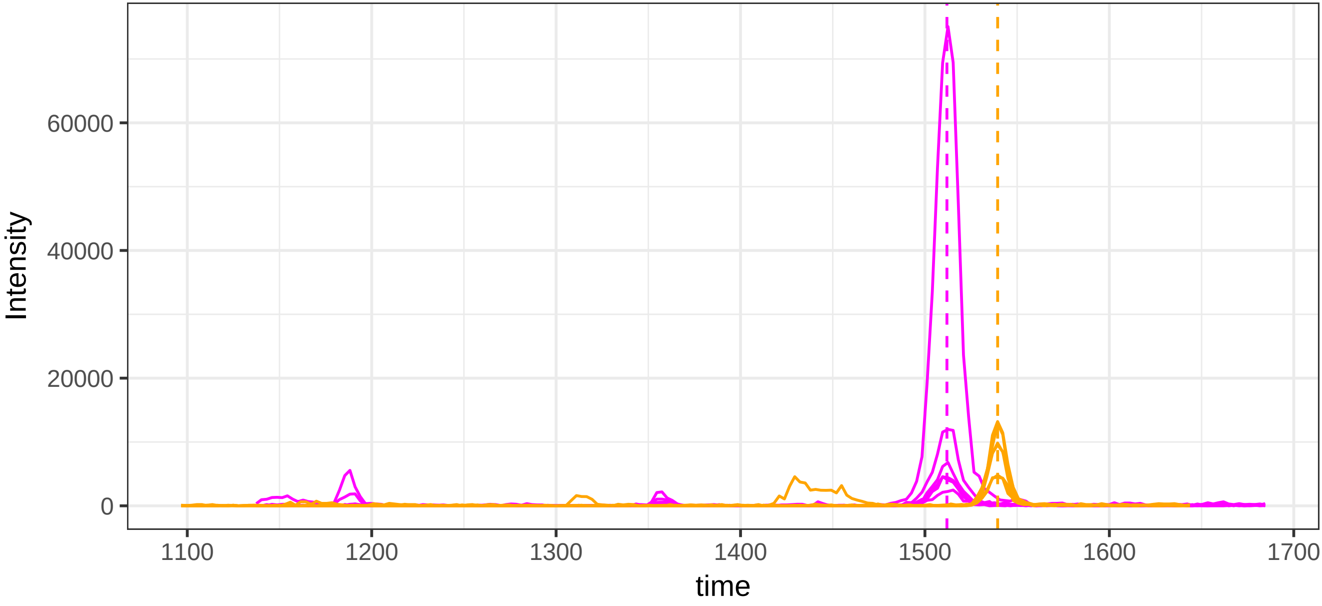

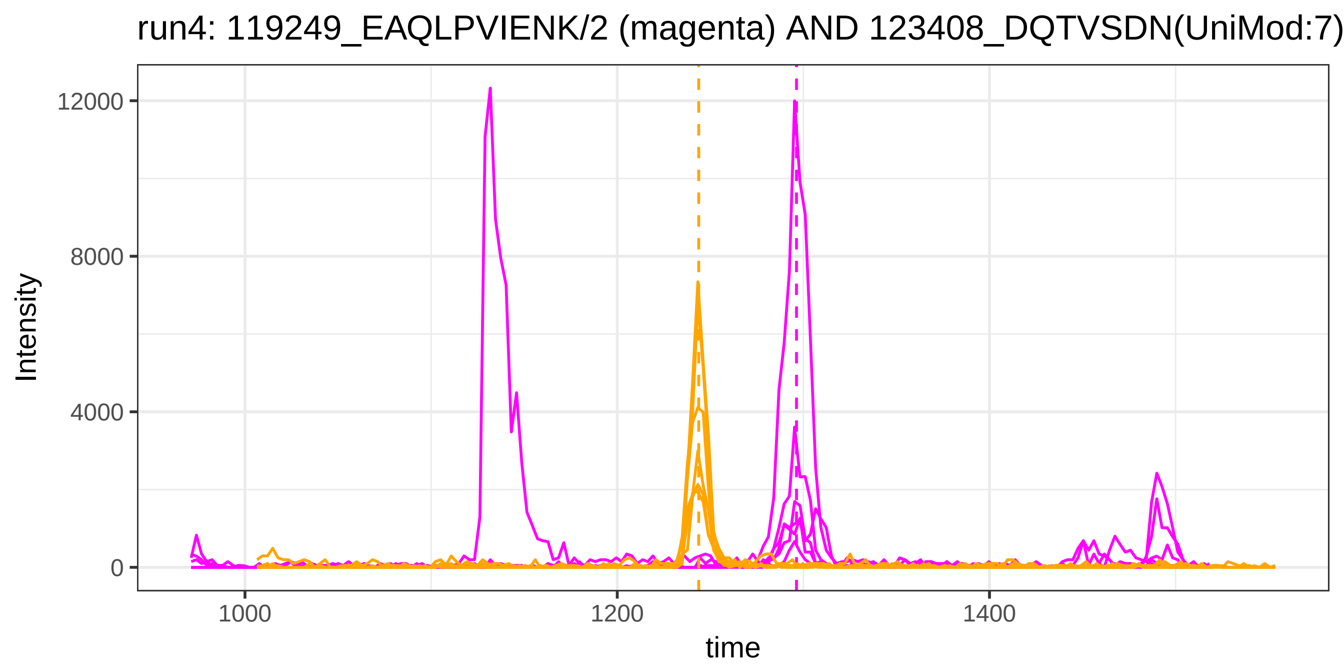
run4: 119249_EAQLPVIENK/2 (magenta) AND 123408_DQTVSDN(UniMod:7)
Intensity
12000
8000
4000
0
1000
1200
1400
time

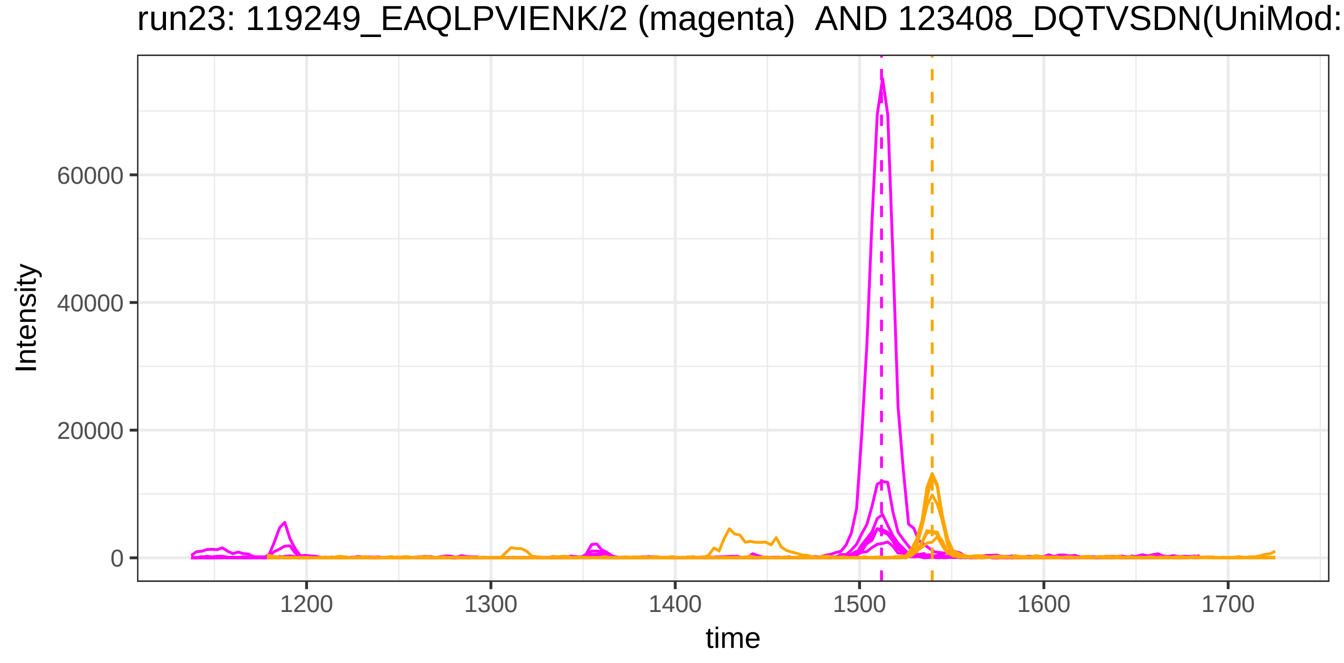
run23: 119249_EAQLPVIENK/2 (magenta) AND 123408_DQTVSDN(UniMod:
Intensity
60000
40000
20000
0
1200
1300
1400
1500
1600
1700
time

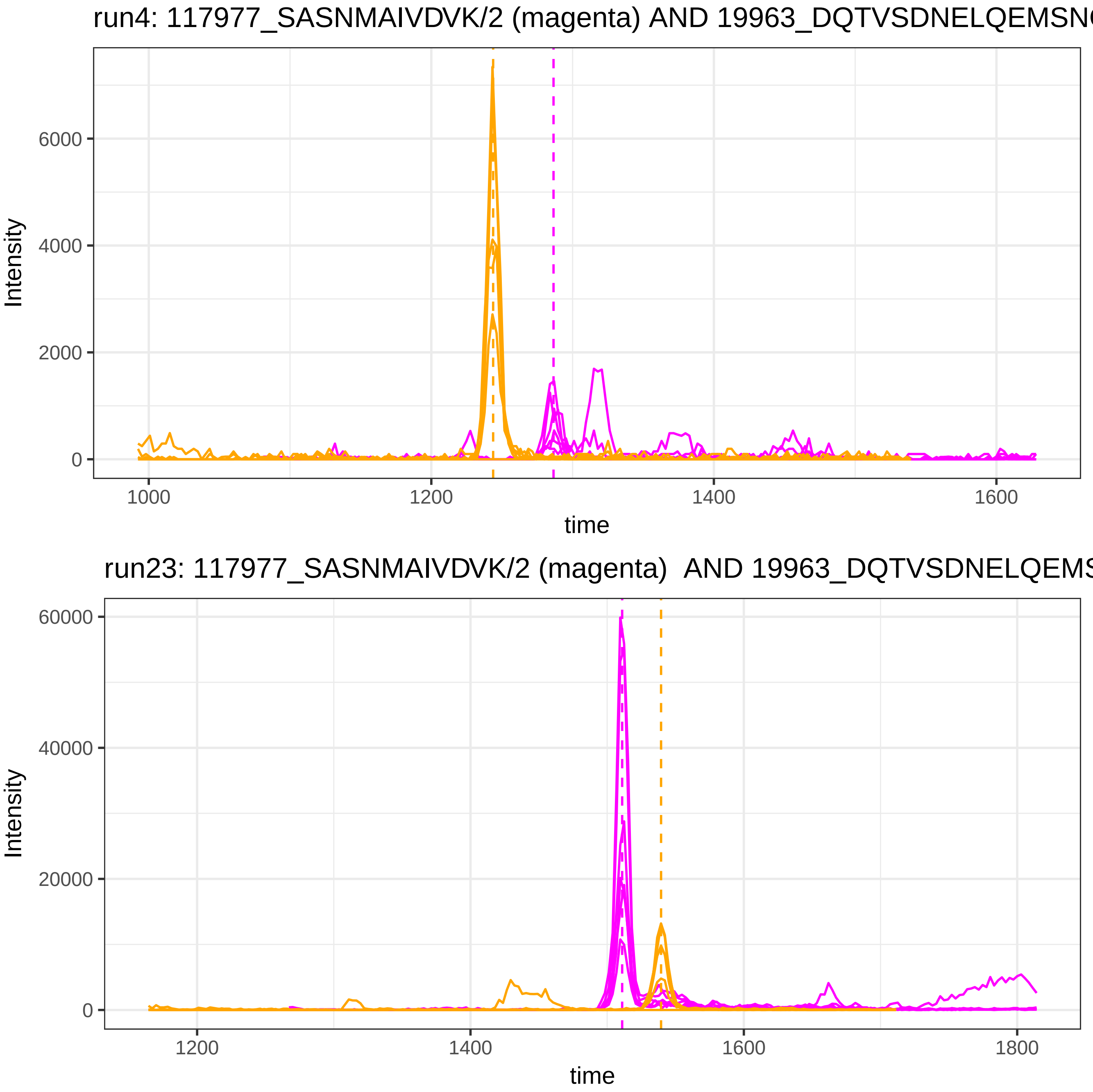

run4: 117977_SASNMAIVDVK/2 (magenta) AND 19963_DQTVSDNELQEMSN
Intensity
6000
4000
2000
0
1000
1200
1400
1600
time
run23: 117977_SASNMAIVDVK/2 (magenta) AND 19963_DQTVSDNELQEMS
Intensity
60000
40000
20000
0
1200
1400
1600
1800
time

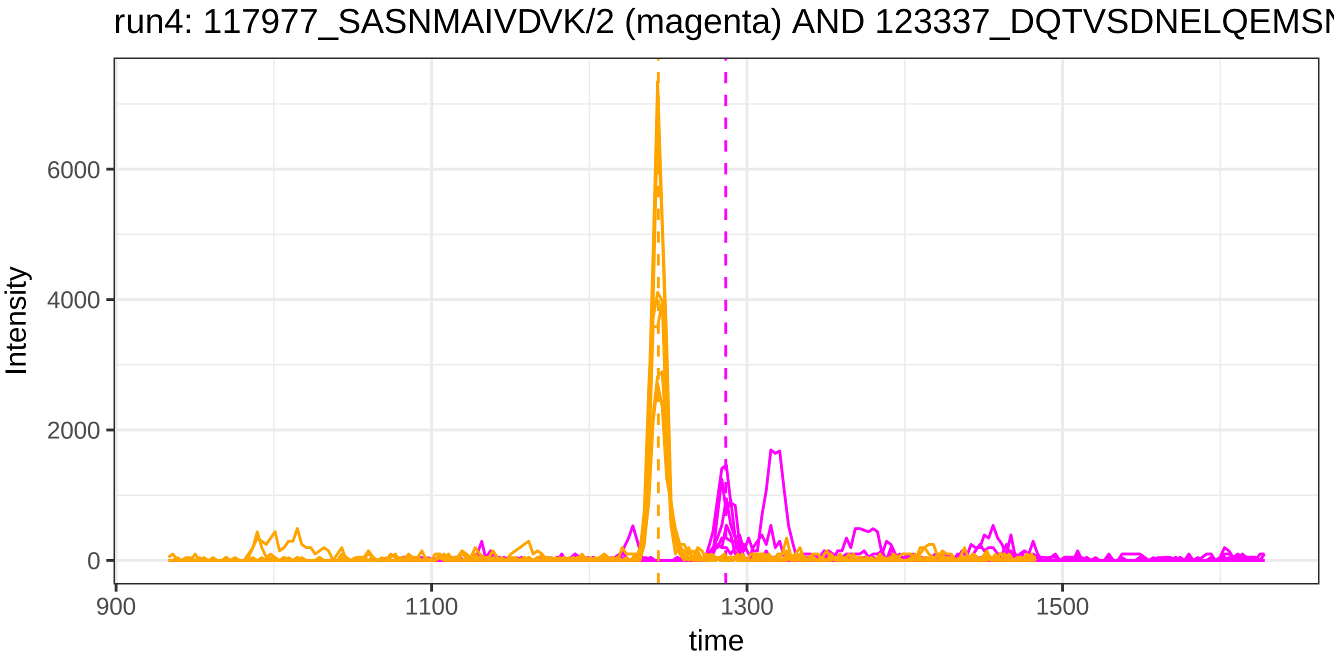
run4: 117977_SASNMAIVDVK/2 (magenta) AND 123337_DQTVSDNELQEMSN
Intensity
6000
4000
2000
0
900
1100
1300
1500
time

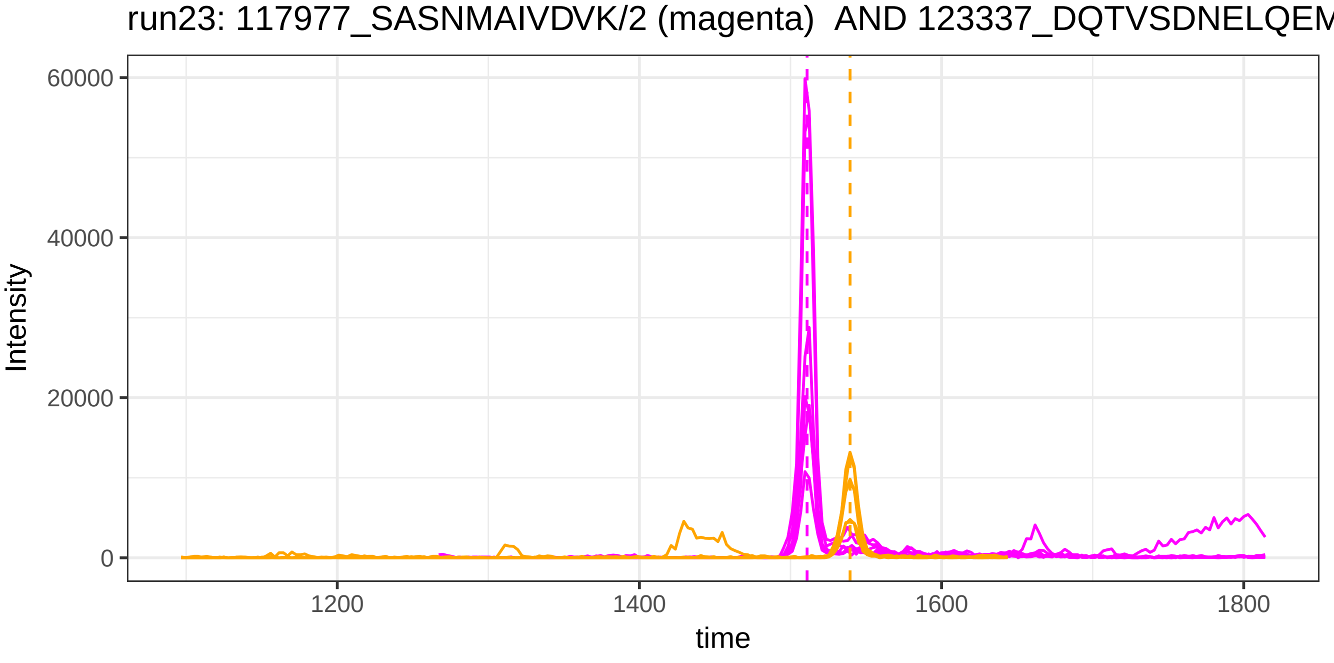
run23: 117977_SASNMAIVDVK/2 (magenta) AND 123337_DQTVSDNELQEM
Intensity
60000
40000
20000
0
1200
1400
1600
1800
time

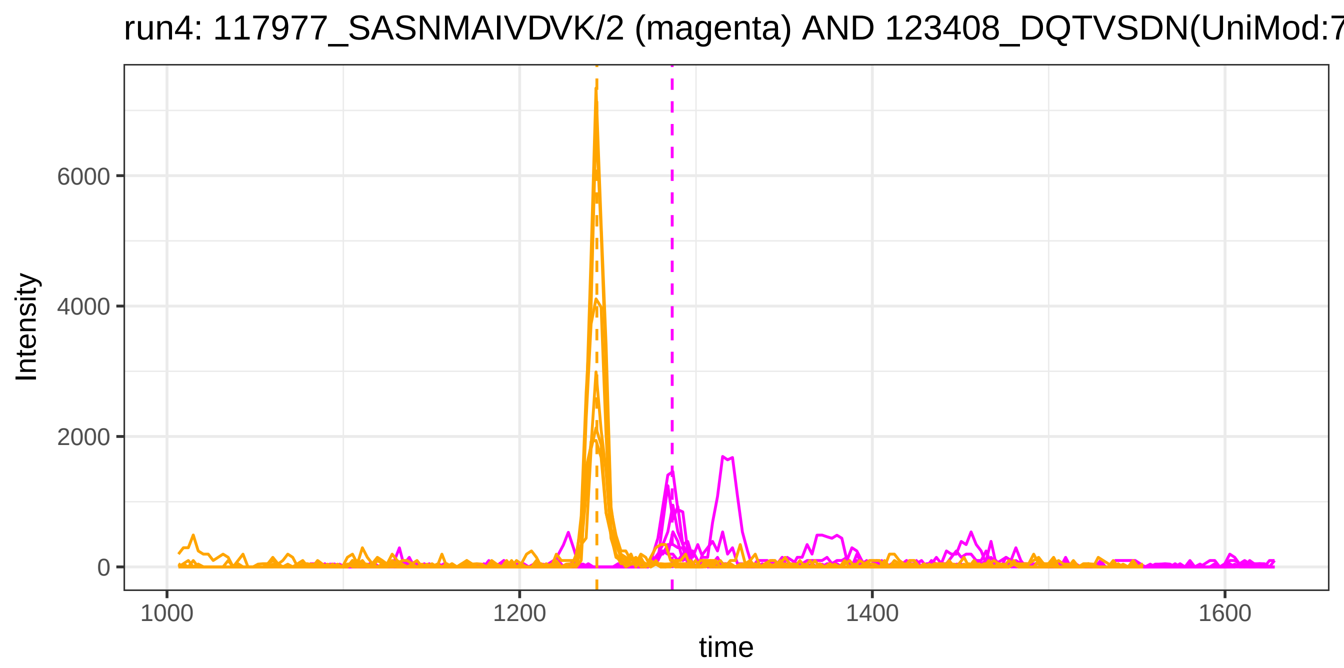
run4: 117977_SASNMAIVDVK/2 (magenta) AND 123408_DQTVSDN(UniMod:7
Intensity
6000
4000
2000
0
1000
1200
1400
1600
time

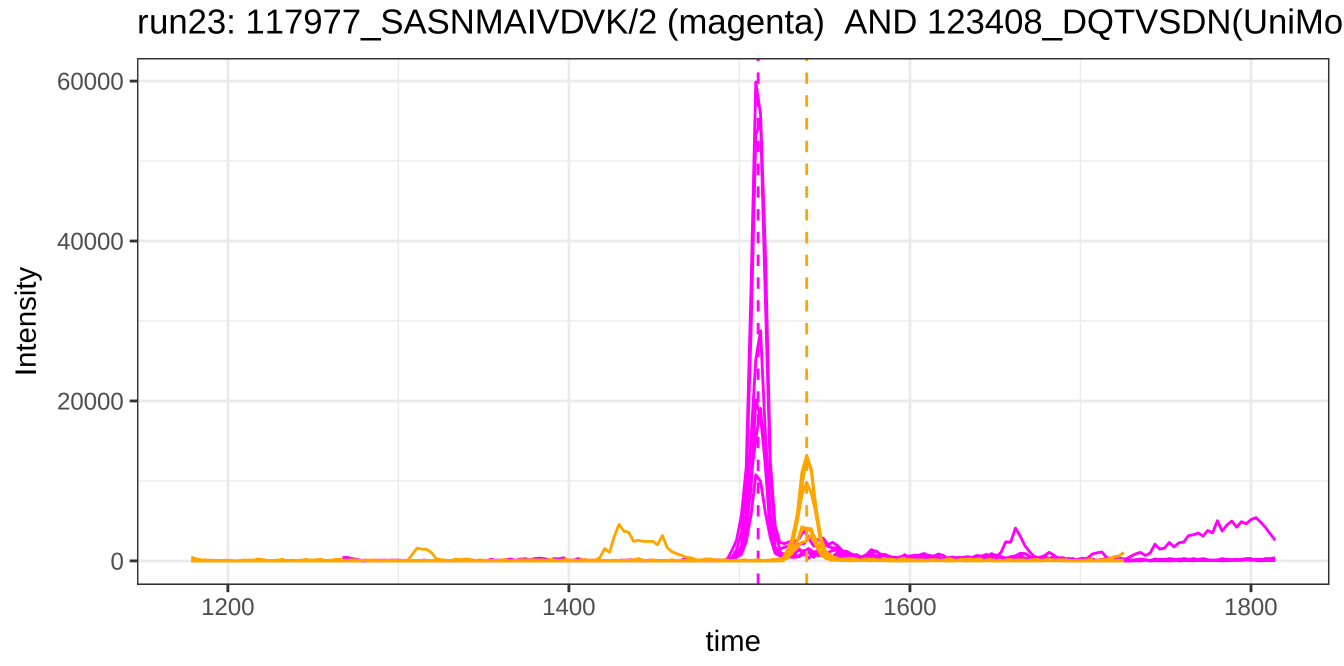
run23: 117977_SASNMAIVDVK/2 (magenta) AND 123408_DQTVSDN(UniMo
Intensity
60000
40000
20000
0
1200
1400
1600
1800
time

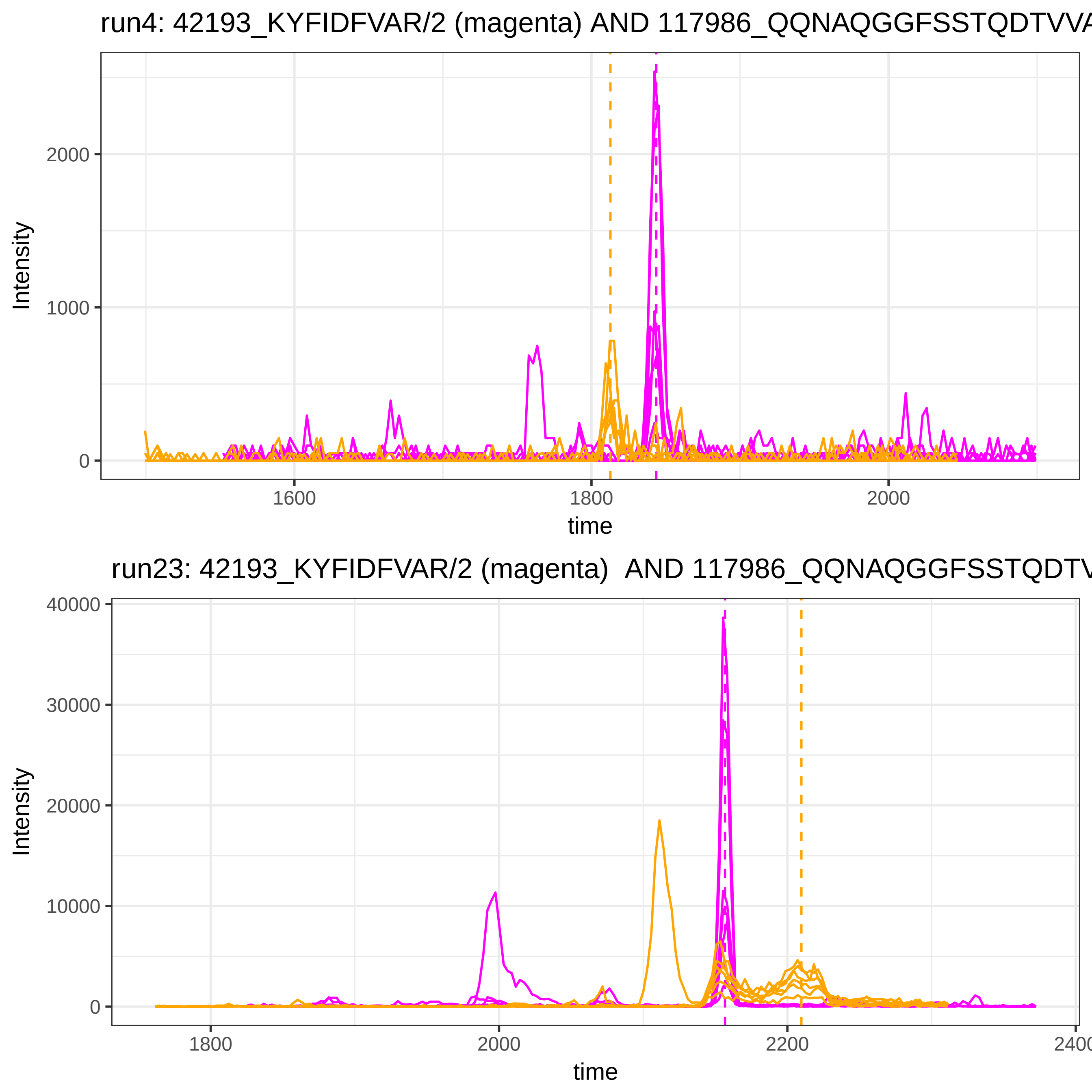
run4: 42193_KYFIDFVAR/2 (magenta) AND 117986_QQNAQGGFSSTQDTVVA
Intensity
2000
1000
0
1600
1800
2000
time
run23: 42193_KYFIDFVAR/2 (magenta) AND 117986_QQNAQGGFSSTQDTV
Intensity
40000
30000
20000
10000
0
1800
2000
2200
time

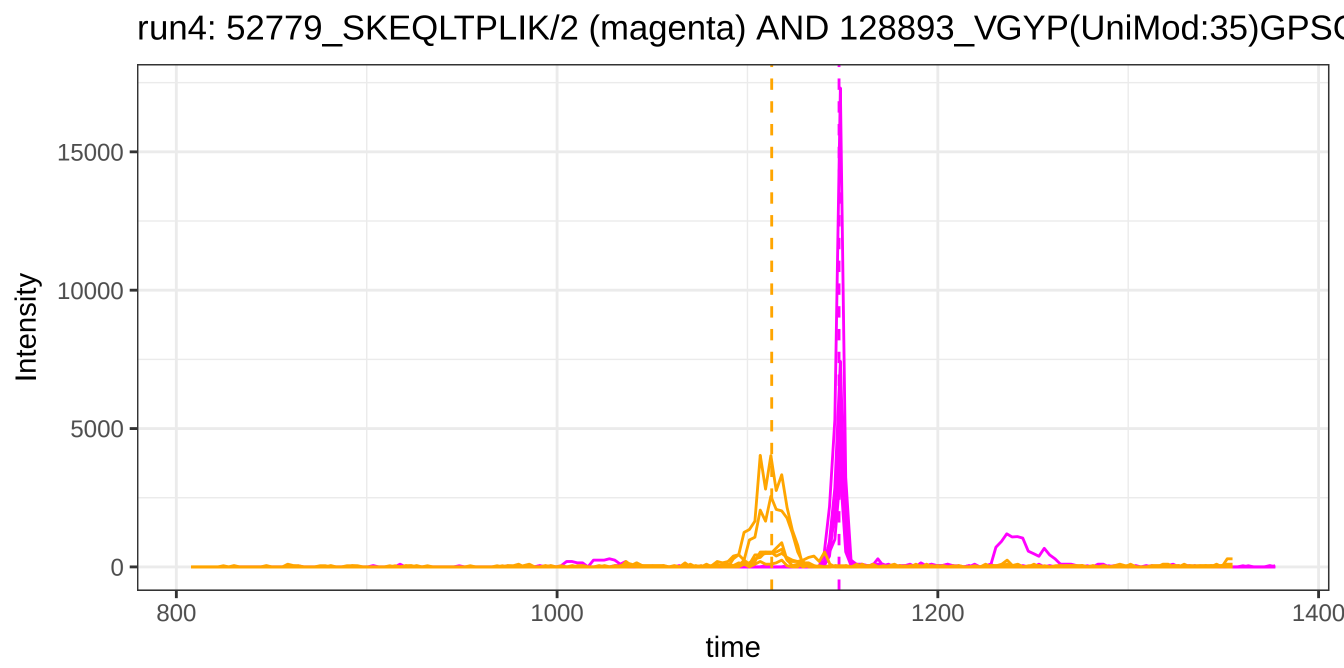
run4: 52779_SKEQLTPLIK/2 (magenta) AND 128893_VGYP(UniMod:35)GPSC
Intensity
15000
10000
5000
0
800
1000
1200
1400
time

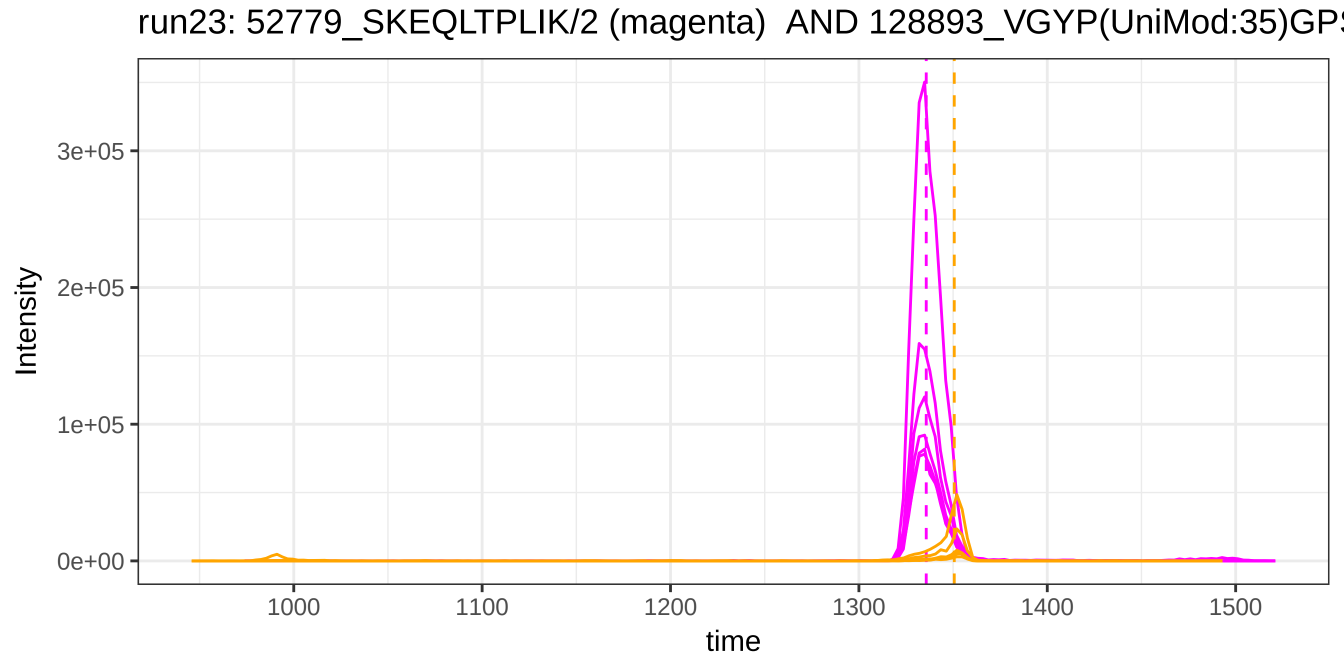
run23: 52779_SKEQLTPLIK/2 (magenta) AND 128893_VGYP(UniMod:35)GPS
Intensity
3e+05
2e+05
1e+05
0e+00
1000
1100
1200
1300
1400
1500
time

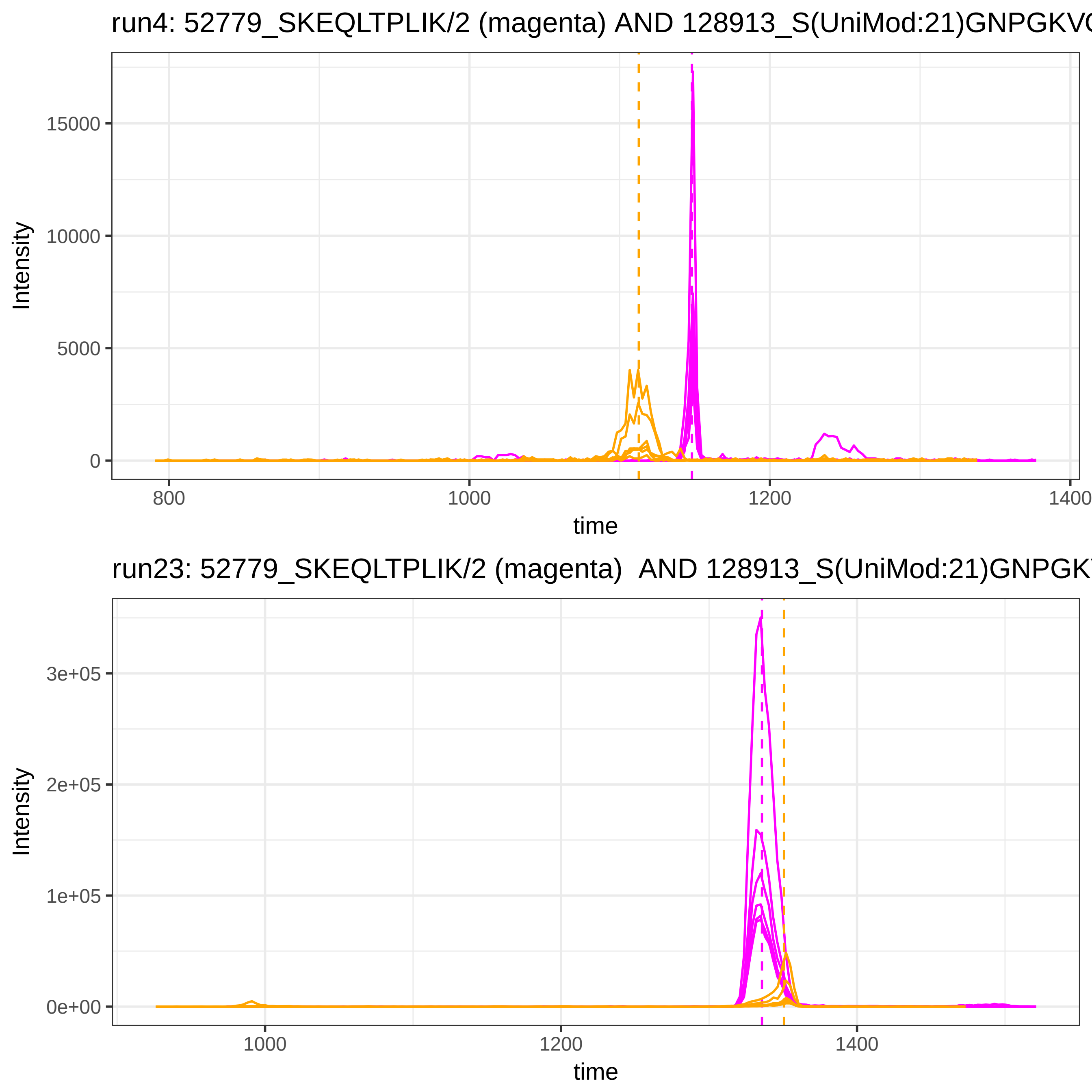
run4: 52779_SKEQLTPLIK/2 (magenta) AND 128913_S(UniMod:21)GNPGKVC
Intensity
15000
10000
5000
0
800
1000
1200
1400
time
run23: 52779_SKEQLTPLIK/2 (magenta) AND 128913_S(UniMod:21)GNPGKV
Intensity
3e+05
2e+05
1e+05
0e+00
1000
1200
1400
time

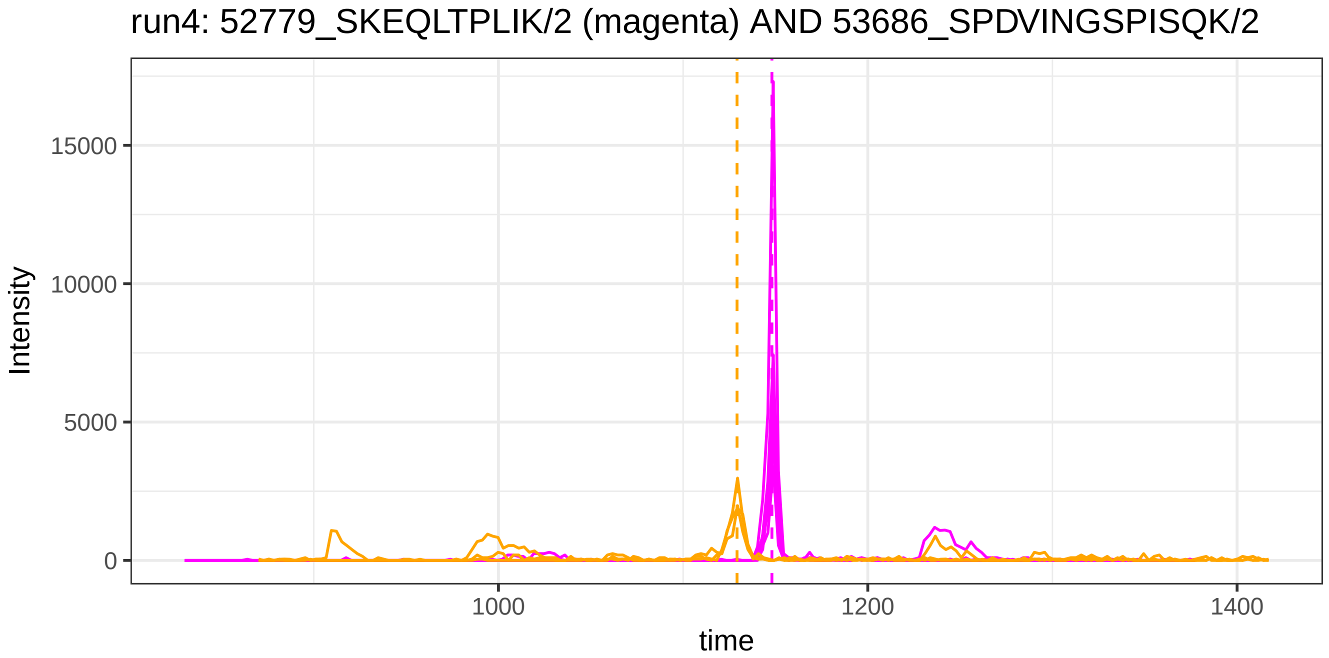
run4: 52779_SKEQLTPLIK/2 (magenta) AND 53686_SPDVINGSPISQK/2
Intensity
15000
10000
5000
0
1000
1200
1400
time

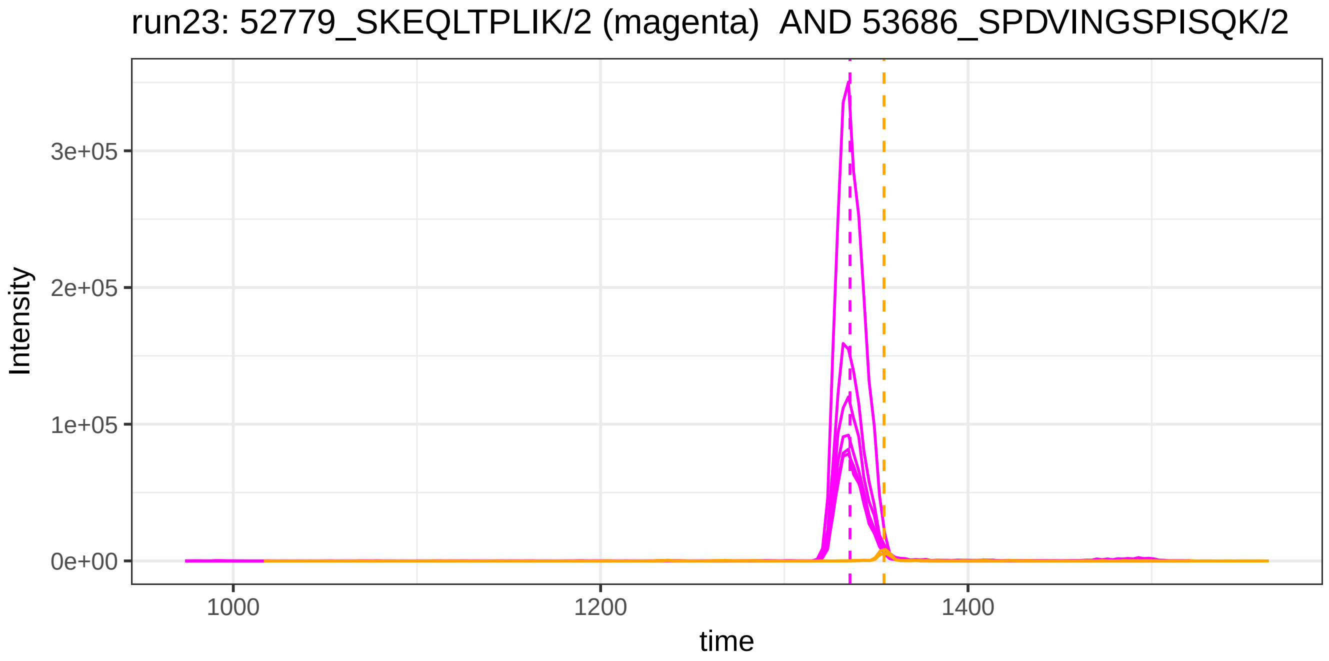
run23: 52779_SKEQLTPLIK/2 (magenta)  AND 53686_SPDVINGSPISQK/2
Intensity
3e+05
2e+05
1e+05
0e+00
1000
1200
1400
time

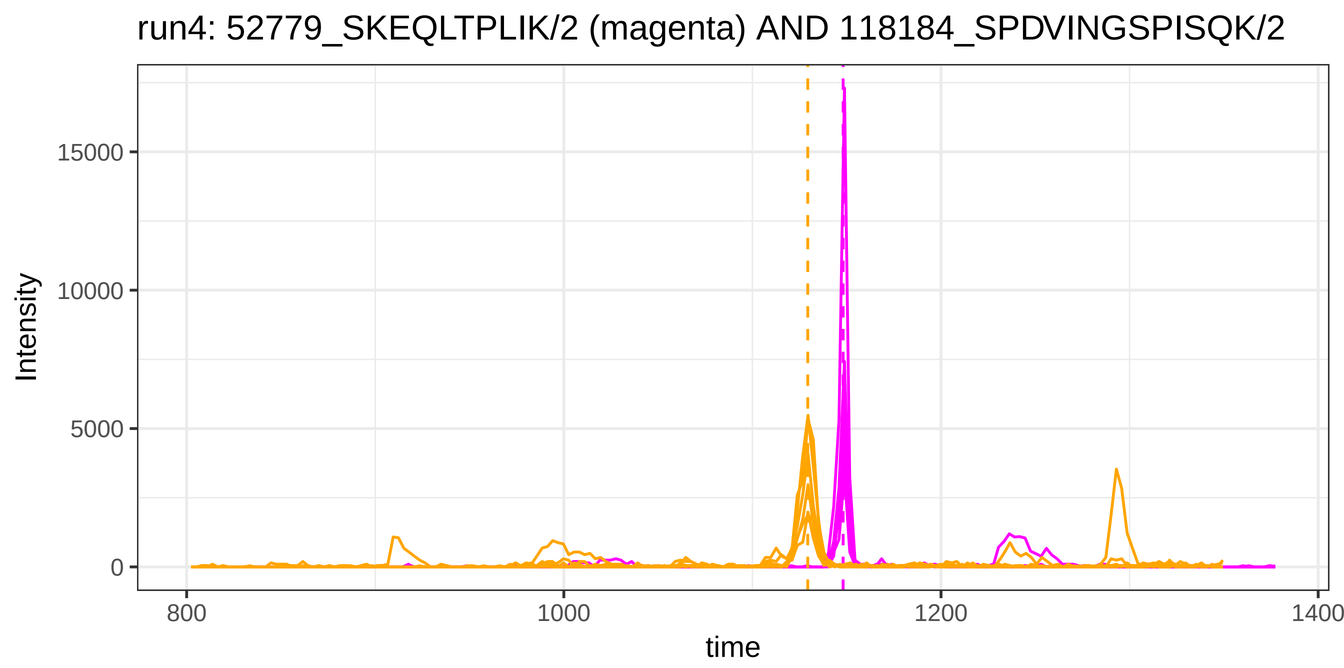
run4: 52779_SKEQLTPLIK/2 (magenta) AND 118184_SPDVINGSPISQK/2
Intensity
15000
10000
5000
0
800
1000
1200
1400
time

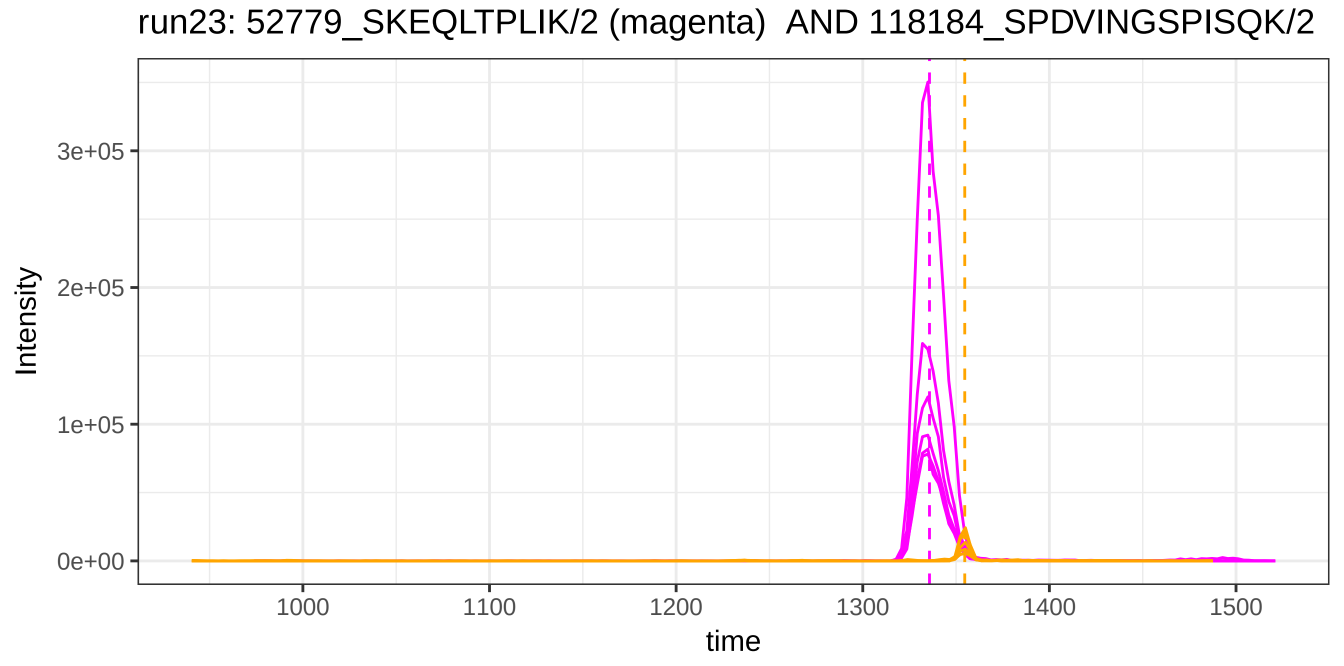
run23: 52779_SKEQLTPLIK/2 (magenta)  AND 118184_SPDVINGSPISQK/2
Intensity
3e+05
2e+05
1e+05
0e+00
1000
1100
1200
1300
1400
1500
time

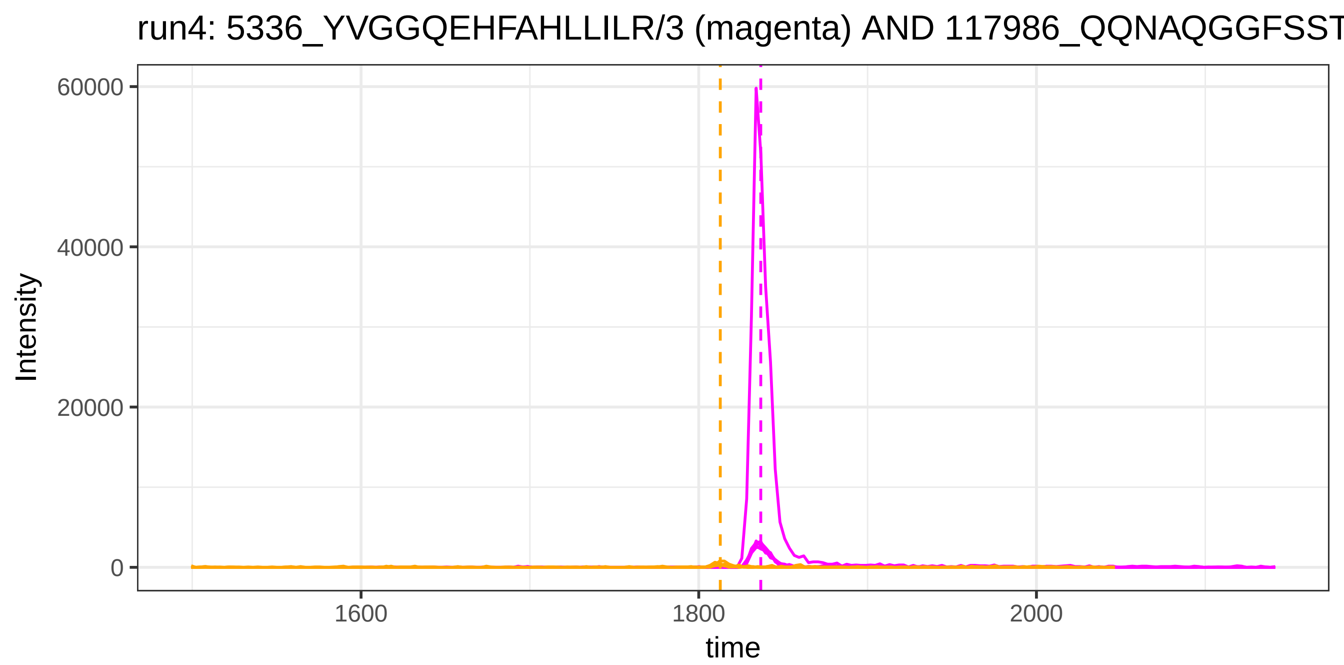
run4: 5336_YVGGQEHFAHLLILR/3 (magenta) AND 117986_QQNAQGGFSST
Intensity
60000
40000
20000
0
1600
1800
2000
time

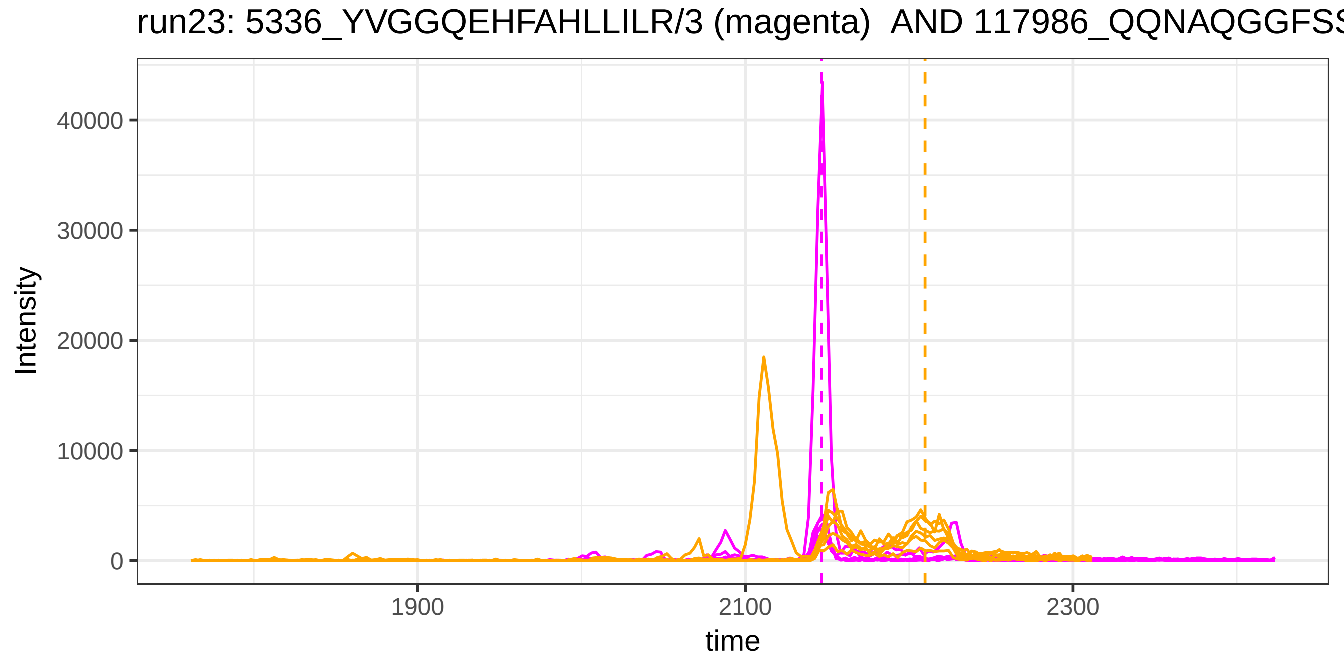
run23: 5336_YVGGQEHFAHLLILR/3 (magenta) AND 117986_QQNAQGGFSS
Intensity
40000
30000
20000
10000
0
1900
2100
2300
time

run4: 54026_SSALDMENFR/2 (magenta) AND 31059_EFQLFSSPHGK/2

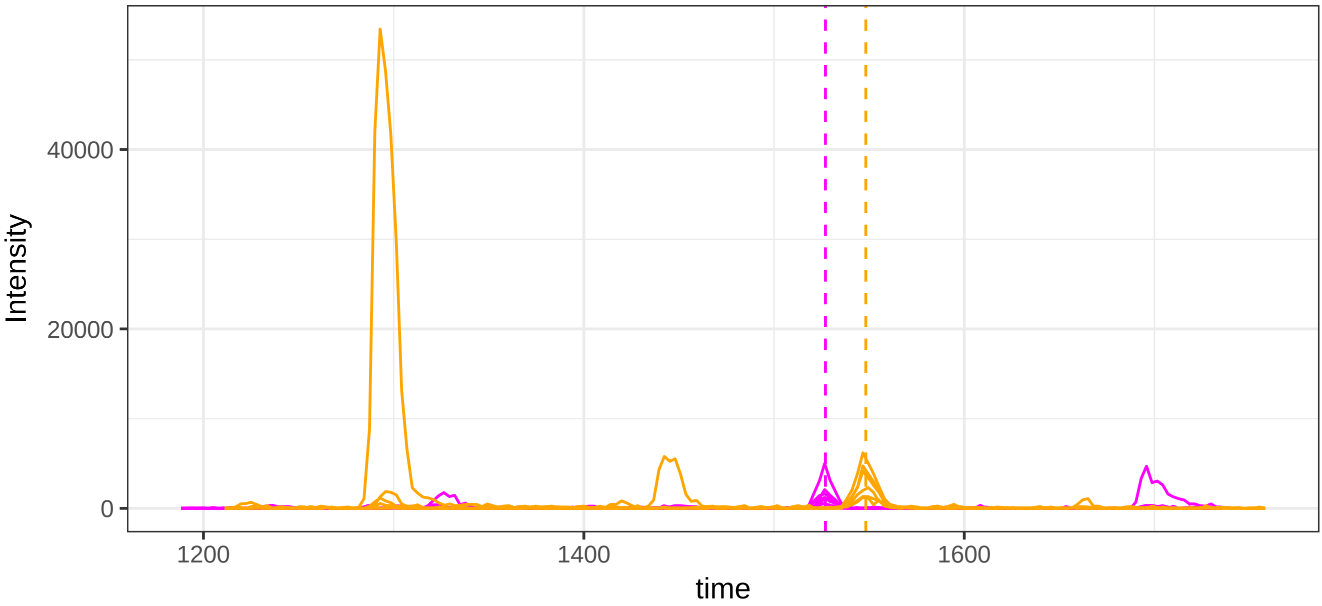


run23: 54026_SSALDMENFR/2 (magenta)  AND 31059_EFQLFSSPHGK/2

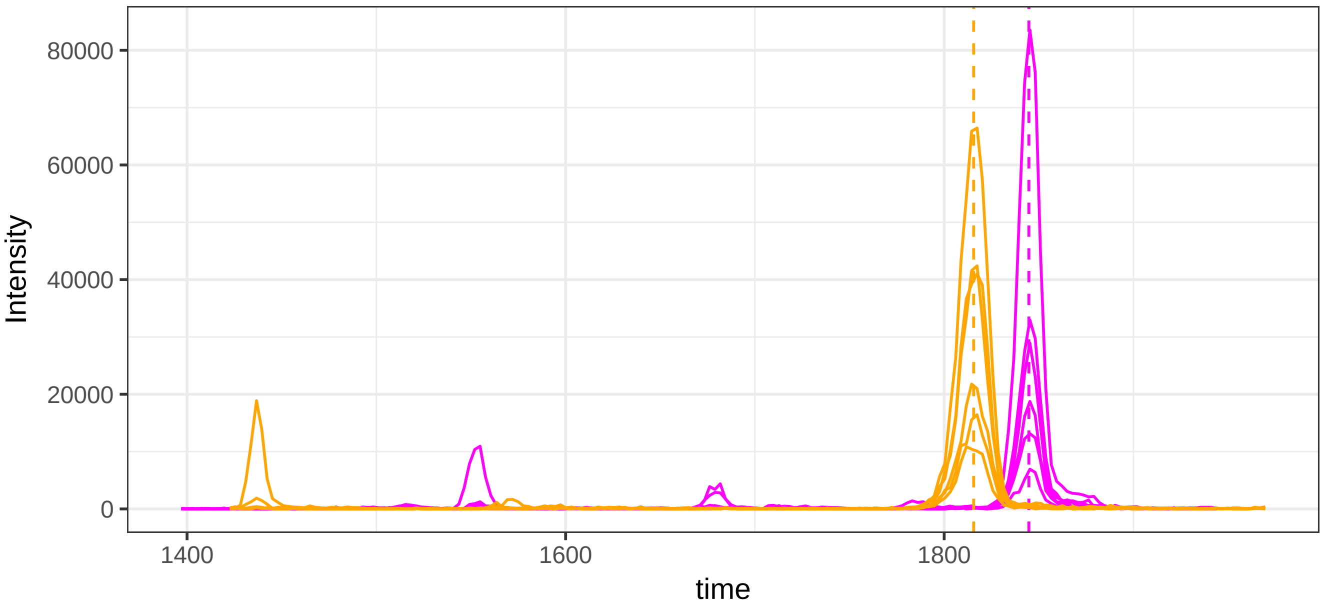

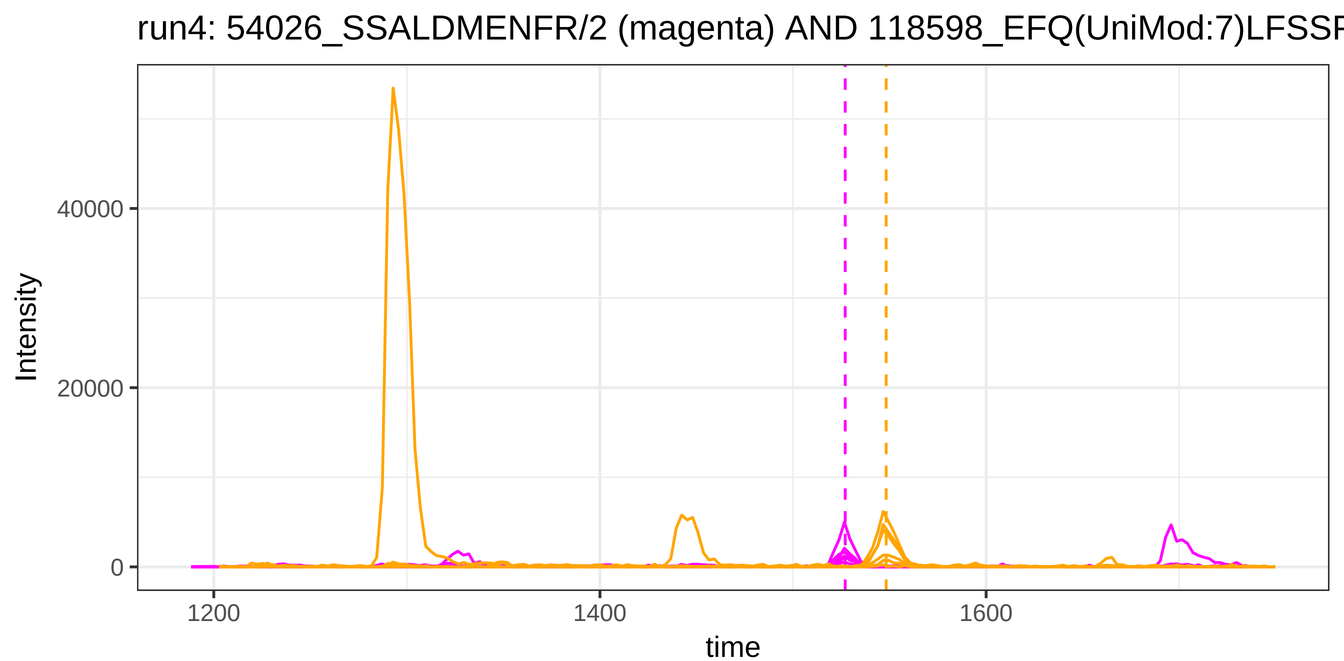
run4: 54026_SSALDMENFR/2 (magenta) AND 118598_EFQ(UniMod:7)LFSSF
Intensity
40000
20000
0
1200
1400
1600
time

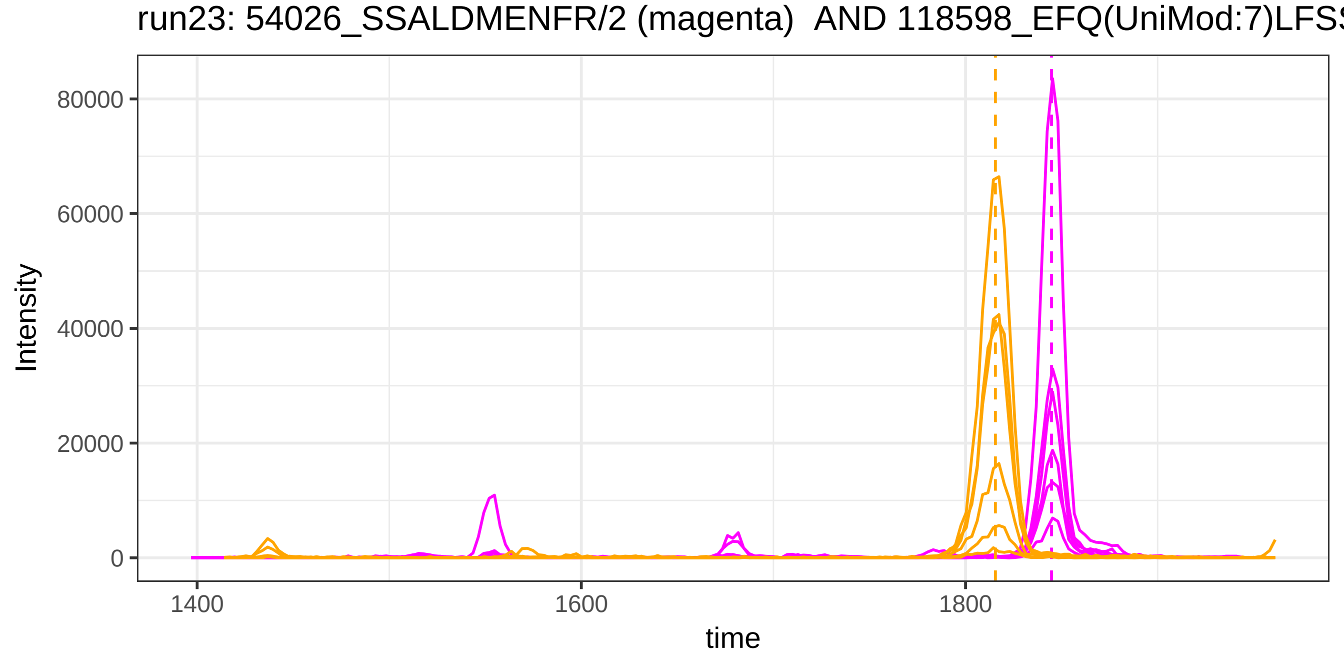
run23: 54026_SSALDMENFR/2 (magenta) AND 118598_EFQ(UniMod:7)LFSS
Intensity
80000
60000
40000
20000
0
1400
1600
1800
time

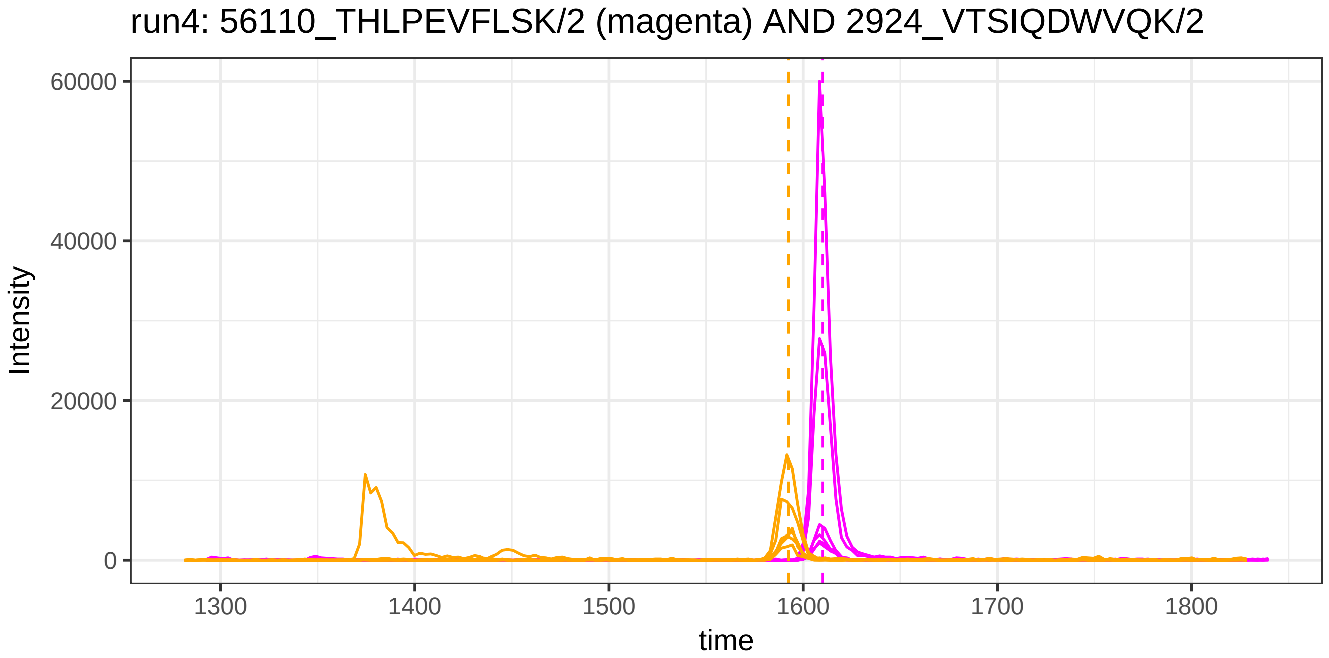
run4: 56110_THLPEVFLSK/2 (magenta) AND 2924_VTSIQDWVQK/2
Intensity
60000
40000
20000
0
1300
1400
1500
1600
1700
1800
time

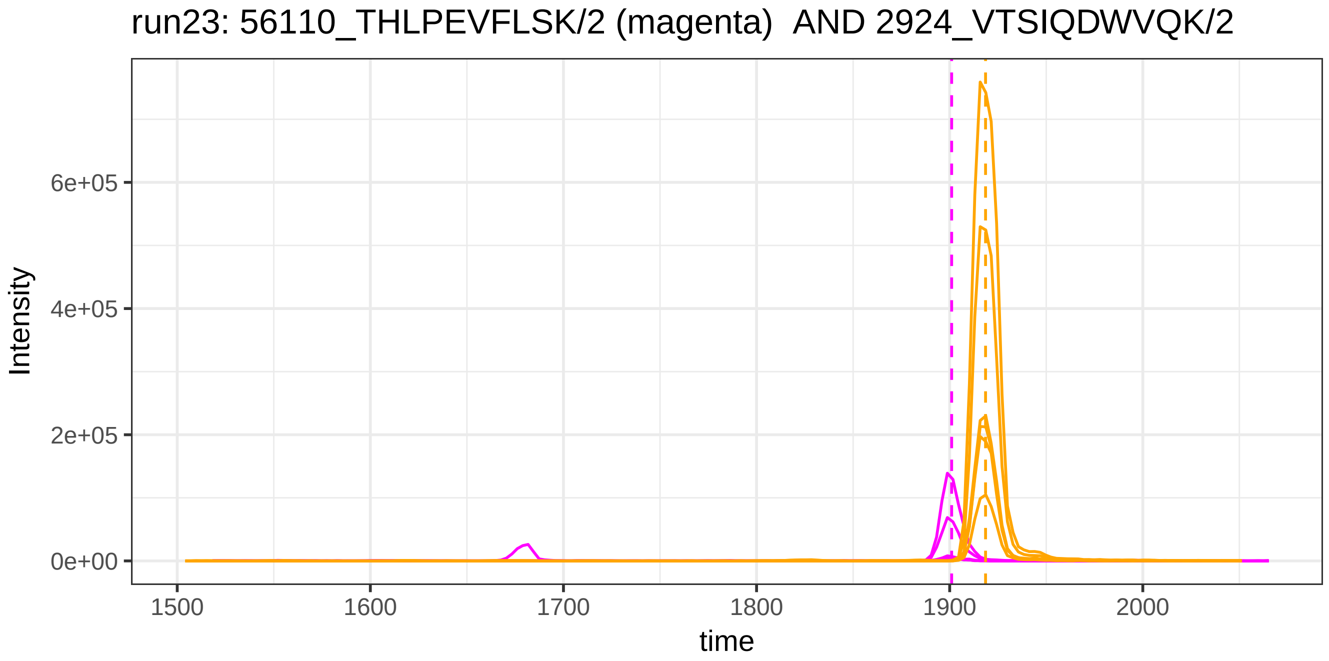
run23: 56110_THLPEVFLSK/2 (magenta)  AND 2924_VTSIQDWVQK/2
Intensity
6e+05
4e+05
2e+05
0e+00
1500
1600
1700
1800
1900
2000
time

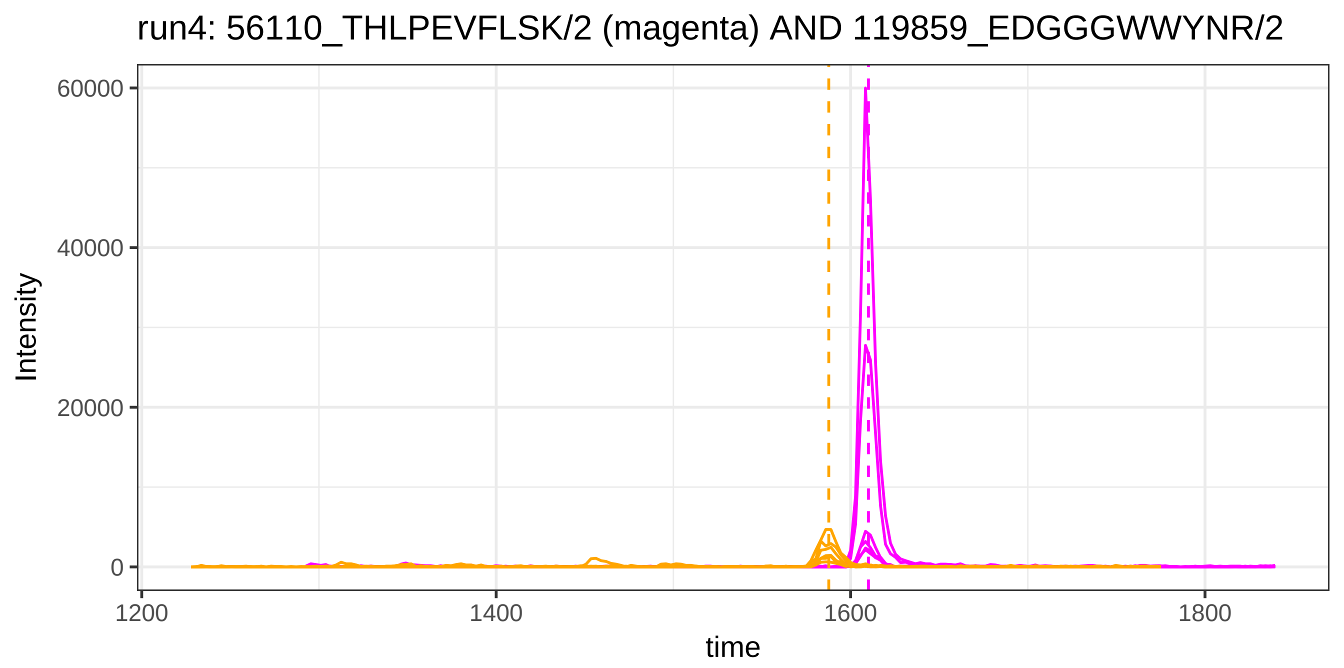

run4: 56110_THLPEVFLSK/2 (magenta) AND 119859_EDGGGWWYNR/2
Intensity
60000
40000
20000
0
1200
1400
1600
1800
time


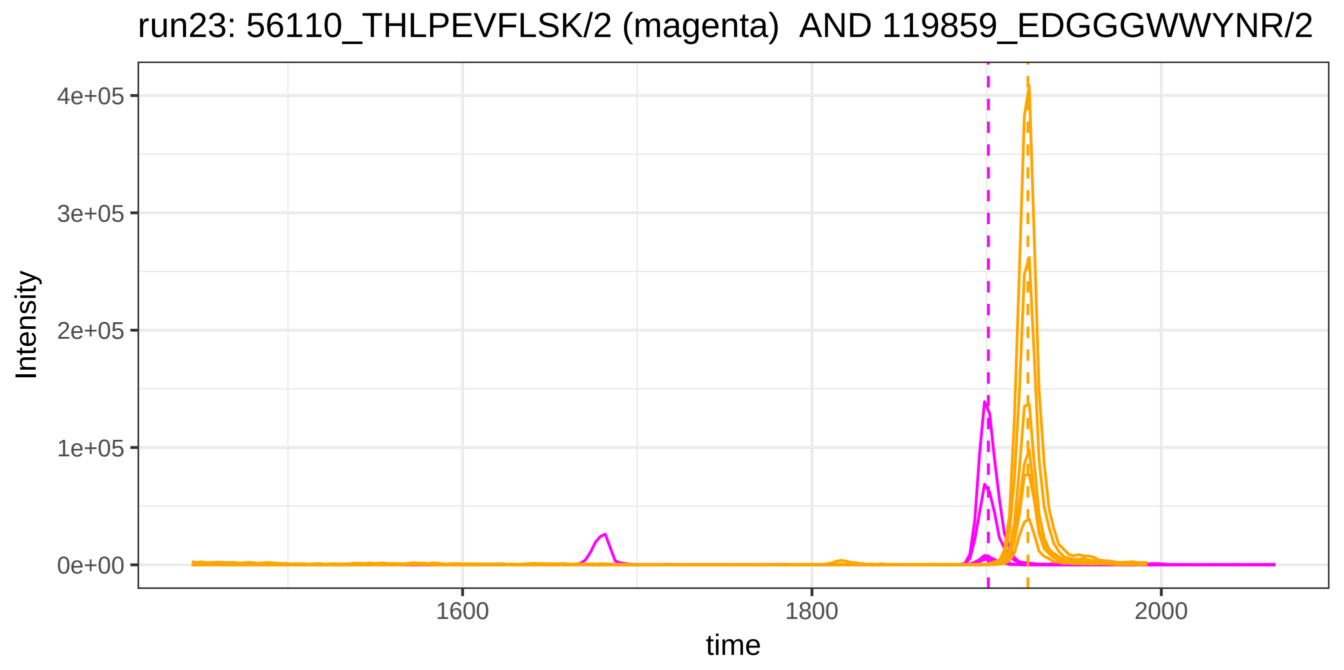

run23: 56110_THLPEVFLSK/2 (magenta) AND 119859_EDGGGWWYNR/2
Intensity
4e+05
3e+05
2e+05
1e+05
0e+00
1600
1800
2000
time

run4: 56110_THLPEVFLSK/2 (magenta) AND 5357_YVLTQPPSVSVAPGETAF

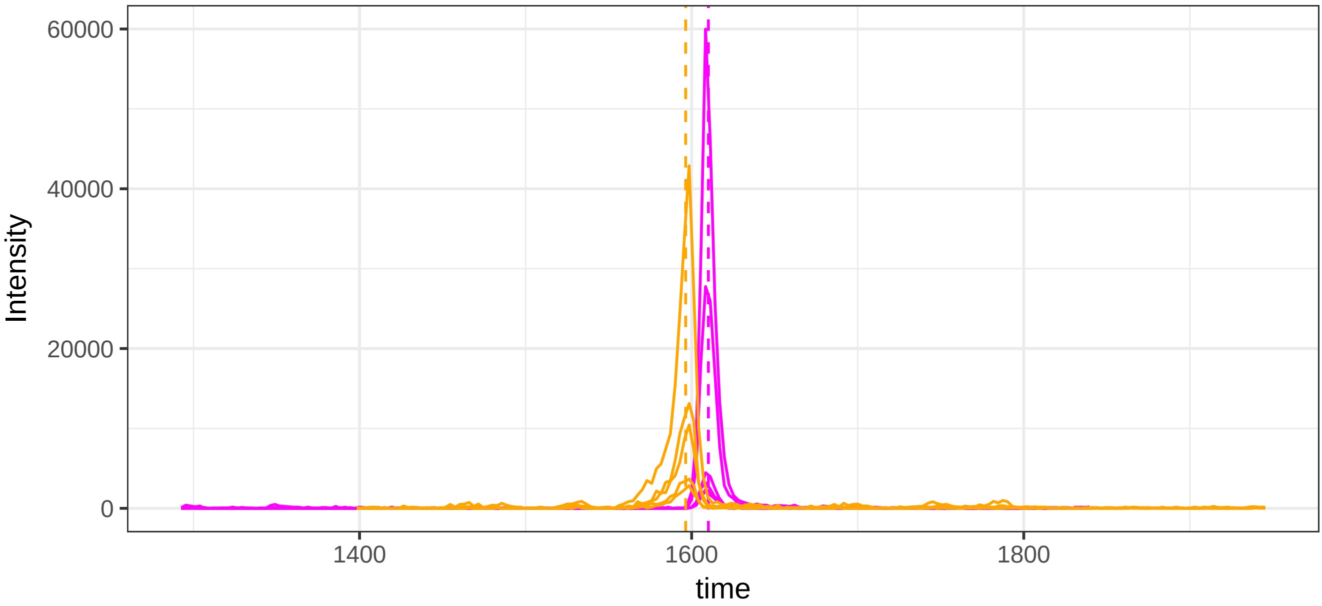


run23: 56110_THLPEVFLSK/2 (magenta)  AND 5357_YVLTQPPSVSVAPGETA

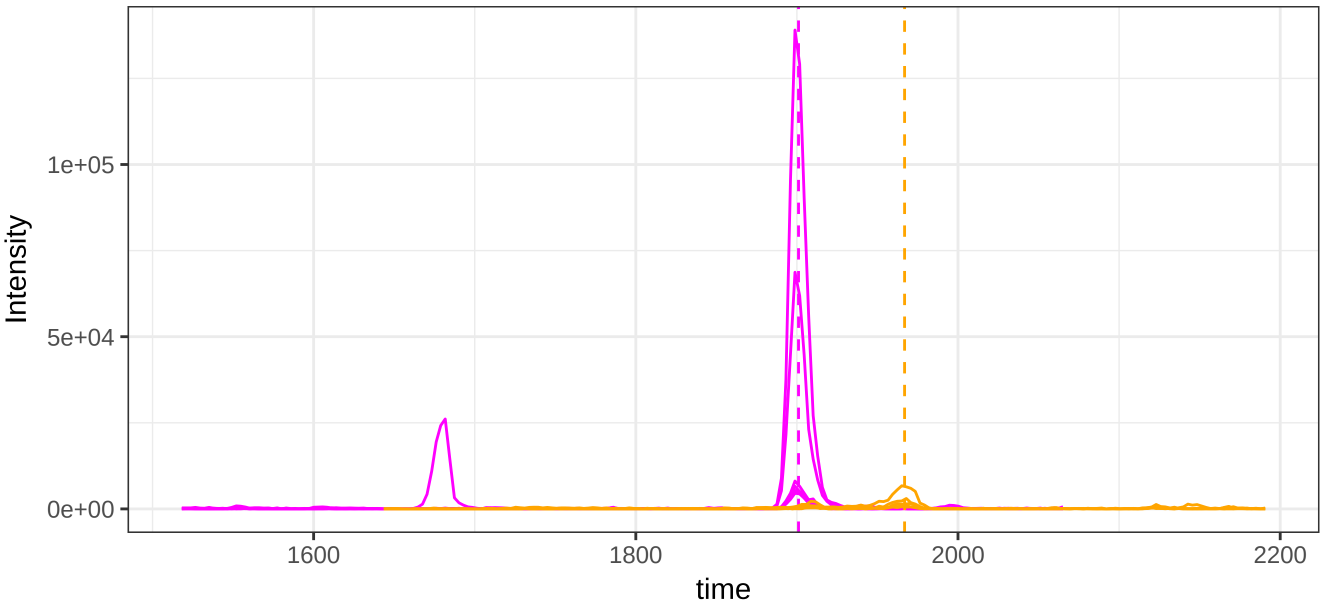

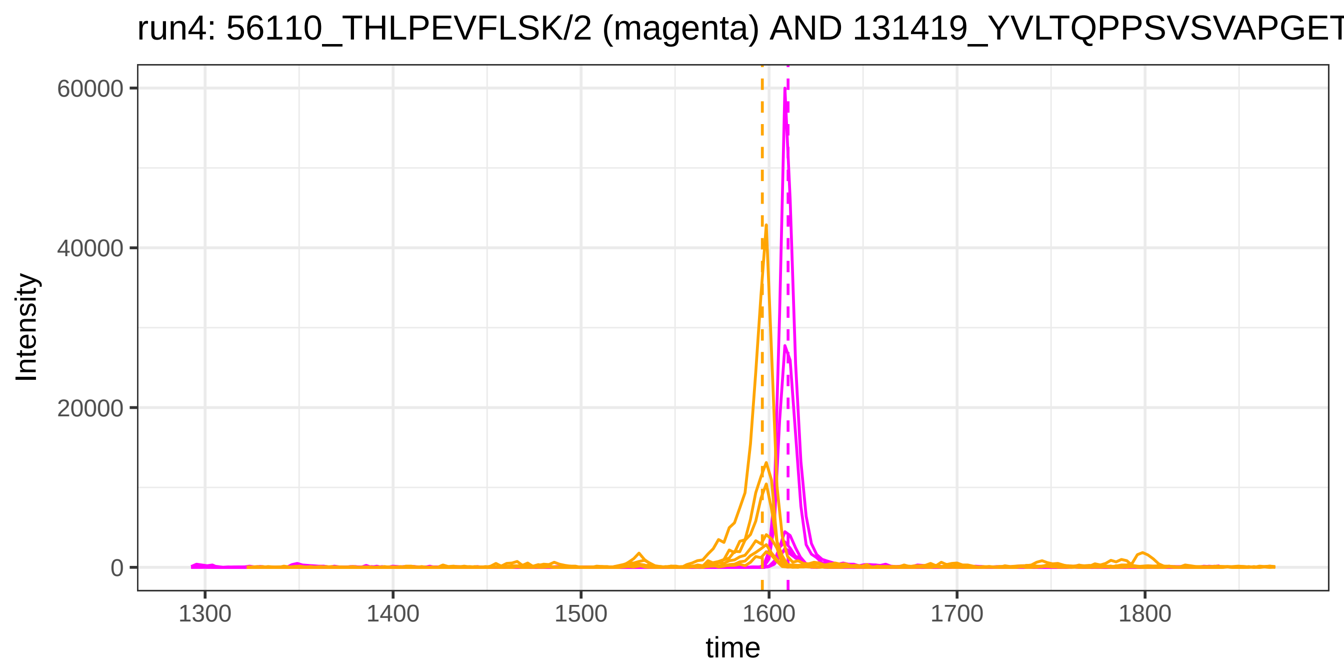
run4: 56110_THLPEVFLSK/2 (magenta) AND 131419_YVLTQPPSVSVAPGET
Intensity
60000
40000
20000
0
1300
1400
1500
1600
1700
1800
time

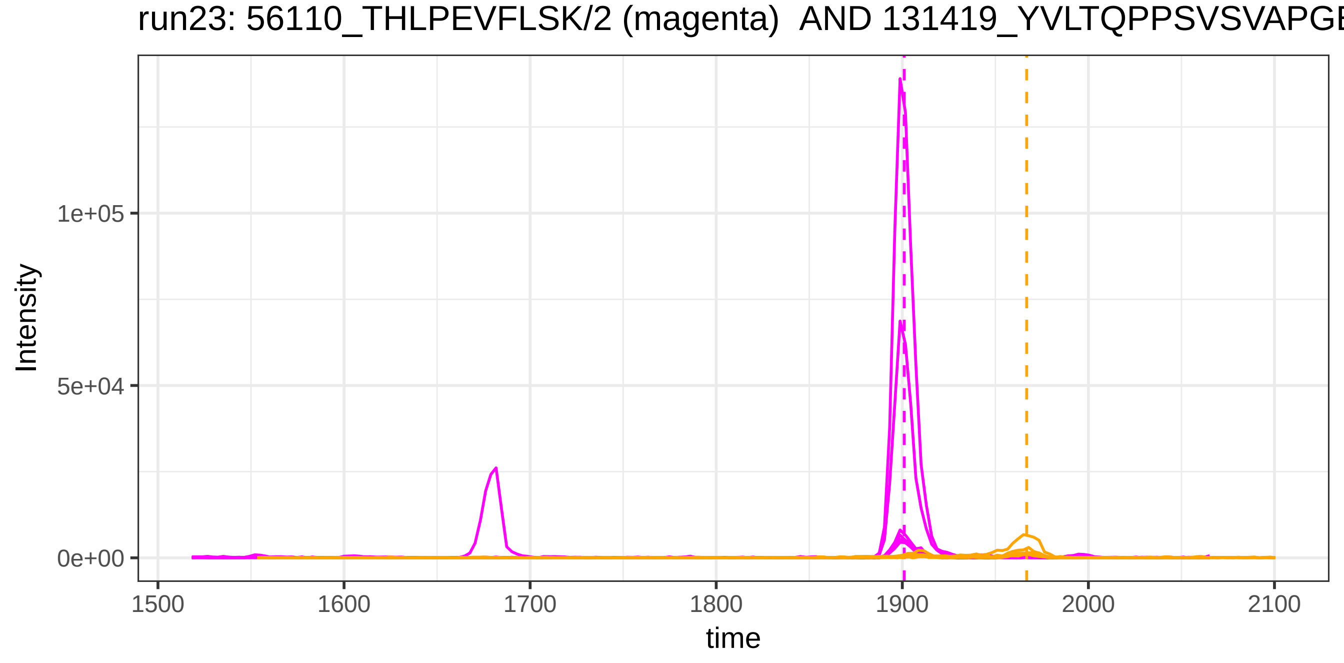
run23: 56110_THLPEVFLSK/2 (magenta) AND 131419_YVLTQPPSVSVAPGE
Intensity
1e+05
5e+04
0e+00
1500
1600
1700
1800
1900
2000
2100
time

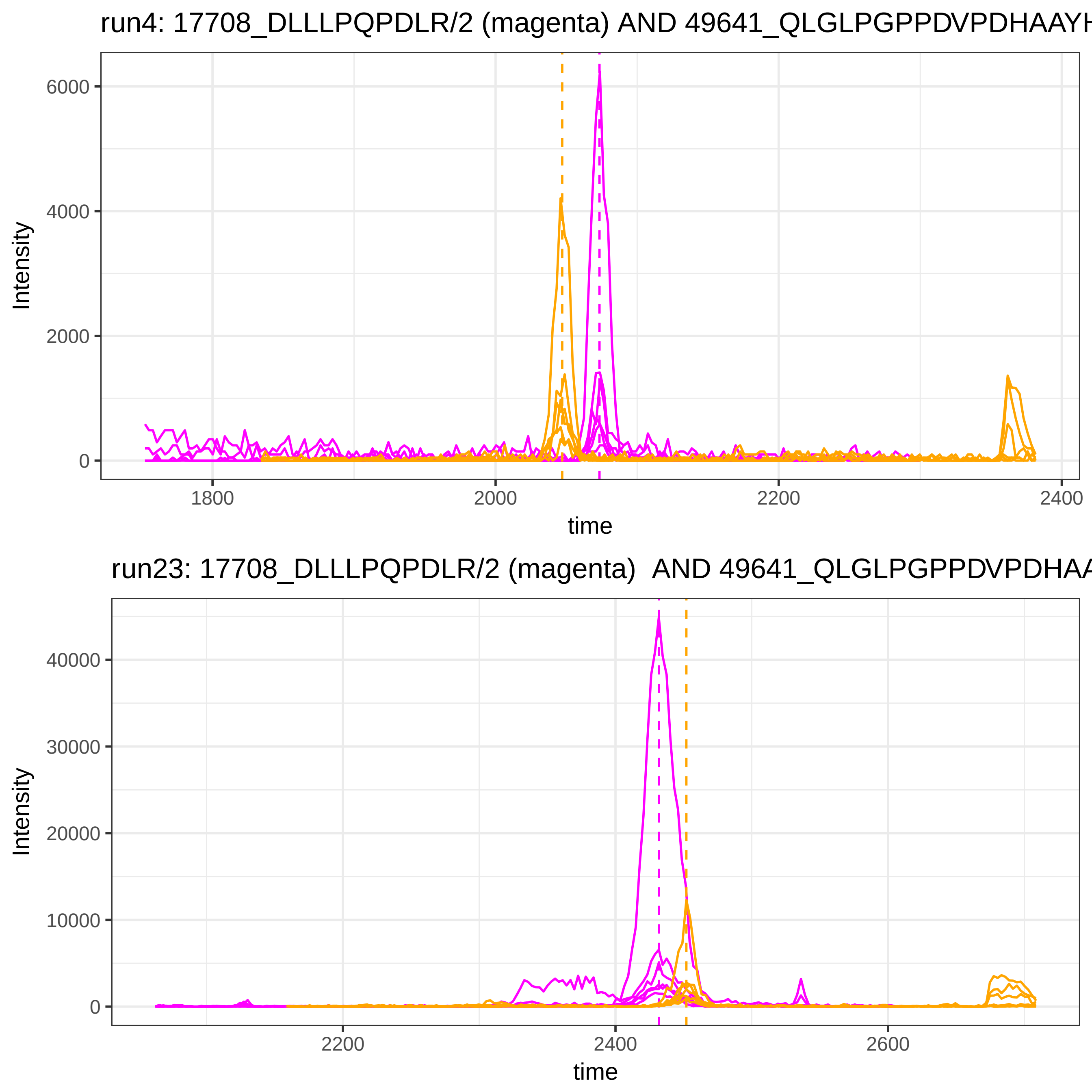

run4: 17708_DLLLPQPDLR/2 (magenta) AND 49641_QLGLPGPPDVPDHAAYH
Intensity
6000
4000
2000
0
1800
2000
2200
2400
time
run23: 17708_DLLLPQPDLR/2 (magenta) AND 49641_QLGLPGPPDVPDHAA
Intensity
40000
30000
20000
10000
0
2200
2400
2600
time

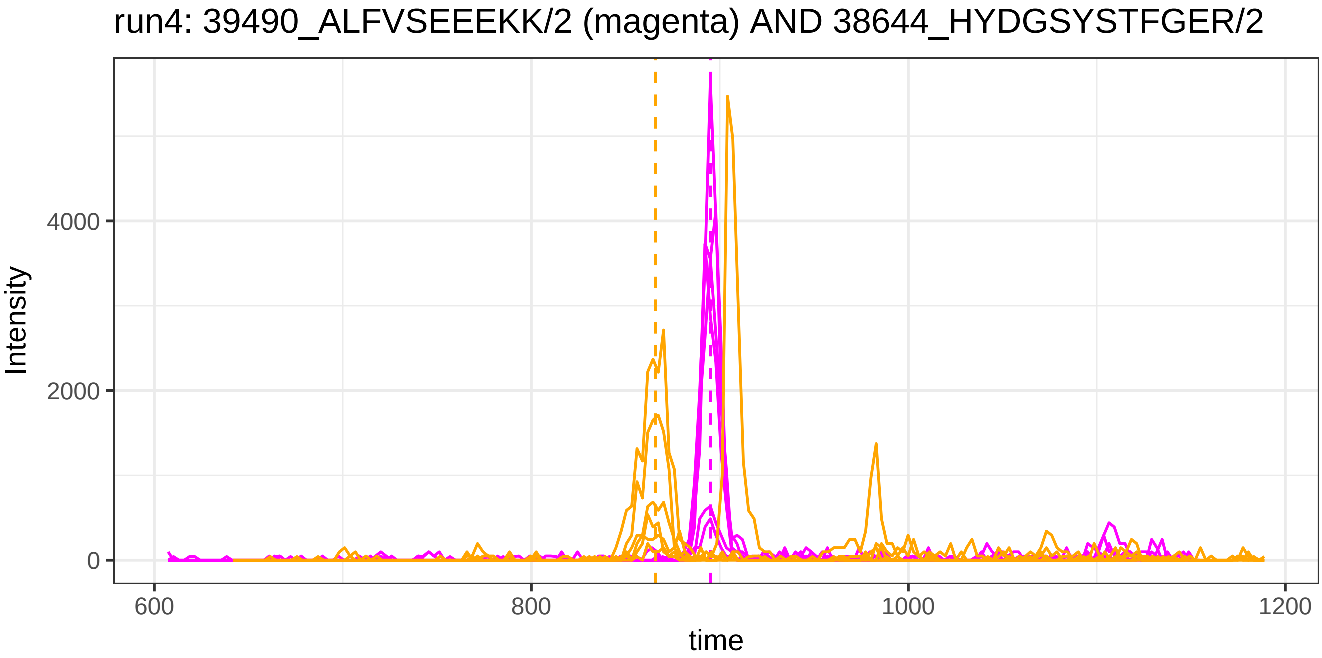
run4: 39490_ALFVSEEEKK/2 (magenta) AND 38644_HYDGSYSTFGER/2
Intensity
4000
2000
0
600
800
1000
1200
time

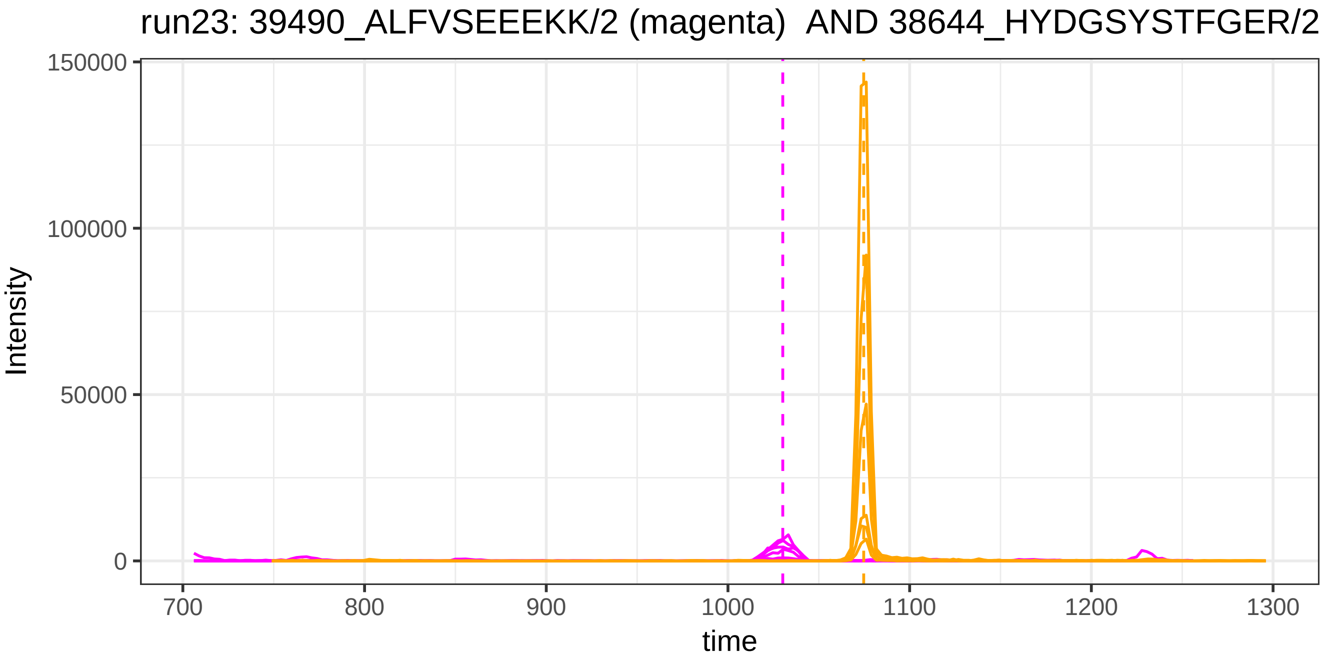
run23: 39490_ALFVSEEEKK/2 (magenta) AND 38644_HYDGSYSTFGER/2
Intensity
150000
100000
50000
0
700
800
900
1000
1100
1200
1300
time

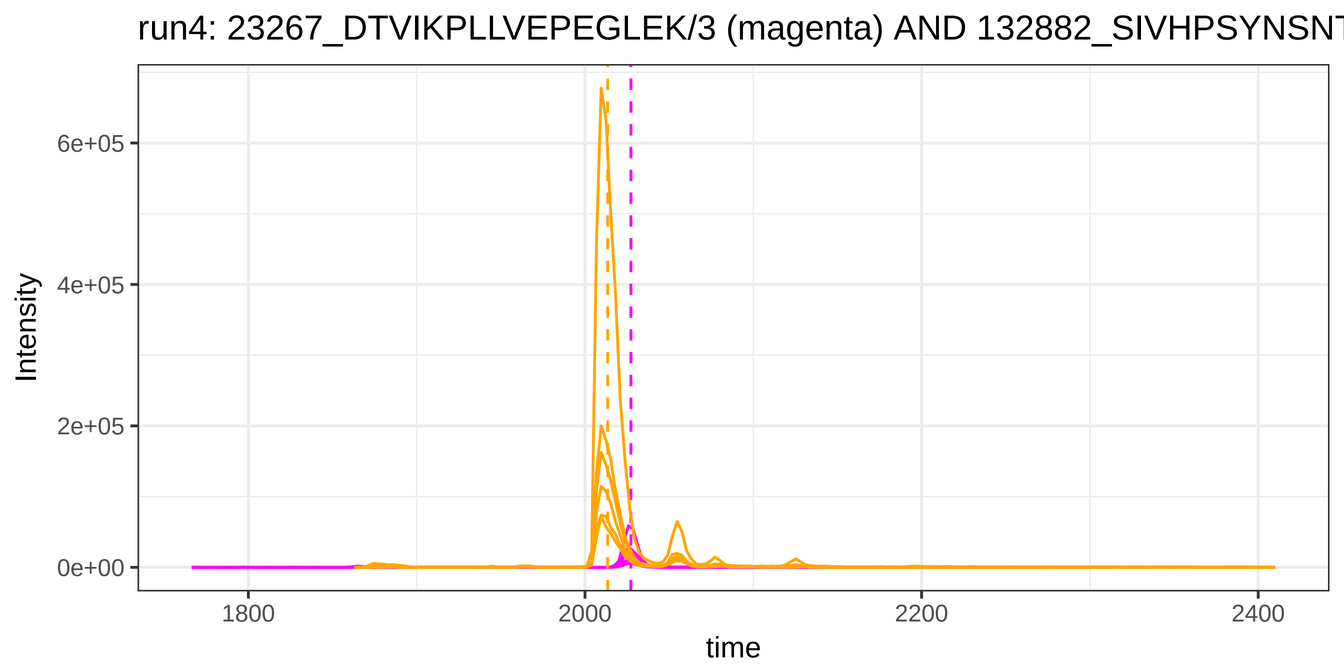

run4: 23267_DTVIKPLLVEPEGLEK/3 (magenta) AND 132882_SIVHPSYNSNT
6e+05
4e+05
2e+05
0e+00
Intensity
1800
2000
2200
2400
time


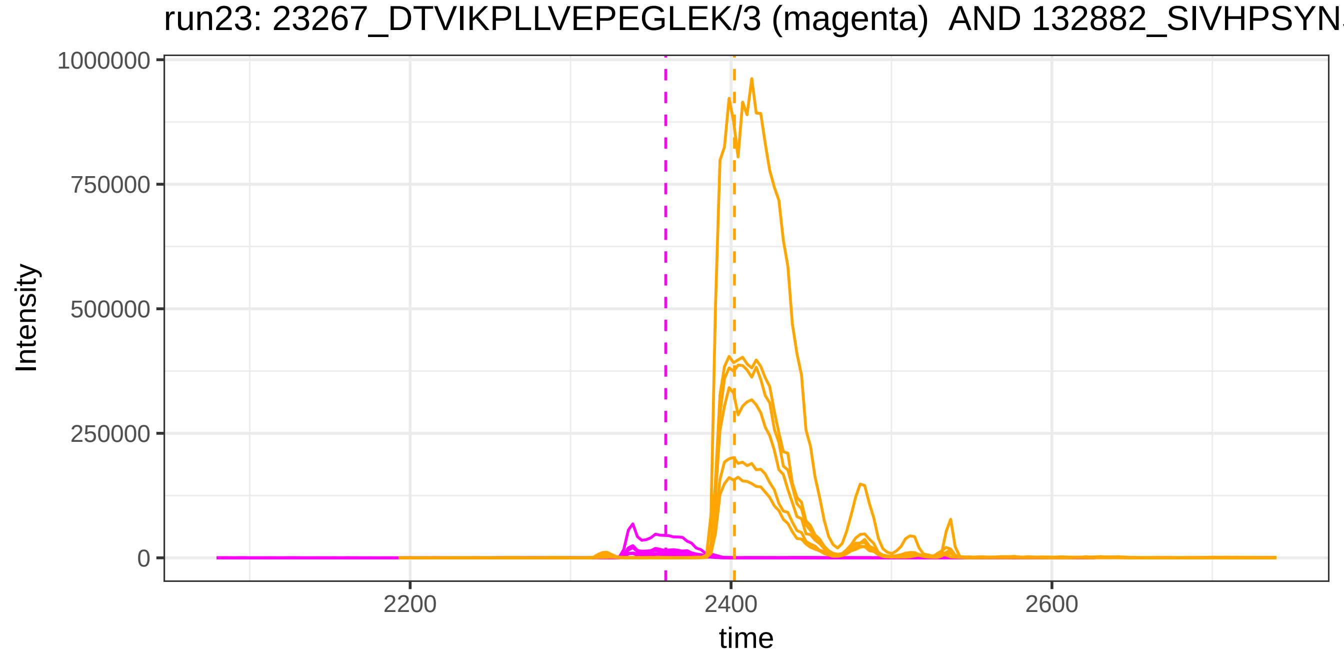

run23: 23267_DTVIKPLLVEPEGLEK/3 (magenta) AND 132882_SIVHPSYN
1000000
750000
500000
250000
0
Intensity
2200
2400
2600
time

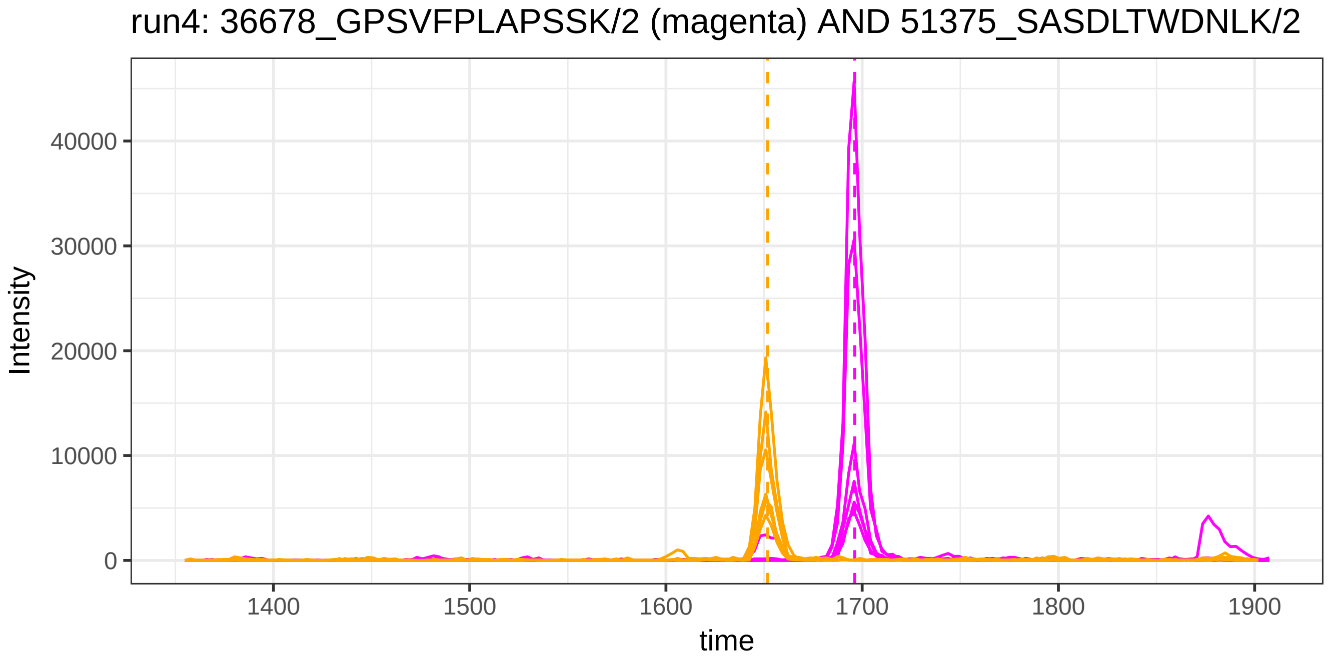

run4: 36678_GPSVFPLAPSSK/2 (magenta) AND 51375_SASDLTWDNLK/2
Intensity
40000
30000
20000
10000
0
1400
1500
1600
1700
1800
1900
time


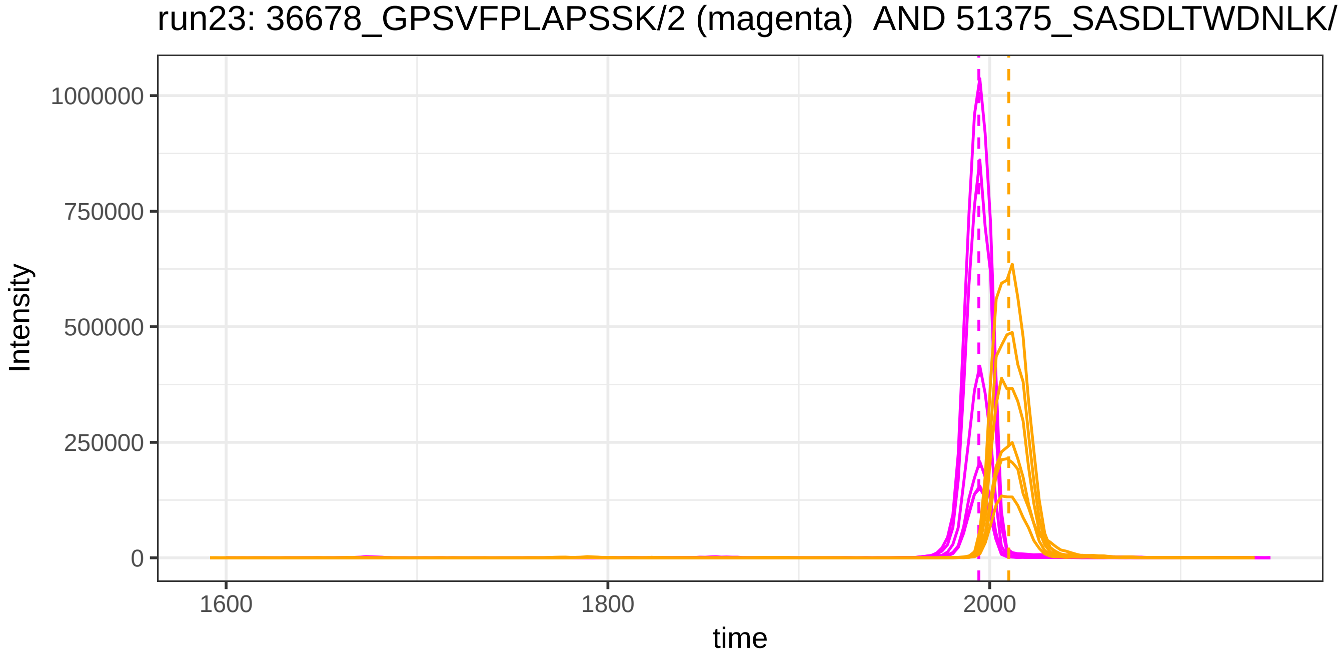

run23: 36678_GPSVFPLAPSSK/2 (magenta) AND 51375_SASDLTWDNLK/
Intensity
1000000
750000
500000
250000
0
1600
1800
2000
time

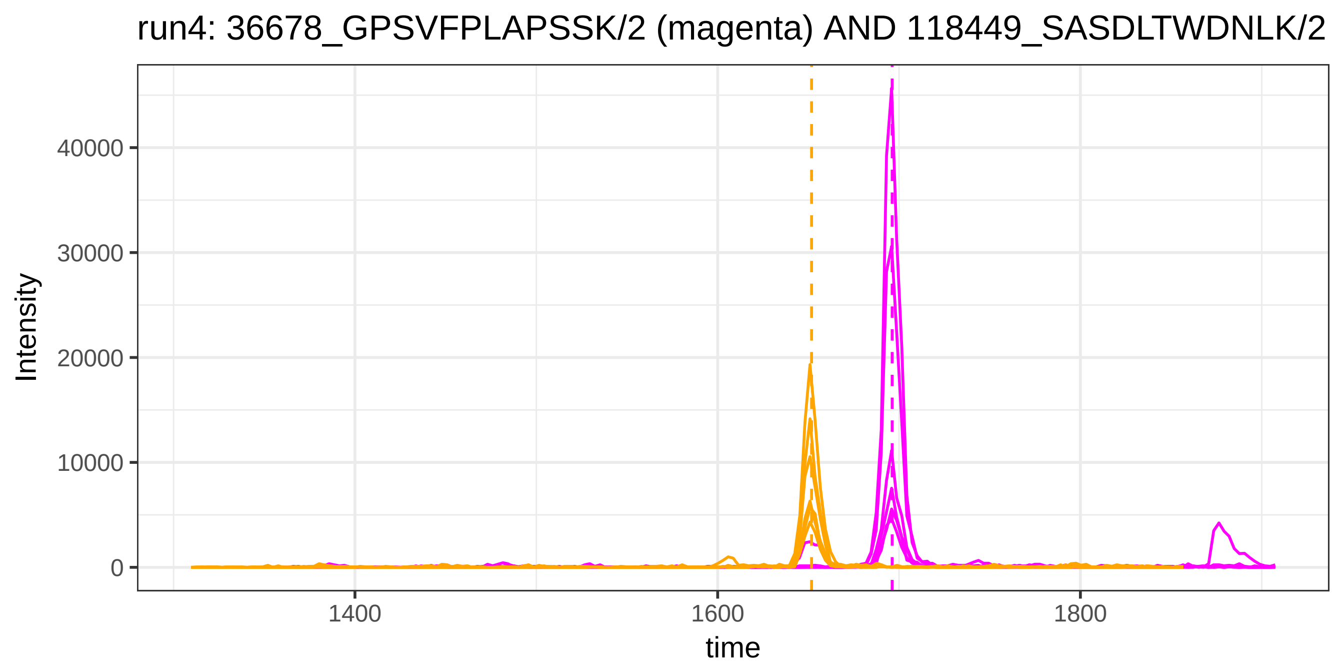
run4: 36678_GPSVFPLAPSSK/2 (magenta) AND 118449_SASDLTWDNLK/2
Intensity
40000
30000
20000
10000
0
1400
1600
1800
time

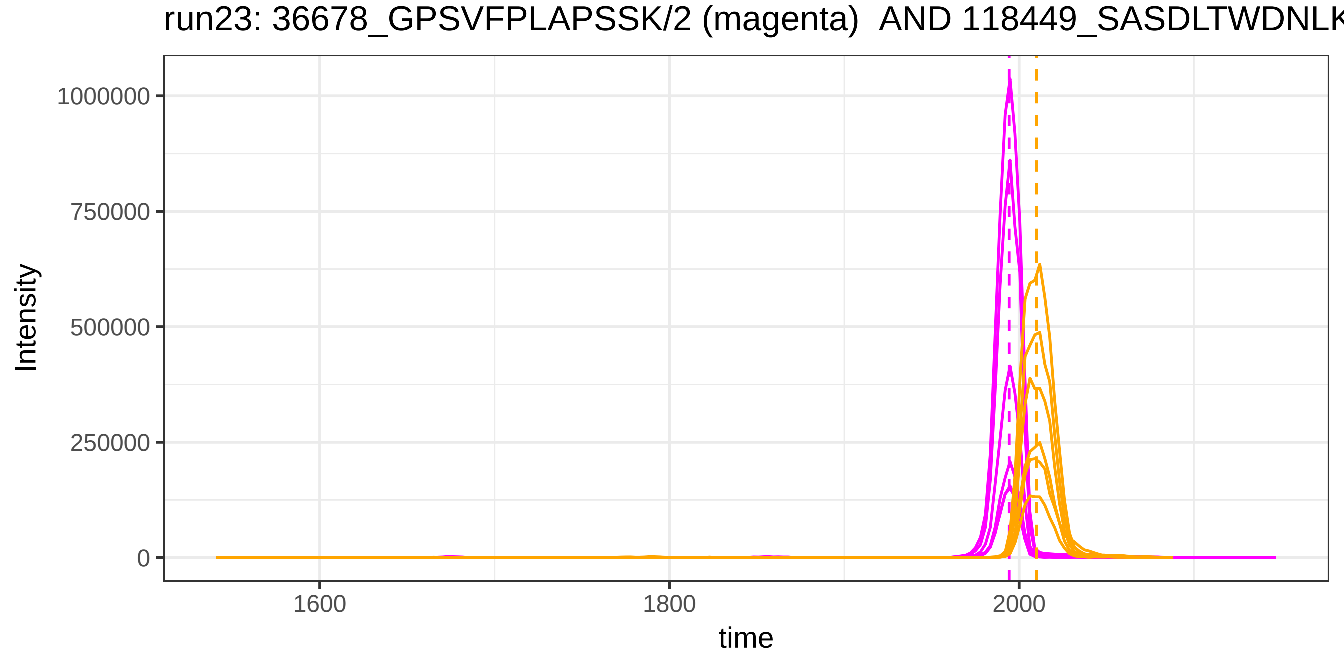
run23: 36678_GPSVFPLAPSSK/2 (magenta) AND 118449_SASDLTWDNLK
Intensity
1000000
750000
500000
250000
0
1600
1800
2000
time

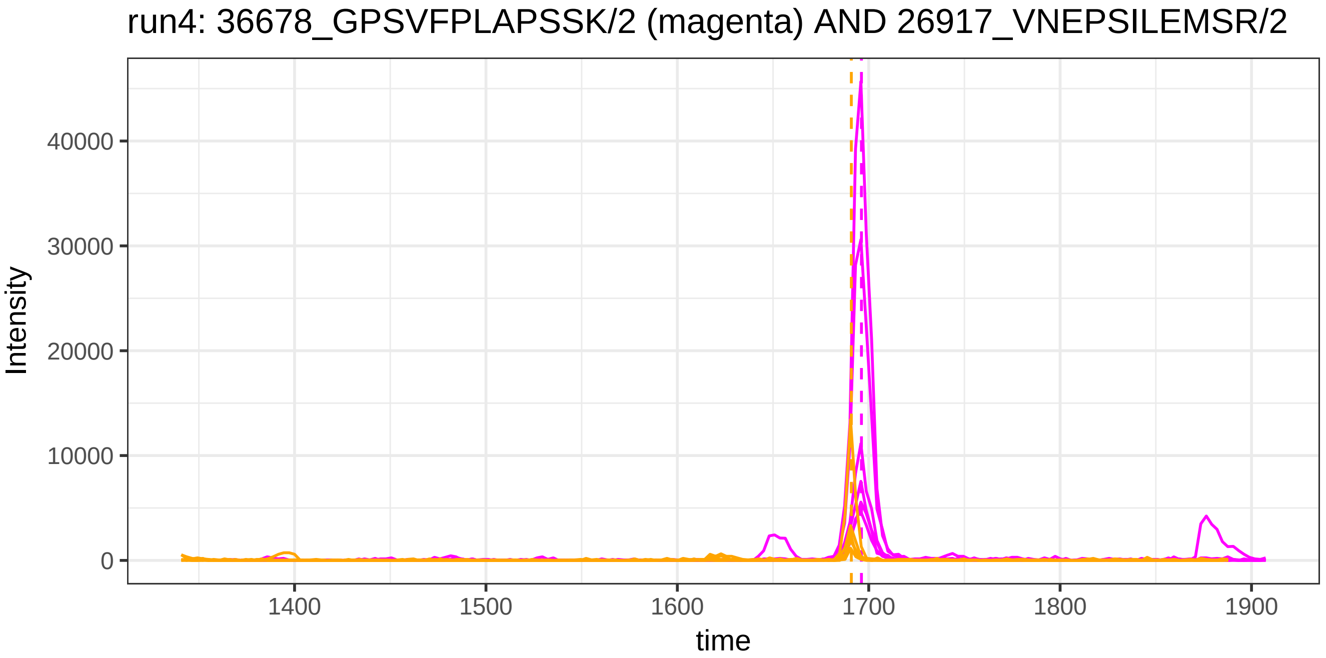
run4: 36678_GPSVFPLAPSSK/2 (magenta) AND 26917_VNEPSILEMSR/2
Intensity
40000
30000
20000
10000
0
1400
1500
1600
1700
1800
1900
time

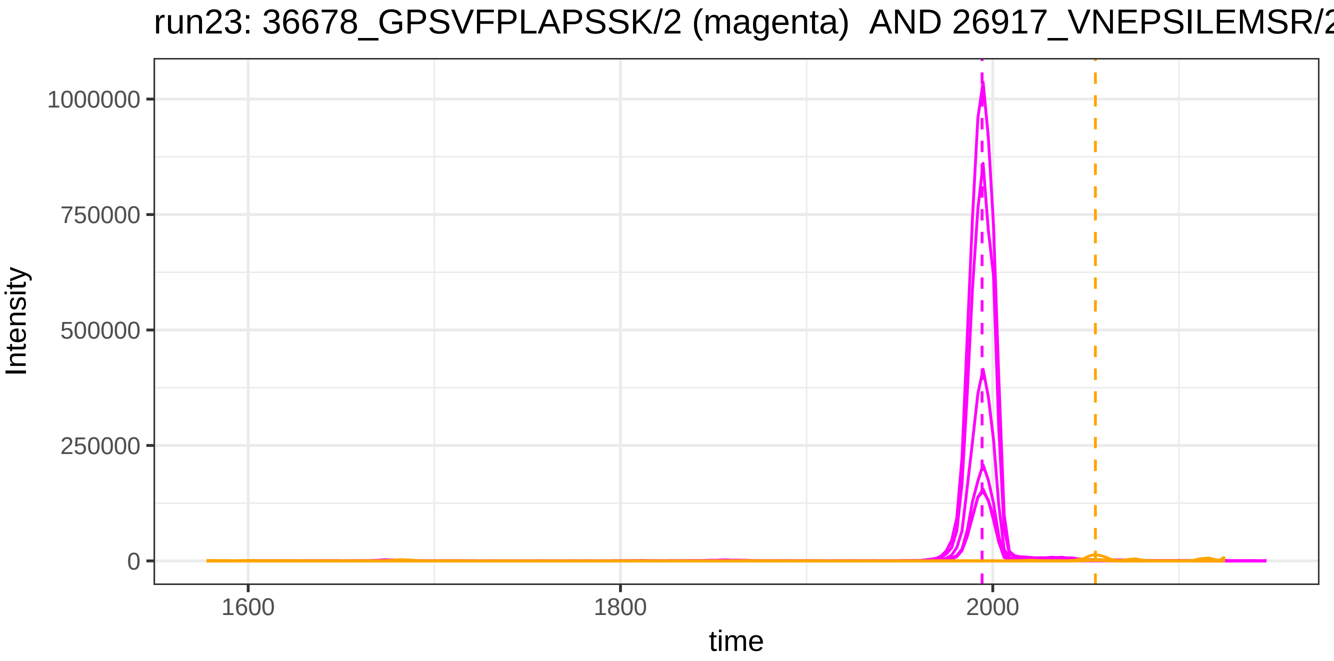
run23: 36678_GPSVFPLAPSSK/2 (magenta) AND 26917_VNEPSILEMSR/2
Intensity
1000000
750000
500000
250000
0
1600
1800
2000
time

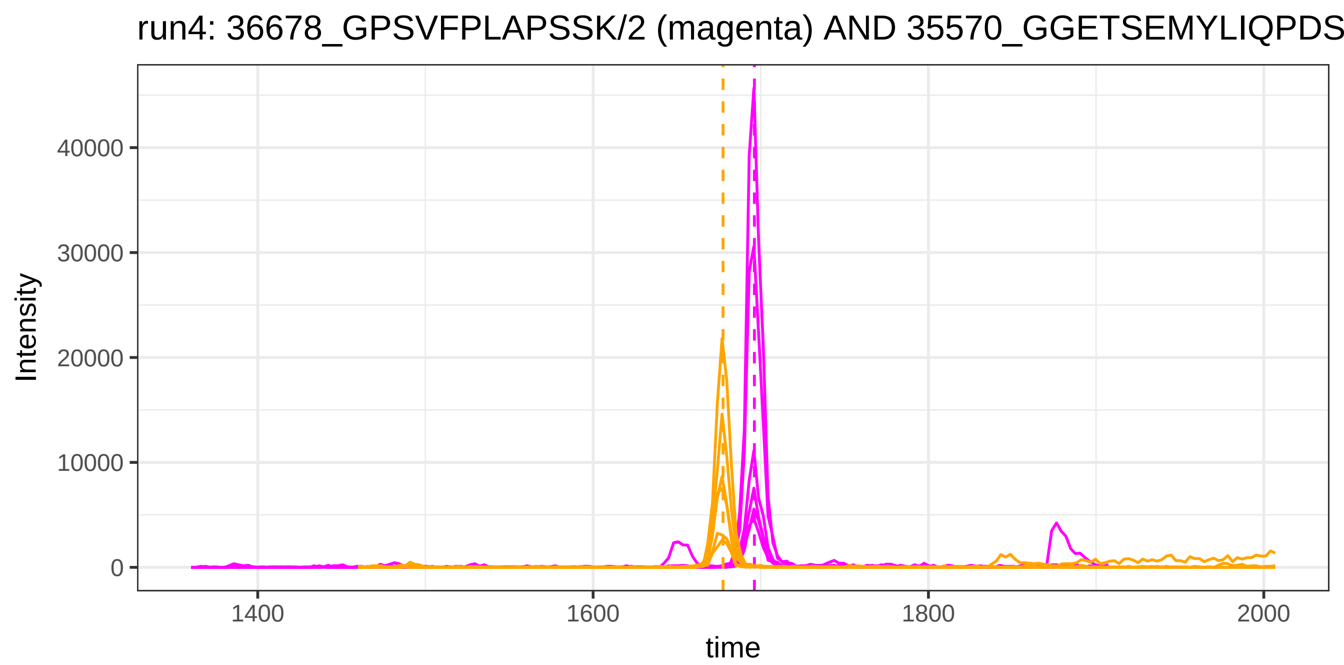
run4: 36678_GPSVFPLAPSSK/2 (magenta) AND 35570_GGETSEMYLIQPDS
Intensity
40000
30000
20000
10000
0
1400
1600
1800
2000
time

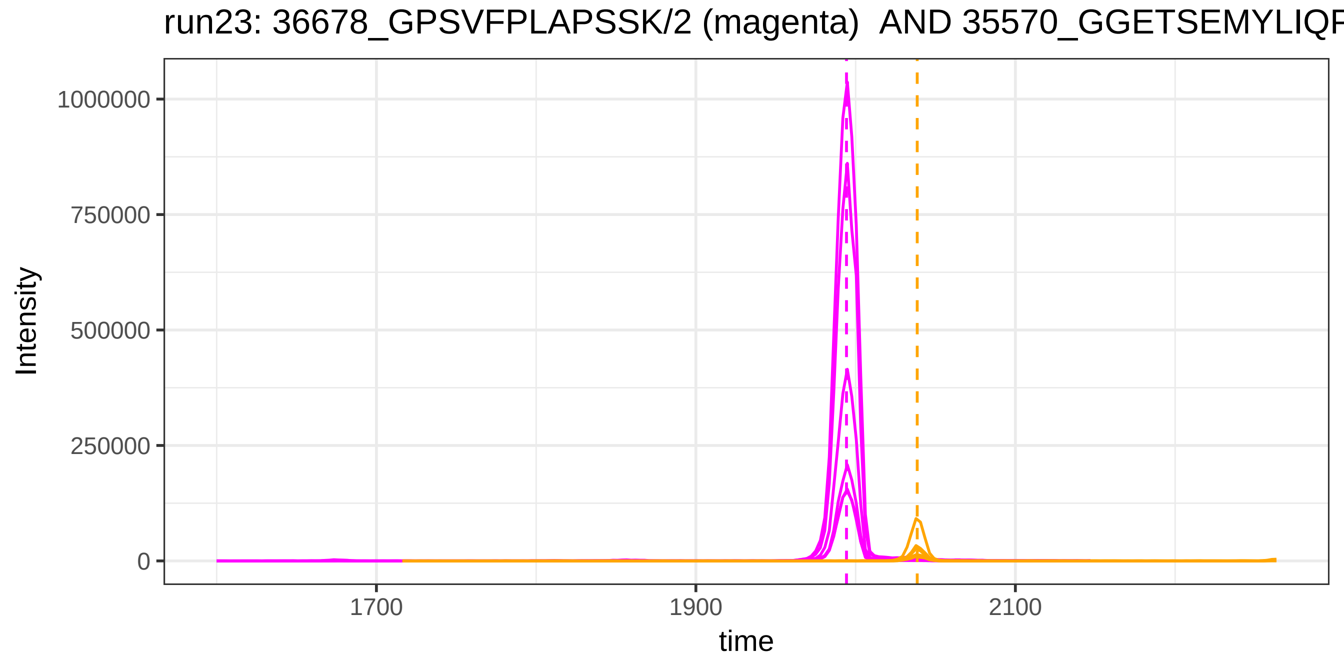
run23: 36678_GPSVFPLAPSSK/2 (magenta) AND 35570_GGETSEMYLIQF
Intensity
1000000
750000
500000
250000
0
1700
1900
2100
time

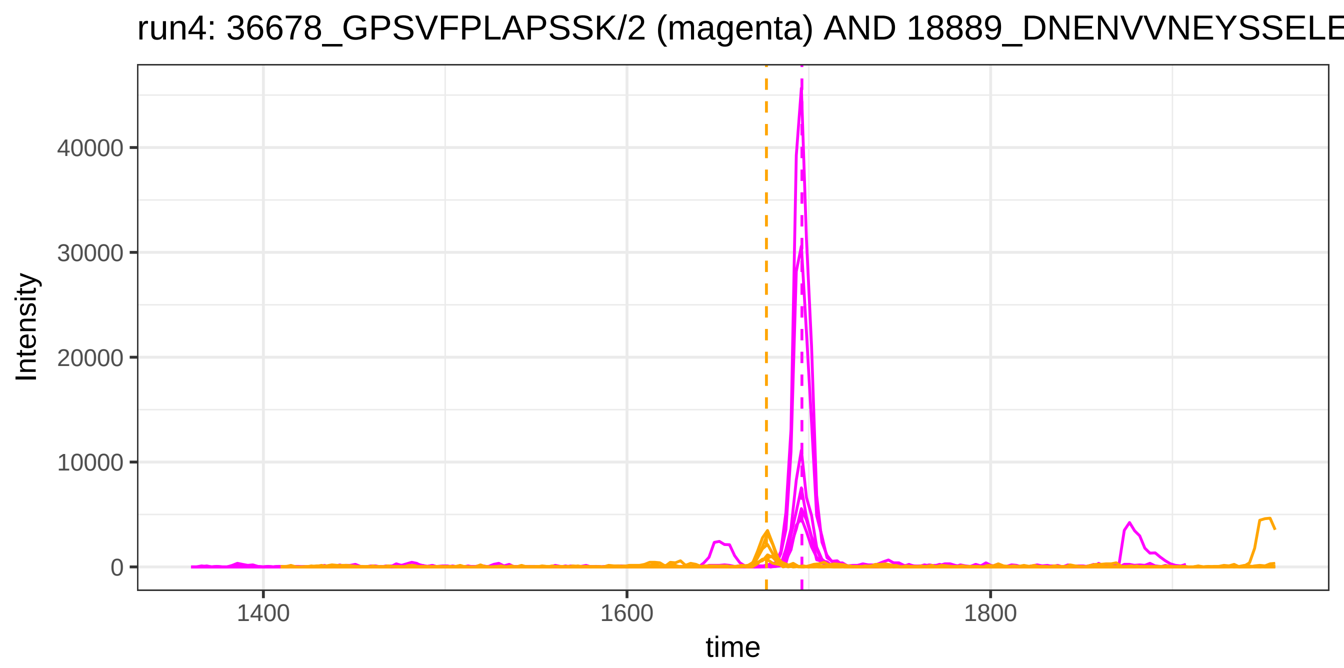

run4: 36678_GPSVFPLAPSSK/2 (magenta) AND 18889_DNENVVNEYSSELE
Intensity
40000
30000
20000
10000
0
1400
1600
1800
time


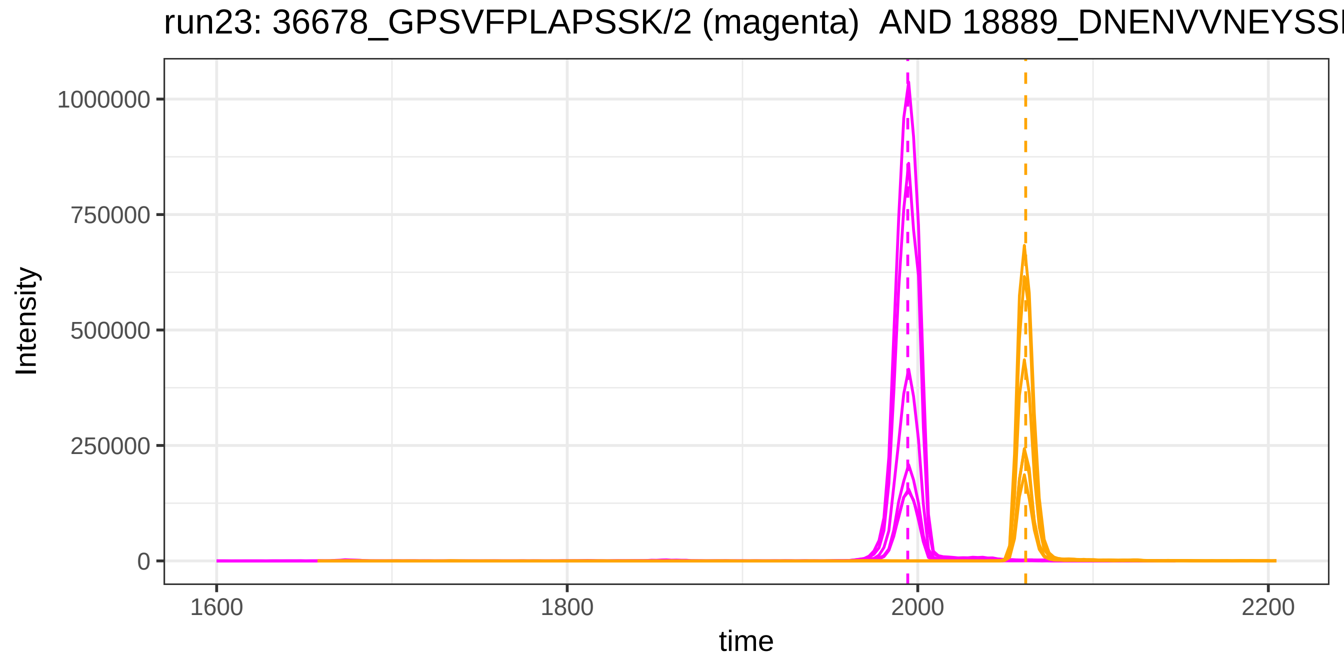

run23: 36678_GPSVFPLAPSSK/2 (magenta) AND 18889_DNENVVNEYSS
Intensity
1000000
750000
500000
250000
0
1600
1800
2000
2200
time

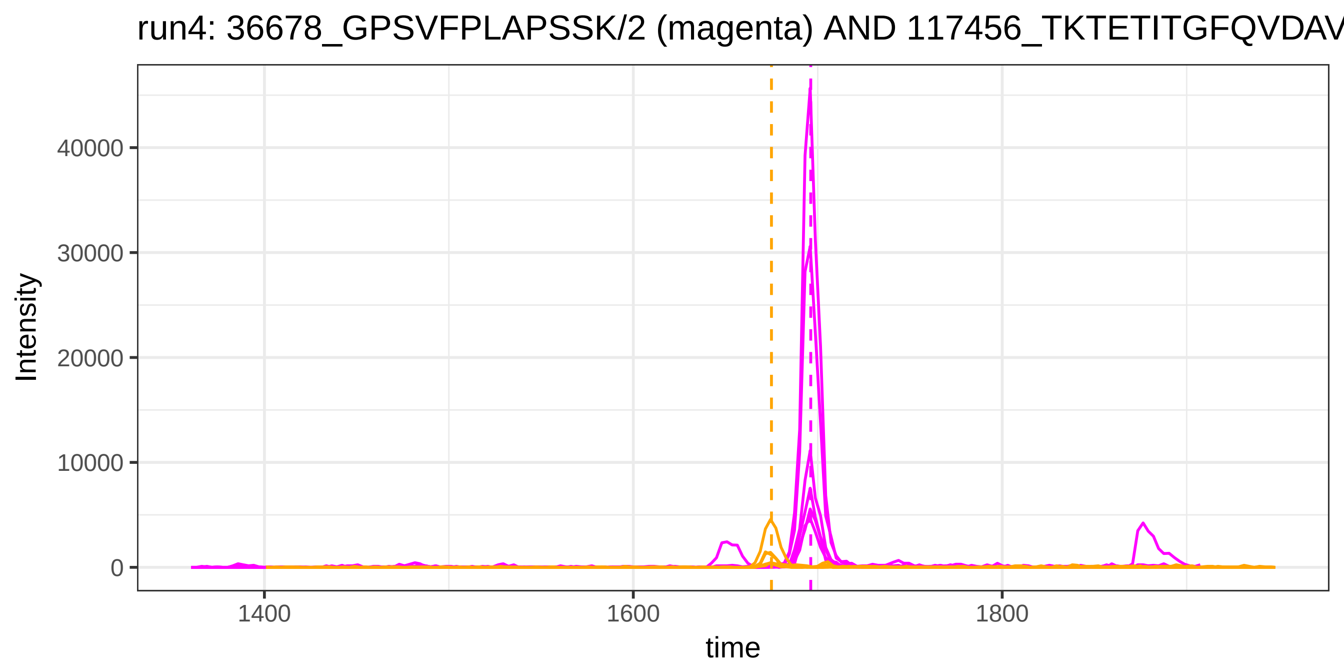

run4: 36678_GPSVFPLAPSSK/2 (magenta) AND 117456_TKTETITGFQVDAV
Intensity
40000
30000
20000
10000
0
1400
1600
1800
time


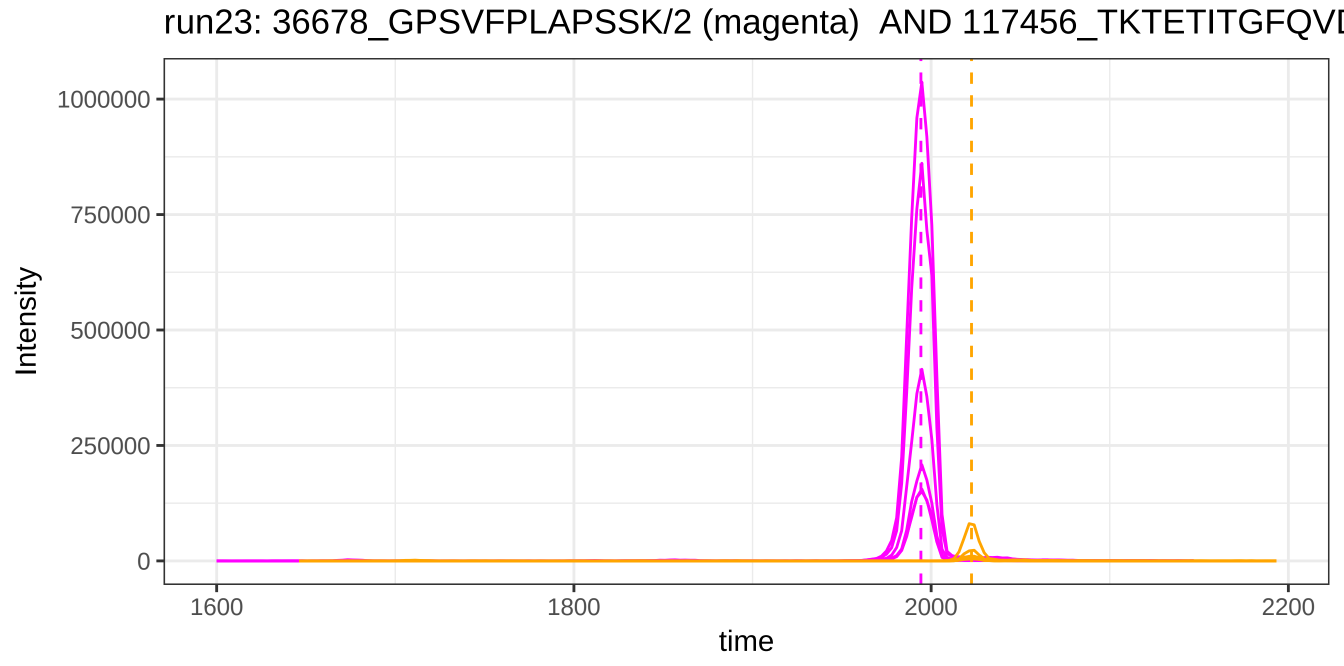

run23: 36678_GPSVFPLAPSSK/2 (magenta) AND 117456_TKTETITGFQVD
Intensity
1000000
750000
500000
250000
0
1600
1800
2000
2200
time

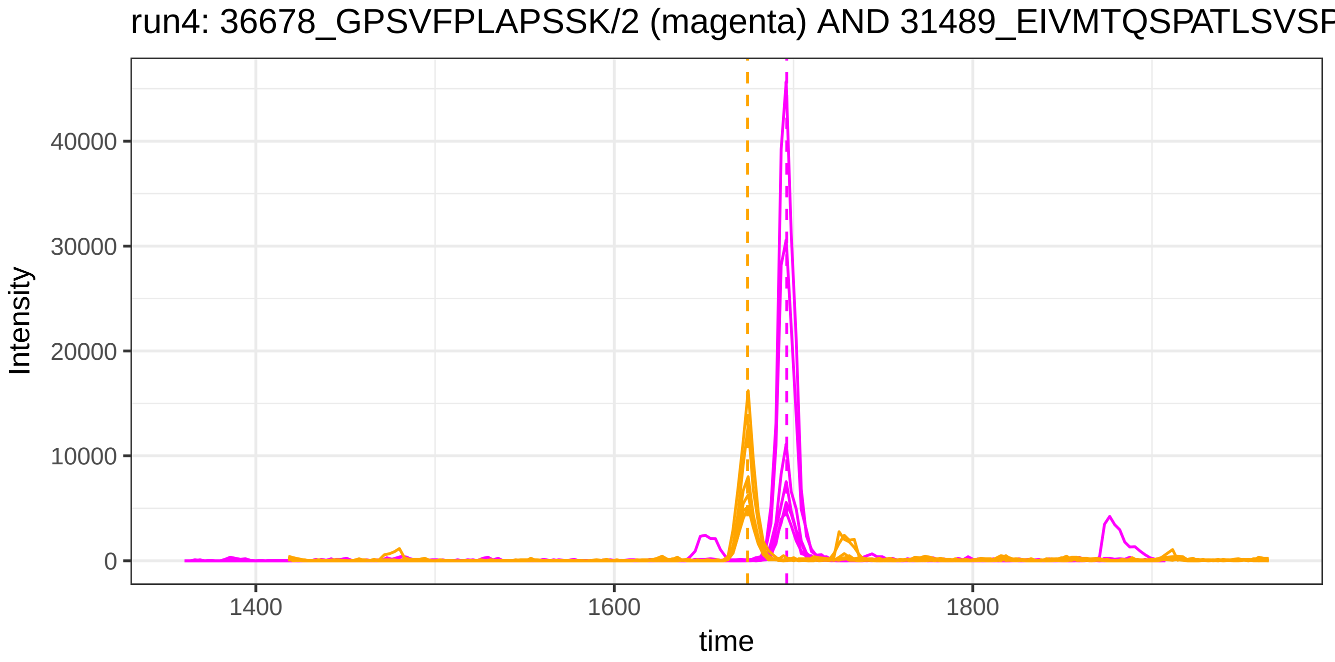
run4: 36678_GPSVFPLAPSSK/2 (magenta) AND 31489_EIVMTQSPATLSVSP
Intensity
0
10000
20000
30000
40000
1400
1600
1800
time

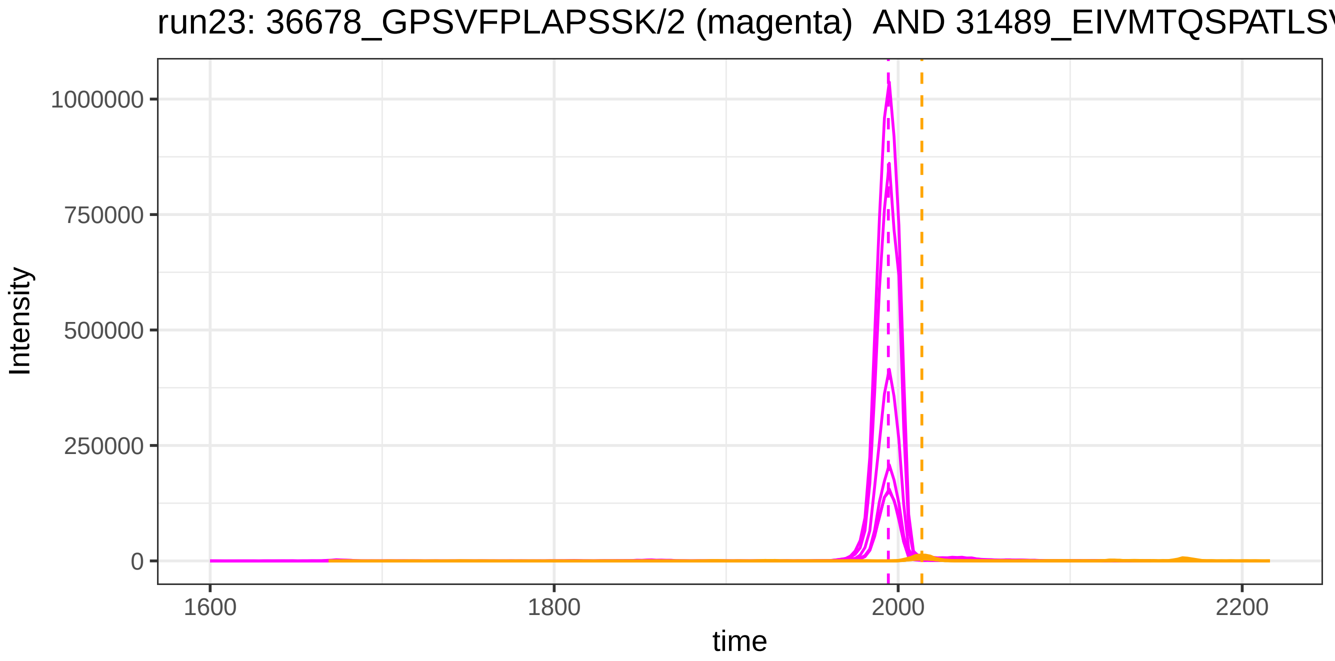
run23: 36678_GPSVFPLAPSSK/2 (magenta) AND 31489_EIVMTQSPATLS
Intensity
0
250000
500000
750000
1000000
1600
1800
2000
2200
time

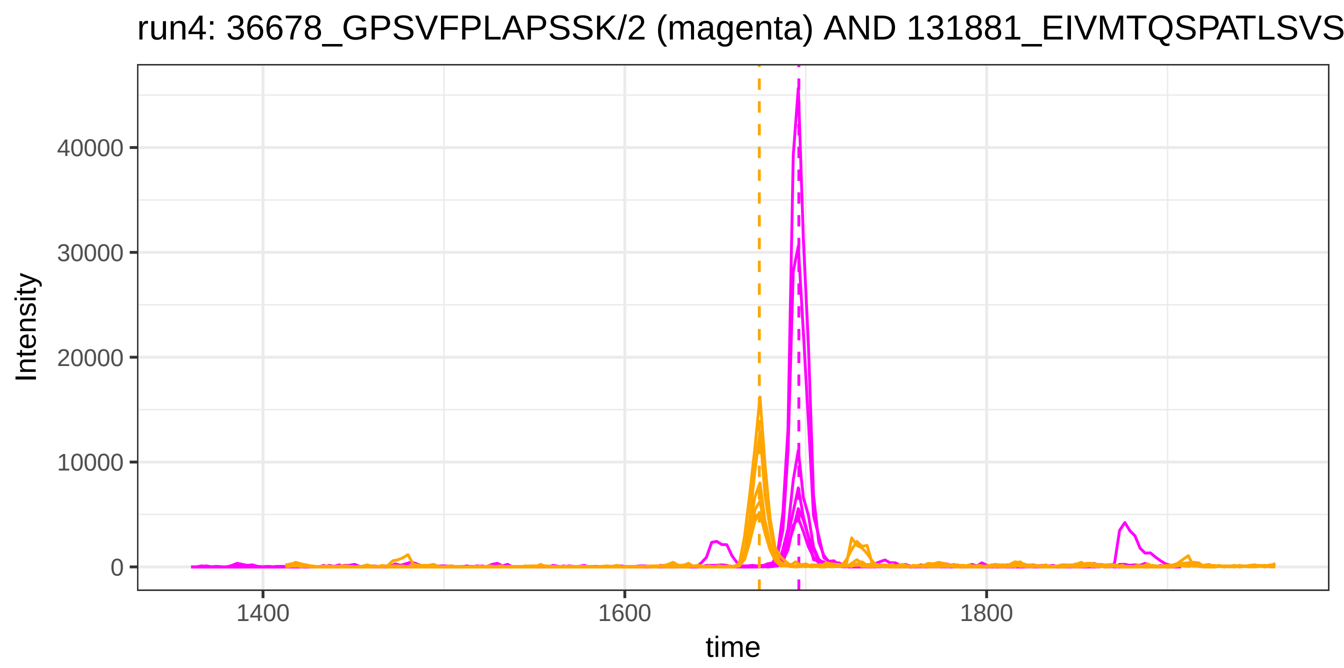
run4: 36678_GPSVFPLAPSSK/2 (magenta) AND 131881_EIVMTQSPATLSVS
Intensity
40000
30000
20000
10000
0
1400
1600
1800
time

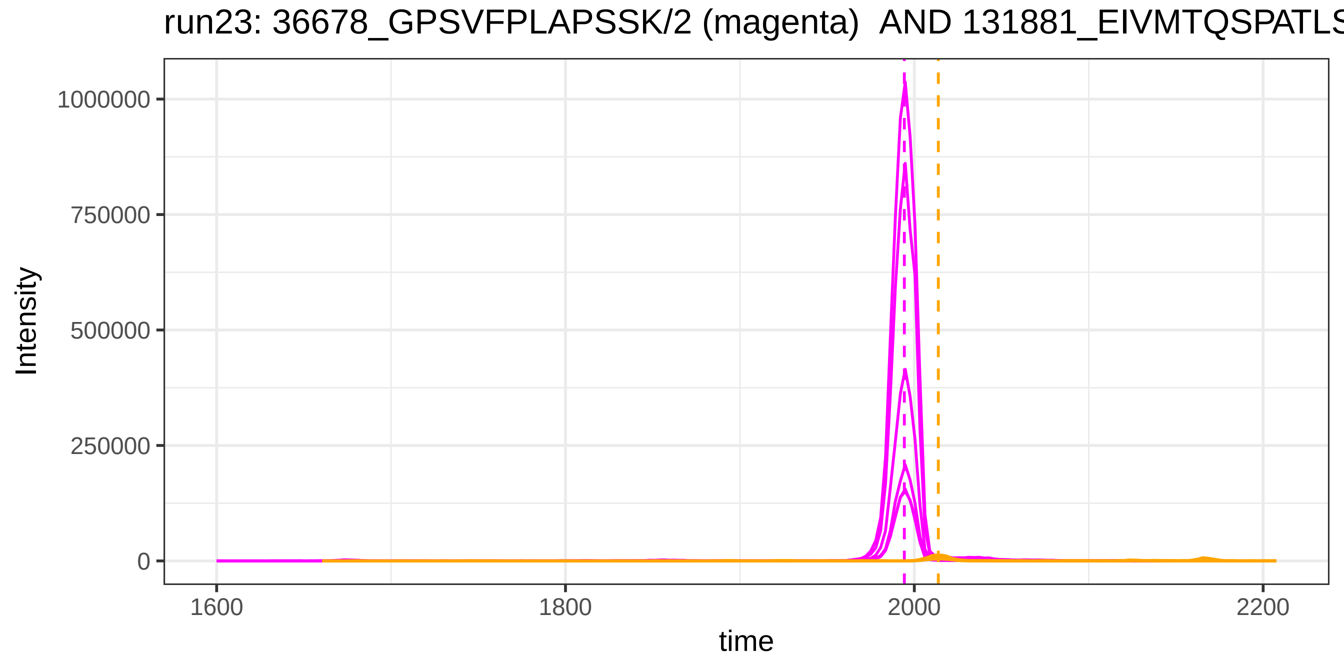
run23: 36678_GPSVFPLAPSSK/2 (magenta) AND 131881_EIVMTQSPATLS
Intensity
1000000
750000
500000
250000
0
1600
1800
2000
2200
time

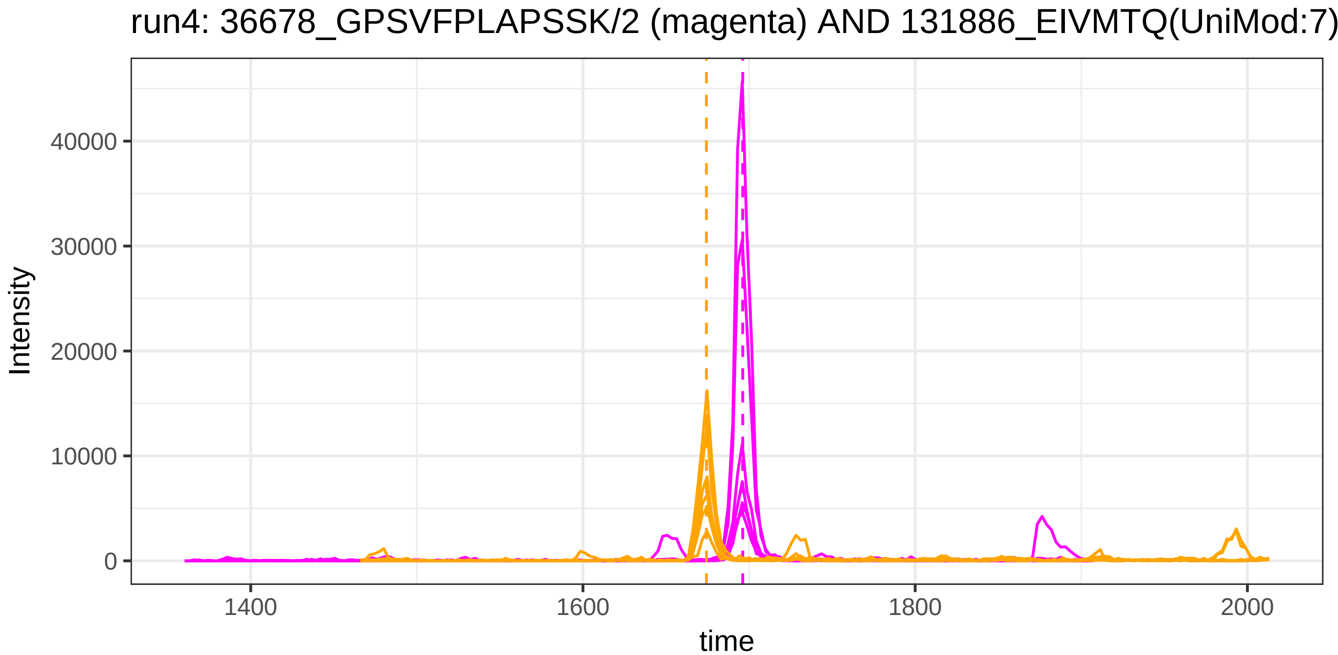
run4: 36678_GPSVFPLAPSSK/2 (magenta) AND 131886_EIVMTQ(UniMod:7)
Intensity
40000
30000
20000
10000
0
1400
1600
1800
2000
time

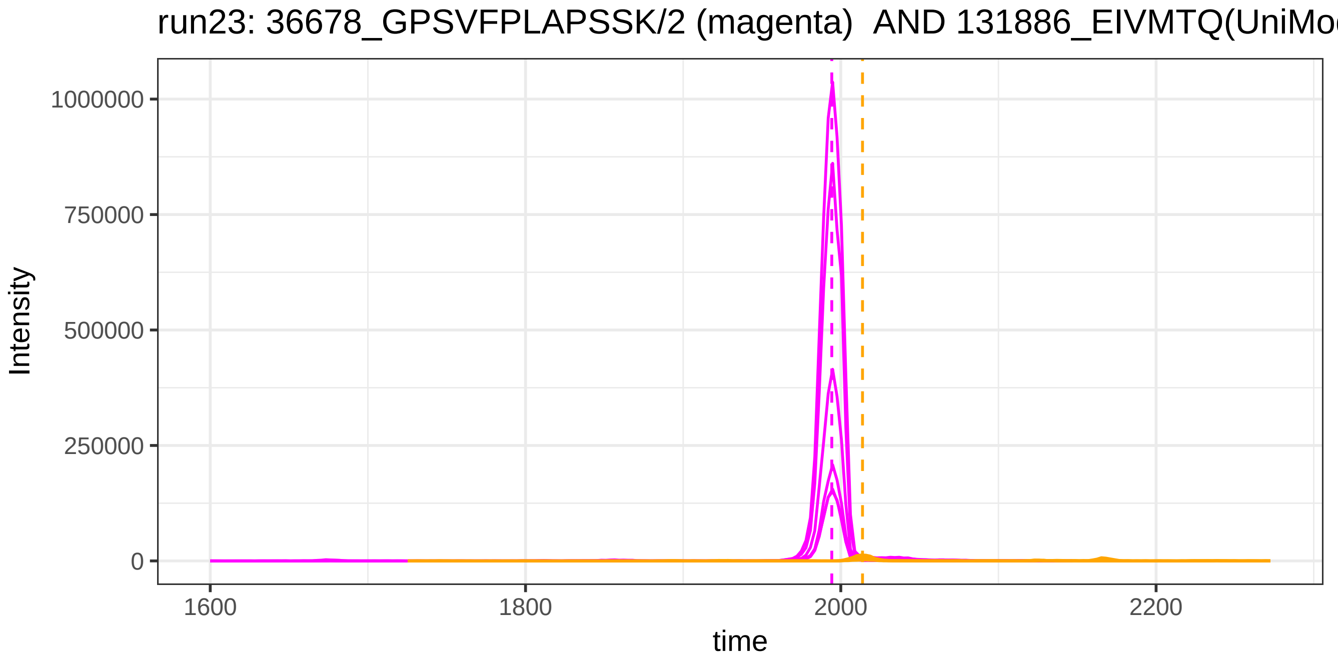
run23: 36678_GPSVFPLAPSSK/2 (magenta) AND 131886_EIVMTQ(UniMo
Intensity
1000000
750000
500000
250000
0
1600
1800
2000
2200
time

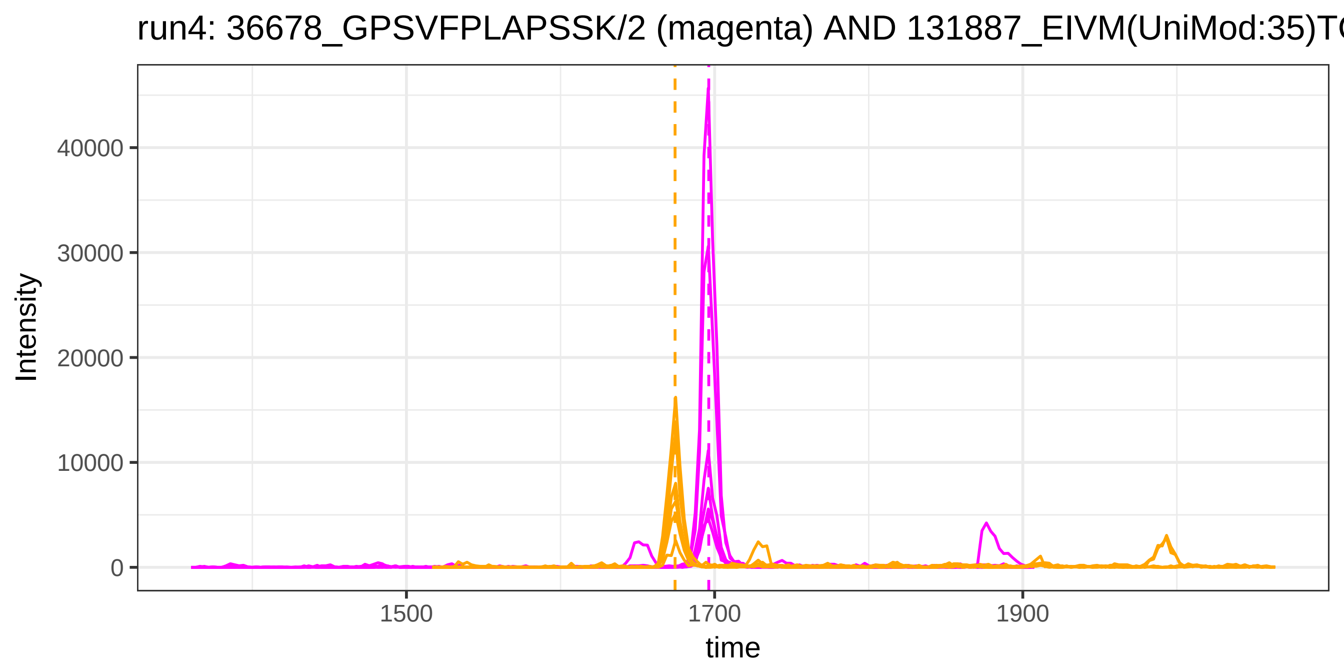

run4: 36678_GPSVFPLAPSSK/2 (magenta) AND 131887_EIVM(UniMod:35)T
Intensity
40000
30000
20000
10000
0
1500
1700
1900
time


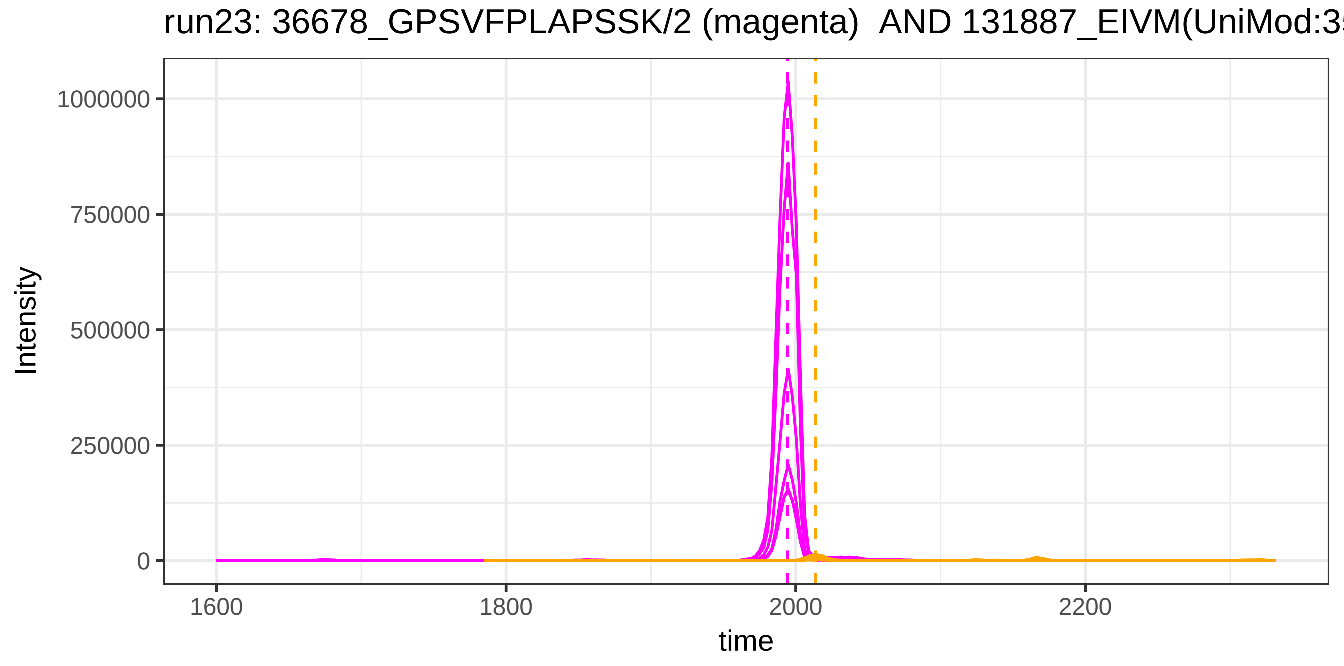

run23: 36678_GPSVFPLAPSSK/2 (magenta) AND 131887_EIVM(UniMod:3
Intensity
1000000
750000
500000
250000
0
1600
1800
2000
2200
time

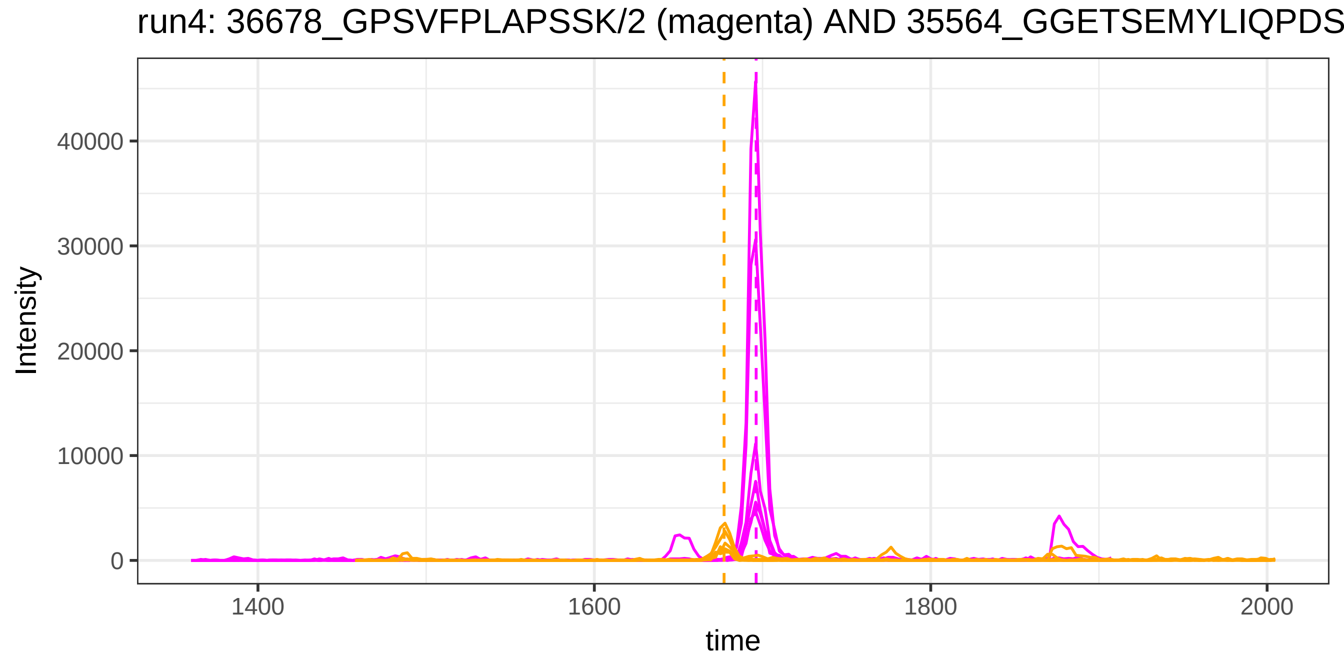
run4: 36678_GPSVFPLAPSSK/2 (magenta) AND 35564_GGETSEMYLIQPDS
Intensity
40000
30000
20000
10000
0
1400
1600
1800
2000
time

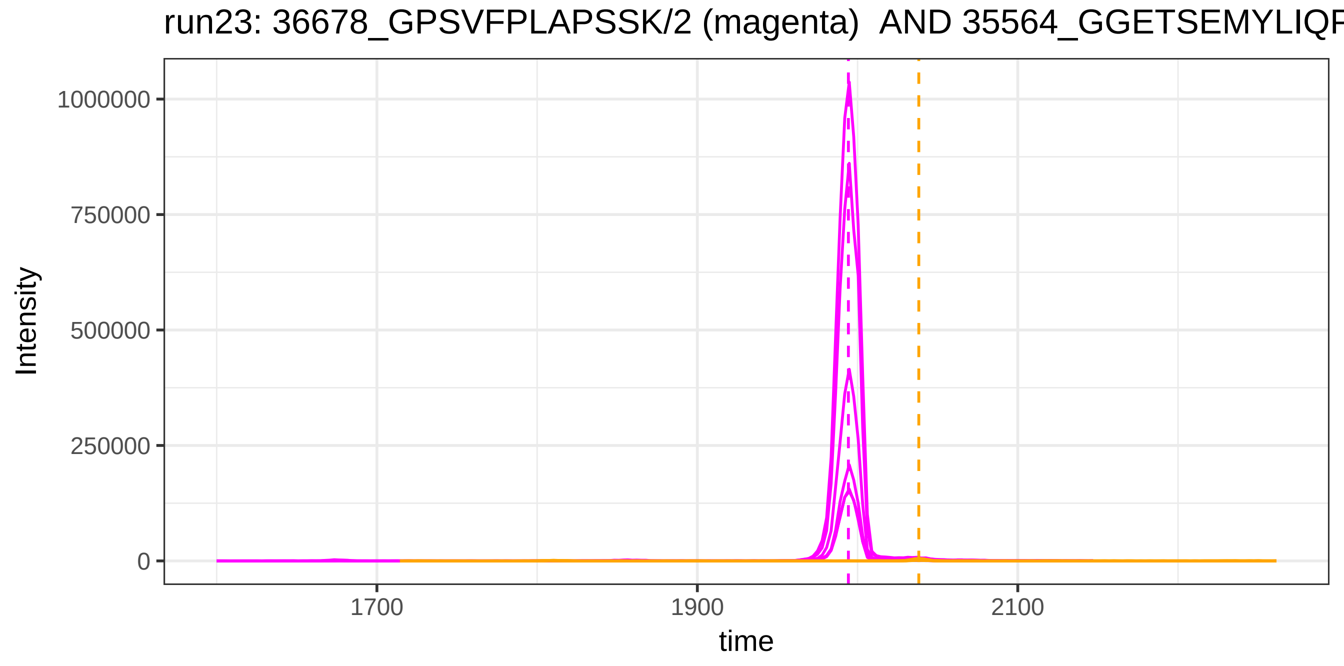
run23: 36678_GPSVFPLAPSSK/2 (magenta) AND 35564_GGETSEMYLIQF
Intensity
1000000
750000
500000
250000
0
1700
1900
2100
time

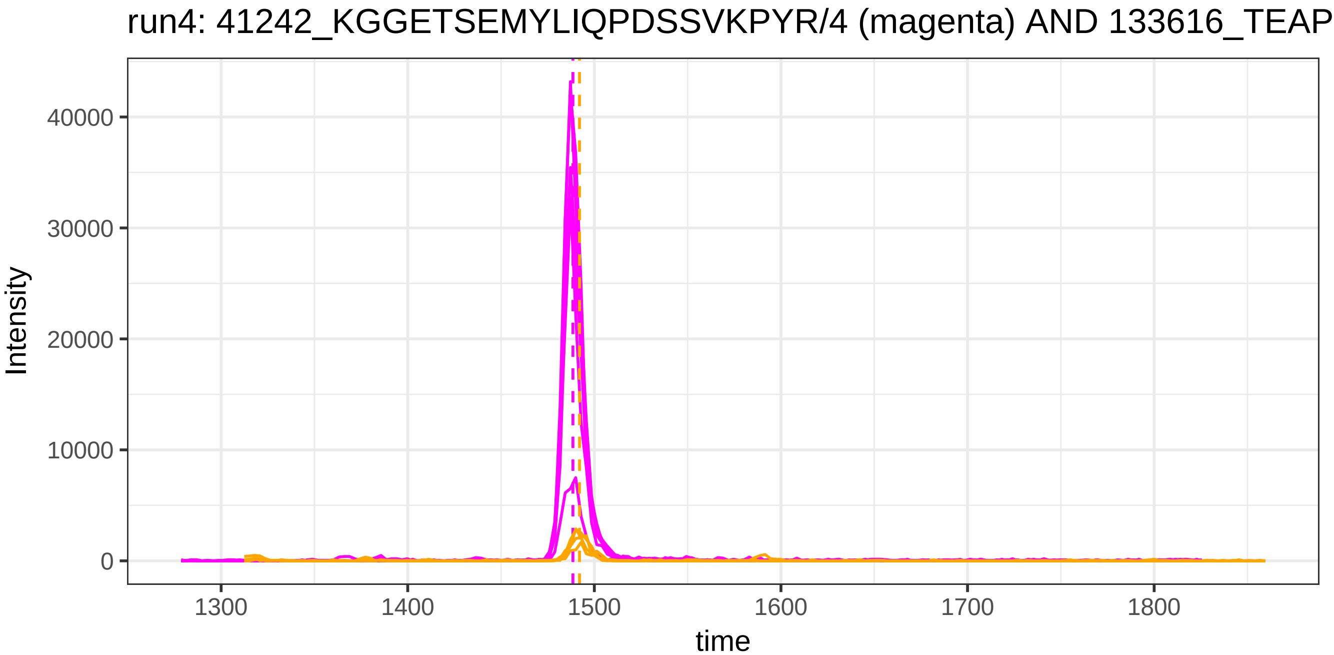

run4: 41242_KGGETSEMYLIQPDSSVKPYR/4 (magenta) AND 133616_TEAP
Intensity
40000
30000
20000
10000
0
1300
1400
1500
1600
1700
1800
time


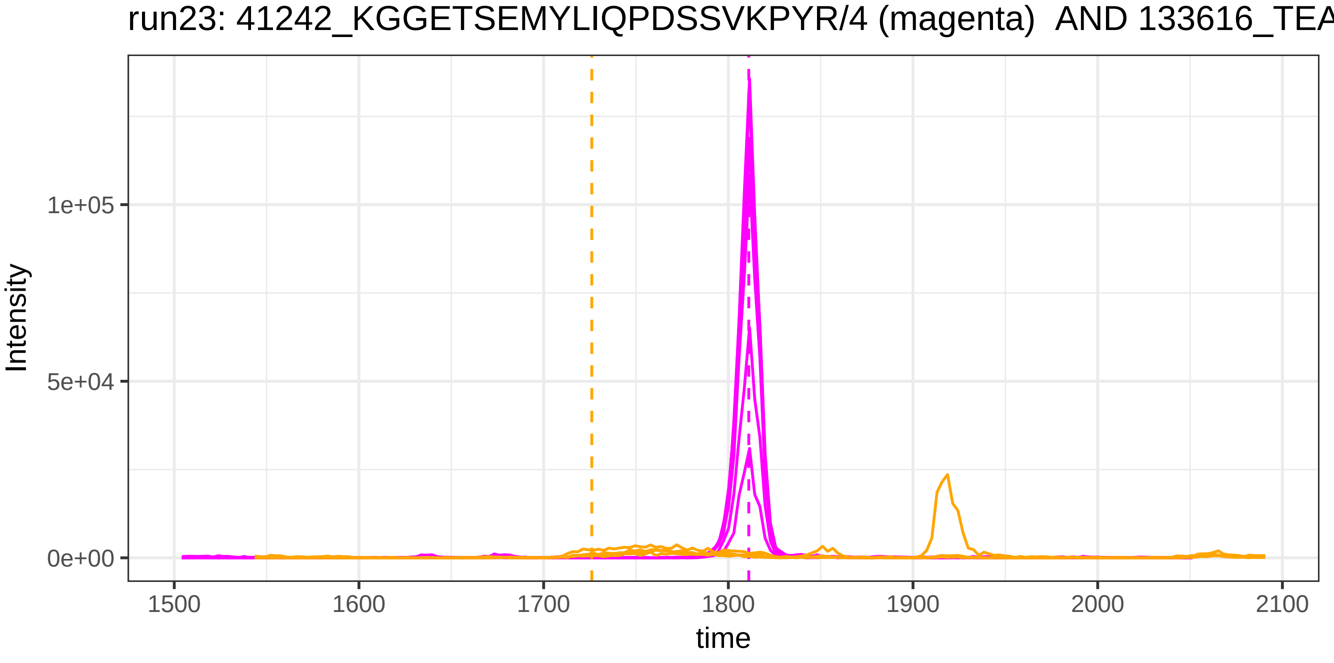

run23: 41242_KGGETSEMYLIQPDSSVKPYR/4 (magenta) AND 133616_TEA
Intensity
1e+05
5e+04
0e+00
1500
1600
1700
1800
1900
2000
2100
time

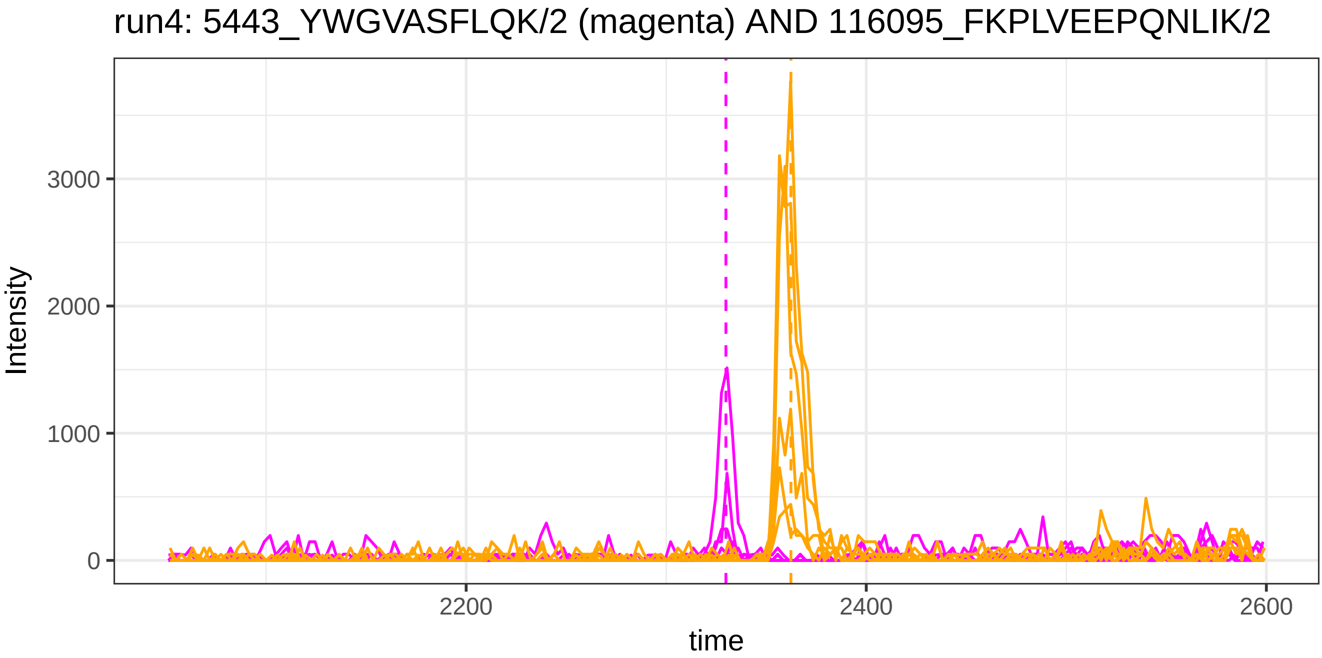
run4: 5443_YWGVASFLQK/2 (magenta) AND 116095_FKPLVEEPQNLIK/2
Intensity
3000
2000
1000
0
2200
2400
2600
time

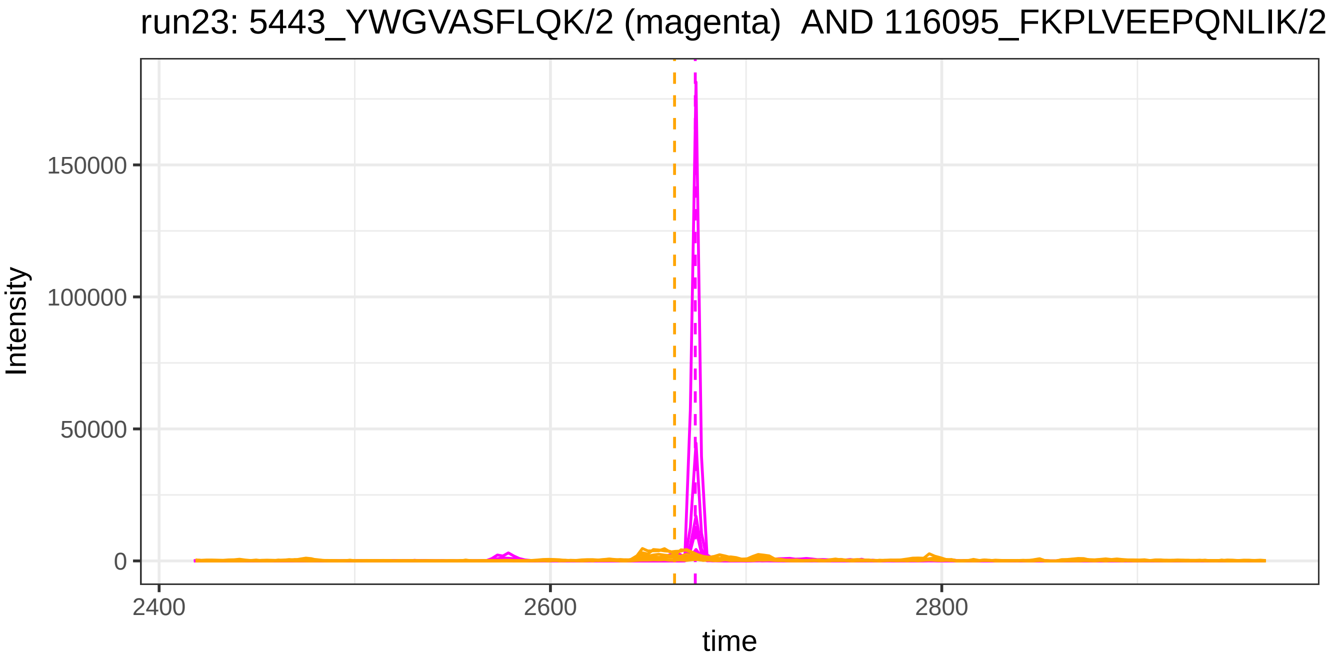
run23: 5443_YWGVASFLQK/2 (magenta)  AND 116095_FKPLVEEPQNLIK/2
Intensity
150000
100000
50000
0
2400
2600
2800
time

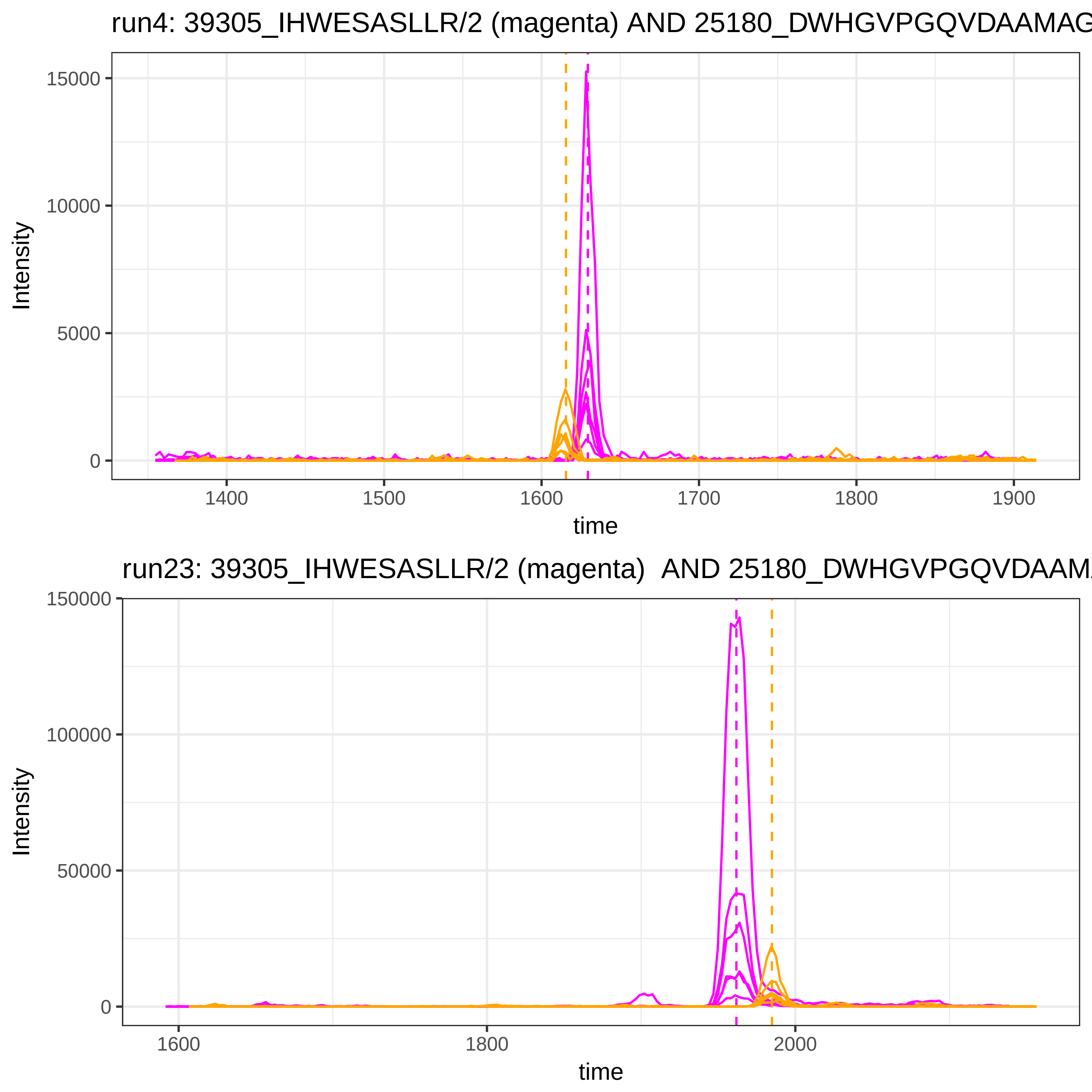
run4: 39305_IHWESASLLR/2 (magenta) AND 25180_DWHGVPGQVDAAMAG
Intensity
15000
10000
5000
0
1400
1500
1600
1700
1800
1900
time
run23: 39305_IHWESASLLR/2 (magenta) AND 25180_DWHGVPGQVDAAM
Intensity
150000
100000
50000
0
1600
1800
2000
time

run4: 49521_ATEHLSTLSEK/2 (magenta) AND 49416_ATEDEGSEQKIPEATN

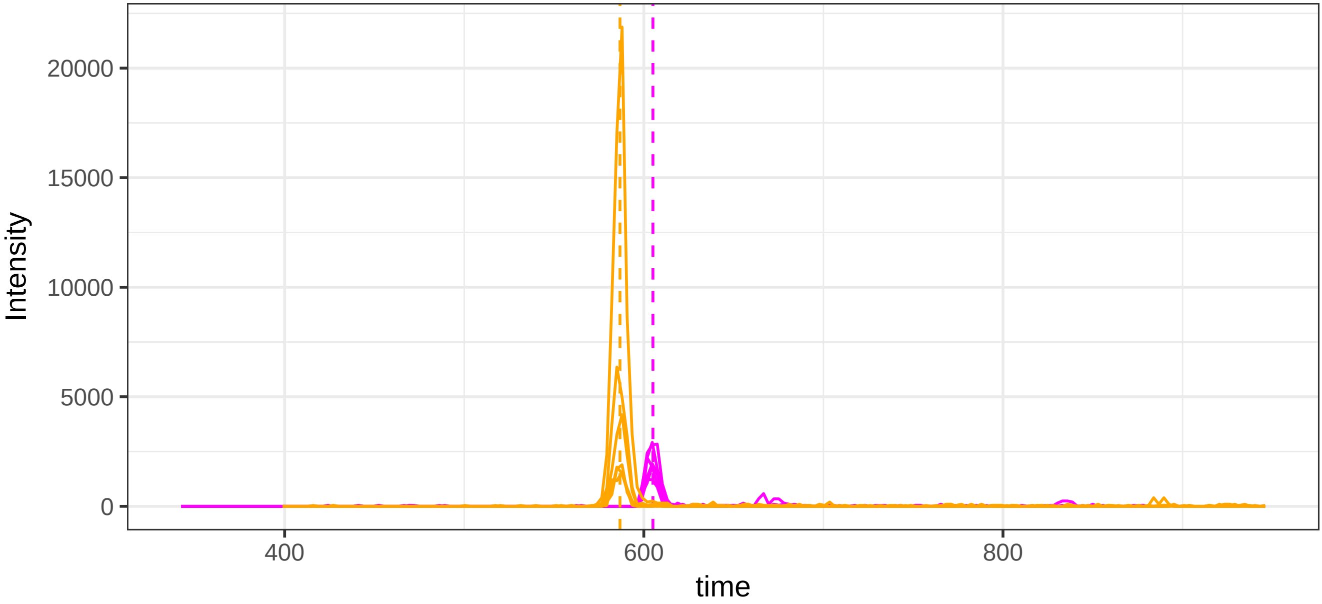


run23: 49521_ATEHLSTLSEK/2 (magenta) AND 49416_ATEDEGSEQKIPEAT

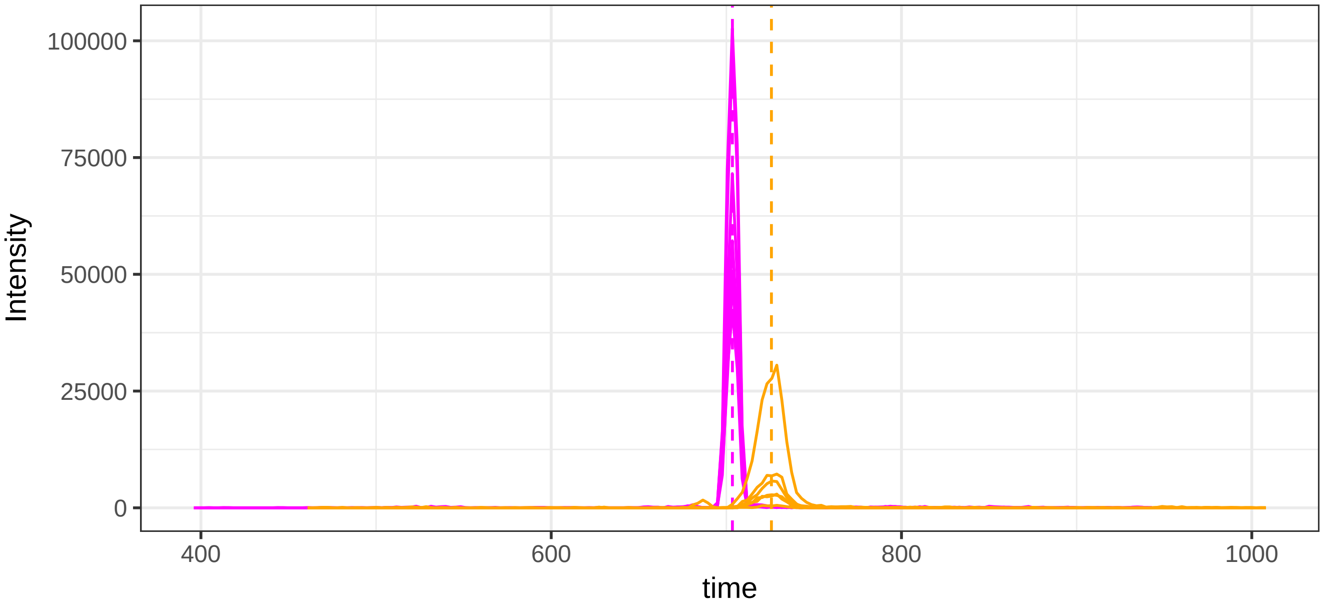

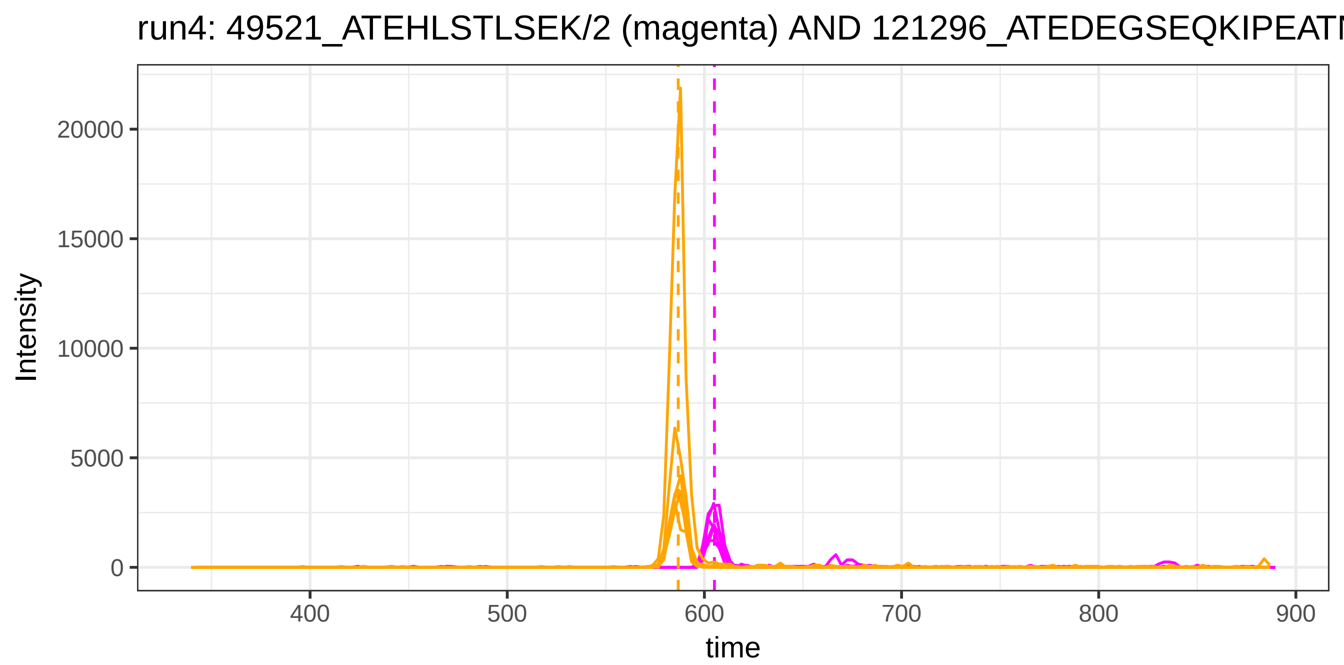
run4: 49521_ATEHLSTLSEK/2 (magenta) AND 121296_ATEDEGSEQKIPEATI
Intensity
20000
15000
10000
5000
0
400
500
600
700
800
900
time

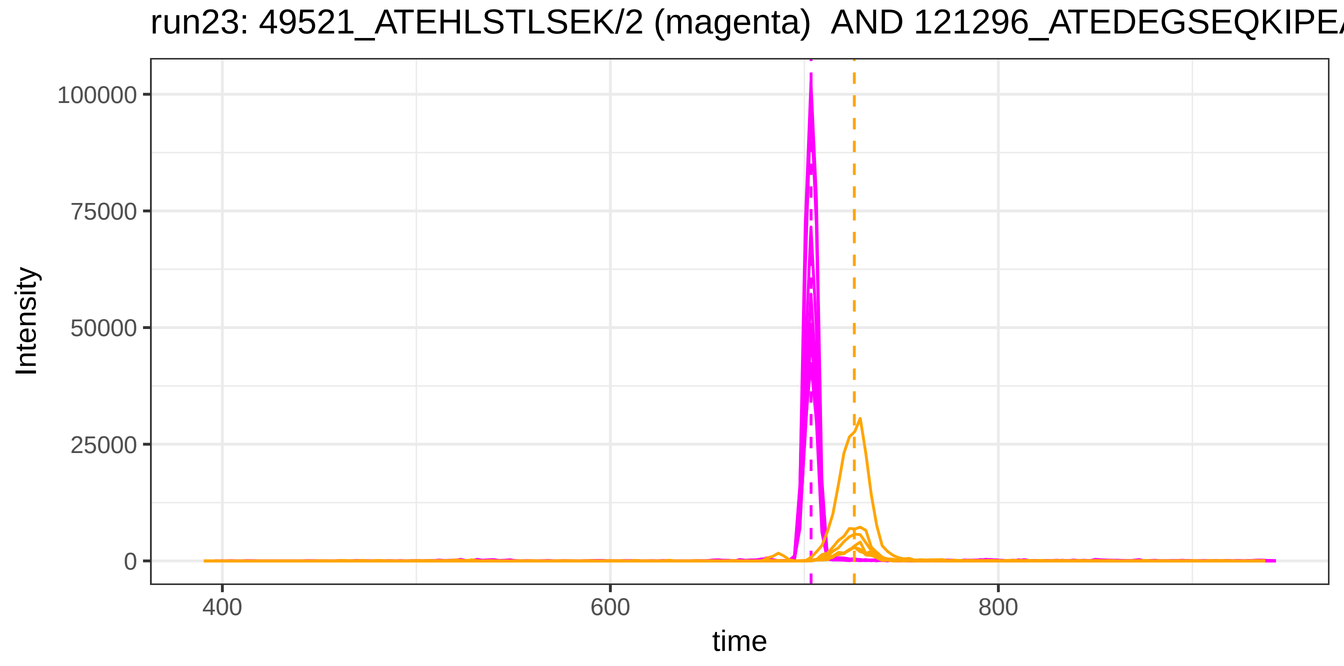
run23: 49521_ATEHLSTLSEK/2 (magenta) AND 121296_ATEDEGSEQKIPEA
Intensity
100000
75000
50000
25000
0
400
600
800
time

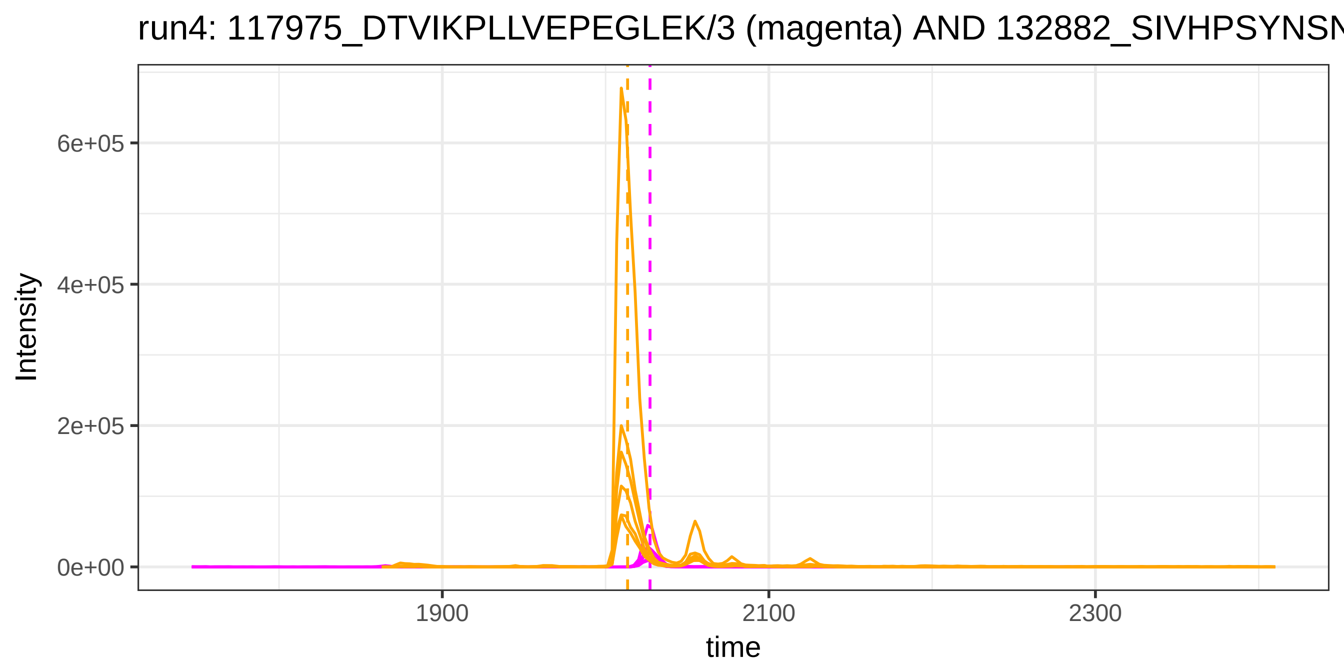
run4: 117975_DTVIKPLLVEPEGLEK/3 (magenta) AND 132882_SIVHPSYNSN
Intensity
6e+05
4e+05
2e+05
0e+00
1900
2100
2300
time

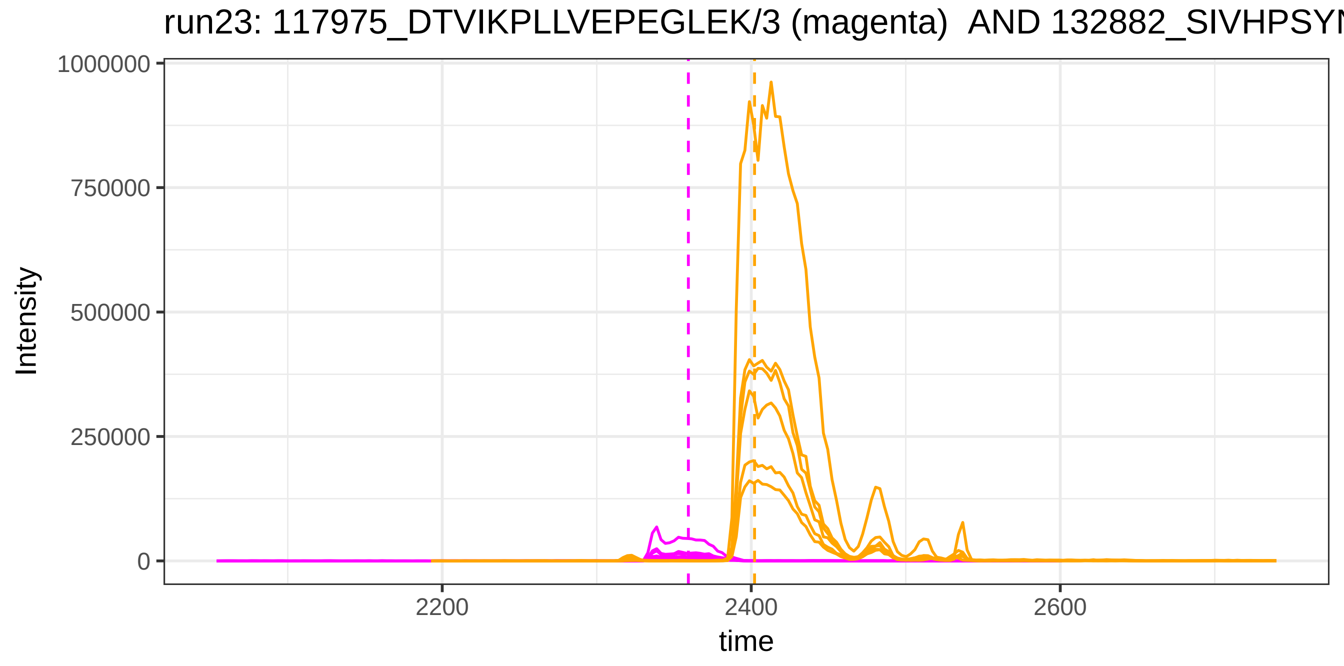
run23: 117975_DTVIKPLLVEPEGLEK/3 (magenta) AND 132882_SIVHPSYN
Intensity
1000000
750000
500000
250000
0
2200
2400
2600
time

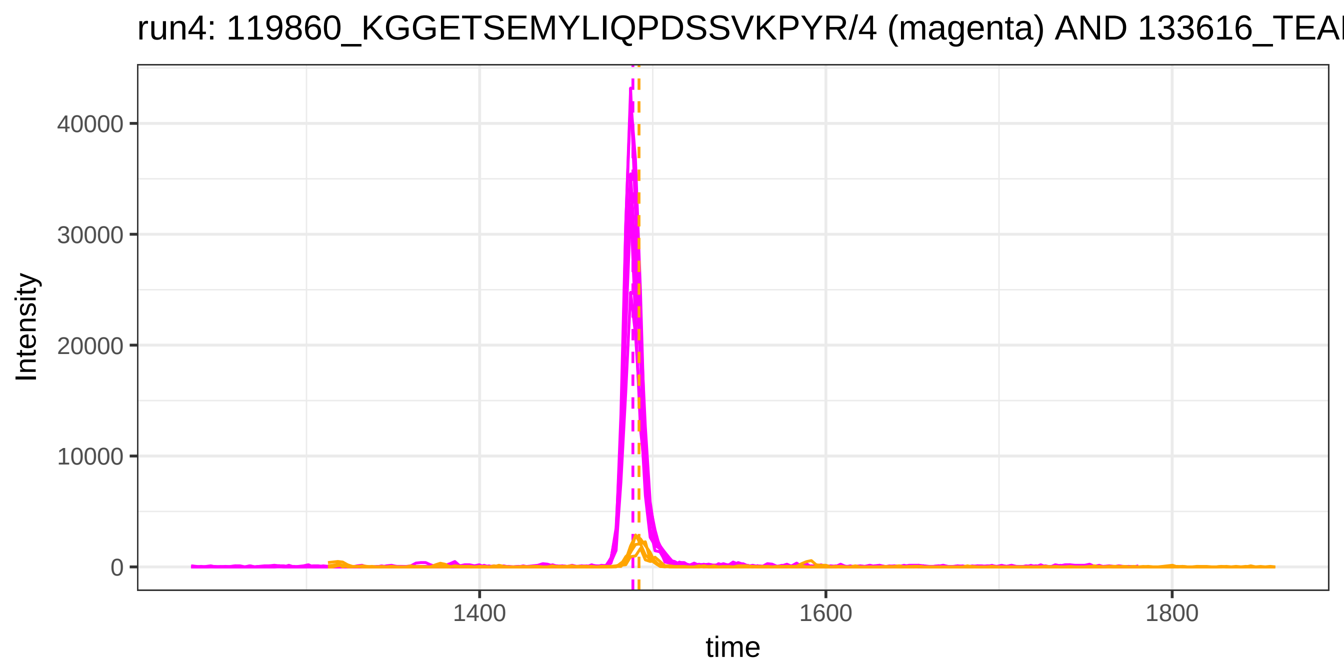
run4: 119860_KGGETSEMYLIQPDSSVKPYR/4 (magenta) AND 133616_TEA
Intensity
40000
30000
20000
10000
0
1400
1600
1800
time

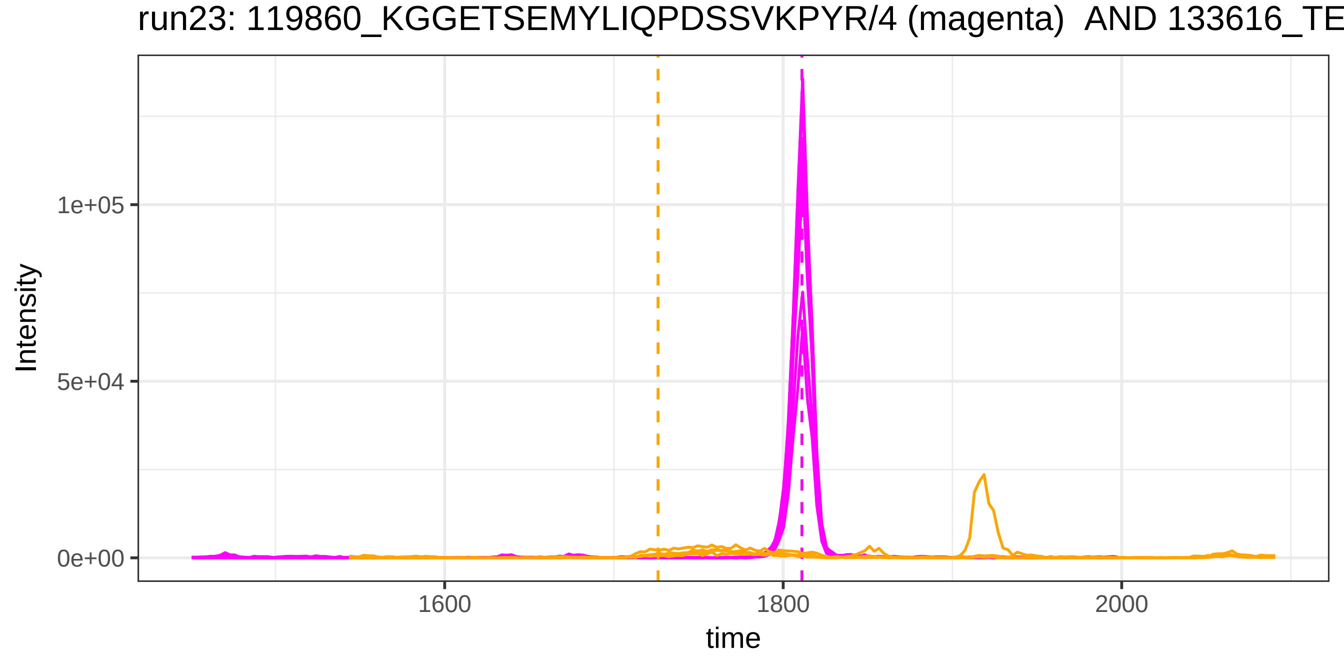
run23: 119860_KGGETSEMYLIQPDSSVKPYR/4 (magenta) AND 133616_TE
Intensity
1e+05
5e+04
0e+00
1600
1800
2000
time

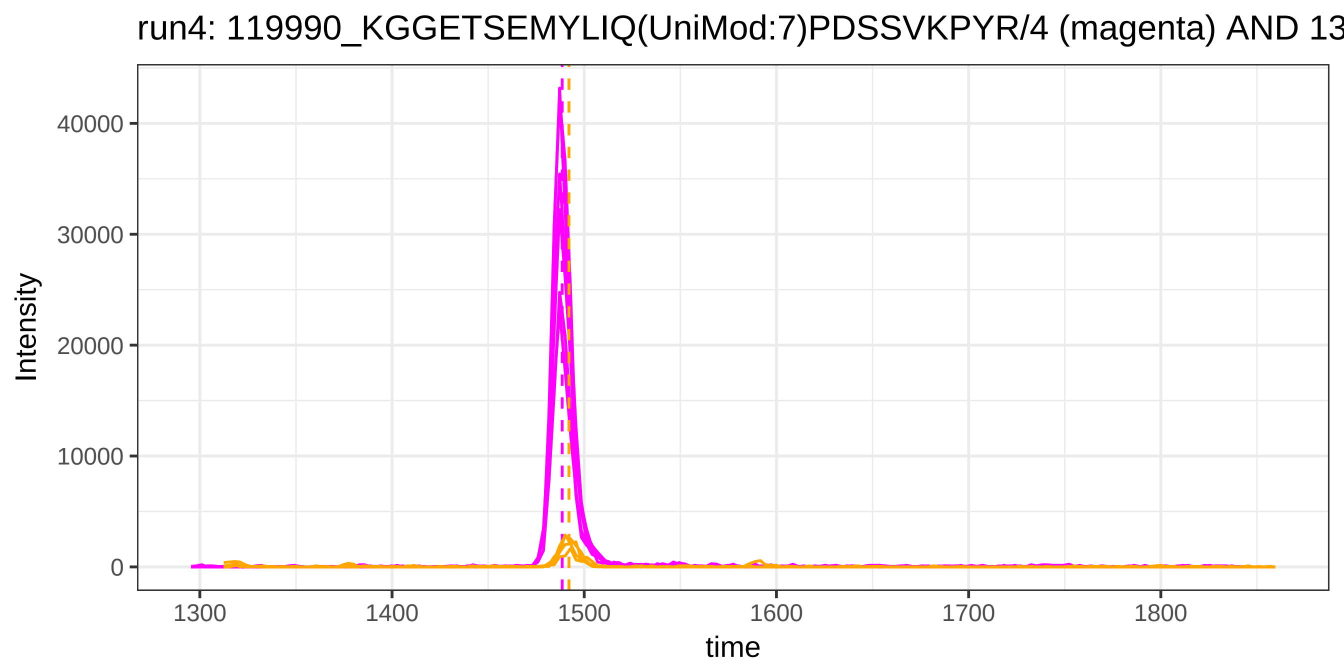
run4: 119990_KGGETSEMYLIQ(UniMod:7)PDSSVKPYR/4 (magenta) AND 13
Intensity
40000
30000
20000
10000
0
1300
1400
1500
1600
1700
1800
time

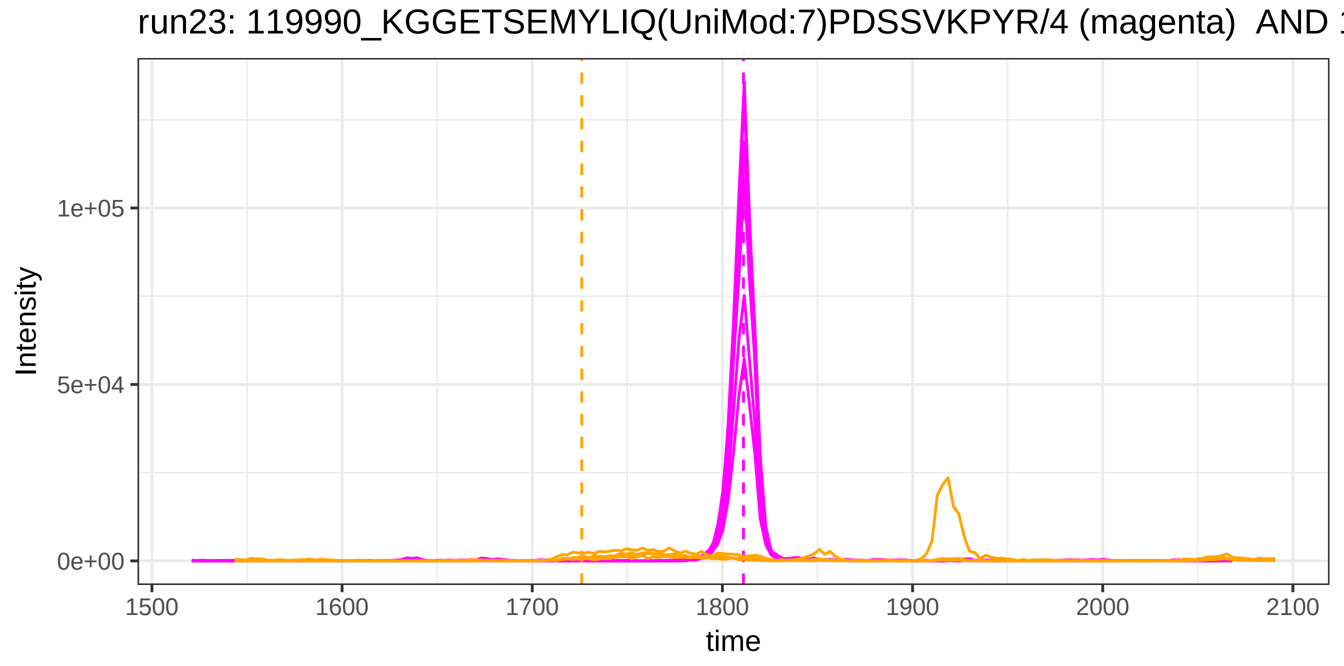
run23: 119990_KGGETSEMYLIQ(UniMod:7)PDSSVKPYR/4 (magenta) AND 1
Intensity
1e+05
5e+04
0e+00
1500
1600
1700
1800
1900
2000
2100
time

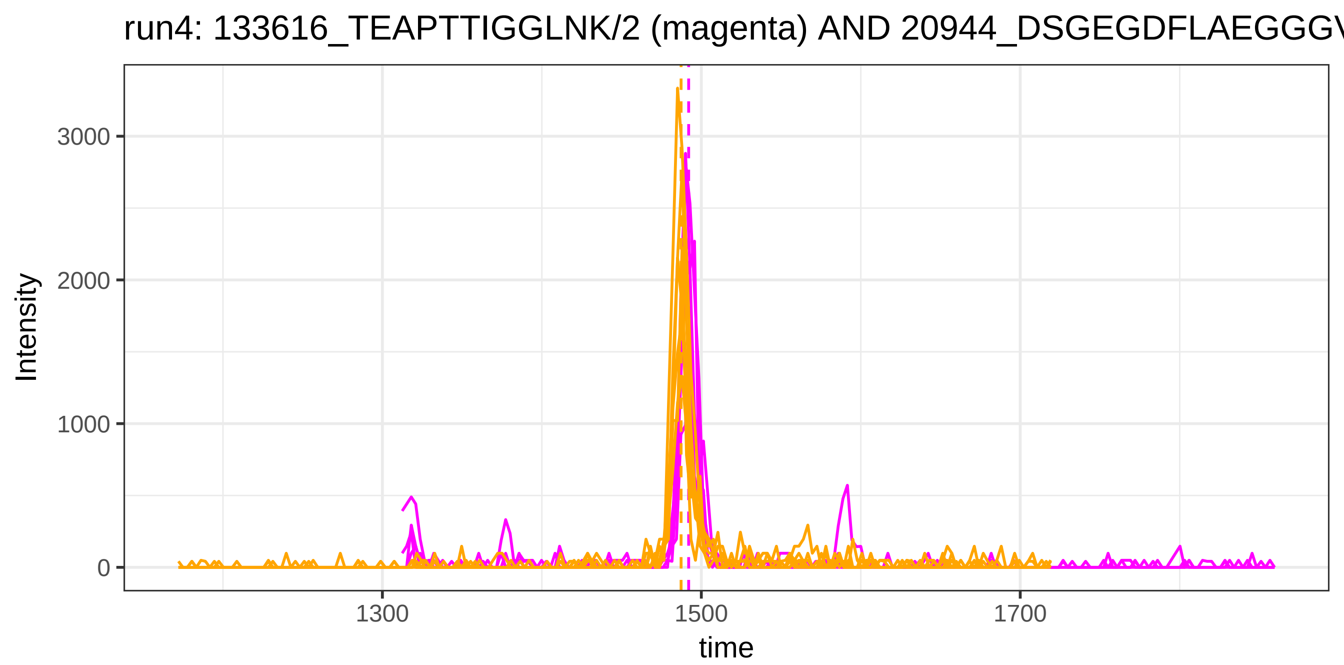
run4: 133616_TEAPTTIGGLNK/2 (magenta) AND 20944_DSGEGDFLAEGGGV
Intensity
3000
2000
1000
0
1300
1500
1700
time

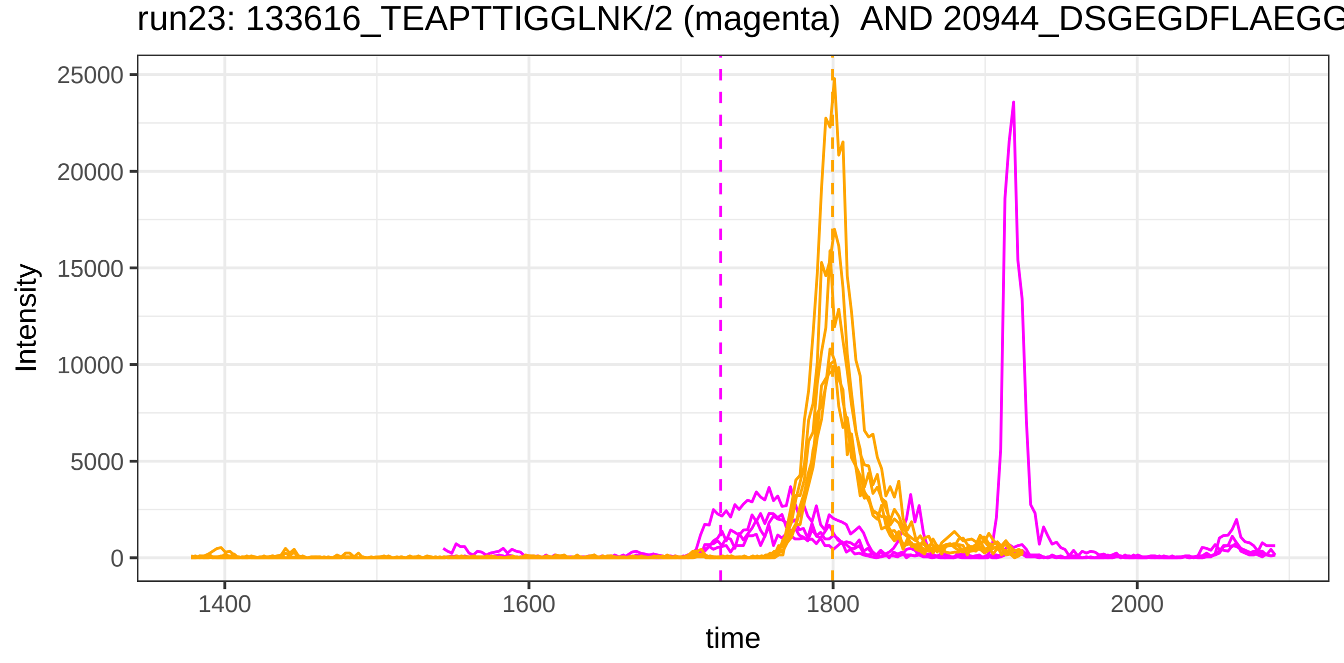
run23: 133616_TEAPTTIGGLNK/2 (magenta) AND 20944_DSGEGDFLAEGG
Intensity
25000
20000
15000
10000
5000
0
1400
1600
1800
2000
time

run4: 133616_TEAPTTIGGLNK/2 (magenta) AND 119499_DSGEGDFLAEGGG

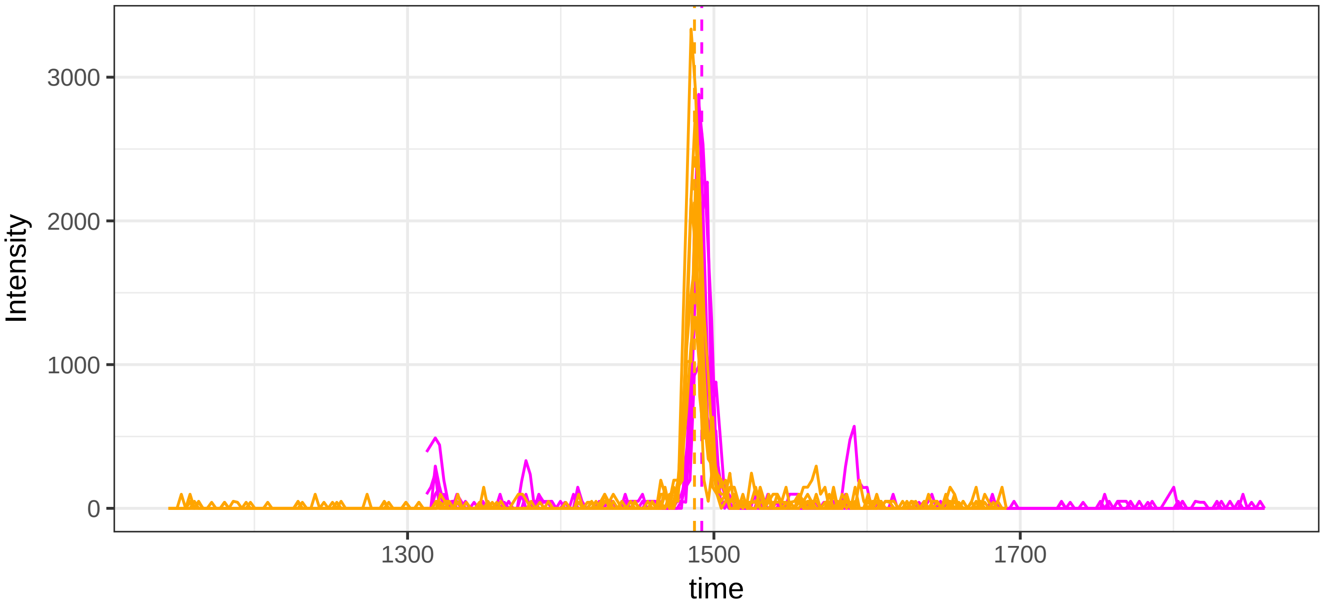


run23: 133616_TEAPTTIGGLNK/2 (magenta)  AND 119499_DSGEGDFLAEG

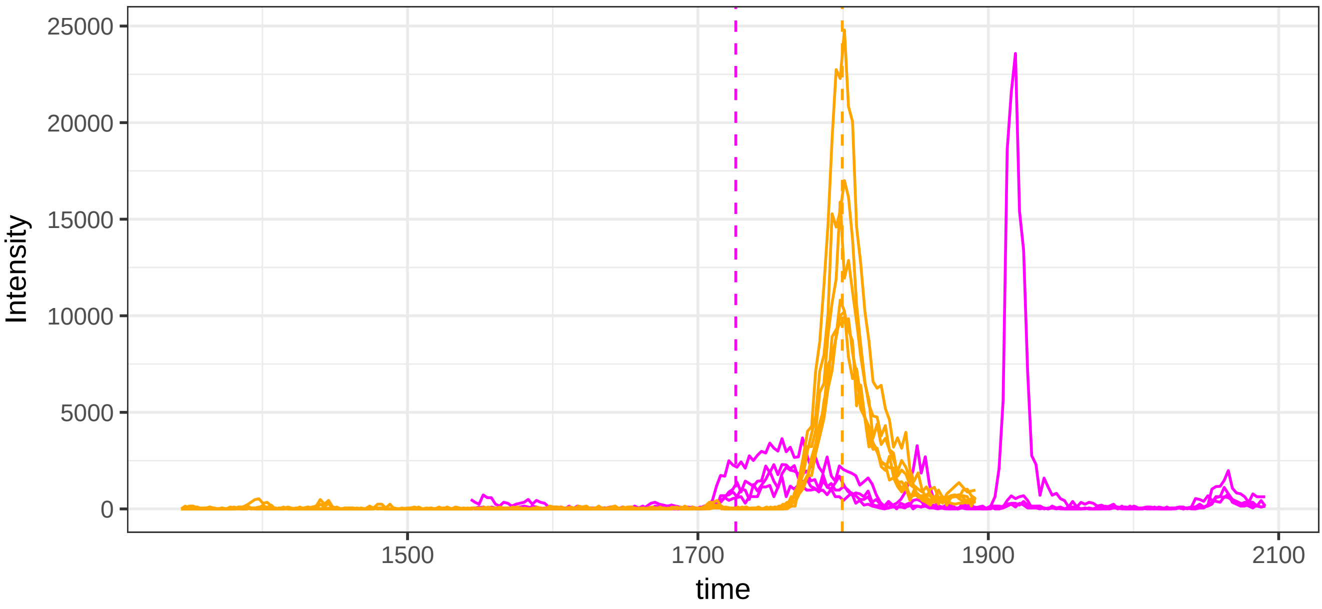

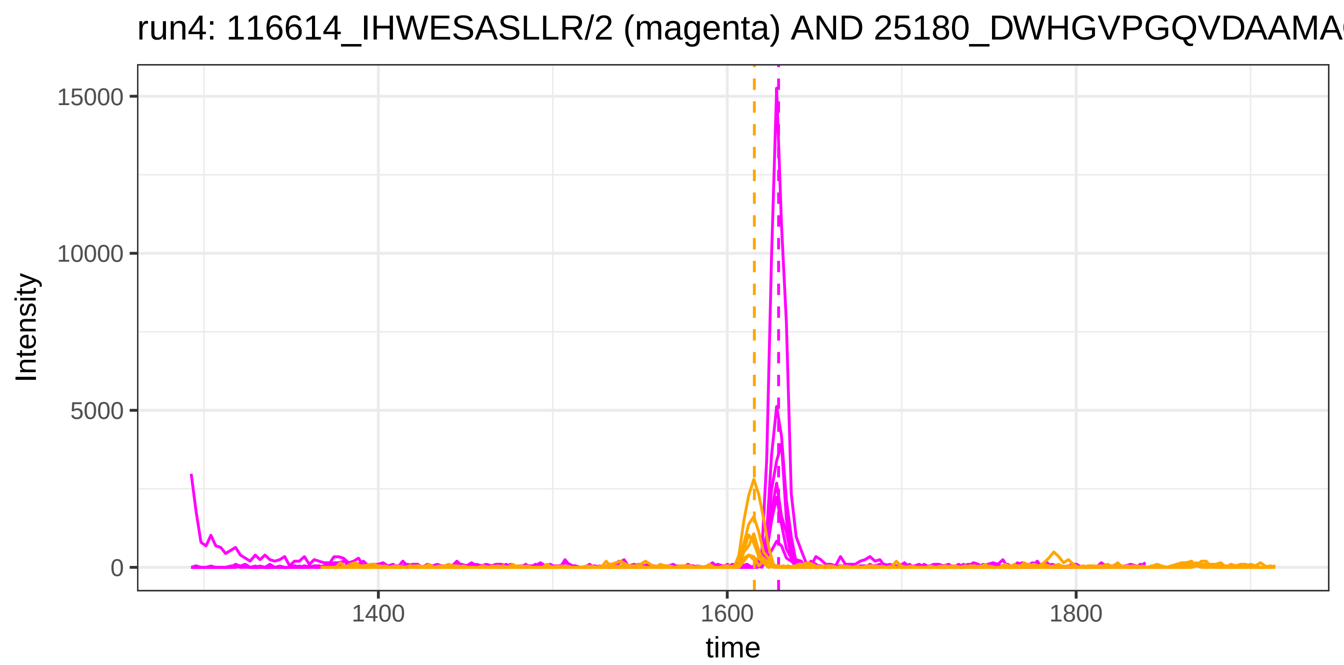

run4: 116614_IHWESASLLR/2 (magenta) AND 25180_DWHGVPGQVDAAMA
Intensity
15000
10000
5000
0
1400
1600
1800
time


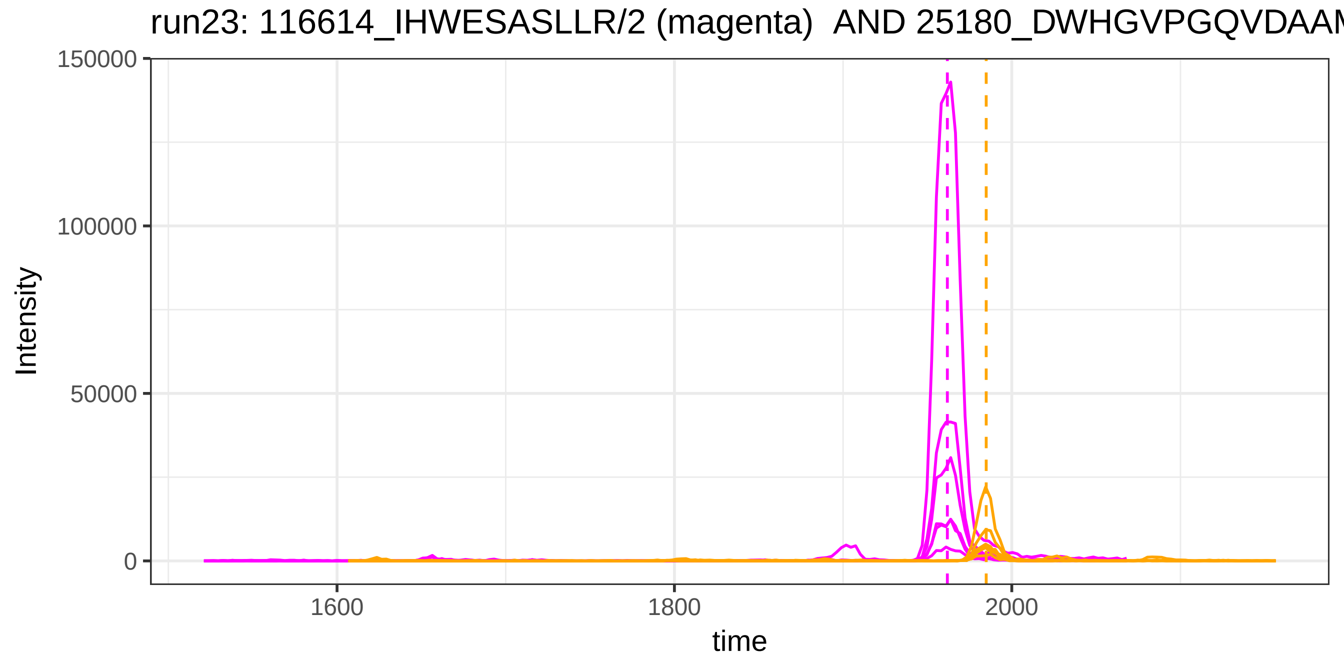

run23: 116614_IHWESASLLR/2 (magenta) AND 25180_DWHGVPGQVDAAM
Intensity
150000
100000
50000
0
1600
1800
2000
time

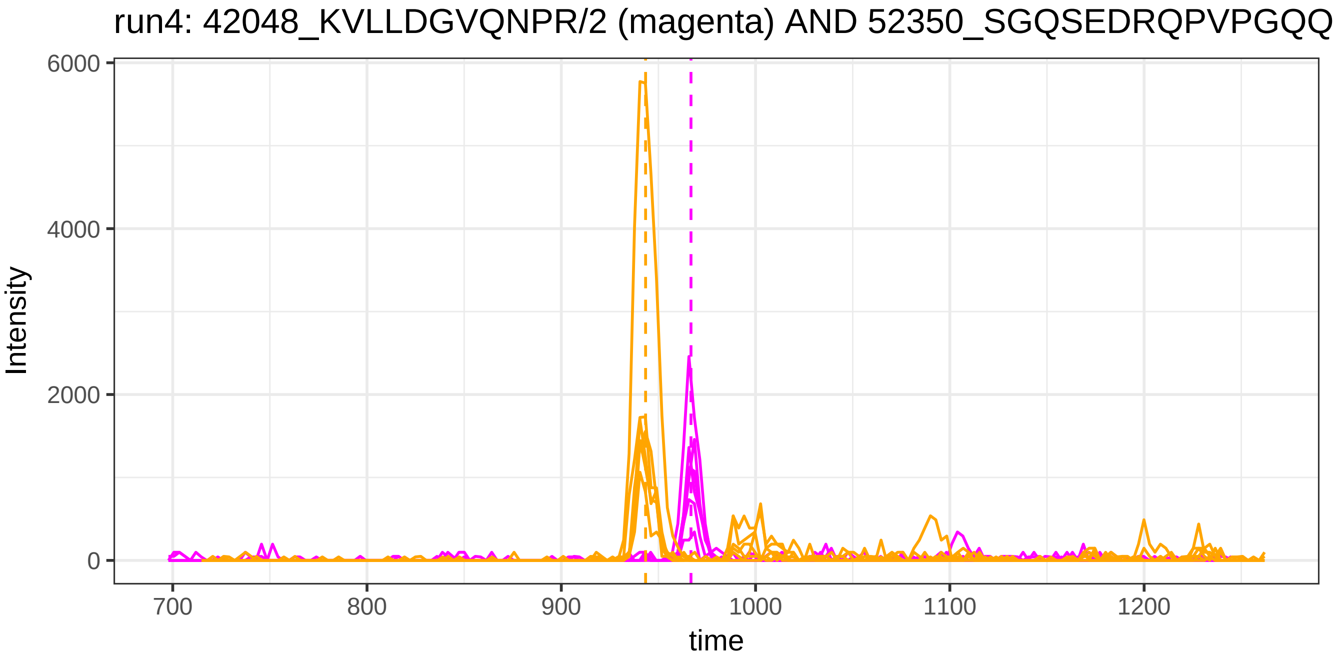

run4: 42048_KVLLDGVQNPR/2 (magenta) AND 52350_SGQSEDRQPVPGQQ
Intensity
6000
4000
2000
0
700
800
900
1000
1100
1200
time


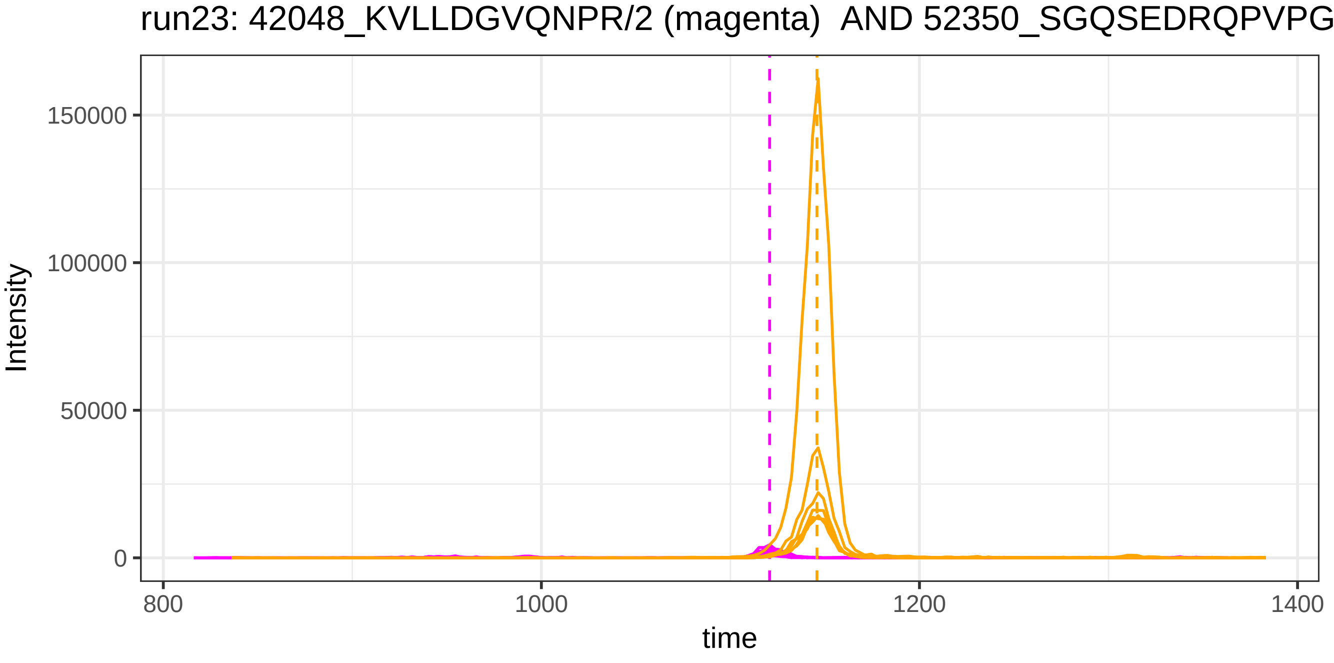

run23: 42048_KVLLDGVQNPR/2 (magenta) AND 52350_SGQSEDRQPVPG
Intensity
150000
100000
50000
0
800
1000
1200
1400
time

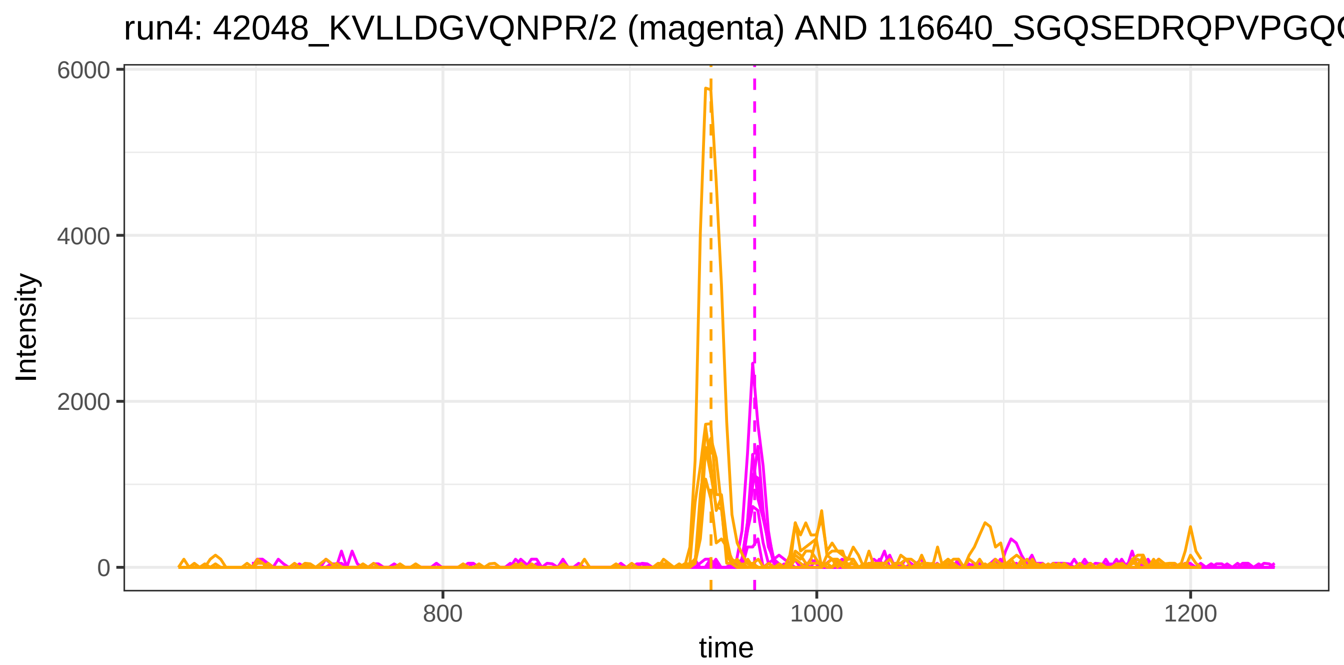
run4: 42048_KVLLDGVQNPR/2 (magenta) AND 116640_SGQSEDRQPVPGQ
Intensity
6000
4000
2000
0
800
1000
1200
time

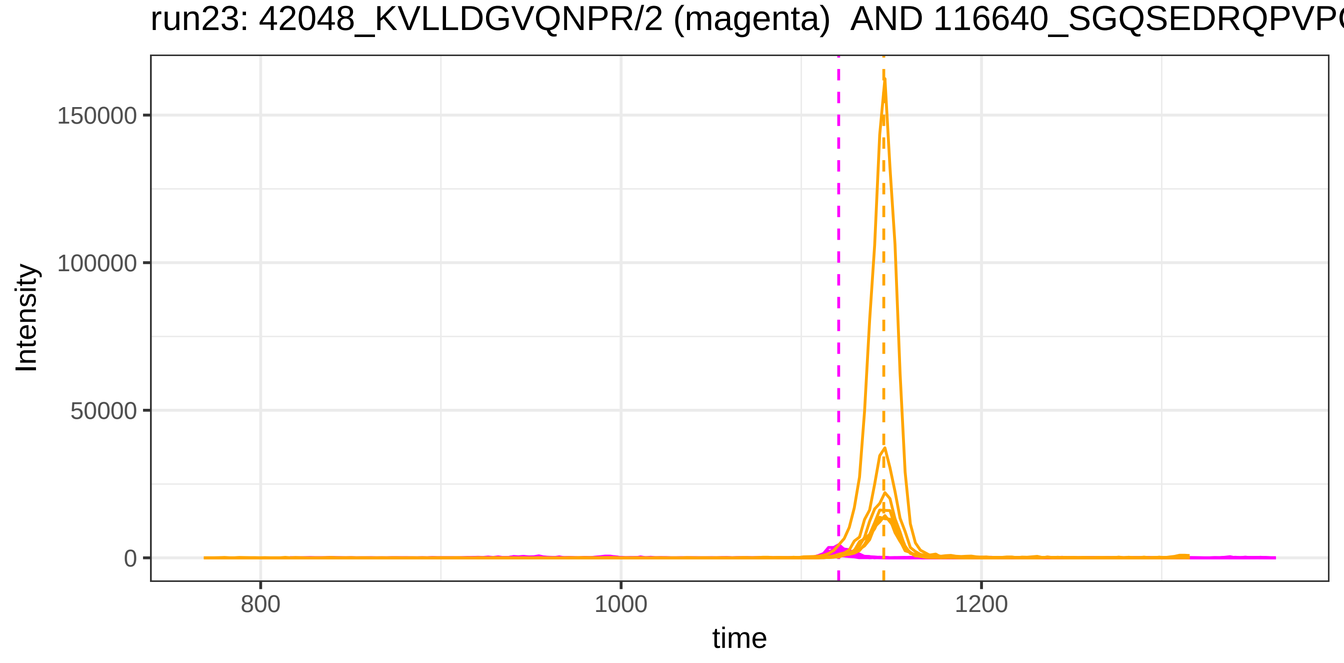
run23: 42048_KVLLDGVQNPR/2 (magenta) AND 116640_SGQSEDRQPVP
Intensity
150000
100000
50000
0
800
1000
1200
time

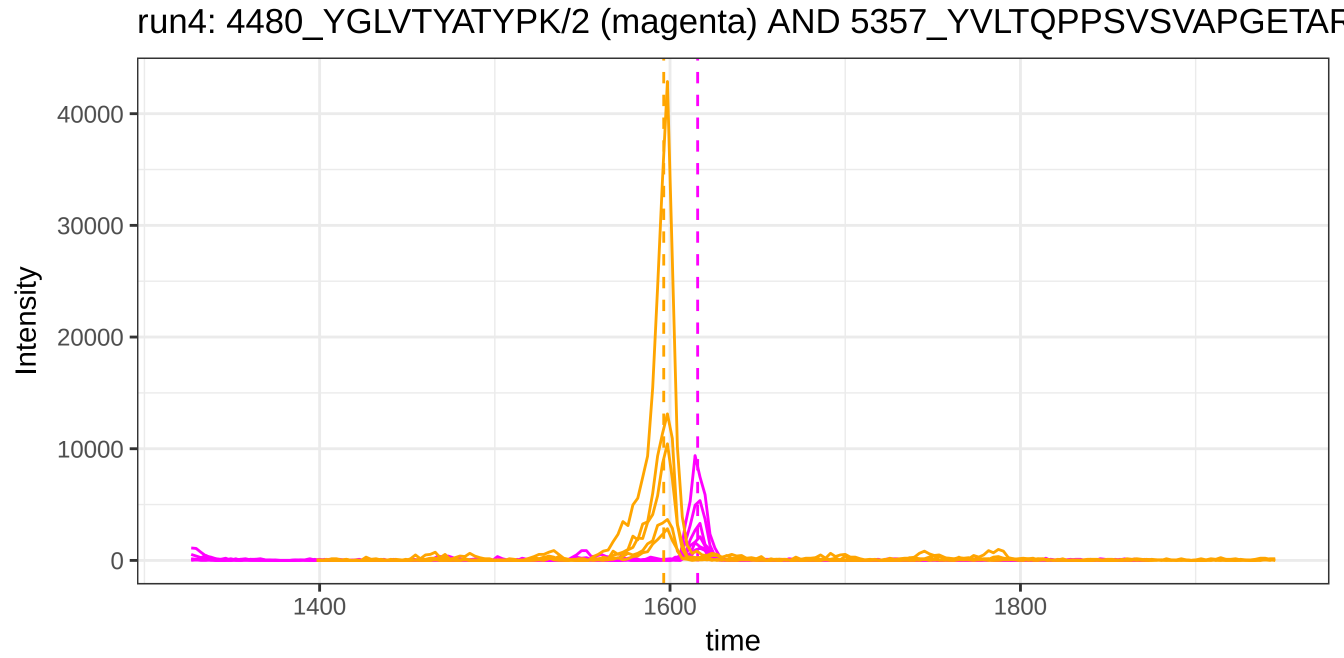
run4: 4480_YGLVTYATYPK/2 (magenta) AND 5357_YVLTQPPSVSVAPGETAR
Intensity
40000
30000
20000
10000
0
1400
1600
1800
time

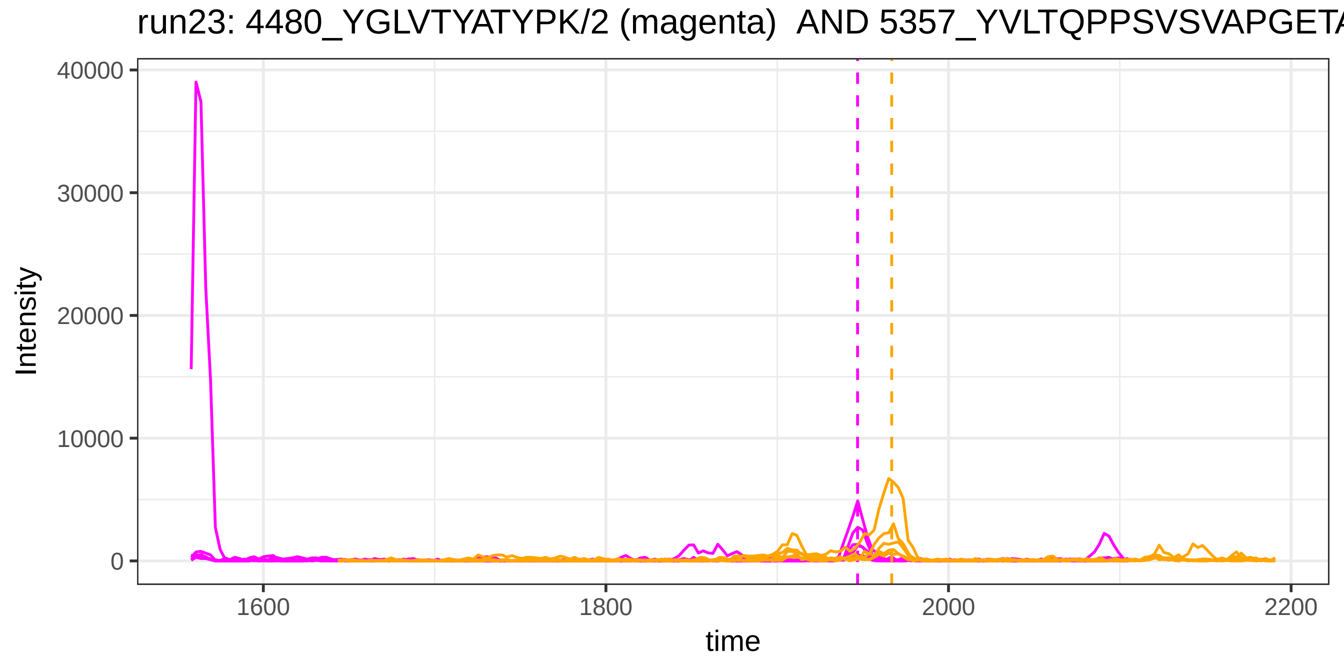
run23: 4480_YGLVTYATYPK/2 (magenta) AND 5357_YVLTQPPSVSVAPGETA
Intensity
40000
30000
20000
10000
0
1600
1800
2000
2200
time

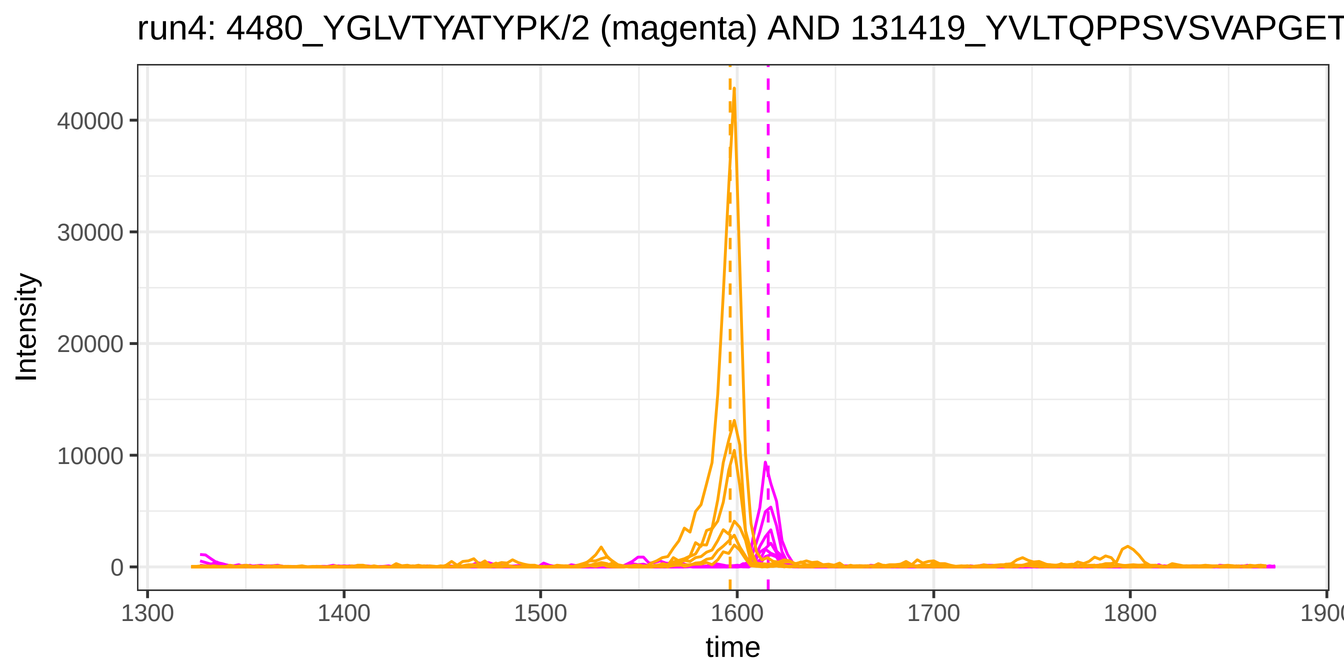
run4: 4480_YGLVTYATYPK/2 (magenta) AND 131419_YVLTQPPSVSVAPGET
Intensity
0
10000
20000
30000
40000
1300
1400
1500
1600
1700
1800
time

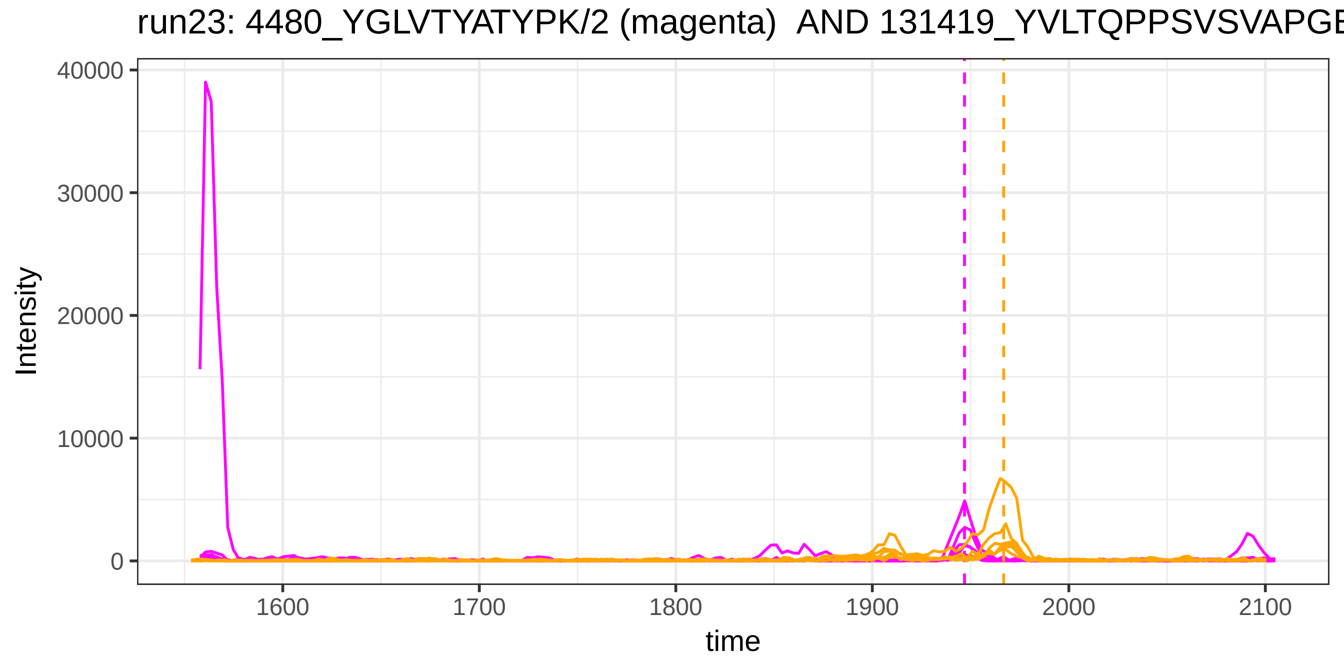
run23: 4480_YGLVTYATYPK/2 (magenta) AND 131419_YVLTQPPSVSVAPGE
Intensity
0
10000
20000
30000
40000
1600
1700
1800
1900
2000
2100
time

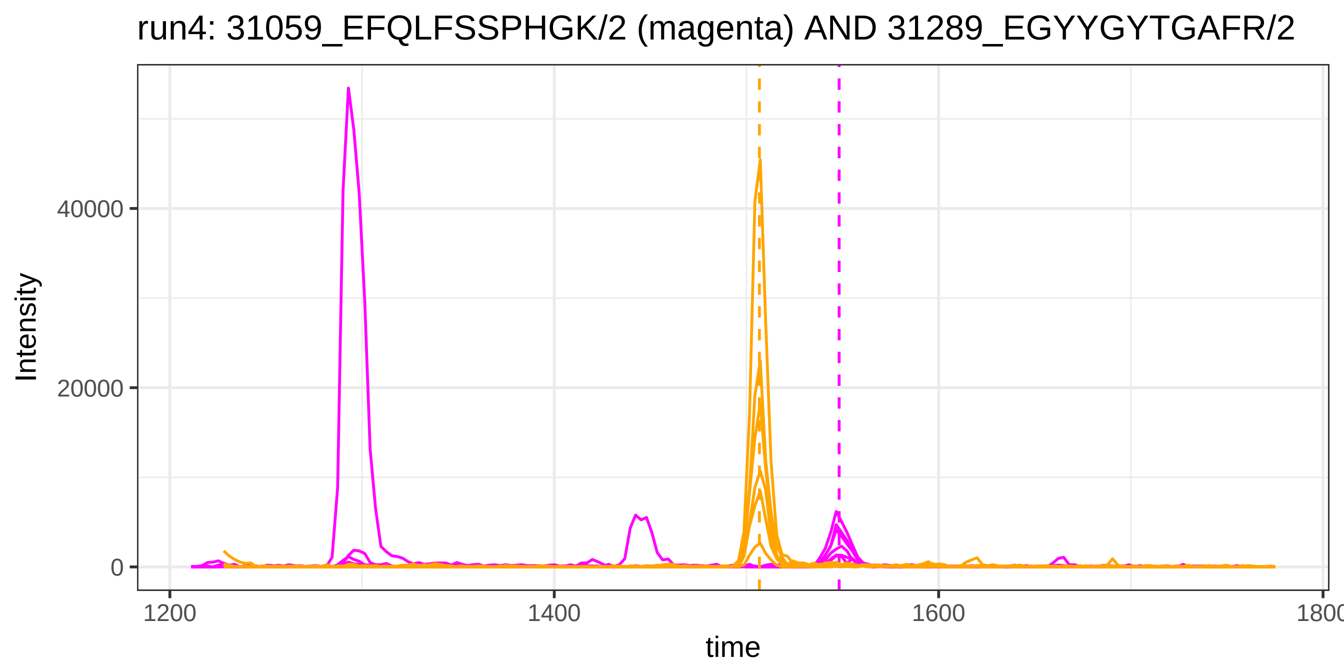
run4: 31059_EFQLFSSPHGK/2 (magenta) AND 31289_EGYYGYTGAFR/2
Intensity
40000
20000
0
1200
1400
1600
1800
time

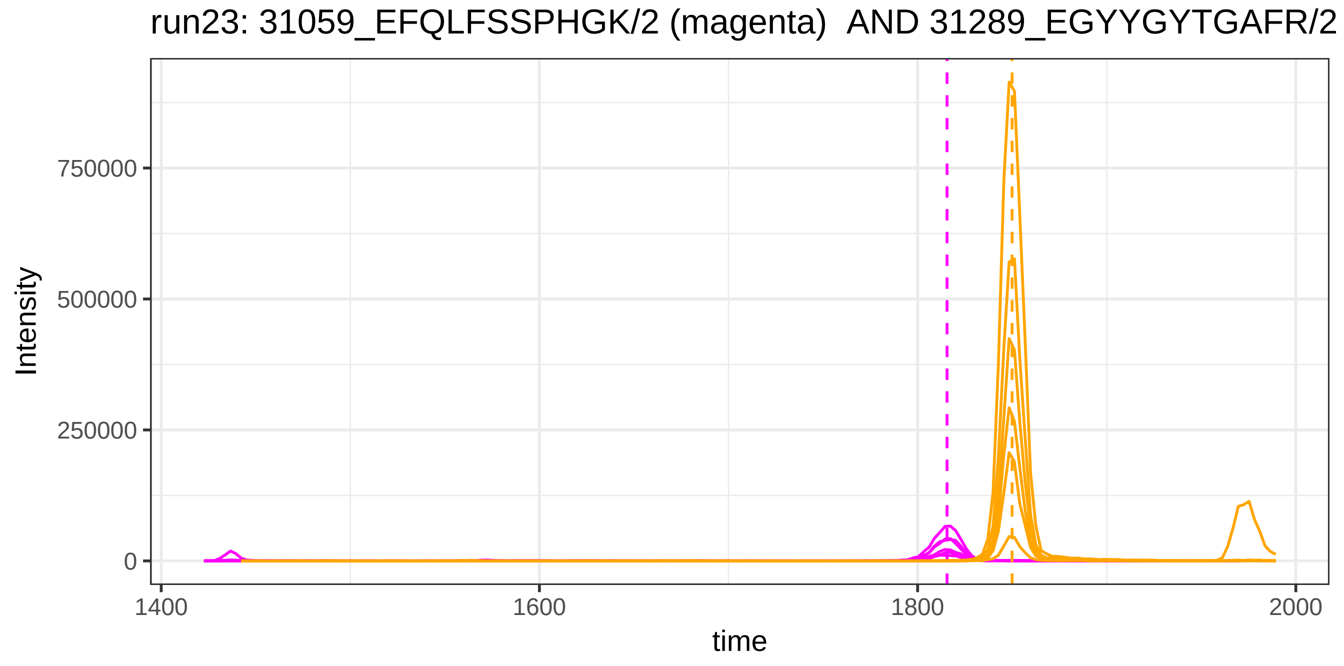
run23: 31059_EFQLFSSPHGK/2 (magenta) AND 31289_EGYYGYTGAFR/2
Intensity
750000
500000
250000
0
1400
1600
1800
2000
time

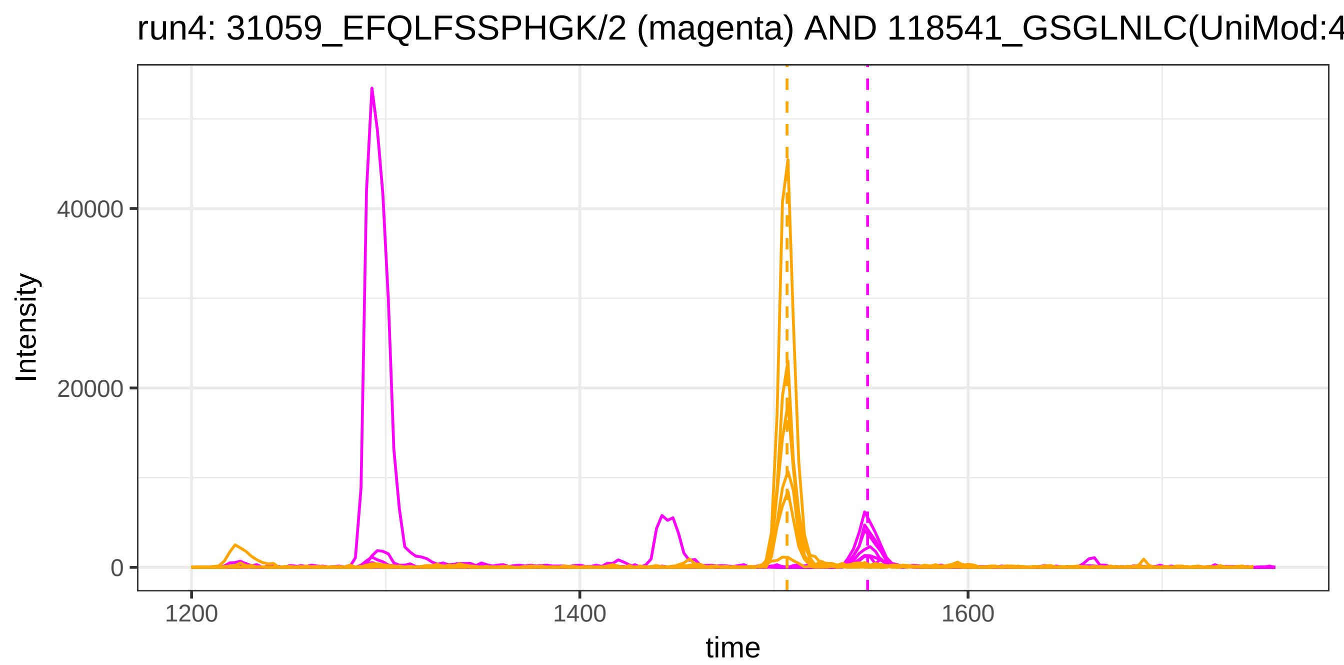

run4: 31059_EFQLFSSPHGK/2 (magenta) AND 118541_GSGLNLC(UniMod:4
Intensity
40000
20000
0
1200
1400
1600
time


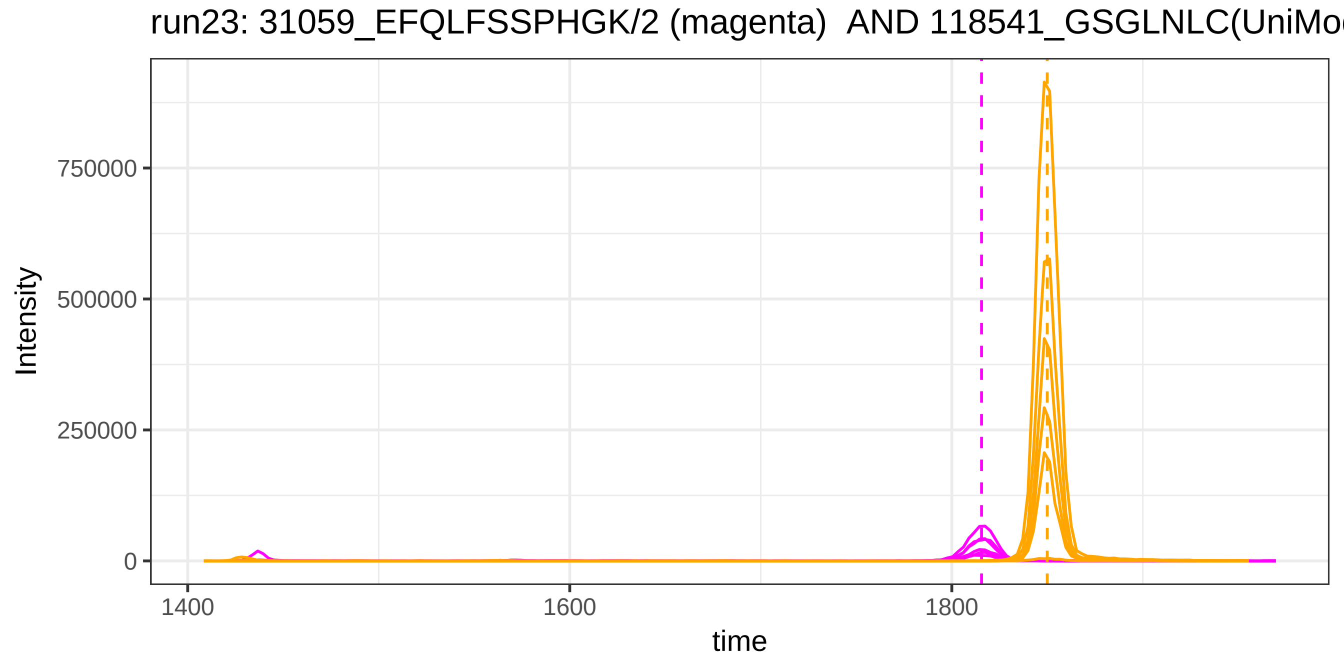

run23: 31059_EFQLFSSPHGK/2 (magenta) AND 118541_GSGLNLC(UniMo
Intensity
750000
500000
250000
0
1400
1600
1800
time

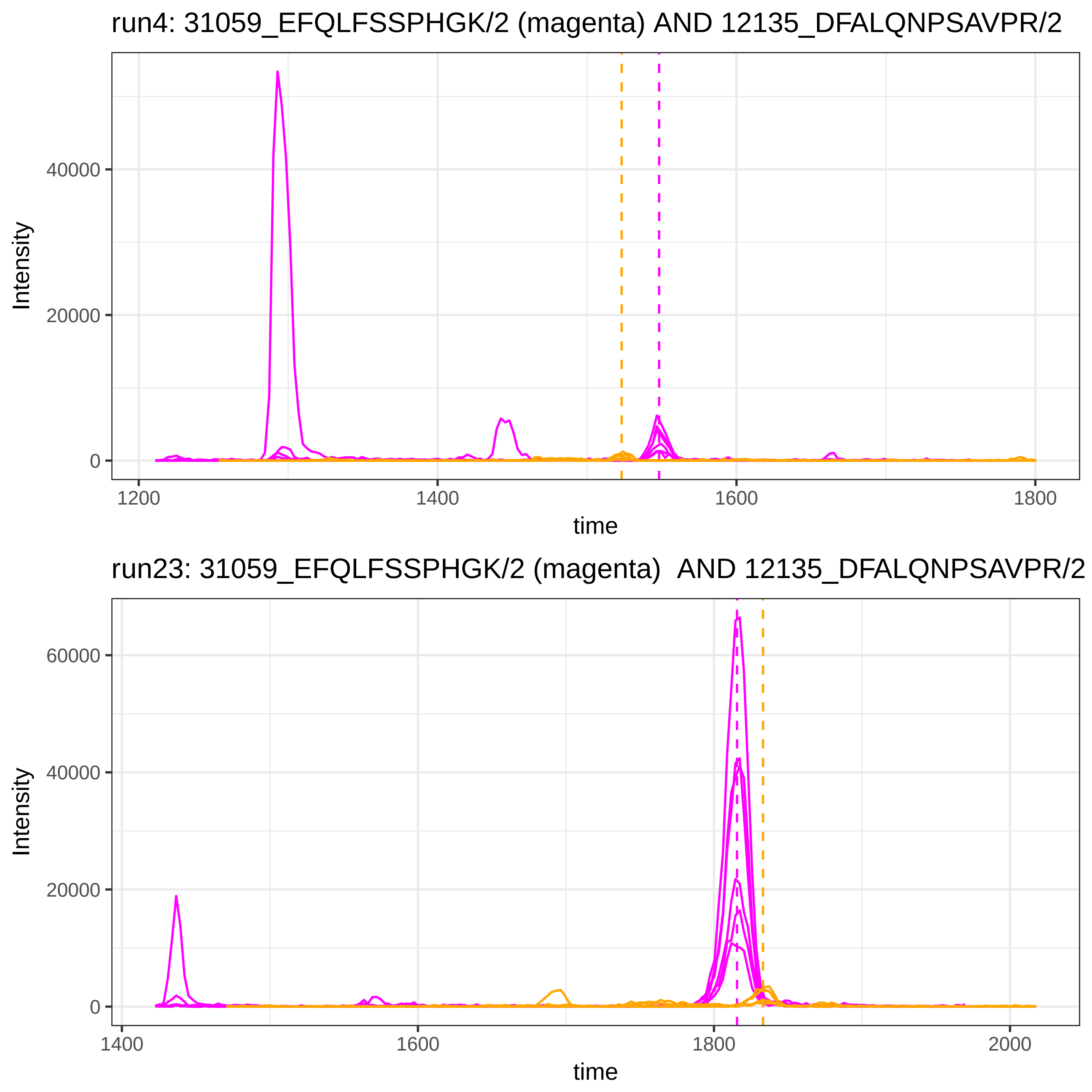

run4: 31059_EFQLFSSPHGK/2 (magenta) AND 12135_DFALQNPSAVPR/2
Intensity
time
run23: 31059_EFQLFSSPHGK/2 (magenta)  AND 12135_DFALQNPSAVPR/2
Intensity
time

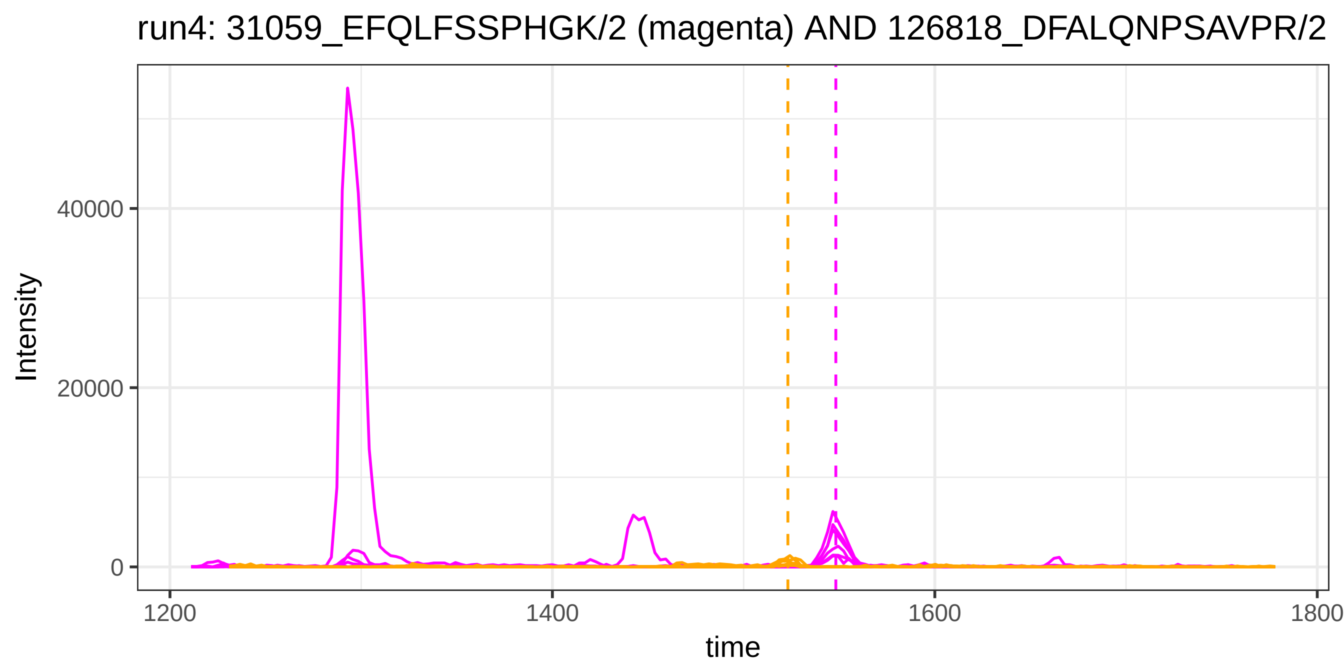
run4: 31059_EFQLFSSPHGK/2 (magenta) AND 126818_DFALQNPSAVPR/2
Intensity
40000
20000
0
1200
1400
1600
1800
time

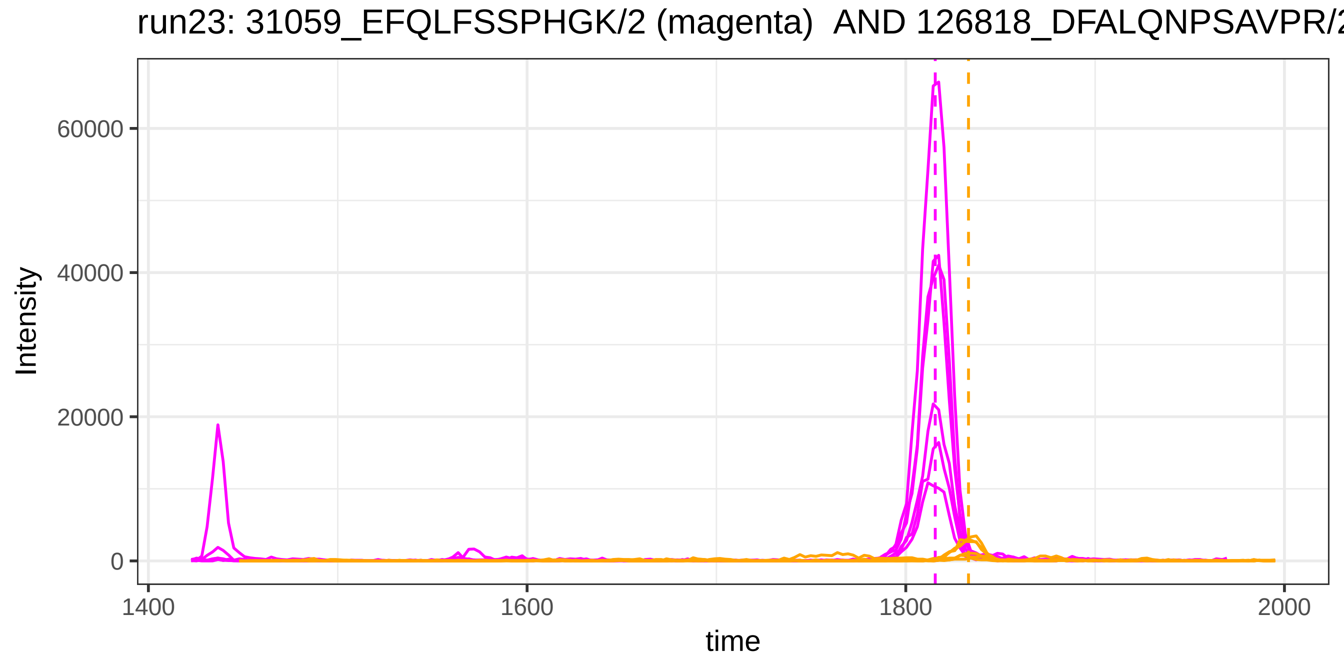
run23: 31059_EFQLFSSPHGK/2 (magenta) AND 126818_DFALQNPSAVPR/2
Intensity
60000
40000
20000
0
1400
1600
1800
2000
time

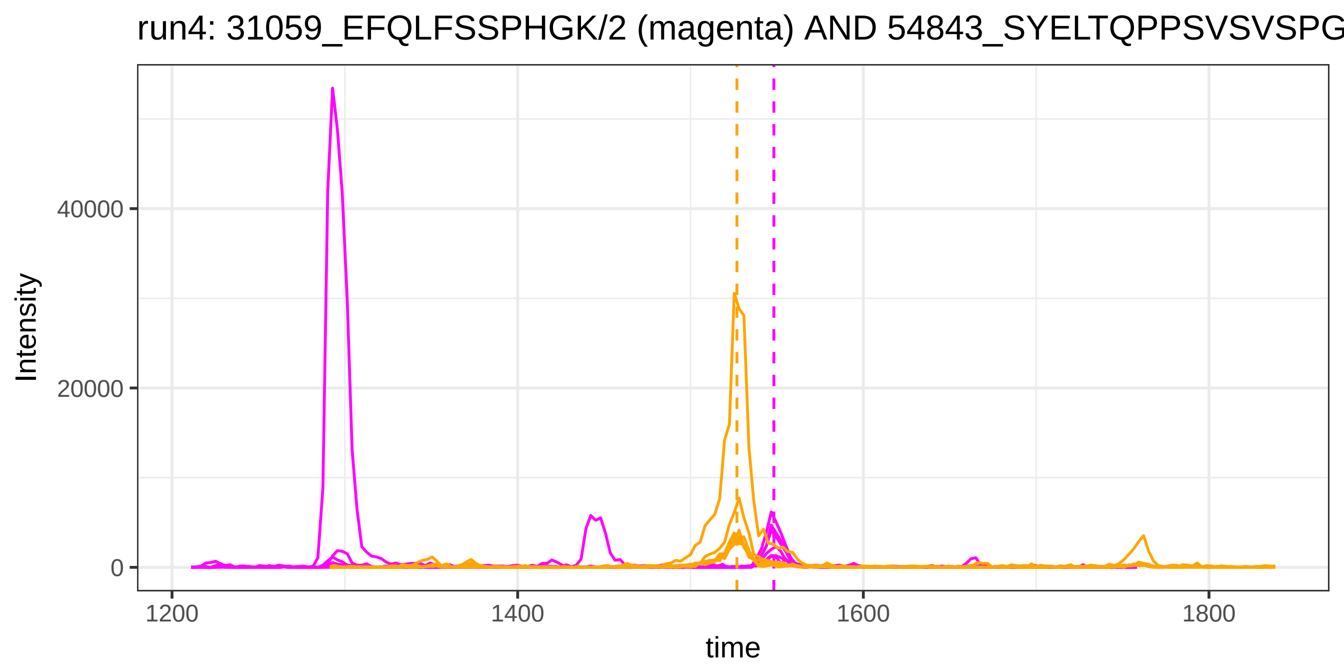

run4: 31059_EFQLFSSPHGK/2 (magenta) AND 54843_SYELTQPPSVSVSPG
Intensity
40000
20000
0
1200
1400
1600
1800
time


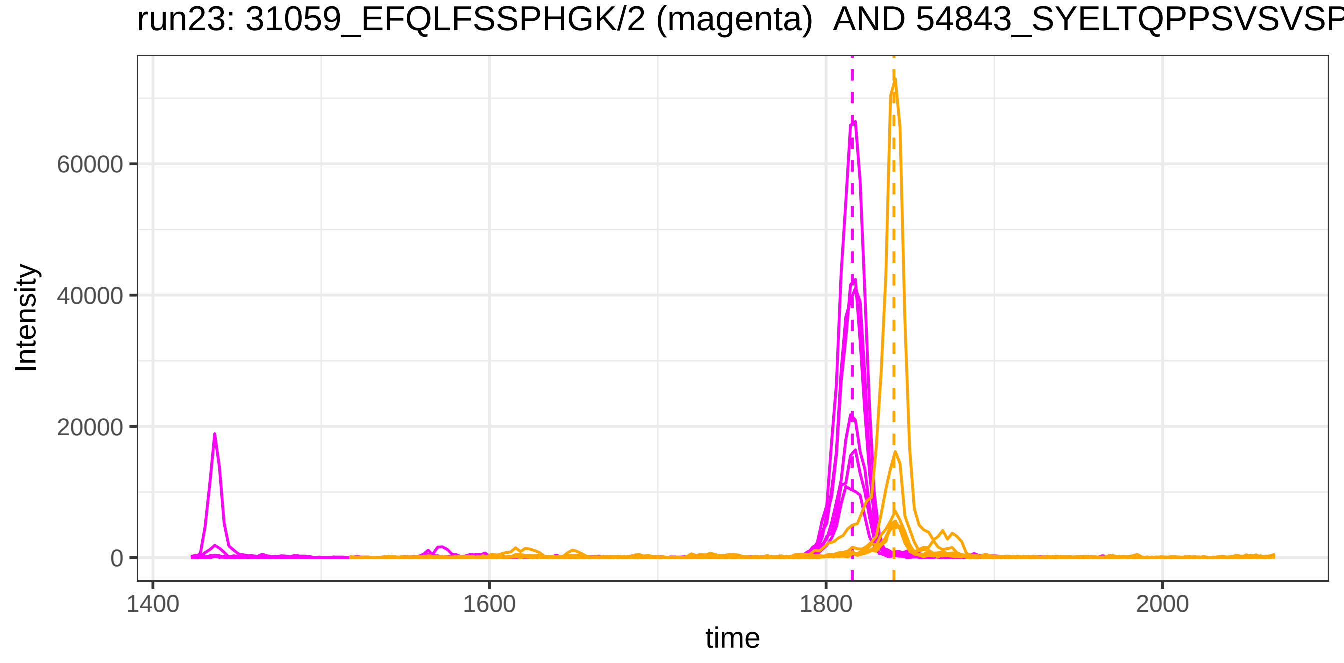

run23: 31059_EFQLFSSPHGK/2 (magenta) AND 54843_SYELTQPPSVSVSP
Intensity
60000
40000
20000
0
1400
1600
1800
2000
time

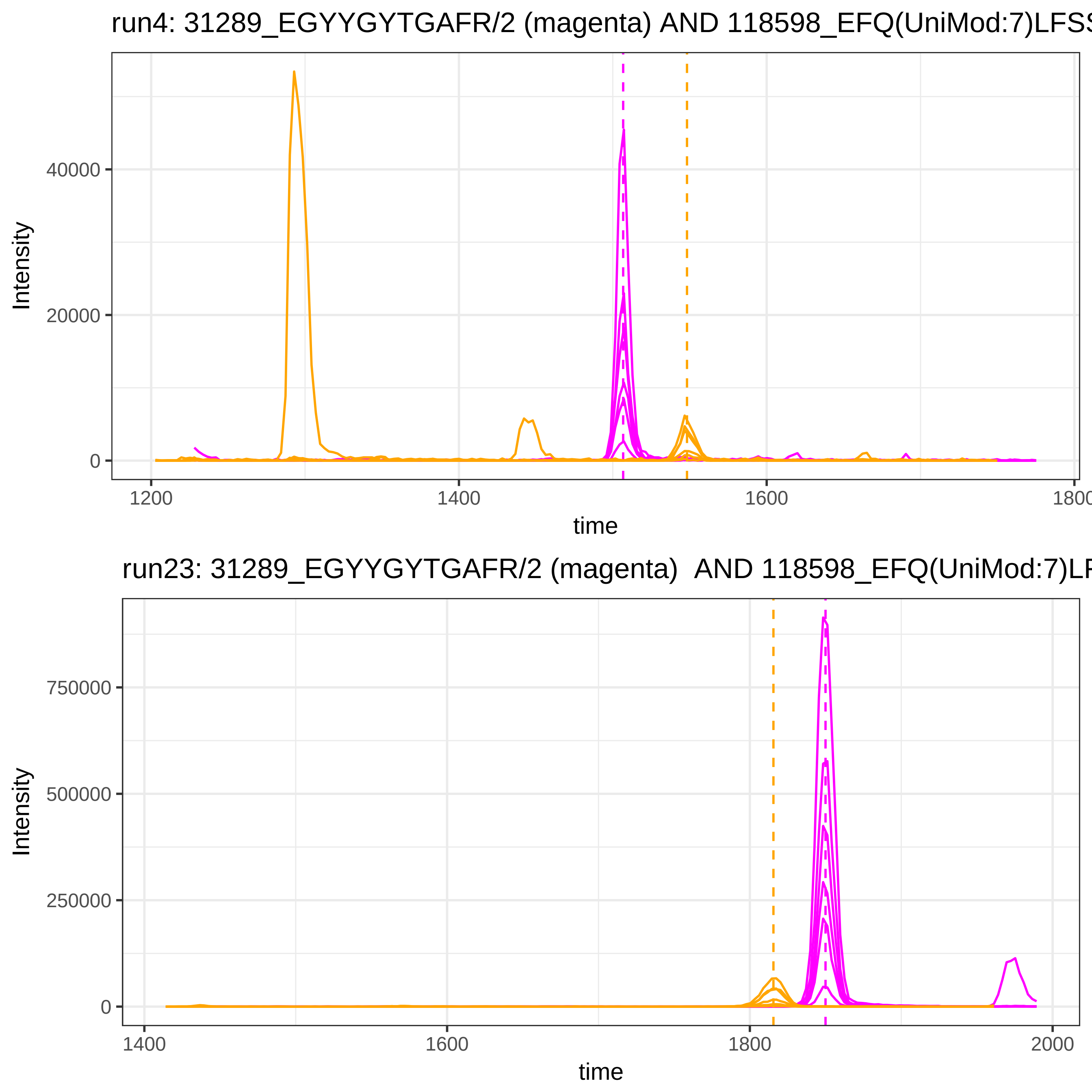
run4: 31289_EGYYGYTGAFR/2 (magenta) AND 118598_EFQ(UniMod:7)LFSS
Intensity
40000
20000
0
1200
1400
1600
1800
time
run23: 31289_EGYYGYTGAFR/2 (magenta) AND 118598_EFQ(UniMod:7)LF
Intensity
750000
500000
250000
0
1400
1600
1800
2000
time

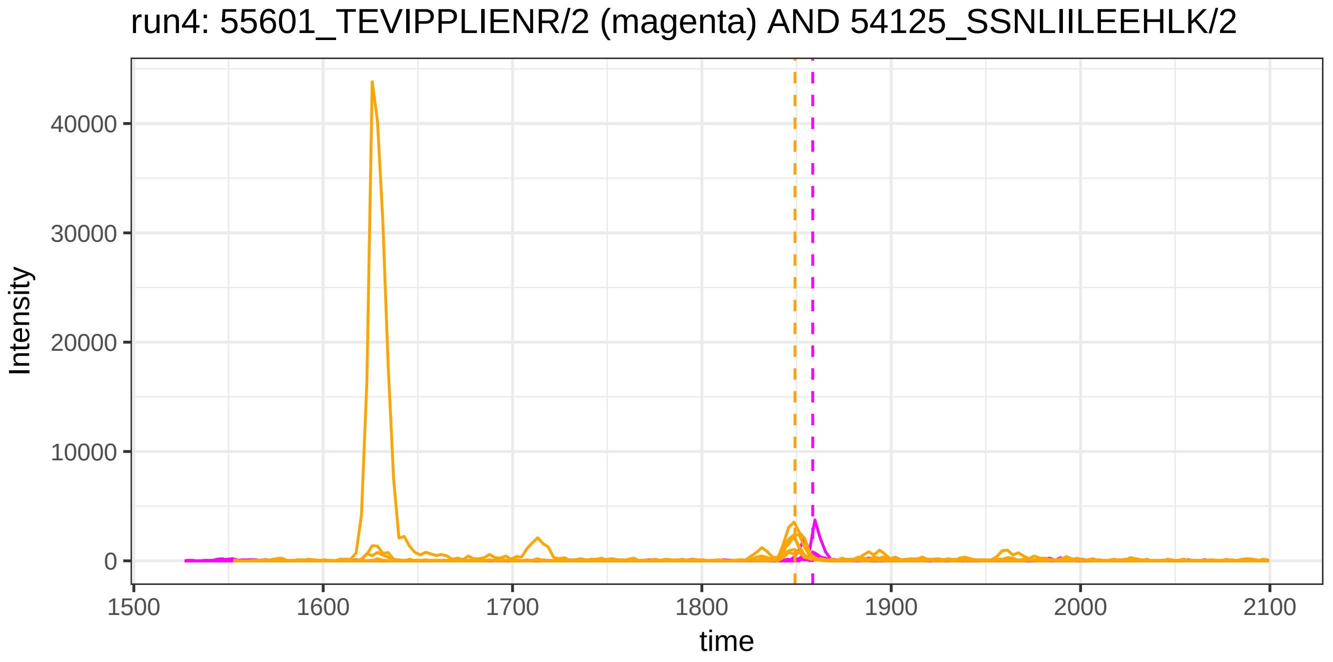
run4: 55601_TEVIPPLIENR/2 (magenta) AND 54125_SSNLIILEEHLK/2
Intensity
40000
30000
20000
10000
0
1500
1600
1700
1800
1900
2000
2100
time

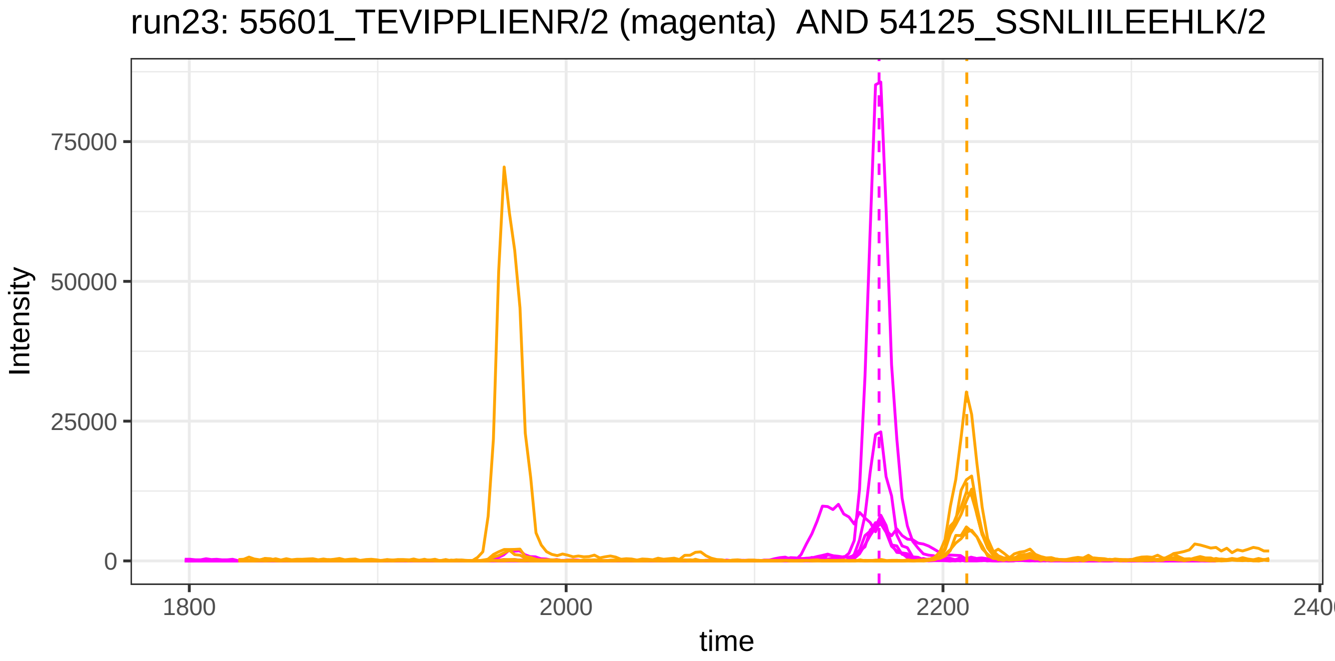
run23: 55601_TEVIPPLIENR/2 (magenta) AND 54125_SSNLIILEEHLK/2
Intensity
75000
50000
25000
0
1800
2000
2200
time

run4: 55601_TEVIPPLIENR/2 (magenta) AND 117986_QQNAQGGFSSTQDTV

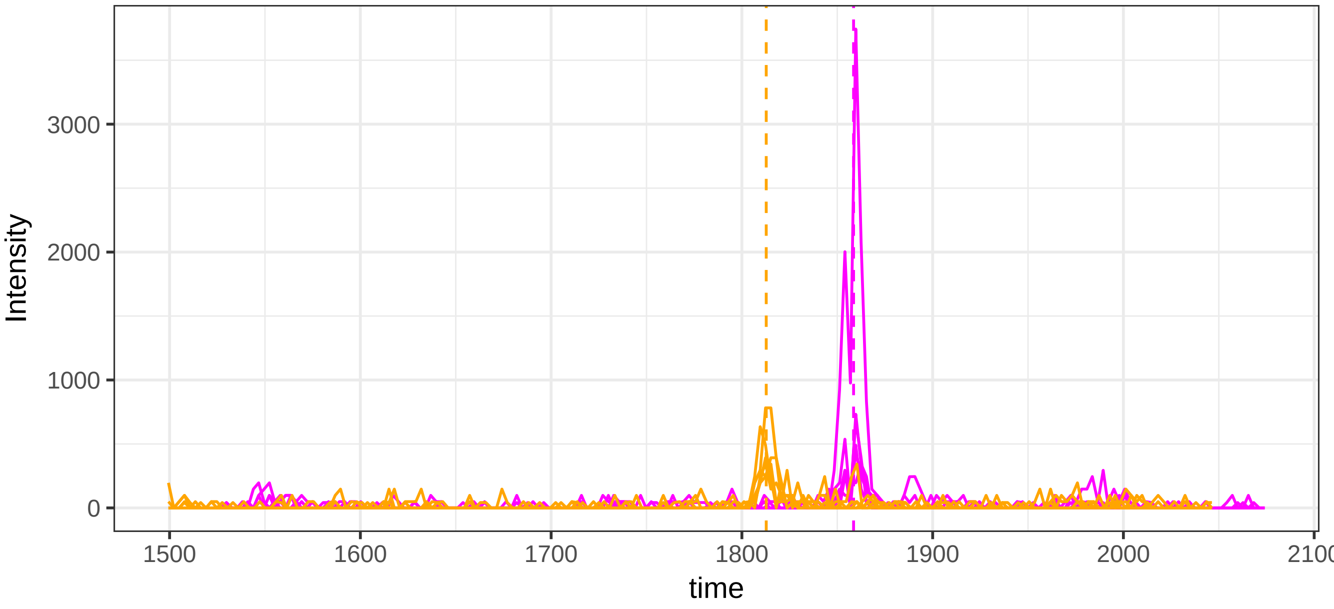


run23: 55601_TEVIPPLIENR/2 (magenta)  AND 117986_QQNAQGGFSSTQD

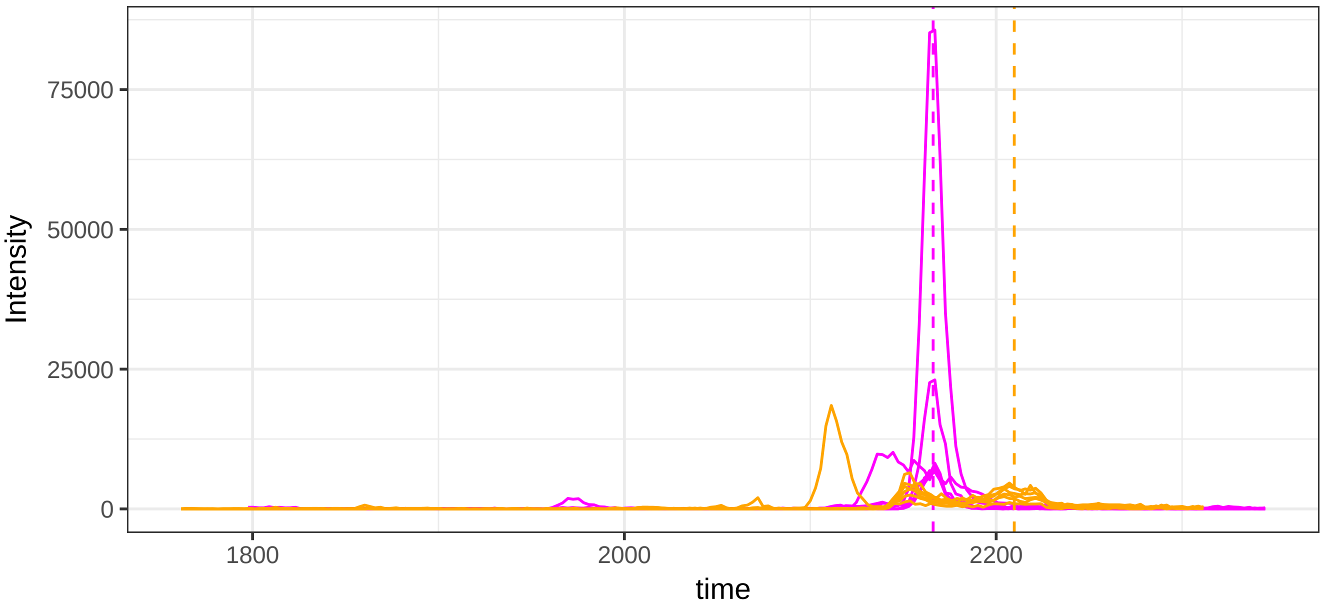

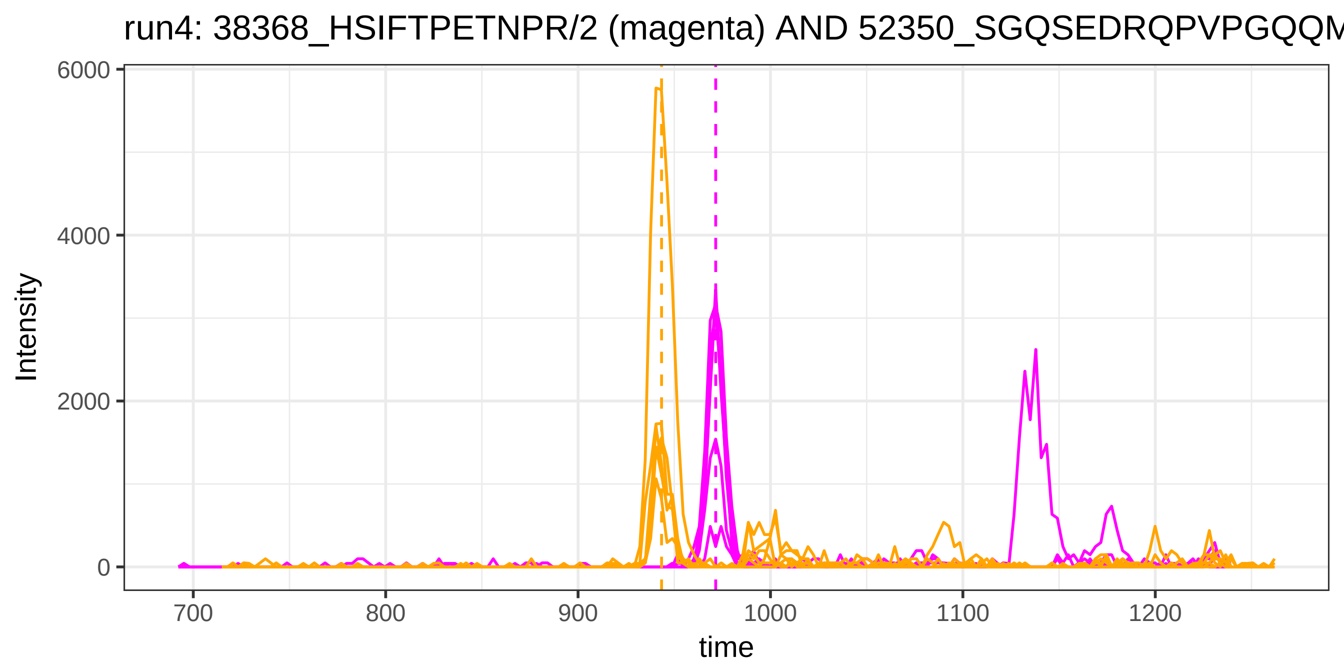

run4: 38368_HSIFTPETNPR/2 (magenta) AND 52350_SGQSEDRQPVPGQQM
Intensity
6000
4000
2000
0
700
800
900
1000
1100
1200
time


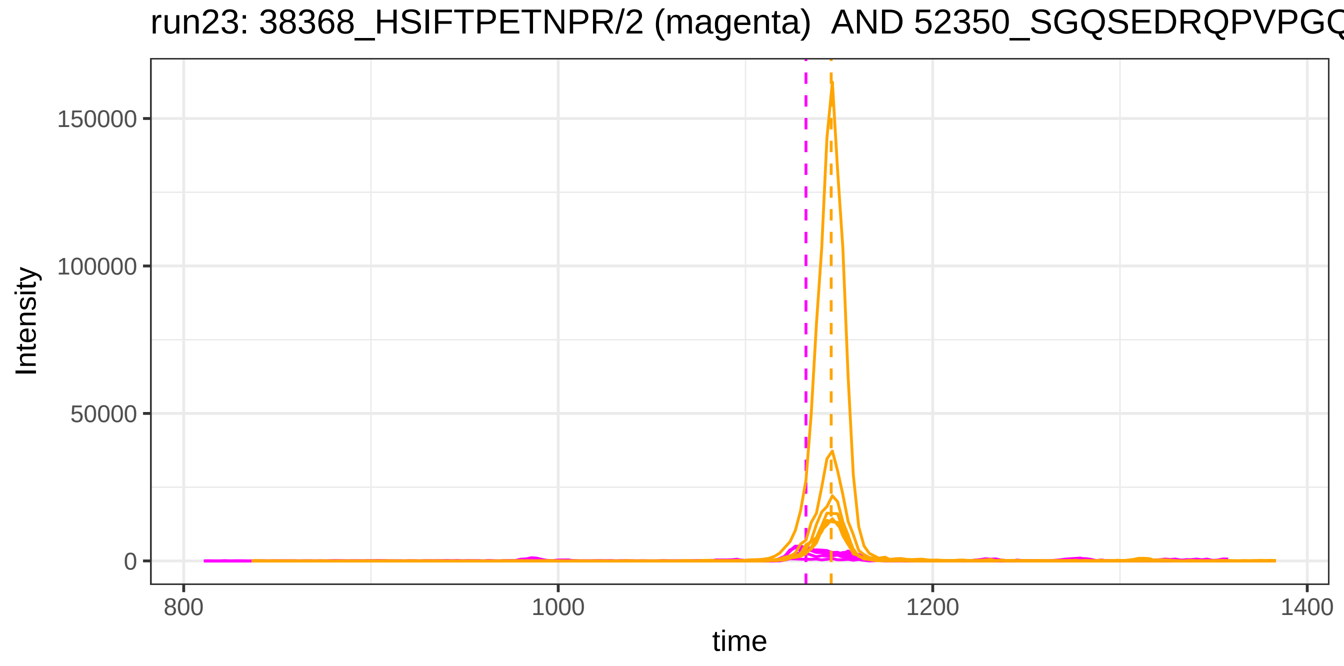

run23: 38368_HSIFTPETNPR/2 (magenta) AND 52350_SGQSEDRQPVPGQ
Intensity
150000
100000
50000
0
800
1000
1200
1400
time

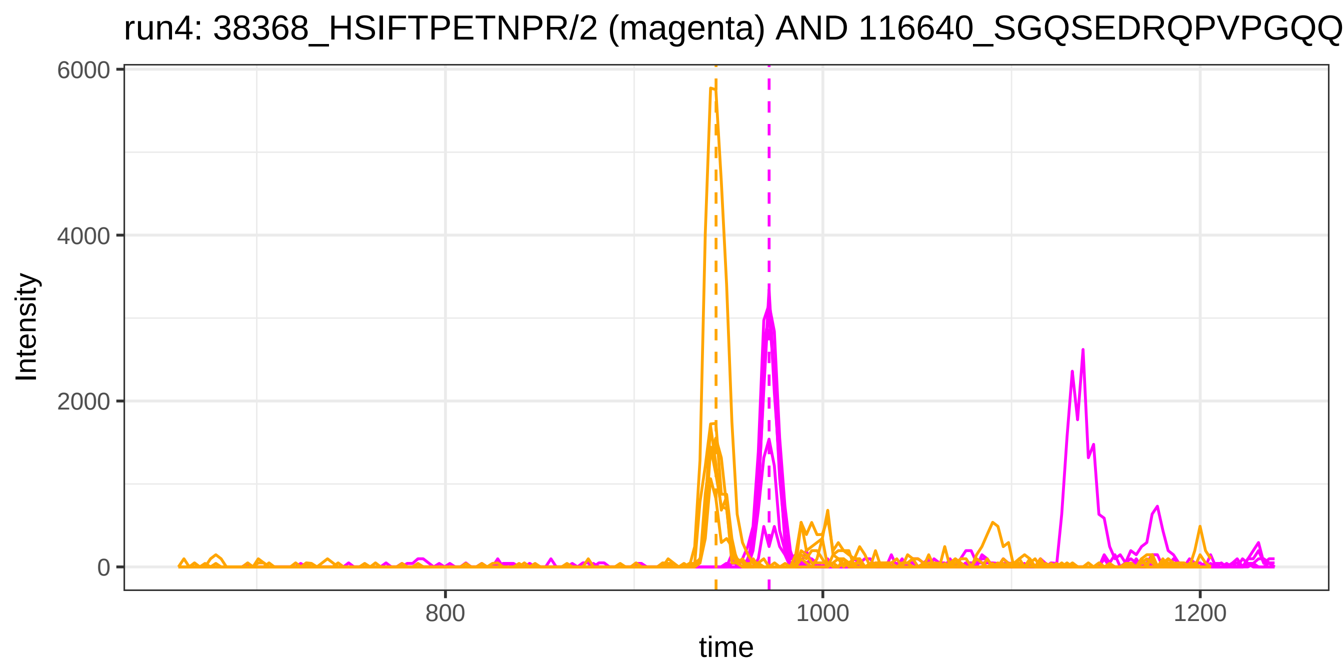
run4: 38368_HSIFTPETNPR/2 (magenta) AND 116640_SGQSEDRQPVPGQQ
Intensity
6000
4000
2000
0
800
1000
1200
time

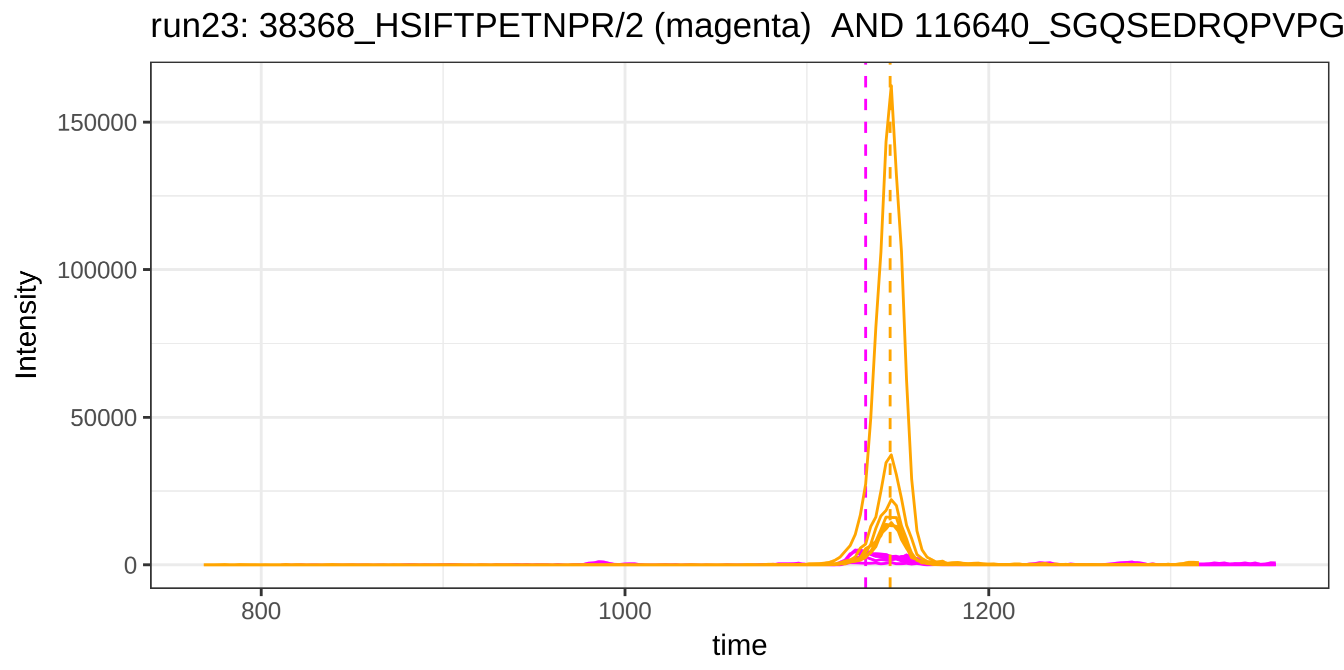
run23: 38368_HSIFTPETNPR/2 (magenta) AND 116640_SGQSEDRQPVPG
Intensity
150000
100000
50000
0
800
1000
1200
time

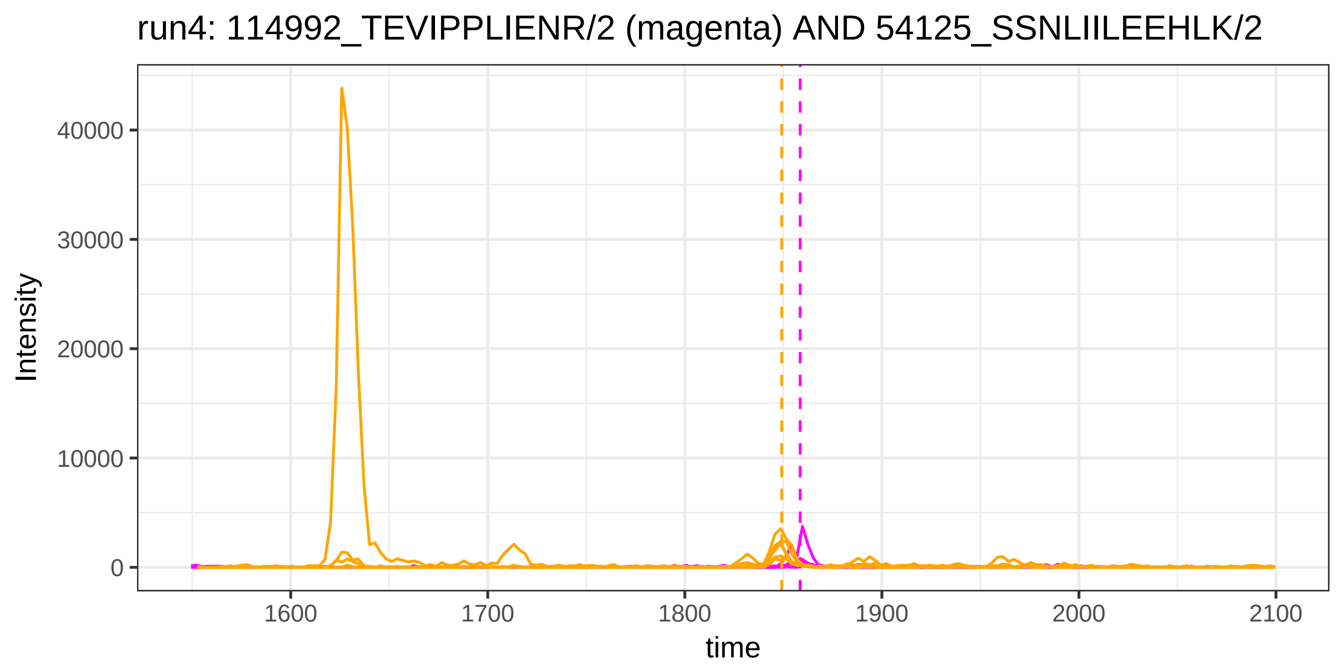
run4: 114992_TEVIPPLIENR/2 (magenta) AND 54125_SSNLIILEEHLK/2
Intensity
40000
30000
20000
10000
0
1600
1700
1800
1900
2000
2100
time

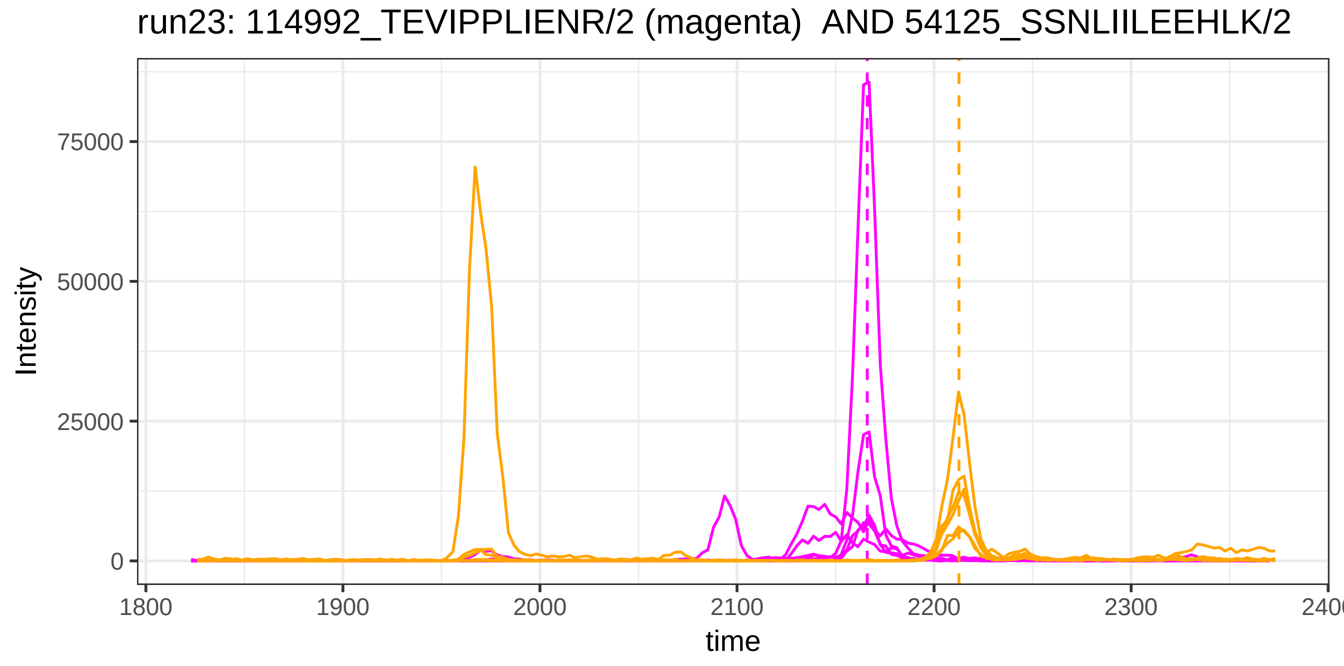
run23: 114992_TEVIPPLIENR/2 (magenta)  AND 54125_SSNLIILEEHLK/2
Intensity
75000
50000
25000
0
1800
1900
2000
2100
2200
2300
time

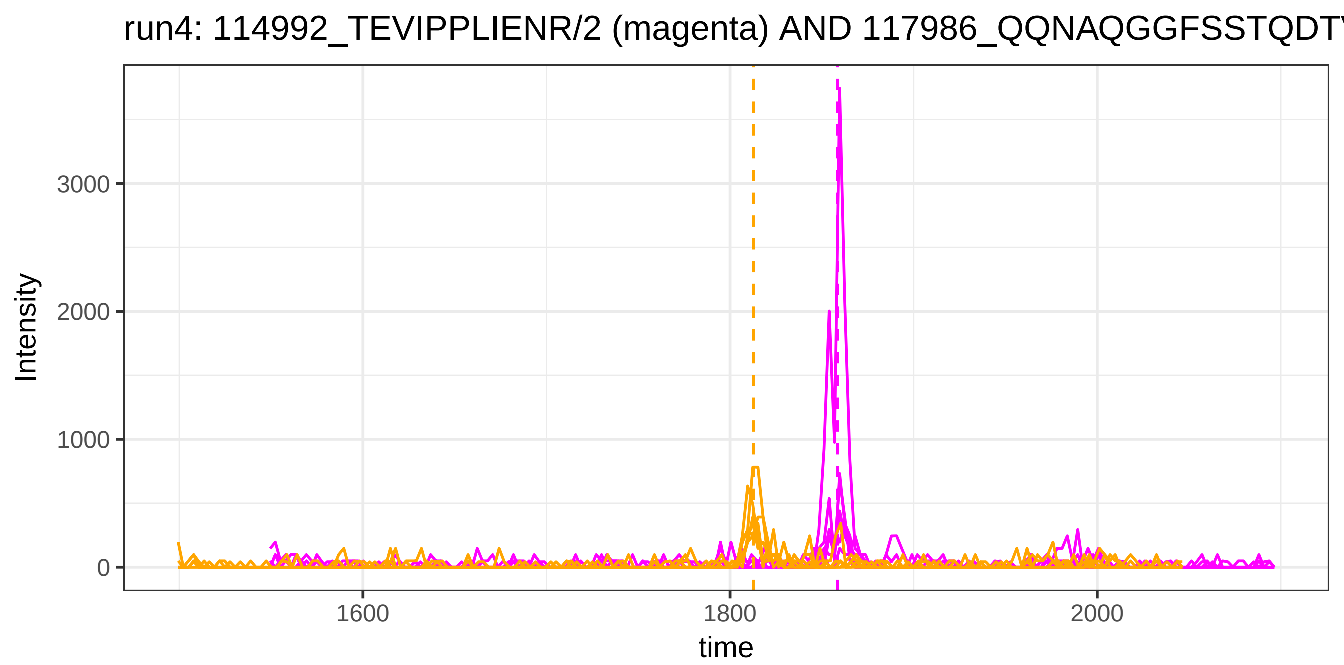
run4: 114992_TEVIPPLIENR/2 (magenta) AND 117986_QQNAQGGFSSTQDT
Intensity
3000
2000
1000
0
1600
1800
2000
time

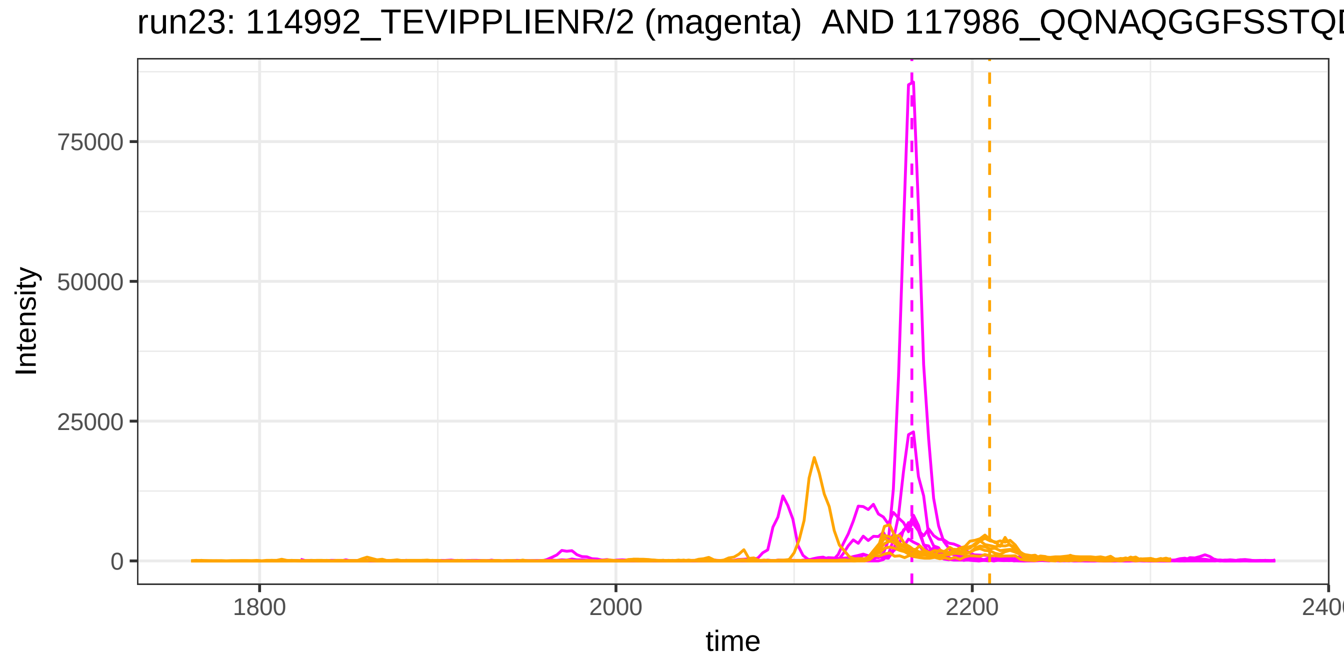
run23: 114992_TEVIPPLIENR/2 (magenta) AND 117986_QQNAQGGFSSTQD
Intensity
75000
50000
25000
0
1800
2000
2200
time

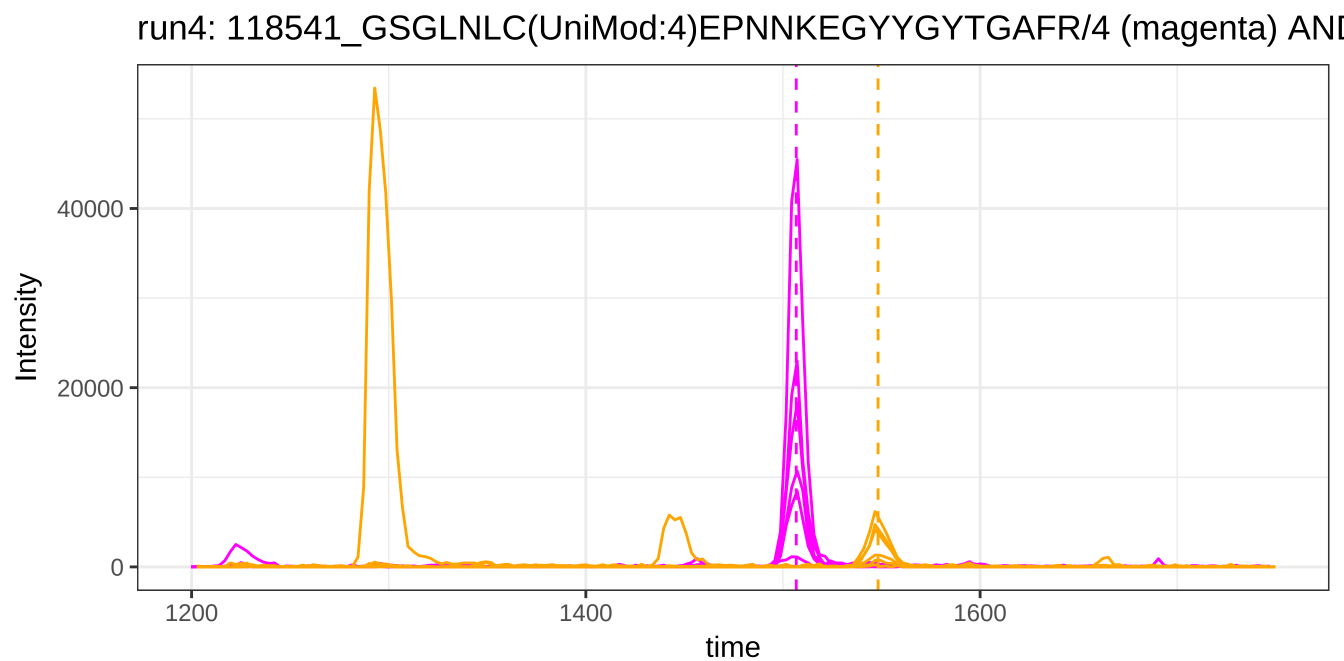
run4: 118541_GSGLNLC(UniMod:4)EPNNKEGYYGYTGAFR/4 (magenta) AND
Intensity
40000
20000
0
1200
1400
1600
time

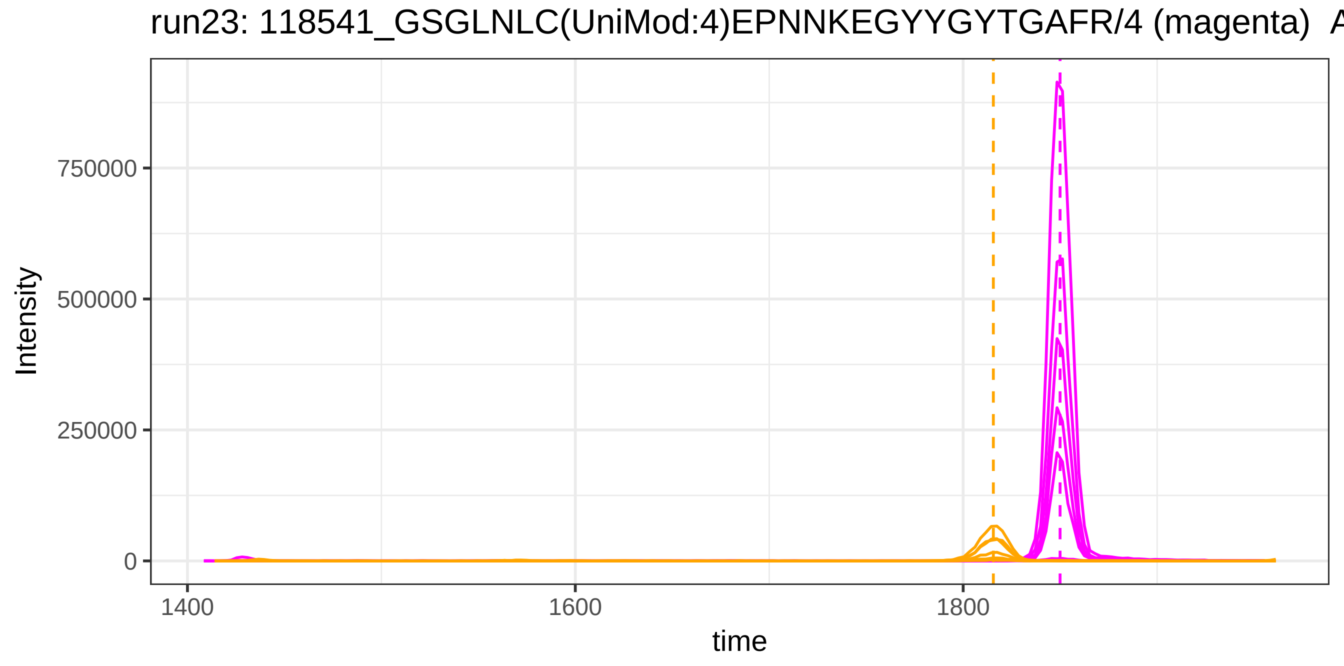
run23: 118541_GSGLNLC(UniMod:4)EPNNKEGYYGYTGAFR/4 (magenta) A
Intensity
750000
500000
250000
0
1400
1600
1800
time

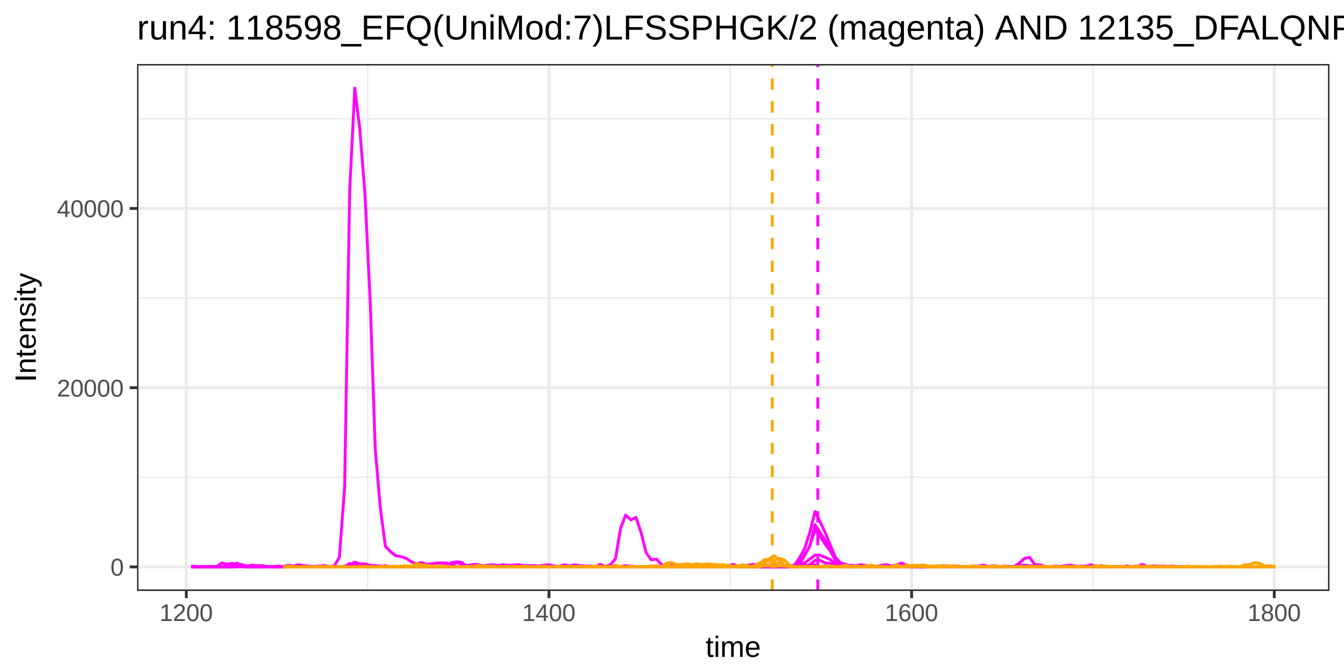
run4: 118598_EFQ(UniMod:7)LFSSPHGK/2 (magenta) AND 12135_DFALQNF
Intensity
40000
20000
0
1200
1400
1600
1800
time

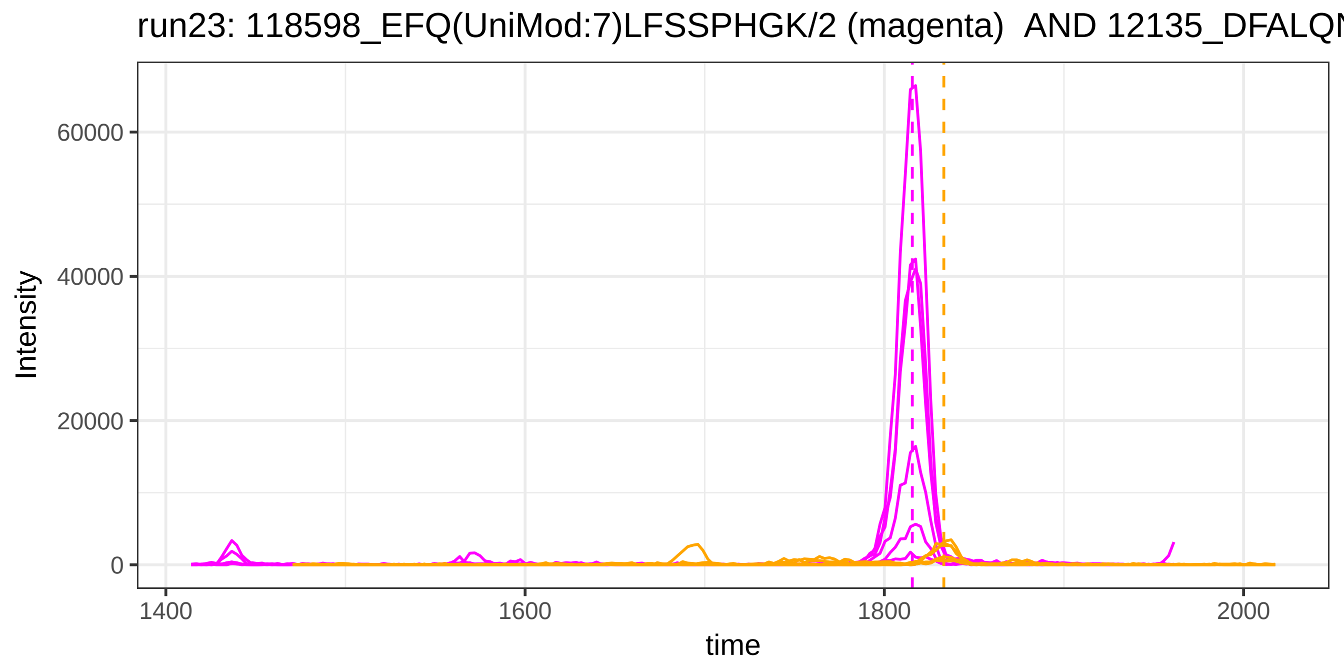
run23: 118598_EFQ(UniMod:7)LFSSPHGK/2 (magenta) AND 12135_DFALQN
Intensity
60000
40000
20000
0
1400
1600
1800
2000
time

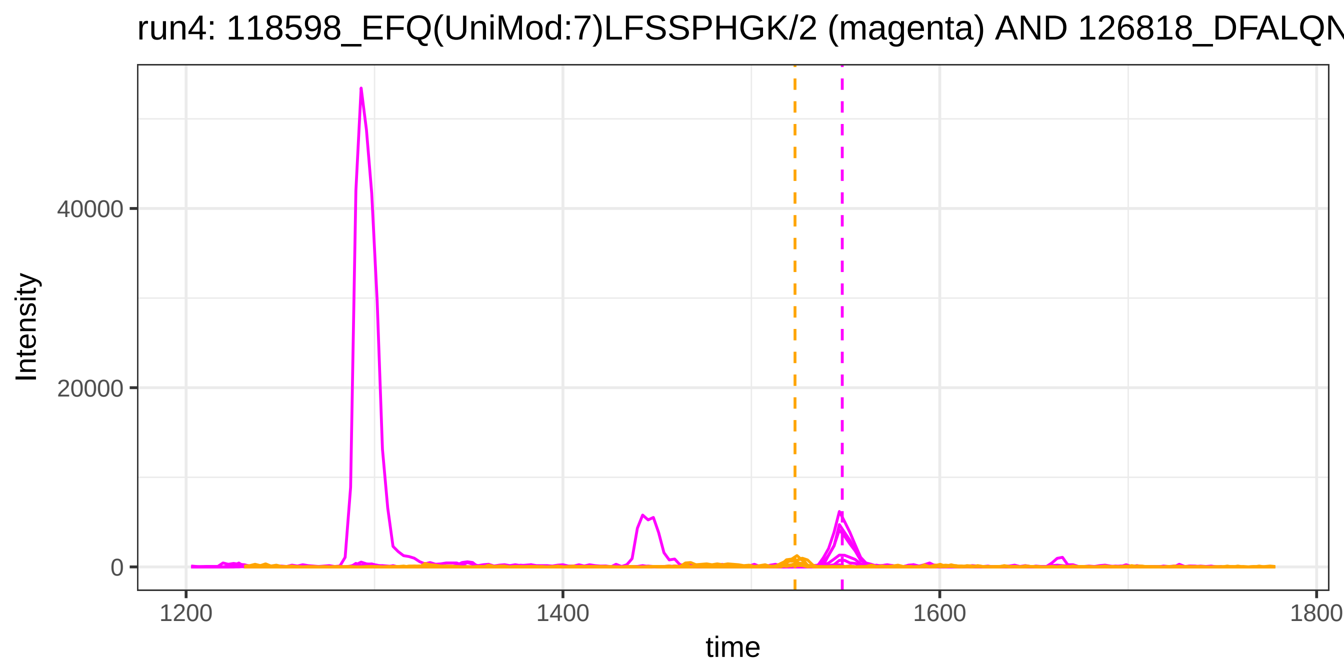
run4: 118598_EFQ(UniMod:7)LFSSPHGK/2 (magenta) AND 126818_DFALQN
Intensity
40000
20000
0
1200
1400
1600
1800
time

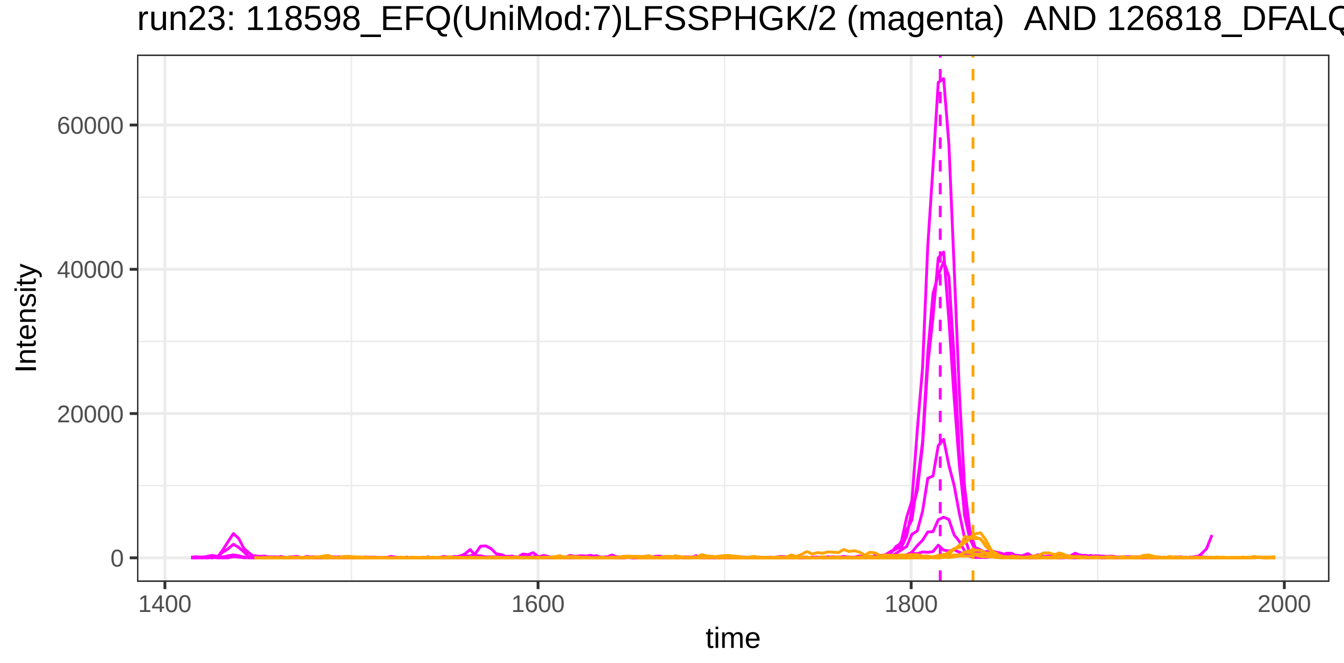
run23: 118598_EFQ(UniMod:7)LFSSPHGK/2 (magenta) AND 126818_DFALQ
Intensity
60000
40000
20000
0
1400
1600
1800
2000
time

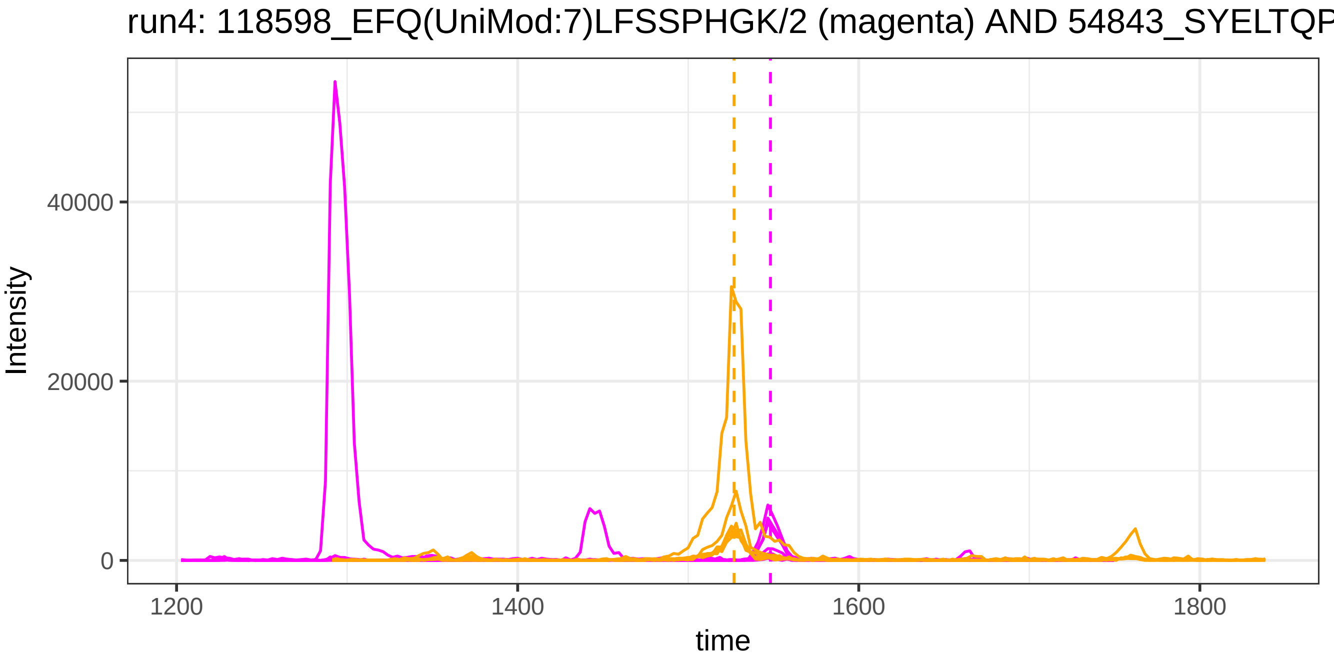
run4: 118598_EFQ(UniMod:7)LFSSPHGK/2 (magenta) AND 54843_SYELTQP
Intensity
40000
20000
0
1200
1400
1600
1800
time

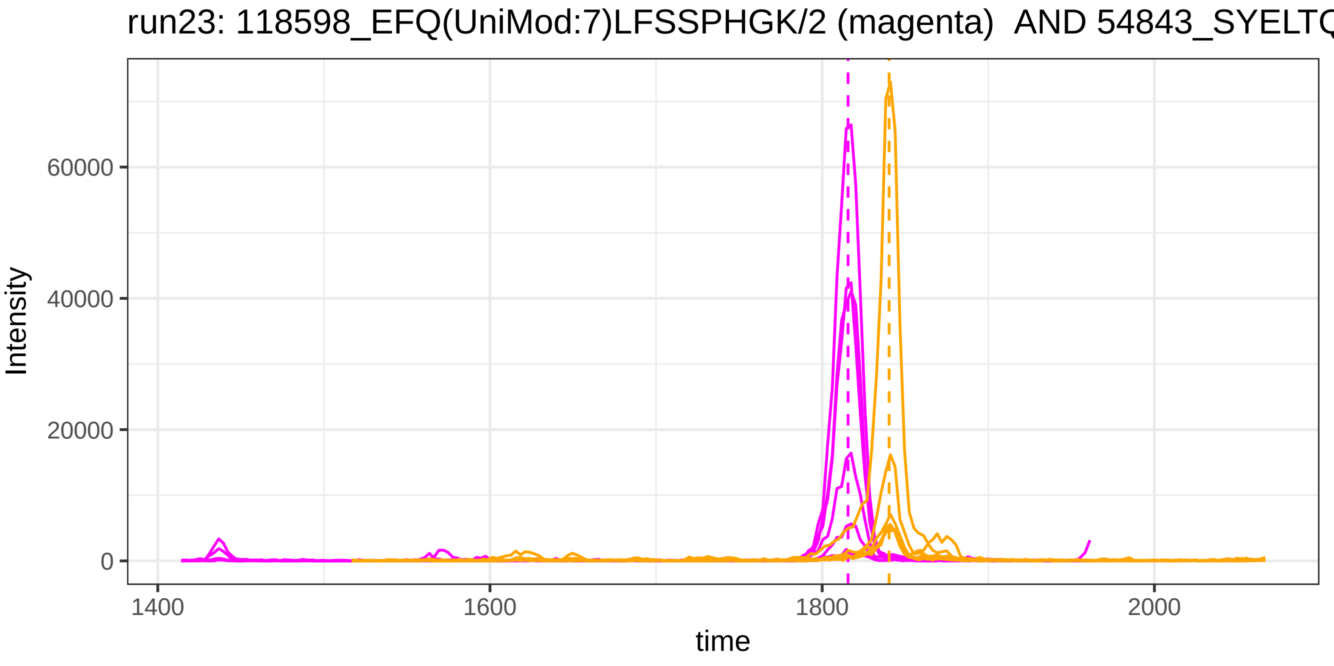
run23: 118598_EFQ(UniMod:7)LFSSPHGK/2 (magenta) AND 54843_SYELTQ
Intensity
60000
40000
20000
0
1400
1600
1800
2000
time

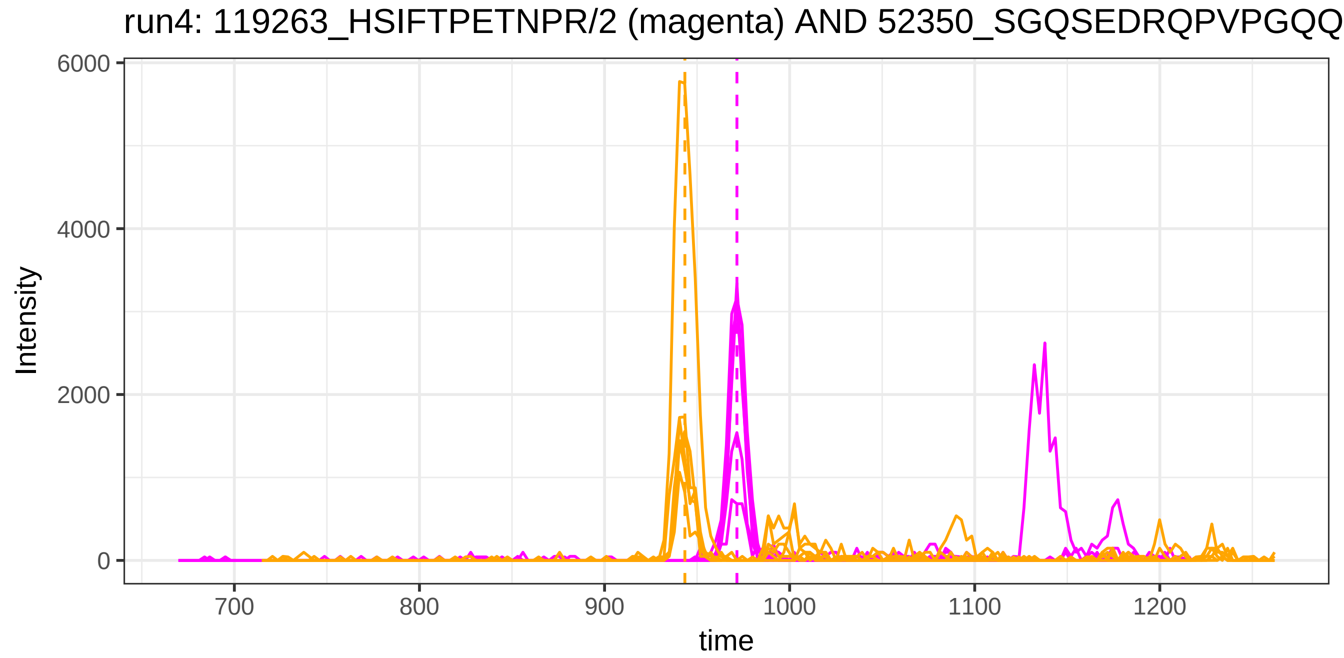

run4: 119263_HSIFTPETNPR/2 (magenta) AND 52350_SGQSEDRQPVPGQQ
Intensity
6000
4000
2000
0
700
800
900
1000
1100
1200
time


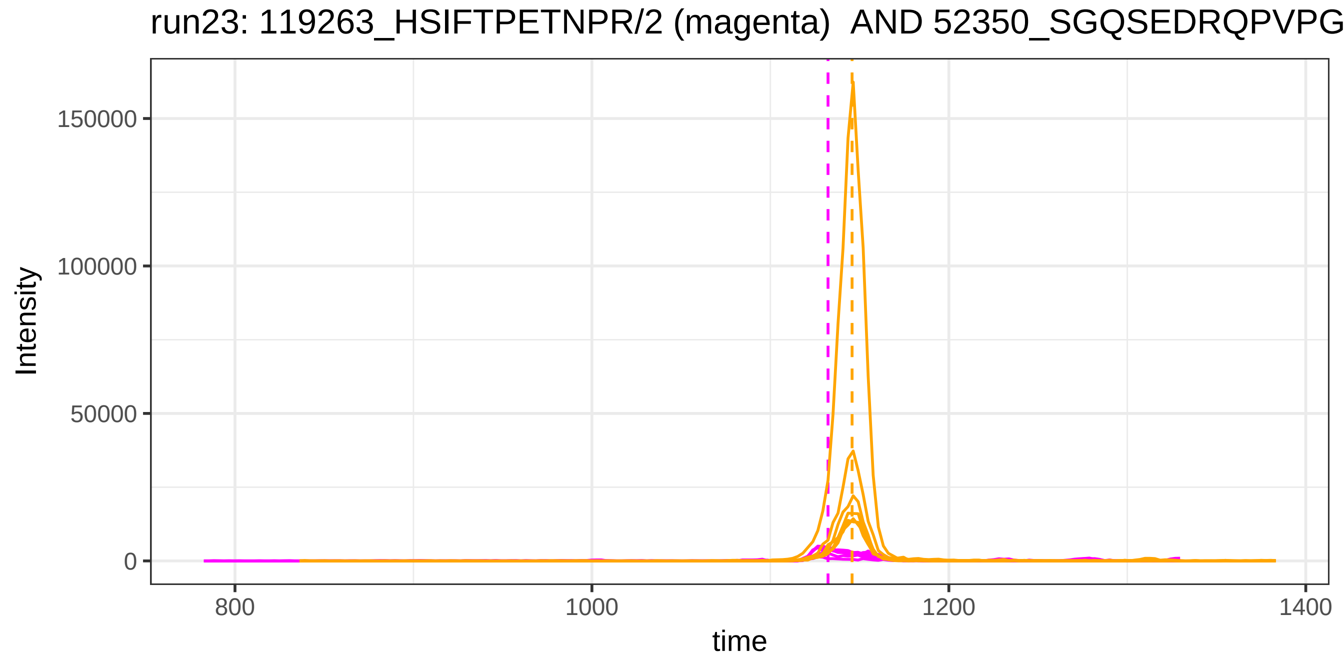

run23: 119263_HSIFTPETNPR/2 (magenta) AND 52350_SGQSEDRQPVPG
Intensity
150000
100000
50000
0
800
1000
1200
1400
time

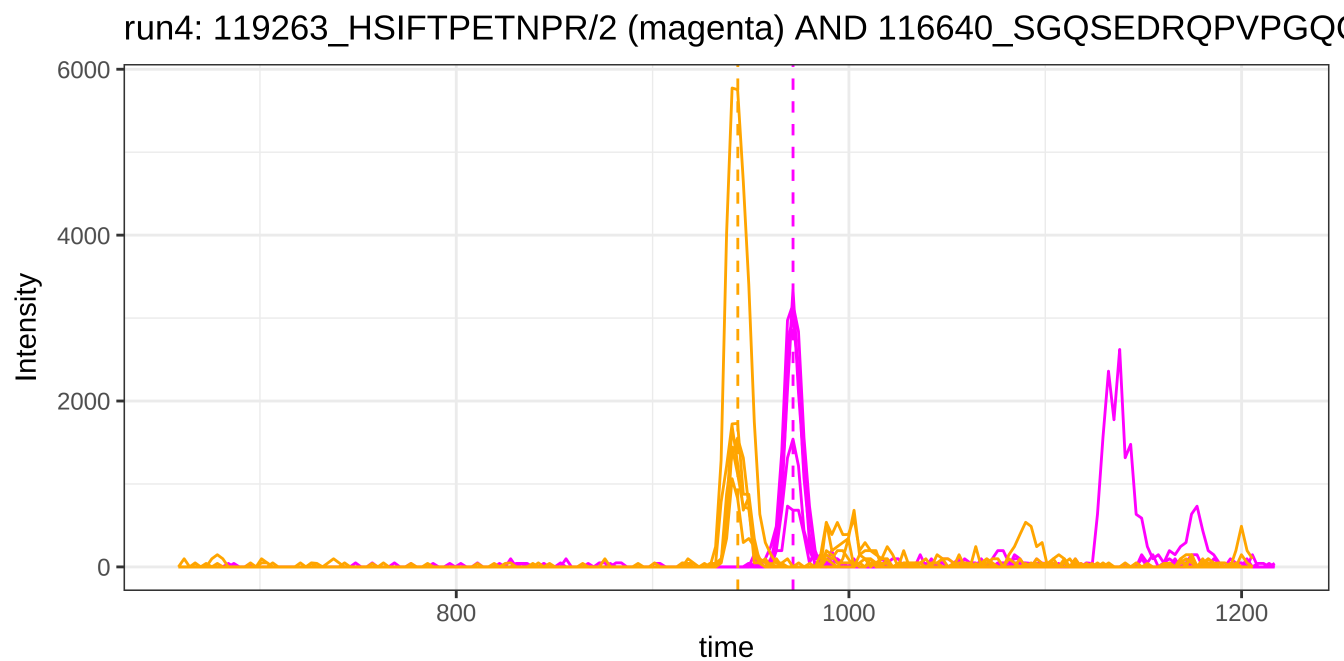
run4: 119263_HSIFTPETNPR/2 (magenta) AND 116640_SGQSEDRQPVPGQQ
Intensity
6000
4000
2000
0
800
1000
1200
time

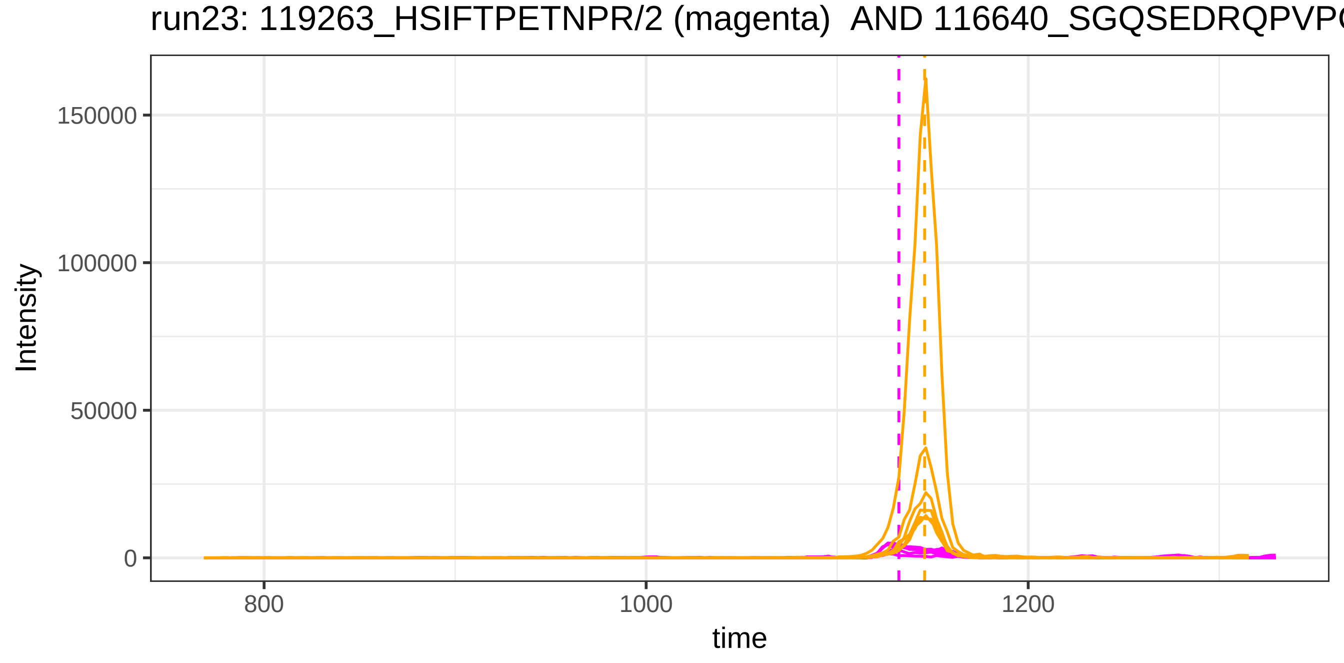
run23: 119263_HSIFTPETNPR/2 (magenta) AND 116640_SGQSEDRQPVPG
Intensity
150000
100000
50000
0
800
1000
1200
time

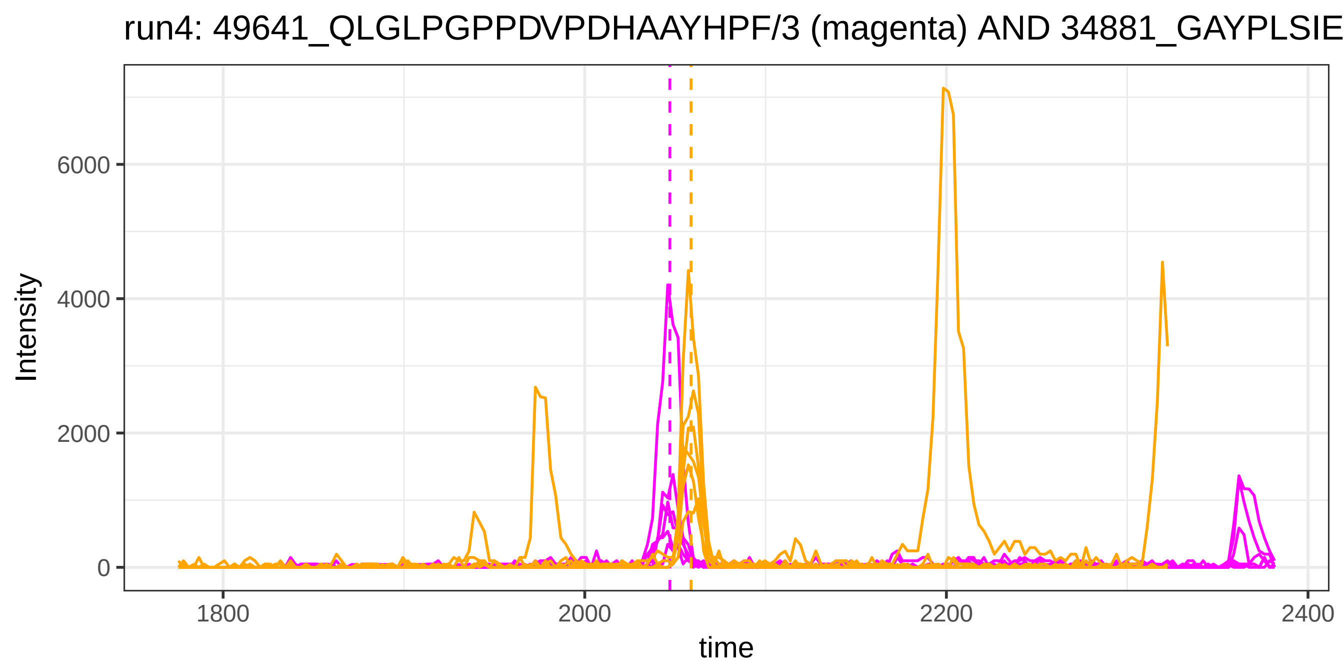
run4: 49641_QLGLPGPPDVPDHAAYHPF/3 (magenta) AND 34881_GAYPLSIE
Intensity
time
0
2000
4000
6000
1800
2000
2200
2400

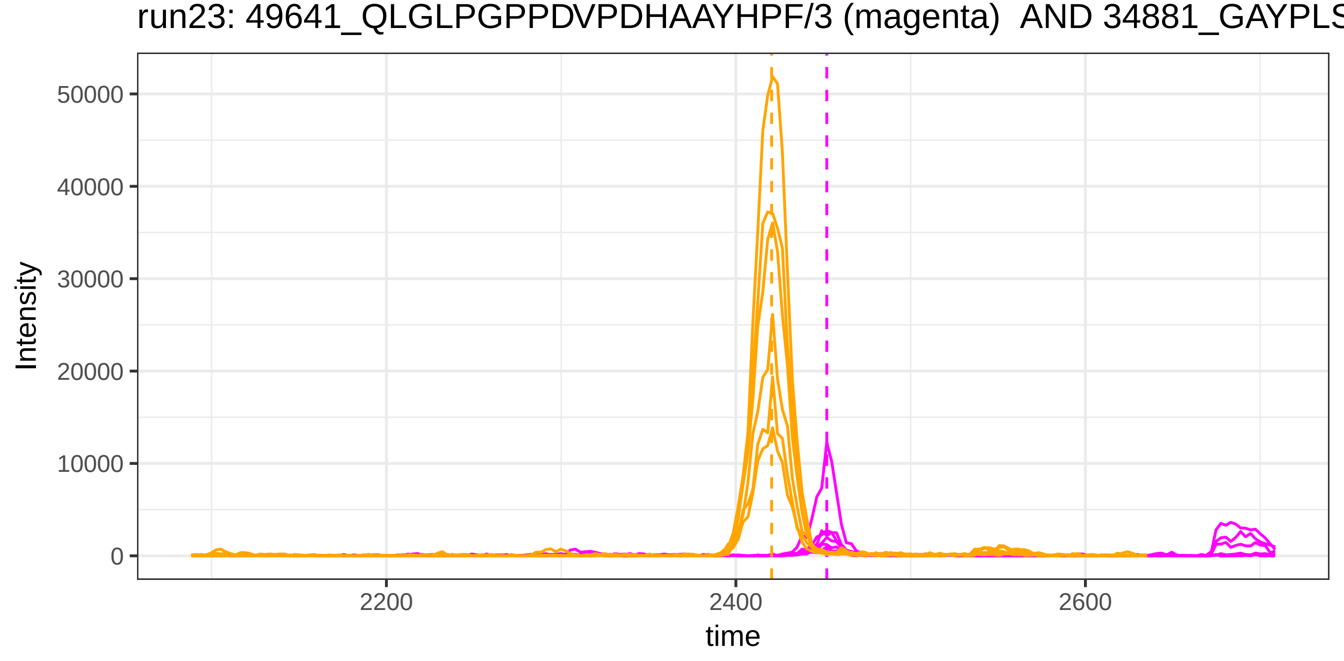
run23: 49641_QLGLPGPPDVPDHAAYHPF/3 (magenta) AND 34881_GAYPLS
Intensity
time
0
10000
20000
30000
40000
50000
2200
2400
2600

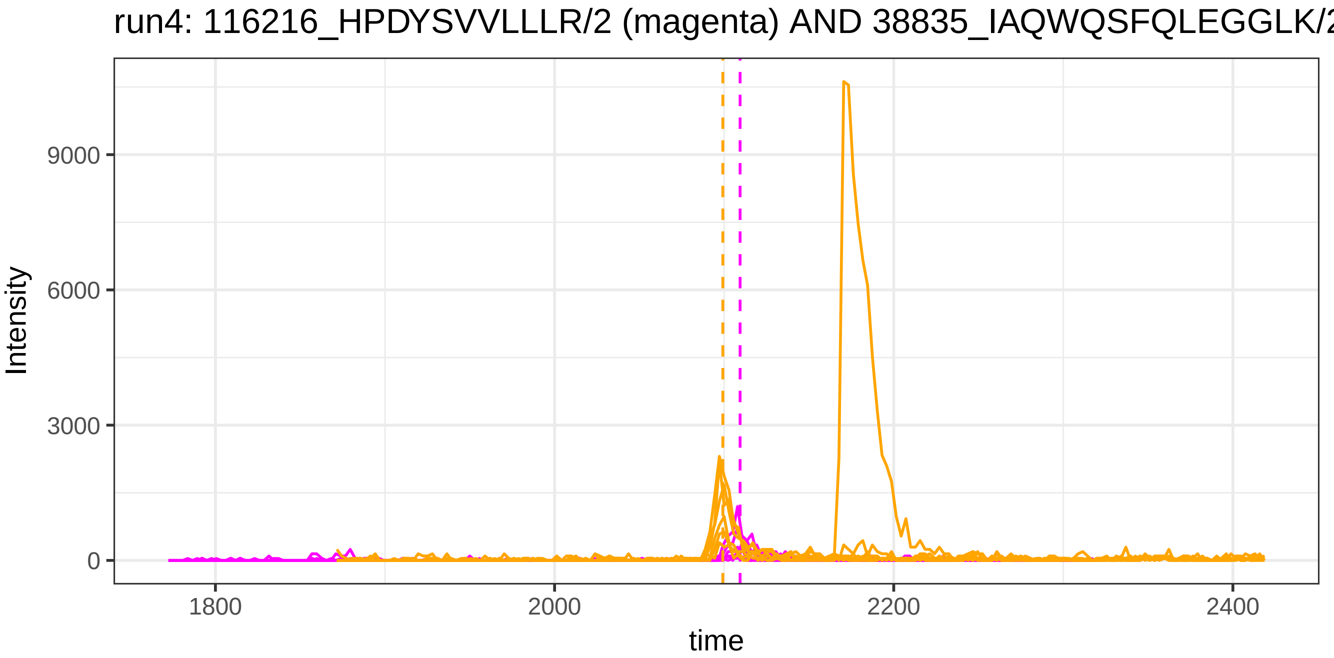
run4: 116216_HPDYSVVLLLR/2 (magenta) AND 38835_IAQWQSFQLEGGLK/2
Intensity
9000
6000
3000
0
1800
2000
2200
2400
time

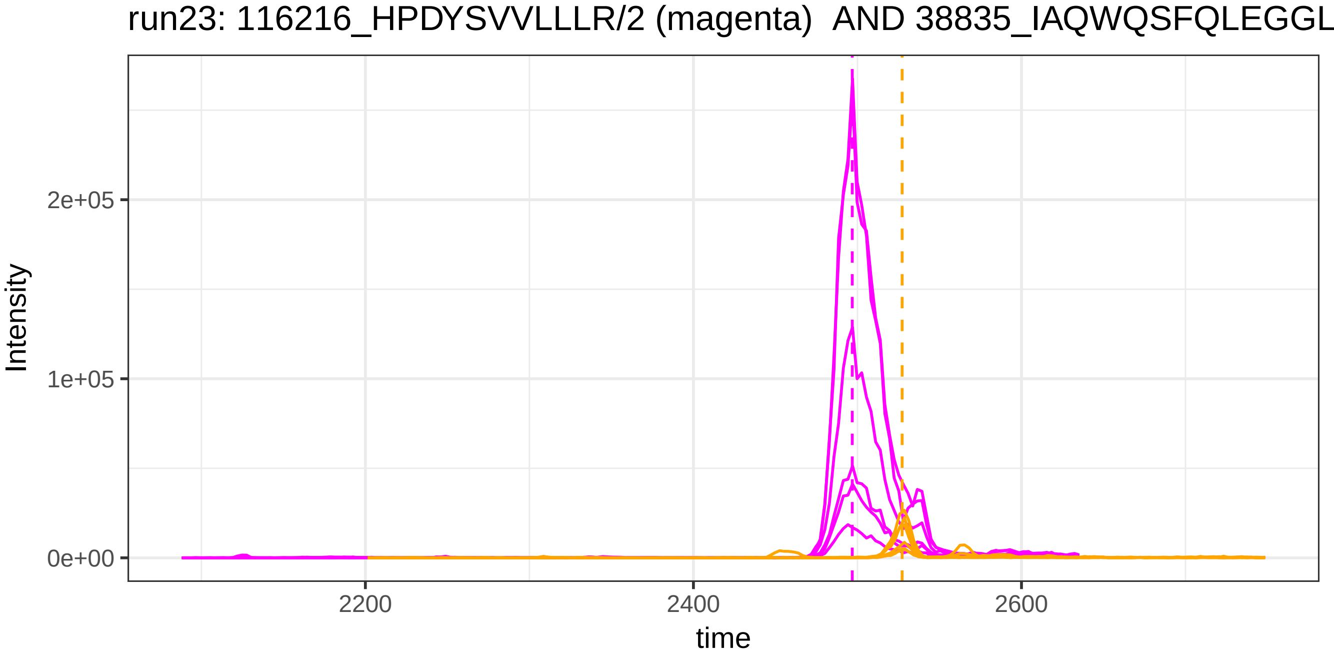
run23: 116216_HPDYSVVLLLR/2 (magenta) AND 38835_IAQWQSFQLEGGL
Intensity
2e+05
1e+05
0e+00
2200
2400
2600
time

run4: 116216_HPDYSVVLLLR/2 (magenta) AND 117978_IAQWQSFQLEGGLK

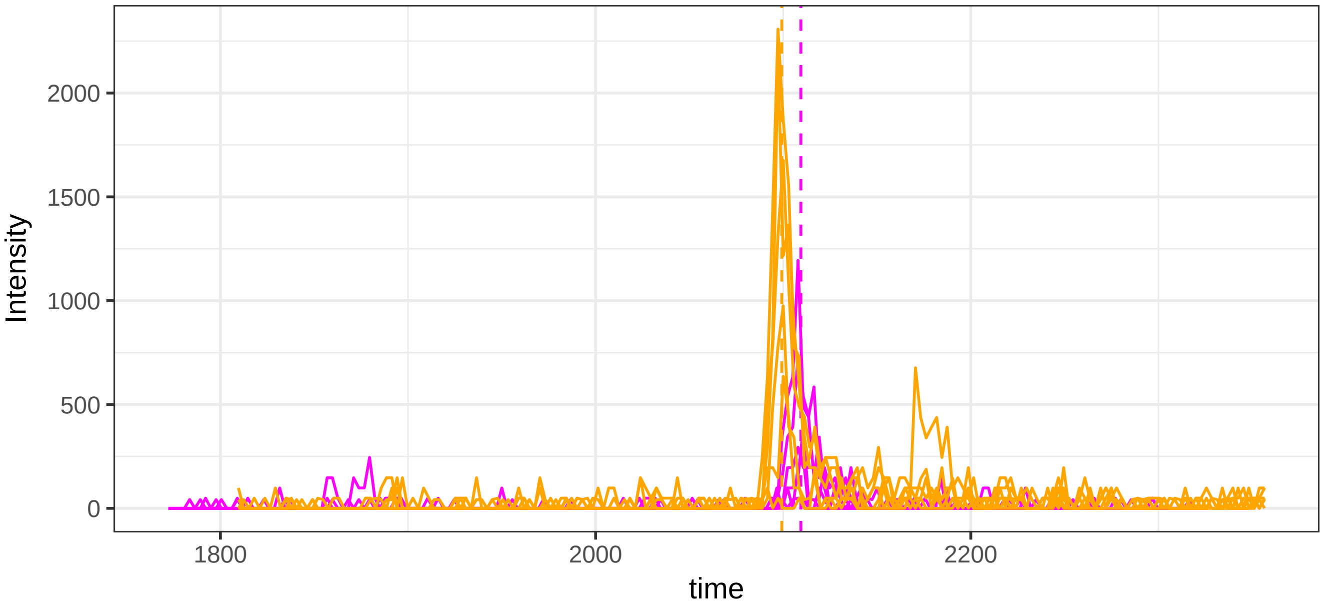


run23: 116216_HPDYSVVLLLR/2 (magenta)  AND 117978_IAQWQSFQLEGG

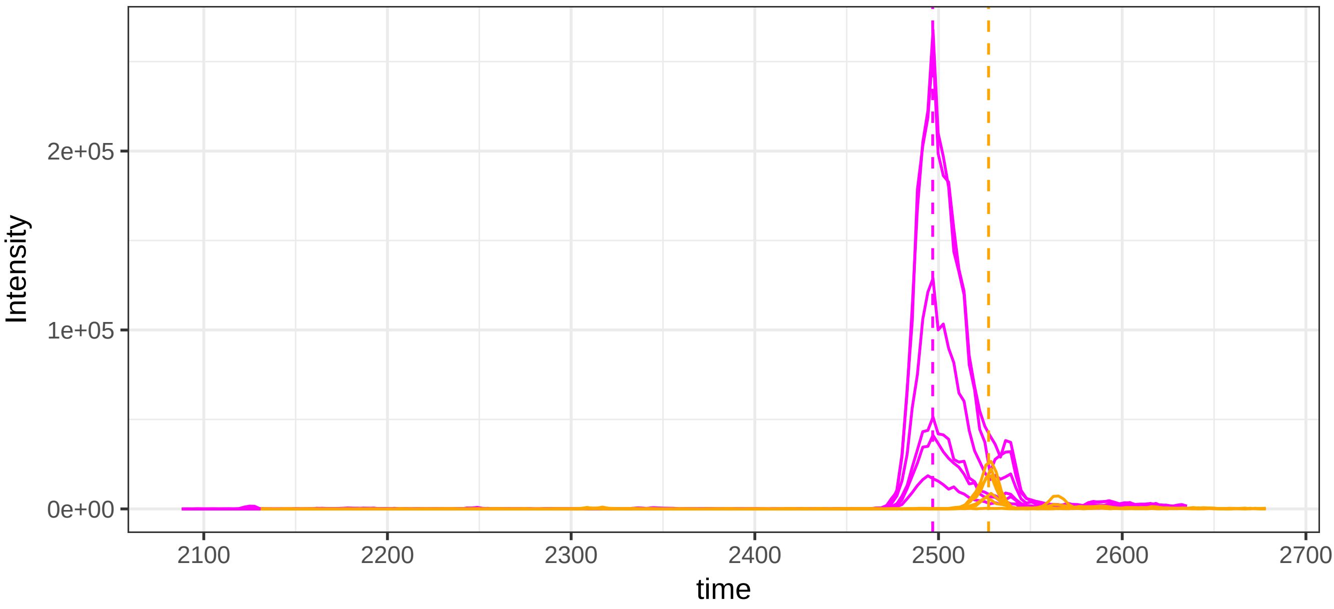

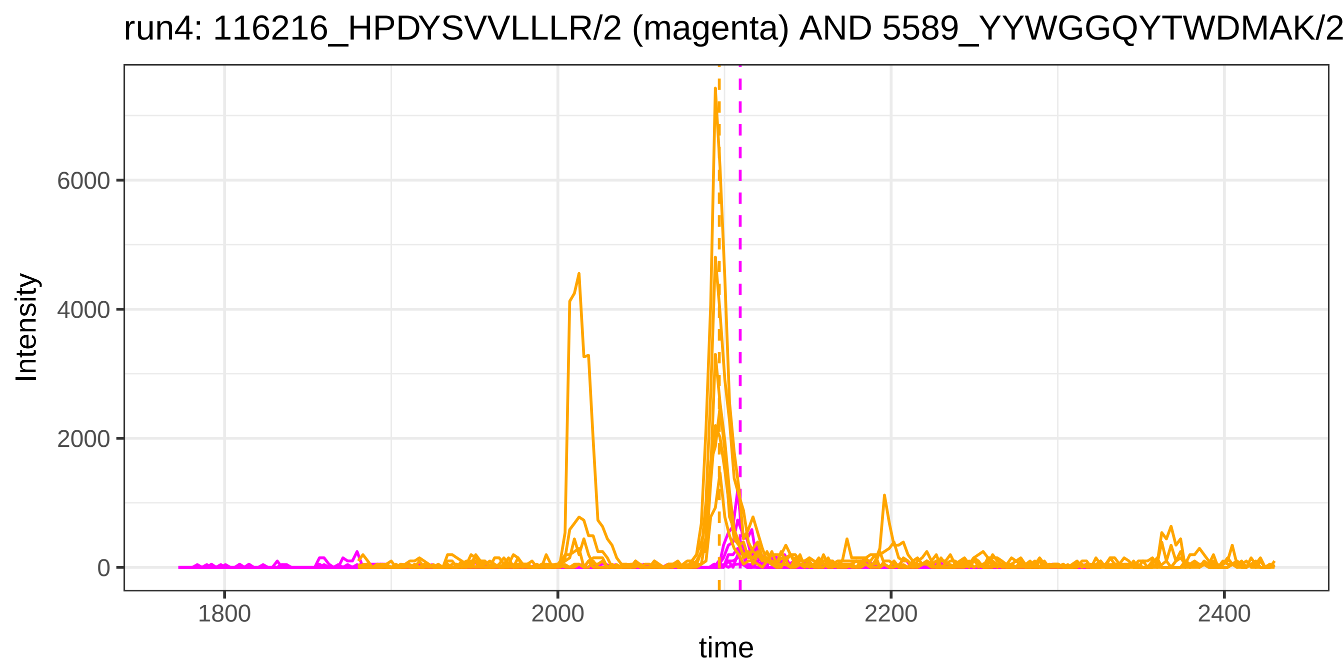
run4: 116216_HPDYSVVLLLR/2 (magenta) AND 5589_YYWGGQYTWDMAK/2
Intensity
6000
4000
2000
0
1800
2000
2200
2400
time

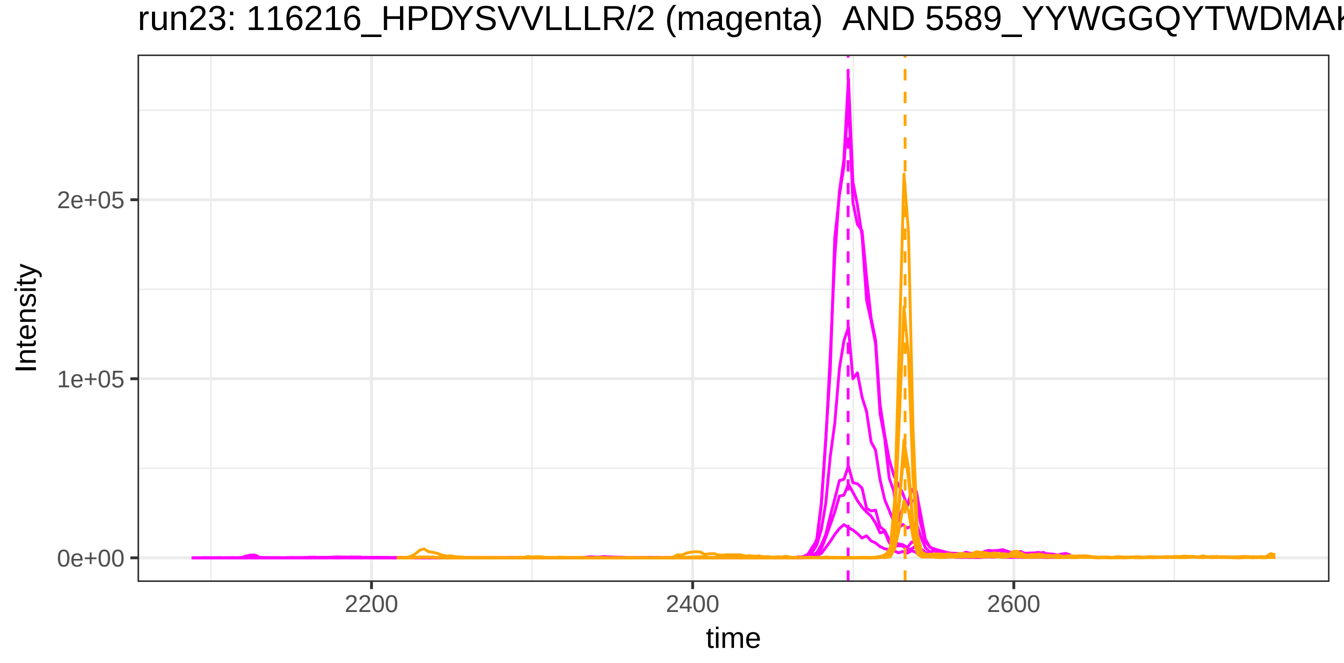
run23: 116216_HPDYSVVLLLR/2 (magenta) AND 5589_YYWGGQYTWDMAK
Intensity
2e+05
1e+05
0e+00
2200
2400
2600
time

run4: 116216_HPDYSVVLLLR/2 (magenta) AND 119947_YYWGGQ(UniMod:7)

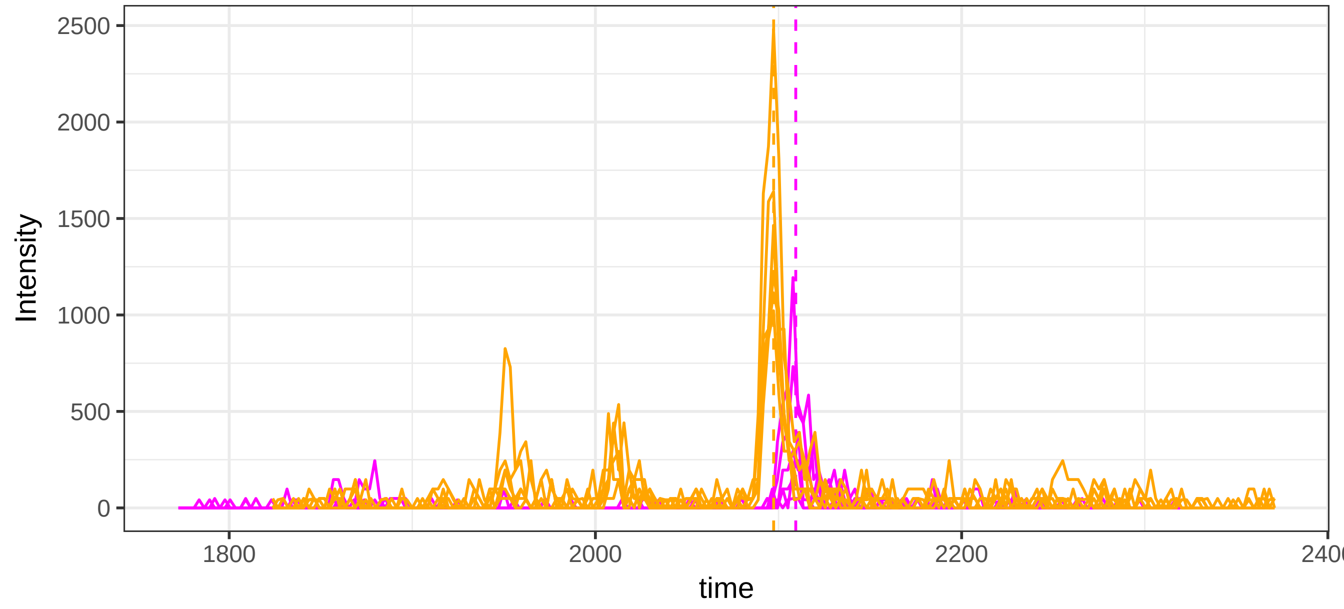


run23: 116216_HPDYSVVLLLR/2 (magenta)  AND 119947_YYWGGQ(UniMod

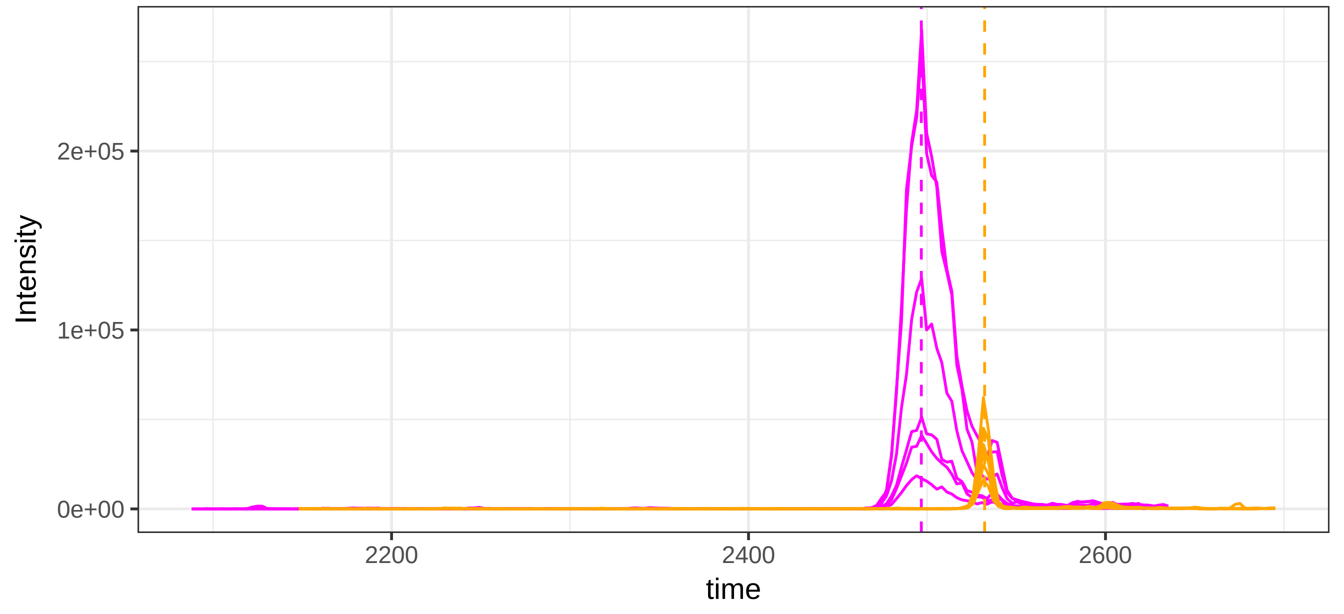

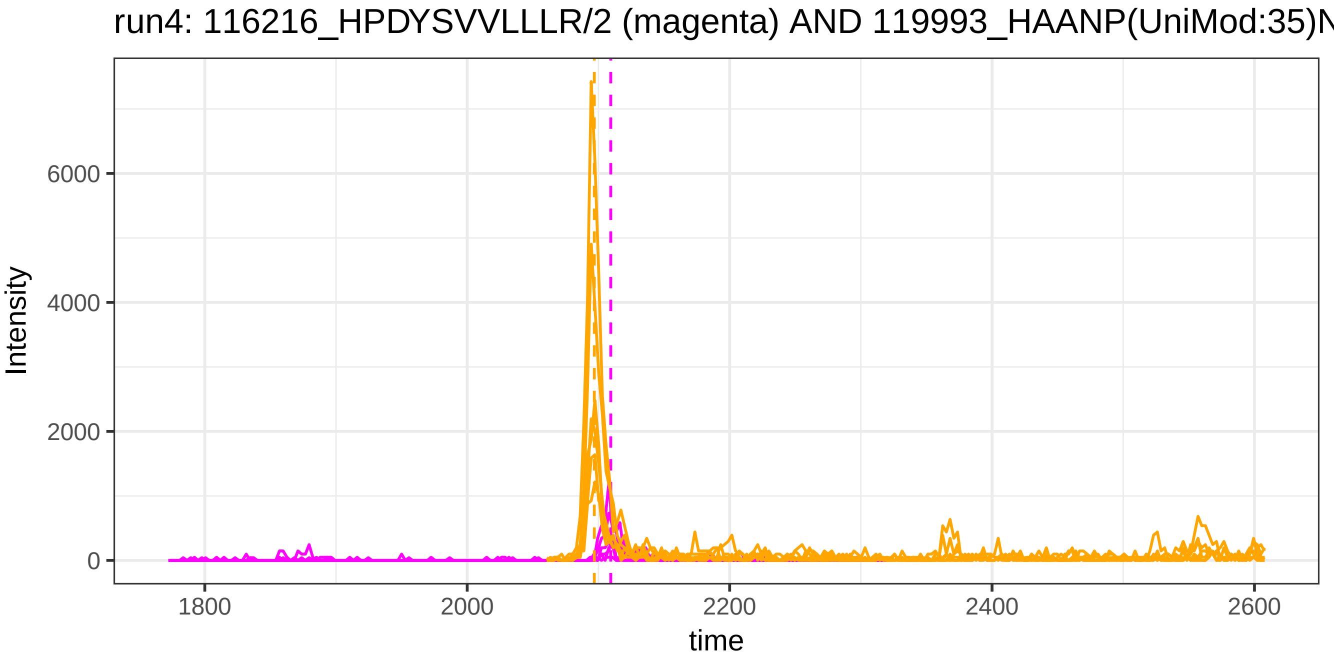

run4: 116216_HPDYSVVLLLR/2 (magenta) AND 119993_HAANP(UniMod:35)N
Intensity
6000
4000
2000
0
1800
2000
2200
2400
2600
time


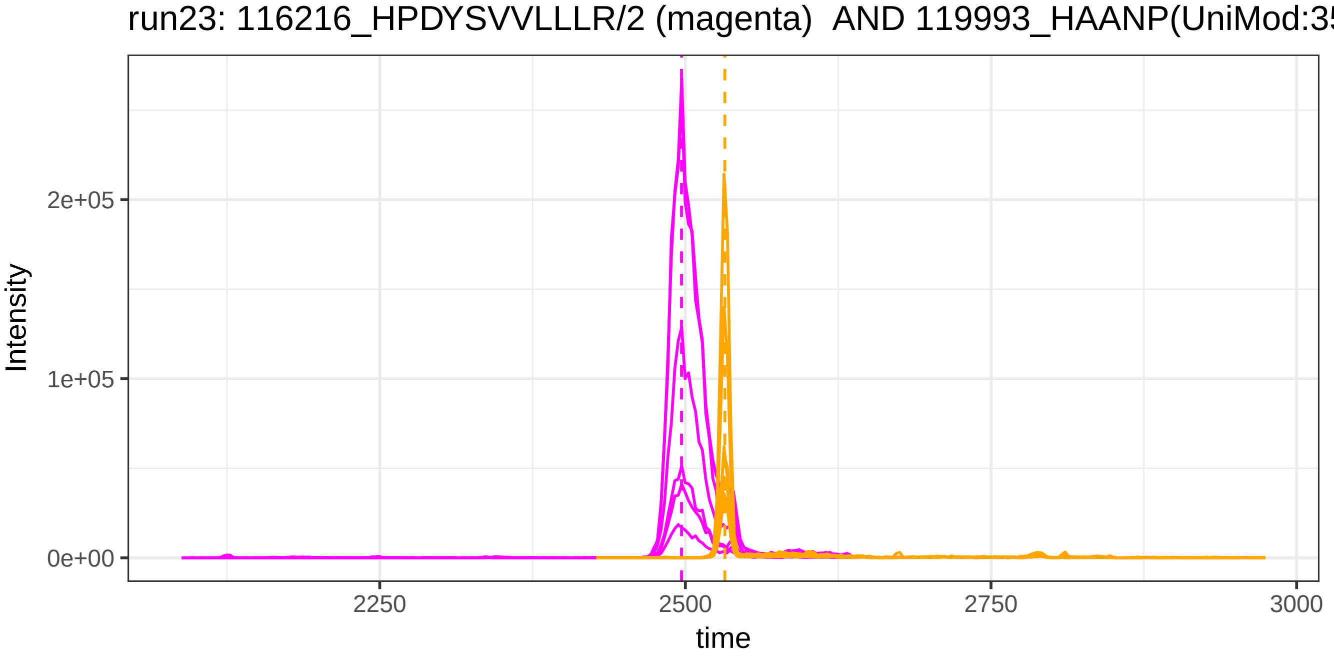

run23: 116216_HPDYSVVLLLR/2 (magenta) AND 119993_HAANP(UniMod:35
Intensity
2e+05
1e+05
0e+00
2250
2500
2750
3000
time

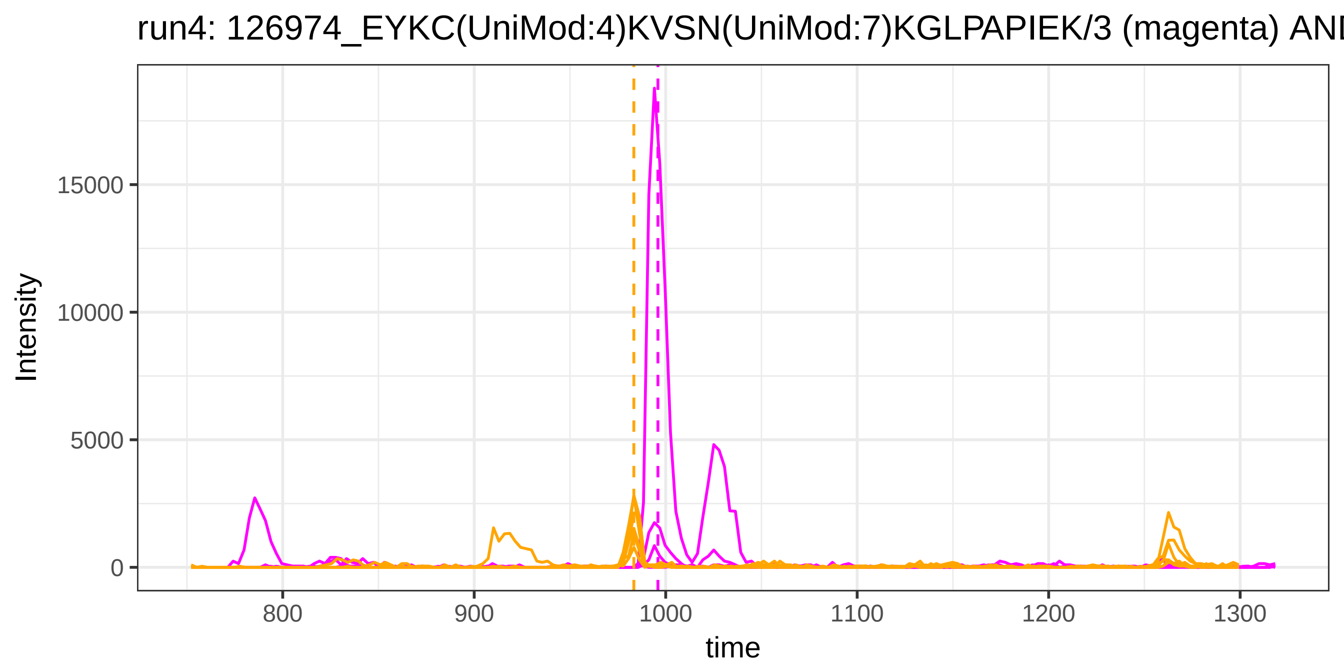

run4: 126974_EYKC(UniMod:4)KVSN(UniMod:7)KGLPAPIEK/3 (magenta) ANI
Intensity
15000
10000
5000
0
800
900
1000
1100
1200
1300
time


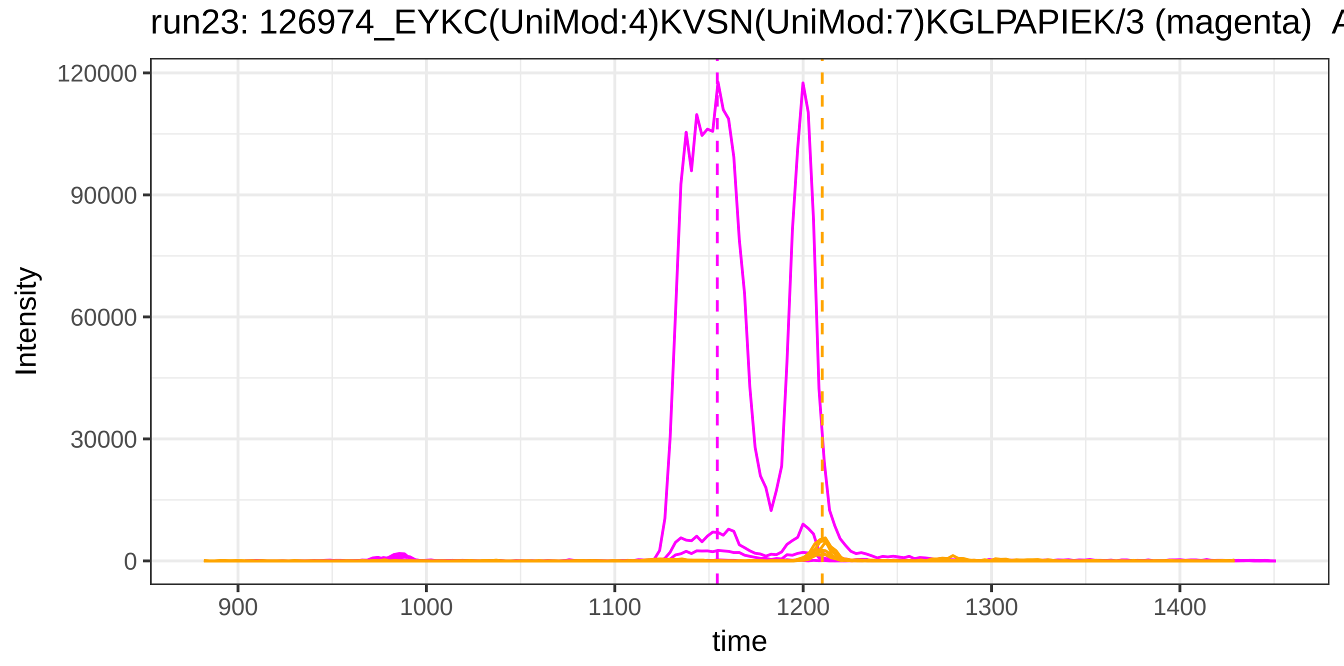

run23: 126974_EYKC(UniMod:4)KVSN(UniMod:7)KGLPAPIEK/3 (magenta) A
Intensity
120000
90000
60000
30000
0
900
1000
1100
1200
1300
1400
time

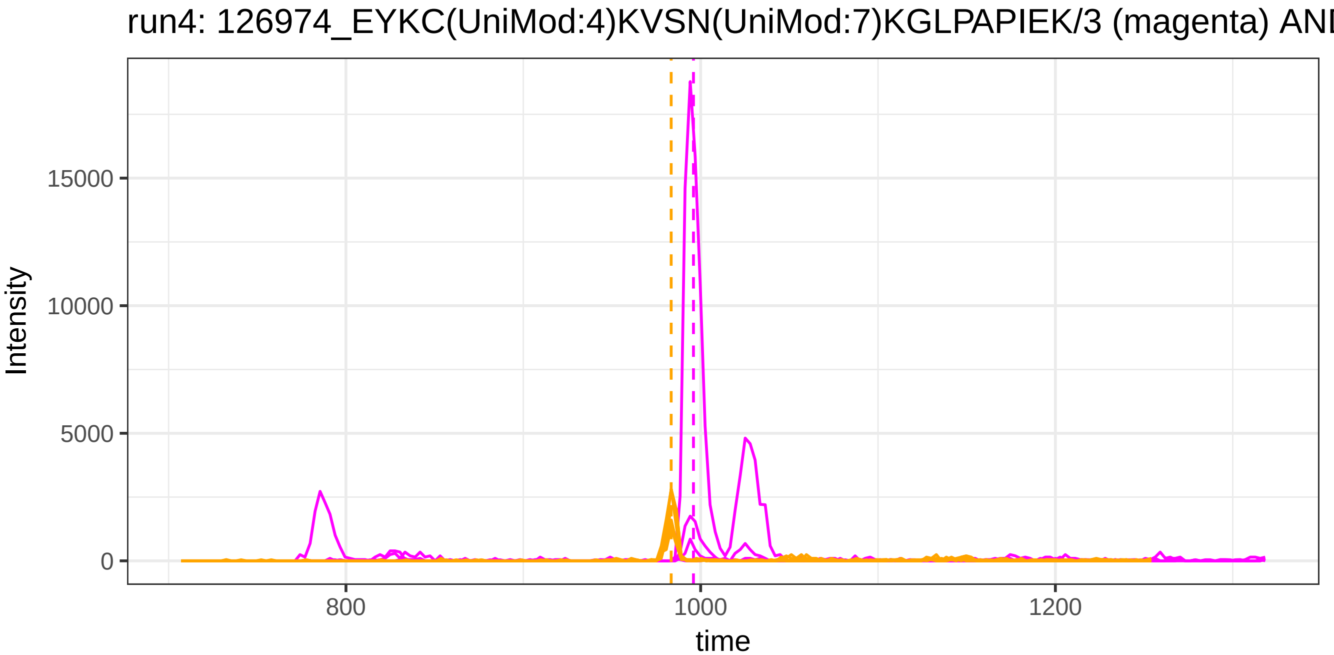

run4: 126974_EYKC(UniMod:4)KVSN(UniMod:7)KGLPAPIEK/3 (magenta) ANI
Intensity
15000
10000
5000
0
800
1000
1200
time


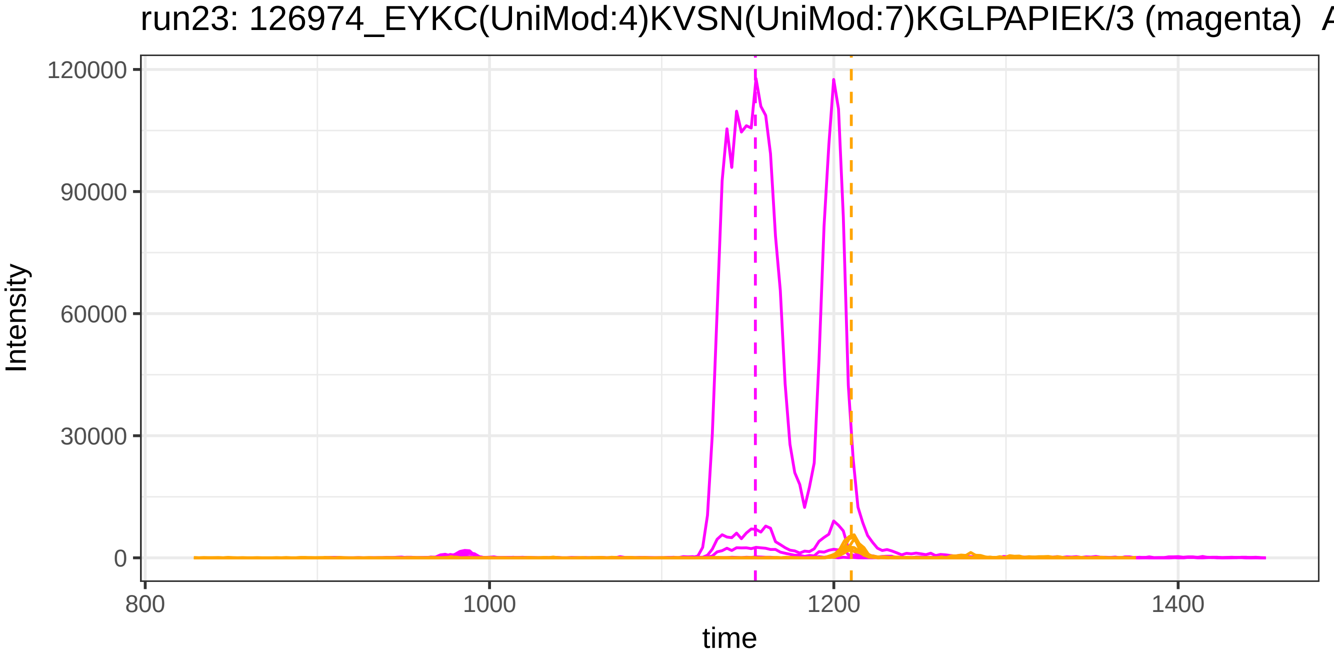

run23: 126974_EYKC(UniMod:4)KVSN(UniMod:7)KGLPAPIEK/3 (magenta) A
Intensity
120000
90000
60000
30000
0
800
1000
1200
1400
time

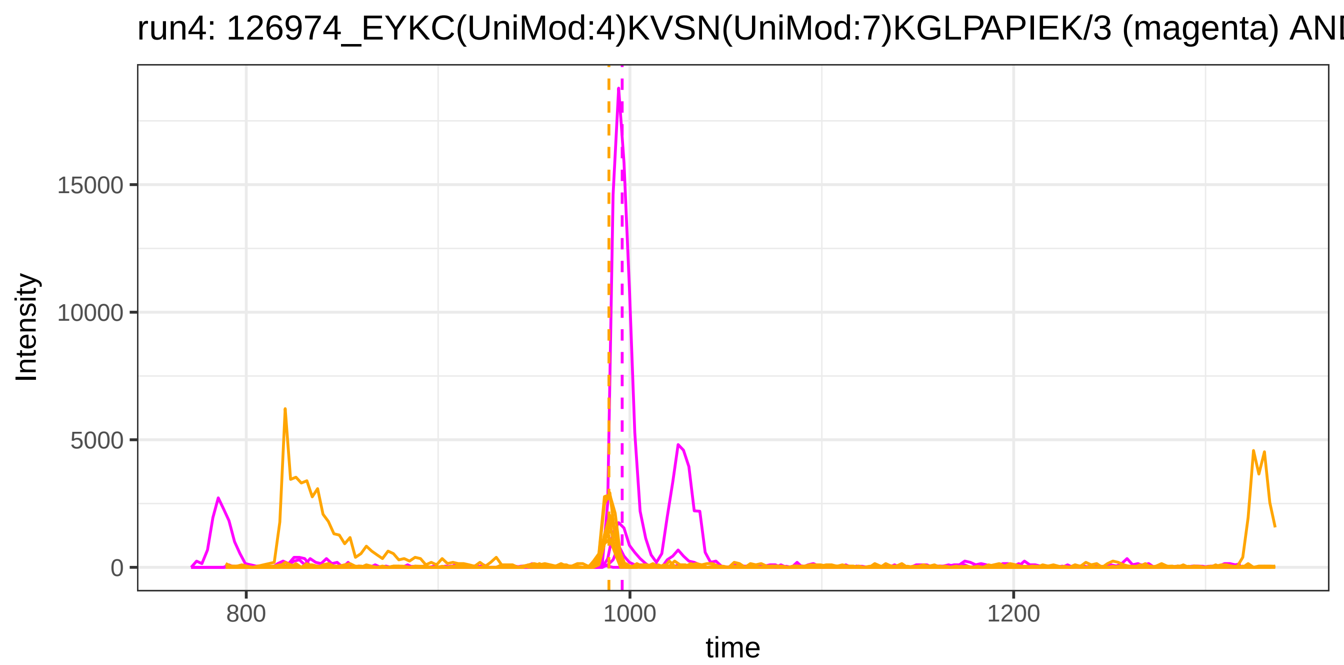
run4: 126974_EYKC(UniMod:4)KVSN(UniMod:7)KGLPAPIEK/3 (magenta) AN
15000
10000
5000
0
800
1000
1200
time
Intensity

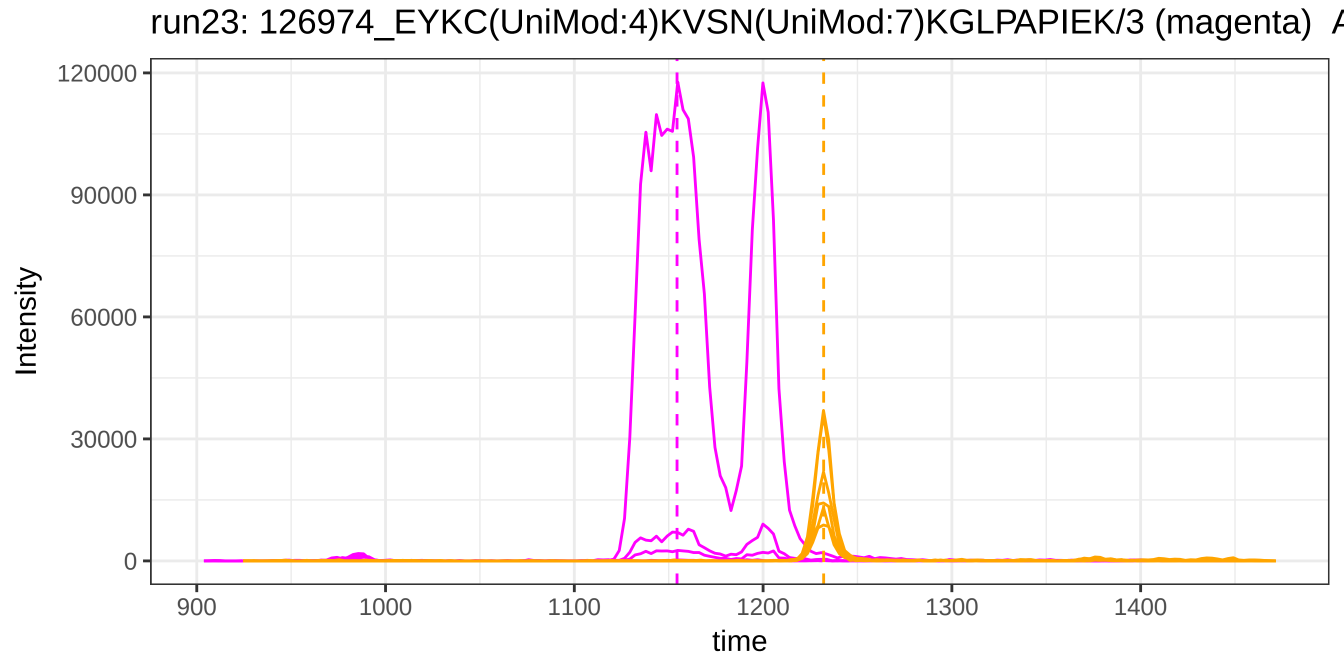
run23: 126974_EYKC(UniMod:4)KVSN(UniMod:7)KGLPAPIEK/3 (magenta) A
120000
90000
60000
30000
0
900
1000
1100
1200
1300
1400
time
Intensity

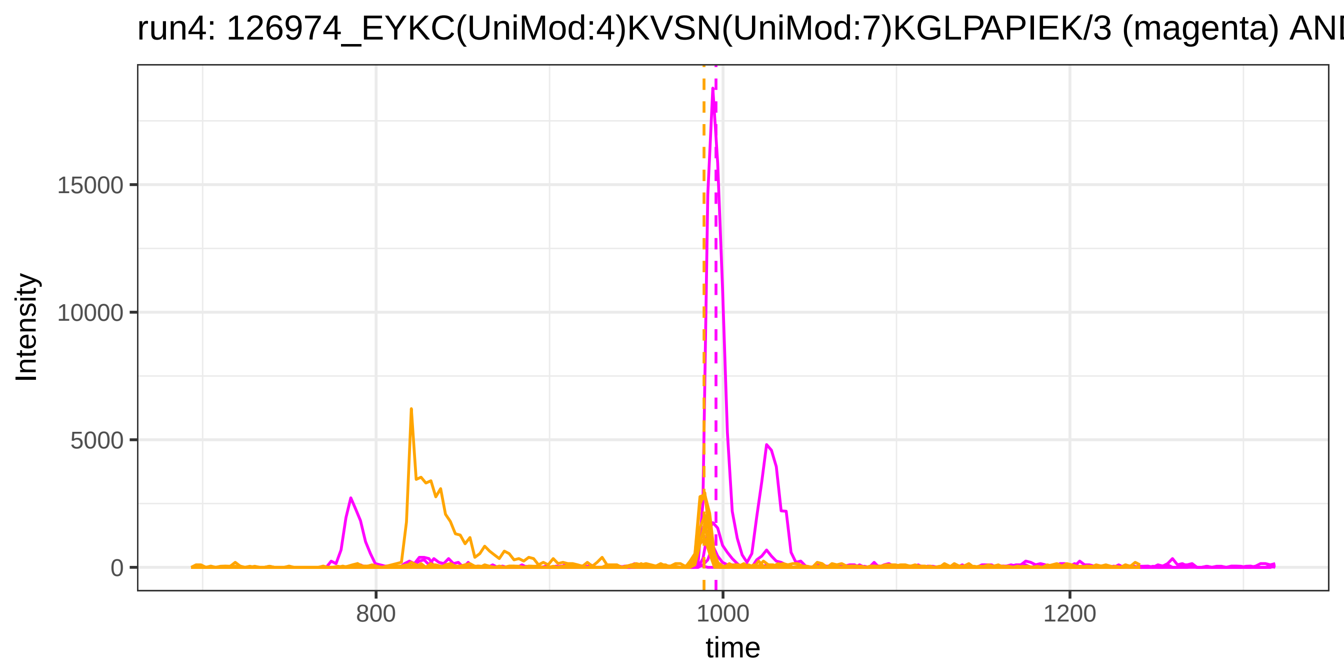

run4: 126974_EYKC(UniMod:4)KVSN(UniMod:7)KGLPAPIEK/3 (magenta) AN
Intensity
15000
10000
5000
0
800
1000
1200
time


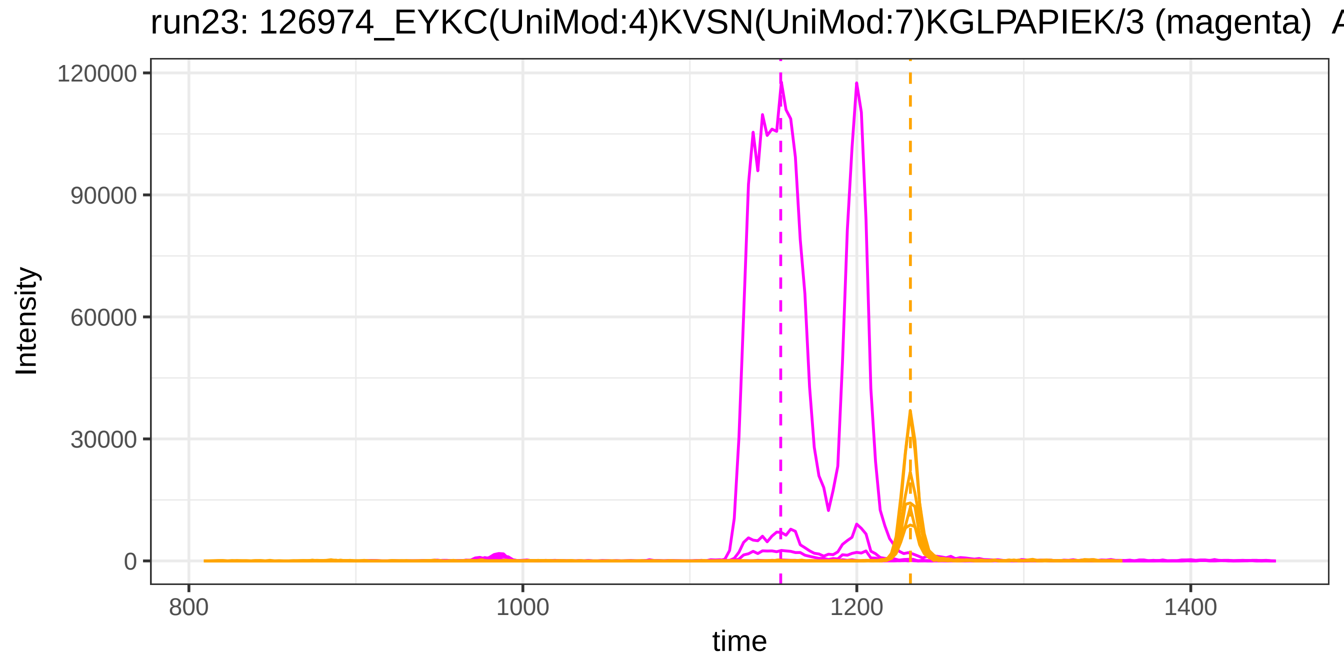

run23: 126974_EYKC(UniMod:4)KVSN(UniMod:7)KGLPAPIEK/3 (magenta) A
Intensity
120000
90000
60000
30000
0
800
1000
1200
1400
time

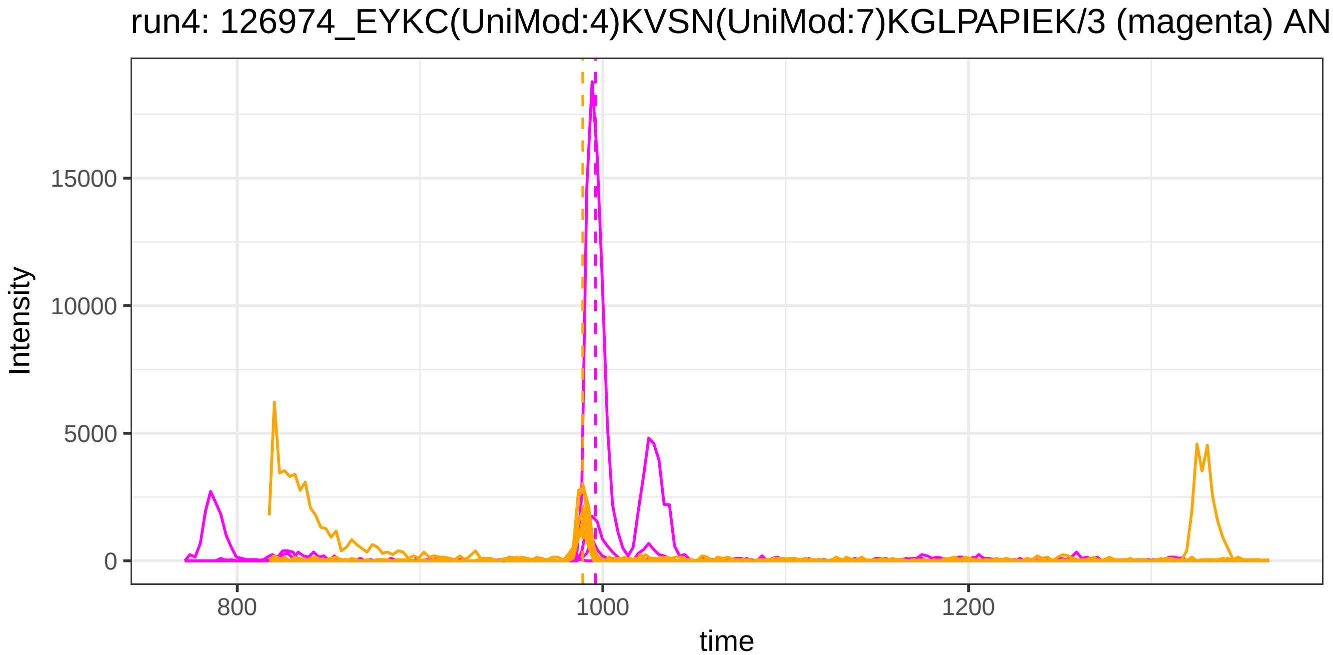
run4: 126974_EYKC(UniMod:4)KVSN(UniMod:7)KGLPAPIEK/3 (magenta) ANI
Intensity
15000
10000
5000
0
800
1000
1200
time

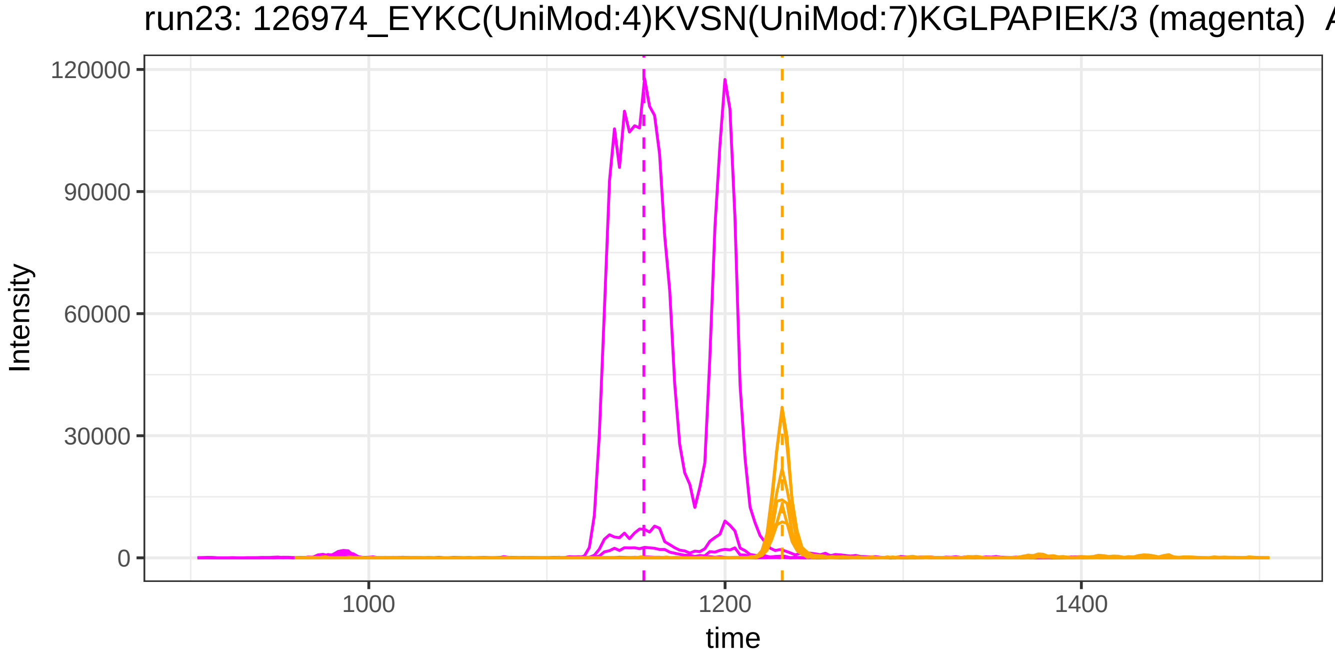
run23: 126974_EYKC(UniMod:4)KVSN(UniMod:7)KGLPAPIEK/3 (magenta) A
Intensity
120000
90000
60000
30000
0
1000
1200
1400
time

run4: 34881_GAYPLSIEPIGVR/2 (magenta) AND 33080_AGDFLEANYMNLQR

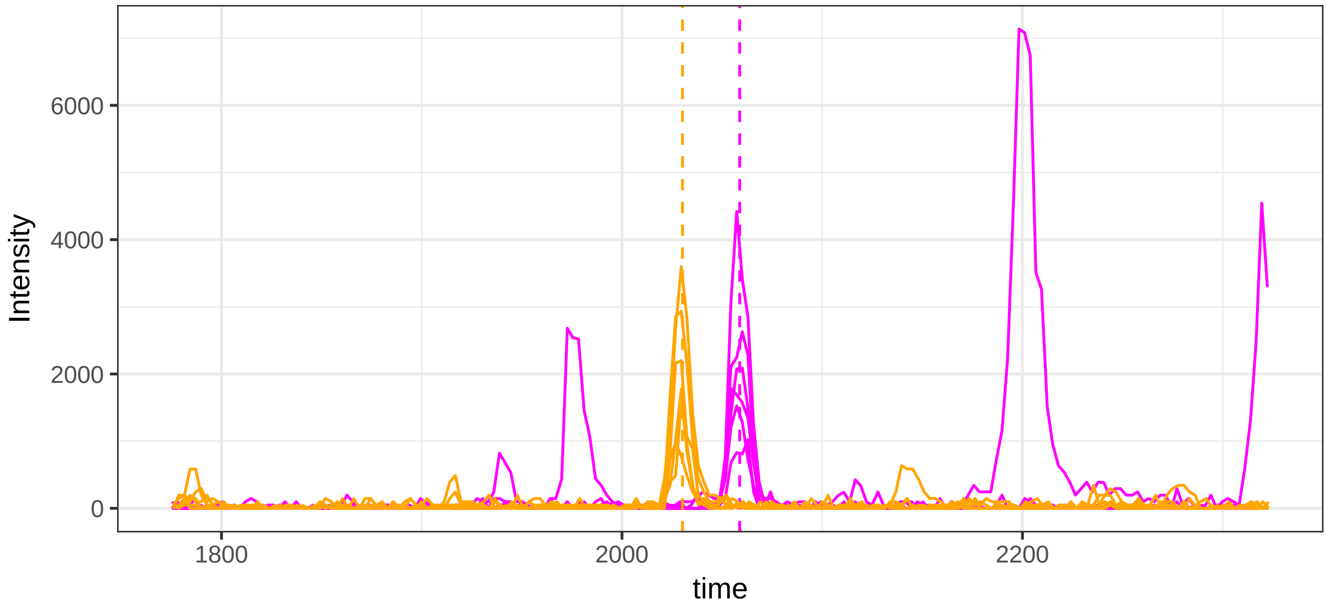


run23: 34881_GAYPLSIEPIGVR/2 (magenta) AND 33080_AGDFLEANYMNLQ

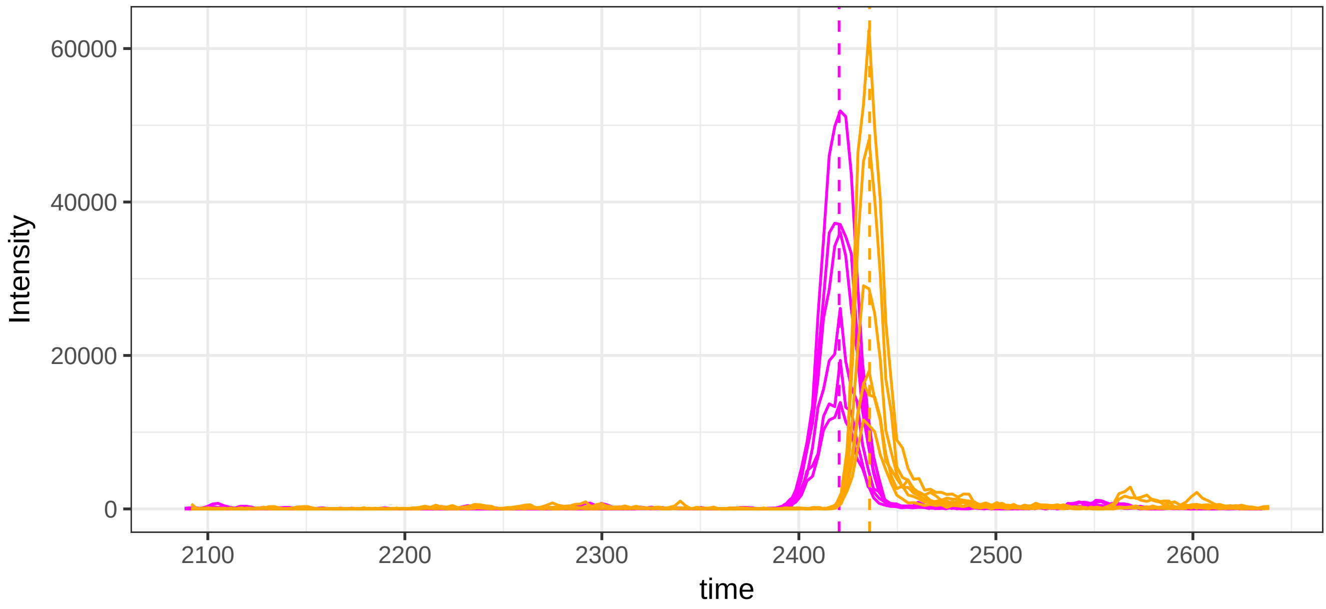

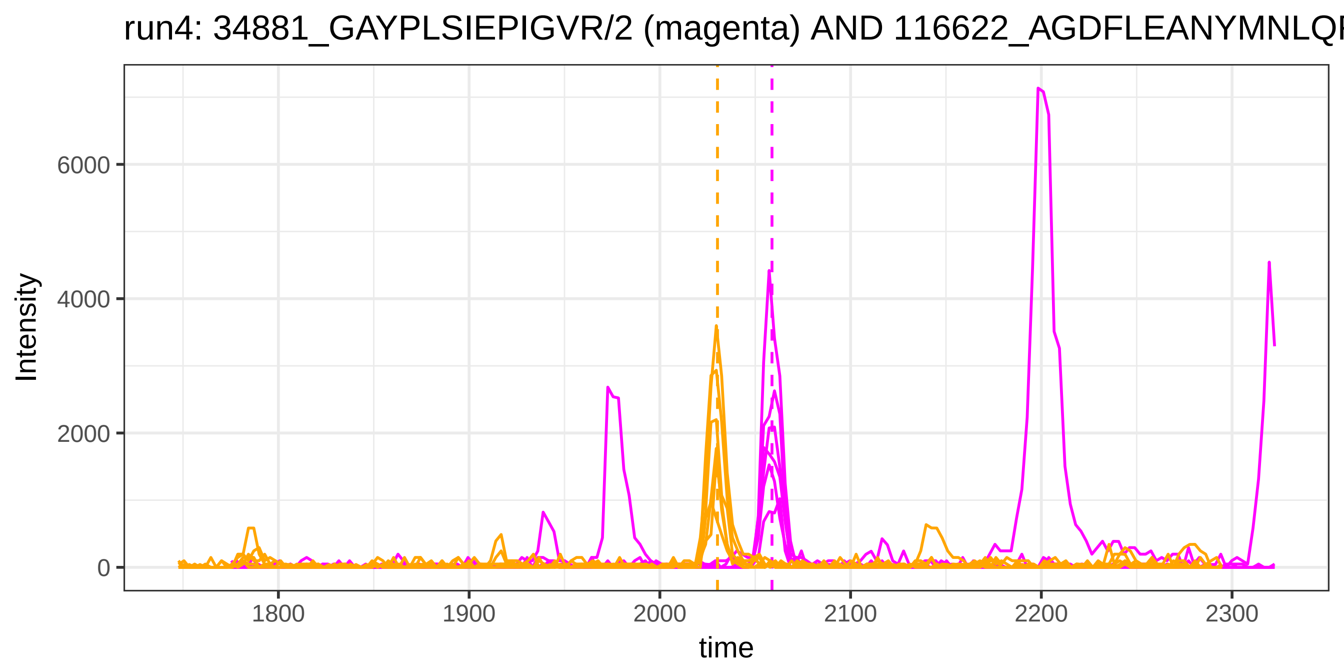

run4: 34881_GAYPLSIEPIGVR/2 (magenta) AND 116622_AGDFLEANYMNLQF
Intensity
6000
4000
2000
0
1800
1900
2000
2100
2200
2300
time


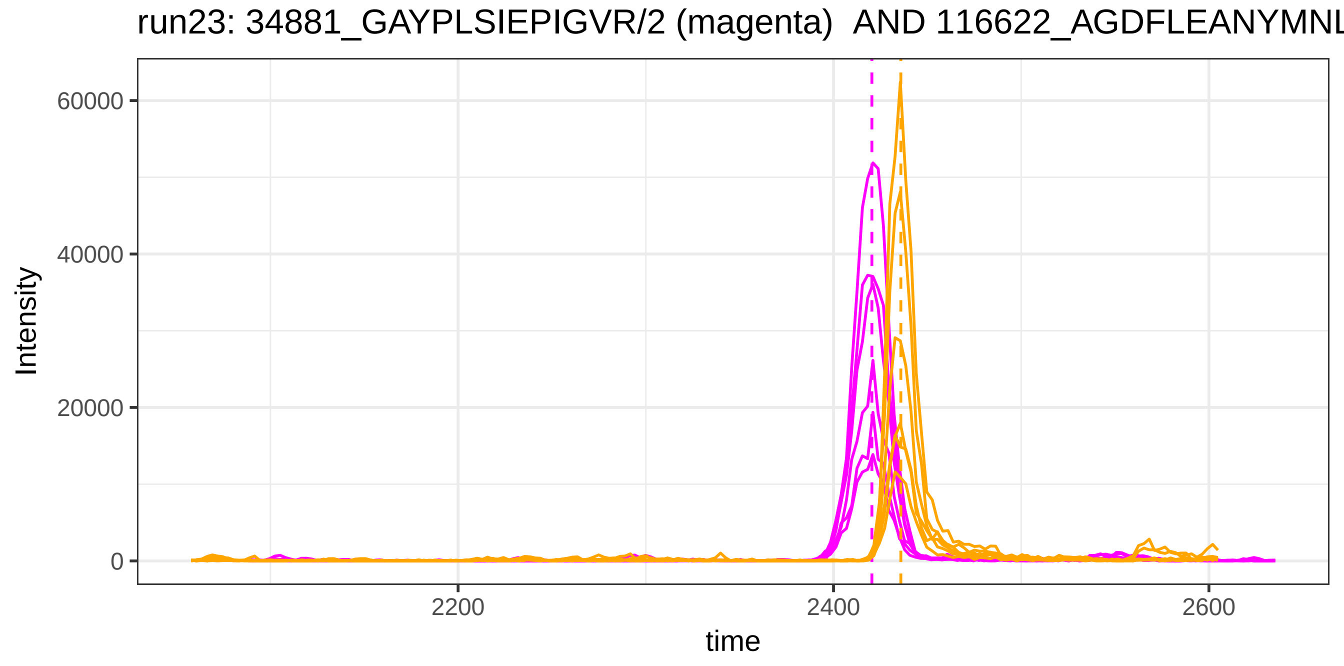

run23: 34881_GAYPLSIEPIGVR/2 (magenta) AND 116622_AGDFLEANYMNL
Intensity
60000
40000
20000
0
2200
2400
2600
time

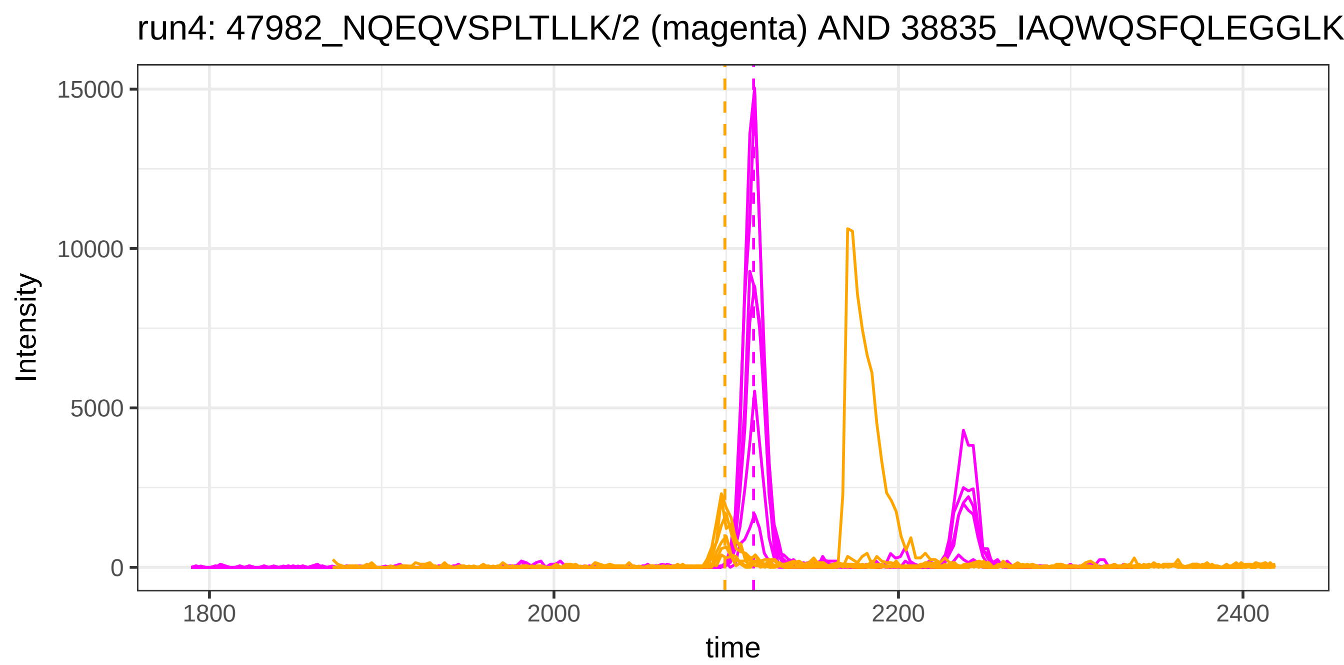
run4: 47982_NQEQVSPLTLLK/2 (magenta) AND 38835_IAQWQSFQLEGGLK
Intensity
15000
10000
5000
0
1800
2000
2200
2400
time

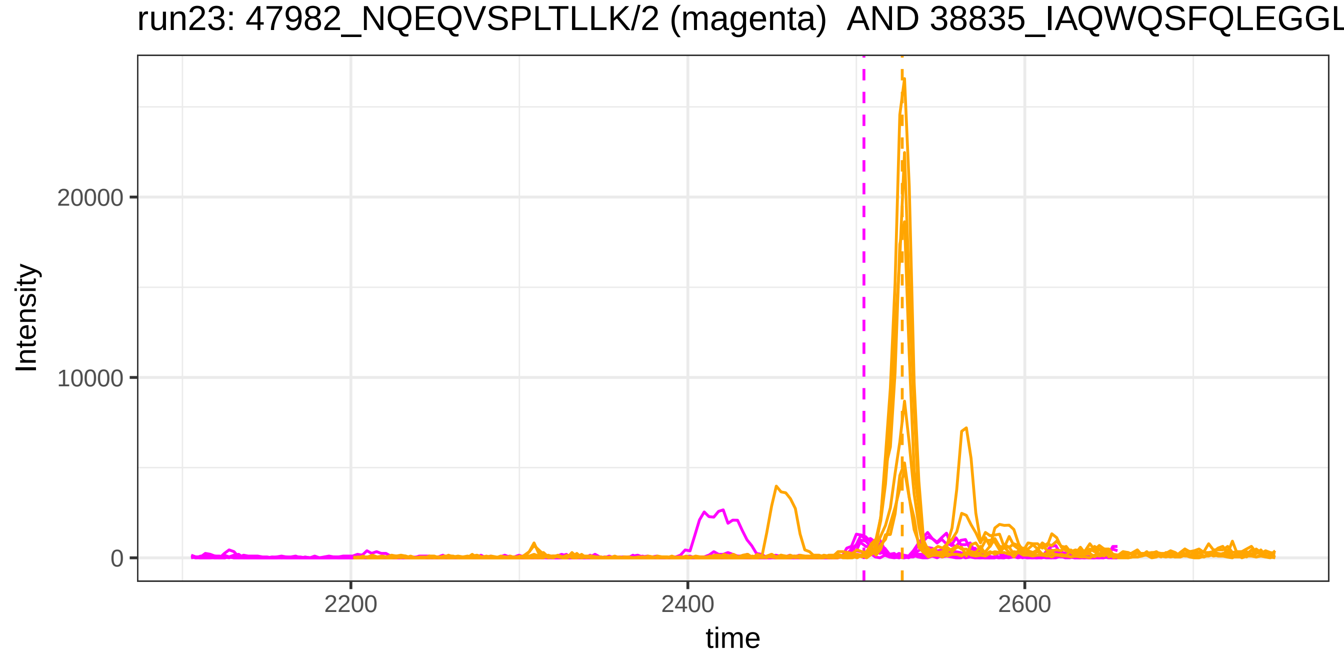
run23: 47982_NQEQVSPLTLLK/2 (magenta) AND 38835_IAQWQSFQLEGGL
Intensity
20000
10000
0
2200
2400
2600
time

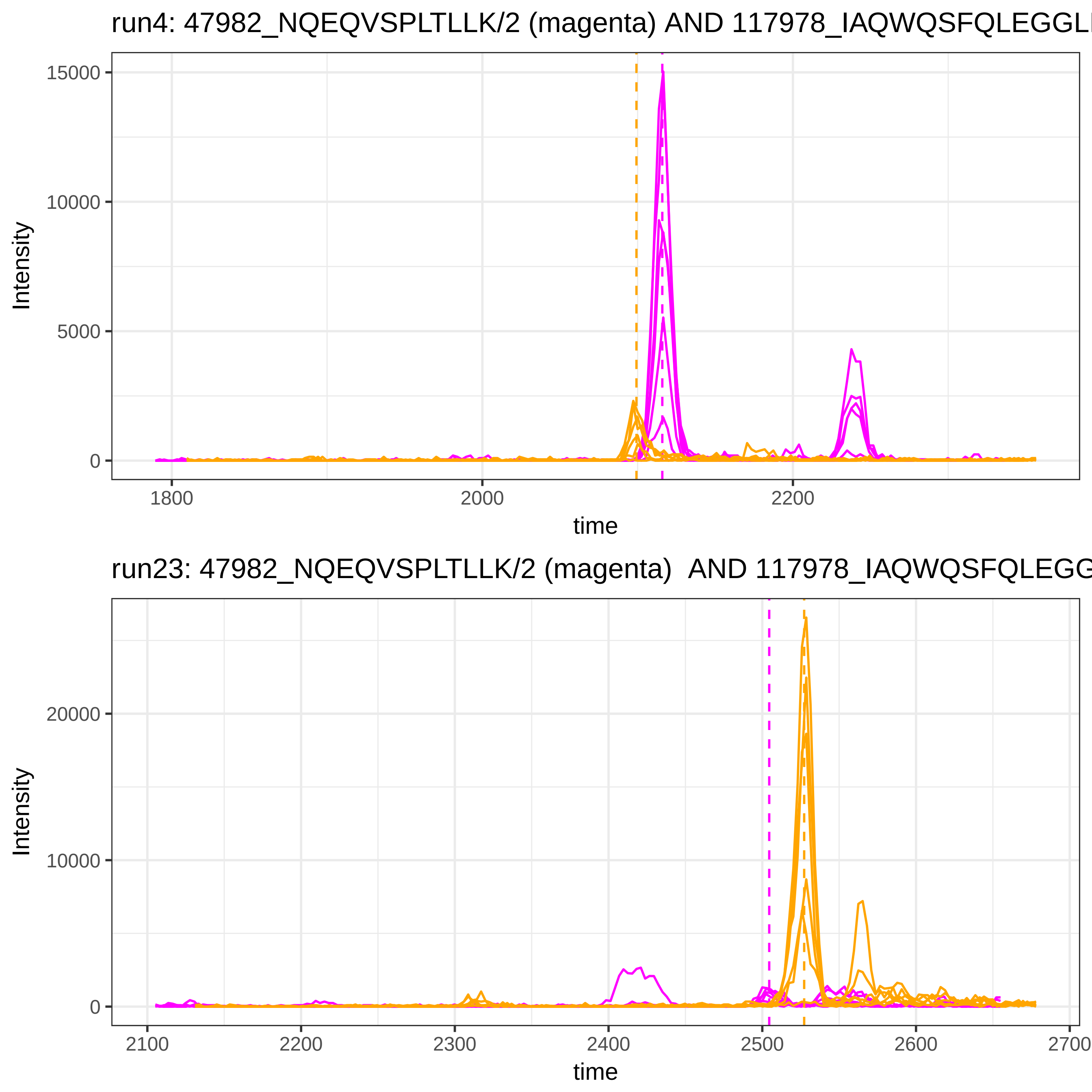

run4: 47982_NQEQVSPLTLLK/2 (magenta) AND 117978_IAQWQSFQLEGGLI
15000
10000
5000
0
Intensity
1800
2000
2200
time
run23: 47982_NQEQVSPLTLLK/2 (magenta)  AND 117978_IAQWQSFQLEGG
20000
10000
0
Intensity
2100
2200
2300
2400
2500
2600
2700
time

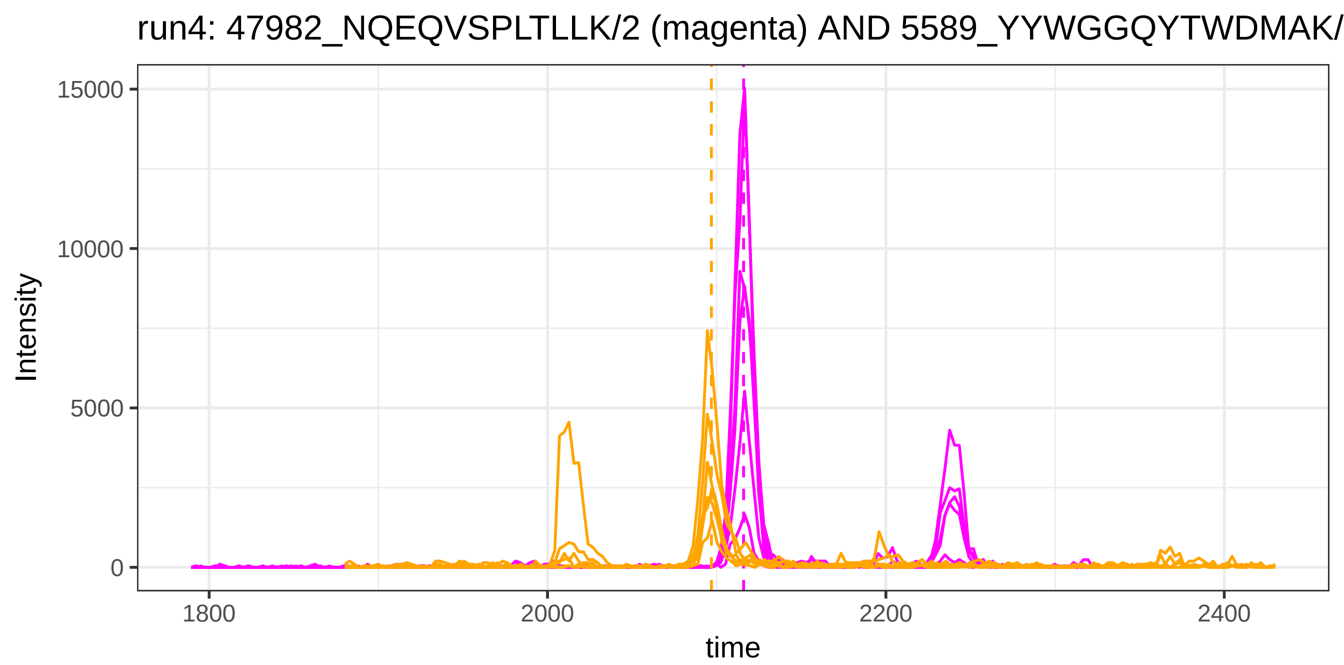
run4: 47982_NQEQVSPLTLLK/2 (magenta) AND 5589_YYWGGQYTWDMAK/
15000
10000
5000
0
Intensity
1800
2000
2200
2400
time

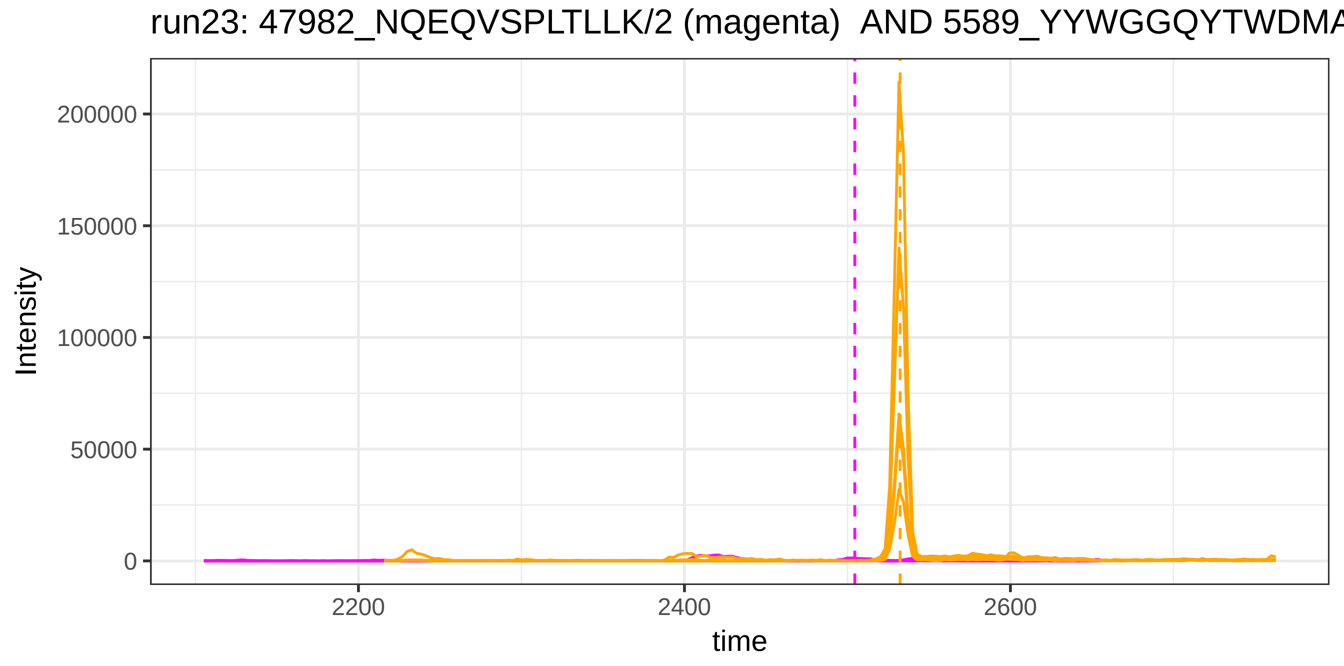
run23: 47982_NQEQVSPLTLLK/2 (magenta) AND 5589_YYWGGQYTWDMA
200000
150000
100000
50000
0
Intensity
2200
2400
2600
time

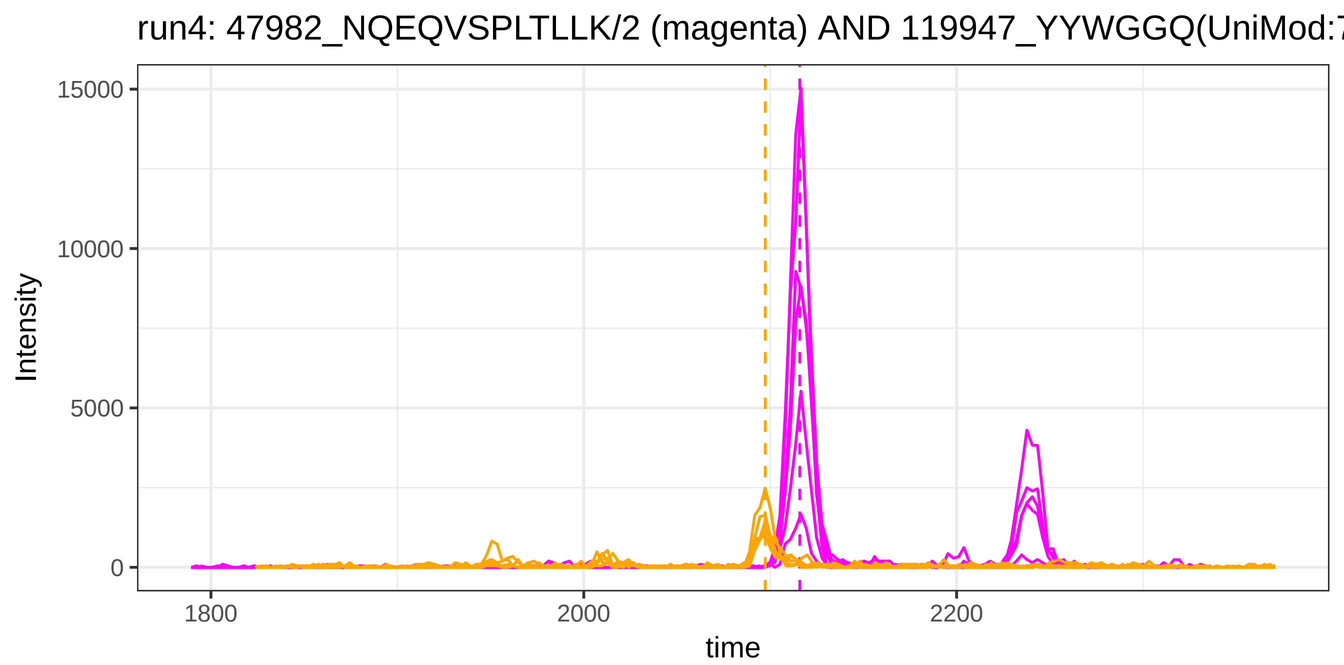
run4: 47982_NQEQVSPLTLLK/2 (magenta) AND 119947_YYWGGQ(UniMod:7
Intensity
15000
10000
5000
0
1800
2000
2200
time

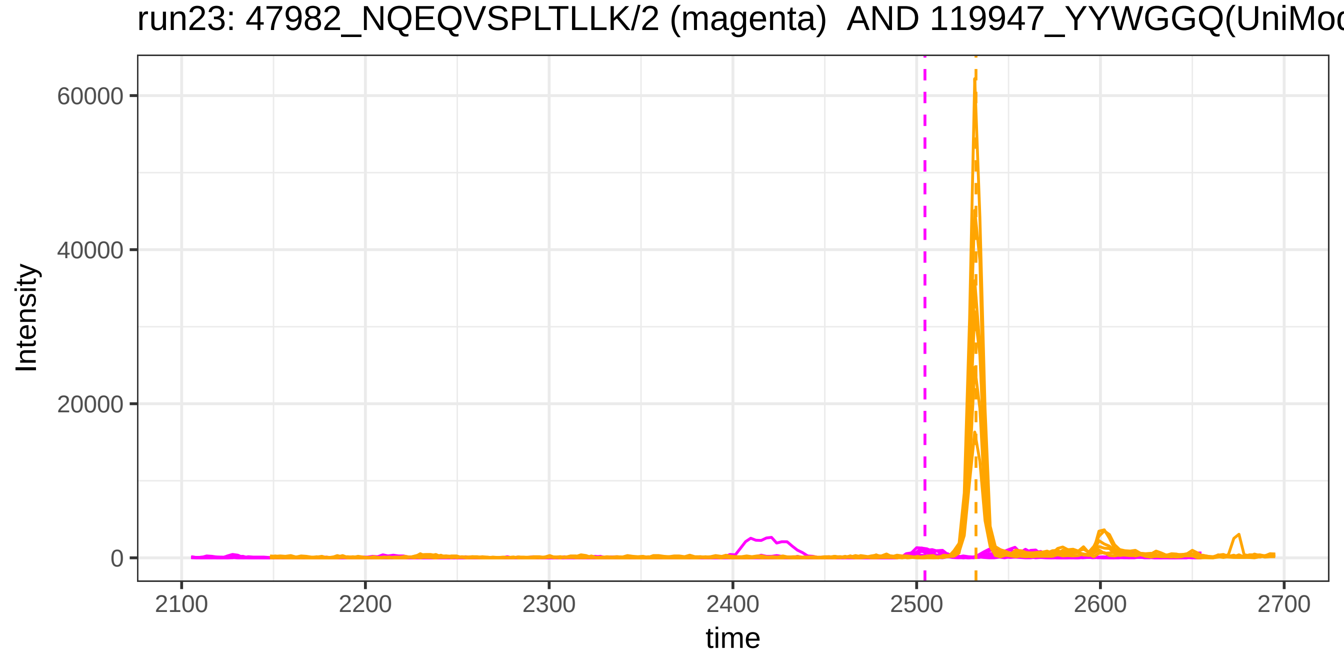
run23: 47982_NQEQVSPLTLLK/2 (magenta)  AND 119947_YYWGGQ(UniMod
Intensity
60000
40000
20000
0
2100
2200
2300
2400
2500
2600
2700
time

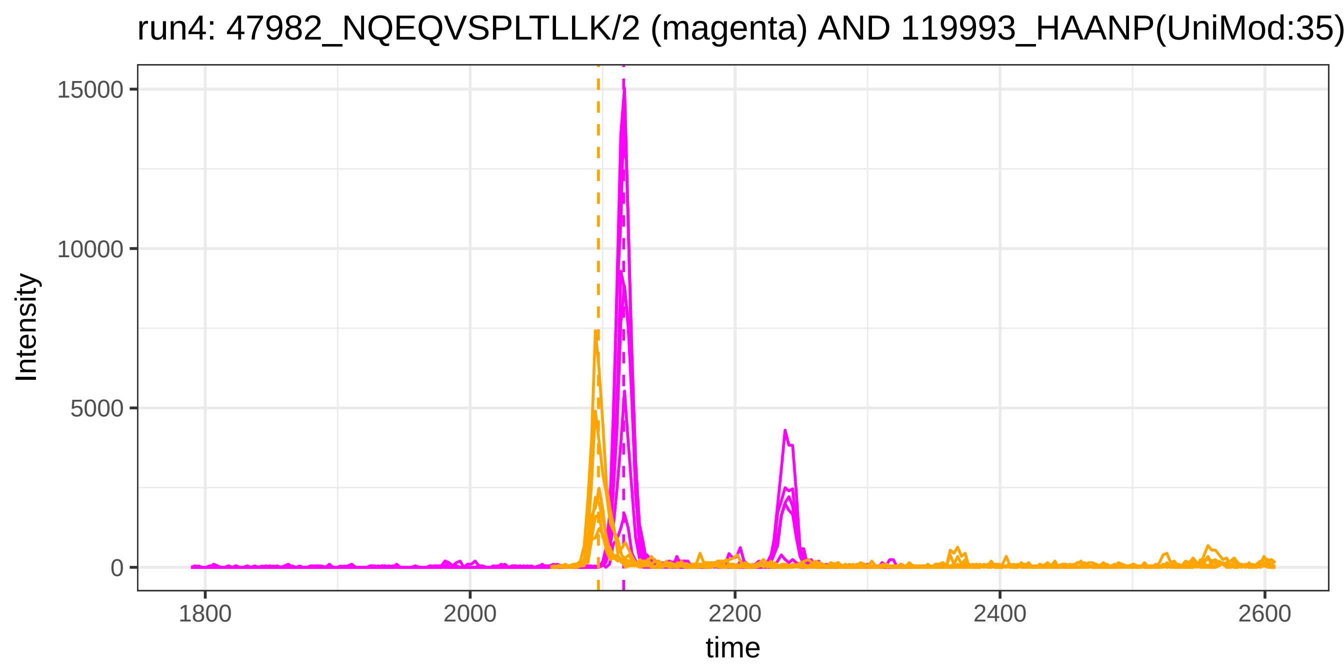
run4: 47982_NQEQVSPLTLLK/2 (magenta) AND 119993_HAANP(UniMod:35)
Intensity
15000
10000
5000
0
1800
2000
2200
2400
2600
time

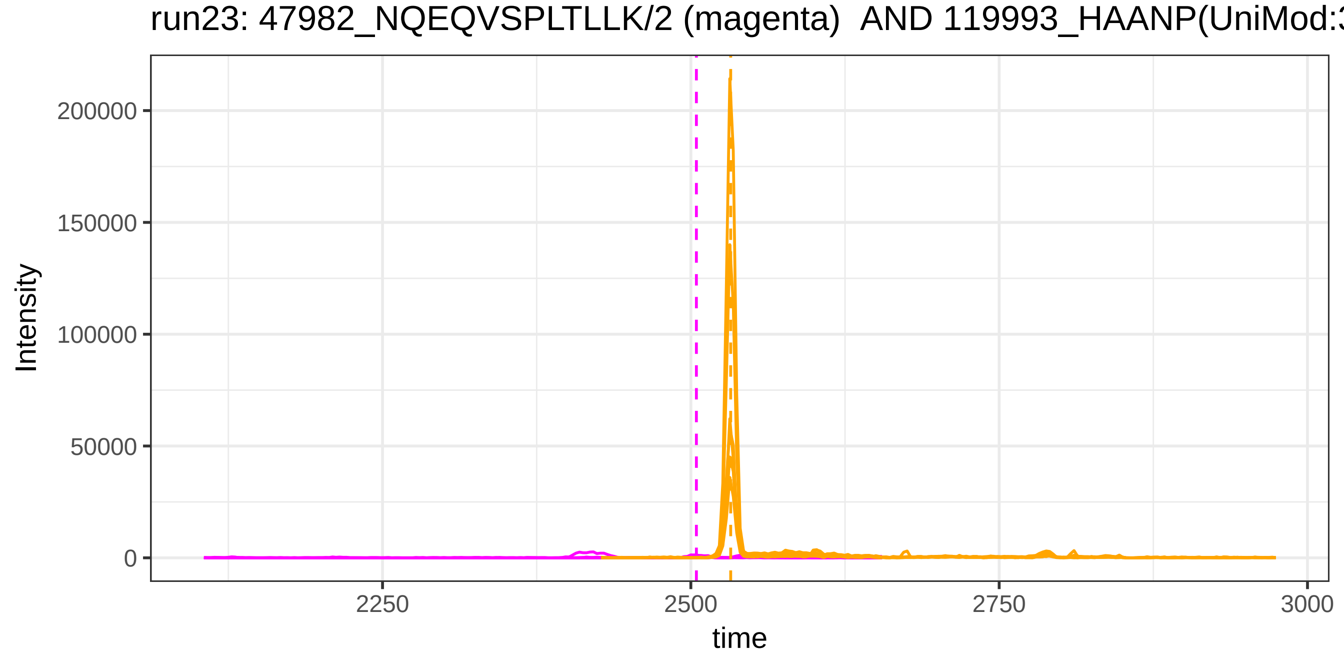
run23: 47982_NQEQVSPLTLLK/2 (magenta) AND 119993_HAANP(UniMod:3
Intensity
200000
150000
100000
50000
0
2250
2500
2750
3000
time

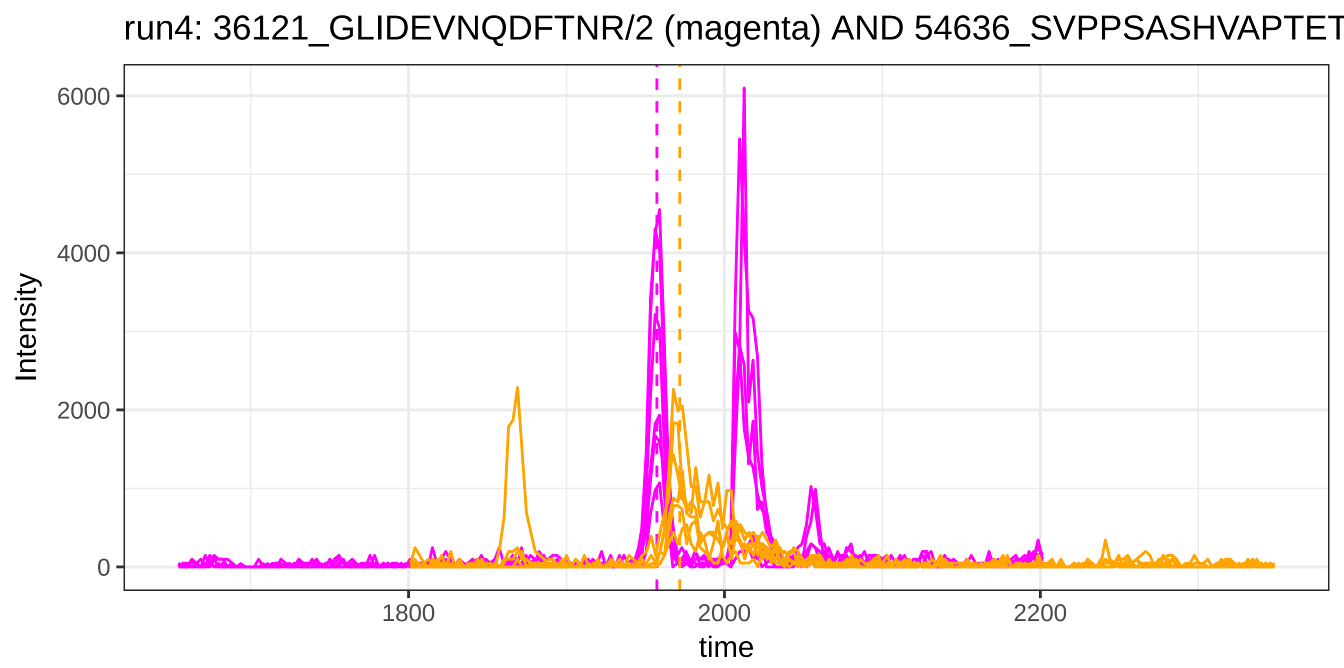
run4: 36121_GLIDEVNQDFTNR/2 (magenta) AND 54636_SVPPSASHVAPTET
Intensity
6000
4000
2000
0
1800
2000
2200
time

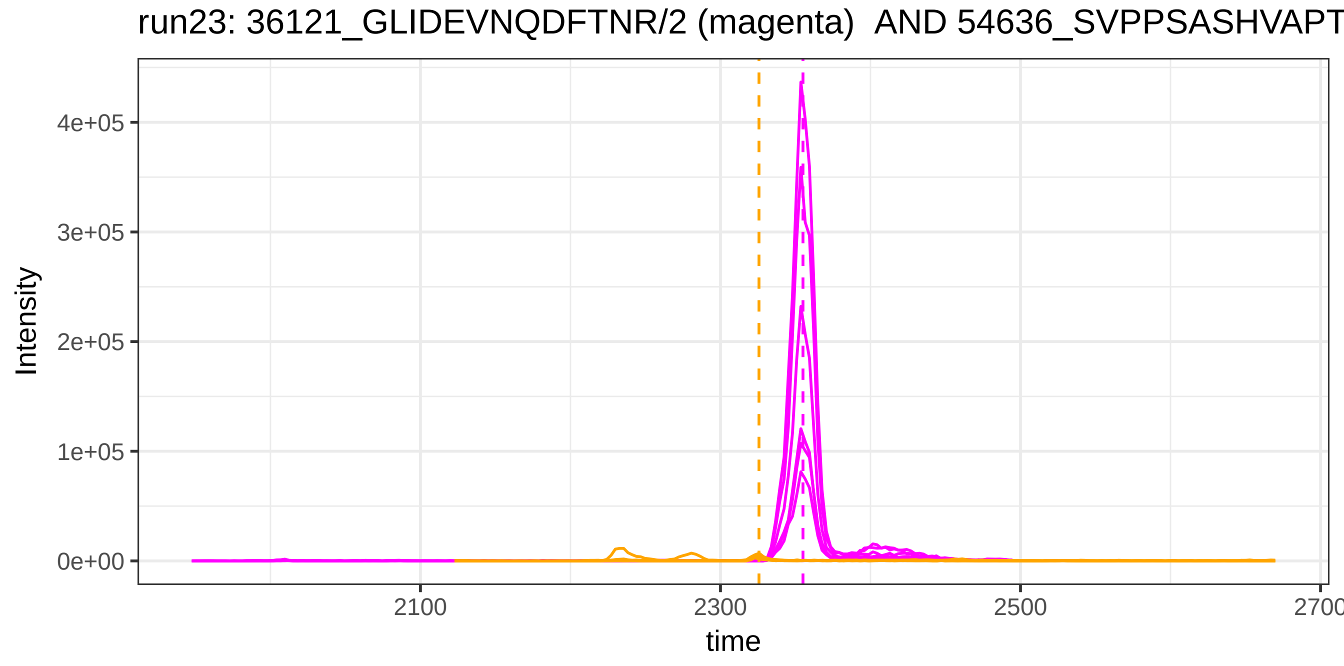
run23: 36121_GLIDEVNQDFTNR/2 (magenta) AND 54636_SVPPSASHVAPT
Intensity
4e+05
3e+05
2e+05
1e+05
0e+00
2100
2300
2500
2700
time
