## Supplemental Figures for "DIAlignR provides precise retention time alignment across distant runs in DIA and targeted proteomics": Supplemental Figure17.pdf

In the figures, blue dashed line indicates RT estimation by LOESS whereas red dashed line indicates RT estimation by chromatogram alignment. Dotted line indicates OpenSWATH annotation.

run4_run23, 128284_AFLLTPR/2

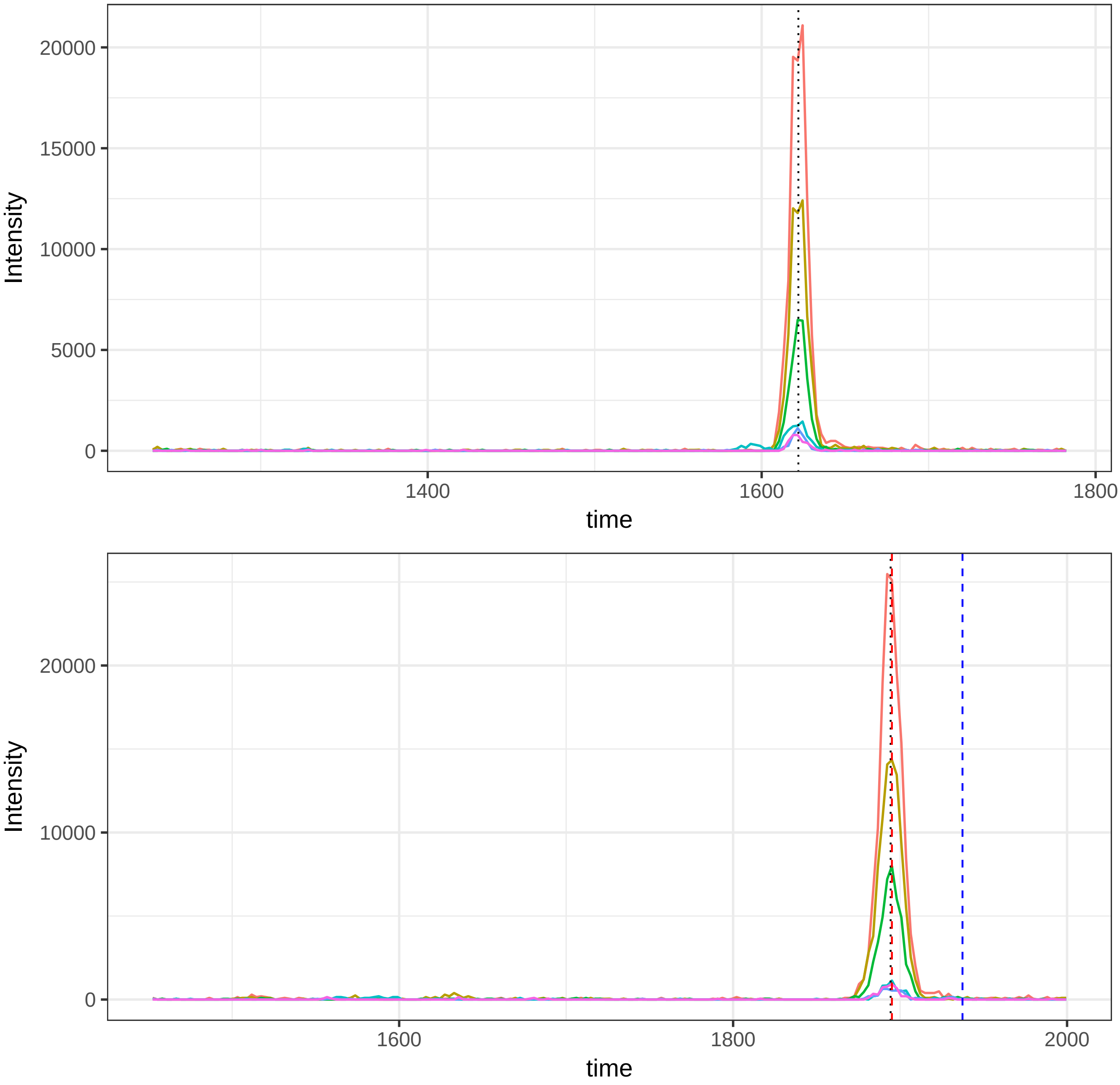

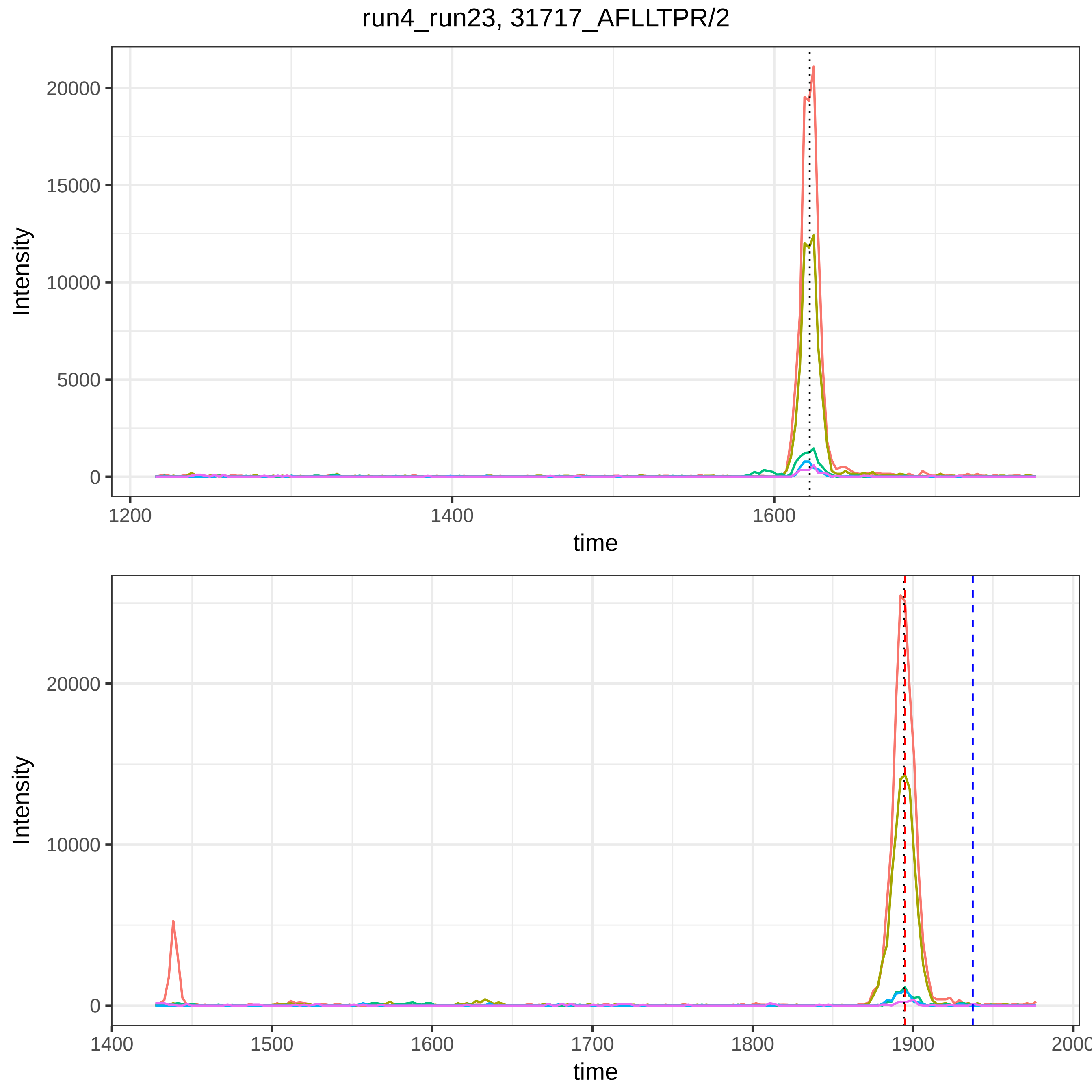

run4_run23, 31717_AFLLTPR/2
Intensity
20000
15000
10000
5000
0
1200
1400
1600
time
Intensity
20000
10000
0
1400
1500
1600
1700
1800
1900
2000
time

run4_run23, 116214_SLHTLFGDKLC(UniMod:4)TVATLR/4

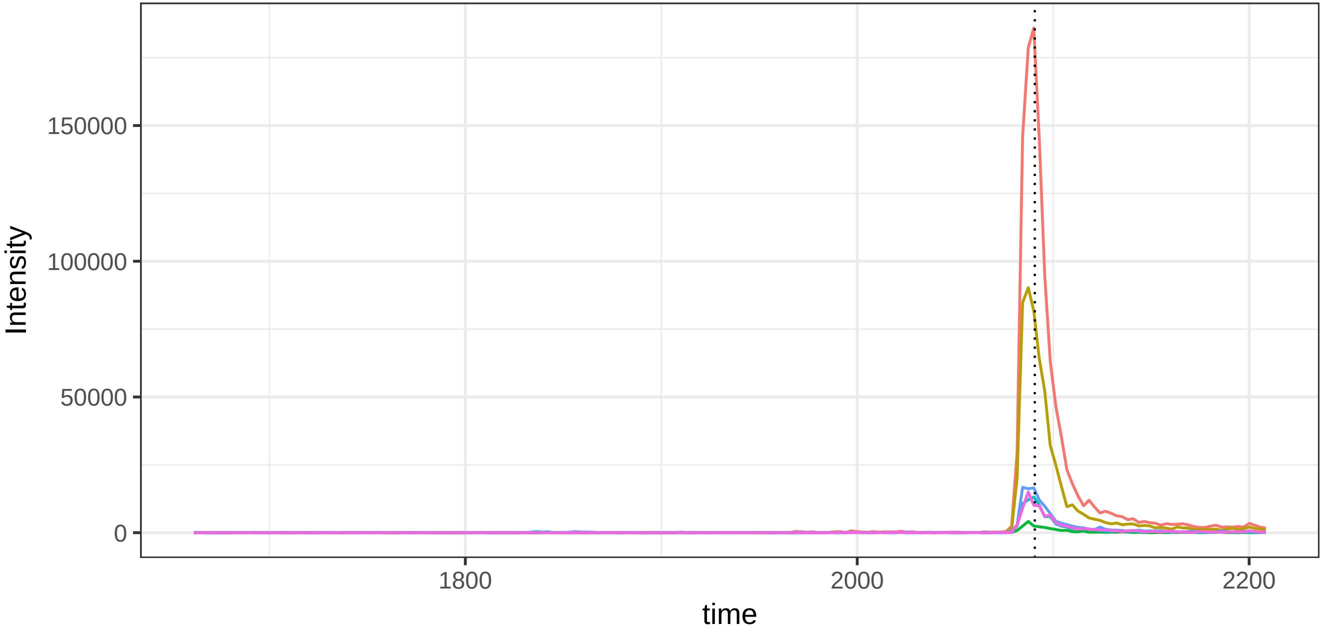


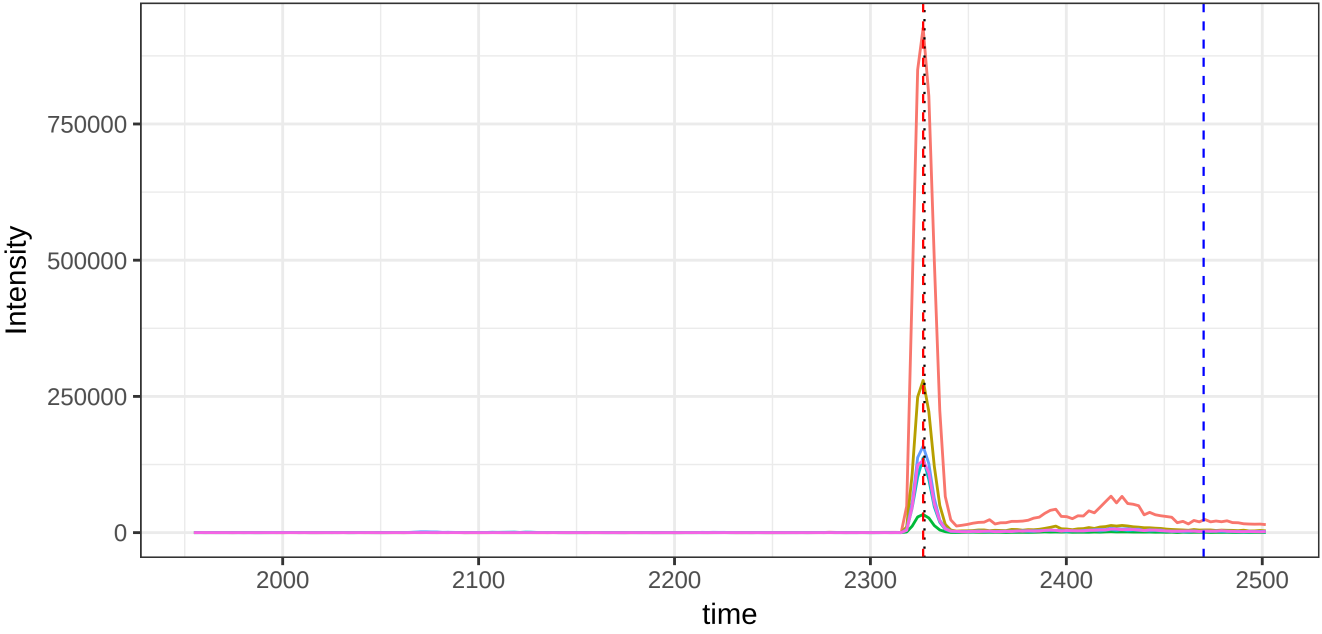

run4_run23, 36678_GPSVFPLAPSSK/2

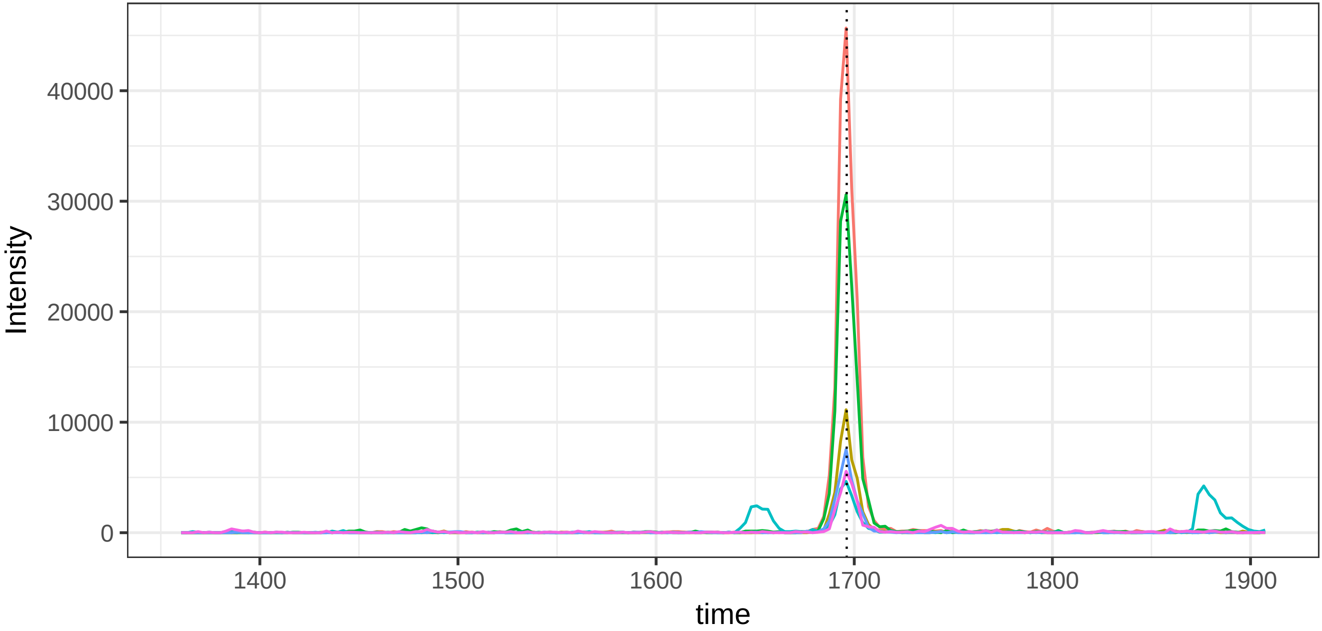


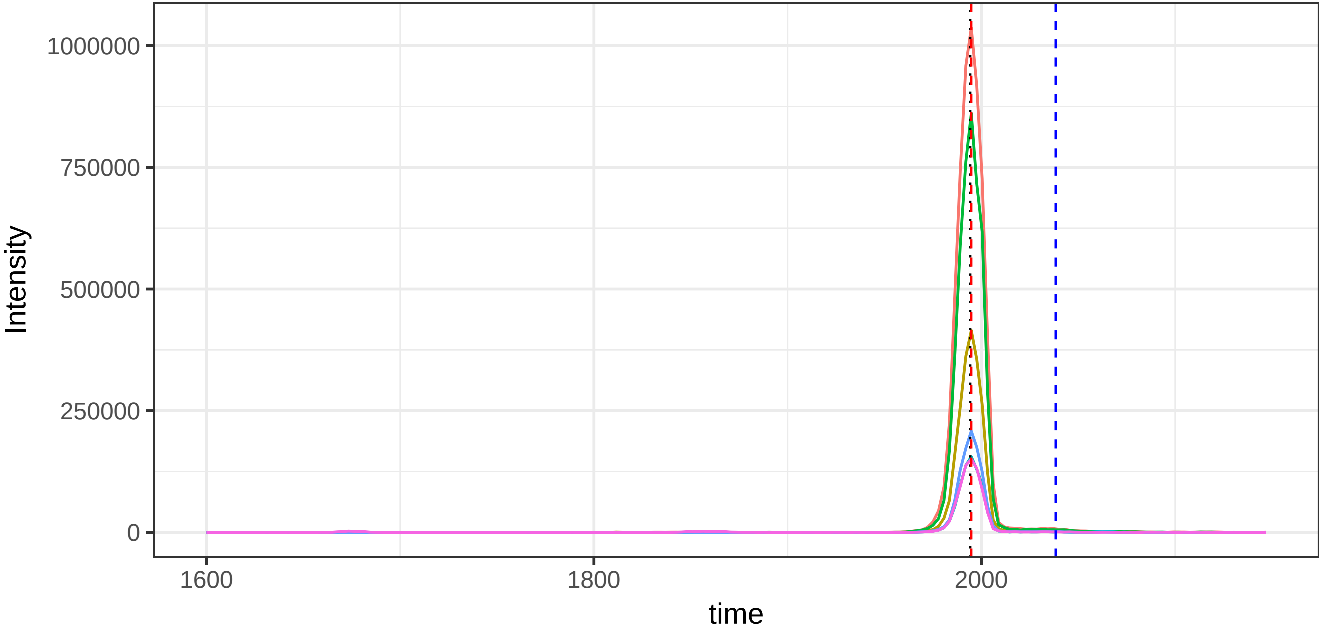

run4_run23, 38644_HYDGSYSTFGER/2

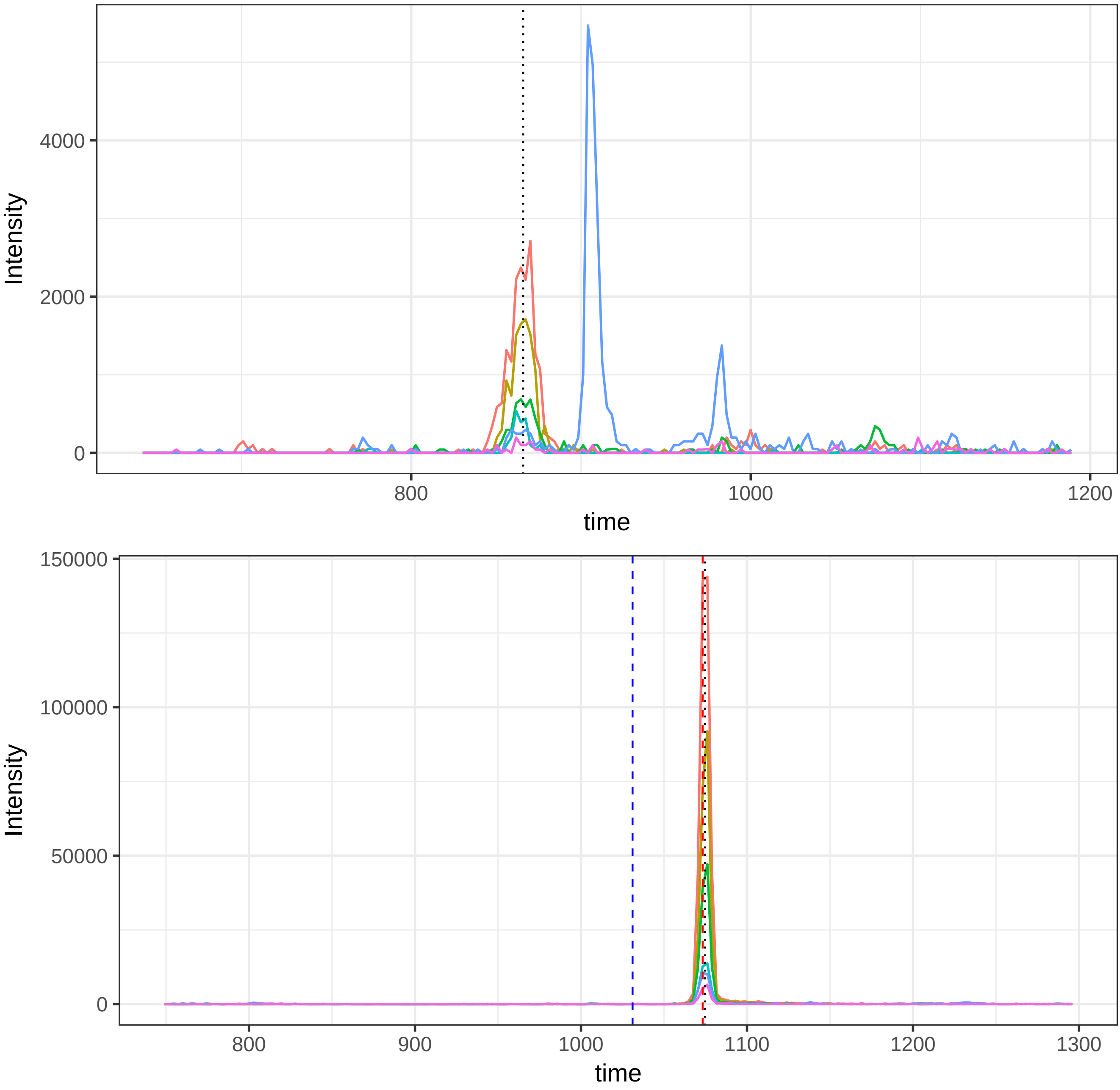

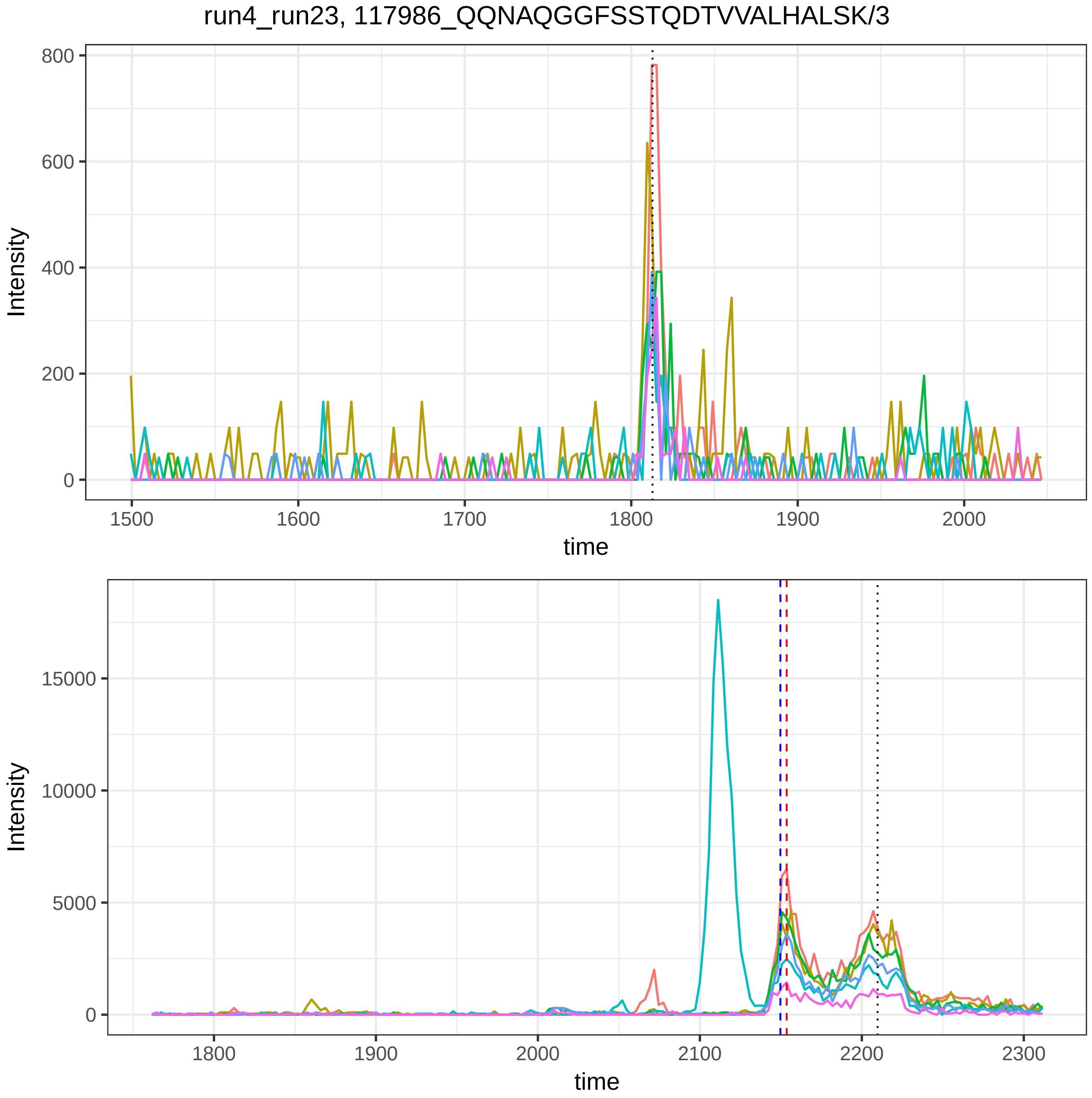
run4_run23, 117986_QQNAQGGFSSTQDTVVALHALSK/3
Intensity
time
800
600
400
200
0
1500
1600
1700
1800
1900
2000
15000
10000
5000
0
1800
1900
2000
2100
2200
2300

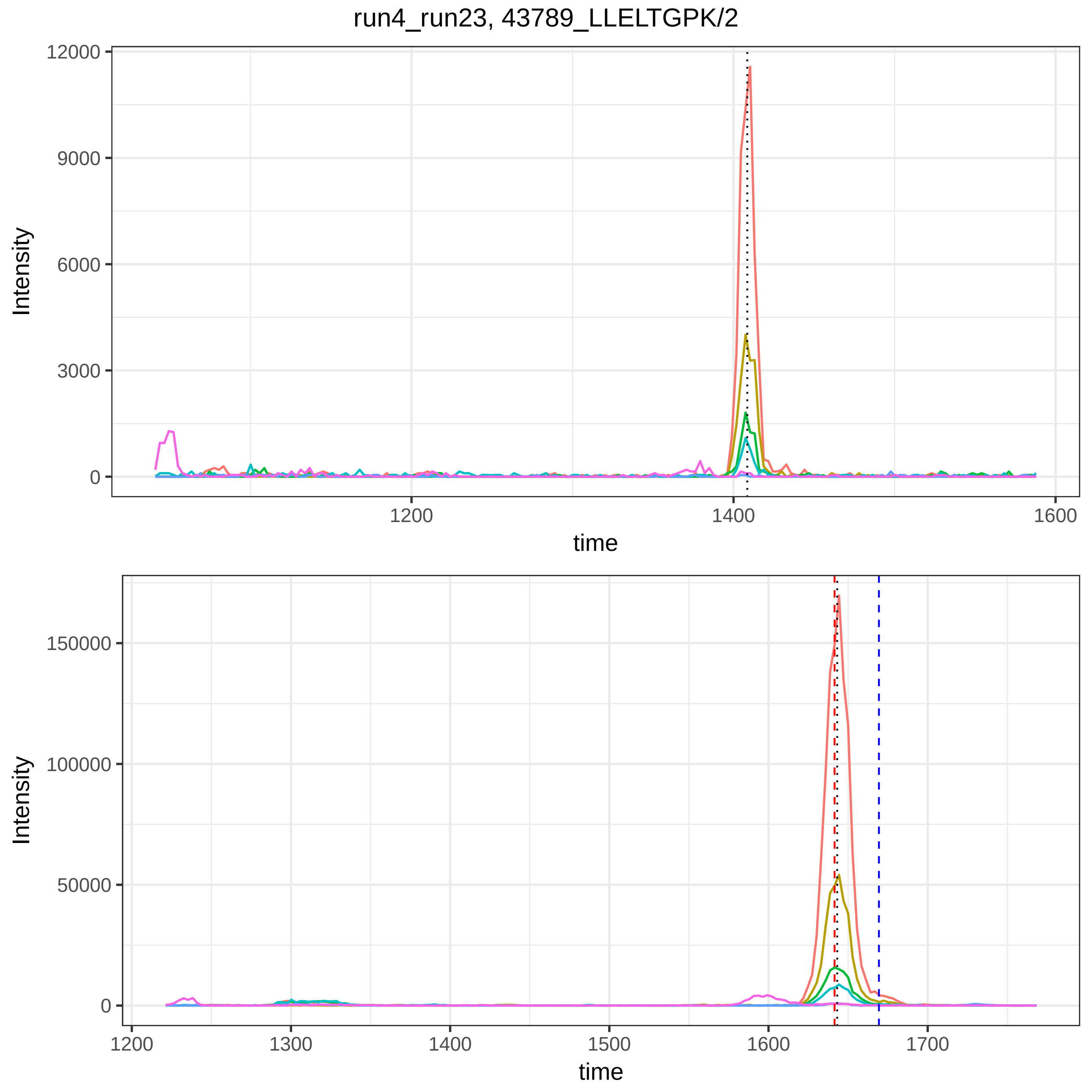

run4_run23, 43789_LLELTGPK/2
Intensity
12000
9000
6000
3000
0
1200
1400
1600
time
Intensity
150000
100000
50000
0
1200
1300
1400
1500
1600
1700
time

run4_run23, 115879_DDNPNLPR/2

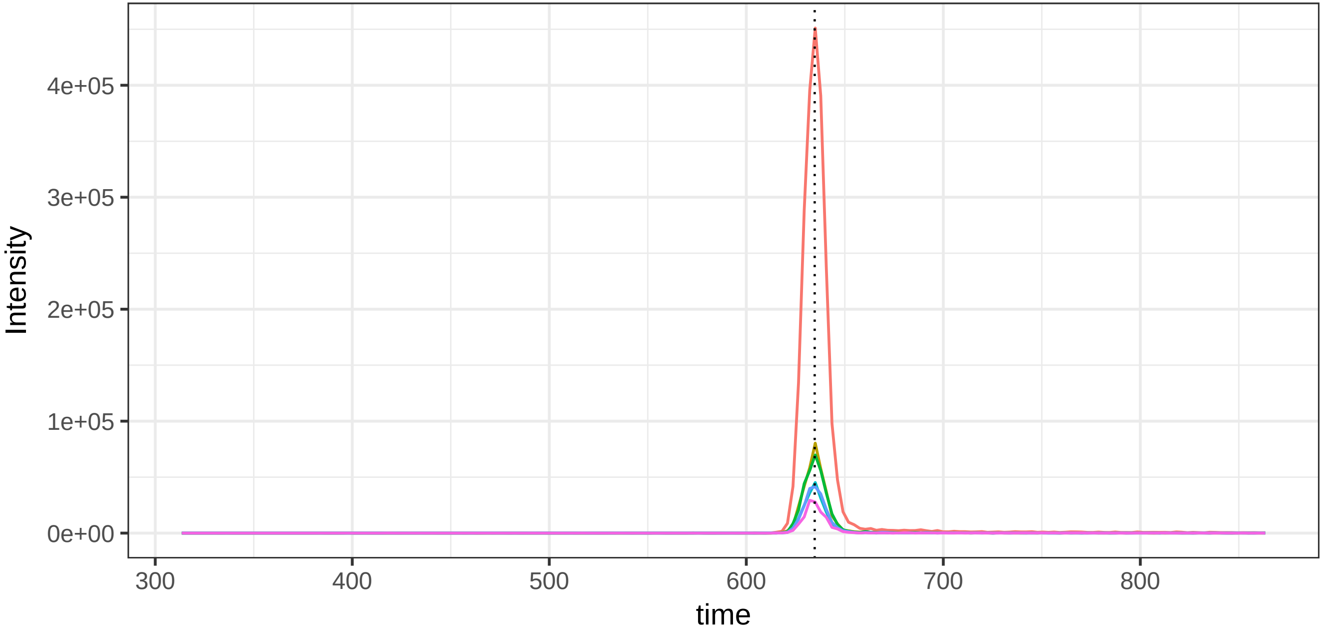


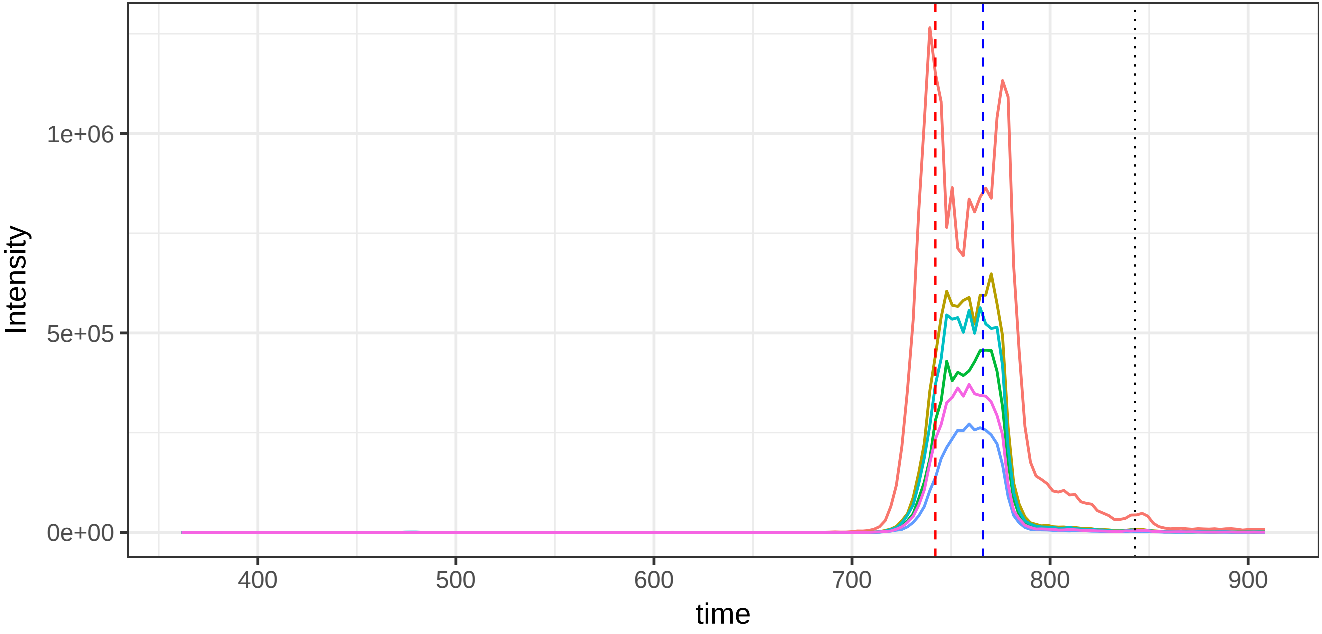

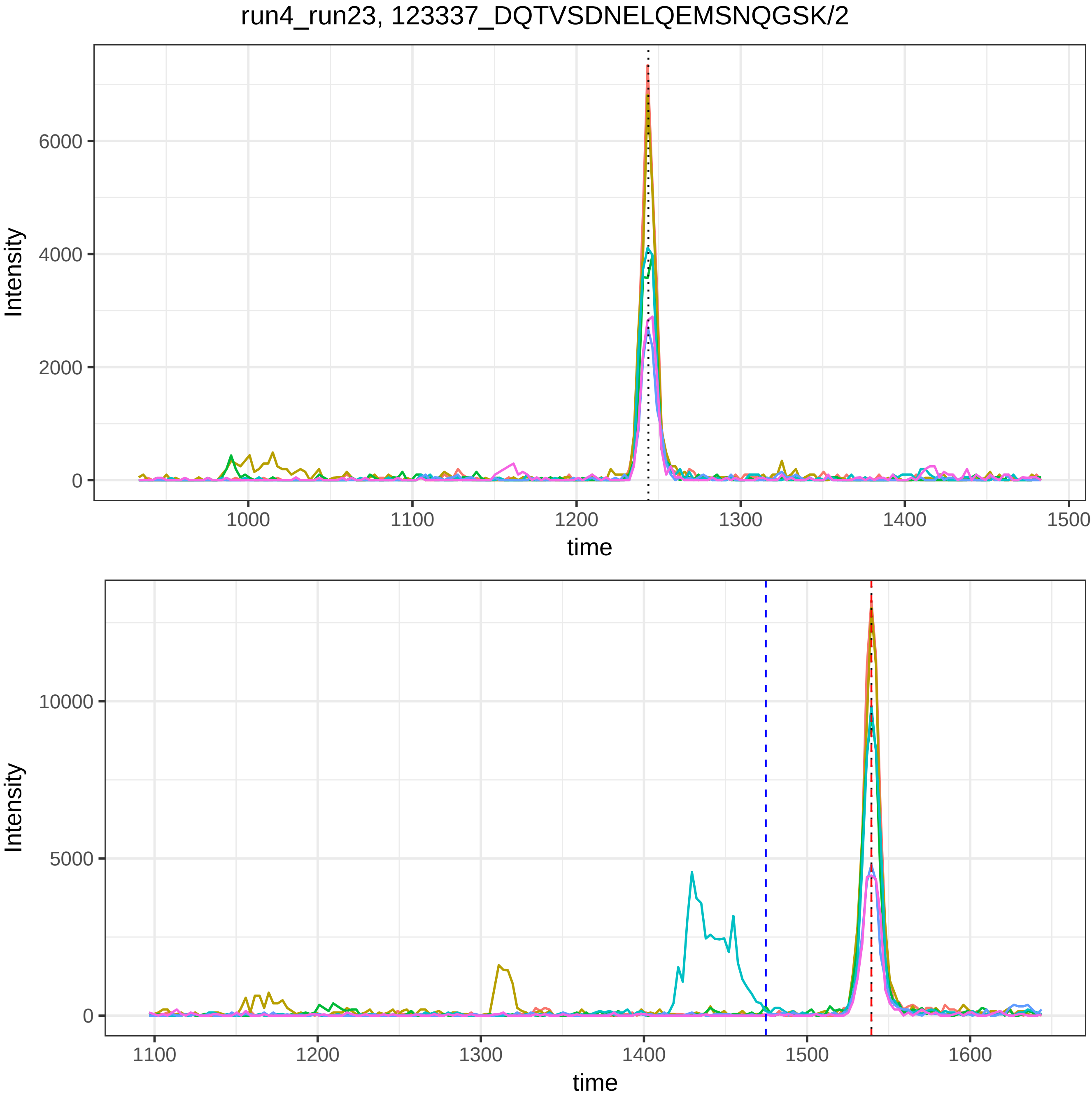
run4_run23, 123337_DQTVSDNELQEMSNQGSK/2
Intensity
6000
4000
2000
0
1000
1100
1200
1300
1400
1500
time
Intensity
10000
5000
0
1100
1200
1300
1400
1500
1600
time

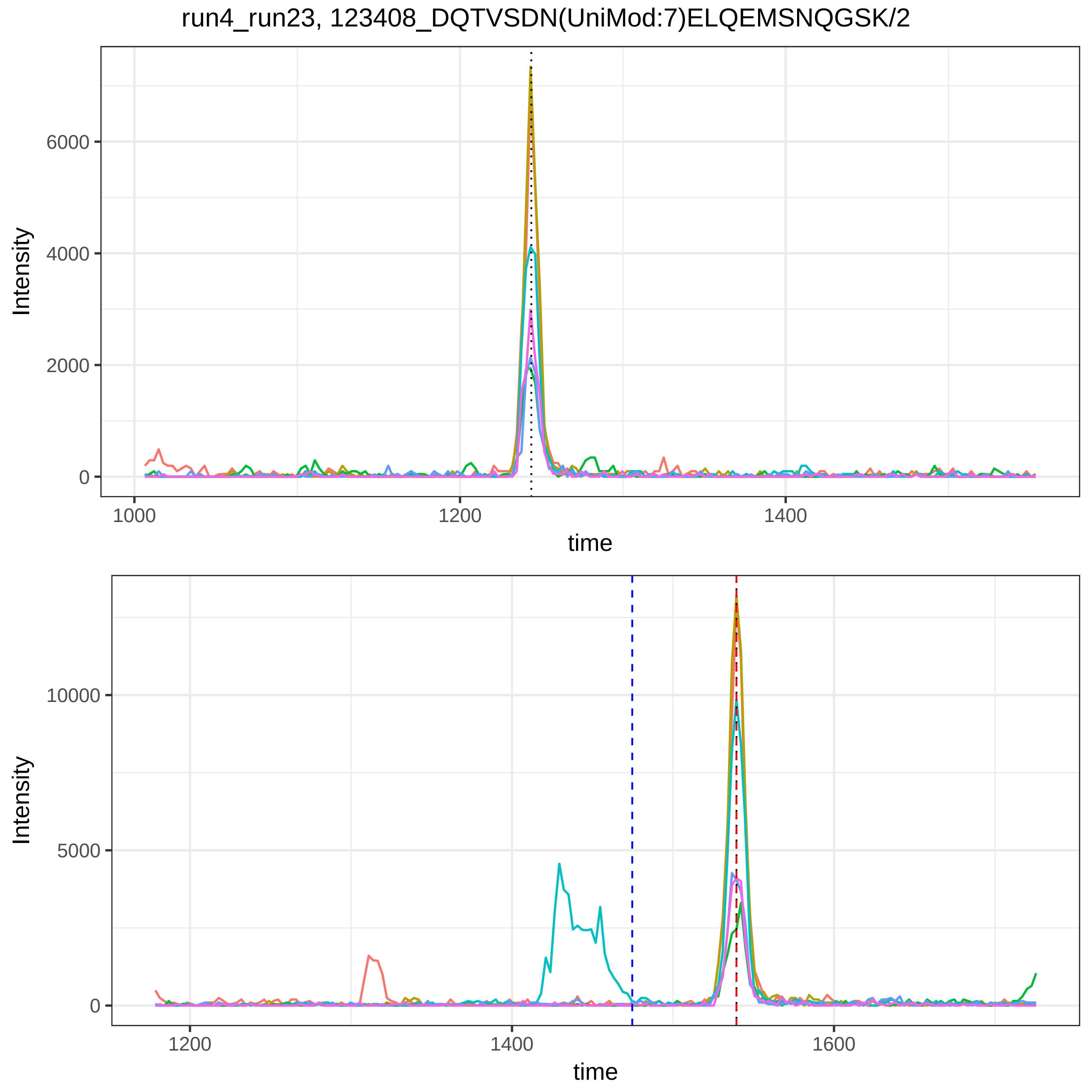

run4_run23, 123408_DQTVSDN(UniMod:7)ELQEMSNQGSK/2
Intensity
6000
4000
2000
0
1000
1200
1400
time
Intensity
10000
5000
0
1200
1400
1600
time

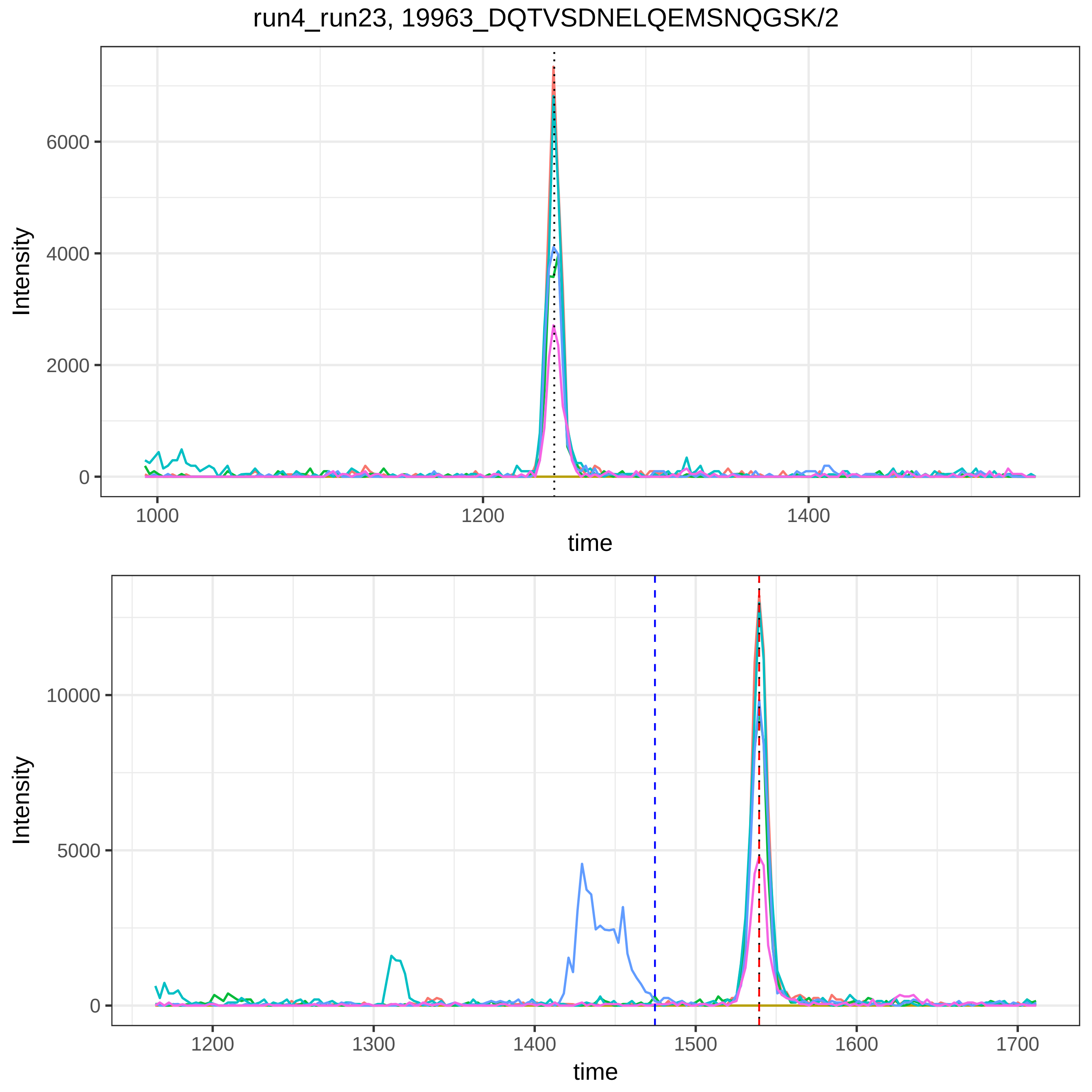

run4_run23, 19963_DQTVSDNELQEMSNQGSK/2
Intensity
time
0
2000
4000
6000
1000
1200
1400
0
5000
10000
1200
1300
1400
1500
1600
1700

run4_run23, 39892_IRPFFPQQ/2

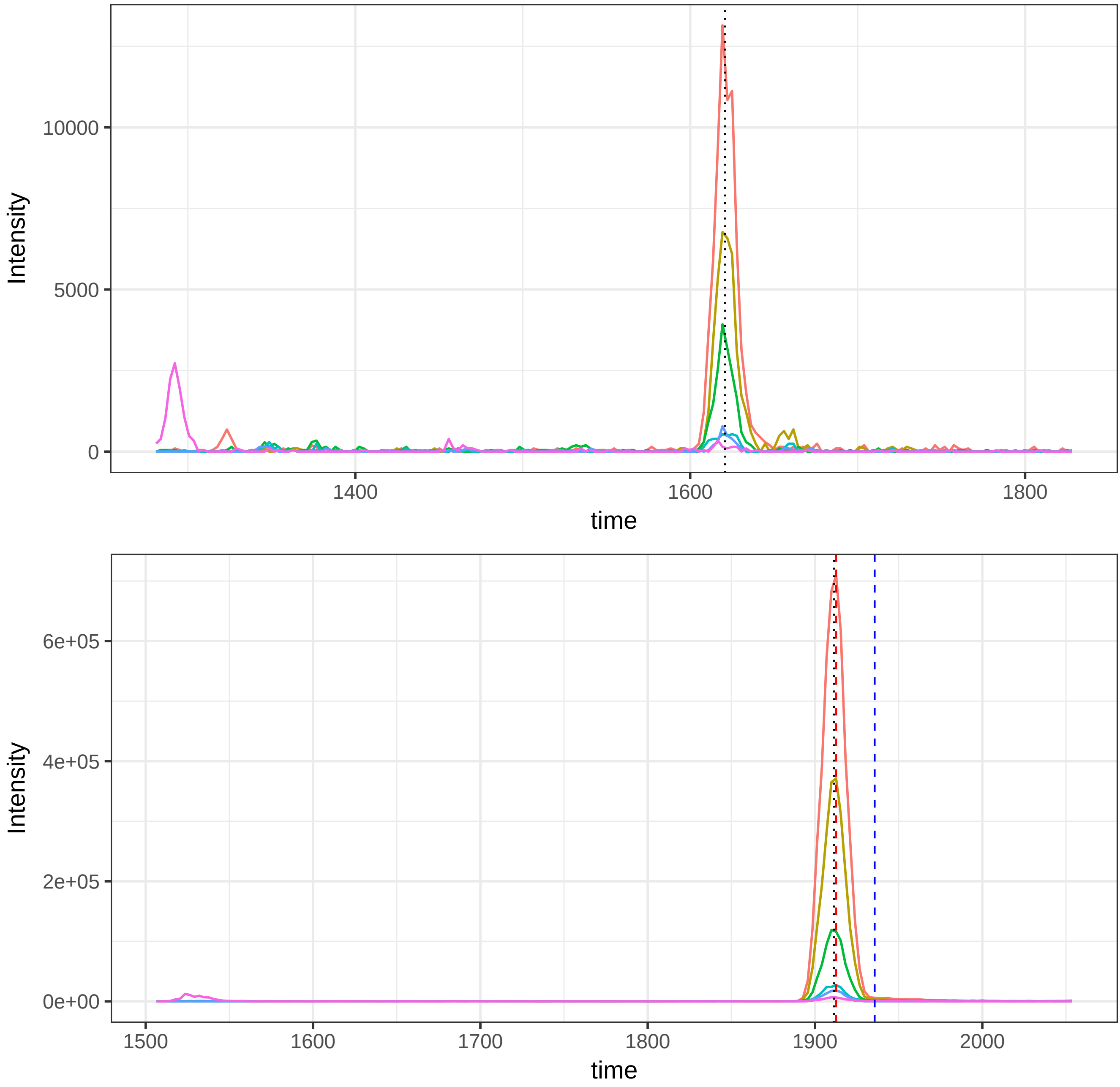

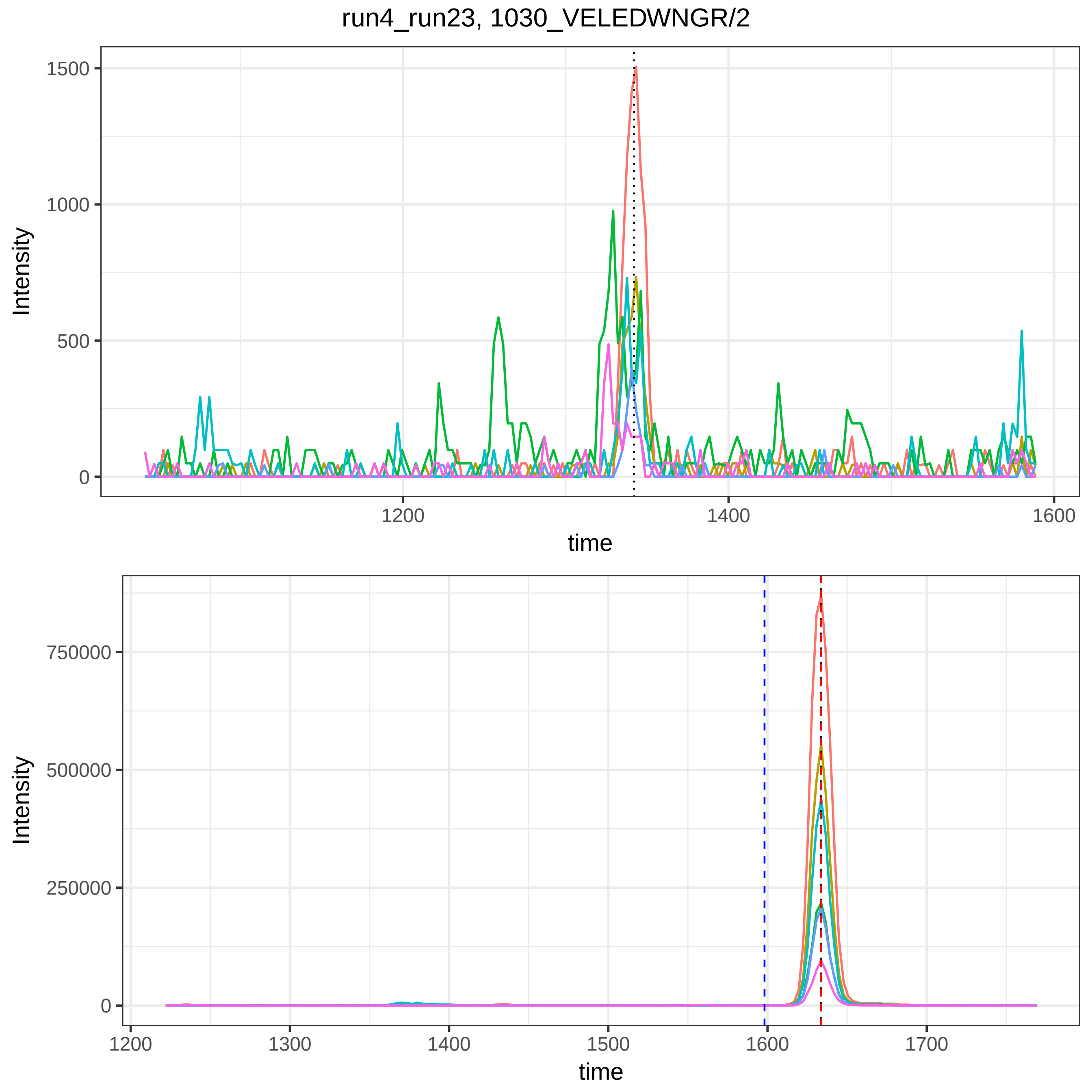
run4_run23, 1030_VELEDWNGR/2
Intensity
1500
1000
500
0
1200
1400
1600
time
Intensity
750000
500000
250000
0
1200
1300
1400
1500
1600
1700
time

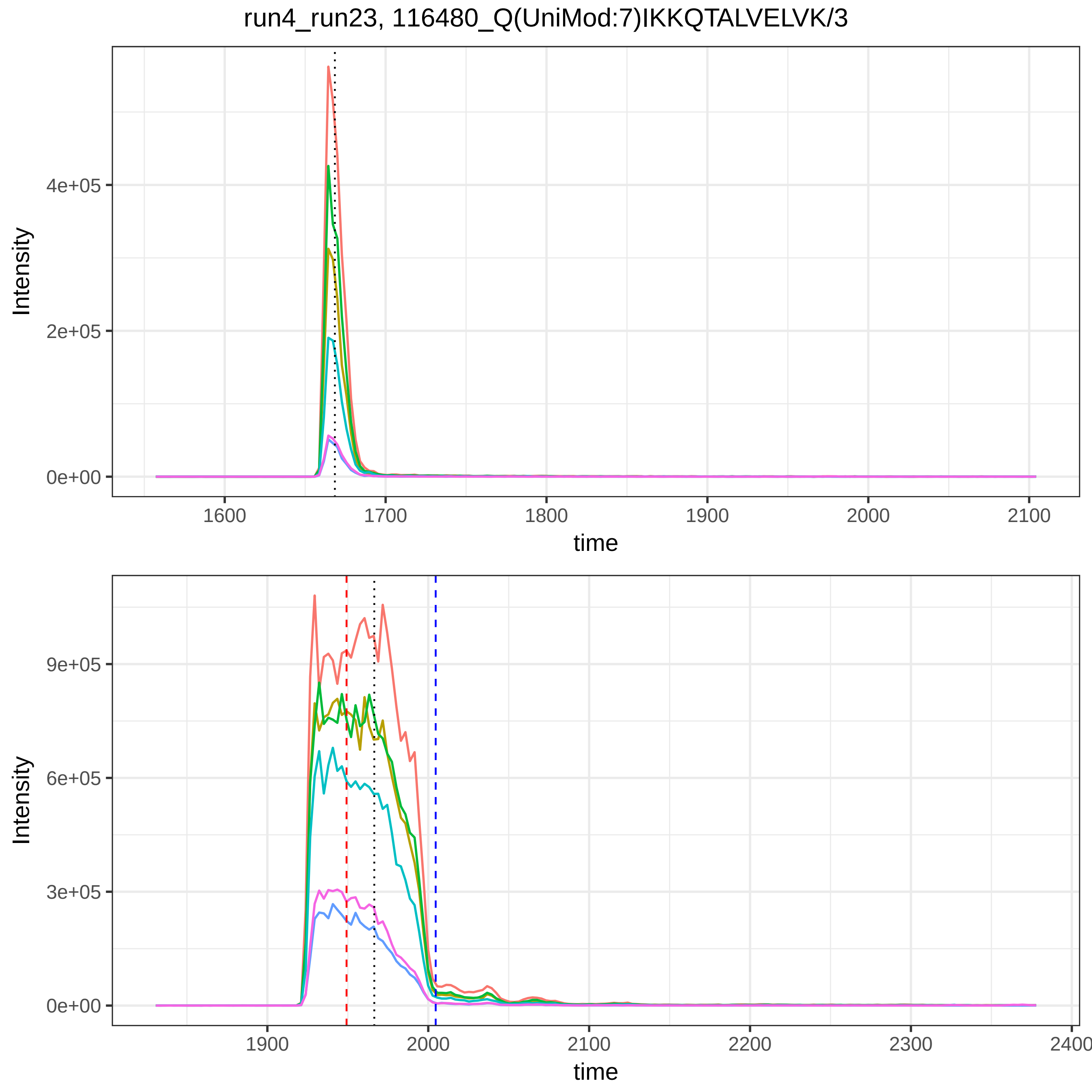

run4_run23, 116480_Q(UniMod:7)IKKQTALVELVK/3
Intensity
4e+05
2e+05
0e+00
1600
1700
1800
1900
2000
2100
time
Intensity
9e+05
6e+05
3e+05
0e+00
1900
2000
2100
2200
2300
2400
time

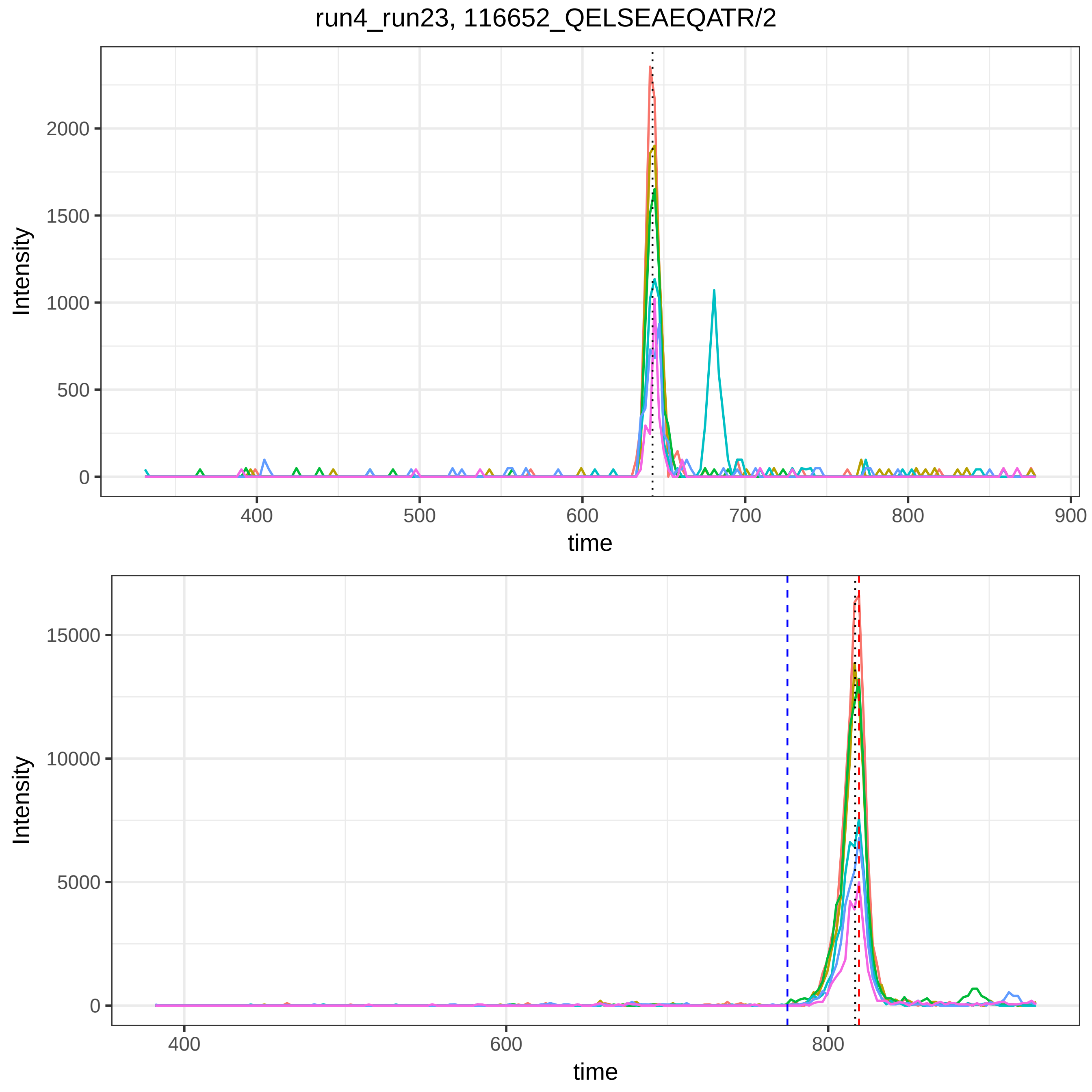

run4_run23, 116652_QELSEAEQATR/2
Intensity
time
Intensity
time

run4_run23, 116798_QGALELIK/2

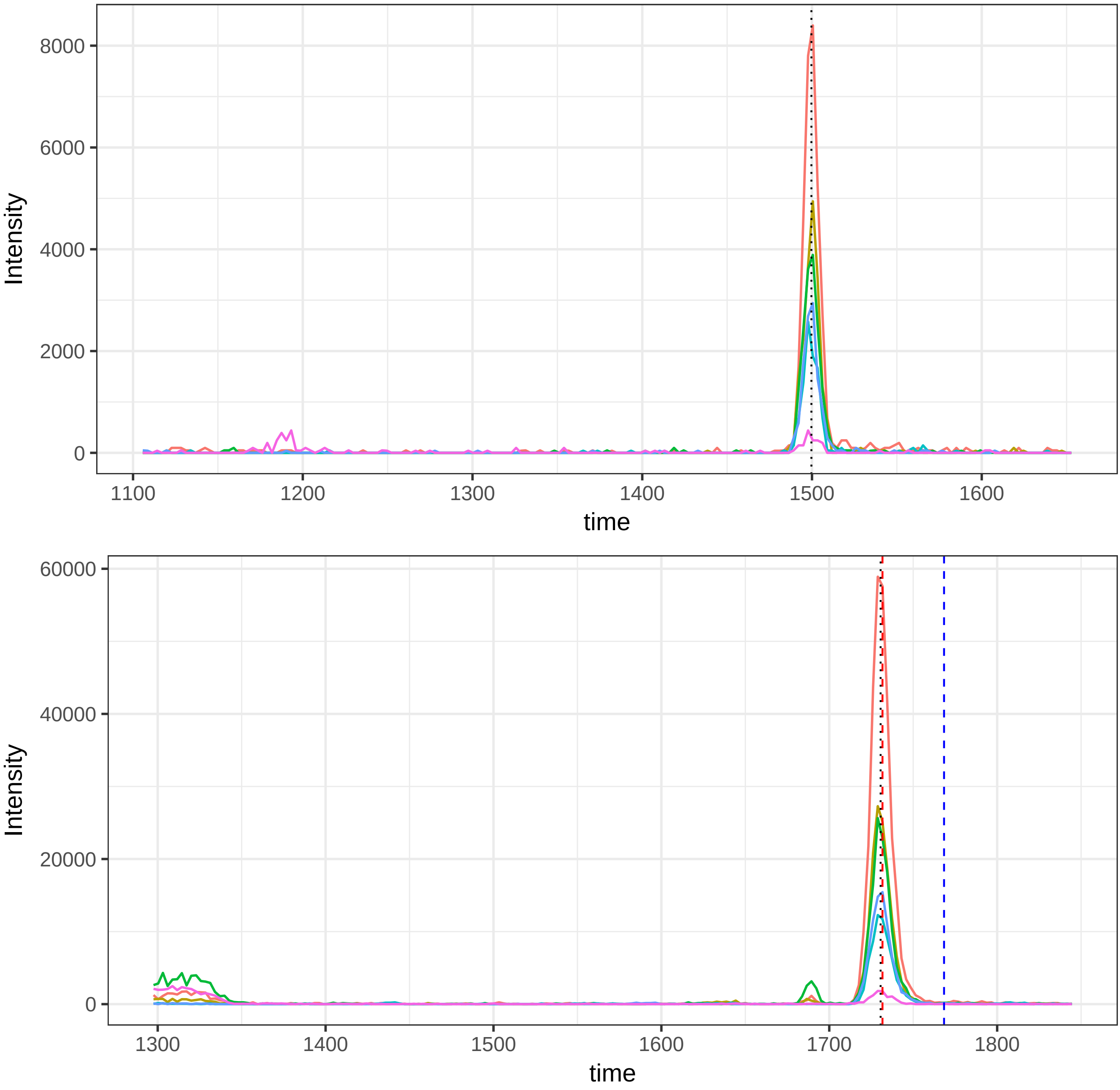

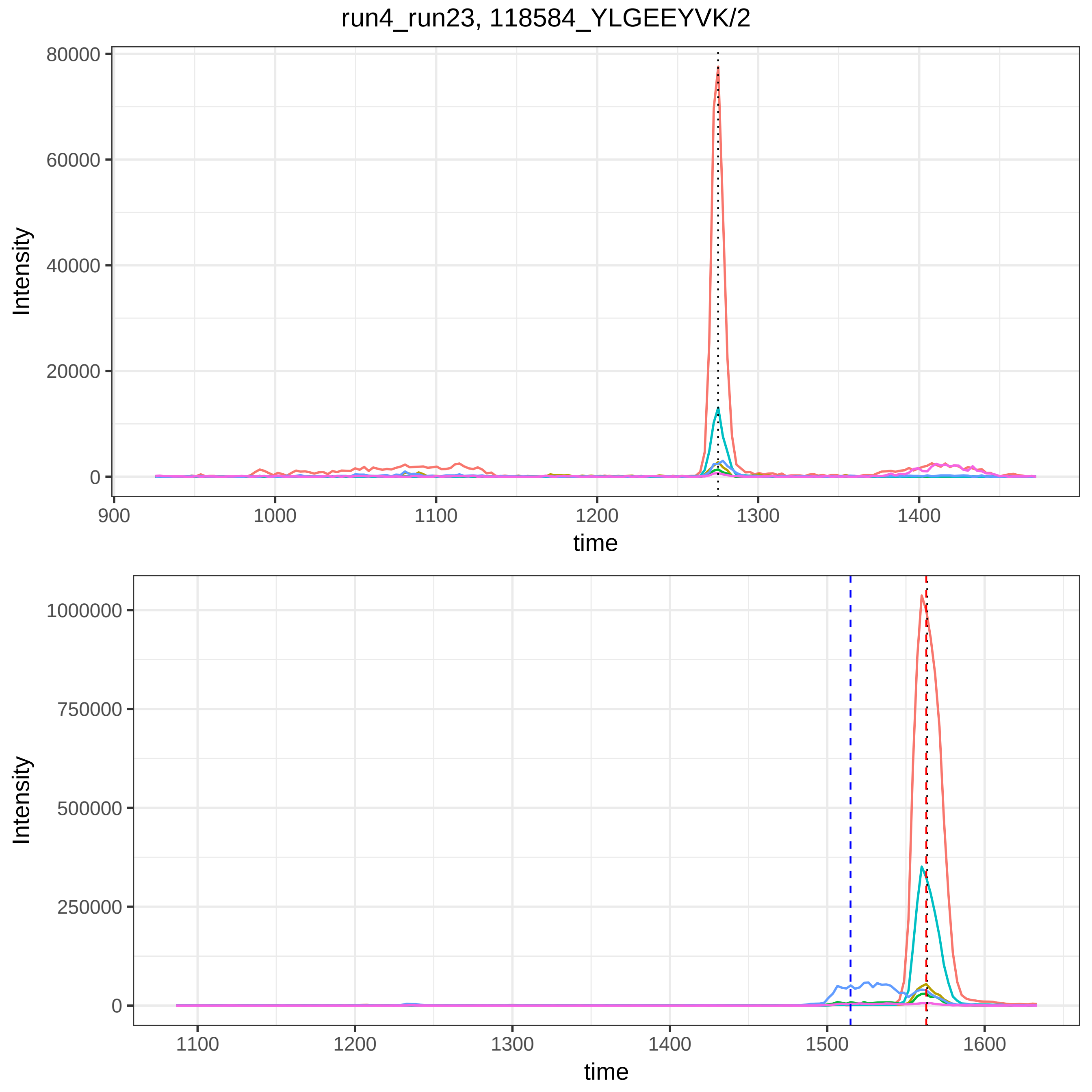

run4_run23, 118584_YLGEEYVK/2
80000
60000
40000
20000
0
Intensity
900
1000
1100
1200
1300
1400
time
1000000
750000
500000
250000
0
Intensity
1100
1200
1300
1400
1500
1600
time

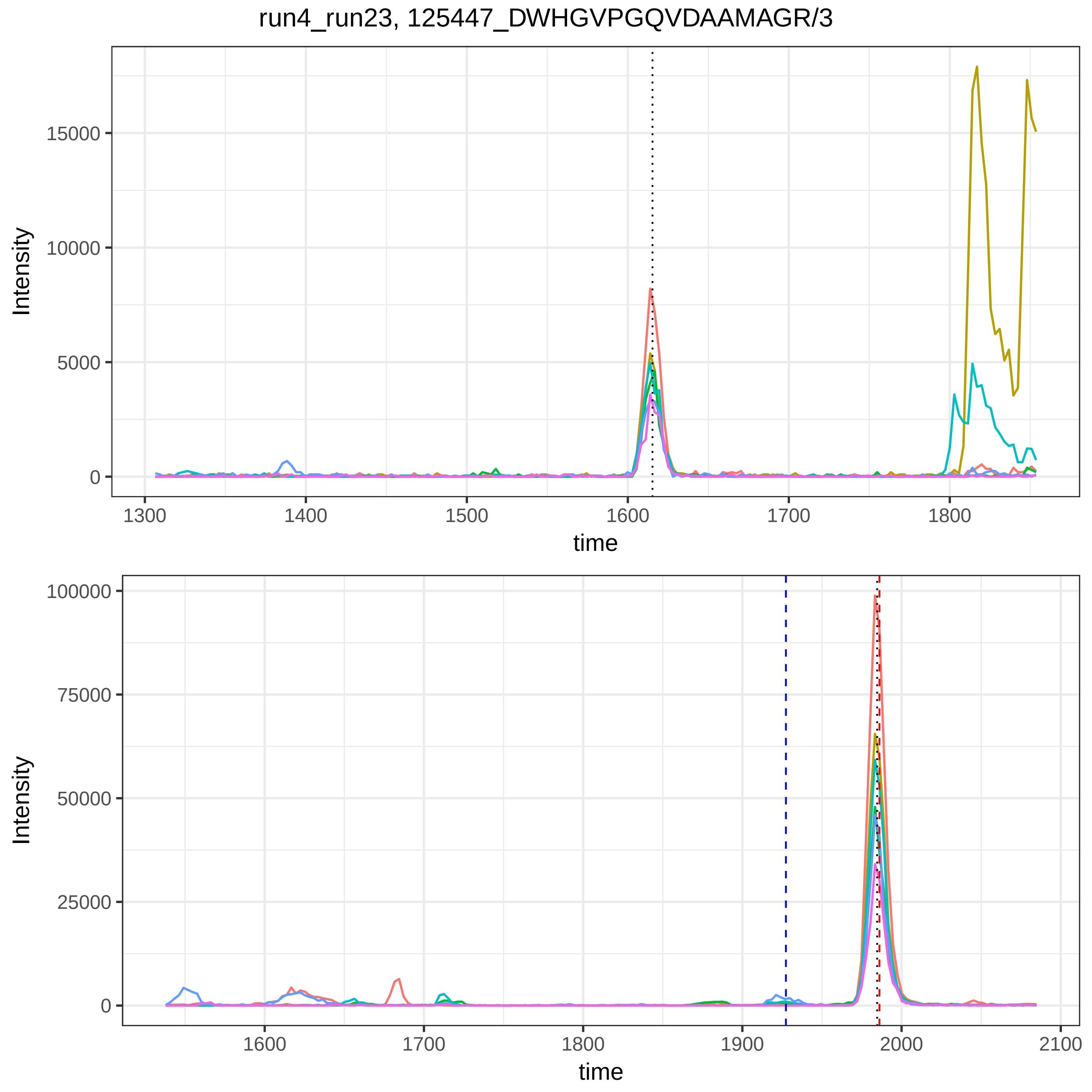

run4_run23, 125447_DWHGVPGQVDAAMAGR/3
Intensity
15000
10000
5000
0
1300
1400
1500
1600
1700
1800
time
Intensity
100000
75000
50000
25000
0
1600
1700
1800
1900
2000
2100
time

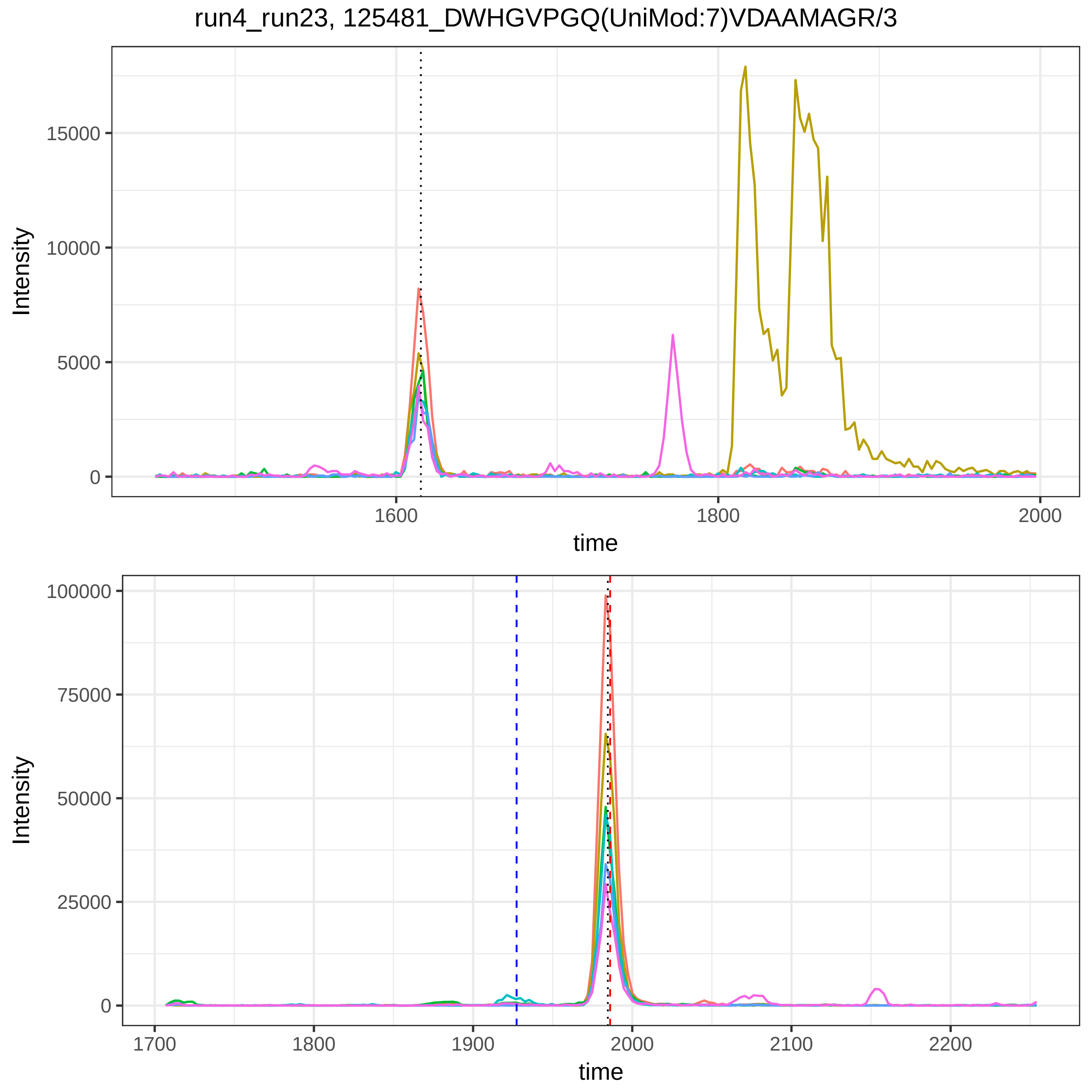
run4_run23, 125481_DWHGVPGQ(UniMod:7)VDAAMAGR/3
Intensity
15000
10000
5000
0
1600
1800
2000
time
Intensity
100000
75000
50000
25000
0
1700
1800
1900
2000
2100
2200
time

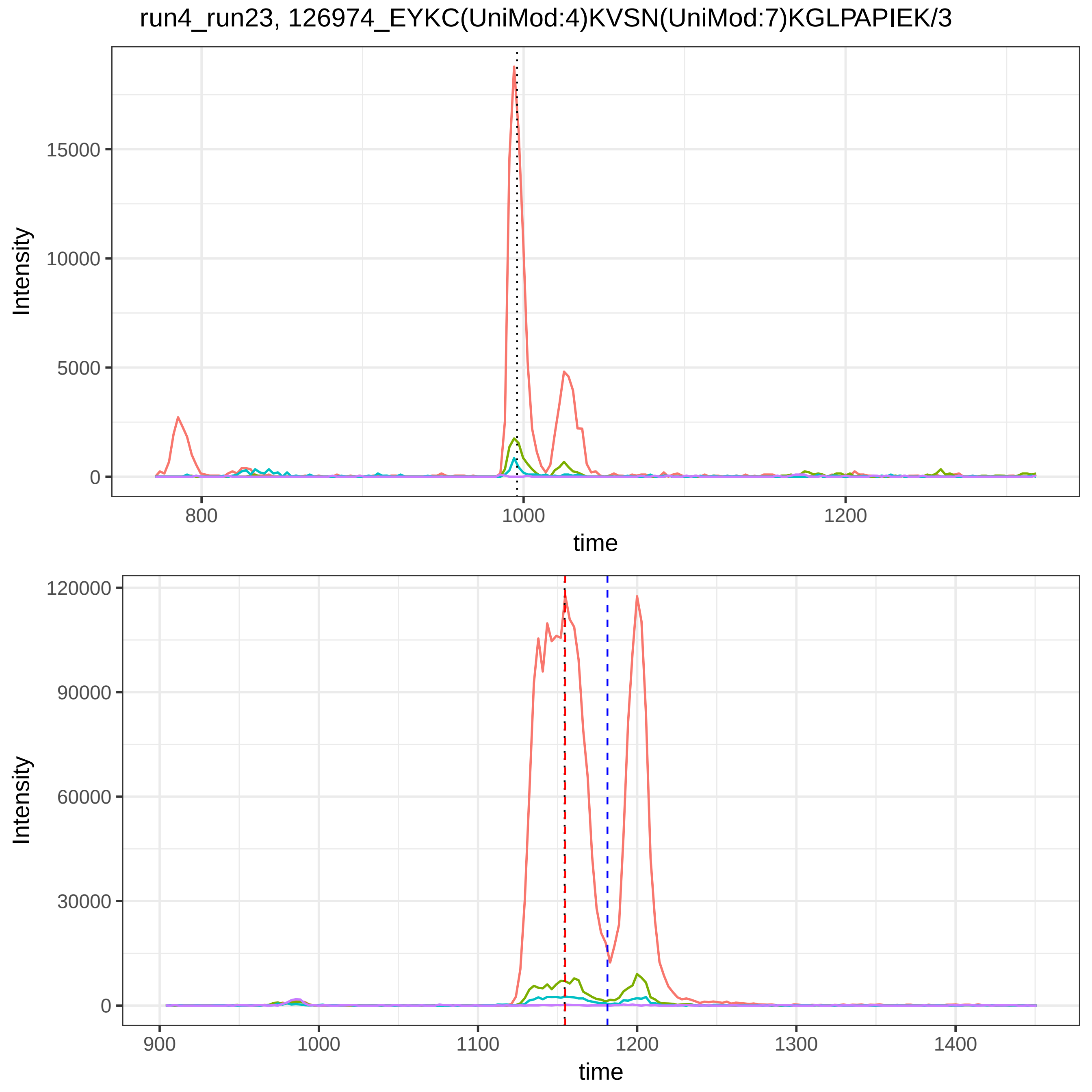

run4_run23, 126974_EYKC(UniMod:4)KVSN(UniMod:7)KGLPAPIEK/3
Intensity
15000
10000
5000
0
800
1000
1200
time
Intensity
120000
90000
60000
30000
0
900
1000
1100
1200
1300
1400
time

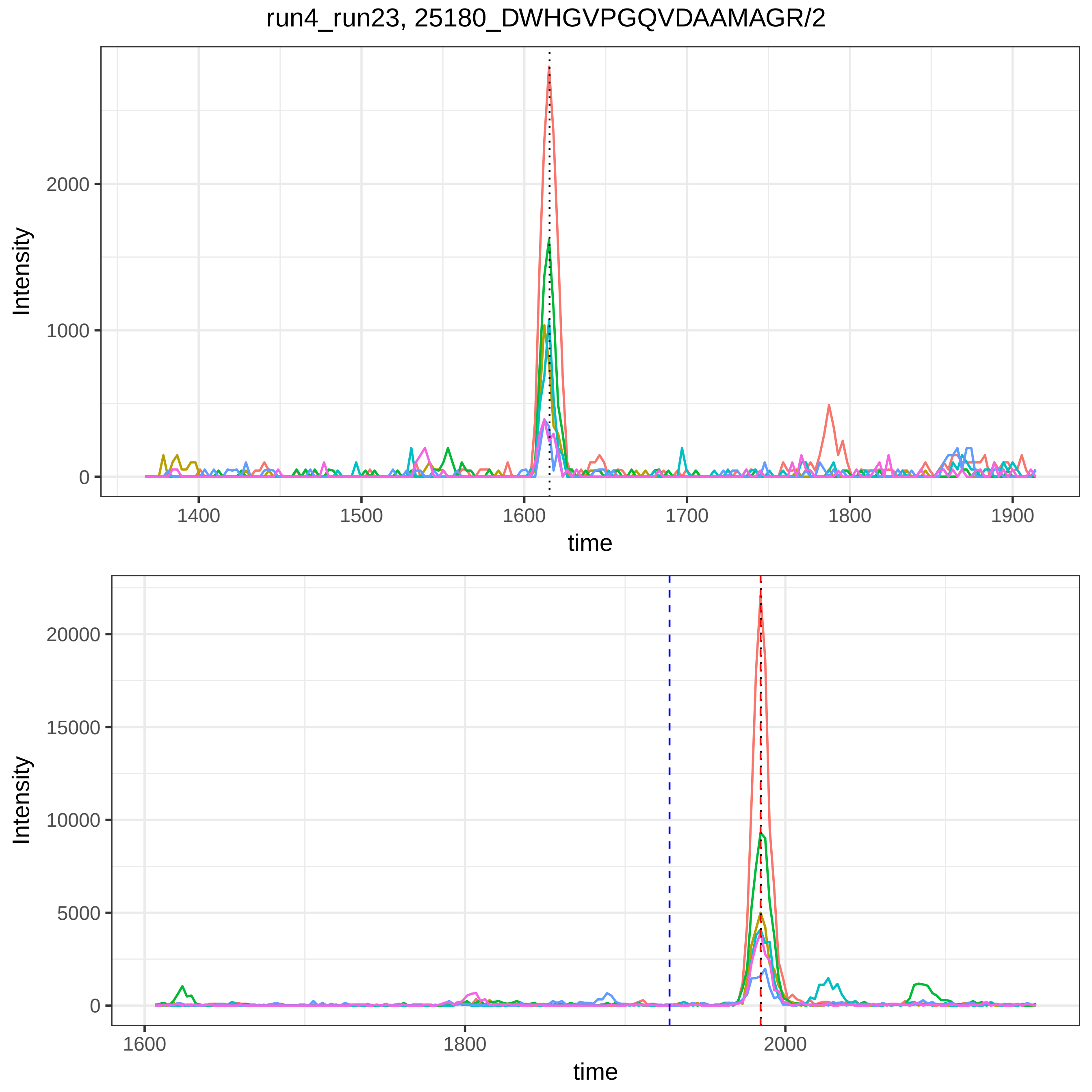
run4_run23, 25180_DWHGVPGQVDAAMAGR/2
Intensity
2000
1000
0
1400
1500
1600
1700
1800
1900
time
Intensity
20000
15000
10000
5000
0
1600
1800
2000
time

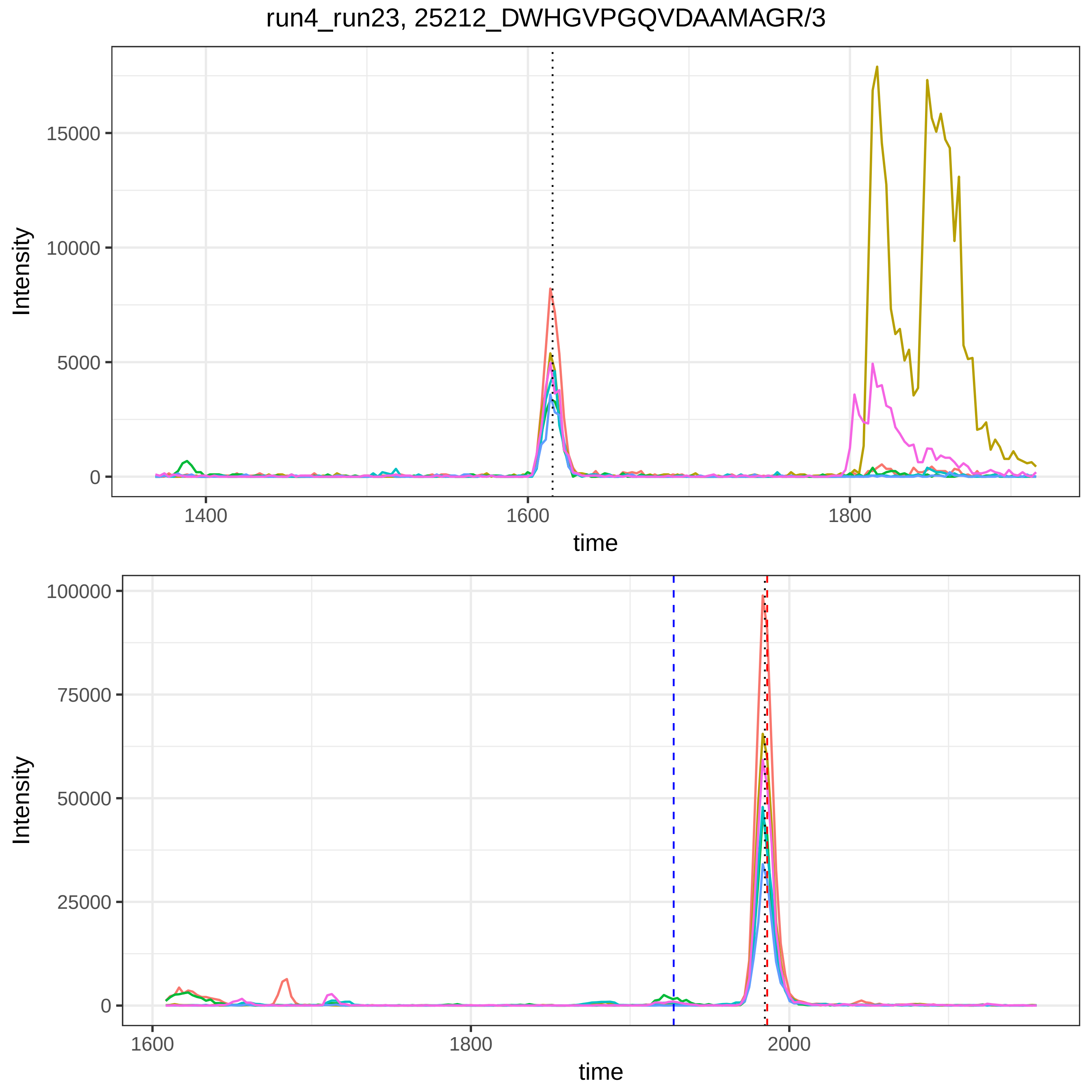
run4_run23, 25212_DWHGVPGQVDAAMAGR/3
Intensity
15000
10000
5000
0
1400
1600
1800
time
Intensity
100000
75000
50000
25000
0
1600
1800
2000
time

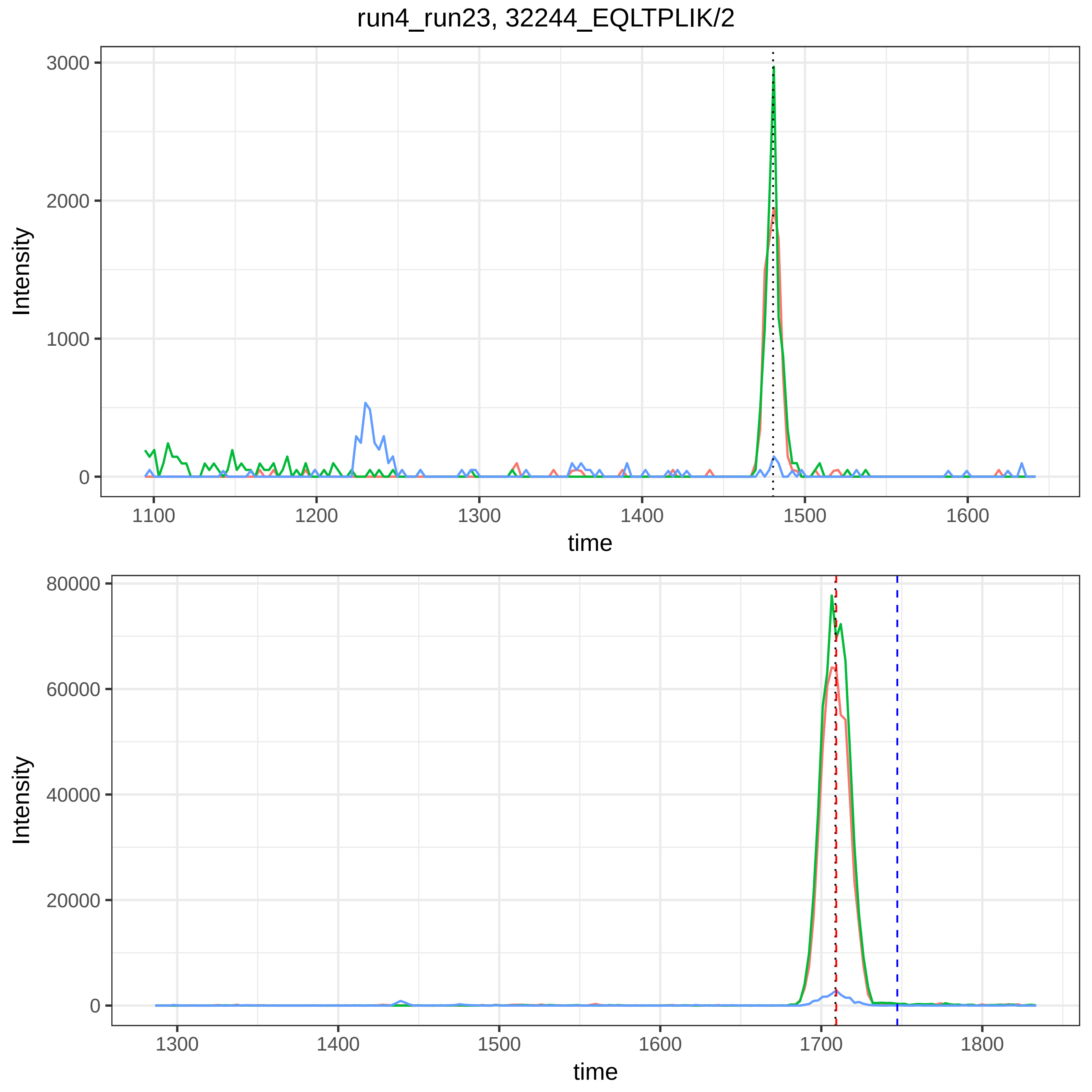

run4_run23, 32244_EQLTPLIK/2
Intensity
time
3000
2000
1000
0
1100
1200
1300
1400
1500
1600
80000
60000
40000
20000
1700
1800

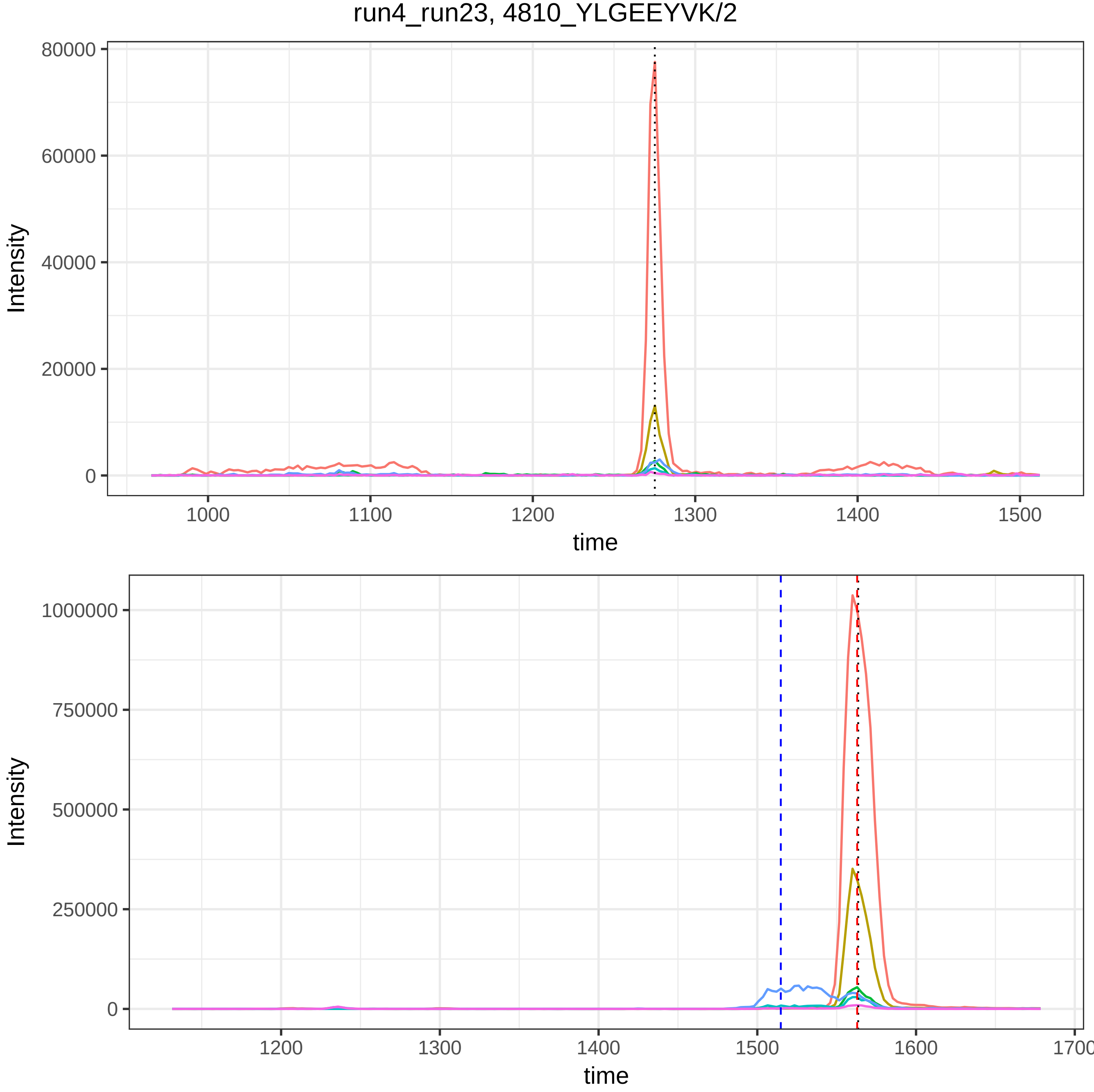

run4_run23, 4810_YLGEEYVK/2
Intensity
80000
60000
40000
20000
0
1000
1100
1200
1300
1400
1500
time
Intensity
1000000
750000
500000
250000
0
1200
1300
1400
1500
1600
1700
time

run4_run23, 49292_QELSEAEQATR/2

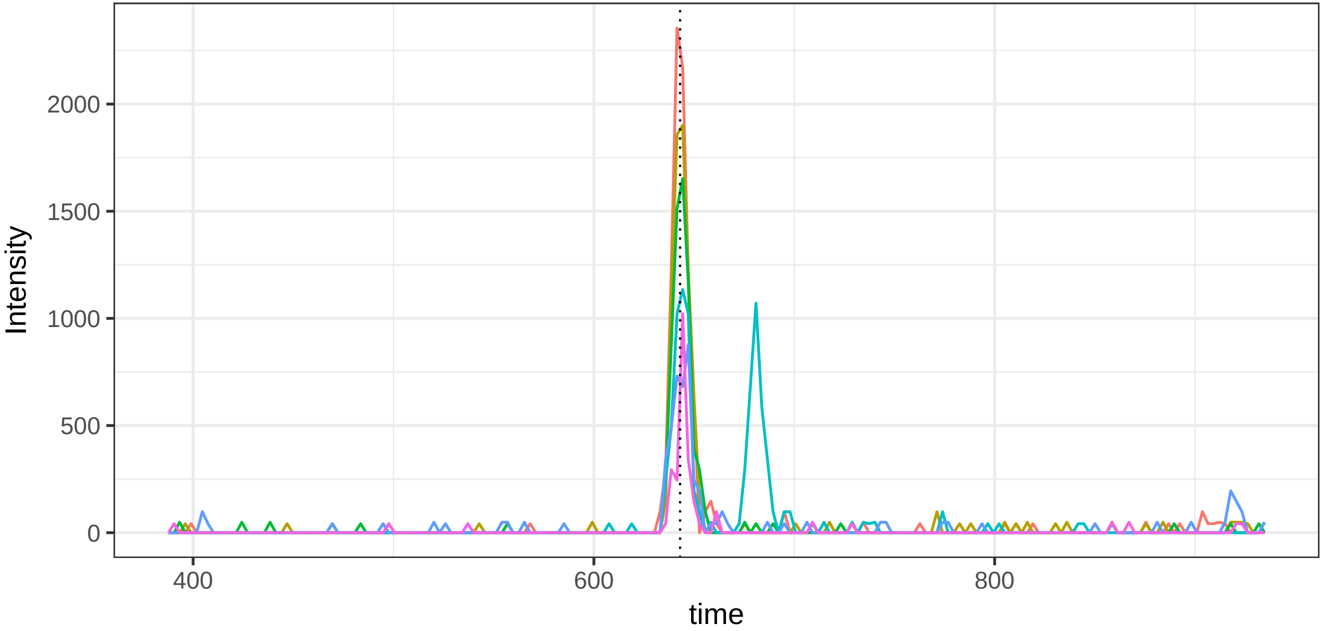


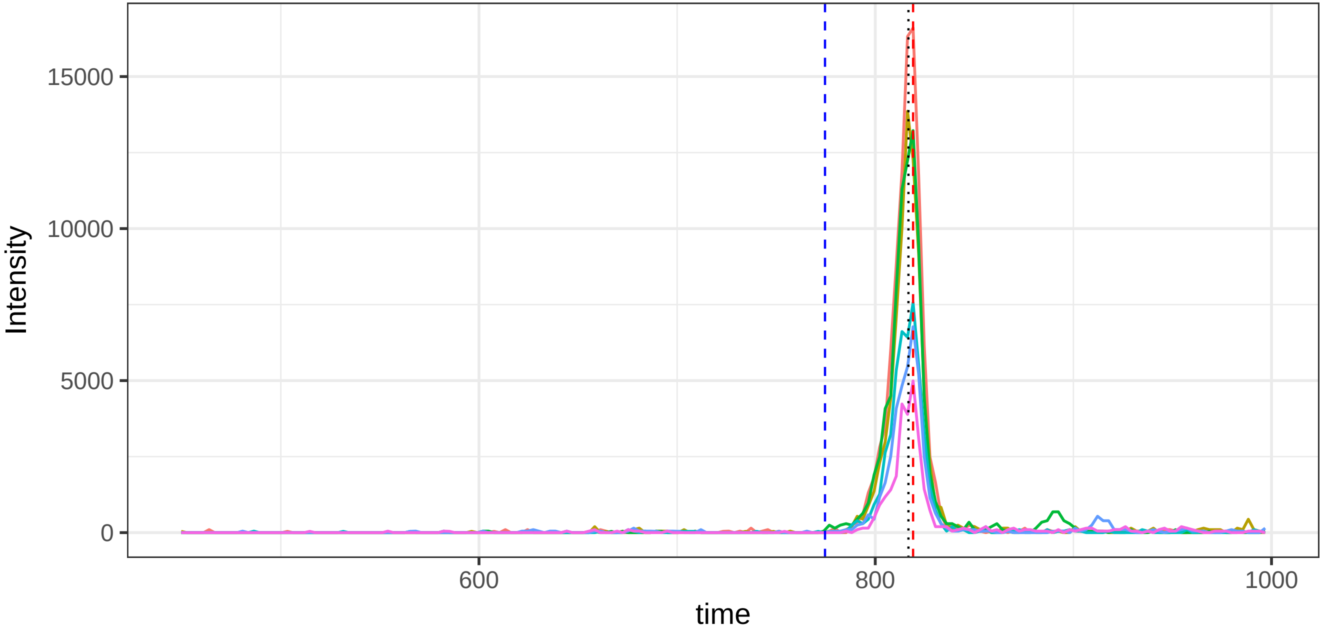

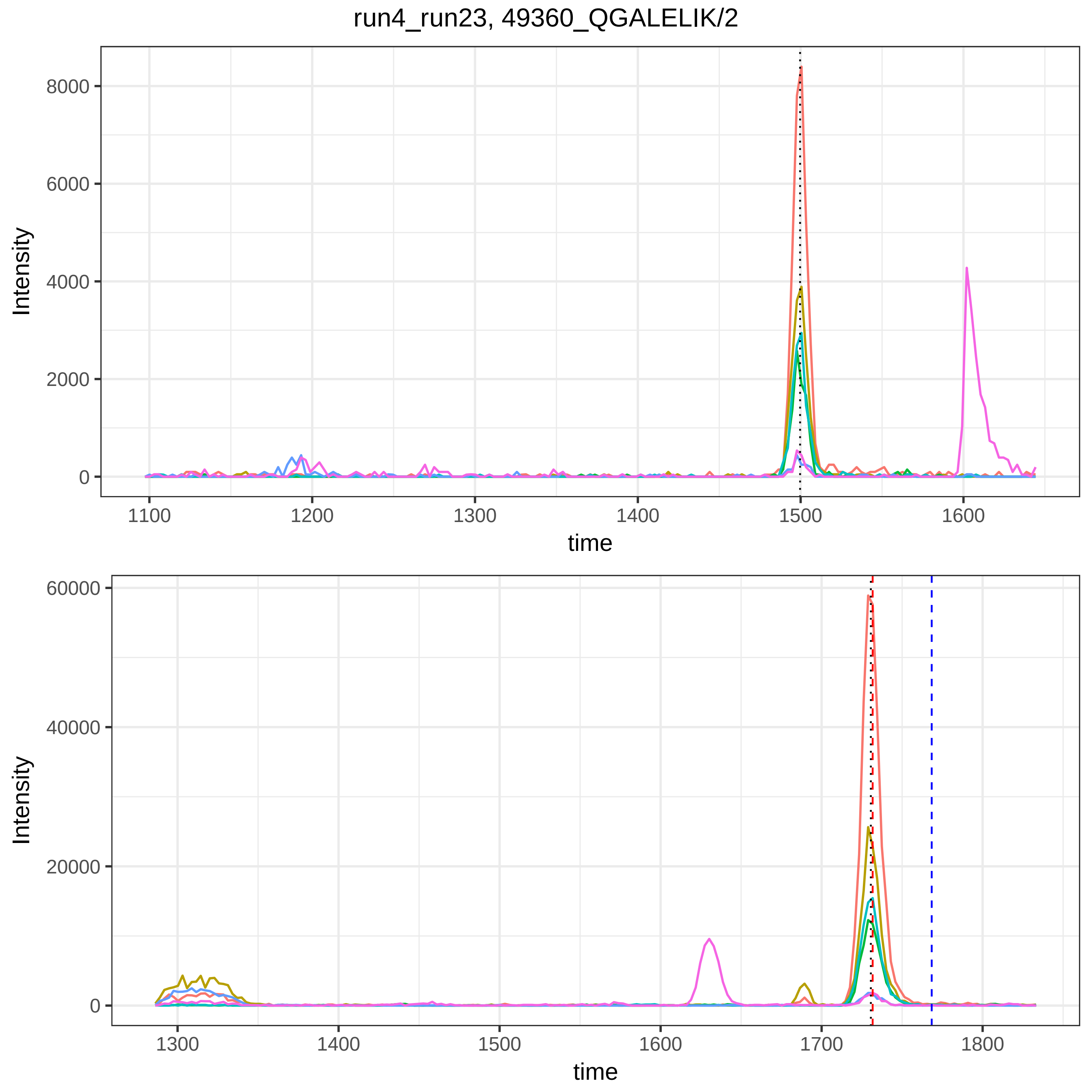
run4_run23, 49360_QGALELIK/2
Intensity
time
8000
6000
4000
2000
0
1100
1200
1300
1400
1500
1600
60000
40000
20000
1700
1800

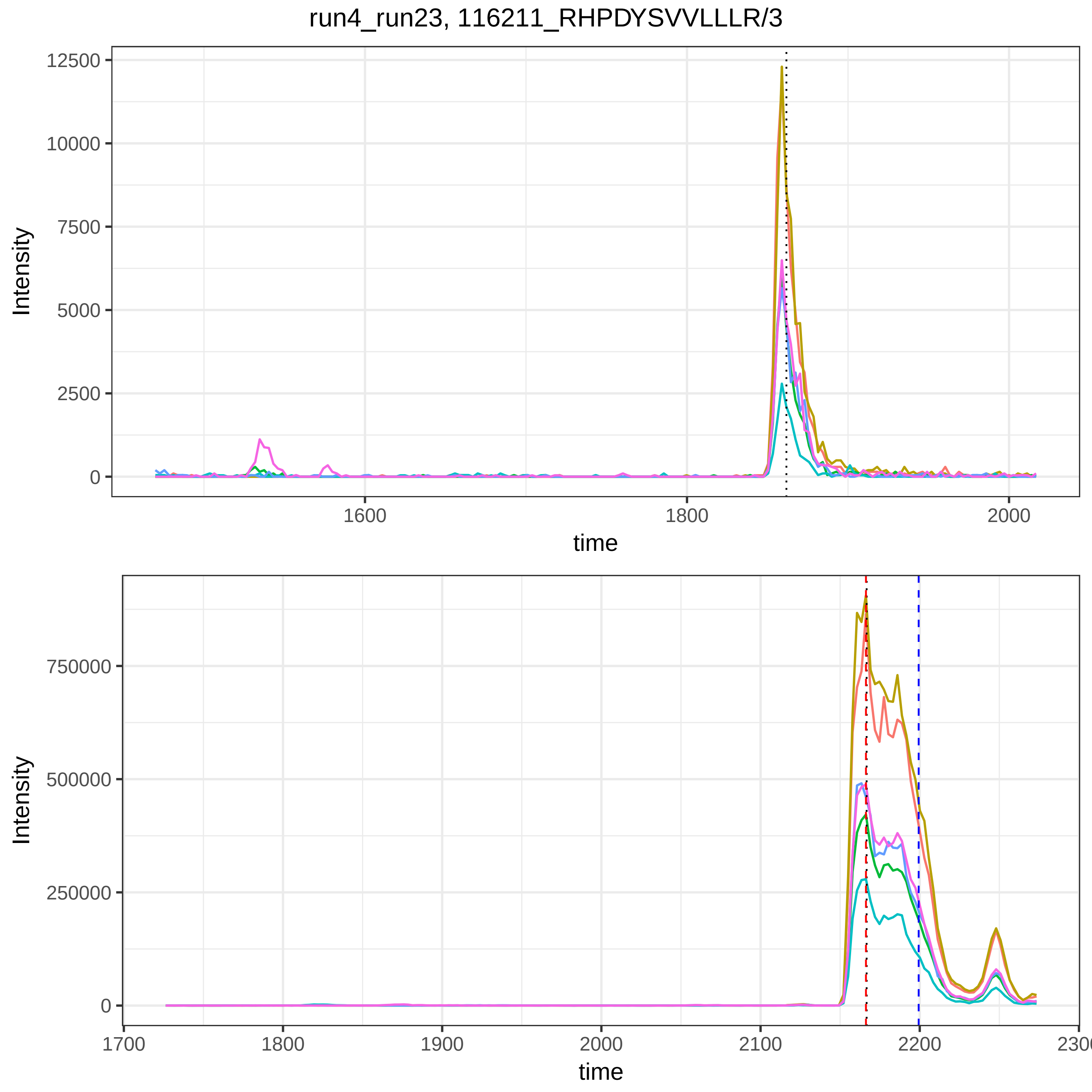

run4_run23, 116211_RHPDYSVVLLLR/3
Intensity
12500
10000
7500
5000
2500
0
1600
1800
2000
time
Intensity
750000
500000
250000
0
1700
1800
1900
2000
2100
2200
2300
time

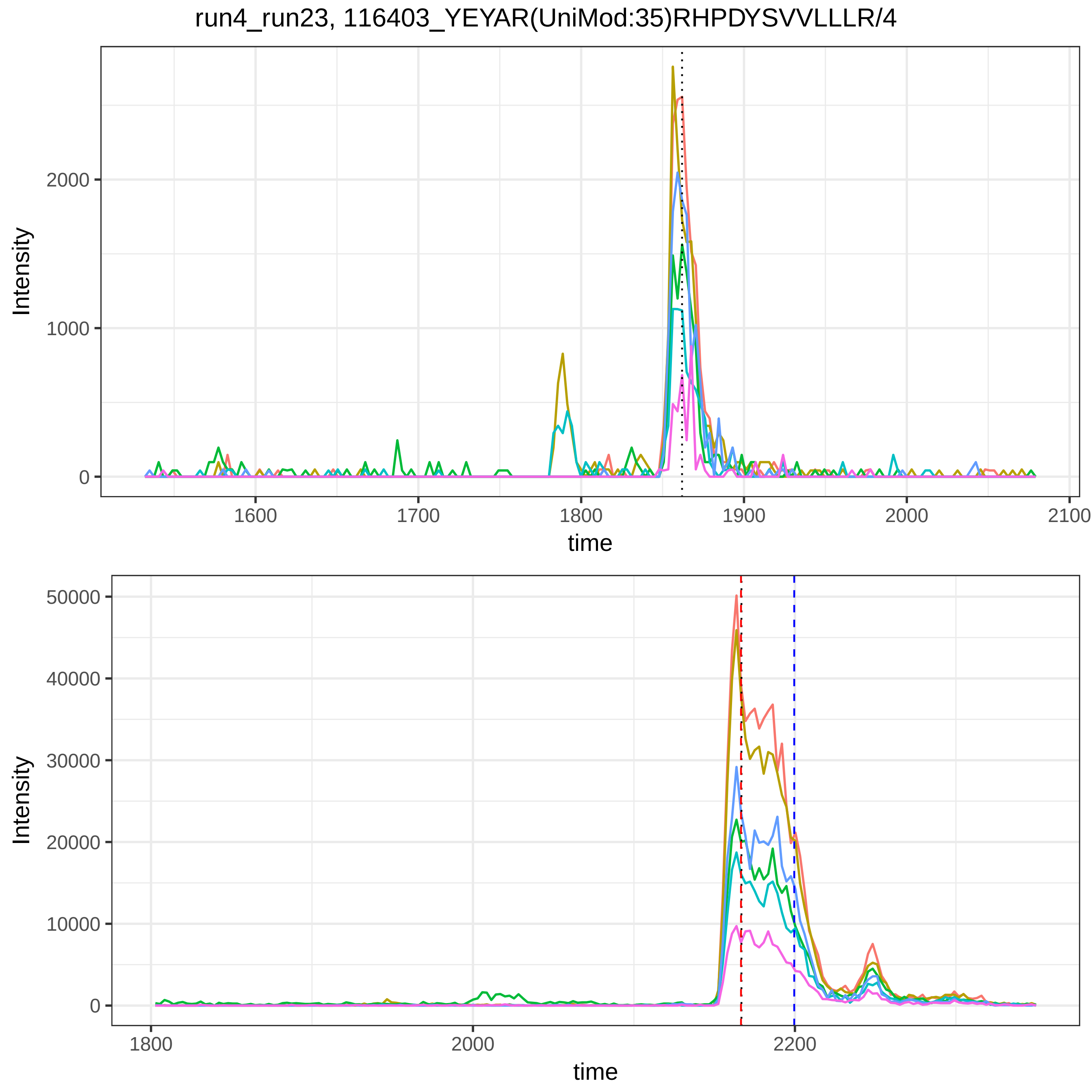
run4_run23, 116403_YEYAR(UniMod:35)RHPDYSVVLLLR/4
Intensity
2000
1000
0
1600
1700
1800
1900
2000
2100
time
Intensity
50000
40000
30000
20000
10000
0
1800
2000
2200
time

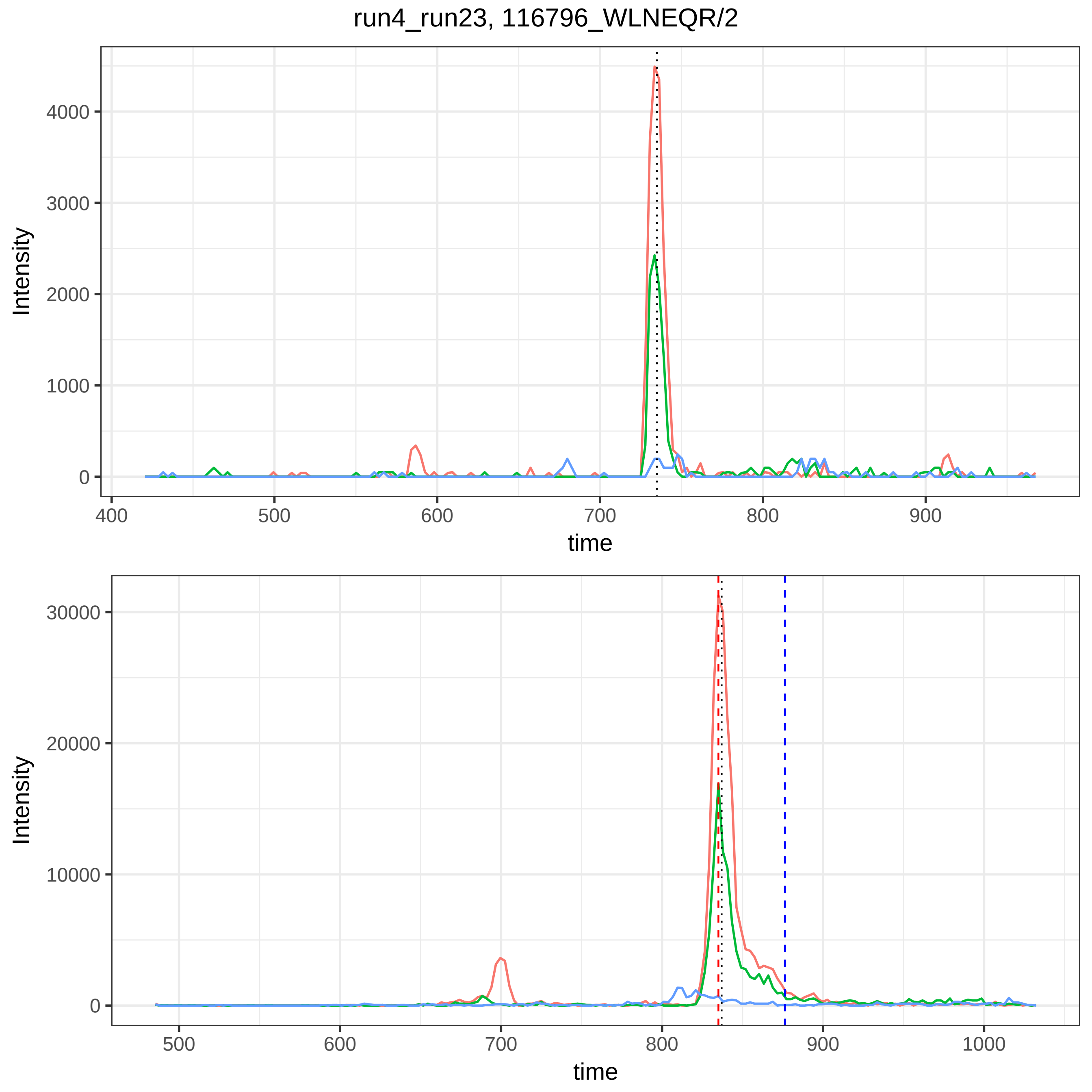

run4_run23, 116796_WLNEQR/2
Intensity
time
0
1000
2000
3000
4000
400
500
600
700
800
900
0
10000
20000
30000
500
600
700
800
900
1000

run4_run23, 118541_GSGLNLC(UniMod:4)EPNNKEGYYGYTGAFR/4

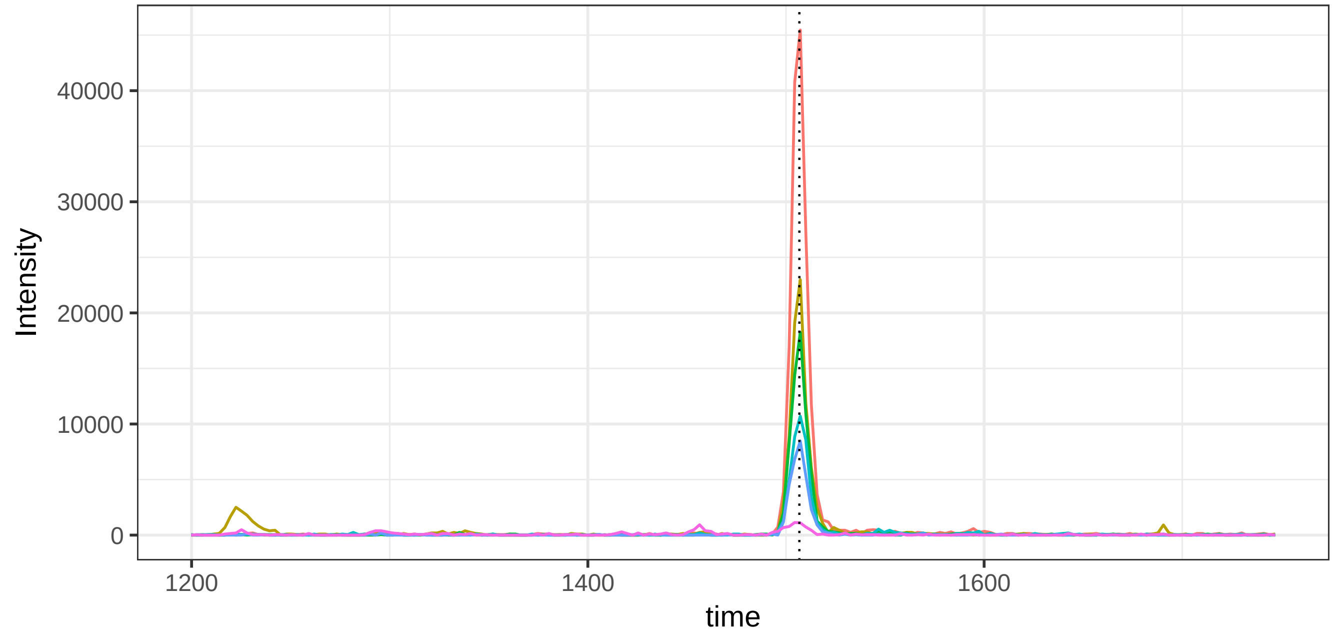


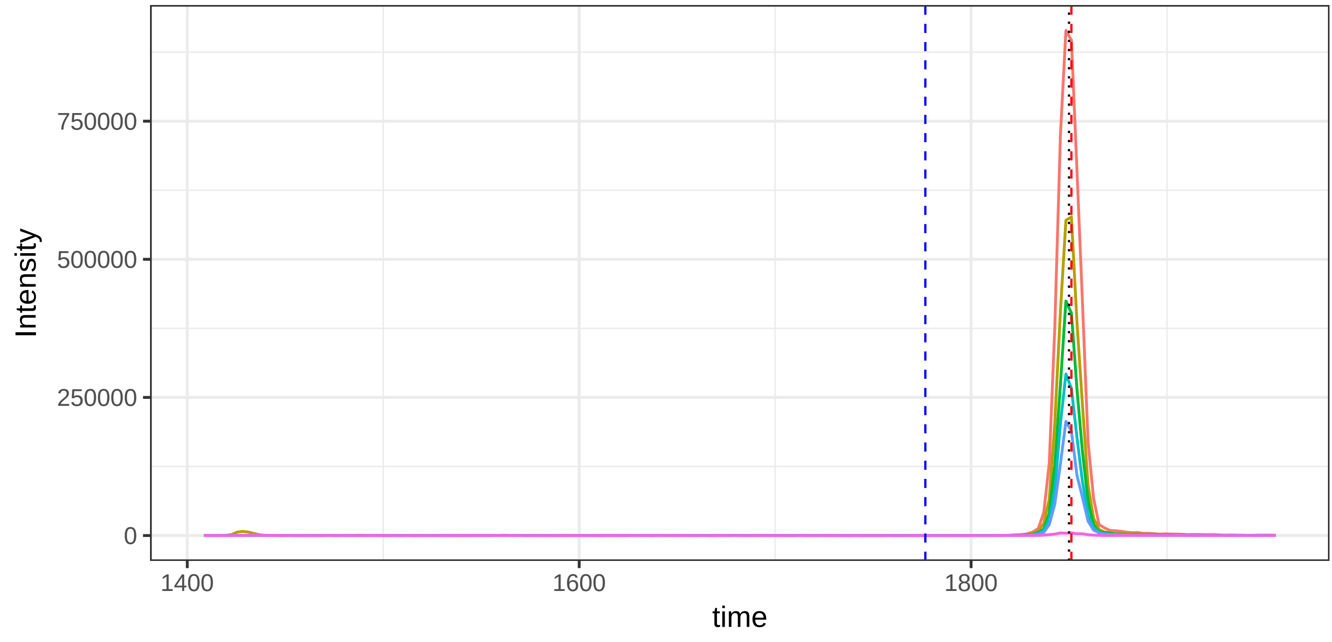

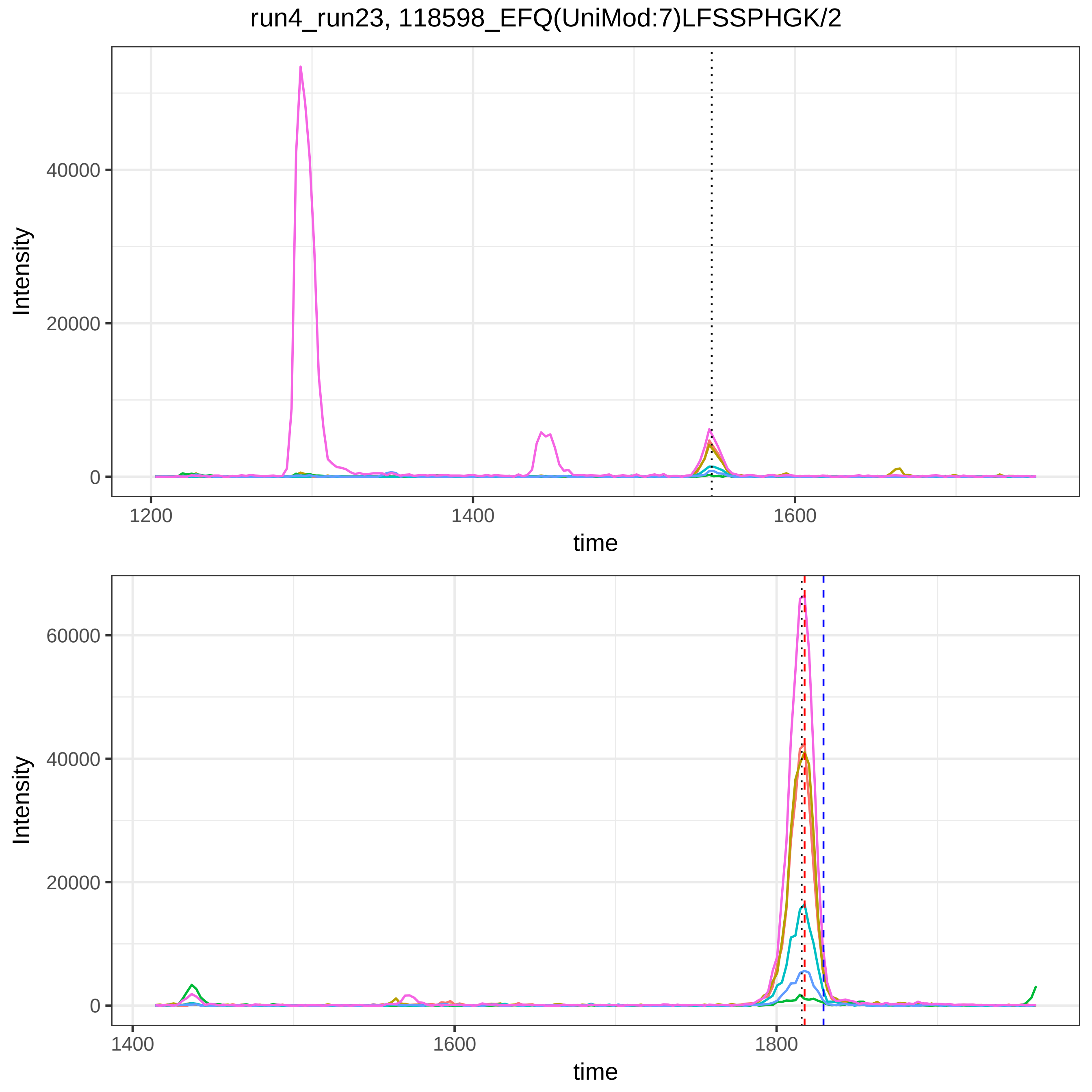
run4_run23, 118598_EFQ(UniMod:7)LFSSPHGK/2
Intensity
40000
20000
0
1200
1400
1600
time
Intensity
60000
40000
20000
0
1400
1600
1800
time

run4_run23, 120092_RTHLPEVFLSK/3

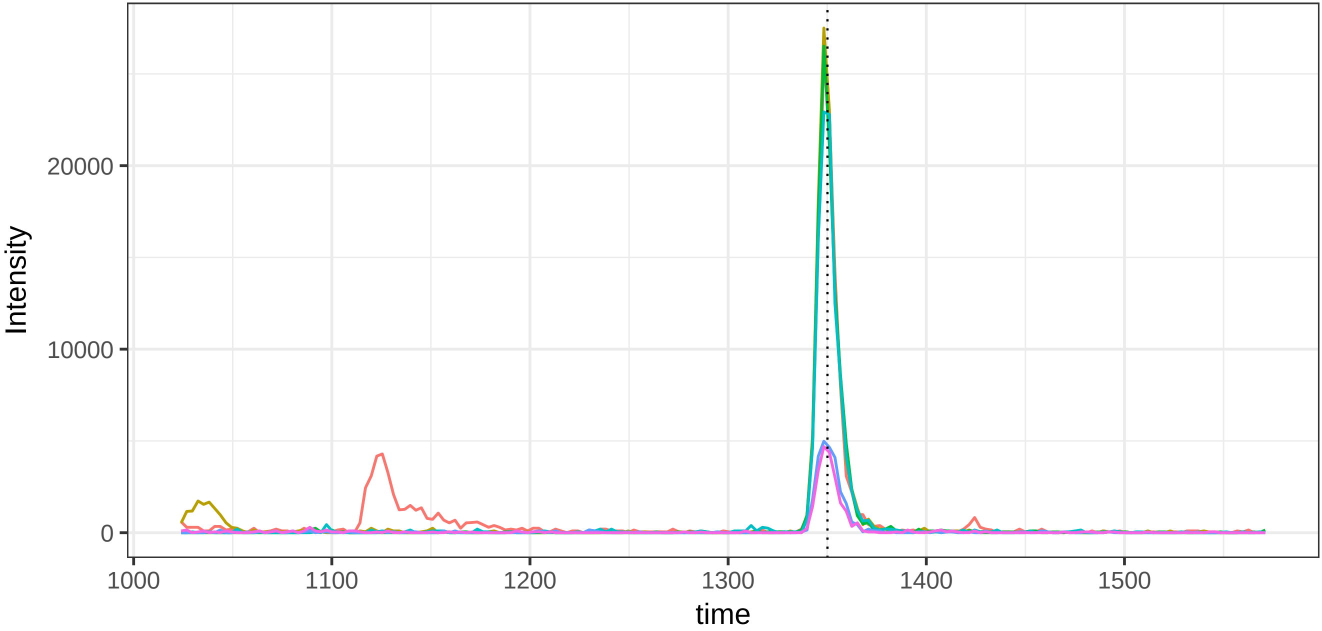


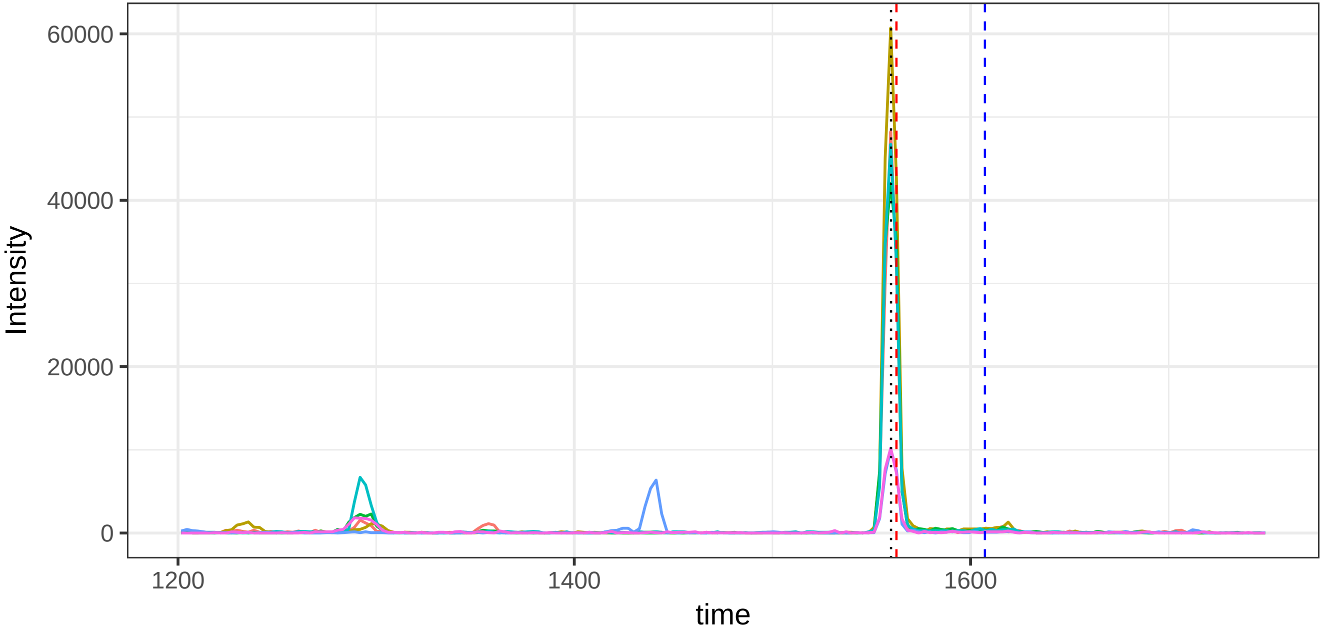

run4_run23, 127018_TPLTATLSK/2

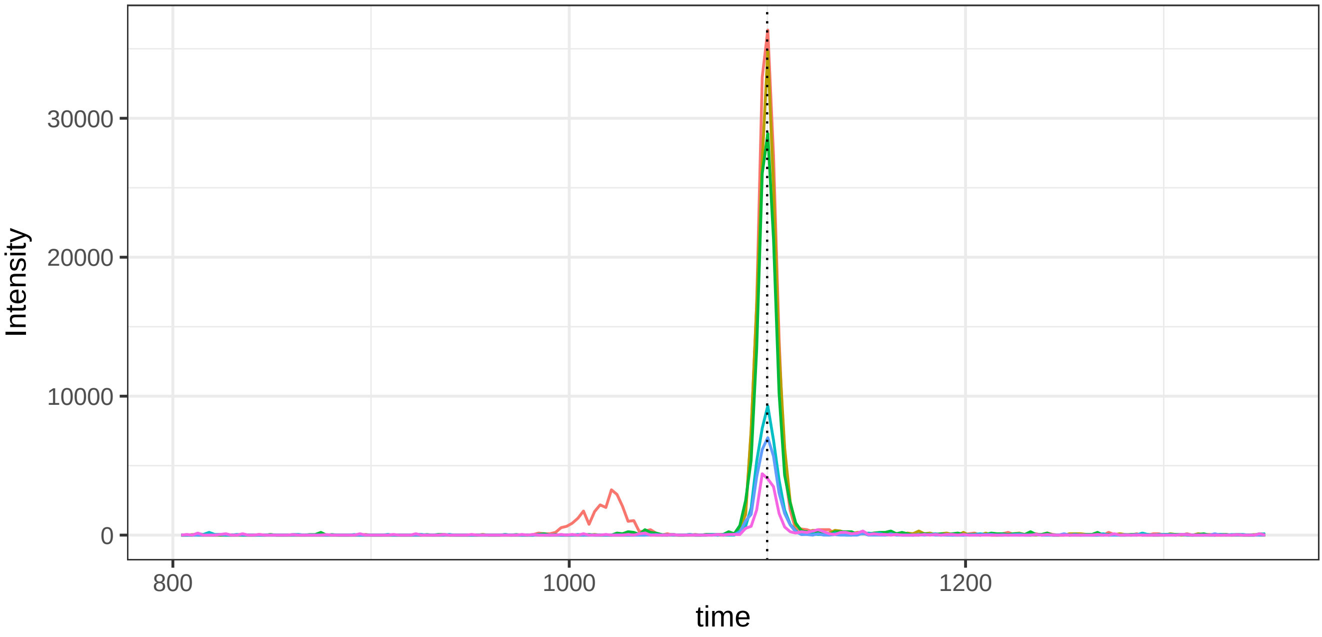


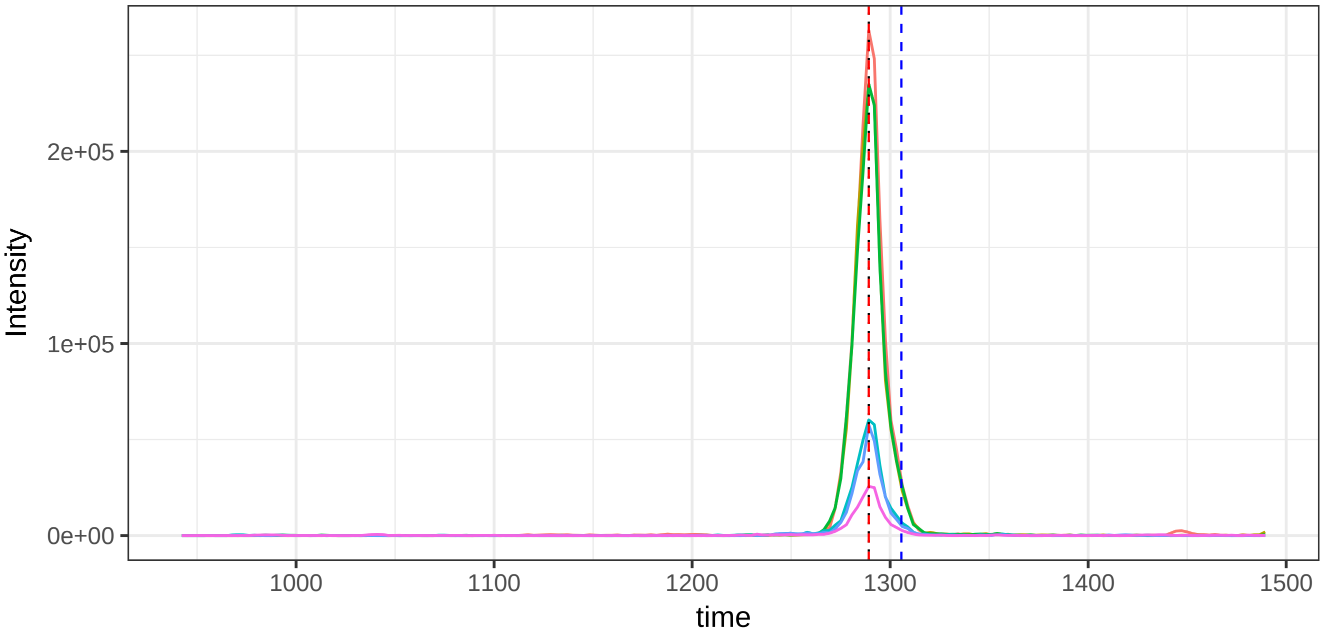

run4_run23, 133616_TEAPTTIGGLNK/2

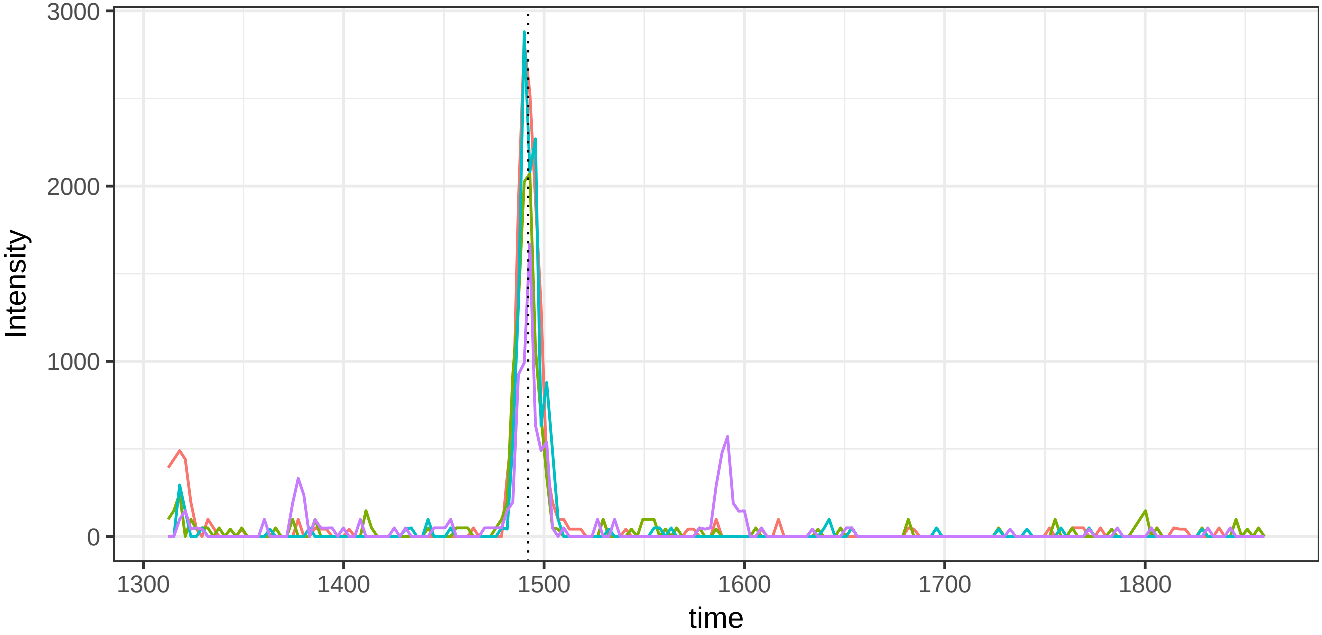


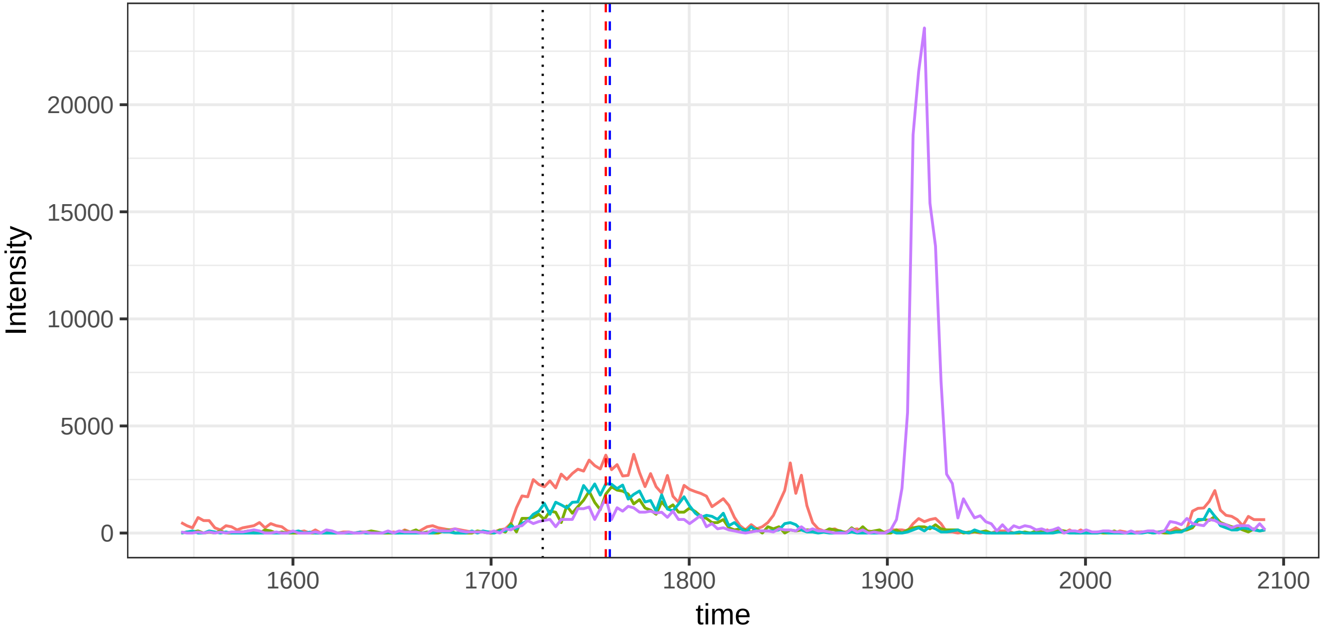

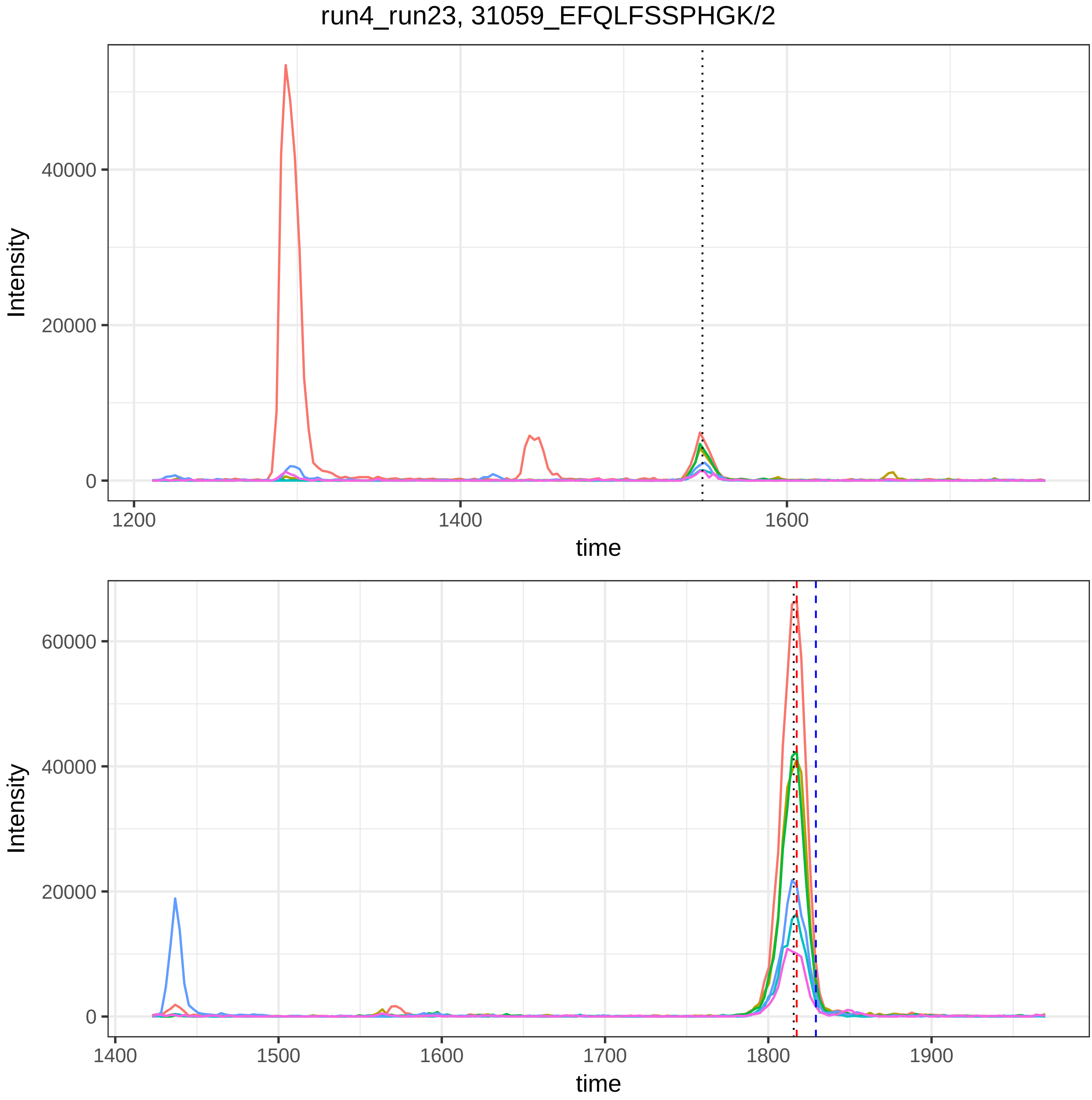

run4_run23, 31059_EFQLFSSPHGK/2
Intensity
40000
20000
0
1200
1400
1600
time
Intensity
60000
40000
20000
0
1400
1500
1600
1700
1800
1900
time

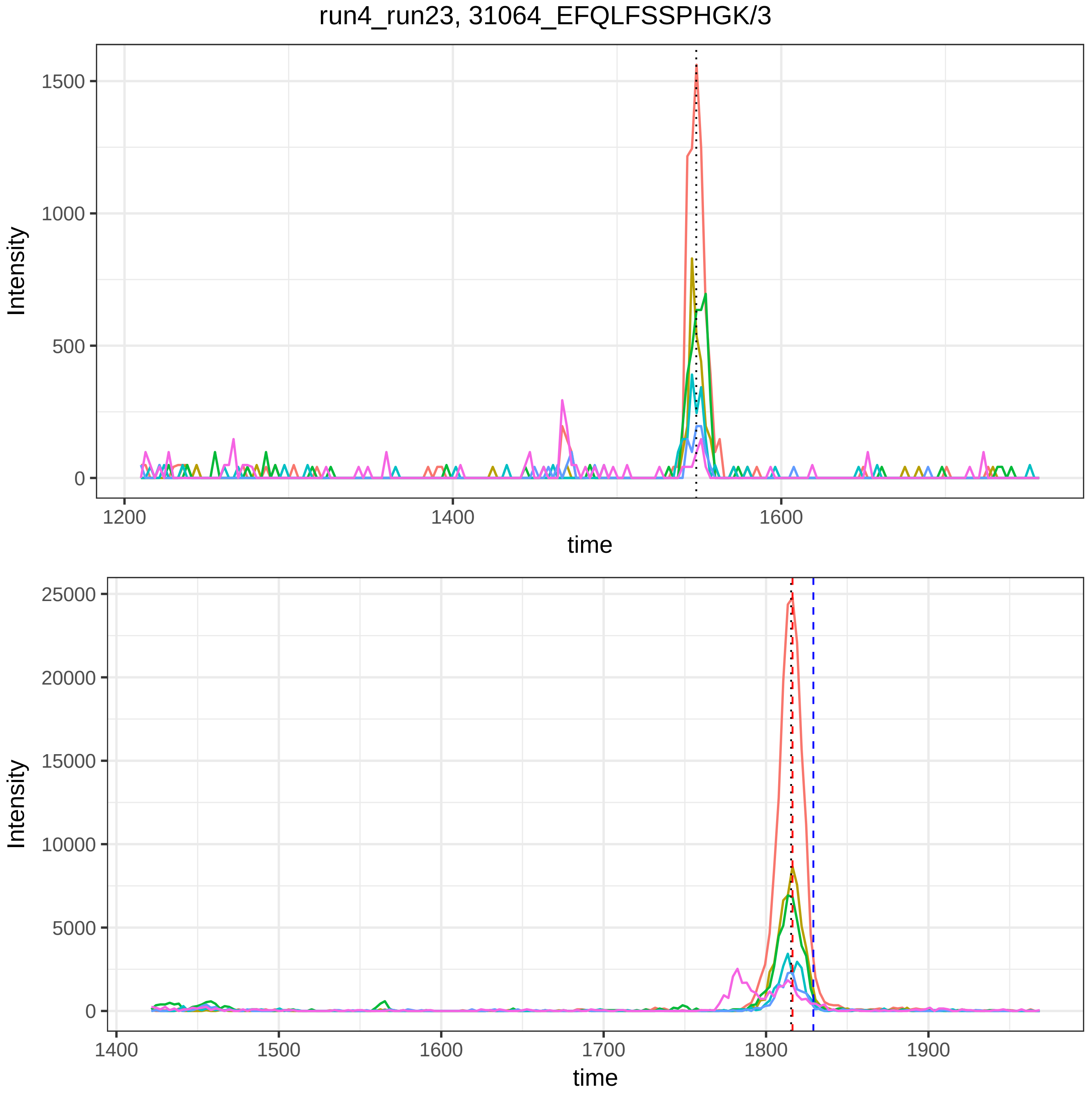

run4_run23, 31064_EFQLFSSPHGK/3
Intensity
time
1500
1000
500
0
1200
1400
1600
25000
20000
15000
10000
5000
1500
1700
1800
1900

run4_run23, 31289_EGYYGYTGAFR/2

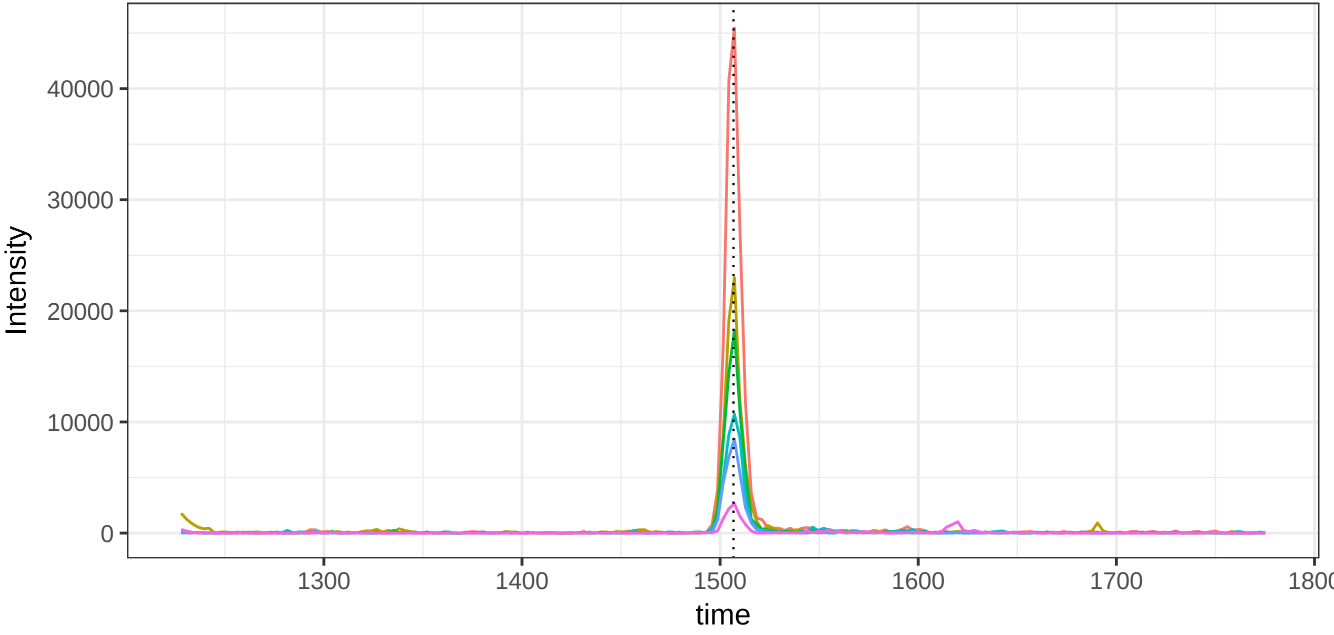


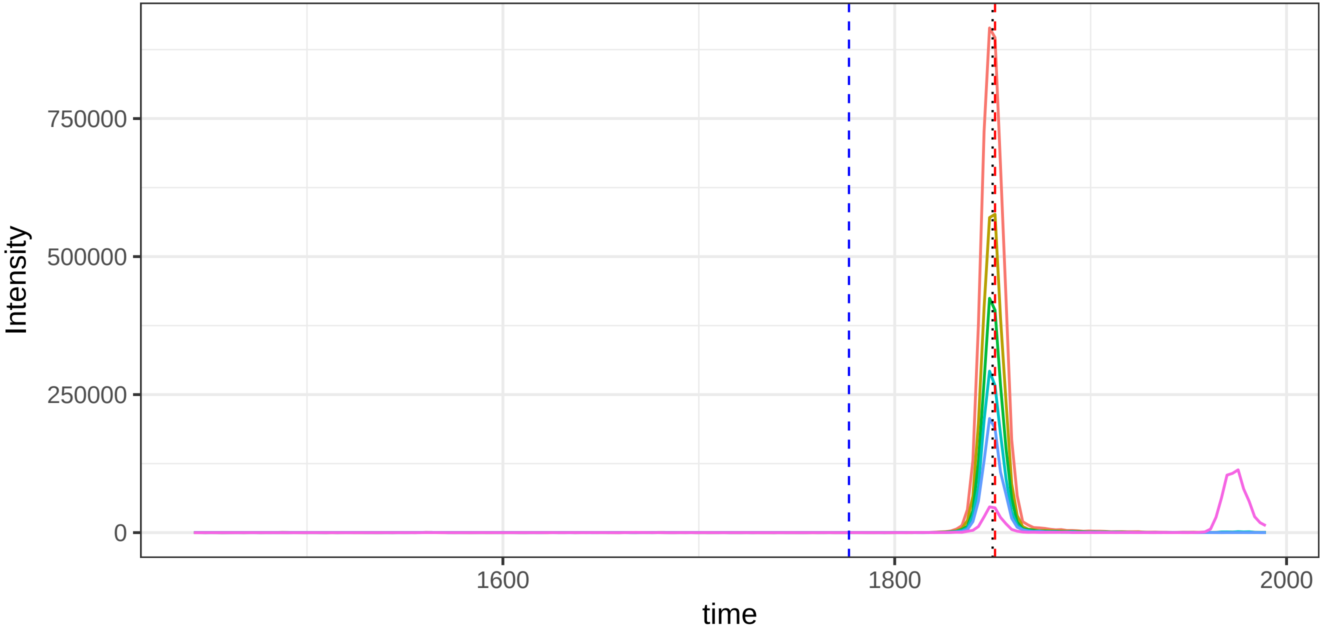

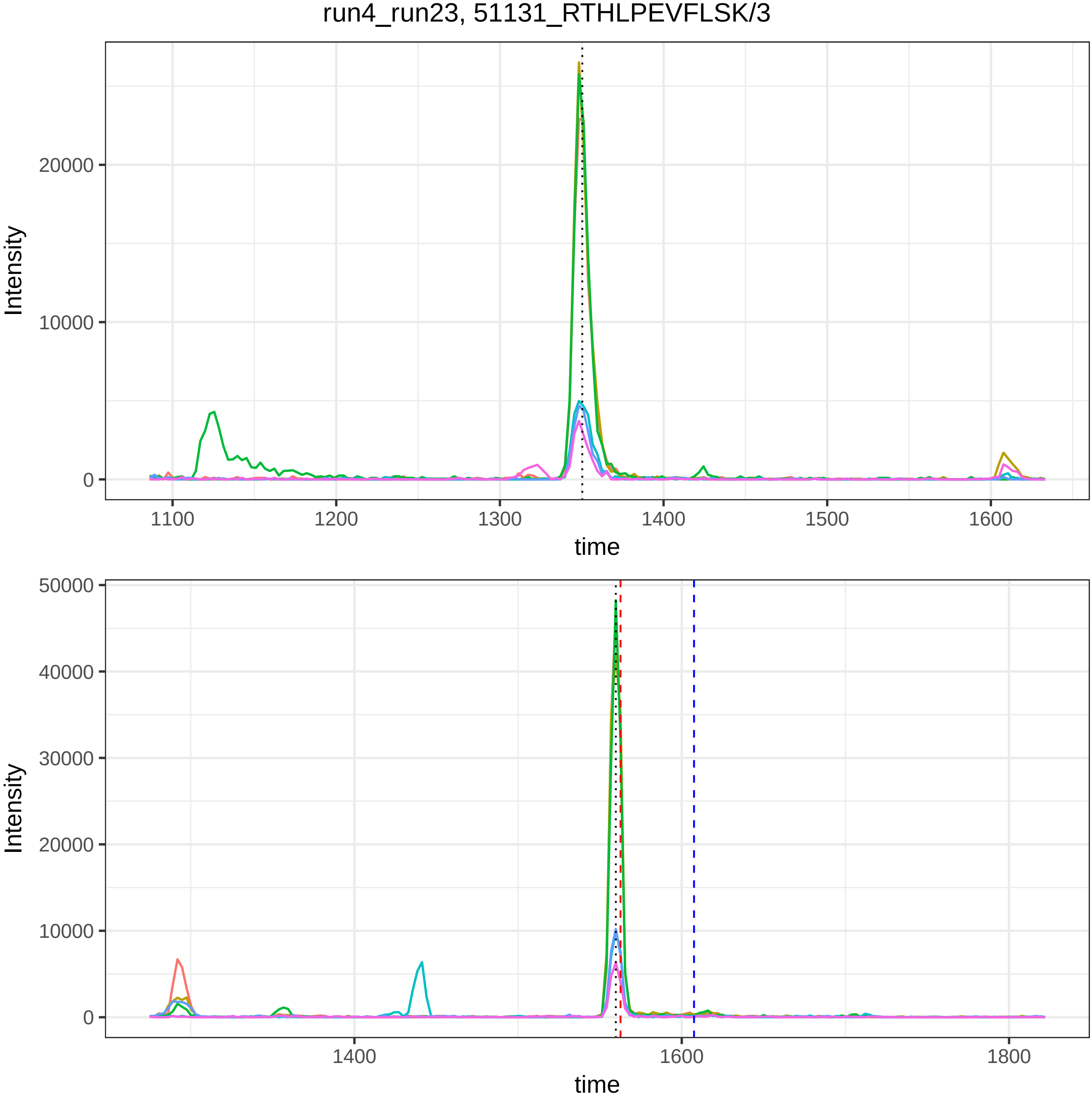
run4_run23, 51131_RTHLPEVFLSK/3
Intensity
20000
10000
0
1100
1200
1300
1400
1500
1600
time
Intensity
50000
40000
30000
20000
10000
0
1400
1600
1800
time

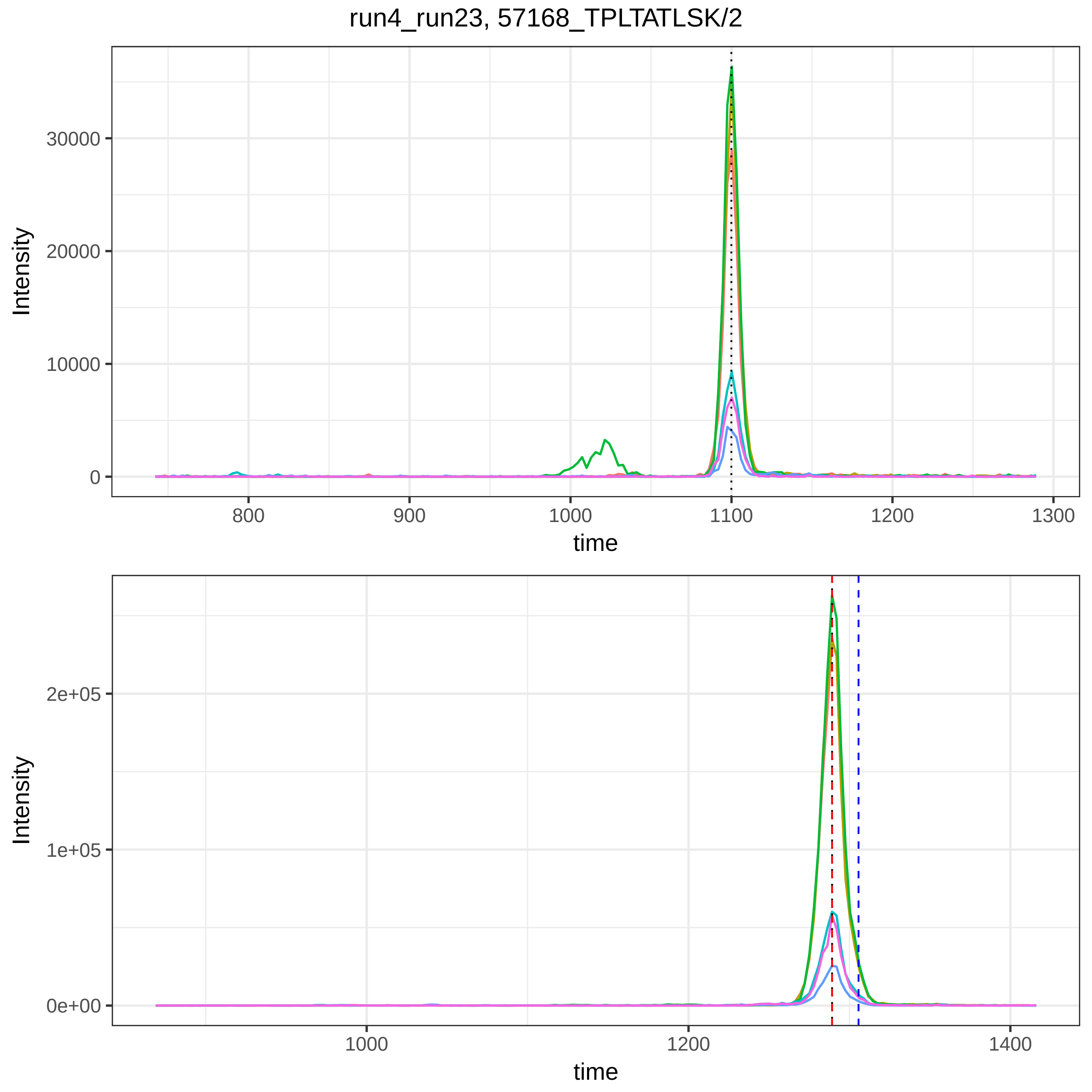

run4_run23, 57168_TPLTATLSK/2
Intensity
30000
20000
10000
0
800
900
1000
1100
1200
1300
time
Intensity
2e+05
1e+05
0e+00
1000
1200
1400
time

run4_run23, 116068_KLVAASQAALG/2

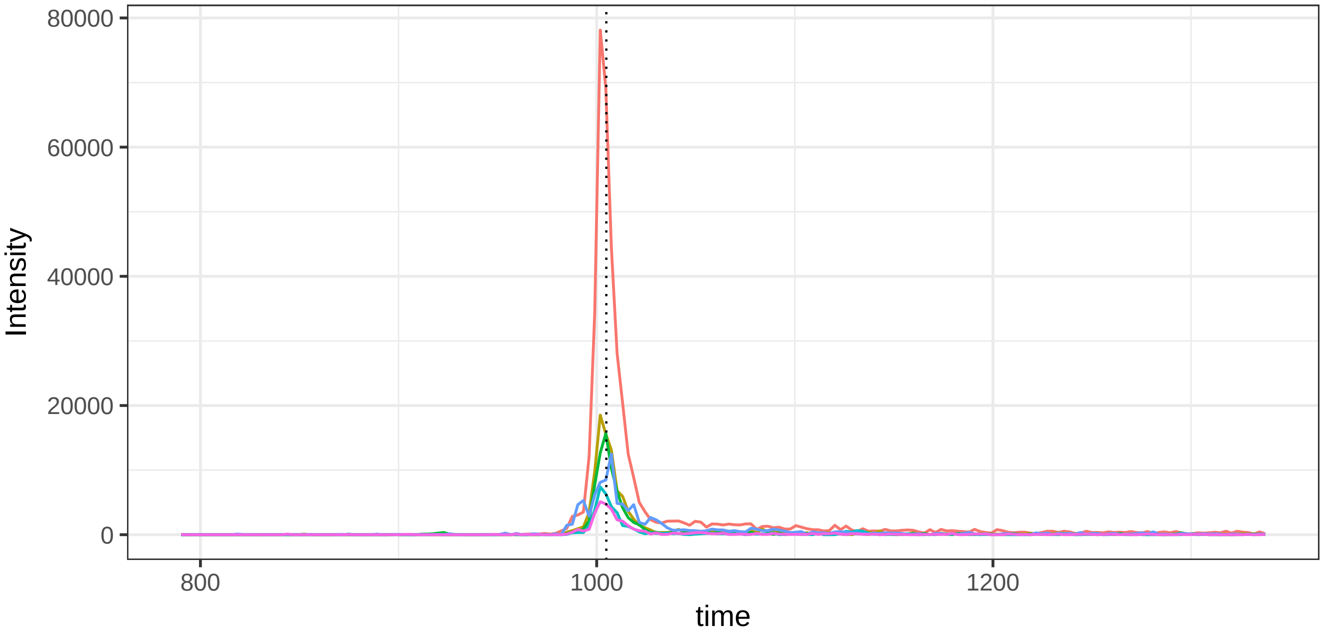


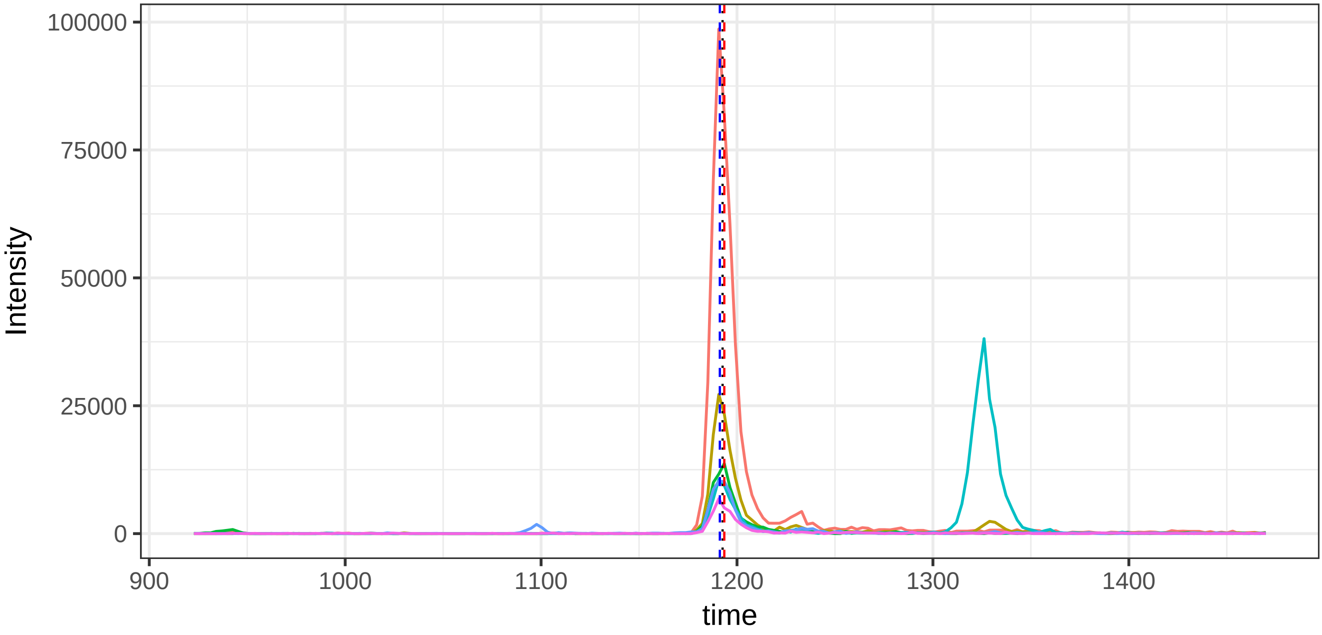

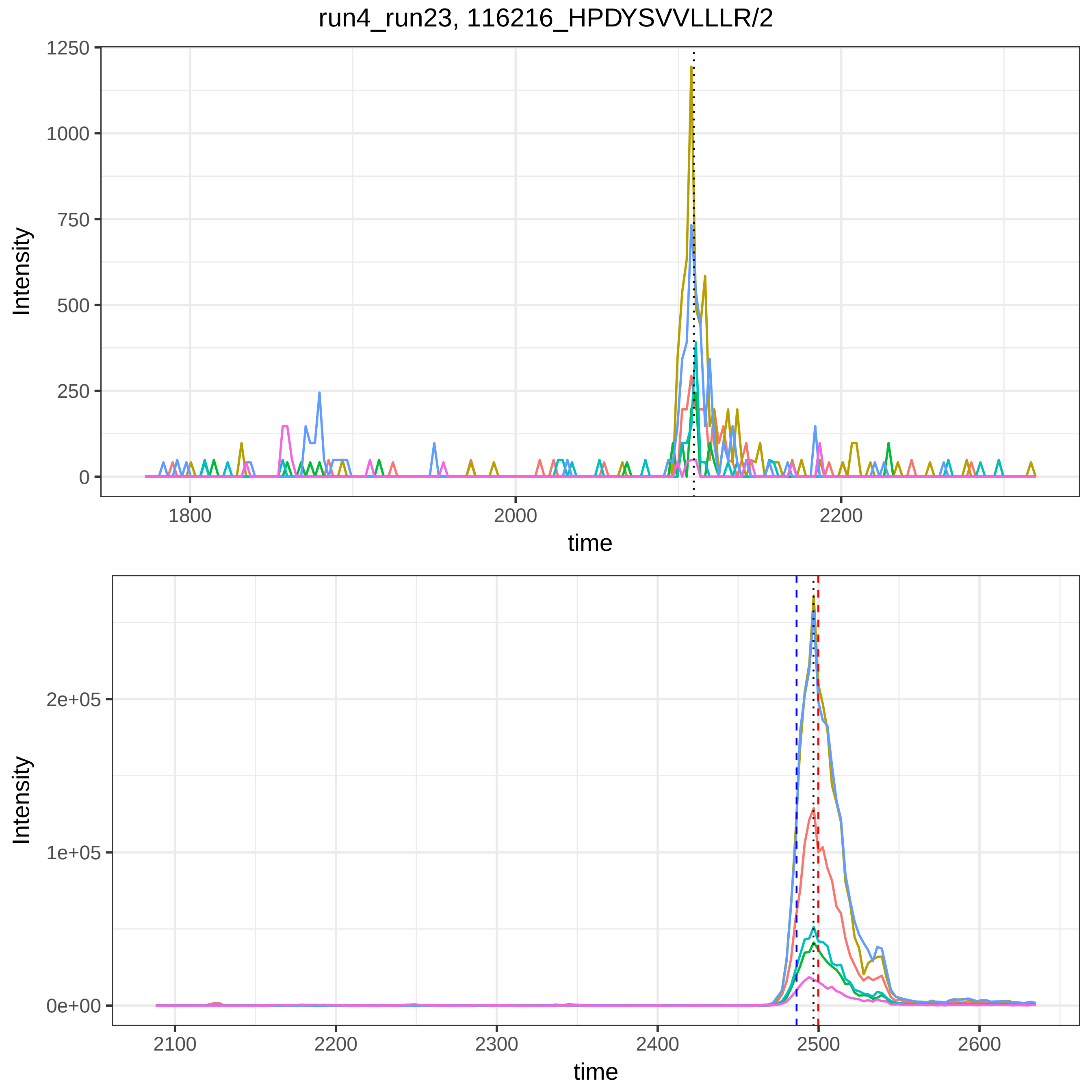
run4_run23, 116216_HPDYSVVLLLR/2
Intensity
1250
1000
750
500
250
0
1800
2000
2200
time
Intensity
2e+05
1e+05
0e+00
2100
2200
2300
2400
2500
2600
time

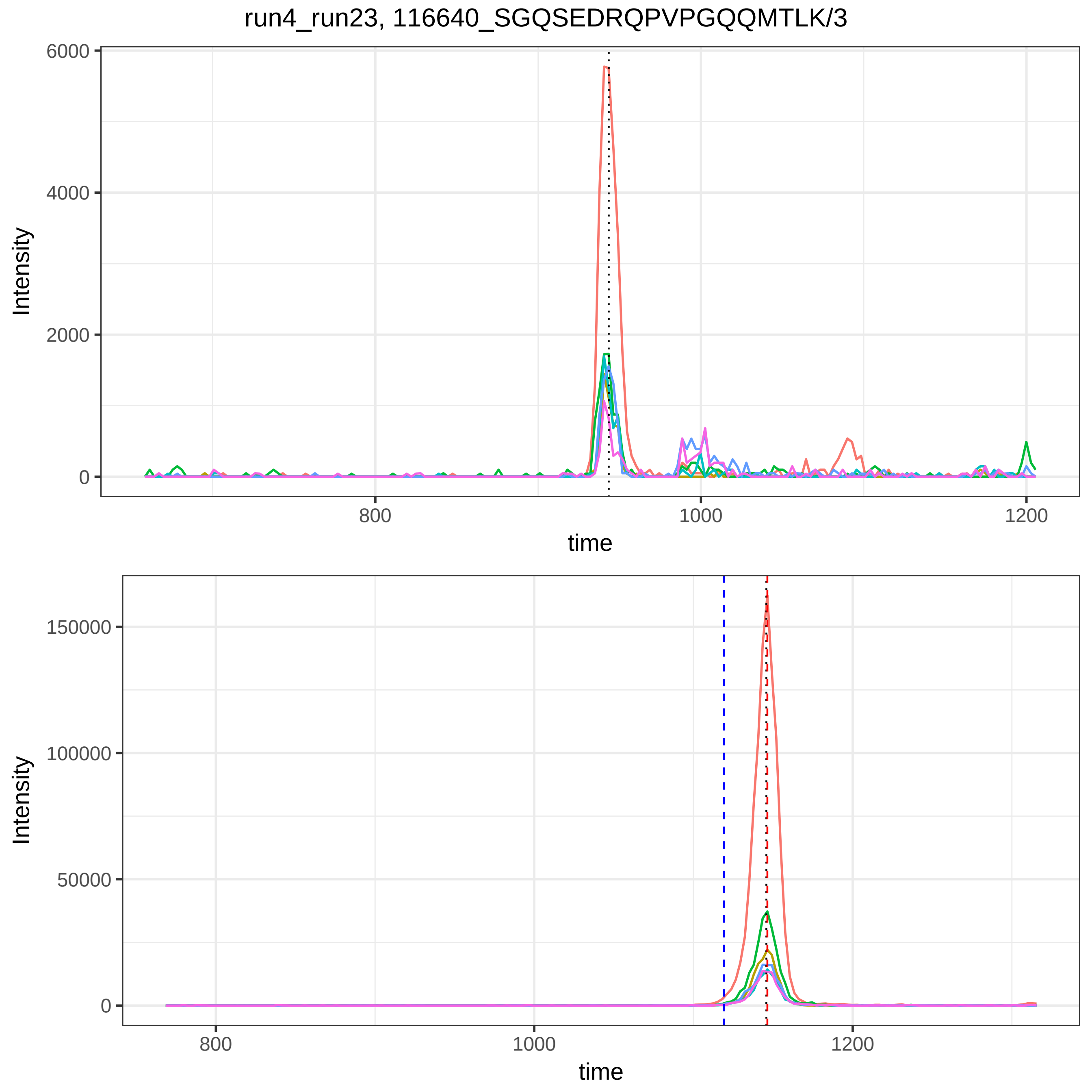
run4_run23, 116640_SGQSEDRQPVPGQQMTLK/3
Intensity
6000
4000
2000
0
800
1000
1200
time
Intensity
150000
100000
50000
0
800
1000
1200
time

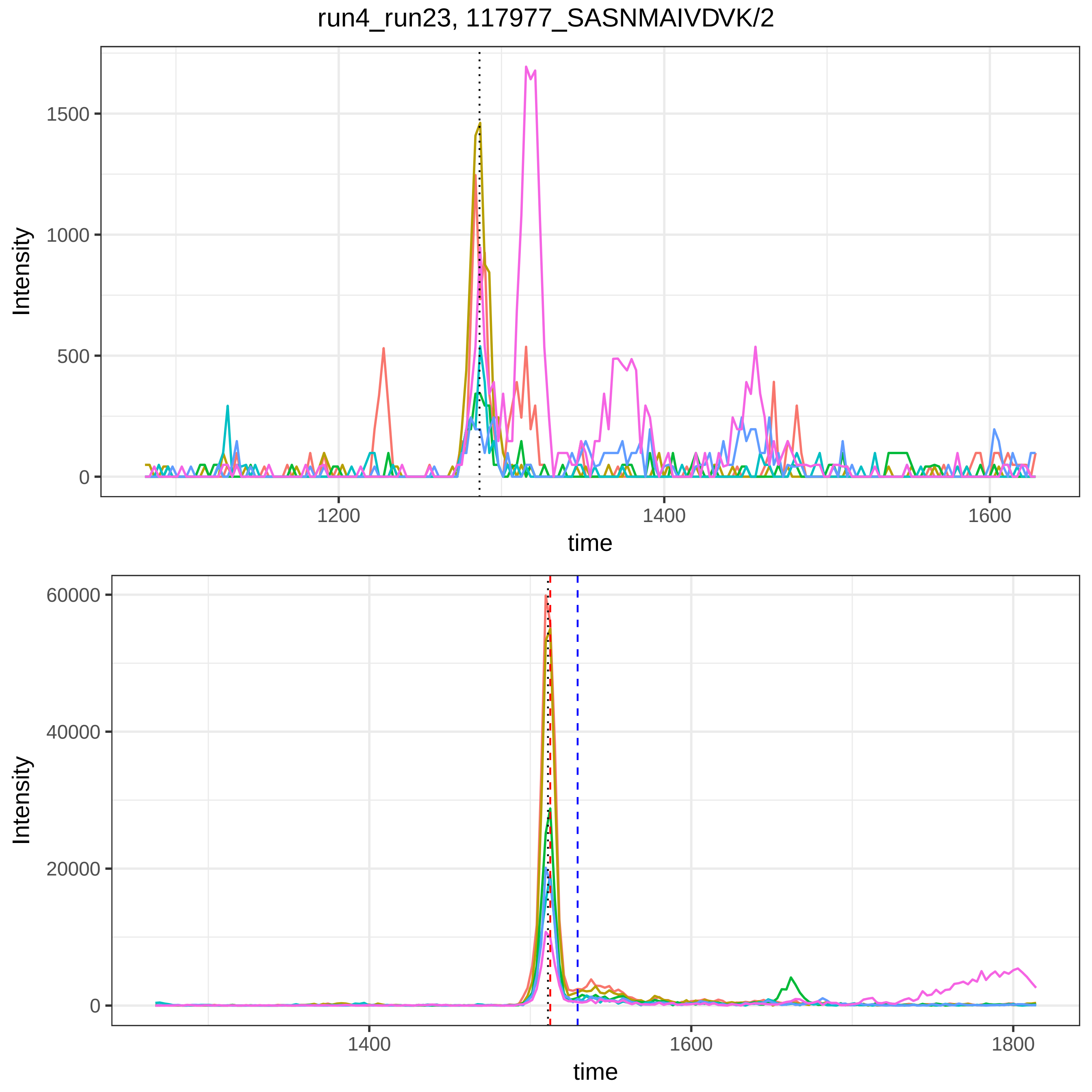

run4_run23, 117977_SASNMAIVDVK/2
Intensity
1500
1000
500
0
1200
1400
1600
time
Intensity
60000
40000
20000
0
1400
1600
1800
time

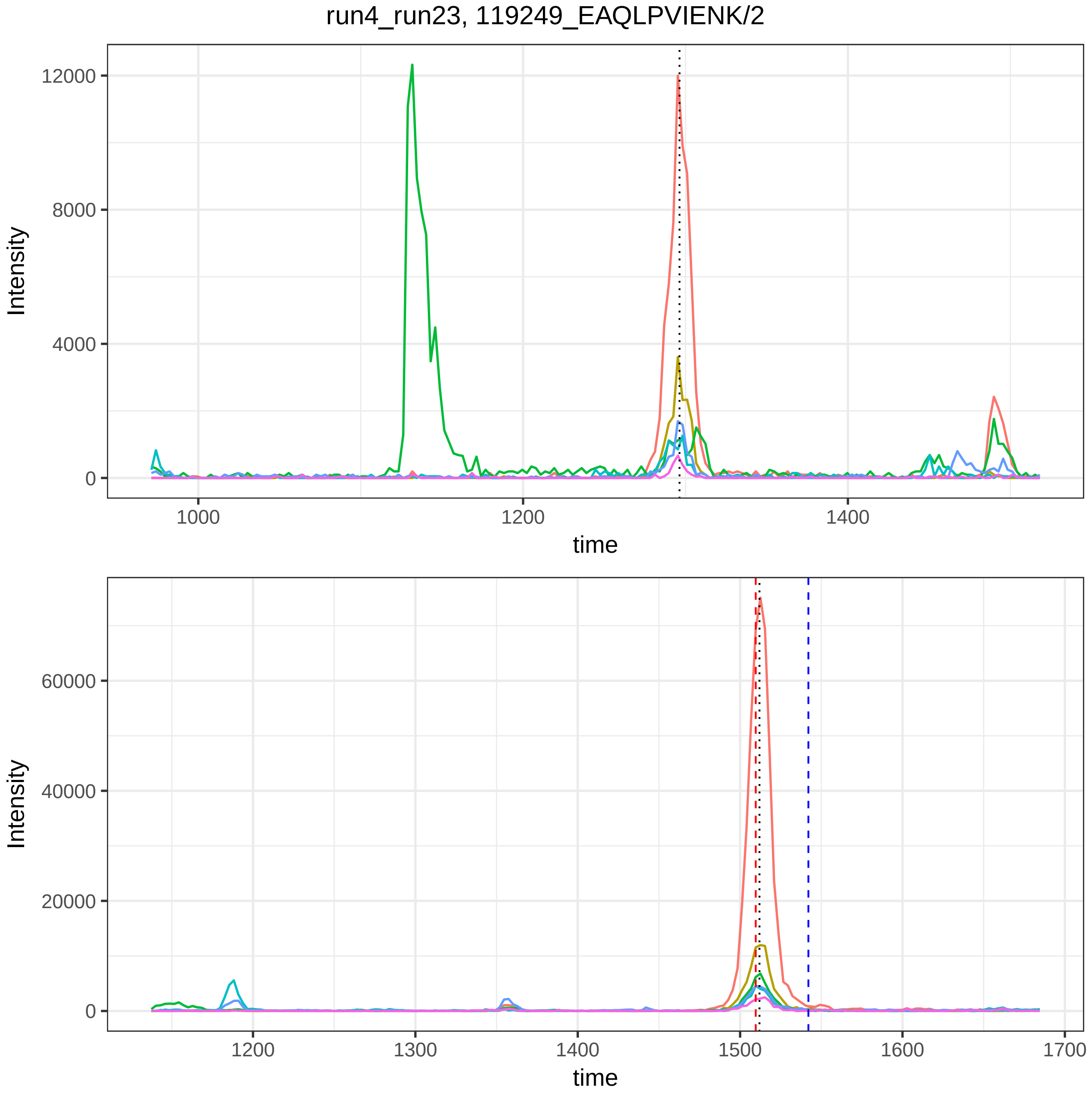
run4_run23, 119249_EAQLPVIENK/2
Intensity
time
Intensity
time

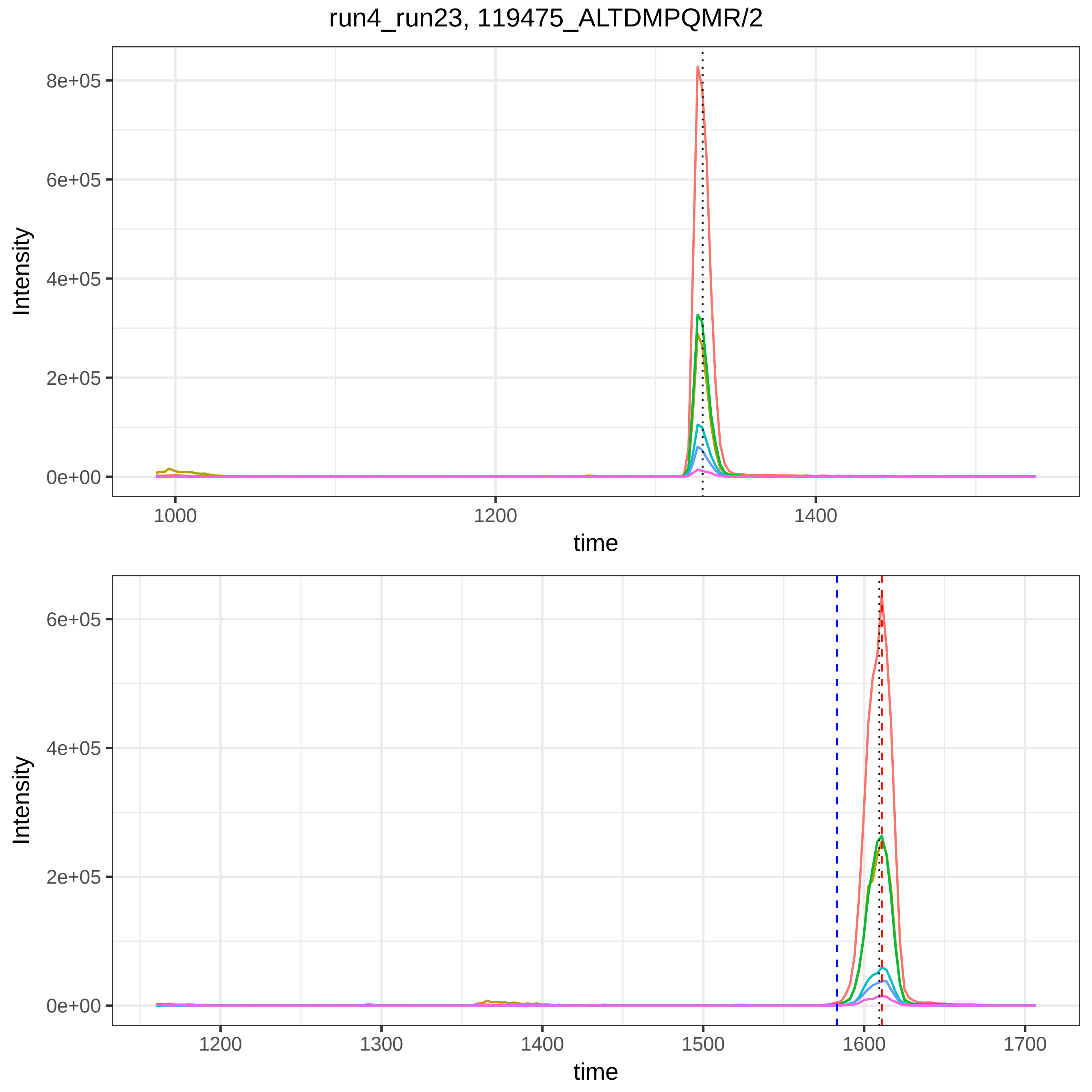

run4_run23, 119475_ALTDMPQMR/2
Intensity
8e+05
6e+05
4e+05
2e+05
0e+00
1000
1200
1400
time
Intensity
6e+05
4e+05
2e+05
0e+00
1200
1300
1400
1500
1600
1700
time

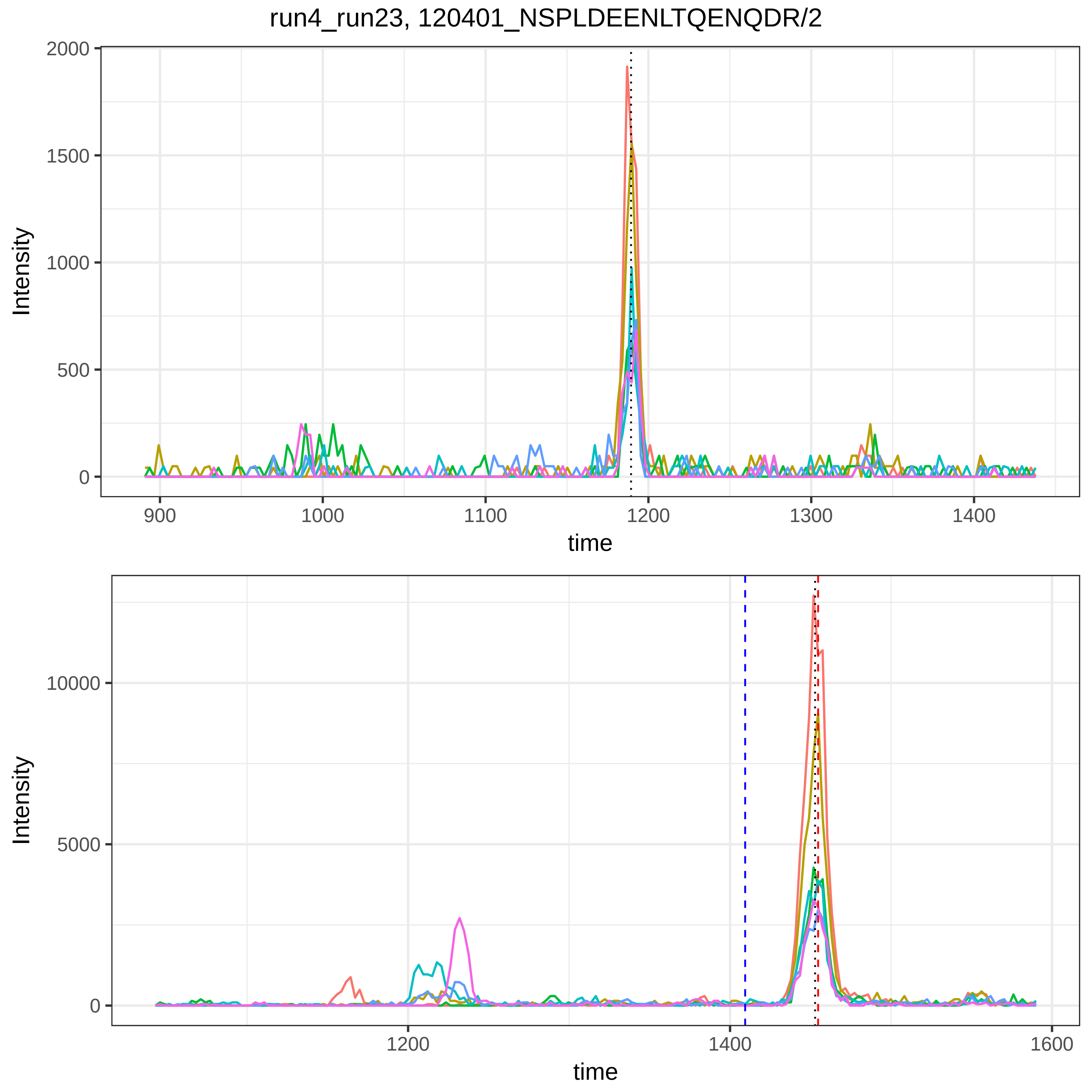
run4_run23, 120401_NSPLDEENLTQENQDR/2
Intensity
2000
1500
1000
500
0
900
1000
1100
1200
1300
1400
time
Intensity
10000
5000
0
1200
1400
1600
time

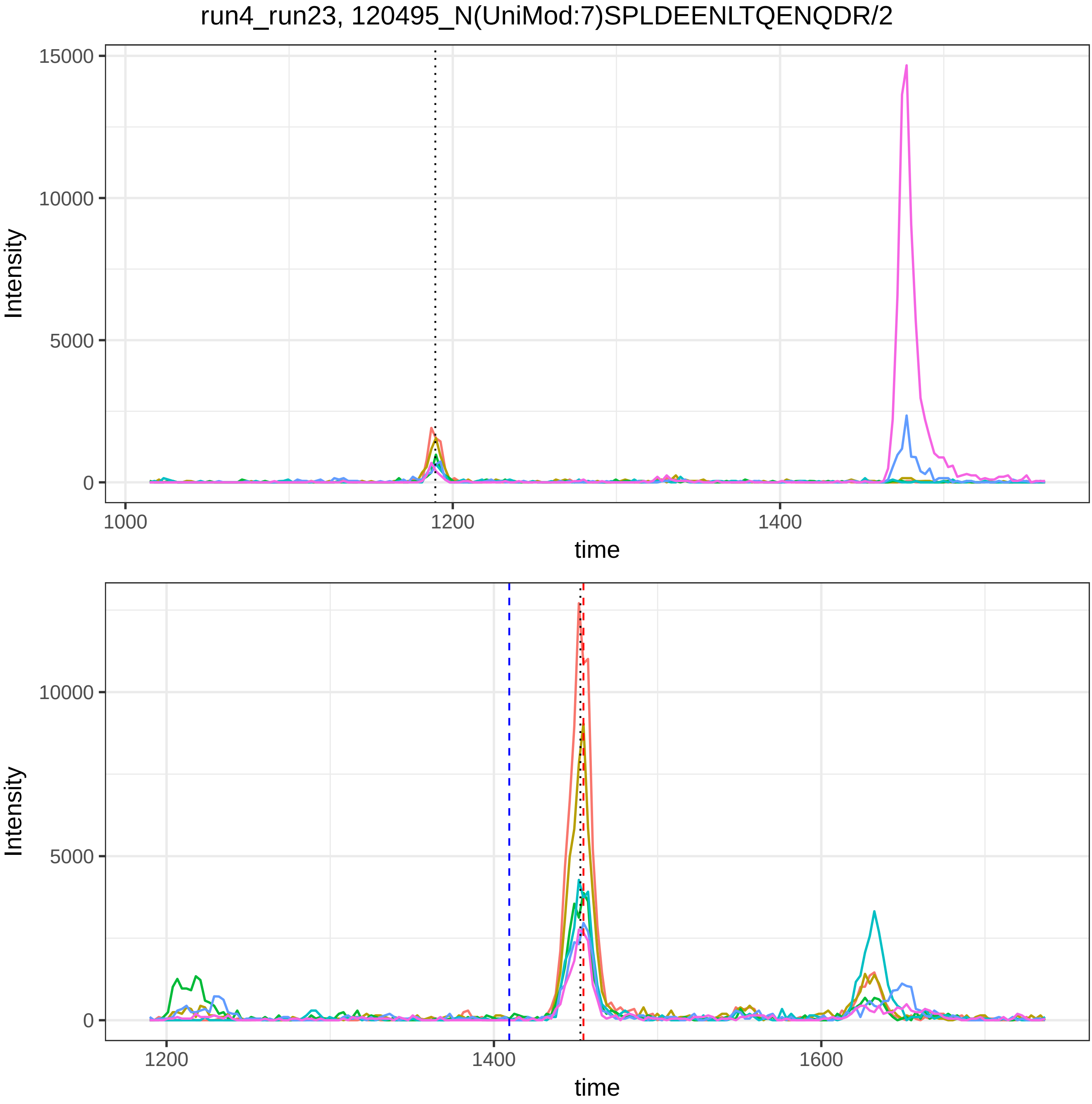
run4_run23, 120495_N(UniMod:7)SPLDEENLTQENQDR/2
Intensity
time
15000
10000
5000
0
1000
1200
1400
1600

run4_run23, 120686_ELAAQTIK/2

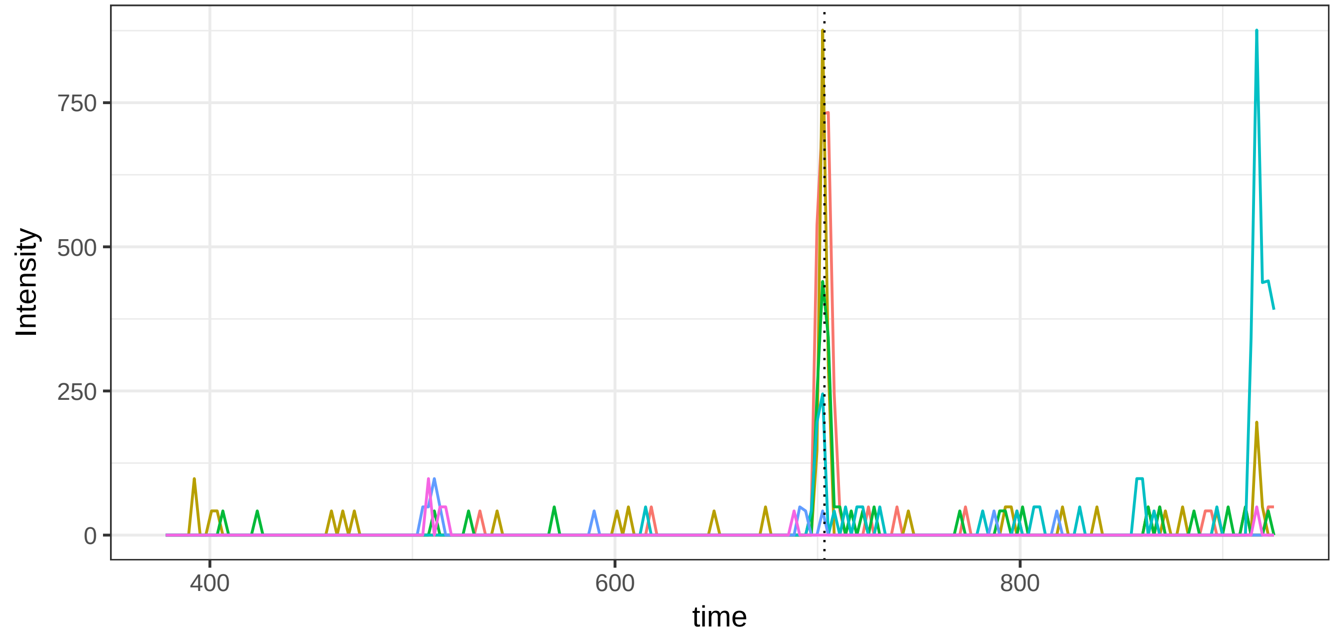


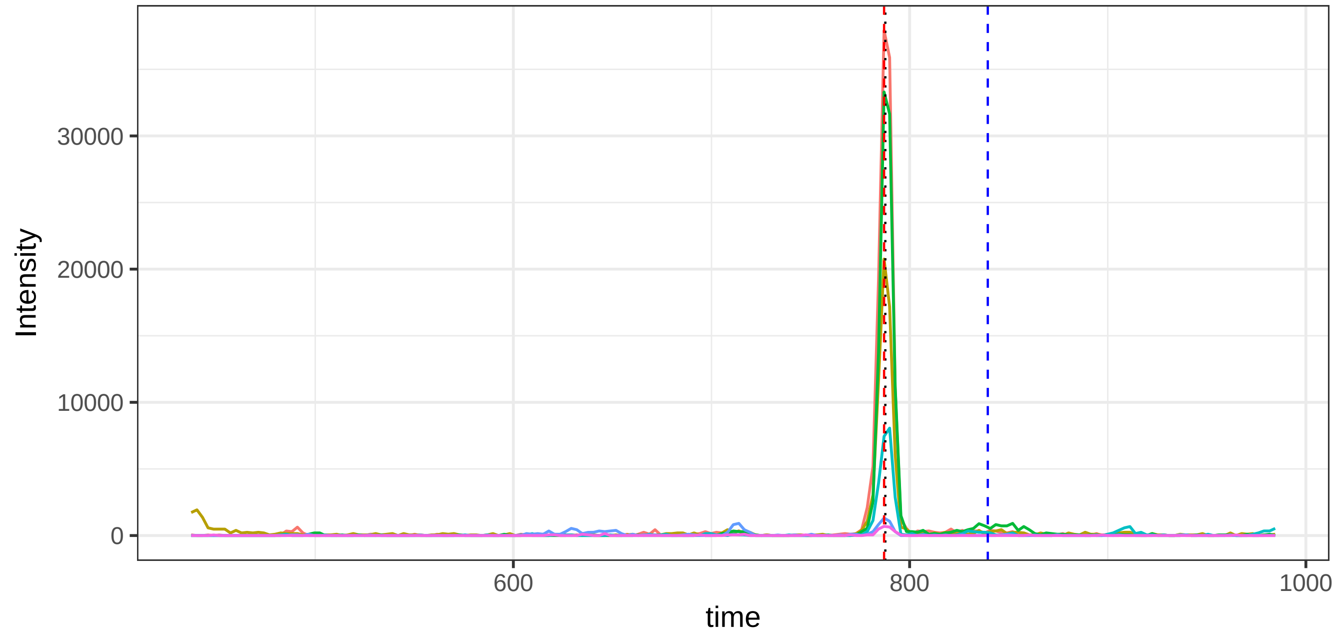

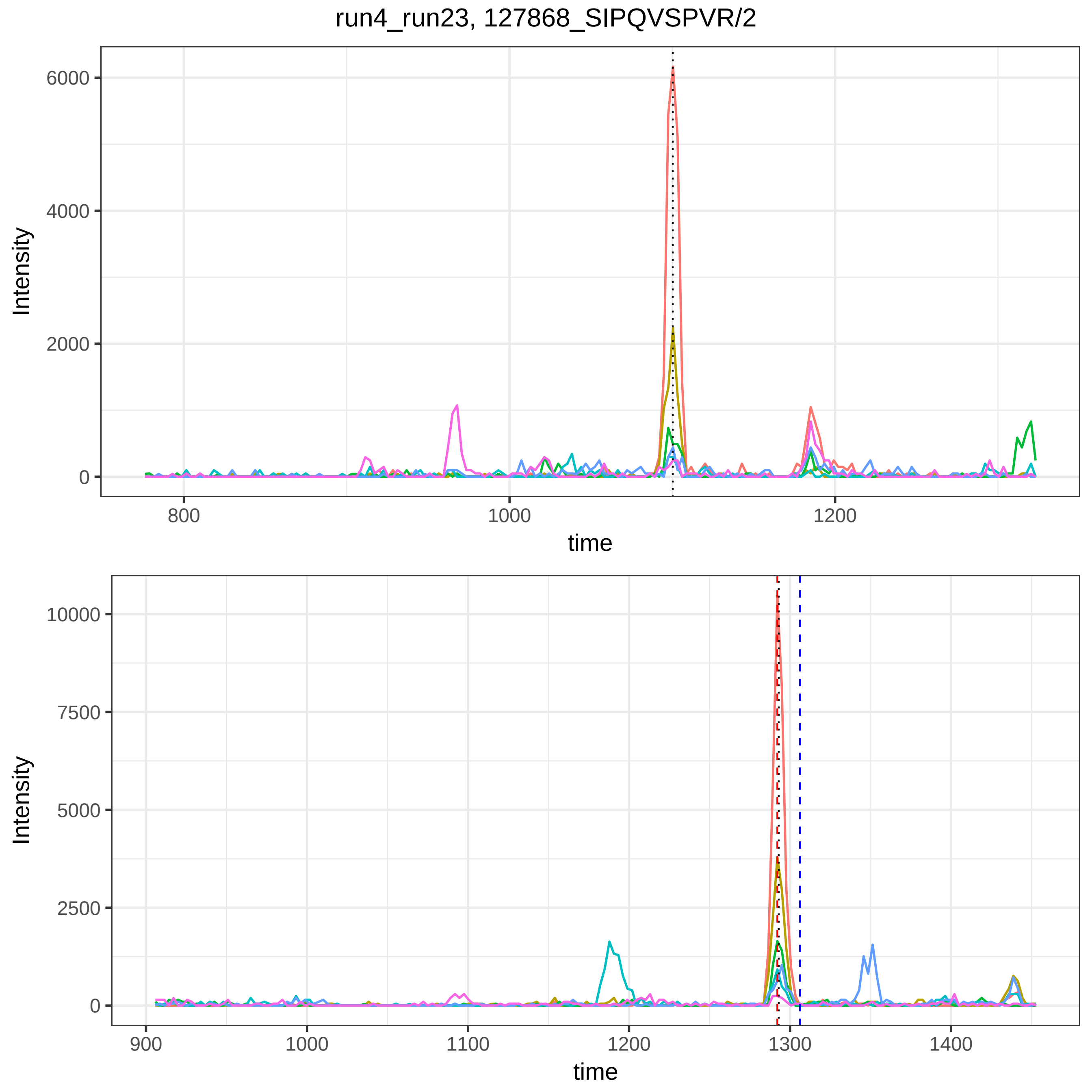
run4_run23, 127868_SIPQVSPVR/2
Intensity
6000
4000
2000
0
800
1000
1200
time
Intensity
10000
7500
5000
2500
0
900
1000
1100
1200
1300
1400
time

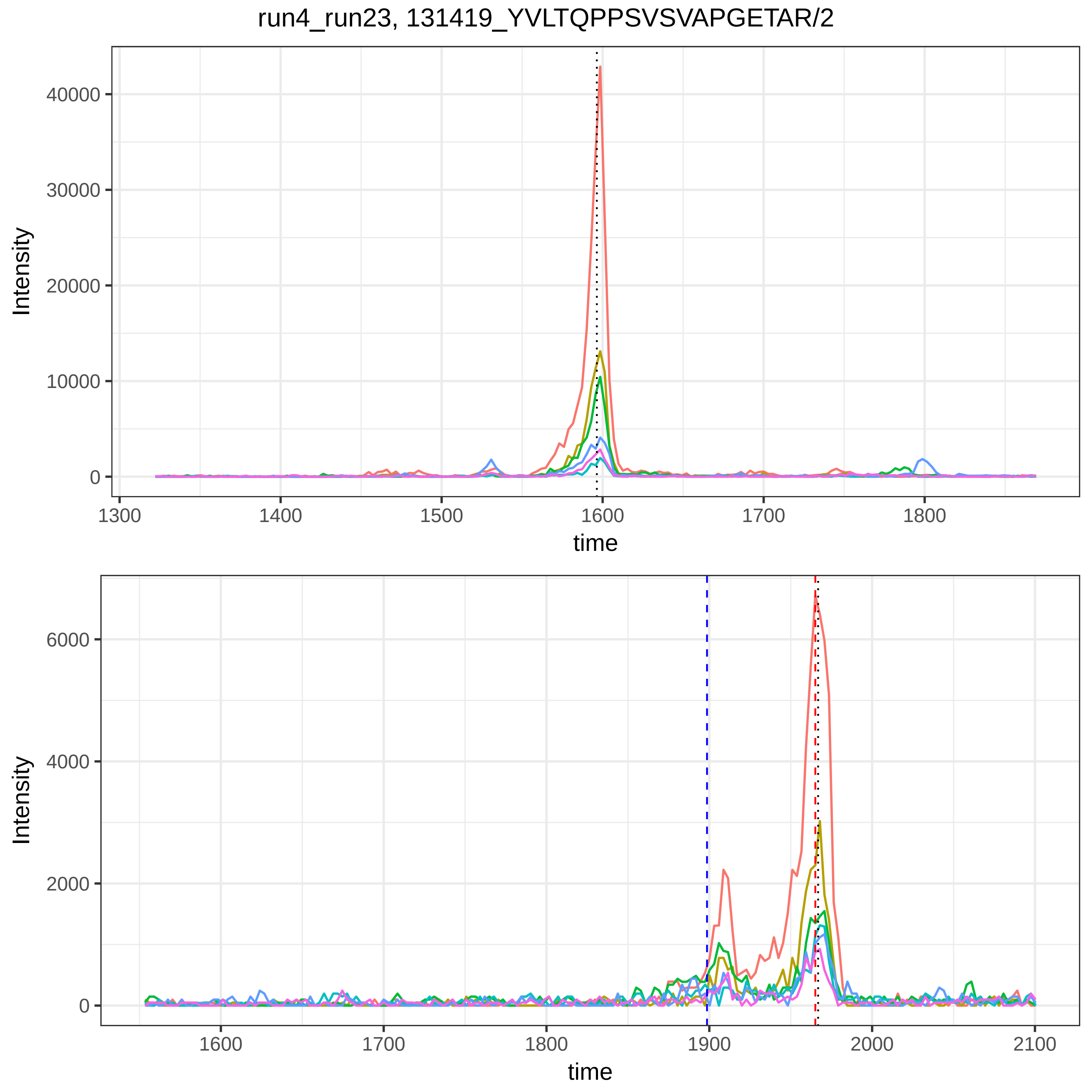
run4_run23, 131419_YVLTQPPSVSVAPGETAR/2
Intensity
time
40000
30000
20000
10000
0
1300
1400
1500
1600
1700
1800
6000
4000
2000
1600
1700
1800
1900
2000
2100

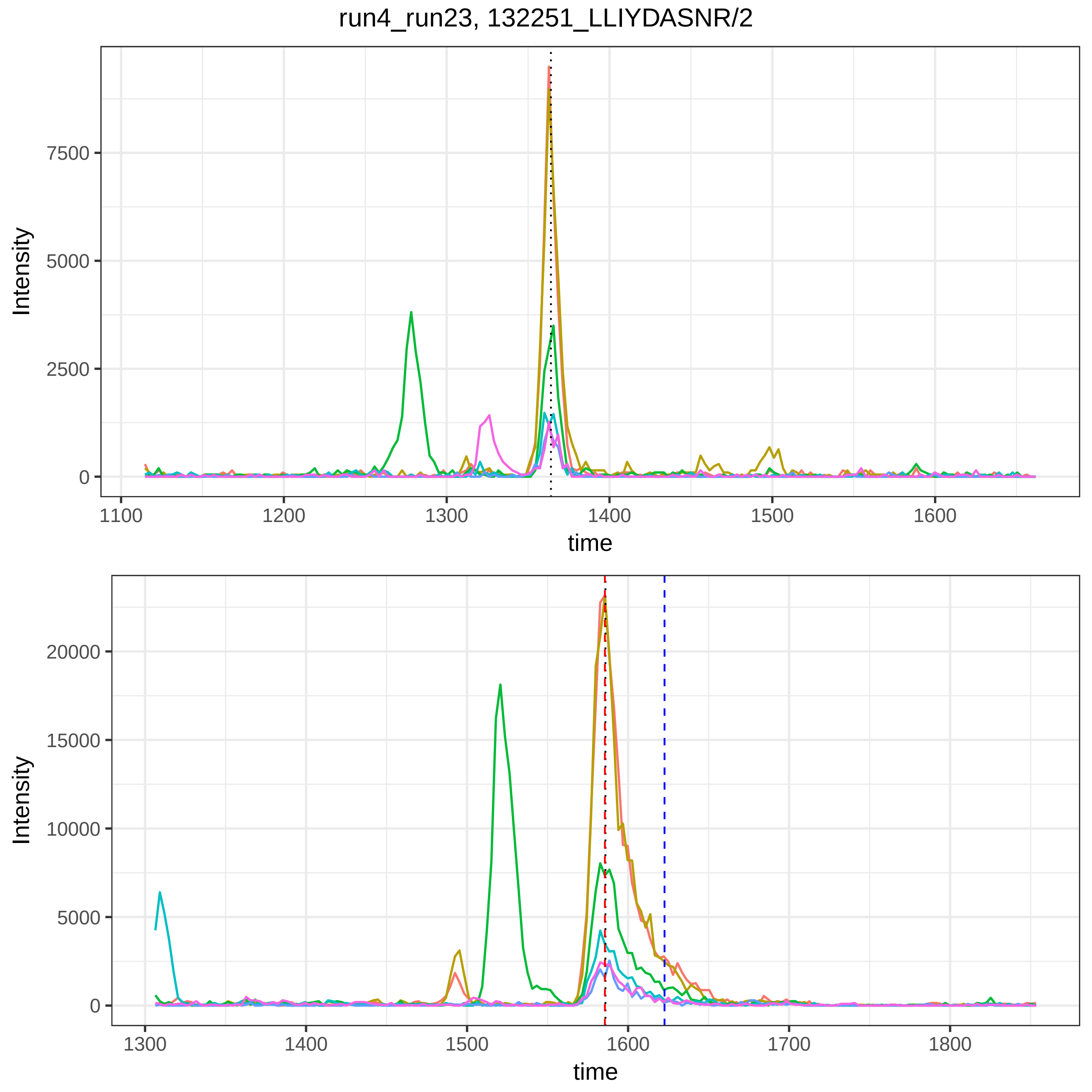
run4_run23, 132251_LLIYDASNR/2
Intensity
time
0
2500
5000
7500
1100
1200
1300
1400
1500
1600
0
5000
10000
15000
20000
1300
1400
1500
1600
1700
1800

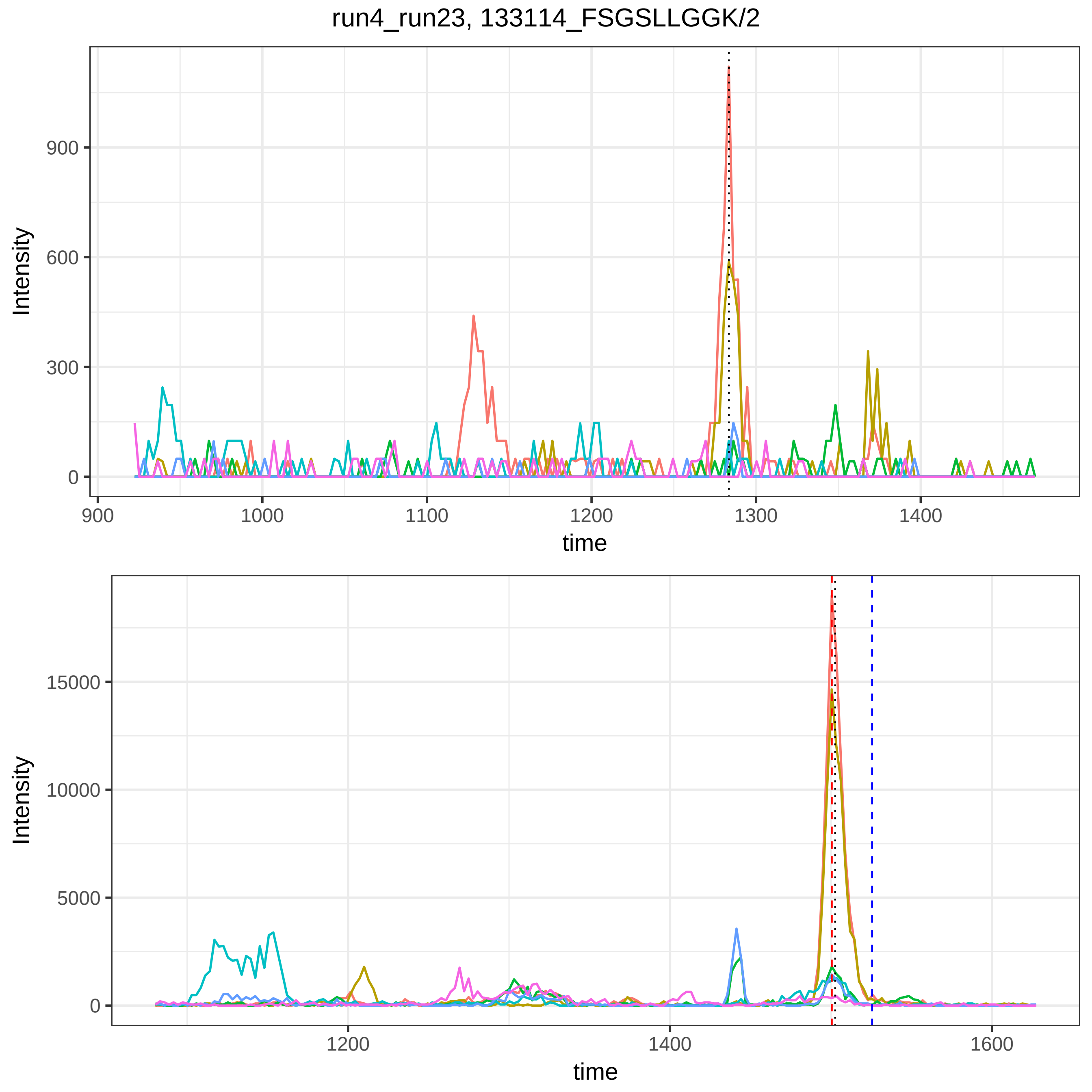

run4_run23, 133114_FSGSLLGGK/2
Intensity
900
600
300
0
900
1000
1100
1200
1300
1400
time
Intensity
15000
10000
5000
0
1200
1400
1600
time

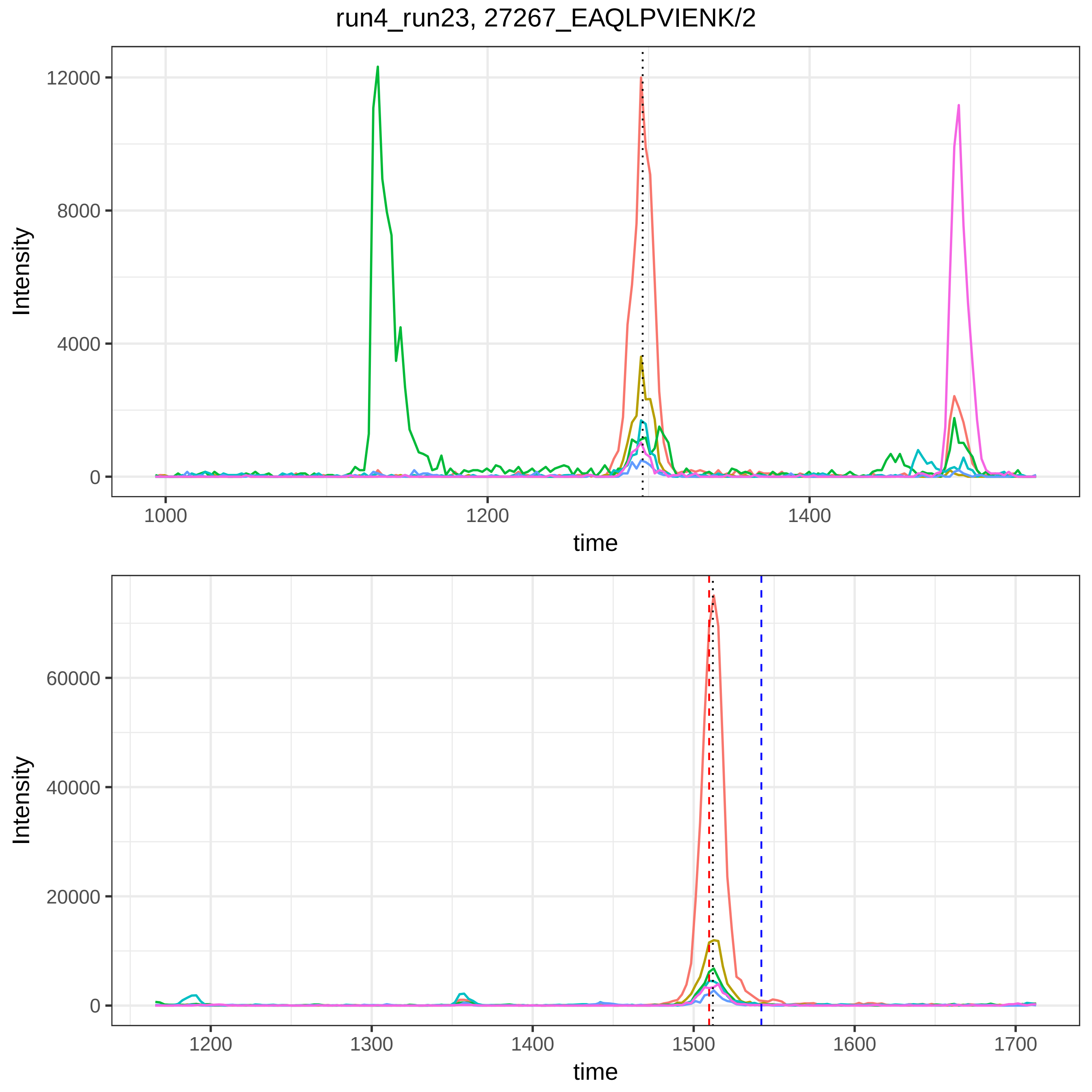

run4_run23, 27267_EAQLPVIENK/2
Intensity
12000
8000
4000
0
1000
1200
1400
time
Intensity
60000
40000
20000
0
1200
1300
1400
1500
1600
1700
time

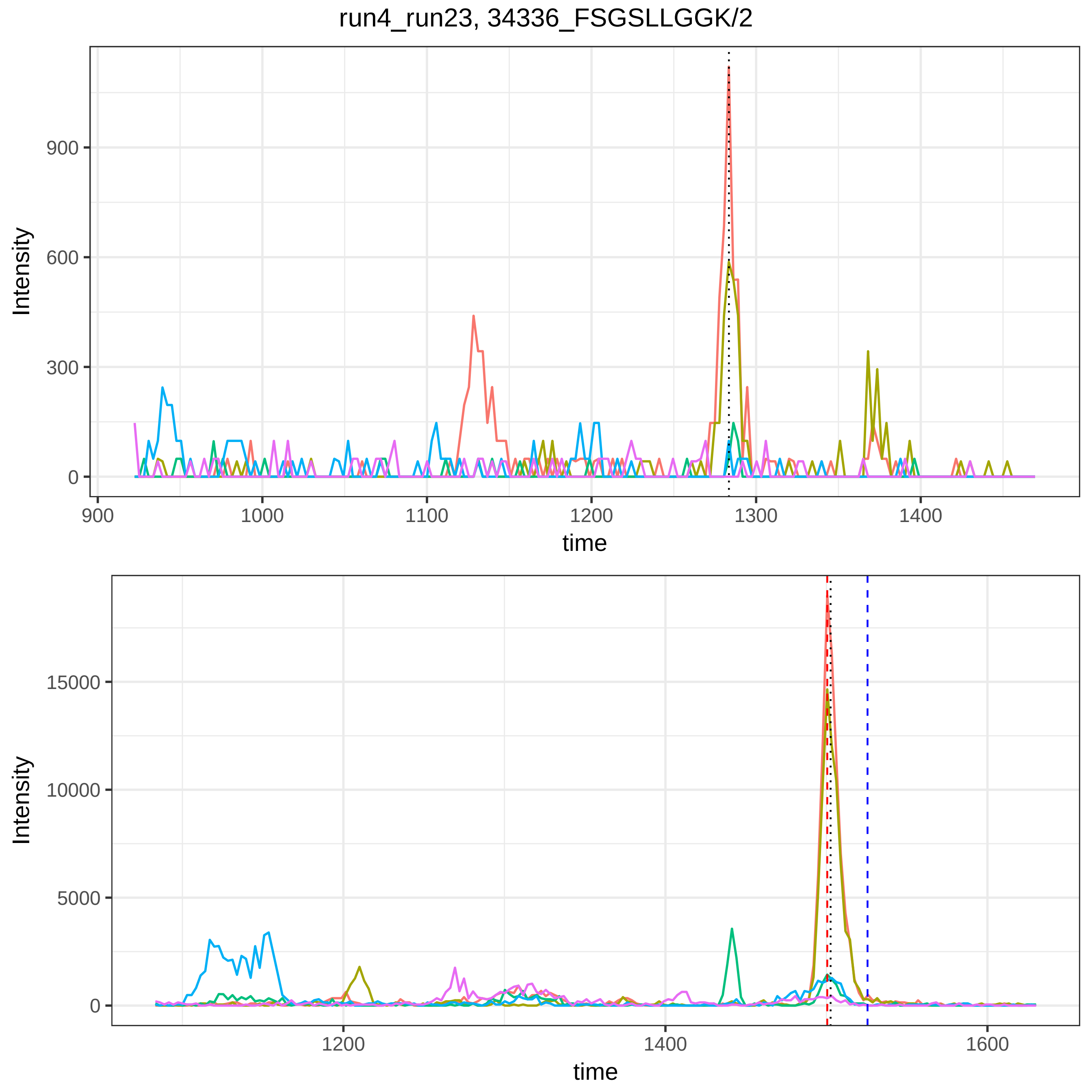
run4_run23, 34336_FSGSLLGGK/2
Intensity
900
600
300
0
900
1000
1100
1200
1300
1400
time
Intensity
15000
10000
5000
0
1200
1400
1600
time

run4_run23, 3842_LLIYGATSR/2

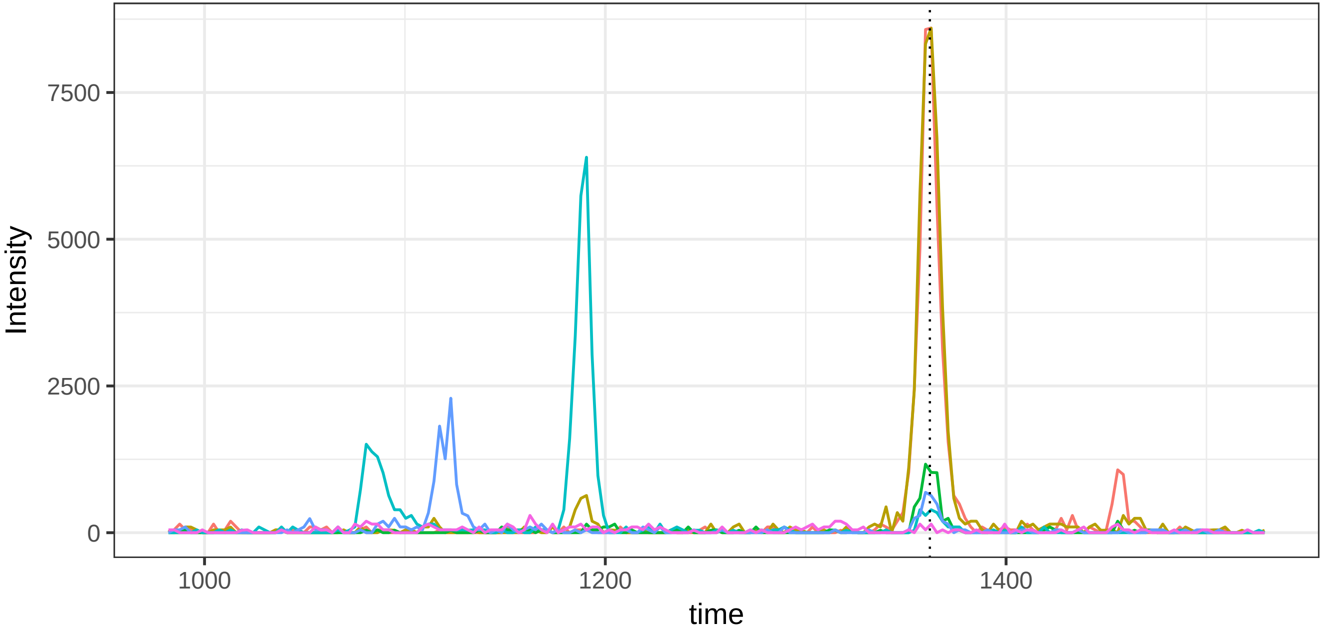


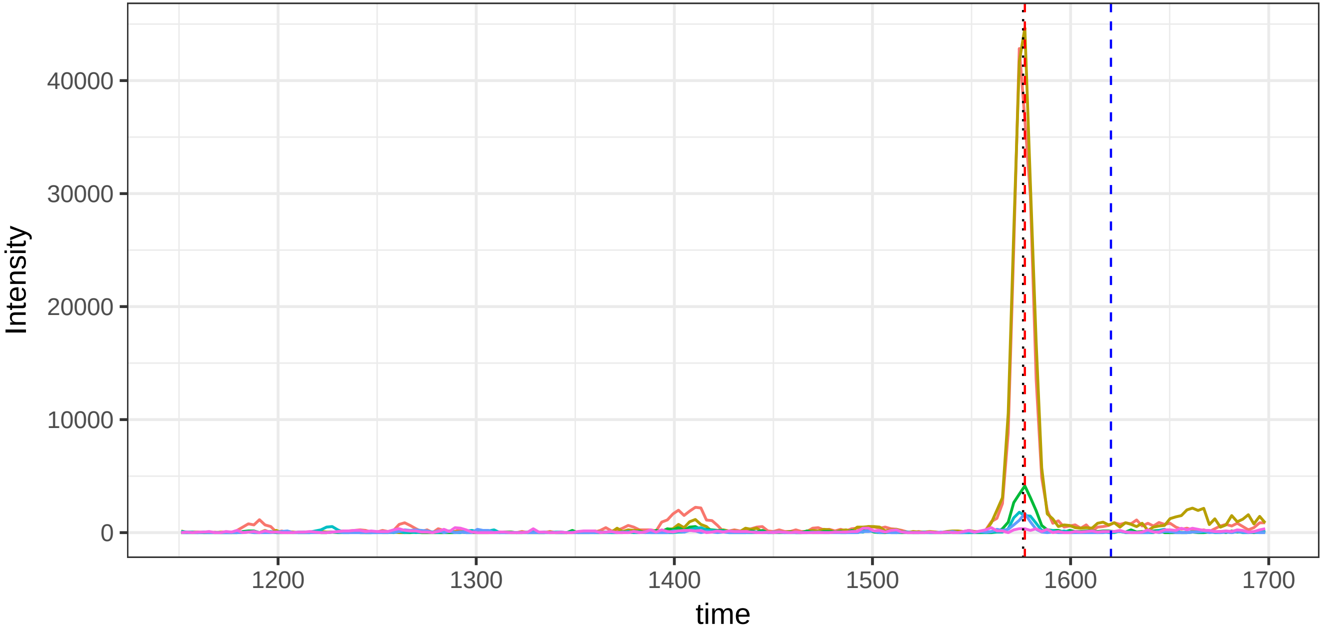

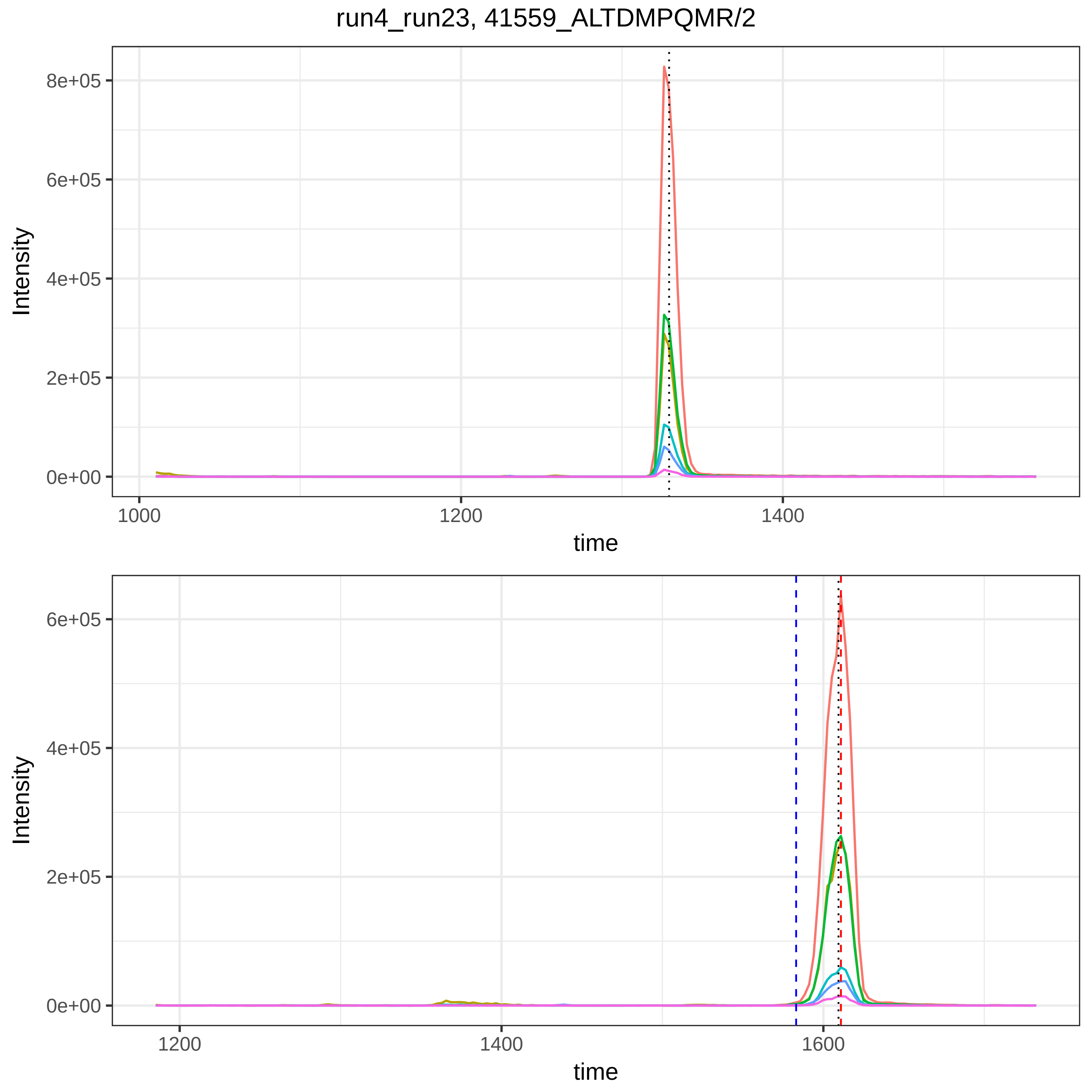

run4_run23, 41559_ALTDMPQMR/2
Intensity
time
8e+05
6e+05
4e+05
2e+05
0e+00
1000
1200
1400
1600

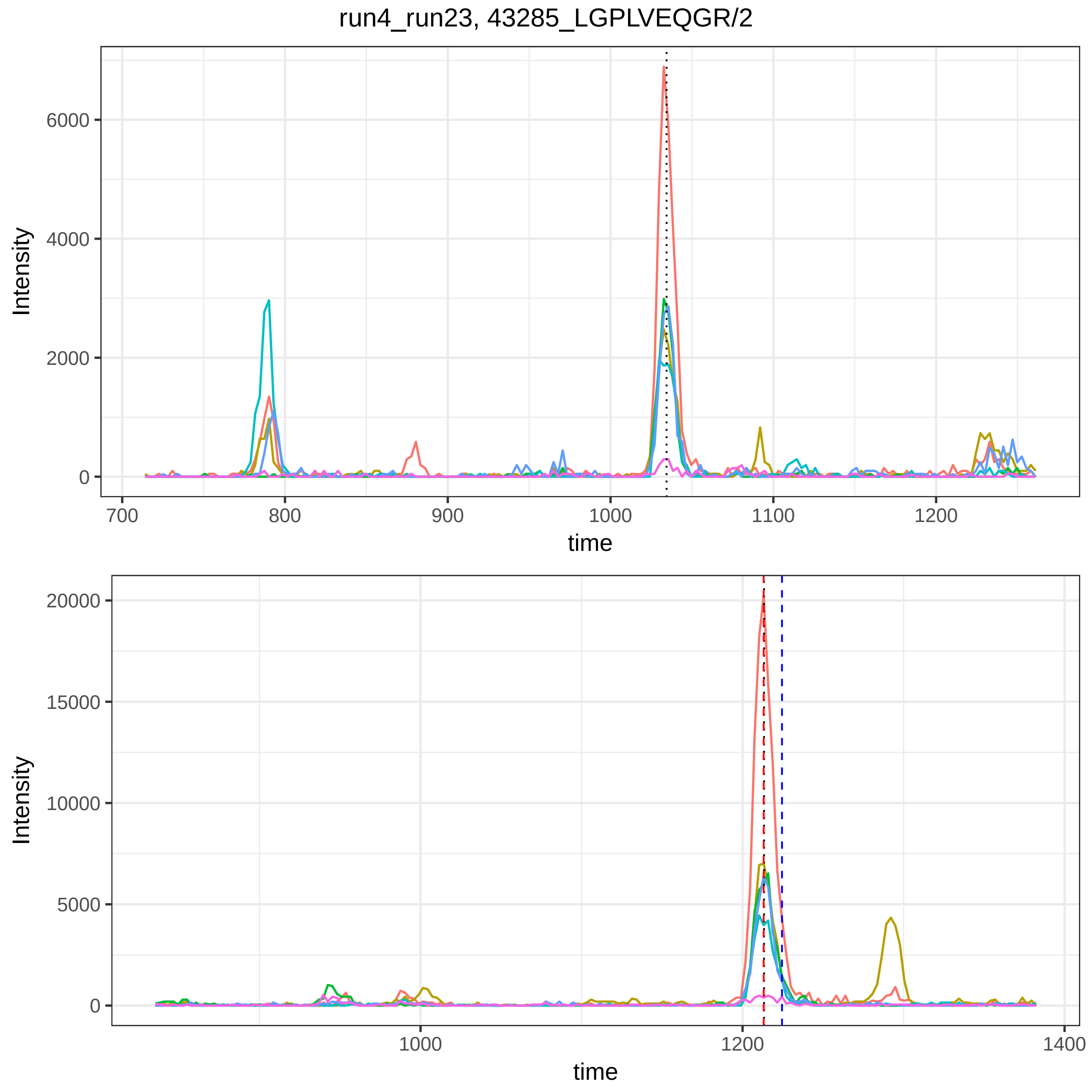

run4_run23, 43285_LGPLVEQGR/2
Intensity
0
2000
4000
6000
700
800
900
1000
1100
1200
time
Intensity
0
5000
10000
15000
20000
1000
1200
1400
time

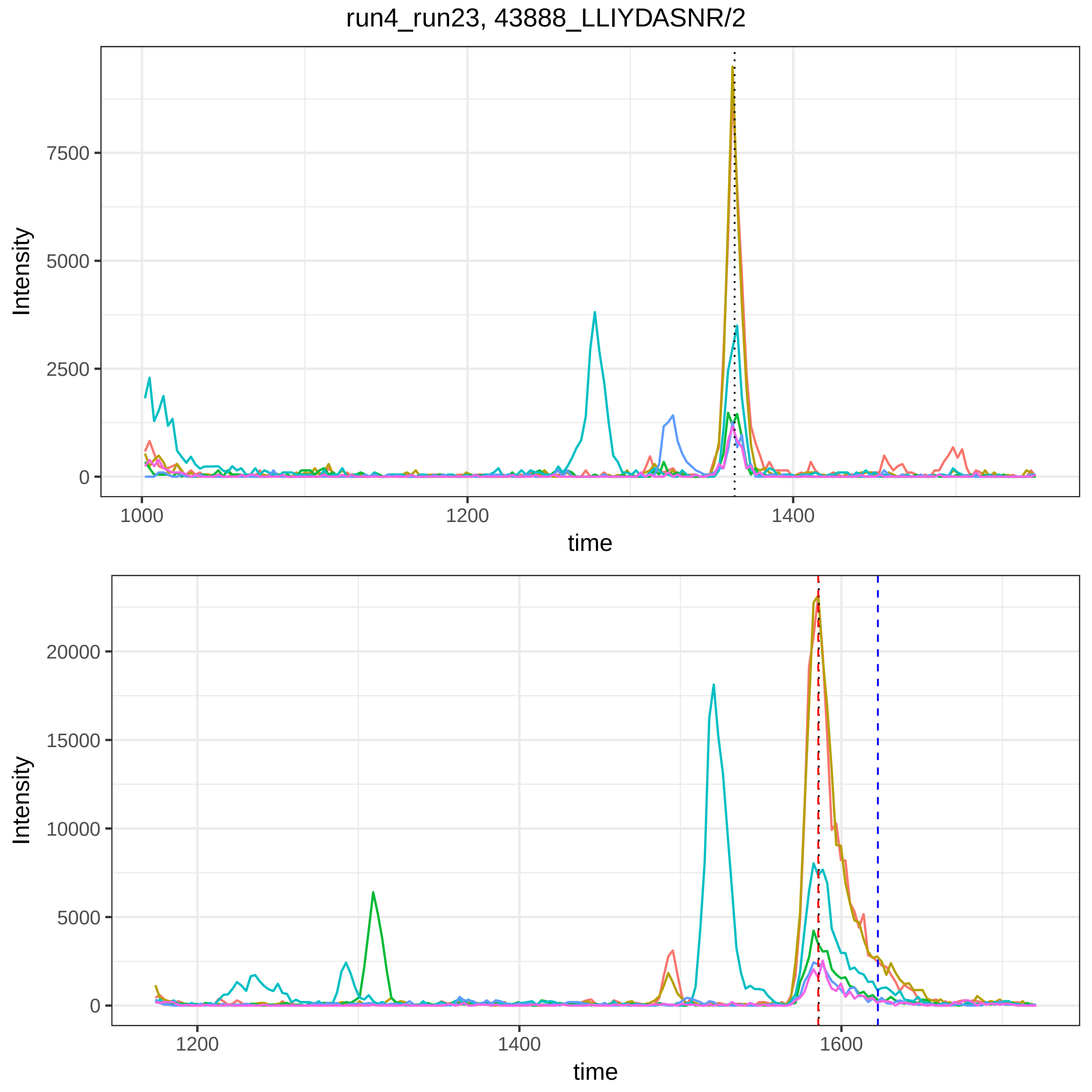

run4_run23, 43888_LLIYDASNR/2
Intensity
7500
5000
2500
0
1000
1200
1400
time
Intensity
20000
15000
10000
5000
0
1200
1400
1600
time

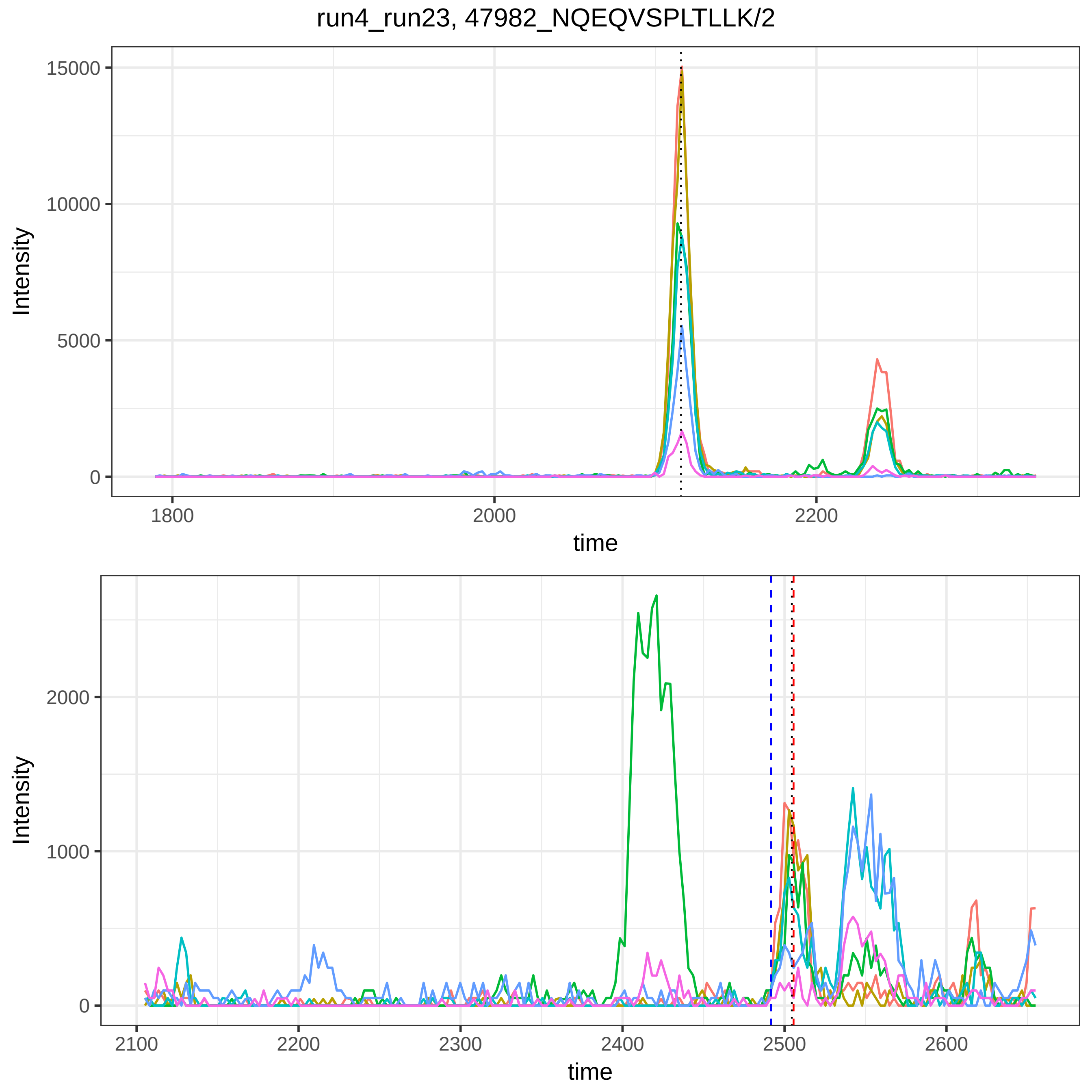
run4_run23, 47982_NQEQVSPLTLLK/2
Intensity
15000
10000
5000
0
1800
2000
2200
time
Intensity
2000
1000
0
2100
2200
2300
2400
2500
2600
time

run4_run23, 4844_YLQEIYNSNNQK/2

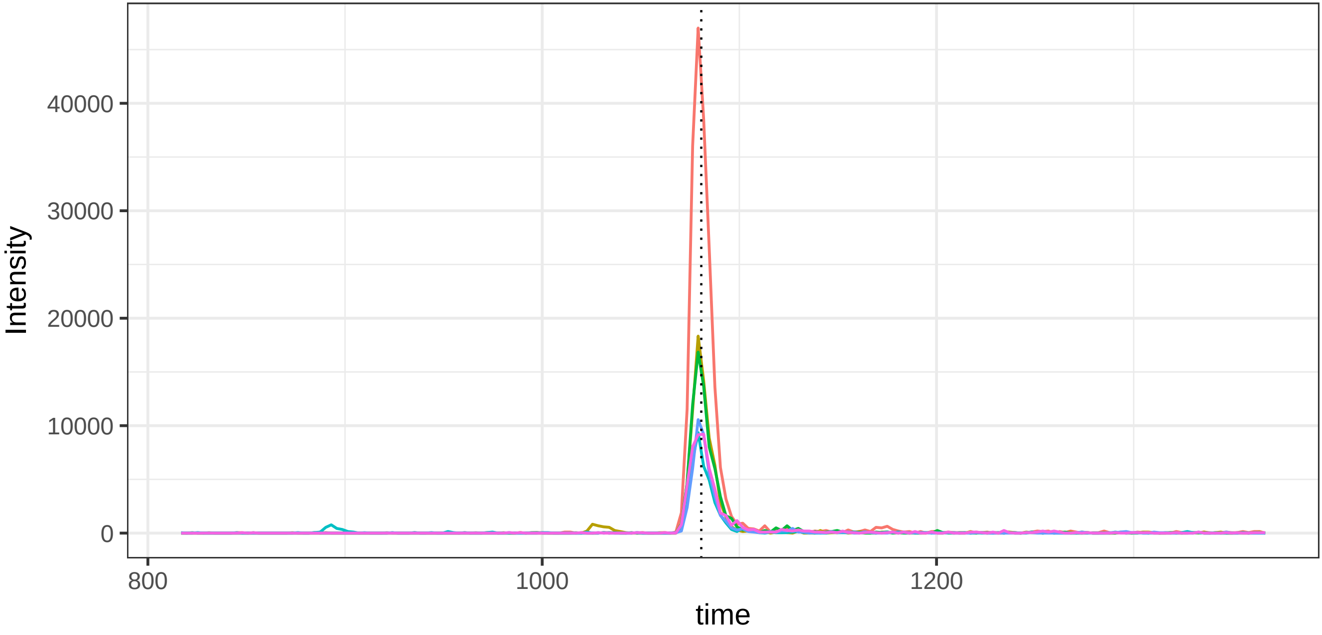


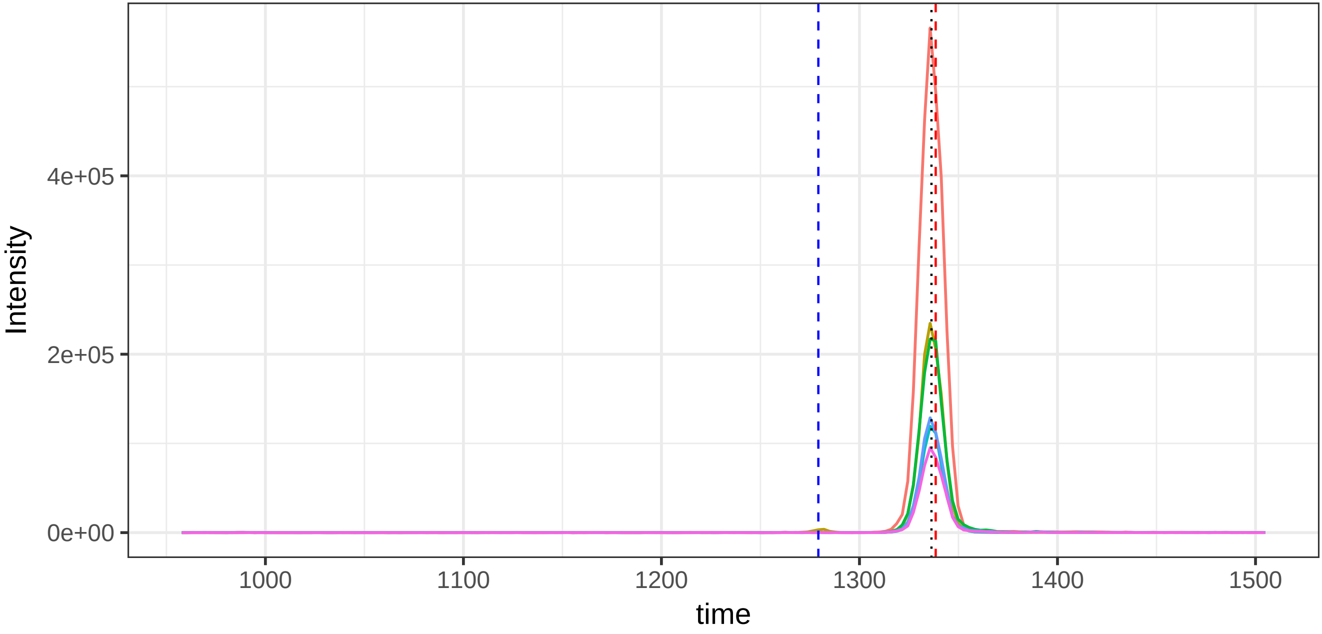

run4_run23, 51391_SASNMAIVDVK/2

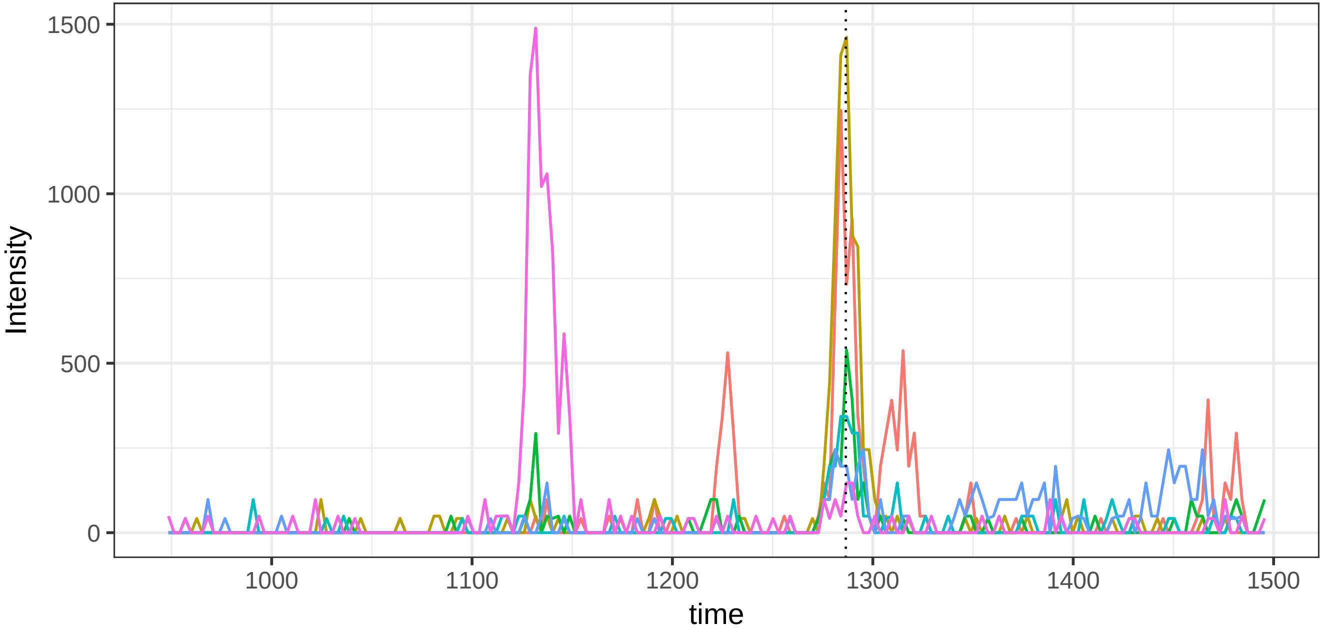


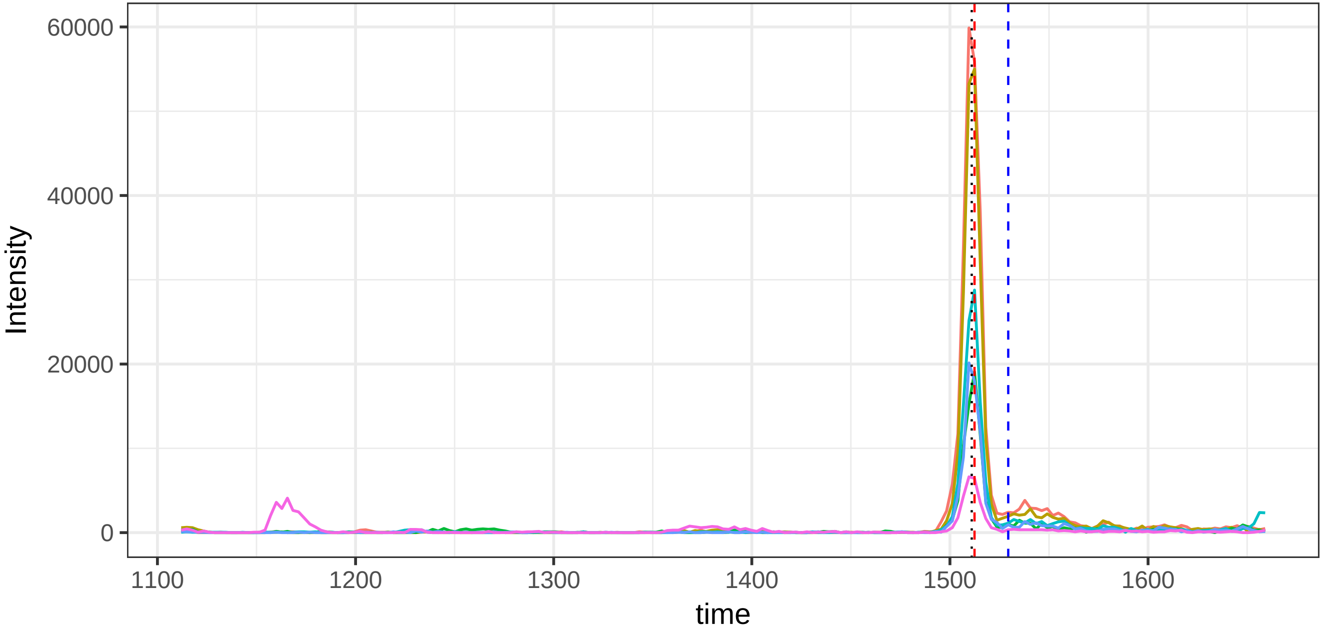

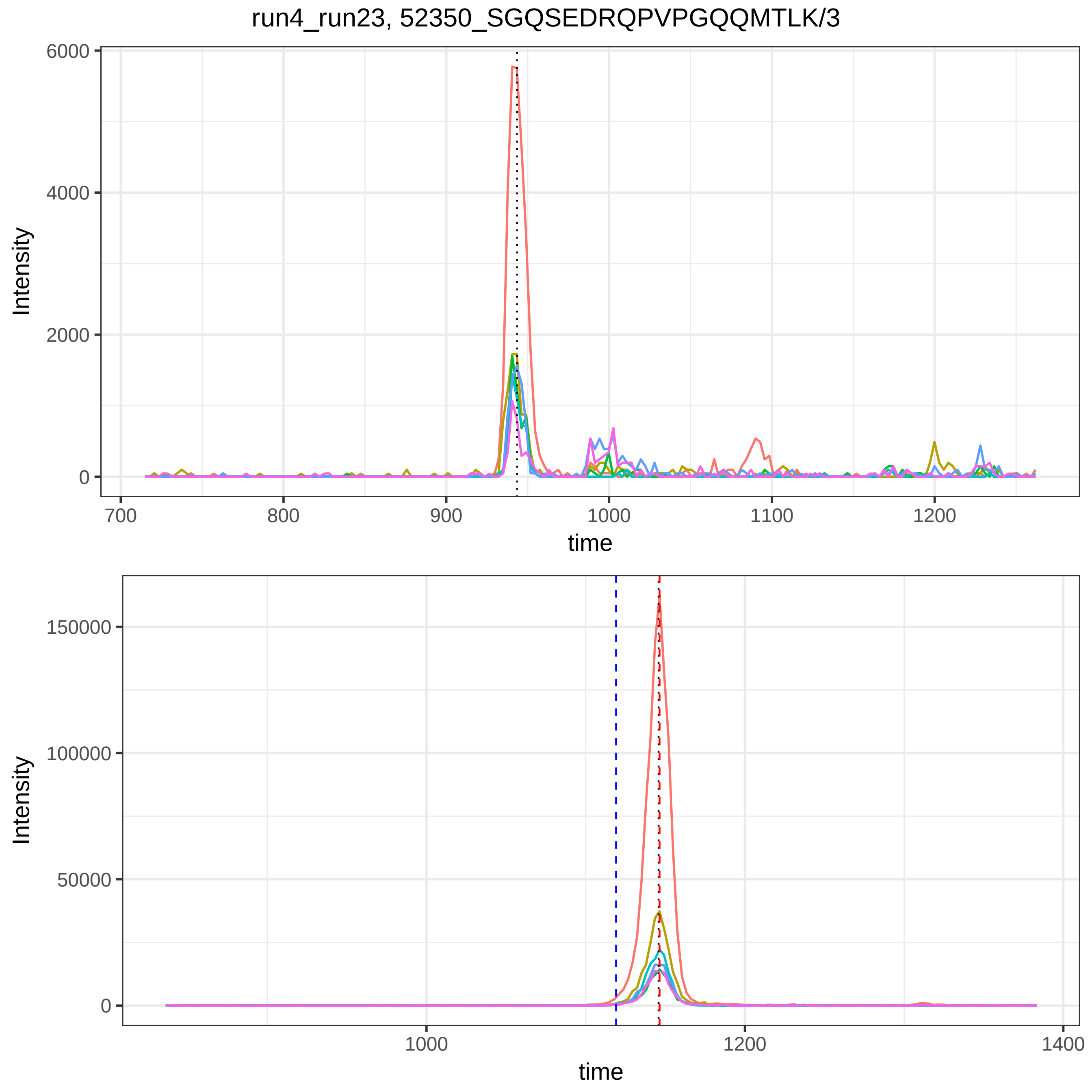

run4_run23, 52350_SGQSEDRQPVPGQQMTLK/3
Intensity
time
Intensity
time

run4_run23, 5357_YVLTQPPSVSVAPGETAR/2

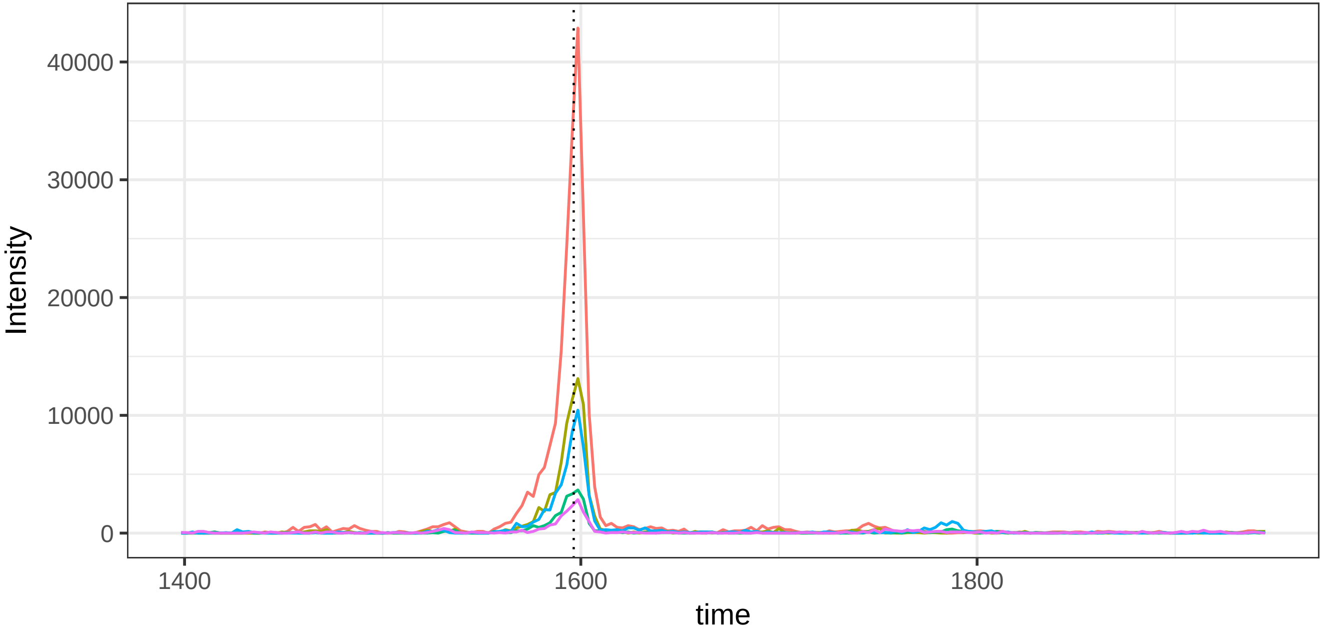


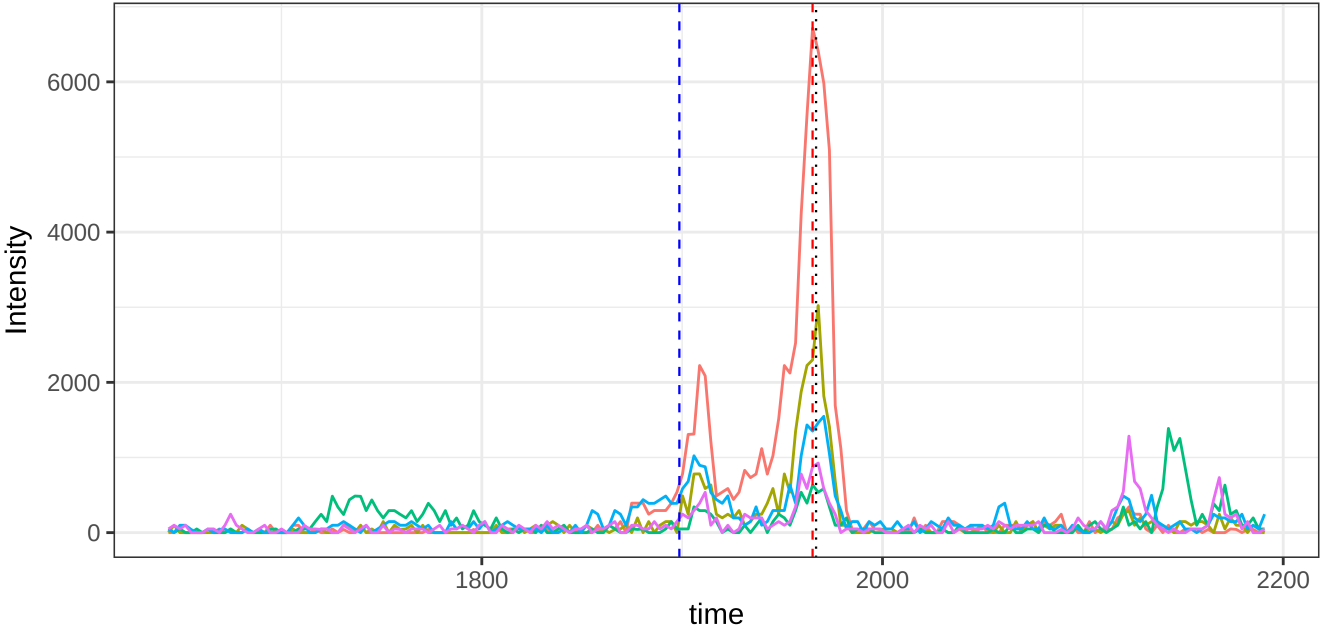

run4_run23, 114992_TEVIPPLIENR/2

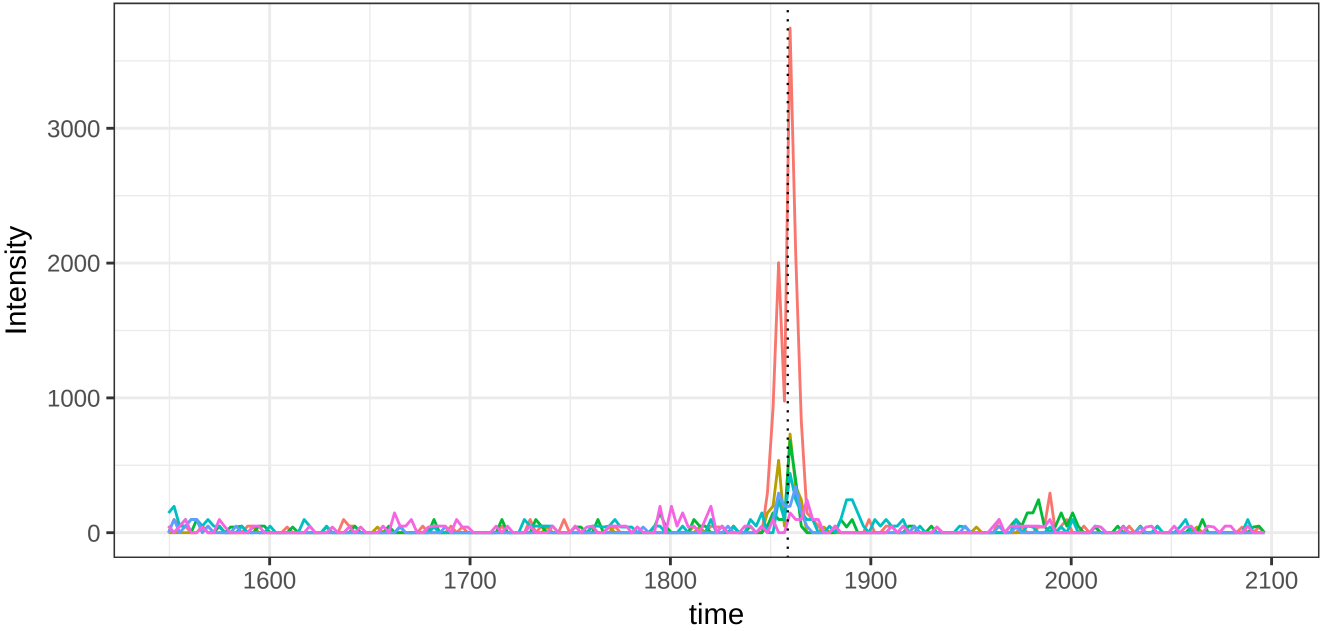


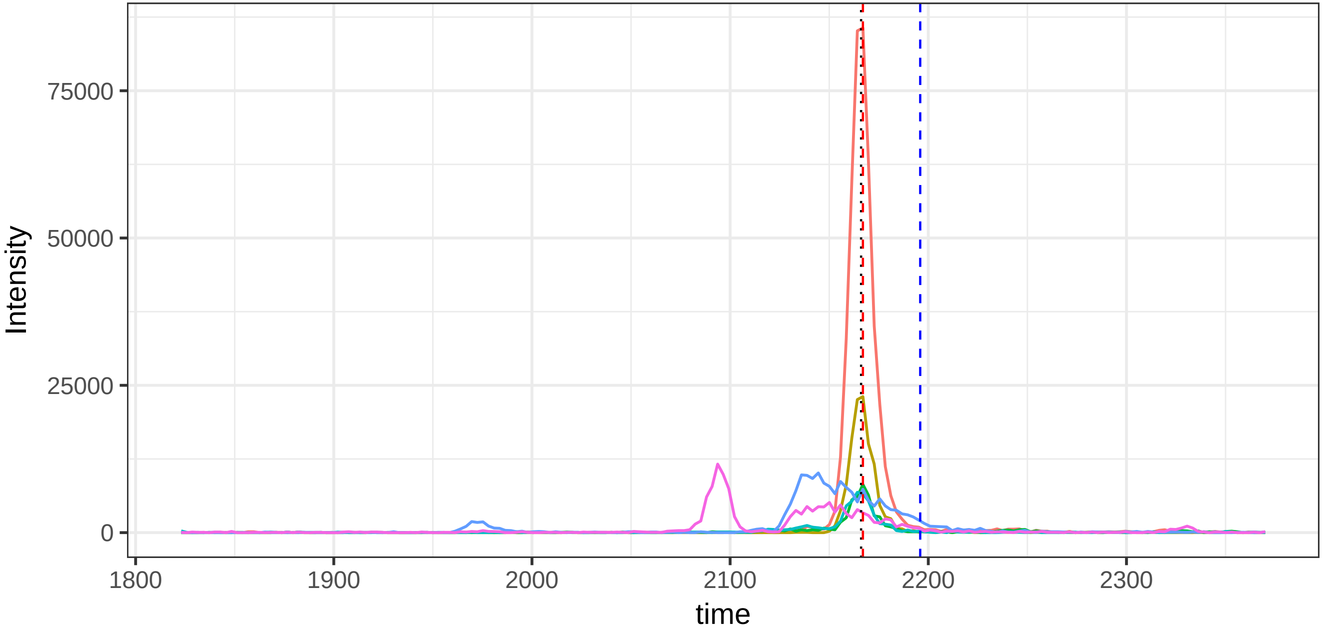

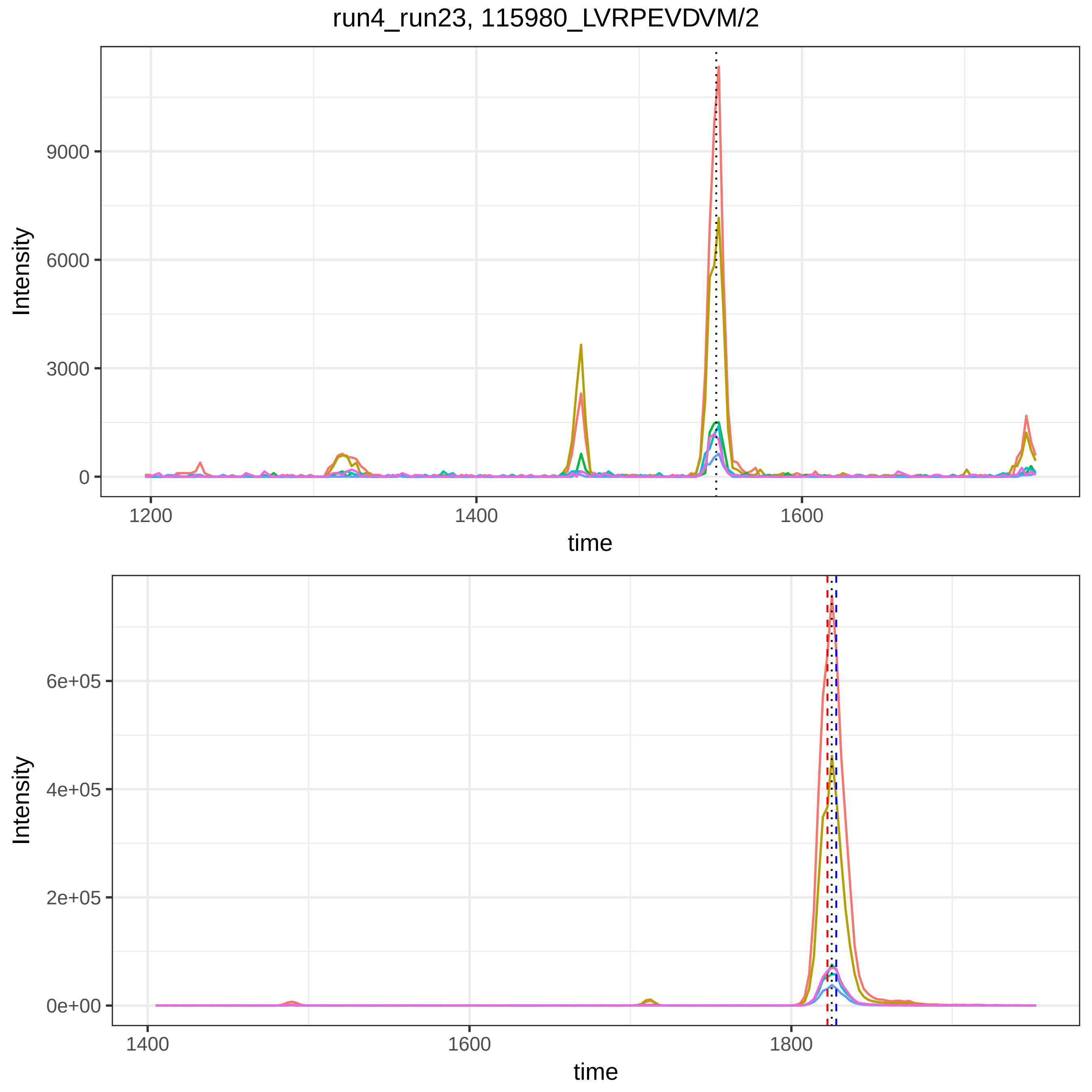
run4_run23, 115980_LVRPEVDVM/2
Intensity
9000
6000
3000
0
1200
1400
1600
time
Intensity
6e+05
4e+05
2e+05
0e+00
1400
1600
1800
time

run4_run23, 116326_EVTEFAK/2

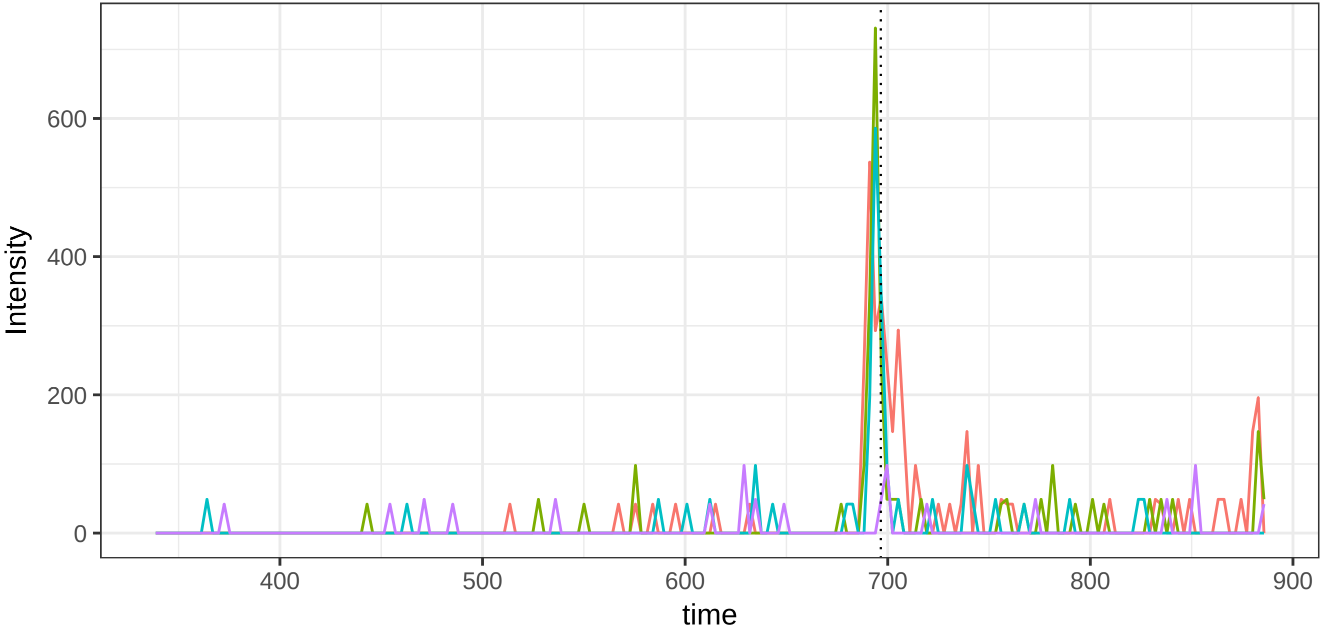


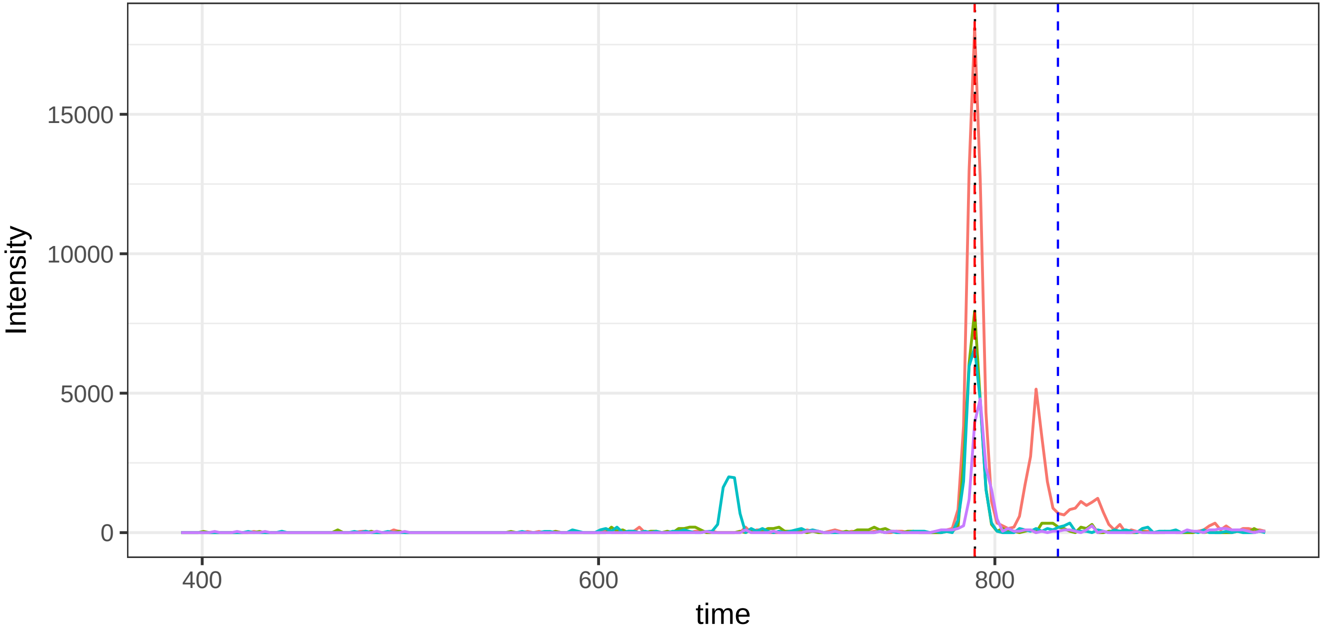

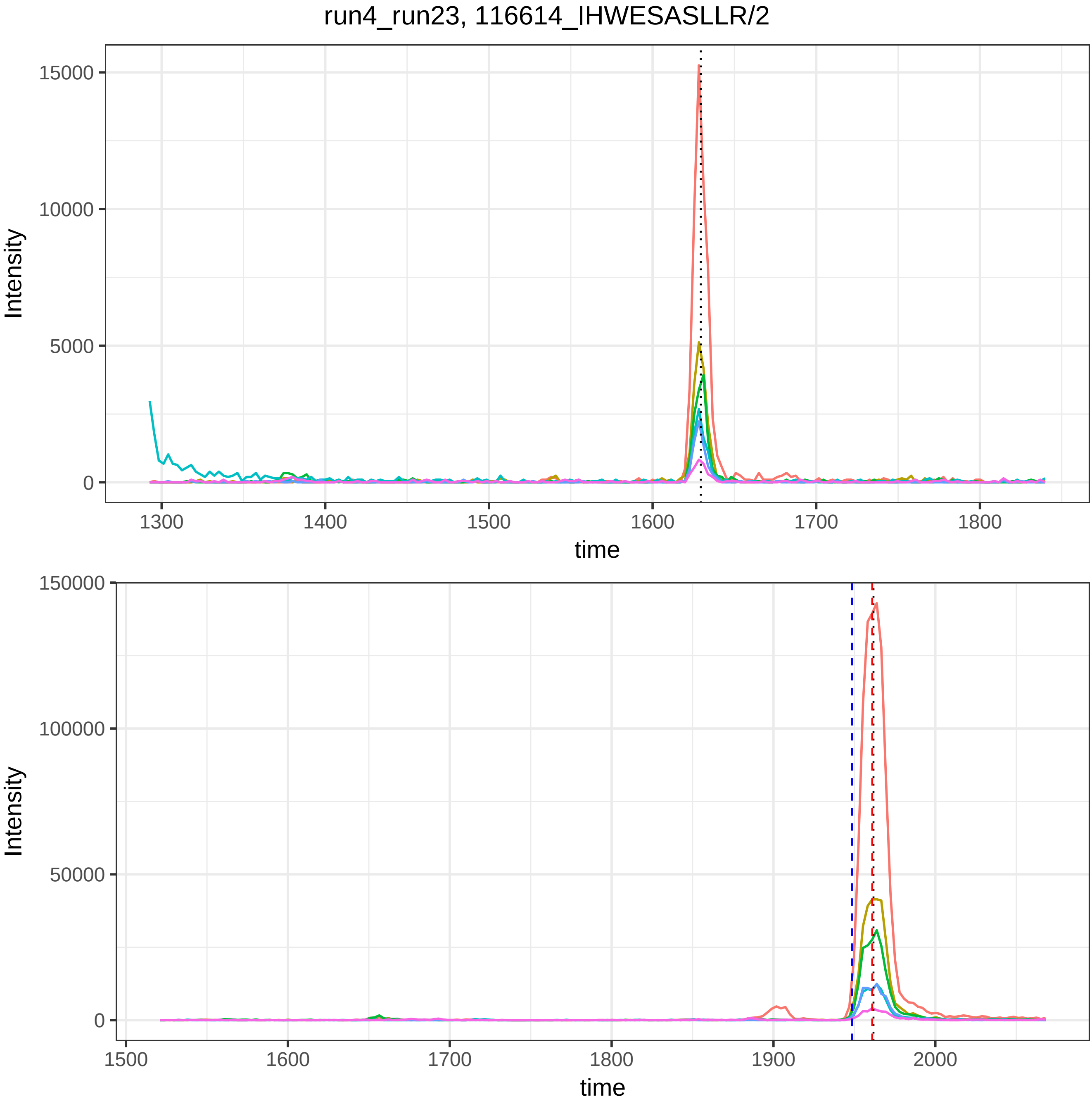

run4_run23, 116614_IHWESASLLR/2
Intensity
time
15000
10000
5000
0
1300
1400
1500
1600
1700
1800
150000
100000
50000
1500
1900
2000

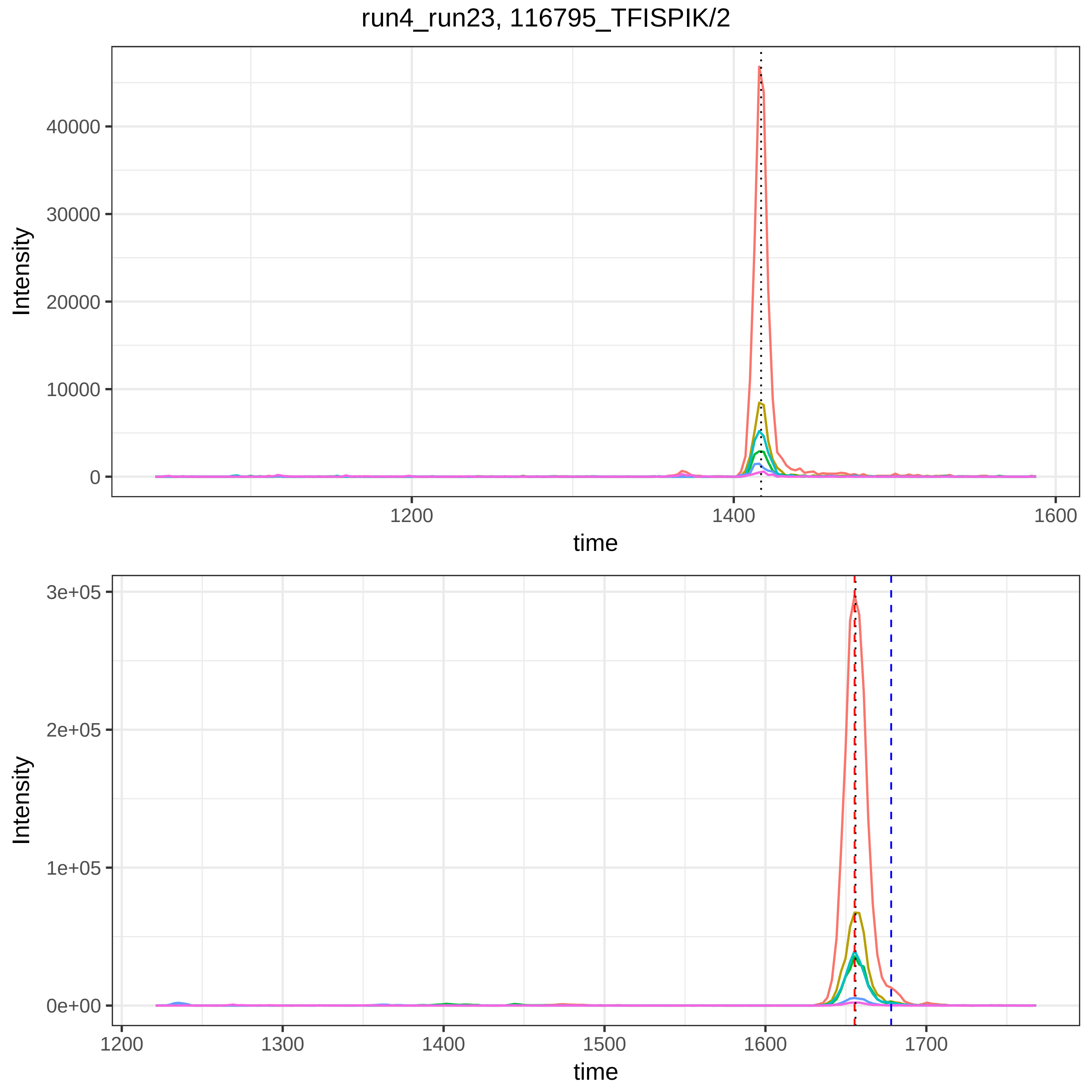
run4_run23, 116795_TFISPIK/2
Intensity
40000
30000
20000
10000
0
1200
1400
1600
time
Intensity
3e+05
2e+05
1e+05
0e+00
1200
1300
1400
1500
1600
1700
time

run4_run23, 119859_EDGGGWWYNR/2

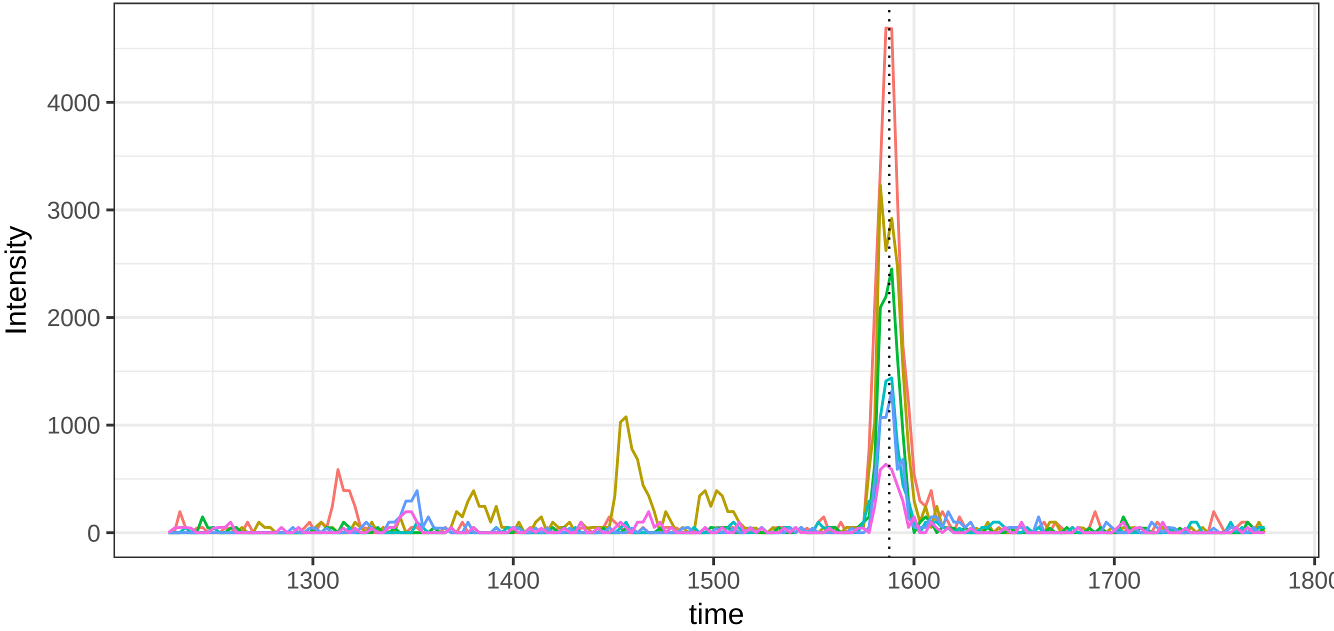


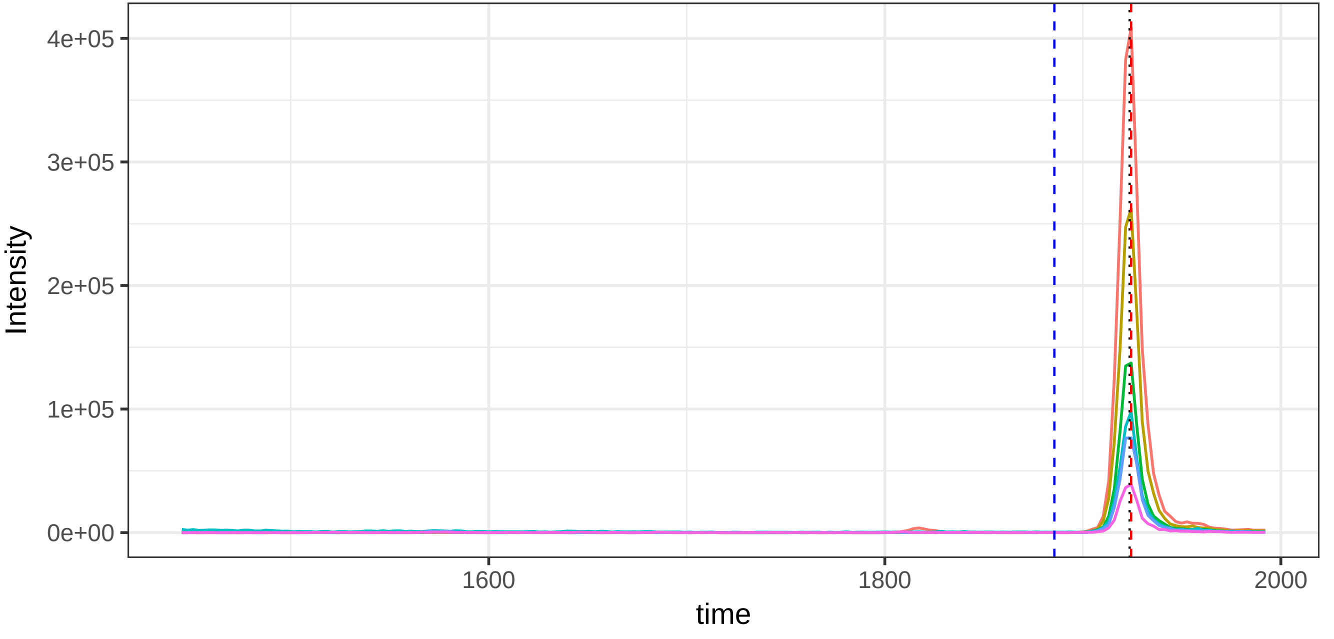

run4_run23, 120396_HPNSPLDEENLTQENQDR/3

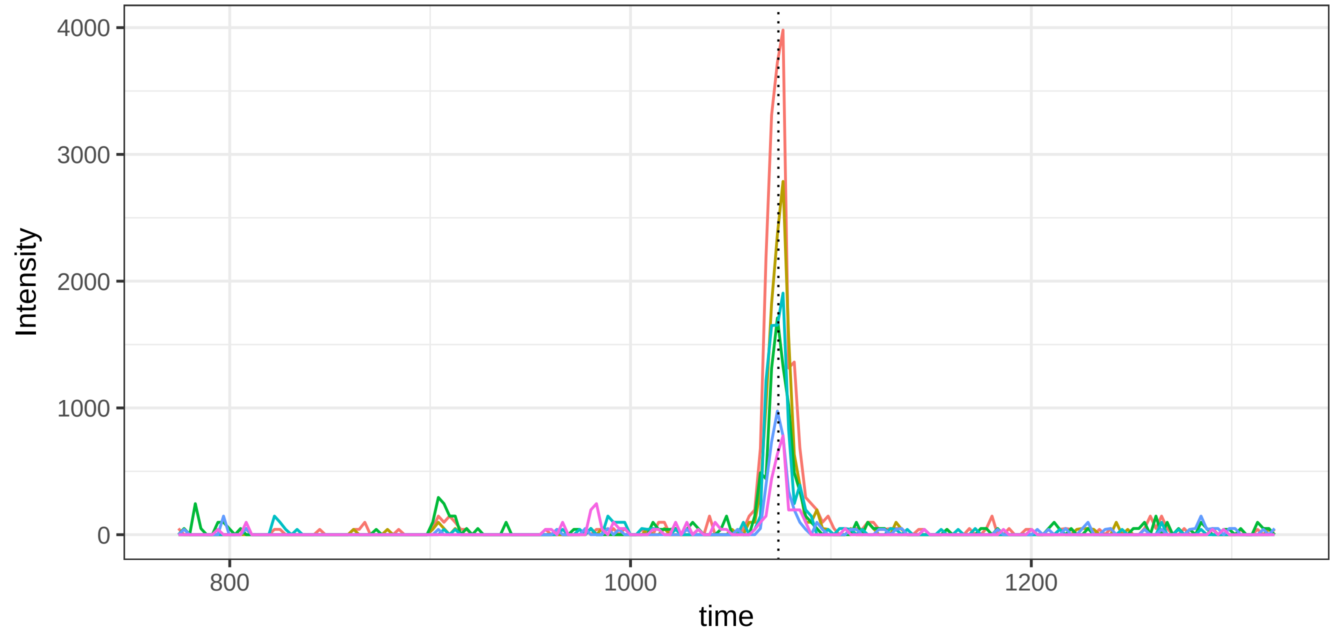


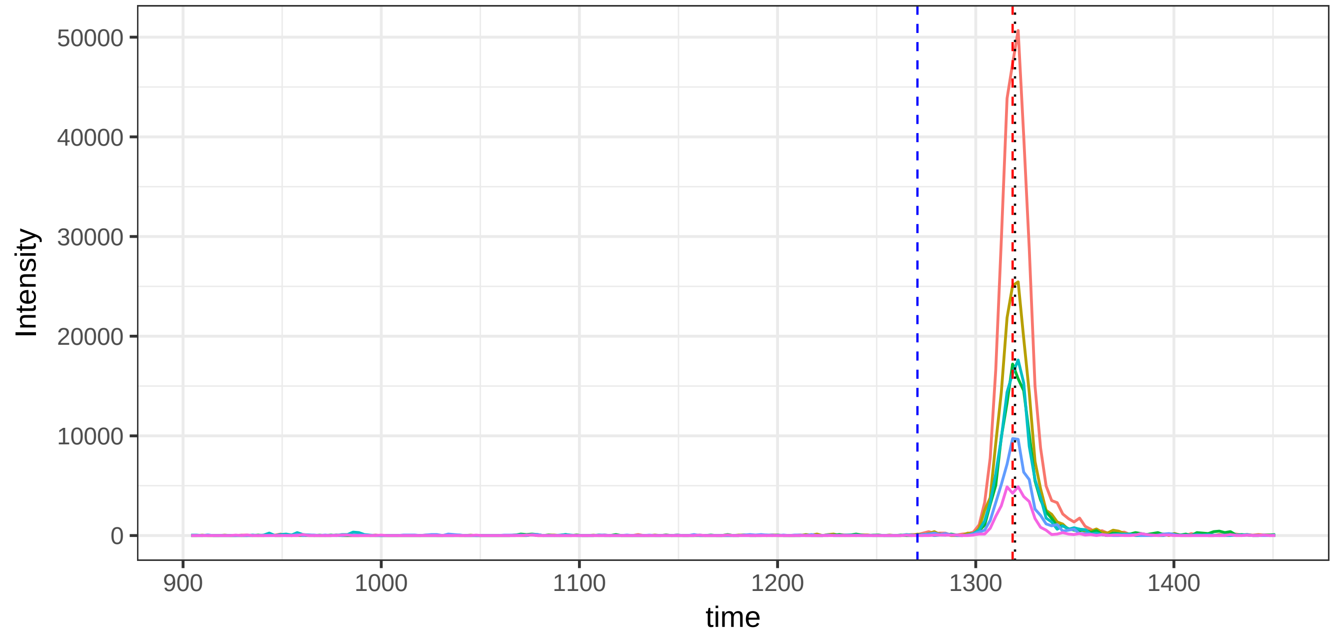

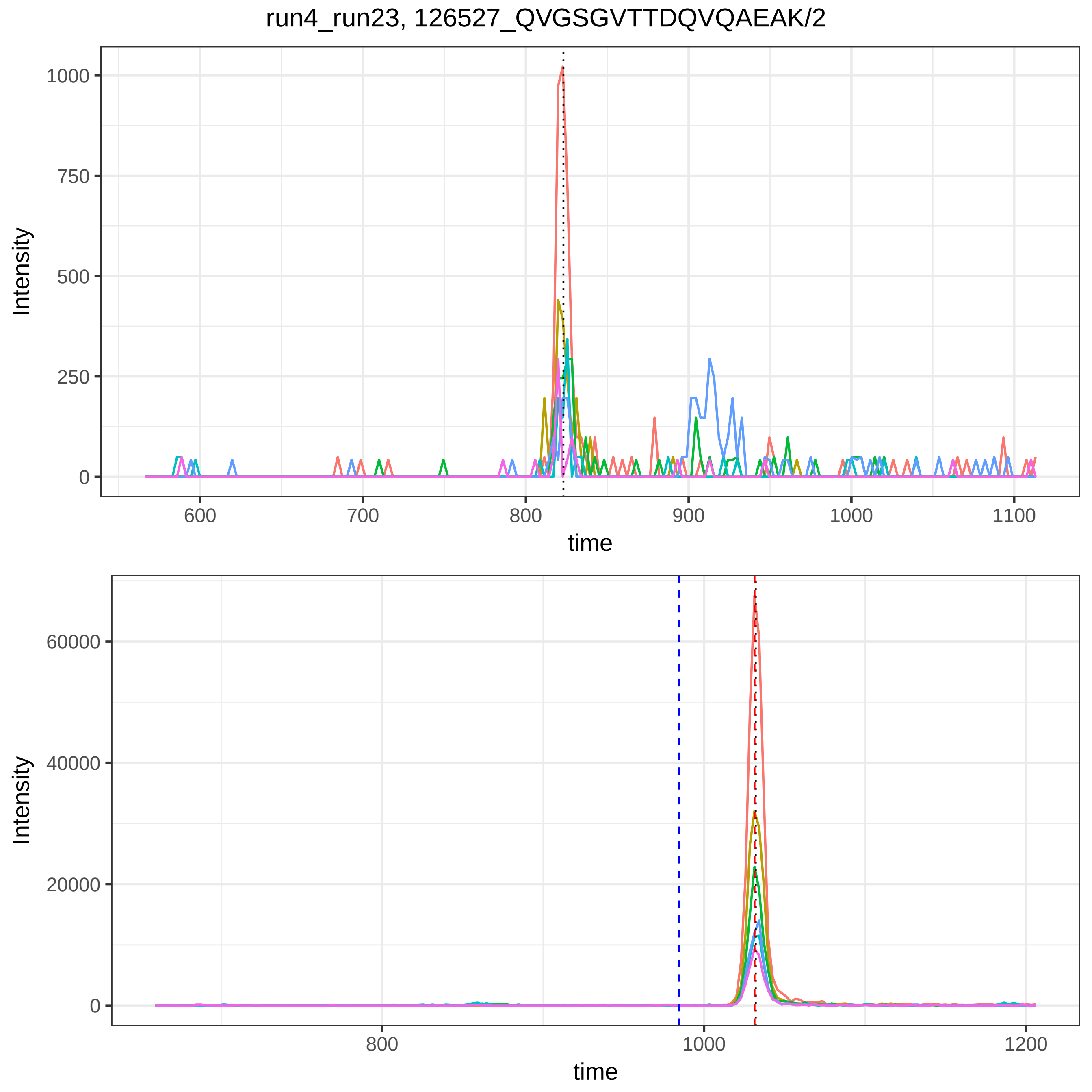
run4_run23, 126527_QVGSGVTTDQVQAEAK/2
Intensity
time
1000
750
500
250
0
600
700
800
900
1000
1100
60000
40000
20000
0
800
1000
1200

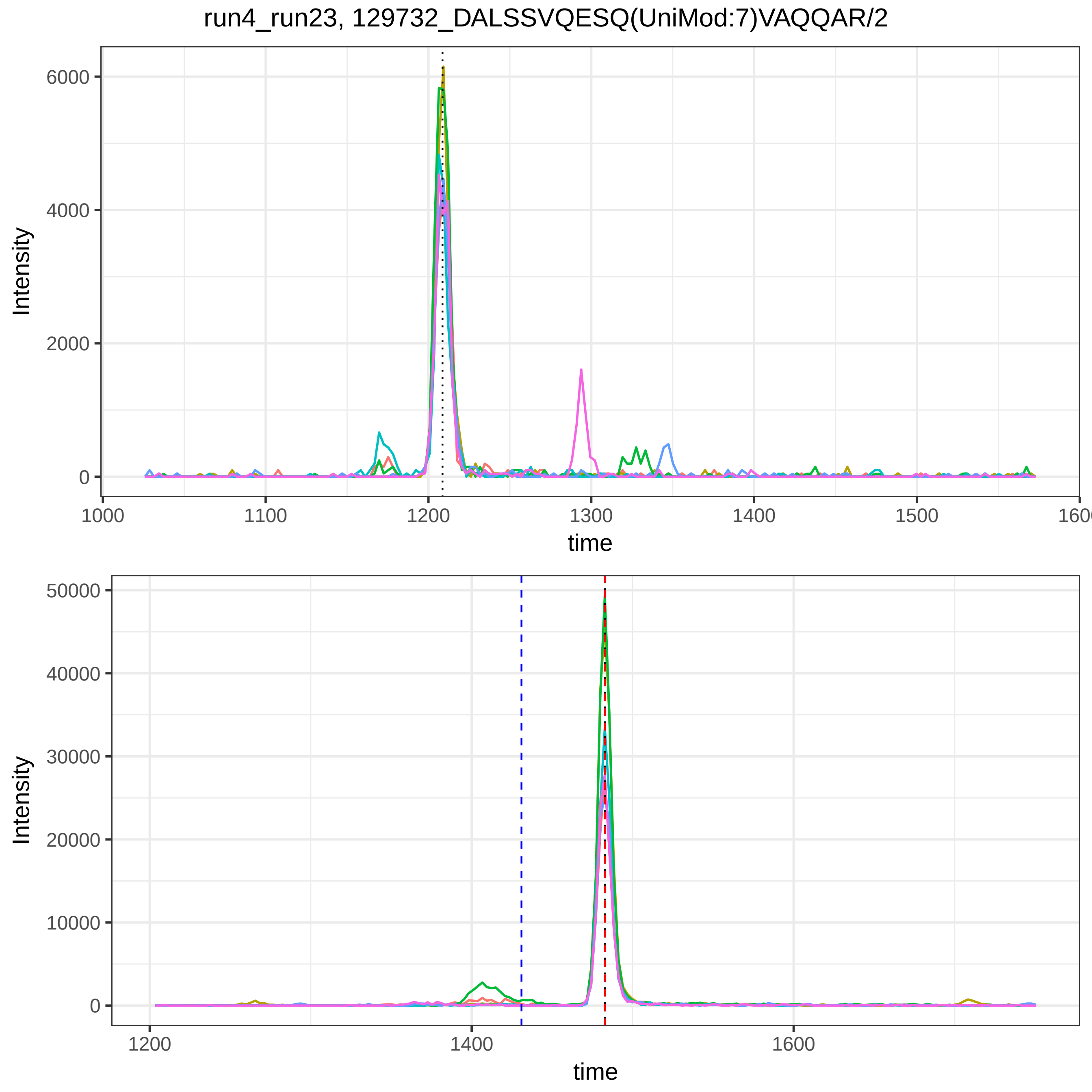

run4_run23, 129732_DALSSVQESQ(UniMod:7)VAQQAR/2
Intensity
6000
4000
2000
0
1000
1100
1200
1300
1400
1500
1600
time
Intensity
50000
40000
30000
20000
10000
0
1200
1400
1600
time

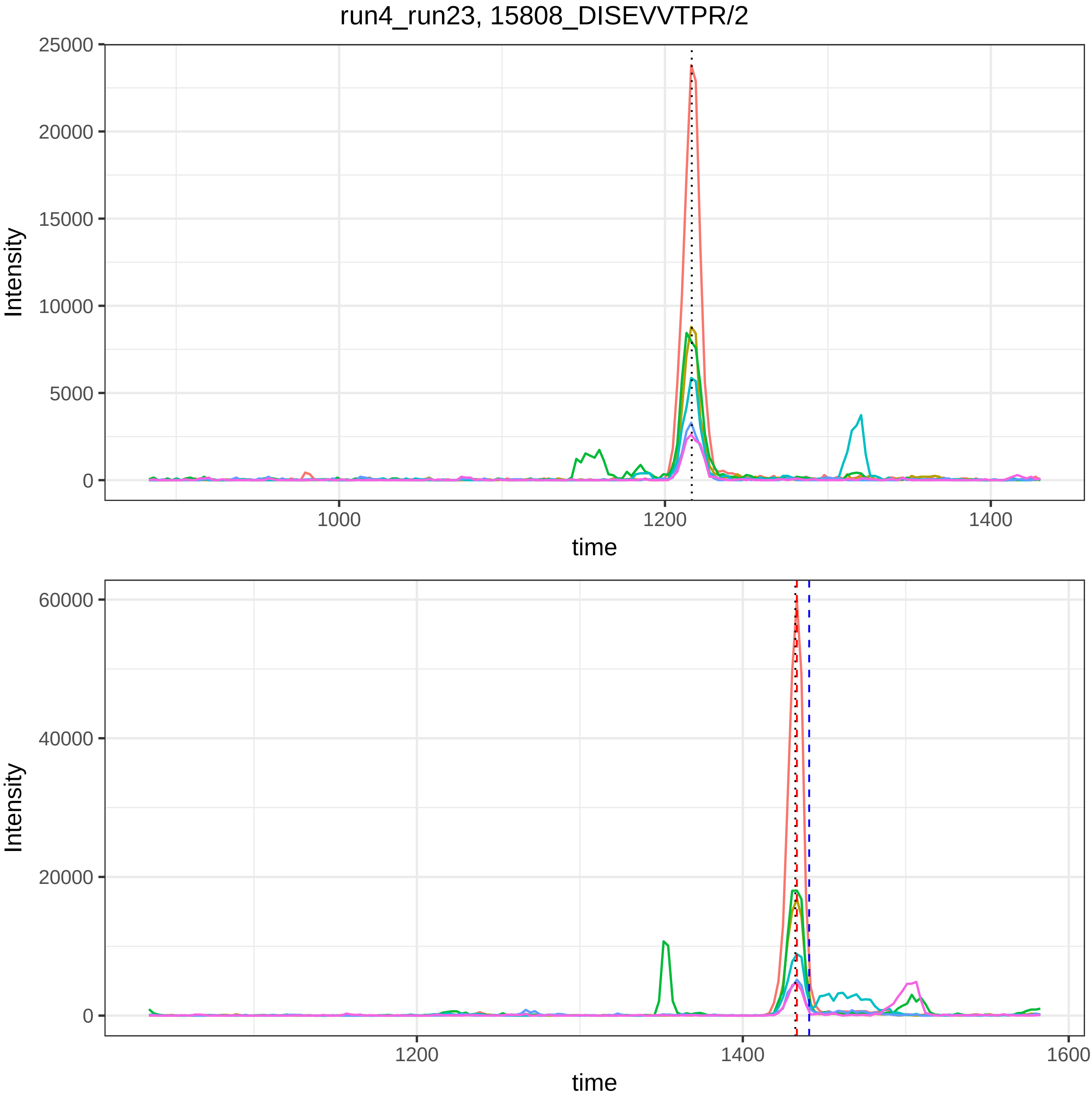
run4_run23, 15808_DISEVVTPR/2
Intensity
time
25000
20000
15000
10000
5000
0
1000
1200
1400
60000
40000
20000
1600

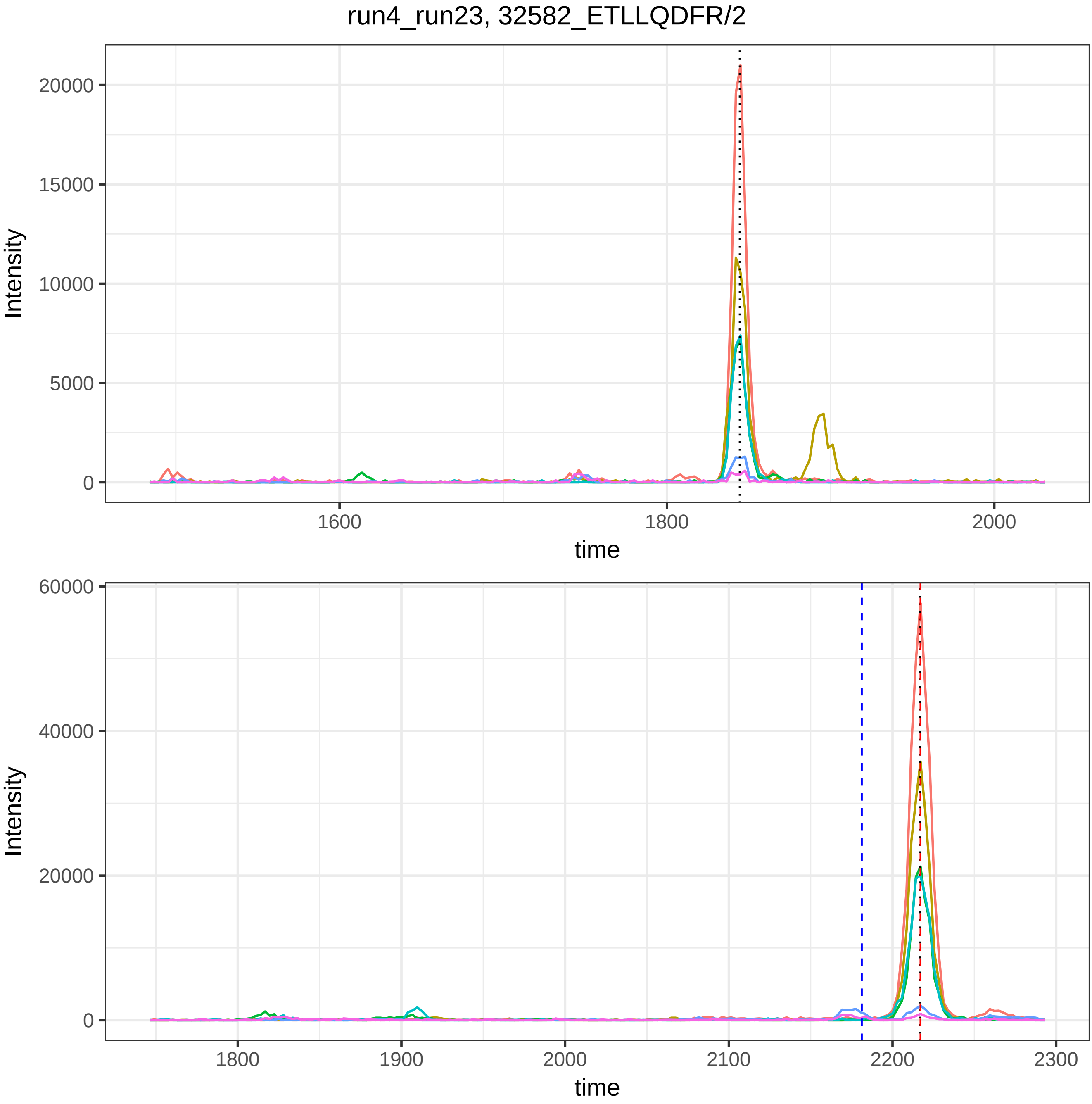
run4_run23, 32582_ETLLQDFR/2
Intensity
20000
15000
10000
5000
0
1600
1800
2000
time
Intensity
60000
40000
20000
0
1800
1900
2000
2100
2200
2300
time

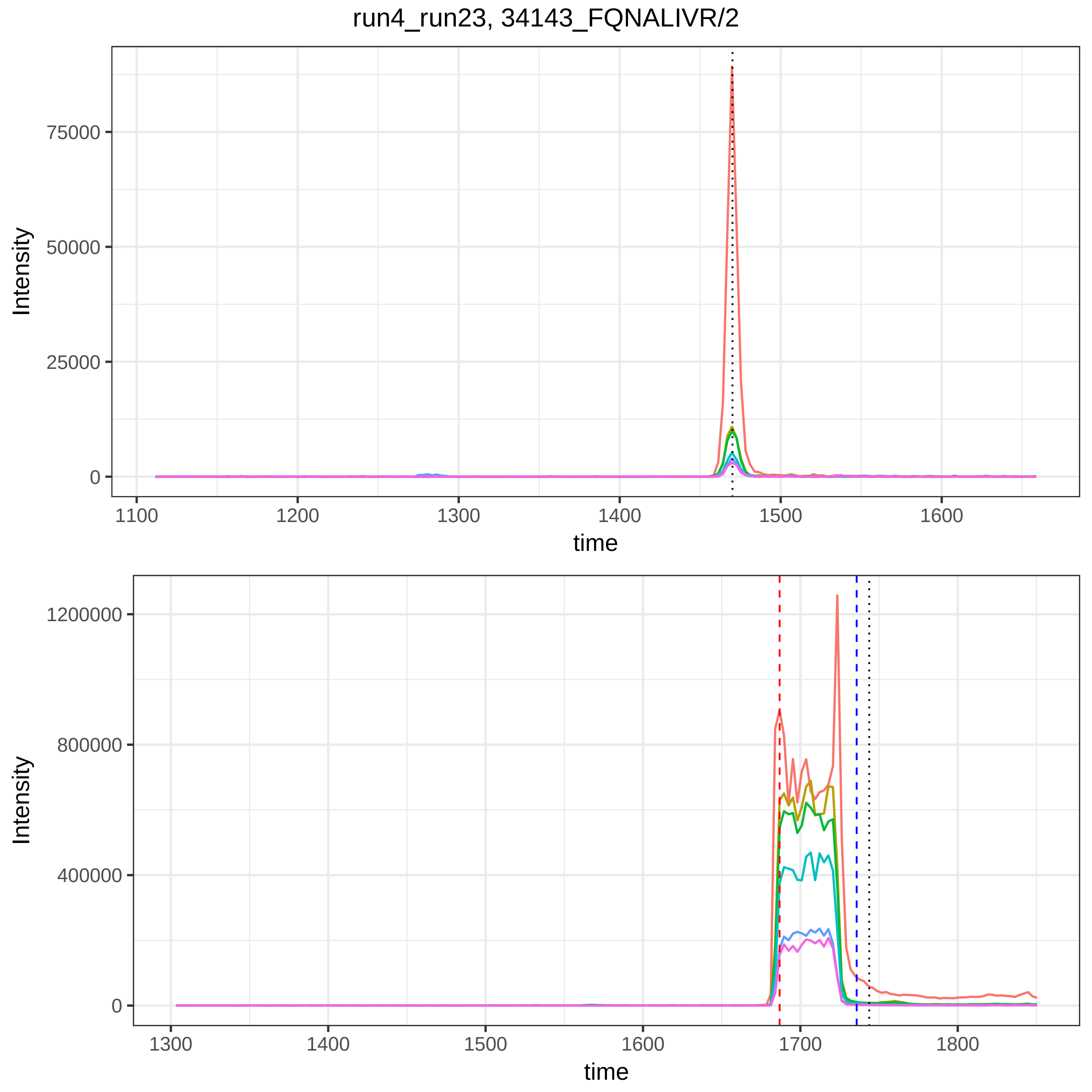

run4_run23, 34143_FQNALIVR/2
Intensity
75000
50000
25000
0
1100
1200
1300
1400
1500
1600
time
Intensity
1200000
800000
400000
0
1300
1400
1500
1600
1700
1800
time

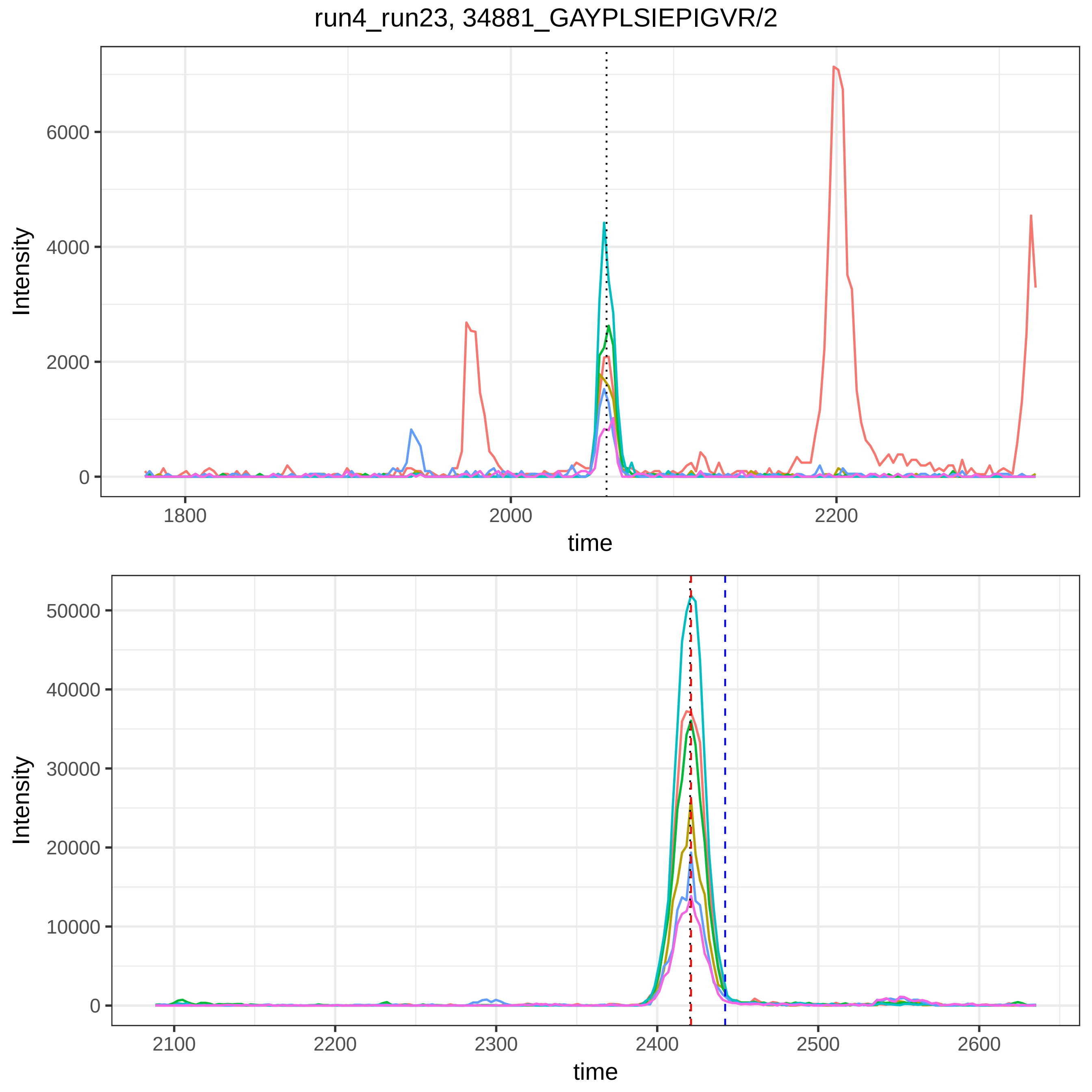

run4_run23, 34881_GAYPLSIEPIGVR/2
Intensity
time
0
2000
4000
6000
1800
2000
2200
0
10000
20000
30000
40000
50000
2100
2200
2300
2400
2500
2600

run4_run23, 38147_HPNSPLDEENLTQENQDR/3

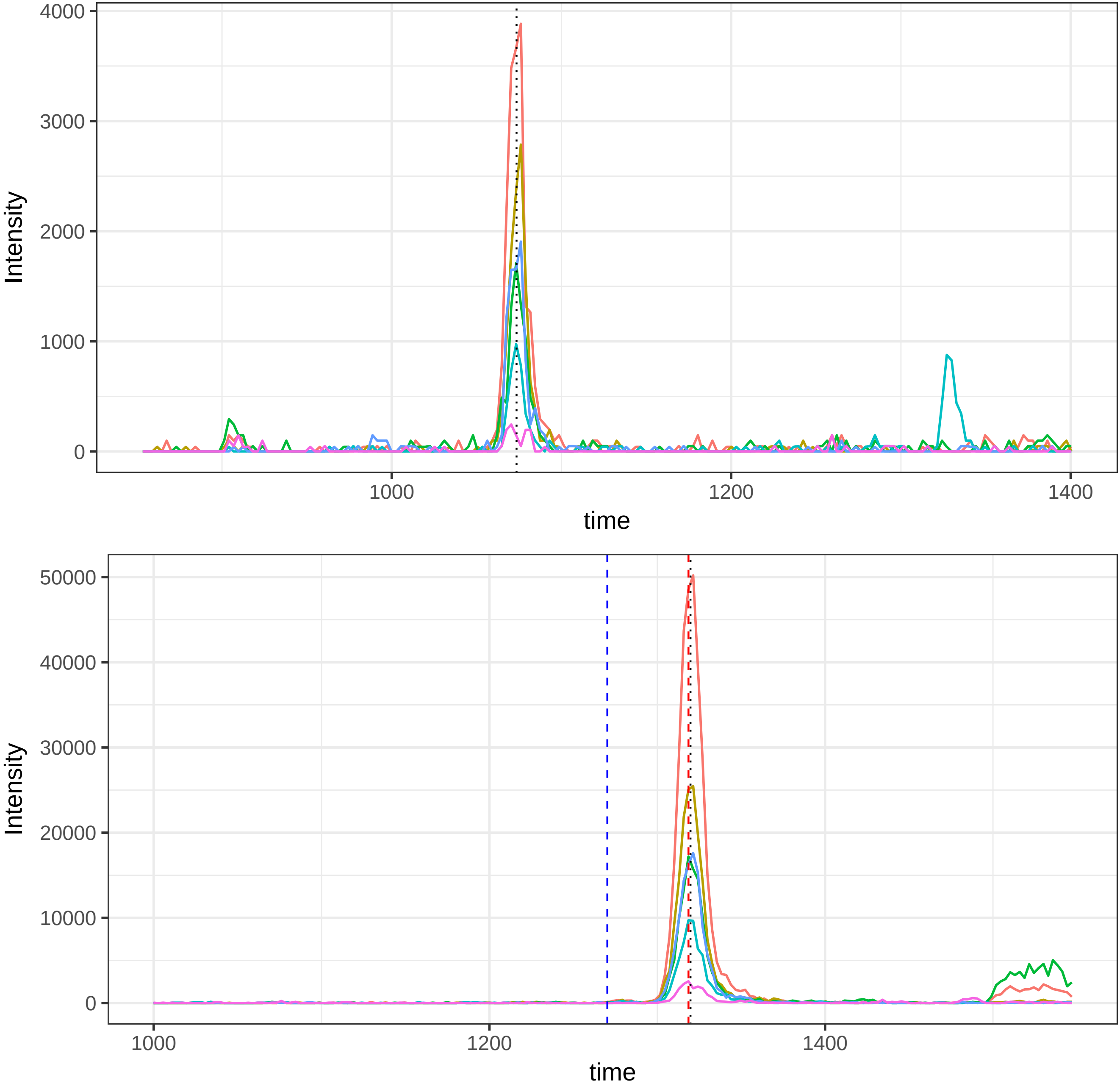

run4_run23, 39305_IHWESASLLR/2

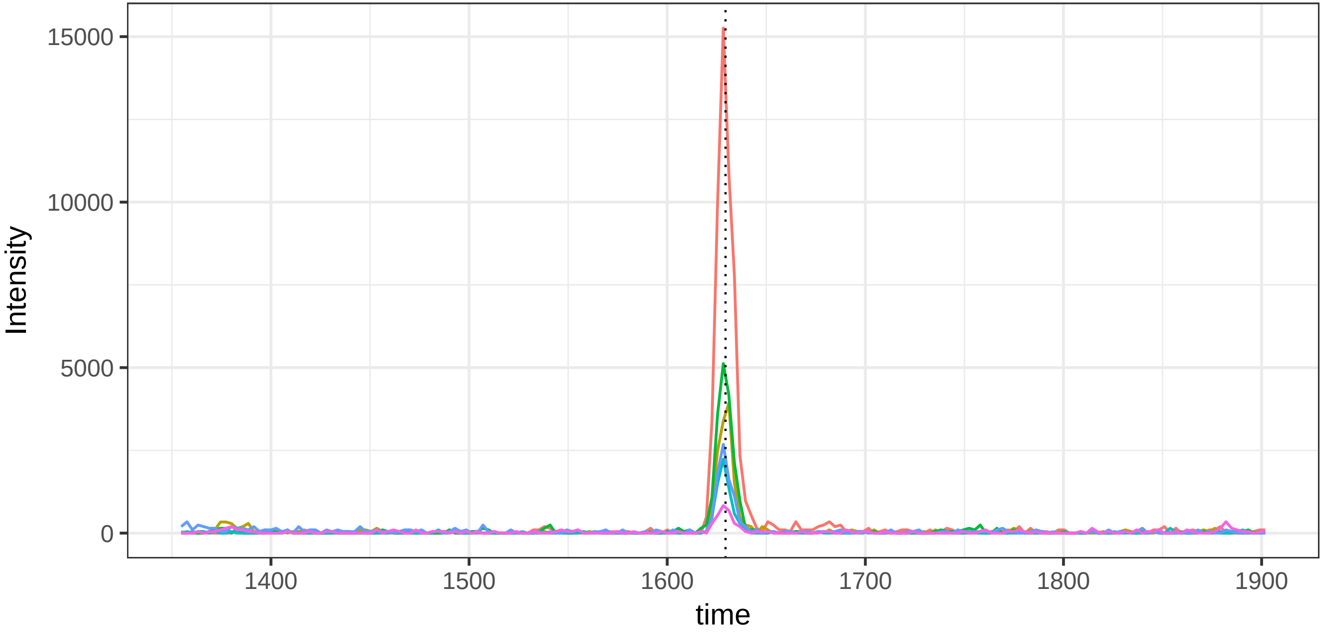


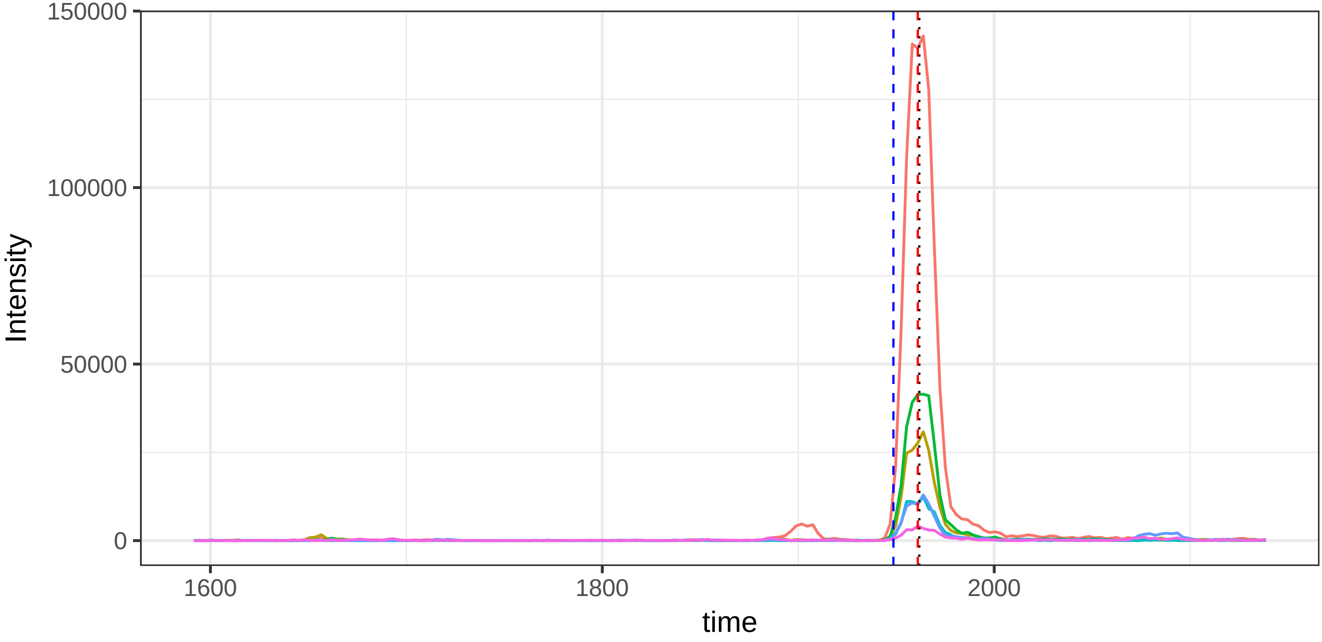

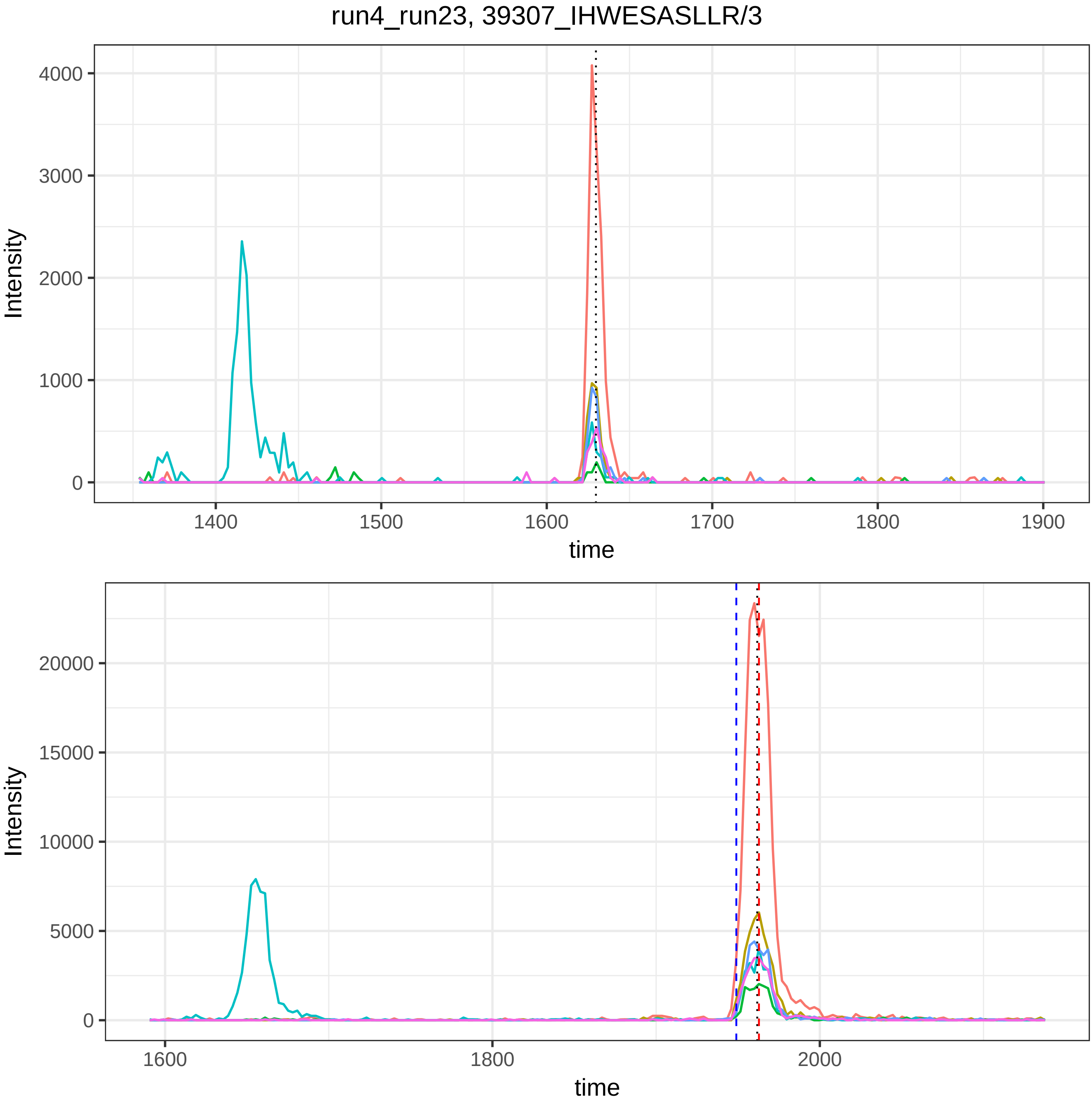

run4_run23, 39307_IHWESASLLR/3
Intensity
time
4000
3000
2000
1000
0
1400
1500
1600
1700
1800
1900
20000
15000
10000
5000
1600
1800
2000

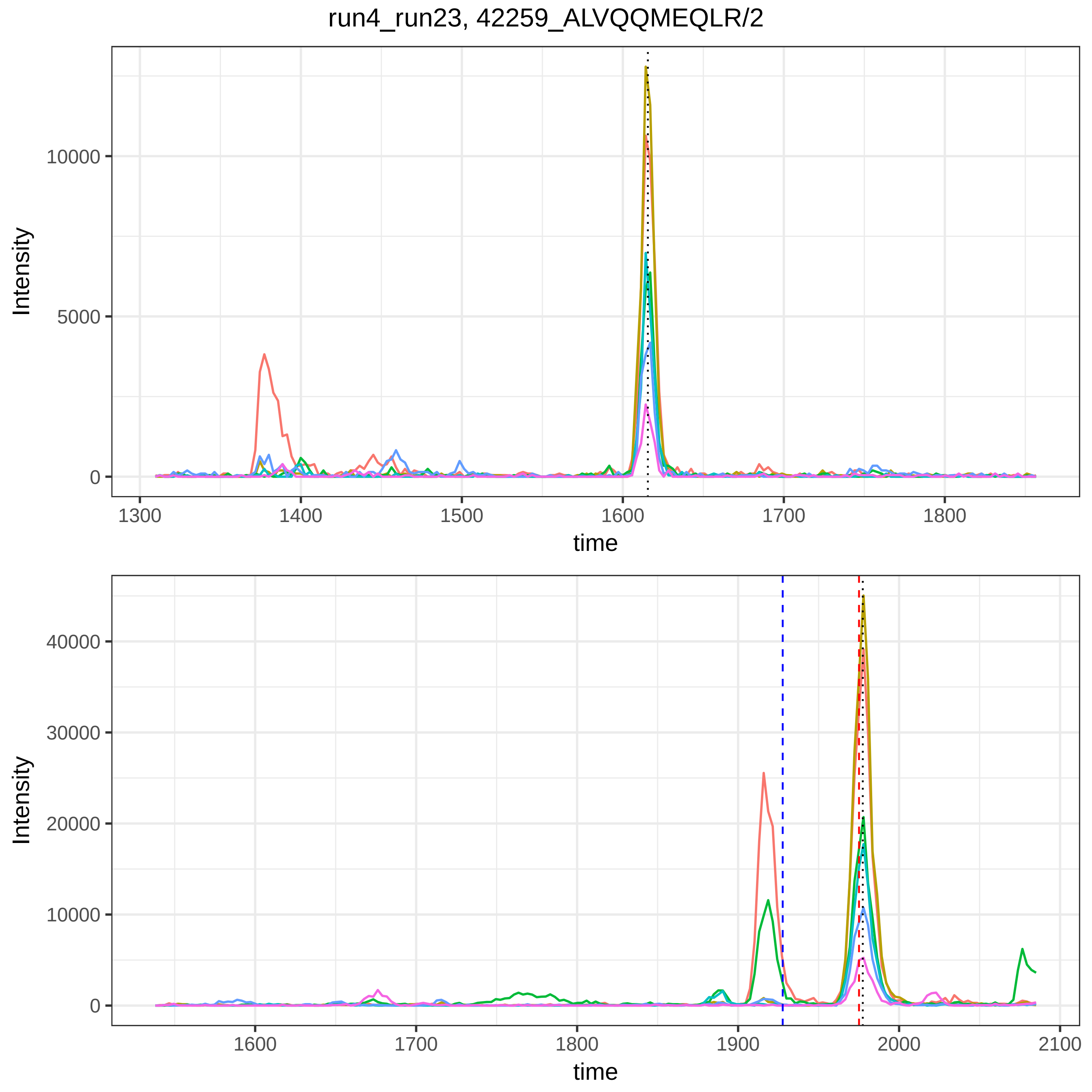
run4_run23, 42259_ALVQQMEQLR/2
Intensity
10000
5000
0
1300
1400
1500
1600
1700
1800
time
Intensity
40000
30000
20000
10000
0
1600
1700
1800
1900
2000
2100
time

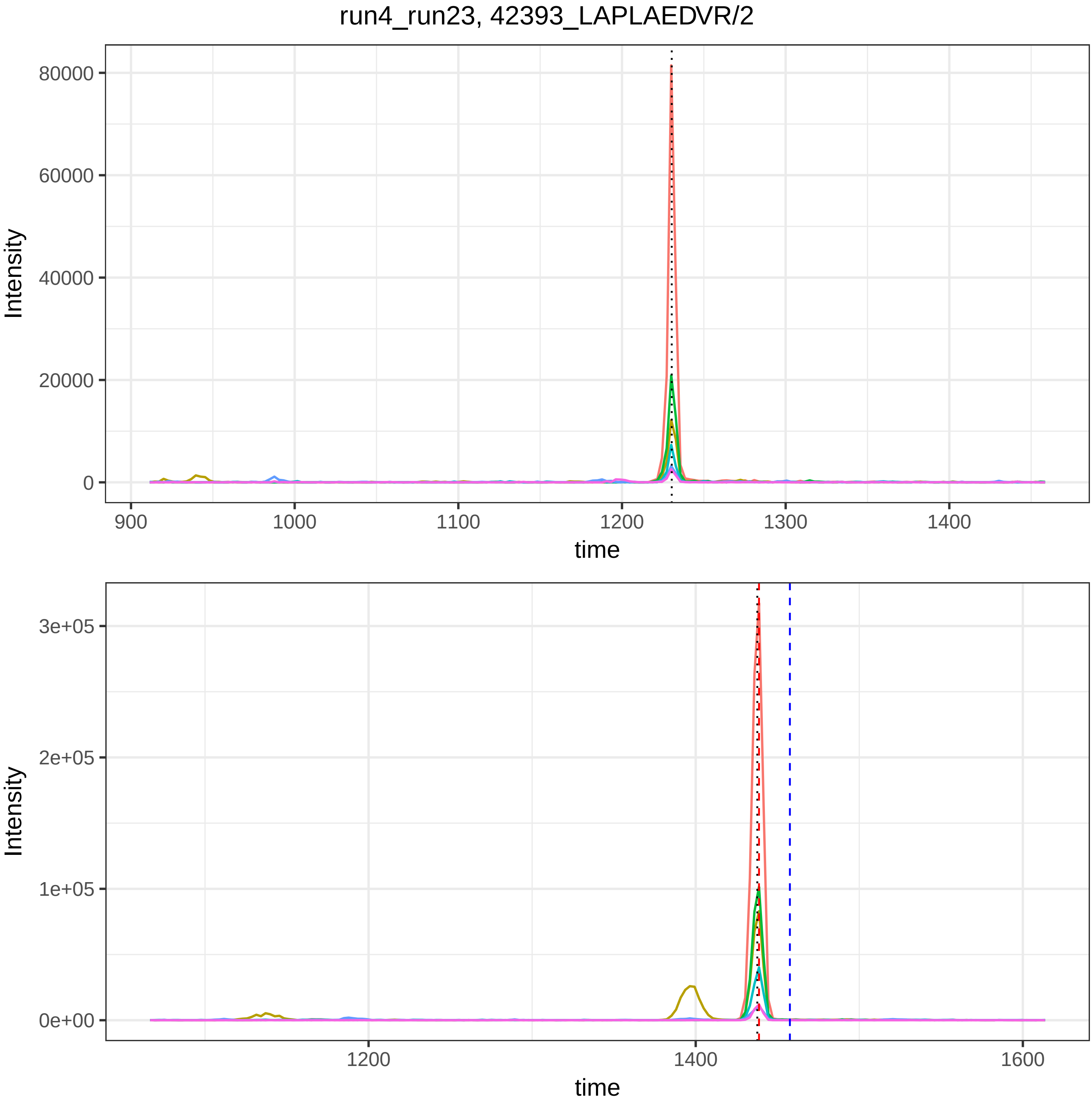
run4_run23, 42393_LAPLAEDVR/2
Intensity
80000
60000
40000
20000
0
900
1000
1100
1200
1300
1400
time
Intensity
3e+05
2e+05
1e+05
0e+00
1200
1400
1600
time

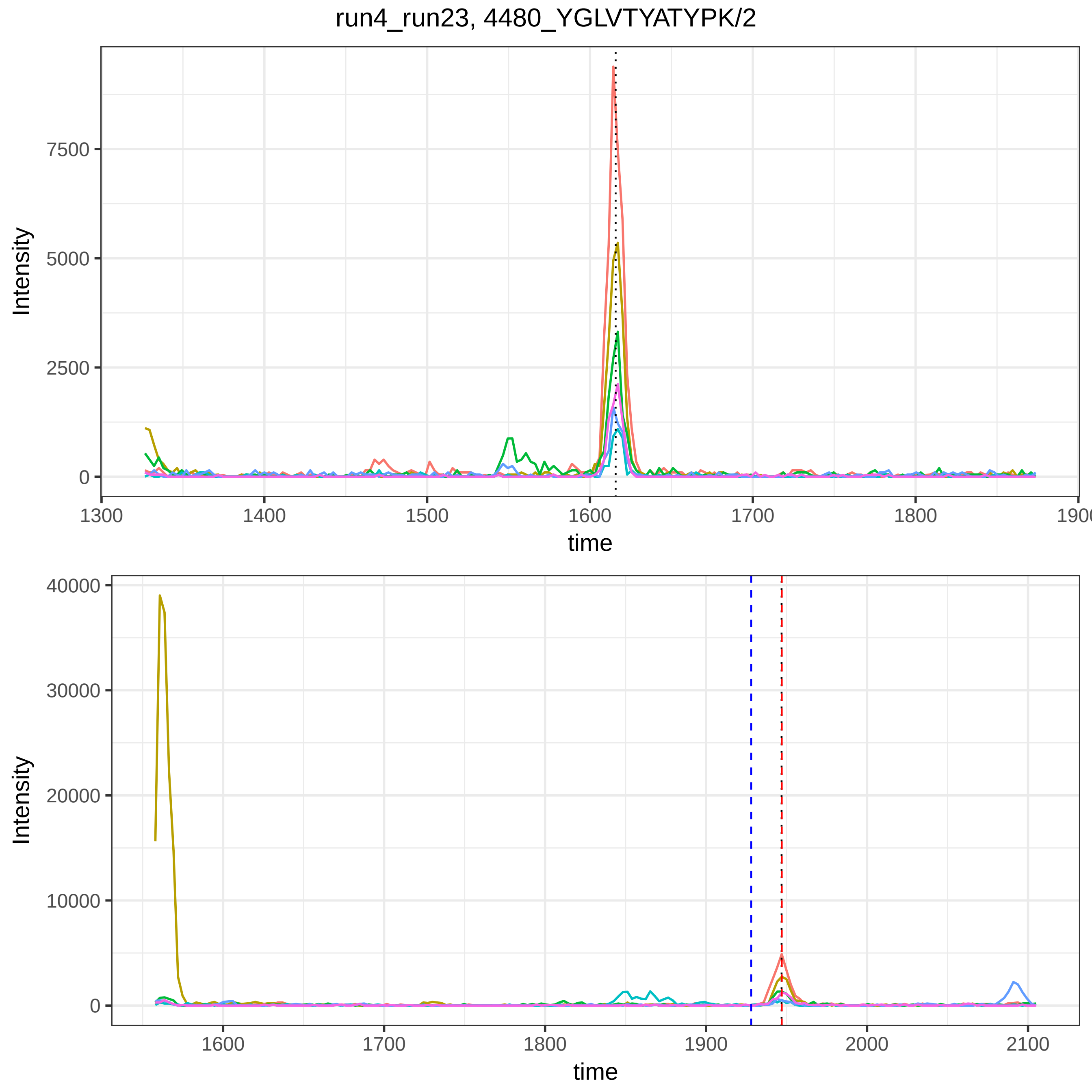
run4_run23, 4480_YGLVTYATYPK/2
Intensity
time
0
2500
5000
7500
1300
1400
1500
1600
1700
1800
1900
0
10000
20000
30000
40000
1600
1700
1800
1900
2000
2100

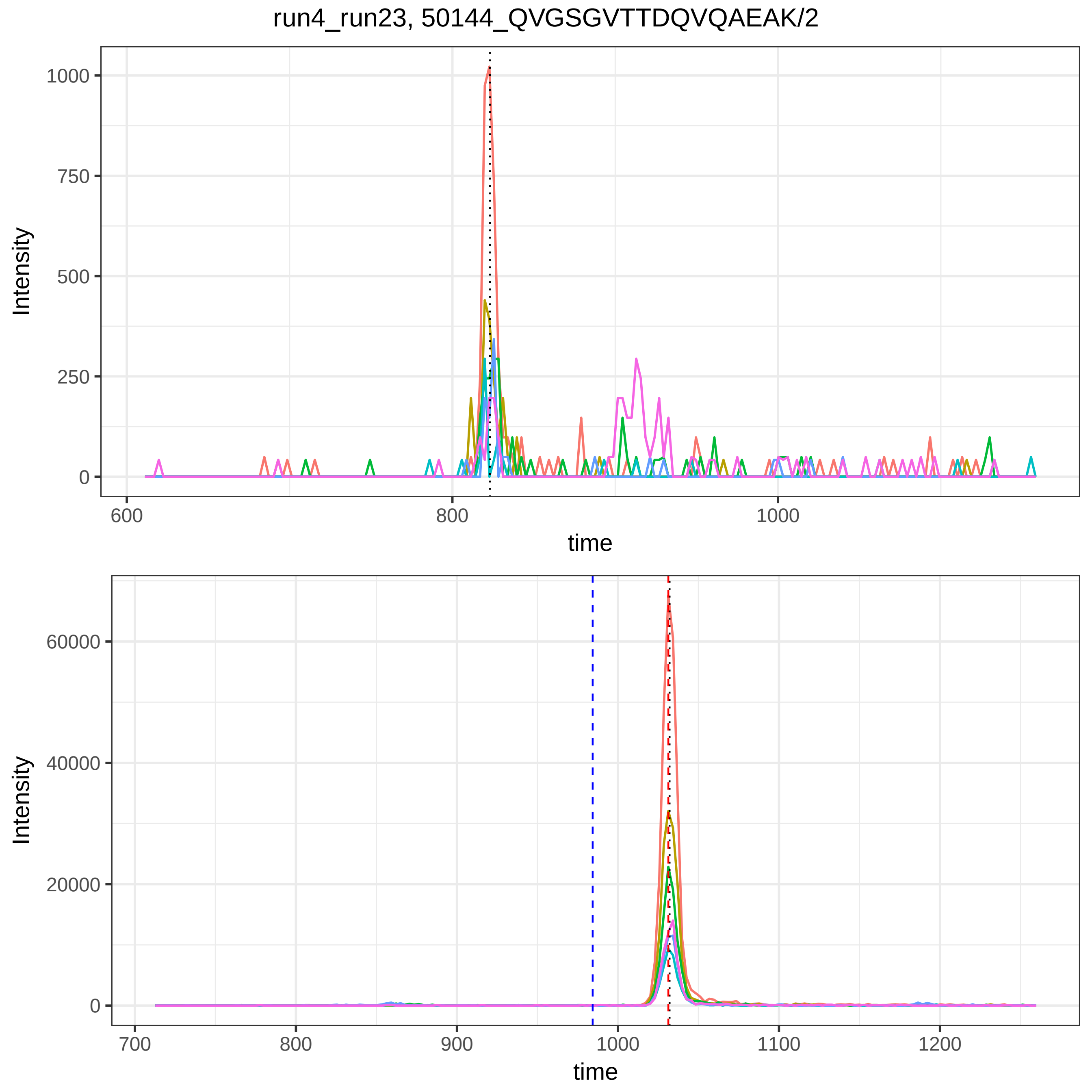

run4_run23, 50144_QVGSGVTTDQVQAEAK/2
Intensity
1000
750
500
250
0
600
800
1000
time
Intensity
60000
40000
20000
0
700
800
900
1000
1100
1200
time

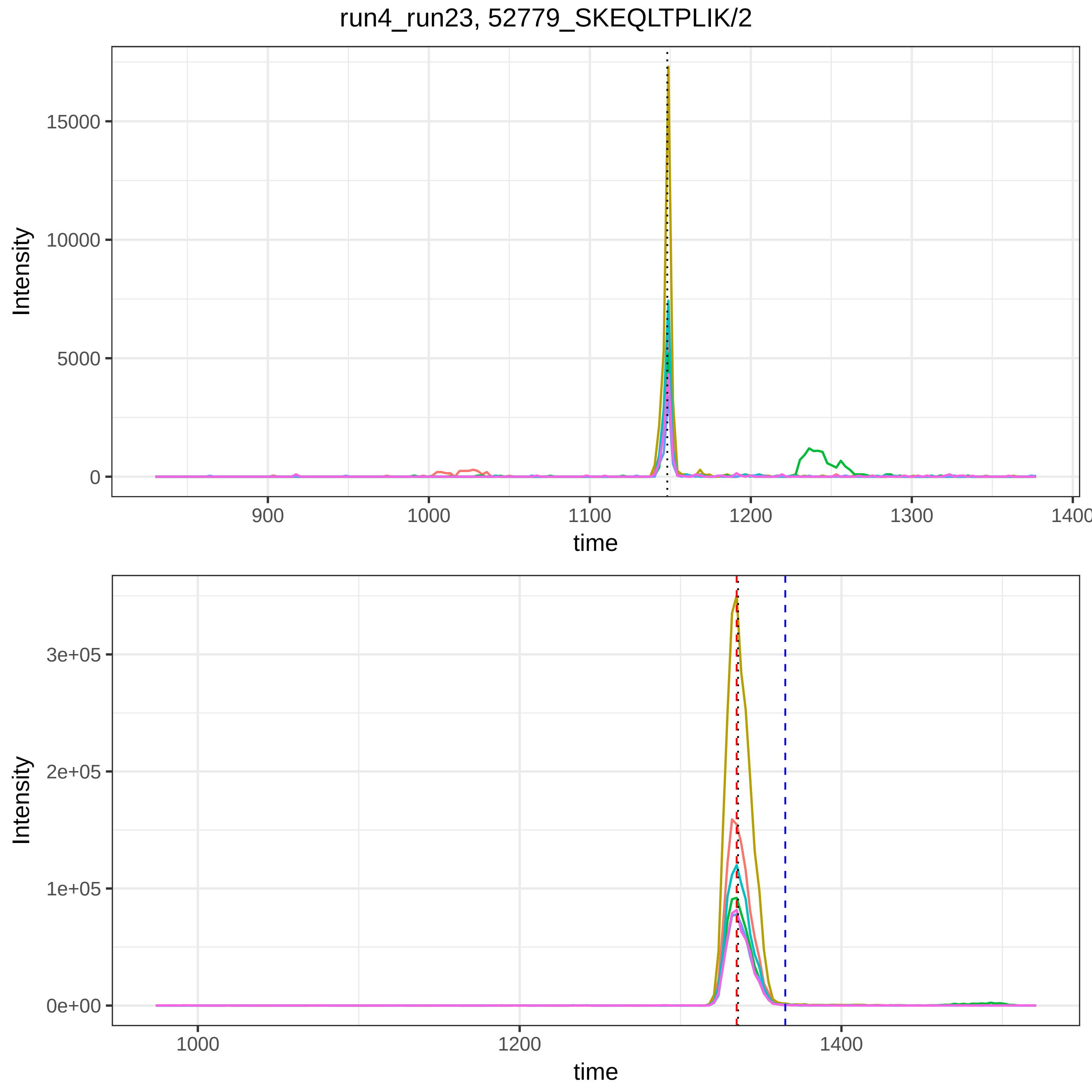
run4_run23, 52779_SKEQLTPLIK/2
Intensity
15000
10000
5000
0
900
1000
1100
1200
1300
1400
time
Intensity
3e+05
2e+05
1e+05
0e+00
1000
1200
1400
time

run4_run23, 54026_SSALDMENFR/2

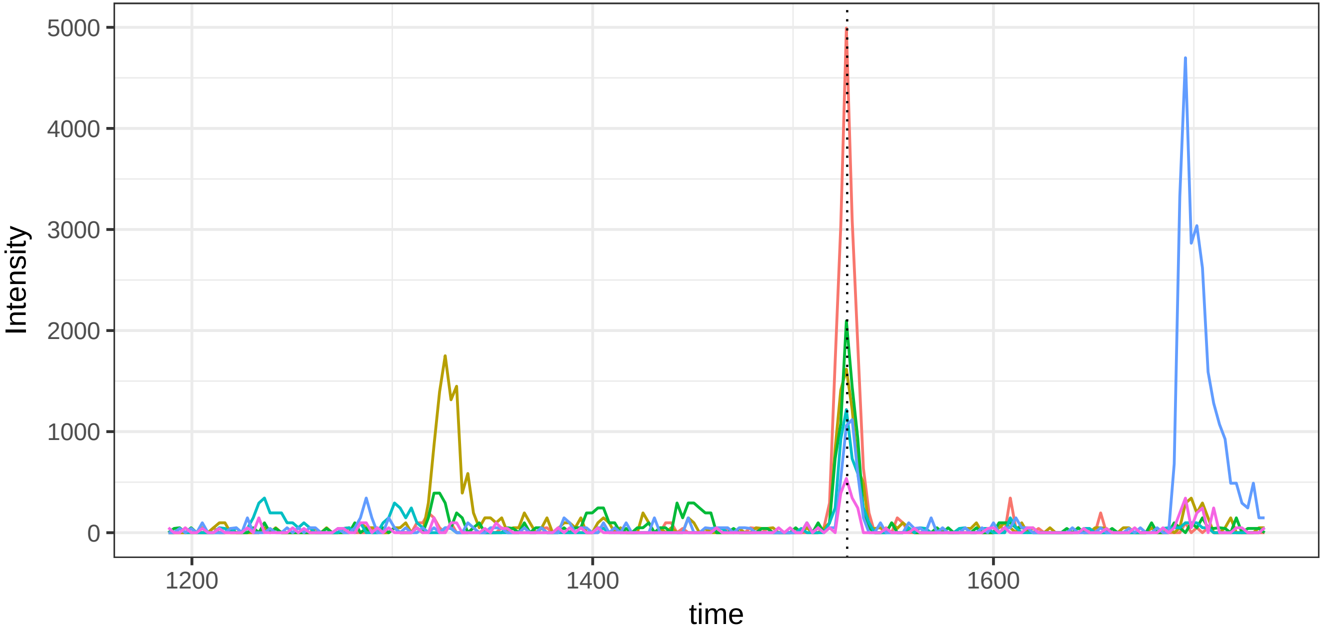


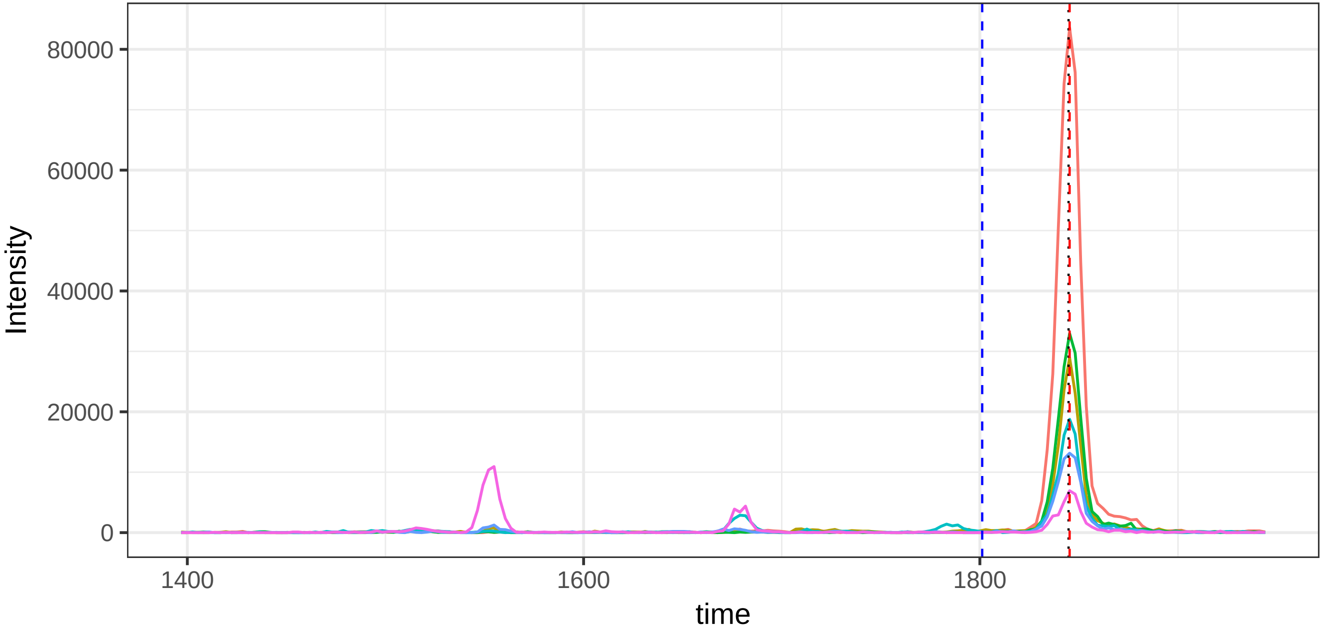

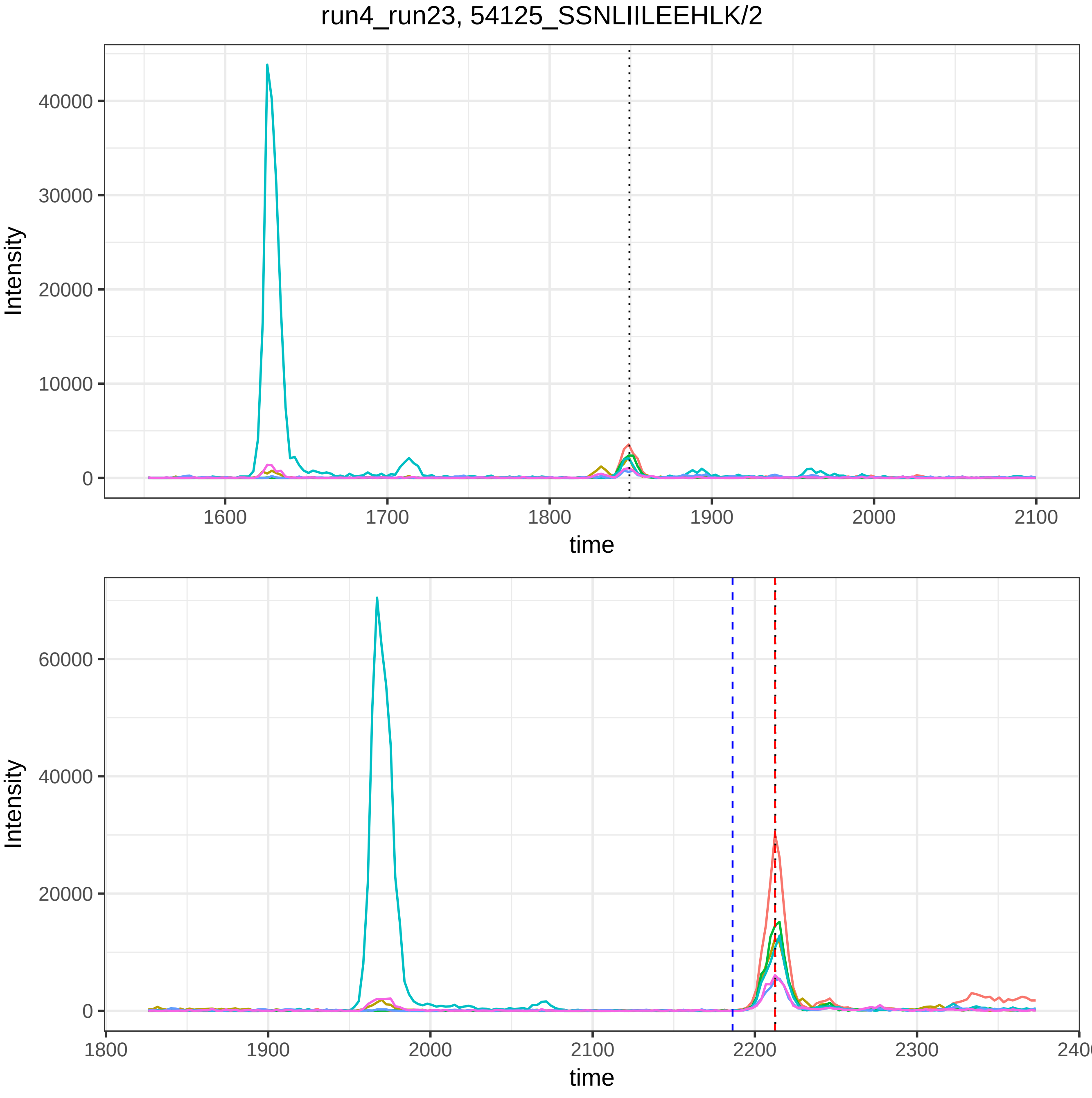
run4_run23, 54125_SSNLIILEEHLK/2
Intensity
time
40000
30000
20000
10000
0
1600
1700
1800
1900
2000
2100
60000
40000
20000
0
1800
1900
2000
2100
2200
2300
2400

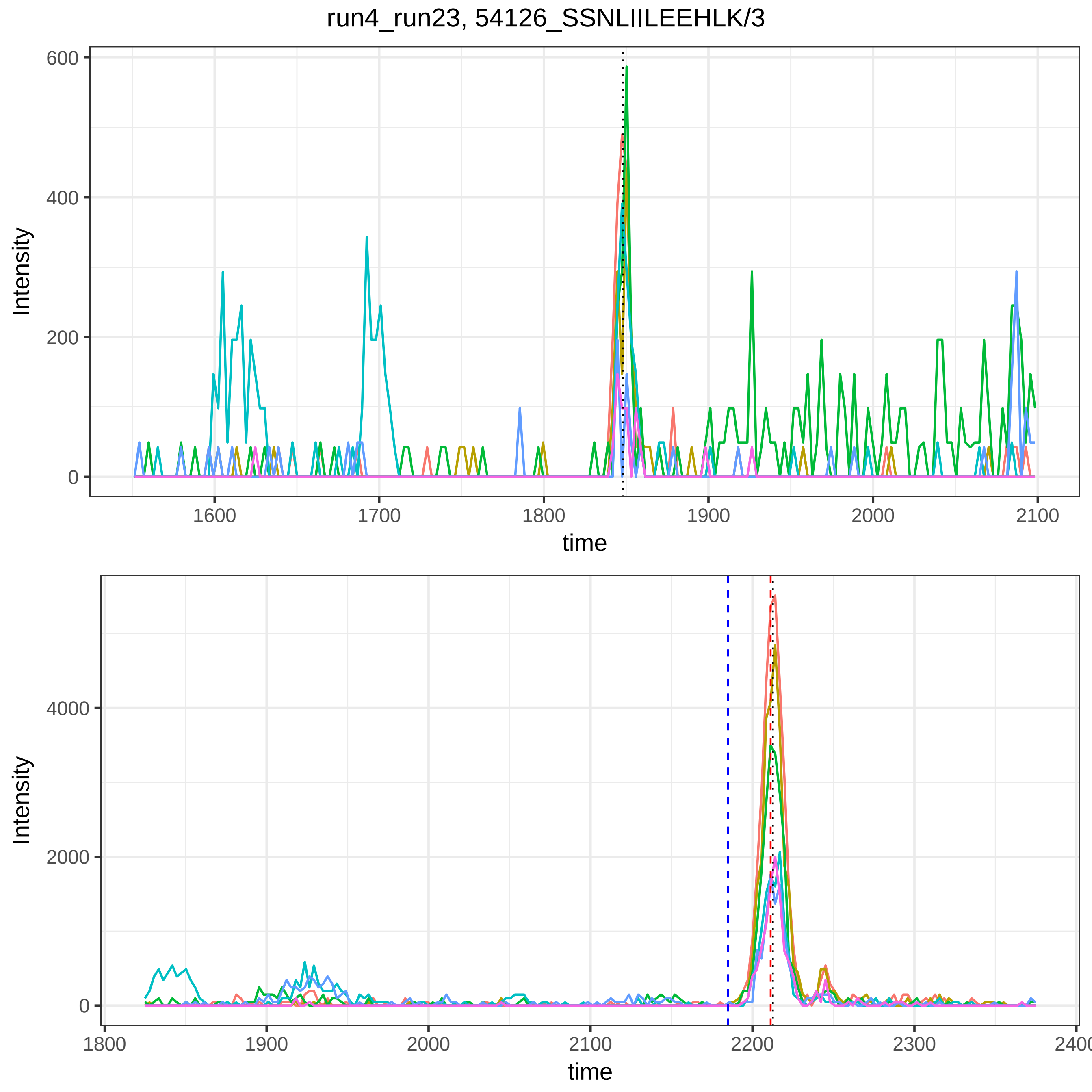
run4_run23, 54126_SSNLIILEEHLK/3
Intensity
600
400
200
0
1600
1700
1800
1900
2000
2100
time
Intensity
4000
2000
0
1800
1900
2000
2100
2200
2300
2400
time

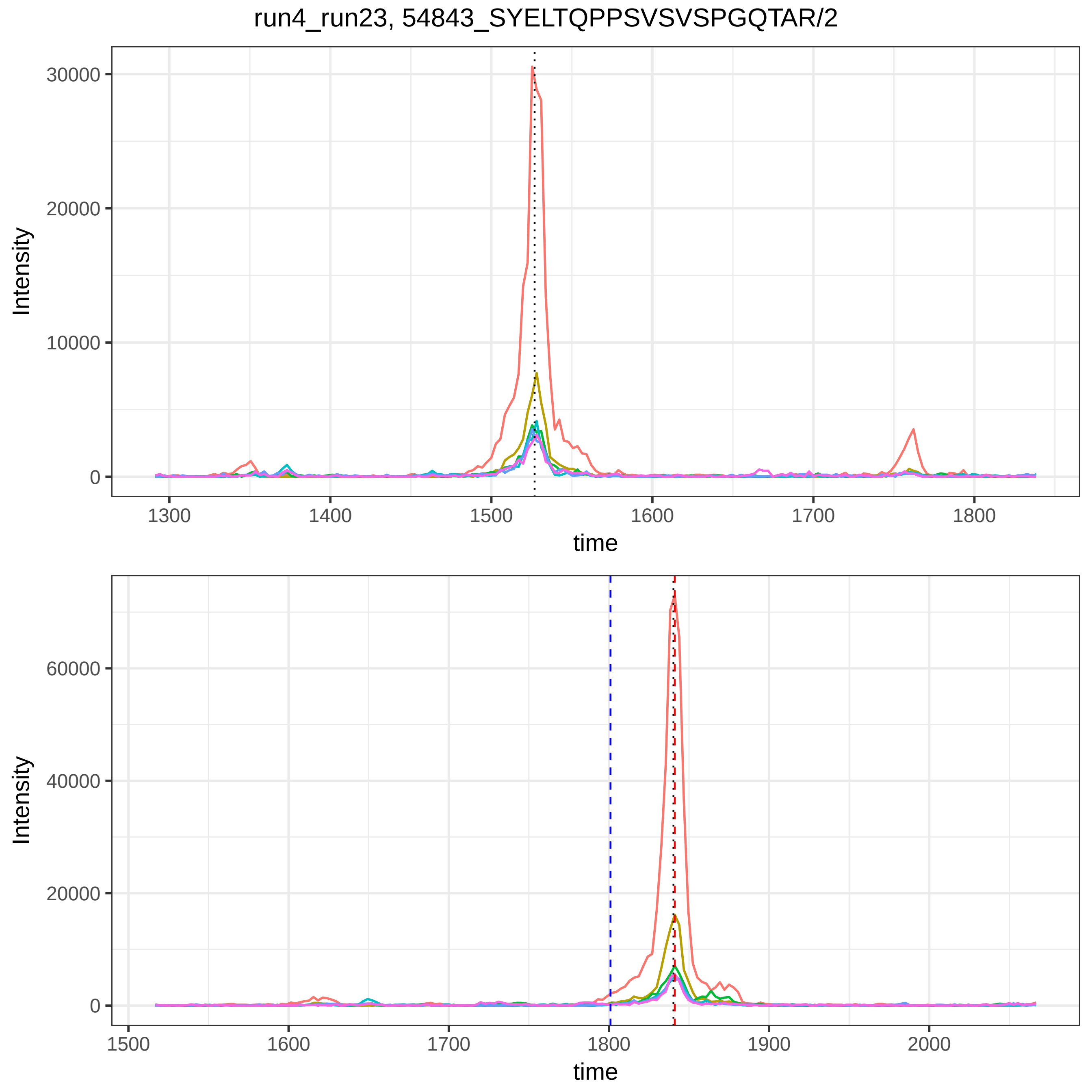

run4_run23, 54843_SYELTQPPSVSVSPGQTAR/2
Intensity
time
30000
20000
10000
0
1300
1400
1500
1600
1700
1800
60000
40000
20000
0
1500
1600
1700
1800
1900
2000

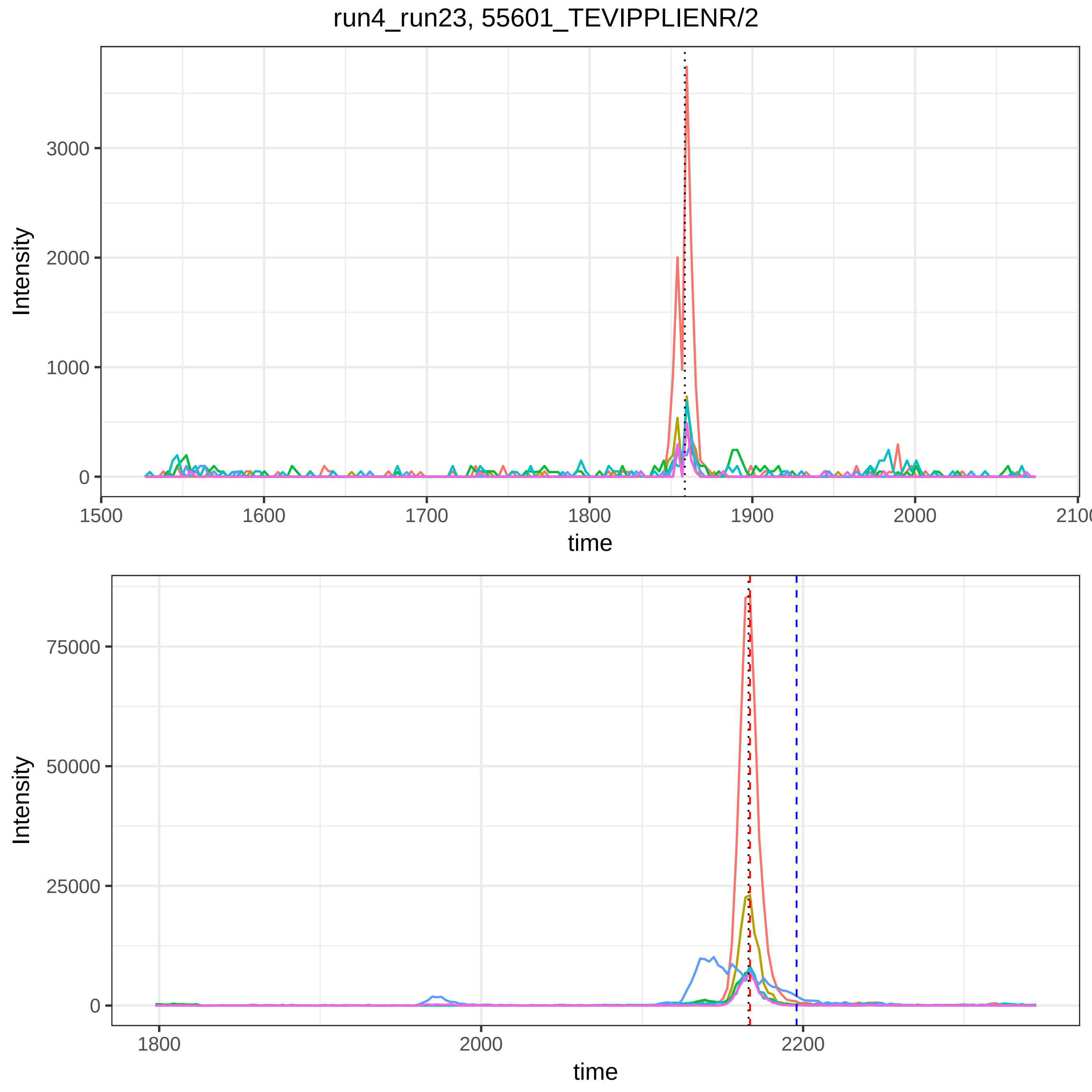
run4_run23, 55601_TEVIPPLIENR/2
Intensity
3000
2000
1000
0
1500
1600
1700
1800
1900
2000
2100
time
Intensity
75000
50000
25000
0
1800
2000
2200
time

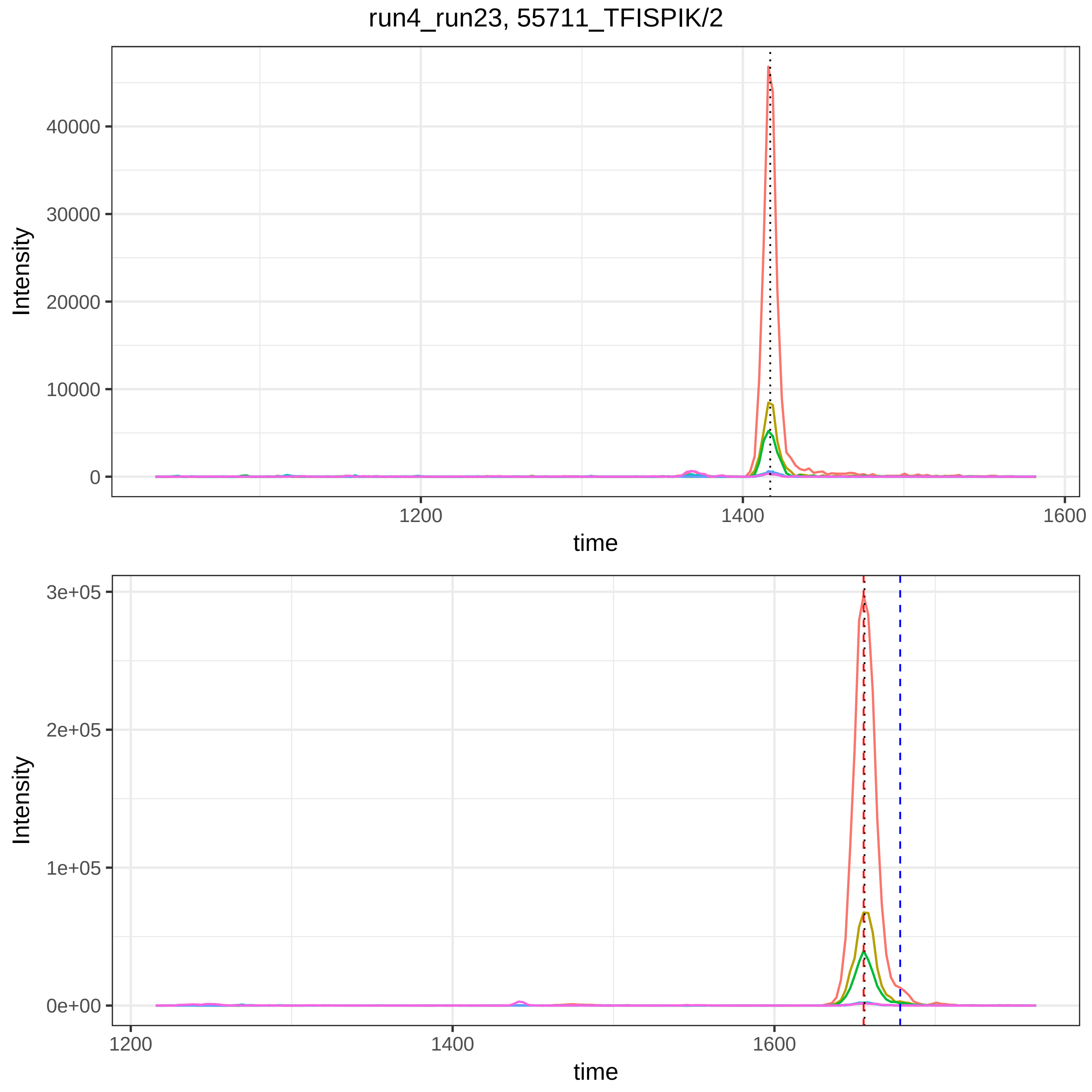

run4_run23, 55711_TFISPIK/2
Intensity
40000
30000
20000
10000
0
1200
1400
1600
time
Intensity
3e+05
2e+05
1e+05
0e+00
1200
1400
1600
time

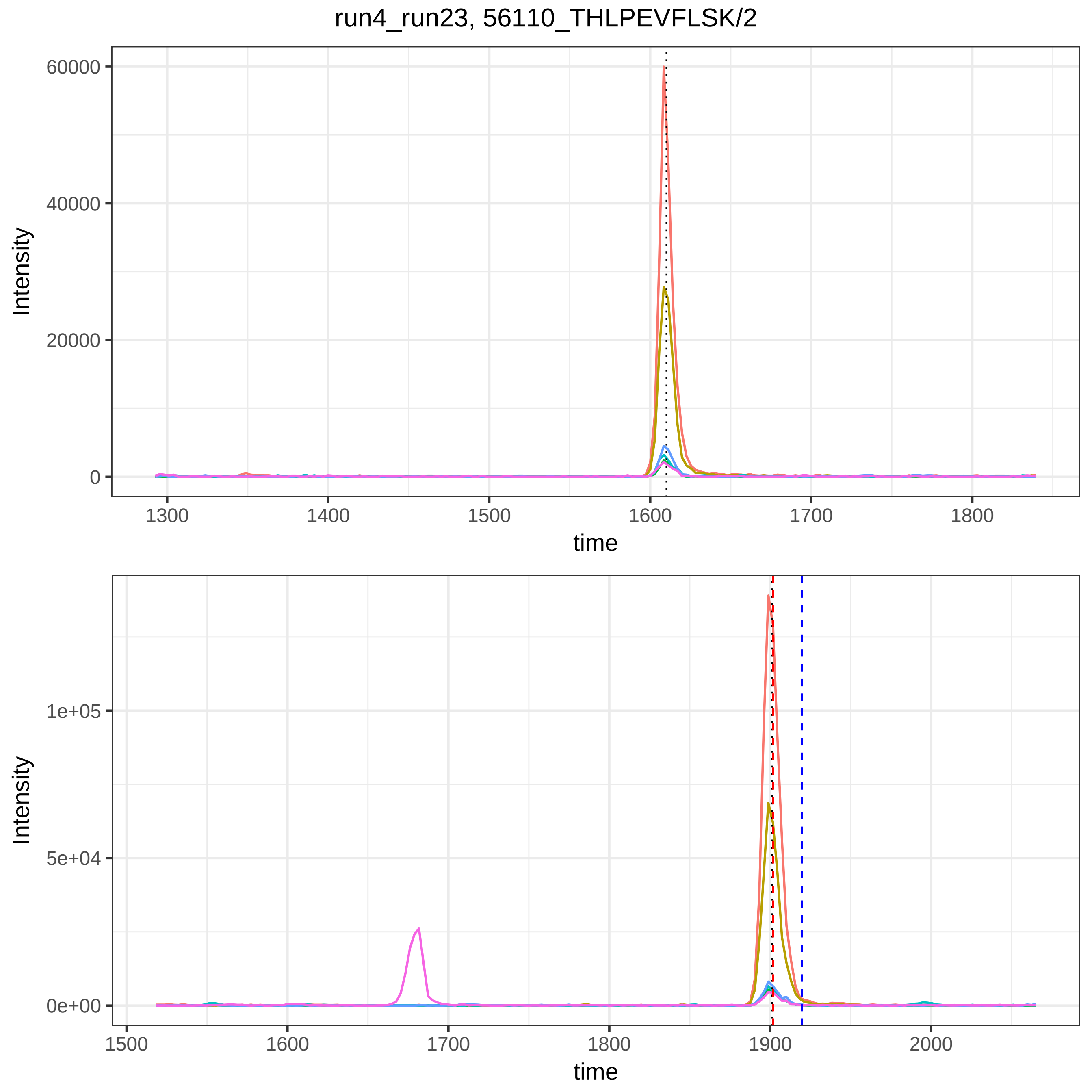

run4_run23, 56110_THLPEVFLSK/2
Intensity
60000
40000
20000
0
1300
1400
1500
1600
1700
1800
time
Intensity
1e+05
5e+04
0e+00
1500
1600
1700
1800
1900
2000
time

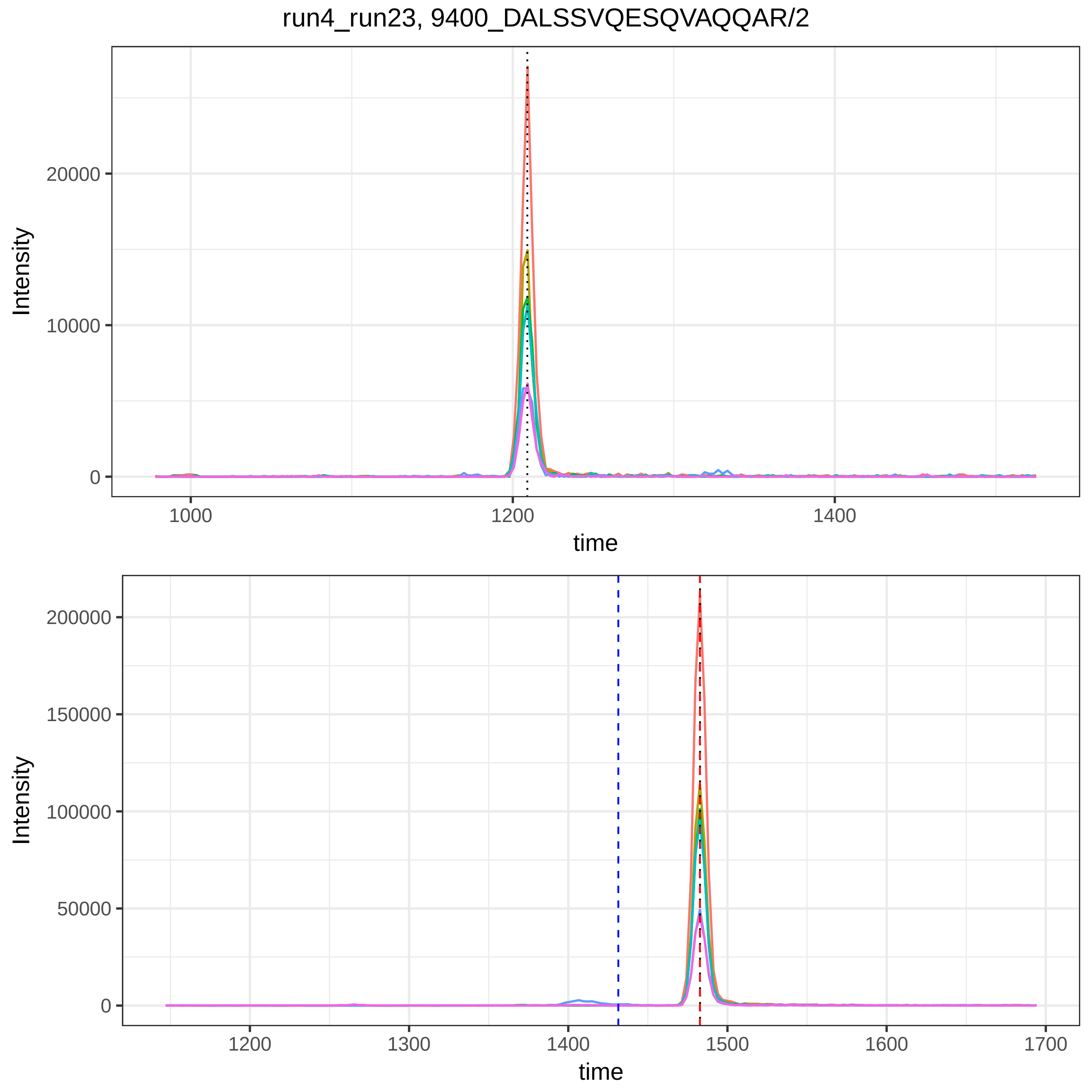
run4_run23, 9400_DALSSVQESQVAQQAR/2
Intensity
20000
10000
0
1000
1200
1400
time
Intensity
200000
150000
100000
50000
0
1200
1300
1400
1500
1600
1700
time

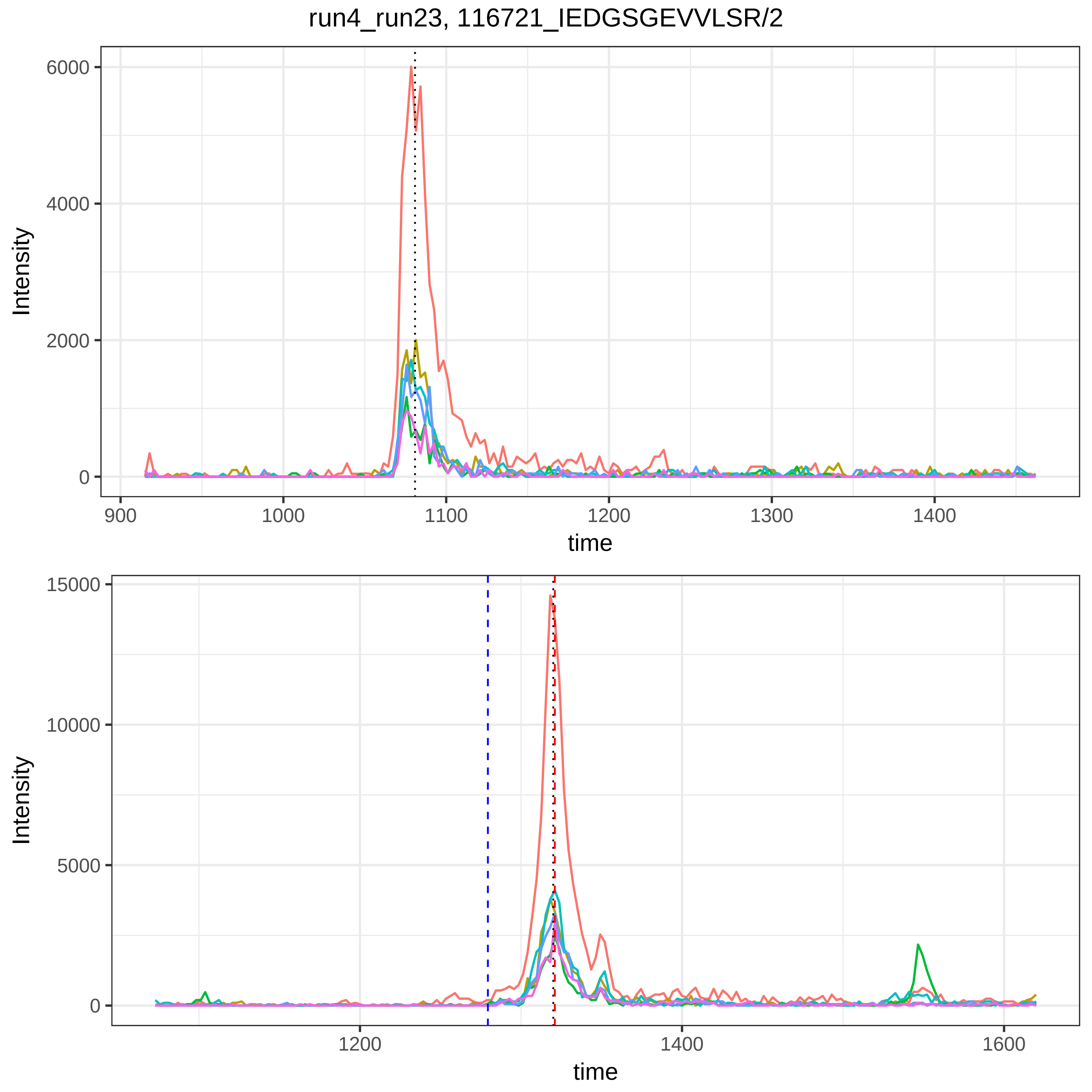
run4_run23, 116721_IEDGSGEVVLSR/2
Intensity
6000
4000
2000
0
900
1000
1100
1200
1300
1400
time
Intensity
15000
10000
5000
0
1200
1400
1600
time

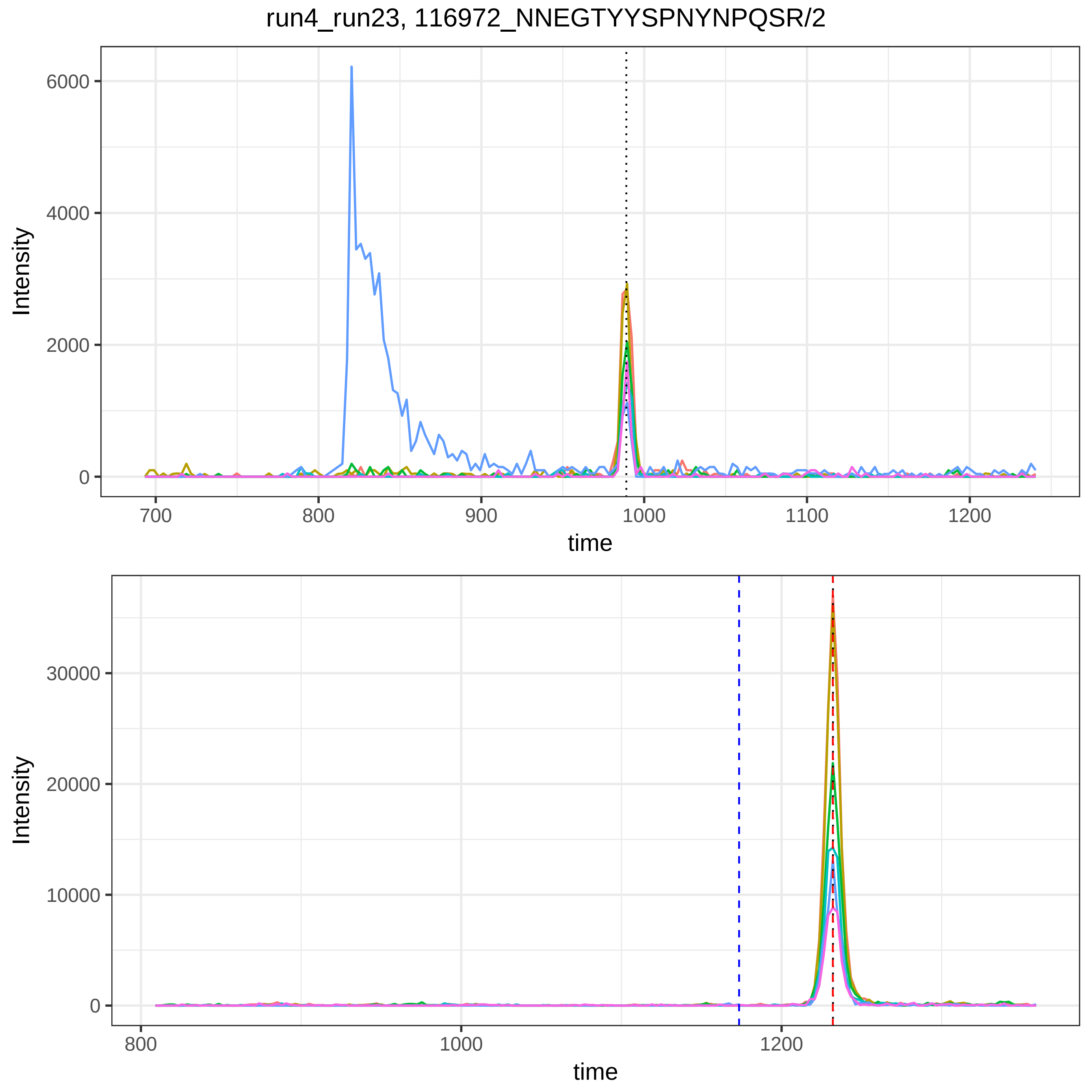

run4_run23, 116972_NNEGTYYSPNYNPQSR/2
Intensity
time
0
2000
4000
6000
700
800
900
1000
1100
1200
0
10000
20000
30000
800
1000
1200

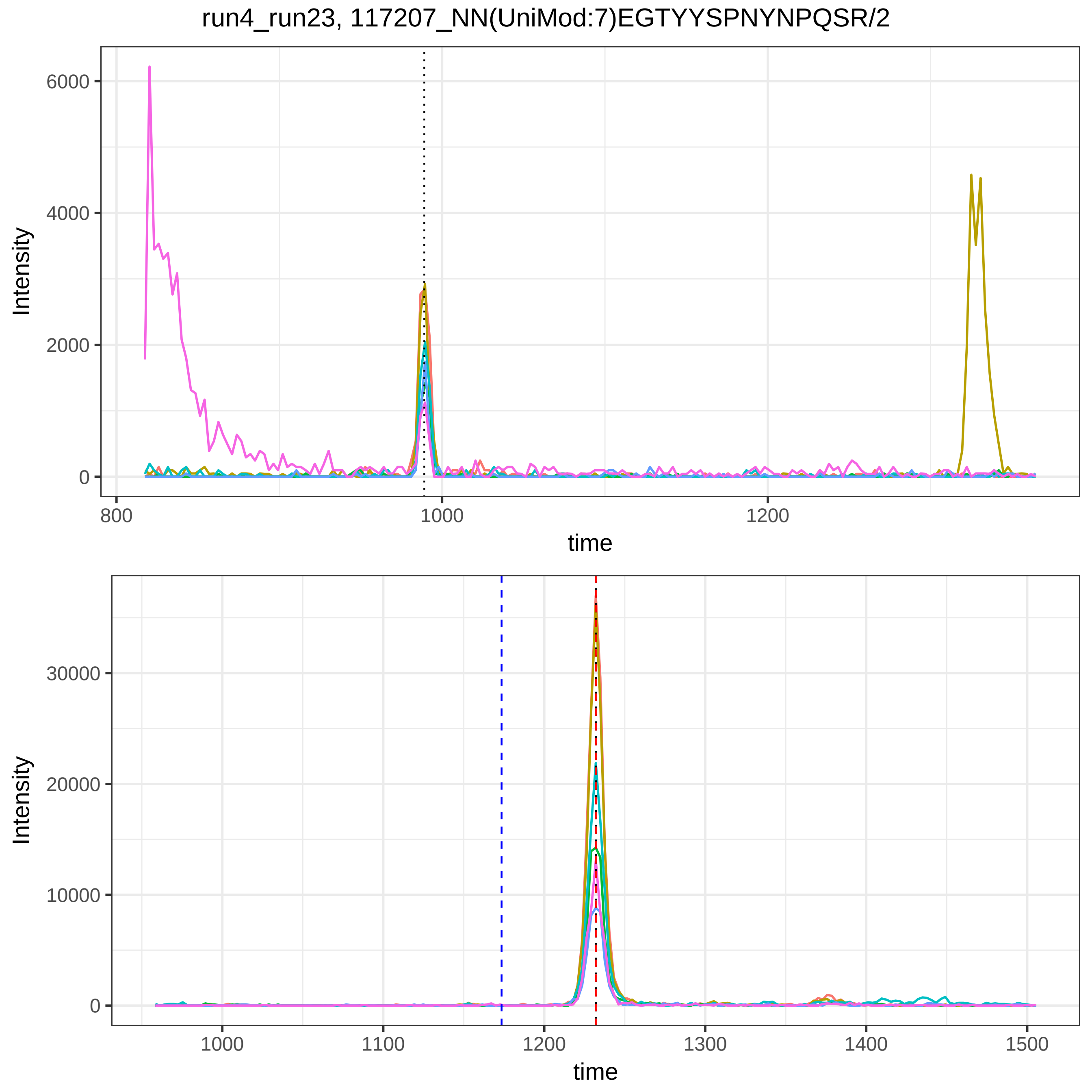
run4_run23, 117207_NN(UniMod:7)EGTYYSPNYNPQSR/2
Intensity
6000
4000
2000
0
800
1000
1200
time
Intensity
30000
20000
10000
0
1000
1100
1200
1300
1400
1500
time

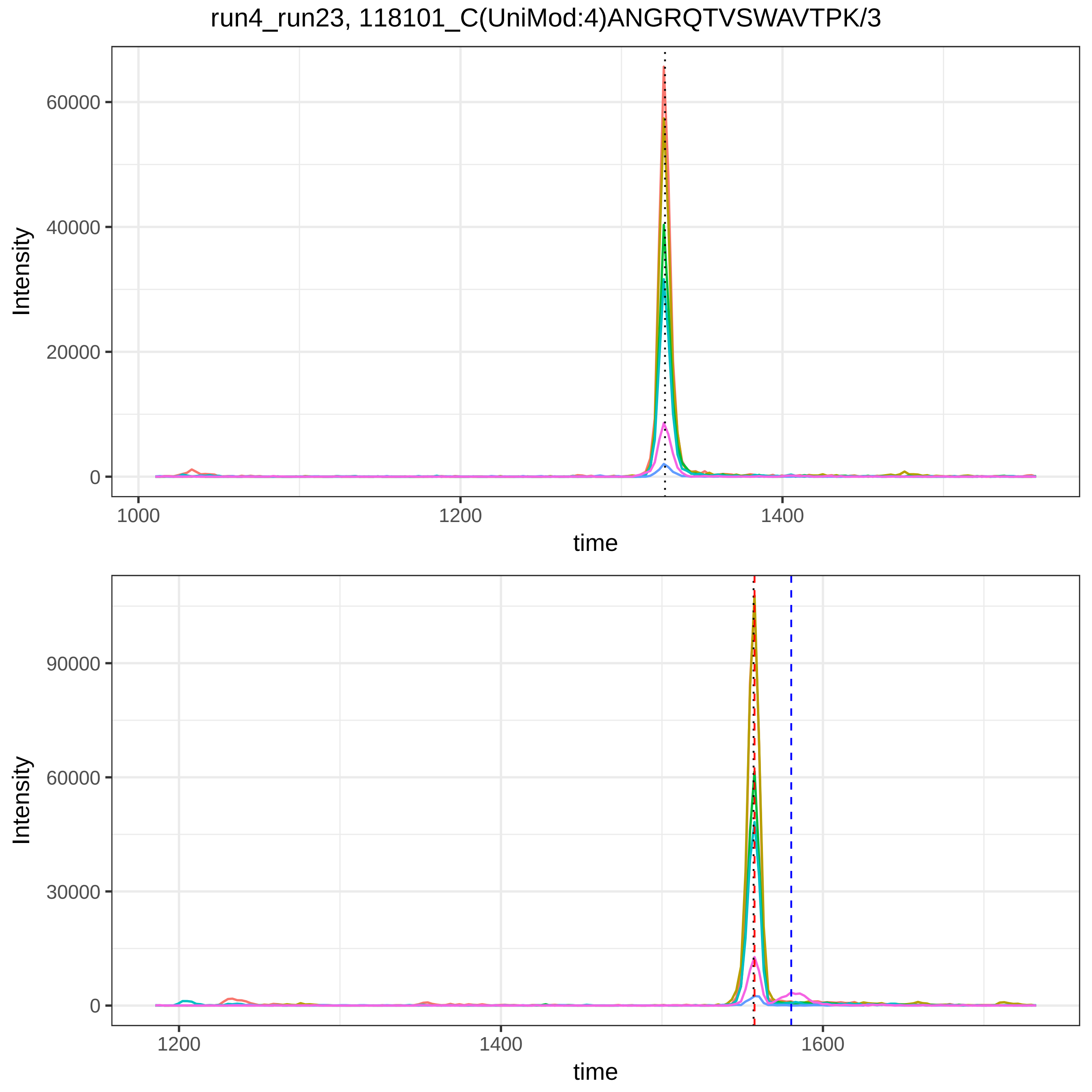
run4_run23, 118101_C(UniMod:4)ANGRQTVSWAVTPK/3
Intensity
60000
40000
20000
0
1000
1200
1400
time
Intensity
90000
60000
30000
0
1200
1400
1600
time

run4_run23, 118454_DSGFQMNQLR/2

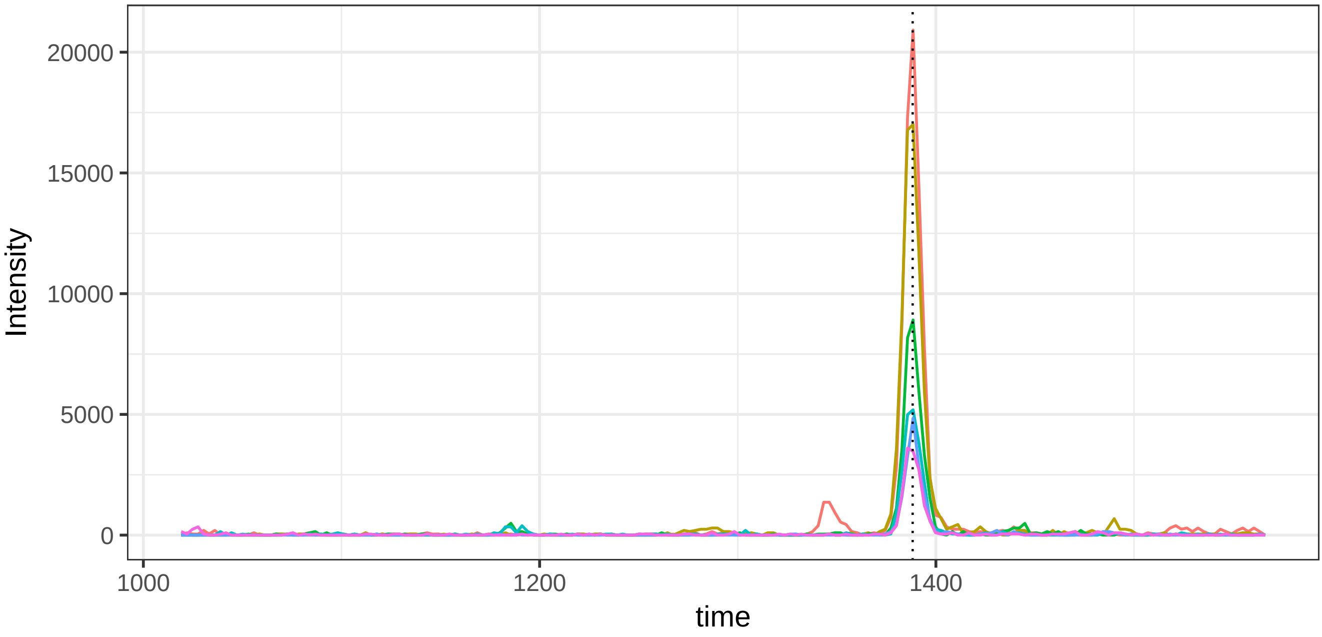


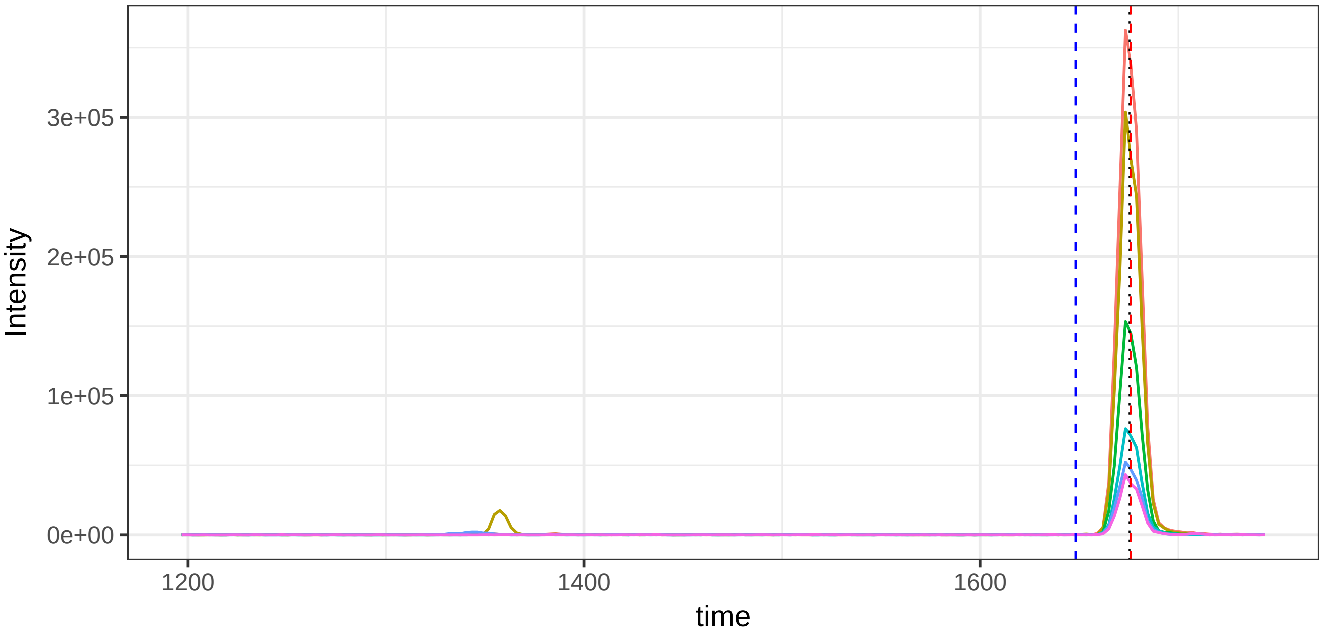

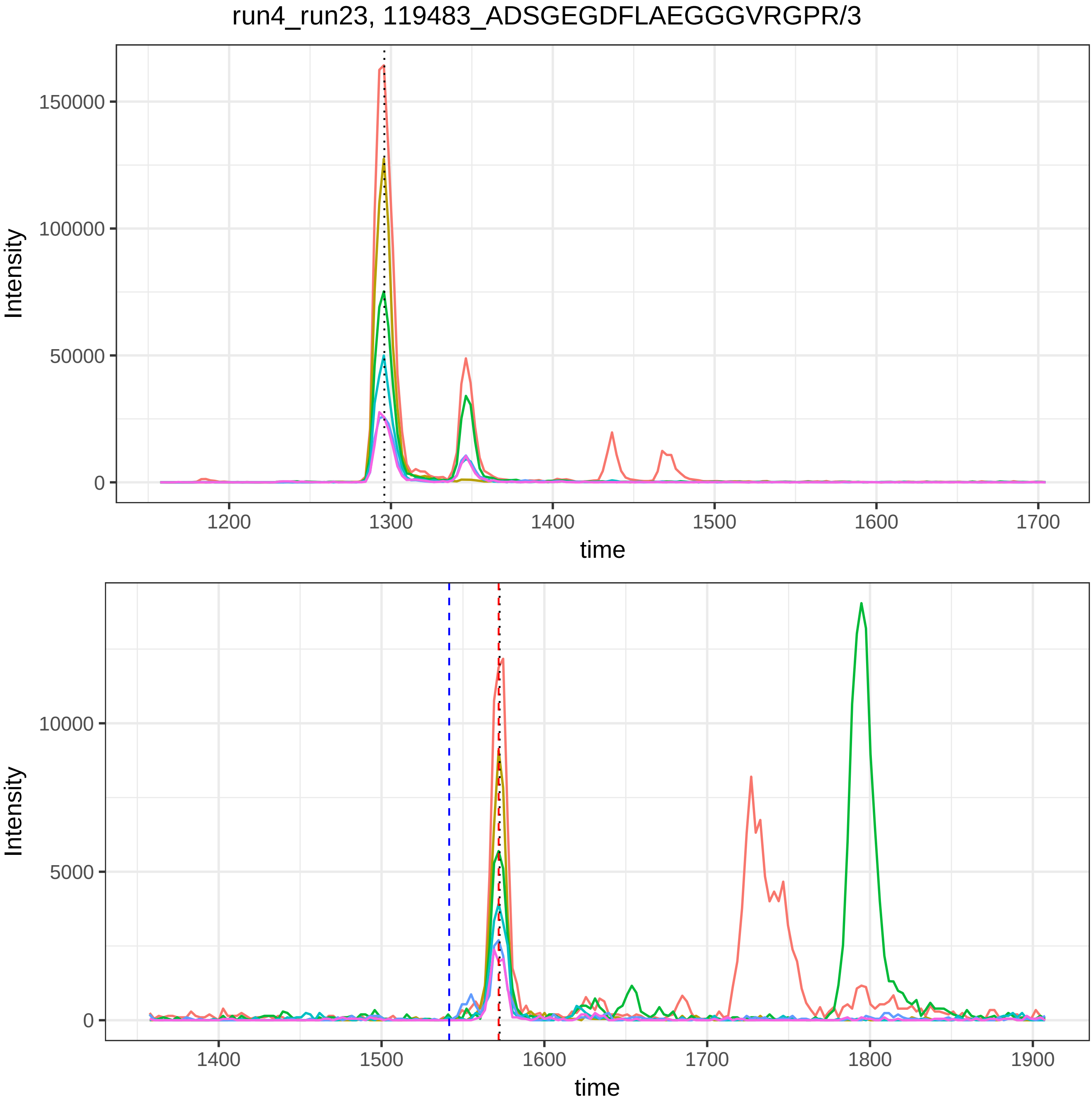

run4_run23, 119483_ADSGEGDFLAEGGGVRGPR/3
Intensity
150000
100000
50000
0
1200
1300
1400
1500
1600
1700
time
Intensity
10000
5000
0
1400
1500
1600
1700
1800
1900
time

run4_run23, 119499_DSGEGDFLAEGGGVR/2

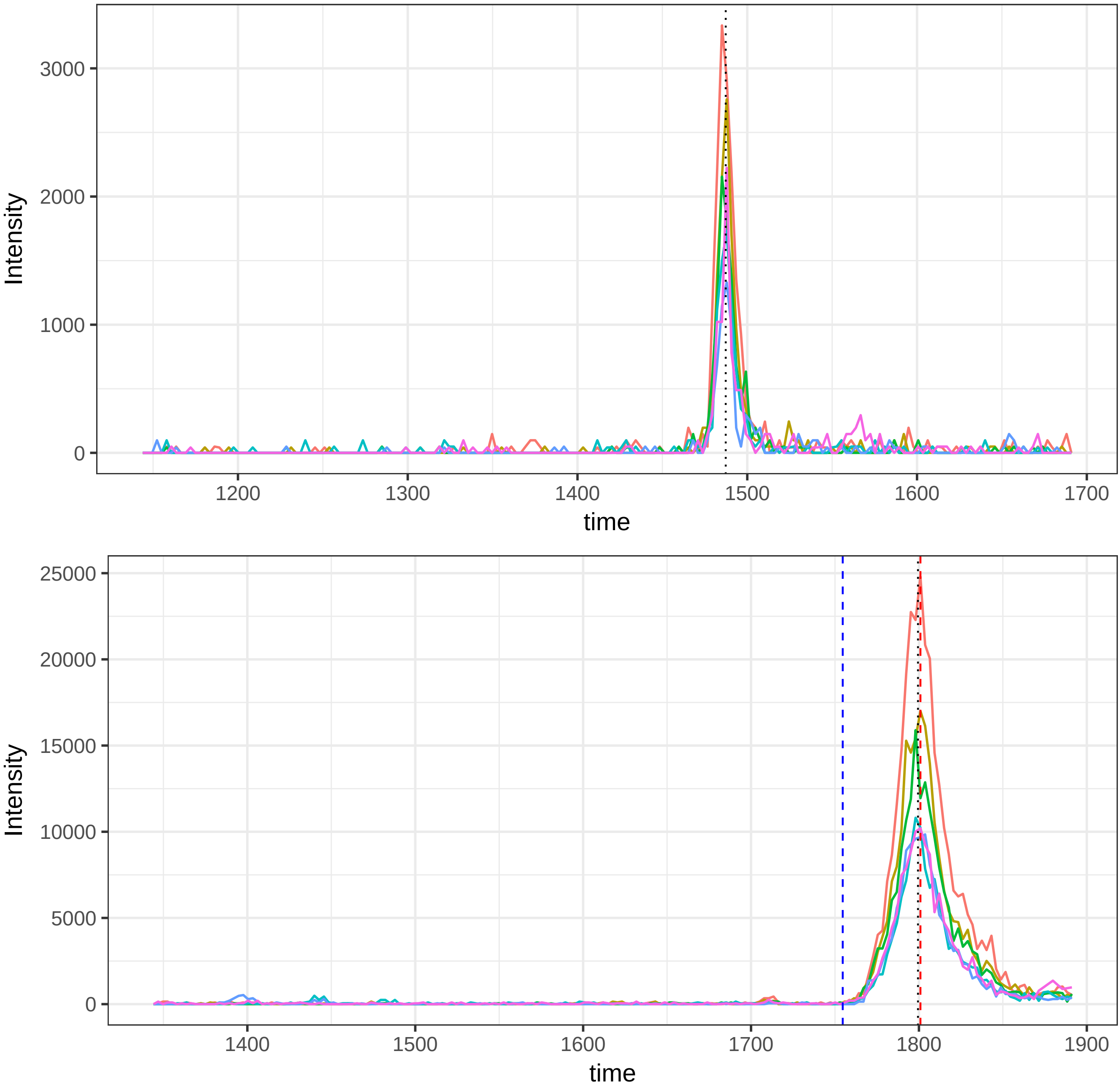

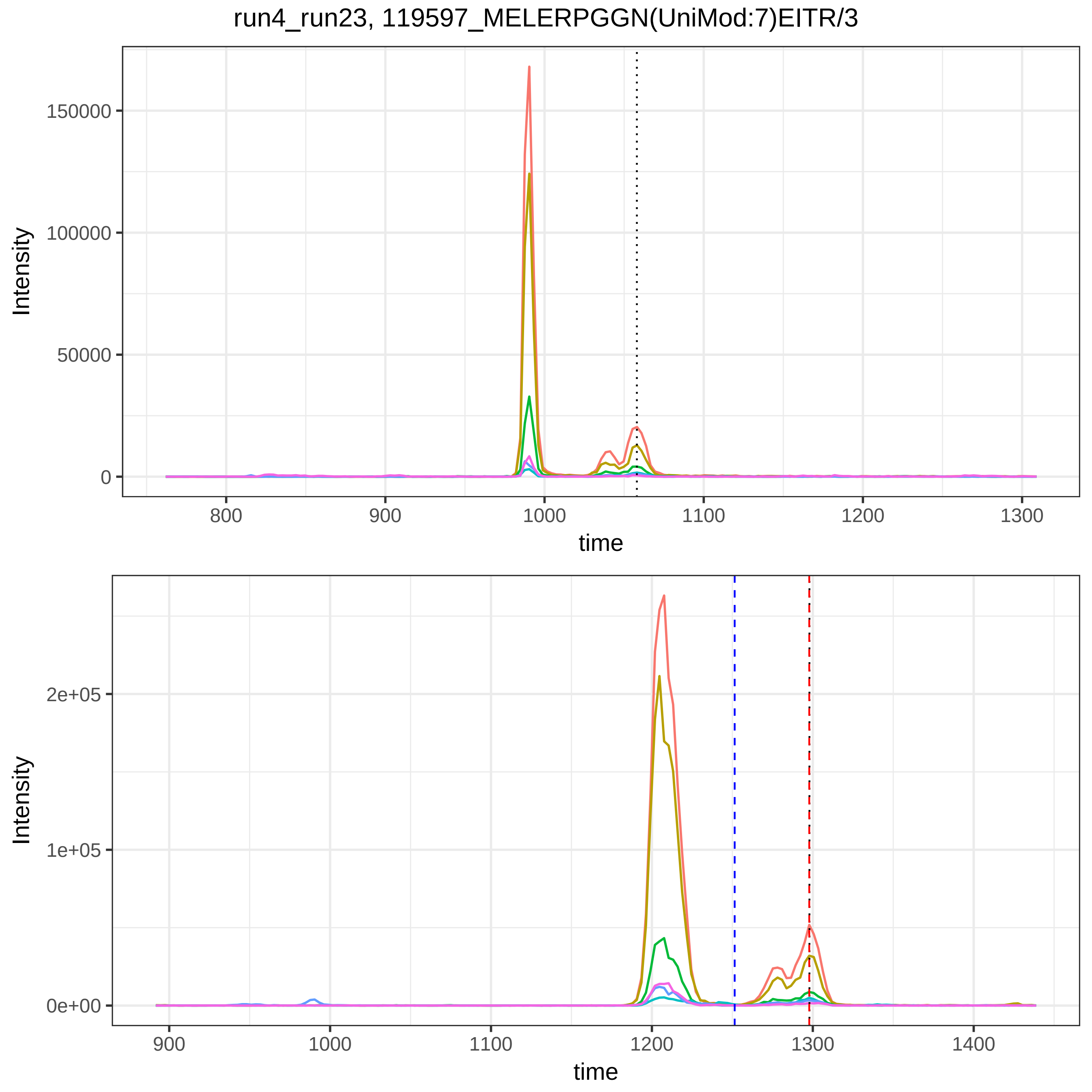
run4_run23, 119597_MELERPGGN(UniMod:7)EITR/3
Intensity
150000
100000
50000
0
800
900
1000
1100
1200
1300
time
Intensity
2e+05
1e+05
0e+00
900
1000
1100
1200
1300
1400
time

run4_run23, 119602_N(UniMod:7)SLFEYQK/2

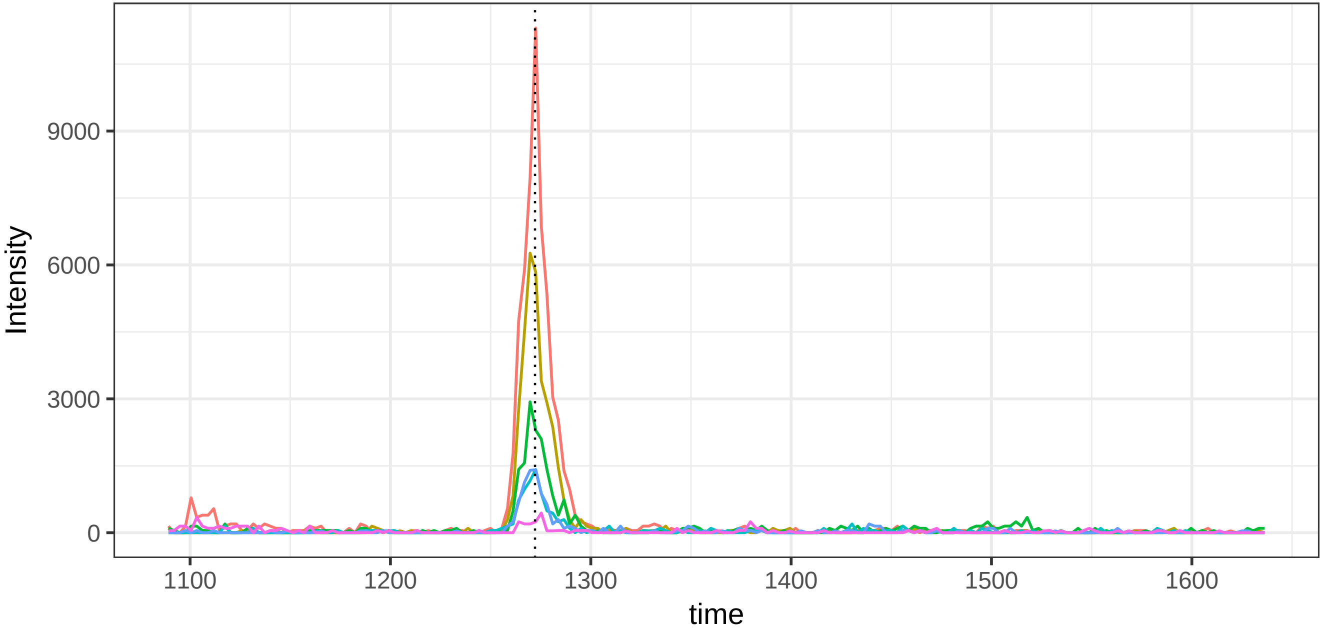


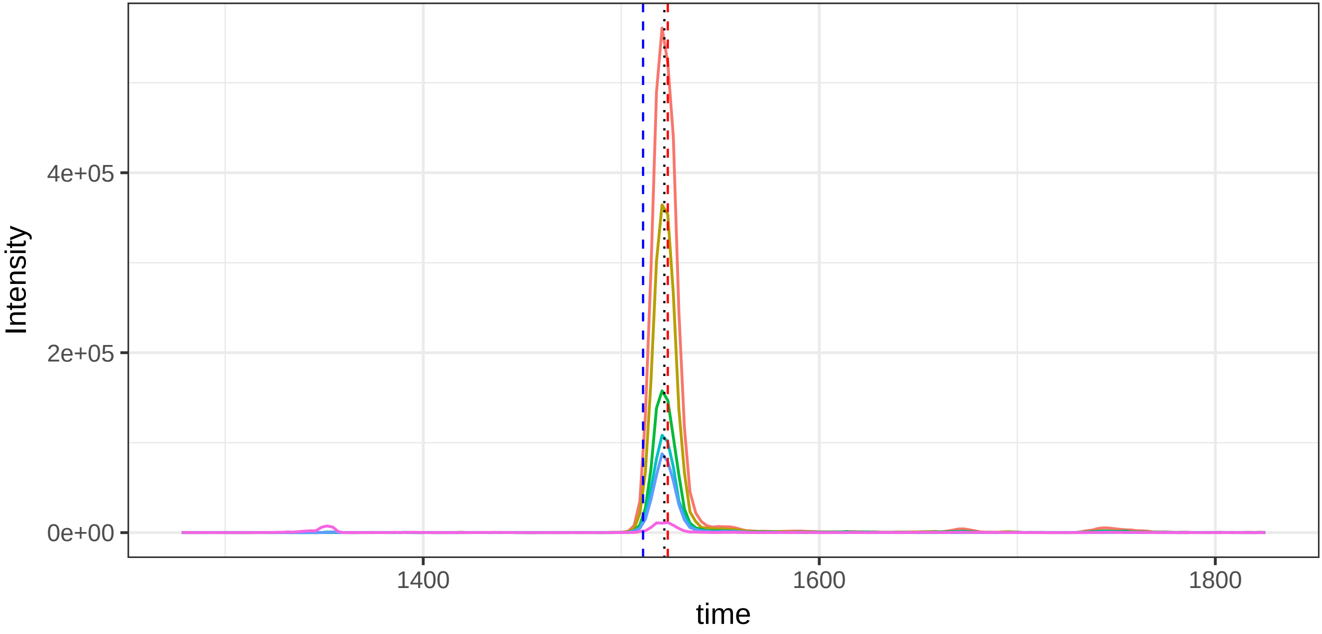

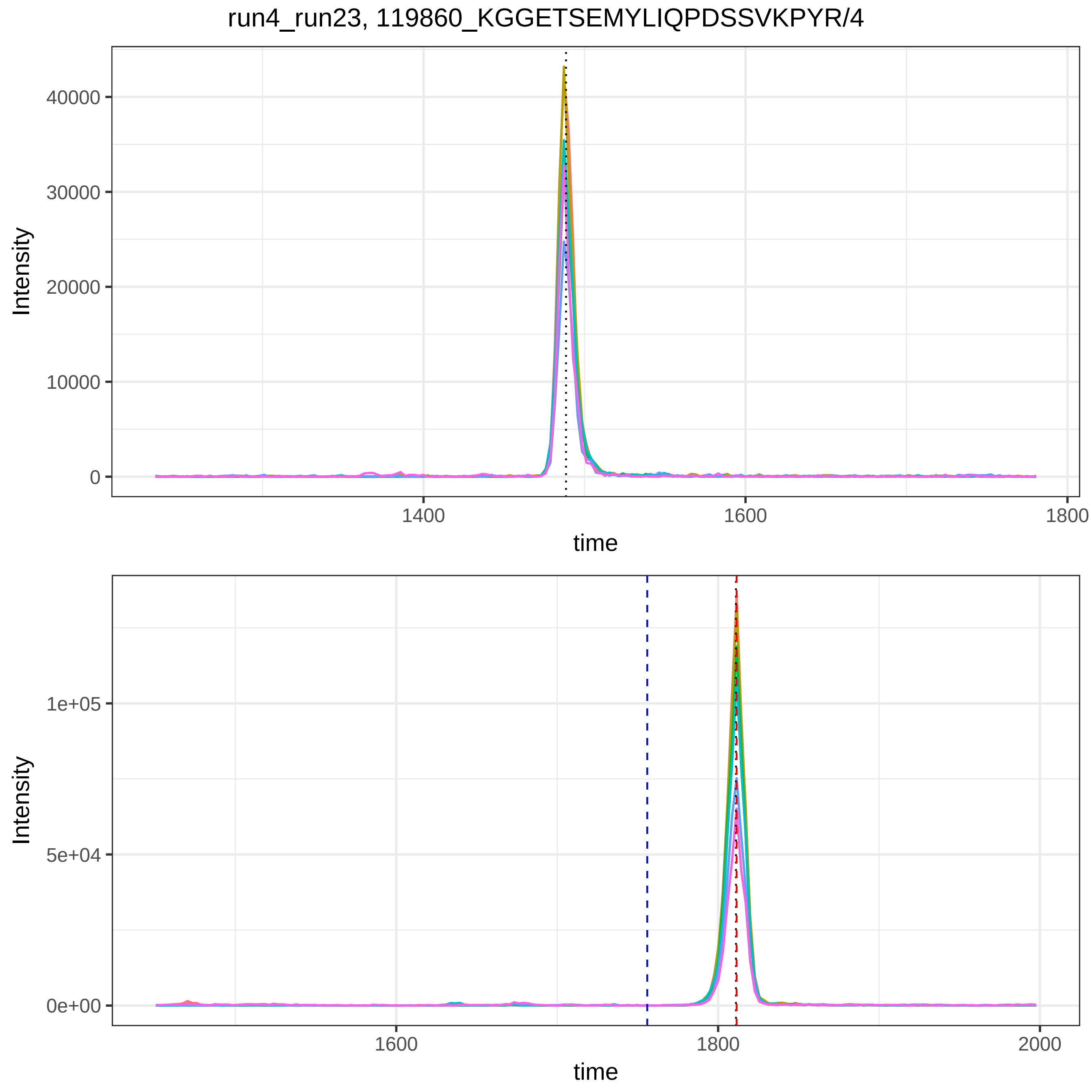

run4_run23, 119860_KGGETSEMYLIQPDSSVKPYR/4
Intensity
time
40000
30000
20000
10000
0
1400
1600
1800
1e+05
5e+04
0e+00
1600
1800
2000

run4_run23, 119990_KGGETSEMYLIQ(UniMod:7)PDSSVKPYR/4

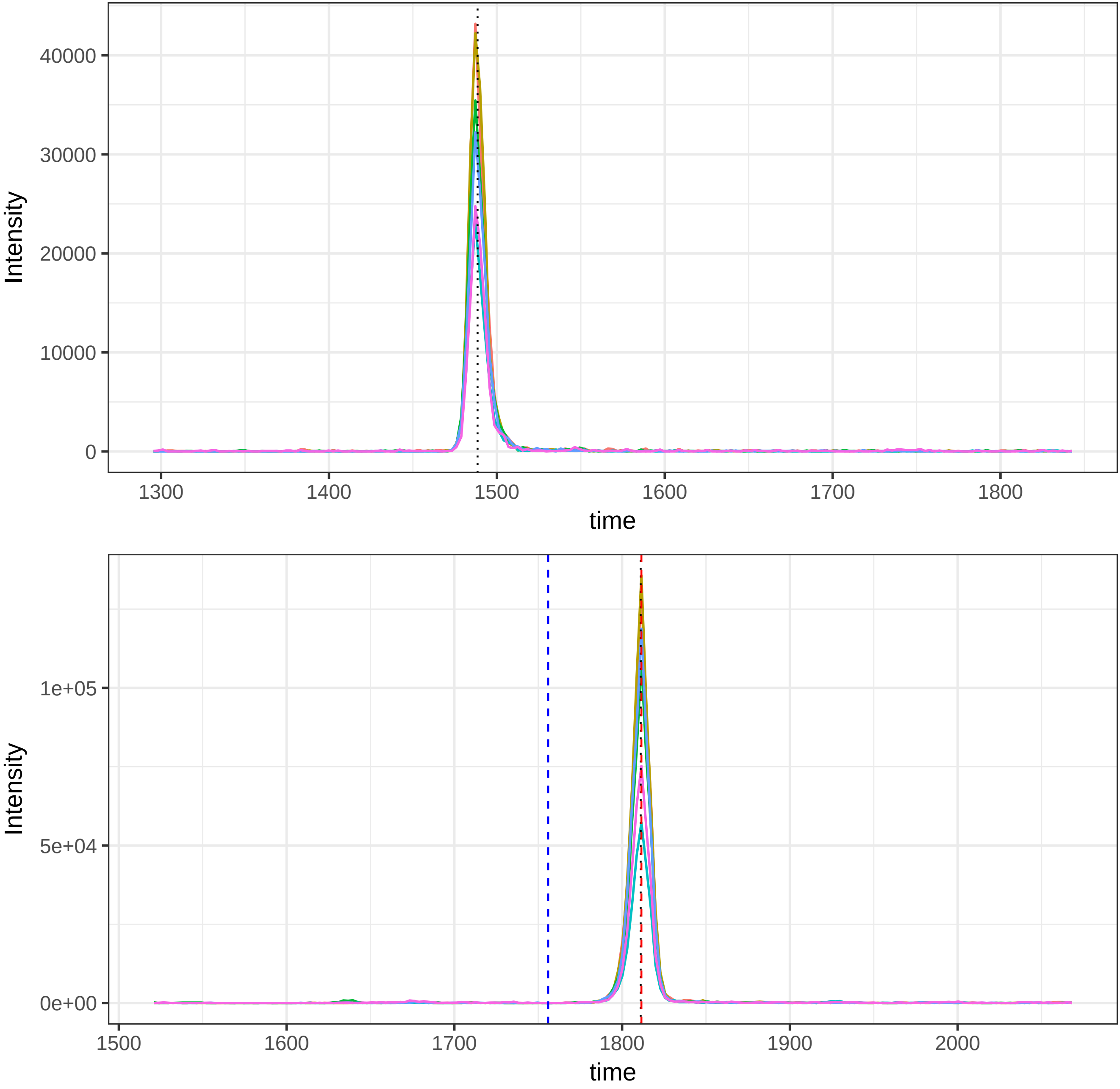

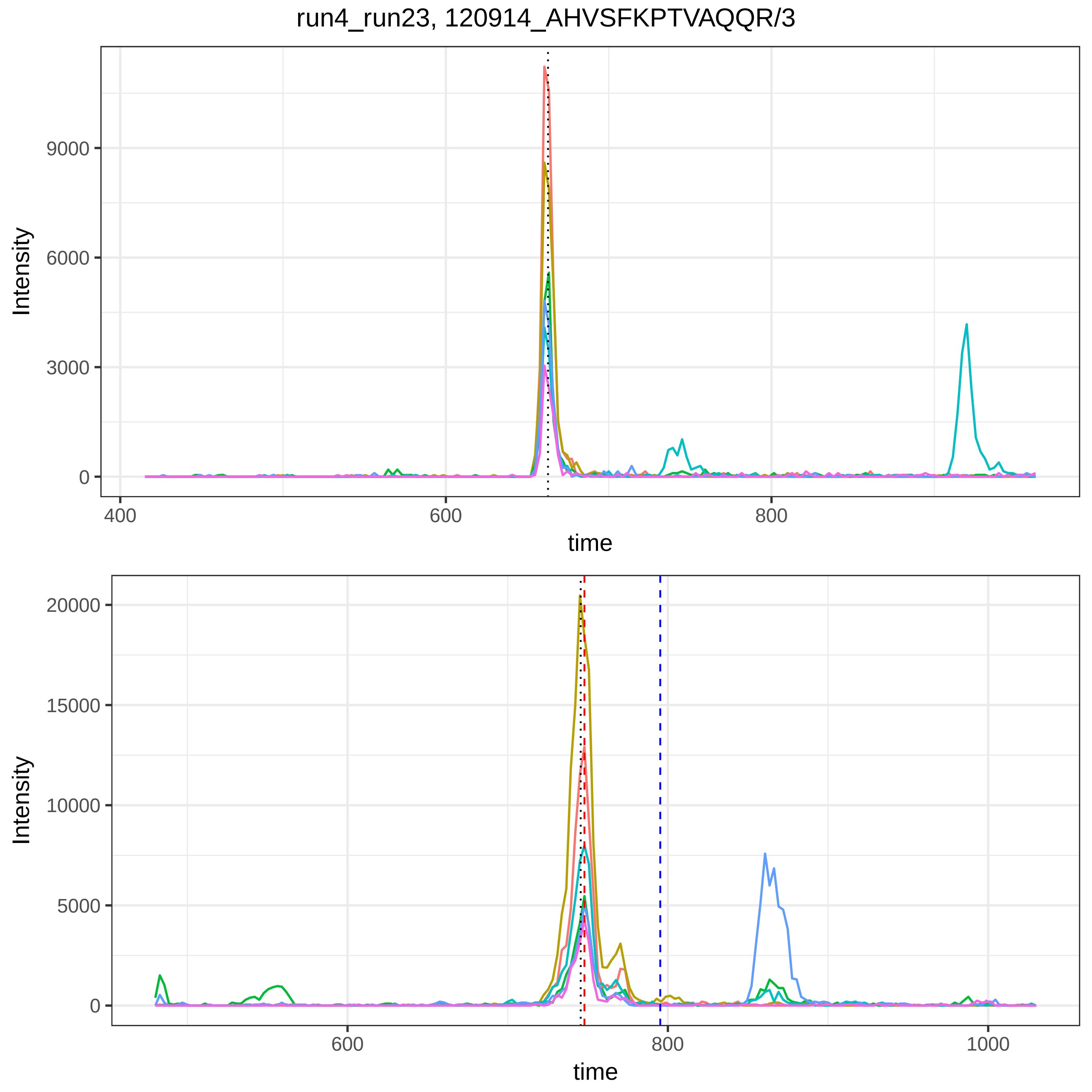

run4_run23, 120914_AHVSFKPTVAQQR/3
Intensity
9000
6000
3000
0
400
600
800
time
Intensity
20000
15000
10000
5000
0
600
800
1000
time

run4_run23, 12135_DFALQNPSAVPR/2

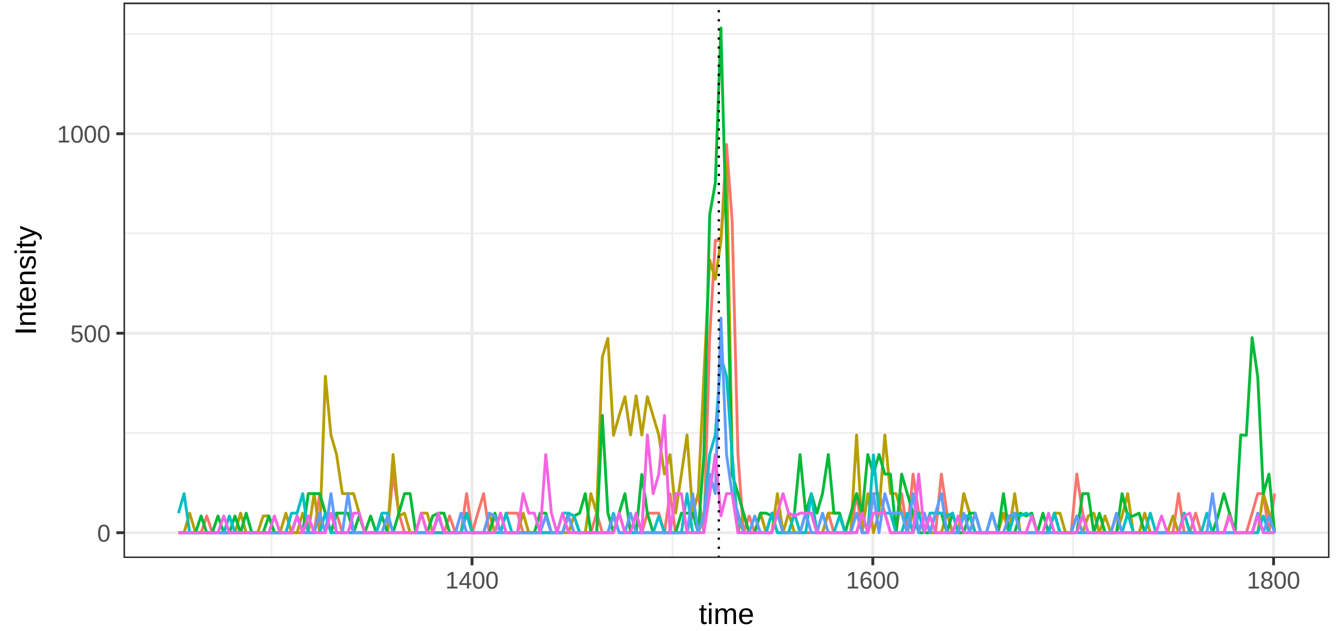


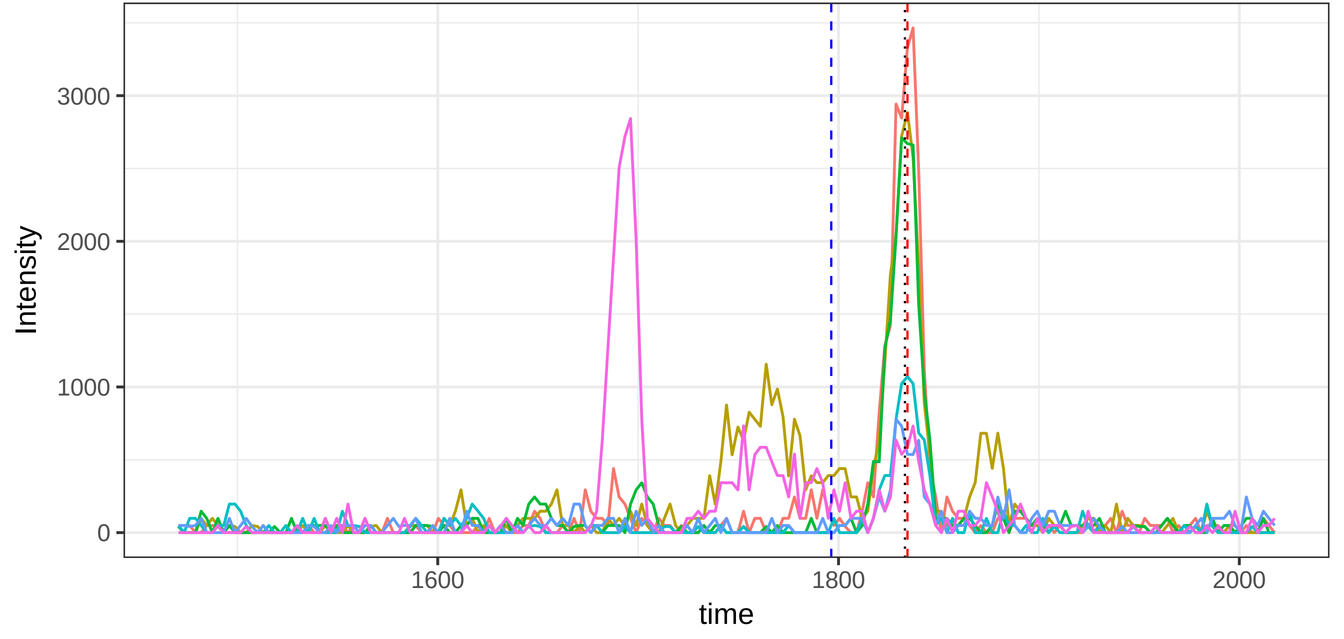

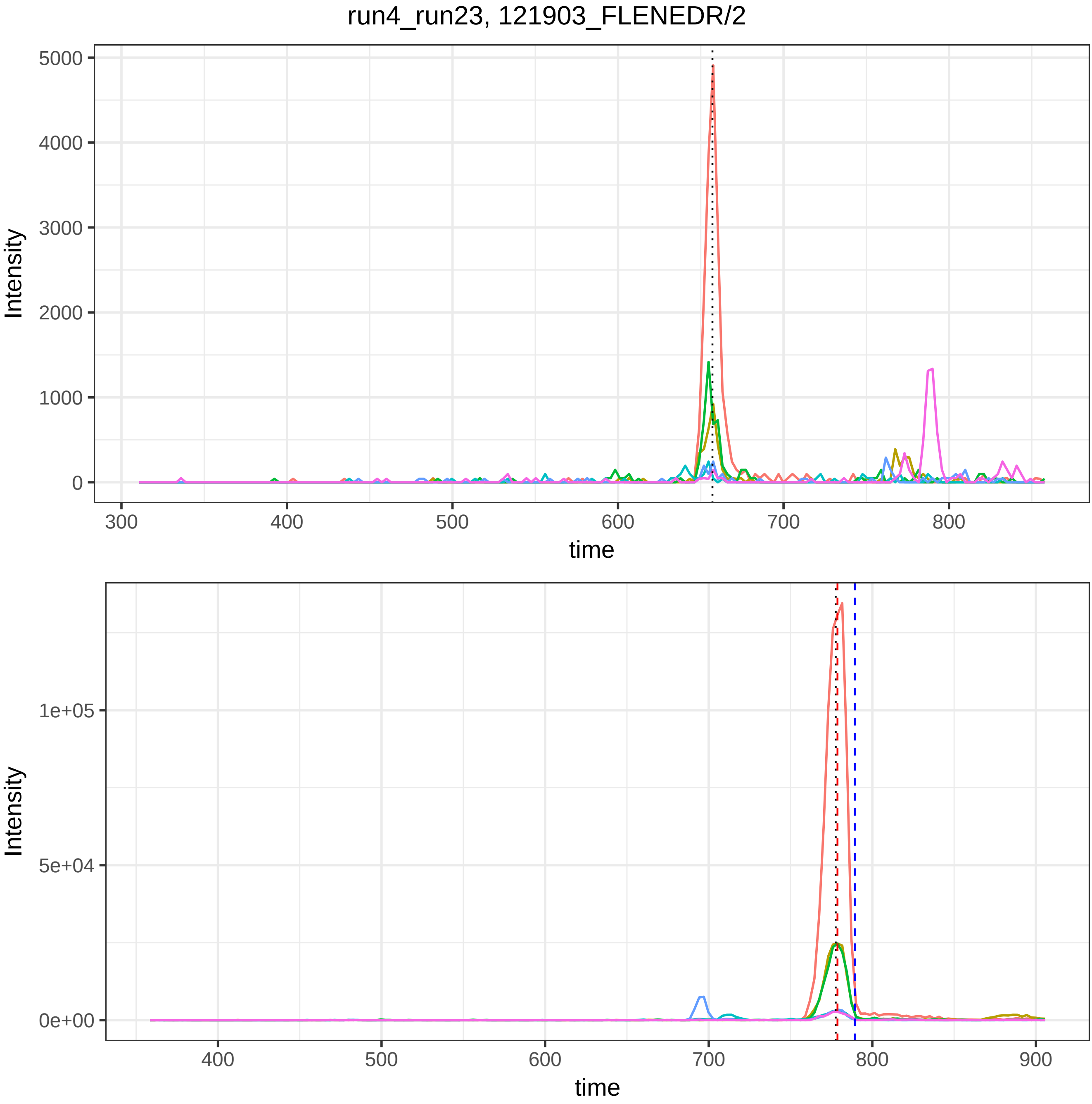

run4_run23, 121903_FLENEDR/2
Intensity
time
5000
4000
3000
2000
1000
0
300
400
500
600
700
800
1e+05
5e+04
0e+00
900

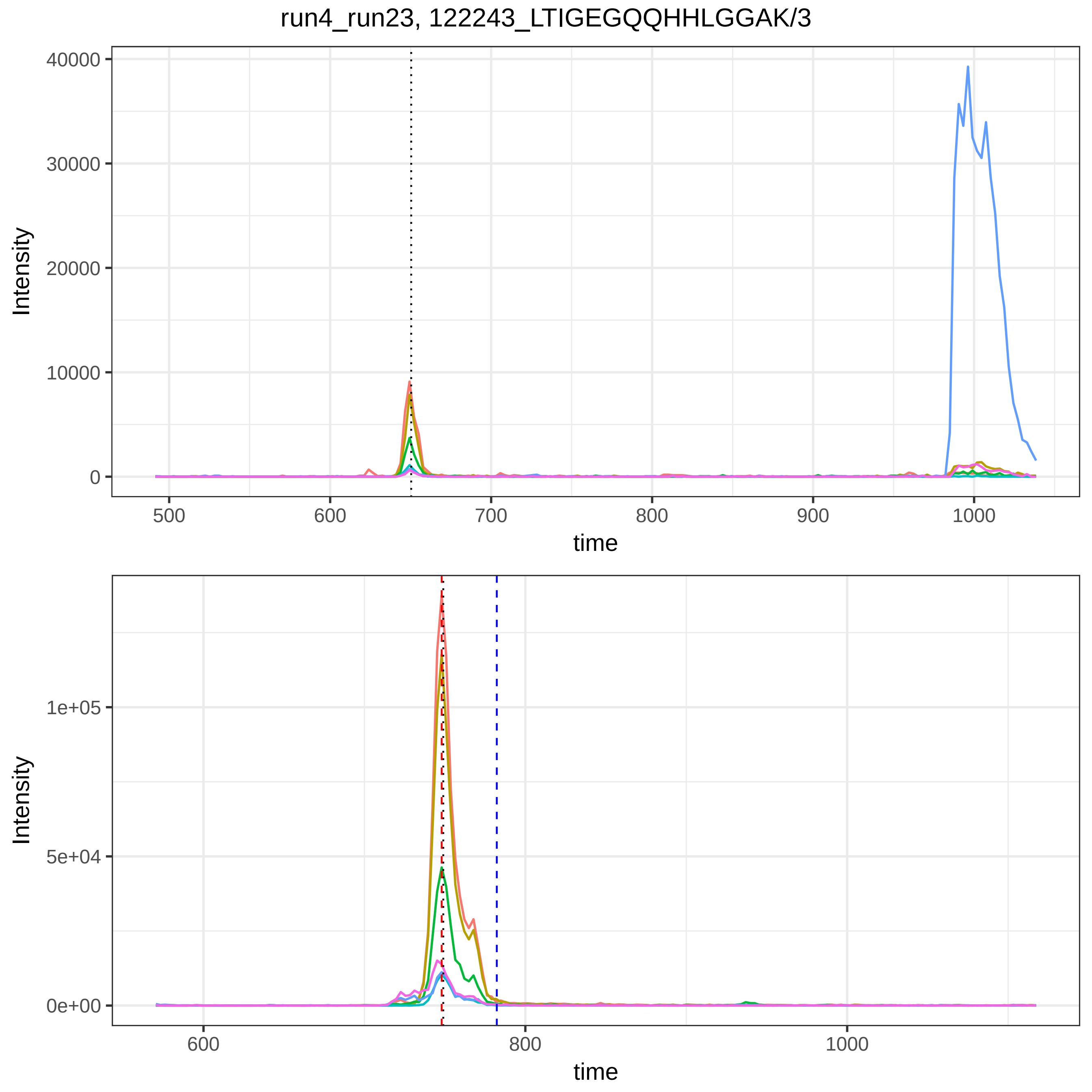
run4_run23, 122243_LTIGEGQQHHLGGAK/3
Intensity
40000
30000
20000
10000
0
500
600
700
800
900
1000
time
Intensity
1e+05
5e+04
0e+00
600
800
1000
time

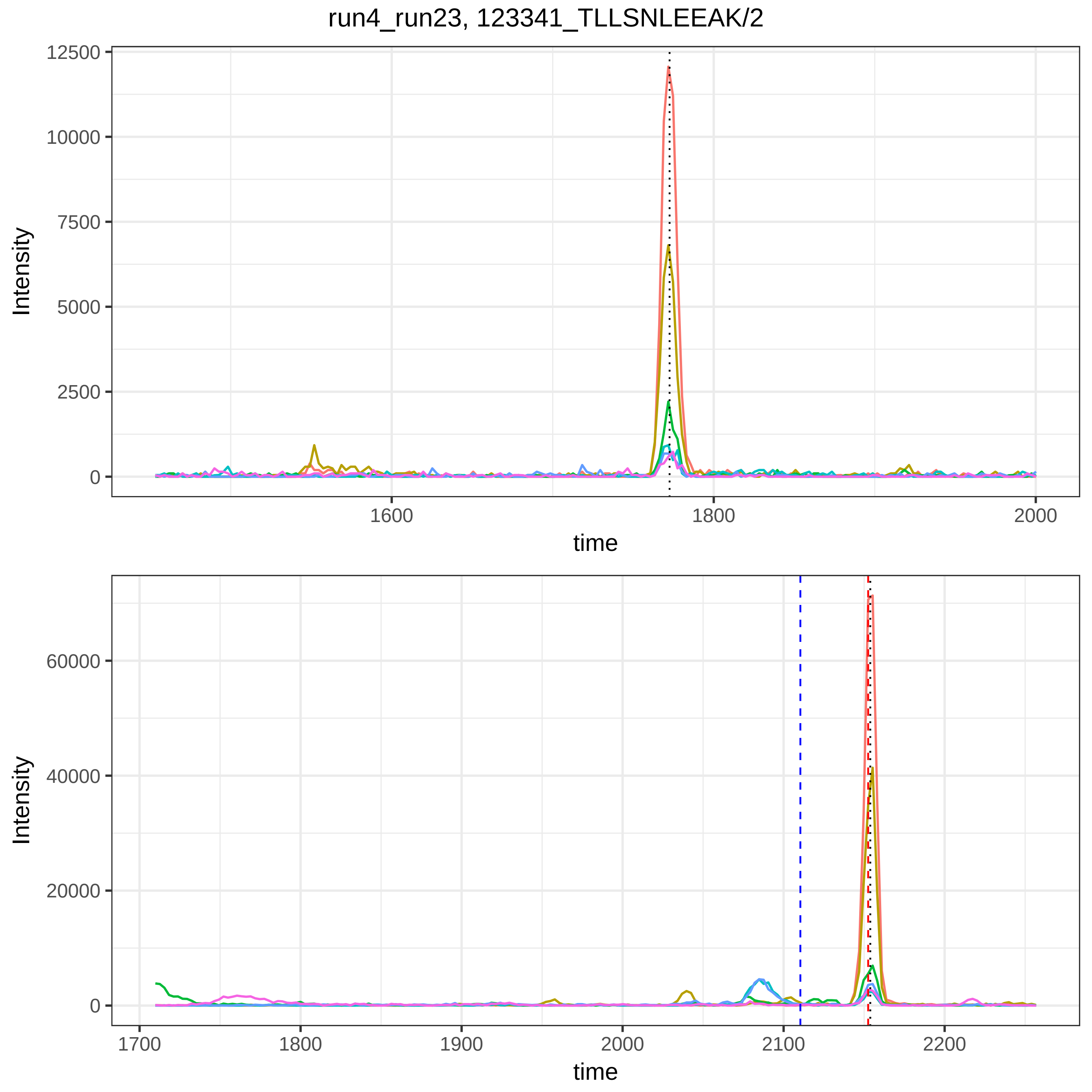

run4_run23, 123341_TLLSNLEEAK/2
Intensity
12500
10000
7500
5000
2500
0
1600
1800
2000
time
Intensity
60000
40000
20000
0
1700
1800
1900
2000
2100
2200
time

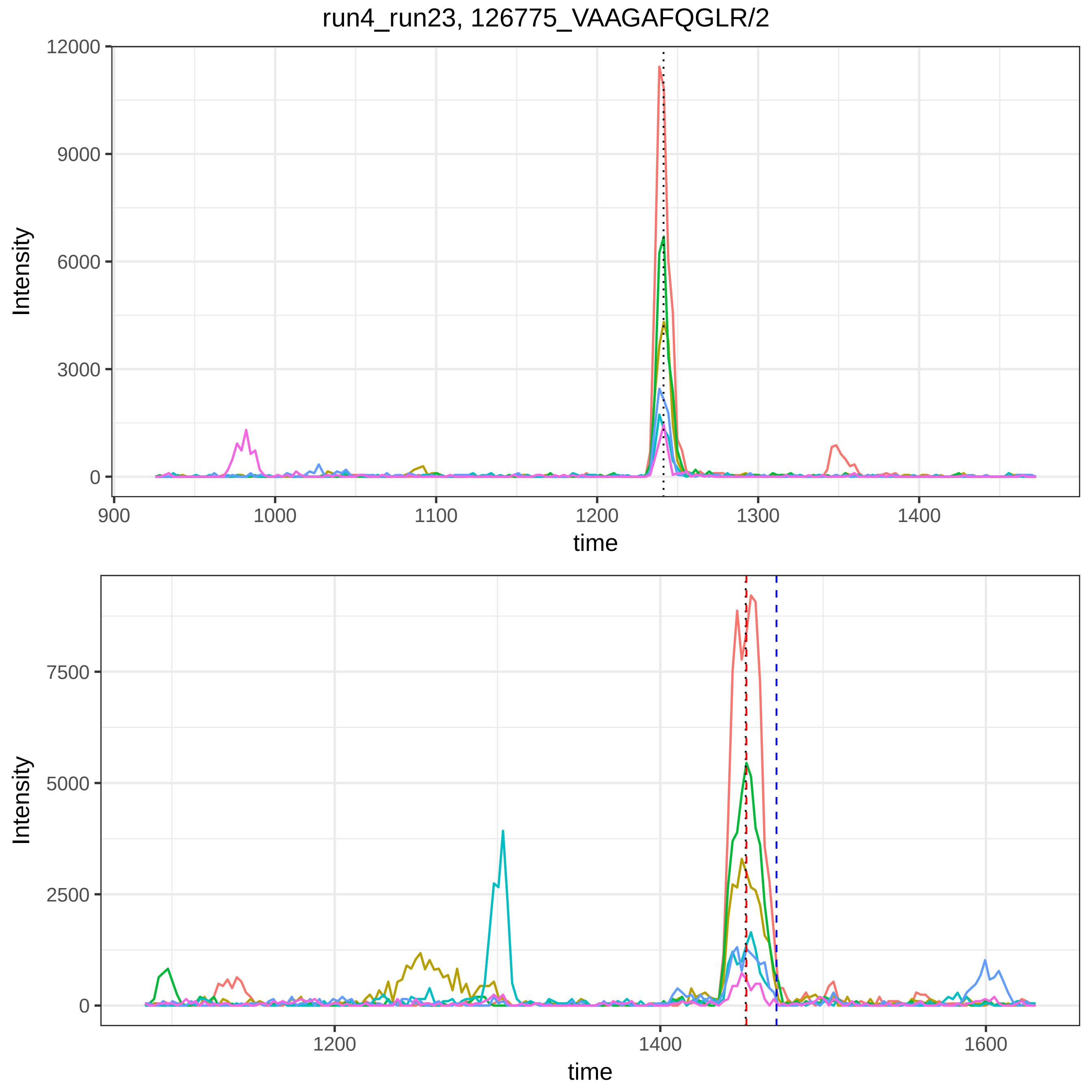
run4_run23, 126775_VAAGAFQGLR/2
Intensity
12000
9000
6000
3000
0
900
1000
1100
1200
1300
1400
time
Intensity
7500
5000
2500
0
1200
1400
1600
time

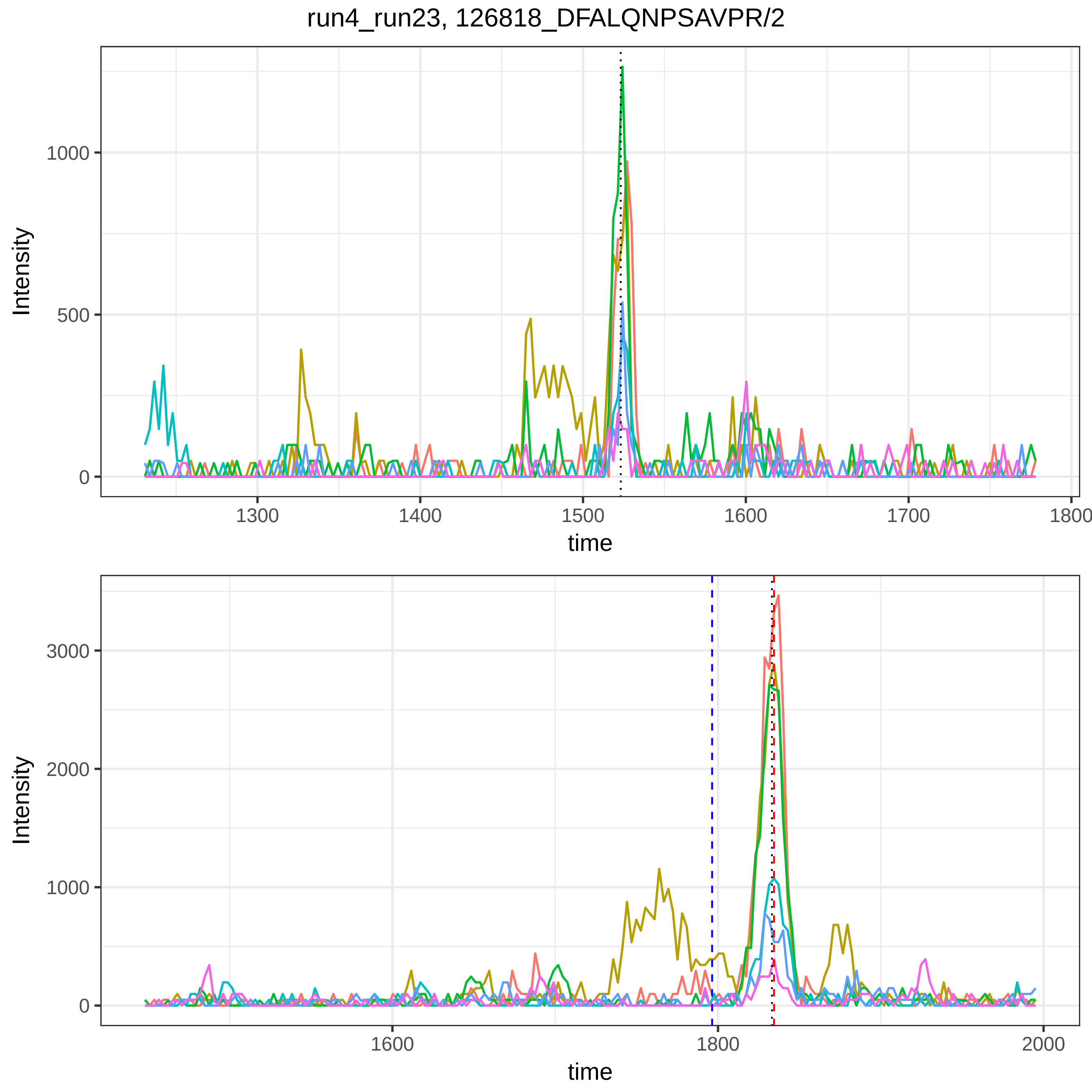
run4_run23, 126818_DFALQNPSAVPR/2
Intensity
1000
500
0
1300
1400
1500
1600
1700
1800
time
Intensity
3000
2000
1000
0
1600
1800
2000
time

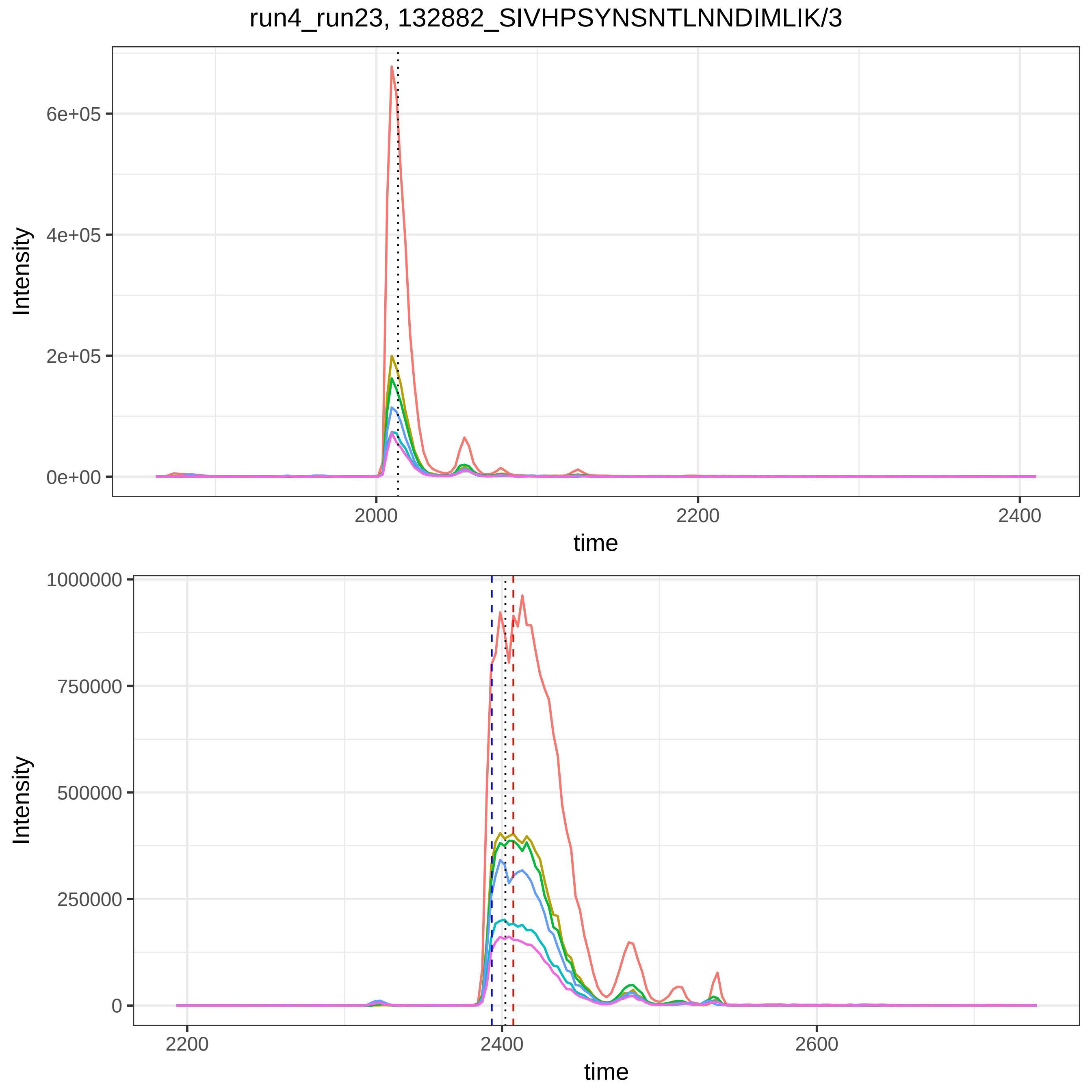

run4_run23, 132882_SIVHPSYNSNTLNNDIMLIK/3
Intensity
6e+05
4e+05
2e+05
0e+00
2000
2200
2400
time
Intensity
1000000
750000
500000
250000
0
2200
2400
2600
time

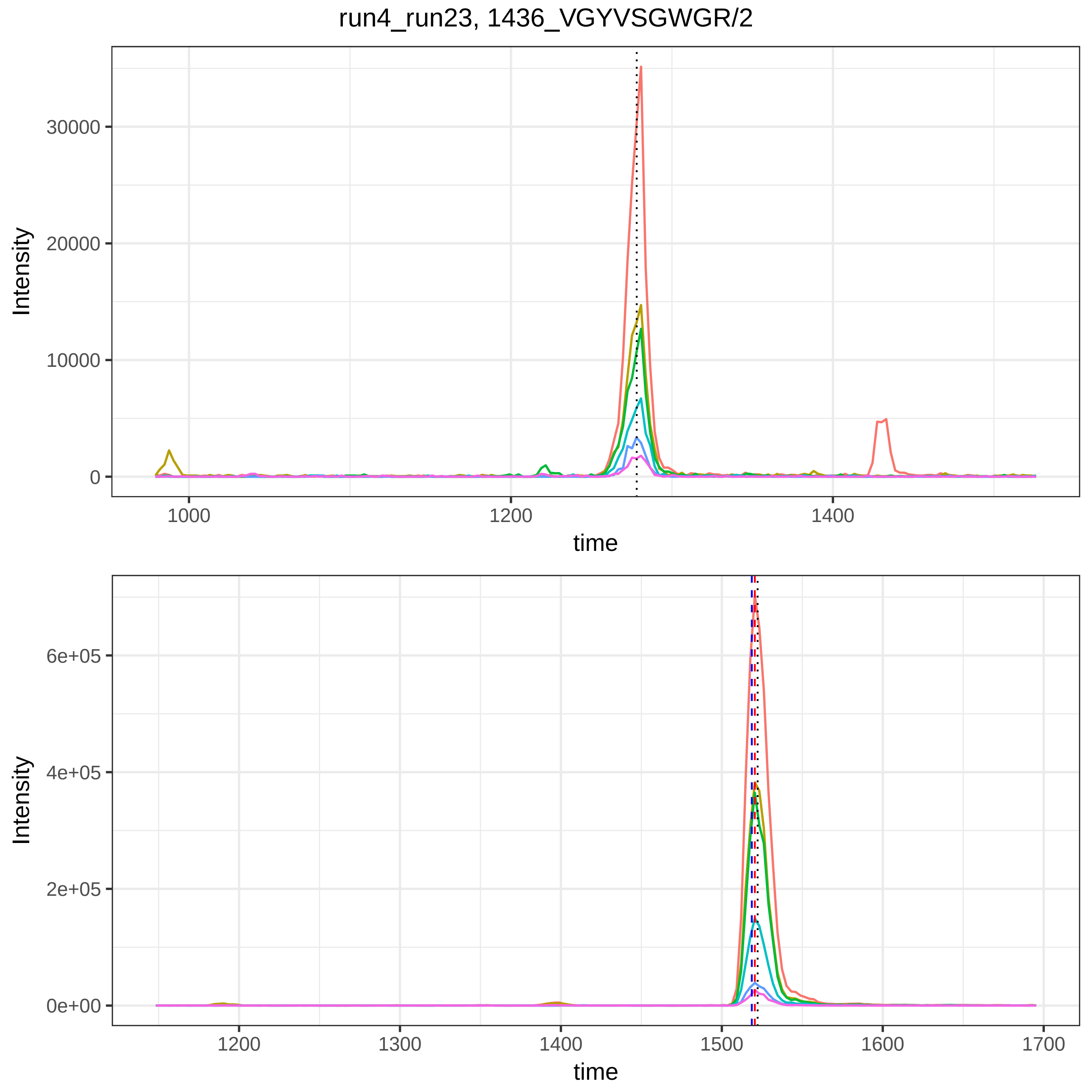
run4_run23, 1436_VGYVSGWGR/2
Intensity
time
30000
20000
10000
0
1000
1200
1400
6e+05
4e+05
2e+05
0e+00
1300
1500
1600
1700

run4_run23, 20944_DSGEGDFLAEGGGVR/2

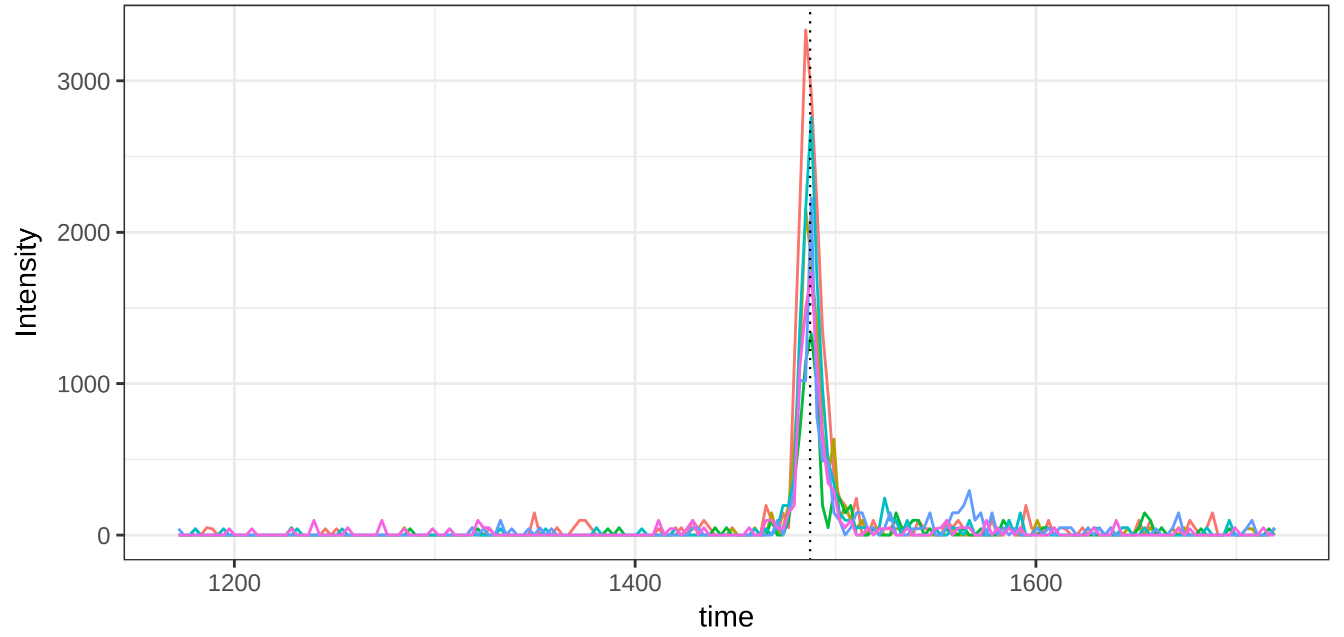


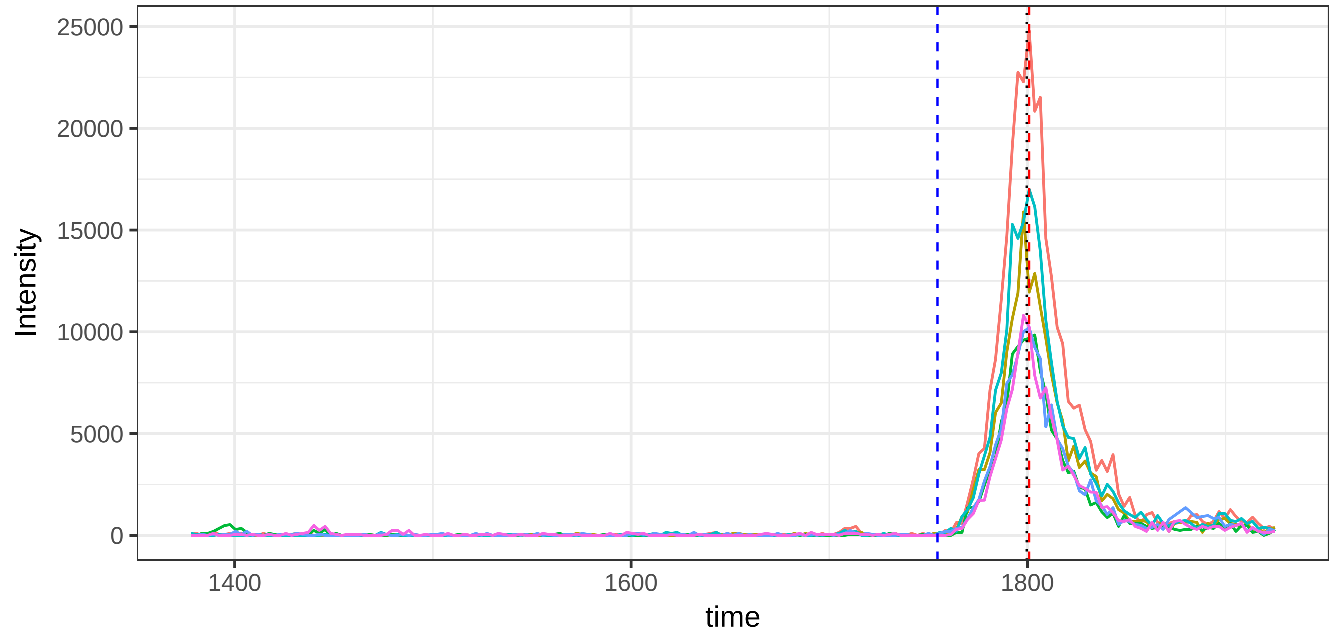

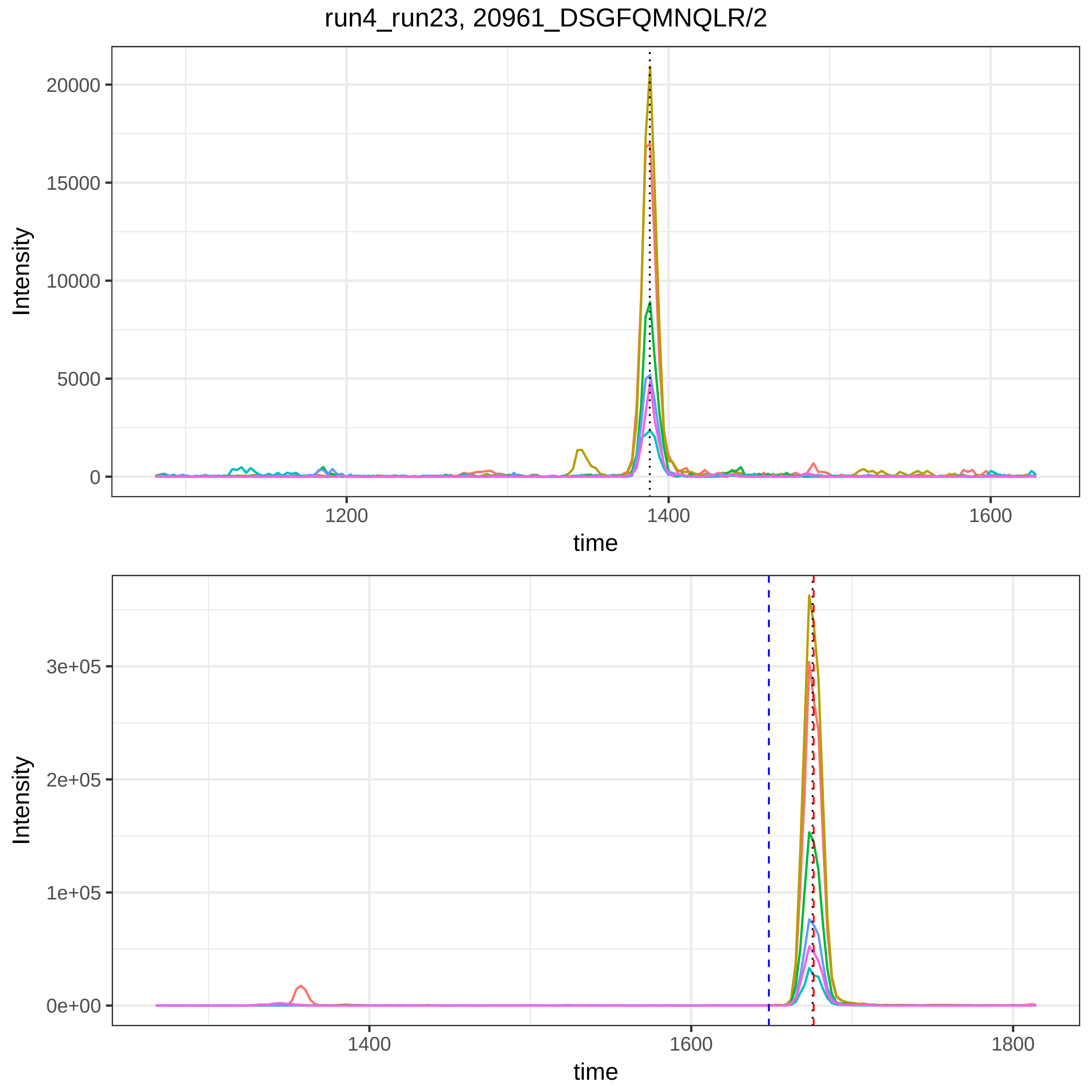
run4_run23, 20961_DSGFQMNQLR/2
Intensity
20000
15000
10000
5000
0
1200
1400
1600
time
Intensity
3e+05
2e+05
1e+05
0e+00
1400
1600
1800
time

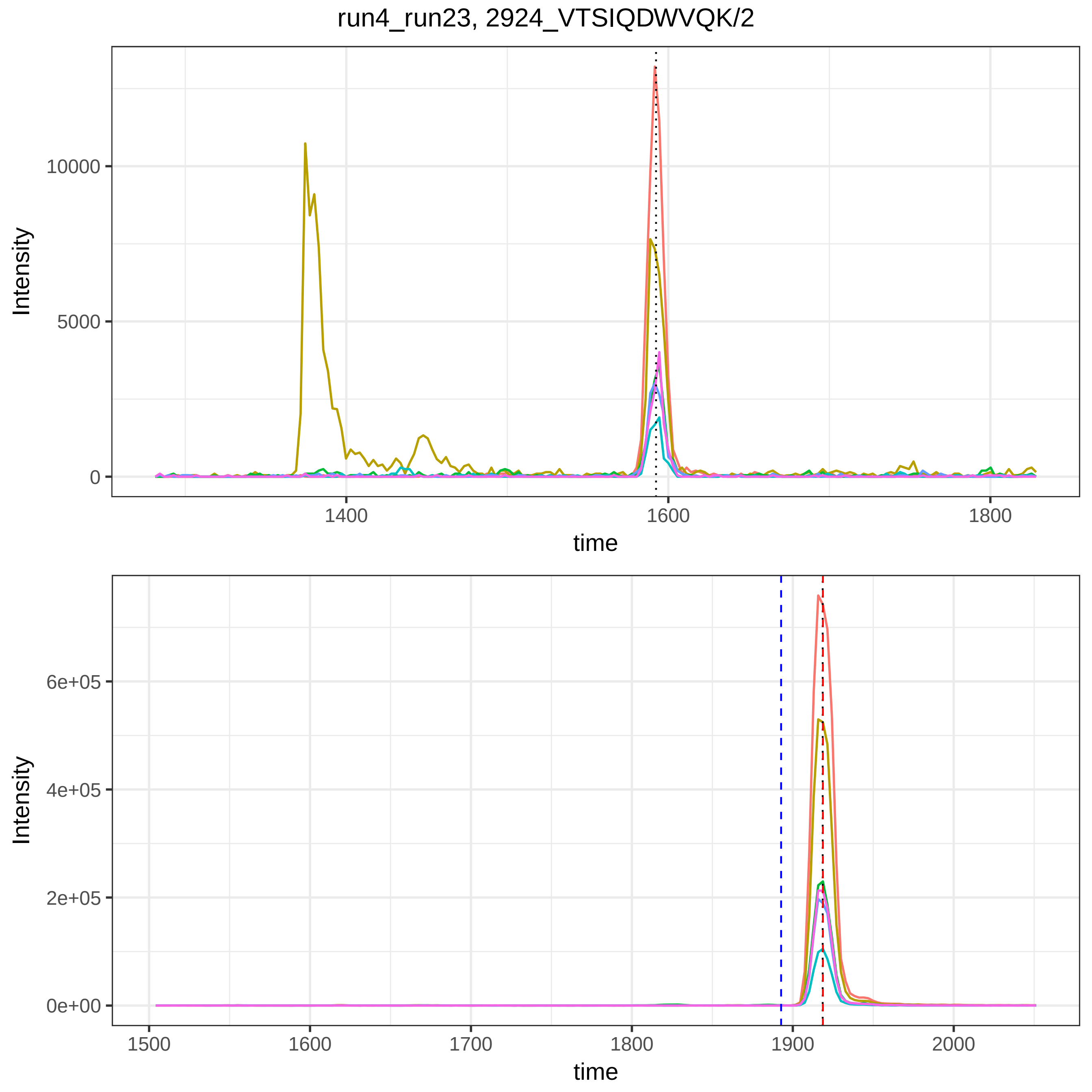
run4_run23, 2924_VTSIQDWVQK/2
Intensity
10000
5000
0
1400
1600
1800
time
Intensity
6e+05
4e+05
2e+05
0e+00
1500
1600
1700
1800
1900
2000
time

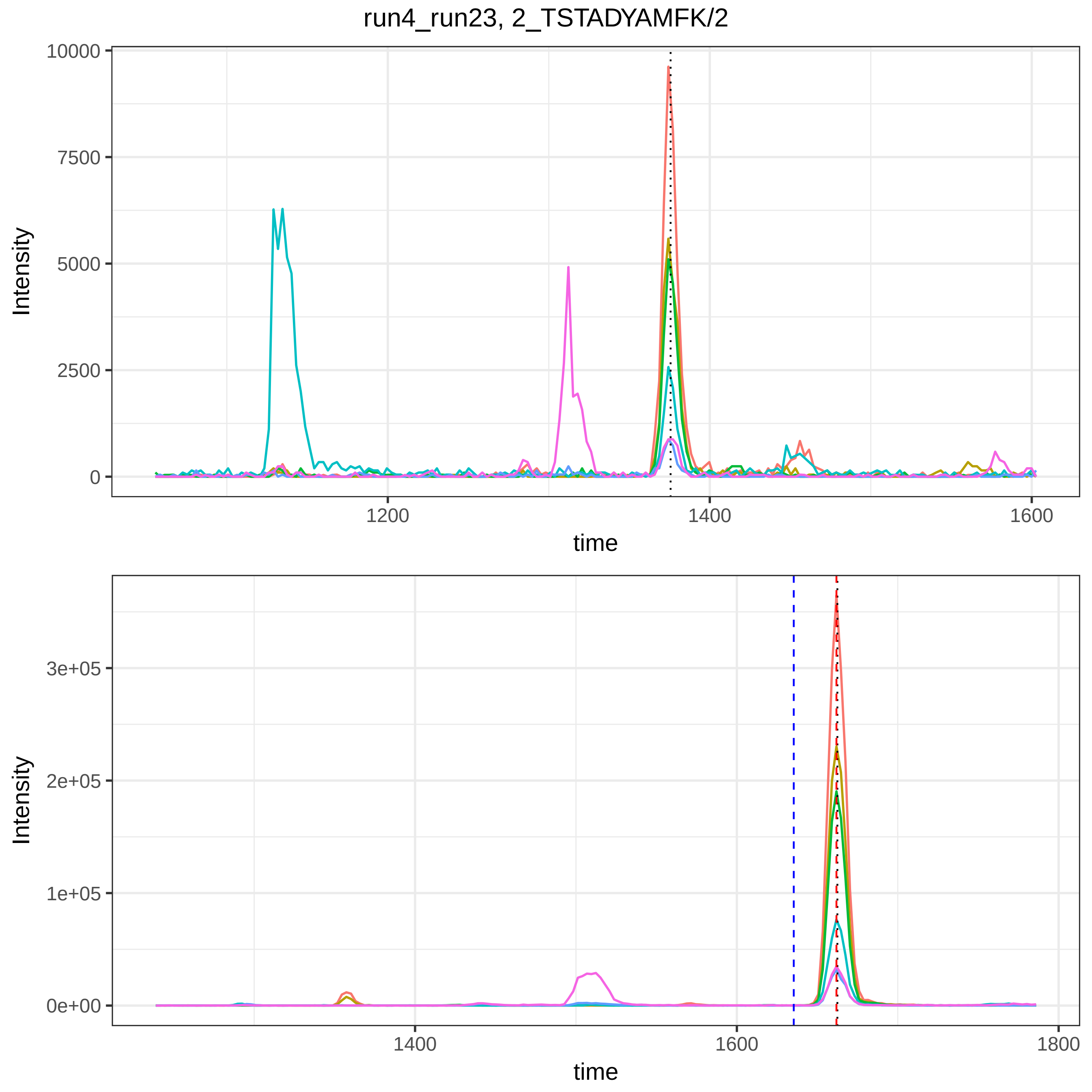

run4_run23, 2_TSTADYAMFK/2
Intensity
10000
7500
5000
2500
0
1200
1400
1600
time
Intensity
3e+05
2e+05
1e+05
0e+00
1400
1600
1800
time

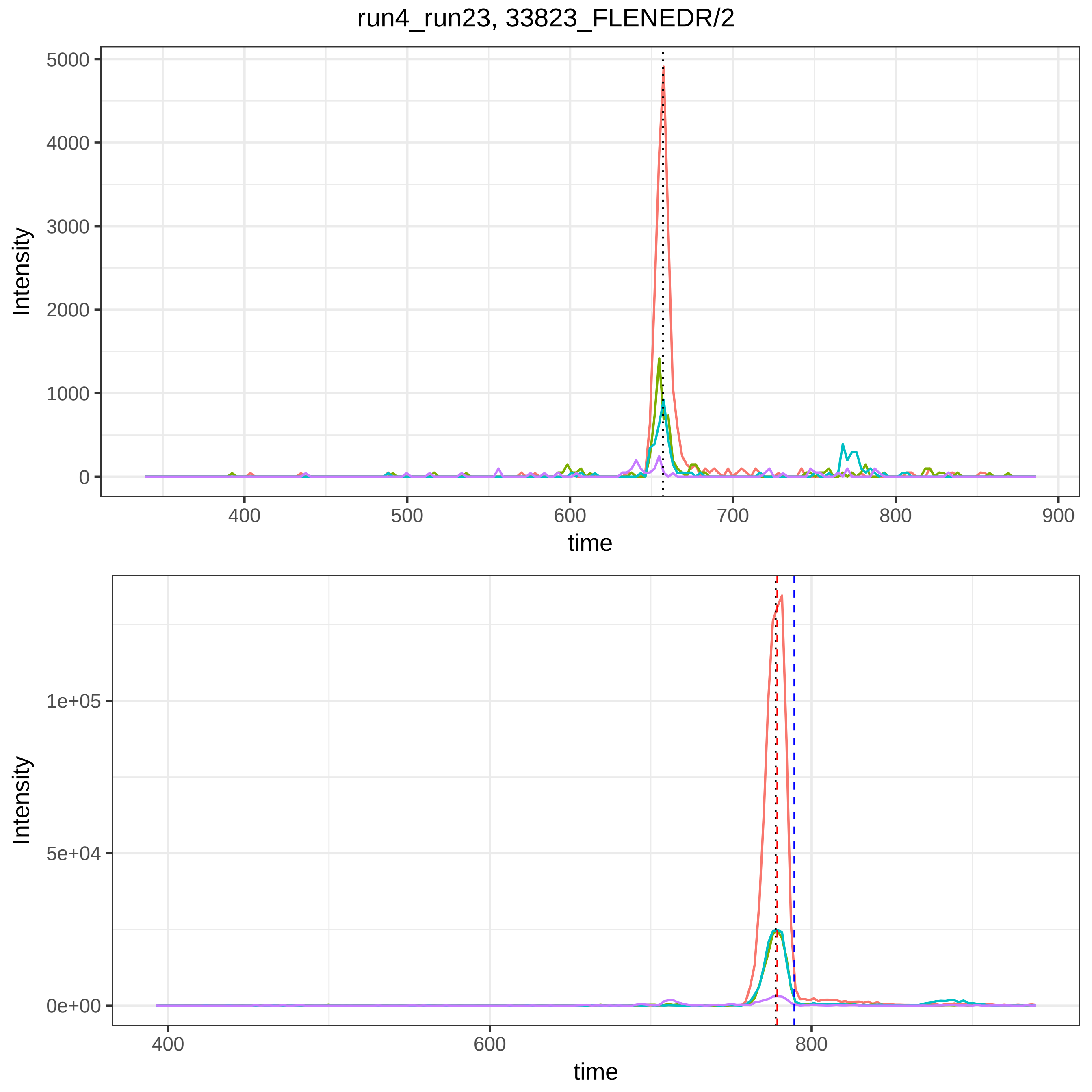
run4_run23, 33823_FLENEDR/2
Intensity
5000
4000
3000
2000
1000
0
400
500
600
700
800
900
time
Intensity
1e+05
5e+04
0e+00
400
600
800
time

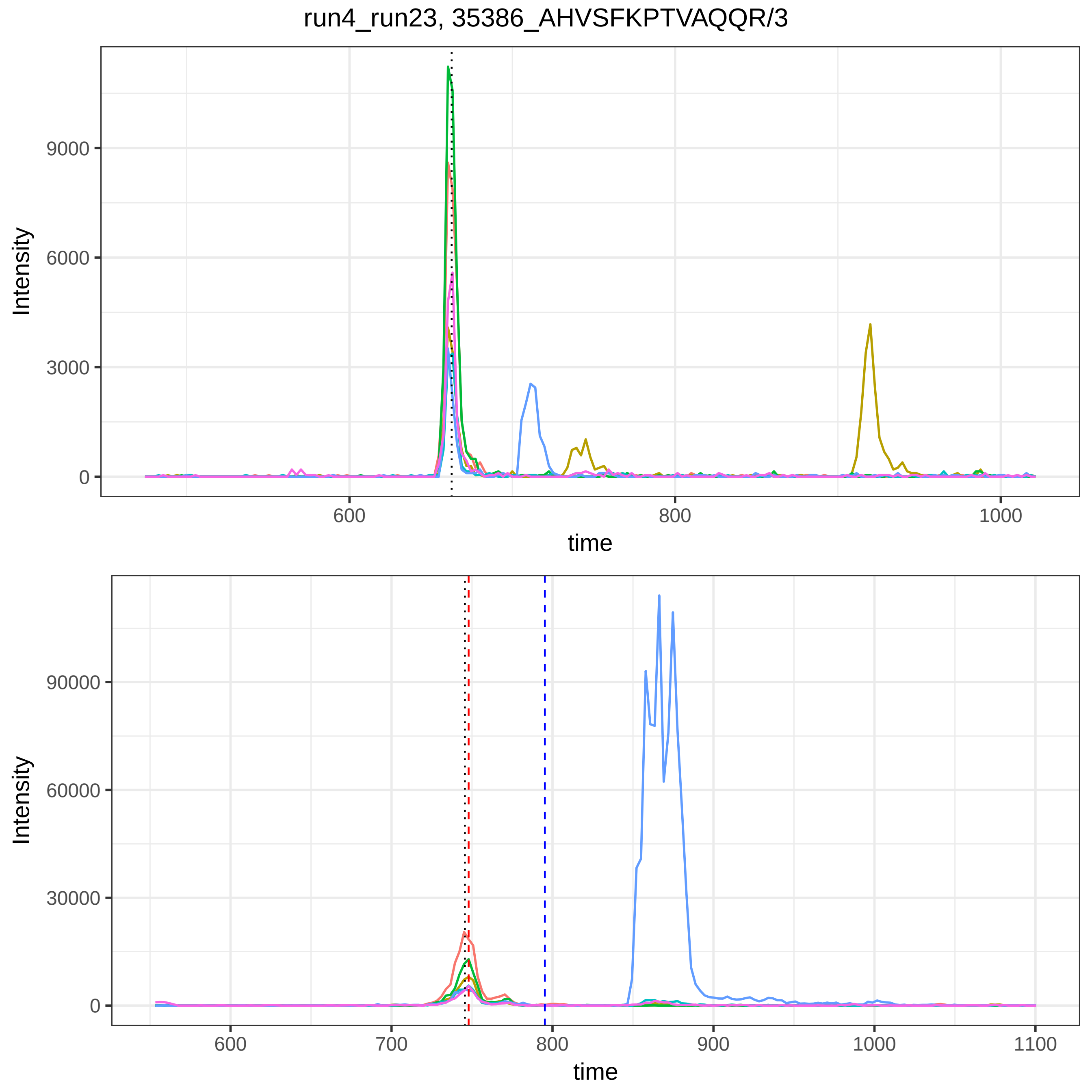

run4_run23, 35386_AHVSFKPTVAQQR/3
Intensity
9000
6000
3000
0
600
800
1000
time
Intensity
90000
60000
30000
0
600
700
800
900
1000
1100
time

run4_run23, 38203_HQFLLTGDTQGR/3

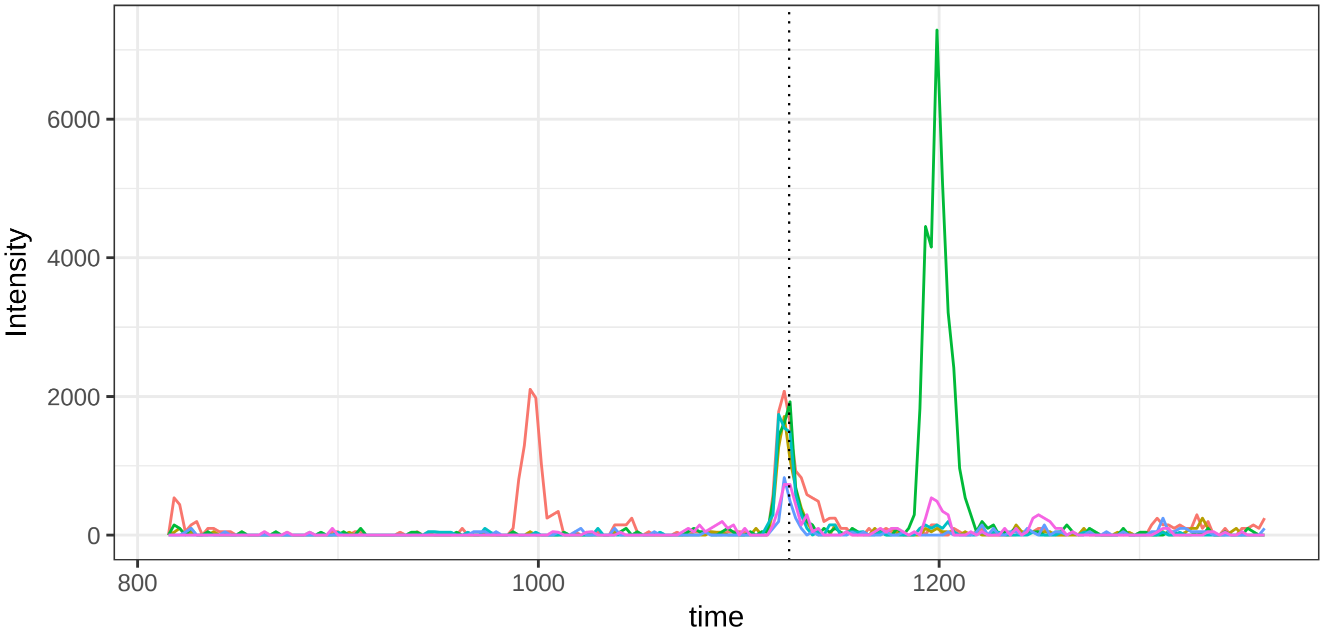


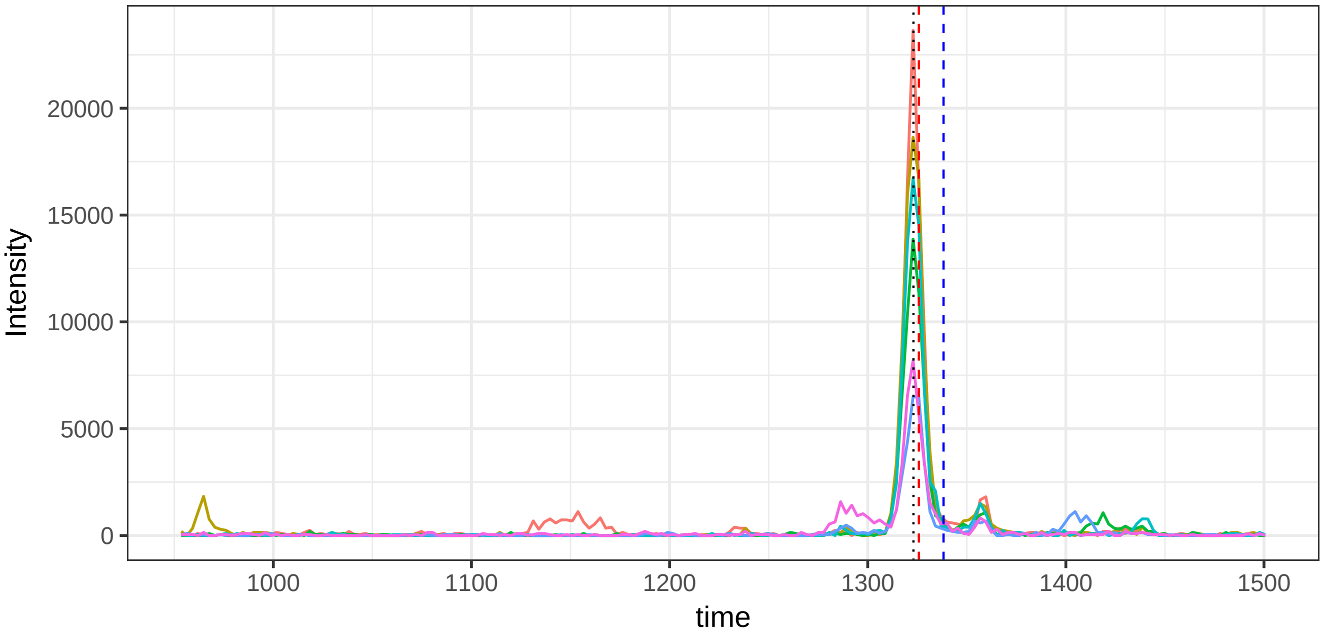

run4_run23, 39035_IEDGSGEVVLSR/2

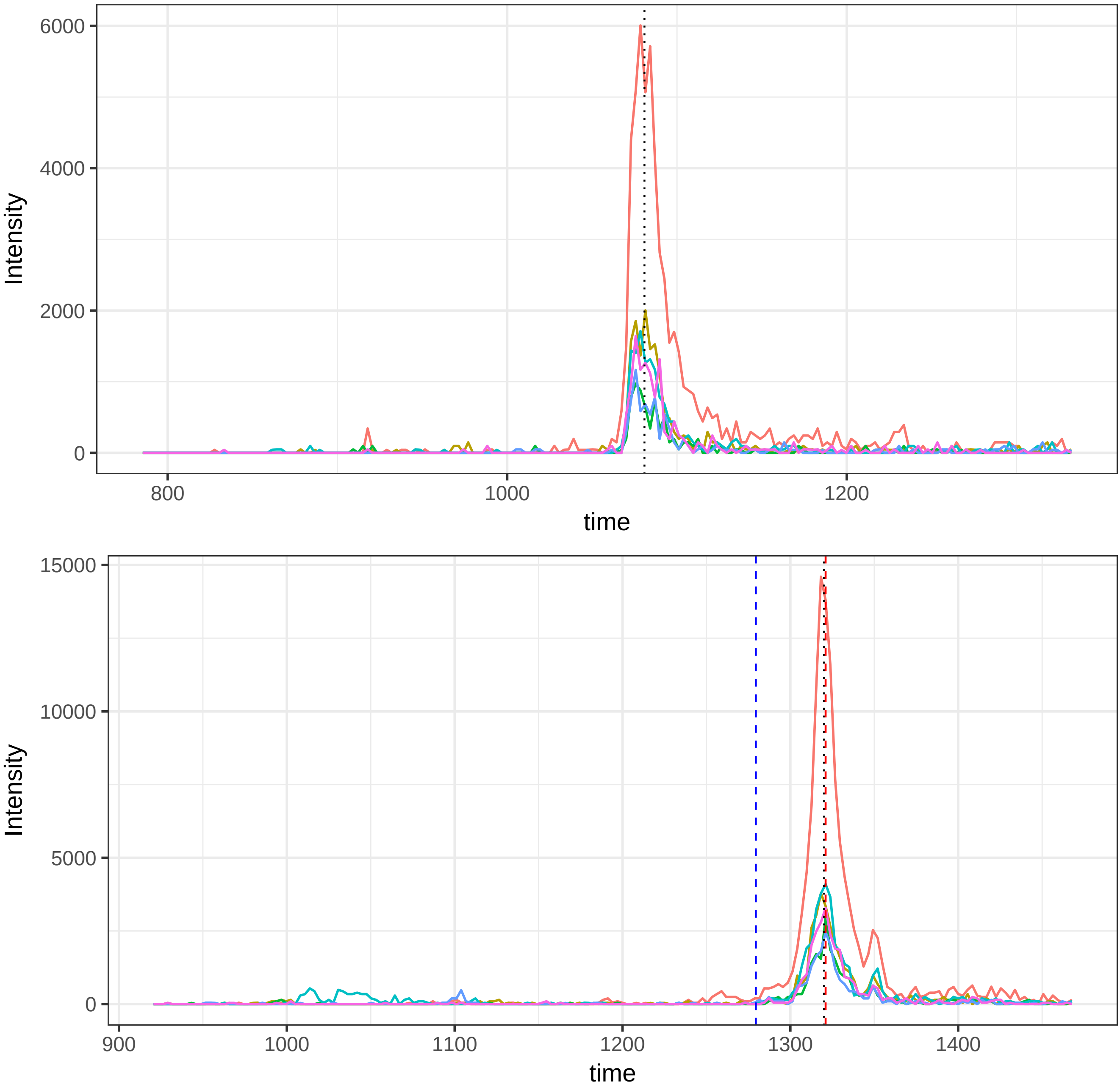

run4_run23, 40510_IYLQPGR/2

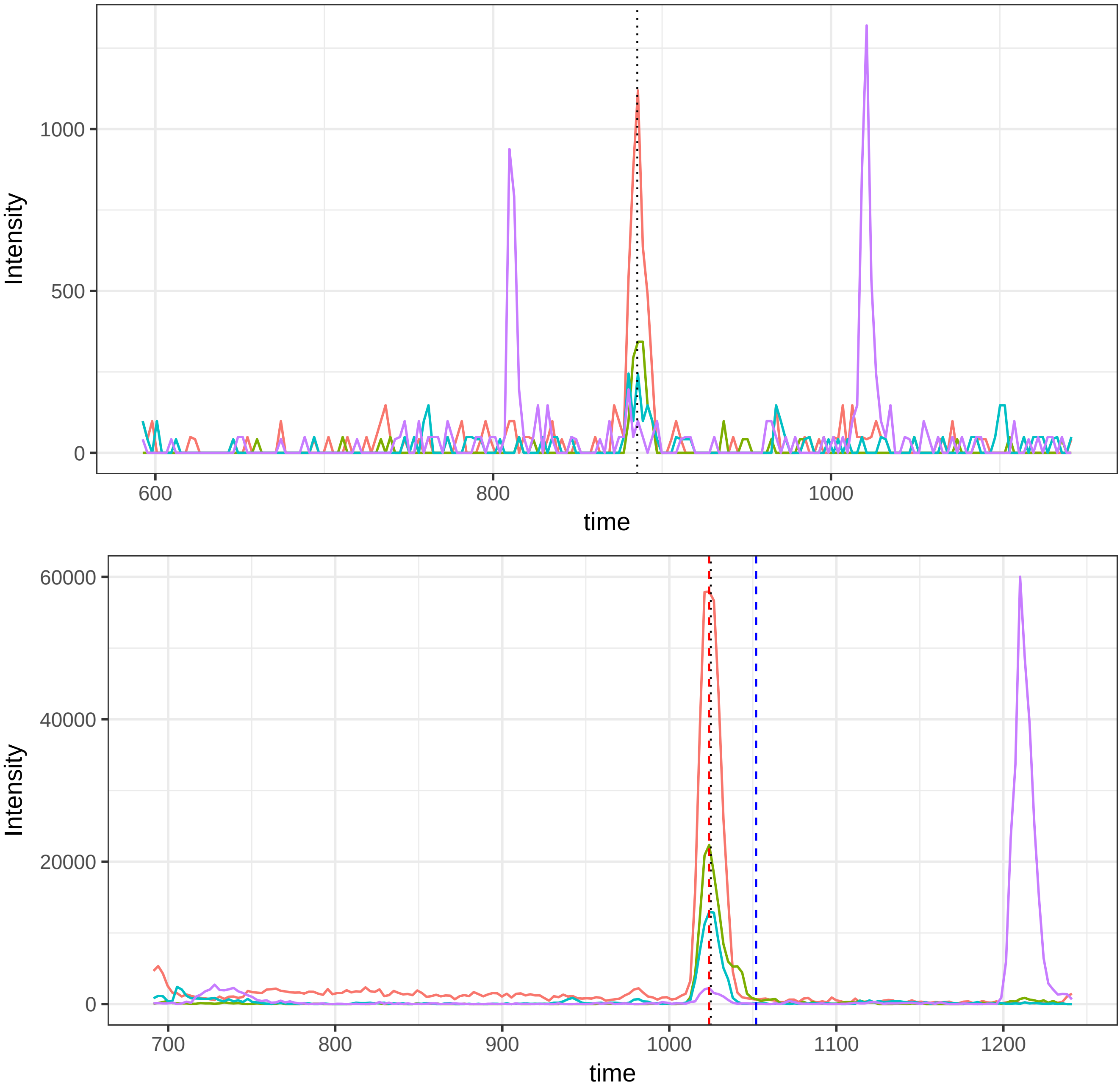

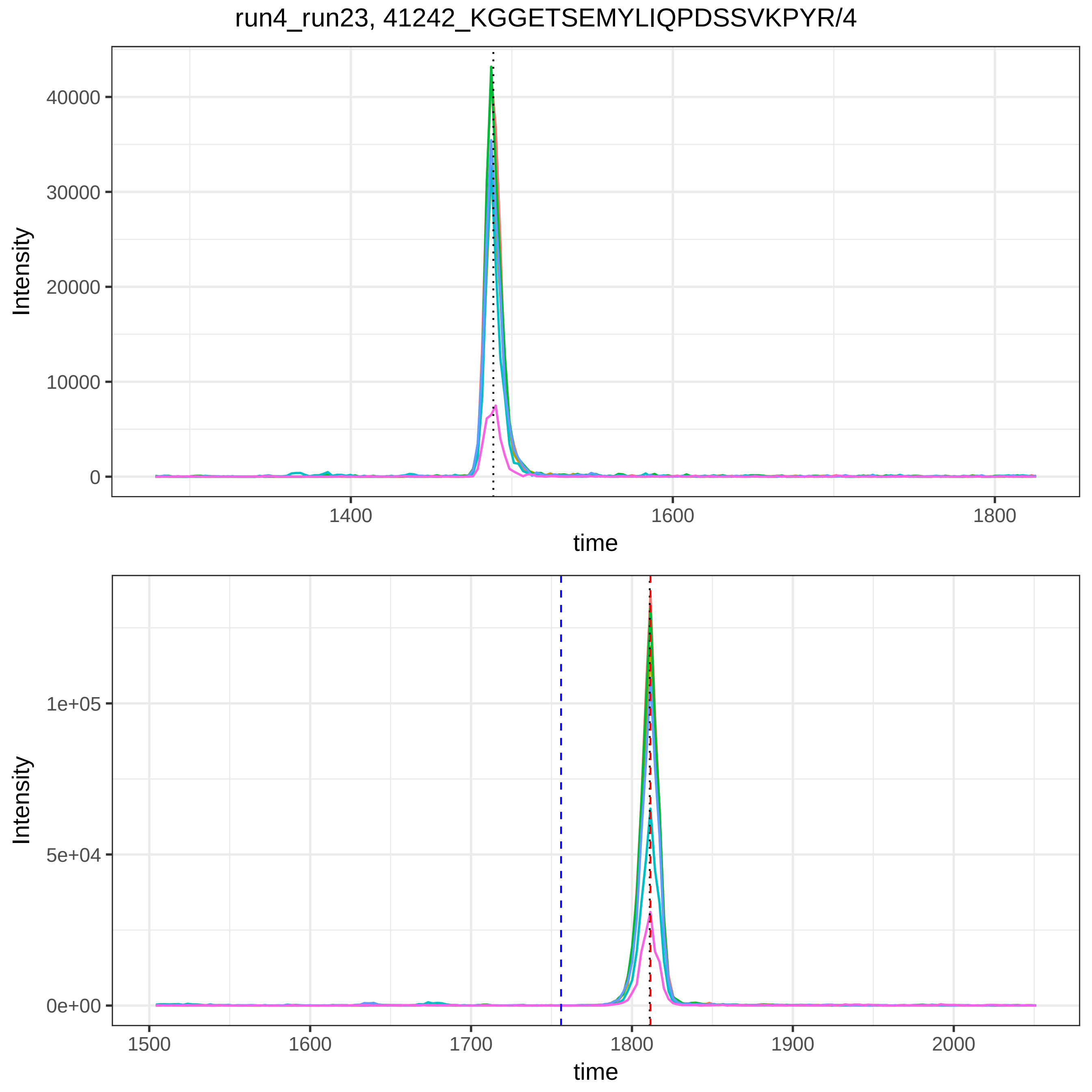

run4_run23, 41242_KGGETSEMYLIQPDSSVKPYR/4
Intensity
40000
30000
20000
10000
0
1400
1600
1800
time
Intensity
1e+05
5e+04
0e+00
1500
1600
1700
1800
1900
2000
time

run4_run23, 41869_KSASDLTWDNLK/2

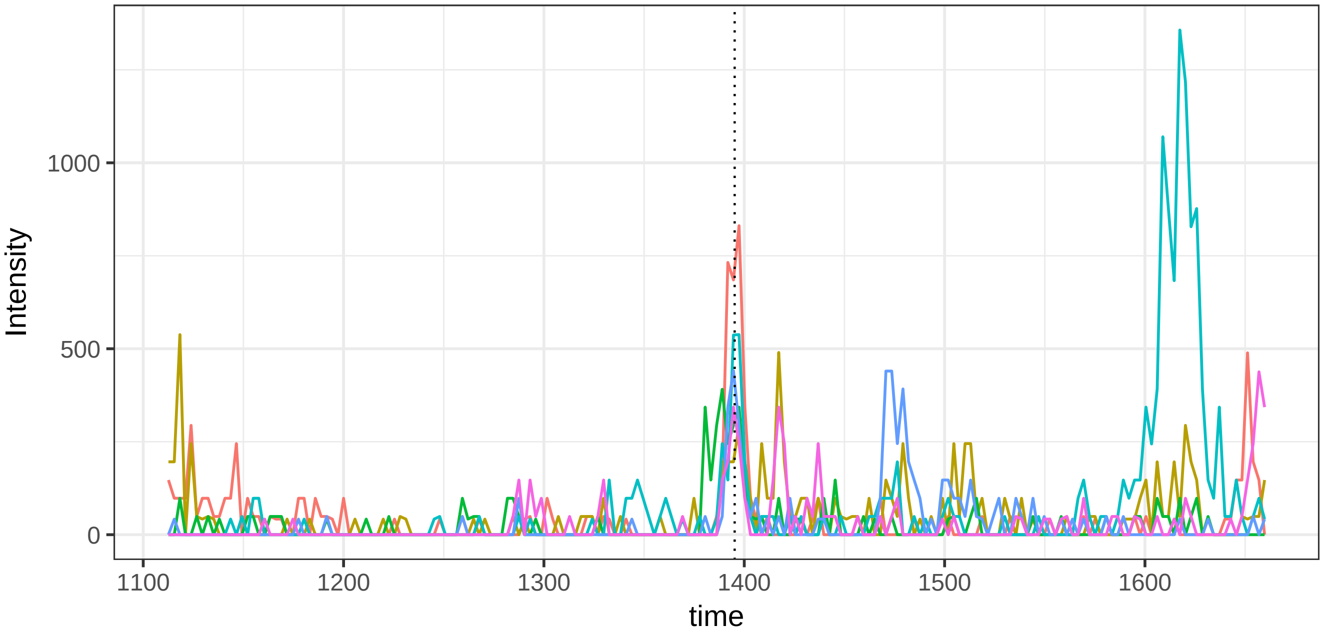


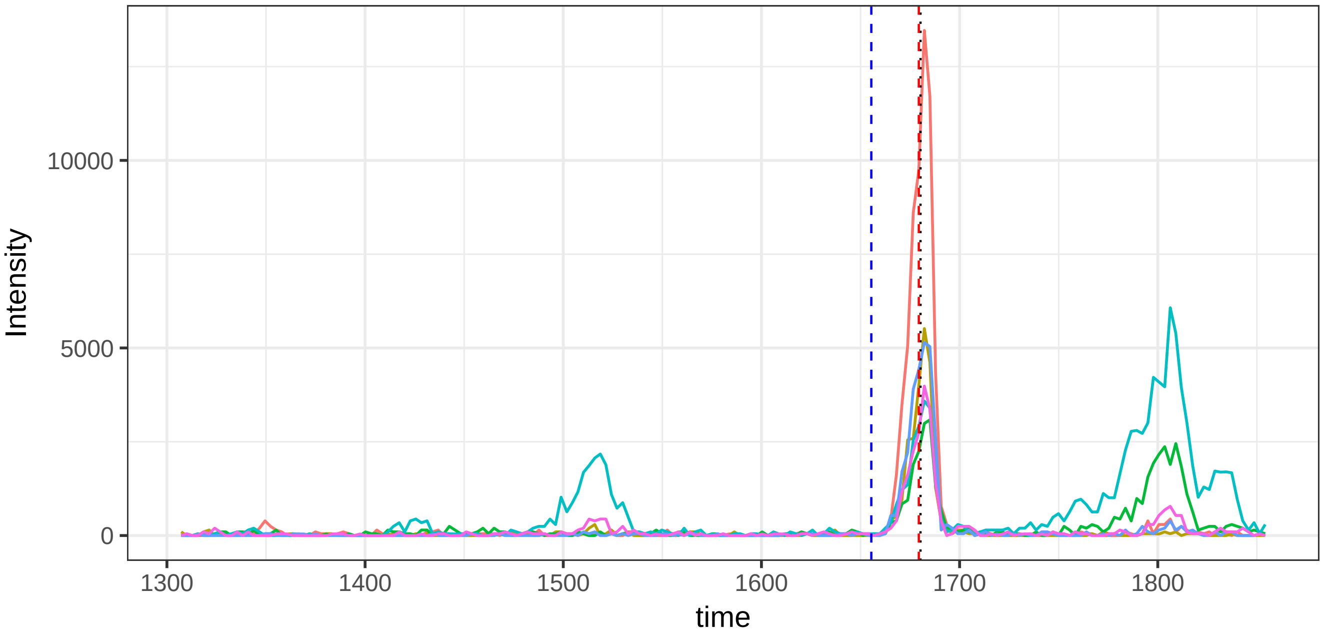

run4_run23, 41874_KSASDLTWDNLK/3

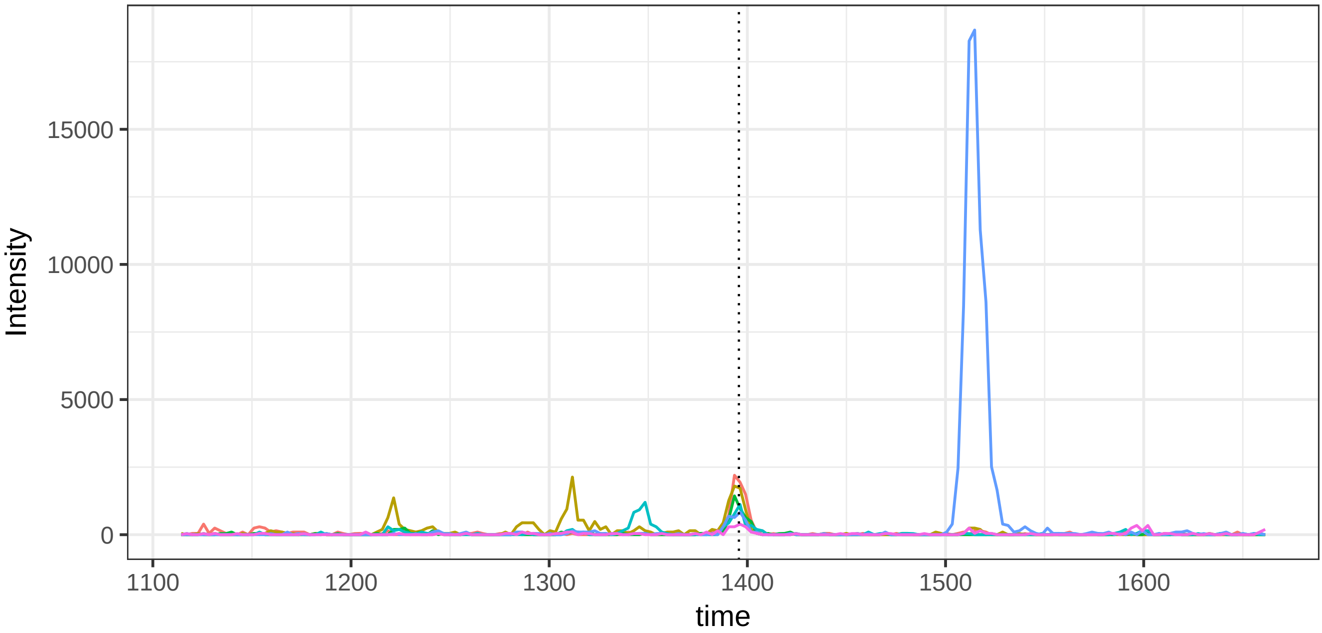


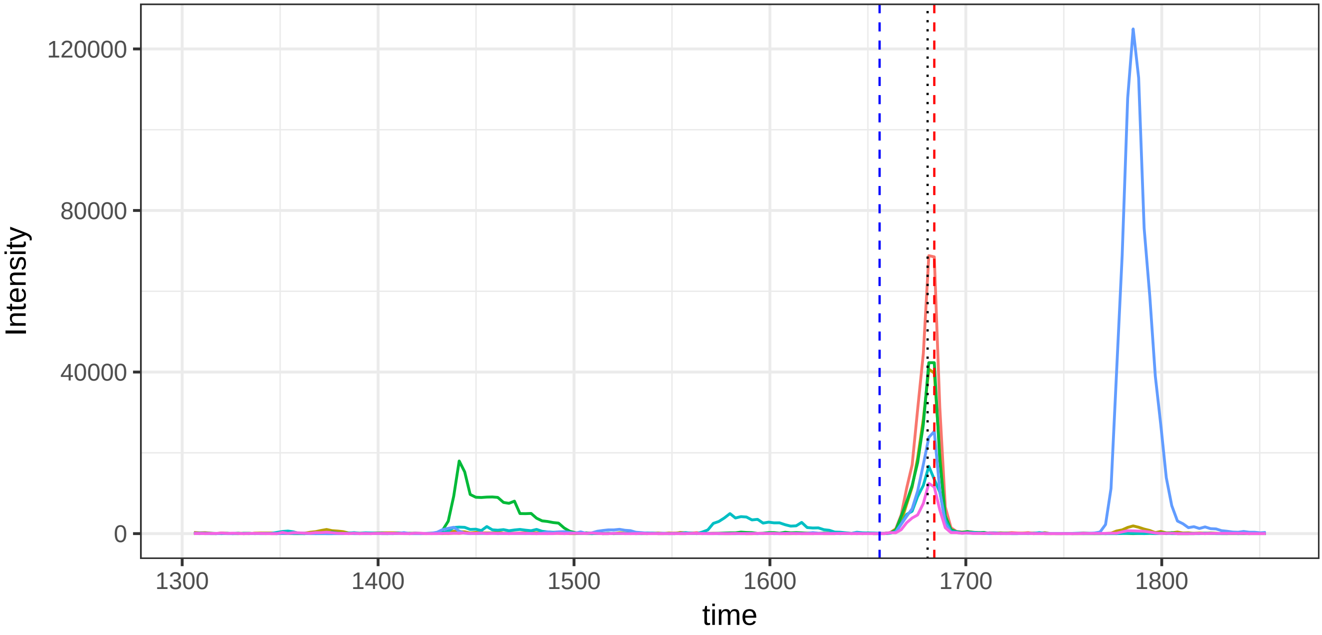

run4_run23, 45181_LTVLGQPK/2

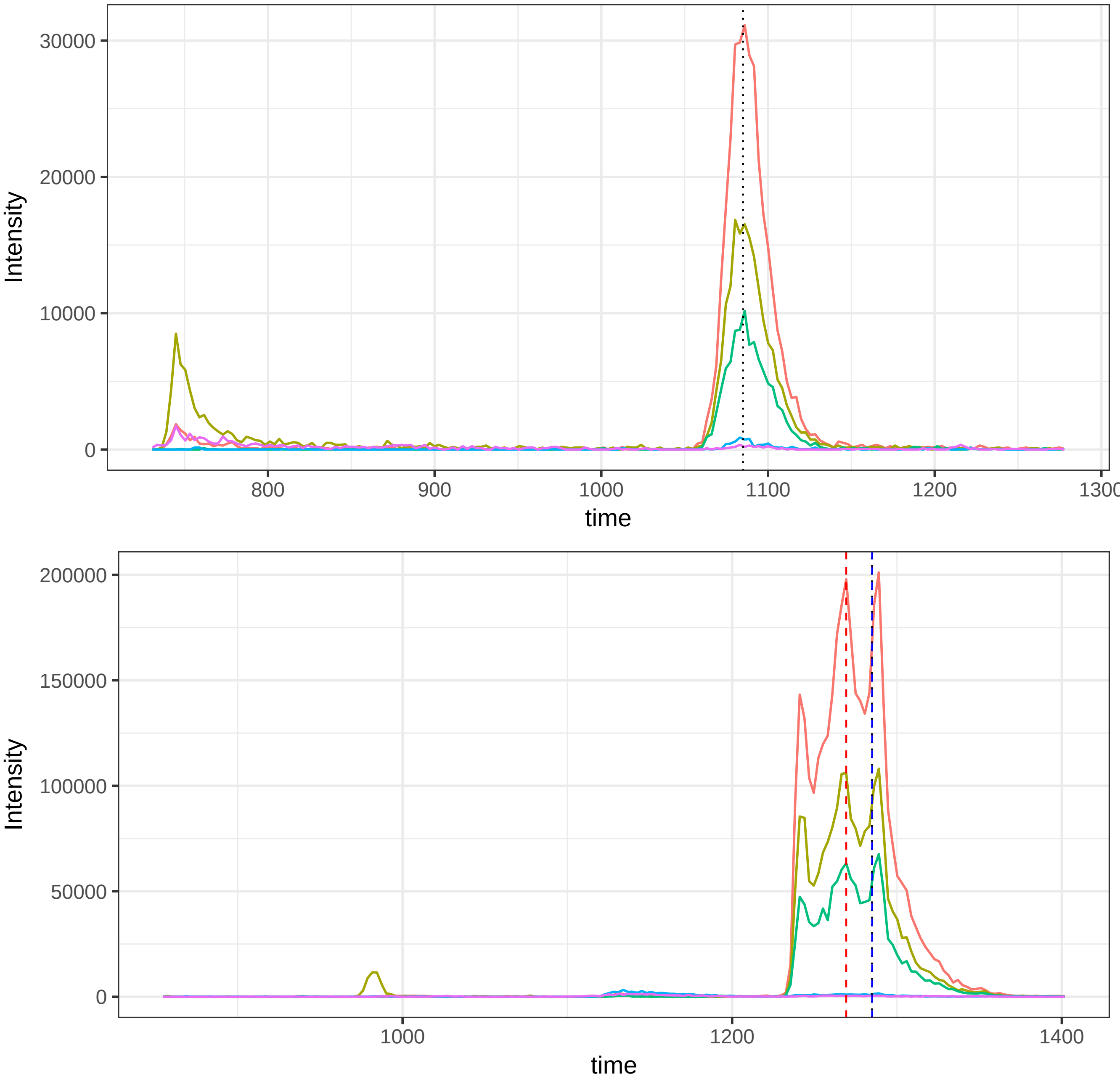

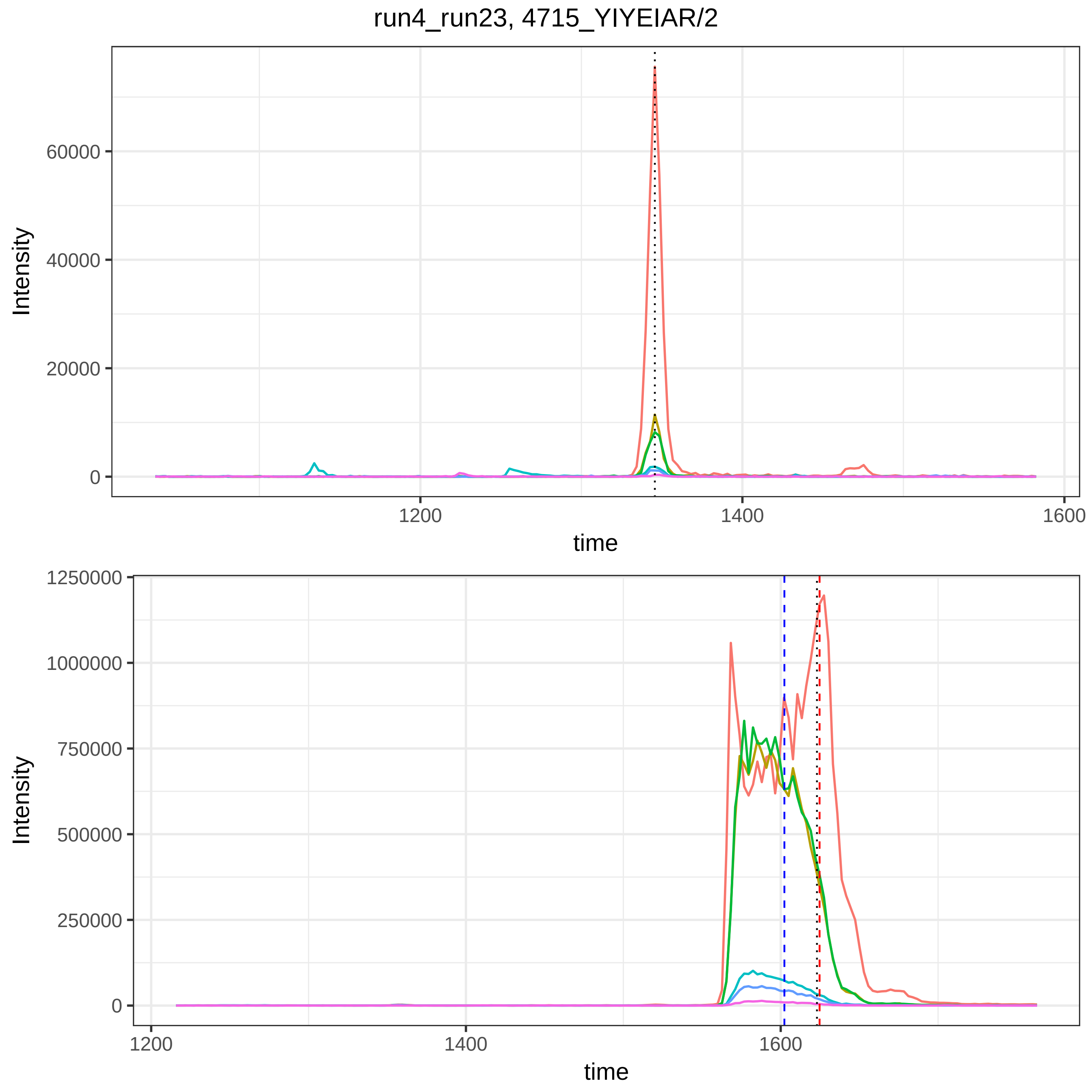

run4_run23, 4715_YIYEIAR/2
Intensity
60000
40000
20000
0
1200
1400
1600
time
Intensity
1250000
1000000
750000
500000
250000
0
1200
1400
1600
time

run4_run23, 47721_NNEGTYYSPNYNPQSR/2

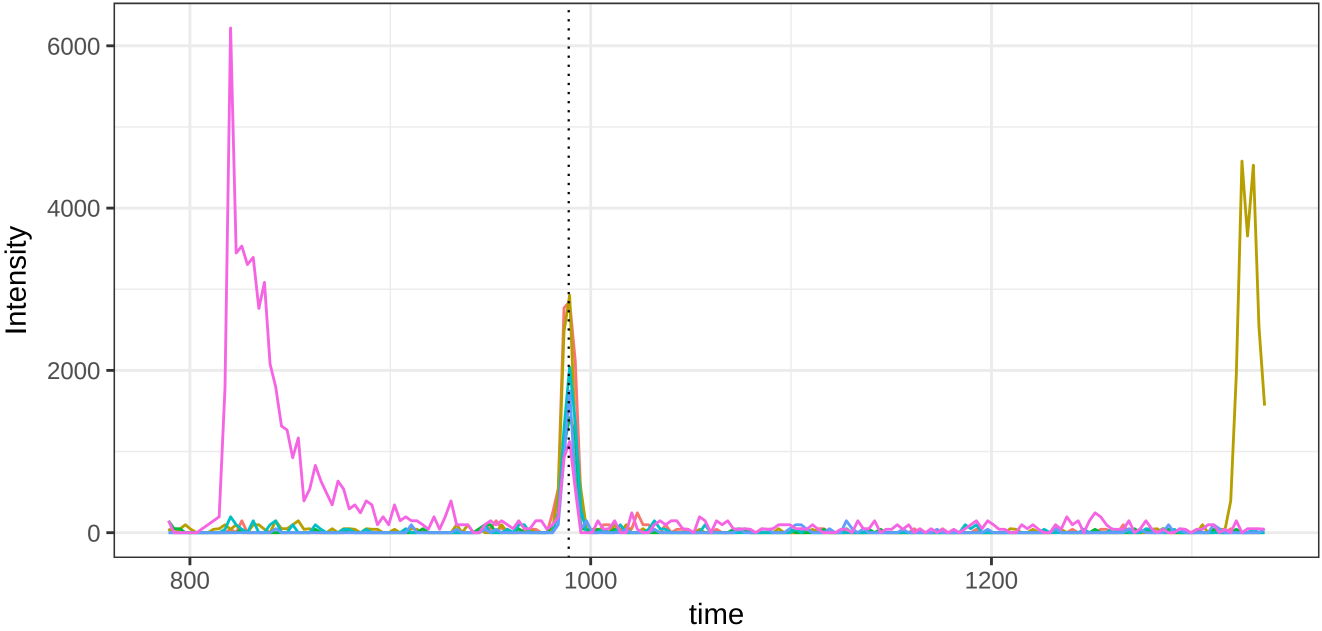


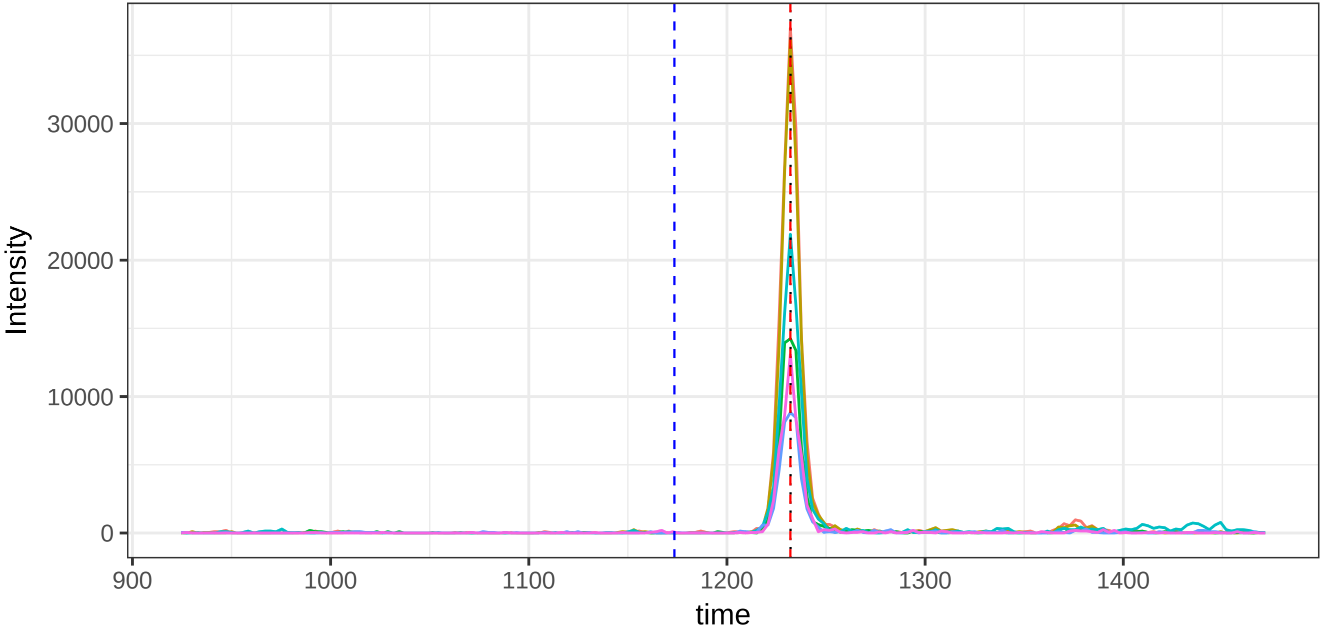

run4_run23, 49641_QLGLPGPPDVPDHAAYHPF/3

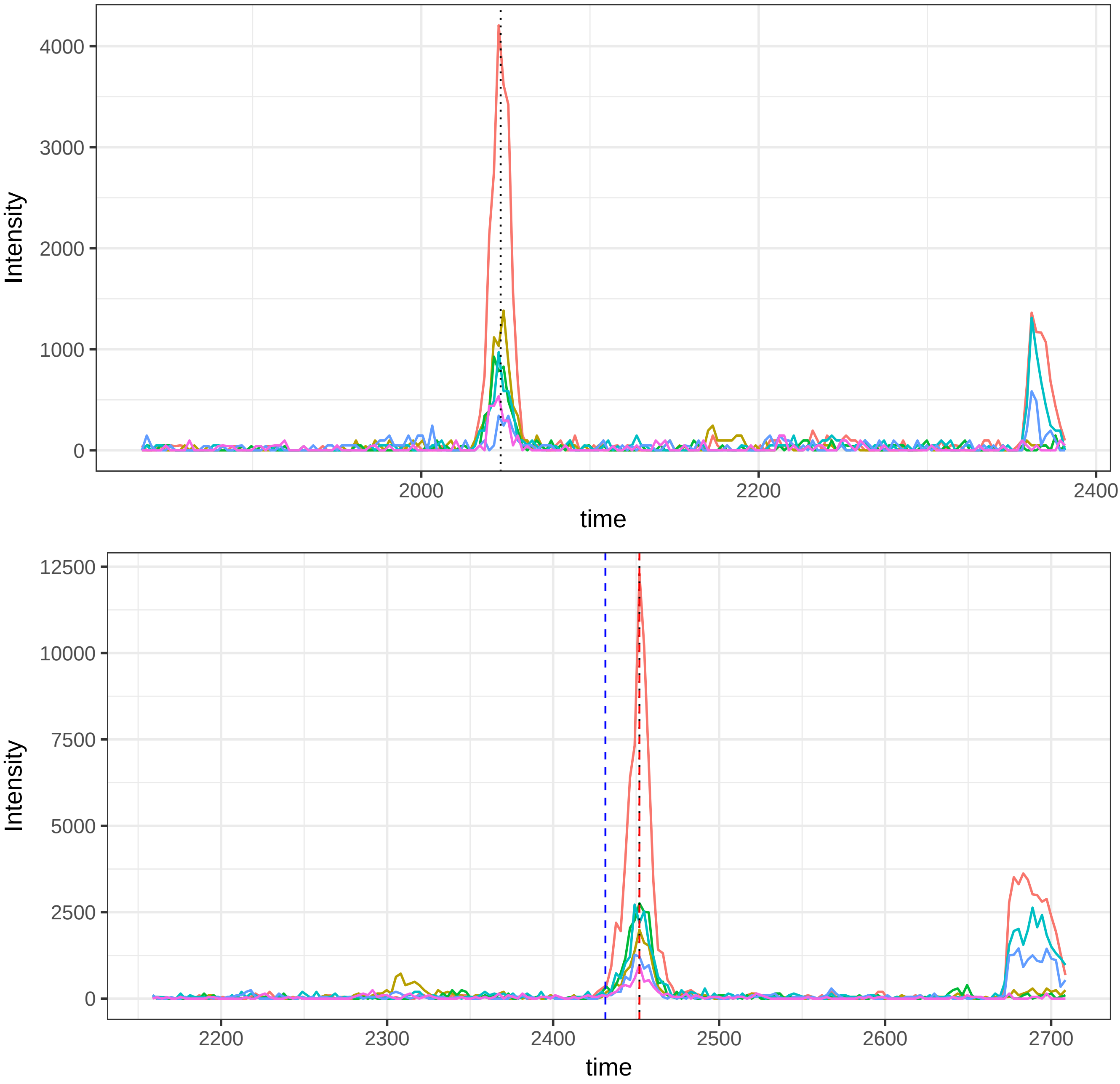

run4_run23, 50302_QYYEGSEIVVAGR/2

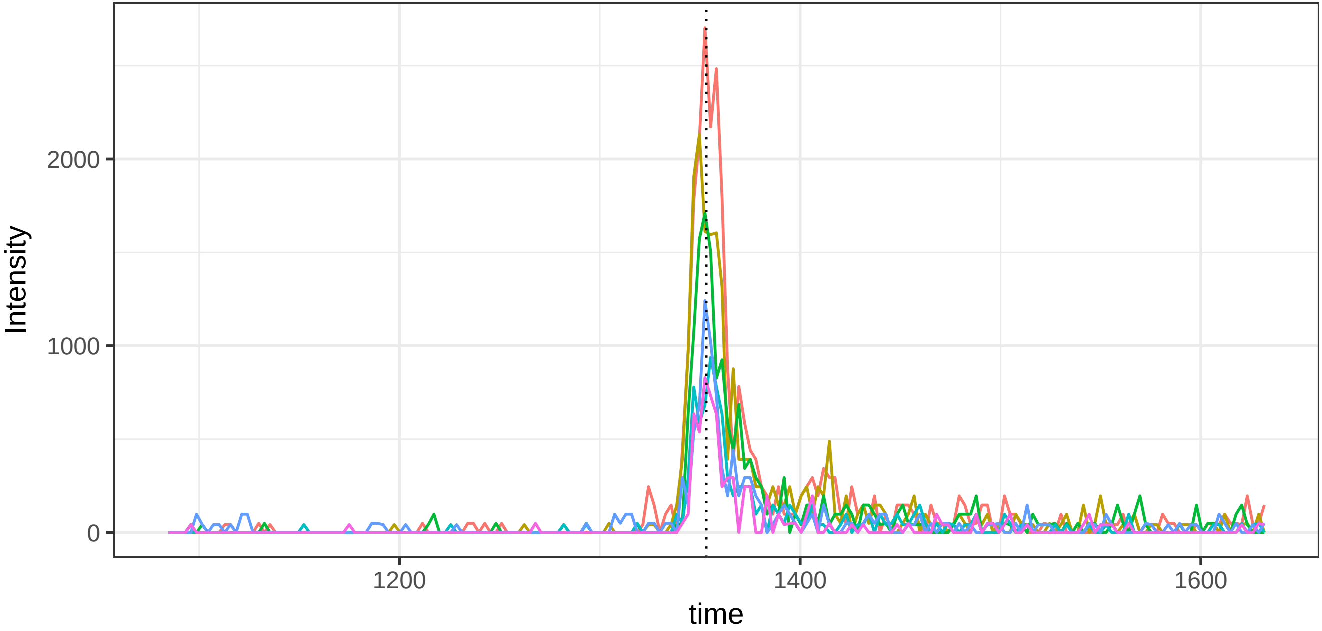


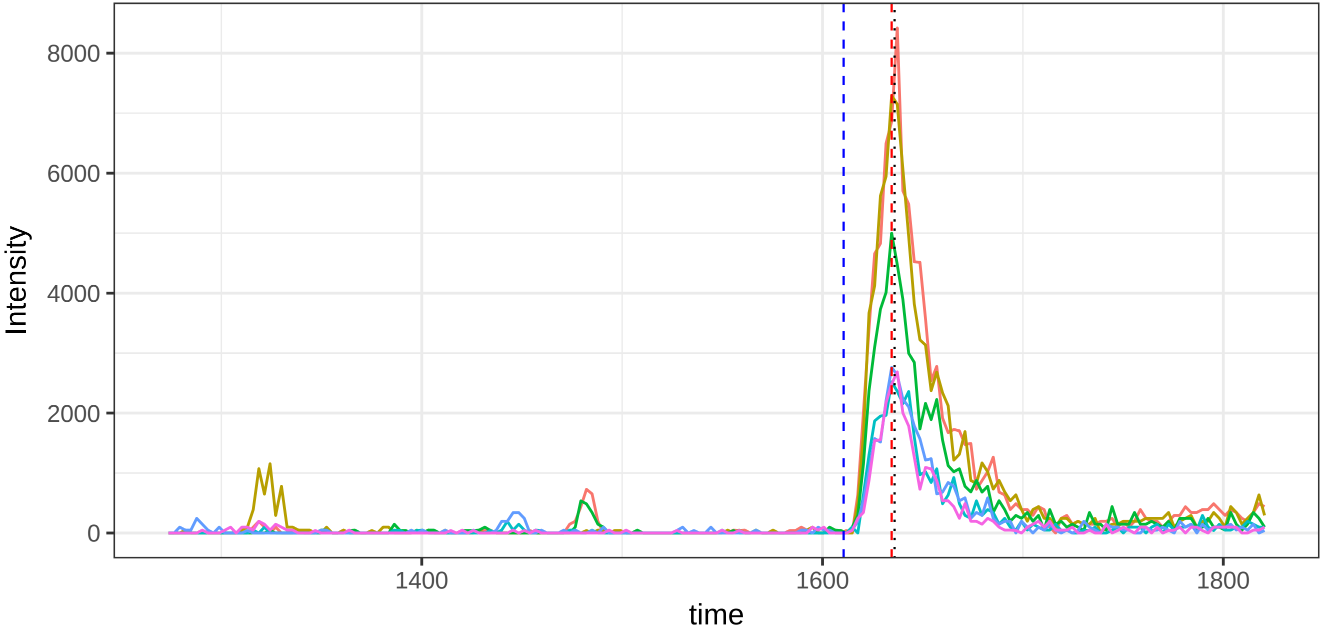

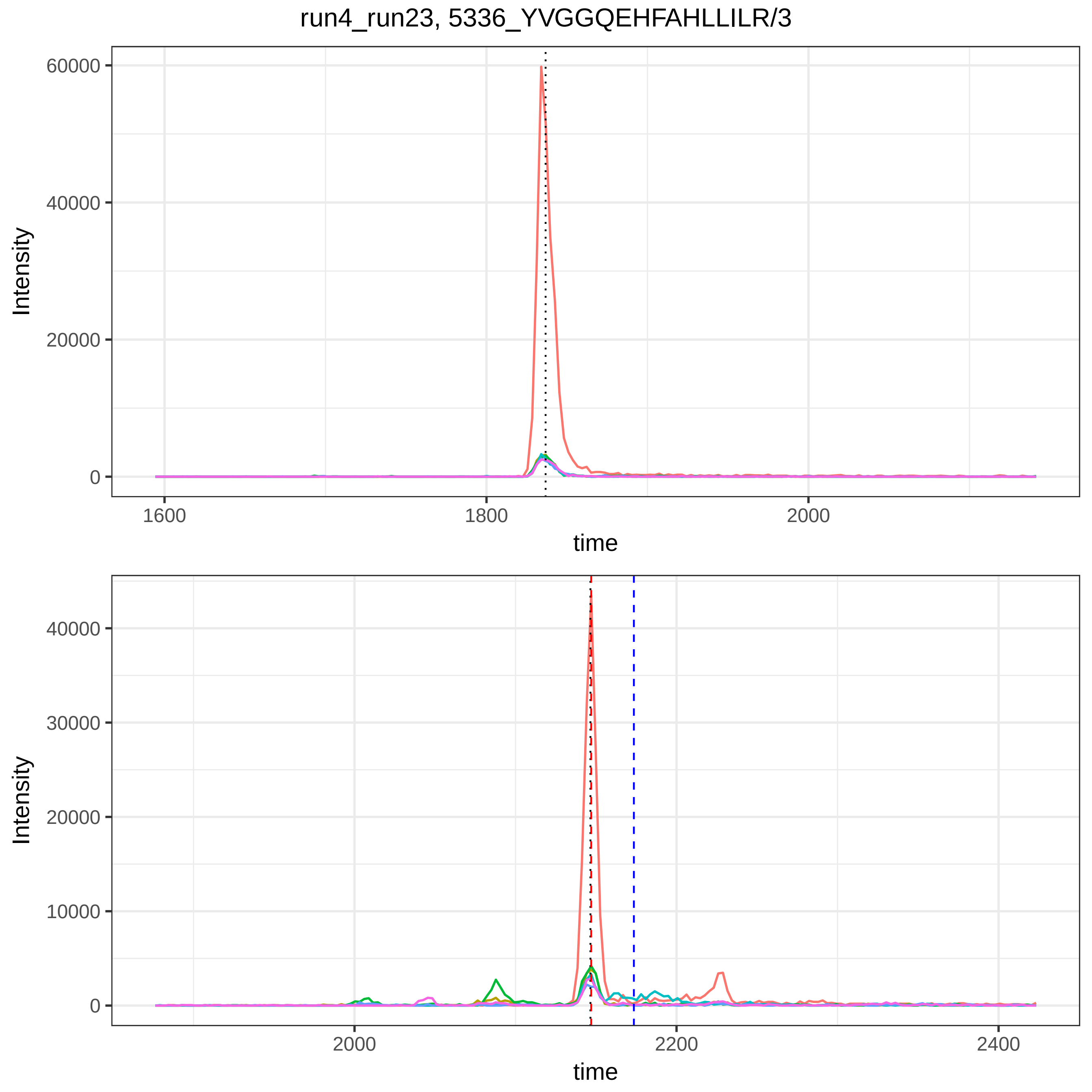

run4_run23, 5336_YVGGQEHFAHLLILR/3
60000
40000
20000
0
Intensity
1600
1800
2000
time
40000
30000
20000
10000
0
Intensity
2000
2200
2400
time

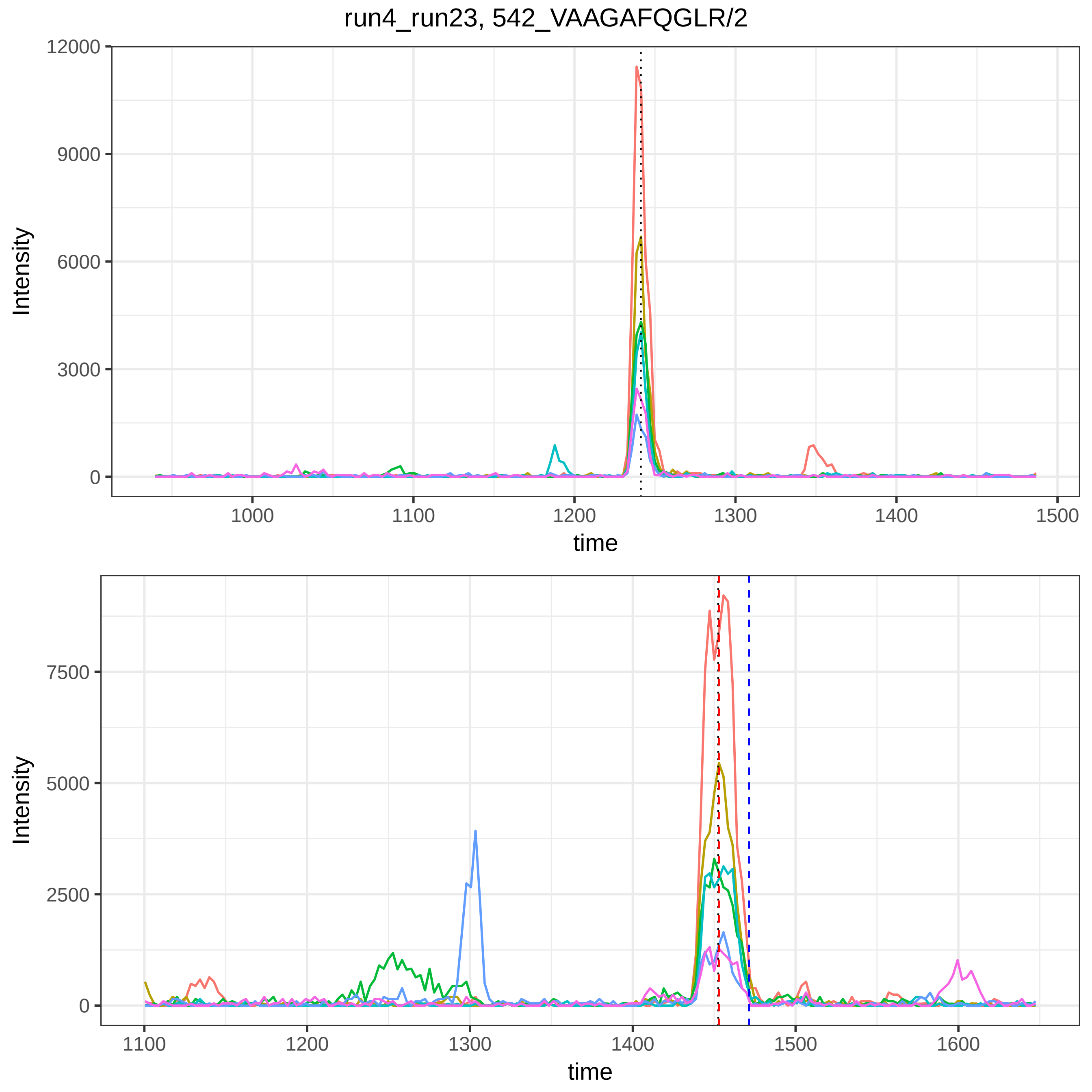
run4_run23, 542_VAAGAFQGLR/2
Intensity
time
12000
9000
6000
3000
0
1000
1100
1200
1300
1400
1500
7500
5000
2500
1600

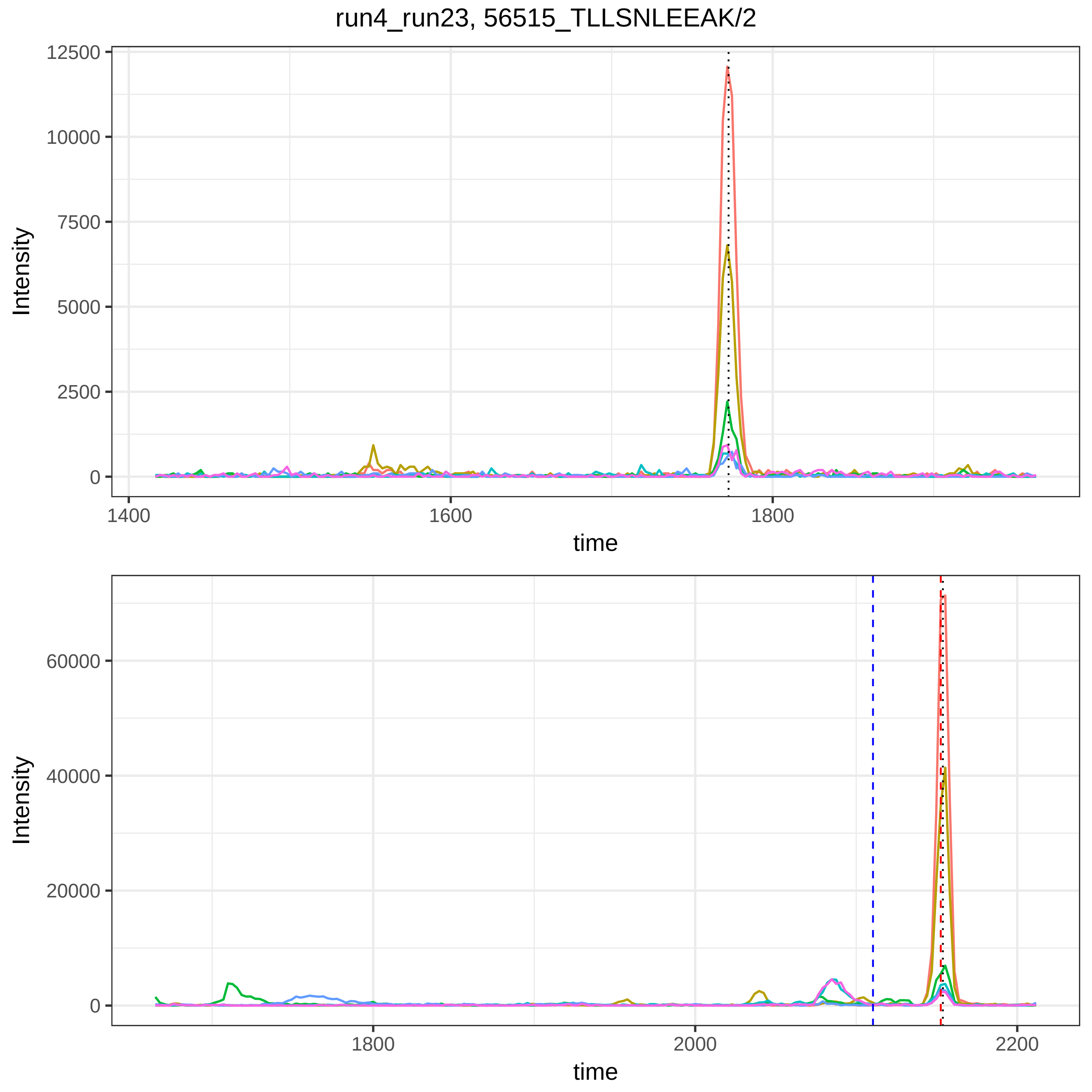

run4_run23, 56515_TLLSNLEEAK/2
Intensity
time
12500
10000
7500
5000
2500
0
1400
1600
1800
60000
40000
20000
2000
2200

run4_run23, 57287_TQVNTQAEQLR/2

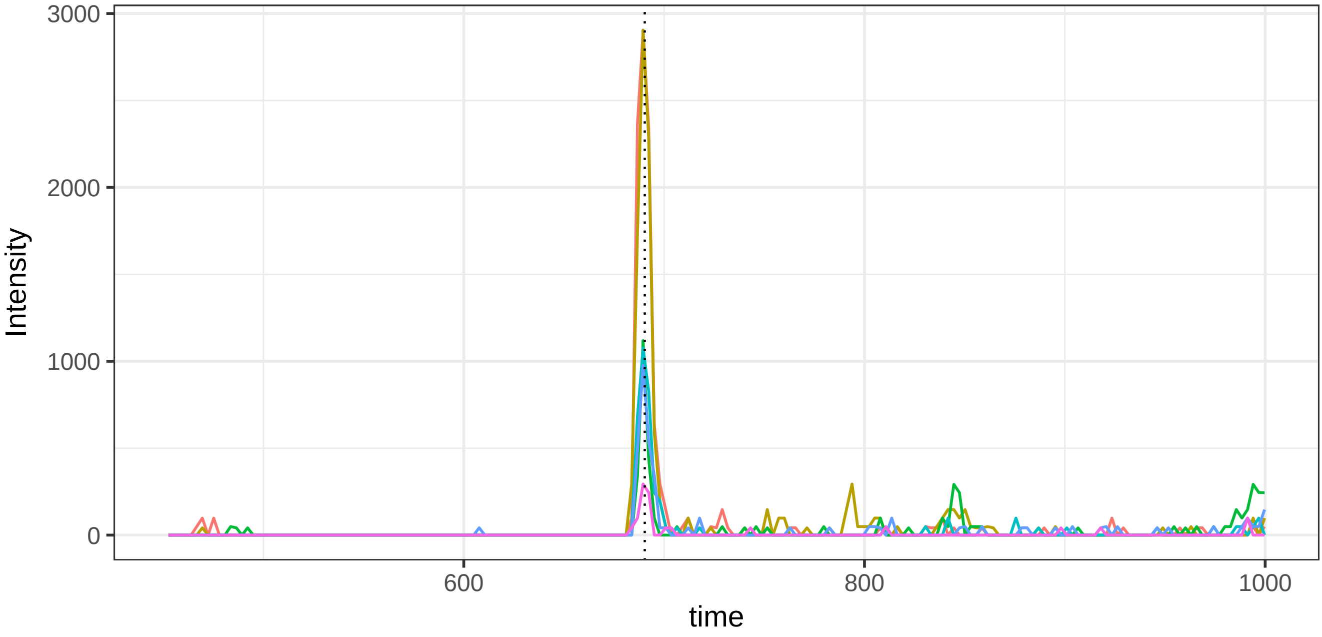


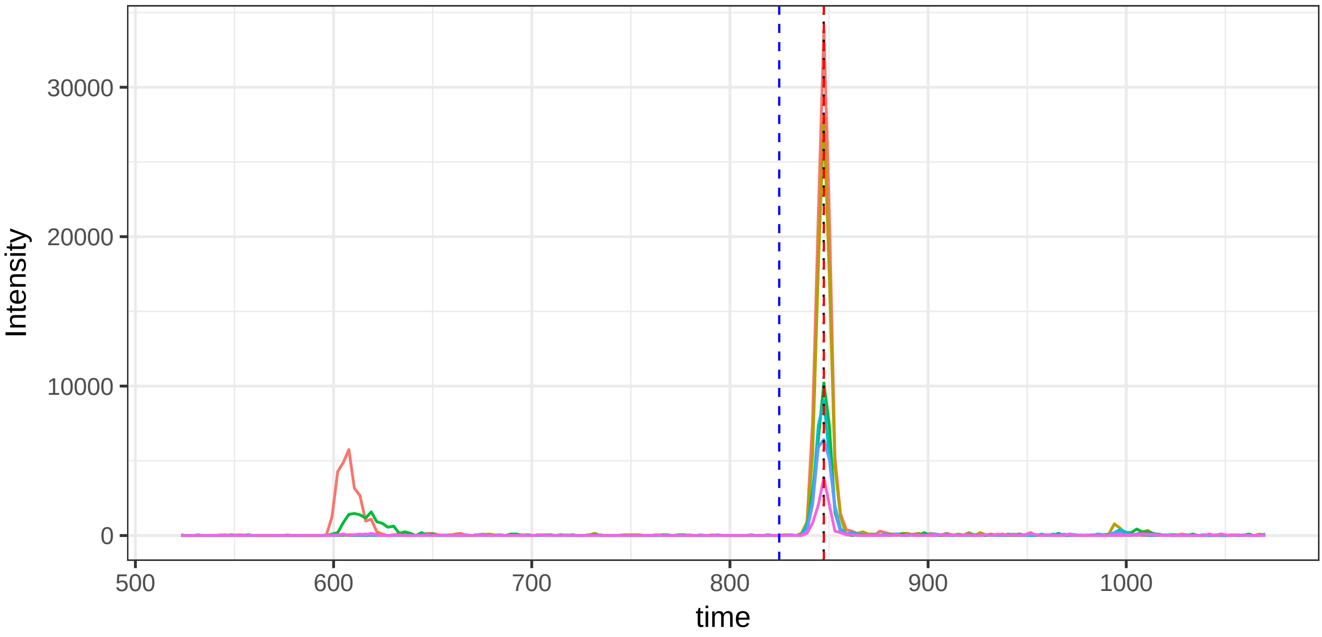

run4_run23, 116622_AGDFLEANYMNLQR/2

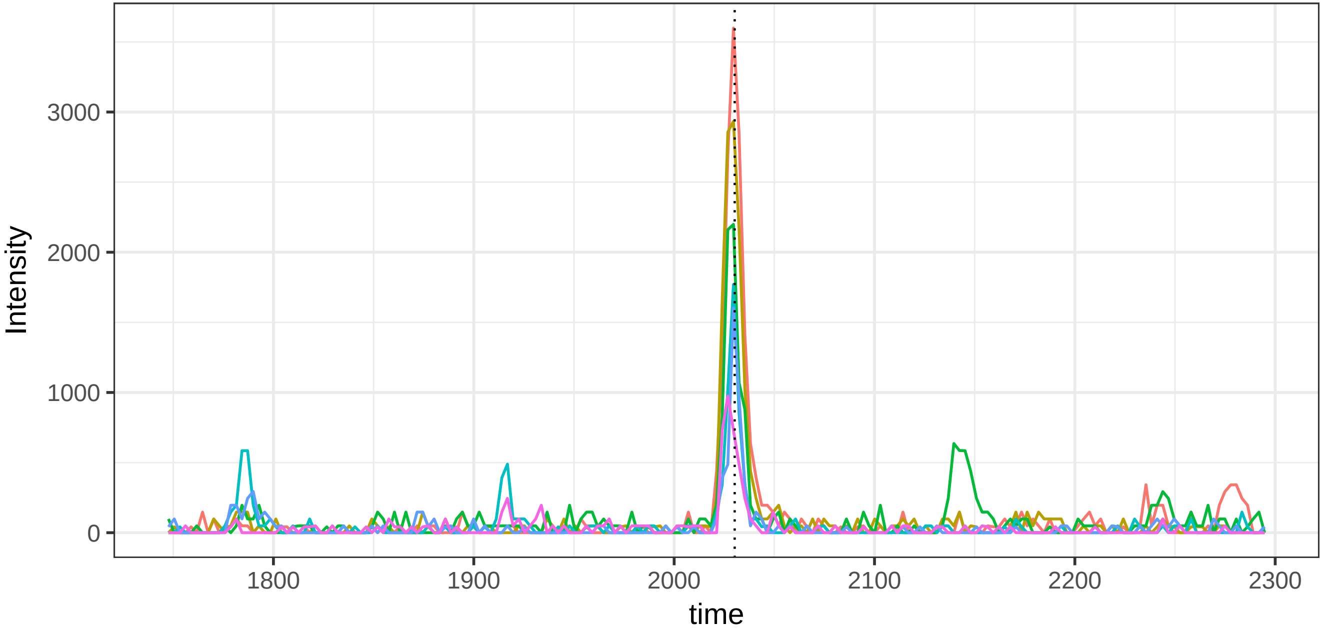


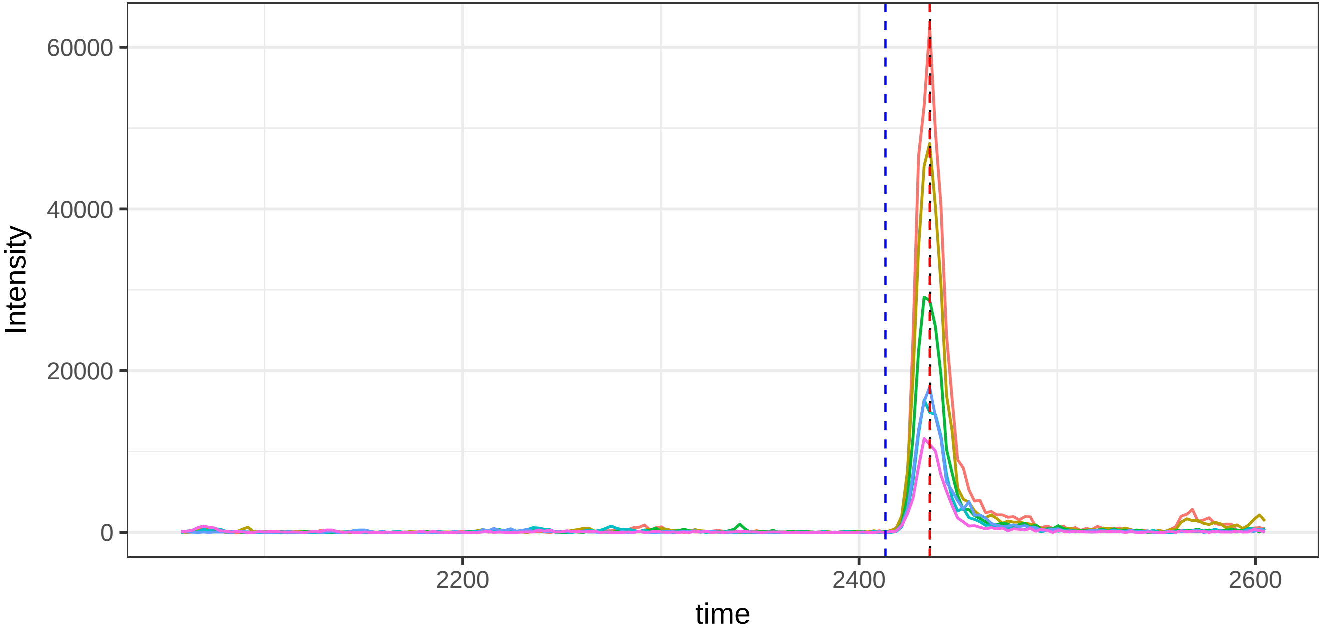

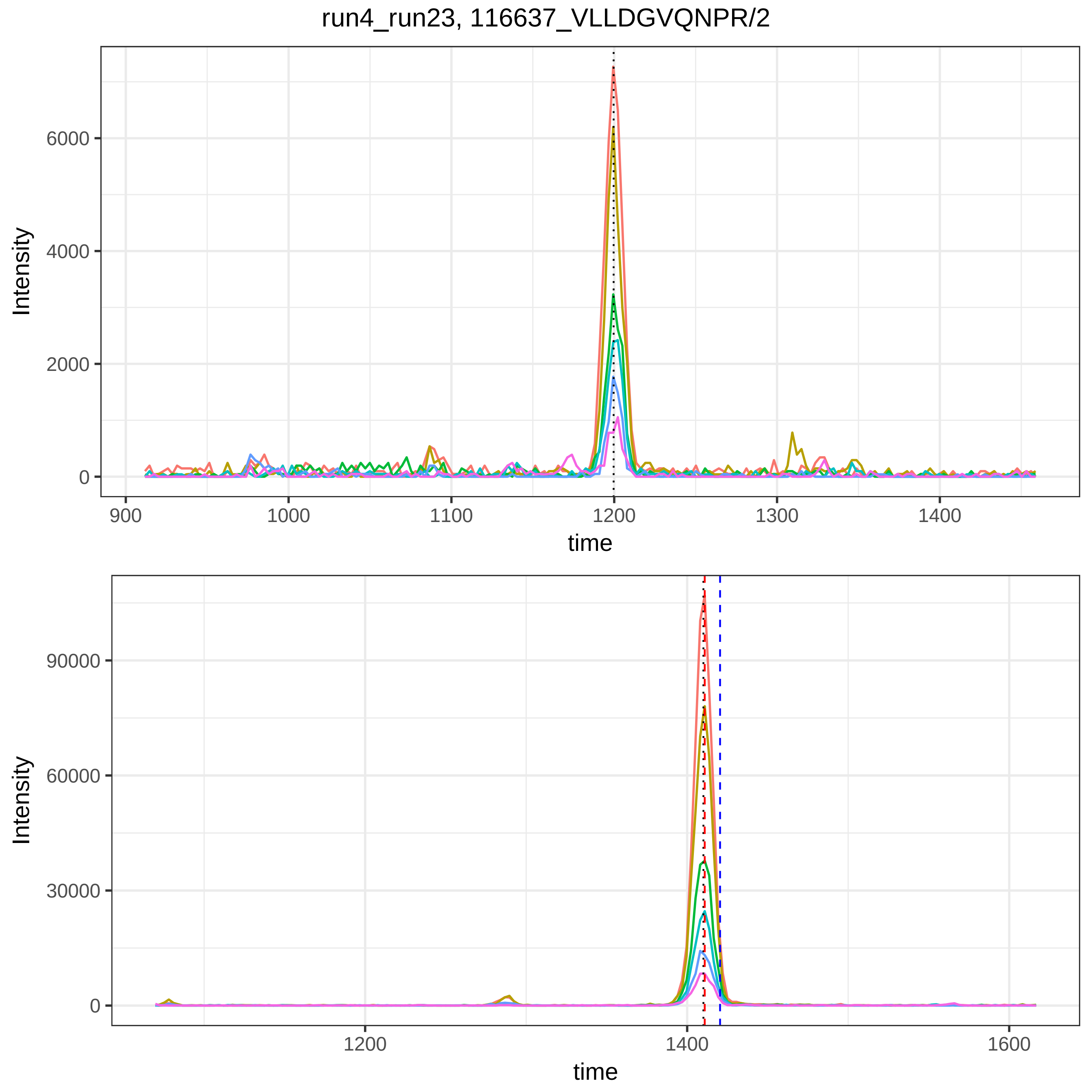

run4_run23, 116637_VLLDGVQNPR/2
Intensity
6000
4000
2000
0
900
1000
1100
1200
1300
1400
time
Intensity
90000
60000
30000
0
1200
1400
1600
time

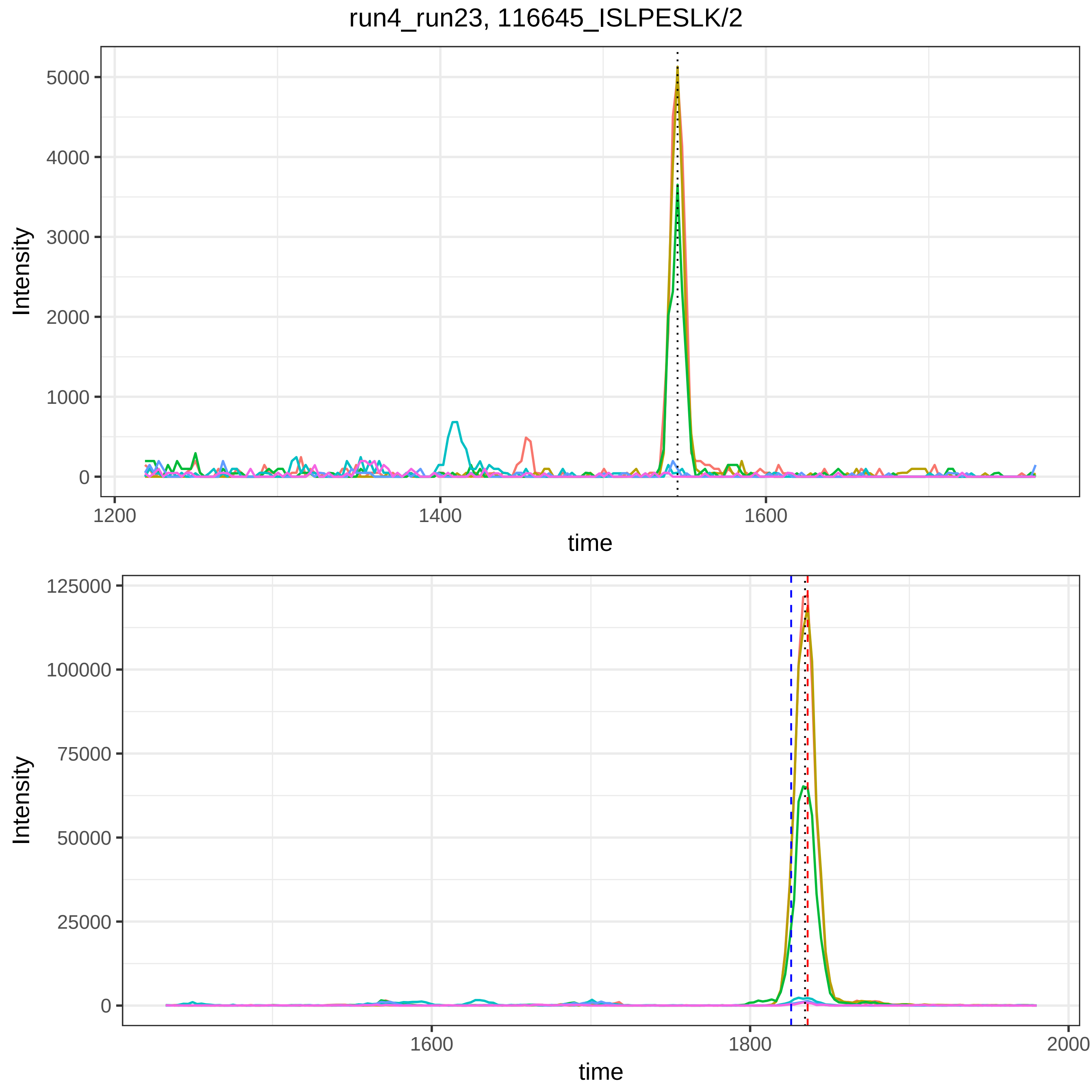

run4_run23, 116645_ISLPESLK/2
Intensity
5000
4000
3000
2000
1000
0
1200
1400
1600
time
Intensity
125000
100000
75000
50000
25000
0
1600
1800
2000
time

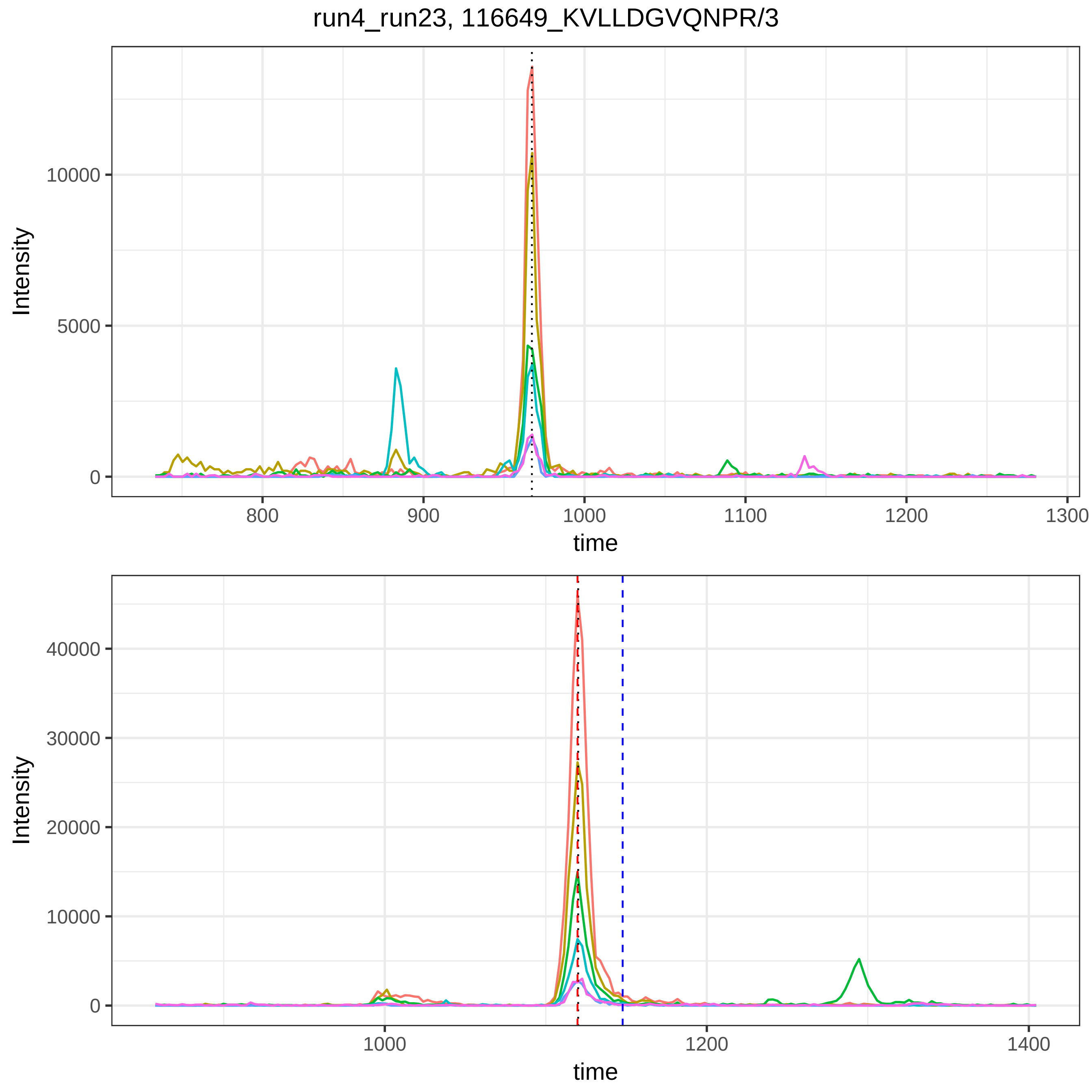
run4_run23, 116649_KVLLDGVQNPR/3
Intensity
10000
5000
0
800
900
1000
1100
1200
1300
time
Intensity
40000
30000
20000
10000
0
1000
1200
1400
time

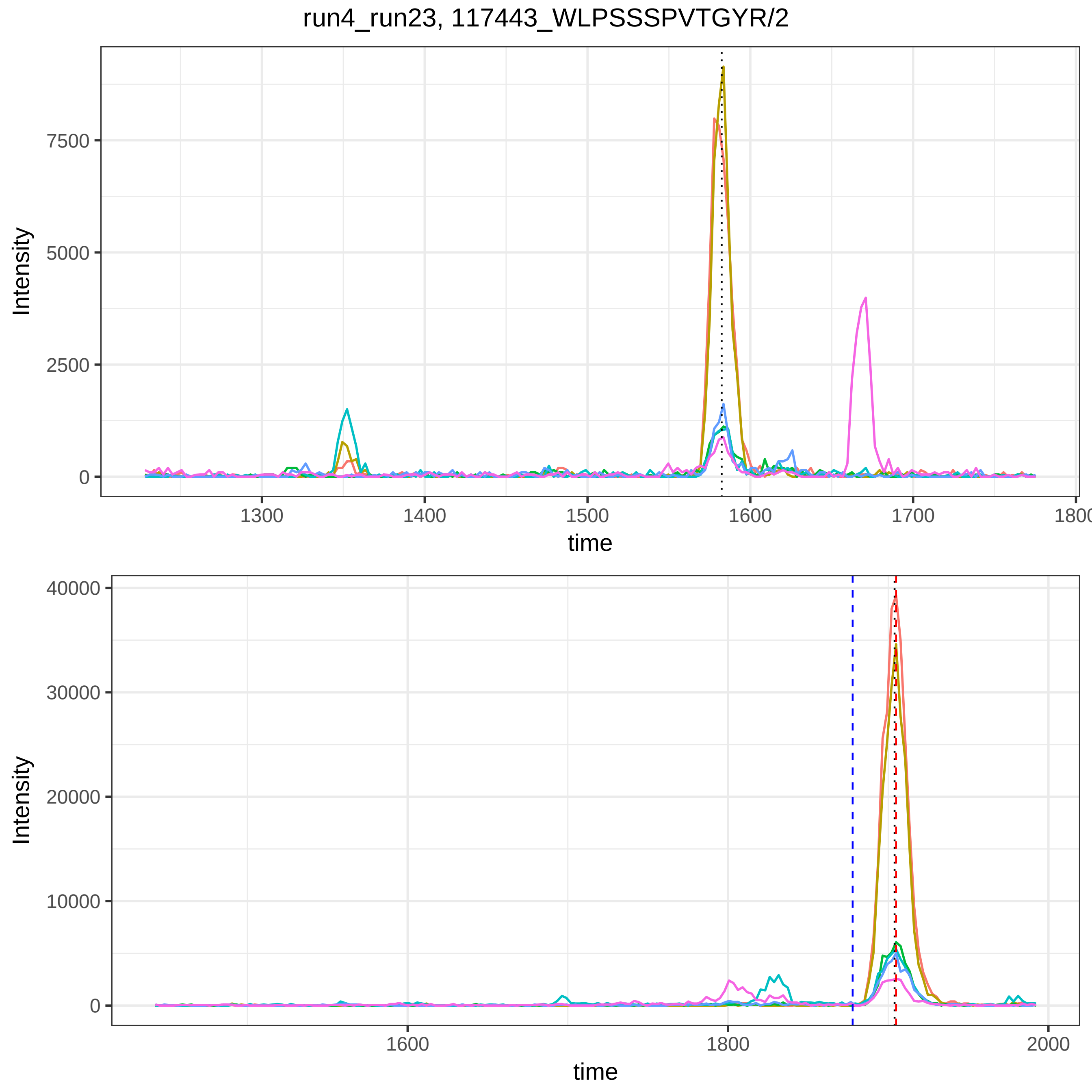
run4_run23, 117443_WLPSSSPVTGYR/2
Intensity
time
0
2500
5000
7500
1300
1400
1500
1600
1700
1800
0
10000
20000
30000
40000
1600
1800
2000

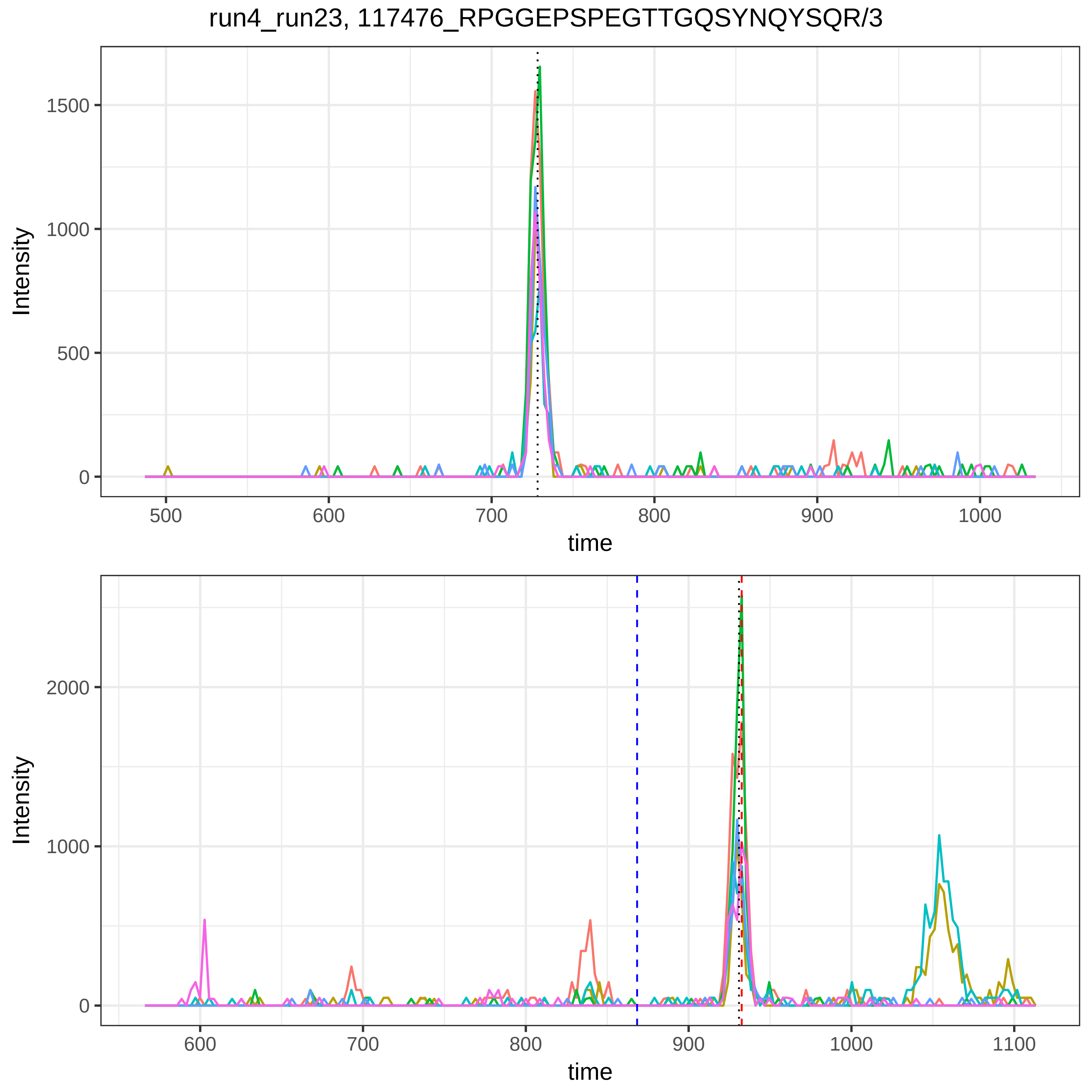
run4_run23, 117476_RPGGEPSPEGTTGQSYNQYSQR/3
Intensity
1500
1000
500
0
500
600
700
800
900
1000
time
Intensity
2000
1000
0
600
700
800
900
1000
1100
time

run4_run23, 117480_TEIDKPSQMQVTDVQDNSISVK/3

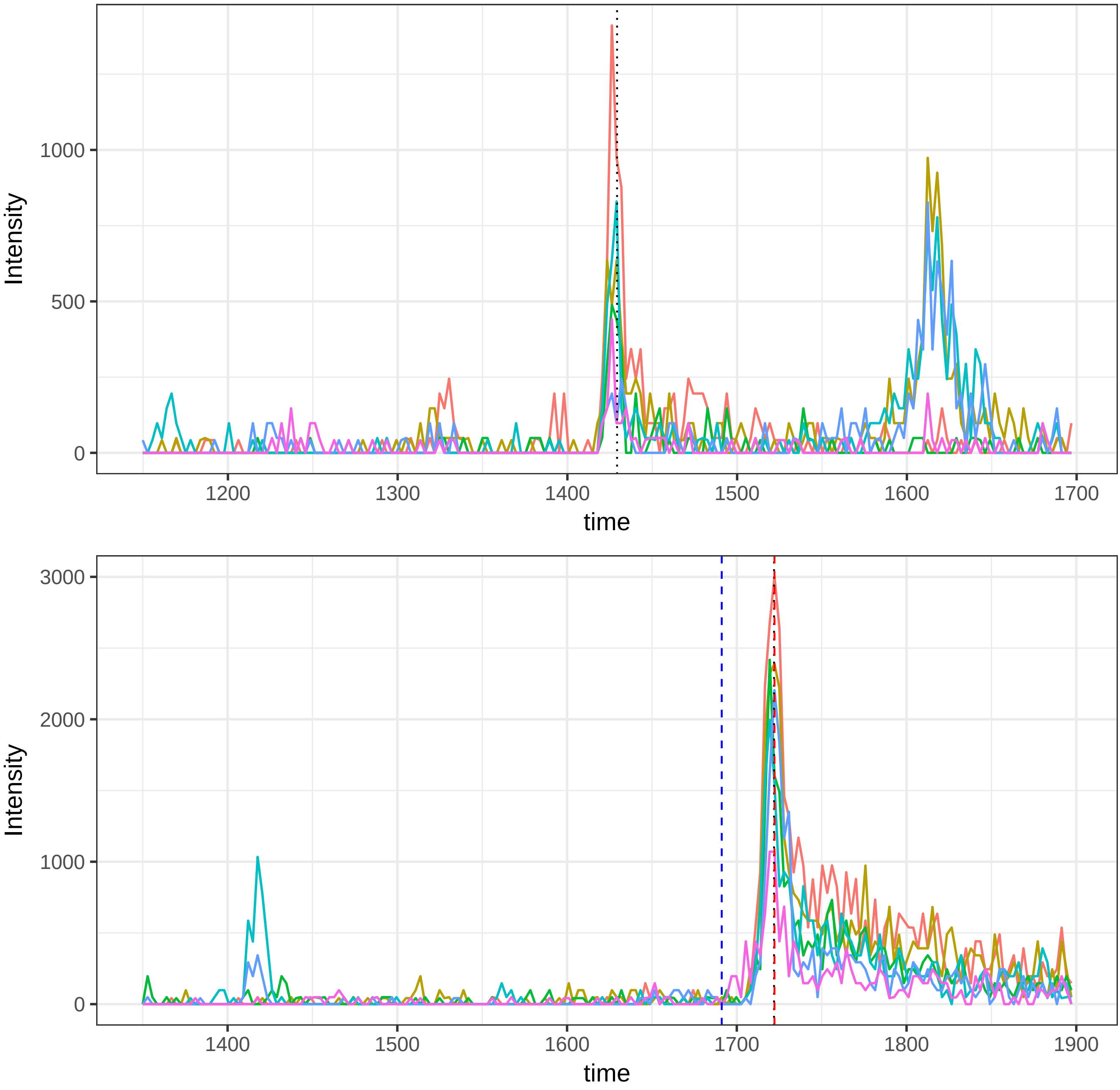

run4_run23, 117683_TEIDKPSQ(UniMod:7)MQ(UniMod:7)VTDVQDNSISVK/3

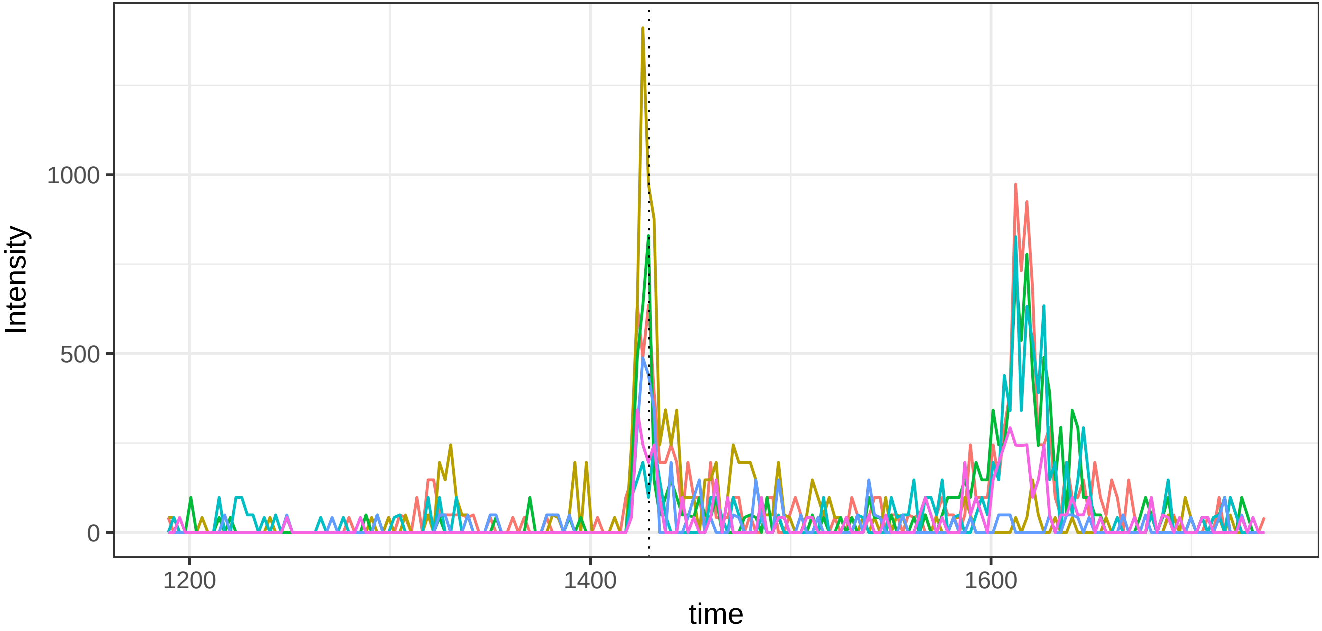


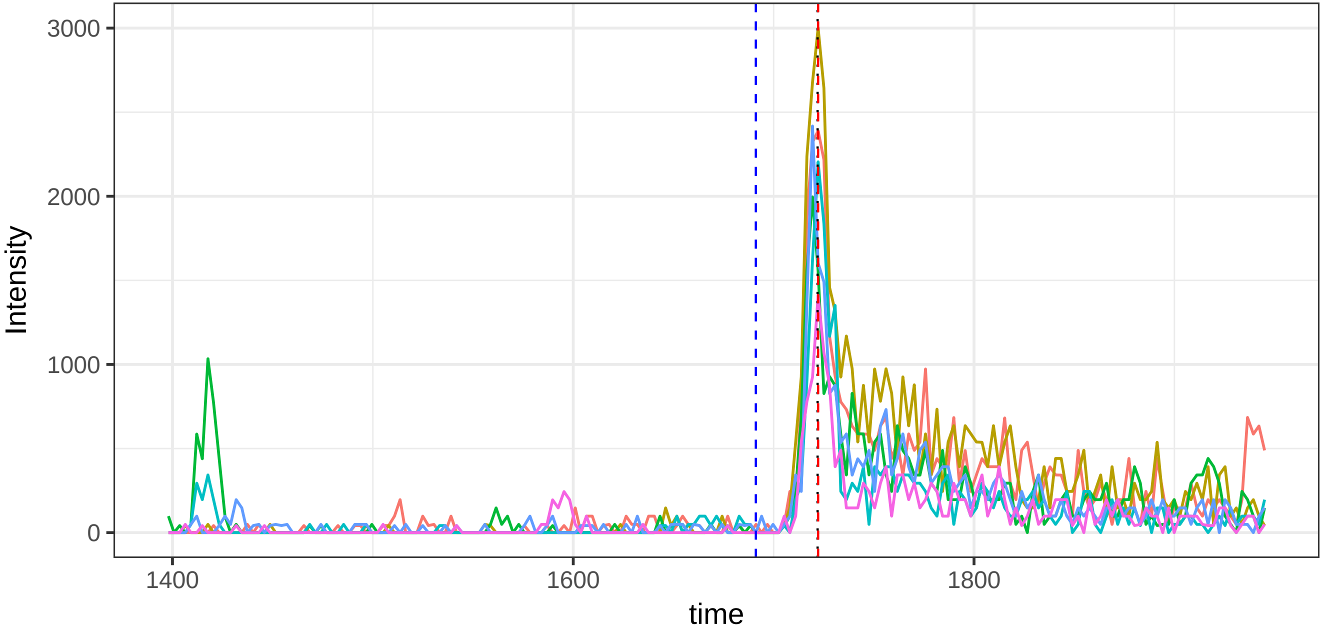

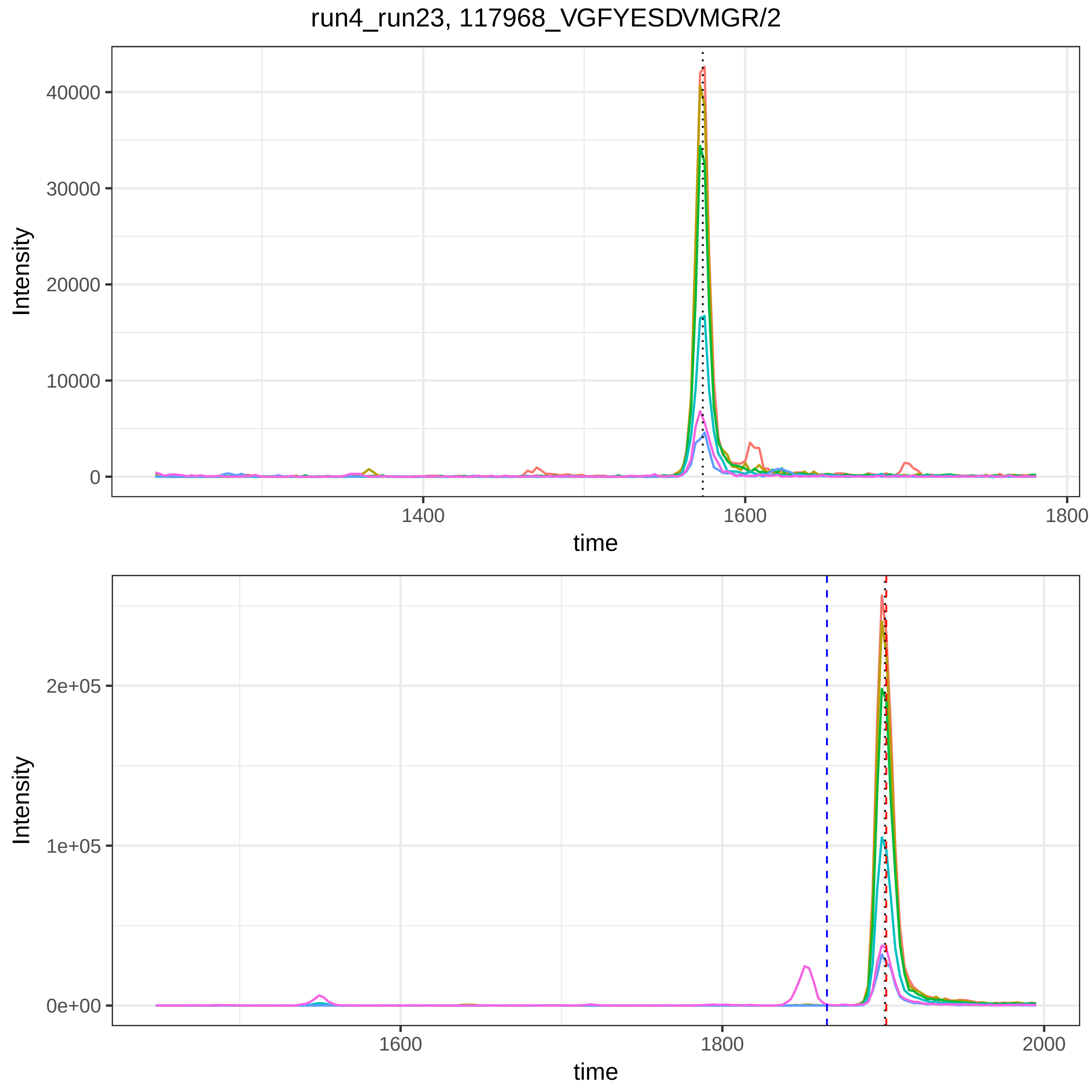

run4_run23, 117968_VGFYESDVMGR/2
Intensity
40000
30000
20000
10000
0
1400
1600
1800
time
Intensity
2e+05
1e+05
0e+00
1600
1800
2000
time

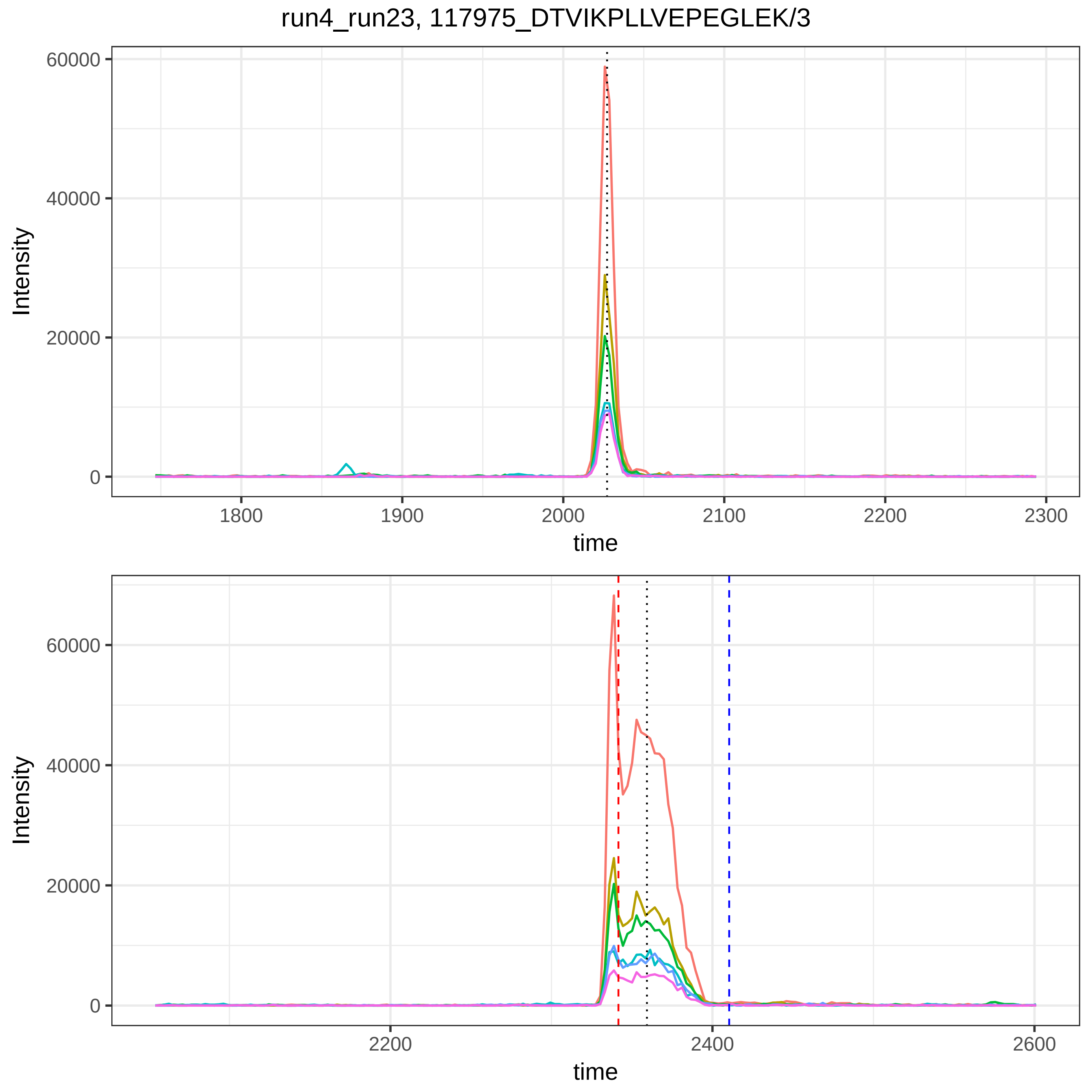

run4_run23, 117975_DTVIKPLLVEPEGLEK/3
Intensity
60000
40000
20000
0
1800
1900
2000
2100
2200
2300
time
Intensity
60000
40000
20000
0
2200
2400
2600
time

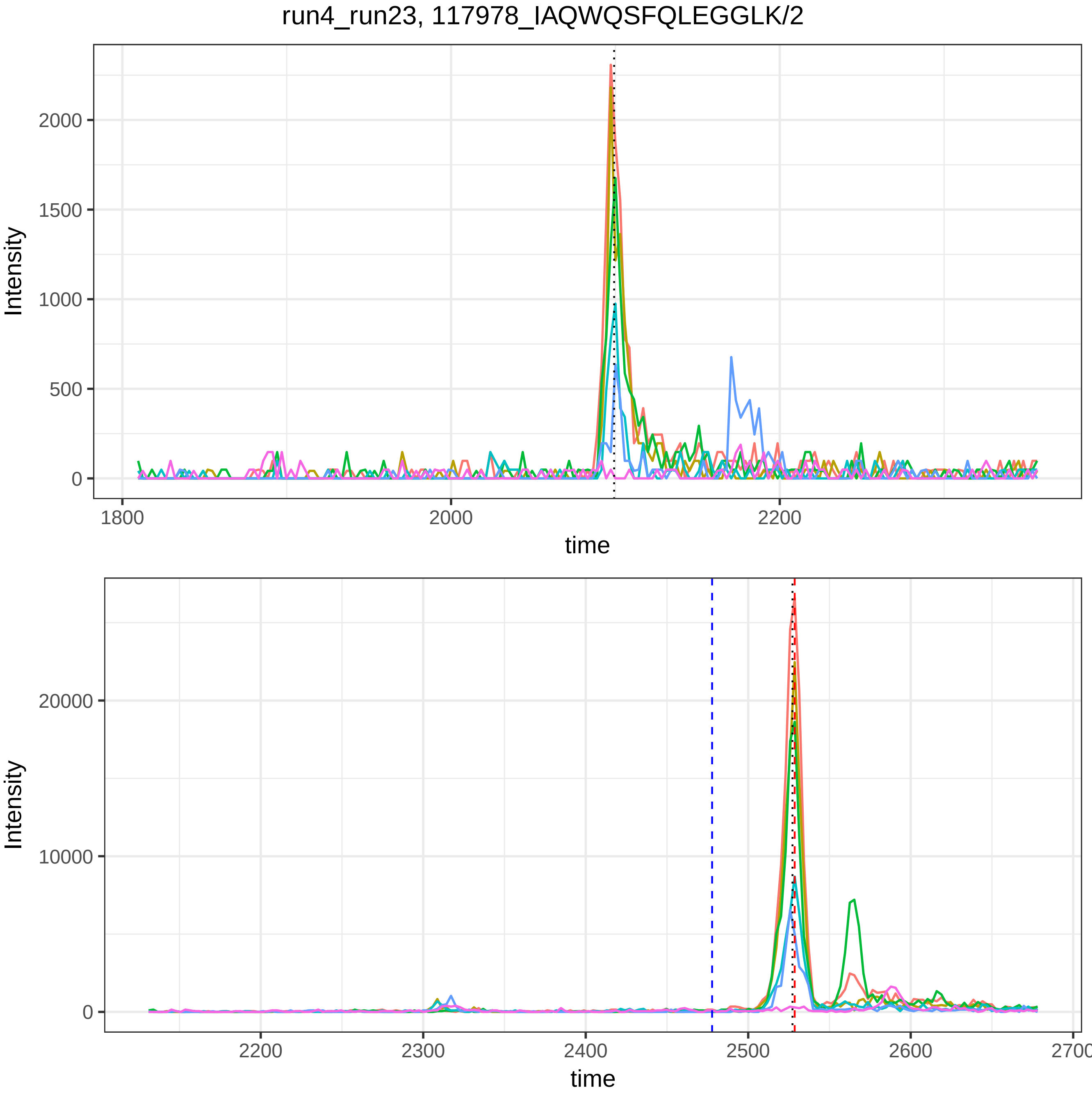
run4_run23, 117978_IAQWQSFQLEGGLK/2
Intensity
time
0
500
1000
1500
2000
1800
2000
2200
Intensity
time
0
10000
20000
2200
2300
2400
2500
2600
2700

run4_run23, 118449_SASDLTWDNLK/2

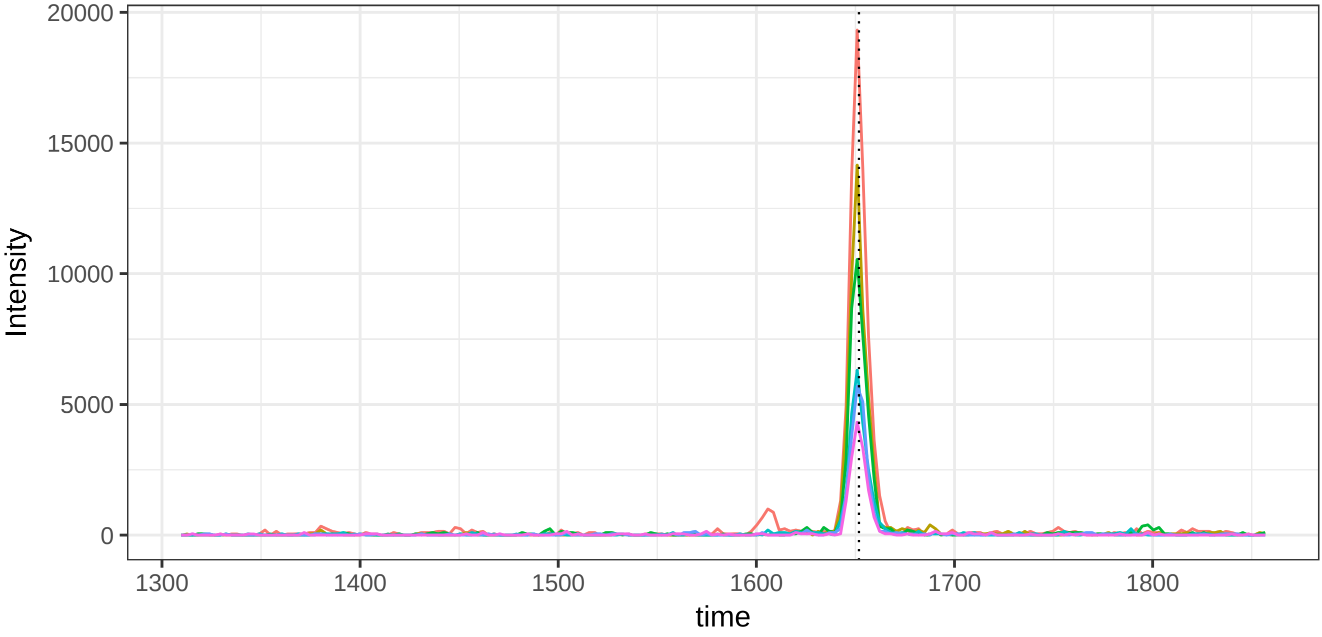


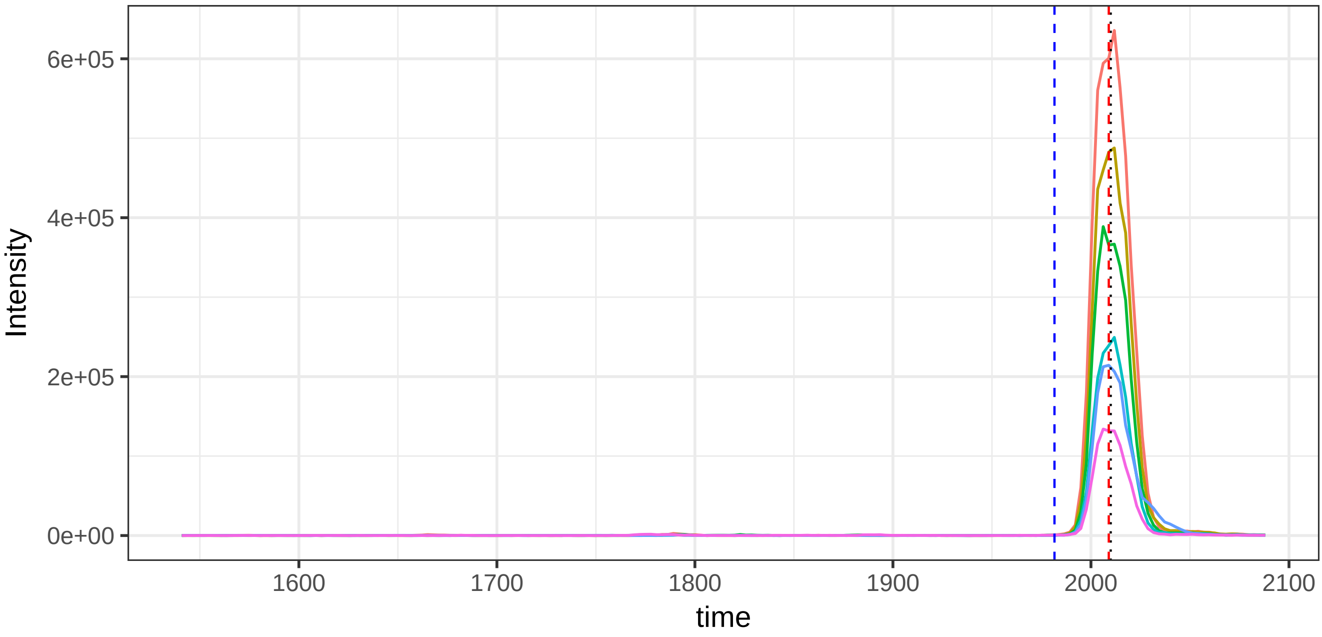

run4_run23, 119263_HSIFTPETNPR/2

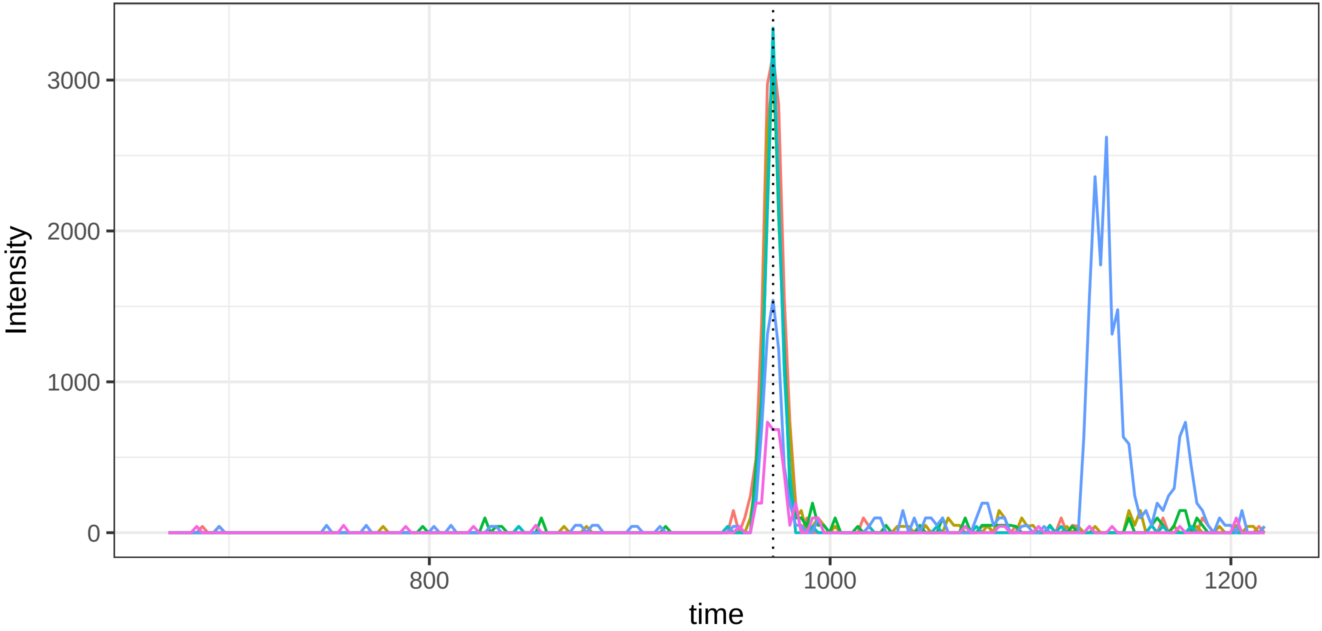


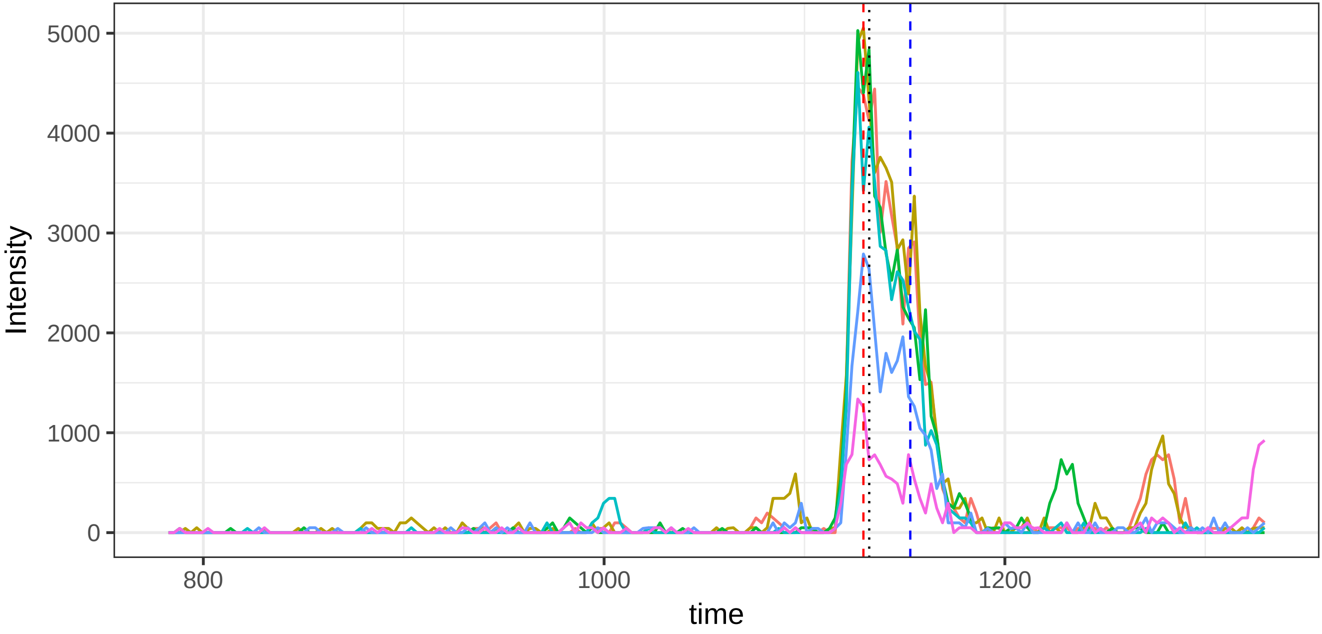

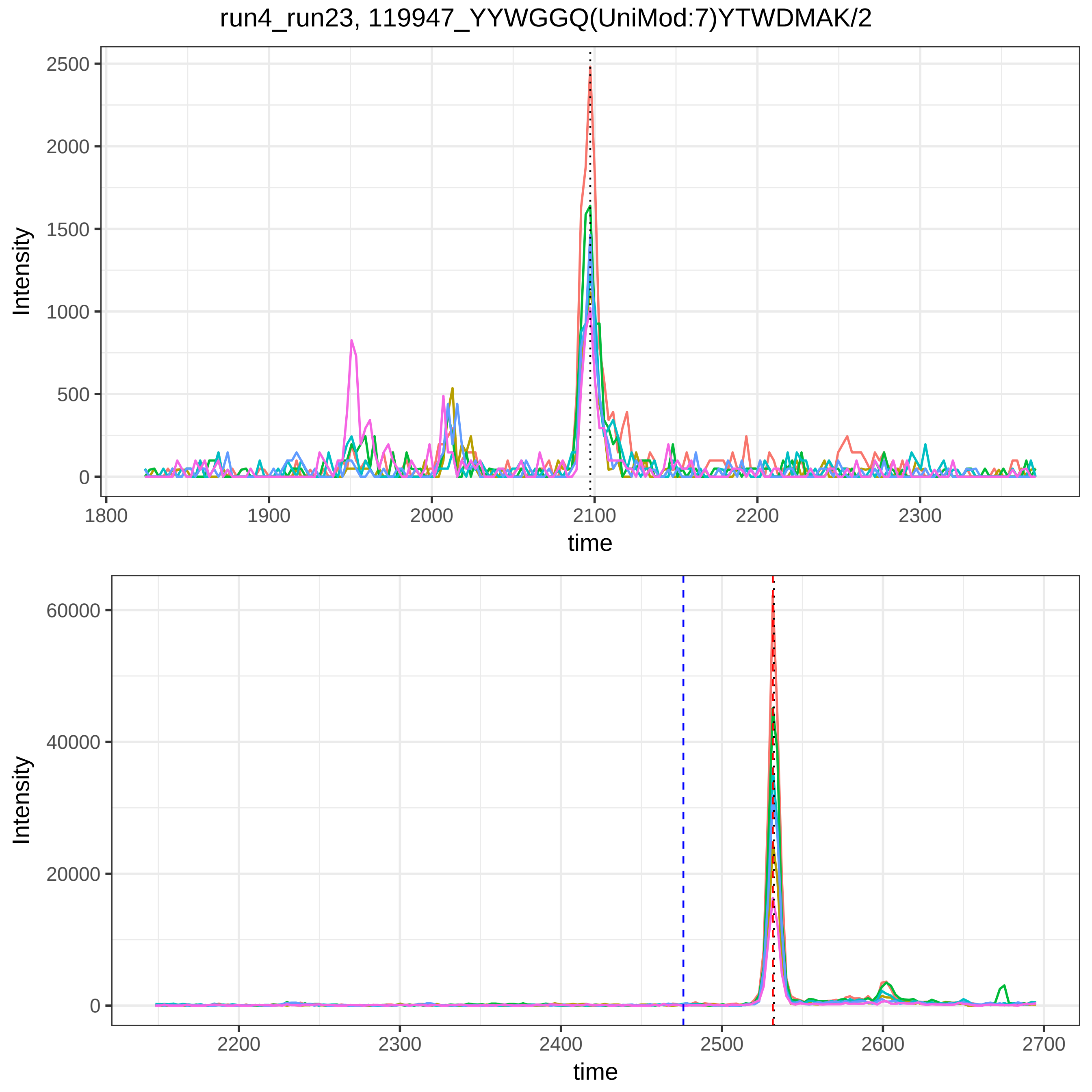
run4_run23, 119947_YYWGGQ(UniMod:7)YTWDMAK/2
Intensity
time
Intensity
time

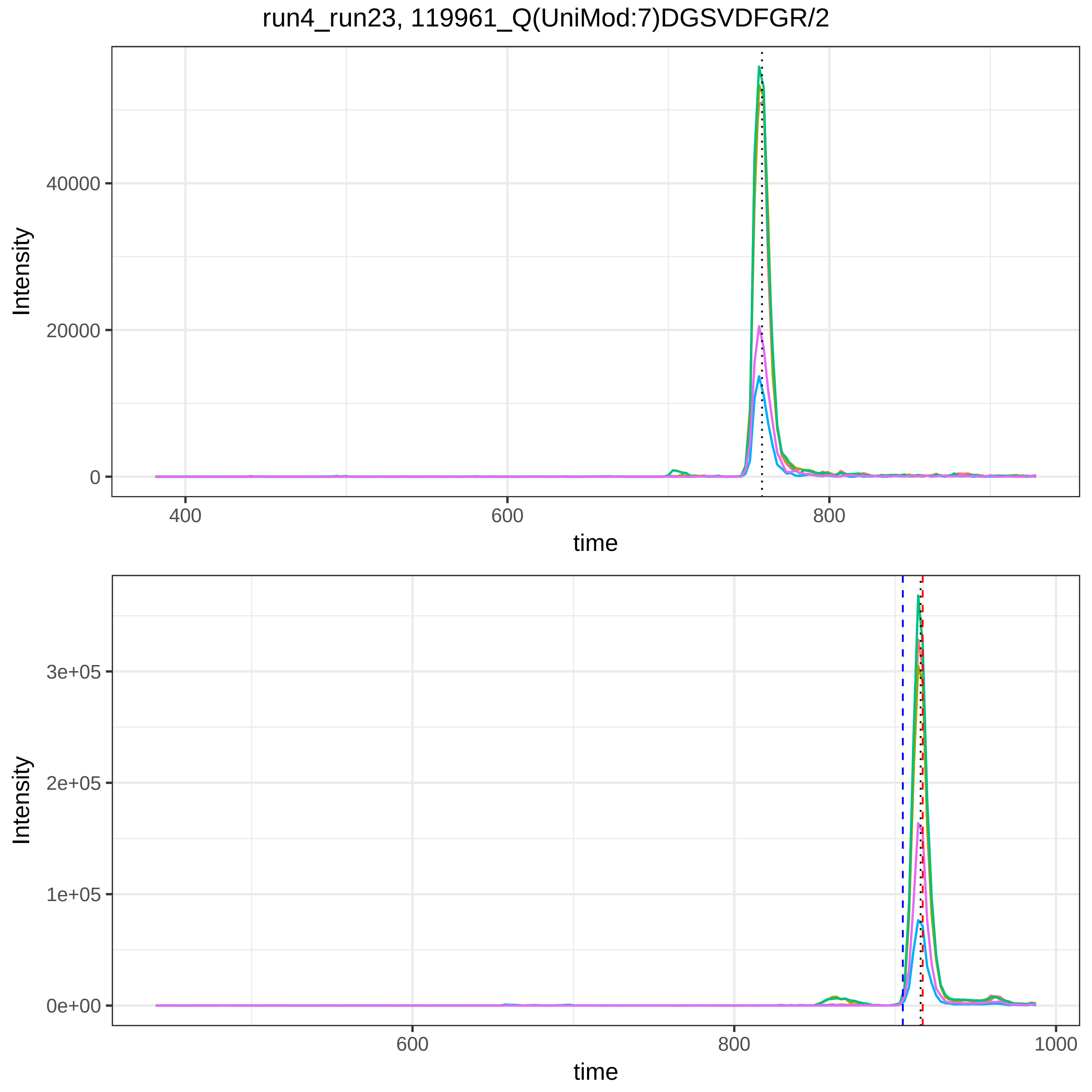
run4_run23, 119961_Q(UniMod:7)DGSVDFGR/2
Intensity
40000
20000
0
400
600
800
time
Intensity
3e+05
2e+05
1e+05
0e+00
600
800
1000
time

run4_run23, 119993_HAANP(UniMod:35)NGRYYWGGQYTWDMAK/3

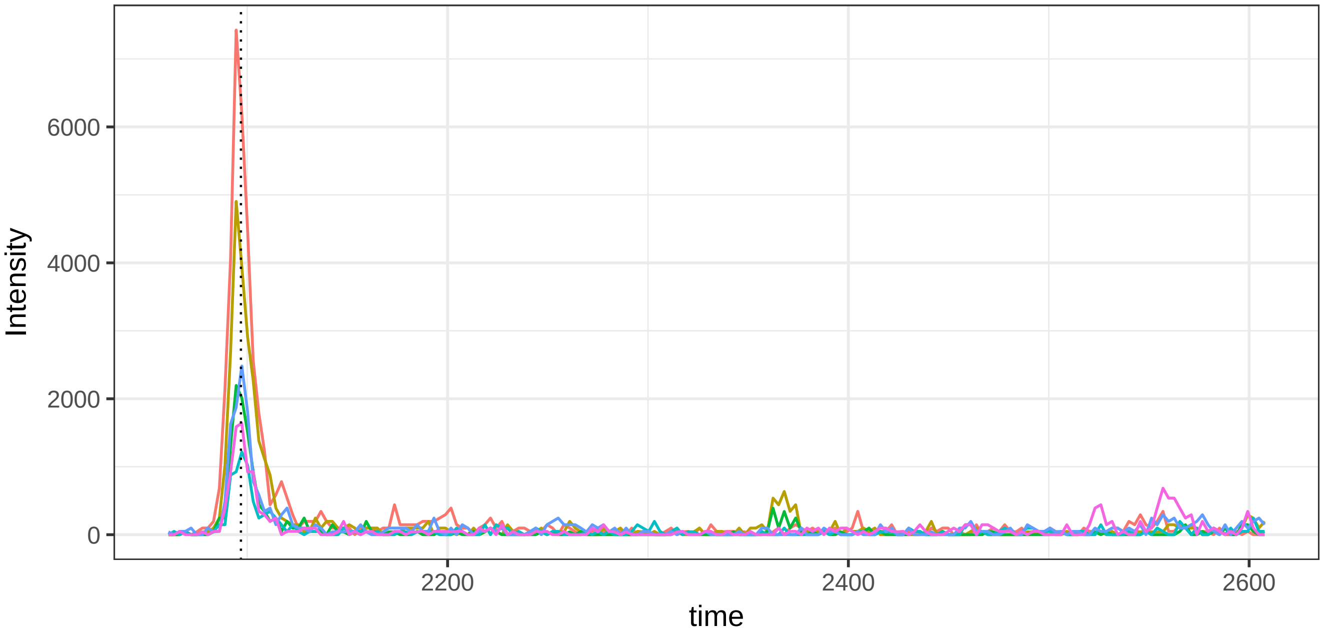


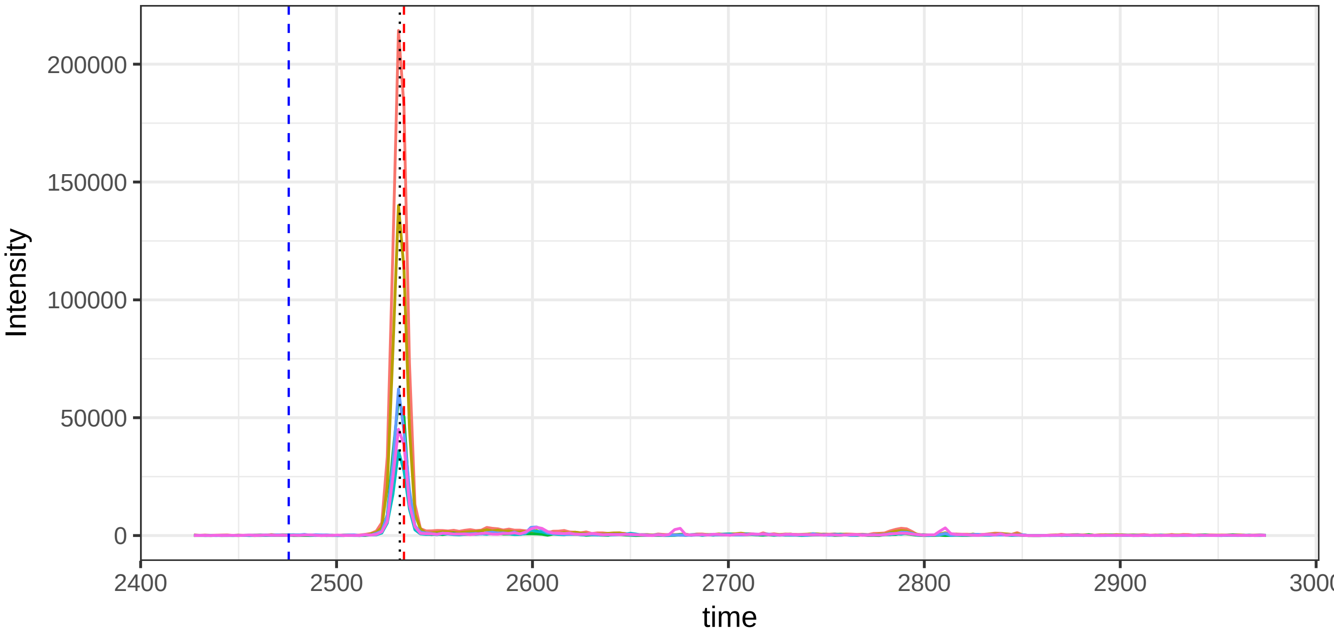

run4_run23, 120588_FTEDAKR(UniMod:35)LYGS(UniMod:21)EAFATDFQDSAAAK/3

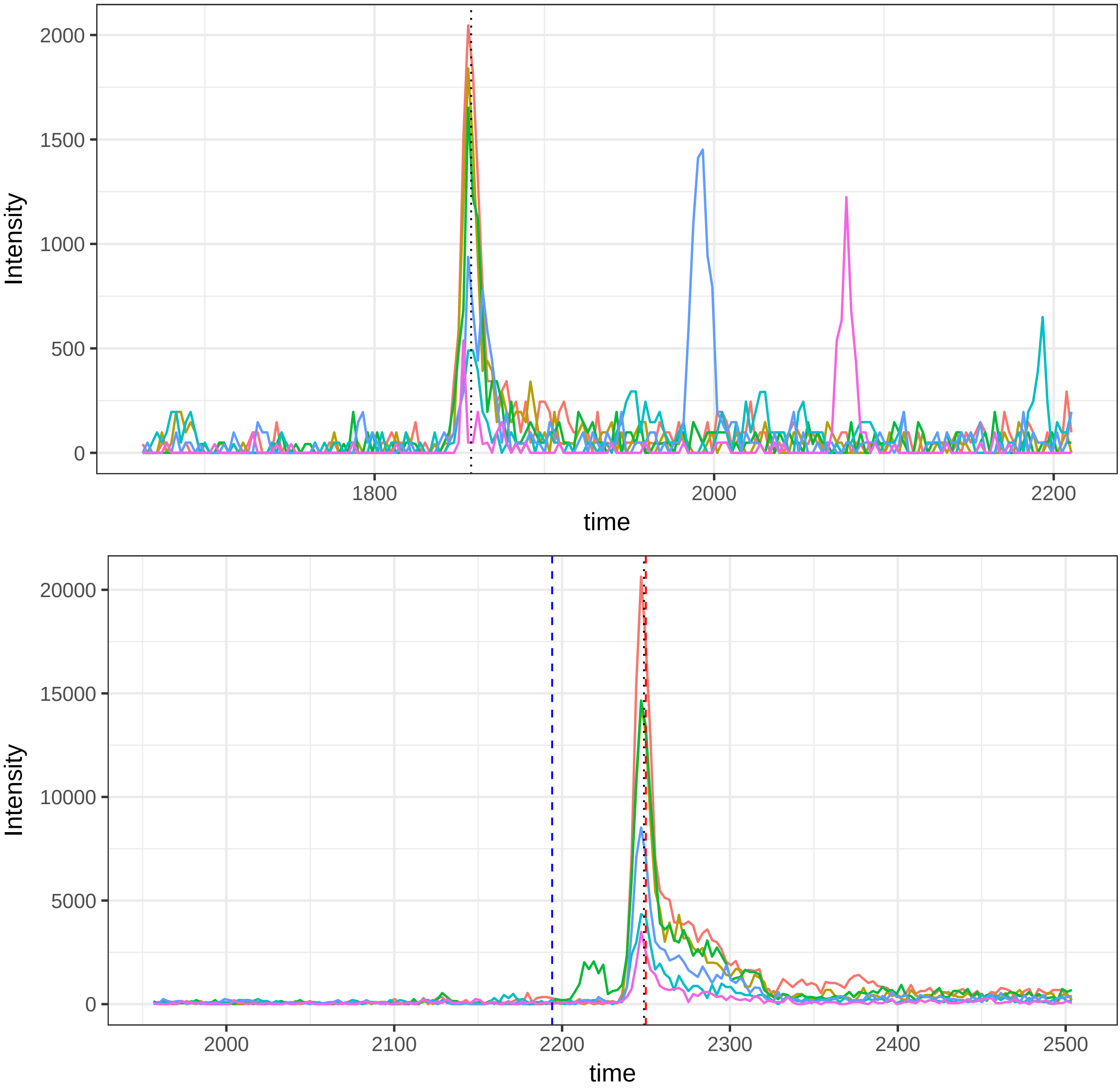

run4_run23, 121296_ATEDEGSEQKIPEATNR/3

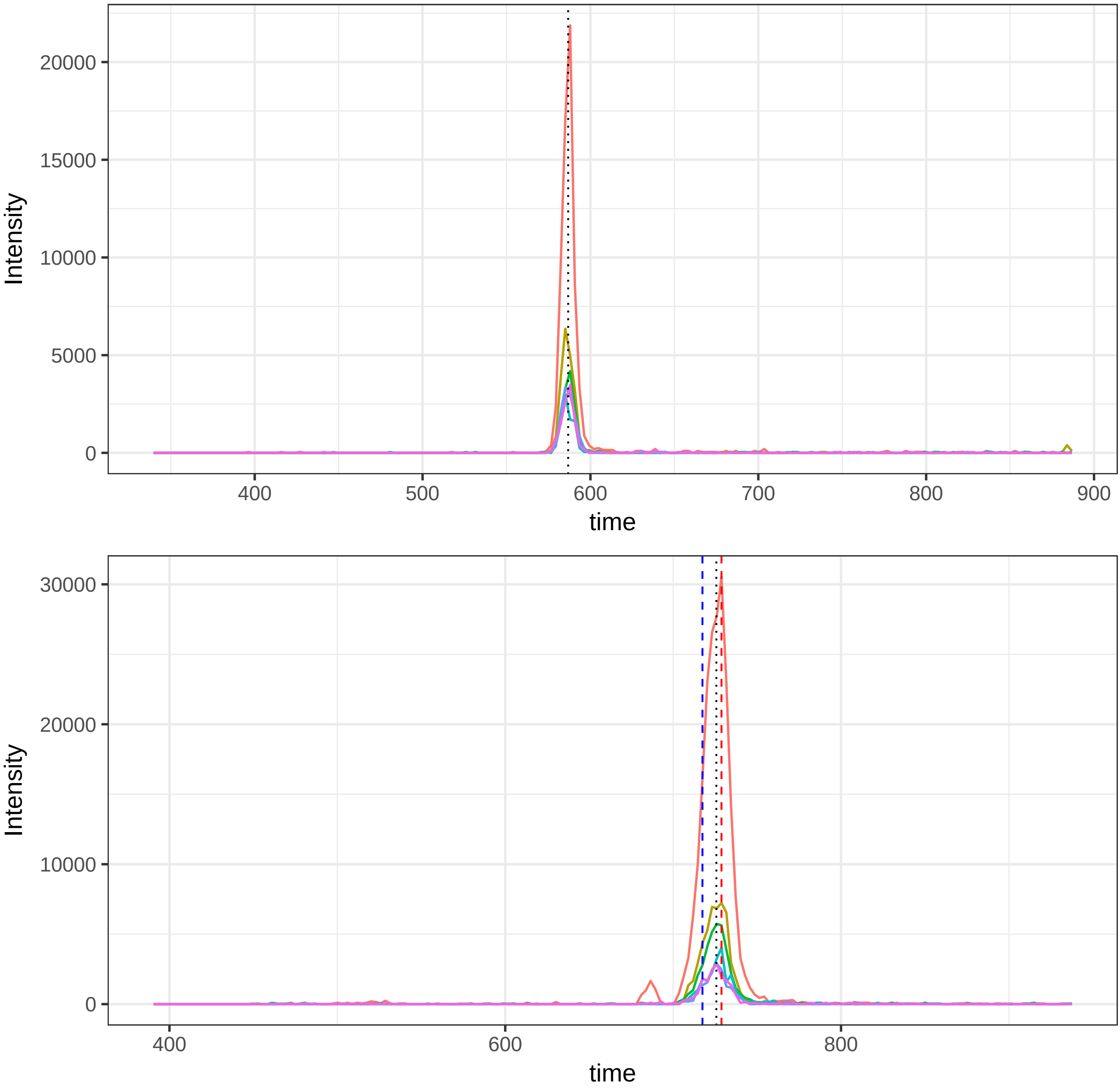

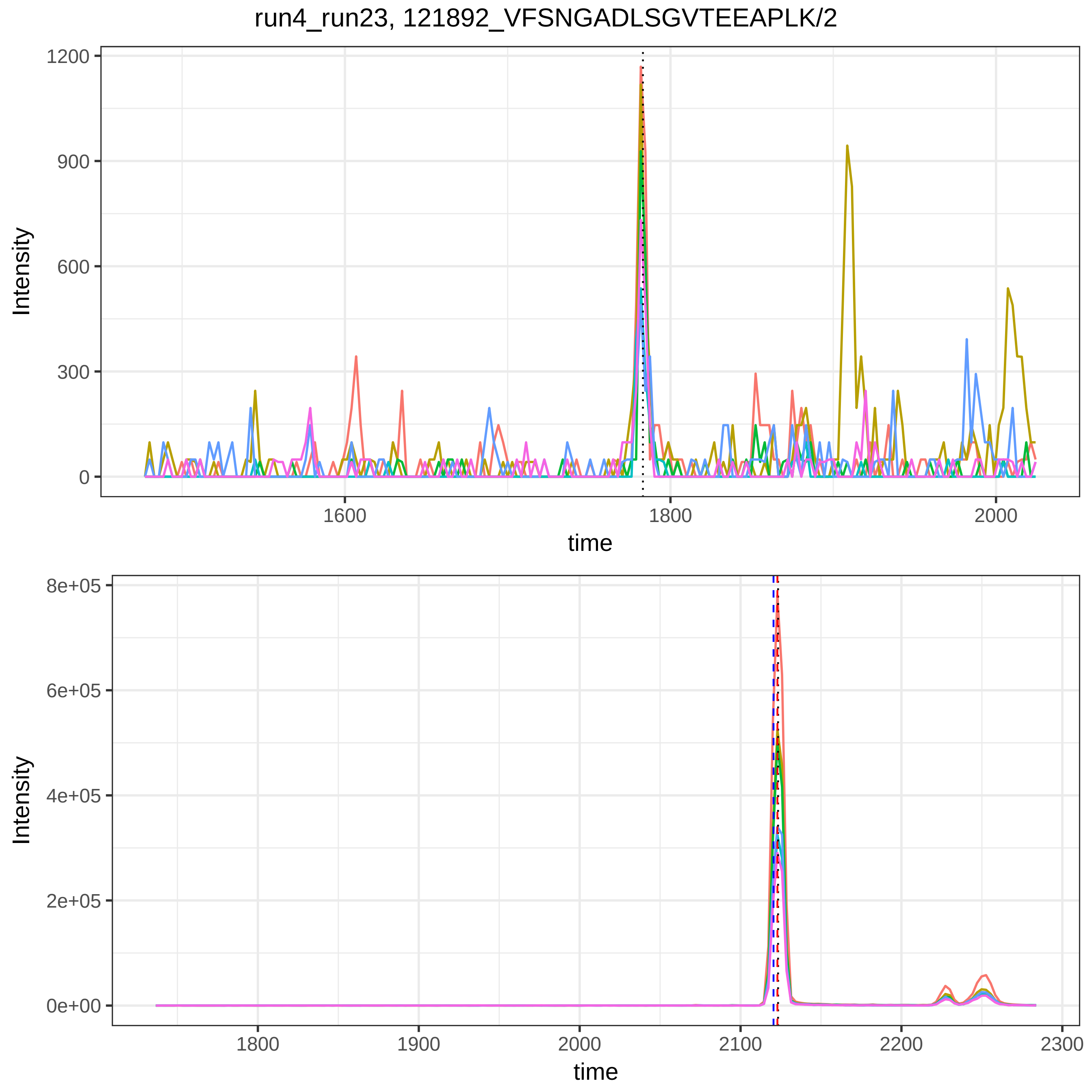
run4_run23, 121892_VFSNGADLSGVTEEAPLK/2
Intensity
1200
900
600
300
0
1600
1800
2000
time
Intensity
8e+05
6e+05
4e+05
2e+05
0e+00
1800
1900
2000
2100
2200
2300
time

run4_run23, 121893_SASLHLPK/2

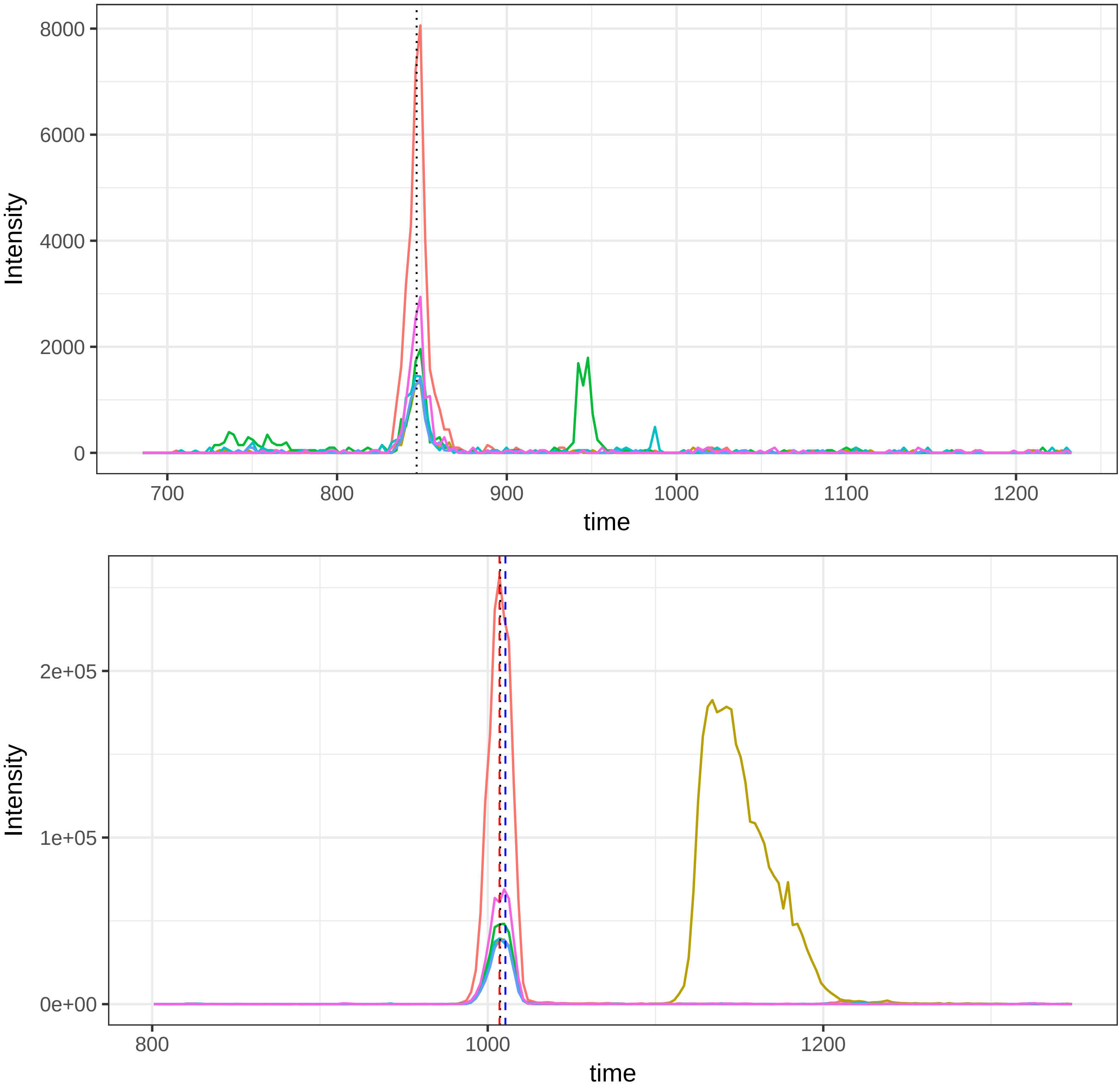

run4_run23, 1256_VFSNGADLSGVTEEAPLK/2

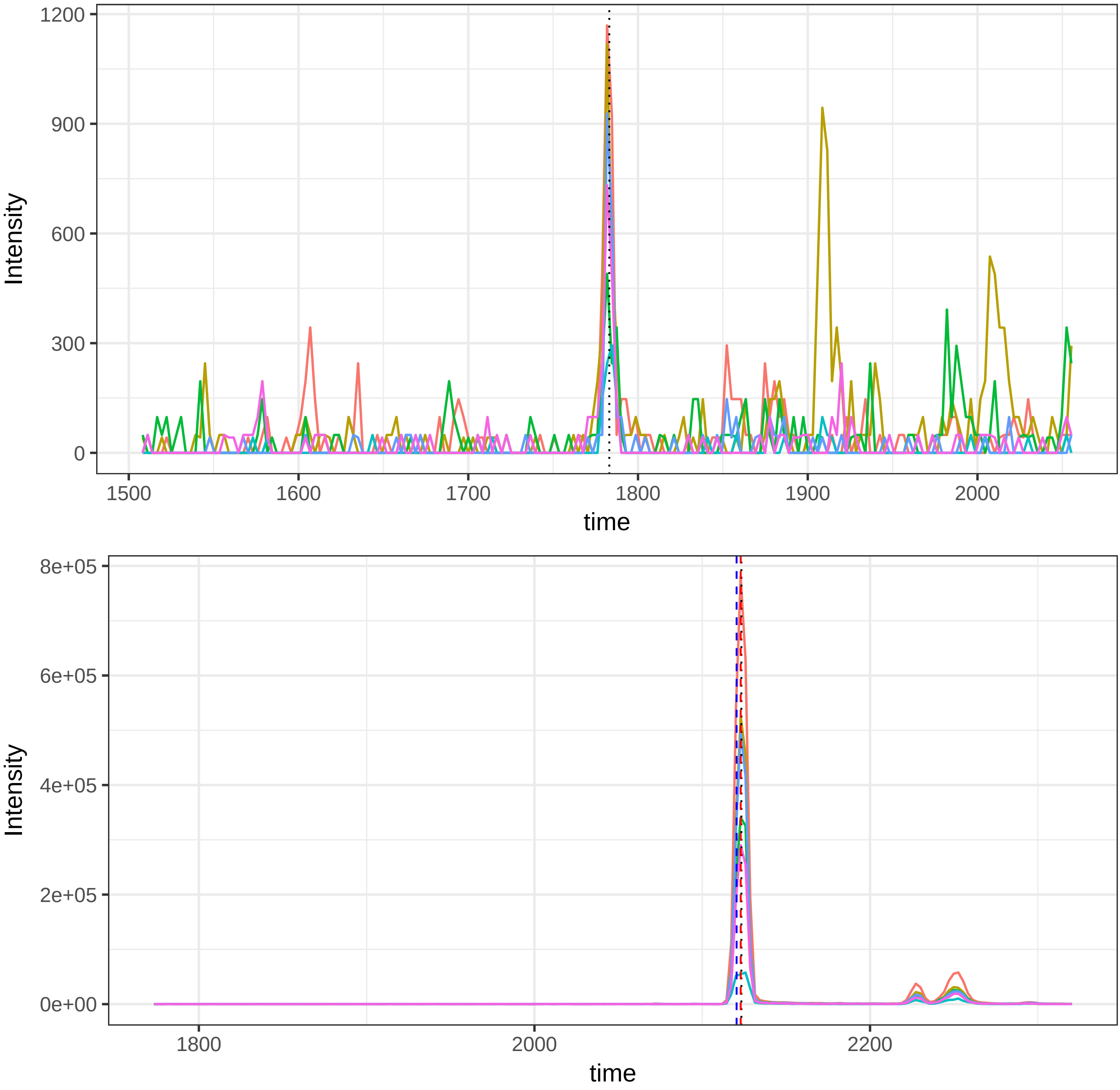

run4_run23, 125820_AATVGSLAGQPLQER/2

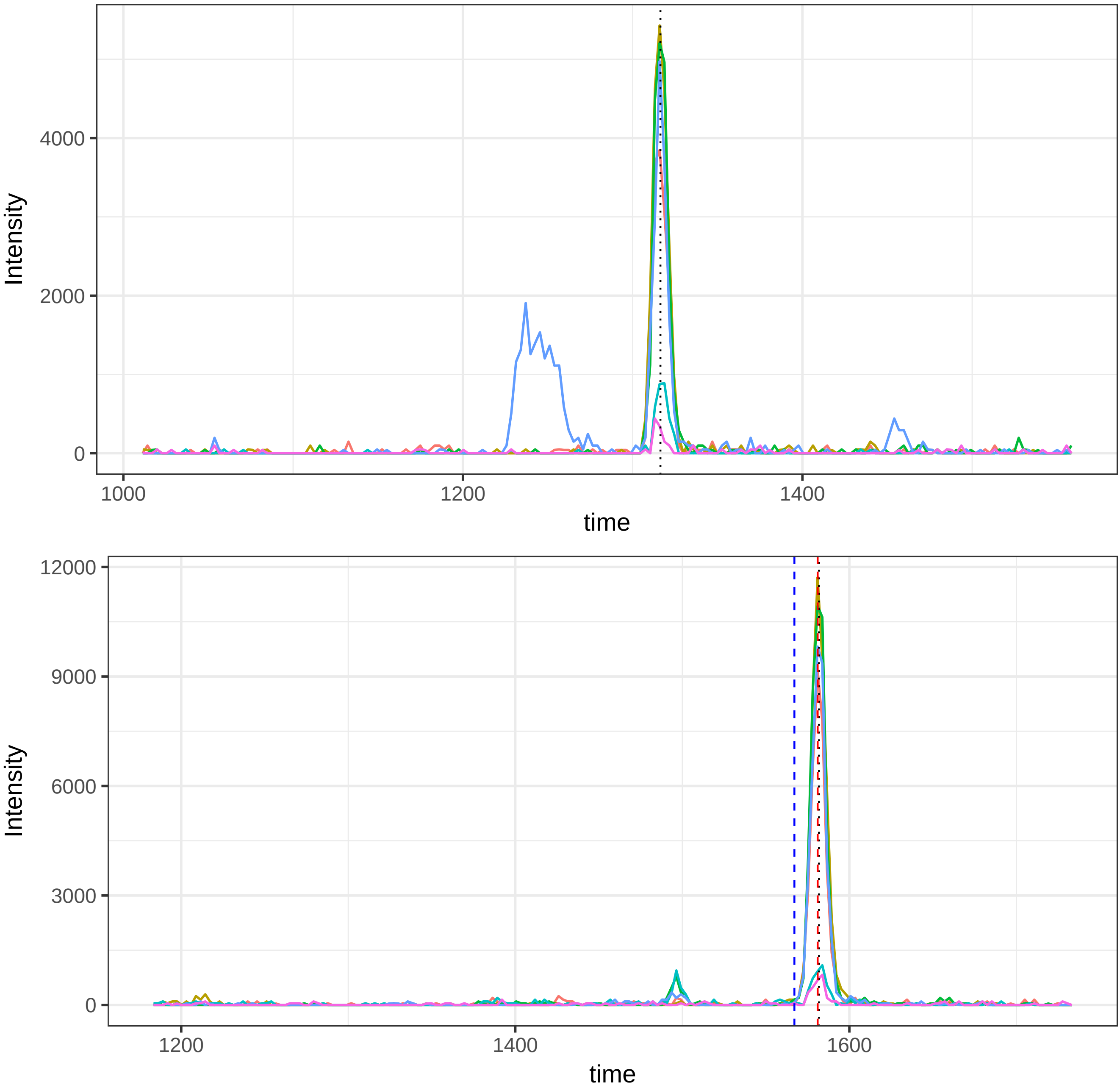

run4_run23, 128343_VTEPISAESGEQVER/2

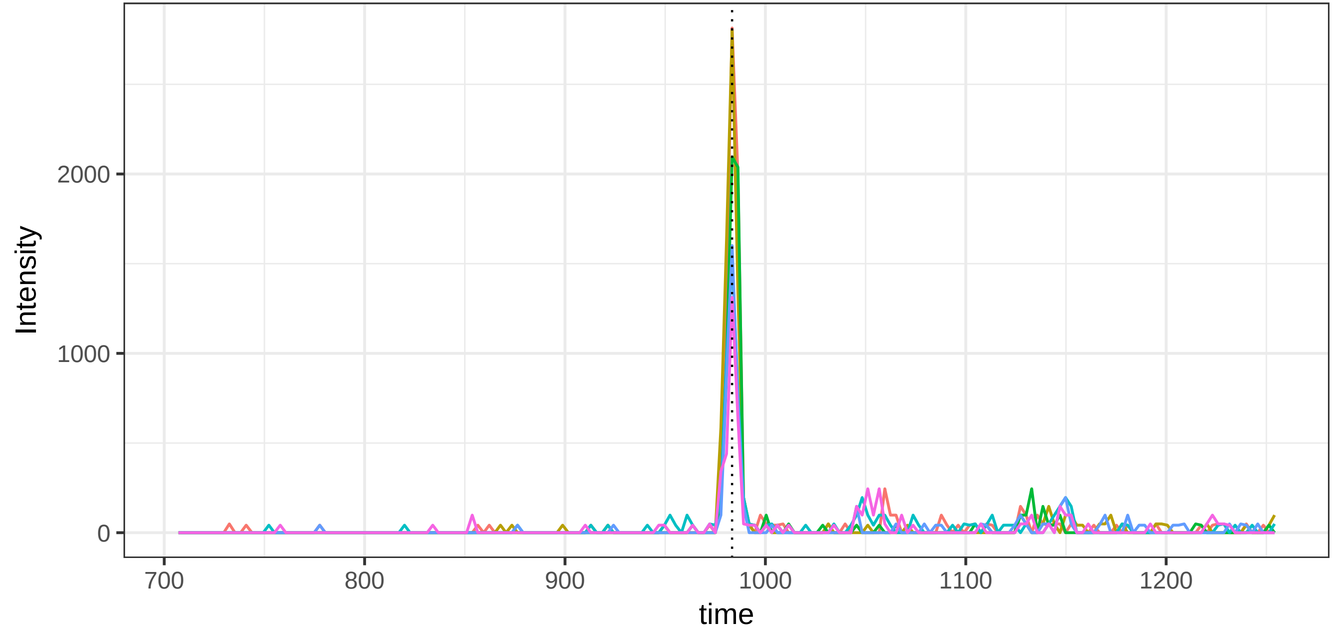


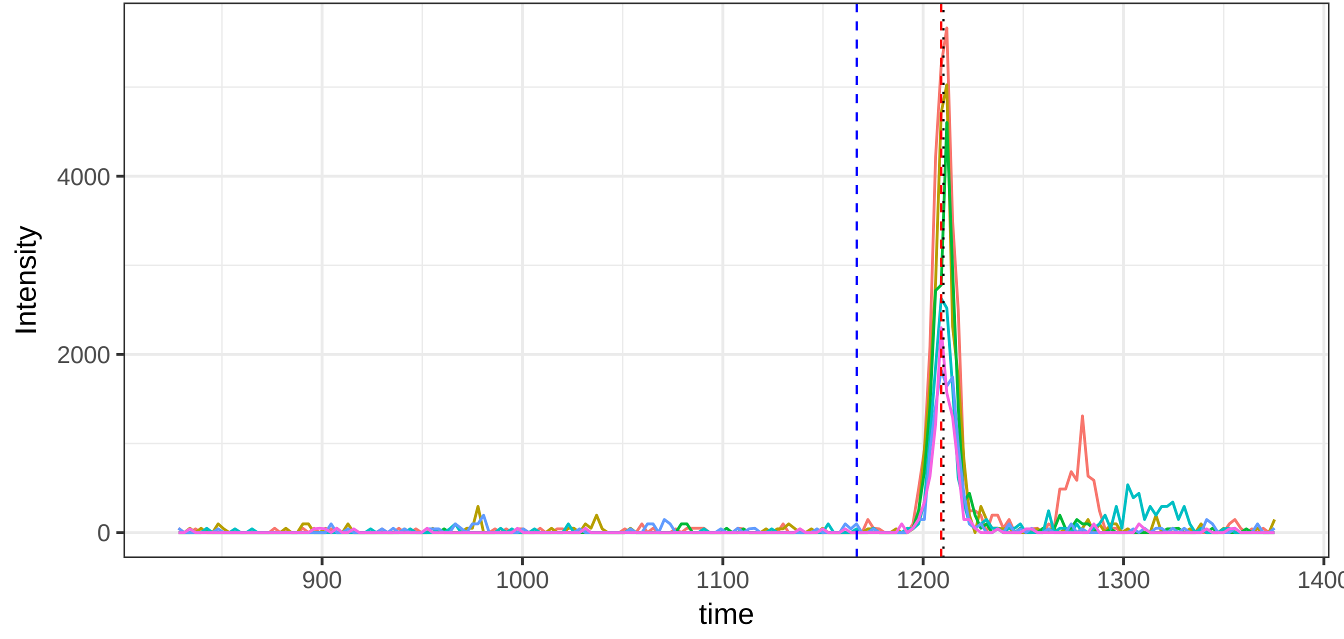

run4_run23, 128893_VGYP(UniMod:35)GPSGPLGAR/2

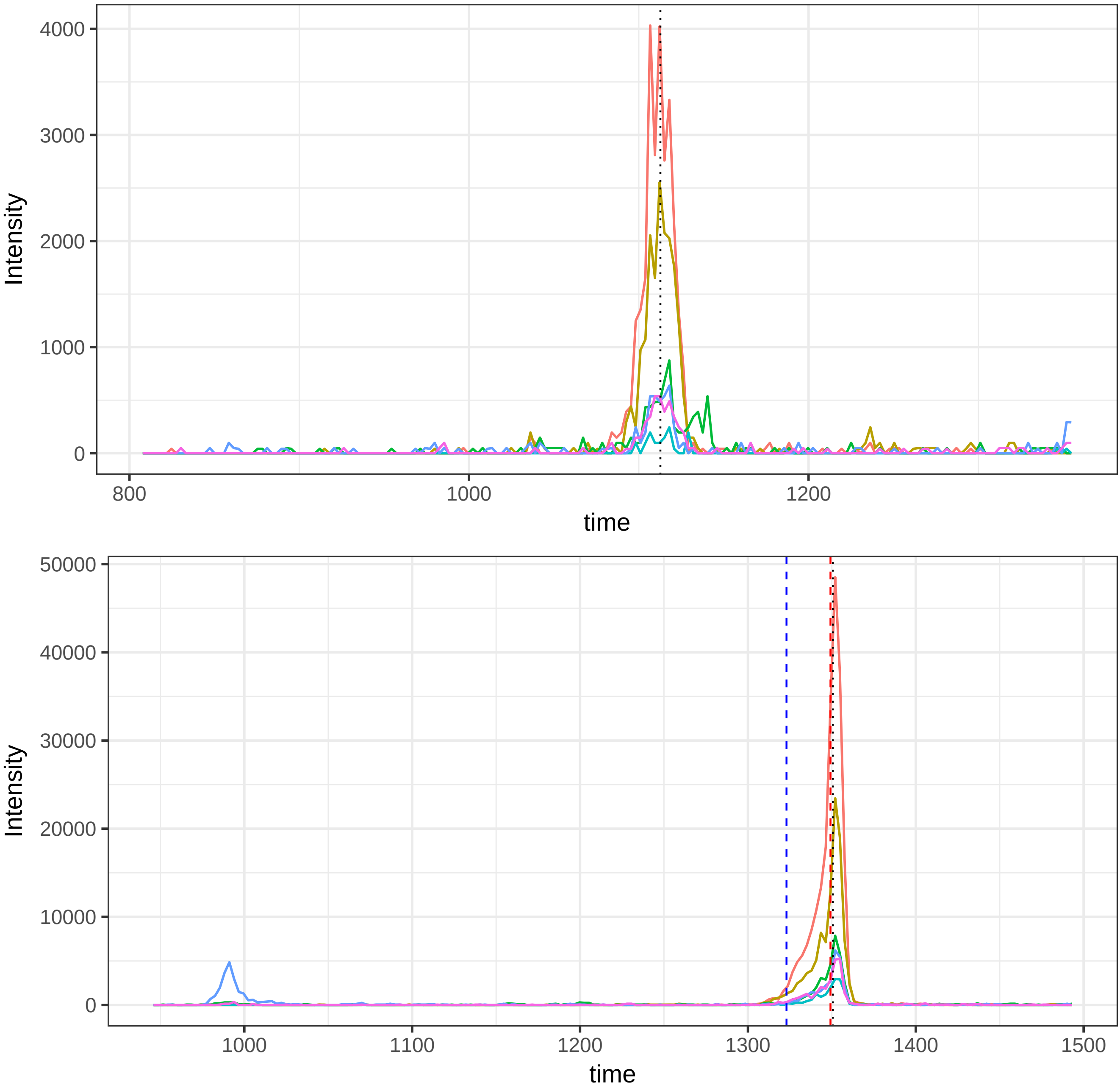

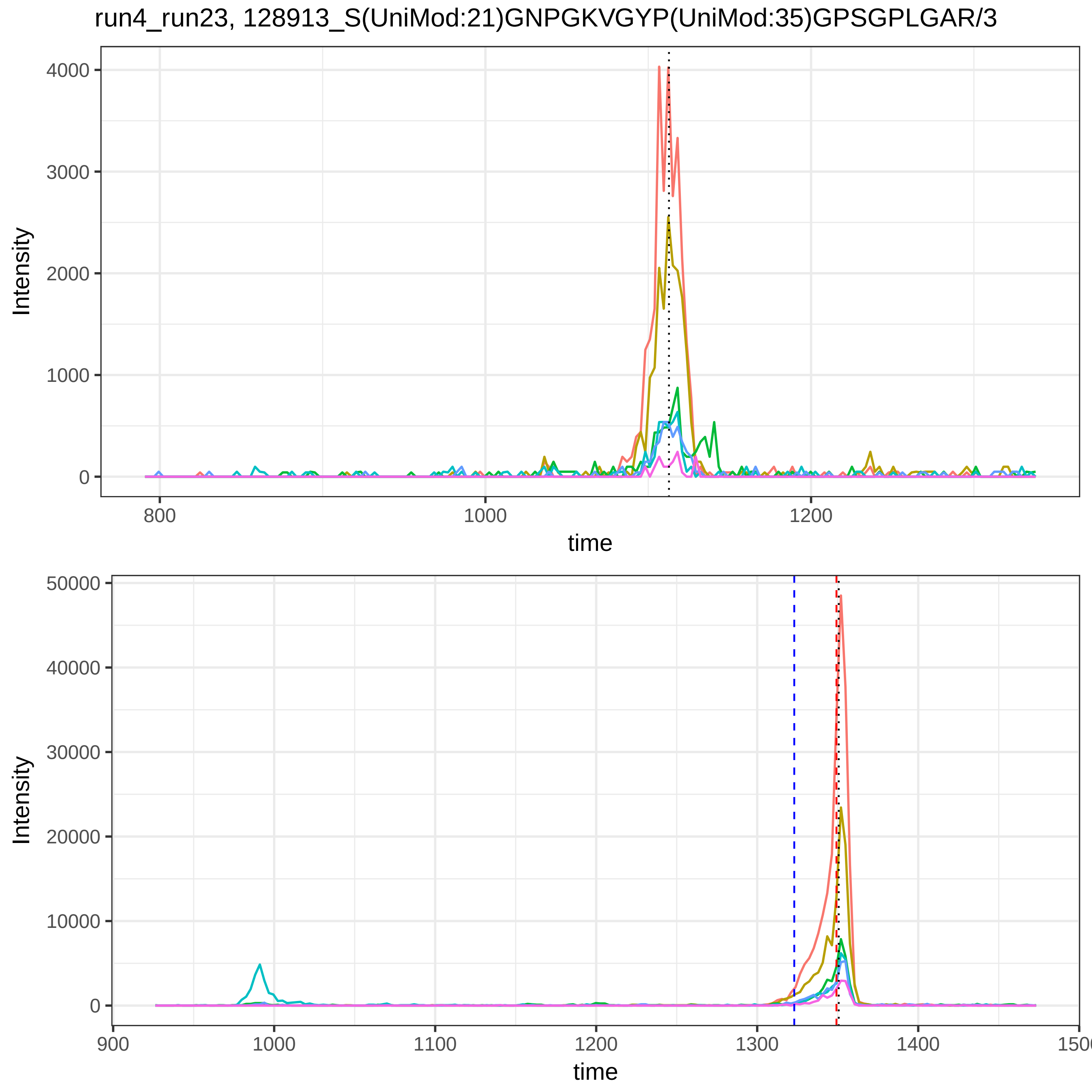

run4_run23, 128913_S(UniMod:21)GNPGKVGYP(UniMod:35)GPSGPLGAR/3
Intensity
4000
3000
2000
1000
0
800
1000
1200
time
Intensity
50000
40000
30000
20000
10000
0
900
1000
1100
1200
1300
1400
time

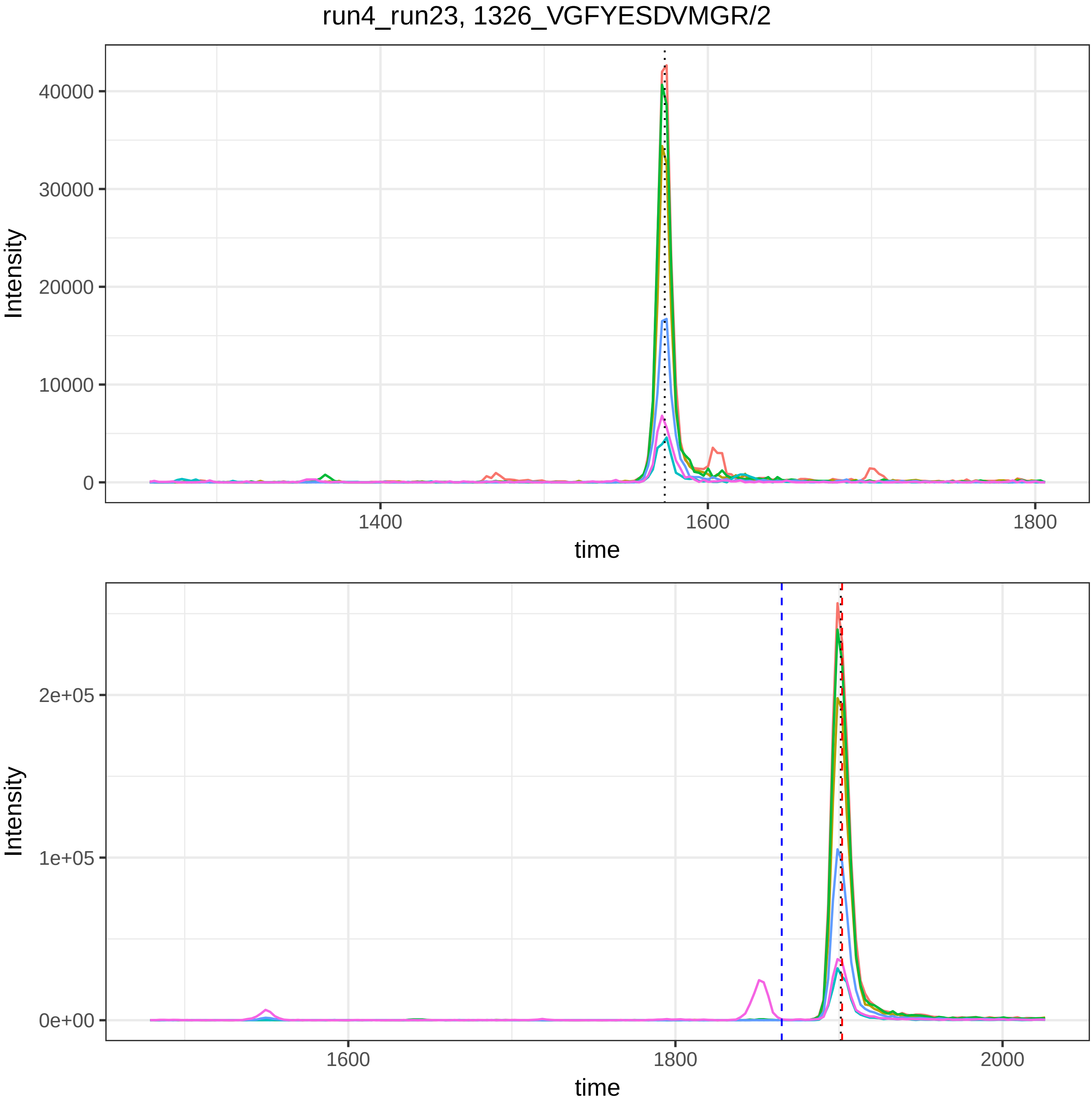

run4_run23, 1326_VGFYESDVMGR/2
Intensity
40000
30000
20000
10000
0
1400
1600
1800
time
Intensity
2e+05
1e+05
0e+00
1600
1800
2000
time

run4_run23, 17708_DLLLPQPDLR/2

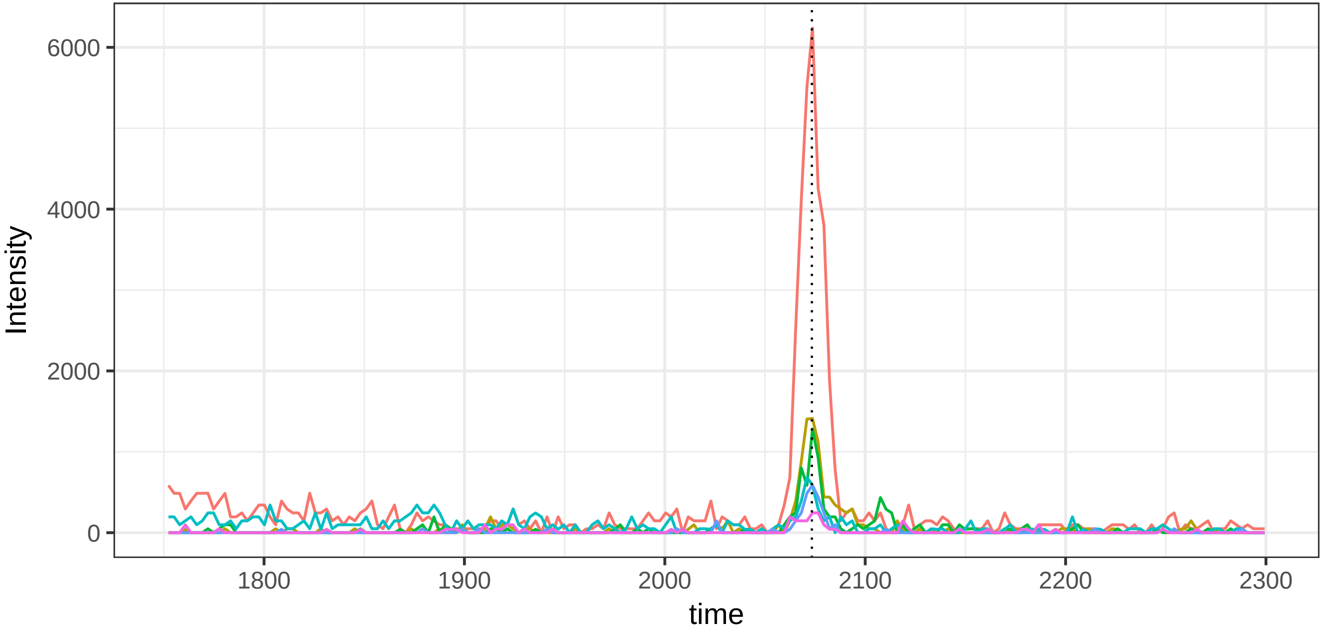


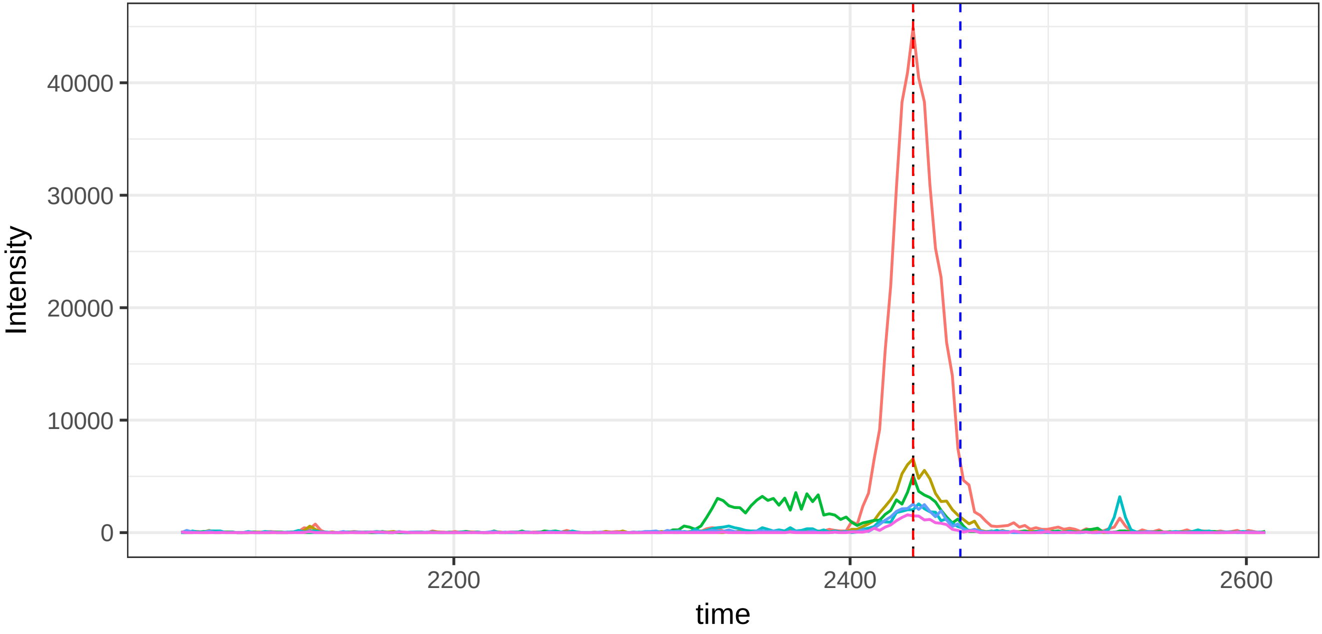

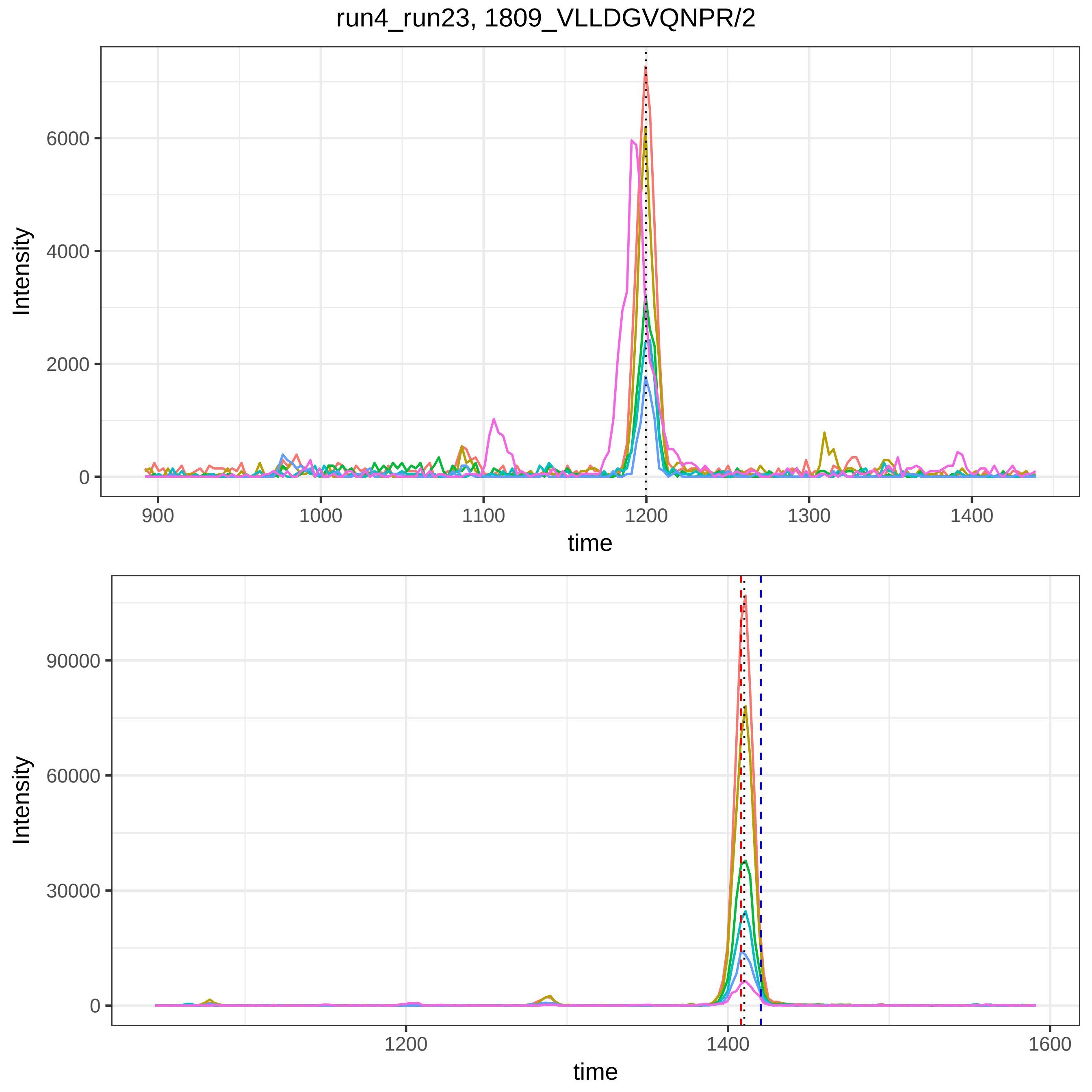
run4_run23, 1809_VLLDGVQNPR/2
Intensity
6000
4000
2000
0
900
1000
1100
1200
1300
1400
time
Intensity
90000
60000
30000
0
1200
1400
1600
time

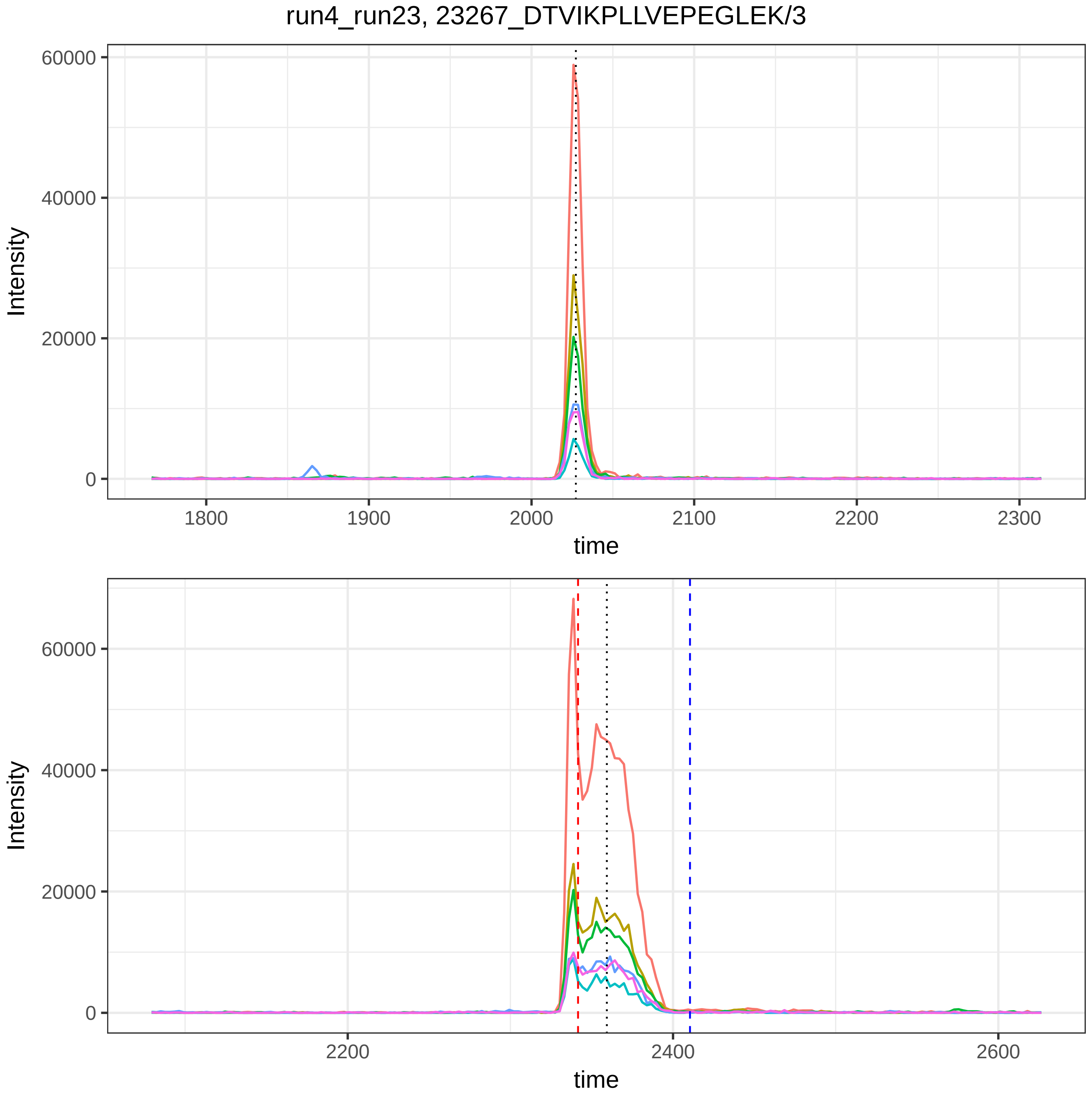

run4_run23, 23267_DTVIKPLLVEPEGLEK/3
Intensity
60000
40000
20000
0
1800
1900
2000
2100
2200
2300
time
Intensity
60000
40000
20000
0
2200
2400
2600
time

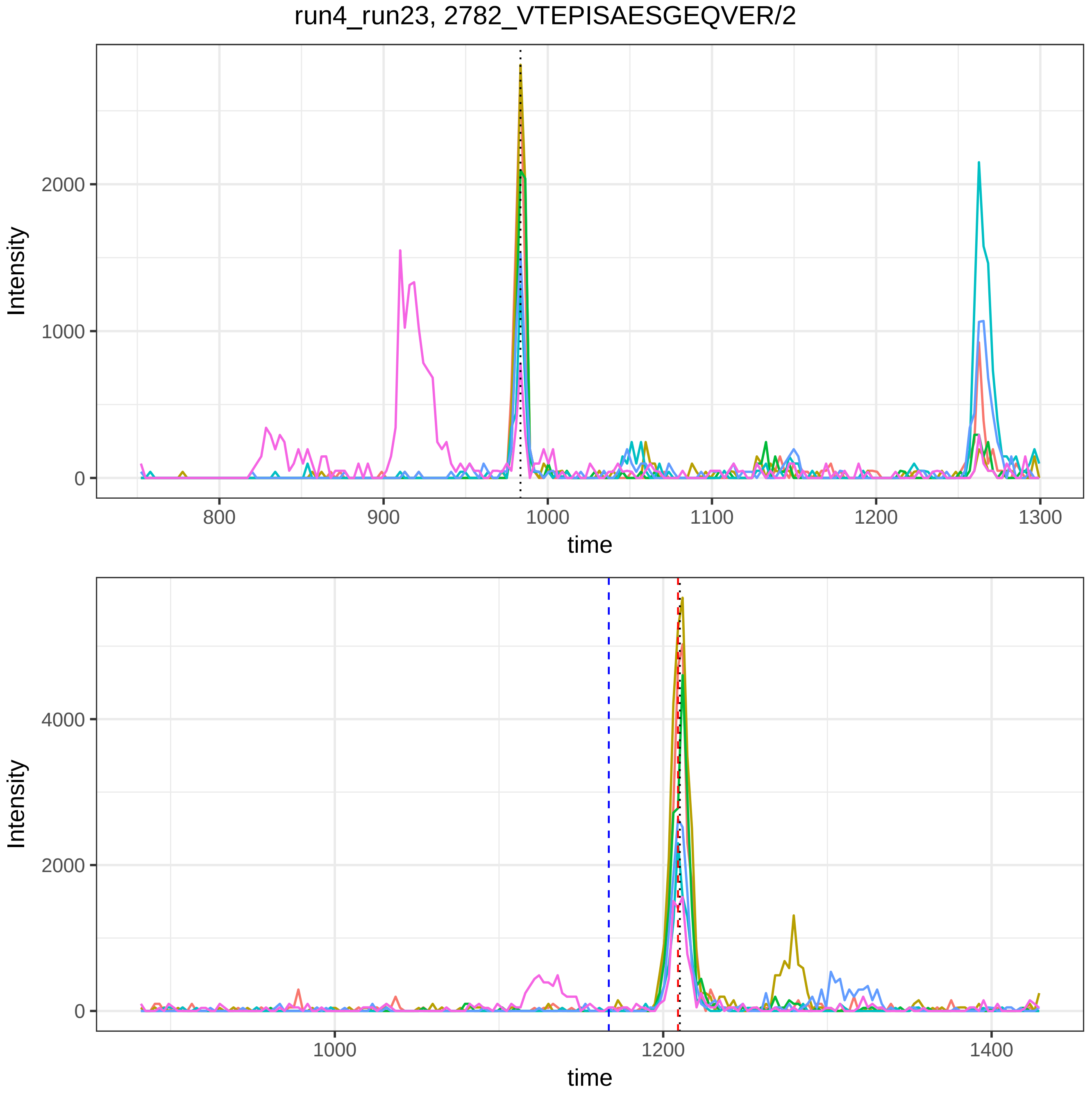
run4_run23, 2782_VTEPISAESGEQVER/2
Intensity
time
0
1000
2000
800
900
1000
1100
1200
1300
0
2000
4000
1000
1200
1400

run4_run23, 32498_ESYSGVTLDPR/2

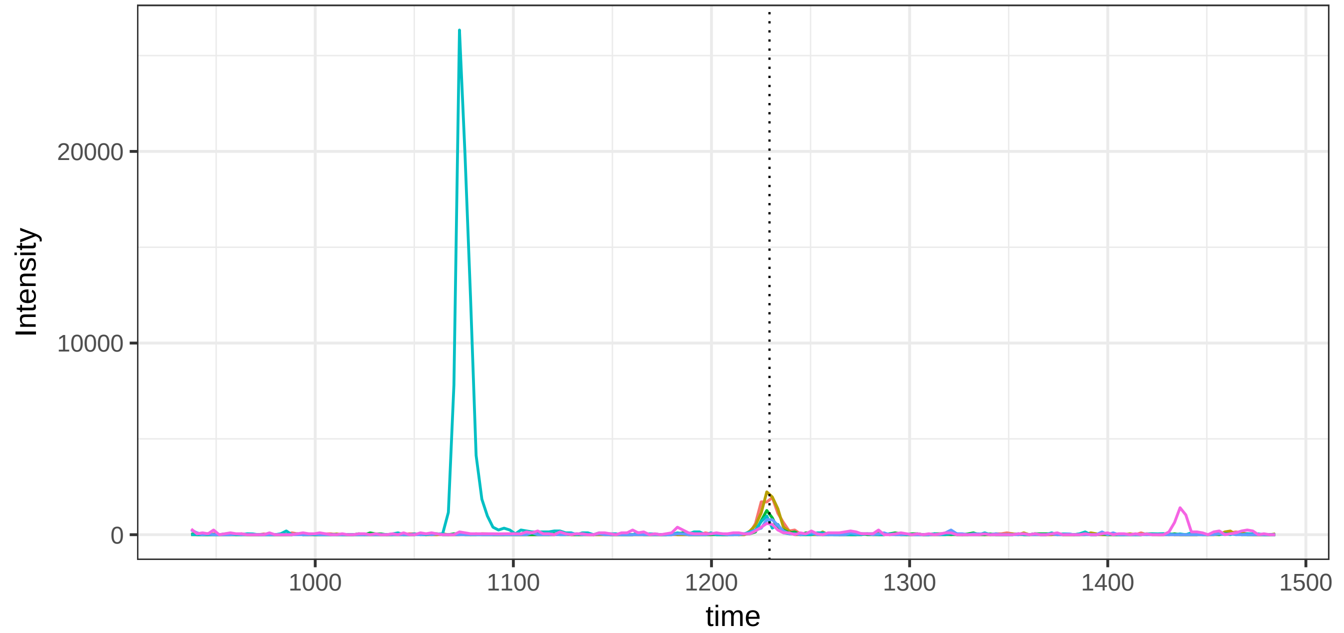


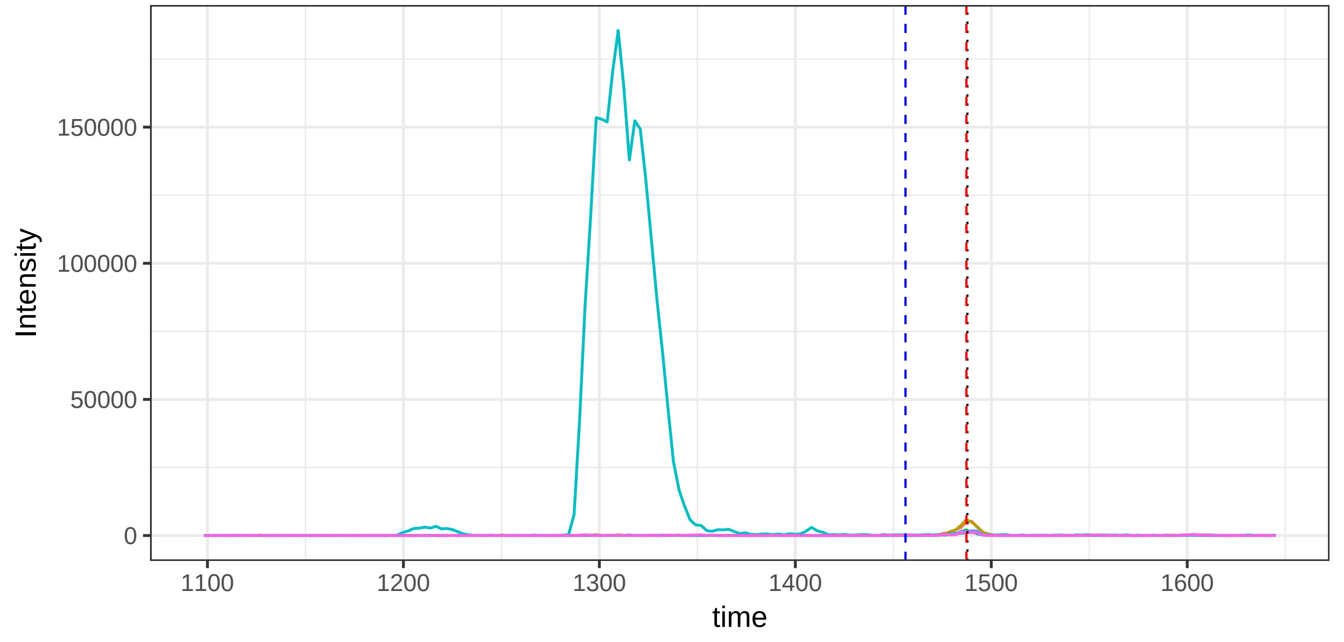

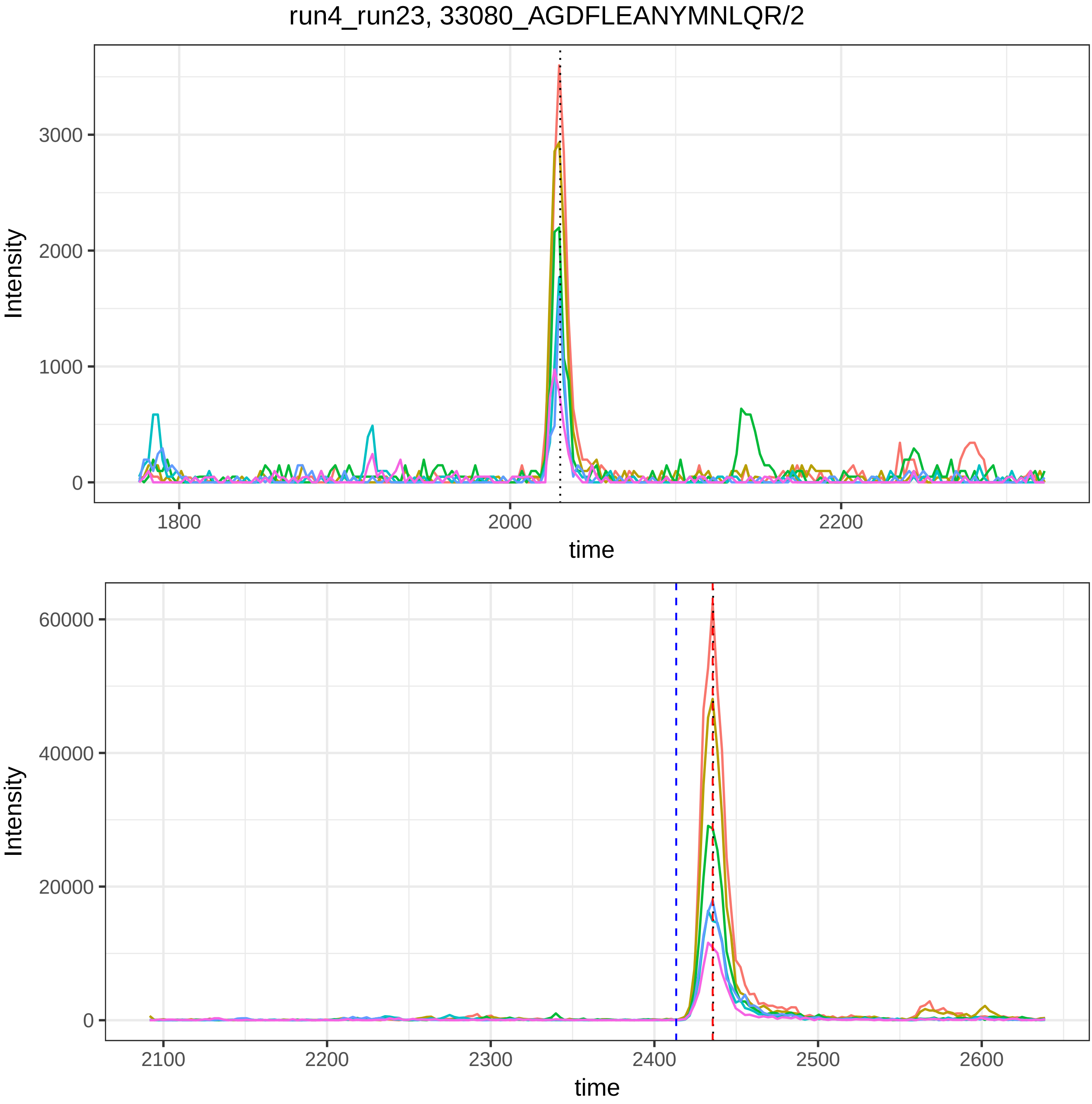

run4_run23, 33080_AGDFLEANYMNLQR/2
Intensity
0
1000
2000
3000
1800
2000
2200
time
Intensity
0
20000
40000
60000
2100
2200
2300
2400
2500
2600
time

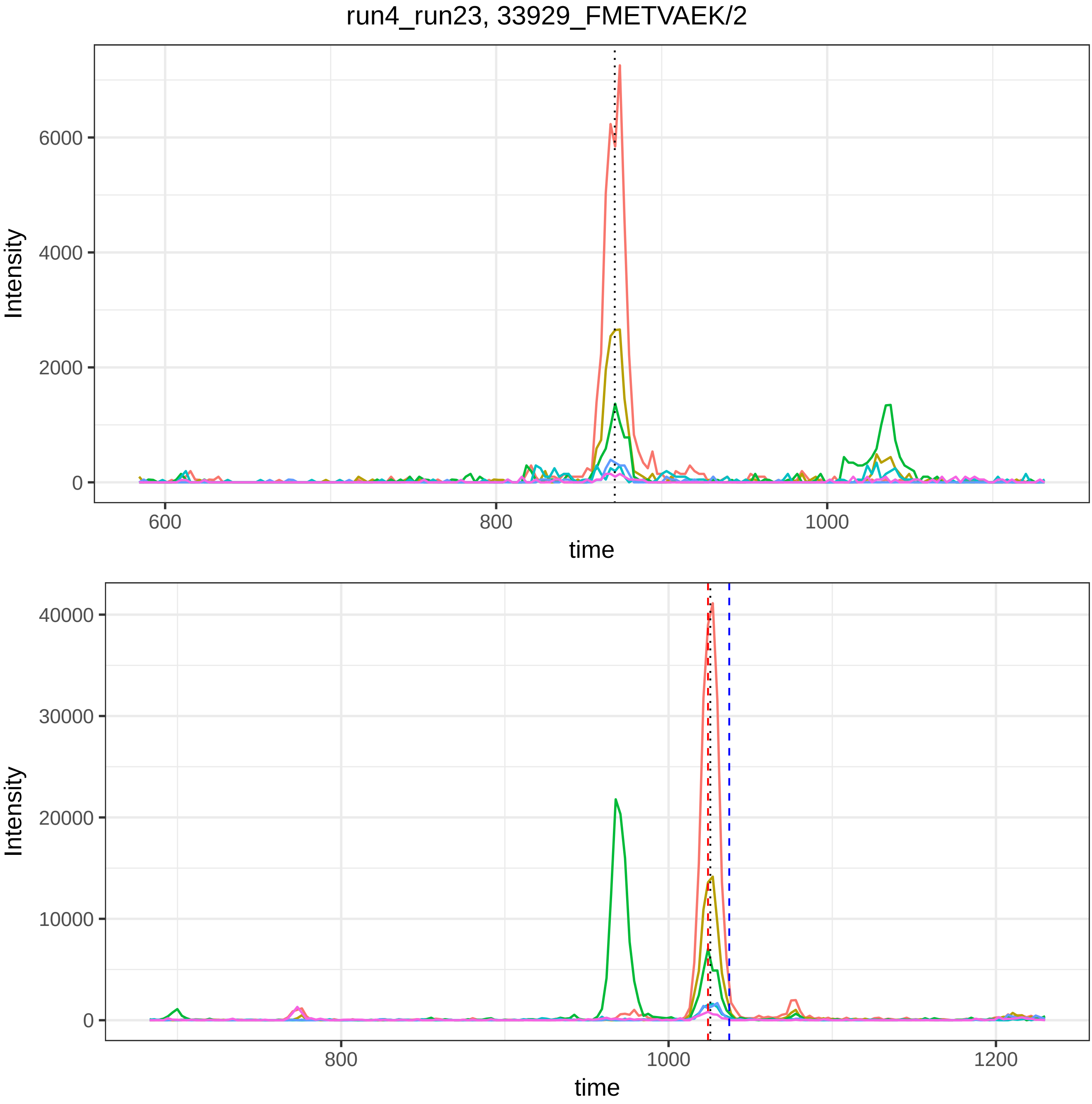

run4_run23, 33929_FMETVAEK/2
Intensity
6000
4000
2000
0
600
800
1000
time
Intensity
40000
30000
20000
10000
0
800
1000
1200
time

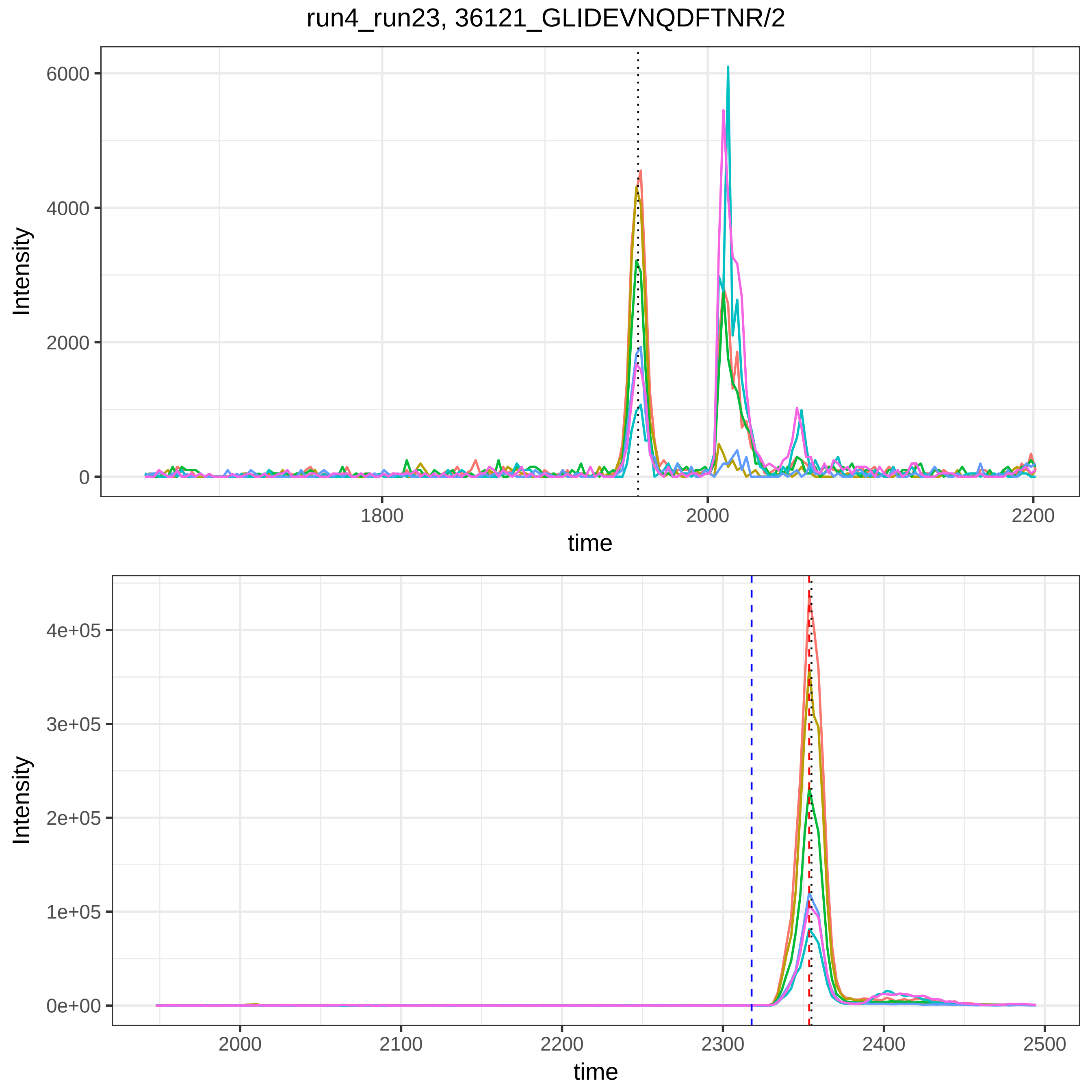
run4_run23, 36121_GLIDEVNQDFTNR/2
Intensity
6000
4000
2000
0
1800
2000
2200
time
Intensity
4e+05
3e+05
2e+05
1e+05
0e+00
2000
2100
2200
2300
2400
2500
time

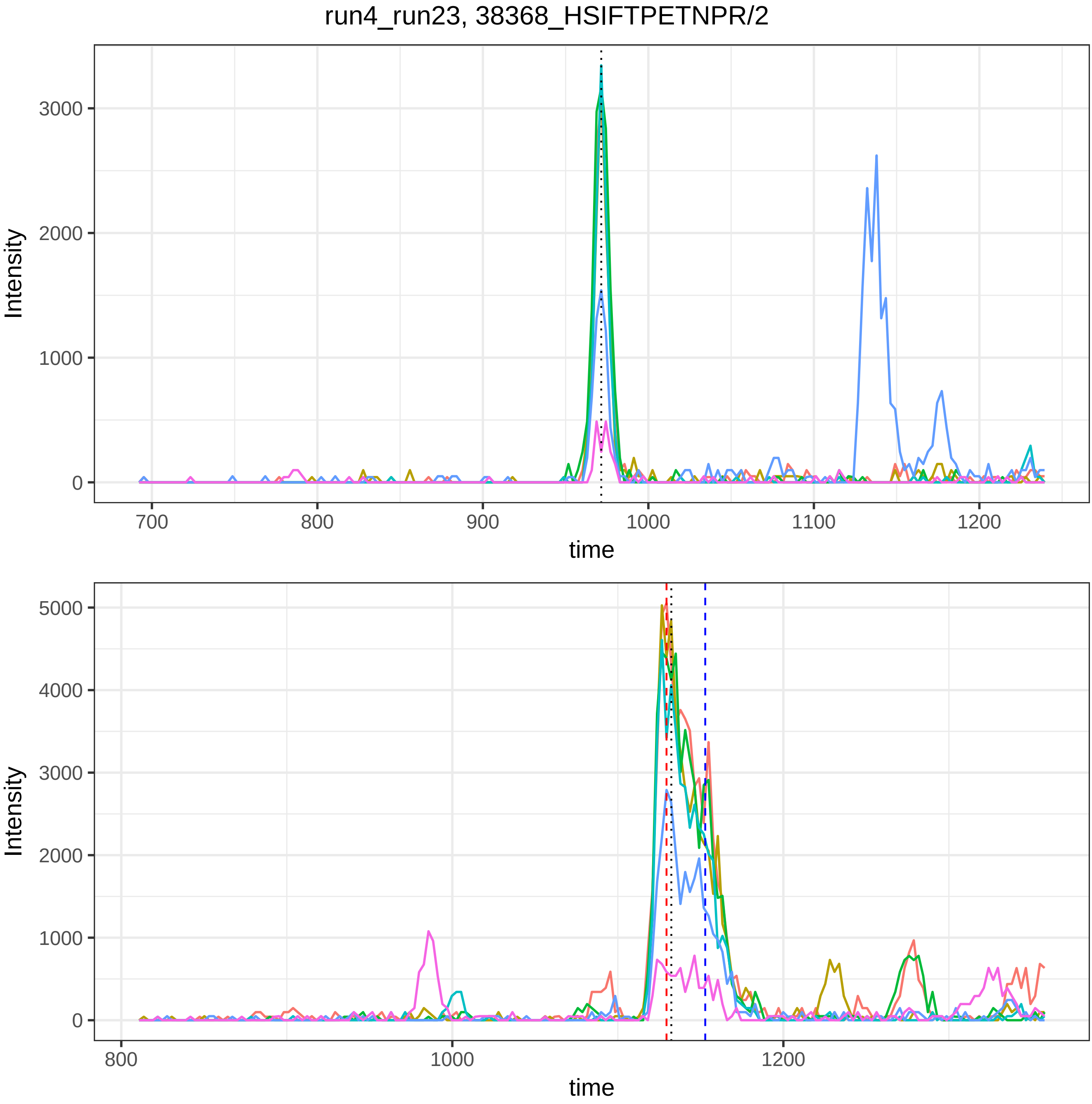
run4_run23, 38368_HSIFTPETNPR/2
Intensity
3000
2000
1000
0
700
800
900
1000
1100
1200
time
Intensity
5000
4000
3000
2000
1000
0
800
1000
1200
time

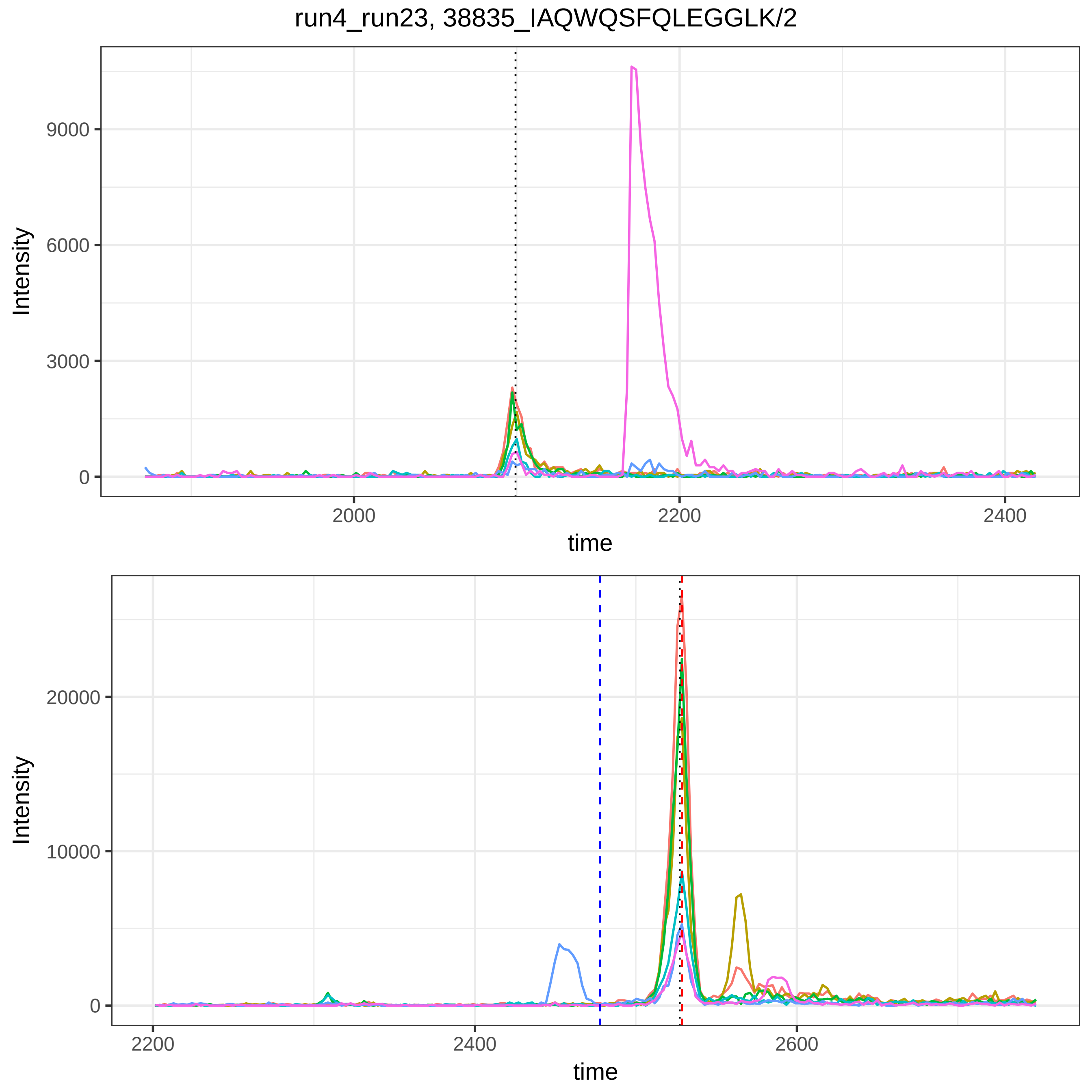

run4_run23, 38835_IAQWQSFQLEGGLK/2
Intensity
9000
6000
3000
0
2000
2200
2400
time
Intensity
20000
10000
0
2200
2400
2600
time

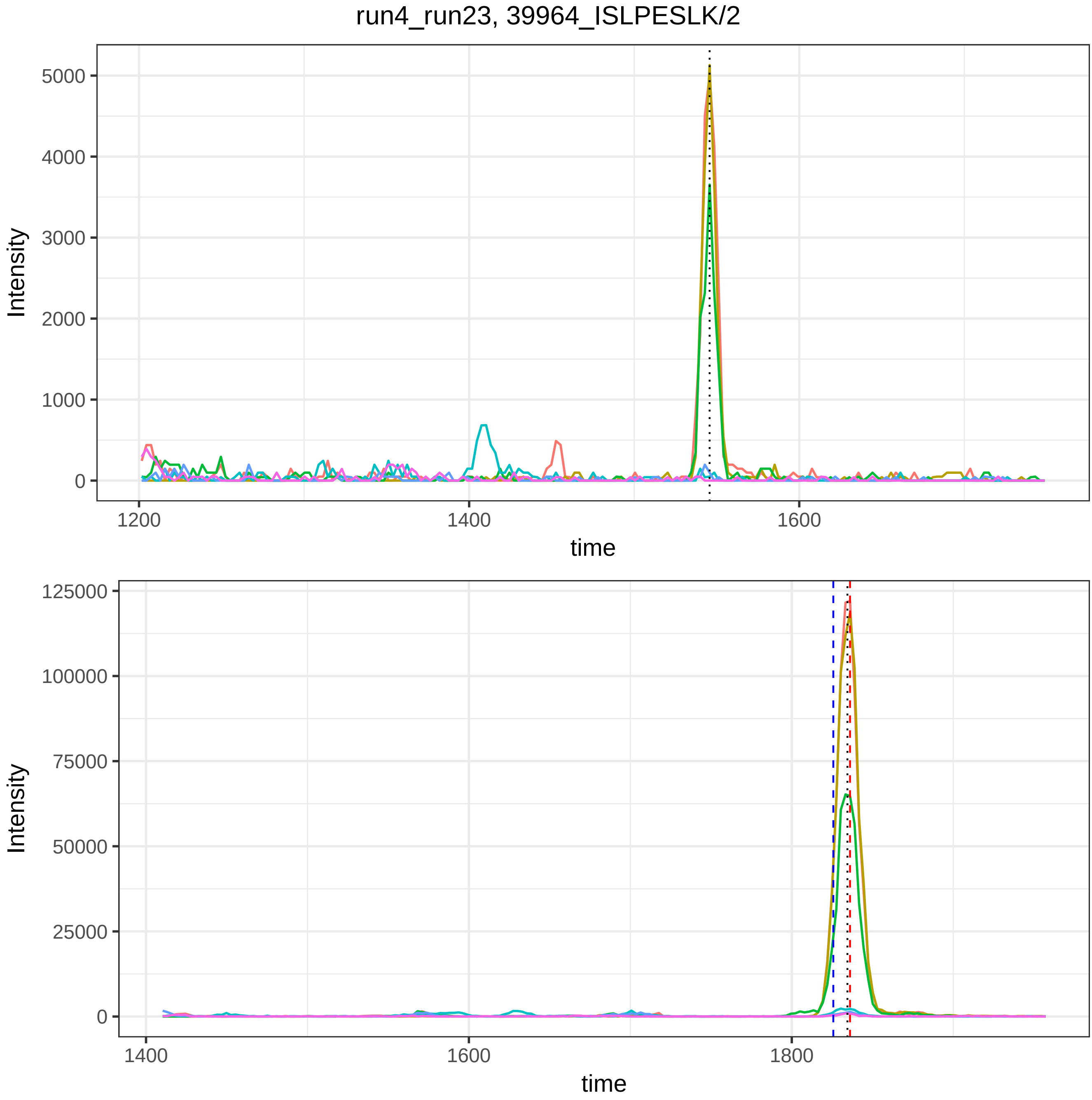

run4_run23, 39964_ISLPESLK/2
Intensity
time
5000
4000
3000
2000
1000
0
1200
1400
1600
125000
100000
75000
50000
25000
0
1400
1600
1800

run4_run23, 40475_IYGNQDTSSQLK/2

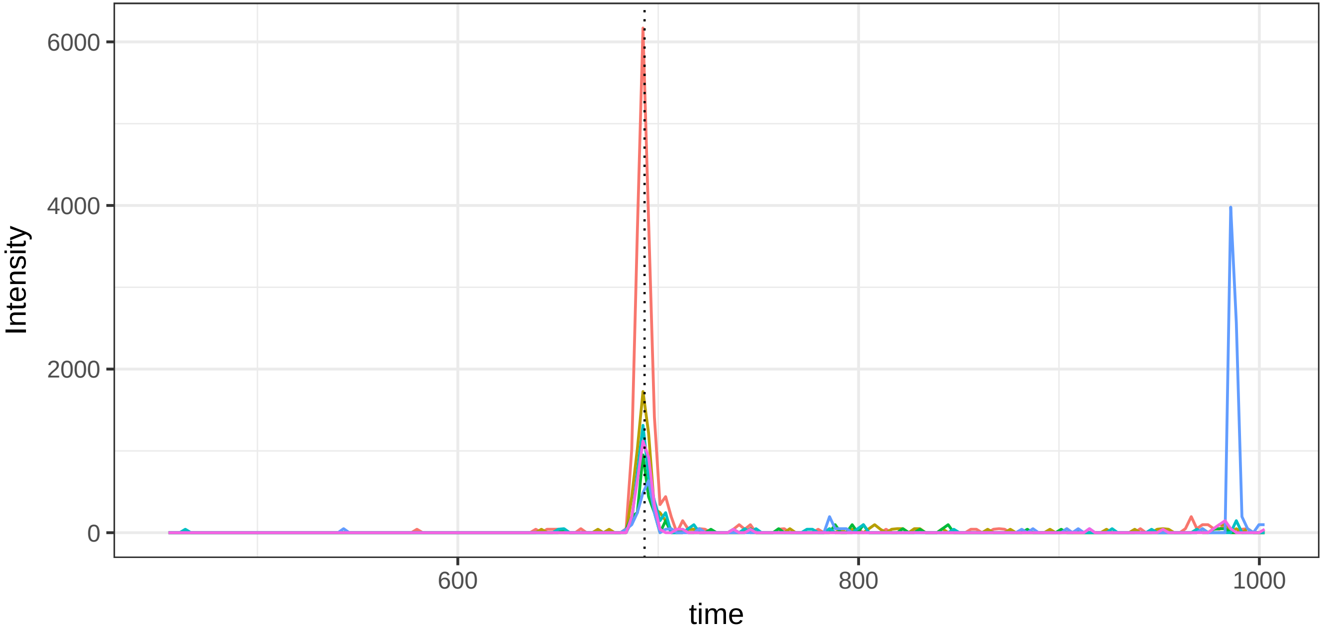


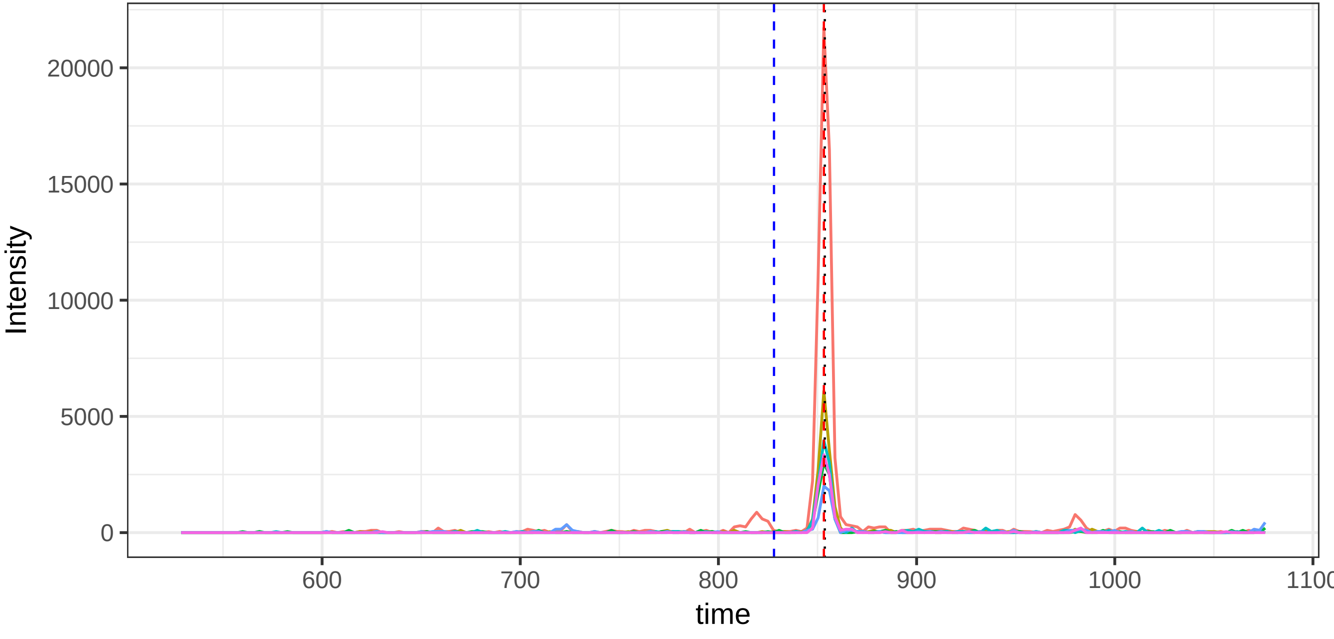

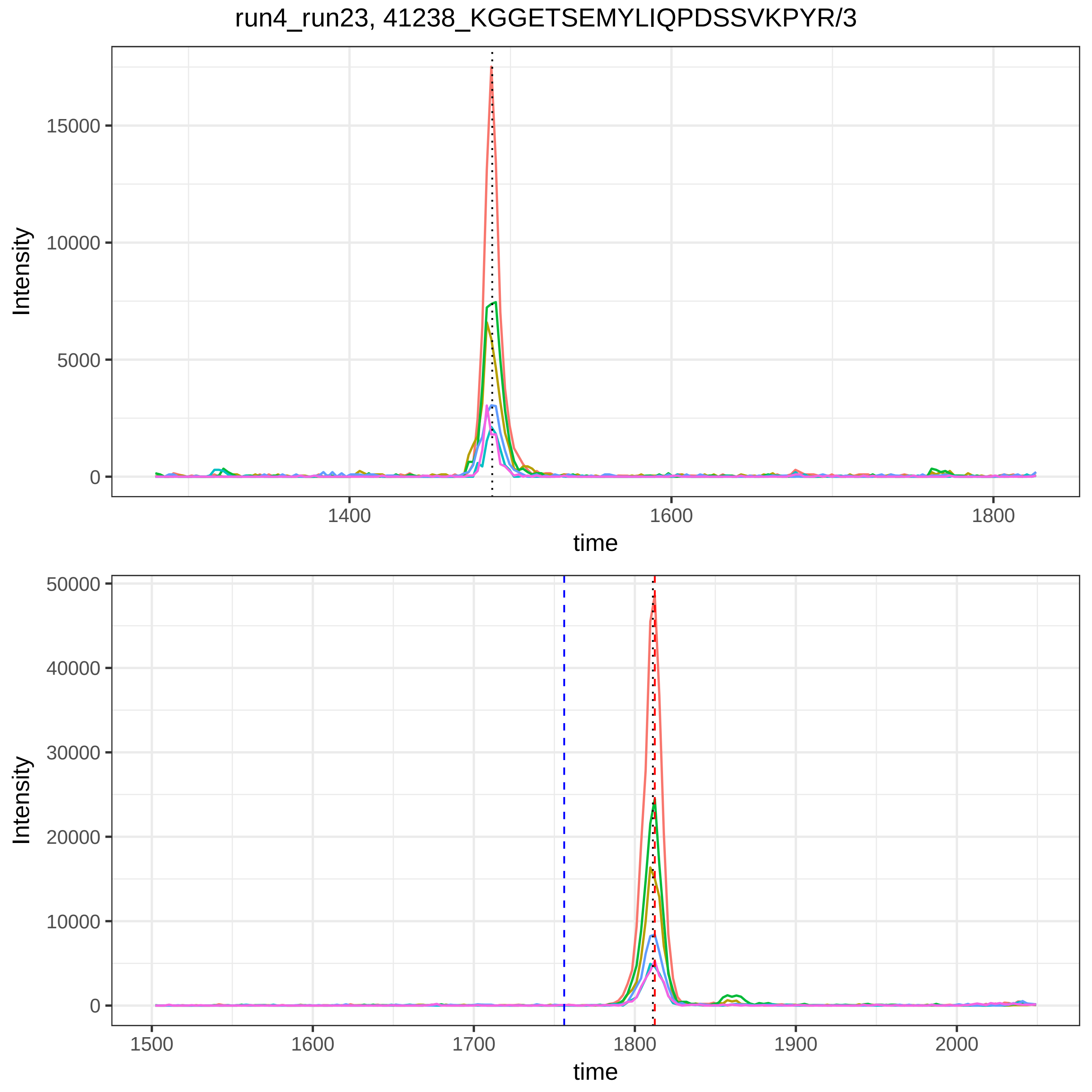

run4_run23, 41238_KGGETSEMYLIQPDSSVKPYR/3
Intensity
15000
10000
5000
0
1400
1600
1800
time
Intensity
50000
40000
30000
20000
10000
0
1500
1600
1700
1800
1900
2000
time

run4_run23, 42048_KVLLDGVQNPR/2

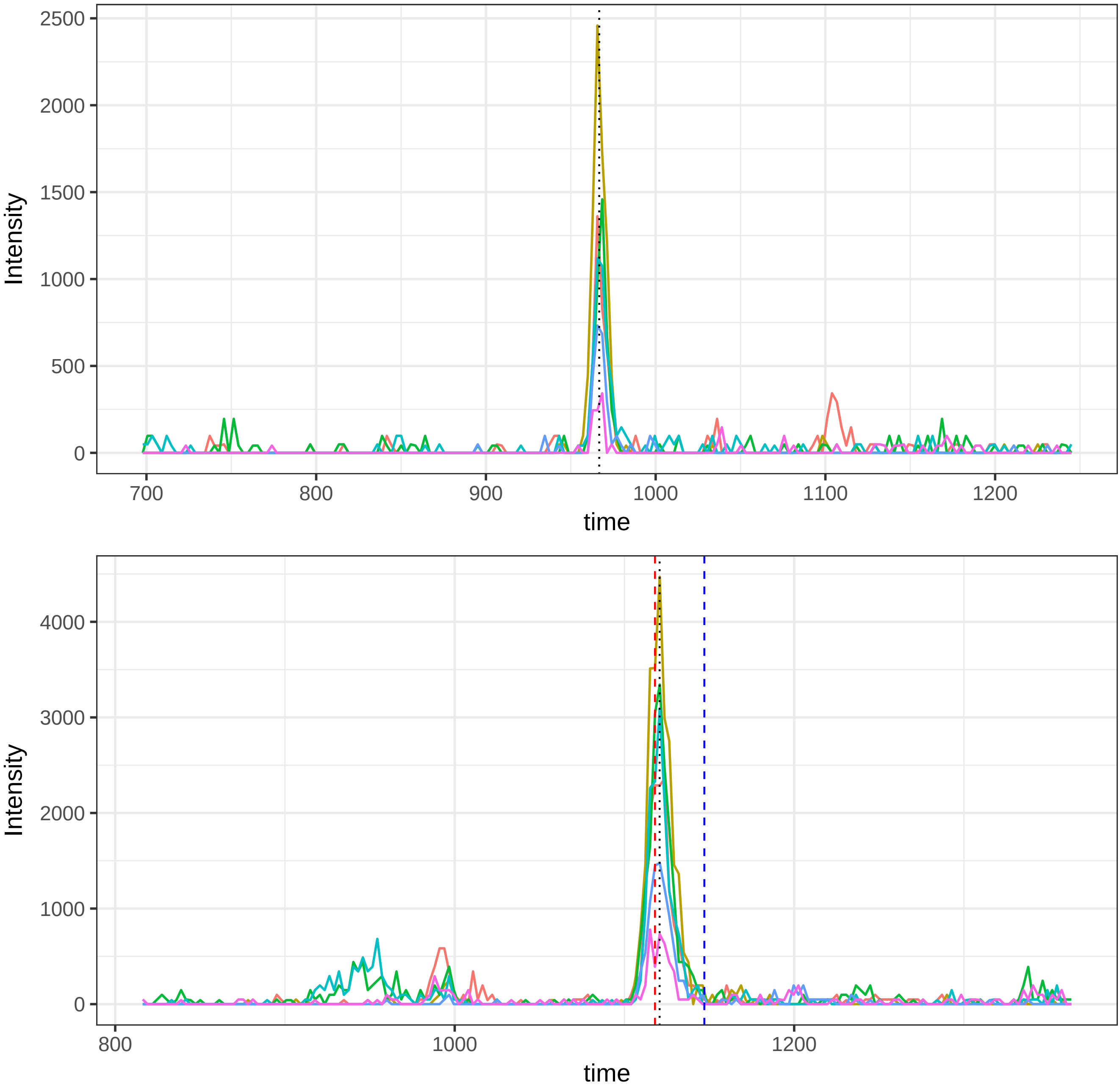

run4_run23, 42052_KVLLDGVQNPR/3

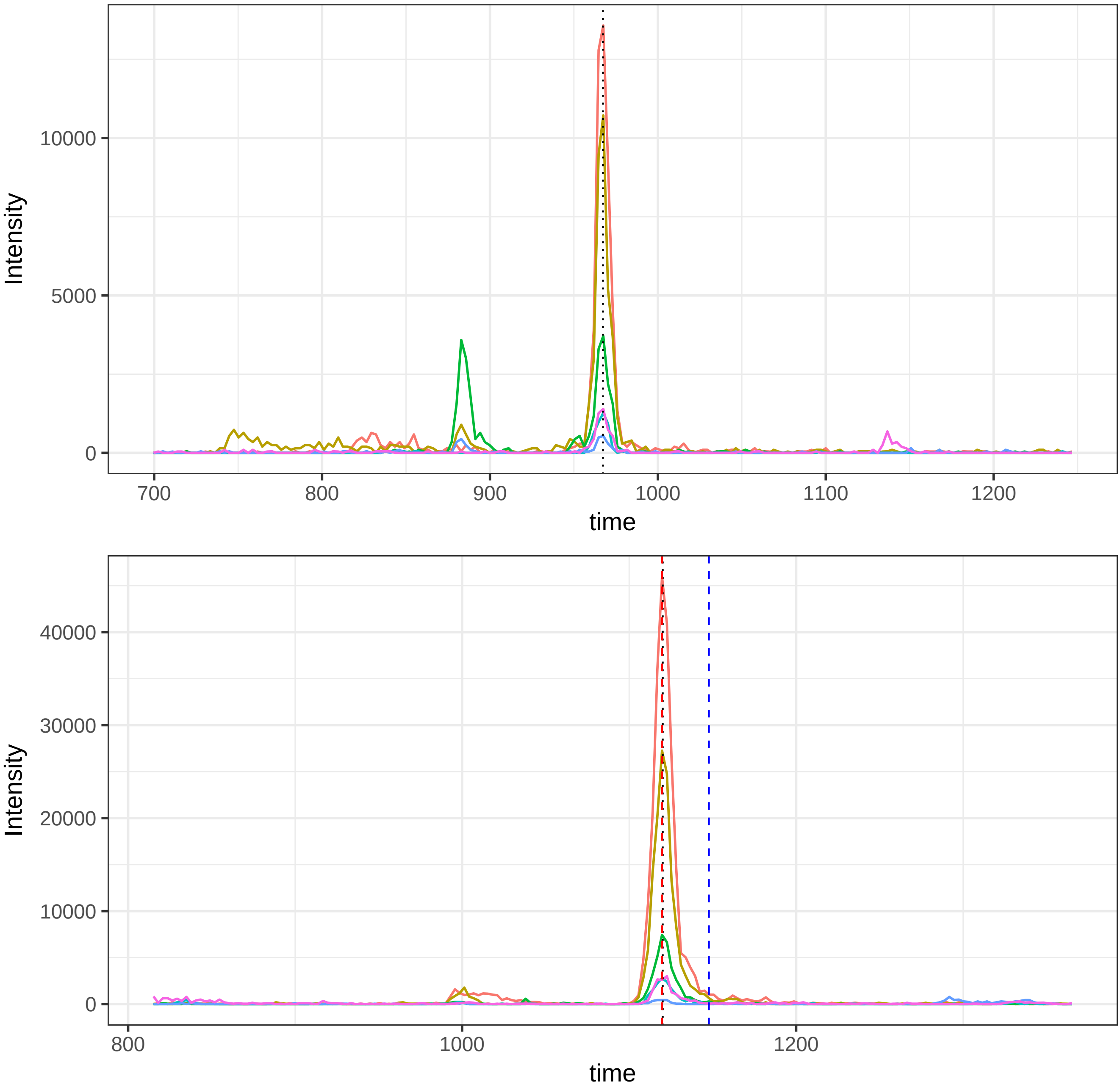

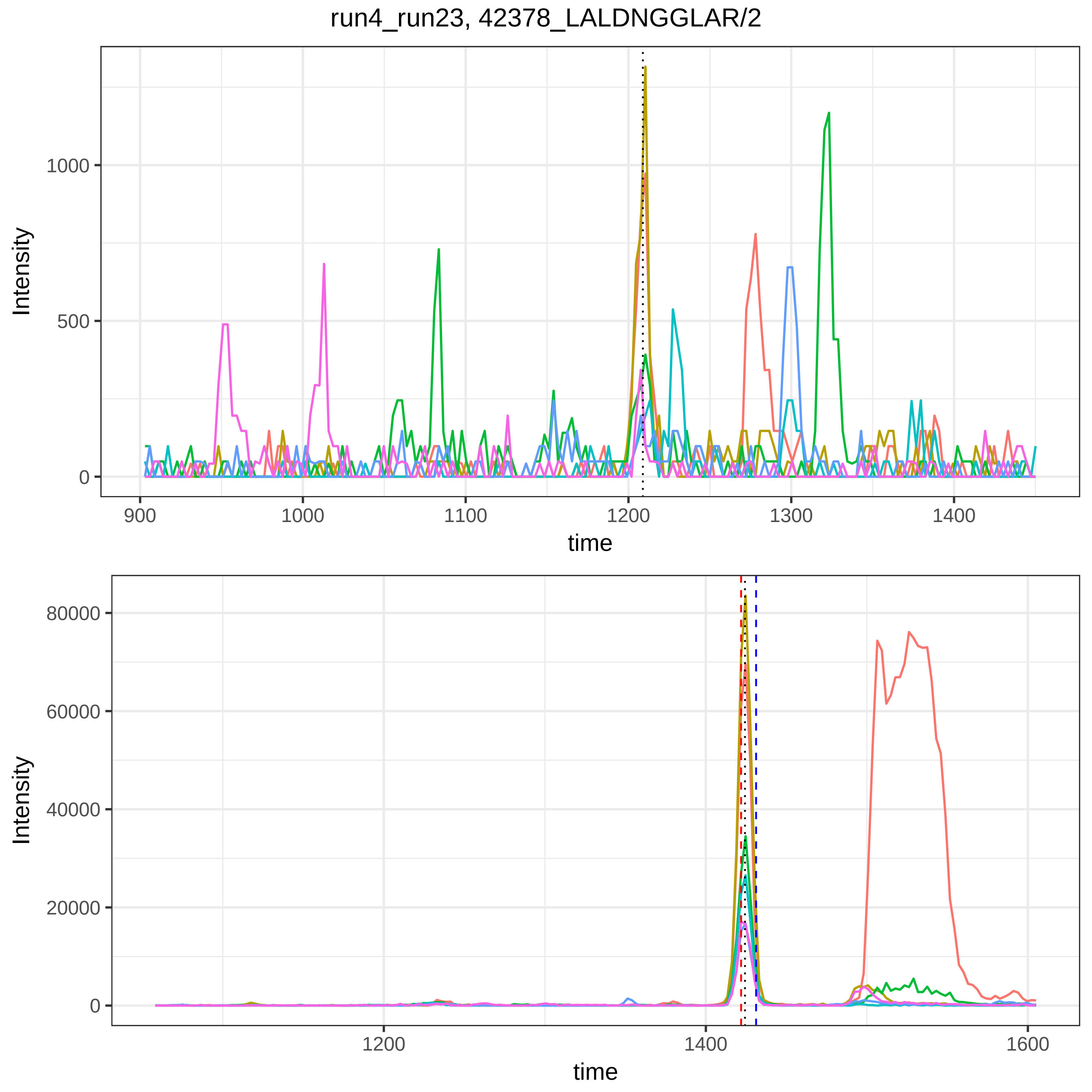
run4_run23, 42378_LALDNGGLAR/2
Intensity
time
1000
500
0
900
1000
1100
1200
1300
1400
80000
60000
40000
20000
0
1200
1400
1600

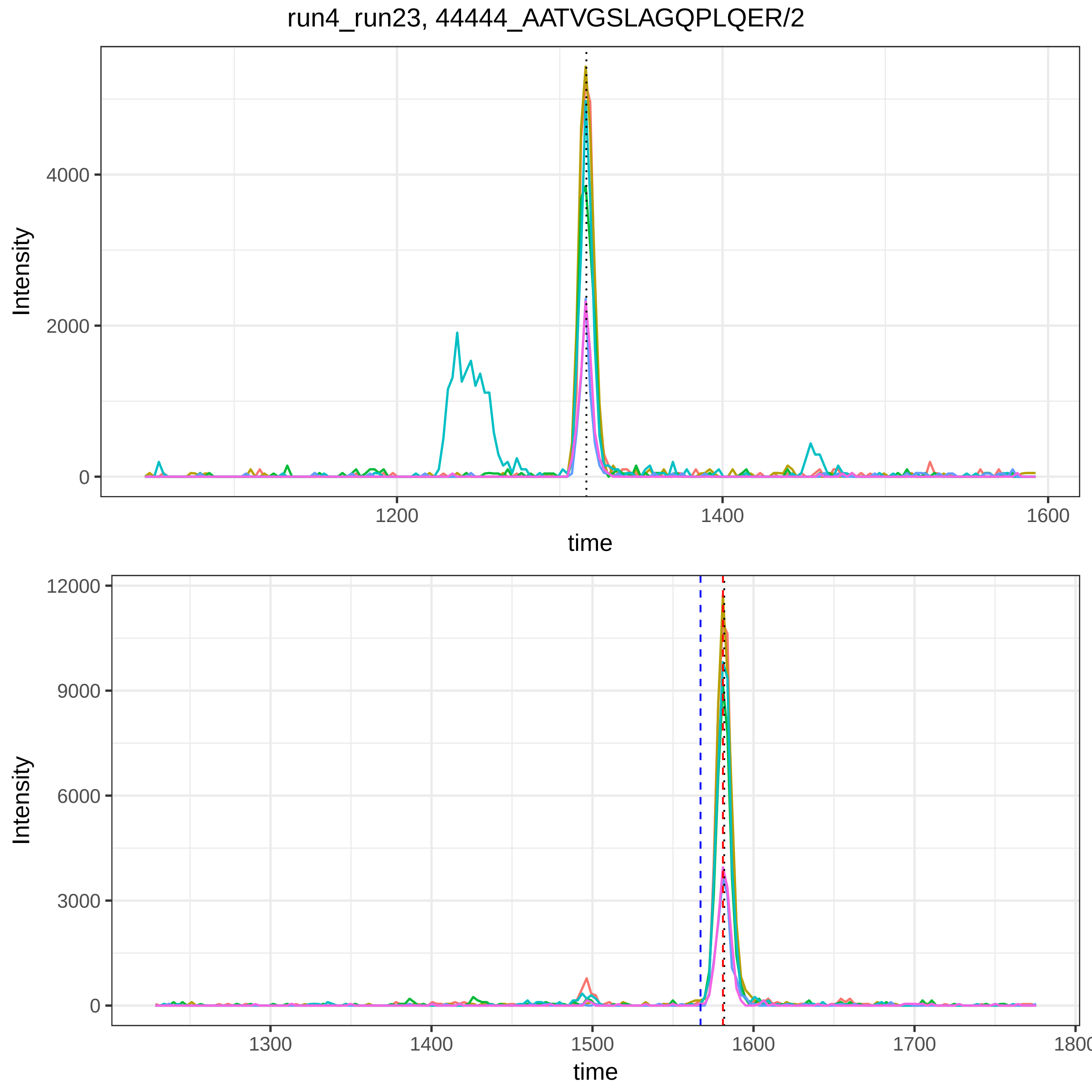

run4_run23, 44444_AATVGSLAGQPLQER/2
Intensity
4000
2000
0
1200
1400
1600
time
Intensity
12000
9000
6000
3000
0
1300
1400
1500
1600
1700
1800
time

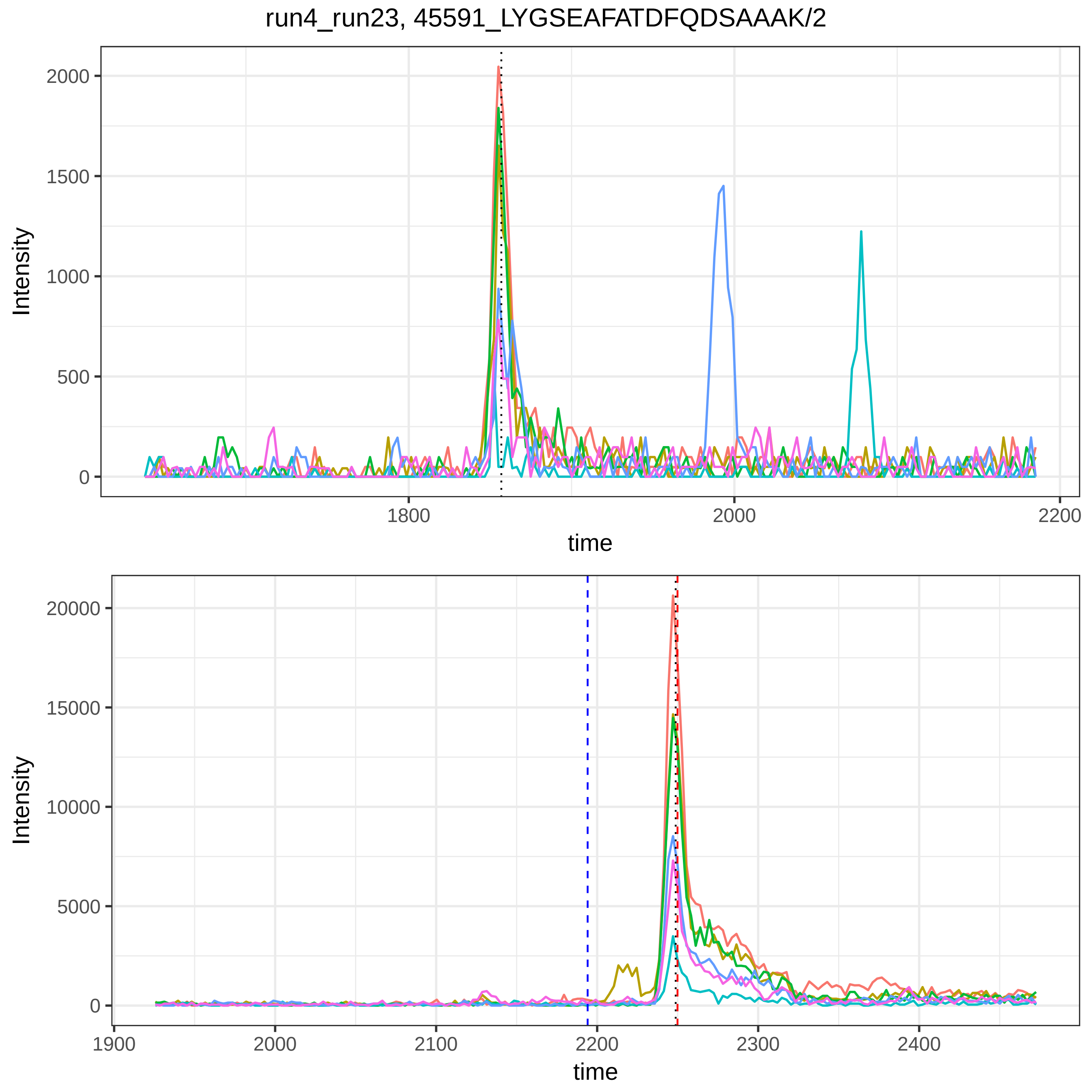
run4_run23, 45591_LYGSEAFATDFQDSAAAK/2
Intensity
0
500
1000
1500
2000
1800
2000
2200
time
Intensity
0
5000
10000
15000
20000
1900
2000
2100
2200
2300
2400
time

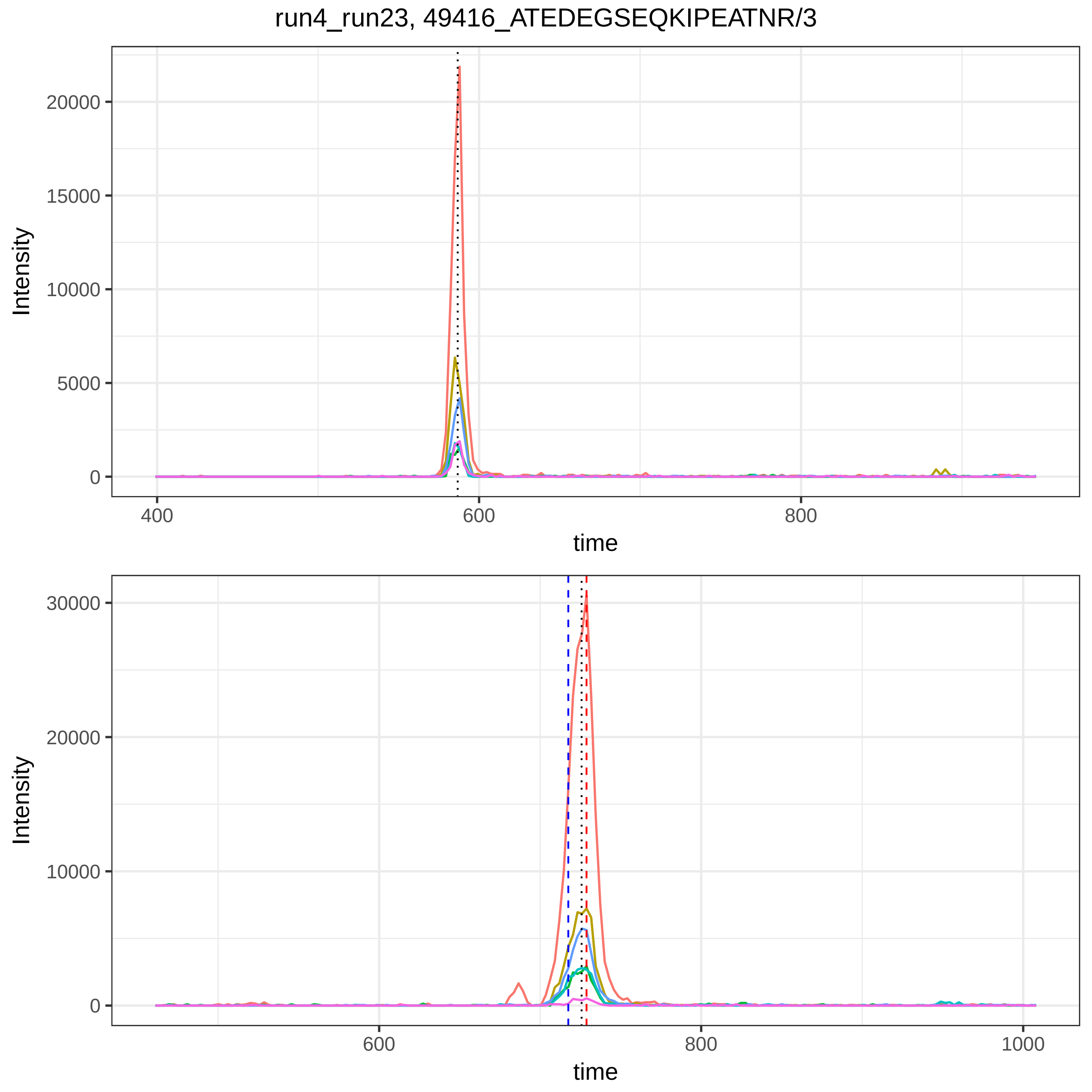
run4_run23, 49416_ATEDEGSEQKIPEATNR/3
Intensity
20000
15000
10000
5000
0
400
600
800
time
Intensity
30000
20000
10000
0
600
800
1000
time

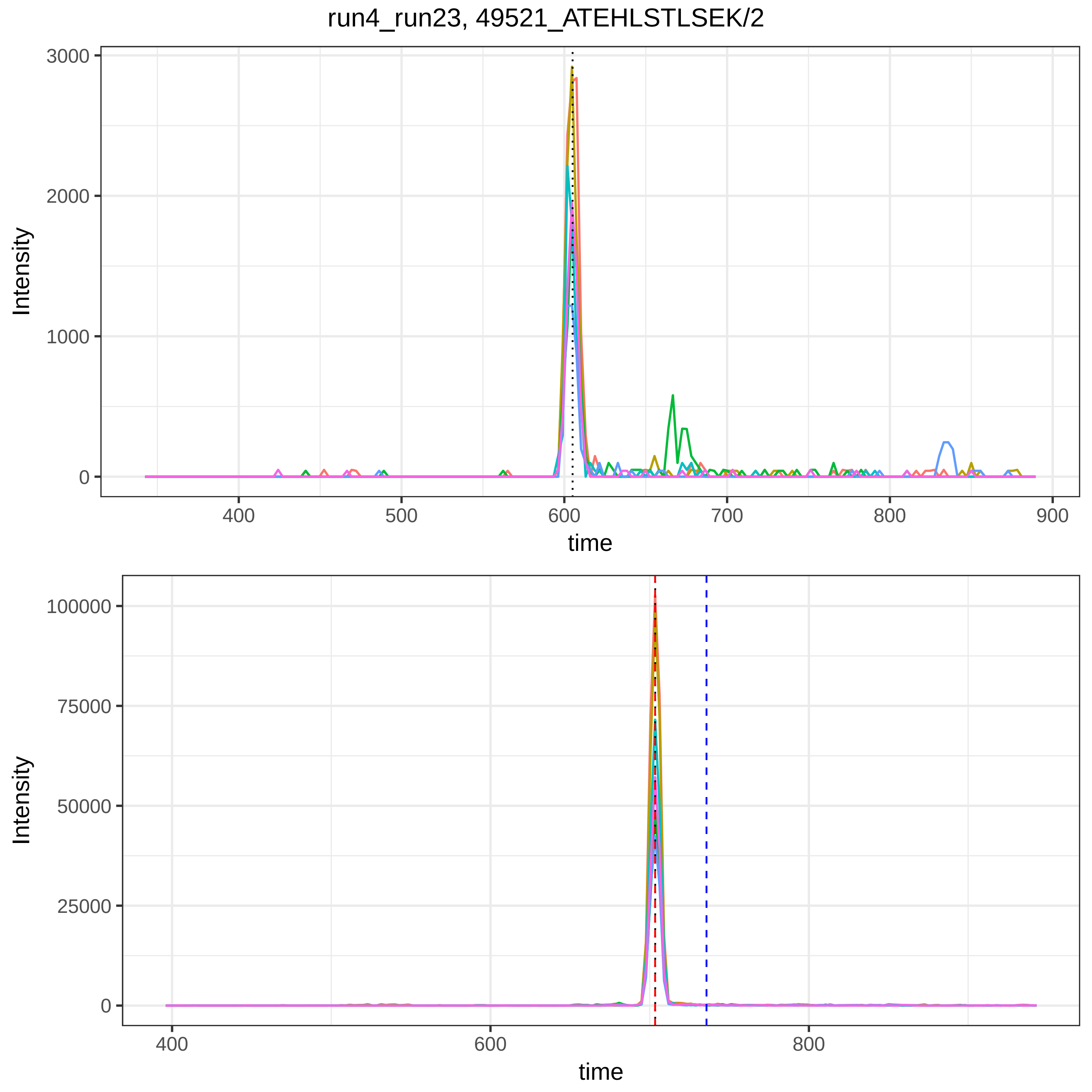

run4_run23, 49521_ATEHLSTLSEK/2
Intensity
time
3000
2000
1000
0
400
500
600
700
800
900
100000
75000
50000
25000
0
400
600
800

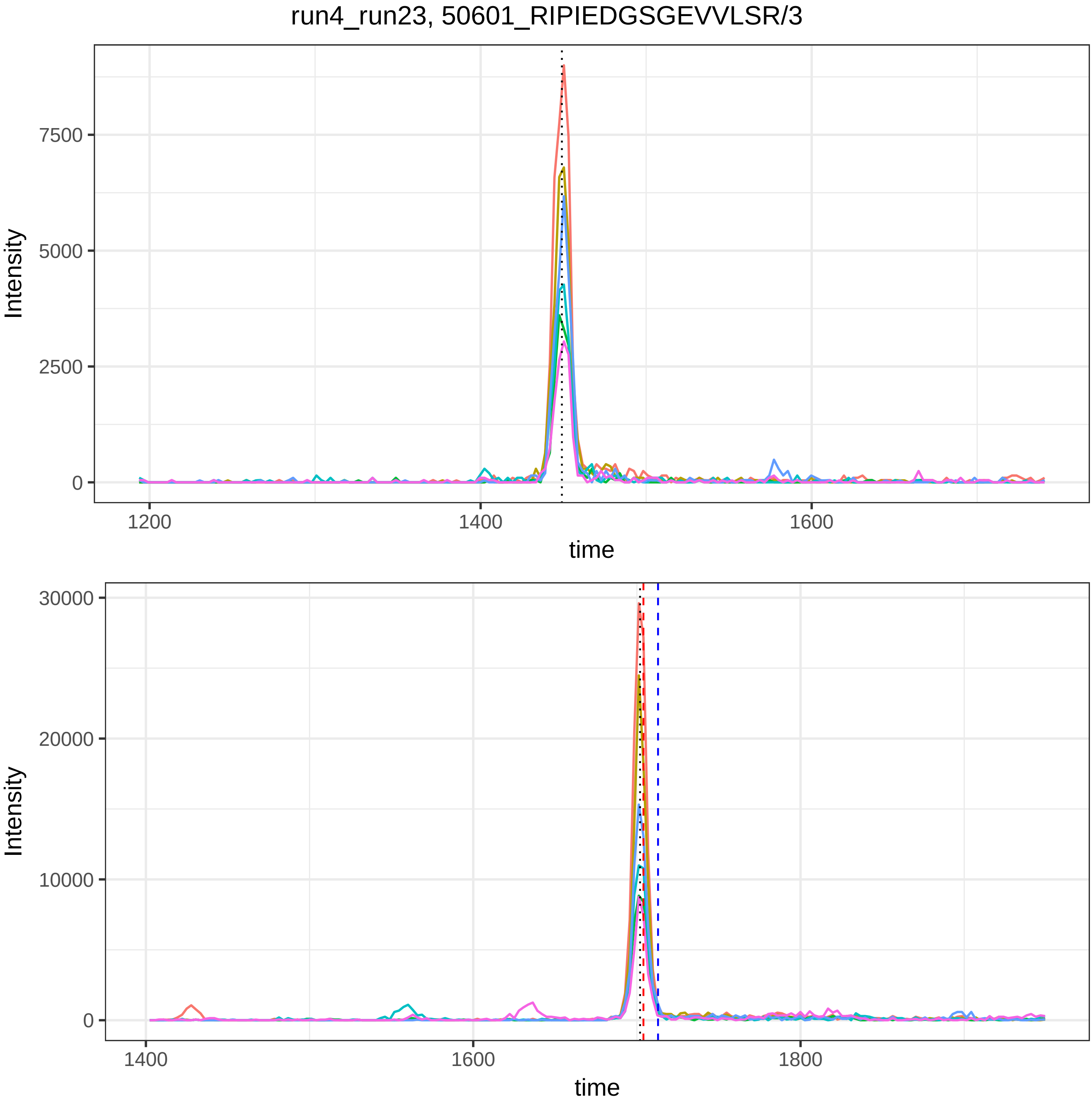
run4_run23, 50601_RIPIEDGSGEVVLSR/3
Intensity
7500
5000
2500
0
1200
1400
1600
time
Intensity
30000
20000
10000
0
1400
1600
1800
time

run4_run23, 50914_RPGGEPSPEGTTGQSYNQYSQR/3

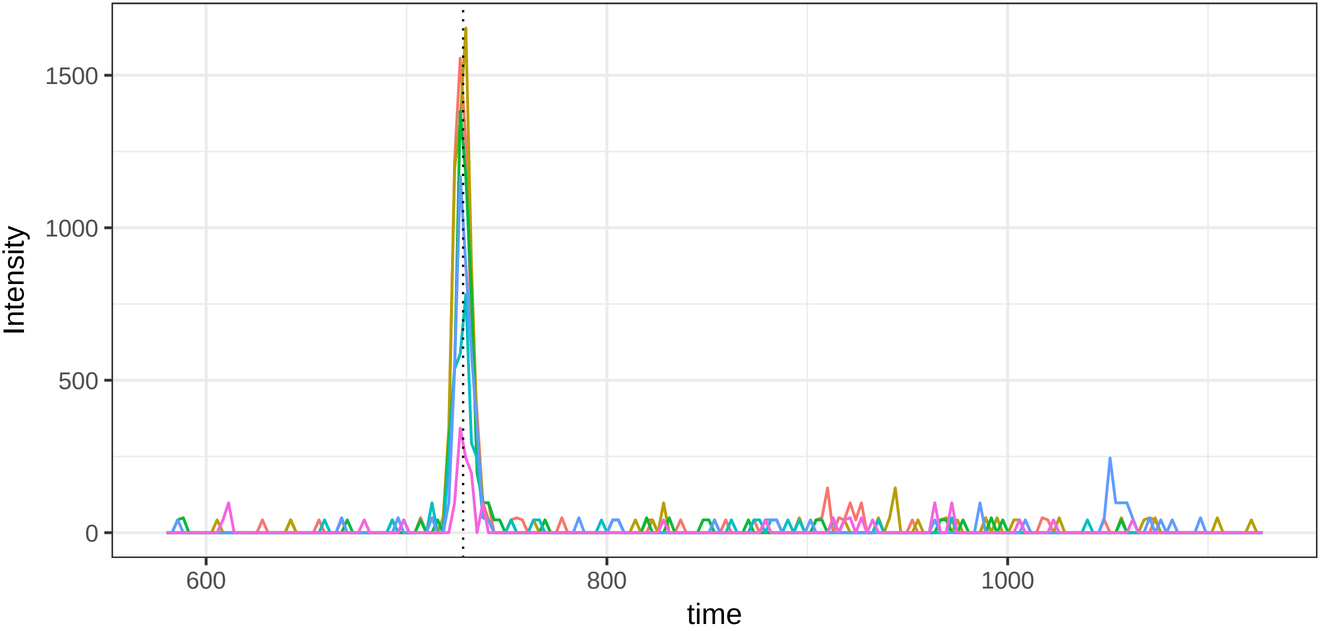


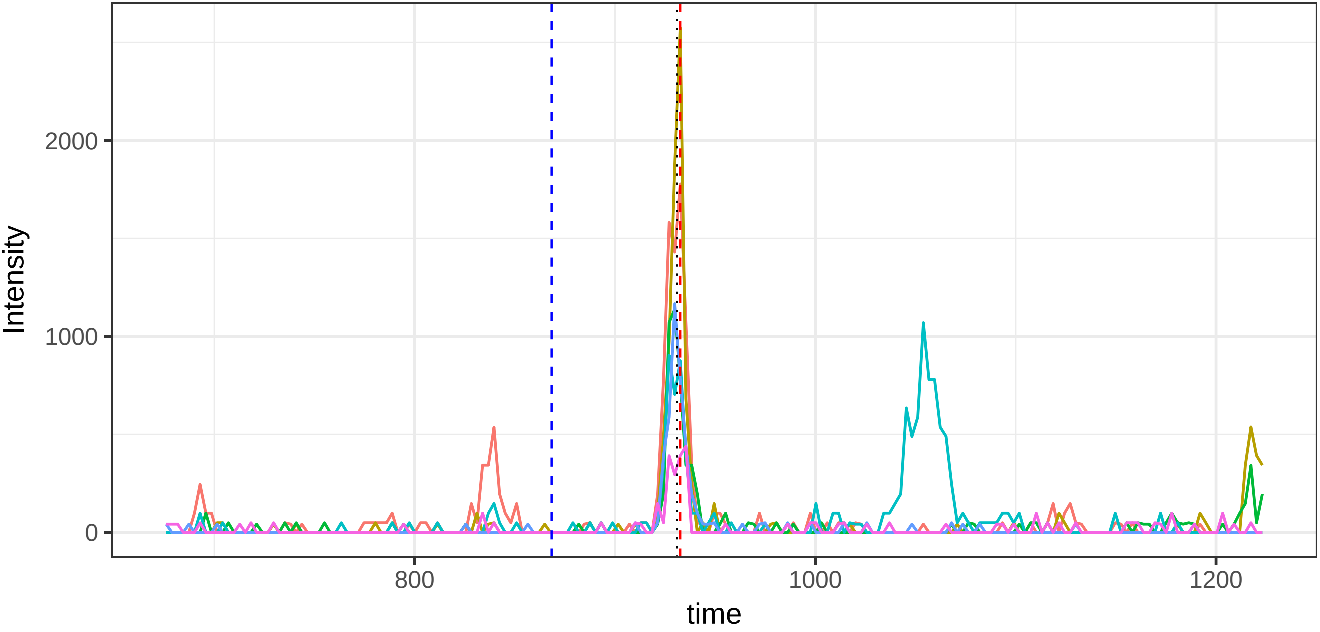

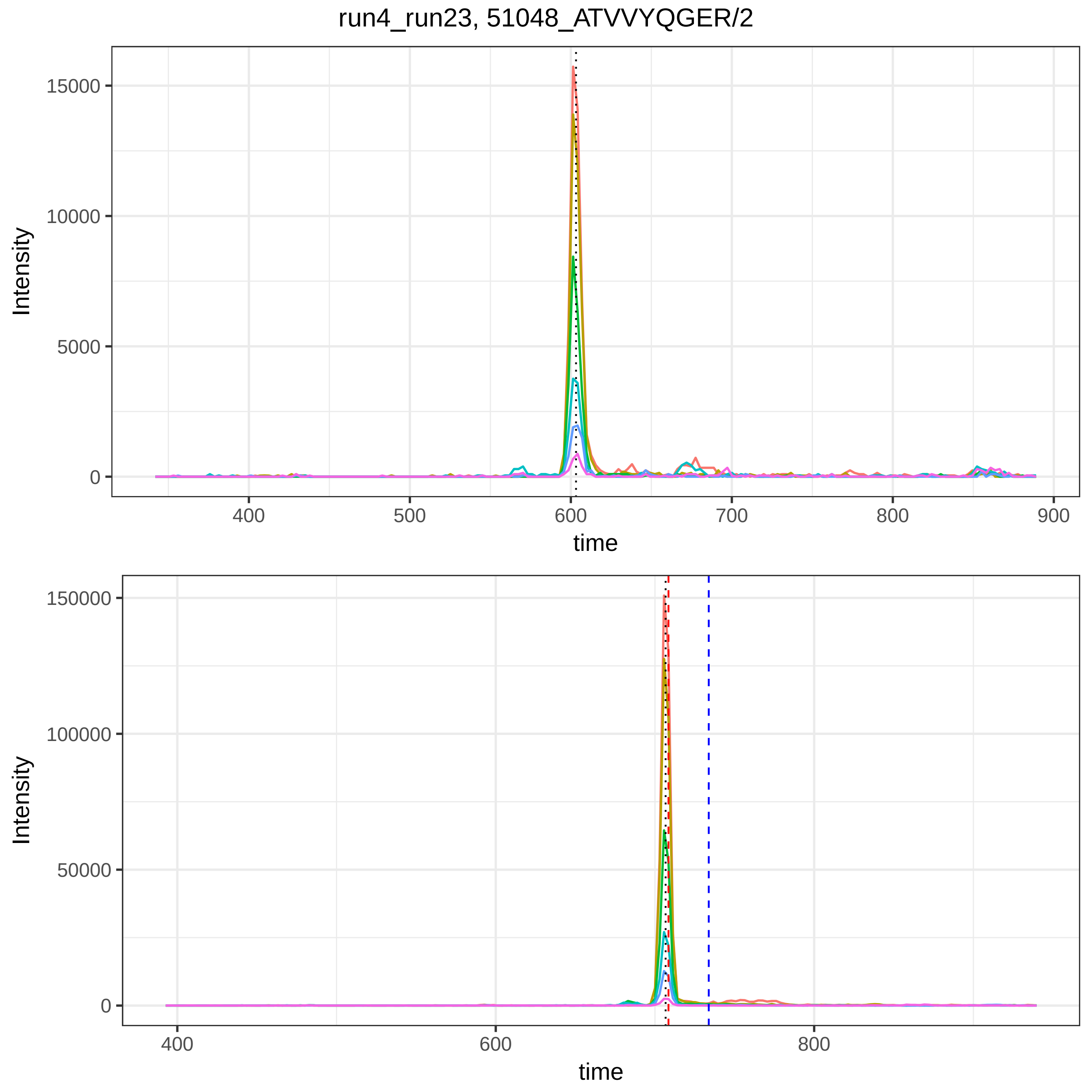

run4_run23, 51048_ATVVYQGER/2
Intensity
15000
10000
5000
0
400
500
600
700
800
900
time
Intensity
150000
100000
50000
0
400
600
800
time

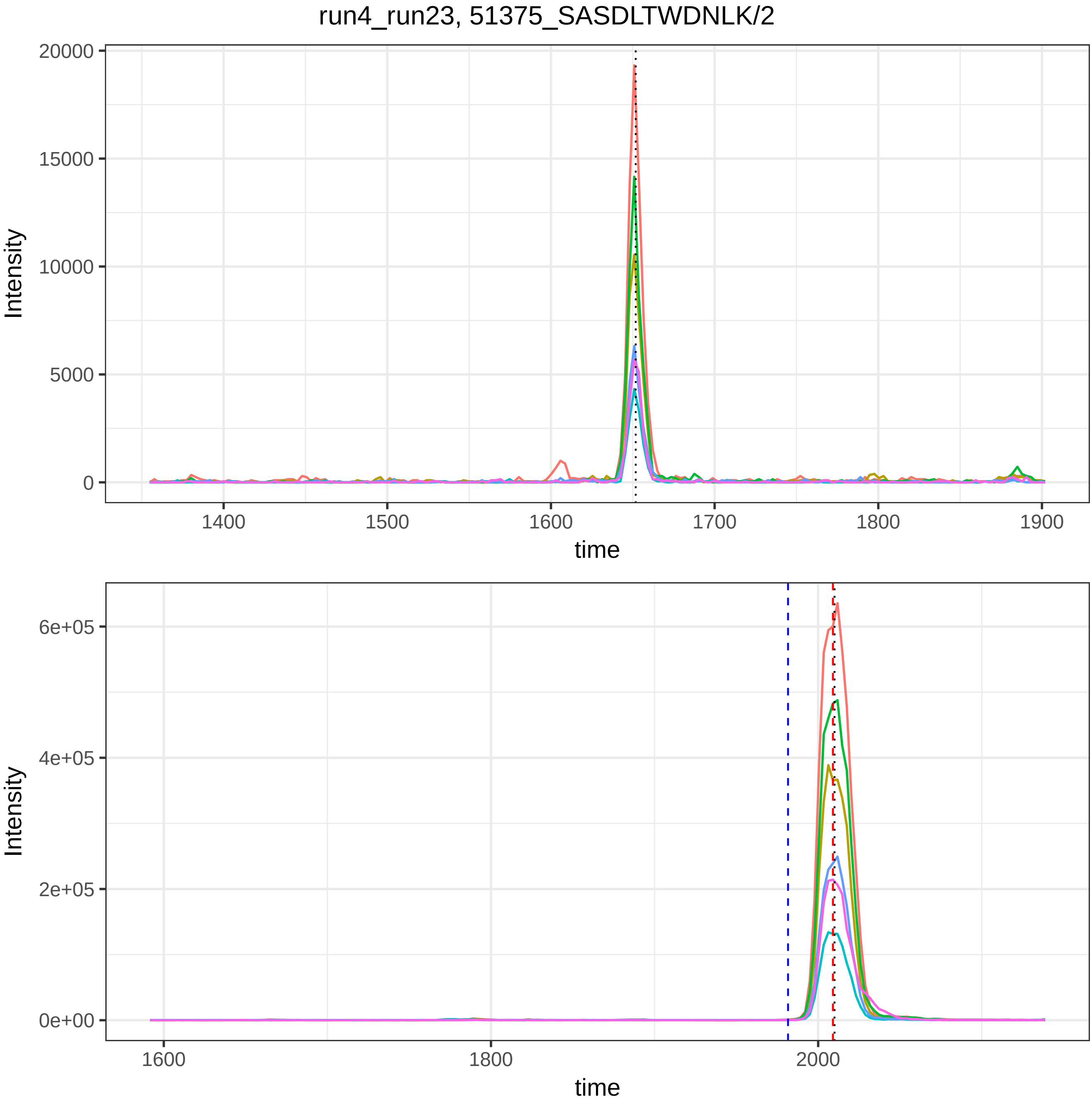
run4_run23, 51375_SASDLTWDNLK/2
20000
15000
10000
5000
0
Intensity
1400
1500
1600
1700
1800
1900
time
6e+05
4e+05
2e+05
0e+00
Intensity
1600
1800
2000
time

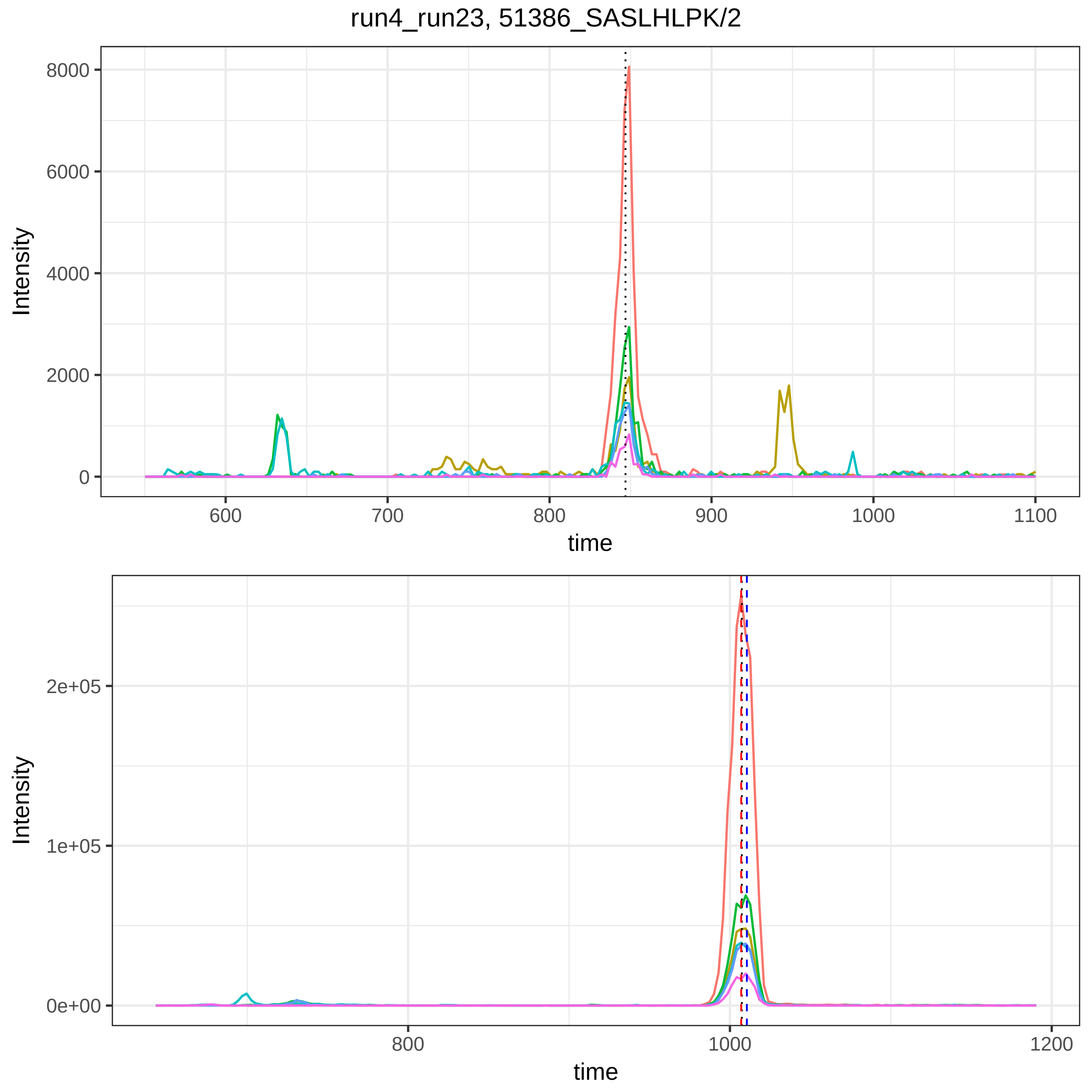
run4_run23, 51386_SASLHLPK/2
Intensity
8000
6000
4000
2000
0
600
700
800
900
1000
1100
time
Intensity
2e+05
1e+05
0e+00
800
1000
1200
time

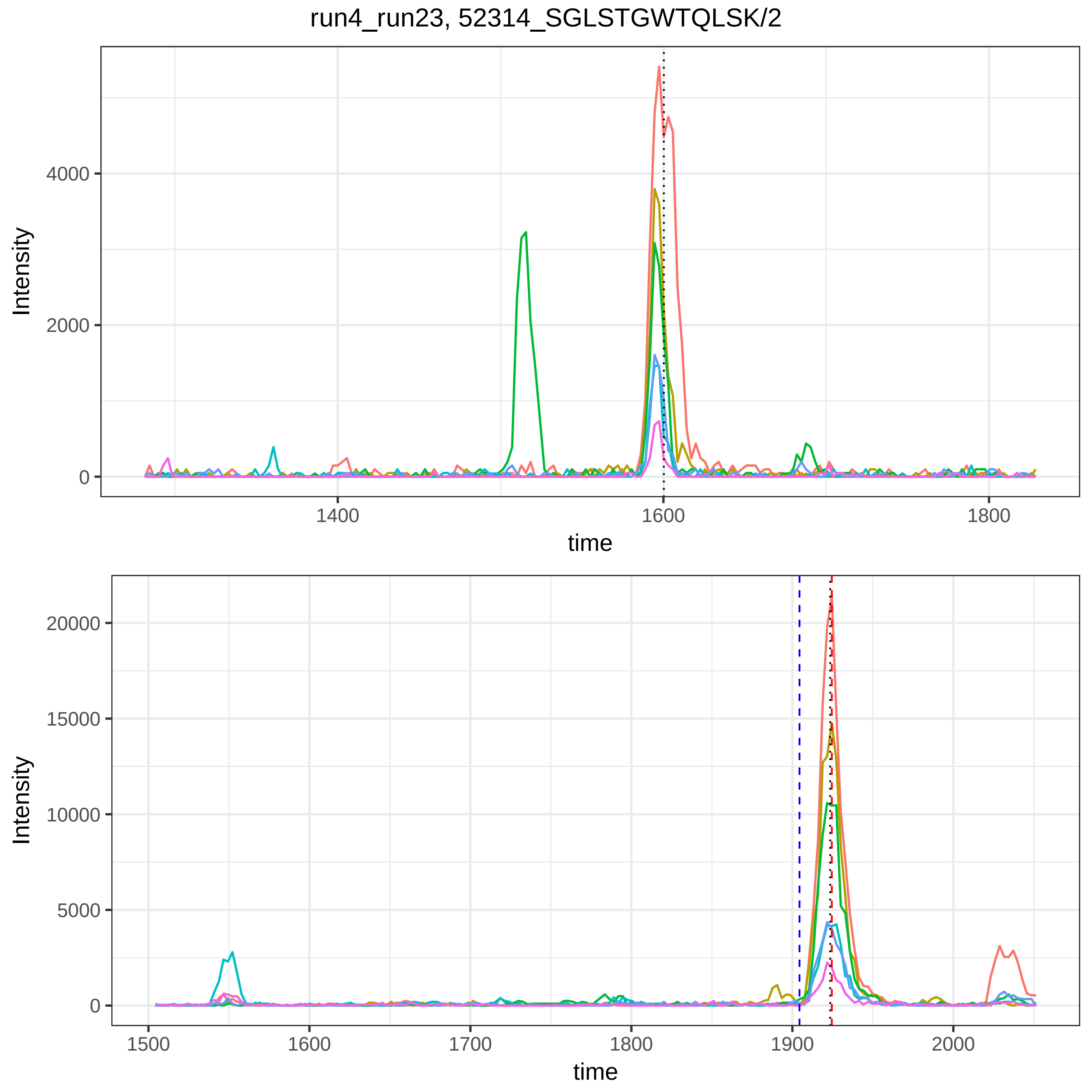

run4_run23, 52314_SGLSTGWTQLSK/2
Intensity
4000
2000
0
1400
1600
1800
time
Intensity
20000
15000
10000
5000
0
1500
1600
1700
1800
1900
2000
time

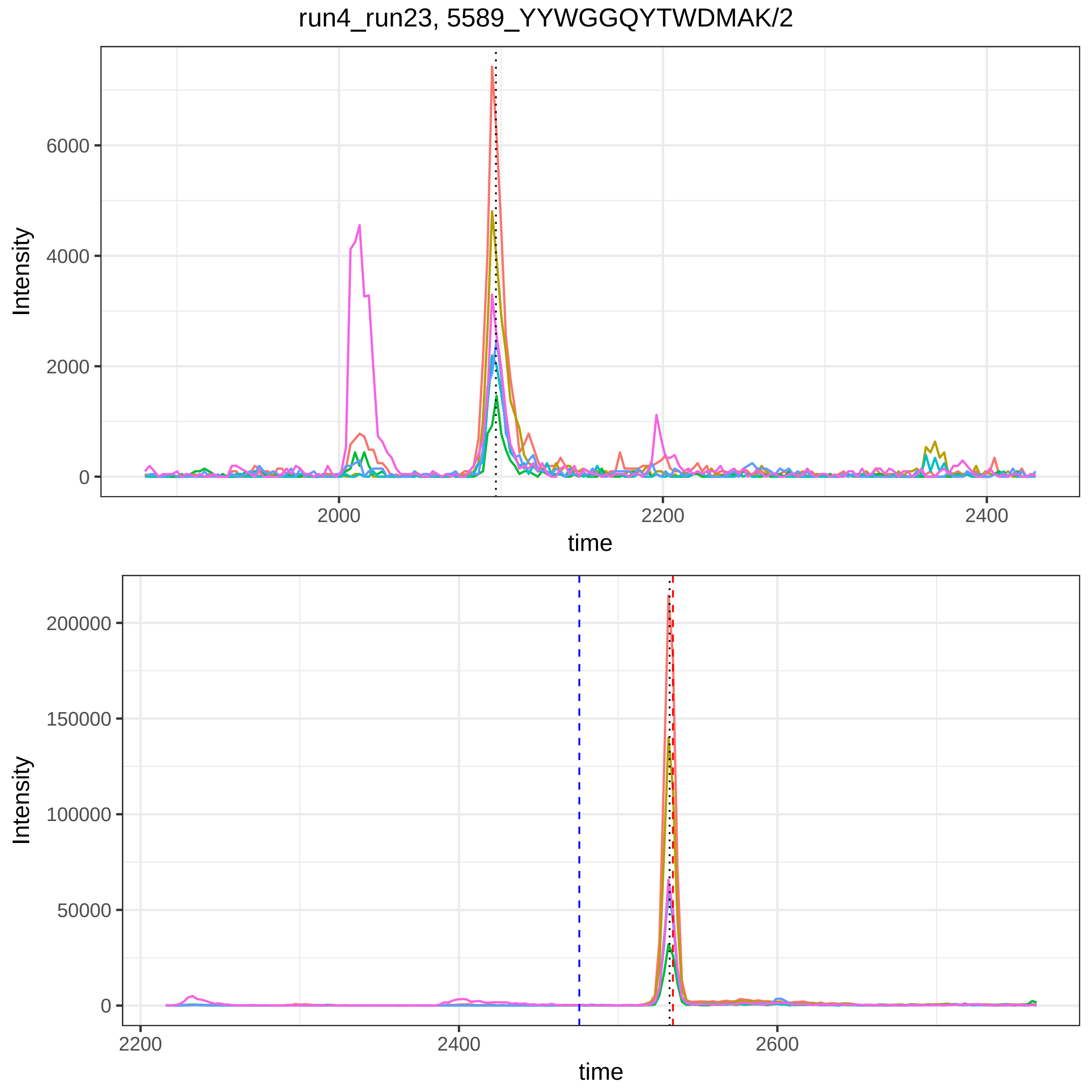

run4_run23, 5589_YYWGGQYTWDMAK/2
Intensity
6000
4000
2000
0
2000
2200
2400
time
Intensity
200000
150000
100000
50000
0
2200
2400
2600
time

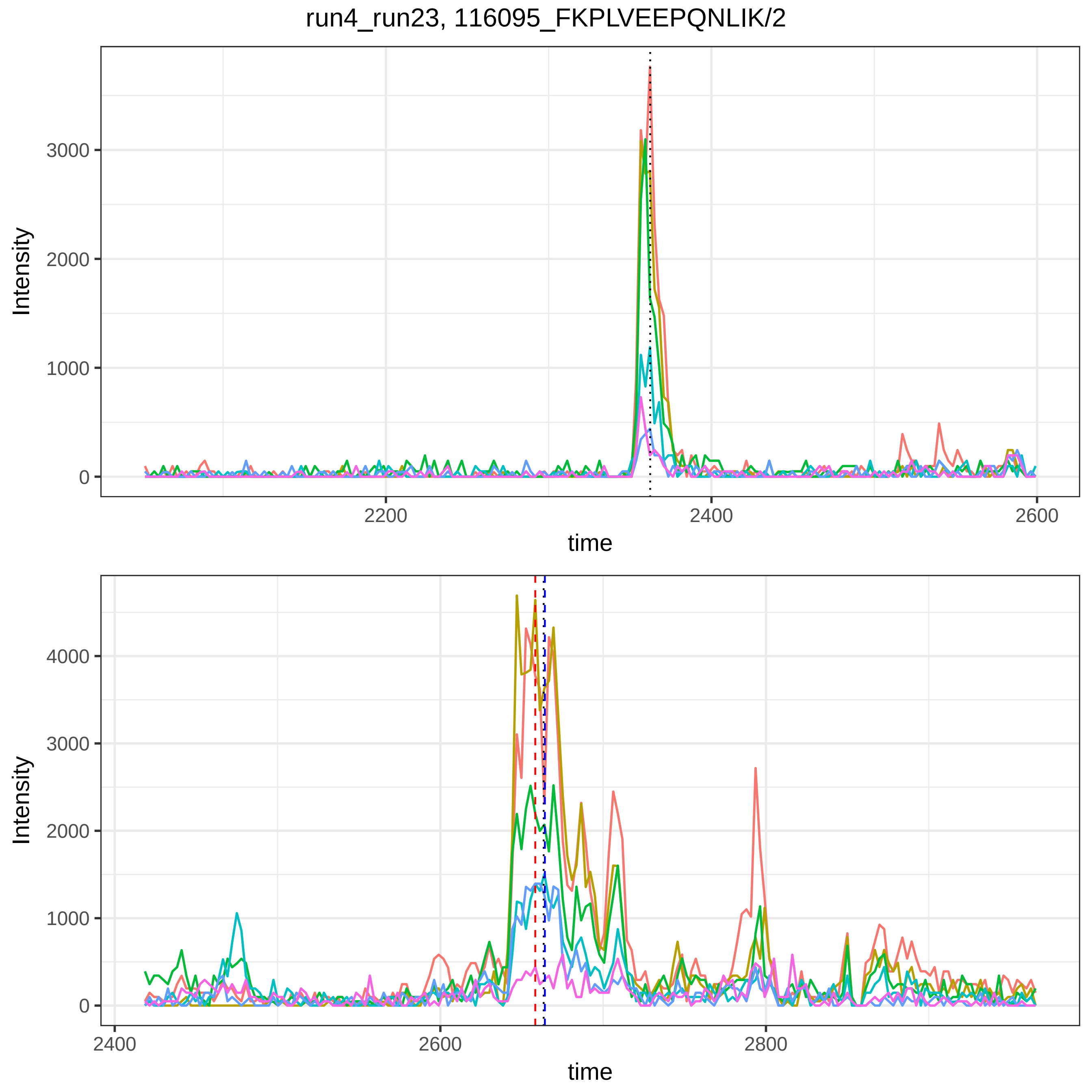

run4_run23, 116095_FKPLVEEPQNLIK/2
Intensity
time
0
1000
2000
3000
2200
2400
2600
4000
2800

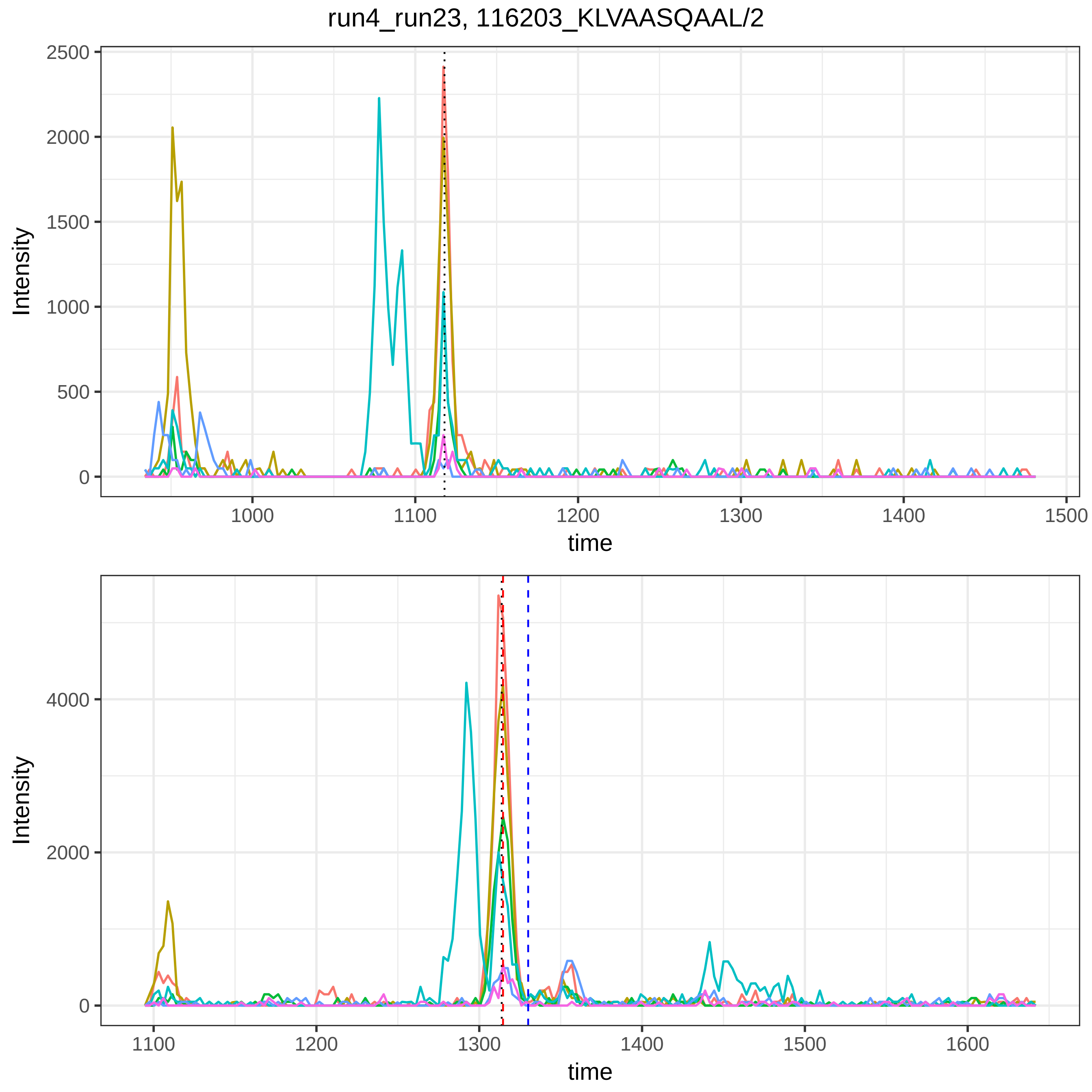

run4_run23, 116203_KLVAASQAAL/2
Intensity
time
2500
2000
1500
1000
500
0
1000
1100
1200
1300
1400
1500
4000
2000
1600

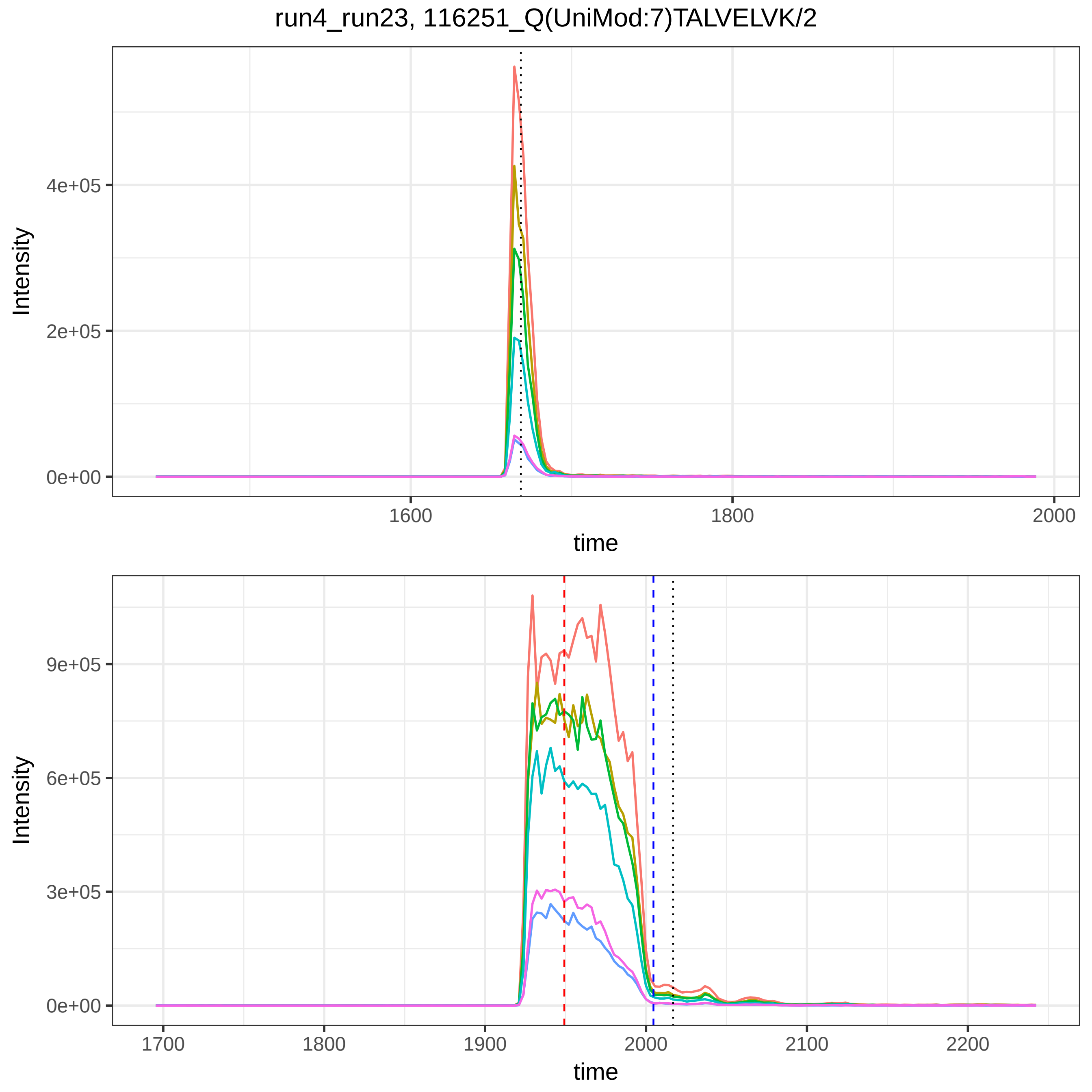

run4_run23, 116251_Q(UniMod:7)TALVELVK/2
Intensity
4e+05
2e+05
0e+00
1600
1800
2000
time
Intensity
9e+05
6e+05
3e+05
0e+00
1700
1800
1900
2000
2100
2200
time

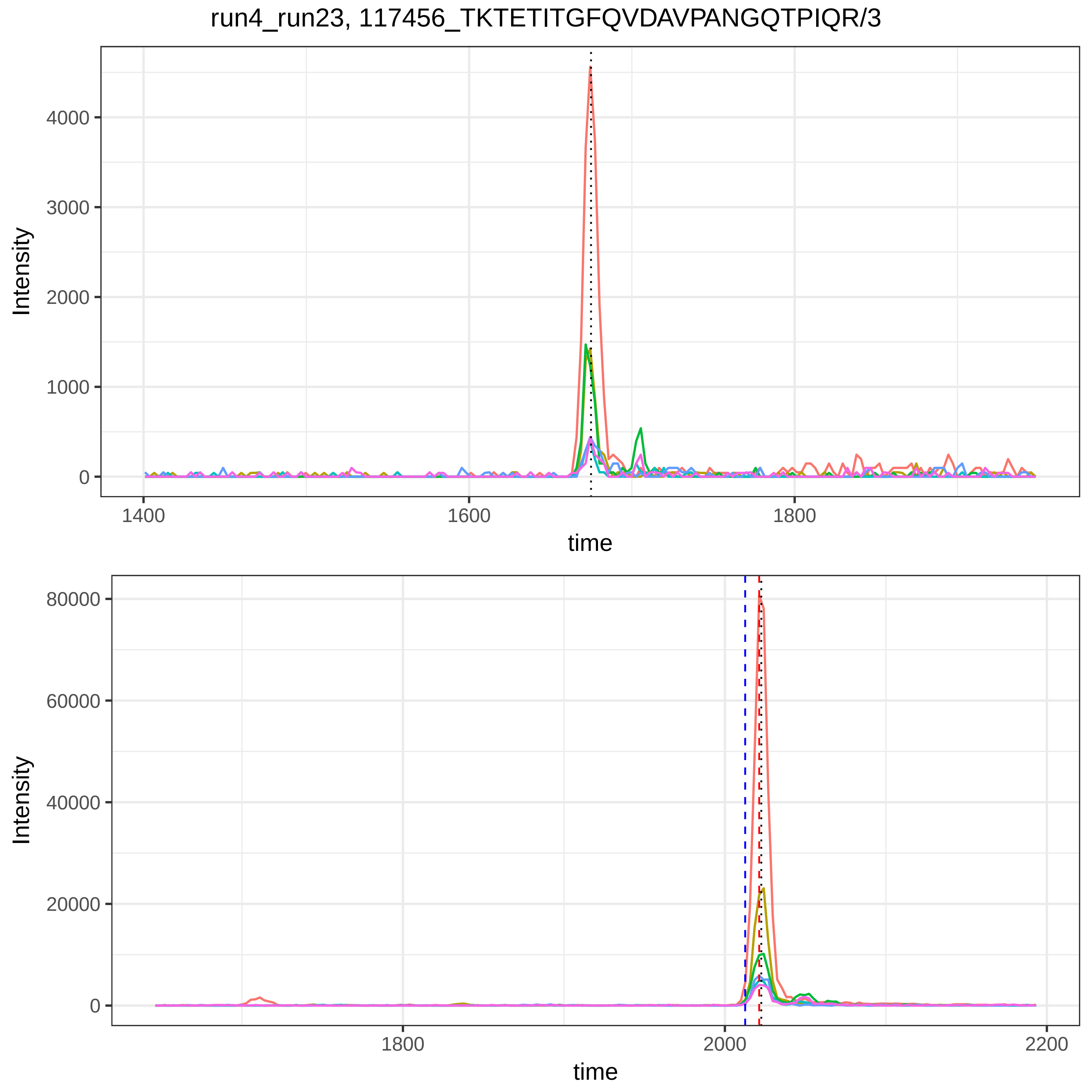

run4_run23, 117456_TKTETITGFQVDAVPANGQTPIQR/3
Intensity
0
1000
2000
3000
4000
time
1400
1600
1800
Intensity
0
20000
40000
60000
80000
time
1800
2000
2200

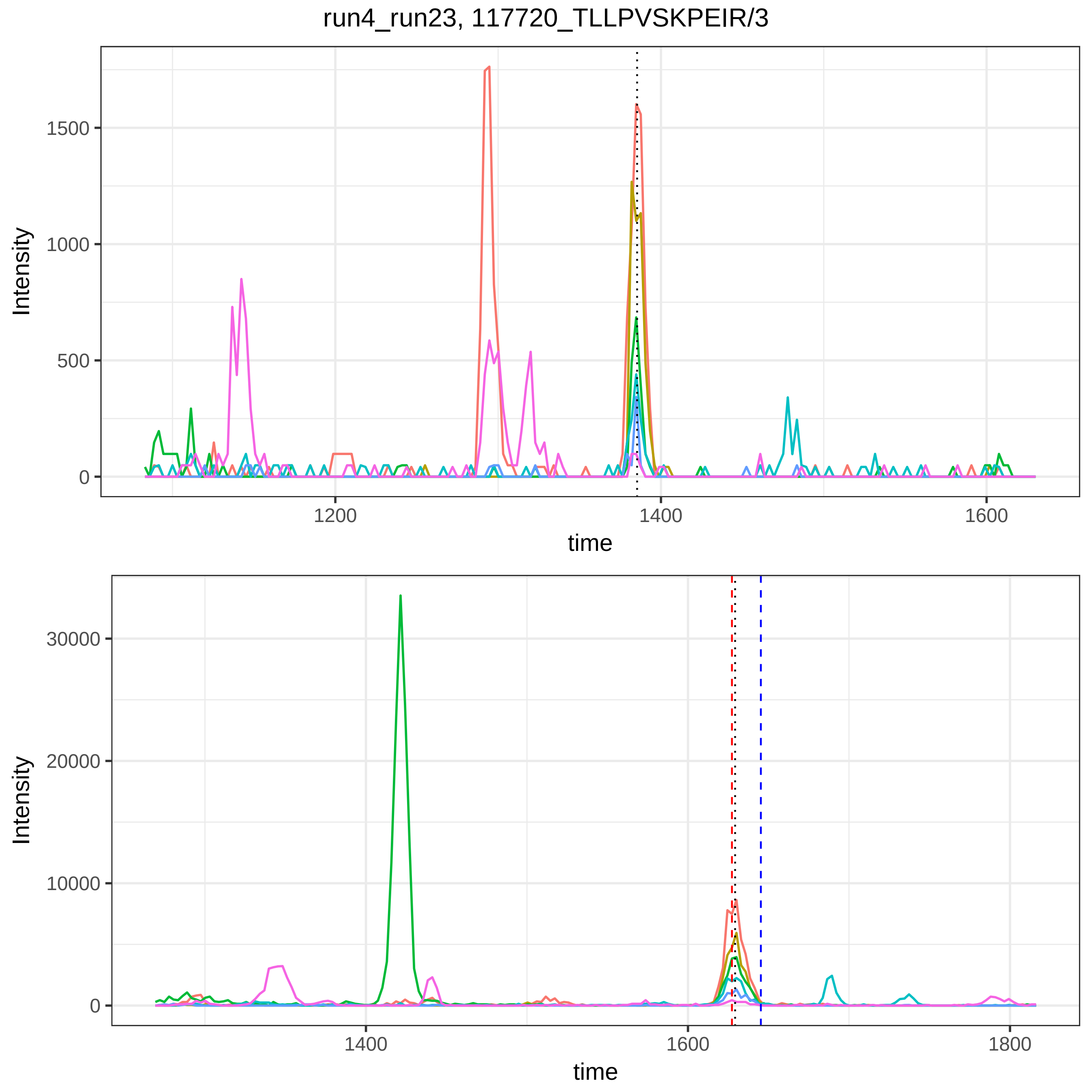

run4_run23, 117720_TLLPVSKPEIR/3
Intensity
time
1500
1000
500
0
1200
1400
1600
30000
20000
10000
1800

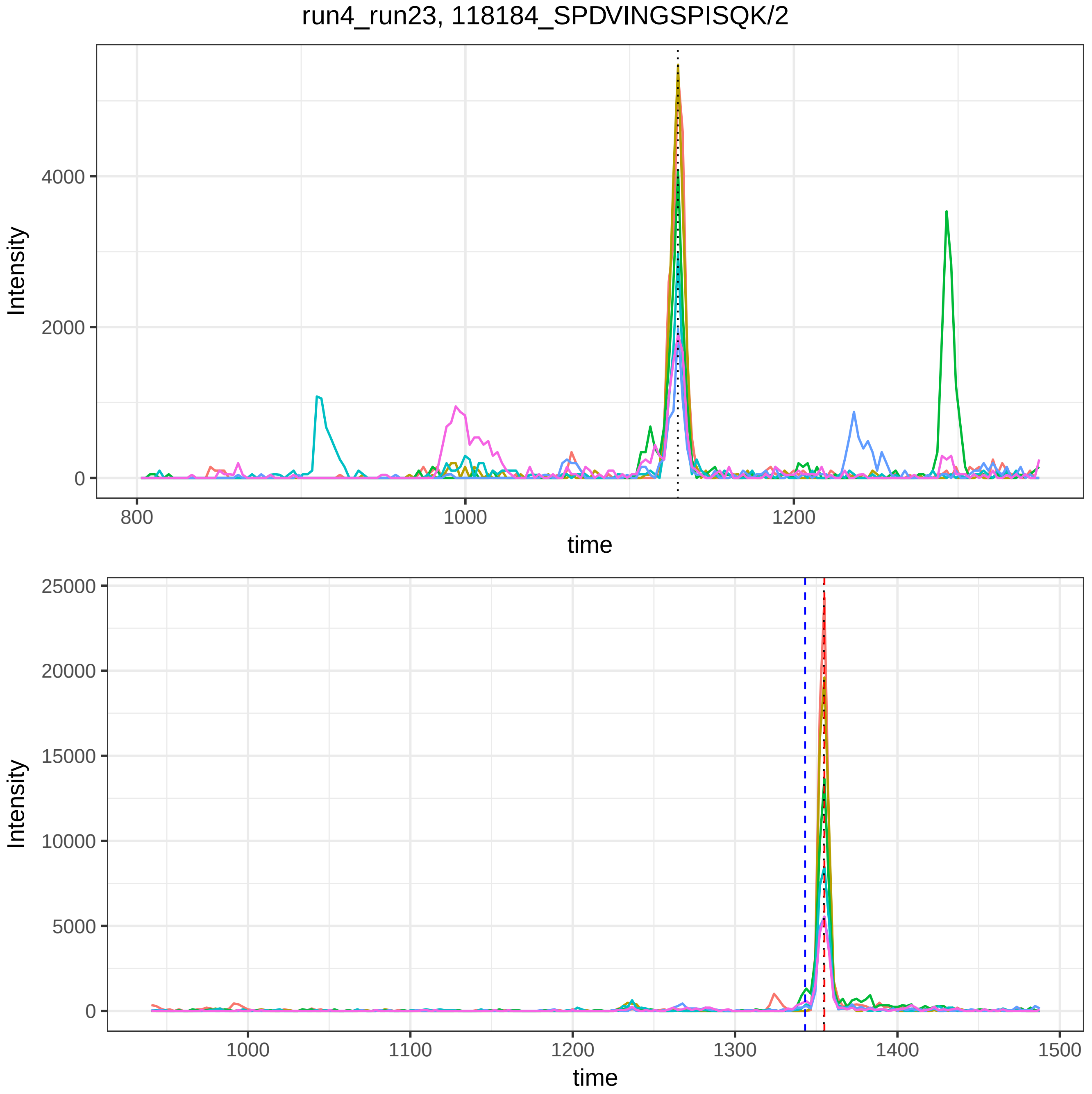
run4_run23, 118184_SPDVINGSPISQK/2
Intensity
time
0
2000
4000
800
1000
1200
0
5000
10000
15000
20000
25000
1100
1300
1400
1500

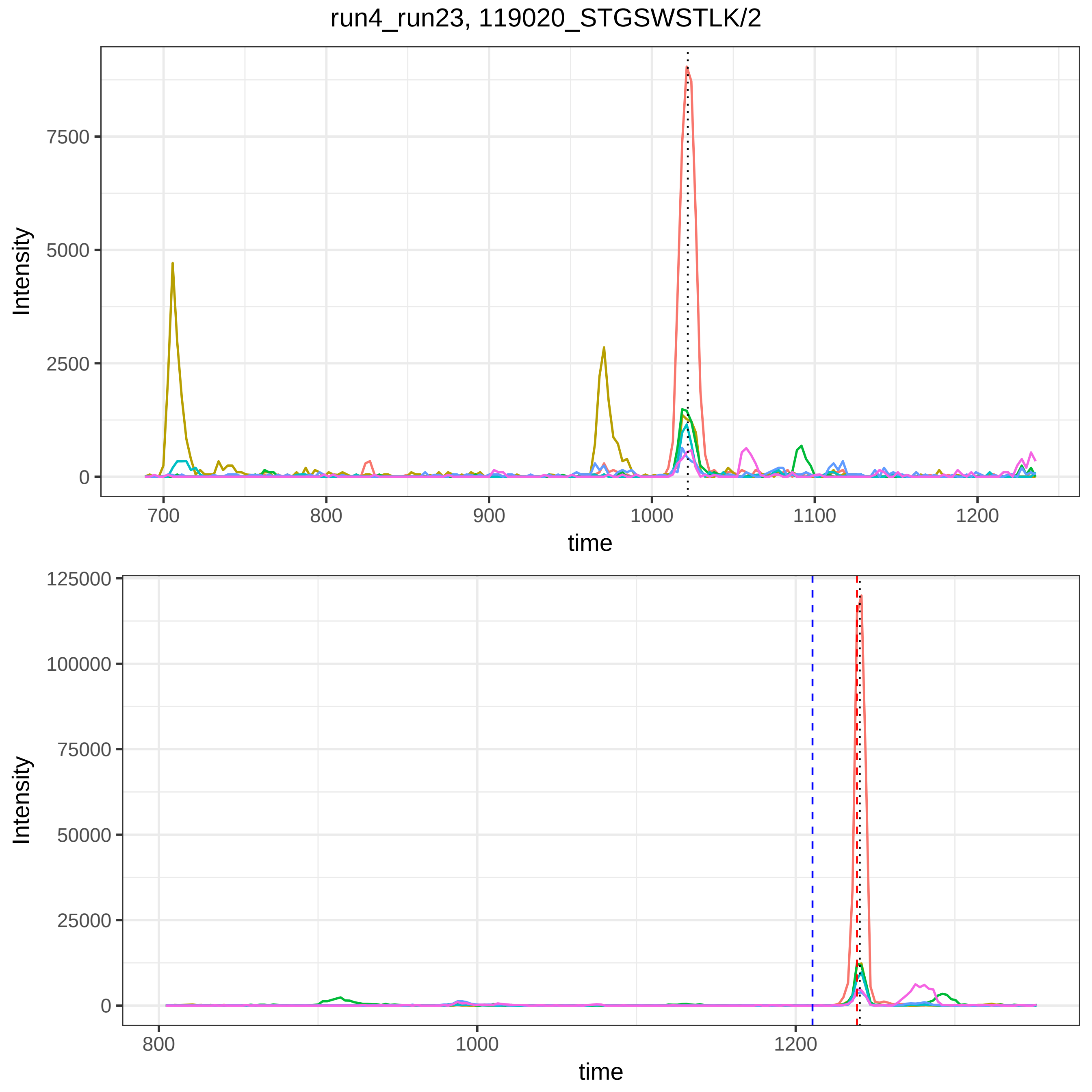

run4_run23, 119020_STGSWSTLK/2
Intensity
7500
5000
2500
0
700
800
900
1000
1100
1200
time
Intensity
125000
100000
75000
50000
25000
0
800
1000
1200
time

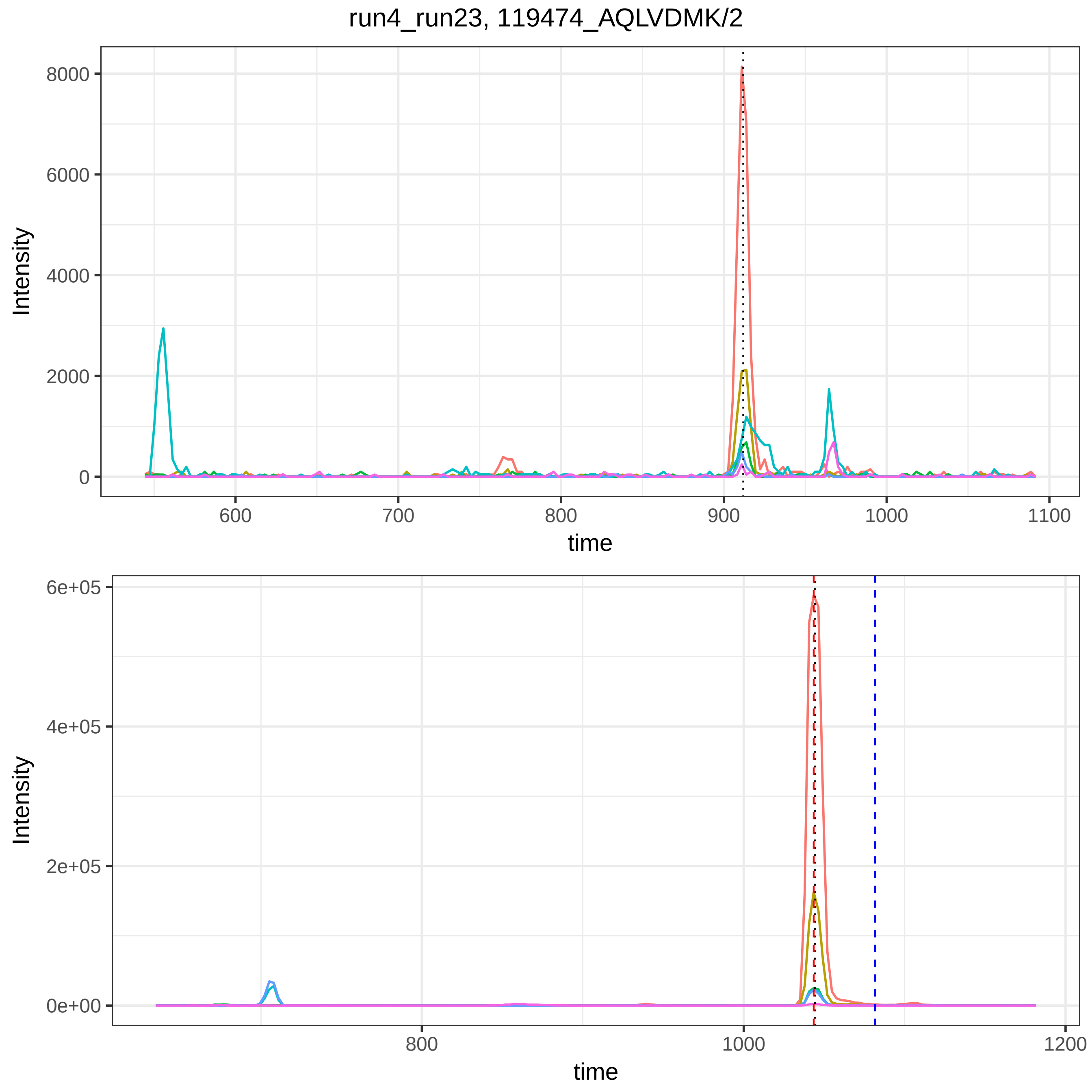
run4_run23, 119474_AQLVDMK/2
Intensity
8000
6000
4000
2000
0
600
700
800
900
1000
1100
time
Intensity
6e+05
4e+05
2e+05
0e+00
800
1000
1200
time

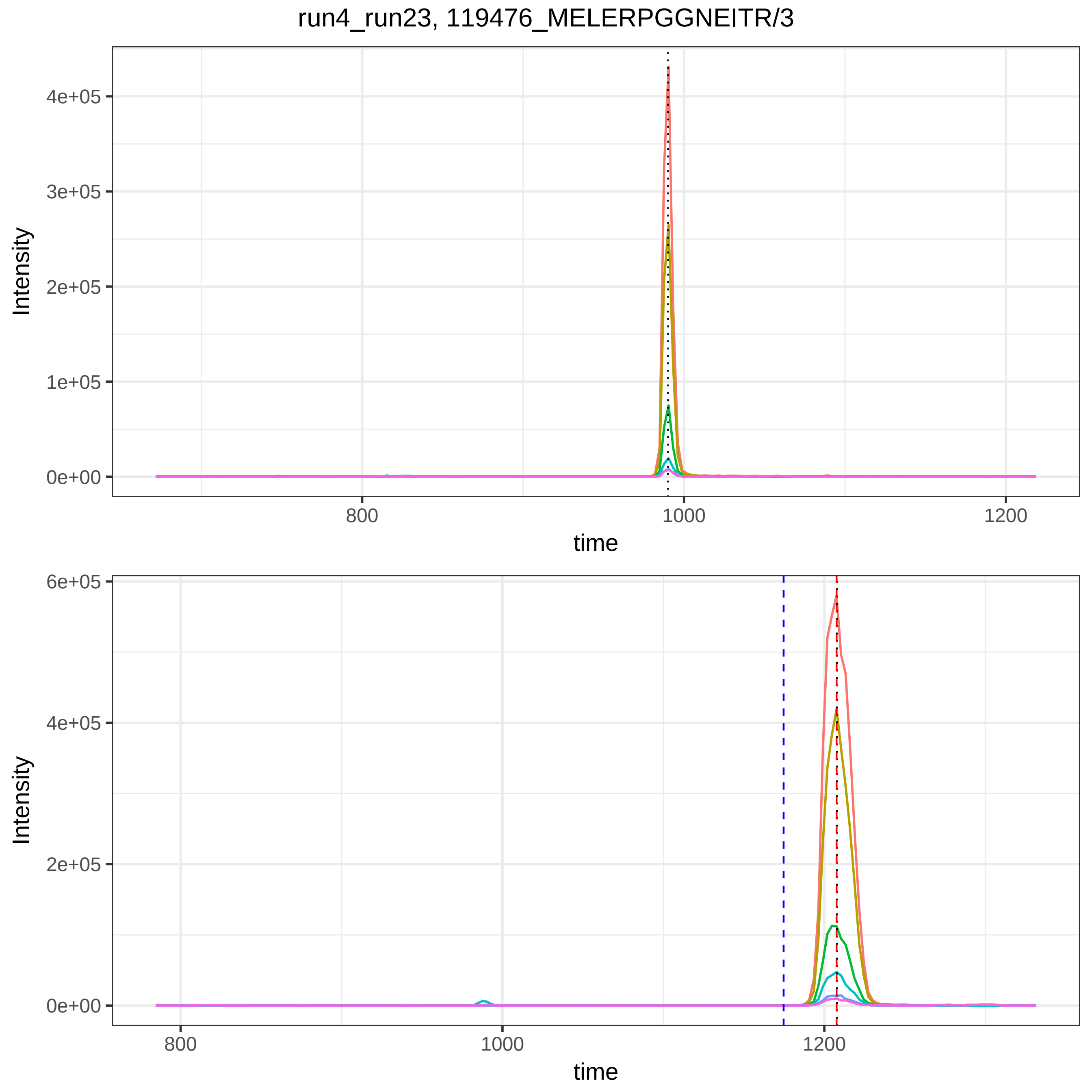

run4_run23, 119476_MELERPGGNEITR/3
Intensity
4e+05
3e+05
2e+05
1e+05
0e+00
800
1000
1200
time
Intensity
6e+05
4e+05
2e+05
0e+00
800
1000
1200
time

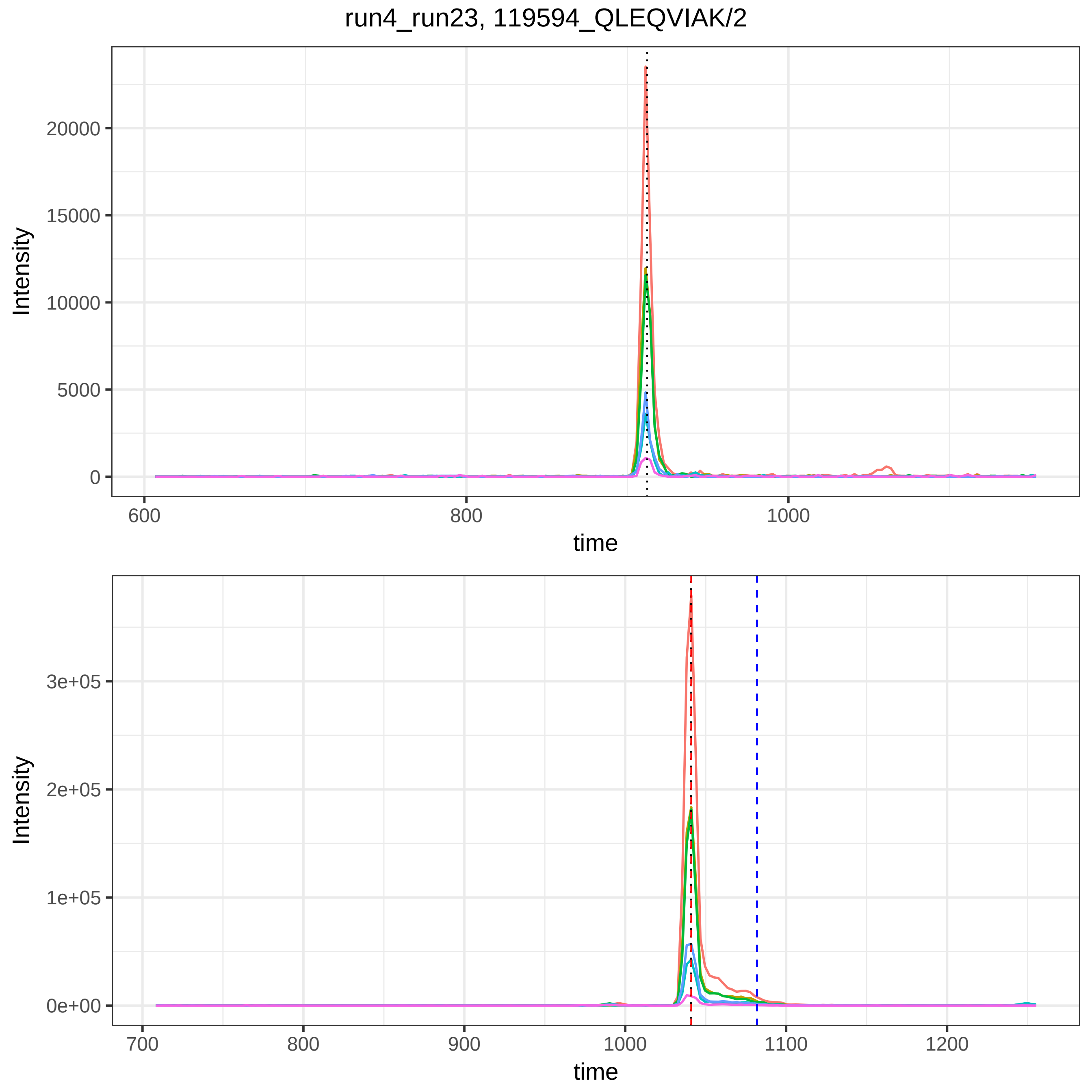
run4_run23, 119594_QLEQVIAK/2
Intensity
20000
15000
10000
5000
0
600
800
1000
time
Intensity
3e+05
2e+05
1e+05
0e+00
700
800
900
1000
1100
1200
time

run4_run23, 119634_AQ(UniMod:7)LVDMK/2

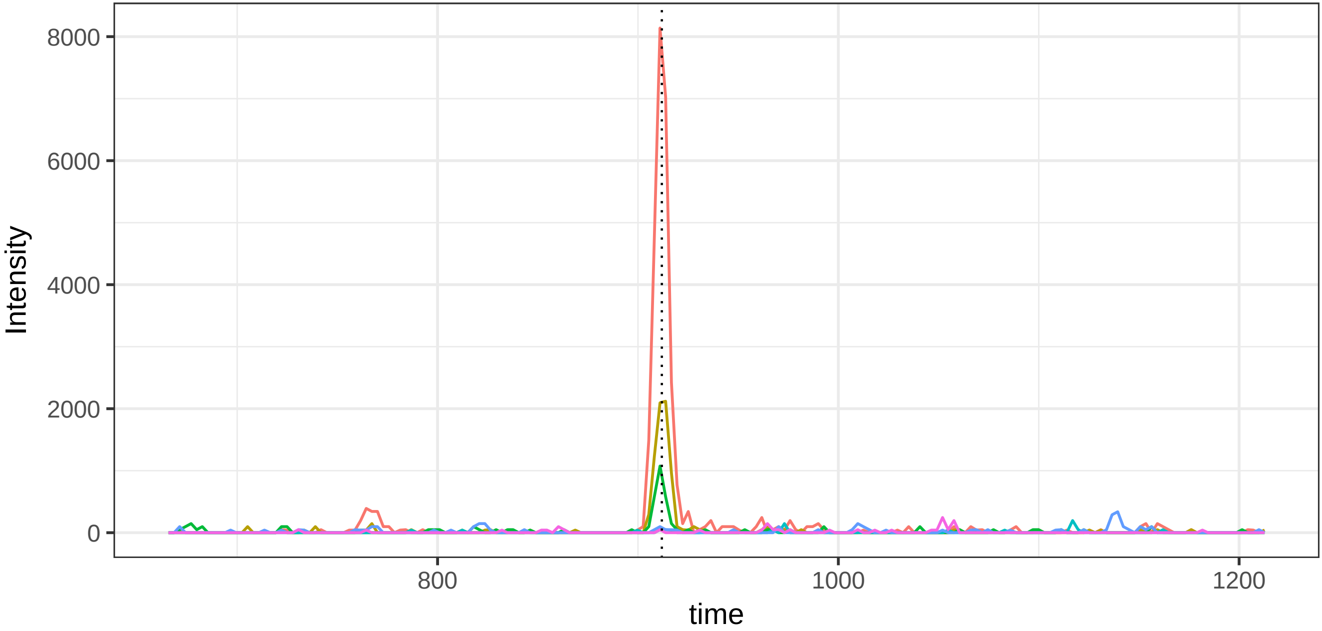


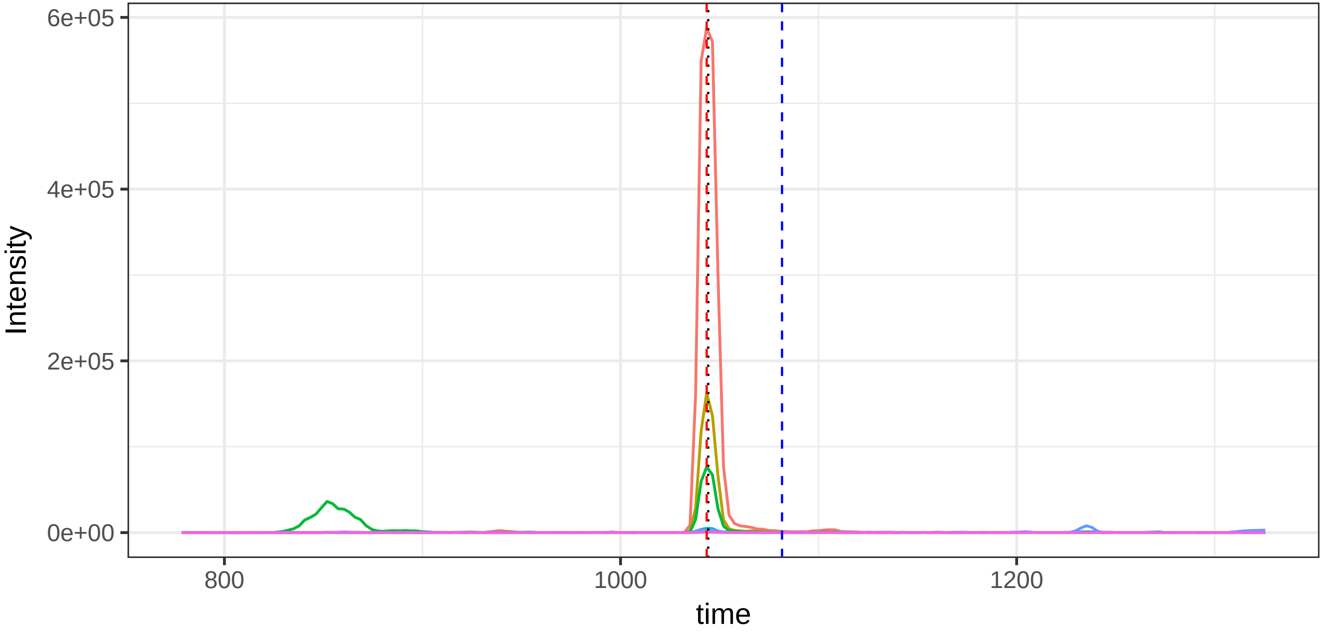

run4_run23, 119655_Q(UniMod:7)LEQVIAK/2

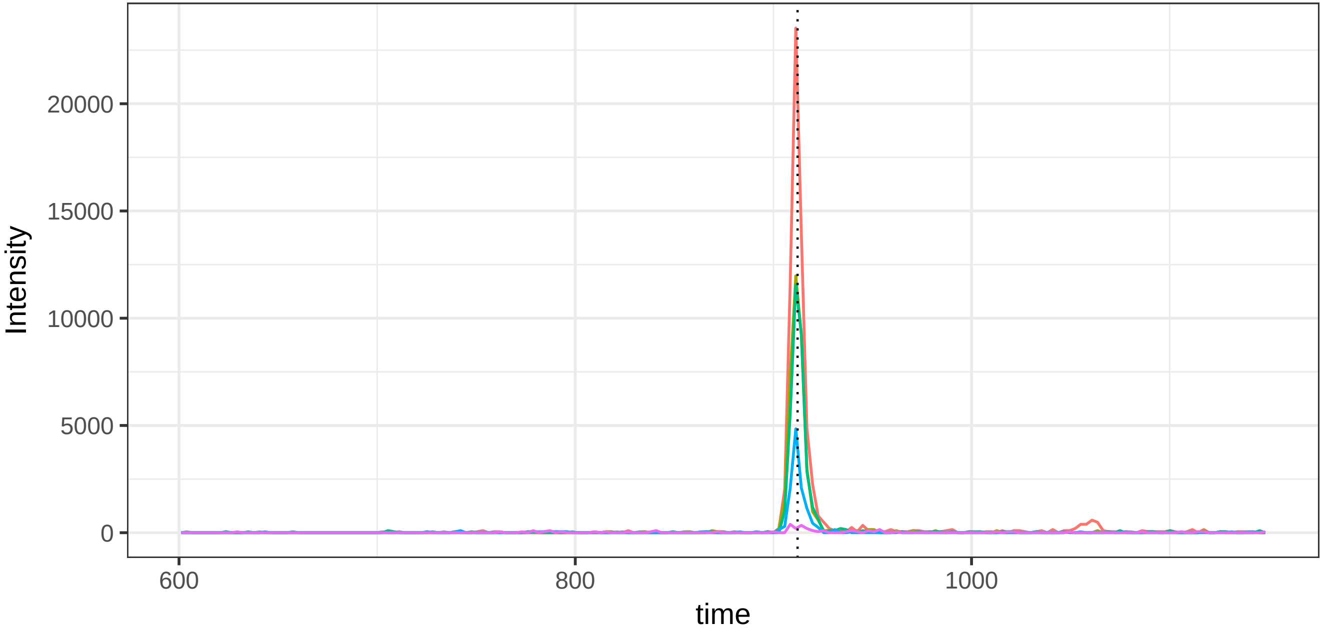


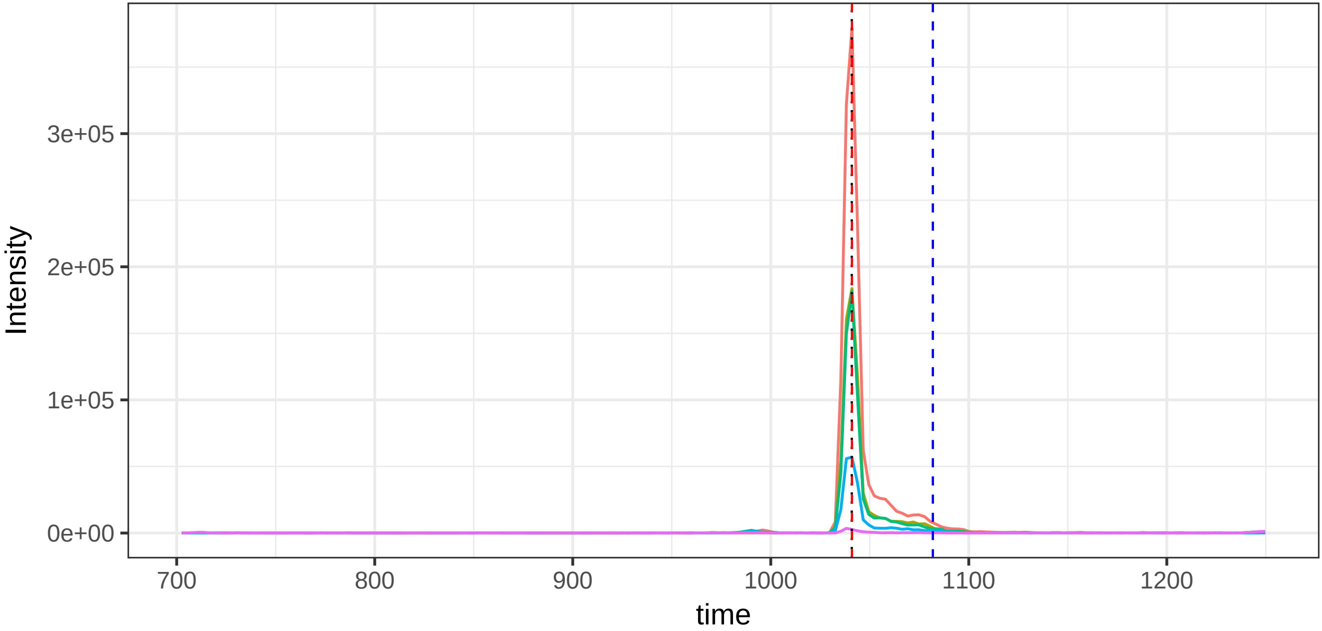

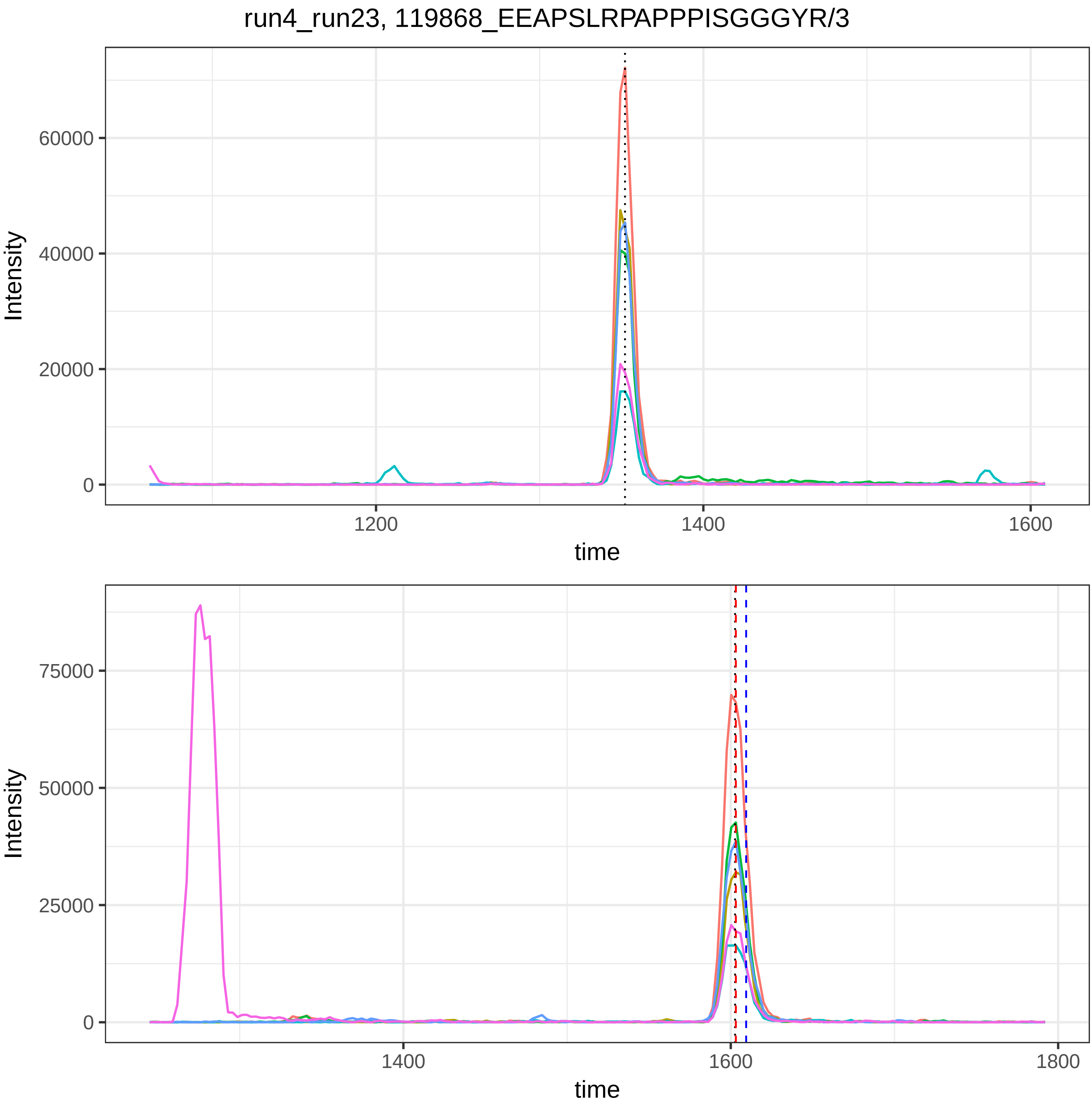
run4_run23, 119868_EEAPSLRPAPPPISGGGYR/3
Intensity
60000
40000
20000
0
1200
1400
1600
time
Intensity
75000
50000
25000
0
1400
1600
1800
time

run4_run23, 119874_VNDNEEGFFSAR/2

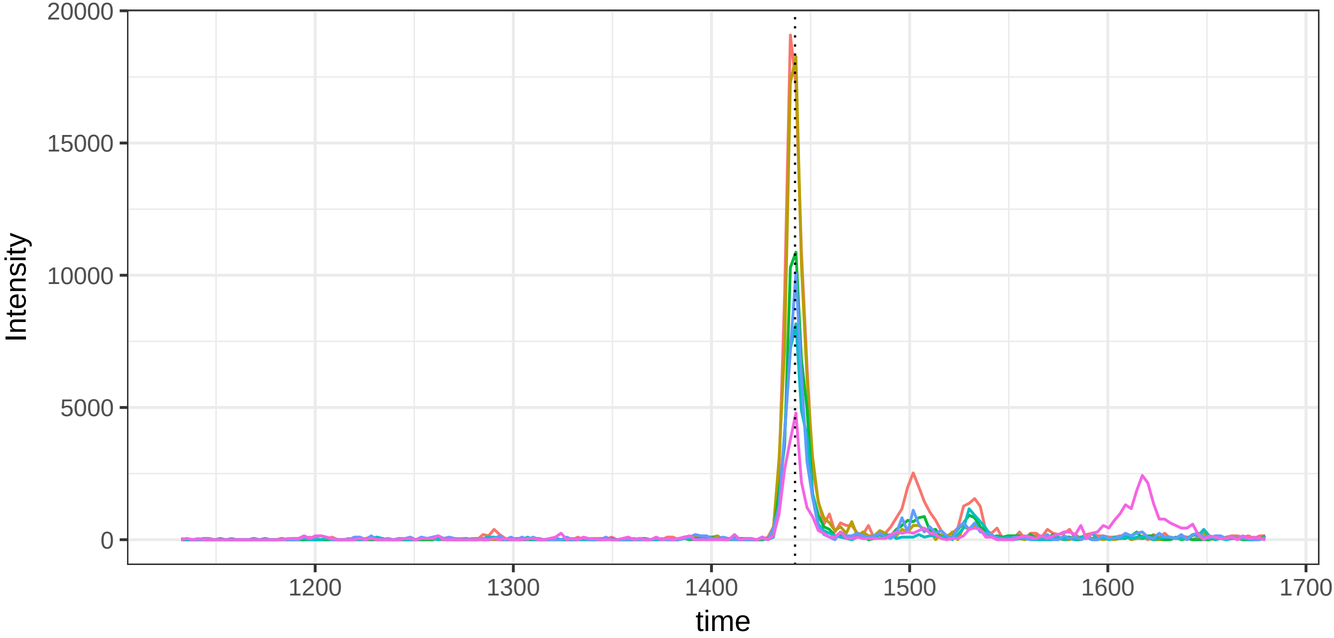


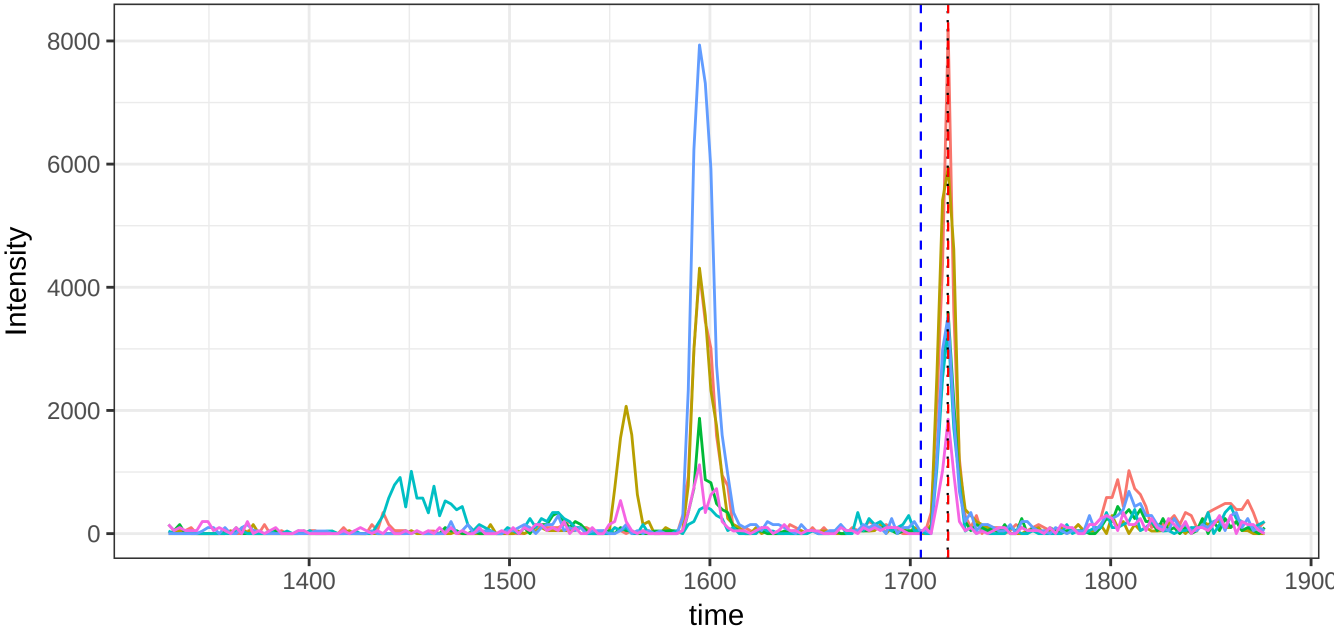

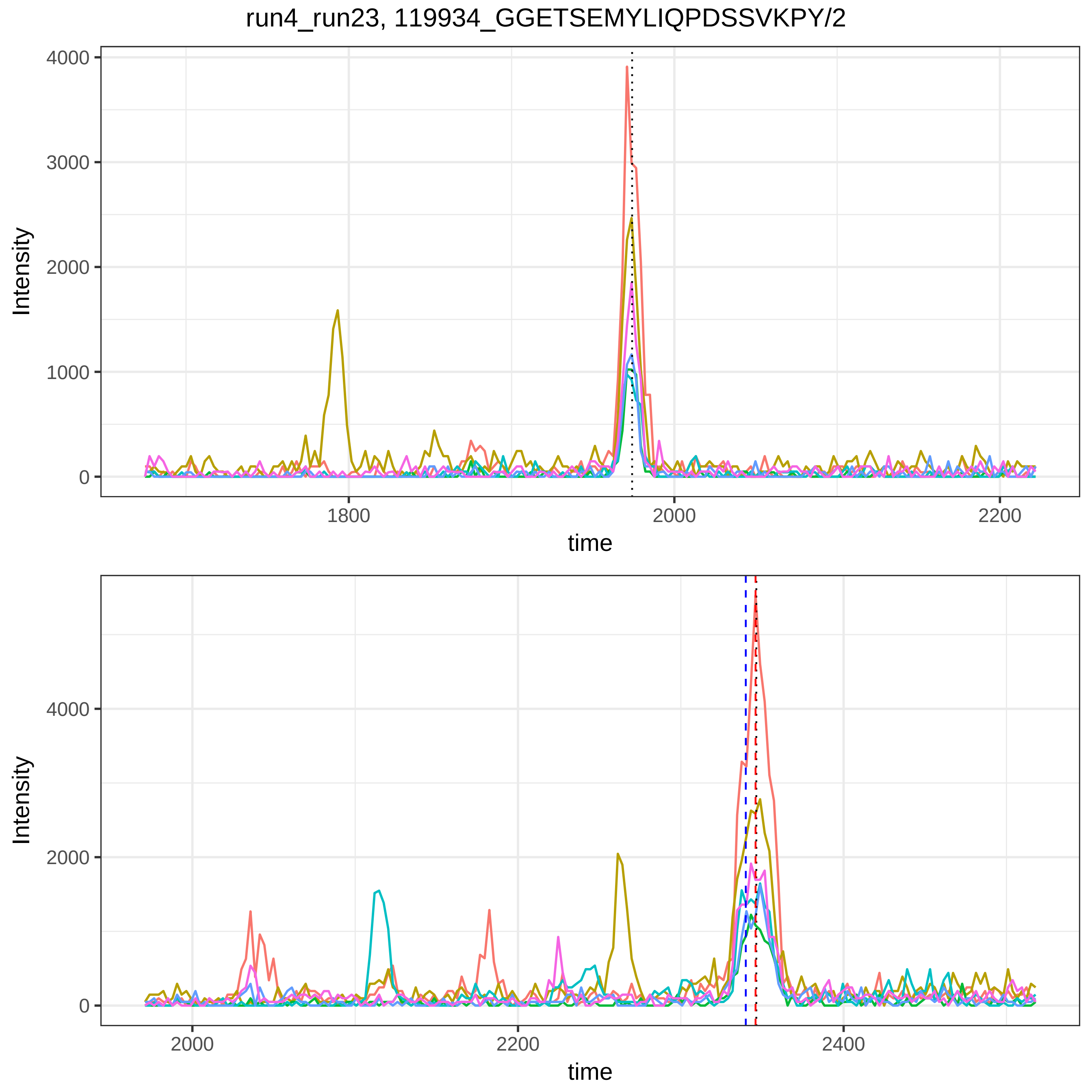
run4_run23, 119934_GGETSEMYLIQPDSSVKPY/2
Intensity
4000
3000
2000
1000
0
1800
2000
2200
time
Intensity
4000
2000
0
2000
2200
2400
time

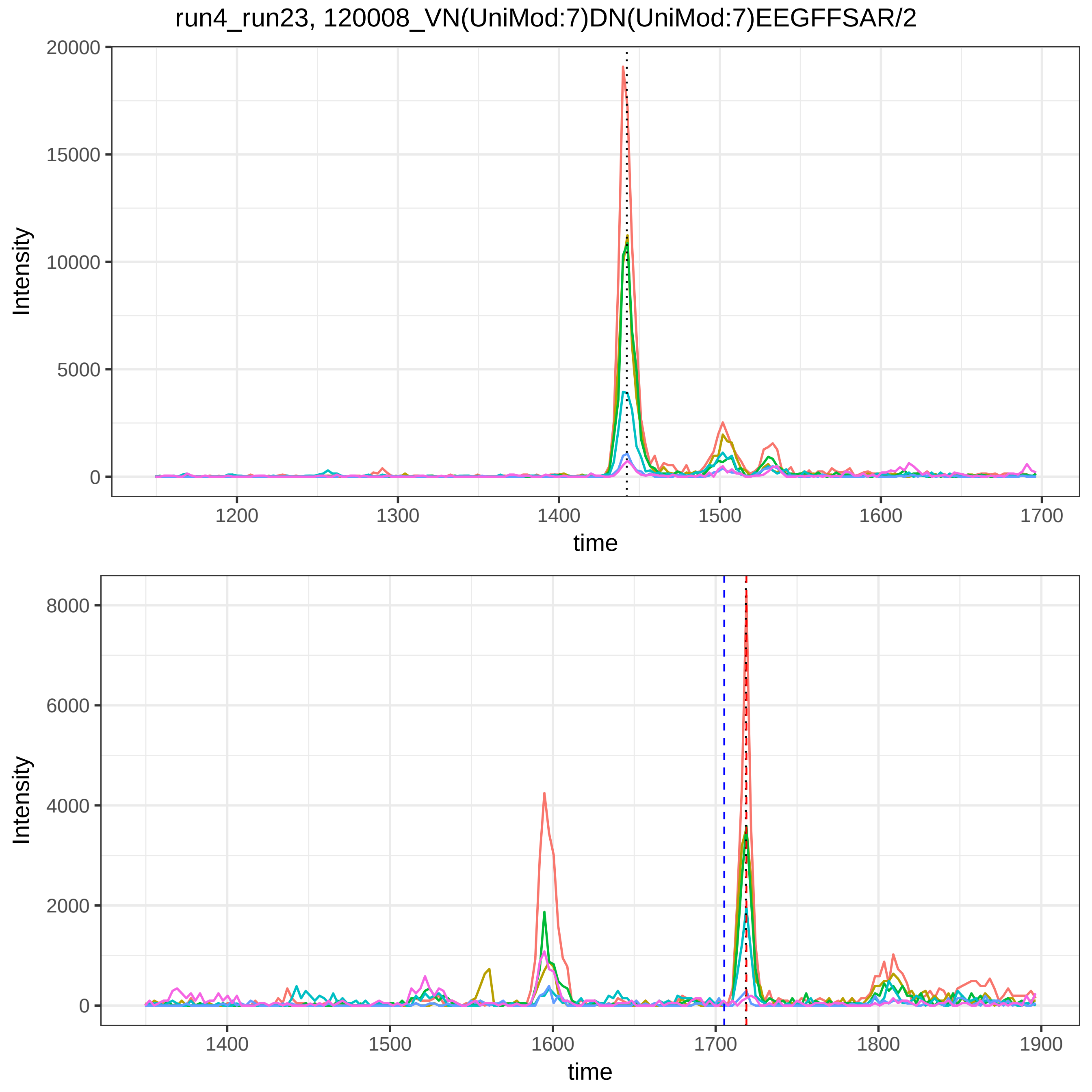
run4_run23, 120008_VN(UniMod:7)DN(UniMod:7)EEGFFSAR/2
Intensity
time
20000
15000
10000
5000
0
1200
1300
1400
1500
1600
1700
8000
6000
4000
2000
1800
1900

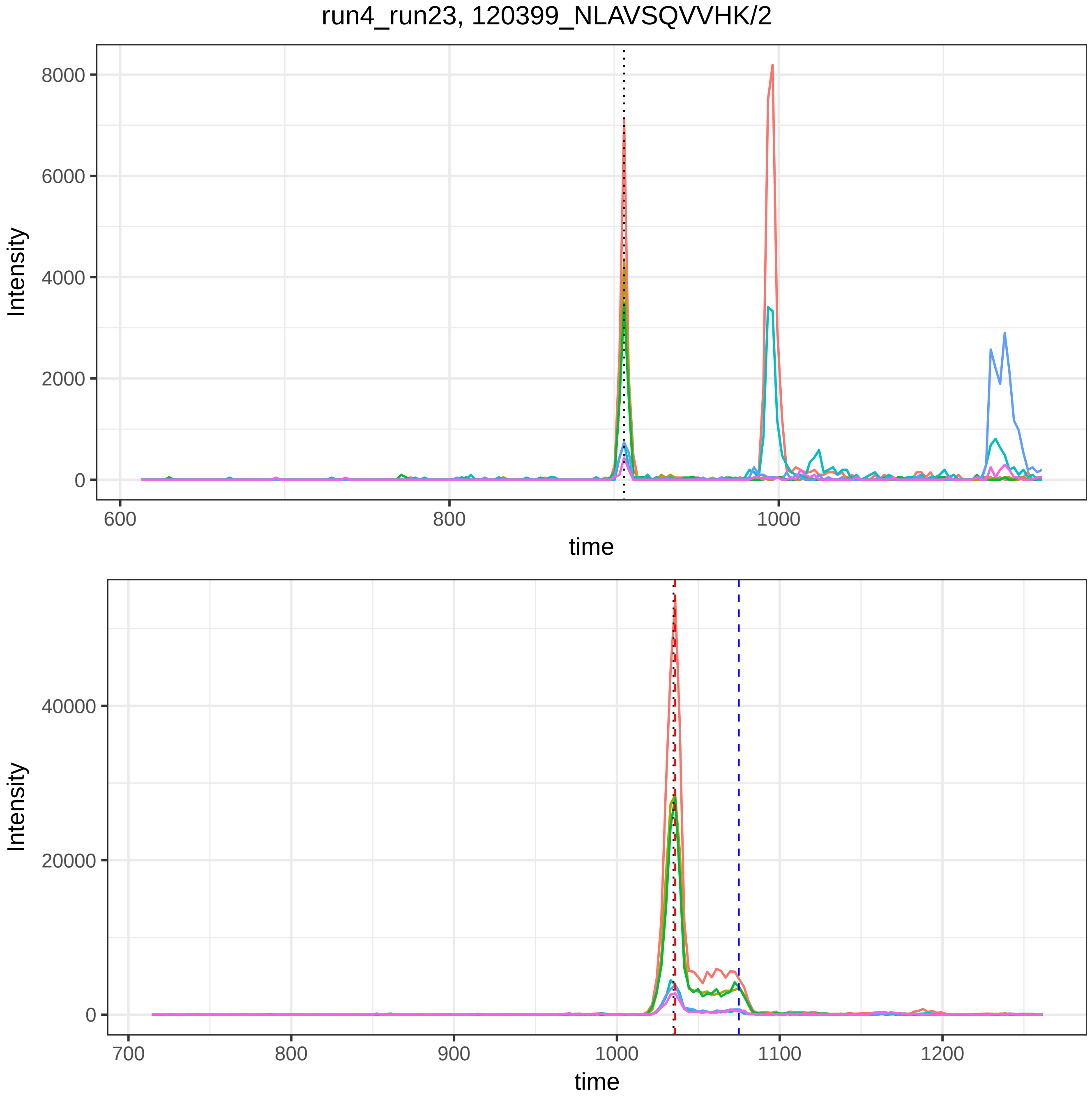

run4_run23, 120399_NLAVSQVVHK/2
Intensity
8000
6000
4000
2000
0
600
800
1000
time
Intensity
40000
20000
0
700
800
900
1000
1100
1200
time

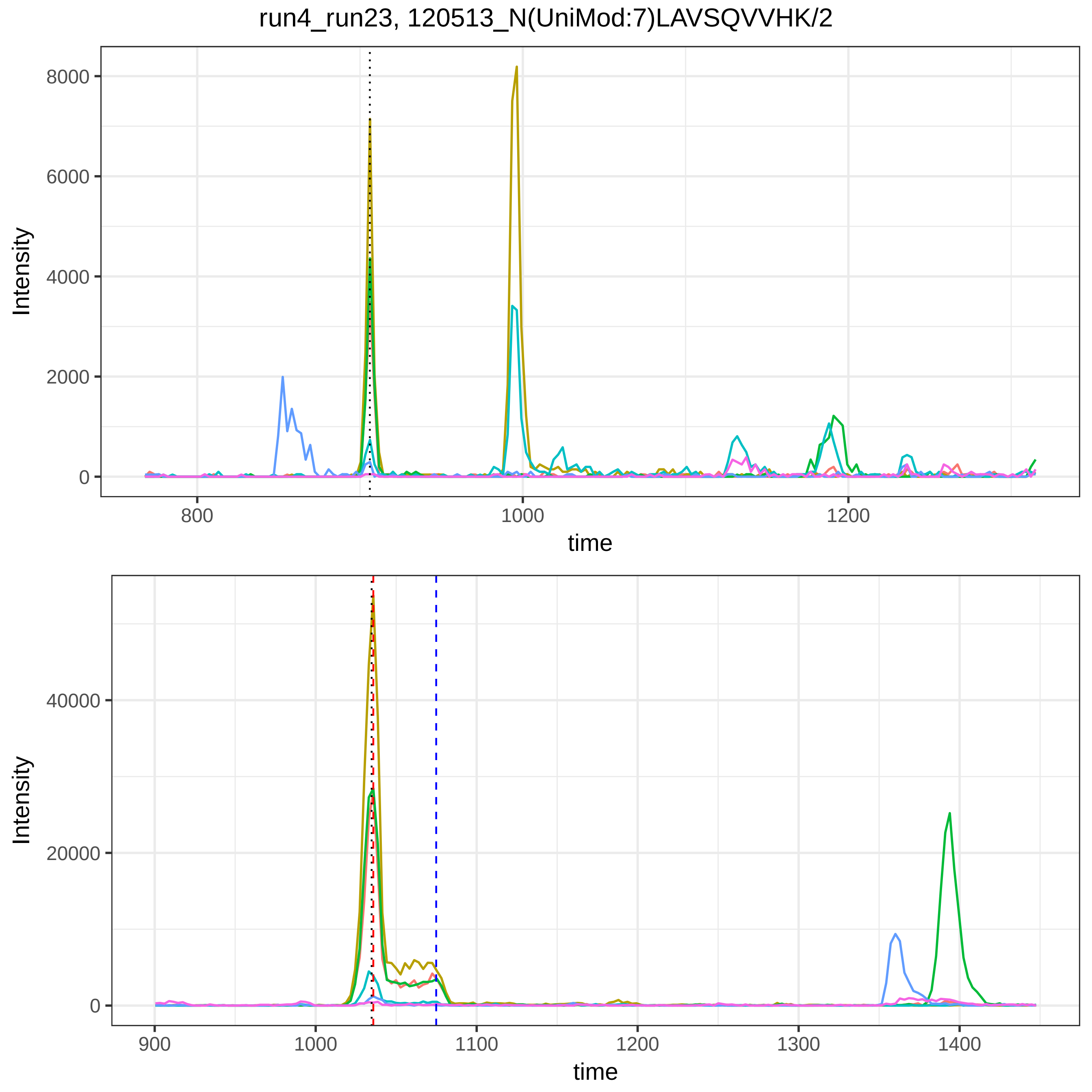
run4_run23, 120513_N(UniMod:7)LAVSQVVHK/2
Intensity
8000
6000
4000
2000
0
800
1000
1200
time
Intensity
40000
20000
0
900
1000
1100
1200
1300
1400
time

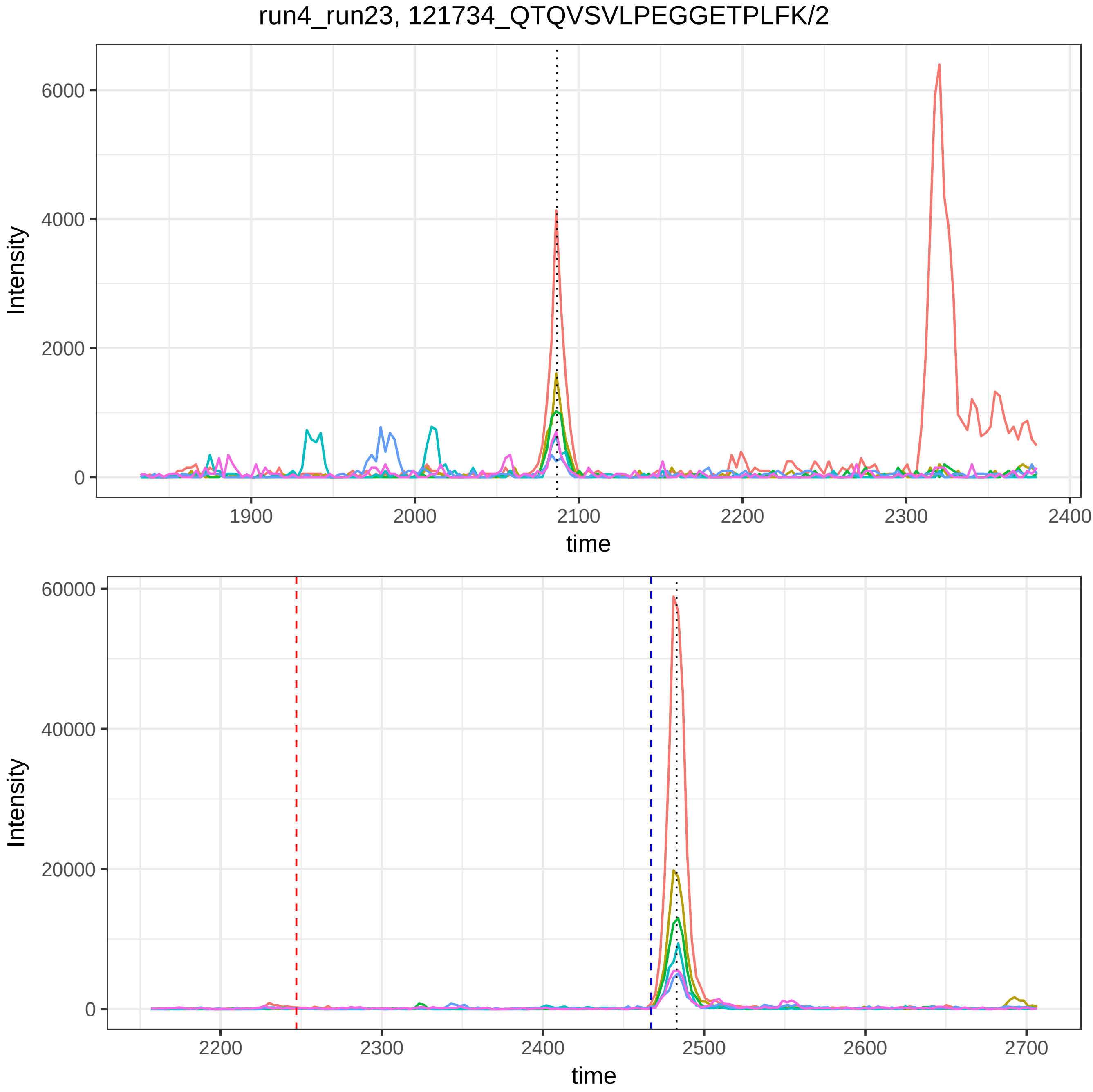
run4_run23, 121734_QTQVSVLPEGGETPLFK/2
Intensity
6000
4000
2000
0
1900
2000
2100
2200
2300
2400
time
Intensity
60000
40000
20000
0
2200
2300
2400
2500
2600
2700
time

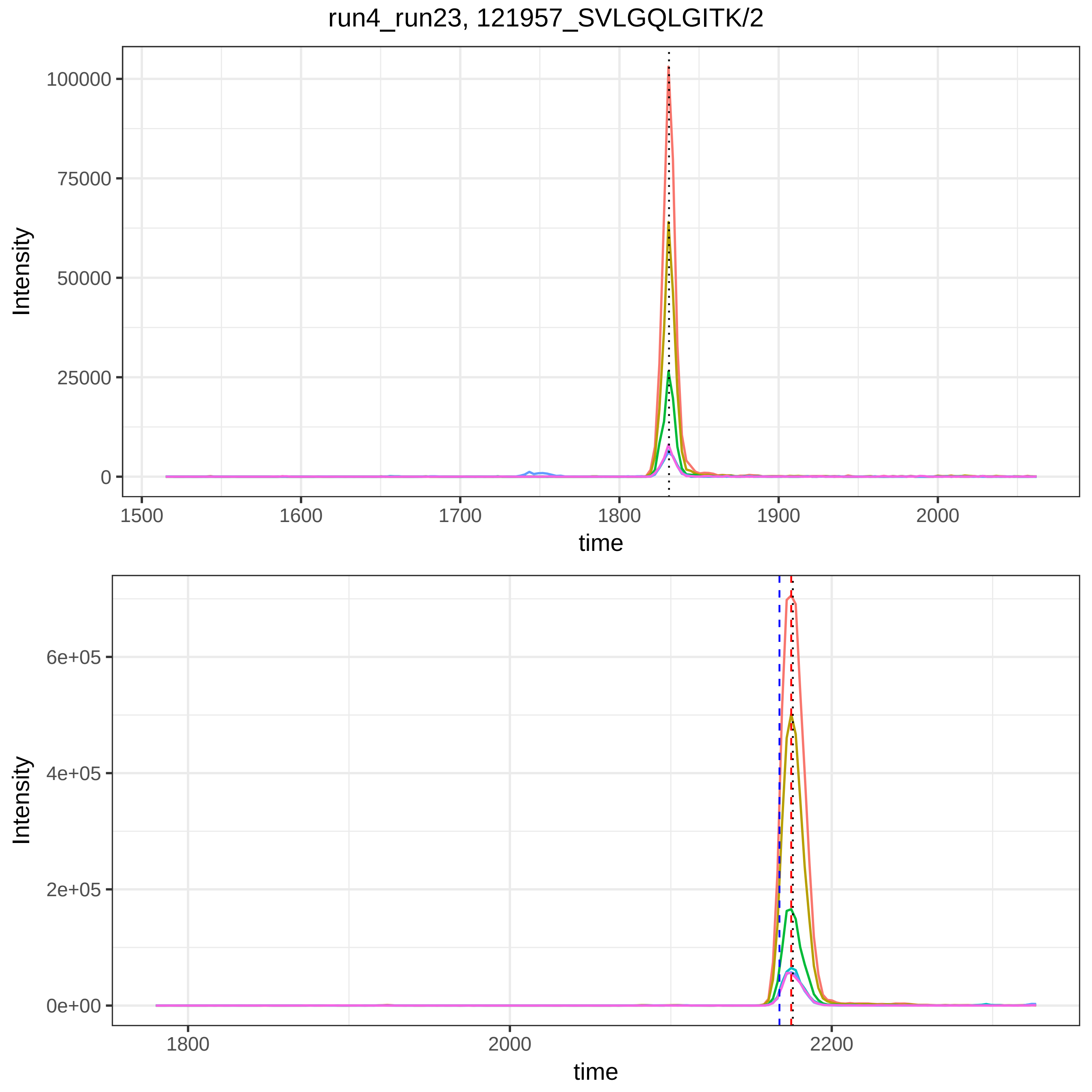
run4_run23, 121957_SVLGQLGITK/2
Intensity
100000
75000
50000
25000
0
1500
1600
1700
1800
1900
2000
time
Intensity
6e+05
4e+05
2e+05
0e+00
1800
2000
2200
time

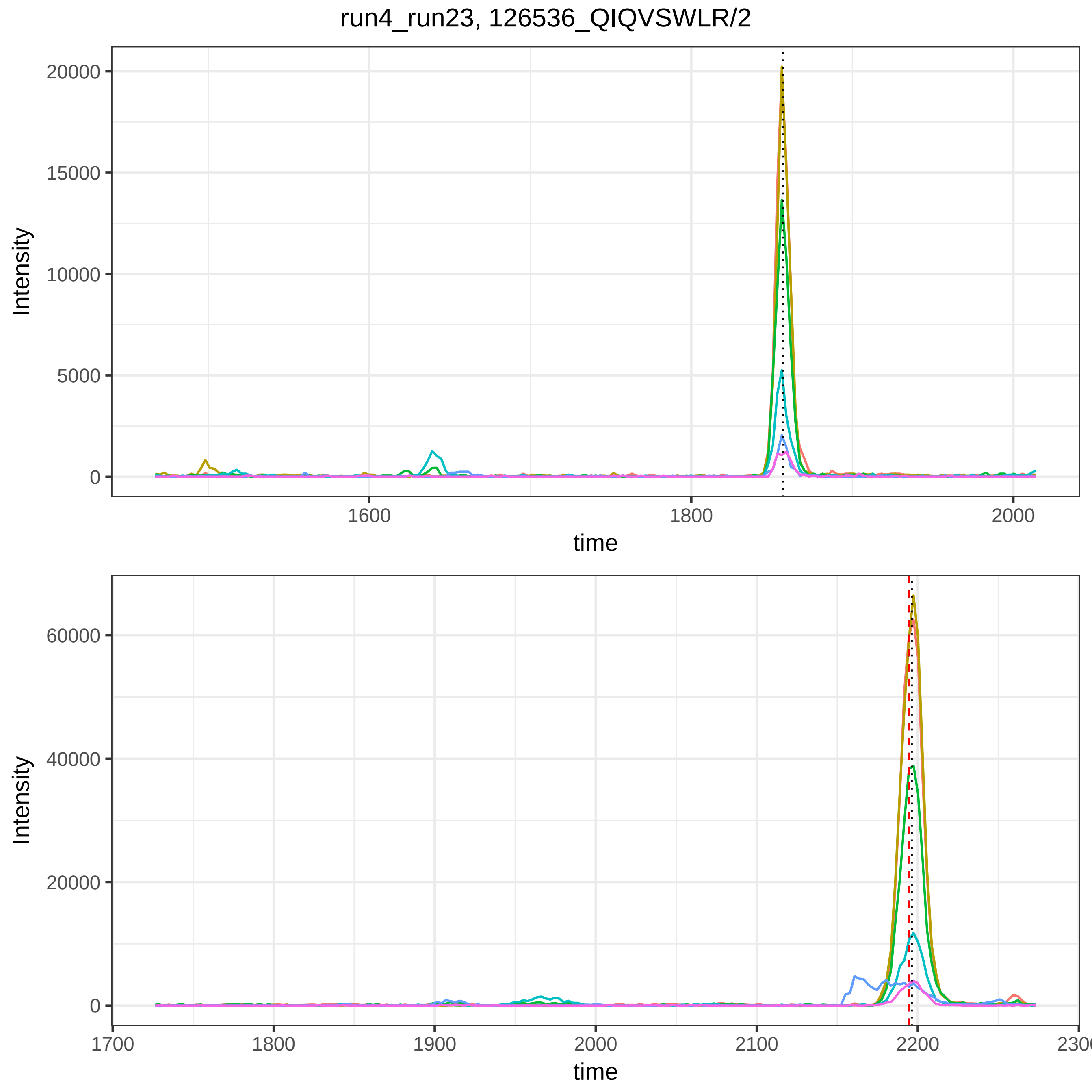
run4_run23, 126536_QIQVSWLR/2
Intensity
20000
15000
10000
5000
0
1600
1800
2000
time
Intensity
60000
40000
20000
0
1700
1800
1900
2000
2100
2200
time

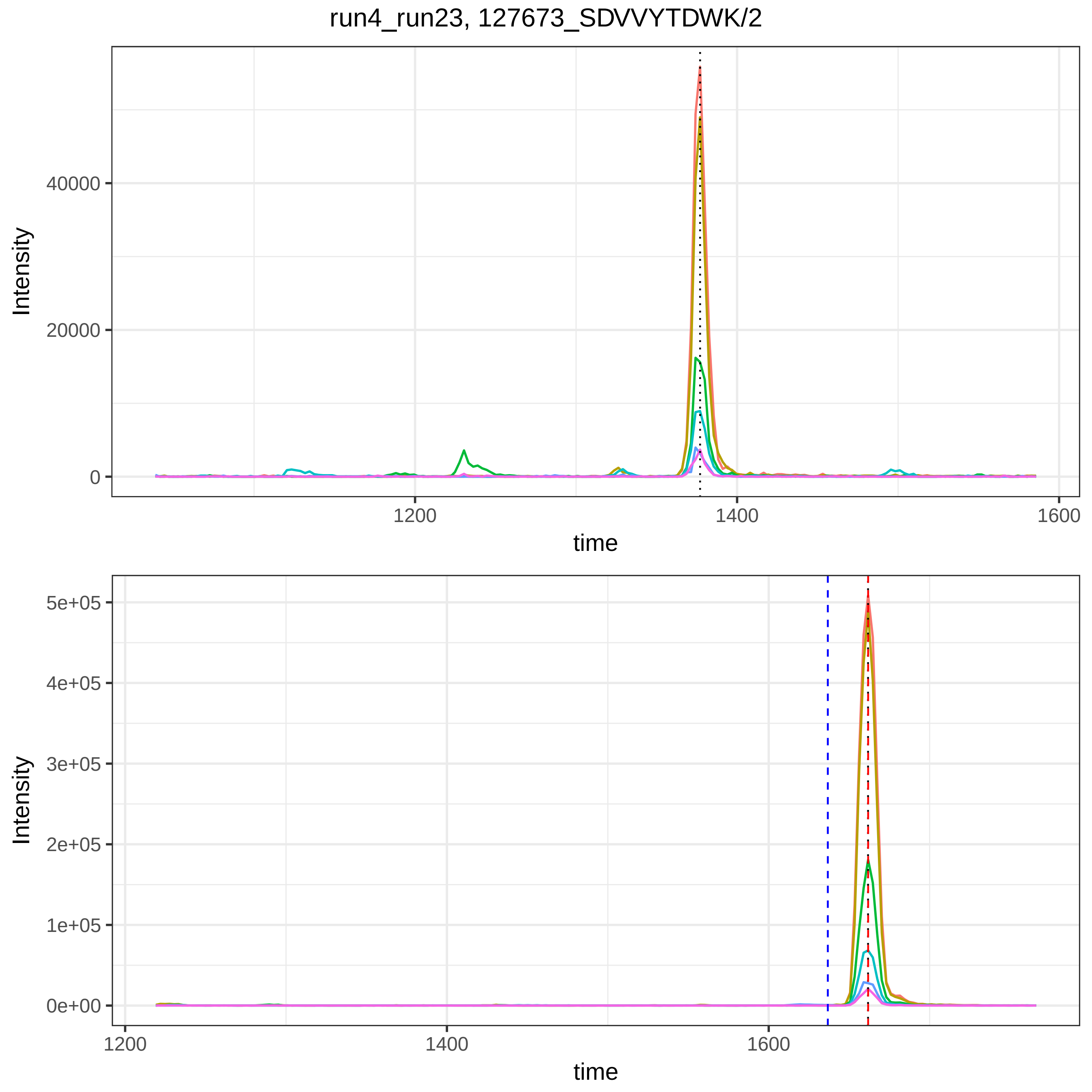
run4_run23, 127673_SDVVYTDWK/2
Intensity
40000
20000
0
1200
1400
1600
time
Intensity
5e+05
4e+05
3e+05
2e+05
1e+05
0e+00
1200
1400
1600
time

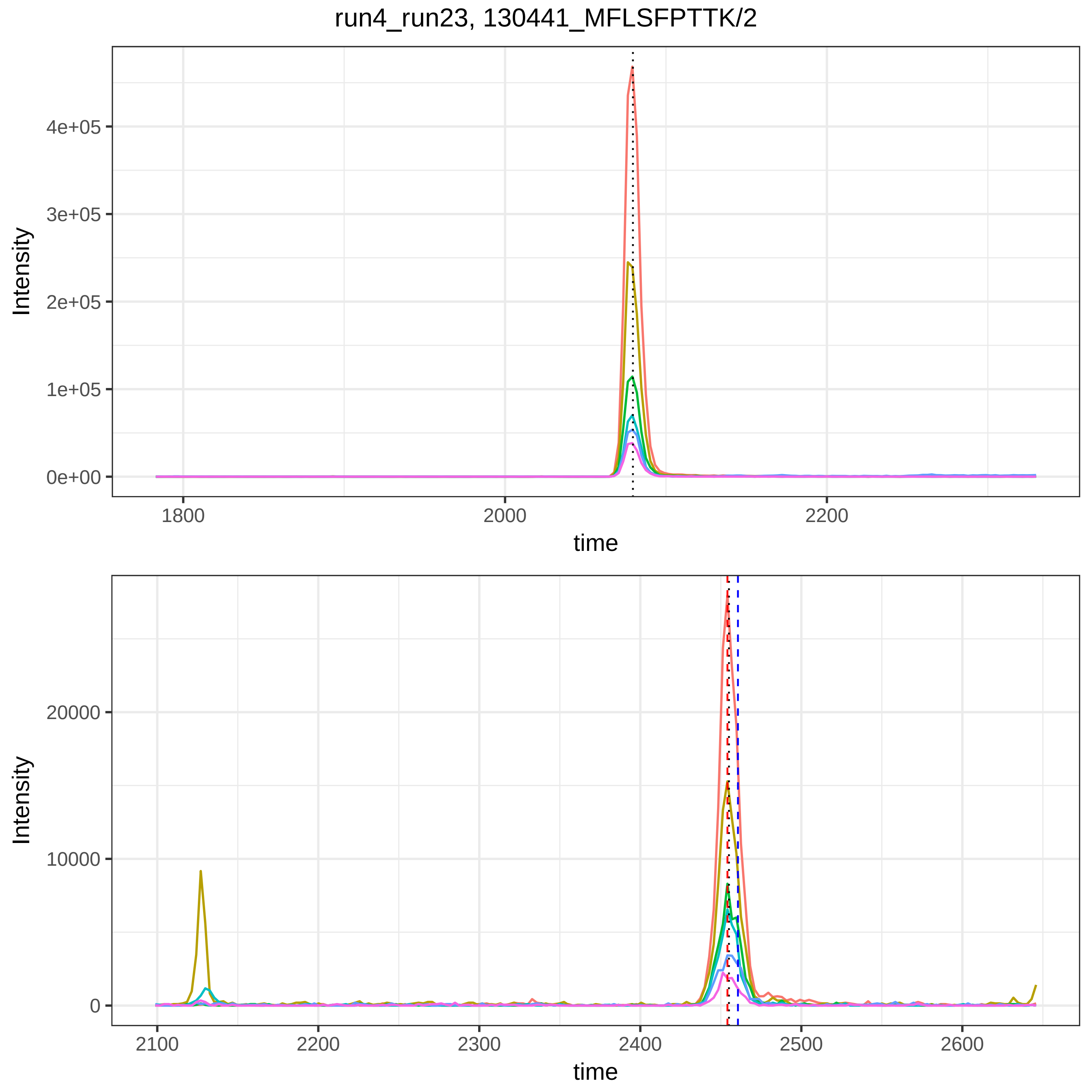
run4_run23, 130441_MFLSFPTTK/2
Intensity
4e+05
3e+05
2e+05
1e+05
0e+00
1800
2000
2200
time
Intensity
20000
10000
0
2100
2200
2300
2400
2500
2600
time

run4_run23, 131881_EIVMTQSPATLSVSPGER/2

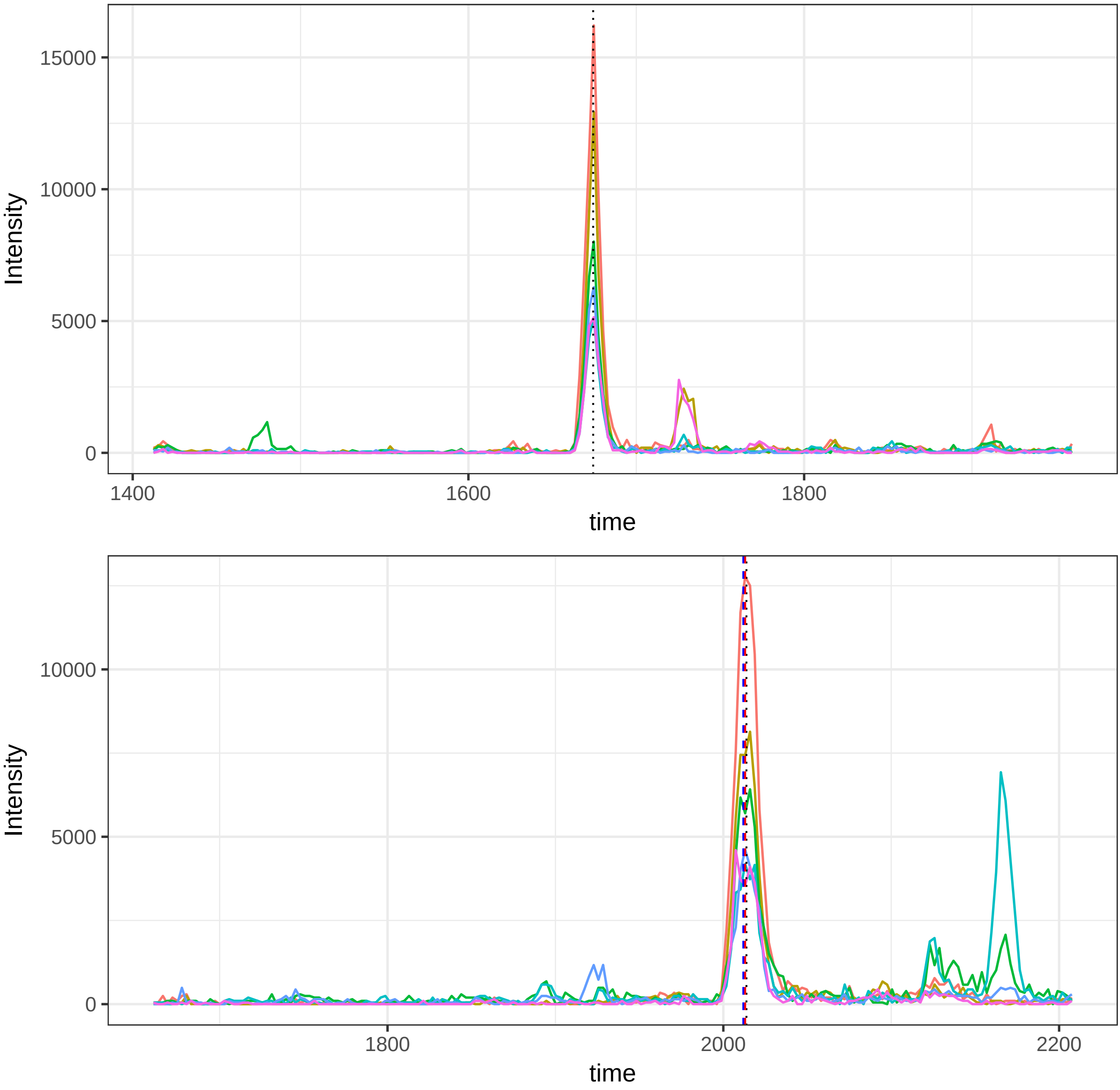

run4_run23, 131886_EIVMTQ(UniMod:7)SPATLSVSPGER/2

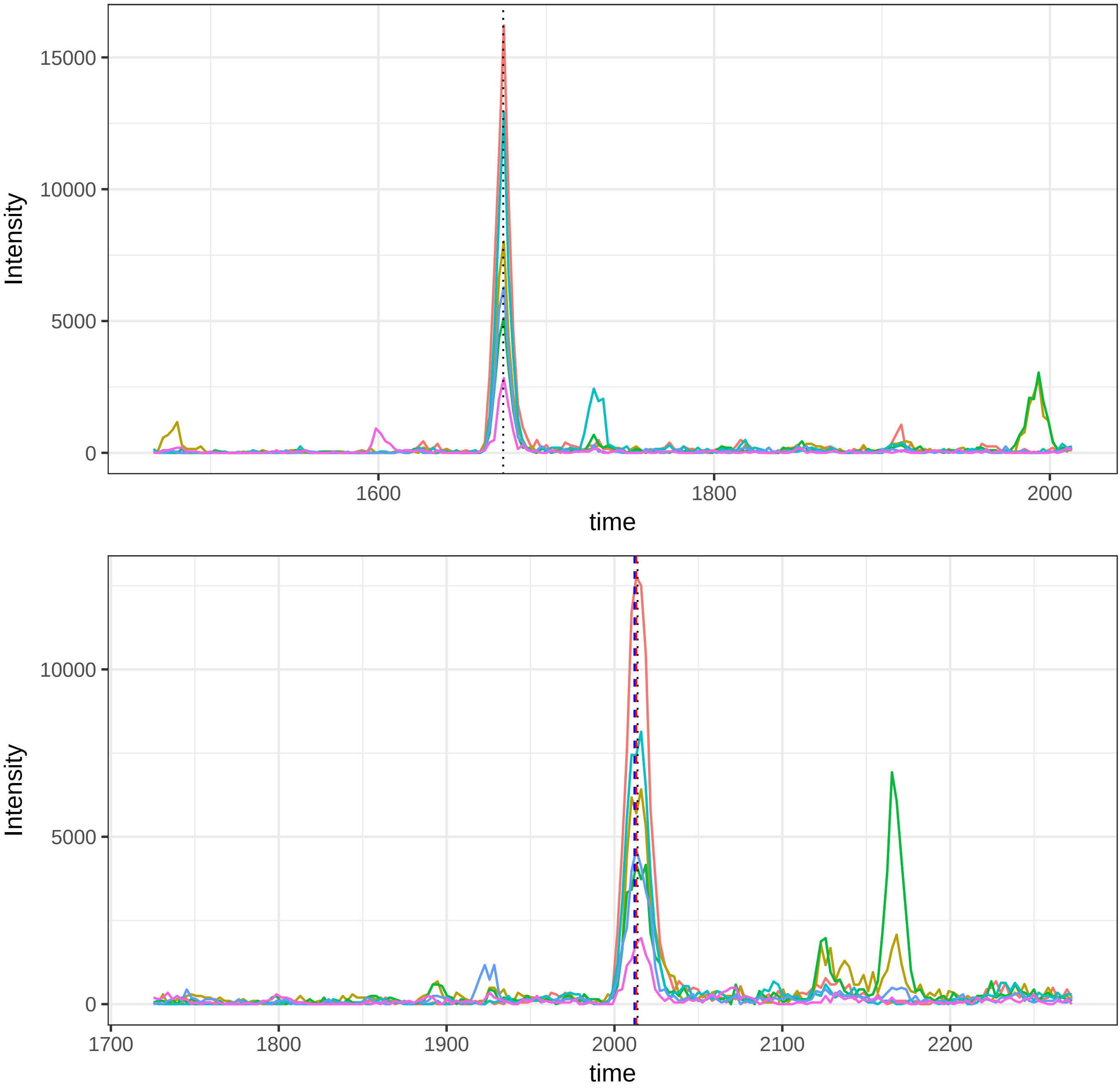

run4_run23, 131887_EIVM(UniMod:35)TQ(UniMod:7)SPATLSVSPGER/2

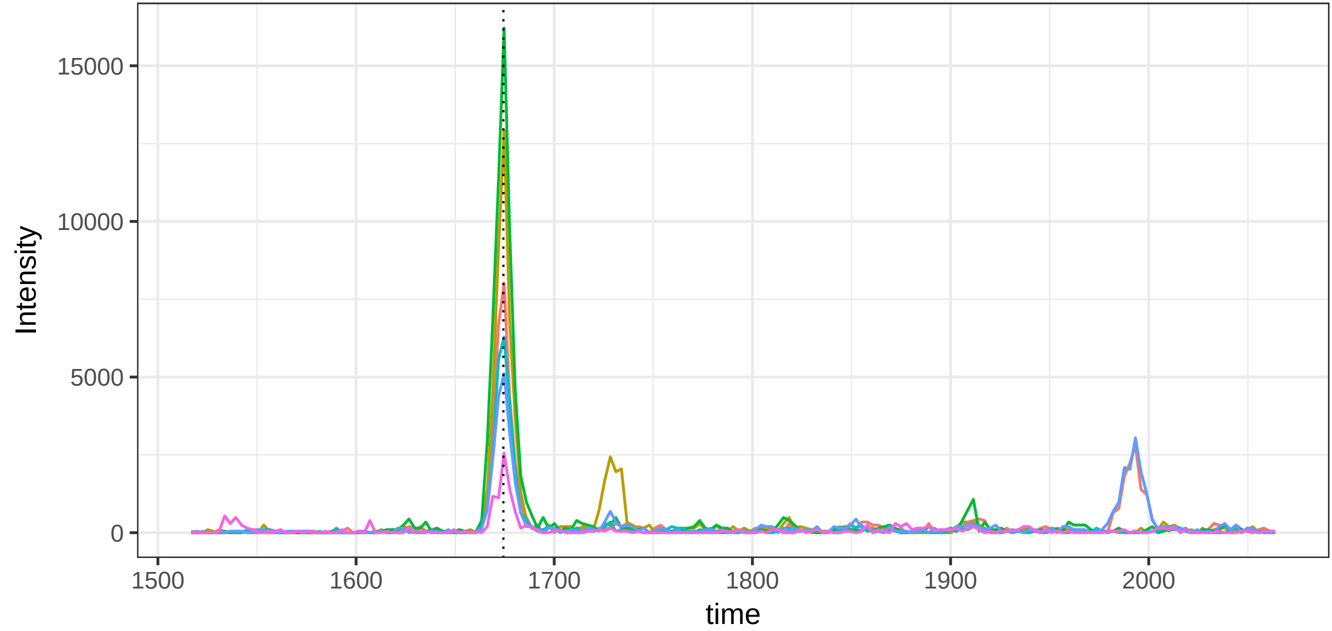


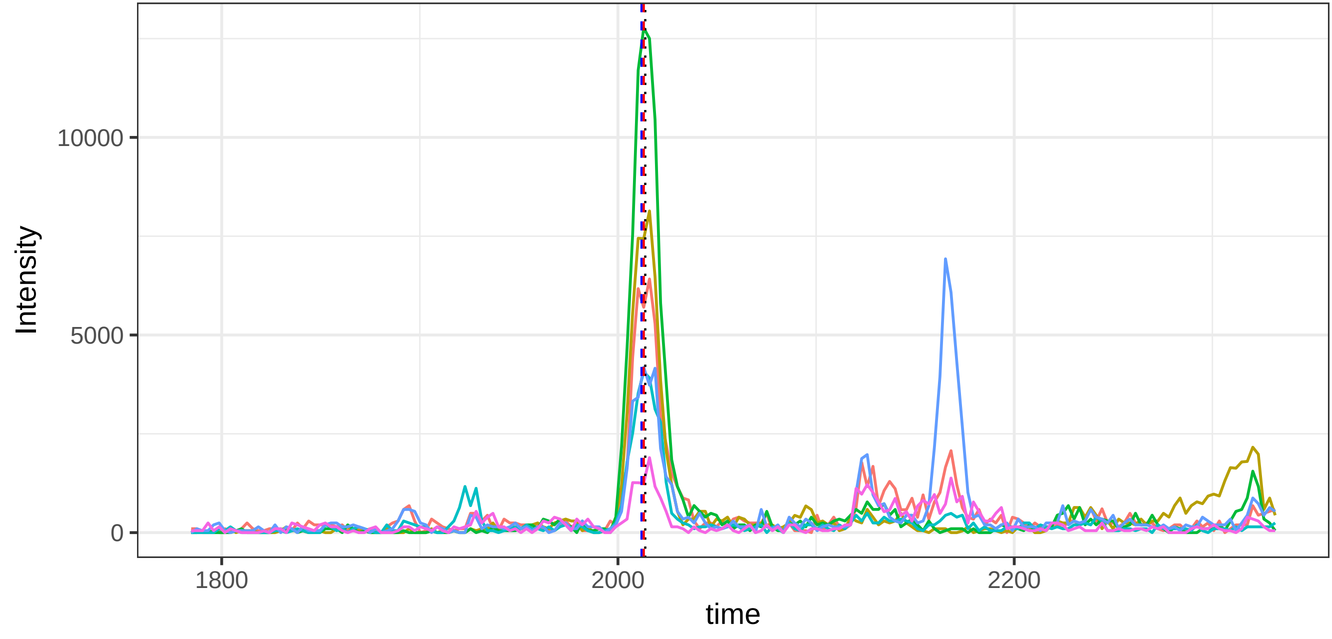

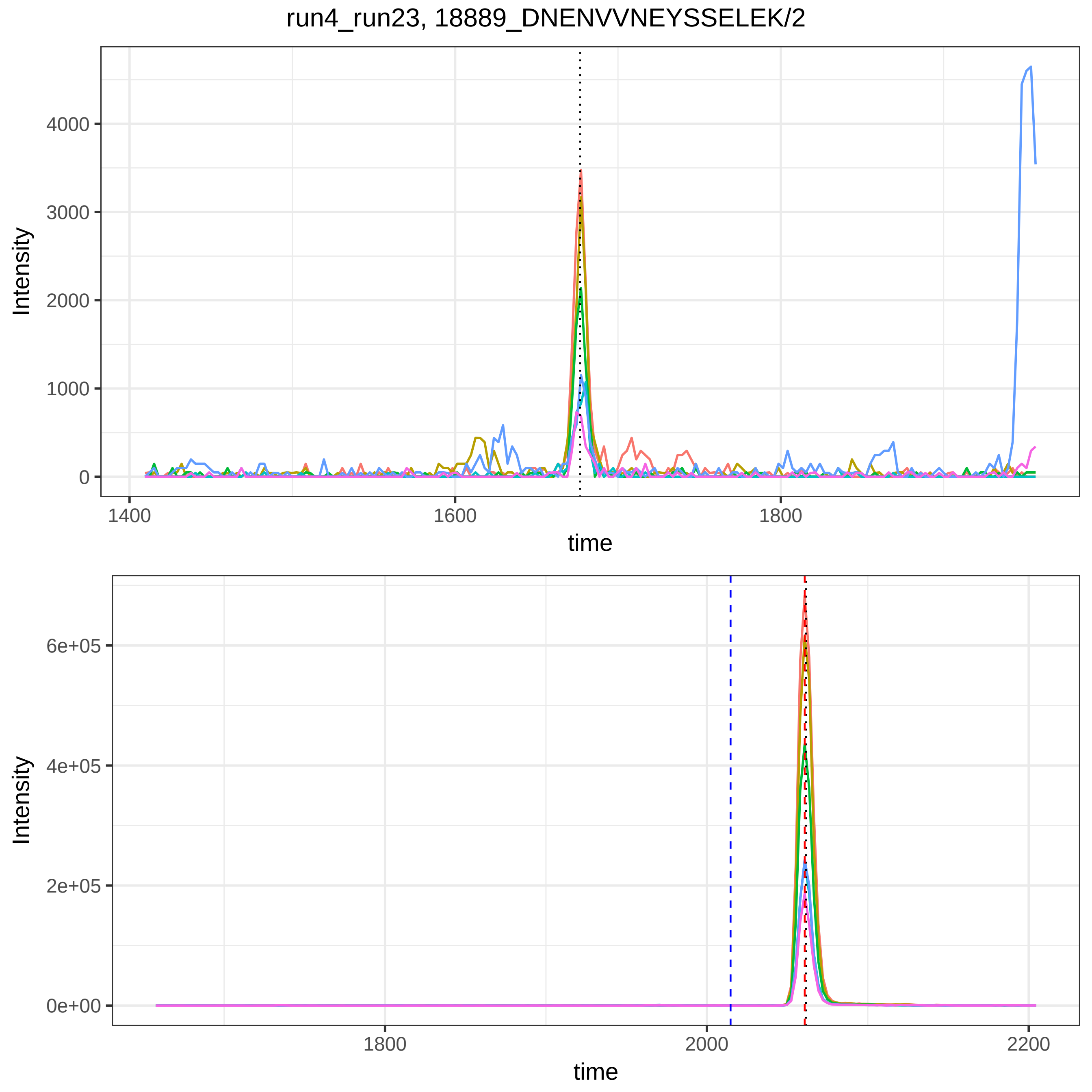

run4_run23, 18889_DNENVVNEYSSELEK/2
Intensity
4000
3000
2000
1000
0
1400
1600
1800
time
Intensity
6e+05
4e+05
2e+05
0e+00
1800
2000
2200
time

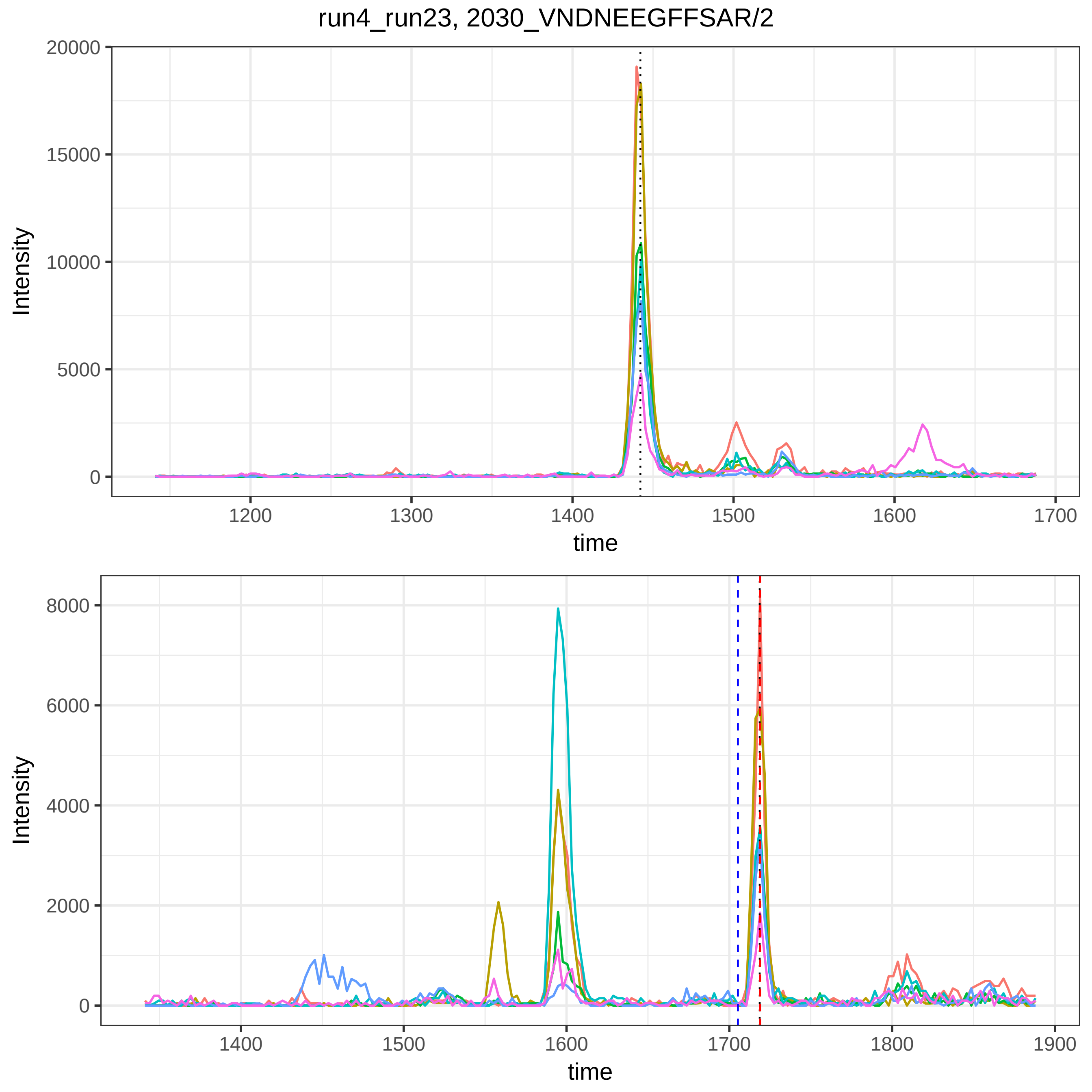

run4_run23, 2030_VNDNEEGFFSAR/2
20000
15000
10000
5000
0
Intensity
1200
1300
1400
1500
1600
1700
time
8000
6000
4000
2000
0
Intensity
1400
1500
1600
1700
1800
1900
time

run4_run23, 26917_VNEPSILEMSR/2

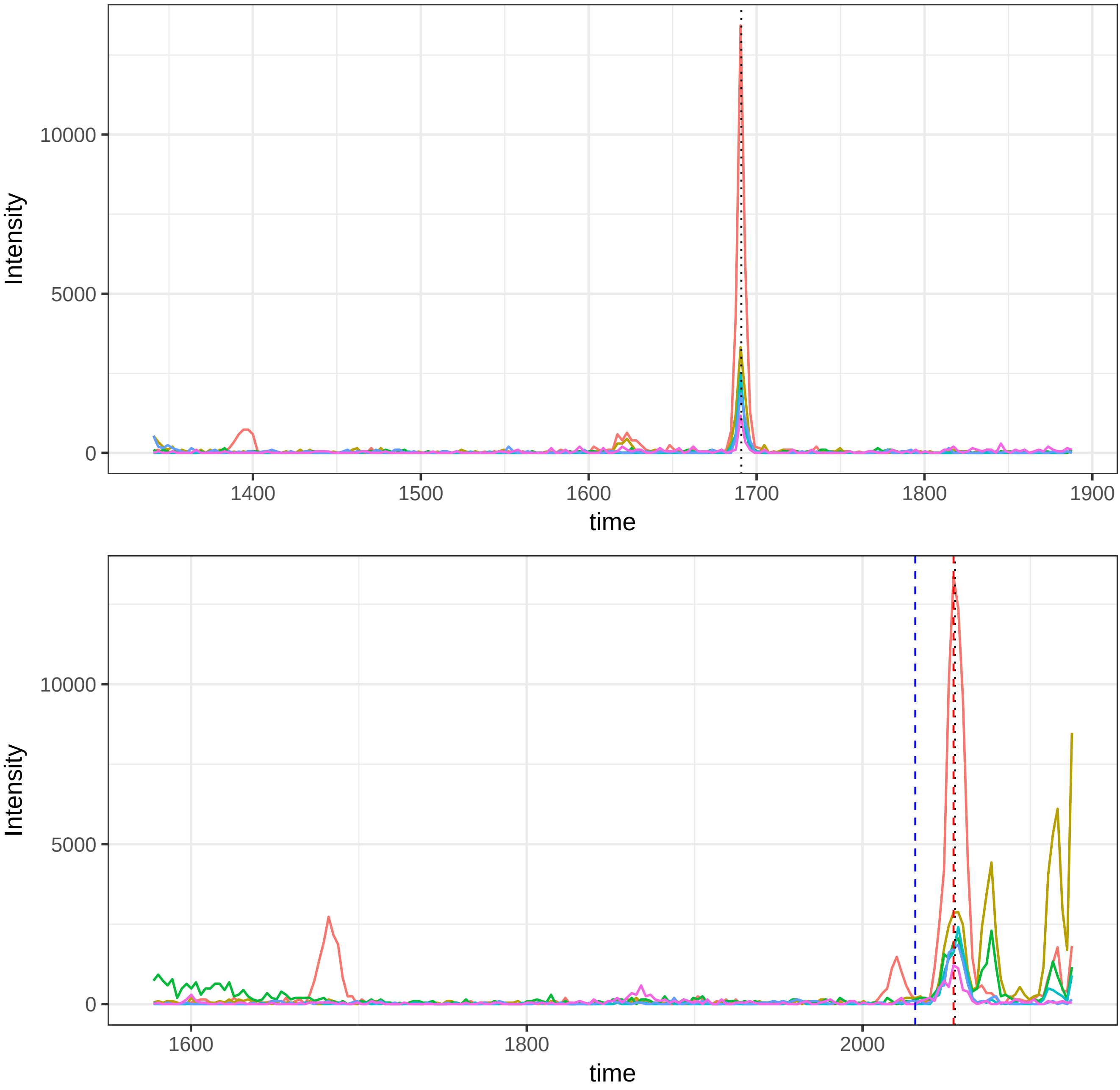

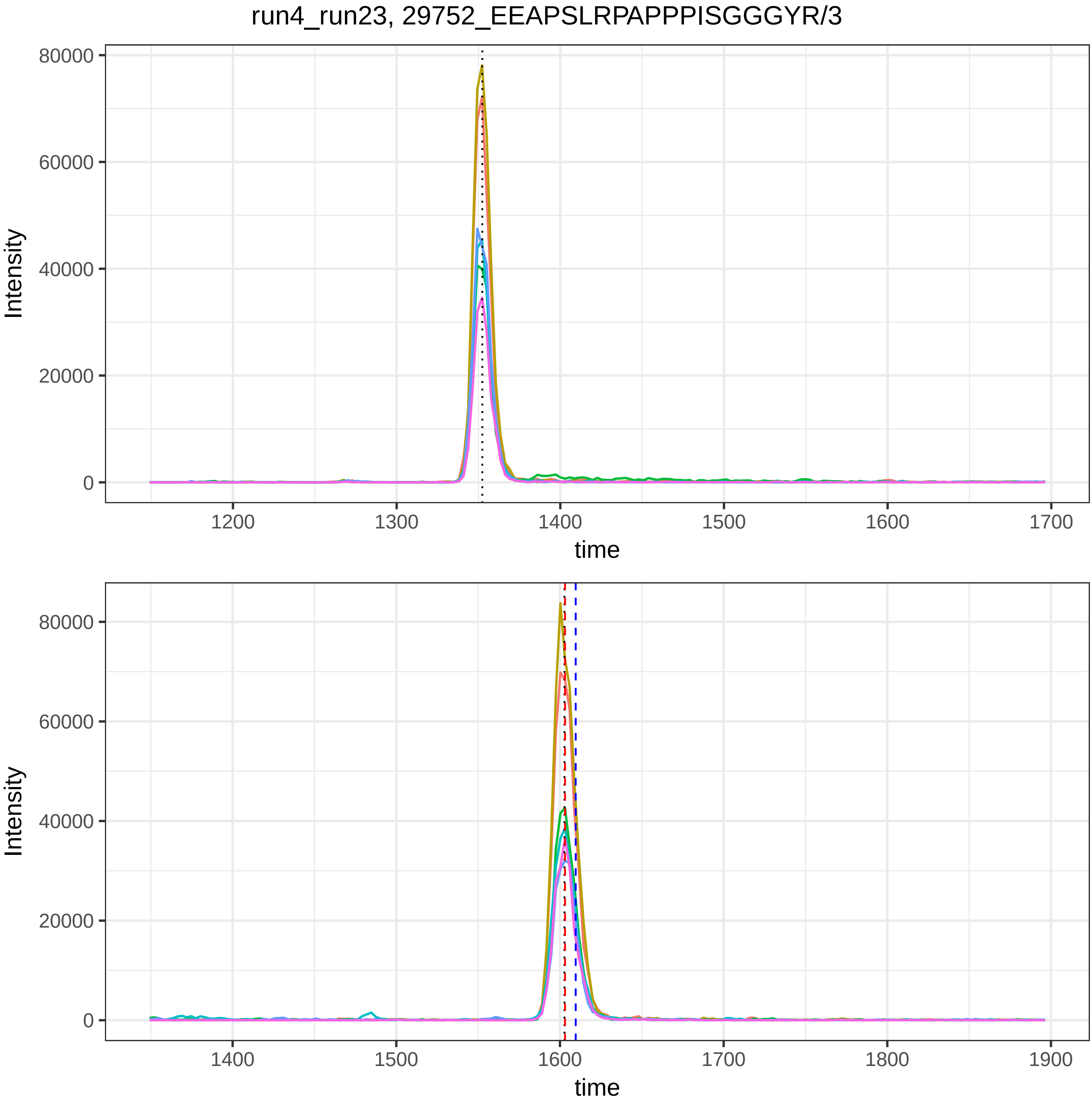

run4_run23, 29752_EEAPSLRPAPPPISGGGYR/3
Intensity
time
80000
60000
40000
20000
0
1200
1300
1400
1500
1600
1700
1800
1900

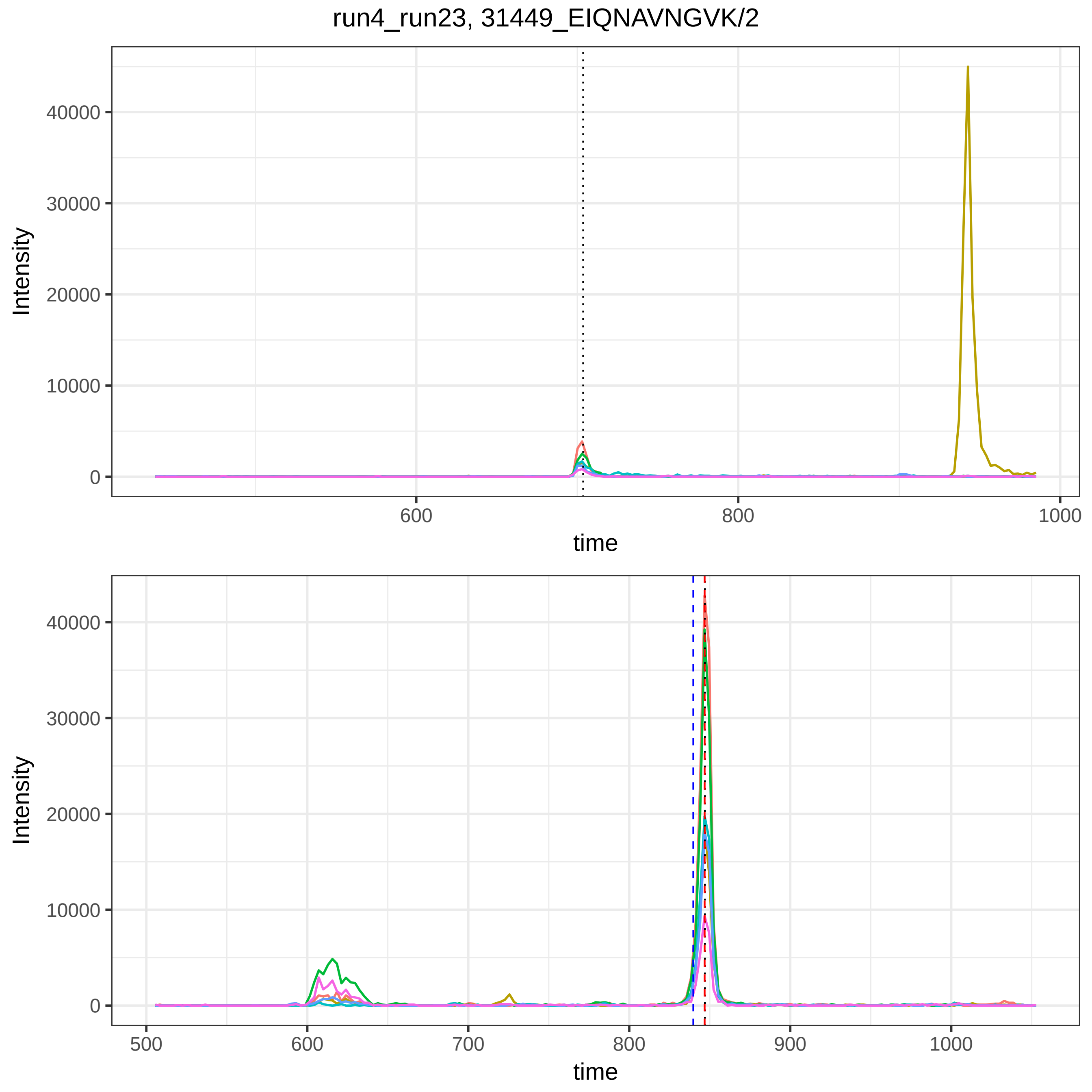

run4_run23, 31449_EIQNAVNGVK/2
Intensity
40000
30000
20000
10000
0
600
800
1000
time
Intensity
40000
30000
20000
10000
0
500
600
700
800
900
1000
time

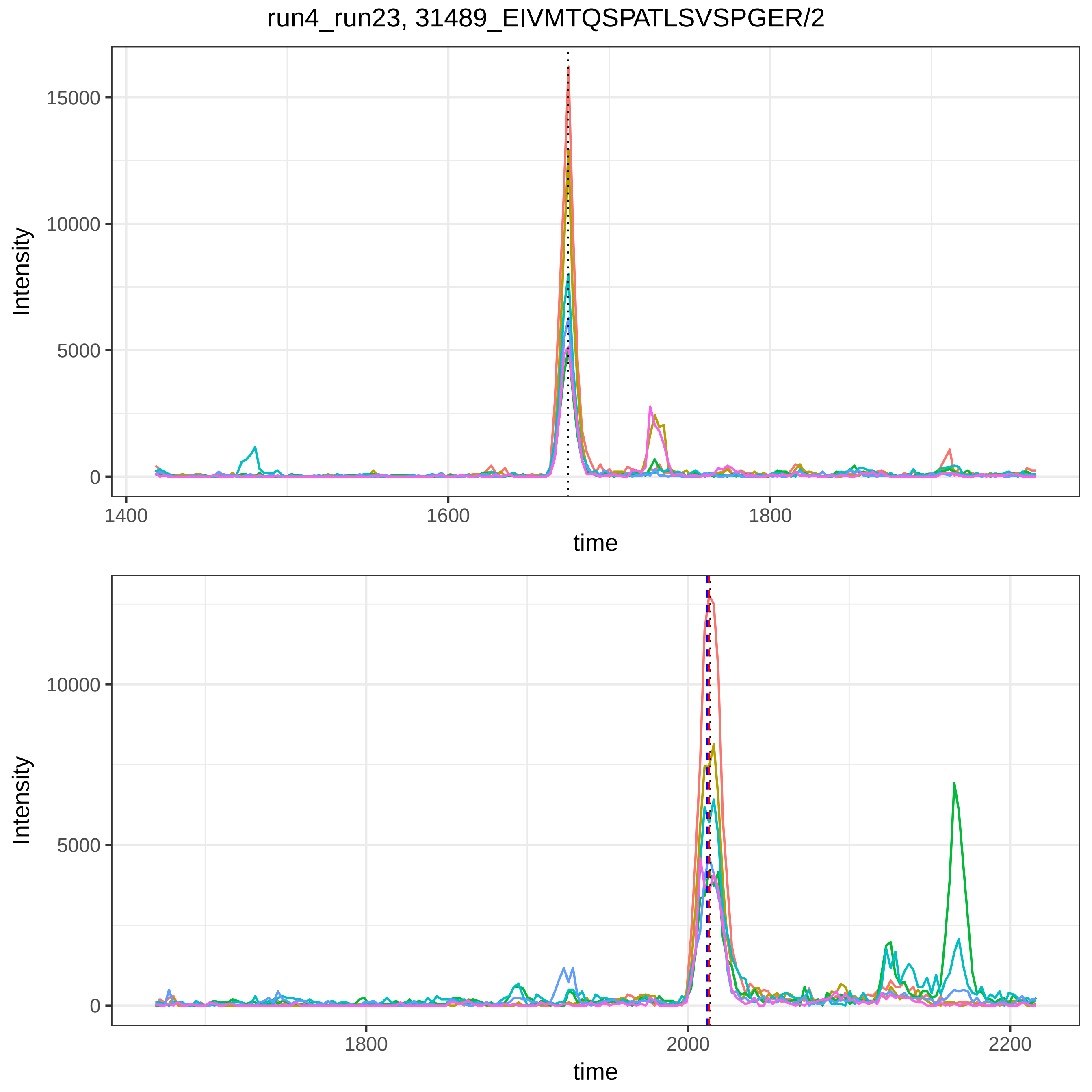

run4_run23, 31489_EIVMTQSPATLSVSPGER/2
Intensity
time
15000
10000
5000
0
1400
1600
1800
2000
2200

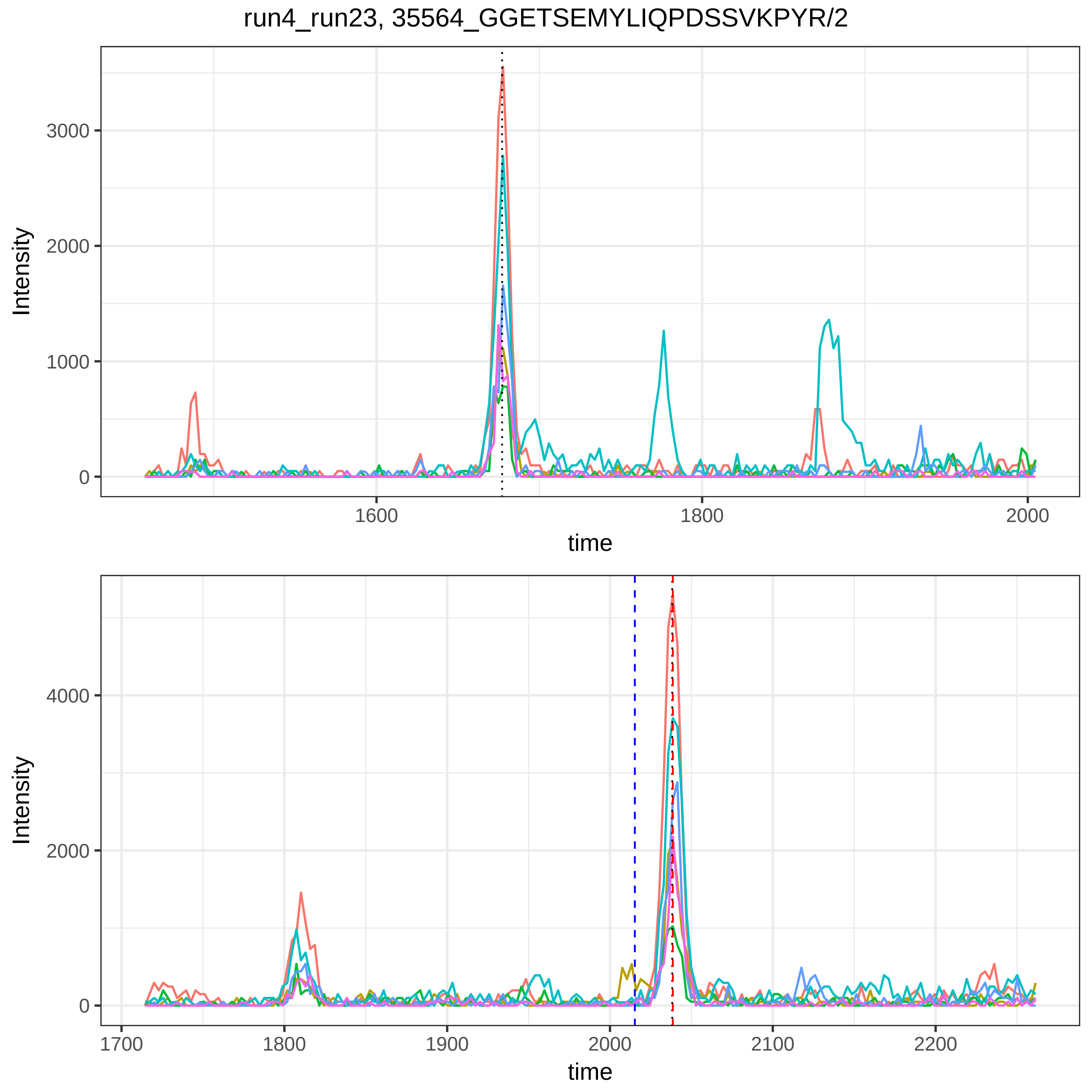
run4_run23, 35564_GGETSEMYLIQPDSSVKPYR/2
Intensity
3000
2000
1000
0
1600
1800
2000
time
Intensity
4000
2000
0
1700
1800
1900
2000
2100
2200
time

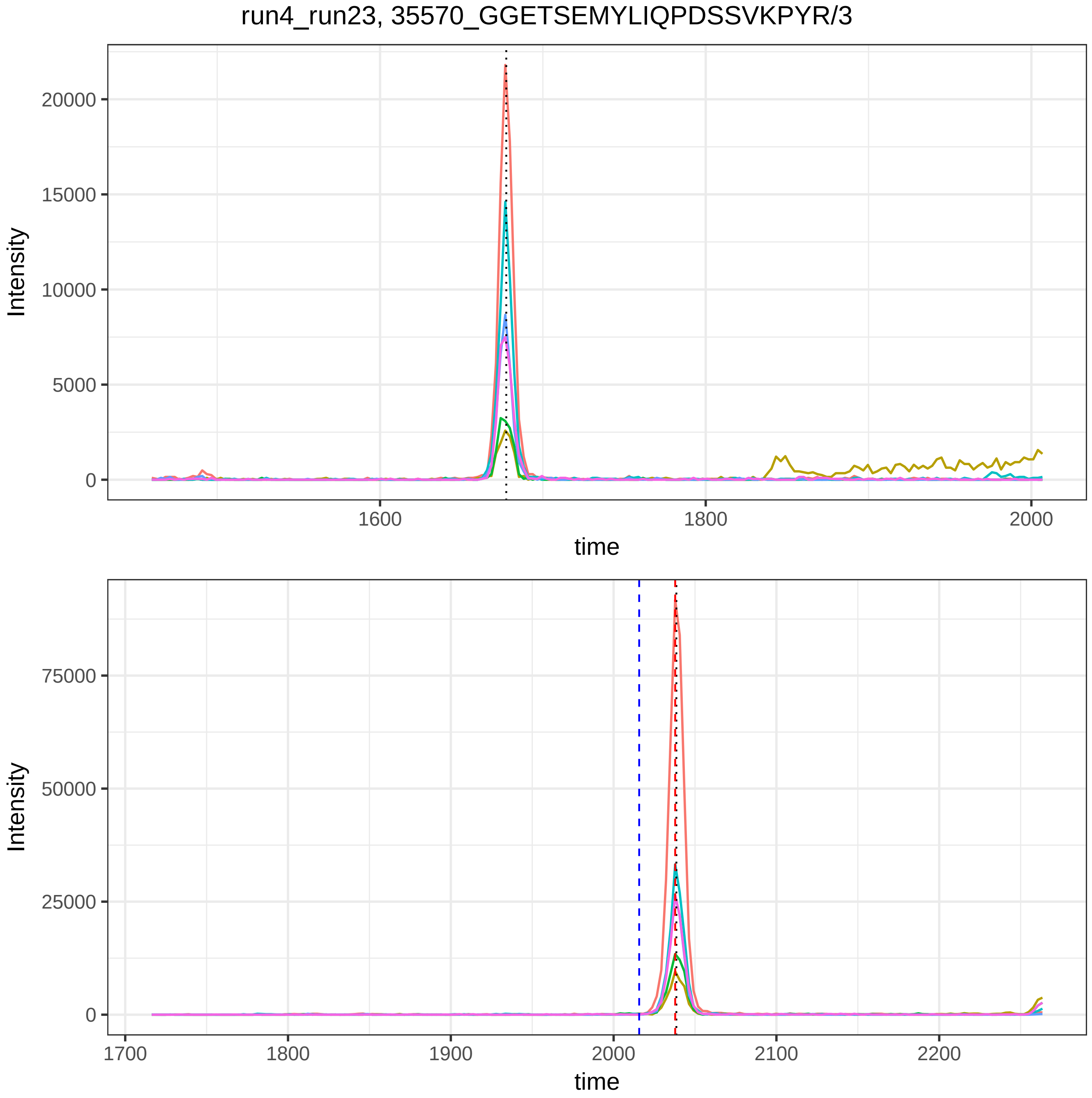

run4_run23, 35570_GGETSEMYLIQPDSSVKPYR/3
Intensity
20000
15000
10000
5000
0
1600
1800
2000
time
Intensity
75000
50000
25000
0
1700
1800
1900
2000
2100
2200
time

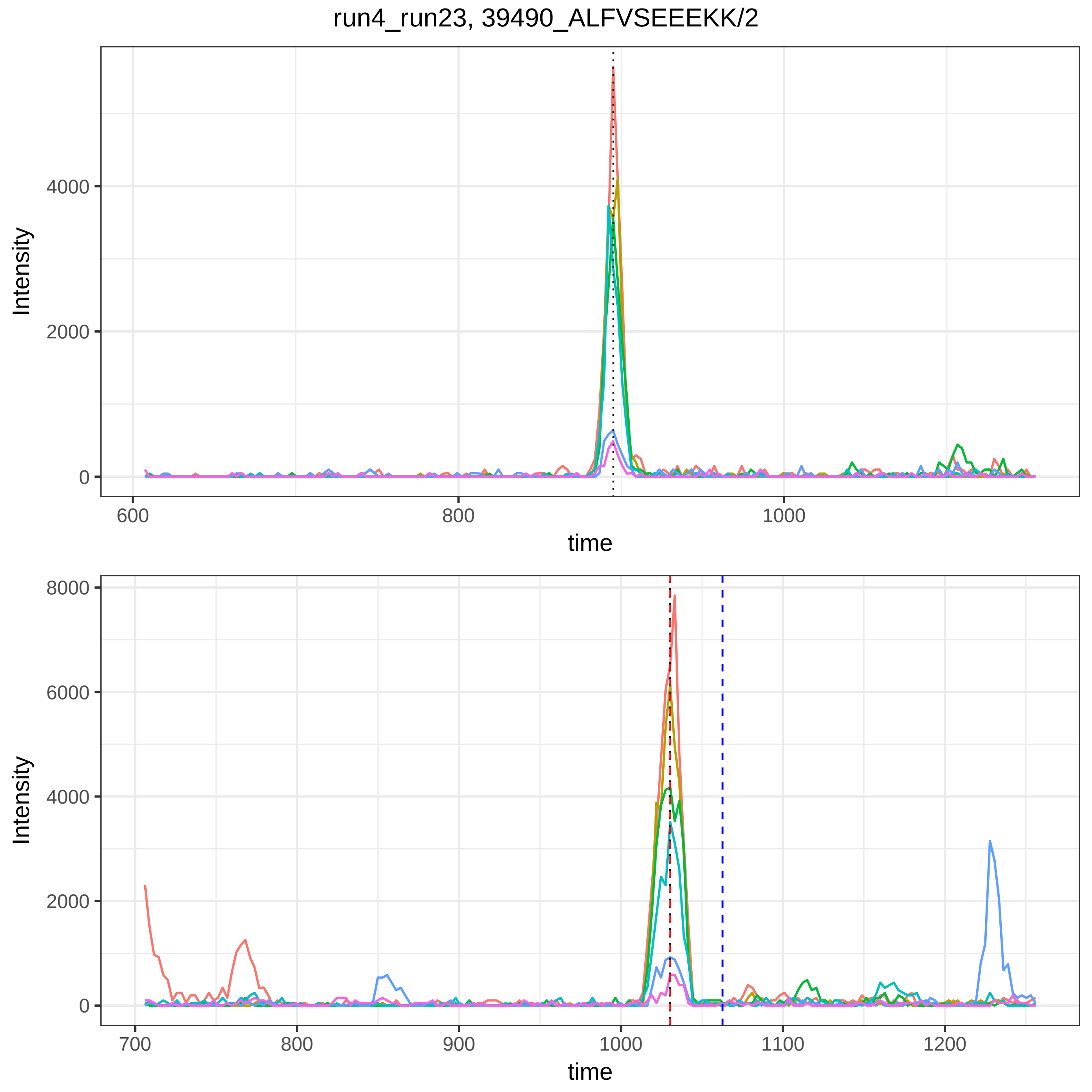
run4_run23, 39490_ALFVSEEEKK/2
Intensity
4000
2000
0
600
800
1000
time
Intensity
8000
6000
4000
2000
0
700
800
900
1000
1100
1200
time

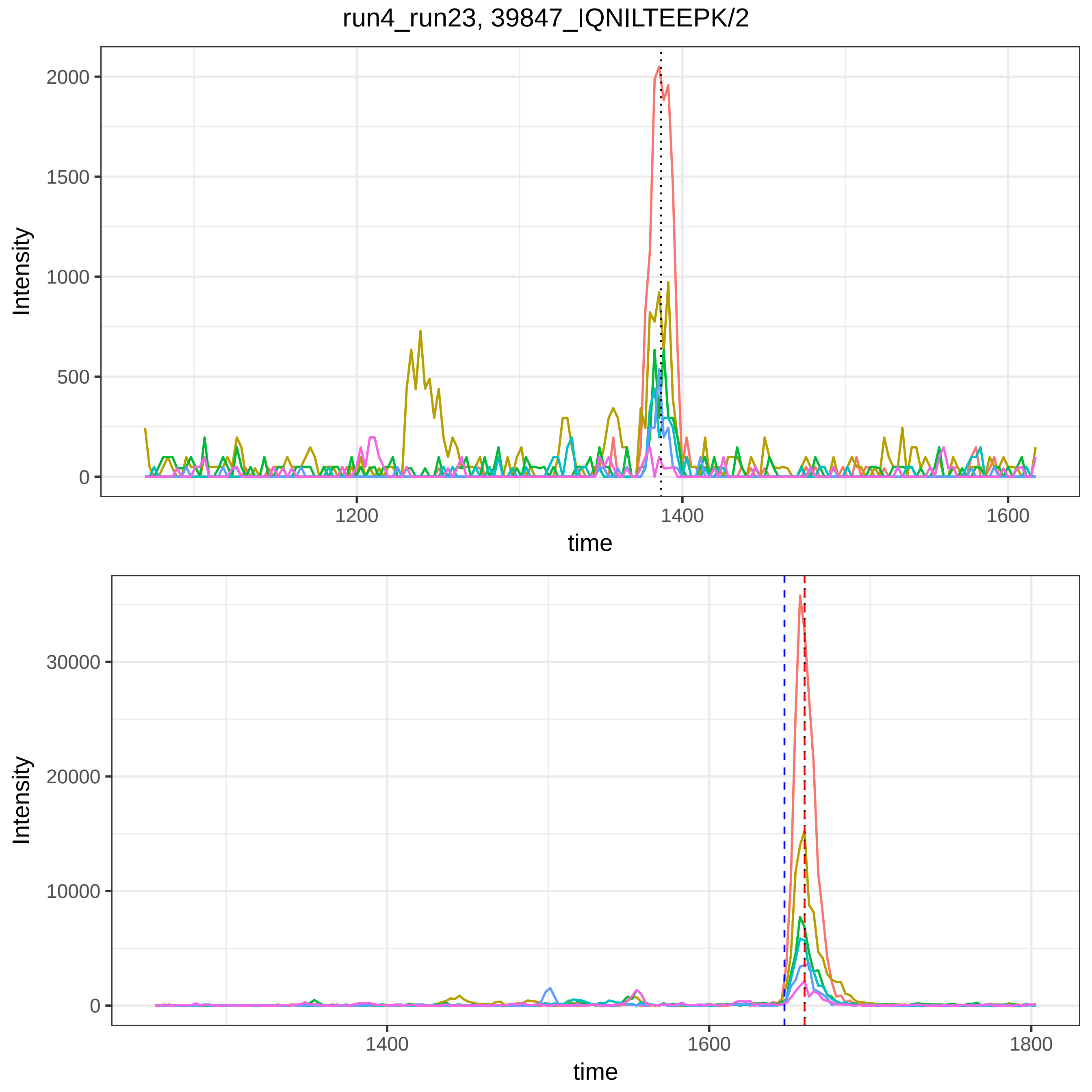
run4_run23, 39847_IQNILTEEPK/2
Intensity
2000
1500
1000
500
0
1200
1400
1600
time
Intensity
30000
20000
10000
0
1400
1600
1800
time

run4_run23, 42193_KYFIDFVAR/2

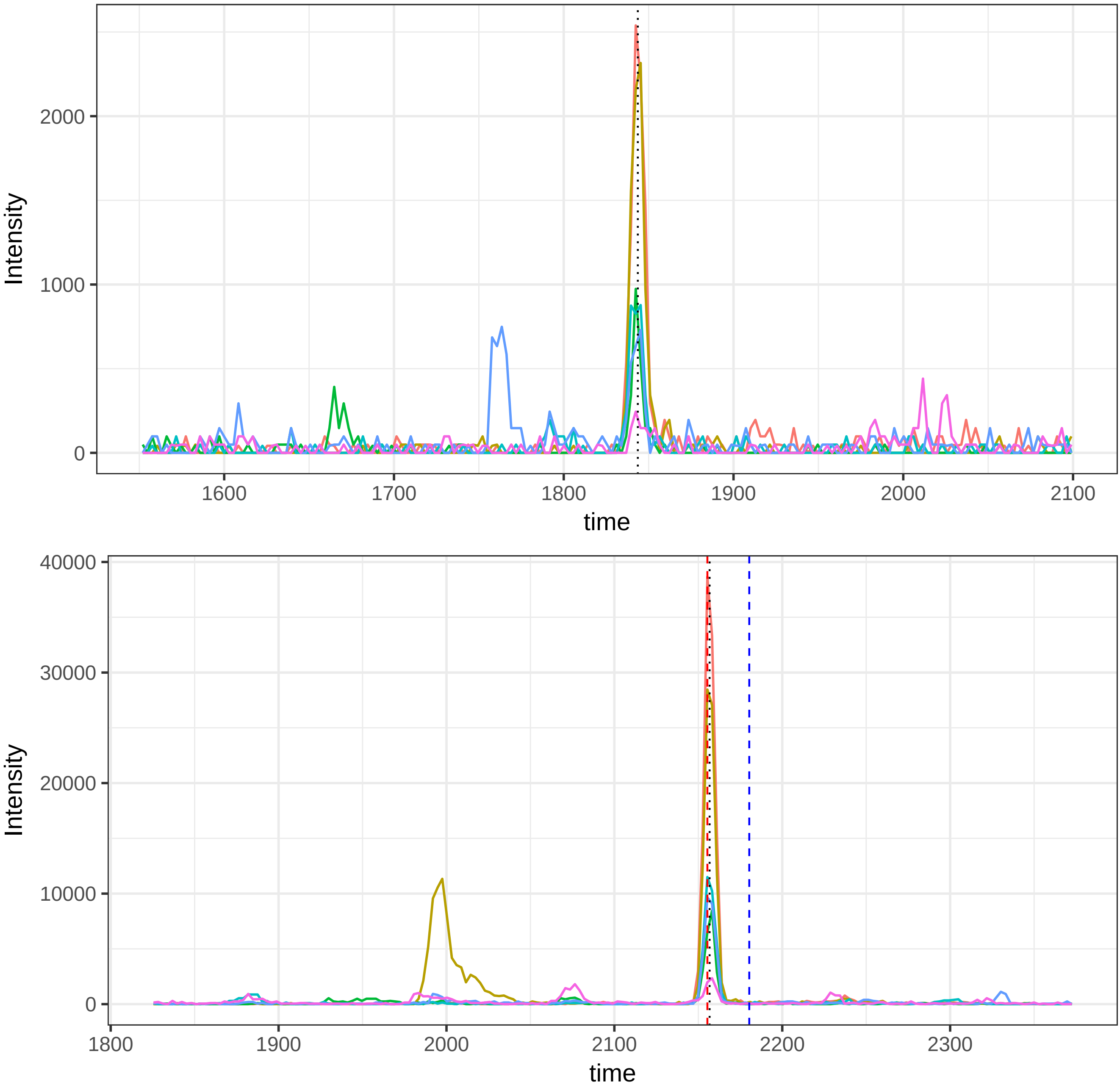

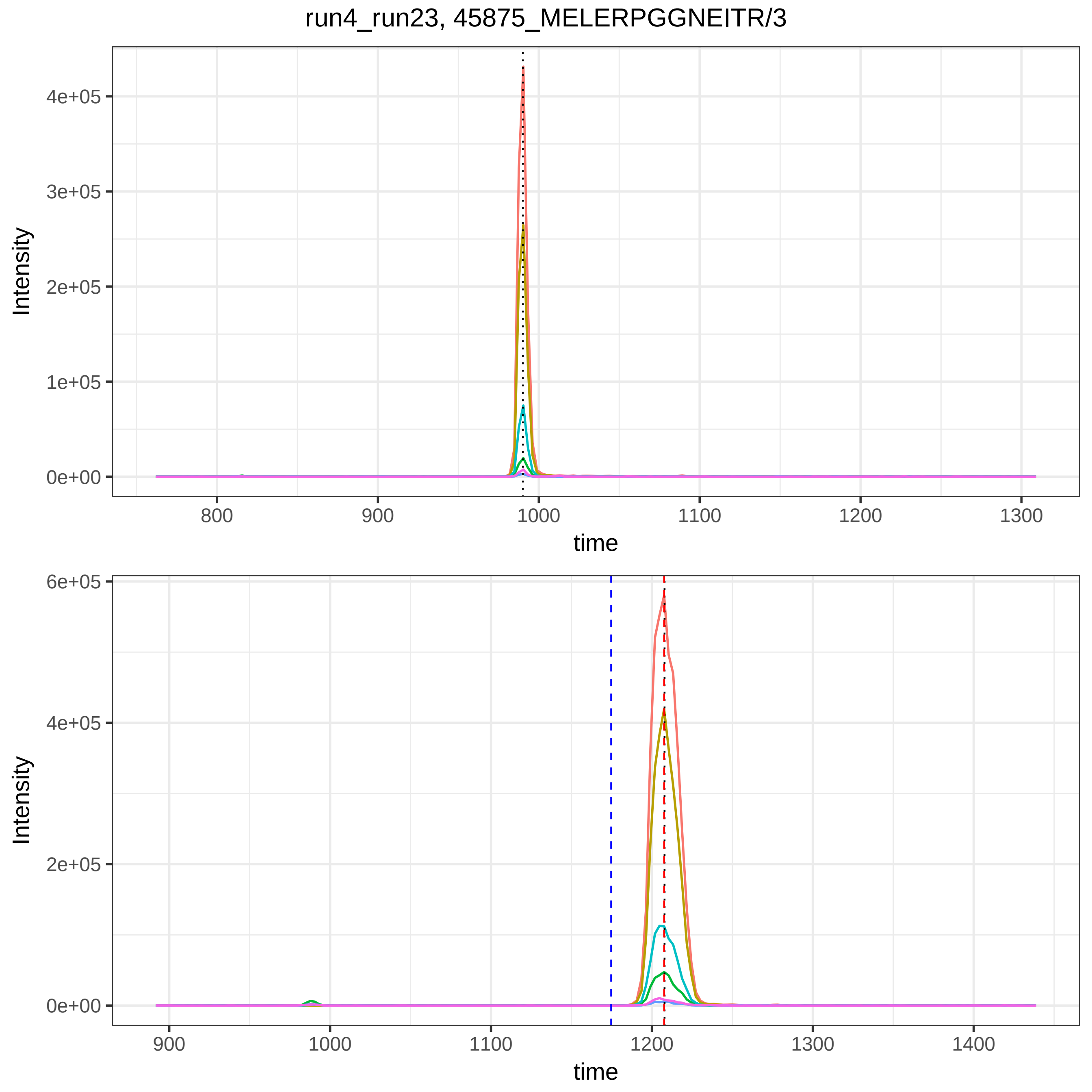
run4_run23, 45875_MELERPGGNEITR/3
Intensity
4e+05
3e+05
2e+05
1e+05
0e+00
800
900
1000
1100
1200
1300
time
Intensity
6e+05
4e+05
2e+05
0e+00
900
1000
1100
1200
1300
1400
time

run4_run23, 46142_AQLVDMK/2

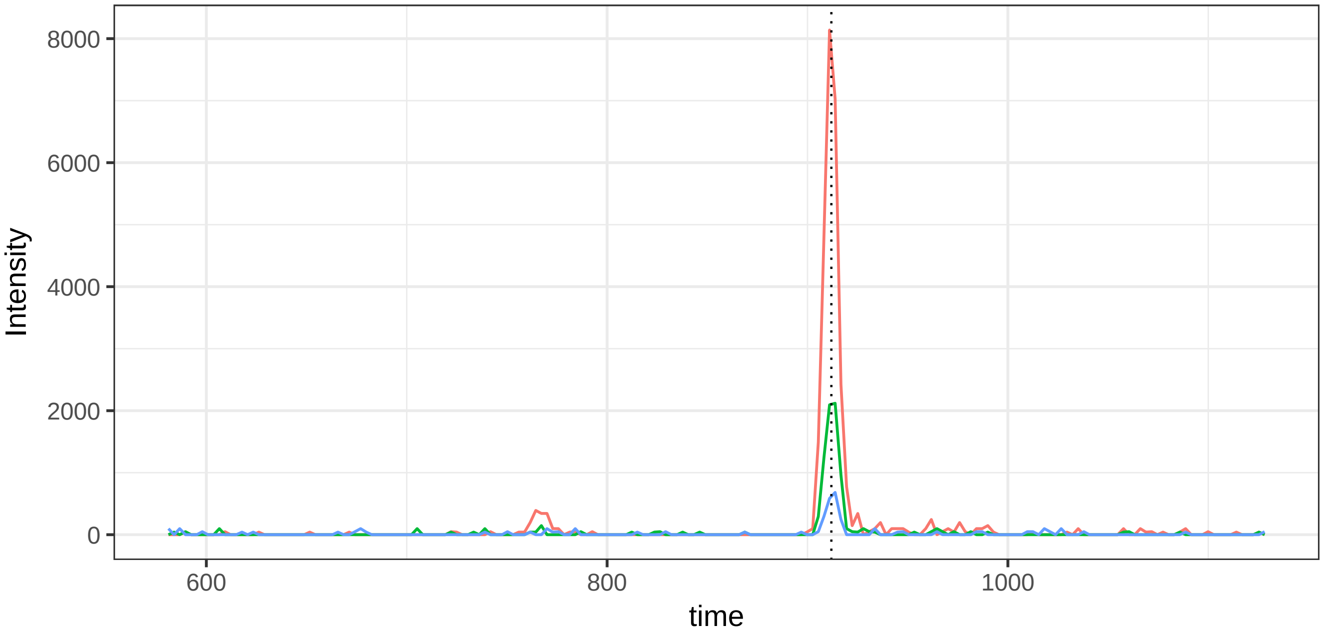


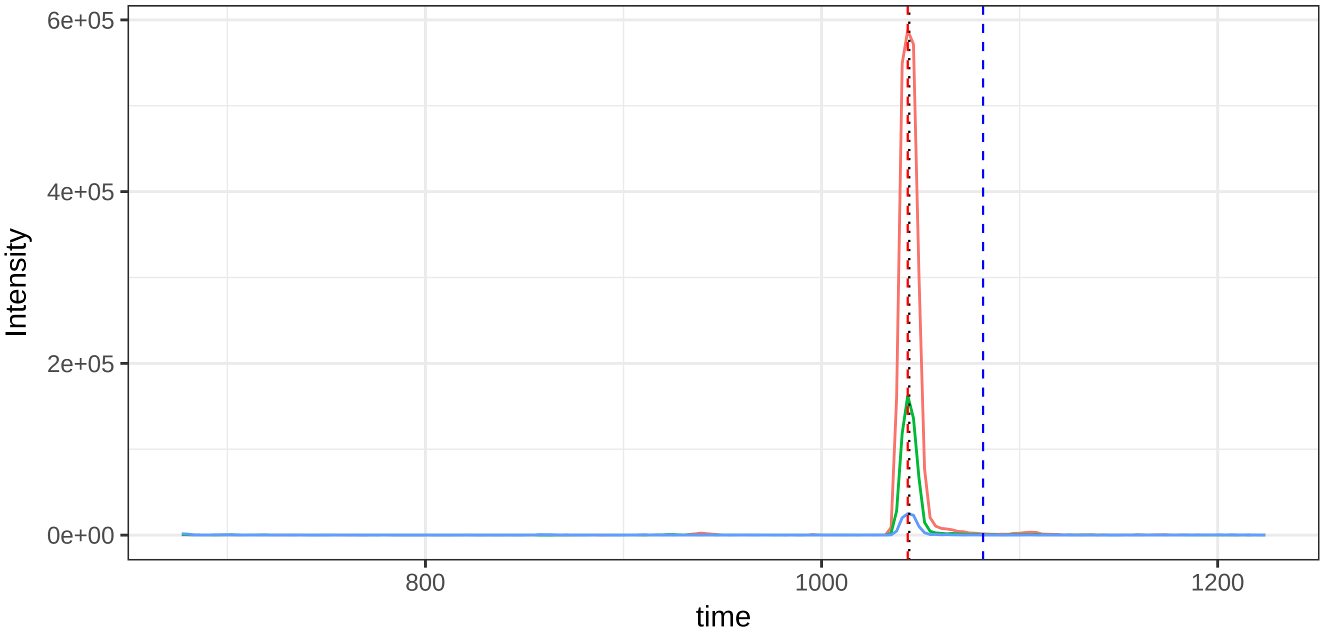

run4_run23, 47477_NLAVSQVVHK/2

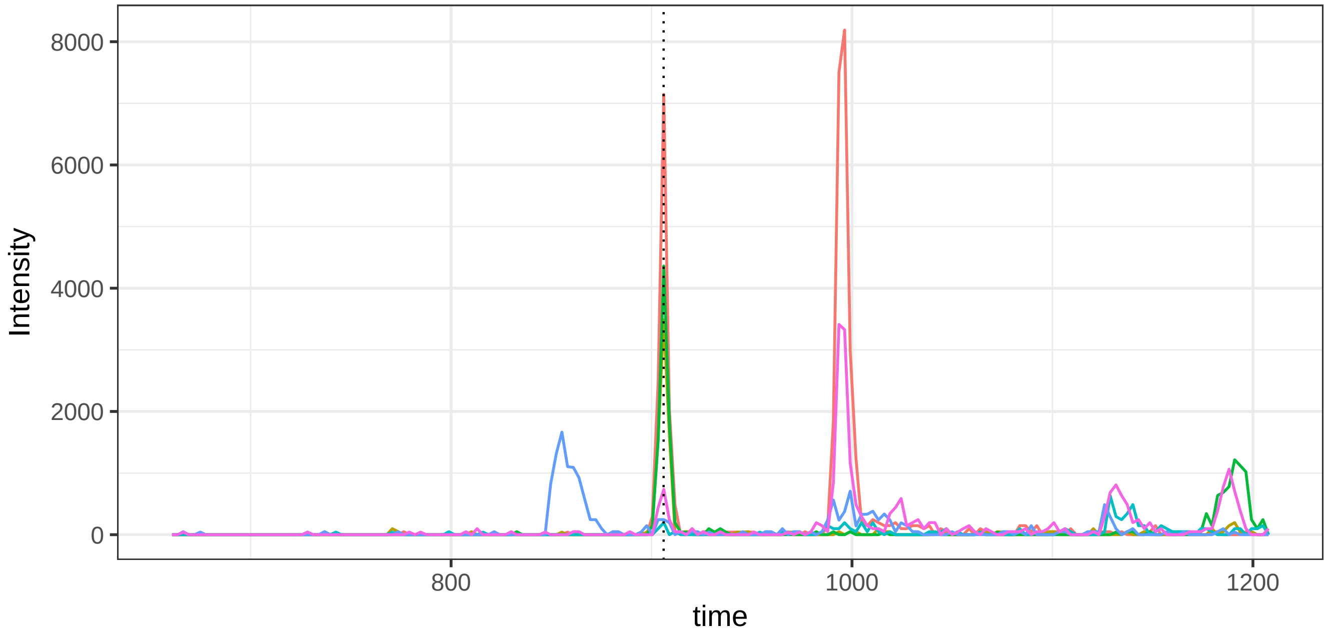


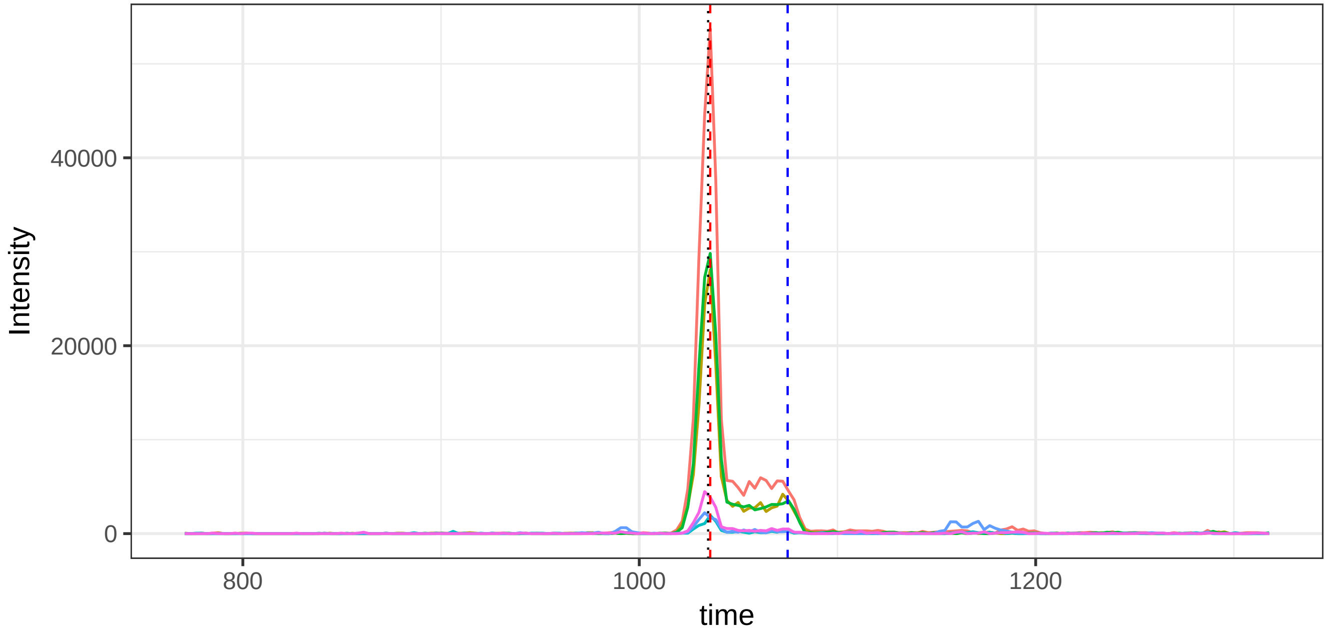

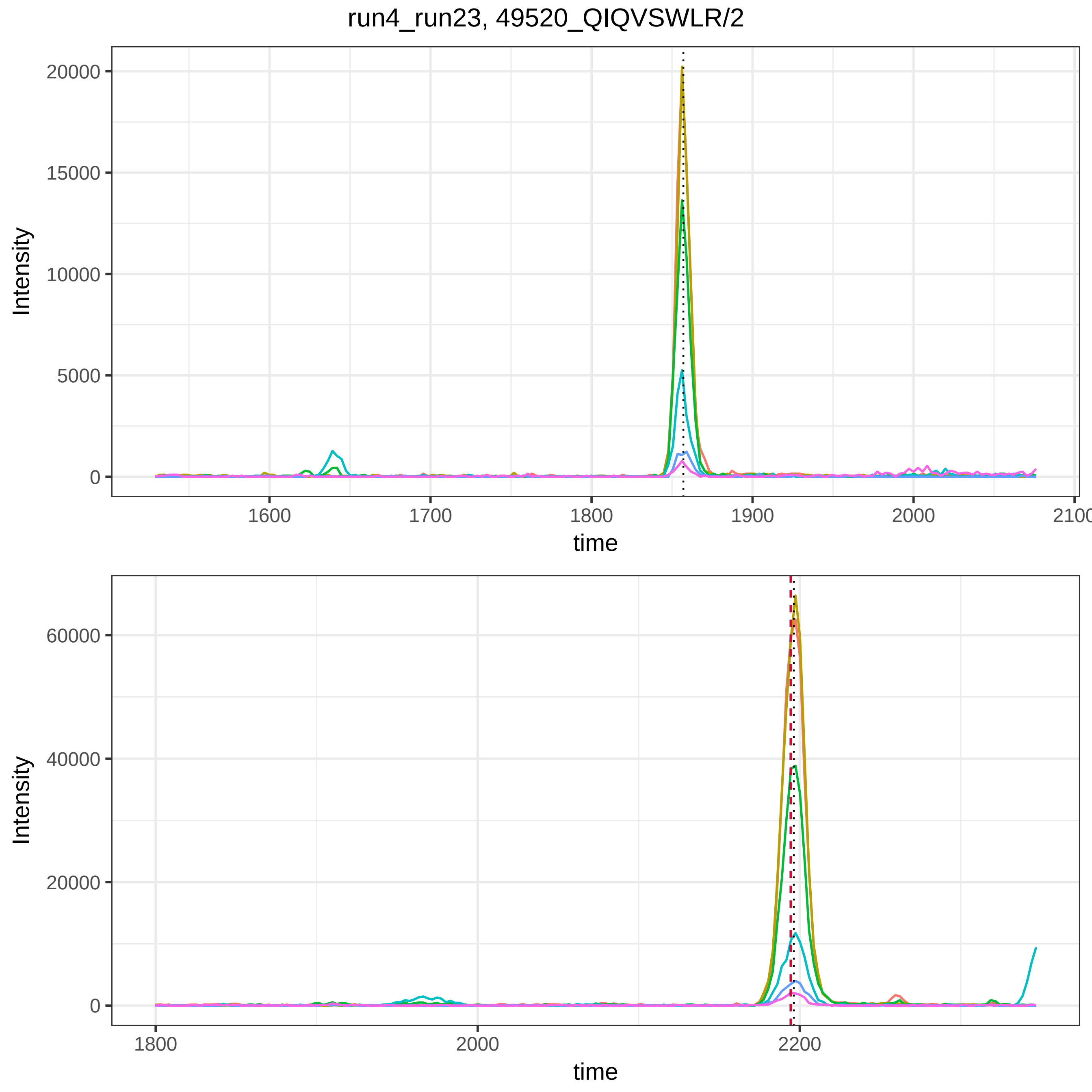

run4_run23, 49520_QIQVSWLR/2
Intensity
time
20000
15000
10000
5000
0
1600
1700
1800
1900
2000
2100
60000
40000
20000
0
1800
2000
2200

run4_run23, 49616_QLEQVIAK/2

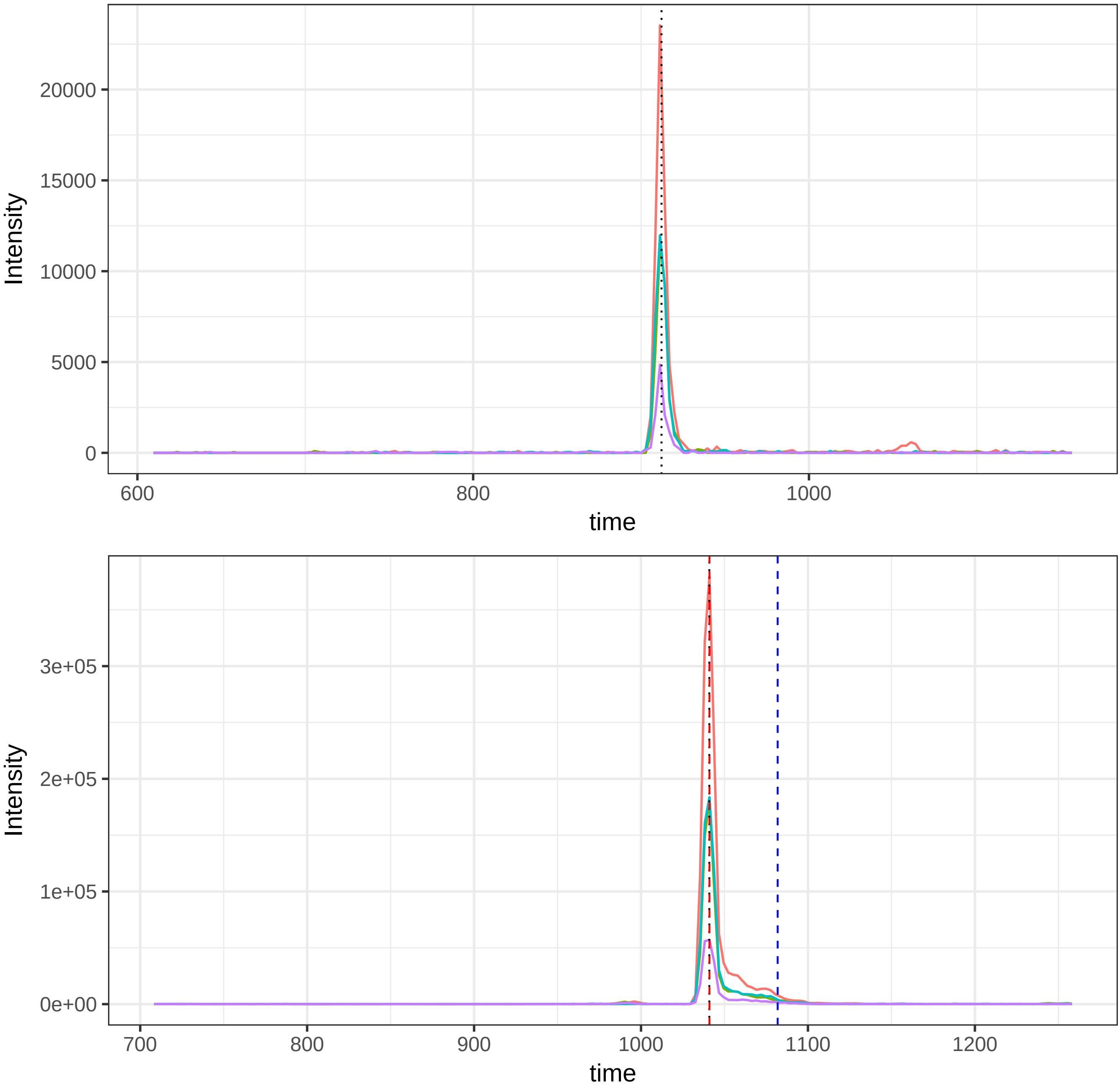

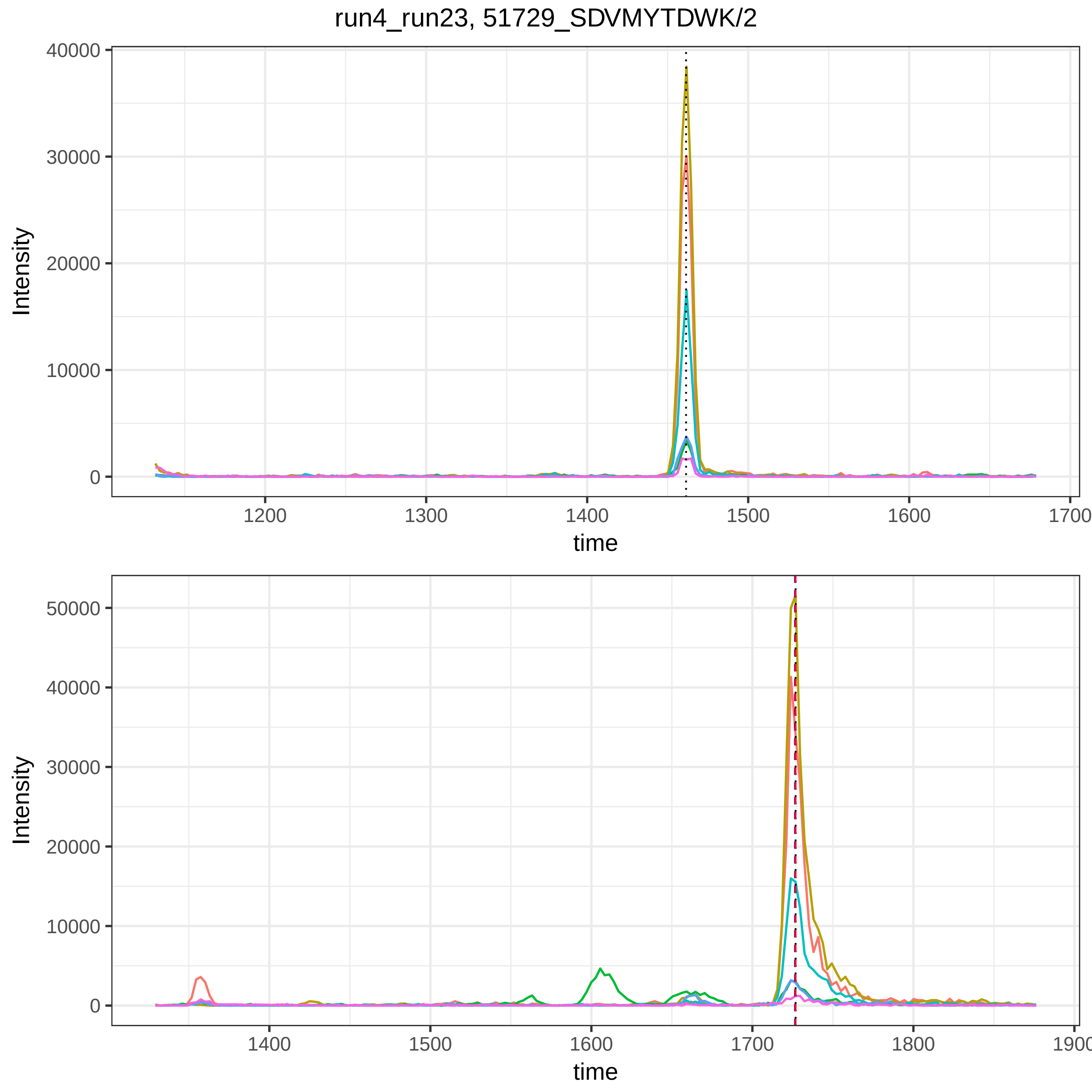
run4_run23, 51729_SDVMYTDWK/2
Intensity
time
0
10000
20000
30000
40000
1200
1300
1400
1500
1600
1700
Intensity
time
0
10000
20000
30000
40000
50000
1400
1500
1600
1700
1800
1900

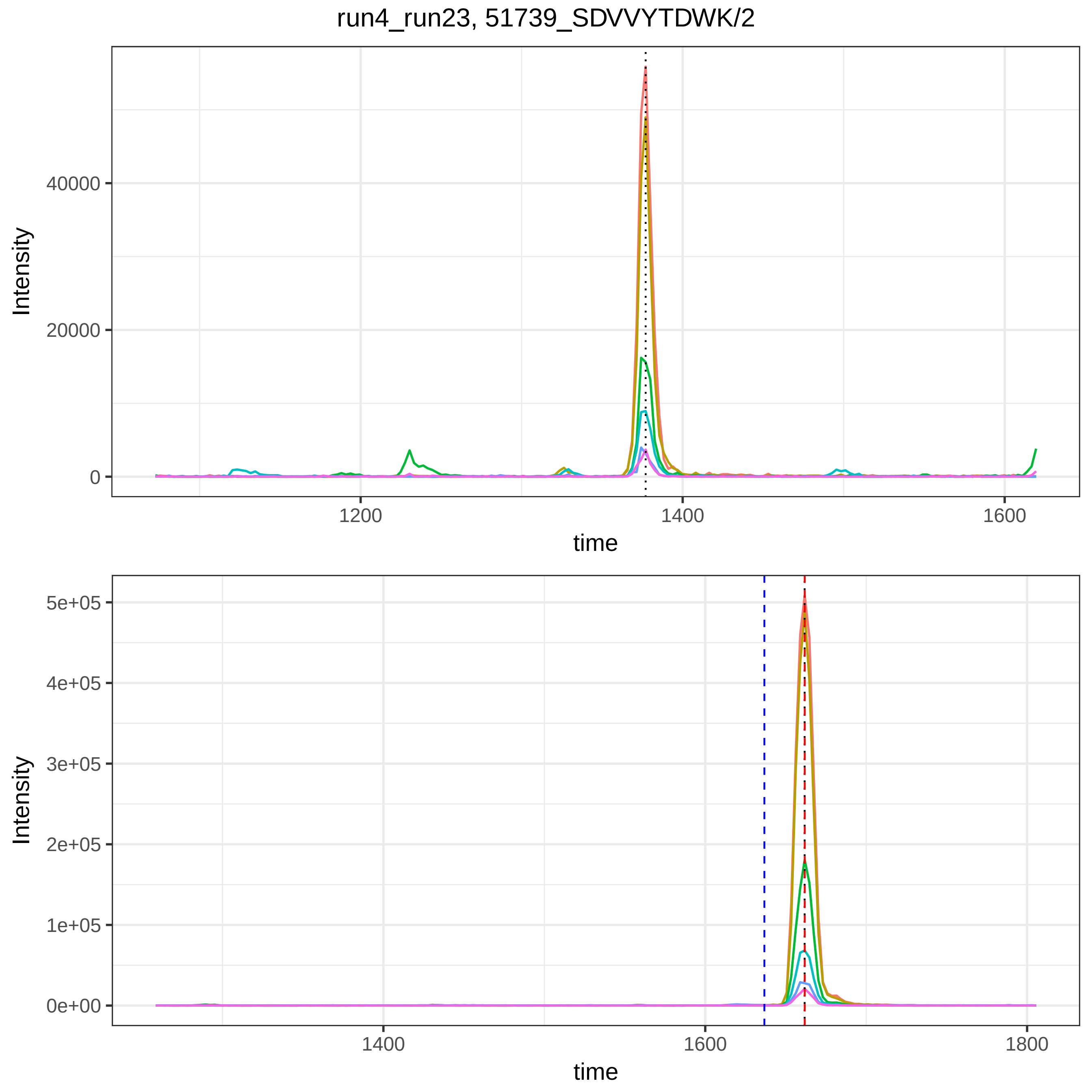
run4_run23, 51739_SDVVYTDWK/2
Intensity
40000
20000
0
1200
1400
1600
time
Intensity
5e+05
4e+05
3e+05
2e+05
1e+05
0e+00
1400
1600
1800
time

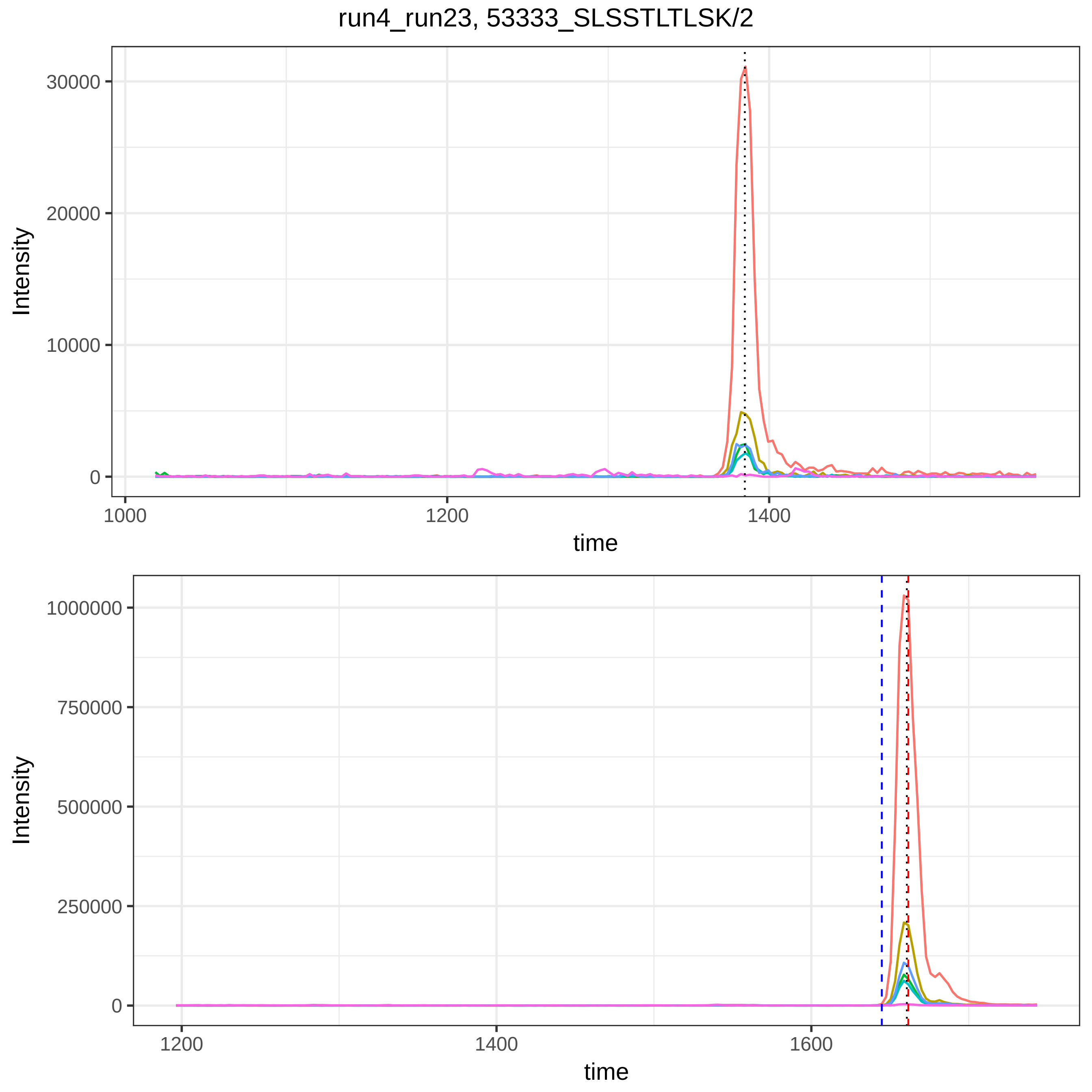
run4_run23, 53333_SLSSTLTLSK/2
Intensity
time
30000
20000
10000
0
1000
1200
1400
1000000
750000
500000
250000
0
1200
1400
1600

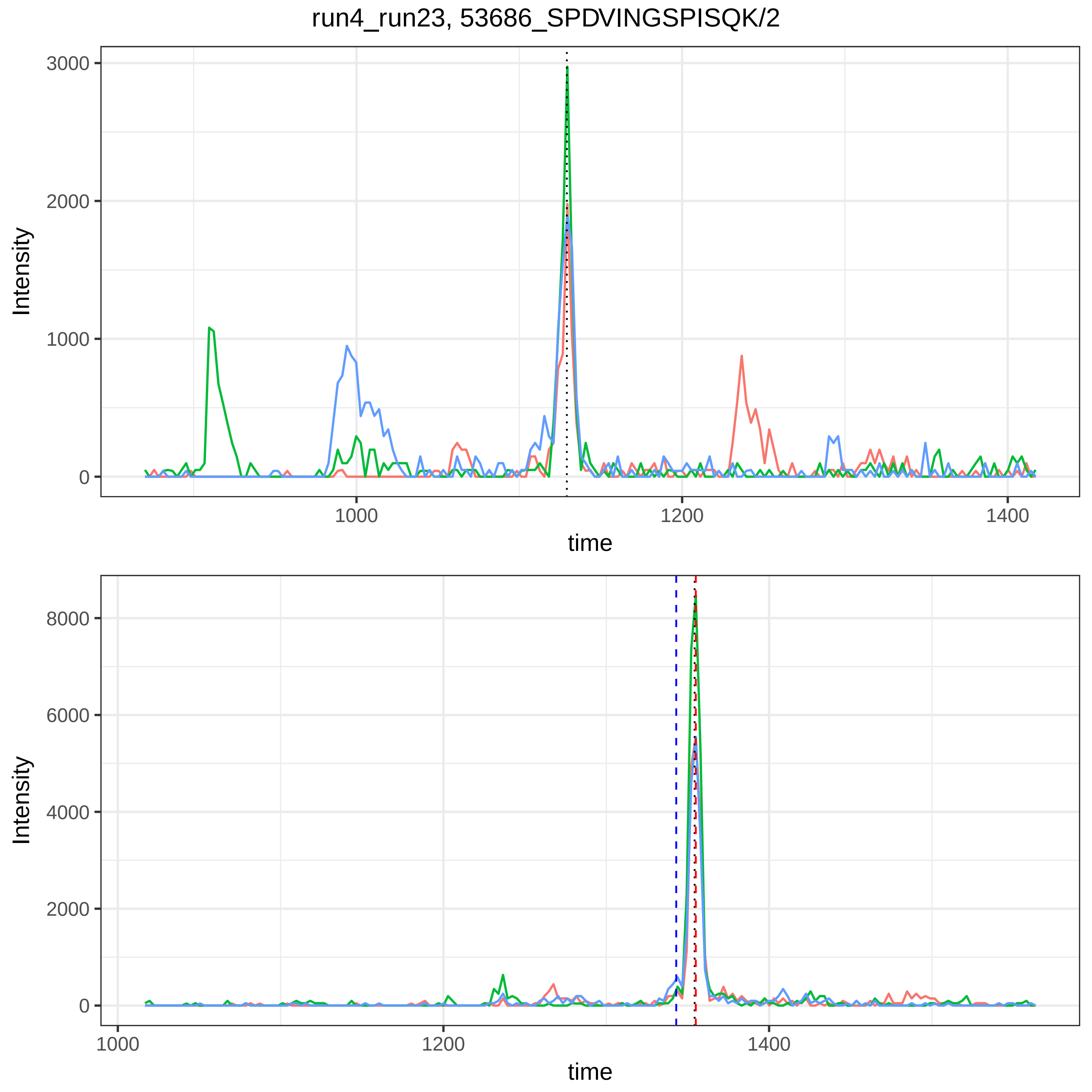

run4_run23, 53686_SPDVINGSPISQK/2
Intensity
0
1000
2000
3000
1000
1200
1400
time
Intensity
0
2000
4000
6000
8000
1000
1200
1400
time

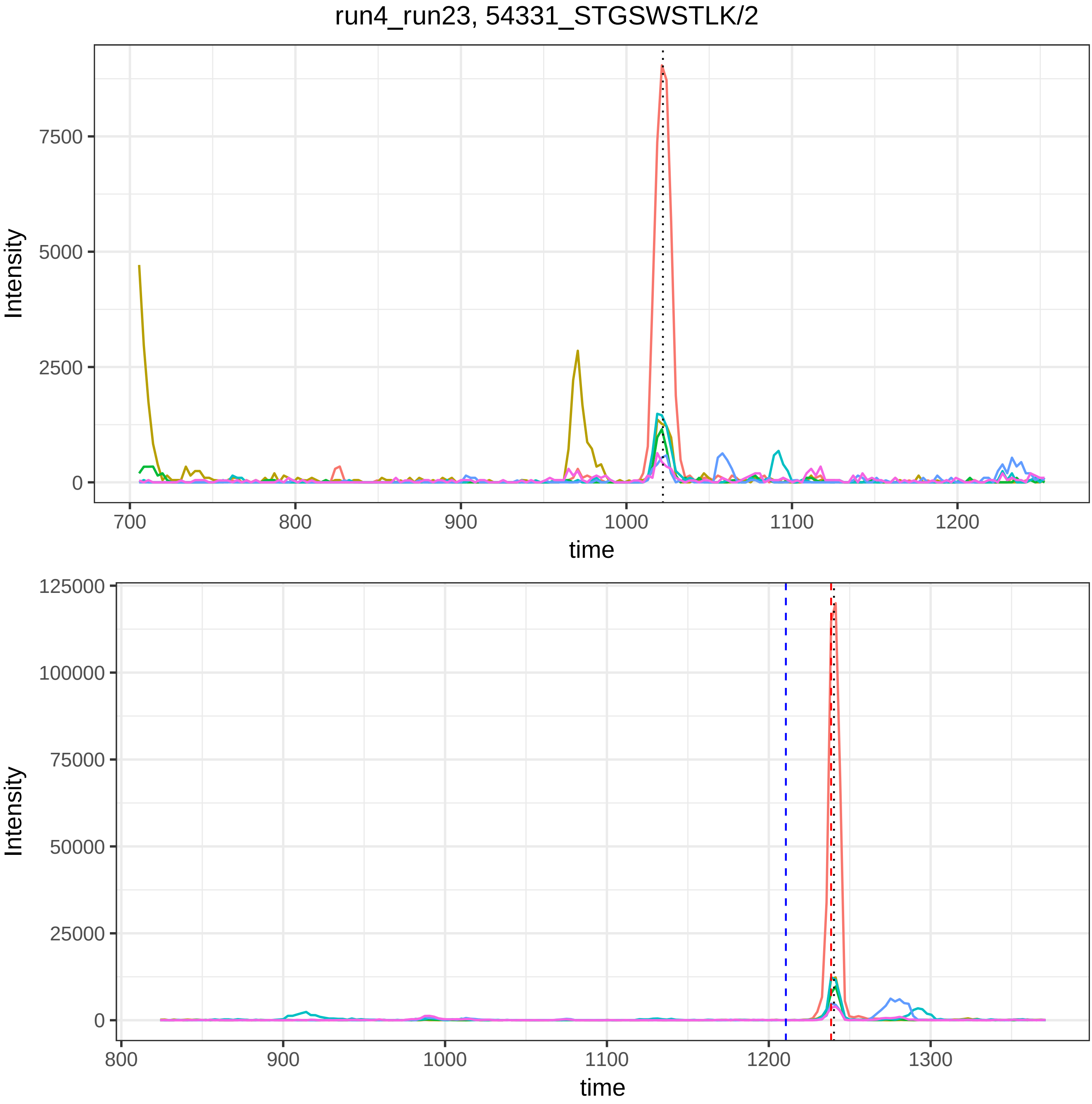

run4_run23, 54331_STGSWSTLK/2
Intensity
7500
5000
2500
0
700
800
900
1000
1100
1200
time
Intensity
125000
100000
75000
50000
25000
0
800
900
1000
1100
1200
1300
time

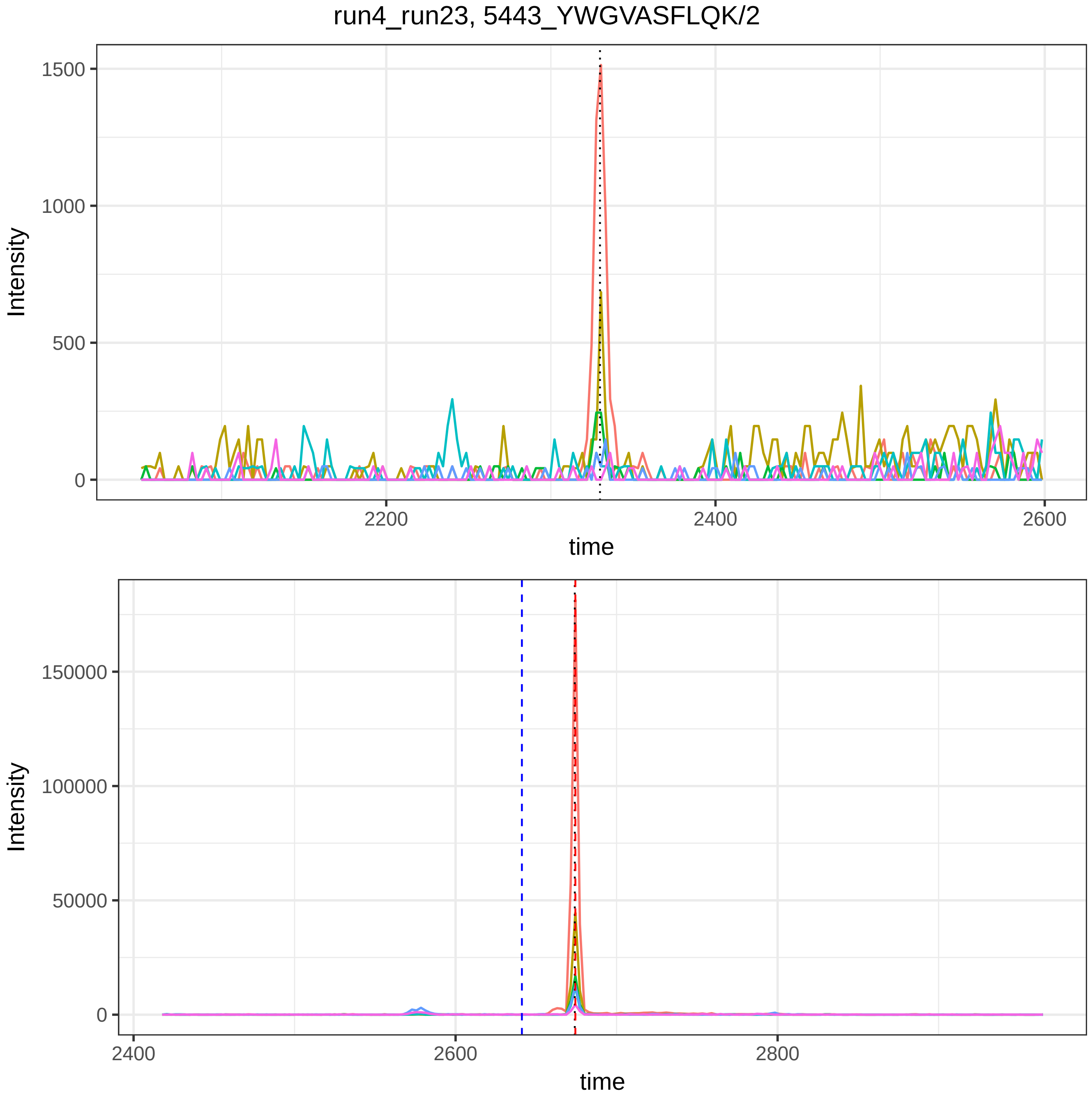
run4_run23, 5443_YWGVASFLQK/2
Intensity
1500
1000
500
0
2200
2400
2600
time
Intensity
150000
100000
50000
0
2400
2600
2800
time

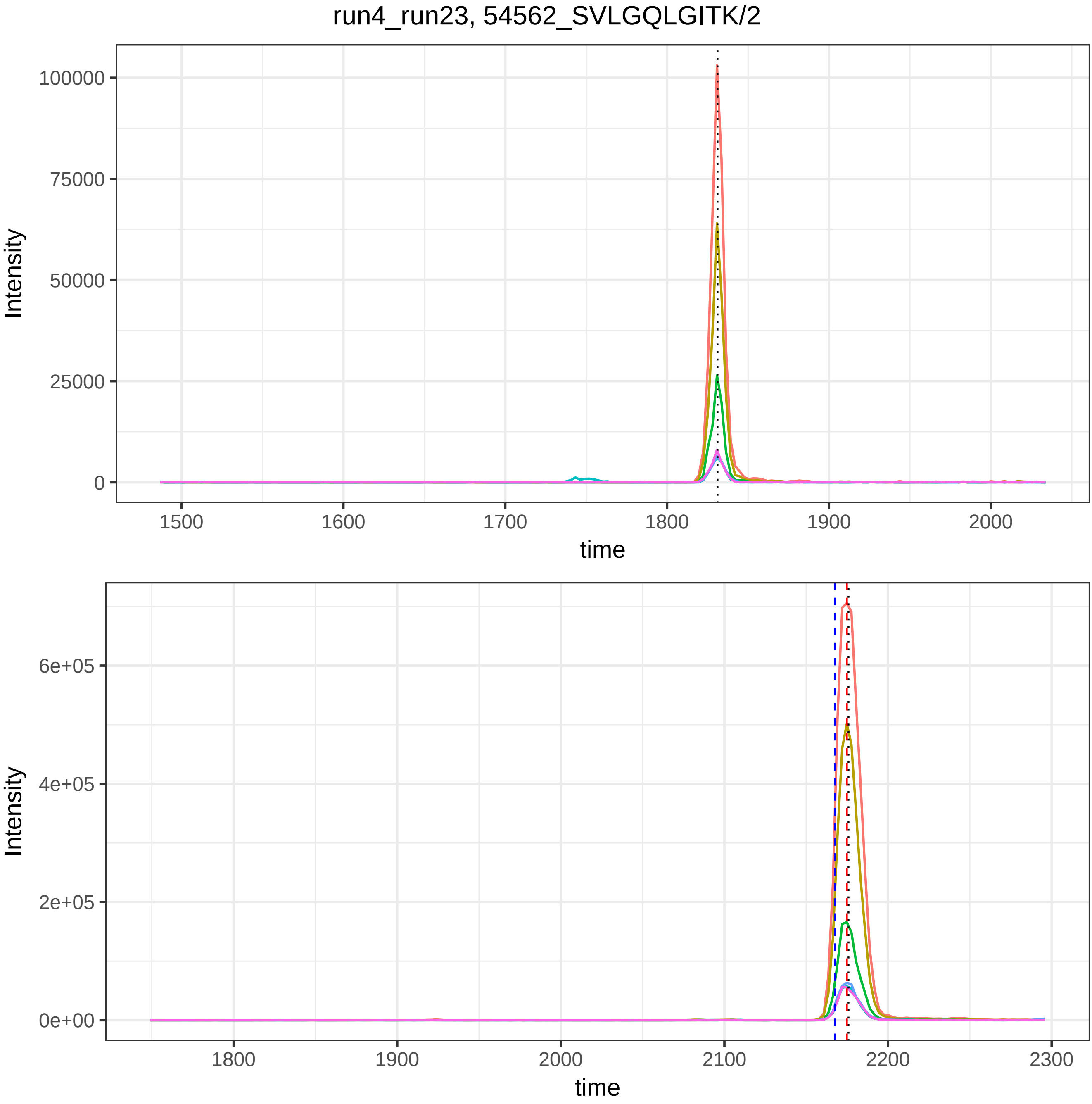
run4_run23, 54562_SVLGQLGITK/2
Intensity
100000
75000
50000
25000
0
1500
1600
1700
1800
1900
2000
time
Intensity
6e+05
4e+05
2e+05
0e+00
1800
1900
2000
2100
2200
2300
time

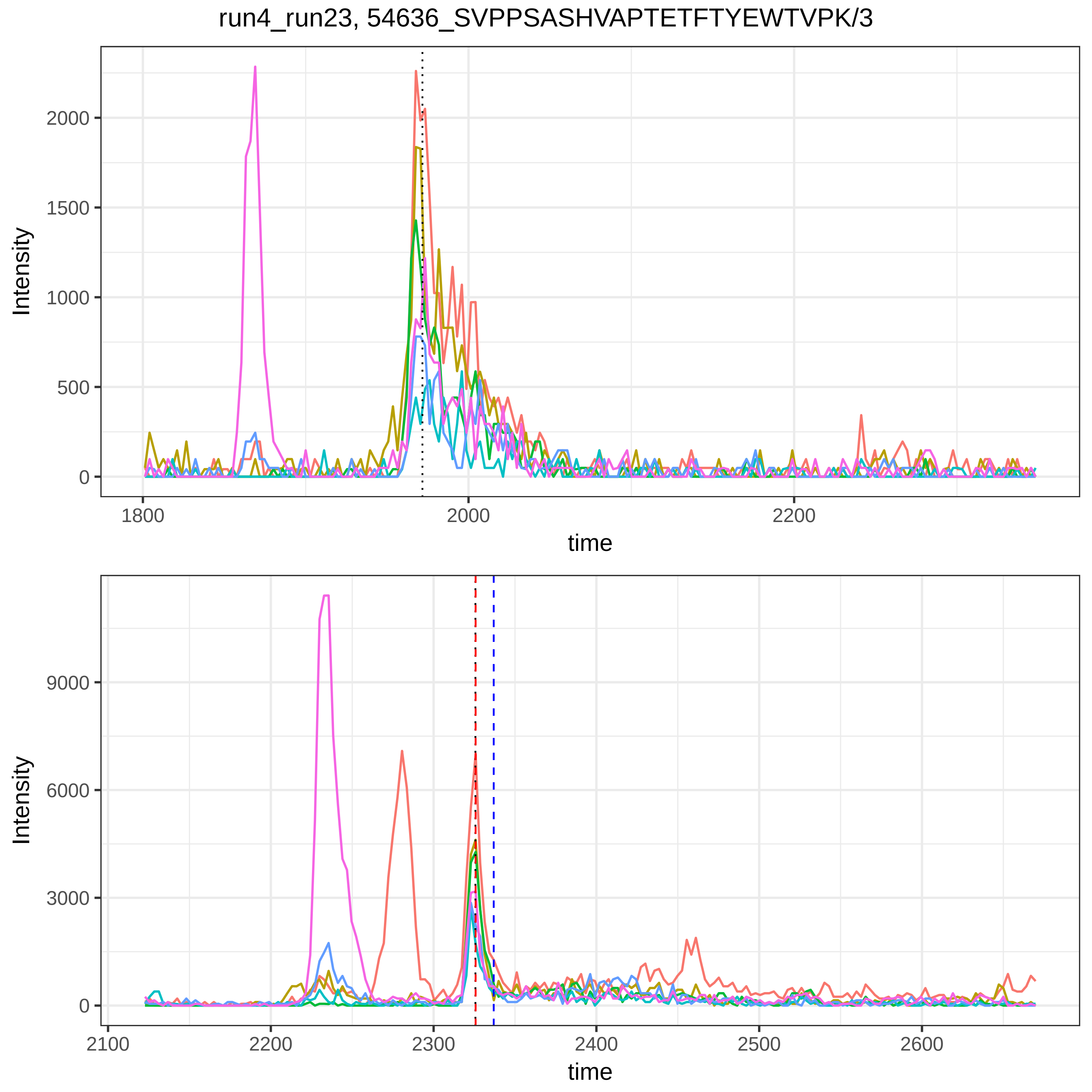

run4_run23, 54636_SVPPSASHVAPTETFTYEWTVPK/3
Intensity
time
0
500
1000
1500
2000
1800
2000
2200
0
3000
6000
9000
2100
2200
2300
2400
2500
2600

run4_run23, 56511_TLLPVSKPEIR/3

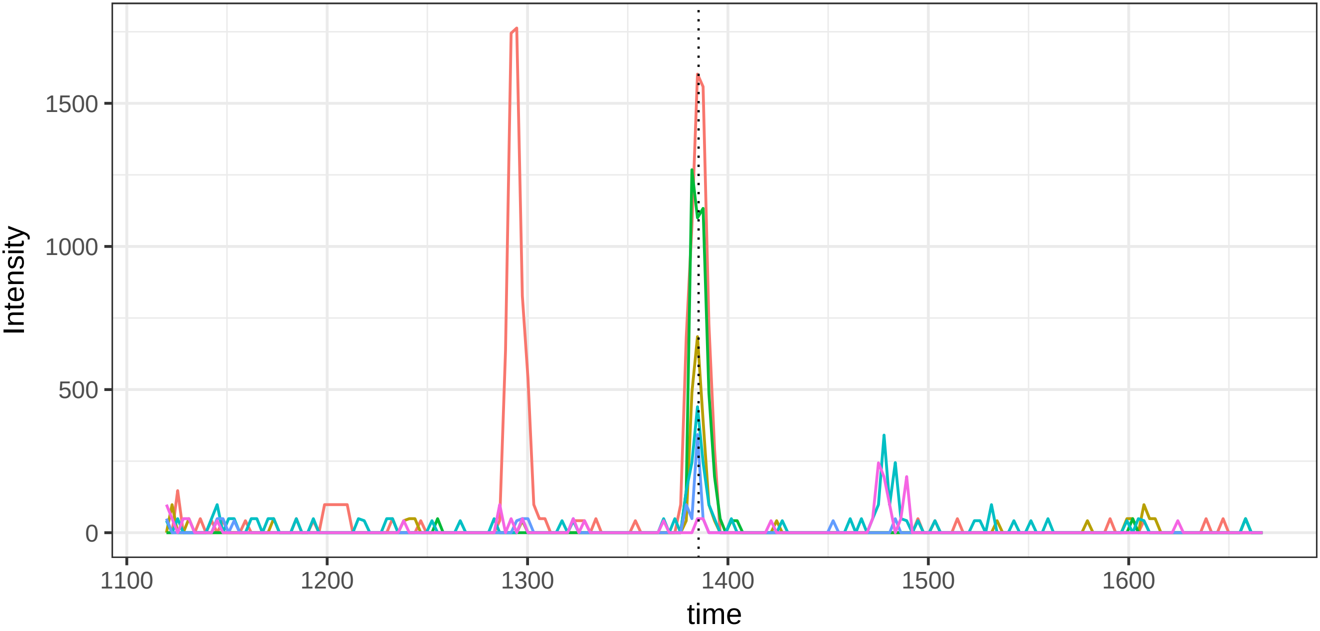


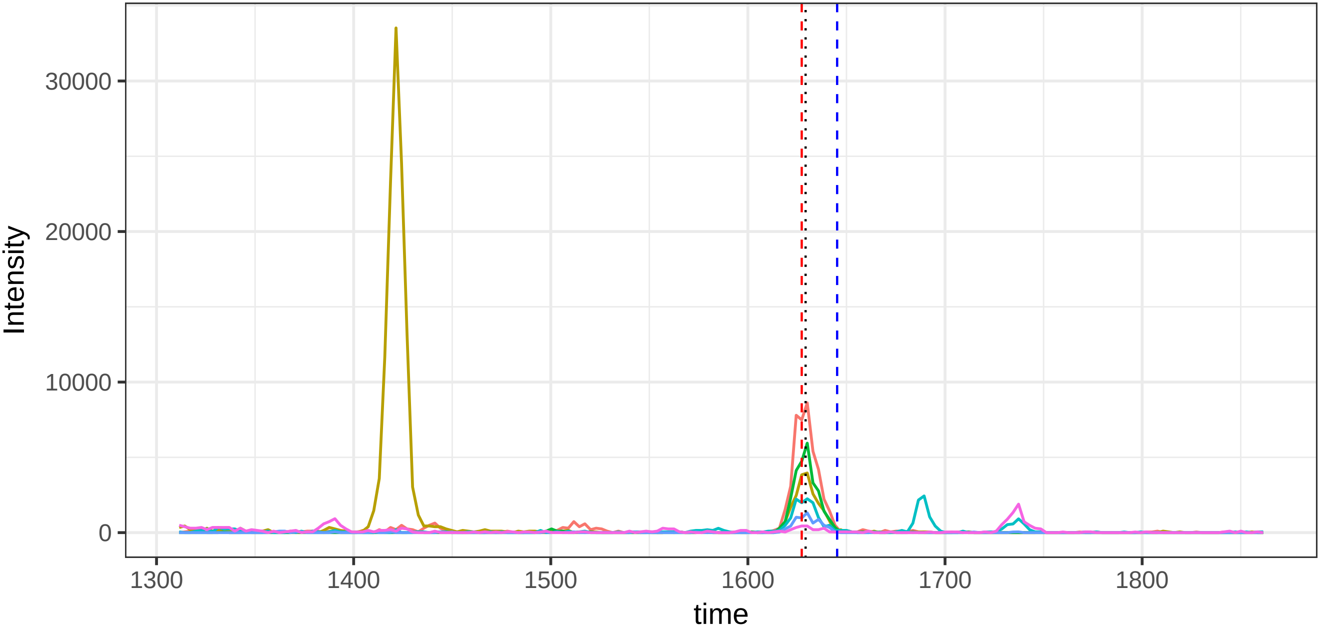
