## Supplemental Figures for "DIAlignR provides precise retention time alignment across distant runs in DIA and targeted proteomics": Supplemental Figure18.pdf

Supplemental Figure 18. In the figures, blue dashed line indicates RT estimation by LOESS whereas red dashed line indicates RT estimation by chromatogram alignment. Dotted line indicates OpenSWATH annotation.

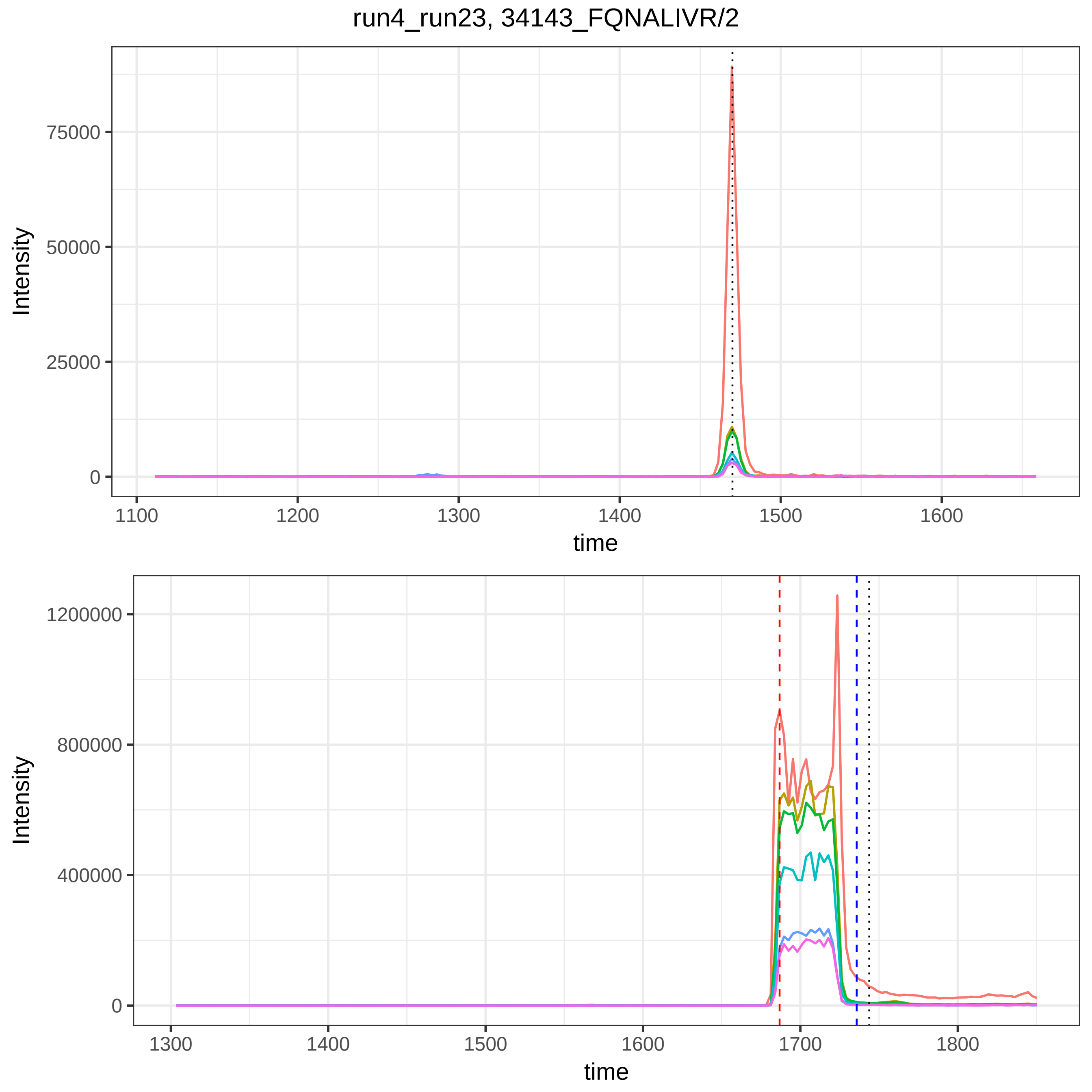
run4_run23, 34143_FQNALIVR/2
Intensity
75000
50000
25000
0
1100
1200
1300
1400
1500
1600
time
Intensity
1200000
800000
400000
0
1300
1400
1500
1600
1700
1800
time

run4_run23, 115879_DDNPNLPR/2

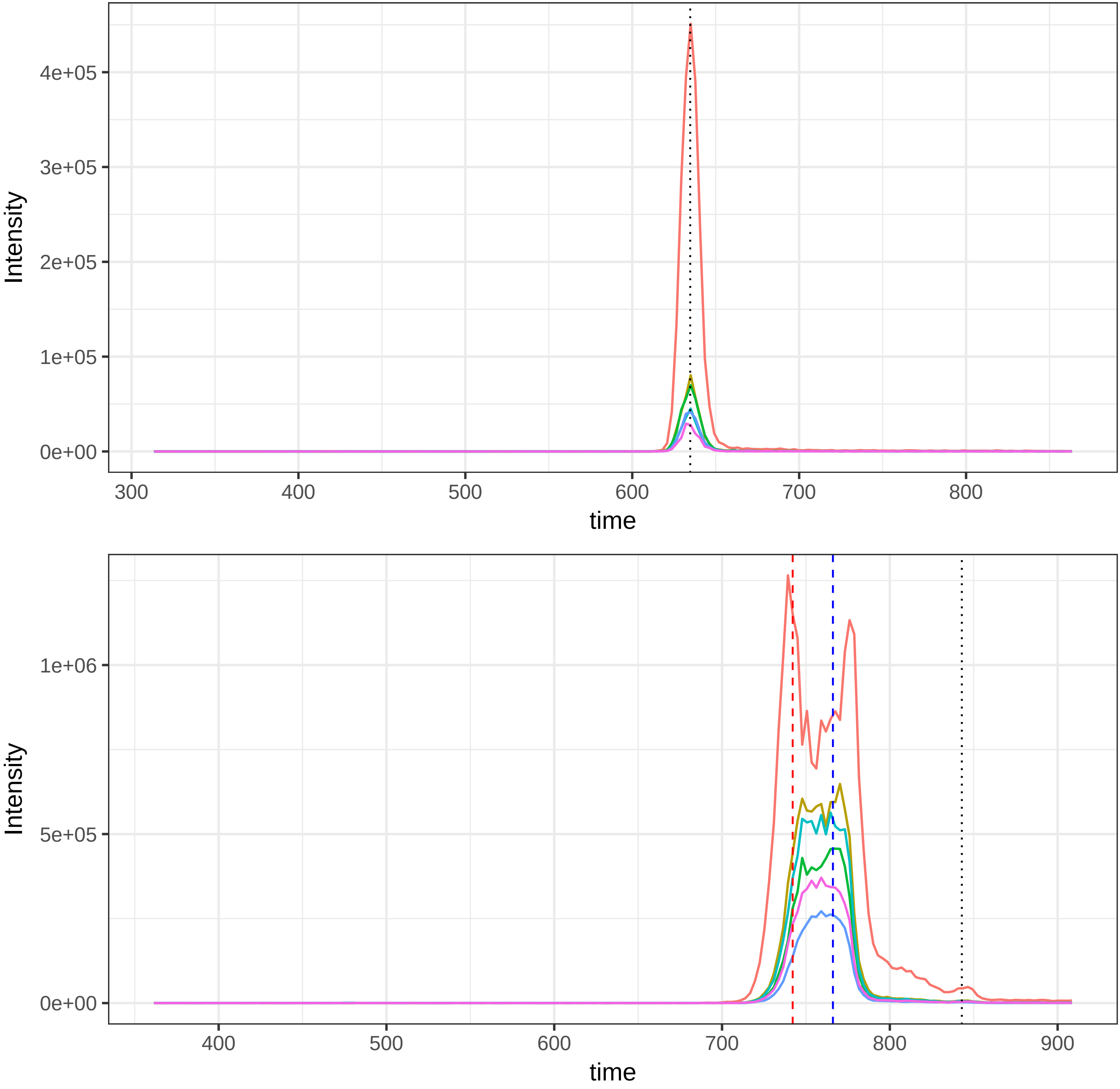

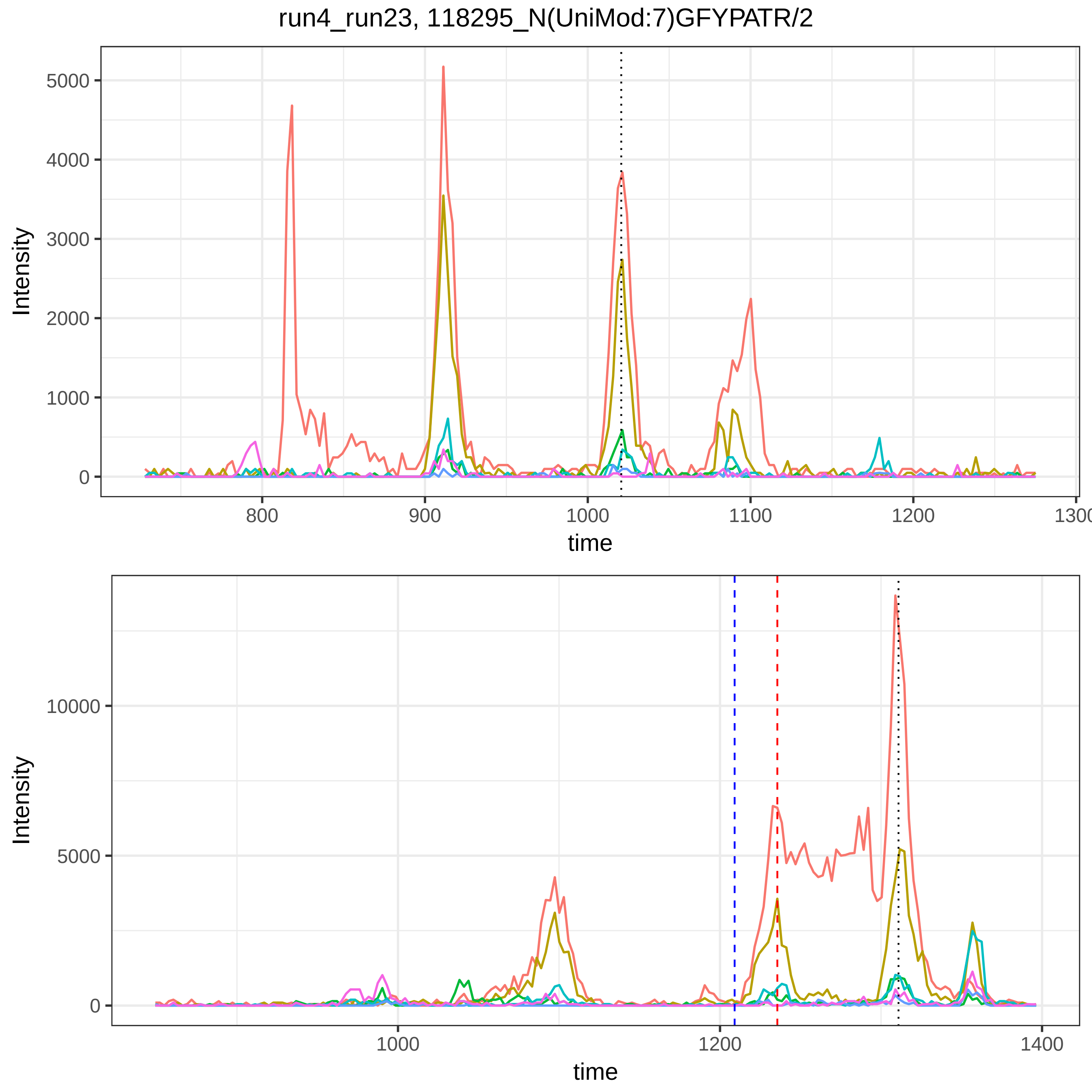

run4_run23, 118295_N(UniMod:7)GFYPATR/2
Intensity
5000
4000
3000
2000
1000
0
800
900
1000
1100
1200
1300
time
Intensity
10000
5000
0
1000
1200
1400
time

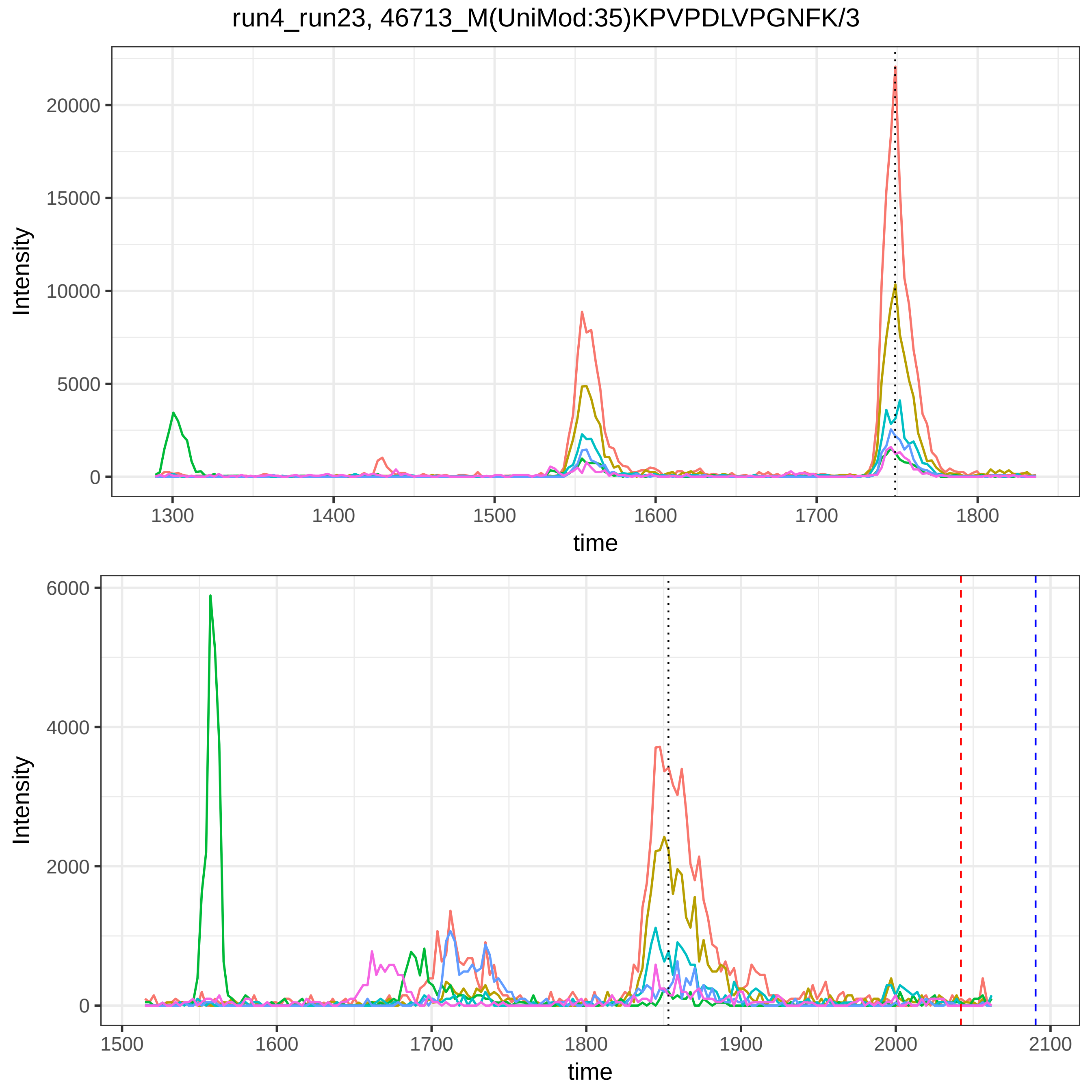
run4_run23, 46713_M(UniMod:35)KPVPDLVPGNFK/3
Intensity
time
0
5000
10000
15000
20000
1300
1400
1500
1600
1700
1800
0
2000
4000
6000
1500
1600
1700
1800
1900
2000
2100

run4_run23, 119603_M(UniMod:35)KPVPDLVPGNFK/3

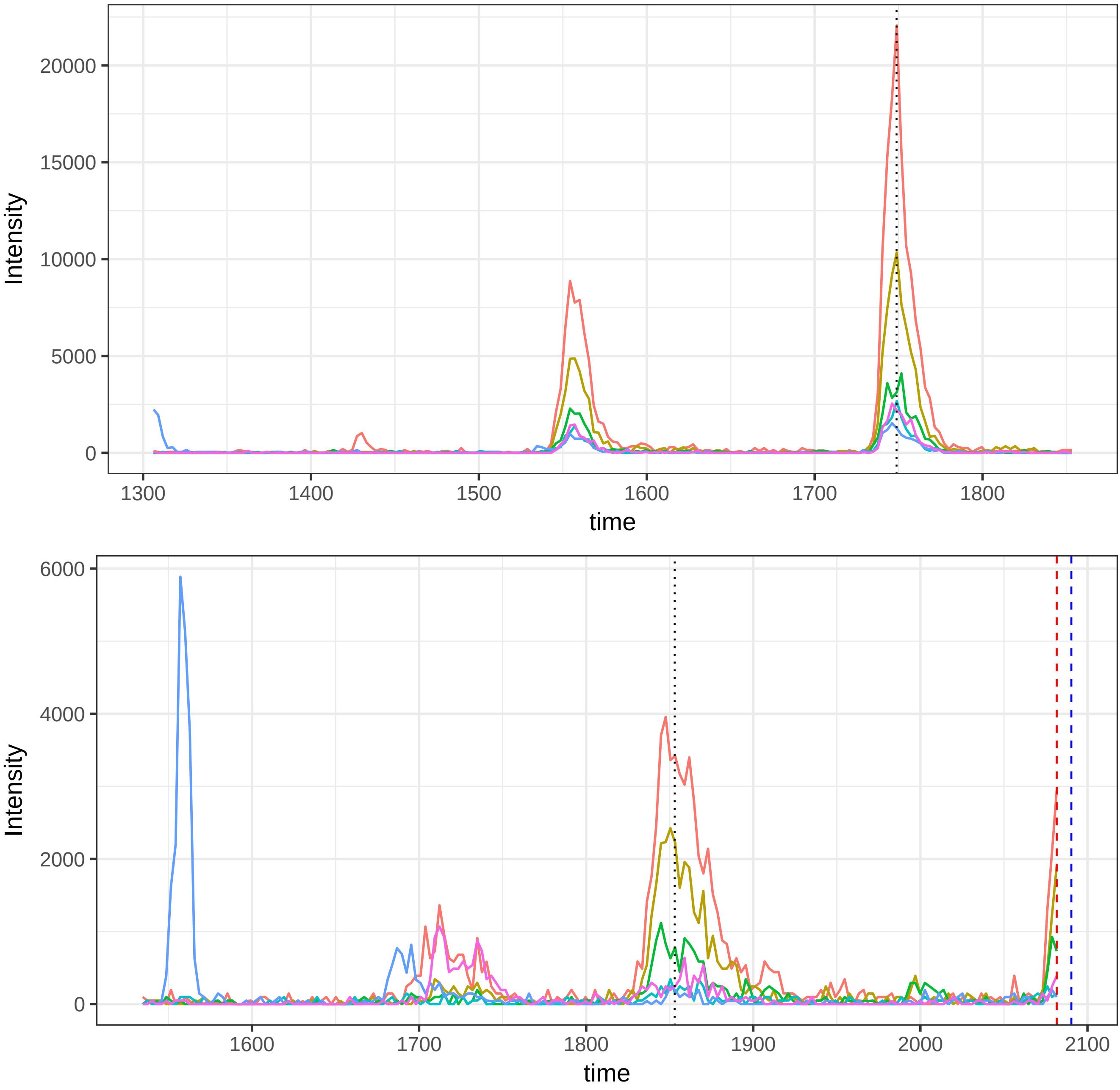

run4_run23, 116251_Q(UniMod:7)TALVELVK/2

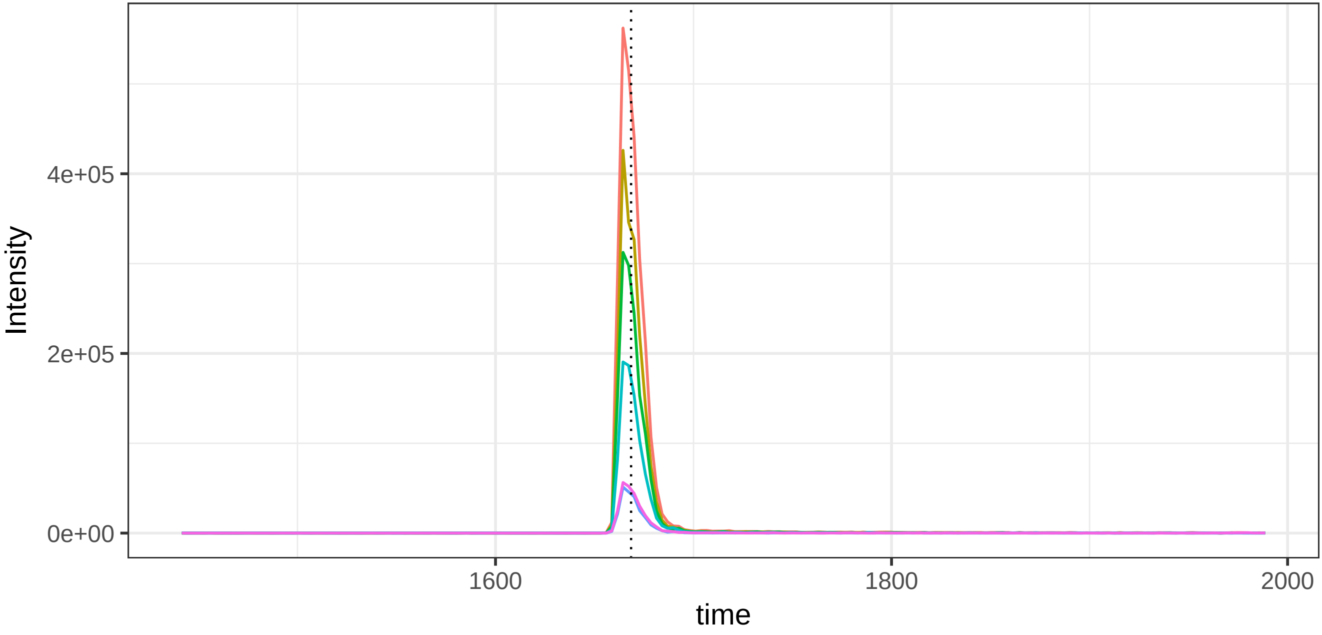


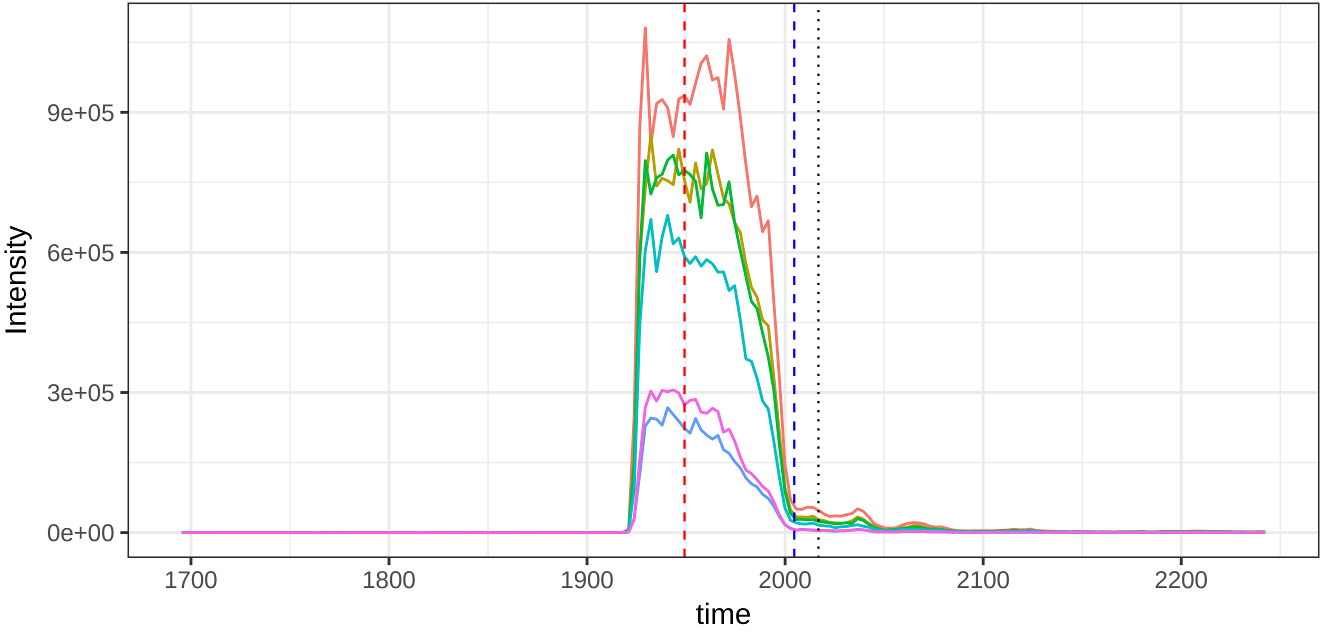

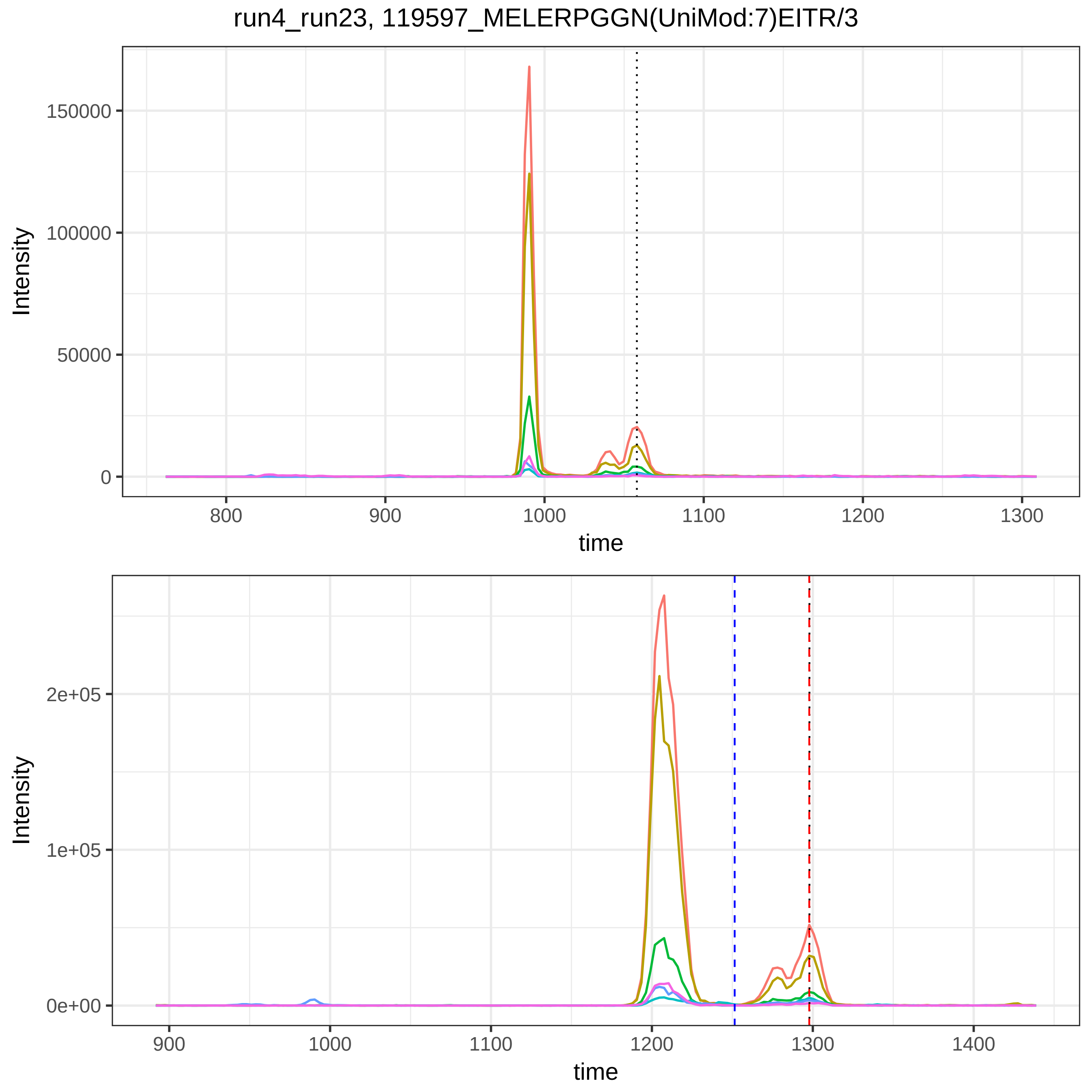

run4_run23, 119597_MELERPGGN(UniMod:7)EITR/3
Intensity
150000
100000
50000
0
800
900
1000
1100
1200
1300
time
Intensity
2e+05
1e+05
0e+00
900
1000
1100
1200
1300
1400
time

run4_run23, 117986_QQNAQGGFSSTQDTVVALHALSK/3

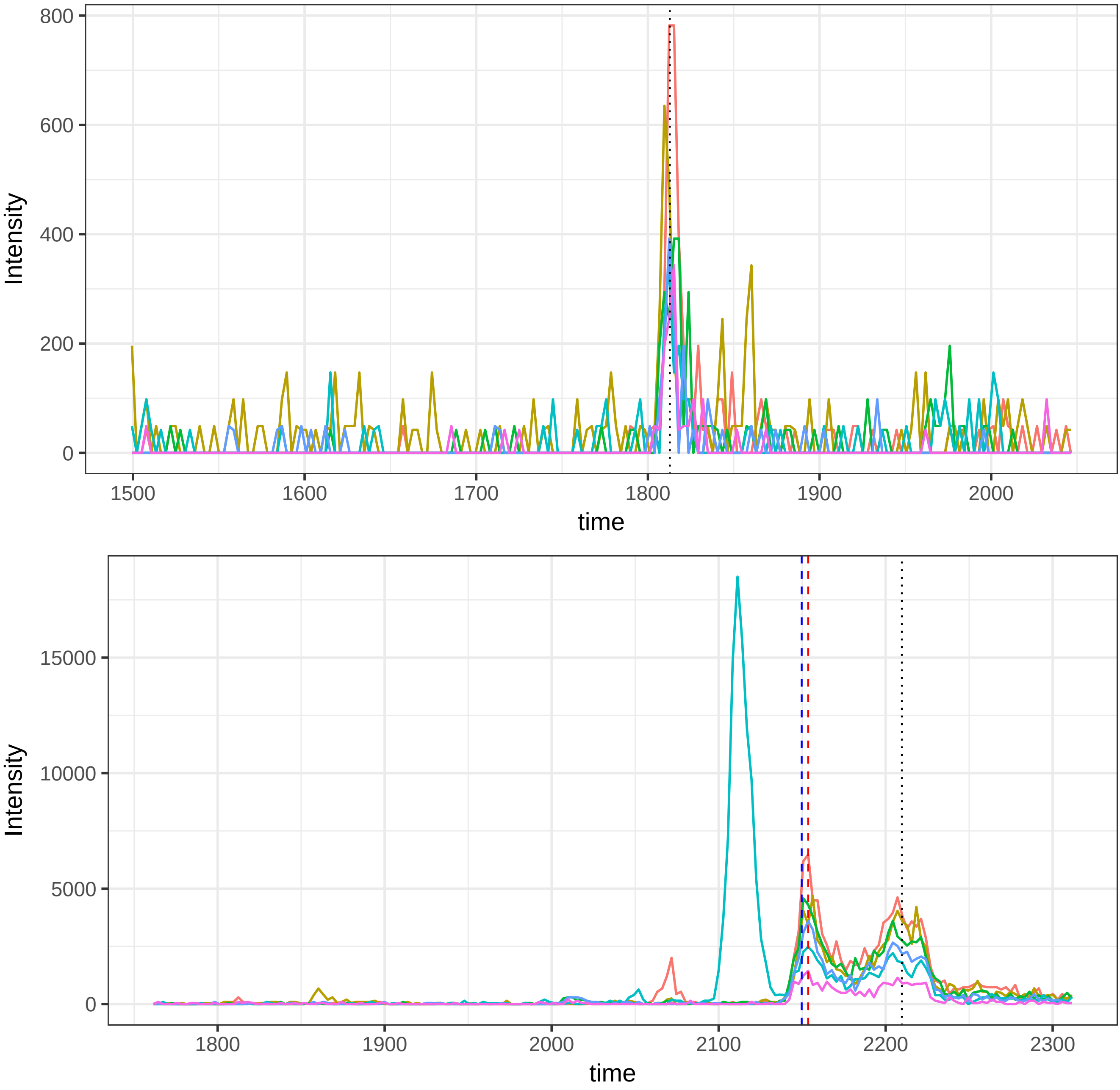

run4_run23, 34143_FQNALIVR/2

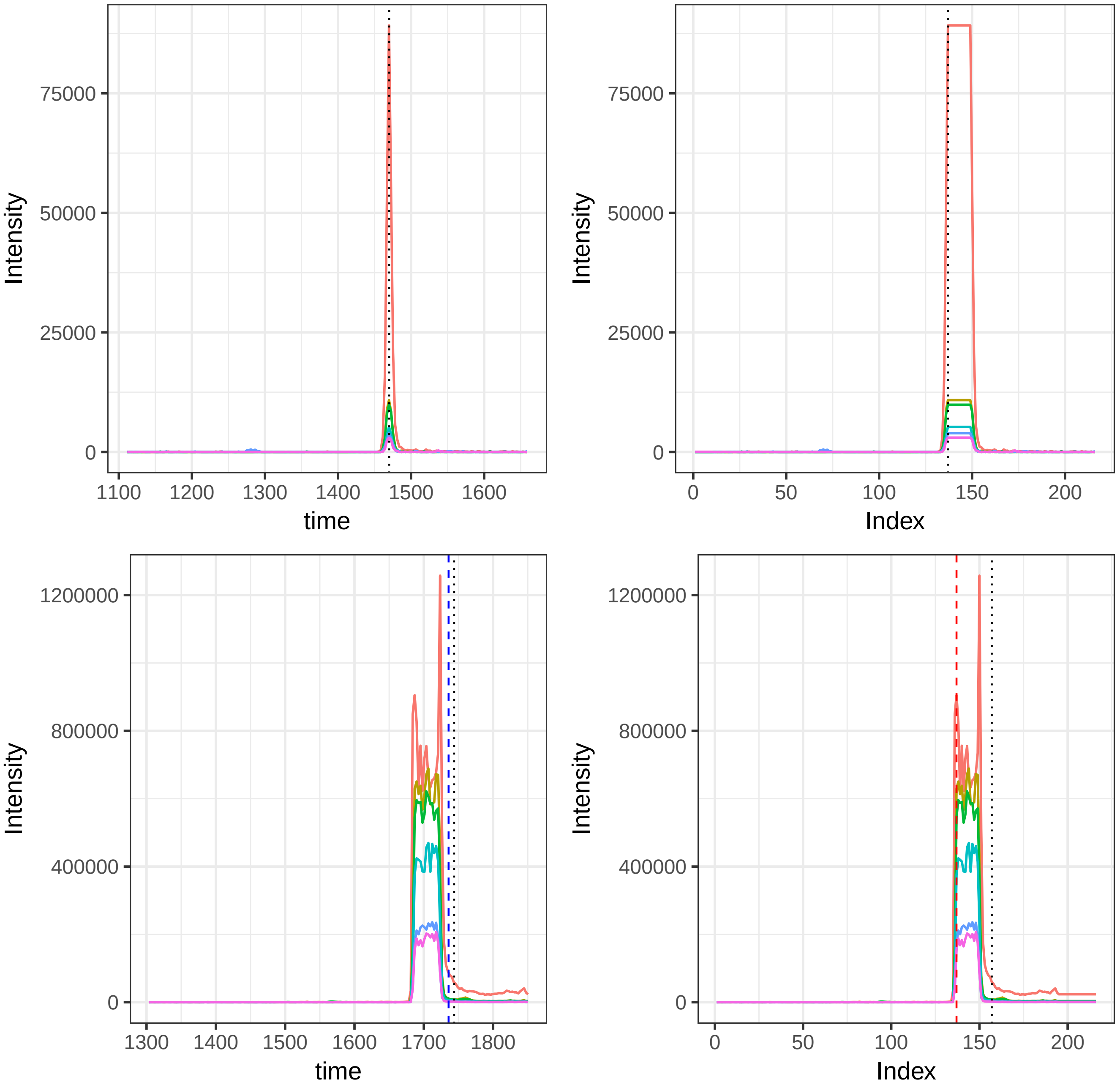

run4_run23, 115879_DDNPNLPR/2

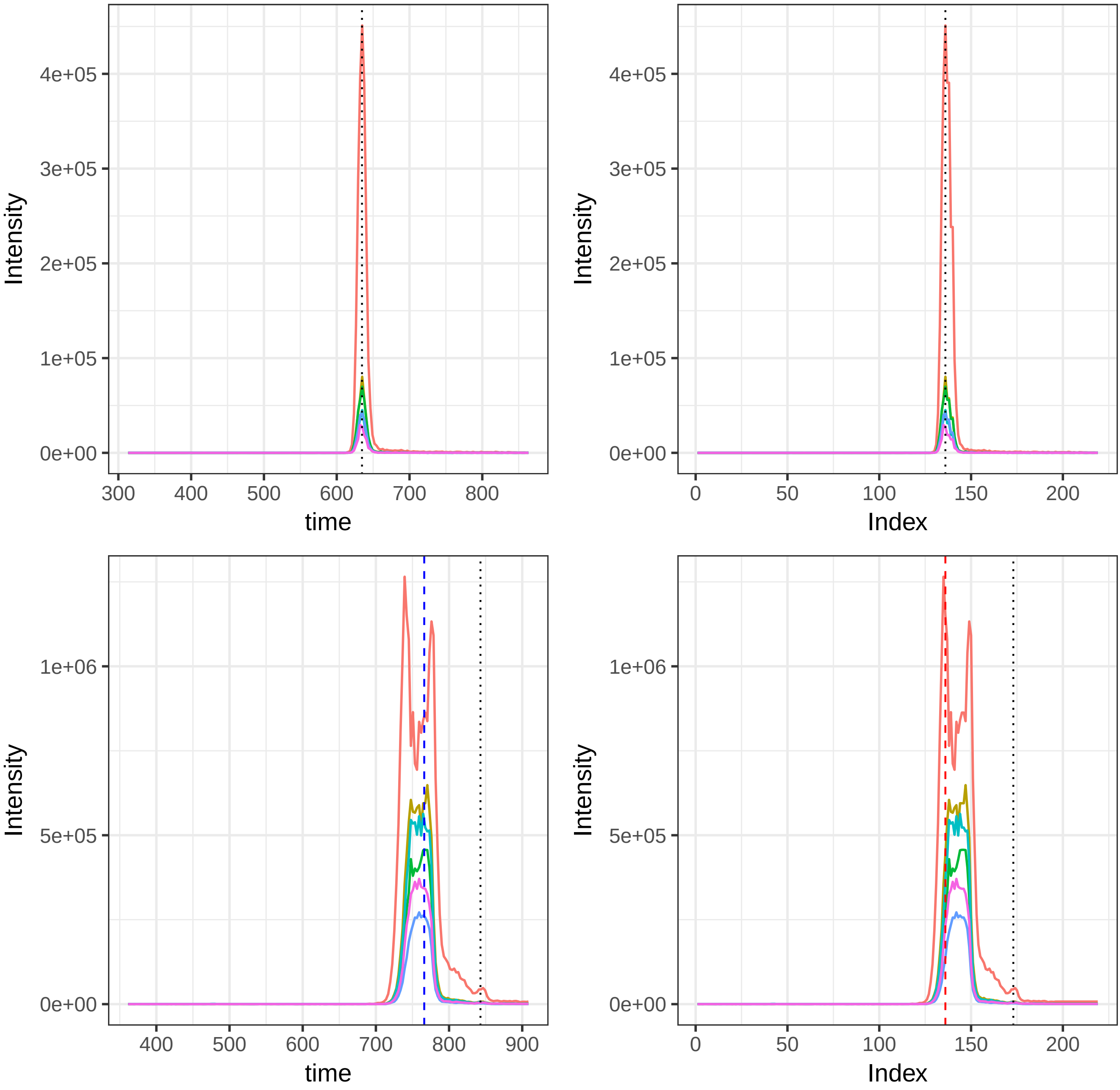

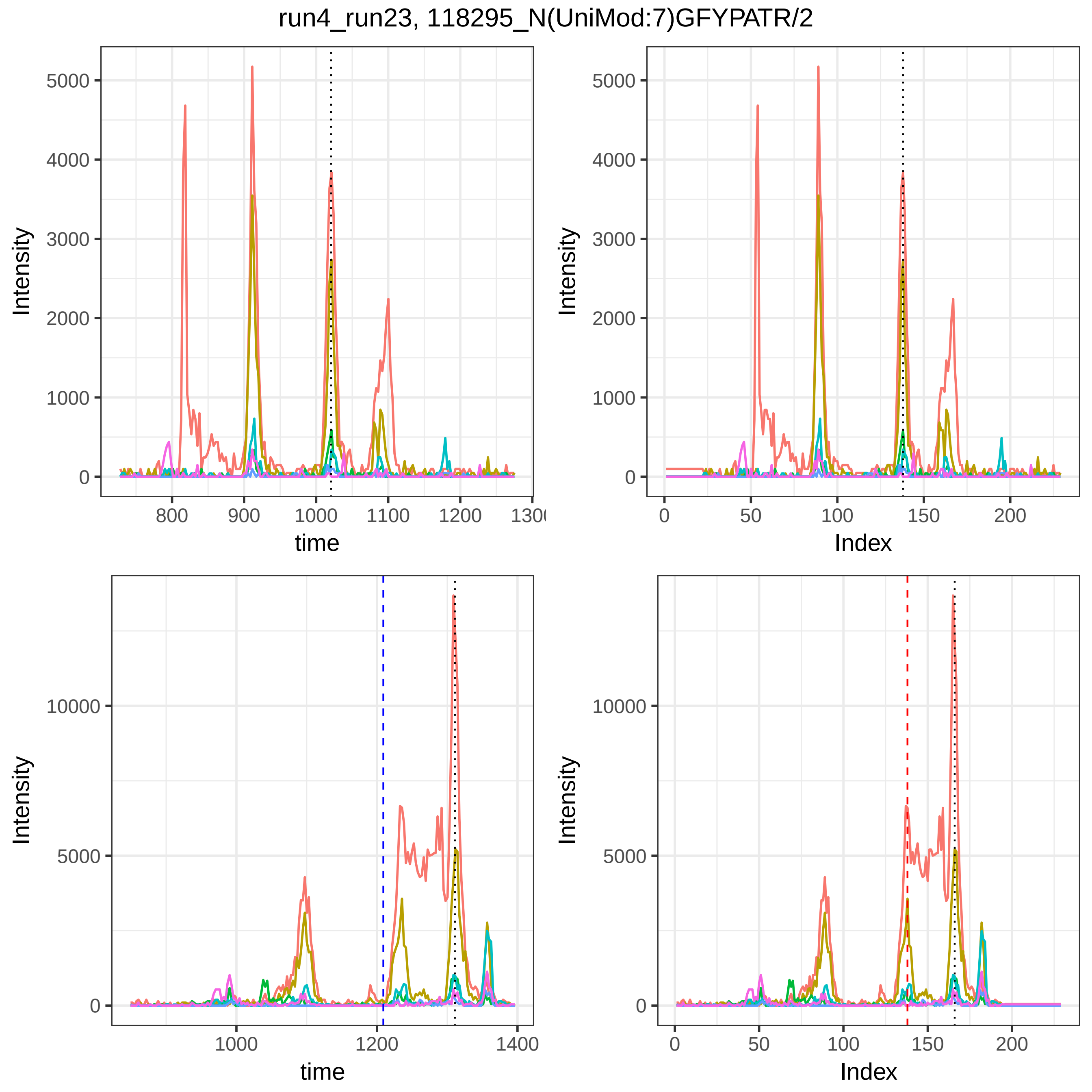

run4_run23, 118295_N(UniMod:7)GFYPATR/2
Intensity
time
Index

run4_run23, 46713_M(UniMod:35)KPVPDLVPGNFK/3

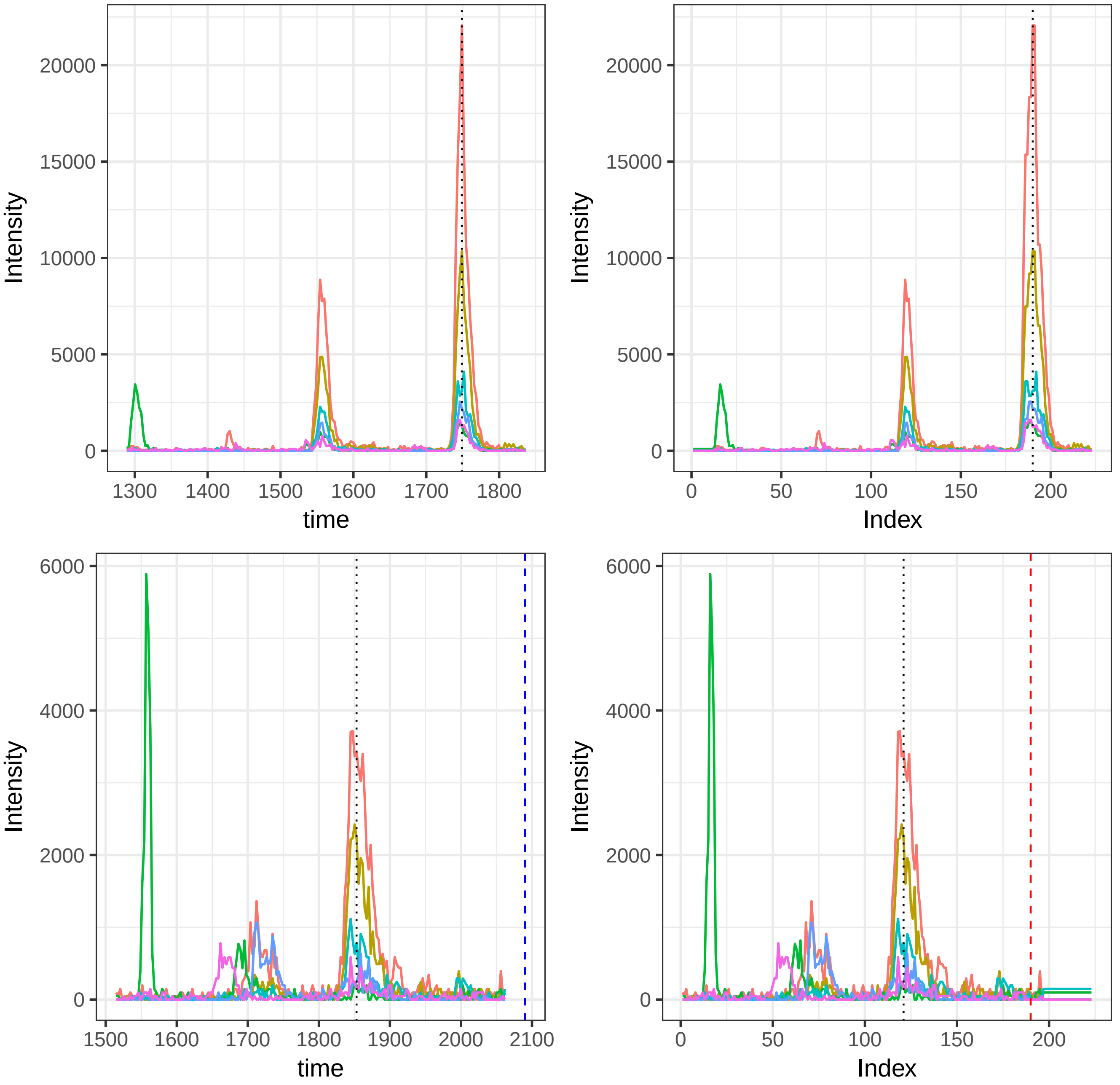

run4_run23, 119603_M(UniMod:35)KPVPDLVPGNFK/3

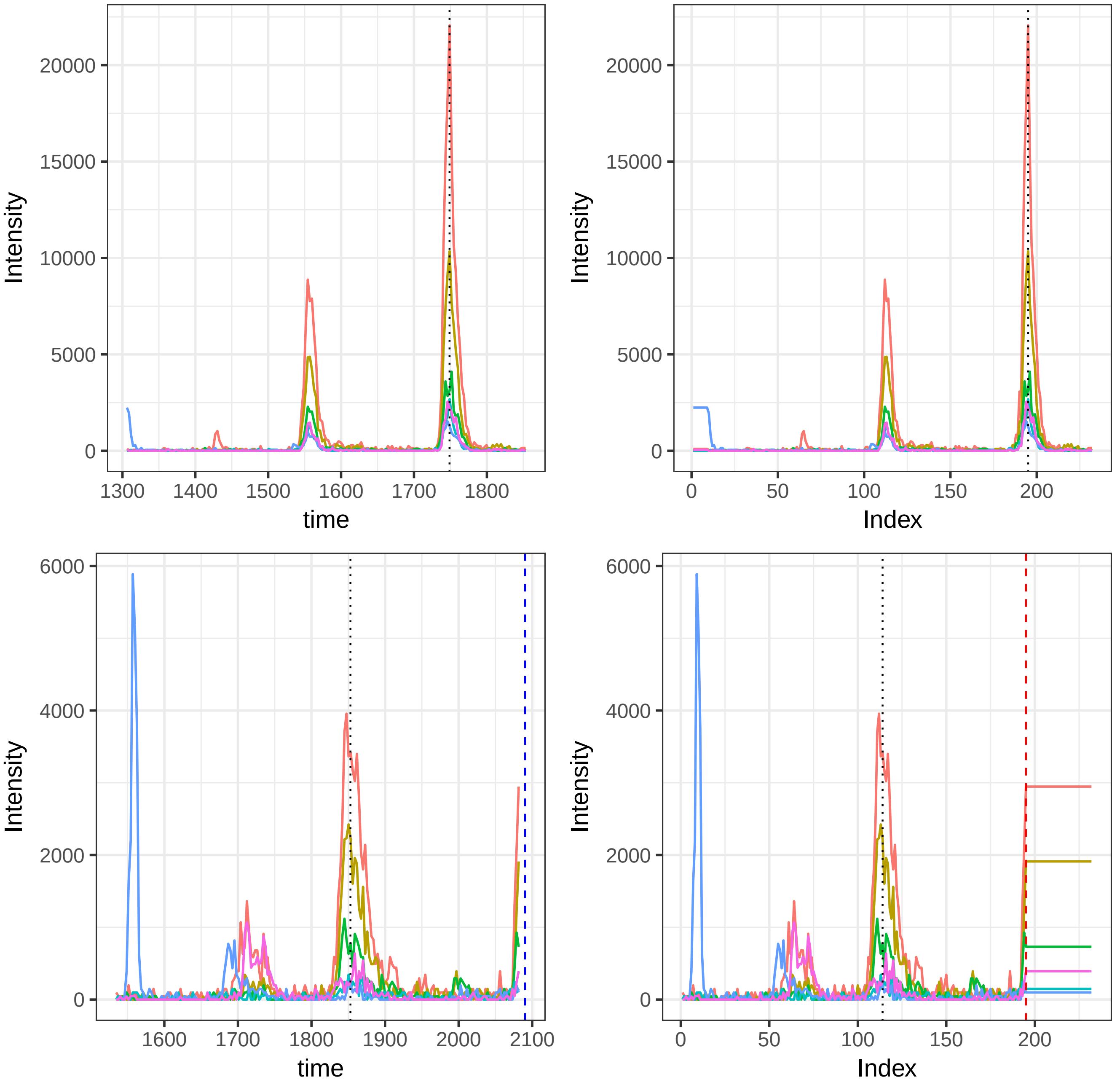

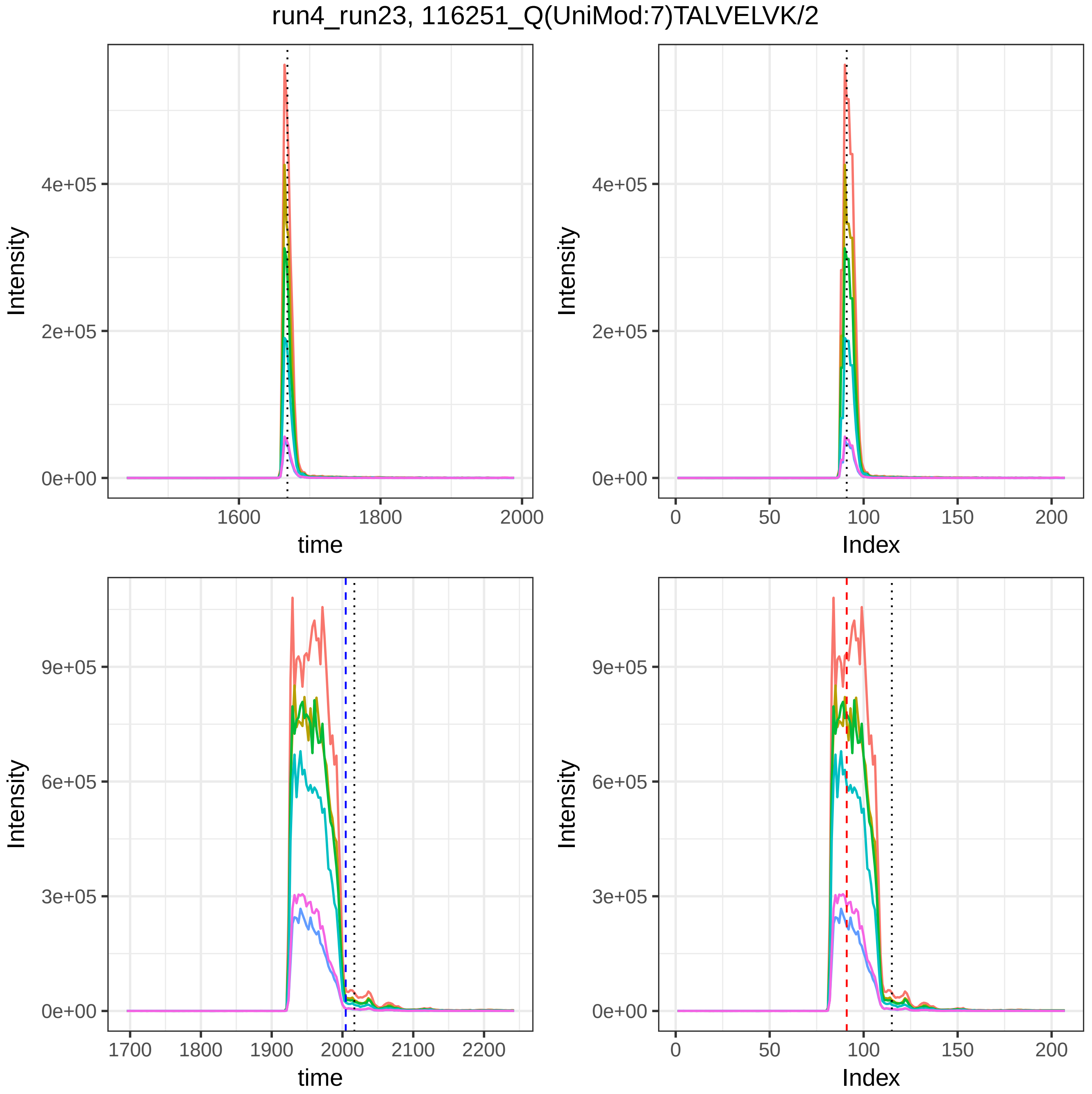
run4_run23, 116251_Q(UniMod:7)TALVELVK/2
Intensity
time
Index
0e+00
2e+05
4e+05
1600
1800
2000
0
50
100
150
200
3e+05
6e+05
9e+05
1700
1900
2100
2200

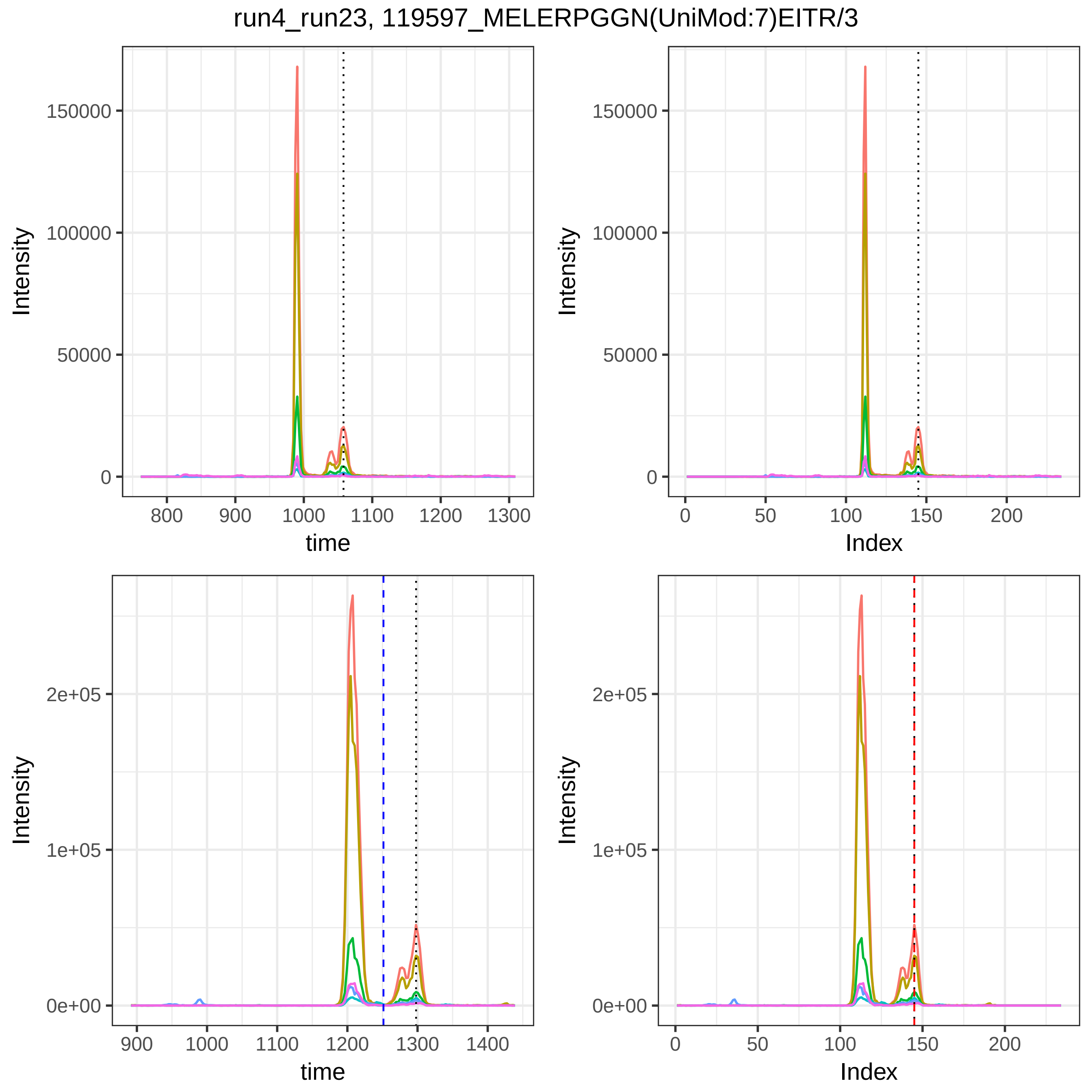
run4_run23, 119597_MELERPGGN(UniMod:7)EITR/3
Intensity
time
Index
150000
100000
50000
0
800
900
1000
1100
1200
1300
0
50
100
150
200
2e+05
1e+05
0e+00
1400

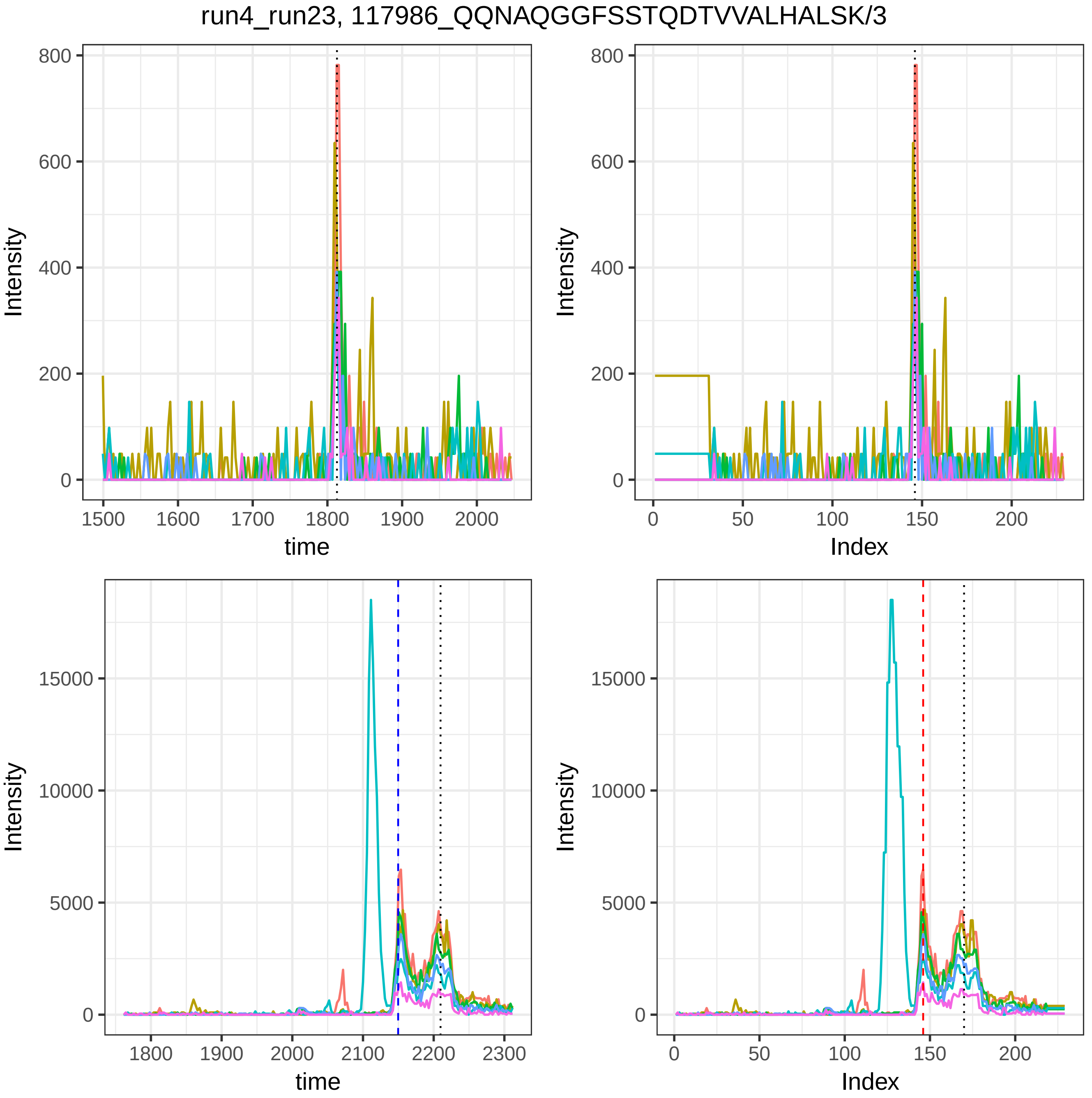
run4_run23, 117986_QQNAQGGFSSTQDTVVALHALSK/3
Intensity
time
Index
0
200
400
600
800
1500
1600
1700
1800
1900
2000
50
100
150
5000
10000
15000
2100
2200
2300
