## Supplemental Figures for "DIAlignR provides precise retention time alignment across distant runs in DIA and targeted proteomics": Supplemental Figure19.pdf

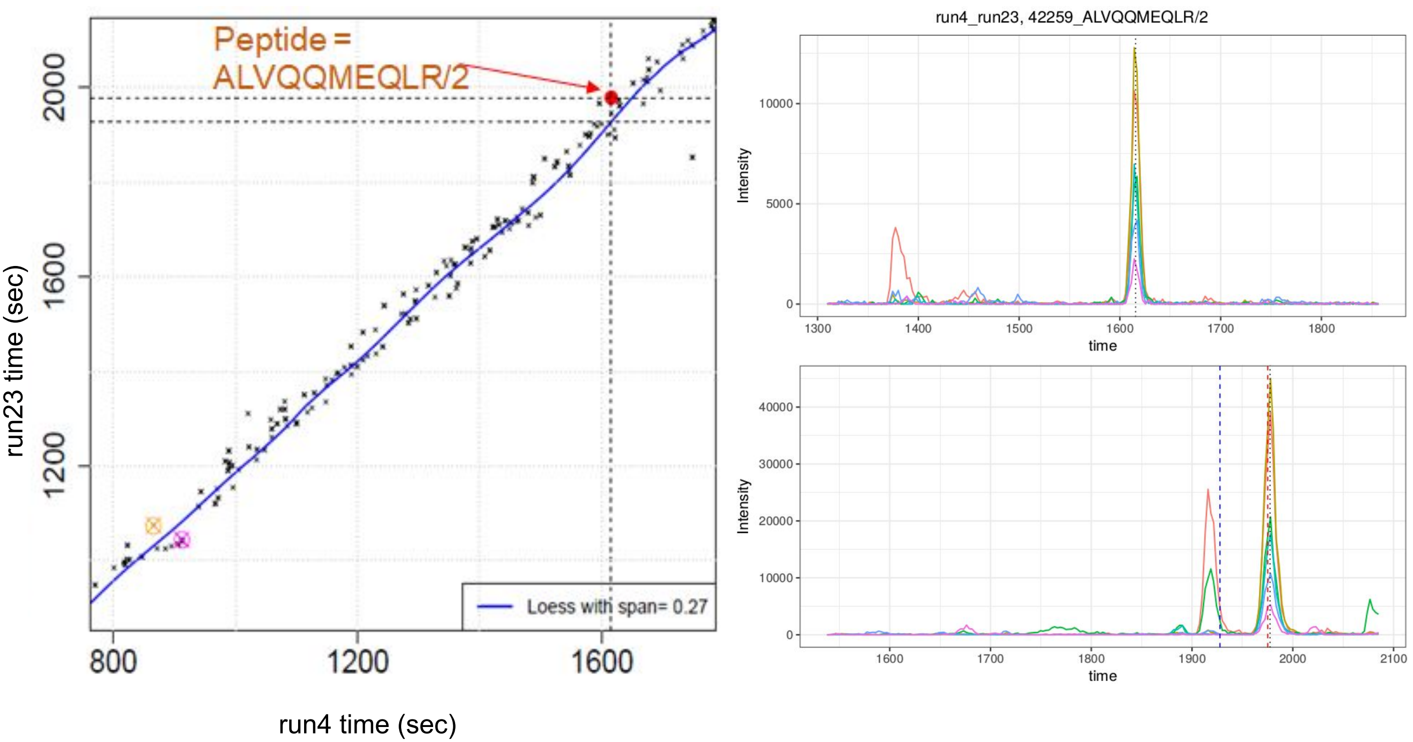

Peptide =
ALVQQMEQLR/2
Loess with span= 0.27
run23 time (sec)
run4 time (sec)
800
1200
1600
2000
run4_run23, 42259_ALVQQMEQLR/2
Intensity
time
1300
1400
1500
1600
1700
1800
0
5000
10000
Intensity
time
1600
1700
1800
1900
2000
2100
0
10000
20000
30000
40000

Supplemental Figure 19. Alignment of peptide 42259_ALVQQMEQLR/2 in run4 and run23 from the clinical plasma dataset. Chromatogram alignment (red-dashed line) maps correct peak-group between upper and lower panel. LOESS (blue-dashed line) maps to wrong peak group.
