## Supplemental Figures for "DIAlignR provides precise retention time alignment across distant runs in DIA and targeted proteomics": Supplemental Figures1-14.pdf

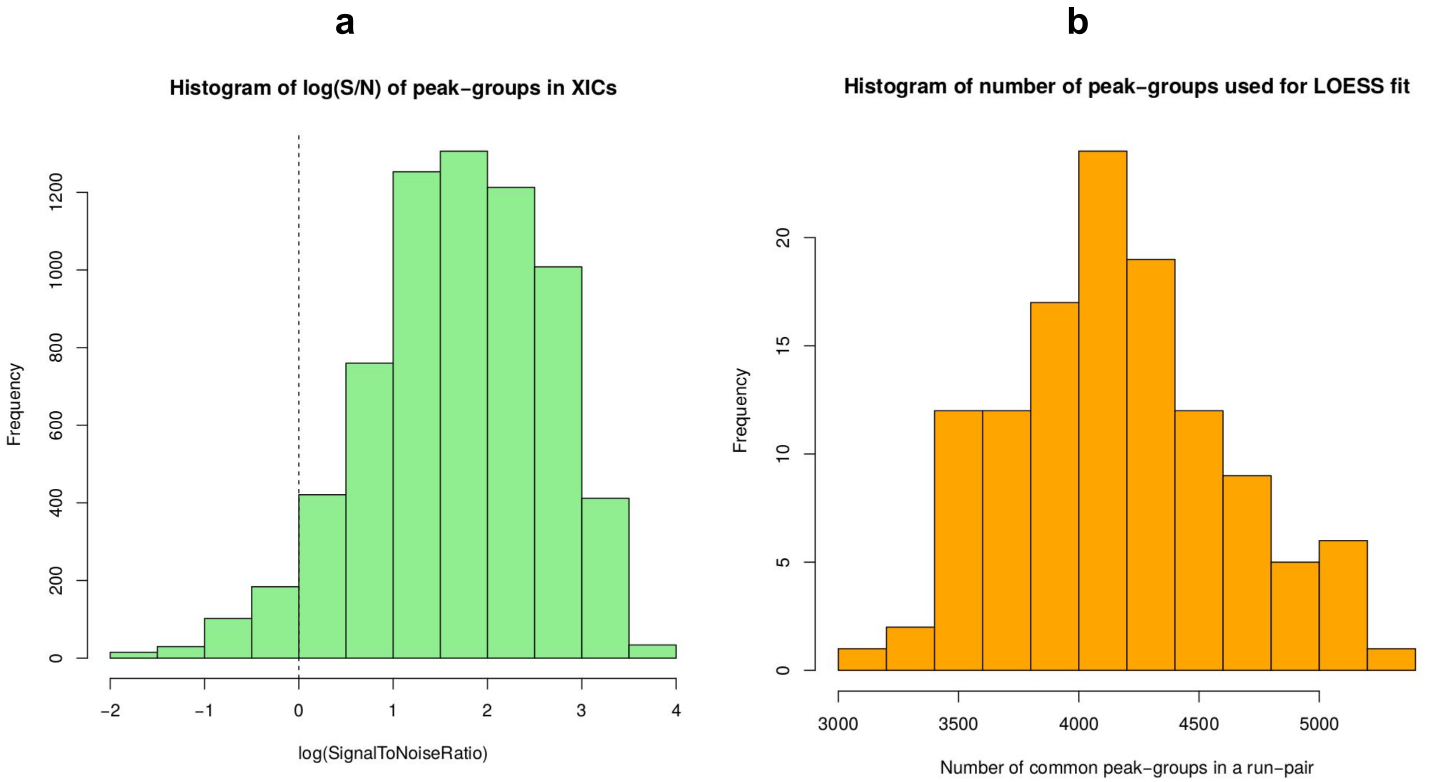

a
Histogram of log(S/N) of peak-groups in XICs
Frequency
0
200
400
600
800
1000
1200
−2
−1
0
1
2
3
4
log(SignalToNoiseRatio)
b
Histogram of number of peak-groups used for LOESS fit
Frequency
0
5
10
15
20
3000
3500
4000
4500
5000
Number of common peak-groups in a run−pair

**Supplemental Figure 1**. *a*, Histogram of log(Signal/Noise) of peptide peak-groups. For each precursor, Signal defines as a sum of fragment-ions' intensities at annotated retention time, whereas, Noise is a sum of average intensities of corresponding fragment-ions' XICs. *b*, Distribution of the number of peak-groups used to obtain global fit is plotted. Minimum 3194 peak-groups and maximum 5260 peak-groups were used to obtain LOESS fit between run-pairs.

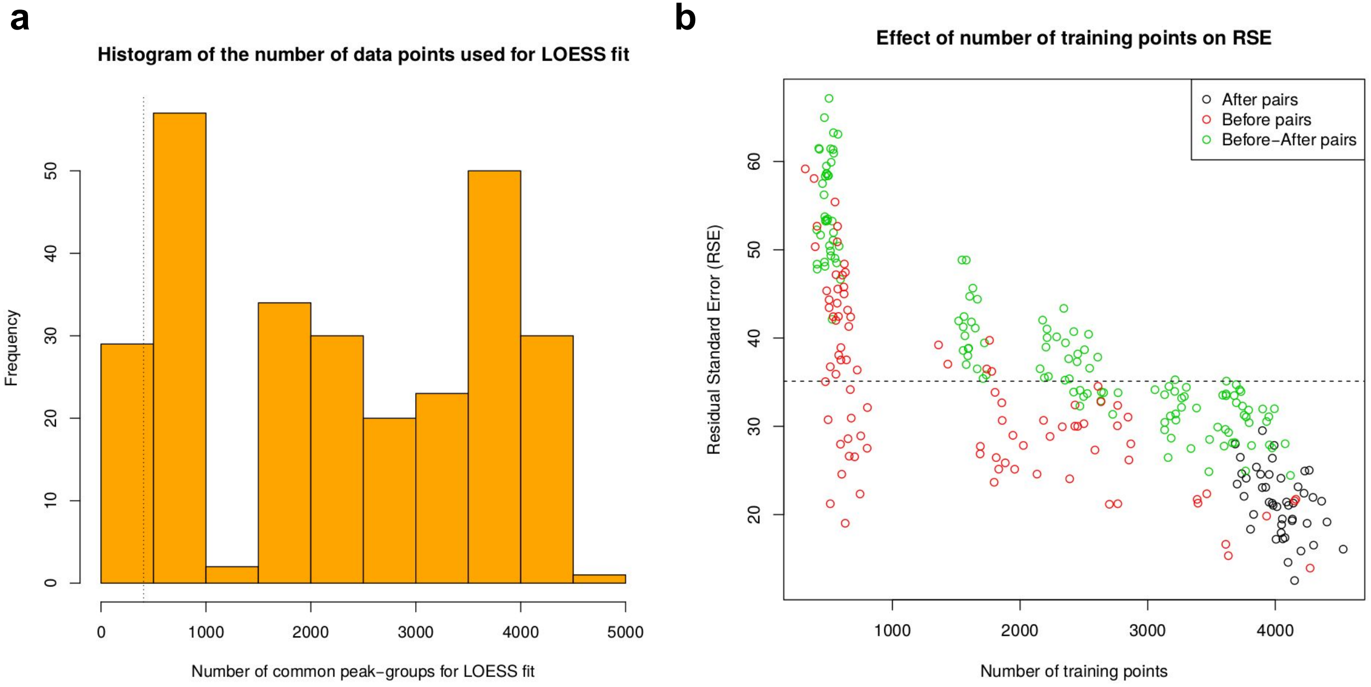
a
Histogram of the number of data points used for LOESS fit
Frequency
0
10
20
30
40
50
0
1000
2000
3000
4000
5000
Number of common peak-groups for LOESS fit
b
Effect of number of training points on RSE
Residual Standard Error (RSE)
20
30
40
50
60
1000
2000
3000
4000
Number of training points
After pairs
Before pairs
Before−After pairs

Supplemental Figure 2. **Number of common peak-groups in a run-pair and its effect on RSE**. *a*, distribution of number of peak-groups used to obtain global fit in clinical plasma dataset is plotted. On average 2262 points are available to obtain a LOESS fit. *b*, effect of number of training points on RSE is plotted.

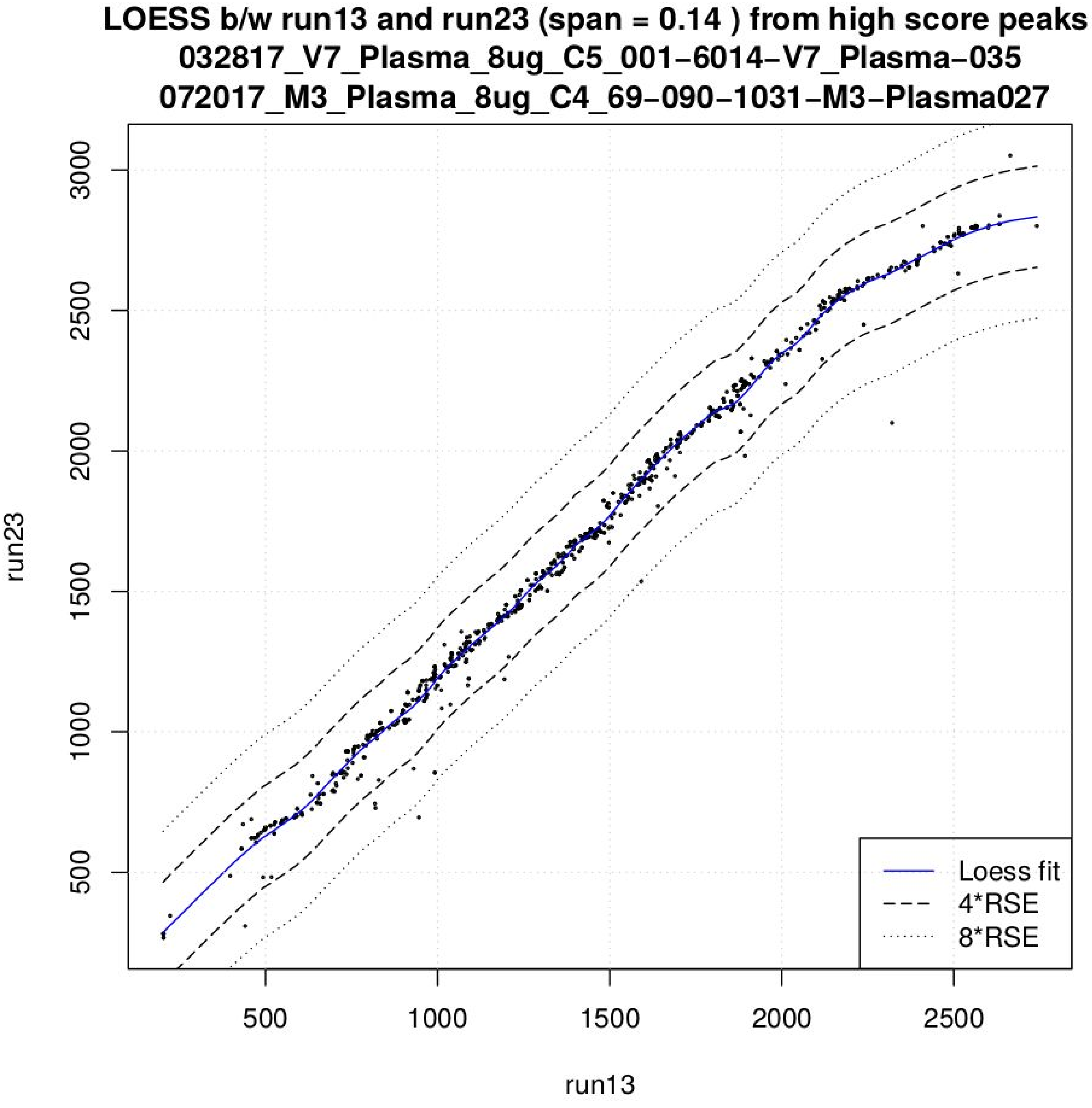


Supplemental Figure 3. LOESS fit between two runs *run13* and *run23* from clinical plasma dataset is plotted in the figure. High scoring peak-groups are shown to be scattered around the fit line.

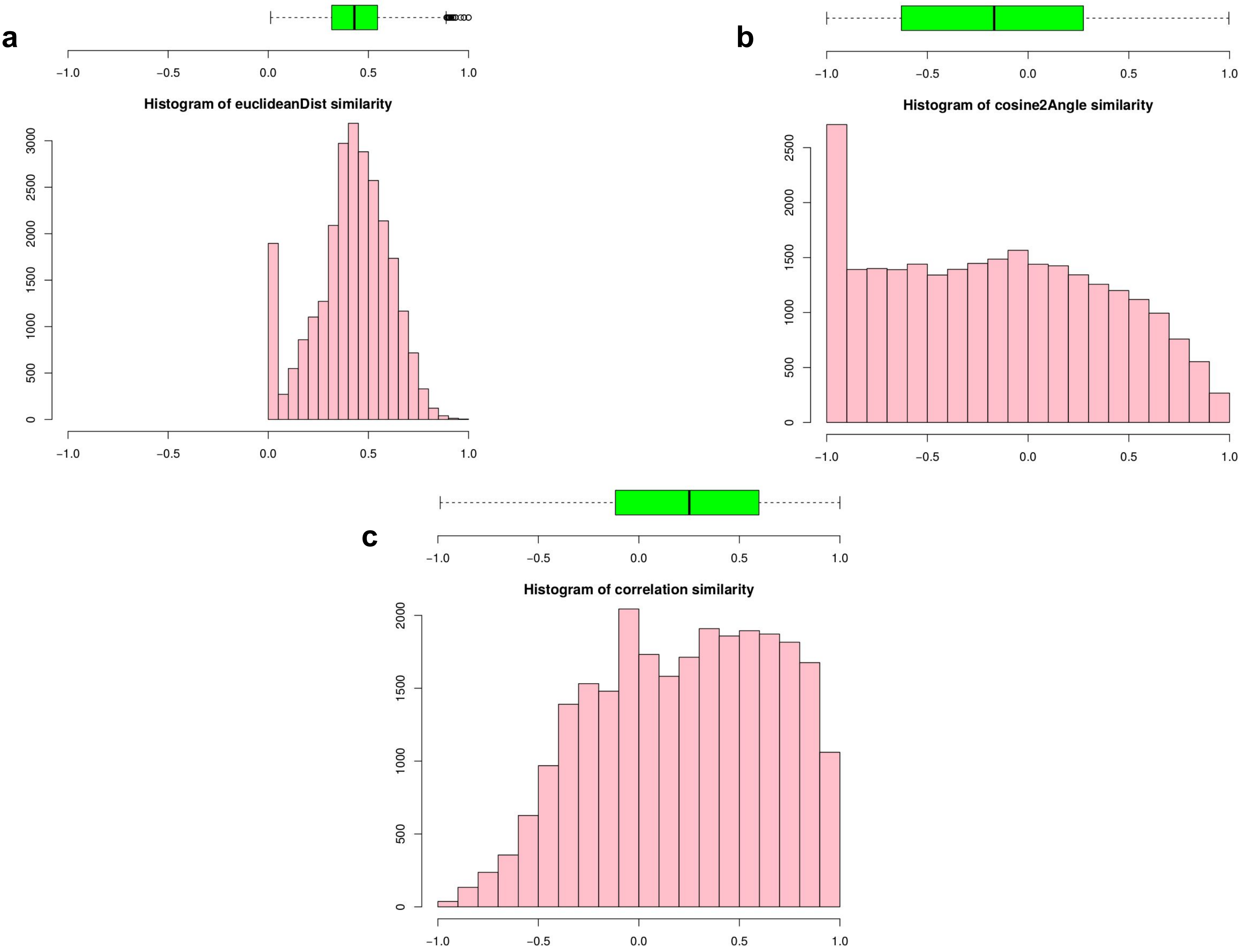

a
Histogram of euclideanDist similarity
b
Histogram of cosine2Angle similarity
c
Histogram of correlation similarity

Supplemental Figure 4. **Distribution of normalized similarity scores for chromatograms of peptide "16372_KALVETDGDMDK/2" from “run11” and “run12” of validation dataset**. *a*, *b* and *c* presents similarity score distribution for Euclidean distance based, cos(2* spectral angle) and pearson’s correlation similarity measures, respectively.

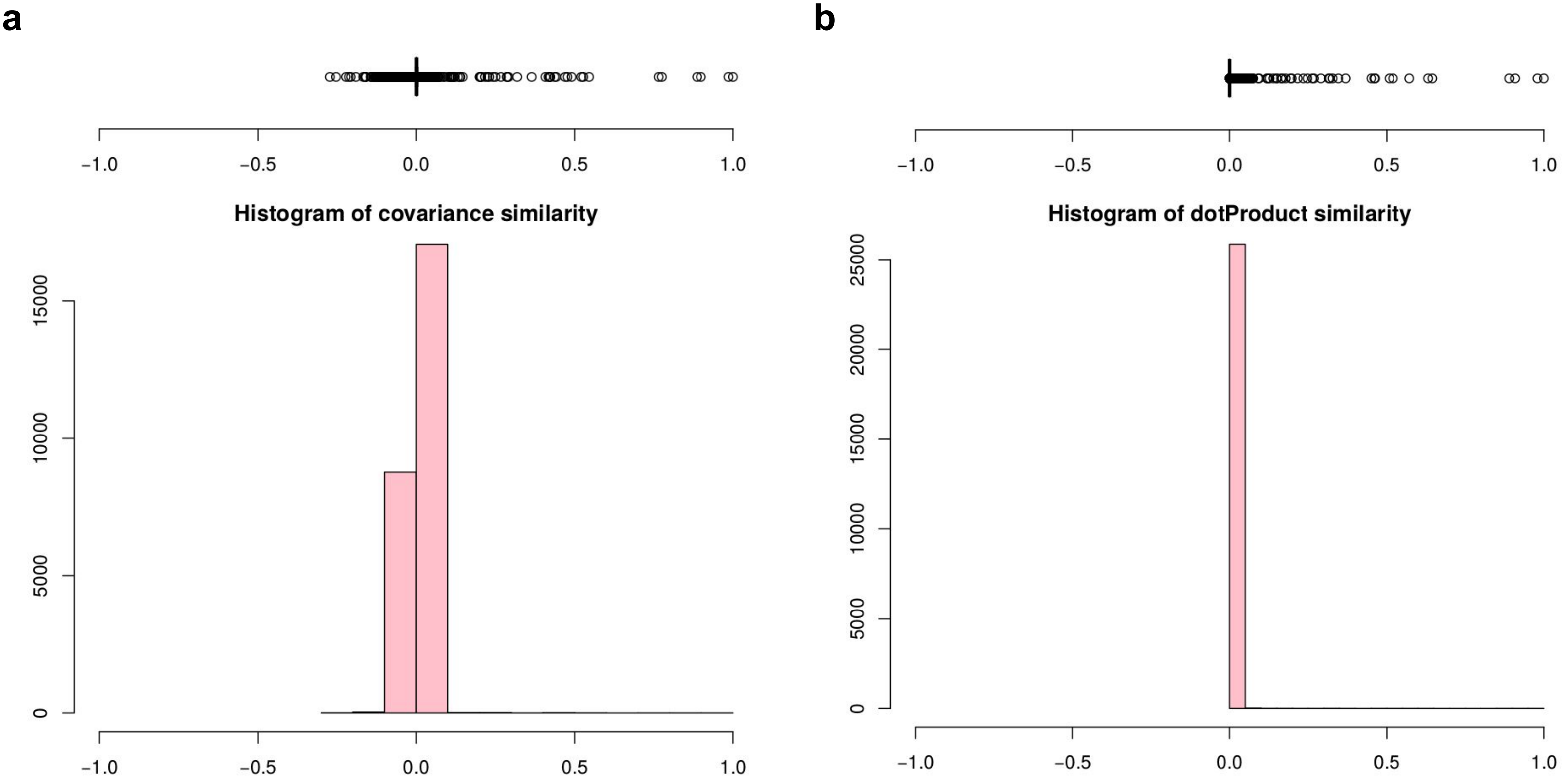
a
b
Histogram of covariance similarity
Histogram of dotProduct similarity
−1.0
−0.5
0.0
0.5
1.0
0
5000
10000
15000
20000
25000

Supplemental Figure 5. **Distribution of normalized similarity scores for chromatograms of peptide "16372_KALVETDGDMDK/2" from “run11” and “run12” of validation dataset**. *a* and *b* presents similarity score distribution for covariance and dot product similarity measures, respectively.

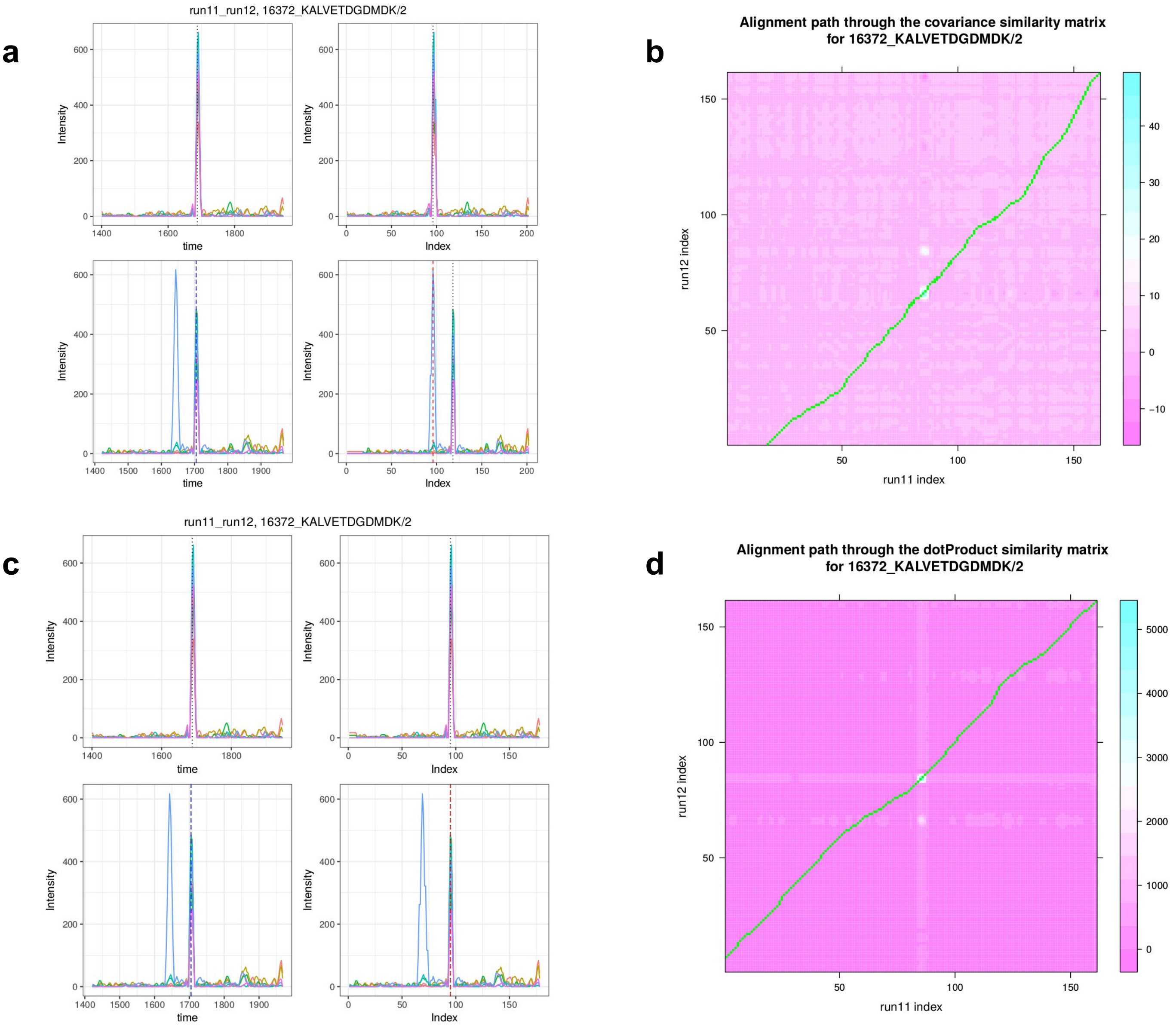

a
run11_run12, 16372_KALVETDGDMDK/2
Intensity
time
Index
b
Alignment path through the covariance similarity matrix
for 16372_KALVETDGDMDK/2
run12 index
run11 index
c
run11_run12, 16372_KALVETDGDMDK/2
Intensity
time
Index
d
Alignment path through the dotProduct similarity matrix
for 16372_KALVETDGDMDK/2
run12 index
run11 index

Supplemental Figure 6. **Alignment of chromatograms of peptide "16372_KALVETDGDMDK/2" from “run11” and “run12” of validation dataset**. *a*, aligned chromatograms resulted using covariance similarity measure. *b*, alignment path in the covariance based similarity matrix passes through lower hot-spot which is due to different precursor in run12. *c*, aligned chromatograms resulted using dot product similarity measure. *d*, alignment path in the dot product based similarity matrix passes through upper hot-spot which is due to co-elution profile of the correct precursor.

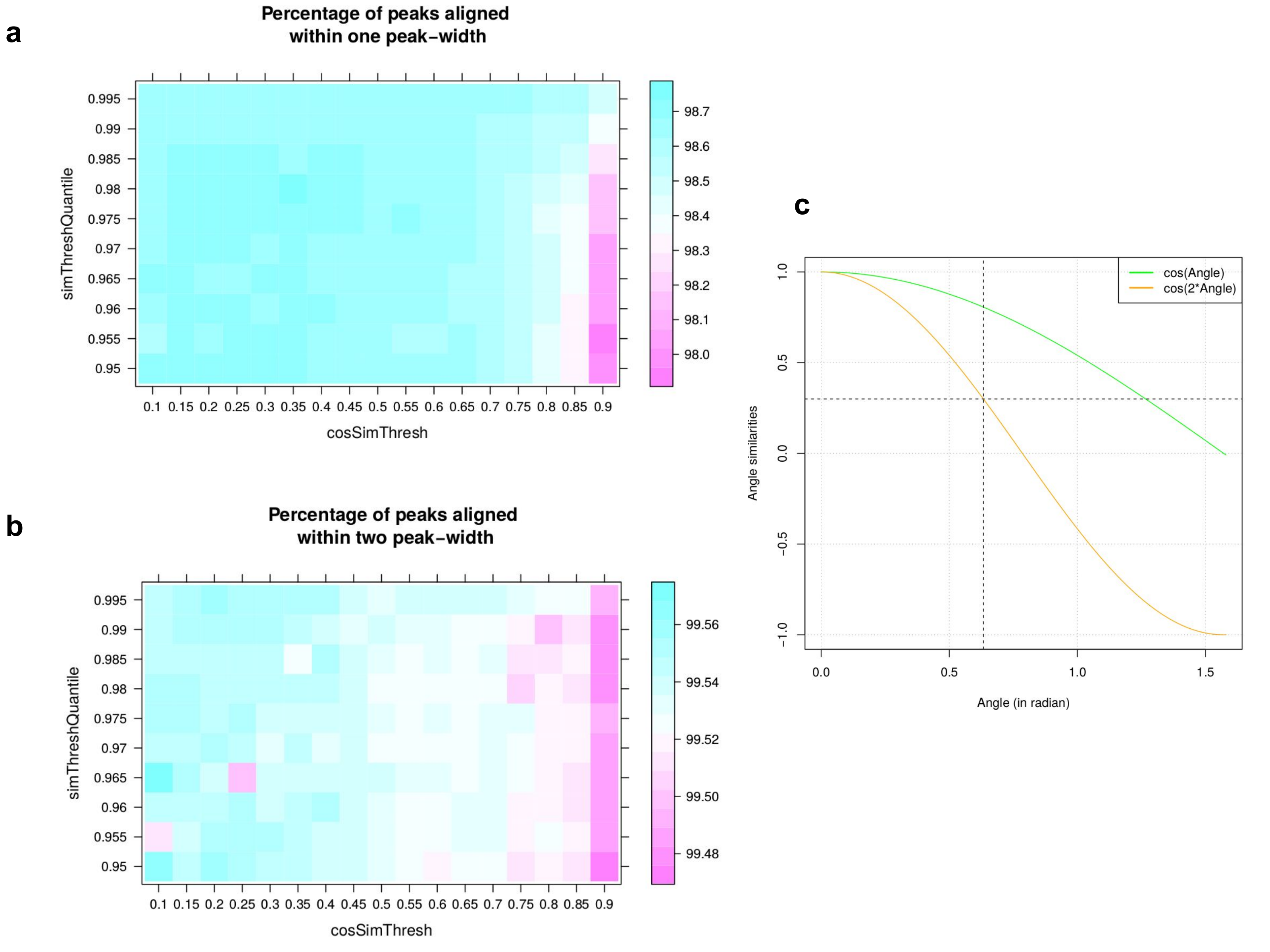

a
Percentage of peaks aligned within one peak-width
simThreshQuantile
cosSimThresh
0.995
0.99
0.985
0.98
0.975
0.97
0.965
0.96
0.955
0.95
0.1 0.15 0.2 0.25 0.3 0.35 0.4 0.45 0.5 0.55 0.6 0.65 0.7 0.75 0.8 0.85 0.9
98.7
98.6
98.5
98.4
98.3
98.2
98.1
98.0
b
Percentage of peaks aligned within two peak-width
simThreshQuantile
cosSimThresh
99.56
99.54
99.52
99.50
99.48
c
cos(Angle)
cos(2*Angle)
Angle similarities
Angle (in radian)
1.0
0.5
0.0
-0.5
-1.0
0.0
0.5
1.0
1.5

Supplemental Figure 7. **Heatmap of percentage of peaks aligned within certain RT tolerance on the gold standard validation dataset for grid search between** $cosSimThresh$ **and** $simThreshQuantile$ **.** *a*, Percentage of peaks aligned within one peak-width. *b*, Percentage of peaks aligned within two peak-width. *c*, similarity score for cos(Angle) and cos(2*Angle). For similarity score > 0.3 with cos(2*Angle), angle between vectors must be less than 0.633 radians.

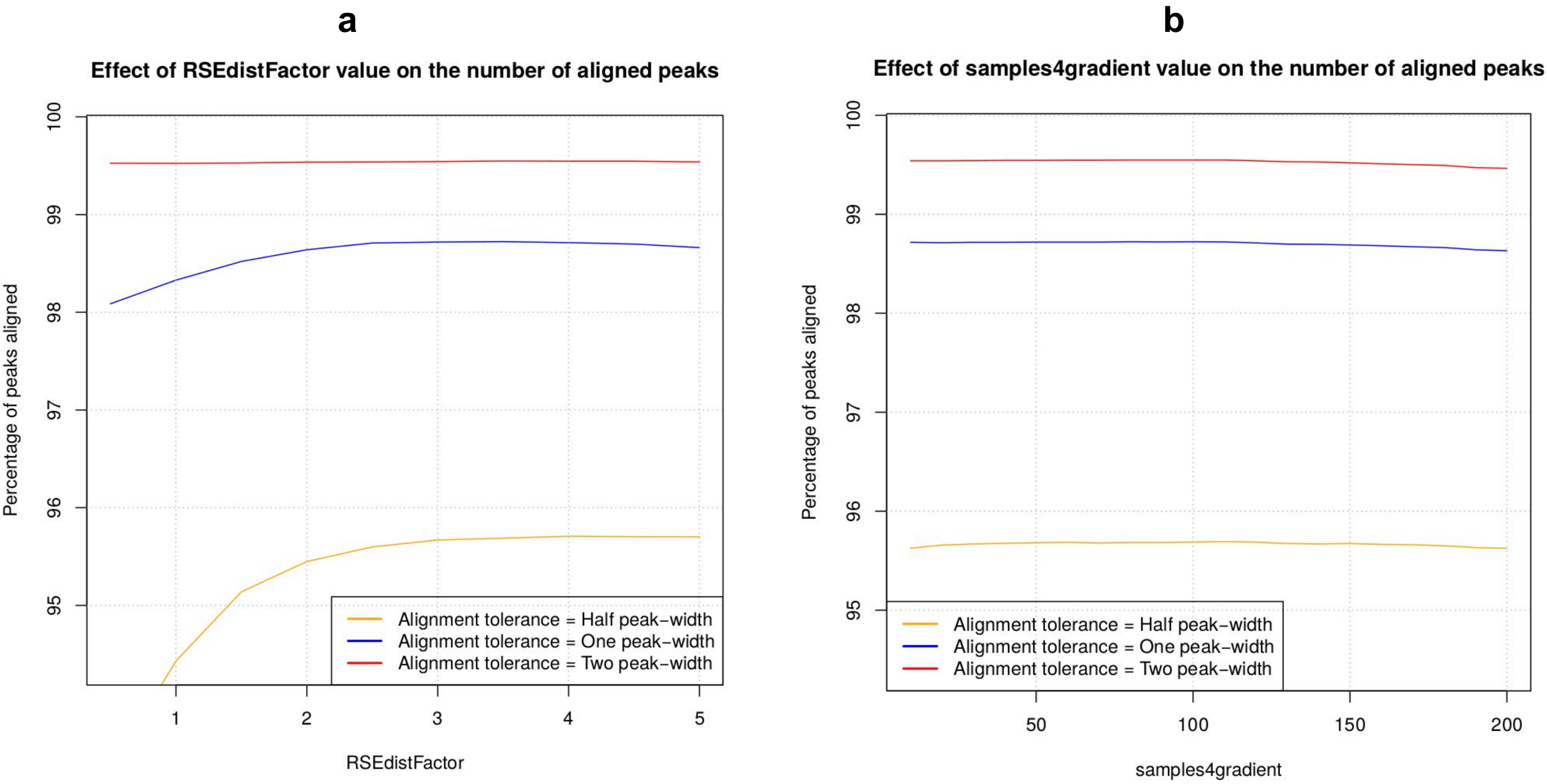
a
Effect of RSEdistFactor value on the number of aligned peaks
Percentage of peaks aligned
100
99
98
97
96
95
1
2
3
4
5
RSEdistFactor
Alignment tolerance = Half peak−width
Alignment tolerance = One peak−width
Alignment tolerance = Two peak−width
b
Effect of samples4gradient value on the number of aligned peaks
Percentage of peaks aligned
100
99
98
97
96
95
50
100
150
200
samples4gradient
Alignment tolerance = Half peak−width
Alignment tolerance = One peak−width
Alignment tolerance = Two peak−width

Supplemental Figure 8. *a*, Effect of RSEdistFactor value on the number of aligned peaks. *b*, Effect of samples4gradient value on the number of aligned peaks.

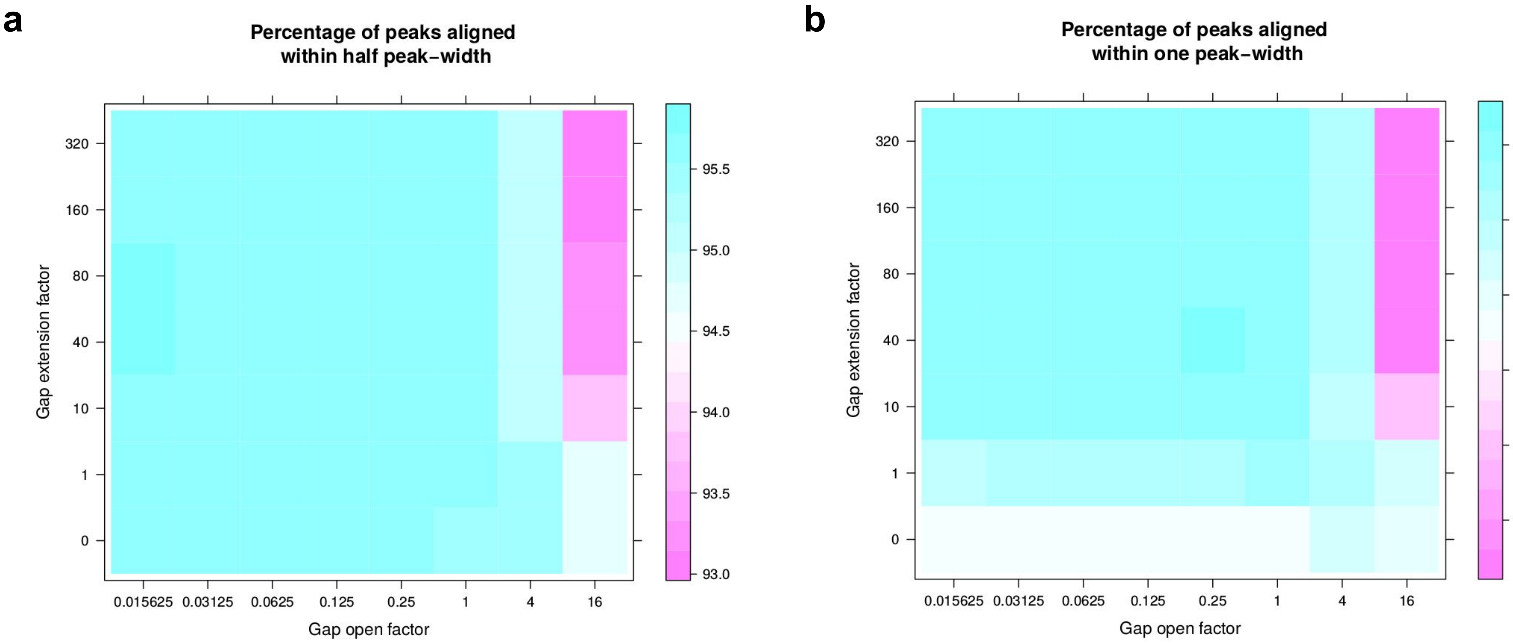
a
Percentage of peaks aligned
within half peak-width
Gap extension factor
320
160
80
40
10
1
0
0.015625
0.03125
0.0625
0.125
0.25
1
4
16
Gap open factor
95.5
95.0
94.5
94.0
93.5
93.0
b
Percentage of peaks aligned
within one peak-width
Gap extension factor
Gap open factor
98.7
98.6
98.5
98.4
98.3
98.2

Supplemental Figure 9. Heatmap of percentage of peaks aligned within RT tolerance of ***a*, half peak-width** and ***b*, one peak-width** on the gold standard validation dataset for grid search between Gap Open Factor and Gap Open Factor.

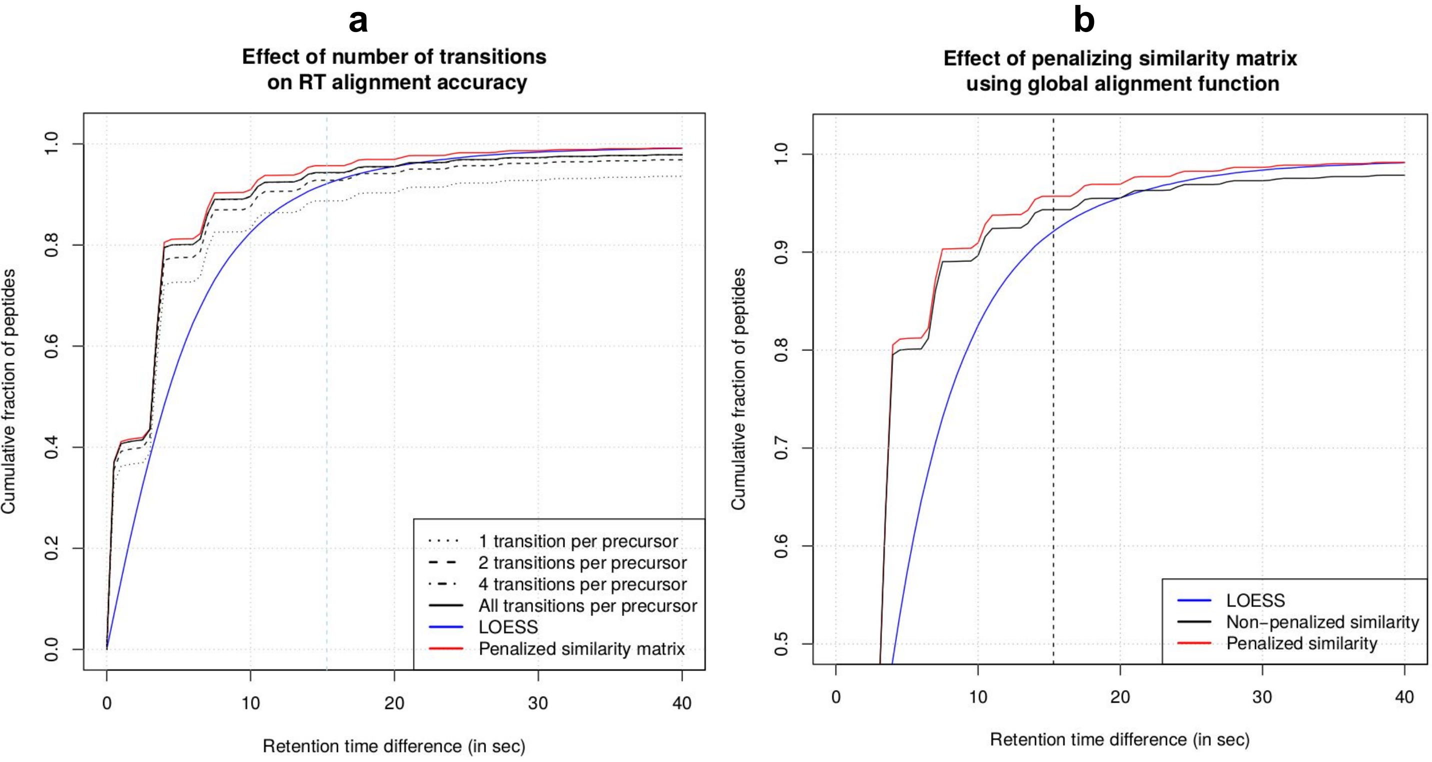
a
Effect of number of transitions
on RT alignment accuracy
Cumulative fraction of peptides
Retention time difference (in sec)
1 transition per precursor
2 transitions per precursor
4 transitions per precursor
All transitions per precursor
LOESS
Penalized similarity matrix
b
Effect of penalizing similarity matrix
using global alignment function
Cumulative fraction of peptides
Retention time difference (in sec)
LOESS
Non−penalized similarity
Penalized similarity

**Supplemental Figure 10**. Effect of *a*, number of transitions and *b*, penalized similarity matrix on the fraction of peptides aligned before certain RT error.

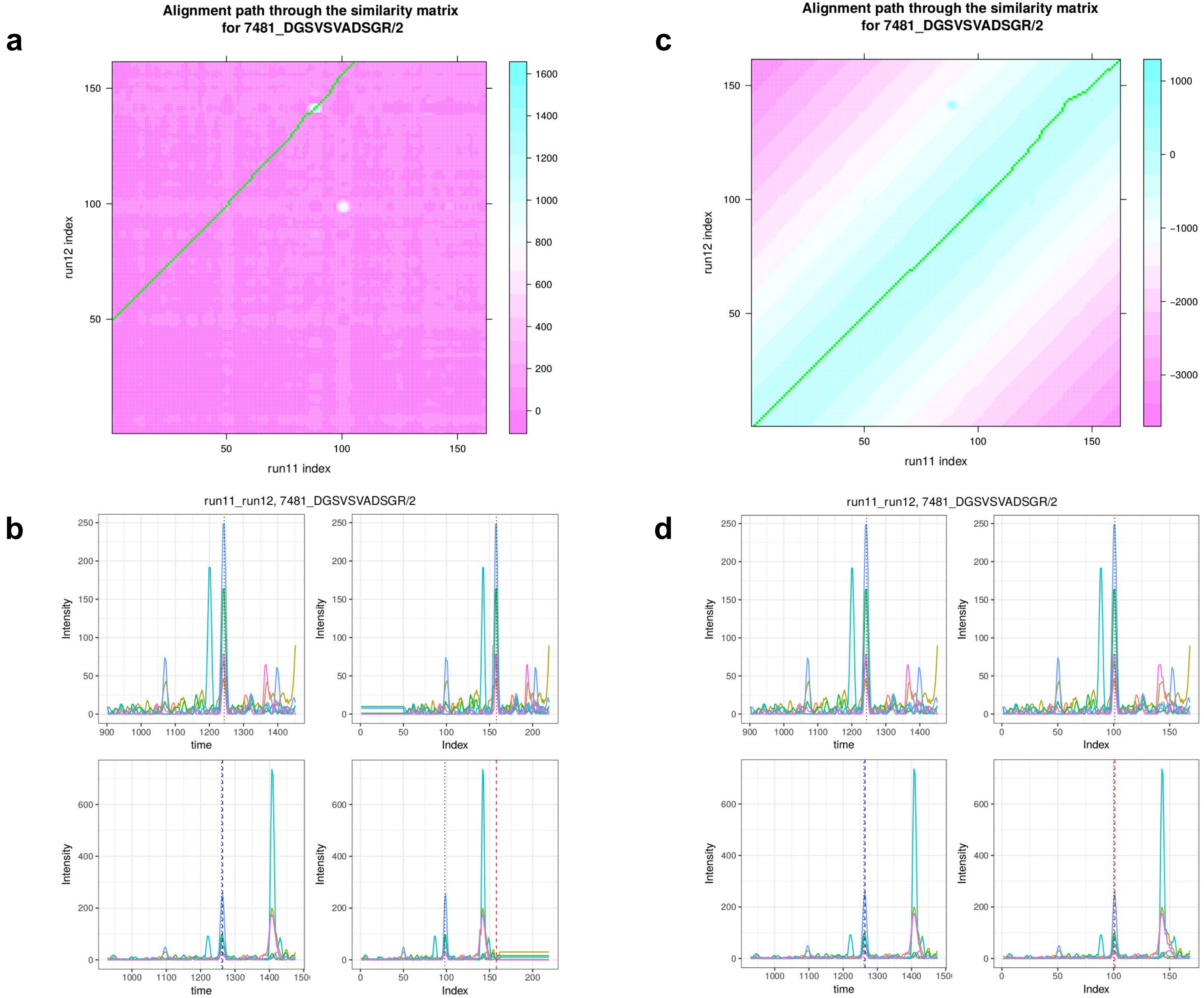
a
Alignment path through the similarity matrix
for 7481_DGSVSVADSGR/2
run12 index
run11 index
b
run11_run12, 7481_DGSVSVADSGR/2
Intensity
time
Index
c
Alignment path through the similarity matrix
for 7481_DGSVSVADSGR/2
run12 index
run11 index
d
run11_run12, 7481_DGSVSVADSGR/2
Intensity
time
Index

Supplemental Figure 11. **Effect of penalizing similarity matrix on the alignment of chromatograms of peptide DGSVSVADSGR/2 for run11 and run12 from the validation dataset**. *a*, similarity matrix derived using masked dot product similarity measure. *b*, chromatograms before and after alignment using the similarity matrix given in *a*. *c*, similarity matrix derived by masked dot product and further penalized with LOESS fit between run11 and run12. *d*, chromatograms before and after alignment using the similarity matrix given in *c*.

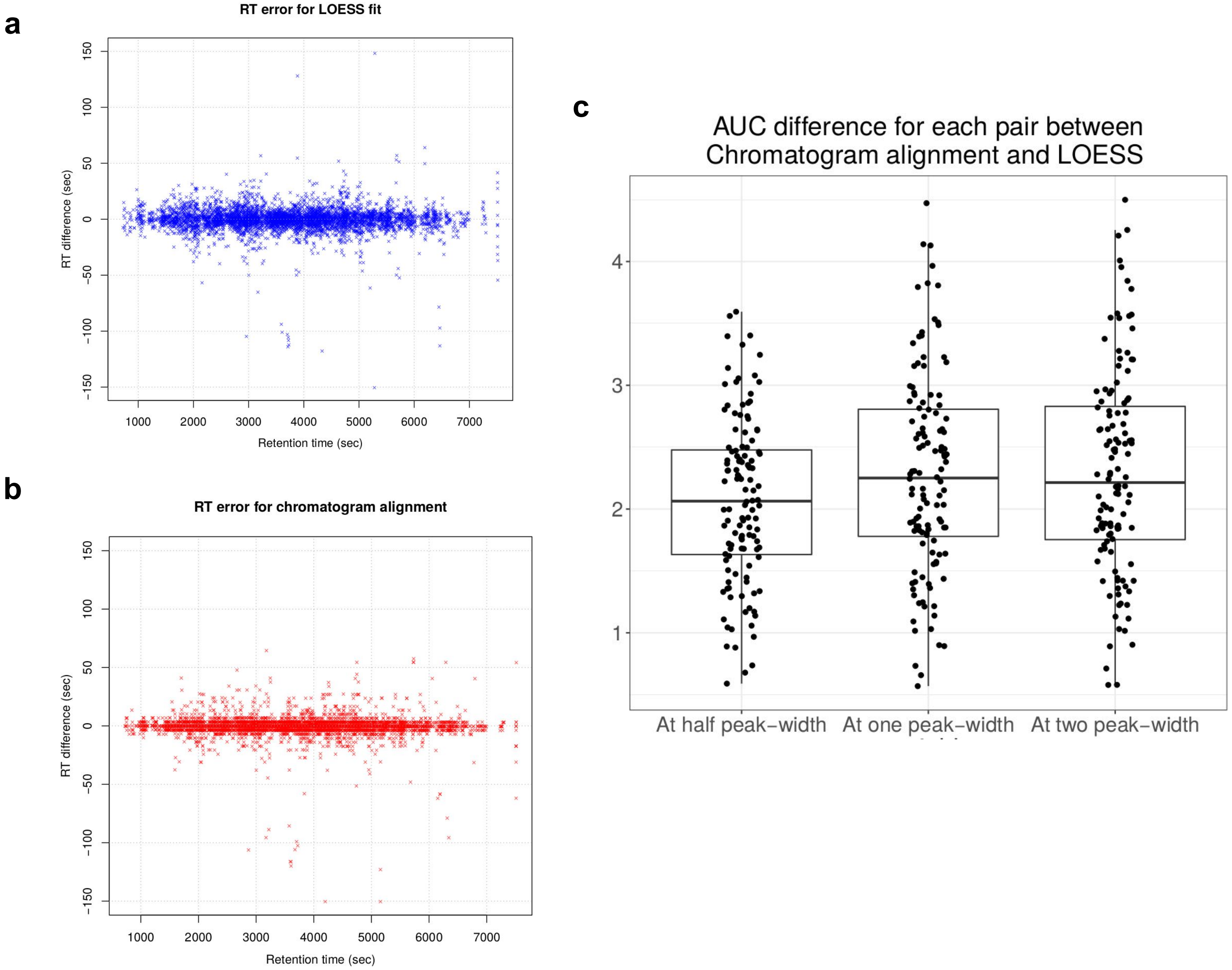

a
RT error for LOESS fit
RT difference (sec)
Retention time (sec)
b
RT error for chromatogram alignment
RT difference (sec)
Retention time (sec)
c
AUC difference for each pair between Chromatogram alignment and LOESS
At half peak-width
At one peak-width
At two peak-width

**Supplemental Figure 12**. Error of reported RTs plotted along the run time for *a*, LOESS and *b*, chromatogram alignment. One point is sampled out of 10 for visualization. *c*, Difference of Area Under the Curve (AUC), represented by fraction of peptides aligned within certain RT error, between chromatogram alignment and LOESS is plotted. Three values of RT error tolerance half peak-width, one peak-width and two peak-width are considered.

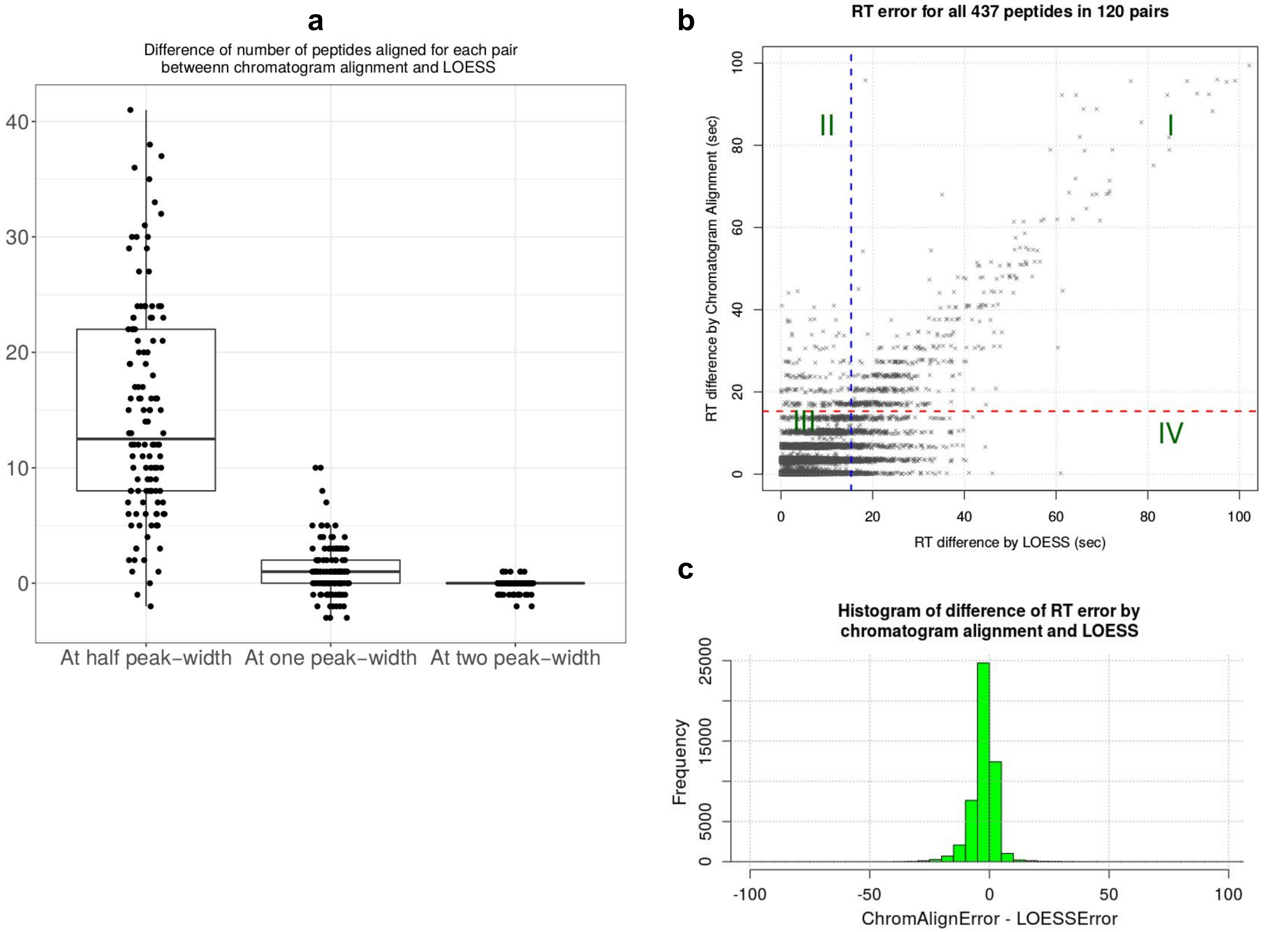

a
Difference of number of peptides aligned for each pair
betweenn chromatogram alignment and LOESS
40
30
20
10
0
At half peak−width
At one peak−width
At two peak−width
b
RT error for all 437 peptides in 120 pairs
RT difference by Chromatogram Alignment (sec)
RT difference by LOESS (sec)
0
20
40
60
80
100
I
II
III
IV
c
Histogram of difference of RT error by
chromatogram alignment and LOESS
Frequency
0
5000
15000
25000
-100
-50
0
50
100
ChromAlignError - LOESSError

Supplemental Figure 13. *a*, Out of 437 annotated peaks, difference of number of peptides aligned within certain RT error, between chromatogram alignment and LOESS is plotted. Three values of RT error tolerance half peak-width, one peak-width and two peak-width are considered. *b*, RT prediction error is plotted for chromatogram alignment and LOESS for each peptide in every pair. For visualization, only one out of three points is plotted. Quadrants I, II, III and IV have 1573, 554, 45055 and 2323 points, respectively. There are total 49505 pairwise alignments. *c*, Histogram of difference of RT error by chromatogram alignment and LOESS.

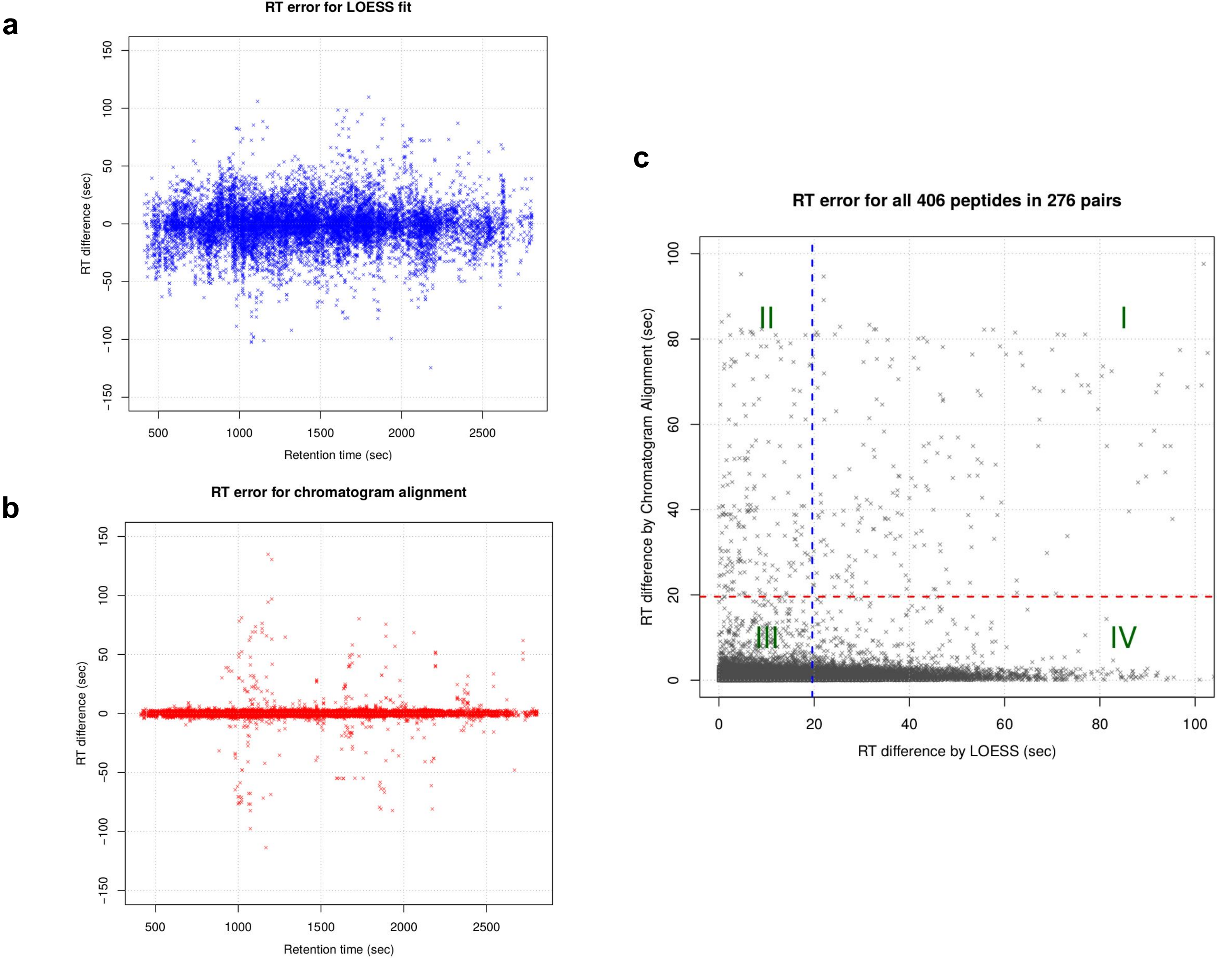
a
RT error for LOESS fit
RT difference (sec)
Retention time (sec)
b
RT error for chromatogram alignment
RT difference (sec)
Retention time (sec)
c
RT error for all 406 peptides in 276 pairs
RT difference by Chromatogram Alignment (sec)
RT difference by LOESS (sec)
I
II
III
IV

Supplemental Figure 14. **From clinical plasma dataset**. Error of reported RTs plotted along the run time for *a*, LOESS and *b*, chromatogram alignment. One point is sampled out of 10 for visualization. *c*, RT prediction error is plotted for chromatogram alignment and LOESS for each peptide in every pair. For visualization, only one out of four points is plotted. Quadrants I, II, III and IV have 1060, 1269, 83926 and 25801 points, respectively.
